## Supplementary Table 3 for "GLYATL1 is associated with metabolic and epigenetic changes and with endocrine resistance in luminal breast cancer"

| ensembl_gene_id | hgnc_symbol | baseMean_KO1_vs_LTED | log2FoldChange_KO1_vs_LTED | lfcSE_KO1_vs_LTED | stat_KO1_vs_LTED | pvalue_KO1_vs_LTED | padj_KO1_vs_LTED |
|---|---|---|---|---|---|---|---|
| ENSG00000000003 | TSPAN6 | 1306.017476 | -0.114700941 | 0.061753672 | -1.857394674 | 0.063255024 | 0.145777601 |
| ENSG00000000419 | DPM1 | 1697.508982 | -0.198170494 | 0.057827507 | -3.426924387 | 0.000610459 | 0.002378385 |
| ENSG00000000457 | SCYL3 | 614.7835509 | -0.087205429 | 0.085173608 | -1.023855058 | 0.30590374 | 0.493512116 |
| ENSG00000000460 | C1orf112 | 428.725597 | -0.427057077 | 0.091615973 | -4.661382309 | 3.14E-06 | 1.72E-05 |
| ENSG00000001036 | FUCA2 | 3167.593799 | 0.635123515 | 0.050047535 | 12.69040554 | 6.68E-37 | 2.43E-35 |
| ENSG00000001084 | GCLC | 1847.371283 | 0.106729098 | 0.061613304 | 1.732241113 | 0.083230642 | 0.182534505 |
| ENSG00000001167 | NFYA | 1761.016949 | 0.083873659 | 0.057832491 | 1.450286137 | 0.146978743 | 0.287050262 |
| ENSG00000001460 | STPG1 | 164.870432 | -0.109426594 | 0.139622355 | -0.783732619 | 0.433197019 | 0.623170067 |
| ENSG00000001461 | NIPAL3 | 1136.408607 | 0.058350887 | 0.066657294 | 0.875386373 | 0.381363712 | 0.573598921 |
| ENSG00000001497 | LAS1L | 2298.65315 | 0.14313611 | 0.052559698 | 2.723305419 | 0.006463228 | 0.02021907 |
| ENSG00000001561 | ENPP4 | 400.8996717 | 0.182959583 | 0.095744348 | 1.910917844 | 0.05601514 | 0.131942428 |
| ENSG00000001617 | SEMA3F | 5496.282674 | 0.361827504 | 0.058164182 | 6.220795905 | 4.95E-10 | 4.05E-09 |
| ENSG00000001629 | ANKIB1 | 3637.046102 | -0.76327449 | 0.050486479 | -15.11839429 | 1.22E-51 | 7.06E-50 |
| ENSG00000001630 | CYP51A1 | 229.7394467 | -1.063554185 | 0.15141268 | -7.02420816 | 2.15E-12 | 2.14E-11 |
| ENSG00000001631 | KRIT1 | 160.8004663 | -0.860730662 | 0.143569113 | -5.995235653 | 2.03E-09 | 1.57E-08 |
| ENSG00000002016 | RAD52 | 236.9662718 | -0.485954005 | 0.120774738 | -4.023639481 | 5.73E-05 | 0.00026595 |
| ENSG00000002330 | BAD | 894.218289 | -0.212176016 | 0.089114853 | -2.380927637 | 0.017269104 | 0.048123572 |
| ENSG00000002549 | LAP3 | 2283.041028 | -0.144782639 | 0.052384862 | -2.763825926 | 0.005712801 | 0.01812108 |
| ENSG00000002586 | CD99 | 1714.770958 | -0.05196366 | 0.056006937 | -0.927807571 | 0.353507396 | 0.545027197 |
| ENSG00000002586 | CD99 | 1714.770958 | -0.05196366 | 0.056006937 | -0.927807571 | 0.353507396 | 0.545027197 |
| ENSG00000002587 | HS3ST1 | 8.608628588 | 0.924943209 | 0.620145076 | 1.491494886 | 0.135831613 | 0.270413712 |
| ENSG00000002822 | MAD1L1 | 4.318136868 | -0.825037833 | 0.877288954 | -0.940440238 | 0.346991791 | 0.537873618 |
| ENSG00000002834 | LASP1 | 10836.71469 | -0.281959716 | 0.044008611 | -6.406921551 | 1.48E-10 | 1.26E-09 |
| ENSG00000002919 | SNX11 | 858.0252452 | 0.183810118 | 0.074002919 | 2.483822525 | 0.012998057 | 0.03743101 |
| ENSG00000003056 | M6PR | 1843.054424 | -0.016817351 | 0.057561743 | -0.292161947 | 0.770162805 | 0.874717039 |
| ENSG00000003096 | KLHL13 | 24.75739597 | -0.45893948 | 0.365287923 | -1.256377369 | 0.208979183 | 0.375688003 |
| ENSG00000003137 | CYP26B1 | 24.3477993 | -0.458414443 | 0.356805255 | -1.28477492 | 0.198870949 | 0.361612204 |
| ENSG00000003147 | ICA1 | 3921.311494 | -1.029823437 | 0.053128978 | -19.38346044 | 1.06E-83 | 1.29E-81 |
| ENSG00000003249 | DBNDD1 | 3022.492955 | 0.612109798 | 0.059528301 | 10.28266877 | 8.44E-25 | 1.85E-23 |
| ENSG00000003393 | ALS2 | 1200.640645 | 0.011908261 | 0.064807918 | 0.183747015 | 0.854211924 | 0.924103822 |
| ENSG00000003400 | CASP10 | 2.352202551 | 0.652228359 | 1.171929256 | 0.556542433 | 0.577840095 | 0.743545771 |

| | | | | | | | |
|---|---|---|---|---|---|---|---|
| ENSG00000003402 | CFLAR | 1087.405088 | 0.058761843 | 0.066014368 | 0.890137173 | 0.373392235 | 0.566156673 |
| ENSG00000003436 | TFPI | 1576.696454 | -1.372189924 | 0.058921719 | -23.28835507 | 5.82E-120 | 1.40E-117 |
| ENSG00000003509 | NDUFAF7 | 362.4092871 | 0.160003292 | 0.103553813 | 1.545122172 | 0.122316715 | 0.249063149 |
| ENSG00000003756 | RBM5 | 2561.808202 | -0.12294613 | 0.059136917 | -2.079008122 | 0.037616604 | 0.094267522 |
| ENSG00000003987 | MTMR7 | 134.0232236 | 0.114904239 | 0.156044951 | 0.736353458 | 0.461515621 | 0.648328558 |
| ENSG00000003989 | SLC7A2 | 4701.147991 | 0.92313444 | 0.050492158 | 18.28272906 | 1.14E-74 | 1.20E-72 |
| ENSG00000004059 | ARF5 | 2585.758732 | -0.015508002 | 0.06123816 | -0.253240824 | 0.800082122 | 0.893437369 |
| ENSG00000004139 | SARM1 | 667.2420576 | 0.101414195 | 0.077384229 | 1.310527938 | 0.190017299 | 0.349195962 |
| ENSG00000004142 | POLDIP2 | 4287.662442 | 0.01787171 | 0.046904655 | 0.381022088 | 0.703186859 | 0.833796972 |
| ENSG00000004399 | PLXND1 | 4296.966577 | 1.768862102 | 0.049624048 | 35.64526032 | 2.79E-278 | 4.92E-275 |
| ENSG00000004455 | AK2 | 3133.343093 | 0.12109116 | 0.048677098 | 2.487641322 | 0.012859335 | 0.037073413 |
| ENSG00000004478 | FKBP4 | 8093.813081 | -0.007255231 | 0.041240915 | -0.175923129 | 0.860354334 | 0.927382816 |
| ENSG00000004487 | KDM1A | 3214.358002 | 0.052591677 | 0.050930676 | 1.032612988 | 0.301785048 | 0.488632764 |
| ENSG00000004534 | RBM6 | 3103.063925 | 0.071623396 | 0.067094956 | 1.067493003 | 0.285749271 | 0.469595837 |
| ENSG00000004660 | CAMKK1 | 225.0451768 | 0.604803706 | 0.12734679 | 4.749265407 | 2.04E-06 | 1.14E-05 |
| ENSG00000004700 | RECQL | 804.1127272 | -0.36003358 | 0.074121759 | -4.857326443 | 1.19E-06 | 6.82E-06 |
| ENSG00000004766 | VPS50 | 739.6291854 | -0.617212385 | 0.076920309 | -8.024049755 | 1.02E-15 | 1.30E-14 |
| ENSG00000004776 | HSPB6 | 8.045029715 | -1.014078296 | 0.6500263 | -1.560057332 | 0.118746333 | 0.243587668 |
| ENSG00000004777 | ARHGAP33 | 663.4429016 | 0.198593973 | 0.095859775 | 2.071713317 | 0.038292185 | 0.095699174 |
| ENSG00000004779 | NDUFAB1 | 1425.751136 | 0.085645622 | 0.05890165 | 1.454044532 | 0.14593396 | 0.285447225 |
| ENSG00000004799 | PDK4 | 4.262723032 | -3.008446205 | 1.093839262 | -2.750354928 | 0.005953075 | 0.018800045 |
| ENSG00000004838 | ZMYND10 | 35.98510254 | -0.422102366 | 0.305525522 | -1.381561719 | 0.167106315 | 0.316661723 |
| ENSG00000004848 | ARX | 75.50177174 | 1.125751645 | 0.222866595 | 5.051235439 | 4.39E-07 | 2.66E-06 |
| ENSG00000004864 | SLC25A13 | 1467.141988 | -0.605486603 | 0.061370386 | -9.866103903 | 5.84E-23 | 1.16E-21 |
| ENSG00000004866 | ST7 | 546.5977635 | -0.592195067 | 0.08317961 | -7.119473965 | 1.08E-12 | 1.11E-11 |
| ENSG00000004897 | CDC27 | 2266.548394 | -0.077395019 | 0.0661851 | -1.16937224 | 0.242253689 | 0.418502871 |
| ENSG00000004939 | SLC4A1 | 8.3944086 | 0.613722441 | 0.608155293 | 1.009154155 | 0.31290071 | 0.501077995 |
| ENSG00000004948 | CALCR | 32.68272532 | -0.751589665 | 0.310909904 | -2.417387334 | 0.015632373 | 0.044109056 |
| ENSG00000004961 | HCCS | 912.5429491 | 0.202407075 | 0.068936696 | 2.936129598 | 0.003323354 | 0.01114913 |
| ENSG00000004975 | DVL2 | 1961.630242 | -0.093508335 | 0.05724212 | -1.633558216 | 0.102351647 | 0.216042215 |
| ENSG00000005001 | PRSS22 | 343.9092416 | -0.066044848 | 0.1094471 | -0.603440821 | 0.546215473 | 0.719191436 |
| ENSG00000005007 | UPF1 | 2644.871556 | 0.030959904 | 0.06369819 | 0.486040552 | 0.626938415 | 0.780060055 |
| ENSG00000005020 | SKAP2 | 245.3726022 | -1.532392919 | 0.126403154 | -12.12305924 | 7.97E-34 | 2.58E-32 |

| | | | | | | | |
|---|---|---|---|---|---|---|---|
| ENSG00000005022 | SLC25A5 | 15216.25919 | 0.023561758 | 0.046023015 | 0.511955983 | 0.608681816 | 0.766911675 |
| ENSG00000005059 | MCUB | 490.0178996 | 0.013774987 | 0.08742135 | 0.157570059 | 0.874795598 | 0.935039187 |
| ENSG00000005073 | HOXA11 | 16.87413001 | -0.946513891 | 0.443621722 | -2.133605828 | 0.032875061 | 0.084037652 |
| ENSG00000005075 | POLR2J | 1345.260871 | -0.376037531 | 0.064376939 | -5.84118381 | 5.18E-09 | 3.85E-08 |
| ENSG00000005100 | DHX33 | 1316.339438 | 0.182469743 | 0.061741589 | 2.955378135 | 0.00312286 | 0.010544365 |
| ENSG00000005108 | THSD7A | 12.43357006 | -1.596923799 | 0.535910733 | -2.979831715 | 0.002884068 | 0.00980737 |
| ENSG00000005156 | LIG3 | 1368.578872 | 0.004366116 | 0.066109944 | 0.06604325 | 0.947343392 | 0.975450308 |
| ENSG00000005175 | RPAP3 | 1016.185196 | 0.089347461 | 0.069311267 | 1.289075575 | 0.197371814 | 0.35945613 |
| ENSG00000005187 | ACSM3 | 10.57457284 | 0.259289752 | 0.541576251 | 0.478768689 | 0.632103194 | 0.783257563 |
| ENSG00000005189 | REXO5 | 269.2915971 | -0.635954379 | 0.117177557 | -5.427271188 | 5.72E-08 | 3.80E-07 |
| ENSG00000005194 | CIAPIN1 | 1351.890241 | -0.0674517 | 0.063072816 | -1.069425847 | 0.284877826 | 0.468768189 |
| ENSG00000005206 | SPPL2B | 1031.071628 | 0.424481712 | 0.073409103 | 5.782412487 | 7.36E-09 | 5.37E-08 |
| ENSG00000005238 | ATOSB | 211.4416114 | -0.003204782 | 0.128833987 | -0.024875281 | 0.980154444 | 0.991041209 |
| ENSG00000005243 | COPZ2 | 101.9008349 | 0.098219722 | 0.181514747 | 0.541111525 | 0.588430715 | 0.751676167 |
| ENSG00000005249 | PRKAR2B | 241.5719737 | -0.761207187 | 0.129365142 | -5.884175416 | 4.00E-09 | 3.01E-08 |
| ENSG00000005302 | MSL3 | 981.1195616 | -0.071058392 | 0.068880003 | -1.031625859 | 0.302247422 | 0.489208767 |
| ENSG00000005339 | CREBBP | 4602.662515 | -0.134382174 | 0.058147383 | -2.311061436 | 0.020829462 | 0.056641519 |
| ENSG00000005379 | TSPOAP1 | 20.72604603 | 3.021513418 | 0.517237928 | 5.841631592 | 5.17E-09 | 3.85E-08 |
| ENSG00000005436 | GCFC2 | 331.1338586 | -0.356575665 | 0.104824751 | -3.401636162 | 0.000669838 | 0.002591841 |
| ENSG00000005448 | WDR54 | 822.7150376 | -0.224279051 | 0.083348372 | -2.690863015 | 0.007126745 | 0.022063095 |
| ENSG00000005469 | CROT | 325.2897933 | -1.132019022 | 0.112248081 | -10.08497437 | 6.44E-24 | 1.34E-22 |
| ENSG00000005471 | ABCB4 | 59.91557504 | -2.311511488 | 0.2636071 | -8.768775522 | 1.81E-18 | 2.75E-17 |
| ENSG00000005483 | KMT2E | 2867.326957 | -0.552029406 | 0.056676581 | -9.739991277 | 2.04E-22 | 3.91E-21 |
| ENSG00000005486 | RHBDD2 | 1749.039648 | 0.184745577 | 0.065419176 | 2.824027866 | 0.004742426 | 0.015365997 |
| ENSG00000005513 | SOX8 | 2.708211761 | 2.0068008 | 1.208740889 | 1.660240684 | 0.096866043 | 0.206652114 |
| ENSG00000005700 | IBTK | 2161.174134 | -0.193644735 | 0.061559245 | -3.145664561 | 0.001657099 | 0.005924854 |
| ENSG00000005801 | ZNF195 | 769.099608 | -0.019683693 | 0.089573356 | -0.219749424 | 0.826066311 | 0.908261176 |
| ENSG00000005810 | MYCBP2 | 1374.6929 | -0.539332644 | 0.146974884 | -3.669556508 | 0.000242972 | 0.001014704 |
| ENSG00000005812 | FBXL3 | 1244.880211 | -0.309516576 | 0.065474932 | -4.727253122 | 2.28E-06 | 1.26E-05 |
| ENSG00000005882 | PDK2 | 1646.151634 | 0.014521219 | 0.06006687 | 0.241750875 | 0.808973206 | 0.898853827 |
| ENSG00000005884 | ITGA3 | 4027.524753 | 0.176486387 | 0.055741623 | 3.16615085 | 0.001544707 | 0.005561138 |
| ENSG00000005889 | ZFX | 1366.86015 | 0.12593698 | 0.065660876 | 1.917991149 | 0.055112133 | 0.130189717 |
| ENSG00000005893 | LAMP2 | 5007.405325 | 0.241064511 | 0.049699455 | 4.85044573 | 1.23E-06 | 7.05E-06 |

| | | | | | | | |
|---|---|---|---|---|---|---|---|
| ENSG00000005961 | ITGA2B | 60.20474491 | -1.150068447 | 0.260828232 | -4.409294343 | 1.04E-05 | 5.35E-05 |
| ENSG00000006007 | GDE1 | 2745.482505 | 0.133185362 | 0.052305065 | 2.54631862 | 0.010886577 | 0.032005393 |
| ENSG00000006015 | REX1BD | 378.2102729 | 0.085895238 | 0.104613273 | 0.821073994 | 0.411604122 | 0.603914203 |
| ENSG00000006016 | CRLF1 | 45.28505183 | -0.744343465 | 0.269900418 | -2.757844803 | 0.005818381 | 0.018420324 |
| ENSG00000006025 | OSBPL7 | 1217.236106 | 0.205468911 | 0.083459483 | 2.461900114 | 0.013820315 | 0.039555587 |
| ENSG00000006042 | TMEM98 | 10.72396756 | 1.893196759 | 0.593167626 | 3.191672432 | 0.001414517 | 0.005134337 |
| ENSG00000006047 | YBX2 | 498.9330601 | -0.63277425 | 0.091434416 | -6.92052597 | 4.50E-12 | 4.36E-11 |
| ENSG00000006062 | MAP3K14 | 343.3039695 | 0.958809585 | 0.107718062 | 8.901103199 | 5.53E-19 | 8.72E-18 |
| ENSG00000006071 | ABCC8 | 391.7804203 | -0.375945664 | 0.103128814 | -3.645398877 | 0.000266977 | 0.001106697 |
| ENSG00000006118 | TMEM132A | 4712.852441 | 0.358107619 | 0.064029054 | 5.592892549 | 2.23E-08 | 1.55E-07 |
| ENSG00000006125 | AP2B1 | 6787.216052 | 0.005163891 | 0.045047124 | 0.114633081 | 0.908735957 | 0.95431011 |
| ENSG00000006194 | ZNF263 | 1474.422458 | 0.228285678 | 0.058453697 | 3.905410427 | 9.41E-05 | 0.000420718 |
| ENSG00000006282 | SPATA20 | 5381.449067 | -0.148973289 | 0.052481808 | -2.838570028 | 0.004531617 | 0.014753804 |
| ENSG00000006283 | CACNA1G | 6.874352331 | -0.022222244 | 0.665176237 | -0.033408055 | 0.973349186 | 0.987970845 |
| ENSG00000006327 | TNFRSF12A | 2399.136784 | 0.488761869 | 0.067867292 | 7.201729307 | 5.95E-13 | 6.18E-12 |
| ENSG00000006432 | MAP3K9 | 1590.340129 | -0.357182865 | 0.080667604 | -4.427835315 | 9.52E-06 | 4.93E-05 |
| ENSG00000006451 | RALA | 2066.951671 | 0.63406151 | 0.061821676 | 10.2562977 | 1.11E-24 | 2.42E-23 |
| ENSG00000006453 | BAIAP2L1 | 666.2206619 | 0.392212047 | 0.077896503 | 5.035040512 | 4.78E-07 | 2.89E-06 |
| ENSG00000006459 | KDM7A | 452.3555424 | -0.020873113 | 0.09537249 | -0.218858841 | 0.826760009 | 0.908541056 |
| ENSG00000006468 | ETV1 | 28.54562858 | -4.90099247 | 0.650266205 | -7.53690171 | 4.81E-14 | 5.37E-13 |
| ENSG00000006530 | AGK | 968.307629 | 0.051016946 | 0.069738819 | 0.731543023 | 0.464447536 | 0.650624508 |
| ENSG00000006534 | ALDH3B1 | 55.07937613 | -0.073504874 | 0.243283752 | -0.302136386 | 0.762548096 | 0.869742236 |
| ENSG00000006555 | TTC22 | 21.3428709 | 0.300666914 | 0.379451224 | 0.792373024 | 0.428143204 | 0.61877975 |
| ENSG00000006576 | PHTF2 | 1292.491998 | 0.708898556 | 0.070612908 | 10.03922059 | 1.02E-23 | 2.11E-22 |
| ENSG00000006607 | FARP2 | 1045.205012 | -0.365605537 | 0.066014753 | -5.53823991 | 3.06E-08 | 2.08E-07 |
| ENSG00000006625 | GGCT | 670.7457554 | -1.160111632 | 0.08110518 | -14.30379217 | 2.07E-46 | 1.04E-44 |
| ENSG00000006634 | DBF4 | 803.2766786 | -0.704116679 | 0.080366568 | -8.761313277 | 1.93E-18 | 2.93E-17 |
| ENSG00000006638 | TBXA2R | 2.946401436 | 1.769363321 | 1.132206826 | 1.562756274 | 0.118109933 | 0.242564177 |
| ENSG00000006652 | IFRD1 | 853.11061 | 0.114602648 | 0.078639786 | 1.4573113 | 0.145030469 | 0.284043282 |
| ENSG00000006695 | COX10 | 588.2432111 | 0.075263978 | 0.08378688 | 0.898278802 | 0.369036931 | 0.561572617 |
| ENSG00000006704 | GTF2IRD1 | 1560.925184 | 0.685612717 | 0.06006506 | 11.41450147 | 3.54E-30 | 9.91E-29 |
| ENSG00000006712 | PAF1 | 3342.420375 | -0.248699374 | 0.050665299 | -4.908672822 | 9.17E-07 | 5.34E-06 |
| ENSG00000006715 | VPS41 | 2334.467073 | 0.778022639 | 0.056521342 | 13.76511254 | 4.13E-43 | 1.87E-41 |

| | | | | | | | |
|---|---|---|---|---|---|---|---|
| ENSG00000006740 | ARHGAP44 | 472.9948924 | 0.63511899 | 0.08919759 | 7.120360404 | 1.08E-12 | 1.10E-11 |
| ENSG00000006744 | ELAC2 | 3358.903088 | 0.078427756 | 0.050216075 | 1.561805776 | 0.118333751 | 0.242958582 |
| ENSG00000006747 | SCIN | 1305.08539 | -2.267845545 | 0.067184645 | -33.75541441 | 8.90E-250 | 1.08E-246 |
| ENSG00000006756 | ARSD | 3976.596818 | 0.011491783 | 0.051362893 | 0.22373707 | 0.822961899 | 0.906383375 |
| ENSG00000006757 | PNPLA4 | 359.6451256 | 0.128801021 | 0.100546115 | 1.281014397 | 0.200188609 | 0.363372189 |
| ENSG00000006831 | ADIPOR2 | 5782.174765 | -0.256375043 | 0.044101852 | -5.8132489 | 6.13E-09 | 4.52E-08 |
| ENSG00000006837 | CDKL3 | 19.93246953 | 0.501441614 | 0.406511294 | 1.233524435 | 0.217380166 | 0.386925901 |
| ENSG00000007047 | MARK4 | 1249.091147 | 0.285921587 | 0.0675538 | 4.232501896 | 2.31E-05 | 0.000113694 |
| ENSG00000007062 | PROM1 | 40.51823542 | -2.568082061 | 0.357467219 | -7.18410507 | 6.76E-13 | 7.02E-12 |
| ENSG00000007080 | CCDC124 | 691.0620672 | -0.07349792 | 0.080718163 | -0.910549961 | 0.362532546 | 0.555227389 |
| ENSG00000007168 | PAFAH1B1 | 2988.303059 | 0.181848921 | 0.050648354 | 3.590421182 | 0.000330144 | 0.001347839 |
| ENSG00000007202 | BLTP2 | 15347.89844 | 0.123771979 | 0.048191236 | 2.568350367 | 0.010218382 | 0.030294597 |
| ENSG00000007255 | TRAPPC6A | 712.9716159 | -0.312912971 | 0.079193788 | -3.951231273 | 7.78E-05 | 0.000353606 |
| ENSG00000007264 | MATK | 486.5728793 | 0.669456106 | 0.097261992 | 6.883018674 | 5.86E-12 | 5.61E-11 |
| ENSG00000007306 | CEACAM7 | 2.265096222 | -0.140465257 | 1.142183629 | -0.122979575 | 0.902123272 | 0.950654947 |
| ENSG00000007341 | ST7L | 371.8714053 | -0.167819568 | 0.103657989 | -1.618973794 | 0.105452902 | 0.22120424 |
| ENSG00000007372 | PAX6 | 2.724263339 | 1.070657142 | 1.089473012 | 0.982729384 | 0.325740645 | 0.516127255 |
| ENSG00000007376 | RPUSD1 | 1633.704532 | 0.413247743 | 0.064813583 | 6.375943475 | 1.82E-10 | 1.54E-09 |
| ENSG00000007384 | RHBDF1 | 1877.673213 | 0.253677006 | 0.06157486 | 4.119814555 | 3.79E-05 | 0.000180276 |
| ENSG00000007392 | LUC7L | 1773.700633 | 0.021907239 | 0.062649828 | 0.349677561 | 0.726580696 | 0.848025671 |
| ENSG00000007402 | CACNA2D2 | 148.4492213 | -0.211890486 | 0.147942126 | -1.432252541 | 0.15207156 | 0.294489413 |
| ENSG00000007516 | BAIAP3 | 548.3937021 | 0.559158055 | 0.090465935 | 6.180868615 | 6.37E-10 | 5.17E-09 |
| ENSG00000007520 | TSR3 | 1070.353612 | 0.035411875 | 0.078030643 | 0.453820115 | 0.649958305 | 0.796349063 |
| ENSG00000007541 | PIGQ | 1355.932773 | 0.289611041 | 0.071055393 | 4.075848825 | 4.58E-05 | 0.000215694 |
| ENSG00000007545 | CRAMP1 | 704.0288518 | -0.032127419 | 0.081192154 | -0.395696105 | 0.692329229 | 0.826457434 |
| ENSG00000007866 | TEAD3 | 1966.442722 | 0.103438242 | 0.061991992 | 1.668574257 | 0.095201784 | 0.203724713 |
| ENSG00000007923 | DNAJC11 | 2110.132429 | -0.006346144 | 0.055804292 | -0.113721423 | 0.90945863 | 0.954587195 |
| ENSG00000007944 | MYLIP | 1229.322089 | -0.341950311 | 0.06471637 | -5.283830212 | 1.27E-07 | 8.14E-07 |
| ENSG00000007968 | E2F2 | 626.4825775 | -0.409484074 | 0.080268318 | -5.101440792 | 3.37E-07 | 2.07E-06 |
| ENSG00000008018 | PSMB1 | 5459.322648 | 0.139926694 | 0.047722937 | 2.932063783 | 0.003367176 | 0.011291188 |
| ENSG00000008056 | SYN1 | 4.379508944 | 0.347689868 | 0.85105422 | 0.408540208 | 0.682877117 | 0.819995869 |
| ENSG00000008083 | JARID2 | 1116.711326 | -0.437058073 | 0.066802018 | -6.542587889 | 6.05E-11 | 5.32E-10 |
| ENSG00000008086 | CDKL5 | 946.5794282 | -0.719375066 | 0.085039876 | -8.459267616 | 2.69E-17 | 3.78E-16 |

| | | | | | | | |
|---|---|---|---|---|---|---|---|
| ENSG00000008128 | CDK11A | 42.54898195 | 0.314416924 | 0.273650611 | 1.148972126 | 0.250567474 | 0.428735728 |
| ENSG00000008130 | NADK | 1561.848801 | 0.101043357 | 0.058861456 | 1.716630264 | 0.086046749 | 0.187467491 |
| ENSG00000008226 | DLEC1 | 26.82502708 | -0.276785984 | 0.339684613 | -0.814832269 | 0.415168327 | 0.60727931 |
| ENSG00000008256 | CYTH3 | 800.4689923 | 0.896825908 | 0.075860607 | 11.82202386 | 3.00E-32 | 9.13E-31 |
| ENSG00000008277 | ADAM22 | 394.5513048 | 0.093161346 | 0.096237076 | 0.968040073 | 0.333024357 | 0.524156346 |
| ENSG00000008282 | SYPL1 | 1945.489011 | -0.049694541 | 0.056318444 | -0.88238485 | 0.377568726 | 0.570318468 |
| ENSG00000008283 | CYB561 | 11812.36311 | -0.073109685 | 0.044623823 | -1.63835549 | 0.101347559 | 0.214159266 |
| ENSG00000008294 | SPAG9 | 3965.032674 | -0.037957149 | 0.050447665 | -0.752406457 | 0.451806665 | 0.639701573 |
| ENSG00000008300 | CELSR3 | 2706.947058 | -0.284525926 | 0.058004759 | -4.905216943 | 9.33E-07 | 5.42E-06 |
| ENSG00000008311 | AASS | 843.0042646 | -1.995435183 | 0.078624983 | -25.37914946 | 4.29E-142 | 1.42E-139 |
| ENSG00000008323 | PLEKHG6 | 87.51823668 | 0.456182733 | 0.1935152 | 2.357348325 | 0.018405973 | 0.050852515 |
| ENSG00000008324 | SS18L2 | 233.2905947 | -0.132486529 | 0.11901827 | -1.113161274 | 0.265639171 | 0.446724037 |
| ENSG00000008382 | MPND | 757.4719346 | 0.119209225 | 0.078205615 | 1.52430519 | 0.127432491 | 0.257332865 |
| ENSG00000008394 | MGST1 | 3053.599509 | -0.705360789 | 0.056295415 | -12.52963123 | 5.14E-36 | 1.81E-34 |
| ENSG00000008405 | CRY1 | 541.7275242 | -0.143342793 | 0.083669614 | -1.713200119 | 0.08667573 | 0.18862928 |
| ENSG00000008441 | NFIX | 962.4980728 | 0.35911856 | 0.070147028 | 5.119512137 | 3.06E-07 | 1.89E-06 |
| ENSG00000008513 | ST3GAL1 | 6091.819972 | -0.301046899 | 0.066118147 | -4.553165979 | 5.28E-06 | 2.81E-05 |
| ENSG00000008516 | MMP25 | 64.58258663 | -1.351453976 | 0.237489884 | -5.690574921 | 1.27E-08 | 9.00E-08 |
| ENSG00000008517 | IL32 | 4.312369623 | 1.074758028 | 0.874451726 | 1.22906502 | 0.219047429 | 0.388837578 |
| ENSG00000008710 | PKD1 | 2771.260419 | 0.229383965 | 0.052178341 | 4.396152852 | 1.10E-05 | 5.66E-05 |
| ENSG00000008735 | MAPK8IP2 | 64.74113426 | 0.09891687 | 0.220486182 | 0.448630697 | 0.653698086 | 0.799053343 |
| ENSG00000008838 | MED24 | 2915.488473 | 0.04247555 | 0.052033854 | 0.816306043 | 0.414325119 | 0.606477249 |
| ENSG00000008853 | RHOBTB2 | 1078.271665 | -0.195985903 | 0.079120065 | -2.477069549 | 0.013246609 | 0.038089396 |
| ENSG00000008869 | HEATR5B | 1460.095961 | 0.091803081 | 0.065311773 | 1.405613057 | 0.159839057 | 0.305660884 |
| ENSG00000008952 | SEC62 | 3199.820156 | 0.0073485 | 0.053671415 | 0.136916452 | 0.891096834 | 0.944150152 |
| ENSG00000008988 | RPS20 | 6316.875863 | 0.179425411 | 0.049624167 | 3.615686099 | 0.000299553 | 0.001233265 |
| ENSG00000009307 | CSDE1 | 100412.113 | 0.2980036 | 0.057990417 | 5.138842205 | 2.76E-07 | 1.71E-06 |
| ENSG00000009335 | UBE3C | 4085.380334 | 0.755037494 | 0.050640354 | 14.90979888 | 2.85E-50 | 1.59E-48 |
| ENSG00000009413 | REV3L | 2194.214939 | -0.032678175 | 0.06491294 | -0.503415423 | 0.61467223 | 0.771450977 |
| ENSG00000009694 | TENM1 | 158.9686021 | 0.658395694 | 0.145485434 | 4.525509381 | 6.03E-06 | 3.19E-05 |
| ENSG00000009724 | MASP2 | 6.147761572 | -1.218417549 | 0.73157405 | -1.665473985 | 0.09581823 | 0.204776429 |
| ENSG00000009765 | IYD | 4.879738413 | -0.508022298 | 0.792320063 | -0.64118318 | 0.521403676 | 0.698452574 |
| ENSG00000009780 | FAM76A | 327.1761992 | -0.326772445 | 0.105618809 | -3.093884966 | 0.001975541 | 0.006956086 |

| | | | | | | | |
|---|---|---|---|---|---|---|---|
| ENSG00000009830 | POMT2 | 1214.515208 | 0.121977789 | 0.064764466 | 1.883406084 | 0.059645356 | 0.138867804 |
| ENSG00000009844 | VTA1 | 2344.985713 | 0.478620949 | 0.052740557 | 9.075007512 | 1.14E-19 | 1.86E-18 |
| ENSG00000009950 | MLXIPL | 269.8882162 | 0.889632356 | 0.130219017 | 6.83181593 | 8.38E-12 | 7.92E-11 |
| ENSG00000009954 | BAZ1B | 5364.072861 | 0.449651642 | 0.045902811 | 9.795732091 | 1.17E-22 | 2.29E-21 |
| ENSG00000010017 | RANBP9 | 2592.573062 | -0.35927599 | 0.050805528 | -7.071592447 | 1.53E-12 | 1.55E-11 |
| ENSG00000010030 | ETV7 | 6.852346516 | -0.076283937 | 0.689451763 | -0.110644343 | 0.911898383 | 0.955697231 |
| ENSG00000010072 | SPRTN | 459.7071949 | -0.103985892 | 0.091413506 | -1.137533131 | 0.255315483 | 0.434762737 |
| ENSG00000010165 | METTL13 | 2708.412824 | -0.22069989 | 0.052169619 | -4.230429406 | 2.33E-05 | 0.0001146 |
| ENSG00000010219 | DYRK4 | 64.88945096 | 0.434204336 | 0.22782823 | 1.905840798 | 0.056670864 | 0.133104292 |
| ENSG00000010244 | ZNF207 | 3492.15941 | 0.167513465 | 0.047665988 | 3.514318539 | 0.000440884 | 0.001762613 |
| ENSG00000010256 | UQCRC1 | 3241.41288 | 0.117993235 | 0.051664488 | 2.283836325 | 0.022381152 | 0.06034251 |
| ENSG00000010270 | STARD3NL | 910.3123163 | 0.288830371 | 0.07030311 | 4.108358374 | 3.98E-05 | 0.000188828 |
| ENSG00000010278 | CD9 | 2975.004258 | -0.167058825 | 0.053120513 | -3.144902295 | 0.001661423 | 0.005939387 |
| ENSG00000010292 | NCAPD2 | 3437.006413 | -0.253715155 | 0.049069692 | -5.170506376 | 2.33E-07 | 1.46E-06 |
| ENSG00000010295 | IFFO1 | 5.174162693 | 1.074368712 | 0.814300018 | 1.319376996 | 0.187043108 | 0.344750836 |
| ENSG00000010310 | GIPR | 68.92296004 | -0.796689032 | 0.228309638 | -3.489511175 | 0.000483905 | 0.001920882 |
| ENSG00000010318 | PHF7 | 123.8075734 | 0.085697961 | 0.162660007 | 0.526853298 | 0.598295467 | 0.758658892 |
| ENSG00000010319 | SEMA3G | 13.98286224 | 0.060465122 | 0.469583388 | 0.128763334 | 0.89754492 | 0.948060798 |
| ENSG00000010322 | NISCH | 3248.646654 | -0.069615947 | 0.050226476 | -1.386040831 | 0.165734436 | 0.314574546 |
| ENSG00000010327 | STAB1 | 8.195441076 | 0.280451163 | 0.609747438 | 0.459946439 | 0.645554666 | 0.792669964 |
| ENSG00000010361 | FUZ | 377.3548943 | -0.569868356 | 0.111636269 | -5.104688295 | 3.31E-07 | 2.03E-06 |
| ENSG00000010404 | IDS | 462.1008021 | 0.447614691 | 0.092391819 | 4.844743789 | 1.27E-06 | 7.25E-06 |
| ENSG00000010539 | ZNF200 | 521.966885 | 0.488906886 | 0.085654222 | 5.707913421 | 1.14E-08 | 8.16E-08 |
| ENSG00000010626 | LRRC23 | 85.18284765 | -0.304119114 | 0.195478818 | -1.555765055 | 0.119763971 | 0.245175196 |
| ENSG00000010671 | BTK | 2.254758436 | -1.931281623 | 1.28248077 | -1.50589519 | 0.132094082 | 0.264672743 |
| ENSG00000010704 | HFE | 63.69486232 | -0.131149649 | 0.228092499 | -0.57498449 | 0.565301787 | 0.733481521 |
| ENSG00000010803 | SCMH1 | 987.0053336 | 0.636425232 | 0.068104846 | 9.344786294 | 9.21E-21 | 1.61E-19 |
| ENSG00000010810 | FYN | 259.0720328 | 0.862553383 | 0.117874258 | 7.317572124 | 2.52E-13 | 2.69E-12 |
| ENSG00000010818 | HIVEP2 | 1209.337103 | 0.708135884 | 0.082450844 | 8.588582573 | 8.80E-18 | 1.28E-16 |
| ENSG00000011007 | ELOA | 2705.646741 | 0.053753151 | 0.051680239 | 1.040110341 | 0.29828864 | 0.48485378 |
| ENSG00000011009 | LYPLA2 | 1207.338469 | 0.250314374 | 0.073537684 | 3.403892528 | 0.000664329 | 0.00257297 |
| ENSG00000011021 | CLCN6 | 897.9813974 | -0.838488399 | 0.079842364 | -10.50179824 | 8.48E-26 | 1.95E-24 |
| ENSG00000011028 | MRC2 | 743.5945889 | 1.872940396 | 0.08377321 | 22.35727143 | 1.03E-110 | 2.03E-108 |

| | | | | | | | |
|---|---|---|---|---|---|---|---|
| ENSG00000011105 | TSPAN9 | 661.6340559 | -0.32190897 | 0.084791649 | -3.796470224 | 0.000146771 | 0.000637193 |
| ENSG00000011114 | BTBD7 | 2799.433811 | -0.567329061 | 0.054730932 | -10.36578468 | 3.55E-25 | 7.94E-24 |
| ENSG00000011132 | APBA3 | 220.1670489 | -0.233165482 | 0.132224686 | -1.763403553 | 0.077832451 | 0.172944249 |
| ENSG00000011143 | MKS1 | 1135.235333 | -0.098823977 | 0.068901996 | -1.434268722 | 0.151495592 | 0.293863146 |
| ENSG00000011198 | ABHD5 | 758.3014034 | -0.039249598 | 0.076186298 | -0.515179226 | 0.606427784 | 0.764953996 |
| ENSG00000011201 | ANOS1 | 540.7868584 | -0.193382058 | 0.087759367 | -2.203548916 | 0.027556083 | 0.072219416 |
| ENSG00000011243 | AKAP8L | 1898.932879 | 0.185258506 | 0.05609316 | 3.302693329 | 0.000957611 | 0.003595938 |
| ENSG00000011258 | MBTD1 | 2144.918157 | 0.133210438 | 0.058825852 | 2.264488028 | 0.023544115 | 0.063088658 |
| ENSG00000011260 | UTP18 | 2237.532631 | 0.182205802 | 0.056895056 | 3.202489222 | 0.001362454 | 0.00496737 |
| ENSG00000011275 | RNF216 | 3377.588422 | 0.316638104 | 0.050147464 | 6.314139954 | 2.72E-10 | 2.27E-09 |
| ENSG00000011295 | TTC19 | 2041.342013 | 0.14171985 | 0.057288537 | 2.473790703 | 0.013368799 | 0.038397427 |
| ENSG00000011304 | PTBP1 | 9520.148794 | -0.104885857 | 0.043216694 | -2.426975458 | 0.015225286 | 0.043082281 |
| ENSG00000011332 | DPF1 | 111.5597275 | -0.579727018 | 0.169881825 | -3.41253113 | 0.000643626 | 0.00249827 |
| ENSG00000011347 | SYT7 | 5092.341148 | 0.471171512 | 0.050896966 | 9.257359478 | 2.10E-20 | 3.58E-19 |
| ENSG00000011376 | LARS2 | 983.7714375 | 0.000693381 | 0.070116264 | 0.009889014 | 0.992109837 | 0.995917623 |
| ENSG00000011405 | PIK3C2A | 2673.411171 | -0.085834297 | 0.05654197 | -1.518063421 | 0.128998411 | 0.259649342 |
| ENSG00000011422 | PLAUR | 33.98778375 | 0.152939209 | 0.301558697 | 0.50716232 | 0.612040931 | 0.769411385 |
| ENSG00000011426 | ANLN | 4435.020321 | 0.489982126 | 0.047603953 | 10.29288729 | 7.59E-25 | 1.68E-23 |
| ENSG00000011451 | WIZ | 2077.261402 | 0.265280412 | 0.060123056 | 4.412290927 | 1.02E-05 | 5.28E-05 |
| ENSG00000011454 | RABGAP1 | 3842.397214 | 0.080858544 | 0.0528231 | 1.530742112 | 0.125833142 | 0.254731051 |
| ENSG00000011478 | QPCTL | 497.2963346 | 0.002695632 | 0.092079893 | 0.029274931 | 0.976645321 | 0.989278333 |
| ENSG00000011485 | PPP5C | 2626.891167 | -0.193319334 | 0.050968401 | -3.792925195 | 0.000148883 | 0.000645629 |
| ENSG00000011523 | CEP68 | 513.6833136 | 0.38536593 | 0.088829645 | 4.338258136 | 1.44E-05 | 7.25E-05 |
| ENSG00000011566 | MAP4K3 | 1051.836883 | -0.053520552 | 0.069026954 | -0.775357297 | 0.438128561 | 0.627656854 |
| ENSG00000011590 | ZBTB32 | 3.47415128 | -0.799098884 | 0.952564721 | -0.838891957 | 0.401529941 | 0.594264313 |
| ENSG00000011638 | LDAF1 | 536.5488318 | 0.167907971 | 0.087753383 | 1.91340738 | 0.05569592 | 0.131379374 |
| ENSG00000011677 | GABRA3 | 116.6869608 | 2.701936263 | 0.207564444 | 13.01733671 | 9.75E-39 | 3.75E-37 |
| ENSG00000012048 | BRCA1 | 1672.540916 | 0.015668458 | 0.055964425 | 0.279971748 | 0.77949918 | 0.880308728 |
| ENSG00000012061 | ERCC1 | 1447.48783 | 0.310043946 | 0.062142174 | 4.989267739 | 6.06E-07 | 3.61E-06 |
| ENSG00000012124 | CD22 | 16.73745176 | 0.391957133 | 0.503691336 | 0.778169298 | 0.436469214 | 0.626296602 |
| ENSG00000012171 | SEMA3B | 292.25724 | -0.536005582 | 0.112994627 | -4.743637802 | 2.10E-06 | 1.17E-05 |
| ENSG00000012174 | MBTPS2 | 1069.920093 | 0.175214094 | 0.071419626 | 2.453304552 | 0.014155044 | 0.040397699 |
| ENSG00000012211 | PRICKLE3 | 467.3923594 | -0.00752876 | 0.094944754 | -0.07929622 | 0.936797013 | 0.969659375 |

| | | | | | | | |
|---|---|---|---|---|---|---|---|
| ENSG00000012223 | LTF | 2.119851123 | -3.598850528 | 1.567570657 | -2.295813916 | 0.021686523 | 0.05865922 |
| ENSG00000012232 | EXTL3 | 1238.28685 | 0.169451225 | 0.07147792 | 2.370679309 | 0.017755429 | 0.049323026 |
| ENSG00000012660 | ELOVL5 | 2734.948873 | 0.372844618 | 0.056535162 | 6.594915531 | 4.25E-11 | 3.78E-10 |
| ENSG00000012822 | CALCOCO1 | 2294.533819 | 0.403147977 | 0.067192753 | 5.999872923 | 1.97E-09 | 1.53E-08 |
| ENSG00000012963 | UBR7 | 2473.957796 | -0.66128908 | 0.051646603 | -12.80411561 | 1.55E-37 | 5.78E-36 |
| ENSG00000012983 | MAP4K5 | 1401.171937 | -0.695159404 | 0.070136806 | -9.911477916 | 3.71E-23 | 7.42E-22 |
| ENSG00000013016 | EHD3 | 630.4903252 | 0.124772357 | 0.081727158 | 1.526693949 | 0.126837133 | 0.256424061 |
| ENSG00000013275 | PSMC4 | 3818.29068 | -0.133315975 | 0.051692595 | -2.579014924 | 0.009908251 | 0.029492055 |
| ENSG00000013288 | MAN2B2 | 1952.09348 | 0.281119482 | 0.056389045 | 4.985356286 | 6.18E-07 | 3.68E-06 |
| ENSG00000013297 | CLDN11 | 6.329254561 | 1.242673415 | 0.728283115 | 1.706305406 | 0.087951232 | 0.190861911 |
| ENSG00000013306 | SLC25A39 | 10828.80309 | 0.245304634 | 0.045099595 | 5.439175994 | 5.35E-08 | 3.56E-07 |
| ENSG00000013364 | MVP | 323.9675091 | 0.924739382 | 0.114453981 | 8.079573783 | 6.50E-16 | 8.33E-15 |
| ENSG00000013374 | NUB1 | 1848.293205 | 0.38785126 | 0.063901651 | 6.069503004 | 1.28E-09 | 1.01E-08 |
| ENSG00000013375 | PGM3 | 2112.565622 | -0.30553554 | 0.053034141 | -5.76111042 | 8.36E-09 | 6.05E-08 |
| ENSG00000013392 | RWDD2A | 710.7618754 | -0.345888144 | 0.079359564 | -4.358493486 | 1.31E-05 | 6.66E-05 |
| ENSG00000013441 | CLK1 | 1163.456711 | -0.326284951 | 0.081902263 | -3.983833171 | 6.78E-05 | 0.000310999 |
| ENSG00000013503 | POLR3B | 586.5307026 | -0.086665966 | 0.081159145 | -1.067852138 | 0.285587215 | 0.469497686 |
| ENSG00000013523 | ANGEL1 | 1679.204339 | -0.472784616 | 0.058896017 | -8.027446272 | 9.95E-16 | 1.26E-14 |
| ENSG00000013561 | RNF14 | 1713.443849 | 0.305621745 | 0.056835695 | 5.377285282 | 7.56E-08 | 4.95E-07 |
| ENSG00000013563 | DNASE1L1 | 1231.209648 | -0.534670558 | 0.064990083 | -8.226956075 | 1.92E-16 | 2.57E-15 |
| ENSG00000013573 | DDX11 | 1108.494496 | -0.253750529 | 0.0807995 | -3.14049627 | 0.001686619 | 0.006018203 |
| ENSG00000013583 | HEBP1 | 4106.708771 | -0.818653864 | 0.049690155 | -16.47517218 | 5.53E-61 | 4.07E-59 |
| ENSG00000013588 | GPRC5A | 3946.984981 | 0.170009671 | 0.053543426 | 3.175173557 | 0.001497469 | 0.005406349 |
| ENSG00000013619 | MAMLD1 | 354.7360792 | -0.730960702 | 0.105254131 | -6.944722198 | 3.79E-12 | 3.69E-11 |
| ENSG00000013725 | CD6 | 8.222287417 | -0.297890647 | 0.601674761 | -0.495102448 | 0.620527806 | 0.775370368 |
| ENSG00000013810 | TACC3 | 3546.800197 | -0.235430495 | 0.050842947 | -4.630543848 | 3.65E-06 | 1.98E-05 |
| ENSG00000014123 | UFL1 | 1399.520771 | -0.017640343 | 0.0690862 | -0.255338159 | 0.79846193 | 0.892453551 |
| ENSG00000014138 | POLA2 | 172.9214606 | -0.094242655 | 0.145060048 | -0.649680294 | 0.515898756 | 0.694607199 |
| ENSG00000014164 | ZC3H3 | 1127.334414 | 0.070011098 | 0.071869433 | 0.974142896 | 0.329985593 | 0.520516366 |
| ENSG00000014216 | CAPN1 | 8669.142768 | -0.140332189 | 0.05167385 | -2.71572932 | 0.006612996 | 0.020634131 |
| ENSG00000014257 | ACP3 | 37.49945279 | -0.243997965 | 0.285343559 | -0.855102411 | 0.392494455 | 0.585424109 |
| ENSG00000014641 | MDH1 | 3405.197629 | 0.04671332 | 0.048352128 | 0.966106796 | 0.333990744 | 0.524957018 |
| ENSG00000014824 | SLC30A9 | 2800.110858 | 0.021471126 | 0.056011412 | 0.383334836 | 0.701471513 | 0.832949765 |

| | | | | | | | |
|---|---|---|---|---|---|---|---|
| ENSG00000014914 | MTMR11 | 722.8167218 | 0.568394464 | 0.087443796 | 6.500111988 | 8.03E-11 | 7.01E-10 |
| ENSG00000014919 | COX15 | 795.7068291 | -0.18318072 | 0.073004344 | -2.509175617 | 0.01210133 | 0.035202196 |
| ENSG00000015133 | CCDC88C | 3238.785075 | -0.180843373 | 0.049998512 | -3.616975089 | 0.000298066 | 0.001227582 |
| ENSG00000015153 | YAF2 | 258.5453295 | 0.130703012 | 0.115076366 | 1.135793705 | 0.256042912 | 0.435502663 |
| ENSG00000015171 | ZMYND11 | 5306.458263 | -1.092458918 | 0.0468223 | -23.33202184 | 2.10E-120 | 5.23E-118 |
| ENSG00000015285 | WAS | 121.4402073 | -0.696820959 | 0.169324799 | -4.115291803 | 3.87E-05 | 0.00018362 |
| ENSG00000015413 | DPEP1 | 7.158591091 | -1.041140853 | 0.666167239 | -1.562882101 | 0.118080328 | 0.242546789 |
| ENSG00000015475 | BID | 1714.529579 | -0.012839748 | 0.06037123 | -0.212679919 | 0.831576631 | 0.910783858 |
| ENSG00000015479 | MATR3 | 27.31570228 | -0.384022481 | 0.338967133 | -1.132919516 | 0.257248054 | 0.436904594 |
| ENSG00000015520 | NPC1L1 | 21.17743821 | 1.475899609 | 0.405233104 | 3.642100295 | 0.000270423 | 0.001120371 |
| ENSG00000015532 | XYLT2 | 1712.303175 | 0.449771094 | 0.062221902 | 7.228501257 | 4.88E-13 | 5.10E-12 |
| ENSG00000015568 | RGPD5 | 6.564779485 | 0.120113351 | 0.728515826 | 0.164874044 | 0.869043122 | 0.931886562 |
| ENSG00000015676 | NUDCD3 | 3966.804428 | 0.580442485 | 0.054456922 | 10.65874575 | 1.59E-26 | 3.82E-25 |
| ENSG00000016082 | ISL1 | 14.87033055 | -2.114743573 | 0.531673082 | -3.977526122 | 6.96E-05 | 0.000318852 |
| ENSG00000016391 | CHDH | 1083.283767 | 0.186818219 | 0.065696242 | 2.843666729 | 0.004459767 | 0.014548785 |
| ENSG00000016402 | IL20RA | 17.21193051 | 2.407047666 | 0.51895363 | 4.638271177 | 3.51E-06 | 1.92E-05 |
| ENSG00000016864 | GLT8D1 | 799.3607004 | -0.043020892 | 0.072205512 | -0.595811747 | 0.551300995 | 0.723235044 |
| ENSG00000017260 | ATP2C1 | 3575.688928 | 0.023335954 | 0.048556634 | 0.480592496 | 0.63080615 | 0.782326489 |
| ENSG00000017483 | SLC38A5 | 3.309077444 | -0.463644943 | 0.960482861 | -0.482720683 | 0.629294075 | 0.7810577 |
| ENSG00000017797 | RALBP1 | 4068.119158 | 0.407704212 | 0.048399281 | 8.423765941 | 3.65E-17 | 5.07E-16 |
| ENSG00000018189 | RUFY3 | 359.6091804 | -0.264559444 | 0.103094201 | -2.566191322 | 0.01028221 | 0.030458922 |
| ENSG00000018236 | CNTN1 | 11.50259075 | -0.723743291 | 0.533075741 | -1.357674408 | 0.174567021 | 0.327545187 |
| ENSG00000018280 | SLC11A1 | 24.37673407 | -0.718761609 | 0.370622793 | -1.939334605 | 0.052460608 | 0.124838832 |
| ENSG00000018408 | WWTR1 | 715.838968 | 0.168420422 | 0.075572069 | 2.22860675 | 0.025840082 | 0.068307383 |
| ENSG00000018610 | STEEP1 | 1311.735695 | -0.118479588 | 0.062841773 | -1.885363532 | 0.059380774 | 0.138307946 |
| ENSG00000018699 | TTC27 | 625.2229626 | 0.114091016 | 0.080114598 | 1.424097724 | 0.154418185 | 0.297737844 |
| ENSG00000019144 | PHLDB1 | 986.4116402 | 0.409089321 | 0.069894875 | 5.852922979 | 4.83E-09 | 3.61E-08 |
| ENSG00000019186 | CYP24A1 | 1.785705404 | -0.424172156 | 1.291848285 | -0.328345178 | 0.742650689 | NA |
| ENSG00000019485 | PRDM11 | 418.8101504 | -0.496880076 | 0.106092003 | -4.683482847 | 2.82E-06 | 1.55E-05 |
| ENSG00000019505 | SYT13 | 2.234002979 | 3.399786847 | 1.521654298 | 2.234270197 | 0.0254653 | 0.067487752 |
| ENSG00000019549 | SNAI2 | 21.67026832 | -2.50072678 | 0.44687652 | -5.596012916 | 2.19E-08 | 1.52E-07 |
| ENSG00000019582 | CD74 | 9.813419292 | 2.567417447 | 0.687869231 | 3.732420831 | 0.000189648 | 0.000808799 |
| ENSG00000019995 | ZRANB1 | 1391.99299 | -0.803279311 | 0.060958913 | -13.17738896 | 1.18E-39 | 4.74E-38 |

| | | | | | | | |
|---|---|---|---|---|---|---|---|
| ENSG00000020129 | NCDN | 2719.177628 | 0.389358531 | 0.06493566 | 5.996066459 | 2.02E-09 | 1.57E-08 |
| ENSG00000020181 | ADGRA2 | 53.0197418 | 0.695115472 | 0.255434813 | 2.721302797 | 0.006502517 | 0.02032258 |
| ENSG00000020256 | ZFP64 | 947.167552 | -0.119092302 | 0.069496853 | -1.71363589 | 0.086595618 | 0.188490699 |
| ENSG00000020426 | MNAT1 | 692.8445987 | -0.776764007 | 0.076536293 | -10.14896295 | 3.35E-24 | 7.08E-23 |
| ENSG00000020577 | SAMD4A | 1074.019472 | 0.16633884 | 0.070428638 | 2.361806867 | 0.018186114 | 0.050324421 |
| ENSG00000020922 | MRE11 | 721.5427579 | -0.034326988 | 0.075897067 | -0.452283464 | 0.651064787 | 0.797236373 |
| ENSG00000021300 | PLEKHB1 | 574.8339118 | 0.253000826 | 0.081624042 | 3.099587087 | 0.001937906 | 0.00683406 |
| ENSG00000021355 | SERPINB1 | 838.8233622 | -0.420352403 | 0.073404992 | -5.726482493 | 1.03E-08 | 7.37E-08 |
| ENSG00000021461 | CYP3A43 | 3.153004788 | -4.171713965 | 1.433068465 | -2.91103605 | 0.003602325 | 0.011995547 |
| ENSG00000021488 | SLC7A9 | 5.479646207 | -0.249725553 | 0.750108351 | -0.332919308 | 0.739195194 | 0.856204886 |
| ENSG00000021574 | SPAST | 788.0572183 | 0.113307371 | 0.077179627 | 1.468099501 | 0.142077197 | 0.279645077 |
| ENSG00000021645 | NRXN3 | 214.5458184 | -1.050140612 | 0.128456636 | -8.175059293 | 2.96E-16 | 3.90E-15 |
| ENSG00000021762 | OSBPL5 | 478.3183939 | 0.078970329 | 0.098802122 | 0.799277664 | 0.424129427 | 0.615019183 |
| ENSG00000021776 | AQR | 2449.475724 | -0.199492576 | 0.057670802 | -3.459160802 | 0.000541861 | 0.002131761 |
| ENSG00000021826 | CPS1 | 284.1688709 | 2.314331133 | 0.131099318 | 17.65326594 | 9.60E-70 | 8.78E-68 |
| ENSG00000022267 | FHL1 | 6.226040844 | -2.024396679 | 0.780369652 | -2.594150956 | 0.009482484 | 0.028390523 |
| ENSG00000022277 | RTF2 | 4240.16082 | -0.073542961 | 0.04480144 | -1.641531172 | 0.100687202 | 0.2129993 |
| ENSG00000022355 | GABRA1 | 37.2933566 | 0.281221297 | 0.29185211 | 0.963574656 | 0.335259218 | 0.526445785 |
| ENSG00000022567 | SLC45A4 | 592.2270631 | 0.456010584 | 0.08194568 | 5.564791045 | 2.62E-08 | 1.80E-07 |
| ENSG00000022840 | RNF10 | 5712.679384 | -0.201842037 | 0.046348299 | -4.354896328 | 1.33E-05 | 6.76E-05 |
| ENSG00000022976 | ZNF839 | 781.0309342 | -0.537475404 | 0.080304697 | -6.692951018 | 2.19E-11 | 1.99E-10 |
| ENSG00000023041 | ZDHHC6 | 1131.631854 | -0.856552978 | 0.070691961 | -12.11669564 | 8.62E-34 | 2.78E-32 |
| ENSG00000023191 | RNH1 | 2672.411633 | 0.036274044 | 0.055013423 | 0.659367157 | 0.509660027 | 0.689658803 |
| ENSG00000023228 | NDUFS1 | 3941.991245 | -0.160886839 | 0.05054018 | -3.183345164 | 0.00145584 | 0.005266828 |
| ENSG00000023287 | RB1CC1 | 2222.630818 | 0.145858106 | 0.060402659 | 2.414762984 | 0.015745452 | 0.044378998 |
| ENSG00000023318 | ERP44 | 1047.192965 | -0.059473542 | 0.066158007 | -0.898962108 | 0.368672844 | 0.561204714 |
| ENSG00000023330 | ALAS1 | 742.6300951 | 0.112143664 | 0.072921574 | 1.537866735 | 0.124081202 | 0.252007329 |
| ENSG00000023445 | BIRC3 | 4.584170557 | 1.230970926 | 0.852532097 | 1.443899802 | 0.148767162 | 0.289655297 |
| ENSG00000023516 | AKAP11 | 2356.433619 | 0.103270479 | 0.059795308 | 1.727066601 | 0.084155703 | 0.184193477 |
| ENSG00000023572 | GLRX2 | 466.7891119 | -0.262754131 | 0.09264238 | -2.836219553 | 0.004565105 | 0.014852292 |
| ENSG00000023608 | SNAPC1 | 118.3893945 | -0.686177163 | 0.171037552 | -4.011850942 | 6.02E-05 | 0.000278687 |
| ENSG00000023697 | DERA | 207.0463249 | -0.362031344 | 0.12915069 | -2.803170039 | 0.005060298 | 0.016287813 |
| ENSG00000023734 | STRAP | 2954.603169 | 0.040594919 | 0.050182841 | 0.808940228 | 0.418549528 | 0.61039531 |

| | | | | | | | |
|---|---|---|---|---|---|---|---|
| ENSG00000023839 | ABCC2 | 28.61403917 | -1.02999295 | 0.333616902 | -3.087352422 | 0.00201948 | 0.007093382 |
| ENSG00000023892 | DEF6 | 119.4884857 | 0.122016774 | 0.182221924 | 0.669605345 | 0.503109406 | 0.684133359 |
| ENSG00000023902 | PLEKHO1 | 468.1287858 | 0.851867083 | 0.092851547 | 9.174506081 | 4.54E-20 | 7.60E-19 |
| ENSG00000023909 | GCLM | 342.3119848 | -0.07305542 | 0.104807891 | -0.697041218 | 0.485776999 | 0.669493804 |
| ENSG00000024048 | UBR2 | 1671.270627 | -0.15134783 | 0.058617048 | -2.581976335 | 0.009823632 | 0.029259169 |
| ENSG00000024422 | EHD2 | 347.8685423 | -2.13667023 | 0.114660522 | -18.63475054 | 1.68E-77 | 1.88E-75 |
| ENSG00000024526 | DEPDC1 | 878.6859317 | -0.136281997 | 0.076292853 | -1.786300961 | 0.074050541 | 0.166231473 |
| ENSG00000024862 | CCDC28A | 307.3033306 | 0.621780673 | 0.110911727 | 5.606085937 | 2.07E-08 | 1.44E-07 |
| ENSG00000025039 | RRAGD | 394.921618 | 1.341129805 | 0.100911523 | 13.29015517 | 2.64E-40 | 1.10E-38 |
| ENSG00000025156 | HSF2 | 394.2746478 | 0.529956803 | 0.0979681 | 5.409483328 | 6.32E-08 | 4.17E-07 |
| ENSG00000025293 | PHF20 | 1200.335217 | -0.025046186 | 0.063203188 | -0.396280417 | 0.691898172 | 0.826372889 |
| ENSG00000025423 | HSD17B6 | 45.15297316 | -0.612801827 | 0.266076216 | -2.303106361 | 0.021272859 | 0.057635455 |
| ENSG00000025434 | NR1H3 | 307.73973 | -0.805358227 | 0.111082773 | -7.250073122 | 4.17E-13 | 4.36E-12 |
| ENSG00000025708 | TYMP | 62.48040169 | -0.950702914 | 0.231441785 | -4.107741021 | 4.00E-05 | 0.000189294 |
| ENSG00000025770 | NCAPH2 | 1185.179561 | -0.362069289 | 0.065853422 | -5.498108951 | 3.84E-08 | 2.60E-07 |
| ENSG00000025772 | TOMM34 | 2415.215594 | -0.148130875 | 0.054419286 | -2.722029009 | 0.006488245 | 0.020280738 |
| ENSG00000025796 | SEC63 | 2996.759752 | -0.012216311 | 0.054474201 | -0.224258651 | 0.822556048 | 0.90619104 |
| ENSG00000025800 | KPNA6 | 3393.035629 | 0.019024708 | 0.053407355 | 0.35621888 | 0.721676654 | 0.845562207 |
| ENSG00000026025 | VIM | 2.597591514 | -0.824638835 | 1.115734736 | -0.739099365 | 0.459846662 | 0.64675789 |
| ENSG00000026036 | RTEL1-TNFRSF | 124.9295902 | -0.156426151 | 0.175460806 | -0.891516204 | 0.372652302 | 0.565349105 |
| ENSG00000026103 | FAS | 77.0612507 | -0.35939426 | 0.212450417 | -1.691661827 | 0.090710471 | 0.195738707 |
| ENSG00000026297 | RNASET2 | 215.0407025 | 0.300165447 | 0.127000026 | 2.363506977 | 0.018102886 | 0.050112248 |
| ENSG00000026508 | CD44 | 1067.590016 | 0.645921634 | 0.067664152 | 9.545994733 | 1.35E-21 | 2.47E-20 |
| ENSG00000026559 | KCNG1 | 74.82947869 | 0.612661969 | 0.209838038 | 2.919689755 | 0.0035038 | 0.011704846 |
| ENSG00000026652 | AGPAT4 | 12.49099575 | -2.985059722 | 0.634044586 | -4.707965005 | 2.50E-06 | 1.38E-05 |
| ENSG00000026950 | BTN3A1 | 5.977633784 | 0.42139768 | 0.726058287 | 0.580390978 | 0.561650988 | 0.730585669 |
| ENSG00000027001 | MIPEP | 371.1216548 | 0.099697682 | 0.103235444 | 0.965731134 | 0.334178736 | 0.525144555 |
| ENSG00000027075 | PRKCH | 1022.667605 | -0.531136995 | 0.066011925 | -8.046076448 | 8.55E-16 | 1.09E-14 |
| ENSG00000027697 | IFNGR1 | 660.8084277 | 0.083505944 | 0.07847436 | 1.064117553 | 0.28727545 | 0.471259927 |
| ENSG00000027847 | B4GALT7 | 637.5255718 | -0.411759266 | 0.083831427 | -4.911753014 | 9.03E-07 | 5.26E-06 |
| ENSG00000028116 | VRK2 | 454.7478639 | 0.240976887 | 0.091385359 | 2.636931022 | 0.008365984 | 0.025405835 |
| ENSG00000028203 | VEZT | 2024.227899 | -0.090017908 | 0.057146246 | -1.575219989 | 0.115205675 | 0.237663505 |
| ENSG00000028277 | POU2F2 | 16.67338414 | -0.267630542 | 0.435225592 | -0.614923724 | 0.538605095 | 0.713330689 |

| | | | | | | | |
|---|---|---|---|---|---|---|---|
| ENSG00000028310 | BRD9 | 1881.941939 | -0.018318329 | 0.061019799 | -0.300203033 | 0.764022292 | 0.870544448 |
| ENSG00000028528 | SNX1 | 4459.483476 | -0.184624344 | 0.047636654 | -3.875678233 | 0.000106328 | 0.000471975 |
| ENSG00000028839 | TBPL1 | 311.581486 | 0.424008317 | 0.115798079 | 3.66161787 | 0.000250628 | 0.001043447 |
| ENSG00000029153 | BMAL2 | 536.7664416 | 0.398642738 | 0.087500872 | 4.555871588 | 5.22E-06 | 2.78E-05 |
| ENSG00000029363 | BCLAF1 | 4893.020849 | 0.236152477 | 0.058927651 | 4.007498561 | 6.14E-05 | 0.000283471 |
| ENSG00000029364 | SLC39A9 | 3642.52706 | -0.664614818 | 0.052301825 | -12.70729689 | 5.39E-37 | 1.96E-35 |
| ENSG00000029534 | ANK1 | 22.08453119 | -0.715241227 | 0.371755703 | -1.923954956 | 0.05436023 | 0.128784847 |
| ENSG00000029559 | IBSP | 30.00059531 | 0.56929883 | 0.324836608 | 1.752569806 | 0.079675878 | 0.17611916 |
| ENSG00000029639 | TFB1M | 166.4928952 | 0.198568422 | 0.141773673 | 1.400601522 | 0.161333266 | 0.308235622 |
| ENSG00000029725 | RABEP1 | 2700.36563 | 0.058160999 | 0.052016972 | 1.118115814 | 0.263517533 | 0.444165482 |
| ENSG00000029993 | HMGB3 | 6971.730268 | -0.824056274 | 0.047573188 | -17.32186343 | 3.22E-67 | 2.71E-65 |
| ENSG00000030066 | NUP160 | 3443.182904 | 0.038345722 | 0.05498736 | 0.6973552 | 0.485580531 | 0.669321308 |
| ENSG00000030110 | BAK1 | 1923.720356 | 0.71321617 | 0.063785821 | 11.18142183 | 5.03E-29 | 1.34E-27 |
| ENSG00000030419 | IKZF2 | 229.9177711 | -0.459594711 | 0.128208413 | -3.584746906 | 0.000337405 | 0.001374792 |
| ENSG00000030582 | GRN | 13934.51179 | 0.111240621 | 0.052264192 | 2.128428962 | 0.033301534 | 0.084938536 |
| ENSG00000031003 | FAM13B | 1824.461108 | -0.17378627 | 0.055703095 | -3.119867376 | 0.001809325 | 0.006415133 |
| ENSG00000031081 | ARHGAP31 | 295.5579276 | 1.308802182 | 0.115739786 | 11.30814416 | 1.20E-29 | 3.28E-28 |
| ENSG00000031691 | CENPQ | 203.9635911 | -0.212083648 | 0.137967787 | -1.537196848 | 0.124245112 | 0.252273222 |
| ENSG00000031698 | SARS1 | 6467.07079 | 0.064500005 | 0.04270688 | 1.510295437 | 0.130968055 | 0.262921781 |
| ENSG00000031823 | RANBP3 | 1452.879027 | 0.079096716 | 0.060856825 | 1.29971807 | 0.193697615 | 0.354453038 |
| ENSG00000032219 | ARID4A | 1815.373549 | -0.569509795 | 0.062666164 | -9.087995165 | 1.01E-19 | 1.66E-18 |
| ENSG00000032389 | EIPR1 | 1286.338528 | -0.062932609 | 0.061858501 | -1.017363952 | 0.308980324 | 0.497118881 |
| ENSG00000032444 | PNPLA6 | 1025.089842 | -0.181593732 | 0.070253469 | -2.584836495 | 0.009742519 | 0.029059079 |
| ENSG00000032742 | IFT88 | 558.8955537 | 0.059046464 | 0.092575791 | 0.637817549 | 0.523592455 | 0.700001783 |
| ENSG00000033011 | ALG1 | 812.4170721 | -0.179150707 | 0.074024052 | -2.420168884 | 0.0155133 | 0.043826984 |
| ENSG00000033030 | ZCCHC8 | 930.1615072 | -0.388679426 | 0.069204313 | -5.616404692 | 1.95E-08 | 1.36E-07 |
| ENSG00000033050 | ABCF2 | 661.2225933 | 0.523462475 | 0.080293481 | 6.519364593 | 7.06E-11 | 6.18E-10 |
| ENSG00000033100 | CHPF2 | 1673.165452 | 0.639858931 | 0.06099366 | 10.49058109 | 9.54E-26 | 2.19E-24 |
| ENSG00000033122 | LRRC7 | 28.57588631 | 3.023795382 | 0.438845161 | 6.890346876 | 5.57E-12 | 5.34E-11 |
| ENSG00000033170 | FUT8 | 6832.896133 | -0.64285019 | 0.048479049 | -13.26037133 | 3.93E-40 | 1.61E-38 |
| ENSG00000033178 | UBA6 | 1646.993964 | -0.236246681 | 0.066000929 | -3.579444791 | 0.000344325 | 0.001399507 |
| ENSG00000033327 | GAB2 | 1434.096414 | 0.162135413 | 0.062639642 | 2.588383476 | 0.009642757 | 0.028810216 |
| ENSG00000033627 | ATP6V0A1 | 2490.892515 | 0.147886646 | 0.051272509 | 2.884326305 | 0.003922521 | 0.012950867 |

| | | | | | | | |
|---|---|---|---|---|---|---|---|
| ENSG00000033800 | PIAS1 | 1536.267801 | 0.004880674 | 0.064304757 | 0.075899117 | 0.939499359 | 0.971233034 |
| ENSG00000033867 | SLC4A7 | 1477.695158 | 0.665785678 | 0.065853943 | 10.11003515 | 4.99E-24 | 1.05E-22 |
| ENSG00000034053 | APBA2 | 78.6622066 | 1.255181683 | 0.21037305 | 5.966456641 | 2.42E-09 | 1.86E-08 |
| ENSG00000034152 | MAP2K3 | 1788.965386 | 0.005707469 | 0.064617253 | 0.088327326 | 0.929616521 | 0.965993019 |
| ENSG00000034510 | TMSB10 | 6982.045955 | -0.463606214 | 0.045917595 | -10.09648292 | 5.73E-24 | 1.20E-22 |
| ENSG00000034533 | ASTE1 | 278.7768011 | -0.316988518 | 0.113744503 | -2.786846928 | 0.005322361 | 0.017021583 |
| ENSG00000034677 | RNF19A | 1980.842713 | -0.487701565 | 0.054147875 | -9.006845838 | 2.12E-19 | 3.43E-18 |
| ENSG00000034693 | PEX3 | 360.7655294 | 0.257742505 | 0.103016015 | 2.501965401 | 0.012350599 | 0.035809999 |
| ENSG00000034713 | GABARAPL2 | 2381.703582 | 0.555213103 | 0.052054917 | 10.66591086 | 1.47E-26 | 3.55E-25 |
| ENSG00000035115 | SH3YL1 | 1629.489082 | 0.257598723 | 0.059281655 | 4.34533622 | 1.39E-05 | 7.04E-05 |
| ENSG00000035141 | FAM136A | 2899.545335 | 0.008992513 | 0.055234283 | 0.162806736 | 0.870670604 | 0.932942429 |
| ENSG00000035403 | VCL | 2244.508639 | -0.536212413 | 0.054957138 | -9.75692025 | 1.72E-22 | 3.32E-21 |
| ENSG00000035499 | DEPDC1B | 735.8976414 | -0.592851015 | 0.07780096 | -7.62009896 | 2.53E-14 | 2.90E-13 |
| ENSG00000035664 | DAPK2 | 217.6316516 | 0.362420342 | 0.127894271 | 2.833749612 | 0.004600536 | 0.014950602 |
| ENSG00000035681 | NSMAF | 1438.098555 | -0.079824226 | 0.067448662 | -1.183481248 | 0.236618474 | 0.411744504 |
| ENSG00000035687 | ADSS2 | 1274.658019 | -0.017124734 | 0.06432191 | -0.266234852 | 0.790058343 | 0.886605444 |
| ENSG00000035862 | TIMP2 | 258.223952 | 1.960327572 | 0.130755657 | 14.99229645 | 8.25E-51 | 4.67E-49 |
| ENSG00000035928 | RFC1 | 2938.690243 | -0.107608342 | 0.05213396 | -2.064073809 | 0.039010719 | 0.097186071 |
| ENSG00000036054 | TBC1D23 | 1028.411689 | -0.040515433 | 0.065755188 | -0.616155679 | 0.537791777 | 0.712782729 |
| ENSG00000036257 | CUL3 | 2497.194959 | -0.050926021 | 0.05104162 | -0.99773521 | 0.318407775 | 0.507523232 |
| ENSG00000036448 | MYOM2 | 5.829078544 | 1.685154121 | 0.788315737 | 2.137663935 | 0.032544029 | 0.083330731 |
| ENSG00000036530 | CYP46A1 | 47.29026862 | 0.222697832 | 0.273552663 | 0.814094913 | 0.41559058 | 0.607741946 |
| ENSG00000036549 | ZZZ3 | 2619.28992 | 0.112671646 | 0.051201696 | 2.200545196 | 0.027768237 | 0.07270068 |
| ENSG00000036672 | USP2 | 239.2584425 | 0.477897302 | 0.124490684 | 3.838819803 | 0.000123627 | 0.00054288 |
| ENSG00000037042 | TUBG2 | 552.4468147 | -0.091626711 | 0.1772809 | -0.516844795 | 0.605264507 | 0.764032785 |
| ENSG00000037241 | RPL26L1 | 588.5713028 | -0.442302436 | 0.081182576 | -5.448243434 | 5.09E-08 | 3.40E-07 |
| ENSG00000037280 | FLT4 | 55.27439312 | -0.90219692 | 0.245456625 | -3.675585937 | 0.000237304 | 0.000993024 |
| ENSG00000037474 | NSUN2 | 4355.902574 | 0.098821407 | 0.048056107 | 2.056375613 | 0.039746325 | 0.098646064 |
| ENSG00000037637 | FBXO42 | 763.0409804 | -0.026455914 | 0.074639699 | -0.354448295 | 0.723002947 | 0.845926239 |
| ENSG00000037749 | MFAP3 | 947.0515061 | -0.018222615 | 0.071418491 | -0.255152625 | 0.79860522 | 0.892570219 |
| ENSG00000037757 | MRI1 | 1062.15225 | 0.220992925 | 0.07943262 | 2.782143208 | 0.005400121 | 0.017236506 |
| ENSG00000037897 | METTL1 | 502.0641454 | 0.117260943 | 0.085424664 | 1.372682511 | 0.169851069 | 0.320412952 |
| ENSG00000037965 | HOXC8 | 57.09520855 | -0.205923894 | 0.242218113 | -0.85015894 | 0.395236726 | 0.588175835 |

| | | | | | | | |
|---|---|---|---|---|---|---|---|
| ENSG00000038002 | AGA | 367.5406742 | -0.247540903 | 0.10542154 | -2.348105552 | 0.018869173 | 0.052007549 |
| ENSG00000038210 | PI4K2B | 975.4426918 | -0.374364849 | 0.076372067 | -4.901855682 | 9.49E-07 | 5.51E-06 |
| ENSG00000038219 | BOD1L1 | 1971.666891 | 0.168741701 | 0.070352376 | 2.3985217 | 0.016461401 | 0.046153085 |
| ENSG00000038274 | MAT2B | 1288.796151 | -0.015567129 | 0.063266235 | -0.246057454 | 0.805637758 | 0.896847196 |
| ENSG00000038358 | EDC4 | 3224.734897 | -0.023220749 | 0.05280622 | -0.439735118 | 0.660128964 | 0.803531701 |
| ENSG00000038382 | TRIO | 4654.842178 | -0.278825059 | 0.051572735 | -5.406443107 | 6.43E-08 | 4.24E-07 |
| ENSG00000038532 | CLEC16A | 1150.99091 | 0.043866857 | 0.064401886 | 0.681142437 | 0.495781367 | 0.677941809 |
| ENSG00000039068 | CDH1 | 22604.64718 | 0.218610954 | 0.043139643 | 5.067518797 | 4.03E-07 | 2.45E-06 |
| ENSG00000039123 | MTREX | 1947.107756 | 0.203306637 | 0.057259117 | 3.55064222 | 0.000384292 | 0.001549054 |
| ENSG00000039139 | DNAH5 | 230.537726 | 0.194972711 | 0.144069136 | 1.353327411 | 0.175951056 | 0.329280243 |
| ENSG00000039319 | ZFYVE16 | 1778.459548 | 0.063468561 | 0.065580073 | 0.967802536 | 0.333142997 | 0.524271136 |
| ENSG00000039523 | RIPOR1 | 1739.750045 | -0.182602634 | 0.065936753 | -2.769360416 | 0.005616647 | 0.017853107 |
| ENSG00000039560 | RAI14 | 1079.171047 | 0.251312697 | 0.064953343 | 3.869126441 | 0.000109226 | 0.000484182 |
| ENSG00000039600 | SOX30 | 6.813711099 | -0.136048872 | 0.70382441 | -0.193299451 | 0.846724456 | 0.91973122 |
| ENSG00000039650 | PNKP | 1204.207303 | 0.221659752 | 0.068286784 | 3.246012444 | 0.001170337 | 0.004328851 |
| ENSG00000040199 | PHLPP2 | 1735.343239 | 0.180059059 | 0.063205014 | 2.848809745 | 0.004388311 | 0.014342194 |
| ENSG00000040275 | SPDL1 | 830.0055734 | -0.518489069 | 0.073243841 | -7.078944251 | 1.45E-12 | 1.47E-11 |
| ENSG00000040341 | STAU2 | 1672.879003 | 0.548841111 | 0.060039157 | 9.141386032 | 6.17E-20 | 1.02E-18 |
| ENSG00000040487 | SLC66A1 | 391.8933764 | 0.389496091 | 0.097837142 | 3.981065689 | 6.86E-05 | 0.000314392 |
| ENSG00000040531 | CTNS | 375.3035988 | -0.015995443 | 0.102455438 | -0.156120976 | 0.875937665 | 0.935616916 |
| ENSG00000040608 | RTN4R | 92.99448215 | -0.284480936 | 0.186838925 | -1.522599944 | 0.127858824 | 0.257973428 |
| ENSG00000040633 | PHF23 | 1292.870327 | -0.043337701 | 0.069826651 | -0.620646994 | 0.534831912 | 0.710342986 |
| ENSG00000040731 | CDH10 | 4.89138479 | -2.13557128 | 0.91938451 | -2.322827128 | 0.020188441 | 0.055126463 |
| ENSG00000040933 | INPP4A | 1042.33574 | 0.320616569 | 0.07049792 | 4.547886959 | 5.42E-06 | 2.88E-05 |
| ENSG00000041353 | RAB27B | 2159.087009 | 0.755624829 | 0.067544993 | 11.18698501 | 4.72E-29 | 1.26E-27 |
| ENSG00000041357 | PSMA4 | 4740.221716 | -0.275916467 | 0.053360661 | -5.17078425 | 2.33E-07 | 1.46E-06 |
| ENSG00000041515 | MYO16 | 21.95559132 | -1.767870805 | 0.412794301 | -4.282691892 | 1.85E-05 | 9.21E-05 |
| ENSG00000041802 | LSG1 | 1997.265482 | -0.086933468 | 0.055103855 | -1.577629499 | 0.114650751 | 0.236731614 |
| ENSG00000041880 | PARP3 | 231.8929931 | -0.021678203 | 0.125426069 | -0.172836504 | 0.862779943 | 0.928773904 |
| ENSG00000041982 | TNC | 3.811217113 | 2.149704593 | 1.026153189 | 2.094915862 | 0.036178473 | 0.091195771 |
| ENSG00000041988 | THAP3 | 278.2176784 | 0.098858332 | 0.111651784 | 0.885416505 | 0.375932034 | 0.568857577 |
| ENSG00000042062 | RIPOR3 | 285.7826217 | -1.597046578 | 0.120923363 | -13.20709698 | 7.98E-40 | 3.22E-38 |
| ENSG00000042088 | TDP1 | 1536.129557 | -0.762906091 | 0.059187195 | -12.88971517 | 5.14E-38 | 1.94E-36 |

| | | | | | | | |
|---|---|---|---|---|---|---|---|
| ENSG00000042286 | AIFM2 | 1088.920182 | -0.696836073 | 0.066735861 | -10.44170354 | 1.60E-25 | 3.64E-24 |
| ENSG00000042317 | SPATA7 | 296.1863724 | -0.347798796 | 0.110938512 | -3.135059152 | 0.001718195 | 0.006122303 |
| ENSG00000042429 | MED17 | 88.46463863 | 0.132186324 | 0.198508925 | 0.665896127 | 0.505477496 | 0.685970999 |
| ENSG00000042445 | RETSAT | 1508.834762 | -0.265482355 | 0.059739277 | -4.444016854 | 8.83E-06 | 4.59E-05 |
| ENSG00000042493 | CAPG | 226.5908221 | 1.475220357 | 0.139229577 | 10.59559604 | 3.12E-26 | 7.39E-25 |
| ENSG00000042753 | AP2S1 | 3057.584463 | -0.049668311 | 0.051185374 | -0.97036139 | 0.331866387 | 0.522871904 |
| ENSG00000042813 | ZPBP | 3.159211338 | 1.418231325 | 1.082891394 | 1.309670881 | 0.190307199 | 0.349587375 |
| ENSG00000042832 | TG | 5.425246069 | 0.726337539 | 0.780945661 | 0.930074364 | 0.352332583 | 0.543727512 |
| ENSG00000042980 | ADAM28 | 2.579108898 | 4.723505697 | 1.505323246 | 3.137868037 | 0.001701815 | 0.006067708 |
| ENSG00000043039 | BARX2 | 304.3436814 | 0.483124806 | 0.109424786 | 4.415131386 | 1.01E-05 | 5.22E-05 |
| ENSG00000043093 | DCUN1D1 | 1293.816008 | -0.150663307 | 0.071639399 | -2.103078875 | 0.035458875 | 0.089584427 |
| ENSG00000043143 | JADE2 | 2043.704083 | -0.185716448 | 0.062687627 | -2.962569453 | 0.00305083 | 0.01031938 |
| ENSG00000043355 | ZIC2 | 152.6239734 | 0.995924723 | 0.158696718 | 6.275647898 | 3.48E-10 | 2.89E-09 |
| ENSG00000043514 | TRIT1 | 500.7089063 | 0.221649814 | 0.088963159 | 2.49147869 | 0.01272126 | 0.036730736 |
| ENSG00000043591 | ADRB1 | 18.42093381 | -0.325098088 | 0.405140987 | -0.802431989 | 0.422303115 | 0.613296624 |
| ENSG00000044090 | CUL7 | 2520.930224 | 0.079669249 | 0.056264192 | 1.415984947 | 0.156779912 | 0.30109432 |
| ENSG00000044115 | CTNNA1 | 16166.08366 | -0.100532736 | 0.043403539 | -2.316233616 | 0.020545517 | 0.055948973 |
| ENSG00000044446 | PHKA2 | 1591.594854 | -0.062271705 | 0.06166879 | -1.009776655 | 0.312602307 | 0.500805847 |
| ENSG00000044459 | CNTLN | 75.25449953 | -0.250546658 | 0.202785879 | -1.235523195 | 0.21663585 | 0.386050368 |
| ENSG00000044524 | EPHA3 | 154.4035927 | -0.232789502 | 0.146796396 | -1.58579849 | 0.112785028 | 0.233573039 |
| ENSG00000044574 | HSPA5 | 10962.5252 | 0.053403495 | 0.041514512 | 1.286381379 | 0.198309994 | 0.360821007 |
| ENSG00000046604 | DSG2 | 1913.83258 | 0.258018472 | 0.059135742 | 4.363156069 | 1.28E-05 | 6.52E-05 |
| ENSG00000046647 | GEMIN8 | 434.7760602 | -0.006405083 | 0.093786795 | -0.068294083 | 0.945551534 | 0.974261906 |
| ENSG00000046651 | OFD1 | 900.8861264 | -0.020176044 | 0.074593787 | -0.270478881 | 0.786791862 | 0.884321672 |
| ENSG00000046653 | GPM6B | 20.94690134 | 0.500355054 | 0.396393511 | 1.262268529 | 0.206852177 | 0.372756397 |
| ENSG00000047056 | WDR37 | 1238.679269 | -1.038571716 | 0.065420521 | -15.87532013 | 9.39E-57 | 6.17E-55 |
| ENSG00000047188 | YTHDC2 | 1715.044256 | 1.000740968 | 0.072367806 | 13.82853814 | 1.71E-43 | 7.84E-42 |
| ENSG00000047230 | CTPS2 | 1386.530244 | -0.345808583 | 0.06062616 | -5.703950007 | 1.17E-08 | 8.34E-08 |
| ENSG00000047249 | ATP6V1H | 1581.575866 | -0.001440971 | 0.057637263 | -0.025000691 | 0.980054413 | 0.991041209 |
| ENSG00000047315 | POLR2B | 4750.754264 | 0.079741318 | 0.052443152 | 1.520528711 | 0.128378148 | 0.258832372 |
| ENSG00000047346 | ATOSA | 7584.906634 | -1.362221422 | 0.0527204 | -25.83860161 | 3.27E-147 | 1.14E-144 |
| ENSG00000047365 | ARAP2 | 448.5169668 | 0.36927655 | 0.093601864 | 3.945183726 | 7.97E-05 | 0.000361933 |
| ENSG00000047410 | TPR | 6666.67432 | -0.149622805 | 0.054348047 | -2.753048397 | 0.005904317 | 0.018664052 |

| | | | | | | | |
|---|---|---|---|---|---|---|---|
| ENSG00000047457 | CP | 8.080419003 | -1.457325038 | 0.651975346 | -2.235245621 | 0.025401228 | 0.067333507 |
| ENSG00000047578 | KATNIP | 1568.066983 | -0.014559092 | 0.05947451 | -0.244795493 | 0.806614784 | 0.897382528 |
| ENSG00000047579 | DTNBP1 | 716.5394219 | -0.365454393 | 0.080775833 | -4.524303599 | 6.06E-06 | 3.21E-05 |
| ENSG00000047597 | XK | 62.81746711 | -0.116351243 | 0.228007748 | -0.510295126 | 0.609844717 | 0.767828639 |
| ENSG00000047621 | C12orf4 | 261.8818065 | 0.157573636 | 0.115963341 | 1.358822833 | 0.174202736 | 0.327022158 |
| ENSG00000047634 | SCML1 | 307.2874309 | -0.319997316 | 0.126919219 | -2.521267612 | 0.011693288 | 0.034145117 |
| ENSG00000047644 | WWC3 | 3638.298244 | 0.034108397 | 0.049470622 | 0.689467725 | 0.490528977 | 0.673681878 |
| ENSG00000047662 | FAM184B | 14.61485947 | -0.773122277 | 0.465416266 | -1.661141509 | 0.09668503 | 0.206321356 |
| ENSG00000047849 | MAP4 | 6336.56847 | 0.015068901 | 0.046168062 | 0.326392322 | 0.744127546 | 0.858493557 |
| ENSG00000047932 | GOPC | 800.1292698 | -0.170137525 | 0.076882713 | -2.212949024 | 0.02690116 | 0.070761681 |
| ENSG00000048028 | USP28 | 565.2114095 | 0.299785029 | 0.083527741 | 3.589047489 | 0.000331888 | 0.001354237 |
| ENSG00000048052 | HDAC9 | 44.60953448 | 0.477816082 | 0.274674697 | 1.7395708 | 0.08193441 | 0.180208568 |
| ENSG00000048140 | TSPAN17 | 2280.075878 | -0.056842377 | 0.05394464 | -1.053716872 | 0.292012562 | 0.476981097 |
| ENSG00000048162 | NOP16 | 1226.832167 | -0.273756438 | 0.065660332 | -4.169281985 | 3.06E-05 | 0.000147881 |
| ENSG00000048342 | CC2D2A | 291.308743 | 0.452178594 | 0.109290331 | 4.137407125 | 3.51E-05 | 0.000167869 |
| ENSG00000048392 | RRM2B | 2350.755648 | -0.182249675 | 0.05343643 | -3.410588531 | 0.000648228 | 0.002514008 |
| ENSG00000048405 | ZNF800 | 633.0573838 | 0.136846686 | 0.078496022 | 1.743358229 | 0.08127106 | 0.178952746 |
| ENSG00000048471 | SNX29 | 1178.978842 | 0.86264358 | 0.087674235 | 9.839191362 | 7.63E-23 | 1.51E-21 |
| ENSG00000048540 | LMO3 | 23.94061397 | 2.638255857 | 0.446886837 | 5.903632946 | 3.56E-09 | 2.69E-08 |
| ENSG00000048544 | MRPS10 | 1542.414096 | -0.013493708 | 0.060815013 | -0.221881204 | 0.824406363 | 0.907523488 |
| ENSG00000048649 | RSF1 | 2818.939421 | 0.044889526 | 0.059931745 | 0.749010834 | 0.453850676 | 0.641526557 |
| ENSG00000048707 | VPS13D | 1285.670191 | -0.090160318 | 0.079054297 | -1.140485984 | 0.254083889 | 0.433037475 |
| ENSG00000048828 | FAM120A | 9399.773649 | 0.353541755 | 0.043415215 | 8.143268569 | 3.85E-16 | 5.01E-15 |
| ENSG00000048991 | R3HDM1 | 2251.608022 | -0.09217596 | 0.056879359 | -1.620552006 | 0.105113752 | 0.220674428 |
| ENSG00000049089 | COL9A2 | 273.4209437 | -2.033371447 | 0.135775545 | -14.97597706 | 1.05E-50 | 5.94E-49 |
| ENSG00000049130 | KITLG | 1247.366948 | -0.194189004 | 0.073876696 | -2.628555617 | 0.008574833 | 0.025988463 |
| ENSG00000049167 | ERCC8 | 260.2533885 | -0.244807462 | 0.117151263 | -2.089669855 | 0.036647467 | 0.092201457 |
| ENSG00000049192 | ADAMTS6 | 26.6741802 | -1.622558128 | 0.368701862 | -4.400732121 | 1.08E-05 | 5.55E-05 |
| ENSG00000049239 | H6PD | 1477.658008 | 0.060398863 | 0.068842237 | 0.877351835 | 0.380295566 | 0.572774111 |
| ENSG00000049245 | VAMP3 | 1937.653304 | -0.013270739 | 0.056189637 | -0.236177691 | 0.81329479 | 0.901448243 |
| ENSG00000049246 | PER3 | 1360.262564 | -0.035888151 | 0.062082811 | -0.578069031 | 0.563217514 | 0.732023217 |
| ENSG00000049247 | UTS2 | 1.853379385 | -0.441804351 | 1.4032538 | -0.314842797 | 0.752881016 | 0.863727271 |
| ENSG00000049283 | EPN3 | 3185.110919 | -0.030086212 | 0.063172213 | -0.47625706 | 0.633891253 | 0.784236938 |

| | | | | | | | |
|---|---|---|---|---|---|---|---|
| ENSG00000049323 | LTBP1 | 2303.455306 | 0.288034105 | 0.064462549 | 4.468239483 | 7.89E-06 | 4.12E-05 |
| ENSG00000049449 | RCN1 | 482.6283428 | 0.236366864 | 0.088789239 | 2.662111601 | 0.007765213 | 0.023821073 |
| ENSG00000049541 | RFC2 | 1345.132516 | 0.291052407 | 0.062166379 | 4.681829787 | 2.84E-06 | 1.56E-05 |
| ENSG00000049618 | ARID1B | 2038.638869 | -0.013188319 | 0.063652412 | -0.207192758 | 0.835859323 | 0.913370821 |
| ENSG00000049656 | CLPTM1L | 4429.694116 | 0.16912716 | 0.047240278 | 3.580147442 | 0.0003434 | 0.001396243 |
| ENSG00000049759 | NEDD4L | 1645.485481 | 0.493774821 | 0.065610476 | 7.525853381 | 5.24E-14 | 5.84E-13 |
| ENSG00000049768 | FOXP3 | 29.91474561 | -0.902790362 | 0.340224685 | -2.653512227 | 0.007965888 | 0.024363365 |
| ENSG00000049769 | PPP1R3F | 218.8039154 | -0.188375722 | 0.132546112 | -1.42120896 | 0.155256021 | 0.298967381 |
| ENSG00000049860 | HEXB | 1639.869255 | 0.582076097 | 0.057501991 | 10.12271202 | 4.38E-24 | 9.23E-23 |
| ENSG00000049883 | PTCD2 | 506.6124152 | -0.10006676 | 0.087527084 | -1.143266236 | 0.252928069 | 0.431735218 |
| ENSG00000050030 | NEXMIF | 130.0366438 | 0.70708872 | 0.178677056 | 3.957356007 | 7.58E-05 | 0.000345211 |
| ENSG00000050130 | JKAMP | 1114.116969 | -0.666695748 | 0.067856141 | -9.825135018 | 8.78E-23 | 1.73E-21 |
| ENSG00000050165 | DKK3 | 4.545784792 | -0.788058776 | 0.863252686 | -0.91289467 | 0.361297941 | 0.553874283 |
| ENSG00000050327 | ARHGEF5 | 507.5519806 | 0.412209353 | 0.093789199 | 4.395062071 | 1.11E-05 | 5.69E-05 |
| ENSG00000050344 | NFE2L3 | 49.32123182 | -0.760843522 | 0.252275397 | -3.015924388 | 0.002561971 | 0.008800782 |
| ENSG00000050393 | MCUR1 | 1664.468386 | -0.049406129 | 0.063206297 | -0.781664662 | 0.434411675 | 0.624359615 |
| ENSG00000050405 | LIMA1 | 1615.429987 | -0.232582944 | 0.061675678 | -3.771064234 | 0.000162553 | 0.000700341 |
| ENSG00000050426 | LETMD1 | 1601.42581 | 0.199272835 | 0.062705457 | 3.177918538 | 0.001483364 | 0.005359643 |
| ENSG00000050438 | SLC4A8 | 330.8644705 | -0.773708242 | 0.108053773 | -7.160400081 | 8.04E-13 | 8.30E-12 |
| ENSG00000050555 | LAMC3 | 4.00030688 | -0.701973182 | 0.89209387 | -0.786882643 | 0.431350569 | 0.621444037 |
| ENSG00000050628 | PTGER3 | 57.84029058 | -1.44679014 | 0.249007668 | -5.810223234 | 6.24E-09 | 4.59E-08 |
| ENSG00000050748 | MAPK9 | 3465.349856 | -0.197722687 | 0.050213514 | -3.937638906 | 8.23E-05 | 0.000372469 |
| ENSG00000050767 | COL23A1 | 36.06781602 | 0.79808236 | 0.3062634 | 2.605869193 | 0.009164145 | 0.027570556 |
| ENSG00000050820 | BCAR1 | 2838.817627 | 0.531437674 | 0.060789209 | 8.742302832 | 2.28E-18 | 3.45E-17 |
| ENSG00000051009 | FHIP1B | 671.1650388 | -0.277771208 | 0.07760799 | -3.579157352 | 0.000344704 | 0.001400798 |
| ENSG00000051108 | HERPUD1 | 1284.188529 | 0.04647663 | 0.062751685 | 0.740643548 | 0.458909596 | 0.646137686 |
| ENSG00000051128 | HOMER3 | 106.8959641 | 0.758479853 | 0.186865046 | 4.058971273 | 4.93E-05 | 0.000230943 |
| ENSG00000051180 | RAD51 | 741.0915071 | -0.37373535 | 0.073378082 | -5.093283141 | 3.52E-07 | 2.15E-06 |
| ENSG00000051341 | POLQ | 773.5006434 | -0.158357347 | 0.074381921 | -2.128976305 | 0.033256221 | 0.084832393 |
| ENSG00000051382 | PIK3CB | 1762.006362 | -0.135802121 | 0.058360889 | -2.326937159 | 0.01996861 | 0.05460483 |
| ENSG00000051523 | CYBA | 673.8369171 | 1.696900507 | 0.091093669 | 18.62808378 | 1.90E-77 | 2.12E-75 |
| ENSG00000051596 | THOC3 | 682.8889408 | -0.471943692 | 0.077578258 | -6.083453102 | 1.18E-09 | 9.30E-09 |
| ENSG00000051620 | HEBP2 | 2086.685476 | 0.076197773 | 0.053877617 | 1.414275114 | 0.157281141 | 0.301851365 |

| | | | | | | | |
|---|---|---|---|---|---|---|---|
| ENSG00000051825 | MPHOSPH9 | 1239.794913 | -0.236084937 | 0.064937438 | -3.635575173 | 0.000277361 | 0.001148495 |
| ENSG00000052126 | PLEKHA5 | 1118.662947 | -0.370668719 | 0.070668843 | -5.245150562 | 1.56E-07 | 9.95E-07 |
| ENSG00000052344 | PRSS8 | 2748.840693 | 0.409786433 | 0.063924431 | 6.410482301 | 1.45E-10 | 1.24E-09 |
| ENSG00000052723 | SIKE1 | 5814.572619 | 0.3954217 | 0.058418586 | 6.768765292 | 1.30E-11 | 1.20E-10 |
| ENSG00000052749 | RRP12 | 2189.693477 | -0.509663315 | 0.05474094 | -9.31045963 | 1.27E-20 | 2.20E-19 |
| ENSG00000052795 | FNIP2 | 268.6016001 | -0.715507966 | 0.121140412 | -5.906434961 | 3.50E-09 | 2.65E-08 |
| ENSG00000052802 | MSMO1 | 1587.586562 | -0.765478529 | 0.062511767 | -12.2453511 | 1.78E-34 | 5.95E-33 |
| ENSG00000052841 | TTC17 | 1916.031086 | 0.108431815 | 0.061508237 | 1.762882849 | 0.07792025 | 0.173091757 |
| ENSG00000053108 | FSTL4 | 34.29097043 | -2.37981859 | 0.351642833 | -6.767715336 | 1.31E-11 | 1.21E-10 |
| ENSG00000053254 | FOXN3 | 2304.783437 | -0.076785566 | 0.069290134 | -1.108174594 | 0.267786421 | 0.449084034 |
| ENSG00000053371 | AKR7A2 | 1218.399257 | 0.091747324 | 0.066778014 | 1.37391514 | 0.169468029 | 0.320018039 |
| ENSG00000053372 | MRTO4 | 1369.221671 | -0.014342674 | 0.061269641 | -0.234091043 | 0.814914299 | 0.902008879 |
| ENSG00000053501 | USE1 | 286.489285 | -0.019804101 | 0.113259875 | -0.174855404 | 0.861193253 | 0.927806003 |
| ENSG00000053524 | MCF2L2 | 66.34253056 | 0.154495143 | 0.219371429 | 0.704262832 | 0.481269109 | 0.665720776 |
| ENSG00000053747 | LAMA3 | 63.44996101 | -0.125917861 | 0.222158585 | -0.566792685 | 0.570855041 | 0.737769839 |
| ENSG00000053770 | AP5M1 | 1285.824067 | -0.54524925 | 0.066097274 | -8.2491942 | 1.59E-16 | 2.15E-15 |
| ENSG00000053900 | ANAPC4 | 784.1493139 | -0.1161759 | 0.077511108 | -1.498829049 | 0.133917987 | 0.267439594 |
| ENSG00000053918 | KCNQ1 | 2.556861587 | -1.240737625 | 1.168534348 | -1.061789606 | 0.288331209 | 0.472316334 |
| ENSG00000054116 | TRAPPC3 | 1637.663078 | 0.098670689 | 0.060473506 | 1.631635 | 0.1027564 | 0.21673702 |
| ENSG00000054118 | THRAP3 | 6259.077782 | -0.352905598 | 0.042257493 | -8.351314169 | 6.75E-17 | 9.27E-16 |
| ENSG00000054148 | PHPT1 | 1895.585786 | -0.215946933 | 0.057651852 | -3.745706769 | 0.000179887 | 0.000769601 |
| ENSG00000054179 | ENTPD2 | 558.0737248 | 0.726011778 | 0.086705383 | 8.373318387 | 5.60E-17 | 7.72E-16 |
| ENSG00000054267 | ARID4B | 1698.309161 | -0.022946463 | 0.061011769 | -0.376098963 | 0.706843325 | 0.836412774 |
| ENSG00000054277 | OPN3 | 225.1447734 | -0.011589434 | 0.124797469 | -0.092865936 | 0.926010068 | 0.964567541 |
| ENSG00000054282 | SDCCAG8 | 407.6600641 | 0.14170777 | 0.10361665 | 1.367615832 | 0.171432358 | 0.322851843 |
| ENSG00000054392 | HHAT | 205.0935103 | 0.262315146 | 0.126521147 | 2.073290923 | 0.038145213 | 0.09540631 |
| ENSG00000054523 | KIF1B | 1703.404908 | -0.363671458 | 0.063602599 | -5.717871039 | 1.08E-08 | 7.73E-08 |
| ENSG00000054598 | FOXC1 | 78.1330105 | 0.709889273 | 0.220782455 | 3.215333717 | 0.00130293 | 0.004774633 |
| ENSG00000054611 | TBC1D22A | 790.7910706 | 0.150846029 | 0.074612128 | 2.021736063 | 0.043203626 | 0.105738951 |
| ENSG00000054654 | SYNE2 | 1802.426411 | -0.805940204 | 0.072331802 | -11.14226634 | 7.81E-29 | 2.07E-27 |
| ENSG00000054690 | PLEKHH1 | 1533.694583 | -0.173665761 | 0.064290115 | -2.701282467 | 0.006907265 | 0.021444398 |
| ENSG00000054793 | ATP9A | 8484.117357 | 0.369260664 | 0.051748492 | 7.135679697 | 9.63E-13 | 9.89E-12 |
| ENSG00000054965 | FAM168A | 3675.966823 | 0.033278379 | 0.054877517 | 0.606411892 | 0.544241274 | 0.717770198 |

| | | | | | | | |
|---|---|---|---|---|---|---|---|
| ENSG00000054967 | RELT | 335.6831366 | 0.299248629 | 0.102999905 | 2.905329173 | 0.00366867 | 0.012189933 |
| ENSG00000054983 | GALC | 911.3455164 | -0.330364329 | 0.073676319 | -4.483996121 | 7.33E-06 | 3.84E-05 |
| ENSG00000055044 | NOP58 | 2370.125232 | 0.050930289 | 0.068298155 | 0.745705187 | 0.455845525 | 0.643117857 |
| ENSG00000055070 | SZRD1 | 4581.487075 | 0.06198434 | 0.055342273 | 1.120017963 | 0.262706108 | 0.443416828 |
| ENSG00000055118 | KCNH2 | 318.5660872 | 1.157971842 | 0.112605143 | 10.28347203 | 8.37E-25 | 1.84E-23 |
| ENSG00000055130 | CUL1 | 2505.000103 | 0.555978434 | 0.051006096 | 10.90023499 | 1.15E-27 | 2.94E-26 |
| ENSG00000055147 | FAM114A2 | 520.6075756 | -0.144329148 | 0.087234624 | -1.654493839 | 0.098027214 | 0.208622719 |
| ENSG00000055163 | CYFIP2 | 1996.836994 | 0.780519416 | 0.060671789 | 12.8646184 | 7.12E-38 | 2.68E-36 |
| ENSG00000055208 | TAB2 | 4290.88924 | 0.204495084 | 0.046323561 | 4.41449405 | 1.01E-05 | 5.23E-05 |
| ENSG00000055211 | GINM1 | 1338.337285 | 0.557545383 | 0.065253993 | 8.544234018 | 1.29E-17 | 1.86E-16 |
| ENSG00000055332 | EIF2AK2 | 1172.947392 | 0.403180631 | 0.069404114 | 5.809174837 | 6.28E-09 | 4.62E-08 |
| ENSG00000055483 | USP36 | 2828.928917 | 0.436432845 | 0.054456471 | 8.014343114 | 1.11E-15 | 1.40E-14 |
| ENSG00000055609 | KMT2C | 2791.540841 | 0.051190316 | 0.143745219 | 0.356118388 | 0.721751907 | 0.845596895 |
| ENSG00000055732 | MCOLN3 | 154.9116882 | -0.15547483 | 0.162364437 | -0.957567024 | 0.338281134 | 0.529559793 |
| ENSG00000055813 | CCDC85A | 100.9165334 | 0.148691372 | 0.176186333 | 0.84394385 | 0.398700783 | 0.591597444 |
| ENSG00000055917 | PUM2 | 4774.778385 | 0.008536323 | 0.049046759 | 0.17404458 | 0.861830427 | 0.928215087 |
| ENSG00000055950 | MRPL43 | 938.2293501 | -0.014864307 | 0.070221077 | -0.211678704 | 0.832357704 | 0.911334752 |
| ENSG00000055955 | ITIH4 | 1.769642685 | 0.071859664 | 1.302146957 | 0.055185525 | 0.955990661 | NA |
| ENSG00000056050 | HPF1 | 242.5078152 | -0.287776095 | 0.126190943 | -2.280481375 | 0.022579154 | 0.060799291 |
| ENSG00000056097 | ZFR | 4554.716793 | -0.094281982 | 0.048403699 | -1.94782599 | 0.051435788 | 0.122708331 |
| ENSG00000056277 | ZNF280C | 285.8540217 | -0.061794024 | 0.110033573 | -0.561592448 | 0.574393727 | 0.740858848 |
| ENSG00000056487 | PHF21B | 116.5729504 | 0.658713561 | 0.174512531 | 3.774591746 | 0.00016027 | 0.000691219 |
| ENSG00000056558 | TRAF1 | 61.98791037 | -0.603235001 | 0.229832651 | -2.624670594 | 0.008673283 | 0.026241772 |
| ENSG00000056586 | RC3H2 | 2395.003556 | -0.076226615 | 0.061694632 | -1.235546957 | 0.216627012 | 0.386050368 |
| ENSG00000056736 | IL17RB | 280.0337541 | -0.118565582 | 0.118343981 | -1.001872517 | 0.316405168 | 0.505194387 |
| ENSG00000056972 | TRAF3IP2 | 941.2090272 | 0.686073429 | 0.076855432 | 8.926804654 | 4.38E-19 | 6.94E-18 |
| ENSG00000056998 | GYG2 | 385.6793084 | -0.164088003 | 0.098241345 | -1.670254035 | 0.094869113 | 0.203107556 |
| ENSG00000057019 | DCBLD2 | 1949.489296 | -0.089967312 | 0.054688484 | -1.645086962 | 0.099951879 | 0.211854024 |
| ENSG00000057252 | SOAT1 | 900.698568 | -0.320772874 | 0.06962206 | -4.607345357 | 4.08E-06 | 2.20E-05 |
| ENSG00000057294 | PKP2 | 617.9819547 | -0.320818797 | 0.082764135 | -3.876302171 | 0.000106056 | 0.000470949 |
| ENSG00000057468 | MSH4 | 4.476380092 | 0.927401694 | 0.847293193 | 1.094546376 | 0.27371542 | 0.455667442 |
| ENSG00000057593 | F7 | 31.62504387 | 0.249356766 | 0.320816362 | 0.777257009 | 0.437007153 | 0.626676508 |
| ENSG00000057608 | GDI2 | 7804.208197 | -1.082422887 | 0.049678693 | -21.78847348 | 2.98E-105 | 5.48E-103 |

| | | | | | | | |
|---|---|---|---|---|---|---|---|
| ENSG00000057663 | ATG5 | 977.610784 | -0.131181571 | 0.069425262 | -1.889536573 | 0.058819967 | 0.137196751 |
| ENSG00000057704 | TMCC3 | 23.72581299 | -0.310698856 | 0.366552756 | -0.847623843 | 0.396647491 | 0.589548623 |
| ENSG00000057757 | PITHD1 | 956.9126757 | 0.046597703 | 0.067059422 | 0.694871821 | 0.487135633 | 0.670520307 |
| ENSG00000057935 | MTA3 | 1583.068818 | 0.15387679 | 0.057453085 | 2.678303349 | 0.007399617 | 0.022804744 |
| ENSG00000058056 | USP13 | 1648.614529 | 0.099502771 | 0.060354984 | 1.648625608 | 0.099224359 | 0.210681858 |
| ENSG00000058063 | ATP11B | 1766.530575 | -0.454549977 | 0.063347254 | -7.175527767 | 7.20E-13 | 7.46E-12 |
| ENSG00000058085 | LAMC2 | 40.84076523 | 1.047823831 | 0.295640212 | 3.544253418 | 0.000393727 | 0.001584577 |
| ENSG00000058091 | CDK14 | 49.34996877 | -1.288877204 | 0.261879724 | -4.921638011 | 8.58E-07 | 5.02E-06 |
| ENSG00000058262 | SEC61A1 | 12317.1709 | 0.175499861 | 0.041479444 | 4.231007987 | 2.33E-05 | 0.000114354 |
| ENSG00000058272 | PPP1R12A | 2522.079849 | -0.177268127 | 0.053434919 | -3.317458513 | 0.000908404 | 0.003427996 |
| ENSG00000058335 | RASGRF1 | 124.3203314 | -9.546804718 | 1.186056234 | -8.049200744 | 8.33E-16 | 1.06E-14 |
| ENSG00000058404 | CAMK2B | 1002.920048 | 1.659648357 | 0.082737772 | 20.05913767 | 1.68E-89 | 2.29E-87 |
| ENSG00000058453 | CROCC | 466.2396573 | 0.077938409 | 0.088639716 | 0.879271868 | 0.379253885 | 0.571734516 |
| ENSG00000058600 | POLR3E | 1497.248205 | 0.235523378 | 0.0617946 | 3.811390928 | 0.000138187 | 0.000602091 |
| ENSG00000058668 | ATP2B4 | 1208.914094 | 1.985712481 | 0.070725899 | 28.07617174 | 1.91E-173 | 8.45E-171 |
| ENSG00000058673 | ZC3H11A | 6.57522017 | 0.378757496 | 0.687122268 | 0.551222851 | 0.581480917 | 0.74646393 |
| ENSG00000058729 | RIOK2 | 757.1228864 | -0.124231265 | 0.092933373 | -1.336777737 | 0.181295198 | 0.336654761 |
| ENSG00000058799 | YIPF1 | 879.6228365 | -0.334821919 | 0.076665402 | -4.367314484 | 1.26E-05 | 6.40E-05 |
| ENSG00000058804 | NDC1 | 1579.6121 | -0.162212337 | 0.063253926 | -2.564462774 | 0.010333566 | 0.030587356 |
| ENSG00000059122 | FLYWCH1 | 890.8294658 | -0.231179719 | 0.071255359 | -3.244383588 | 0.001177051 | 0.004352983 |
| ENSG00000059145 | UNKL | 726.5051308 | 0.462146377 | 0.077916296 | 5.931318593 | 3.01E-09 | 2.29E-08 |
| ENSG00000059377 | TBXAS1 | 35.24855568 | 0.154356363 | 0.307919473 | 0.501288085 | 0.61616839 | 0.772610159 |
| ENSG00000059378 | PARP12 | 180.3290312 | -0.454331315 | 0.134370738 | -3.381177495 | 0.000721759 | 0.002775252 |
| ENSG00000059573 | ALDH18A1 | 5659.286305 | -0.507841923 | 0.047268617 | -10.74374396 | 6.34E-27 | 1.57E-25 |
| ENSG00000059588 | TARBP1 | 1342.898398 | -0.02544722 | 0.076385603 | -0.333141573 | 0.739027419 | 0.856140238 |
| ENSG00000059691 | GATB | 636.3742232 | -0.308708978 | 0.077990418 | -3.95829366 | 7.55E-05 | 0.000343996 |
| ENSG00000059728 | MXD1 | 531.9699344 | 0.304662274 | 0.088812695 | 3.430391058 | 0.000602712 | 0.002350197 |
| ENSG00000059758 | CDK17 | 794.2466013 | -0.246824341 | 0.077737932 | -3.175082418 | 0.001497939 | 0.005407196 |
| ENSG00000059769 | DNAJC25 | 154.2215834 | 0.330377668 | 0.156200803 | 2.115083038 | 0.034422871 | 0.08738084 |
| ENSG00000059915 | PSD | 186.8828938 | -0.140684549 | 0.135932402 | -1.034959629 | 0.300687765 | 0.487482358 |
| ENSG00000060069 | CTDP1 | 697.4479394 | 0.633965722 | 0.078358606 | 8.090569124 | 5.94E-16 | 7.64E-15 |
| ENSG00000060138 | YBX3 | 2349.358102 | 0.140102278 | 0.05357029 | 2.61529811 | 0.008914961 | 0.026898487 |
| ENSG00000060140 | STYK1 | 10.56758232 | -1.038834713 | 0.576070738 | -1.803311024 | 0.071339382 | 0.161186391 |

| | | | | | | | |
|---|---|---|---|---|---|---|---|
| ENSG00000060237 | WNK1 | 4361.647526 | -0.141327855 | 0.056048227 | -2.521540159 | 0.011684234 | 0.03412302 |
| ENSG00000060303 | RPS17P5 | 13.29261706 | 0.287444171 | 0.484012581 | 0.593877478 | 0.552594064 | 0.724232303 |
| ENSG00000060339 | CCAR1 | 3199.992561 | -0.547816648 | 0.057889402 | -9.46315955 | 2.99E-21 | 5.38E-20 |
| ENSG00000060491 | OGFR | 2841.586736 | -0.013634008 | 0.05852416 | -0.232963757 | 0.815789549 | 0.902352305 |
| ENSG00000060558 | GNA15 | 1.823552423 | -0.970692121 | 1.34969815 | -0.719192006 | 0.472022624 | 0.657714066 |
| ENSG00000060642 | PIGV | 352.0699952 | -0.680262076 | 0.107172892 | -6.347333396 | 2.19E-10 | 1.85E-09 |
| ENSG00000060656 | PTPRU | 2177.364189 | -0.147958626 | 0.063111137 | -2.344413876 | 0.019057011 | 0.052437068 |
| ENSG00000060688 | SNRNP40 | 1495.226934 | 0.045920874 | 0.059559736 | 0.771005335 | 0.44070377 | 0.630020584 |
| ENSG00000060749 | QSER1 | 1795.651733 | 0.015972012 | 0.059639612 | 0.267808776 | 0.788846515 | 0.885806405 |
| ENSG00000060762 | MPC1 | 304.6263432 | -0.244412591 | 0.113649448 | -2.150583182 | 0.031509114 | 0.081033528 |
| ENSG00000060971 | ACAA1 | 1867.193924 | 0.071232679 | 0.058617125 | 1.215219604 | 0.224282345 | 0.395808478 |
| ENSG00000061273 | HDAC7 | 2588.75519 | 0.228839885 | 0.058216529 | 3.930840437 | 8.46E-05 | 0.000382406 |
| ENSG00000061455 | PRDM6 | 287.7468742 | 0.098850074 | 0.11238152 | 0.879593673 | 0.379079468 | 0.571584264 |
| ENSG00000061656 | SPAG4 | 56.69761161 | -0.495562159 | 0.236890996 | -2.091941727 | 0.036443729 | 0.091794398 |
| ENSG00000061676 | NCKAP1 | 5146.864533 | 0.087751715 | 0.054639197 | 1.606021314 | 0.108269227 | 0.22597544 |
| ENSG00000061794 | MRPS35 | 1234.814401 | 0.152351147 | 0.063977851 | 2.381310791 | 0.01725115 | 0.048085224 |
| ENSG00000061918 | GUCY1B1 | 265.2219176 | 0.060587588 | 0.11453249 | 0.528999137 | 0.596806048 | 0.757649737 |
| ENSG00000061936 | SFSWAP | 1817.111229 | -0.094626345 | 0.05715706 | -1.655549539 | 0.097813076 | 0.208224942 |
| ENSG00000061938 | TNK2 | 2655.832776 | -0.384188901 | 0.058288347 | -6.59117854 | 4.36E-11 | 3.87E-10 |
| ENSG00000061987 | MON2 | 2311.591885 | -0.063577596 | 0.063449162 | -1.002024205 | 0.316331902 | 0.505126955 |
| ENSG00000062038 | CDH3 | 2596.420788 | 0.265212134 | 0.061302487 | 4.326286708 | 1.52E-05 | 7.64E-05 |
| ENSG00000062194 | GPBP1 | 3133.887029 | 0.026554604 | 0.054836349 | 0.484251863 | 0.628207134 | 0.780689734 |
| ENSG00000062282 | DGAT2 | 377.2260363 | 0.427158686 | 0.099219325 | 4.305196461 | 1.67E-05 | 8.37E-05 |
| ENSG00000062370 | ZNF112 | 146.0280614 | 0.502659876 | 0.149351485 | 3.36561685 | 0.000763727 | 0.002923394 |
| ENSG00000062485 | CS | 7025.637939 | -0.053436219 | 0.046659392 | -1.145240363 | 0.252109602 | 0.430783992 |
| ENSG00000062524 | LTK | 7.545456083 | 0.355103229 | 0.662609479 | 0.535916313 | 0.59201639 | 0.754408176 |
| ENSG00000062582 | MRPS24 | 7.769006111 | 0.516111723 | 0.654392866 | 0.788687881 | 0.430294453 | 0.620532741 |
| ENSG00000062598 | ELMO2 | 1631.405275 | -0.106186481 | 0.061762834 | -1.719261792 | 0.085566714 | 0.186658465 |
| ENSG00000062650 | WAPL | 3423.487435 | -0.45476397 | 0.051551487 | -8.821549 | 1.13E-18 | 1.75E-17 |
| ENSG00000062716 | VMP1 | 10928.09242 | 0.199229993 | 0.053847355 | 3.699903054 | 0.000215682 | 0.000908916 |
| ENSG00000062725 | APPBP2 | 19882.40005 | 0.337964181 | 0.049611698 | 6.812187381 | 9.61E-12 | 9.01E-11 |
| ENSG00000062822 | POLD1 | 1377.798171 | -0.085543906 | 0.064206588 | -1.332322879 | 0.182754112 | 0.33865195 |
| ENSG00000063046 | EIF4B | 23922.33922 | 0.16538737 | 0.047070579 | 3.513603918 | 0.000442071 | 0.001767053 |

| | | | | | | | |
|---|---|---|---|---|---|---|---|
| ENSG00000063127 | SLC6A16 | 12.06758953 | 0.464372308 | 0.513330222 | 0.904626863 | 0.365663099 | 0.558395881 |
| ENSG00000063169 | BICRA | 569.0033567 | -0.033980443 | 0.083556461 | -0.406676426 | 0.684245657 | 0.820994579 |
| ENSG00000063176 | SPHK2 | 339.1353916 | -0.062354978 | 0.106919894 | -0.583193419 | 0.559763101 | 0.729268829 |
| ENSG00000063177 | RPL18 | 11022.63421 | 0.227589369 | 0.04336148 | 5.248653192 | 1.53E-07 | 9.76E-07 |
| ENSG00000063180 | CA11 | 255.4735466 | 1.51879154 | 0.123874845 | 12.26069375 | 1.47E-34 | 4.95E-33 |
| ENSG00000063241 | ISOC2 | 1231.092573 | -0.072147355 | 0.070546064 | -1.022698507 | 0.306450417 | 0.494094221 |
| ENSG00000063244 | U2AF2 | 8447.503017 | -0.163276908 | 0.045433518 | -3.593754483 | 0.000325947 | 0.001332364 |
| ENSG00000063245 | EPN1 | 4108.786247 | 0.014252838 | 0.059399252 | 0.239949783 | 0.810369186 | 0.899645977 |
| ENSG00000063322 | MED29 | 2763.540187 | 0.067304786 | 0.06137862 | 1.096550986 | 0.272837724 | 0.454862706 |
| ENSG00000063438 | AHRR | 7.997843685 | -1.267082497 | 0.650170738 | -1.948845778 | 0.051313845 | 0.122478369 |
| ENSG00000063587 | ZNF275 | 859.3281736 | -0.316573409 | 0.071961868 | -4.399182765 | 1.09E-05 | 5.59E-05 |
| ENSG00000063601 | MTMR1 | 2471.69972 | -0.598810172 | 0.051738162 | -11.57385871 | 5.59E-31 | 1.61E-29 |
| ENSG00000063660 | GPC1 | 1209.284464 | 0.291008044 | 0.068775835 | 4.231254251 | 2.32E-05 | 0.000114278 |
| ENSG00000063761 | ADCK1 | 355.036681 | -0.502671782 | 0.103501988 | -4.856638893 | 1.19E-06 | 6.84E-06 |
| ENSG00000063854 | HAGH | 1262.141029 | -0.052259878 | 0.068831981 | -0.759238328 | 0.447710002 | 0.636060411 |
| ENSG00000063978 | RNF4 | 3036.286243 | 0.185621629 | 0.049306398 | 3.764656038 | 0.000166778 | 0.00071767 |
| ENSG00000064012 | CASP8 | 383.3764754 | 0.768315804 | 0.099374291 | 7.731534965 | 1.06E-14 | 1.24E-13 |
| ENSG00000064042 | LIMCH1 | 149.2689893 | -2.24620541 | 0.171600294 | -13.08975272 | 3.77E-39 | 1.48E-37 |
| ENSG00000064102 | INTS13 | 701.3451256 | -0.15545512 | 0.081282828 | -1.912521059 | 0.055809394 | 0.131565872 |
| ENSG00000064115 | TM7SF3 | 1811.728116 | -0.445365 | 0.05709003 | -7.801099413 | 6.14E-15 | 7.34E-14 |
| ENSG00000064195 | DLX3 | 503.0867793 | -0.480676315 | 0.090544042 | -5.308756998 | 1.10E-07 | 7.13E-07 |
| ENSG00000064199 | SPA17 | 204.9150854 | 0.302710078 | 0.137498026 | 2.20155945 | 0.027696443 | 0.07253755 |
| ENSG00000064205 | CCN5 | 41.13617616 | -2.117208457 | 0.317429601 | -6.66985199 | 2.56E-11 | 2.32E-10 |
| ENSG00000064225 | ST3GAL6 | 4.971831051 | -0.973565764 | 0.793598287 | -1.226774024 | 0.219907537 | 0.390032387 |
| ENSG00000064270 | ATP2C2 | 3033.527596 | 0.158503053 | 0.05135218 | 3.086588604 | 0.002024676 | 0.007109455 |
| ENSG00000064300 | NGFR | 18.77877025 | -0.415643921 | 0.403057457 | -1.031227468 | 0.302434162 | 0.489372906 |
| ENSG00000064309 | CDON | 443.3030513 | -0.011755024 | 0.095005855 | -0.123729469 | 0.901529478 | 0.950373891 |
| ENSG00000064313 | TAF2 | 1542.303528 | -0.160718232 | 0.067579147 | -2.378222273 | 0.017396336 | 0.04844282 |
| ENSG00000064393 | HIPK2 | 3255.753572 | -0.105791791 | 0.178221053 | -0.593598734 | 0.552780528 | 0.724326573 |
| ENSG00000064419 | TNPO3 | 3115.508507 | -0.051162043 | 0.047781911 | -1.070740826 | 0.28428598 | 0.468096496 |
| ENSG00000064490 | RFXANK | 527.6803863 | -0.294820163 | 0.092396257 | -3.190823652 | 0.001418678 | 0.005146186 |
| ENSG00000064545 | TMEM161A | 193.7211235 | -0.187244054 | 0.132897583 | -1.408934985 | 0.158854393 | 0.304183621 |
| ENSG00000064547 | LPAR2 | 396.496537 | 0.311766856 | 0.0952692 | 3.272483207 | 0.001066072 | 0.003969436 |

| | | | | | | | |
|---|---|---|---|---|---|---|---|
| ENSG00000064601 | CTSA | 4750.124013 | 0.412175059 | 0.054613601 | 7.547113808 | 4.45E-14 | 4.97E-13 |
| ENSG00000064607 | SUGP2 | 1302.4487 | 0.242357187 | 0.07122008 | 3.402933354 | 0.000666665 | 0.002581148 |
| ENSG00000064651 | SLC12A2 | 2989.373841 | -0.401710447 | 0.050345589 | -7.979059501 | 1.47E-15 | 1.84E-14 |
| ENSG00000064652 | SNX24 | 348.8162992 | 0.199640911 | 0.100319121 | 1.990058423 | 0.0465845 | 0.112726628 |
| ENSG00000064655 | EYA2 | 896.8918492 | -0.322993552 | 0.072495685 | -4.455348619 | 8.38E-06 | 4.36E-05 |
| ENSG00000064666 | CNN2 | 6548.956167 | -0.150562999 | 0.047011831 | -3.202661855 | 0.001361638 | 0.004965972 |
| ENSG00000064687 | ABCA7 | 630.6485052 | -0.127578453 | 0.085296067 | -1.495713201 | 0.134728409 | 0.268637088 |
| ENSG00000064692 | SNCAIP | 23.97212017 | 0.974484599 | 0.369360249 | 2.638303932 | 0.008332187 | 0.0253099 |
| ENSG00000064703 | DDX20 | 939.0436643 | -0.186495494 | 0.072499854 | -2.572356805 | 0.010100873 | 0.029995344 |
| ENSG00000064726 | BTBD1 | 3240.034101 | 0.286087797 | 0.055345181 | 5.169154602 | 2.35E-07 | 1.47E-06 |
| ENSG00000064763 | FAR2 | 22.07795501 | -1.615181425 | 0.400173063 | -4.036207271 | 5.43E-05 | 0.00025282 |
| ENSG00000064787 | BCAS1 | 3797.560051 | -0.651943957 | 0.062479454 | -10.43453351 | 1.72E-25 | 3.91E-24 |
| ENSG00000064932 | SBNO2 | 1888.963853 | 0.529975025 | 0.062466563 | 8.484139319 | 2.17E-17 | 3.06E-16 |
| ENSG00000064933 | PMS1 | 390.6862429 | -0.128873235 | 0.098191466 | -1.312468794 | 0.189362009 | 0.348223801 |
| ENSG00000064961 | HMG20B | 4988.373663 | -0.353832462 | 0.049317191 | -7.174627212 | 7.25E-13 | 7.50E-12 |
| ENSG00000064989 | CALCRL | 17.50487187 | 0.013440493 | 0.442672533 | 0.030362158 | 0.975778225 | 0.988882862 |
| ENSG00000064995 | TAF11 | 1889.58012 | 0.002391553 | 0.054505869 | 0.043876977 | 0.965002467 | 0.984148726 |
| ENSG00000064999 | ANKS1A | 1859.078364 | 0.23023309 | 0.062883419 | 3.661268614 | 0.000250969 | 0.001044681 |
| ENSG00000065000 | AP3D1 | 6420.207028 | 0.264730141 | 0.045803327 | 5.779714295 | 7.48E-09 | 5.45E-08 |
| ENSG00000065029 | ZNF76 | 1624.770161 | 0.067706777 | 0.06178062 | 1.095922579 | 0.273112657 | 0.455038495 |
| ENSG00000065054 | NHERF2 | 2802.438291 | -0.608181716 | 0.068708332 | -8.851644257 | 8.62E-19 | 1.35E-17 |
| ENSG00000065057 | NTHL1 | 545.9011238 | -0.04800078 | 0.08968176 | -0.535234586 | 0.592487653 | 0.754736754 |
| ENSG00000065060 | BLTP3A | 2229.224688 | 0.080924491 | 0.055064889 | 1.469620535 | 0.141664555 | 0.279071267 |
| ENSG00000065135 | GNAI3 | 3952.795731 | -0.241827818 | 0.050736317 | -4.766365212 | 1.88E-06 | 1.05E-05 |
| ENSG00000065150 | IPO5 | 4105.150831 | 0.083155346 | 0.054342249 | 1.530215396 | 0.125963422 | 0.254972284 |
| ENSG00000065154 | OAT | 2744.321847 | -0.529204433 | 0.050751965 | -10.42726972 | 1.86E-25 | 4.22E-24 |
| ENSG00000065183 | WDR3 | 1789.27024 | 0.076954385 | 0.063179698 | 1.218023961 | 0.223214868 | 0.394495305 |
| ENSG00000065243 | PKN2 | 1295.318768 | -0.207077163 | 0.065586489 | -3.157314373 | 0.001592296 | 0.005721689 |
| ENSG00000065268 | WDR18 | 1293.853031 | 0.069150362 | 0.064675143 | 1.069195337 | 0.284981659 | 0.468871783 |
| ENSG00000065308 | TRAM2 | 2171.423388 | -0.363784542 | 0.052028621 | -6.992008193 | 2.71E-12 | 2.68E-11 |
| ENSG00000065320 | NTN1 | 94.81405501 | 0.30081038 | 0.189367448 | 1.588501 | 0.112173095 | 0.232578706 |
| ENSG00000065328 | MCM10 | 895.8705684 | -0.716873384 | 0.070372896 | -10.18678242 | 2.27E-24 | 4.87E-23 |
| ENSG00000065357 | DGKA | 32.06437889 | -0.453445613 | 0.31722331 | -1.429420848 | 0.15288331 | 0.295699219 |

| | | | | | | | |
|---|---|---|---|---|---|---|---|
| ENSG00000065361 | ERBB3 | 11853.02093 | 0.032886716 | 0.046238081 | 0.711247427 | 0.476930924 | 0.661968041 |
| ENSG00000065413 | ANKRD44 | 139.290786 | 0.202322151 | 0.155589021 | 1.300362651 | 0.193476705 | 0.354218324 |
| ENSG00000065427 | KARS1 | 7456.497219 | 0.411877936 | 0.044567126 | 9.241743176 | 2.43E-20 | 4.13E-19 |
| ENSG00000065457 | ADAT1 | 2006.389697 | 0.333725467 | 0.057143904 | 5.84008864 | 5.22E-09 | 3.88E-08 |
| ENSG00000065485 | PDIA5 | 551.2811051 | -0.152203261 | 0.083935353 | -1.813339133 | 0.069779531 | 0.158191438 |
| ENSG00000065491 | TBC1D22B | 713.4413615 | 0.009866871 | 0.075381643 | 0.13089222 | 0.895860569 | 0.947292908 |
| ENSG00000065518 | NDUFB4 | 1538.651147 | 0.056967797 | 0.057637268 | 0.988384754 | 0.322964251 | 0.512862991 |
| ENSG00000065526 | SPEN | 2403.342886 | -0.051510336 | 0.076001093 | -0.677757842 | 0.497925245 | 0.679791539 |
| ENSG00000065534 | MYLK | 149.069113 | 1.994692879 | 0.172142812 | 11.58743054 | 4.77E-31 | 1.38E-29 |
| ENSG00000065548 | ZC3H15 | 2513.980107 | -0.021564093 | 0.055236656 | -0.39039462 | 0.696244765 | 0.829343624 |
| ENSG00000065559 | MAP2K4 | 1277.623492 | -0.124903933 | 0.062075404 | -2.012132434 | 0.044205979 | 0.10785845 |
| ENSG00000065600 | PACC1 | 186.8606915 | 0.197005955 | 0.132723002 | 1.484339206 | 0.137718954 | 0.27322259 |
| ENSG00000065609 | SNAP91 | 30.02390078 | -0.633354232 | 0.331772293 | -1.909002788 | 0.056261732 | 0.132387335 |
| ENSG00000065613 | SLK | 1519.965619 | -0.109245066 | 0.063577548 | -1.718296295 | 0.085742585 | 0.186970998 |
| ENSG00000065615 | CYB5R4 | 551.2425105 | 0.076095902 | 0.086998589 | 0.874679723 | 0.381748196 | 0.574061201 |
| ENSG00000065621 | GSTO2 | 316.9393148 | 0.248866006 | 0.109271101 | 2.277509824 | 0.022755799 | 0.061169208 |
| ENSG00000065665 | SEC61A2 | 173.1783208 | 0.079385588 | 0.143597982 | 0.552832194 | 0.580378321 | 0.745630889 |
| ENSG00000065717 | TLE2 | 1620.656831 | -0.305838549 | 0.067629813 | -4.522244498 | 6.12E-06 | 3.24E-05 |
| ENSG00000065802 | ASB1 | 1079.886759 | -0.347278396 | 0.068829871 | -5.04546052 | 4.52E-07 | 2.74E-06 |
| ENSG00000065809 | FAM107B | 2520.698959 | -0.949027854 | 0.056205697 | -16.88490495 | 5.81E-64 | 4.58E-62 |
| ENSG00000065833 | ME1 | 891.3870877 | 0.191215069 | 0.074635915 | 2.561971246 | 0.010407994 | 0.030775892 |
| ENSG00000065882 | TBC1D1 | 1041.046765 | 0.820054066 | 0.067632213 | 12.12519936 | 7.77E-34 | 2.51E-32 |
| ENSG00000065883 | CDK13 | 2630.169623 | 0.498603837 | 0.050446648 | 9.883785223 | 4.89E-23 | 9.76E-22 |
| ENSG00000065911 | MTHFD2 | 5153.663906 | 0.058588915 | 0.064771891 | 0.904542288 | 0.365707922 | 0.558395881 |
| ENSG00000065923 | SLC9A7 | 2720.956989 | 0.134771288 | 0.052603766 | 2.562008361 | 0.010406881 | 0.030775892 |
| ENSG00000065970 | FOXJ2 | 781.2254905 | -0.138015322 | 0.072250633 | -1.910229946 | 0.056103613 | 0.13209657 |
| ENSG00000065978 | YBX1 | 9512.110334 | 0.043047167 | 0.052113711 | 0.826023825 | 0.408790577 | 0.601170321 |
| ENSG00000065989 | PDE4A | 63.90449681 | -0.671743012 | 0.221387875 | -3.034235775 | 0.002411459 | 0.008348757 |
| ENSG00000066027 | PPP2R5A | 1051.772203 | -0.111691427 | 0.070027921 | -1.594955641 | 0.110722156 | 0.229965256 |
| ENSG00000066032 | CTNNA2 | 2.60934274 | -0.202311991 | 1.125882649 | -0.179691899 | 0.857394454 | 0.926087485 |
| ENSG00000066044 | ELAVL1 | 3235.100445 | 0.039284145 | 0.048600017 | 0.80831546 | 0.418909005 | 0.610638152 |
| ENSG00000066084 | DIP2B | 2592.764673 | -0.148779347 | 0.05690056 | -2.614725518 | 0.008929919 | 0.026929451 |
| ENSG00000066117 | SMARCD1 | 3844.215073 | -0.114261831 | 0.050202416 | -2.27602255 | 0.02284466 | 0.061357745 |

| | | | | | | | |
|---|---|---|---|---|---|---|---|
| ENSG00000066135 | KDM4A | 2282.798423 | -0.021139097 | 0.054898604 | -0.385057093 | 0.700195117 | 0.831778571 |
| ENSG00000066136 | NFYC | 1681.47184 | -0.071169673 | 0.062223295 | -1.143778601 | 0.252715467 | 0.431538842 |
| ENSG00000066185 | ZMYND12 | 15.1396134 | -0.658298272 | 0.450895638 | -1.459979241 | 0.144295779 | 0.282918391 |
| ENSG00000066230 | SLC9A3 | 28.30085368 | 0.434884613 | 0.347088809 | 1.252949107 | 0.210224229 | 0.377439447 |
| ENSG00000066248 | NGEF | 95.35096353 | 0.739425332 | 0.196869318 | 3.755919603 | 0.000172706 | 0.000741231 |
| ENSG00000066279 | ASPM | 2426.429819 | -0.111332936 | 0.069513062 | -1.601611752 | 0.109241491 | 0.2274047 |
| ENSG00000066322 | ELOVL1 | 2707.004636 | -0.14332796 | 0.057751962 | -2.481785117 | 0.013072608 | 0.037626804 |
| ENSG00000066336 | SPI1 | 2.123329606 | 0.993286608 | 1.292444472 | 0.768533294 | 0.442170418 | 0.631436093 |
| ENSG00000066379 | POLR1H | 297.0173231 | -0.189189794 | 0.114520734 | -1.652013459 | 0.098531807 | 0.209444001 |
| ENSG00000066382 | MPPED2 | 51.17661777 | 0.017256057 | 0.24437145 | 0.070614045 | 0.943704932 | 0.973189082 |
| ENSG00000066422 | ZBTB11 | 1665.169137 | -0.006987582 | 0.057659364 | -0.121187292 | 0.90354269 | 0.951246259 |
| ENSG00000066427 | ATXN3 | 631.0775849 | -0.494020034 | 0.085814723 | -5.756821401 | 8.57E-09 | 6.20E-08 |
| ENSG00000066455 | GOLGA5 | 2451.273562 | -0.529448015 | 0.052468826 | -10.09071588 | 6.07E-24 | 1.27E-22 |
| ENSG00000066468 | FGFR2 | 518.4967361 | -1.548892102 | 0.091819448 | -16.86888933 | 7.62E-64 | 5.99E-62 |
| ENSG00000066557 | LRRC40 | 367.2235938 | -0.555308131 | 0.105248044 | -5.276184813 | 1.32E-07 | 8.47E-07 |
| ENSG00000066583 | ISOC1 | 3124.181723 | 0.04923437 | 0.051748141 | 0.951422966 | 0.341389707 | 0.532463956 |
| ENSG00000066629 | EML1 | 581.7892992 | 0.169987483 | 0.083576752 | 2.033908704 | 0.041960801 | 0.10314839 |
| ENSG00000066651 | TRMT11 | 271.5713494 | 0.148703801 | 0.113800466 | 1.306706431 | 0.191312434 | 0.351214476 |
| ENSG00000066654 | THUMPD1 | 1729.989053 | 0.101995825 | 0.057994887 | 1.75870376 | 0.078627838 | 0.174272716 |
| ENSG00000066697 | MSANTD3 | 317.0088332 | 0.03378651 | 0.105656664 | 0.319776418 | 0.749137825 | 0.861285106 |
| ENSG00000066735 | KIF26A | 388.4298508 | -1.36198858 | 0.111270137 | -12.24037843 | 1.89E-34 | 6.32E-33 |
| ENSG00000066739 | ATG2B | 1749.926822 | -0.660756551 | 0.066026431 | -10.00745528 | 1.41E-23 | 2.88E-22 |
| ENSG00000066777 | ARFGEF1 | 3548.594103 | 0.58164285 | 0.054571445 | 10.65837354 | 1.59E-26 | 3.84E-25 |
| ENSG00000066827 | ZFAT | 600.056514 | 0.01436942 | 0.082552434 | 0.174064153 | 0.861815045 | 0.928215087 |
| ENSG00000066855 | MTFR1 | 2060.015731 | 0.207404835 | 0.055108625 | 3.763563971 | 0.000167509 | 0.000720678 |
| ENSG00000066923 | STAG3 | 29.73036381 | -0.56724285 | 0.323649044 | -1.752648003 | 0.079662446 | 0.176106439 |
| ENSG00000066926 | FECH | 1115.871075 | 0.386979676 | 0.065922347 | 5.870235101 | 4.35E-09 | 3.27E-08 |
| ENSG00000066933 | MYO9A | 1500.366318 | -0.310673648 | 0.0641438 | -4.84339329 | 1.28E-06 | 7.29E-06 |
| ENSG00000067048 | DDX3Y | 1.595678251 | -0.190901637 | 1.549737695 | -0.123183192 | 0.901962035 | NA |
| ENSG00000067057 | PFKP | 3978.918865 | -1.097856654 | 0.048008296 | -22.86806148 | 9.66E-116 | 2.09E-113 |
| ENSG00000067064 | IDI1 | 6248.352726 | -1.646339117 | 0.060256228 | -27.32230635 | 2.30E-164 | 9.28E-162 |
| ENSG00000067066 | SP100 | 3.725120698 | 2.178338423 | 1.050747187 | 2.073132766 | 0.038159925 | 0.095431014 |
| ENSG00000067082 | KLF6 | 1880.418708 | -1.247822298 | 0.058044943 | -21.49751947 | 1.64E-102 | 2.77E-100 |

| | | | | | | | |
|---|---|---|---|---|---|---|---|
| ENSG00000067113 | PLPP1 | 158.5524751 | 0.853432033 | 0.151446496 | 5.635204862 | 1.75E-08 | 1.23E-07 |
| ENSG00000067141 | NEO1 | 4046.305136 | -0.058394853 | 0.052301252 | -1.116509671 | 0.264204033 | 0.444900296 |
| ENSG00000067167 | TRAM1 | 3704.880186 | 0.382524704 | 0.062857621 | 6.085574004 | 1.16E-09 | 9.18E-09 |
| ENSG00000067177 | PHKA1 | 968.6977489 | 0.539062622 | 0.069605036 | 7.744592281 | 9.59E-15 | 1.12E-13 |
| ENSG00000067182 | TNFRSF1A | 1479.910299 | -0.219463493 | 0.059574777 | -3.683832386 | 0.000229753 | 0.000964065 |
| ENSG00000067191 | CACNB1 | 224.6533683 | 0.145302691 | 0.132487875 | 1.096724449 | 0.272761866 | 0.454862706 |
| ENSG00000067208 | EVI5 | 302.8997445 | 0.079128299 | 0.108303567 | 0.730615824 | 0.465013844 | 0.65100803 |
| ENSG00000067221 | STOML1 | 199.6016522 | -0.414405851 | 0.138015869 | -3.002595669 | 0.002676878 | 0.009168048 |
| ENSG00000067225 | PKM | 36074.27064 | -0.224297939 | 0.041109601 | -5.456096223 | 4.87E-08 | 3.25E-07 |
| ENSG00000067248 | DHX29 | 1291.51533 | 0.046264824 | 0.062256463 | 0.743132878 | 0.457401236 | 0.644622241 |
| ENSG00000067334 | DNTTIP2 | 1388.587269 | 0.040711841 | 0.062815745 | 0.648115231 | 0.516910424 | 0.695234822 |
| ENSG00000067365 | METTL22 | 455.1329075 | -0.11866454 | 0.101411393 | -1.170130254 | 0.241948555 | 0.418133182 |
| ENSG00000067369 | TP53BP1 | 2885.326358 | -0.193259543 | 0.056489648 | -3.421149691 | 0.00062357 | 0.002424827 |
| ENSG00000067533 | RRP15 | 1265.70785 | 0.384231975 | 0.065716543 | 5.846807485 | 5.01E-09 | 3.73E-08 |
| ENSG00000067560 | RHOA | 7670.527252 | -0.089402345 | 0.042830472 | -2.087353711 | 0.036856173 | 0.092624957 |
| ENSG00000067596 | DHX8 | 2961.975287 | 0.150191941 | 0.05038776 | 2.980722693 | 0.002875691 | 0.009784685 |
| ENSG00000067601 | PMS2P4 | 62.28583478 | 0.536737092 | 0.230604645 | 2.32752073 | 0.019937567 | 0.054552455 |
| ENSG00000067606 | PRKCZ | 1568.43319 | -0.179895519 | 0.065617429 | -2.741581345 | 0.006114422 | 0.01928039 |
| ENSG00000067704 | IARS2 | 4842.99516 | -0.09650695 | 0.045739537 | -2.109924087 | 0.034864895 | 0.088317215 |
| ENSG00000067715 | SYT1 | 931.1370849 | 0.294207836 | 0.073357877 | 4.010582762 | 6.06E-05 | 0.000280075 |
| ENSG00000067798 | NAV3 | 44.66215803 | -0.746903302 | 0.263069773 | -2.839183286 | 0.004522917 | 0.014729658 |
| ENSG00000067829 | IDH3G | 2639.203074 | -0.455982033 | 0.057446558 | -7.937499581 | 2.06E-15 | 2.55E-14 |
| ENSG00000067836 | ROGDI | 329.9600701 | -0.054620676 | 0.108044301 | -0.505539633 | 0.613179869 | 0.770125715 |
| ENSG00000067840 | PDZD4 | 791.2870544 | -0.757247124 | 0.075436428 | -10.03821553 | 1.04E-23 | 2.13E-22 |
| ENSG00000067900 | ROCK1 | 3705.717685 | 0.00222949 | 0.055882351 | 0.03989613 | 0.968175936 | 0.985838518 |
| ENSG00000067955 | CBFB | 1086.372272 | -0.000504668 | 0.068625477 | -0.007353941 | 0.994132457 | 0.996533822 |
| ENSG00000067992 | PDK3 | 1722.850242 | 0.222350482 | 0.056221854 | 3.954876362 | 7.66E-05 | 0.000348465 |
| ENSG00000068001 | HYAL2 | 367.323954 | 0.474255299 | 0.10290009 | 4.608891005 | 4.05E-06 | 2.19E-05 |
| ENSG00000068024 | HDAC4 | 423.6275882 | -0.387290287 | 0.100379281 | -3.858269185 | 0.000114193 | 0.000504737 |
| ENSG00000068028 | RASSF1 | 282.1453583 | -0.474059894 | 0.114109599 | -4.154426082 | 3.26E-05 | 0.000157029 |
| ENSG00000068078 | FGFR3 | 2645.800221 | -1.664503766 | 0.055006987 | -30.25986078 | 3.87E-201 | 2.40E-198 |
| ENSG00000068079 | IFI35 | 47.4921043 | 1.403782016 | 0.278240884 | 5.045203991 | 4.53E-07 | 2.75E-06 |
| ENSG00000068097 | HEATR6 | 11962.37574 | 0.431603985 | 0.045229606 | 9.542510319 | 1.39E-21 | 2.55E-20 |

| | | | | | | | |
|---|---|---|---|---|---|---|---|
| ENSG00000068120 | COASY | 1812.916315 | -0.28689527 | 0.059815986 | -4.796297593 | 1.62E-06 | 9.12E-06 |
| ENSG00000068137 | PLEKHH3 | 1729.921899 | 0.220375998 | 0.063910532 | 3.448195354 | 0.000564346 | 0.002211117 |
| ENSG00000068305 | MEF2A | 1403.667364 | 0.219095691 | 0.061390133 | 3.568907264 | 0.000358473 | 0.001451867 |
| ENSG00000068308 | OTUD5 | 2648.113734 | -0.062591796 | 0.05075841 | -1.23313155 | 0.217526688 | 0.387066571 |
| ENSG00000068323 | TFE3 | 2515.741215 | 0.030735937 | 0.055597286 | 0.552831616 | 0.580378717 | 0.745630889 |
| ENSG00000068354 | TBC1D25 | 645.7733148 | -0.06673954 | 0.078623677 | -0.84884786 | 0.395965955 | 0.588993581 |
| ENSG00000068366 | ACSL4 | 236.8798134 | -1.563272021 | 0.126902752 | -12.31866129 | 7.19E-35 | 2.44E-33 |
| ENSG00000068383 | INPP5A | 733.9646181 | -0.787693837 | 0.084177398 | -9.35754556 | 8.16E-21 | 1.43E-19 |
| ENSG00000068394 | GPKOW | 978.2451611 | -0.13156388 | 0.075712105 | -1.737686189 | 0.082266124 | 0.180747522 |
| ENSG00000068400 | GRIPAP1 | 2636.926312 | -0.080200274 | 0.052218612 | -1.535856101 | 0.124573677 | 0.252749278 |
| ENSG00000068438 | FTSJ1 | 1551.647013 | -0.108082006 | 0.057899235 | -1.866725984 | 0.061939863 | 0.14319263 |
| ENSG00000068489 | PRR11 | 4945.955658 | 0.025452706 | 0.048224077 | 0.527800788 | 0.59763761 | 0.758106883 |
| ENSG00000068615 | REEP1 | 300.7937228 | -0.996476118 | 0.113071217 | -8.812818565 | 1.22E-18 | 1.88E-17 |
| ENSG00000068650 | ATP11A | 1731.946781 | 0.226986941 | 0.06377351 | 3.559266831 | 0.000371892 | 0.00150197 |
| ENSG00000068654 | POLR1A | 3175.506678 | 0.014648756 | 0.06035976 | 0.242690767 | 0.80824496 | 0.898401971 |
| ENSG00000068697 | LAPTM4A | 3399.066633 | 0.294678086 | 0.047109983 | 6.255109154 | 3.97E-10 | 3.29E-09 |
| ENSG00000068724 | TTC7A | 725.1764456 | 0.098123515 | 0.07489404 | 1.310164526 | 0.190140184 | 0.349392487 |
| ENSG00000068745 | IP6K2 | 2396.786345 | 0.113991151 | 0.061533484 | 1.852506025 | 0.063953193 | 0.147081516 |
| ENSG00000068784 | SRBD1 | 860.1167957 | -0.031882334 | 0.072831632 | -0.43775394 | 0.661564664 | 0.804425601 |
| ENSG00000068796 | KIF2A | 2158.754729 | 0.100399082 | 0.057208625 | 1.754964096 | 0.079265449 | 0.175482475 |
| ENSG00000068831 | RASGRP2 | 2.510880146 | -0.095191671 | 1.132529293 | -0.084052281 | 0.933014864 | 0.967820981 |
| ENSG00000068878 | PSME4 | 2291.718954 | 0.072100759 | 0.060352938 | 1.194652018 | 0.232223017 | 0.406253031 |
| ENSG00000068885 | IFT80 | 274.4068579 | 0.214987151 | 0.123646372 | 1.738725911 | 0.082082986 | 0.180448845 |
| ENSG00000068903 | SIRT2 | 1092.556922 | -0.260823814 | 0.078031092 | -3.342562648 | 0.000830086 | 0.003154245 |
| ENSG00000068912 | ERLEC1 | 1470.316971 | 0.153752508 | 0.059274825 | 2.59389224 | 0.009489622 | 0.028404775 |
| ENSG00000068971 | PPP2R5B | 354.4404907 | 0.140595847 | 0.103736637 | 1.355315262 | 0.175317134 | 0.328520536 |
| ENSG00000068976 | PYGM | 36.9268484 | -0.295702892 | 0.296339991 | -0.997850109 | 0.318352048 | 0.507469667 |
| ENSG00000069011 | PITX1 | 311.4566658 | 0.526847022 | 0.110559711 | 4.765271325 | 1.89E-06 | 1.06E-05 |
| ENSG00000069020 | MAST4 | 403.1524817 | 0.455135741 | 0.095981197 | 4.741926085 | 2.12E-06 | 1.18E-05 |
| ENSG00000069188 | SDK2 | 7.675749041 | 1.13210573 | 0.686985432 | 1.64793266 | 0.099366491 | 0.210925076 |
| ENSG00000069248 | NUP133 | 2134.491654 | -0.107344052 | 0.056687954 | -1.893595446 | 0.058278728 | 0.136239074 |
| ENSG00000069275 | NUCKS1 | 36531.99956 | 0.070031328 | 0.052291128 | 1.339258327 | 0.18048659 | 0.335659698 |
| ENSG00000069329 | VPS35 | 3508.414845 | -0.123523996 | 0.049518634 | -2.494495227 | 0.012613642 | 0.036465905 |

| | | | | | | | |
|---|---|---|---|---|---|---|---|
| ENSG00000069345 | DNAJA2 | 2028.796982 | 0.149671693 | 0.054080165 | 2.767589414 | 0.005647255 | 0.017923837 |
| ENSG00000069399 | BCL3 | 709.9442109 | -0.469908609 | 0.081169199 | -5.789247815 | 7.07E-09 | 5.17E-08 |
| ENSG00000069424 | KCNAB2 | 794.3844917 | 0.237481759 | 0.073225064 | 3.243175841 | 0.001182052 | 0.004369365 |
| ENSG00000069431 | ABCC9 | 2.175864211 | 0.935662836 | 1.239741013 | 0.754724435 | 0.450414342 | 0.638625693 |
| ENSG00000069482 | GAL | 52.92200503 | -0.179828378 | 0.24310081 | -0.739727598 | 0.459465299 | 0.646475341 |
| ENSG00000069493 | CLEC2D | 159.7238592 | 0.16590516 | 0.14131218 | 1.174032984 | 0.240381814 | 0.416178024 |
| ENSG00000069509 | FUNDC1 | 612.2476116 | -0.187108224 | 0.081940703 | -2.283458859 | 0.022403354 | 0.060391649 |
| ENSG00000069535 | MAOB | 123.051368 | 3.525418519 | 0.236251207 | 14.92233019 | 2.36E-50 | 1.32E-48 |
| ENSG00000069667 | RORA | 12.02016233 | -0.438275722 | 0.516293288 | -0.848889057 | 0.395943028 | 0.588993581 |
| ENSG00000069696 | DRD4 | 2.957297265 | 0.160483226 | 1.0107469 | 0.158776867 | 0.873844675 | 0.934676032 |
| ENSG00000069702 | TGFBR3 | 32.66417194 | 1.352102193 | 0.324576902 | 4.165737565 | 3.10E-05 | 0.000150134 |
| ENSG00000069812 | HES2 | 39.64090774 | -0.209164569 | 0.289029704 | -0.723678452 | 0.469263165 | 0.655237507 |
| ENSG00000069849 | ATP1B3 | 2016.647993 | 0.125329763 | 0.053888936 | 2.325704911 | 0.020034298 | 0.054751823 |
| ENSG00000069869 | NEDD4 | 2109.574622 | -0.645216401 | 0.055445497 | -11.63694861 | 2.67E-31 | 7.84E-30 |
| ENSG00000069943 | PIGB | 410.8367186 | -0.200463222 | 0.097769167 | -2.050372611 | 0.040328084 | 0.099776342 |
| ENSG00000069956 | MAPK6 | 12465.47305 | -1.111325349 | 0.057160103 | -19.44232587 | 3.38E-84 | 4.15E-82 |
| ENSG00000069966 | GNB5 | 4526.56121 | -0.820965969 | 0.047618688 | -17.24041563 | 1.32E-66 | 1.09E-64 |
| ENSG00000069974 | RAB27A | 764.9190983 | -0.672520841 | 0.076426453 | -8.799582002 | 1.37E-18 | 2.11E-17 |
| ENSG00000069998 | HDHD5 | 1490.89115 | -0.528681326 | 0.06090167 | -8.680900276 | 3.93E-18 | 5.85E-17 |
| ENSG00000070010 | UFD1 | 2051.166096 | -0.729754057 | 0.058428958 | -12.48959568 | 8.51E-36 | 2.96E-34 |
| ENSG00000070018 | LRP6 | 964.5115452 | -0.002206621 | 0.071456919 | -0.030880439 | 0.97536489 | 0.988740606 |
| ENSG00000070047 | PHRF1 | 1889.028107 | 0.108911535 | 0.058294917 | 1.868285261 | 0.06172232 | 0.142833655 |
| ENSG00000070061 | ELP1 | 2625.628937 | -0.048983049 | 0.055508572 | -0.882441172 | 0.377538279 | 0.570310031 |
| ENSG00000070081 | NUCB2 | 1746.697173 | -0.527573765 | 0.05776595 | -9.132953974 | 6.67E-20 | 1.10E-18 |
| ENSG00000070087 | PFN2 | 5181.977276 | -0.183240498 | 0.051769253 | -3.539562357 | 0.000400791 | 0.001610748 |
| ENSG00000070159 | PTPN3 | 1736.686463 | 0.03877695 | 0.058987877 | 0.657371515 | 0.510942063 | 0.69065033 |
| ENSG00000070182 | SPTB | 208.828704 | -0.420982338 | 0.131522137 | -3.200847756 | 0.001370239 | 0.004993373 |
| ENSG00000070214 | SLC44A1 | 3405.503851 | -0.236016918 | 0.050658921 | -4.658940918 | 3.18E-06 | 1.74E-05 |
| ENSG00000070269 | TMEM260 | 800.5927109 | -0.251318387 | 0.07374599 | -3.40789222 | 0.000654667 | 0.002537694 |
| ENSG00000070366 | SMG6 | 1611.286153 | -0.107919422 | 0.066133411 | -1.631844176 | 0.102712317 | 0.216683883 |
| ENSG00000070367 | EXOC5 | 2130.069062 | -0.727860545 | 0.056978404 | -12.77432318 | 2.28E-37 | 8.45E-36 |
| ENSG00000070371 | CLTCL1 | 142.1231461 | -2.453650382 | 0.184984294 | -13.26410113 | 3.74E-40 | 1.54E-38 |
| ENSG00000070388 | FGF22 | 10.43839925 | 0.740200315 | 0.546162876 | 1.355273944 | 0.175330293 | 0.328520536 |

| | | | | | | | |
|---|---|---|---|---|---|---|---|
| ENSG00000070404 | FSTL3 | 685.3586618 | 0.861263625 | 0.091117203 | 9.452261414 | 3.32E-21 | 5.96E-20 |
| ENSG00000070413 | DGCR2 | 4373.48232 | -0.413689223 | 0.057672947 | -7.1730204 | 7.34E-13 | 7.59E-12 |
| ENSG00000070423 | RNF126 | 765.6425163 | 0.112358066 | 0.089860192 | 1.250365312 | 0.21116613 | 0.378677286 |
| ENSG00000070444 | MNT | 966.3619268 | 0.222025035 | 0.070130691 | 3.165875466 | 0.00154617 | 0.005564659 |
| ENSG00000070476 | ZXDC | 1770.300802 | 0.049449069 | 0.067292671 | 0.734835881 | 0.462439452 | 0.6490279 |
| ENSG00000070495 | JMJD6 | 749.9943457 | 0.053603423 | 0.072880613 | 0.735496316 | 0.462037283 | 0.648626151 |
| ENSG00000070501 | POLB | 316.2376119 | 0.291700744 | 0.106546223 | 2.737785882 | 0.006185433 | 0.019477535 |
| ENSG00000070540 | WIPI1 | 846.7222591 | 0.339100467 | 0.071475827 | 4.744267831 | 2.09E-06 | 1.17E-05 |
| ENSG00000070610 | GBA2 | 1717.758479 | -0.051504422 | 0.061803817 | -0.83335342 | 0.404645437 | 0.597525994 |
| ENSG00000070614 | NDST1 | 7339.24158 | 0.185710435 | 0.056170996 | 3.306162387 | 0.000945833 | 0.003556951 |
| ENSG00000070669 | ASNS | 34.84266986 | -0.801453093 | 0.307792141 | -2.603877702 | 0.009217564 | 0.027702204 |
| ENSG00000070718 | AP3M2 | 569.7996323 | 0.497900058 | 0.085636633 | 5.814101298 | 6.10E-09 | 4.49E-08 |
| ENSG00000070731 | ST6GALNAC2 | 259.3068235 | -0.654107484 | 0.118366748 | -5.526108424 | 3.27E-08 | 2.23E-07 |
| ENSG00000070756 | PABPC1 | 56516.97036 | 0.121737413 | 0.060521705 | 2.011467004 | 0.044276152 | 0.107995208 |
| ENSG00000070759 | TESK2 | 808.0804871 | 1.3023578 | 0.084556753 | 15.40217373 | 1.58E-53 | 9.55E-52 |
| ENSG00000070761 | CFAP20 | 681.6801968 | -0.006992438 | 0.075604558 | -0.092486989 | 0.926311128 | 0.964662124 |
| ENSG00000070770 | CSNK2A2 | 1976.244953 | 0.095576975 | 0.059344391 | 1.610547746 | 0.107278331 | 0.224322275 |
| ENSG00000070778 | PTPN21 | 472.0575735 | 0.048959078 | 0.09104435 | 0.537749771 | 0.590749813 | 0.753338191 |
| ENSG00000070785 | EIF2B3 | 553.6093646 | -0.118392613 | 0.084701165 | -1.397768411 | 0.162182623 | 0.30929166 |
| ENSG00000070814 | TCOF1 | 3119.00295 | 0.039433668 | 0.05198479 | 0.758561655 | 0.448114818 | 0.636320267 |
| ENSG00000070831 | CDC42 | 3991.045163 | -0.024256185 | 0.049095128 | -0.494065007 | 0.62126027 | 0.775691207 |
| ENSG00000070882 | OSBPL3 | 637.9348451 | -2.314952038 | 0.103481258 | -22.3707373 | 7.59E-111 | 1.53E-108 |
| ENSG00000070950 | RAD18 | 1248.661427 | -0.005284916 | 0.070641106 | -0.074813605 | 0.940363017 | 0.971621655 |
| ENSG00000070961 | ATP2B1 | 5770.269548 | -0.001639621 | 0.049980991 | -0.032804891 | 0.973830178 | 0.988106737 |
| ENSG00000071051 | NCK2 | 872.4683481 | -0.1138576 | 0.071384246 | -1.594996185 | 0.110713089 | 0.229965256 |
| ENSG00000071054 | MAP4K4 | 1842.758934 | 0.685466541 | 0.058288945 | 11.75980354 | 6.29E-32 | 1.89E-30 |
| ENSG00000071073 | MGAT4A | 2323.704948 | -1.22598805 | 0.056705757 | -21.62016917 | 1.16E-103 | 2.05E-101 |
| ENSG00000071082 | RPL31 | 6860.362421 | 0.216926772 | 0.048728577 | 4.451736254 | 8.52E-06 | 4.43E-05 |
| ENSG00000071127 | WDR1 | 7738.038513 | -0.13454551 | 0.044790677 | -3.003873107 | 0.002665665 | 0.00913237 |
| ENSG00000071189 | SNX13 | 2987.897103 | -1.698295049 | 0.059282055 | -28.64770882 | 1.71E-180 | 8.36E-178 |
| ENSG00000071205 | ARHGAP10 | 1226.916126 | -0.445313108 | 0.062589227 | -7.114852317 | 1.12E-12 | 1.14E-11 |
| ENSG00000071242 | RPS6KA2 | 231.6758305 | -0.932212213 | 0.128135895 | -7.275184027 | 3.46E-13 | 3.64E-12 |
| ENSG00000071243 | ING3 | 358.7681239 | -0.057183603 | 0.09923565 | -0.576240528 | 0.564452615 | 0.732969584 |

| | | | | | | | |
|---|---|---|---|---|---|---|---|
| ENSG00000071246 | VASH1 | 804.1290278 | 0.365499544 | 0.080602012 | 4.534620588 | 5.77E-06 | 3.07E-05 |
| ENSG00000071282 | LMCD1 | 1525.487063 | -1.231833213 | 0.068135456 | -18.07918048 | 4.65E-73 | 4.64E-71 |
| ENSG00000071462 | BUD23 | 1880.60145 | 0.558911892 | 0.059945578 | 9.323655078 | 1.12E-20 | 1.95E-19 |
| ENSG00000071537 | SEL1L | 4245.482458 | -0.490859372 | 0.048494114 | -10.12204018 | 4.41E-24 | 9.28E-23 |
| ENSG00000071539 | TRIP13 | 1315.113142 | -0.099890237 | 0.06868029 | -1.454423635 | 0.145828891 | 0.285279492 |
| ENSG00000071553 | ATP6AP1 | 10911.67176 | -0.373039533 | 0.046098957 | -8.092147002 | 5.86E-16 | 7.55E-15 |
| ENSG00000071564 | TCF3 | 5338.981335 | -0.072595071 | 0.053107913 | -1.366935118 | 0.171645643 | 0.323226979 |
| ENSG00000071575 | TRIB2 | 276.1129465 | 0.798235968 | 0.116525846 | 6.850291136 | 7.37E-12 | 6.99E-11 |
| ENSG00000071626 | DAZAP1 | 3965.086073 | 0.019844152 | 0.046867939 | 0.423405689 | 0.671999295 | 0.812095037 |
| ENSG00000071655 | MBD3 | 1756.735279 | -0.12828167 | 0.063823062 | -2.009957925 | 0.044435642 | 0.108303616 |
| ENSG00000071794 | HLTF | 3112.481403 | -0.555931017 | 0.053258264 | -10.43839913 | 1.66E-25 | 3.76E-24 |
| ENSG00000071859 | FAM50A | 4418.833947 | -0.432559646 | 0.047281688 | -9.148566073 | 5.77E-20 | 9.61E-19 |
| ENSG00000071889 | FAM3A | 1495.534727 | -0.515488262 | 0.070021636 | -7.361842549 | 1.81E-13 | 1.95E-12 |
| ENSG00000071894 | CPSF1 | 7824.787843 | 0.009923094 | 0.050344505 | 0.197103816 | 0.843746297 | 0.917672106 |
| ENSG00000071967 | CYBRD1 | 8.711218227 | -4.137790025 | 0.971754497 | -4.258061102 | 2.06E-05 | 0.000102213 |
| ENSG00000071994 | PDCD2 | 2790.68645 | 0.261892241 | 0.053082048 | 4.933725285 | 8.07E-07 | 4.73E-06 |
| ENSG00000072042 | RDH11 | 2602.679857 | -0.830153638 | 0.053041257 | -15.65109298 | 3.27E-55 | 2.06E-53 |
| ENSG00000072062 | PRKACA | 3317.709086 | -0.183155553 | 0.055435816 | -3.303920949 | 0.000953427 | 0.003582575 |
| ENSG00000072071 | ADGRL1 | 2658.456328 | 0.507191663 | 0.065392664 | 7.756094173 | 8.76E-15 | 1.03E-13 |
| ENSG00000072110 | ACTN1 | 6273.647734 | -0.636654064 | 0.049706385 | -12.80829549 | 1.47E-37 | 5.50E-36 |
| ENSG00000072121 | ZFYVE26 | 1621.010769 | -0.600265656 | 0.062431364 | -9.614809287 | 6.92E-22 | 1.29E-20 |
| ENSG00000072133 | RPS6KA6 | 664.8561554 | 0.155863127 | 0.085327393 | 1.826648173 | 0.067752685 | 0.154451615 |
| ENSG00000072134 | EPN2 | 1645.548263 | 0.190794838 | 0.057942331 | 3.292840238 | 0.000991808 | 0.003714019 |
| ENSG00000072135 | PTPN18 | 2541.568866 | -0.170969372 | 0.057934834 | -2.951063457 | 0.003166818 | 0.010675506 |
| ENSG00000072163 | LIMS2 | 2.012154995 | -0.604198482 | 1.36070873 | -0.444032193 | 0.657019308 | 0.801277136 |
| ENSG00000072182 | ASIC4 | 4.134516044 | -0.984051577 | 0.888973702 | -1.106952405 | 0.268314506 | 0.449838104 |
| ENSG00000072195 | SPEG | 72.99293934 | 0.007338664 | 0.210679588 | 0.034833293 | 0.972212673 | 0.987410366 |
| ENSG00000072201 | LNX1 | 643.0401703 | -0.02720197 | 0.080264355 | -0.338904738 | 0.734681495 | 0.853692306 |
| ENSG00000072210 | ALDH3A2 | 2793.506116 | 0.304885264 | 0.049736038 | 6.130067383 | 8.78E-10 | 7.02E-09 |
| ENSG00000072274 | TFRC | 7574.754108 | 0.371107423 | 0.055175482 | 6.725948043 | 1.74E-11 | 1.60E-10 |
| ENSG00000072310 | SREBF1 | 27495.29551 | 0.030693399 | 0.043172187 | 0.710953064 | 0.477113322 | 0.661968041 |
| ENSG00000072364 | AFF4 | 4704.605516 | 0.070112794 | 0.054571883 | 1.284778722 | 0.19886962 | 0.361612204 |
| ENSG00000072401 | UBE2D1 | 195.441092 | -0.306752922 | 0.132400116 | -2.316862947 | 0.020511199 | 0.055872261 |

| | | | | | | | |
|---|---|---|---|---|---|---|---|
| ENSG00000072415 | PALS1 | 1648.815588 | -0.525454147 | 0.060536913 | -8.679896595 | 3.96E-18 | 5.90E-17 |
| ENSG00000072422 | RHOBTB1 | 219.5979258 | -0.655134514 | 0.124614964 | -5.25727003 | 1.46E-07 | 9.35E-07 |
| ENSG00000072501 | SMC1A | 8137.050164 | -0.18721265 | 0.045223739 | -4.13969865 | 3.48E-05 | 0.00016634 |
| ENSG00000072506 | HSD17B10 | 3385.485717 | -0.246867484 | 0.054424082 | -4.535997188 | 5.73E-06 | 3.05E-05 |
| ENSG00000072518 | MARK2 | 3187.060815 | 0.154066304 | 0.058197005 | 2.647323582 | 0.008113168 | 0.024742897 |
| ENSG00000072571 | HMMR | 806.5255783 | -0.372760136 | 0.083206107 | -4.479961248 | 7.47E-06 | 3.91E-05 |
| ENSG00000072609 | CHFR | 1310.901705 | -0.030994714 | 0.065937029 | -0.47006538 | 0.638308307 | 0.786942719 |
| ENSG00000072682 | P4HA2 | 524.7545516 | -0.028081985 | 0.086206323 | -0.325753187 | 0.7446111 | 0.858792289 |
| ENSG00000072694 | FCGR2B | 447.1804327 | -1.247124329 | 0.098795131 | -12.62333797 | 1.57E-36 | 5.62E-35 |
| ENSG00000072736 | NFATC3 | 1216.315678 | 0.022460338 | 0.061647643 | 0.36433409 | 0.715608546 | 0.84172279 |
| ENSG00000072756 | TRNT1 | 1009.175031 | -0.085092237 | 0.06942856 | -1.225608553 | 0.220346018 | 0.390628875 |
| ENSG00000072778 | ACADVL | 5174.583303 | 0.222206119 | 0.051197743 | 4.34015457 | 1.42E-05 | 7.19E-05 |
| ENSG00000072786 | STK10 | 1754.591602 | -0.3988719 | 0.06043841 | -6.59964253 | 4.12E-11 | 3.66E-10 |
| ENSG00000072803 | FBXW11 | 2552.138309 | -0.25368216 | 0.051586646 | -4.917593569 | 8.76E-07 | 5.12E-06 |
| ENSG00000072818 | ACAP1 | 11.55721438 | 0.0272563 | 0.559277478 | 0.048734842 | 0.961130609 | 0.982116705 |
| ENSG00000072849 | DERL2 | 624.3672534 | -0.276679253 | 0.078567506 | -3.521548137 | 0.000429035 | 0.001719137 |
| ENSG00000072858 | SIDT1 | 1880.953718 | -0.574556979 | 0.056217148 | -10.2203154 | 1.61E-24 | 3.49E-23 |
| ENSG00000072864 | NDE1 | 1170.36209 | -0.25400472 | 0.067311575 | -3.773566732 | 0.00016093 | 0.000693936 |
| ENSG00000072954 | TMEM38A | 213.9033837 | 0.186014677 | 0.131739754 | 1.41198591 | 0.15795411 | 0.3028642 |
| ENSG00000072958 | AP1M1 | 957.3688796 | -0.411527792 | 0.07622362 | -5.398953687 | 6.70E-08 | 4.41E-07 |
| ENSG00000073008 | PVR | 1670.130494 | 0.014360829 | 0.057580132 | 0.249405968 | 0.80304677 | 0.895135709 |
| ENSG00000073050 | XRCC1 | 1748.00757 | -0.329773834 | 0.060712421 | -5.431735846 | 5.58E-08 | 3.71E-07 |
| ENSG00000073060 | SCARB1 | 3028.046775 | 0.476251071 | 0.053532124 | 8.89654725 | 5.76E-19 | 9.07E-18 |
| ENSG00000073067 | CYP2W1 | 2.936056189 | -1.460694848 | 1.060990386 | -1.376727695 | 0.168596453 | 0.318767216 |
| ENSG00000073111 | MCM2 | 5817.242273 | -0.163623775 | 0.043431073 | -3.767435683 | 0.000164933 | 0.000710128 |
| ENSG00000073146 | MOV10L1 | 4.974756271 | -1.377162469 | 0.825798556 | -1.667673623 | 0.095380534 | 0.204012071 |
| ENSG00000073150 | PANX2 | 347.0998689 | 0.080411772 | 0.105933598 | 0.759077141 | 0.447806413 | 0.636157983 |
| ENSG00000073169 | SELENOO | 633.6627326 | -0.226919191 | 0.086776533 | -2.614983362 | 0.00892318 | 0.026912668 |
| ENSG00000073282 | TP63 | 300.4224157 | 0.527047982 | 0.114791281 | 4.591358997 | 4.40E-06 | 2.36E-05 |
| ENSG00000073331 | ALPK1 | 248.2348987 | 0.077278094 | 0.119512863 | 0.646609008 | 0.517885027 | 0.695770572 |
| ENSG00000073350 | LLGL2 | 8340.371148 | 0.150254783 | 0.044824963 | 3.35203363 | 0.000802203 | 0.003055884 |
| ENSG00000073417 | PDE8A | 1860.715362 | -1.190458778 | 0.060789773 | -19.58320811 | 2.15E-85 | 2.71E-83 |
| ENSG00000073464 | CLCN4 | 35.77902549 | -0.335240901 | 0.292277596 | -1.14699486 | 0.251383745 | 0.429840721 |

| | | | | | | | |
|---|---|---|---|---|---|---|---|
| ENSG00000073536 | NLE1 | 743.231575 | 0.237271254 | 0.075457726 | 3.144426222 | 0.001664129 | 0.005946279 |
| ENSG00000073578 | SDHA | 388.9927947 | -0.078981772 | 0.102231267 | -0.772579414 | 0.439771332 | 0.629163256 |
| ENSG00000073584 | SMARCE1 | 397.8033718 | 0.412305181 | 0.104715486 | 3.93738496 | 8.24E-05 | 0.000372789 |
| ENSG00000073598 | FNDC8 | 1.734180714 | 0.676835232 | 1.355734938 | 0.499238614 | 0.617611295 | NA |
| ENSG00000073605 | GSDMB | 196.6648432 | -0.789777502 | 0.160746221 | -4.913194824 | 8.96E-07 | 5.23E-06 |
| ENSG00000073614 | KDM5A | 1610.307081 | -0.044712569 | 0.063175411 | -0.707752718 | 0.479098833 | 0.663653377 |
| ENSG00000073670 | ADAM11 | 211.1463441 | 0.274349329 | 0.126217894 | 2.173616752 | 0.029733929 | 0.077073032 |
| ENSG00000073711 | PPP2R3A | 428.35825 | 0.978216284 | 0.094181604 | 10.38648995 | 2.86E-25 | 6.44E-24 |
| ENSG00000073712 | FERMT2 | 1578.144544 | -1.113494755 | 0.060891615 | -18.28650403 | 1.06E-74 | 1.12E-72 |
| ENSG00000073792 | IGF2BP2 | 38.55841221 | -1.689657487 | 0.307410072 | -5.49642852 | 3.88E-08 | 2.62E-07 |
| ENSG00000073803 | MAP3K13 | 547.0708114 | -0.277433406 | 0.083216589 | -3.33387142 | 0.000856462 | 0.003247479 |
| ENSG00000073849 | ST6GAL1 | 7.027211708 | -0.074904511 | 0.658016811 | -0.113833734 | 0.909369597 | 0.954587195 |
| ENSG00000073905 | VDAC1P1 | 535.0886652 | -0.052807262 | 0.084113445 | -0.62781 | 0.530128415 | 0.705746726 |
| ENSG00000073910 | FRY | 73.53708412 | 0.111473617 | 0.209813105 | 0.53129959 | 0.59521119 | 0.756588059 |
| ENSG00000073921 | PICALM | 4600.194993 | 0.049112878 | 0.05299447 | 0.926754765 | 0.354053876 | 0.545582595 |
| ENSG00000073969 | NSF | 1958.902529 | 0.041383768 | 0.054797929 | 0.755206793 | 0.450124913 | 0.638403176 |
| ENSG00000074047 | GLI2 | 2.41732918 | -0.346474534 | 1.104577334 | -0.313671595 | 0.753770477 | 0.864272242 |
| ENSG00000074054 | CLASP1 | 2040.964169 | -0.055953933 | 0.055352398 | -1.010867383 | 0.312079907 | 0.500357357 |
| ENSG00000074071 | MRPS34 | 2332.985568 | -0.108423085 | 0.064763986 | -1.674126193 | 0.094105801 | 0.20169925 |
| ENSG00000074181 | NOTCH3 | 7026.39048 | 0.69571566 | 0.058854217 | 11.82099941 | 3.04E-32 | 9.23E-31 |
| ENSG00000074201 | CLNS1A | 3495.274075 | 0.240956316 | 0.050881193 | 4.735665606 | 2.18E-06 | 1.21E-05 |
| ENSG00000074211 | PPP2R2C | 2171.942319 | 0.17785586 | 0.058073024 | 3.062624419 | 0.002194052 | 0.007650336 |
| ENSG00000074219 | TEAD2 | 1471.901671 | 1.384862753 | 0.068275741 | 20.28337934 | 1.80E-91 | 2.52E-89 |
| ENSG00000074266 | EED | 700.3112261 | -0.108574536 | 0.079207364 | -1.370763151 | 0.170448801 | 0.321342497 |
| ENSG00000074276 | CDHR2 | 11.38534418 | -0.769861703 | 0.535741222 | -1.437002926 | 0.150717162 | 0.292706714 |
| ENSG00000074317 | SNCB | 3.969035632 | 0.944073411 | 0.913351034 | 1.033636988 | 0.301305901 | 0.488132583 |
| ENSG00000074319 | TSG101 | 2700.972478 | -0.009882634 | 0.050583361 | -0.195373208 | 0.845100789 | 0.918622636 |
| ENSG00000074356 | NCBP3 | 1458.137436 | -0.071180043 | 0.05968553 | -1.192584581 | 0.233032107 | 0.40735781 |
| ENSG00000074370 | ATP2A3 | 5785.86668 | -0.048750023 | 0.065652238 | -0.742549286 | 0.457754602 | 0.64497823 |
| ENSG00000074410 | CA12 | 1933.299819 | -0.045298651 | 0.061969343 | -0.730984856 | 0.464788403 | 0.65089103 |
| ENSG00000074416 | MGLL | 130.4609564 | -0.367793753 | 0.172826733 | -2.128106838 | 0.033328227 | 0.084997167 |
| ENSG00000074527 | NTN4 | 2.853419654 | -0.98084092 | 1.089933486 | -0.899908969 | 0.368168697 | 0.560734956 |
| ENSG00000074582 | BCS1L | 780.1490272 | -0.012768063 | 0.079147421 | -0.161320016 | 0.87184136 | 0.933577967 |

| | | | | | | | |
|---|---|---|---|---|---|---|---|
| ENSG00000074590 | NUAK1 | 3353.649008 | 0.186617694 | 0.057846466 | 3.226086343 | 0.001254955 | 0.00461209 |
| ENSG00000074603 | DPP8 | 2035.812405 | 0.068860483 | 0.056162216 | 1.226099813 | 0.220161116 | 0.39039159 |
| ENSG00000074621 | SLC24A1 | 1037.801256 | -0.335985347 | 0.074632738 | -4.501849399 | 6.74E-06 | 3.55E-05 |
| ENSG00000074657 | ZNF532 | 178.6916975 | 0.115973146 | 0.136687266 | 0.848456106 | 0.396184008 | 0.589145937 |
| ENSG00000074660 | SCARF1 | 17.0262006 | 0.257059741 | 0.434429662 | 0.591717748 | 0.55403961 | 0.725423029 |
| ENSG00000074695 | LMAN1 | 2357.108768 | 0.594682142 | 0.058405672 | 10.18192454 | 2.39E-24 | 5.11E-23 |
| ENSG00000074696 | HACD3 | 6198.871522 | -0.29601832 | 0.042476214 | -6.969037284 | 3.19E-12 | 3.14E-11 |
| ENSG00000074755 | ZZEF1 | 2361.822039 | 0.019575717 | 0.066239017 | 0.295531521 | 0.767587877 | 0.873089344 |
| ENSG00000074800 | ENO1 | 33544.9328 | -0.342352694 | 0.044570679 | -7.681119109 | 1.58E-14 | 1.82E-13 |
| ENSG00000074803 | SLC12A1 | 5.136527427 | 2.204454606 | 0.911589183 | 2.418254457 | 0.015595167 | 0.044014903 |
| ENSG00000074842 | MYDGF | 995.6904405 | 0.087943853 | 0.071032164 | 1.238084944 | 0.215684566 | 0.384923288 |
| ENSG00000074855 | ANO8 | 1027.147563 | -0.034724003 | 0.069833943 | -0.497236752 | 0.619022107 | 0.77417746 |
| ENSG00000074935 | TUBE1 | 161.5678454 | -0.190931132 | 0.147797732 | -1.291840743 | 0.196412302 | 0.358049763 |
| ENSG00000074964 | ARHGEF10L | 708.0760994 | 0.555608187 | 0.07803362 | 7.120112924 | 1.08E-12 | 1.10E-11 |
| ENSG00000074966 | TXK | 6.197856576 | 1.356661345 | 0.736714771 | 1.841501486 | 0.065548103 | 0.150277182 |
| ENSG00000075043 | KCNQ2 | 6.824346558 | 0.979916831 | 0.7057737 | 1.388429223 | 0.165006383 | 0.313477965 |
| ENSG00000075073 | TACR2 | 8.9888134 | 1.893614113 | 0.660927159 | 2.865087458 | 0.004168941 | 0.013675891 |
| ENSG00000075089 | ACTR6 | 544.5163752 | -0.028573697 | 0.08633423 | -0.330966026 | 0.740670148 | 0.856961376 |
| ENSG00000075131 | TIPIN | 388.5641066 | -0.130305275 | 0.102710936 | -1.268660182 | 0.20456229 | 0.369443167 |
| ENSG00000075142 | SRI | 1111.232933 | 0.272959532 | 0.066760373 | 4.088646018 | 4.34E-05 | 0.000204639 |
| ENSG00000075151 | EIF4G3 | 4268.801584 | 0.074406872 | 0.051156888 | 1.454483947 | 0.145812181 | 0.285279492 |
| ENSG00000075188 | NUP37 | 702.1082049 | -0.238394751 | 0.078571789 | -3.034101073 | 0.002412536 | 0.008349966 |
| ENSG00000075213 | SEMA3A | 18.21169618 | 1.131866879 | 0.433652581 | 2.610077578 | 0.009052169 | 0.027262277 |
| ENSG00000075218 | GTSE1 | 917.5330003 | -0.583256441 | 0.069376814 | -8.407080177 | 4.20E-17 | 5.82E-16 |
| ENSG00000075223 | SEMA3C | 5915.021418 | 1.195658095 | 0.049921372 | 23.95082604 | 9.06E-127 | 2.47E-124 |
| ENSG00000075234 | TTC38 | 615.9138898 | 0.256557516 | 0.086151605 | 2.977977186 | 0.002901576 | 0.00986252 |
| ENSG00000075239 | ACAT1 | 1320.253337 | -0.268767759 | 0.069564626 | -3.863569378 | 0.000111742 | 0.000494858 |
| ENSG00000075240 | GRAMD4 | 1634.845769 | -0.201844345 | 0.060260863 | -3.349509709 | 0.000809547 | 0.003081814 |
| ENSG00000075275 | CELSR1 | 1959.251798 | 0.779711477 | 0.071484911 | 10.90735749 | 1.06E-27 | 2.72E-26 |
| ENSG00000075290 | WNT8B | 23.16077655 | 0.309451034 | 0.371537332 | 0.832893514 | 0.404904788 | 0.597789441 |
| ENSG00000075292 | ZNF638 | 4618.31206 | -0.027209409 | 0.049511929 | -0.549552601 | 0.582626276 | 0.747090374 |
| ENSG00000075303 | SLC25A40 | 702.9395475 | -0.566896438 | 0.080421401 | -7.049074417 | 1.80E-12 | 1.81E-11 |
| ENSG00000075336 | TIMM21 | 503.1496087 | 0.336754871 | 0.092258554 | 3.650120837 | 0.000262117 | 0.001088516 |

| | | | | | | | |
|---|---|---|---|---|---|---|---|
| ENSG00000075391 | RASAL2 | 440.1528708 | 0.029809474 | 0.093805902 | 0.317778231 | 0.750653164 | 0.862253188 |
| ENSG00000075399 | VPS9D1 | 894.4735082 | -0.160820842 | 0.075499467 | -2.130092422 | 0.033163984 | 0.084644169 |
| ENSG00000075407 | ZNF37A | 647.8507282 | -0.18432437 | 0.087399388 | -2.108989267 | 0.034945508 | 0.088482335 |
| ENSG00000075413 | MARK3 | 3925.676581 | -0.406410777 | 0.049235505 | -8.254424837 | 1.53E-16 | 2.06E-15 |
| ENSG00000075415 | SLC25A3 | 11235.36679 | -0.007872618 | 0.048530637 | -0.162219546 | 0.871132968 | 0.933087382 |
| ENSG00000075420 | FNDC3B | 1001.477684 | 0.422751655 | 0.069005863 | 6.126315023 | 8.99E-10 | 7.17E-09 |
| ENSG00000075426 | FOSL2 | 2013.334369 | 0.571607029 | 0.061978074 | 9.222729778 | 2.90E-20 | 4.90E-19 |
| ENSG00000075461 | CACNG4 | 11987.43628 | 1.030440504 | 0.048655315 | 21.17837514 | 1.51E-99 | 2.42E-97 |
| ENSG00000075539 | FRYL | 1808.528076 | -0.040267577 | 0.075992071 | -0.529891828 | 0.596186932 | 0.757115159 |
| ENSG00000075568 | TMEM131 | 2580.497975 | 0.138540581 | 0.051760147 | 2.676587858 | 0.007437606 | 0.022904912 |
| ENSG00000075618 | FSCN1 | 300.5325024 | 1.003815802 | 0.118689226 | 8.457514087 | 2.73E-17 | 3.83E-16 |
| ENSG00000075624 | ACTB | 46055.44535 | -0.268866722 | 0.049444755 | -5.437719805 | 5.40E-08 | 3.59E-07 |
| ENSG00000075643 | MOCOS | 309.6603141 | 0.622833154 | 0.10832148 | 5.749858236 | 8.93E-09 | 6.45E-08 |
| ENSG00000075651 | PLD1 | 30.17999762 | -0.755007234 | 0.335665433 | -2.249285031 | 0.024494368 | 0.0653449 |
| ENSG00000075673 | ATP12A | 1.681262288 | -0.804643375 | 1.375965979 | -0.584784353 | 0.558692726 | NA |
| ENSG00000075702 | WDR62 | 1420.97745 | -0.463536139 | 0.068940683 | -6.723695218 | 1.77E-11 | 1.62E-10 |
| ENSG00000075711 | DLG1 | 2272.331297 | -0.043557797 | 0.056751577 | -0.767516953 | 0.442774218 | 0.631641123 |
| ENSG00000075785 | RAB7A | 9227.927743 | 0.141983823 | 0.0444452 | 3.194581679 | 0.001400337 | 0.005087697 |
| ENSG00000075790 | BCAP29 | 13.51589168 | -0.292521841 | 0.484961019 | -0.603186296 | 0.546384763 | 0.719356431 |
| ENSG00000075826 | SEC31B | 116.8286034 | 0.324961456 | 0.18310241 | 1.774752475 | 0.075938767 | 0.169620949 |
| ENSG00000075856 | SART3 | 2605.471912 | -0.089014438 | 0.050480138 | -1.763355689 | 0.077840518 | 0.172945423 |
| ENSG00000075884 | ARHGAP15 | 2.719002112 | -2.989499168 | 1.374506156 | -2.174962371 | 0.029632936 | 0.076828612 |
| ENSG00000075914 | EXOSC7 | 405.1236201 | -0.015349749 | 0.094460454 | -0.162499206 | 0.870912754 | 0.932975557 |
| ENSG00000075945 | KIFAP3 | 736.2686028 | -0.109440052 | 0.077738816 | -1.407791587 | 0.159192792 | 0.304704435 |
| ENSG00000075975 | MKRN2 | 1401.926688 | 0.0822887 | 0.058828545 | 1.398788629 | 0.161876377 | 0.308913077 |
| ENSG00000076003 | MCM6 | 4187.049023 | -0.0820056 | 0.049362578 | -1.661290872 | 0.096655042 | 0.206276553 |
| ENSG00000076043 | REXO2 | 364.215972 | -0.114895239 | 0.097674006 | -1.176313358 | 0.23946968 | 0.41508372 |
| ENSG00000076053 | RBM7 | 183.4989301 | -0.057637872 | 0.136077513 | -0.423566468 | 0.671882014 | 0.812021148 |
| ENSG00000076067 | RBMS2 | 99.81231683 | -0.512984683 | 0.187095212 | -2.741837585 | 0.006109654 | 0.019270654 |
| ENSG00000076108 | BAZ2A | 5251.624206 | -0.081264022 | 0.047556377 | -1.708793363 | 0.087489233 | 0.190039106 |
| ENSG00000076201 | PTPN23 | 1837.604341 | 0.060146474 | 0.060749558 | 0.990072614 | 0.322138628 | 0.511949389 |
| ENSG00000076242 | MLH1 | 1488.042428 | -0.335859615 | 0.063527206 | -5.286862656 | 1.24E-07 | 8.02E-07 |
| ENSG00000076248 | UNG | 3274.999126 | -0.636736416 | 0.048356404 | -13.16757155 | 1.35E-39 | 5.37E-38 |

| | | | | | | | |
|---|---|---|---|---|---|---|---|
| ENSG00000076258 | FMO4 | 52.75665365 | -0.934985568 | 0.250240416 | -3.736349166 | 0.000186711 | 0.000796718 |
| ENSG00000076321 | KLHL20 | 781.5795781 | -0.242602486 | 0.072275434 | -3.356638217 | 0.000788963 | 0.003010447 |
| ENSG00000076344 | RGS11 | 91.10116143 | 0.724675121 | 0.203725489 | 3.557115635 | 0.000374949 | 0.00151352 |
| ENSG00000076351 | SLC46A1 | 1087.831757 | 0.32596052 | 0.066213106 | 4.922900316 | 8.53E-07 | 4.99E-06 |
| ENSG00000076356 | PLXNA2 | 75.43820077 | 0.968045122 | 0.227918407 | 4.247331897 | 2.16E-05 | 0.000106815 |
| ENSG00000076382 | SPAG5 | 2289.306041 | 0.036774769 | 0.055052833 | 0.667990499 | 0.504139657 | 0.685087598 |
| ENSG00000076513 | ANKRD13A | 716.0212754 | -0.040444871 | 0.077083498 | -0.524689096 | 0.599799339 | 0.759867528 |
| ENSG00000076554 | TPD52 | 1203.068158 | 0.037259265 | 0.083434794 | 0.446567469 | 0.655187388 | 0.799808359 |
| ENSG00000076555 | ACACB | 299.6367073 | -0.561762385 | 0.107954373 | -5.203701976 | 1.95E-07 | 1.23E-06 |
| ENSG00000076604 | TRAF4 | 3520.665122 | -0.023926419 | 0.053640582 | -0.446050703 | 0.655560621 | 0.800136241 |
| ENSG00000076641 | PAG1 | 53.7555839 | 0.792398737 | 0.251337757 | 3.152724634 | 0.001617543 | 0.005803322 |
| ENSG00000076650 | GPATCH1 | 552.7068751 | -0.284449279 | 0.083802872 | -3.394266489 | 0.000688127 | 0.002656172 |
| ENSG00000076662 | ICAM3 | 399.8631694 | -0.403161415 | 0.095922593 | -4.202987025 | 2.63E-05 | 0.000128569 |
| ENSG00000076685 | NT5C2 | 1349.185997 | 0.348629255 | 0.060392842 | 5.772691638 | 7.80E-09 | 5.67E-08 |
| ENSG00000076706 | MCAM | 4.879790438 | 1.513396415 | 0.849456026 | 1.781606545 | 0.074813416 | 0.167615956 |
| ENSG00000076716 | GPC4 | 1071.72959 | -0.440856687 | 0.065606928 | -6.719666665 | 1.82E-11 | 1.67E-10 |
| ENSG00000076770 | MBNL3 | 84.1519249 | 1.247778478 | 0.204309113 | 6.107306999 | 1.01E-09 | 8.05E-09 |
| ENSG00000076826 | CAMSAP3 | 1454.003629 | 0.141427993 | 0.064586637 | 2.189740776 | 0.028543042 | 0.074398067 |
| ENSG00000076864 | RAP1GAP | 846.0288465 | 0.090508801 | 0.081868568 | 1.105537853 | 0.268926602 | 0.450503558 |
| ENSG00000076924 | XAB2 | 1284.599223 | -0.055662645 | 0.06792044 | -0.819527149 | 0.412485719 | 0.604588997 |
| ENSG00000076928 | ARHGEF1 | 2911.993329 | -0.049596021 | 0.058354498 | -0.849909133 | 0.395375608 | 0.588306184 |
| ENSG00000076944 | STXBP2 | 908.0344074 | 0.000897775 | 0.075159596 | 0.011944914 | 0.990469564 | 0.994958761 |
| ENSG00000076984 | MAP2K7 | 1315.185204 | 0.058004093 | 0.063041003 | 0.920101041 | 0.35751996 | 0.549809472 |
| ENSG00000077044 | DGKD | 1331.558969 | -0.252653685 | 0.065826403 | -3.838181551 | 0.000123949 | 0.000544189 |
| ENSG00000077063 | CTTNBP2 | 7.626080945 | -5.485477747 | 1.290065224 | -4.252093341 | 2.12E-05 | 0.000104817 |
| ENSG00000077092 | RARB | 2.954467482 | -1.906907898 | 1.120884223 | -1.701253224 | 0.088895447 | 0.192546649 |
| ENSG00000077097 | TOP2B | 11001.51583 | -0.221064941 | 0.046510569 | -4.753004464 | 2.00E-06 | 1.12E-05 |
| ENSG00000077147 | TM9SF3 | 6057.655356 | -0.320549573 | 0.054835948 | -5.845610104 | 5.05E-09 | 3.76E-08 |
| ENSG00000077150 | NFKB2 | 957.2324586 | 0.046402787 | 0.069281299 | 0.66977363 | 0.503002106 | 0.684068551 |
| ENSG00000077152 | UBE2T | 1413.14458 | 0.04227692 | 0.062101194 | 0.680774679 | 0.496014074 | 0.677981461 |
| ENSG00000077157 | PPP1R12B | 1521.757524 | 0.015090607 | 0.064273789 | 0.234786335 | 0.814374574 | 0.901759557 |
| ENSG00000077232 | DNAJC10 | 2477.069567 | -0.190730493 | 0.058690716 | -3.249755768 | 0.001155042 | 0.004279867 |
| ENSG00000077235 | GTF3C1 | 5974.945495 | 0.07591165 | 0.048685031 | 1.559240048 | 0.118939575 | 0.243875032 |

| | | | | | | | |
|---|---|---|---|---|---|---|---|
| ENSG00000077238 | IL4R | 461.722512 | 0.923046523 | 0.102078437 | 9.042522082 | 1.53E-19 | 2.50E-18 |
| ENSG00000077254 | USP33 | 1922.523137 | 0.050019262 | 0.055936562 | 0.894214092 | 0.371207354 | 0.563924535 |
| ENSG00000077264 | PAK3 | 37.20675037 | -0.399598576 | 0.291424741 | -1.37118961 | 0.170315856 | 0.321144644 |
| ENSG00000077274 | CAPN6 | 2.802772642 | 3.784091304 | 1.461225427 | 2.589669762 | 0.009606804 | 0.028725246 |
| ENSG00000077312 | SNRPA | 2276.892463 | 0.005071571 | 0.051303461 | 0.098854361 | 0.921253906 | 0.9619656 |
| ENSG00000077348 | EXOSC5 | 687.2979081 | 0.20422632 | 0.08382824 | 2.436247266 | 0.014840536 | 0.042083008 |
| ENSG00000077380 | DYNC1I2 | 1370.930378 | -0.213597958 | 0.06258652 | -3.412842835 | 0.00064289 | 0.002495837 |
| ENSG00000077454 | LRCH4 | 599.8160634 | -0.526802362 | 0.083169031 | -6.334116841 | 2.39E-10 | 2.01E-09 |
| ENSG00000077458 | FAM76B | 348.7079362 | -0.207313967 | 0.114183702 | -1.815617856 | 0.069429013 | 0.157599599 |
| ENSG00000077463 | SIRT6 | 465.7193871 | -0.013879109 | 0.095123725 | -0.145905865 | 0.883995702 | 0.939910479 |
| ENSG00000077514 | POLD3 | 1095.899771 | 0.148467963 | 0.068863464 | 2.155975827 | 0.031085553 | 0.080105884 |
| ENSG00000077549 | CAPZB | 4323.335441 | -0.211443689 | 0.048473674 | -4.36203142 | 1.29E-05 | 6.55E-05 |
| ENSG00000077585 | GPR137B | 749.1090758 | 0.418809851 | 0.074614849 | 5.612955809 | 1.99E-08 | 1.39E-07 |
| ENSG00000077684 | JADE1 | 669.0723127 | -0.510841211 | 0.077313249 | -6.607421332 | 3.91E-11 | 3.49E-10 |
| ENSG00000077713 | SLC25A43 | 1788.455196 | -0.08122649 | 0.05726313 | -1.418478009 | 0.156051255 | 0.300219371 |
| ENSG00000077721 | UBE2A | 3214.243875 | -0.065275572 | 0.04867346 | -1.341091678 | 0.179890689 | 0.334858834 |
| ENSG00000077782 | FGFR1 | 1841.739672 | -0.059691275 | 0.060129568 | -0.992710868 | 0.320850877 | 0.510355035 |
| ENSG00000077935 | SMC1B | 13.24022836 | -2.340595068 | 0.564048082 | -4.149637492 | 3.33E-05 | 0.000159947 |
| ENSG00000077942 | FBLN1 | 180.8857652 | -1.188644185 | 0.162194487 | -7.328511639 | 2.33E-13 | 2.48E-12 |
| ENSG00000077984 | CST7 | 62.64090977 | 9.319608858 | 1.1925246 | 7.81502441 | 5.50E-15 | 6.60E-14 |
| ENSG00000078018 | MAP2 | 677.1094068 | 0.551728492 | 0.079796317 | 6.914210014 | 4.70E-12 | 4.54E-11 |
| ENSG00000078043 | PIAS2 | 646.548658 | 0.750973572 | 0.080700852 | 9.305646094 | 1.33E-20 | 2.29E-19 |
| ENSG00000078053 | AMPH | 28.72634605 | 0.121958287 | 0.334746195 | 0.364330615 | 0.715611141 | 0.84172279 |
| ENSG00000078061 | ARAF | 2901.534653 | -0.055488925 | 0.057695642 | -0.961752459 | 0.336173964 | 0.527274737 |
| ENSG00000078070 | MCCC1 | 1819.262649 | -0.046888825 | 0.05559438 | -0.843409442 | 0.398999493 | 0.591855698 |
| ENSG00000078081 | LAMP3 | 7.167994474 | 0.962852623 | 0.672861204 | 1.430982522 | 0.152435224 | 0.295019324 |
| ENSG00000078114 | NEBL | 2845.717805 | 0.128316316 | 0.057063983 | 2.248639301 | 0.024535454 | 0.065424074 |
| ENSG00000078124 | ACER3 | 690.3725092 | 0.240617584 | 0.077845171 | 3.090976369 | 0.001994995 | 0.007013822 |
| ENSG00000078140 | UBE2K | 3212.041522 | 0.011599754 | 0.054411721 | 0.213184837 | 0.831182795 | 0.910670869 |
| ENSG00000078142 | PIK3C3 | 734.6670096 | 0.620249273 | 0.075092815 | 8.259768586 | 1.46E-16 | 1.97E-15 |
| ENSG00000078177 | N4BP2 | 1157.174734 | -0.553849463 | 0.074664723 | -7.417819848 | 1.19E-13 | 1.29E-12 |
| ENSG00000078237 | TIGAR | 71.74061642 | 0.006750555 | 0.211010108 | 0.03199162 | 0.974478734 | 0.988514222 |
| ENSG00000078246 | TULP3 | 938.9560426 | -0.050046732 | 0.067551171 | -0.740871423 | 0.458771404 | 0.646014119 |

| | | | | | | | |
|---|---|---|---|---|---|---|---|
| ENSG00000078269 | SYNJ2 | 1063.447291 | -0.163258641 | 0.068487549 | -2.38377112 | 0.017136257 | 0.047811448 |
| ENSG00000078304 | PPP2R5C | 3012.065399 | -0.0731976 | 0.048865131 | -1.497951572 | 0.134145833 | 0.267684883 |
| ENSG00000078319 | PMS2P1 | 549.2240209 | -0.283180677 | 0.090915852 | -3.11475579 | 0.001840973 | 0.006519282 |
| ENSG00000078369 | GNB1 | 10260.24584 | -0.088159812 | 0.042933089 | -2.053423445 | 0.040031527 | 0.099181687 |
| ENSG00000078401 | EDN1 | 285.0731137 | -2.378665202 | 0.133574994 | -17.80771337 | 6.16E-71 | 5.81E-69 |
| ENSG00000078403 | MLLT10 | 997.3050515 | -0.375987005 | 0.068860486 | -5.460127129 | 4.76E-08 | 3.19E-07 |
| ENSG00000078487 | ZCWPW1 | 271.5693784 | -1.008419599 | 0.118618159 | -8.501393112 | 1.87E-17 | 2.66E-16 |
| ENSG00000078589 | P2RY10 | 2.197105287 | 4.479828152 | 1.554157494 | 2.882480167 | 0.00394558 | 0.013015478 |
| ENSG00000078618 | NRDC | 8940.611094 | -0.066274935 | 0.051030972 | -1.298719827 | 0.194040097 | 0.354796734 |
| ENSG00000078668 | VDAC3 | 2414.028288 | -0.085392299 | 0.059573638 | -1.43339069 | 0.151746218 | 0.294083095 |
| ENSG00000078674 | PCM1 | 3213.438515 | 0.026516724 | 0.062891244 | 0.421628228 | 0.673296399 | 0.81300239 |
| ENSG00000078687 | TNRC6C | 1383.715919 | -0.421747002 | 0.068262084 | -6.178349331 | 6.48E-10 | 5.25E-09 |
| ENSG00000078699 | CBFA2T2 | 951.8544862 | -0.379221062 | 0.068638869 | -5.524873404 | 3.30E-08 | 2.24E-07 |
| ENSG00000078747 | ITCH | 1866.447827 | -0.695486414 | 0.058784811 | -11.83105649 | 2.70E-32 | 8.25E-31 |
| ENSG00000078795 | PKD2L2 | 4.476783618 | 0.061429742 | 0.829631387 | 0.074044622 | 0.94097488 | 0.971771654 |
| ENSG00000078804 | TP53INP2 | 298.9590269 | -0.07397358 | 0.115658286 | -0.639587377 | 0.522440891 | 0.699229524 |
| ENSG00000078808 | SDF4 | 2986.808019 | 0.148400387 | 0.055276925 | 2.68467155 | 0.007260111 | 0.022445681 |
| ENSG00000078814 | MYH7B | 5.675274946 | -1.24135627 | 0.761826333 | -1.629447835 | 0.103218249 | 0.217491195 |
| ENSG00000078898 | BPIFB2 | 1.859014394 | -1.63727904 | 1.414511175 | -1.157487526 | 0.247073233 | 0.424302071 |
| ENSG00000078900 | TP73 | 565.5107119 | -0.043282613 | 0.089888333 | -0.481515364 | 0.630150263 | 0.781766645 |
| ENSG00000078902 | TOLLIP | 1116.858265 | 0.321184249 | 0.072142655 | 4.452071383 | 8.50E-06 | 4.42E-05 |
| ENSG00000078967 | UBE2D4 | 661.0671611 | 1.149348838 | 0.09091754 | 12.64166224 | 1.24E-36 | 4.47E-35 |
| ENSG00000079101 | CLUL1 | 1.900242183 | -0.022322064 | 1.307758901 | -0.017068945 | 0.986381614 | 0.993497695 |
| ENSG00000079102 | RUNX1T1 | 153.7785378 | -1.016447263 | 0.152276005 | -6.675032362 | 2.47E-11 | 2.24E-10 |
| ENSG00000079134 | THOC1 | 841.272655 | 0.139377041 | 0.071025283 | 1.962358112 | 0.049720806 | 0.11923481 |
| ENSG00000079150 | FKBP7 | 14.86842264 | 0.292601121 | 0.448351508 | 0.652615451 | 0.514004222 | 0.693073347 |
| ENSG00000079156 | OSBPL6 | 347.8182201 | 0.018454835 | 0.10927157 | 0.168889627 | 0.865883459 | 0.930281311 |
| ENSG00000079215 | SLC1A3 | 12.79530337 | -4.738719925 | 0.926845666 | -5.112738937 | 3.18E-07 | 1.96E-06 |
| ENSG00000079246 | XRCC5 | 11267.24983 | -0.051083802 | 0.043836996 | -1.165312557 | 0.243892498 | 0.420637088 |
| ENSG00000079257 | LXN | 1873.307974 | -0.852228707 | 0.056888348 | -14.98072519 | 9.81E-51 | 5.55E-49 |
| ENSG00000079263 | SP140 | 2.306290588 | -0.213298245 | 1.138790659 | -0.187302419 | 0.851423526 | 0.922260215 |
| ENSG00000079277 | MKNK1 | 914.4739659 | -0.164139699 | 0.069746836 | -2.353364096 | 0.018604407 | 0.051333306 |
| ENSG00000079308 | TNS1 | 70.63852093 | 0.796264898 | 0.221811843 | 3.589821386 | 0.000330905 | 0.001350703 |

| | | | | | | | |
|---|---|---|---|---|---|---|---|
| ENSG00000079313 | REXO1 | 767.8786763 | 0.072136605 | 0.077745462 | 0.927856147 | 0.353482194 | 0.545024972 |
| ENSG00000079332 | SAR1A | 2895.688959 | -0.551921804 | 0.0527255 | -10.46783448 | 1.21E-25 | 2.77E-24 |
| ENSG00000079335 | CDC14A | 176.2150798 | 0.14986878 | 0.136750613 | 1.095927667 | 0.27311043 | 0.455038495 |
| ENSG00000079337 | RAPGEF3 | 61.44463066 | 0.059281595 | 0.224884881 | 0.26360863 | 0.792081508 | 0.888134796 |
| ENSG00000079385 | CEACAM1 | 1.883187827 | 3.134543813 | 1.604498916 | 1.953596716 | 0.050748929 | 0.121382592 |
| ENSG00000079387 | SENP1 | 1152.134576 | -0.381110173 | 0.064668688 | -5.893272077 | 3.79E-09 | 2.86E-08 |
| ENSG00000079393 | DUSP13 | 7.536927024 | 0.233654739 | 0.639413444 | 0.365420436 | 0.714797589 | 0.841413008 |
| ENSG00000079432 | CIC | 3961.003115 | -0.144115455 | 0.050171375 | -2.872463718 | 0.004072848 | 0.013397067 |
| ENSG00000079459 | FDFT1 | 6710.766079 | -0.594695142 | 0.046619184 | -12.75644682 | 2.87E-37 | 1.06E-35 |
| ENSG00000079462 | PAFAH1B3 | 1609.36826 | -0.078652283 | 0.064182896 | -1.22543992 | 0.220409514 | 0.390650873 |
| ENSG00000079482 | OPHN1 | 871.3819022 | -0.053239035 | 0.069506704 | -0.765955396 | 0.443702842 | 0.632561258 |
| ENSG00000079616 | KIF22 | 5315.972235 | -0.253226572 | 0.047444711 | -5.337298193 | 9.43E-08 | 6.13E-07 |
| ENSG00000079691 | CARMIL1 | 1202.790885 | -0.302402635 | 0.066690314 | -4.534431105 | 5.78E-06 | 3.07E-05 |
| ENSG00000079739 | PGM1 | 984.797756 | 0.459967342 | 0.071306894 | 6.450531176 | 1.11E-10 | 9.59E-10 |
| ENSG00000079785 | DDX1 | 4043.922843 | -0.249699494 | 0.049871791 | -5.006828281 | 5.53E-07 | 3.31E-06 |
| ENSG00000079805 | DNM2 | 6808.470269 | -0.075227603 | 0.052673427 | -1.428188876 | 0.153237502 | 0.296122339 |
| ENSG00000079819 | EPB41L2 | 2963.098168 | 0.470908166 | 0.051929752 | 9.068176708 | 1.21E-19 | 1.98E-18 |
| ENSG00000079841 | RIMS1 | 155.7917304 | -3.023068439 | 0.186073711 | -16.2466177 | 2.36E-59 | 1.66E-57 |
| ENSG00000079950 | STX7 | 1091.092918 | 0.244288643 | 0.067912604 | 3.597103173 | 0.000321781 | 0.001317211 |
| ENSG00000079974 | RABL2B | 523.3370743 | -0.017584008 | 0.09297726 | -0.189121603 | 0.84999751 | 0.921503648 |
| ENSG00000079999 | KEAP1 | 3149.391127 | -0.104966141 | 0.059965702 | -1.750436283 | 0.08004306 | 0.17670944 |
| ENSG00000080007 | DDX43 | 344.0306616 | 0.241103412 | 0.104570467 | 2.305654925 | 0.021129922 | 0.057309104 |
| ENSG00000080031 | PTPRH | 2.4774695 | 0.367821913 | 1.128701685 | 0.325880539 | 0.74451474 | 0.858724327 |
| ENSG00000080189 | SLC35C2 | 2387.444117 | -0.031492285 | 0.063680339 | -0.494537023 | 0.620926966 | 0.775560477 |
| ENSG00000080200 | CRYBG3 | 69.91174761 | 1.601249479 | 0.232121297 | 6.898330733 | 5.26E-12 | 5.06E-11 |
| ENSG00000080224 | EPHA6 | 104.3614165 | 1.000630685 | 0.180158648 | 5.554164033 | 2.79E-08 | 1.91E-07 |
| ENSG00000080298 | RFX3 | 228.8151296 | -0.245340431 | 0.124057931 | -1.977627939 | 0.047970695 | 0.115653993 |
| ENSG00000080345 | RIF1 | 2120.457465 | -0.172291778 | 0.070357255 | -2.448813265 | 0.014332773 | 0.040813533 |
| ENSG00000080371 | RAB21 | 2256.931834 | 0.055729387 | 0.057650783 | 0.966671812 | 0.333708123 | 0.524728517 |
| ENSG00000080503 | SMARCA2 | 3277.061236 | -0.097417458 | 0.051734721 | -1.883018927 | 0.059697803 | 0.138961705 |
| ENSG00000080546 | SESN1 | 561.8980936 | -0.184220685 | 0.08238251 | -2.236162558 | 0.025341125 | 0.067215527 |
| ENSG00000080561 | MID2 | 1548.399704 | -0.276942801 | 0.068050724 | -4.069652561 | 4.71E-05 | 0.000221058 |
| ENSG00000080603 | SRCAP | 237.8071613 | -0.173434963 | 0.119454648 | -1.451889624 | 0.146532298 | 0.286397795 |

| | | | | | | | |
|---|---|---|---|---|---|---|---|
| ENSG00000080608 | PUM3 | 781.0523237 | -0.008371054 | 0.074719013 | -0.112033786 | 0.910796618 | 0.955226736 |
| ENSG00000080644 | CHRNA3 | 23.85643423 | 0.260443198 | 0.374131244 | 0.696127902 | 0.486348734 | 0.669947245 |
| ENSG00000080709 | KCNN2 | 85.15974477 | 1.592425453 | 0.21441404 | 7.426871175 | 1.11E-13 | 1.21E-12 |
| ENSG00000080802 | CNOT4 | 1060.424947 | -0.071466506 | 0.066536037 | -1.074102236 | 0.282776863 | 0.466348039 |
| ENSG00000080815 | PSEN1 | 3186.291548 | -0.53143172 | 0.050930588 | -10.43443119 | 1.73E-25 | 3.91E-24 |
| ENSG00000080819 | CPOX | 292.8267658 | -0.082373975 | 0.112258262 | -0.733789865 | 0.463076819 | 0.64960449 |
| ENSG00000080822 | CLDND1 | 707.6949563 | -0.28052257 | 0.076756492 | -3.654708052 | 0.000257475 | 0.001069937 |
| ENSG00000080823 | MOK | 207.9176513 | -0.085339965 | 0.130203237 | -0.655436588 | 0.512186701 | 0.691842603 |
| ENSG00000080824 | HSP90AA1 | 46685.47028 | -0.070966265 | 0.048379093 | -1.466878767 | 0.142409038 | 0.280128908 |
| ENSG00000080839 | RBL1 | 520.9430497 | -0.499571673 | 0.101111389 | -4.940805154 | 7.78E-07 | 4.58E-06 |
| ENSG00000080845 | DLGAP4 | 2150.358245 | -0.319308556 | 0.055381231 | -5.76564574 | 8.13E-09 | 5.90E-08 |
| ENSG00000080854 | IGSF9B | 257.1928134 | -0.070296282 | 0.121539088 | -0.578384148 | 0.563004794 | 0.731788202 |
| ENSG00000080947 | CROCCP3 | 108.7746416 | 0.695825836 | 0.173080131 | 4.020252544 | 5.81E-05 | 0.00026953 |
| ENSG00000080986 | NDC80 | 776.2608408 | 0.055020336 | 0.072638092 | 0.757458443 | 0.448775254 | 0.636942667 |
| ENSG00000081014 | AP4E1 | 6509.130328 | -1.607068403 | 0.051357361 | -31.2918805 | 6.02E-215 | 4.60E-212 |
| ENSG00000081019 | RSBN1 | 1928.546869 | 0.271829445 | 0.05966287 | 4.556090646 | 5.21E-06 | 2.78E-05 |
| ENSG00000081026 | MAGI3 | 1393.511319 | -0.136643134 | 0.064922528 | -2.104710616 | 0.035316506 | 0.089313267 |
| ENSG00000081052 | COL4A4 | 13.96284587 | 0.406129563 | 0.47180965 | 0.860791133 | 0.38935309 | 0.582063205 |
| ENSG00000081059 | TCF7 | 47.78889518 | 2.927344169 | 0.336460069 | 8.700420754 | 3.31E-18 | 4.95E-17 |
| ENSG00000081087 | OSTM1 | 494.246744 | 0.554670282 | 0.089155089 | 6.221409133 | 4.93E-10 | 4.03E-09 |
| ENSG00000081138 | CDH7 | 118.2160033 | -1.752157338 | 0.182731399 | -9.588704241 | 8.92E-22 | 1.65E-20 |
| ENSG00000081148 | IMPG2 | 11.26585878 | -0.343257677 | 0.528670717 | -0.649284452 | 0.516154534 | 0.694735284 |
| ENSG00000081154 | PCNP | 2354.858113 | 0.01027962 | 0.059116943 | 0.173886194 | 0.861954903 | 0.928215087 |
| ENSG00000081177 | EXD2 | 1863.283531 | -0.554887091 | 0.055813955 | -9.941726754 | 2.74E-23 | 5.54E-22 |
| ENSG00000081181 | ARG2 | 216.8481319 | -0.473052354 | 0.124717191 | -3.7930004 | 0.000148838 | 0.000645555 |
| ENSG00000081189 | MEF2C | 154.452536 | -5.014849955 | 0.290785194 | -17.24589169 | 1.20E-66 | 9.99E-65 |
| ENSG00000081277 | PKP1 | 885.8025417 | -0.285277732 | 0.074131561 | -3.848262822 | 0.000118958 | 0.000523883 |
| ENSG00000081307 | UBA5 | 1650.788289 | -0.323932702 | 0.061645066 | -5.254803417 | 1.48E-07 | 9.46E-07 |
| ENSG00000081320 | STK17B | 274.9021381 | -0.368190074 | 0.111160955 | -3.312224824 | 0.000925571 | 0.003488762 |
| ENSG00000081377 | CDC14B | 253.5483041 | 0.094272652 | 0.11770861 | 0.800898526 | 0.423190394 | 0.614118651 |
| ENSG00000081386 | ZNF510 | 739.8638631 | -0.086511999 | 0.076369763 | -1.132804336 | 0.25729643 | 0.436954405 |
| ENSG00000081479 | LRP2 | 41.6895085 | -1.696113694 | 0.320883637 | -5.285759383 | 1.25E-07 | 8.06E-07 |
| ENSG00000081692 | JMJD4 | 839.3987448 | 0.353736068 | 0.072738948 | 4.863090218 | 1.16E-06 | 6.64E-06 |

| | | | | | | | |
|---|---|---|---|---|---|---|---|
| ENSG00000081721 | DUSP12 | 721.7353297 | 0.151903538 | 0.078211179 | 1.942222834 | 0.05211013 | 0.124094922 |
| ENSG00000081760 | AACS | 4072.806585 | -0.334382241 | 0.046330308 | -7.217354208 | 5.30E-13 | 5.52E-12 |
| ENSG00000081791 | DELE1 | 1999.450861 | 0.379881297 | 0.057025106 | 6.661649945 | 2.71E-11 | 2.45E-10 |
| ENSG00000081803 | CADPS2 | 1021.652199 | 0.226813923 | 0.067599537 | 3.35525851 | 0.000792909 | 0.003023491 |
| ENSG00000081818 | PCDHB4 | 18.24254683 | 0.917398595 | 0.431970454 | 2.123753109 | 0.033690794 | 0.085750229 |
| ENSG00000081842 | PCDHA6 | 145.5517907 | 0.135710297 | 0.155633749 | 0.871985016 | 0.383216552 | 0.575627796 |
| ENSG00000081853 | PCDHGA2 | 63.42408186 | -2.234557316 | 0.255929802 | -8.731133672 | 2.52E-18 | 3.80E-17 |
| ENSG00000081870 | IFT25 | 455.5678601 | -0.507448811 | 0.093156285 | -5.447284731 | 5.11E-08 | 3.41E-07 |
| ENSG00000081913 | PHLPP1 | 657.2757598 | 0.568443569 | 0.082234149 | 6.912500208 | 4.76E-12 | 4.60E-11 |
| ENSG00000081923 | ATP8B1 | 1459.422856 | 0.829179083 | 0.065876544 | 12.5868637 | 2.49E-36 | 8.90E-35 |
| ENSG00000082014 | SMARCD3 | 723.2477241 | 1.443212339 | 0.080172483 | 18.00134262 | 1.90E-72 | 1.86E-70 |
| ENSG00000082068 | WDR70 | 1108.034042 | 0.017920662 | 0.069679841 | 0.257185751 | 0.797035384 | 0.891799572 |
| ENSG00000082126 | MPP4 | 1.661437555 | 1.023653201 | 1.431360752 | 0.71516087 | 0.474509645 | NA |
| ENSG00000082146 | STRADB | 316.4308319 | 0.15501018 | 0.109931394 | 1.410062893 | 0.158521113 | 0.303748273 |
| ENSG00000082153 | BZW1 | 4823.794955 | -0.408529428 | 0.057161992 | -7.146871788 | 8.88E-13 | 9.13E-12 |
| ENSG00000082175 | PGR | 493.0098047 | 4.775559768 | 0.302715285 | 15.77574706 | 4.57E-56 | 2.95E-54 |
| ENSG00000082196 | C1QTNF3 | 50.53709753 | -0.12125952 | 0.262806193 | -0.461402825 | 0.644509625 | 0.791770208 |
| ENSG00000082212 | ME2 | 670.0084655 | 0.835215338 | 0.080332923 | 10.39692451 | 2.56E-25 | 5.79E-24 |
| ENSG00000082213 | C5orf22 | 1060.397347 | -0.089922668 | 0.068118211 | -1.320097322 | 0.186802527 | 0.34452886 |
| ENSG00000082258 | CCNT2 | 1221.96073 | -0.007215792 | 0.065556427 | -0.110069944 | 0.912353905 | 0.955898542 |
| ENSG00000082269 | FAM135A | 78.38843729 | -0.376554137 | 0.202024883 | -1.863899789 | 0.062335778 | 0.143977321 |
| ENSG00000082438 | COBLL1 | 884.2794798 | 0.909322275 | 0.074135112 | 12.26574363 | 1.38E-34 | 4.66E-33 |
| ENSG00000082458 | DLG3 | 2835.434969 | 0.144243855 | 0.051857854 | 2.781523773 | 0.005410437 | 0.017262228 |
| ENSG00000082482 | KCNK2 | 933.6530794 | 2.551257429 | 0.080252858 | 31.79023772 | 8.83E-222 | 7.50E-219 |
| ENSG00000082497 | SERTAD4 | 501.5100883 | -0.387624894 | 0.089665483 | -4.323011251 | 1.54E-05 | 7.74E-05 |
| ENSG00000082512 | TRAF5 | 451.9712007 | -1.327559426 | 0.101645087 | -13.06073383 | 5.52E-39 | 2.15E-37 |
| ENSG00000082515 | MRPL22 | 540.2045648 | -0.09644305 | 0.083049157 | -1.16127669 | 0.245529396 | 0.422634077 |
| ENSG00000082516 | GEMIN5 | 1750.249125 | 0.177159904 | 0.059533325 | 2.975810671 | 0.002922152 | 0.009925106 |
| ENSG00000082641 | NFE2L1 | 9464.692671 | -0.056145815 | 0.04617648 | -1.215896372 | 0.2240244 | 0.395469007 |
| ENSG00000082684 | SEMA5B | 8.466581761 | -1.394374575 | 0.683394993 | -2.040364048 | 0.04131408 | 0.101853304 |
| ENSG00000082701 | GSK3B | 4947.476732 | -0.231909551 | 0.051660104 | -4.489142205 | 7.15E-06 | 3.76E-05 |
| ENSG00000082781 | ITGB5 | 2858.732823 | 0.076770505 | 0.056002729 | 1.370835077 | 0.170426374 | 0.321326622 |
| ENSG00000082805 | ERC1 | 1089.976548 | -0.135106258 | 0.079470049 | -1.700090277 | 0.089113946 | 0.192922949 |

| | | | | | | | |
|---|---|---|---|---|---|---|---|
| ENSG00000082898 | XPO1 | 8005.988346 | -0.202646104 | 0.05863073 | -3.456312121 | 0.000547621 | 0.002152574 |
| ENSG00000082929 | LINC01587 | 5.755667756 | 0.026433394 | 0.755531457 | 0.03498649 | 0.972090514 | 0.987370219 |
| ENSG00000082996 | RNF13 | 1405.02429 | -0.14561496 | 0.063510507 | -2.292769583 | 0.021861274 | 0.059090094 |
| ENSG00000083093 | PALB2 | 747.8478469 | -0.066919642 | 0.077670108 | -0.861588122 | 0.38891421 | 0.581558697 |
| ENSG00000083097 | DOP1A | 715.1269276 | -0.129795005 | 0.076098569 | -1.705616901 | 0.088079431 | 0.191085884 |
| ENSG00000083099 | LYRM2 | 1931.160892 | -0.0151862 | 0.054831228 | -0.276962611 | 0.781808816 | 0.881574709 |
| ENSG00000083123 | BCKDHB | 405.4098055 | 0.082896615 | 0.095230344 | 0.8704853 | 0.384035249 | 0.576480082 |
| ENSG00000083168 | KAT6A | 2322.542128 | -0.141306716 | 0.068572828 | -2.060680877 | 0.039333495 | 0.097832634 |
| ENSG00000083223 | TUT7 | 1057.558333 | 0.065810537 | 0.066578348 | 0.988467555 | 0.322923717 | 0.512862991 |
| ENSG00000083290 | ULK2 | 1368.386527 | -0.031090771 | 0.061929869 | -0.502031918 | 0.615645069 | 0.772264951 |
| ENSG00000083307 | GRHL2 | 5560.884047 | -0.449268378 | 0.044502715 | -10.09530277 | 5.80E-24 | 1.21E-22 |
| ENSG00000083312 | TNPO1 | 3934.893262 | 0.134315708 | 0.051496329 | 2.608257954 | 0.009100435 | 0.027393252 |
| ENSG00000083444 | PLOD1 | 1853.724582 | 0.215224515 | 0.068802073 | 3.12816903 | 0.00175899 | 0.006255035 |
| ENSG00000083454 | P2RX5 | 5.910437333 | 2.546887854 | 0.890568364 | 2.859845417 | 0.004238476 | 0.013888106 |
| ENSG00000083457 | ITGAE | 291.4830354 | 0.120370841 | 0.109684946 | 1.09742353 | 0.272456292 | 0.454523114 |
| ENSG00000083520 | DIS3 | 1383.049061 | 0.043091238 | 0.066003386 | 0.652864052 | 0.513843926 | 0.692979404 |
| ENSG00000083535 | PIBF1 | 280.3388634 | 0.14573242 | 0.110402487 | 1.320010302 | 0.186831579 | 0.344541361 |
| ENSG00000083544 | TDRD3 | 162.1095372 | -0.213167731 | 0.144013843 | -1.480189177 | 0.138822768 | 0.274770863 |
| ENSG00000083635 | NUFIP1 | 472.9548882 | 0.062452282 | 0.088266428 | 0.70754287 | 0.479229182 | 0.663653377 |
| ENSG00000083642 | PDS5B | 3340.88452 | -0.748977682 | 0.048954127 | -15.29958218 | 7.69E-53 | 4.55E-51 |
| ENSG00000083720 | OXCT1 | 4059.446454 | 0.089292644 | 0.05022471 | 1.777862832 | 0.075426386 | 0.168643401 |
| ENSG00000083750 | RRAGB | 500.5027948 | 0.254729688 | 0.091704311 | 2.777728602 | 0.005474032 | 0.017440874 |
| ENSG00000083799 | CYLD | 538.0786488 | 0.656093383 | 0.088045943 | 7.451716192 | 9.21E-14 | 1.01E-12 |
| ENSG00000083807 | SLC27A5 | 277.3259139 | 0.141491398 | 0.113221464 | 1.249687053 | 0.211413889 | 0.378974706 |
| ENSG00000083812 | ZNF324 | 581.384153 | 0.017405356 | 0.085376913 | 0.203864901 | 0.838459074 | 0.91500172 |
| ENSG00000083817 | ZNF416 | 245.6587661 | 0.591952318 | 0.122710565 | 4.823971889 | 1.41E-06 | 7.98E-06 |
| ENSG00000083828 | ZNF586 | 328.0834504 | 0.016379255 | 0.103649695 | 0.158025111 | 0.874437012 | 0.934873703 |
| ENSG00000083838 | ZNF446 | 547.4108412 | 0.084887066 | 0.091324288 | 0.92951249 | 0.352623556 | 0.544055279 |
| ENSG00000083845 | RPS5 | 11451.69924 | 0.27424759 | 0.044550122 | 6.155933576 | 7.46E-10 | 6.01E-09 |
| ENSG00000083857 | FAT1 | 1463.644747 | -0.477015334 | 0.075548337 | -6.314041476 | 2.72E-10 | 2.28E-09 |
| ENSG00000083896 | YTHDC1 | 3218.815301 | -0.137517322 | 0.051336415 | -2.678748055 | 0.007389797 | 0.022782146 |
| ENSG00000083937 | CHMP2B | 1305.689749 | -0.129591709 | 0.068609452 | -1.888831723 | 0.058914381 | 0.137386285 |
| ENSG00000084070 | SMAP2 | 2012.56014 | -0.742699005 | 0.056274777 | -13.1977246 | 9.04E-40 | 3.64E-38 |

| | | | | | | | |
|---|---|---|---|---|---|---|---|
| ENSG00000084072 | PPIE | 1543.713302 | -0.089984623 | 0.058849632 | -1.529060087 | 0.126249549 | 0.255393708 |
| ENSG00000084073 | ZMPSTE24 | 1939.316851 | -0.261854065 | 0.064977337 | -4.029929161 | 5.58E-05 | 0.000259143 |
| ENSG00000084090 | STARD7 | 6192.260784 | -0.101989185 | 0.045713742 | -2.231039959 | 0.025678482 | 0.067927166 |
| ENSG00000084092 | NOA1 | 1777.599003 | 0.40288443 | 0.057562108 | 6.999125716 | 2.58E-12 | 2.55E-11 |
| ENSG00000084093 | REST | 2496.479019 | -0.035186207 | 0.052542728 | -0.669668445 | 0.503069171 | 0.6841192 |
| ENSG00000084110 | HAL | 8.319708926 | -0.00858604 | 0.603224549 | -0.014233572 | 0.988643636 | 0.994082067 |
| ENSG00000084112 | SSH1 | 2279.23576 | -0.587942542 | 0.058287537 | -10.08693398 | 6.31E-24 | 1.32E-22 |
| ENSG00000084234 | APLP2 | 12391.89501 | 0.377036586 | 0.043320355 | 8.703450869 | 3.22E-18 | 4.83E-17 |
| ENSG00000084444 | FAM234B | 2078.779201 | -0.80461656 | 0.059798432 | -13.45547934 | 2.86E-41 | 1.23E-39 |
| ENSG00000084453 | SLCO1A2 | 2.777510185 | 0.219912842 | 1.10951496 | 0.198206288 | 0.842883668 | 0.917308531 |
| ENSG00000084463 | WBP11 | 2597.162622 | -0.032573989 | 0.050544978 | -0.644455499 | 0.519280106 | 0.696787882 |
| ENSG00000084623 | EIF3I | 6836.567341 | 0.151670251 | 0.041350494 | 3.667918677 | 0.000244533 | 0.001019925 |
| ENSG00000084628 | NKAIN1 | 578.2130426 | 1.741900391 | 0.089196734 | 19.52874623 | 6.26E-85 | 7.80E-83 |
| ENSG00000084636 | COL16A1 | 82.8819728 | 0.557227248 | 0.197931827 | 2.81524834 | 0.004873955 | 0.015756558 |
| ENSG00000084652 | TXLNA | 3455.160312 | -0.0794242 | 0.047953041 | -1.656291206 | 0.097662861 | 0.207967775 |
| ENSG00000084676 | NCOA1 | 1969.92729 | -0.178922617 | 0.059596968 | -3.002210065 | 0.002680272 | 0.009177483 |
| ENSG00000084693 | AGBL5 | 1072.462328 | 0.020691845 | 0.066021837 | 0.313409101 | 0.753969871 | 0.864328061 |
| ENSG00000084710 | EFR3B | 1323.560311 | 0.620159128 | 0.065859663 | 9.416372678 | 4.67E-21 | 8.30E-20 |
| ENSG00000084731 | KIF3C | 1355.093274 | 0.028082968 | 0.06273278 | 0.447660192 | 0.654398455 | 0.799387444 |
| ENSG00000084733 | RAB10 | 5233.326655 | -0.035183377 | 0.050949257 | -0.69055723 | 0.489843833 | 0.672996199 |
| ENSG00000084754 | HADHA | 7585.877489 | 0.011272306 | 0.043666816 | 0.258143531 | 0.796296137 | 0.891280831 |
| ENSG00000084764 | MAPRE3 | 513.5778026 | 0.667041563 | 0.086871963 | 7.678444684 | 1.61E-14 | 1.86E-13 |
| ENSG00000084774 | CAD | 4302.688863 | -0.146996315 | 0.052366627 | -2.807060994 | 0.004999577 | 0.016108189 |
| ENSG00000085063 | CD59 | 1168.573107 | 0.399210878 | 0.064956647 | 6.145804859 | 7.96E-10 | 6.38E-09 |
| ENSG00000085117 | CD82 | 83.85151711 | 1.360834608 | 0.218212082 | 6.236293594 | 4.48E-10 | 3.69E-09 |
| ENSG00000085185 | BCORL1 | 660.5698016 | -0.450801287 | 0.090353758 | -4.989292029 | 6.06E-07 | 3.61E-06 |
| ENSG00000085224 | ATRX | 4242.04273 | -0.260190326 | 0.048960047 | -5.314339734 | 1.07E-07 | 6.92E-07 |
| ENSG00000085231 | AK6 | 327.590494 | 0.03013975 | 0.104167759 | 0.289338568 | 0.772322301 | 0.875955383 |
| ENSG00000085274 | MYNN | 1156.71591 | -0.102418489 | 0.064111193 | -1.597513387 | 0.110151316 | 0.22900772 |
| ENSG00000085276 | MECOM | 443.3397011 | 1.326484465 | 0.101288797 | 13.09606303 | 3.47E-39 | 1.36E-37 |
| ENSG00000085365 | SCAMP1 | 1720.745984 | -0.059889647 | 0.07573452 | -0.790783998 | 0.429070049 | 0.619398838 |
| ENSG00000085377 | PREP | 1483.700839 | 0.093957427 | 0.060096542 | 1.563441474 | 0.117948791 | 0.242363425 |
| ENSG00000085382 | HACE1 | 378.2326722 | 0.102632127 | 0.097696321 | 1.050521927 | 0.293478215 | 0.479003219 |

| | | | | | | | |
|---|---|---|---|---|---|---|---|
| ENSG00000085415 | SEH1L | 1849.972178 | 0.356123592 | 0.057185758 | 6.227487481 | 4.74E-10 | 3.89E-09 |
| ENSG00000085433 | WDR47 | 502.2597166 | -0.004870205 | 0.087157644 | -0.055878113 | 0.955438907 | 0.979458252 |
| ENSG00000085449 | WDFY1 | 835.7374518 | 0.081487769 | 0.072863618 | 1.118360192 | 0.263413189 | 0.44412799 |
| ENSG00000085465 | OVGP1 | 29.68968443 | 0.626117466 | 0.324351144 | 1.930369224 | 0.053561105 | 0.127167435 |
| ENSG00000085491 | SLC25A24 | 2911.721573 | -0.060974798 | 0.051869871 | -1.175534013 | 0.239781138 | 0.415482989 |
| ENSG00000085511 | MAP3K4 | 800.6815436 | -0.468743477 | 0.073043187 | -6.417346997 | 1.39E-10 | 1.18E-09 |
| ENSG00000085514 | PILRA | 13.44068363 | -1.052612459 | 0.512771915 | -2.05278883 | 0.040093062 | 0.09930197 |
| ENSG00000085552 | IGSF9 | 1457.83435 | 0.714255485 | 0.063033039 | 11.33144614 | 9.17E-30 | 2.53E-28 |
| ENSG00000085644 | ZNF213 | 1181.29837 | 0.054415126 | 0.072840339 | 0.747046585 | 0.455035441 | 0.642488646 |
| ENSG00000085719 | CPNE3 | 10591.68703 | 0.361000829 | 0.048186515 | 7.491739743 | 6.80E-14 | 7.51E-13 |
| ENSG00000085721 | RRN3 | 2079.101818 | 0.069979478 | 0.059903473 | 1.168204016 | 0.242724481 | 0.419006978 |
| ENSG00000085733 | CTTN | 10305.54063 | 0.288143398 | 0.043883796 | 6.566054527 | 5.17E-11 | 4.57E-10 |
| ENSG00000085741 | WNT11 | 14.89818932 | 0.072461637 | 0.458084326 | 0.158184057 | 0.874311766 | 0.934789447 |
| ENSG00000085760 | MTIF2 | 1434.413137 | -0.089393 | 0.061257797 | -1.459291793 | 0.144484813 | 0.283143825 |
| ENSG00000085788 | DDHD2 | 1248.046918 | 0.071991357 | 0.062972329 | 1.143222076 | 0.252946399 | 0.431735218 |
| ENSG00000085831 | TTC39A | 2434.713683 | -0.089618773 | 0.052556387 | -1.705192819 | 0.088158469 | 0.191239269 |
| ENSG00000085832 | EPS15 | 2450.315167 | 0.110672419 | 0.053433879 | 2.07120315 | 0.038339815 | 0.095807774 |
| ENSG00000085840 | ORC1 | 698.311765 | -0.136939474 | 0.07636399 | -1.793246702 | 0.072933486 | 0.164205924 |
| ENSG00000085871 | MGST2 | 362.709883 | -0.052409309 | 0.100430772 | -0.521845128 | 0.601778166 | 0.761366787 |
| ENSG00000085872 | CHERP | 1926.236736 | 0.122363194 | 0.059904555 | 2.042635871 | 0.041088497 | 0.101358648 |
| ENSG00000085978 | ATG16L1 | 1043.887787 | -0.380654256 | 0.066836881 | -5.695272545 | 1.23E-08 | 8.76E-08 |
| ENSG00000085982 | USP40 | 1015.561712 | -0.077809529 | 0.067685014 | -1.149582819 | 0.250315738 | 0.42846165 |
| ENSG00000085998 | POMGNT1 | 2086.056124 | 0.372316285 | 0.053733373 | 6.928957934 | 4.24E-12 | 4.11E-11 |
| ENSG00000085999 | RAD54L | 806.7608297 | -0.011848696 | 0.074907905 | -0.15817684 | 0.874317453 | 0.934789447 |
| ENSG00000086015 | MAST2 | 2848.847006 | -0.16281381 | 0.054424727 | -2.991541148 | 0.002775731 | 0.009479719 |
| ENSG00000086061 | DNAJA1 | 5821.903374 | 0.060765744 | 0.042721787 | 1.422359609 | 0.154921883 | 0.298472158 |
| ENSG00000086062 | B4GALT1 | 2215.688638 | 0.376105154 | 0.054956388 | 6.843702179 | 7.72E-12 | 7.30E-11 |
| ENSG00000086065 | CHMP5 | 1347.664599 | 0.186594609 | 0.061173246 | 3.050264956 | 0.002286396 | 0.007946957 |
| ENSG00000086102 | NFX1 | 1503.517669 | -0.144603663 | 0.063026029 | -2.294348317 | 0.021770499 | 0.058865542 |
| ENSG00000086159 | AQP6 | 8.848483608 | 0.552208174 | 0.623107235 | 0.88621692 | 0.375500648 | 0.56836558 |
| ENSG00000086189 | DIMT1 | 599.5553785 | 0.136913093 | 0.082810213 | 1.653335842 | 0.098262532 | 0.208949063 |
| ENSG00000086200 | IPO11 | 890.3841341 | -0.021716633 | 0.070153822 | -0.309557371 | 0.756897578 | 0.865910156 |
| ENSG00000086205 | FOLH1 | 62.15610039 | -3.684801079 | 0.326396047 | -11.28935571 | 1.48E-29 | 4.05E-28 |

| | | | | | | | |
|---|---|---|---|---|---|---|---|
| ENSG00000086232 | EIF2AK1 | 11436.77379 | 0.238036562 | 0.042821043 | 5.55886881 | 2.72E-08 | 1.86E-07 |
| ENSG00000086300 | SNX10 | 201.1187155 | -1.328221684 | 0.141347316 | -9.39686526 | 5.62E-21 | 9.94E-20 |
| ENSG00000086475 | SEPHS1 | 2675.20432 | -0.605368239 | 0.051079443 | -11.85150422 | 2.11E-32 | 6.49E-31 |
| ENSG00000086504 | MRPL28 | 2003.796677 | -0.098760547 | 0.059312708 | -1.665082414 | 0.095896315 | 0.204897612 |
| ENSG00000086544 | ITPKC | 1129.146057 | 0.155251368 | 0.069864013 | 2.222193686 | 0.026270219 | 0.069276793 |
| ENSG00000086548 | CEACAM6 | 319.8041561 | 0.214900294 | 0.121090021 | 1.774715145 | 0.075944933 | 0.169620949 |
| ENSG00000086570 | FAT2 | 3.419155569 | -0.867637498 | 0.957940427 | -0.905732208 | 0.36507761 | 0.557837983 |
| ENSG00000086589 | RBM22 | 2052.627835 | 0.013321586 | 0.052901641 | 0.251818014 | 0.801181733 | 0.894273499 |
| ENSG00000086598 | TMED2 | 5798.708876 | -0.028252625 | 0.047311137 | -0.59716647 | 0.550396243 | 0.722615811 |
| ENSG00000086619 | ERO1B | 83.52595124 | -1.541034787 | 0.209010699 | -7.372994756 | 1.67E-13 | 1.80E-12 |
| ENSG00000086666 | ZFAND6 | 1355.296038 | -0.029816128 | 0.062698379 | -0.475548624 | 0.634395985 | 0.784488377 |
| ENSG00000086712 | TXLNG | 1521.696301 | 0.223575091 | 0.061253428 | 3.650001302 | 0.000262239 | 0.001088826 |
| ENSG00000086717 | PPEF1 | 28.27604589 | -1.921599143 | 0.364020296 | -5.278824188 | 1.30E-07 | 8.36E-07 |
| ENSG00000086730 | LAT2 | 9.776537846 | 0.651253254 | 0.582828271 | 1.117401619 | 0.263822643 | 0.444499989 |
| ENSG00000086758 | HUWE1 | 16126.70375 | 0.54064621 | 0.064347377 | 8.401993028 | 4.39E-17 | 6.07E-16 |
| ENSG00000086827 | ZW10 | 705.2966097 | -0.06437582 | 0.076254633 | -0.844221747 | 0.398545504 | 0.591564793 |
| ENSG00000086848 | ALG9 | 121.9983504 | -0.035313093 | 0.168950423 | -0.20901453 | 0.834436896 | 0.912652922 |
| ENSG00000087008 | ACOX3 | 780.497471 | 0.211961508 | 0.072182999 | 2.936446404 | 0.003319962 | 0.011139378 |
| ENSG00000087053 | MTMR2 | 825.6089339 | -0.085132943 | 0.076079017 | -1.119006869 | 0.263137209 | 0.443948486 |
| ENSG00000087074 | PPP1R15A | 737.249112 | 0.613714209 | 0.078704536 | 7.797698077 | 6.30E-15 | 7.52E-14 |
| ENSG00000087076 | HSD17B14 | 15.99536546 | 0.142678893 | 0.453963095 | 0.314296239 | 0.753296055 | 0.864030575 |
| ENSG00000087077 | TRIP6 | 90.72812117 | -0.191856979 | 0.193850356 | -0.989716927 | 0.322312499 | 0.512145787 |
| ENSG00000087085 | ACHE | 770.028906 | -2.229307561 | 0.091246336 | -24.43174879 | 7.87E-132 | 2.31E-129 |
| ENSG00000087086 | FTL | 8003.496492 | 0.534329516 | 0.043731456 | 12.21842494 | 2.48E-34 | 8.22E-33 |
| ENSG00000087087 | SRRT | 5835.596571 | -0.561824914 | 0.047252707 | -11.88979328 | 1.34E-32 | 4.12E-31 |
| ENSG00000087088 | BAX | 1670.00035 | -0.126288542 | 0.061414568 | -2.056328763 | 0.039750838 | 0.098646064 |
| ENSG00000087095 | NLK | 630.0638385 | -0.006361338 | 0.078786763 | -0.0807412 | 0.935647771 | 0.969248877 |
| ENSG00000087111 | PIGS | 3183.9873 | 0.040399488 | 0.053661991 | 0.752851095 | 0.451539399 | 0.639481035 |
| ENSG00000087128 | TMPRSS11E | 3.683606475 | 1.304048858 | 1.015936112 | 1.28359337 | 0.199284271 | 0.362162811 |
| ENSG00000087152 | ATXN7L3 | 3781.935053 | -0.093679808 | 0.052782756 | -1.774818422 | 0.075927874 | 0.169615875 |
| ENSG00000087157 | PGS1 | 1408.107917 | 0.166828751 | 0.068040493 | 2.451903911 | 0.01421026 | 0.040519996 |
| ENSG00000087191 | PSMC5 | 9522.654034 | 0.022560279 | 0.041328998 | 0.54587045 | 0.585154992 | 0.74932765 |
| ENSG00000087206 | UIMC1 | 1438.646047 | -0.577312296 | 0.06342576 | -9.102173929 | 8.85E-20 | 1.46E-18 |

| | | | | | | | |
|---|---|---|---|---|---|---|---|
| ENSG00000087253 | LPCAT2 | 241.0918007 | 1.990976328 | 0.134556652 | 14.79656565 | 1.54E-49 | 8.44E-48 |
| ENSG00000087258 | GNAO1 | 8.03947327 | 0.463938723 | 0.627971731 | 0.738789184 | 0.460035021 | 0.646880187 |
| ENSG00000087263 | OGFOD1 | 1128.560527 | 0.119046687 | 0.065323596 | 1.822414804 | 0.06839207 | 0.155661681 |
| ENSG00000087266 | SH3BP2 | 2015.742113 | 0.794581087 | 0.063635654 | 12.48641355 | 8.86E-36 | 3.07E-34 |
| ENSG00000087269 | NOP14 | 3540.296422 | 0.014083136 | 0.046454173 | 0.303161919 | 0.761766469 | 0.869297509 |
| ENSG00000087274 | ADD1 | 6937.566629 | -0.118354611 | 0.045064195 | -2.626355836 | 0.008630454 | 0.026137427 |
| ENSG00000087299 | L2HGDH | 922.8270224 | -0.506745721 | 0.072002484 | -7.037892157 | 1.95E-12 | 1.95E-11 |
| ENSG00000087301 | TXNDC16 | 834.7799718 | -0.009650663 | 0.072289679 | -0.133499867 | 0.893798069 | 0.945700277 |
| ENSG00000087302 | RTRAF | 2903.878677 | -0.63843913 | 0.056809348 | -11.23827601 | 2.64E-29 | 7.15E-28 |
| ENSG00000087338 | GMCL1 | 1189.712875 | 0.071900754 | 0.068998237 | 1.042066526 | 0.297380837 | 0.483809983 |
| ENSG00000087365 | SF3B2 | 12679.74729 | -0.041089143 | 0.041884392 | -0.981013246 | 0.326586209 | 0.517145554 |
| ENSG00000087448 | KLHL42 | 1197.112245 | 0.291811222 | 0.070110881 | 4.162138836 | 3.15E-05 | 0.000152392 |
| ENSG00000087460 | GNAS | 49666.53989 | -0.323923389 | 0.052571455 | -6.161583059 | 7.20E-10 | 5.81E-09 |
| ENSG00000087470 | DNM1L | 1925.528672 | 0.240530274 | 0.058111026 | 4.139150332 | 3.49E-05 | 0.000166703 |
| ENSG00000087494 | PTHLH | 2.980992866 | 0.092566329 | 1.016003477 | 0.09110828 | 0.927406554 | 0.965137362 |
| ENSG00000087495 | PHACTR3 | 2.638200672 | 3.698907947 | 1.471728347 | 2.513308896 | 0.011960456 | 0.034840998 |
| ENSG00000087502 | ERGIC2 | 872.7602064 | 0.176769263 | 0.073802782 | 2.395157169 | 0.016613242 | 0.046531247 |
| ENSG00000087510 | TFAP2C | 2620.645293 | 0.155406006 | 0.051428458 | 3.021790087 | 0.002512847 | 0.008654064 |
| ENSG00000087586 | AURKA | 3953.851211 | -0.275554926 | 0.047349548 | -5.819589369 | 5.90E-09 | 4.36E-08 |
| ENSG00000087589 | CASS4 | 4.479335056 | 2.849510867 | 1.074785949 | 2.651235691 | 0.008019786 | 0.024497413 |
| ENSG00000087842 | PIR | 394.096927 | -0.200462487 | 0.097578103 | -2.054379836 | 0.039938943 | 0.099016465 |
| ENSG00000087884 | AAMDC | 274.1594812 | -0.151701426 | 0.112325191 | -1.350555686 | 0.176837804 | 0.330589131 |
| ENSG00000087903 | RFX2 | 234.5750665 | -0.936494813 | 0.121737919 | -7.692712487 | 1.44E-14 | 1.67E-13 |
| ENSG00000087995 | METTL2A | 2240.082677 | 0.146288412 | 0.059017449 | 2.478731518 | 0.013185051 | 0.037926655 |
| ENSG00000088002 | SULT2B1 | 1014.481712 | 0.180083448 | 0.089346076 | 2.015571991 | 0.043844751 | 0.107113801 |
| ENSG00000088035 | ALG6 | 346.3713884 | -0.084096745 | 0.101969055 | -0.824728101 | 0.409525974 | 0.602065433 |
| ENSG00000088038 | CNOT3 | 3048.83621 | -0.156765464 | 0.056182767 | -2.790276662 | 0.005266302 | 0.016853929 |
| ENSG00000088053 | GP6 | 3.59988492 | 0.894144572 | 0.940245245 | 0.950969522 | 0.341619848 | 0.532714248 |
| ENSG00000088179 | PTPN4 | 683.7843877 | -0.101640477 | 0.077411546 | -1.312988591 | 0.189186792 | 0.348002968 |
| ENSG00000088205 | DDX18 | 3193.000035 | 0.068588491 | 0.053673995 | 1.277871911 | 0.201294594 | 0.364934116 |
| ENSG00000088247 | KHSRP | 6058.594398 | -0.002230609 | 0.046462111 | -0.048009214 | 0.9617089 | 0.982361474 |
| ENSG00000088256 | GNA11 | 2847.784487 | 0.056070614 | 0.057968235 | 0.967264473 | 0.333411839 | 0.52456184 |
| ENSG00000088280 | ASAP3 | 653.737558 | 0.146728662 | 0.084395659 | 1.738580683 | 0.082108547 | 0.180487741 |

| | | | | | | | |
|---|---|---|---|---|---|---|---|
| ENSG00000088298 | EDEM2 | 425.3492861 | -0.737744768 | 0.102321073 | -7.210096055 | 5.59E-13 | 5.82E-12 |
| ENSG00000088305 | DNMT3B | 968.6115047 | -1.973351674 | 0.075295359 | -26.20814472 | 2.15E-151 | 7.69E-149 |
| ENSG00000088325 | TPX2 | 4829.307227 | -0.737757331 | 0.047517001 | -15.5261761 | 2.31E-54 | 1.42E-52 |
| ENSG00000088340 | FER1L4 | 236.7653587 | 0.423979839 | 0.13021526 | 3.255991961 | 0.00112997 | 0.004191891 |
| ENSG00000088356 | PDRG1 | 1172.172863 | -0.126217198 | 0.066619025 | -1.894611915 | 0.058143835 | 0.135937584 |
| ENSG00000088367 | EPB41L1 | 1906.881734 | 0.290465439 | 0.055228725 | 5.259318202 | 1.45E-07 | 9.25E-07 |
| ENSG00000088387 | DOCK9 | 1044.376661 | -0.111735424 | 0.075575524 | -1.478460457 | 0.139284574 | 0.275447252 |
| ENSG00000088448 | ANKRD10 | 1116.138473 | 0.043843083 | 0.063839936 | 0.686765773 | 0.492230342 | 0.675102478 |
| ENSG00000088451 | TGDS | 190.0879751 | -0.081509572 | 0.131420817 | -0.620218117 | 0.535114196 | 0.71059448 |
| ENSG00000088538 | DOCK3 | 111.8412242 | 0.40383979 | 0.169728498 | 2.379328127 | 0.017344229 | 0.048315315 |
| ENSG00000088543 | C3orf18 | 87.17435201 | -0.152912138 | 0.187719582 | -0.814577447 | 0.415314224 | 0.607398286 |
| ENSG00000088682 | COQ9 | 1012.007861 | 0.092057489 | 0.072467936 | 1.270320282 | 0.203970567 | 0.368731414 |
| ENSG00000088726 | TMEM40 | 31.92207599 | 0.407199327 | 0.31412685 | 1.296289466 | 0.194875776 | 0.355975658 |
| ENSG00000088727 | KIF9 | 127.543397 | 0.346752195 | 0.160179105 | 2.164777956 | 0.030404694 | 0.078616284 |
| ENSG00000088756 | ARHGAP28 | 43.48567597 | 2.75621842 | 0.340398194 | 8.097041852 | 5.63E-16 | 7.26E-15 |
| ENSG00000088766 | CRLS1 | 10.27480712 | 0.085056478 | 0.542943769 | 0.156657987 | 0.8755144 | 0.935458795 |
| ENSG00000088808 | PPP1R13B | 1410.226932 | -0.201247619 | 0.060688625 | -3.316068188 | 0.000912936 | 0.003443397 |
| ENSG00000088812 | ATRN | 1901.294876 | 0.168450874 | 0.062157362 | 2.710071167 | 0.006726877 | 0.020949574 |
| ENSG00000088826 | SMOX | 72.87904185 | 1.250046052 | 0.217193935 | 5.755437187 | 8.64E-09 | 6.25E-08 |
| ENSG00000088832 | FKBP1A | 2179.173576 | 0.1516419 | 0.057841797 | 2.621666457 | 0.008750102 | 0.026449775 |
| ENSG00000088833 | NSFL1C | 1735.535438 | -0.156530108 | 0.063673924 | -2.458307866 | 0.013959345 | 0.039903722 |
| ENSG00000088836 | SLC4A11 | 251.2373687 | 0.282768379 | 0.119807038 | 2.360198396 | 0.018265164 | 0.050518792 |
| ENSG00000088854 | DNAAF9 | 599.9535904 | -0.664008878 | 0.080627357 | -8.235528316 | 1.79E-16 | 2.39E-15 |
| ENSG00000088876 | ZNF343 | 155.3379237 | -0.26121046 | 0.143718248 | -1.81751771 | 0.069137879 | 0.157063076 |
| ENSG00000088881 | EBF4 | 67.30988565 | -1.567390803 | 0.23287565 | -6.730591222 | 1.69E-11 | 1.55E-10 |
| ENSG00000088882 | CPXM1 | 1.569619885 | 1.141925957 | 1.512342092 | 0.755071199 | 0.450206263 | NA |
| ENSG00000088888 | MAVS | 9517.83983 | 0.152516922 | 0.047271956 | 3.226372142 | 0.001253702 | 0.004608225 |
| ENSG00000088899 | LZTS3 | 69.34570313 | 0.580676149 | 0.217892792 | 2.664962632 | 0.007699688 | 0.023632705 |
| ENSG00000088930 | XRN2 | 4894.242472 | -0.38790811 | 0.04565523 | -8.496466062 | 1.95E-17 | 2.77E-16 |
| ENSG00000088970 | KIZ | 221.2595441 | -0.058485646 | 0.126265038 | -0.463197468 | 0.643222831 | 0.79088448 |
| ENSG00000088986 | DYNLL1 | 3364.814675 | -0.365950414 | 0.050965346 | -7.18037737 | 6.95E-13 | 7.21E-12 |
| ENSG00000088992 | TESC | 1062.939706 | 3.539230996 | 0.089776181 | 39.42282847 | 0 | 0 |
| ENSG00000089006 | SNX5 | 2672.792802 | -0.22149787 | 0.06861081 | -3.228323183 | 0.001245182 | 0.004579108 |

| | | | | | | | |
|---|---|---|---|---|---|---|---|
| ENSG00000089009 | RPL6 | 6573.89137 | 0.184803655 | 0.054542867 | 3.388227717 | 0.000703458 | 0.00270897 |
| ENSG00000089022 | MAPKAPK5 | 1569.975085 | 0.120152872 | 0.058767863 | 2.044533623 | 0.040900859 | 0.101019133 |
| ENSG00000089041 | P2RX7 | 3.101563294 | -0.549729644 | 1.013177571 | -0.542579761 | 0.587419177 | 0.751053167 |
| ENSG00000089048 | ESF1 | 837.2394527 | -0.205811944 | 0.075397051 | -2.729708134 | 0.006339042 | 0.019892972 |
| ENSG00000089050 | RBBP9 | 793.9340274 | -0.338362923 | 0.073383164 | -4.610906735 | 4.01E-06 | 2.17E-05 |
| ENSG00000089053 | ANAPC5 | 4604.483543 | -0.117671947 | 0.047187399 | -2.493715459 | 0.012641384 | 0.0365415 |
| ENSG00000089057 | SLC23A2 | 1037.795416 | 0.257853675 | 0.067791994 | 3.803600713 | 0.000142608 | 0.000620176 |
| ENSG00000089060 | SLC8B1 | 288.4468884 | 0.578707492 | 0.112416523 | 5.147886425 | 2.63E-07 | 1.64E-06 |
| ENSG00000089063 | TMEM230 | 1575.668415 | 0.039934565 | 0.059451872 | 0.671712495 | 0.501766745 | 0.682833786 |
| ENSG00000089091 | DZANK1 | 168.5776416 | -0.272413149 | 0.14864979 | -1.832583469 | 0.066864538 | 0.152867501 |
| ENSG00000089094 | KDM2B | 1984.298189 | 0.004168802 | 0.056708494 | 0.073512834 | 0.941398032 | 0.972164889 |
| ENSG00000089101 | CFAP61 | 3.090162142 | -1.566820125 | 1.042392531 | -1.503099915 | 0.132813281 | 0.265904753 |
| ENSG00000089123 | TASP1 | 413.8872886 | -0.423301186 | 0.09628831 | -4.396184599 | 1.10E-05 | 5.66E-05 |
| ENSG00000089127 | OAS1 | 25.13032591 | 4.787913109 | 0.71082915 | 6.735673553 | 1.63E-11 | 1.50E-10 |
| ENSG00000089154 | GCN1 | 8956.29095 | 0.017707301 | 0.048986041 | 0.361476476 | 0.717743287 | 0.843020378 |
| ENSG00000089157 | RPLP0 | 17718.89922 | 0.059365918 | 0.05467643 | 1.085768001 | 0.277581647 | 0.460354019 |
| ENSG00000089159 | PXN | 5944.756846 | -0.109361657 | 0.050342597 | -2.172348348 | 0.029829398 | 0.077294294 |
| ENSG00000089163 | SIRT4 | 10.72801181 | -0.480637106 | 0.53202749 | -0.903406525 | 0.366310178 | 0.559017794 |
| ENSG00000089177 | KIF16B | 574.229719 | -0.06944467 | 0.08572599 | -0.810077199 | 0.417895808 | 0.609908631 |
| ENSG00000089195 | TRMT6 | 587.7283537 | 0.032884201 | 0.083254716 | 0.394983048 | 0.692855399 | 0.826827386 |
| ENSG00000089220 | PEBP1 | 8907.317159 | 0.166625404 | 0.041795172 | 3.986714184 | 6.70E-05 | 0.000307371 |
| ENSG00000089234 | BRAP | 994.2195213 | 0.12699903 | 0.06660854 | 1.906647865 | 0.056566202 | 0.132899292 |
| ENSG00000089248 | ERP29 | 3739.256202 | 0.15141359 | 0.04868033 | 3.1103649 | 0.001868564 | 0.006607807 |
| ENSG00000089280 | FUS | 13054.69894 | -0.290471319 | 0.053699966 | -5.409152706 | 6.33E-08 | 4.18E-07 |
| ENSG00000089289 | IGBP1 | 2754.314035 | 0.192719916 | 0.052024064 | 3.704437953 | 0.00021186 | 0.000894217 |
| ENSG00000089327 | FXYD5 | 50.74228926 | 2.012940401 | 0.284748601 | 7.069184521 | 1.56E-12 | 1.57E-11 |
| ENSG00000089335 | ZNF302 | 785.6459971 | 0.265381609 | 0.083612379 | 3.173951186 | 0.00150379 | 0.005426607 |
| ENSG00000089351 | GRAMD1A | 2497.260528 | 0.856094784 | 0.059956483 | 14.27860244 | 2.98E-46 | 1.47E-44 |
| ENSG00000089356 | FXYD3 | 22.12382124 | 0.035167757 | 0.381132872 | 0.092271645 | 0.926482216 | 0.964663921 |
| ENSG00000089486 | CDIP1 | 2.351614765 | 0.87406727 | 1.194884858 | 0.731507529 | 0.464469207 | 0.650624508 |
| ENSG00000089505 | CMTM1 | 42.16378133 | 0.110700868 | 0.271653205 | 0.40750805 | 0.683634883 | 0.820502629 |
| ENSG00000089597 | GANAB | 17665.21037 | -0.117764177 | 0.04159911 | -2.830930234 | 0.004641284 | 0.015076616 |
| ENSG00000089639 | GMIP | 438.5837329 | 0.522077457 | 0.092309316 | 5.6557396 | 1.55E-08 | 1.09E-07 |

| | | | | | | | |
|---|---|---|---|---|---|---|---|
| ENSG00000089682 | RBM41 | 588.1628654 | 0.272913699 | 0.088140612 | 3.096344508 | 0.001959226 | 0.006901818 |
| ENSG00000089685 | BIRC5 | 2840.421918 | -0.08944542 | 0.050996481 | -1.753952804 | 0.079438595 | 0.175733747 |
| ENSG00000089692 | LAG3 | 77.5449258 | -0.963560697 | 0.215851506 | -4.463998019 | 8.04E-06 | 4.19E-05 |
| ENSG00000089693 | MLF2 | 3828.384861 | 0.049381687 | 0.046854456 | 1.053937898 | 0.291911351 | 0.476936684 |
| ENSG00000089723 | OTUB2 | 131.4340219 | -0.504080633 | 0.159856032 | -3.153341325 | 0.00161413 | 0.005791981 |
| ENSG00000089737 | DDX24 | 5443.995809 | -0.609827767 | 0.045459201 | -13.41483697 | 4.95E-41 | 2.11E-39 |
| ENSG00000089775 | ZBTB25 | 550.668728 | -0.716107465 | 0.087808098 | -8.155369285 | 3.48E-16 | 4.55E-15 |
| ENSG00000089818 | NECAP1 | 641.2998111 | 0.029387612 | 0.080500346 | 0.365061937 | 0.715065173 | 0.841598433 |
| ENSG00000089820 | ARHGAP4 | 428.8472067 | -0.87830842 | 0.093805254 | -9.363104785 | 7.74E-21 | 1.36E-19 |
| ENSG00000089847 | ANKRD24 | 73.48236453 | -0.553947621 | 0.20648123 | -2.682798919 | 0.007300887 | 0.022550472 |
| ENSG00000089876 | DHX32 | 1224.777045 | -0.336679163 | 0.062167563 | -5.415672557 | 6.11E-08 | 4.03E-07 |
| ENSG00000089902 | RCOR1 | 2732.369434 | -0.377499247 | 0.05547839 | -6.804437713 | 1.01E-11 | 9.49E-11 |
| ENSG00000089916 | GPATCH2L | 1785.461407 | -0.460507242 | 0.066237202 | -6.952395725 | 3.59E-12 | 3.51E-11 |
| ENSG00000090006 | LTBP4 | 1262.41565 | 0.314513933 | 0.072378809 | 4.345386995 | 1.39E-05 | 7.04E-05 |
| ENSG00000090013 | BLVRB | 1227.411048 | -0.473071706 | 0.06503625 | -7.273969659 | 3.49E-13 | 3.67E-12 |
| ENSG00000090020 | SLC9A1 | 4905.197269 | 0.279773641 | 0.05663421 | 4.940011395 | 7.81E-07 | 4.59E-06 |
| ENSG00000090054 | SPTLC1 | 1650.750747 | -0.016896776 | 0.057947476 | -0.291587785 | 0.770601816 | 0.874999042 |
| ENSG00000090060 | PAPOLA | 10673.18585 | -0.219446997 | 0.047381385 | -4.631502388 | 3.63E-06 | 1.97E-05 |
| ENSG00000090061 | CCNK | 2608.89176 | -0.5259146 | 0.055760175 | -9.431724361 | 4.03E-21 | 7.21E-20 |
| ENSG00000090097 | PCBP4 | 616.9889445 | 0.101397406 | 0.079853256 | 1.269796756 | 0.204157037 | 0.368972065 |
| ENSG00000090238 | YPEL3 | 1378.569622 | 0.25100093 | 0.063110927 | 3.977139036 | 6.97E-05 | 0.000319308 |
| ENSG00000090263 | MRPS33 | 1313.037687 | -0.252914171 | 0.062510028 | -4.045977583 | 5.21E-05 | 0.000243045 |
| ENSG00000090266 | NDUFB2 | 1600.923362 | -0.196910225 | 0.058088321 | -3.389841927 | 0.000699329 | 0.002694879 |
| ENSG00000090273 | NUDC | 2515.491733 | 0.052276605 | 0.050661988 | 1.031870399 | 0.302132835 | 0.489092317 |
| ENSG00000090316 | MAEA | 2378.942768 | 0.11799489 | 0.053529614 | 2.204291816 | 0.027503828 | 0.072107179 |
| ENSG00000090339 | ICAM1 | 360.0696638 | 0.255731699 | 0.100649942 | 2.540803259 | 0.011059813 | 0.032464763 |
| ENSG00000090372 | STRN4 | 4246.981333 | 0.032606613 | 0.051326547 | 0.635277756 | 0.525247279 | 0.701327047 |
| ENSG00000090376 | IRAK3 | 1.925621632 | -0.14805094 | 1.252276012 | -0.118225486 | 0.905888996 | 0.95262805 |
| ENSG00000090382 | LYZ | 7.601517567 | 1.113886091 | 0.681846583 | 1.633631551 | 0.102336239 | 0.216029568 |
| ENSG00000090432 | MUL1 | 616.7238781 | -0.102136289 | 0.089767724 | -1.137784102 | 0.255210645 | 0.434699451 |
| ENSG00000090447 | TFAP4 | 432.6677065 | -0.066759279 | 0.095997726 | -0.69542563 | 0.486788603 | 0.670243716 |
| ENSG00000090470 | PDCD7 | 1086.049349 | 0.057918012 | 0.064320263 | 0.900462923 | 0.367873948 | 0.56039766 |
| ENSG00000090487 | SPG21 | 3337.179891 | 0.182171662 | 0.055132422 | 3.304256448 | 0.000952287 | 0.003579462 |

| | | | | | | | |
|---|---|---|---|---|---|---|---|
| ENSG00000090520 | DNAJB11 | 776.3017004 | -0.133251259 | 0.075407588 | -1.767080236 | 0.077214792 | 0.171971585 |
| ENSG00000090530 | P3H2 | 1.611251199 | 0.43414744 | 1.404493436 | 0.309113186 | 0.757235429 | NA |
| ENSG00000090534 | THPO | 19.05457607 | 1.076234533 | 0.437203608 | 2.46163232 | 0.013830637 | 0.039580191 |
| ENSG00000090539 | CHRD | 424.9799119 | 1.187559186 | 0.102782174 | 11.55413571 | 7.04E-31 | 2.03E-29 |
| ENSG00000090554 | FLT3LG | 4.442427324 | -0.673569757 | 0.833110012 | -0.808500374 | 0.418802591 | 0.610531992 |
| ENSG00000090565 | RAB11FIP3 | 2728.932396 | 0.220029081 | 0.057376453 | 3.834832397 | 0.00012565 | 0.00055113 |
| ENSG00000090581 | GNPTG | 1109.875498 | 0.262745022 | 0.074328571 | 3.534912846 | 0.00040791 | 0.001638209 |
| ENSG00000090612 | ZNF268 | 1006.151872 | 0.19962164 | 0.071039086 | 2.810025474 | 0.004953758 | 0.01597627 |
| ENSG00000090615 | GOLGA3 | 3568.659532 | -0.166854609 | 0.053413969 | -3.123800972 | 0.001785312 | 0.006340773 |
| ENSG00000090621 | PABPC4 | 6259.600326 | 0.249184737 | 0.048130521 | 5.177270761 | 2.25E-07 | 1.41E-06 |
| ENSG00000090661 | CERS4 | 2178.90959 | 0.246136139 | 0.071419866 | 3.446325985 | 0.000568264 | 0.00222571 |
| ENSG00000090674 | MCOLN1 | 277.6688354 | 0.32864635 | 0.110135975 | 2.984005442 | 0.002845017 | 0.009694696 |
| ENSG00000090686 | USP48 | 1736.366831 | 0.001870776 | 0.05656502 | 0.033073026 | 0.973616353 | 0.988001378 |
| ENSG00000090776 | EFNB1 | 227.1912125 | -0.321671293 | 0.136462051 | -2.357221588 | 0.018412256 | 0.050858177 |
| ENSG00000090857 | PDPR | 3330.253925 | -0.007579598 | 0.054018113 | -0.140315869 | 0.888410427 | 0.94247758 |
| ENSG00000090861 | AARS1 | 9662.921009 | -0.413541705 | 0.042943147 | -9.629981343 | 5.97E-22 | 1.11E-20 |
| ENSG00000090863 | GLG1 | 8112.853423 | 0.035987705 | 0.049337928 | 0.729412577 | 0.465749327 | 0.651726716 |
| ENSG00000090889 | KIF4A | 2402.313773 | -0.313071324 | 0.0516019 | -6.06705032 | 1.30E-09 | 1.03E-08 |
| ENSG00000090905 | TNRC6A | 2272.087168 | -0.016773742 | 0.054474773 | -0.307917607 | 0.758145028 | 0.866725209 |
| ENSG00000090924 | PLEKHG2 | 2407.942511 | -0.579809099 | 0.05993759 | -9.673547164 | 3.91E-22 | 7.36E-21 |
| ENSG00000090971 | NAT14 | 574.0942825 | -0.091476527 | 0.086656855 | -1.055617894 | 0.291142822 | 0.476106098 |
| ENSG00000090975 | PITPNM2 | 216.9257871 | 0.995697939 | 0.135387314 | 7.354440431 | 1.92E-13 | 2.05E-12 |
| ENSG00000090989 | EXOC1 | 1279.521547 | 0.008247546 | 0.065613394 | 0.125699125 | 0.899970094 | 0.949471761 |
| ENSG00000091009 | RBM27 | 361.9752671 | -0.24731837 | 0.10886651 | -2.271758047 | 0.023101128 | 0.062003028 |
| ENSG00000091010 | POU4F3 | 3.960598007 | -0.224031998 | 0.891613356 | -0.251265861 | 0.801608567 | 0.894413497 |
| ENSG00000091039 | OSBPL8 | 2546.311835 | 0.026785806 | 0.063339176 | 0.422894761 | 0.672372046 | 0.812271037 |
| ENSG00000091073 | DTX2 | 269.830913 | 0.62276485 | 0.126203977 | 4.934589741 | 8.03E-07 | 4.71E-06 |
| ENSG00000091127 | PUS7 | 1051.091789 | 0.047001941 | 0.068838977 | 0.682780927 | 0.494745283 | 0.67711556 |
| ENSG00000091128 | LAMB4 | 3.225280155 | -0.722035124 | 0.996619575 | -0.724484188 | 0.468768531 | 0.654666389 |
| ENSG00000091129 | NRCAM | 2618.641109 | 0.770606417 | 0.052860226 | 14.57819007 | 3.87E-48 | 2.01E-46 |
| ENSG00000091136 | LAMB1 | 1115.869414 | -2.241628945 | 0.072144188 | -31.07151132 | 5.84E-212 | 4.19E-209 |
| ENSG00000091137 | SLC26A4 | 2.762973438 | 0.017562975 | 1.041833468 | 0.016857756 | 0.986550094 | 0.993497695 |
| ENSG00000091140 | DLD | 2471.767598 | -0.238410967 | 0.053973624 | -4.417175478 | 1.00E-05 | 5.18E-05 |

| | | | | | | | |
|---|---|---|---|---|---|---|---|
| ENSG00000091157 | WDR7 | 656.2303704 | 0.28485636 | 0.084424973 | 3.374077003 | 0.000740636 | 0.002841646 |
| ENSG00000091164 | TXNL1 | 2192.692984 | 0.564960518 | 0.055635954 | 10.15459392 | 3.16E-24 | 6.70E-23 |
| ENSG00000091181 | IL5RA | 2.270817477 | -0.179658018 | 1.159395468 | -0.154958358 | 0.876854146 | 0.935974786 |
| ENSG00000091262 | ABCC6 | 10.75583833 | -1.532654882 | 0.561423967 | -2.729942026 | 0.006334546 | 0.019887025 |
| ENSG00000091317 | CMTM6 | 2374.773547 | 0.137023081 | 0.052303109 | 2.61978847 | 0.008798432 | 0.026581857 |
| ENSG00000091409 | ITGA6 | 101.1175127 | -0.897669144 | 0.181858945 | -4.936073647 | 7.97E-07 | 4.68E-06 |
| ENSG00000091428 | RAPGEF4 | 637.8509549 | -0.206341098 | 0.080281184 | -2.570229883 | 0.010163105 | 0.03016843 |
| ENSG00000091436 | MAP3K20 | 1391.617129 | -0.127239503 | 0.062817029 | -2.025557474 | 0.042810153 | 0.104999991 |
| ENSG00000091483 | FH | 1410.439551 | -0.072679974 | 0.061701449 | -1.177929766 | 0.238824607 | 0.414357623 |
| ENSG00000091490 | SEL1L3 | 337.69203 | -0.051286528 | 0.100833336 | -0.508626706 | 0.611013906 | 0.768826384 |
| ENSG00000091513 | TF | 6.845118365 | -0.566016386 | 0.675266938 | -0.838211312 | 0.401912035 | 0.594561301 |
| ENSG00000091527 | CDV3 | 12994.17314 | -0.038998351 | 0.063649197 | -0.612707664 | 0.540069654 | 0.714414141 |
| ENSG00000091536 | MYO15A | 16.7237206 | -0.969705947 | 0.458392919 | -2.115447046 | 0.034391864 | 0.087311792 |
| ENSG00000091542 | ALKBH5 | 3771.792628 | -0.151972208 | 0.050547784 | -3.00650588 | 0.002642689 | 0.00905907 |
| ENSG00000091622 | PITPNM3 | 126.0752305 | 1.704359768 | 0.180724575 | 9.430702854 | 4.07E-21 | 7.27E-20 |
| ENSG00000091640 | SPAG7 | 1297.68694 | 0.156101055 | 0.064062262 | 2.436708459 | 0.014821624 | 0.04204898 |
| ENSG00000091651 | ORC6 | 439.8063395 | -0.001699191 | 0.104498419 | -0.016260444 | 0.987026615 | 0.99369967 |
| ENSG00000091732 | ZC3HC1 | 374.8849135 | -0.607563841 | 0.101235617 | -6.001483036 | 1.96E-09 | 1.52E-08 |
| ENSG00000091831 | ESR1 | 28639.75848 | 0.085045866 | 0.042398471 | 2.00587108 | 0.044870004 | 0.109176697 |
| ENSG00000091844 | RGS17 | 195.060508 | -0.01398478 | 0.130747402 | -0.106960293 | 0.91482048 | 0.957703529 |
| ENSG00000091947 | TMEM101 | 918.1170544 | 0.056974279 | 0.070452366 | 0.808692198 | 0.418692217 | 0.61040988 |
| ENSG00000091986 | CCDC80 | 7.358906361 | 1.106411201 | 0.695177249 | 1.591552662 | 0.111485254 | 0.231277964 |
| ENSG00000092010 | PSME1 | 3910.965171 | -0.109408293 | 0.051073736 | -2.142163516 | 0.032180326 | 0.082546873 |
| ENSG00000092020 | PPP2R3C | 611.2614714 | -0.638361199 | 0.08970967 | -7.115857163 | 1.11E-12 | 1.13E-11 |
| ENSG00000092036 | HAUS4 | 495.173993 | -0.088213873 | 0.087550847 | -1.00757303 | 0.313659482 | 0.501980571 |
| ENSG00000092068 | SLC7A8 | 329.8763089 | 0.888697542 | 0.111206048 | 7.99144973 | 1.33E-15 | 1.67E-14 |
| ENSG00000092094 | OSGEP | 892.3816444 | -0.510355281 | 0.094789663 | -5.384081574 | 7.28E-08 | 4.77E-07 |
| ENSG00000092096 | SLC22A17 | 399.7881327 | 0.110728963 | 0.102723914 | 1.077927807 | 0.281065975 | 0.464494882 |
| ENSG00000092098 | RNF31 | 33.16261312 | -0.678833471 | 0.310244063 | -2.188062729 | 0.028665033 | 0.074658792 |
| ENSG00000092108 | SCFD1 | 1884.178482 | -0.657879768 | 0.059504947 | -11.05588369 | 2.05E-28 | 5.35E-27 |
| ENSG00000092140 | G2E3 | 854.54033 | -0.473259822 | 0.072390862 | -6.537563043 | 6.25E-11 | 5.50E-10 |
| ENSG00000092148 | HECTD1 | 9059.165346 | -0.422691565 | 0.054425956 | -7.766359919 | 8.08E-15 | 9.54E-14 |
| ENSG00000092199 | HNRNPC | 12098.21213 | -0.102649089 | 0.047985791 | -2.139155935 | 0.032423042 | 0.083104422 |

| | | | | | | | |
|---|---|---|---|---|---|---|---|
| ENSG00000092200 | RPGRIP1 | 33.61576681 | -1.207284417 | 0.32644523 | -3.698275568 | 0.000217069 | 0.000914521 |
| ENSG00000092201 | SUPT16H | 9590.913468 | -0.44193084 | 0.045752084 | -9.659250579 | 4.49E-22 | 8.42E-21 |
| ENSG00000092203 | TOX4 | 3569.183465 | -0.359209409 | 0.047681172 | -7.533569184 | 4.94E-14 | 5.51E-13 |
| ENSG00000092208 | GEMIN2 | 537.4555173 | -0.396976663 | 0.086105081 | -4.610374446 | 4.02E-06 | 2.17E-05 |
| ENSG00000092295 | TGM1 | 81.39416643 | 0.233062668 | 0.205913629 | 1.131846733 | 0.257698876 | 0.437443556 |
| ENSG00000092330 | TINF2 | 1605.073036 | -0.407012751 | 0.058705083 | -6.933177353 | 4.11E-12 | 3.99E-11 |
| ENSG00000092421 | SEMA6A | 382.1561149 | 1.095218937 | 0.107141527 | 10.22217034 | 1.58E-24 | 3.43E-23 |
| ENSG00000092439 | TRPM7 | 22467.03681 | -1.316890373 | 0.055014258 | -23.9372558 | 1.25E-126 | 3.39E-124 |
| ENSG00000092445 | TYRO3 | 1351.270996 | -0.712358223 | 0.07249159 | -9.826770557 | 8.63E-23 | 1.70E-21 |
| ENSG00000092470 | WDR76 | 712.826761 | -0.458897192 | 0.080266834 | -5.717145757 | 1.08E-08 | 7.76E-08 |
| ENSG00000092529 | CAPN3 | 2.714768929 | 1.583760892 | 1.180389944 | 1.341726859 | 0.179684575 | 0.334588775 |
| ENSG00000092531 | SNAP23 | 1366.213386 | -0.351566271 | 0.063561516 | -5.531118402 | 3.18E-08 | 2.17E-07 |
| ENSG00000092607 | TBX15 | 1.719990989 | 1.284343266 | 1.388086563 | 0.925261651 | 0.354829821 | NA |
| ENSG00000092621 | PHGDH | 5125.059244 | 0.152930821 | 0.047068552 | 3.249108293 | 0.001157674 | 0.004288235 |
| ENSG00000092758 | COL9A3 | 869.7827759 | -0.453119832 | 0.077245845 | -5.865944373 | 4.47E-09 | 3.34E-08 |
| ENSG00000092820 | EZR | 6482.199296 | -0.104698913 | 0.048305562 | -2.167429752 | 0.030202101 | 0.078145296 |
| ENSG00000092841 | MYL6 | 7690.123177 | -0.366620654 | 0.042023217 | -8.72424049 | 2.68E-18 | 4.03E-17 |
| ENSG00000092847 | AGO1 | 2830.359164 | 0.026771033 | 0.061006029 | 0.438826028 | 0.660787599 | 0.803906848 |
| ENSG00000092853 | CLSPN | 977.6996016 | -0.360582832 | 0.066785985 | -5.399079334 | 6.70E-08 | 4.41E-07 |
| ENSG00000092871 | RFFL | 351.52075 | -0.008577136 | 0.102889541 | -0.083362564 | 0.933563254 | 0.968127167 |
| ENSG00000092929 | UNC13D | 1752.352502 | -0.276873692 | 0.069201891 | -4.00095559 | 6.31E-05 | 0.000290643 |
| ENSG00000092931 | MFSD11 | 669.7423503 | -0.193046313 | 0.083564834 | -2.310138188 | 0.020880505 | 0.056766862 |
| ENSG00000092964 | DPYSL2 | 1314.806133 | -1.023884786 | 0.061385666 | -16.67954195 | 1.85E-62 | 1.39E-60 |
| ENSG00000092969 | TGFB2 | 307.6938975 | -1.280333435 | 0.119485506 | -10.71538695 | 8.62E-27 | 2.11E-25 |
| ENSG00000092978 | GPATCH2 | 1729.8976 | 0.619927639 | 0.058210889 | 10.64968504 | 1.75E-26 | 4.20E-25 |
| ENSG00000093000 | NUP50 | 1952.258493 | -0.07662009 | 0.058903716 | -1.300768355 | 0.193337757 | 0.353992189 |
| ENSG00000093009 | CDC45 | 1016.907382 | -0.716193282 | 0.066792732 | -10.722623 | 7.97E-27 | 1.96E-25 |
| ENSG00000093010 | COMT | 2734.27226 | -0.684560973 | 0.068341108 | -10.01682574 | 1.29E-23 | 2.63E-22 |
| ENSG00000093072 | ADA2 | 6.950813162 | -5.358140863 | 1.300819081 | -4.119051558 | 3.80E-05 | 0.000180799 |
| ENSG00000093144 | ECHDC1 | 1977.103739 | 0.40825408 | 0.057657465 | 7.08068034 | 1.43E-12 | 1.45E-11 |
| ENSG00000093167 | LRRFIP2 | 908.6888177 | 0.019290002 | 0.070334842 | 0.274259551 | 0.783885167 | 0.882792721 |
| ENSG00000093183 | SEC22C | 949.2704914 | -0.473750537 | 0.070843046 | -6.687325871 | 2.27E-11 | 2.07E-10 |
| ENSG00000093217 | XYLB | 285.3837654 | 0.389027713 | 0.1180178 | 3.296347787 | 0.000979507 | 0.00367215 |

| | | | | | | | |
|---|---|---|---|---|---|---|---|
| ENSG00000094631 | HDAC6 | 1692.68347 | -0.279475601 | 0.067331808 | -4.150721765 | 3.31E-05 | 0.000159325 |
| ENSG00000094755 | GABRP | 10.96690677 | 0.196162566 | 0.534032983 | 0.367322941 | 0.713378144 | 0.84060482 |
| ENSG00000094804 | CDC6 | 1503.912577 | -0.202311918 | 0.064762571 | -3.123901903 | 0.001784699 | 0.00633958 |
| ENSG00000094841 | UPRT | 487.4309066 | 0.123201549 | 0.086191199 | 1.429398245 | 0.152889803 | 0.295699219 |
| ENSG00000094880 | CDC23 | 1650.45743 | 0.082751715 | 0.056321085 | 1.469284809 | 0.141755555 | 0.279226553 |
| ENSG00000094914 | AAAS | 1685.907073 | -0.323161872 | 0.05789553 | -5.581810377 | 2.38E-08 | 1.64E-07 |
| ENSG00000094916 | CBX5 | 9291.072695 | -0.2452929 | 0.052290905 | -4.6909286 | 2.72E-06 | 1.50E-05 |
| ENSG00000094975 | SUCO | 2259.681695 | -0.236355588 | 0.058971513 | -4.007962054 | 6.12E-05 | 0.00028303 |
| ENSG00000095002 | MSH2 | 2718.554282 | -0.090988885 | 0.049384768 | -1.842448352 | 0.065409595 | 0.150004609 |
| ENSG00000095015 | MAP3K1 | 1852.792631 | 0.917629712 | 0.062681906 | 14.63946727 | 1.57E-48 | 8.28E-47 |
| ENSG00000095059 | DHPS | 82.46500103 | -0.002495957 | 0.198812651 | -0.012554318 | 0.989983366 | 0.994622626 |
| ENSG00000095066 | HOOK2 | 2166.143474 | -0.059158487 | 0.059272399 | -0.998078155 | 0.318241462 | 0.507399162 |
| ENSG00000095139 | ARCN1 | 7419.468962 | -0.104796358 | 0.04650248 | -2.253564936 | 0.024223551 | 0.064697668 |
| ENSG00000095203 | EPB41L4B | 1024.491889 | 0.610795233 | 0.066926411 | 9.126370616 | 7.08E-20 | 1.17E-18 |
| ENSG00000095209 | TMEM38B | 380.3038388 | 0.144924695 | 0.098421265 | 1.472493722 | 0.140887601 | 0.277898673 |
| ENSG00000095261 | PSMD5 | 1933.451893 | 0.191549184 | 0.053682085 | 3.568214306 | 0.000359422 | 0.001455455 |
| ENSG00000095303 | PTGS1 | 3.763454437 | 2.134911242 | 1.027755027 | 2.07725692 | 0.037777853 | 0.094619931 |
| ENSG00000095319 | NUP188 | 4421.067623 | -0.071672445 | 0.049438922 | -1.449716983 | 0.147137458 | 0.28733577 |
| ENSG00000095321 | CRAT | 67.19514181 | -0.999456299 | 0.228255809 | -4.378667538 | 1.19E-05 | 6.09E-05 |
| ENSG00000095380 | NANS | 1381.266753 | 0.062985217 | 0.060669918 | 1.038162223 | 0.299194537 | 0.485913109 |
| ENSG00000095383 | TBC1D2 | 446.9323977 | -0.063909021 | 0.094673786 | -0.67504453 | 0.499647474 | 0.681005336 |
| ENSG00000095397 | WHRN | 483.6042787 | -0.122810419 | 0.094827031 | -1.295099269 | 0.195285987 | 0.356488274 |
| ENSG00000095485 | CWF19L1 | 768.7608397 | -0.127834683 | 0.073582899 | -1.737287947 | 0.082336358 | 0.18088451 |
| ENSG00000095539 | SEMA4G | 45.37837957 | 0.303354972 | 0.266664152 | 1.137591872 | 0.255290942 | 0.434762737 |
| ENSG00000095564 | BTAF1 | 1950.271325 | -0.134853935 | 0.064070442 | -2.104776116 | 0.035310801 | 0.089313025 |
| ENSG00000095574 | IKZF5 | 428.8057131 | -0.603501859 | 0.098088408 | -6.152631812 | 7.62E-10 | 6.13E-09 |
| ENSG00000095585 | BLNK | 364.148696 | -0.775119327 | 0.099522311 | -7.788397562 | 6.79E-15 | 8.08E-14 |
| ENSG00000095587 | TLL2 | 6.472864428 | -0.292870937 | 0.690465573 | -0.424164431 | 0.671445899 | 0.811665117 |
| ENSG00000095596 | CYP26A1 | 5.210232154 | -0.500867159 | 0.773828962 | -0.647258223 | 0.517464835 | 0.695665852 |
| ENSG00000095637 | SORBS1 | 52.34915015 | 0.295753799 | 0.251764049 | 1.174726099 | 0.240104314 | 0.415917312 |
| ENSG00000095739 | BAMBI | 5579.813771 | -0.617097562 | 0.050824437 | -12.14174899 | 6.35E-34 | 2.06E-32 |
| ENSG00000095752 | IL11 | 92.12548863 | 0.607844691 | 0.189405917 | 3.209217012 | 0.00133097 | 0.004865729 |
| ENSG00000095787 | WAC | 5120.864996 | -0.534800717 | 0.04749654 | -11.2597827 | 2.07E-29 | 5.63E-28 |

| | | | | | | | |
|---|---|---|---|---|---|---|---|
| ENSG00000095794 | CREM | 292.0473783 | -0.793500003 | 0.111857916 | -7.093820753 | 1.30E-12 | 1.32E-11 |
| ENSG00000095906 | NUBP2 | 1727.053811 | 0.080525676 | 0.070796598 | 1.13742296 | 0.255361513 | 0.434762737 |
| ENSG00000095951 | HIVEP1 | 550.4707381 | -0.637896876 | 0.087229405 | -7.312865197 | 2.62E-13 | 2.78E-12 |
| ENSG00000096006 | CRISP3 | 237.626332 | -1.772938427 | 0.139853373 | -12.67712307 | 7.92E-37 | 2.86E-35 |
| ENSG00000096060 | FKBP5 | 1467.535656 | 0.316725671 | 0.065234187 | 4.855209895 | 1.20E-06 | 6.89E-06 |
| ENSG00000096063 | SRPK1 | 2043.10876 | 0.1998382 | 0.055788203 | 3.582087057 | 0.00034086 | 0.001387144 |
| ENSG00000096070 | BRPF3 | 2425.795799 | -0.200945125 | 0.058674484 | -3.424744641 | 0.000615378 | 0.002395513 |
| ENSG00000096080 | MRPS18A | 953.843212 | -0.077727451 | 0.076850588 | -1.011409981 | 0.311820246 | 0.500045889 |
| ENSG00000096092 | TMEM14A | 484.6130328 | -0.29225449 | 0.088996993 | -3.283869252 | 0.001023924 | 0.003825541 |
| ENSG00000096093 | EFHC1 | 663.1362929 | -0.311508571 | 0.087884918 | -3.544505459 | 0.00039335 | 0.001583374 |
| ENSG00000096384 | HSP90AB1 | 48662.9506 | -0.026666388 | 0.052235576 | -0.510502411 | 0.609699526 | 0.767761904 |
| ENSG00000096401 | CDC5L | 2519.969463 | 0.030225901 | 0.051944922 | 0.581883655 | 0.560645051 | 0.729860281 |
| ENSG00000096433 | ITPR3 | 7173.851866 | 0.343631887 | 0.048777562 | 7.044876235 | 1.86E-12 | 1.86E-11 |
| ENSG00000096654 | ZNF184 | 270.5641323 | -0.167016556 | 0.11499163 | -1.452423597 | 0.146383859 | 0.286180819 |
| ENSG00000096696 | DSP | 10254.1005 | -0.55231415 | 0.04872548 | -11.33522228 | 8.78E-30 | 2.43E-28 |
| ENSG00000096717 | SIRT1 | 998.3969401 | -0.481440456 | 0.068345841 | -7.044180749 | 1.87E-12 | 1.87E-11 |
| ENSG00000096746 | HNRNPH3 | 5325.14844 | -0.317432428 | 0.055439649 | -5.72572941 | 1.03E-08 | 7.40E-08 |
| ENSG00000096872 | IFT74 | 268.7407817 | 0.05318457 | 0.11264099 | 0.472160001 | 0.636812588 | 0.786213586 |
| ENSG00000096968 | JAK2 | 238.2958943 | -0.190615947 | 0.120128142 | -1.586771791 | 0.11256434 | 0.233234333 |
| ENSG00000097007 | ABL1 | 3924.708805 | 0.0350054 | 0.059351158 | 0.58980147 | 0.555323757 | 0.726455691 |
| ENSG00000097021 | ACOT7 | 1100.483679 | -0.049455307 | 0.064703343 | -0.764339287 | 0.444665079 | 0.633380605 |
| ENSG00000097033 | SH3GLB1 | 1325.466594 | 0.222520495 | 0.064710764 | 3.438693684 | 0.000584528 | 0.002283951 |
| ENSG00000097046 | CDC7 | 536.5441643 | -0.117491859 | 0.08634182 | -1.360775798 | 0.173584552 | 0.326021748 |
| ENSG00000097096 | SYDE2 | 547.8972278 | -0.109628565 | 0.084392784 | -1.299027704 | 0.193934422 | 0.354631776 |
| ENSG00000099194 | SCD | 109564.5876 | -0.558369531 | 0.055271516 | -10.10230175 | 5.40E-24 | 1.13E-22 |
| ENSG00000099203 | TMED1 | 651.4085887 | -0.040268356 | 0.084452185 | -0.476818401 | 0.633491441 | 0.78408836 |
| ENSG00000099204 | ABLIM1 | 622.1938564 | -1.518562061 | 0.084712073 | -17.92615865 | 7.37E-72 | 7.07E-70 |
| ENSG00000099219 | ERMP1 | 1725.357942 | -0.115995171 | 0.057656222 | -2.011841329 | 0.044236666 | 0.107921845 |
| ENSG00000099246 | RAB18 | 1811.478699 | -0.45584273 | 0.056726749 | -8.035763333 | 9.30E-16 | 1.18E-14 |
| ENSG00000099250 | NRP1 | 1579.062403 | -0.630691768 | 0.061664077 | -10.22786367 | 1.49E-24 | 3.24E-23 |
| ENSG00000099251 | HSD17B7P2 | 73.87832467 | -1.226574353 | 0.224099633 | -5.473343862 | 4.42E-08 | 2.97E-07 |
| ENSG00000099256 | PRTFDC1 | 8.427530153 | -1.989484251 | 0.679822841 | -2.926474561 | 0.003428275 | 0.011475943 |
| ENSG00000099260 | PALMD | 7.162774826 | -3.852422959 | 1.001810041 | -3.845462514 | 0.000120325 | 0.000529394 |

| | | | | | | | |
|---|---|---|---|---|---|---|---|
| ENSG00000099282 | TSPAN15 | 1083.10049 | -0.963433991 | 0.069977138 | -13.76783924 | 3.98E-43 | 1.80E-41 |
| ENSG00000099284 | MACROH2A2 | 704.1661685 | -1.449936998 | 0.085858678 | -16.88748343 | 5.56E-64 | 4.40E-62 |
| ENSG00000099290 | WASHC2A | 1897.389395 | -0.701234088 | 0.061181739 | -11.46149316 | 2.06E-30 | 5.81E-29 |
| ENSG00000099308 | MAST3 | 510.7178046 | 0.401707778 | 0.085837234 | 4.679877962 | 2.87E-06 | 1.58E-05 |
| ENSG00000099326 | MZF1 | 996.2996307 | 0.582301481 | 0.078180879 | 7.448131689 | 9.47E-14 | 1.03E-12 |
| ENSG00000099330 | OCEL1 | 233.7583354 | -0.126744967 | 0.12745645 | -0.994417834 | 0.320019489 | 0.509515996 |
| ENSG00000099331 | MYO9B | 2670.548491 | 0.081735717 | 0.056027722 | 1.458844209 | 0.144607992 | 0.283316647 |
| ENSG00000099337 | KCNK6 | 1775.80499 | 0.620352387 | 0.058412294 | 10.6202366 | 2.40E-26 | 5.71E-25 |
| ENSG00000099338 | CATSPERG | 377.3731829 | 0.702244701 | 0.112784225 | 6.226444364 | 4.77E-10 | 3.91E-09 |
| ENSG00000099341 | PSMD8 | 4493.879709 | -0.010422626 | 0.048450939 | -0.215117119 | 0.829676007 | 0.909788127 |
| ENSG00000099364 | FBXL19 | 1333.350323 | 0.062474707 | 0.064167882 | 0.973613366 | 0.330248547 | 0.52089533 |
| ENSG00000099365 | STX1B | 32.95407243 | 0.159847519 | 0.324281687 | 0.492927986 | 0.622063472 | 0.776353086 |
| ENSG00000099377 | HSD3B7 | 871.9561458 | -0.212726234 | 0.078523136 | -2.709089906 | 0.006746805 | 0.021003083 |
| ENSG00000099381 | SETD1A | 2107.350392 | -0.089568038 | 0.055055576 | -1.62686587 | 0.103765588 | 0.218403761 |
| ENSG00000099385 | BCL7C | 1233.476892 | 0.032641464 | 0.069659048 | 0.468589012 | 0.639363433 | 0.787619975 |
| ENSG00000099617 | EFNA2 | 25.41279075 | 0.967634382 | 0.362009143 | 2.672955644 | 0.007518619 | 0.023135764 |
| ENSG00000099622 | CIRBP | 5096.244967 | -0.157552089 | 0.051971159 | -3.031529238 | 0.002433184 | 0.008410009 |
| ENSG00000099624 | ATP5F1D | 1873.336093 | 0.191591681 | 0.059790765 | 3.204369131 | 0.001353588 | 0.004941329 |
| ENSG00000099625 | CBARP | 361.1388643 | 0.225550026 | 0.101512088 | 2.221903137 | 0.026289852 | 0.069318451 |
| ENSG00000099725 | PRKY | 1.694897974 | 1.312718236 | 1.542508219 | 0.851028358 | 0.3947536 | NA |
| ENSG00000099769 | IGFALS | 10.60040191 | 0.435717161 | 0.553978057 | 0.786524223 | 0.431560434 | 0.62160049 |
| ENSG00000099783 | HNRNPM | 7589.549096 | -0.31088609 | 0.04297784 | -7.233636916 | 4.70E-13 | 4.91E-12 |
| ENSG00000099785 | MARCHF2 | 296.7359116 | -0.054603963 | 0.109691354 | -0.497796418 | 0.618627541 | 0.773916986 |
| ENSG00000099795 | NDUFB7 | 907.2987874 | 0.100950798 | 0.083895971 | 1.203285404 | 0.228865895 | 0.40191255 |
| ENSG00000099797 | TECR | 1576.342801 | -0.036484683 | 0.059688484 | -0.611251627 | 0.54103301 | 0.715266093 |
| ENSG00000099800 | TIMM13 | 573.581999 | 0.241527599 | 0.082831556 | 2.915888714 | 0.00354677 | 0.011834604 |
| ENSG00000099804 | CDC34 | 1268.602507 | 0.042364192 | 0.074466432 | 0.568903216 | 0.569421823 | 0.736830811 |
| ENSG00000099810 | MTAP | 3.112812979 | 0.737490416 | 1.035657094 | 0.712099033 | 0.476403453 | 0.661463213 |
| ENSG00000099812 | MISP | 2345.316701 | -0.0907833 | 0.067673491 | -1.341489841 | 0.179761466 | 0.334691018 |
| ENSG00000099814 | CEP170B | 4926.685615 | -0.20814465 | 0.054792451 | -3.798783379 | 0.000145408 | 0.000631635 |
| ENSG00000099817 | POLR2E | 3341.660619 | 0.070073656 | 0.052151725 | 1.343649825 | 0.17906165 | 0.333821556 |
| ENSG00000099821 | POLRMT | 925.8086015 | 0.119205099 | 0.070644605 | 1.687391406 | 0.091528115 | 0.197169214 |
| ENSG00000099822 | HCN2 | 850.0152279 | 0.760750395 | 0.075811968 | 10.03470054 | 1.07E-23 | 2.21E-22 |

| | | | | | | | |
|---|---|---|---|---|---|---|---|
| ENSG00000099840 | IZUMO4 | 98.25864039 | 0.423928104 | 0.182783766 | 2.319287505 | 0.020379452 | 0.055542906 |
| ENSG00000099849 | RASSF7 | 1017.114119 | -0.042611226 | 0.082067427 | -0.51922215 | 0.603605834 | 0.762610445 |
| ENSG00000099860 | GADD45B | 524.6301852 | -0.18620199 | 0.085327363 | -2.182207258 | 0.02909424 | 0.075577156 |
| ENSG00000099864 | PALM | 1501.417275 | 0.424317588 | 0.076400482 | 5.553860068 | 2.79E-08 | 1.91E-07 |
| ENSG00000099866 | MADCAM1 | 13.15113798 | -0.513693308 | 0.477563068 | -1.075655433 | 0.282081386 | 0.465502266 |
| ENSG00000099875 | MKNK2 | 3130.250728 | -0.205986494 | 0.055408472 | -3.71759928 | 0.000201125 | 0.000852672 |
| ENSG00000099889 | ARVCF | 764.4748315 | -0.576321213 | 0.074000712 | -7.788049568 | 6.81E-15 | 8.09E-14 |
| ENSG00000099899 | TRMT2A | 1304.559124 | -0.25587065 | 0.068608018 | -3.729456963 | 0.000191893 | 0.00081735 |
| ENSG00000099901 | RANBP1 | 2084.360059 | -0.521148919 | 0.055165826 | -9.446952139 | 3.49E-21 | 6.25E-20 |
| ENSG00000099904 | ZDHHC8 | 1242.217247 | -0.434372999 | 0.075233117 | -5.773693999 | 7.76E-09 | 5.64E-08 |
| ENSG00000099910 | KLHL22 | 645.9526838 | -0.280527049 | 0.078553299 | -3.571168283 | 0.000355392 | 0.001440916 |
| ENSG00000099917 | MED15 | 1641.276064 | -0.422968734 | 0.060975655 | -6.936682114 | 4.01E-12 | 3.90E-11 |
| ENSG00000099940 | SNAP29 | 1575.23384 | -0.759381262 | 0.057102574 | -13.29854687 | 2.36E-40 | 9.84E-39 |
| ENSG00000099942 | CRKL | 3154.567426 | -0.762037462 | 0.052630763 | -14.47893624 | 1.65E-47 | 8.43E-46 |
| ENSG00000099949 | LZTR1 | 373.1739533 | -0.494416348 | 0.099504267 | -4.968795422 | 6.74E-07 | 3.99E-06 |
| ENSG00000099953 | MMP11 | 53.92869927 | 0.5309646 | 0.24647086 | 2.154269272 | 0.031219061 | 0.080413796 |
| ENSG00000099954 | CECR2 | 349.4736155 | -1.796685034 | 0.109742475 | -16.37182898 | 3.04E-60 | 2.19E-58 |
| ENSG00000099956 | SMARCB1 | 3454.788143 | -0.556601707 | 0.050476246 | -11.0270028 | 2.83E-28 | 7.35E-27 |
| ENSG00000099957 | P2RX6 | 1.755953873 | 0.080515168 | 1.445230894 | 0.055710937 | 0.955572086 | NA |
| ENSG00000099968 | BCL2L13 | 2162.740208 | -0.574296585 | 0.053958322 | -10.6433367 | 1.87E-26 | 4.49E-25 |
| ENSG00000099974 | DDTL | 38.4799934 | -0.195621638 | 0.28395915 | -0.688907674 | 0.49088137 | 0.673915141 |
| ENSG00000099991 | CABIN1 | 3175.541644 | -0.609862323 | 0.058803188 | -10.37124583 | 3.35E-25 | 7.51E-24 |
| ENSG00000099992 | TBC1D10A | 269.3020399 | -0.513704337 | 0.12693092 | -4.047117419 | 5.19E-05 | 0.00024216 |
| ENSG00000099994 | SUSD2 | 475.1535617 | 3.788001499 | 0.132727661 | 28.53965376 | 3.78E-179 | 1.77E-176 |
| ENSG00000099995 | SF3A1 | 4794.135798 | -0.326863003 | 0.044842505 | -7.289133456 | 3.12E-13 | 3.29E-12 |
| ENSG00000099998 | GGT5 | 11.71022254 | -2.527340397 | 0.619233755 | -4.081399596 | 4.48E-05 | 0.000210779 |
| ENSG00000099999 | RNF215 | 481.7634569 | -0.396790981 | 0.093530297 | -4.242379136 | 2.21E-05 | 0.000109013 |
| ENSG00000100003 | SEC14L2 | 61.9807961 | 0.009543429 | 0.23236498 | 0.041070858 | 0.967239407 | 0.985409779 |
| ENSG00000100014 | SPECC1L | 1000.46943 | -0.591061846 | 0.071853518 | -8.225927768 | 1.94E-16 | 2.59E-15 |
| ENSG00000100023 | PPIL2 | 1951.745824 | -0.417402411 | 0.064217025 | -6.499871464 | 8.04E-11 | 7.02E-10 |
| ENSG00000100027 | YPEL1 | 119.2835995 | 0.592105773 | 0.168019367 | 3.524032864 | 0.000425032 | 0.001703393 |
| ENSG00000100028 | SNRPD3 | 1602.401074 | -0.403955462 | 0.060035296 | -6.728632848 | 1.71E-11 | 1.57E-10 |
| ENSG00000100029 | PES1 | 2955.849351 | -0.395863778 | 0.049293034 | -8.030825977 | 9.68E-16 | 1.23E-14 |

| | | | | | | | |
|---|---|---|---|---|---|---|---|
| ENSG00000100030 | MAPK1 | 3869.119203 | -0.535389925 | 0.048141968 | -11.12106445 | 9.91E-29 | 2.61E-27 |
| ENSG00000100033 | PRODH | 28.256387 | -0.853384632 | 0.331059488 | -2.577738026 | 0.009944936 | 0.029578951 |
| ENSG00000100034 | PPM1F | 1029.474217 | -0.60358333 | 0.071015729 | -8.499290776 | 1.91E-17 | 2.70E-16 |
| ENSG00000100036 | SLC35E4 | 53.94815115 | -0.639071979 | 0.264915409 | -2.412362425 | 0.015849518 | 0.044628444 |
| ENSG00000100038 | TOP3B | 883.0253768 | -0.674184221 | 0.074392997 | -9.062468911 | 1.28E-19 | 2.08E-18 |
| ENSG00000100053 | CRYBB3 | 1.984314765 | -0.21952511 | 1.233373107 | -0.177987592 | 0.85873272 | 0.926723937 |
| ENSG00000100056 | ESS2 | 694.946509 | -0.645220451 | 0.081990366 | -7.869466633 | 3.56E-15 | 4.33E-14 |
| ENSG00000100058 | CRYBB2P1 | 281.9750382 | -0.519923889 | 0.113045515 | -4.599243828 | 4.24E-06 | 2.28E-05 |
| ENSG00000100060 | MFNG | 4.980340406 | 0.515279564 | 0.796797034 | 0.646688607 | 0.517833498 | 0.69574209 |
| ENSG00000100065 | CARD10 | 270.8999236 | 0.327503788 | 0.117907883 | 2.777624177 | 0.005475791 | 0.017444057 |
| ENSG00000100068 | LRP5L | 123.2703576 | -0.789891126 | 0.169145308 | -4.669896764 | 3.01E-06 | 1.65E-05 |
| ENSG00000100075 | SLC25A1 | 5653.020406 | -0.749361049 | 0.045200661 | -16.57854175 | 9.96E-62 | 7.47E-60 |
| ENSG00000100077 | GRK3 | 491.8618839 | -0.789802539 | 0.090749906 | -8.703067307 | 3.23E-18 | 4.85E-17 |
| ENSG00000100078 | PLA2G3 | 433.1681185 | -1.668360889 | 0.101330886 | -16.46448542 | 6.60E-61 | 4.82E-59 |
| ENSG00000100083 | GGA1 | 1787.025068 | -0.191989657 | 0.062754111 | -3.059395689 | 0.00221784 | 0.007730725 |
| ENSG00000100084 | HIRA | 315.204629 | -0.568951288 | 0.104864042 | -5.425608959 | 5.78E-08 | 3.83E-07 |
| ENSG00000100092 | SH3BP1 | 89.46392445 | -0.089554975 | 0.188480577 | -0.475141665 | 0.634686003 | 0.784593248 |
| ENSG00000100097 | LGALS1 | 739.102273 | -0.636985555 | 0.079217908 | -8.040928772 | 8.92E-16 | 1.14E-14 |
| ENSG00000100099 | HPS4 | 891.8268193 | -0.518203717 | 0.071814645 | -7.215850151 | 5.36E-13 | 5.58E-12 |
| ENSG00000100100 | PIK3IP1 | 667.6288687 | -0.35965956 | 0.08653183 | -4.156384537 | 3.23E-05 | 0.000155787 |
| ENSG00000100104 | SRRD | 759.3589496 | -0.598259151 | 0.078190835 | -7.651269474 | 1.99E-14 | 2.29E-13 |
| ENSG00000100105 | PATZ1 | 4240.303116 | -0.729513618 | 0.049681184 | -14.68390173 | 8.17E-49 | 4.34E-47 |
| ENSG00000100106 | TRIOBP | 1329.298127 | -0.23794217 | 0.06344223 | -3.75053291 | 0.000176459 | 0.000756348 |
| ENSG00000100109 | TFIP11 | 1325.691483 | -0.675783213 | 0.062573674 | -10.79980085 | 3.45E-27 | 8.63E-26 |
| ENSG00000100116 | GCAT | 480.057378 | 0.228999408 | 0.087857164 | 2.606496703 | 0.00914737 | 0.027527309 |
| ENSG00000100124 | ANKRD54 | 889.616779 | -0.06231228 | 0.077430739 | -0.804748616 | 0.420964759 | 0.612069605 |
| ENSG00000100129 | EIF3L | 4512.015875 | 0.405099736 | 0.047477478 | 8.532461163 | 1.43E-17 | 2.05E-16 |
| ENSG00000100138 | SNU13 | 2829.800106 | 0.057721229 | 0.052756668 | 1.094103008 | 0.273909803 | 0.455920105 |
| ENSG00000100139 | MICALL1 | 1261.808811 | -0.087888915 | 0.067744148 | -1.297365423 | 0.194505484 | 0.355505999 |
| ENSG00000100142 | POLR2F | 726.8653843 | 0.165021848 | 0.077858277 | 2.1195158 | 0.0340469 | 0.086503033 |
| ENSG00000100147 | CCDC134 | 134.3238914 | -0.430960666 | 0.169813503 | -2.537846859 | 0.011153678 | 0.03270681 |
| ENSG00000100150 | DEPDC5 | 369.4431837 | -0.433307721 | 0.097993074 | -4.421819851 | 9.79E-06 | 5.07E-05 |
| ENSG00000100151 | PICK1 | 667.1581211 | 0.068581071 | 0.080317582 | 0.853873695 | 0.393174976 | 0.586058087 |

| | | | | | | | |
|---|---|---|---|---|---|---|---|
| ENSG00000100154 | TTC28 | 363.0346658 | -0.152806181 | 0.0978847 | -1.561083413 | 0.118504071 | 0.24317769 |
| ENSG00000100156 | SLC16A8 | 3.547387113 | 1.286549124 | 1.06327696 | 1.209984955 | 0.226284666 | 0.398447209 |
| ENSG00000100162 | CENPM | 434.3246546 | 0.085283774 | 0.105644652 | 0.807270148 | 0.419510859 | 0.610929766 |
| ENSG00000100167 | SEPTIN3 | 395.2237096 | -0.18875946 | 0.095081784 | -1.985232638 | 0.047118595 | 0.113790984 |
| ENSG00000100181 | TPTEP1 | 2.310025436 | -0.591095651 | 1.140307751 | -0.518365021 | 0.604203615 | 0.763029194 |
| ENSG00000100196 | KDELR3 | 552.5928826 | 0.295467153 | 0.091181687 | 3.240422095 | 0.001193529 | 0.004405398 |
| ENSG00000100197 | CYP2D6 | 14.49973189 | 1.253709886 | 0.511201956 | 2.452474742 | 0.014187734 | 0.040472073 |
| ENSG00000100201 | DDX17 | 12587.49288 | 0.054288068 | 0.04945283 | 1.097774754 | 0.272302858 | 0.454366276 |
| ENSG00000100206 | DMC1 | 113.1431365 | 0.882405393 | 0.176118524 | 5.010292926 | 5.43E-07 | 3.26E-06 |
| ENSG00000100207 | TCF20 | 2258.712194 | -0.067885535 | 0.058051369 | -1.169404536 | 0.242240683 | 0.418502871 |
| ENSG00000100209 | HSCB | 732.0935017 | 0.204389142 | 0.074754125 | 2.734152038 | 0.006254116 | 0.01966412 |
| ENSG00000100211 | CBY1 | 561.8171789 | -0.012380212 | 0.083652518 | -0.147995684 | 0.882346174 | 0.939121523 |
| ENSG00000100216 | TOMM22 | 1744.311573 | 0.056679652 | 0.057042268 | 0.993643031 | 0.320396687 | 0.509913303 |
| ENSG00000100218 | RSPH14 | 2.361658654 | 0.223096065 | 1.194567425 | 0.186758872 | 0.851849695 | 0.922505405 |
| ENSG00000100219 | XBP1 | 24546.76632 | -0.322516032 | 0.057376524 | -5.621045163 | 1.90E-08 | 1.33E-07 |
| ENSG00000100220 | RTCB | 2317.291743 | -0.645332622 | 0.052307254 | -12.33734459 | 5.70E-35 | 1.94E-33 |
| ENSG00000100221 | JOSD1 | 2114.188668 | -0.180444776 | 0.052723465 | -3.422475655 | 0.000620537 | 0.002414367 |
| ENSG00000100225 | FBXO7 | 2270.371696 | -0.616146043 | 0.059577386 | -10.34194483 | 4.55E-25 | 1.01E-23 |
| ENSG00000100226 | GTPBP1 | 1157.642539 | 0.20252133 | 0.067100853 | 3.018163263 | 0.002543119 | 0.008746498 |
| ENSG00000100227 | POLDIP3 | 2914.043854 | -0.047361895 | 0.052284188 | -0.905855034 | 0.365012587 | 0.557779693 |
| ENSG00000100228 | RAB36 | 277.4914684 | -0.171051492 | 0.112544532 | -1.519856085 | 0.12854715 | 0.259013756 |
| ENSG00000100234 | TIMP3 | 90.57271554 | 1.563287206 | 0.203151461 | 7.695180704 | 1.41E-14 | 1.64E-13 |
| ENSG00000100239 | PPP6R2 | 2005.261097 | 0.271022761 | 0.056701902 | 4.779782551 | 1.75E-06 | 9.87E-06 |
| ENSG00000100241 | SBF1 | 3336.605551 | 0.252927829 | 0.060644009 | 4.170697667 | 3.04E-05 | 0.000147058 |
| ENSG00000100242 | SUN2 | 1576.161064 | 0.031775901 | 0.072059363 | 0.440968389 | 0.659235882 | 0.802870627 |
| ENSG00000100243 | CYB5R3 | 3436.445635 | -0.145709529 | 0.056329584 | -2.586731835 | 0.009689097 | 0.028924281 |
| ENSG00000100246 | DNAL4 | 469.2119856 | 0.060637513 | 0.09172838 | 0.661055099 | 0.508576979 | 0.688791823 |
| ENSG00000100253 | MIOX | 7.939874358 | -0.835299171 | 0.664337346 | -1.257341885 | 0.208629863 | 0.375149882 |
| ENSG00000100258 | LMF2 | 1057.242807 | -0.234880736 | 0.070332365 | -3.339582496 | 0.000839044 | 0.003185646 |
| ENSG00000100263 | RHBDD3 | 586.2626159 | -0.981557435 | 0.088005258 | -11.15339529 | 6.89E-29 | 1.83E-27 |
| ENSG00000100266 | PACSIN2 | 1747.290619 | -0.671999845 | 0.058724579 | -11.44324663 | 2.54E-30 | 7.15E-29 |
| ENSG00000100271 | TTLL1 | 212.6801957 | -0.024808132 | 0.125674916 | -0.19739923 | 0.843515132 | 0.917551194 |
| ENSG00000100276 | RASL10A | 74.39147779 | -1.648771335 | 0.2207953 | -7.467420444 | 8.18E-14 | 8.96E-13 |

| | | | | | | | |
|---|---|---|---|---|---|---|---|
| ENSG00000100280 | AP1B1 | 5800.706067 | -0.771857512 | 0.052694314 | -14.64783302 | 1.39E-48 | 7.33E-47 |
| ENSG00000100281 | HMGXB4 | 1151.424865 | -0.048320736 | 0.063416005 | -0.761964359 | 0.446081276 | 0.634689852 |
| ENSG00000100284 | TOM1 | 764.8232168 | 0.360988255 | 0.080727217 | 4.471704473 | 7.76E-06 | 4.05E-05 |
| ENSG00000100288 | CHKB | 18.02105129 | -0.807610466 | 0.419807813 | -1.923762352 | 0.054384378 | 0.128815451 |
| ENSG00000100290 | BIK | 152.0017014 | -0.10836501 | 0.148786254 | -0.728326759 | 0.466413585 | 0.652370444 |
| ENSG00000100292 | HMOX1 | 91.15057139 | 2.048590998 | 0.215817473 | 9.492238837 | 2.26E-21 | 4.10E-20 |
| ENSG00000100294 | MCAT | 511.2073123 | 0.079562369 | 0.094554336 | 0.841446015 | 0.400098116 | 0.592910258 |
| ENSG00000100296 | THOC5 | 1599.99559 | -0.613671974 | 0.058122641 | -10.55822599 | 4.65E-26 | 1.09E-24 |
| ENSG00000100297 | MCM5 | 2320.873042 | -0.059400349 | 0.053484387 | -1.110611019 | 0.266735818 | 0.447845982 |
| ENSG00000100298 | APOBEC3H | 2.7057658 | 0.704979493 | 1.061088389 | 0.664392807 | 0.506438935 | 0.686747601 |
| ENSG00000100299 | ARSA | 373.1187175 | 0.437372798 | 0.105632053 | 4.14053108 | 3.47E-05 | 0.000165841 |
| ENSG00000100300 | TSPO | 971.3133032 | 0.537277134 | 0.091318665 | 5.883541258 | 4.02E-09 | 3.02E-08 |
| ENSG00000100302 | RASD2 | 11.04968679 | 0.89446357 | 0.545923486 | 1.63844127 | 0.101329676 | 0.214152094 |
| ENSG00000100304 | TTLL12 | 2665.162452 | 0.379986699 | 0.057600307 | 6.596956136 | 4.20E-11 | 3.73E-10 |
| ENSG00000100307 | CBX7 | 492.2383851 | 0.227613532 | 0.090622832 | 2.511657696 | 0.012016558 | 0.034973338 |
| ENSG00000100311 | PDGFB | 988.439097 | 0.190918751 | 0.071578703 | 2.667256372 | 0.007647331 | 0.023481432 |
| ENSG00000100314 | CABP7 | 3.167285171 | 0.220086056 | 0.994256654 | 0.221357388 | 0.824814169 | 0.907715593 |
| ENSG00000100316 | RPL3 | 20749.1264 | 0.158649765 | 0.046619615 | 3.403068934 | 0.000666335 | 0.002580303 |
| ENSG00000100319 | ZMAT5 | 299.7745264 | -0.848576987 | 0.112144274 | -7.566832951 | 3.82E-14 | 4.30E-13 |
| ENSG00000100320 | RBFOX2 | 1834.683555 | 0.137987687 | 0.055833003 | 2.471435866 | 0.013457169 | 0.038616145 |
| ENSG00000100321 | SYNGR1 | 1542.912685 | -0.045247801 | 0.079397949 | -0.569886272 | 0.568754837 | 0.736073456 |
| ENSG00000100324 | TAB1 | 954.4695942 | -0.021718601 | 0.076025107 | -0.285676698 | 0.775125754 | 0.877702986 |
| ENSG00000100325 | ASCC2 | 1998.494298 | -0.695744899 | 0.055547359 | -12.52525605 | 5.43E-36 | 1.91E-34 |
| ENSG00000100330 | MTMR3 | 2322.469938 | -0.574575087 | 0.058242659 | -9.865193304 | 5.89E-23 | 1.17E-21 |
| ENSG00000100335 | MIEF1 | 1568.621956 | 0.097404349 | 0.062798276 | 1.551067255 | 0.120885567 | 0.246955687 |
| ENSG00000100336 | APOL4 | 64.62063881 | -0.120724831 | 0.226500575 | -0.53300011 | 0.594033504 | 0.755928806 |
| ENSG00000100342 | APOL1 | 3.844379958 | 2.089651249 | 1.032465677 | 2.023942583 | 0.04297606 | 0.105271843 |
| ENSG00000100344 | PNPLA3 | 41.30764535 | -0.321140213 | 0.273591647 | -1.173793924 | 0.240477578 | 0.416312403 |
| ENSG00000100345 | MYH9 | 28089.74614 | 0.197330556 | 0.054397685 | 3.627554275 | 0.000286119 | 0.001180921 |
| ENSG00000100346 | CACNA1I | 11.53348649 | 0.949988741 | 0.534296238 | 1.778018772 | 0.075400772 | 0.168602564 |
| ENSG00000100347 | SAMM50 | 1103.160845 | -0.038467323 | 0.070456446 | -0.54597308 | 0.585084442 | 0.749279132 |
| ENSG00000100348 | TXN2 | 1916.451382 | -0.113006119 | 0.057804694 | -1.954964391 | 0.050587273 | 0.12105905 |
| ENSG00000100350 | FOXRED2 | 2991.386916 | 0.189765701 | 0.054639494 | 3.473050153 | 0.000514579 | 0.002031397 |

| | | | | | | | |
|---|---|---|---|---|---|---|---|
| ENSG00000100353 | EIF3D | 5307.4796 | 0.337916557 | 0.049515842 | 6.824412962 | 8.83E-12 | 8.31E-11 |
| ENSG00000100354 | TNRC6B | 794.2707722 | -0.26506106 | 0.086277027 | -3.072208998 | 0.002124809 | 0.007425825 |
| ENSG00000100359 | SGSM3 | 22.00195373 | -0.513413336 | 0.388282081 | -1.322268941 | 0.186078611 | 0.343442219 |
| ENSG00000100360 | IFT27 | 476.3151676 | -0.180944807 | 0.088819293 | -2.037224136 | 0.041627586 | 0.102461033 |
| ENSG00000100362 | PVALB | 51.34604086 | -2.399550238 | 0.286420403 | -8.377721056 | 5.40E-17 | 7.44E-16 |
| ENSG00000100364 | KIAA0930 | 1827.887072 | -0.034986042 | 0.065553551 | -0.533701703 | 0.59354793 | 0.755562373 |
| ENSG00000100365 | NCF4 | 2.032126889 | 0.215946557 | 1.237771398 | 0.174464007 | 0.861500816 | 0.928006607 |
| ENSG00000100368 | CSF2RB | 2.785692546 | 0.33255341 | 1.035155549 | 0.321259361 | 0.748013851 | 0.860436606 |
| ENSG00000100372 | SLC25A17 | 586.6238858 | 0.011462995 | 0.088371979 | 0.12971301 | 0.896793489 | 0.947893263 |
| ENSG00000100373 | UPK3A | 80.27918614 | -1.431918845 | 0.211576658 | -6.767848852 | 1.31E-11 | 1.21E-10 |
| ENSG00000100376 | FAM118A | 810.4867717 | -0.597079876 | 0.078738002 | -7.583122029 | 3.37E-14 | 3.82E-13 |
| ENSG00000100379 | KCTD17 | 295.8957686 | 0.478433103 | 0.113734165 | 4.20659088 | 2.59E-05 | 0.000126672 |
| ENSG00000100380 | ST13 | 4387.058746 | 0.198861638 | 0.045644772 | 4.356723183 | 1.32E-05 | 6.71E-05 |
| ENSG00000100385 | IL2RB | 5.839990618 | -2.104126233 | 0.835990574 | -2.516925787 | 0.011838376 | 0.034520446 |
| ENSG00000100387 | RBX1 | 753.9244617 | -0.238656637 | 0.075678922 | -3.153541696 | 0.001613022 | 0.005788911 |
| ENSG00000100393 | EP300 | 3044.953808 | -0.095855717 | 0.059897575 | -1.600327182 | 0.10952602 | 0.227976308 |
| ENSG00000100395 | L3MBTL2 | 1267.224967 | -0.053745768 | 0.071078248 | -0.756149311 | 0.44955968 | 0.637858675 |
| ENSG00000100399 | CHADL | 17.18335124 | 1.460069711 | 0.467664853 | 3.122042852 | 0.001796008 | 0.006374814 |
| ENSG00000100401 | RANGAP1 | 4504.470859 | -0.20842613 | 0.050485183 | -4.128461397 | 3.65E-05 | 0.000174027 |
| ENSG00000100403 | ZC3H7B | 4600.804999 | 0.015012704 | 0.055954878 | 0.268300184 | 0.788468264 | 0.8855117 |
| ENSG00000100410 | PHF5A | 1034.400543 | -0.191661504 | 0.066254132 | -2.892823419 | 0.003817959 | 0.012638013 |
| ENSG00000100412 | ACO2 | 5974.739844 | 0.112675691 | 0.048417693 | 2.327159439 | 0.019956781 | 0.054588511 |
| ENSG00000100413 | POLR3H | 3773.530004 | 0.151729897 | 0.055144561 | 2.75149342 | 0.005932421 | 0.018747726 |
| ENSG00000100416 | TRMU | 1015.599715 | -0.12059788 | 0.068778723 | -1.753418417 | 0.079530212 | 0.175881911 |
| ENSG00000100417 | PMM1 | 638.2257478 | -0.064139359 | 0.083361764 | -0.769409814 | 0.441650064 | 0.631007254 |
| ENSG00000100418 | DESI1 | 935.2929562 | 0.428590427 | 0.072265767 | 5.930753163 | 3.02E-09 | 2.30E-08 |
| ENSG00000100422 | CERK | 2133.233506 | 0.808851721 | 0.055568198 | 14.55601849 | 5.35E-48 | 2.77E-46 |
| ENSG00000100425 | BRD1 | 1208.479622 | -0.024753711 | 0.066840619 | -0.37033934 | 0.711129665 | 0.839248612 |
| ENSG00000100426 | ZBED4 | 1156.367442 | 0.050729337 | 0.066861815 | 0.758719115 | 0.4480206 | 0.636304641 |
| ENSG00000100429 | HDAC10 | 59.07631629 | 0.374508854 | 0.241737824 | 1.549235644 | 0.121325084 | 0.247571378 |
| ENSG00000100439 | ABHD4 | 817.3331071 | 0.045454302 | 0.085874929 | 0.529308172 | 0.596591688 | 0.757503366 |
| ENSG00000100441 | KHNYN | 4090.726787 | -0.894213452 | 0.04836311 | -18.48957727 | 2.51E-76 | 2.72E-74 |
| ENSG00000100442 | FKBP3 | 3109.72424 | -0.814185425 | 0.052507958 | -15.50594326 | 3.16E-54 | 1.94E-52 |

| | | | | | | | |
|---|---|---|---|---|---|---|---|
| ENSG00000100445 | SDR39U1 | 1392.241971 | -0.474684773 | 0.063379325 | -7.489583865 | 6.91E-14 | 7.63E-13 |
| ENSG00000100461 | RBM23 | 3140.837393 | -0.361842823 | 0.054179243 | -6.678624547 | 2.41E-11 | 2.19E-10 |
| ENSG00000100462 | PRMT5 | 4325.795116 | -0.460029605 | 0.046387835 | -9.917031267 | 3.51E-23 | 7.05E-22 |
| ENSG00000100473 | COCH | 360.5426797 | -0.764500848 | 0.101327545 | -7.544847256 | 4.53E-14 | 5.06E-13 |
| ENSG00000100478 | AP4S1 | 509.6957766 | -0.711125166 | 0.086000828 | -8.268817663 | 1.35E-16 | 1.83E-15 |
| ENSG00000100479 | POLE2 | 280.2274448 | -1.253338773 | 0.11765917 | -10.652283 | 1.70E-26 | 4.09E-25 |
| ENSG00000100483 | VCPKMT | 266.5839715 | -0.604113454 | 0.126041359 | -4.79297795 | 1.64E-06 | 9.26E-06 |
| ENSG00000100485 | SOS2 | 1426.689133 | -0.693942441 | 0.064564233 | -10.74809397 | 6.05E-27 | 1.50E-25 |
| ENSG00000100490 | CDKL1 | 9.536889842 | 0.343926363 | 0.574698524 | 0.598446574 | 0.549541998 | 0.721915345 |
| ENSG00000100503 | NIN | 2775.173477 | -0.664373393 | 0.057596249 | -11.53501155 | 8.79E-31 | 2.51E-29 |
| ENSG00000100504 | PYGL | 600.3859802 | -1.515293674 | 0.085133688 | -17.79899016 | 7.20E-71 | 6.76E-69 |
| ENSG00000100505 | TRIM9 | 4.74754562 | -1.461815079 | 0.90121404 | -1.622050938 | 0.104792442 | 0.22016106 |
| ENSG00000100519 | PSMC6 | 1799.919044 | -0.689069275 | 0.057595406 | -11.96396247 | 5.49E-33 | 1.71E-31 |
| ENSG00000100522 | GNPNAT1 | 3983.388073 | -0.7943224 | 0.053818246 | -14.75935129 | 2.68E-49 | 1.45E-47 |
| ENSG00000100523 | DDHD1 | 1085.096817 | -0.537443421 | 0.06807711 | -7.894627476 | 2.91E-15 | 3.56E-14 |
| ENSG00000100526 | CDKN3 | 715.8136578 | -0.878590321 | 0.077302408 | -11.36562685 | 6.20E-30 | 1.73E-28 |
| ENSG00000100528 | CNIH1 | 2229.149166 | -0.524507607 | 0.054337244 | -9.652819388 | 4.78E-22 | 8.95E-21 |
| ENSG00000100532 | CGRRF1 | 314.1779324 | -0.537192126 | 0.109509028 | -4.905459704 | 9.32E-07 | 5.42E-06 |
| ENSG00000100554 | ATP6V1D | 1635.109693 | -0.502933842 | 0.058801726 | -8.55304556 | 1.20E-17 | 1.72E-16 |
| ENSG00000100558 | PLEK2 | 308.1689683 | -0.743049009 | 0.107193063 | -6.931875908 | 4.15E-12 | 4.03E-11 |
| ENSG00000100564 | PIGH | 395.6296659 | -0.730001376 | 0.100289404 | -7.278948176 | 3.36E-13 | 3.55E-12 |
| ENSG00000100567 | PSMA3 | 3238.699211 | -0.458229654 | 0.057001942 | -8.038842813 | 9.07E-16 | 1.15E-14 |
| ENSG00000100568 | VTI1B | 3087.509658 | -0.364489293 | 0.048740858 | -7.478105756 | 7.54E-14 | 8.28E-13 |
| ENSG00000100575 | TIMM9 | 909.0501529 | -0.493128638 | 0.069917023 | -7.053055444 | 1.75E-12 | 1.76E-11 |
| ENSG00000100577 | GSTZ1 | 1615.091358 | -0.548522697 | 0.059187239 | -9.267583757 | 1.90E-20 | 3.26E-19 |
| ENSG00000100578 | KIAA0586 | 985.1543832 | -0.748752255 | 0.071400757 | -10.48661507 | 9.95E-26 | 2.28E-24 |
| ENSG00000100580 | TMED8 | 2059.648521 | -0.266625475 | 0.060533005 | -4.404629776 | 1.06E-05 | 5.46E-05 |
| ENSG00000100583 | SAMD15 | 178.9042837 | 0.029820162 | 0.14166963 | 0.210490859 | 0.833284586 | 0.911870839 |
| ENSG00000100591 | AHSA1 | 4749.582881 | -0.439372615 | 0.047841327 | -9.18395538 | 4.16E-20 | 6.97E-19 |
| ENSG00000100592 | DAAM1 | 2241.650014 | -0.65631196 | 0.055773954 | -11.76735583 | 5.75E-32 | 1.73E-30 |
| ENSG00000100593 | ISM2 | 3.155902622 | 1.397437569 | 1.033046448 | 1.3527345 | 0.176140465 | 0.329554054 |
| ENSG00000100596 | SPTLC2 | 3614.834932 | -0.604612758 | 0.049948606 | -12.10469722 | 9.97E-34 | 3.20E-32 |
| ENSG00000100599 | RIN3 | 204.2534075 | 0.163969484 | 0.130202294 | 1.259344046 | 0.207906091 | 0.374332134 |

| | | | | | | | |
|---|---|---|---|---|---|---|---|
| ENSG00000100600 | LGMN | 681.116377 | -0.763864866 | 0.079712243 | -9.582779725 | 9.45E-22 | 1.74E-20 |
| ENSG00000100601 | ALKBH1 | 528.0659953 | -0.691643495 | 0.08523373 | -8.114668856 | 4.87E-16 | 6.31E-15 |
| ENSG00000100603 | SNW1 | 3142.147891 | -0.473306502 | 0.048665603 | -9.725688721 | 2.34E-22 | 4.47E-21 |
| ENSG00000100604 | CHGA | 6.01714715 | -6.136368015 | 1.31944375 | -4.650723469 | 3.31E-06 | 1.81E-05 |
| ENSG00000100605 | ITPK1 | 4957.335239 | -0.240750035 | 0.046591989 | -5.167198107 | 2.38E-07 | 1.48E-06 |
| ENSG00000100612 | DHRS7 | 663.9079374 | -0.614387196 | 0.080515749 | -7.630646264 | 2.34E-14 | 2.68E-13 |
| ENSG00000100614 | PPM1A | 2144.466974 | -0.579278776 | 0.052065648 | -11.12593039 | 9.38E-29 | 2.47E-27 |
| ENSG00000100625 | SIX4 | 780.7112671 | -0.129545207 | 0.078170869 | -1.657205666 | 0.097477901 | 0.207665589 |
| ENSG00000100626 | GALNT16 | 172.8061377 | -0.43244021 | 0.145591147 | -2.970236976 | 0.002975701 | 0.01008906 |
| ENSG00000100629 | CEP128 | 306.2078602 | -1.226137334 | 0.111393956 | -11.00721599 | 3.53E-28 | 9.12E-27 |
| ENSG00000100632 | ERH | 3169.889914 | -0.662290225 | 0.053637626 | -12.34749331 | 5.03E-35 | 1.72E-33 |
| ENSG00000100644 | HIF1A | 5067.438027 | -0.482508452 | 0.048958854 | -9.855387002 | 6.50E-23 | 1.29E-21 |
| ENSG00000100647 | SUSD6 | 2221.135898 | -0.715069991 | 0.059741714 | -11.96935844 | 5.14E-33 | 1.61E-31 |
| ENSG00000100650 | SRSF5 | 4853.385576 | -0.487081894 | 0.06332141 | -7.692214843 | 1.45E-14 | 1.68E-13 |
| ENSG00000100664 | EIF5 | 11717.25039 | -0.301530308 | 0.052819315 | -5.708712968 | 1.14E-08 | 8.13E-08 |
| ENSG00000100665 | SERPINA4 | 2.044880892 | -0.263948043 | 1.239302802 | -0.212981075 | 0.831341724 | 0.910700499 |
| ENSG00000100697 | DICER1 | 3070.328438 | -0.828240217 | 0.053417956 | -15.50490274 | 3.21E-54 | 1.97E-52 |
| ENSG00000100711 | ZFYVE21 | 1484.610471 | -0.420998399 | 0.061833358 | -6.808596716 | 9.86E-12 | 9.24E-11 |
| ENSG00000100714 | MTHFD1 | 6756.813441 | -0.975296139 | 0.046949972 | -20.77309302 | 7.58E-96 | 1.14E-93 |
| ENSG00000100722 | ZC3H14 | 3356.880939 | -0.405651168 | 0.049072812 | -8.266311821 | 1.38E-16 | 1.87E-15 |
| ENSG00000100726 | TELO2 | 1943.183879 | 0.049070294 | 0.063114093 | 0.777485531 | 0.436872367 | 0.626600732 |
| ENSG00000100731 | PCNX1 | 2571.819399 | -0.661663923 | 0.057067891 | -11.59432932 | 4.40E-31 | 1.28E-29 |
| ENSG00000100744 | GSKIP | 653.688872 | -0.516855999 | 0.079265442 | -6.520571718 | 7.00E-11 | 6.13E-10 |
| ENSG00000100749 | VRK1 | 1026.951496 | -0.593482252 | 0.069923639 | -8.487576746 | 2.11E-17 | 2.98E-16 |
| ENSG00000100764 | PSMC1 | 490.2142083 | -0.787485697 | 0.093139174 | -8.454935413 | 2.79E-17 | 3.91E-16 |
| ENSG00000100767 | PAPLN | 100.5186768 | -0.313638277 | 0.177827265 | -1.763724345 | 0.0777784 | 0.17284089 |
| ENSG00000100784 | RPS6KA5 | 467.3459792 | -0.947025159 | 0.103903034 | -9.114509181 | 7.90E-20 | 1.30E-18 |
| ENSG00000100796 | PPP4R3A | 4453.279103 | -0.612416665 | 0.051493487 | -11.89308985 | 1.29E-32 | 3.96E-31 |
| ENSG00000100802 | C14orf93 | 281.2806915 | -0.304386643 | 0.113497139 | -2.681888247 | 0.007320791 | 0.022605862 |
| ENSG00000100804 | PSMB5 | 4784.481916 | -0.425920821 | 0.045883656 | -9.28262606 | 1.65E-20 | 2.84E-19 |
| ENSG00000100811 | YY1 | 3686.491677 | -0.100108545 | 0.047930523 | -2.088617813 | 0.03674214 | 0.09242951 |
| ENSG00000100813 | ACIN1 | 8535.852175 | -0.521105578 | 0.044166781 | -11.79858619 | 3.97E-32 | 1.20E-30 |
| ENSG00000100814 | CCNB1IP1 | 1031.623031 | -0.036445903 | 0.065364348 | -0.55758076 | 0.577130698 | 0.743117322 |

| | | | | | | | |
|---|---|---|---|---|---|---|---|
| ENSG00000100815 | TRIP11 | 2489.400255 | -0.517902605 | 0.060544167 | -8.554128846 | 1.19E-17 | 1.71E-16 |
| ENSG00000100823 | APEX1 | 6630.248207 | -0.344556608 | 0.046058049 | -7.480920533 | 7.38E-14 | 8.12E-13 |
| ENSG00000100836 | PABPN1 | 1470.090984 | -0.400798186 | 0.071510068 | -5.60477985 | 2.09E-08 | 1.45E-07 |
| ENSG00000100852 | ARHGAP5 | 3692.295311 | -0.554619516 | 0.052155484 | -10.63396347 | 2.07E-26 | 4.95E-25 |
| ENSG00000100865 | CINP | 923.0602579 | -0.338551002 | 0.07130006 | -4.748256912 | 2.05E-06 | 1.14E-05 |
| ENSG00000100867 | DHRS2 | 1978.992544 | 0.456213255 | 0.056158778 | 8.123632141 | 4.52E-16 | 5.87E-15 |
| ENSG00000100883 | SRP54 | 3506.430829 | -0.553270219 | 0.052133772 | -10.61251091 | 2.61E-26 | 6.19E-25 |
| ENSG00000100888 | CHD8 | 4952.543061 | -0.387054184 | 0.057359858 | -6.747823257 | 1.50E-11 | 1.39E-10 |
| ENSG00000100889 | PCK2 | 3337.839483 | -0.512348765 | 0.060278546 | -8.499686802 | 1.90E-17 | 2.70E-16 |
| ENSG00000100890 | PRORP | 160.440215 | -0.437032717 | 0.149463365 | -2.924012298 | 0.003455511 | 0.011555309 |
| ENSG00000100897 | DCAF11 | 4222.944678 | -0.063022716 | 0.054121494 | -1.164467407 | 0.244234644 | 0.421005616 |
| ENSG00000100902 | PSMA6 | 218.6306616 | -0.411254036 | 0.126107029 | -3.261150789 | 0.00110961 | 0.004120845 |
| ENSG00000100906 | NFKBIA | 2580.41411 | -0.510725556 | 0.052341908 | -9.757488362 | 1.71E-22 | 3.30E-21 |
| ENSG00000100908 | EMC9 | 751.0016661 | -0.538254513 | 0.076467245 | -7.039020619 | 1.94E-12 | 1.93E-11 |
| ENSG00000100911 | PSME2 | 1810.185637 | -0.345460171 | 0.058579622 | -5.897275534 | 3.70E-09 | 2.79E-08 |
| ENSG00000100916 | BRMS1L | 594.9632244 | -0.483409332 | 0.086292982 | -5.601954163 | 2.12E-08 | 1.47E-07 |
| ENSG00000100918 | REC8 | 70.04153951 | 0.848377772 | 0.229797507 | 3.691849322 | 0.000222629 | 0.00093657 |
| ENSG00000100926 | TM9SF1 | 125.6268382 | 0.035139004 | 0.17598359 | 0.199672046 | 0.841737078 | 0.916877899 |
| ENSG00000100934 | SEC23A | 3372.844337 | -0.409750385 | 0.04775816 | -8.579693725 | 9.51E-18 | 1.38E-16 |
| ENSG00000100938 | GMPR2 | 2171.88867 | -0.69508856 | 0.054945473 | -12.65051565 | 1.11E-36 | 4.01E-35 |
| ENSG00000100941 | PNN | 6689.090236 | -0.085869985 | 0.143386664 | -0.598870095 | 0.549259515 | 0.721774261 |
| ENSG00000100949 | RABGGTA | 1039.141181 | -0.118645439 | 0.070630917 | -1.679794689 | 0.092997269 | 0.199733036 |
| ENSG00000100968 | NFATC4 | 340.283486 | -0.044992941 | 0.103511301 | -0.434666943 | 0.663804196 | 0.805985984 |
| ENSG00000100979 | PLTP | 54.34351971 | -0.523067642 | 0.239513842 | -2.18387229 | 0.028971634 | 0.0753013 |
| ENSG00000100982 | PCIF1 | 2173.264136 | -0.486885012 | 0.059116788 | -8.235985593 | 1.78E-16 | 2.39E-15 |
| ENSG00000100983 | GSS | 2135.435896 | -0.8653899 | 0.056496934 | -15.3174665 | 5.85E-53 | 3.46E-51 |
| ENSG00000100985 | MMP9 | 91.62452202 | -1.180310463 | 0.205769013 | -5.736094293 | 9.69E-09 | 6.98E-08 |
| ENSG00000100987 | VSX1 | 3.742059064 | -1.945171689 | 1.019862266 | -1.907288616 | 0.056483224 | 0.13278272 |
| ENSG00000100991 | TRPC4AP | 2533.489791 | -0.525572298 | 0.055223756 | -9.517141384 | 1.78E-21 | 3.25E-20 |
| ENSG00000100994 | PYGB | 4672.854271 | -0.285357691 | 0.049099259 | -5.811853367 | 6.18E-09 | 4.55E-08 |
| ENSG00000100997 | ABHD12 | 2285.74283 | -0.371602064 | 0.057824481 | -6.426379616 | 1.31E-10 | 1.12E-09 |
| ENSG00000101000 | PROCR | 61.41702104 | 0.682561638 | 0.234589731 | 2.90959725 | 0.003618948 | 0.012042162 |
| ENSG00000101003 | GINS1 | 1370.066866 | -0.859359565 | 0.060957852 | -14.09760243 | 3.93E-45 | 1.91E-43 |

| | | | | | | | |
|---|---|---|---|---|---|---|---|
| ENSG00000101004 | NINL | 951.4228448 | -0.413751766 | 0.070088004 | -5.903317891 | 3.56E-09 | 2.69E-08 |
| ENSG00000101019 | UQCC1 | 1226.626538 | -0.446209761 | 0.064492967 | -6.918735176 | 4.56E-12 | 4.41E-11 |
| ENSG00000101040 | ZMYND8 | 5113.814878 | -0.400708972 | 0.046990195 | -8.527501757 | 1.50E-17 | 2.14E-16 |
| ENSG00000101049 | SGK2 | 3.726538142 | 0.464887431 | 0.894350622 | 0.519804448 | 0.603199879 | 0.762307472 |
| ENSG00000101052 | IFT52 | 768.1014401 | -0.494927375 | 0.073170165 | -6.764059828 | 1.34E-11 | 1.24E-10 |
| ENSG00000101057 | MYBL2 | 7737.30575 | -0.501826873 | 0.044591633 | -11.2538348 | 2.22E-29 | 6.01E-28 |
| ENSG00000101079 | NDRG3 | 1122.942091 | -0.114227271 | 0.068230022 | -1.6741497 | 0.094101182 | 0.20169925 |
| ENSG00000101082 | SLA2 | 2.441008755 | 0.443455629 | 1.122308186 | 0.395128214 | 0.692748268 | 0.826785561 |
| ENSG00000101084 | RAB5IF | 49.7453746 | -0.410723744 | 0.272395746 | -1.50781996 | 0.131600612 | 0.263919952 |
| ENSG00000101096 | NFATC2 | 267.4546167 | 1.408001302 | 0.119277617 | 11.80440508 | 3.70E-32 | 1.12E-30 |
| ENSG00000101098 | RIMS4 | 825.9016284 | 1.462127403 | 0.080464221 | 18.1711497 | 8.73E-74 | 8.87E-72 |
| ENSG00000101104 | PABPC1L | 2379.490492 | -0.30759063 | 0.20242309 | -1.519543199 | 0.128625823 | 0.259126756 |
| ENSG00000101109 | STK4 | 1998.151146 | -0.276604134 | 0.056045562 | -4.935344057 | 8.00E-07 | 4.70E-06 |
| ENSG00000101115 | SALL4 | 1314.424718 | 1.440560752 | 0.065833643 | 21.88183268 | 3.87E-106 | 7.34E-104 |
| ENSG00000101126 | ADNP | 7632.145833 | -0.118280517 | 0.045863545 | -2.578965883 | 0.009909657 | 0.029492416 |
| ENSG00000101132 | PFDN4 | 3490.019786 | 0.330028477 | 0.067807363 | 4.867148121 | 1.13E-06 | 6.51E-06 |
| ENSG00000101138 | CSTF1 | 2355.541576 | -0.09055646 | 0.05194481 | -1.743320638 | 0.081277622 | 0.178952746 |
| ENSG00000101144 | BMP7 | 6529.207397 | -0.458829801 | 0.059261126 | -7.742508945 | 9.75E-15 | 1.14E-13 |
| ENSG00000101146 | RAE1 | 5067.639409 | -0.055182247 | 0.04572373 | -1.206862342 | 0.227485162 | 0.400223148 |
| ENSG00000101150 | TPD52L2 | 6689.346653 | -0.292831595 | 0.045215505 | -6.476353488 | 9.40E-11 | 8.14E-10 |
| ENSG00000101152 | DNAJC5 | 6486.307982 | -0.125343805 | 0.052096693 | -2.405983921 | 0.016128975 | 0.045342975 |
| ENSG00000101158 | NELFCD | 9477.045722 | -0.269477026 | 0.047056969 | -5.7266125 | 1.02E-08 | 7.37E-08 |
| ENSG00000101160 | CTSZ | 570.0216473 | 0.107108534 | 0.097002801 | 1.10417981 | 0.269515146 | 0.451059856 |
| ENSG00000101161 | PRPF6 | 7446.539331 | -0.124785293 | 0.046826526 | -2.66484199 | 0.00770245 | 0.023638022 |
| ENSG00000101162 | TUBB1 | 1.855750241 | 0.589371034 | 1.347938342 | 0.437238867 | 0.661938125 | 0.804709093 |
| ENSG00000101166 | PRELID3B | 5787.488009 | 0.08042272 | 0.053106864 | 1.5143564 | 0.129935471 | 0.261168817 |
| ENSG00000101180 | HRH3 | 6.926306287 | 0.666375497 | 0.709575265 | 0.939118837 | 0.347669734 | 0.538596954 |
| ENSG00000101181 | MTG2 | 1076.559 | -0.515157711 | 0.0644477 | -7.993422791 | 1.31E-15 | 1.65E-14 |
| ENSG00000101182 | PSMA7 | 5050.36647 | -0.414792058 | 0.043767336 | -9.477205986 | 2.61E-21 | 4.71E-20 |
| ENSG00000101187 | SLCO4A1 | 1182.610504 | -0.538079243 | 0.064427596 | -8.351688915 | 6.73E-17 | 9.24E-16 |
| ENSG00000101188 | NTSR1 | 19.3724459 | -2.767139151 | 0.489749393 | -5.650112469 | 1.60E-08 | 1.13E-07 |
| ENSG00000101189 | MRGBP | 1699.572322 | -0.37697698 | 0.056030823 | -6.728028656 | 1.72E-11 | 1.58E-10 |
| ENSG00000101190 | TCFL5 | 1547.654063 | 0.069439628 | 0.059401251 | 1.168992689 | 0.242406577 | 0.418672404 |

| | | | | | | | |
|---|---|---|---|---|---|---|---|
| ENSG00000101191 | DIDO1 | 4684.312234 | -0.210666161 | 0.051423235 | -4.096711538 | 4.19E-05 | 0.000198048 |
| ENSG00000101193 | GID8 | 6979.630359 | -0.261161175 | 0.041476868 | -6.296550046 | 3.04E-10 | 2.54E-09 |
| ENSG00000101194 | SLC17A9 | 2.174876578 | 0.985466433 | 1.281356459 | 0.76908063 | 0.441845446 | 0.631129174 |
| ENSG00000101197 | BIRC7 | 2.11429931 | 0.542163885 | 1.269611255 | 0.42703141 | 0.66935645 | 0.810007484 |
| ENSG00000101199 | ARFGAP1 | 3710.857086 | -0.347076314 | 0.051661977 | -6.718215941 | 1.84E-11 | 1.68E-10 |
| ENSG00000101203 | COL20A1 | 3.849320497 | -0.593754035 | 0.938514747 | -0.632652856 | 0.526960362 | 0.703022432 |
| ENSG00000101210 | EEF1A2 | 22124.14872 | 0.064829935 | 0.039677374 | 1.633927045 | 0.10227417 | 0.215938285 |
| ENSG00000101213 | PTK6 | 555.4597836 | -0.455608781 | 0.088062027 | -5.173725817 | 2.29E-07 | 1.44E-06 |
| ENSG00000101216 | GMEB2 | 1124.773756 | -0.225207562 | 0.067310495 | -3.345801586 | 0.000820451 | 0.003121768 |
| ENSG00000101220 | ADISSP | 2449.588794 | 0.162463399 | 0.056896798 | 2.85540495 | 0.004298199 | 0.014069731 |
| ENSG00000101222 | SPEF1 | 22.12384961 | 0.033221588 | 0.391847965 | 0.084781832 | 0.932434837 | 0.967598417 |
| ENSG00000101224 | CDC25B | 5955.111556 | -0.217018652 | 0.049543863 | -4.380333657 | 1.18E-05 | 6.05E-05 |
| ENSG00000101230 | ISM1 | 11.15231253 | 0.417043204 | 0.52880505 | 0.788652083 | 0.430315381 | 0.620532741 |
| ENSG00000101236 | RNF24 | 726.1684923 | -0.294038078 | 0.080680822 | -3.644460631 | 0.000267953 | 0.001110541 |
| ENSG00000101246 | ARFRP1 | 1096.092644 | 0.059307689 | 0.070533719 | 0.840841655 | 0.400436648 | 0.593258635 |
| ENSG00000101247 | NDUFAF5 | 275.4429915 | -0.200619126 | 0.124106411 | -1.616508962 | 0.105984317 | 0.222136157 |
| ENSG00000101255 | TRIB3 | 2431.776252 | -0.110393605 | 0.053001121 | -2.082854171 | 0.037264518 | 0.093507771 |
| ENSG00000101265 | RASSF2 | 37.52711019 | -0.669218305 | 0.287494116 | -2.327763486 | 0.019924666 | 0.054530164 |
| ENSG00000101266 | CSNK2A1 | 2508.560008 | 0.137785092 | 0.061768088 | 2.230684107 | 0.025702061 | 0.067973862 |
| ENSG00000101276 | SLC52A3 | 248.3968475 | -1.542969754 | 0.135089736 | -11.42181338 | 3.25E-30 | 9.14E-29 |
| ENSG00000101278 | RPS10P5 | 21.79665875 | 0.212086048 | 0.376277999 | 0.563641904 | 0.572997863 | 0.739747367 |
| ENSG00000101290 | CDS2 | 1341.258317 | -0.147891738 | 0.061675313 | -2.397908196 | 0.016488997 | 0.046213512 |
| ENSG00000101294 | HM13 | 4383.73133 | -0.320457707 | 0.051257961 | -6.251862155 | 4.06E-10 | 3.35E-09 |
| ENSG00000101306 | MYLK2 | 12.04471363 | -0.173294407 | 0.504936937 | -0.3432001 | 0.73144793 | 0.851271651 |
| ENSG00000101310 | SEC23B | 3193.150633 | -0.353489678 | 0.050217839 | -7.039125687 | 1.93E-12 | 1.93E-11 |
| ENSG00000101311 | FERMT1 | 317.5545411 | 1.167141179 | 0.114645642 | 10.18042341 | 2.43E-24 | 5.19E-23 |
| ENSG00000101331 | CCM2L | 19.77814925 | -0.047260895 | 0.397824514 | -0.118798347 | 0.905435118 | 0.95262805 |
| ENSG00000101333 | PLCB4 | 1177.619297 | -1.137291263 | 0.06936171 | -16.39652868 | 2.02E-60 | 1.47E-58 |
| ENSG00000101336 | HCK | 3.869891643 | 0.162271637 | 0.886070238 | 0.183136313 | 0.854691064 | 0.924486249 |
| ENSG00000101337 | TM9SF4 | 3044.839804 | -0.401783006 | 0.054863949 | -7.323260709 | 2.42E-13 | 2.58E-12 |
| ENSG00000101342 | TLDC2 | 1.778111155 | -0.403384177 | 1.299758677 | -0.310353133 | 0.75629243 | NA |
| ENSG00000101343 | CRNKL1 | 1642.459982 | -0.351773003 | 0.056078562 | -6.272860614 | 3.54E-10 | 2.94E-09 |
| ENSG00000101346 | POFUT1 | 1977.7642 | -0.213452273 | 0.055870636 | -3.82047331 | 0.000133196 | 0.000581449 |

| | | | | | | | |
|---|---|---|---|---|---|---|---|
| ENSG00000101347 | SAMHD1 | 641.4984115 | -0.246271661 | 0.082092276 | -2.999937042 | 0.002700354 | 0.009240174 |
| ENSG00000101349 | PAK5 | 3.042952728 | 1.702475763 | 1.111935336 | 1.531092418 | 0.125746553 | 0.254641967 |
| ENSG00000101350 | KIF3B | 2824.369762 | -0.24708856 | 0.050708079 | -4.872765175 | 1.10E-06 | 6.34E-06 |
| ENSG00000101353 | MROH8 | 59.18909819 | -0.520068237 | 0.246174206 | -2.112602478 | 0.034634806 | 0.087831357 |
| ENSG00000101361 | NOP56 | 3514.076073 | 0.050874907 | 0.049285727 | 1.032244229 | 0.30195772 | 0.488877839 |
| ENSG00000101363 | MANBAL | 1024.122976 | -0.688434312 | 0.06838043 | -10.06770968 | 7.67E-24 | 1.59E-22 |
| ENSG00000101365 | IDH3B | 1405.5673 | -0.229619516 | 0.065670786 | -3.496524577 | 0.000471361 | 0.001874007 |
| ENSG00000101367 | MAPRE1 | 3428.956947 | -0.740860991 | 0.049481611 | -14.97245097 | 1.11E-50 | 6.25E-49 |
| ENSG00000101384 | JAG1 | 467.7444421 | 0.423359531 | 0.097369384 | 4.347973814 | 1.37E-05 | 6.95E-05 |
| ENSG00000101391 | CDK5RAP1 | 786.9185949 | -0.697030616 | 0.074233749 | -9.389672781 | 6.02E-21 | 1.06E-19 |
| ENSG00000101400 | SNTA1 | 108.9087065 | 0.546305417 | 0.174509729 | 3.130515528 | 0.001744998 | 0.006210091 |
| ENSG00000101407 | TTI1 | 1203.098978 | -0.227511508 | 0.066191852 | -3.437152788 | 0.000587864 | 0.002296203 |
| ENSG00000101412 | E2F1 | 1396.487798 | -0.420292672 | 0.059937384 | -7.012195819 | 2.35E-12 | 2.33E-11 |
| ENSG00000101413 | RPRD1B | 1660.871731 | -0.198343555 | 0.058987931 | -3.362442999 | 0.000772561 | 0.002953757 |
| ENSG00000101417 | PXMP4 | 949.853019 | -1.089110775 | 0.069711008 | -15.62322506 | 5.06E-55 | 3.18E-53 |
| ENSG00000101421 | CHMP4B | 2870.112392 | -0.494163218 | 0.049767966 | -9.929343364 | 3.10E-23 | 6.24E-22 |
| ENSG00000101439 | CST3 | 2616.398845 | -0.095292754 | 0.061720741 | -1.54393406 | 0.122604308 | 0.249515864 |
| ENSG00000101442 | ACTR5 | 293.1523305 | -0.518810822 | 0.115181589 | -4.504286038 | 6.66E-06 | 3.51E-05 |
| ENSG00000101443 | WFDC2 | 452.9968407 | -0.846214925 | 0.096055631 | -8.809633746 | 1.26E-18 | 1.93E-17 |
| ENSG00000101444 | AHCY | 5544.060088 | -0.070459381 | 0.04519116 | -1.559140776 | 0.118963064 | 0.243879598 |
| ENSG00000101447 | FAM83D | 738.0152775 | -0.538686805 | 0.074718204 | -7.209579111 | 5.61E-13 | 5.84E-12 |
| ENSG00000101452 | DHX35 | 301.8886184 | -0.270426661 | 0.110421165 | -2.449047353 | 0.014323462 | 0.040792081 |
| ENSG00000101457 | DNTTIP1 | 1235.796188 | -0.371355454 | 0.063930554 | -5.808732009 | 6.29E-09 | 4.63E-08 |
| ENSG00000101463 | SYNDIG1 | 11.2470754 | -6.058533365 | 1.254809095 | -4.828251076 | 1.38E-06 | 7.82E-06 |
| ENSG00000101464 | PIGU | 667.9910639 | -0.409363984 | 0.079189375 | -5.169430703 | 2.35E-07 | 1.47E-06 |
| ENSG00000101470 | TNNC2 | 8.171754039 | -0.603025108 | 0.611739012 | -0.98575552 | 0.324253097 | 0.514445401 |
| ENSG00000101473 | ACOT8 | 667.0944752 | -0.019089755 | 0.092671678 | -0.205993409 | 0.83679606 | 0.913925996 |
| ENSG00000101474 | APMAP | 4420.958691 | -0.187729054 | 0.052123843 | -3.601596601 | 0.000316269 | 0.001297426 |
| ENSG00000101489 | CELF4 | 5.094451377 | 1.073583344 | 0.802498836 | 1.3378005 | 0.180961479 | 0.33637925 |
| ENSG00000101493 | ZNF516 | 743.164027 | 1.8882283 | 0.082726475 | 22.82495793 | 2.59E-115 | 5.56E-113 |
| ENSG00000101544 | ADNP2 | 1997.53184 | 0.426228523 | 0.055383718 | 7.695917492 | 1.40E-14 | 1.63E-13 |
| ENSG00000101546 | RBFA | 347.3277566 | 0.485153194 | 0.100786637 | 4.813665877 | 1.48E-06 | 8.39E-06 |
| ENSG00000101557 | USP14 | 3590.188208 | 0.175264471 | 0.056033886 | 3.127829997 | 0.00176102 | 0.006260313 |

| | | | | | | | |
|---|---|---|---|---|---|---|---|
| ENSG00000101558 | VAPA | 3137.299477 | 0.136458392 | 0.054419806 | 2.507513411 | 0.012158397 | 0.035345765 |
| ENSG00000101574 | METTL4 | 306.8516581 | 0.408429188 | 0.108137994 | 3.776925876 | 0.000158776 | 0.000685162 |
| ENSG00000101577 | LPIN2 | 1550.17024 | 0.708746195 | 0.061188859 | 11.58292868 | 5.03E-31 | 1.45E-29 |
| ENSG00000101596 | SMCHD1 | 2160.804288 | 0.598429559 | 0.062156425 | 9.627798823 | 6.10E-22 | 1.14E-20 |
| ENSG00000101605 | MYOM1 | 16.91308064 | -0.007399421 | 0.423905867 | -0.01745534 | 0.986073361 | 0.993350589 |
| ENSG00000101608 | MYL12A | 6559.672129 | -0.158907056 | 0.045530468 | -3.490125699 | 0.000482793 | 0.001916802 |
| ENSG00000101624 | CEP76 | 354.5236072 | 0.352328996 | 0.100685332 | 3.499308068 | 0.000466467 | 0.001856482 |
| ENSG00000101638 | ST8SIA5 | 6.409095061 | 0.737609027 | 0.721910415 | 1.021745928 | 0.306901167 | 0.494681898 |
| ENSG00000101639 | CEP192 | 1629.005824 | 0.168837775 | 0.064348314 | 2.623810394 | 0.008695218 | 0.026301731 |
| ENSG00000101654 | RNMT | 2223.851429 | 0.449374616 | 0.054644741 | 8.22356566 | 1.98E-16 | 2.64E-15 |
| ENSG00000101665 | SMAD7 | 755.8671021 | 0.373778337 | 0.078199865 | 4.779782354 | 1.75E-06 | 9.87E-06 |
| ENSG00000101695 | RNF125 | 83.89279856 | 2.085193125 | 0.220431586 | 9.459593166 | 3.09E-21 | 5.56E-20 |
| ENSG00000101745 | ANKRD12 | 1287.635908 | 0.22556447 | 0.076039129 | 2.966426266 | 0.003012826 | 0.010202868 |
| ENSG00000101746 | NOL4 | 4.575746975 | 3.565671807 | 1.247821678 | 2.857517119 | 0.004269696 | 0.013982416 |
| ENSG00000101751 | POLI | 1074.012051 | 0.2074833 | 0.075692754 | 2.741125007 | 0.006122921 | 0.019301883 |
| ENSG00000101752 | MIB1 | 2216.456096 | 0.20540312 | 0.063929863 | 3.212944759 | 0.001313816 | 0.004809915 |
| ENSG00000101773 | RBBP8 | 1823.232804 | 0.53835944 | 0.059301579 | 9.078332302 | 1.10E-19 | 1.81E-18 |
| ENSG00000101782 | RIOK3 | 1489.28631 | 0.086387448 | 0.062001231 | 1.393318273 | 0.163523567 | 0.311158882 |
| ENSG00000101811 | CSTF2 | 1463.06143 | -0.154587253 | 0.062060924 | -2.490895133 | 0.012742172 | 0.036777231 |
| ENSG00000101825 | MXRA5 | 6.471201759 | 2.36947032 | 0.81871755 | 2.894124258 | 0.003802177 | 0.012593406 |
| ENSG00000101842 | VSIG1 | 10.87048742 | -0.109003174 | 0.5344989 | -0.203935263 | 0.838404088 | 0.91499129 |
| ENSG00000101843 | PSMD10 | 1708.770373 | -0.143189468 | 0.056022957 | -2.555907 | 0.010591142 | 0.031257018 |
| ENSG00000101844 | ATG4A | 555.5746662 | 0.098254735 | 0.083841932 | 1.171904473 | 0.241235411 | 0.417215043 |
| ENSG00000101846 | STS | 558.0325119 | -0.048035597 | 0.082630952 | -0.581326922 | 0.561020139 | 0.730103564 |
| ENSG00000101849 | TBL1X | 5199.613763 | -0.253568019 | 0.048414189 | -5.237473205 | 1.63E-07 | 1.04E-06 |
| ENSG00000101850 | GPR143 | 204.4357752 | 0.610099574 | 0.136817732 | 4.459214198 | 8.23E-06 | 4.28E-05 |
| ENSG00000101856 | PGRMC1 | 2468.664402 | -0.239780483 | 0.056670312 | -4.231148064 | 2.33E-05 | 0.000114307 |
| ENSG00000101868 | POLA1 | 1354.801399 | -0.055499524 | 0.064482402 | -0.860692568 | 0.389407387 | 0.582084259 |
| ENSG00000101871 | MID1 | 115.7767383 | -1.236711717 | 0.187898121 | -6.58182055 | 4.65E-11 | 4.12E-10 |
| ENSG00000101882 | NKAP | 1272.934434 | 0.069319562 | 0.061733176 | 1.122889932 | 0.261484244 | 0.441733324 |
| ENSG00000101888 | NXT2 | 560.0250328 | -0.163924576 | 0.084847361 | -1.931993799 | 0.053360271 | 0.12679559 |
| ENSG00000101901 | ALG13 | 1442.347252 | 0.061099573 | 0.063711969 | 0.958996774 | 0.33756037 | 0.528785427 |
| ENSG00000101911 | PRPS2 | 1444.84935 | 0.186116058 | 0.061356667 | 3.033346927 | 0.002418574 | 0.008365815 |

| | | | | | | | |
|---|---|---|---|---|---|---|---|
| ENSG00000101928 | MOSPD1 | 713.5621061 | 0.178048112 | 0.075351118 | 2.362912681 | 0.018131941 | 0.050186623 |
| ENSG00000101935 | AMMECR1 | 1308.556391 | -0.143338985 | 0.07127942 | -2.010944884 | 0.044331279 | 0.108095189 |
| ENSG00000101940 | WDR13 | 1389.012172 | 0.006040199 | 0.070549552 | 0.085616401 | 0.93177136 | 0.967362192 |
| ENSG00000101945 | SUV39H1 | 924.0618309 | -0.215522793 | 0.070201342 | -3.070066589 | 0.002140111 | 0.007472471 |
| ENSG00000101955 | SRPX | 9.6288048 | 0.283685004 | 0.596666483 | 0.475449875 | 0.634466353 | 0.784490817 |
| ENSG00000101966 | XIAP | 2965.155279 | -0.223373045 | 0.057489791 | -3.885438447 | 0.000102145 | 0.000454817 |
| ENSG00000101972 | STAG2 | 5504.250456 | -0.124218128 | 0.051769606 | -2.399441253 | 0.016420114 | 0.046054215 |
| ENSG00000101974 | ATP11C | 2014.11145 | -0.543501414 | 0.062583962 | -8.684356084 | 3.81E-18 | 5.68E-17 |
| ENSG00000101977 | MCF2 | 4.610703297 | 0.453138104 | 0.816142323 | 0.555219464 | 0.578744557 | 0.744317998 |
| ENSG00000101986 | ABCD1 | 645.8201031 | -0.373132723 | 0.086857861 | -4.295900416 | 1.74E-05 | 8.70E-05 |
| ENSG00000101997 | CCDC22 | 990.4431307 | -0.167876867 | 0.06738739 | -2.491220804 | 0.012730498 | 0.036752783 |
| ENSG00000102003 | SYP | 31.30635796 | -0.610025511 | 0.313964659 | -1.942975089 | 0.052019168 | 0.123904031 |
| ENSG00000102007 | PLP2 | 3001.321066 | -0.116951782 | 0.049507687 | -2.36229541 | 0.018162163 | 0.050264208 |
| ENSG00000102024 | PLS3 | 1229.612287 | -0.131951266 | 0.06107045 | -2.160640139 | 0.030723148 | 0.079332398 |
| ENSG00000102030 | NAA10 | 1010.356515 | -0.509549544 | 0.067810941 | -7.514267403 | 5.72E-14 | 6.36E-13 |
| ENSG00000102032 | RENBP | 58.78801106 | 1.954850124 | 0.261161675 | 7.485210548 | 7.14E-14 | 7.87E-13 |
| ENSG00000102034 | ELF4 | 2052.953174 | 0.072658496 | 0.056596058 | 1.283808431 | 0.199208993 | 0.36205469 |
| ENSG00000102038 | SMARCA1 | 1424.988545 | -0.447431254 | 0.062909765 | -7.11227031 | 1.14E-12 | 1.16E-11 |
| ENSG00000102048 | ASB9 | 315.7912515 | 1.718816021 | 0.1170539 | 14.6839705 | 8.17E-49 | 4.34E-47 |
| ENSG00000102054 | RBBP7 | 5746.979978 | 0.146525606 | 0.045070248 | 3.251049484 | 0.001149798 | 0.004262504 |
| ENSG00000102057 | KCND1 | 177.4824418 | 0.037875868 | 0.148838729 | 0.25447589 | 0.799127928 | 0.892849925 |
| ENSG00000102078 | SLC25A14 | 502.1705193 | 0.056435657 | 0.090431919 | 0.624067895 | 0.532582997 | 0.708011935 |
| ENSG00000102081 | FMR1 | 2874.845404 | -0.061005823 | 0.059018405 | -1.033674534 | 0.301288342 | 0.488132583 |
| ENSG00000102096 | PIM2 | 550.2198616 | 0.040201925 | 0.087655677 | 0.458634583 | 0.646496598 | 0.793463804 |
| ENSG00000102098 | SCML2 | 320.8801034 | 0.150119604 | 0.108389837 | 1.384997044 | 0.166053371 | 0.31510169 |
| ENSG00000102100 | SLC35A2 | 1492.46401 | -0.048836568 | 0.0617317 | -0.791110046 | 0.428879777 | 0.61920798 |
| ENSG00000102103 | PQBP1 | 2286.468718 | -0.007907335 | 0.060615291 | -0.13045116 | 0.896209492 | 0.947508791 |
| ENSG00000102109 | PCSK1N | 314.2101624 | -0.011173922 | 0.120470211 | -0.092752568 | 0.926100133 | 0.96457376 |
| ENSG00000102119 | EMD | 2964.970182 | -0.667007217 | 0.049961416 | -13.35044655 | 1.18E-40 | 4.96E-39 |
| ENSG00000102125 | TAFAZZIN | 738.5138041 | -1.096539232 | 0.077664412 | -14.11894068 | 2.90E-45 | 1.42E-43 |
| ENSG00000102128 | RAB40AL | 2.033935291 | 1.027535087 | 1.271259219 | 0.808281326 | 0.41892865 | 0.610638152 |
| ENSG00000102144 | PGK1 | 13595.48582 | -0.079564826 | 0.044312748 | -1.79552904 | 0.07256945 | 0.163530765 |
| ENSG00000102158 | MAGT1 | 2896.43512 | -0.019193744 | 0.049961063 | -0.384174047 | 0.700849452 | 0.832426553 |

| | | | | | | | |
|---|---|---|---|---|---|---|---|
| ENSG00000102172 | SMS | 2734.238097 | -0.38508892 | 0.050323151 | -7.652321253 | 1.97E-14 | 2.27E-13 |
| ENSG00000102174 | PHEX | 135.8308546 | 1.175293818 | 0.167512432 | 7.016158768 | 2.28E-12 | 2.26E-11 |
| ENSG00000102178 | UBL4A | 3884.891189 | -0.66741688 | 0.052321754 | -12.75601129 | 2.89E-37 | 1.06E-35 |
| ENSG00000102181 | CD99L2 | 1275.412667 | -0.093480432 | 0.062176644 | -1.503465386 | 0.132719077 | 0.26573934 |
| ENSG00000102189 | EEA1 | 984.1276246 | -0.045270244 | 0.077659655 | -0.582931297 | 0.559939551 | 0.729374442 |
| ENSG00000102195 | GPR50 | 66.97685629 | 4.422884219 | 0.393589327 | 11.23730731 | 2.67E-29 | 7.22E-28 |
| ENSG00000102218 | RP2 | 452.158847 | -0.19208394 | 0.089514967 | -2.145830435 | 0.031886509 | 0.081912265 |
| ENSG00000102221 | JADE3 | 1361.117399 | 0.03226959 | 0.059580471 | 0.541613551 | 0.588084754 | 0.751401597 |
| ENSG00000102225 | CDK16 | 5360.901092 | 0.078196573 | 0.048912752 | 1.598695019 | 0.109888385 | 0.228594597 |
| ENSG00000102226 | USP11 | 4737.787667 | -0.310133152 | 0.047980709 | -6.463705053 | 1.02E-10 | 8.81E-10 |
| ENSG00000102241 | HTATSF1 | 5629.287693 | -0.045396837 | 0.043452111 | -1.044755604 | 0.296135939 | 0.482345813 |
| ENSG00000102243 | VGLL1 | 5.806883257 | 1.410325834 | 0.786618323 | 1.792897258 | 0.072989354 | 0.164267221 |
| ENSG00000102265 | TIMP1 | 255.3516762 | 3.140996093 | 0.159997046 | 19.63158798 | 8.31E-86 | 1.06E-83 |
| ENSG00000102287 | GABRE | 54.86577764 | 4.401873528 | 0.433079488 | 10.16412378 | 2.87E-24 | 6.10E-23 |
| ENSG00000102290 | PCDH11X | 67.13992392 | 1.051656795 | 0.228602838 | 4.600366317 | 4.22E-06 | 2.27E-05 |
| ENSG00000102302 | FGD1 | 232.0022205 | -1.269984699 | 0.126041613 | -10.07591599 | 7.06E-24 | 1.46E-22 |
| ENSG00000102309 | PIN4 | 559.9904369 | 0.09250927 | 0.08301343 | 1.114389207 | 0.265112251 | 0.446099541 |
| ENSG00000102312 | PORCN | 346.8173902 | 0.024195282 | 0.100620057 | 0.240461817 | 0.809972261 | 0.899408839 |
| ENSG00000102313 | ITIH6 | 2.395684579 | -0.627309868 | 1.128642561 | -0.555809155 | 0.578341327 | 0.743966244 |
| ENSG00000102316 | MAGED2 | 6828.78807 | -0.607266615 | 0.048039974 | -12.64086049 | 1.26E-36 | 4.51E-35 |
| ENSG00000102317 | RBM3 | 6927.438314 | 0.248247699 | 0.063150896 | 3.93102417 | 8.46E-05 | 0.000382189 |
| ENSG00000102349 | KLF8 | 138.4334194 | -0.335084778 | 0.154312283 | -2.171471848 | 0.029895523 | 0.077430653 |
| ENSG00000102362 | SYTL4 | 166.7689397 | 0.313622174 | 0.145744159 | 2.151867872 | 0.031407762 | 0.080809115 |
| ENSG00000102384 | CENPI | 713.708803 | -0.396092674 | 0.079838546 | -4.961170964 | 7.01E-07 | 4.14E-06 |
| ENSG00000102385 | DRP2 | 47.17474666 | -0.207579424 | 0.262970003 | -0.78936541 | 0.429898466 | 0.620204441 |
| ENSG00000102387 | TAF7L | 28.5608374 | 2.06944294 | 0.380371877 | 5.440578194 | 5.31E-08 | 3.54E-07 |
| ENSG00000102390 | PBDC1 | 1014.257782 | -0.227625045 | 0.070195054 | -3.242750466 | 0.001183818 | 0.004375189 |
| ENSG00000102393 | GLA | 848.8723723 | 0.107035344 | 0.071120277 | 1.504990545 | 0.132326508 | 0.265068991 |
| ENSG00000102401 | ARMCX3 | 1660.475009 | -0.055423391 | 0.061487569 | -0.901375545 | 0.36738868 | 0.559881506 |
| ENSG00000102409 | BEX4 | 28.76695458 | -0.822691646 | 0.33820901 | -2.432494766 | 0.014995208 | 0.042499392 |
| ENSG00000102471 | NDFIP2 | 664.557494 | -0.072842796 | 0.07938842 | -0.917549393 | 0.358854823 | 0.551455629 |
| ENSG00000102524 | TNFSF13B | 7.984002932 | 0.367880026 | 0.624776374 | 0.588818722 | 0.555982885 | 0.726719554 |
| ENSG00000102531 | FNDC3A | 1048.098991 | 0.028741757 | 0.066475373 | 0.432366989 | 0.665474706 | 0.807126455 |

| | | | | | | | |
|---|---|---|---|---|---|---|---|
| ENSG00000102543 | CDADC1 | 108.906192 | -0.249151512 | 0.169141588 | -1.473035198 | 0.140741545 | 0.277706094 |
| ENSG00000102547 | CAB39L | 487.7079947 | -0.502073433 | 0.10133423 | -4.954628173 | 7.25E-07 | 4.28E-06 |
| ENSG00000102554 | KLF5 | 616.2646392 | 1.361040086 | 0.084855048 | 16.0395889 | 6.76E-58 | 4.57E-56 |
| ENSG00000102572 | STK24 | 2742.077213 | 0.100815632 | 0.052976025 | 1.903042597 | 0.057034987 | 0.133822503 |
| ENSG00000102575 | ACP5 | 3.312427369 | 3.099141265 | 1.301694626 | 2.38085124 | 0.017272686 | 0.048127707 |
| ENSG00000102580 | DNAJC3 | 1263.13466 | 0.2200056 | 0.066774022 | 3.294778306 | 0.000984994 | 0.003690307 |
| ENSG00000102595 | UGGT2 | 333.2372904 | 0.02802809 | 0.107402261 | 0.260963688 | 0.79412051 | 0.88974732 |
| ENSG00000102606 | ARHGEF7 | 1454.431649 | -0.117292034 | 0.062219675 | -1.885127721 | 0.059412596 | 0.138368016 |
| ENSG00000102683 | SGCG | 2122.612664 | 0.235206019 | 0.055161573 | 4.263946909 | 2.01E-05 | 9.97E-05 |
| ENSG00000102699 | PARP4 | 3901.955486 | 0.164380223 | 0.048412464 | 3.395411184 | 0.000685256 | 0.002645981 |
| ENSG00000102710 | SUPT20H | 1779.518128 | -0.175378168 | 0.059556936 | -2.944714399 | 0.003232529 | 0.010876242 |
| ENSG00000102738 | MRPS31 | 609.9609005 | -0.238818629 | 0.080119006 | -2.980798695 | 0.002874977 | 0.009784685 |
| ENSG00000102743 | SLC25A15 | 1835.820352 | 0.039907111 | 0.055818719 | 0.714941367 | 0.474645274 | 0.659944021 |
| ENSG00000102753 | KPNA3 | 1354.234971 | 0.027990967 | 0.059617033 | 0.469512918 | 0.638703053 | 0.787290808 |
| ENSG00000102755 | FLT1 | 11.20458542 | -2.200222266 | 0.627561363 | -3.505987456 | 0.000454917 | 0.001813658 |
| ENSG00000102760 | RGCC | 5.521296124 | 0.69960331 | 0.750830269 | 0.931772918 | 0.351453893 | 0.542773146 |
| ENSG00000102763 | VWA8 | 1110.865413 | 0.070436834 | 0.065278465 | 1.079020993 | 0.280578369 | 0.463989893 |
| ENSG00000102780 | DGKH | 880.423281 | -0.293911235 | 0.071729003 | -4.097522904 | 4.18E-05 | 0.000197437 |
| ENSG00000102781 | KATNAL1 | 329.0231963 | -0.515548635 | 0.119059262 | -4.330185056 | 1.49E-05 | 7.51E-05 |
| ENSG00000102786 | INTS6 | 939.2305425 | -0.192128789 | 0.072202286 | -2.660979308 | 0.007791375 | 0.023885358 |
| ENSG00000102796 | DHRS12 | 80.04890605 | -0.154562426 | 0.19884666 | -0.777294553 | 0.436985008 | 0.626676508 |
| ENSG00000102804 | TSC22D1 | 5518.83107 | -0.023316944 | 0.043650337 | -0.534175571 | 0.593220069 | 0.755354595 |
| ENSG00000102805 | CLN5 | 221.1231121 | -0.124406908 | 0.124189751 | -1.001748596 | 0.316465029 | 0.50523403 |
| ENSG00000102854 | MSLN | 5.693466672 | 0.313072886 | 0.759825847 | 0.412032424 | 0.680315658 | 0.818419487 |
| ENSG00000102858 | MGRN1 | 2396.979832 | 0.094392351 | 0.05769156 | 1.63615529 | 0.101807091 | 0.215070878 |
| ENSG00000102870 | ZNF629 | 3034.687425 | 0.079392294 | 0.058821578 | 1.349713757 | 0.177107818 | 0.331012982 |
| ENSG00000102871 | TRADD | 418.9188725 | -0.147873721 | 0.09693078 | -1.525560003 | 0.127119479 | 0.256859056 |
| ENSG00000102878 | HSF4 | 717.8979788 | 0.116607606 | 0.084275408 | 1.383649262 | 0.166465877 | 0.315675557 |
| ENSG00000102879 | CORO1A | 142.1339355 | -0.344572079 | 0.157398359 | -2.189171999 | 0.028584341 | 0.074488786 |
| ENSG00000102882 | MAPK3 | 3133.911005 | -0.159076656 | 0.0548264 | -2.9014609 | 0.003714271 | 0.012325385 |
| ENSG00000102886 | GDPD3 | 192.5295578 | 0.749492433 | 0.140862382 | 5.320742275 | 1.03E-07 | 6.69E-07 |
| ENSG00000102890 | ELMO3 | 1455.560032 | 0.897927921 | 0.066790511 | 13.44394444 | 3.34E-41 | 1.44E-39 |
| ENSG00000102893 | PHKB | 1455.343813 | -0.254516107 | 0.064810103 | -3.927105441 | 8.60E-05 | 0.000387857 |

| | | | | | | | |
|---|---|---|---|---|---|---|---|
| ENSG00000102897 | LYRM1 | 416.1836691 | -0.472038519 | 0.093295421 | -5.059610797 | 4.20E-07 | 2.56E-06 |
| ENSG00000102898 | NUTF2 | 2994.196553 | 0.041968871 | 0.049050026 | 0.855634038 | 0.392200236 | 0.585061348 |
| ENSG00000102900 | NUP93 | 1294.764601 | -0.050199752 | 0.061254444 | -0.819528321 | 0.412485051 | 0.604588997 |
| ENSG00000102901 | CENPT | 1285.085864 | -0.032747863 | 0.068876554 | -0.475457337 | 0.634461035 | 0.784490817 |
| ENSG00000102904 | TSNAXIP1 | 47.56081383 | -0.112552642 | 0.257843699 | -0.436515 | 0.662463119 | 0.804963395 |
| ENSG00000102908 | NFAT5 | 994.0388343 | 0.006917813 | 0.070753134 | 0.097773945 | 0.922111797 | 0.962216621 |
| ENSG00000102910 | LONP2 | 2586.828647 | -0.218921374 | 0.050947465 | -4.297002252 | 1.73E-05 | 8.67E-05 |
| ENSG00000102921 | N4BP1 | 900.7075341 | -0.06764581 | 0.069063349 | -0.979474801 | 0.327345434 | 0.517679904 |
| ENSG00000102931 | ARL2BP | 1087.946093 | 0.037438865 | 0.066864426 | 0.559922035 | 0.575532618 | 0.741934153 |
| ENSG00000102934 | PLLP | 103.2610489 | 0.603279929 | 0.179805723 | 3.355176464 | 0.000793144 | 0.003023885 |
| ENSG00000102935 | ZNF423 | 7.981558072 | -1.732066869 | 0.665051978 | -2.604408266 | 0.009203305 | 0.0276666 |
| ENSG00000102962 | CCL22 | 4.768182663 | 5.605163385 | 1.354340413 | 4.138666564 | 3.49E-05 | 0.00016702 |
| ENSG00000102967 | DHODH | 512.2114579 | 0.178154046 | 0.086150216 | 2.067946602 | 0.038645045 | 0.096392011 |
| ENSG00000102974 | CTCF | 4351.60983 | -0.136253943 | 0.047184664 | -2.887674345 | 0.003881015 | 0.012824903 |
| ENSG00000102977 | ACD | 775.9717213 | 0.241571284 | 0.077179111 | 3.130008618 | 0.001748012 | 0.006218853 |
| ENSG00000102978 | POLR2C | 1790.089963 | -0.108427197 | 0.058399607 | -1.856642578 | 0.063362022 | 0.145965534 |
| ENSG00000102981 | PARD6A | 205.5482759 | 0.374036166 | 0.135129728 | 2.767978385 | 0.00564052 | 0.017906659 |
| ENSG00000102984 | ZNF821 | 415.0041201 | 0.310942461 | 0.097403794 | 3.192303386 | 0.00141143 | 0.005124755 |
| ENSG00000102996 | MMP15 | 1375.545715 | 0.4471676 | 0.071832325 | 6.225158403 | 4.81E-10 | 3.94E-09 |
| ENSG00000103005 | USB1 | 830.1963536 | -0.066746607 | 0.070361449 | -0.948624673 | 0.342811538 | 0.533778686 |
| ENSG00000103018 | CYB5B | 3017.633022 | -0.247572158 | 0.04938084 | -5.013526702 | 5.34E-07 | 3.21E-06 |
| ENSG00000103021 | CCDC113 | 308.4811874 | 0.549555709 | 0.110393463 | 4.978154446 | 6.42E-07 | 3.81E-06 |
| ENSG00000103024 | NME3 | 985.5182249 | 0.105489986 | 0.074020675 | 1.425142171 | 0.154116109 | 0.297344283 |
| ENSG00000103034 | NDRG4 | 556.7558494 | -0.345521638 | 0.086666969 | -3.986774208 | 6.70E-05 | 0.000307355 |
| ENSG00000103035 | PSMD7 | 6021.836056 | 0.34796083 | 0.043669852 | 7.967987377 | 1.61E-15 | 2.01E-14 |
| ENSG00000103037 | SETD6 | 557.0617791 | 0.044206244 | 0.089944219 | 0.491485111 | 0.62308338 | 0.777075826 |
| ENSG00000103042 | SLC38A7 | 528.2739066 | 0.329371002 | 0.085992275 | 3.830239423 | 0.000128019 | 0.000560876 |
| ENSG00000103043 | VAC14 | 1701.647036 | -0.017195413 | 0.057267838 | -0.300263 | 0.763976553 | 0.870544448 |
| ENSG00000103044 | HAS3 | 78.05884837 | 0.419348578 | 0.202602627 | 2.069808202 | 0.038470309 | 0.096029259 |
| ENSG00000103047 | TANGO6 | 256.4431119 | 0.005401356 | 0.118556324 | 0.045559413 | 0.963661419 | 0.983511721 |
| ENSG00000103051 | COG4 | 3281.24444 | -0.277540123 | 0.053227843 | -5.214190676 | 1.85E-07 | 1.17E-06 |
| ENSG00000103056 | SMPD3 | 95.66758106 | -0.907193032 | 0.19116908 | -4.74550086 | 2.08E-06 | 1.16E-05 |
| ENSG00000103061 | SLC7A6OS | 867.0264362 | 0.005225437 | 0.07463144 | 0.07001657 | 0.944180471 | 0.973504427 |

| | | | | | | | |
|---|---|---|---|---|---|---|---|
| ENSG00000103064 | SLC7A6 | 2897.585368 | 0.394729505 | 0.052202219 | 7.561546513 | 3.98E-14 | 4.48E-13 |
| ENSG00000103066 | PLA2G15 | 549.7583889 | -0.107294243 | 0.086119577 | -1.245875169 | 0.212810228 | 0.38088996 |
| ENSG00000103067 | ESRP2 | 4973.960568 | -0.007874414 | 0.049874495 | -0.157884594 | 0.874547738 | 0.934948509 |
| ENSG00000103089 | FA2H | 282.7633342 | 0.220699899 | 0.111882191 | 1.972609728 | 0.048540044 | 0.116805686 |
| ENSG00000103091 | WDR59 | 3218.572998 | 0.537804575 | 0.053646842 | 10.02490658 | 1.18E-23 | 2.43E-22 |
| ENSG00000103111 | MON1B | 6690.251677 | -0.652938976 | 0.046439621 | -14.0599548 | 6.69E-45 | 3.21E-43 |
| ENSG00000103121 | CMC2 | 845.8984692 | 0.168154364 | 0.070694933 | 2.378591452 | 0.017378925 | 0.048406089 |
| ENSG00000103126 | AXIN1 | 1600.506889 | 0.157734216 | 0.061893133 | 2.548492998 | 0.010818946 | 0.03183104 |
| ENSG00000103145 | HCFC1R1 | 694.6304244 | -0.399520524 | 0.092784274 | -4.305907743 | 1.66E-05 | 8.34E-05 |
| ENSG00000103148 | NPRL3 | 1550.123415 | -0.216547324 | 0.06100823 | -3.549477226 | 0.000385997 | 0.001555105 |
| ENSG00000103150 | MLYCD | 833.6172366 | 0.462191727 | 0.07345445 | 6.292222326 | 3.13E-10 | 2.61E-09 |
| ENSG00000103152 | MPG | 1193.349092 | 0.142842571 | 0.071563478 | 1.996026123 | 0.045931079 | 0.111381115 |
| ENSG00000103154 | NECAB2 | 124.5448101 | 1.199938663 | 0.170017264 | 7.057745966 | 1.69E-12 | 1.70E-11 |
| ENSG00000103160 | HSDL1 | 2438.511505 | 0.676009812 | 0.055828217 | 12.10874807 | 9.49E-34 | 3.05E-32 |
| ENSG00000103168 | TAF1C | 3585.625558 | 0.372434682 | 0.062048857 | 6.002281113 | 1.95E-09 | 1.51E-08 |
| ENSG00000103174 | NAGPA | 313.9472826 | -0.09238004 | 0.106516387 | -0.867284766 | 0.385786002 | 0.578389052 |
| ENSG00000103184 | SEC14L5 | 5.059868512 | 0.453363438 | 0.774776234 | 0.585154033 | 0.55844415 | 0.728502064 |
| ENSG00000103187 | COTL1 | 1154.743418 | 0.43681311 | 0.066515575 | 6.567080075 | 5.13E-11 | 4.54E-10 |
| ENSG00000103194 | USP10 | 5792.67378 | 0.256523847 | 0.046943417 | 5.464532894 | 4.64E-08 | 3.11E-07 |
| ENSG00000103196 | CRISPLD2 | 216.6021942 | -1.124155029 | 0.127001985 | -8.851476078 | 8.64E-19 | 1.35E-17 |
| ENSG00000103197 | TSC2 | 4645.383757 | 0.201610734 | 0.05199694 | 3.877357685 | 0.000105597 | 0.000469184 |
| ENSG00000103199 | ZNF500 | 365.2391542 | 0.063503403 | 0.100532013 | 0.631673449 | 0.527600281 | 0.703507931 |
| ENSG00000103202 | NME4 | 3837.072381 | 0.341184682 | 0.053231788 | 6.409416125 | 1.46E-10 | 1.24E-09 |
| ENSG00000103222 | ABCC1 | 2476.079306 | 0.615372621 | 0.05640658 | 10.90958919 | 1.04E-27 | 2.66E-26 |
| ENSG00000103226 | NOMO3 | 384.8455864 | 0.121534576 | 0.095914324 | 1.267116011 | 0.205113811 | 0.370205416 |
| ENSG00000103227 | LMF1 | 508.3467797 | 0.029797809 | 0.092199287 | 0.323189152 | 0.746551996 | 0.859604617 |
| ENSG00000103245 | CIAO3 | 1007.23366 | 0.07962658 | 0.070623774 | 1.127475572 | 0.259541463 | 0.439484833 |
| ENSG00000103248 | MTHFSD | 701.6965742 | 0.213661447 | 0.077294477 | 2.764252445 | 0.005705338 | 0.018099911 |
| ENSG00000103249 | CLCN7 | 2504.943515 | 0.522892751 | 0.061833237 | 8.456499721 | 2.76E-17 | 3.87E-16 |
| ENSG00000103253 | HAGHL | 1343.296912 | 0.017144216 | 0.0617146 | 0.277798373 | 0.78116714 | 0.881300566 |
| ENSG00000103254 | ANTKMT | 331.782335 | 0.061761853 | 0.105529713 | 0.585255572 | 0.558375883 | 0.728502064 |
| ENSG00000103257 | SLC7A5 | 17533.09791 | 0.56643739 | 0.045356819 | 12.48847247 | 8.63E-36 | 3.00E-34 |
| ENSG00000103260 | METRN | 1436.138268 | 0.525703679 | 0.071308024 | 7.372293489 | 1.68E-13 | 1.80E-12 |

| | | | | | | | |
|---|---|---|---|---|---|---|---|
| ENSG00000103264 | FBXO31 | 1581.389778 | 0.577236284 | 0.063894783 | 9.034169297 | 1.65E-19 | 2.69E-18 |
| ENSG00000103266 | STUB1 | 3950.572837 | 0.08062053 | 0.051820796 | 1.555756302 | 0.119766053 | 0.245175196 |
| ENSG00000103269 | RHBDL1 | 256.7918654 | -0.193773899 | 0.122355102 | -1.583701017 | 0.113261773 | 0.234454528 |
| ENSG00000103274 | NUBP1 | 813.1966429 | 0.081881697 | 0.074779601 | 1.094973705 | 0.273528156 | 0.455516218 |
| ENSG00000103275 | UBE2I | 4081.054422 | 0.158159513 | 0.046506444 | 3.400808551 | 0.000671869 | 0.002598662 |
| ENSG00000103313 | MEFV | 4.270249638 | -1.295449195 | 0.876530324 | -1.477928556 | 0.139426902 | 0.275629863 |
| ENSG00000103316 | CRYM | 19.46961142 | -3.131716722 | 0.533545046 | -5.869638839 | 4.37E-09 | 3.28E-08 |
| ENSG00000103319 | EEF2K | 1650.213258 | -0.12296225 | 0.05889285 | -2.087897782 | 0.036807056 | 0.092529606 |
| ENSG00000103326 | CAPN15 | 1702.837608 | 0.181547296 | 0.064182923 | 2.828591875 | 0.004675328 | 0.01517431 |
| ENSG00000103335 | PIEZO1 | 16565.86228 | 0.666462748 | 0.047025644 | 14.17232594 | 1.36E-45 | 6.68E-44 |
| ENSG00000103342 | GSPT1 | 8464.537531 | 0.046165957 | 0.044093512 | 1.047001133 | 0.295099056 | 0.481018631 |
| ENSG00000103343 | ZNF174 | 342.179935 | 0.15520615 | 0.104050001 | 1.491649673 | 0.135791009 | 0.27035634 |
| ENSG00000103351 | CLUAP1 | 584.1538379 | 0.138702674 | 0.085054831 | 1.630744213 | 0.102944303 | 0.217053521 |
| ENSG00000103353 | UBFD1 | 4749.847879 | 0.058466875 | 0.046447199 | 1.2587815 | 0.208109264 | 0.374550959 |
| ENSG00000103356 | EARS2 | 1445.358613 | 0.265268389 | 0.059378728 | 4.467397673 | 7.92E-06 | 4.13E-05 |
| ENSG00000103363 | ELOB | 2567.695665 | 0.227756841 | 0.062999117 | 3.615238617 | 0.000300071 | 0.001234954 |
| ENSG00000103365 | GGA2 | 3088.167851 | 0.250260366 | 0.051184435 | 4.889384175 | 1.01E-06 | 5.85E-06 |
| ENSG00000103381 | CPPED1 | 242.6485814 | 0.687503767 | 0.120142612 | 5.722397379 | 1.05E-08 | 7.54E-08 |
| ENSG00000103404 | USP31 | 467.5614298 | 0.111756909 | 0.09393828 | 1.189684427 | 0.234170448 | 0.408911478 |
| ENSG00000103415 | HMOX2 | 1812.375602 | -0.08775954 | 0.062746396 | -1.398638734 | 0.161921345 | 0.308947488 |
| ENSG00000103423 | DNAJA3 | 2917.386 | 0.174775857 | 0.053054297 | 3.294282756 | 0.000986732 | 0.003696216 |
| ENSG00000103429 | BFAR | 2100.395523 | 0.141163042 | 0.056855738 | 2.48282843 | 0.013034385 | 0.037526204 |
| ENSG00000103460 | TOX3 | 2.201838909 | 4.482030325 | 1.543621992 | 2.903580247 | 0.003689224 | 0.012249356 |
| ENSG00000103479 | RBL2 | 4713.663679 | 0.19092252 | 0.049617661 | 3.847874282 | 0.000119147 | 0.000524513 |
| ENSG00000103485 | QPRT | 606.1307971 | -0.953662365 | 0.090471933 | -10.54097481 | 5.59E-26 | 1.30E-24 |
| ENSG00000103489 | XYLT1 | 492.4573435 | 0.609008717 | 0.098987494 | 6.15238041 | 7.63E-10 | 6.14E-09 |
| ENSG00000103490 | PYCARD | 414.3934956 | -0.057289768 | 0.095825752 | -0.597853567 | 0.549937645 | 0.722293002 |
| ENSG00000103494 | RPGRIP1L | 294.1604762 | 0.172235126 | 0.10896263 | 1.580680691 | 0.113951066 | 0.235652865 |
| ENSG00000103495 | MAZ | 14721.8561 | -0.132358823 | 0.041840296 | -3.163429391 | 0.001559222 | 0.005609872 |
| ENSG00000103496 | STX4 | 1426.167752 | 0.059745968 | 0.059420682 | 1.005474286 | 0.314668525 | 0.503189213 |
| ENSG00000103502 | CDIPT | 2673.936975 | 0.196003332 | 0.056602954 | 3.462775651 | 0.000534634 | 0.00210441 |
| ENSG00000103507 | BCKDK | 1531.677026 | 0.102332304 | 0.059474434 | 1.720609951 | 0.085321627 | 0.186245998 |
| ENSG00000103510 | KAT8 | 905.7918925 | -0.023497604 | 0.06928006 | -0.339168347 | 0.734482913 | 0.853578385 |

| | | | | | | | |
|---|---|---|---|---|---|---|---|
| ENSG00000103512 | NOMO1 | 2086.930802 | 0.108110207 | 0.055962798 | 1.931822745 | 0.053381388 | 0.126819494 |
| ENSG00000103522 | IL21R | 6.53115914 | -2.883616607 | 0.85826288 | -3.35982911 | 0.000779907 | 0.002978865 |
| ENSG00000103528 | SYT17 | 444.1874699 | -0.815550519 | 0.097475996 | -8.366680542 | 5.93E-17 | 8.16E-16 |
| ENSG00000103534 | TMC5 | 70.17965625 | -2.800431015 | 0.266572203 | -10.5053377 | 8.16E-26 | 1.88E-24 |
| ENSG00000103540 | CCP110 | 866.9151459 | -0.364543037 | 0.079489402 | -4.586058374 | 4.52E-06 | 2.42E-05 |
| ENSG00000103544 | VPS35L | 252.6278194 | -1.230579953 | 0.11961327 | -10.28798852 | 7.98E-25 | 1.76E-23 |
| ENSG00000103549 | RNF40 | 5747.264067 | 0.018007082 | 0.049679697 | 0.362463607 | 0.717005615 | 0.842671694 |
| ENSG00000103550 | KNOP1 | 1509.920466 | 0.017700106 | 0.057697969 | 0.306771726 | 0.759017131 | 0.867296921 |
| ENSG00000103591 | AAGAB | 3148.686723 | -0.118885498 | 0.047770024 | -2.488705005 | 0.01282093 | 0.036976632 |
| ENSG00000103599 | IQCH | 154.3593307 | 0.045648024 | 0.145976967 | 0.312707035 | 0.754503251 | 0.864498202 |
| ENSG00000103642 | LACTB | 505.1965424 | 1.286874238 | 0.089897176 | 14.31495739 | 1.76E-46 | 8.84E-45 |
| ENSG00000103647 | CORO2B | 76.11845962 | -0.772260622 | 0.20713657 | -3.728267894 | 0.0001928 | 0.000821021 |
| ENSG00000103653 | CSK | 3617.824924 | 0.015811827 | 0.053368182 | 0.296278155 | 0.767017667 | 0.872778717 |
| ENSG00000103657 | HERC1 | 1898.208288 | -0.137609816 | 0.129101513 | -1.06590398 | 0.28646705 | 0.470236431 |
| ENSG00000103671 | TRIP4 | 1085.224689 | -0.323637678 | 0.066538475 | -4.863917906 | 1.15E-06 | 6.61E-06 |
| ENSG00000103707 | MTFMT | 503.3678276 | 0.00097698 | 0.087095973 | 0.011217278 | 0.991050094 | 0.995176993 |
| ENSG00000103723 | AP3B2 | 109.461894 | -0.512576987 | 0.171756894 | -2.984316819 | 0.002842123 | 0.009687713 |
| ENSG00000103740 | ACSBG1 | 24.50977189 | -0.924102271 | 0.358170776 | -2.580060498 | 0.009878301 | 0.029406725 |
| ENSG00000103742 | IGDCC4 | 23.25730251 | 0.33810054 | 0.370003747 | 0.91377599 | 0.360834565 | 0.553385808 |
| ENSG00000103769 | RAB11A | 5867.586975 | -0.030730255 | 0.049159238 | -0.625116598 | 0.531894535 | 0.707424666 |
| ENSG00000103811 | CTSH | 1026.355139 | 0.475158843 | 0.084665473 | 5.612191443 | 2.00E-08 | 1.39E-07 |
| ENSG00000103832 | GOLGA8UP | 2.71894561 | -0.320019342 | 1.052347338 | -0.304100491 | 0.761051333 | 0.868799083 |
| ENSG00000103852 | TTC23 | 392.6850642 | 0.048474126 | 0.099247345 | 0.488417362 | 0.625254247 | 0.778594473 |
| ENSG00000103855 | CD276 | 4097.214583 | 0.120952923 | 0.051107489 | 2.366637975 | 0.017950482 | 0.049768438 |
| ENSG00000103876 | FAH | 560.1831116 | -0.269905962 | 0.085529415 | -3.155709217 | 0.001601084 | 0.005749666 |
| ENSG00000103888 | CEMIP | 495.3451485 | -0.878674682 | 0.094203034 | -9.327456278 | 1.08E-20 | 1.88E-19 |
| ENSG00000103932 | RPAP1 | 1151.487297 | -0.149793089 | 0.067578628 | -2.216574876 | 0.026652156 | 0.070179116 |
| ENSG00000103942 | HOMER2 | 1775.046368 | 0.528349727 | 0.057210273 | 9.235224697 | 2.58E-20 | 4.38E-19 |
| ENSG00000103966 | EHD4 | 1564.063747 | -0.757274129 | 0.064389188 | -11.7608895 | 6.21E-32 | 1.86E-30 |
| ENSG00000103978 | TMEM87A | 1671.919912 | -0.095607734 | 0.057207769 | -1.671236885 | 0.094674897 | 0.202729594 |
| ENSG00000103994 | ZNF106 | 2391.837547 | -0.409364384 | 0.058837516 | -6.957540263 | 3.46E-12 | 3.39E-11 |
| ENSG00000103995 | CEP152 | 372.9756813 | -0.477869361 | 0.105515777 | -4.528890126 | 5.93E-06 | 3.15E-05 |
| ENSG00000104043 | ATP8B4 | 24.15053221 | -0.700338633 | 0.374511705 | -1.870004658 | 0.061483171 | 0.142452676 |

| | | | | | | | |
|---|---|---|---|---|---|---|---|
| ENSG00000104047 | DTWD1 | 3400.366277 | -1.27274265 | 0.063249393 | -20.12260662 | 4.68E-90 | 6.43E-88 |
| ENSG00000104064 | GABPB1 | 2764.520784 | -1.529951927 | 0.056886588 | -26.89477393 | 2.53E-159 | 9.51E-157 |
| ENSG00000104067 | TJP1 | 2089.293403 | -0.161947072 | 0.054077235 | -2.994736549 | 0.00274682 | 0.009392167 |
| ENSG00000104081 | BMF | 3880.400606 | 0.491243461 | 0.057263229 | 8.578689449 | 9.60E-18 | 1.39E-16 |
| ENSG00000104093 | DMXL2 | 7497.315498 | -1.435354995 | 0.046409978 | -30.92772384 | 5.06E-210 | 3.32E-207 |
| ENSG00000104129 | DNAJC17 | 367.4067722 | -0.434748459 | 0.101831251 | -4.269302924 | 1.96E-05 | 9.74E-05 |
| ENSG00000104131 | EIF3J | 2649.576893 | -0.026431963 | 0.067400611 | -0.392162072 | 0.694938466 | 0.828271093 |
| ENSG00000104133 | SPG11 | 2690.269975 | -0.172541032 | 0.062851059 | -2.745236671 | 0.006046727 | 0.019082669 |
| ENSG00000104140 | RHOV | 392.7167611 | 0.838962938 | 0.097265351 | 8.625506734 | 6.38E-18 | 9.34E-17 |
| ENSG00000104142 | VPS18 | 1420.682277 | 0.178138877 | 0.069377316 | 2.567681891 | 0.010238106 | 0.030340028 |
| ENSG00000104147 | OIP5 | 261.1629561 | -0.25676766 | 0.114176546 | -2.248865188 | 0.024521074 | 0.065393333 |
| ENSG00000104154 | SLC30A4 | 150.0375531 | -0.649925136 | 0.155770782 | -4.17231734 | 3.02E-05 | 0.000146047 |
| ENSG00000104164 | BLOC1S6 | 1295.541496 | -0.174654778 | 0.065799808 | -2.654335675 | 0.007946473 | 0.024321828 |
| ENSG00000104177 | MYEF2 | 1985.979915 | 0.2478209 | 0.065916117 | 3.759640433 | 0.000170158 | 0.000731251 |
| ENSG00000104205 | SGK3 | 52.5278447 | 0.713395673 | 0.266538889 | 2.676516267 | 0.007439195 | 0.022906731 |
| ENSG00000104213 | PDGFRL | 131.5561628 | 0.746762907 | 0.168494181 | 4.431980403 | 9.34E-06 | 4.84E-05 |
| ENSG00000104218 | CSPP1 | 669.3839386 | 0.186900923 | 0.078946447 | 2.367439329 | 0.017911657 | 0.049690821 |
| ENSG00000104219 | ZDHHC2 | 299.6197237 | 0.115699109 | 0.109787354 | 1.053847319 | 0.291952825 | 0.476936684 |
| ENSG00000104221 | BRF2 | 362.1186666 | -0.160173926 | 0.100589651 | -1.592349962 | 0.111306093 | 0.230989847 |
| ENSG00000104228 | TRIM35 | 870.9355448 | -0.099697433 | 0.070540346 | -1.413339162 | 0.157556024 | 0.302227209 |
| ENSG00000104231 | ZFAND1 | 1288.1942 | 0.106333685 | 0.073534431 | 1.446039403 | 0.14816615 | 0.288803763 |
| ENSG00000104267 | CA2 | 111.4446543 | 1.491409721 | 0.185207848 | 8.052626996 | 8.10E-16 | 1.03E-14 |
| ENSG00000104290 | FZD3 | 530.8880732 | 0.013400109 | 0.089293664 | 0.150067863 | 0.880711075 | 0.938252557 |
| ENSG00000104299 | INTS9 | 462.9994646 | -0.103450691 | 0.092592469 | -1.117268953 | 0.263879346 | 0.44451404 |
| ENSG00000104312 | RIPK2 | 1376.95225 | 0.315709824 | 0.06037082 | 5.229510285 | 1.70E-07 | 1.08E-06 |
| ENSG00000104313 | EYA1 | 2.933203023 | 0.466202025 | 1.016158726 | 0.458788585 | 0.646385993 | 0.793412953 |
| ENSG00000104320 | NBN | 7100.716932 | 0.443649403 | 0.047446121 | 9.350593797 | 8.72E-21 | 1.53E-19 |
| ENSG00000104324 | CPQ | 96.43055831 | 0.93017872 | 0.187635625 | 4.957367336 | 7.15E-07 | 4.22E-06 |
| ENSG00000104325 | DECR1 | 3124.874095 | 0.361981243 | 0.051420259 | 7.039662017 | 1.93E-12 | 1.93E-11 |
| ENSG00000104327 | CALB1 | 2.355496448 | 2.525135429 | 1.405372681 | 1.796772815 | 0.072371694 | 0.163181311 |
| ENSG00000104331 | BPNT2 | 3024.191387 | 0.196815553 | 0.052533778 | 3.746457238 | 0.00017935 | 0.000767446 |
| ENSG00000104341 | LAPTM4B | 10909.99368 | 0.324067852 | 0.050018765 | 6.478925473 | 9.24E-11 | 8.01E-10 |
| ENSG00000104343 | UBE2W | 500.5849713 | 0.618110045 | 0.096487223 | 6.406133648 | 1.49E-10 | 1.27E-09 |

| | | | | | | | |
|---|---|---|---|---|---|---|---|
| ENSG00000104356 | POP1 | 918.0208862 | -0.284567466 | 0.070211433 | -4.053007498 | 5.06E-05 | 0.000236478 |
| ENSG00000104361 | NIPAL2 | 836.9762883 | 0.021687149 | 0.073320079 | 0.295787312 | 0.767392513 | 0.873024431 |
| ENSG00000104365 | IKBKB | 1242.739778 | 0.263367598 | 0.066580044 | 3.95565371 | 7.63E-05 | 0.000347408 |
| ENSG00000104368 | PLAT | 1.74179757 | 3.033522379 | 1.612242835 | 1.881554263 | 0.059896561 | NA |
| ENSG00000104369 | JPH1 | 1378.174842 | 1.744273177 | 0.074159063 | 23.52070144 | 2.50E-122 | 6.38E-120 |
| ENSG00000104375 | STK3 | 1134.918278 | 0.141040709 | 0.064554263 | 2.184839585 | 0.028900611 | 0.075142242 |
| ENSG00000104381 | GDAP1 | 3321.872386 | 1.466609011 | 0.166722312 | 8.796717081 | 1.41E-18 | 2.16E-17 |
| ENSG00000104388 | RAB2A | 2355.036456 | 0.469961944 | 0.056653875 | 8.295318617 | 1.08E-16 | 1.47E-15 |
| ENSG00000104408 | EIF3E | 9740.448417 | 0.033599941 | 0.05884825 | 0.570959056 | 0.568027397 | 0.735690408 |
| ENSG00000104412 | EMC2 | 1361.524402 | 0.02185491 | 0.063522451 | 0.344050173 | 0.730808554 | 0.850978918 |
| ENSG00000104413 | ESRP1 | 5963.345024 | -0.31269442 | 0.048262006 | -6.479101184 | 9.23E-11 | 8.00E-10 |
| ENSG00000104419 | NDRG1 | 639.6320446 | -0.538609751 | 0.08324465 | -6.470202566 | 9.79E-11 | 8.46E-10 |
| ENSG00000104427 | ZC2HC1A | 290.5103204 | -0.07152635 | 0.111804364 | -0.639745603 | 0.522338002 | 0.69916229 |
| ENSG00000104432 | IL7 | 2.414468552 | -0.32640054 | 1.126394009 | -0.289774748 | 0.771988568 | 0.875706757 |
| ENSG00000104442 | ARMC1 | 1378.981209 | 0.441963116 | 0.065165055 | 6.782210426 | 1.18E-11 | 1.10E-10 |
| ENSG00000104447 | TRPS1 | 6573.338071 | -0.06698864 | 0.051015447 | -1.313105023 | 0.189147562 | 0.347958706 |
| ENSG00000104450 | SPAG1 | 501.696182 | -0.370795369 | 0.08739983 | -4.242518183 | 2.21E-05 | 0.000108969 |
| ENSG00000104472 | CHRAC1 | 1059.676429 | -0.143517051 | 0.071636253 | -2.003413695 | 0.045132903 | 0.10970002 |
| ENSG00000104490 | NCALD | 57.44782939 | -1.237094461 | 0.249599621 | -4.956315454 | 7.18E-07 | 4.25E-06 |
| ENSG00000104497 | SNX16 | 333.2379438 | -0.022776721 | 0.109105697 | -0.20875831 | 0.834636918 | 0.912770412 |
| ENSG00000104517 | UBR5 | 8606.277458 | -0.300760749 | 0.048628744 | -6.184834747 | 6.22E-10 | 5.05E-09 |
| ENSG00000104522 | GFUS | 2371.59654 | 0.037085728 | 0.06043545 | 0.613641965 | 0.539451946 | 0.714049685 |
| ENSG00000104524 | PYCR3 | 866.1050327 | 0.282148481 | 0.083240199 | 3.389570004 | 0.000700023 | 0.0026971 |
| ENSG00000104529 | EEF1D | 11990.24006 | 0.311357585 | 0.043719194 | 7.12175953 | 1.07E-12 | 1.09E-11 |
| ENSG00000104549 | SQLE | 12842.62526 | -1.258252353 | 0.050226022 | -25.05180179 | 1.67E-138 | 5.24E-136 |
| ENSG00000104611 | SH2D4A | 580.039158 | -0.27108966 | 0.08634265 | -3.139695858 | 0.001691233 | 0.006033731 |
| ENSG00000104613 | INTS10 | 1265.5092 | 0.167440713 | 0.067988333 | 2.462785959 | 0.013786219 | 0.039467847 |
| ENSG00000104626 | ERI1 | 460.2920384 | -0.02125431 | 0.089975604 | -0.236223035 | 0.813259606 | 0.901448243 |
| ENSG00000104635 | SLC39A14 | 1413.885245 | 0.075983952 | 0.058991911 | 1.288040198 | 0.19773197 | 0.35999773 |
| ENSG00000104643 | MTMR9 | 425.1061441 | 0.027076837 | 0.096016841 | 0.282000916 | 0.777942808 | 0.879540484 |
| ENSG00000104660 | LEPROTL1 | 818.1980055 | 0.183428233 | 0.073457447 | 2.497067896 | 0.012522498 | 0.036243515 |
| ENSG00000104671 | DCTN6 | 261.0659676 | -0.089069676 | 0.119302494 | -0.746586875 | 0.455312974 | 0.64272227 |
| ENSG00000104679 | R3HCC1 | 695.5901703 | 0.145087866 | 0.079458903 | 1.82594852 | 0.067858017 | 0.154676361 |

| | | | | | | | |
|---|---|---|---|---|---|---|---|
| ENSG00000104687 | GSR | 3709.662604 | -0.292553521 | 0.048713153 | -6.005637208 | 1.91E-09 | 1.48E-08 |
| ENSG00000104689 | TNFRSF10A | 372.2592047 | -0.318483578 | 0.098432735 | -3.235545346 | 0.001214106 | 0.004476308 |
| ENSG00000104691 | UBXN8 | 186.4626546 | 0.39085378 | 0.138359354 | 2.824917638 | 0.004729277 | 0.015325933 |
| ENSG00000104695 | PPP2CB | 1262.77528 | -0.010004527 | 0.063174387 | -0.158363659 | 0.874170248 | 0.934789447 |
| ENSG00000104714 | ERICH1 | 338.3659958 | -0.065986062 | 0.103237218 | -0.639169317 | 0.52271279 | 0.699409032 |
| ENSG00000104723 | TUSC3 | 1874.257104 | -0.056593827 | 0.057904598 | -0.977363274 | 0.328389338 | 0.518747128 |
| ENSG00000104728 | ARHGEF10 | 14.61896022 | 0.841987082 | 0.48912036 | 1.72143127 | 0.085172594 | 0.186046401 |
| ENSG00000104731 | KLHDC4 | 1876.94777 | 0.312045058 | 0.066851531 | 4.667732405 | 3.05E-06 | 1.67E-05 |
| ENSG00000104738 | MCM4 | 6533.644599 | 0.133504841 | 0.046698013 | 2.858897667 | 0.004251159 | 0.013927676 |
| ENSG00000104756 | KCTD9 | 488.2280805 | -0.437082458 | 0.089192084 | -4.900462436 | 9.56E-07 | 5.54E-06 |
| ENSG00000104763 | ASAH1 | 2118.934058 | 0.477556104 | 0.053358592 | 8.94993826 | 3.56E-19 | 5.67E-18 |
| ENSG00000104765 | BNIP3L | 1595.031183 | 0.137692377 | 0.061269038 | 2.247340292 | 0.024618288 | 0.065614444 |
| ENSG00000104774 | MAN2B1 | 836.8633832 | 0.243722862 | 0.077461115 | 3.146389818 | 0.001652995 | 0.005913867 |
| ENSG00000104783 | KCNN4 | 20.70958704 | 1.572712752 | 0.428717316 | 3.668414348 | 0.000244059 | 0.00101832 |
| ENSG00000104805 | NUCB1 | 4376.962312 | 0.133396595 | 0.058074531 | 2.296989609 | 0.021619363 | 0.058518959 |
| ENSG00000104808 | DHDH | 4.633848213 | 0.238208044 | 0.824626615 | 0.288867761 | 0.772682576 | 0.876166615 |
| ENSG00000104812 | GYS1 | 2874.690752 | -0.213748142 | 0.05402346 | -3.956579972 | 7.60E-05 | 0.000346265 |
| ENSG00000104814 | MAP4K1 | 59.71362234 | -0.603245531 | 0.23770765 | -2.537762381 | 0.01115637 | 0.032710524 |
| ENSG00000104823 | ECH1 | 1460.899005 | -0.74717908 | 0.077299155 | -9.666070533 | 4.20E-22 | 7.89E-21 |
| ENSG00000104824 | HNRNPL | 3908.428472 | -0.080487321 | 0.047897186 | -1.680418589 | 0.092875902 | 0.199547924 |
| ENSG00000104825 | NFKBIB | 538.6272415 | -0.136214096 | 0.090372072 | -1.507258749 | 0.131744347 | 0.264133483 |
| ENSG00000104826 | LHB | 5.10043646 | 0.374875946 | 0.783236526 | 0.478624188 | 0.632206008 | 0.783300341 |
| ENSG00000104833 | TUBB4A | 33.05646916 | -0.381815041 | 0.307252702 | -1.242674312 | 0.213987874 | 0.382430805 |
| ENSG00000104835 | SARS2 | 108.8409231 | -0.095631004 | 0.177504946 | -0.538751211 | 0.590058533 | 0.752932545 |
| ENSG00000104852 | SNRNP70 | 6300.480312 | -0.002540748 | 0.048915797 | -0.051941256 | 0.958575501 | 0.981140547 |
| ENSG00000104853 | CLPTM1 | 5107.298466 | 0.145500027 | 0.050029562 | 2.908281049 | 0.003634216 | 0.012087706 |
| ENSG00000104856 | RELB | 78.13904432 | 1.913577467 | 0.226800565 | 8.437269393 | 3.25E-17 | 4.53E-16 |
| ENSG00000104859 | CLASRP | 1235.473558 | -0.023867765 | 0.069590162 | -0.342976138 | 0.731616412 | 0.851381347 |
| ENSG00000104863 | LIN7B | 119.94994 | -0.419734236 | 0.163407822 | -2.568630012 | 0.01021014 | 0.030280623 |
| ENSG00000104866 | PPP1R37 | 2331.557522 | 0.215581145 | 0.061598278 | 3.499791769 | 0.000465622 | 0.00185376 |
| ENSG00000104870 | FCGRT | 15.93984618 | 0.995566225 | 0.459947148 | 2.164523097 | 0.030424226 | 0.078643039 |
| ENSG00000104872 | PIH1D1 | 1697.945571 | -0.243568093 | 0.063297071 | -3.848015217 | 0.000119079 | 0.000524312 |
| ENSG00000104880 | ARHGEF18 | 418.5936081 | -0.022200193 | 0.103127938 | -0.215268462 | 0.829558017 | 0.909783073 |

| | | | | | | | |
|---|---|---|---|---|---|---|---|
| ENSG00000104881 | PPP1R13L | 1330.854135 | -0.012456735 | 0.070798282 | -0.175946856 | 0.860335694 | 0.927382816 |
| ENSG00000104883 | PEX11G | 60.48013065 | -0.653280979 | 0.234642066 | -2.784159679 | 0.005366661 | 0.017153679 |
| ENSG00000104884 | ERCC2 | 1039.661628 | -0.26535866 | 0.065592041 | -4.045592385 | 5.22E-05 | 0.000243395 |
| ENSG00000104885 | DOT1L | 1741.436104 | 0.211320419 | 0.057309377 | 3.687362022 | 0.000226591 | 0.000951491 |
| ENSG00000104886 | PLEKHJ1 | 755.5131558 | -0.114816871 | 0.081531901 | -1.408244747 | 0.15905861 | 0.304498416 |
| ENSG00000104888 | SLC17A7 | 19.74426886 | -0.170497851 | 0.405345207 | -0.420623824 | 0.674029794 | 0.813630998 |
| ENSG00000104889 | RNASEH2A | 2.874913416 | -0.785160415 | 1.061102511 | -0.739947749 | 0.4593317 | 0.646406306 |
| ENSG00000104892 | KLC3 | 98.80767669 | 0.504757023 | 0.186317562 | 2.709122084 | 0.006746151 | 0.021003083 |
| ENSG00000104894 | CD37 | 3.561103092 | 0.902334862 | 0.945023932 | 0.954827524 | 0.339664926 | 0.530711987 |
| ENSG00000104897 | SF3A2 | 1959.835597 | -0.039800784 | 0.056102976 | -0.709423748 | 0.478061555 | 0.662842676 |
| ENSG00000104899 | AMH | 565.0312936 | 0.03809676 | 0.094029441 | 0.405157779 | 0.685361538 | 0.821903585 |
| ENSG00000104901 | DKKL1 | 6.963377425 | 1.010010698 | 0.677885721 | 1.48994243 | 0.136239374 | 0.270896354 |
| ENSG00000104904 | OAZ1 | 6064.16661 | -0.040984741 | 0.044537482 | -0.920230302 | 0.357452423 | 0.549742463 |
| ENSG00000104907 | TRMT1 | 963.1725236 | 0.119219964 | 0.069049379 | 1.726589953 | 0.084241332 | 0.184310553 |
| ENSG00000104915 | STX10 | 699.2196593 | -0.086145313 | 0.078765205 | -1.09369757 | 0.27408764 | 0.456149918 |
| ENSG00000104936 | DMPK | 1046.846707 | 0.280906009 | 0.071129712 | 3.949207751 | 7.84E-05 | 0.000356062 |
| ENSG00000104946 | TBC1D17 | 2310.317244 | -0.087669603 | 0.056282002 | -1.55768452 | 0.119308052 | 0.244390277 |
| ENSG00000104951 | IL4I1 | 5.427330491 | 1.194392077 | 0.797071076 | 1.49847625 | 0.134009559 | 0.267506028 |
| ENSG00000104953 | TLE6 | 33.27051616 | -1.112113841 | 0.316706377 | -3.511498096 | 0.000445589 | 0.001779562 |
| ENSG00000104957 | YJU2B | 406.4734139 | -0.069160163 | 0.102296551 | -0.676075214 | 0.498992893 | 0.680567986 |
| ENSG00000104960 | PTOV1 | 5073.464369 | -0.242020388 | 0.058866267 | -4.111359513 | 3.93E-05 | 0.000186583 |
| ENSG00000104964 | TLE5 | 5772.352686 | -0.015311274 | 0.049068452 | -0.312039074 | 0.755010828 | 0.864705683 |
| ENSG00000104967 | NOVA2 | 2.059454438 | 0.541245474 | 1.229335769 | 0.44027473 | 0.659738141 | 0.803183832 |
| ENSG00000104969 | SGTA | 2200.789796 | 0.010838118 | 0.057895476 | 0.187201468 | 0.851502674 | 0.922260215 |
| ENSG00000104973 | MED25 | 1559.415333 | 0.178325419 | 0.057940293 | 3.07774451 | 0.002085737 | 0.00730151 |
| ENSG00000104976 | SNAPC2 | 476.7230826 | 0.020383058 | 0.093882151 | 0.217113236 | 0.828120099 | 0.909124955 |
| ENSG00000104979 | C19orf53 | 1434.009895 | -0.027315086 | 0.066731348 | -0.409329143 | 0.682298131 | 0.819815584 |
| ENSG00000104980 | TIMM44 | 1128.115642 | 0.103949857 | 0.065880362 | 1.577858025 | 0.11459823 | 0.23666408 |
| ENSG00000104983 | CCDC61 | 184.3507899 | -0.014999918 | 0.134931095 | -0.111167244 | 0.911483726 | 0.955510518 |
| ENSG00000104998 | IL27RA | 424.8397239 | 0.216240086 | 0.101791286 | 2.124347716 | 0.033641078 | 0.085671219 |
| ENSG00000105011 | ASF1B | 1685.545766 | -0.47077836 | 0.060256952 | -7.812847264 | 5.59E-15 | 6.70E-14 |
| ENSG00000105048 | TNNT1 | 343.1196267 | -1.176898735 | 0.115715011 | -10.17066603 | 2.68E-24 | 5.72E-23 |
| ENSG00000105053 | VRK3 | 1056.469511 | -0.053798449 | 0.070672942 | -0.761231211 | 0.446518979 | 0.634997544 |

| | | | | | | | |
|---|---|---|---|---|---|---|---|
| ENSG00000105058 | FAM32A | 1855.520198 | -0.06964118 | 0.058822078 | -1.183929273 | 0.236441061 | 0.411560661 |
| ENSG00000105063 | PPP6R1 | 4798.780617 | 0.081991606 | 0.051060057 | 1.605787605 | 0.108320585 | 0.22605043 |
| ENSG00000105072 | C19orf44 | 92.62895753 | -0.609071477 | 0.187266649 | -3.252428988 | 0.001144232 | 0.004243239 |
| ENSG00000105085 | MED26 | 207.4650228 | 0.268262454 | 0.132498037 | 2.024652296 | 0.04290308 | 0.105171688 |
| ENSG00000105088 | OLFM2 | 271.286105 | 0.726300643 | 0.115801367 | 6.271952224 | 3.57E-10 | 2.96E-09 |
| ENSG00000105127 | AKAP8 | 980.3732136 | -0.033341745 | 0.067906325 | -0.490996162 | 0.623429162 | 0.777266998 |
| ENSG00000105131 | EPHX3 | 4.040569317 | 0.515372652 | 0.94801739 | 0.543632066 | 0.586694689 | 0.750419636 |
| ENSG00000105135 | ILVBL | 1443.561004 | -0.147634281 | 0.077196912 | -1.912437641 | 0.055820084 | 0.131577551 |
| ENSG00000105136 | ZNF419 | 315.8763763 | 0.438752326 | 0.105778946 | 4.147822832 | 3.36E-05 | 0.000161119 |
| ENSG00000105137 | SYDE1 | 71.76359401 | -0.547882386 | 0.211725682 | -2.587699243 | 0.00966193 | 0.028860164 |
| ENSG00000105141 | CASP14 | 3.600292124 | -1.507796841 | 1.004903507 | -1.500439426 | 0.133500613 | 0.266768647 |
| ENSG00000105171 | POP4 | 1313.750074 | -0.153183999 | 0.061533802 | -2.489428476 | 0.012794866 | 0.036910743 |
| ENSG00000105173 | CCNE1 | 402.0055055 | 0.541532035 | 0.099823734 | 5.42488257 | 5.80E-08 | 3.84E-07 |
| ENSG00000105176 | URI1 | 2662.880051 | 0.239579095 | 0.053635052 | 4.466838097 | 7.94E-06 | 4.14E-05 |
| ENSG00000105185 | PDCD5 | 1119.716981 | -0.039530468 | 0.065257061 | -0.605765372 | 0.544670567 | 0.717913747 |
| ENSG00000105186 | ANKRD27 | 2112.435652 | 0.054234859 | 0.055403391 | 0.978908666 | 0.32762511 | 0.51794656 |
| ENSG00000105193 | RPS16 | 9735.1588 | 0.258227334 | 0.050666478 | 5.096611074 | 3.46E-07 | 2.12E-06 |
| ENSG00000105197 | TIMM50 | 1704.438898 | 0.130987843 | 0.061066182 | 2.145014464 | 0.031951691 | 0.082061328 |
| ENSG00000105202 | FBL | 4599.603368 | 0.134422271 | 0.044847376 | 2.99732747 | 0.00272358 | 0.009318258 |
| ENSG00000105204 | DYRK1B | 1334.508671 | -0.039340239 | 0.064469365 | -0.610216016 | 0.541718722 | 0.715677694 |
| ENSG00000105219 | CCNP | 35.09048591 | 0.231848244 | 0.297923554 | 0.778213878 | 0.436442937 | 0.626296602 |
| ENSG00000105220 | GPI | 11504.7128 | 0.141718741 | 0.043416838 | 3.264142366 | 0.00109796 | 0.004080881 |
| ENSG00000105221 | AKT2 | 6078.537912 | 0.120959457 | 0.049336013 | 2.451747743 | 0.014216429 | 0.040532547 |
| ENSG00000105223 | PLD3 | 2291.585408 | -0.090376666 | 0.068318284 | -1.322876697 | 0.185876386 | 0.343151802 |
| ENSG00000105227 | PRX | 252.0215307 | 0.238312064 | 0.117488415 | 2.028387769 | 0.042520687 | 0.104379301 |
| ENSG00000105229 | PIAS4 | 946.6180935 | 0.017489613 | 0.080056574 | 0.218465671 | 0.827066303 | 0.90866384 |
| ENSG00000105245 | NUMBL | 596.4850921 | 0.354339911 | 0.085841902 | 4.127819895 | 3.66E-05 | 0.000174477 |
| ENSG00000105248 | YJU2 | 720.8740832 | -0.107392051 | 0.077068091 | -1.393469716 | 0.163477797 | 0.311117928 |
| ENSG00000105254 | TBCB | 2384.89882 | -0.099753259 | 0.055142711 | -1.809001712 | 0.070450737 | 0.159445532 |
| ENSG00000105255 | FSD1 | 219.6114279 | -0.133425989 | 0.123690455 | -1.07870885 | 0.280717539 | 0.464119665 |
| ENSG00000105258 | POLR2I | 1029.604815 | 0.155110027 | 0.071227968 | 2.17765622 | 0.029431641 | 0.076367135 |
| ENSG00000105270 | CLIP3 | 217.2461607 | -0.091731282 | 0.131559298 | -0.697261867 | 0.485638928 | 0.669343728 |
| ENSG00000105281 | SLC1A5 | 11129.77474 | -0.127764892 | 0.046670574 | -2.737589878 | 0.006189121 | 0.019486471 |

| | | | | | | | |
|---|---|---|---|---|---|---|---|
| ENSG00000105287 | PRKD2 | 1435.383246 | -0.019909125 | 0.067457584 | -0.295135457 | 0.767890405 | 0.873216967 |
| ENSG00000105289 | TJP3 | 1210.379276 | 0.469905409 | 0.076938988 | 6.107507032 | 1.01E-09 | 8.04E-09 |
| ENSG00000105290 | APLP1 | 241.5934133 | 0.779676725 | 0.126231353 | 6.176569497 | 6.55E-10 | 5.30E-09 |
| ENSG00000105298 | CACTIN | 821.6379811 | 0.010815121 | 0.072570771 | 0.149028611 | 0.881531056 | 0.938595484 |
| ENSG00000105321 | CCDC9 | 721.9662174 | -0.126745996 | 0.085663598 | -1.479578241 | 0.138985837 | 0.274951285 |
| ENSG00000105323 | HNRNPUL1 | 14084.49654 | -0.075507717 | 0.038837632 | -1.944189509 | 0.051872602 | 0.123606264 |
| ENSG00000105325 | FZR1 | 1547.236414 | -0.012430643 | 0.065237965 | -0.190543077 | 0.848883591 | 0.920956268 |
| ENSG00000105327 | BBC3 | 419.8130075 | 0.51230433 | 0.101331228 | 5.055739878 | 4.29E-07 | 2.61E-06 |
| ENSG00000105329 | TGFB1 | 1803.737696 | 0.34691232 | 0.067709128 | 5.123567967 | 3.00E-07 | 1.85E-06 |
| ENSG00000105339 | DENND3 | 341.5960495 | -0.59218913 | 0.10364135 | -5.713830729 | 1.10E-08 | 7.91E-08 |
| ENSG00000105341 | DMAC2 | 1460.859064 | 0.18262988 | 0.06030392 | 3.028491026 | 0.002457784 | 0.008487364 |
| ENSG00000105355 | PLIN3 | 1676.284538 | -0.213545457 | 0.073456726 | -2.907091961 | 0.003648059 | 0.012126718 |
| ENSG00000105357 | MYH14 | 5055.246778 | 0.774415961 | 0.061615179 | 12.56859063 | 3.14E-36 | 1.12E-34 |
| ENSG00000105364 | MRPL4 | 2237.403381 | 0.103457751 | 0.064668364 | 1.599820151 | 0.109638487 | 0.228189702 |
| ENSG00000105369 | CD79A | 6.035700498 | 0.359580719 | 0.702928246 | 0.511546835 | 0.608968202 | 0.767188235 |
| ENSG00000105372 | RPS19 | 9976.786831 | 0.199456882 | 0.042787241 | 4.66159716 | 3.14E-06 | 1.72E-05 |
| ENSG00000105373 | NOP53 | 4587.38284 | 0.258462429 | 0.050537913 | 5.114228408 | 3.15E-07 | 1.94E-06 |
| ENSG00000105376 | ICAM5 | 305.1030763 | 0.253878229 | 0.107572869 | 2.360058178 | 0.018272069 | 0.05052519 |
| ENSG00000105379 | ETFB | 386.7701888 | -0.134485418 | 0.100786585 | -1.334358309 | 0.182086456 | 0.337922766 |
| ENSG00000105383 | CD33 | 1.913451737 | 3.153071073 | 1.571098991 | 2.006920691 | 0.044758108 | 0.108927541 |
| ENSG00000105388 | CEACAM5 | 86.87161591 | 0.896083142 | 0.199486191 | 4.491955743 | 7.06E-06 | 3.71E-05 |
| ENSG00000105393 | BABAM1 | 409.4205918 | -0.182157093 | 0.096970321 | -1.878482929 | 0.06031513 | 0.140137279 |
| ENSG00000105397 | TYK2 | 3580.590726 | -0.06928489 | 0.053574669 | -1.293239713 | 0.195928164 | 0.357394409 |
| ENSG00000105401 | CDC37 | 5527.387341 | -0.07948076 | 0.048610081 | -1.635067406 | 0.102034918 | 0.215492636 |
| ENSG00000105402 | NAPA | 2109.890276 | 0.065623084 | 0.058740355 | 1.117172082 | 0.263920755 | 0.444524801 |
| ENSG00000105404 | RABAC1 | 62.42051839 | -0.278508536 | 0.228528556 | -1.218703432 | 0.222956775 | 0.394130262 |
| ENSG00000105409 | ATP1A3 | 6.14500988 | 1.451076325 | 0.774216365 | 1.874251682 | 0.060895744 | 0.14127714 |
| ENSG00000105419 | MEIS3 | 857.8022151 | -0.501938455 | 0.072940099 | -6.881515955 | 5.92E-12 | 5.66E-11 |
| ENSG00000105426 | PTPRS | 3087.781459 | 0.594620068 | 0.061467185 | 9.673780729 | 3.90E-22 | 7.35E-21 |
| ENSG00000105427 | CNFN | 43.30963784 | 0.5013943 | 0.273838098 | 1.830988105 | 0.067102319 | 0.153319441 |
| ENSG00000105429 | MEGF8 | 1441.353463 | -0.162537326 | 0.071188791 | -2.283187058 | 0.022419352 | 0.060413587 |
| ENSG00000105438 | KDELR1 | 6711.489541 | -0.049505462 | 0.045626066 | -1.085025885 | 0.277910191 | 0.460673443 |
| ENSG00000105443 | CYTH2 | 3457.193093 | 0.051226482 | 0.058310167 | 0.878517148 | 0.379663136 | 0.572075049 |

| | | | | | | | |
|---|---|---|---|---|---|---|---|
| ENSG00000105447 | GRWD1 | 1959.473506 | 0.129068034 | 0.061705627 | 2.091673634 | 0.036467721 | 0.091839885 |
| ENSG00000105464 | GRIN2D | 28.88957872 | -0.782082094 | 0.339575913 | -2.30311416 | 0.021272421 | 0.057635455 |
| ENSG00000105467 | SYNGR4 | 2.627704585 | -0.885980672 | 1.095212976 | -0.808957427 | 0.418539635 | 0.61039531 |
| ENSG00000105472 | CLEC11A | 94.84651038 | 0.459566832 | 0.187505568 | 2.450950325 | 0.014247961 | 0.04060731 |
| ENSG00000105479 | ODAD1 | 173.9535343 | -0.008106094 | 0.146965794 | -0.055156329 | 0.95601392 | 0.979848062 |
| ENSG00000105483 | CARD8 | 428.1560711 | -0.940116133 | 0.096347138 | -9.757592766 | 1.71E-22 | 3.30E-21 |
| ENSG00000105486 | LIG1 | 2724.862874 | -0.043384389 | 0.053753266 | -0.807102384 | 0.419607499 | 0.611006476 |
| ENSG00000105499 | PLA2G4C | 43.78996424 | 0.216415186 | 0.2650752 | 0.8164294 | 0.414254587 | 0.606477249 |
| ENSG00000105514 | RAB3D | 3908.604548 | 0.123036127 | 0.051101028 | 2.407703581 | 0.016053209 | 0.045157648 |
| ENSG00000105516 | DBP | 458.5111485 | 0.19889828 | 0.097180751 | 2.046683921 | 0.040689129 | 0.10055032 |
| ENSG00000105518 | TMEM205 | 1662.825465 | 0.021285165 | 0.065202912 | 0.326445004 | 0.744087693 | 0.858490754 |
| ENSG00000105519 | CAPS | 121.9148498 | -0.152428293 | 0.166492894 | -0.915524318 | 0.359916442 | 0.55249486 |
| ENSG00000105520 | PLPPR2 | 2215.985146 | 0.593415958 | 0.063507396 | 9.344044903 | 9.27E-21 | 1.62E-19 |
| ENSG00000105523 | FAM83E | 6.87769968 | 0.091976862 | 0.654205807 | 0.140593161 | 0.888191351 | 0.942392571 |
| ENSG00000105538 | RASIP1 | 33.43798584 | -1.412372376 | 0.337999112 | -4.178627469 | 2.93E-05 | 0.000142265 |
| ENSG00000105549 | THEG | 79.24055445 | 0.026913078 | 0.196830745 | 0.136732083 | 0.89124257 | 0.944238 |
| ENSG00000105550 | FGF21 | 1.687431957 | -0.231660533 | 1.321653312 | -0.175280863 | 0.860858948 | NA |
| ENSG00000105552 | BCAT2 | 1704.246324 | 0.042433846 | 0.059139203 | 0.717524828 | 0.473050322 | 0.658601347 |
| ENSG00000105556 | MIER2 | 587.3050867 | 0.189204966 | 0.093502039 | 2.023538388 | 0.04301767 | 0.105340024 |
| ENSG00000105559 | PLEKHA4 | 71.88155184 | 0.519023971 | 0.224321504 | 2.313750403 | 0.020681417 | 0.056285646 |
| ENSG00000105568 | PPP2R1A | 10966.55163 | 0.146542914 | 0.049664879 | 2.950634697 | 0.003171217 | 0.010685624 |
| ENSG00000105576 | TNPO2 | 3472.309155 | -0.130647201 | 0.05117843 | -2.552778606 | 0.01068674 | 0.031490536 |
| ENSG00000105607 | GCDH | 488.855689 | 0.180993758 | 0.091584098 | 1.976257476 | 0.048125623 | 0.115966575 |
| ENSG00000105610 | KLF1 | 3.393647887 | -0.035450607 | 0.944244873 | -0.03754387 | 0.970051362 | 0.986563569 |
| ENSG00000105612 | DNASE2 | 1548.171802 | 0.312845706 | 0.061423373 | 5.093268065 | 3.52E-07 | 2.15E-06 |
| ENSG00000105613 | MAST1 | 86.52363486 | -0.208002601 | 0.203705532 | -1.021094512 | 0.307209663 | 0.495040016 |
| ENSG00000105617 | LENG1 | 236.9689147 | -0.304472345 | 0.121957468 | -2.496545306 | 0.012540965 | 0.036278655 |
| ENSG00000105618 | PRPF31 | 2605.460928 | -0.003965161 | 0.051945833 | -0.076332613 | 0.93915448 | 0.971039083 |
| ENSG00000105619 | TFPT | 1077.670984 | -0.267673923 | 0.075346757 | -3.552560639 | 0.000381501 | 0.001538884 |
| ENSG00000105639 | JAK3 | 88.44908054 | 0.937517379 | 0.198079859 | 4.733027292 | 2.21E-06 | 1.23E-05 |
| ENSG00000105640 | RPL18A | 1380.547371 | 0.371652877 | 0.064996945 | 5.718005342 | 1.08E-08 | 7.73E-08 |
| ENSG00000105641 | SLC5A5 | 1.658117373 | -0.212182739 | 1.394067509 | -0.152204063 | 0.87902599 | NA |
| ENSG00000105642 | KCNN1 | 8.053346718 | -0.935043881 | 0.629813204 | -1.484636833 | 0.137640053 | 0.273136922 |

| | | | | | | | |
|---|---|---|---|---|---|---|---|
| ENSG00000105643 | ARRDC2 | 495.6168724 | 0.176715017 | 0.087802166 | 2.012649856 | 0.044151478 | 0.107748394 |
| ENSG00000105647 | PIK3R2 | 3.508188345 | -1.118250712 | 0.96307252 | -1.161128252 | 0.245589747 | 0.422701956 |
| ENSG00000105649 | RAB3A | 118.0578066 | 0.826576678 | 0.167458818 | 4.935999707 | 7.97E-07 | 4.68E-06 |
| ENSG00000105655 | ISYNA1 | 1742.240336 | 0.672367529 | 0.068652978 | 9.793712448 | 1.20E-22 | 2.33E-21 |
| ENSG00000105656 | ELL | 503.8999031 | 0.15774181 | 0.093510959 | 1.686880469 | 0.091626338 | 0.197325216 |
| ENSG00000105662 | CRTC1 | 755.8624679 | 0.574008243 | 0.074311968 | 7.72430416 | 1.12E-14 | 1.31E-13 |
| ENSG00000105668 | UPK1A | 638.0205686 | 1.305834816 | 0.107678302 | 12.12718618 | 7.58E-34 | 2.46E-32 |
| ENSG00000105669 | COPE | 755.326832 | 0.092026002 | 0.076719478 | 1.199512874 | 0.230328582 | 0.403677719 |
| ENSG00000105671 | DDX49 | 751.0464417 | 0.212059438 | 0.079982671 | 2.65131727 | 0.008017849 | 0.024497413 |
| ENSG00000105672 | ETV2 | 25.49988316 | -0.025827872 | 0.345507746 | -0.074753381 | 0.940410935 | 0.971621655 |
| ENSG00000105676 | ARMC6 | 430.1918935 | 0.215762179 | 0.093539506 | 2.306642265 | 0.021074771 | 0.057206869 |
| ENSG00000105677 | TMEM147 | 1765.586393 | 0.353272381 | 0.062123381 | 5.686625111 | 1.30E-08 | 9.19E-08 |
| ENSG00000105679 | GAPDHS | 4.821618247 | 0.086495651 | 0.807628477 | 0.107098318 | 0.914710981 | 0.957703529 |
| ENSG00000105695 | MAG | 1.711515057 | -1.419327597 | 1.372170185 | -1.034367028 | 0.300964612 | NA |
| ENSG00000105696 | TMEM59L | 16.26588792 | -0.268830494 | 0.433758439 | -0.619770061 | 0.535409182 | 0.710903898 |
| ENSG00000105698 | USF2 | 4132.384283 | 0.343197765 | 0.049976268 | 6.867214719 | 6.55E-12 | 6.24E-11 |
| ENSG00000105699 | LSR | 4143.52655 | 0.412296804 | 0.065480646 | 6.296468176 | 3.05E-10 | 2.54E-09 |
| ENSG00000105700 | KXD1 | 1506.962722 | -0.048668672 | 0.065029232 | -0.748412217 | 0.454211556 | 0.641838792 |
| ENSG00000105701 | FKBP8 | 5207.278116 | -0.135033566 | 0.06522664 | -2.070221109 | 0.038431643 | 0.095964069 |
| ENSG00000105705 | SUGP1 | 535.9555498 | -0.075972273 | 0.091203854 | -0.832994105 | 0.404848054 | 0.597786712 |
| ENSG00000105707 | HPN | 5.967095712 | -0.347681472 | 0.715274554 | -0.486081142 | 0.626909637 | 0.780060055 |
| ENSG00000105708 | ZNF14 | 152.62582 | 0.810612913 | 0.149049292 | 5.438555934 | 5.37E-08 | 3.58E-07 |
| ENSG00000105711 | SCN1B | 295.4282703 | 1.523377644 | 0.121250439 | 12.56389381 | 3.34E-36 | 1.18E-34 |
| ENSG00000105717 | PBX4 | 13.13628804 | 0.424080346 | 0.492262734 | 0.861491877 | 0.388967193 | 0.581600014 |
| ENSG00000105722 | ERF | 2271.322394 | 0.104615788 | 0.052555669 | 1.990570945 | 0.046528077 | 0.112625735 |
| ENSG00000105723 | GSK3A | 1714.79751 | 0.021609938 | 0.05941853 | 0.36369021 | 0.716089355 | 0.842069397 |
| ENSG00000105726 | ATP13A1 | 1200.306823 | 0.340480962 | 0.067881042 | 5.015847611 | 5.28E-07 | 3.17E-06 |
| ENSG00000105732 | ZNF574 | 894.8677229 | 0.00825047 | 0.081619435 | 0.101084619 | 0.919483288 | 0.960822968 |
| ENSG00000105737 | GRIK5 | 9.378356081 | -0.418753226 | 0.582938531 | -0.718348855 | 0.472542214 | 0.658173551 |
| ENSG00000105738 | SIPA1L3 | 1653.754914 | -0.136862825 | 0.072191449 | -1.89583154 | 0.057982324 | 0.135656758 |
| ENSG00000105750 | ZNF85 | 101.0089934 | 0.175477467 | 0.182807395 | 0.959903543 | 0.337103763 | 0.528329598 |
| ENSG00000105755 | ETHE1 | 396.4381444 | 0.020599767 | 0.096792288 | 0.212824466 | 0.83146388 | 0.91071463 |
| ENSG00000105767 | CADM4 | 140.1130694 | 0.419482409 | 0.159096483 | 2.636654194 | 0.008372814 | 0.025416481 |

| | | | | | | | |
|---|---|---|---|---|---|---|---|
| ENSG00000105771 | SMG9 | 1571.635684 | -0.360216464 | 0.061790711 | -5.829621615 | 5.56E-09 | 4.12E-08 |
| ENSG00000105778 | AVL9 | 5967.359278 | 0.041057998 | 0.044422612 | 0.92425898 | 0.355351494 | 0.547061491 |
| ENSG00000105784 | RUNDC3B | 60.45627185 | 0.608724237 | 0.22791351 | 2.670856319 | 0.007565802 | 0.023255997 |
| ENSG00000105792 | CFAP69 | 22.30233628 | 0.994072882 | 0.394341551 | 2.520842349 | 0.011707429 | 0.034178377 |
| ENSG00000105793 | GTPBP10 | 467.9541728 | -0.35587583 | 0.091057599 | -3.908249648 | 9.30E-05 | 0.000416537 |
| ENSG00000105808 | RASA4 | 32.47465853 | 0.061151054 | 0.31063605 | 0.196857557 | 0.843939008 | 0.917708773 |
| ENSG00000105810 | CDK6 | 722.7083825 | -1.449760553 | 0.08292574 | -17.4826363 | 1.94E-68 | 1.69E-66 |
| ENSG00000105819 | PMPCB | 2221.703472 | -0.392334983 | 0.057339881 | -6.842270679 | 7.79E-12 | 7.37E-11 |
| ENSG00000105821 | DNAJC2 | 1230.200602 | -0.359475508 | 0.077227663 | -4.654750563 | 3.24E-06 | 1.77E-05 |
| ENSG00000105825 | TFPI2 | 17.79739233 | -1.996550086 | 0.468563822 | -4.260999229 | 2.04E-05 | 0.000100965 |
| ENSG00000105829 | BET1 | 263.1712419 | -0.449228321 | 0.113727945 | -3.95002583 | 7.81E-05 | 0.00035511 |
| ENSG00000105835 | NAMPT | 1023.812638 | 0.052339358 | 0.070376852 | 0.743701331 | 0.457057184 | 0.644312162 |
| ENSG00000105849 | POLR1F | 1089.504641 | -1.706834169 | 0.079456657 | -21.48132369 | 2.33E-102 | 3.90E-100 |
| ENSG00000105854 | PON2 | 352.4130536 | -0.167406199 | 0.108272074 | -1.5461623 | 0.122065376 | 0.248772186 |
| ENSG00000105855 | ITGB8 | 127.8973548 | -0.69637198 | 0.162172769 | -4.294013006 | 1.75E-05 | 8.77E-05 |
| ENSG00000105856 | HBP1 | 1742.8638 | 0.351331346 | 0.058506693 | 6.004977039 | 1.91E-09 | 1.49E-08 |
| ENSG00000105865 | DUS4L | 99.2950557 | -0.148155323 | 0.193016503 | -0.767578523 | 0.442737626 | 0.631641123 |
| ENSG00000105866 | SP4 | 326.8385811 | -1.703796721 | 0.110656797 | -15.39712662 | 1.71E-53 | 1.03E-51 |
| ENSG00000105875 | WDR91 | 475.1219366 | 0.005365579 | 0.090597991 | 0.059224042 | 0.95277366 | 0.978056463 |
| ENSG00000105877 | DNAH11 | 23.0692467 | -2.046581663 | 0.417883467 | -4.897493735 | 9.71E-07 | 5.62E-06 |
| ENSG00000105879 | CBLL1 | 1026.639235 | -0.0005696 | 0.069388155 | -0.008208888 | 0.993450328 | 0.996403923 |
| ENSG00000105880 | DLX5 | 15.34266054 | -1.807671742 | 0.506849203 | -3.566488281 | 0.000361797 | 0.001464812 |
| ENSG00000105887 | MTPN | 5641.092697 | -0.073741585 | 0.050478651 | -1.460846974 | 0.144057441 | 0.282571843 |
| ENSG00000105894 | PTN | 1.741839556 | -0.264987793 | 1.395008185 | -0.189954292 | 0.849344948 | NA |
| ENSG00000105926 | PALS2 | 992.141685 | -1.585367782 | 0.071902496 | -22.04885599 | 9.80E-108 | 1.87E-105 |
| ENSG00000105928 | GSDME | 2.20981808 | -1.908665661 | 1.295978777 | -1.472759967 | 0.14081577 | 0.277804762 |
| ENSG00000105929 | ATP6V0A4 | 60.72315178 | 1.750555036 | 0.264966449 | 6.60670451 | 3.93E-11 | 3.50E-10 |
| ENSG00000105939 | ZC3HAV1 | 2043.784925 | -0.223320063 | 0.055885749 | -3.996010887 | 6.44E-05 | 0.000296264 |
| ENSG00000105948 | TTC26 | 277.8367168 | -0.204197955 | 0.113844785 | -1.793652246 | 0.072868692 | 0.164076148 |
| ENSG00000105953 | OGDH | 7167.381401 | 0.716365238 | 0.054463345 | 13.15316264 | 1.63E-39 | 6.45E-38 |
| ENSG00000105963 | ADAP1 | 153.6719251 | 0.394693932 | 0.172058918 | 2.293946381 | 0.021793579 | 0.058921002 |
| ENSG00000105967 | TFEC | 1.785769997 | -1.470465202 | 1.425622032 | -1.031455161 | 0.302327425 | NA |
| ENSG00000105968 | H2AZ2 | 7634.885123 | 0.712825361 | 0.045516133 | 15.66093837 | 2.80E-55 | 1.78E-53 |

| | | | | | | | |
|---|---|---|---|---|---|---|---|
| ENSG00000105971 | CAV2 | 149.9495176 | -0.102147985 | 0.153838495 | -0.66399496 | 0.506693537 | 0.686971025 |
| ENSG00000105974 | CAV1 | 495.9252464 | -0.957990386 | 0.104771066 | -9.143654086 | 6.04E-20 | 1.00E-18 |
| ENSG00000105976 | MET | 1562.932109 | -0.079867606 | 0.058328542 | -1.369271429 | 0.170914445 | 0.322023475 |
| ENSG00000105982 | RNF32 | 54.59527653 | 0.169869746 | 0.239595977 | 0.70898413 | 0.478334325 | 0.66295052 |
| ENSG00000105983 | LMBR1 | 2171.873259 | 0.775211582 | 0.055225 | 14.03733055 | 9.21E-45 | 4.38E-43 |
| ENSG00000105988 | NHP2P1 | 461.4259392 | -0.265870877 | 0.088788534 | -2.994428049 | 0.002749599 | 0.009400268 |
| ENSG00000105993 | DNAJB6 | 1787.026572 | 0.396359909 | 0.058509754 | 6.774253585 | 1.25E-11 | 1.16E-10 |
| ENSG00000105997 | HOXA3 | 3.360613275 | -0.763366187 | 0.962273089 | -0.793294747 | 0.427606117 | 0.618297152 |
| ENSG00000106003 | LFNG | 1071.348439 | 2.722858478 | 0.086370173 | 31.52544887 | 3.89E-218 | 3.19E-215 |
| ENSG00000106004 | HOXA5 | 21.33666535 | -1.706938196 | 0.41904416 | -4.073408863 | 4.63E-05 | 0.000217655 |
| ENSG00000106006 | HOXA6 | 11.20137376 | -1.936778792 | 0.571126312 | -3.391156654 | 0.000695983 | 0.00268469 |
| ENSG00000106009 | BRAT1 | 2604.480249 | 0.32881385 | 0.058558423 | 5.61514184 | 1.96E-08 | 1.37E-07 |
| ENSG00000106012 | IQCE | 3915.645368 | 0.59441925 | 0.05105818 | 11.6419985 | 2.52E-31 | 7.41E-30 |
| ENSG00000106018 | VIPR2 | 4.17151521 | 1.241715919 | 0.912020308 | 1.361500296 | 0.17335564 | 0.325725152 |
| ENSG00000106025 | TSPAN12 | 161.5824178 | 0.730427839 | 0.152408912 | 4.792553322 | 1.65E-06 | 9.28E-06 |
| ENSG00000106028 | SSBP1 | 1977.904093 | 0.002338988 | 0.053741622 | 0.043522836 | 0.965284761 | 0.984336434 |
| ENSG00000106031 | HOXA13 | 15.37602448 | -1.989346884 | 0.493565093 | -4.030566403 | 5.56E-05 | 0.000258546 |
| ENSG00000106034 | CPED1 | 3.49817725 | 0.660889763 | 0.981234571 | 0.67352882 | 0.50061092 | 0.681732072 |
| ENSG00000106038 | EVX1 | 2.124584745 | -3.598892829 | 1.566999719 | -2.296677392 | 0.02163718 | 0.058553369 |
| ENSG00000106049 | HIBADH | 666.4735733 | -1.246295553 | 0.079847569 | -15.60843448 | 6.38E-55 | 3.99E-53 |
| ENSG00000106052 | TAX1BP1 | 2007.188238 | -1.273987213 | 0.056023449 | -22.74024956 | 1.79E-114 | 3.77E-112 |
| ENSG00000106070 | GRB10 | 1258.692132 | 1.5161514 | 0.065080047 | 23.29671631 | 4.79E-120 | 1.17E-117 |
| ENSG00000106077 | ABHD11 | 1132.590269 | 1.032413882 | 0.070139002 | 14.71954053 | 4.83E-49 | 2.59E-47 |
| ENSG00000106078 | COBL | 1284.314776 | 1.373335884 | 0.069045734 | 19.89023513 | 4.94E-88 | 6.67E-86 |
| ENSG00000106080 | FKBP14 | 61.73332588 | -1.089859788 | 0.232512168 | -4.687323672 | 2.77E-06 | 1.52E-05 |
| ENSG00000106086 | PLEKHA8 | 387.0867088 | -1.57351916 | 0.10303805 | -15.27124358 | 1.19E-52 | 6.99E-51 |
| ENSG00000106089 | STX1A | 252.8673363 | 0.586534973 | 0.119393582 | 4.912617267 | 8.99E-07 | 5.24E-06 |
| ENSG00000106100 | NOD1 | 113.1578222 | -1.618791687 | 0.179711887 | -9.007705115 | 2.10E-19 | 3.40E-18 |
| ENSG00000106105 | GARS1 | 3406.910238 | -1.394420623 | 0.063919716 | -21.81518803 | 1.66E-105 | 3.11E-103 |
| ENSG00000106123 | EPHB6 | 14.45420592 | 1.058650867 | 0.484023094 | 2.18719082 | 0.028728597 | 0.074785642 |
| ENSG00000106125 | MINDY4 | 2.033415775 | -1.759414035 | 1.374109219 | -1.280403341 | 0.20040332 | 0.363598513 |
| ENSG00000106133 | NSUN5P2 | 375.5834542 | 0.611027637 | 0.10281143 | 5.943187823 | 2.80E-09 | 2.13E-08 |
| ENSG00000106144 | CASP2 | 2175.496641 | -0.124175146 | 0.056318809 | -2.204861012 | 0.027463849 | 0.072010595 |

| | | | | | | | |
|---|---|---|---|---|---|---|---|
| ENSG00000106153 | CHCHD2 | 3839.459558 | 0.520165071 | 0.049275932 | 10.5561691 | 4.76E-26 | 1.11E-24 |
| ENSG00000106211 | HSPB1 | 14477.47708 | 0.756240292 | 0.046453472 | 16.27952138 | 1.38E-59 | 9.74E-58 |
| ENSG00000106244 | PDAP1 | 4136.170982 | -0.46505513 | 0.046812492 | -9.934423791 | 2.95E-23 | 5.95E-22 |
| ENSG00000106245 | BUD31 | 2215.535558 | -0.3427746 | 0.055045567 | -6.227106397 | 4.75E-10 | 3.90E-09 |
| ENSG00000106246 | PTCD1 | 209.0569086 | -0.293303284 | 0.135372095 | -2.166645079 | 0.030261927 | 0.078282433 |
| ENSG00000106258 | CYP3A5 | 2.93416349 | 0.848554585 | 1.058042533 | 0.802004228 | 0.422550512 | 0.613489389 |
| ENSG00000106261 | ZKSCAN1 | 4676.567036 | -0.299032369 | 0.048759452 | -6.1328083 | 8.63E-10 | 6.90E-09 |
| ENSG00000106263 | EIF3B | 15238.09302 | 0.323258818 | 0.042339419 | 7.634937419 | 2.26E-14 | 2.59E-13 |
| ENSG00000106266 | SNX8 | 1483.092346 | 0.442338032 | 0.062222853 | 7.108932025 | 1.17E-12 | 1.19E-11 |
| ENSG00000106268 | NUDT1 | 441.0804444 | -0.132277381 | 0.094789706 | -1.395482556 | 0.162870371 | 0.3102164 |
| ENSG00000106290 | TAF6 | 2804.068355 | -0.737028052 | 0.052988567 | -13.90919007 | 5.57E-44 | 2.60E-42 |
| ENSG00000106299 | WASL | 2396.309957 | 0.958805538 | 0.055362191 | 17.3187788 | 3.39E-67 | 2.85E-65 |
| ENSG00000106302 | HYAL4 | 2.92331711 | 3.853704551 | 1.447703908 | 2.661942493 | 0.007769115 | 0.023829857 |
| ENSG00000106304 | SPAM1 | 18.73278526 | 2.588313128 | 0.503357573 | 5.142096323 | 2.72E-07 | 1.69E-06 |
| ENSG00000106305 | AIMP2 | 1320.133856 | 0.243220645 | 0.065598999 | 3.70768838 | 0.00020916 | 0.000883958 |
| ENSG00000106328 | FSCN3 | 3.929315294 | 0.946220599 | 0.937551761 | 1.00924625 | 0.312856551 | 0.501042256 |
| ENSG00000106330 | MOSPD3 | 618.5375133 | -0.481318044 | 0.087519405 | -5.49955802 | 3.81E-08 | 2.58E-07 |
| ENSG00000106333 | PCOLCE | 269.0747475 | -0.647799697 | 0.120588284 | -5.371995316 | 7.79E-08 | 5.09E-07 |
| ENSG00000106336 | FBXO24 | 23.78332446 | -0.822647015 | 0.384622605 | -2.138842086 | 0.03244846 | 0.08316028 |
| ENSG00000106344 | RBM28 | 1518.027016 | 0.067010218 | 0.059583659 | 1.124640874 | 0.260741246 | 0.441084878 |
| ENSG00000106346 | USP42 | 1120.339292 | 0.32402444 | 0.063432618 | 5.108167545 | 3.25E-07 | 2.00E-06 |
| ENSG00000106348 | IMPDH1 | 3830.18078 | -0.123752265 | 0.051153374 | -2.419239557 | 0.015552994 | 0.043928303 |
| ENSG00000106351 | AGFG2 | 518.4217667 | -0.52156323 | 0.085280849 | -6.11583065 | 9.61E-10 | 7.65E-09 |
| ENSG00000106355 | LSM5 | 1790.853163 | -0.054586119 | 0.067013714 | -0.814551471 | 0.415329099 | 0.607398286 |
| ENSG00000106366 | SERPINE1 | 27.81769825 | 0.076520524 | 0.333666241 | 0.229332532 | 0.81861047 | 0.903986346 |
| ENSG00000106367 | AP1S1 | 3136.412235 | -0.241659191 | 0.05372236 | -4.49829808 | 6.85E-06 | 3.60E-05 |
| ENSG00000106392 | C1GALT1 | 176.1195641 | 0.264462938 | 0.138133725 | 1.914542877 | 0.055550826 | 0.131077551 |
| ENSG00000106397 | PLOD3 | 2350.771859 | -0.345660391 | 0.063964149 | -5.40397077 | 6.52E-08 | 4.29E-07 |
| ENSG00000106399 | RPA3 | 1288.8178 | -1.230355208 | 0.065529441 | -18.77560956 | 1.20E-78 | 1.34E-76 |
| ENSG00000106400 | ZNHIT1 | 1501.677562 | -0.218962799 | 0.06499878 | -3.368721673 | 0.000755176 | 0.002892111 |
| ENSG00000106404 | CLDN15 | 266.1428375 | -0.591301995 | 0.114443848 | -5.166743364 | 2.38E-07 | 1.49E-06 |
| ENSG00000106415 | GLCCI1 | 981.0148094 | -0.9016875 | 0.06889031 | -13.08874211 | 3.82E-39 | 1.50E-37 |
| ENSG00000106443 | PHF14 | 4232.111515 | -1.688022197 | 0.053924881 | -31.30321628 | 4.22E-215 | 3.34E-212 |

| | | | | | | | |
|---|---|---|---|---|---|---|---|
| ENSG00000106459 | NRF1 | 696.752751 | -0.184001425 | 0.080713649 | -2.279681668 | 0.022626575 | 0.0608932 |
| ENSG00000106460 | TMEM106B | 2395.551517 | -1.275124527 | 0.059635658 | -21.3819143 | 1.97E-101 | 3.25E-99 |
| ENSG00000106462 | EZH2 | 1787.659518 | 0.49560831 | 0.072705696 | 6.816636621 | 9.32E-12 | 8.74E-11 |
| ENSG00000106477 | CEP41 | 340.9965003 | -0.009707712 | 0.101596046 | -0.095552067 | 0.923876336 | 0.963279648 |
| ENSG00000106479 | ZNF862 | 960.5918283 | 0.414894996 | 0.077441855 | 5.357503305 | 8.44E-08 | 5.50E-07 |
| ENSG00000106484 | MEST | 2193.121043 | -0.194365213 | 0.05165968 | -3.762416137 | 0.00016828 | 0.000723858 |
| ENSG00000106524 | ANKMY2 | 662.0703928 | -2.060389195 | 0.084915837 | -24.26389785 | 4.72E-130 | 1.35E-127 |
| ENSG00000106526 | ACTR3C | 101.0730537 | 0.593796224 | 0.180687303 | 3.286319598 | 0.001015058 | 0.003797362 |
| ENSG00000106537 | TSPAN13 | 5116.817069 | -1.614544773 | 0.048956711 | -32.97902871 | 1.62E-238 | 1.77E-235 |
| ENSG00000106541 | AGR2 | 3629.099495 | -2.418775555 | 0.0509493 | -47.47416676 | 0 | 0 |
| ENSG00000106546 | AHR | 253.3229872 | -0.502973583 | 0.120490056 | -4.174399103 | 2.99E-05 | 0.00014478 |
| ENSG00000106554 | CHCHD3 | 2268.949053 | -0.087723978 | 0.052133556 | -1.682677808 | 0.09243748 | 0.198792143 |
| ENSG00000106565 | TMEM176B | 3.246167154 | 2.448992546 | 1.183697219 | 2.06893495 | 0.038552191 | 0.096212714 |
| ENSG00000106571 | GLI3 | 803.0448886 | -0.036791579 | 0.072593242 | -0.506818244 | 0.612282354 | 0.769537956 |
| ENSG00000106588 | PSMA2 | 110.3060496 | 0.656328202 | 0.177922546 | 3.688842238 | 0.000225277 | 0.000946666 |
| ENSG00000106591 | MRPL32 | 1021.60123 | 0.611547116 | 0.069250268 | 8.830971158 | 1.04E-18 | 1.61E-17 |
| ENSG00000106603 | COA1 | 1976.632691 | 0.572349574 | 0.056523681 | 10.1258369 | 4.24E-24 | 8.95E-23 |
| ENSG00000106605 | BLVRA | 2056.644317 | 0.099072374 | 0.05286416 | 1.874093403 | 0.060917552 | 0.141299156 |
| ENSG00000106608 | URGCP | 1656.531136 | 0.760211484 | 0.065102044 | 11.6772291 | 1.67E-31 | 4.95E-30 |
| ENSG00000106609 | TMEM248 | 2240.27402 | 0.731372487 | 0.053783925 | 13.59834723 | 4.10E-42 | 1.82E-40 |
| ENSG00000106610 | STAG3L4 | 292.7645256 | 0.781066087 | 0.11041766 | 7.073742413 | 1.51E-12 | 1.52E-11 |
| ENSG00000106615 | RHEB | 1079.611834 | 0.191142735 | 0.065270237 | 2.928482318 | 0.003406212 | 0.011410412 |
| ENSG00000106617 | PRKAG2 | 356.9096096 | 0.872591048 | 0.101477907 | 8.598827837 | 8.05E-18 | 1.17E-16 |
| ENSG00000106624 | AEBP1 | 111.4602632 | -0.038578436 | 0.174656447 | -0.220881834 | 0.825184442 | 0.907770158 |
| ENSG00000106628 | POLD2 | 5008.550074 | 0.956375438 | 0.048558459 | 19.69534169 | 2.36E-86 | 3.05E-84 |
| ENSG00000106635 | BCL7B | 1606.934939 | 0.39029612 | 0.059937609 | 6.511706479 | 7.43E-11 | 6.50E-10 |
| ENSG00000106636 | YKT6 | 4569.378271 | 0.6388534 | 0.047290738 | 13.50905968 | 1.38E-41 | 6.04E-40 |
| ENSG00000106638 | TBL2 | 935.1197611 | 0.381985329 | 0.069538868 | 5.493119762 | 3.95E-08 | 2.67E-07 |
| ENSG00000106665 | CLIP2 | 120.4260667 | 1.789313316 | 0.188078428 | 9.513655204 | 1.84E-21 | 3.36E-20 |
| ENSG00000106682 | EIF4H | 6595.946035 | 0.47490414 | 0.045762832 | 10.37750772 | 3.14E-25 | 7.06E-24 |
| ENSG00000106683 | LIMK1 | 962.7140742 | 1.038134049 | 0.074603764 | 13.91530397 | 5.11E-44 | 2.39E-42 |
| ENSG00000106686 | SPATA6L | 12.70536956 | -0.423358016 | 0.48620888 | -0.870732793 | 0.383900069 | 0.576314873 |
| ENSG00000106688 | SLC1A1 | 36.15448943 | 0.243379679 | 0.290896115 | 0.836654965 | 0.40278654 | 0.59550936 |

| | | | | | | | |
|---|---|---|---|---|---|---|---|
| ENSG00000106689 | LHX2 | 77.30414901 | -0.87310115 | 0.20431661 | -4.273275432 | 1.93E-05 | 9.58E-05 |
| ENSG00000106692 | FKTN | 684.4481667 | 0.302017548 | 0.077769304 | 3.883505878 | 0.000102961 | 0.000458093 |
| ENSG00000106701 | FSD1L | 77.56574848 | -0.249612666 | 0.203431723 | -1.227009543 | 0.219819004 | 0.38993566 |
| ENSG00000106723 | SPIN1 | 3121.057006 | -0.137390801 | 0.049717191 | -2.763446545 | 0.005719446 | 0.018134636 |
| ENSG00000106733 | NMRK1 | 295.8405456 | -0.075023349 | 0.112359437 | -0.667708478 | 0.504319697 | 0.685175989 |
| ENSG00000106771 | TMEM245 | 3105.183413 | 0.400827105 | 0.056737227 | 7.064622749 | 1.61E-12 | 1.62E-11 |
| ENSG00000106772 | PRUNE2 | 2.904314246 | -0.151201712 | 1.009254302 | -0.149815276 | 0.880910357 | 0.938252557 |
| ENSG00000106780 | MEGF9 | 3023.073164 | 1.085993752 | 0.055572268 | 19.54200872 | 4.82E-85 | 6.05E-83 |
| ENSG00000106785 | TRIM14 | 412.2775124 | -0.374265956 | 0.096082744 | -3.895246347 | 9.81E-05 | 0.000437819 |
| ENSG00000106789 | CORO2A | 674.8346685 | -0.298917928 | 0.08118284 | -3.682033402 | 0.000231381 | 0.000970717 |
| ENSG00000106799 | TGFBR1 | 1430.147218 | -0.926203793 | 0.06334479 | -14.62162535 | 2.04E-48 | 1.07E-46 |
| ENSG00000106803 | SEC61B | 1358.176922 | 0.040467025 | 0.061079735 | 0.662527841 | 0.507632998 | 0.687838215 |
| ENSG00000106804 | C5 | 292.273211 | 0.144284322 | 0.11404753 | 1.265124486 | 0.205826707 | 0.37129134 |
| ENSG00000106829 | TLE4 | 124.6534383 | 1.682784257 | 0.179069766 | 9.397366711 | 5.59E-21 | 9.90E-20 |
| ENSG00000106852 | LHX6 | 55.1285547 | -0.142010065 | 0.237094344 | -0.598960154 | 0.549199456 | 0.721774261 |
| ENSG00000106853 | PTGR1 | 664.3871603 | 0.897179778 | 0.08058215 | 11.13372843 | 8.60E-29 | 2.27E-27 |
| ENSG00000106868 | SUSD1 | 310.0097558 | -0.333816843 | 0.106790588 | -3.125901357 | 0.00177261 | 0.006298586 |
| ENSG00000106927 | AMBP | 4.092053123 | -2.471311094 | 1.025027579 | -2.410970342 | 0.015910142 | 0.04479365 |
| ENSG00000106948 | AKNA | 254.2021632 | 0.27553016 | 0.118925008 | 2.316839525 | 0.020512475 | 0.055872261 |
| ENSG00000106976 | DNM1 | 1678.268063 | -0.098765645 | 0.062808305 | -1.5724934 | 0.115836169 | 0.238792389 |
| ENSG00000106991 | ENG | 1.709076558 | 0.125653341 | 1.358793872 | 0.092474174 | 0.926321309 | NA |
| ENSG00000106992 | AK1 | 102.805644 | 0.426614576 | 0.176069524 | 2.422989321 | 0.015393379 | 0.04353108 |
| ENSG00000106993 | CDC37L1 | 252.9918813 | 0.029481927 | 0.120467669 | 0.244728953 | 0.806666308 | 0.897382528 |
| ENSG00000107014 | RLN2 | 22.01507736 | -0.352314957 | 0.386113428 | -0.912464917 | 0.361524029 | 0.554099627 |
| ENSG00000107018 | RLN1 | 6.298641679 | -0.634761735 | 0.721180923 | -0.880169891 | 0.378767282 | 0.571354252 |
| ENSG00000107020 | PLGRKT | 437.0670674 | -1.225370728 | 0.095414851 | -12.84255771 | 9.47E-38 | 3.55E-36 |
| ENSG00000107021 | TBC1D13 | 1533.244493 | 0.260620558 | 0.068569865 | 3.800803143 | 0.000144228 | 0.000626864 |
| ENSG00000107036 | RIC1 | 724.2968309 | 0.038121961 | 0.080366159 | 0.4743534 | 0.635247918 | 0.784991303 |
| ENSG00000107077 | KDM4C | 658.4345191 | 0.155919262 | 0.077756863 | 2.00521543 | 0.044940021 | 0.109323869 |
| ENSG00000107099 | DOCK8 | 228.8391155 | 1.285110919 | 0.133419954 | 9.632074343 | 5.85E-22 | 1.09E-20 |
| ENSG00000107104 | KANK1 | 610.8509227 | 1.210944905 | 0.086094162 | 14.0653545 | 6.20E-45 | 2.99E-43 |
| ENSG00000107105 | ELAVL2 | 59.57219435 | -0.749232196 | 0.235355095 | -3.183411844 | 0.001455504 | 0.005266446 |
| ENSG00000107130 | NCS1 | 1912.077017 | 0.734585393 | 0.058687763 | 12.51684087 | 6.04E-36 | 2.11E-34 |

| | | | | | | | |
|---|---|---|---|---|---|---|---|
| ENSG00000107140 | TESK1 | 962.2435387 | -0.343251971 | 0.073491036 | -4.670664455 | 3.00E-06 | 1.65E-05 |
| ENSG00000107159 | CA9 | 1.775834635 | 1.975186965 | 1.479717934 | 1.334840188 | 0.181928657 | NA |
| ENSG00000107164 | FUBP3 | 3030.547944 | 0.045891161 | 0.049689109 | 0.923565787 | 0.355712434 | 0.547543665 |
| ENSG00000107165 | TYRP1 | 291.9859987 | 0.261661531 | 0.118891571 | 2.200841728 | 0.02774723 | 0.072662268 |
| ENSG00000107175 | CREB3 | 912.9381985 | -0.117100695 | 0.071192243 | -1.644851889 | 0.100000358 | 0.211937197 |
| ENSG00000107185 | RGP1 | 1763.263443 | -0.346483616 | 0.061915395 | -5.596081844 | 2.19E-08 | 1.52E-07 |
| ENSG00000107186 | MPDZ | 1033.605553 | 0.230819771 | 0.069170489 | 3.336968924 | 0.000846974 | 0.003213627 |
| ENSG00000107187 | LHX3 | 2.622545157 | -0.174159901 | 1.07295332 | -0.162318246 | 0.871055247 | 0.93308463 |
| ENSG00000107201 | RIGI | 129.6321248 | 0.777924159 | 0.165495852 | 4.700565895 | 2.59E-06 | 1.43E-05 |
| ENSG00000107223 | EDF1 | 4630.252232 | 0.19112436 | 0.050570098 | 3.779394667 | 0.00015721 | 0.000678916 |
| ENSG00000107242 | PIP5K1B | 5.20251681 | 0.842906953 | 0.810385198 | 1.04013123 | 0.298278936 | 0.48485378 |
| ENSG00000107249 | GLIS3 | 15.48627246 | -0.667319472 | 0.449539081 | -1.484452634 | 0.13768888 | 0.273201132 |
| ENSG00000107262 | BAG1 | 3691.919492 | -0.03390533 | 0.049360146 | -0.686896875 | 0.492147716 | 0.675029514 |
| ENSG00000107263 | RAPGEF1 | 5191.13048 | 0.211442617 | 0.054032604 | 3.913241313 | 9.11E-05 | 0.000408574 |
| ENSG00000107281 | NPDC1 | 3599.604882 | 0.059408684 | 0.055211471 | 1.076020678 | 0.281918009 | 0.465366562 |
| ENSG00000107282 | APBA1 | 54.16560661 | -0.256639182 | 0.247079252 | -1.038691753 | 0.298948118 | 0.485654092 |
| ENSG00000107290 | SETX | 2667.984746 | -0.098997257 | 0.06354768 | -1.557842178 | 0.119270665 | 0.244355331 |
| ENSG00000107331 | ABCA2 | 7806.301817 | 0.721154021 | 0.052243432 | 13.80372597 | 2.42E-43 | 1.10E-41 |
| ENSG00000107338 | SHB | 220.2578546 | -0.049519698 | 0.127888944 | -0.387208595 | 0.698601788 | 0.830862695 |
| ENSG00000107341 | UBE2R2 | 1849.802964 | 0.084199554 | 0.057470697 | 1.465086706 | 0.142897263 | 0.280776293 |
| ENSG00000107362 | ABHD17B | 260.6940237 | -0.215406873 | 0.117471551 | -1.833693959 | 0.066699436 | 0.152535645 |
| ENSG00000107371 | EXOSC3 | 292.2632125 | -0.102667339 | 0.108090279 | -0.949829527 | 0.34219888 | 0.53329254 |
| ENSG00000107372 | ZFAND5 | 1935.944632 | 0.19276494 | 0.061974736 | 3.110379369 | 0.001868472 | 0.006607807 |
| ENSG00000107404 | DVL1 | 1698.23731 | 0.20815877 | 0.057808023 | 3.60086299 | 0.000317163 | 0.00130035 |
| ENSG00000107438 | PDLIM1 | 829.2970547 | 0.333301728 | 0.080275513 | 4.15197256 | 3.30E-05 | 0.000158545 |
| ENSG00000107443 | CCNJ | 582.2925405 | -0.537122992 | 0.089323339 | -6.013243556 | 1.82E-09 | 1.41E-08 |
| ENSG00000107485 | GATA3 | 26195.67384 | -0.311820161 | 0.041249254 | -7.559413331 | 4.05E-14 | 4.55E-13 |
| ENSG00000107518 | ATRNL1 | 32.04177583 | 0.852363412 | 0.34493014 | 2.471118967 | 0.0134691 | 0.038637134 |
| ENSG00000107521 | HPS1 | 2293.899834 | -0.541311341 | 0.058924028 | -9.186597706 | 4.05E-20 | 6.81E-19 |
| ENSG00000107537 | PHYH | 493.0083556 | 0.037912872 | 0.089177686 | 0.425138548 | 0.670735678 | 0.811118793 |
| ENSG00000107551 | RASSF4 | 12.70790923 | 0.361708202 | 0.498811183 | 0.725140523 | 0.468365827 | 0.654303152 |
| ENSG00000107554 | DNMBP | 596.1351899 | -0.408190539 | 0.082152867 | -4.968670672 | 6.74E-07 | 4.00E-06 |
| ENSG00000107560 | RAB11FIP2 | 378.3996369 | -0.465458934 | 0.099014013 | -4.70094 | 2.59E-06 | 1.43E-05 |

| | | | | | | | |
|---|---|---|---|---|---|---|---|
| ENSG00000107562 | CXCL12 | 300.9734678 | 0.456735491 | 0.129451949 | 3.528224145 | 0.000418358 | 0.001678406 |
| ENSG00000107566 | ERLIN1 | 876.5355241 | 0.227029336 | 0.070855503 | 3.204117194 | 0.001354773 | 0.004944868 |
| ENSG00000107581 | EIF3A | 10496.52348 | -0.266989252 | 0.044444206 | -6.007290437 | 1.89E-09 | 1.47E-08 |
| ENSG00000107614 | TRDMT1 | 39.27825812 | -0.007760836 | 0.283379356 | -0.027386738 | 0.978151276 | 0.989932072 |
| ENSG00000107625 | DDX50 | 1177.817329 | -0.732163711 | 0.064951051 | -11.27254596 | 1.79E-29 | 4.89E-28 |
| ENSG00000107643 | MAPK8 | 725.5484834 | -0.381850502 | 0.082861687 | -4.608287796 | 4.06E-06 | 2.19E-05 |
| ENSG00000107651 | SEC23IP | 2245.686376 | -0.544259295 | 0.052474878 | -10.37180685 | 3.33E-25 | 7.47E-24 |
| ENSG00000107669 | ATE1 | 999.3012709 | -0.621631994 | 0.067308408 | -9.235577178 | 2.57E-20 | 4.37E-19 |
| ENSG00000107672 | NSMCE4A | 855.9355719 | -0.568277529 | 0.073167023 | -7.76685323 | 8.05E-15 | 9.50E-14 |
| ENSG00000107679 | PLEKHA1 | 582.3740503 | -0.549376033 | 0.08745961 | -6.281482794 | 3.35E-10 | 2.79E-09 |
| ENSG00000107719 | PALD1 | 63.86697694 | 0.133335137 | 0.221418187 | 0.602186925 | 0.547049718 | 0.719888603 |
| ENSG00000107731 | UNC5B | 119.5607989 | 0.020308857 | 0.164485862 | 0.123468711 | 0.901735949 | 0.950420022 |
| ENSG00000107742 | SPOCK2 | 48.42846043 | -0.689971815 | 0.267837288 | -2.576085729 | 0.009992587 | 0.02970839 |
| ENSG00000107745 | MICU1 | 1514.903461 | -0.62961811 | 0.0579791 | -10.85939769 | 1.80E-27 | 4.57E-26 |
| ENSG00000107758 | PPP3CB | 1408.999204 | -0.60994292 | 0.065522003 | -9.308978531 | 1.29E-20 | 2.22E-19 |
| ENSG00000107771 | CCSER2 | 1493.804886 | -0.456516831 | 0.062930475 | -7.254304523 | 4.04E-13 | 4.23E-12 |
| ENSG00000107779 | BMPR1A | 1324.223715 | -0.246729539 | 0.065439592 | -3.770340417 | 0.000163025 | 0.000702177 |
| ENSG00000107789 | MINPP1 | 643.558337 | -0.392524483 | 0.079354497 | -4.946468026 | 7.56E-07 | 4.45E-06 |
| ENSG00000107796 | ACTA2 | 77.87367646 | -0.002894169 | 0.199405407 | -0.014513995 | 0.988419914 | 0.994043197 |
| ENSG00000107798 | LIPA | 1461.790432 | -0.301400589 | 0.059015766 | -5.107119791 | 3.27E-07 | 2.01E-06 |
| ENSG00000107815 | TWNK | 781.0296698 | 0.325534513 | 0.073023808 | 4.457922997 | 8.28E-06 | 4.31E-05 |
| ENSG00000107816 | LZTS2 | 1442.581441 | 0.125632115 | 0.067780582 | 1.853511896 | 0.063809023 | 0.146833081 |
| ENSG00000107819 | SFXN3 | 441.4417114 | 0.007751404 | 0.090836355 | 0.085333719 | 0.931996086 | 0.967452912 |
| ENSG00000107821 | KAZALD1 | 288.9329711 | -1.121460988 | 0.115272748 | -9.728760757 | 2.27E-22 | 4.34E-21 |
| ENSG00000107829 | FBXW4 | 967.2333796 | 0.040280882 | 0.07096067 | 0.567650812 | 0.570272097 | 0.737339443 |
| ENSG00000107833 | NPM3 | 442.5837274 | -0.153361974 | 0.092071638 | -1.665680959 | 0.095776977 | 0.204707336 |
| ENSG00000107854 | TNKS2 | 3174.885936 | -0.287867015 | 0.052369001 | -5.496897202 | 3.87E-08 | 2.61E-07 |
| ENSG00000107862 | GBF1 | 2665.315208 | 0.016416595 | 0.058068824 | 0.28270928 | 0.777399702 | 0.879192919 |
| ENSG00000107863 | ARHGAP21 | 1339.661918 | -0.510943832 | 0.064972937 | -7.863948541 | 3.72E-15 | 4.52E-14 |
| ENSG00000107864 | CPEB3 | 168.5317034 | -0.499672818 | 0.143277078 | -3.487458196 | 0.000487635 | 0.001934016 |
| ENSG00000107872 | FBXL15 | 212.4791168 | 0.155910095 | 0.142064038 | 1.097463487 | 0.272438834 | 0.454523114 |
| ENSG00000107874 | CUEDC2 | 1036.148973 | -0.149782247 | 0.07137836 | -2.098426565 | 0.035867481 | 0.090507096 |
| ENSG00000107882 | SUFU | 767.2719198 | 0.168489806 | 0.074994623 | 2.246691809 | 0.02465973 | 0.065705069 |

| | | | | | | | |
|---|---|---|---|---|---|---|---|
| ENSG00000107890 | ANKRD26 | 377.7499608 | -0.581673844 | 0.098511888 | -5.904605598 | 3.53E-09 | 2.68E-08 |
| ENSG00000107897 | ACBD5 | 961.8764424 | -0.548820635 | 0.06850207 | -8.011737946 | 1.13E-15 | 1.43E-14 |
| ENSG00000107902 | LHPP | 450.13942 | -0.43328433 | 0.093299863 | -4.643997508 | 3.42E-06 | 1.87E-05 |
| ENSG00000107929 | LARP4B | 3489.679902 | -1.199149878 | 0.054312189 | -22.07883529 | 5.05E-108 | 9.73E-106 |
| ENSG00000107937 | GTPBP4 | 3532.784881 | -1.063210684 | 0.053456572 | -19.88924175 | 5.04E-88 | 6.77E-86 |
| ENSG00000107938 | EDRF1 | 853.2988968 | -0.459140115 | 0.080571749 | -5.698524838 | 1.21E-08 | 8.60E-08 |
| ENSG00000107949 | BCCIP | 1395.305843 | -0.531964595 | 0.068319155 | -7.786463283 | 6.89E-15 | 8.19E-14 |
| ENSG00000107951 | MTPAP | 645.0663948 | -0.693558821 | 0.081022535 | -8.560073063 | 1.13E-17 | 1.63E-16 |
| ENSG00000107954 | NEURL1 | 20.89466695 | -0.140240189 | 0.394533684 | -0.355458087 | 0.722246439 | 0.845625137 |
| ENSG00000107957 | SH3PXD2A | 38.0676469 | 0.895481795 | 0.295489282 | 3.030505162 | 0.00244145 | 0.008436039 |
| ENSG00000107959 | PITRM1 | 4549.984288 | -1.173567395 | 0.049625253 | -23.64859282 | 1.22E-123 | 3.18E-121 |
| ENSG00000107960 | STN1 | 597.1346892 | 0.046331856 | 0.087059284 | 0.532187425 | 0.594596192 | 0.756355993 |
| ENSG00000107968 | MAP3K8 | 77.73024332 | -0.380842821 | 0.208851728 | -1.82350812 | 0.068226468 | 0.155392691 |
| ENSG00000107984 | DKK1 | 5066.146076 | -0.671101627 | 0.049770994 | -13.48378992 | 1.95E-41 | 8.47E-40 |
| ENSG00000108001 | EBF3 | 10.81117665 | -2.414280164 | 0.621100083 | -3.887103269 | 0.000101448 | 0.000451973 |
| ENSG00000108010 | GLRX3 | 1390.706939 | -0.533856644 | 0.062998884 | -8.474065145 | 2.37E-17 | 3.34E-16 |
| ENSG00000108018 | SORCS1 | 29.16866281 | -4.494319027 | 0.578780912 | -7.765147282 | 8.16E-15 | 9.62E-14 |
| ENSG00000108021 | TASOR2 | 3982.198375 | -0.882179341 | 0.054160595 | -16.28821355 | 1.20E-59 | 8.47E-58 |
| ENSG00000108039 | XPNPEP1 | 1462.077416 | -0.317376632 | 0.058755318 | -5.40166645 | 6.60E-08 | 4.34E-07 |
| ENSG00000108055 | SMC3 | 2237.995893 | -0.645708198 | 0.061204563 | -10.55000095 | 5.08E-26 | 1.18E-24 |
| ENSG00000108061 | SHOC2 | 1054.916392 | -0.364671329 | 0.066570479 | -5.477973671 | 4.30E-08 | 2.90E-07 |
| ENSG00000108064 | TFAM | 1373.195038 | -0.489248579 | 0.063081149 | -7.755860341 | 8.77E-15 | 1.03E-13 |
| ENSG00000108091 | CCDC6 | 4171.850534 | -0.605783131 | 0.049061918 | -12.34731861 | 5.04E-35 | 1.72E-33 |
| ENSG00000108094 | CUL2 | 1244.132167 | -0.525895272 | 0.064575289 | -8.143908951 | 3.83E-16 | 4.99E-15 |
| ENSG00000108100 | CCNY | 2144.613993 | -0.496806109 | 0.054190387 | -9.167790341 | 4.83E-20 | 8.08E-19 |
| ENSG00000108106 | UBE2S | 988.1860183 | -0.000798619 | 0.07107092 | -0.011236932 | 0.991034414 | 0.995176993 |
| ENSG00000108107 | RPL28 | 10533.18294 | 0.253875462 | 0.044352811 | 5.723999399 | 1.04E-08 | 7.48E-08 |
| ENSG00000108175 | ZMIZ1 | 4047.945352 | -0.419828959 | 0.053905683 | -7.788213371 | 6.80E-15 | 8.09E-14 |
| ENSG00000108176 | DNAJC12 | 166.9891779 | 0.258563284 | 0.140654073 | 1.838292194 | 0.066019361 | 0.151227493 |
| ENSG00000108179 | PPIF | 2349.768355 | -0.315279933 | 0.055757951 | -5.654439027 | 1.56E-08 | 1.10E-07 |
| ENSG00000108187 | PBLD | 226.3699017 | -0.4895005 | 0.124499673 | -3.931741266 | 8.43E-05 | 0.000381276 |
| ENSG00000108219 | TSPAN14 | 3410.258988 | -0.056481182 | 0.054975246 | -1.02739298 | 0.304235456 | 0.491594094 |
| ENSG00000108239 | TBC1D12 | 580.4722684 | -0.696335772 | 0.084524306 | -8.23829036 | 1.75E-16 | 2.34E-15 |

| | | | | | | | |
|---|---|---|---|---|---|---|---|
| ENSG00000108244 | KRT23 | 75.67033329 | 4.208435448 | 0.351773608 | 11.96347696 | 5.52E-33 | 1.72E-31 |
| ENSG00000108256 | NUFIP2 | 3648.540881 | 0.053724258 | 0.051223986 | 1.048810571 | 0.294265311 | 0.480117087 |
| ENSG00000108262 | GIT1 | 4130.856288 | 0.220579381 | 0.053839457 | 4.096983768 | 4.19E-05 | 0.000197856 |
| ENSG00000108298 | RPL19 | 12836.62195 | 0.285794815 | 0.047858077 | 5.971715422 | 2.35E-09 | 1.81E-08 |
| ENSG00000108306 | FBXL20 | 565.5441739 | 0.345242475 | 0.083029755 | 4.158057228 | 3.21E-05 | 0.000154912 |
| ENSG00000108309 | RUNDC3A | 73.18088016 | 0.701421668 | 0.210766858 | 3.327950481 | 0.000874874 | 0.003310179 |
| ENSG00000108312 | UBTF | 7616.542793 | -0.236463988 | 0.0434603 | -5.440919322 | 5.30E-08 | 3.53E-07 |
| ENSG00000108344 | PSMD3 | 3932.306776 | -0.139698182 | 0.045444068 | -3.074068577 | 0.002111609 | 0.007384193 |
| ENSG00000108349 | CASC3 | 4634.97687 | 0.057183308 | 0.046477878 | 1.230333897 | 0.218572096 | 0.388287106 |
| ENSG00000108352 | RAPGEFL1 | 1031.443281 | 2.717319583 | 0.083620136 | 32.4959959 | 1.21E-231 | 1.21E-228 |
| ENSG00000108370 | RGS9 | 25.78529046 | 0.026166436 | 0.351761548 | 0.074386858 | 0.940702565 | 0.971736581 |
| ENSG00000108375 | RNF43 | 1546.499299 | 0.333720619 | 0.061008854 | 5.47003583 | 4.50E-08 | 3.02E-07 |
| ENSG00000108379 | WNT3 | 171.058909 | 1.010525878 | 0.150344094 | 6.721420533 | 1.80E-11 | 1.65E-10 |
| ENSG00000108384 | RAD51C | 3374.429983 | -0.168127823 | 0.053888373 | -3.119927609 | 0.001808955 | 0.006414813 |
| ENSG00000108387 | SEPTIN4 | 30.10257629 | -1.022421965 | 0.327617861 | -3.120776024 | 0.001803752 | 0.00639834 |
| ENSG00000108389 | MTMR4 | 5085.709878 | 0.278961183 | 0.051636849 | 5.402366518 | 6.58E-08 | 4.33E-07 |
| ENSG00000108395 | TRIM37 | 33388.8481 | 0.380060278 | 0.046563716 | 8.162155169 | 3.29E-16 | 4.32E-15 |
| ENSG00000108406 | DHX40 | 7024.166906 | 0.377379176 | 0.049020128 | 7.69845346 | 1.38E-14 | 1.60E-13 |
| ENSG00000108417 | KRT37 | 4.876354267 | -0.914841267 | 0.804737649 | -1.13681927 | 0.255613846 | 0.434934106 |
| ENSG00000108423 | TUBD1 | 2496.500939 | 0.251945648 | 0.051684776 | 4.874658839 | 1.09E-06 | 6.28E-06 |
| ENSG00000108424 | KPNB1 | 17588.36978 | 0.000893201 | 0.04398128 | 0.020308671 | 0.983797139 | 0.992410257 |
| ENSG00000108433 | GOSR2 | 1301.934754 | 0.199205182 | 0.066240586 | 3.007298008 | 0.002635812 | 0.009036846 |
| ENSG00000108439 | PNPO | 1692.86054 | 0.0099478 | 0.059541516 | 0.167073346 | 0.867312338 | 0.93096599 |
| ENSG00000108443 | RPS6KB1 | 19701.45286 | 0.234597871 | 0.047226713 | 4.967482548 | 6.78E-07 | 4.02E-06 |
| ENSG00000108448 | TRIM16L | 325.5670426 | 0.008466209 | 0.105101223 | 0.080552905 | 0.935797521 | 0.969261113 |
| ENSG00000108465 | CDK5RAP3 | 893.2044651 | -0.00214727 | 0.093262518 | -0.02302393 | 0.981631185 | 0.991616512 |
| ENSG00000108468 | CBX1 | 7147.40673 | -0.081985286 | 0.04339217 | -1.889402779 | 0.058837879 | 0.137224577 |
| ENSG00000108469 | RECQL5 | 1894.752013 | -0.037968958 | 0.061480235 | -0.617579907 | 0.536852293 | 0.711908785 |
| ENSG00000108474 | PIGL | 302.3746479 | -0.173766474 | 0.118458321 | -1.466899689 | 0.142403345 | 0.280128908 |
| ENSG00000108479 | GALK1 | 1011.565936 | 0.671480337 | 0.075932004 | 8.843179419 | 9.30E-19 | 1.45E-17 |
| ENSG00000108506 | INTS2 | 2444.346434 | -0.131849822 | 0.056236431 | -2.344562414 | 0.019049422 | 0.052422473 |
| ENSG00000108509 | CAMTA2 | 1354.354067 | -0.073903808 | 0.067052474 | -1.102178682 | 0.270383998 | 0.452151109 |
| ENSG00000108510 | MED13 | 11130.4398 | -0.00704558 | 0.05592369 | -0.125985601 | 0.899743323 | 0.949347215 |

| | | | | | | | |
|---|---|---|---|---|---|---|---|
| ENSG00000108515 | ENO3 | 368.8937707 | -0.713234851 | 0.098788527 | -7.219814654 | 5.21E-13 | 5.43E-12 |
| ENSG00000108518 | PFN1 | 9056.090371 | -0.326047316 | 0.042389791 | -7.691647247 | 1.45E-14 | 1.68E-13 |
| ENSG00000108523 | RNF167 | 2080.124366 | 0.011491174 | 0.057301175 | 0.200539932 | 0.841058331 | 0.916573782 |
| ENSG00000108528 | SLC25A11 | 1296.059902 | 0.048117768 | 0.067675618 | 0.711005964 | 0.47708054 | 0.661968041 |
| ENSG00000108551 | RASD1 | 696.165506 | -1.591094973 | 0.098163857 | -16.20856207 | 4.39E-59 | 3.05E-57 |
| ENSG00000108556 | CHRNE | 10.67248139 | 0.660092726 | 0.541999866 | 1.21788356 | 0.223268225 | 0.394498427 |
| ENSG00000108557 | RAI1 | 4053.993405 | -0.041186375 | 0.057180976 | -0.720281089 | 0.471351946 | 0.657149904 |
| ENSG00000108559 | NUP88 | 1525.561267 | 0.079561785 | 0.062627072 | 1.270405635 | 0.203940177 | 0.368725383 |
| ENSG00000108561 | C1QBP | 2698.820939 | 0.15220574 | 0.052039743 | 2.92479806 | 0.003446798 | 0.011529535 |
| ENSG00000108578 | BLMH | 1043.906217 | -0.076991402 | 0.064578733 | -1.192209859 | 0.233178969 | 0.407490329 |
| ENSG00000108582 | CPD | 1212.469982 | 0.376574637 | 0.068847543 | 5.46968884 | 4.51E-08 | 3.03E-07 |
| ENSG00000108587 | GOSR1 | 1283.006214 | 0.158660826 | 0.060413202 | 2.626260834 | 0.008632864 | 0.02613935 |
| ENSG00000108588 | CCDC47 | 8444.321064 | 0.187847456 | 0.048926401 | 3.839388414 | 0.000123341 | 0.000541832 |
| ENSG00000108590 | MED31 | 114.564168 | 0.131430882 | 0.16561662 | 0.793585096 | 0.427437012 | 0.618217808 |
| ENSG00000108591 | DRG2 | 1349.633341 | -0.205950637 | 0.063206459 | -3.258379592 | 0.001120504 | 0.004158609 |
| ENSG00000108592 | FTSJ3 | 7956.182259 | -0.055598452 | 0.041468892 | -1.340726729 | 0.180009193 | 0.33498912 |
| ENSG00000108599 | AKAP10 | 1041.474165 | -0.154162479 | 0.069214374 | -2.227318849 | 0.025925972 | 0.06850658 |
| ENSG00000108602 | ALDH3A1 | 2.336584242 | 4.57124348 | 1.513338562 | 3.020635034 | 0.002522452 | 0.008683231 |
| ENSG00000108604 | SMARCD2 | 12570.40877 | 0.164367566 | 0.04113363 | 3.995941178 | 6.44E-05 | 0.000296292 |
| ENSG00000108639 | SYNGR2 | 7569.695543 | 0.065129076 | 0.052693341 | 1.236002032 | 0.216457808 | 0.385888244 |
| ENSG00000108641 | B9D1 | 417.0040722 | 0.259855813 | 0.096286472 | 2.698778013 | 0.006959458 | 0.021585988 |
| ENSG00000108651 | UTP6 | 923.2038798 | 0.067985854 | 0.06931494 | 0.980825411 | 0.326678845 | 0.517185142 |
| ENSG00000108654 | DDX5 | 29328.93495 | 0.086165801 | 0.112062094 | 0.768911215 | 0.441946019 | 0.631233528 |
| ENSG00000108666 | C17orf75 | 365.1938313 | 0.05846218 | 0.098913316 | 0.59104458 | 0.554490551 | 0.72573819 |
| ENSG00000108669 | CYTH1 | 1009.616213 | 0.266131914 | 0.068032168 | 3.911854068 | 9.16E-05 | 0.000410849 |
| ENSG00000108671 | PSMD11 | 3145.232355 | -0.126561578 | 0.050788963 | -2.491911055 | 0.012705785 | 0.036695293 |
| ENSG00000108679 | LGALS3BP | 597.293692 | -0.964025731 | 0.089335144 | -10.79111411 | 3.79E-27 | 9.45E-26 |
| ENSG00000108733 | PEX12 | 149.7616446 | -0.059434519 | 0.145309836 | -0.409019243 | 0.682525539 | 0.819871514 |
| ENSG00000108759 | KRT32 | 4.083915022 | -0.50148334 | 0.859337913 | -0.583569434 | 0.55951003 | 0.72907705 |
| ENSG00000108771 | DHX58 | 9.801872407 | 0.112504937 | 0.554786909 | 0.20278946 | 0.839299597 | 0.915309377 |
| ENSG00000108773 | KAT2A | 2538.510836 | 0.183701665 | 0.060248559 | 3.049063223 | 0.002295562 | 0.007973983 |
| ENSG00000108774 | RAB5C | 1090.037491 | 0.03884465 | 0.066263465 | 0.586215187 | 0.557730917 | 0.727915708 |
| ENSG00000108784 | NAGLU | 768.7038195 | 0.059089579 | 0.074621833 | 0.79185377 | 0.428445945 | 0.619042354 |

| | | | | | | | |
|---|---|---|---|---|---|---|---|
| ENSG00000108786 | HSD17B1 | 34.63485468 | 0.312145979 | 0.328784586 | 0.949393591 | 0.342420469 | 0.533424321 |
| ENSG00000108788 | MLX | 1975.914116 | -0.066068064 | 0.055620937 | -1.187827222 | 0.234901487 | 0.409876026 |
| ENSG00000108797 | CNTNAP1 | 94.03799963 | -0.834996534 | 0.186621721 | -4.474273023 | 7.67E-06 | 4.01E-05 |
| ENSG00000108798 | ABI3 | 2.114216114 | -1.798415109 | 1.426175493 | -1.261005478 | 0.207306873 | 0.37337047 |
| ENSG00000108799 | EZH1 | 1261.932855 | 0.186349102 | 0.067865213 | 2.745870729 | 0.006035054 | 0.01904845 |
| ENSG00000108813 | DLX4 | 188.6714195 | -0.449159354 | 0.132755505 | -3.383357656 | 0.000716053 | 0.002754697 |
| ENSG00000108819 | PPP1R9B | 4194.558124 | -0.069017587 | 0.051796908 | -1.332465384 | 0.182707309 | 0.33864132 |
| ENSG00000108821 | COL1A1 | 1323.203182 | -0.927113304 | 0.087985003 | -10.53717418 | 5.82E-26 | 1.35E-24 |
| ENSG00000108826 | MRPL27 | 1490.075926 | -0.156657406 | 0.058508547 | -2.677513164 | 0.007417094 | 0.022854014 |
| ENSG00000108828 | VAT1 | 5521.19466 | 0.479919935 | 0.054054266 | 8.878484035 | 6.78E-19 | 1.06E-17 |
| ENSG00000108829 | LRRC59 | 8323.019968 | -0.069841932 | 0.043116133 | -1.619856109 | 0.10526319 | 0.220911002 |
| ENSG00000108830 | RND2 | 27.70796909 | 0.673350686 | 0.352791864 | 1.908634394 | 0.056309272 | 0.13248561 |
| ENSG00000108839 | ALOX12 | 1.633450164 | -1.398839893 | 1.427059346 | -0.980225452 | 0.326974843 | NA |
| ENSG00000108840 | HDAC5 | 1095.398049 | 0.785792263 | 0.071378238 | 11.0088492 | 3.46E-28 | 8.97E-27 |
| ENSG00000108846 | ABCC3 | 1705.924036 | -0.208206344 | 0.065379738 | -3.18456987 | 0.001449693 | 0.005246247 |
| ENSG00000108848 | LUC7L3 | 5209.107085 | -0.027895551 | 0.068418656 | -0.407718495 | 0.683480357 | 0.820419622 |
| ENSG00000108852 | MPP2 | 405.5839396 | -0.249840897 | 0.094385909 | -2.64701479 | 0.00812058 | 0.02476221 |
| ENSG00000108854 | SMURF2 | 1918.337202 | 0.023702933 | 0.05371335 | 0.441285685 | 0.659006188 | 0.802779253 |
| ENSG00000108861 | DUSP3 | 2291.247212 | 0.585629383 | 0.057610849 | 10.16526213 | 2.83E-24 | 6.03E-23 |
| ENSG00000108878 | CACNG1 | 42.9781321 | 2.240276541 | 0.315865697 | 7.09249711 | 1.32E-12 | 1.33E-11 |
| ENSG00000108883 | EFTUD2 | 5202.664571 | -0.070654151 | 0.04703435 | -1.502181929 | 0.13305013 | 0.266193092 |
| ENSG00000108924 | HLF | 97.71847687 | -0.695865602 | 0.185398613 | -3.753348476 | 0.000174488 | 0.000748318 |
| ENSG00000108932 | SLC16A6 | 624.1258113 | -0.31275566 | 0.0867891 | -3.603628344 | 0.000313806 | 0.001288013 |
| ENSG00000108946 | PRKAR1A | 9686.815075 | 0.027903202 | 0.047375576 | 0.588978637 | 0.555875605 | 0.726680327 |
| ENSG00000108947 | EFNB3 | 501.3746308 | 0.201071807 | 0.089183818 | 2.254577235 | 0.024159878 | 0.064556939 |
| ENSG00000108953 | YWHAE | 10592.48255 | -0.051172113 | 0.04314341 | -1.186093382 | 0.235585424 | 0.41056975 |
| ENSG00000108958 | ENSG0000001C | 125.3411183 | -0.208879957 | 0.158430283 | -1.318434536 | 0.187358226 | 0.345220699 |
| ENSG00000108960 | MMD | 469.5032187 | -1.701715575 | 0.097096254 | -17.52606828 | 9.06E-69 | 8.06E-67 |
| ENSG00000108961 | RANGRF | 50.81884274 | -0.713093794 | 0.252424179 | -2.824982125 | 0.004728325 | 0.015325933 |
| ENSG00000108963 | DPH1 | 843.9492878 | 0.144791972 | 0.071674697 | 2.020126753 | 0.043370241 | 0.106112771 |
| ENSG00000108984 | MAP2K6 | 267.6515997 | 1.277279307 | 0.119706581 | 10.67008429 | 1.40E-26 | 3.40E-25 |
| ENSG00000109016 | DHRS7B | 442.2256788 | -0.330728818 | 0.100152899 | -3.302239089 | 0.000959163 | 0.003601177 |
| ENSG00000109046 | WSB1 | 1339.547414 | -0.490367686 | 0.17957612 | -2.73069541 | 0.006320085 | 0.019847058 |

| | | | | | | | |
|---|---|---|---|---|---|---|---|
| ENSG00000109062 | NHERF1 | 10362.75563 | 0.168271704 | 0.040887051 | 4.115525538 | 3.86E-05 | 0.000183472 |
| ENSG00000109063 | MYH3 | 23.46259784 | -1.146500994 | 0.378319024 | -3.030513725 | 0.002441381 | 0.008436039 |
| ENSG00000109065 | NAT9 | 1193.941311 | 0.310185949 | 0.064381114 | 4.817964947 | 1.45E-06 | 8.22E-06 |
| ENSG00000109066 | TMEM104 | 756.3276286 | 0.228233667 | 0.085171062 | 2.679709054 | 0.007368618 | 0.022732126 |
| ENSG00000109072 | VTN | 60.94600849 | -1.055901793 | 0.24534962 | -4.303661816 | 1.68E-05 | 8.42E-05 |
| ENSG00000109079 | TNFAIP1 | 2092.526595 | 0.160175904 | 0.055896569 | 2.865576653 | 0.004162505 | 0.013659696 |
| ENSG00000109083 | IFT20 | 524.5486143 | 0.00369457 | 0.086462662 | 0.042730232 | 0.96591658 | 0.984632112 |
| ENSG00000109084 | TMEM97 | 2171.14446 | -0.432649337 | 0.053652204 | -8.063962104 | 7.39E-16 | 9.45E-15 |
| ENSG00000109089 | CDR2L | 1263.190184 | 0.114271404 | 0.081257622 | 1.406285351 | 0.159639408 | 0.305306589 |
| ENSG00000109099 | PMP22 | 15714.15998 | 0.084559217 | 0.04223409 | 2.002155536 | 0.045268006 | 0.10998179 |
| ENSG00000109101 | FOXN1 | 13.48999853 | 1.746262759 | 0.54755287 | 3.189213049 | 0.001426607 | 0.005170042 |
| ENSG00000109103 | UNC119 | 1045.773716 | 0.684493632 | 0.074193098 | 9.225839793 | 2.81E-20 | 4.77E-19 |
| ENSG00000109107 | ALDOC | 353.591228 | -0.138109341 | 0.104228405 | -1.325064334 | 0.185149814 | 0.342140868 |
| ENSG00000109111 | SUPT6H | 6956.859033 | -0.066377432 | 0.046811115 | -1.417984434 | 0.15619531 | 0.300364147 |
| ENSG00000109113 | RAB34 | 3.029127897 | 0.351250143 | 1.058739474 | 0.331762584 | 0.740068539 | 0.856567724 |
| ENSG00000109118 | PHF12 | 2341.910758 | -0.207256518 | 0.059294929 | -3.495349799 | 0.000473441 | 0.001881624 |
| ENSG00000109133 | TMEM33 | 1303.675713 | -0.073020373 | 0.065267373 | -1.118788302 | 0.263230464 | 0.444040506 |
| ENSG00000109171 | SLAIN2 | 1713.556175 | -0.165774013 | 0.058245728 | -2.846114526 | 0.004425627 | 0.0144518 |
| ENSG00000109180 | OCIAD1 | 2694.917874 | -0.020433565 | 0.054182868 | -0.377122244 | 0.706082762 | 0.835743179 |
| ENSG00000109181 | UGT2B10 | 3.781703344 | -1.984398364 | 0.998073007 | -1.98822967 | 0.046786295 | 0.11312616 |
| ENSG00000109184 | DCUN1D4 | 1028.962322 | 0.053313104 | 0.070852558 | 0.752451369 | 0.451779665 | 0.639701573 |
| ENSG00000109189 | USP46 | 1135.389716 | 0.240987512 | 0.066555041 | 3.620875435 | 0.000293608 | 0.001210308 |
| ENSG00000109205 | ODAM | 15.10159557 | 1.909341134 | 0.510688389 | 3.738759633 | 0.00018493 | 0.000789854 |
| ENSG00000109220 | CHIC2 | 188.0038023 | -0.184600987 | 0.134290106 | -1.374643242 | 0.169242076 | 0.319698058 |
| ENSG00000109255 | NMU | 33.92335217 | -1.443822587 | 0.329820187 | -4.377605272 | 1.20E-05 | 6.12E-05 |
| ENSG00000109265 | CRACD | 330.3395044 | 0.850773019 | 0.114723524 | 7.415855005 | 1.21E-13 | 1.31E-12 |
| ENSG00000109270 | LAMTOR3 | 1043.95843 | 0.099604454 | 0.065770049 | 1.514434845 | 0.129915587 | 0.261151732 |
| ENSG00000109320 | NFKB1 | 1334.785213 | -0.185061291 | 0.063938669 | -2.894356329 | 0.003799367 | 0.012585919 |
| ENSG00000109321 | AREG | 38.16714646 | 1.265305408 | 0.310460091 | 4.075581509 | 4.59E-05 | 0.000215854 |
| ENSG00000109323 | MANBA | 1173.069473 | 0.104750737 | 0.067910108 | 1.542491096 | 0.1229543 | 0.25009502 |
| ENSG00000109332 | UBE2D3 | 4674.89648 | -0.043205887 | 0.051269892 | -0.842714597 | 0.399388081 | 0.592163734 |
| ENSG00000109339 | MAPK10 | 61.00526636 | -0.481281773 | 0.224936699 | -2.139632059 | 0.032384514 | 0.083024201 |
| ENSG00000109381 | ELF2 | 1321.241001 | -0.185512228 | 0.061094741 | -3.03646801 | 0.002393675 | 0.008293446 |

| | | | | | | | |
|---|---|---|---|---|---|---|---|
| ENSG00000109390 | NDUFC1 | 742.2523471 | -0.474804286 | 0.074715899 | -6.354795831 | 2.09E-10 | 1.76E-09 |
| ENSG00000109436 | TBC1D9 | 9046.086218 | 1.14970537 | 0.04997126 | 23.00733224 | 3.94E-117 | 8.94E-115 |
| ENSG00000109445 | ZNF330 | 775.7412549 | -0.059371603 | 0.077365333 | -0.767418705 | 0.442832611 | 0.631673927 |
| ENSG00000109452 | INPP4B | 5658.488935 | 0.683249187 | 0.0500944 | 13.63923275 | 2.34E-42 | 1.05E-40 |
| ENSG00000109458 | GAB1 | 780.4385112 | -0.088809757 | 0.076302154 | -1.163922017 | 0.244455616 | 0.421228261 |
| ENSG00000109466 | KLHL2 | 330.4276241 | 0.244867427 | 0.103962642 | 2.355340559 | 0.018505737 | 0.05107334 |
| ENSG00000109472 | CPE | 1269.328131 | -1.058704595 | 0.067612722 | -15.65836374 | 2.91E-55 | 1.85E-53 |
| ENSG00000109475 | RPL34 | 4480.655143 | 0.341310385 | 0.053233626 | 6.41155617 | 1.44E-10 | 1.23E-09 |
| ENSG00000109501 | WFS1 | 1569.725398 | 0.678984385 | 0.062125152 | 10.92929932 | 8.35E-28 | 2.14E-26 |
| ENSG00000109519 | GRPEL1 | 1731.355391 | 0.120791601 | 0.055265824 | 2.185647355 | 0.028841415 | 0.075026819 |
| ENSG00000109534 | GAR1 | 670.9571969 | -0.173725062 | 0.081268233 | -2.137674891 | 0.032543139 | 0.083330731 |
| ENSG00000109536 | FRG1 | 257.8237893 | -0.0635057 | 0.115007959 | -0.552185262 | 0.580821431 | 0.745901787 |
| ENSG00000109572 | CLCN3 | 2096.17639 | 0.152873166 | 0.058685127 | 2.604972922 | 0.009188152 | 0.02763191 |
| ENSG00000109576 | AADAT | 291.4677273 | -0.12558412 | 0.113672297 | -1.104790911 | 0.2692502 | 0.450879596 |
| ENSG00000109586 | GALNT7 | 1274.837025 | -0.424849194 | 0.064003552 | -6.637900208 | 3.18E-11 | 2.86E-10 |
| ENSG00000109606 | DHX15 | 8623.363366 | -0.421972899 | 0.046198507 | -9.133907605 | 6.61E-20 | 1.10E-18 |
| ENSG00000109618 | SEPSECS | 608.6313482 | -0.065673149 | 0.087505202 | -0.750505658 | 0.452950217 | 0.640609 |
| ENSG00000109654 | TRIM2 | 646.0426261 | 1.094562033 | 0.084883964 | 12.89480348 | 4.81E-38 | 1.82E-36 |
| ENSG00000109670 | FBXW7 | 829.6358293 | -0.177608208 | 0.075881466 | -2.340600636 | 0.019252749 | 0.052899516 |
| ENSG00000109674 | NEIL3 | 153.0801566 | -0.340067233 | 0.146000095 | -2.329226104 | 0.019847091 | 0.054343789 |
| ENSG00000109680 | TBC1D19 | 57.79122801 | 0.270345819 | 0.237621577 | 1.137715784 | 0.255239181 | 0.434715777 |
| ENSG00000109685 | NSD2 | 7808.358226 | -0.281626769 | 0.04516125 | -6.236026887 | 4.49E-10 | 3.69E-09 |
| ENSG00000109686 | SH3D19 | 1141.434885 | 0.350099006 | 0.064167053 | 5.456055526 | 4.87E-08 | 3.25E-07 |
| ENSG00000109689 | STIM2 | 1037.886325 | -0.014209264 | 0.071766759 | -0.197992269 | 0.843051112 | 0.917338822 |
| ENSG00000109705 | NKX3-2 | 42.47108958 | -0.119411826 | 0.272588616 | -0.438066079 | 0.661338383 | 0.804235714 |
| ENSG00000109736 | MFSD10 | 3168.450529 | 0.038158408 | 0.057199398 | 0.667112058 | 0.504700559 | 0.685322056 |
| ENSG00000109738 | GLRB | 63.00549572 | 0.268443673 | 0.221458404 | 1.212162957 | 0.225450008 | 0.39749621 |
| ENSG00000109743 | BST1 | 4.776560476 | 0.648181214 | 0.812746985 | 0.797519063 | 0.425149636 | 0.616060327 |
| ENSG00000109756 | RAPGEF2 | 1020.739584 | -0.010677127 | 0.06928594 | -0.154102356 | 0.877529031 | 0.936390175 |
| ENSG00000109758 | HGFAC | 50.2762484 | 0.78312285 | 0.258795867 | 3.026025331 | 0.002477915 | 0.008545307 |
| ENSG00000109762 | SNX25 | 215.7086724 | -0.308772307 | 0.126209993 | -2.446496505 | 0.01442522 | 0.041056394 |
| ENSG00000109771 | LRP2BP | 5.108095302 | 0.06168895 | 0.787136856 | 0.078371314 | 0.937532691 | 0.970153346 |
| ENSG00000109775 | UFSP2 | 155.7607209 | -0.161275264 | 0.146522985 | -1.10068235 | 0.271034931 | 0.45297547 |

| | | | | | | | |
|---|---|---|---|---|---|---|---|
| ENSG00000109787 | KLF3 | 952.4734823 | 0.195852903 | 0.069589843 | 2.814389204 | 0.004887002 | 0.015789835 |
| ENSG00000109790 | KLHL5 | 745.4092102 | 0.989920572 | 0.078898867 | 12.54670205 | 4.14E-36 | 1.46E-34 |
| ENSG00000109794 | FAM149A | 3.371920091 | -1.76283599 | 1.027331763 | -1.715936422 | 0.08617368 | 0.187696943 |
| ENSG00000109805 | NCAPG | 1083.333866 | -0.32454995 | 0.06794528 | -4.77663718 | 1.78E-06 | 1.00E-05 |
| ENSG00000109814 | UGDH | 3328.847517 | -0.863789703 | 0.052138516 | -16.56720908 | 1.20E-61 | 8.96E-60 |
| ENSG00000109819 | PPARGC1A | 25.83100766 | -2.649230112 | 0.418063785 | -6.336904101 | 2.34E-10 | 1.97E-09 |
| ENSG00000109854 | HTATIP2 | 1197.860975 | -0.067225448 | 0.065589319 | -1.024945046 | 0.305389118 | 0.492902721 |
| ENSG00000109861 | CTSC | 194.5127149 | -0.933538516 | 0.133035601 | -7.017208263 | 2.26E-12 | 2.25E-11 |
| ENSG00000109881 | CCDC34 | 396.7844102 | 0.145852235 | 0.101130478 | 1.442218381 | 0.149240777 | 0.290478824 |
| ENSG00000109911 | ELP4 | 555.3510744 | -0.071690128 | 0.082431805 | -0.869690142 | 0.384469761 | 0.576981311 |
| ENSG00000109917 | ZPR1 | 1252.139611 | 0.150519262 | 0.061783505 | 2.436237029 | 0.014840956 | 0.042083008 |
| ENSG00000109919 | MTCH2 | 3511.611813 | 0.015920547 | 0.047490854 | 0.335233966 | 0.737448602 | 0.855304663 |
| ENSG00000109920 | FNBP4 | 3392.118231 | -0.130489722 | 0.07168746 | -1.820258685 | 0.06871962 | 0.156252164 |
| ENSG00000109927 | TECTA | 8.400133534 | 0.090344847 | 0.594864633 | 0.151874631 | 0.879285817 | 0.93743629 |
| ENSG00000109929 | SC5D | 1341.96014 | -0.125102752 | 0.066918171 | -1.869488524 | 0.061554879 | 0.142575618 |
| ENSG00000109944 | JHY | 152.1813557 | 0.037388189 | 0.149059512 | 0.250827258 | 0.801947666 | 0.894633256 |
| ENSG00000109971 | HSPA8 | 32747.03671 | 0.069312197 | 0.050351872 | 1.376556507 | 0.168649406 | 0.318841055 |
| ENSG00000110002 | VWA5A | 8.1792498 | 0.591506527 | 0.614559035 | 0.962489351 | 0.33580385 | 0.526940305 |
| ENSG00000110011 | DNAJC4 | 549.7099439 | -0.132951858 | 0.088717584 | -1.498596464 | 0.133978351 | 0.267469008 |
| ENSG00000110013 | SIAE | 499.4939569 | -0.137123122 | 0.085577326 | -1.602330057 | 0.109082642 | 0.227177106 |
| ENSG00000110025 | SNX15 | 31.72795981 | 0.161300594 | 0.311309332 | 0.51813607 | 0.604363335 | 0.763147185 |
| ENSG00000110031 | LPXN | 80.98211296 | -0.114082589 | 0.206760486 | -0.551762048 | 0.581111392 | 0.746190615 |
| ENSG00000110042 | DTX4 | 551.2845002 | -0.045481872 | 0.085266489 | -0.53340852 | 0.593750821 | 0.755710631 |
| ENSG00000110046 | ATG2A | 1937.255418 | 0.168145978 | 0.060901667 | 2.760942146 | 0.005763489 | 0.018259139 |
| ENSG00000110047 | EHD1 | 1738.042257 | 0.029862646 | 0.05966247 | 0.50052647 | 0.616704423 | 0.772907138 |
| ENSG00000110048 | OSBP | 2745.730992 | 0.005141823 | 0.051764357 | 0.099331332 | 0.920875202 | 0.961795535 |
| ENSG00000110057 | UNC93B1 | 820.5660224 | 0.276969051 | 0.088951857 | 3.113696127 | 0.001847597 | 0.00654072 |
| ENSG00000110060 | PUS3 | 199.1655029 | -0.009568662 | 0.133435071 | -0.071710249 | 0.9428325 | 0.972907279 |
| ENSG00000110063 | DCPS | 603.472776 | 0.126508406 | 0.086381775 | 1.464526593 | 0.143050122 | 0.281004436 |
| ENSG00000110066 | KMT5B | 2328.372738 | -0.081684346 | 0.053698768 | -1.521158682 | 0.128220021 | 0.258596158 |
| ENSG00000110074 | FOXRED1 | 712.9428711 | -0.277573484 | 0.078829024 | -3.521209197 | 0.000429584 | 0.001720735 |
| ENSG00000110075 | PPP6R3 | 6167.273202 | -0.13130314 | 0.044802182 | -2.930730891 | 0.003381656 | 0.011334773 |
| ENSG00000110076 | NRXN2 | 1.98000695 | -1.197387889 | 1.293869049 | -0.925432052 | 0.354741213 | 0.546426756 |

| | | | | | | | |
|---|---|---|---|---|---|---|---|
| ENSG00000110080 | ST3GAL4 | 955.607099 | 0.023519115 | 0.072098353 | 0.326208773 | 0.744266404 | 0.858567399 |
| ENSG00000110090 | CPT1A | 4340.861295 | -0.653063606 | 0.048127042 | -13.56957707 | 6.07E-42 | 2.68E-40 |
| ENSG00000110092 | CCND1 | 13773.87325 | -0.189798578 | 0.041727612 | -4.54851286 | 5.40E-06 | 2.87E-05 |
| ENSG00000110104 | CCDC86 | 1270.273798 | -0.067134378 | 0.066250564 | -1.013340473 | 0.310897562 | 0.498917791 |
| ENSG00000110107 | PRPF19 | 5252.046773 | -0.095511486 | 0.04399164 | -2.171128124 | 0.029921489 | 0.077489157 |
| ENSG00000110108 | TMEM109 | 2371.942128 | -0.272450816 | 0.056567527 | -4.81638196 | 1.46E-06 | 8.28E-06 |
| ENSG00000110169 | HPX | 43.99536888 | -1.170501638 | 0.281341247 | -4.160433817 | 3.18E-05 | 0.00015347 |
| ENSG00000110171 | TRIM3 | 676.1824207 | -0.666914677 | 0.081284058 | -8.204741449 | 2.31E-16 | 3.06E-15 |
| ENSG00000110172 | CHORDC1 | 594.8964629 | 0.062474771 | 0.082090061 | 0.761051584 | 0.446626258 | 0.635071366 |
| ENSG00000110200 | ANAPC15 | 1011.222722 | 0.111859818 | 0.066582966 | 1.680006542 | 0.092956043 | 0.199682706 |
| ENSG00000110218 | PANX1 | 760.4067305 | 0.357954396 | 0.074131645 | 4.828631485 | 1.37E-06 | 7.81E-06 |
| ENSG00000110237 | ARHGEF17 | 2428.054815 | 0.109570491 | 0.059119166 | 1.853383588 | 0.063827398 | 0.146845904 |
| ENSG00000110274 | CEP164 | 1132.994305 | 0.018387801 | 0.067060444 | 0.274197429 | 0.783932904 | 0.882792721 |
| ENSG00000110315 | RNF141 | 638.3480456 | 0.216939779 | 0.086841475 | 2.498112549 | 0.012485655 | 0.036155127 |
| ENSG00000110318 | CEP126 | 112.6351767 | -0.079845035 | 0.170698565 | -0.467754575 | 0.639960109 | 0.78809775 |
| ENSG00000110321 | EIF4G2 | 27477.21235 | -0.012386362 | 0.047136054 | -0.262778942 | 0.792720967 | 0.888678053 |
| ENSG00000110324 | IL10RA | 6.086806844 | 1.274122148 | 0.75633539 | 1.684599405 | 0.092065886 | 0.198141611 |
| ENSG00000110328 | GALNT18 | 110.3382038 | 0.390354732 | 0.173381081 | 2.251426337 | 0.024358548 | 0.065035508 |
| ENSG00000110330 | BIRC2 | 919.2297929 | -0.069943223 | 0.074268096 | -0.941766739 | 0.346312078 | 0.537146657 |
| ENSG00000110344 | UBE4A | 4113.266984 | -0.160380716 | 0.050199232 | -3.194883868 | 0.001398871 | 0.005083984 |
| ENSG00000110367 | DDX6 | 7422.397314 | -0.150754274 | 0.0444424 | -3.392127205 | 0.000693522 | 0.002676098 |
| ENSG00000110375 | UPK2 | 1535.7647 | 1.189878838 | 0.142287696 | 8.362485826 | 6.14E-17 | 8.45E-16 |
| ENSG00000110395 | CBL | 1114.706512 | -0.311696104 | 0.082864345 | -3.761522544 | 0.000168882 | 0.000726314 |
| ENSG00000110400 | NECTIN1 | 1974.484293 | -0.09000544 | 0.060679879 | -1.48328313 | 0.137999203 | 0.273620515 |
| ENSG00000110422 | HIPK3 | 2057.787003 | -0.062157406 | 0.0576843 | -1.077544593 | 0.28123704 | 0.464677161 |
| ENSG00000110427 | KIAA1549L | 8.322133233 | -0.092031212 | 0.607651888 | -0.15145384 | 0.879617721 | 0.937615842 |
| ENSG00000110429 | FBXO3 | 802.5638633 | 0.024678921 | 0.071448007 | 0.345410902 | 0.729785483 | 0.850415471 |
| ENSG00000110435 | PDHX | 1069.418153 | 0.109877878 | 0.065120374 | 1.687304154 | 0.091544883 | 0.197186818 |
| ENSG00000110436 | SLC1A2 | 104.5559975 | 0.7366547 | 0.178893685 | 4.117835118 | 3.82E-05 | 0.000181681 |
| ENSG00000110442 | COMMD9 | 908.3940269 | -0.16832056 | 0.071075484 | -2.368194351 | 0.017875143 | 0.049613525 |
| ENSG00000110446 | SLC15A3 | 6.321160501 | -0.655699686 | 0.721771016 | -0.908459429 | 0.363635539 | 0.556351473 |
| ENSG00000110448 | CD5 | 1.921411204 | 0.370650385 | 1.340774704 | 0.276444942 | 0.782206342 | 0.881889885 |
| ENSG00000110455 | ACCS | 374.1526261 | 0.139444273 | 0.108540266 | 1.284723886 | 0.198888788 | 0.361615979 |

| | | | | | | | |
|---|---|---|---|---|---|---|---|
| ENSG00000110492 | MDK | 3431.392427 | 0.847793785 | 0.048588468 | 17.44845698 | 3.54E-68 | 3.06E-66 |
| ENSG00000110497 | AMBRA1 | 2135.372416 | 0.139446856 | 0.055735092 | 2.501957949 | 0.012350859 | 0.035809999 |
| ENSG00000110514 | MADD | 4554.418844 | -0.251348481 | 0.048504664 | -5.18194455 | 2.20E-07 | 1.38E-06 |
| ENSG00000110536 | PTPMT1 | 469.9069253 | 0.139979147 | 0.09282027 | 1.508066569 | 0.13153749 | 0.263857121 |
| ENSG00000110583 | NAA40 | 1284.309136 | 0.423935868 | 0.063895218 | 6.634860645 | 3.25E-11 | 2.92E-10 |
| ENSG00000110619 | CARS1 | 3018.519064 | 0.195013522 | 0.052471633 | 3.716551423 | 0.000201961 | 0.00085574 |
| ENSG00000110628 | SLC22A18 | 486.1090003 | 0.053928703 | 0.088652775 | 0.608313764 | 0.542979394 | 0.71662275 |
| ENSG00000110651 | CD81 | 4481.052284 | 0.37739232 | 0.055002184 | 6.861406109 | 6.82E-12 | 6.48E-11 |
| ENSG00000110660 | SLC35F2 | 780.1259698 | 0.449959328 | 0.074563569 | 6.034573404 | 1.59E-09 | 1.25E-08 |
| ENSG00000110665 | C11orf21 | 2.96160508 | 0.814176763 | 1.036931904 | 0.785178621 | 0.432348848 | 0.622408697 |
| ENSG00000110693 | SOX6 | 14.57632801 | -0.759504973 | 0.473362463 | -1.604489228 | 0.108606257 | 0.226438937 |
| ENSG00000110696 | C11orf58 | 3629.656104 | 0.003521458 | 0.054201632 | 0.064969585 | 0.948198217 | 0.976023831 |
| ENSG00000110697 | PITPNM1 | 917.4566355 | 0.407985035 | 0.082325365 | 4.955763434 | 7.20E-07 | 4.26E-06 |
| ENSG00000110700 | RPS13 | 5659.856523 | 0.306522602 | 0.052143877 | 5.878400634 | 4.14E-09 | 3.11E-08 |
| ENSG00000110711 | AIP | 1203.372595 | -0.398182802 | 0.071449485 | -5.572927515 | 2.50E-08 | 1.72E-07 |
| ENSG00000110713 | NUP98 | 3408.442198 | 0.023047948 | 0.053675152 | 0.429396975 | 0.667634355 | 0.808785741 |
| ENSG00000110717 | NDUFS8 | 1760.729122 | 0.064692461 | 0.063133445 | 1.024693978 | 0.305507605 | 0.49301985 |
| ENSG00000110719 | TCIRG1 | 519.0197482 | 0.531067826 | 0.094674333 | 5.609417134 | 2.03E-08 | 1.41E-07 |
| ENSG00000110721 | CHKA | 1375.715947 | 0.093755736 | 0.060695431 | 1.544691837 | 0.12242082 | 0.249186657 |
| ENSG00000110723 | EXPH5 | 182.4832835 | -0.187986019 | 0.137279607 | -1.369365941 | 0.170884915 | 0.322023475 |
| ENSG00000110756 | HPS5 | 683.2194247 | -0.108770403 | 0.075833149 | -1.434338459 | 0.1514757 | 0.293856453 |
| ENSG00000110768 | GTF2H1 | 1845.47538 | 0.341205799 | 0.061503981 | 5.547702665 | 2.89E-08 | 1.98E-07 |
| ENSG00000110799 | VWF | 6.822016912 | 2.456395455 | 0.812499844 | 3.023256524 | 0.002500702 | 0.008618704 |
| ENSG00000110801 | PSMD9 | 181.7569664 | 0.020512559 | 0.133473656 | 0.153682452 | 0.877860122 | 0.936395015 |
| ENSG00000110811 | P3H3 | 8.450960076 | -0.47734416 | 0.611179663 | -0.781021014 | 0.434790135 | 0.624786125 |
| ENSG00000110841 | PPFIBP1 | 969.9680692 | 0.853411765 | 0.070433714 | 12.11652374 | 8.63E-34 | 2.78E-32 |
| ENSG00000110844 | PRPF40B | 746.5283902 | -0.089053046 | 0.07472375 | -1.191763613 | 0.233353947 | 0.407702936 |
| ENSG00000110851 | PRDM4 | 1958.837187 | -0.404622797 | 0.056951039 | -7.104748289 | 1.21E-12 | 1.22E-11 |
| ENSG00000110871 | COQ5 | 936.5892336 | -0.171079902 | 0.072140619 | -2.37147814 | 0.017717095 | 0.049244366 |
| ENSG00000110876 | SELPLG | 2.562039618 | 0.541748714 | 1.148388602 | 0.471746857 | 0.637107487 | 0.786385275 |
| ENSG00000110880 | CORO1C | 2178.1954 | 0.207657161 | 0.055762252 | 3.72397372 | 0.000196111 | 0.00083359 |
| ENSG00000110881 | ASIC1 | 755.5404961 | 1.097344183 | 0.077998282 | 14.06882504 | 5.90E-45 | 2.86E-43 |
| ENSG00000110888 | CAPRIN2 | 196.5260814 | -0.424595356 | 0.140979616 | -3.01174999 | 0.002597464 | 0.008914696 |

| | | | | | | | |
|---|---|---|---|---|---|---|---|
| ENSG00000110906 | KCTD10 | 1840.924015 | -0.024466939 | 0.054938892 | -0.445348245 | 0.656068108 | 0.800627854 |
| ENSG00000110911 | SLC11A2 | 1816.141714 | 0.004565936 | 0.060638432 | 0.075297729 | 0.939977828 | 0.971354177 |
| ENSG00000110917 | MLEC | 9291.109062 | 0.065036056 | 0.044556545 | 1.459629686 | 0.144391876 | 0.283010051 |
| ENSG00000110921 | MVK | 2058.139784 | -0.949258387 | 0.055221118 | -17.19013336 | 3.15E-66 | 2.58E-64 |
| ENSG00000110925 | CSRNP2 | 1212.146026 | -0.028298451 | 0.066277923 | -0.426966478 | 0.669403744 | 0.810007484 |
| ENSG00000110931 | CAMKK2 | 2330.95006 | 0.333768518 | 0.059118788 | 5.64572666 | 1.64E-08 | 1.15E-07 |
| ENSG00000110934 | BIN2 | 4.188487103 | 2.326108965 | 1.016689313 | 2.287925067 | 0.022141885 | 0.059785175 |
| ENSG00000110944 | IL23A | 8.969238996 | -0.82393273 | 0.594834355 | -1.38514651 | 0.166007673 | 0.315041034 |
| ENSG00000110955 | ATP5F1B | 17451.45823 | 0.002319977 | 0.04476294 | 0.051828077 | 0.958665683 | 0.981163147 |
| ENSG00000110958 | PTGES3 | 7020.32389 | -0.112504239 | 0.059439462 | -1.892753314 | 0.058390683 | 0.136459073 |
| ENSG00000110975 | SYT10 | 140.2849929 | -0.10374928 | 0.154690807 | -0.670688081 | 0.502419258 | 0.683507467 |
| ENSG00000110987 | BCL7A | 1027.725078 | 0.301489295 | 0.073551048 | 4.099048232 | 4.15E-05 | 0.000196181 |
| ENSG00000111011 | RSRC2 | 2009.709174 | -0.373845966 | 0.066525426 | -5.61959523 | 1.91E-08 | 1.34E-07 |
| ENSG00000111012 | CYP27B1 | 187.2850028 | 0.117629397 | 0.142286851 | 0.826706026 | 0.408403705 | 0.600793959 |
| ENSG00000111052 | LIN7A | 1088.321452 | 0.710938256 | 0.066797903 | 10.64312235 | 1.88E-26 | 4.50E-25 |
| ENSG00000111057 | KRT18 | 43806.85288 | 0.229499858 | 0.042514172 | 5.398196606 | 6.73E-08 | 4.42E-07 |
| ENSG00000111058 | ACSS3 | 1867.846094 | 0.684527476 | 0.058347375 | 11.73193264 | 8.74E-32 | 2.62E-30 |
| ENSG00000111077 | TNS2 | 1392.129382 | 0.028150256 | 0.068575304 | 0.410501368 | 0.6814382 | 0.819131835 |
| ENSG00000111087 | GLI1 | 2.545045769 | -1.210574513 | 1.122718722 | -1.078252717 | 0.280920992 | 0.464389101 |
| ENSG00000111110 | PPM1H | 2638.139478 | 0.304928612 | 0.052365949 | 5.823032308 | 5.78E-09 | 4.27E-08 |
| ENSG00000111142 | METAP2 | 3298.279178 | -0.014578764 | 0.052581837 | -0.277258555 | 0.781581581 | 0.881489169 |
| ENSG00000111144 | LTA4H | 2897.371931 | -0.137255529 | 0.053448254 | -2.568007722 | 0.010228488 | 0.03031544 |
| ENSG00000111145 | ELK3 | 343.3193713 | 0.354514751 | 0.105403256 | 3.363413662 | 0.000769849 | 0.00294388 |
| ENSG00000111181 | SLC6A12 | 3.55682502 | 3.239120826 | 1.287237729 | 2.516334593 | 0.011858255 | 0.034569623 |
| ENSG00000111186 | WNT5B | 5.642885355 | -0.495675233 | 0.741190897 | -0.668755155 | 0.503651678 | 0.684667822 |
| ENSG00000111196 | MAGOHB | 204.2791307 | -0.312129904 | 0.129023997 | -2.41916164 | 0.015556326 | 0.043932305 |
| ENSG00000111199 | TRPV4 | 19.00183852 | -0.000390124 | 0.400875555 | -0.000973179 | 0.999223515 | 0.999528515 |
| ENSG00000111203 | ITFG2 | 506.1797924 | 0.112769588 | 0.088680749 | 1.271635496 | 0.203502659 | 0.368079397 |
| ENSG00000111206 | FOXM1 | 4380.979184 | -0.264074079 | 0.049539184 | -5.330610227 | 9.79E-08 | 6.35E-07 |
| ENSG00000111215 | PRR4 | 3.505251984 | -0.876606222 | 0.931106883 | -0.941466805 | 0.346465693 | 0.537239623 |
| ENSG00000111224 | PARP11 | 137.7773191 | 1.165423657 | 0.162089904 | 7.189983019 | 6.48E-13 | 6.73E-12 |
| ENSG00000111229 | ARPC3 | 2046.946424 | -0.188289444 | 0.056873693 | -3.310659726 | 0.000930763 | 0.003506027 |
| ENSG00000111231 | GPN3 | 315.0456803 | -0.209185381 | 0.105250909 | -1.98749239 | 0.046867858 | 0.1132928 |

| | | | | | | | |
|---|---|---|---|---|---|---|---|
| ENSG00000111237 | VPS29 | 985.6960517 | -0.239917161 | 0.070999429 | -3.379142132 | 0.000727124 | 0.002794008 |
| ENSG00000111247 | RAD51AP1 | 267.9862933 | -0.151623221 | 0.112745886 | -1.344822658 | 0.178682511 | 0.333358556 |
| ENSG00000111249 | CUX2 | 314.4296454 | 2.019010103 | 0.125110293 | 16.13784172 | 1.38E-58 | 9.47E-57 |
| ENSG00000111252 | SH2B3 | 212.0170981 | 0.159450598 | 0.127175218 | 1.253786709 | 0.209919541 | 0.377039953 |
| ENSG00000111254 | AKAP3 | 5.80633337 | -0.583782729 | 0.724492804 | -0.805781265 | 0.420368982 | 0.611649401 |
| ENSG00000111261 | MANSC1 | 743.837452 | 0.221514663 | 0.082119369 | 2.697471573 | 0.006986825 | 0.021659157 |
| ENSG00000111266 | DUSP16 | 2711.710488 | -0.476279629 | 0.055156147 | -8.635114266 | 5.87E-18 | 8.61E-17 |
| ENSG00000111269 | CREBL2 | 849.6813052 | -0.121386889 | 0.073449305 | -1.652662197 | 0.098399632 | 0.209182426 |
| ENSG00000111271 | ACAD10 | 912.9420515 | 0.279487945 | 0.072658811 | 3.846580215 | 0.000119778 | 0.000527087 |
| ENSG00000111275 | ALDH2 | 12.24400449 | 0.389967032 | 0.501788189 | 0.777154665 | 0.437067526 | 0.626723906 |
| ENSG00000111276 | CDKN1B | 1639.567302 | -0.217774226 | 0.059204207 | -3.678357261 | 0.000234741 | 0.000983734 |
| ENSG00000111300 | NAA25 | 1846.380284 | 0.103255478 | 0.060736373 | 1.700060014 | 0.089119638 | 0.192922949 |
| ENSG00000111319 | SCNN1A | 122.50714 | 1.348348962 | 0.174092496 | 7.745014823 | 9.56E-15 | 1.12E-13 |
| ENSG00000111321 | LTBR | 1762.475929 | 0.171641471 | 0.062950981 | 2.726589306 | 0.006399263 | 0.020054522 |
| ENSG00000111325 | OGFOD2 | 74.83013742 | -0.269802032 | 0.209012966 | -1.290838736 | 0.196759603 | 0.358540415 |
| ENSG00000111328 | CDK2AP1 | 3244.209391 | -0.247435812 | 0.050701232 | -4.880272144 | 1.06E-06 | 6.12E-06 |
| ENSG00000111331 | OAS3 | 856.8966471 | 1.201028544 | 0.073819814 | 16.2697313 | 1.62E-59 | 1.14E-57 |
| ENSG00000111335 | OAS2 | 16.87239607 | 3.456305769 | 0.638391218 | 5.414087273 | 6.16E-08 | 4.07E-07 |
| ENSG00000111341 | MGP | 65.40855636 | 3.259396102 | 0.305088314 | 10.68345115 | 1.22E-26 | 2.95E-25 |
| ENSG00000111344 | RASAL1 | 74.11447339 | 1.248703398 | 0.221176205 | 5.645740239 | 1.64E-08 | 1.15E-07 |
| ENSG00000111348 | ARHGDIB | 15.18111857 | 4.797006622 | 0.912514473 | 5.256910178 | 1.46E-07 | 9.37E-07 |
| ENSG00000111358 | GTF2H3 | 1173.147142 | -0.143638234 | 0.06962059 | -2.063157369 | 0.039097679 | 0.097341085 |
| ENSG00000111361 | EIF2B1 | 1745.331928 | -0.06043492 | 0.060283378 | -1.002513818 | 0.316095496 | 0.504854883 |
| ENSG00000111364 | DDX55 | 1111.306947 | -0.014656979 | 0.076437394 | -0.191751423 | 0.847936924 | 0.920424395 |
| ENSG00000111371 | SLC38A1 | 38180.20579 | -0.41124519 | 0.049608727 | -8.289775068 | 1.13E-16 | 1.54E-15 |
| ENSG00000111404 | RERGL | 14.80042256 | -2.804079387 | 0.572902609 | -4.894513209 | 9.85E-07 | 5.70E-06 |
| ENSG00000111412 | SPRING1 | 4244.442445 | 0.367939237 | 0.048370588 | 7.606672829 | 2.81E-14 | 3.20E-13 |
| ENSG00000111424 | VDR | 1313.996784 | -0.719040322 | 0.072980246 | -9.852533586 | 6.68E-23 | 1.32E-21 |
| ENSG00000111445 | RFC5 | 1193.489591 | -0.224677747 | 0.062486986 | -3.595592618 | 0.000323654 | 0.001323934 |
| ENSG00000111450 | STX2 | 152.6757315 | -1.073188991 | 0.150950407 | -7.109546847 | 1.16E-12 | 1.18E-11 |
| ENSG00000111452 | ADGRD1 | 10.90516835 | -1.158862568 | 0.546911059 | -2.118923266 | 0.034096952 | 0.08659184 |
| ENSG00000111481 | COPZ1 | 4150.455207 | -0.003414952 | 0.048144422 | -0.070931424 | 0.943452334 | 0.973031555 |
| ENSG00000111490 | TBC1D30 | 513.3990464 | -0.174894013 | 0.088928499 | -1.966681274 | 0.049219967 | 0.11819387 |

| | | | | | | | |
|---|---|---|---|---|---|---|---|
| ENSG00000111530 | CAND1 | 6751.125741 | -0.101663437 | 0.052972516 | -1.919173257 | 0.054962411 | 0.129929689 |
| ENSG00000111540 | RAB5B | 6853.102532 | 0.197925504 | 0.044516053 | 4.446160243 | 8.74E-06 | 4.54E-05 |
| ENSG00000111554 | MDM1 | 296.6550869 | -0.011990668 | 0.106546108 | -0.112539707 | 0.910395488 | 0.95506154 |
| ENSG00000111581 | NUP107 | 2097.524879 | -0.183024844 | 0.064465428 | -2.839116256 | 0.004523867 | 0.014730662 |
| ENSG00000111596 | CNOT2 | 2452.307995 | -0.04643839 | 0.051020464 | -0.910191441 | 0.362721557 | 0.555462785 |
| ENSG00000111602 | TIMELESS | 4209.772823 | -0.012513956 | 0.046979975 | -0.266367878 | 0.789955902 | 0.886574774 |
| ENSG00000111605 | CPSF6 | 3402.357067 | 0.009617337 | 0.053326801 | 0.180347165 | 0.85688003 | 0.925855018 |
| ENSG00000111615 | KRR1 | 2394.471952 | 0.425231304 | 0.057730503 | 7.365799396 | 1.76E-13 | 1.89E-12 |
| ENSG00000111639 | MRPL51 | 1807.213362 | -0.162947118 | 0.05498458 | -2.963505739 | 0.003041564 | 0.010291948 |
| ENSG00000111640 | GAPDH | 35614.52994 | -0.020536008 | 0.049918232 | -0.411392926 | 0.68078444 | 0.818841625 |
| ENSG00000111641 | NOP2 | 14.05389439 | 0.117309195 | 0.475009089 | 0.246962 | 0.804937633 | 0.896404708 |
| ENSG00000111642 | CHD4 | 455.7632151 | 0.226761476 | 0.095776759 | 2.367604397 | 0.017903668 | 0.049680676 |
| ENSG00000111644 | ACRBP | 20.34680687 | 0.183743884 | 0.407697267 | 0.450687064 | 0.652215107 | 0.798091138 |
| ENSG00000111647 | BLTP3B | 824.9600058 | -0.167600434 | 0.073791323 | -2.271275639 | 0.023130297 | 0.062066793 |
| ENSG00000111652 | COPS7A | 1243.165969 | 0.041203539 | 0.068695826 | 0.599796834 | 0.548641643 | 0.721458257 |
| ENSG00000111653 | ING4 | 455.4817659 | 0.354394749 | 0.091996077 | 3.852281088 | 0.000117023 | 0.00051615 |
| ENSG00000111664 | GNB3 | 4.280089562 | -0.83466602 | 0.851460347 | -0.980275855 | 0.32694997 | 0.517435827 |
| ENSG00000111665 | CDCA3 | 817.9823372 | -0.434581656 | 0.077342092 | -5.618953995 | 1.92E-08 | 1.34E-07 |
| ENSG00000111666 | CHPT1 | 594.529267 | -0.265487 | 0.088734831 | -2.991914197 | 0.002772342 | 0.009469553 |
| ENSG00000111667 | USP5 | 3342.591709 | -0.026472593 | 0.052046836 | -0.508630202 | 0.611011455 | 0.768826384 |
| ENSG00000111669 | TPI1 | 11056.58168 | 0.016904065 | 0.044424654 | 0.380510911 | 0.703566199 | 0.834098636 |
| ENSG00000111670 | GNPTAB | 3249.43221 | -0.195485908 | 0.052017298 | -3.758094237 | 0.000171212 | 0.000735095 |
| ENSG00000111671 | SPSB2 | 107.5383893 | -0.028666166 | 0.171180837 | -0.167461305 | 0.867007091 | 0.930846618 |
| ENSG00000111674 | ENO2 | 1748.068725 | -0.539536934 | 0.057116721 | -9.446216957 | 3.51E-21 | 6.29E-20 |
| ENSG00000111676 | ATN1 | 6681.98187 | -0.030048515 | 0.054654322 | -0.549792108 | 0.582461972 | 0.747046721 |
| ENSG00000111678 | C12orf57 | 1482.69257 | -0.12035575 | 0.069815499 | -1.723911618 | 0.0847238 | 0.185171871 |
| ENSG00000111679 | PTPN6 | 875.4739426 | 0.01184597 | 0.07681841 | 0.154207436 | 0.877446179 | 0.936345321 |
| ENSG00000111684 | LPCAT3 | 1259.588402 | -0.464071765 | 0.066717852 | -6.955736049 | 3.51E-12 | 3.43E-11 |
| ENSG00000111696 | NT5DC3 | 218.9958217 | 0.404018444 | 0.12791851 | 3.158404855 | 0.001586351 | 0.005703007 |
| ENSG00000111707 | SUDS3 | 2961.309718 | -0.106946196 | 0.05120571 | -2.088559955 | 0.036747353 | 0.092432487 |
| ENSG00000111711 | GOLT1B | 418.1942838 | -0.168700903 | 0.109203302 | -1.544833347 | 0.122386579 | 0.249161175 |
| ENSG00000111725 | PRKAB1 | 1267.2867 | -0.064132438 | 0.063328948 | -1.012687569 | 0.311209418 | 0.499310675 |
| ENSG00000111726 | CMAS | 780.8114761 | 0.085602021 | 0.073656853 | 1.162173214 | 0.245165113 | 0.42213373 |

| | | | | | | | |
|---|---|---|---|---|---|---|---|
| ENSG00000111727 | HCFC2 | 247.5505862 | -0.15425894 | 0.118711017 | -1.299449238 | 0.193789803 | 0.354483437 |
| ENSG00000111728 | ST8SIA1 | 93.14315252 | 1.845665192 | 0.208144016 | 8.867250781 | 7.50E-19 | 1.17E-17 |
| ENSG00000111731 | C2CD5 | 777.1994442 | -0.237630746 | 0.073834462 | -3.218425933 | 0.001288963 | 0.004728738 |
| ENSG00000111737 | RAB35 | 2523.557108 | -0.045578253 | 0.061440135 | -0.741831911 | 0.458189184 | 0.64527349 |
| ENSG00000111752 | PHC1 | 318.4185998 | -0.134593438 | 0.105071261 | -1.280972896 | 0.200203186 | 0.363372189 |
| ENSG00000111775 | COX6A1 | 965.2203696 | 0.002832646 | 0.069079912 | 0.041005348 | 0.967291632 | 0.985419222 |
| ENSG00000111785 | RIC8B | 387.731979 | 0.061693181 | 0.095787055 | 0.644065957 | 0.519532664 | 0.696979701 |
| ENSG00000111786 | SRSF9 | 5408.047544 | -0.161399402 | 0.04760249 | -3.39056638 | 0.000697484 | 0.002689122 |
| ENSG00000111788 | DDX12B | 162.3660497 | -0.354073836 | 0.157482752 | -2.248334066 | 0.024554896 | 0.065468307 |
| ENSG00000111790 | FGFR1OP2 | 426.0705104 | -0.372960181 | 0.093831342 | -3.974793227 | 7.04E-05 | 0.000322214 |
| ENSG00000111799 | COL12A1 | 3762.303134 | 2.366161584 | 0.055795449 | 42.40778829 | 0 | 0 |
| ENSG00000111801 | BTN3A3 | 5.057444832 | 0.286508062 | 0.784732126 | 0.365103012 | 0.715034513 | 0.841598433 |
| ENSG00000111802 | TDP2 | 1129.635948 | -0.090566391 | 0.067520113 | -1.341324645 | 0.179815072 | 0.334763654 |
| ENSG00000111816 | FRK | 26.05197936 | 0.678512605 | 0.361948161 | 1.874612662 | 0.06084603 | 0.141204646 |
| ENSG00000111817 | DSE | 312.9755359 | 0.046817313 | 0.1136174 | 0.412061115 | 0.680294629 | 0.818419487 |
| ENSG00000111832 | RWDD1 | 1019.384505 | 0.055049695 | 0.067418489 | 0.816537056 | 0.414193039 | 0.606432749 |
| ENSG00000111834 | RSPH4A | 11.24309919 | -0.138045793 | 0.548822612 | -0.251530805 | 0.801403749 | 0.894398618 |
| ENSG00000111843 | TMEM14C | 1338.330776 | -0.770256589 | 0.060754659 | -12.6781486 | 7.82E-37 | 2.83E-35 |
| ENSG00000111845 | PAK1IP1 | 462.0001945 | -0.578278613 | 0.09286178 | -6.22730488 | 4.75E-10 | 3.89E-09 |
| ENSG00000111846 | GCNT2 | 3.418221387 | -1.773456362 | 1.058401754 | -1.675598472 | 0.093816868 | 0.201192761 |
| ENSG00000111850 | SMIM8 | 119.7666721 | 0.193952675 | 0.168191603 | 1.153165033 | 0.248842653 | 0.42673431 |
| ENSG00000111859 | NEDD9 | 776.4290412 | 0.711738538 | 0.076451224 | 9.3097075 | 1.28E-20 | 2.21E-19 |
| ENSG00000111860 | CEP85L | 120.6600698 | 0.26576224 | 0.162000868 | 1.640498864 | 0.100901485 | 0.213373899 |
| ENSG00000111875 | ASF1A | 1043.243338 | 0.113737108 | 0.066041114 | 1.722216678 | 0.085030275 | 0.185770905 |
| ENSG00000111877 | MCM9 | 869.1378906 | 0.450616286 | 0.072105634 | 6.24939083 | 4.12E-10 | 3.40E-09 |
| ENSG00000111879 | FAM184A | 175.6650839 | 0.735196155 | 0.142855983 | 5.146414884 | 2.66E-07 | 1.65E-06 |
| ENSG00000111880 | RNGTT | 1087.280299 | 0.061894632 | 0.065493459 | 0.945050586 | 0.344633051 | 0.535542116 |
| ENSG00000111885 | MAN1A1 | 736.6049911 | -0.453555328 | 0.079838757 | -5.680891649 | 1.34E-08 | 9.50E-08 |
| ENSG00000111886 | GABRR2 | 32.88853792 | -0.363215858 | 0.321288572 | -1.130497282 | 0.258266743 | 0.437889067 |
| ENSG00000111897 | SERINC1 | 4685.774035 | 0.283581864 | 0.054700521 | 5.184262548 | 2.17E-07 | 1.36E-06 |
| ENSG00000111906 | HDDC2 | 2091.719062 | 0.465922048 | 0.054442626 | 8.558037675 | 1.15E-17 | 1.65E-16 |
| ENSG00000111907 | TPD52L1 | 1924.407601 | 0.803319004 | 0.060455081 | 13.28786576 | 2.72E-40 | 1.13E-38 |
| ENSG00000111911 | HINT3 | 1292.818629 | 0.573154758 | 0.063366867 | 9.045022753 | 1.50E-19 | 2.44E-18 |

| | | | | | | | |
|---|---|---|---|---|---|---|---|
| ENSG00000111912 | NCOA7 | 353.1185361 | 0.734176291 | 0.11446264 | 6.414112889 | 1.42E-10 | 1.21E-09 |
| ENSG00000111913 | RIPOR2 | 3.198958435 | -4.18550383 | 1.47083803 | -2.845659241 | 0.004431959 | 0.014468357 |
| ENSG00000111961 | SASH1 | 661.4427364 | -0.708501478 | 0.090544439 | -7.824903322 | 5.08E-15 | 6.12E-14 |
| ENSG00000111962 | UST | 257.1035663 | -0.31440855 | 0.115486725 | -2.722464848 | 0.006479693 | 0.020256766 |
| ENSG00000111981 | ULBP1 | 206.2511911 | 0.441595845 | 0.128685981 | 3.431576933 | 0.000600083 | 0.002340344 |
| ENSG00000112029 | FBXO5 | 686.7592145 | -0.07234748 | 0.079885015 | -0.905645186 | 0.365123684 | 0.557837983 |
| ENSG00000112031 | MTRF1L | 554.1788155 | 0.191310527 | 0.088401767 | 2.164102976 | 0.030456447 | 0.078714612 |
| ENSG00000112033 | PPARD | 1129.740581 | 0.650596873 | 0.07064021 | 9.210007643 | 3.26E-20 | 5.51E-19 |
| ENSG00000112039 | FANCE | 285.9282634 | 0.034004605 | 0.110380171 | 0.308068053 | 0.75803055 | 0.866644446 |
| ENSG00000112062 | MAPK14 | 2132.659357 | -0.079890329 | 0.053932656 | -1.481297893 | 0.13852721 | 0.274470046 |
| ENSG00000112077 | RHAG | 1.940832738 | 1.394229938 | 1.33156105 | 1.047064224 | 0.295069958 | 0.481018631 |
| ENSG00000112078 | KCTD20 | 3223.755093 | -0.142287711 | 0.048169887 | -2.953872647 | 0.003138135 | 0.010588147 |
| ENSG00000112079 | STK38 | 1969.519302 | 0.25572777 | 0.055503851 | 4.607387875 | 4.08E-06 | 2.20E-05 |
| ENSG00000112081 | SRSF3 | 6928.761616 | -0.175034056 | 0.046734212 | -3.745308847 | 0.000180172 | 0.000770534 |
| ENSG00000112110 | MRPL18 | 1082.736449 | -0.076988863 | 0.066434311 | -1.15887201 | 0.24650836 | 0.423702607 |
| ENSG00000112118 | MCM3 | 5151.557125 | -0.051824054 | 0.045053398 | -1.150280692 | 0.250028281 | 0.428097385 |
| ENSG00000112130 | RNF8 | 670.7754163 | 0.159130865 | 0.080655996 | 1.972957647 | 0.048500388 | 0.116746999 |
| ENSG00000112137 | PHACTR1 | 4.367670046 | 2.924132049 | 1.109011904 | 2.636700326 | 0.008371675 | 0.025416481 |
| ENSG00000112139 | MDGA1 | 47.31423708 | -0.15841464 | 0.262078935 | -0.604453924 | 0.545541894 | 0.718666765 |
| ENSG00000112144 | CILK1 | 919.0836973 | -0.001239333 | 0.073921792 | -0.016765468 | 0.986623718 | 0.993497695 |
| ENSG00000112146 | FBXO9 | 2387.795763 | 0.153144401 | 0.056037118 | 2.732910015 | 0.006277748 | 0.019719504 |
| ENSG00000112149 | CD83 | 56.26809414 | 0.1683103 | 0.23793037 | 0.707393092 | 0.479322229 | 0.663653377 |
| ENSG00000112159 | MDN1 | 1536.991671 | -0.076223831 | 0.082914465 | -0.919306789 | 0.357935128 | 0.550300371 |
| ENSG00000112167 | SAYSD1 | 617.1466903 | -0.347240435 | 0.078622034 | -4.416579125 | 1.00E-05 | 5.19E-05 |
| ENSG00000112182 | BACH2 | 18.60260355 | -1.537714407 | 0.452675656 | -3.396945225 | 0.000681426 | 0.002632965 |
| ENSG00000112183 | RBM24 | 19.81933683 | 1.130854445 | 0.442819183 | 2.553761191 | 0.010656632 | 0.031421997 |
| ENSG00000112186 | CAP2 | 601.5705475 | -0.741036071 | 0.084877085 | -8.73069654 | 2.53E-18 | 3.81E-17 |
| ENSG00000112200 | ZNF451 | 964.9955517 | -0.185475557 | 0.070481814 | -2.631537791 | 0.008499941 | 0.025771696 |
| ENSG00000112208 | BAG2 | 204.6662173 | -0.741592264 | 0.130066252 | -5.701650127 | 1.19E-08 | 8.45E-08 |
| ENSG00000112210 | RAB23 | 303.8628692 | -0.291595295 | 0.108826884 | -2.679441752 | 0.007374503 | 0.022744166 |
| ENSG00000112218 | GPR63 | 74.62366162 | 0.093914531 | 0.20632691 | 0.45517345 | 0.64898446 | 0.795495779 |
| ENSG00000112234 | FBXL4 | 753.8217324 | 0.104073178 | 0.079033187 | 1.316828816 | 0.187896012 | 0.346044838 |
| ENSG00000112237 | CCNC | 1411.172194 | 0.064290174 | 0.074480122 | 0.863185672 | 0.38803539 | 0.580736682 |

| | | | | | | | |
|---|---|---|---|---|---|---|---|
| ENSG00000112242 | E2F3 | 969.3412286 | -0.437604575 | 0.071617584 | -6.110295137 | 9.94E-10 | 7.90E-09 |
| ENSG00000112245 | PTP4A1 | 1.952587642 | -0.12139534 | 1.293107601 | -0.093878761 | 0.925205465 | 0.964342468 |
| ENSG00000112249 | ASCC3 | 2812.524088 | 0.050256698 | 0.056915885 | 0.8829995 | 0.377236542 | 0.570022519 |
| ENSG00000112276 | BVES | 10.81261615 | -1.144505701 | 0.546327692 | -2.094906988 | 0.036179262 | 0.091195771 |
| ENSG00000112282 | MED23 | 1477.469532 | 0.172182416 | 0.058622721 | 2.937127653 | 0.003312677 | 0.011122626 |
| ENSG00000112290 | WASF1 | 686.8219044 | -0.112765812 | 0.077029263 | -1.463934725 | 0.143211784 | 0.281297914 |
| ENSG00000112293 | GPLD1 | 105.2395026 | 0.980927376 | 0.178572123 | 5.493171958 | 3.95E-08 | 2.67E-07 |
| ENSG00000112294 | ALDH5A1 | 2217.783102 | 0.760429698 | 0.058317153 | 13.03955458 | 7.29E-39 | 2.82E-37 |
| ENSG00000112297 | CRYBG1 | 2162.019237 | -0.518572947 | 0.057378884 | -9.037696694 | 1.60E-19 | 2.61E-18 |
| ENSG00000112304 | ACOT13 | 591.4386006 | -0.03670963 | 0.083396389 | -0.440182493 | 0.659804938 | 0.80321925 |
| ENSG00000112305 | SMAP1 | 1455.980124 | 0.164213258 | 0.059461989 | 2.761650922 | 0.005750993 | 0.018224586 |
| ENSG00000112306 | RPS12 | 7013.433549 | 0.434756268 | 0.049579606 | 8.768852794 | 1.80E-18 | 2.75E-17 |
| ENSG00000112308 | C6orf62 | 3452.882049 | 0.054195535 | 0.058224333 | 0.930805595 | 0.351954135 | 0.543362801 |
| ENSG00000112309 | B3GAT2 | 17.88132801 | 0.118383638 | 0.442581601 | 0.267484319 | 0.789096286 | 0.885944625 |
| ENSG00000112312 | GMNN | 848.7864134 | -0.375216219 | 0.072699213 | -5.161214341 | 2.45E-07 | 1.53E-06 |
| ENSG00000112319 | EYA4 | 21.46062676 | -1.284265387 | 0.397420488 | -3.231502722 | 0.001231412 | 0.004533555 |
| ENSG00000112320 | SOBP | 102.2247007 | -0.923323107 | 0.181501191 | -5.087146259 | 3.63E-07 | 2.22E-06 |
| ENSG00000112335 | SNX3 | 2577.799892 | 0.051424345 | 0.05255225 | 0.978537446 | 0.32780858 | 0.518115532 |
| ENSG00000112339 | HBS1L | 2100.135544 | 0.380765152 | 0.055267184 | 6.889534191 | 5.60E-12 | 5.37E-11 |
| ENSG00000112357 | PEX7 | 406.7011139 | 0.436666477 | 0.097274385 | 4.489018139 | 7.16E-06 | 3.76E-05 |
| ENSG00000112365 | ZBTB24 | 555.2363842 | 0.090265696 | 0.085670317 | 1.05364028 | 0.29204764 | 0.476981097 |
| ENSG00000112367 | FIG4 | 684.6383979 | 0.398894028 | 0.078129498 | 5.105549603 | 3.30E-07 | 2.03E-06 |
| ENSG00000112378 | PERP | 5354.665354 | 0.600193974 | 0.049450478 | 12.13727342 | 6.70E-34 | 2.18E-32 |
| ENSG00000112379 | ARFGEF3 | 4711.752535 | -0.448567977 | 0.055694963 | -8.054013332 | 8.01E-16 | 1.02E-14 |
| ENSG00000112394 | SLC16A10 | 49.03379673 | -0.122458443 | 0.24840532 | -0.492978343 | 0.62202789 | 0.776353086 |
| ENSG00000112406 | HECA | 710.4490033 | 0.632311128 | 0.076986817 | 8.213239044 | 2.15E-16 | 2.86E-15 |
| ENSG00000112414 | ADGRG6 | 3834.777494 | -0.153989244 | 0.054544309 | -2.823195432 | 0.004754758 | 0.015403777 |
| ENSG00000112419 | PHACTR2 | 1214.315293 | 0.593138385 | 0.068546476 | 8.653083574 | 5.01E-18 | 7.40E-17 |
| ENSG00000112425 | EPM2A | 268.1026848 | 0.228588062 | 0.114072924 | 2.003876585 | 0.045083282 | 0.109602639 |
| ENSG00000112473 | SLC39A7 | 4133.023777 | -0.118864224 | 0.048258937 | -2.463051028 | 0.013776031 | 0.039443603 |
| ENSG00000112511 | PHF1 | 1150.90969 | -0.278978484 | 0.06934616 | -4.02298385 | 5.75E-05 | 0.000266638 |
| ENSG00000112514 | CUTA | 2436.540212 | 0.426471656 | 0.053120517 | 8.02837917 | 9.88E-16 | 1.25E-14 |
| ENSG00000112530 | PACRG | 57.31368466 | 0.114906768 | 0.241481928 | 0.47584003 | 0.634188349 | 0.784362944 |

| | | | | | | | |
|---|---|---|---|---|---|---|---|
| ENSG00000112531 | QKI | 1606.362425 | 0.441929962 | 0.062484445 | 7.072639601 | 1.52E-12 | 1.53E-11 |
| ENSG00000112541 | PDE10A | 234.5717687 | -3.504694674 | 0.166142488 | -21.09451181 | 8.93E-99 | 1.41E-96 |
| ENSG00000112561 | TFEB | 118.6539898 | -0.60405809 | 0.182058688 | -3.317930586 | 0.00090687 | 0.003422771 |
| ENSG00000112562 | SMOC2 | 18.62878822 | -2.091959057 | 0.454513399 | -4.602634511 | 4.17E-06 | 2.25E-05 |
| ENSG00000112576 | CCND3 | 1699.545026 | 0.195263556 | 0.067029398 | 2.913103234 | 0.003578563 | 0.011923345 |
| ENSG00000112578 | BYSL | 981.6818428 | 0.104477833 | 0.068173464 | 1.53252933 | 0.125391866 | 0.254126967 |
| ENSG00000112584 | FAM120B | 2536.060639 | 0.321272458 | 0.053292894 | 6.028429607 | 1.66E-09 | 1.29E-08 |
| ENSG00000112592 | TBP | 784.8653432 | 0.234177846 | 0.073066191 | 3.20500962 | 0.00135058 | 0.004931916 |
| ENSG00000112599 | GUCA1B | 17.04879044 | -0.116085433 | 0.421086735 | -0.275680574 | 0.782793418 | 0.882248256 |
| ENSG00000112624 | BICRAL | 981.5551403 | -0.252472243 | 0.076062539 | -3.319271828 | 0.000902525 | 0.003408054 |
| ENSG00000112640 | PPP2R5D | 2383.914137 | -0.226247475 | 0.051943576 | -4.355639167 | 1.33E-05 | 6.74E-05 |
| ENSG00000112651 | MRPL2 | 79.81005834 | 0.035624517 | 0.208519587 | 0.170844945 | 0.864345688 | 0.929719103 |
| ENSG00000112655 | PTK7 | 1307.579183 | 0.586951781 | 0.072233456 | 8.125760699 | 4.45E-16 | 5.77E-15 |
| ENSG00000112658 | SRF | 2415.439858 | -0.192651013 | 0.059381936 | -3.244269678 | 0.001177522 | 0.004354023 |
| ENSG00000112659 | CUL9 | 1922.680618 | 0.065569272 | 0.063782201 | 1.02801832 | 0.30394121 | 0.491257053 |
| ENSG00000112667 | DNPH1 | 737.2374038 | 0.39872686 | 0.083131252 | 4.796353371 | 1.62E-06 | 9.12E-06 |
| ENSG00000112679 | DUSP22 | 400.0957518 | -0.298430676 | 0.095337075 | -3.130268838 | 0.001746464 | 0.006214345 |
| ENSG00000112685 | EXOC2 | 933.3378021 | -0.540294022 | 0.069817787 | -7.738630041 | 1.00E-14 | 1.18E-13 |
| ENSG00000112695 | COX7A2 | 2131.635028 | 0.023472194 | 0.052621802 | 0.446054549 | 0.655557843 | 0.800136241 |
| ENSG00000112697 | TMEM30A | 8864.048839 | 0.53622231 | 0.052649858 | 10.18468664 | 2.32E-24 | 4.98E-23 |
| ENSG00000112699 | GMDS | 501.2019609 | -0.065958887 | 0.086904049 | -0.758985204 | 0.447861408 | 0.636196712 |
| ENSG00000112701 | SENP6 | 3111.552643 | 0.095428398 | 0.05311467 | 1.796648623 | 0.07239142 | 0.163195761 |
| ENSG00000112715 | VEGFA | 2031.074418 | 0.271799173 | 0.068155763 | 3.987911824 | 6.67E-05 | 0.000306069 |
| ENSG00000112739 | PRPF4B | 2526.983353 | -0.511747206 | 0.064807996 | -7.896359062 | 2.87E-15 | 3.52E-14 |
| ENSG00000112742 | TTK | 775.3771067 | -0.033451548 | 0.087548491 | -0.382091653 | 0.702393383 | 0.833224956 |
| ENSG00000112759 | SLC29A1 | 1274.970306 | 0.330984473 | 0.063623339 | 5.202249347 | 1.97E-07 | 1.24E-06 |
| ENSG00000112763 | BTN2A1 | 497.0533147 | 0.156116803 | 0.090878291 | 1.717866849 | 0.085820905 | 0.187088431 |
| ENSG00000112773 | TENT5A | 1306.754282 | -0.302130065 | 0.062539829 | -4.831002434 | 1.36E-06 | 7.73E-06 |
| ENSG00000112782 | CLIC5 | 5.668200212 | -0.243518425 | 0.72767714 | -0.334651746 | 0.737887806 | 0.855511284 |
| ENSG00000112787 | FBRSL1 | 2277.766884 | 0.033731148 | 0.059379404 | 0.56806142 | 0.569993263 | 0.737196 |
| ENSG00000112796 | ENPP5 | 197.5178029 | 0.257002947 | 0.129835549 | 1.979449764 | 0.04776539 | 0.115219564 |
| ENSG00000112812 | PRSS16 | 50.40547144 | 1.3917772 | 0.268505121 | 5.183428894 | 2.18E-07 | 1.37E-06 |
| ENSG00000112851 | ERBIN | 3294.441424 | -0.06423862 | 0.051339987 | -1.251239513 | 0.210847106 | 0.37826172 |

| | | | | | | | |
|---|---|---|---|---|---|---|---|
| ENSG00000112852 | PCDHB2 | 607.1715548 | 0.51790907 | 0.081191668 | 6.378845023 | 1.78E-10 | 1.51E-09 |
| ENSG00000112855 | HARS2 | 1843.607654 | 0.148796935 | 0.054447569 | 2.732848093 | 0.006278928 | 0.019720512 |
| ENSG00000112874 | NUDT12 | 460.2229788 | -0.230364318 | 0.097221064 | -2.369489791 | 0.017812647 | 0.049464002 |
| ENSG00000112877 | CEP72 | 355.0878771 | -0.043560757 | 0.101662843 | -0.428482571 | 0.66829982 | 0.809226597 |
| ENSG00000112893 | MAN2A1 | 1561.596274 | 0.217879202 | 0.06284285 | 3.467048391 | 0.000526207 | 0.002073732 |
| ENSG00000112902 | SEMA5A | 183.3435956 | 4.142698353 | 0.228325888 | 18.14379611 | 1.44E-73 | 1.45E-71 |
| ENSG00000112941 | TENT4A | 1677.477046 | -0.084875764 | 0.056763155 | -1.495261565 | 0.134846192 | 0.268825205 |
| ENSG00000112964 | GHR | 1464.34464 | 0.351022892 | 0.064553484 | 5.437706348 | 5.40E-08 | 3.59E-07 |
| ENSG00000112972 | HMGCS1 | 8047.64631 | -0.577806735 | 0.060204315 | -9.597430644 | 8.20E-22 | 1.52E-20 |
| ENSG00000112977 | DAP | 7871.963734 | 0.086489202 | 0.044976033 | 1.923006468 | 0.054479238 | 0.128946937 |
| ENSG00000112981 | NME5 | 6.787332655 | 1.820922064 | 0.771990019 | 2.358737832 | 0.018337205 | 0.050681384 |
| ENSG00000112983 | BRD8 | 3273.068205 | 0.267084583 | 0.057699163 | 4.62891606 | 3.68E-06 | 1.99E-05 |
| ENSG00000112984 | KIF20A | 2200.176465 | -0.015470381 | 0.055882712 | -0.27683662 | 0.781905561 | 0.881637432 |
| ENSG00000112992 | NNT | 3307.161892 | -0.064415084 | 0.051025988 | -1.262397595 | 0.206805754 | 0.372702019 |
| ENSG00000112996 | MRPS30 | 1365.925406 | 0.138176975 | 0.062520314 | 2.210113265 | 0.027097302 | 0.071187828 |
| ENSG00000113013 | HSPA9 | 13608.13598 | 0.10233385 | 0.047069909 | 2.174082143 | 0.029698967 | 0.076991106 |
| ENSG00000113048 | MRPS27 | 3054.755452 | 0.223313006 | 0.053047165 | 4.209706694 | 2.56E-05 | 0.000125044 |
| ENSG00000113068 | PFDN1 | 1118.339068 | -0.194364698 | 0.063679673 | -3.052225132 | 0.002271517 | 0.007901227 |
| ENSG00000113070 | HBEGF | 51.54655449 | 0.364620405 | 0.249839312 | 1.459419667 | 0.144449636 | 0.283099074 |
| ENSG00000113083 | LOX | 12.53461237 | -1.331643928 | 0.512765135 | -2.596986098 | 0.009404573 | 0.02819037 |
| ENSG00000113108 | APBB3 | 514.3658255 | -0.060202447 | 0.086794511 | -0.693620443 | 0.48792027 | 0.671365247 |
| ENSG00000113119 | TMCO6 | 285.7242322 | -0.028079117 | 0.112766725 | -0.249001792 | 0.803359395 | 0.89531017 |
| ENSG00000113140 | SPARC | 5.849852603 | 1.074472168 | 0.75206731 | 1.428691494 | 0.153092924 | 0.295942671 |
| ENSG00000113141 | IK | 2450.773517 | -0.092456411 | 0.052304616 | -1.767653019 | 0.077118929 | 0.171808121 |
| ENSG00000113161 | HMGCR | 6916.598595 | -0.648094106 | 0.053185263 | -12.1855955 | 3.71E-34 | 1.22E-32 |
| ENSG00000113163 | CERT1 | 1806.032978 | -0.18067826 | 0.065731343 | -2.748738289 | 0.005982513 | 0.018887814 |
| ENSG00000113194 | FAF2 | 4373.383146 | -0.406734567 | 0.049461494 | -8.223256769 | 1.98E-16 | 2.64E-15 |
| ENSG00000113205 | PCDHB3 | 252.7761493 | -0.192587504 | 0.124127548 | -1.551529102 | 0.120774939 | 0.246808329 |
| ENSG00000113209 | PCDHB5 | 8.702258663 | 0.689996708 | 0.617438419 | 1.117515021 | 0.263774181 | 0.444467438 |
| ENSG00000113211 | PCDHB6 | 5.515299901 | 1.18708673 | 0.815491478 | 1.455670307 | 0.145483782 | 0.284736623 |
| ENSG00000113212 | PCDHB7 | 3.839974756 | 0.540102717 | 0.960733888 | 0.562177231 | 0.573995273 | 0.740618232 |
| ENSG00000113240 | CLK4 | 487.8380709 | -0.729605665 | 0.094659207 | -7.70770948 | 1.28E-14 | 1.49E-13 |
| ENSG00000113248 | PCDHB15 | 2.218282872 | -0.13434663 | 1.18218428 | -0.113642714 | 0.909521026 | 0.954587195 |

| | | | | | | | |
|---|---|---|---|---|---|---|---|
| ENSG00000113269 | RNF130 | 2224.831177 | -0.289211951 | 0.051871387 | -5.575558503 | 2.47E-08 | 1.70E-07 |
| ENSG00000113272 | THG1L | 835.5276201 | 0.499824146 | 0.071155615 | 7.024380947 | 2.15E-12 | 2.14E-11 |
| ENSG00000113273 | ARSB | 153.8404581 | -0.576436072 | 0.147122947 | -3.918056863 | 8.93E-05 | 0.000401473 |
| ENSG00000113282 | CLINT1 | 3126.895385 | -0.06972715 | 0.051036259 | -1.366227698 | 0.171867506 | 0.323565088 |
| ENSG00000113296 | THBS4 | 4.664948917 | 0.157966058 | 0.814868417 | 0.193854683 | 0.846289668 | 0.919479255 |
| ENSG00000113300 | CNOT6 | 4098.427101 | -0.328065238 | 0.048659002 | -6.742128382 | 1.56E-11 | 1.44E-10 |
| ENSG00000113312 | TTC1 | 1537.343882 | -0.027852774 | 0.057198407 | -0.486950172 | 0.626293643 | 0.779556993 |
| ENSG00000113318 | MSH3 | 971.6563177 | 0.192350369 | 0.068928726 | 2.790569055 | 0.005261548 | 0.016841063 |
| ENSG00000113319 | RASGRF2 | 6.897121213 | 0.354064475 | 0.676266485 | 0.523557626 | 0.60058626 | 0.760636889 |
| ENSG00000113327 | GABRG2 | 72.52434553 | 0.477789599 | 0.207154445 | 2.306441457 | 0.021085978 | 0.057216978 |
| ENSG00000113328 | CCNG1 | 1888.911527 | -0.375306155 | 0.062896266 | -5.967065715 | 2.42E-09 | 1.86E-08 |
| ENSG00000113356 | POLR3G | 103.3859744 | 0.721138063 | 0.184155765 | 3.915913592 | 9.01E-05 | 0.000404629 |
| ENSG00000113360 | DROSHA | 3925.56594 | -0.146396195 | 0.055014318 | -2.661056254 | 0.007789594 | 0.023883091 |
| ENSG00000113368 | LMNB1 | 5266.557284 | -0.399978118 | 0.045312129 | -8.827175584 | 1.07E-18 | 1.66E-17 |
| ENSG00000113369 | ARRDC3 | 2008.528367 | 0.093431913 | 0.147057332 | 0.635343448 | 0.525204442 | 0.701327047 |
| ENSG00000113384 | GOLPH3 | 3844.827976 | -0.019930467 | 0.048780328 | -0.408575906 | 0.682850916 | 0.819995869 |
| ENSG00000113387 | SUB1 | 3287.860696 | 0.015776227 | 0.056443855 | 0.279502997 | 0.779858838 | 0.880534036 |
| ENSG00000113389 | NPR3 | 205.1997347 | 1.37050597 | 0.134774303 | 10.16889672 | 2.73E-24 | 5.82E-23 |
| ENSG00000113391 | FAM172A | 869.8002727 | 0.607269291 | 0.081352165 | 7.464697393 | 8.35E-14 | 9.14E-13 |
| ENSG00000113396 | SLC27A6 | 116.7515156 | 0.603372998 | 0.171808226 | 3.511898185 | 0.000444918 | 0.001777195 |
| ENSG00000113407 | TARS1 | 5753.457083 | -0.10222176 | 0.047326579 | -2.159922856 | 0.030778642 | 0.079430988 |
| ENSG00000113430 | IRX4 | 203.5365129 | 0.103417146 | 0.138423266 | 0.747108121 | 0.454998299 | 0.642475746 |
| ENSG00000113441 | LNPEP | 2088.642736 | 0.298219123 | 0.069084642 | 4.31672095 | 1.58E-05 | 7.95E-05 |
| ENSG00000113448 | PDE4D | 136.5024018 | -2.423213718 | 0.181117374 | -13.37924499 | 8.00E-41 | 3.38E-39 |
| ENSG00000113456 | RAD1 | 1435.323526 | -0.098702711 | 0.061603135 | -1.602235188 | 0.109103612 | 0.22720015 |
| ENSG00000113460 | BRIX1 | 1226.14578 | -0.065611128 | 0.064084231 | -1.023826407 | 0.305917275 | 0.493512116 |
| ENSG00000113492 | AGXT2 | 5.34766011 | -0.246290665 | 0.759133407 | -0.324436605 | 0.74560751 | 0.859379806 |
| ENSG00000113494 | PRLR | 6051.401151 | -0.61640822 | 0.063029033 | -9.779750585 | 1.38E-22 | 2.67E-21 |
| ENSG00000113504 | SLC12A7 | 2879.976108 | 0.25925593 | 0.053577747 | 4.838873301 | 1.31E-06 | 7.44E-06 |
| ENSG00000113522 | RAD50 | 157.1804214 | 0.109172923 | 0.149706665 | 0.729245579 | 0.465851455 | 0.651744797 |
| ENSG00000113532 | ST8SIA4 | 2225.753313 | 0.127853962 | 0.052486193 | 2.435954215 | 0.014852564 | 0.042110719 |
| ENSG00000113552 | GNPDA1 | 1582.537547 | 0.286173286 | 0.060403347 | 4.737705761 | 2.16E-06 | 1.20E-05 |
| ENSG00000113558 | SKP1 | 3558.831119 | 0.058662848 | 0.05216211 | 1.12462567 | 0.260747691 | 0.441084878 |

| | | | | | | | |
|---|---|---|---|---|---|---|---|
| ENSG00000113569 | NUP155 | 2899.560896 | -0.118907431 | 0.052825105 | -2.250964408 | 0.024387792 | 0.065091622 |
| ENSG00000113575 | PPP2CA | 3967.902848 | 0.011819409 | 0.055024243 | 0.214803671 | 0.829920391 | 0.909838651 |
| ENSG00000113580 | NR3C1 | 2977.106936 | -0.810820138 | 0.056728146 | -14.29308364 | 2.42E-46 | 1.20E-44 |
| ENSG00000113583 | C5orf15 | 1380.239234 | -0.024783685 | 0.061464545 | -0.4032192 | 0.686786976 | 0.823053664 |
| ENSG00000113593 | PPWD1 | 707.6394011 | -0.081327643 | 0.079038862 | -1.028957678 | 0.303499563 | 0.490679798 |
| ENSG00000113594 | LIFR | 1271.619308 | -0.278173412 | 0.074097177 | -3.754170202 | 0.000173917 | 0.000746147 |
| ENSG00000113595 | TRIM23 | 540.5814885 | 0.028633204 | 0.089788857 | 0.318894848 | 0.749806251 | 0.861664934 |
| ENSG00000113597 | TRAPPC13 | 503.7714577 | 0.02006805 | 0.09041392 | 0.22195753 | 0.824346945 | 0.907501628 |
| ENSG00000113615 | SEC24A | 1075.886246 | -0.257691528 | 0.06440797 | -4.000926116 | 6.31E-05 | 0.000290643 |
| ENSG00000113621 | TXNDC15 | 536.5174542 | -0.18131614 | 0.085500789 | -2.120637033 | 0.033952359 | 0.086310629 |
| ENSG00000113638 | TTC33 | 301.5162106 | -0.731110538 | 0.108645798 | -6.729303401 | 1.70E-11 | 1.57E-10 |
| ENSG00000113643 | RARS1 | 2446.926047 | -0.258300625 | 0.055784087 | -4.630363929 | 3.65E-06 | 1.98E-05 |
| ENSG00000113645 | WWC1 | 2157.93234 | -0.151328803 | 0.05647507 | -2.67956823 | 0.007371718 | 0.022738632 |
| ENSG00000113648 | MACROH2A1 | 8706.094769 | 0.026961909 | 0.039887666 | 0.675946027 | 0.499074914 | 0.680567986 |
| ENSG00000113649 | TCERG1 | 2703.716353 | 0.127727286 | 0.062036092 | 2.05891894 | 0.039502004 | 0.09816396 |
| ENSG00000113657 | DPYSL3 | 24.94179121 | -1.157340348 | 0.365236296 | -3.168744072 | 0.001530991 | 0.005516956 |
| ENSG00000113658 | SMAD5 | 1529.685971 | 0.09170838 | 0.067602787 | 1.356576925 | 0.174915678 | 0.328015068 |
| ENSG00000113712 | CSNK1A1 | 3772.953728 | -0.009983692 | 0.046604909 | -0.214219743 | 0.830375701 | 0.910033374 |
| ENSG00000113716 | HMGXB3 | 2790.735594 | 0.293909663 | 0.053777136 | 5.465327534 | 4.62E-08 | 3.10E-07 |
| ENSG00000113719 | ERGIC1 | 9593.781103 | -0.949777163 | 0.049231861 | -19.2919208 | 6.28E-83 | 7.46E-81 |
| ENSG00000113732 | ATP6V0E1 | 3025.005707 | -0.323151593 | 0.0505882 | -6.387884761 | 1.68E-10 | 1.43E-09 |
| ENSG00000113734 | BNIP1 | 381.306248 | -0.44168234 | 0.097000486 | -4.55340338 | 5.28E-06 | 2.81E-05 |
| ENSG00000113739 | STC2 | 9719.646817 | 0.682535776 | 0.058314301 | 11.704432 | 1.21E-31 | 3.59E-30 |
| ENSG00000113742 | CPEB4 | 550.8439588 | -0.111434994 | 0.083314708 | -1.337518882 | 0.181053323 | 0.336455523 |
| ENSG00000113749 | HRH2 | 19.44048353 | -0.529290215 | 0.417102509 | -1.268969145 | 0.204452069 | 0.369388868 |
| ENSG00000113758 | DBN1 | 6008.228414 | 0.044569364 | 0.055251581 | 0.806662245 | 0.419861102 | 0.61118178 |
| ENSG00000113761 | ZNF346 | 947.5452683 | -0.142763128 | 0.072586513 | -1.966799664 | 0.049206311 | 0.118173449 |
| ENSG00000113763 | UNC5A | 32.27615635 | -1.004495817 | 0.317411669 | -3.164646787 | 0.001552713 | 0.005587332 |
| ENSG00000113790 | EHHADH | 192.3324824 | -0.772854763 | 0.136652495 | -5.655621293 | 1.55E-08 | 1.09E-07 |
| ENSG00000113810 | SMC4 | 5711.311095 | -0.13648378 | 0.052710162 | -2.589325744 | 0.009616408 | 0.028746468 |
| ENSG00000113811 | SELENOK | 526.7547786 | 0.011921455 | 0.084193227 | 0.141596371 | 0.887398834 | 0.941998797 |
| ENSG00000113812 | ACTR8 | 1045.306726 | 0.017396189 | 0.067649105 | 0.257153284 | 0.797060445 | 0.891799572 |
| ENSG00000113838 | TBCCD1 | 318.9333306 | -0.297188248 | 0.106292576 | -2.79594548 | 0.005174813 | 0.016602826 |

| | | | | | | | |
|---|---|---|---|---|---|---|---|
| ENSG00000113845 | TIMMDC1 | 2037.717828 | -0.232562737 | 0.05291068 | -4.395383629 | 1.11E-05 | 5.68E-05 |
| ENSG00000113851 | CRBN | 1287.013598 | -0.004290783 | 0.065470699 | -0.065537456 | 0.947746085 | 0.975733561 |
| ENSG00000113905 | HRG | 2.16540533 | 3.395862665 | 1.524667611 | 2.227280649 | 0.025928523 | 0.06850658 |
| ENSG00000113916 | BCL6 | 774.9043575 | 0.301229298 | 0.086694954 | 3.474588613 | 0.000511637 | 0.002020131 |
| ENSG00000113924 | HGD | 34.59652988 | 0.009585761 | 0.31168801 | 0.030754348 | 0.975465448 | 0.988740606 |
| ENSG00000113966 | ARL6 | 83.00794904 | 0.59849047 | 0.196943896 | 3.038888144 | 0.00237453 | 0.008230843 |
| ENSG00000113971 | NPHP3 | 100.8874629 | -0.138452934 | 0.186619369 | -0.741900128 | 0.458147848 | 0.64527349 |
| ENSG00000114019 | AMOTL2 | 2033.739018 | -0.232781554 | 0.06567402 | -3.544499851 | 0.000393359 | 0.001583374 |
| ENSG00000114021 | NIT2 | 1696.103977 | 0.206833603 | 0.05780269 | 3.578269481 | 0.000345877 | 0.001404569 |
| ENSG00000114023 | FAM162A | 1289.133108 | -0.531052269 | 0.061259808 | -8.668852998 | 4.36E-18 | 6.47E-17 |
| ENSG00000114026 | OGG1 | 886.8576744 | 0.029906311 | 0.072217423 | 0.414114893 | 0.678789966 | 0.817269817 |
| ENSG00000114030 | KPNA1 | 2918.146669 | -0.173675298 | 0.052469352 | -3.310033201 | 0.000932849 | 0.003512731 |
| ENSG00000114054 | PCCB | 2564.087322 | 0.255111125 | 0.052410356 | 4.867570893 | 1.13E-06 | 6.50E-06 |
| ENSG00000114062 | UBE3A | 1869.267732 | 0.117972998 | 0.062776626 | 1.879250392 | 0.060210312 | 0.139941697 |
| ENSG00000114098 | ARMC8 | 1636.291238 | -0.20730574 | 0.057803115 | -3.586411244 | 0.00033526 | 0.001366779 |
| ENSG00000114107 | CEP70 | 578.3985747 | -0.051383315 | 0.085231952 | -0.602864468 | 0.546598855 | 0.719597001 |
| ENSG00000114120 | SLC25A36 | 2277.295335 | -0.439961429 | 0.078742598 | -5.587336967 | 2.31E-08 | 1.59E-07 |
| ENSG00000114125 | RNF7 | 1280.150764 | 0.014206413 | 0.061016199 | 0.232830194 | 0.815893265 | 0.902352305 |
| ENSG00000114126 | TFDP2 | 1447.890416 | 0.71927487 | 0.064062947 | 11.22762691 | 2.98E-29 | 8.03E-28 |
| ENSG00000114127 | XRN1 | 1200.100129 | -0.044053123 | 0.067767 | -0.650067481 | 0.515648633 | 0.6944375 |
| ENSG00000114166 | KAT2B | 794.067884 | -0.482329548 | 0.073489006 | -6.563288475 | 5.26E-11 | 4.65E-10 |
| ENSG00000114200 | BCHE | 8.779952027 | 5.489446577 | 1.28243585 | 4.280484345 | 1.86E-05 | 9.30E-05 |
| ENSG00000114209 | PDCD10 | 1562.860314 | -0.345516803 | 0.060491374 | -5.71183594 | 1.12E-08 | 7.99E-08 |
| ENSG00000114251 | WNT5A | 15.56220531 | -1.754052547 | 0.508213689 | -3.451407517 | 0.000557671 | 0.002186832 |
| ENSG00000114268 | PFKFB4 | 356.0331345 | -0.004659552 | 0.10151664 | -0.045899389 | 0.963390441 | 0.983294741 |
| ENSG00000114270 | COL7A1 | 628.7700828 | 0.011109434 | 0.08968199 | 0.123875864 | 0.901413564 | 0.950295365 |
| ENSG00000114279 | FGF12 | 567.8412374 | 0.061265606 | 0.083000267 | 0.738137457 | 0.460430924 | 0.647248857 |
| ENSG00000114302 | PRKAR2A | 2743.536231 | -0.10319564 | 0.053925292 | -1.913677906 | 0.055661324 | 0.131324773 |
| ENSG00000114315 | HES1 | 1143.165047 | 0.049536361 | 0.071481958 | 0.692991107 | 0.488315133 | 0.67162154 |
| ENSG00000114316 | USP4 | 1265.34676 | -0.051900204 | 0.064351893 | -0.806506261 | 0.419951 | 0.611273853 |
| ENSG00000114331 | ACAP2 | 1189.199885 | -0.202029099 | 0.067859588 | -2.977163656 | 0.002909287 | 0.009884336 |
| ENSG00000114346 | ECT2 | 3273.928075 | -0.442744358 | 0.054466578 | -8.128734605 | 4.34E-16 | 5.64E-15 |
| ENSG00000114349 | GNAT1 | 22.82288588 | 0.98385286 | 0.379104478 | 2.595202423 | 0.009453522 | 0.028322294 |

| | | | | | | | |
|---|---|---|---|---|---|---|---|
| ENSG00000114353 | GNAI2 | 5624.223578 | -0.092018668 | 0.04787343 | -1.922123969 | 0.054590161 | 0.129196151 |
| ENSG00000114354 | TFG | 3671.762225 | -0.035337562 | 0.047060202 | -0.7509012 | 0.452712118 | 0.6403907 |
| ENSG00000114378 | HYAL1 | 4.985689305 | 0.361447117 | 0.781966508 | 0.462228386 | 0.643917549 | 0.791314543 |
| ENSG00000114383 | TUSC2 | 719.2093757 | 0.196402638 | 0.083697528 | 2.346576342 | 0.018946784 | 0.052171316 |
| ENSG00000114388 | NPRL2 | 553.8020184 | -0.261081252 | 0.091929478 | -2.840016678 | 0.004511118 | 0.014699579 |
| ENSG00000114391 | RPL24 | 6325.79381 | 0.137423168 | 0.056517665 | 2.431508219 | 0.015036107 | 0.042599518 |
| ENSG00000114395 | CYB561D2 | 159.9632683 | 0.125142826 | 0.149028801 | 0.839722424 | 0.401064038 | 0.593886889 |
| ENSG00000114405 | C3orf14 | 3449.659187 | -0.220250872 | 0.056105414 | -3.92566161 | 8.65E-05 | 0.000389961 |
| ENSG00000114416 | FXR1 | 3386.11507 | 0.061813661 | 0.049360388 | 1.252292844 | 0.210463176 | 0.377779753 |
| ENSG00000114423 | CBLB | 808.8288555 | 0.268032613 | 0.072114754 | 3.716751414 | 0.000201801 | 0.000855221 |
| ENSG00000114439 | BBX | 1813.287383 | 0.396079566 | 0.069159111 | 5.72707714 | 1.02E-08 | 7.35E-08 |
| ENSG00000114446 | IFT57 | 1452.804928 | 0.686080984 | 0.061798179 | 11.10196123 | 1.23E-28 | 3.22E-27 |
| ENSG00000114450 | GNB4 | 11.17195669 | -0.517248273 | 0.520557596 | -0.993642733 | 0.320396832 | 0.509913303 |
| ENSG00000114473 | IQCG | 111.4786971 | -0.8688036 | 0.17245364 | -5.037896565 | 4.71E-07 | 2.85E-06 |
| ENSG00000114480 | GBE1 | 871.8808037 | -0.083280718 | 0.075692911 | -1.10024462 | 0.271225554 | 0.453228017 |
| ENSG00000114491 | UMPS | 2160.338998 | -0.023081746 | 0.053184789 | -0.433991502 | 0.664294612 | 0.806241836 |
| ENSG00000114503 | NCBP2 | 3185.477573 | -0.188720823 | 0.050497875 | -3.737203296 | 0.000186078 | 0.000794166 |
| ENSG00000114520 | SNX4 | 1527.960038 | 0.064256664 | 0.060926592 | 1.054657124 | 0.291582168 | 0.476552788 |
| ENSG00000114529 | C3orf52 | 117.3056219 | -0.11813639 | 0.171212286 | -0.689999489 | 0.490194509 | 0.67331669 |
| ENSG00000114541 | FRMD4B | 128.9321521 | -0.209386815 | 0.157154307 | -1.332364464 | 0.182740453 | 0.33865195 |
| ENSG00000114544 | SLC41A3 | 2305.549632 | 0.338850497 | 0.057850198 | 5.857378306 | 4.70E-09 | 3.51E-08 |
| ENSG00000114554 | PLXNA1 | 7307.081187 | -0.050932607 | 0.057276221 | -0.889245233 | 0.373871298 | 0.566636335 |
| ENSG00000114573 | ATP6V1A | 3777.287818 | -0.249493967 | 0.048204759 | -5.175712406 | 2.27E-07 | 1.42E-06 |
| ENSG00000114626 | ABTB1 | 643.2173652 | 0.187950517 | 0.080679944 | 2.32958164 | 0.019828273 | 0.05431159 |
| ENSG00000114631 | PODXL2 | 2756.634357 | 0.713979719 | 0.054762706 | 13.03769984 | 7.47E-39 | 2.89E-37 |
| ENSG00000114646 | CSPG5 | 84.31154938 | 0.800955489 | 0.19556644 | 4.095567149 | 4.21E-05 | 0.000198988 |
| ENSG00000114648 | KLHL18 | 680.0290308 | -0.013808557 | 0.07708634 | -0.179131055 | 0.857834798 | 0.926188651 |
| ENSG00000114650 | SCAP | 2514.875591 | 0.116029901 | 0.059215583 | 1.959448753 | 0.050060256 | 0.119960542 |
| ENSG00000114654 | EFCC1 | 107.1067661 | 0.846061657 | 0.177622452 | 4.763258514 | 1.90E-06 | 1.07E-05 |
| ENSG00000114656 | CFAP92 | 170.5202892 | 1.241878671 | 0.144111042 | 8.617512263 | 6.84E-18 | 1.00E-16 |
| ENSG00000114670 | NEK11 | 300.6976816 | -0.694358913 | 0.108172398 | -6.419002691 | 1.37E-10 | 1.17E-09 |
| ENSG00000114686 | MRPL3 | 3240.171986 | 0.115794444 | 0.051442251 | 2.250959899 | 0.024388077 | 0.065091622 |
| ENSG00000114698 | PLSCR4 | 30.25463719 | 1.417154712 | 0.340736325 | 4.159094901 | 3.20E-05 | 0.000154275 |

| | | | | | | | |
|---|---|---|---|---|---|---|---|
| ENSG00000114735 | HEMK1 | 719.5676882 | -0.452158284 | 0.089886559 | -5.030321451 | 4.90E-07 | 2.95E-06 |
| ENSG00000114737 | CISH | 205.2871483 | 0.82264505 | 0.133666439 | 6.15446224 | 7.53E-10 | 6.06E-09 |
| ENSG00000114738 | MAPKAPK3 | 1101.030064 | 0.515761946 | 0.067804967 | 7.606551039 | 2.82E-14 | 3.20E-13 |
| ENSG00000114739 | ACVR2B | 731.7801888 | -0.204484247 | 0.081901678 | -2.496703987 | 0.012535355 | 0.036267 |
| ENSG00000114742 | WDR48 | 1304.530634 | 0.200935993 | 0.063455998 | 3.166540587 | 0.001542638 | 0.005554563 |
| ENSG00000114744 | COMMD2 | 1290.065395 | 0.003210629 | 0.060272274 | 0.053268748 | 0.957517781 | 0.980775937 |
| ENSG00000114745 | GORASP1 | 1916.137814 | -0.008249972 | 0.069300181 | -0.119046899 | 0.9052382 | 0.952576345 |
| ENSG00000114767 | RRP9 | 647.8332303 | 0.134510295 | 0.079152322 | 1.69938533 | 0.089246604 | 0.193088475 |
| ENSG00000114770 | ABCC5 | 10841.85588 | -0.941411568 | 0.051741451 | -18.1945335 | 5.70E-74 | 5.81E-72 |
| ENSG00000114771 | AADAC | 1.962526787 | 2.301252531 | 1.513192844 | 1.520792634 | 0.128311883 | 0.258721509 |
| ENSG00000114779 | ABHD14B | 1006.550515 | 0.189844829 | 0.079365318 | 2.392037648 | 0.016755124 | 0.046877648 |
| ENSG00000114784 | EIF1B | 702.7329706 | -0.484747551 | 0.075936779 | -6.383567482 | 1.73E-10 | 1.47E-09 |
| ENSG00000114790 | ARHGEF26 | 1550.772551 | 0.207898025 | 0.057216895 | 3.633507669 | 0.000279594 | 0.001156698 |
| ENSG00000114796 | KLHL24 | 713.8848673 | 0.853831411 | 0.092800035 | 9.200766014 | 3.55E-20 | 5.99E-19 |
| ENSG00000114805 | PLCH1 | 991.2666328 | 0.642785596 | 0.076791811 | 8.37049666 | 5.74E-17 | 7.90E-16 |
| ENSG00000114812 | VIPR1 | 725.850258 | 0.208609529 | 0.080353022 | 2.596162842 | 0.009427137 | 0.028250625 |
| ENSG00000114841 | DNAH1 | 290.8354287 | 0.157866426 | 0.110177478 | 1.43283754 | 0.15190427 | 0.294264817 |
| ENSG00000114850 | SSR3 | 3492.420857 | -0.217012014 | 0.056371186 | -3.8496975 | 0.000118264 | 0.000521324 |
| ENSG00000114853 | ZBTB47 | 386.2876898 | 0.810355062 | 0.101548865 | 7.979951895 | 1.46E-15 | 1.83E-14 |
| ENSG00000114854 | TNNC1 | 10.73792347 | -0.070149928 | 0.536517589 | -0.130750473 | 0.895972704 | 0.947302844 |
| ENSG00000114857 | NKTR | 1687.1766 | -0.223834375 | 0.070572395 | -3.171698737 | 0.001515501 | 0.005464571 |
| ENSG00000114859 | CLCN2 | 594.5479943 | 0.217543709 | 0.086838845 | 2.505142818 | 0.012240196 | 0.03554305 |
| ENSG00000114861 | FOXP1 | 419.600622 | 0.201193854 | 0.097638625 | 2.060596962 | 0.039341507 | 0.097841952 |
| ENSG00000114867 | EIF4G1 | 25699.78318 | -0.026123935 | 0.041351668 | -0.631750455 | 0.527549953 | 0.703507931 |
| ENSG00000114902 | SPCS1 | 1260.772128 | -0.117724532 | 0.061376361 | -1.918076118 | 0.05510136 | 0.130177673 |
| ENSG00000114904 | NEK4 | 1169.775692 | -0.073655758 | 0.06343988 | -1.161032429 | 0.245628712 | 0.422701956 |
| ENSG00000114923 | SLC4A3 | 287.4505045 | 0.019391449 | 0.110612113 | 0.175310352 | 0.860835778 | 0.927524107 |
| ENSG00000114933 | INO80D | 431.288508 | -0.437216923 | 0.095923765 | -4.557962504 | 5.17E-06 | 2.76E-05 |
| ENSG00000114942 | EEF1B2 | 4194.685734 | 0.180486284 | 0.051296602 | 3.51848417 | 0.00043402 | 0.001737897 |
| ENSG00000114948 | ADAM23 | 18.08345526 | -0.161689505 | 0.415353137 | -0.389282011 | 0.697067542 | 0.829782025 |
| ENSG00000114956 | DGUOK | 1210.418183 | -0.265078984 | 0.061685425 | -4.297270929 | 1.73E-05 | 8.66E-05 |
| ENSG00000114978 | MOB1A | 4139.100246 | -0.006264782 | 0.053121884 | -0.117932228 | 0.906121357 | 0.952631711 |
| ENSG00000114982 | KANSL3 | 2361.632855 | 0.058523636 | 0.057830896 | 1.011978721 | 0.311548228 | 0.49971447 |

| | | | | | | | |
|---|---|---|---|---|---|---|---|
| ENSG00000114988 | LMAN2L | 933.2664633 | 0.22830323 | 0.072288758 | 3.158212085 | 0.0015874 | 0.005705886 |
| ENSG00000114993 | RTKN | 1450.10587 | 0.091050209 | 0.061527453 | 1.479830633 | 0.138918451 | 0.274881021 |
| ENSG00000114999 | TTL | 1576.088879 | -0.142037127 | 0.058102039 | -2.444615202 | 0.014500676 | 0.041245569 |
| ENSG00000115020 | PIKFYVE | 1366.388062 | -0.160438995 | 0.070901621 | -2.262839605 | 0.023645578 | 0.063308773 |
| ENSG00000115041 | KCNIP3 | 118.8025342 | 0.337716198 | 0.164028652 | 2.058885411 | 0.039505217 | 0.09816396 |
| ENSG00000115042 | FAHD2A | 610.9649739 | 0.137280883 | 0.089905037 | 1.5269543 | 0.126772376 | 0.256315732 |
| ENSG00000115053 | NCL | 28717.02333 | -0.038428585 | 0.048091111 | -0.799078752 | 0.424244749 | 0.615065066 |
| ENSG00000115073 | ACTR1B | 2862.735577 | 0.00335628 | 0.056625986 | 0.059271021 | 0.952736242 | 0.978056463 |
| ENSG00000115084 | SLC35F5 | 1682.731143 | 0.00394778 | 0.069034348 | 0.057185735 | 0.954397241 | 0.979088978 |
| ENSG00000115085 | ZAP70 | 6.235218603 | -1.398516105 | 0.743160839 | -1.881848493 | 0.05985659 | 0.139274791 |
| ENSG00000115091 | ACTR3 | 3517.407385 | -0.245024203 | 0.049873425 | -4.912921066 | 8.97E-07 | 5.23E-06 |
| ENSG00000115107 | STEAP3 | 969.7254499 | 0.63798133 | 0.075833075 | 8.412969262 | 4.00E-17 | 5.54E-16 |
| ENSG00000115109 | EPB41L5 | 2209.841706 | 0.30351143 | 0.060212817 | 5.040644886 | 4.64E-07 | 2.81E-06 |
| ENSG00000115128 | SF3B6 | 1799.148909 | -0.104509042 | 0.057344205 | -1.822486533 | 0.068381195 | 0.155652373 |
| ENSG00000115129 | TP53I3 | 685.9719535 | -0.908733082 | 0.081143876 | -11.19903467 | 4.12E-29 | 1.10E-27 |
| ENSG00000115137 | DNAJC27 | 182.1726308 | -0.008715171 | 0.137599239 | -0.063337349 | 0.949497875 | 0.976618218 |
| ENSG00000115145 | STAM2 | 1025.401139 | 0.049338976 | 0.065650996 | 0.751534312 | 0.452331159 | 0.640066851 |
| ENSG00000115159 | GPD2 | 1659.586403 | -0.362613991 | 0.059292308 | -6.115700419 | 9.61E-10 | 7.65E-09 |
| ENSG00000115163 | CENPA | 400.3397877 | -0.369686511 | 0.100290459 | -3.686158329 | 0.000227665 | 0.00095565 |
| ENSG00000115170 | ACVR1 | 601.7623499 | 0.4937885 | 0.084634525 | 5.834362492 | 5.40E-09 | 4.00E-08 |
| ENSG00000115183 | TANC1 | 1150.415082 | 0.410007537 | 0.066104389 | 6.202425349 | 5.56E-10 | 4.53E-09 |
| ENSG00000115194 | SLC30A3 | 148.3711277 | -0.120133815 | 0.150903976 | -0.796094432 | 0.42597715 | 0.616835995 |
| ENSG00000115204 | MPV17 | 1106.497214 | -0.024842309 | 0.071728016 | -0.346340395 | 0.729086914 | 0.849990029 |
| ENSG00000115207 | GTF3C2 | 3440.804545 | 0.113034208 | 0.0515682 | 2.191936299 | 0.028384107 | 0.074051109 |
| ENSG00000115211 | EIF2B4 | 1704.291548 | -0.029185892 | 0.057598591 | -0.506711908 | 0.612356974 | 0.769537956 |
| ENSG00000115216 | NRBP1 | 6274.634674 | 0.147937588 | 0.04576273 | 3.23270894 | 0.001226224 | 0.004515907 |
| ENSG00000115221 | ITGB6 | 271.2327818 | 0.831565392 | 0.116946125 | 7.110670782 | 1.15E-12 | 1.18E-11 |
| ENSG00000115226 | FNDC4 | 6.796744415 | -0.970796663 | 0.679419495 | -1.428861947 | 0.153043916 | 0.295872867 |
| ENSG00000115233 | PSMD14 | 2814.476881 | -0.06487625 | 0.05269513 | -1.231162174 | 0.218262215 | 0.387918951 |
| ENSG00000115234 | SNX17 | 2954.221347 | -0.419308086 | 0.052353258 | -8.009207123 | 1.15E-15 | 1.46E-14 |
| ENSG00000115239 | ASB3 | 411.5714292 | -0.122652218 | 0.104075541 | -1.178492252 | 0.23860042 | 0.414125265 |
| ENSG00000115241 | PPM1G | 5860.04521 | -0.050922265 | 0.042951946 | -1.185563632 | 0.235794672 | 0.410790661 |
| ENSG00000115252 | PDE1A | 229.5454561 | 0.344480993 | 0.135159251 | 2.548704515 | 0.010812387 | 0.031815823 |

| | | | | | | | |
|---|---|---|---|---|---|---|---|
| ENSG00000115255 | REEP6 | 1330.983602 | 0.574482739 | 0.070619914 | 8.134854707 | 4.12E-16 | 5.37E-15 |
| ENSG00000115257 | PCSK4 | 99.35599995 | 0.193057441 | 0.185672293 | 1.039775176 | 0.298444365 | 0.485007829 |
| ENSG00000115266 | APC2 | 35.02225727 | 0.402722354 | 0.309783399 | 1.300012705 | 0.193596615 | 0.354353016 |
| ENSG00000115267 | IFIH1 | 89.63243239 | 0.807105461 | 0.19972442 | 4.041095537 | 5.32E-05 | 0.000247859 |
| ENSG00000115268 | RPS15 | 4175.721929 | 0.107516005 | 0.046832852 | 2.295739004 | 0.021690809 | 0.058663895 |
| ENSG00000115271 | GCA | 496.0699702 | -0.047888529 | 0.085810449 | -0.558073398 | 0.576794265 | 0.742934331 |
| ENSG00000115274 | INO80B | 20.48273161 | -0.133545768 | 0.388186002 | -0.344025204 | 0.730827332 | 0.850978918 |
| ENSG00000115275 | MOGS | 2093.835405 | 0.156414459 | 0.062475928 | 2.503595608 | 0.012293846 | 0.035679083 |
| ENSG00000115282 | TTC31 | 1047.447789 | 0.027828472 | 0.069159842 | 0.402379054 | 0.687405083 | 0.823383061 |
| ENSG00000115286 | NDUFS7 | 581.4229178 | 0.023735505 | 0.086554175 | 0.274227158 | 0.783910059 | 0.882792721 |
| ENSG00000115289 | PCGF1 | 384.2103103 | -0.080770404 | 0.09986424 | -0.808802068 | 0.418629007 | 0.61039531 |
| ENSG00000115290 | GRB14 | 265.9691454 | 0.500235752 | 0.113974724 | 4.389006051 | 1.14E-05 | 5.83E-05 |
| ENSG00000115295 | CLIP4 | 77.35479599 | 3.160233573 | 0.274612443 | 11.50797663 | 1.20E-30 | 3.42E-29 |
| ENSG00000115306 | SPTBN1 | 4985.841071 | -0.049269561 | 0.051582283 | -0.955164415 | 0.339494558 | 0.530665194 |
| ENSG00000115307 | AUP1 | 3799.781732 | -0.043267837 | 0.048356238 | -0.894772595 | 0.370908663 | 0.563598379 |
| ENSG00000115310 | RTN4 | 2781.971596 | 0.078689824 | 0.048548983 | 1.620833606 | 0.105053329 | 0.220567762 |
| ENSG00000115317 | HTRA2 | 773.4187489 | -0.147586272 | 0.077583589 | -1.902287248 | 0.057133612 | 0.134026491 |
| ENSG00000115318 | LOXL3 | 212.2631287 | -0.496247102 | 0.125066708 | -3.967859314 | 7.25E-05 | 0.000331072 |
| ENSG00000115325 | DOK1 | 329.6103709 | -0.077510996 | 0.103163047 | -0.751344573 | 0.452445311 | 0.640092225 |
| ENSG00000115339 | GALNT3 | 1376.844317 | 0.10926877 | 0.059685025 | 1.830756887 | 0.067136838 | 0.153367762 |
| ENSG00000115350 | POLE4 | 766.1550011 | -0.195448853 | 0.07393478 | -2.643530612 | 0.008204635 | 0.024975361 |
| ENSG00000115355 | CCDC88A | 4.882835702 | -0.917616892 | 0.815860004 | -1.124723467 | 0.260706234 | 0.4410798 |
| ENSG00000115361 | ACADL | 21.06559128 | -1.118705704 | 0.402521766 | -2.779242757 | 0.005448579 | 0.017371842 |
| ENSG00000115363 | EVA1A | 2.918417725 | -1.146735723 | 1.066360761 | -1.075373143 | 0.282207701 | 0.465675823 |
| ENSG00000115364 | MRPL19 | 1522.999572 | -0.003496661 | 0.061877295 | -0.056509602 | 0.954935846 | 0.979311919 |
| ENSG00000115365 | LANCL1 | 1947.991747 | 0.338636007 | 0.058014908 | 5.837051507 | 5.31E-09 | 3.94E-08 |
| ENSG00000115368 | WDR75 | 1296.050812 | 0.022938904 | 0.070379229 | 0.325932863 | 0.744475151 | 0.858721841 |
| ENSG00000115380 | EFEMP1 | 1049.761395 | 0.013216179 | 0.069145399 | 0.191136063 | 0.848418994 | 0.920714588 |
| ENSG00000115386 | REG1A | 1.888297808 | 0.902986038 | 1.391011653 | 0.649157781 | 0.516236397 | 0.694776354 |
| ENSG00000115392 | FANCL | 398.2144196 | 0.034853977 | 0.093855652 | 0.371357247 | 0.710371464 | 0.838742158 |
| ENSG00000115414 | FN1 | 1997.551471 | 0.795599796 | 0.056061342 | 14.19159391 | 1.03E-45 | 5.08E-44 |
| ENSG00000115415 | STAT1 | 3566.062756 | 0.368207376 | 0.047354728 | 7.775514472 | 7.51E-15 | 8.90E-14 |
| ENSG00000115419 | GLS | 765.1889701 | 0.097451513 | 0.081804174 | 1.191278 | 0.233544467 | 0.408004728 |

| | | | | | | | |
|---|---|---|---|---|---|---|---|
| ENSG00000115421 | PAPOLG | 433.4007549 | -0.062531997 | 0.097900435 | -0.638730533 | 0.522998246 | 0.699689728 |
| ENSG00000115423 | DNAH6 | 66.2631318 | 0.776931751 | 0.238266426 | 3.260768901 | 0.001111106 | 0.00412573 |
| ENSG00000115425 | PECR | 234.6454796 | 0.199367 | 0.129345598 | 1.541351256 | 0.123231321 | 0.250547416 |
| ENSG00000115446 | UNC50 | 557.2033156 | 0.000728778 | 0.084033328 | 0.008672491 | 0.99308044 | 0.996294293 |
| ENSG00000115457 | IGFBP2 | 3025.420748 | -0.520166855 | 0.05627925 | -9.242604625 | 2.41E-20 | 4.10E-19 |
| ENSG00000115459 | ELMOD3 | 439.5347065 | -0.144146875 | 0.091391526 | -1.57724551 | 0.114739045 | 0.236892601 |
| ENSG00000115461 | IGFBP5 | 2643.079211 | -0.59386882 | 0.050877863 | -11.67244027 | 1.76E-31 | 5.23E-30 |
| ENSG00000115464 | USP34 | 4471.099821 | -0.130109649 | 0.051301507 | -2.536175959 | 0.011207041 | 0.032845836 |
| ENSG00000115468 | EFHD1 | 399.7898691 | -0.733635007 | 0.096474356 | -7.604456143 | 2.86E-14 | 3.25E-13 |
| ENSG00000115474 | KCNJ13 | 23.27781265 | -2.404517907 | 0.437353733 | -5.497879007 | 3.84E-08 | 2.60E-07 |
| ENSG00000115484 | CCT4 | 6373.432434 | -0.052220729 | 0.045364729 | -1.151130628 | 0.249678499 | 0.427721961 |
| ENSG00000115486 | GGCX | 1563.819469 | 0.103255929 | 0.06032992 | 1.71152106 | 0.086984967 | 0.189176634 |
| ENSG00000115504 | EHBP1 | 1026.995692 | 0.109267178 | 0.06923313 | 1.578249868 | 0.114508218 | 0.236501172 |
| ENSG00000115507 | OTX1 | 101.3446734 | 0.172463404 | 0.17959533 | 0.960288915 | 0.336909828 | 0.528123295 |
| ENSG00000115514 | TXNDC9 | 714.1824356 | -0.040207952 | 0.081893111 | -0.49098089 | 0.623439964 | 0.777266998 |
| ENSG00000115520 | COQ10B | 700.1711071 | 0.186135087 | 0.075315563 | 2.471402709 | 0.013458417 | 0.038616145 |
| ENSG00000115524 | SF3B1 | 11156.77298 | -0.070612483 | 0.042859151 | -1.64754741 | 0.09944558 | 0.211034376 |
| ENSG00000115525 | ST3GAL5 | 1161.193083 | -1.738168009 | 0.072479548 | -23.98149633 | 4.34E-127 | 1.20E-124 |
| ENSG00000115526 | CHST10 | 519.3315013 | 0.212599102 | 0.089011195 | 2.388453505 | 0.016919449 | 0.047275537 |
| ENSG00000115539 | PDCL3 | 765.4193359 | -0.002764659 | 0.075992474 | -0.036380693 | 0.970978809 | 0.98680905 |
| ENSG00000115540 | MOB4 | 405.737969 | 0.26718554 | 0.108252905 | 2.468160457 | 0.013580943 | 0.038915338 |
| ENSG00000115541 | HSPE1 | 306.3047404 | 0.193946476 | 0.108905255 | 1.780873433 | 0.07493313 | 0.167758045 |
| ENSG00000115548 | KDM3A | 1327.27278 | 0.431684021 | 0.062487331 | 6.908344716 | 4.90E-12 | 4.73E-11 |
| ENSG00000115556 | PLCD4 | 17.03966096 | 1.272338053 | 0.467221762 | 2.72319947 | 0.006465301 | 0.020222798 |
| ENSG00000115561 | CHMP3 | 2039.341149 | 0.199585398 | 0.054158698 | 3.685195639 | 0.000228527 | 0.000959094 |
| ENSG00000115568 | ZNF142 | 1442.154044 | 0.09909978 | 0.068137969 | 1.454398804 | 0.145835771 | 0.285279492 |
| ENSG00000115590 | IL1R2 | 3.235320366 | 0.116485228 | 0.97935498 | 0.118940763 | 0.905322287 | 0.95262113 |
| ENSG00000115594 | IL1R1 | 2595.002058 | 0.174926073 | 0.06759913 | 2.587697096 | 0.00966199 | 0.028860164 |
| ENSG00000115596 | WNT6 | 6.666291345 | -2.620770419 | 0.839466398 | -3.121947973 | 0.001796587 | 0.006375882 |
| ENSG00000115616 | SLC9A2 | 935.0282352 | 0.753696646 | 0.073406912 | 10.26737979 | 9.89E-25 | 2.17E-23 |
| ENSG00000115641 | FHL2 | 40.22349213 | 1.436929309 | 0.299508943 | 4.797617376 | 1.61E-06 | 9.06E-06 |
| ENSG00000115648 | MLPH | 3595.309259 | -0.735294844 | 0.057198546 | -12.85513173 | 8.05E-38 | 3.02E-36 |
| ENSG00000115649 | CNPPD1 | 2637.245941 | -0.236459166 | 0.053364342 | -4.431033067 | 9.38E-06 | 4.86E-05 |

| | | | | | | | |
|---|---|---|---|---|---|---|---|
| ENSG00000115652 | UXS1 | 990.9305342 | 0.283231512 | 0.069985745 | 4.046988608 | 5.19E-05 | 0.000242195 |
| ENSG00000115657 | ABCB6 | 45.30916196 | -0.044938485 | 0.266308732 | -0.16874582 | 0.865996577 | 0.930359269 |
| ENSG00000115661 | STK16 | 969.0982346 | 0.161993438 | 0.072809456 | 2.224895596 | 0.026088248 | 0.068860247 |
| ENSG00000115677 | HDLBP | 19233.6204 | -0.469224226 | 0.041425554 | -11.32692709 | 9.65E-30 | 2.66E-28 |
| ENSG00000115685 | PPP1R7 | 2235.143154 | -0.690693258 | 0.054617685 | -12.6459636 | 1.18E-36 | 4.24E-35 |
| ENSG00000115687 | PASK | 995.9189715 | -1.332055588 | 0.07371713 | -18.06982439 | 5.51E-73 | 5.45E-71 |
| ENSG00000115694 | STK25 | 3433.535263 | -0.695676052 | 0.054728344 | -12.71143978 | 5.11E-37 | 1.87E-35 |
| ENSG00000115705 | TPO | 3.168234497 | 0.248923008 | 0.970704779 | 0.256435338 | 0.797614704 | 0.891941177 |
| ENSG00000115718 | PROC | 2.964307406 | -1.133805349 | 1.047117896 | -1.082786717 | 0.278903103 | 0.461918791 |
| ENSG00000115738 | ID2 | 2197.140775 | -1.254167512 | 0.056680343 | -22.12702758 | 1.74E-108 | 3.40E-106 |
| ENSG00000115750 | TAF1B | 692.3259964 | 0.140434398 | 0.0836014 | 1.679809165 | 0.092994451 | 0.199733036 |
| ENSG00000115756 | HPCAL1 | 1107.058114 | 0.207668069 | 0.064736229 | 3.207911112 | 0.001337028 | 0.004887098 |
| ENSG00000115758 | ODC1 | 4248.275074 | -0.09964807 | 0.051919153 | -1.919293076 | 0.054947254 | 0.129907245 |
| ENSG00000115760 | BIRC6 | 2975.88737 | 0.024376685 | 0.118592647 | 0.205549716 | 0.837142659 | 0.914148686 |
| ENSG00000115761 | NOL10 | 1526.732585 | 0.132735155 | 0.058981345 | 2.250459948 | 0.024419763 | 0.06516861 |
| ENSG00000115762 | PLEKHB2 | 3527.653324 | 0.144154898 | 0.052582022 | 2.74152442 | 0.006115481 | 0.019281081 |
| ENSG00000115806 | GORASP2 | 3905.015115 | -0.023308163 | 0.04847664 | -0.480812261 | 0.630649935 | 0.78221733 |
| ENSG00000115808 | STRN | 1341.715571 | -0.059278973 | 0.067050279 | -0.884097331 | 0.376643674 | 0.569446113 |
| ENSG00000115816 | CEBPZ | 1589.322696 | 0.016152784 | 0.059506461 | 0.271445879 | 0.78604812 | 0.883788848 |
| ENSG00000115825 | PRKD3 | 466.3662342 | -0.021963414 | 0.088603064 | -0.247885494 | 0.804223004 | 0.895880917 |
| ENSG00000115827 | DCAF17 | 558.9170524 | 0.040344045 | 0.087189003 | 0.462719422 | 0.643565495 | 0.791178589 |
| ENSG00000115839 | RAB3GAP1 | 2121.126169 | 0.053002285 | 0.05434183 | 0.975349653 | 0.329386848 | 0.519786359 |
| ENSG00000115840 | SLC25A12 | 306.4425305 | 0.088612597 | 0.107139089 | 0.827079989 | 0.408191725 | 0.600670309 |
| ENSG00000115841 | RMDN2 | 32.27883392 | 0.440396226 | 0.314450559 | 1.400526136 | 0.161355823 | 0.308253045 |
| ENSG00000115844 | DLX2 | 157.8696046 | -1.102468548 | 0.151577535 | -7.273297774 | 3.51E-13 | 3.68E-12 |
| ENSG00000115850 | LCT | 2.362529196 | 1.873090063 | 1.277204108 | 1.466554994 | 0.142497151 | 0.280182107 |
| ENSG00000115866 | DARS1 | 2273.205737 | -0.020578672 | 0.05652583 | -0.364057848 | 0.715814812 | 0.841891334 |
| ENSG00000115875 | SRSF7 | 3896.778001 | -0.173901088 | 0.058330706 | -2.981295763 | 0.002870314 | 0.009772189 |
| ENSG00000115884 | SDC1 | 1869.119117 | 1.609684132 | 0.070551457 | 22.81574621 | 3.20E-115 | 6.80E-113 |
| ENSG00000115896 | PLCL1 | 15.90018493 | 1.218969679 | 0.456783048 | 2.668596575 | 0.007616888 | 0.02339648 |
| ENSG00000115902 | SLC1A4 | 1163.730104 | -0.203566718 | 0.063460183 | -3.207786506 | 0.001337608 | 0.004888437 |
| ENSG00000115904 | SOS1 | 1843.305975 | -0.043760925 | 0.0708107 | -0.617998757 | 0.536576158 | 0.711682859 |
| ENSG00000115919 | KYNU | 2596.281321 | 0.887858488 | 0.055867509 | 15.89221529 | 7.17E-57 | 4.73E-55 |

| | | | | | | | |
|---|---|---|---|---|---|---|---|
| ENSG00000115935 | WIPF1 | 245.5422377 | 1.390470446 | 0.124735009 | 11.14739526 | 7.37E-29 | 1.96E-27 |
| ENSG00000115942 | ORC2 | 1670.982938 | 0.111981914 | 0.058298507 | 1.920836731 | 0.054752294 | 0.129486353 |
| ENSG00000115944 | COX7A2L | 2875.150402 | 0.196819032 | 0.050263164 | 3.915770816 | 9.01E-05 | 0.000404789 |
| ENSG00000115946 | PNO1 | 741.7128752 | 0.032819909 | 0.077700349 | 0.422390755 | 0.672739825 | 0.812672542 |
| ENSG00000115947 | ORC4 | 944.5887302 | 0.130299694 | 0.072201551 | 1.804666147 | 0.07112694 | 0.160753893 |
| ENSG00000115963 | RND3 | 1129.997983 | 1.803875133 | 0.073019793 | 24.70392002 | 9.71E-135 | 2.93E-132 |
| ENSG00000115966 | ATF2 | 1024.288776 | -0.394923255 | 0.066773207 | -5.914397009 | 3.33E-09 | 2.53E-08 |
| ENSG00000115970 | THADA | 1287.381601 | -0.326969823 | 0.064876067 | -5.039914421 | 4.66E-07 | 2.82E-06 |
| ENSG00000115977 | AAK1 | 589.2007434 | 0.064906132 | 0.082904569 | 0.782901746 | 0.433684812 | 0.623549332 |
| ENSG00000115993 | TRAK2 | 2764.547725 | 0.227185029 | 0.049029184 | 4.63366935 | 3.59E-06 | 1.96E-05 |
| ENSG00000115998 | C2orf42 | 252.6893807 | 0.013582641 | 0.12079635 | 0.112442483 | 0.910472572 | 0.95506154 |
| ENSG00000116001 | TIA1 | 1609.17341 | -0.046125047 | 0.067357827 | -0.684776346 | 0.493485064 | 0.67625731 |
| ENSG00000116005 | PCYOX1 | 1985.900495 | 0.155932153 | 0.057467797 | 2.713383144 | 0.006660005 | 0.020761044 |
| ENSG00000116014 | KISS1R | 65.67442864 | 0.650951209 | 0.225464058 | 2.887161766 | 0.003887343 | 0.012843965 |
| ENSG00000116016 | EPAS1 | 466.2184204 | 1.807035465 | 0.108049677 | 16.72411729 | 8.75E-63 | 6.69E-61 |
| ENSG00000116017 | ARID3A | 1356.681921 | -0.06107506 | 0.062751785 | -0.973280046 | 0.330414137 | 0.521084855 |
| ENSG00000116030 | SUMO1 | 2277.837089 | -0.156126361 | 0.054329103 | -2.873715049 | 0.004056748 | 0.013349849 |
| ENSG00000116031 | CD207 | 12.98078362 | 0.238626885 | 0.494353813 | 0.48270465 | 0.629305461 | 0.7810577 |
| ENSG00000116032 | GRIN3B | 92.01721715 | 0.766302174 | 0.190835378 | 4.015514219 | 5.93E-05 | 0.000274504 |
| ENSG00000116035 | VAX2 | 127.7528774 | 0.08786863 | 0.15941883 | 0.551181002 | 0.581509602 | 0.74646393 |
| ENSG00000116039 | ATP6V1B1 | 10.03791379 | 0.237270388 | 0.547856148 | 0.433088848 | 0.664950225 | 0.806706252 |
| ENSG00000116044 | NFE2L2 | 2330.177269 | -0.047481211 | 0.051704696 | -0.918315241 | 0.35845385 | 0.550913261 |
| ENSG00000116062 | MSH6 | 3572.112896 | 0.127105756 | 0.047036575 | 2.702274892 | 0.00688668 | 0.021386278 |
| ENSG00000116095 | PLEKHA3 | 493.1242846 | 0.341239692 | 0.087757066 | 3.888458305 | 0.000100883 | 0.000449545 |
| ENSG00000116096 | SPR | 1479.680193 | 0.331446943 | 0.064022994 | 5.176998496 | 2.25E-07 | 1.41E-06 |
| ENSG00000116106 | EPHA4 | 1316.151202 | 0.972625489 | 0.06411895 | 15.1690801 | 5.67E-52 | 3.29E-50 |
| ENSG00000116117 | PARD3B | 456.571274 | 0.30801005 | 0.090905006 | 3.388262806 | 0.000703368 | 0.00270897 |
| ENSG00000116120 | FARSB | 2229.301347 | -1.128829094 | 0.061867626 | -18.24587698 | 2.23E-74 | 2.32E-72 |
| ENSG00000116127 | ALMS1 | 1093.601066 | 0.171088564 | 0.084017562 | 2.036342864 | 0.041715938 | 0.102656471 |
| ENSG00000116128 | BCL9 | 2036.711426 | 0.499661503 | 0.061220873 | 8.161620087 | 3.31E-16 | 4.34E-15 |
| ENSG00000116133 | DHCR24 | 33474.93388 | -0.417851271 | 0.041545664 | -10.05763851 | 8.50E-24 | 1.76E-22 |
| ENSG00000116138 | DNAJC16 | 598.3752655 | 0.15989712 | 0.082735813 | 1.932622826 | 0.053282678 | 0.126624329 |
| ENSG00000116141 | MARK1 | 129.1299593 | 1.234805469 | 0.169984668 | 7.264216715 | 3.75E-13 | 3.93E-12 |

| | | | | | | | |
|---|---|---|---|---|---|---|---|
| ENSG00000116151 | MORN1 | 201.446062 | -0.362945477 | 0.130740695 | -2.776071188 | 0.005502016 | 0.017513008 |
| ENSG00000116161 | CACYBP | 2509.856014 | -0.06584482 | 0.053567496 | -1.229193544 | 0.218999249 | 0.388782137 |
| ENSG00000116171 | SCP2 | 2349.089781 | -0.320171182 | 0.051391435 | -6.230049485 | 4.66E-10 | 3.83E-09 |
| ENSG00000116183 | PAPPA2 | 2.828133489 | 0.226682513 | 1.077654536 | 0.210348034 | 0.833396048 | 0.91194931 |
| ENSG00000116191 | RALGPS2 | 1420.394721 | 0.051229044 | 0.062498457 | 0.819684939 | 0.412395738 | 0.604534362 |
| ENSG00000116198 | CEP104 | 1291.42501 | -0.096781934 | 0.06448741 | -1.500788046 | 0.133410391 | 0.266681281 |
| ENSG00000116199 | FAM20B | 2633.931768 | -0.042684802 | 0.053039799 | -0.804769311 | 0.420952814 | 0.612069605 |
| ENSG00000116205 | TCEANC2 | 454.8814683 | 0.219122864 | 0.091355521 | 2.398572767 | 0.016459106 | 0.046152291 |
| ENSG00000116209 | TMEM59 | 4177.54763 | 0.147727823 | 0.047136866 | 3.134018795 | 0.001724298 | 0.006141189 |
| ENSG00000116212 | LRRC42 | 849.3087775 | -0.501250752 | 0.078056353 | -6.421652218 | 1.35E-10 | 1.15E-09 |
| ENSG00000116213 | WRAP73 | 764.0250524 | 0.021523214 | 0.074596737 | 0.288527553 | 0.772942942 | 0.876182887 |
| ENSG00000116221 | MRPL37 | 3083.596486 | 0.069592319 | 0.050214941 | 1.385888698 | 0.165780892 | 0.31463669 |
| ENSG00000116237 | ICMT | 3288.015466 | 0.122525043 | 0.048272017 | 2.53822094 | 0.011141762 | 0.032681246 |
| ENSG00000116251 | RPL22 | 3168.067012 | 0.592349546 | 0.050047375 | 11.83577663 | 2.55E-32 | 7.81E-31 |
| ENSG00000116254 | CHD5 | 299.3005056 | -1.131451035 | 0.110757018 | -10.21561483 | 1.69E-24 | 3.66E-23 |
| ENSG00000116260 | QSOX1 | 6183.194437 | 1.257296078 | 0.051992631 | 24.1821977 | 3.42E-129 | 9.58E-127 |
| ENSG00000116266 | STXBP3 | 1279.85235 | 0.13206159 | 0.063994455 | 2.063641142 | 0.039051754 | 0.097237893 |
| ENSG00000116273 | PHF13 | 963.9193633 | -0.004841464 | 0.066808082 | -0.072468231 | 0.942229287 | 0.972585486 |
| ENSG00000116285 | ERRFI1 | 827.5691389 | 0.300686914 | 0.073175952 | 4.109094667 | 3.97E-05 | 0.000188305 |
| ENSG00000116288 | PARK7 | 3491.983051 | 0.065862787 | 0.046545462 | 1.415020587 | 0.15706246 | 0.30150735 |
| ENSG00000116299 | ELAPOR1 | 5298.975682 | -0.407990758 | 0.046594968 | -8.756111901 | 2.02E-18 | 3.06E-17 |
| ENSG00000116329 | OPRD1 | 3.707787916 | 1.502391603 | 1.044676417 | 1.438140632 | 0.150394156 | 0.292203094 |
| ENSG00000116337 | AMPD2 | 1975.66856 | 0.65610852 | 0.065691146 | 9.987777069 | 1.72E-23 | 3.50E-22 |
| ENSG00000116350 | SRSF4 | 2432.025791 | -0.209325729 | 0.05279271 | -3.965049926 | 7.34E-05 | 0.000334795 |
| ENSG00000116353 | MECR | 516.0737197 | 0.015799912 | 0.087800438 | 0.179952538 | 0.857189829 | 0.92601529 |
| ENSG00000116396 | KCNC4 | 28.1919632 | -1.478075641 | 0.35293511 | -4.187952968 | 2.81E-05 | 0.000136863 |
| ENSG00000116406 | EDEM3 | 1302.725535 | -0.156673067 | 0.07115536 | -2.20184489 | 0.027676267 | 0.072492985 |
| ENSG00000116455 | WDR77 | 789.6554505 | 0.129819369 | 0.072891316 | 1.780999112 | 0.074912596 | 0.167739868 |
| ENSG00000116459 | ATP5PB | 3907.488427 | 0.051885763 | 0.045646653 | 1.136682745 | 0.255670935 | 0.43496672 |
| ENSG00000116473 | RAP1A | 1201.279058 | 0.062540257 | 0.063151573 | 0.990319868 | 0.322017799 | 0.511819324 |
| ENSG00000116478 | HDAC1 | 6997.483365 | -0.216083805 | 0.042195865 | -5.120971039 | 3.04E-07 | 1.88E-06 |
| ENSG00000116489 | CAPZA1 | 4972.816763 | -0.015238038 | 0.054604661 | -0.279061125 | 0.780197917 | 0.880664282 |
| ENSG00000116497 | S100PBP | 890.2961438 | -0.153327168 | 0.069627338 | -2.202111593 | 0.027657427 | 0.072451909 |

| | | | | | | | |
|---|---|---|---|---|---|---|---|
| ENSG00000116514 | RNF19B | 553.7860876 | -0.192432395 | 0.084740844 | -2.270834073 | 0.023157024 | 0.062116714 |
| ENSG00000116521 | SCAMP3 | 3023.678312 | 0.213278263 | 0.048530974 | 4.39468332 | 1.11E-05 | 5.69E-05 |
| ENSG00000116525 | TRIM62 | 654.6654984 | 0.021374496 | 0.089130611 | 0.23981094 | 0.810476825 | 0.899645977 |
| ENSG00000116539 | ASH1L | 3499.156472 | 0.285508408 | 0.059568119 | 4.792973369 | 1.64E-06 | 9.26E-06 |
| ENSG00000116544 | DLGAP3 | 28.81470943 | 0.271384461 | 0.334603005 | 0.811064028 | 0.417328902 | 0.609511434 |
| ENSG00000116560 | SFPQ | 10042.29259 | -0.174367363 | 0.056608643 | -3.080225113 | 0.002068442 | 0.007247603 |
| ENSG00000116574 | RHOU | 641.8198328 | -0.434439016 | 0.078789233 | -5.513938873 | 3.51E-08 | 2.38E-07 |
| ENSG00000116580 | GON4L | 2300.967267 | 0.241748782 | 0.055733252 | 4.337604132 | 1.44E-05 | 7.27E-05 |
| ENSG00000116584 | ARHGEF2 | 1269.956328 | 0.579093523 | 0.070346306 | 8.232038782 | 1.84E-16 | 2.46E-15 |
| ENSG00000116586 | LAMTOR2 | 1024.574846 | 0.111474603 | 0.070568866 | 1.579657006 | 0.114185437 | 0.236017414 |
| ENSG00000116604 | MEF2D | 1818.237051 | 0.077855107 | 0.060099269 | 1.295441827 | 0.195167857 | 0.356397365 |
| ENSG00000116641 | DOCK7 | 1653.214251 | 0.075792187 | 0.06578223 | 1.152168104 | 0.249252002 | 0.427278502 |
| ENSG00000116649 | SRM | 3326.159659 | 0.284441085 | 0.047868162 | 5.942176841 | 2.81E-09 | 2.15E-08 |
| ENSG00000116652 | DLEU2L | 7.169949932 | -0.116059128 | 0.657136288 | -0.176613481 | 0.859812004 | 0.92728725 |
| ENSG00000116661 | FBXO2 | 298.0165231 | 0.88638525 | 0.111733476 | 7.933032119 | 2.14E-15 | 2.64E-14 |
| ENSG00000116663 | FBXO6 | 240.0091097 | 0.656461069 | 0.120040505 | 5.468663007 | 4.53E-08 | 3.04E-07 |
| ENSG00000116667 | C1orf21 | 344.6987873 | -0.019269777 | 0.109243013 | -0.17639368 | 0.859984669 | 0.927319435 |
| ENSG00000116668 | SWT1 | 93.31847356 | -0.191725368 | 0.18697402 | -1.02541181 | 0.305168918 | 0.492616634 |
| ENSG00000116670 | MAD2L2 | 653.8180912 | 0.026003285 | 0.080184215 | 0.324294314 | 0.745715224 | 0.859386985 |
| ENSG00000116675 | DNAJC6 | 1257.051989 | -0.803149755 | 0.064936413 | -12.36824954 | 3.88E-35 | 1.33E-33 |
| ENSG00000116678 | LEPR | 180.7953625 | -0.69210079 | 0.138880856 | -4.983413905 | 6.25E-07 | 3.71E-06 |
| ENSG00000116679 | IVNS1ABP | 4414.343828 | -0.560644195 | 0.046820608 | -11.9743041 | 4.84E-33 | 1.52E-31 |
| ENSG00000116685 | KIAA2013 | 1390.76815 | 0.129337592 | 0.062918612 | 2.055633262 | 0.039817879 | 0.098791061 |
| ENSG00000116688 | MFN2 | 3189.763715 | 0.095321846 | 0.054348552 | 1.75389854 | 0.079447894 | 0.175733747 |
| ENSG00000116690 | PRG4 | 7.212945563 | -0.516664039 | 0.661465144 | -0.781090346 | 0.434749359 | 0.624766667 |
| ENSG00000116691 | MIIP | 342.5735645 | 0.111235671 | 0.101267112 | 1.098438258 | 0.272013164 | 0.454041488 |
| ENSG00000116698 | SMG7 | 6578.295374 | -0.088067114 | 0.044059371 | -1.998828251 | 0.04562694 | 0.11076 |
| ENSG00000116701 | NCF2 | 19.74792002 | 1.842872685 | 0.452150195 | 4.075797614 | 4.59E-05 | 0.000215698 |
| ENSG00000116704 | SLC35D1 | 711.6337675 | 0.09405827 | 0.078049285 | 1.205113788 | 0.228159378 | 0.40104462 |
| ENSG00000116717 | GADD45A | 225.1285663 | 0.133806051 | 0.123986658 | 1.07919718 | 0.280499836 | 0.463926911 |
| ENSG00000116729 | WLS | 1076.763192 | 1.510953611 | 0.070677405 | 21.37817053 | 2.13E-101 | 3.50E-99 |
| ENSG00000116731 | PRDM2 | 1111.168023 | 0.049389952 | 0.066569056 | 0.741935584 | 0.458126365 | 0.64527349 |
| ENSG00000116741 | RGS2 | 22.68410534 | -0.17729714 | 0.373082759 | -0.475222013 | 0.634628739 | 0.784564738 |

| | | | | | | | |
|---|---|---|---|---|---|---|---|
| ENSG00000116747 | RO60 | 1306.078624 | -0.207461952 | 0.067924508 | -3.054301856 | 0.00225585 | 0.007854945 |
| ENSG00000116748 | AMPD1 | 29.65390213 | 0.976916626 | 0.337178976 | 2.897323661 | 0.003763613 | 0.012474682 |
| ENSG00000116750 | UCHL5 | 1308.266667 | -0.31054856 | 0.066667196 | -4.658191434 | 3.19E-06 | 1.74E-05 |
| ENSG00000116752 | BCAS2 | 7240.304356 | 0.320669326 | 0.048038308 | 6.675283572 | 2.47E-11 | 2.23E-10 |
| ENSG00000116754 | SRSF11 | 3771.980543 | -0.107350564 | 0.063796852 | -1.682693729 | 0.092434396 | 0.198792143 |
| ENSG00000116761 | CTH | 48.34967062 | -0.713927912 | 0.274650021 | -2.599409637 | 0.009338426 | 0.028028718 |
| ENSG00000116771 | AGMAT | 279.3409787 | 0.357072853 | 0.116714958 | 3.059358109 | 0.002218118 | 0.007730725 |
| ENSG00000116774 | OLFML3 | 2000.176879 | 4.517281362 | 0.093290935 | 48.42143957 | 0 | 0 |
| ENSG00000116785 | CFHR3 | 18.03814285 | -0.337385025 | 0.415232795 | -0.81252018 | 0.416493212 | 0.608557597 |
| ENSG00000116786 | PLEKHM2 | 1850.29295 | 0.151307521 | 0.060035052 | 2.520319648 | 0.011724831 | 0.034224152 |
| ENSG00000116791 | CRYZ | 889.9733957 | -0.160693257 | 0.072016676 | -2.231333981 | 0.025659014 | 0.067891325 |
| ENSG00000116793 | PHTF1 | 1985.593914 | 0.430229992 | 0.053836799 | 7.991373884 | 1.33E-15 | 1.67E-14 |
| ENSG00000116809 | ZBTB17 | 589.6590695 | 0.048532423 | 0.094447196 | 0.513857746 | 0.607351451 | 0.765740164 |
| ENSG00000116815 | CD58 | 201.2854878 | 0.259303149 | 0.139390742 | 1.860260917 | 0.06284862 | 0.144972081 |
| ENSG00000116819 | TFAP2E | 23.78234409 | 0.2729105 | 0.370996177 | 0.735615399 | 0.461964789 | 0.648626151 |
| ENSG00000116830 | TTF2 | 3503.231925 | 0.026582447 | 0.053469233 | 0.497154073 | 0.619080406 | 0.774187991 |
| ENSG00000116833 | NR5A2 | 34.56928048 | -3.428828426 | 0.423212438 | -8.101908448 | 5.41E-16 | 6.98E-15 |
| ENSG00000116852 | KIF21B | 56.51278605 | 1.506651605 | 0.259261836 | 5.811312709 | 6.20E-09 | 4.56E-08 |
| ENSG00000116857 | TMEM9 | 3647.366581 | 0.317801756 | 0.051558262 | 6.163934685 | 7.10E-10 | 5.73E-09 |
| ENSG00000116863 | ADPRS | 1035.89825 | -0.234344152 | 0.072295701 | -3.241467293 | 0.001189161 | 0.004389982 |
| ENSG00000116871 | MAP7D1 | 1712.194517 | 0.206246767 | 0.060727336 | 3.396275582 | 0.000683095 | 0.002638083 |
| ENSG00000116874 | WARS2 | 536.594588 | 0.477299413 | 0.085976129 | 5.55153411 | 2.83E-08 | 1.94E-07 |
| ENSG00000116885 | OSCP1 | 158.4140053 | 0.268913806 | 0.148606041 | 1.809575204 | 0.070361688 | 0.159259706 |
| ENSG00000116898 | MRPS15 | 1874.63007 | -0.196215857 | 0.056982788 | -3.443423229 | 0.0005744 | 0.002247438 |
| ENSG00000116903 | EXOC8 | 676.8263277 | -0.00729141 | 0.076076617 | -0.095842989 | 0.923645275 | 0.963279648 |
| ENSG00000116906 | GNPAT | 1528.507537 | -0.188530574 | 0.060964478 | -3.092465962 | 0.00198501 | 0.00698514 |
| ENSG00000116918 | TSNAX | 518.1391926 | -0.094080365 | 0.087913631 | -1.07014537 | 0.28455388 | 0.468335916 |
| ENSG00000116922 | C1orf109 | 747.9393357 | -0.016260108 | 0.073551578 | -0.221070819 | 0.825037291 | 0.90772273 |
| ENSG00000116954 | RRAGC | 550.5522807 | 0.188556895 | 0.084716817 | 2.225731587 | 0.026032166 | 0.068743856 |
| ENSG00000116977 | LGALS8 | 3053.767637 | -0.222432798 | 0.049801239 | -4.466410943 | 7.95E-06 | 4.15E-05 |
| ENSG00000116984 | MTR | 1932.367293 | -0.106477533 | 0.055890103 | -1.905123223 | 0.056764055 | 0.133282234 |
| ENSG00000116985 | BMP8B | 636.3880743 | 0.500345091 | 0.082845879 | 6.039468645 | 1.55E-09 | 1.21E-08 |
| ENSG00000116990 | MYCL | 235.3912263 | 0.968131184 | 0.124634402 | 7.767768516 | 7.99E-15 | 9.45E-14 |

| | | | | | | | |
|---|---|---|---|---|---|---|---|
| ENSG00000116991 | SIPA1L2 | 2549.532681 | -0.060172616 | 0.054714735 | -1.099751577 | 0.271440376 | 0.453421846 |
| ENSG00000117000 | RLF | 1017.931431 | -0.415624056 | 0.070889948 | -5.862947721 | 4.55E-09 | 3.40E-08 |
| ENSG00000117009 | KMO | 8.893951536 | -1.40705872 | 0.628793149 | -2.237713186 | 0.025239765 | 0.067006157 |
| ENSG00000117010 | ZNF684 | 53.66464232 | 0.066743056 | 0.24507172 | 0.272340915 | 0.785359899 | 0.883215101 |
| ENSG00000117013 | KCNQ4 | 73.97975302 | 0.725329558 | 0.210816403 | 3.440574574 | 0.00058048 | 0.002268908 |
| ENSG00000117016 | RIMS3 | 351.1664377 | -0.059898631 | 0.102167816 | -0.586276902 | 0.557689451 | 0.727915708 |
| ENSG00000117020 | AKT3 | 12.14702361 | -0.228228197 | 0.500636175 | -0.455876359 | 0.648478891 | 0.795128387 |
| ENSG00000117036 | ETV3 | 1612.259183 | -0.023539643 | 0.066438866 | -0.354305317 | 0.723110083 | 0.8459455 |
| ENSG00000117054 | ACADM | 998.2777272 | -0.208304436 | 0.069883564 | -2.980735737 | 0.002875568 | 0.009784685 |
| ENSG00000117069 | ST6GALNAC5 | 107.3236225 | 0.872852195 | 0.177626165 | 4.91398434 | 8.92E-07 | 5.21E-06 |
| ENSG00000117114 | ADGRL2 | 366.6610016 | 1.260147936 | 0.105089107 | 11.99123271 | 3.95E-33 | 1.24E-31 |
| ENSG00000117115 | PADI2 | 66.1879558 | -0.118223717 | 0.220100644 | -0.537134807 | 0.5911745 | 0.753709593 |
| ENSG00000117118 | SDHB | 1189.135886 | 0.16953642 | 0.062692956 | 2.70423393 | 0.006846207 | 0.02126923 |
| ENSG00000117122 | MFAP2 | 149.9443852 | -0.64009682 | 0.150624771 | -4.2496119 | 2.14E-05 | 0.000105848 |
| ENSG00000117133 | RPF1 | 1008.128364 | 0.07613656 | 0.066627482 | 1.14272005 | 0.253154842 | 0.431939753 |
| ENSG00000117139 | KDM5B | 12245.08977 | 0.648129022 | 0.043786117 | 14.80215785 | 1.42E-49 | 7.79E-48 |
| ENSG00000117143 | UAP1 | 1593.638544 | 0.122097126 | 0.057586022 | 2.120256312 | 0.033984436 | 0.086373029 |
| ENSG00000117148 | ACTL8 | 6.054586964 | -1.955165804 | 0.787438851 | -2.482943029 | 0.013030193 | 0.037518842 |
| ENSG00000117151 | CTBS | 120.3188888 | -0.063794853 | 0.161411478 | -0.395231208 | 0.692672263 | 0.826737862 |
| ENSG00000117153 | KLHL12 | 2978.507991 | 0.347203279 | 0.051932513 | 6.68566298 | 2.30E-11 | 2.09E-10 |
| ENSG00000117155 | SSX2IP | 759.238359 | 0.25616427 | 0.077420081 | 3.30875744 | 0.00093711 | 0.003525303 |
| ENSG00000117174 | ZNHIT6 | 400.1351028 | -0.291626107 | 0.107428459 | -2.714607553 | 0.006635435 | 0.020692887 |
| ENSG00000117222 | RBBP5 | 2260.848156 | 0.309949448 | 0.053692444 | 5.772682789 | 7.80E-09 | 5.67E-08 |
| ENSG00000117226 | GBP3 | 32.12603038 | 2.142585576 | 0.358045053 | 5.984122823 | 2.18E-09 | 1.68E-08 |
| ENSG00000117228 | GBP1 | 3.78940307 | -0.03812972 | 0.90426298 | -0.042166627 | 0.966365867 | 0.985001021 |
| ENSG00000117242 | PINK1-AS | 222.8782927 | 0.35598111 | 0.126668418 | 2.810338333 | 0.004948945 | 0.015967481 |
| ENSG00000117245 | KIF17 | 6.088053419 | 1.217842642 | 0.761820159 | 1.598595979 | 0.109910404 | 0.228600729 |
| ENSG00000117262 | GPR89A | 533.4442886 | 0.295387139 | 0.08417059 | 3.509386578 | 0.000449142 | 0.001792567 |
| ENSG00000117266 | CDK18 | 810.8192547 | 0.825709675 | 0.084535849 | 9.76756823 | 1.55E-22 | 3.00E-21 |
| ENSG00000117280 | RAB29 | 1503.083122 | 0.647207393 | 0.060575873 | 10.68424384 | 1.21E-26 | 2.93E-25 |
| ENSG00000117298 | ECE1 | 2658.556919 | -0.276582643 | 0.05744135 | -4.815044251 | 1.47E-06 | 8.33E-06 |
| ENSG00000117305 | HMGCL | 791.1622855 | -0.347931304 | 0.074708381 | -4.657192397 | 3.21E-06 | 1.75E-05 |
| ENSG00000117308 | GALE | 390.5649338 | -0.151022331 | 0.098234725 | -1.537361974 | 0.124204693 | 0.252213477 |

| | | | | | | | |
|---|---|---|---|---|---|---|---|
| ENSG00000117318 | ID3 | 1626.061942 | -2.052443565 | 0.07186443 | -28.55993655 | 2.11E-179 | 1.01E-176 |
| ENSG00000117335 | CD46 | 3252.251752 | -0.125856296 | 0.056707129 | -2.219408718 | 0.026458931 | 0.069710334 |
| ENSG00000117360 | PRPF3 | 5997.221397 | 0.503120504 | 0.058266241 | 8.634854304 | 5.88E-18 | 8.62E-17 |
| ENSG00000117362 | APH1A | 11980.0718 | 0.355521449 | 0.043249484 | 8.220247226 | 2.03E-16 | 2.71E-15 |
| ENSG00000117385 | P3H1 | 657.3850842 | -0.38880492 | 0.089278163 | -4.354983441 | 1.33E-05 | 6.76E-05 |
| ENSG00000117394 | SLC2A1 | 2873.845455 | -0.095916576 | 0.050533729 | -1.898070407 | 0.057686808 | 0.135061785 |
| ENSG00000117395 | EBNA1BP2 | 2490.420971 | 0.116082022 | 0.050343014 | 2.305821853 | 0.021120589 | 0.05729734 |
| ENSG00000117399 | CDC20 | 2270.933434 | -0.295720427 | 0.05481809 | -5.394577324 | 6.87E-08 | 4.51E-07 |
| ENSG00000117400 | MPL | 16.08573179 | 0.318707439 | 0.459902419 | 0.692989265 | 0.488316289 | 0.67162154 |
| ENSG00000117407 | ARTN | 262.4391399 | -0.08991768 | 0.1195029 | -0.752430943 | 0.451791944 | 0.639701573 |
| ENSG00000117408 | IPO13 | 2162.199649 | -0.029393895 | 0.05531473 | -0.531393622 | 0.59514604 | 0.756588059 |
| ENSG00000117410 | ATP6V0B | 2953.404948 | 0.071686269 | 0.054614499 | 1.312586768 | 0.189322231 | 0.348196262 |
| ENSG00000117411 | B4GALT2 | 2291.943299 | 0.210206334 | 0.060898809 | 3.451731451 | 0.000557002 | 0.002184867 |
| ENSG00000117419 | ERI3 | 1695.95895 | 0.004280163 | 0.062196583 | 0.068816685 | 0.945135536 | 0.973920097 |
| ENSG00000117425 | PTCH2 | 18.65052505 | 0.628818476 | 0.447799477 | 1.404241203 | 0.160247036 | 0.306389982 |
| ENSG00000117448 | AKR1A1 | 1929.854313 | -0.032692463 | 0.06305765 | -0.518453558 | 0.604141855 | 0.763029194 |
| ENSG00000117450 | PRDX1 | 13372.24272 | -0.096294947 | 0.048342553 | -1.991929287 | 0.046378817 | 0.112288133 |
| ENSG00000117461 | PIK3R3 | 6493.524006 | -0.337506175 | 0.045429475 | -7.429233404 | 1.09E-13 | 1.19E-12 |
| ENSG00000117472 | TSPAN1 | 209.7084948 | 0.225366829 | 0.150524133 | 1.497213933 | 0.134337601 | 0.267974311 |
| ENSG00000117475 | BLZF1 | 931.0357031 | -0.153230376 | 0.069629309 | -2.20065914 | 0.027760164 | 0.072687838 |
| ENSG00000117477 | CCDC181 | 2.27126589 | 0.266324628 | 1.157418269 | 0.230102319 | 0.818012263 | 0.90364036 |
| ENSG00000117479 | SLC19A2 | 854.633176 | 0.112146529 | 0.072379532 | 1.549423239 | 0.121280011 | 0.247545462 |
| ENSG00000117480 | FAAH | 1266.975477 | 0.001663385 | 0.073633504 | 0.022590062 | 0.981977271 | 0.991627389 |
| ENSG00000117481 | NSUN4 | 1489.614109 | 0.022688619 | 0.06400878 | 0.354461041 | 0.722993396 | 0.845926239 |
| ENSG00000117500 | TMED5 | 692.9692279 | -0.012890835 | 0.078538744 | -0.164133451 | 0.869626089 | 0.932287011 |
| ENSG00000117505 | DR1 | 995.9863434 | -0.187347733 | 0.068158932 | -2.748689392 | 0.005983406 | 0.018888032 |
| ENSG00000117519 | CNN3 | 1.893031755 | 1.472344068 | 1.432531776 | 1.02779156 | 0.304047887 | 0.491394853 |
| ENSG00000117523 | PRRC2C | 8303.937495 | -0.031113368 | 0.062522181 | -0.497637282 | 0.618739721 | 0.773972911 |
| ENSG00000117525 | F3 | 15.50761523 | -0.573589122 | 0.455320683 | -1.259747565 | 0.207760441 | 0.374099255 |
| ENSG00000117528 | ABCD3 | 777.2303547 | -0.261934972 | 0.083360134 | -3.142209101 | 0.001676782 | 0.005986831 |
| ENSG00000117533 | VAMP4 | 197.5443718 | 0.0548823 | 0.140267349 | 0.39126925 | 0.695598226 | 0.82880711 |
| ENSG00000117543 | DPH5 | 489.210359 | 0.376737599 | 0.0881347 | 4.27456607 | 1.92E-05 | 9.53E-05 |
| ENSG00000117569 | PTBP2 | 380.0224893 | 0.72530584 | 0.108478882 | 6.686147834 | 2.29E-11 | 2.08E-10 |

| | | | | | | | |
|---|---|---|---|---|---|---|---|
| ENSG00000117586 | TNFSF4 | 6.112405078 | 0.093892883 | 0.731939906 | 0.128279497 | 0.89792779 | 0.948320221 |
| ENSG00000117592 | PRDX6 | 7763.010767 | -0.165123841 | 0.043656259 | -3.782363492 | 0.000155346 | 0.000671625 |
| ENSG00000117593 | DARS2 | 2215.967604 | -0.556922925 | 0.060207888 | -9.249999372 | 2.24E-20 | 3.84E-19 |
| ENSG00000117595 | IRF6 | 1047.242655 | -0.031103683 | 0.067856272 | -0.45837595 | 0.646682367 | 0.793519085 |
| ENSG00000117597 | UTP25 | 736.5940213 | 0.268188328 | 0.075290352 | 3.562054374 | 0.000367964 | 0.00148768 |
| ENSG00000117598 | PLPPR5 | 123.4984551 | 0.387678152 | 0.163414032 | 2.372367578 | 0.017674497 | 0.049139858 |
| ENSG00000117601 | SERPINC1 | 3.116182609 | 1.801208258 | 1.100816599 | 1.636247363 | 0.101787827 | 0.215049987 |
| ENSG00000117602 | RCAN3 | 518.8721398 | -0.775372193 | 0.087496724 | -8.861728239 | 7.88E-19 | 1.23E-17 |
| ENSG00000117614 | SYF2 | 921.5390135 | 0.177652532 | 0.068992731 | 2.574945633 | 0.010025584 | 0.029794909 |
| ENSG00000117616 | RSRP1 | 647.2532643 | 0.069560807 | 0.098962255 | 0.7029024 | 0.482116575 | 0.666287182 |
| ENSG00000117620 | SLC35A3 | 562.3413547 | 0.02617012 | 0.09088574 | 0.28794528 | 0.773388626 | 0.876471449 |
| ENSG00000117625 | RCOR3 | 675.1406632 | -0.173768282 | 0.080440985 | -2.160195853 | 0.030757511 | 0.079403253 |
| ENSG00000117632 | STMN1 | 6579.082688 | -0.506995375 | 0.046852175 | -10.82117045 | 2.73E-27 | 6.87E-26 |
| ENSG00000117640 | MTFR1L | 1666.933696 | -0.036050525 | 0.061366563 | -0.587462024 | 0.556893448 | 0.72734774 |
| ENSG00000117643 | MAN1C1 | 171.4755671 | 0.3100287 | 0.145580942 | 2.129596739 | 0.033204921 | 0.084710952 |
| ENSG00000117650 | NEK2 | 501.160195 | -0.137782535 | 0.088626025 | -1.554650965 | 0.120029219 | 0.245626252 |
| ENSG00000117676 | RPS6KA1 | 1683.98904 | 0.001456286 | 0.066804866 | 0.021799094 | 0.982608217 | 0.991991925 |
| ENSG00000117682 | DHDDS | 828.8003361 | -0.348208578 | 0.072361768 | -4.812051831 | 1.49E-06 | 8.45E-06 |
| ENSG00000117691 | NENF | 545.4339378 | 0.308109264 | 0.092157886 | 3.343276153 | 0.000827955 | 0.003147188 |
| ENSG00000117697 | NSL1 | 353.3442658 | -0.280566925 | 0.099667274 | -2.815035608 | 0.004877183 | 0.015764771 |
| ENSG00000117713 | ARID1A | 3514.8748 | -0.059486808 | 0.055804466 | -1.065986509 | 0.286429741 | 0.470208834 |
| ENSG00000117724 | CENPF | 2562.721224 | -0.197110375 | 0.055408879 | -3.557378848 | 0.000374574 | 0.001512271 |
| ENSG00000117748 | RPA2 | 2251.577078 | -0.38044063 | 0.055569326 | -6.84623438 | 7.58E-12 | 7.18E-11 |
| ENSG00000117751 | PPP1R8 | 2145.342309 | -0.18675608 | 0.05256564 | -3.552816641 | 0.00038113 | 0.001537658 |
| ENSG00000117758 | STX12 | 1615.734419 | 0.107199372 | 0.060284679 | 1.778219157 | 0.075367868 | 0.168578285 |
| ENSG00000117791 | MTARC2 | 416.8381928 | -0.339969877 | 0.092671529 | -3.668547203 | 0.000243933 | 0.001017976 |
| ENSG00000117859 | OSBPL9 | 3139.169172 | 0.367733739 | 0.050597826 | 7.267777478 | 3.65E-13 | 3.83E-12 |
| ENSG00000117862 | TXNDC12 | 98.57166168 | -0.082883669 | 0.178321394 | -0.464799358 | 0.642075147 | 0.790034632 |
| ENSG00000117868 | ESYT2 | 3347.538612 | 0.8674985 | 0.048895649 | 17.74183426 | 1.99E-70 | 1.86E-68 |
| ENSG00000117877 | POLR1G | 1079.26024 | 0.408269406 | 0.066781605 | 6.113500957 | 9.75E-10 | 7.76E-09 |
| ENSG00000117899 | MESD | 2242.284158 | -0.186875425 | 0.05212152 | -3.585379402 | 0.000336589 | 0.001371707 |
| ENSG00000117906 | RCN2 | 2212.144621 | 0.187617797 | 0.05717208 | 3.281633235 | 0.001032078 | 0.003852239 |
| ENSG00000117971 | CHRNB4 | 26.19545596 | -1.338052125 | 0.35702206 | -3.747813583 | 0.000178383 | 0.000763452 |

| | | | | | | | |
|---|---|---|---|---|---|---|---|
| ENSG00000117983 | MUC5B | 8.840690801 | -1.242267306 | 0.618836894 | -2.007422826 | 0.04470466 | 0.108832103 |
| ENSG00000117984 | CTSD | 2228.575814 | 0.537390997 | 0.055767716 | 9.636238294 | 5.62E-22 | 1.05E-20 |
| ENSG00000118007 | STAG1 | 2826.517029 | -0.650818297 | 0.050290033 | -12.94129796 | 2.63E-38 | 1.01E-36 |
| ENSG00000118046 | STK11 | 2563.043699 | 0.09797573 | 0.052264204 | 1.874623989 | 0.060844471 | 0.141204646 |
| ENSG00000118058 | KMT2A | 2437.595477 | -0.18621164 | 0.066982333 | -2.780011276 | 0.005435701 | 0.017335602 |
| ENSG00000118094 | TREH | 2.660015816 | 0.457826493 | 1.142138456 | 0.400850257 | 0.688530375 | 0.823987156 |
| ENSG00000118096 | IFT46 | 1201.376125 | -0.308909875 | 0.063318472 | -4.878669105 | 1.07E-06 | 6.16E-06 |
| ENSG00000118156 | ZNF541 | 72.88878707 | -0.189589431 | 0.206636956 | -0.917500116 | 0.358880632 | 0.55145835 |
| ENSG00000118160 | SLC8A2 | 5.668362191 | -0.023047183 | 0.724962269 | -0.031790872 | 0.974638826 | 0.988514222 |
| ENSG00000118162 | KPTN | 309.3746498 | -0.249018915 | 0.109237104 | -2.279618431 | 0.022630329 | 0.060896158 |
| ENSG00000118181 | RPS25 | 5797.402961 | 0.331822489 | 0.055198837 | 6.011403628 | 1.84E-09 | 1.43E-08 |
| ENSG00000118193 | KIF14 | 1852.275653 | 0.122640908 | 0.064902609 | 1.889614462 | 0.058809542 | 0.137186382 |
| ENSG00000118194 | TNNT2 | 37.30825125 | 1.062054765 | 0.299021665 | 3.551765269 | 0.000382656 | 0.001543272 |
| ENSG00000118197 | DDX59 | 975.4765426 | 0.388825741 | 0.067981045 | 5.71961994 | 1.07E-08 | 7.66E-08 |
| ENSG00000118200 | CAMSAP2 | 4028.088675 | 0.42790761 | 0.053340072 | 8.022254124 | 1.04E-15 | 1.31E-14 |
| ENSG00000118217 | ATF6 | 2085.921174 | 0.410641175 | 0.055492188 | 7.399981736 | 1.36E-13 | 1.47E-12 |
| ENSG00000118242 | MREG | 1918.003961 | 0.056240836 | 0.060021405 | 0.937012985 | 0.348751878 | 0.539945204 |
| ENSG00000118246 | FASTKD2 | 794.6930785 | 0.388629905 | 0.07717267 | 5.035848863 | 4.76E-07 | 2.88E-06 |
| ENSG00000118257 | NRP2 | 14.32215633 | 0.25866025 | 0.471296372 | 0.548827162 | 0.583124068 | 0.74747799 |
| ENSG00000118260 | CREB1 | 1322.407965 | -0.19157977 | 0.06654316 | -2.87903024 | 0.003989001 | 0.013145766 |
| ENSG00000118263 | KLF7 | 80.55475206 | -1.104791328 | 0.210216603 | -5.255490352 | 1.48E-07 | 9.43E-07 |
| ENSG00000118271 | TTR | 3.157255473 | 0.138979677 | 1.02059474 | 0.136175185 | 0.891682792 | 0.94446409 |
| ENSG00000118276 | B4GALT6 | 362.4449984 | 0.342962059 | 0.098561726 | 3.47966776 | 0.000502036 | 0.001985709 |
| ENSG00000118292 | C1orf54 | 49.66894634 | 0.848320755 | 0.258861959 | 3.277116331 | 0.001048732 | 0.003909947 |
| ENSG00000118298 | CA14 | 5.258389792 | 0.206632384 | 0.772574721 | 0.267459416 | 0.789115457 | 0.885944625 |
| ENSG00000118307 | DNAI7 | 8.015449877 | 0.932115169 | 0.678650271 | 1.373483825 | 0.169601986 | 0.320165542 |
| ENSG00000118322 | ATP10B | 6.388889965 | 2.669415881 | 0.860124745 | 3.103521781 | 0.001912322 | 0.00675318 |
| ENSG00000118363 | SPCS2 | 741.5504888 | 0.042091271 | 0.073810774 | 0.570259177 | 0.568501925 | 0.736037682 |
| ENSG00000118369 | USP35 | 346.3369761 | -0.29443245 | 0.110307384 | -2.669199831 | 0.00760322 | 0.023358493 |
| ENSG00000118402 | ELOVL4 | 164.9686237 | -0.032592657 | 0.155342523 | -0.209811561 | 0.83381475 | 0.912258208 |
| ENSG00000118407 | FILIP1 | 33.34183562 | 3.072539541 | 0.413263971 | 7.434811052 | 1.05E-13 | 1.14E-12 |
| ENSG00000118412 | CASP8AP2 | 941.025007 | -0.321375253 | 0.067477946 | -4.762670928 | 1.91E-06 | 1.07E-05 |
| ENSG00000118418 | HMGN3 | 1040.379353 | -0.345979843 | 0.067646517 | -5.114525593 | 3.15E-07 | 1.94E-06 |

| | | | | | | | |
|---|---|---|---|---|---|---|---|
| ENSG00000118420 | UBE3D | 134.65641 | 0.140643419 | 0.15565635 | 0.903550797 | 0.366233641 | 0.558975365 |
| ENSG00000118432 | CNR1 | 24.33065973 | 1.191300709 | 0.383037271 | 3.110143055 | 0.001869968 | 0.006611753 |
| ENSG00000118454 | ANKRD13C | 964.9533226 | 0.191288957 | 0.068511797 | 2.792058653 | 0.005237386 | 0.016766067 |
| ENSG00000118482 | PHF3 | 2777.640715 | -0.015056318 | 0.055160796 | -0.272953244 | 0.784889157 | 0.883106023 |
| ENSG00000118495 | PLAGL1 | 111.2699707 | 0.241762191 | 0.174680841 | 1.384022369 | 0.166351606 | 0.315511024 |
| ENSG00000118496 | FBXO30 | 350.0257643 | 0.248490703 | 0.099477598 | 2.497956397 | 0.012491156 | 0.036161927 |
| ENSG00000118503 | TNFAIP3 | 40.15502727 | 1.240563626 | 0.290026241 | 4.277418557 | 1.89E-05 | 9.42E-05 |
| ENSG00000118507 | AKAP7 | 65.92571647 | 0.926828061 | 0.223349315 | 4.14967945 | 3.33E-05 | 0.000159947 |
| ENSG00000118508 | RAB32 | 464.5438143 | -0.141668632 | 0.093116499 | -1.521412794 | 0.12815628 | 0.258498643 |
| ENSG00000118513 | MYB | 1684.874662 | 0.776634658 | 0.076487404 | 10.15375884 | 3.19E-24 | 6.75E-23 |
| ENSG00000118514 | ALDH8A1 | 2.864583093 | -0.172404465 | 1.050656365 | -0.164092153 | 0.869658599 | 0.932287011 |
| ENSG00000118515 | SGK1 | 60.29014503 | 0.144059781 | 0.234624282 | 0.61400201 | 0.539213999 | 0.713848634 |
| ENSG00000118518 | RNF146 | 877.183235 | 0.40789666 | 0.072376551 | 5.635757082 | 1.74E-08 | 1.22E-07 |
| ENSG00000118523 | CCN2 | 94.89188643 | -0.945788719 | 0.194786939 | -4.855503786 | 1.20E-06 | 6.88E-06 |
| ENSG00000118557 | PMFBP1 | 16.875182 | 0.177597075 | 0.451686391 | 0.393186686 | 0.694181603 | 0.827678065 |
| ENSG00000118564 | FBXL5 | 1389.462716 | -0.220202273 | 0.061949741 | -3.554530998 | 0.000378654 | 0.001528205 |
| ENSG00000118579 | MED28 | 1770.026427 | 0.062544643 | 0.059433545 | 1.052345827 | 0.292640913 | 0.477894251 |
| ENSG00000118596 | SLC16A7 | 505.6786669 | 0.005017767 | 0.093455308 | 0.053691617 | 0.957180862 | 0.980624826 |
| ENSG00000118600 | RXYLT1 | 745.4774658 | 0.126818118 | 0.076324378 | 1.661567653 | 0.096599494 | 0.206177185 |
| ENSG00000118620 | ZNF430 | 127.1232121 | -0.018217579 | 0.162523413 | -0.112092026 | 0.91075044 | 0.955221978 |
| ENSG00000118640 | VAMP8 | 1864.98771 | -0.168430441 | 0.059117116 | -2.84909771 | 0.004384341 | 0.014331261 |
| ENSG00000118655 | DCLRE1B | 6246.759226 | 0.060551643 | 0.047154112 | 1.284122218 | 0.199099195 | 0.361883808 |
| ENSG00000118680 | MYL12B | 12332.64925 | -0.050210436 | 0.048269765 | -1.040204706 | 0.298244806 | 0.484851239 |
| ENSG00000118689 | FOXO3 | 2327.452034 | 0.233218449 | 0.057524141 | 4.054270894 | 5.03E-05 | 0.0002353 |
| ENSG00000118690 | ARMC2 | 47.89384119 | -0.110500069 | 0.255566187 | -0.432373587 | 0.665469912 | 0.807126455 |
| ENSG00000118705 | RPN2 | 8351.654149 | -0.337495249 | 0.042687517 | -7.906181297 | 2.65E-15 | 3.26E-14 |
| ENSG00000118707 | TGIF2 | 1412.796489 | -0.505184068 | 0.060627069 | -8.332648653 | 7.91E-17 | 1.08E-15 |
| ENSG00000118729 | CASQ2 | 2.101572473 | 0.438330009 | 1.253913721 | 0.349569513 | 0.726661794 | 0.848067845 |
| ENSG00000118762 | PKD2 | 644.5437933 | 0.037239498 | 0.081265571 | 0.458244464 | 0.646776818 | 0.793552988 |
| ENSG00000118777 | ABCG2 | 472.5204985 | -1.48582605 | 0.094512982 | -15.72086734 | 1.09E-55 | 6.95E-54 |
| ENSG00000118804 | STBD1 | 28.45022397 | -0.021742546 | 0.340886946 | -0.063782277 | 0.94914359 | 0.976558753 |
| ENSG00000118816 | CCNI | 10943.90579 | 0.088744237 | 0.049776456 | 1.782855661 | 0.074609802 | 0.167274127 |
| ENSG00000118855 | MFSD1 | 1620.529136 | 0.115067922 | 0.058060939 | 1.981847398 | 0.047496323 | 0.114630788 |

| | | | | | | | |
|---|---|---|---|---|---|---|---|
| ENSG00000118873 | RAB3GAP2 | 1556.342068 | -0.175277076 | 0.060046958 | -2.91900012 | 0.003511561 | 0.011725649 |
| ENSG00000118894 | EEF2KMT | 485.9626646 | -0.164229448 | 0.089523732 | -1.834479474 | 0.066582852 | 0.152299394 |
| ENSG00000118898 | PPL | 1942.850938 | 0.387990358 | 0.067121583 | 5.7804113 | 7.45E-09 | 5.43E-08 |
| ENSG00000118900 | UBN1 | 3342.644817 | 0.06182143 | 0.050081133 | 1.234425545 | 0.217044375 | 0.386508148 |
| ENSG00000118922 | KLF12 | 285.5930965 | -0.253292846 | 0.108415011 | -2.336326342 | 0.01947424 | 0.05344996 |
| ENSG00000118939 | UCHL3 | 42.32298665 | -0.348206703 | 0.271630839 | -1.281911526 | 0.199873684 | 0.362946435 |
| ENSG00000118946 | PCDH17 | 18.66290337 | -0.759719577 | 0.406475196 | -1.869042893 | 0.061616847 | 0.142690336 |
| ENSG00000118960 | HS1BP3 | 891.9322479 | 0.469171867 | 0.082195748 | 5.707982223 | 1.14E-08 | 8.16E-08 |
| ENSG00000118961 | LDAH | 681.1063369 | 0.546021261 | 0.080375187 | 6.793405761 | 1.10E-11 | 1.02E-10 |
| ENSG00000118965 | WDR35 | 360.5971097 | 0.080090831 | 0.098868021 | 0.810078234 | 0.417895213 | 0.609908631 |
| ENSG00000118976 | OTUD4P1 | 92.6118058 | 0.050743668 | 0.194534577 | 0.260846525 | 0.794210864 | 0.88974732 |
| ENSG00000118985 | ELL2 | 342.7093149 | 0.061913598 | 0.100635627 | 0.615225438 | 0.538405851 | 0.713216899 |
| ENSG00000118997 | DNAH7 | 24.38415582 | 0.65911835 | 0.374366195 | 1.760624645 | 0.078301954 | 0.173651086 |
| ENSG00000119004 | CYP20A1 | 508.2829933 | -0.122552165 | 0.085888476 | -1.426875531 | 0.153615775 | 0.296685344 |
| ENSG00000119013 | NDUFB3 | 787.6640854 | -0.193754832 | 0.077342558 | -2.505151599 | 0.012239892 | 0.03554305 |
| ENSG00000119041 | GTF3C3 | 1419.552627 | 0.031516849 | 0.066534588 | 0.473691201 | 0.635720129 | 0.785317815 |
| ENSG00000119042 | SATB2 | 974.0793188 | 0.072942369 | 0.075006734 | 0.972477603 | 0.330813002 | 0.521570465 |
| ENSG00000119048 | UBE2B | 984.8670517 | 0.197507125 | 0.069538762 | 2.840245044 | 0.004507889 | 0.014691146 |
| ENSG00000119121 | TRPM6 | 1.869951107 | 0.601553592 | 1.346478087 | 0.446760774 | 0.655047796 | 0.799765631 |
| ENSG00000119138 | KLF9 | 227.6461378 | -0.212386686 | 0.124684605 | -1.703391411 | 0.088494844 | 0.191841969 |
| ENSG00000119139 | TJP2 | 44.32050621 | -0.024194154 | 0.274284548 | -0.088208229 | 0.929711177 | 0.966010255 |
| ENSG00000119185 | ITGB1BP1 | 1676.664312 | 0.063158684 | 0.055737988 | 1.133135351 | 0.257157418 | 0.43681534 |
| ENSG00000119203 | CPSF3 | 1713.322316 | -0.179986934 | 0.062553176 | -2.877342836 | 0.004010396 | 0.013206789 |
| ENSG00000119227 | PIGZ | 269.6950055 | 0.41988494 | 0.115766627 | 3.626994681 | 0.000286739 | 0.001183057 |
| ENSG00000119231 | SENP5 | 1243.760233 | -0.015980254 | 0.062786316 | -0.254518098 | 0.799095324 | 0.892849925 |
| ENSG00000119242 | CCDC92 | 1325.61458 | -0.210764109 | 0.067901088 | -3.103987207 | 0.001909316 | 0.006743604 |
| ENSG00000119280 | C1orf198 | 1514.007585 | -0.113929502 | 0.057717371 | -1.973920488 | 0.048390785 | 0.116544315 |
| ENSG00000119283 | TRIM67 | 4.293713782 | -0.440447735 | 0.866352006 | -0.508393508 | 0.611177405 | 0.768877202 |
| ENSG00000119285 | HEATR1 | 2146.909992 | -0.027359916 | 0.056639313 | -0.483055234 | 0.629056518 | 0.7810577 |
| ENSG00000119314 | PTBP3 | 3611.594998 | -0.109874016 | 0.055304151 | -1.986722754 | 0.046953129 | 0.113475008 |
| ENSG00000119318 | RAD23B | 4957.624525 | -0.168410024 | 0.045567466 | -3.695839148 | 0.000219162 | 0.000922659 |
| ENSG00000119321 | FKBP15 | 1675.578684 | 0.030054844 | 0.059939862 | 0.501416637 | 0.616077933 | 0.772581194 |
| ENSG00000119326 | CTNNAL1 | 812.3399281 | -0.042920453 | 0.071099585 | -0.603666714 | 0.546065249 | 0.719092449 |

| | | | | | | | |
|---|---|---|---|---|---|---|---|
| ENSG00000119328 | ABITRAM | 650.9930648 | 0.158868194 | 0.078921768 | 2.012983213 | 0.044116395 | 0.107685689 |
| ENSG00000119333 | DYNC2I2 | 3275.540429 | -0.225320406 | 0.058533117 | -3.849451685 | 0.000118383 | 0.000521547 |
| ENSG00000119335 | SET | 13268.37203 | -0.044619876 | 0.05420939 | -0.823102347 | 0.410449794 | 0.602953271 |
| ENSG00000119383 | PTPA | 8038.71847 | -0.062823877 | 0.048790715 | -1.287619531 | 0.197878437 | 0.360235803 |
| ENSG00000119392 | GLE1 | 2814.166816 | 0.075252558 | 0.049152234 | 1.531009945 | 0.125766935 | 0.254641967 |
| ENSG00000119396 | RAB14 | 3556.520396 | 0.086529375 | 0.051744047 | 1.672257577 | 0.094473541 | 0.202361003 |
| ENSG00000119397 | CNTRL | 334.5929705 | -0.430381821 | 0.101956258 | -4.221239882 | 2.43E-05 | 0.000119093 |
| ENSG00000119401 | TRIM32 | 798.9148722 | 0.066075937 | 0.076999502 | 0.858134598 | 0.390818145 | 0.583644831 |
| ENSG00000119402 | FBXW2 | 3288.74608 | 0.042025502 | 0.049530728 | 0.848473329 | 0.39617442 | 0.589145937 |
| ENSG00000119403 | PHF19 | 1966.629964 | -0.40553474 | 0.059659133 | -6.797529886 | 1.06E-11 | 9.93E-11 |
| ENSG00000119408 | NEK6 | 1655.95687 | 1.278302878 | 0.064673669 | 19.7654301 | 5.91E-87 | 7.70E-85 |
| ENSG00000119411 | BSPRY | 1051.309273 | -0.034006187 | 0.071806273 | -0.473582403 | 0.635797726 | 0.785317815 |
| ENSG00000119414 | PPP6C | 2343.46231 | 0.141366922 | 0.055690838 | 2.53842333 | 0.01113532 | 0.032669681 |
| ENSG00000119421 | NDUFA8 | 1018.388935 | 0.120839556 | 0.068602072 | 1.761456367 | 0.078161192 | 0.173489864 |
| ENSG00000119431 | HDHD3 | 634.0527116 | 0.548963228 | 0.079818327 | 6.877658904 | 6.08E-12 | 5.81E-11 |
| ENSG00000119446 | RBM18 | 919.1093879 | 0.236376731 | 0.069365783 | 3.407684875 | 0.000655165 | 0.002539194 |
| ENSG00000119471 | HSDL2 | 994.9083081 | 0.379680752 | 0.066456017 | 5.713263766 | 1.11E-08 | 7.93E-08 |
| ENSG00000119487 | MAPKAP1 | 3036.785209 | 0.106652426 | 0.051088824 | 2.087588202 | 0.036834997 | 0.092581882 |
| ENSG00000119508 | NR4A3 | 59.72241982 | 1.219196354 | 0.241486624 | 5.048711747 | 4.45E-07 | 2.70E-06 |
| ENSG00000119509 | INVS | 561.315666 | -0.17114755 | 0.087802857 | -1.949225301 | 0.051268525 | 0.122395666 |
| ENSG00000119514 | GALNT12 | 645.1118861 | -0.834823569 | 0.089302882 | -9.348226476 | 8.91E-21 | 1.56E-19 |
| ENSG00000119522 | DENND1A | 1277.321363 | 0.247967925 | 0.061230978 | 4.049713636 | 5.13E-05 | 0.000239587 |
| ENSG00000119523 | ALG2 | 765.0447218 | 0.016926362 | 0.073973498 | 0.228816563 | 0.819011493 | 0.904057533 |
| ENSG00000119535 | CSF3R | 2.258482143 | -3.658176412 | 1.521337672 | -2.404578864 | 0.016191112 | 0.04549536 |
| ENSG00000119537 | KDSR | 983.6622777 | 1.002948819 | 0.068359927 | 14.67158987 | 9.80E-49 | 5.19E-47 |
| ENSG00000119541 | VPS4B | 1645.716613 | 0.487736066 | 0.06087619 | 8.011934854 | 1.13E-15 | 1.43E-14 |
| ENSG00000119547 | ONECUT2 | 189.2931179 | 0.341274265 | 0.137059065 | 2.489979516 | 0.012775046 | 0.036862836 |
| ENSG00000119559 | C19orf25 | 660.4003302 | 0.198843076 | 0.088684437 | 2.242141707 | 0.024952215 | 0.066380223 |
| ENSG00000119574 | ZBTB45 | 497.1998944 | -0.242987284 | 0.09665812 | -2.51388383 | 0.011940976 | 0.034793089 |
| ENSG00000119596 | YLPM1 | 3923.818877 | -0.590175394 | 0.049321169 | -11.96596537 | 5.36E-33 | 1.67E-31 |
| ENSG00000119599 | DCAF4 | 1972.12212 | -1.240207919 | 0.056093837 | -22.10952199 | 2.56E-108 | 4.98E-106 |
| ENSG00000119616 | FCF1 | 1480.084527 | -0.543858033 | 0.061349317 | -8.864940272 | 7.65E-19 | 1.20E-17 |
| ENSG00000119630 | PGF | 6.059746801 | -0.129909651 | 0.739279289 | -0.175724727 | 0.860510208 | 0.927426758 |

| | | | | | | | |
|---|---|---|---|---|---|---|---|
| ENSG00000119632 | IFI27L2 | 188.2515374 | -0.098224914 | 0.138899387 | -0.707165932 | 0.479463367 | 0.66373081 |
| ENSG00000119636 | BBOF1 | 275.8443859 | -1.252575982 | 0.118323382 | -10.58603942 | 3.46E-26 | 8.14E-25 |
| ENSG00000119638 | NEK9 | 54.47538807 | -0.489758486 | 0.246608472 | -1.98597592 | 0.047035998 | 0.113639072 |
| ENSG00000119640 | ACYP1 | 174.8820217 | -0.454664895 | 0.137012199 | -3.318426369 | 0.000905262 | 0.003417263 |
| ENSG00000119650 | IFT43 | 721.0488524 | -0.586072331 | 0.076918612 | -7.619382558 | 2.55E-14 | 2.91E-13 |
| ENSG00000119655 | NPC2 | 1342.214219 | -0.366356864 | 0.059692172 | -6.13743565 | 8.39E-10 | 6.71E-09 |
| ENSG00000119661 | DNAL1 | 621.8380168 | -0.438046082 | 0.084317413 | -5.195203059 | 2.04E-07 | 1.29E-06 |
| ENSG00000119669 | IRF2BPL | 2101.227476 | 0.148279143 | 0.064066622 | 2.314452332 | 0.020642923 | 0.056194216 |
| ENSG00000119673 | ACOT2 | 161.9458539 | -1.319929011 | 0.151983872 | -8.684665004 | 3.80E-18 | 5.67E-17 |
| ENSG00000119681 | LTBP2 | 128.2778597 | -0.134185471 | 0.156714981 | -0.856238953 | 0.39186562 | 0.584770973 |
| ENSG00000119682 | AREL1 | 2966.083135 | -0.612038174 | 0.049686437 | -12.31801283 | 7.25E-35 | 2.46E-33 |
| ENSG00000119684 | MLH3 | 861.0231487 | -0.364445893 | 0.07179064 | -5.07650991 | 3.84E-07 | 2.34E-06 |
| ENSG00000119685 | TTLL5 | 1642.972439 | -0.773592875 | 0.058753557 | -13.16674122 | 1.36E-39 | 5.42E-38 |
| ENSG00000119686 | FLVCR2 | 542.574974 | -0.352598918 | 0.08774009 | -4.018675123 | 5.85E-05 | 0.000271231 |
| ENSG00000119688 | ABCD4 | 1103.578186 | -0.958106887 | 0.076282154 | -12.56003984 | 3.50E-36 | 1.24E-34 |
| ENSG00000119689 | DLST | 5135.86736 | -0.621541925 | 0.047171107 | -13.17632694 | 1.20E-39 | 4.80E-38 |
| ENSG00000119699 | TGFB3 | 557.4912016 | -0.824759936 | 0.093557025 | -8.815585316 | 1.19E-18 | 1.84E-17 |
| ENSG00000119703 | ZC2HC1C | 39.50805385 | 0.239805645 | 0.305291132 | 0.785498234 | 0.432161505 | 0.622217092 |
| ENSG00000119705 | SLIRP | 991.6867878 | -0.39517833 | 0.072884828 | -5.421956024 | 5.90E-08 | 3.90E-07 |
| ENSG00000119707 | RBM25 | 4507.755129 | -0.529959609 | 0.064740326 | -8.18592747 | 2.70E-16 | 3.57E-15 |
| ENSG00000119711 | ALDH6A1 | 2574.196229 | -1.043540962 | 0.0561556 | -18.58302584 | 4.41E-77 | 4.84E-75 |
| ENSG00000119714 | GPR68 | 220.3107546 | -0.192113555 | 0.127288914 | -1.509271694 | 0.131229365 | 0.263377221 |
| ENSG00000119715 | ESRRB | 2.425786509 | -1.127809366 | 1.151459611 | -0.979460639 | 0.327352428 | 0.517679904 |
| ENSG00000119718 | EIF2B2 | 2150.164132 | -0.790209471 | 0.053218607 | -14.84836825 | 7.13E-50 | 3.92E-48 |
| ENSG00000119720 | NRDE2 | 1151.793747 | -0.740383705 | 0.068633602 | -10.78748142 | 3.94E-27 | 9.82E-26 |
| ENSG00000119723 | COQ6 | 1075.358385 | -1.252777963 | 0.071603625 | -17.49601314 | 1.54E-68 | 1.34E-66 |
| ENSG00000119725 | ZNF410 | 21.70707326 | -0.730613482 | 0.379635014 | -1.924515536 | 0.054289995 | 0.128634341 |
| ENSG00000119729 | RHOQ | 2216.13024 | 0.325852453 | 0.065135973 | 5.002649648 | 5.65E-07 | 3.38E-06 |
| ENSG00000119737 | GPR75 | 17.01790517 | -0.174592455 | 0.432760849 | -0.403438656 | 0.686625554 | 0.822946197 |
| ENSG00000119760 | SUPT7L | 1671.152391 | -0.219649159 | 0.065566846 | -3.350003432 | 0.000808106 | 0.003076837 |
| ENSG00000119771 | KLHL29 | 191.3785422 | 0.57224107 | 0.135456465 | 4.224538625 | 2.39E-05 | 0.000117437 |
| ENSG00000119772 | DNMT3A | 2614.902707 | -0.402832744 | 0.058696803 | -6.862941776 | 6.75E-12 | 6.42E-11 |
| ENSG00000119777 | TMEM214 | 2306.098241 | 0.112213822 | 0.059184147 | 1.896011489 | 0.057958525 | 0.135614909 |

| | | | | | | | |
|---|---|---|---|---|---|---|---|
| ENSG00000119778 | ATAD2B | 993.058642 | -0.107095399 | 0.070440716 | -1.520362164 | 0.128419978 | 0.258848458 |
| ENSG00000119782 | FKBP1B | 3.091884297 | -1.604737602 | 1.084993696 | -1.479029426 | 0.139132451 | 0.275217592 |
| ENSG00000119787 | ATL2 | 1701.246794 | 0.082595362 | 0.06285517 | 1.314058379 | 0.188826559 | 0.347563289 |
| ENSG00000119801 | YPEL5 | 1499.765885 | 0.230371288 | 0.060694935 | 3.795560327 | 0.00014731 | 0.000639413 |
| ENSG00000119812 | FAM98A | 1527.926999 | -0.009319851 | 0.060603807 | -0.15378326 | 0.877780634 | 0.936395015 |
| ENSG00000119820 | YIPF4 | 916.950064 | 0.26414069 | 0.071245285 | 3.707483101 | 0.000209329 | 0.000884512 |
| ENSG00000119844 | AFTPH | 1789.03581 | -0.040331214 | 0.059746538 | -0.675038509 | 0.499651299 | 0.681005336 |
| ENSG00000119862 | LGALSL | 420.0051875 | -0.317045734 | 0.092248176 | -3.436878067 | 0.00058846 | 0.00229775 |
| ENSG00000119866 | BCL11A | 3.465976381 | 0.650824726 | 0.94974576 | 0.68526205 | 0.493178575 | 0.675918061 |
| ENSG00000119878 | CRIPT | 300.1455543 | 0.152056474 | 0.111852546 | 1.359436861 | 0.174008197 | 0.326710431 |
| ENSG00000119888 | EPCAM | 5502.743878 | -0.120473042 | 0.047082138 | -2.558784407 | 0.010503886 | 0.031023451 |
| ENSG00000119899 | SLC17A5 | 409.9480855 | 0.383175819 | 0.100172948 | 3.825142679 | 0.000130696 | 0.000571952 |
| ENSG00000119900 | OGFRL1 | 1332.039991 | -0.008587762 | 0.06997325 | -0.122729214 | 0.902321529 | 0.950707258 |
| ENSG00000119906 | SLF2 | 1179.606068 | -0.176507501 | 0.067446759 | -2.616990124 | 0.008870891 | 0.026783132 |
| ENSG00000119912 | IDE | 2926.039465 | -0.577470034 | 0.058318154 | -9.902063037 | 4.08E-23 | 8.15E-22 |
| ENSG00000119915 | ELOVL3 | 4.023824475 | -0.514051616 | 0.890907837 | -0.576997524 | 0.563941128 | 0.73250719 |
| ENSG00000119917 | IFIT3 | 31.61607242 | 3.263564441 | 0.434838014 | 7.505241809 | 6.13E-14 | 6.80E-13 |
| ENSG00000119919 | NKX2-3 | 1.799469322 | 0.070166394 | 1.310920481 | 0.053524523 | 0.957313992 | 0.980698565 |
| ENSG00000119922 | IFIT2 | 43.78179069 | 3.283596484 | 0.373384648 | 8.794138981 | 1.44E-18 | 2.21E-17 |
| ENSG00000119927 | GPAM | 643.7457839 | -0.550361204 | 0.079214796 | -6.947707104 | 3.71E-12 | 3.62E-11 |
| ENSG00000119929 | CUTC | 422.0789137 | -0.099274222 | 0.096241375 | -1.031512919 | 0.302300353 | 0.489241117 |
| ENSG00000119938 | PPP1R3C | 1800.013449 | 0.240719955 | 0.058148876 | 4.139718144 | 3.48E-05 | 0.00016634 |
| ENSG00000119943 | PYROXD2 | 685.3716782 | -1.028317813 | 0.079459429 | -12.94141959 | 2.63E-38 | 1.01E-36 |
| ENSG00000119946 | CNNM1 | 800.6519536 | -0.211240628 | 0.075532829 | -2.79667304 | 0.005163175 | 0.016577081 |
| ENSG00000119950 | MXI1 | 797.7806775 | 0.172782583 | 0.078289602 | 2.206967175 | 0.027316353 | 0.071681209 |
| ENSG00000119953 | SMNDC1 | 686.1422662 | -0.448146589 | 0.081270408 | -5.514265314 | 3.50E-08 | 2.38E-07 |
| ENSG00000119965 | C10orf88 | 243.0867386 | -0.483832843 | 0.117966905 | -4.101428641 | 4.11E-05 | 0.000194213 |
| ENSG00000119969 | HELLS | 1099.310839 | -0.612207969 | 0.070703551 | -8.658800855 | 4.77E-18 | 7.05E-17 |
| ENSG00000119977 | TCTN3 | 1914.455603 | -0.290398295 | 0.059070178 | -4.916157429 | 8.83E-07 | 5.15E-06 |
| ENSG00000119979 | DENND10 | 378.9830994 | -0.313754464 | 0.098327807 | -3.190902707 | 0.00141829 | 0.005145592 |
| ENSG00000119986 | AVPI1 | 559.2456124 | -0.838862445 | 0.089134066 | -9.411243986 | 4.90E-21 | 8.70E-20 |
| ENSG00000120008 | WDR11 | 1846.291362 | -0.483413248 | 0.056960431 | -8.486825665 | 2.12E-17 | 3.00E-16 |
| ENSG00000120029 | ARMH3 | 1012.699895 | 0.038440739 | 0.069184992 | 0.555622519 | 0.578468934 | 0.74408867 |

| | | | | | | | |
|---|---|---|---|---|---|---|---|
| ENSG00000120049 | KCNIP2 | 12.045901 | -0.037551211 | 0.513475111 | -0.073131512 | 0.941701466 | 0.972259401 |
| ENSG00000120053 | GOT1 | 1869.894183 | -0.197143456 | 0.058037629 | -3.39682136 | 0.000681735 | 0.002633714 |
| ENSG00000120055 | C10orf95 | 39.29479106 | 0.328361371 | 0.283106949 | 1.15984921 | 0.246110205 | 0.423284457 |
| ENSG00000120063 | GNA13 | 5137.227059 | -0.056638116 | 0.051854456 | -1.092251675 | 0.274722495 | 0.45683264 |
| ENSG00000120071 | KANSL1 | 2006.60864 | -0.142787308 | 0.054741545 | -2.60839017 | 0.00909692 | 0.027386265 |
| ENSG00000120129 | DUSP1 | 363.7767574 | -0.804832285 | 0.106566313 | -7.552408061 | 4.27E-14 | 4.79E-13 |
| ENSG00000120137 | PANK3 | 5258.939747 | -0.792751626 | 0.05669777 | -13.98206008 | 2.01E-44 | 9.47E-43 |
| ENSG00000120149 | MSX2 | 1043.138818 | -0.853398536 | 0.072054911 | -11.8437248 | 2.32E-32 | 7.11E-31 |
| ENSG00000120158 | RCL1 | 662.5434447 | 0.144858462 | 0.081080858 | 1.786592605 | 0.074003358 | 0.166190605 |
| ENSG00000120159 | CAAP1 | 528.5324594 | -0.048035791 | 0.091262453 | -0.526347795 | 0.59864658 | 0.75887321 |
| ENSG00000120162 | MOB3B | 119.6997748 | -0.558629212 | 0.168546248 | -3.314397193 | 0.000918409 | 0.003462905 |
| ENSG00000120211 | INSL4 | 5.298754502 | 1.135166747 | 0.855283773 | 1.327239897 | 0.184429339 | 0.341111749 |
| ENSG00000120217 | CD274 | 4.1930475 | -2.16822891 | 1.048282632 | -2.068362905 | 0.038605911 | 0.096336298 |
| ENSG00000120251 | GRIA2 | 2.89299629 | 0.49163437 | 1.017687861 | 0.483089549 | 0.629032155 | 0.7810577 |
| ENSG00000120253 | NUP43 | 1288.851665 | 0.783681851 | 0.06555199 | 11.95511908 | 6.10E-33 | 1.89E-31 |
| ENSG00000120254 | MTHFD1L | 2052.611065 | 0.590889863 | 0.058002846 | 10.18725636 | 2.26E-24 | 4.86E-23 |
| ENSG00000120256 | LRP11 | 59.88525137 | 0.386084213 | 0.230144926 | 1.677569955 | 0.093431082 | 0.200477881 |
| ENSG00000120262 | CCDC170 | 4339.604163 | -0.223997847 | 0.046443845 | -4.822982376 | 1.41E-06 | 8.02E-06 |
| ENSG00000120265 | PCMT1 | 3306.272664 | 0.09795643 | 0.053935241 | 1.816186024 | 0.069341841 | 0.15746405 |
| ENSG00000120278 | PLEKHG1 | 25.42832153 | -1.899955109 | 0.384095067 | -4.946575136 | 7.55E-07 | 4.45E-06 |
| ENSG00000120306 | CYSTM1 | 975.6280444 | 0.014534547 | 0.077900565 | 0.186578199 | 0.851991361 | 0.922571717 |
| ENSG00000120314 | WDR55 | 1932.03367 | 0.062167772 | 0.057698244 | 1.07746385 | 0.281273092 | 0.464703258 |
| ENSG00000120318 | ARAP3 | 10.67854448 | 1.382567248 | 0.570972931 | 2.421423457 | 0.015459856 | 0.043702909 |
| ENSG00000120322 | PCDHB8 | 46.40856448 | 0.173785249 | 0.259803755 | 0.668909691 | 0.503553088 | 0.684589655 |
| ENSG00000120324 | PCDHB10 | 13.73710556 | 0.743907855 | 0.488581628 | 1.522586631 | 0.127862157 | 0.257973428 |
| ENSG00000120327 | PCDHB14 | 35.30668212 | 0.283989983 | 0.295370983 | 0.961468794 | 0.336316509 | 0.527420065 |
| ENSG00000120328 | PCDHB12 | 9.610973831 | -0.221212332 | 0.602842977 | -0.366948509 | 0.713657427 | 0.840800958 |
| ENSG00000120333 | MRPS14 | 706.4435487 | -0.283054397 | 0.079301502 | -3.5693447 | 0.000357875 | 0.001449957 |
| ENSG00000120334 | CENPL | 649.1334217 | -0.151673454 | 0.079597251 | -1.905511217 | 0.056713651 | 0.133191149 |
| ENSG00000120370 | GORAB | 321.3963537 | 0.073937269 | 0.109255701 | 0.676736024 | 0.498573456 | 0.680263789 |
| ENSG00000120437 | ACAT2 | 2252.267956 | -0.679254692 | 0.054014004 | -12.57552935 | 2.88E-36 | 1.03E-34 |
| ENSG00000120438 | TCP1 | 5941.390375 | 0.070341126 | 0.048974817 | 1.436271332 | 0.150925148 | 0.293036217 |
| ENSG00000120451 | SNX19 | 1916.297675 | 0.060143393 | 0.062888 | 0.956357225 | 0.338891788 | 0.530045518 |

| | | | | | | | |
|---|---|---|---|---|---|---|---|
| ENSG00000120457 | KCNJ5 | 4.019656358 | -1.207382959 | 0.925974697 | -1.303904915 | 0.192266004 | 0.352395505 |
| ENSG00000120458 | MSANTD2 | 149.1482333 | -0.578173161 | 0.151205844 | -3.823748767 | 0.000131438 | 0.000574759 |
| ENSG00000120471 | TP53AIP1 | 4.13404976 | 0.244253349 | 0.936737806 | 0.260748897 | 0.794286156 | 0.889780468 |
| ENSG00000120509 | PDZD11 | 1408.090856 | -0.146607879 | 0.061180465 | -2.396318469 | 0.016560694 | 0.046403118 |
| ENSG00000120519 | SLC10A7 | 151.0492169 | -0.373852119 | 0.154460746 | -2.420369769 | 0.015504732 | 0.043808172 |
| ENSG00000120526 | NUDCD1 | 1550.090369 | -0.387333711 | 0.060910982 | -6.359012752 | 2.03E-10 | 1.72E-09 |
| ENSG00000120533 | ENY2 | 2228.287136 | -0.330899592 | 0.054300705 | -6.093835988 | 1.10E-09 | 8.74E-09 |
| ENSG00000120539 | MASTL | 811.7205714 | -0.301537658 | 0.074560712 | -4.044189623 | 5.25E-05 | 0.000244757 |
| ENSG00000120549 | KIAA1217 | 534.2175465 | -0.696144421 | 0.085130199 | -8.177408607 | 2.90E-16 | 3.83E-15 |
| ENSG00000120555 | SEPTIN7P9 | 7.420011928 | -0.84681039 | 0.655066707 | -1.292708638 | 0.196111849 | 0.357615725 |
| ENSG00000120594 | PLXDC2 | 1178.019531 | -0.624140451 | 0.065845198 | -9.478906117 | 2.57E-21 | 4.64E-20 |
| ENSG00000120616 | EPC1 | 1099.413392 | -0.54922623 | 0.066193237 | -8.297316348 | 1.06E-16 | 1.45E-15 |
| ENSG00000120647 | CCDC77 | 165.5624444 | 0.249506888 | 0.141689966 | 1.760935478 | 0.078249324 | 0.173578744 |
| ENSG00000120656 | TAF12 | 847.155472 | -0.289971894 | 0.071542447 | -4.053144749 | 5.05E-05 | 0.000236387 |
| ENSG00000120658 | ENOX1 | 29.23376397 | -0.256664851 | 0.320586684 | -0.800609831 | 0.423357558 | 0.614254144 |
| ENSG00000120662 | MTRF1 | 147.4905932 | 0.30409641 | 0.149383495 | 2.035676102 | 0.04178289 | 0.102777129 |
| ENSG00000120664 | SPART-AS1 | 4.302340553 | 0.62997709 | 0.856580584 | 0.73545572 | 0.462061998 | 0.648626151 |
| ENSG00000120669 | SOHLH2 | 4.228756596 | 1.877754368 | 0.962406931 | 1.951102291 | 0.051044879 | 0.121975992 |
| ENSG00000120675 | DNAJC15 | 1098.218638 | 0.048431275 | 0.063677758 | 0.760568154 | 0.446915047 | 0.635403234 |
| ENSG00000120685 | PROSER1 | 2974.463267 | -0.111669091 | 0.059010977 | -1.892344382 | 0.058445111 | 0.136558449 |
| ENSG00000120686 | UFM1 | 1747.528412 | 0.133955601 | 0.063685194 | 2.103402554 | 0.035430595 | 0.089542198 |
| ENSG00000120688 | WBP4 | 778.4438525 | 0.056552932 | 0.07552057 | 0.748841434 | 0.453952783 | 0.641552294 |
| ENSG00000120690 | ELF1 | 3922.499809 | 0.095973273 | 0.051601666 | 1.859887099 | 0.062901501 | 0.145079472 |
| ENSG00000120693 | SMAD9 | 140.7529192 | -0.942420694 | 0.155435712 | -6.06308988 | 1.34E-09 | 1.05E-08 |
| ENSG00000120694 | HSPH1 | 7176.379849 | 0.253890317 | 0.045376405 | 5.595205634 | 2.20E-08 | 1.53E-07 |
| ENSG00000120696 | KBTBD7 | 771.4740103 | -0.030660048 | 0.076810953 | -0.399162454 | 0.689773506 | 0.824651044 |
| ENSG00000120697 | ALG5 | 811.7672609 | -0.233479455 | 0.072619899 | -3.215089229 | 0.00130404 | 0.004777939 |
| ENSG00000120699 | EXOSC8 | 1042.903448 | -0.51121187 | 0.150754505 | -3.391022186 | 0.000696325 | 0.002685556 |
| ENSG00000120705 | ETF1 | 5314.489041 | 0.171829039 | 0.045792199 | 3.752364871 | 0.000175174 | 0.000751121 |
| ENSG00000120708 | TGFBI | 29.87033654 | -0.733455978 | 0.328011109 | -2.23607054 | 0.025347151 | 0.06722123 |
| ENSG00000120709 | FAM53C | 2343.735939 | 0.132481934 | 0.056577924 | 2.341583501 | 0.01920213 | 0.05279205 |
| ENSG00000120725 | SIL1 | 806.0965058 | 0.085618541 | 0.074759432 | 1.145254036 | 0.25210394 | 0.430783992 |
| ENSG00000120727 | PAIP2 | 3281.872794 | -0.13268528 | 0.049940481 | -2.656868286 | 0.007887025 | 0.024152764 |

| | | | | | | | |
|---|---|---|---|---|---|---|---|
| ENSG00000120733 | KDM3B | 4709.046825 | 0.101266342 | 0.047469619 | 2.133287444 | 0.032901154 | 0.084076246 |
| ENSG00000120738 | EGR1 | 1929.083699 | 0.417217449 | 0.064331018 | 6.485478749 | 8.85E-11 | 7.70E-10 |
| ENSG00000120742 | SERP1 | 5374.363758 | -0.060589661 | 0.058002877 | -1.04459752 | 0.296209026 | 0.482430605 |
| ENSG00000120756 | PLS1 | 863.8561551 | 0.211661927 | 0.070091013 | 3.019815481 | 0.002529287 | 0.008704148 |
| ENSG00000120784 | ZFP30 | 509.4730624 | 0.012707077 | 0.087542903 | 0.145152569 | 0.884590414 | 0.940249752 |
| ENSG00000120798 | NR2C1 | 641.7321562 | 0.2113083 | 0.0869209 | 2.431041319 | 0.015055498 | 0.042649187 |
| ENSG00000120800 | UTP20 | 1097.637908 | 0.03348093 | 0.071028587 | 0.471372608 | 0.637374673 | 0.78651829 |
| ENSG00000120802 | TMPO | 7769.723141 | -0.351785998 | 0.049454162 | -7.113374937 | 1.13E-12 | 1.15E-11 |
| ENSG00000120805 | ARL1 | 2011.814957 | -0.140407075 | 0.05319111 | -2.639671866 | 0.008298633 | 0.025224679 |
| ENSG00000120820 | GLT8D2 | 1.831655675 | 0.56712904 | 1.39286944 | 0.407165973 | 0.683886091 | 0.82069193 |
| ENSG00000120832 | MTERF2 | 339.3194457 | 0.208347052 | 0.105355607 | 1.977560174 | 0.047978346 | 0.115660283 |
| ENSG00000120833 | SOCS2 | 301.1265016 | -0.675772131 | 0.11225011 | -6.020235824 | 1.74E-09 | 1.36E-08 |
| ENSG00000120837 | NFYB | 1299.362695 | -0.501160695 | 0.069053584 | -7.257562397 | 3.94E-13 | 4.13E-12 |
| ENSG00000120860 | WASHC3 | 427.4241414 | -0.349917233 | 0.095494707 | -3.66425789 | 0.000248057 | 0.001033307 |
| ENSG00000120868 | APAF1 | 624.8748234 | -0.79968115 | 0.084777804 | -9.432671186 | 4.00E-21 | 7.15E-20 |
| ENSG00000120875 | DUSP4 | 1299.264133 | -0.270370048 | 0.063493293 | -4.258245802 | 2.06E-05 | 0.00010215 |
| ENSG00000120885 | CLU | 11045.23504 | -0.925332594 | 0.064340394 | -14.38182977 | 6.73E-47 | 3.42E-45 |
| ENSG00000120889 | TNFRSF10B | 1479.764373 | 0.18299837 | 0.0658492 | 2.779052303 | 0.005451775 | 0.017379615 |
| ENSG00000120896 | SORBS3 | 1537.127759 | -0.56867066 | 0.066159145 | -8.595495855 | 8.29E-18 | 1.21E-16 |
| ENSG00000120899 | PTK2B | 282.2850419 | -0.946318453 | 0.111935517 | -8.454139321 | 2.81E-17 | 3.94E-16 |
| ENSG00000120910 | PPP3CC | 249.2823059 | -0.224552273 | 0.117275503 | -1.9147415 | 0.055525478 | 0.131031218 |
| ENSG00000120913 | PDLIM2 | 113.6832163 | -0.138457731 | 0.173366214 | -0.798643101 | 0.424497388 | 0.615236927 |
| ENSG00000120915 | EPHX2 | 129.9388105 | -0.674798879 | 0.158298066 | -4.262837153 | 2.02E-05 | 0.000100225 |
| ENSG00000120925 | RNF170 | 341.1979496 | 0.013765652 | 0.101763984 | 0.135270368 | 0.892398112 | 0.944983167 |
| ENSG00000120942 | UBIAD1 | 848.8297959 | 0.122796051 | 0.074713654 | 1.643555698 | 0.100268011 | 0.21242595 |
| ENSG00000120948 | TARDBP | 2479.024952 | -0.125241073 | 0.057423599 | -2.181003537 | 0.029183155 | 0.075799544 |
| ENSG00000120963 | ZNF706 | 1906.725192 | 0.039171302 | 0.055731339 | 0.702859526 | 0.482143297 | 0.666287182 |
| ENSG00000120992 | LYPLA1 | 1672.099447 | -0.066754082 | 0.06845416 | -0.975164736 | 0.329478551 | 0.519863226 |
| ENSG00000121005 | CRISPLD1 | 432.8748189 | 1.182568675 | 0.098596854 | 11.99397983 | 3.82E-33 | 1.20E-31 |
| ENSG00000121022 | COPS5 | 1563.176039 | 0.151149128 | 0.061017234 | 2.477154706 | 0.013243448 | 0.038085083 |
| ENSG00000121039 | RDH10 | 570.9891302 | 0.72684238 | 0.085637106 | 8.487470142 | 2.11E-17 | 2.98E-16 |
| ENSG00000121057 | AKAP1 | 4589.346097 | 0.26284273 | 0.045471604 | 5.780370707 | 7.45E-09 | 5.43E-08 |
| ENSG00000121058 | COIL | 1562.981281 | 0.064468537 | 0.057184403 | 1.127379735 | 0.259581963 | 0.439501826 |

| | | | | | | | |
|---|---|---|---|---|---|---|---|
| ENSG00000121060 | TRIM25 | 6419.709586 | 0.45091751 | 0.043013421 | 10.4831817 | 1.03E-25 | 2.37E-24 |
| ENSG00000121064 | SCPEP1 | 1088.198409 | 0.84029183 | 0.065685744 | 12.79260571 | 1.80E-37 | 6.69E-36 |
| ENSG00000121067 | SPOP | 2295.014154 | -0.111131353 | 0.055409678 | -2.005630719 | 0.044895662 | 0.109227541 |
| ENSG00000121068 | TBX2 | 6569.528325 | 0.008222971 | 0.049365727 | 0.166572459 | 0.867706466 | 0.930975461 |
| ENSG00000121073 | SLC35B1 | 2393.169192 | -0.01933416 | 0.060150669 | -0.321428839 | 0.747885432 | 0.860419264 |
| ENSG00000121075 | TBX4 | 49.78796203 | 0.075249133 | 0.2629737 | 0.286146992 | 0.774765541 | 0.877499635 |
| ENSG00000121089 | NACA3P | 498.8704991 | 0.249241778 | 0.08573317 | 2.907180262 | 0.00364703 | 0.012125052 |
| ENSG00000121101 | TEX14 | 243.7437835 | -1.575821164 | 0.145181851 | -10.85411956 | 1.91E-27 | 4.83E-26 |
| ENSG00000121104 | FAM117A | 610.1895244 | 0.048122228 | 0.080770746 | 0.595787842 | 0.551316967 | 0.723235044 |
| ENSG00000121152 | NCAPH | 1511.934735 | -0.295093189 | 0.061863856 | -4.770041968 | 1.84E-06 | 1.03E-05 |
| ENSG00000121207 | LRAT | 12.39237058 | -0.147904258 | 0.488820797 | -0.30257358 | 0.762214852 | 0.869521542 |
| ENSG00000121210 | TMEM131L | 1464.122569 | -0.635879264 | 0.06128792 | -10.37527896 | 3.21E-25 | 7.22E-24 |
| ENSG00000121211 | MND1 | 206.6159576 | 0.086811237 | 0.133682912 | 0.649381702 | 0.516091688 | 0.694703869 |
| ENSG00000121236 | TRIM6 | 1.927107127 | 0.743242928 | 1.379342243 | 0.538838661 | 0.589998184 | 0.752932545 |
| ENSG00000121270 | ABCC11 | 32.87878821 | -1.319445169 | 0.349108311 | -3.779472234 | 0.000157161 | 0.000678832 |
| ENSG00000121274 | TENT4B | 435.7587915 | 0.242115942 | 0.100446701 | 2.410392163 | 0.015935381 | 0.044848196 |
| ENSG00000121281 | ADCY7 | 140.3326263 | 0.366681602 | 0.158317317 | 2.316118099 | 0.020551821 | 0.055959499 |
| ENSG00000121289 | CEP89 | 1317.61549 | -0.242114042 | 0.062181753 | -3.893650962 | 9.87E-05 | 0.000440367 |
| ENSG00000121297 | TSHZ3 | 6.741646857 | -0.35339538 | 0.672687475 | -0.525348536 | 0.599340923 | 0.759438841 |
| ENSG00000121310 | ECHDC2 | 781.2720735 | 0.021269494 | 0.082254245 | 0.258582323 | 0.795957524 | 0.891000127 |
| ENSG00000121316 | PLBD1 | 119.7668215 | 1.500547199 | 0.182584323 | 8.218379165 | 2.06E-16 | 2.74E-15 |
| ENSG00000121318 | TAS2R10 | 2.573892235 | 0.966929468 | 1.100636723 | 0.878518268 | 0.379662528 | 0.572075049 |
| ENSG00000121350 | PYROXD1 | 254.0282095 | 0.61767602 | 0.123804447 | 4.989126287 | 6.07E-07 | 3.61E-06 |
| ENSG00000121361 | KCNJ8 | 65.7034678 | 0.457903257 | 0.224399941 | 2.040567639 | 0.041293821 | 0.101814302 |
| ENSG00000121390 | PSPC1 | 2825.540331 | 0.287241583 | 0.049903229 | 5.755971873 | 8.61E-09 | 6.23E-08 |
| ENSG00000121410 | A1BG | 9.953822935 | -0.255947986 | 0.554751044 | -0.461374502 | 0.644529942 | 0.791770208 |
| ENSG00000121413 | ZSCAN18 | 426.1523392 | 0.87649616 | 0.095172217 | 9.209580143 | 3.27E-20 | 5.53E-19 |
| ENSG00000121417 | ZNF211 | 546.2401519 | 0.317453331 | 0.085306298 | 3.721335228 | 0.000198172 | 0.000841699 |
| ENSG00000121440 | PDZRN3 | 295.6875738 | 1.001226835 | 0.115469769 | 8.67090016 | 4.29E-18 | 6.36E-17 |
| ENSG00000121454 | LHX4 | 56.95080849 | 1.233575821 | 0.245037378 | 5.034235311 | 4.80E-07 | 2.90E-06 |
| ENSG00000121481 | RNF2 | 574.2136437 | -0.221782862 | 0.083503392 | -2.655974277 | 0.007907965 | 0.024210424 |
| ENSG00000121486 | TRMT1L | 590.2376052 | -0.297297402 | 0.086130812 | -3.451696279 | 0.000557074 | 0.002184867 |
| ENSG00000121542 | SEC22A | 360.4727892 | -0.21909884 | 0.099832654 | -2.194661072 | 0.028187919 | 0.07363139 |

| | | | | | | | |
|---|---|---|---|---|---|---|---|
| ENSG00000121552 | CSTA | 89.31926976 | -3.286331511 | 0.264371416 | -12.4307369 | 1.78E-35 | 6.15E-34 |
| ENSG00000121577 | POPDC2 | 13.66279772 | 0.05634812 | 0.467332732 | 0.120573878 | 0.90402856 | 0.951652678 |
| ENSG00000121578 | B4GALT4 | 620.3210501 | 0.494549626 | 0.080411814 | 6.150211069 | 7.74E-10 | 6.21E-09 |
| ENSG00000121579 | NAA50 | 5412.028958 | -0.127429718 | 0.05105296 | -2.496029973 | 0.012559199 | 0.036322242 |
| ENSG00000121621 | KIF18A | 418.3395856 | -0.439440218 | 0.094924716 | -4.629355084 | 3.67E-06 | 1.99E-05 |
| ENSG00000121644 | DESI2 | 1189.720311 | -0.169362792 | 0.068426112 | -2.475119329 | 0.013319167 | 0.038278838 |
| ENSG00000121653 | MAPK8IP1 | 911.6050943 | 1.019343151 | 0.07694705 | 13.24733245 | 4.68E-40 | 1.92E-38 |
| ENSG00000121671 | CRY2 | 1361.949391 | -0.007049138 | 0.072081163 | -0.097794447 | 0.922095517 | 0.962216621 |
| ENSG00000121680 | PEX16 | 896.7805709 | 0.445896527 | 0.072607174 | 6.14121856 | 8.19E-10 | 6.56E-09 |
| ENSG00000121691 | CAT | 1570.309118 | 0.071428359 | 0.061098883 | 1.169061615 | 0.242378808 | 0.418655963 |
| ENSG00000121716 | PILRB | 159.4769777 | -0.388786345 | 0.15041857 | -2.584696457 | 0.009746476 | 0.029067104 |
| ENSG00000121741 | ZMYM2 | 4953.245421 | 0.524435589 | 0.051977945 | 10.08957912 | 6.14E-24 | 1.28E-22 |
| ENSG00000121743 | GJA3 | 42.93111697 | -0.245391652 | 0.277079891 | -0.885635009 | 0.37581424 | 0.568791889 |
| ENSG00000121749 | TBC1D15 | 957.8403117 | 0.104656633 | 0.072348303 | 1.446566512 | 0.148018371 | 0.288564752 |
| ENSG00000121753 | ADGRB2 | 539.6568304 | -0.017054158 | 0.08460507 | -0.201573717 | 0.840249994 | 0.915953947 |
| ENSG00000121764 | HCRTR1 | 3.903917567 | -0.354849257 | 0.881671607 | -0.402473273 | 0.687335754 | 0.823367217 |
| ENSG00000121766 | ZCCHC17 | 1335.211935 | 0.037137519 | 0.061984998 | 0.599137211 | 0.549081389 | 0.72174691 |
| ENSG00000121769 | FABP3 | 2.301556966 | -0.217735118 | 1.161297459 | -0.187492977 | 0.851274129 | 0.9222591 |
| ENSG00000121774 | KHDRBS1 | 6799.689964 | -0.063882804 | 0.042552335 | -1.501276113 | 0.133284161 | 0.266452169 |
| ENSG00000121775 | TMEM39B | 426.2317456 | -0.101807251 | 0.09328593 | -1.091346253 | 0.275120553 | 0.457338078 |
| ENSG00000121807 | CCR2 | 1.809373839 | 0.10605002 | 1.27799841 | 0.08298134 | 0.933866377 | 0.968266427 |
| ENSG00000121851 | POLR3GL | 286.7212105 | -0.263895604 | 0.113571487 | -2.323607905 | 0.020146518 | 0.055032285 |
| ENSG00000121864 | ZNF639 | 886.0286774 | -0.327392161 | 0.074830027 | -4.375144237 | 1.21E-05 | 6.18E-05 |
| ENSG00000121871 | SLITRK3 | 2.897315246 | 1.626444473 | 1.106305535 | 1.470158488 | 0.141518835 | 0.278903958 |
| ENSG00000121879 | PIK3CA | 814.8602068 | -0.171910969 | 0.071767294 | -2.395394324 | 0.016602499 | 0.046508894 |
| ENSG00000121892 | PDS5A | 7189.507769 | -0.006508759 | 0.053680892 | -0.12124909 | 0.903493743 | 0.951246259 |
| ENSG00000121897 | LIAS | 266.9352018 | 0.02139418 | 0.116369693 | 0.183846662 | 0.854133749 | 0.924103822 |
| ENSG00000121898 | CPXM2 | 73.01480931 | -0.823114449 | 0.219498713 | -3.749973914 | 0.000176853 | 0.000757753 |
| ENSG00000121900 | TMEM54 | 2050.805128 | -0.240275123 | 0.061312918 | -3.918833592 | 8.90E-05 | 0.000400307 |
| ENSG00000121903 | ZSCAN20 | 162.1751295 | -0.067249926 | 0.154499994 | -0.435274618 | 0.663363106 | 0.805636147 |
| ENSG00000121905 | HPCA | 38.67547367 | 0.020606356 | 0.285629667 | 0.072143614 | 0.942487618 | 0.972720833 |
| ENSG00000121931 | LRIF1 | 700.3183335 | 0.238966515 | 0.075568178 | 3.162263829 | 0.001565477 | 0.005629728 |
| ENSG00000121940 | CLCC1 | 1302.938527 | 0.348932666 | 0.061543876 | 5.669656959 | 1.43E-08 | 1.01E-07 |

| | | | | | | | |
|---|---|---|---|---|---|---|---|
| ENSG00000121957 | GPSM2 | 1586.267022 | -0.089590693 | 0.057768271 | -1.550863347 | 0.120934436 | 0.246971954 |
| ENSG00000121964 | GTDC1 | 498.0448507 | 0.433226268 | 0.089695389 | 4.829972574 | 1.37E-06 | 7.76E-06 |
| ENSG00000121966 | CXCR4 | 2888.440141 | -0.558451231 | 0.052524242 | -10.63225694 | 2.11E-26 | 5.04E-25 |
| ENSG00000121988 | ZRANB3 | 362.8481676 | 0.000325688 | 0.099808464 | 0.003263131 | 0.997396403 | 0.998440979 |
| ENSG00000121989 | ACVR2A | 260.0308248 | -0.138993287 | 0.11777399 | -1.180169642 | 0.237932751 | 0.413417262 |
| ENSG00000122008 | POLK | 288.5052406 | -0.227756521 | 0.112133519 | -2.031119 | 0.042242925 | 0.103775186 |
| ENSG00000122012 | SV2C | 2.732762979 | -0.97916908 | 1.127887116 | -0.868144574 | 0.385315192 | 0.577892122 |
| ENSG00000122025 | FLT3 | 2.445731236 | 0.04638494 | 1.104664002 | 0.041990089 | 0.966506599 | 0.98505693 |
| ENSG00000122026 | RPL21 | 204.8991136 | 0.022106682 | 0.134443996 | 0.164430415 | 0.869392322 | 0.932217438 |
| ENSG00000122033 | MTIF3 | 1012.369634 | 0.078797887 | 0.069586185 | 1.132378312 | 0.257475419 | 0.437161284 |
| ENSG00000122034 | GTF3A | 3771.74527 | -0.040165667 | 0.051376328 | -0.781793264 | 0.434336081 | 0.624290081 |
| ENSG00000122035 | RASL11A | 15.95697066 | 0.076169573 | 0.458617267 | 0.16608527 | 0.868089846 | 0.931081965 |
| ENSG00000122042 | UBL3 | 5013.816518 | -0.456635069 | 0.053439596 | -8.544882554 | 1.29E-17 | 1.85E-16 |
| ENSG00000122068 | FYTTD1 | 3009.23228 | -0.260943385 | 0.051892503 | -5.028537252 | 4.94E-07 | 2.98E-06 |
| ENSG00000122085 | MTERF4 | 1440.327866 | -1.440021806 | 0.069366619 | -20.75957906 | 1.00E-95 | 1.50E-93 |
| ENSG00000122126 | OCRL | 3814.13212 | 0.236003507 | 0.046983504 | 5.023114172 | 5.08E-07 | 3.06E-06 |
| ENSG00000122140 | MRPS2 | 1886.102328 | 0.094096631 | 0.057906307 | 1.624980706 | 0.10416667 | 0.219107226 |
| ENSG00000122203 | KIAA1191 | 4060.611132 | -0.279650627 | 0.045812131 | -6.104292099 | 1.03E-09 | 8.20E-09 |
| ENSG00000122218 | COPA | 7326.369476 | -0.072764371 | 0.045761546 | -1.590076773 | 0.111817501 | 0.231925268 |
| ENSG00000122257 | RBBP6 | 3067.505949 | -0.081466129 | 0.052265234 | -1.558705906 | 0.119066003 | 0.244031299 |
| ENSG00000122299 | ZC3H7A | 1969.683099 | -0.293857865 | 0.056684245 | -5.184118886 | 2.17E-07 | 1.36E-06 |
| ENSG00000122335 | SERAC1 | 201.2653649 | -0.018315194 | 0.133106526 | -0.137598015 | 0.890558124 | 0.943913993 |
| ENSG00000122359 | ANXA11 | 3267.489157 | -0.497571029 | 0.048882911 | -10.17883383 | 2.46E-24 | 5.27E-23 |
| ENSG00000122367 | LDB3 | 66.74970258 | -0.184879346 | 0.219448587 | -0.842472252 | 0.399523665 | 0.59228854 |
| ENSG00000122376 | SHLD2 | 1623.354339 | -0.422779997 | 0.064073303 | -6.598379941 | 4.16E-11 | 3.69E-10 |
| ENSG00000122378 | PRXL2A | 3769.167775 | -0.217509622 | 0.046845443 | -4.643133014 | 3.43E-06 | 1.87E-05 |
| ENSG00000122386 | ZNF205 | 377.4258954 | -0.090697294 | 0.09812111 | -0.924340274 | 0.355309181 | 0.54703306 |
| ENSG00000122390 | NAA60 | 2365.873655 | 0.091346697 | 0.062827503 | 1.453928498 | 0.145966131 | 0.285485808 |
| ENSG00000122406 | RPL5 | 11668.23903 | 0.273660252 | 0.046996955 | 5.822935844 | 5.78E-09 | 4.27E-08 |
| ENSG00000122417 | ODF2L | 311.2362824 | -0.192717703 | 0.104440966 | -1.84523095 | 0.06500395 | 0.149163811 |
| ENSG00000122432 | SPATA1 | 10.222627 | 0.763252113 | 0.568867702 | 1.341704072 | 0.179691966 | 0.334588775 |
| ENSG00000122435 | TRMT13 | 244.1624478 | 0.177412903 | 0.131989673 | 1.344142306 | 0.178902374 | 0.333714462 |
| ENSG00000122477 | LRRC39 | 10.693206 | 0.305351014 | 0.530158608 | 0.575961627 | 0.564641119 | 0.733084344 |

| | | | | | | | |
|---|---|---|---|---|---|---|---|
| ENSG00000122481 | RWDD3 | 35.0614648 | -0.146286137 | 0.298878903 | -0.489449526 | 0.62452348 | 0.778068139 |
| ENSG00000122482 | ZNF644 | 1125.627827 | -0.07537795 | 0.070945216 | -1.062481085 | 0.28801734 | 0.472004414 |
| ENSG00000122483 | CCDC18 | 242.2044727 | 0.041948357 | 0.124290431 | 0.337502709 | 0.73573797 | 0.854314084 |
| ENSG00000122484 | RPAP2 | 574.7985191 | 0.058207452 | 0.090088492 | 0.646114183 | 0.518205411 | 0.695932225 |
| ENSG00000122490 | SLC66A2 | 803.0408601 | 0.701446247 | 0.085500393 | 8.204011993 | 2.32E-16 | 3.08E-15 |
| ENSG00000122507 | BBS9 | 495.3725492 | -0.265854323 | 0.08713712 | -3.050988188 | 0.002280895 | 0.007929041 |
| ENSG00000122512 | PMS2 | 1161.463457 | 0.386124655 | 0.063336138 | 6.096435095 | 1.08E-09 | 8.60E-09 |
| ENSG00000122515 | ZMIZ2 | 5364.018525 | 0.838233856 | 0.063115877 | 13.28087157 | 2.99E-40 | 1.24E-38 |
| ENSG00000122543 | OCM | 1.828772765 | -0.486769186 | 1.303693807 | -0.373376926 | 0.70886793 | 0.837745175 |
| ENSG00000122545 | SEPTIN7 | 5189.350297 | 0.357777552 | 0.052735446 | 6.784384655 | 1.17E-11 | 1.08E-10 |
| ENSG00000122547 | EEPD1 | 114.2314954 | 0.973058911 | 0.173171958 | 5.619032781 | 1.92E-08 | 1.34E-07 |
| ENSG00000122550 | KLHL7 | 1312.430861 | -2.248808134 | 0.068594642 | -32.78402011 | 9.95E-236 | 1.04E-232 |
| ENSG00000122557 | HERPUD2 | 1105.685382 | 0.546176644 | 0.064631969 | 8.450564844 | 2.90E-17 | 4.06E-16 |
| ENSG00000122565 | CBX3 | 3855.227406 | -1.805110668 | 0.060447796 | -29.8623075 | 6.08E-196 | 3.40E-193 |
| ENSG00000122566 | HNRNPA2B1 | 20365.59874 | -1.041449449 | 0.058963991 | -17.66246547 | 8.16E-70 | 7.52E-68 |
| ENSG00000122584 | NXPH1 | 15.18450909 | -2.140922001 | 0.515691391 | -4.151556603 | 3.30E-05 | 0.000158778 |
| ENSG00000122591 | HYCC1 | 98.81691096 | -1.913496643 | 0.19675011 | -9.725517521 | 2.35E-22 | 4.47E-21 |
| ENSG00000122592 | HOXA7 | 19.56358564 | -1.64325484 | 0.443047771 | -3.708978911 | 0.000208097 | 0.00087979 |
| ENSG00000122641 | INHBA | 28.51997725 | -1.601457943 | 0.356827236 | -4.488048503 | 7.19E-06 | 3.77E-05 |
| ENSG00000122642 | FKBP9 | 3157.256997 | 0.315625564 | 0.048800102 | 6.467723441 | 9.95E-11 | 8.59E-10 |
| ENSG00000122643 | NT5C3A | 914.4030647 | 0.060985806 | 0.073961746 | 0.824558759 | 0.409622143 | 0.602084447 |
| ENSG00000122644 | ARL4A | 1362.784435 | -0.916946204 | 0.061654979 | -14.87221661 | 4.99E-50 | 2.77E-48 |
| ENSG00000122674 | CCZ1 | 491.0545787 | 0.260604152 | 0.08650215 | 3.01268989 | 0.002589434 | 0.008891126 |
| ENSG00000122678 | POLM | 973.1568732 | 0.670576034 | 0.074368266 | 9.016964724 | 1.93E-19 | 3.13E-18 |
| ENSG00000122687 | MRM2 | 11.10124144 | 0.185561281 | 0.529982855 | 0.35012695 | 0.726243426 | 0.847853251 |
| ENSG00000122692 | SMU1 | 1634.534653 | -0.07648704 | 0.05916567 | -1.292760481 | 0.196093912 | 0.357611444 |
| ENSG00000122694 | GLIPR2 | 27.68844938 | 0.618782799 | 0.339552653 | 1.822347118 | 0.068402333 | 0.155669595 |
| ENSG00000122696 | SLC25A51 | 246.0900858 | 0.09459088 | 0.118256571 | 0.799878424 | 0.42378124 | 0.614665 |
| ENSG00000122705 | CLTA | 2400.583869 | 0.134925166 | 0.051074613 | 2.641726671 | 0.008248459 | 0.025102103 |
| ENSG00000122707 | RECK | 2.647638172 | -0.17780279 | 1.049775693 | -0.169372173 | 0.86550391 | 0.930134906 |
| ENSG00000122711 | SPINK4 | 2.338831018 | -2.038854494 | 1.270649892 | -1.604576136 | 0.108587117 | 0.226432911 |
| ENSG00000122728 | TAF1L | 77.11994947 | -0.001959022 | 0.202291219 | -0.009684167 | 0.992273273 | 0.99592006 |
| ENSG00000122729 | ACO1 | 986.5730843 | 0.571375647 | 0.073227106 | 7.802788847 | 6.06E-15 | 7.24E-14 |

| | | | | | | | |
|---|---|---|---|---|---|---|---|
| ENSG00000122735 | DNAI1 | 6.234727102 | 1.373209346 | 0.740817256 | 1.853641143 | 0.063790517 | 0.146805224 |
| ENSG00000122741 | DCAF10 | 832.976968 | -0.079829131 | 0.079465873 | -1.004571242 | 0.31510335 | 0.503784196 |
| ENSG00000122756 | CNTFR | 68.09274571 | 4.114997432 | 0.361296598 | 11.38952721 | 4.72E-30 | 1.32E-28 |
| ENSG00000122778 | KIAA1549 | 883.324537 | 0.194055558 | 0.074713182 | 2.597340306 | 0.009394879 | 0.028168675 |
| ENSG00000122779 | TRIM24 | 5633.047259 | -0.253045654 | 0.049953498 | -5.065624354 | 4.07E-07 | 2.48E-06 |
| ENSG00000122783 | CYREN | 1409.220859 | -0.297334065 | 0.066061821 | -4.500845738 | 6.77E-06 | 3.57E-05 |
| ENSG00000122786 | CALD1 | 12.97219475 | -1.027509259 | 0.505022332 | -2.034581825 | 0.041892968 | 0.103014758 |
| ENSG00000122787 | AKR1D1 | 13.65772994 | -1.079840482 | 0.493764523 | -2.186954373 | 0.028745855 | 0.074816192 |
| ENSG00000122861 | PLAU | 36.63093335 | -8.748149184 | 1.205034296 | -7.259668221 | 3.88E-13 | 4.07E-12 |
| ENSG00000122863 | CHST3 | 542.6585184 | -0.624028134 | 0.087170985 | -7.158667915 | 8.15E-13 | 8.40E-12 |
| ENSG00000122873 | CISD1 | 395.5923379 | -0.751913567 | 0.095900788 | -7.840535897 | 4.49E-15 | 5.42E-14 |
| ENSG00000122877 | EGR2 | 46.79893002 | 0.535010956 | 0.294775344 | 1.814978649 | 0.069527191 | 0.157728813 |
| ENSG00000122882 | ECD | 1210.763455 | -0.474470737 | 0.063776062 | -7.439636832 | 1.01E-13 | 1.10E-12 |
| ENSG00000122884 | P4HA1 | 724.4722437 | -1.038627124 | 0.079717801 | -13.02879797 | 8.39E-39 | 3.24E-37 |
| ENSG00000122912 | SLC25A16 | 348.174688 | -0.589064189 | 0.101317276 | -5.814054717 | 6.10E-09 | 4.49E-08 |
| ENSG00000122952 | ZWINT | 2307.964493 | -0.926390747 | 0.054261386 | -17.07274393 | 2.37E-65 | 1.91E-63 |
| ENSG00000122958 | VPS26A | 1689.916155 | -0.410423966 | 0.06174736 | -6.646826107 | 2.99E-11 | 2.70E-10 |
| ENSG00000122965 | RBM19 | 1122.400111 | -0.095872859 | 0.06868427 | -1.395848849 | 0.162760017 | 0.310042036 |
| ENSG00000122966 | CIT | 1713.486544 | -0.342944007 | 0.063279149 | -5.419541986 | 5.98E-08 | 3.95E-07 |
| ENSG00000122970 | IFT81 | 288.7317075 | 0.248079825 | 0.10947361 | 2.266115318 | 0.023444324 | 0.062828596 |
| ENSG00000122971 | ACADS | 238.5148058 | 0.346195187 | 0.129465452 | 2.67403529 | 0.007494456 | 0.023067601 |
| ENSG00000123009 | NME2P1 | 191.8579063 | 0.215826932 | 0.142815051 | 1.511233796 | 0.130728895 | 0.262602526 |
| ENSG00000123064 | DDX54 | 3880.885226 | 0.245926158 | 0.052118538 | 4.7185928 | 2.37E-06 | 1.32E-05 |
| ENSG00000123066 | MED13L | 7912.452338 | 0.522272204 | 0.052154941 | 10.01385852 | 1.32E-23 | 2.70E-22 |
| ENSG00000123080 | CDKN2C | 618.1789213 | 0.016905974 | 0.083134434 | 0.20335706 | 0.83885596 | 0.915260676 |
| ENSG00000123091 | RNF11 | 2448.942472 | 0.129996086 | 0.055647943 | 2.336044778 | 0.019488908 | 0.05347154 |
| ENSG00000123094 | RASSF8 | 2081.190172 | 0.929650487 | 0.057897796 | 16.056751 | 5.13E-58 | 3.48E-56 |
| ENSG00000123095 | BHLHE41 | 127.1396735 | 2.339342678 | 0.192271132 | 12.166895 | 4.66E-34 | 1.53E-32 |
| ENSG00000123096 | SSPN | 65.22428009 | 0.6193474 | 0.227022376 | 2.728133725 | 0.006369378 | 0.019971779 |
| ENSG00000123104 | ITPR2 | 462.4563888 | 0.065366987 | 0.095500062 | 0.684470623 | 0.493678033 | 0.676438565 |
| ENSG00000123106 | CCDC91 | 367.5073976 | 0.241605078 | 0.101188205 | 2.387680247 | 0.016955086 | 0.047369343 |
| ENSG00000123119 | NECAB1 | 189.4913623 | 4.816463467 | 0.264510972 | 18.20893643 | 4.38E-74 | 4.49E-72 |
| ENSG00000123124 | WWP1 | 6499.700059 | 0.33081551 | 0.050891932 | 6.500352704 | 8.01E-11 | 7.00E-10 |

| | | | | | | | |
|---|---|---|---|---|---|---|---|
| ENSG00000123130 | ACOT9 | 754.9904427 | -0.487070441 | 0.081413968 | -5.982639766 | 2.20E-09 | 1.69E-08 |
| ENSG00000123131 | PRDX4 | 1258.754563 | 0.021994224 | 0.061017465 | 0.360457843 | 0.718504777 | 0.843432038 |
| ENSG00000123136 | DDX39A | 2460.827107 | -0.4152272 | 0.053523864 | -7.75779573 | 8.64E-15 | 1.02E-13 |
| ENSG00000123143 | PKN1 | 2840.482595 | -0.009546272 | 0.054831792 | -0.17410104 | 0.861786056 | 0.928215087 |
| ENSG00000123144 | TRIR | 2283.555004 | -0.019562109 | 0.057403427 | -0.34078295 | 0.733266987 | 0.852610101 |
| ENSG00000123146 | ADGRE5 | 303.0461093 | 0.198501483 | 0.112320518 | 1.76727713 | 0.077181828 | 0.171914859 |
| ENSG00000123154 | WDR83 | 279.8230014 | -0.114979619 | 0.111829643 | -1.028167629 | 0.303870983 | 0.491178153 |
| ENSG00000123159 | GIPC1 | 3084.971163 | 0.016978265 | 0.062282496 | 0.272600912 | 0.785160011 | 0.883118404 |
| ENSG00000123178 | SPRYD7 | 245.5602728 | -0.0354089 | 0.115814676 | -0.305737591 | 0.759804449 | 0.867980383 |
| ENSG00000123179 | EBPL | 411.0731938 | 0.301731408 | 0.094123762 | 3.205687928 | 0.0013474 | 0.004922657 |
| ENSG00000123191 | ATP7B | 938.0238846 | -0.459529341 | 0.068806874 | -6.678538269 | 2.41E-11 | 2.19E-10 |
| ENSG00000123200 | ZC3H13 | 3444.78238 | -0.018706708 | 0.053084068 | -0.352397773 | 0.724539972 | 0.846625252 |
| ENSG00000123201 | GUCY1B2 | 206.3897218 | -0.192092975 | 0.131551655 | -1.460209493 | 0.144232507 | 0.282839939 |
| ENSG00000123213 | NLN | 1337.102113 | -0.058782323 | 0.063853199 | -0.920585412 | 0.357266922 | 0.549567705 |
| ENSG00000123219 | CENPK | 281.904601 | -0.270033489 | 0.114213859 | -2.364279536 | 0.018065176 | 0.050014691 |
| ENSG00000123240 | OPTN | 731.7232392 | -0.135466934 | 0.074843181 | -1.810010383 | 0.070294177 | 0.159122599 |
| ENSG00000123268 | ATF1 | 1013.036459 | -0.075537338 | 0.068740302 | -1.098879933 | 0.271820441 | 0.453891462 |
| ENSG00000123297 | TSFM | 28.37088456 | 0.503741235 | 0.328759388 | 1.532248976 | 0.125461007 | 0.25420204 |
| ENSG00000123329 | ARHGAP9 | 5.039380154 | -1.596576386 | 0.861854131 | -1.852490263 | 0.063955454 | 0.147081516 |
| ENSG00000123342 | MMP19 | 145.4544286 | 0.218833447 | 0.158758185 | 1.378407344 | 0.168077557 | 0.318022036 |
| ENSG00000123349 | PFDN5 | 4659.449569 | 0.10027171 | 0.04590161 | 2.184492211 | 0.028926099 | 0.07519999 |
| ENSG00000123352 | SPATS2 | 1943.349626 | -0.271339805 | 0.058628396 | -4.628129444 | 3.69E-06 | 2.00E-05 |
| ENSG00000123353 | ORMDL2 | 797.3147261 | -0.06591953 | 0.072638676 | -0.907499057 | 0.364142948 | 0.556807655 |
| ENSG00000123358 | NR4A1 | 960.2927522 | 0.731230918 | 0.074710182 | 9.787567111 | 1.27E-22 | 2.47E-21 |
| ENSG00000123364 | HOXC13 | 1436.650816 | -0.050573518 | 0.063851933 | -0.792043641 | 0.42833523 | 0.618922284 |
| ENSG00000123374 | CDK2 | 2201.650758 | -0.114220946 | 0.052493921 | -2.175889021 | 0.02956356 | 0.076674739 |
| ENSG00000123384 | LRP1 | 384.5854715 | -0.376684267 | 0.100998538 | -3.729601169 | 0.000191783 | 0.000817144 |
| ENSG00000123388 | HOXC11 | 504.1156459 | -0.005658153 | 0.092950212 | -0.060872941 | 0.951460399 | 0.977495703 |
| ENSG00000123395 | ATG101 | 1492.163518 | -0.043970054 | 0.06476693 | -0.678896682 | 0.497203328 | 0.679040564 |
| ENSG00000123405 | NFE2 | 1.542642409 | -0.110867108 | 1.501209692 | -0.073851847 | 0.941128272 | NA |
| ENSG00000123407 | HOXC12 | 41.84541416 | 0.64282029 | 0.276306916 | 2.326471955 | 0.019993387 | 0.054666066 |
| ENSG00000123411 | IKZF4 | 398.0419339 | 0.05619415 | 0.099298464 | 0.565911568 | 0.571453898 | 0.738377403 |
| ENSG00000123415 | SMUG1 | 808.9540548 | -0.278580099 | 0.078185803 | -3.563052225 | 0.000366568 | 0.001482557 |

| | | | | | | | |
|---|---|---|---|---|---|---|---|
| ENSG00000123416 | TUBA1B | 22836.30583 | -0.327336882 | 0.056122424 | -5.83255067 | 5.46E-09 | 4.05E-08 |
| ENSG00000123427 | EEF1AKMT3 | 269.792469 | -0.077960426 | 0.11327251 | -0.688255487 | 0.491291908 | 0.674258831 |
| ENSG00000123444 | KBTBD4 | 1385.331423 | -0.1001244 | 0.064043947 | -1.563370223 | 0.11796554 | 0.242376129 |
| ENSG00000123453 | SARDH | 13.51373153 | -0.397948006 | 0.474609111 | -0.838475277 | 0.401763827 | 0.594410925 |
| ENSG00000123472 | ATPAF1 | 2411.471854 | -0.115249611 | 0.050472971 | -2.283392634 | 0.022407251 | 0.060395057 |
| ENSG00000123473 | STIL | 1556.541739 | -0.413721286 | 0.062386157 | -6.631619981 | 3.32E-11 | 2.98E-10 |
| ENSG00000123485 | HJURP | 882.5120649 | -0.28873991 | 0.072061064 | -4.006878255 | 6.15E-05 | 0.000284159 |
| ENSG00000123505 | AMD1 | 2523.829756 | -0.083952033 | 0.058129966 | -1.444212659 | 0.148679165 | 0.289573458 |
| ENSG00000123545 | NDUFAF4 | 391.3762054 | 0.265827095 | 0.095754445 | 2.776133218 | 0.005500966 | 0.017512097 |
| ENSG00000123552 | USP45 | 268.4197009 | -0.064013446 | 0.131458882 | -0.48694653 | 0.626296224 | 0.779556993 |
| ENSG00000123560 | PLP1 | 1.956873175 | -0.174088671 | 1.22964451 | -0.141576423 | 0.887414591 | 0.941998797 |
| ENSG00000123562 | MORF4L2 | 9692.197225 | -0.229531502 | 0.045471955 | -5.047759755 | 4.47E-07 | 2.71E-06 |
| ENSG00000123570 | RAB9B | 429.138886 | 0.793132373 | 0.100706447 | 7.875686175 | 3.39E-15 | 4.13E-14 |
| ENSG00000123572 | NRK | 25.34541787 | -1.074502737 | 0.369662791 | -2.906710558 | 0.00365251 | 0.012139753 |
| ENSG00000123575 | FAM199X | 3051.945406 | 0.06600113 | 0.05811351 | 1.135727824 | 0.256070492 | 0.435517281 |
| ENSG00000123595 | RAB9A | 679.6756939 | 0.208528873 | 0.076812505 | 2.714777675 | 0.006632028 | 0.020685073 |
| ENSG00000123600 | METTL8 | 414.6325248 | -0.165726149 | 0.09283036 | -1.785258062 | 0.074219468 | 0.166512919 |
| ENSG00000123607 | TTC21B | 598.9091114 | -0.139806593 | 0.086364689 | -1.618793456 | 0.105491711 | 0.221265415 |
| ENSG00000123609 | NMI | 74.20748476 | -0.63853971 | 0.212526305 | -3.004520825 | 0.002659995 | 0.009114309 |
| ENSG00000123612 | ACVR1C | 19.80594585 | 1.260436587 | 0.426234188 | 2.957145677 | 0.003105014 | 0.010488738 |
| ENSG00000123636 | BAZ2B | 1082.404499 | -0.439013111 | 0.066447802 | -6.606886846 | 3.92E-11 | 3.50E-10 |
| ENSG00000123643 | SLC36A1 | 586.8779748 | 0.12771198 | 0.087668972 | 1.456752334 | 0.145184758 | 0.284272648 |
| ENSG00000123684 | LPGAT1 | 1384.360445 | 0.231908764 | 0.060243443 | 3.849527036 | 0.000118346 | 0.000521547 |
| ENSG00000123728 | RAP2C | 2683.710808 | -0.086040351 | 0.060496647 | -1.422233379 | 0.154958512 | 0.29847233 |
| ENSG00000123737 | EXOSC9 | 684.2579446 | -0.291167749 | 0.080861387 | -3.600825559 | 0.000317208 | 0.00130035 |
| ENSG00000123739 | PLA2G12A | 844.8770549 | 0.403310442 | 0.071322136 | 5.654772315 | 1.56E-08 | 1.10E-07 |
| ENSG00000123810 | B9D2 | 87.95522362 | 0.151778087 | 0.192255639 | 0.789459742 | 0.42984335 | 0.620163928 |
| ENSG00000123815 | COQ8B | 691.2402944 | -0.229711501 | 0.086196745 | -2.664967231 | 0.007699583 | 0.023632705 |
| ENSG00000123836 | PFKFB2 | 746.6532202 | 0.66003068 | 0.080924414 | 8.156137909 | 3.46E-16 | 4.52E-15 |
| ENSG00000123870 | ZNF137P | 94.0195842 | -0.054673349 | 0.182621304 | -0.299381001 | 0.764649357 | 0.871042619 |
| ENSG00000123901 | GPR83 | 3.93355138 | 0.704724733 | 0.913716082 | 0.771273207 | 0.44054501 | 0.629859927 |
| ENSG00000123908 | AGO2 | 3349.355203 | -0.072179531 | 0.057125667 | -1.263521905 | 0.20640168 | 0.372090729 |
| ENSG00000123933 | MXD4 | 4960.620249 | 0.001002033 | 0.057306615 | 0.017485461 | 0.986049332 | 0.993350589 |

| | | | | | | | |
|---|---|---|---|---|---|---|---|
| ENSG00000123965 | PMS2P5 | 28.42291384 | 0.248683285 | 0.325633152 | 0.763691545 | 0.445051081 | 0.633676984 |
| ENSG00000123975 | CKS2 | 1053.650044 | -0.18270962 | 0.066963555 | -2.728493434 | 0.006362436 | 0.019955465 |
| ENSG00000123983 | ACSL3 | 9751.05543 | -1.26037942 | 0.048656292 | -25.90372923 | 6.05E-148 | 2.13E-145 |
| ENSG00000123989 | CHPF | 3111.95275 | 0.297744706 | 0.069532214 | 4.282111681 | 1.85E-05 | 9.23E-05 |
| ENSG00000123992 | DNPEP | 4110.795382 | 0.157568522 | 0.052963472 | 2.975041398 | 0.00292949 | 0.009944141 |
| ENSG00000123999 | INHA | 68.67041009 | 0.56088051 | 0.21996759 | 2.549832497 | 0.010777469 | 0.031725283 |
| ENSG00000124006 | OBSL1 | 6891.805556 | 0.054769112 | 0.05056412 | 1.083161579 | 0.27873671 | 0.46170988 |
| ENSG00000124067 | SLC12A4 | 30.85582729 | -1.01009912 | 0.337663246 | -2.991439349 | 0.002776657 | 0.009481469 |
| ENSG00000124074 | ENKD1 | 25.39634371 | 0.29735231 | 0.359972649 | 0.826041398 | 0.408780609 | 0.601170321 |
| ENSG00000124097 | HMGB1P1 | 101.866763 | -0.029200668 | 0.190415544 | -0.153352331 | 0.878120434 | 0.936542041 |
| ENSG00000124098 | FAM210B | 2131.599296 | 0.385082539 | 0.054254025 | 7.097769036 | 1.27E-12 | 1.29E-11 |
| ENSG00000124103 | FAM209A | 10.10742029 | 0.556378058 | 0.55613559 | 1.000435987 | 0.317099562 | 0.506106168 |
| ENSG00000124104 | SNX21 | 825.2168466 | 0.269364303 | 0.080174804 | 3.359712627 | 0.000780236 | 0.002979626 |
| ENSG00000124116 | WFDC3 | 5.222797012 | -0.696215939 | 0.771625781 | -0.902271485 | 0.366912669 | 0.559399478 |
| ENSG00000124120 | TTPAL | 1565.117259 | -0.475271667 | 0.06035914 | -7.874062916 | 3.43E-15 | 4.18E-14 |
| ENSG00000124126 | PREX1 | 94628.88707 | -1.444878624 | 0.049135646 | -29.40591497 | 4.61E-190 | 2.46E-187 |
| ENSG00000124140 | SLC12A5 | 675.24053 | -0.79686198 | 0.084101235 | -9.475033055 | 2.67E-21 | 4.81E-20 |
| ENSG00000124143 | ARHGAP40 | 4.600595917 | -1.759097986 | 0.912495722 | -1.927787653 | 0.053881544 | 0.127848844 |
| ENSG00000124145 | SDC4 | 6928.429583 | 0.047921169 | 0.04668058 | 1.026576121 | 0.304620103 | 0.492007686 |
| ENSG00000124151 | NCOA3 | 63815.48915 | 0.001561149 | 0.049197908 | 0.03173201 | 0.974685767 | 0.988518127 |
| ENSG00000124155 | PIGT | 3698.665416 | -0.309234298 | 0.051619076 | -5.990698053 | 2.09E-09 | 1.62E-08 |
| ENSG00000124159 | MATN4 | 3.950658863 | -1.376754595 | 0.942059535 | -1.461430561 | 0.143897319 | 0.282378485 |
| ENSG00000124160 | NCOA5 | 3579.164249 | -0.2553467 | 0.051088437 | -4.998130943 | 5.79E-07 | 3.45E-06 |
| ENSG00000124164 | VAPB | 7786.104504 | -0.047512502 | 0.042930846 | -1.10672178 | 0.268414236 | 0.449874922 |
| ENSG00000124171 | PARD6B | 6756.706471 | 0.490054348 | 0.055567597 | 8.819066833 | 1.15E-18 | 1.78E-17 |
| ENSG00000124172 | ATP5F1E | 5528.16314 | -0.18934532 | 0.050085669 | -3.780429106 | 0.000156558 | 0.000676355 |
| ENSG00000124177 | CHD6 | 6129.905402 | -0.011342427 | 0.052174228 | -0.217395207 | 0.827900366 | 0.909057744 |
| ENSG00000124181 | PLCG1 | 11545.93735 | -0.44482476 | 0.048015881 | -9.264117466 | 1.97E-20 | 3.37E-19 |
| ENSG00000124191 | TOX2 | 22.22937618 | -1.098647281 | 0.388526237 | -2.827729958 | 0.004687933 | 0.015208767 |
| ENSG00000124193 | SRSF6 | 4052.554184 | -0.065041321 | 0.058775326 | -1.10660928 | 0.268462893 | 0.44992247 |
| ENSG00000124198 | ARFGEF2 | 3635.742221 | -0.106552684 | 0.051475135 | -2.069983585 | 0.038453882 | 0.095999376 |
| ENSG00000124201 | ZNFX1 | 2437.397499 | 0.401721778 | 0.059133706 | 6.793448314 | 1.09E-11 | 1.02E-10 |
| ENSG00000124207 | CSE1L | 7615.095715 | -0.259565773 | 0.057596373 | -4.506634019 | 6.59E-06 | 3.48E-05 |

| | | | | | | | |
|---|---|---|---|---|---|---|---|
| ENSG00000124209 | RAB22A | 4019.395601 | 0.114468756 | 0.054682797 | 2.093323008 | 0.03632033 | 0.091518934 |
| ENSG00000124214 | STAU1 | 6778.380877 | -0.055944474 | 0.045182792 | -1.238180975 | 0.215648964 | 0.384917388 |
| ENSG00000124216 | SNAI1 | 46.13184568 | 0.236178648 | 0.27035516 | 0.873586611 | 0.38234342 | 0.574617273 |
| ENSG00000124217 | MOCS3 | 848.8885661 | 0.10519158 | 0.077754261 | 1.352872228 | 0.176096453 | 0.329498583 |
| ENSG00000124222 | STX16 | 4140.606938 | -0.039110826 | 0.06701184 | -0.583640539 | 0.55946218 | 0.72907705 |
| ENSG00000124224 | PPP4R1L | 413.8231982 | -0.734362813 | 0.102198048 | -7.18568337 | 6.69E-13 | 6.94E-12 |
| ENSG00000124225 | PMEPA1 | 3026.307288 | 0.656724021 | 0.061775025 | 10.63089852 | 2.14E-26 | 5.10E-25 |
| ENSG00000124226 | RNF114 | 4596.455378 | -0.065658348 | 0.046110969 | -1.423920349 | 0.154469531 | 0.297800742 |
| ENSG00000124228 | DDX27 | 4041.712578 | -0.265476859 | 0.04776207 | -5.558319762 | 2.72E-08 | 1.87E-07 |
| ENSG00000124243 | BCAS4 | 828.9748889 | 0.291268485 | 0.087162926 | 3.341655648 | 0.000832803 | 0.003164045 |
| ENSG00000124249 | KCNK15 | 378.5632043 | 0.552765278 | 0.097597603 | 5.663717749 | 1.48E-08 | 1.05E-07 |
| ENSG00000124251 | TP53TG5 | 3.100545289 | -3.17611042 | 1.322428266 | -2.401726053 | 0.016317924 | 0.045801195 |
| ENSG00000124257 | NEURL2 | 36.11616271 | 0.707601872 | 0.321618155 | 2.200130375 | 0.027797646 | 0.072761069 |
| ENSG00000124275 | MTRR | 994.6828456 | 0.167948632 | 0.067739663 | 2.479324888 | 0.013163135 | 0.037868361 |
| ENSG00000124279 | FASTKD3 | 417.2062085 | 0.009135076 | 0.098429179 | 0.092808623 | 0.926055601 | 0.964571171 |
| ENSG00000124299 | PEPD | 686.6190734 | 0.197327312 | 0.083369871 | 2.366889992 | 0.017938264 | 0.049746588 |
| ENSG00000124302 | CHST8 | 10.53234628 | -0.085130209 | 0.533489946 | -0.159572283 | 0.873218014 | 0.934604743 |
| ENSG00000124313 | IQSEC2 | 1722.038643 | -1.446790279 | 0.070973607 | -20.38490547 | 2.28E-92 | 3.24E-90 |
| ENSG00000124333 | VAMP7 | 2012.158395 | -0.398934704 | 0.066702558 | -5.980800686 | 2.22E-09 | 1.71E-08 |
| ENSG00000124333 | VAMP7 | 2012.158395 | -0.398934704 | 0.066702558 | -5.980800686 | 2.22E-09 | 1.71E-08 |
| ENSG00000124343 | XG | 129.2249109 | 1.40682105 | 0.173512182 | 8.107909382 | 5.15E-16 | 6.67E-15 |
| ENSG00000124356 | STAMBP | 1519.756043 | -0.095563185 | 0.060404067 | -1.58206541 | 0.113634639 | 0.235120287 |
| ENSG00000124357 | NAGK | 1442.350538 | -0.241744043 | 0.06324686 | -3.822229944 | 0.00013225 | 0.000577761 |
| ENSG00000124370 | MCEE | 124.6383478 | -0.383806915 | 0.160418732 | -2.392531788 | 0.016732579 | 0.046833254 |
| ENSG00000124374 | PAIP2B | 460.5612411 | -0.655663771 | 0.09011882 | -7.275547692 | 3.45E-13 | 3.63E-12 |
| ENSG00000124380 | SNRNP27 | 320.515395 | -0.23218427 | 0.106528191 | -2.179557053 | 0.029290311 | 0.076053007 |
| ENSG00000124383 | MPHOSPH10 | 1346.715462 | -0.037057376 | 0.061390177 | -0.603636893 | 0.546085079 | 0.719092449 |
| ENSG00000124391 | IL17C | 12.69923342 | 0.837612883 | 0.498981411 | 1.678645468 | 0.09322116 | 0.200121037 |
| ENSG00000124399 | NDUFB4P12 | 134.6392246 | 0.146900562 | 0.152931571 | 0.960564006 | 0.336771434 | 0.52795303 |
| ENSG00000124406 | ATP8A1 | 558.3442471 | -0.594262965 | 0.084318997 | -7.047794485 | 1.82E-12 | 1.82E-11 |
| ENSG00000124422 | USP22 | 13036.69709 | 0.152274421 | 0.042447035 | 3.587398293 | 0.000333994 | 0.001362102 |
| ENSG00000124429 | POF1B | 189.0966418 | 0.977250059 | 0.136312522 | 7.1691877 | 7.54E-13 | 7.80E-12 |
| ENSG00000124444 | ZNF576 | 206.4188641 | 0.081524443 | 0.132518038 | 0.615195067 | 0.538425905 | 0.713216899 |

| | | | | | | | |
|---|---|---|---|---|---|---|---|
| ENSG00000124459 | ZNF45 | 729.9470322 | -0.045507272 | 0.074177449 | -0.613492005 | 0.539551068 | 0.714103677 |
| ENSG00000124466 | LYPD3 | 815.7847455 | -1.476273477 | 0.091161542 | -16.19403809 | 5.56E-59 | 3.85E-57 |
| ENSG00000124479 | NDP | 78.13522959 | -1.776453246 | 0.220099499 | -8.071137175 | 6.96E-16 | 8.92E-15 |
| ENSG00000124486 | USP9X | 4958.992296 | -0.117622799 | 0.052404807 | -2.244503984 | 0.024799993 | 0.06601437 |
| ENSG00000124493 | GRM4 | 1338.500523 | -0.094743077 | 0.074927418 | -1.26446473 | 0.206063272 | 0.371601248 |
| ENSG00000124496 | TRERF1 | 2366.454602 | -0.284493408 | 0.058349649 | -4.875666166 | 1.08E-06 | 6.25E-06 |
| ENSG00000124507 | PACSIN1 | 855.2071726 | 0.54675658 | 0.082449881 | 6.63138108 | 3.33E-11 | 2.98E-10 |
| ENSG00000124508 | BTN2A2 | 264.4424007 | 1.138707227 | 0.116687481 | 9.758606609 | 1.69E-22 | 3.27E-21 |
| ENSG00000124523 | SIRT5 | 420.6818106 | -0.613831715 | 0.094655749 | -6.484885714 | 8.88E-11 | 7.72E-10 |
| ENSG00000124532 | MRS2 | 1155.953397 | 0.195582476 | 0.065146565 | 3.002191712 | 0.002680433 | 0.009177483 |
| ENSG00000124535 | WRNIP1 | 1680.67558 | -0.51531751 | 0.059393132 | -8.676382141 | 4.09E-18 | 6.08E-17 |
| ENSG00000124541 | RRP36 | 1722.892933 | 0.019530985 | 0.05792165 | 0.337196624 | 0.735968682 | 0.854386347 |
| ENSG00000124549 | BTN2A3P | 39.73292647 | -0.255540312 | 0.294794452 | -0.866842337 | 0.386028402 | 0.578639018 |
| ENSG00000124562 | SNRPC | 2232.234088 | -0.068459907 | 0.052668864 | -1.299817434 | 0.193663549 | 0.354418971 |
| ENSG00000124570 | SERPINB6 | 1370.104373 | -0.745555787 | 0.062834885 | -11.86531637 | 1.79E-32 | 5.51E-31 |
| ENSG00000124571 | XPO5 | 3651.838597 | -0.013899892 | 0.051648201 | -0.269126362 | 0.787832445 | 0.885231009 |
| ENSG00000124574 | ABCC10 | 1138.611249 | 0.216909759 | 0.068932508 | 3.146697618 | 0.001651256 | 0.005909437 |
| ENSG00000124575 | H1-3 | 32.96463772 | -0.200567683 | 0.30741917 | -0.652424124 | 0.514127606 | 0.69315823 |
| ENSG00000124587 | PEX6 | 559.7992787 | 0.689362909 | 0.090623697 | 7.606872507 | 2.81E-14 | 3.20E-13 |
| ENSG00000124588 | NQO2 | 700.7104512 | -0.777944788 | 0.081599831 | -9.533656885 | 1.52E-21 | 2.78E-20 |
| ENSG00000124593 | ENSG0000012 | 35.96807993 | -0.050487952 | 0.296762281 | -0.170129275 | 0.864908471 | 0.929799912 |
| ENSG00000124596 | OARD1 | 834.4392872 | -0.220687075 | 0.07123069 | -3.098202132 | 0.001946986 | 0.00686397 |
| ENSG00000124602 | UNC5CL | 10.74373192 | 0.039615454 | 0.534035138 | 0.074181362 | 0.940866076 | 0.971736581 |
| ENSG00000124608 | AARS2 | 923.0173876 | 0.119620375 | 0.070943281 | 1.686141016 | 0.091768642 | 0.197594578 |
| ENSG00000124613 | ZNF391 | 169.706863 | 0.184350815 | 0.149341192 | 1.234427102 | 0.217043795 | 0.386508148 |
| ENSG00000124614 | RPS10 | 50.64568198 | 0.047106226 | 0.260754501 | 0.180653549 | 0.856639521 | 0.925725957 |
| ENSG00000124615 | MOCS1 | 329.595593 | 0.392776349 | 0.109431365 | 3.589248363 | 0.000331633 | 0.001353435 |
| ENSG00000124635 | H2BC11 | 188.9739452 | 0.574382344 | 0.13677231 | 4.199551369 | 2.67E-05 | 0.000130369 |
| ENSG00000124641 | MED20 | 943.0362772 | 0.082331597 | 0.068933343 | 1.194365362 | 0.23233508 | 0.406370643 |
| ENSG00000124657 | OR2B6 | 18.35909998 | 0.681419023 | 0.41052703 | 1.659863963 | 0.096941823 | 0.206773167 |
| ENSG00000124659 | TBCC | 648.1492004 | -0.069848469 | 0.083950386 | -0.83202082 | 0.405397193 | 0.598135555 |
| ENSG00000124664 | SPDEF | 7104.104936 | -0.795397752 | 0.054726436 | -14.5340683 | 7.37E-48 | 3.78E-46 |
| ENSG00000124688 | MAD2L1BP | 755.2445108 | -0.198636747 | 0.077580875 | -2.560382905 | 0.010455689 | 0.030889055 |

| | | | | | | | |
|---|---|---|---|---|---|---|---|
| ENSG00000124702 | KLHDC3 | 3756.616552 | 0.164010817 | 0.053699017 | 3.054261055 | 0.002256157 | 0.007854945 |
| ENSG00000124713 | GNMT | 12.54651484 | -0.198745798 | 0.493445457 | -0.402771563 | 0.687116283 | 0.823269645 |
| ENSG00000124721 | DNAH8 | 8.57907378 | -1.668887644 | 0.668565715 | -2.4962208 | 0.012552444 | 0.036307284 |
| ENSG00000124733 | MEA1 | 2393.452216 | -0.132759997 | 0.065798268 | -2.017682254 | 0.043624364 | 0.106700385 |
| ENSG00000124743 | KLHL31 | 36.41246899 | 0.404192634 | 0.295141869 | 1.36948592 | 0.170847433 | 0.321988179 |
| ENSG00000124749 | COL21A1 | 2.012727962 | 0.786913497 | 1.253011094 | 0.628017981 | 0.529992161 | 0.705630887 |
| ENSG00000124762 | CDKN1A | 5270.893719 | 0.258523953 | 0.049688658 | 5.202876519 | 1.96E-07 | 1.24E-06 |
| ENSG00000124766 | SOX4 | 3426.646787 | 0.576218734 | 0.053171658 | 10.83695249 | 2.30E-27 | 5.81E-26 |
| ENSG00000124767 | GLO1 | 5013.771204 | -0.910763839 | 0.052413776 | -17.37642106 | 1.24E-67 | 1.06E-65 |
| ENSG00000124782 | RREB1 | 1472.428773 | -0.279432057 | 0.059498947 | -4.696420168 | 2.65E-06 | 1.46E-05 |
| ENSG00000124783 | SSR1 | 3567.047559 | -0.552561324 | 0.054881639 | -10.06823659 | 7.63E-24 | 1.58E-22 |
| ENSG00000124784 | RIOK1 | 555.6043179 | -0.721555963 | 0.090502916 | -7.972737165 | 1.55E-15 | 1.94E-14 |
| ENSG00000124786 | SLC35B3 | 381.8429409 | -0.463871867 | 0.098394773 | -4.714395396 | 2.42E-06 | 1.34E-05 |
| ENSG00000124787 | RPP40 | 205.1039777 | -0.059097887 | 0.130781196 | -0.451883669 | 0.65135279 | 0.797503896 |
| ENSG00000124788 | ATXN1 | 342.7562102 | -0.215221167 | 0.101800465 | -2.114147182 | 0.034502698 | 0.0875641 |
| ENSG00000124789 | NUP153 | 2977.153659 | -0.82676861 | 0.054475319 | -15.17693923 | 5.03E-52 | 2.93E-50 |
| ENSG00000124795 | DEK | 3504.694426 | -0.711810449 | 0.063166244 | -11.26884249 | 1.87E-29 | 5.09E-28 |
| ENSG00000124802 | EEF1E1 | 69.68717552 | -0.118242146 | 0.213970562 | -0.552609411 | 0.580530896 | 0.745653906 |
| ENSG00000124812 | CRISP1 | 2.776635965 | 0.505436431 | 1.147353232 | 0.440523822 | 0.659557763 | 0.803092096 |
| ENSG00000124813 | RUNX2 | 167.607529 | 0.525713172 | 0.15393936 | 3.415066624 | 0.000637664 | 0.002476386 |
| ENSG00000124831 | LRRFIP1 | 3772.82755 | 0.038872004 | 0.047415341 | 0.819819136 | 0.41231922 | 0.604534362 |
| ENSG00000124839 | RAB17 | 984.5256949 | -0.327852324 | 0.076949359 | -4.260624509 | 2.04E-05 | 0.000101113 |
| ENSG00000124920 | MYRF | 29.60993464 | 0.533476328 | 0.327742246 | 1.627731348 | 0.103581863 | 0.218097103 |
| ENSG00000124942 | AHNAK | 14191.23223 | -0.242917147 | 0.121913654 | -1.99253438 | 0.046312457 | 0.11214055 |
| ENSG00000125037 | EMC3 | 2882.132594 | -0.016765383 | 0.052477586 | -0.319477033 | 0.749364805 | 0.861330224 |
| ENSG00000125046 | SSUH2 | 9.683672428 | -0.67346746 | 0.57069874 | -1.180075254 | 0.237970286 | 0.413438227 |
| ENSG00000125107 | CNOT1 | 8397.126412 | -0.038992527 | 0.051400919 | -0.758595905 | 0.448094323 | 0.636320267 |
| ENSG00000125122 | FBXL9P | 25.04577373 | 0.502931886 | 0.351949449 | 1.428988984 | 0.1530074 | 0.295852137 |
| ENSG00000125124 | BBS2 | 877.517876 | 0.196585521 | 0.070335519 | 2.794967943 | 0.005190486 | 0.016639149 |
| ENSG00000125148 | MT2A | 1369.558553 | -0.424566987 | 0.063510753 | -6.684962263 | 2.31E-11 | 2.10E-10 |
| ENSG00000125149 | PHAF1 | 565.4881066 | -0.193257834 | 0.086048585 | -2.245915296 | 0.024709434 | 0.065803856 |
| ENSG00000125166 | GOT2 | 4305.135456 | -0.011318364 | 0.048454261 | -0.233588611 | 0.815304371 | 0.902135938 |
| ENSG00000125170 | DOK4 | 588.8667589 | -0.096258108 | 0.088055199 | -1.093156442 | 0.274325118 | 0.456373753 |

| | | | | | | | |
|---|---|---|---|---|---|---|---|
| ENSG00000125245 | GPR18 | 29.2412608 | 3.966142472 | 0.529039504 | 7.496873945 | 6.54E-14 | 7.23E-13 |
| ENSG00000125246 | CLYBL | 40.86952143 | 0.662281718 | 0.279982749 | 2.365437585 | 0.018008779 | 0.049888048 |
| ENSG00000125247 | TMTC4 | 812.8015074 | 0.276080375 | 0.07165517 | 3.852902399 | 0.000116726 | 0.000515038 |
| ENSG00000125249 | RAP2A | 963.8424206 | -0.16347207 | 0.070201181 | -2.32862279 | 0.019879058 | 0.054411834 |
| ENSG00000125257 | ABCC4 | 389.4188776 | -0.601332572 | 0.097372043 | -6.175618304 | 6.59E-10 | 5.33E-09 |
| ENSG00000125266 | EFNB2 | 2051.501863 | 0.290944534 | 0.058565539 | 4.967845272 | 6.77E-07 | 4.01E-06 |
| ENSG00000125304 | TM9SF2 | 5004.922643 | -0.045434984 | 0.048348858 | -0.9397323 | 0.347354893 | 0.538290971 |
| ENSG00000125319 | HROB | 388.3721958 | -0.114075636 | 0.096704617 | -1.17962968 | 0.238147533 | 0.413526941 |
| ENSG00000125347 | IRF1 | 370.7782284 | -0.076210955 | 0.100969885 | -0.754788966 | 0.450375615 | 0.638625693 |
| ENSG00000125351 | UPF3B | 1504.646304 | -0.106922348 | 0.059999465 | -1.782055033 | 0.074740258 | 0.16751749 |
| ENSG00000125352 | RNF113A | 731.2334875 | -0.062340706 | 0.078889869 | -0.790224481 | 0.429396682 | 0.619714368 |
| ENSG00000125354 | SEPTIN6 | 18.14078213 | -0.628358222 | 0.425431776 | -1.476989396 | 0.13967848 | 0.275940779 |
| ENSG00000125356 | NDUFA1 | 2228.923229 | -0.058818981 | 0.051733197 | -1.136967832 | 0.255551733 | 0.434925199 |
| ENSG00000125375 | DMAC2L | 485.1966296 | -0.066176295 | 0.090274649 | -0.733055133 | 0.463524804 | 0.649941774 |
| ENSG00000125378 | BMP4 | 349.4913109 | -1.904242587 | 0.121661393 | -15.65198732 | 3.22E-55 | 2.04E-53 |
| ENSG00000125384 | PTGER2 | 16.25880399 | -1.905378641 | 0.47861867 | -3.980995228 | 6.86E-05 | 0.000314422 |
| ENSG00000125386 | FAM193A | 1464.422262 | -0.094533754 | 0.059577194 | -1.586743975 | 0.112570642 | 0.233234333 |
| ENSG00000125388 | GRK4 | 140.0335162 | -0.054322022 | 0.151318056 | -0.358992336 | 0.719600823 | 0.844168774 |
| ENSG00000125398 | SOX9 | 489.7993823 | 1.804754814 | 0.098746299 | 18.27668316 | 1.27E-74 | 1.33E-72 |
| ENSG00000125409 | TEKT3 | 18.67583112 | -0.782526534 | 0.417330143 | -1.875077915 | 0.060782005 | 0.14108461 |
| ENSG00000125430 | HS3ST3B1 | 8.388306575 | -0.017794393 | 0.594840137 | -0.029914581 | 0.976135177 | 0.989157212 |
| ENSG00000125434 | SLC25A35 | 90.93528361 | -0.195163168 | 0.193077321 | -1.010803168 | 0.312110646 | 0.500364186 |
| ENSG00000125445 | MRPS7 | 2949.770282 | 0.013949805 | 0.050867885 | 0.274235985 | 0.783903276 | 0.882792721 |
| ENSG00000125447 | GGA3 | 2295.771059 | 0.264575337 | 0.056436695 | 4.688001949 | 2.76E-06 | 1.52E-05 |
| ENSG00000125449 | ARMC7 | 906.4714909 | -0.811170908 | 0.0798098 | -10.16380074 | 2.88E-24 | 6.11E-23 |
| ENSG00000125450 | NUP85 | 2497.619414 | -0.165985898 | 0.052080649 | -3.18709349 | 0.001437103 | 0.005203969 |
| ENSG00000125454 | SLC25A19 | 493.967583 | 0.121331122 | 0.089150557 | 1.360968749 | 0.173523566 | 0.32596058 |
| ENSG00000125457 | MIF4GD | 1315.036522 | 0.016667395 | 0.061122958 | 0.272686331 | 0.785094343 | 0.883106023 |
| ENSG00000125458 | NT5C | 827.6223542 | 0.08118921 | 0.070712617 | 1.148157339 | 0.250903617 | 0.429243715 |
| ENSG00000125459 | MSTO1 | 449.9564377 | 0.178486687 | 0.09976802 | 1.78901703 | 0.073612073 | 0.165425558 |
| ENSG00000125462 | MIR9-1HG | 15.7107245 | -0.806088098 | 0.456379687 | -1.766266381 | 0.077351169 | 0.172174899 |
| ENSG00000125482 | TTF1 | 616.052161 | 0.148612094 | 0.082273842 | 1.806310381 | 0.070869871 | 0.160251833 |
| ENSG00000125484 | GTF3C4 | 2880.663599 | -0.046354258 | 0.062800974 | -0.738113674 | 0.460445375 | 0.647248857 |

| | | | | | | | |
|---|---|---|---|---|---|---|---|
| ENSG00000125485 | DDX31 | 741.9893492 | 0.046310822 | 0.074768374 | 0.61939052 | 0.535659125 | 0.711031295 |
| ENSG00000125503 | PPP1R12C | 1951.24855 | 0.104525748 | 0.055427918 | 1.885796021 | 0.059322447 | 0.138186123 |
| ENSG00000125505 | MBOAT7 | 6460.164127 | 0.304425447 | 0.057060618 | 5.335123598 | 9.55E-08 | 6.20E-07 |
| ENSG00000125510 | OPRL1 | 178.6919488 | -0.31756257 | 0.141809691 | -2.239357326 | 0.025132676 | 0.06676822 |
| ENSG00000125520 | SLC2A4RG | 4209.20739 | -0.154847711 | 0.054864895 | -2.822345908 | 0.004767373 | 0.015442464 |
| ENSG00000125531 | FNDC11 | 3.347336078 | -0.7821586 | 0.966412413 | -0.80934246 | 0.418318189 | 0.610291228 |
| ENSG00000125534 | PPDPF | 7328.266823 | -0.091126901 | 0.05903251 | -1.543673155 | 0.122667533 | 0.249600248 |
| ENSG00000125551 | PLGLB2 | 10.20017582 | 0.355632507 | 0.557241236 | 0.638202064 | 0.523342154 | 0.699904915 |
| ENSG00000125611 | CHCHD5 | 441.1044802 | -0.21132804 | 0.091681196 | -2.305031458 | 0.021164812 | 0.057396948 |
| ENSG00000125618 | PAX8 | 129.5642401 | 0.196082306 | 0.158408306 | 1.237828444 | 0.215779679 | 0.385021679 |
| ENSG00000125629 | INSIG2 | 760.3227802 | 0.540824735 | 0.074250282 | 7.283807184 | 3.25E-13 | 3.42E-12 |
| ENSG00000125630 | POLR1B | 1716.354077 | 0.094734179 | 0.057177274 | 1.656850205 | 0.097549764 | 0.207780091 |
| ENSG00000125633 | CCDC93 | 1937.417057 | 0.183701242 | 0.058396348 | 3.145765901 | 0.001656525 | 0.005923724 |
| ENSG00000125637 | PSD4 | 3740.953083 | -0.462285548 | 0.053219954 | -8.686319947 | 3.74E-18 | 5.59E-17 |
| ENSG00000125648 | SLC25A23 | 3525.043383 | -0.232668453 | 0.047904061 | -4.856967239 | 1.19E-06 | 6.83E-06 |
| ENSG00000125650 | PSPN | 76.52641655 | 0.352806944 | 0.201472558 | 1.751141435 | 0.07992155 | 0.176577132 |
| ENSG00000125651 | GTF2F1 | 3419.784466 | -0.029318187 | 0.051674393 | -0.567363932 | 0.57046695 | 0.737372898 |
| ENSG00000125652 | ALKBH7 | 305.9889575 | 0.214934639 | 0.119446028 | 1.799428933 | 0.071950856 | 0.162400574 |
| ENSG00000125656 | CLPP | 992.3878839 | -0.133637484 | 0.070142394 | -1.90523129 | 0.056750013 | 0.133262902 |
| ENSG00000125657 | TNFSF9 | 10.99808291 | 0.328242709 | 0.530483315 | 0.618761608 | 0.536073417 | 0.711332959 |
| ENSG00000125676 | THOC2 | 7034.170547 | -0.242754737 | 0.05279397 | -4.598152722 | 4.26E-06 | 2.29E-05 |
| ENSG00000125686 | MED1 | 2469.648908 | -0.082499574 | 0.06079502 | -1.357012043 | 0.174777384 | 0.327859448 |
| ENSG00000125691 | RPL23 | 9747.392631 | 0.202297594 | 0.052505875 | 3.852856352 | 0.000116748 | 0.000515038 |
| ENSG00000125703 | ATG4C | 245.4781715 | -0.279564332 | 0.118072971 | -2.367725062 | 0.017897831 | 0.049670486 |
| ENSG00000125730 | C3 | 39.57771035 | -0.330088103 | 0.288958745 | -1.142336438 | 0.253314201 | 0.431990496 |
| ENSG00000125731 | SH2D3A | 306.0198984 | -0.677411022 | 0.11002072 | -6.157122237 | 7.41E-10 | 5.97E-09 |
| ENSG00000125733 | TRIP10 | 864.9452146 | -0.085503497 | 0.071992252 | -1.187676366 | 0.234960938 | 0.40994858 |
| ENSG00000125734 | GPR108 | 1295.159458 | -0.182436951 | 0.072590231 | -2.513243826 | 0.011962662 | 0.034843001 |
| ENSG00000125740 | FOSB | 94.88923913 | 0.531117155 | 0.182299693 | 2.913428677 | 0.003574835 | 0.011912654 |
| ENSG00000125741 | OPA3 | 1280.109494 | -0.139935427 | 0.064258553 | -2.177693396 | 0.029428871 | 0.076367135 |
| ENSG00000125743 | SNRPD2 | 3604.621614 | 0.184865544 | 0.048309038 | 3.826727894 | 0.000129858 | 0.000568717 |
| ENSG00000125744 | RTN2 | 294.2857285 | 0.196631633 | 0.115052379 | 1.709061861 | 0.087439491 | 0.189967036 |
| ENSG00000125746 | EML2 | 920.0659841 | -0.028435334 | 0.07352963 | -0.386719397 | 0.698963955 | 0.831090713 |

| | | | | | | | |
|---|---|---|---|---|---|---|---|
| ENSG00000125753 | VASP | 3232.65559 | -0.347710171 | 0.059237641 | -5.869750448 | 4.36E-09 | 3.27E-08 |
| ENSG00000125755 | SYMPK | 3839.843289 | -0.105440359 | 0.051753436 | -2.037359597 | 0.04161402 | 0.102438632 |
| ENSG00000125772 | GPCPD1 | 467.8124341 | 0.033736919 | 0.098463494 | 0.342633783 | 0.731873984 | 0.851637881 |
| ENSG00000125775 | SDCBP2 | 11.4720482 | -0.158849663 | 0.542603163 | -0.292754768 | 0.769709604 | 0.874359188 |
| ENSG00000125779 | PANK2 | 885.9174143 | 0.004045087 | 0.07083586 | 0.057105079 | 0.954461491 | 0.979088978 |
| ENSG00000125787 | GNRH2 | 5.22761015 | 1.033503869 | 0.809926366 | 1.276046702 | 0.20193901 | 0.365799187 |
| ENSG00000125788 | DEFB126 | 6.742992653 | 0.005058126 | 0.670763161 | 0.007540852 | 0.993983328 | 0.996533822 |
| ENSG00000125812 | GZF1 | 936.6935407 | -0.359534806 | 0.071435478 | -5.033000663 | 4.83E-07 | 2.91E-06 |
| ENSG00000125814 | NAPB | 246.0985178 | -0.457490058 | 0.144061856 | -3.175650175 | 0.001495011 | 0.005399174 |
| ENSG00000125817 | CENPB | 5341.144805 | -0.033085199 | 0.052974851 | -0.624545396 | 0.532269467 | 0.707691769 |
| ENSG00000125818 | PSMF1 | 3069.081994 | 0.037456444 | 0.053804233 | 0.696161657 | 0.486327597 | 0.669947245 |
| ENSG00000125821 | DTD1 | 775.9698133 | -0.05358879 | 0.076361569 | -0.701776966 | 0.482818266 | 0.666938702 |
| ENSG00000125823 | CSTL1 | 2.345467378 | 1.369656443 | 1.268261384 | 1.079948077 | 0.280165302 | 0.463535885 |
| ENSG00000125826 | RBCK1 | 2562.430039 | 0.328383316 | 0.056458021 | 5.816415603 | 6.01E-09 | 4.44E-08 |
| ENSG00000125827 | TMX4 | 1113.490452 | -0.141821448 | 0.066860201 | -2.12116395 | 0.033908008 | 0.086216992 |
| ENSG00000125834 | STK35 | 1505.145566 | -0.156960235 | 0.061842419 | -2.538067519 | 0.011146648 | 0.032690373 |
| ENSG00000125835 | SNRPB | 2507.490602 | -0.221444777 | 0.058637848 | -3.776481978 | 0.000159059 | 0.000686254 |
| ENSG00000125841 | NRSN2 | 1400.424408 | -0.163798468 | 0.061509194 | -2.662991602 | 0.007744935 | 0.023762044 |
| ENSG00000125843 | AP5S1 | 744.2268979 | -0.01295544 | 0.082777975 | -0.156508295 | 0.875632382 | 0.93549771 |
| ENSG00000125844 | RRBP1 | 4026.047496 | -0.335607763 | 0.048807459 | -6.876157256 | 6.15E-12 | 5.87E-11 |
| ENSG00000125845 | BMP2 | 7.281422117 | 0.429263829 | 0.663290043 | 0.647173636 | 0.517519572 | 0.695665852 |
| ENSG00000125846 | ZNF133 | 252.1747468 | -0.676633648 | 0.116783778 | -5.793901002 | 6.88E-09 | 5.04E-08 |
| ENSG00000125848 | FLRT3 | 123.0682214 | -0.349838147 | 0.162859156 | -2.148102415 | 0.03170562 | 0.081493214 |
| ENSG00000125850 | OVOL2 | 186.9725565 | -0.59895217 | 0.133664388 | -4.481015307 | 7.43E-06 | 3.89E-05 |
| ENSG00000125863 | MKKS | 936.1485472 | -0.018077871 | 0.070951424 | -0.254792225 | 0.798883581 | 0.892721951 |
| ENSG00000125864 | BFSP1 | 92.15742541 | 0.29465867 | 0.18985483 | 1.552020931 | 0.120657215 | 0.24662537 |
| ENSG00000125868 | DSTN | 7059.516831 | -0.19162784 | 0.049863008 | -3.84308625 | 0.000121497 | 0.000534139 |
| ENSG00000125870 | SNRPB2 | 1416.362189 | -0.142102373 | 0.059833567 | -2.374960746 | 0.017550817 | 0.04884335 |
| ENSG00000125871 | MGME1 | 1176.302229 | -0.542716965 | 0.06410276 | -8.466358806 | 2.53E-17 | 3.56E-16 |
| ENSG00000125875 | TBC1D20 | 1189.842673 | 0.223884048 | 0.067902581 | 3.297136036 | 0.000976762 | 0.003664212 |
| ENSG00000125877 | ITPA | 809.2857803 | 0.076077417 | 0.076389441 | 0.995915353 | 0.319291275 | 0.508689685 |
| ENSG00000125885 | MCM8 | 211.2827934 | -0.0460955 | 0.126723364 | -0.363749028 | 0.716045429 | 0.842060908 |
| ENSG00000125898 | FAM110A | 337.9123755 | 0.167332793 | 0.103594026 | 1.615274542 | 0.106251254 | 0.222593952 |

| | | | | | | | |
|---|---|---|---|---|---|---|---|
| ENSG00000125901 | MRPS26 | 1521.958168 | -0.035657572 | 0.065517457 | -0.544245357 | 0.586272644 | 0.750100097 |
| ENSG00000125910 | S1PR4 | 146.6008207 | 1.021019558 | 0.155695029 | 6.557817326 | 5.46E-11 | 4.82E-10 |
| ENSG00000125912 | NCLN | 1960.208989 | 0.341065946 | 0.059934912 | 5.690605614 | 1.27E-08 | 9.00E-08 |
| ENSG00000125944 | HNRNPR | 5227.42097 | -0.086583435 | 0.044936377 | -1.926800525 | 0.054004494 | 0.127994947 |
| ENSG00000125945 | ZNF436 | 479.6964904 | -0.018446946 | 0.087424881 | -0.211003383 | 0.832884632 | 0.911607112 |
| ENSG00000125952 | MAX | 1617.626307 | -0.496148513 | 0.060942373 | -8.141273301 | 3.91E-16 | 5.09E-15 |
| ENSG00000125962 | ARMCX5 | 438.9709751 | 0.139860313 | 0.090889469 | 1.538795578 | 0.12385421 | 0.251655176 |
| ENSG00000125966 | MMP24 | 10.80993611 | -0.407599976 | 0.526767548 | -0.773775791 | 0.439063391 | 0.628529149 |
| ENSG00000125967 | NECAB3 | 935.3153377 | -0.443534665 | 0.075669198 | -5.861495552 | 4.59E-09 | 3.43E-08 |
| ENSG00000125968 | ID1 | 1456.347949 | -2.179112981 | 0.070347489 | -30.9764147 | 1.12E-210 | 7.56E-208 |
| ENSG00000125970 | RALY | 3542.933364 | -0.640687648 | 0.053782944 | -11.91246884 | 1.02E-32 | 3.15E-31 |
| ENSG00000125971 | DYNLRB1 | 142.5911938 | -0.731920471 | 0.161712514 | -4.526059555 | 6.01E-06 | 3.19E-05 |
| ENSG00000125977 | EIF2S2 | 1922.939371 | -0.3755736 | 0.060903781 | -6.166671331 | 6.97E-10 | 5.63E-09 |
| ENSG00000125991 | ERGIC3 | 4576.749899 | -0.238434922 | 0.051020522 | -4.67331403 | 2.96E-06 | 1.63E-05 |
| ENSG00000125995 | ROMO1 | 864.4152976 | -0.509666609 | 0.077308318 | -6.592649081 | 4.32E-11 | 3.83E-10 |
| ENSG00000125999 | BPIFB1 | 17.77334849 | 0.415090418 | 0.428369117 | 0.969001734 | 0.332544324 | 0.523610549 |
| ENSG00000126001 | CEP250 | 1261.69986 | -0.805800328 | 0.063162519 | -12.75757114 | 2.83E-37 | 1.04E-35 |
| ENSG00000126003 | PLAGL2 | 1453.473929 | -0.42215875 | 0.062655258 | -6.737802457 | 1.61E-11 | 1.48E-10 |
| ENSG00000126005 | MMP24OS | 511.261619 | 0.001981661 | 0.096190999 | 0.020601311 | 0.983563695 | 0.9923499 |
| ENSG00000126010 | GRPR | 18.09533002 | 1.369954224 | 0.448428627 | 3.055010634 | 0.002250525 | 0.007837719 |
| ENSG00000126012 | KDM5C | 6799.259174 | 0.140415356 | 0.054310687 | 2.585409329 | 0.009726345 | 0.029018384 |
| ENSG00000126016 | AMOT | 1097.846632 | 2.450308508 | 0.084386262 | 29.03681786 | 2.26E-185 | 1.13E-182 |
| ENSG00000126062 | TMEM115 | 1138.870062 | 0.140162138 | 0.071439153 | 1.961979288 | 0.049764896 | 0.119327554 |
| ENSG00000126067 | PSMB2 | 3419.765318 | -0.056556076 | 0.04661391 | -1.213287543 | 0.225019904 | 0.396890404 |
| ENSG00000126070 | AGO3 | 564.3902662 | -0.437242075 | 0.086634829 | -5.046954882 | 4.49E-07 | 2.72E-06 |
| ENSG00000126088 | UROD | 933.8684772 | -0.432502664 | 0.072777643 | -5.942795747 | 2.80E-09 | 2.14E-08 |
| ENSG00000126091 | ST3GAL3 | 28.0245455 | 0.186503219 | 0.338952641 | 0.550233856 | 0.582158985 | 0.746825138 |
| ENSG00000126106 | TMEM53 | 219.285002 | -0.27965902 | 0.126806765 | -2.205395109 | 0.027426381 | 0.071928796 |
| ENSG00000126107 | HECTD3 | 1635.348689 | 0.064870077 | 0.071398213 | 0.908567232 | 0.36357861 | 0.556345471 |
| ENSG00000126214 | KLC1 | 572.6888354 | -0.337147688 | 0.082015137 | -4.110798304 | 3.94E-05 | 0.000186998 |
| ENSG00000126215 | XRCC3 | 830.3252359 | -0.633857268 | 0.079237111 | -7.999499953 | 1.25E-15 | 1.57E-14 |
| ENSG00000126216 | TUBGCP3 | 810.1682829 | -0.044787142 | 0.075835159 | -0.59058546 | 0.554798209 | 0.726016596 |
| ENSG00000126217 | MCF2L | 2109.201722 | -0.246829497 | 0.058593537 | -4.212572039 | 2.52E-05 | 0.000123573 |

| | | | | | | | |
|---|---|---|---|---|---|---|---|
| ENSG00000126226 | PCID2 | 1019.833411 | -0.261947191 | 0.066696074 | -3.927475408 | 8.58E-05 | 0.000387413 |
| ENSG00000126233 | SLURP1 | 3.500110833 | 1.237144192 | 0.977588811 | 1.26550568 | 0.205690114 | 0.371128772 |
| ENSG00000126243 | LRFN3 | 450.0208472 | 0.333477488 | 0.10020135 | 3.328073799 | 0.000874487 | 0.003310179 |
| ENSG00000126246 | IGFLR1 | 14.27060222 | -0.03931913 | 0.463222739 | -0.084881693 | 0.932355446 | 0.967574483 |
| ENSG00000126247 | CAPNS1 | 10005.14369 | -0.412828157 | 0.043684694 | -9.450178601 | 3.38E-21 | 6.07E-20 |
| ENSG00000126249 | PDCD2L | 134.8988373 | 0.290682091 | 0.15949646 | 1.8224987 | 0.068379351 | 0.155652373 |
| ENSG00000126254 | RBM42 | 2077.105332 | -0.235351259 | 0.070779122 | -3.325150881 | 0.000883707 | 0.003341945 |
| ENSG00000126261 | UBA2 | 4897.16005 | -0.025603151 | 0.051568187 | -0.496491205 | 0.619547891 | 0.774603696 |
| ENSG00000126262 | FFAR2 | 245.1648774 | -0.362314575 | 0.120117336 | -3.016338758 | 0.002558473 | 0.008790079 |
| ENSG00000126264 | HCST | 9.516285274 | -0.278557338 | 0.558816716 | -0.498477103 | 0.618147804 | 0.773612145 |
| ENSG00000126267 | COX6B1 | 3338.263277 | 0.046741154 | 0.048545845 | 0.962825019 | 0.335635343 | 0.526875672 |
| ENSG00000126351 | THRA | 1860.192119 | 0.206081995 | 0.055036508 | 3.744459865 | 0.000180782 | 0.000772856 |
| ENSG00000126353 | CCR7 | 12.35086082 | -0.944252623 | 0.544212247 | -1.735081538 | 0.082726364 | 0.181550061 |
| ENSG00000126368 | NR1D1 | 389.0033566 | 0.165237027 | 0.116504581 | 1.418287806 | 0.156106755 | 0.300275782 |
| ENSG00000126391 | FRMD8 | 1189.564619 | 0.143033164 | 0.070217064 | 2.037014318 | 0.041648607 | 0.102501775 |
| ENSG00000126432 | PRDX5 | 3838.346527 | -0.336539536 | 0.046986713 | -7.162440545 | 7.93E-13 | 8.18E-12 |
| ENSG00000126453 | BCL2L12 | 695.5237889 | -0.151002177 | 0.080873354 | -1.867143752 | 0.061881516 | 0.143086582 |
| ENSG00000126456 | IRF3 | 2352.895318 | -0.075044941 | 0.055003208 | -1.364373888 | 0.172449922 | 0.324421946 |
| ENSG00000126457 | PRMT1 | 5757.848308 | 0.080650057 | 0.043311548 | 1.862091298 | 0.062590222 | 0.144463196 |
| ENSG00000126458 | RRAS | 133.489987 | -0.403942569 | 0.156343905 | -2.583679667 | 0.009775254 | 0.029141562 |
| ENSG00000126460 | PRRG2 | 238.9557231 | 0.034017195 | 0.119345293 | 0.285031732 | 0.775619831 | 0.878132412 |
| ENSG00000126461 | SCAF1 | 2179.166931 | -0.058428469 | 0.069449521 | -0.841308444 | 0.400175162 | 0.592982712 |
| ENSG00000126464 | PRR12 | 1974.978042 | -0.081459792 | 0.056829537 | -1.433405874 | 0.151741881 | 0.294083095 |
| ENSG00000126522 | ASL | 549.2248544 | 0.50333914 | 0.087002211 | 5.785360304 | 7.24E-09 | 5.28E-08 |
| ENSG00000126524 | SBDS | 1616.557551 | 0.298202591 | 0.059616198 | 5.002039753 | 5.67E-07 | 3.39E-06 |
| ENSG00000126561 | STAT5A | 79.48842578 | 0.936435427 | 0.209088114 | 4.478664083 | 7.51E-06 | 3.93E-05 |
| ENSG00000126562 | WNK4 | 42.00844885 | -1.015868379 | 0.288872982 | -3.516661095 | 0.000437011 | 0.001747836 |
| ENSG00000126581 | BECN1 | 2850.162011 | 0.055970549 | 0.050912142 | 1.099355613 | 0.271612984 | 0.453578063 |
| ENSG00000126583 | PRKCG | 5.507567055 | -1.130397058 | 0.775697903 | -1.457264553 | 0.145043368 | 0.284044293 |
| ENSG00000126602 | TRAP1 | 4510.027571 | 0.255433419 | 0.047170311 | 5.415131142 | 6.12E-08 | 4.04E-07 |
| ENSG00000126603 | GLIS2 | 756.4900942 | 0.41500471 | 0.090628902 | 4.579165166 | 4.67E-06 | 2.50E-05 |
| ENSG00000126653 | NSRP1 | 711.6817358 | -0.14392213 | 0.078641325 | -1.830108159 | 0.067233767 | 0.153548477 |
| ENSG00000126698 | DNAJC8 | 3799.413218 | -0.241318856 | 0.047064451 | -5.127412541 | 2.94E-07 | 1.81E-06 |

| | | | | | | | |
|---|---|---|---|---|---|---|---|
| ENSG00000126705 | AHDC1 | 609.0625071 | 0.466591708 | 0.087610104 | 5.325775072 | 1.01E-07 | 6.51E-07 |
| ENSG00000126709 | IFI6 | 140.6372966 | 1.613688724 | 0.32382795 | 4.983166912 | 6.26E-07 | 3.72E-06 |
| ENSG00000126733 | DACH2 | 3.239986344 | 1.542522786 | 1.058759707 | 1.456914895 | 0.145139874 | 0.284209024 |
| ENSG00000126746 | ZNF384 | 1741.25681 | -0.17735867 | 0.063377102 | -2.798466056 | 0.005134597 | 0.016494559 |
| ENSG00000126749 | EMG1 | 877.1531923 | -0.07176525 | 0.071874174 | -0.998484529 | 0.318044463 | 0.507207853 |
| ENSG00000126756 | UXT | 1259.792596 | 0.076342514 | 0.062952499 | 1.212700302 | 0.225244424 | 0.39722533 |
| ENSG00000126759 | CFP | 12.59657922 | 0.404014288 | 0.509059988 | 0.793647699 | 0.427400556 | 0.618217808 |
| ENSG00000126767 | ELK1 | 2029.627106 | 0.093272561 | 0.058175496 | 1.603296359 | 0.10886924 | 0.226815036 |
| ENSG00000126768 | TIMM17B | 1428.506959 | -0.103252805 | 0.0697812 | -1.479665091 | 0.138962646 | 0.274929116 |
| ENSG00000126773 | PCNX4 | 1462.993028 | -0.730085033 | 0.069363525 | -10.52548924 | 6.59E-26 | 1.53E-24 |
| ENSG00000126775 | ATG14 | 717.8799529 | -0.388796154 | 0.082390601 | -4.718938193 | 2.37E-06 | 1.31E-05 |
| ENSG00000126777 | KTN1 | 10948.18765 | -0.438736338 | 0.045714313 | -9.597351572 | 8.20E-22 | 1.52E-20 |
| ENSG00000126778 | SIX1 | 332.34123 | -0.177448577 | 0.104161095 | -1.70359746 | 0.088456316 | 0.191776571 |
| ENSG00000126787 | DLGAP5 | 1643.653457 | -0.761782831 | 0.063711896 | -11.95668127 | 5.99E-33 | 1.86E-31 |
| ENSG00000126790 | L3HYPDH | 165.0239663 | -0.442652724 | 0.145515964 | -3.041953016 | 0.002350486 | 0.008152426 |
| ENSG00000126803 | HSPA2 | 193.1806231 | -1.139186512 | 0.134842075 | -8.448301583 | 2.96E-17 | 4.13E-16 |
| ENSG00000126804 | ZBTB1 | 2337.61573 | -0.213464618 | 0.055248746 | -3.863700718 | 0.000111682 | 0.000494687 |
| ENSG00000126814 | TRMT5 | 737.0571868 | -0.239382231 | 0.077917534 | -3.072251103 | 0.002124509 | 0.007425825 |
| ENSG00000126821 | SGPP1 | 541.9910001 | -2.916315631 | 0.108587353 | -26.85686264 | 7.01E-159 | 2.59E-156 |
| ENSG00000126822 | PLEKHG3 | 2110.761583 | -0.556892271 | 0.056696674 | -9.822309294 | 9.03E-23 | 1.77E-21 |
| ENSG00000126858 | RHOT1 | 958.7259872 | -0.077870068 | 0.067407817 | -1.155208277 | 0.248005146 | 0.425586329 |
| ENSG00000126870 | DYNC2I1 | 557.7779307 | 0.607292613 | 0.086337994 | 7.033897622 | 2.01E-12 | 2.00E-11 |
| ENSG00000126878 | AIF1L | 4799.243285 | 0.346149044 | 0.047563358 | 7.277640984 | 3.40E-13 | 3.58E-12 |
| ENSG00000126882 | FAM78A | 237.9246804 | -0.006302885 | 0.124773512 | -0.050514611 | 0.959712306 | 0.981621201 |
| ENSG00000126883 | NUP214 | 2434.567901 | -0.032252333 | 0.055412939 | -0.582036133 | 0.560542343 | 0.729849205 |
| ENSG00000126903 | SLC10A3 | 1503.212162 | -0.642128227 | 0.061364968 | -10.46408476 | 1.26E-25 | 2.88E-24 |
| ENSG00000126934 | MAP2K2 | 3141.744554 | 0.027846525 | 0.05486265 | 0.507567989 | 0.611756345 | 0.76921553 |
| ENSG00000126945 | HNRNPH2 | 3849.118475 | -0.01546962 | 0.049490784 | -0.312575759 | 0.754602998 | 0.864498202 |
| ENSG00000126947 | ARMCX1 | 439.9751991 | 0.160532291 | 0.091675791 | 1.75108705 | 0.079930916 | 0.176580819 |
| ENSG00000126952 | NXF5 | 1.798470548 | -0.41280271 | 1.277720184 | -0.323077553 | 0.74663651 | 0.859607712 |
| ENSG00000126953 | TIMM8A | 698.6918553 | -0.017531423 | 0.075547364 | -0.232058695 | 0.816492426 | 0.902797332 |
| ENSG00000126970 | ZC4H2 | 156.6840528 | -0.726575824 | 0.148303053 | -4.89926409 | 9.62E-07 | 5.57E-06 |
| ENSG00000127022 | CANX | 25097.589 | -0.234925576 | 0.051498171 | -4.561823637 | 5.07E-06 | 2.71E-05 |

| | | | | | | | |
|---|---|---|---|---|---|---|---|
| ENSG00000127054 | INTS11 | 2336.04906 | 0.061961527 | 0.064382693 | 0.962394152 | 0.33585165 | 0.526943222 |
| ENSG00000127080 | IPPK | 695.4611198 | 0.349812148 | 0.07877573 | 4.440608149 | 8.97E-06 | 4.66E-05 |
| ENSG00000127081 | ZNF484 | 148.5720448 | -0.321750357 | 0.151647806 | -2.121694768 | 0.033863378 | 0.086132154 |
| ENSG00000127084 | FGD3 | 234.8210474 | 0.453699055 | 0.121024615 | 3.748816348 | 0.000177671 | 0.000760832 |
| ENSG00000127124 | HIVEP3 | 474.7908465 | -0.618197037 | 0.096224804 | -6.42450812 | 1.32E-10 | 1.13E-09 |
| ENSG00000127125 | PPCS | 1372.07764 | -0.08238891 | 0.060212627 | -1.368299534 | 0.171218337 | 0.322528218 |
| ENSG00000127129 | EDN2 | 24.79978578 | -0.532018109 | 0.370863037 | -1.434540668 | 0.151418032 | 0.293769423 |
| ENSG00000127152 | BCL11B | 289.0925326 | -0.332212101 | 0.11069715 | -3.001089927 | 0.002690151 | 0.009208007 |
| ENSG00000127184 | COX7C | 3492.236488 | 0.039314639 | 0.050684911 | 0.775667527 | 0.437945318 | 0.627472712 |
| ENSG00000127191 | TRAF2 | 902.8240162 | -0.187235612 | 0.072208358 | -2.592990845 | 0.00951453 | 0.02847532 |
| ENSG00000127220 | ABHD8 | 35.64375049 | -0.227397254 | 0.301929208 | -0.753147586 | 0.451361232 | 0.639390671 |
| ENSG00000127249 | ATP13A4 | 5.759030249 | -0.24974401 | 0.721355609 | -0.34621483 | 0.72918127 | 0.850013636 |
| ENSG00000127311 | HELB | 66.61463838 | -0.523128244 | 0.218130115 | -2.398239438 | 0.016474093 | 0.046183024 |
| ENSG00000127314 | RAP1B | 730.4084201 | 0.138176875 | 0.081777312 | 1.68967249 | 0.091090629 | 0.196411226 |
| ENSG00000127325 | BEST3 | 2.012716821 | 0.269836493 | 1.228946641 | 0.219567298 | 0.826208163 | 0.908362226 |
| ENSG00000127328 | RAB3IP | 1032.978654 | 0.156051571 | 0.066440542 | 2.348740186 | 0.018837046 | 0.051925238 |
| ENSG00000127329 | PTPRB | 11.07493124 | -1.707480573 | 0.576426699 | -2.962181618 | 0.003054675 | 0.010330865 |
| ENSG00000127334 | DYRK2 | 1289.247965 | 0.236240686 | 0.067420427 | 3.503992707 | 0.000458338 | 0.001826663 |
| ENSG00000127337 | YEATS4 | 627.4739315 | 0.153309515 | 0.080246088 | 1.910492086 | 0.056069884 | 0.132053329 |
| ENSG00000127364 | TAS2R4 | 8.198867836 | -0.223395307 | 0.624726112 | -0.357589195 | 0.720650765 | 0.844878037 |
| ENSG00000127366 | TAS2R5 | 6.637490175 | -0.477251469 | 0.683927262 | -0.697810273 | 0.485295854 | 0.669112086 |
| ENSG00000127399 | LRRC61 | 396.7459452 | -0.636538608 | 0.10342415 | -6.154641885 | 7.52E-10 | 6.06E-09 |
| ENSG00000127415 | IDUA | 876.1267692 | 0.236386744 | 0.078364086 | 3.016518862 | 0.002556953 | 0.008786175 |
| ENSG00000127418 | FGFRL1 | 1662.329625 | 0.892003899 | 0.066611586 | 13.39112245 | 6.81E-41 | 2.89E-39 |
| ENSG00000127419 | TMEM175 | 548.6992111 | -0.144000868 | 0.083698283 | -1.720475776 | 0.085345993 | 0.186245998 |
| ENSG00000127423 | AUNIP | 335.6157642 | 0.191321389 | 0.106899619 | 1.789729376 | 0.073497428 | 0.165232359 |
| ENSG00000127445 | PIN1 | 834.0680144 | -0.045785544 | 0.070770372 | -0.646959216 | 0.517658339 | 0.695710487 |
| ENSG00000127452 | FBXL12 | 447.3030638 | -0.14526272 | 0.092000751 | -1.57892972 | 0.114352178 | 0.236253263 |
| ENSG00000127463 | EMC1 | 1710.508103 | -0.049455661 | 0.059564346 | -0.830289657 | 0.406375035 | 0.599115893 |
| ENSG00000127481 | UBR4 | 3099.076388 | -0.019994136 | 0.063975034 | -0.312530286 | 0.75463755 | 0.864498202 |
| ENSG00000127483 | HP1BP3 | 4750.357455 | -0.052835134 | 0.045635489 | -1.157764163 | 0.246960292 | 0.424205847 |
| ENSG00000127511 | SIN3B | 1166.625895 | -0.034834738 | 0.069156633 | -0.503707838 | 0.614466699 | 0.771235218 |
| ENSG00000127526 | SLC35E1 | 2743.784074 | 0.116915827 | 0.048683574 | 2.401545697 | 0.01632597 | 0.045818173 |

| | | | | | | | |
|---|---|---|---|---|---|---|---|
| ENSG00000127527 | EPS15L1 | 1389.9492 | -0.234179588 | 0.059277992 | -3.95053169 | 7.80E-05 | 0.000354571 |
| ENSG00000127528 | KLF2 | 19.77975216 | -0.094073295 | 0.38764579 | -0.242678489 | 0.808254473 | 0.898401971 |
| ENSG00000127530 | OR7C1 | 2.856091135 | -0.996482929 | 1.11189388 | -0.896203268 | 0.370144211 | 0.562809584 |
| ENSG00000127540 | UQCR11 | 318.6720154 | 0.129843686 | 0.105287628 | 1.233228326 | 0.21749059 | 0.387062384 |
| ENSG00000127554 | GFER | 1089.01295 | -0.015815812 | 0.06671188 | -0.237076389 | 0.812597529 | 0.901009586 |
| ENSG00000127561 | SYNGR3 | 280.648844 | 0.812672679 | 0.115756983 | 7.020506721 | 2.21E-12 | 2.20E-11 |
| ENSG00000127564 | PKMYT1 | 1434.414937 | -0.479601542 | 0.062700659 | -7.649066991 | 2.02E-14 | 2.33E-13 |
| ENSG00000127578 | WFIKKN1 | 65.66275203 | 0.937070199 | 0.231898852 | 4.04085743 | 5.33E-05 | 0.00024806 |
| ENSG00000127580 | WDR24 | 668.7610546 | -0.152712509 | 0.081979126 | -1.862821878 | 0.062487331 | 0.144254742 |
| ENSG00000127585 | FBXL16 | 1692.039368 | 0.35207938 | 0.065799825 | 5.350764645 | 8.76E-08 | 5.70E-07 |
| ENSG00000127586 | CHTF18 | 1447.10721 | 0.034622709 | 0.066569831 | 0.520096084 | 0.602996608 | 0.762148806 |
| ENSG00000127588 | GNG13 | 32.75891832 | 0.669565324 | 0.317103544 | 2.111503757 | 0.034729035 | 0.088021663 |
| ENSG00000127589 | TUBBP1 | 1383.890197 | -0.552691874 | 0.061496347 | -8.987393523 | 2.53E-19 | 4.08E-18 |
| ENSG00000127603 | MACF1 | 4459.307447 | -0.035103587 | 0.070191559 | -0.500111226 | 0.616996762 | 0.772972842 |
| ENSG00000127616 | SMARCA4 | 9103.586683 | 0.06496153 | 0.043083384 | 1.507809354 | 0.131603328 | 0.263919952 |
| ENSG00000127663 | KDM4B | 1272.458694 | 0.307865053 | 0.062741351 | 4.906892319 | 9.25E-07 | 5.38E-06 |
| ENSG00000127666 | TICAM1 | 237.4734137 | 0.014049145 | 0.119353575 | 0.117710298 | 0.906297207 | 0.952693306 |
| ENSG00000127720 | METTL25 | 158.9484085 | -0.049551909 | 0.146582749 | -0.338047344 | 0.735327511 | 0.854010383 |
| ENSG00000127774 | EMC6 | 471.0278599 | -0.127686718 | 0.092939267 | -1.373872659 | 0.169481219 | 0.320018039 |
| ENSG00000127804 | METTL16 | 1070.877589 | 0.158579766 | 0.065538245 | 2.419652307 | 0.015535353 | 0.043883882 |
| ENSG00000127824 | TUBA4A | 210.016705 | 1.395151798 | 0.136018863 | 10.25704647 | 1.10E-24 | 2.41E-23 |
| ENSG00000127831 | VIL1 | 87.59016487 | -1.171696252 | 0.201085605 | -5.826852967 | 5.65E-09 | 4.18E-08 |
| ENSG00000127837 | AAMP | 4716.898209 | -0.041158613 | 0.049531907 | -0.830951516 | 0.406001019 | 0.598679911 |
| ENSG00000127838 | PNKD | 1553.375576 | -0.421408659 | 0.06414705 | -6.569416041 | 5.05E-11 | 4.47E-10 |
| ENSG00000127870 | RNF6 | 2153.512633 | 0.050199854 | 0.053275799 | 0.942263756 | 0.34605762 | 0.5369335 |
| ENSG00000127884 | ECHS1 | 4067.351756 | -0.420542057 | 0.047703729 | -8.815706183 | 1.19E-18 | 1.83E-17 |
| ENSG00000127914 | AKAP9 | 2272.699105 | -0.707140509 | 0.062260283 | -11.35781079 | 6.78E-30 | 1.88E-28 |
| ENSG00000127922 | SEM1 | 1693.013593 | -0.362708062 | 0.060059303 | -6.039165312 | 1.55E-09 | 1.21E-08 |
| ENSG00000127928 | GNGT1 | 131.1576904 | -0.897342886 | 0.167534022 | -5.356183042 | 8.50E-08 | 5.54E-07 |
| ENSG00000127946 | HIP1 | 1439.704688 | 0.492405907 | 0.066755306 | 7.376281117 | 1.63E-13 | 1.75E-12 |
| ENSG00000127947 | PTPN12 | 2294.600302 | 0.214990724 | 0.05382909 | 3.993950531 | 6.50E-05 | 0.000298672 |
| ENSG00000127948 | POR | 2160.600184 | 0.471641496 | 0.067242671 | 7.014020868 | 2.32E-12 | 2.30E-11 |
| ENSG00000127952 | STYXL1 | 780.7005039 | 0.223729441 | 0.074724837 | 2.994043899 | 0.002753063 | 0.00941071 |

| | | | | | | | |
|---|---|---|---|---|---|---|---|
| ENSG00000127954 | STEAP4 | 17.92257191 | 0.736573257 | 0.43015228 | 1.712354649 | 0.086831331 | 0.188914145 |
| ENSG00000127955 | GNAI1 | 11.07742624 | -2.998339737 | 0.674763445 | -4.443542046 | 8.85E-06 | 4.59E-05 |
| ENSG00000127957 | PMS2P3 | 103.523066 | 0.654535193 | 0.180332387 | 3.629604214 | 0.000283856 | 0.001172005 |
| ENSG00000127980 | PEX1 | 1290.009601 | -0.331078742 | 0.068327909 | -4.845439405 | 1.26E-06 | 7.23E-06 |
| ENSG00000127989 | MTERF1 | 378.1015931 | -0.67850293 | 0.103626335 | -6.547591721 | 5.85E-11 | 5.16E-10 |
| ENSG00000127990 | SGCE | 88.37069308 | -7.567539622 | 0.848172633 | -8.922169057 | 4.57E-19 | 7.23E-18 |
| ENSG00000127993 | RBM48 | 419.5130838 | -0.840211407 | 0.09754991 | -8.613143817 | 7.11E-18 | 1.04E-16 |
| ENSG00000127995 | CASD1 | 516.5651512 | -0.541422961 | 0.10225059 | -5.295059541 | 1.19E-07 | 7.67E-07 |
| ENSG00000128000 | ZNF780B | 235.9086335 | -0.171686388 | 0.120788211 | -1.421383649 | 0.155205257 | 0.29891769 |
| ENSG00000128011 | LRFN1 | 51.59162472 | -0.166276944 | 0.25560139 | -0.650532237 | 0.515348485 | 0.694378703 |
| ENSG00000128016 | ZFP36 | 667.1090241 | -0.092368273 | 0.082013645 | -1.126254937 | 0.260057626 | 0.440209706 |
| ENSG00000128039 | SRD5A3 | 625.3344066 | -0.016702609 | 0.080262415 | -0.208100001 | 0.835150886 | 0.913085565 |
| ENSG00000128040 | SPINK2 | 8.440317986 | -0.149350097 | 0.598631834 | -0.249485725 | 0.802985082 | 0.895135709 |
| ENSG00000128045 | RASL11B | 4.838208572 | 2.095900738 | 0.916754232 | 2.286218776 | 0.022241463 | 0.060025783 |
| ENSG00000128050 | PAICS | 7012.612199 | 0.045947023 | 0.05106465 | 0.899781416 | 0.368236586 | 0.560801121 |
| ENSG00000128052 | KDR | 6.330642564 | -1.242296324 | 0.77307371 | -1.606957147 | 0.108063769 | 0.225608196 |
| ENSG00000128059 | PPAT | 1202.718249 | 0.133863835 | 0.066883496 | 2.001447943 | 0.045344138 | 0.110143428 |
| ENSG00000128159 | TUBGCP6 | 1597.936753 | -0.171608028 | 0.06699838 | -2.561375773 | 0.010425852 | 0.030820753 |
| ENSG00000128165 | ADM2 | 422.3937692 | -0.242764785 | 0.093720898 | -2.590295126 | 0.009589368 | 0.028676849 |
| ENSG00000128185 | DGCR6L | 1243.002251 | -0.484424597 | 0.074719914 | -6.48320605 | 8.98E-11 | 7.81E-10 |
| ENSG00000128191 | DGCR8 | 1609.763012 | -0.161491693 | 0.061713673 | -2.616789524 | 0.008876105 | 0.026795349 |
| ENSG00000128203 | ASPHD2 | 8.364494356 | -0.907199341 | 0.627713221 | -1.445244916 | 0.148389104 | 0.289090959 |
| ENSG00000128218 | VPREB3 | 5.975734725 | -1.707336103 | 0.776657708 | -2.198312184 | 0.027926867 | 0.073057626 |
| ENSG00000128228 | SDF2L1 | 349.3543565 | -0.613835916 | 0.111388084 | -5.510786191 | 3.57E-08 | 2.42E-07 |
| ENSG00000128245 | YWHAH | 1741.606666 | -0.85904731 | 0.058492792 | -14.68637897 | 7.88E-49 | 4.20E-47 |
| ENSG00000128254 | YWHAH-AS1 | 7.589581672 | -1.066887074 | 0.648436279 | -1.645322923 | 0.099903235 | 0.211770487 |
| ENSG00000128262 | POM121L9P | 3.672183669 | -0.465107471 | 0.907405934 | -0.512568249 | 0.608253368 | 0.766456043 |
| ENSG00000128266 | GNAZ | 199.616953 | 0.132972883 | 0.135154433 | 0.983858836 | 0.325184926 | 0.51535352 |
| ENSG00000128271 | ADORA2A | 7.767406135 | -1.093870577 | 0.66832296 | -1.636739484 | 0.101684915 | 0.214852349 |
| ENSG00000128272 | ATF4 | 9656.856047 | 0.057928888 | 0.047168461 | 1.228127581 | 0.219399079 | 0.389341291 |
| ENSG00000128274 | A4GALT | 196.7509358 | -0.362101145 | 0.12921052 | -2.802412254 | 0.005072201 | 0.016321545 |
| ENSG00000128283 | CDC42EP1 | 639.8859166 | 0.145171661 | 0.085262203 | 1.702649652 | 0.088633652 | 0.192088425 |
| ENSG00000128284 | APOL3 | 3.82259198 | 5.288631957 | 1.395889351 | 3.788718607 | 0.000151426 | 0.000656038 |

| | | | | | | | |
|---|---|---|---|---|---|---|---|
| ENSG00000128294 | TPST2 | 174.8356444 | 0.148356531 | 0.142729872 | 1.039421735 | 0.29860864 | 0.485167661 |
| ENSG00000128298 | BAIAP2L2 | 8.835085536 | -0.134851375 | 0.596233145 | -0.226172222 | 0.821067475 | 0.905064046 |
| ENSG00000128309 | MPST | 926.3465358 | -0.43631337 | 0.073496612 | -5.936509963 | 2.91E-09 | 2.22E-08 |
| ENSG00000128310 | GALR3 | 2.173418249 | -0.397361761 | 1.170105319 | -0.339594867 | 0.734161644 | 0.853390834 |
| ENSG00000128311 | TST | 1195.460443 | -0.471758734 | 0.07638689 | -6.175912314 | 6.58E-10 | 5.32E-09 |
| ENSG00000128335 | APOL2 | 341.4519568 | 0.143337137 | 0.102280586 | 1.401410983 | 0.16109121 | 0.307829774 |
| ENSG00000128342 | LIF | 28.69678081 | -0.314101346 | 0.333188296 | -0.942714225 | 0.345827095 | 0.536750313 |
| ENSG00000128346 | C22orf23 | 11.58879539 | -0.166043606 | 0.530025255 | -0.313274894 | 0.754071822 | 0.864401739 |
| ENSG00000128383 | APOBEC3A | 2.007911145 | -1.717406203 | 1.3468701 | -1.275109012 | 0.20227066 | 0.366284255 |
| ENSG00000128394 | APOBEC3F | 2.333234316 | -1.061155056 | 1.193935009 | -0.888787956 | 0.37411705 | 0.566896435 |
| ENSG00000128408 | RIBC2 | 50.31999334 | -1.061653525 | 0.256852232 | -4.133324119 | 3.58E-05 | 0.000170703 |
| ENSG00000128422 | KRT17 | 64.11644444 | 2.444253365 | 0.269448272 | 9.071326921 | 1.18E-19 | 1.93E-18 |
| ENSG00000128463 | EMC4 | 1780.696254 | -0.198409521 | 0.058903514 | -3.368381768 | 0.000756108 | 0.002895196 |
| ENSG00000128482 | RNF112 | 3.098204805 | -1.175502842 | 1.036623641 | -1.133972636 | 0.256806027 | 0.436347696 |
| ENSG00000128487 | SPECC1 | 2893.639359 | 0.245863877 | 0.052639983 | 4.670667844 | 3.00E-06 | 1.65E-05 |
| ENSG00000128512 | DOCK4 | 82.36727496 | 0.704345041 | 0.197336147 | 3.569265197 | 0.000357984 | 0.001450141 |
| ENSG00000128513 | POT1 | 596.7336563 | 0.115971436 | 0.082102035 | 1.412528149 | 0.157794508 | 0.302634063 |
| ENSG00000128524 | ATP6V1F | 2288.77241 | 0.095028922 | 0.052347315 | 1.815354283 | 0.069469482 | 0.157626974 |
| ENSG00000128534 | LSM8 | 744.1064257 | -0.075664613 | 0.077969066 | -0.970444012 | 0.33182522 | 0.522842952 |
| ENSG00000128536 | CDHR3 | 40.03763551 | 0.46905771 | 0.281188221 | 1.668127163 | 0.095290486 | 0.203857479 |
| ENSG00000128563 | PRKRIP1 | 1395.273274 | -0.447302253 | 0.061358573 | -7.289971586 | 3.10E-13 | 3.28E-12 |
| ENSG00000128564 | VGF | 155.4167352 | -0.614202807 | 0.161430837 | -3.804742759 | 0.000141952 | 0.000617556 |
| ENSG00000128567 | PODXL | 1622.170851 | -0.274667402 | 0.061490729 | -4.466810015 | 7.94E-06 | 4.14E-05 |
| ENSG00000128573 | FOXP2 | 132.5021031 | 0.414553534 | 0.158605601 | 2.613738297 | 0.008955761 | 0.026993187 |
| ENSG00000128578 | STRIP2 | 142.2057232 | 0.137968976 | 0.154889576 | 0.890757018 | 0.373059539 | 0.565817244 |
| ENSG00000128581 | IFT22 | 1256.807252 | -0.272818751 | 0.0654346 | -4.169334723 | 3.05E-05 | 0.000147878 |
| ENSG00000128585 | MKLN1 | 1207.226553 | -0.001036233 | 0.066702758 | -0.015535086 | 0.987605293 | 0.993927018 |
| ENSG00000128590 | DNAJB9 | 285.3800551 | -0.196068694 | 0.11363379 | -1.725443594 | 0.084447561 | 0.184673694 |
| ENSG00000128591 | FLNC | 3.286694217 | -0.542188371 | 1.024917152 | -0.529007023 | 0.596800578 | 0.757649737 |
| ENSG00000128594 | LRRC4 | 42.14564482 | 1.035684229 | 0.290432646 | 3.56600487 | 0.000362465 | 0.001467256 |
| ENSG00000128595 | CALU | 3415.98797 | -0.073139847 | 0.053277884 | -1.372799418 | 0.169814712 | 0.320412952 |
| ENSG00000128596 | CCDC136 | 102.5935029 | -0.360547097 | 0.176223903 | -2.045960228 | 0.040760284 | 0.10070446 |
| ENSG00000128602 | SMO | 597.4398795 | 0.481508206 | 0.088495904 | 5.441022517 | 5.30E-08 | 3.53E-07 |

| | | | | | | | |
|---|---|---|---|---|---|---|---|
| ENSG00000128604 | IRF5 | 510.603929 | 0.420253708 | 0.089459199 | 4.697713736 | 2.63E-06 | 1.45E-05 |
| ENSG00000128606 | LRRC17 | 13.23085035 | -0.291460659 | 0.487706468 | -0.597614914 | 0.550096911 | 0.72241946 |
| ENSG00000128607 | KLHDC10 | 1861.085572 | -0.00058809 | 0.05690665 | -0.010334285 | 0.99175458 | 0.995660834 |
| ENSG00000128609 | NDUFA5 | 1002.079207 | 0.210839365 | 0.075250885 | 2.80181909 | 0.005081536 | 0.016342414 |
| ENSG00000128610 | FEZF1 | 20.40522699 | -1.775589177 | 0.432122856 | -4.108991578 | 3.97E-05 | 0.00018835 |
| ENSG00000128626 | MRPS12 | 1039.103432 | -0.025438433 | 0.069167324 | -0.367781077 | 0.71303648 | 0.84037489 |
| ENSG00000128641 | MYO1B | 10953.74829 | -0.524541126 | 0.045713693 | -11.47448585 | 1.77E-30 | 5.01E-29 |
| ENSG00000128654 | MTX2 | 834.9681226 | -0.011785281 | 0.070717045 | -0.166654031 | 0.867642277 | 0.930973171 |
| ENSG00000128655 | PDE11A | 8.796209372 | -0.780998647 | 0.588356367 | -1.327424486 | 0.184368304 | 0.341053859 |
| ENSG00000128656 | CHN1 | 499.9605547 | 0.545098286 | 0.088366079 | 6.168637247 | 6.89E-10 | 5.57E-09 |
| ENSG00000128683 | GAD1 | 24.32341132 | 1.37163802 | 0.388707129 | 3.528718454 | 0.000417577 | 0.001675568 |
| ENSG00000128692 | EIF2S2P4 | 1012.074193 | -0.457106378 | 0.074583238 | -6.128808409 | 8.85E-10 | 7.07E-09 |
| ENSG00000128694 | OSGEPL1 | 147.3704649 | -0.002619719 | 0.149002698 | -0.01758169 | 0.985972564 | 0.993350589 |
| ENSG00000128699 | ORMDL1 | 838.6222054 | 0.129716926 | 0.078818973 | 1.645757631 | 0.099813668 | 0.21166083 |
| ENSG00000128708 | HAT1 | 2180.439059 | -0.161451454 | 0.054259722 | -2.975530412 | 0.002924823 | 0.009931239 |
| ENSG00000128731 | HERC2 | 1654.839534 | -0.170026997 | 0.069334157 | -2.452283338 | 0.014195284 | 0.040487356 |
| ENSG00000128789 | PSMG2 | 1281.678903 | 0.173212329 | 0.061484759 | 2.817158785 | 0.004845057 | 0.015669759 |
| ENSG00000128791 | TWSG1 | 776.2961392 | 0.519444113 | 0.076301283 | 6.807803138 | 9.91E-12 | 9.28E-11 |
| ENSG00000128829 | EIF2AK4 | 1654.637602 | -0.016826648 | 0.059138144 | -0.284531217 | 0.776003314 | 0.878332993 |
| ENSG00000128833 | MYO5C | 18075.2786 | -1.504081998 | 0.047152306 | -31.89837621 | 2.81E-223 | 2.69E-220 |
| ENSG00000128849 | CGNL1 | 134.1975467 | -1.251600049 | 0.183871725 | -6.806919588 | 9.97E-12 | 9.34E-11 |
| ENSG00000128872 | TMOD2 | 5614.124982 | -1.017829747 | 0.053674065 | -18.9631576 | 3.44E-80 | 3.92E-78 |
| ENSG00000128881 | TTBK2 | 246.7213152 | -0.820034202 | 0.121946639 | -6.724533012 | 1.76E-11 | 1.61E-10 |
| ENSG00000128886 | ELL3 | 11.92906888 | 0.784708371 | 0.512979251 | 1.529707818 | 0.126089068 | 0.25509394 |
| ENSG00000128891 | CCDC32 | 636.56923 | 0.030219631 | 0.079319619 | 0.380985584 | 0.703213946 | 0.833796972 |
| ENSG00000128908 | INO80 | 2070.850598 | -0.326807736 | 0.056514059 | -5.782768736 | 7.35E-09 | 5.36E-08 |
| ENSG00000128915 | ICE2 | 1083.363102 | -0.184716683 | 0.065487366 | -2.820646104 | 0.004792704 | 0.0155151 |
| ENSG00000128917 | DLL4 | 39.59107958 | 0.475026304 | 0.286304053 | 1.659167234 | 0.097082098 | 0.207033869 |
| ENSG00000128918 | ALDH1A2 | 11.36542622 | -2.664083285 | 0.630111161 | -4.227957619 | 2.36E-05 | 0.000115692 |
| ENSG00000128923 | MINDY2 | 546.5206318 | -0.289298774 | 0.086854766 | -3.330833605 | 0.000865863 | 0.003279872 |
| ENSG00000128928 | IVD | 4393.551916 | -0.030228757 | 0.051558661 | -0.586298334 | 0.557675051 | 0.727915708 |
| ENSG00000128944 | KNSTRN | 1160.956205 | -0.408243058 | 0.067030832 | -6.090377272 | 1.13E-09 | 8.92E-09 |
| ENSG00000128951 | DUT | 1478.202503 | -0.267436493 | 0.066887892 | -3.998279581 | 6.38E-05 | 0.000293734 |

| | | | | | | | |
|---|---|---|---|---|---|---|---|
| ENSG00000128965 | CHAC1 | 220.5304247 | -0.058306366 | 0.134710464 | -0.432827295 | 0.665140242 | 0.806891442 |
| ENSG00000128973 | CLN6 | 980.0478062 | 0.025337533 | 0.070527082 | 0.359259631 | 0.719400871 | 0.844059737 |
| ENSG00000128989 | ARPP19 | 32083.78968 | -1.387195631 | 0.058073187 | -23.88702423 | 4.18E-126 | 1.10E-123 |
| ENSG00000129003 | VPS13C | 1348.032436 | 0.069472009 | 0.070373525 | 0.987189555 | 0.323549721 | 0.513619561 |
| ENSG00000129007 | CALML4 | 13.97251026 | -0.060312287 | 0.471452405 | -0.127928687 | 0.898205408 | 0.948554546 |
| ENSG00000129028 | THAP10 | 171.2063259 | 0.080035972 | 0.142829469 | 0.560360358 | 0.575233665 | 0.741632118 |
| ENSG00000129038 | LOXL1 | 78.79722378 | -0.098127855 | 0.207043868 | -0.473947168 | 0.635537582 | 0.785180533 |
| ENSG00000129048 | ACKR4 | 1.794746841 | 0.618444196 | 1.313204955 | 0.47094263 | 0.637681704 | 0.78659182 |
| ENSG00000129055 | ANAPC13 | 1422.758893 | -0.137328089 | 0.065331319 | -2.102025364 | 0.035551054 | 0.089787645 |
| ENSG00000129071 | MBD4 | 1439.845908 | 0.17059113 | 0.061623604 | 2.768275753 | 0.005635375 | 0.017897695 |
| ENSG00000129083 | COPB1 | 4149.581135 | -0.078376934 | 0.050464295 | -1.553116594 | 0.120395282 | 0.246243561 |
| ENSG00000129084 | PSMA1 | 287.9797042 | 0.080959743 | 0.119768251 | 0.675969992 | 0.499059698 | 0.680567986 |
| ENSG00000129103 | SUMF2 | 3451.853928 | 0.085229317 | 0.048393657 | 1.761167107 | 0.078210123 | 0.173564887 |
| ENSG00000129116 | PALLD | 915.8883302 | -0.533906423 | 0.073196425 | -7.29415983 | 3.01E-13 | 3.18E-12 |
| ENSG00000129128 | SPCS3 | 1321.675315 | 0.111187975 | 0.064164549 | 1.732856799 | 0.083121124 | 0.182329183 |
| ENSG00000129158 | SERGEF | 261.1116992 | 0.024351315 | 0.116273674 | 0.209431024 | 0.834111776 | 0.912384329 |
| ENSG00000129159 | KCNC1 | 33.66661579 | -1.336090018 | 0.317504337 | -4.208100056 | 2.58E-05 | 0.000125856 |
| ENSG00000129167 | TPH1 | 4.597607784 | -0.719805544 | 0.857201496 | -0.839715688 | 0.401067816 | 0.593886889 |
| ENSG00000129173 | E2F8 | 680.1651585 | -0.725516148 | 0.081108181 | -8.945042768 | 3.72E-19 | 5.92E-18 |
| ENSG00000129187 | DCTD | 1662.208267 | 1.430346372 | 0.058458518 | 24.46771519 | 3.26E-132 | 9.71E-130 |
| ENSG00000129194 | SOX15 | 4.213587801 | 0.13461921 | 0.859829078 | 0.156565082 | 0.875587624 | 0.93549346 |
| ENSG00000129195 | PIMREG | 420.6323105 | -0.149421329 | 0.096852633 | -1.54276993 | 0.122886608 | 0.249979496 |
| ENSG00000129197 | RPAIN | 835.7462805 | 0.053776364 | 0.072983814 | 0.736825895 | 0.461228234 | 0.648052463 |
| ENSG00000129204 | USP6 | 27.77291982 | 0.2747332 | 0.377896045 | 0.727007346 | 0.467221456 | 0.653190484 |
| ENSG00000129214 | SHBG | 6.2418104 | -0.76543553 | 0.710913146 | -1.076693453 | 0.281617238 | 0.465070869 |
| ENSG00000129219 | PLD2 | 798.622768 | 0.460888605 | 0.07666154 | 6.01199253 | 1.83E-09 | 1.42E-08 |
| ENSG00000129235 | TXNDC17 | 692.8167868 | -0.053213502 | 0.078884669 | -0.674573437 | 0.499946814 | 0.681205745 |
| ENSG00000129244 | ATP1B2 | 2.080304231 | -1.193113467 | 1.262497547 | -0.945042206 | 0.344637329 | 0.535542116 |
| ENSG00000129245 | FXR2 | 1480.340927 | -0.084428717 | 0.063124519 | -1.337494801 | 0.181061178 | 0.336455523 |
| ENSG00000129250 | KIF1C | 7704.933978 | 0.175417653 | 0.050548487 | 3.470284954 | 0.000519906 | 0.002050663 |
| ENSG00000129255 | MPDU1 | 1056.885382 | 0.05275147 | 0.069731116 | 0.756498292 | 0.449350496 | 0.637640751 |
| ENSG00000129292 | PHF20L1 | 1808.966819 | -0.092126455 | 0.062581301 | -1.472108339 | 0.140991624 | 0.27805604 |
| ENSG00000129295 | DNAAF11 | 99.20999138 | 0.535375321 | 0.181406604 | 2.951244935 | 0.003164958 | 0.010672371 |

| | | | | | | | |
|---|---|---|---|---|---|---|---|
| ENSG00000129315 | CCNT1 | 1996.782476 | 0.006684826 | 0.05599811 | 0.119375925 | 0.904977534 | 0.952389422 |
| ENSG00000129317 | PUS7L | 655.1641922 | 0.021097512 | 0.087269035 | 0.241752552 | 0.808971906 | 0.898853827 |
| ENSG00000129347 | KRI1 | 1990.148619 | -0.028586335 | 0.057960265 | -0.493205727 | 0.621867232 | 0.776234318 |
| ENSG00000129351 | ILF3 | 15886.81559 | 0.049173617 | 0.040889764 | 1.202589893 | 0.229135061 | 0.402216177 |
| ENSG00000129353 | SLC44A2 | 11355.83218 | 0.185734578 | 0.04761908 | 3.900423474 | 9.60E-05 | 0.000428978 |
| ENSG00000129354 | AP1M2 | 5490.65922 | -0.092331733 | 0.048466721 | -1.905054245 | 0.05677302 | 0.133289641 |
| ENSG00000129355 | CDKN2D | 149.3780924 | -0.499160274 | 0.149239896 | -3.344683872 | 0.000823764 | 0.003132816 |
| ENSG00000129422 | MTUS1 | 973.5180171 | 0.409672931 | 0.072017001 | 5.6885586 | 1.28E-08 | 9.10E-08 |
| ENSG00000129437 | KLK14 | 8.159546569 | 1.662010926 | 0.668174542 | 2.487390379 | 0.012868411 | 0.037094915 |
| ENSG00000129460 | NGDN | 1078.79436 | -0.457406482 | 0.065560393 | -6.976872127 | 3.02E-12 | 2.97E-11 |
| ENSG00000129467 | ADCY4 | 9.56881642 | -1.266735143 | 0.588645301 | -2.151949811 | 0.031401307 | 0.080809115 |
| ENSG00000129472 | RAB2B | 399.7965884 | -0.291573978 | 0.101303443 | -2.87822377 | 0.003999213 | 0.013173745 |
| ENSG00000129473 | BCL2L2 | 1000.237597 | 0.043160403 | 0.071117883 | 0.606885377 | 0.543926983 | 0.717456866 |
| ENSG00000129474 | AJUBA | 1675.768503 | -0.693131753 | 0.058164503 | -11.91674847 | 9.68E-33 | 2.99E-31 |
| ENSG00000129480 | DTD2 | 503.9794039 | -0.526584837 | 0.092166085 | -5.713433922 | 1.11E-08 | 7.92E-08 |
| ENSG00000129484 | PARP2 | 908.0606017 | -0.63184686 | 0.081548751 | -7.748087496 | 9.33E-15 | 1.10E-13 |
| ENSG00000129493 | HEATR5A | 616.3837071 | -0.38138323 | 0.082743848 | -4.609203477 | 4.04E-06 | 2.18E-05 |
| ENSG00000129514 | FOXA1 | 7711.126015 | -0.403918282 | 0.047279836 | -8.543140509 | 1.31E-17 | 1.87E-16 |
| ENSG00000129515 | SNX6 | 2560.511873 | -0.327850823 | 0.055057974 | -5.954647448 | 2.61E-09 | 2.00E-08 |
| ENSG00000129518 | EAPP | 1780.836471 | -0.521931272 | 0.055502818 | -9.403689561 | 5.27E-21 | 9.34E-20 |
| ENSG00000129521 | EGLN3 | 3694.268399 | -1.070778509 | 0.066165019 | -16.1834534 | 6.60E-59 | 4.56E-57 |
| ENSG00000129534 | MIS18BP1 | 1859.248312 | -0.476419615 | 0.068529809 | -6.952005614 | 3.60E-12 | 3.52E-11 |
| ENSG00000129535 | NRL | 69.71098573 | -0.041035568 | 0.21207378 | -0.193496658 | 0.846570023 | 0.919696738 |
| ENSG00000129559 | NEDD8 | 995.6378996 | -0.580897947 | 0.070443627 | -8.24628107 | 1.63E-16 | 2.20E-15 |
| ENSG00000129562 | DAD1 | 1770.828492 | -0.54186349 | 0.057866457 | -9.364034299 | 7.67E-21 | 1.35E-19 |
| ENSG00000129566 | TEP1 | 999.8534695 | -0.884774688 | 0.066877583 | -13.22976464 | 5.91E-40 | 2.41E-38 |
| ENSG00000129595 | EPB41L4A | 1490.397032 | -0.010086599 | 0.061790392 | -0.163238963 | 0.87033029 | 0.932790446 |
| ENSG00000129596 | CDO1 | 4.051397199 | 1.174589878 | 0.927047295 | 1.267022603 | 0.205147208 | 0.370236563 |
| ENSG00000129625 | REEP5 | 9175.138093 | 0.762216798 | 0.041306822 | 18.45256453 | 4.97E-76 | 5.38E-74 |
| ENSG00000129636 | ITFG1 | 831.887442 | 0.10910845 | 0.072827835 | 1.498169629 | 0.134089184 | 0.267600657 |
| ENSG00000129646 | QRICH2 | 147.1247061 | 0.815930275 | 0.156625468 | 5.209435516 | 1.89E-07 | 1.20E-06 |
| ENSG00000129654 | FOXJ1 | 2.139403466 | 1.031969597 | 1.300627951 | 0.793439505 | 0.427521802 | 0.618217808 |
| ENSG00000129657 | SEC14L1 | 4142.77977 | 0.387051966 | 0.054590525 | 7.09009424 | 1.34E-12 | 1.36E-11 |

| | | | | | | | |
|---|---|---|---|---|---|---|---|
| ENSG00000129667 | RHBDF2 | 690.7993998 | 0.601215686 | 0.080192735 | 7.49713408 | 6.52E-14 | 7.21E-13 |
| ENSG00000129673 | AANAT | 4.633851996 | 3.005080233 | 1.075669328 | 2.793684038 | 0.005211137 | 0.016689024 |
| ENSG00000129675 | ARHGEF6 | 243.8434841 | 0.927762436 | 0.127380231 | 7.283409911 | 3.25E-13 | 3.43E-12 |
| ENSG00000129680 | MAP7D3 | 979.1428156 | -0.084236273 | 0.074264038 | -1.134280809 | 0.256676777 | 0.436202619 |
| ENSG00000129682 | FGF13 | 254.958593 | 0.306763589 | 0.115787159 | 2.649374873 | 0.008064083 | 0.024614712 |
| ENSG00000129691 | ASH2L | 1410.643338 | -0.011849317 | 0.062321728 | -0.1901314 | 0.849206165 | 0.921106747 |
| ENSG00000129696 | TTI2 | 421.9054757 | -0.353749838 | 0.092893336 | -3.80812932 | 0.000140022 | 0.000609739 |
| ENSG00000129749 | CHRNA10 | 20.7446642 | 0.035860704 | 0.392539053 | 0.091355762 | 0.927209911 | 0.965137362 |
| ENSG00000129757 | CDKN1C | 8.424334679 | -0.645082359 | 0.631146103 | -1.022080872 | 0.306742626 | 0.494495842 |
| ENSG00000129810 | SGO1 | 617.4386834 | -0.260195144 | 0.084517009 | -3.078612776 | 0.002079668 | 0.007282471 |
| ENSG00000129910 | CDH15 | 7.056435414 | 0.33809149 | 0.695854472 | 0.485865226 | 0.627062725 | 0.780131681 |
| ENSG00000129911 | KLF16 | 721.4707766 | 0.164161395 | 0.08460137 | 1.94041059 | 0.052329811 | 0.124553419 |
| ENSG00000129925 | PGAP6 | 1789.419877 | 0.336864994 | 0.063954843 | 5.267231993 | 1.38E-07 | 8.87E-07 |
| ENSG00000129932 | DOHH | 255.7316669 | -0.13715234 | 0.116533132 | -1.176938582 | 0.239220021 | 0.414824045 |
| ENSG00000129933 | MAU2 | 959.7165936 | 0.232065566 | 0.071430585 | 3.248826365 | 0.001158822 | 0.004291794 |
| ENSG00000129946 | SHC2 | 226.0731285 | -1.363026033 | 0.126279439 | -10.79372888 | 3.69E-27 | 9.20E-26 |
| ENSG00000129951 | PLPPR3 | 45.17705821 | 0.884208854 | 0.282498028 | 3.129964694 | 0.001748273 | 0.006218853 |
| ENSG00000129968 | ABHD17A | 1014.748888 | 0.054768043 | 0.076384791 | 0.717001928 | 0.473372907 | 0.658890509 |
| ENSG00000129988 | LBP | 2.709127967 | -0.224856141 | 1.108013803 | -0.202936228 | 0.839184878 | 0.915309377 |
| ENSG00000129990 | SYT5 | 2.363988626 | -3.705440878 | 1.51655943 | -2.443320588 | 0.014552803 | 0.041368192 |
| ENSG00000129993 | CBFA2T3 | 1827.522063 | -0.04222539 | 0.058731096 | -0.718961385 | 0.472164712 | 0.657847464 |
| ENSG00000130005 | GAMT | 419.2944777 | 0.385576432 | 0.099237758 | 3.885380315 | 0.00010217 | 0.000454837 |
| ENSG00000130021 | PUDP | 401.3439801 | -0.584489624 | 0.097748354 | -5.979534189 | 2.24E-09 | 1.72E-08 |
| ENSG00000130023 | ERMARD | 869.2244416 | 0.003079437 | 0.081902935 | 0.037598617 | 0.970007711 | 0.986563569 |
| ENSG00000130024 | PHF10 | 2554.889941 | 0.294461818 | 0.055268595 | 5.327832554 | 9.94E-08 | 6.44E-07 |
| ENSG00000130038 | CRACR2A | 253.5598263 | 0.195658073 | 0.124944016 | 1.565965937 | 0.117356592 | 0.2413844 |
| ENSG00000130045 | NXNL2 | 35.88653345 | 1.350989106 | 0.31125565 | 4.340448462 | 1.42E-05 | 7.18E-05 |
| ENSG00000130052 | STARD8 | 65.56721756 | 0.143246997 | 0.217177995 | 0.659583385 | 0.509521219 | 0.689541429 |
| ENSG00000130054 | NALF2 | 594.8225464 | 0.20238282 | 0.087092318 | 2.323773486 | 0.020137638 | 0.055014577 |
| ENSG00000130055 | GDPD2 | 4.00179863 | 0.790303349 | 0.911809808 | 0.866741443 | 0.386083693 | 0.578684084 |
| ENSG00000130066 | SAT1 | 1176.029335 | -0.573987978 | 0.06605585 | -8.6894345 | 3.64E-18 | 5.45E-17 |
| ENSG00000130119 | GNL3L | 3071.995321 | 0.024592198 | 0.050638954 | 0.485637968 | 0.627223872 | 0.780154165 |
| ENSG00000130147 | SH3BP4 | 2406.747827 | 0.245927723 | 0.065778928 | 3.738700662 | 0.000184974 | 0.000789892 |

| | | | | | | | |
|---|---|---|---|---|---|---|---|
| ENSG00000130150 | MOSPD2 | 545.1139307 | 0.286036327 | 0.087153186 | 3.281995091 | 0.001030754 | 0.003848848 |
| ENSG00000130158 | DOCK6 | 1670.698175 | 0.652781301 | 0.068621518 | 9.512778548 | 1.86E-21 | 3.38E-20 |
| ENSG00000130159 | ECSIT | 553.6486789 | 0.581028308 | 0.084866782 | 6.846357255 | 7.58E-12 | 7.18E-11 |
| ENSG00000130164 | LDLR | 8043.448177 | -0.334509225 | 0.046968908 | -7.121928962 | 1.06E-12 | 1.09E-11 |
| ENSG00000130165 | ELOF1 | 1321.876376 | 0.028195365 | 0.067933485 | 0.415043704 | 0.67810991 | 0.81682581 |
| ENSG00000130173 | ANGPTL8 | 1.690740998 | 0.357904194 | 1.388516323 | 0.257760163 | 0.796592011 | NA |
| ENSG00000130175 | PRKCSH | 8607.629591 | 0.145447681 | 0.049755732 | 2.923234684 | 0.003464153 | 0.011580832 |
| ENSG00000130177 | CDC16 | 1474.649735 | -0.075861614 | 0.060474107 | -1.254447861 | 0.209679266 | 0.376711163 |
| ENSG00000130193 | THEM6 | 3728.849376 | -0.08154021 | 0.065921244 | -1.236933715 | 0.216111692 | 0.385420717 |
| ENSG00000130202 | NECTIN2 | 6557.853377 | -0.089652215 | 0.061111045 | -1.467037846 | 0.14236576 | 0.280091813 |
| ENSG00000130203 | APOE | 233.9154195 | -1.156135065 | 0.12826046 | -9.013963216 | 1.99E-19 | 3.22E-18 |
| ENSG00000130204 | TOMM40 | 3585.08063 | 0.128593452 | 0.055400718 | 2.321151353 | 0.020278677 | 0.055295052 |
| ENSG00000130208 | APOC1 | 8.963212705 | -1.626609716 | 0.62873037 | -2.587133998 | 0.009677795 | 0.028899846 |
| ENSG00000130222 | GADD45G | 138.4774392 | -0.663192225 | 0.152677971 | -4.34373225 | 1.40E-05 | 7.08E-05 |
| ENSG00000130227 | XPO7 | 3492.03878 | 0.133845421 | 0.047289314 | 2.830352347 | 0.004649677 | 0.015099601 |
| ENSG00000130244 | FAM98C | 544.4677429 | -0.163922603 | 0.083457538 | -1.964143763 | 0.049513422 | 0.118811497 |
| ENSG00000130254 | SAFB2 | 2019.976771 | -0.07938996 | 0.053109128 | -1.49484585 | 0.134954677 | 0.2690181 |
| ENSG00000130255 | RPL36 | 6059.785191 | -0.123214513 | 0.047285524 | -2.60575545 | 0.009167189 | 0.027576096 |
| ENSG00000130270 | ATP8B3 | 56.43360342 | 0.155748585 | 0.234905078 | 0.663027748 | 0.507312783 | 0.687526156 |
| ENSG00000130294 | KIF1A | 1398.718538 | -0.331120538 | 0.071905315 | -4.604952179 | 4.13E-06 | 2.22E-05 |
| ENSG00000130299 | GTPBP3 | 528.6337946 | -0.053377418 | 0.091324419 | -0.58448133 | 0.558896523 | 0.728758429 |
| ENSG00000130303 | BST2 | 9.236029913 | 0.831589757 | 0.598643562 | 1.38912336 | 0.164795241 | 0.313102769 |
| ENSG00000130304 | SLC27A1 | 255.2859357 | 0.464485736 | 0.116028998 | 4.003186666 | 6.25E-05 | 0.000288168 |
| ENSG00000130305 | NSUN5 | 761.325298 | 0.200653076 | 0.073286078 | 2.737942631 | 0.006182486 | 0.019471062 |
| ENSG00000130309 | COLGALT1 | 1727.558772 | 0.289071822 | 0.056754088 | 5.093409672 | 3.52E-07 | 2.15E-06 |
| ENSG00000130311 | DDA1 | 981.1485783 | 0.266811897 | 0.07207719 | 3.701752201 | 0.000214116 | 0.000902907 |
| ENSG00000130312 | MRPL34 | 623.5418322 | -0.027962858 | 0.081526648 | -0.342990398 | 0.731605684 | 0.851381347 |
| ENSG00000130313 | PGLS | 1134.85111 | 0.010368377 | 0.075521376 | 0.137290622 | 0.890801083 | 0.944056489 |
| ENSG00000130332 | LSM7 | 821.5314012 | 0.133152474 | 0.072940582 | 1.825492346 | 0.067926765 | 0.154802304 |
| ENSG00000130338 | TULP4 | 1779.778995 | -1.048190752 | 0.057980168 | -18.07843583 | 4.71E-73 | 4.68E-71 |
| ENSG00000130340 | SNX9 | 1113.144629 | -0.120490094 | 0.063566737 | -1.895489674 | 0.058027558 | 0.135748745 |
| ENSG00000130347 | RTN4IP1 | 145.8170338 | -0.071793003 | 0.151605525 | -0.47355136 | 0.635819868 | 0.785317815 |
| ENSG00000130348 | QRSL1 | 624.0828204 | -0.010282587 | 0.082055369 | -0.125312788 | 0.900275929 | 0.949707063 |

| | | | | | | | |
|---|---|---|---|---|---|---|---|
| ENSG00000130349 | MTRES1 | 338.0110233 | 0.023811024 | 0.104313863 | 0.228263278 | 0.819441571 | 0.904358204 |
| ENSG00000130363 | RSPH3 | 372.3328647 | 0.145308991 | 0.106403692 | 1.365638615 | 0.17205242 | 0.323833485 |
| ENSG00000130377 | ACSBG2 | 6.44303779 | -0.292590407 | 0.695701128 | -0.420569114 | 0.674069751 | 0.813636416 |
| ENSG00000130382 | MLLT1 | 2934.139746 | 0.097080928 | 0.056767516 | 1.710149289 | 0.08723827 | 0.189601698 |
| ENSG00000130396 | AFDN | 2996.381726 | -0.273929098 | 0.050865112 | -5.385402449 | 7.23E-08 | 4.74E-07 |
| ENSG00000130402 | ACTN4 | 13097.76934 | -0.013259324 | 0.052748087 | -0.251370711 | 0.80152751 | 0.894398618 |
| ENSG00000130414 | NDUFA10 | 3090.517704 | -0.001318793 | 0.051089595 | -0.025813331 | 0.979406229 | 0.990765043 |
| ENSG00000130427 | EPO | 1.99091024 | -0.683812348 | 1.230130179 | -0.555886165 | 0.578288676 | 0.743940233 |
| ENSG00000130429 | ARPC1B | 30.29329781 | 0.007648643 | 0.321533043 | 0.023788048 | 0.981021674 | 0.99123506 |
| ENSG00000130433 | CACNG6 | 47.26586265 | -5.225907797 | 0.562338295 | -9.293174306 | 1.50E-20 | 2.58E-19 |
| ENSG00000130449 | ZSWIM6 | 555.8047081 | 0.391559081 | 0.086901309 | 4.505790365 | 6.61E-06 | 3.49E-05 |
| ENSG00000130475 | FCHO1 | 266.6364 | 0.310329908 | 0.115001852 | 2.698477484 | 0.006965745 | 0.021602535 |
| ENSG00000130477 | UNC13A | 10.7497549 | -1.422634012 | 0.556967931 | -2.554247621 | 0.010641755 | 0.031382164 |
| ENSG00000130479 | MAP1S | 574.1486151 | 0.179697555 | 0.084748327 | 2.120366983 | 0.033975109 | 0.086358892 |
| ENSG00000130487 | KLHDC7B | 39.54053601 | 0.374765183 | 0.292137585 | 1.282837959 | 0.199548852 | 0.362528762 |
| ENSG00000130508 | PXDN | 3729.07413 | 0.777158584 | 0.055879919 | 13.90765415 | 5.69E-44 | 2.65E-42 |
| ENSG00000130511 | SSBP4 | 856.200274 | 0.015401727 | 0.070977042 | 0.216995899 | 0.828211541 | 0.909138327 |
| ENSG00000130513 | GDF15 | 169.1102409 | 1.130486857 | 0.147204824 | 7.679686219 | 1.59E-14 | 1.84E-13 |
| ENSG00000130517 | PGPEP1 | 1162.881158 | 0.241616169 | 0.066496262 | 3.633530107 | 0.00027957 | 0.001156698 |
| ENSG00000130518 | IQCN | 635.154731 | -0.22739217 | 0.084057893 | -2.705185206 | 0.006826632 | 0.021217034 |
| ENSG00000130520 | LSM4 | 2101.986171 | -0.342167141 | 0.059613196 | -5.739788537 | 9.48E-09 | 6.84E-08 |
| ENSG00000130522 | JUND | 1348.871893 | 0.068033308 | 0.071784551 | 0.947743024 | 0.343260292 | 0.534255452 |
| ENSG00000130528 | HRC | 6.234577365 | -0.788846121 | 0.709170929 | -1.112349772 | 0.265987792 | 0.446949893 |
| ENSG00000130529 | TRPM4 | 1625.017451 | -0.044771992 | 0.059389183 | -0.753874522 | 0.450924571 | 0.639083755 |
| ENSG00000130544 | ZNF557 | 400.9967283 | -0.082890344 | 0.100878186 | -0.821687492 | 0.41125478 | 0.603671145 |
| ENSG00000130545 | CRB3 | 442.2652047 | 0.050862771 | 0.101724868 | 0.500003307 | 0.617072749 | 0.773024159 |
| ENSG00000130558 | OLFM1 | 859.7840263 | 0.12718953 | 0.072037604 | 1.765599128 | 0.077463126 | 0.172323906 |
| ENSG00000130559 | CAMSAP1 | 2718.781388 | 0.167342774 | 0.052343839 | 3.196990861 | 0.001388693 | 0.005053398 |
| ENSG00000130560 | UBAC1 | 1521.580289 | 0.224941007 | 0.059493057 | 3.780962319 | 0.000156223 | 0.000675163 |
| ENSG00000130584 | ZBTB46 | 278.18232 | -0.226678131 | 0.117202956 | -1.93406496 | 0.053105141 | 0.126241652 |
| ENSG00000130589 | HELZ2 | 1240.680189 | 0.257635581 | 0.069498413 | 3.707071414 | 0.00020967 | 0.000885787 |
| ENSG00000130590 | SAMD10 | 648.7278091 | 0.369502464 | 0.084945828 | 4.349860043 | 1.36E-05 | 6.90E-05 |
| ENSG00000130600 | H19 | 60.77933474 | -2.659088335 | 0.281794981 | -9.436251582 | 3.86E-21 | 6.91E-20 |

| | | | | | | | |
|---|---|---|---|---|---|---|---|
| ENSG00000130612 | CYP2G1P | 1.564023183 | 2.866626186 | 1.637985246 | 1.7500928 | 0.080102302 | NA |
| ENSG00000130635 | COL5A1 | 846.028439 | -2.221967493 | 0.087526019 | -25.38636532 | 3.57E-142 | 1.20E-139 |
| ENSG00000130638 | ATXN10 | 2572.803451 | 0.0630837 | 0.051308727 | 1.2294926 | 0.218887171 | 0.388613241 |
| ENSG00000130640 | TUBGCP2 | 2590.937066 | -0.55933785 | 0.054071094 | -10.34448928 | 4.43E-25 | 9.87E-24 |
| ENSG00000130649 | CYP2E1 | 12.4183044 | -0.450964593 | 0.489907343 | -0.920509968 | 0.357306327 | 0.549591467 |
| ENSG00000130653 | PNPLA7 | 414.2650879 | -0.350618624 | 0.102265865 | -3.428501036 | 0.000606924 | 0.002366221 |
| ENSG00000130656 | HBZ | 1.591464773 | 2.854039207 | 1.638389885 | 1.741978044 | 0.081512286 | NA |
| ENSG00000130669 | PAK4 | 3576.017491 | -0.102433587 | 0.060959092 | -1.680366018 | 0.092886124 | 0.199551196 |
| ENSG00000130675 | MNX1 | 146.1532927 | 0.605713767 | 0.151532072 | 3.99726447 | 6.41E-05 | 0.000294819 |
| ENSG00000130684 | ZNF337 | 804.6800903 | -0.584798486 | 0.072974097 | -8.013781793 | 1.11E-15 | 1.41E-14 |
| ENSG00000130695 | CEP85 | 810.3332286 | 0.025872967 | 0.073760649 | 0.350769245 | 0.725761472 | 0.847784533 |
| ENSG00000130699 | TAF4 | 1175.651587 | -0.362473017 | 0.063593196 | -5.699871034 | 1.20E-08 | 8.54E-08 |
| ENSG00000130701 | RBBP8NL | 613.7839844 | 0.340959484 | 0.085141654 | 4.004614269 | 6.21E-05 | 0.000286606 |
| ENSG00000130702 | LAMA5 | 7175.340231 | -0.047975306 | 0.051090134 | -0.939032701 | 0.347713955 | 0.538629086 |
| ENSG00000130703 | OSBPL2 | 2880.020791 | -0.018944393 | 0.053969687 | -0.351019137 | 0.725573992 | 0.847660795 |
| ENSG00000130706 | ADRM1 | 4493.38732 | -0.238831601 | 0.053104075 | -4.497425142 | 6.88E-06 | 3.62E-05 |
| ENSG00000130707 | ASS1 | 5030.851508 | 0.930948067 | 0.061761709 | 15.07322384 | 2.43E-51 | 1.39E-49 |
| ENSG00000130711 | PRDM12 | 15.88858827 | -0.148341039 | 0.441420598 | -0.336053731 | 0.736830349 | 0.854938089 |
| ENSG00000130713 | EXOSC2 | 1290.193976 | -0.018190999 | 0.063592994 | -0.286053514 | 0.774837135 | 0.877499635 |
| ENSG00000130714 | POMT1 | 1993.129043 | -0.111633571 | 0.0566062 | -1.972108564 | 0.048597214 | 0.116930994 |
| ENSG00000130717 | UCK1 | 1861.00384 | -0.096195553 | 0.070802662 | -1.358643172 | 0.174259688 | 0.327102302 |
| ENSG00000130720 | FIBCD1 | 377.6688175 | 0.070902291 | 0.102277084 | 0.69323731 | 0.488160637 | 0.671528246 |
| ENSG00000130724 | CHMP2A | 1593.788565 | -0.170388575 | 0.058554332 | -2.909922614 | 0.003615183 | 0.012031378 |
| ENSG00000130725 | UBE2M | 1794.041934 | -0.122164008 | 0.055837866 | -2.187834484 | 0.028681661 | 0.074683008 |
| ENSG00000130726 | TRIM28 | 16296.89224 | -0.06402735 | 0.04246567 | -1.507743783 | 0.131620115 | 0.263930546 |
| ENSG00000130731 | METTL26 | 1343.276369 | 0.085276979 | 0.063631834 | 1.340162217 | 0.180192612 | 0.335194496 |
| ENSG00000130733 | YIPF2 | 1821.861659 | 0.175276479 | 0.065484724 | 2.676601005 | 0.007437314 | 0.022904912 |
| ENSG00000130734 | ATG4D | 1089.581508 | 0.019598719 | 0.068031754 | 0.288081936 | 0.773284019 | 0.876416754 |
| ENSG00000130741 | EIF2S3 | 5765.215395 | 0.223010664 | 0.053571983 | 4.162822667 | 3.14E-05 | 0.000151968 |
| ENSG00000130748 | TMEM160 | 145.2323182 | 0.84615879 | 0.156353858 | 5.411819074 | 6.24E-08 | 4.12E-07 |
| ENSG00000130749 | ZC3H4 | 2880.745689 | 0.387943755 | 0.057300572 | 6.770329564 | 1.28E-11 | 1.19E-10 |
| ENSG00000130751 | NPAS1 | 37.92666064 | 0.473301301 | 0.288081024 | 1.64294508 | 0.100394297 | 0.212634583 |
| ENSG00000130755 | GMFG | 2.731771343 | 0.321933938 | 1.073367686 | 0.299928852 | 0.764231426 | 0.870644539 |

| | | | | | | | |
|---|---|---|---|---|---|---|---|
| ENSG00000130758 | MAP3K10 | 335.2929422 | -0.058070915 | 0.107821563 | -0.538583504 | 0.590174272 | 0.753022849 |
| ENSG00000130762 | ARHGEF16 | 1287.474177 | -0.057369232 | 0.064179687 | -0.893884576 | 0.371383651 | 0.564017991 |
| ENSG00000130764 | LRRC47 | 1579.776296 | 0.123166537 | 0.059515537 | 2.069485426 | 0.038500558 | 0.096094309 |
| ENSG00000130766 | SESN2 | 1551.451071 | -0.106965008 | 0.058941371 | -1.814769594 | 0.069559325 | 0.157770508 |
| ENSG00000130768 | SMPDL3B | 230.6291337 | 0.683682251 | 0.122032631 | 5.602454419 | 2.11E-08 | 1.47E-07 |
| ENSG00000130770 | ATP5IF1 | 2795.051595 | -0.027096245 | 0.049848797 | -0.543568677 | 0.586738319 | 0.7504336 |
| ENSG00000130772 | MED18 | 541.2780127 | -0.225661916 | 0.087648249 | -2.574631183 | 0.010034703 | 0.029818145 |
| ENSG00000130775 | THEMIS2 | 3.007966337 | 2.197728922 | 1.176993035 | 1.86724038 | 0.061868027 | 0.143069812 |
| ENSG00000130779 | CLIP1 | 2413.442662 | 0.044639629 | 0.055507989 | 0.80420188 | 0.421280393 | 0.61231529 |
| ENSG00000130783 | CCDC62 | 20.83510847 | -0.573714834 | 0.382917695 | -1.498271929 | 0.134062614 | 0.267588651 |
| ENSG00000130787 | HIP1R | 3422.590756 | 0.063761341 | 0.051640746 | 1.234709914 | 0.216938485 | 0.386409569 |
| ENSG00000130803 | ZNF317 | 1217.396894 | 0.005094806 | 0.067408625 | 0.075580925 | 0.939752513 | 0.971233034 |
| ENSG00000130810 | PPAN | 228.3838614 | 0.180519392 | 0.130349994 | 1.384882237 | 0.166088479 | 0.315116179 |
| ENSG00000130811 | EIF3G | 4273.503689 | 0.18884388 | 0.049181187 | 3.839758482 | 0.000123155 | 0.00054112 |
| ENSG00000130812 | ANGPTL6 | 12.86451355 | 0.128817778 | 0.495619616 | 0.259912589 | 0.7949312 | 0.890293445 |
| ENSG00000130813 | SHFL | 343.6749357 | 0.966224082 | 0.107647742 | 8.975795164 | 2.81E-19 | 4.52E-18 |
| ENSG00000130816 | DNMT1 | 5997.087006 | -0.224670852 | 0.043316661 | -5.186707582 | 2.14E-07 | 1.35E-06 |
| ENSG00000130818 | ZNF426 | 20.81965177 | 0.983149175 | 0.406841593 | 2.41654047 | 0.015668784 | 0.044190056 |
| ENSG00000130821 | SLC6A8 | 817.8570371 | 0.282815579 | 0.071494285 | 3.955778851 | 7.63E-05 | 0.000347359 |
| ENSG00000130822 | PNCK | 4.760400369 | 0.239723115 | 0.855437742 | 0.280234438 | 0.779297647 | 0.88025447 |
| ENSG00000130826 | DKC1 | 5521.664155 | -0.451148534 | 0.049388999 | -9.134595607 | 6.57E-20 | 1.09E-18 |
| ENSG00000130827 | PLXNA3 | 8318.052141 | -0.369847415 | 0.052347894 | -7.065182369 | 1.60E-12 | 1.61E-11 |
| ENSG00000130829 | DUSP9 | 14.74674119 | -2.547548942 | 0.554752018 | -4.592230147 | 4.39E-06 | 2.35E-05 |
| ENSG00000130844 | ZNF331 | 1066.591583 | 0.113749431 | 0.065194421 | 1.744772454 | 0.081024486 | 0.178515338 |
| ENSG00000130856 | ZNF236 | 678.2794146 | 0.378835146 | 0.07798191 | 4.857987549 | 1.19E-06 | 6.80E-06 |
| ENSG00000130881 | LRP3 | 1232.285238 | 0.203407975 | 0.075726098 | 2.686101382 | 0.007229115 | 0.022357152 |
| ENSG00000130921 | MTRFR | 450.8293824 | -0.308938928 | 0.092210871 | -3.350352561 | 0.000807088 | 0.003073471 |
| ENSG00000130935 | NOL11 | 3829.961139 | 0.107326441 | 0.052839454 | 2.031179986 | 0.04223674 | 0.103771106 |
| ENSG00000130939 | UBE4B | 2757.09131 | 0.20993777 | 0.057017798 | 3.681969113 | 0.000231439 | 0.000970785 |
| ENSG00000130940 | CASZ1 | 1282.450254 | -0.026615815 | 0.063527218 | -0.418967113 | 0.675240168 | 0.814492031 |
| ENSG00000130950 | NUTM2F | 4.247138145 | -0.308025436 | 0.857522411 | -0.359203949 | 0.719442523 | 0.844065436 |
| ENSG00000130956 | HABP4 | 406.3981701 | -0.023577753 | 0.095677421 | -0.246429646 | 0.80534966 | 0.896657018 |
| ENSG00000130957 | FBP2 | 9.901360385 | 0.124911414 | 0.585975172 | 0.213168441 | 0.831195583 | 0.910670869 |

| | | | | | | | |
|---|---|---|---|---|---|---|---|
| ENSG00000130958 | SLC35D2 | 92.93697899 | 0.098655063 | 0.185508332 | 0.531809335 | 0.594858056 | 0.756439984 |
| ENSG00000130962 | PRRG1 | 267.8703993 | 0.033289753 | 0.113896181 | 0.292281554 | 0.770071361 | 0.874656485 |
| ENSG00000130985 | UBA1 | 14499.84954 | -0.226449553 | 0.044088756 | -5.136220054 | 2.80E-07 | 1.74E-06 |
| ENSG00000130997 | POLN | 11.69359525 | 1.145639237 | 0.535924331 | 2.137688419 | 0.03254204 | 0.083330731 |
| ENSG00000131013 | PPIL4 | 1201.389543 | -0.11587654 | 0.065498409 | -1.769150446 | 0.076868771 | 0.171350657 |
| ENSG00000131015 | ULBP2 | 89.4746078 | 0.391808847 | 0.191394836 | 2.047123398 | 0.04064597 | 0.100497797 |
| ENSG00000131016 | AKAP12 | 336.783747 | 1.337770776 | 0.1205032 | 11.10153739 | 1.23E-28 | 3.24E-27 |
| ENSG00000131018 | SYNE1 | 444.0568352 | 1.312694586 | 0.106674949 | 12.30555626 | 8.46E-35 | 2.86E-33 |
| ENSG00000131019 | ULBP3 | 122.6905921 | 0.323365917 | 0.162810967 | 1.986143335 | 0.047017411 | 0.113618393 |
| ENSG00000131023 | LATS1 | 1667.640936 | 0.250751832 | 0.062847635 | 3.989837192 | 6.61E-05 | 0.000303717 |
| ENSG00000131037 | EPS8L1 | 740.6417525 | -0.329873883 | 0.082013722 | -4.022179138 | 5.77E-05 | 0.000267442 |
| ENSG00000131043 | AAR2 | 1398.609879 | -0.697715491 | 0.065932365 | -10.58229123 | 3.60E-26 | 8.45E-25 |
| ENSG00000131044 | TTLL9 | 34.95265228 | 2.211086624 | 0.355308396 | 6.223006977 | 4.88E-10 | 3.99E-09 |
| ENSG00000131051 | RBM39 | 6688.555114 | -0.367992996 | 0.065114677 | -5.651460085 | 1.59E-08 | 1.12E-07 |
| ENSG00000131061 | ZNF341 | 271.7167481 | -0.049266888 | 0.11144892 | -0.442058006 | 0.658447232 | 0.802336104 |
| ENSG00000131067 | GGT7 | 758.7888277 | -0.999217916 | 0.076717992 | -13.02455768 | 8.87E-39 | 3.42E-37 |
| ENSG00000131069 | ACSS2 | 1692.747966 | -1.087845232 | 0.060568959 | -17.96044115 | 3.98E-72 | 3.83E-70 |
| ENSG00000131080 | EDA2R | 216.8906134 | 0.034575733 | 0.124728463 | 0.277208041 | 0.781620366 | 0.881489169 |
| ENSG00000131089 | ARHGEF9 | 1021.828779 | -0.106370373 | 0.066657755 | -1.595768906 | 0.110540398 | 0.229670929 |
| ENSG00000131094 | C1QL1 | 24.37462539 | 0.100961042 | 0.355547113 | 0.283959673 | 0.776441285 | 0.878628802 |
| ENSG00000131095 | GFAP | 2.762499692 | -0.069141925 | 1.160740301 | -0.059567093 | 0.952500428 | 0.977951028 |
| ENSG00000131096 | PYY | 4.212282889 | 1.750499639 | 1.024764067 | 1.708197715 | 0.087599662 | 0.190235581 |
| ENSG00000131100 | ATP6V1E1 | 2539.843757 | -0.477226674 | 0.05205788 | -9.167232113 | 4.85E-20 | 8.11E-19 |
| ENSG00000131115 | ZNF227 | 700.6326142 | 0.296529287 | 0.076027611 | 3.900284141 | 9.61E-05 | 0.000429142 |
| ENSG00000131116 | ZNF428 | 658.5523964 | 0.118940755 | 0.079892559 | 1.488758857 | 0.13655088 | 0.271374616 |
| ENSG00000131127 | ZNF141 | 576.151521 | 0.228694707 | 0.096714033 | 2.364648645 | 0.018047183 | 0.049976145 |
| ENSG00000131142 | CCL25 | 1.905359626 | 1.443528994 | 1.348732586 | 1.070285547 | 0.284490798 | 0.468265689 |
| ENSG00000131143 | COX4I1 | 7666.413827 | 0.528724136 | 0.043370003 | 12.19100985 | 3.47E-34 | 1.14E-32 |
| ENSG00000131148 | EMC8 | 1340.345078 | 0.224823237 | 0.061450569 | 3.658603014 | 0.000253594 | 0.00105484 |
| ENSG00000131149 | GSE1 | 10289.04135 | 0.200101689 | 0.042091218 | 4.754000944 | 1.99E-06 | 1.11E-05 |
| ENSG00000131153 | GINS2 | 1902.903565 | 0.393767868 | 0.057733573 | 6.820431324 | 9.08E-12 | 8.54E-11 |
| ENSG00000131165 | CHMP1A | 5088.784312 | 0.161004976 | 0.05573635 | 2.888688915 | 0.003868516 | 0.01279097 |
| ENSG00000131171 | SH3BGRL | 5344.613108 | 0.705383582 | 0.054510818 | 12.94024942 | 2.67E-38 | 1.02E-36 |

| | | | | | | | |
|---|---|---|---|---|---|---|---|
| ENSG00000131174 | COX7B | 1865.119879 | 0.183019371 | 0.053812189 | 3.4010765 | 0.00067121 | 0.002596554 |
| ENSG00000131187 | F12 | 459.2710641 | -0.350941724 | 0.104066618 | -3.372279513 | 0.000745487 | 0.002858824 |
| ENSG00000131188 | PRR7 | 326.5206479 | -0.007040885 | 0.120863821 | -0.058254693 | 0.953545756 | 0.978498754 |
| ENSG00000131196 | NFATC1 | 516.2879458 | 0.865256932 | 0.087263635 | 9.915435364 | 3.57E-23 | 7.15E-22 |
| ENSG00000131236 | CAP1 | 7108.125414 | -0.275965055 | 0.04371676 | -6.312568857 | 2.74E-10 | 2.30E-09 |
| ENSG00000131238 | PPT1 | 7648.829784 | -0.982876722 | 0.046959611 | -20.9302571 | 2.84E-97 | 4.31E-95 |
| ENSG00000131242 | RAB11FIP4 | 1846.014996 | 0.938186781 | 0.060924437 | 15.39918664 | 1.66E-53 | 9.98E-52 |
| ENSG00000131263 | RLIM | 2309.657798 | 0.081569539 | 0.061523212 | 1.325833557 | 0.184894835 | 0.341752453 |
| ENSG00000131269 | ABCB7 | 747.4860129 | -0.013614548 | 0.075594286 | -0.180100224 | 0.857073887 | 0.925991782 |
| ENSG00000131323 | TRAF3 | 1525.512942 | 0.006285389 | 0.063647015 | 0.098753864 | 0.921333701 | 0.9619656 |
| ENSG00000131351 | HAUS8 | 306.7762089 | -0.217789044 | 0.109044022 | -1.997257994 | 0.045797164 | 0.11110268 |
| ENSG00000131368 | MRPS25 | 2759.772315 | 0.220929776 | 0.064372389 | 3.43205802 | 0.000599019 | 0.002336991 |
| ENSG00000131370 | SH3BP5 | 3855.67994 | -0.827205448 | 0.047997475 | -17.23435351 | 1.47E-66 | 1.21E-64 |
| ENSG00000131373 | HACL1 | 701.6422491 | 0.181319982 | 0.07952052 | 2.280165932 | 0.022597849 | 0.060830164 |
| ENSG00000131374 | TBC1D5 | 5587.830454 | -0.000710959 | 0.048777089 | -0.014575668 | 0.988370711 | 0.994043197 |
| ENSG00000131375 | CAPN7 | 2757.161501 | 0.322945395 | 0.052697086 | 6.128334967 | 8.88E-10 | 7.09E-09 |
| ENSG00000131378 | RFTN1 | 1745.864849 | -3.597746003 | 0.074067829 | -48.57366615 | 0 | 0 |
| ENSG00000131379 | C3orf20 | 13.62109586 | -0.227705465 | 0.476314319 | -0.478057149 | 0.63260953 | 0.783513891 |
| ENSG00000131381 | RBSN | 3296.52503 | -0.183300856 | 0.052779303 | -3.472968508 | 0.000514736 | 0.002031665 |
| ENSG00000131386 | GALNT15 | 7.221991086 | 0.7486051 | 0.662851661 | 1.129370482 | 0.25874158 | 0.438528839 |
| ENSG00000131389 | SLC6A6 | 5271.337206 | 0.004967185 | 0.044173965 | 0.112445981 | 0.910469799 | 0.95506154 |
| ENSG00000131398 | KCNC3 | 1247.329191 | 1.560916826 | 0.072167355 | 21.62912631 | 9.56E-104 | 1.71E-101 |
| ENSG00000131400 | NAPSA | 61.65099413 | -1.184889949 | 0.241569889 | -4.904957135 | 9.34E-07 | 5.43E-06 |
| ENSG00000131408 | NR1H2 | 1629.448736 | 0.084513956 | 0.068209573 | 1.239033652 | 0.215333036 | 0.384448225 |
| ENSG00000131409 | LRRC4B | 3.774722946 | 1.152227827 | 0.950724509 | 1.211947117 | 0.225532624 | 0.397545342 |
| ENSG00000131437 | KIF3A | 533.8510448 | -0.052183423 | 0.089389404 | -0.583776385 | 0.559370768 | 0.729047521 |
| ENSG00000131446 | MGAT1 | 2123.732437 | -0.189816481 | 0.058771636 | -3.229729431 | 0.001239074 | 0.004558108 |
| ENSG00000131459 | GFPT2 | 79.3389261 | -0.447370907 | 0.199895909 | -2.238019325 | 0.025219795 | 0.066976394 |
| ENSG00000131462 | TUBG1 | 3002.843939 | -0.259193464 | 0.05148454 | -5.034394118 | 4.79E-07 | 2.90E-06 |
| ENSG00000131467 | PSME3 | 5326.278019 | 0.029329767 | 0.045396631 | 0.646078048 | 0.518228811 | 0.695932225 |
| ENSG00000131469 | RPL27 | 6739.154813 | 0.183037017 | 0.047360238 | 3.864782448 | 0.000111188 | 0.00049269 |
| ENSG00000131470 | PSMC3IP | 485.3196247 | -0.259720462 | 0.099053937 | -2.622010498 | 0.008741274 | 0.026430054 |
| ENSG00000131471 | AOC3 | 12.30901024 | -0.284487693 | 0.497573418 | -0.571750184 | 0.567491231 | 0.735328109 |

| | | | | | | | |
|---|---|---|---|---|---|---|---|
| ENSG00000131473 | ACLY | 28223.18783 | -0.451212898 | 0.043457204 | -10.3829253 | 2.97E-25 | 6.68E-24 |
| ENSG00000131475 | VPS25 | 1763.422178 | -0.172202174 | 0.055148783 | -3.122501804 | 0.00179321 | 0.006365868 |
| ENSG00000131477 | RAMP2 | 7.696085827 | 0.713271015 | 0.64551012 | 1.10497263 | 0.269171449 | 0.450813539 |
| ENSG00000131480 | AOC2 | 31.24956216 | 0.220869662 | 0.315151113 | 0.700837321 | 0.483404544 | 0.667507389 |
| ENSG00000131495 | NDUFA2 | 1194.06354 | -0.004890395 | 0.065424742 | -0.074748402 | 0.940414896 | 0.971621655 |
| ENSG00000131503 | ANKHD1 | 43.50336999 | -0.013330752 | 0.266468887 | -0.050027424 | 0.960100534 | 0.981799414 |
| ENSG00000131504 | DIAPH1 | 7164.246738 | -0.102411237 | 0.04345861 | -2.356523565 | 0.018446897 | 0.050947727 |
| ENSG00000131507 | NDFIP1 | 2617.480505 | 0.140335256 | 0.058294591 | 2.407346092 | 0.016068934 | 0.045196339 |
| ENSG00000131508 | UBE2D2 | 2817.013093 | 0.010912811 | 0.054401555 | 0.200597407 | 0.841013386 | 0.916568399 |
| ENSG00000131558 | EXOC4 | 2468.042399 | 0.041428493 | 0.052695329 | 0.786189095 | 0.431756714 | 0.621776495 |
| ENSG00000131584 | ACAP3 | 2126.061568 | 0.299816416 | 0.068039394 | 4.406512143 | 1.05E-05 | 5.41E-05 |
| ENSG00000131591 | C1orf159 | 359.0429681 | 0.314892461 | 0.108486512 | 2.902595491 | 0.003700843 | 0.012284379 |
| ENSG00000131620 | ANO1 | 42.90076516 | 0.282238088 | 0.273310827 | 1.032663401 | 0.301761447 | 0.488629038 |
| ENSG00000131626 | PPFIA1 | 3493.321401 | 0.266919248 | 0.048146977 | 5.543842314 | 2.96E-08 | 2.02E-07 |
| ENSG00000131650 | KREMEN2 | 240.159716 | -0.015370744 | 0.123626217 | -0.124332396 | 0.901052098 | 0.950176295 |
| ENSG00000131652 | THOC6 | 803.8624995 | -0.368129149 | 0.073231084 | -5.026952072 | 4.98E-07 | 3.00E-06 |
| ENSG00000131653 | TRAF7 | 6376.753763 | 0.545228115 | 0.050411457 | 10.81555959 | 2.91E-27 | 7.29E-26 |
| ENSG00000131669 | NINJ1 | 2756.257487 | 0.550982697 | 0.06785534 | 8.119960742 | 4.66E-16 | 6.05E-15 |
| ENSG00000131697 | NPHP4 | 673.2355263 | 0.215523828 | 0.088445558 | 2.436796513 | 0.014818015 | 0.042043942 |
| ENSG00000131711 | MAP1B | 242.0254033 | -0.50206867 | 0.123245873 | -4.073715881 | 4.63E-05 | 0.000217413 |
| ENSG00000131724 | IL13RA1 | 3217.592699 | 0.508149411 | 0.048889449 | 10.39384609 | 2.64E-25 | 5.97E-24 |
| ENSG00000131725 | WDR44 | 1062.307457 | -0.48697244 | 0.071337776 | -6.826291267 | 8.71E-12 | 8.21E-11 |
| ENSG00000131732 | ZCCHC9 | 428.5915015 | -0.195196677 | 0.093853709 | -2.079797151 | 0.037544143 | 0.094106494 |
| ENSG00000131746 | TNS4 | 21.03122177 | -0.291549294 | 0.404142776 | -0.721401721 | 0.470662388 | 0.656511929 |
| ENSG00000131747 | TOP2A | 10901.79394 | -0.159282753 | 0.051184253 | -3.111948382 | 0.00185857 | 0.006576522 |
| ENSG00000131748 | STARD3 | 1069.250437 | -0.007314758 | 0.066091975 | -0.110675435 | 0.911873726 | 0.955697231 |
| ENSG00000131759 | RARA | 2641.669373 | 0.22538596 | 0.053410144 | 4.219909224 | 2.44E-05 | 0.000119773 |
| ENSG00000131773 | KHDRBS3 | 35.94604688 | -0.00512308 | 0.292491513 | -0.017515311 | 0.986025518 | 0.993350589 |
| ENSG00000131778 | CHD1L | 51.49232267 | 0.272273428 | 0.265361993 | 1.026045309 | 0.304870228 | 0.492307689 |
| ENSG00000131779 | PEX11B | 1282.459738 | 0.274934428 | 0.062531536 | 4.39673236 | 1.10E-05 | 5.65E-05 |
| ENSG00000131781 | FMO5 | 247.5672464 | 0.831299915 | 0.120497552 | 6.898894643 | 5.24E-12 | 5.04E-11 |
| ENSG00000131788 | PIAS3 | 6630.532141 | 0.225405361 | 0.0529765 | 4.254817901 | 2.09E-05 | 0.000103638 |
| ENSG00000131791 | PRKAB2 | 2063.816971 | 0.265850001 | 0.058690953 | 4.529658948 | 5.91E-06 | 3.14E-05 |

| | | | | | | | |
|---|---|---|---|---|---|---|---|
| ENSG00000131797 | CLUHP3 | 799.9358459 | 0.53505323 | 0.082212298 | 6.508189665 | 7.61E-11 | 6.65E-10 |
| ENSG00000131828 | PDHA1 | 3617.511116 | 0.131391385 | 0.046840858 | 2.805059337 | 0.005030732 | 0.016197191 |
| ENSG00000131831 | RAI2 | 292.177068 | -2.809387522 | 0.137828317 | -20.38323901 | 2.36E-92 | 3.34E-90 |
| ENSG00000131844 | MCCC2 | 6706.773888 | -0.056575963 | 0.044585964 | -1.268918699 | 0.204470062 | 0.369392285 |
| ENSG00000131845 | ZNF304 | 362.3947463 | 0.750066976 | 0.100095924 | 7.493481725 | 6.71E-14 | 7.41E-13 |
| ENSG00000131848 | ZSCAN5A | 156.6418096 | 0.250856127 | 0.146513728 | 1.712168065 | 0.086865701 | 0.188961514 |
| ENSG00000131849 | ZNF132 | 4.47018068 | -0.925160039 | 0.832135672 | -1.111789905 | 0.266228493 | 0.447190572 |
| ENSG00000131871 | SELENOS | 1606.976264 | -0.182944715 | 0.056603207 | -3.232055659 | 0.001229031 | 0.004525518 |
| ENSG00000131873 | CHSY1 | 1838.166742 | 0.063021014 | 0.055129098 | 1.143153377 | 0.252974916 | 0.431735218 |
| ENSG00000131876 | SNRPA1 | 988.2026445 | 0.17865641 | 0.068825033 | 2.595805667 | 0.009436942 | 0.028276313 |
| ENSG00000131899 | LLGL1 | 1763.186795 | 0.175011087 | 0.062437132 | 2.80299688 | 0.005063016 | 0.016294274 |
| ENSG00000131931 | THAP1 | 315.582979 | -0.070848595 | 0.107214403 | -0.660812284 | 0.508732704 | 0.688880717 |
| ENSG00000131941 | RHPN2 | 1209.714775 | -0.551348433 | 0.066401338 | -8.303272924 | 1.01E-16 | 1.38E-15 |
| ENSG00000131943 | C19orf12 | 679.2049198 | -0.178763443 | 0.079087902 | -2.260313384 | 0.023801808 | 0.063675041 |
| ENSG00000131944 | FAAP24 | 143.2404159 | -0.334269584 | 0.154717482 | -2.16051592 | 0.030732753 | 0.079348266 |
| ENSG00000131966 | ACTR10 | 1654.30489 | -0.353662393 | 0.056662972 | -6.241507964 | 4.33E-10 | 3.57E-09 |
| ENSG00000131979 | GCH1 | 935.1290347 | -0.371509971 | 0.072951483 | -5.092562254 | 3.53E-07 | 2.16E-06 |
| ENSG00000131981 | LGALS3 | 2649.97089 | 0.527366439 | 0.054002683 | 9.765559929 | 1.58E-22 | 3.06E-21 |
| ENSG00000132000 | PODNL1 | 14.09671752 | -0.352781577 | 0.472474905 | -0.746667333 | 0.455264393 | 0.642693242 |
| ENSG00000132002 | DNAJB1 | 2905.942644 | 0.233963625 | 0.058398705 | 4.006315261 | 6.17E-05 | 0.000284722 |
| ENSG00000132003 | ZSWIM4 | 203.5220091 | 0.875913305 | 0.142000816 | 6.168368111 | 6.90E-10 | 5.58E-09 |
| ENSG00000132004 | FBXW9 | 359.4350602 | 0.312473226 | 0.103237677 | 3.026736315 | 0.002472095 | 0.00852847 |
| ENSG00000132005 | RFX1 | 539.4710817 | 0.062124533 | 0.090310129 | 0.687902158 | 0.491514397 | 0.674428199 |
| ENSG00000132010 | ZNF20 | 7.812434687 | 0.458792354 | 0.661798821 | 0.693250485 | 0.488152371 | 0.671528246 |
| ENSG00000132016 | BRME1 | 144.7084515 | -0.556793115 | 0.154848282 | -3.595733231 | 0.000323479 | 0.001323455 |
| ENSG00000132017 | DCAF15 | 754.4248636 | -0.01000168 | 0.081884124 | -0.122144316 | 0.902784726 | 0.950907167 |
| ENSG00000132024 | CC2D1A | 1851.118906 | -0.055475123 | 0.057662457 | -0.962066583 | 0.336016159 | 0.527129227 |
| ENSG00000132026 | RTBDN | 4.084880374 | -1.0222054 | 0.953761857 | -1.07176167 | 0.283827094 | 0.467676593 |
| ENSG00000132031 | MATN3 | 26.95580214 | 0.998662278 | 0.358752837 | 2.783705592 | 0.005374179 | 0.017168036 |
| ENSG00000132109 | TRIM21 | 99.02812738 | 0.185118212 | 0.190288876 | 0.972827296 | 0.330639144 | 0.521332185 |
| ENSG00000132122 | SPATA6 | 142.5176837 | 0.77447731 | 0.154911215 | 4.999491542 | 5.75E-07 | 3.43E-06 |
| ENSG00000132128 | LRRC41 | 3221.23088 | -0.050233467 | 0.051940972 | -0.967126064 | 0.333481017 | 0.524587159 |
| ENSG00000132141 | CCT6B | 45.14572305 | 0.358838451 | 0.261598179 | 1.371716167 | 0.170151814 | 0.320914462 |

| | | | | | | | |
|---|---|---|---|---|---|---|---|
| ENSG00000132153 | DHX30 | 2662.30925 | -0.005762189 | 0.063230951 | -0.091129241 | 0.927389899 | 0.965137362 |
| ENSG00000132155 | RAF1 | 6474.798654 | -0.053927401 | 0.04471859 | -1.20592801 | 0.227845249 | 0.40067229 |
| ENSG00000132182 | NUP210 | 8930.587956 | 0.302173618 | 0.044698067 | 6.760328497 | 1.38E-11 | 1.27E-10 |
| ENSG00000132185 | FCRLA | 4.419648034 | 1.393892399 | 0.904095624 | 1.541753286 | 0.123133558 | 0.250393034 |
| ENSG00000132196 | HSD17B7 | 590.8009716 | -0.458251845 | 0.081102176 | -5.650302696 | 1.60E-08 | 1.13E-07 |
| ENSG00000132199 | ENOSF1 | 1788.157266 | 0.214754054 | 0.079695399 | 2.694685723 | 0.007045505 | 0.021823371 |
| ENSG00000132204 | LINC00470 | 5.569025377 | 0.170825335 | 0.737928182 | 0.231493172 | 0.81693169 | 0.903062617 |
| ENSG00000132205 | EMILIN2 | 15.14655122 | 0.731983036 | 0.454694908 | 1.609833369 | 0.107434238 | 0.224580046 |
| ENSG00000132254 | ARFIP2 | 2633.926231 | -0.474832612 | 0.054813293 | -8.662727395 | 4.61E-18 | 6.82E-17 |
| ENSG00000132256 | TRIM5 | 69.76233678 | 1.248273629 | 0.225758757 | 5.52923681 | 3.22E-08 | 2.19E-07 |
| ENSG00000132259 | CNGA4 | 4.064576685 | 0.190570254 | 0.874407493 | 0.217942155 | 0.827474181 | 0.908850796 |
| ENSG00000132275 | RRP8 | 426.2746554 | 0.131656294 | 0.098434509 | 1.337501398 | 0.181059026 | 0.336455523 |
| ENSG00000132286 | TIMM10B | 630.9266209 | 0.123872827 | 0.0789854 | 1.568300292 | 0.116811068 | 0.240413233 |
| ENSG00000132294 | EFR3A | 2196.614144 | 0.135743889 | 0.053930591 | 2.517010957 | 0.011835515 | 0.034516982 |
| ENSG00000132300 | PTCD3 | 1998.919421 | 0.052304602 | 0.064773262 | 0.807502981 | 0.419376758 | 0.610892467 |
| ENSG00000132305 | IMMT | 3979.707234 | -0.207031844 | 0.048035454 | -4.30997999 | 1.63E-05 | 8.19E-05 |
| ENSG00000132313 | MRPL35 | 1124.286104 | 0.123411537 | 0.066531887 | 1.854923152 | 0.063607201 | 0.146427416 |
| ENSG00000132321 | IQCA1 | 2.803176168 | 1.220764306 | 1.113474359 | 1.09635601 | 0.272923007 | 0.454904728 |
| ENSG00000132323 | ILKAP | 477.0063569 | -0.013400603 | 0.088536015 | -0.151357644 | 0.8796936 | 0.937619587 |
| ENSG00000132326 | PER2 | 824.8162941 | -0.134507304 | 0.075837994 | -1.773613686 | 0.076127072 | 0.169945021 |
| ENSG00000132329 | RAMP1 | 19.24636064 | -0.407890179 | 0.416370966 | -0.979631656 | 0.327267973 | 0.517679904 |
| ENSG00000132330 | SCLY | 29.14087469 | 0.057011125 | 0.330332827 | 0.172586918 | 0.862976135 | 0.92885434 |
| ENSG00000132334 | PTPRE | 61.50117886 | 0.162520184 | 0.227129425 | 0.715539978 | 0.474275447 | 0.659706449 |
| ENSG00000132341 | RAN | 10357.0708 | -0.062775502 | 0.050789484 | -1.235994081 | 0.216460764 | 0.385888244 |
| ENSG00000132356 | PRKAA1 | 2257.740777 | -0.079297301 | 0.059512059 | -1.332457692 | 0.182709835 | 0.33864132 |
| ENSG00000132357 | CARD6 | 2.211265268 | -0.852716396 | 1.191435368 | -0.715705123 | 0.474173447 | 0.659616385 |
| ENSG00000132359 | RAP1GAP2 | 930.6601449 | 0.542453976 | 0.073045708 | 7.426226596 | 1.12E-13 | 1.22E-12 |
| ENSG00000132361 | CLUH | 4470.135292 | 0.318750911 | 0.055090646 | 5.785935308 | 7.21E-09 | 5.26E-08 |
| ENSG00000132376 | INPP5K | 1057.421536 | 0.281468502 | 0.071320505 | 3.946529825 | 7.93E-05 | 0.000359976 |
| ENSG00000132382 | MYBBP1A | 2288.641717 | 0.289152952 | 0.057814013 | 5.001433676 | 5.69E-07 | 3.40E-06 |
| ENSG00000132383 | RPA1 | 3661.281432 | 0.046171156 | 0.046360861 | 0.995908093 | 0.319294803 | 0.508689685 |
| ENSG00000132386 | SERPINF1 | 18.64882916 | 0.385894231 | 0.411200307 | 0.93845803 | 0.348009076 | 0.538977059 |
| ENSG00000132388 | UBE2G1 | 1161.003112 | 0.116253004 | 0.070912988 | 1.639375347 | 0.101135113 | 0.213808818 |

| | | | | | | | |
|---|---|---|---|---|---|---|---|
| ENSG00000132394 | EEFSEC | 911.3491246 | -0.365736796 | 0.072926791 | -5.015122567 | 5.30E-07 | 3.18E-06 |
| ENSG00000132405 | TBC1D14 | 3590.604588 | 0.09565823 | 0.05057254 | 1.891505368 | 0.058556915 | 0.136763962 |
| ENSG00000132406 | TMEM128 | 409.3474479 | -0.271567325 | 0.093629209 | -2.900455181 | 0.003726211 | 0.012363217 |
| ENSG00000132423 | COQ3 | 202.2539865 | -0.351633749 | 0.132939409 | -2.645067787 | 0.008167455 | 0.024881995 |
| ENSG00000132424 | PNISR | 3010.095667 | 0.069847631 | 0.076796842 | 0.909511762 | 0.363080052 | 0.555826253 |
| ENSG00000132429 | POPDC3 | 2.632513313 | -0.521032951 | 1.156560937 | -0.450501944 | 0.652348552 | 0.798160975 |
| ENSG00000132432 | SEC61G | 1015.033025 | 0.413912794 | 0.069987957 | 5.914057346 | 3.34E-09 | 2.53E-08 |
| ENSG00000132434 | LANCL2 | 1202.841665 | 0.977608653 | 0.066162701 | 14.77582749 | 2.10E-49 | 1.15E-47 |
| ENSG00000132436 | FIGNL1 | 1010.477061 | 0.546126431 | 0.07330351 | 7.450208477 | 9.32E-14 | 1.02E-12 |
| ENSG00000132437 | DDC | 4.240071975 | -0.813772348 | 0.877891761 | -0.926962053 | 0.353946237 | 0.54552047 |
| ENSG00000132463 | GRSF1 | 4664.767586 | 0.077739719 | 0.051453344 | 1.510877874 | 0.130819569 | 0.262715654 |
| ENSG00000132466 | ANKRD17 | 4023.039804 | -0.045185533 | 0.063733843 | -0.708972358 | 0.478341631 | 0.66295052 |
| ENSG00000132467 | UTP3 | 1485.471847 | -0.08446702 | 0.059734974 | -1.414029582 | 0.157353217 | 0.301920349 |
| ENSG00000132470 | ITGB4 | 8131.522224 | 0.139389434 | 0.057295489 | 2.432816904 | 0.014981875 | 0.042466849 |
| ENSG00000132471 | WBP2 | 5104.931394 | 0.120908021 | 0.051516104 | 2.346994658 | 0.018925526 | 0.052119036 |
| ENSG00000132475 | H3-3B | 20861.46691 | 0.208107147 | 0.050711862 | 4.103717358 | 4.07E-05 | 0.00019238 |
| ENSG00000132478 | UNK | 1782.549137 | 0.235489537 | 0.057028888 | 4.129302604 | 3.64E-05 | 0.0001735 |
| ENSG00000132481 | TRIM47 | 33.49940376 | 0.076018317 | 0.303805921 | 0.250219999 | 0.80241722 | 0.89482529 |
| ENSG00000132485 | ZRANB2 | 1445.397223 | 0.196261305 | 0.081755462 | 2.400589515 | 0.016368687 | 0.045915587 |
| ENSG00000132507 | EIF5A | 6525.991868 | 0.049766826 | 0.044899753 | 1.108398665 | 0.267689681 | 0.44895462 |
| ENSG00000132510 | KDM6B | 1500.304004 | 0.119213214 | 0.063348851 | 1.881852833 | 0.059856 | 0.139274791 |
| ENSG00000132522 | GPS2 | 57.18330438 | -0.184881506 | 0.247926426 | -0.745711173 | 0.455841908 | 0.643117857 |
| ENSG00000132530 | XAF1 | 5.106658154 | 3.172776382 | 1.071432069 | 2.96124829 | 0.003063948 | 0.010360698 |
| ENSG00000132535 | DLG4 | 129.2679292 | -0.43007071 | 0.159377948 | -2.69843298 | 0.006966676 | 0.021602535 |
| ENSG00000132541 | RIDA | 757.8627455 | -0.557192858 | 0.074641413 | -7.464929179 | 8.33E-14 | 9.13E-13 |
| ENSG00000132549 | VPS13B | 2402.0681 | -0.474351074 | 0.072483715 | -6.544243375 | 5.98E-11 | 5.27E-10 |
| ENSG00000132561 | MATN2 | 4302.809267 | 1.904248524 | 0.053634776 | 35.50398928 | 4.27E-276 | 6.99E-273 |
| ENSG00000132563 | REEP2 | 105.308689 | -0.0075028 | 0.174657737 | -0.042957159 | 0.965735684 | 0.984577423 |
| ENSG00000132570 | PCBD2 | 596.4423374 | 0.038447647 | 0.082588196 | 0.465534409 | 0.641548802 | 0.789631802 |
| ENSG00000132581 | SDF2 | 1167.235274 | -0.087482722 | 0.068305665 | -1.280753538 | 0.200280248 | 0.363454544 |
| ENSG00000132589 | FLOT2 | 4938.540093 | -0.234778377 | 0.053339932 | -4.401549977 | 1.07E-05 | 5.53E-05 |
| ENSG00000132591 | ERAL1 | 1928.898758 | -0.171275316 | 0.061021274 | -2.806813176 | 0.005003425 | 0.016116059 |
| ENSG00000132600 | PRMT7 | 855.1555653 | -0.175517751 | 0.076825047 | -2.284642299 | 0.022333811 | 0.060225416 |

| | | | | | | | |
|---|---|---|---|---|---|---|---|
| ENSG00000132603 | NIP7 | 1284.210052 | 0.074276627 | 0.062842399 | 1.18195084 | 0.237225207 | 0.412400055 |
| ENSG00000132604 | TERF2 | 1092.556488 | -0.062231419 | 0.064291915 | -0.967950938 | 0.333068873 | 0.524190447 |
| ENSG00000132612 | VPS4A | 1426.873688 | 0.206831723 | 0.064867757 | 3.188513581 | 0.001430063 | 0.005180929 |
| ENSG00000132613 | MTSS2 | 5270.569704 | 0.032958121 | 0.062076277 | 0.530929407 | 0.595467699 | 0.756704466 |
| ENSG00000132623 | ANKEF1 | 700.3076779 | 0.090505105 | 0.078098044 | 1.158865201 | 0.246511136 | 0.423702607 |
| ENSG00000132635 | PCED1A | 924.9524592 | -0.086812996 | 0.077636247 | -1.118201859 | 0.263480791 | 0.444136184 |
| ENSG00000132639 | SNAP25 | 11.59292142 | -0.771562594 | 0.570566764 | -1.35227399 | 0.176287682 | 0.329802595 |
| ENSG00000132640 | BTBD3 | 717.6199663 | -0.583762249 | 0.079451621 | -7.347392569 | 2.02E-13 | 2.16E-12 |
| ENSG00000132646 | PCNA | 2361.63227 | -0.24424885 | 0.054790249 | -4.457888999 | 8.28E-06 | 4.31E-05 |
| ENSG00000132661 | NXT1 | 454.6690771 | -0.31786988 | 0.097425639 | -3.262692284 | 0.001103593 | 0.004099825 |
| ENSG00000132664 | POLR3F | 593.349972 | -0.432881599 | 0.083680518 | -5.173027228 | 2.30E-07 | 1.44E-06 |
| ENSG00000132669 | RIN2 | 411.1095281 | -0.677959422 | 0.095503822 | -7.098767447 | 1.26E-12 | 1.28E-11 |
| ENSG00000132670 | PTPRA | 2189.383523 | -0.134478437 | 0.061460362 | -2.188051491 | 0.028665852 | 0.074658792 |
| ENSG00000132676 | DAP3 | 4312.949475 | 0.273056999 | 0.045857631 | 5.95445068 | 2.61E-09 | 2.00E-08 |
| ENSG00000132677 | RHBG | 225.8629677 | -0.985620458 | 0.132722227 | -7.426189864 | 1.12E-13 | 1.22E-12 |
| ENSG00000132680 | KHDC4 | 1498.40034 | 0.460322082 | 0.082268579 | 5.595357151 | 2.20E-08 | 1.52E-07 |
| ENSG00000132688 | NES | 296.4823406 | -1.543443078 | 0.114347557 | -13.49782292 | 1.61E-41 | 7.02E-40 |
| ENSG00000132692 | BCAN | 1.720567635 | 0.35172336 | 1.423190706 | 0.247137195 | 0.804802049 | NA |
| ENSG00000132694 | ARHGEF11 | 2673.178393 | 0.140184153 | 0.059044913 | 2.374195275 | 0.017587246 | 0.04893286 |
| ENSG00000132698 | RAB25 | 3429.178981 | 0.170019243 | 0.052543684 | 3.235769377 | 0.001213154 | 0.004473589 |
| ENSG00000132716 | DCAF8 | 1048.805169 | 0.404026204 | 0.070159021 | 5.758720672 | 8.48E-09 | 6.14E-08 |
| ENSG00000132718 | SYT11 | 3.518046871 | 1.258595077 | 0.971825697 | 1.295083143 | 0.19529155 | 0.356488274 |
| ENSG00000132740 | IGHMBP2 | 866.8865738 | -0.137164078 | 0.076013462 | -1.804470867 | 0.071157522 | 0.160807168 |
| ENSG00000132746 | ALDH3B2 | 10576.25559 | -0.199252831 | 0.047684761 | -4.178543154 | 2.93E-05 | 0.000142287 |
| ENSG00000132749 | TESMIN | 874.0590581 | 0.17635969 | 0.070837924 | 2.489622519 | 0.012787883 | 0.036895239 |
| ENSG00000132763 | MMACHC | 526.5563033 | -0.343738221 | 0.087216343 | -3.941213422 | 8.11E-05 | 0.000367468 |
| ENSG00000132768 | DPH2 | 1559.238935 | 0.036933828 | 0.065353449 | 0.565139696 | 0.57197875 | 0.738847487 |
| ENSG00000132773 | TOE1 | 449.2228601 | -0.198650698 | 0.094148225 | -2.109978162 | 0.034860237 | 0.088315167 |
| ENSG00000132780 | NASP | 4009.094912 | -0.141255747 | 0.057406035 | -2.460642815 | 0.013868836 | 0.039667335 |
| ENSG00000132781 | MUTYH | 43.49326539 | -0.503460002 | 0.271410324 | -1.854977346 | 0.063599462 | 0.146424293 |
| ENSG00000132792 | CTNNBL1 | 1122.000185 | -0.281935079 | 0.06323306 | -4.458665726 | 8.25E-06 | 4.29E-05 |
| ENSG00000132793 | LPIN3 | 2258.260719 | -0.005057551 | 0.063276028 | -0.079928391 | 0.936294209 | 0.969525239 |
| ENSG00000132801 | ZSWIM3 | 303.2215348 | -0.479780307 | 0.118447208 | -4.050583496 | 5.11E-05 | 0.000238746 |

| | | | | | | | |
|---|---|---|---|---|---|---|---|
| ENSG00000132819 | RBM38 | 1710.397744 | 0.161860408 | 0.05678424 | 2.850445962 | 0.004365797 | 0.014272678 |
| ENSG00000132823 | OSER1 | 1399.793997 | -0.012345735 | 0.059203717 | -0.208529726 | 0.834815375 | 0.912892778 |
| ENSG00000132824 | SERINC3 | 5692.687653 | 0.073734094 | 0.047536573 | 1.551102426 | 0.12087714 | 0.246955687 |
| ENSG00000132825 | PPP1R3D | 1208.17669 | -0.176125697 | 0.063030072 | -2.794312139 | 0.005201025 | 0.016668275 |
| ENSG00000132837 | DMGDH | 13.26679739 | 1.7386396 | 0.534916365 | 3.250301756 | 0.001152826 | 0.004272348 |
| ENSG00000132842 | AP3B1 | 2018.514478 | 0.002356538 | 0.058247541 | 0.040457292 | 0.967728555 | 0.985650827 |
| ENSG00000132846 | ZBED3 | 871.4097505 | 0.186394032 | 0.084606678 | 2.203065245 | 0.02759015 | 0.072292182 |
| ENSG00000132849 | PATJ | 2366.271374 | 0.288793661 | 0.059862255 | 4.824303048 | 1.40E-06 | 7.97E-06 |
| ENSG00000132874 | SLC14A2 | 3.250037696 | 4.012735445 | 1.423394467 | 2.819130985 | 0.004815386 | 0.015582588 |
| ENSG00000132879 | FBXO44 | 561.1898629 | 1.073973083 | 0.091300049 | 11.76311617 | 6.05E-32 | 1.82E-30 |
| ENSG00000132881 | CPLANE2 | 134.7232614 | -0.035752212 | 0.157283222 | -0.227311035 | 0.820181893 | 0.904627311 |
| ENSG00000132906 | CASP9 | 263.9357656 | 0.485650531 | 0.125581318 | 3.867219557 | 0.000110083 | 0.000487889 |
| ENSG00000132911 | NMUR2 | 2.225368747 | -2.621872161 | 1.413134125 | -1.855359739 | 0.063544873 | 0.146327984 |
| ENSG00000132912 | DCTN4 | 3146.756065 | 0.283188761 | 0.050075119 | 5.655278817 | 1.56E-08 | 1.10E-07 |
| ENSG00000132932 | ATP8A2 | 874.2230599 | 0.994526655 | 0.074229957 | 13.39791499 | 6.22E-41 | 2.65E-39 |
| ENSG00000132938 | MTUS2 | 26.59613014 | 1.074189741 | 0.356974999 | 3.009145579 | 0.002619835 | 0.008987442 |
| ENSG00000132950 | ZMYM5 | 459.8631732 | 0.616621123 | 0.093031643 | 6.628079474 | 3.40E-11 | 3.05E-10 |
| ENSG00000132952 | USPL1 | 829.6793048 | -0.20691937 | 0.074484668 | -2.77801292 | 0.005469244 | 0.017430462 |
| ENSG00000132953 | XPO4 | 1701.685592 | 0.023549663 | 0.058584249 | 0.40197942 | 0.687699172 | 0.823383061 |
| ENSG00000132958 | TPTE2 | 1.687876367 | 3.008605088 | 1.618513434 | 1.858869396 | 0.063045653 | NA |
| ENSG00000132963 | POMP | 1980.452706 | -0.013385812 | 0.061891089 | -0.216280119 | 0.828769405 | 0.909446073 |
| ENSG00000132964 | CDK8 | 615.644104 | -0.033952805 | 0.080026845 | -0.4242677 | 0.671370593 | 0.811616853 |
| ENSG00000132967 | HMGB1P5 | 2919.339758 | -0.360490455 | 0.062272605 | -5.788909173 | 7.08E-09 | 5.18E-08 |
| ENSG00000132970 | WASF3 | 114.0867904 | 0.243791182 | 0.167278432 | 1.457397578 | 0.145006665 | 0.28402091 |
| ENSG00000133019 | CHRM3 | 119.1038928 | -0.708032649 | 0.163938309 | -4.318896859 | 1.57E-05 | 7.88E-05 |
| ENSG00000133026 | MYH10 | 3114.458975 | -0.06398159 | 0.05109915 | -1.252106733 | 0.210530975 | 0.377812764 |
| ENSG00000133027 | PEMT | 530.8451922 | 0.034510252 | 0.085560469 | 0.403343417 | 0.686695606 | 0.822987159 |
| ENSG00000133028 | SCO1 | 1333.937926 | 0.039851995 | 0.060627897 | 0.657321089 | 0.510974479 | 0.690652518 |
| ENSG00000133030 | MPRIP | 4526.306834 | 0.15685585 | 0.054981572 | 2.85288042 | 0.004332492 | 0.014173897 |
| ENSG00000133056 | PIK3C2B | 3989.223736 | 0.599554369 | 0.055977203 | 10.71068816 | 9.07E-27 | 2.22E-25 |
| ENSG00000133059 | DSTYK | 2920.80685 | 0.440343958 | 0.058136894 | 7.574260107 | 3.61E-14 | 4.07E-13 |
| ENSG00000133065 | SLC41A1 | 3634.066318 | 0.233060119 | 0.052294135 | 4.456716168 | 8.32E-06 | 4.33E-05 |
| ENSG00000133067 | LGR6 | 2.746993589 | 1.991834125 | 1.204489849 | 1.65367448 | 0.098193671 | 0.208880082 |

| | | | | | | | |
|---|---|---|---|---|---|---|---|
| ENSG00000133069 | TMCC2 | 53.46953981 | 0.269258123 | 0.240833874 | 1.118024298 | 0.263556616 | 0.44419872 |
| ENSG00000133083 | DCLK1 | 1446.035035 | 0.078348052 | 0.062957384 | 1.244461685 | 0.213329693 | 0.381522035 |
| ENSG00000133101 | CCNA1 | 16.8230396 | -1.685183485 | 0.465724707 | -3.618411176 | 0.000296417 | 0.001221669 |
| ENSG00000133103 | COG6 | 1403.384676 | 0.055296361 | 0.060441618 | 0.91487228 | 0.360258684 | 0.552798275 |
| ENSG00000133104 | SPART | 588.9447812 | 0.51715073 | 0.081631024 | 6.335222901 | 2.37E-10 | 1.99E-09 |
| ENSG00000133107 | TRPC4 | 4.373809971 | 0.83528524 | 0.847841944 | 0.985189806 | 0.324530847 | 0.514743665 |
| ENSG00000133111 | RFXAP | 179.4925474 | 0.273167909 | 0.141722584 | 1.927483264 | 0.053919432 | 0.127897452 |
| ENSG00000133112 | TPT1 | 33629.98496 | 0.184214323 | 0.057400634 | 3.209273333 | 0.001330709 | 0.004865551 |
| ENSG00000133114 | GPALPP1 | 1087.244358 | 0.197099987 | 0.072653524 | 2.712875788 | 0.00667021 | 0.020790031 |
| ENSG00000133116 | KL | 19.23918806 | -3.240655061 | 0.549980821 | -5.892305576 | 3.81E-09 | 2.87E-08 |
| ENSG00000133119 | RFC3 | 1392.095603 | -0.052480439 | 0.067824255 | -0.77377097 | 0.439066243 | 0.628529149 |
| ENSG00000133121 | STARD13 | 599.9209187 | -0.825043708 | 0.084414255 | -9.773748646 | 1.46E-22 | 2.83E-21 |
| ENSG00000133124 | IRS4 | 85.77980733 | 1.512677992 | 0.205739577 | 7.352391863 | 1.95E-13 | 2.08E-12 |
| ENSG00000133131 | MORC4 | 785.7188564 | -0.442508294 | 0.074894418 | -5.908428259 | 3.45E-09 | 2.62E-08 |
| ENSG00000133134 | BEX2 | 75.85643524 | 0.349752484 | 0.203286519 | 1.720490303 | 0.085343355 | 0.186245998 |
| ENSG00000133135 | RNF128 | 7.998876255 | -3.601340508 | 0.89986638 | -4.002083631 | 6.28E-05 | 0.000289398 |
| ENSG00000133136 | GNG5B | 55.55693827 | 0.108106247 | 0.241528207 | 0.447592636 | 0.654447219 | 0.799387444 |
| ENSG00000133138 | TBC1D8B | 129.9920002 | -1.480169831 | 0.167832662 | -8.819319288 | 1.15E-18 | 1.78E-17 |
| ENSG00000133142 | TCEAL4 | 4595.803568 | -0.110408773 | 0.045985439 | -2.400950756 | 0.016352537 | 0.045875895 |
| ENSG00000133193 | FAM104A | 1339.672032 | 0.232505223 | 0.060816855 | 3.823039254 | 0.000131817 | 0.000576086 |
| ENSG00000133195 | SLC39A11 | 1106.115728 | 0.328760723 | 0.069546283 | 4.727222035 | 2.28E-06 | 1.26E-05 |
| ENSG00000133216 | EPHB2 | 586.2406847 | 0.588038501 | 0.09244612 | 6.360878094 | 2.01E-10 | 1.69E-09 |
| ENSG00000133226 | SRRM1 | 2766.298794 | -0.207600292 | 0.053090744 | -3.910291658 | 9.22E-05 | 0.000413354 |
| ENSG00000133243 | BTBD2 | 2324.590113 | 0.035781181 | 0.064918995 | 0.55116659 | 0.581519481 | 0.74646393 |
| ENSG00000133246 | PRAM1 | 12.00464314 | -0.977934808 | 0.510331954 | -1.91627195 | 0.055330489 | 0.130638269 |
| ENSG00000133247 | KMT5C | 546.0208528 | 0.071263241 | 0.084916117 | 0.839219256 | 0.401346284 | 0.594137714 |
| ENSG00000133250 | ZNF414 | 202.8324517 | 0.054656905 | 0.134503581 | 0.40636022 | 0.684477944 | 0.821187388 |
| ENSG00000133256 | PDE6B | 287.9171221 | -0.993660089 | 0.112270194 | -8.850613425 | 8.70E-19 | 1.36E-17 |
| ENSG00000133265 | HSPBP1 | 968.4274405 | -0.047338442 | 0.073034249 | -0.648167705 | 0.516876487 | 0.69523431 |
| ENSG00000133275 | CSNK1G2 | 2026.00678 | 0.242437178 | 0.057674295 | 4.203556866 | 2.63E-05 | 0.000128273 |
| ENSG00000133302 | SLF1 | 434.5485542 | 0.241363032 | 0.09634452 | 2.50520767 | 0.012237952 | 0.03554305 |
| ENSG00000133313 | CNDP2 | 3065.455243 | 0.264209539 | 0.05018114 | 5.265116342 | 1.40E-07 | 8.97E-07 |
| ENSG00000133315 | MACROD1 | 748.7243302 | 0.275163456 | 0.074139641 | 3.711421505 | 0.000206099 | 0.000872146 |

| | | | | | | | |
|---|---|---|---|---|---|---|---|
| ENSG00000133316 | WDR74 | 1792.284637 | 0.099837249 | 0.058833222 | 1.69695361 | 0.089705431 | 0.193916566 |
| ENSG00000133318 | RTN3 | 4472.969279 | -0.094012028 | 0.048013436 | -1.958035839 | 0.050225807 | 0.120294427 |
| ENSG00000133321 | PLAAT4 | 5.984205552 | 2.236286831 | 0.8295167 | 2.695891271 | 0.007020058 | 0.02175336 |
| ENSG00000133328 | PLAAT2 | 8.77506609 | 0.343068051 | 0.596700583 | 0.574941706 | 0.565330722 | 0.733481521 |
| ENSG00000133392 | MYH11 | 6.321132382 | -0.359482348 | 0.74243896 | -0.484191116 | 0.628250242 | 0.780689734 |
| ENSG00000133393 | CEP20 | 1622.38422 | 0.148065362 | 0.061611039 | 2.403227823 | 0.016251059 | 0.045641443 |
| ENSG00000133398 | MED10 | 932.6787751 | 0.255025497 | 0.07069897 | 3.60720246 | 0.000309516 | 0.001271772 |
| ENSG00000133401 | PDZD2 | 53.59129648 | 0.229300188 | 0.243008242 | 0.943590169 | 0.345379118 | 0.536243449 |
| ENSG00000133422 | MORC2 | 3154.776918 | -0.468457003 | 0.051224909 | -9.145101708 | 5.96E-20 | 9.92E-19 |
| ENSG00000133424 | LARGE1 | 1028.434531 | 0.104055476 | 0.073616508 | 1.413480197 | 0.15751458 | 0.302177286 |
| ENSG00000133433 | GSTT2B | 21.41560056 | -0.775503025 | 0.384485702 | -2.016987944 | 0.043696771 | 0.106809157 |
| ENSG00000133460 | SLC2A11 | 573.098494 | -0.563531594 | 0.24284299 | -2.320559449 | 0.020310633 | 0.055368499 |
| ENSG00000133466 | C1QTNF6 | 600.9426825 | 0.194913172 | 0.091777107 | 2.123766799 | 0.033689649 | 0.085750229 |
| ENSG00000133477 | FAM83F | 6.975730805 | 5.139303889 | 1.319309807 | 3.895448865 | 9.80E-05 | 0.000437539 |
| ENSG00000133519 | ZDHHC8BP | 282.7418915 | -2.230877972 | 0.133396124 | -16.72370903 | 8.81E-63 | 6.71E-61 |
| ENSG00000133597 | ADCK2 | 838.7322817 | -0.155594985 | 0.070212779 | -2.216049366 | 0.026688121 | 0.070265753 |
| ENSG00000133606 | MKRN1 | 4728.204821 | 0.040663753 | 0.043897509 | 0.926333936 | 0.354272465 | 0.545839905 |
| ENSG00000133612 | AGAP3 | 3628.811688 | 0.567231104 | 0.055591239 | 10.20360606 | 1.91E-24 | 4.13E-23 |
| ENSG00000133619 | KRBA1 | 512.9589547 | 0.569940495 | 0.093877492 | 6.071109102 | 1.27E-09 | 1.00E-08 |
| ENSG00000133624 | ZNF767P | 420.541955 | 0.815959004 | 0.114795878 | 7.107912037 | 1.18E-12 | 1.20E-11 |
| ENSG00000133627 | ACTR3B | 247.0598696 | 0.750783392 | 0.124763056 | 6.017673958 | 1.77E-09 | 1.38E-08 |
| ENSG00000133636 | NTS | 8.039609592 | 0.622510326 | 0.654168167 | 0.951605959 | 0.341296858 | 0.532356356 |
| ENSG00000133639 | BTG1 | 1894.704642 | -0.051632538 | 0.055945621 | -0.922905797 | 0.356056299 | 0.54799943 |
| ENSG00000133640 | LRRIQ1 | 4.412154851 | 1.054601993 | 0.857668772 | 1.229614541 | 0.218841482 | 0.388562198 |
| ENSG00000133641 | C12orf29 | 363.574065 | 0.238359812 | 0.101441062 | 2.349736947 | 0.018786683 | 0.051792633 |
| ENSG00000133657 | ATP13A3 | 2472.577397 | -0.241094287 | 0.063336413 | -3.806566792 | 0.000140909 | 0.00061337 |
| ENSG00000133661 | SFTPD | 56.66528276 | -0.416206591 | 0.241394419 | -1.724176525 | 0.084675981 | 0.185137914 |
| ENSG00000133678 | TMEM254 | 1109.209358 | -0.29226935 | 0.064002724 | -4.566514242 | 4.96E-06 | 2.65E-05 |
| ENSG00000133687 | TMTC1 | 1628.318946 | 3.455170316 | 0.077128005 | 44.79786966 | 0 | 0 |
| ENSG00000133703 | KRAS | 949.2683377 | 0.194061365 | 0.078408796 | 2.474994839 | 0.01332381 | 0.038287386 |
| ENSG00000133704 | IPO8 | 1427.783142 | -0.281762677 | 0.065778153 | -4.28352974 | 1.84E-05 | 9.18E-05 |
| ENSG00000133706 | LARS1 | 4558.329291 | 0.222415689 | 0.055378864 | 4.01625588 | 5.91E-05 | 0.000273697 |
| ENSG00000133710 | SPINK5 | 52.60858492 | -0.673417566 | 0.244779435 | -2.751119864 | 0.005939191 | 0.018763949 |

| | | | | | | | |
|---|---|---|---|---|---|---|---|
| ENSG00000133731 | IMPA1 | 187.5872432 | -0.222321456 | 0.135410332 | -1.641835244 | 0.100624153 | 0.21295227 |
| ENSG00000133739 | LRRCC1 | 686.6641877 | 0.148755566 | 0.08177802 | 1.819016487 | 0.068908915 | 0.156651522 |
| ENSG00000133740 | E2F5 | 462.8618443 | 0.005938951 | 0.087993753 | 0.067492872 | 0.946189337 | 0.974743281 |
| ENSG00000133773 | CCDC59 | 847.8823 | -0.146325692 | 0.071337288 | -2.051181026 | 0.040249321 | 0.099624452 |
| ENSG00000133789 | SWAP70 | 567.5532205 | -0.00608523 | 0.092450599 | -0.065821419 | 0.947520003 | 0.975544584 |
| ENSG00000133794 | BMAL1 | 398.0849248 | -0.213523765 | 0.095279385 | -2.241027944 | 0.025024265 | 0.066511023 |
| ENSG00000133805 | AMPD3 | 86.17247742 | 0.604057664 | 0.199678463 | 3.025151807 | 0.002485084 | 0.008566163 |
| ENSG00000133812 | SBF2 | 657.9775629 | 0.434417309 | 0.086091691 | 5.04598415 | 4.51E-07 | 2.74E-06 |
| ENSG00000133816 | MICAL2 | 394.6774503 | 2.102709959 | 0.109537843 | 19.19619652 | 3.98E-82 | 4.64E-80 |
| ENSG00000133818 | RRAS2 | 497.8326095 | -0.089580798 | 0.088719859 | -1.009704028 | 0.312637112 | 0.500805847 |
| ENSG00000133835 | HSD17B4 | 3683.392516 | -0.239195375 | 0.051298251 | -4.662836824 | 3.12E-06 | 1.71E-05 |
| ENSG00000133858 | ZFC3H1 | 1264.269984 | 0.338094983 | 0.066223411 | 5.105369501 | 3.30E-07 | 2.03E-06 |
| ENSG00000133872 | SARAF | 3540.477235 | 0.173479148 | 0.0470058 | 3.690590253 | 0.000223734 | 0.000940701 |
| ENSG00000133874 | RNF122 | 75.43980735 | 0.24344981 | 0.219561848 | 1.108798325 | 0.267517193 | 0.448763759 |
| ENSG00000133884 | DPF2 | 2463.354898 | -0.110367798 | 0.053867508 | -2.048875154 | 0.040474324 | 0.100116563 |
| ENSG00000133895 | MEN1 | 2036.518424 | -0.184105831 | 0.058003695 | -3.174036224 | 0.001503349 | 0.00542587 |
| ENSG00000133935 | ERG28 | 1919.474139 | -0.856576341 | 0.05765184 | -14.85774498 | 6.20E-50 | 3.43E-48 |
| ENSG00000133937 | GSC | 48.94013171 | -0.807222632 | 0.268172088 | -3.010091912 | 0.002611686 | 0.008960827 |
| ENSG00000133943 | DGLUCY | 185.2882634 | -0.577177858 | 0.133503247 | -4.323324504 | 1.54E-05 | 7.74E-05 |
| ENSG00000133958 | UNC79 | 14.81135124 | -0.20837846 | 0.458557497 | -0.454421662 | 0.649525364 | 0.796021186 |
| ENSG00000133961 | NUMB | 2317.550474 | -0.784976907 | 0.055821354 | -14.06230492 | 6.47E-45 | 3.11E-43 |
| ENSG00000133962 | CATSPERB | 23.04936733 | -0.321732855 | 0.374630218 | -0.858801132 | 0.390450243 | 0.583343425 |
| ENSG00000133983 | COX16 | 73.83400801 | 0.300063782 | 0.205178945 | 1.462449191 | 0.143618158 | 0.281975398 |
| ENSG00000133985 | TTC9 | 729.8358029 | -1.191004396 | 0.083377042 | -14.28456049 | 2.73E-46 | 1.36E-44 |
| ENSG00000133997 | MED6 | 680.0953855 | -0.726848284 | 0.078051391 | -9.312432158 | 1.25E-20 | 2.16E-19 |
| ENSG00000134001 | EIF2S1 | 4490.65637 | -0.519802857 | 0.054718539 | -9.499574902 | 2.11E-21 | 3.83E-20 |
| ENSG00000134013 | LOXL2 | 25.55831225 | 0.597400055 | 0.350877621 | 1.702588082 | 0.088645182 | 0.192095265 |
| ENSG00000134014 | ELP3 | 764.9638138 | -0.17422885 | 0.072518168 | -2.402554483 | 0.016281009 | 0.045714363 |
| ENSG00000134020 | PEBP4 | 2.898631299 | -0.182811191 | 1.029488209 | -0.177574828 | 0.859056895 | 0.926792302 |
| ENSG00000134030 | CTIF | 1015.739514 | 1.067633629 | 0.074410939 | 14.34780483 | 1.10E-46 | 5.54E-45 |
| ENSG00000134046 | MBD2 | 1870.390847 | 0.424834275 | 0.056030171 | 7.582241313 | 3.40E-14 | 3.84E-13 |
| ENSG00000134049 | IER3IP1 | 282.5447105 | 0.640592021 | 0.24785523 | 2.584541071 | 0.009750869 | 0.029072646 |
| ENSG00000134056 | MRPS36 | 424.3489451 | -0.145129306 | 0.100500087 | -1.444071436 | 0.148718882 | 0.289610454 |

| | | | | | | | |
|---|---|---|---|---|---|---|---|
| ENSG00000134057 | CCNB1 | 3608.066241 | -0.250875381 | 0.052796779 | -4.751717583 | 2.02E-06 | 1.13E-05 |
| ENSG00000134058 | CDK7 | 738.187691 | 0.014992193 | 0.075240621 | 0.199256637 | 0.842061998 | 0.917009895 |
| ENSG00000134070 | IRAK2 | 5.391221654 | 0.868330669 | 0.794366698 | 1.093110614 | 0.274345236 | 0.456373753 |
| ENSG00000134072 | CAMK1 | 170.1979058 | 0.049819007 | 0.143162452 | 0.347989342 | 0.727848188 | 0.848977344 |
| ENSG00000134077 | THUMPD3 | 2555.383189 | -0.135393893 | 0.052058938 | -2.600780923 | 0.009301183 | 0.027931554 |
| ENSG00000134086 | VHL | 4368.833942 | 0.031655651 | 0.046675843 | 0.678202018 | 0.497643613 | 0.679520506 |
| ENSG00000134107 | BHLHE40 | 3395.879093 | 0.056276696 | 0.054183714 | 1.038627508 | 0.298978007 | 0.4856646 |
| ENSG00000134108 | ARL8B | 2734.742234 | 0.151321444 | 0.056931031 | 2.657978274 | 0.007861096 | 0.024079789 |
| ENSG00000134109 | EDEM1 | 1788.033888 | 0.312063417 | 0.058177837 | 5.363957057 | 8.14E-08 | 5.31E-07 |
| ENSG00000134115 | CNTN6 | 3.170518478 | 0.717763294 | 0.993685544 | 0.722324379 | 0.470095067 | 0.655959905 |
| ENSG00000134138 | MEIS2 | 319.5518547 | -0.023864571 | 0.104758739 | -0.227805065 | 0.819797789 | 0.904443968 |
| ENSG00000134146 | DPH6 | 136.9243145 | -0.125201339 | 0.152252495 | -0.822327012 | 0.410890807 | 0.603369294 |
| ENSG00000134152 | KATNBL1 | 438.197542 | -0.29403865 | 0.10104156 | -2.910076318 | 0.003613405 | 0.012027207 |
| ENSG00000134153 | EMC7 | 1241.281897 | -0.193757103 | 0.062775326 | -3.086516884 | 0.002025164 | 0.007110081 |
| ENSG00000134183 | GNAT2 | 6.024235382 | -0.084454037 | 0.713795576 | -0.11831684 | 0.905816615 | 0.95262805 |
| ENSG00000134184 | GSTM1 | 10.09738681 | -0.415885588 | 0.544590031 | -0.763667282 | 0.445065543 | 0.633676984 |
| ENSG00000134186 | PRPF38B | 2265.311283 | -0.049769518 | 0.067324666 | -0.739246411 | 0.459757384 | 0.646687987 |
| ENSG00000134201 | GSTM5 | 10.56798184 | -1.073541045 | 0.568777172 | -1.887454521 | 0.059099218 | 0.137736062 |
| ENSG00000134202 | GSTM3 | 6397.02339 | -0.297405946 | 0.042843374 | -6.941702226 | 3.87E-12 | 3.77E-11 |
| ENSG00000134215 | VAV3 | 451.517414 | 1.714984669 | 0.098625687 | 17.38882351 | 1.00E-67 | 8.58E-66 |
| ENSG00000134222 | PSRC1 | 740.6160389 | -0.461554312 | 0.0750758 | -6.14784411 | 7.85E-10 | 6.30E-09 |
| ENSG00000134240 | HMGCS2 | 1.606932243 | -1.366263942 | 1.463639558 | -0.933470221 | 0.350577239 | NA |
| ENSG00000134242 | PTPN22 | 17.75190559 | -0.010800985 | 0.42232596 | -0.025574997 | 0.979596329 | 0.990826269 |
| ENSG00000134243 | SORT1 | 4957.94186 | 0.129775534 | 0.049169969 | 2.6393251 | 0.008307128 | 0.025247152 |
| ENSG00000134245 | WNT2B | 780.2583763 | -0.748079001 | 0.080486125 | -9.294508821 | 1.48E-20 | 2.55E-19 |
| ENSG00000134247 | PTGFRN | 4358.031679 | 0.453077642 | 0.056386984 | 8.035145859 | 9.35E-16 | 1.19E-14 |
| ENSG00000134248 | LAMTOR5 | 1257.075149 | -0.088338275 | 0.062196557 | -1.420308113 | 0.155518 | 0.299394338 |
| ENSG00000134250 | NOTCH2 | 7361.19204 | -0.214950167 | 0.059257482 | -3.627392851 | 0.000286298 | 0.001181447 |
| ENSG00000134253 | TRIM45 | 1090.228338 | -0.567168109 | 0.077170044 | -7.349589061 | 1.99E-13 | 2.13E-12 |
| ENSG00000134255 | CEPT1 | 696.2039161 | 0.04441193 | 0.0782497 | 0.567566785 | 0.570329166 | 0.737339443 |
| ENSG00000134256 | CD101 | 92.08848562 | 0.781414197 | 0.187594444 | 4.165444245 | 3.11E-05 | 0.000150264 |
| ENSG00000134258 | VTCN1 | 385.7440027 | -1.131399311 | 0.109238529 | -10.35714524 | 3.88E-25 | 8.68E-24 |
| ENSG00000134262 | AP4B1 | 2593.055927 | 0.135844248 | 0.066665027 | 2.037713851 | 0.041578558 | 0.10237331 |

| | | | | | | | |
|---|---|---|---|---|---|---|---|
| ENSG00000134265 | NAPG | 1296.745472 | 0.313969193 | 0.063921638 | 4.91178269 | 9.03E-07 | 5.26E-06 |
| ENSG00000134278 | SPIRE1 | 1903.968239 | 0.312253081 | 0.059615075 | 5.237820859 | 1.62E-07 | 1.03E-06 |
| ENSG00000134283 | PPHLN1 | 1574.725417 | -0.041093318 | 0.069146689 | -0.594291914 | 0.552316886 | 0.724008535 |
| ENSG00000134285 | FKBP11 | 245.9334991 | 0.206555 | 0.12381846 | 1.668208433 | 0.095274357 | 0.203857479 |
| ENSG00000134287 | ARF3 | 7511.667958 | 0.173904645 | 0.04254302 | 4.08773628 | 4.36E-05 | 0.000205358 |
| ENSG00000134291 | TMEM106C | 6538.221247 | -0.773491102 | 0.045482295 | -17.00642184 | 7.36E-65 | 5.92E-63 |
| ENSG00000134294 | SLC38A2 | 20009.80518 | -0.361007531 | 0.053837032 | -6.705561574 | 2.01E-11 | 1.83E-10 |
| ENSG00000134297 | PLEKHA8P1 | 81.94980363 | -0.260826504 | 0.199878969 | -1.304922204 | 0.19191934 | 0.351928829 |
| ENSG00000134308 | YWHAQ | 7432.278208 | -0.053664544 | 0.049403783 | -1.086243618 | 0.277371224 | 0.460157474 |
| ENSG00000134313 | KIDINS220 | 2823.823119 | 0.04903835 | 0.059560326 | 0.823339182 | 0.410315138 | 0.602871278 |
| ENSG00000134317 | GRHL1 | 969.4747387 | 0.464043097 | 0.066586114 | 6.96906709 | 3.19E-12 | 3.14E-11 |
| ENSG00000134318 | ROCK2 | 2177.428612 | -0.150214026 | 0.057910745 | -2.593888672 | 0.009489721 | 0.028404775 |
| ENSG00000134321 | RSAD2 | 4.952545064 | 1.876781815 | 0.877744161 | 2.138187752 | 0.032501508 | 0.083259028 |
| ENSG00000134323 | MYCN | 34.587929 | -0.16762865 | 0.298331449 | -0.561887293 | 0.574192812 | 0.740748122 |
| ENSG00000134324 | LPIN1 | 3841.125424 | -0.437522507 | 0.056251533 | -7.77796588 | 7.37E-15 | 8.73E-14 |
| ENSG00000134326 | CMPK2 | 13.66430517 | 0.257624447 | 0.48308625 | 0.533288718 | 0.593833736 | 0.755758441 |
| ENSG00000134330 | IAH1 | 1065.046351 | 0.038908902 | 0.064986711 | 0.598720902 | 0.549359016 | 0.721787983 |
| ENSG00000134333 | LDHA | 8797.464195 | -0.22836853 | 0.052555295 | -4.345300134 | 1.39E-05 | 7.04E-05 |
| ENSG00000134352 | IL6ST | 1309.445564 | 0.183166188 | 0.06708925 | 2.730186847 | 0.006329844 | 0.019874982 |
| ENSG00000134363 | FST | 47.13156579 | -0.600487679 | 0.258828439 | -2.320022023 | 0.020339686 | 0.055441112 |
| ENSG00000134369 | NAV1 | 216.9216107 | 1.006514339 | 0.135872017 | 7.407811844 | 1.28E-13 | 1.39E-12 |
| ENSG00000134371 | CDC73 | 1918.109137 | -0.325170343 | 0.055635751 | -5.844629326 | 5.08E-09 | 3.78E-08 |
| ENSG00000134375 | TIMM17A | 1961.30794 | 0.005761063 | 0.057427543 | 0.100318812 | 0.920091223 | 0.961195531 |
| ENSG00000134376 | CRB1 | 1.898191183 | 0.404976064 | 1.315812841 | 0.307776343 | 0.758252524 | 0.866725443 |
| ENSG00000134389 | CFHR5 | 3.245726423 | -0.635680523 | 0.999729185 | -0.635852722 | 0.52487242 | 0.701099828 |
| ENSG00000134419 | RPS15A | 1427.926267 | 0.218855311 | 0.062431456 | 3.505529489 | 0.0004557 | 0.001816465 |
| ENSG00000134440 | NARS1 | 4663.424959 | 0.68763172 | 0.046686423 | 14.72873005 | 4.22E-49 | 2.27E-47 |
| ENSG00000134444 | RELCH | 1157.404323 | 0.513290253 | 0.065291564 | 7.861509588 | 3.80E-15 | 4.61E-14 |
| ENSG00000134452 | FBH1 | 2912.062124 | -1.065586474 | 0.057993004 | -18.37439707 | 2.11E-75 | 2.26E-73 |
| ENSG00000134453 | RBM17 | 3063.736464 | -1.251175274 | 0.058750863 | -21.29628776 | 1.23E-100 | 2.00E-98 |
| ENSG00000134461 | ANKRD16 | 258.1516252 | -0.901998438 | 0.123423396 | -7.308164124 | 2.71E-13 | 2.87E-12 |
| ENSG00000134463 | ECHDC3 | 53.76956261 | -0.234875793 | 0.240786577 | -0.975452189 | 0.329336007 | 0.51977764 |
| ENSG00000134470 | IL15RA | 2.745456069 | -2.980185175 | 1.340184599 | -2.223712448 | 0.026167797 | 0.069022567 |

| | | | | | | | |
|---|---|---|---|---|---|---|---|
| ENSG00000134480 | CCNH | 717.7681317 | -0.063858363 | 0.076105887 | -0.839072589 | 0.401428577 | 0.594190964 |
| ENSG00000134490 | TMEM241 | 570.9641226 | -0.232826523 | 0.082407673 | -2.825301516 | 0.004723614 | 0.015313696 |
| ENSG00000134504 | KCTD1 | 368.202143 | 0.285179061 | 0.100006913 | 2.851593467 | 0.00435007 | 0.014223289 |
| ENSG00000134508 | CABLES1 | 2103.538291 | -0.473694663 | 0.055425748 | -8.546473094 | 1.27E-17 | 1.82E-16 |
| ENSG00000134531 | EMP1 | 22.15467019 | -0.467890788 | 0.384571733 | -1.216654131 | 0.223735838 | 0.395135581 |
| ENSG00000134532 | SOX5 | 50.26971787 | -1.194276737 | 0.264378408 | -4.51730058 | 6.26E-06 | 3.31E-05 |
| ENSG00000134533 | RERG | 296.4773381 | 0.76314058 | 0.115574393 | 6.603024796 | 4.03E-11 | 3.59E-10 |
| ENSG00000134569 | LRP4 | 954.3969092 | 0.019455038 | 0.074262326 | 0.26197723 | 0.793338997 | 0.88906676 |
| ENSG00000134574 | DDB2 | 1286.797434 | -0.74076184 | 0.066391854 | -11.15742059 | 6.59E-29 | 1.75E-27 |
| ENSG00000134575 | ACP2 | 1003.265167 | -0.542560237 | 0.071137351 | -7.626939021 | 2.40E-14 | 2.75E-13 |
| ENSG00000134590 | RTL8C | 2778.464262 | -0.067159376 | 0.057846225 | -1.160998435 | 0.245642536 | 0.422701956 |
| ENSG00000134594 | RAB33A | 9.006700768 | 0.094924795 | 0.591857445 | 0.160384558 | 0.872578153 | 0.934146135 |
| ENSG00000134595 | SOX3 | 70.93461694 | -1.131088931 | 0.217839167 | -5.192312043 | 2.08E-07 | 1.31E-06 |
| ENSG00000134597 | RBMX2 | 1418.994796 | 0.032721024 | 0.06037462 | 0.541966541 | 0.587841555 | 0.751216382 |
| ENSG00000134602 | STK26 | 1782.185144 | 0.252710647 | 0.064561041 | 3.914290121 | 9.07E-05 | 0.000406962 |
| ENSG00000134612 | FOLH1B | 4.66261291 | -2.700975806 | 0.979422331 | -2.757723325 | 0.005820544 | 0.018424628 |
| ENSG00000134627 | PIWIL4 | 10.24368013 | 0.256047524 | 0.56183822 | 0.455731765 | 0.648582877 | 0.795130996 |
| ENSG00000134644 | PUM1 | 3473.441676 | 0.102773103 | 0.048941815 | 2.099903786 | 0.035737306 | 0.090208384 |
| ENSG00000134668 | SPOCD1 | 105.4727178 | -1.213904114 | 0.183237306 | -6.624765119 | 3.48E-11 | 3.11E-10 |
| ENSG00000134684 | YARS1 | 5087.689108 | 0.135586973 | 0.047403438 | 2.860277191 | 0.004232709 | 0.013871191 |
| ENSG00000134686 | PHC2 | 1910.410836 | 0.149244235 | 0.063439896 | 2.352529637 | 0.018646204 | 0.051442443 |
| ENSG00000134690 | CDCA8 | 1838.839212 | -0.1586612 | 0.055466625 | -2.860480513 | 0.004229995 | 0.01386428 |
| ENSG00000134697 | GNL2 | 1950.38573 | 0.09422163 | 0.057637571 | 1.634725899 | 0.102106521 | 0.215624007 |
| ENSG00000134698 | AGO4 | 683.8465277 | -0.080761449 | 0.082541089 | -0.978439339 | 0.32785708 | 0.518156487 |
| ENSG00000134709 | HOOK1 | 1620.559275 | 0.036123043 | 0.064855935 | 0.556973587 | 0.577545475 | 0.743451272 |
| ENSG00000134716 | CYP2J2 | 433.5691005 | -0.869075288 | 0.092909247 | -9.354023603 | 8.44E-21 | 1.48E-19 |
| ENSG00000134717 | BTF3L4 | 1605.213141 | 0.203772233 | 0.063022552 | 3.233322473 | 0.001223594 | 0.00450839 |
| ENSG00000134744 | TUT4 | 423.9793845 | -0.33915758 | 0.094677668 | -3.582234204 | 0.000340668 | 0.001386609 |
| ENSG00000134748 | PRPF38A | 2069.962482 | -0.003179376 | 0.052797598 | -0.060218184 | 0.951981863 | 0.977646532 |
| ENSG00000134755 | DSC2 | 389.2195934 | 0.048053891 | 0.100095768 | 0.480079143 | 0.631171119 | 0.782652187 |
| ENSG00000134758 | RNF138 | 352.636422 | 0.709263065 | 0.100996584 | 7.022644092 | 2.18E-12 | 2.17E-11 |
| ENSG00000134759 | ELP2 | 3357.399335 | 0.320411411 | 0.058305269 | 5.495410934 | 3.90E-08 | 2.63E-07 |
| ENSG00000134765 | DSC1 | 7.327987665 | 0.572465459 | 0.691418051 | 0.82795851 | 0.407693997 | 0.600443008 |

| | | | | | | | |
|---|---|---|---|---|---|---|---|
| ENSG00000134769 | DTNA | 2229.649413 | 0.794647296 | 0.062558096 | 12.70254927 | 5.72E-37 | 2.08E-35 |
| ENSG00000134775 | FHOD3 | 20.69946506 | 0.508876936 | 0.394193034 | 1.290933357 | 0.196726788 | 0.358509096 |
| ENSG00000134779 | TPGS2 | 3426.249297 | 0.644194837 | 0.050624015 | 12.72508397 | 4.29E-37 | 1.57E-35 |
| ENSG00000134780 | DAGLA | 330.7355503 | 0.98970818 | 0.108479482 | 9.123459658 | 7.28E-20 | 1.20E-18 |
| ENSG00000134802 | SLC43A3 | 2.432612015 | 2.428240285 | 1.390933254 | 1.745763341 | 0.080852083 | 0.178221081 |
| ENSG00000134809 | TIMM10 | 768.6577156 | -0.156075982 | 0.079094128 | -1.973294172 | 0.048462057 | 0.116679217 |
| ENSG00000134815 | DHX34 | 1148.106364 | -0.070387818 | 0.070623686 | -0.99666022 | 0.318929465 | 0.508248831 |
| ENSG00000134824 | FADS2 | 899.8458939 | 0.141409501 | 0.075370453 | 1.876192793 | 0.060628811 | 0.140771754 |
| ENSG00000134825 | TMEM258 | 777.8178982 | 0.03809592 | 0.073425608 | 0.518836974 | 0.603874432 | 0.762847209 |
| ENSG00000134827 | TCN1 | 15.90645529 | 4.848182694 | 0.906274506 | 5.349574176 | 8.82E-08 | 5.74E-07 |
| ENSG00000134830 | C5AR2 | 107.134166 | 1.225067567 | 0.188798775 | 6.488747437 | 8.66E-11 | 7.54E-10 |
| ENSG00000134851 | TMEM165 | 1259.676552 | -0.149526854 | 0.061823798 | -2.418597034 | 0.015580489 | 0.043984301 |
| ENSG00000134852 | CLOCK | 1961.294298 | -0.115642651 | 0.063553451 | -1.819612472 | 0.068818041 | 0.156460442 |
| ENSG00000134864 | GGACT | 71.76843309 | 0.111113996 | 0.208992825 | 0.531664168 | 0.594958614 | 0.756476588 |
| ENSG00000134871 | COL4A2 | 1.713406655 | 3.025186661 | 1.605290236 | 1.884510721 | 0.059495926 | NA |
| ENSG00000134882 | UBAC2 | 1660.32271 | 0.168509392 | 0.05752231 | 2.929461489 | 0.003395499 | 0.011377847 |
| ENSG00000134884 | ARGLU1 | 2037.615373 | 0.207805721 | 0.141820203 | 1.465275865 | 0.142845668 | 0.280698958 |
| ENSG00000134897 | BIVM | 217.064909 | 0.119535234 | 0.128295967 | 0.931714667 | 0.351484004 | 0.542783108 |
| ENSG00000134899 | ERCC5 | 44.69049174 | -0.317740411 | 0.27194009 | -1.168420631 | 0.242637137 | 0.418944447 |
| ENSG00000134900 | TPP2 | 1222.377234 | -0.142906873 | 0.069298617 | -2.062189411 | 0.039189706 | 0.097527866 |
| ENSG00000134901 | POGLUT2 | 142.7853894 | -0.471327096 | 0.155634226 | -3.02842831 | 0.002458294 | 0.008487848 |
| ENSG00000134905 | CARS2 | 720.2463331 | -0.195821855 | 0.074158527 | -2.640584485 | 0.008276315 | 0.025180196 |
| ENSG00000134909 | ARHGAP32 | 2052.889716 | 0.030912459 | 0.054093875 | 0.571459505 | 0.567688203 | 0.735500247 |
| ENSG00000134910 | STT3A | 3227.077558 | -0.272926329 | 0.050327296 | -5.423027826 | 5.86E-08 | 3.88E-07 |
| ENSG00000134940 | ACRV1 | 2.469917236 | -1.589989023 | 1.217436073 | -1.306014385 | 0.191547666 | 0.351528276 |
| ENSG00000134955 | SLC37A2 | 15.81388845 | 0.28102628 | 0.445363705 | 0.631004001 | 0.528037909 | 0.704009626 |
| ENSG00000134962 | KLB | 40.39048311 | -0.065401765 | 0.276973266 | -0.236130246 | 0.813331605 | 0.901448243 |
| ENSG00000134970 | TMED7 | 781.3094078 | 0.079494595 | 0.095045032 | 0.836388742 | 0.402936244 | 0.595590718 |
| ENSG00000134982 | APC | 1843.711085 | 0.488009284 | 0.05990262 | 8.146710142 | 3.74E-16 | 4.88E-15 |
| ENSG00000134986 | NREP | 679.67713 | 0.346663035 | 0.080952117 | 4.282322047 | 1.85E-05 | 9.22E-05 |
| ENSG00000134987 | WDR36 | 1117.914531 | -0.142909513 | 0.065292954 | -2.188743271 | 0.028615505 | 0.074561527 |
| ENSG00000134996 | OSTF1 | 705.7998318 | -0.048776374 | 0.076546509 | -0.637212266 | 0.523986589 | 0.700317662 |
| ENSG00000135002 | RFK | 386.7088164 | 0.116877963 | 0.100999251 | 1.157216138 | 0.247184066 | 0.424399556 |

| | | | | | | | |
|---|---|---|---|---|---|---|---|
| ENSG00000135018 | UBQLN1 | 4830.144474 | -0.026530398 | 0.050198005 | -0.528514978 | 0.597141954 | 0.75769879 |
| ENSG00000135040 | NAA35 | 1339.588913 | 0.021544422 | 0.062684608 | 0.343695566 | 0.731075246 | 0.851119615 |
| ENSG00000135045 | C9orf40 | 218.0123104 | -0.050476649 | 0.126453132 | -0.399172789 | 0.689765892 | 0.824651044 |
| ENSG00000135046 | ANXA1 | 8.662375465 | 0.509640297 | 0.614513656 | 0.829339254 | 0.406912465 | 0.599725179 |
| ENSG00000135047 | CTSL | 1167.08546 | -0.301973543 | 0.067102806 | -4.500162646 | 6.79E-06 | 3.58E-05 |
| ENSG00000135048 | CEMIP2 | 869.4548705 | 0.005762426 | 0.076192553 | 0.075629777 | 0.939713646 | 0.971233034 |
| ENSG00000135049 | AGTPBP1 | 525.4821003 | -0.700476192 | 0.087108468 | -8.041424754 | 8.88E-16 | 1.13E-14 |
| ENSG00000135052 | GOLM1 | 584.4412513 | 0.238465712 | 0.08704181 | 2.739668579 | 0.006150117 | 0.019379626 |
| ENSG00000135063 | ENTREP1 | 473.2068472 | 1.051413494 | 0.100050268 | 10.50885233 | 7.86E-26 | 1.82E-24 |
| ENSG00000135070 | ISCA1 | 895.6698925 | 0.391725022 | 0.069804882 | 5.611713854 | 2.00E-08 | 1.39E-07 |
| ENSG00000135074 | ADAM19 | 18.37030132 | 0.155341106 | 0.411192838 | 0.377781644 | 0.705592811 | 0.83533566 |
| ENSG00000135083 | CCNJL | 234.3206524 | 1.014331631 | 0.127369598 | 7.963687129 | 1.67E-15 | 2.08E-14 |
| ENSG00000135090 | TAOK3 | 1650.30186 | 0.2863718 | 0.060630212 | 4.723252513 | 2.32E-06 | 1.29E-05 |
| ENSG00000135093 | USP30 | 588.3392662 | -0.159330099 | 0.083910305 | -1.898814437 | 0.057588878 | 0.1348738 |
| ENSG00000135097 | MSI1 | 175.9425889 | -1.470269494 | 0.142649362 | -10.30687744 | 6.56E-25 | 1.45E-23 |
| ENSG00000135100 | HNF1A | 2.676462356 | 1.117695549 | 1.109301608 | 1.00756687 | 0.313662441 | 0.501980571 |
| ENSG00000135108 | FBXO21 | 2909.74142 | 0.030062135 | 0.053215583 | 0.56491225 | 0.572133451 | 0.738964099 |
| ENSG00000135111 | TBX3 | 571.2616349 | 0.239373371 | 0.094566266 | 2.531276542 | 0.011364821 | 0.033262182 |
| ENSG00000135114 | OASL | 52.39999049 | 3.043073313 | 0.326184023 | 9.329314437 | 1.07E-20 | 1.85E-19 |
| ENSG00000135116 | HRK | 65.60174626 | 1.300510038 | 0.2378142 | 5.468597072 | 4.54E-08 | 3.04E-07 |
| ENSG00000135119 | RNFT2 | 187.9018963 | 0.370700997 | 0.136072176 | 2.724296829 | 0.006443856 | 0.020169473 |
| ENSG00000135124 | P2RX4 | 945.2936976 | 0.119927857 | 0.070908898 | 1.691294893 | 0.090780495 | 0.195852963 |
| ENSG00000135127 | BICDL1 | 950.6681389 | -0.137559876 | 0.067204519 | -2.046884317 | 0.040669444 | 0.100523333 |
| ENSG00000135148 | TRAFD1 | 2301.577143 | -0.339344208 | 0.056029209 | -6.056558997 | 1.39E-09 | 1.09E-08 |
| ENSG00000135164 | DMTF1 | 1528.063093 | -0.551916972 | 0.078606298 | -7.02128179 | 2.20E-12 | 2.19E-11 |
| ENSG00000135185 | TMEM243 | 389.5304154 | -0.443668506 | 0.098844425 | -4.488553649 | 7.17E-06 | 3.77E-05 |
| ENSG00000135205 | CCDC146 | 107.3088585 | 1.584321995 | 0.193203202 | 8.200288496 | 2.40E-16 | 3.17E-15 |
| ENSG00000135211 | TMEM60 | 263.3676351 | 0.207574381 | 0.113776331 | 1.824407403 | 0.068090504 | 0.155144632 |
| ENSG00000135241 | PNPLA8 | 877.0856558 | -0.019236773 | 0.070514484 | -0.272805981 | 0.785002362 | 0.883106023 |
| ENSG00000135245 | HILPDA | 1206.126542 | -0.694034389 | 0.068022763 | -10.20297266 | 1.92E-24 | 4.15E-23 |
| ENSG00000135249 | RINT1 | 726.7022291 | -0.240631216 | 0.077347052 | -3.111058683 | 0.001864179 | 0.006595352 |
| ENSG00000135250 | SRPK2 | 2970.211582 | -0.396964344 | 0.048602516 | -8.167567715 | 3.15E-16 | 4.13E-15 |
| ENSG00000135253 | KCP | 31.59515574 | -0.035818786 | 0.317185717 | -0.112926855 | 0.910088545 | 0.954920698 |

| | | | | | | | |
|---|---|---|---|---|---|---|---|
| ENSG00000135269 | TES | 1398.102617 | 0.078538993 | 0.060122316 | 1.306320154 | 0.191443706 | 0.351421831 |
| ENSG00000135297 | MTO1 | 1077.00841 | -0.273287761 | 0.067539992 | -4.046310238 | 5.20E-05 | 0.000242798 |
| ENSG00000135298 | ADGRB3 | 4.068493541 | -1.141490738 | 0.917170958 | -1.244577936 | 0.213286935 | 0.381505052 |
| ENSG00000135299 | ANKRD6 | 76.63164077 | -1.164270982 | 0.213268982 | -5.459166973 | 4.78E-08 | 3.20E-07 |
| ENSG00000135312 | HTR1B | 2.437255304 | -2.052493656 | 1.25624802 | -1.633828371 | 0.102294893 | 0.215962162 |
| ENSG00000135314 | KHDC1 | 187.7153391 | -0.063698549 | 0.148261224 | -0.429637282 | 0.667459513 | 0.808691146 |
| ENSG00000135315 | CEP162 | 200.6192853 | 0.131056707 | 0.128919656 | 1.016576616 | 0.309354883 | 0.497378821 |
| ENSG00000135316 | SYNCRIP | 7892.177242 | -0.0250507 | 0.051893412 | -0.482733724 | 0.629284815 | 0.7810577 |
| ENSG00000135317 | SNX14 | 1694.36343 | -0.24374581 | 0.063252672 | -3.853525927 | 0.000116429 | 0.000513828 |
| ENSG00000135318 | NT5E | 6.429435852 | -0.24055511 | 0.687955035 | -0.349666909 | 0.726588691 | 0.848025671 |
| ENSG00000135333 | EPHA7 | 837.3582839 | 0.987861121 | 0.074719208 | 13.22097953 | 6.64E-40 | 2.70E-38 |
| ENSG00000135334 | AKIRIN2 | 1446.875229 | -0.10444788 | 0.059396241 | -1.758493096 | 0.078663645 | 0.174318395 |
| ENSG00000135336 | ORC3 | 984.5557519 | -0.143131356 | 0.068003763 | -2.104756416 | 0.035312517 | 0.089313025 |
| ENSG00000135338 | LCA5 | 58.72880885 | -0.007679706 | 0.229827103 | -0.033415146 | 0.973343532 | 0.987970845 |
| ENSG00000135341 | MAP3K7 | 1644.862 | 0.07850425 | 0.057533309 | 1.364500878 | 0.172409978 | 0.324392851 |
| ENSG00000135346 | CGA | 4.863972944 | -0.35302083 | 0.790419329 | -0.446624744 | 0.655146027 | 0.799808359 |
| ENSG00000135362 | PRR5L | 98.25086741 | 0.414112795 | 0.184322618 | 2.246673792 | 0.024660882 | 0.065705069 |
| ENSG00000135363 | LMO2 | 30.34436754 | 0.962248509 | 0.335062782 | 2.871845403 | 0.004080826 | 0.013419458 |
| ENSG00000135365 | PHF21A | 873.0998363 | -0.431682584 | 0.068937425 | -6.261948155 | 3.80E-10 | 3.15E-09 |
| ENSG00000135372 | NAT10 | 2838.61396 | -0.062175681 | 0.052223761 | -1.190563064 | 0.233825159 | 0.4083707 |
| ENSG00000135373 | EHF | 130.2742895 | 1.887234426 | 0.174254 | 10.83036504 | 2.47E-27 | 6.24E-26 |
| ENSG00000135374 | ELF5 | 207.7474407 | 0.621289894 | 0.126932229 | 4.894658343 | 9.85E-07 | 5.70E-06 |
| ENSG00000135378 | PRRG4 | 253.530984 | 0.373346505 | 0.115683613 | 3.22730675 | 0.001249614 | 0.004593934 |
| ENSG00000135387 | CAPRIN1 | 11448.42922 | 0.149756682 | 0.044226047 | 3.386164766 | 0.000708768 | 0.002728502 |
| ENSG00000135390 | ATP5MC2 | 5433.414724 | 0.221171263 | 0.046513021 | 4.755039705 | 1.98E-06 | 1.11E-05 |
| ENSG00000135392 | DNAJC14 | 1556.61431 | -0.054051442 | 0.061557681 | -0.878061693 | 0.37991024 | 0.572385453 |
| ENSG00000135404 | CD63 | 6172.101259 | 0.181257548 | 0.043960477 | 4.123193347 | 3.74E-05 | 0.000177762 |
| ENSG00000135406 | PRPH | 134.9947617 | -0.253252031 | 0.155087174 | -1.632965666 | 0.102476218 | 0.216245464 |
| ENSG00000135407 | AVIL | 160.7533017 | 0.771630479 | 0.156516077 | 4.930039744 | 8.22E-07 | 4.82E-06 |
| ENSG00000135414 | GDF11 | 1666.261282 | 0.90912438 | 0.059261976 | 15.3407707 | 4.08E-53 | 2.43E-51 |
| ENSG00000135423 | GLS2 | 91.45457332 | 0.738136782 | 0.193091945 | 3.822721767 | 0.000131987 | 0.000576719 |
| ENSG00000135424 | ITGA7 | 69.16493094 | -0.853485056 | 0.217148161 | -3.930427283 | 8.48E-05 | 0.000382988 |
| ENSG00000135436 | FAM186B | 36.70246981 | 0.148031525 | 0.287510813 | 0.514872894 | 0.606641843 | 0.7650978 |

| | | | | | | | |
|---|---|---|---|---|---|---|---|
| ENSG00000135437 | RDH5 | 23.73710116 | -0.054324085 | 0.362170408 | -0.149995923 | 0.880767832 | 0.938252557 |
| ENSG00000135439 | AGAP2 | 296.5524581 | -0.173503117 | 0.108879942 | -1.593526901 | 0.111042038 | 0.230512312 |
| ENSG00000135441 | BLOC1S1 | 130.6046634 | 0.19286025 | 0.156979895 | 1.228566563 | 0.219234358 | 0.389109182 |
| ENSG00000135446 | CDK4 | 7220.847184 | 0.028411491 | 0.042675813 | 0.665751605 | 0.505569883 | 0.686015208 |
| ENSG00000135451 | TROAP | 1812.412889 | -0.344240034 | 0.068248003 | -5.043957603 | 4.56E-07 | 2.76E-06 |
| ENSG00000135452 | TSPAN31 | 1058.625242 | -0.039963873 | 0.066349559 | -0.602323112 | 0.546959079 | 0.719888603 |
| ENSG00000135454 | B4GALNT1 | 945.2562912 | -0.541541929 | 0.069373405 | -7.806189311 | 5.89E-15 | 7.05E-14 |
| ENSG00000135469 | COQ10A | 256.7471239 | 0.098930603 | 0.116707737 | 0.847678188 | 0.396617217 | 0.589541823 |
| ENSG00000135473 | PAN2 | 2224.016217 | -0.510981843 | 0.064185548 | -7.96101087 | 1.71E-15 | 2.12E-14 |
| ENSG00000135476 | ESPL1 | 2663.807144 | -0.477838859 | 0.052514667 | -9.099150592 | 9.10E-20 | 1.50E-18 |
| ENSG00000135477 | KRT87P | 1.777259216 | 1.218264088 | 1.374806206 | 0.886135139 | 0.37554471 | NA |
| ENSG00000135480 | KRT7 | 159.9842273 | -0.226189179 | 0.162176298 | -1.394711691 | 0.163102798 | 0.310540451 |
| ENSG00000135482 | ZC3H10 | 565.3446718 | -0.2312219 | 0.084472982 | -2.737229048 | 0.006195914 | 0.01949983 |
| ENSG00000135486 | HNRNPA1 | 7029.412793 | 0.339357069 | 0.066093855 | 5.134472355 | 2.83E-07 | 1.75E-06 |
| ENSG00000135502 | SLC26A10P | 10.03120265 | -0.490483459 | 0.590247225 | -0.830979694 | 0.405985101 | 0.598679911 |
| ENSG00000135503 | ACVR1B | 1465.488715 | 0.047051844 | 0.063552046 | 0.740367097 | 0.459077278 | 0.646246487 |
| ENSG00000135506 | OS9 | 7336.118694 | 0.024619582 | 0.045808401 | 0.537446883 | 0.590958966 | 0.753563018 |
| ENSG00000135517 | MIP | 2.768759285 | -0.608474818 | 1.067669916 | -0.569909116 | 0.568739342 | 0.736073456 |
| ENSG00000135519 | KCNH3 | 41.12243359 | 0.084618062 | 0.27436991 | 0.30840868 | 0.757771378 | 0.866391318 |
| ENSG00000135521 | LTV1 | 531.0638634 | 0.375903211 | 0.090597196 | 4.149170477 | 3.34E-05 | 0.00016024 |
| ENSG00000135525 | MAP7 | 2921.869702 | -0.205548201 | 0.052717958 | -3.899016775 | 9.66E-05 | 0.00043131 |
| ENSG00000135535 | CD164 | 5692.157125 | -0.101688475 | 0.058724222 | -1.731627468 | 0.083339912 | 0.182721742 |
| ENSG00000135537 | AFG1L | 109.8367364 | -0.044423769 | 0.17594375 | -0.252488475 | 0.800663521 | 0.893869047 |
| ENSG00000135540 | NHSL1 | 487.9703166 | 0.483295965 | 0.090998992 | 5.311003495 | 1.09E-07 | 7.05E-07 |
| ENSG00000135541 | AHI1 | 553.3532489 | 0.35823487 | 0.085284798 | 4.200453993 | 2.66E-05 | 0.000129906 |
| ENSG00000135547 | HEY2 | 20.83140141 | -0.669668681 | 0.40201036 | -1.665799562 | 0.095753343 | 0.204703291 |
| ENSG00000135549 | PKIB | 186.8256551 | 1.757627932 | 0.15358093 | 11.44431103 | 2.51E-30 | 7.07E-29 |
| ENSG00000135587 | SMPD2 | 462.0257496 | 0.603474393 | 0.089771778 | 6.72231748 | 1.79E-11 | 1.64E-10 |
| ENSG00000135596 | MICAL1 | 1442.578437 | 0.14047977 | 0.078028281 | 1.800369924 | 0.071802246 | 0.162104471 |
| ENSG00000135597 | REPS1 | 1706.007648 | 0.35493116 | 0.059541723 | 5.961049515 | 2.51E-09 | 1.92E-08 |
| ENSG00000135604 | STX11 | 2.117198244 | 0.423955436 | 1.252078855 | 0.338601226 | 0.734910159 | 0.853785022 |
| ENSG00000135605 | TEC | 96.01749551 | 0.472217759 | 0.188093415 | 2.510549132 | 0.012054355 | 0.035069993 |
| ENSG00000135617 | PRADC1 | 448.5688084 | -0.073179778 | 0.10583212 | -0.691470401 | 0.489269975 | 0.672489708 |

| | | | | | | | |
|---|---|---|---|---|---|---|---|
| ENSG00000135622 | SEMA4F | 235.8441771 | -0.309834852 | 0.125142206 | -2.475862151 | 0.013291489 | 0.038208866 |
| ENSG00000135624 | CCT7 | 9860.080863 | 0.001347284 | 0.043913012 | 0.030680738 | 0.975524153 | 0.988756421 |
| ENSG00000135625 | EGR4 | 9.187769783 | 2.12824506 | 0.669440407 | 3.179140426 | 0.001477125 | 0.005339623 |
| ENSG00000135631 | RAB11FIP5 | 137.2435405 | 0.221133097 | 0.159493465 | 1.386471203 | 0.165603067 | 0.31435121 |
| ENSG00000135632 | SMYD5 | 1237.841079 | 0.265172846 | 0.062668263 | 4.231373806 | 2.32E-05 | 0.000114242 |
| ENSG00000135636 | DYSF | 5.457102052 | -0.068985439 | 0.770790845 | -0.089499557 | 0.928684906 | 0.965737636 |
| ENSG00000135637 | CCDC142 | 650.759568 | -0.358474337 | 0.084521495 | -4.241220958 | 2.22E-05 | 0.00010953 |
| ENSG00000135643 | KCNMB4 | 540.9044536 | 0.39777778 | 0.08819219 | 4.510351526 | 6.47E-06 | 3.42E-05 |
| ENSG00000135655 | USP15 | 1619.160018 | -0.199826132 | 0.065028997 | -3.072877372 | 0.002120056 | 0.007411472 |
| ENSG00000135677 | GNS | 5896.242532 | 0.320353611 | 0.045193593 | 7.08847412 | 1.36E-12 | 1.37E-11 |
| ENSG00000135678 | CPM | 79.96049638 | 1.958494188 | 0.221752298 | 8.831900311 | 1.03E-18 | 1.60E-17 |
| ENSG00000135679 | MDM2 | 3276.749414 | -0.125753304 | 0.053028947 | -2.371408671 | 0.017720426 | 0.049244366 |
| ENSG00000135686 | KLHL36 | 3197.928039 | 0.414825613 | 0.051177944 | 8.105554396 | 5.25E-16 | 6.79E-15 |
| ENSG00000135697 | BCO1 | 2.008420147 | -0.173719444 | 1.224150905 | -0.141910155 | 0.887150973 | 0.941925545 |
| ENSG00000135698 | MPHOSPH6 | 729.4320673 | -0.231577583 | 0.081032726 | -2.857827871 | 0.004265517 | 0.013970726 |
| ENSG00000135702 | CHST5 | 2.798940407 | 2.122505604 | 1.217640057 | 1.743130567 | 0.08131081 | 0.179008636 |
| ENSG00000135709 | KIAA0513 | 2976.601416 | 0.468632022 | 0.11923629 | 3.930280143 | 8.48E-05 | 0.000383147 |
| ENSG00000135720 | DYNC1LI2 | 3726.076849 | 0.103244362 | 0.059433898 | 1.737129231 | 0.082364363 | 0.180928707 |
| ENSG00000135722 | FBXL8 | 299.3615022 | -0.098526702 | 0.119061667 | -0.82752664 | 0.407938629 | 0.600533412 |
| ENSG00000135723 | FHOD1 | 1794.811335 | 0.117667425 | 0.058407947 | 2.014579003 | 0.043948779 | 0.107322227 |
| ENSG00000135736 | CCDC102A | 117.7784981 | 0.092211028 | 0.167628655 | 0.550091082 | 0.582256903 | 0.746908984 |
| ENSG00000135740 | SLC9A5 | 195.0493975 | -0.857825632 | 0.15120118 | -5.673405678 | 1.40E-08 | 9.92E-08 |
| ENSG00000135747 | ZNF670-ZNF69 | 17.43617895 | 0.561145818 | 0.419628906 | 1.337243002 | 0.181143329 | 0.336558397 |
| ENSG00000135749 | PCNX2 | 753.6723625 | 0.356213955 | 0.077040002 | 4.623753179 | 3.77E-06 | 2.04E-05 |
| ENSG00000135750 | KCNK1 | 606.3530692 | 0.685848158 | 0.082767557 | 8.286437185 | 1.17E-16 | 1.58E-15 |
| ENSG00000135763 | URB2 | 878.4154548 | -0.196959472 | 0.070495574 | -2.793926789 | 0.005207227 | 0.016681159 |
| ENSG00000135766 | EGLN1 | 111.6930786 | -0.251940244 | 0.174233374 | -1.445993029 | 0.148179157 | 0.288804576 |
| ENSG00000135773 | CAPN9 | 116.5252564 | -0.633833801 | 0.165726589 | -3.824575192 | 0.000130998 | 0.000573053 |
| ENSG00000135775 | COG2 | 1558.760452 | -0.066330682 | 0.0576011 | -1.151552344 | 0.249505074 | 0.427488714 |
| ENSG00000135776 | ABCB10 | 772.8564356 | -0.331072773 | 0.073870555 | -4.481796226 | 7.40E-06 | 3.88E-05 |
| ENSG00000135778 | NTPCR | 1023.408304 | -0.007158926 | 0.067414804 | -0.10619219 | 0.915429867 | 0.958033237 |
| ENSG00000135801 | TAF5L | 1183.315277 | -0.029939154 | 0.06311533 | -0.47435629 | 0.635245857 | 0.784991303 |
| ENSG00000135821 | GLUL | 6392.189317 | 0.214856975 | 0.042210474 | 5.090134126 | 3.58E-07 | 2.19E-06 |

| | | | | | | | |
|---|---|---|---|---|---|---|---|
| ENSG00000135823 | STX6 | 2382.793814 | 0.186514735 | 0.053815593 | 3.465812139 | 0.000528633 | 0.002082574 |
| ENSG00000135828 | RNASEL | 273.3858696 | 0.302285918 | 0.119146353 | 2.537097518 | 0.011177581 | 0.032764339 |
| ENSG00000135829 | DHX9 | 11916.85852 | -0.22300299 | 0.046060709 | -4.841501418 | 1.29E-06 | 7.35E-06 |
| ENSG00000135835 | KIAA1614 | 8.606298837 | 0.144366538 | 0.589288082 | 0.244984656 | 0.806468312 | 0.897336329 |
| ENSG00000135837 | CEP350 | 2270.677516 | -0.259732124 | 0.128216047 | -2.025738034 | 0.042791637 | 0.104977024 |
| ENSG00000135838 | NPL | 27.93554107 | -0.306659501 | 0.342921998 | -0.894254389 | 0.371185798 | 0.563924535 |
| ENSG00000135842 | NIBAN1 | 7.172724313 | -0.386534075 | 0.650176825 | -0.59450608 | 0.552173677 | 0.723944914 |
| ENSG00000135845 | PIGC | 895.8084838 | -0.033118258 | 0.069796619 | -0.474496598 | 0.635145824 | 0.784950177 |
| ENSG00000135862 | LAMC1 | 3292.457437 | 0.273289854 | 0.052576933 | 5.197904109 | 2.02E-07 | 1.27E-06 |
| ENSG00000135870 | RC3H1 | 1309.030731 | 0.061341863 | 0.067767591 | 0.905179925 | 0.365370075 | 0.558177246 |
| ENSG00000135899 | SP110 | 321.1693435 | 0.803100284 | 0.109377978 | 7.342431257 | 2.10E-13 | 2.24E-12 |
| ENSG00000135900 | MRPL44 | 701.0675285 | 0.001952628 | 0.079498224 | 0.024561906 | 0.980404405 | 0.991070363 |
| ENSG00000135905 | DOCK10 | 144.4712559 | 1.115821518 | 0.163544946 | 6.822720867 | 8.93E-12 | 8.41E-11 |
| ENSG00000135912 | TTLL4 | 728.4397099 | -0.283300887 | 0.078829288 | -3.593853156 | 0.000325824 | 0.001332096 |
| ENSG00000135913 | USP37 | 1220.789664 | -0.213692429 | 0.064668395 | -3.304433777 | 0.000951685 | 0.003577966 |
| ENSG00000135916 | ITM2C | 628.7750816 | -1.229478475 | 0.089995136 | -13.66161027 | 1.72E-42 | 7.73E-41 |
| ENSG00000135919 | SERPINE2 | 297.1447394 | -1.396083917 | 0.111669765 | -12.50189716 | 7.29E-36 | 2.55E-34 |
| ENSG00000135924 | DNAJB2 | 1648.853675 | 0.134532494 | 0.06572095 | 2.047025991 | 0.040655533 | 0.100499776 |
| ENSG00000135926 | TMBIM1 | 95.72811677 | -0.399822529 | 0.182982126 | -2.185035986 | 0.028886208 | 0.075113309 |
| ENSG00000135929 | CYP27A1 | 7.961443633 | 0.815902268 | 0.653884694 | 1.247776979 | 0.212112741 | 0.379908361 |
| ENSG00000135930 | EIF4E2 | 2011.037296 | 0.224770795 | 0.053605514 | 4.193053631 | 2.75E-05 | 0.000133962 |
| ENSG00000135931 | ARMC9 | 959.1414085 | 0.007167017 | 0.068422188 | 0.104746976 | 0.916576585 | 0.958874843 |
| ENSG00000135932 | CAB39 | 1445.140806 | 0.102848054 | 0.063923237 | 1.608930632 | 0.10763151 | 0.224910444 |
| ENSG00000135940 | COX5B | 1693.663976 | 0.204225216 | 0.059597019 | 3.426768984 | 0.000610808 | 0.002379342 |
| ENSG00000135945 | REV1 | 1123.028086 | -0.029670427 | 0.067505866 | -0.439523693 | 0.660282118 | 0.80367002 |
| ENSG00000135951 | TSGA10 | 269.6934178 | 0.414300941 | 0.114150219 | 3.629436247 | 0.000284041 | 0.001172557 |
| ENSG00000135953 | MFSD9 | 599.7490427 | 0.020182235 | 0.08095866 | 0.249290632 | 0.803135977 | 0.895191649 |
| ENSG00000135956 | TMEM127 | 3657.408401 | 0.194910941 | 0.054284283 | 3.590559359 | 0.000329969 | 0.001347364 |
| ENSG00000135966 | TGFBRAP1 | 1543.101228 | 0.170096039 | 0.060146295 | 2.828038515 | 0.004683417 | 0.015196264 |
| ENSG00000135968 | GCC2 | 931.9110567 | 0.023444787 | 0.082400396 | 0.284522753 | 0.7760098 | 0.878332993 |
| ENSG00000135972 | MRPS9 | 984.4211113 | -0.007796407 | 0.066718648 | -0.11685499 | 0.906974966 | 0.95304868 |
| ENSG00000135973 | GPR45 | 2.211676255 | -0.160250219 | 1.241668499 | -0.129060388 | 0.897309865 | 0.948060798 |
| ENSG00000135974 | C2orf49 | 669.1711413 | -0.022164729 | 0.080711314 | -0.274617369 | 0.783610221 | 0.882691054 |

| | | | | | | | |
|---|---|---|---|---|---|---|---|
| ENSG00000135976 | ANKRD36 | 112.9943856 | -0.511037753 | 0.171189837 | -2.985210809 | 0.00283383 | 0.009662314 |
| ENSG00000135999 | EPC2 | 1309.492998 | -0.066027658 | 0.064648239 | -1.021337292 | 0.307094664 | 0.494924237 |
| ENSG00000136002 | ARHGEF4 | 56.65795449 | 0.198501836 | 0.238131389 | 0.833581143 | 0.404517055 | 0.597451793 |
| ENSG00000136003 | ISCU | 1260.014318 | -0.026986251 | 0.06154344 | -0.438491101 | 0.66103032 | 0.80407421 |
| ENSG00000136010 | ALDH1L2 | 372.6493695 | 0.157098573 | 0.108780174 | 1.44418387 | 0.148687261 | 0.289573458 |
| ENSG00000136021 | SCYL2 | 2040.983724 | -0.022308771 | 0.057112615 | -0.390610219 | 0.696085371 | 0.829258331 |
| ENSG00000136026 | CKAP4 | 1283.159061 | 0.093243356 | 0.065691361 | 1.419415798 | 0.155777829 | 0.299819062 |
| ENSG00000136040 | PLXNC1 | 13.43606637 | -1.49132866 | 0.502675912 | -2.966779636 | 0.003009366 | 0.010192654 |
| ENSG00000136044 | APPL2 | 1616.561463 | -0.10175055 | 0.058611178 | -1.736026364 | 0.082559173 | 0.181258536 |
| ENSG00000136045 | PWP1 | 1928.537807 | -0.287738405 | 0.057550321 | -4.99977064 | 5.74E-07 | 3.43E-06 |
| ENSG00000136048 | DRAM1 | 3184.237929 | -0.949193976 | 0.058012593 | -16.36186099 | 3.58E-60 | 2.57E-58 |
| ENSG00000136051 | WASHC4 | 1281.146672 | 0.030680903 | 0.065567705 | 0.467927057 | 0.639836754 | 0.787988143 |
| ENSG00000136052 | SLC41A2 | 340.688253 | 0.431736663 | 0.102341294 | 4.21859686 | 2.46E-05 | 0.000120446 |
| ENSG00000136059 | VILL | 7.404544864 | 1.622894326 | 0.704233422 | 2.304483534 | 0.021195516 | 0.057461341 |
| ENSG00000136068 | FLNB | 7686.249608 | -0.333060426 | 0.045690005 | -7.289568647 | 3.11E-13 | 3.29E-12 |
| ENSG00000136098 | NEK3 | 207.5997913 | -0.17755478 | 0.141992548 | -1.250451401 | 0.211134698 | 0.378659211 |
| ENSG00000136099 | PCDH8 | 27.94252864 | -0.693880332 | 0.348712859 | -1.989832935 | 0.046609342 | 0.112774845 |
| ENSG00000136100 | VPS36 | 849.4535041 | -0.004468411 | 0.069672167 | -0.064134805 | 0.948862889 | 0.976445122 |
| ENSG00000136104 | RNASEH2B | 422.7407372 | 0.055259614 | 0.096751722 | 0.571148638 | 0.567898891 | 0.735607034 |
| ENSG00000136108 | CKAP2 | 2264.849951 | -0.073911169 | 0.057948397 | -1.275465299 | 0.202144599 | 0.366084874 |
| ENSG00000136111 | TBC1D4 | 765.1335023 | 0.156091624 | 0.076381453 | 2.043580203 | 0.040995037 | 0.101209424 |
| ENSG00000136114 | THSD1 | 19.21799508 | 1.323525558 | 0.443210959 | 2.986220288 | 0.002824491 | 0.009633337 |
| ENSG00000136122 | BORA | 283.112872 | -0.134431001 | 0.111669639 | -1.203827664 | 0.228656195 | 0.401605781 |
| ENSG00000136141 | LRCH1 | 2466.046468 | -0.011638305 | 0.05333253 | -0.218221503 | 0.827256532 | 0.908785789 |
| ENSG00000136143 | SUCLA2 | 978.2002901 | 0.022287969 | 0.067142015 | 0.331952642 | 0.739925019 | 0.856567724 |
| ENSG00000136144 | RCBTB1 | 420.5143608 | 0.602737792 | 0.100685426 | 5.986345933 | 2.15E-09 | 1.66E-08 |
| ENSG00000136146 | MED4 | 805.5358368 | -0.086880382 | 0.076898821 | -1.129801232 | 0.258559989 | 0.438289082 |
| ENSG00000136147 | PHF11 | 277.2731904 | 0.082793199 | 0.115589998 | 0.716266118 | 0.473827044 | 0.6593226 |
| ENSG00000136149 | RPL13AP25 | 431.1504122 | 0.356949005 | 0.097354244 | 3.666496588 | 0.000245896 | 0.001025238 |
| ENSG00000136152 | COG3 | 1745.337245 | -0.029513783 | 0.055632719 | -0.530511245 | 0.595757515 | 0.756946961 |
| ENSG00000136153 | LMO7 | 574.578577 | 0.038824878 | 0.086744782 | 0.447575945 | 0.654459267 | 0.799387444 |
| ENSG00000136156 | ITM2B | 2306.076484 | 0.150303537 | 0.053591348 | 2.804623182 | 0.005037543 | 0.016216846 |
| ENSG00000136158 | SPRY2 | 194.8030502 | -0.019359864 | 0.138526616 | -0.139755552 | 0.888853133 | 0.942772834 |

| | | | | | | | |
|---|---|---|---|---|---|---|---|
| ENSG00000136159 | NUDT15 | 563.3458847 | 0.105302983 | 0.087577492 | 1.20239779 | 0.229209445 | 0.402238729 |
| ENSG00000136161 | RCBTB2 | 41.4931454 | 0.242423884 | 0.287049688 | 0.844536309 | 0.398369783 | 0.591380497 |
| ENSG00000136167 | LCP1 | 389.7920944 | -0.749072899 | 0.097520957 | -7.681147957 | 1.58E-14 | 1.82E-13 |
| ENSG00000136169 | SETDB2 | 246.8471523 | -0.19302869 | 0.116357681 | -1.6589252 | 0.097130866 | 0.207118615 |
| ENSG00000136193 | SCRN1 | 4115.060569 | -1.302035496 | 0.047705354 | -27.29327828 | 5.10E-164 | 2.02E-161 |
| ENSG00000136197 | C7orf25 | 6.329304009 | 1.401007389 | 0.733334281 | 1.910462152 | 0.056073735 | 0.132053329 |
| ENSG00000136205 | TNS3 | 3608.348463 | 0.177830167 | 0.07069781 | 2.515356093 | 0.011891221 | 0.034661323 |
| ENSG00000136213 | CHST12 | 594.717063 | 0.544351514 | 0.083190595 | 6.543426143 | 6.01E-11 | 5.29E-10 |
| ENSG00000136235 | GPNMB | 35.78787986 | 1.147688483 | 0.30394742 | 3.775944147 | 0.000159403 | 0.000687608 |
| ENSG00000136237 | RAPGEF5 | 397.666715 | -1.669965329 | 0.104864931 | -15.924917 | 4.26E-57 | 2.83E-55 |
| ENSG00000136238 | RAC1 | 7969.109517 | 0.150096199 | 0.047032735 | 3.191313457 | 0.001416275 | 0.005139095 |
| ENSG00000136240 | KDELR2 | 5980.449288 | 0.219929875 | 0.045642921 | 4.818488202 | 1.45E-06 | 8.20E-06 |
| ENSG00000136243 | NUP42 | 965.1123335 | -1.797888067 | 0.077764427 | -23.11967237 | 2.94E-118 | 6.94E-116 |
| ENSG00000136247 | ZDHHC4 | 2180.897929 | 0.020796766 | 0.055956133 | 0.371661951 | 0.710144557 | 0.838646906 |
| ENSG00000136250 | AOAH | 3.29253209 | 1.9158555 | 1.092746705 | 1.753247565 | 0.079559522 | 0.175910395 |
| ENSG00000136261 | BZW2 | 5726.697162 | -1.465788603 | 0.046066992 | -31.81863031 | 3.58E-222 | 3.16E-219 |
| ENSG00000136270 | TBRG4 | 3073.992061 | 0.588879738 | 0.054108779 | 10.88325689 | 1.39E-27 | 3.53E-26 |
| ENSG00000136271 | DDX56 | 4268.790092 | 0.682054215 | 0.046870982 | 14.55173728 | 5.69E-48 | 2.94E-46 |
| ENSG00000136273 | HUS1 | 635.7009468 | 0.631087351 | 0.082195565 | 7.677875899 | 1.62E-14 | 1.87E-13 |
| ENSG00000136279 | DBNL | 3917.227192 | 0.796423336 | 0.05420182 | 14.69366405 | 7.08E-49 | 3.78E-47 |
| ENSG00000136280 | CCM2 | 1529.885873 | 0.598563626 | 0.06090876 | 9.827217327 | 8.60E-23 | 1.69E-21 |
| ENSG00000136286 | MYO1G | 8.848224461 | -1.489855262 | 0.615869892 | -2.419107152 | 0.015558656 | 0.043933478 |
| ENSG00000136295 | TTYH3 | 3286.544686 | 0.334053153 | 0.063398303 | 5.269118217 | 1.37E-07 | 8.78E-07 |
| ENSG00000136305 | CIDEB | 39.66377456 | -0.462672191 | 0.313313748 | -1.476705677 | 0.13975455 | 0.276043514 |
| ENSG00000136319 | TTC5 | 766.4209965 | -0.463933166 | 0.073782605 | -6.287839309 | 3.22E-10 | 2.68E-09 |
| ENSG00000136367 | ZFHX2 | 237.9855239 | -0.238282684 | 0.124182921 | -1.918803994 | 0.055009144 | 0.129999976 |
| ENSG00000136371 | MTHFS | 28.57972212 | 0.035707302 | 0.326711865 | 0.109292946 | 0.912970141 | 0.956500504 |
| ENSG00000136378 | ADAMTS7 | 12.89651669 | 0.151535749 | 0.500068233 | 0.303030145 | 0.761866889 | 0.869297509 |
| ENSG00000136379 | ABHD17C | 1577.244228 | -1.224834472 | 0.059276529 | -20.66305977 | 7.45E-95 | 1.10E-92 |
| ENSG00000136381 | IREB2 | 4178.60341 | -0.158072053 | 0.053824555 | -2.936801876 | 0.003316159 | 0.011128245 |
| ENSG00000136383 | ALPK3 | 51.23232292 | 0.069827148 | 0.24337877 | 0.286907308 | 0.774183295 | 0.87711205 |
| ENSG00000136404 | TM6SF1 | 2.52851861 | -0.365223778 | 1.104559169 | -0.330651167 | 0.740907992 | 0.857108928 |
| ENSG00000136425 | CIB2 | 6.673731705 | -1.949930526 | 0.743969543 | -2.620981657 | 0.008767698 | 0.026495981 |

| | | | | | | | |
|---|---|---|---|---|---|---|---|
| ENSG00000136436 | CALCOCO2 | 2387.557162 | 0.143436485 | 0.054799378 | 2.617483788 | 0.00885807 | 0.026747942 |
| ENSG00000136444 | RSAD1 | 1938.519125 | 0.024831244 | 0.061310057 | 0.405010942 | 0.685469468 | 0.821959935 |
| ENSG00000136448 | NMT1 | 4137.544753 | -0.127080796 | 0.054845622 | -2.317063652 | 0.020500265 | 0.055852265 |
| ENSG00000136449 | MYCBPAP | 3.141272166 | -1.211805939 | 1.021757387 | -1.186001641 | 0.235621651 | 0.410570504 |
| ENSG00000136450 | SRSF1 | 1626.551378 | -0.379786892 | 0.079098408 | -4.80144799 | 1.58E-06 | 8.90E-06 |
| ENSG00000136451 | VEZF1 | 3500.537053 | 0.207229304 | 0.049551242 | 4.182121309 | 2.89E-05 | 0.000140214 |
| ENSG00000136457 | CHAD | 4.832333493 | -3.243174205 | 1.064786383 | -3.04584493 | 0.002320275 | 0.00805373 |
| ENSG00000136463 | TACO1 | 2192.896692 | -0.062381032 | 0.053223729 | -1.172053006 | 0.241175776 | 0.417149317 |
| ENSG00000136478 | TEX2 | 3675.058748 | 0.132842762 | 0.047717009 | 2.783970858 | 0.005369786 | 0.017158781 |
| ENSG00000136485 | DCAF7 | 13783.39485 | 0.033997958 | 0.042796535 | 0.794409128 | 0.426957292 | 0.617748362 |
| ENSG00000136490 | LIMD2 | 132.3615133 | -1.396598121 | 0.166314947 | -8.397309715 | 4.57E-17 | 6.31E-16 |
| ENSG00000136492 | BRIP1 | 9291.822616 | 0.000173984 | 0.049163847 | 0.003538858 | 0.997176406 | 0.99830788 |
| ENSG00000136504 | KAT7 | 3218.294855 | 0.059790712 | 0.052926391 | 1.129695613 | 0.258604506 | 0.438332154 |
| ENSG00000136518 | ACTL6A | 1977.703787 | -0.027668185 | 0.063271145 | -0.437295463 | 0.661897085 | 0.804701846 |
| ENSG00000136521 | NDUFB5 | 1468.203901 | -0.082363153 | 0.058851698 | -1.399503429 | 0.161662071 | 0.308658169 |
| ENSG00000136522 | MRPL47 | 856.7902535 | -0.130621888 | 0.070488824 | -1.853086501 | 0.063869961 | 0.14691436 |
| ENSG00000136527 | TRA2B | 3918.90145 | -0.297149314 | 0.047714458 | -6.22765774 | 4.73E-10 | 3.89E-09 |
| ENSG00000136535 | TBR1 | 5.549062047 | -1.579304019 | 0.783545778 | -2.015586151 | 0.043843269 | 0.107113801 |
| ENSG00000136536 | MARCHF7 | 2526.421318 | -0.059728585 | 0.055081504 | -1.084367353 | 0.278201953 | 0.461023824 |
| ENSG00000136542 | GALNT5 | 1.572065846 | 3.996101089 | 1.665395765 | 2.399490364 | 0.016417912 | NA |
| ENSG00000136560 | TANK | 1044.927514 | 0.435740929 | 0.066055397 | 6.596598484 | 4.21E-11 | 3.73E-10 |
| ENSG00000136603 | SKIL | 1751.699886 | -0.102562529 | 0.061095541 | -1.678723642 | 0.093205916 | 0.20010704 |
| ENSG00000136628 | EPRS1 | 7921.486784 | 0.0661396 | 0.049339217 | 1.340507709 | 0.18008034 | 0.335094338 |
| ENSG00000136631 | VPS45 | 2520.386957 | 0.062355803 | 0.051531141 | 1.210060579 | 0.226255648 | 0.398426696 |
| ENSG00000136636 | KCTD3 | 6297.486477 | 0.776948432 | 0.045439716 | 17.09844382 | 1.52E-65 | 1.24E-63 |
| ENSG00000136643 | RPS6KC1 | 555.3440228 | -0.025701412 | 0.085339005 | -0.301168399 | 0.763286086 | 0.87031501 |
| ENSG00000136682 | ZNG1B | 638.6815384 | -0.125253123 | 0.080729709 | -1.551512134 | 0.120779002 | 0.246808329 |
| ENSG00000136689 | IL1RN | 4.756381665 | 2.987346718 | 1.059112066 | 2.820614374 | 0.004793178 | 0.0155151 |
| ENSG00000136695 | IL36RN | 2.537025387 | -0.792871797 | 1.112041952 | -0.712987307 | 0.475853612 | 0.661019853 |
| ENSG00000136699 | SMPD4 | 3825.038335 | -0.210092309 | 0.047448094 | -4.427834532 | 9.52E-06 | 4.93E-05 |
| ENSG00000136709 | WDR33 | 2675.284154 | -0.130306016 | 0.053178482 | -2.450352304 | 0.01427165 | 0.040659668 |
| ENSG00000136710 | CCDC115 | 836.1653903 | -0.181977413 | 0.071916396 | -2.530402289 | 0.011393181 | 0.033336681 |
| ENSG00000136715 | SAP130 | 1927.448122 | -0.14861903 | 0.054455813 | -2.729167406 | 0.006349446 | 0.019920172 |

| | | | | | | | |
|---|---|---|---|---|---|---|---|
| ENSG00000136717 | BIN1 | 4.490927981 | 1.29947008 | 0.95693388 | 1.357951794 | 0.174478981 | 0.327433559 |
| ENSG00000136718 | IMP4 | 2253.546122 | 0.051123992 | 0.05480124 | 0.932898463 | 0.350872396 | 0.542203622 |
| ENSG00000136720 | HS6ST1 | 2247.591357 | -0.20813655 | 0.068982548 | -3.017234888 | 0.002550921 | 0.008768393 |
| ENSG00000136731 | UGGT1 | 2509.525869 | 0.015041595 | 0.065124038 | 0.230968411 | 0.817339343 | 0.903254867 |
| ENSG00000136738 | STAM | 731.1656285 | -0.802253442 | 0.075365595 | -10.64482337 | 1.84E-26 | 4.42E-25 |
| ENSG00000136754 | ABI1 | 1272.808209 | -0.39191546 | 0.061855988 | -6.335934013 | 2.36E-10 | 1.98E-09 |
| ENSG00000136758 | YME1L1 | 5303.713564 | -0.568749903 | 0.04654142 | -12.22029556 | 2.42E-34 | 8.05E-33 |
| ENSG00000136770 | DNAJC1 | 878.6788855 | -0.586474435 | 0.071610872 | -8.18974021 | 2.62E-16 | 3.46E-15 |
| ENSG00000136783 | NIPSNAP3A | 321.0568761 | -0.029217809 | 0.105495028 | -0.276959112 | 0.781811503 | 0.881574709 |
| ENSG00000136802 | LRRC8A | 1595.916339 | 0.545897584 | 0.063759365 | 8.56184159 | 1.11E-17 | 1.60E-16 |
| ENSG00000136807 | CDK9 | 1594.335328 | 0.189938769 | 0.06216506 | 3.055394279 | 0.002247648 | 0.007830076 |
| ENSG00000136810 | TXN | 2669.682189 | 0.070970057 | 0.05480891 | 1.294863495 | 0.195367324 | 0.356598218 |
| ENSG00000136811 | ODF2 | 3237.458056 | -0.00307466 | 0.050188257 | -0.061262542 | 0.951150122 | 0.977483371 |
| ENSG00000136813 | ECPAS | 4522.509498 | 0.123525647 | 0.053098993 | 2.326327477 | 0.020001087 | 0.054680603 |
| ENSG00000136816 | TOR1B | 1172.161828 | -0.010449021 | 0.066956755 | -0.156056265 | 0.875988672 | 0.935616916 |
| ENSG00000136819 | C9orf78 | 2416.826953 | -0.040104321 | 0.052133883 | -0.769256366 | 0.441741135 | 0.631058764 |
| ENSG00000136824 | SMC2 | 1291.094505 | -0.327174304 | 0.065015055 | -5.03228527 | 4.85E-07 | 2.92E-06 |
| ENSG00000136826 | KLF4 | 184.4514783 | 0.057310713 | 0.135507777 | 0.422933019 | 0.672344131 | 0.812271037 |
| ENSG00000136827 | TOR1A | 1818.073679 | 0.117452223 | 0.060255709 | 1.949229793 | 0.051267989 | 0.122395666 |
| ENSG00000136828 | RALGPS1 | 574.0933417 | 0.57558909 | 0.083681232 | 6.878353469 | 6.05E-12 | 5.79E-11 |
| ENSG00000136830 | NIBAN2 | 6600.801705 | 0.485286167 | 0.056930048 | 8.524253612 | 1.54E-17 | 2.20E-16 |
| ENSG00000136840 | ST6GALNAC4 | 169.3312291 | 2.735026568 | 0.176119642 | 15.52936703 | 2.20E-54 | 1.36E-52 |
| ENSG00000136842 | TMOD1 | 51.94845712 | -0.019481201 | 0.255258689 | -0.076319443 | 0.939164958 | 0.971039083 |
| ENSG00000136848 | DAB2IP | 2631.628909 | 0.047013902 | 0.05646023 | 0.832690574 | 0.405019262 | 0.597811154 |
| ENSG00000136854 | STXBP1 | 1586.1217 | 0.134481888 | 0.057589966 | 2.335161796 | 0.01953497 | 0.053578696 |
| ENSG00000136856 | SLC2A8 | 659.6815608 | -0.146020453 | 0.082179168 | -1.776854846 | 0.075592125 | 0.168898739 |
| ENSG00000136861 | CDK5RAP2 | 2063.615725 | -0.127731673 | 0.056052753 | -2.278776092 | 0.022680379 | 0.061004698 |
| ENSG00000136867 | SLC31A2 | 287.330384 | 0.51341731 | 0.118401254 | 4.336248934 | 1.45E-05 | 7.31E-05 |
| ENSG00000136868 | SLC31A1 | 1837.27562 | 0.1749454 | 0.061294087 | 2.854197 | 0.004314577 | 0.014119314 |
| ENSG00000136870 | ZNF189 | 405.8526624 | -0.35844397 | 0.096843732 | -3.701261416 | 0.00021453 | 0.000904489 |
| ENSG00000136874 | STX17 | 1355.904421 | 0.20237398 | 0.06386715 | 3.168670936 | 0.001531377 | 0.005517478 |
| ENSG00000136875 | PRPF4 | 1852.596271 | -0.15906504 | 0.055193009 | -2.881978041 | 0.003951873 | 0.013030898 |
| ENSG00000136877 | FPGS | 1451.140939 | 0.159068618 | 0.068134319 | 2.334632836 | 0.01956261 | 0.053648089 |

| | | | | | | | |
|---|---|---|---|---|---|---|---|
| ENSG00000136878 | USP20 | 1275.385867 | 0.251958227 | 0.065070518 | 3.872079633 | 0.000107911 | 0.000478537 |
| ENSG00000136881 | BAAT | 3.063153821 | -0.292419134 | 0.993921056 | -0.294207605 | 0.76859927 | 0.873806485 |
| ENSG00000136883 | KIF12 | 380.1063461 | -0.864351395 | 0.113088301 | -7.643154847 | 2.12E-14 | 2.43E-13 |
| ENSG00000136888 | ATP6V1G1 | 4395.407548 | -0.115675683 | 0.046798239 | -2.471795614 | 0.013443635 | 0.038583384 |
| ENSG00000136891 | TEX10 | 1058.988176 | 0.181727529 | 0.072932296 | 2.49172918 | 0.012712293 | 0.036709465 |
| ENSG00000136895 | GARNL3 | 132.5182448 | 1.261202292 | 0.17105479 | 7.373089586 | 1.67E-13 | 1.80E-12 |
| ENSG00000136897 | MRPL50 | 655.093683 | -0.137976567 | 0.083601212 | -1.650413481 | 0.098858396 | 0.2100409 |
| ENSG00000136908 | DPM2 | 1270.95661 | -0.053883465 | 0.075178957 | -0.716736005 | 0.473537005 | 0.658998963 |
| ENSG00000136918 | WDR38 | 12.43629367 | 0.382517975 | 0.507903566 | 0.753131106 | 0.451371134 | 0.639390671 |
| ENSG00000136925 | TSTD2 | 909.7140785 | -0.001338155 | 0.071220257 | -0.018788962 | 0.985009459 | 0.993193404 |
| ENSG00000136928 | GABBR2 | 225.0613972 | 1.017086172 | 0.135308086 | 7.516817377 | 5.61E-14 | 6.25E-13 |
| ENSG00000136930 | PSMB7 | 4029.236118 | 0.057805457 | 0.048138488 | 1.200815801 | 0.229822661 | 0.403037371 |
| ENSG00000136931 | NR5A1 | 2.542214885 | 1.939225396 | 1.235005036 | 1.57021659 | 0.116364731 | 0.239688151 |
| ENSG00000136932 | TRMO | 389.1240042 | -0.086306573 | 0.098998058 | -0.871800667 | 0.383317131 | 0.575703482 |
| ENSG00000136933 | RABEPK | 463.4777753 | 0.122250152 | 0.089060746 | 1.37266031 | 0.169857973 | 0.320412952 |
| ENSG00000136935 | GOLGA1 | 1246.590336 | -0.11503389 | 0.063878406 | -1.800825926 | 0.071730321 | 0.161958028 |
| ENSG00000136936 | XPA | 220.2034166 | -0.209107197 | 0.13097191 | -1.596580493 | 0.11035925 | 0.229377657 |
| ENSG00000136937 | NCBP1 | 2439.229635 | 0.197121538 | 0.053567326 | 3.679883859 | 0.00023334 | 0.000978221 |
| ENSG00000136938 | ANP32B | 5836.885019 | -0.126481205 | 0.048823401 | -2.590585699 | 0.009581276 | 0.028660122 |
| ENSG00000136940 | PDCL | 808.7044962 | 0.102181077 | 0.070946455 | 1.440256274 | 0.149794907 | 0.291340928 |
| ENSG00000136942 | RPL35 | 7183.260515 | 0.254319919 | 0.045904453 | 5.540201534 | 3.02E-08 | 2.06E-07 |
| ENSG00000136943 | CTSV | 93.69686358 | -0.141003702 | 0.183795703 | -0.767176273 | 0.442976718 | 0.631832054 |
| ENSG00000136944 | LMX1B | 1773.590018 | 0.435696497 | 0.05692074 | 7.654441886 | 1.94E-14 | 2.24E-13 |
| ENSG00000136950 | ARPC5L | 1485.173128 | -0.010333886 | 0.06064667 | -0.170394944 | 0.864699548 | 0.929799912 |
| ENSG00000136982 | DSCC1 | 494.8582301 | -0.195733381 | 0.089613601 | -2.184192803 | 0.028948084 | 0.075248616 |
| ENSG00000136986 | DERL1 | 4539.311746 | -0.206071103 | 0.047428609 | -4.344869217 | 1.39E-05 | 7.05E-05 |
| ENSG00000136997 | MYC | 5311.632521 | 0.213426907 | 0.049266447 | 4.332094519 | 1.48E-05 | 7.45E-05 |
| ENSG00000137038 | DMAC1 | 692.9591962 | 0.25498078 | 0.076995281 | 3.311641657 | 0.000927503 | 0.003495468 |
| ENSG00000137040 | RANBP6 | 574.1014837 | -0.060884869 | 0.083349242 | -0.730478976 | 0.46509746 | 0.651045626 |
| ENSG00000137054 | POLR1E | 946.9166046 | 0.298033919 | 0.068974628 | 4.320920992 | 1.55E-05 | 7.81E-05 |
| ENSG00000137055 | PLAA | 985.6318076 | -0.090492945 | 0.076857525 | -1.177411645 | 0.239031244 | 0.414653414 |
| ENSG00000137070 | IL11RA | 83.4152311 | -0.866083116 | 0.197929493 | -4.375715325 | 1.21E-05 | 6.17E-05 |
| ENSG00000137073 | UBAP2 | 1663.076426 | -0.053951412 | 0.056772546 | -0.950308125 | 0.341955712 | 0.533056811 |

| | | | | | | | |
|---|---|---|---|---|---|---|---|
| ENSG00000137074 | APTX | 782.1096594 | 0.057327192 | 0.074884369 | 0.76554284 | 0.443948367 | 0.632753994 |
| ENSG00000137075 | RNF38 | 670.7412369 | -0.043447902 | 0.082283405 | -0.528027516 | 0.597480238 | 0.757960331 |
| ENSG00000137076 | TLN1 | 4937.313915 | -0.19371797 | 0.054365603 | -3.563245164 | 0.000366298 | 0.001481729 |
| ENSG00000137094 | DNAJB5 | 92.70329638 | -0.867917421 | 0.201478747 | -4.307736846 | 1.65E-05 | 8.27E-05 |
| ENSG00000137098 | SPAG8 | 11.71581759 | -0.060627168 | 0.510421192 | -0.118778705 | 0.90545068 | 0.95262805 |
| ENSG00000137100 | DCTN3 | 1000.742026 | -0.001662643 | 0.068407781 | -0.024304879 | 0.980609421 | 0.991070363 |
| ENSG00000137101 | CD72 | 12.96791468 | -0.185166914 | 0.496719235 | -0.372779834 | 0.709312313 | 0.838047653 |
| ENSG00000137103 | TMEM8B | 165.9138946 | 0.56389929 | 0.147093704 | 3.8336059 | 0.000126278 | 0.000553463 |
| ENSG00000137106 | GRHPR | 1736.148249 | -0.105065989 | 0.058843436 | -1.78551758 | 0.074177402 | 0.166451102 |
| ENSG00000137124 | ALDH1B1 | 721.010863 | 0.14047205 | 0.078613175 | 1.786876692 | 0.07395742 | 0.166103703 |
| ENSG00000137133 | HINT2 | 26.80270204 | -0.453673926 | 0.338323198 | -1.340948325 | 0.179937231 | 0.33490954 |
| ENSG00000137135 | ARHGEF39 | 286.3049667 | -0.543975478 | 0.118442262 | -4.592748122 | 4.37E-06 | 2.35E-05 |
| ENSG00000137142 | IGFBPL1 | 95.87593441 | -1.312159778 | 0.196176735 | -6.688661522 | 2.25E-11 | 2.05E-10 |
| ENSG00000137145 | DENND4C | 1395.574923 | 0.086975655 | 0.06693296 | 1.29944432 | 0.19379149 | 0.354483437 |
| ENSG00000137154 | RPS6 | 14308.92838 | 0.259615663 | 0.048940552 | 5.304714652 | 1.13E-07 | 7.29E-07 |
| ENSG00000137161 | CNPY3 | 1698.509758 | -0.279767704 | 0.067535411 | -4.142533526 | 3.43E-05 | 0.000164605 |
| ENSG00000137166 | FOXP4 | 2714.21598 | -0.063309124 | 0.059845518 | -1.057875786 | 0.290112076 | 0.474555834 |
| ENSG00000137168 | PPIL1 | 1393.520625 | -0.270785847 | 0.064689908 | -4.185905602 | 2.84E-05 | 0.000137986 |
| ENSG00000137171 | KLC4 | 816.0819249 | 0.137895583 | 0.073258487 | 1.882315455 | 0.059793197 | 0.139157583 |
| ENSG00000137177 | KIF13A | 1877.09287 | -0.031470754 | 0.061048792 | -0.515501675 | 0.606202499 | 0.764860977 |
| ENSG00000137185 | ZSCAN9 | 422.08489 | -0.066970393 | 0.095365305 | -0.702251128 | 0.482522566 | 0.666610518 |
| ENSG00000137193 | PIM1 | 175.3807022 | 0.438585689 | 0.138220003 | 3.173098538 | 0.001508213 | 0.005440002 |
| ENSG00000137198 | GMPR | 60.75942917 | -1.119020793 | 0.234399352 | -4.773992691 | 1.81E-06 | 1.01E-05 |
| ENSG00000137200 | CMTR1 | 3069.127097 | -0.089907054 | 0.049381495 | -1.82066285 | 0.068658123 | 0.156143287 |
| ENSG00000137203 | TFAP2A | 1145.58702 | -0.805132983 | 0.066087128 | -12.18290162 | 3.83E-34 | 1.26E-32 |
| ENSG00000137207 | YIPF3 | 3288.853369 | 0.012671343 | 0.052838987 | 0.239810493 | 0.810477172 | 0.899645977 |
| ENSG00000137210 | TMEM14B | 687.8730769 | -0.729984938 | 0.077766897 | -9.386833829 | 6.18E-21 | 1.09E-19 |
| ENSG00000137216 | TMEM63B | 2256.420615 | 0.135954643 | 0.063266387 | 2.148923781 | 0.031640442 | 0.081353031 |
| ENSG00000137218 | FRS3 | 123.1282128 | -0.374224571 | 0.166846306 | -2.242929924 | 0.024901334 | 0.06626106 |
| ENSG00000137221 | TJAP1 | 969.1162429 | 0.166428727 | 0.074661309 | 2.229116119 | 0.02580618 | 0.068225627 |
| ENSG00000137261 | KIAA0319 | 72.67435142 | 0.681213936 | 0.214244688 | 3.17960713 | 0.001474748 | 0.005332653 |
| ENSG00000137266 | SLC22A23 | 744.0249681 | -0.423260531 | 0.084345325 | -5.018186039 | 5.22E-07 | 3.13E-06 |
| ENSG00000137267 | TUBB2A | 74.76805444 | -1.432250288 | 0.227527008 | -6.294858365 | 3.08E-10 | 2.57E-09 |

| | | | | | | | |
|---|---|---|---|---|---|---|---|
| ENSG00000137269 | LRRC1 | 1766.410963 | -0.181702501 | 0.057146974 | -3.179564713 | 0.001474964 | 0.005332653 |
| ENSG00000137274 | BPHL | 441.8601025 | -0.623475066 | 0.091858835 | -6.787317344 | 1.14E-11 | 1.06E-10 |
| ENSG00000137275 | RIPK1 | 980.3494821 | -0.479031652 | 0.069672517 | -6.875475037 | 6.18E-12 | 5.90E-11 |
| ENSG00000137285 | TUBB2B | 8.318086212 | -1.344585198 | 0.660342165 | -2.036194671 | 0.041730811 | 0.102673291 |
| ENSG00000137288 | UQCC2 | 1007.459572 | 0.060135515 | 0.072033941 | 0.834821946 | 0.403817967 | 0.596617498 |
| ENSG00000137309 | HMGA1 | 4659.549322 | 0.643589158 | 0.05047877 | 12.74969971 | 3.13E-37 | 1.15E-35 |
| ENSG00000137310 | TCF19 | 1592.004048 | -0.338693336 | 0.05906782 | -5.733973844 | 9.81E-09 | 7.06E-08 |
| ENSG00000137312 | FLOT1 | 3793.260366 | -0.304321506 | 0.059766222 | -5.091864483 | 3.55E-07 | 2.17E-06 |
| ENSG00000137331 | IER3 | 2023.062909 | -1.048116143 | 0.059819159 | -17.5214121 | 9.84E-69 | 8.71E-67 |
| ENSG00000137337 | MDC1 | 2606.925489 | -0.201251985 | 0.056361773 | -3.570717796 | 0.000356004 | 0.001442887 |
| ENSG00000137338 | PGBD1 | 171.0547266 | -1.247403055 | 0.151756329 | -8.219776132 | 2.04E-16 | 2.71E-15 |
| ENSG00000137343 | ATAT1 | 373.0631046 | 0.133454816 | 0.1111554 | 1.200614774 | 0.229900667 | 0.403062894 |
| ENSG00000137364 | TPMT | 344.304208 | -0.008663113 | 0.101298626 | -0.085520542 | 0.931847565 | 0.967397526 |
| ENSG00000137393 | RNF144B | 735.2117462 | -1.471429245 | 0.084361825 | -17.44188497 | 3.97E-68 | 3.42E-66 |
| ENSG00000137404 | NRM | 442.3501966 | -0.346214039 | 0.091166674 | -3.797594297 | 0.000146107 | 0.000634551 |
| ENSG00000137409 | MTCH1 | 5331.979363 | -0.182928822 | 0.04738167 | -3.860750846 | 0.000113039 | 0.000500119 |
| ENSG00000137411 | VARS2 | 25.19031622 | -0.613213093 | 0.401943677 | -1.525619455 | 0.127104664 | 0.256851744 |
| ENSG00000137413 | TAF8 | 728.3208974 | 0.069040517 | 0.077665215 | 0.888950314 | 0.374029783 | 0.56680164 |
| ENSG00000137414 | FAM8A1 | 878.8695543 | -0.492000259 | 0.077571643 | -6.342527275 | 2.26E-10 | 1.90E-09 |
| ENSG00000137434 | C6orf52 | 66.09760088 | -1.994444349 | 0.262548669 | -7.596474794 | 3.04E-14 | 3.45E-13 |
| ENSG00000137441 | FGFBP2 | 15.06465133 | -1.767988457 | 0.487913334 | -3.62357069 | 0.000290564 | 0.001198406 |
| ENSG00000137449 | CPEB2 | 439.0985587 | 0.82796299 | 0.095117703 | 8.704615068 | 3.19E-18 | 4.79E-17 |
| ENSG00000137460 | FHDC1 | 617.5112591 | 0.322511446 | 0.079165997 | 4.073863268 | 4.62E-05 | 0.00021732 |
| ENSG00000137462 | TLR2 | 11.61793694 | 0.537762157 | 0.52970458 | 1.015211456 | 0.310005041 | 0.497921249 |
| ENSG00000137474 | MYO7A | 99.67609676 | -0.497580292 | 0.179193021 | -2.776783883 | 0.005489967 | 0.01748436 |
| ENSG00000137478 | FCHSD2 | 1386.389586 | 0.037721078 | 0.061338638 | 0.614964392 | 0.538578236 | 0.713330689 |
| ENSG00000137486 | ARRB1 | 1939.510168 | 0.496658302 | 0.063609675 | 7.807905092 | 5.81E-15 | 6.96E-14 |
| ENSG00000137492 | THAP12 | 2541.468438 | -0.135385217 | 0.063188263 | -2.142569068 | 0.032147717 | 0.082490898 |
| ENSG00000137494 | ANKRD42 | 557.7252233 | 0.422784212 | 0.084371235 | 5.010999458 | 5.41E-07 | 3.25E-06 |
| ENSG00000137496 | IL18BP | 282.8361786 | -0.066607809 | 0.116089386 | -0.573763128 | 0.566128103 | 0.734059387 |
| ENSG00000137497 | NUMA1 | 22373.36221 | -0.294860431 | 0.049979123 | -5.899671965 | 3.64E-09 | 2.75E-08 |
| ENSG00000137500 | CCDC90B | 1123.557028 | -0.06896574 | 0.071132941 | -0.969533087 | 0.33227928 | 0.523342718 |
| ENSG00000137501 | SYTL2 | 11849.79168 | 0.616833732 | 0.052562103 | 11.73533208 | 8.40E-32 | 2.52E-30 |

| | | | | | | | |
|---|---|---|---|---|---|---|---|
| ENSG00000137502 | RAB30 | 967.1917336 | 0.199936461 | 0.071688776 | 2.788950684 | 0.005287912 | 0.016920727 |
| ENSG00000137504 | CREBZF | 2149.50369 | -0.090153837 | 0.130156641 | -0.692656453 | 0.488525173 | 0.671800345 |
| ENSG00000137509 | PRCP | 640.3805921 | -0.565476294 | 0.082603899 | -6.845636841 | 7.61E-12 | 7.21E-11 |
| ENSG00000137513 | NARS2 | 797.4804139 | -0.229951326 | 0.073078033 | -3.14665456 | 0.001651499 | 0.005909437 |
| ENSG00000137522 | RNF121 | 1046.969373 | 0.129028906 | 0.071611996 | 1.801777827 | 0.071580368 | 0.161667188 |
| ENSG00000137547 | MRPL15 | 1205.420502 | -0.147826986 | 0.064678848 | -2.285553773 | 0.022280377 | 0.060102523 |
| ENSG00000137558 | PI15 | 30.55841495 | 2.7062455 | 0.399134049 | 6.780292258 | 1.20E-11 | 1.11E-10 |
| ENSG00000137561 | TTPA | 5.417700943 | 0.707732274 | 0.766777772 | 0.922995293 | 0.356009658 | 0.547964409 |
| ENSG00000137563 | GGH | 402.2137535 | 0.225540537 | 0.100823631 | 2.236980909 | 0.025287588 | 0.067109816 |
| ENSG00000137571 | SLCO5A1 | 1.920024931 | 0.789731354 | 1.27525228 | 0.619274607 | 0.53573547 | 0.711031295 |
| ENSG00000137573 | SULF1 | 73.84431837 | 3.240036298 | 0.287085322 | 11.28596987 | 1.54E-29 | 4.20E-28 |
| ENSG00000137574 | TGS1 | 643.3712551 | -0.013111897 | 0.077797613 | -0.168538557 | 0.866159613 | 0.930447274 |
| ENSG00000137575 | SDCBP | 1523.836873 | -0.217944274 | 0.063075615 | -3.455285745 | 0.00054971 | 0.002159919 |
| ENSG00000137601 | NEK1 | 373.0574602 | 0.00148229 | 0.101378927 | 0.014621279 | 0.988334323 | 0.994043197 |
| ENSG00000137628 | DDX60 | 25.37832686 | 2.348663333 | 0.41737055 | 5.627285718 | 1.83E-08 | 1.28E-07 |
| ENSG00000137642 | SORL1 | 968.6887013 | 0.205547462 | 0.081463546 | 2.523183354 | 0.011629775 | 0.033978055 |
| ENSG00000137648 | TMPRSS4 | 111.1206332 | 1.874854695 | 0.193349629 | 9.696706954 | 3.11E-22 | 5.90E-21 |
| ENSG00000137656 | BUD13 | 504.5223312 | -0.091127693 | 0.087377051 | -1.042924791 | 0.296983127 | 0.483279629 |
| ENSG00000137672 | TRPC6 | 13.48333023 | 3.295798378 | 0.668992321 | 4.926511521 | 8.37E-07 | 4.90E-06 |
| ENSG00000137692 | DCUN1D5 | 671.553635 | -0.158264425 | 0.076589672 | -2.066393833 | 0.038791308 | 0.096672743 |
| ENSG00000137693 | YAP1 | 2025.516867 | -0.309469715 | 0.058617422 | -5.27948352 | 1.30E-07 | 8.33E-07 |
| ENSG00000137699 | TRIM29 | 10.26126099 | 2.396451426 | 0.662779896 | 3.61575757 | 0.000299471 | 0.001233146 |
| ENSG00000137700 | SLC37A4 | 2542.554447 | 0.433014899 | 0.056363704 | 7.682513185 | 1.56E-14 | 1.81E-13 |
| ENSG00000137709 | POU2F3 | 102.5638054 | -0.608881307 | 0.182147714 | -3.342788632 | 0.00082941 | 0.0031522 |
| ENSG00000137710 | RDX | 1452.096639 | 0.685467136 | 0.065477551 | 10.46873509 | 1.20E-25 | 2.75E-24 |
| ENSG00000137713 | PPP2R1B | 730.1234443 | 0.176928088 | 0.078669106 | 2.249016114 | 0.024511471 | 0.065375322 |
| ENSG00000137714 | FDX1 | 307.4377077 | 0.00246744 | 0.108351992 | 0.022772449 | 0.981831785 | 0.991627389 |
| ENSG00000137720 | CFAP68 | 99.44090688 | -0.053571194 | 0.180401054 | -0.296956103 | 0.766500023 | 0.872468208 |
| ENSG00000137726 | FXYD6 | 22.86512634 | -0.172199465 | 0.374843931 | -0.459389764 | 0.645954298 | 0.793095247 |
| ENSG00000137731 | FXYD2 | 4.686337153 | 2.545212583 | 0.990827316 | 2.568775144 | 0.010205866 | 0.030271859 |
| ENSG00000137745 | MMP13 | 1.740350383 | 1.268877157 | 1.383336305 | 0.917258625 | 0.359007133 | NA |
| ENSG00000137747 | TMPRSS13 | 861.1918469 | 1.78022055 | 0.077666986 | 22.92120047 | 2.86E-116 | 6.24E-114 |
| ENSG00000137760 | ALKBH8 | 187.2480328 | -0.020111981 | 0.135931669 | -0.147956551 | 0.882377058 | 0.939121523 |

| | | | | | | | |
|---|---|---|---|---|---|---|---|
| ENSG00000137764 | MAP2K5 | 741.1336251 | 0.112108773 | 0.075209223 | 1.490625337 | 0.136059888 | 0.270680239 |
| ENSG00000137767 | SQOR | 260.8075729 | 0.149117691 | 0.116116236 | 1.284210515 | 0.199068307 | 0.361856336 |
| ENSG00000137770 | CTDSPL2 | 1308.410478 | -0.135637346 | 0.066392666 | -2.042956759 | 0.041056719 | 0.101327717 |
| ENSG00000137776 | SLTM | 4808.282477 | -0.163951186 | 0.048585009 | -3.374522115 | 0.00073944 | 0.00283773 |
| ENSG00000137801 | THBS1 | 19741.88427 | 0.828953787 | 0.045362614 | 18.27394214 | 1.33E-74 | 1.39E-72 |
| ENSG00000137802 | MAPKBP1 | 586.7012017 | -0.358803604 | 0.085203166 | -4.211153433 | 2.54E-05 | 0.000124299 |
| ENSG00000137804 | NUSAP1 | 10.87578317 | -0.174038839 | 0.53791743 | -0.323541921 | 0.746284865 | 0.859604617 |
| ENSG00000137806 | NDUFAF1 | 274.5034894 | 0.116820406 | 0.120915393 | 0.966133455 | 0.333977406 | 0.524957018 |
| ENSG00000137807 | KIF23 | 2748.228354 | -0.492621669 | 0.055759334 | -8.834783883 | 1.00E-18 | 1.56E-17 |
| ENSG00000137812 | KNL1 | 1129.789939 | -0.399023602 | 0.06696666 | -5.958541205 | 2.54E-09 | 1.95E-08 |
| ENSG00000137814 | HAUS2 | 779.0194429 | -0.297293783 | 0.073212619 | -4.060690437 | 4.89E-05 | 0.000229296 |
| ENSG00000137815 | RTF1 | 3607.251846 | 0.039727132 | 0.048428426 | 0.820326707 | 0.412029885 | 0.604345625 |
| ENSG00000137817 | PARP6 | 2936.769921 | 0.175433799 | 0.069034127 | 2.541261929 | 0.011045314 | 0.03242635 |
| ENSG00000137818 | RPLP1 | 10547.13451 | -0.084748165 | 0.054580371 | -1.552722402 | 0.120489468 | 0.246348342 |
| ENSG00000137819 | PAQR5 | 294.0970142 | -0.751574356 | 0.112489677 | -6.68127403 | 2.37E-11 | 2.15E-10 |
| ENSG00000137821 | LRRC49 | 68.75759368 | -0.194028832 | 0.215198664 | -0.901626564 | 0.367255275 | 0.559826966 |
| ENSG00000137822 | TUBGCP4 | 1606.422013 | 0.158623464 | 0.058493981 | 2.711791239 | 0.006692072 | 0.020849673 |
| ENSG00000137824 | RMDN3 | 1839.543597 | -0.151394171 | 0.057669269 | -2.625213965 | 0.008659453 | 0.026203384 |
| ENSG00000137825 | ITPKA | 68.1424782 | 1.171432527 | 0.224838015 | 5.210117727 | 1.89E-07 | 1.19E-06 |
| ENSG00000137831 | UACA | 1137.719547 | 0.048984702 | 0.067054312 | 0.73052277 | 0.4650707 | 0.651045626 |
| ENSG00000137834 | SMAD6 | 1111.895562 | -0.734164699 | 0.065498919 | -11.2088064 | 3.69E-29 | 9.89E-28 |
| ENSG00000137841 | PLCB2 | 18.73344667 | 1.207458541 | 0.443913253 | 2.720032646 | 0.006527547 | 0.020395251 |
| ENSG00000137842 | TMEM62 | 439.8267134 | -0.120285203 | 0.091617207 | -1.312910608 | 0.189213072 | 0.348023402 |
| ENSG00000137843 | PAK6 | 464.2969328 | 0.052983845 | 0.093093889 | 0.569144183 | 0.569258296 | 0.736660762 |
| ENSG00000137845 | ADAM10 | 3193.289841 | -0.301420365 | 0.055333275 | -5.447361756 | 5.11E-08 | 3.41E-07 |
| ENSG00000137868 | STRA6 | 34.38541423 | -2.464997784 | 0.351404261 | -7.014706592 | 2.30E-12 | 2.29E-11 |
| ENSG00000137869 | CYP19A1 | 7.366052417 | -0.82491736 | 0.68692701 | -1.200880659 | 0.229797498 | 0.403024052 |
| ENSG00000137871 | ZNF280D | 507.6773035 | -0.567102265 | 0.087740941 | -6.46337114 | 1.02E-10 | 8.83E-10 |
| ENSG00000137872 | SEMA6D | 156.5087661 | -0.96648214 | 0.1527788 | -6.32602261 | 2.52E-10 | 2.11E-09 |
| ENSG00000137875 | BCL2L10 | 3.137921139 | -0.018800389 | 0.993313982 | -0.018926935 | 0.984899392 | 0.993193404 |
| ENSG00000137876 | RSL24D1 | 3432.832198 | -0.3751709 | 0.062702713 | -5.983327994 | 2.19E-09 | 1.69E-08 |
| ENSG00000137877 | SPTBN5 | 76.6689348 | -0.597820405 | 0.21277766 | -2.809601367 | 0.00496029 | 0.015995086 |
| ENSG00000137878 | GCOM1 | 8.10385744 | 0.958929662 | 0.643788077 | 1.489511371 | 0.136352761 | 0.271074819 |

| | | | | | | | |
|---|---|---|---|---|---|---|---|
| ENSG00000137880 | GCHFR | 292.0693061 | -1.090424204 | 0.11803451 | -9.238181311 | 2.51E-20 | 4.27E-19 |
| ENSG00000137936 | BCAR3 | 389.1001169 | -1.051642763 | 0.108760887 | -9.66931027 | 4.07E-22 | 7.66E-21 |
| ENSG00000137941 | TTLL7 | 147.5379677 | -0.871877845 | 0.169824829 | -5.133983355 | 2.84E-07 | 1.76E-06 |
| ENSG00000137942 | FNBP1L | 1840.803179 | -0.08245113 | 0.063746677 | -1.29341849 | 0.195866358 | 0.357338497 |
| ENSG00000137944 | KYAT3 | 468.8449622 | 0.118291116 | 0.092591176 | 1.2775636 | 0.201403342 | 0.365059471 |
| ENSG00000137947 | GTF2B | 522.6637438 | 0.052771964 | 0.085913135 | 0.614247915 | 0.539051515 | 0.713716751 |
| ENSG00000137948 | BRDT | 1.687876367 | 3.008605088 | 1.618513434 | 1.858869396 | 0.063045653 | NA |
| ENSG00000137955 | RABGGTB | 992.0230348 | 0.233439213 | 0.089386943 | 2.611558289 | 0.009013062 | 0.027155194 |
| ENSG00000137959 | IFI44L | 3.356903611 | 5.086110344 | 1.469921442 | 3.460123921 | 0.000539927 | 0.002124515 |
| ENSG00000137962 | ARHGAP29 | 866.6787595 | 0.557695448 | 0.072688802 | 7.672370868 | 1.69E-14 | 1.95E-13 |
| ENSG00000137965 | IFI44 | 5.954946997 | 5.918815252 | 1.325266444 | 4.466132285 | 7.96E-06 | 4.15E-05 |
| ENSG00000137968 | SLC44A5 | 25.34399611 | 1.585011029 | 0.39965998 | 3.965898787 | 7.31E-05 | 0.000333739 |
| ENSG00000137970 | RPL7P9 | 5767.612772 | 0.440833195 | 0.056801865 | 7.760892972 | 8.43E-15 | 9.94E-14 |
| ENSG00000137975 | CLCA2 | 3.978133654 | 4.327274584 | 1.400117024 | 3.090652074 | 0.001997175 | 0.00702041 |
| ENSG00000137976 | DNASE2B | 7.053468511 | 0.864613995 | 0.709877859 | 1.217975717 | 0.223233201 | 0.394497313 |
| ENSG00000137992 | DBT | 584.5466235 | 0.120542698 | 0.083099967 | 1.450574555 | 0.146898365 | 0.286917709 |
| ENSG00000137996 | RTCA | 452.5128835 | 0.182142875 | 0.09210926 | 1.977465408 | 0.047989047 | 0.115673924 |
| ENSG00000138002 | IFT172 | 734.6428067 | 0.099172339 | 0.082158075 | 1.207091798 | 0.227396793 | 0.400129051 |
| ENSG00000138018 | SELENOI | 2602.285869 | 0.045224872 | 0.055325445 | 0.817433506 | 0.413680734 | 0.605914701 |
| ENSG00000138028 | CGREF1 | 938.170066 | 0.311129578 | 0.073694848 | 4.221863326 | 2.42E-05 | 0.000118789 |
| ENSG00000138029 | HADHB | 2130.250178 | -0.048899939 | 0.055262303 | -0.884869732 | 0.376226895 | 0.569228662 |
| ENSG00000138030 | KHK | 242.74294 | -0.279211934 | 0.12180006 | -2.292379284 | 0.021883766 | 0.059143921 |
| ENSG00000138031 | ADCY3 | 2377.144758 | 0.679110455 | 0.054898105 | 12.37038058 | 3.78E-35 | 1.30E-33 |
| ENSG00000138032 | PPM1B | 316.095326 | -0.012053295 | 0.105082353 | -0.114703322 | 0.908680281 | 0.95431011 |
| ENSG00000138035 | PNPT1 | 1169.071046 | 0.093279045 | 0.067773966 | 1.3763256 | 0.168720851 | 0.318930758 |
| ENSG00000138036 | DYNC2LI1 | 284.9233391 | -0.036242113 | 0.108706438 | -0.33339436 | 0.73883662 | 0.856040341 |
| ENSG00000138050 | THUMPD2 | 259.5364908 | 0.16317422 | 0.126041489 | 1.294607206 | 0.195455765 | 0.356702884 |
| ENSG00000138061 | CYP1B1 | 1405.937878 | 0.283376142 | 0.066607702 | 4.254405029 | 2.10E-05 | 0.000103807 |
| ENSG00000138069 | RAB1A | 3682.546954 | 0.029177185 | 0.052846687 | 0.552110006 | 0.580872987 | 0.745926238 |
| ENSG00000138071 | ACTR2 | 6650.802715 | -0.060460278 | 0.049048082 | -1.232673651 | 0.217697546 | 0.387214171 |
| ENSG00000138073 | PREB | 2099.878035 | -0.078427803 | 0.066798652 | -1.174092611 | 0.240357933 | 0.416168827 |
| ENSG00000138074 | SLC5A6 | 3678.31975 | -0.035910062 | 0.051587817 | -0.696095779 | 0.48636885 | 0.669947245 |
| ENSG00000138078 | PREPL | 2035.987219 | 0.260101046 | 0.059482433 | 4.372737157 | 1.23E-05 | 6.25E-05 |

| | | | | | | | |
|---|---|---|---|---|---|---|---|
| ENSG00000138079 | SLC3A1 | 7.444562045 | -0.096535974 | 0.651239576 | -0.148234194 | 0.882157947 | 0.939050334 |
| ENSG00000138080 | EMILIN1 | 24.48562074 | -1.163258325 | 0.3623295 | -3.210498524 | 0.001325049 | 0.004847541 |
| ENSG00000138081 | FBXO11 | 1739.524528 | 0.094700928 | 0.061549951 | 1.538602817 | 0.123901291 | 0.251686496 |
| ENSG00000138085 | ATRAID | 1893.387669 | -0.221634056 | 0.055848544 | -3.968484029 | 7.23E-05 | 0.000330358 |
| ENSG00000138092 | CENPO | 1534.057442 | -0.140690845 | 0.058218384 | -2.416605111 | 0.015666002 | 0.044187642 |
| ENSG00000138095 | LRPPRC | 8631.641392 | 0.032824297 | 0.049932725 | 0.657370442 | 0.510942753 | 0.69065033 |
| ENSG00000138100 | TRIM54 | 31.78597182 | 1.047823181 | 0.330465653 | 3.170747611 | 0.001520472 | 0.005481632 |
| ENSG00000138101 | DTNB | 629.5217867 | 0.56142749 | 0.081582537 | 6.881711552 | 5.91E-12 | 5.66E-11 |
| ENSG00000138107 | ACTR1A | 2372.748486 | -0.210687699 | 0.054284515 | -3.881174925 | 0.000103953 | 0.000462058 |
| ENSG00000138111 | MFSD13A | 81.51997421 | -0.635485923 | 0.204451827 | -3.108242809 | 0.001882034 | 0.006651341 |
| ENSG00000138115 | CYP2C8 | 30.66415552 | 2.051901677 | 0.362934956 | 5.65363474 | 1.57E-08 | 1.11E-07 |
| ENSG00000138119 | MYOF | 5593.987404 | -0.137495033 | 0.054931476 | -2.503028188 | 0.012313574 | 0.035724469 |
| ENSG00000138134 | STAMBPL1 | 53.82148164 | 0.237017269 | 0.24188391 | 0.979880263 | 0.327145227 | 0.517600364 |
| ENSG00000138138 | ATAD1 | 1438.405783 | -0.453274079 | 0.073106273 | -6.20020776 | 5.64E-10 | 4.60E-09 |
| ENSG00000138160 | KIF11 | 2660.490922 | -0.809538646 | 0.0612193 | -13.22358557 | 6.41E-40 | 2.61E-38 |
| ENSG00000138162 | TACC2 | 1039.7086 | -0.546454029 | 0.069142615 | -7.903288435 | 2.72E-15 | 3.33E-14 |
| ENSG00000138166 | DUSP5 | 130.4274168 | 0.774642204 | 0.159698892 | 4.850642327 | 1.23E-06 | 7.05E-06 |
| ENSG00000138172 | CALHM2 | 371.9978153 | 0.362348827 | 0.101048398 | 3.585893826 | 0.000335926 | 0.001369249 |
| ENSG00000138175 | ARL3 | 654.1223281 | 0.389172338 | 0.078182779 | 4.977724551 | 6.43E-07 | 3.82E-06 |
| ENSG00000138180 | CEP55 | 1580.69778 | -0.808745071 | 0.062562913 | -12.92690875 | 3.17E-38 | 1.21E-36 |
| ENSG00000138182 | KIF20B | 1472.037929 | -0.278764622 | 0.073556996 | -3.789777121 | 0.000150783 | 0.000653495 |
| ENSG00000138185 | ENTPD1 | 24.95725543 | -1.033526642 | 0.357874447 | -2.887958751 | 0.003877507 | 0.012815158 |
| ENSG00000138190 | EXOC6 | 899.2032668 | -0.294069021 | 0.075185152 | -3.911264555 | 9.18E-05 | 0.000411772 |
| ENSG00000138193 | PLCE1 | 234.5363825 | 0.139623692 | 0.126530085 | 1.103482164 | 0.269817833 | 0.451434694 |
| ENSG00000138231 | DBR1 | 744.1760444 | -0.016919408 | 0.077580294 | -0.218088991 | 0.827359775 | 0.908805627 |
| ENSG00000138246 | DNAJC13 | 3099.906927 | -0.28146029 | 0.053622164 | -5.248954372 | 1.53E-07 | 9.75E-07 |
| ENSG00000138271 | GPR87 | 44.24715652 | 1.94759031 | 0.301235619 | 6.465338718 | 1.01E-10 | 8.73E-10 |
| ENSG00000138279 | ANXA7 | 2832.471594 | -0.73445778 | 0.05267532 | -13.94310993 | 3.47E-44 | 1.63E-42 |
| ENSG00000138286 | FAM149B1 | 694.2394635 | -0.970842508 | 0.07765737 | -12.5016146 | 7.32E-36 | 2.55E-34 |
| ENSG00000138303 | ASCC1 | 936.1189689 | -1.101231069 | 0.069028363 | -15.95331287 | 2.70E-57 | 1.81E-55 |
| ENSG00000138311 | ZNF365 | 19.15287542 | 0.524060991 | 0.410085795 | 1.277930124 | 0.201274065 | 0.364934116 |
| ENSG00000138326 | RPS24 | 9613.248109 | -0.211371114 | 0.059639328 | -3.544156559 | 0.000393871 | 0.001584775 |
| ENSG00000138336 | TET1 | 115.703527 | -1.291426659 | 0.185396493 | -6.965755604 | 3.27E-12 | 3.21E-11 |

| | | | | | | | |
|---|---|---|---|---|---|---|---|
| ENSG00000138346 | DNA2 | 454.2369446 | -0.87021477 | 0.092853194 | -9.371942258 | 7.12E-21 | 1.26E-19 |
| ENSG00000138356 | AOX1 | 76.61807024 | -1.085010261 | 0.208570853 | -5.202118342 | 1.97E-07 | 1.24E-06 |
| ENSG00000138363 | ATIC | 6736.701703 | -0.146404675 | 0.047763847 | -3.065177632 | 0.002175408 | 0.007588785 |
| ENSG00000138375 | SMARCAL1 | 557.6380566 | -0.321503852 | 0.088191369 | -3.645525143 | 0.000266846 | 0.001106354 |
| ENSG00000138376 | BARD1 | 999.34729 | 0.254573803 | 0.069754443 | 3.649571131 | 0.000262678 | 0.001090082 |
| ENSG00000138378 | STAT4 | 45.87581549 | -1.199799569 | 0.267324927 | -4.488169446 | 7.18E-06 | 3.77E-05 |
| ENSG00000138379 | MSTN | 6.314164551 | -0.339898163 | 0.687967927 | -0.49406106 | 0.621263058 | 0.775691207 |
| ENSG00000138380 | CARF | 251.40254 | 0.070563758 | 0.117827186 | 0.598875019 | 0.549256231 | 0.721774261 |
| ENSG00000138381 | ASNSD1 | 486.5287724 | -0.062673502 | 0.08915132 | -0.703001391 | 0.482054883 | 0.666287182 |
| ENSG00000138382 | METTL5 | 545.6479754 | 0.044702525 | 0.083921827 | 0.532668631 | 0.594262984 | 0.756068903 |
| ENSG00000138385 | SSB | 2926.619361 | 0.039934332 | 0.058363064 | 0.684239818 | 0.493823742 | 0.676438565 |
| ENSG00000138386 | NAB1 | 192.6078819 | -0.130831725 | 0.12986796 | -1.007421115 | 0.313732449 | 0.501980571 |
| ENSG00000138398 | PPIG | 2241.641805 | 0.010123711 | 0.055593988 | 0.182100821 | 0.855503605 | 0.925108546 |
| ENSG00000138399 | FASTKD1 | 415.6121541 | -0.1205594 | 0.097361025 | -1.238271676 | 0.215615342 | 0.384889577 |
| ENSG00000138400 | MDH1B | 36.40938318 | 0.417192073 | 0.303758791 | 1.373432099 | 0.169618057 | 0.320165542 |
| ENSG00000138411 | HECW2 | 292.2636947 | 0.302774259 | 0.113592439 | 2.665443784 | 0.007688679 | 0.023605231 |
| ENSG00000138413 | IDH1 | 3323.409428 | -0.127021236 | 0.054202467 | -2.343458589 | 0.019105883 | 0.052558935 |
| ENSG00000138430 | OLA1 | 3106.221617 | 0.376320086 | 0.054764143 | 6.871651163 | 6.35E-12 | 6.05E-11 |
| ENSG00000138433 | CIR1 | 650.0179953 | 0.036007452 | 0.078899604 | 0.45637051 | 0.648123568 | 0.794865258 |
| ENSG00000138434 | ITPRID2 | 1856.206036 | -0.079446212 | 0.057237381 | -1.388012717 | 0.165133173 | 0.313666886 |
| ENSG00000138439 | FAM117B | 539.0038374 | 0.334356897 | 0.08789202 | 3.804178078 | 0.000142276 | 0.000618848 |
| ENSG00000138442 | WDR12 | 860.6644889 | 0.1302882 | 0.072466678 | 1.797904973 | 0.072192068 | 0.162856332 |
| ENSG00000138443 | ABI2 | 2342.977756 | 0.165061822 | 0.055233238 | 2.988450922 | 0.002803955 | 0.009568988 |
| ENSG00000138448 | ITGAV | 2633.582277 | 0.233165819 | 0.056094001 | 4.156697961 | 3.23E-05 | 0.000155639 |
| ENSG00000138459 | SLC35A5 | 519.4612341 | 0.089879939 | 0.085210306 | 1.05480127 | 0.291516223 | 0.476478957 |
| ENSG00000138463 | SLC49A4 | 523.8912339 | 0.028228607 | 0.089094396 | 0.316839307 | 0.751365537 | 0.862482741 |
| ENSG00000138468 | SENP7 | 597.022931 | -0.460670361 | 0.083123506 | -5.541998661 | 2.99E-08 | 2.04E-07 |
| ENSG00000138495 | COX17 | 725.8417307 | -0.159772101 | 0.077292019 | -2.067122878 | 0.038722577 | 0.096553904 |
| ENSG00000138496 | PARP9 | 198.9750076 | 0.337584566 | 0.13122746 | 2.572514671 | 0.010096268 | 0.029985549 |
| ENSG00000138587 | MNS1 | 58.21427688 | -1.390000182 | 0.253103224 | -5.491831198 | 3.98E-08 | 2.69E-07 |
| ENSG00000138592 | USP8 | 18432.65921 | -1.259024226 | 0.050167725 | -25.09629897 | 5.46E-139 | 1.74E-136 |
| ENSG00000138593 | SECISBP2L | 1322.059058 | -0.030753231 | 0.06207434 | -0.495425818 | 0.620299574 | 0.775247738 |
| ENSG00000138594 | TMOD3 | 14766.20144 | -1.294988002 | 0.047128893 | -27.47758131 | 3.25E-166 | 1.33E-163 |

| | | | | | | | |
|---|---|---|---|---|---|---|---|
| ENSG00000138600 | SPPL2A | 13023.48852 | -1.435117737 | 0.052971869 | -27.09207305 | 1.22E-161 | 4.67E-159 |
| ENSG00000138604 | GLCE | 1710.3577 | -0.308849373 | 0.059922372 | -5.154158001 | 2.55E-07 | 1.59E-06 |
| ENSG00000138606 | SHF | 65.91613059 | -0.646921964 | 0.232041494 | -2.787958108 | 0.00530414 | 0.016967923 |
| ENSG00000138613 | APH1B | 358.1270272 | -0.808841363 | 0.101604323 | -7.960698334 | 1.71E-15 | 2.12E-14 |
| ENSG00000138614 | INTS14 | 2186.399531 | -0.188465701 | 0.054555533 | -3.454566213 | 0.000551179 | 0.002164705 |
| ENSG00000138615 | CILP | 13.78782831 | -0.023492452 | 0.48975372 | -0.047967889 | 0.961741834 | 0.982361474 |
| ENSG00000138617 | PARP16 | 595.8306204 | 0.192910835 | 0.083072556 | 2.322196947 | 0.020222333 | 0.05518681 |
| ENSG00000138621 | PPCDC | 519.553707 | -0.420867043 | 0.089672225 | -4.693393541 | 2.69E-06 | 1.48E-05 |
| ENSG00000138622 | HCN4 | 13.23825924 | 0.423065796 | 0.479135549 | 0.882977264 | 0.377248556 | 0.570022519 |
| ENSG00000138623 | SEMA7A | 94.86550371 | 0.5959455 | 0.189217316 | 3.149529405 | 0.001635337 | 0.005862575 |
| ENSG00000138629 | UBL7 | 1176.272583 | -0.322044437 | 0.069658796 | -4.623169735 | 3.78E-06 | 2.05E-05 |
| ENSG00000138639 | ARHGAP24 | 55.72140526 | 1.568081838 | 0.254304057 | 6.166169188 | 7.00E-10 | 5.65E-09 |
| ENSG00000138640 | FAM13A | 136.7459614 | -0.7284577 | 0.155672256 | -4.679431765 | 2.88E-06 | 1.58E-05 |
| ENSG00000138641 | HERC3 | 7.343073892 | 0.600838407 | 0.667614173 | 0.899978508 | 0.368131688 | 0.560715818 |
| ENSG00000138642 | HERC6 | 107.3125355 | 0.935152608 | 0.176922766 | 5.285654471 | 1.25E-07 | 8.06E-07 |
| ENSG00000138646 | HERC5 | 19.35886796 | 2.126054582 | 0.476230967 | 4.464335013 | 8.03E-06 | 4.19E-05 |
| ENSG00000138650 | PCDH10 | 418.0740735 | 1.412331483 | 0.100789709 | 14.01265569 | 1.30E-44 | 6.18E-43 |
| ENSG00000138653 | NDST4 | 214.6693094 | 3.382474329 | 0.176141324 | 19.20318444 | 3.48E-82 | 4.09E-80 |
| ENSG00000138658 | ZGRF1 | 437.7358405 | -0.672703723 | 0.113052205 | -5.950381245 | 2.68E-09 | 2.05E-08 |
| ENSG00000138660 | AP1AR | 991.6528184 | 0.008052127 | 0.068111959 | 0.118218994 | 0.905894141 | 0.95262805 |
| ENSG00000138663 | COPS4 | 1315.698727 | -0.208784893 | 0.060485045 | -3.451843256 | 0.000556771 | 0.002184798 |
| ENSG00000138668 | HNRNPD | 6149.384891 | 0.034394167 | 0.049980296 | 0.68815453 | 0.491355474 | 0.67430573 |
| ENSG00000138669 | PRKG2 | 29.90311744 | -0.092944803 | 0.344574274 | -0.269738079 | 0.787361763 | 0.884875507 |
| ENSG00000138670 | RASGEF1B | 34.00376224 | 0.339320677 | 0.315880134 | 1.074207083 | 0.282729879 | 0.466304077 |
| ENSG00000138674 | SEC31A | 5220.914972 | -0.029730981 | 0.048437196 | -0.613804756 | 0.539344354 | 0.713980004 |
| ENSG00000138678 | GPAT3 | 99.49267917 | -0.02640278 | 0.177519463 | -0.148731745 | 0.881765311 | 0.938683755 |
| ENSG00000138685 | FGF2 | 2.754071699 | -0.947325492 | 1.134483113 | -0.835028289 | 0.403701781 | 0.596554938 |
| ENSG00000138686 | BBS7 | 397.5914175 | 0.263769552 | 0.096368789 | 2.737084855 | 0.00619863 | 0.019505703 |
| ENSG00000138688 | BLTP1 | 2090.993896 | -0.043984114 | 0.078249047 | -0.562104149 | 0.574045062 | 0.740640817 |
| ENSG00000138696 | BMPR1B | 87.73456817 | -0.129223357 | 0.189597505 | -0.681566757 | 0.495512941 | 0.677781102 |
| ENSG00000138698 | RAP1GDS1 | 1215.007765 | -0.066423684 | 0.064044382 | -1.037150834 | 0.299665567 | 0.486368198 |
| ENSG00000138709 | LARP1B | 799.0097647 | 0.266710139 | 0.073479498 | 3.629721859 | 0.000283727 | 0.001171893 |
| ENSG00000138735 | PDE5A | 208.8467594 | 0.979252371 | 0.132834647 | 7.371965046 | 1.68E-13 | 1.81E-12 |

| | | | | | | | |
|---|---|---|---|---|---|---|---|
| ENSG00000138738 | PRDM5 | 123.5227201 | 0.39628248 | 0.161992658 | 2.446299011 | 0.014433125 | 0.041073797 |
| ENSG00000138744 | NAAA | 956.3406167 | 0.150529451 | 0.0705257 | 2.134391446 | 0.032810751 | 0.08390456 |
| ENSG00000138750 | NUP54 | 1197.234655 | -0.139851319 | 0.065483102 | -2.135685614 | 0.032705048 | 0.083689274 |
| ENSG00000138756 | BMP2K | 547.3033889 | -0.110688759 | 0.087493007 | -1.2651155 | 0.205829927 | 0.37129134 |
| ENSG00000138757 | G3BP2 | 4227.202312 | -0.092476475 | 0.051494021 | -1.795868183 | 0.072515484 | 0.163441265 |
| ENSG00000138758 | SEPTIN11 | 6346.01677 | 0.059894654 | 0.046538045 | 1.287004094 | 0.198092861 | 0.360483122 |
| ENSG00000138759 | FRAS1 | 744.5512937 | 1.079860538 | 0.098622085 | 10.94947994 | 6.68E-28 | 1.72E-26 |
| ENSG00000138760 | SCARB2 | 5113.794552 | 0.107490492 | 0.044944409 | 2.391632122 | 0.016773646 | 0.046919574 |
| ENSG00000138764 | CCNG2 | 2783.370297 | -0.393696642 | 0.056870105 | -6.922734545 | 4.43E-12 | 4.29E-11 |
| ENSG00000138767 | CNOT6L | 1117.08179 | 0.145147885 | 0.069608735 | 2.085196417 | 0.037051475 | 0.093015966 |
| ENSG00000138768 | USO1 | 2299.280102 | 0.068880578 | 0.061943384 | 1.111992496 | 0.266141377 | 0.447089532 |
| ENSG00000138769 | CDKL2 | 12.37158954 | -1.32044237 | 0.536562486 | -2.460929353 | 0.013857765 | 0.039642989 |
| ENSG00000138771 | SHROOM3 | 3086.441124 | 0.929880589 | 0.069060528 | 13.46471876 | 2.52E-41 | 1.09E-39 |
| ENSG00000138772 | ANXA3 | 908.5343046 | -0.027258162 | 0.077442431 | -0.351979676 | 0.724853504 | 0.846905337 |
| ENSG00000138777 | PPA2 | 1821.177482 | 0.163389 | 0.055525312 | 2.942603918 | 0.003254645 | 0.010944234 |
| ENSG00000138778 | CENPE | 1933.437028 | 0.05329571 | 0.065090327 | 0.818796156 | 0.412902725 | 0.604968289 |
| ENSG00000138780 | GSTCD | 948.7885374 | -0.104912002 | 0.068027956 | -1.542189527 | 0.123027544 | 0.250221817 |
| ENSG00000138785 | INTS12 | 697.9690983 | -0.074103697 | 0.075720972 | -0.978641647 | 0.327757074 | 0.518069818 |
| ENSG00000138792 | ENPEP | 286.8360814 | -0.237282709 | 0.108310145 | -2.190770854 | 0.028468379 | 0.074220322 |
| ENSG00000138794 | CASP6 | 711.048108 | -0.119039016 | 0.074910171 | -1.58909016 | 0.112040039 | 0.232323827 |
| ENSG00000138795 | LEF1 | 75.84486884 | 2.370242817 | 0.241392336 | 9.819047496 | 9.32E-23 | 1.83E-21 |
| ENSG00000138796 | HADH | 1933.735029 | -0.111008319 | 0.058534704 | -1.896453054 | 0.057900162 | 0.135500234 |
| ENSG00000138798 | EGF | 21.3345688 | 3.206487893 | 0.525619209 | 6.100400898 | 1.06E-09 | 8.40E-09 |
| ENSG00000138801 | PAPSS1 | 2966.92651 | 0.155535823 | 0.051456541 | 3.022663772 | 0.002505605 | 0.008634305 |
| ENSG00000138802 | SEC24B | 1362.672485 | -0.013786538 | 0.061356038 | -0.224697328 | 0.822214743 | 0.906023932 |
| ENSG00000138814 | PPP3CA | 4346.064964 | -0.181366266 | 0.05451292 | -3.327032697 | 0.000877761 | 0.003320005 |
| ENSG00000138821 | SLC39A8 | 1301.833178 | -0.084678002 | 0.068713438 | -1.232335413 | 0.217823817 | 0.387354911 |
| ENSG00000138829 | FBN2 | 253.2722602 | 0.493344318 | 0.126255989 | 3.907492394 | 9.33E-05 | 0.000417436 |
| ENSG00000138834 | MAPK8IP3 | 3009.646329 | 0.026940645 | 0.065629774 | 0.410494258 | 0.681443415 | 0.819131835 |
| ENSG00000138835 | RGS3 | 1391.929355 | -0.437258347 | 0.06431845 | -6.798334639 | 1.06E-11 | 9.88E-11 |
| ENSG00000138867 | GUCD1 | 3445.948077 | -0.967623717 | 0.054059141 | -17.89935422 | 1.19E-71 | 1.14E-69 |
| ENSG00000138942 | RNF185 | 1431.850856 | -0.776976185 | 0.060969509 | -12.74368453 | 3.38E-37 | 1.24E-35 |
| ENSG00000139044 | B4GALNT3 | 37.21259574 | 0.939251961 | 0.3037034 | 3.09266199 | 0.001983699 | 0.00698267 |

| | | | | | | | |
|---|---|---|---|---|---|---|---|
| ENSG00000139055 | ERP27 | 29.2956421 | 0.575743563 | 0.337542456 | 1.705692286 | 0.088065387 | 0.191073486 |
| ENSG00000139083 | ETV6 | 333.0687449 | 0.138591549 | 0.103171867 | 1.343307562 | 0.179172404 | 0.333903242 |
| ENSG00000139112 | GABARAPL1 | 10.05346674 | 1.282960426 | 0.596151963 | 2.152069449 | 0.031391885 | 0.080809115 |
| ENSG00000139116 | KIF21A | 1351.803853 | 0.016627344 | 0.066569023 | 0.24977599 | 0.802760589 | 0.895077666 |
| ENSG00000139131 | YARS2 | 329.8227018 | -0.106885592 | 0.106141501 | -1.007010365 | 0.313929793 | 0.502165083 |
| ENSG00000139132 | FGD4 | 321.6447889 | 0.075708428 | 0.105024383 | 0.720865248 | 0.470992426 | 0.656892404 |
| ENSG00000139133 | ALG10 | 194.9795064 | -0.648465967 | 0.135755162 | -4.77673158 | 1.78E-06 | 1.00E-05 |
| ENSG00000139144 | PIK3C2G | 16.42480801 | -1.479794095 | 0.457324378 | -3.235764738 | 0.001213173 | 0.004473589 |
| ENSG00000139146 | SINHCAF | 578.7474196 | -0.196294078 | 0.081476158 | -2.409221092 | 0.01598661 | 0.04497582 |
| ENSG00000139154 | AEBP2 | 533.7018614 | -0.034054469 | 0.094124252 | -0.361803342 | 0.717498995 | 0.842913738 |
| ENSG00000139160 | ETFBKMT | 22.07638936 | 0.458142965 | 0.386067958 | 1.18668995 | 0.235349941 | 0.410346404 |
| ENSG00000139163 | ETNK1 | 702.6452117 | -0.169564502 | 0.079343462 | -2.137094824 | 0.03259028 | 0.083430535 |
| ENSG00000139168 | ZCRB1 | 1043.74943 | 0.083429291 | 0.067834861 | 1.229888133 | 0.218738999 | 0.388470436 |
| ENSG00000139173 | TMEM117 | 117.5300122 | 0.56014176 | 0.167879852 | 3.336563334 | 0.000848211 | 0.003217787 |
| ENSG00000139174 | PRICKLE1 | 228.3656957 | 2.670679107 | 0.152490825 | 17.51370362 | 1.13E-68 | 9.90E-67 |
| ENSG00000139178 | C1RL | 3.436322864 | 1.453367412 | 1.028848277 | 1.412615878 | 0.157768698 | 0.302609861 |
| ENSG00000139180 | NDUFA9 | 95.72986208 | 0.003602875 | 0.19077832 | 0.018885139 | 0.984932734 | 0.993193404 |
| ENSG00000139182 | CLSTN3 | 595.1730673 | 0.442435217 | 0.09659645 | 4.580243048 | 4.64E-06 | 2.49E-05 |
| ENSG00000139187 | KLRG1 | 2.28027358 | -0.597377939 | 1.236023792 | -0.483306181 | 0.628878352 | 0.781033479 |
| ENSG00000139190 | VAMP1 | 119.1209043 | -0.080373603 | 0.168593295 | -0.476730719 | 0.633553885 | 0.784123341 |
| ENSG00000139192 | TAPBPL | 2.325247682 | 1.110972332 | 1.19448286 | 0.930086457 | 0.352326322 | 0.543727512 |
| ENSG00000139194 | RBP5 | 3.964928104 | 0.764883376 | 0.894865997 | 0.854746273 | 0.392691627 | 0.58568012 |
| ENSG00000139197 | PEX5 | 964.91779 | 0.462436297 | 0.070848945 | 6.527073852 | 6.71E-11 | 5.88E-10 |
| ENSG00000139211 | AMIGO2 | 917.6462713 | 1.161178593 | 0.070591532 | 16.44926193 | 8.49E-61 | 6.18E-59 |
| ENSG00000139218 | SCAF11 | 4729.774138 | 0.01555164 | 0.049595137 | 0.313571876 | 0.753846222 | 0.864272694 |
| ENSG00000139219 | COL2A1 | 157.3890729 | -1.812135504 | 0.159281571 | -11.3769314 | 5.45E-30 | 1.52E-28 |
| ENSG00000139223 | ANP32D | 8.446369203 | -0.904807205 | 0.650473375 | -1.390998062 | 0.164226014 | 0.312279905 |
| ENSG00000139233 | LLPH | 624.3480499 | 0.155536972 | 0.080517209 | 1.931723349 | 0.053393661 | 0.126835517 |
| ENSG00000139239 | RPL14P1 | 5039.742084 | 0.201134064 | 0.052325665 | 3.843889304 | 0.0001211 | 0.000532699 |
| ENSG00000139263 | LRIG3 | 212.3308188 | 0.133111577 | 0.125803226 | 1.058093516 | 0.29001281 | 0.474427288 |
| ENSG00000139266 | MARCHF9 | 1238.736089 | -0.281585924 | 0.067723463 | -4.157878413 | 3.21E-05 | 0.000154968 |
| ENSG00000139269 | INHBE | 23.27606715 | -2.526120453 | 0.445008316 | -5.676569088 | 1.37E-08 | 9.74E-08 |
| ENSG00000139278 | GLIPR1 | 47.50037563 | -0.113888059 | 0.267165309 | -0.426283111 | 0.669901563 | 0.810353399 |

| | | | | | | | |
|---|---|---|---|---|---|---|---|
| ENSG00000139289 | PHLDA1 | 1180.437939 | 2.572693378 | 0.073940637 | 34.79403838 | 2.99E-265 | 3.92E-262 |
| ENSG00000139291 | TMEM19 | 421.6927145 | 0.240855393 | 0.096244076 | 2.502547716 | 0.0123303 | 0.035760457 |
| ENSG00000139304 | PTPRQ | 110.25749 | 0.955792522 | 0.185379051 | 5.155882063 | 2.52E-07 | 1.57E-06 |
| ENSG00000139318 | DUSP6 | 128.3647552 | -0.034386053 | 0.159593723 | -0.215459931 | 0.829408749 | 0.909755986 |
| ENSG00000139323 | POC1B | 599.2508902 | 0.235249018 | 0.084759769 | 2.775479708 | 0.005512034 | 0.0175303 |
| ENSG00000139324 | TMTC3 | 983.2486203 | -0.300006465 | 0.078957613 | -3.799588861 | 0.000144936 | 0.000629704 |
| ENSG00000139343 | SNRPF | 1121.950626 | -0.027876765 | 0.063860565 | -0.436525496 | 0.662455505 | 0.804963395 |
| ENSG00000139344 | AMDHD1 | 8.968842933 | 0.196020131 | 0.591477373 | 0.331407659 | 0.74033658 | 0.856705062 |
| ENSG00000139350 | NEDD1 | 897.4307091 | -0.327475849 | 0.074431634 | -4.399686392 | 1.08E-05 | 5.58E-05 |
| ENSG00000139351 | SYCP3 | 5.710848903 | 0.819732466 | 0.753061438 | 1.088533317 | 0.276359737 | 0.458899193 |
| ENSG00000139352 | ASCL1 | 229.165468 | 7.334804649 | 0.523204452 | 14.01900275 | 1.19E-44 | 5.67E-43 |
| ENSG00000139354 | GAS2L3 | 642.3764807 | -0.089020455 | 0.086402374 | -1.030301039 | 0.302868711 | 0.489937821 |
| ENSG00000139364 | TMEM132B | 64.57390824 | 0.399016219 | 0.224463057 | 1.777647621 | 0.075461747 | 0.168689581 |
| ENSG00000139370 | SLC15A4 | 571.6705849 | -0.174337529 | 0.084505689 | -2.063027136 | 0.03911005 | 0.097361318 |
| ENSG00000139372 | TDG | 1956.063854 | 0.023991182 | 0.06028881 | 0.397937556 | 0.690676216 | 0.825601187 |
| ENSG00000139405 | RITA1 | 1310.022892 | 0.009977142 | 0.065410099 | 0.152532132 | 0.878767249 | 0.937101185 |
| ENSG00000139410 | SDSL | 311.9063677 | -0.166216134 | 0.11001927 | -1.510791099 | 0.130841683 | 0.262737064 |
| ENSG00000139428 | MMAB | 1354.149825 | -0.739156291 | 0.062660158 | -11.79627231 | 4.08E-32 | 1.23E-30 |
| ENSG00000139433 | GLTP | 1660.110147 | -0.020863226 | 0.05650124 | -0.369252536 | 0.7119395 | 0.839858692 |
| ENSG00000139436 | GIT2 | 1102.21112 | -0.067561192 | 0.070390599 | -0.959804199 | 0.337153769 | 0.528371871 |
| ENSG00000139437 | TCHP | 779.9505291 | 0.192607662 | 0.075200301 | 2.561261842 | 0.010429272 | 0.030822919 |
| ENSG00000139438 | FAM222A | 256.8191797 | -0.153107133 | 0.115889926 | -1.321142731 | 0.186453778 | 0.343996273 |
| ENSG00000139496 | NUP58 | 2486.044233 | 0.109078053 | 0.052680088 | 2.070574622 | 0.038398566 | 0.095902352 |
| ENSG00000139505 | MTMR6 | 1060.38938 | 0.088076489 | 0.066752155 | 1.319455357 | 0.187016925 | 0.344730275 |
| ENSG00000139508 | SLC46A3 | 298.64388 | 0.733673041 | 0.110175832 | 6.6591105 | 2.75E-11 | 2.49E-10 |
| ENSG00000139514 | SLC7A1 | 8214.885278 | -0.037042887 | 0.044426647 | -0.833798837 | 0.404394351 | 0.597309021 |
| ENSG00000139517 | LNX2 | 1266.58469 | 0.018590635 | 0.061972632 | 0.299981376 | 0.764191361 | 0.870644539 |
| ENSG00000139531 | SUOX | 835.0789943 | -0.495747192 | 0.080587612 | -6.151655045 | 7.67E-10 | 6.16E-09 |
| ENSG00000139537 | CCDC65 | 3.061721778 | -1.931593528 | 1.157923762 | -1.668152595 | 0.095285439 | 0.203857479 |
| ENSG00000139540 | SLC39A5 | 5.044307333 | 0.141183148 | 0.815167375 | 0.173195288 | 0.862497927 | 0.928557464 |
| ENSG00000139546 | TARBP2 | 1130.46386 | -0.166655592 | 0.065526984 | -2.543312419 | 0.0109807 | 0.0322573 |
| ENSG00000139547 | RDH16 | 16.13443874 | 0.368624007 | 0.456450118 | 0.807588809 | 0.419327331 | 0.610870102 |
| ENSG00000139572 | GPR84 | 1.787152592 | 0.640383186 | 1.29624544 | 0.494029268 | 0.621285509 | NA |

| | | | | | | | |
|---|---|---|---|---|---|---|---|
| ENSG00000139579 | NABP2 | 1569.462204 | -0.261838719 | 0.059290997 | -4.416163225 | 1.00E-05 | 5.20E-05 |
| ENSG00000139597 | N4BP2L1 | 127.1730512 | 0.944516564 | 0.164835714 | 5.730048061 | 1.00E-08 | 7.23E-08 |
| ENSG00000139613 | SMARCC2 | 6450.33461 | 0.191102999 | 0.048472394 | 3.942512055 | 8.06E-05 | 0.000365628 |
| ENSG00000139618 | BRCA2 | 709.0779388 | -0.279957737 | 0.088513004 | -3.162899509 | 0.001562062 | 0.005619212 |
| ENSG00000139620 | KANSL2 | 1084.453025 | 0.114822521 | 0.067666948 | 1.696877489 | 0.089719825 | 0.193929405 |
| ENSG00000139624 | CERS5 | 1033.898544 | 0.077145861 | 0.066970802 | 1.151932755 | 0.249348707 | 0.427335695 |
| ENSG00000139625 | MAP3K12 | 887.832003 | 0.412345535 | 0.078547884 | 5.249607141 | 1.52E-07 | 9.72E-07 |
| ENSG00000139626 | ITGB7 | 67.3416576 | 1.147681848 | 0.229098266 | 5.009561474 | 5.46E-07 | 3.27E-06 |
| ENSG00000139629 | GALNT6 | 2559.391848 | -0.373907892 | 0.060568126 | -6.173344257 | 6.69E-10 | 5.41E-09 |
| ENSG00000139631 | CSAD | 776.0167314 | -0.0114192 | 0.167370381 | -0.068227128 | 0.945604832 | 0.974261906 |
| ENSG00000139636 | LMBR1L | 391.4081008 | 0.07770409 | 0.097474316 | 0.79717502 | 0.425349391 | 0.61626573 |
| ENSG00000139637 | MYG1 | 1469.571676 | 0.259031175 | 0.065671921 | 3.944321561 | 8.00E-05 | 0.000363094 |
| ENSG00000139641 | ESYT1 | 9731.942268 | -0.221775296 | 0.046911466 | -4.72752857 | 2.27E-06 | 1.26E-05 |
| ENSG00000139644 | TMBIM6 | 30905.16315 | 0.006250592 | 0.044111826 | 0.14169878 | 0.887317938 | 0.941998797 |
| ENSG00000139645 | ANKRD52 | 7345.6262 | 0.670595758 | 0.045092278 | 14.87163173 | 5.04E-50 | 2.79E-48 |
| ENSG00000139651 | ZNF740 | 2769.168355 | -0.101349336 | 0.051236547 | -1.978067246 | 0.047921121 | 0.115570912 |
| ENSG00000139668 | WDFY2 | 427.913983 | -0.333026509 | 0.100648463 | -3.308808697 | 0.000936938 | 0.003525236 |
| ENSG00000139675 | HNRNPA1L2 | 491.6083557 | 0.287479612 | 0.095419083 | 3.012810471 | 0.002588405 | 0.008888924 |
| ENSG00000139684 | ESD | 5069.967851 | 0.093630862 | 0.046189564 | 2.027099935 | 0.042652194 | 0.104679718 |
| ENSG00000139687 | RB1 | 1606.579914 | -0.126218427 | 0.061417615 | -2.055085138 | 0.039870782 | 0.09887954 |
| ENSG00000139697 | SBNO1 | 3433.514382 | -0.072011257 | 0.049240173 | -1.462449305 | 0.143618127 | 0.281975398 |
| ENSG00000139714 | MORN3 | 44.30034282 | -0.048297457 | 0.268195402 | -0.180083091 | 0.857087337 | 0.925991782 |
| ENSG00000139718 | SETD1B | 2661.39338 | 0.105988529 | 0.055395529 | 1.913304734 | 0.055709052 | 0.131393527 |
| ENSG00000139719 | VPS33A | 380.8850733 | -0.006378578 | 0.097661873 | -0.065312874 | 0.947924892 | 0.975873851 |
| ENSG00000139722 | VPS37B | 734.6518978 | -0.088231023 | 0.077537241 | -1.137918016 | 0.255154718 | 0.434668018 |
| ENSG00000139725 | RHOF | 1376.262124 | 0.171820766 | 0.064341744 | 2.670439969 | 0.007575192 | 0.023281738 |
| ENSG00000139726 | DENR | 4676.962904 | 0.063211401 | 0.053284234 | 1.186305916 | 0.235501511 | 0.410506577 |
| ENSG00000139734 | DIAPH3 | 395.2713401 | -0.156479921 | 0.097224142 | -1.60947597 | 0.107512305 | 0.224702285 |
| ENSG00000139737 | SLAIN1 | 345.2778565 | 0.50907667 | 0.105278231 | 4.835535962 | 1.33E-06 | 7.56E-06 |
| ENSG00000139746 | RBM26 | 1268.279749 | -0.152622566 | 0.063799239 | -2.392231785 | 0.016746263 | 0.046865838 |
| ENSG00000139793 | MBNL2 | 793.4571836 | 0.251250651 | 0.076395328 | 3.288822202 | 0.001006076 | 0.003764988 |
| ENSG00000139800 | ZIC5 | 30.69250176 | 0.886304192 | 0.358385777 | 2.473045108 | 0.013396723 | 0.038463183 |
| ENSG00000139826 | ABHD13 | 233.5015241 | -0.088963093 | 0.131618288 | -0.675917417 | 0.499093079 | 0.680567986 |

| | | | | | | | |
|---|---|---|---|---|---|---|---|
| ENSG00000139832 | RAB20 | 335.7467009 | -0.088530081 | 0.109622745 | -0.807588618 | 0.419327441 | 0.610870102 |
| ENSG00000139835 | GRTP1 | 767.0074109 | -0.186032152 | 0.074470924 | -2.498050802 | 0.01248783 | 0.036156861 |
| ENSG00000139842 | CUL4A | 2678.971399 | -0.014170747 | 0.051374646 | -0.275831521 | 0.782677472 | 0.88216092 |
| ENSG00000139865 | TTC6 | 233.9086171 | -0.865934501 | 0.126194154 | -6.861922499 | 6.79E-12 | 6.46E-11 |
| ENSG00000139880 | CDH24 | 904.4943402 | 0.254114623 | 0.072861798 | 3.48762493 | 0.000487331 | 0.001933145 |
| ENSG00000139890 | REM2 | 18.42410988 | -0.62064369 | 0.40740724 | -1.523398776 | 0.127658968 | 0.257676791 |
| ENSG00000139899 | CBLN3 | 97.48263852 | -0.950382236 | 0.184494408 | -5.151279351 | 2.59E-07 | 1.61E-06 |
| ENSG00000139908 | TSSK4 | 3.730687656 | -0.516718632 | 0.909255797 | -0.568287421 | 0.569839819 | 0.737080657 |
| ENSG00000139910 | NOVA1 | 294.3573421 | -0.30126235 | 0.113942613 | -2.643983161 | 0.008193673 | 0.024945304 |
| ENSG00000139914 | FITM1 | 5.599241348 | -0.189045111 | 0.736208033 | -0.25678219 | 0.797346919 | 0.89183367 |
| ENSG00000139915 | MDGA2 | 47.0297915 | 0.24870169 | 0.256682652 | 0.968907278 | 0.332591454 | 0.523618692 |
| ENSG00000139921 | TMX1 | 1700.553884 | -0.66132117 | 0.05913781 | -11.18271319 | 4.96E-29 | 1.32E-27 |
| ENSG00000139926 | FRMD6 | 1065.268755 | -0.505781672 | 0.072533276 | -6.973098396 | 3.10E-12 | 3.05E-11 |
| ENSG00000139946 | PELI2 | 2.138413255 | -4.640367063 | 1.550729084 | -2.992377656 | 0.002768136 | 0.009459413 |
| ENSG00000139970 | RTN1 | 39.65860474 | -0.510563862 | 0.308777903 | -1.653498699 | 0.098229411 | 0.208928542 |
| ENSG00000139971 | ARMH4 | 20.23056339 | -0.624925255 | 0.395796005 | -1.578907436 | 0.11435729 | 0.236253263 |
| ENSG00000139974 | SLC38A6 | 117.4738121 | -0.603100218 | 0.174871643 | -3.448816553 | 0.000563049 | 0.002206791 |
| ENSG00000139977 | NAA30 | 993.9560939 | -0.576664228 | 0.074043279 | -7.788204869 | 6.80E-15 | 8.09E-14 |
| ENSG00000139990 | DCAF5 | 3727.370644 | -0.644678617 | 0.048809316 | -13.20810588 | 7.88E-40 | 3.18E-38 |
| ENSG00000139998 | RAB15 | 2653.009555 | 0.278320606 | 0.056211076 | 4.951348163 | 7.37E-07 | 4.35E-06 |
| ENSG00000140006 | WDR89 | 669.3972954 | -1.180072008 | 0.085854849 | -13.74496625 | 5.46E-43 | 2.46E-41 |
| ENSG00000140009 | ESR2 | 2.421124617 | 0.018508447 | 1.183705991 | 0.015636017 | 0.987524772 | 0.993927018 |
| ENSG00000140015 | KCNH5 | 2.726136334 | 1.090671324 | 1.081172653 | 1.008785526 | 0.313077504 | 0.501291124 |
| ENSG00000140022 | STON2 | 120.4667964 | -1.90773928 | 0.178685146 | -10.67654094 | 1.31E-26 | 3.18E-25 |
| ENSG00000140025 | EFCAB11 | 324.6279727 | -0.551936455 | 0.105007923 | -5.256141073 | 1.47E-07 | 9.40E-07 |
| ENSG00000140030 | GPR65 | 4.290692227 | -0.139344191 | 0.882985683 | -0.157810249 | 0.874606322 | 0.934962079 |
| ENSG00000140043 | PTGR2 | 223.7315592 | -0.788700864 | 0.124765236 | -6.32147938 | 2.59E-10 | 2.17E-09 |
| ENSG00000140044 | JDP2 | 700.0818205 | -0.276383327 | 0.079604444 | -3.471958532 | 0.000516676 | 0.002038622 |
| ENSG00000140057 | AK7 | 1030.296453 | 0.196566833 | 0.066590544 | 2.951873071 | 0.003158527 | 0.010653818 |
| ENSG00000140104 | CLBA1 | 960.8657234 | -0.163697452 | 0.069723412 | -2.347811848 | 0.018884058 | 0.05203607 |
| ENSG00000140105 | WARS1 | 4172.772835 | -0.579764189 | 0.049792235 | -11.64366663 | 2.47E-31 | 7.28E-30 |
| ENSG00000140153 | WDR20 | 822.9029335 | -0.371970116 | 0.077001832 | -4.830665803 | 1.36E-06 | 7.73E-06 |
| ENSG00000140157 | NIPA2 | 1106.486924 | -0.13641151 | 0.065887647 | -2.070365476 | 0.038418132 | 0.095940775 |

| | | | | | | | |
|---|---|---|---|---|---|---|---|
| ENSG00000140199 | SLC12A6 | 252.0233259 | 0.218079125 | 0.117612758 | 1.854213167 | 0.063708669 | 0.146631571 |
| ENSG00000140254 | DUOXA1 | 7.947113803 | -0.426858325 | 0.6182771 | -0.690399702 | 0.489942864 | 0.673011336 |
| ENSG00000140259 | MFAP1 | 1395.410113 | -0.344323621 | 0.062638894 | -5.496962042 | 3.86E-08 | 2.61E-07 |
| ENSG00000140262 | TCF12 | 1579.420834 | -0.575286892 | 0.057925458 | -9.931503539 | 3.04E-23 | 6.12E-22 |
| ENSG00000140263 | SORD | 1922.533053 | -0.424832887 | 0.055811948 | -7.611862724 | 2.70E-14 | 3.08E-13 |
| ENSG00000140264 | SERF2 | 6233.600134 | -0.168873166 | 0.047189892 | -3.57858767 | 0.000345456 | 0.001403358 |
| ENSG00000140265 | ZSCAN29 | 1363.692455 | -0.206633644 | 0.059114572 | -3.495477286 | 0.000473215 | 0.001881051 |
| ENSG00000140279 | DUOX2 | 5.154361342 | -0.445892117 | 0.775866064 | -0.574702436 | 0.56549256 | 0.733608512 |
| ENSG00000140280 | LYSMD2 | 1589.246419 | -1.248170573 | 0.062952796 | -19.82708703 | 1.74E-87 | 2.32E-85 |
| ENSG00000140284 | SLC27A2 | 1570.910526 | -0.594940363 | 0.061134796 | -9.731615996 | 2.21E-22 | 4.23E-21 |
| ENSG00000140285 | FGF7 | 16.26148709 | -3.100586353 | 0.565062075 | -5.487160598 | 4.08E-08 | 2.75E-07 |
| ENSG00000140287 | HDC | 7.858834659 | 3.01223513 | 0.848348257 | 3.550705864 | 0.0003842 | 0.001549054 |
| ENSG00000140299 | BNIP2 | 1573.26441 | -0.156984391 | 0.062805961 | -2.499514174 | 0.012436372 | 0.036030612 |
| ENSG00000140307 | GTF2A2 | 864.6813933 | -0.233913181 | 0.072162231 | -3.241490438 | 0.001189064 | 0.004389982 |
| ENSG00000140319 | SRP14 | 5382.975362 | -0.190596733 | 0.046200826 | -4.125396624 | 3.70E-05 | 0.000176214 |
| ENSG00000140320 | BAHD1 | 2573.854734 | -0.026400442 | 0.063680074 | -0.414579316 | 0.678449892 | 0.816989004 |
| ENSG00000140323 | DISP2 | 140.1561496 | -0.30475949 | 0.153319321 | -1.987743546 | 0.04684006 | 0.113237536 |
| ENSG00000140326 | CDAN1 | 787.227387 | 0.086298738 | 0.079368926 | 1.087311403 | 0.276899211 | 0.459495652 |
| ENSG00000140332 | TLE3 | 5643.221329 | -0.314029043 | 0.048454262 | -6.480937544 | 9.12E-11 | 7.92E-10 |
| ENSG00000140350 | ANP32A | 6223.909268 | -0.344619539 | 0.043909263 | -7.848447403 | 4.21E-15 | 5.09E-14 |
| ENSG00000140365 | COMMD4 | 1837.143787 | -0.321392166 | 0.062794276 | -5.118176159 | 3.09E-07 | 1.90E-06 |
| ENSG00000140367 | UBE2Q2 | 1833.805417 | -0.088504118 | 0.067060856 | -1.319758252 | 0.186915744 | 0.344626842 |
| ENSG00000140374 | ETFA | 4775.473254 | -0.453401625 | 0.044490221 | -10.19104007 | 2.17E-24 | 4.68E-23 |
| ENSG00000140382 | HMG20A | 1818.065166 | 0.070434146 | 0.060064307 | 1.172645618 | 0.240937948 | 0.416857721 |
| ENSG00000140386 | SCAPER | 465.5702267 | -0.079325373 | 0.091296593 | -0.868875506 | 0.384915228 | 0.577574263 |
| ENSG00000140391 | TSPAN3 | 4437.278062 | 0.378546631 | 0.045444349 | 8.329894416 | 8.09E-17 | 1.10E-15 |
| ENSG00000140395 | SKIC8 | 2277.30197 | -0.280972913 | 0.052692983 | -5.332264294 | 9.70E-08 | 6.30E-07 |
| ENSG00000140396 | NCOA2 | 3609.206799 | 0.224008215 | 0.060461879 | 3.704949624 | 0.000211433 | 0.000892578 |
| ENSG00000140398 | NEIL1 | 404.669 | -0.773517636 | 0.112992622 | -6.845735782 | 7.61E-12 | 7.21E-11 |
| ENSG00000140400 | MAN2C1 | 3083.236285 | 0.146609131 | 0.065120921 | 2.251336885 | 0.024364208 | 0.065043051 |
| ENSG00000140403 | DNAJA4 | 747.7806013 | -0.323872208 | 0.074813532 | -4.32905916 | 1.50E-05 | 7.54E-05 |
| ENSG00000140406 | TLNRD1 | 656.8908752 | -0.19691151 | 0.082174488 | -2.396260857 | 0.016563298 | 0.046404745 |
| ENSG00000140416 | TPM1 | 6644.860596 | 0.140731796 | 0.042364676 | 3.321913637 | 0.000894024 | 0.003378732 |

| | | | | | | | |
|---|---|---|---|---|---|---|---|
| ENSG00000140443 | IGF1R | 2561.948488 | 1.001716167 | 0.062962609 | 15.90969905 | 5.43E-57 | 3.60E-55 |
| ENSG00000140450 | ARRDC4 | 549.7454017 | -0.046232871 | 0.090744232 | -0.509485508 | 0.610411954 | 0.768331974 |
| ENSG00000140451 | PIF1 | 319.9470205 | -0.468757365 | 0.107601816 | -4.356407547 | 1.32E-05 | 6.72E-05 |
| ENSG00000140455 | USP3 | 1625.470845 | -0.064897462 | 0.057719837 | -1.124352825 | 0.260863377 | 0.441163976 |
| ENSG00000140463 | BBS4 | 940.7679442 | -0.177618097 | 0.067439566 | -2.633737273 | 0.008445081 | 0.025615516 |
| ENSG00000140464 | PML | 717.0964703 | -0.122488023 | 0.078975113 | -1.550969903 | 0.120908897 | 0.246956175 |
| ENSG00000140465 | CYP1A1 | 779.7038244 | -0.033902617 | 0.080974094 | -0.418684728 | 0.675446558 | 0.814669479 |
| ENSG00000140470 | ADAMTS17 | 48.64938133 | -0.979778683 | 0.261399439 | -3.748204992 | 0.000178105 | 0.000762546 |
| ENSG00000140471 | LINS1 | 427.75227 | 0.087639725 | 0.092269946 | 0.94981876 | 0.342204352 | 0.53329254 |
| ENSG00000140474 | ULK3 | 3769.768311 | 0.082280464 | 0.059188048 | 1.390153351 | 0.164482316 | 0.3126636 |
| ENSG00000140479 | PCSK6 | 2425.125678 | 1.055440968 | 0.060167476 | 17.54171915 | 6.88E-69 | 6.17E-67 |
| ENSG00000140481 | CCDC33 | 12.81778093 | -3.012388715 | 0.64434908 | -4.675088099 | 2.94E-06 | 1.61E-05 |
| ENSG00000140497 | SCAMP2 | 3162.533868 | -0.278234826 | 0.055217156 | -5.038919903 | 4.68E-07 | 2.83E-06 |
| ENSG00000140505 | CYP1A2 | 4.130724694 | 0.524912272 | 0.865025766 | 0.60681692 | 0.543972418 | 0.717456866 |
| ENSG00000140511 | HAPLN3 | 17.3098204 | 0.143552618 | 0.423254464 | 0.33916386 | 0.734486293 | 0.853578385 |
| ENSG00000140519 | RHCG | 43.11529917 | 0.491515914 | 0.270647417 | 1.816074656 | 0.069358921 | 0.157487246 |
| ENSG00000140521 | POLG | 4522.196214 | 0.12469967 | 0.045702529 | 2.728506999 | 0.006362174 | 0.019955465 |
| ENSG00000140525 | FANCI | 6789.136383 | 0.000486712 | 0.050248559 | 0.009686093 | 0.992271737 | 0.99592006 |
| ENSG00000140526 | ABHD2 | 2250.802437 | 0.14233684 | 0.060121511 | 2.367486078 | 0.017909394 | 0.049690553 |
| ENSG00000140527 | WDR93 | 11.68210192 | 0.428983034 | 0.543825557 | 0.788824704 | 0.430214469 | 0.620481249 |
| ENSG00000140534 | TICRR | 1529.246161 | -0.171427107 | 0.061210381 | -2.800621459 | 0.005100431 | 0.01639169 |
| ENSG00000140538 | NTRK3 | 28.74536142 | 7.225677389 | 1.208520557 | 5.97894454 | 2.25E-09 | 1.73E-08 |
| ENSG00000140543 | DET1 | 27.47507519 | 0.645980053 | 0.346163784 | 1.866111023 | 0.062025834 | 0.143362482 |
| ENSG00000140545 | MFGE8 | 430.4882423 | 0.973278197 | 0.095916856 | 10.14710278 | 3.41E-24 | 7.21E-23 |
| ENSG00000140548 | ZNF710 | 1191.933171 | -0.136916436 | 0.073222046 | -1.869879952 | 0.06150049 | 0.142464026 |
| ENSG00000140553 | UNC45A | 5780.5018 | 0.267035799 | 0.055154813 | 4.84156839 | 1.29E-06 | 7.35E-06 |
| ENSG00000140563 | MCTP2 | 62.93870266 | 1.643673336 | 0.241915149 | 6.794420868 | 1.09E-11 | 1.01E-10 |
| ENSG00000140564 | FURIN | 3980.707222 | 0.017007302 | 0.063371882 | 0.268373002 | 0.788412219 | 0.885492107 |
| ENSG00000140575 | IQGAP1 | 11790.33551 | -0.101717511 | 0.048896395 | -2.080266063 | 0.037501136 | 0.094029517 |
| ENSG00000140577 | CRTC3 | 2125.041997 | 0.481633997 | 0.05841387 | 8.245199305 | 1.65E-16 | 2.22E-15 |
| ENSG00000140598 | EFL1 | 1616.396918 | -0.084096791 | 0.059466251 | -1.414193607 | 0.157305064 | 0.301872024 |
| ENSG00000140600 | SH3GL3 | 21.65913369 | -1.80776268 | 0.422139004 | -4.282387234 | 1.85E-05 | 9.22E-05 |
| ENSG00000140612 | SEC11A | 5118.303317 | 0.045443125 | 0.048999273 | 0.927424481 | 0.353706185 | 0.545223752 |

| | | | | | | | |
|---|---|---|---|---|---|---|---|
| ENSG00000140632 | GLYR1 | 4250.313961 | 0.093511651 | 0.055213539 | 1.69363623 | 0.090334429 | 0.195018991 |
| ENSG00000140650 | PMM2 | 920.5979081 | 0.107553787 | 0.071405892 | 1.506231252 | 0.132007819 | 0.264546115 |
| ENSG00000140675 | SLC5A2 | 28.74643052 | -0.328265204 | 0.34495964 | -0.95160467 | 0.341297512 | 0.532356356 |
| ENSG00000140682 | TGFB1I1 | 354.6359655 | -1.214772771 | 0.11277014 | -10.77211371 | 4.66E-27 | 1.16E-25 |
| ENSG00000140688 | RUSF1 | 2996.437717 | 0.108850459 | 0.058043827 | 1.875314974 | 0.060749404 | 0.141023208 |
| ENSG00000140691 | ARMC5 | 481.0828214 | 0.172204293 | 0.09624744 | 1.789183106 | 0.073585331 | 0.165397561 |
| ENSG00000140694 | PARN | 1464.352524 | 0.144250006 | 0.060962402 | 2.366212648 | 0.017971119 | 0.049819634 |
| ENSG00000140718 | FTO | 1536.916696 | 0.181471428 | 0.062032191 | 2.92543958 | 0.0034397 | 0.011510826 |
| ENSG00000140740 | UQCRC2 | 3169.33169 | -0.077912533 | 0.05135392 | -1.517168161 | 0.129224233 | 0.259989818 |
| ENSG00000140743 | CDR2 | 389.779299 | -0.053243866 | 0.097718852 | -0.544867901 | 0.585844374 | 0.749814489 |
| ENSG00000140750 | ARHGAP17 | 1104.038271 | -0.083729485 | 0.065778921 | -1.272892354 | 0.203056243 | 0.36735885 |
| ENSG00000140795 | MYLK3 | 4.196918367 | -1.284509239 | 0.887063315 | -1.44804685 | 0.147603949 | 0.288050581 |
| ENSG00000140798 | ABCC12 | 2.428719191 | 0.379972552 | 1.145966718 | 0.331573811 | 0.740211098 | 0.856646281 |
| ENSG00000140807 | NKD1 | 43.93649474 | 2.410406992 | 0.320743234 | 7.515067308 | 5.69E-14 | 6.32E-13 |
| ENSG00000140829 | DHX38 | 4830.484027 | 0.32087423 | 0.046847457 | 6.849341541 | 7.42E-12 | 7.04E-11 |
| ENSG00000140830 | TXNL4B | 939.9816032 | 0.329094684 | 0.069863889 | 4.710511927 | 2.47E-06 | 1.37E-05 |
| ENSG00000140832 | MARVELD3 | 1218.647529 | 0.235994212 | 0.064912132 | 3.63559482 | 0.00027734 | 0.001148495 |
| ENSG00000140836 | ZFHX3 | 3302.329558 | -0.282525994 | 0.14784703 | -1.910934521 | 0.056012996 | 0.131942428 |
| ENSG00000140839 | CLEC18B | 19.36151115 | -0.144065526 | 0.406915871 | -0.354042533 | 0.723307008 | 0.846082679 |
| ENSG00000140848 | CPNE2 | 626.4519683 | 0.003701494 | 0.087137803 | 0.042478627 | 0.96611715 | 0.984791265 |
| ENSG00000140853 | NLRC5 | 7.872047809 | 1.002824082 | 0.639756152 | 1.567509867 | 0.116995562 | 0.240771345 |
| ENSG00000140854 | KATNB1 | 689.7526947 | 0.177588096 | 0.077300027 | 2.297387238 | 0.021596689 | 0.058464485 |
| ENSG00000140859 | KIFC3 | 147.6149767 | -0.585597787 | 0.153438512 | -3.816498078 | 0.000135359 | 0.00059033 |
| ENSG00000140873 | ADAMTS18 | 28.46421746 | 0.828087035 | 0.33484029 | 2.47308063 | 0.013395392 | 0.038463183 |
| ENSG00000140876 | NUDT7 | 464.2689676 | -0.601478348 | 0.08955009 | -6.716669373 | 1.86E-11 | 1.70E-10 |
| ENSG00000140905 | GCSH | 27.10950869 | -0.158095025 | 0.359194332 | -0.440137863 | 0.659837261 | 0.80321925 |
| ENSG00000140931 | CMTM3 | 433.5774141 | -0.280464607 | 0.093935398 | -2.985717997 | 0.002829134 | 0.009647738 |
| ENSG00000140932 | CMTM2 | 1.743264462 | -0.293746188 | 1.372476023 | -0.214026462 | 0.830526421 | NA |
| ENSG00000140937 | CDH11 | 98.22492769 | -0.964092837 | 0.192942418 | -4.996790467 | 5.83E-07 | 3.47E-06 |
| ENSG00000140939 | NOL3 | 1205.443244 | -0.01869744 | 0.06797839 | -0.275049761 | 0.783278008 | 0.882490914 |
| ENSG00000140941 | MAP1LC3B | 3067.21609 | 0.574113214 | 0.051027173 | 11.25112731 | 2.29E-29 | 6.19E-28 |
| ENSG00000140943 | MBTPS1 | 10995.23319 | 0.657173819 | 0.044513691 | 14.76340871 | 2.52E-49 | 1.37E-47 |
| ENSG00000140945 | CDH13 | 2.346436408 | -1.487958983 | 1.217014305 | -1.222630644 | 0.221469234 | 0.392226065 |

| | | | | | | | |
|---|---|---|---|---|---|---|---|
| ENSG00000140948 | ZCCHC14 | 2927.617115 | 0.364935133 | 0.052481915 | 6.953540769 | 3.56E-12 | 3.48E-11 |
| ENSG00000140950 | MEAK7 | 750.5381036 | 0.199587661 | 0.074792156 | 2.66856408 | 0.007617625 | 0.02339648 |
| ENSG00000140955 | ADAD2 | 1.546961365 | 1.880190412 | 1.607046532 | 1.169966379 | 0.242014499 | NA |
| ENSG00000140961 | OSGIN1 | 412.289706 | 0.117268835 | 0.095256545 | 1.231084278 | 0.218291345 | 0.387918951 |
| ENSG00000140983 | RHOT2 | 3742.414219 | 0.069452413 | 0.056194 | 1.235940009 | 0.216480864 | 0.385894088 |
| ENSG00000140986 | RPL3L | 9.45847004 | -0.814967479 | 0.610258667 | -1.335445972 | 0.181730427 | 0.3373834 |
| ENSG00000140987 | ZSCAN32 | 780.7209451 | 0.444231222 | 0.077817489 | 5.708629596 | 1.14E-08 | 8.13E-08 |
| ENSG00000140988 | RPS2 | 13208.73259 | 0.112148037 | 0.057834332 | 1.939125653 | 0.05248604 | 0.124886397 |
| ENSG00000140990 | NDUFB10 | 2439.415658 | 0.100749931 | 0.0553324 | 1.820812583 | 0.068635351 | 0.156106975 |
| ENSG00000140992 | PDPK1 | 2974.403137 | -0.010923754 | 0.0511096 | -0.21373194 | 0.830756102 | 0.910363284 |
| ENSG00000140993 | TIGD7 | 452.5055495 | 0.498102358 | 0.091938466 | 5.417779731 | 6.03E-08 | 3.99E-07 |
| ENSG00000140995 | DEF8 | 1891.753579 | 0.603120787 | 0.059786031 | 10.08798843 | 6.24E-24 | 1.30E-22 |
| ENSG00000141002 | TCF25 | 4449.957497 | 0.598076801 | 0.050586224 | 11.82291844 | 2.97E-32 | 9.04E-31 |
| ENSG00000141012 | GALNS | 1770.693852 | 0.345559107 | 0.065475329 | 5.277699468 | 1.31E-07 | 8.40E-07 |
| ENSG00000141013 | GAS8 | 1831.13526 | 0.685920231 | 0.060192866 | 11.39537411 | 4.41E-30 | 1.23E-28 |
| ENSG00000141026 | MED9 | 538.7471586 | -0.203638606 | 0.087660197 | -2.323045276 | 0.02017672 | 0.055101662 |
| ENSG00000141027 | NCOR1 | 6623.785445 | -0.152806714 | 0.055293858 | -2.763538646 | 0.005717832 | 0.018132025 |
| ENSG00000141030 | COPS3 | 2328.539457 | 0.074592203 | 0.054178037 | 1.376797804 | 0.16857477 | 0.318752491 |
| ENSG00000141034 | GID4 | 549.7222131 | 0.224406518 | 0.08316725 | 2.698255827 | 0.006970385 | 0.021611114 |
| ENSG00000141040 | ZNF287 | 252.400287 | 0.033675626 | 0.120263221 | 0.280016004 | 0.779465227 | 0.880308728 |
| ENSG00000141068 | KSR1 | 130.1724504 | -0.303413071 | 0.16012963 | -1.894796554 | 0.05811936 | 0.135908065 |
| ENSG00000141076 | UTP4 | 1779.180785 | -0.052752858 | 0.067466673 | -0.781909888 | 0.434267534 | 0.624230668 |
| ENSG00000141084 | RANBP10 | 1554.20832 | 0.191840431 | 0.063019683 | 3.044135121 | 0.002333503 | 0.008097196 |
| ENSG00000141086 | CTRL | 4.466042307 | 0.136798031 | 0.823518918 | 0.166114012 | 0.868067228 | 0.931081965 |
| ENSG00000141098 | GFOD2 | 839.51068 | 0.474880329 | 0.074761641 | 6.351924931 | 2.13E-10 | 1.79E-09 |
| ENSG00000141101 | NOB1 | 2536.111317 | 0.272690255 | 0.052722172 | 5.172212084 | 2.31E-07 | 1.45E-06 |
| ENSG00000141127 | PRPSAP2 | 886.0131931 | -0.116801031 | 0.070913992 | -1.647080187 | 0.099541565 | 0.21117946 |
| ENSG00000141179 | PCTP | 642.3762436 | 0.078002634 | 0.079788129 | 0.97762204 | 0.328261292 | 0.518686516 |
| ENSG00000141198 | TOM1L1 | 2610.053961 | -0.110570524 | 0.052855996 | -2.091920174 | 0.036445657 | 0.091794398 |
| ENSG00000141219 | C17orf80 | 1031.978641 | -0.210521925 | 0.070142128 | -3.001362121 | 0.002687747 | 0.009201152 |
| ENSG00000141232 | TOB1 | 5355.215282 | -0.592628698 | 0.046713475 | -12.68646148 | 7.03E-37 | 2.55E-35 |
| ENSG00000141252 | VPS53 | 1279.138911 | -0.047179232 | 0.06150391 | -0.767093218 | 0.443026094 | 0.631832054 |
| ENSG00000141258 | SGSM2 | 6289.57769 | -0.165735186 | 0.05424545 | -3.055282734 | 0.002248484 | 0.0078318 |

| | | | | | | | |
|---|---|---|---|---|---|---|---|
| ENSG00000141279 | NPEPPS | 5603.431507 | 0.361977358 | 0.048505281 | 7.46263812 | 8.48E-14 | 9.28E-13 |
| ENSG00000141294 | LRRC46 | 49.34457224 | -0.401954096 | 0.250664253 | -1.603555715 | 0.108812019 | 0.226775369 |
| ENSG00000141295 | SCRN2 | 871.6430506 | 0.143231279 | 0.079066488 | 1.81152954 | 0.070058921 | 0.158668342 |
| ENSG00000141298 | SSH2 | 609.0445151 | 0.110818175 | 0.081303463 | 1.363019124 | 0.172876484 | 0.325037828 |
| ENSG00000141314 | RHBDL3 | 96.11084214 | 1.181667922 | 0.19614144 | 6.024570457 | 1.70E-09 | 1.32E-08 |
| ENSG00000141337 | ARSG | 736.7886181 | 0.248257984 | 0.077922075 | 3.185977591 | 0.001442658 | 0.005222435 |
| ENSG00000141349 | G6PC3 | 1849.984713 | 0.050210442 | 0.057275389 | 0.876649528 | 0.380677029 | 0.573201906 |
| ENSG00000141367 | CLTC | 42013.48251 | 0.037403863 | 0.050938195 | 0.73429895 | 0.462766558 | 0.649288369 |
| ENSG00000141371 | CHCT1 | 4.400792006 | 0.336127522 | 0.83255542 | 0.403729906 | 0.686411346 | 0.822742745 |
| ENSG00000141376 | BCAS3 | 20167.37943 | -0.22747096 | 0.043570906 | -5.220707573 | 1.78E-07 | 1.13E-06 |
| ENSG00000141378 | PTRH2 | 1663.860734 | 0.250819094 | 0.056802004 | 4.415673304 | 1.01E-05 | 5.21E-05 |
| ENSG00000141380 | SS18 | 3016.108834 | 0.141935723 | 0.050476092 | 2.811939642 | 0.004924374 | 0.015890441 |
| ENSG00000141384 | TAF4B | 377.1429254 | 0.285045534 | 0.100184255 | 2.845212889 | 0.004438175 | 0.014484526 |
| ENSG00000141385 | AFG3L2 | 4273.002954 | 0.281474382 | 0.04507967 | 6.243931698 | 4.27E-10 | 3.52E-09 |
| ENSG00000141391 | PRELID3A | 183.8948348 | 0.284006309 | 0.134372153 | 2.11358011 | 0.034551146 | 0.08765679 |
| ENSG00000141401 | IMPA2 | 696.5658686 | 0.190998627 | 0.076728189 | 2.489288883 | 0.012799891 | 0.036920597 |
| ENSG00000141404 | GNAL | 6.793615957 | 0.009805464 | 0.662143595 | 0.014808668 | 0.988184825 | 0.994030148 |
| ENSG00000141424 | SLC39A6 | 15116.21991 | 0.184746953 | 0.049736167 | 3.714539437 | 0.000203574 | 0.000862392 |
| ENSG00000141425 | RPRD1A | 2375.746843 | 0.670382793 | 0.060220228 | 11.13218622 | 8.75E-29 | 2.31E-27 |
| ENSG00000141428 | C18orf21 | 259.7558889 | 0.665825417 | 0.117458851 | 5.668584437 | 1.44E-08 | 1.02E-07 |
| ENSG00000141429 | GALNT1 | 2117.12426 | 0.867013256 | 0.054843254 | 15.80893169 | 2.70E-56 | 1.75E-54 |
| ENSG00000141441 | GAREM1 | 386.7803989 | 0.580940849 | 0.104019103 | 5.584943843 | 2.34E-08 | 1.61E-07 |
| ENSG00000141446 | ESCO1 | 1245.711261 | 0.082518159 | 0.064390717 | 1.281522597 | 0.200010168 | 0.363136783 |
| ENSG00000141447 | OSBPL1A | 639.2673169 | 0.485831775 | 0.080964713 | 6.000537215 | 1.97E-09 | 1.52E-08 |
| ENSG00000141448 | GATA6 | 120.3708021 | 0.34908561 | 0.167941307 | 2.078616721 | 0.037652593 | 0.094337099 |
| ENSG00000141449 | GREB1L | 235.8912315 | 1.570965205 | 0.134078933 | 11.71671921 | 1.05E-31 | 3.12E-30 |
| ENSG00000141452 | RMC1 | 650.1150889 | 0.048890905 | 0.081886613 | 0.597056145 | 0.550469896 | 0.722629536 |
| ENSG00000141456 | PELP1 | 2065.215369 | 0.064893219 | 0.05807871 | 1.11733231 | 0.263852266 | 0.444501063 |
| ENSG00000141458 | NPC1 | 1918.022777 | -0.075576202 | 0.054885602 | -1.376976814 | 0.168519417 | 0.318674092 |
| ENSG00000141469 | SLC14A1 | 4.60650184 | 1.698792096 | 0.933085637 | 1.82061756 | 0.068665012 | 0.156143474 |
| ENSG00000141480 | ARRB2 | 903.9266269 | -0.049018207 | 0.070467062 | -0.695618711 | 0.486667645 | 0.670117394 |
| ENSG00000141497 | ZMYND15 | 33.99674857 | 0.219044645 | 0.298352776 | 0.734180014 | 0.462839032 | 0.64935034 |
| ENSG00000141499 | WRAP53 | 423.0412336 | -0.268710084 | 0.094383901 | -2.84699067 | 0.004413466 | 0.01441619 |

| | | | | | | | |
|---|---|---|---|---|---|---|---|
| ENSG00000141503 | MINK1 | 2813.003333 | 0.08675855 | 0.066229556 | 1.309967254 | 0.190206914 | 0.349487112 |
| ENSG00000141504 | SAT2 | 159.8729134 | 0.42918399 | 0.144997779 | 2.959934926 | 0.00307704 | 0.010401902 |
| ENSG00000141505 | ASGR1 | 6.085288229 | 0.03742623 | 0.723534108 | 0.051726974 | 0.958746244 | 0.981197069 |
| ENSG00000141506 | PIK3R5 | 8.042164863 | 1.08266943 | 0.653682065 | 1.656263018 | 0.097668566 | 0.207967775 |
| ENSG00000141510 | TP53 | 3781.794179 | 0.261455251 | 0.048673151 | 5.37165248 | 7.80E-08 | 5.10E-07 |
| ENSG00000141519 | CCDC40 | 127.1189119 | 0.73023531 | 0.167665443 | 4.35531197 | 1.33E-05 | 6.75E-05 |
| ENSG00000141522 | ARHGDIA | 12677.48138 | 0.037094391 | 0.047665202 | 0.778227917 | 0.436434662 | 0.626296602 |
| ENSG00000141524 | TMC6 | 1412.904894 | 0.11150189 | 0.062549691 | 1.782612982 | 0.074649325 | 0.167346381 |
| ENSG00000141526 | SLC16A3 | 404.9197084 | -0.397135965 | 0.101141439 | -3.926540568 | 8.62E-05 | 0.000388615 |
| ENSG00000141527 | CARD14 | 338.3746563 | 0.0346612 | 0.104859502 | 0.330548968 | 0.740985198 | 0.857152965 |
| ENSG00000141540 | TTYH2 | 42.52564386 | 0.248809612 | 0.273191377 | 0.910752067 | 0.362426023 | 0.555140503 |
| ENSG00000141542 | RAB40B | 605.4352737 | 0.4244418 | 0.084317137 | 5.03387348 | 4.81E-07 | 2.90E-06 |
| ENSG00000141543 | EIF4A3 | 4425.986649 | -0.002621129 | 0.044872545 | -0.058412762 | 0.953419849 | 0.978413319 |
| ENSG00000141551 | CSNK1D | 8959.388962 | 0.202028699 | 0.047229466 | 4.277598644 | 1.89E-05 | 9.41E-05 |
| ENSG00000141552 | ANAPC11 | 1795.060543 | 0.152359717 | 0.058250135 | 2.615611385 | 0.008906787 | 0.026877359 |
| ENSG00000141556 | TBCD | 4631.344962 | 0.466176089 | 0.048082604 | 9.695317022 | 3.16E-22 | 5.97E-21 |
| ENSG00000141560 | FN3KRP | 1587.037102 | 0.232727434 | 0.05655524 | 4.115046381 | 3.87E-05 | 0.000183777 |
| ENSG00000141562 | NARF | 1685.564991 | 0.596710692 | 0.063824634 | 9.349222244 | 8.83E-21 | 1.54E-19 |
| ENSG00000141564 | RPTOR | 1916.910999 | -0.030330715 | 0.059953815 | -0.505901334 | 0.612925916 | 0.769891065 |
| ENSG00000141568 | FOXK2 | 5175.88069 | 0.172571378 | 0.0460428 | 3.748064396 | 0.000178205 | 0.000762831 |
| ENSG00000141569 | TRIM65 | 1463.32395 | 0.27975476 | 0.061173011 | 4.573172965 | 4.80E-06 | 2.57E-05 |
| ENSG00000141570 | CBX8 | 1063.684717 | 0.18670378 | 0.071566816 | 2.608803767 | 0.009085933 | 0.027357481 |
| ENSG00000141574 | SECTM1 | 9.895178149 | 1.181538391 | 0.582062576 | 2.02991644 | 0.042365034 | 0.104043065 |
| ENSG00000141576 | RNF157 | 513.5990555 | 0.4504999 | 0.09248033 | 4.871305084 | 1.11E-06 | 6.38E-06 |
| ENSG00000141577 | CEP131 | 1076.775758 | 0.164484702 | 0.066640837 | 2.468226836 | 0.013578424 | 0.038915338 |
| ENSG00000141579 | ZNF750 | 93.40957385 | 0.000511332 | 0.182785499 | 0.002797443 | 0.997767967 | 0.998725768 |
| ENSG00000141580 | WDR45B | 4510.572166 | 0.47300784 | 0.047219283 | 10.01726019 | 1.28E-23 | 2.62E-22 |
| ENSG00000141582 | CBX4 | 1809.528741 | 0.301397321 | 0.059708679 | 5.047797496 | 4.47E-07 | 2.71E-06 |
| ENSG00000141622 | RNF165 | 149.0163268 | 0.813493639 | 0.15775488 | 5.156693982 | 2.51E-07 | 1.57E-06 |
| ENSG00000141627 | DYM | 1765.208808 | 0.675896996 | 0.060895857 | 11.09922788 | 1.27E-28 | 3.32E-27 |
| ENSG00000141639 | MAPK4 | 373.6382459 | -1.515424174 | 0.10577625 | -14.32669602 | 1.49E-46 | 7.48E-45 |
| ENSG00000141642 | ELAC1 | 53.46084905 | 0.537152351 | 0.250003451 | 2.148579743 | 0.031667729 | 0.081404942 |
| ENSG00000141644 | MBD1 | 1983.207659 | 0.452458166 | 0.053657188 | 8.432386811 | 3.39E-17 | 4.72E-16 |

| | | | | | | | |
|---|---|---|---|---|---|---|---|
| ENSG00000141646 | SMAD4 | 765.6944302 | 0.253453499 | 0.073189981 | 3.46295349 | 0.000534281 | 0.002103381 |
| ENSG00000141655 | TNFRSF11A | 271.7793974 | 1.047979977 | 0.117050477 | 8.953231167 | 3.45E-19 | 5.51E-18 |
| ENSG00000141664 | ZCCHC2 | 933.1199124 | 0.541528401 | 0.073445294 | 7.373221277 | 1.67E-13 | 1.79E-12 |
| ENSG00000141665 | FBXO15 | 40.6197052 | -1.260089038 | 0.299678622 | -4.204801233 | 2.61E-05 | 0.000127624 |
| ENSG00000141668 | CBLN2 | 361.4200601 | -0.710040632 | 0.10574462 | -6.714673826 | 1.88E-11 | 1.72E-10 |
| ENSG00000141682 | PMAIP1 | 86.05555184 | 0.795402664 | 0.1954647 | 4.069290593 | 4.72E-05 | 0.000221357 |
| ENSG00000141696 | P3H4 | 2095.406485 | -0.071589747 | 0.056616904 | -1.264458895 | 0.206065365 | 0.371601248 |
| ENSG00000141698 | NT5C3B | 1392.586427 | 0.394009923 | 0.059530771 | 6.618592679 | 3.63E-11 | 3.24E-10 |
| ENSG00000141699 | RETREG3 | 2066.183967 | 0.13469364 | 0.061304938 | 2.197109138 | 0.028012653 | 0.073240284 |
| ENSG00000141736 | ERBB2 | 9261.896212 | -0.432814769 | 0.061561875 | -7.030565108 | 2.06E-12 | 2.05E-11 |
| ENSG00000141738 | GRB7 | 599.5020715 | 0.105017346 | 0.08883837 | 1.182116983 | 0.237159287 | 0.41238001 |
| ENSG00000141741 | MIEN1 | 614.6832089 | 0.203018301 | 0.08747148 | 2.320965648 | 0.020288698 | 0.055315276 |
| ENSG00000141744 | PNMT | 2.158780111 | -0.431888908 | 1.17158427 | -0.368636657 | 0.712398569 | 0.839972059 |
| ENSG00000141753 | IGFBP4 | 7050.996458 | 0.058790628 | 0.047074487 | 1.24888515 | 0.211707084 | 0.379270637 |
| ENSG00000141756 | FKBP10 | 5751.459118 | 0.214776309 | 0.048891104 | 4.392952741 | 1.12E-05 | 5.73E-05 |
| ENSG00000141759 | TXNL4A | 1680.525139 | 0.526357183 | 0.056334872 | 9.343363437 | 9.33E-21 | 1.62E-19 |
| ENSG00000141854 | MISP3 | 550.0318945 | 0.328322468 | 0.097087616 | 3.381713137 | 0.000720353 | 0.002770311 |
| ENSG00000141858 | SAMD1 | 2228.153214 | -0.380749764 | 0.05892136 | -6.461998898 | 1.03E-10 | 8.90E-10 |
| ENSG00000141867 | BRD4 | 2107.744799 | -0.052074049 | 0.059450404 | -0.875924219 | 0.381071232 | 0.573376841 |
| ENSG00000141873 | SLC39A3 | 251.3152943 | 0.389257078 | 0.128925915 | 3.019230676 | 0.002534175 | 0.008718352 |
| ENSG00000141905 | NFIC | 2932.856229 | 0.472262498 | 0.055187146 | 8.557472753 | 1.15E-17 | 1.66E-16 |
| ENSG00000141933 | TPGS1 | 76.28798409 | 0.205298742 | 0.208481912 | 0.984731672 | 0.324755891 | 0.51488701 |
| ENSG00000141934 | PLPP2 | 1583.752289 | 0.391292377 | 0.067570947 | 5.790837498 | 7.00E-09 | 5.12E-08 |
| ENSG00000141956 | PRDM15 | 858.888731 | 0.014039632 | 0.069276304 | 0.202661388 | 0.839399705 | 0.915367 |
| ENSG00000141959 | PFKL | 7476.265248 | -0.082033664 | 0.052800352 | -1.553657532 | 0.120266128 | 0.246023273 |
| ENSG00000141965 | FEM1A | 1465.592979 | 0.16978406 | 0.063430475 | 2.676695384 | 0.00743522 | 0.022903712 |
| ENSG00000141968 | VAV1 | 255.5506515 | 0.125699182 | 0.120066712 | 1.046911168 | 0.295140551 | 0.481018631 |
| ENSG00000141971 | MVB12A | 592.3270911 | -0.373876089 | 0.082809503 | -4.514893506 | 6.33E-06 | 3.35E-05 |
| ENSG00000141977 | CIB3 | 6.736170826 | -1.951204738 | 0.737899181 | -2.644270095 | 0.00818673 | 0.024927474 |
| ENSG00000141985 | SH3GL1 | 2606.732985 | 0.024566748 | 0.061790619 | 0.397580542 | 0.690939406 | 0.825657653 |
| ENSG00000141994 | DUS3L | 580.5085373 | 0.192623754 | 0.090957423 | 2.1177354 | 0.034197483 | 0.08682792 |
| ENSG00000142002 | DPP9 | 1976.681951 | 0.093462717 | 0.069554746 | 1.343728818 | 0.179036095 | 0.333812913 |
| ENSG00000142039 | CCDC97 | 1774.143818 | -0.18532133 | 0.061430143 | -3.016781672 | 0.002554737 | 0.008779877 |

| | | | | | | | |
|---|---|---|---|---|---|---|---|
| ENSG00000142046 | TMEM91 | 56.18217965 | -0.024661778 | 0.237786114 | -0.103714123 | 0.917396219 | 0.959384996 |
| ENSG00000142065 | ZFP14 | 338.3112756 | 0.21722778 | 0.106493403 | 2.039823824 | 0.041367876 | 0.101923035 |
| ENSG00000142082 | SIRT3 | 436.7934946 | 0.026532154 | 0.092331934 | 0.287356202 | 0.773839593 | 0.876765954 |
| ENSG00000142089 | IFITM3 | 2403.598482 | -0.027742599 | 0.06001791 | -0.462238675 | 0.643910172 | 0.791314543 |
| ENSG00000142102 | PGGHG | 464.5350071 | 0.190005398 | 0.092476266 | 2.054639605 | 0.039913827 | 0.098964894 |
| ENSG00000142149 | HUNK | 83.16438413 | 0.375250222 | 0.225514997 | 1.663970141 | 0.096118397 | 0.205245836 |
| ENSG00000142156 | COL6A1 | 34.78311355 | -0.835149241 | 0.298764139 | -2.795346337 | 0.005184414 | 0.016622007 |
| ENSG00000142166 | IFNAR1 | 1791.4457 | -0.010020329 | 0.058375613 | -0.171652657 | 0.863710608 | 0.929172842 |
| ENSG00000142168 | SOD1 | 6324.098964 | 0.018382579 | 0.044795337 | 0.410368148 | 0.681535908 | 0.819200091 |
| ENSG00000142173 | COL6A2 | 131.2073222 | -0.237469623 | 0.160663951 | -1.478051684 | 0.139393945 | 0.275616023 |
| ENSG00000142178 | SIK1 | 4.514680084 | 0.022031292 | 0.835837675 | 0.026358338 | 0.978971524 | 0.990493497 |
| ENSG00000142186 | SCYL1 | 4753.982259 | 0.09840743 | 0.053125537 | 1.852356444 | 0.063974655 | 0.147110926 |
| ENSG00000142188 | TMEM50B | 1312.622956 | 0.076242256 | 0.061791439 | 1.233864385 | 0.217253443 | 0.386840952 |
| ENSG00000142192 | APP | 14451.71156 | 0.462394566 | 0.046325256 | 9.98147889 | 1.84E-23 | 3.72E-22 |
| ENSG00000142197 | DOP1B | 3780.630755 | -0.346803827 | 0.050355321 | -6.887133656 | 5.69E-12 | 5.46E-11 |
| ENSG00000142207 | URB1 | 1904.509054 | 0.082354387 | 0.056058637 | 1.469075799 | 0.141812231 | 0.279314208 |
| ENSG00000142208 | AKT1 | 17504.907 | -0.351282224 | 0.040672726 | -8.636800606 | 5.78E-18 | 8.49E-17 |
| ENSG00000142227 | EMP3 | 120.6488545 | -0.54631231 | 0.167117789 | -3.269025481 | 0.001079186 | 0.004015657 |
| ENSG00000142230 | SAE1 | 6763.792501 | 0.187711449 | 0.042762502 | 4.389627325 | 1.14E-05 | 5.81E-05 |
| ENSG00000142233 | NTN5 | 53.78044681 | 0.852801071 | 0.271961263 | 3.135744633 | 0.001714184 | 0.00610991 |
| ENSG00000142235 | LMTK3 | 676.3837733 | 0.117193755 | 0.090077366 | 1.301034432 | 0.193246669 | 0.353910154 |
| ENSG00000142252 | GEMIN7 | 520.0504033 | -0.113309477 | 0.091698142 | -1.23567909 | 0.216577873 | 0.386016447 |
| ENSG00000142273 | CBLC | 4.278378548 | -0.554902959 | 0.852192088 | -0.651147748 | 0.514951116 | 0.693942232 |
| ENSG00000142279 | WTIP | 291.6293899 | 0.550355968 | 0.11249772 | 4.89215221 | 9.97E-07 | 5.77E-06 |
| ENSG00000142319 | SLC6A3 | 3.598448873 | 0.612934351 | 0.94231546 | 0.650455582 | 0.515397984 | 0.694381263 |
| ENSG00000142327 | RNPEPL1 | 1887.670156 | 0.091787298 | 0.063997322 | 1.434236549 | 0.15150477 | 0.293863146 |
| ENSG00000142330 | CAPN10 | 472.3996612 | 0.114482702 | 0.090178 | 1.269519202 | 0.204255946 | 0.369121744 |
| ENSG00000142347 | MYO1F | 19.48087301 | -0.123300203 | 0.410425807 | -0.3004202 | 0.763856658 | 0.870485431 |
| ENSG00000142396 | ERVK3-1 | 452.1649911 | 0.346089002 | 0.109250949 | 3.167835211 | 0.001535785 | 0.005531625 |
| ENSG00000142408 | CACNG8 | 48.12681771 | -2.377236468 | 0.297707532 | -7.985140478 | 1.40E-15 | 1.76E-14 |
| ENSG00000142409 | ZNF787 | 585.950754 | 0.023798168 | 0.083648902 | 0.284500663 | 0.776026726 | 0.878332993 |
| ENSG00000142444 | TIMM29 | 627.8761747 | -0.181907374 | 0.088225378 | -2.061848621 | 0.03922215 | 0.097598018 |
| ENSG00000142449 | FBN3 | 3.243842287 | -0.976919289 | 0.983704351 | -0.993102539 | 0.320659987 | 0.510226822 |

| | | | | | | | |
|---|---|---|---|---|---|---|---|
| ENSG00000142453 | CARM1 | 3979.621246 | -0.021012917 | 0.051725583 | -0.406238384 | 0.684567453 | 0.821251824 |
| ENSG00000142459 | EVI5L | 735.7770249 | -0.152495467 | 0.07964134 | -1.914777766 | 0.055520851 | 0.131031218 |
| ENSG00000142494 | SLC47A1 | 19.169501 | 1.130623795 | 0.417490492 | 2.708142624 | 0.006766094 | 0.021048846 |
| ENSG00000142507 | PSMB6 | 1902.741631 | -0.217706527 | 0.056775562 | -3.834511161 | 0.000125814 | 0.000551663 |
| ENSG00000142528 | ZNF473 | 854.3611654 | 0.130908605 | 0.070439558 | 1.858452962 | 0.063104717 | 0.14548967 |
| ENSG00000142530 | GARIN5A | 76.00739315 | 0.157770164 | 0.224157091 | 0.703837489 | 0.481533984 | 0.665926549 |
| ENSG00000142534 | RPS11 | 11627.84451 | 0.2596305 | 0.041691156 | 6.227471791 | 4.74E-10 | 3.89E-09 |
| ENSG00000142541 | RPL13A | 13227.78831 | 0.373193277 | 0.045780864 | 8.151730702 | 3.59E-16 | 4.69E-15 |
| ENSG00000142544 | CTU1 | 74.80622161 | 0.419618102 | 0.211053775 | 1.988204669 | 0.046789059 | 0.11312616 |
| ENSG00000142546 | NOSIP | 1099.888132 | 0.173435783 | 0.08188047 | 2.118158111 | 0.034161679 | 0.086746616 |
| ENSG00000142552 | RCN3 | 4.308913747 | -0.029952393 | 0.850783714 | -0.035205649 | 0.971915758 | 0.987280143 |
| ENSG00000142556 | ZNF614 | 518.1895075 | 0.321896172 | 0.085844202 | 3.749771827 | 0.000176996 | 0.000758222 |
| ENSG00000142599 | RERE | 2958.035391 | -0.053459725 | 0.051235539 | -1.043410998 | 0.296757982 | 0.483068171 |
| ENSG00000142606 | MMEL1 | 62.24802127 | -0.006564529 | 0.225031159 | -0.029171643 | 0.976727697 | 0.989278333 |
| ENSG00000142609 | CFAP74 | 4.916936731 | -1.153783057 | 0.826155911 | -1.39656818 | 0.162543465 | 0.309748054 |
| ENSG00000142611 | PRDM16 | 252.9209972 | 1.938002147 | 0.131223278 | 14.76873753 | 2.33E-49 | 1.27E-47 |
| ENSG00000142619 | PADI3 | 11.63186648 | -2.147000543 | 0.579591365 | -3.704334936 | 0.000211946 | 0.000894416 |
| ENSG00000142621 | FHAD1 | 2.802693451 | -0.302985152 | 1.070233103 | -0.283102019 | 0.777098634 | 0.87898243 |
| ENSG00000142623 | PADI1 | 1.528026877 | 1.864545912 | 1.60982242 | 1.158230802 | 0.246769862 | NA |
| ENSG00000142627 | EPHA2 | 214.9230175 | 0.016827168 | 0.132881744 | 0.126632653 | 0.899231152 | 0.949051063 |
| ENSG00000142632 | ARHGEF19 | 1689.466897 | 0.66988916 | 0.06224335 | 10.7624214 | 5.18E-27 | 1.28E-25 |
| ENSG00000142634 | EFHD2 | 637.1462802 | 0.415129599 | 0.089613994 | 4.632419321 | 3.61E-06 | 1.97E-05 |
| ENSG00000142655 | PEX14 | 436.5748954 | -0.004361536 | 0.100443843 | -0.043422633 | 0.965364637 | 0.984374128 |
| ENSG00000142657 | PGD | 7987.584366 | -0.235625362 | 0.042522384 | -5.541207722 | 3.00E-08 | 2.05E-07 |
| ENSG00000142669 | SH3BGRL3 | 2145.726643 | -0.160991571 | 0.056343296 | -2.857333232 | 0.004272171 | 0.013988523 |
| ENSG00000142675 | CNKSR1 | 781.6453694 | -0.01378271 | 0.084434722 | -0.163235095 | 0.870333335 | 0.932790446 |
| ENSG00000142676 | RPL11 | 9951.1487 | 0.239437662 | 0.049067332 | 4.879777518 | 1.06E-06 | 6.13E-06 |
| ENSG00000142677 | IL22RA1 | 2.242460308 | 1.704894421 | 1.288370453 | 1.323295188 | 0.185737231 | 0.342977709 |
| ENSG00000142684 | ZNF593 | 345.7319473 | -0.144419228 | 0.102300426 | -1.411716783 | 0.15803337 | 0.30294021 |
| ENSG00000142686 | C1orf216 | 810.8159431 | 0.246891688 | 0.077065986 | 3.203640172 | 0.00135702 | 0.004951492 |
| ENSG00000142687 | KIAA0319L | 4007.107685 | -0.024846808 | 0.058686002 | -0.423385595 | 0.672013953 | 0.812095037 |
| ENSG00000142694 | EVA1B | 111.8033109 | 0.285279351 | 0.172937364 | 1.649610843 | 0.099022556 | 0.210331244 |
| ENSG00000142731 | PLK4 | 898.568456 | -0.249162183 | 0.075255493 | -3.31088369 | 0.000930019 | 0.003504373 |

| | | | | | | | |
|---|---|---|---|---|---|---|---|
| ENSG00000142733 | MAP3K6 | 557.7743241 | 0.112840513 | 0.084024151 | 1.342953327 | 0.179287087 | 0.334051802 |
| ENSG00000142751 | GPN2 | 987.1083916 | 0.024747117 | 0.066567035 | 0.371762341 | 0.710069805 | 0.838644973 |
| ENSG00000142765 | SYTL1 | 1128.574168 | 0.058438959 | 0.070723998 | 0.826296034 | 0.408636183 | 0.601020327 |
| ENSG00000142784 | WDTC1 | 2219.083163 | 0.379341884 | 0.065554502 | 5.786664135 | 7.18E-09 | 5.24E-08 |
| ENSG00000142794 | NBPF3 | 709.4431869 | -0.068416728 | 0.076410004 | -0.895389674 | 0.370578819 | 0.563283733 |
| ENSG00000142798 | HSPG2 | 312.5380651 | 1.87593459 | 0.12902821 | 14.53894921 | 6.86E-48 | 3.53E-46 |
| ENSG00000142856 | ITGB3BP | 317.2832158 | -0.10953153 | 0.120411999 | -0.909639661 | 0.363012575 | 0.555760042 |
| ENSG00000142864 | SERBP1 | 8890.456775 | 0.072751271 | 0.047457574 | 1.532974937 | 0.12528203 | 0.253951557 |
| ENSG00000142867 | BCL10 | 499.1487584 | -0.028318576 | 0.090617294 | -0.31250741 | 0.754654933 | 0.864498202 |
| ENSG00000142871 | CCN1 | 299.2119143 | -0.481872998 | 0.114075583 | -4.22415547 | 2.40E-05 | 0.000117612 |
| ENSG00000142875 | PRKACB | 1130.999322 | -0.108145789 | 0.066775466 | -1.619543759 | 0.105330319 | 0.220987518 |
| ENSG00000142892 | PIGK | 385.565478 | -0.180381205 | 0.098903313 | -1.823813567 | 0.068180262 | 0.155318293 |
| ENSG00000142910 | TINAGL1 | 6.008613614 | -0.644956369 | 0.759327028 | -0.849378918 | 0.39567048 | 0.588668579 |
| ENSG00000142920 | AZIN2 | 23.14755678 | -0.092916461 | 0.368778625 | -0.251957285 | 0.80107408 | 0.894196847 |
| ENSG00000142937 | RPS8 | 15235.96 | 0.162492077 | 0.052649348 | 3.086307476 | 0.002026591 | 0.007114002 |
| ENSG00000142945 | KIF2C | 2053.573829 | -0.352972082 | 0.053963518 | -6.540938996 | 6.11E-11 | 5.38E-10 |
| ENSG00000142949 | PTPRF | 25508.42982 | 0.313507302 | 0.058464677 | 5.362337004 | 8.22E-08 | 5.36E-07 |
| ENSG00000142959 | BEST4 | 2.204164876 | 1.683468569 | 1.290274687 | 1.30473657 | 0.191982564 | 0.351988493 |
| ENSG00000142961 | MOB3C | 331.6956793 | 0.027854404 | 0.110172673 | 0.252824978 | 0.800403465 | 0.893665701 |
| ENSG00000142973 | CYP4B1 | 888.7929775 | 0.62677782 | 0.083462986 | 7.509650113 | 5.93E-14 | 6.58E-13 |
| ENSG00000143001 | TMEM61 | 5.048967799 | 2.281720475 | 0.933096595 | 2.445320759 | 0.014472337 | 0.04117517 |
| ENSG00000143013 | LMO4 | 618.879189 | 0.651486536 | 0.082698415 | 7.87785998 | 3.33E-15 | 4.06E-14 |
| ENSG00000143028 | SYPL2 | 3.438781067 | -1.41876085 | 1.065288765 | -1.331808704 | 0.182923057 | 0.338899606 |
| ENSG00000143033 | MTF2 | 538.9662233 | -0.248259882 | 0.092337691 | -2.688608282 | 0.007175056 | 0.022200673 |
| ENSG00000143036 | SLC44A3 | 72.73466475 | 0.543999855 | 0.207827283 | 2.617557456 | 0.008856158 | 0.02674569 |
| ENSG00000143061 | IGSF3 | 11823.72317 | -0.708126123 | 0.05216263 | -13.57535308 | 5.61E-42 | 2.48E-40 |
| ENSG00000143067 | ZNF697 | 132.5457426 | 0.453295989 | 0.158138452 | 2.866450141 | 0.004151035 | 0.013628847 |
| ENSG00000143079 | CTTNBP2NL | 716.5650321 | 0.021335633 | 0.078669418 | 0.271206183 | 0.786232458 | 0.883866146 |
| ENSG00000143093 | STRIP1 | 1429.539851 | -0.005510033 | 0.062366484 | -0.088349266 | 0.929599083 | 0.965993019 |
| ENSG00000143106 | PSMA5 | 3877.050418 | 0.037349687 | 0.04801746 | 0.777835539 | 0.436665974 | 0.626500559 |
| ENSG00000143110 | C1orf162 | 9.411826132 | -0.114742833 | 0.574554406 | -0.199707516 | 0.841709335 | 0.916877899 |
| ENSG00000143126 | CELSR2 | 3982.326534 | 1.117271713 | 0.056399236 | 19.81005055 | 2.44E-87 | 3.21E-85 |
| ENSG00000143127 | ITGA10 | 82.19799747 | -0.44744753 | 0.211833045 | -2.112265016 | 0.034663725 | 0.087894976 |

| | | | | | | | |
|---|---|---|---|---|---|---|---|
| ENSG00000143147 | GPR161 | 275.7831462 | 0.082856012 | 0.114445601 | 0.723977251 | 0.469079702 | 0.65506108 |
| ENSG00000143149 | ALDH9A1 | 1742.79813 | 0.176204504 | 0.057029723 | 3.089695963 | 0.002003615 | 0.007040889 |
| ENSG00000143153 | ATP1B1 | 2518.960563 | 0.377696096 | 0.050764691 | 7.440133924 | 1.01E-13 | 1.10E-12 |
| ENSG00000143155 | TIPRL | 1399.737311 | 0.316218635 | 0.062015128 | 5.099056359 | 3.41E-07 | 2.09E-06 |
| ENSG00000143156 | NME7 | 287.0239122 | -0.54838452 | 0.112564677 | -4.871728296 | 1.11E-06 | 6.37E-06 |
| ENSG00000143157 | POGK | 3684.19304 | -0.076568478 | 0.047890872 | -1.598811511 | 0.109862491 | 0.228572967 |
| ENSG00000143158 | MPC2 | 867.8957524 | -0.044158076 | 0.070706246 | -0.624528649 | 0.532280462 | 0.707691769 |
| ENSG00000143162 | CREG1 | 1935.179186 | 0.156949997 | 0.053855467 | 2.914281583 | 0.003565082 | 0.011885333 |
| ENSG00000143164 | DCAF6 | 1949.467298 | 0.386377606 | 0.05392161 | 7.165542889 | 7.75E-13 | 8.00E-12 |
| ENSG00000143171 | RXRG | 2.081751418 | -0.20678355 | 1.235680109 | -0.167343918 | 0.86709945 | 0.930846618 |
| ENSG00000143178 | TBX19 | 38.42614733 | 0.500204091 | 0.28773608 | 1.738412824 | 0.082138099 | 0.180518106 |
| ENSG00000143179 | UCK2 | 1723.172238 | 0.253059419 | 0.057757422 | 4.381418205 | 1.18E-05 | 6.02E-05 |
| ENSG00000143183 | TMCO1 | 2336.28593 | 0.261454656 | 0.052992212 | 4.9338317 | 8.06E-07 | 4.73E-06 |
| ENSG00000143190 | POU2F1 | 708.7839096 | 0.105670085 | 0.081829077 | 1.291351305 | 0.196581888 | 0.358301963 |
| ENSG00000143195 | ILDR2 | 4.69424427 | -0.846575049 | 0.859684144 | -0.984751266 | 0.324746264 | 0.51488701 |
| ENSG00000143198 | MGST3 | 535.0051034 | 0.220693195 | 0.083600987 | 2.639839572 | 0.008294528 | 0.025215541 |
| ENSG00000143199 | ADCY10 | 3.898181168 | 0.280722597 | 0.883705565 | 0.317665304 | 0.750738832 | 0.862304667 |
| ENSG00000143207 | COP1 | 1765.029698 | -0.147266214 | 0.056076316 | -2.62617492 | 0.008635043 | 0.02613935 |
| ENSG00000143217 | NECTIN4 | 1259.707694 | 0.92544021 | 0.068209003 | 13.56771345 | 6.22E-42 | 2.75E-40 |
| ENSG00000143222 | UFC1 | 2126.40911 | 0.457764291 | 0.055855869 | 8.195455516 | 2.50E-16 | 3.30E-15 |
| ENSG00000143224 | PPOX | 356.4469545 | 0.284932181 | 0.101731171 | 2.800834569 | 0.005097064 | 0.016385461 |
| ENSG00000143228 | NUF2 | 707.9980024 | -0.132260543 | 0.085351336 | -1.549601312 | 0.121237238 | 0.247480178 |
| ENSG00000143248 | RGS5 | 14.60795154 | 1.050567235 | 0.47197218 | 2.225909235 | 0.026020262 | 0.068720332 |
| ENSG00000143252 | SDHC | 1501.447489 | 0.071003748 | 0.057744317 | 1.229623141 | 0.21883826 | 0.388562198 |
| ENSG00000143256 | PFDN2 | 763.47531 | 0.285044255 | 0.076430761 | 3.729444152 | 0.000191903 | 0.00081735 |
| ENSG00000143257 | NR1I3 | 3.933582225 | -0.67100382 | 0.889110532 | -0.754691117 | 0.450434338 | 0.638625693 |
| ENSG00000143258 | USP21 | 1003.457301 | 0.079506516 | 0.073539832 | 1.081135413 | 0.279636883 | 0.462933542 |
| ENSG00000143294 | PRCC | 3788.668007 | 0.132474391 | 0.049375432 | 2.683002197 | 0.007296451 | 0.02254284 |
| ENSG00000143303 | METTL25B | 1001.795691 | -0.016091165 | 0.07319581 | -0.219837238 | 0.825997917 | 0.908261176 |
| ENSG00000143314 | MRPL24 | 1674.215272 | 0.172216707 | 0.060069545 | 2.866955418 | 0.004144414 | 0.013611003 |
| ENSG00000143315 | PIGM | 1119.634736 | 0.158690282 | 0.067476494 | 2.35178612 | 0.018683514 | 0.051520594 |
| ENSG00000143319 | ISG20L2 | 2919.465125 | 0.182851467 | 0.049729415 | 3.676927811 | 0.00023606 | 0.000988538 |
| ENSG00000143320 | CRABP2 | 7221.022539 | 0.363329691 | 0.045791596 | 7.934418667 | 2.11E-15 | 2.61E-14 |

| | | | | | | | |
|---|---|---|---|---|---|---|---|
| ENSG00000143321 | HDGF | 14684.60534 | -0.053964552 | 0.039106352 | -1.37994342 | 0.167604066 | 0.317361724 |
| ENSG00000143322 | ABL2 | 1218.26939 | -0.179766488 | 0.080800524 | -2.224818348 | 0.026093435 | 0.068866015 |
| ENSG00000143324 | XPR1 | 1574.680567 | -0.3979263 | 0.057951653 | -6.866521977 | 6.58E-12 | 6.26E-11 |
| ENSG00000143333 | RGS16 | 86.87604194 | -0.398495985 | 0.189120026 | -2.107106237 | 0.035108372 | 0.088855478 |
| ENSG00000143337 | TOR1AIP1 | 2210.664099 | 0.039801797 | 0.058059431 | 0.685535423 | 0.493006115 | 0.675789422 |
| ENSG00000143340 | FAM163A | 2.108129641 | 0.095085449 | 1.214629842 | 0.078283479 | 0.937602559 | 0.970153346 |
| ENSG00000143341 | HMCN1 | 7.614867837 | -1.621395399 | 0.678719115 | -2.388904869 | 0.016898677 | 0.047229003 |
| ENSG00000143344 | RGL1 | 55.36653962 | 0.710655457 | 0.254189974 | 2.795765094 | 0.005177702 | 0.016608159 |
| ENSG00000143353 | LYPLAL1 | 260.107725 | 0.724505227 | 0.115284225 | 6.284513117 | 3.29E-10 | 2.74E-09 |
| ENSG00000143355 | LHX9 | 2.532782165 | -2.830491447 | 1.426593481 | -1.984091113 | 0.047245684 | 0.114073885 |
| ENSG00000143363 | PRUNE1 | 3006.524617 | 0.305508434 | 0.050628689 | 6.034294803 | 1.60E-09 | 1.25E-08 |
| ENSG00000143365 | RORC | 232.8661971 | 1.946081234 | 0.13833186 | 14.06820698 | 5.96E-45 | 2.88E-43 |
| ENSG00000143367 | TUFT1 | 2981.970549 | 0.693454892 | 0.059179787 | 11.71776584 | 1.03E-31 | 3.09E-30 |
| ENSG00000143368 | SF3B4 | 7099.624197 | 0.105456739 | 0.044363826 | 2.377088451 | 0.017449903 | 0.048580192 |
| ENSG00000143369 | ECM1 | 65.70168149 | 1.023707609 | 0.240444771 | 4.257558211 | 2.07E-05 | 0.000102421 |
| ENSG00000143373 | ZNF687 | 5774.617658 | 0.278508363 | 0.057724262 | 4.824805992 | 1.40E-06 | 7.95E-06 |
| ENSG00000143374 | TARS2 | 2593.421528 | 0.278660918 | 0.05051373 | 5.51653816 | 3.46E-08 | 2.35E-07 |
| ENSG00000143375 | CGN | 4727.410318 | 0.6548173 | 0.047336459 | 13.83325484 | 1.61E-43 | 7.35E-42 |
| ENSG00000143376 | SNX27 | 2643.262189 | 0.411952561 | 0.051706032 | 7.967205122 | 1.62E-15 | 2.02E-14 |
| ENSG00000143379 | SETDB1 | 5596.649564 | 0.373991522 | 0.045519803 | 8.216017995 | 2.10E-16 | 2.80E-15 |
| ENSG00000143382 | ADAMTSL4 | 13.94900727 | 0.588570224 | 0.47631865 | 1.235664873 | 0.21658316 | 0.386016447 |
| ENSG00000143384 | MCL1 | 13270.12679 | 0.387372895 | 0.041510757 | 9.331867727 | 1.04E-20 | 1.81E-19 |
| ENSG00000143387 | CTSK | 328.798457 | 0.814564855 | 0.118890196 | 6.851404751 | 7.31E-12 | 6.94E-11 |
| ENSG00000143390 | RFX5 | 4291.988685 | -0.380039887 | 0.048835232 | -7.782084206 | 7.13E-15 | 8.48E-14 |
| ENSG00000143393 | PI4KB | 8479.697807 | 0.405114468 | 0.045754653 | 8.854060451 | 8.44E-19 | 1.32E-17 |
| ENSG00000143398 | PIP5K1A | 6815.822334 | 0.274724208 | 0.04609704 | 5.959693011 | 2.53E-09 | 1.94E-08 |
| ENSG00000143401 | ANP32E | 2585.568801 | 0.326191452 | 0.06485998 | 5.029163636 | 4.93E-07 | 2.97E-06 |
| ENSG00000143409 | MINDY1 | 1387.630535 | 0.389068412 | 0.069734197 | 5.579305803 | 2.41E-08 | 1.66E-07 |
| ENSG00000143412 | ANXA9 | 1445.770788 | 1.281687721 | 0.0729228 | 17.57595331 | 3.77E-69 | 3.40E-67 |
| ENSG00000143416 | SELENBP1 | 5120.328233 | 0.255901742 | 0.051893702 | 4.931267826 | 8.17E-07 | 4.79E-06 |
| ENSG00000143418 | CERS2 | 21648.27291 | 0.362625196 | 0.042895672 | 8.453654709 | 2.82E-17 | 3.95E-16 |
| ENSG00000143420 | ENSA | 13080.02011 | 0.172014286 | 0.038822971 | 4.430734778 | 9.39E-06 | 4.87E-05 |
| ENSG00000143429 | LSP1P4 | 6.191668305 | 0.123268467 | 0.689568141 | 0.178761836 | 0.858124713 | 0.926370866 |

| | | | | | | | |
|---|---|---|---|---|---|---|---|
| ENSG00000143434 | SEMA6C | 150.1170674 | 0.982106223 | 0.15127574 | 6.492159417 | 8.46E-11 | 7.38E-10 |
| ENSG00000143436 | MRPL9 | 4160.258457 | 0.2330362 | 0.047333992 | 4.923231496 | 8.51E-07 | 4.98E-06 |
| ENSG00000143437 | ARNT | 4284.506082 | 0.275752458 | 0.046569659 | 5.921290061 | 3.19E-09 | 2.43E-08 |
| ENSG00000143442 | POGZ | 7141.902394 | 0.163306921 | 0.05010975 | 3.258984965 | 0.001118116 | 0.004151089 |
| ENSG00000143443 | C1orf56 | 471.6458021 | 0.615104926 | 0.095414311 | 6.446673629 | 1.14E-10 | 9.83E-10 |
| ENSG00000143450 | OAZ3 | 110.7666091 | 0.133533813 | 0.18936991 | 0.705147999 | 0.480718138 | 0.665078951 |
| ENSG00000143457 | GOLPH3L | 1989.437341 | 0.540886496 | 0.055031642 | 9.828645401 | 8.48E-23 | 1.67E-21 |
| ENSG00000143458 | GABPB2 | 1116.516495 | 0.34558115 | 0.066186115 | 5.221354227 | 1.78E-07 | 1.12E-06 |
| ENSG00000143476 | DTL | 631.1717895 | -0.216318338 | 0.081787295 | -2.644889274 | 0.008171765 | 0.02489182 |
| ENSG00000143479 | DYRK3 | 141.2549623 | 0.763907242 | 0.152749196 | 5.001055729 | 5.70E-07 | 3.41E-06 |
| ENSG00000143486 | EIF2D | 3544.593044 | 0.608545375 | 0.049632554 | 12.2610127 | 1.47E-34 | 4.93E-33 |
| ENSG00000143493 | INTS7 | 783.798024 | -0.002196132 | 0.073845131 | -0.029739704 | 0.976274647 | 0.989211149 |
| ENSG00000143494 | VASH2 | 82.78089915 | 0.329677552 | 0.19536812 | 1.687468516 | 0.091513299 | 0.197169214 |
| ENSG00000143498 | TAF1A | 123.3452001 | -0.312875869 | 0.164602051 | -1.900801766 | 0.057327983 | 0.134413729 |
| ENSG00000143499 | SMYD2 | 1069.542898 | -0.054695324 | 0.06735877 | -0.812000038 | 0.41679161 | 0.608838482 |
| ENSG00000143502 | SUSD4 | 709.5089425 | 1.694439065 | 0.080668193 | 21.00504559 | 5.90E-98 | 9.14E-96 |
| ENSG00000143507 | DUSP10 | 143.6024271 | 0.313620691 | 0.157141502 | 1.995785253 | 0.045957303 | 0.111420474 |
| ENSG00000143512 | HHIPL2 | 2.670281547 | 0.369881068 | 1.087746637 | 0.340043403 | 0.733823842 | 0.853084627 |
| ENSG00000143514 | TP53BP2 | 1168.270175 | 0.094328735 | 0.068678798 | 1.373476783 | 0.169604174 | 0.320165542 |
| ENSG00000143515 | ATP8B2 | 3284.298157 | -0.180084453 | 0.055824736 | -3.225889936 | 0.001255816 | 0.004614517 |
| ENSG00000143537 | ADAM15 | 4267.198448 | 1.435356796 | 0.054602681 | 26.28729506 | 2.68E-152 | 9.76E-150 |
| ENSG00000143543 | JTB | 342.4320542 | 0.485110191 | 0.112896283 | 4.296954487 | 1.73E-05 | 8.67E-05 |
| ENSG00000143545 | RAB13 | 446.1207562 | 0.23460243 | 0.099772986 | 2.351362222 | 0.018704815 | 0.051573133 |
| ENSG00000143549 | TPM3 | 8752.277085 | -0.021131102 | 0.044615291 | -0.473629132 | 0.635764398 | 0.785317815 |
| ENSG00000143552 | NUP210L | 692.7148178 | 0.23316446 | 0.07527876 | 3.097347224 | 0.00195261 | 0.006881683 |
| ENSG00000143553 | SNAPIN | 936.0593642 | 0.335176774 | 0.068213848 | 4.913617732 | 8.94E-07 | 5.22E-06 |
| ENSG00000143554 | SLC27A3 | 1271.433419 | 0.242524049 | 0.068143121 | 3.559039355 | 0.000372214 | 0.001503007 |
| ENSG00000143569 | UBAP2L | 8940.89212 | 0.482905064 | 0.043021744 | 11.22467441 | 3.09E-29 | 8.30E-28 |
| ENSG00000143570 | SLC39A1 | 114.4442032 | 0.023787054 | 0.17292004 | 0.137561001 | 0.890587379 | 0.943913993 |
| ENSG00000143575 | HAX1 | 2732.376604 | 0.116814775 | 0.049992682 | 2.336637495 | 0.019458041 | 0.053421174 |
| ENSG00000143578 | CREB3L4 | 2300.433083 | -0.504068621 | 0.059957656 | -8.407076859 | 4.20E-17 | 5.82E-16 |
| ENSG00000143590 | EFNA3 | 313.677864 | 2.201541667 | 0.122546522 | 17.9649462 | 3.67E-72 | 3.55E-70 |
| ENSG00000143612 | C1orf43 | 8231.997854 | 0.3639961 | 0.041902311 | 8.686778613 | 3.73E-18 | 5.57E-17 |

| | | | | | | | |
|---|---|---|---|---|---|---|---|
| ENSG00000143614 | GATAD2B | 2324.484439 | 0.317596502 | 0.054646798 | 5.811804378 | 6.18E-09 | 4.55E-08 |
| ENSG00000143621 | ILF2 | 7845.564417 | 0.146275828 | 0.044900415 | 3.257783447 | 0.001122861 | 0.004166681 |
| ENSG00000143622 | RIT1 | 930.7552118 | 0.248426644 | 0.067717906 | 3.66855177 | 0.000243928 | 0.001017976 |
| ENSG00000143624 | INTS3 | 4974.798732 | 0.121207475 | 0.055447694 | 2.185978647 | 0.028817168 | 0.074976276 |
| ENSG00000143627 | PKLR | 3.199357483 | 0.746407849 | 0.987653568 | 0.755738523 | 0.449805982 | 0.638126966 |
| ENSG00000143630 | HCN3 | 474.9587363 | 0.005949187 | 0.105407414 | 0.056439932 | 0.954991347 | 0.979325056 |
| ENSG00000143632 | ACTA1 | 7.669764107 | -0.061352024 | 0.678624321 | -0.090406463 | 0.927964221 | 0.965331908 |
| ENSG00000143633 | C1orf131 | 614.1495068 | 0.02518964 | 0.083089667 | 0.303162128 | 0.761766309 | 0.869297509 |
| ENSG00000143641 | GALNT2 | 2828.2911 | 0.373038181 | 0.054629289 | 6.828538136 | 8.58E-12 | 8.09E-11 |
| ENSG00000143643 | TTC13 | 570.5067673 | -0.136284444 | 0.082991408 | -1.642151242 | 0.100558664 | 0.212884437 |
| ENSG00000143653 | SCCPDH | 3334.279075 | 0.270473185 | 0.050888993 | 5.314964366 | 1.07E-07 | 6.90E-07 |
| ENSG00000143669 | LYST | 195.3268692 | 0.266271199 | 0.136659705 | 1.94842509 | 0.05136412 | 0.122572861 |
| ENSG00000143674 | MAP3K21 | 426.7950534 | 0.55294488 | 0.093718837 | 5.900039941 | 3.63E-09 | 2.75E-08 |
| ENSG00000143702 | CEP170 | 631.4105938 | 0.000110995 | 0.080387731 | 0.001380749 | 0.998898322 | 0.999333951 |
| ENSG00000143727 | ACP1 | 2576.727967 | 0.180740246 | 0.050480511 | 3.580396501 | 0.000343073 | 0.00139516 |
| ENSG00000143740 | SNAP47 | 1512.02971 | 0.055236265 | 0.069156615 | 0.798712676 | 0.424457035 | 0.615226766 |
| ENSG00000143742 | SRP9 | 2610.570058 | 0.016547435 | 0.072227264 | 0.229102338 | 0.818789376 | 0.903986346 |
| ENSG00000143748 | NVL | 1798.555435 | -1.374341299 | 0.058961543 | -23.30911338 | 3.58E-120 | 8.84E-118 |
| ENSG00000143751 | SDE2 | 765.7669023 | 0.068992124 | 0.077228546 | 0.893350029 | 0.371669754 | 0.564268972 |
| ENSG00000143753 | DEGS1 | 1046.835429 | -1.084112796 | 0.073457706 | -14.75832629 | 2.72E-49 | 1.47E-47 |
| ENSG00000143756 | FBXO28 | 1610.586144 | -0.060501229 | 0.065946052 | -0.917435196 | 0.358914636 | 0.551473661 |
| ENSG00000143761 | ARF1 | 11892.08042 | -0.059824725 | 0.042223127 | -1.416871018 | 0.156520641 | 0.300794464 |
| ENSG00000143771 | CNIH4 | 1113.416408 | -0.251304954 | 0.06519775 | -3.854503494 | 0.000115965 | 0.000511977 |
| ENSG00000143772 | ITPKB | 994.4509576 | -0.516444633 | 0.067952332 | -7.600101636 | 2.96E-14 | 3.36E-13 |
| ENSG00000143774 | GUK1 | 3240.779556 | 0.16965857 | 0.059123545 | 2.869560165 | 0.004110431 | 0.013511003 |
| ENSG00000143776 | CDC42BPA | 2284.390207 | -0.006387808 | 0.069203483 | -0.09230472 | 0.926455938 | 0.964663921 |
| ENSG00000143786 | CNIH3 | 101.1564165 | 1.146618578 | 0.189581291 | 6.04816316 | 1.47E-09 | 1.15E-08 |
| ENSG00000143793 | C1orf35 | 740.732727 | -0.369706469 | 0.075010999 | -4.928696754 | 8.28E-07 | 4.85E-06 |
| ENSG00000143797 | MBOAT2 | 1832.513658 | -0.210298012 | 0.056427886 | -3.726845457 | 0.000193891 | 0.00082498 |
| ENSG00000143799 | PARP1 | 7430.215954 | -0.082185857 | 0.04234054 | -1.94106775 | 0.05225006 | 0.124389413 |
| ENSG00000143801 | PSEN2 | 536.4839134 | 0.025394828 | 0.088268715 | 0.287699082 | 0.773577093 | 0.876598425 |
| ENSG00000143811 | PYCR2 | 751.5755463 | -0.148342879 | 0.076094394 | -1.949458703 | 0.05124067 | 0.122367371 |
| ENSG00000143815 | LBR | 1478.58256 | -0.790181165 | 0.060028035 | -13.16353552 | 1.42E-39 | 5.65E-38 |

| | | | | | | | |
|---|---|---|---|---|---|---|---|
| ENSG00000143816 | WNT9A | 182.0364162 | -0.345901072 | 0.133711294 | -2.586924875 | 0.00968367 | 0.028913627 |
| ENSG00000143819 | EPHX1 | 636.882872 | 0.290402986 | 0.086947199 | 3.339992415 | 0.000837807 | 0.003181475 |
| ENSG00000143839 | REN | 5.201205862 | -1.429039306 | 0.817087902 | -1.748941948 | 0.080301055 | 0.177159665 |
| ENSG00000143842 | SOX13 | 3579.097573 | 0.79582209 | 0.057478718 | 13.84550877 | 1.35E-43 | 6.24E-42 |
| ENSG00000143845 | ETNK2 | 1244.838101 | 0.632890108 | 0.065990733 | 9.590590642 | 8.76E-22 | 1.62E-20 |
| ENSG00000143847 | PPFIA4 | 26.96536059 | 0.922782712 | 0.348522424 | 2.647699682 | 0.008104148 | 0.024721963 |
| ENSG00000143850 | PLEKHA6 | 4551.152538 | -0.053952929 | 0.056917813 | -0.947909386 | 0.343175586 | 0.534159855 |
| ENSG00000143862 | ARL8A | 2230.272966 | 0.361728553 | 0.062179669 | 5.817473164 | 5.97E-09 | 4.41E-08 |
| ENSG00000143867 | OSR1 | 4.473094759 | -1.64833099 | 0.884081018 | -1.86445694 | 0.062257563 | 0.14382061 |
| ENSG00000143870 | PDIA6 | 7198.831312 | 0.092148709 | 0.041337165 | 2.229197608 | 0.02580076 | 0.068219159 |
| ENSG00000143878 | RHOB | 8904.770791 | 0.59790256 | 0.045022931 | 13.27995647 | 3.03E-40 | 1.25E-38 |
| ENSG00000143882 | ATP6V1C2 | 323.2423946 | 0.334659058 | 0.10642073 | 3.144679213 | 0.00166269 | 0.005942065 |
| ENSG00000143889 | HNRNPLL | 1689.708148 | 0.066081853 | 0.057835749 | 1.142577978 | 0.253213853 | 0.431939753 |
| ENSG00000143891 | GALM | 779.7218639 | 0.460530722 | 0.076366643 | 6.030522027 | 1.63E-09 | 1.28E-08 |
| ENSG00000143919 | CAMKMT | 178.430624 | 0.005199384 | 0.137522248 | 0.037807581 | 0.9698411 | 0.986499047 |
| ENSG00000143921 | ABCG8 | 1.870343491 | -2.362905761 | 1.467185656 | -1.610502224 | 0.10728826 | 0.224322275 |
| ENSG00000143924 | EML4 | 1881.120891 | 0.02871098 | 0.059830547 | 0.479871589 | 0.631318704 | 0.782730991 |
| ENSG00000143933 | CALM2 | 5658.282613 | -0.107578111 | 0.054216569 | -1.98422943 | 0.047230269 | 0.114048671 |
| ENSG00000143942 | CHAC2 | 73.79364164 | -0.169758157 | 0.219099081 | -0.774800863 | 0.438457339 | 0.627947553 |
| ENSG00000143947 | RPS27A | 4645.882648 | 0.175856757 | 0.061036285 | 2.881183832 | 0.003961845 | 0.013061903 |
| ENSG00000143951 | WDPCP | 148.9062458 | 0.122851244 | 0.150142951 | 0.818228514 | 0.413226717 | 0.605404322 |
| ENSG00000143952 | VPS54 | 1378.783802 | 0.23407273 | 0.060445095 | 3.872485088 | 0.000107731 | 0.000477926 |
| ENSG00000143970 | ASXL2 | 1732.0059 | -0.09352278 | 0.069676424 | -1.34224426 | 0.17951681 | 0.334425501 |
| ENSG00000143971 | ETAA1 | 309.5622578 | -0.179948555 | 0.109613371 | -1.641666095 | 0.100659222 | 0.212985769 |
| ENSG00000143977 | SNRPG | 595.4518236 | -0.194544209 | 0.082856441 | -2.347967231 | 0.018876182 | 0.052020617 |
| ENSG00000143994 | ABHD1 | 19.0787525 | -1.098926627 | 0.418121271 | -2.628248556 | 0.008582578 | 0.026005064 |
| ENSG00000143995 | MEIS1 | 43.38570533 | -0.679324835 | 0.270913563 | -2.507533503 | 0.012157706 | 0.035345765 |
| ENSG00000144021 | CIAO1 | 3665.65228 | 0.186547906 | 0.051349839 | 3.632881978 | 0.000280273 | 0.001159089 |
| ENSG00000144026 | ZNF514 | 542.3402169 | -0.053234018 | 0.089553029 | -0.594441288 | 0.552217001 | 0.723957622 |
| ENSG00000144028 | SNRNP200 | 16918.20775 | -0.037242136 | 0.046089111 | -0.80804631 | 0.419063923 | 0.610728065 |
| ENSG00000144029 | MRPS5 | 1705.23971 | -0.04735053 | 0.057174446 | -0.828176446 | 0.407570581 | 0.600369311 |
| ENSG00000144031 | ANKRD53 | 3.405522785 | 0.858771653 | 0.982976824 | 0.873643845 | 0.382312241 | 0.574617273 |
| ENSG00000144034 | TPRKB | 338.8900718 | 0.04992252 | 0.106374333 | 0.469309833 | 0.638848188 | 0.787290808 |

| | | | | | | | |
|---|---|---|---|---|---|---|---|
| ENSG00000144036 | EXOC6B | 1364.812592 | -0.24559886 | 0.059856362 | -4.103137124 | 4.08E-05 | 0.000192824 |
| ENSG00000144040 | SFXN5 | 814.9381135 | 0.085541211 | 0.074198942 | 1.152862946 | 0.248966643 | 0.426853124 |
| ENSG00000144043 | TEX261 | 4058.031595 | -0.008439272 | 0.050324114 | -0.167698364 | 0.866820583 | 0.930846618 |
| ENSG00000144045 | DQX1 | 8.043932601 | -1.451999701 | 0.645403125 | -2.24975623 | 0.024464424 | 0.065272608 |
| ENSG00000144048 | DUSP11 | 554.7650357 | 0.030619138 | 0.084540461 | 0.362183238 | 0.717215105 | 0.842788367 |
| ENSG00000144061 | NPHP1 | 68.05420722 | 0.348657366 | 0.232357234 | 1.50052297 | 0.133478988 | 0.266756797 |
| ENSG00000144063 | MALL | 2.269392896 | 0.242666763 | 1.157034906 | 0.209731584 | 0.833877173 | 0.912258208 |
| ENSG00000144115 | THNSL2 | 609.6488484 | -0.414318378 | 0.085410313 | -4.850917474 | 1.23E-06 | 7.04E-06 |
| ENSG00000144118 | RALB | 2287.400292 | -0.244215492 | 0.054524207 | -4.479028749 | 7.50E-06 | 3.92E-05 |
| ENSG00000144120 | TMEM177 | 231.2087065 | 0.181713085 | 0.126301374 | 1.43872611 | 0.150228139 | 0.291929987 |
| ENSG00000144130 | NT5DC4 | 4.539222739 | 0.052178342 | 0.811962042 | 0.064262046 | 0.948761574 | 0.976428466 |
| ENSG00000144134 | RABL2A | 231.6099565 | 0.25718449 | 0.124625515 | 2.063658389 | 0.039050117 | 0.097237893 |
| ENSG00000144136 | SLC20A1 | 2220.963786 | -1.262356821 | 0.054885273 | -22.99991865 | 4.67E-117 | 1.05E-114 |
| ENSG00000144152 | FBLN7 | 21.92258452 | 1.485359984 | 0.398943529 | 3.72323368 | 0.000196687 | 0.000835712 |
| ENSG00000144158 | SNRPA1P1 | 342.9107785 | 0.098767008 | 0.101215946 | 0.975804819 | 0.329161196 | 0.519573236 |
| ENSG00000144161 | ZC3H8 | 535.71691 | 0.028129686 | 0.089215206 | 0.315301469 | 0.75253277 | 0.863414111 |
| ENSG00000144182 | LIPT1 | 61.98896028 | 0.040505492 | 0.235855414 | 0.171738655 | 0.863642995 | 0.929172842 |
| ENSG00000144191 | CNGA3 | 2.021656867 | 1.526703892 | 1.397858117 | 1.092173715 | 0.274756754 | 0.45683264 |
| ENSG00000144199 | FAHD2B | 89.92872721 | 0.27382339 | 0.204219168 | 1.340830995 | 0.179975331 | 0.334953276 |
| ENSG00000144214 | LYG1 | 18.06409269 | -0.672992329 | 0.419518536 | -1.604201654 | 0.10866961 | 0.226543153 |
| ENSG00000144218 | AFF3 | 135.3293905 | -1.702456525 | 0.169507586 | -10.0435418 | 9.81E-24 | 2.02E-22 |
| ENSG00000144224 | UBXN4 | 4451.274888 | 0.012788405 | 0.048394418 | 0.264253727 | 0.791584414 | 0.887763734 |
| ENSG00000144228 | SPOPL | 1942.284302 | 0.076353213 | 0.058293761 | 1.309800774 | 0.190263241 | 0.349534617 |
| ENSG00000144229 | THSD7B | 1.687431957 | -0.231660533 | 1.321653312 | -0.175280863 | 0.860858948 | NA |
| ENSG00000144231 | POLR2D | 1683.107906 | 0.030490464 | 0.058935843 | 0.517350088 | 0.604911795 | 0.763691421 |
| ENSG00000144233 | AMMECR1L | 1816.689227 | 0.119844833 | 0.056742233 | 2.112092287 | 0.034678534 | 0.087918612 |
| ENSG00000144278 | GALNT13 | 31.87724171 | -0.578152456 | 0.315572826 | -1.832073008 | 0.066940544 | 0.153010769 |
| ENSG00000144283 | PKP4 | 4914.04145 | 0.001151858 | 0.044749536 | 0.025740101 | 0.979464639 | 0.990775111 |
| ENSG00000144290 | SLC4A10 | 32.43290339 | 0.094182527 | 0.308445482 | 0.305345783 | 0.76010281 | 0.868106598 |
| ENSG00000144306 | SCRN3 | 516.7724556 | 0.178176146 | 0.090972113 | 1.958579832 | 0.050162013 | 0.120179278 |
| ENSG00000144320 | LNPK | 812.4899386 | -0.225317162 | 0.077745576 | -2.898134821 | 0.003753892 | 0.012446058 |
| ENSG00000144331 | ZNF385B | 9.096532438 | 1.314261387 | 0.622263142 | 2.112066903 | 0.034680711 | 0.087918612 |
| ENSG00000144354 | CDCA7 | 137.3645572 | -0.979387449 | 0.157139504 | -6.232598597 | 4.59E-10 | 3.77E-09 |

| | | | | | | | |
|---|---|---|---|---|---|---|---|
| ENSG00000144355 | DLX1 | 984.4501544 | -0.621534074 | 0.070856367 | -8.771746267 | 1.76E-18 | 2.68E-17 |
| ENSG00000144357 | UBR3 | 2102.303564 | -0.128712703 | 0.057577308 | -2.23547623 | 0.0253861 | 0.067308963 |
| ENSG00000144362 | PHOSPHO2 | 50.17676276 | -0.493701695 | 0.26805593 | -1.841786131 | 0.065506439 | 0.150196674 |
| ENSG00000144366 | GULP1 | 140.4932891 | 0.788353909 | 0.154833299 | 5.091630246 | 3.55E-07 | 2.17E-06 |
| ENSG00000144369 | FAM171B | 587.9797064 | 0.533435173 | 0.088125611 | 6.053123085 | 1.42E-09 | 1.11E-08 |
| ENSG00000144381 | HSPD1 | 14510.0651 | 0.136887409 | 0.055369058 | 2.472272667 | 0.013425707 | 0.038536752 |
| ENSG00000144395 | CCDC150 | 101.2218357 | -0.485744527 | 0.195532029 | -2.484219749 | 0.012983566 | 0.03739746 |
| ENSG00000144401 | METTL21A | 345.4227966 | -0.102635533 | 0.104305775 | -0.98398706 | 0.325121876 | 0.515289196 |
| ENSG00000144426 | NBEAL1 | 451.9088633 | -0.318799169 | 0.104911846 | -3.038733754 | 0.002375747 | 0.008233818 |
| ENSG00000144445 | KANSL1L | 251.6174855 | 0.553587727 | 0.117982537 | 4.692115813 | 2.70E-06 | 1.49E-05 |
| ENSG00000144452 | ABCA12 | 1610.762758 | 1.517069061 | 0.064390709 | 23.56037202 | 9.83E-123 | 2.53E-120 |
| ENSG00000144455 | SUMF1 | 1357.580305 | 0.051693546 | 0.062083807 | 0.83264136 | 0.405047025 | 0.597811154 |
| ENSG00000144468 | RHBDD1 | 565.5258371 | 0.10957613 | 0.084410866 | 1.298128259 | 0.194243265 | 0.355083314 |
| ENSG00000144476 | ACKR3 | 5284.172414 | 1.52358071 | 0.062680709 | 24.30701145 | 1.65E-130 | 4.80E-128 |
| ENSG00000144485 | HES6 | 569.1615796 | 0.045735109 | 0.087561398 | 0.522320454 | 0.60144723 | 0.761283686 |
| ENSG00000144504 | ANKMY1 | 434.1699584 | 0.17570319 | 0.091441734 | 1.921477018 | 0.054671597 | 0.129322173 |
| ENSG00000144524 | COPS7B | 914.8354358 | 0.198545675 | 0.068505034 | 2.898264014 | 0.003752346 | 0.012442731 |
| ENSG00000144535 | DIS3L2 | 560.8362968 | 0.128129146 | 0.085557248 | 1.497583764 | 0.134241428 | 0.267828571 |
| ENSG00000144550 | CPNE9 | 1.611240058 | -0.166246268 | 1.346420489 | -0.12347277 | 0.901732735 | NA |
| ENSG00000144554 | FANCD2 | 2434.141374 | -0.377404376 | 0.062773567 | -6.012154415 | 1.83E-09 | 1.42E-08 |
| ENSG00000144559 | TAMM41 | 605.6251555 | 0.023387138 | 0.087056785 | 0.268642336 | 0.78820493 | 0.885432696 |
| ENSG00000144560 | VGLL4 | 2193.740675 | 0.507435168 | 0.055369113 | 9.164589021 | 4.97E-20 | 8.31E-19 |
| ENSG00000144566 | RAB5A | 2602.693732 | 0.007062696 | 0.050527141 | 0.139780241 | 0.888833625 | 0.942772834 |
| ENSG00000144567 | RETREG2 | 4255.892696 | 0.180308603 | 0.049390641 | 3.650663382 | 0.000261564 | 0.001086416 |
| ENSG00000144579 | CTDSP1 | 4756.559168 | -0.332365103 | 0.060285025 | -5.513228215 | 3.52E-08 | 2.39E-07 |
| ENSG00000144580 | CNOT9 | 2404.091018 | -0.010594045 | 0.050970647 | -0.207845995 | 0.835349218 | 0.913215357 |
| ENSG00000144589 | STK11IP | 1289.979975 | 0.130827765 | 0.065606984 | 1.994113398 | 0.046139663 | 0.111803515 |
| ENSG00000144591 | GMPPA | 1179.284879 | -0.290555993 | 0.069299063 | -4.192783846 | 2.76E-05 | 0.000134065 |
| ENSG00000144597 | EAF1 | 3744.632905 | 0.179916932 | 0.047283665 | 3.805054745 | 0.000141773 | 0.000616895 |
| ENSG00000144619 | CNTN4 | 171.4839705 | -0.584389304 | 0.14088043 | -4.148122653 | 3.35E-05 | 0.000160941 |
| ENSG00000144635 | DYNC1LI1 | 1394.304778 | -0.268031823 | 0.06546053 | -4.094556278 | 4.23E-05 | 0.000199817 |
| ENSG00000144645 | OSBPL10 | 1493.845543 | -0.134984253 | 0.060154371 | -2.243964145 | 0.024834708 | 0.066099108 |
| ENSG00000144647 | POMGNT2 | 278.8895476 | 0.137485815 | 0.114626527 | 1.199424066 | 0.230363094 | 0.403707363 |

| | | | | | | | |
|---|---|---|---|---|---|---|---|
| ENSG00000144648 | ACKR2 | 13.94025296 | -1.385717603 | 0.489192279 | -2.832664501 | 0.004616181 | 0.014999318 |
| ENSG00000144649 | GASK1A | 24.5532334 | 0.891385732 | 0.374011369 | 2.383311861 | 0.017157652 | 0.047853683 |
| ENSG00000144655 | CSRNP1 | 887.8647612 | 0.185202485 | 0.072225872 | 2.564212509 | 0.010341021 | 0.030605473 |
| ENSG00000144659 | SLC25A38 | 1982.535395 | 0.114514133 | 0.054143894 | 2.114996261 | 0.034430266 | 0.087389943 |
| ENSG00000144668 | ITGA9 | 25.47855796 | -1.898025408 | 0.383441802 | -4.949969976 | 7.42E-07 | 4.38E-06 |
| ENSG00000144671 | SLC22A14 | 2.678900856 | 0.170555482 | 1.090204718 | 0.156443537 | 0.875683422 | 0.935508672 |
| ENSG00000144674 | GOLGA4 | 4485.319582 | -0.006652556 | 0.056374456 | -0.118006567 | 0.906062454 | 0.95262805 |
| ENSG00000144677 | CTDSPL | 326.9808133 | 0.641618114 | 0.105213481 | 6.098250031 | 1.07E-09 | 8.51E-09 |
| ENSG00000144711 | IQSEC1 | 3713.390937 | 0.204840595 | 0.057528996 | 3.560649557 | 0.000369939 | 0.001495136 |
| ENSG00000144712 | CAND2 | 3.727902134 | -0.262104535 | 0.91189078 | -0.287429745 | 0.773783288 | 0.876745462 |
| ENSG00000144713 | RPL32 | 13555.08001 | 0.2268564 | 0.051470524 | 4.407501258 | 1.05E-05 | 5.39E-05 |
| ENSG00000144724 | PTPRG | 1142.742264 | 0.595490788 | 0.068437767 | 8.701201348 | 3.28E-18 | 4.92E-17 |
| ENSG00000144730 | IL17RD | 84.17105968 | -0.159385054 | 0.204233361 | -0.780406557 | 0.43515161 | 0.625148918 |
| ENSG00000144736 | SHQ1 | 705.1093921 | -0.005184602 | 0.075107443 | -0.069029143 | 0.944966422 | 0.973833321 |
| ENSG00000144741 | SLC25A26 | 474.2439676 | 0.200864731 | 0.093188581 | 2.155465061 | 0.03112546 | 0.080190706 |
| ENSG00000144744 | UBA3 | 664.0562232 | 0.165659216 | 0.07839605 | 2.113106677 | 0.034591638 | 0.087741285 |
| ENSG00000144746 | ARL6IP5 | 1271.206038 | 0.006529089 | 0.062670318 | 0.104181527 | 0.917025293 | 0.959128265 |
| ENSG00000144747 | TMF1 | 1652.46017 | 0.086671513 | 0.05705001 | 1.519219925 | 0.128707148 | 0.259176788 |
| ENSG00000144749 | LRIG1 | 1484.490344 | 0.297729729 | 0.060955982 | 4.884339772 | 1.04E-06 | 6.00E-06 |
| ENSG00000144791 | LIMD1 | 1428.106431 | 0.144671467 | 0.05906693 | 2.449280275 | 0.014314202 | 0.04077077 |
| ENSG00000144802 | NFKBIZ | 354.3103988 | -0.973344025 | 0.113168289 | -8.600854852 | 7.91E-18 | 1.15E-16 |
| ENSG00000144815 | NXPE3 | 1341.258208 | -0.419034129 | 0.061453856 | -6.818679223 | 9.19E-12 | 8.63E-11 |
| ENSG00000144821 | MYH15 | 2.900218184 | 0.233443563 | 1.106981405 | 0.210883003 | 0.832978567 | 0.911666428 |
| ENSG00000144824 | PHLDB2 | 54.73538863 | -0.763565955 | 0.246257183 | -3.100685011 | 0.001930736 | 0.006810868 |
| ENSG00000144827 | ABHD10 | 596.4578716 | 0.004111441 | 0.08428687 | 0.048779138 | 0.961095308 | 0.982116705 |
| ENSG00000144840 | RABL3 | 1215.552397 | 0.139032477 | 0.063388341 | 2.193344638 | 0.028282558 | 0.073819761 |
| ENSG00000144843 | ADPRH | 47.93042555 | -0.699572292 | 0.257929723 | -2.712259311 | 0.006682629 | 0.02082308 |
| ENSG00000144847 | IGSF11 | 9.005773723 | 0.30260428 | 0.605037469 | 0.500141388 | 0.616975525 | 0.772972842 |
| ENSG00000144848 | ATG3 | 1608.734653 | -0.051723771 | 0.059720264 | -0.866100831 | 0.386434869 | 0.578997258 |
| ENSG00000144852 | NR1I2 | 2.172404331 | 0.2858244 | 1.211093653 | 0.2360052 | 0.813428635 | 0.901448243 |
| ENSG00000144857 | BOC | 68.99484897 | 0.940374449 | 0.221361805 | 4.248133271 | 2.16E-05 | 0.000106457 |
| ENSG00000144867 | SRPRB | 2296.445814 | 0.169414495 | 0.052816631 | 3.207597512 | 0.001338487 | 0.004890871 |
| ENSG00000144891 | AGTR1 | 16.30695851 | 2.01321858 | 0.517946091 | 3.88692687 | 0.000101521 | 0.000452213 |

| | | | | | | | |
|---|---|---|---|---|---|---|---|
| ENSG00000144893 | MED12L | 960.413377 | 0.188908647 | 0.069758423 | 2.70804067 | 0.006768173 | 0.021052459 |
| ENSG00000144895 | EIF2A | 4264.725769 | 0.145661409 | 0.058896382 | 2.473180952 | 0.013391632 | 0.038458191 |
| ENSG00000144908 | ALDH1L1 | 3.595573426 | 1.668572365 | 1.004704352 | 1.660759567 | 0.096761745 | 0.206465857 |
| ENSG00000144909 | OSBPL11 | 1154.421602 | -0.069900858 | 0.065262721 | -1.071068703 | 0.284138539 | 0.46795449 |
| ENSG00000144935 | TRPC1 | 185.9263546 | 0.325548102 | 0.13784603 | 2.361679204 | 0.018192377 | 0.050335681 |
| ENSG00000144959 | NCEH1 | 434.1107594 | -0.073460447 | 0.091908921 | -0.799274394 | 0.424131323 | 0.615019183 |
| ENSG00000145002 | FAM86B2 | 3.032539837 | 0.091971365 | 1.006634522 | 0.0913652 | 0.927202412 | 0.965137362 |
| ENSG00000145012 | LPP | 1506.835557 | 0.017610651 | 0.081466986 | 0.216169171 | 0.828855883 | 0.909497463 |
| ENSG00000145014 | TMEM44 | 600.1410522 | -0.27852874 | 0.081354112 | -3.423659017 | 0.000617841 | 0.002404287 |
| ENSG00000145016 | RUBCN | 1401.193726 | -0.105365982 | 0.061995045 | -1.69958717 | 0.089208605 | 0.193070799 |
| ENSG00000145022 | TCTA | 902.7215972 | 0.466401428 | 0.079517301 | 5.865408164 | 4.48E-09 | 3.35E-08 |
| ENSG00000145040 | UCN2 | 6.166635878 | 0.412449292 | 0.715785113 | 0.576219432 | 0.564466872 | 0.732969584 |
| ENSG00000145041 | DCAF1 | 1156.145403 | 0.182749171 | 0.064415349 | 2.837043853 | 0.004553336 | 0.014818204 |
| ENSG00000145050 | MANF | 904.2659052 | -0.115518582 | 0.069715941 | -1.656989502 | 0.097521598 | 0.207739386 |
| ENSG00000145063 | ENSG0000014 | 7.832384926 | 0.224154819 | 0.625331061 | 0.358457835 | 0.720000717 | 0.844366665 |
| ENSG00000145087 | STXBP5L | 4.057981209 | 0.416167271 | 0.867785519 | 0.479573884 | 0.63153042 | 0.782886131 |
| ENSG00000145088 | EAF2 | 30.9345611 | -0.663945713 | 0.326229548 | -2.035210227 | 0.041829724 | 0.102881299 |
| ENSG00000145103 | ILDR1 | 413.1394135 | -0.811981073 | 0.102191976 | -7.945644145 | 1.93E-15 | 2.39E-14 |
| ENSG00000145113 | MUC4 | 40.18663028 | -0.714008961 | 0.287403135 | -2.484346462 | 0.012978946 | 0.037390058 |
| ENSG00000145147 | SLIT2 | 514.5502788 | 0.152280259 | 0.088592075 | 1.718892569 | 0.085633936 | 0.186751828 |
| ENSG00000145191 | EIF2B5 | 2739.297645 | -0.028932363 | 0.049875276 | -0.580094286 | 0.561851037 | 0.730744007 |
| ENSG00000145198 | VWA5B2 | 36.06229614 | 0.567557644 | 0.298060348 | 1.904170238 | 0.056888017 | 0.133518631 |
| ENSG00000145214 | DGKQ | 868.1764192 | 0.229900739 | 0.073774005 | 3.11628384 | 0.001831459 | 0.006489588 |
| ENSG00000145216 | FIP1L1 | 772.4818918 | -0.106250468 | 0.072756356 | -1.460359959 | 0.144191171 | 0.282785797 |
| ENSG00000145217 | SLC26A1 | 20.73933195 | -0.100527047 | 0.384536436 | -0.261423984 | 0.793765563 | 0.889457895 |
| ENSG00000145220 | LYAR | 668.9101389 | -0.083662762 | 0.078727619 | -1.062686303 | 0.287924234 | 0.471919258 |
| ENSG00000145241 | CENPC | 585.9502662 | -0.040025691 | 0.089834006 | -0.445551661 | 0.655921135 | 0.80049108 |
| ENSG00000145242 | EPHA5 | 5.097689465 | -2.472258524 | 0.933547122 | -2.648241813 | 0.008091163 | 0.024688916 |
| ENSG00000145246 | ATP10D | 67.52036928 | 0.271268496 | 0.215627029 | 1.258044956 | 0.208375498 | 0.374853665 |
| ENSG00000145247 | OCIAD2 | 1091.041513 | 0.341805573 | 0.065852971 | 5.190435149 | 2.10E-07 | 1.32E-06 |
| ENSG00000145248 | SLC10A4 | 1.838677282 | -0.449060575 | 1.272796557 | -0.352814102 | 0.724227811 | 0.846509142 |
| ENSG00000145284 | SCD5 | 621.0941179 | 0.46706571 | 0.080584801 | 5.795952859 | 6.79E-09 | 4.98E-08 |
| ENSG00000145287 | PLAC8 | 4.875030752 | -1.040996383 | 0.860138285 | -1.210266303 | 0.226176724 | 0.398342014 |

| | | | | | | | |
|---|---|---|---|---|---|---|---|
| ENSG00000145293 | ENOPH1 | 2042.963295 | 0.076519676 | 0.055330281 | 1.382962004 | 0.166676517 | 0.315995619 |
| ENSG00000145331 | TRMT10A | 289.0326089 | 0.080832294 | 0.11055116 | 0.731175446 | 0.464671996 | 0.650767741 |
| ENSG00000145332 | KLHL8 | 890.7527207 | -0.035929225 | 0.069706684 | -0.515434433 | 0.606249477 | 0.764860977 |
| ENSG00000145335 | SNCA | 1.725192729 | -2.223435253 | 1.551125186 | -1.433433789 | 0.151733908 | NA |
| ENSG00000145337 | PYURF | 1164.171704 | 0.130056644 | 0.064215025 | 2.02533044 | 0.042833445 | 0.105026294 |
| ENSG00000145348 | TBCK | 939.2535182 | -0.246035391 | 0.068786838 | -3.576780083 | 0.000347853 | 0.001411843 |
| ENSG00000145349 | CAMK2D | 1292.543211 | -0.066067624 | 0.061886203 | -1.067566279 | 0.285716202 | 0.469595837 |
| ENSG00000145354 | CISD2 | 826.2275402 | 0.14975973 | 0.073920213 | 2.025964535 | 0.042768419 | 0.104931287 |
| ENSG00000145358 | DDIT4L | 34.58816602 | 0.546486863 | 0.298673551 | 1.829712948 | 0.067292874 | 0.153588121 |
| ENSG00000145362 | ANK2 | 42.88457922 | 1.099047891 | 0.281369639 | 3.906064268 | 9.38E-05 | 0.000419746 |
| ENSG00000145365 | TIFA | 125.2369136 | 0.183294279 | 0.158742543 | 1.15466387 | 0.248228101 | 0.425881047 |
| ENSG00000145375 | SPATA5 | 284.3613097 | -0.3101854 | 0.112482021 | -2.757644272 | 0.005821952 | 0.018426541 |
| ENSG00000145386 | CCNA2 | 1811.470592 | -0.303970155 | 0.05960415 | -5.0998153 | 3.40E-07 | 2.08E-06 |
| ENSG00000145388 | METTL14 | 773.6353178 | -0.051076836 | 0.074742662 | -0.683369238 | 0.494373553 | 0.676868356 |
| ENSG00000145390 | USP53 | 894.6863281 | -0.210611819 | 0.070468026 | -2.988757168 | 0.002801147 | 0.009562247 |
| ENSG00000145391 | SETD7 | 5952.587404 | 0.224490872 | 0.045363668 | 4.948693078 | 7.47E-07 | 4.40E-06 |
| ENSG00000145414 | NAF1 | 238.9900302 | -0.119147386 | 0.118854946 | -1.002460477 | 0.316121246 | 0.504860859 |
| ENSG00000145416 | MARCHF1 | 51.05088654 | 0.022033862 | 0.249549821 | 0.08829444 | 0.929642658 | 0.965993019 |
| ENSG00000145425 | RPS3A | 2640.298409 | 0.187763894 | 0.056235229 | 3.338901589 | 0.000841104 | 0.003192408 |
| ENSG00000145431 | PDGFC | 343.8930978 | -0.14948028 | 0.100324068 | -1.489974272 | 0.136231001 | 0.270896354 |
| ENSG00000145439 | CBR4 | 337.8618853 | 0.349797235 | 0.102832566 | 3.401619226 | 0.000669879 | 0.002591841 |
| ENSG00000145451 | GLRA3 | 15.78505658 | 0.549988724 | 0.446164557 | 1.232703755 | 0.21768631 | 0.387214171 |
| ENSG00000145476 | CYP4V2 | 305.9710047 | -0.130662097 | 0.107230188 | -1.218519711 | 0.223026539 | 0.394223209 |
| ENSG00000145491 | ROPN1L | 4.339170195 | -1.293614539 | 0.912081495 | -1.418310257 | 0.156100203 | 0.300275782 |
| ENSG00000145494 | NDUFS6 | 1378.321237 | 0.204021198 | 0.061792019 | 3.301740288 | 0.00096087 | 0.003606997 |
| ENSG00000145495 | MARCHF6 | 5755.240297 | -0.00023501 | 0.052853782 | -0.004446419 | 0.996452282 | 0.997800662 |
| ENSG00000145506 | NKD2 | 114.2177244 | -0.263213418 | 0.179232901 | -1.46855525 | 0.14195346 | 0.279535433 |
| ENSG00000145526 | CDH18 | 96.46948228 | 1.970928019 | 0.203981271 | 9.662298927 | 4.36E-22 | 8.18E-21 |
| ENSG00000145545 | SRD5A1 | 1014.748417 | 0.185512116 | 0.071642699 | 2.589407135 | 0.009614135 | 0.028743419 |
| ENSG00000145555 | MYO10 | 1176.020614 | 0.658739591 | 0.067241688 | 9.796595131 | 1.16E-22 | 2.27E-21 |
| ENSG00000145569 | OTULINL | 193.6099007 | -0.412587008 | 0.135185535 | -3.052005596 | 0.002273179 | 0.007905809 |
| ENSG00000145592 | RPL37 | 12504.91668 | 0.265835368 | 0.049778594 | 5.340355106 | 9.28E-08 | 6.03E-07 |
| ENSG00000145604 | SKP2 | 714.7844719 | -0.417001647 | 0.07899578 | -5.278783842 | 1.30E-07 | 8.36E-07 |

| | | | | | | | |
|---|---|---|---|---|---|---|---|
| ENSG00000145623 | OSMR | 620.2277883 | 0.444017078 | 0.084135611 | 5.277397656 | 1.31E-07 | 8.42E-07 |
| ENSG00000145632 | PLK2 | 7010.988453 | 0.45176517 | 0.045443171 | 9.941321439 | 2.75E-23 | 5.56E-22 |
| ENSG00000145675 | PIK3R1 | 3943.666075 | 1.158417316 | 0.054868533 | 21.11259849 | 6.09E-99 | 9.71E-97 |
| ENSG00000145681 | HAPLN1 | 2.127030706 | -1.318367133 | 1.234319036 | -1.068092685 | 0.285478705 | 0.469386575 |
| ENSG00000145685 | LHFPL2 | 647.5219257 | 0.228854483 | 0.077236192 | 2.963047228 | 0.003046098 | 0.010304895 |
| ENSG00000145687 | SSBP2 | 1433.800127 | 0.502215239 | 0.060877733 | 8.249571961 | 1.59E-16 | 2.14E-15 |
| ENSG00000145700 | ANKRD31 | 23.45708712 | 0.26139214 | 0.382283889 | 0.68376447 | 0.494123906 | 0.676647708 |
| ENSG00000145703 | IQGAP2 | 10.91317421 | -1.169013062 | 0.559366977 | -2.089885729 | 0.036628066 | 0.092162754 |
| ENSG00000145715 | RASA1 | 1627.30811 | 0.041798128 | 0.056761501 | 0.736381662 | 0.461498461 | 0.648328558 |
| ENSG00000145721 | LIX1 | 390.7116736 | 1.260233988 | 0.100317348 | 12.56247307 | 3.40E-36 | 1.20E-34 |
| ENSG00000145723 | GIN1 | 181.4898071 | -0.102417377 | 0.133811303 | -0.765386594 | 0.444041374 | 0.632807922 |
| ENSG00000145725 | PPIP5K2 | 1185.199999 | 0.024025508 | 0.067517242 | 0.35584256 | 0.721958474 | 0.845596895 |
| ENSG00000145730 | PAM | 1247.197651 | 1.509077857 | 0.065703712 | 22.96792396 | 9.76E-117 | 2.17E-114 |
| ENSG00000145734 | BDP1 | 1610.162335 | -0.130957953 | 0.063265302 | -2.069980687 | 0.038454153 | 0.095999376 |
| ENSG00000145736 | GTF2H2 | 145.7475431 | -0.419905078 | 0.148137838 | -2.834556549 | 0.004588934 | 0.014917123 |
| ENSG00000145740 | SLC30A5 | 1904.617336 | 0.022662503 | 0.057709462 | 0.392699946 | 0.694541111 | 0.827977606 |
| ENSG00000145741 | BTF3 | 10126.53462 | 0.239662634 | 0.043127794 | 5.557034389 | 2.74E-08 | 1.88E-07 |
| ENSG00000145743 | FBXL17 | 828.2624122 | 0.199916106 | 0.072664078 | 2.75123707 | 0.005937066 | 0.01875982 |
| ENSG00000145757 | SPATA9 | 10.07925044 | 1.038329745 | 0.591158253 | 1.756432799 | 0.079014536 | 0.175028338 |
| ENSG00000145779 | TNFAIP8 | 80.7937828 | 0.253568313 | 0.202634787 | 1.25135628 | 0.210804521 | 0.378214899 |
| ENSG00000145780 | FEM1C | 541.2120727 | 0.413640148 | 0.092830963 | 4.455842473 | 8.36E-06 | 4.35E-05 |
| ENSG00000145781 | COMMD10 | 624.1153379 | 0.26710539 | 0.085679094 | 3.117509525 | 0.001823861 | 0.006463674 |
| ENSG00000145782 | ATG12 | 1093.158558 | 0.122809174 | 0.068579039 | 1.790768375 | 0.073330472 | 0.164905503 |
| ENSG00000145794 | MEGF10 | 10.1547643 | 1.23410556 | 0.571471865 | 2.159521115 | 0.030809761 | 0.079484471 |
| ENSG00000145808 | ADAMTS19 | 620.3353554 | 0.766736462 | 0.086209946 | 8.893828366 | 5.90E-19 | 9.28E-18 |
| ENSG00000145817 | YIPF5 | 1226.862354 | -0.073790714 | 0.062307971 | -1.184290106 | 0.236298244 | 0.411406195 |
| ENSG00000145819 | ARHGAP26 | 391.2652694 | -0.023458733 | 0.103066929 | -0.22760679 | 0.819951941 | 0.904529311 |
| ENSG00000145832 | SLC25A48 | 2.554577605 | -2.130979155 | 1.247196821 | -1.708614966 | 0.087522295 | 0.190092922 |
| ENSG00000145833 | DDX46 | 4387.298417 | 0.058193157 | 0.049330327 | 1.179662896 | 0.238134317 | 0.413526941 |
| ENSG00000145860 | RNF145 | 2324.823009 | 0.059866269 | 0.057777195 | 1.036157409 | 0.300128713 | 0.48680172 |
| ENSG00000145861 | C1QTNF2 | 1.622143349 | 0.410567706 | 1.341968565 | 0.305944354 | 0.759647014 | NA |
| ENSG00000145864 | GABRB2 | 106.2983512 | -0.01123715 | 0.186229608 | -0.060340297 | 0.951884608 | 0.977646532 |
| ENSG00000145868 | FBXO38 | 1068.974833 | -0.093149986 | 0.065776505 | -1.416158944 | 0.156728974 | 0.301077953 |

| | | | | | | | |
|---|---|---|---|---|---|---|---|
| ENSG00000145882 | PCYOX1L | 498.2187914 | -0.272595745 | 0.086083598 | -3.166639785 | 0.001542112 | 0.00555354 |
| ENSG00000145901 | TNIP1 | 2313.3082 | 0.229466153 | 0.066180087 | 3.467299053 | 0.000525717 | 0.002072155 |
| ENSG00000145907 | G3BP1 | 6501.172519 | 0.203023973 | 0.047524416 | 4.271993025 | 1.94E-05 | 9.63E-05 |
| ENSG00000145911 | N4BP3 | 1809.84307 | -0.644643555 | 0.073232861 | -8.802654245 | 1.34E-18 | 2.05E-17 |
| ENSG00000145912 | NHP2 | 3778.354844 | -0.359169126 | 0.045689925 | -7.861013662 | 3.81E-15 | 4.62E-14 |
| ENSG00000145916 | RMND5B | 3589.118916 | -0.584791485 | 0.055693133 | -10.50024395 | 8.62E-26 | 1.98E-24 |
| ENSG00000145919 | BOD1 | 3311.839764 | -0.141306694 | 0.04976784 | -2.839317418 | 0.004521016 | 0.014725558 |
| ENSG00000145920 | CPLX2 | 4.278273071 | -2.5776228 | 1.022223949 | -2.521583262 | 0.011682802 | 0.03412302 |
| ENSG00000145945 | FAM50B | 105.8465709 | -10.27827881 | 1.187261433 | -8.657131884 | 4.84E-18 | 7.15E-17 |
| ENSG00000145949 | MYLK4 | 28.97862942 | -0.000617022 | 0.327771823 | -0.001882475 | 0.998498003 | 0.999161772 |
| ENSG00000145979 | TBC1D7 | 503.5527638 | -0.431896322 | 0.086995065 | -4.964607138 | 6.88E-07 | 4.07E-06 |
| ENSG00000145982 | FARS2 | 238.2818705 | -0.009203198 | 0.123823228 | -0.0743253 | 0.940751547 | 0.971736581 |
| ENSG00000145990 | GFOD1 | 247.2915821 | -0.372888211 | 0.119465613 | -3.121301628 | 0.001800535 | 0.006388906 |
| ENSG00000145996 | CDKAL1 | 295.3166368 | -0.828235222 | 0.109625819 | -7.555110935 | 4.19E-14 | 4.69E-13 |
| ENSG00000146005 | PSD2 | 8.011032989 | 1.864606351 | 0.702805139 | 2.653091514 | 0.007975824 | 0.024385634 |
| ENSG00000146007 | ZMAT2 | 3278.475411 | 0.146855505 | 0.049955537 | 2.939724265 | 0.003285044 | 0.011039982 |
| ENSG00000146013 | GFRA3 | 36.05790126 | -0.070805983 | 0.29781872 | -0.237748598 | 0.812076089 | 0.900518464 |
| ENSG00000146021 | KLHL3 | 2.750706155 | 0.679425134 | 1.057918025 | 0.642228526 | 0.520724814 | 0.697950759 |
| ENSG00000146038 | DCDC2 | 1061.34997 | 0.148323944 | 0.067323112 | 2.203165298 | 0.0275831 | 0.072281965 |
| ENSG00000146049 | KAAG1 | 7.087000578 | -0.147464762 | 0.65915019 | -0.223719516 | 0.822975559 | 0.906383375 |
| ENSG00000146054 | TRIM7 | 137.09523 | -1.459614959 | 0.161768421 | -9.022867062 | 1.83E-19 | 2.97E-18 |
| ENSG00000146063 | TRIM41 | 2470.45959 | -0.39701752 | 0.057400582 | -6.916611431 | 4.63E-12 | 4.47E-11 |
| ENSG00000146066 | HIGD2A | 1653.623904 | -0.236903801 | 0.057694805 | -4.106154828 | 4.02E-05 | 0.000190559 |
| ENSG00000146067 | FAM193B | 1614.11733 | -0.276491963 | 0.07047958 | -3.923008094 | 8.75E-05 | 0.000393818 |
| ENSG00000146072 | TNFRSF21 | 531.3261867 | 0.645808158 | 0.092433491 | 6.986733388 | 2.81E-12 | 2.77E-11 |
| ENSG00000146083 | RNF44 | 2532.619163 | 0.12147089 | 0.057199871 | 2.123621743 | 0.033701786 | 0.08576869 |
| ENSG00000146085 | MMUT | 1078.908008 | -0.270763651 | 0.065572755 | -4.12920964 | 3.64E-05 | 0.000173534 |
| ENSG00000146090 | RASGEF1C | 21.52087542 | 1.460866732 | 0.412585816 | 3.540758495 | 0.000398979 | 0.001604025 |
| ENSG00000146094 | DOK3 | 381.2552982 | 0.272307074 | 0.108186902 | 2.517005935 | 0.011835684 | 0.034516982 |
| ENSG00000146109 | ABT1 | 925.525564 | -0.168465101 | 0.06883124 | -2.447509321 | 0.01438474 | 0.040951346 |
| ENSG00000146112 | PPP1R18 | 753.4778146 | 0.020914999 | 0.076482502 | 0.273461232 | 0.784498689 | 0.883042195 |
| ENSG00000146122 | DAAM2 | 16.98149678 | 0.43805971 | 0.455506074 | 0.961698943 | 0.336200853 | 0.527274737 |
| ENSG00000146143 | PRIM2 | 464.6445037 | -0.410846283 | 0.088673877 | -4.633227939 | 3.60E-06 | 1.96E-05 |

| | | | | | | | |
|---|---|---|---|---|---|---|---|
| ENSG00000146151 | HMGCLL1 | 3.23214051 | -1.283103149 | 1.043218702 | -1.22994646 | 0.218717155 | 0.388461717 |
| ENSG00000146197 | SCUBE3 | 34.65144931 | 0.37954544 | 0.299492395 | 1.267295755 | 0.205049558 | 0.370120491 |
| ENSG00000146205 | ANO7 | 24.2136511 | -0.947180445 | 0.373066773 | -2.538903256 | 0.011120057 | 0.032629074 |
| ENSG00000146216 | TTBK1 | 34.30902013 | -0.504168046 | 0.301441571 | -1.672523279 | 0.094421181 | 0.202280715 |
| ENSG00000146221 | TCTE1 | 6.849757177 | 0.242909404 | 0.678834364 | 0.357833099 | 0.720468219 | 0.844793547 |
| ENSG00000146223 | RPL7L1 | 6803.107424 | -0.099668063 | 0.045505202 | -2.190256471 | 0.028505642 | 0.074309025 |
| ENSG00000146232 | NFKBIE | 279.6862154 | 0.030653926 | 0.114872999 | 0.266850576 | 0.789584212 | 0.886238142 |
| ENSG00000146242 | TPBG | 2325.652352 | 0.675438245 | 0.064125066 | 10.53313916 | 6.08E-26 | 1.41E-24 |
| ENSG00000146243 | IRAK1BP1 | 91.70103394 | 0.419173912 | 0.186860916 | 2.243240159 | 0.024881332 | 0.066215517 |
| ENSG00000146247 | PHIP | 1844.897846 | 0.020756511 | 0.05804408 | 0.357599097 | 0.720643354 | 0.844878037 |
| ENSG00000146263 | MMS22L | 749.2876832 | -0.14890546 | 0.077274492 | -1.926967834 | 0.053983639 | 0.127971962 |
| ENSG00000146267 | FAXC | 198.5839787 | 0.304262282 | 0.137799561 | 2.208006175 | 0.027243843 | 0.071515477 |
| ENSG00000146278 | PNRC1 | 1337.920526 | -0.239941833 | 0.062090208 | -3.864406941 | 0.000111359 | 0.000493354 |
| ENSG00000146281 | PM20D2 | 690.0419763 | -0.090158506 | 0.077867112 | -1.157850894 | 0.24692489 | 0.424176799 |
| ENSG00000146282 | RARS2 | 1444.425771 | 0.070292268 | 0.063369284 | 1.109248252 | 0.267323102 | 0.448602192 |
| ENSG00000146350 | TBC1D32 | 148.4472304 | 0.232077206 | 0.165811105 | 1.399648144 | 0.16161871 | 0.308626754 |
| ENSG00000146376 | ARHGAP18 | 308.6226493 | -0.409247588 | 0.111858132 | -3.658630628 | 0.000253566 | 0.00105484 |
| ENSG00000146386 | ABRACL | 772.0768486 | 0.580163983 | 0.074173962 | 7.82166631 | 5.21E-15 | 6.28E-14 |
| ENSG00000146409 | SLC18B1 | 257.3349784 | -0.290073972 | 0.117424427 | -2.47030349 | 0.013499847 | 0.038705972 |
| ENSG00000146410 | MTFR2 | 196.8895077 | 0.054475385 | 0.129122609 | 0.421888823 | 0.673106169 | 0.81285826 |
| ENSG00000146411 | SLC2A12 | 6.924076082 | 0.217225135 | 0.660509819 | 0.328874952 | 0.742250207 | 0.85777393 |
| ENSG00000146414 | SHPRH | 600.4233742 | -0.221842138 | 0.083864685 | -2.645239037 | 0.008163323 | 0.024873258 |
| ENSG00000146416 | AIG1 | 1121.492299 | 0.593491272 | 0.067140769 | 8.839506591 | 9.61E-19 | 1.50E-17 |
| ENSG00000146425 | DYNLT1 | 1407.956643 | -0.274853037 | 0.064729909 | -4.246152072 | 2.17E-05 | 0.000107333 |
| ENSG00000146426 | TIAM2 | 126.1444656 | 0.173282224 | 0.162800221 | 1.06438568 | 0.287154018 | 0.471161803 |
| ENSG00000146433 | TMEM181 | 3274.224592 | -1.971444278 | 0.051201171 | -38.50389017 | 0 | 0 |
| ENSG00000146453 | PNLDC1 | 2.13226219 | -0.406235983 | 1.191560132 | -0.340927808 | 0.73315793 | 0.852569723 |
| ENSG00000146457 | WTAP | 1501.42825 | -0.14640632 | 0.06558792 | -2.232214728 | 0.025600774 | 0.067776315 |
| ENSG00000146463 | ZMYM4 | 3231.067178 | -0.043610302 | 0.052538662 | -0.830061152 | 0.40650421 | 0.599267821 |
| ENSG00000146476 | ARMT1 | 12278.20303 | 0.150187876 | 0.056231131 | 2.670902635 | 0.007564759 | 0.023255905 |
| ENSG00000146530 | VWDE | 817.4065516 | -0.521832948 | 0.078588604 | -6.64005873 | 3.14E-11 | 2.83E-10 |
| ENSG00000146535 | GNA12 | 4004.670557 | -0.296516342 | 0.054679228 | -5.422833367 | 5.87E-08 | 3.88E-07 |
| ENSG00000146540 | C7orf50 | 5393.575573 | -0.071140592 | 0.047237191 | -1.506029269 | 0.13205966 | 0.264626887 |

| | | | | | | | |
|---|---|---|---|---|---|---|---|
| ENSG00000146555 | SDK1 | 2562.958105 | 1.594174688 | 0.068214199 | 23.37012993 | 8.60E-121 | 2.17E-118 |
| ENSG00000146556 | WASH2P | 169.0288001 | -0.33896676 | 0.148928375 | -2.2760388 | 0.022843687 | 0.061357745 |
| ENSG00000146574 | CCZ1B | 1070.980856 | 0.227159885 | 0.083431065 | 2.722725453 | 0.006474584 | 0.020243554 |
| ENSG00000146576 | INTS15 | 1396.868228 | 0.248981984 | 0.062419963 | 3.988819779 | 6.64E-05 | 0.000304961 |
| ENSG00000146587 | RBAK | 1557.416291 | 0.220028944 | 0.063800446 | 3.448705407 | 0.000563281 | 0.002207322 |
| ENSG00000146648 | EGFR | 136.1834315 | -0.152727101 | 0.164639546 | -0.9276453 | 0.353591591 | 0.545083738 |
| ENSG00000146670 | CDCA5 | 2183.788234 | -0.438351805 | 0.054095019 | -8.103367326 | 5.35E-16 | 6.90E-15 |
| ENSG00000146674 | IGFBP3 | 2159.666378 | 0.986784863 | 0.062168294 | 15.87279948 | 9.78E-57 | 6.41E-55 |
| ENSG00000146676 | PURB | 3581.379276 | 0.619654701 | 0.053136234 | 11.66162241 | 2.00E-31 | 5.92E-30 |
| ENSG00000146677 | RPL32P18 | 1416.044774 | 0.25927886 | 0.063708586 | 4.069763244 | 4.71E-05 | 0.000220999 |
| ENSG00000146700 | SSC4D | 22.08933208 | 0.226578524 | 0.372189761 | 0.608771513 | 0.542675898 | 0.716405887 |
| ENSG00000146701 | MDH2 | 6516.530698 | 0.557489056 | 0.04512219 | 12.35509759 | 4.57E-35 | 1.56E-33 |
| ENSG00000146707 | POMZP3 | 89.60161725 | 0.88880942 | 0.192326997 | 4.621345076 | 3.81E-06 | 2.06E-05 |
| ENSG00000146729 | NIPSNAP2 | 1493.327732 | 0.443114378 | 0.060351334 | 7.342246615 | 2.10E-13 | 2.24E-12 |
| ENSG00000146731 | CCT6A | 6274.977767 | 0.579905535 | 0.048179709 | 12.03630226 | 2.29E-33 | 7.24E-32 |
| ENSG00000146733 | PSPH | 781.6686868 | 0.446381057 | 0.074207123 | 6.015339792 | 1.80E-09 | 1.40E-08 |
| ENSG00000146755 | TRIM50 | 2.772014875 | 0.701904585 | 1.073972571 | 0.653559135 | 0.513395879 | 0.69261947 |
| ENSG00000146757 | ZNF92 | 386.8939737 | 0.52397374 | 0.097943232 | 5.349769724 | 8.81E-08 | 5.73E-07 |
| ENSG00000146776 | ATXN7L1 | 334.9860725 | -0.092683197 | 0.103029393 | -0.899580156 | 0.368343722 | 0.560889809 |
| ENSG00000146802 | TMEM168 | 483.8098061 | 0.11237785 | 0.092201422 | 1.218829902 | 0.22290876 | 0.394075751 |
| ENSG00000146826 | TRAPPC14 | 768.5578053 | -0.211266463 | 0.081304989 | -2.598444043 | 0.009364731 | 0.028096642 |
| ENSG00000146828 | SLC12A9 | 2053.080497 | -0.459292301 | 0.062012222 | -7.406480356 | 1.30E-13 | 1.40E-12 |
| ENSG00000146830 | GIGYF1 | 5283.467505 | -0.615218654 | 0.059506421 | -10.33869365 | 4.71E-25 | 1.05E-23 |
| ENSG00000146833 | TRIM4 | 2019.858749 | -0.154460353 | 0.0527353 | -2.928974563 | 0.003400823 | 0.011394022 |
| ENSG00000146834 | MEPCE | 3552.376464 | -0.625582848 | 0.051129078 | -12.23536339 | 2.01E-34 | 6.70E-33 |
| ENSG00000146842 | TMEM209 | 632.9530051 | 0.019671699 | 0.078452909 | 0.250745313 | 0.802011025 | 0.894633256 |
| ENSG00000146858 | ZC3HAV1L | 227.0541067 | 0.154426726 | 0.120981075 | 1.276453571 | 0.201795229 | 0.365567603 |
| ENSG00000146859 | TMEM140 | 52.2178462 | -0.390263568 | 0.245925655 | -1.586916858 | 0.112531476 | 0.233195309 |
| ENSG00000146872 | TLK2 | 3383.339225 | 0.084491892 | 0.053063295 | 1.592285073 | 0.111320665 | 0.230999192 |
| ENSG00000146904 | EPHA1 | 721.417899 | 0.119524894 | 0.080744528 | 1.480284754 | 0.13879727 | 0.27476781 |
| ENSG00000146909 | NOM1 | 604.6743507 | 0.300991025 | 0.080655248 | 3.731821941 | 0.0001901 | 0.000810574 |
| ENSG00000146918 | NCAPG2 | 1826.172462 | 0.394748365 | 0.056955744 | 6.930791163 | 4.18E-12 | 4.06E-11 |
| ENSG00000146938 | NLGN4X | 9.381491594 | -2.024142635 | 0.62906238 | -3.217713697 | 0.001292167 | 0.00473898 |

| | | | | | | | |
|---|---|---|---|---|---|---|---|
| ENSG00000146950 | SHROOM2 | 817.1751388 | 0.104486247 | 0.078479308 | 1.331385938 | 0.183062054 | 0.339074971 |
| ENSG00000146955 | RAB19 | 31.91409443 | -1.137408528 | 0.326535296 | -3.48326365 | 0.00049534 | 0.001961862 |
| ENSG00000146963 | LUC7L2 | 613.5176437 | -0.195529747 | 0.081830635 | -2.38944433 | 0.016873881 | 0.047171195 |
| ENSG00000146966 | DENND2A | 2.059880245 | -0.332394595 | 1.238162264 | -0.268458024 | 0.788346781 | 0.885492107 |
| ENSG00000147003 | CLTRN | 4.426873282 | -0.040918611 | 0.889252194 | -0.04601463 | 0.963298589 | 0.983272654 |
| ENSG00000147010 | SH3KBP1 | 1259.811474 | 0.537836608 | 0.06484159 | 8.294623988 | 1.09E-16 | 1.48E-15 |
| ENSG00000147027 | TMEM47 | 498.8334824 | -1.879321577 | 0.097301424 | -19.31443043 | 4.06E-83 | 4.88E-81 |
| ENSG00000147041 | SYTL5 | 118.1038749 | 0.728012136 | 0.174216253 | 4.178784249 | 2.93E-05 | 0.000142197 |
| ENSG00000147044 | CASK | 2280.355316 | 0.297157942 | 0.054272521 | 5.475292744 | 4.37E-08 | 2.94E-07 |
| ENSG00000147050 | KDM6A | 1270.735918 | -0.108933642 | 0.064199695 | -1.696793751 | 0.089735661 | 0.19394536 |
| ENSG00000147082 | CCNB3 | 9.731836305 | -0.625485075 | 0.62974452 | -0.993236234 | 0.320594844 | 0.510158555 |
| ENSG00000147099 | HDAC8 | 159.631932 | 0.081445094 | 0.140963978 | 0.577772388 | 0.563417799 | 0.732154597 |
| ENSG00000147100 | SLC16A2 | 1105.849386 | 0.332624367 | 0.067237719 | 4.94699062 | 7.54E-07 | 4.44E-06 |
| ENSG00000147117 | ZNF157 | 2.822431938 | -2.382583549 | 1.233433795 | -1.931667154 | 0.053400601 | 0.126838869 |
| ENSG00000147118 | ZNF182 | 231.7982276 | -0.61982897 | 0.121163055 | -5.11565978 | 3.13E-07 | 1.93E-06 |
| ENSG00000147119 | CHST7 | 66.30751332 | 0.083256222 | 0.224003285 | 0.371674112 | 0.710135502 | 0.838646906 |
| ENSG00000147121 | KRBOX4 | 350.3500747 | -0.07633234 | 0.10214281 | -0.747309967 | 0.454876477 | 0.642397145 |
| ENSG00000147123 | NDUFB11 | 2391.653329 | 0.043113487 | 0.060910411 | 0.707818039 | 0.479058263 | 0.663653377 |
| ENSG00000147124 | ZNF41 | 771.5679172 | -0.189443075 | 0.073332937 | -2.583328616 | 0.009785207 | 0.029167444 |
| ENSG00000147130 | ZMYM3 | 3611.252358 | 0.093868508 | 0.051755194 | 1.813702188 | 0.069723588 | 0.158096186 |
| ENSG00000147133 | TAF1 | 2374.780725 | -0.012444438 | 0.054687207 | -0.227556663 | 0.819990914 | 0.904529311 |
| ENSG00000147140 | NONO | 20568.7415 | 0.127817615 | 0.042535876 | 3.004936713 | 0.002656361 | 0.009104576 |
| ENSG00000147144 | CCDC120 | 1169.019618 | 0.206889075 | 0.066682012 | 3.102621954 | 0.001918145 | 0.006770618 |
| ENSG00000147155 | EBP | 1159.096165 | -0.53032656 | 0.067282294 | -7.882111747 | 3.22E-15 | 3.93E-14 |
| ENSG00000147162 | OGT | 6942.614297 | 0.059219952 | 0.07048396 | 0.840190485 | 0.400801593 | 0.593722623 |
| ENSG00000147164 | SNX12 | 2391.487663 | -0.042800805 | 0.05307634 | -0.806400842 | 0.420011762 | 0.611284724 |
| ENSG00000147166 | ITGB1BP2 | 22.4534487 | -0.19540857 | 0.371301895 | -0.526279485 | 0.598694035 | 0.75887321 |
| ENSG00000147174 | GCNA | 84.3685891 | 0.146646318 | 0.202199199 | 0.725256672 | 0.468294581 | 0.654243465 |
| ENSG00000147180 | ZNF711 | 334.8635159 | 0.276592917 | 0.104045763 | 2.658377519 | 0.007851788 | 0.024054491 |
| ENSG00000147202 | DIAPH2 | 861.3759976 | 0.365506099 | 0.073304917 | 4.986106196 | 6.16E-07 | 3.66E-06 |
| ENSG00000147224 | PRPS1 | 1241.993129 | -0.620793692 | 0.063984376 | -9.702270074 | 2.95E-22 | 5.60E-21 |
| ENSG00000147231 | RADX | 39.71774447 | -4.786205509 | 0.53827575 | -8.891735336 | 6.02E-19 | 9.45E-18 |
| ENSG00000147234 | FRMPD3 | 58.75739006 | -0.680731506 | 0.231183635 | -2.944548843 | 0.003234259 | 0.010880466 |

| | | | | | | | |
|---|---|---|---|---|---|---|---|
| ENSG00000147246 | HTR2C | 287.0076947 | -0.033260339 | 0.109050774 | -0.304998654 | 0.760367177 | 0.868320542 |
| ENSG00000147251 | DOCK11 | 11.73043603 | 0.495265809 | 0.531593802 | 0.931662121 | 0.351511168 | 0.542788516 |
| ENSG00000147255 | IGSF1 | 37.91683279 | 4.203400451 | 0.497843542 | 8.443215779 | 3.09E-17 | 4.31E-16 |
| ENSG00000147256 | ARHGAP36 | 4.18944757 | 2.826709326 | 1.093130607 | 2.585884348 | 0.009712951 | 0.028989735 |
| ENSG00000147257 | GPC3 | 18.2711888 | -0.394029793 | 0.41115933 | -0.958338444 | 0.337892124 | 0.529131362 |
| ENSG00000147274 | RBMX | 11231.60607 | -0.109390669 | 0.057371623 | -1.906703399 | 0.056559007 | 0.132895996 |
| ENSG00000147316 | MCPH1 | 417.1232166 | -0.056636038 | 0.098328186 | -0.575989865 | 0.564622032 | 0.733084344 |
| ENSG00000147324 | MFHAS1 | 660.3451489 | 0.007470181 | 0.079773616 | 0.093642253 | 0.925393344 | 0.964370625 |
| ENSG00000147364 | FBXO25 | 717.7535325 | 0.112450739 | 0.07581251 | 1.483274183 | 0.138001579 | 0.273620515 |
| ENSG00000147378 | FATE1 | 154.4962187 | 4.43269209 | 0.264440905 | 16.76250536 | 4.59E-63 | 3.54E-61 |
| ENSG00000147381 | MAGEA4 | 4.471714521 | 3.576772563 | 1.227019661 | 2.915008355 | 0.00355679 | 0.011861138 |
| ENSG00000147383 | NSDHL | 2316.614821 | -0.742343927 | 0.053755052 | -13.80975183 | 2.23E-43 | 1.01E-41 |
| ENSG00000147394 | ZNF185 | 1571.372807 | 0.151248193 | 0.066100667 | 2.288149278 | 0.022128829 | 0.059756956 |
| ENSG00000147400 | CETN2 | 2044.223427 | -0.403048423 | 0.054206413 | -7.435438033 | 1.04E-13 | 1.14E-12 |
| ENSG00000147403 | RPL10 | 9033.740158 | -0.319602921 | 0.051032905 | -6.262683297 | 3.78E-10 | 3.13E-09 |
| ENSG00000147408 | CSGALNACT1 | 177.3806755 | -0.944174121 | 0.143929717 | -6.559966502 | 5.38E-11 | 4.75E-10 |
| ENSG00000147416 | ATP6V1B2 | 1428.351523 | 0.147851374 | 0.059416044 | 2.488408228 | 0.012831635 | 0.037002854 |
| ENSG00000147419 | CCDC25 | 1306.217419 | 0.028783817 | 0.061099924 | 0.47109415 | 0.637573503 | 0.78659182 |
| ENSG00000147421 | HMBOX1 | 330.2980657 | -0.330368438 | 0.105857861 | -3.120868261 | 0.001803187 | 0.006397325 |
| ENSG00000147437 | GNRH1 | 12.88108527 | 0.273441317 | 0.489269897 | 0.558876234 | 0.57624619 | 0.742571005 |
| ENSG00000147439 | BIN3 | 397.0491095 | 0.05370188 | 0.095492732 | 0.562366154 | 0.573866574 | 0.740493824 |
| ENSG00000147454 | SLC25A37 | 709.5438231 | 0.320993635 | 0.087406492 | 3.672423254 | 0.000240261 | 0.001004116 |
| ENSG00000147457 | CHMP7 | 1518.896484 | 0.061085311 | 0.059441149 | 1.02766033 | 0.304109634 | 0.491460022 |
| ENSG00000147459 | DOCK5 | 1044.379223 | -0.742072948 | 0.066875136 | -11.09639531 | 1.31E-28 | 3.42E-27 |
| ENSG00000147471 | PLPBP | 1093.965829 | -0.125817928 | 0.065795085 | -1.912269417 | 0.055841646 | 0.131614854 |
| ENSG00000147475 | ERLIN2 | 1499.35901 | -0.125723075 | 0.057379566 | -2.191077488 | 0.028446185 | 0.074179322 |
| ENSG00000147485 | PXDNL | 28.14764205 | 0.224919036 | 0.341839831 | 0.657966145 | 0.510559886 | 0.690498984 |
| ENSG00000147509 | RGS20 | 14.00878642 | -0.36758358 | 0.475564238 | -0.772942014 | 0.439556699 | 0.629034976 |
| ENSG00000147526 | TACC1 | 6938.487155 | 0.40678992 | 0.05096391 | 7.981921342 | 1.44E-15 | 1.80E-14 |
| ENSG00000147533 | GOLGA7 | 1086.087435 | -0.060936379 | 0.068854395 | -0.885003476 | 0.376154757 | 0.569157056 |
| ENSG00000147535 | PLPP5 | 604.8762666 | 0.054575333 | 0.086748768 | 0.629119399 | 0.529270892 | 0.705079806 |
| ENSG00000147536 | GINS4 | 522.6592753 | -0.232362812 | 0.084089636 | -2.763275271 | 0.005722448 | 0.018141648 |
| ENSG00000147548 | NSD3 | 2246.132685 | -0.151672913 | 0.056592843 | -2.680072319 | 0.007360626 | 0.022710525 |

| | | | | | | | |
|---|---|---|---|---|---|---|---|
| ENSG00000147586 | MRPS28 | 75.21743226 | -0.354102968 | 0.212073136 | -1.669720998 | 0.094974577 | 0.203276433 |
| ENSG00000147592 | LACTB2 | 429.2821368 | 0.951074636 | 0.095156186 | 9.994879724 | 1.60E-23 | 3.26E-22 |
| ENSG00000147601 | TERF1 | 754.5851993 | 0.306746322 | 0.076354341 | 4.017405121 | 5.88E-05 | 0.000272531 |
| ENSG00000147604 | RPL7 | 1236.610026 | 0.391090795 | 0.076082319 | 5.14036377 | 2.74E-07 | 1.70E-06 |
| ENSG00000147606 | SLC26A7 | 32.28309823 | -3.088395853 | 0.407833072 | -7.572695959 | 3.66E-14 | 4.12E-13 |
| ENSG00000147614 | ATP6V0D2 | 2.386175024 | 0.872262318 | 1.192193474 | 0.731644936 | 0.464385314 | 0.650604562 |
| ENSG00000147642 | SYBU | 112.4623943 | 0.489084877 | 0.169433791 | 2.886584039 | 0.003894487 | 0.012865717 |
| ENSG00000147649 | MTDH | 7526.465671 | -0.192048658 | 0.04898951 | -3.920199626 | 8.85E-05 | 0.000398279 |
| ENSG00000147650 | LRP12 | 573.1321758 | 0.326296896 | 0.081947703 | 3.981769894 | 6.84E-05 | 0.000313524 |
| ENSG00000147654 | EBAG9 | 1039.053974 | -0.549647412 | 0.069948352 | -7.857903611 | 3.91E-15 | 4.73E-14 |
| ENSG00000147669 | POLR2K | 1331.72642 | -0.527609527 | 0.061353729 | -8.599469648 | 8.01E-18 | 1.17E-16 |
| ENSG00000147676 | MAL2 | 6439.239321 | -0.217845232 | 0.055603826 | -3.917810104 | 8.94E-05 | 0.000401774 |
| ENSG00000147677 | EIF3H | 7233.937536 | 0.455413939 | 0.04545576 | 10.01883886 | 1.26E-23 | 2.58E-22 |
| ENSG00000147679 | UTP23 | 1265.639898 | 0.474751761 | 0.072238693 | 6.571987089 | 4.96E-11 | 4.40E-10 |
| ENSG00000147684 | NDUFB9 | 3071.470049 | -0.235163562 | 0.049409297 | -4.759500276 | 1.94E-06 | 1.08E-05 |
| ENSG00000147687 | TATDN1 | 1516.265962 | -0.122877019 | 0.067100496 | -1.831238611 | 0.067064936 | 0.153249291 |
| ENSG00000147689 | FAM83A | 26.61848891 | -1.358649437 | 0.363720732 | -3.735419283 | 0.000187403 | 0.00079952 |
| ENSG00000147789 | ZNF7 | 1167.807052 | 0.066651768 | 0.067949594 | 0.980900174 | 0.326641972 | 0.517185142 |
| ENSG00000147799 | ARHGAP39 | 1321.994101 | -0.041888701 | 0.060821279 | -0.688717862 | 0.491000833 | 0.674019811 |
| ENSG00000147804 | SLC39A4 | 686.9088169 | 0.323741753 | 0.077973607 | 4.15194021 | 3.30E-05 | 0.000158545 |
| ENSG00000147813 | NAPRT | 771.5111964 | 0.322340462 | 0.077543774 | 4.156883882 | 3.23E-05 | 0.000155545 |
| ENSG00000147852 | VLDLR | 547.2244013 | 0.859636134 | 0.087532805 | 9.820731039 | 9.17E-23 | 1.80E-21 |
| ENSG00000147853 | AK3 | 1200.646272 | -0.035547758 | 0.065032128 | -0.546618406 | 0.584640921 | 0.748878371 |
| ENSG00000147854 | UHRF2 | 616.3451857 | -0.211153714 | 0.090260051 | -2.339392811 | 0.019315113 | 0.053064515 |
| ENSG00000147862 | NFIB | 1432.014613 | 0.963665698 | 0.062473045 | 15.42530387 | 1.11E-53 | 6.71E-52 |
| ENSG00000147872 | PLIN2 | 248.6645419 | -0.266899495 | 0.119039867 | -2.24210175 | 0.024954797 | 0.066380223 |
| ENSG00000147874 | HAUS6 | 557.633413 | 0.01574034 | 0.083102003 | 0.189409874 | 0.849771586 | 0.921342291 |
| ENSG00000147883 | CDKN2B | 744.8239173 | 0.619951941 | 0.077900044 | 7.958300316 | 1.74E-15 | 2.16E-14 |
| ENSG00000147889 | CDKN2A | 40.14107808 | -0.0056272 | 0.283330905 | -0.019860875 | 0.984154356 | 0.992635461 |
| ENSG00000147894 | C9orf72 | 162.4132839 | 0.189686534 | 0.151072613 | 1.255598422 | 0.209261604 | 0.376093795 |
| ENSG00000147905 | ZCCHC7 | 437.7870092 | 0.200832708 | 0.094741958 | 2.119786344 | 0.034024067 | 0.086454598 |
| ENSG00000147912 | FBXO10 | 97.23396213 | 0.277405771 | 0.188612492 | 1.470770932 | 0.141353076 | 0.278625156 |
| ENSG00000147955 | SIGMAR1 | 851.8425225 | -0.125846772 | 0.071491629 | -1.760300794 | 0.07835682 | 0.173755963 |

| | | | | | | | |
|---|---|---|---|---|---|---|---|
| ENSG00000147996 | ZNG1E | 204.4742955 | -0.222552858 | 0.131920916 | -1.687017233 | 0.091600038 | 0.197287098 |
| ENSG00000148019 | CEP78 | 1090.921768 | -0.173388696 | 0.066371808 | -2.612384723 | 0.008991301 | 0.027093188 |
| ENSG00000148057 | IDNK | 212.2168988 | -0.454411402 | 0.126505831 | -3.592019417 | 0.000328125 | 0.00134079 |
| ENSG00000148082 | SHC3 | 100.0173719 | -0.727681501 | 0.178983703 | -4.065629945 | 4.79E-05 | 0.000224722 |
| ENSG00000148090 | AUH | 273.8651354 | 0.448618126 | 0.11937145 | 3.758169369 | 0.000171161 | 0.000735012 |
| ENSG00000148110 | MFSD14B | 1424.107133 | -0.033165369 | 0.060944934 | -0.544185832 | 0.586313601 | 0.750100097 |
| ENSG00000148120 | AOPEP | 1112.515361 | 0.364193312 | 0.067072607 | 5.4298368 | 5.64E-08 | 3.75E-07 |
| ENSG00000148123 | PLPPR1 | 178.9021363 | -0.743390612 | 0.14015895 | -5.303911125 | 1.13E-07 | 7.32E-07 |
| ENSG00000148143 | ZNF462 | 25.08003742 | -0.159069886 | 0.348592561 | -0.456320368 | 0.648159619 | 0.794866977 |
| ENSG00000148153 | INIP | 1305.245062 | 0.062067079 | 0.061829304 | 1.003845664 | 0.31545301 | 0.504207396 |
| ENSG00000148154 | UGCG | 1162.180308 | 0.823953052 | 0.067154 | 12.269605 | 1.32E-34 | 4.45E-33 |
| ENSG00000148158 | SNX30 | 1207.060964 | 0.332330493 | 0.062852967 | 5.287427256 | 1.24E-07 | 7.99E-07 |
| ENSG00000148175 | STOM | 688.7399326 | -0.83113583 | 0.076600979 | -10.85019857 | 1.99E-27 | 5.03E-26 |
| ENSG00000148180 | GSN | 891.7462651 | 0.914433712 | 0.079179259 | 11.54890458 | 7.48E-31 | 2.15E-29 |
| ENSG00000148187 | MRRF | 1264.315172 | -0.100028252 | 0.070837285 | -1.412084792 | 0.157924996 | 0.302833688 |
| ENSG00000148200 | NR6A1 | 383.3488568 | 0.233752774 | 0.108517062 | 2.1540647 | 0.031235098 | 0.080446071 |
| ENSG00000148204 | CRB2 | 1.780982923 | 0.110153164 | 1.294164325 | 0.085115284 | 0.93216974 | NA |
| ENSG00000148218 | ALAD | 1953.900793 | 0.461633138 | 0.063958152 | 7.217737246 | 5.29E-13 | 5.51E-12 |
| ENSG00000148219 | ASTN2 | 408.7902583 | -0.003238275 | 0.09741232 | -0.033242969 | 0.973480833 | 0.98799497 |
| ENSG00000148225 | WDR31 | 149.4164542 | -0.155365262 | 0.152155163 | -1.021097532 | 0.307208233 | 0.495040016 |
| ENSG00000148229 | POLE3 | 4001.481386 | -0.08000371 | 0.048366993 | -1.654097235 | 0.098107758 | 0.2087554 |
| ENSG00000148248 | SURF4 | 10219.45312 | 0.089107417 | 0.043384409 | 2.053904163 | 0.039984968 | 0.099087736 |
| ENSG00000148290 | SURF1 | 1678.067482 | -0.006264724 | 0.057887769 | -0.1082219 | 0.913819673 | 0.957171969 |
| ENSG00000148291 | SURF2 | 902.660843 | 0.287863647 | 0.074894075 | 3.843610399 | 0.000121237 | 0.000533203 |
| ENSG00000148296 | SURF6 | 2416.752047 | 0.041336814 | 0.05350935 | 0.772515712 | 0.439809045 | 0.629163256 |
| ENSG00000148297 | MED22 | 2205.548966 | 0.047862565 | 0.055899205 | 0.856229796 | 0.391870685 | 0.584770973 |
| ENSG00000148300 | REXO4 | 2546.941271 | 0.139032019 | 0.05535432 | 2.511674246 | 0.012015995 | 0.034973338 |
| ENSG00000148303 | RPL7A | 17755.99603 | 0.139750452 | 0.053302813 | 2.621821348 | 0.008746126 | 0.026441242 |
| ENSG00000148308 | GTF3C5 | 2590.900985 | -0.145016437 | 0.05955794 | -2.434879998 | 0.014896729 | 0.042230718 |
| ENSG00000148331 | ASB6 | 1291.431993 | -0.006690901 | 0.070019353 | -0.095557875 | 0.923871723 | 0.963279648 |
| ENSG00000148334 | PTGES2 | 1739.196942 | 0.087445452 | 0.057164985 | 1.52970307 | 0.126090243 | 0.25509394 |
| ENSG00000148335 | NTMT1 | 944.7352549 | 0.050991946 | 0.084167839 | 0.605836461 | 0.544623356 | 0.717913747 |
| ENSG00000148337 | CIZ1 | 6771.639973 | 0.161259222 | 0.046629449 | 3.458312835 | 0.00054357 | 0.002138116 |

| | | | | | | | |
|---|---|---|---|---|---|---|---|
| ENSG00000148339 | SLC25A25 | 840.1703356 | -0.065664596 | 0.071711939 | -0.915671746 | 0.359839088 | 0.552413082 |
| ENSG00000148341 | SH3GLB2 | 6065.471569 | 0.44211634 | 0.054652781 | 8.089548797 | 5.99E-16 | 7.70E-15 |
| ENSG00000148343 | MIGA2 | 1350.081078 | 0.108914515 | 0.066932132 | 1.627238093 | 0.10368654 | 0.218277437 |
| ENSG00000148344 | PTGES | 13.10552222 | 3.78737952 | 0.763342209 | 4.961574868 | 6.99E-07 | 4.14E-06 |
| ENSG00000148356 | LRSAM1 | 1253.771267 | -0.374992128 | 0.06274476 | -5.976469201 | 2.28E-09 | 1.76E-08 |
| ENSG00000148357 | HMCN2 | 1.676961935 | 1.257595055 | 1.473618083 | 0.853406367 | 0.393433992 | NA |
| ENSG00000148358 | GPR107 | 4593.872716 | 0.121909401 | 0.049046373 | 2.485594595 | 0.012933521 | 0.03727683 |
| ENSG00000148362 | PAXX | 1146.299247 | 0.212620225 | 0.07917396 | 2.685481745 | 0.007242533 | 0.022394354 |
| ENSG00000148377 | IDI2 | 3.618266469 | -1.540120597 | 0.988987434 | -1.557270138 | 0.119406362 | 0.244547982 |
| ENSG00000148384 | INPP5E | 975.635725 | 0.082872781 | 0.072417342 | 1.144377566 | 0.252467088 | 0.431178901 |
| ENSG00000148396 | SEC16A | 8698.060871 | -0.159369051 | 0.051392369 | -3.101025618 | 0.001928516 | 0.006804085 |
| ENSG00000148399 | DPH7 | 1154.270686 | -0.062126653 | 0.06823139 | -0.910528918 | 0.362543638 | 0.555227389 |
| ENSG00000148400 | NOTCH1 | 910.9473525 | 0.527390674 | 0.074777706 | 7.0527795 | 1.75E-12 | 1.76E-11 |
| ENSG00000148408 | CACNA1B | 5.318832247 | -0.085702713 | 0.756086701 | -0.113350377 | 0.909752779 | 0.954699394 |
| ENSG00000148411 | NACC2 | 1874.160253 | 0.295221529 | 0.059717443 | 4.943639854 | 7.67E-07 | 4.52E-06 |
| ENSG00000148426 | PROSER2 | 290.2038485 | -0.409080696 | 0.111175194 | -3.679604064 | 0.000233596 | 0.000979116 |
| ENSG00000148429 | USP6NL | 961.3287916 | -0.765086469 | 0.068643617 | -11.14577734 | 7.51E-29 | 1.99E-27 |
| ENSG00000148444 | COMMD3 | 141.4743178 | -0.437295383 | 0.154100243 | -2.837733255 | 0.004543514 | 0.014788337 |
| ENSG00000148450 | MSRB2 | 448.6690827 | -0.640727131 | 0.091480985 | -7.003937802 | 2.49E-12 | 2.46E-11 |
| ENSG00000148459 | PDSS1 | 244.9655876 | -0.617479778 | 0.128507525 | -4.805008726 | 1.55E-06 | 8.75E-06 |
| ENSG00000148468 | FAM171A1 | 10.65743264 | -0.848041902 | 0.556405266 | -1.524144278 | 0.127472674 | 0.257368697 |
| ENSG00000148481 | MINDY3 | 494.5494494 | -0.537032078 | 0.093218986 | -5.760973173 | 8.36E-09 | 6.06E-08 |
| ENSG00000148484 | RSU1 | 1236.706581 | -0.976972725 | 0.062547073 | -15.61979937 | 5.34E-55 | 3.35E-53 |
| ENSG00000148488 | ST8SIA6 | 737.4341026 | -2.656212045 | 0.088016581 | -30.17854144 | 4.53E-200 | 2.73E-197 |
| ENSG00000148498 | PARD3 | 2163.523864 | -0.927322058 | 0.055152161 | -16.8138843 | 1.93E-63 | 1.50E-61 |
| ENSG00000148516 | ZEB1 | 4.718091338 | -1.236466671 | 0.839491396 | -1.47287593 | 0.140784493 | 0.277766946 |
| ENSG00000148541 | FAM13C | 6.046157953 | 1.979838429 | 0.817824237 | 2.420860547 | 0.015483816 | 0.043759854 |
| ENSG00000148572 | NRBF2 | 437.1855388 | -0.401306036 | 0.091041326 | -4.40795465 | 1.04E-05 | 5.38E-05 |
| ENSG00000148606 | POLR3A | 1402.102808 | -0.592395612 | 0.059847606 | -9.898401122 | 4.23E-23 | 8.44E-22 |
| ENSG00000148634 | HERC4 | 1201.511441 | -0.640899135 | 0.065419523 | -9.79675648 | 1.16E-22 | 2.27E-21 |
| ENSG00000148655 | LRMDA | 71.67629373 | 0.081255173 | 0.211197308 | 0.384735838 | 0.70043314 | 0.83200432 |
| ENSG00000148660 | CAMK2G | 1695.904003 | -0.383489389 | 0.063087284 | -6.0787113 | 1.21E-09 | 9.57E-09 |
| ENSG00000148671 | ADIRF | 118.9900613 | -1.262075563 | 0.180533501 | -6.990810892 | 2.73E-12 | 2.70E-11 |

| | | | | | | | |
|---|---|---|---|---|---|---|---|
| ENSG00000148672 | GLUD1 | 4235.672392 | -0.698670528 | 0.046496849 | -15.02619102 | 4.95E-51 | 2.82E-49 |
| ENSG00000148677 | ANKRD1 | 493.8435224 | 0.79224665 | 0.104477117 | 7.582968115 | 3.38E-14 | 3.82E-13 |
| ENSG00000148680 | HTR7 | 39.90804332 | 3.534974999 | 0.412055124 | 8.578888577 | 9.58E-18 | 1.39E-16 |
| ENSG00000148688 | RPP30 | 706.3148599 | -0.356959938 | 0.085888432 | -4.156088668 | 3.24E-05 | 0.000155956 |
| ENSG00000148690 | FRA10AC1 | 444.7115211 | -0.454265247 | 0.095883546 | -4.737676729 | 2.16E-06 | 1.20E-05 |
| ENSG00000148700 | ADD3 | 537.1391431 | -0.806117622 | 0.085507915 | -9.427403594 | 4.20E-21 | 7.49E-20 |
| ENSG00000148702 | HABP2 | 33.56879082 | -2.239588788 | 0.347265824 | -6.449205853 | 1.12E-10 | 9.67E-10 |
| ENSG00000148719 | DNAJB12 | 1753.366872 | -0.502502651 | 0.0609823 | -8.240139408 | 1.72E-16 | 2.31E-15 |
| ENSG00000148730 | EIF4EBP2 | 6002.012072 | -0.422496902 | 0.053058464 | -7.962855847 | 1.68E-15 | 2.09E-14 |
| ENSG00000148735 | PLEKHS1 | 667.6675181 | -3.092269717 | 0.102744083 | -30.09681562 | 5.33E-199 | 3.14E-196 |
| ENSG00000148737 | TCF7L2 | 182.6549795 | -11.06020806 | 1.183739598 | -9.343446889 | 9.32E-21 | 1.62E-19 |
| ENSG00000148773 | MKI67 | 6304.595079 | -0.868287652 | 0.055567382 | -15.62585132 | 4.85E-55 | 3.06E-53 |
| ENSG00000148814 | LRRC27 | 832.7689033 | 0.126346136 | 0.070211536 | 1.799506792 | 0.071938551 | 0.162396216 |
| ENSG00000148824 | MTG1 | 256.7746444 | -0.537044919 | 0.124351302 | -4.318771978 | 1.57E-05 | 7.89E-05 |
| ENSG00000148826 | NKX6-2 | 1.740829641 | -1.437443056 | 1.380549177 | -1.041211048 | 0.297777609 | NA |
| ENSG00000148834 | GSTO1 | 695.809212 | -0.16621372 | 0.07999591 | -2.077777726 | 0.037729836 | 0.094509985 |
| ENSG00000148835 | TAF5 | 217.9773316 | -0.112925381 | 0.123490143 | -0.914448543 | 0.360481204 | 0.553036645 |
| ENSG00000148840 | PPRC1 | 1676.944234 | 0.091725918 | 0.059354063 | 1.545402521 | 0.122248931 | 0.248960929 |
| ENSG00000148841 | ITPRIP | 70.32904072 | -0.75851114 | 0.234519021 | -3.234326743 | 0.001219299 | 0.004494731 |
| ENSG00000148842 | CNNM2 | 342.7175735 | 0.263160978 | 0.102345885 | 2.571290255 | 0.010132037 | 0.030083992 |
| ENSG00000148843 | PDCD11 | 2018.577801 | 0.136219131 | 0.053076027 | 2.566490735 | 0.010273337 | 0.030436569 |
| ENSG00000148908 | RGS10 | 265.5920696 | -0.174072804 | 0.11456595 | -1.519411341 | 0.128658989 | 0.259148056 |
| ENSG00000148925 | BTBD10 | 982.2969161 | -0.173986404 | 0.066692249 | -2.608794984 | 0.009086166 | 0.027357481 |
| ENSG00000148926 | ADM | 68.61501433 | -0.070767883 | 0.214682419 | -0.329639863 | 0.741672099 | 0.857515143 |
| ENSG00000148935 | GAS2 | 17.67211766 | -0.00518334 | 0.454786533 | -0.011397303 | 0.990906465 | 0.995176993 |
| ENSG00000148942 | SLC5A12 | 113.3953637 | 0.592024724 | 0.175703909 | 3.369445382 | 0.000753196 | 0.002885491 |
| ENSG00000148943 | LIN7C | 679.2829944 | -0.183298468 | 0.086968914 | -2.107632006 | 0.035062833 | 0.088759809 |
| ENSG00000148948 | LRRC4C | 85.18012381 | 2.927668051 | 0.25242753 | 11.5980537 | 4.22E-31 | 1.23E-29 |
| ENSG00000148950 | IMMP1L | 246.3482543 | -0.111507703 | 0.116782469 | -0.954832553 | 0.339662382 | 0.530711987 |
| ENSG00000148985 | PGAP2 | 591.388788 | -0.020468963 | 0.089336989 | -0.229120802 | 0.818775026 | 0.903986346 |
| ENSG00000149016 | TUT1 | 268.7317792 | -0.222084401 | 0.115782318 | -1.918120186 | 0.055095773 | 0.130177673 |
| ENSG00000149043 | SYT8 | 8.841815929 | 1.80911358 | 0.655459461 | 2.760069368 | 0.005778909 | 0.018305464 |
| ENSG00000149084 | HSD17B12 | 484.4056179 | -0.426220619 | 0.090770282 | -4.695596487 | 2.66E-06 | 1.47E-05 |

| | | | | | | | |
|---|---|---|---|---|---|---|---|
| ENSG00000149089 | APIP | 611.8877465 | 0.159273795 | 0.08828379 | 1.804111435 | 0.071213839 | 0.160918585 |
| ENSG00000149090 | PAMR1 | 5.829482069 | 0.894100736 | 0.75313965 | 1.187164606 | 0.235162699 | 0.410175814 |
| ENSG00000149091 | DGKZ | 2188.53581 | -0.017424704 | 0.054212747 | -0.321413402 | 0.747897128 | 0.860419264 |
| ENSG00000149100 | EIF3M | 3484.503295 | 0.239890839 | 0.057261148 | 4.189417233 | 2.80E-05 | 0.000136041 |
| ENSG00000149115 | TNKS1BP1 | 3384.910515 | 0.321924984 | 0.055153572 | 5.836883664 | 5.32E-09 | 3.95E-08 |
| ENSG00000149131 | SERPING1 | 2.877796325 | -0.129503575 | 1.033921924 | -0.125254694 | 0.900321918 | 0.949711905 |
| ENSG00000149136 | SSRP1 | 6560.738108 | -0.195608375 | 0.04317146 | -4.53096501 | 5.87E-06 | 3.12E-05 |
| ENSG00000149150 | SLC43A1 | 138.0685755 | -0.269040319 | 0.157570766 | -1.707425346 | 0.087743023 | 0.190482156 |
| ENSG00000149177 | PTPRJ | 2251.023509 | -0.611487184 | 0.064900673 | -9.421892836 | 4.43E-21 | 7.88E-20 |
| ENSG00000149179 | CSTPP1 | 1141.074003 | 0.142916878 | 0.069145153 | 2.066910996 | 0.038742541 | 0.096593185 |
| ENSG00000149182 | ARFGAP2 | 5191.172162 | -0.098716896 | 0.04744669 | -2.080585511 | 0.037471861 | 0.093966386 |
| ENSG00000149187 | CELF1 | 5330.07158 | -0.187383617 | 0.048840748 | -3.836624665 | 0.000124737 | 0.000547334 |
| ENSG00000149196 | HIKESHI | 677.4941514 | -0.153755581 | 0.076435723 | -2.011567045 | 0.044265596 | 0.107980942 |
| ENSG00000149201 | CCDC81 | 14.68546308 | -0.330636851 | 0.463863378 | -0.712789296 | 0.47597615 | 0.661069981 |
| ENSG00000149212 | SESN3 | 1006.967253 | -0.467586777 | 0.069807648 | -6.698217076 | 2.11E-11 | 1.92E-10 |
| ENSG00000149218 | ENDOD1 | 370.1394454 | 0.321945856 | 0.104878845 | 3.069692996 | 0.002142789 | 0.007480685 |
| ENSG00000149231 | CCDC82 | 82.18624154 | 0.459207477 | 0.200968875 | 2.284968144 | 0.022314696 | 0.060188021 |
| ENSG00000149243 | KLHL35 | 168.313279 | 0.681382636 | 0.143588372 | 4.745388705 | 2.08E-06 | 1.16E-05 |
| ENSG00000149256 | TENM4 | 205.7922494 | 0.784097766 | 0.12940308 | 6.059343915 | 1.37E-09 | 1.07E-08 |
| ENSG00000149257 | SERPINH1 | 4126.365718 | -0.326176498 | 0.059534314 | -5.478798313 | 4.28E-08 | 2.88E-07 |
| ENSG00000149260 | CAPN5 | 635.4976888 | 0.993788116 | 0.089952707 | 11.04789558 | 2.24E-28 | 5.84E-27 |
| ENSG00000149262 | INTS4 | 878.3774025 | -0.209477611 | 0.071900216 | -2.913448989 | 0.003574602 | 0.011912654 |
| ENSG00000149269 | PAK1 | 1349.764986 | 0.533986067 | 0.062820701 | 8.500160858 | 1.89E-17 | 2.69E-16 |
| ENSG00000149273 | RPS3 | 24414.78205 | 0.187086078 | 0.051002682 | 3.668161597 | 0.000244301 | 0.001019142 |
| ENSG00000149289 | ZC3H12C | 8.838033142 | 1.62092607 | 0.659597522 | 2.45744718 | 0.013992839 | 0.039994483 |
| ENSG00000149292 | TTC12 | 406.9769376 | -0.092903661 | 0.093046721 | -0.998462495 | 0.318055142 | 0.507207853 |
| ENSG00000149295 | DRD2 | 3.392298087 | 0.243329803 | 0.968710113 | 0.251189494 | 0.801667607 | 0.894424147 |
| ENSG00000149308 | NPAT | 455.5043006 | -0.096859297 | 0.099208698 | -0.976318598 | 0.328906606 | 0.519350052 |
| ENSG00000149311 | ATM | 691.2320258 | -0.377086446 | 0.083625475 | -4.509229352 | 6.51E-06 | 3.44E-05 |
| ENSG00000149313 | AASDHPPT | 1270.761432 | 0.05546843 | 0.066255749 | 0.837186675 | 0.402487646 | 0.5952209 |
| ENSG00000149328 | GLB1L2 | 479.0695812 | 1.124490075 | 0.092044272 | 12.21683921 | 2.53E-34 | 8.35E-33 |
| ENSG00000149346 | SLX4IP | 565.9290053 | -0.483524668 | 0.081766339 | -5.913492956 | 3.35E-09 | 2.54E-08 |
| ENSG00000149357 | LAMTOR1 | 3418.104585 | 0.020426199 | 0.054355671 | 0.375787808 | 0.707074653 | 0.836461297 |

| | | | | | | | |
|---|---|---|---|---|---|---|---|
| ENSG00000149380 | P4HA3 | 19.61224746 | 0.485551626 | 0.412547252 | 1.176960028 | 0.239211461 | 0.414824045 |
| ENSG00000149418 | ST14 | 3417.22267 | 0.329213171 | 0.066368535 | 4.960380243 | 7.04E-07 | 4.16E-06 |
| ENSG00000149428 | HYOU1 | 12356.28612 | 0.3774202 | 0.043634523 | 8.649577828 | 5.17E-18 | 7.62E-17 |
| ENSG00000149474 | KAT14 | 1087.555708 | -0.155244208 | 0.06371609 | -2.436499274 | 0.014830199 | 0.042068106 |
| ENSG00000149476 | TKFC | 1993.97657 | -0.217368507 | 0.058425336 | -3.720449411 | 0.000198869 | 0.000844198 |
| ENSG00000149480 | MTA2 | 7533.321037 | -0.175016399 | 0.041838017 | -4.183190613 | 2.87E-05 | 0.000139615 |
| ENSG00000149483 | TMEM138 | 591.7511715 | -0.138153404 | 0.082838969 | -1.66773447 | 0.095368449 | 0.204005243 |
| ENSG00000149485 | FADS1 | 6405.974953 | -0.53889631 | 0.045557231 | -11.82899628 | 2.76E-32 | 8.43E-31 |
| ENSG00000149489 | ROM1 | 94.04494352 | 0.478210652 | 0.190700564 | 2.507652002 | 0.01215363 | 0.035340888 |
| ENSG00000149499 | EML3 | 2765.976945 | -0.287780757 | 0.057164952 | -5.034216747 | 4.80E-07 | 2.90E-06 |
| ENSG00000149503 | INCENP | 2014.903493 | -0.300172774 | 0.054747377 | -5.482870468 | 4.18E-08 | 2.82E-07 |
| ENSG00000149527 | PLCH2 | 25.69467999 | -0.338459055 | 0.353317044 | -0.95794715 | 0.338089409 | 0.52937814 |
| ENSG00000149531 | FRG1BP | 559.869056 | -0.263160363 | 0.084879999 | -3.100381315 | 0.001932717 | 0.006816808 |
| ENSG00000149532 | CPSF7 | 4033.770263 | -0.099234901 | 0.047986636 | -2.067969525 | 0.038642889 | 0.096392011 |
| ENSG00000149541 | B3GAT3 | 1149.723061 | 0.601505877 | 0.067152438 | 8.957320058 | 3.33E-19 | 5.32E-18 |
| ENSG00000149547 | EI24 | 3369.3655 | 0.00930358 | 0.04882994 | 0.190530241 | 0.848893648 | 0.920956268 |
| ENSG00000149548 | CCDC15 | 86.43774232 | -1.038350924 | 0.202164558 | -5.136166977 | 2.80E-07 | 1.74E-06 |
| ENSG00000149554 | CHEK1 | 649.790526 | -0.326352759 | 0.079775321 | -4.090898734 | 4.30E-05 | 0.000202703 |
| ENSG00000149557 | FEZ1 | 224.2432852 | -0.244267577 | 0.121667863 | -2.007658977 | 0.044679543 | 0.108794046 |
| ENSG00000149564 | ESAM | 27.94097028 | -0.384687984 | 0.352951906 | -1.089916153 | 0.27575008 | 0.458019466 |
| ENSG00000149573 | MPZL2 | 338.238033 | 1.393234637 | 0.108269945 | 12.86815685 | 6.80E-38 | 2.56E-36 |
| ENSG00000149577 | SIDT2 | 1506.302137 | 0.033564732 | 0.067083232 | 0.500344588 | 0.616832463 | 0.772960336 |
| ENSG00000149582 | TMEM25 | 788.4811576 | 0.365360532 | 0.076978414 | 4.746272512 | 2.07E-06 | 1.15E-05 |
| ENSG00000149591 | TAGLN | 730.9645934 | -0.626669567 | 0.101096879 | -6.198703382 | 5.69E-10 | 4.64E-09 |
| ENSG00000149599 | DUSP15 | 9.263712724 | 0.151265467 | 0.584392719 | 0.258842149 | 0.795757035 | 0.890871567 |
| ENSG00000149600 | COMMD7 | 779.3067124 | -1.141676391 | 0.078151262 | -14.60854701 | 2.48E-48 | 1.29E-46 |
| ENSG00000149609 | C20orf144 | 11.6305122 | -0.520619743 | 0.54247001 | -0.959720783 | 0.337195761 | 0.528401581 |
| ENSG00000149634 | SPATA25 | 27.4117373 | 0.2050976 | 0.335257532 | 0.611761349 | 0.540695666 | 0.714868747 |
| ENSG00000149636 | DSN1 | 696.8316643 | -0.954892861 | 0.084250639 | -11.33395399 | 8.91E-30 | 2.46E-28 |
| ENSG00000149639 | SOGA1 | 4264.238317 | -0.478558208 | 0.115485049 | -4.143897519 | 3.41E-05 | 0.000163693 |
| ENSG00000149646 | CNBD2 | 46.74857502 | -0.88415237 | 0.259917809 | -3.401661363 | 0.000669776 | 0.002591841 |
| ENSG00000149654 | CDH22 | 4.223488314 | 0.978264192 | 0.877563089 | 1.114750842 | 0.264957207 | 0.445969501 |
| ENSG00000149657 | LSM14B | 2449.796748 | -0.399737047 | 0.054165264 | -7.37995192 | 1.58E-13 | 1.71E-12 |

| | | | | | | | |
|---|---|---|---|---|---|---|---|
| ENSG00000149658 | YTHDF1 | 3326.488488 | -0.118057034 | 0.048877727 | -2.415354418 | 0.015719905 | 0.044317884 |
| ENSG00000149679 | CABLES2 | 1906.360534 | 0.068081648 | 0.056177192 | 1.21190906 | 0.225547194 | 0.397545342 |
| ENSG00000149716 | LTO1 | 937.6341685 | -0.202260057 | 0.071804146 | -2.816829778 | 0.004850022 | 0.015683608 |
| ENSG00000149743 | TRPT1 | 653.5454474 | -0.175071091 | 0.079130924 | -2.212423188 | 0.026937438 | 0.070832741 |
| ENSG00000149761 | NUDT22 | 840.5959613 | -0.22957638 | 0.073397128 | -3.127865988 | 0.001760804 | 0.006260313 |
| ENSG00000149781 | FERMT3 | 40.39517851 | 0.195966527 | 0.284437174 | 0.688962431 | 0.49084691 | 0.673915141 |
| ENSG00000149782 | PLCB3 | 2173.580244 | 0.136309433 | 0.059345596 | 2.296875302 | 0.021625884 | 0.058529706 |
| ENSG00000149792 | MRPL49 | 3388.12535 | -0.14245324 | 0.047840352 | -2.977679622 | 0.002904394 | 0.009870637 |
| ENSG00000149806 | FAU | 2402.407224 | 0.117836423 | 0.054151347 | 2.176057093 | 0.029550992 | 0.076650808 |
| ENSG00000149809 | TM7SF2 | 2425.467225 | -0.364262807 | 0.059905546 | -6.080619058 | 1.20E-09 | 9.46E-09 |
| ENSG00000149823 | VPS51 | 2575.537222 | 0.126490644 | 0.057496767 | 2.199961002 | 0.027809662 | 0.072775912 |
| ENSG00000149922 | TBX6 | 83.43151902 | -0.97827547 | 0.201172166 | -4.862876862 | 1.16E-06 | 6.65E-06 |
| ENSG00000149923 | PPP4C | 3482.334662 | -0.188807752 | 0.056839409 | -3.32177542 | 0.000894467 | 0.003379849 |
| ENSG00000149925 | ALDOA | 498.547883 | 0.170511944 | 0.105082692 | 1.622645375 | 0.104665235 | 0.219994547 |
| ENSG00000149926 | TLCD3B | 55.56933154 | 0.15133299 | 0.240803917 | 0.628449039 | 0.52970982 | 0.705336852 |
| ENSG00000149927 | DOC2A | 408.8055541 | 0.354034422 | 0.102029503 | 3.469922051 | 0.000520609 | 0.00205273 |
| ENSG00000149929 | HIRIP3 | 511.1547208 | -0.332121119 | 0.090296441 | -3.678119713 | 0.00023496 | 0.00098447 |
| ENSG00000149930 | TAOK2 | 4087.272951 | -0.259969232 | 0.055770941 | -4.661374303 | 3.14E-06 | 1.72E-05 |
| ENSG00000149932 | TMEM219 | 1424.067869 | 0.100509653 | 0.075478706 | 1.331629262 | 0.182982044 | 0.338981515 |
| ENSG00000149972 | CNTN5 | 8.634177071 | 0.313488307 | 0.621143884 | 0.504695152 | 0.613772966 | 0.770659652 |
| ENSG00000150051 | MKX | 34.67237687 | 2.978098339 | 0.404751642 | 7.357841271 | 1.87E-13 | 2.00E-12 |
| ENSG00000150054 | MPP7 | 894.941405 | -0.538650532 | 0.072842206 | -7.394758642 | 1.42E-13 | 1.53E-12 |
| ENSG00000150093 | ITGB1 | 3863.155417 | -0.318084119 | 0.050127242 | -6.345534034 | 2.22E-10 | 1.87E-09 |
| ENSG00000150281 | CTF1 | 217.9190775 | -0.584927266 | 0.134783644 | -4.339749617 | 1.43E-05 | 7.20E-05 |
| ENSG00000150316 | CWC15 | 782.8214854 | -0.218861808 | 0.071998384 | -3.03981554 | 0.002367231 | 0.008206782 |
| ENSG00000150347 | ARID5B | 1201.187424 | -0.364422709 | 0.070950736 | -5.136278085 | 2.80E-07 | 1.74E-06 |
| ENSG00000150401 | DCUN1D2 | 283.4195093 | -0.144833153 | 0.110230095 | -1.313916602 | 0.188874271 | 0.347567446 |
| ENSG00000150403 | TMCO3 | 1028.695297 | -0.199078198 | 0.066562347 | -2.990853025 | 0.002781994 | 0.009498278 |
| ENSG00000150433 | TMEM218 | 279.7739615 | -0.422732908 | 0.123101585 | -3.434016786 | 0.000594707 | 0.002320563 |
| ENSG00000150455 | TIRAP | 275.9024504 | -0.439064424 | 0.114415933 | -3.837441262 | 0.000124323 | 0.000545622 |
| ENSG00000150456 | EEF1AKMT1 | 370.7888648 | -0.207191854 | 0.098333413 | -2.107034102 | 0.035114624 | 0.088861497 |
| ENSG00000150457 | LATS2 | 749.8934803 | -0.241441325 | 0.078316833 | -3.08287907 | 0.002050084 | 0.007187672 |
| ENSG00000150459 | SAP18 | 4563.798282 | 0.001725304 | 0.044940678 | 0.038390703 | 0.969376173 | 0.986272649 |

| | | | | | | | |
|---|---|---|---|---|---|---|---|
| ENSG00000150471 | ADGRL3 | 128.5662288 | -0.728446151 | 0.158321494 | -4.601056559 | 4.20E-06 | 2.26E-05 |
| ENSG00000150477 | KIAA1328 | 164.7324025 | 0.21227543 | 0.140574863 | 1.510052546 | 0.131030017 | 0.263023152 |
| ENSG00000150510 | FAM124A | 18.38387609 | 0.198120473 | 0.422124133 | 0.469341735 | 0.638825388 | 0.787290808 |
| ENSG00000150527 | MIA2 | 1157.751488 | -0.459687179 | 0.064675098 | -7.107637925 | 1.18E-12 | 1.20E-11 |
| ENSG00000150540 | HNMT | 26.66767172 | 1.961778932 | 0.383563994 | 5.114606592 | 3.14E-07 | 1.94E-06 |
| ENSG00000150551 | LYPD1 | 895.3897016 | 1.164306783 | 0.08186518 | 14.22224664 | 6.67E-46 | 3.29E-44 |
| ENSG00000150556 | LYPD6B | 484.0105568 | 1.16226512 | 0.095120062 | 12.21892733 | 2.46E-34 | 8.18E-33 |
| ENSG00000150593 | PDCD4 | 2974.006646 | -0.17960768 | 0.051961037 | -3.456583817 | 0.000547069 | 0.002150774 |
| ENSG00000150627 | WDR17 | 82.25800473 | 0.319453315 | 0.201840135 | 1.582704625 | 0.113488804 | 0.234860907 |
| ENSG00000150630 | VEGFC | 62.86676623 | 0.313034646 | 0.227270654 | 1.377365008 | 0.168399426 | 0.318506788 |
| ENSG00000150636 | CCDC102B | 19.13936749 | 0.810365555 | 0.418734768 | 1.93527172 | 0.052956961 | 0.125928546 |
| ENSG00000150637 | CD226 | 3.293361747 | 0.260843053 | 1.003672249 | 0.259888677 | 0.794949645 | 0.890293445 |
| ENSG00000150667 | FSIP1 | 92.69754413 | -0.346453698 | 0.200373666 | -1.729038077 | 0.08380228 | 0.183612637 |
| ENSG00000150672 | DLG2 | 42.36510601 | -0.082072374 | 0.273003701 | -0.30062733 | 0.763698689 | 0.870479014 |
| ENSG00000150676 | CCDC83 | 178.4049061 | 0.795869673 | 0.137484536 | 5.788794108 | 7.09E-09 | 5.18E-08 |
| ENSG00000150687 | PRSS23 | 443.2357406 | 0.866601663 | 0.098059701 | 8.837490337 | 9.79E-19 | 1.52E-17 |
| ENSG00000150712 | MTMR12 | 2658.593427 | -0.278754579 | 0.05188251 | -5.372804436 | 7.75E-08 | 5.07E-07 |
| ENSG00000150722 | PPP1R1C | 8.265799188 | 0.214245101 | 0.602174774 | 0.355785579 | 0.72200115 | 0.845596895 |
| ENSG00000150753 | CCT5 | 15786.88866 | -0.025106865 | 0.044924667 | -0.558865924 | 0.576253227 | 0.742571005 |
| ENSG00000150756 | ATPSCKMT | 741.8636908 | -0.111872528 | 0.075548832 | -1.48079758 | 0.138660522 | 0.274591882 |
| ENSG00000150760 | DOCK1 | 1183.004958 | 0.127641566 | 0.067578276 | 1.888795825 | 0.058919192 | 0.137386285 |
| ENSG00000150764 | DIXDC1 | 314.2098594 | -0.120137429 | 0.105860211 | -1.134868601 | 0.256430377 | 0.435903139 |
| ENSG00000150768 | DLAT | 1501.227699 | -0.321115863 | 0.060877878 | -5.27475451 | 1.33E-07 | 8.53E-07 |
| ENSG00000150773 | PIH1D2 | 11.8613803 | -0.026514493 | 0.510325803 | -0.051956012 | 0.958563744 | 0.981140547 |
| ENSG00000150776 | NKAPD1 | 978.1445837 | 0.04825924 | 0.068209482 | 0.707515116 | 0.479246422 | 0.663653377 |
| ENSG00000150779 | TIMM8B | 753.0663867 | 0.06278269 | 0.079904914 | 0.785717504 | 0.432033006 | 0.62208207 |
| ENSG00000150782 | IL18 | 1.718140277 | 3.028410241 | 1.611424033 | 1.879337889 | 0.060198371 | NA |
| ENSG00000150787 | PTS | 229.9437017 | 0.102708177 | 0.129668988 | 0.792079731 | 0.428314188 | 0.618922284 |
| ENSG00000150867 | PIP4K2A | 1159.993154 | -0.281248709 | 0.064667126 | -4.349175949 | 1.37E-05 | 6.92E-05 |
| ENSG00000150873 | C2orf50 | 12.9105386 | 0.265576459 | 0.519858417 | 0.510863055 | 0.609446952 | 0.767622747 |
| ENSG00000150893 | FREM2 | 2193.565502 | -0.284844551 | 0.07153453 | -3.981916848 | 6.84E-05 | 0.000313393 |
| ENSG00000150907 | FOXO1 | 171.3949301 | 0.045177068 | 0.143784767 | 0.314199265 | 0.753369701 | 0.864071843 |
| ENSG00000150938 | CRIM1 | 1694.086007 | 0.78286769 | 0.064877058 | 12.06694193 | 1.58E-33 | 5.02E-32 |

| | | | | | | | |
|---|---|---|---|---|---|---|---|
| ENSG00000150961 | SEC24D | 1276.779616 | 0.074667539 | 0.061680997 | 1.210543651 | 0.226070351 | 0.398222672 |
| ENSG00000150967 | ABCB9 | 690.2008249 | 0.195382357 | 0.08037888 | 2.430767361 | 0.015066885 | 0.042676176 |
| ENSG00000150977 | RILPL2 | 753.2077905 | 0.340744768 | 0.079763592 | 4.271933615 | 1.94E-05 | 9.63E-05 |
| ENSG00000150990 | DHX37 | 1386.057063 | 0.216874481 | 0.06459411 | 3.357496214 | 0.000786518 | 0.003001618 |
| ENSG00000150991 | UBC | 19326.75549 | -0.135520207 | 0.038986243 | -3.476103296 | 0.000508756 | 0.002009793 |
| ENSG00000150995 | ITPR1 | 2028.052522 | -2.014881258 | 0.067131481 | -30.01395496 | 6.45E-198 | 3.70E-195 |
| ENSG00000151006 | PRSS53 | 73.96886003 | -0.655616404 | 0.242055745 | -2.708534777 | 0.006758103 | 0.021026839 |
| ENSG00000151012 | SLC7A11 | 617.0458514 | -0.563451792 | 0.188823027 | -2.984020544 | 0.002844877 | 0.009694696 |
| ENSG00000151014 | NOCT | 230.7828944 | 0.365323386 | 0.12162718 | 3.003632791 | 0.002667771 | 0.009138221 |
| ENSG00000151023 | ENKUR | 5.36477538 | -0.566516079 | 0.756130673 | -0.749230389 | 0.453718356 | 0.641379041 |
| ENSG00000151025 | GPR158 | 94.53519662 | 0.576784464 | 0.183351235 | 3.145789912 | 0.001656389 | 0.005923724 |
| ENSG00000151062 | CACNA2D4 | 3.318545013 | -0.178366867 | 1.026194901 | -0.173813831 | 0.862011774 | 0.928215087 |
| ENSG00000151065 | DCP1B | 306.3696655 | 0.225342802 | 0.108827437 | 2.070643294 | 0.038392143 | 0.095902352 |
| ENSG00000151067 | CACNA1C | 3.014561813 | 1.710388873 | 1.089123507 | 1.570426918 | 0.116315824 | 0.239661995 |
| ENSG00000151090 | THRB | 547.2097927 | 0.355293233 | 0.087149118 | 4.076842552 | 4.57E-05 | 0.000214863 |
| ENSG00000151092 | NGLY1 | 1010.123138 | 0.153000008 | 0.068904721 | 2.220457539 | 0.026387725 | 0.069546692 |
| ENSG00000151093 | OXSM | 392.2924287 | -0.215801097 | 0.099348749 | -2.172157176 | 0.02984381 | 0.077322904 |
| ENSG00000151116 | UEVLD | 1067.410692 | 0.191822763 | 0.070180791 | 2.73326591 | 0.006270968 | 0.019706303 |
| ENSG00000151117 | TMEM86A | 328.6019425 | 0.068068248 | 0.108246458 | 0.628826558 | 0.529462611 | 0.705157753 |
| ENSG00000151131 | NOPCHAP1 | 293.3359525 | 0.475956196 | 0.11479716 | 4.146062473 | 3.38E-05 | 0.000162294 |
| ENSG00000151135 | TMEM263 | 743.1461916 | -0.434071406 | 0.076555237 | -5.670041985 | 1.43E-08 | 1.01E-07 |
| ENSG00000151136 | ABTB3 | 14.82785437 | 0.255043898 | 0.458623159 | 0.556107761 | 0.57813719 | 0.743870499 |
| ENSG00000151148 | UBE3B | 2217.726512 | -0.126708904 | 0.054569356 | -2.321979087 | 0.020234062 | 0.055212249 |
| ENSG00000151150 | ANK3 | 2568.432736 | -1.133283334 | 0.063812646 | -17.75954162 | 1.45E-70 | 1.36E-68 |
| ENSG00000151151 | IPMK | 262.0561237 | -0.529960576 | 0.129808063 | -4.082647593 | 4.45E-05 | 0.000209736 |
| ENSG00000151164 | RAD9B | 10.85426261 | -1.536643119 | 0.555099151 | -2.768231795 | 0.005636135 | 0.017897695 |
| ENSG00000151176 | PLBD2 | 1740.649937 | 0.442793317 | 0.05683324 | 7.791097491 | 6.64E-15 | 7.92E-14 |
| ENSG00000151208 | DLG5 | 4974.786592 | -0.78844442 | 0.04907797 | -16.06513936 | 4.48E-58 | 3.05E-56 |
| ENSG00000151229 | SLC2A13 | 124.7422142 | 0.321907836 | 0.179544196 | 1.792916969 | 0.072986202 | 0.164267221 |
| ENSG00000151233 | GXYLT1 | 1142.963192 | -0.047055474 | 0.065396182 | -0.719544659 | 0.471805396 | 0.657466637 |
| ENSG00000151239 | TWF1 | 1589.09078 | 0.151537782 | 0.068939906 | 2.198114135 | 0.027940973 | 0.073086195 |
| ENSG00000151240 | DIP2C | 2327.512237 | -0.880889403 | 0.05539749 | -15.9012512 | 6.21E-57 | 4.11E-55 |
| ENSG00000151247 | EIF4E | 619.5935822 | 0.0923141 | 0.087897974 | 1.050241489 | 0.293607099 | 0.479179485 |

| | | | | | | | |
|---|---|---|---|---|---|---|---|
| ENSG00000151276 | MAGI1 | 1556.688125 | -0.073127723 | 0.066061816 | -1.106959023 | 0.268311645 | 0.449838104 |
| ENSG00000151287 | TEX30 | 203.8927265 | -0.601706253 | 0.139923296 | -4.300257855 | 1.71E-05 | 8.55E-05 |
| ENSG00000151292 | CSNK1G3 | 1227.098 | -0.144743473 | 0.0670698 | -2.158102062 | 0.030919898 | 0.079750669 |
| ENSG00000151304 | SRFBP1 | 344.972665 | 0.096071187 | 0.10798758 | 0.889650334 | 0.37365367 | 0.566456198 |
| ENSG00000151320 | AKAP6 | 148.4480685 | -2.253313283 | 0.19240219 | -11.7114742 | 1.11E-31 | 3.32E-30 |
| ENSG00000151322 | NPAS3 | 445.1082801 | 0.236499778 | 0.091359426 | 2.588674088 | 0.009634623 | 0.028789665 |
| ENSG00000151327 | FAM177A1 | 1829.177159 | -0.540855356 | 0.058828435 | -9.193774273 | 3.79E-20 | 6.37E-19 |
| ENSG00000151332 | MBIP | 574.6608634 | -0.486897979 | 0.084678695 | -5.749946674 | 8.93E-09 | 6.45E-08 |
| ENSG00000151338 | MIPOL1 | 170.847732 | -0.089970912 | 0.143509022 | -0.626935582 | 0.530701462 | 0.706329284 |
| ENSG00000151348 | EXT2 | 1705.928251 | 0.22599336 | 0.057657864 | 3.919558346 | 8.87E-05 | 0.000399262 |
| ENSG00000151353 | TMEM18 | 1641.911382 | 0.155397068 | 0.057215399 | 2.716000759 | 0.006607577 | 0.020622831 |
| ENSG00000151364 | KCTD14 | 6.520763123 | 0.264592751 | 0.690420313 | 0.383234309 | 0.701546042 | 0.832995145 |
| ENSG00000151366 | NDUFC2 | 542.6312232 | 0.520677203 | 0.085835458 | 6.065991991 | 1.31E-09 | 1.03E-08 |
| ENSG00000151388 | ADAMTS12 | 2.046285768 | 3.291239487 | 1.547765251 | 2.126446168 | 0.033466127 | 0.085283567 |
| ENSG00000151413 | NUBPL | 358.8619199 | -0.525639074 | 0.100613774 | -5.224325199 | 1.75E-07 | 1.11E-06 |
| ENSG00000151414 | NEK7 | 1637.92665 | 0.20677773 | 0.057729866 | 3.581815546 | 0.000341215 | 0.001388341 |
| ENSG00000151422 | FER | 397.9467906 | -0.523639062 | 0.094582504 | -5.53632055 | 3.09E-08 | 2.11E-07 |
| ENSG00000151445 | VIPAS39 | 1438.58682 | -0.602685523 | 0.058707439 | -10.26591403 | 1.00E-24 | 2.20E-23 |
| ENSG00000151458 | ANKRD50 | 3003.092275 | 0.635351475 | 0.059216604 | 10.7292791 | 7.42E-27 | 1.83E-25 |
| ENSG00000151461 | UPF2 | 1938.351463 | -0.52620835 | 0.059252707 | -8.880747867 | 6.64E-19 | 1.04E-17 |
| ENSG00000151465 | CDC123 | 1659.8874 | -0.557992626 | 0.057299916 | -9.738105457 | 2.07E-22 | 3.98E-21 |
| ENSG00000151466 | SCLT1 | 342.1811137 | -0.017643799 | 0.102725793 | -0.171756269 | 0.863629147 | 0.929172842 |
| ENSG00000151468 | CCDC3 | 2.951531122 | -1.532370375 | 1.067202503 | -1.435875918 | 0.151037652 | 0.293155406 |
| ENSG00000151470 | C4orf33 | 50.56851338 | -0.704308669 | 0.251856098 | -2.79647257 | 0.00516638 | 0.016582727 |
| ENSG00000151474 | FRMD4A | 146.3586036 | -0.899148682 | 0.153289754 | -5.865680252 | 4.47E-09 | 3.35E-08 |
| ENSG00000151491 | EPS8 | 1949.637012 | -0.235624541 | 0.054328492 | -4.337034412 | 1.44E-05 | 7.29E-05 |
| ENSG00000151498 | ACAD8 | 746.5467894 | -0.174525081 | 0.074481803 | -2.343190885 | 0.019119598 | 0.052584053 |
| ENSG00000151500 | THYN1 | 460.0037483 | -0.310692284 | 0.097323774 | -3.192357535 | 0.001411165 | 0.005124606 |
| ENSG00000151502 | VPS26B | 2179.902654 | 0.08803632 | 0.054582577 | 1.61290151 | 0.106765907 | 0.223386529 |
| ENSG00000151503 | NCAPD3 | 1704.85463 | -0.143697981 | 0.064286964 | -2.235258464 | 0.025400385 | 0.067333507 |
| ENSG00000151532 | VTI1A | 895.9192952 | -0.950728797 | 0.078155789 | -12.16453455 | 4.80E-34 | 1.57E-32 |
| ENSG00000151552 | QDPR | 435.3288329 | -0.817212286 | 0.095634926 | -8.545123842 | 1.28E-17 | 1.84E-16 |
| ENSG00000151553 | FHIP2A | 983.2416031 | -0.93033754 | 0.076325426 | -12.18909068 | 3.55E-34 | 1.17E-32 |

| | | | | | | | |
|---|---|---|---|---|---|---|---|
| ENSG00000151575 | TEX9 | 91.48905534 | 0.639383324 | 0.19511911 | 3.276887256 | 0.001049583 | 0.003911848 |
| ENSG00000151576 | QTRT2 | 1465.984512 | -0.089135171 | 0.060793711 | -1.466190648 | 0.142596356 | 0.280353137 |
| ENSG00000151611 | MMAA | 141.6924876 | 0.061531101 | 0.152361435 | 0.403849575 | 0.686323339 | 0.822712932 |
| ENSG00000151612 | ZNF827 | 249.2770504 | -0.80613754 | 0.120776772 | -6.674607435 | 2.48E-11 | 2.24E-10 |
| ENSG00000151617 | EDNRA | 36.84402369 | -0.482516631 | 0.293647797 | -1.643181509 | 0.100345384 | 0.212550611 |
| ENSG00000151623 | NR3C2 | 281.5811771 | -0.872453581 | 0.112140278 | -7.780019753 | 7.25E-15 | 8.61E-14 |
| ENSG00000151632 | AKR1C2 | 64.67203918 | 2.456058591 | 0.265634997 | 9.245990262 | 2.33E-20 | 3.98E-19 |
| ENSG00000151640 | DPYSL4 | 8.780138089 | 0.171136744 | 0.605051221 | 0.282846704 | 0.777294351 | 0.879138067 |
| ENSG00000151651 | ADAM8 | 247.1476311 | -0.890855921 | 0.122156885 | -7.292719715 | 3.04E-13 | 3.22E-12 |
| ENSG00000151657 | KIN | 423.266049 | -1.140882503 | 0.096496688 | -11.82302235 | 2.97E-32 | 9.04E-31 |
| ENSG00000151665 | PIGF | 338.2506088 | -0.134617964 | 0.104887572 | -1.283450094 | 0.199334433 | 0.362196586 |
| ENSG00000151687 | ANKAR | 33.64763329 | 0.126662258 | 0.300353782 | 0.421710216 | 0.673236547 | 0.812972911 |
| ENSG00000151689 | INPP1 | 137.7702738 | 0.275864197 | 0.156853143 | 1.758741912 | 0.078621355 | 0.174272716 |
| ENSG00000151690 | MFSD6 | 1491.588812 | 0.508168794 | 0.061898007 | 8.20977642 | 2.22E-16 | 2.94E-15 |
| ENSG00000151692 | RNF144A | 206.2571118 | -0.328711716 | 0.127955181 | -2.568959797 | 0.010200429 | 0.030260415 |
| ENSG00000151693 | ASAP2 | 2432.085223 | 0.13295298 | 0.054391461 | 2.444372294 | 0.014510444 | 0.041268236 |
| ENSG00000151694 | ADAM17 | 1222.72402 | 0.459596219 | 0.064240982 | 7.154252752 | 8.41E-13 | 8.66E-12 |
| ENSG00000151715 | TMEM45B | 583.8912114 | 0.107513808 | 0.085687263 | 1.254723332 | 0.209579213 | 0.376605604 |
| ENSG00000151718 | WWC2 | 363.5039904 | -0.55147442 | 0.111796053 | -4.932861278 | 8.10E-07 | 4.75E-06 |
| ENSG00000151725 | CENPU | 756.1082718 | -0.262826422 | 0.092601008 | -2.838267393 | 0.004535917 | 0.014765706 |
| ENSG00000151726 | ACSL1 | 650.9888556 | 0.465228409 | 0.078114658 | 5.955712064 | 2.59E-09 | 1.98E-08 |
| ENSG00000151729 | SLC25A4 | 442.0624924 | -0.2373859 | 0.091267827 | -2.600981178 | 0.009295755 | 0.027918909 |
| ENSG00000151743 | AMN1 | 131.6605372 | 0.198346846 | 0.159430737 | 1.244094137 | 0.213464919 | 0.381674609 |
| ENSG00000151746 | BICD1 | 261.8127034 | 0.033985927 | 0.118645076 | 0.286450382 | 0.774533191 | 0.877335147 |
| ENSG00000151748 | SAV1 | 795.536235 | -0.591287248 | 0.074225161 | -7.966129597 | 1.64E-15 | 2.04E-14 |
| ENSG00000151773 | CCDC122 | 292.4829554 | 0.29183005 | 0.110042943 | 2.651965147 | 0.00800248 | 0.024460612 |
| ENSG00000151779 | NBAS | 1122.649148 | 0.058708113 | 0.06612842 | 0.887789447 | 0.374654021 | 0.567485192 |
| ENSG00000151790 | TDO2 | 8.268335482 | 1.856633333 | 0.682489809 | 2.720382498 | 0.006520644 | 0.020376458 |
| ENSG00000151806 | GUF1 | 1262.149582 | 0.325442512 | 0.068706902 | 4.736678616 | 2.17E-06 | 1.21E-05 |
| ENSG00000151834 | GABRA2 | 2.939844268 | 2.188639582 | 1.218182864 | 1.796642891 | 0.072392331 | 0.163195761 |
| ENSG00000151835 | SACS | 232.7639621 | 0.108391543 | 0.121084547 | 0.895172388 | 0.370694943 | 0.563385583 |
| ENSG00000151846 | PABPC3 | 893.4897578 | 0.125952046 | 0.084488956 | 1.490751598 | 0.136026724 | 0.270648295 |
| ENSG00000151849 | CENPJ | 747.4032867 | -0.431143281 | 0.077745055 | -5.545603931 | 2.93E-08 | 2.00E-07 |

| | | | | | | | |
|---|---|---|---|---|---|---|---|
| ENSG00000151876 | FBXO4 | 625.4299024 | 0.026846938 | 0.08870702 | 0.302647275 | 0.762158683 | 0.869504919 |
| ENSG00000151881 | TMEM267 | 330.6129384 | 0.243523868 | 0.1056744 | 2.304473632 | 0.021196071 | 0.057461341 |
| ENSG00000151882 | CCL28 | 109.3420465 | 1.330519754 | 0.190066588 | 7.000282217 | 2.55E-12 | 2.53E-11 |
| ENSG00000151883 | PARP8 | 343.3625779 | 0.139734034 | 0.102015149 | 1.36973808 | 0.170768677 | 0.321866195 |
| ENSG00000151892 | GFRA1 | 4442.826926 | 2.332127482 | 0.066277726 | 35.18719821 | 3.14E-271 | 4.80E-268 |
| ENSG00000151893 | CACUL1 | 1895.676466 | -0.410312389 | 0.057324346 | -7.157733396 | 8.20E-13 | 8.45E-12 |
| ENSG00000151914 | DST | 1866.950603 | -0.005426977 | 0.067226179 | -0.080727139 | 0.935658953 | 0.969248877 |
| ENSG00000151917 | BEND6 | 260.7545331 | 0.731800271 | 0.118222171 | 6.190042587 | 6.01E-10 | 4.89E-09 |
| ENSG00000151923 | TIAL1 | 2602.184492 | -0.588462771 | 0.056791996 | -10.3617202 | 3.70E-25 | 8.28E-24 |
| ENSG00000151929 | BAG3 | 2215.364362 | -0.989033403 | 0.059184783 | -16.71094081 | 1.09E-62 | 8.29E-61 |
| ENSG00000151967 | SCHIP1 | 4.133163193 | -0.066038168 | 0.843954263 | -0.078248515 | 0.937630371 | 0.970153346 |
| ENSG00000152049 | KCNE4 | 2.627704585 | -0.885980672 | 1.095212976 | -0.808957427 | 0.418539635 | 0.61039531 |
| ENSG00000152056 | AP1S3 | 130.5044991 | 1.714552365 | 0.172905259 | 9.916137753 | 3.54E-23 | 7.11E-22 |
| ENSG00000152061 | RABGAP1L | 834.7579953 | 0.02655185 | 0.074297786 | 0.357370677 | 0.720814325 | 0.845026606 |
| ENSG00000152076 | CCDC74B | 87.5731545 | 1.032166479 | 0.216075605 | 4.776876503 | 1.78E-06 | 1.00E-05 |
| ENSG00000152078 | TLCD4 | 164.4366228 | 0.361220171 | 0.147562133 | 2.447919148 | 0.014368389 | 0.040909874 |
| ENSG00000152082 | MZT2B | 1591.17948 | -0.107472932 | 0.067088435 | -1.601959142 | 0.109164645 | 0.227285982 |
| ENSG00000152092 | ASTN1 | 9.13909936 | 0.107599976 | 0.584631553 | 0.1840475 | 0.853976192 | 0.923979523 |
| ENSG00000152102 | FAM168B | 6404.719095 | -0.001368232 | 0.048082151 | -0.028456139 | 0.97729835 | 0.989387067 |
| ENSG00000152104 | PTPN14 | 678.5943385 | -0.089470803 | 0.083405393 | -1.072722036 | 0.28339585 | 0.46720092 |
| ENSG00000152117 | SMPD4BP | 1220.467342 | 0.038674251 | 0.072504716 | 0.533403252 | 0.593754467 | 0.755710631 |
| ENSG00000152127 | MGAT5 | 3196.081827 | 0.133801136 | 0.052569064 | 2.545244759 | 0.010920117 | 0.032091657 |
| ENSG00000152128 | TMEM163 | 7.015803502 | 0.403466793 | 0.667071273 | 0.60483311 | 0.545289891 | 0.718492252 |
| ENSG00000152133 | GPATCH11 | 297.4308125 | -0.21202117 | 0.110628025 | -1.916523136 | 0.055298541 | 0.130576278 |
| ENSG00000152137 | HSPB8 | 778.3793644 | 1.323132047 | 0.078449453 | 16.86604549 | 8.00E-64 | 6.26E-62 |
| ENSG00000152147 | GEMIN6 | 544.9653711 | -0.149636603 | 0.084682716 | -1.767026509 | 0.077223789 | 0.171974926 |
| ENSG00000152154 | TMEM178A | 9.895852182 | 1.361594517 | 0.640316128 | 2.126441076 | 0.033466551 | 0.085283567 |
| ENSG00000152193 | OBI1 | 334.0822635 | -0.02464478 | 0.110671538 | -0.22268399 | 0.823781462 | 0.906966152 |
| ENSG00000152213 | ARL11 | 24.10770197 | 7.93897716 | 1.215798177 | 6.529847887 | 6.58E-11 | 5.78E-10 |
| ENSG00000152217 | SETBP1 | 430.6058193 | 0.651292642 | 0.09580173 | 6.798339034 | 1.06E-11 | 9.88E-11 |
| ENSG00000152219 | ARL14EP | 419.6898465 | 0.106587774 | 0.092763049 | 1.149032674 | 0.250542508 | 0.428735728 |
| ENSG00000152223 | EPG5 | 1124.259631 | 0.590663517 | 0.081168498 | 7.277004366 | 3.41E-13 | 3.59E-12 |
| ENSG00000152229 | PSTPIP2 | 132.6869935 | 0.050680872 | 0.157582293 | 0.321615272 | 0.747744173 | 0.86037274 |

| | | | | | | | |
|---|---|---|---|---|---|---|---|
| ENSG00000152234 | ATP5F1A | 12031.85472 | 0.409519975 | 0.047755923 | 8.575270818 | 9.89E-18 | 1.43E-16 |
| ENSG00000152240 | HAUS1 | 571.1661149 | 0.363838165 | 0.083031191 | 4.381945636 | 1.18E-05 | 6.01E-05 |
| ENSG00000152242 | C18orf25 | 1099.394509 | 0.358607315 | 0.064142534 | 5.590788097 | 2.26E-08 | 1.56E-07 |
| ENSG00000152253 | SPC25 | 328.4669983 | -0.261396298 | 0.119015879 | -2.196314476 | 0.028069442 | 0.073372038 |
| ENSG00000152256 | PDK1 | 259.7704574 | 0.24053844 | 0.123674565 | 1.944930553 | 0.051783337 | 0.123444846 |
| ENSG00000152270 | PDE3B | 723.0560164 | -0.074835767 | 0.076742036 | -0.975160035 | 0.329480883 | 0.519863226 |
| ENSG00000152284 | TCF7L1 | 449.4453282 | -0.618825473 | 0.093333693 | -6.630247354 | 3.35E-11 | 3.01E-10 |
| ENSG00000152291 | TGOLN2 | 6594.850956 | 0.293898629 | 0.045648323 | 6.438322643 | 1.21E-10 | 1.04E-09 |
| ENSG00000152292 | SH2D6 | 2.570052538 | -2.207592759 | 1.249627314 | -1.766600916 | 0.077295087 | 0.172100292 |
| ENSG00000152315 | KCNK13 | 13.52707277 | 0.367385957 | 0.501605614 | 0.732419947 | 0.463912288 | 0.650169158 |
| ENSG00000152332 | UHMK1 | 5292.737011 | 0.077800898 | 0.054691543 | 1.42253982 | 0.1548696 | 0.298411239 |
| ENSG00000152348 | ATG10 | 372.1451196 | -0.250705962 | 0.097614172 | -2.568335701 | 0.010218814 | 0.030294597 |
| ENSG00000152359 | POC5 | 371.2503739 | -0.070602997 | 0.101164858 | -0.697900422 | 0.48523947 | 0.669074559 |
| ENSG00000152377 | SPOCK1 | 3.482197401 | 2.452554361 | 1.135367198 | 2.160141992 | 0.030761679 | 0.079405077 |
| ENSG00000152380 | FAM151B | 67.90436112 | -0.411641677 | 0.235730524 | -1.746238333 | 0.080769546 | 0.178056255 |
| ENSG00000152382 | TADA1 | 607.4435227 | -0.039682688 | 0.084575372 | -0.469199097 | 0.638927331 | 0.787290808 |
| ENSG00000152402 | GUCY1A2 | 368.7861132 | 0.165514571 | 0.121621242 | 1.360901828 | 0.173544716 | 0.325973616 |
| ENSG00000152404 | CWF19L2 | 287.8282653 | -0.293496512 | 0.10922724 | -2.687026724 | 0.007209118 | 0.022303058 |
| ENSG00000152409 | JMY | 888.9460201 | 0.362585115 | 0.07389566 | 4.906717325 | 9.26E-07 | 5.39E-06 |
| ENSG00000152413 | HOMER1 | 746.1418227 | 0.284774478 | 0.073322863 | 3.883842882 | 0.000102818 | 0.000457547 |
| ENSG00000152422 | XRCC4 | 180.9019863 | -0.131526664 | 0.137832384 | -0.954250814 | 0.339956699 | 0.530878603 |
| ENSG00000152430 | BOLL | 1.722418348 | -0.789003607 | 1.337439249 | -0.589936034 | 0.555233535 | NA |
| ENSG00000152433 | ZNF547 | 71.07360388 | 1.252064644 | 0.22762646 | 5.5005233 | 3.79E-08 | 2.57E-07 |
| ENSG00000152439 | ZNF773 | 86.54400437 | 0.374002036 | 0.191050854 | 1.957604629 | 0.050276424 | 0.120377952 |
| ENSG00000152443 | ZNF776 | 800.6975031 | 0.384054005 | 0.073484772 | 5.226307364 | 1.73E-07 | 1.10E-06 |
| ENSG00000152454 | ZNF256 | 154.8584181 | 0.877154248 | 0.150970585 | 5.810100329 | 6.24E-09 | 4.59E-08 |
| ENSG00000152455 | SUV39H2 | 531.6105511 | -0.681601115 | 0.093026965 | -7.326919805 | 2.36E-13 | 2.51E-12 |
| ENSG00000152457 | DCLRE1C | 423.8203253 | -0.654808626 | 0.092636568 | -7.06857603 | 1.57E-12 | 1.58E-11 |
| ENSG00000152464 | RPP38 | 380.1714203 | -0.541544127 | 0.102563594 | -5.280081416 | 1.29E-07 | 8.30E-07 |
| ENSG00000152465 | NMT2 | 354.9275824 | -0.258406906 | 0.102048443 | -2.53219841 | 0.011334983 | 0.033183322 |
| ENSG00000152475 | ZNF837 | 76.26446446 | -0.067824338 | 0.207566612 | -0.326759383 | 0.743849882 | 0.858490754 |
| ENSG00000152484 | USP12 | 1520.472222 | -0.033345224 | 0.06800607 | -0.490327167 | 0.623902406 | 0.77763223 |
| ENSG00000152492 | CCDC50 | 1101.227745 | 0.139266343 | 0.068329055 | 2.038171662 | 0.041532768 | 0.102271544 |

| | | | | | | | |
|---|---|---|---|---|---|---|---|
| ENSG00000152495 | CAMK4 | 1.598474914 | -0.178247836 | 1.39679048 | -0.127612436 | 0.898455689 | NA |
| ENSG00000152503 | TRIM36 | 297.2524329 | -0.528214843 | 0.109084286 | -4.842263387 | 1.28E-06 | 7.33E-06 |
| ENSG00000152518 | ZFP36L2 | 5242.835305 | -0.443644823 | 0.050539511 | -8.778178018 | 1.66E-18 | 2.53E-17 |
| ENSG00000152520 | PAN3 | 1527.024233 | -0.123490997 | 0.066509638 | -1.856738366 | 0.063348386 | 0.145948778 |
| ENSG00000152527 | PLEKHH2 | 351.593552 | -0.678646483 | 0.113871706 | -5.95974636 | 2.53E-09 | 1.94E-08 |
| ENSG00000152556 | PFKM | 1023.7673 | -0.757133885 | 0.069053884 | -10.96439248 | 5.67E-28 | 1.46E-26 |
| ENSG00000152558 | TMEM123 | 2091.49436 | -0.047684386 | 0.060571769 | -0.787237799 | 0.431142674 | 0.621257094 |
| ENSG00000152580 | IGSF10 | 41.50228583 | -0.408838343 | 0.274779058 | -1.487880282 | 0.136782469 | 0.271740697 |
| ENSG00000152582 | SPEF2 | 247.3034024 | 0.508405381 | 0.122666087 | 4.144628665 | 3.40E-05 | 0.000163244 |
| ENSG00000152583 | SPARCL1 | 11.12805294 | 0.545976604 | 0.536863002 | 1.016975657 | 0.30916501 | 0.497224342 |
| ENSG00000152592 | DMP1 | 8.64442861 | 0.495085574 | 0.626656092 | 0.790043503 | 0.429502363 | 0.619827894 |
| ENSG00000152595 | MEPE | 1.686444324 | -0.172489906 | 1.381534262 | -0.124853875 | 0.900639235 | NA |
| ENSG00000152601 | MBNL1 | 2630.082368 | -0.210497797 | 0.055995247 | -3.759208302 | 0.000170452 | 0.000732378 |
| ENSG00000152620 | NADK2 | 964.5103189 | 0.027506151 | 0.07267506 | 0.378481289 | 0.705073091 | 0.834997903 |
| ENSG00000152642 | GPD1L | 3512.562345 | -0.500717062 | 0.047557795 | -10.52860124 | 6.38E-26 | 1.48E-24 |
| ENSG00000152661 | GJA1 | 9.968542947 | 5.673372811 | 1.265471865 | 4.483207384 | 7.35E-06 | 3.85E-05 |
| ENSG00000152669 | CCNO | 117.0344757 | 1.975939552 | 0.194577476 | 10.15502717 | 3.15E-24 | 6.67E-23 |
| ENSG00000152672 | CLEC4F | 8.293293023 | -0.246998781 | 0.657972282 | -0.375393899 | 0.707367541 | 0.836573253 |
| ENSG00000152683 | SLC30A6 | 1038.364916 | -0.129470811 | 0.066632739 | -1.943051014 | 0.052009995 | 0.123895045 |
| ENSG00000152684 | PELO | 569.8024271 | -0.124581603 | 0.083382686 | -1.494094381 | 0.135150952 | 0.269305262 |
| ENSG00000152689 | RASGRP3 | 22.55673869 | -2.211616309 | 0.434802494 | -5.086484872 | 3.65E-07 | 2.23E-06 |
| ENSG00000152700 | SAR1B | 1919.296812 | -0.095273891 | 0.058597184 | -1.62591247 | 0.103968277 | 0.218730032 |
| ENSG00000152705 | CATSPER3 | 11.36504019 | -0.103892894 | 0.522193018 | -0.198954965 | 0.842297973 | 0.917009895 |
| ENSG00000152749 | GPR180 | 338.4604376 | 0.141885205 | 0.10363251 | 1.36911867 | 0.170962183 | 0.322072143 |
| ENSG00000152763 | DNAI4 | 176.3246957 | 0.477899169 | 0.146852304 | 3.254284438 | 0.001136784 | 0.004216301 |
| ENSG00000152766 | ANKRD22 | 16.33932066 | 0.334060267 | 0.430163985 | 0.776588183 | 0.437401776 | 0.626920679 |
| ENSG00000152767 | FARP1 | 2015.425654 | -0.015610775 | 0.057234345 | -0.272751875 | 0.785043955 | 0.883106023 |
| ENSG00000152778 | IFIT5 | 342.5168807 | -0.022158794 | 0.101598963 | -0.218100597 | 0.827350733 | 0.908805627 |
| ENSG00000152782 | PANK1 | 748.3877966 | -0.481684931 | 0.074219134 | -6.490037057 | 8.58E-11 | 7.48E-10 |
| ENSG00000152784 | PRDM8 | 5.406281513 | -0.784924674 | 0.761546328 | -1.030698521 | 0.302682218 | 0.489670669 |
| ENSG00000152795 | HNRNPDL | 5813.669491 | -0.038107447 | 0.066271152 | -0.575023166 | 0.56527563 | 0.733481521 |
| ENSG00000152804 | HHEX | 79.53901737 | -0.326414294 | 0.198176447 | -1.647089244 | 0.099539704 | 0.21117946 |
| ENSG00000152818 | UTRN | 943.8033217 | 0.49252336 | 0.080850535 | 6.091776144 | 1.12E-09 | 8.85E-09 |

| | | | | | | | |
|---|---|---|---|---|---|---|---|
| ENSG00000152822 | GRM1 | 1.801406909 | -0.841809647 | 1.418048245 | -0.593639638 | 0.552753164 | 0.724326573 |
| ENSG00000152894 | PTPRK | 4726.355449 | 0.992307313 | 0.047982409 | 20.68064783 | 5.17E-95 | 7.66E-93 |
| ENSG00000152904 | GGPS1 | 496.4169383 | -0.391862067 | 0.087418787 | -4.482584118 | 7.37E-06 | 3.86E-05 |
| ENSG00000152926 | ZNF117 | 887.3809407 | 1.319200794 | 0.085135844 | 15.49524531 | 3.74E-54 | 2.28E-52 |
| ENSG00000152931 | PART1 | 2.308163583 | -0.19004019 | 1.121704177 | -0.169420952 | 0.865465544 | 0.930134906 |
| ENSG00000152936 | LMNTD1 | 7.788457388 | 1.922156124 | 0.723499287 | 2.656749162 | 0.007889812 | 0.024158075 |
| ENSG00000152939 | MARVELD2 | 1482.483005 | -0.027537968 | 0.05930796 | -0.464321621 | 0.642417336 | 0.790314651 |
| ENSG00000152942 | RAD17 | 972.4948278 | 0.005377872 | 0.076736018 | 0.070082756 | 0.944127792 | 0.973493866 |
| ENSG00000152944 | MED21 | 449.5789766 | 0.002589181 | 0.089376472 | 0.02896938 | 0.976889011 | 0.989309285 |
| ENSG00000152952 | PLOD2 | 1411.406244 | 0.68043153 | 0.068632364 | 9.914149613 | 3.61E-23 | 7.24E-22 |
| ENSG00000152977 | ZIC1 | 608.8735601 | -0.321663362 | 0.081549902 | -3.944374593 | 8.00E-05 | 0.000363085 |
| ENSG00000152990 | ADGRA3 | 1545.87775 | 0.246963896 | 0.058256088 | 4.239280438 | 2.24E-05 | 0.000110434 |
| ENSG00000153006 | SREK1IP1 | 440.518749 | 0.019277222 | 0.090409191 | 0.21322193 | 0.831153864 | 0.910670869 |
| ENSG00000153012 | LGI2 | 19.38597917 | -0.763186661 | 0.411370137 | -1.855231071 | 0.063563236 | 0.14635558 |
| ENSG00000153015 | CWC27 | 968.3661159 | -0.023241977 | 0.070690674 | -0.328784201 | 0.742318806 | 0.85777393 |
| ENSG00000153029 | MR1 | 392.500677 | 0.036478486 | 0.095096912 | 0.383592751 | 0.701280313 | 0.832852595 |
| ENSG00000153037 | SRP19 | 328.2314172 | 0.821818387 | 0.107646382 | 7.63442648 | 2.27E-14 | 2.60E-13 |
| ENSG00000153044 | CENPH | 366.6031415 | -0.031409617 | 0.106349695 | -0.295342801 | 0.767732024 | 0.87314707 |
| ENSG00000153046 | CDYL | 1351.440288 | -0.84456643 | 0.063189587 | -13.3655951 | 9.61E-41 | 4.05E-39 |
| ENSG00000153048 | CARHSP1 | 1581.923899 | 0.125706478 | 0.064456295 | 1.950259151 | 0.051145239 | 0.122164908 |
| ENSG00000153060 | TEKT5 | 11.97228367 | -1.263264386 | 0.536501582 | -2.354633106 | 0.018541002 | 0.051164512 |
| ENSG00000153064 | BANK1 | 82.26416082 | 0.061572884 | 0.196674965 | 0.313069253 | 0.754228048 | 0.864498202 |
| ENSG00000153066 | TXNDC11 | 1062.935966 | -0.013022558 | 0.065438409 | -0.199004804 | 0.842258987 | 0.917009895 |
| ENSG00000153071 | DAB2 | 250.2036262 | -2.477961153 | 0.137176346 | -18.06405563 | 6.12E-73 | 6.02E-71 |
| ENSG00000153093 | ACOXL | 5.144359438 | -1.042687294 | 0.78762455 | -1.323837982 | 0.185556859 | 0.342755 |
| ENSG00000153094 | BCL2L11 | 1886.291807 | 0.376030129 | 0.062013648 | 6.06366729 | 1.33E-09 | 1.05E-08 |
| ENSG00000153107 | ANAPC1 | 1599.732934 | 0.127538661 | 0.062137869 | 2.05251103 | 0.040120024 | 0.099358022 |
| ENSG00000153113 | CAST | 3134.189601 | 0.846535123 | 0.049222393 | 17.19817079 | 2.74E-66 | 2.25E-64 |
| ENSG00000153130 | SCOC | 999.2285023 | 0.035935843 | 0.079783881 | 0.450414831 | 0.652411353 | 0.798160975 |
| ENSG00000153132 | CLGN | 140.7835308 | -1.68513513 | 0.174862635 | -9.63690801 | 5.58E-22 | 1.04E-20 |
| ENSG00000153140 | CETN3 | 517.5915312 | -0.004798157 | 0.086852844 | -0.055244673 | 0.95594354 | 0.979819704 |
| ENSG00000153147 | SMARCA5 | 5278.346358 | -0.059877275 | 0.047585451 | -1.25831055 | 0.208279467 | 0.374751426 |
| ENSG00000153162 | BMP6 | 6.983974906 | -1.019933983 | 0.668936356 | -1.524710048 | 0.127331434 | 0.257151431 |

| | | | | | | | |
|---|---|---|---|---|---|---|---|
| ENSG00000153165 | RGPD3 | 9.310752018 | 0.280152455 | 0.578930137 | 0.483914098 | 0.628446835 | 0.780792125 |
| ENSG00000153179 | RASSF3 | 1188.887219 | -0.095580998 | 0.063042234 | -1.516142302 | 0.129483374 | 0.260442665 |
| ENSG00000153187 | HNRNPU | 15713.91293 | -0.173500295 | 0.0471803 | -3.677388544 | 0.000235634 | 0.000986935 |
| ENSG00000153201 | RANBP2 | 4227.371556 | -0.007925449 | 0.067656197 | -0.117142979 | 0.906746752 | 0.952939824 |
| ENSG00000153207 | AHCTF1 | 2260.880424 | -0.018105404 | 0.057903983 | -0.312679763 | 0.754523972 | 0.864498202 |
| ENSG00000153208 | MERTK | 52.5404002 | 0.537897985 | 0.245687687 | 2.189356706 | 0.028570924 | 0.07446228 |
| ENSG00000153214 | TMEM87B | 1113.675713 | -0.276164512 | 0.071960149 | -3.837742345 | 0.000124171 | 0.000545058 |
| ENSG00000153234 | NR4A2 | 257.7255904 | -0.341852963 | 0.119010191 | -2.872467978 | 0.004072794 | 0.013397067 |
| ENSG00000153237 | CCDC148 | 23.4585128 | 0.130105415 | 0.367331399 | 0.354190834 | 0.723195872 | 0.846002719 |
| ENSG00000153246 | PLA2R1 | 106.834047 | -0.076505149 | 0.177928243 | -0.429977545 | 0.667211975 | 0.808556374 |
| ENSG00000153250 | RBMS1 | 808.2895906 | 0.599575459 | 0.082419848 | 7.274648895 | 3.47E-13 | 3.65E-12 |
| ENSG00000153283 | CD96 | 1.900675452 | 2.075098982 | 1.469186865 | 1.412413241 | 0.15782832 | 0.302673605 |
| ENSG00000153291 | SLC25A27 | 7.004406354 | -0.471343414 | 0.66056505 | -0.713545795 | 0.475508086 | 0.660696059 |
| ENSG00000153294 | ADGRF4 | 85.32395044 | 1.20377898 | 0.206083581 | 5.841217297 | 5.18E-09 | 3.85E-08 |
| ENSG00000153310 | CYRIB | 2812.779829 | -0.006287275 | 0.050267228 | -0.12507701 | 0.900462584 | 0.949808511 |
| ENSG00000153317 | ASAP1 | 4171.731964 | -0.024393018 | 0.048356807 | -0.504438137 | 0.613953524 | 0.770801984 |
| ENSG00000153339 | TRAPPC8 | 1430.323802 | 0.506286465 | 0.063248173 | 8.004760309 | 1.20E-15 | 1.51E-14 |
| ENSG00000153363 | LINC00467 | 255.6481395 | -0.468366158 | 0.128366197 | -3.648672079 | 0.000263599 | 0.001093682 |
| ENSG00000153391 | INO80C | 194.5053546 | 0.526570056 | 0.131683173 | 3.998764957 | 6.37E-05 | 0.000293191 |
| ENSG00000153395 | LPCAT1 | 10619.60275 | -0.329144246 | 0.043675331 | -7.536159213 | 4.84E-14 | 5.40E-13 |
| ENSG00000153404 | PLEKHG4B | 3.818378094 | 0.981440477 | 0.937843269 | 1.046486667 | 0.295336397 | 0.481282727 |
| ENSG00000153406 | NMRAL1 | 2093.975464 | 0.032881234 | 0.056641496 | 0.580514924 | 0.561567426 | 0.730540816 |
| ENSG00000153443 | UBALD1 | 697.3316342 | -0.061180367 | 0.080359228 | -0.761335913 | 0.446456455 | 0.634987357 |
| ENSG00000153485 | LYSET | 349.3325209 | -0.585457623 | 0.105964232 | -5.525049459 | 3.29E-08 | 2.24E-07 |
| ENSG00000153487 | ING1 | 634.4262401 | -0.63686817 | 0.081855584 | -7.780387603 | 7.23E-15 | 8.59E-14 |
| ENSG00000153495 | TEX29 | 1.911955101 | 0.883518893 | 1.309646559 | 0.674623918 | 0.499914733 | 0.681205745 |
| ENSG00000153531 | ADPRHL1 | 510.4824099 | -0.589656536 | 0.09597286 | -6.143992564 | 8.05E-10 | 6.45E-09 |
| ENSG00000153551 | CMTM7 | 7.978791153 | -0.669519062 | 0.614037968 | -1.090354501 | 0.275557016 | 0.457842682 |
| ENSG00000153558 | FBXL2 | 79.65945635 | 0.289215662 | 0.19973832 | 1.447972839 | 0.147624647 | 0.288066468 |
| ENSG00000153560 | UBP1 | 5391.202165 | 0.111195464 | 0.045924601 | 2.421261406 | 0.015466751 | 0.043717011 |
| ENSG00000153561 | RMND5A | 3258.844767 | 0.192229592 | 0.052146056 | 3.686368762 | 0.000227477 | 0.000955036 |
| ENSG00000153563 | CD8A | 104.4475268 | -0.402988042 | 0.181076391 | -2.225513989 | 0.026046753 | 0.068774461 |
| ENSG00000153574 | RPIA | 673.9539272 | -0.024948157 | 0.077061499 | -0.32374347 | 0.746132258 | 0.859596562 |

| | | | | | | | |
|---|---|---|---|---|---|---|---|
| ENSG00000153707 | PTPRD | 43.81678024 | 0.569567387 | 0.271772047 | 2.095754117 | 0.036104008 | 0.091043856 |
| ENSG00000153714 | LURAP1L | 359.6533225 | 1.058696486 | 0.103607531 | 10.2183352 | 1.64E-24 | 3.56E-23 |
| ENSG00000153721 | CNKSR3 | 66.49791166 | 1.430895254 | 0.244762855 | 5.846047409 | 5.03E-09 | 3.75E-08 |
| ENSG00000153767 | GTF2E1 | 777.0371735 | -0.206315672 | 0.075946313 | -2.71659892 | 0.006595648 | 0.020588403 |
| ENSG00000153774 | CFDP1 | 1313.308345 | -0.10452671 | 0.062079587 | -1.683753315 | 0.092229353 | 0.198418959 |
| ENSG00000153786 | ZDHHC7 | 5813.292356 | 0.511924732 | 0.049839156 | 10.271537 | 9.47E-25 | 2.08E-23 |
| ENSG00000153814 | JAZF1 | 17.8034283 | -2.640153297 | 0.504244515 | -5.235859228 | 1.64E-07 | 1.04E-06 |
| ENSG00000153815 | CMIP | 1580.569453 | 0.494350952 | 0.061087017 | 8.09256985 | 5.84E-16 | 7.53E-15 |
| ENSG00000153823 | PID1 | 76.27012168 | -0.057424545 | 0.204203225 | -0.281212721 | 0.778547248 | 0.879753405 |
| ENSG00000153827 | TRIP12 | 5528.781749 | 0.072611339 | 0.049386017 | 1.470281326 | 0.141485577 | 0.27886237 |
| ENSG00000153832 | FBXO36 | 156.380232 | 0.149979828 | 0.147362945 | 1.017758083 | 0.308792937 | 0.496900138 |
| ENSG00000153879 | CEBPG | 1539.438961 | 0.087261579 | 0.065422791 | 1.333810102 | 0.1822661 | 0.338201434 |
| ENSG00000153885 | KCTD15 | 2256.048728 | -0.251984874 | 0.055363015 | -4.551501993 | 5.33E-06 | 2.83E-05 |
| ENSG00000153896 | ZNF599 | 151.3756285 | 0.238449564 | 0.147308977 | 1.618703545 | 0.105511064 | 0.221285775 |
| ENSG00000153898 | MCOLN2 | 66.02974864 | 1.37103138 | 0.236826664 | 5.789176593 | 7.07E-09 | 5.17E-08 |
| ENSG00000153904 | DDAH1 | 1469.173525 | 0.924500074 | 0.060259045 | 15.34209643 | 4.00E-53 | 2.40E-51 |
| ENSG00000153914 | SREK1 | 2292.365843 | -0.108406936 | 0.070684435 | -1.533674775 | 0.125109683 | 0.253669446 |
| ENSG00000153922 | CHD1 | 1699.236172 | 0.069945722 | 0.06401684 | 1.092614409 | 0.274563134 | 0.456643101 |
| ENSG00000153933 | DGKE | 803.7370716 | 0.291683342 | 0.08157185 | 3.575784306 | 0.000349179 | 0.001416978 |
| ENSG00000153936 | HS2ST1 | 870.2869601 | 0.198205524 | 0.072430476 | 2.736493469 | 0.006209783 | 0.019538118 |
| ENSG00000153944 | MSI2 | 6039.706887 | 0.039310488 | 0.047207131 | 0.832723523 | 0.405000675 | 0.597811154 |
| ENSG00000153956 | CACNA2D1 | 123.6185109 | 3.093612067 | 0.220351829 | 14.03942085 | 8.94E-45 | 4.27E-43 |
| ENSG00000153975 | ZUP1 | 390.0564247 | -0.0643825 | 0.099497867 | -0.647074175 | 0.517583938 | 0.695665852 |
| ENSG00000153982 | GDPD1 | 64.00802474 | 0.050063884 | 0.228050571 | 0.219529746 | 0.826237411 | 0.908362226 |
| ENSG00000153989 | NUS1 | 1205.484862 | 0.149251412 | 0.06667183 | 2.238597789 | 0.025182098 | 0.066891771 |
| ENSG00000153993 | SEMA3D | 53.6296845 | 0.47171354 | 0.24469892 | 1.927730368 | 0.053888673 | 0.127852534 |
| ENSG00000154001 | PPP2R5E | 2804.193555 | -0.801737765 | 0.051793713 | -15.47944181 | 4.78E-54 | 2.91E-52 |
| ENSG00000154016 | GRAP | 4.151978012 | -1.271855036 | 0.886940236 | -1.433980537 | 0.151577817 | 0.293948007 |
| ENSG00000154025 | SLC5A10 | 22.0260826 | -0.363615944 | 0.375963608 | -0.967157291 | 0.333465409 | 0.524587159 |
| ENSG00000154040 | CABYR | 127.520571 | 0.063711512 | 0.161979703 | 0.39333022 | 0.694075601 | 0.827653275 |
| ENSG00000154059 | IMPACT | 1095.628618 | 0.203018025 | 0.065828591 | 3.084040264 | 0.0020421 | 0.007164056 |
| ENSG00000154065 | ANKRD29 | 2.610345517 | -1.235308579 | 1.116158731 | -1.106749913 | 0.268402069 | 0.449874922 |
| ENSG00000154079 | SDHAF4 | 215.9588419 | 0.082593959 | 0.123806984 | 0.667118742 | 0.504696289 | 0.685322056 |

| | | | | | | | |
|---|---|---|---|---|---|---|---|
| ENSG00000154080 | CHST9 | 2.557418528 | 0.774935304 | 1.148726344 | 0.674603928 | 0.499927437 | 0.681205745 |
| ENSG00000154099 | DNAAF1 | 56.53541626 | 0.396007867 | 0.253604725 | 1.561516127 | 0.118402022 | 0.243076998 |
| ENSG00000154102 | C16orf74 | 264.4400625 | 1.882989411 | 0.141485867 | 13.30867496 | 2.06E-40 | 8.63E-39 |
| ENSG00000154114 | TBCEL | 298.0117383 | 0.094805445 | 0.107207046 | 0.884321026 | 0.376522941 | 0.569338624 |
| ENSG00000154118 | JPH3 | 1472.588313 | 0.894521105 | 0.078336064 | 11.41902036 | 3.36E-30 | 9.42E-29 |
| ENSG00000154122 | ANKH | 349.9777472 | 0.459438195 | 0.111526869 | 4.119529212 | 3.80E-05 | 0.000180462 |
| ENSG00000154124 | OTULIN | 1367.147133 | 0.016915551 | 0.063767489 | 0.265269199 | 0.79080209 | 0.887220617 |
| ENSG00000154127 | UBASH3B | 4.912700971 | -0.758747922 | 0.807652239 | -0.939448794 | 0.347500371 | 0.53848004 |
| ENSG00000154134 | ROBO3 | 19.05746833 | -0.38117718 | 0.412092373 | -0.924979944 | 0.35497634 | 0.546644725 |
| ENSG00000154143 | PANX3 | 12.1056453 | -0.399745738 | 0.519922386 | -0.768856562 | 0.441978467 | 0.63124057 |
| ENSG00000154144 | TBRG1 | 500.4587696 | 0.033387965 | 0.091082892 | 0.366566802 | 0.713942175 | 0.840918855 |
| ENSG00000154146 | NRGN | 124.9119527 | -0.574392179 | 0.166846592 | -3.442636576 | 0.000576073 | 0.0022536 |
| ENSG00000154153 | RETREG1 | 1074.523874 | 1.643099199 | 0.072457255 | 22.6768071 | 7.59E-114 | 1.55E-111 |
| ENSG00000154174 | TOMM70 | 4047.700687 | 0.053220143 | 0.048515018 | 1.096982854 | 0.272648888 | 0.454731892 |
| ENSG00000154188 | ANGPT1 | 38.23472607 | -2.742559093 | 0.353432237 | -7.759787611 | 8.51E-15 | 1.00E-13 |
| ENSG00000154198 | CYP4Z2P | 16.36094431 | -1.405418419 | 0.4494307 | -3.127108185 | 0.00176535 | 0.00627376 |
| ENSG00000154217 | PITPNC1 | 2101.501013 | 0.252273302 | 0.055335121 | 4.559008787 | 5.14E-06 | 2.74E-05 |
| ENSG00000154222 | CC2D1B | 1319.315474 | -0.01732605 | 0.065966754 | -0.262648211 | 0.792821737 | 0.88871611 |
| ENSG00000154229 | PRKCA | 628.359013 | 0.437838272 | 0.079976674 | 5.474574671 | 4.39E-08 | 2.95E-07 |
| ENSG00000154237 | LRRK1 | 144.5899307 | -0.114631504 | 0.15751453 | -0.727751935 | 0.466765452 | 0.652743201 |
| ENSG00000154240 | CEP112 | 21.43652957 | -0.393319973 | 0.403207216 | -0.975478507 | 0.329322958 | 0.51977764 |
| ENSG00000154258 | ABCA9 | 3.588087704 | 4.159219018 | 1.398732575 | 2.973562704 | 0.002943643 | 0.00998627 |
| ENSG00000154262 | ABCA6 | 6.12037947 | 1.291290831 | 0.735780769 | 1.754993994 | 0.079260335 | 0.175482475 |
| ENSG00000154263 | ABCA10 | 26.07358974 | 0.999810377 | 0.357864216 | 2.793826076 | 0.005208849 | 0.016684025 |
| ENSG00000154265 | ABCA5 | 782.1792146 | 0.401505922 | 0.078285136 | 5.128763156 | 2.92E-07 | 1.80E-06 |
| ENSG00000154269 | ENPP3 | 7.34089286 | -2.317644997 | 0.751772948 | -3.082905555 | 0.002049902 | 0.007187672 |
| ENSG00000154274 | C4orf19 | 21.00959966 | 0.575593279 | 0.395968328 | 1.453634643 | 0.146047627 | 0.285620849 |
| ENSG00000154305 | MIA3 | 2462.286992 | -0.231402226 | 0.056157511 | -4.12059266 | 3.78E-05 | 0.000179706 |
| ENSG00000154309 | DISP1 | 197.9222109 | 0.155193205 | 0.143132931 | 1.08425926 | 0.278249863 | 0.461069912 |
| ENSG00000154310 | TNIK | 63.38171027 | -0.673784424 | 0.230953882 | -2.91739813 | 0.003529649 | 0.01177919 |
| ENSG00000154316 | TDH | 21.08262601 | -0.479644378 | 0.381668645 | -1.256703646 | 0.208860968 | 0.375520855 |
| ENSG00000154319 | FAM167A | 25.19771199 | 0.697507041 | 0.359986931 | 1.937589899 | 0.052673276 | 0.125305916 |
| ENSG00000154328 | NEIL2 | 190.1191772 | 0.697212438 | 0.135852676 | 5.132121501 | 2.86E-07 | 1.77E-06 |

| | | | | | | | |
|---|---|---|---|---|---|---|---|
| ENSG00000154330 | PGM5 | 84.01081762 | -1.374468101 | 0.219667747 | -6.257031892 | 3.92E-10 | 3.25E-09 |
| ENSG00000154342 | WNT3A | 3.959440933 | -2.096063738 | 1.031816634 | -2.031430459 | 0.042211347 | 0.103730938 |
| ENSG00000154358 | OBSCN | 1351.924457 | -0.496204226 | 0.082305592 | -6.02880332 | 1.65E-09 | 1.29E-08 |
| ENSG00000154359 | LONRF1 | 797.9852541 | -0.053615864 | 0.083282605 | -0.643782264 | 0.519716636 | 0.69707672 |
| ENSG00000154370 | TRIM11 | 1562.235147 | -0.196230976 | 0.062325426 | -3.148489902 | 0.001641164 | 0.005879791 |
| ENSG00000154380 | ENAH | 3487.517066 | 0.445277084 | 0.052373267 | 8.501991823 | 1.86E-17 | 2.65E-16 |
| ENSG00000154429 | CCSAP | 433.78458 | 0.259306944 | 0.099425698 | 2.608047499 | 0.009106032 | 0.027406503 |
| ENSG00000154447 | SH3RF1 | 658.7857994 | 0.159708345 | 0.08373786 | 1.907241778 | 0.056489286 | 0.13278272 |
| ENSG00000154473 | BUB3 | 3517.85766 | -0.515408225 | 0.048840187 | -10.55295347 | 4.92E-26 | 1.15E-24 |
| ENSG00000154478 | GPR26 | 2.80606308 | 0.689058847 | 1.05262431 | 0.654610425 | 0.51271861 | 0.692235002 |
| ENSG00000154479 | CFAP210 | 9.769528806 | -0.284892963 | 0.578875815 | -0.492148671 | 0.622614247 | 0.776597778 |
| ENSG00000154485 | MMP21 | 2.136547723 | -0.453446665 | 1.243244476 | -0.364728478 | 0.715314099 | 0.841656152 |
| ENSG00000154511 | DIPK1A | 44.49705003 | 0.103696191 | 0.263591103 | 0.393397915 | 0.694025609 | 0.827653275 |
| ENSG00000154518 | ATP5MC3 | 3039.482682 | 0.111275317 | 0.049314617 | 2.256436805 | 0.024043289 | 0.064283572 |
| ENSG00000154545 | MAGED4 | 6.008676951 | -0.386323272 | 0.788712549 | -0.489815045 | 0.624264784 | 0.777957092 |
| ENSG00000154582 | ELOC | 1211.110508 | 0.428337521 | 0.06223827 | 6.882220866 | 5.89E-12 | 5.64E-11 |
| ENSG00000154608 | CEP170P1 | 6.860385175 | 0.575463497 | 0.694973762 | 0.828036293 | 0.407649947 | 0.600416679 |
| ENSG00000154639 | CXADR | 591.7664399 | 0.717086357 | 0.087018693 | 8.24060128 | 1.71E-16 | 2.30E-15 |
| ENSG00000154640 | BTG3 | 294.2315113 | -0.133479581 | 0.10765377 | -1.239896956 | 0.21501351 | 0.383994544 |
| ENSG00000154642 | C21orf91 | 558.1663512 | 0.1221131 | 0.095241691 | 1.282139135 | 0.199793843 | 0.362858899 |
| ENSG00000154645 | CHODL | 11.59578183 | 0.355594118 | 0.524614342 | 0.677820048 | 0.497885798 | 0.679791539 |
| ENSG00000154646 | TMPRSS15 | 26.48698696 | -1.728588842 | 0.37312006 | -4.632795245 | 3.61E-06 | 1.96E-05 |
| ENSG00000154654 | NCAM2 | 3227.813903 | 1.015120652 | 0.055404257 | 18.32206966 | 5.52E-75 | 5.86E-73 |
| ENSG00000154710 | RABGEF1 | 68.84299436 | 0.494875266 | 0.217168611 | 2.278760569 | 0.022681302 | 0.061004698 |
| ENSG00000154719 | MRPL39 | 587.2176277 | -0.189427997 | 0.08204186 | -2.308918843 | 0.020948085 | 0.056916863 |
| ENSG00000154721 | JAM2 | 6.088058525 | -1.566966415 | 0.761557134 | -2.057582215 | 0.039630255 | 0.098421355 |
| ENSG00000154723 | ATP5PF | 1714.708506 | -0.053427797 | 0.055853494 | -0.956570369 | 0.338784151 | 0.530045518 |
| ENSG00000154727 | GABPA | 567.3661701 | 0.112126203 | 0.087518974 | 1.281164505 | 0.200135891 | 0.363307536 |
| ENSG00000154734 | ADAMTS1 | 69.3722914 | -2.299746343 | 0.256055317 | -8.981443422 | 2.67E-19 | 4.30E-18 |
| ENSG00000154743 | TSEN2 | 560.9193279 | 0.251062603 | 0.086295313 | 2.909342286 | 0.003621901 | 0.01205024 |
| ENSG00000154767 | XPC | 3268.939032 | 0.0421985 | 0.05226018 | 0.807469477 | 0.419396053 | 0.610892467 |
| ENSG00000154781 | CCDC174 | 1080.407779 | -0.235779482 | 0.065214883 | -3.61542444 | 0.000299856 | 0.001234289 |
| ENSG00000154783 | FGD5 | 17.63668944 | -3.544550067 | 0.592303652 | -5.984346123 | 2.17E-09 | 1.68E-08 |

| | | | | | | | |
|---|---|---|---|---|---|---|---|
| ENSG00000154803 | FLCN | 953.0452575 | 0.344426884 | 0.074341454 | 4.633039387 | 3.60E-06 | 1.96E-05 |
| ENSG00000154813 | DPH3 | 825.3516265 | -0.076820768 | 0.074151511 | -1.035997346 | 0.300203381 | 0.486862181 |
| ENSG00000154814 | OXNAD1 | 598.4008163 | 0.102464271 | 0.080187736 | 1.27780476 | 0.201318275 | 0.364934116 |
| ENSG00000154822 | PLCL2 | 1.826861464 | -0.413175538 | 1.286372949 | -0.321194206 | 0.748063223 | 0.860436606 |
| ENSG00000154832 | CXXC1 | 1736.011777 | 0.241348932 | 0.059352948 | 4.066334343 | 4.78E-05 | 0.00022409 |
| ENSG00000154839 | SKA1 | 338.0150598 | 0.348186176 | 0.104137532 | 3.34352246 | 0.00082722 | 0.003144917 |
| ENSG00000154845 | PPP4R1 | 2477.941925 | -0.011367009 | 0.051352615 | -0.221352097 | 0.824818288 | 0.907715593 |
| ENSG00000154856 | APCDD1 | 121.615079 | 0.532889228 | 0.172812026 | 3.083635097 | 0.002044882 | 0.007172722 |
| ENSG00000154874 | CCDC144BP | 27.93538808 | -0.473490973 | 0.334275564 | -1.416468998 | 0.156638235 | 0.300995234 |
| ENSG00000154889 | MPPE1 | 497.0032237 | 0.111278972 | 0.089483999 | 1.243562787 | 0.213660519 | 0.381905276 |
| ENSG00000154898 | CCDC144CP | 15.56936344 | 0.40875901 | 0.449558732 | 0.909244957 | 0.363220838 | 0.556004673 |
| ENSG00000154914 | USP43 | 401.5358322 | -0.262254684 | 0.102443644 | -2.559989798 | 0.010467524 | 0.030920035 |
| ENSG00000154917 | RAB6B | 392.8202368 | 1.113499968 | 0.102900038 | 10.82118131 | 2.73E-27 | 6.87E-26 |
| ENSG00000154920 | EME1 | 389.6129785 | -0.547740506 | 0.107318002 | -5.103901473 | 3.33E-07 | 2.04E-06 |
| ENSG00000154928 | EPHB1 | 15.4031992 | -1.484885367 | 0.480292182 | -3.091629265 | 0.001990613 | 0.007001634 |
| ENSG00000154930 | ACSS1 | 1059.235519 | 1.151819228 | 0.072673701 | 15.84918904 | 1.42E-56 | 9.25E-55 |
| ENSG00000154945 | ANKRD40 | 2793.46791 | -0.10245479 | 0.049063193 | -2.088221003 | 0.036777904 | 0.092489049 |
| ENSG00000154957 | ZNF18 | 251.947871 | 0.009083631 | 0.123918433 | 0.073303308 | 0.94156476 | 0.972202592 |
| ENSG00000154978 | VOPP1 | 935.4290686 | 0.68102308 | 0.071130281 | 9.574306027 | 1.03E-21 | 1.89E-20 |
| ENSG00000155008 | APOOL | 1219.65334 | 0.249445151 | 0.069645546 | 3.581638238 | 0.000341446 | 0.001389037 |
| ENSG00000155016 | CYP2U1 | 298.5201046 | 0.25514901 | 0.110449785 | 2.310090604 | 0.020883139 | 0.056767294 |
| ENSG00000155026 | RSPH10B | 4.30297065 | -0.330630814 | 0.855541023 | -0.386458165 | 0.69915738 | 0.831105311 |
| ENSG00000155034 | FBXL18 | 2527.285524 | 0.040712692 | 0.063634842 | 0.639786171 | 0.522311625 | 0.69916229 |
| ENSG00000155052 | CNTNAP5 | 2.878350689 | -1.452326364 | 1.075894634 | -1.349877877 | 0.177055159 | 0.330941526 |
| ENSG00000155066 | PROM2 | 1816.674145 | 1.086500001 | 0.060788748 | 17.87337352 | 1.90E-71 | 1.81E-69 |
| ENSG00000155085 | AK9 | 256.7787167 | 0.077175247 | 0.117046647 | 0.659354618 | 0.509668077 | 0.689658803 |
| ENSG00000155090 | KLF10 | 1304.561939 | -0.055128502 | 0.060659019 | -0.908826143 | 0.363441905 | 0.556231723 |
| ENSG00000155093 | PTPRN2 | 2.312886064 | -0.610472577 | 1.134016767 | -0.538327646 | 0.590350868 | 0.753103247 |
| ENSG00000155096 | AZIN1 | 8136.405161 | -0.189599973 | 0.044823063 | -4.229964692 | 2.34E-05 | 0.000114812 |
| ENSG00000155097 | ATP6V1C1 | 3763.350109 | -0.120196749 | 0.050799251 | -2.366112608 | 0.017975976 | 0.049827077 |
| ENSG00000155099 | PIP4P2 | 153.3357828 | 0.145010677 | 0.149023575 | 0.973072064 | 0.330517487 | 0.521212013 |
| ENSG00000155100 | OTUD6B | 828.5269146 | 0.185643843 | 0.075622077 | 2.454889512 | 0.01409279 | 0.040240054 |
| ENSG00000155111 | CDK19 | 1830.561029 | 0.164177668 | 0.06350499 | 2.58527192 | 0.009730223 | 0.029026178 |

| | | | | | | | |
|---|---|---|---|---|---|---|---|
| ENSG00000155115 | GTF3C6 | 842.386369 | -0.241101949 | 0.074359172 | -3.242396917 | 0.001185288 | 0.004379209 |
| ENSG00000155158 | TTC39B | 512.1985597 | 0.675498073 | 0.089467595 | 7.550198168 | 4.35E-14 | 4.86E-13 |
| ENSG00000155189 | AGPAT5 | 1024.985629 | 0.103877465 | 0.069923634 | 1.48558447 | 0.137389065 | 0.272686032 |
| ENSG00000155229 | MMS19 | 3615.238461 | -0.571661925 | 0.0503181 | -11.36096007 | 6.54E-30 | 1.82E-28 |
| ENSG00000155252 | PI4K2A | 1190.261545 | -0.730384152 | 0.068262485 | -10.69964201 | 1.02E-26 | 2.50E-25 |
| ENSG00000155254 | MARVELD1 | 2970.786777 | -0.559253484 | 0.053855938 | -10.38424919 | 2.92E-25 | 6.59E-24 |
| ENSG00000155256 | ZFYVE27 | 1351.44246 | -0.326422304 | 0.065927397 | -4.951239059 | 7.37E-07 | 4.35E-06 |
| ENSG00000155265 | GOLGA7B | 101.3971935 | -1.149987549 | 0.18535611 | -6.204206328 | 5.50E-10 | 4.49E-09 |
| ENSG00000155269 | GPR78 | 3.063651683 | 0.289938718 | 0.998183146 | 0.290466453 | 0.771459411 | 0.87553945 |
| ENSG00000155275 | TRMT44 | 374.8541128 | -0.013835742 | 0.100779869 | -0.137286765 | 0.890804131 | 0.944056489 |
| ENSG00000155287 | SLC25A28 | 737.0471969 | -0.050764036 | 0.077694302 | -0.653381707 | 0.513510229 | 0.692692264 |
| ENSG00000155304 | HSPA13 | 639.1069717 | 0.014285904 | 0.093007244 | 0.153599907 | 0.87792521 | 0.936420901 |
| ENSG00000155313 | USP25 | 1187.976659 | 0.244080598 | 0.067658023 | 3.607563254 | 0.000309086 | 0.00127046 |
| ENSG00000155324 | GRAMD2B | 309.4012033 | 0.310152977 | 0.106034269 | 2.92502583 | 0.003444276 | 0.011522779 |
| ENSG00000155329 | ZCCHC10 | 447.7680784 | 0.156110799 | 0.0903528 | 1.727791486 | 0.084025614 | 0.183978963 |
| ENSG00000155330 | C16orf87 | 230.1882051 | -0.222772235 | 0.122997652 | -1.811190954 | 0.070111298 | 0.158755619 |
| ENSG00000155363 | MOV10 | 2621.056073 | 0.045937921 | 0.057976684 | 0.792351646 | 0.428155666 | 0.61877975 |
| ENSG00000155366 | RHOC | 3289.411002 | 0.127358886 | 0.061114602 | 2.083935464 | 0.037166038 | 0.093281062 |
| ENSG00000155367 | PPM1J | 54.28810488 | 1.046143523 | 0.255538969 | 4.093870795 | 4.24E-05 | 0.000200326 |
| ENSG00000155368 | DBI | 4826.783311 | -0.159544536 | 0.050662022 | -3.149194032 | 0.001637215 | 0.005867474 |
| ENSG00000155380 | SLC16A1 | 483.0135395 | 0.005809176 | 0.091720777 | 0.063335441 | 0.949499394 | 0.976618218 |
| ENSG00000155393 | HEATR3 | 579.954187 | 0.473234653 | 0.085389236 | 5.542087946 | 2.99E-08 | 2.04E-07 |
| ENSG00000155428 | TRIM74 | 5.468301965 | 0.344669918 | 0.751801318 | 0.458458785 | 0.646622867 | 0.793491418 |
| ENSG00000155438 | NIFK | 1444.894102 | 0.063483634 | 0.059714532 | 1.063118678 | 0.287728135 | 0.471631541 |
| ENSG00000155463 | OXA1L | 3665.463522 | -0.220456103 | 0.052713937 | -4.18212177 | 2.89E-05 | 0.000140214 |
| ENSG00000155465 | SLC7A7 | 2.694855047 | -1.75828211 | 1.137404415 | -1.545872415 | 0.122135384 | 0.248855268 |
| ENSG00000155506 | LARP1 | 13524.30534 | 0.114924302 | 0.043129349 | 2.664642597 | 0.007707018 | 0.023648877 |
| ENSG00000155508 | CNOT8 | 1489.417304 | -0.33885049 | 0.065524555 | -5.171351221 | 2.32E-07 | 1.45E-06 |
| ENSG00000155530 | LRGUK | 8.26700483 | 1.134660766 | 0.653576481 | 1.736079555 | 0.082549769 | 0.181258536 |
| ENSG00000155542 | SETD9 | 115.0243058 | 0.307389113 | 0.177857708 | 1.728286708 | 0.083936835 | 0.183854769 |
| ENSG00000155545 | MIER3 | 1041.044004 | -0.025525544 | 0.069587044 | -0.3668146 | 0.713757316 | 0.840814785 |
| ENSG00000155561 | NUP205 | 4265.96022 | -0.384984583 | 0.049290065 | -7.810591952 | 5.69E-15 | 6.82E-14 |
| ENSG00000155592 | ZKSCAN2 | 622.0246566 | -0.023099586 | 0.08274733 | -0.279158087 | 0.780123507 | 0.880664282 |

| | | | | | | | |
|---|---|---|---|---|---|---|---|
| ENSG00000155621 | C9orf85 | 271.9393362 | 0.076475888 | 0.114785668 | 0.666249454 | 0.505251668 | 0.685867404 |
| ENSG00000155629 | PIK3AP1 | 10.67179927 | 0.170319829 | 0.53637744 | 0.317537271 | 0.750835963 | 0.862329879 |
| ENSG00000155636 | RBM45 | 434.2118285 | -0.122641886 | 0.094379443 | -1.299455491 | 0.193787659 | 0.354483437 |
| ENSG00000155657 | TTN | 6.189421121 | 0.060692376 | 0.751351144 | 0.080777645 | 0.935618786 | 0.969248877 |
| ENSG00000155660 | PDIA4 | 5806.101986 | 0.308465423 | 0.045883625 | 6.722778021 | 1.78E-11 | 1.63E-10 |
| ENSG00000155666 | KDM8 | 149.4298984 | -0.03235756 | 0.148720446 | -0.217573042 | 0.827761791 | 0.908992603 |
| ENSG00000155729 | KCTD18 | 750.1565413 | 0.061683217 | 0.075078145 | 0.821586856 | 0.411312073 | 0.603671145 |
| ENSG00000155744 | HYCC2 | 524.872154 | -0.495014441 | 0.085532367 | -5.787451659 | 7.15E-09 | 5.22E-08 |
| ENSG00000155749 | FLACC1 | 10.46191762 | -0.51206515 | 0.555133129 | -0.922418648 | 0.356310246 | 0.548353485 |
| ENSG00000155755 | TMEM237 | 495.1204489 | 0.021041621 | 0.088231823 | 0.238481088 | 0.811507982 | 0.900221351 |
| ENSG00000155760 | FZD7 | 295.3017245 | 0.359533027 | 0.108826143 | 3.303737649 | 0.000954051 | 0.003584331 |
| ENSG00000155761 | SPAG17 | 13.78372255 | -0.348134924 | 0.511765951 | -0.680261989 | 0.496338588 | 0.678222969 |
| ENSG00000155792 | DEPTOR | 383.5173072 | 0.775131962 | 0.099050566 | 7.825618706 | 5.05E-15 | 6.09E-14 |
| ENSG00000155816 | FMN2 | 1.829137984 | -3.349666097 | 1.609297752 | -2.081445832 | 0.037393117 | 0.093799684 |
| ENSG00000155827 | RNF20 | 2175.273902 | 0.006285093 | 0.056045551 | 0.112142587 | 0.910710351 | 0.955221978 |
| ENSG00000155846 | PPARGC1B | 212.3134423 | 1.113884985 | 0.13269786 | 8.394144287 | 4.69E-17 | 6.48E-16 |
| ENSG00000155849 | ELMO1 | 39.76215247 | 2.156923631 | 0.325739467 | 6.621622023 | 3.55E-11 | 3.18E-10 |
| ENSG00000155850 | SLC26A2 | 542.1968761 | -0.618708735 | 0.094505537 | -6.546798815 | 5.88E-11 | 5.18E-10 |
| ENSG00000155858 | LSM11 | 490.0166423 | 0.088414631 | 0.086636713 | 1.020521536 | 0.307481181 | 0.495407943 |
| ENSG00000155868 | MED7 | 376.6250065 | 0.113887427 | 0.097914104 | 1.163136084 | 0.244774294 | 0.421650695 |
| ENSG00000155875 | SAXO1 | 3.077830267 | -0.328428052 | 0.997363536 | -0.32929623 | 0.741931793 | 0.857590509 |
| ENSG00000155876 | RRAGA | 1355.826738 | -0.004487913 | 0.064676004 | -0.069390697 | 0.944678633 | 0.973711185 |
| ENSG00000155893 | PXYLP1 | 403.5943976 | 0.566920608 | 0.099210908 | 5.714297141 | 1.10E-08 | 7.89E-08 |
| ENSG00000155903 | RASA2 | 415.2943264 | -0.288937907 | 0.1043608 | -2.768644055 | 0.005629009 | 0.017887447 |
| ENSG00000155906 | RMND1 | 5888.655355 | 0.102695525 | 0.049325151 | 2.082011361 | 0.037341432 | 0.093680277 |
| ENSG00000155957 | TMBIM4 | 223.2477987 | 0.363535907 | 0.139006991 | 2.615234702 | 0.008916616 | 0.026899944 |
| ENSG00000155959 | VBP1 | 1956.377033 | -0.67775735 | 0.064835822 | -10.45343962 | 1.41E-25 | 3.22E-24 |
| ENSG00000155961 | RAB39B | 605.0175645 | -0.488940332 | 0.082995674 | -5.891154414 | 3.84E-09 | 2.89E-08 |
| ENSG00000155962 | CLIC2 | 37.7237733 | -0.793910841 | 0.294553631 | -2.695301491 | 0.007032497 | 0.02178602 |
| ENSG00000155966 | AFF2 | 157.3971475 | 0.036615894 | 0.151624152 | 0.241491169 | 0.80917446 | 0.898908577 |
| ENSG00000155974 | GRIP1 | 564.7213217 | 0.726369641 | 0.08296171 | 8.755480622 | 2.03E-18 | 3.08E-17 |
| ENSG00000155975 | VPS37A | 597.6940391 | 0.15907985 | 0.08089924 | 1.96639487 | 0.049253016 | 0.118236101 |
| ENSG00000155980 | KIF5A | 32.82205494 | -4.671971772 | 0.57198375 | -8.168014865 | 3.14E-16 | 4.12E-15 |

| | | | | | | | |
|---|---|---|---|---|---|---|---|
| ENSG00000156006 | NAT2 | 2.537032849 | 1.380000078 | 1.155878378 | 1.193897303 | 0.232518143 | 0.406552301 |
| ENSG00000156011 | PSD3 | 1096.285808 | -0.831857514 | 0.079020798 | -10.5270705 | 6.48E-26 | 1.50E-24 |
| ENSG00000156017 | CARNMT1 | 352.6067447 | 0.283696664 | 0.100951217 | 2.810235211 | 0.004950531 | 0.015970352 |
| ENSG00000156026 | MCU | 660.2097202 | -0.533486301 | 0.081517553 | -6.54443467 | 5.97E-11 | 5.26E-10 |
| ENSG00000156030 | MIDEAS | 4050.155521 | -0.555721247 | 0.048980087 | -11.34586065 | 7.78E-30 | 2.15E-28 |
| ENSG00000156042 | CFAP70 | 11.09617577 | -0.371404805 | 0.549111548 | -0.676374056 | 0.498803185 | 0.68045582 |
| ENSG00000156049 | GNA14 | 265.5865414 | -0.254008009 | 0.115627244 | -2.19678339 | 0.02803592 | 0.073292764 |
| ENSG00000156050 | FAM161B | 590.3329902 | -1.004228456 | 0.088822962 | -11.30595554 | 1.23E-29 | 3.36E-28 |
| ENSG00000156052 | GNAQ | 2273.118009 | -0.216182884 | 0.054448285 | -3.970425979 | 7.17E-05 | 0.000327851 |
| ENSG00000156103 | MMP16 | 2154.290512 | 0.823874802 | 0.063859033 | 12.90146061 | 4.42E-38 | 1.67E-36 |
| ENSG00000156110 | ADK | 1481.086195 | -0.691687908 | 0.062346001 | -11.09434285 | 1.34E-28 | 3.50E-27 |
| ENSG00000156113 | KCNMA1 | 455.5547366 | -4.731459616 | 0.160134505 | -29.54678395 | 7.22E-192 | 3.94E-189 |
| ENSG00000156127 | BATF | 528.6452388 | -0.266130545 | 0.092229509 | -2.88552491 | 0.003907615 | 0.01290537 |
| ENSG00000156136 | DCK | 427.7605724 | -0.122626712 | 0.101199657 | -1.211730509 | 0.225615557 | 0.397621757 |
| ENSG00000156140 | ADAMTS3 | 3.795135466 | 0.194361732 | 0.926559338 | 0.209767172 | 0.833849396 | 0.912258208 |
| ENSG00000156162 | DPY19L4 | 2205.978536 | -0.441912341 | 0.064724124 | -6.827629567 | 8.63E-12 | 8.14E-11 |
| ENSG00000156170 | NDUFAF6 | 1150.878236 | -0.201157215 | 0.065195197 | -3.08546066 | 0.002032371 | 0.0071332 |
| ENSG00000156171 | DRAM2 | 585.8957462 | 0.318225668 | 0.085087858 | 3.739965654 | 0.000184045 | 0.000786366 |
| ENSG00000156172 | CFAP418 | 141.1559463 | -0.478084824 | 0.152533489 | -3.134294165 | 0.001722681 | 0.006136381 |
| ENSG00000156206 | CFAP161 | 14.57850909 | -0.515555833 | 0.457244931 | -1.127526623 | 0.259519891 | 0.439484833 |
| ENSG00000156218 | ADAMTSL3 | 22.34838026 | -0.808646784 | 0.391787742 | -2.063992044 | 0.039018471 | 0.097186071 |
| ENSG00000156232 | WHAMM | 653.1519447 | -0.224046673 | 0.078904381 | -2.839470646 | 0.004518845 | 0.014722669 |
| ENSG00000156239 | N6AMT1 | 461.1501503 | 0.186359496 | 0.091391616 | 2.039131204 | 0.041436935 | 0.102057471 |
| ENSG00000156253 | RWDD2B | 1075.296123 | -0.275106868 | 0.070137686 | -3.922383042 | 8.77E-05 | 0.000394764 |
| ENSG00000156256 | USP16 | 1485.634506 | 0.031198769 | 0.062463129 | 0.499474958 | 0.617444824 | 0.773221455 |
| ENSG00000156261 | CCT8 | 5924.152477 | -0.026467643 | 0.04547536 | -0.582021615 | 0.560552122 | 0.729849205 |
| ENSG00000156273 | BACH1 | 1102.173555 | -0.96673241 | 0.067361691 | -14.35136789 | 1.04E-46 | 5.28E-45 |
| ENSG00000156298 | TSPAN7 | 1.679837707 | -0.211068775 | 1.368167696 | -0.154271129 | 0.87739596 | NA |
| ENSG00000156299 | TIAM1 | 2256.640108 | -0.700065798 | 0.058161602 | -12.0365631 | 2.28E-33 | 7.22E-32 |
| ENSG00000156304 | SCAF4 | 2138.115424 | -0.052244731 | 0.057571822 | -0.907470521 | 0.364158032 | 0.556807655 |
| ENSG00000156313 | RPGR | 174.7058097 | 0.125532107 | 0.138182994 | 0.90844831 | 0.363641412 | 0.556351473 |
| ENSG00000156345 | CDK20 | 370.9833331 | 0.359887117 | 0.097995534 | 3.672484865 | 0.000240203 | 0.001004057 |
| ENSG00000156374 | PCGF6 | 188.1300283 | -0.078493435 | 0.134509337 | -0.583553798 | 0.559520552 | 0.72907705 |

| | | | | | | | |
|---|---|---|---|---|---|---|---|
| ENSG00000156381 | ANKRD9 | 1805.218518 | -0.306004358 | 0.062966792 | -4.859773647 | 1.18E-06 | 6.75E-06 |
| ENSG00000156384 | SFR1 | 55.39003553 | -0.468991966 | 0.245827839 | -1.907806565 | 0.056416222 | 0.13268281 |
| ENSG00000156398 | SFXN2 | 459.0774814 | 0.159733313 | 0.09812882 | 1.627792047 | 0.103568987 | 0.21809001 |
| ENSG00000156411 | ATP5MJ | 2369.249347 | 0.125317085 | 0.05291191 | 2.368409775 | 0.017864737 | 0.049590642 |
| ENSG00000156427 | FGF18 | 5.928864873 | -0.641781155 | 0.732266092 | -0.876431617 | 0.380795436 | 0.573267313 |
| ENSG00000156453 | PCDH1 | 1414.389228 | 0.206482215 | 0.071796455 | 2.875938846 | 0.004028277 | 0.01326377 |
| ENSG00000156463 | SH3RF2 | 28.40569847 | -1.59714076 | 0.361632246 | -4.416477729 | 1.00E-05 | 5.19E-05 |
| ENSG00000156467 | UQCRB | 4813.668382 | -0.05951171 | 0.049769975 | -1.19573519 | 0.231799914 | 0.405746011 |
| ENSG00000156469 | MTERF3 | 457.6051478 | -0.247408003 | 0.102848511 | -2.40555745 | 0.016147813 | 0.045390372 |
| ENSG00000156471 | PTDSS1 | 4337.73373 | 0.161080468 | 0.046337195 | 3.476267113 | 0.000508446 | 0.002009258 |
| ENSG00000156475 | PPP2R2B | 58.24393665 | 1.955674612 | 0.277133165 | 7.056804673 | 1.70E-12 | 1.71E-11 |
| ENSG00000156482 | RPL30 | 12358.11613 | -0.136781986 | 0.05504009 | -2.485133758 | 0.012950277 | 0.037316838 |
| ENSG00000156500 | PABIR3 | 146.4312666 | -0.202284599 | 0.150355148 | -1.345378607 | 0.178503 | 0.333156579 |
| ENSG00000156502 | SUPV3L1 | 825.5718923 | -0.539810796 | 0.077405124 | -6.973838037 | 3.08E-12 | 3.03E-11 |
| ENSG00000156504 | PABIR2 | 3205.259658 | 0.60933505 | 0.05838906 | 10.43577435 | 1.70E-25 | 3.87E-24 |
| ENSG00000156508 | EEF1A1 | 27373.44366 | 0.106225988 | 0.112277489 | 0.946102277 | 0.344096423 | 0.535012331 |
| ENSG00000156509 | FBXO43 | 10.95647089 | -1.25818901 | 0.563193704 | -2.234025345 | 0.025481405 | 0.067514835 |
| ENSG00000156515 | HK1 | 7298.922424 | -0.729361441 | 0.042011528 | -17.36098352 | 1.63E-67 | 1.38E-65 |
| ENSG00000156521 | TYSND1 | 1265.836019 | -0.773175567 | 0.066030881 | -11.70930264 | 1.14E-31 | 3.40E-30 |
| ENSG00000156531 | PHF6 | 1753.491417 | 0.035873034 | 0.060608042 | 0.591885708 | 0.553927125 | 0.725418397 |
| ENSG00000156535 | CD109 | 56.0391643 | -0.864781798 | 0.238174612 | -3.630873129 | 0.000282464 | 0.001166898 |
| ENSG00000156564 | LRFN2 | 4.798888301 | -0.839454488 | 0.849320986 | -0.988383076 | 0.322965073 | 0.512862991 |
| ENSG00000156587 | UBE2L6 | 343.0881343 | -0.553133185 | 0.105039556 | -5.265951294 | 1.39E-07 | 8.93E-07 |
| ENSG00000156599 | ZDHHC5 | 3460.464431 | -0.042924511 | 0.052842692 | -0.812307435 | 0.416615245 | 0.60862208 |
| ENSG00000156603 | MED19 | 324.8272128 | -0.056582457 | 0.105433188 | -0.536666476 | 0.591498016 | 0.753962137 |
| ENSG00000156639 | ZFAND3 | 3622.980777 | 0.175534126 | 0.048642271 | 3.608674532 | 0.000307765 | 0.001265712 |
| ENSG00000156642 | NPTN | 2949.403101 | 0.314265385 | 0.048616072 | 6.464228176 | 1.02E-10 | 8.79E-10 |
| ENSG00000156650 | KAT6B | 986.6025115 | -0.339034691 | 0.069027238 | -4.911607348 | 9.03E-07 | 5.26E-06 |
| ENSG00000156671 | SAMD8 | 527.3594721 | -0.784535063 | 0.089243471 | -8.790951891 | 1.48E-18 | 2.26E-17 |
| ENSG00000156675 | RAB11FIP1 | 1255.612472 | 0.32493024 | 0.063922294 | 5.08320681 | 3.71E-07 | 2.26E-06 |
| ENSG00000156697 | UTP14A | 1865.062953 | -0.349214542 | 0.055269282 | -6.318420077 | 2.64E-10 | 2.21E-09 |
| ENSG00000156709 | AIFM1 | 3692.510469 | -0.158409741 | 0.049457225 | -3.202964613 | 0.001360207 | 0.004961544 |
| ENSG00000156711 | MAPK13 | 1550.922993 | -0.522503386 | 0.062578013 | -8.349632041 | 6.85E-17 | 9.39E-16 |

| | | | | | | | |
|---|---|---|---|---|---|---|---|
| ENSG00000156735 | BAG4 | 481.084949 | -0.348830763 | 0.087148268 | -4.002727429 | 6.26E-05 | 0.00028867 |
| ENSG00000156787 | TBC1D31 | 375.6496324 | -0.442327563 | 0.110731923 | -3.994580336 | 6.48E-05 | 0.000297939 |
| ENSG00000156795 | NTAQ1 | 512.5316657 | -0.189674213 | 0.092307608 | -2.05480586 | 0.03989776 | 0.098935749 |
| ENSG00000156802 | ATAD2 | 6303.989582 | -0.389805053 | 0.046918861 | -8.308067228 | 9.73E-17 | 1.32E-15 |
| ENSG00000156804 | FBXO32 | 266.222767 | 0.792039725 | 0.117199927 | 6.758022345 | 1.40E-11 | 1.29E-10 |
| ENSG00000156831 | NSMCE2 | 1073.176131 | -0.65295254 | 0.066442121 | -9.827388616 | 8.58E-23 | 1.69E-21 |
| ENSG00000156853 | ZNF689 | 615.4379183 | -0.339579768 | 0.08627309 | -3.936102977 | 8.28E-05 | 0.000374711 |
| ENSG00000156858 | PRR14 | 1376.899205 | -0.183158804 | 0.06556765 | -2.793432496 | 0.005215191 | 0.016699677 |
| ENSG00000156860 | FBRS | 3930.76586 | -0.03535312 | 0.057248472 | -0.617538231 | 0.536879772 | 0.711908785 |
| ENSG00000156869 | FRRS1 | 23.22885446 | 0.636649169 | 0.378554904 | 1.681788197 | 0.092609918 | 0.199125647 |
| ENSG00000156873 | PHKG2 | 1453.823029 | -0.090223205 | 0.07217961 | -1.249981885 | 0.211306165 | 0.378848255 |
| ENSG00000156875 | MFSD14A | 42.75876205 | 0.265974385 | 0.280333941 | 0.948776962 | 0.342734061 | 0.533726113 |
| ENSG00000156876 | SASS6 | 206.7586662 | -0.202888007 | 0.12568341 | -1.614278353 | 0.106467063 | 0.222980747 |
| ENSG00000156925 | ZIC3 | 225.7429325 | -2.255765637 | 0.14134621 | -15.95915193 | 2.46E-57 | 1.65E-55 |
| ENSG00000156928 | MALSU1 | 681.90553 | -1.725028731 | 0.08114795 | -21.25782277 | 2.79E-100 | 4.51E-98 |
| ENSG00000156931 | VPS8 | 1480.965957 | -0.270491561 | 0.066517456 | -4.066474828 | 4.77E-05 | 0.000224001 |
| ENSG00000156958 | GALK2 | 683.5522793 | -0.327009785 | 0.077776382 | -4.204486942 | 2.62E-05 | 0.000127774 |
| ENSG00000156959 | LHFPL4 | 1.949855572 | -1.021636489 | 1.326864176 | -0.769963126 | 0.441321766 | 0.630616751 |
| ENSG00000156966 | B3GNT7 | 18.44510011 | -0.174460191 | 0.410083178 | -0.425426353 | 0.670525898 | 0.81098034 |
| ENSG00000156970 | BUB1B | 1545.431136 | -0.078346629 | 0.058064184 | -1.349310764 | 0.17723717 | 0.331225701 |
| ENSG00000156973 | PDE6D | 341.853129 | -0.150625114 | 0.108602366 | -1.386941363 | 0.165459644 | 0.314130947 |
| ENSG00000156976 | EIF4A2 | 7624.670953 | 0.443711274 | 0.049697236 | 8.928288705 | 4.33E-19 | 6.86E-18 |
| ENSG00000156983 | BRPF1 | 1980.127586 | -0.129789317 | 0.057468309 | -2.258450269 | 0.023917602 | 0.063962437 |
| ENSG00000156990 | RPUSD3 | 1133.775194 | -0.270960973 | 0.064799004 | -4.181560786 | 2.90E-05 | 0.00014053 |
| ENSG00000157014 | TATDN2 | 1555.598861 | 0.156914324 | 0.060432194 | 2.596535271 | 0.009416924 | 0.028223704 |
| ENSG00000157017 | GHRL | 11.76114956 | 0.686216034 | 0.526648907 | 1.302985774 | 0.192579618 | 0.352829362 |
| ENSG00000157020 | SEC13 | 4910.380723 | -0.281027013 | 0.047700212 | -5.891525423 | 3.83E-09 | 2.88E-08 |
| ENSG00000157021 | CIBAR1P1 | 103.853275 | 0.80855268 | 0.183210499 | 4.413244253 | 1.02E-05 | 5.26E-05 |
| ENSG00000157036 | EXOG | 269.9737658 | -0.001298957 | 0.115872574 | -0.011210218 | 0.991055728 | 0.995176993 |
| ENSG00000157045 | NTAN1 | 539.506027 | -0.228056472 | 0.084165005 | -2.709635325 | 0.006735722 | 0.020974273 |
| ENSG00000157064 | NMNAT2 | 701.4406808 | 1.010712709 | 0.08248752 | 12.25291668 | 1.62E-34 | 5.43E-33 |
| ENSG00000157077 | ZFYVE9 | 1231.401795 | 0.119752096 | 0.06205694 | 1.929713185 | 0.053642385 | 0.127334056 |
| ENSG00000157106 | SMG1 | 3081.528849 | -0.172730166 | 0.063766866 | -2.708776177 | 0.006753188 | 0.0210201 |

| | | | | | | | |
|---|---|---|---|---|---|---|---|
| ENSG00000157107 | FCHO2 | 510.187028 | 0.185799052 | 0.090285044 | 2.057916161 | 0.039598182 | 0.098352349 |
| ENSG00000157110 | RBPMS | 618.1191179 | -0.017467797 | 0.080336132 | -0.217433882 | 0.827870229 | 0.909057744 |
| ENSG00000157111 | TMEM171 | 2.853298559 | 1.647193535 | 1.133468461 | 1.453232791 | 0.146159131 | 0.285790186 |
| ENSG00000157152 | SYN2 | 6.756653672 | 0.396230004 | 0.683477327 | 0.579726625 | 0.562098985 | 0.730942164 |
| ENSG00000157181 | ODR4 | 829.8167424 | -0.159414112 | 0.080281866 | -1.98568019 | 0.047068846 | 0.11368281 |
| ENSG00000157184 | CPT2 | 1111.659692 | -0.328693317 | 0.069241946 | -4.747025981 | 2.06E-06 | 1.15E-05 |
| ENSG00000157191 | NECAP2 | 1203.978309 | -0.289072164 | 0.067827736 | -4.261857791 | 2.03E-05 | 0.000100631 |
| ENSG00000157193 | LRP8 | 3469.954082 | 0.027189994 | 0.053045193 | 0.512581682 | 0.60824397 | 0.766456043 |
| ENSG00000157212 | PAXIP1 | 1112.96812 | 0.515477706 | 0.066541711 | 7.746685424 | 9.43E-15 | 1.11E-13 |
| ENSG00000157214 | STEAP2 | 13.49512376 | 2.889416019 | 0.623018107 | 4.637772139 | 3.52E-06 | 1.92E-05 |
| ENSG00000157216 | SSBP3 | 1789.230422 | 0.003249454 | 0.057722785 | 0.056294135 | 0.955107491 | 0.979400378 |
| ENSG00000157224 | CLDN12 | 536.9123625 | -0.691004763 | 0.093173788 | -7.416300019 | 1.20E-13 | 1.31E-12 |
| ENSG00000157227 | MMP14 | 28.11333418 | -1.228681535 | 0.371525781 | -3.307123219 | 0.000942594 | 0.003545354 |
| ENSG00000157240 | FZD1 | 550.7717546 | -0.79432896 | 0.083217012 | -9.545271351 | 1.36E-21 | 2.49E-20 |
| ENSG00000157259 | GATAD1 | 1759.456 | -0.728483083 | 0.060466869 | -12.0476403 | 2.00E-33 | 6.32E-32 |
| ENSG00000157303 | SUSD3 | 20.73888658 | 2.879594016 | 0.497225735 | 5.791321352 | 6.98E-09 | 5.11E-08 |
| ENSG00000157306 | ZFHX2-AS1 | 44.70008809 | -0.634826883 | 0.278981975 | -2.27551218 | 0.022875223 | 0.061425449 |
| ENSG00000157322 | CLEC18A | 5.763782473 | 0.840751685 | 0.738516318 | 1.138433457 | 0.25493953 | 0.434366669 |
| ENSG00000157326 | DHRS4 | 459.6833648 | -0.195738673 | 0.095462506 | -2.050424629 | 0.040323012 | 0.099774554 |
| ENSG00000157343 | ARMC12 | 33.99927221 | 0.580685768 | 0.309309472 | 1.877361735 | 0.060468532 | 0.140442253 |
| ENSG00000157349 | DDX19B | 638.3339824 | 0.017479336 | 0.079096305 | 0.220988022 | 0.82510176 | 0.90772273 |
| ENSG00000157350 | ST3GAL2 | 85.96906224 | -0.206268817 | 0.208436294 | -0.989601249 | 0.322369059 | 0.512200181 |
| ENSG00000157353 | FCSK | 966.3201037 | 0.482735637 | 0.080170878 | 6.021334076 | 1.73E-09 | 1.35E-08 |
| ENSG00000157379 | DHRS1 | 765.8401341 | -0.177518675 | 0.075711052 | -2.3446864 | 0.019043089 | 0.052411333 |
| ENSG00000157388 | CACNA1D | 713.2794997 | -0.960634881 | 0.079311646 | -12.11215416 | 9.11E-34 | 2.93E-32 |
| ENSG00000157404 | KIT | 2.628721962 | 2.070265746 | 1.231497437 | 1.681096268 | 0.092744217 | 0.199377035 |
| ENSG00000157426 | AASDH | 438.2197846 | -0.365496407 | 0.093074615 | -3.926918256 | 8.60E-05 | 0.000388082 |
| ENSG00000157429 | ZNF19 | 118.5889676 | 0.57324945 | 0.177729697 | 3.225400479 | 0.001257966 | 0.004621674 |
| ENSG00000157445 | CACNA2D3 | 17.42576008 | -1.688259478 | 0.459418802 | -3.674772276 | 0.000238062 | 0.000996012 |
| ENSG00000157450 | RNF111 | 902.4545032 | -0.121438934 | 0.075160627 | -1.615725404 | 0.106153696 | 0.222470837 |
| ENSG00000157456 | CCNB2 | 1743.258107 | -0.402571085 | 0.058412885 | -6.891819913 | 5.51E-12 | 5.29E-11 |
| ENSG00000157470 | FAM81A | 79.21631183 | 1.128473028 | 0.207359605 | 5.442106369 | 5.27E-08 | 3.51E-07 |
| ENSG00000157483 | MYO1E | 451.1586523 | 0.256857503 | 0.093960848 | 2.733665242 | 0.006263369 | 0.019687816 |

| | | | | | | | |
|---|---|---|---|---|---|---|---|
| ENSG00000157500 | APPL1 | 1445.503667 | -0.136793724 | 0.061227778 | -2.234177506 | 0.025471396 | 0.06749611 |
| ENSG00000157502 | PWWP3B | 183.0934511 | -4.203510866 | 0.218257212 | -19.25943627 | 1.18E-82 | 1.39E-80 |
| ENSG00000157514 | TSC22D3 | 3699.438356 | -0.528883274 | 0.051648341 | -10.2400825 | 1.31E-24 | 2.86E-23 |
| ENSG00000157538 | VPS26C | 1476.529031 | 0.098601945 | 0.058191151 | 1.694449126 | 0.090179971 | 0.194722189 |
| ENSG00000157540 | DYRK1A | 2088.299193 | -0.180775866 | 0.054121279 | -3.340199452 | 0.000837182 | 0.003180007 |
| ENSG00000157551 | KCNJ15 | 3.008373541 | -0.849094727 | 1.014379548 | -0.837058208 | 0.40255985 | 0.595250932 |
| ENSG00000157557 | ETS2 | 1216.619348 | -0.53257692 | 0.066594999 | -7.997251022 | 1.27E-15 | 1.60E-14 |
| ENSG00000157578 | LCA5L | 175.0578739 | 0.418941772 | 0.144849516 | 2.892255253 | 0.003824871 | 0.012657567 |
| ENSG00000157593 | SLC35B2 | 2962.970793 | 0.048755319 | 0.056429017 | 0.864011488 | 0.38758158 | 0.580322528 |
| ENSG00000157600 | TMEM164 | 3341.247164 | 0.944175008 | 0.052387826 | 18.02279421 | 1.29E-72 | 1.27E-70 |
| ENSG00000157601 | MX1 | 55.62601583 | 0.126703432 | 0.238475697 | 0.531305428 | 0.595207145 | 0.756588059 |
| ENSG00000157613 | CREB3L1 | 94.63508906 | 0.373145327 | 0.185099667 | 2.015915715 | 0.04380879 | 0.107048748 |
| ENSG00000157617 | C2CD2 | 637.6927502 | 0.118641372 | 0.077867235 | 1.523636641 | 0.127599505 | 0.257581428 |
| ENSG00000157625 | TAB3 | 1912.543743 | -0.118571254 | 0.063259756 | -1.874355236 | 0.060881479 | 0.141266215 |
| ENSG00000157637 | SLC38A10 | 3399.215831 | 0.438282276 | 0.062005961 | 7.068389409 | 1.57E-12 | 1.58E-11 |
| ENSG00000157653 | C9orf43 | 45.58510371 | -0.078658568 | 0.268712708 | -0.292723661 | 0.769733383 | 0.874359188 |
| ENSG00000157654 | PALM2AKAP2 | 139.6151641 | -0.174573984 | 0.160522198 | -1.087537962 | 0.276799131 | 0.459362806 |
| ENSG00000157657 | ZNF618 | 456.8556179 | -0.194859099 | 0.097023697 | -2.008366054 | 0.044604407 | 0.10864569 |
| ENSG00000157680 | DGKI | 3.453791886 | -0.795485318 | 0.96101122 | -0.827758616 | 0.407807216 | 0.600511443 |
| ENSG00000157693 | TMEM268 | 1420.25096 | -0.537808235 | 0.059844397 | -8.986776665 | 2.55E-19 | 4.10E-18 |
| ENSG00000157703 | SVOPL | 28.5739417 | -0.431350723 | 0.329101546 | -1.310691879 | 0.189961884 | 0.349148755 |
| ENSG00000157734 | SNX22 | 260.8680659 | -0.349176435 | 0.141140383 | -2.473965471 | 0.013362261 | 0.038383455 |
| ENSG00000157741 | UBN2 | 845.6538438 | -0.142597135 | 0.077749741 | -1.834052872 | 0.066646146 | 0.152428972 |
| ENSG00000157764 | BRAF | 990.2850252 | -0.470207413 | 0.07646919 | -6.148978609 | 7.80E-10 | 6.26E-09 |
| ENSG00000157778 | PSMG3 | 1199.517333 | -1.47E-05 | 0.062594735 | -0.000234249 | 0.999813097 | 0.999914377 |
| ENSG00000157782 | CABP1 | 3.519517442 | -1.430360763 | 1.011359467 | -1.414295124 | 0.157275268 | 0.301851365 |
| ENSG00000157796 | WDR19 | 536.5013425 | 0.622972226 | 0.092700007 | 6.720304012 | 1.81E-11 | 1.66E-10 |
| ENSG00000157800 | SLC37A3 | 913.0059789 | -0.620445217 | 0.069494409 | -8.927987537 | 4.34E-19 | 6.87E-18 |
| ENSG00000157823 | AP3S2 | 684.7525634 | 0.249019049 | 0.080919484 | 3.077368242 | 0.002088371 | 0.007308503 |
| ENSG00000157827 | FMNL2 | 623.2037165 | 0.304087552 | 0.081752251 | 3.71962297 | 0.00019952 | 0.000846495 |
| ENSG00000157833 | GAREM2 | 36.29283107 | 0.714702337 | 0.305521311 | 2.339288005 | 0.019320533 | 0.053066694 |
| ENSG00000157837 | SPPL3 | 2210.515031 | 0.117948603 | 0.054843893 | 2.150624188 | 0.031505875 | 0.081033528 |
| ENSG00000157851 | DPYSL5 | 245.8275924 | -0.914734991 | 0.120362433 | -7.599837969 | 2.97E-14 | 3.37E-13 |

| | | | | | | | |
|---|---|---|---|---|---|---|---|
| ENSG00000157869 | RAB28 | 384.3437038 | 0.063058617 | 0.09888439 | 0.637700421 | 0.523668712 | 0.700010847 |
| ENSG00000157870 | PRXL2B | 1203.341384 | -0.147746555 | 0.067911388 | -2.175578495 | 0.029586793 | 0.07672632 |
| ENSG00000157881 | PANK4 | 585.7183098 | 0.036761095 | 0.089131519 | 0.412436539 | 0.680019486 | 0.818289926 |
| ENSG00000157895 | C12orf43 | 693.7172652 | 0.002944726 | 0.081721325 | 0.036033752 | 0.971255446 | 0.986933608 |
| ENSG00000157911 | PEX10 | 885.6517883 | 0.048957907 | 0.074201022 | 0.65980097 | 0.509381561 | 0.689403699 |
| ENSG00000157916 | RER1 | 2458.665355 | 0.050860676 | 0.050560448 | 1.005938004 | 0.314445393 | 0.502910429 |
| ENSG00000157927 | RADIL | 54.61795297 | -0.233114916 | 0.241809642 | -0.964043094 | 0.335024321 | 0.526221015 |
| ENSG00000157933 | SKI | 1602.383399 | 0.158603596 | 0.0600755 | 2.640071181 | 0.008288862 | 0.025204995 |
| ENSG00000157954 | WIPI2 | 3853.280378 | 0.543581762 | 0.054407441 | 9.990945237 | 1.67E-23 | 3.39E-22 |
| ENSG00000157978 | LDLRAP1 | 814.4091413 | 0.303638437 | 0.074564402 | 4.072163539 | 4.66E-05 | 0.000218778 |
| ENSG00000157985 | AGAP1 | 2345.809611 | -0.328336174 | 0.063083731 | -5.204767821 | 1.94E-07 | 1.23E-06 |
| ENSG00000157992 | KRTCAP3 | 77.79043391 | 0.473298528 | 0.219892695 | 2.152406786 | 0.03136533 | 0.080763348 |
| ENSG00000157999 | ANKRD61 | 6.732096825 | 0.646773116 | 0.691626835 | 0.935147516 | 0.349712282 | 0.540957502 |
| ENSG00000158006 | PAFAH2 | 574.9881536 | -0.180050086 | 0.081826031 | -2.200401053 | 0.027778453 | 0.072719128 |
| ENSG00000158008 | EXTL1 | 11.38523481 | 0.175322352 | 0.561076307 | 0.312475059 | 0.754679515 | 0.864498202 |
| ENSG00000158019 | BABAM2 | 642.2176915 | -0.388612817 | 0.080266325 | -4.841542435 | 1.29E-06 | 7.35E-06 |
| ENSG00000158023 | CFAP251 | 45.31746977 | 0.158336669 | 0.261398296 | 0.605729537 | 0.544694367 | 0.717913747 |
| ENSG00000158042 | MRPL17 | 957.5771809 | -0.131220175 | 0.067323118 | -1.949110193 | 0.051282267 | 0.122415733 |
| ENSG00000158050 | DUSP2 | 1021.327624 | -0.411281093 | 0.074711141 | -5.504950003 | 3.69E-08 | 2.50E-07 |
| ENSG00000158055 | GRHL3 | 1402.524043 | 1.237507218 | 0.07342706 | 16.85355816 | 9.88E-64 | 7.71E-62 |
| ENSG00000158062 | UBXN11 | 514.2892177 | -0.005827034 | 0.094803923 | -0.061464059 | 0.950989637 | 0.977409778 |
| ENSG00000158079 | PTPDC1 | 258.541774 | -0.041619511 | 0.116971636 | -0.355808578 | 0.721983925 | 0.845596895 |
| ENSG00000158089 | GALNT14 | 509.5094619 | -0.630646411 | 0.09858205 | -6.397172837 | 1.58E-10 | 1.35E-09 |
| ENSG00000158092 | NCK1 | 698.6893794 | -0.036483442 | 0.084993509 | -0.429249744 | 0.667741486 | 0.808785741 |
| ENSG00000158106 | RHPN1 | 4372.395007 | 0.506657315 | 0.059981372 | 8.446911075 | 2.99E-17 | 4.18E-16 |
| ENSG00000158109 | TPRG1L | 1366.560942 | 0.270090284 | 0.060478731 | 4.465872226 | 7.97E-06 | 4.16E-05 |
| ENSG00000158122 | PRXL2C | 251.4221221 | -0.153157211 | 0.120609839 | -1.26985669 | 0.204135683 | 0.36896254 |
| ENSG00000158156 | XKR8 | 313.9845012 | 0.200598278 | 0.109634016 | 1.829708379 | 0.067293557 | 0.153588121 |
| ENSG00000158158 | CNNM4 | 2594.415716 | 0.041341287 | 0.065283379 | 0.633258998 | 0.526564524 | 0.702576059 |
| ENSG00000158161 | EYA3 | 1215.055475 | 0.065562397 | 0.06749726 | 0.971334199 | 0.331381884 | 0.522252021 |
| ENSG00000158163 | DZIP1L | 14.12368721 | -0.662787279 | 0.475259074 | -1.394581008 | 0.163142225 | 0.310579473 |
| ENSG00000158164 | TMSB15A | 12.16060326 | -0.416884358 | 0.49918583 | -0.835128589 | 0.403645312 | 0.596512767 |
| ENSG00000158169 | FANCC | 673.3696832 | -0.094675636 | 0.078912248 | -1.19975844 | 0.230233169 | 0.403541328 |

| | | | | | | | |
|---|---|---|---|---|---|---|---|
| ENSG00000158186 | MRAS | 137.3713969 | 0.685070316 | 0.155903756 | 4.394187368 | 1.11E-05 | 5.70E-05 |
| ENSG00000158195 | WASF2 | 5921.444077 | -0.136663938 | 0.046507881 | -2.938511383 | 0.003297925 | 0.011080025 |
| ENSG00000158201 | ABHD3 | 300.5638589 | 0.347599246 | 0.108688576 | 3.198121254 | 0.001383261 | 0.005035228 |
| ENSG00000158220 | ESYT3 | 149.4793894 | 0.552838795 | 0.151229606 | 3.655625433 | 0.000256556 | 0.001066773 |
| ENSG00000158234 | FAIM | 501.3029227 | -0.433565104 | 0.086704816 | -5.000473119 | 5.72E-07 | 3.42E-06 |
| ENSG00000158246 | TENT5B | 121.0191993 | 0.12090681 | 0.163927229 | 0.737563922 | 0.460779486 | 0.647569773 |
| ENSG00000158258 | CLSTN2 | 7021.555413 | 0.859777411 | 0.061142081 | 14.06195856 | 6.51E-45 | 3.12E-43 |
| ENSG00000158286 | RNF207 | 124.3635153 | 0.340539693 | 0.174486673 | 1.951665917 | 0.050977882 | 0.121828587 |
| ENSG00000158290 | CUL4B | 3499.381149 | -0.17334026 | 0.053992586 | -3.210445615 | 0.001325293 | 0.004847541 |
| ENSG00000158292 | GPR153 | 399.9556244 | 0.357418256 | 0.107747657 | 3.317178906 | 0.000909314 | 0.0034303 |
| ENSG00000158296 | SLC13A3 | 57.36540313 | 1.622204751 | 0.250704605 | 6.470582198 | 9.76E-11 | 8.44E-10 |
| ENSG00000158301 | GPRASP2 | 576.5610742 | 0.069844427 | 0.0827239 | 0.844307708 | 0.39849748 | 0.591531784 |
| ENSG00000158321 | AUTS2 | 140.839318 | 4.072564282 | 0.25548366 | 15.94060567 | 3.31E-57 | 2.21E-55 |
| ENSG00000158352 | SHROOM4 | 32.3834042 | 1.744500241 | 0.33724262 | 5.172834443 | 2.31E-07 | 1.44E-06 |
| ENSG00000158373 | H2BC5 | 1156.766636 | -0.292084247 | 0.072172879 | -4.047008413 | 5.19E-05 | 0.000242195 |
| ENSG00000158402 | CDC25C | 438.2735916 | -0.215418345 | 0.091089484 | -2.364909059 | 0.018034499 | 0.049947048 |
| ENSG00000158406 | H4C8 | 618.8357725 | 0.197368117 | 0.078927122 | 2.500637461 | 0.012397001 | 0.035921082 |
| ENSG00000158411 | MITD1 | 281.0718881 | -0.229594721 | 0.117067608 | -1.961214754 | 0.049853977 | 0.11952866 |
| ENSG00000158417 | EIF5B | 4677.303481 | 0.012377162 | 0.051159568 | 0.241932491 | 0.808832474 | 0.898836887 |
| ENSG00000158423 | RIBC1 | 137.8004277 | -0.859975927 | 0.159356496 | -5.396553963 | 6.79E-08 | 4.46E-07 |
| ENSG00000158427 | TMSB15B | 31.15402614 | -0.167143727 | 0.329226085 | -0.507686769 | 0.611673029 | 0.769201803 |
| ENSG00000158435 | CNOT11 | 1828.354654 | -0.025013198 | 0.057572133 | -0.43446711 | 0.663949274 | 0.806043413 |
| ENSG00000158445 | KCNB1 | 25.23226978 | 0.925942524 | 0.359547627 | 2.575298668 | 0.010015356 | 0.029768369 |
| ENSG00000158457 | TSPAN33 | 2.489657673 | 0.609588728 | 1.135965655 | 0.53662602 | 0.591525967 | 0.753962137 |
| ENSG00000158458 | NRG2 | 3.986712403 | 0.739141023 | 0.939266759 | 0.786934081 | 0.431320456 | 0.621444037 |
| ENSG00000158467 | AHCYL2 | 945.7232659 | -0.03842709 | 0.075957145 | -0.505904871 | 0.612923433 | 0.769891065 |
| ENSG00000158470 | B4GALT5 | 3028.271996 | 0.239559192 | 0.051130823 | 4.685220728 | 2.80E-06 | 1.54E-05 |
| ENSG00000158480 | SPATA2 | 1329.828733 | -0.077284366 | 0.060847982 | -1.27012208 | 0.204041148 | 0.368820733 |
| ENSG00000158483 | FAM86C1P | 414.3676314 | -0.096872613 | 0.09406164 | -1.029884378 | 0.303064285 | 0.490150501 |
| ENSG00000158486 | DNAH3 | 90.99450547 | 0.662292954 | 0.199784337 | 3.315039429 | 0.000916302 | 0.003455526 |
| ENSG00000158526 | TSR2 | 2340.40983 | -0.151749068 | 0.053375197 | -2.843063403 | 0.004468218 | 0.014574282 |
| ENSG00000158528 | PPP1R9A | 650.9274696 | -0.205071544 | 0.083684725 | -2.450525398 | 0.01426479 | 0.040650221 |
| ENSG00000158545 | ZC3H18 | 3796.852129 | 0.420905794 | 0.048066783 | 8.756687492 | 2.01E-18 | 3.05E-17 |

| | | | | | | | |
|---|---|---|---|---|---|---|---|
| ENSG00000158552 | ZFAND2B | 759.215857 | 0.385859397 | 0.077177024 | 4.999666735 | 5.74E-07 | 3.43E-06 |
| ENSG00000158555 | GDPD5 | 163.6008468 | 0.518660395 | 0.149042771 | 3.479943323 | 0.00050152 | 0.001984282 |
| ENSG00000158560 | DYNC1I1 | 24.8637977 | -0.290151934 | 0.349201776 | -0.83090051 | 0.406029836 | 0.59868392 |
| ENSG00000158571 | PFKFB1 | 15.85621555 | -0.989974098 | 0.471279928 | -2.100607389 | 0.035675445 | 0.090062144 |
| ENSG00000158604 | TMED4 | 2373.509541 | 0.696271991 | 0.051979167 | 13.39521245 | 6.45E-41 | 2.74E-39 |
| ENSG00000158615 | PPP1R15B | 6060.170373 | 0.407043317 | 0.045168415 | 9.011680269 | 2.03E-19 | 3.29E-18 |
| ENSG00000158623 | COPG2 | 804.4070145 | 0.343333102 | 0.074080026 | 4.6346245 | 3.58E-06 | 1.95E-05 |
| ENSG00000158636 | EMSY | 1784.455325 | -0.192294581 | 0.057298326 | -3.35602443 | 0.000790716 | 0.003016133 |
| ENSG00000158639 | PAGE5 | 1.639630974 | -0.169059384 | 1.329067842 | -0.12720147 | 0.898780942 | NA |
| ENSG00000158669 | GPAT4 | 2310.367156 | 0.183540143 | 0.054130619 | 3.3906899 | 0.000697169 | 0.002688362 |
| ENSG00000158683 | PKD1L1 | 6.403426305 | 0.433763084 | 0.694341258 | 0.624711666 | 0.532160315 | 0.707691769 |
| ENSG00000158691 | ZSCAN12 | 85.26655029 | -0.732293597 | 0.19688755 | -3.719349423 | 0.000199737 | 0.000847172 |
| ENSG00000158710 | TAGLN2 | 5920.265767 | 0.54414167 | 0.047480182 | 11.46039568 | 2.09E-30 | 5.88E-29 |
| ENSG00000158711 | ELK4 | 3667.392633 | 0.207231818 | 0.057021802 | 3.634255833 | 0.000278784 | 0.001153763 |
| ENSG00000158715 | SLC45A3 | 162.3762376 | 1.228297992 | 0.156273299 | 7.859935133 | 3.84E-15 | 4.66E-14 |
| ENSG00000158716 | DUSP23 | 300.5047513 | 0.599808726 | 0.110945186 | 5.406351975 | 6.43E-08 | 4.24E-07 |
| ENSG00000158717 | RNF166 | 906.1389003 | 0.416735977 | 0.07556027 | 5.515279115 | 3.48E-08 | 2.37E-07 |
| ENSG00000158747 | NBL1 | 293.0719677 | 0.083387142 | 0.110971446 | 0.751428808 | 0.452394631 | 0.640066851 |
| ENSG00000158748 | HTR6 | 8.034800237 | 0.694696225 | 0.639905173 | 1.085623705 | 0.277645508 | 0.460401037 |
| ENSG00000158769 | F11R | 2378.164522 | 0.098448953 | 0.052361147 | 1.880190933 | 0.060082061 | 0.139671917 |
| ENSG00000158773 | USF1 | 1295.93064 | 0.296299143 | 0.063294454 | 4.681281294 | 2.85E-06 | 1.57E-05 |
| ENSG00000158786 | PLA2G2F | 2.077876872 | 0.540146277 | 1.247203779 | 0.433085825 | 0.664952421 | 0.806706252 |
| ENSG00000158792 | SPATA2L | 848.4210251 | 0.63606115 | 0.08947623 | 7.108716438 | 1.17E-12 | 1.19E-11 |
| ENSG00000158793 | NIT1 | 543.201496 | 0.060178494 | 0.086869484 | 0.692746071 | 0.488468921 | 0.67179119 |
| ENSG00000158796 | DEDD | 1257.688735 | 0.027078752 | 0.060905423 | 0.444603305 | 0.656606459 | 0.801029152 |
| ENSG00000158805 | ZNF276 | 2486.05274 | 0.597940548 | 0.066569335 | 8.982221988 | 2.65E-19 | 4.27E-18 |
| ENSG00000158813 | EDA | 124.1530493 | 0.003473526 | 0.161531975 | 0.021503643 | 0.982843898 | 0.992098874 |
| ENSG00000158828 | PINK1 | 1199.392463 | 0.092301652 | 0.069617843 | 1.32583327 | 0.18489493 | 0.341752453 |
| ENSG00000158850 | B4GALT3 | 1418.285676 | 0.551134994 | 0.068104792 | 8.092455377 | 5.85E-16 | 7.53E-15 |
| ENSG00000158856 | DMTN | 750.6118047 | 0.243124516 | 0.077007738 | 3.157143982 | 0.001593227 | 0.005724137 |
| ENSG00000158863 | FHIP2B | 1444.112347 | 0.209337177 | 0.067600448 | 3.096683272 | 0.001956988 | 0.006894995 |
| ENSG00000158864 | NDUFS2 | 2706.67435 | 0.17553251 | 0.050802714 | 3.455179811 | 0.000549926 | 0.002160154 |
| ENSG00000158869 | FCER1G | 3.596526431 | 0.368863823 | 0.916901197 | 0.402293971 | 0.687467691 | 0.823383061 |

| | | | | | | | |
|---|---|---|---|---|---|---|---|
| ENSG00000158882 | TOMM40L | 428.9444929 | -0.101250993 | 0.09169742 | -1.104185847 | 0.269512528 | 0.451059856 |
| ENSG00000158941 | CCAR2 | 4239.296852 | 0.019777203 | 0.050224093 | 0.393779209 | 0.693744055 | 0.827371387 |
| ENSG00000158966 | CACHD1 | 310.1324883 | 0.990813201 | 0.114544091 | 8.650059472 | 5.15E-18 | 7.59E-17 |
| ENSG00000158985 | CDC42SE2 | 1314.342219 | 0.240574517 | 0.066258077 | 3.630870798 | 0.000282467 | 0.001166898 |
| ENSG00000158987 | RAPGEF6 | 125.1389732 | -0.686520047 | 0.175274525 | -3.916827309 | 8.97E-05 | 0.000403336 |
| ENSG00000159023 | EPB41 | 2953.971414 | 0.49656258 | 0.056779229 | 8.745497013 | 2.22E-18 | 3.36E-17 |
| ENSG00000159055 | MIS18A | 477.4729372 | -0.027493803 | 0.087845086 | -0.312980541 | 0.754295446 | 0.864498202 |
| ENSG00000159063 | ALG8 | 1238.96733 | -0.332501269 | 0.063236124 | -5.258090566 | 1.46E-07 | 9.31E-07 |
| ENSG00000159069 | FBXW5 | 3547.6857 | 0.107882477 | 0.063357583 | 1.702755565 | 0.088613822 | 0.192063593 |
| ENSG00000159079 | CFAP298 | 210.4416094 | -0.043139975 | 0.129130695 | -0.334079943 | 0.738319235 | 0.855807824 |
| ENSG00000159082 | SYNJ1 | 445.6090855 | -0.22364846 | 0.091161194 | -2.453329644 | 0.014154056 | 0.040397699 |
| ENSG00000159086 | PAXBP1 | 1378.219103 | 0.120694893 | 0.084924328 | 1.421205166 | 0.155257123 | 0.298967381 |
| ENSG00000159110 | IFNAR2 | 112.3478752 | 0.537764423 | 0.171361086 | 3.138194534 | 0.00169992 | 0.006061895 |
| ENSG00000159111 | MRPL10 | 2870.015206 | 0.071684211 | 0.051591897 | 1.389447094 | 0.164696838 | 0.31296765 |
| ENSG00000159128 | IFNGR2 | 733.1469135 | 0.392737151 | 0.082093087 | 4.784046541 | 1.72E-06 | 9.67E-06 |
| ENSG00000159131 | GART | 2588.300233 | -0.201922242 | 0.052302106 | -3.860690493 | 0.000113067 | 0.000500146 |
| ENSG00000159140 | SON | 11606.99957 | -0.093276026 | 0.045384039 | -2.055260553 | 0.039853845 | 0.098863118 |
| ENSG00000159147 | DONSON | 461.165602 | 0.07170372 | 0.096922024 | 0.739808323 | 0.459416308 | 0.646458326 |
| ENSG00000159164 | SV2A | 9.290351822 | 0.523723265 | 0.604094428 | 0.866955961 | 0.38596614 | 0.578583497 |
| ENSG00000159166 | LAD1 | 2590.134884 | 0.841430075 | 0.060097396 | 14.00110696 | 1.53E-44 | 7.26E-43 |
| ENSG00000159167 | STC1 | 1776.512683 | 3.383355687 | 0.071845841 | 47.09187975 | 0 | 0 |
| ENSG00000159173 | TNNI1 | 1.764018817 | 2.017844007 | 1.529139629 | 1.319594345 | 0.186970492 | NA |
| ENSG00000159176 | CSRP1 | 16002.81092 | 0.143167801 | 0.041501458 | 3.449705316 | 0.000561199 | 0.002199933 |
| ENSG00000159184 | HOXB13 | 18.30144184 | -2.39032154 | 0.47608829 | -5.020752641 | 5.15E-07 | 3.09E-06 |
| ENSG00000159197 | KCNE2 | 1.941258545 | 0.334476381 | 1.238171841 | 0.270137286 | 0.787054638 | 0.88457368 |
| ENSG00000159199 | ATP5MC1 | 1495.135115 | 0.120373775 | 0.060902471 | 1.976500692 | 0.048098097 | 0.115912423 |
| ENSG00000159200 | RCAN1 | 232.4299562 | -0.339707662 | 0.11900197 | -2.854638969 | 0.004308578 | 0.014101694 |
| ENSG00000159202 | UBE2Z | 7659.402287 | -0.174762653 | 0.04220098 | -4.141198916 | 3.45E-05 | 0.000165463 |
| ENSG00000159208 | CIART | 757.6697266 | 0.471831632 | 0.074152524 | 6.362988144 | 1.98E-10 | 1.67E-09 |
| ENSG00000159210 | SNF8 | 2471.35313 | -0.090510535 | 0.054995693 | -1.64577497 | 0.099810097 | 0.21166083 |
| ENSG00000159214 | CCDC24 | 409.6038542 | -0.1028462 | 0.10130788 | -1.015184604 | 0.310017838 | 0.497921249 |
| ENSG00000159216 | RUNX1 | 752.4103324 | 1.757423514 | 0.081085796 | 21.67362955 | 3.64E-104 | 6.57E-102 |
| ENSG00000159217 | IGF2BP1 | 18.69394011 | -0.77036202 | 0.41228773 | -1.868505811 | 0.061691601 | 0.142820196 |

| | | | | | | | |
|---|---|---|---|---|---|---|---|
| ENSG00000159228 | CBR1 | 1368.774589 | -0.046195603 | 0.065281177 | -0.707640481 | 0.479168548 | 0.663653377 |
| ENSG00000159231 | CBR3 | 121.1097348 | -0.028085233 | 0.163308232 | -0.171976835 | 0.86345574 | 0.929172842 |
| ENSG00000159239 | ENSG0000015 | 2.412180892 | 0.826850912 | 1.152765097 | 0.717276151 | 0.47320372 | 0.658734955 |
| ENSG00000159247 | TUBBP5 | 6.168591661 | -0.687655718 | 0.713450072 | -0.963845607 | 0.335123337 | 0.526340501 |
| ENSG00000159256 | MORC3 | 882.8677629 | 0.028725763 | 0.071603301 | 0.401179306 | 0.688288115 | 0.823819752 |
| ENSG00000159259 | CHAF1B | 767.0687007 | 0.106486557 | 0.073553914 | 1.447734742 | 0.14769125 | 0.288147412 |
| ENSG00000159263 | SIM2 | 701.0091 | -0.437963086 | 0.074865055 | -5.850033586 | 4.91E-09 | 3.67E-08 |
| ENSG00000159267 | HLCS | 1422.489495 | 0.134429897 | 0.062332882 | 2.156644994 | 0.031033336 | 0.079998284 |
| ENSG00000159289 | GOLGA6A | 2.218773271 | -1.919223133 | 1.273906936 | -1.506564631 | 0.131922289 | 0.264420909 |
| ENSG00000159314 | ARHGAP27 | 992.4810032 | -0.061521947 | 0.069399637 | -0.886488016 | 0.375354609 | 0.568358728 |
| ENSG00000159322 | ADPGK | 1557.076735 | -0.14699551 | 0.062794452 | -2.340899639 | 0.019237338 | 0.052869836 |
| ENSG00000159335 | PTMS | 6081.877816 | -0.622534627 | 0.044863523 | -13.87618685 | 8.83E-44 | 4.09E-42 |
| ENSG00000159346 | ADIPOR1 | 7814.318224 | 0.467977698 | 0.042990982 | 10.88548526 | 1.35E-27 | 3.45E-26 |
| ENSG00000159348 | CYB5R1 | 1644.117123 | 0.481620477 | 0.065752134 | 7.324788514 | 2.39E-13 | 2.55E-12 |
| ENSG00000159352 | PSMD4 | 8260.095108 | 0.05690486 | 0.043516381 | 1.307665285 | 0.190986863 | 0.35069548 |
| ENSG00000159363 | ATP13A2 | 1958.061065 | 0.12532896 | 0.063938509 | 1.960148304 | 0.049978459 | 0.119802074 |
| ENSG00000159377 | PSMB4 | 10475.39933 | 0.187510796 | 0.043347912 | 4.325716866 | 1.52E-05 | 7.66E-05 |
| ENSG00000159388 | BTG2 | 2161.470748 | 0.085115604 | 0.062018544 | 1.372421826 | 0.16993216 | 0.320526537 |
| ENSG00000159399 | HK2 | 2659.361937 | 0.180775376 | 0.049464079 | 3.654679896 | 0.000257503 | 0.001069937 |
| ENSG00000159403 | C1R | 6.062493388 | -0.004663482 | 0.717128021 | -0.006502997 | 0.994811395 | 0.99681023 |
| ENSG00000159409 | CELF3 | 46.40669299 | -1.28401722 | 0.270303091 | -4.750286848 | 2.03E-06 | 1.13E-05 |
| ENSG00000159423 | ALDH4A1 | 1980.985199 | -0.387795217 | 0.068559326 | -5.656345263 | 1.55E-08 | 1.09E-07 |
| ENSG00000159433 | STARD9 | 24.88464724 | -0.96330364 | 0.365445238 | -2.635972618 | 0.00838965 | 0.02546422 |
| ENSG00000159445 | THEM4 | 965.3613517 | 0.68007485 | 0.071425219 | 9.521494776 | 1.71E-21 | 3.12E-20 |
| ENSG00000159450 | TCHH | 262.2429741 | 0.715933924 | 0.131322287 | 5.451732084 | 4.99E-08 | 3.33E-07 |
| ENSG00000159459 | UBR1 | 1198.805071 | -0.440010099 | 0.068186483 | -6.45303997 | 1.10E-10 | 9.44E-10 |
| ENSG00000159461 | AMFR | 2602.669967 | 0.526089393 | 0.052430322 | 10.03406755 | 1.08E-23 | 2.22E-22 |
| ENSG00000159479 | MED8 | 801.4128744 | 0.109890698 | 0.075932637 | 1.44721298 | 0.147837282 | 0.288340802 |
| ENSG00000159496 | RGL4 | 5.288172494 | -0.849944735 | 0.789981134 | -1.075905106 | 0.281969698 | 0.465393058 |
| ENSG00000159556 | ISL2 | 23.93881742 | -0.331740218 | 0.382486653 | -0.867324953 | 0.385763988 | 0.578389052 |
| ENSG00000159579 | RSPRY1 | 825.3888061 | 0.039494549 | 0.071760016 | 0.550369842 | 0.582065729 | 0.746825138 |
| ENSG00000159588 | CCDC17 | 37.45275661 | -0.363787081 | 0.296290816 | -1.227804109 | 0.219520513 | 0.389516399 |
| ENSG00000159592 | GPBP1L1 | 3760.896575 | 0.063545972 | 0.053157903 | 1.195419091 | 0.23192333 | 0.40588433 |

| | | | | | | | |
|---|---|---|---|---|---|---|---|
| ENSG00000159593 | NAE1 | 1061.422184 | 0.259912702 | 0.070997072 | 3.660893229 | 0.000251337 | 0.001046024 |
| ENSG00000159596 | TMEM69 | 1041.417896 | 0.263888254 | 0.070007 | 3.769455281 | 0.000163604 | 0.000704539 |
| ENSG00000159618 | ADGRG5 | 129.3650985 | -0.466623864 | 0.158175913 | -2.950031107 | 0.003177419 | 0.010704949 |
| ENSG00000159625 | DRC7 | 2.269818703 | -0.562790205 | 1.143061878 | -0.492353227 | 0.622469658 | 0.776597778 |
| ENSG00000159640 | ACE | 83.72769049 | -0.175641687 | 0.21178805 | -0.829327658 | 0.406919024 | 0.599725179 |
| ENSG00000159658 | EFCAB14 | 4934.659277 | 0.027031182 | 0.050013832 | 0.540474117 | 0.588870106 | 0.752153688 |
| ENSG00000159674 | SPON2 | 15.48816784 | 0.356265715 | 0.458064247 | 0.777763637 | 0.436708369 | 0.6265222 |
| ENSG00000159685 | CHCHD6 | 288.3094115 | 0.286437127 | 0.109563275 | 2.614353468 | 0.00893965 | 0.026951711 |
| ENSG00000159692 | CTBP1 | 4314.563731 | -0.068797517 | 0.05270343 | -1.30537077 | 0.191766627 | 0.351761268 |
| ENSG00000159712 | ANKRD18CP | 7.678162356 | -0.706225237 | 0.636169803 | -1.110120653 | 0.266947039 | 0.448069442 |
| ENSG00000159714 | ZDHHC1 | 3.832466753 | 1.084262473 | 0.916512012 | 1.183031383 | 0.23679671 | 0.412023402 |
| ENSG00000159720 | ATP6V0D1 | 3359.369136 | 0.06179627 | 0.052001927 | 1.188345757 | 0.234697217 | 0.409617481 |
| ENSG00000159733 | ZFYVE28 | 427.3952157 | 0.014098792 | 0.096360378 | 0.146313165 | 0.883674174 | 0.939712848 |
| ENSG00000159753 | CARMIL2 | 79.47843578 | 0.430558526 | 0.199729076 | 2.155712801 | 0.031106098 | 0.080149825 |
| ENSG00000159761 | C16orf86 | 18.81124659 | 0.248268045 | 0.41452272 | 0.59892506 | 0.549222859 | 0.721774261 |
| ENSG00000159784 | FAM131B | 8.248699063 | 0.129054564 | 0.609391984 | 0.211775946 | 0.832281835 | 0.911334752 |
| ENSG00000159788 | RGS12 | 914.4468904 | 0.298522306 | 0.068614049 | 4.350746119 | 1.36E-05 | 6.88E-05 |
| ENSG00000159792 | PSKH1 | 1874.329522 | 0.387856776 | 0.063238627 | 6.133225749 | 8.61E-10 | 6.89E-09 |
| ENSG00000159840 | ZYX | 2345.273752 | 0.133296165 | 0.062640685 | 2.127948703 | 0.033341337 | 0.0850117 |
| ENSG00000159842 | ABR | 3950.740275 | 0.303477262 | 0.053107729 | 5.714370949 | 1.10E-08 | 7.89E-08 |
| ENSG00000159873 | CCDC117 | 2518.589862 | 0.065806365 | 0.057988716 | 1.134813281 | 0.25645356 | 0.435910246 |
| ENSG00000159882 | ZNF230 | 113.1623157 | 0.258937349 | 0.167058666 | 1.549978554 | 0.121146663 | 0.247362062 |
| ENSG00000159884 | CCDC107 | 103.8795017 | -0.108667777 | 0.175814213 | -0.618083003 | 0.536520626 | 0.711679378 |
| ENSG00000159885 | ZNF222 | 61.15426664 | 0.065114184 | 0.229715753 | 0.283455458 | 0.776827721 | 0.878849333 |
| ENSG00000159899 | NPR2 | 217.8572949 | -0.80415546 | 0.132292652 | -6.078610188 | 1.21E-09 | 9.57E-09 |
| ENSG00000159904 | ZNF890P | 6.554641887 | -0.736724383 | 0.687622893 | -1.071407586 | 0.283986203 | 0.467822586 |
| ENSG00000159905 | ZNF221 | 18.48576922 | 0.478127135 | 0.411882252 | 1.160834518 | 0.245709203 | 0.422753252 |
| ENSG00000159915 | ZNF233 | 183.1774505 | 0.069610271 | 0.13807687 | 0.504141433 | 0.614161993 | 0.770979321 |
| ENSG00000159917 | ZNF235 | 295.5785142 | 0.195123941 | 0.110314755 | 1.768792772 | 0.076928463 | 0.171450398 |
| ENSG00000159921 | GNE | 237.2323519 | -0.398400451 | 0.129887361 | -3.067276494 | 0.002160189 | 0.00753799 |
| ENSG00000159958 | TNFRSF13C | 23.04913587 | -0.168632615 | 0.360910832 | -0.467241767 | 0.640326914 | 0.78846481 |
| ENSG00000160007 | ARHGAP35 | 10456.90401 | -0.025759041 | 0.059150814 | -0.43548076 | 0.663213501 | 0.805549688 |
| ENSG00000160014 | CALM3 | 10618.89114 | -0.326402512 | 0.044144387 | -7.393975357 | 1.43E-13 | 1.54E-12 |

| | | | | | | | |
|---|---|---|---|---|---|---|---|
| ENSG00000160049 | DFFA | 1978.302026 | 0.106526059 | 0.058432021 | 1.82307673 | 0.068291771 | 0.155495108 |
| ENSG00000160050 | CCDC28B | 63.96033407 | 0.571349191 | 0.22404523 | 2.550151101 | 0.010767624 | 0.031700371 |
| ENSG00000160051 | IQCC | 314.0149003 | -0.089364886 | 0.116610925 | -0.766350886 | 0.443467548 | 0.632343707 |
| ENSG00000160055 | TMEM234 | 210.332541 | 0.255306662 | 0.128044596 | 1.993888619 | 0.046164228 | 0.111851224 |
| ENSG00000160058 | BSDC1 | 2865.149042 | 0.373721383 | 0.055435646 | 6.741535625 | 1.57E-11 | 1.44E-10 |
| ENSG00000160062 | ZBTB8A | 175.3772738 | -0.115993761 | 0.144627261 | -0.80201865 | 0.42254217 | 0.613489389 |
| ENSG00000160072 | ATAD3B | 799.4274285 | 0.446603971 | 0.079370646 | 5.626815386 | 1.84E-08 | 1.28E-07 |
| ENSG00000160075 | SSU72 | 2776.518893 | 0.022036111 | 0.050359916 | 0.437572426 | 0.661696263 | 0.804542974 |
| ENSG00000160087 | UBE2J2 | 1144.63422 | 0.105224036 | 0.066830855 | 1.574482872 | 0.115375858 | 0.237907612 |
| ENSG00000160094 | ZNF362 | 1532.375153 | -0.119895645 | 0.060903861 | -1.968605001 | 0.048998468 | 0.117748256 |
| ENSG00000160097 | FNDC5 | 57.89577382 | 0.530804526 | 0.239155531 | 2.219495092 | 0.026453061 | 0.069702873 |
| ENSG00000160111 | CPAMD8 | 311.6370449 | -0.050050256 | 0.107802584 | -0.464276959 | 0.642449329 | 0.790314651 |
| ENSG00000160113 | NR2F6 | 924.554329 | -0.04276794 | 0.070394332 | -0.607548055 | 0.543487262 | 0.716981872 |
| ENSG00000160117 | ANKLE1 | 14.32791155 | -0.374873816 | 0.474086575 | -0.790728605 | 0.42910238 | 0.619406532 |
| ENSG00000160124 | MIX23 | 426.5406153 | -0.238219831 | 0.094089529 | -2.531842112 | 0.011346507 | 0.03321282 |
| ENSG00000160131 | VMA21 | 1433.439037 | -0.33975649 | 0.064189339 | -5.293036113 | 1.20E-07 | 7.75E-07 |
| ENSG00000160145 | KALRN | 248.5584375 | 1.305058653 | 0.124075247 | 10.51828377 | 7.12E-26 | 1.64E-24 |
| ENSG00000160161 | CILP2 | 4.81774738 | -0.480843003 | 0.789597054 | -0.60897264 | 0.542542574 | 0.71631532 |
| ENSG00000160172 | FAM86C2P | 282.5714398 | -0.086104128 | 0.109353477 | -0.78739269 | 0.431052024 | 0.62116549 |
| ENSG00000160179 | ABCG1 | 5434.275383 | -0.155293919 | 0.047807472 | -3.248318971 | 0.001160891 | 0.004298068 |
| ENSG00000160180 | TFF3 | 256.150551 | 1.444972619 | 0.133425434 | 10.82981387 | 2.49E-27 | 6.27E-26 |
| ENSG00000160182 | TFF1 | 2280.296782 | -0.755387108 | 0.071293482 | -10.59545824 | 3.13E-26 | 7.39E-25 |
| ENSG00000160183 | TMPRSS3 | 34.93342865 | -0.5694152 | 0.32551169 | -1.749292626 | 0.08024045 | 0.177060384 |
| ENSG00000160188 | RSPH1 | 160.2648542 | 1.389734854 | 0.153209669 | 9.070803868 | 1.18E-19 | 1.93E-18 |
| ENSG00000160190 | SLC37A1 | 1564.183331 | 0.488344585 | 0.064466906 | 7.575120569 | 3.59E-14 | 4.05E-13 |
| ENSG00000160191 | PDE9A | 93.38791583 | 0.141776203 | 0.183952244 | 0.77072288 | 0.440871207 | 0.630129937 |
| ENSG00000160193 | WDR4 | 400.1635215 | 0.19207032 | 0.094328495 | 2.03618557 | 0.041731725 | 0.102673291 |
| ENSG00000160194 | NDUFV3 | 1149.187206 | 0.115816265 | 0.068144867 | 1.699559622 | 0.089213791 | 0.193070799 |
| ENSG00000160199 | PKNOX1 | 741.8176859 | 0.15039773 | 0.075289522 | 1.997591774 | 0.045760936 | 0.111038277 |
| ENSG00000160200 | CBS | 694.7432452 | 0.155539648 | 0.082268567 | 1.890632768 | 0.058673383 | 0.136994138 |
| ENSG00000160201 | U2AF1 | 60.09910428 | -0.43140303 | 0.251285061 | -1.716787413 | 0.086018022 | 0.187429086 |
| ENSG00000160207 | HSF2BP | 43.69017771 | -0.207260104 | 0.271509629 | -0.763361894 | 0.445247599 | 0.633857511 |
| ENSG00000160208 | RRP1B | 2804.178058 | 0.089473328 | 0.050855929 | 1.759349001 | 0.078518249 | 0.174097102 |

| | | | | | | | |
|---|---|---|---|---|---|---|---|
| ENSG00000160209 | PDXK | 7154.189849 | 0.512874806 | 0.045541718 | 11.26164818 | 2.03E-29 | 5.52E-28 |
| ENSG00000160211 | G6PD | 9609.555842 | -0.440953964 | 0.045478328 | -9.695914124 | 3.14E-22 | 5.94E-21 |
| ENSG00000160213 | CSTB | 3094.444695 | -0.034058171 | 0.047893835 | -0.711118066 | 0.477011076 | 0.661968041 |
| ENSG00000160214 | RRP1 | 2076.647061 | 0.134310738 | 0.053569204 | 2.507237902 | 0.012167878 | 0.03536439 |
| ENSG00000160216 | AGPAT3 | 5141.605384 | 0.173739405 | 0.048426714 | 3.587676926 | 0.000333637 | 0.001360889 |
| ENSG00000160218 | TRAPPC10 | 1698.294498 | 0.107270449 | 0.05573919 | 1.924506763 | 0.054291094 | 0.128634341 |
| ENSG00000160219 | GAB3 | 1.809362698 | -0.430248475 | 1.276450142 | -0.337066417 | 0.736066833 | 0.854386347 |
| ENSG00000160221 | GATD3 | 36.07318178 | -0.197643827 | 0.309654805 | -0.638271469 | 0.523296981 | 0.699904915 |
| ENSG00000160223 | ICOSLG | 3.991324523 | -0.39736499 | 0.904827407 | -0.439161089 | 0.660544817 | 0.803707149 |
| ENSG00000160226 | CFAP410 | 653.6477802 | 0.095522096 | 0.081032432 | 1.178813148 | 0.238472588 | 0.413937895 |
| ENSG00000160229 | ZNF66 | 119.7636267 | 0.425131352 | 0.167209336 | 2.542509666 | 0.011005956 | 0.032323214 |
| ENSG00000160233 | LRRC3 | 172.9297705 | -0.357511051 | 0.142848892 | -2.50272191 | 0.012324234 | 0.035750875 |
| ENSG00000160255 | ITGB2 | 24.63398484 | 0.340193435 | 0.359806909 | 0.945488889 | 0.344409341 | 0.535390003 |
| ENSG00000160256 | SLX9 | 516.8677062 | 0.155174642 | 0.088829115 | 1.746889418 | 0.080656522 | 0.177892569 |
| ENSG00000160271 | RALGDS | 342.7412389 | 0.682157188 | 0.106104703 | 6.429094767 | 1.28E-10 | 1.10E-09 |
| ENSG00000160282 | FTCD | 48.67683364 | -0.108367896 | 0.253308663 | -0.427809674 | 0.668789693 | 0.809734289 |
| ENSG00000160284 | SPATC1L | 204.4884343 | 0.436877462 | 0.135338825 | 3.228027589 | 0.00124647 | 0.004583108 |
| ENSG00000160285 | LSS | 19761.56497 | -0.63086205 | 0.045590155 | -13.8376815 | 1.51E-43 | 6.93E-42 |
| ENSG00000160293 | VAV2 | 3853.526908 | 0.075981241 | 0.058131965 | 1.307047527 | 0.191196571 | 0.351052452 |
| ENSG00000160294 | MCM3AP | 5060.914054 | -0.130976408 | 0.047515661 | -2.756489238 | 0.005842554 | 0.018484096 |
| ENSG00000160298 | C21orf58 | 774.8344694 | -0.125984812 | 0.092482264 | -1.362259162 | 0.173116111 | 0.325355037 |
| ENSG00000160299 | PCNT | 3410.128875 | -0.110109105 | 0.046905506 | -2.347466535 | 0.018901571 | 0.052065567 |
| ENSG00000160305 | DIP2A | 2577.713418 | -0.356099157 | 0.056426163 | -6.310887289 | 2.77E-10 | 2.32E-09 |
| ENSG00000160310 | PRMT2 | 3401.54506 | -0.204081962 | 0.049100379 | -4.156423373 | 3.23E-05 | 0.000155787 |
| ENSG00000160318 | CLDND2 | 6.772371202 | -0.012516812 | 0.693157922 | -0.018057662 | 0.985592853 | 0.993350589 |
| ENSG00000160323 | ADAMTS13 | 360.4637525 | -0.425780955 | 0.100684961 | -4.22884364 | 2.35E-05 | 0.000115311 |
| ENSG00000160325 | CACFD1 | 1349.275106 | 0.357661477 | 0.078208216 | 4.573195706 | 4.80E-06 | 2.57E-05 |
| ENSG00000160326 | SLC2A6 | 1031.972248 | -0.503945024 | 0.072755009 | -6.926602434 | 4.31E-12 | 4.18E-11 |
| ENSG00000160336 | ZNF761 | 1284.746272 | 0.136373289 | 0.063002938 | 2.164554419 | 0.030421825 | 0.078643039 |
| ENSG00000160345 | PIERCE1 | 189.8165992 | 0.20740994 | 0.136559816 | 1.518821176 | 0.128807514 | 0.259333366 |
| ENSG00000160352 | ZNF714 | 351.0833891 | -0.221241518 | 0.099385954 | -2.226084354 | 0.026008532 | 0.068697262 |
| ENSG00000160360 | GPSM1 | 937.001761 | 0.392891094 | 0.084931793 | 4.625960188 | 3.73E-06 | 2.02E-05 |
| ENSG00000160392 | C19orf47 | 448.9864257 | -0.238488445 | 0.09365327 | -2.546504191 | 0.01088079 | 0.031992481 |

| | | | | | | | |
|---|---|---|---|---|---|---|---|
| ENSG00000160396 | HIPK4 | 4.603591765 | 1.469484958 | 0.867998153 | 1.692958624 | 0.090463343 | 0.195242176 |
| ENSG00000160401 | CFAP157 | 13.92017572 | 0.984923344 | 0.505793989 | 1.947281632 | 0.05150098 | 0.122848339 |
| ENSG00000160404 | TOR2A | 468.6309659 | -0.090075231 | 0.091439828 | -0.985076554 | 0.324586469 | 0.514760705 |
| ENSG00000160408 | ST6GALNAC6 | 3.200378863 | 2.319920902 | 1.160312781 | 1.999392698 | 0.045565881 | 0.110623486 |
| ENSG00000160410 | SHKBP1 | 1534.84213 | 0.058072547 | 0.072820164 | 0.797478933 | 0.425172933 | 0.616060327 |
| ENSG00000160439 | RDH13 | 577.9954523 | 0.068570374 | 0.081300575 | 0.843418076 | 0.398994666 | 0.591855698 |
| ENSG00000160445 | ZER1 | 2997.755 | 0.136321412 | 0.066193089 | 2.059450837 | 0.03945107 | 0.098071906 |
| ENSG00000160446 | ZDHHC12 | 1220.329049 | 0.024148849 | 0.079466824 | 0.303885915 | 0.761214809 | 0.868899224 |
| ENSG00000160447 | PKN3 | 868.2907665 | -0.285635341 | 0.074762606 | -3.820564268 | 0.000133147 | 0.000581345 |
| ENSG00000160460 | SPTBN4 | 114.1703393 | 1.512712204 | 0.180086902 | 8.399901293 | 4.47E-17 | 6.18E-16 |
| ENSG00000160469 | BRSK1 | 197.455709 | -0.105409869 | 0.13741147 | -0.76711114 | 0.443015439 | 0.631832054 |
| ENSG00000160472 | TMEM190 | 4.128565893 | 0.209631325 | 0.935432034 | 0.224101076 | 0.822678655 | 0.906215744 |
| ENSG00000160539 | PLPP7 | 1.69688806 | -0.81891525 | 1.387869336 | -0.590052124 | 0.555155705 | NA |
| ENSG00000160551 | TAOK1 | 2891.631705 | -0.29733699 | 0.07281375 | -4.083528044 | 4.44E-05 | 0.000208986 |
| ENSG00000160563 | MED27 | 345.7603976 | 0.003704901 | 0.102882013 | 0.036011163 | 0.971273458 | 0.986933608 |
| ENSG00000160570 | DEDD2 | 790.5854372 | 0.31290777 | 0.078758292 | 3.973013637 | 7.10E-05 | 0.000324502 |
| ENSG00000160584 | SIK3 | 631.5691652 | 0.05388269 | 0.08234901 | 0.654321044 | 0.512904991 | 0.692356657 |
| ENSG00000160588 | MPZL3 | 439.021191 | 0.604904839 | 0.095708221 | 6.32030178 | 2.61E-10 | 2.19E-09 |
| ENSG00000160602 | NEK8 | 765.2447796 | -1.147986087 | 0.087754787 | -13.08174888 | 4.19E-39 | 1.64E-37 |
| ENSG00000160606 | TLCD1 | 340.093375 | -0.333278741 | 0.105870515 | -3.147984506 | 0.001644004 | 0.005888127 |
| ENSG00000160613 | PCSK7 | 947.6605139 | 0.144975954 | 0.070753743 | 2.04902169 | 0.040459994 | 0.100091908 |
| ENSG00000160633 | SAFB | 3356.989034 | -0.10359778 | 0.047230089 | -2.193469934 | 0.028273538 | 0.073804617 |
| ENSG00000160678 | S100A1 | 13.82131744 | -0.173079127 | 0.48485639 | -0.356969878 | 0.721114356 | 0.845162383 |
| ENSG00000160679 | CHTOP | 3467.404775 | 0.138346495 | 0.049278904 | 2.80741824 | 0.004994036 | 0.016092594 |
| ENSG00000160683 | CXCR5 | 28.265039 | -0.820660804 | 0.341793211 | -2.401044776 | 0.016348336 | 0.04587533 |
| ENSG00000160685 | ZBTB7B | 2307.614484 | 0.50652037 | 0.065822523 | 7.695243941 | 1.41E-14 | 1.64E-13 |
| ENSG00000160688 | FLAD1 | 1317.085086 | 0.278487352 | 0.06651491 | 4.186841016 | 2.83E-05 | 0.000137506 |
| ENSG00000160691 | SHC1 | 2207.559057 | 0.483344414 | 0.05562288 | 8.689668933 | 3.63E-18 | 5.44E-17 |
| ENSG00000160695 | VPS11 | 3704.96481 | 0.440103281 | 0.053646468 | 8.203769923 | 2.33E-16 | 3.09E-15 |
| ENSG00000160703 | NLRX1 | 550.9995632 | -0.376903822 | 0.087298714 | -4.317404051 | 1.58E-05 | 7.93E-05 |
| ENSG00000160710 | ADAR | 14040.16866 | 0.399572124 | 0.044897653 | 8.899621614 | 5.60E-19 | 8.83E-18 |
| ENSG00000160712 | IL6R | 211.1519414 | 1.224261883 | 0.148457897 | 8.246525835 | 1.63E-16 | 2.19E-15 |
| ENSG00000160714 | UBE2Q1 | 4089.68817 | 0.347152234 | 0.046374579 | 7.485830461 | 7.11E-14 | 7.83E-13 |

| | | | | | | | |
|---|---|---|---|---|---|---|---|
| ENSG00000160716 | CHRNB2 | 9.654774239 | -3.515882745 | 0.79892287 | -4.400778696 | 1.08E-05 | 5.55E-05 |
| ENSG00000160741 | CRTC2 | 1834.289338 | 0.400195051 | 0.064091324 | 6.244137721 | 4.26E-10 | 3.51E-09 |
| ENSG00000160746 | ANO10 | 709.605875 | -0.021261737 | 0.077342111 | -0.274905058 | 0.783389181 | 0.882572823 |
| ENSG00000160752 | FDPS | 6674.449752 | -0.481764705 | 0.046732606 | -10.30896304 | 6.42E-25 | 1.42E-23 |
| ENSG00000160753 | RUSC1 | 4088.781414 | -0.134579828 | 0.055169198 | -2.439401558 | 0.01471161 | 0.041773034 |
| ENSG00000160766 | GBAP1 | 515.8367702 | 0.242789797 | 0.092187271 | 2.633658586 | 0.008447038 | 0.025618066 |
| ENSG00000160767 | ENTREP3 | 1926.295768 | 0.109402314 | 0.05638321 | 1.940334972 | 0.052338994 | 0.124562353 |
| ENSG00000160781 | PAQR6 | 824.4548304 | 0.883390411 | 0.088889516 | 9.938072036 | 2.84E-23 | 5.74E-22 |
| ENSG00000160783 | PMF1 | 645.5636411 | 0.115889719 | 0.084956586 | 1.364105192 | 0.172534461 | 0.324527758 |
| ENSG00000160785 | SLC25A44 | 2916.992858 | 0.264394078 | 0.055101374 | 4.798320944 | 1.60E-06 | 9.03E-06 |
| ENSG00000160789 | LMNA | 5355.227789 | -0.120770711 | 0.052331878 | -2.307784778 | 0.02101111 | 0.057047563 |
| ENSG00000160796 | NBEAL2 | 7280.119844 | -0.200603664 | 0.058530381 | -3.427342521 | 0.00060952 | 0.002375129 |
| ENSG00000160799 | CCDC12 | 661.7602793 | -0.181012307 | 0.080945988 | -2.23621097 | 0.025337955 | 0.067215527 |
| ENSG00000160803 | UBQLN4 | 2821.103687 | 0.234718752 | 0.053548299 | 4.383309169 | 1.17E-05 | 5.97E-05 |
| ENSG00000160813 | PPP1R35 | 558.22637 | -0.588484578 | 0.098375739 | -5.982009207 | 2.20E-09 | 1.70E-08 |
| ENSG00000160818 | GPATCH4 | 1949.617119 | 0.206301723 | 0.055392398 | 3.724368844 | 0.000195804 | 0.000832732 |
| ENSG00000160867 | FGFR4 | 457.2990177 | -0.748245008 | 0.097854435 | -7.646510952 | 2.07E-14 | 2.37E-13 |
| ENSG00000160877 | NACC1 | 3782.802329 | 0.165670562 | 0.052327278 | 3.166045875 | 0.001545264 | 0.005562272 |
| ENSG00000160883 | HK3 | 2.893550654 | -0.73405905 | 1.054908276 | -0.695851068 | 0.486522104 | 0.670074159 |
| ENSG00000160888 | IER2 | 1201.209962 | 0.153202664 | 0.066988855 | 2.286987354 | 0.022196561 | 0.05991165 |
| ENSG00000160908 | ZNF394 | 625.0168274 | -0.325607666 | 0.081372701 | -4.001436118 | 6.30E-05 | 0.000290133 |
| ENSG00000160917 | CPSF4 | 1425.349993 | -0.614003645 | 0.060616221 | -10.12936204 | 4.09E-24 | 8.64E-23 |
| ENSG00000160932 | LY6E | 3736.763817 | 0.183209852 | 0.060007083 | 3.053137117 | 0.002264624 | 0.007880838 |
| ENSG00000160948 | VPS28 | 3551.022702 | 0.140707612 | 0.058284695 | 2.414143396 | 0.015772254 | 0.044443619 |
| ENSG00000160949 | TONSL | 2032.848493 | 0.119343077 | 0.05922323 | 2.015139628 | 0.043890021 | 0.107209541 |
| ENSG00000160953 | PWWP3A | 1313.967731 | 0.200764974 | 0.060905038 | 3.2963607 | 0.000979462 | 0.00367215 |
| ENSG00000160957 | RECQL4 | 3613.088631 | -0.087129669 | 0.057555086 | -1.513848299 | 0.130064319 | 0.2613591 |
| ENSG00000160959 | LRRC14 | 2133.094268 | -0.019165649 | 0.061975723 | -0.309244452 | 0.757135582 | 0.866096054 |
| ENSG00000160961 | ZNF333 | 165.8176871 | 0.089902051 | 0.146722466 | 0.612735411 | 0.540051304 | 0.714414141 |
| ENSG00000160963 | COL26A1 | 5.537390013 | -0.291057453 | 0.732723391 | -0.397226915 | 0.691200136 | 0.825797152 |
| ENSG00000160972 | PPP1R16A | 1974.20977 | 0.193969983 | 0.059664666 | 3.251002571 | 0.001149988 | 0.004262519 |
| ENSG00000160973 | FOXH1 | 29.38265962 | 0.162202081 | 0.324487985 | 0.499870839 | 0.617166027 | 0.773024159 |
| ENSG00000160991 | ORAI2 | 1720.469701 | 0.173668434 | 0.06011609 | 2.888884392 | 0.003866112 | 0.012786708 |

| | | | | | | | |
|---|---|---|---|---|---|---|---|
| ENSG00000160993 | ALKBH4 | 478.5543608 | -0.560913442 | 0.098206316 | -5.711582153 | 1.12E-08 | 8.00E-08 |
| ENSG00000160999 | SH2B2 | 98.34790614 | 0.210318283 | 0.180605293 | 1.164518937 | 0.244213774 | 0.421001275 |
| ENSG00000161010 | MRNIP | 1257.635265 | -0.486417119 | 0.159693319 | -3.04594533 | 0.0023195 | 0.00805226 |
| ENSG00000161011 | SQSTM1 | 11329.04514 | 0.418801149 | 0.04678672 | 8.951282558 | 3.51E-19 | 5.61E-18 |
| ENSG00000161013 | MGAT4B | 6129.363439 | -0.052300857 | 0.049727631 | -1.051746394 | 0.29291592 | 0.478255601 |
| ENSG00000161016 | RPL8 | 22530.04837 | 0.160419989 | 0.046058737 | 3.482943722 | 0.000495933 | 0.001963869 |
| ENSG00000161021 | MAML1 | 4038.157186 | -0.274186836 | 0.056681958 | -4.837285866 | 1.32E-06 | 7.50E-06 |
| ENSG00000161031 | PGLYRP2 | 9.621786651 | 1.45535685 | 0.602340727 | 2.416168764 | 0.01568479 | 0.044224321 |
| ENSG00000161036 | LRWD1 | 774.5405076 | -0.513845861 | 0.082776351 | -6.207640923 | 5.38E-10 | 4.39E-09 |
| ENSG00000161040 | FBXL13 | 11.68055623 | 0.202826024 | 0.521651636 | 0.388815082 | 0.697412943 | 0.830107037 |
| ENSG00000161048 | NAPEPLD | 175.938685 | -0.217209609 | 0.135465674 | -1.603429136 | 0.108839943 | 0.226775369 |
| ENSG00000161057 | PSMC2 | 3274.52336 | -0.550519588 | 0.0479745 | -11.47525425 | 1.76E-30 | 4.97E-29 |
| ENSG00000161082 | CELF5 | 39.12396074 | -0.273344114 | 0.278412454 | -0.981795571 | 0.32620057 | 0.516713236 |
| ENSG00000161091 | MFSD12 | 1390.10925 | 0.22760387 | 0.066518843 | 3.421645047 | 0.000622435 | 0.002420933 |
| ENSG00000161133 | USP41 | 1.912509465 | -1.012078951 | 1.388050292 | -0.72913709 | 0.465917808 | 0.651756481 |
| ENSG00000161179 | YDJC | 697.8805265 | -0.332880645 | 0.084588792 | -3.935280738 | 8.31E-05 | 0.000375922 |
| ENSG00000161180 | CCDC116 | 8.058250883 | -1.907589333 | 0.677883038 | -2.81403904 | 0.004892329 | 0.015798145 |
| ENSG00000161202 | DVL3 | 5411.788418 | 0.066758513 | 0.050115853 | 1.332083737 | 0.182832674 | 0.338759514 |
| ENSG00000161203 | AP2M1 | 13223.55703 | -0.275869953 | 0.045548752 | -6.056586361 | 1.39E-09 | 1.09E-08 |
| ENSG00000161204 | ABCF3 | 2580.560507 | -0.155901597 | 0.053248683 | -2.927801902 | 0.003413675 | 0.011432073 |
| ENSG00000161217 | PCYT1A | 2899.214343 | -0.343228985 | 0.049765379 | -6.896942985 | 5.31E-12 | 5.11E-11 |
| ENSG00000161243 | FBXO27 | 862.6573777 | -0.095574547 | 0.071955346 | -1.328248031 | 0.184096182 | 0.34062183 |
| ENSG00000161249 | DMKN | 2668.644756 | -0.83855001 | 0.052881136 | -15.85726172 | 1.25E-56 | 8.18E-55 |
| ENSG00000161265 | U2AF1L4 | 73.94108603 | 0.11556544 | 0.207623845 | 0.556609671 | 0.577794145 | 0.743545771 |
| ENSG00000161267 | BDH1 | 1837.430965 | 0.169392952 | 0.058327262 | 2.904181452 | 0.003682147 | 0.01223294 |
| ENSG00000161277 | THAP8 | 190.7394392 | 0.025692647 | 0.140971814 | 0.182253788 | 0.855383563 | 0.925022342 |
| ENSG00000161298 | ZNF382 | 41.28356302 | 1.690176843 | 0.299246997 | 5.648099591 | 1.62E-08 | 1.14E-07 |
| ENSG00000161328 | LRRC56 | 338.3768113 | 0.218753878 | 0.101171862 | 2.162200766 | 0.030602705 | 0.079048085 |
| ENSG00000161381 | PLXDC1 | 2.978060184 | -1.161192134 | 1.02639531 | -1.13133032 | 0.257916087 | 0.4375795 |
| ENSG00000161395 | PGAP3 | 595.8266999 | 0.170548032 | 0.081772799 | 2.085632813 | 0.037011897 | 0.092945037 |
| ENSG00000161405 | IKZF3 | 9.055772833 | -2.67695389 | 0.720870288 | -3.713502881 | 0.00020441 | 0.00086548 |
| ENSG00000161509 | GRIN2C | 27.68472878 | 0.023873221 | 0.336657147 | 0.070912564 | 0.943467344 | 0.973031555 |
| ENSG00000161513 | FDXR | 594.4694077 | -0.007289127 | 0.081725834 | -0.089189994 | 0.928930917 | 0.965899521 |

| | | | | | | | |
|---|---|---|---|---|---|---|---|
| ENSG00000161526 | SAP30BP | 2933.697757 | -0.042009475 | 0.049532936 | -0.848111938 | 0.396375634 | 0.58929728 |
| ENSG00000161533 | ACOX1 | 2673.268726 | 0.180004746 | 0.053416934 | 3.369806783 | 0.000752209 | 0.002882191 |
| ENSG00000161542 | PRPSAP1 | 2026.181741 | 0.226057896 | 0.053085121 | 4.258404114 | 2.06E-05 | 0.0001021 |
| ENSG00000161544 | CYGB | 2.254192931 | -0.552105323 | 1.154957377 | -0.478030907 | 0.632628207 | 0.783513891 |
| ENSG00000161547 | SRSF2 | 5557.525507 | -0.164619248 | 0.053600912 | -3.071202383 | 0.002131986 | 0.00744637 |
| ENSG00000161558 | TMEM143 | 153.2638743 | -0.712722263 | 0.149133336 | -4.779094223 | 1.76E-06 | 9.91E-06 |
| ENSG00000161594 | KLHL10 | 5.425660411 | 0.037101683 | 0.788577748 | 0.047048859 | 0.962474287 | 0.982786597 |
| ENSG00000161618 | ALDH16A1 | 680.6716027 | -0.11277924 | 0.080797187 | -1.395831271 | 0.162765311 | 0.310042036 |
| ENSG00000161638 | ITGA5 | 146.4900144 | 0.776696438 | 0.160596129 | 4.83633351 | 1.32E-06 | 7.53E-06 |
| ENSG00000161642 | ZNF385A | 1627.31053 | -0.370173661 | 0.064049438 | -5.779498998 | 7.49E-09 | 5.45E-08 |
| ENSG00000161647 | MPP3 | 216.4653263 | 0.179346921 | 0.135258947 | 1.325952371 | 0.184855474 | 0.341752453 |
| ENSG00000161653 | NAGS | 41.14519484 | -0.03544508 | 0.290095419 | -0.122184211 | 0.90275313 | 0.950907167 |
| ENSG00000161654 | LSM12 | 268.3790239 | 0.108689218 | 0.113850235 | 0.954668363 | 0.339745434 | 0.530782032 |
| ENSG00000161664 | ASB16 | 6.459281092 | 0.730899918 | 0.700373821 | 1.043585434 | 0.296677235 | 0.482987422 |
| ENSG00000161671 | EMC10 | 3073.041967 | 0.460166243 | 0.059269192 | 7.764004027 | 8.23E-15 | 9.71E-14 |
| ENSG00000161677 | JOSD2 | 429.8541902 | -0.117147406 | 0.101206975 | -1.157503286 | 0.247066798 | 0.424302071 |
| ENSG00000161682 | FAM171A2 | 461.4942935 | 0.118378879 | 0.097309727 | 1.216516398 | 0.223788268 | 0.395173803 |
| ENSG00000161692 | DBF4B | 585.5760686 | -0.017240691 | 0.082782163 | -0.208265771 | 0.835021457 | 0.913001315 |
| ENSG00000161714 | PLCD3 | 973.3141458 | 0.523161572 | 0.071743922 | 7.292068223 | 3.05E-13 | 3.23E-12 |
| ENSG00000161791 | FMNL3 | 341.661778 | -0.61510316 | 0.105663608 | -5.821334038 | 5.84E-09 | 4.31E-08 |
| ENSG00000161800 | RACGAP1 | 3410.231181 | -0.2074603 | 0.048101412 | -4.312977288 | 1.61E-05 | 8.09E-05 |
| ENSG00000161813 | LARP4 | 2420.269547 | 0.16834824 | 0.061711881 | 2.727971292 | 0.006372515 | 0.019976155 |
| ENSG00000161835 | TAMALIN | 12.04073356 | 0.683656885 | 0.508201428 | 1.345247863 | 0.178545204 | 0.333156579 |
| ENSG00000161847 | RAVER1 | 1423.526349 | 0.087965689 | 0.060411912 | 1.456098423 | 0.145365414 | 0.28455351 |
| ENSG00000161860 | SYCE2 | 25.90808638 | -1.759259992 | 0.37224421 | -4.726090943 | 2.29E-06 | 1.27E-05 |
| ENSG00000161888 | SPC24 | 747.5642993 | -0.260768555 | 0.075717278 | -3.443976888 | 0.000573225 | 0.002243606 |
| ENSG00000161904 | LEMD2 | 1642.548366 | -0.055319616 | 0.0613713 | -0.901392285 | 0.367379782 | 0.559881506 |
| ENSG00000161905 | ALOX15 | 496.4319 | -0.778728682 | 0.101131996 | -7.700121763 | 1.36E-14 | 1.58E-13 |
| ENSG00000161912 | ADCY10P1 | 72.83485395 | 0.334530292 | 0.22487582 | 1.487622333 | 0.13685052 | 0.271828811 |
| ENSG00000161914 | ZNF653 | 124.2440238 | -0.052907154 | 0.164269288 | -0.322075748 | 0.74739531 | 0.860057608 |
| ENSG00000161920 | MED11 | 372.8306681 | -0.071602827 | 0.104308184 | -0.686454541 | 0.492426521 | 0.675169708 |
| ENSG00000161921 | CXCL16 | 774.9714314 | 0.881234863 | 0.0758415 | 11.61942819 | 3.28E-31 | 9.58E-30 |
| ENSG00000161955 | TNFSF13 | 38.391446 | 0.418145498 | 0.293540199 | 1.424491429 | 0.154304264 | 0.297632206 |

| | | | | | | | |
|---|---|---|---|---|---|---|---|
| ENSG00000161956 | SENP3 | 267.6371489 | 0.049573448 | 0.113001376 | 0.438697734 | 0.660880569 | 0.803977317 |
| ENSG00000161958 | FGF11 | 14.76621373 | 0.241391764 | 0.4575076 | 0.527623506 | 0.597760675 | 0.758176522 |
| ENSG00000161960 | EIF4A1 | 140.5059935 | 0.169564352 | 0.168159823 | 1.008352346 | 0.313285342 | 0.501553893 |
| ENSG00000161970 | RPL26 | 572.11502 | 0.2928751 | 0.082271263 | 3.559871206 | 0.000371037 | 0.001499046 |
| ENSG00000161980 | POLR3K | 469.2065413 | -0.009466795 | 0.088515551 | -0.106950644 | 0.914828135 | 0.957703529 |
| ENSG00000161981 | SNRNP25 | 1019.156744 | -0.14661546 | 0.070141367 | -2.090285175 | 0.03659219 | 0.092102792 |
| ENSG00000161996 | WDR90 | 2023.555295 | 0.683358235 | 0.069285809 | 9.862888947 | 6.03E-23 | 1.20E-21 |
| ENSG00000161999 | JMJD8 | 2074.926135 | 0.247813943 | 0.062414688 | 3.970442716 | 7.17E-05 | 0.000327851 |
| ENSG00000162004 | CCDC78 | 1057.034936 | 0.425407285 | 0.067708059 | 6.282963805 | 3.32E-10 | 2.76E-09 |
| ENSG00000162006 | MSLNL | 11.73194391 | 2.372823449 | 0.609517671 | 3.892952674 | 9.90E-05 | 0.000441551 |
| ENSG00000162009 | SSTR5 | 1.719990989 | 1.284343266 | 1.388086563 | 0.925261651 | 0.354829821 | NA |
| ENSG00000162032 | SPSB3 | 896.2088373 | -0.038196246 | 0.081433438 | -0.469048668 | 0.639034849 | 0.787361694 |
| ENSG00000162039 | MEIOB | 3.775330437 | -0.753669857 | 0.936925046 | -0.804407845 | 0.421161472 | 0.612220022 |
| ENSG00000162062 | TEDC2 | 505.093366 | -0.063015254 | 0.086111621 | -0.731785715 | 0.46429937 | 0.650563617 |
| ENSG00000162063 | CCNF | 1738.613133 | -0.108567441 | 0.063497703 | -1.709785334 | 0.087305576 | 0.189730003 |
| ENSG00000162065 | TBC1D24 | 802.2033949 | -0.014861253 | 0.075462556 | -0.196935463 | 0.843878042 | 0.917708773 |
| ENSG00000162066 | AMDHD2 | 660.5669652 | 0.550447971 | 0.080632619 | 6.826616555 | 8.69E-12 | 8.19E-11 |
| ENSG00000162068 | NTN3 | 7.141566803 | 0.763347591 | 0.676143721 | 1.128972387 | 0.258909482 | 0.438739453 |
| ENSG00000162069 | BICDL2 | 1142.617495 | 0.071329189 | 0.065974593 | 1.081161486 | 0.279625287 | 0.462933542 |
| ENSG00000162073 | PAQR4 | 1790.453958 | -0.019211456 | 0.057741536 | -0.33271467 | 0.739349675 | 0.856254116 |
| ENSG00000162076 | FLYWCH2 | 883.3651765 | -0.337490358 | 0.086466836 | -3.903119112 | 9.50E-05 | 0.000424475 |
| ENSG00000162078 | ZG16B | 440.3975975 | -0.519149343 | 0.095122843 | -5.457672685 | 4.82E-08 | 3.23E-07 |
| ENSG00000162086 | ZNF75A | 1158.562326 | 0.10030135 | 0.064357671 | 1.558498763 | 0.119115061 | 0.244103939 |
| ENSG00000162104 | ADCY9 | 1538.535308 | -0.217041171 | 0.067616593 | -3.209880332 | 0.001327902 | 0.004856063 |
| ENSG00000162105 | SHANK2 | 2531.926671 | -0.254889761 | 0.057201439 | -4.45600257 | 8.35E-06 | 4.34E-05 |
| ENSG00000162129 | CLPB | 915.191863 | -0.223343522 | 0.072537582 | -3.079004237 | 0.002076937 | 0.007275148 |
| ENSG00000162139 | NEU3 | 771.1427054 | 0.129704602 | 0.077173712 | 1.680683729 | 0.092824363 | 0.199493244 |
| ENSG00000162144 | CYB561A3 | 1635.576007 | 0.190693066 | 0.064008344 | 2.979190718 | 0.002890108 | 0.009826454 |
| ENSG00000162148 | PPP1R32 | 39.23211348 | 1.409524533 | 0.30336855 | 4.646244748 | 3.38E-06 | 1.85E-05 |
| ENSG00000162174 | ASRGL1 | 68.34341548 | -3.944974203 | 0.333803303 | -11.81825994 | 3.14E-32 | 9.52E-31 |
| ENSG00000162188 | GNG3 | 2.798468831 | 0.703939719 | 1.069999377 | 0.657887971 | 0.510610121 | 0.690526214 |
| ENSG00000162191 | UBXN1 | 2862.770666 | 0.101580802 | 0.055422197 | 1.832854118 | 0.066824269 | 0.152790663 |
| ENSG00000162194 | LBHD1 | 647.1811932 | -1.189785739 | 0.10227068 | -11.63369346 | 2.78E-31 | 8.14E-30 |

| | | | | | | | |
|---|---|---|---|---|---|---|---|
| ENSG00000162222 | TTC9C | 648.283653 | -0.185687822 | 0.084300737 | -2.202683263 | 0.027617081 | 0.072354481 |
| ENSG00000162227 | TAF6L | 775.8487019 | 0.015598347 | 0.078201052 | 0.199464674 | 0.841899274 | 0.917009895 |
| ENSG00000162231 | NXF1 | 3392.713364 | -0.164899351 | 0.059001111 | -2.794851609 | 0.005192354 | 0.016642812 |
| ENSG00000162236 | STX5 | 1844.414595 | -0.030576228 | 0.064051393 | -0.477370222 | 0.633098515 | 0.783728883 |
| ENSG00000162241 | SLC25A45 | 215.2424032 | -1.20424431 | 0.138874861 | -8.67143483 | 4.27E-18 | 6.33E-17 |
| ENSG00000162244 | RPL29 | 5513.384095 | 0.175535364 | 0.047100557 | 3.726821386 | 0.00019391 | 0.00082498 |
| ENSG00000162298 | SYVN1 | 2280.599101 | 0.000196376 | 0.059142094 | 0.003320416 | 0.997350697 | 0.998438795 |
| ENSG00000162300 | ZFPL1 | 943.3380069 | -0.048534774 | 0.070966819 | -0.683907989 | 0.494033269 | 0.67660437 |
| ENSG00000162302 | RPS6KA4 | 990.3947876 | 0.016716662 | 0.083334416 | 0.200597342 | 0.841013437 | 0.916568399 |
| ENSG00000162337 | LRP5 | 4357.821783 | 0.006896253 | 0.060406864 | 0.114163404 | 0.909108262 | 0.95454703 |
| ENSG00000162341 | TPCN2 | 415.3894359 | 0.077573818 | 0.094423763 | 0.821549743 | 0.411333203 | 0.603671145 |
| ENSG00000162368 | CMPK1 | 2324.323198 | 0.088145338 | 0.061298782 | 1.437962305 | 0.15044475 | 0.292251889 |
| ENSG00000162373 | BEND5 | 7.964688082 | 1.165437106 | 0.648506746 | 1.797108686 | 0.072318367 | 0.163093132 |
| ENSG00000162377 | COA7 | 1664.754685 | -0.057077678 | 0.060036936 | -0.950709387 | 0.341751922 | 0.53284776 |
| ENSG00000162378 | ZYG11B | 1156.567872 | -0.115170357 | 0.068175671 | -1.689317539 | 0.091158594 | 0.19652083 |
| ENSG00000162383 | SLC1A7 | 2.473154223 | -3.790398937 | 1.495172893 | -2.535090727 | 0.011241821 | 0.032935807 |
| ENSG00000162384 | CZIB | 957.5092292 | -0.231578842 | 0.068538042 | -3.378836555 | 0.000727933 | 0.002796647 |
| ENSG00000162385 | MAGOH | 687.0547531 | -0.371148591 | 0.077687692 | -4.777443929 | 1.78E-06 | 9.98E-06 |
| ENSG00000162390 | ACOT11 | 26.12140297 | 0.522100579 | 0.344669617 | 1.514785618 | 0.129826704 | 0.260995933 |
| ENSG00000162396 | PARS2 | 400.8195361 | 0.441171149 | 0.100635382 | 4.383857234 | 1.17E-05 | 5.96E-05 |
| ENSG00000162402 | USP24 | 2476.209393 | -0.242479015 | 0.055432145 | -4.374339349 | 1.22E-05 | 6.21E-05 |
| ENSG00000162407 | PLPP3 | 460.4786069 | 0.263812252 | 0.0900914 | 2.928273419 | 0.003408502 | 0.011416415 |
| ENSG00000162408 | NOL9 | 678.2657586 | -0.290020067 | 0.081911272 | -3.540661242 | 0.000399126 | 0.001604336 |
| ENSG00000162409 | PRKAA2 | 23.59249833 | 1.746903414 | 0.396201139 | 4.409132741 | 1.04E-05 | 5.35E-05 |
| ENSG00000162413 | KLHL21 | 1132.155304 | 0.570447727 | 0.064852888 | 8.796026552 | 1.42E-18 | 2.17E-17 |
| ENSG00000162415 | ZSWIM5 | 1195.52922 | -0.095691962 | 0.063346753 | -1.510605649 | 0.130888954 | 0.26278598 |
| ENSG00000162419 | GMEB1 | 842.1223925 | -0.067320516 | 0.07088633 | -0.949696743 | 0.342266365 | 0.53329254 |
| ENSG00000162426 | SLC45A1 | 7.393593227 | -0.487234414 | 0.654153401 | -0.744832042 | 0.456373262 | 0.643478245 |
| ENSG00000162430 | SELENON | 2241.53703 | 0.630200268 | 0.058471155 | 10.77796845 | 4.37E-27 | 1.09E-25 |
| ENSG00000162433 | AK4 | 1878.112242 | 0.381436483 | 0.060903414 | 6.262973743 | 3.78E-10 | 3.13E-09 |
| ENSG00000162434 | JAK1 | 3918.328918 | -0.074252842 | 0.047506692 | -1.562997521 | 0.118053178 | 0.242512752 |
| ENSG00000162437 | RAVER2 | 989.7687832 | 0.348071055 | 0.070414954 | 4.943141136 | 7.69E-07 | 4.53E-06 |
| ENSG00000162438 | CTRC | 1.618419641 | 1.767506494 | 1.515495449 | 1.166289542 | 0.2434974 | NA |

| | | | | | | | |
|---|---|---|---|---|---|---|---|
| ENSG00000162441 | LZIC | 1021.335043 | -0.057946457 | 0.066964796 | -0.865327171 | 0.386859241 | 0.579392244 |
| ENSG00000162444 | RBP7 | 5.208863278 | 0.647553675 | 0.872653829 | 0.742051033 | 0.458056416 | 0.64527349 |
| ENSG00000162458 | FBLIM1 | 305.6157827 | 0.837318479 | 0.11250058 | 7.44279258 | 9.86E-14 | 1.08E-12 |
| ENSG00000162461 | SLC25A34 | 34.25400795 | 0.632957896 | 0.309772319 | 2.04330038 | 0.041022712 | 0.101265578 |
| ENSG00000162482 | AKR7A3 | 8.850198301 | -0.073081477 | 0.599070781 | -0.121991389 | 0.902905838 | 0.950907167 |
| ENSG00000162490 | DRAXIN | 5.917462619 | 1.07500384 | 0.742863492 | 1.447108186 | 0.147866626 | 0.288366947 |
| ENSG00000162496 | DHRS3 | 43.58283308 | 1.452534627 | 0.316901964 | 4.583545673 | 4.57E-06 | 2.45E-05 |
| ENSG00000162510 | MATN1 | 6.592814698 | -0.569913852 | 0.722502198 | -0.7888057 | 0.430225578 | 0.620481249 |
| ENSG00000162511 | LAPTM5 | 9.816390249 | -0.207369567 | 0.584590356 | -0.354726287 | 0.722794656 | 0.845926239 |
| ENSG00000162512 | SDC3 | 1678.17965 | 0.508645696 | 0.067124307 | 7.577667802 | 3.52E-14 | 3.97E-13 |
| ENSG00000162517 | PEF1 | 2248.049618 | -0.202117694 | 0.058794706 | -3.437685249 | 0.000586709 | 0.002292083 |
| ENSG00000162520 | SYNC | 61.67707234 | 0.148151963 | 0.229266632 | 0.646199411 | 0.518150221 | 0.695932225 |
| ENSG00000162521 | RBBP4 | 5548.801868 | 0.144201413 | 0.051341672 | 2.808662193 | 0.004974782 | 0.016039563 |
| ENSG00000162522 | KIAA1522 | 13253.24006 | 0.20808409 | 0.051042 | 4.076722917 | 4.57E-05 | 0.00021493 |
| ENSG00000162526 | TSSK3 | 6.904492569 | -0.220686222 | 0.670255608 | -0.329256807 | 0.741961588 | 0.857590509 |
| ENSG00000162542 | TMCO4 | 336.542985 | -0.210247269 | 0.107777879 | -1.95074602 | 0.051087266 | 0.122064564 |
| ENSG00000162545 | CAMK2N1 | 704.7444095 | -0.334753574 | 0.076964269 | -4.349467313 | 1.36E-05 | 6.91E-05 |
| ENSG00000162552 | WNT4 | 45.52044909 | 0.961095501 | 0.277065496 | 3.468838652 | 0.000522713 | 0.002060671 |
| ENSG00000162571 | TTLL10 | 5.554420986 | 0.880677982 | 0.7842329 | 1.122980154 | 0.261445923 | 0.44171229 |
| ENSG00000162572 | SCNN1D | 109.8250427 | -0.31468096 | 0.177036005 | -1.777496954 | 0.075486512 | 0.168728498 |
| ENSG00000162576 | MXRA8 | 6.24079755 | 1.083758991 | 0.824341961 | 1.314695893 | 0.188612127 | 0.347196453 |
| ENSG00000162585 | FAAP20 | 642.3191825 | 0.009713161 | 0.085416085 | 0.11371583 | 0.909463064 | 0.954587195 |
| ENSG00000162591 | MEGF6 | 628.9985747 | -0.451079837 | 0.079636346 | -5.664245764 | 1.48E-08 | 1.04E-07 |
| ENSG00000162595 | DIRAS3 | 2.510005926 | 0.216706442 | 1.139605583 | 0.190159161 | 0.849184412 | 0.921106747 |
| ENSG00000162599 | NFIA | 279.6610775 | -1.221552818 | 0.114473645 | -10.67103973 | 1.39E-26 | 3.37E-25 |
| ENSG00000162600 | OMA1 | 300.719384 | -0.209190681 | 0.106456415 | -1.965035935 | 0.049410079 | 0.118588324 |
| ENSG00000162601 | MYSM1 | 1178.538347 | -0.29261975 | 0.077011434 | -3.799692285 | 0.000144876 | 0.000629561 |
| ENSG00000162604 | TM2D1 | 367.7797956 | 0.093941935 | 0.099205145 | 0.9469462 | 0.343666194 | 0.53463329 |
| ENSG00000162607 | USP1 | 2840.558498 | -0.270202754 | 0.065440131 | -4.129006908 | 3.64E-05 | 0.000173651 |
| ENSG00000162613 | FUBP1 | 4489.373954 | -0.082119477 | 0.047325763 | -1.735196048 | 0.082706086 | 0.181522926 |
| ENSG00000162614 | NEXN | 254.9436269 | 6.234267399 | 0.346482438 | 17.99302564 | 2.21E-72 | 2.15E-70 |
| ENSG00000162616 | DNAJB4 | 146.6920737 | 0.746827129 | 0.165094847 | 4.523624699 | 6.08E-06 | 3.22E-05 |
| ENSG00000162620 | LRRIQ3 | 24.24829693 | -0.156888686 | 0.361500748 | -0.4339927 | 0.664293742 | 0.806241836 |

| | | | | | | | |
|---|---|---|---|---|---|---|---|
| ENSG00000162621 | LRRC53 | 5.202127801 | 1.499482445 | 0.841999103 | 1.780859908 | 0.07493534 | 0.167758045 |
| ENSG00000162623 | TYW3 | 1044.150501 | 0.35093542 | 0.068988288 | 5.086884033 | 3.64E-07 | 2.22E-06 |
| ENSG00000162627 | SNX7 | 789.5878008 | 0.275522854 | 0.073470142 | 3.750133674 | 0.00017674 | 0.000757411 |
| ENSG00000162631 | NTNG1 | 4.162827851 | -0.360760831 | 0.871636315 | -0.41388917 | 0.678955275 | 0.817353392 |
| ENSG00000162636 | EEIG2 | 5774.481602 | -0.773913003 | 0.055907552 | -13.84272752 | 1.41E-43 | 6.47E-42 |
| ENSG00000162639 | HENMT1 | 1311.301298 | -0.242756361 | 0.062946397 | -3.856556897 | 0.000114995 | 0.000508088 |
| ENSG00000162641 | AKNAD1 | 1.497762967 | 1.846018595 | 1.694416448 | 1.089471598 | 0.275945973 | NA |
| ENSG00000162642 | C1orf52 | 466.643893 | 0.076836485 | 0.097667027 | 0.786718785 | 0.431446505 | 0.621507498 |
| ENSG00000162643 | DNAI3 | 12.05856821 | 0.725845749 | 0.512104422 | 1.417378404 | 0.156372323 | 0.30055979 |
| ENSG00000162645 | GBP2 | 576.2813997 | -1.674901536 | 0.088347304 | -18.958151 | 3.78E-80 | 4.30E-78 |
| ENSG00000162650 | ATXN7L2 | 566.1651791 | 0.526459027 | 0.088300469 | 5.962131716 | 2.49E-09 | 1.91E-08 |
| ENSG00000162664 | ZNF326 | 918.4946385 | 0.043397448 | 0.074292182 | 0.584145556 | 0.559122387 | 0.728807456 |
| ENSG00000162669 | HFM1 | 2.576319593 | -0.407797999 | 1.106300692 | -0.368614069 | 0.712415408 | 0.839972059 |
| ENSG00000162670 | BRINP3 | 198.3157255 | 2.306499745 | 0.155316108 | 14.85035761 | 6.92E-50 | 3.82E-48 |
| ENSG00000162676 | GFI1 | 43.6485311 | 0.147872174 | 0.27010931 | 0.547453079 | 0.584067498 | 0.748394593 |
| ENSG00000162687 | KCNT2 | 2.095918861 | -1.797922202 | 1.295777021 | -1.387524376 | 0.165281924 | 0.313819511 |
| ENSG00000162688 | AGL | 1005.391407 | 0.177221608 | 0.073098601 | 2.424418605 | 0.01533292 | 0.043376148 |
| ENSG00000162694 | EXTL2 | 97.31231947 | 0.001588257 | 0.185495852 | 0.008562221 | 0.993168419 | 0.996338984 |
| ENSG00000162695 | SLC30A7 | 463.7814953 | -0.138783015 | 0.092064051 | -1.507461522 | 0.1316924 | 0.264052412 |
| ENSG00000162702 | ZNF281 | 1589.888757 | 0.03044276 | 0.06065432 | 0.501905879 | 0.615733729 | 0.772318429 |
| ENSG00000162704 | ARPC5 | 3133.305488 | -0.277063375 | 0.053478159 | -5.180869701 | 2.21E-07 | 1.39E-06 |
| ENSG00000162711 | NLRP3 | 1.889733854 | 1.461831553 | 1.352702832 | 1.080674571 | 0.279841898 | 0.463190856 |
| ENSG00000162714 | ZNF496 | 2623.982324 | 0.234757238 | 0.054719227 | 4.290214789 | 1.79E-05 | 8.92E-05 |
| ENSG00000162729 | IGSF8 | 770.7950296 | 0.658207016 | 0.079450902 | 8.28444984 | 1.19E-16 | 1.60E-15 |
| ENSG00000162733 | DDR2 | 16.61299529 | -1.10463239 | 0.439630193 | -2.51263996 | 0.011983156 | 0.034893831 |
| ENSG00000162734 | PEA15 | 2980.914334 | 0.284367374 | 0.05517944 | 5.153502365 | 2.56E-07 | 1.59E-06 |
| ENSG00000162735 | PEX19 | 1006.779509 | 0.498765018 | 0.06676491 | 7.470466371 | 7.99E-14 | 8.76E-13 |
| ENSG00000162736 | NCSTN | 2403.203087 | 0.388560798 | 0.051211753 | 7.587336376 | 3.27E-14 | 3.70E-13 |
| ENSG00000162745 | OLFML2B | 2.217879346 | 1.729508648 | 1.287037476 | 1.343790434 | 0.179016164 | 0.333812913 |
| ENSG00000162746 | FCRLB | 32.73528897 | 0.211686563 | 0.306459601 | 0.690748674 | 0.489723495 | 0.672936425 |
| ENSG00000162755 | KLHDC9 | 121.1730722 | 0.534853128 | 0.166271969 | 3.216736609 | 0.001296576 | 0.004752868 |
| ENSG00000162757 | C1orf74 | 143.3563676 | -0.119142749 | 0.150350147 | -0.792435205 | 0.428106959 | 0.61877975 |
| ENSG00000162769 | FLVCR1 | 331.3094816 | -0.174358709 | 0.103884708 | -1.678386669 | 0.093271638 | 0.200210666 |

| | | | | | | | |
|---|---|---|---|---|---|---|---|
| ENSG00000162772 | ATF3 | 194.1971952 | 0.40398027 | 0.130859753 | 3.087123884 | 0.002021033 | 0.007097751 |
| ENSG00000162775 | RBM15 | 627.387643 | -0.247845299 | 0.081925558 | -3.025250036 | 0.002484277 | 0.008564669 |
| ENSG00000162777 | DENND2D | 1557.977279 | 0.218843294 | 0.061225755 | 3.574366622 | 0.000351077 | 0.001424425 |
| ENSG00000162779 | AXDND1 | 3.791808147 | -1.615701583 | 0.944344645 | -1.710923646 | 0.087095209 | 0.189380483 |
| ENSG00000162782 | TDRD5 | 429.6934286 | -0.03233934 | 0.095455514 | -0.338789651 | 0.734768198 | 0.853727216 |
| ENSG00000162783 | IER5 | 1003.872723 | -0.167219914 | 0.068368867 | -2.445848833 | 0.014451158 | 0.041120015 |
| ENSG00000162804 | SNED1 | 38.72461699 | -0.142080666 | 0.284061234 | -0.500176191 | 0.616951021 | 0.772972842 |
| ENSG00000162813 | BPNT1 | 826.679838 | -0.441676745 | 0.076580852 | -5.767456692 | 8.05E-09 | 5.84E-08 |
| ENSG00000162814 | SPATA17 | 374.9967645 | 0.222796679 | 0.099396006 | 2.241505346 | 0.02499336 | 0.066451979 |
| ENSG00000162817 | C1orf115 | 541.2702937 | -0.741376103 | 0.08896441 | -8.33340099 | 7.86E-17 | 1.07E-15 |
| ENSG00000162819 | BROX | 1289.571047 | 0.02280666 | 0.069239943 | 0.329385881 | 0.741864039 | 0.857590509 |
| ENSG00000162825 | NBPF20 | 256.0338 | 0.261695113 | 0.119734743 | 2.185623866 | 0.028843135 | 0.075026819 |
| ENSG00000162836 | ACP6 | 885.9964633 | 0.17112815 | 0.072638979 | 2.355872185 | 0.018479275 | 0.051014695 |
| ENSG00000162840 | MT2P1 | 24.04738497 | -0.190453068 | 0.362364447 | -0.52558431 | 0.59917706 | 0.75931509 |
| ENSG00000162851 | TFB2M | 751.6372565 | 0.131362526 | 0.077337169 | 1.69856911 | 0.089400401 | 0.193330052 |
| ENSG00000162852 | CNST | 1163.184081 | -0.008396932 | 0.070845184 | -0.118525088 | 0.905651618 | 0.95262805 |
| ENSG00000162869 | PPP1R21 | 801.7917671 | -0.27404195 | 0.074314548 | -3.687594895 | 0.000226384 | 0.000950969 |
| ENSG00000162873 | KLHDC8A | 2.045314161 | 1.515013457 | 1.316955544 | 1.150390736 | 0.249982974 | 0.42805176 |
| ENSG00000162878 | PKDCC | 5.430029139 | 0.883828454 | 0.76029238 | 1.162484957 | 0.245038532 | 0.421979124 |
| ENSG00000162885 | B3GALNT2 | 1219.205456 | 0.104863513 | 0.064632995 | 1.622445509 | 0.104707992 | 0.220044094 |
| ENSG00000162889 | MAPKAPK2 | 6836.894591 | 0.390924519 | 0.049315892 | 7.926948207 | 2.25E-15 | 2.77E-14 |
| ENSG00000162891 | IL20 | 46.41874524 | 2.023196352 | 0.313395182 | 6.455735337 | 1.08E-10 | 9.28E-10 |
| ENSG00000162892 | IL24 | 1.667083704 | 0.369517592 | 1.364546011 | 0.270798924 | 0.786545687 | NA |
| ENSG00000162894 | FCMR | 58.68474633 | 1.978129418 | 0.278682711 | 7.098141876 | 1.26E-12 | 1.28E-11 |
| ENSG00000162909 | CAPN2 | 376.518153 | 0.105644913 | 0.097258534 | 1.086227685 | 0.277378272 | 0.460157474 |
| ENSG00000162910 | MRPL55 | 586.0044906 | -0.040683242 | 0.085592585 | -0.475312693 | 0.634564113 | 0.784558953 |
| ENSG00000162913 | OBSCN-AS1 | 140.1573539 | 0.642917869 | 0.159013579 | 4.043163316 | 5.27E-05 | 0.000245732 |
| ENSG00000162923 | WDR26 | 4426.359545 | -0.005210841 | 0.050293635 | -0.103608351 | 0.91748016 | 0.959385307 |
| ENSG00000162924 | REL | 363.8673835 | 0.817602532 | 0.109047927 | 7.497643941 | 6.50E-14 | 7.19E-13 |
| ENSG00000162927 | PUS10 | 191.9029172 | -0.120141126 | 0.138742091 | -0.865931344 | 0.386527813 | 0.579047086 |
| ENSG00000162928 | PEX13 | 883.1843099 | 0.323783501 | 0.070992142 | 4.560835764 | 5.10E-06 | 2.72E-05 |
| ENSG00000162929 | SANBR | 313.7750206 | 0.208729491 | 0.108100394 | 1.930885565 | 0.053497206 | 0.127055171 |
| ENSG00000162946 | DISC1 | 144.1463222 | 0.749245577 | 0.155272804 | 4.825349692 | 1.40E-06 | 7.93E-06 |

| | | | | | | | |
|---|---|---|---|---|---|---|---|
| ENSG00000162949 | CAPN13 | 120.8305413 | -0.181518751 | 0.164880294 | -1.100912343 | 0.270934809 | 0.452841131 |
| ENSG00000162959 | MEMO1 | 32.05069927 | 0.425103639 | 0.309688289 | 1.372682322 | 0.169851127 | 0.320412952 |
| ENSG00000162961 | DPY30 | 1287.669923 | -0.300429122 | 0.062874396 | -4.778242706 | 1.77E-06 | 9.95E-06 |
| ENSG00000162971 | TYW5 | 424.0341534 | 0.032226521 | 0.101239319 | 0.318320206 | 0.750242058 | 0.862079384 |
| ENSG00000162972 | MAIP1 | 421.7803527 | -0.174422075 | 0.092164929 | -1.892499431 | 0.058424469 | 0.136524124 |
| ENSG00000162975 | KCNF1 | 19.52228263 | 2.693867348 | 0.500231045 | 5.385246232 | 7.23E-08 | 4.74E-07 |
| ENSG00000162976 | SLC66A3 | 465.2241229 | 0.342316642 | 0.091840701 | 3.727286906 | 0.000193552 | 0.000823916 |
| ENSG00000162980 | ARL5A | 327.7908531 | 0.335006584 | 0.106629356 | 3.141785691 | 0.001679209 | 0.005993629 |
| ENSG00000162981 | LRATD1 | 1815.241 | -1.024069117 | 0.061196967 | -16.73398475 | 7.41E-63 | 5.69E-61 |
| ENSG00000162989 | KCNJ3 | 85.79087454 | -0.306725041 | 0.1943984 | -1.577816696 | 0.114607727 | 0.23666408 |
| ENSG00000162994 | CLHC1 | 70.58478205 | 0.646382632 | 0.213902272 | 3.021859586 | 0.002512271 | 0.008653376 |
| ENSG00000162997 | PRORSD1P | 42.81172191 | 0.237130471 | 0.278839532 | 0.850419125 | 0.395092107 | 0.588013388 |
| ENSG00000162998 | FRZB | 2.102521799 | 0.481614948 | 1.204096128 | 0.399980481 | 0.689170894 | 0.82436022 |
| ENSG00000162999 | DUSP19 | 18.44029075 | -0.140848447 | 0.409654722 | -0.343822345 | 0.730979895 | 0.851072364 |
| ENSG00000163001 | CFAP36 | 968.5527116 | 0.251232571 | 0.067724372 | 3.709633077 | 0.00020756 | 0.000877682 |
| ENSG00000163002 | NUP35 | 373.4988743 | -0.161095696 | 0.098076035 | -1.642559214 | 0.100474165 | 0.212764456 |
| ENSG00000163006 | CCDC138 | 186.4657656 | 0.115642047 | 0.135384854 | 0.854172707 | 0.393009303 | 0.585963435 |
| ENSG00000163013 | FBXO41 | 1497.498595 | 0.121241679 | 0.06911527 | 1.754195266 | 0.079397054 | 0.175689054 |
| ENSG00000163016 | ALMS1P1 | 15.04731657 | -0.141792856 | 0.476797069 | -0.297386174 | 0.766171698 | 0.87225701 |
| ENSG00000163017 | ACTG2 | 21.89229473 | -0.924055281 | 0.384319006 | -2.404396521 | 0.016199192 | 0.045512487 |
| ENSG00000163026 | WDCP | 236.7486267 | -0.096948864 | 0.123703253 | -0.783721221 | 0.433203708 | 0.623170067 |
| ENSG00000163029 | SMC6 | 1020.447505 | 0.142911143 | 0.075974897 | 1.881031082 | 0.059967691 | 0.139505002 |
| ENSG00000163040 | CCDC74A | 267.5892831 | 1.293621784 | 0.121337768 | 10.66132831 | 1.54E-26 | 3.72E-25 |
| ENSG00000163041 | H3-3A | 367.7109262 | -0.042012982 | 0.103599285 | -0.405533512 | 0.68508539 | 0.821615373 |
| ENSG00000163050 | COQ8A | 1269.873854 | 0.051422077 | 0.063642029 | 0.807989277 | 0.419096755 | 0.610728065 |
| ENSG00000163053 | SLC16A14 | 639.8658618 | 0.26054521 | 0.080648003 | 3.230646754 | 0.001235105 | 0.004545693 |
| ENSG00000163060 | TEKT4 | 10.94846206 | 0.019180497 | 0.52989784 | 0.036196594 | 0.971125602 | 0.986914513 |
| ENSG00000163069 | SGCB | 620.3422582 | -0.005105799 | 0.079306211 | -0.064380826 | 0.948666997 | 0.97640107 |
| ENSG00000163071 | SPATA18 | 253.584014 | -0.439354987 | 0.115761156 | -3.795357638 | 0.000147431 | 0.000639694 |
| ENSG00000163072 | NOSTRIN | 1.962481018 | -0.579728676 | 1.327949591 | -0.436559249 | 0.662431022 | 0.804963395 |
| ENSG00000163075 | CFAP221 | 6.992492823 | -1.030103685 | 0.677605833 | -1.520210768 | 0.128458012 | 0.258879627 |
| ENSG00000163082 | SGPP2 | 94.51616048 | 1.17121924 | 0.198526084 | 5.899573572 | 3.64E-09 | 2.75E-08 |
| ENSG00000163083 | INHBB | 363.4974793 | 0.059203356 | 0.100825851 | 0.587184295 | 0.557079939 | 0.727416829 |

| | | | | | | | |
|---|---|---|---|---|---|---|---|
| ENSG00000163093 | BBS5 | 149.8293269 | 0.249902245 | 0.149649302 | 1.669919216 | 0.094935348 | 0.203230392 |
| ENSG00000163104 | SMARCAD1 | 1327.801914 | -0.001457104 | 0.064841556 | -0.022471757 | 0.982071641 | 0.991627389 |
| ENSG00000163110 | PDLIM5 | 1540.090164 | -0.049496808 | 0.059120479 | -0.837219341 | 0.402469288 | 0.5952209 |
| ENSG00000163116 | STPG2 | 2.13843932 | 3.381866788 | 1.540921669 | 2.194703895 | 0.028184845 | 0.07363139 |
| ENSG00000163125 | RPRD2 | 5638.996892 | 0.204073385 | 0.050041868 | 4.078052943 | 4.54E-05 | 0.000213792 |
| ENSG00000163126 | ANKRD23 | 38.20033374 | -0.188054649 | 0.286249273 | -0.65696114 | 0.511205904 | 0.690883907 |
| ENSG00000163131 | CTSS | 5.142720478 | -0.070042289 | 0.796017769 | -0.087990861 | 0.92988394 | 0.966032078 |
| ENSG00000163132 | MSX1 | 40.29251507 | -0.52819146 | 0.276043688 | -1.913434295 | 0.055692477 | 0.131379374 |
| ENSG00000163138 | PACRGL | 319.8778735 | 0.188683809 | 0.105926636 | 1.781268773 | 0.074868553 | 0.16767962 |
| ENSG00000163141 | BNIPL | 345.810734 | 2.253532241 | 0.123547762 | 18.24017043 | 2.48E-74 | 2.55E-72 |
| ENSG00000163155 | LYSMD1 | 254.9506414 | -0.178025955 | 0.118786729 | -1.498702383 | 0.133950858 | 0.267469008 |
| ENSG00000163156 | SCNM1 | 1617.223961 | 0.323352437 | 0.060335973 | 5.359198218 | 8.36E-08 | 5.45E-07 |
| ENSG00000163157 | TMOD4 | 69.71706489 | 0.721530123 | 0.229628637 | 3.142160889 | 0.001677059 | 0.005986885 |
| ENSG00000163159 | VPS72 | 3608.427002 | 0.26852367 | 0.049107222 | 5.468109563 | 4.55E-08 | 3.05E-07 |
| ENSG00000163161 | ERCC3 | 3301.978429 | -0.048794702 | 0.05079466 | -0.960626616 | 0.336739941 | 0.527939738 |
| ENSG00000163162 | RNF149 | 681.6575589 | 0.457601295 | 0.078116529 | 5.857931745 | 4.69E-09 | 3.50E-08 |
| ENSG00000163166 | IWS1 | 1662.710576 | 0.008165705 | 0.058088596 | 0.140573288 | 0.888207052 | 0.942392571 |
| ENSG00000163170 | BOLA3 | 487.3100069 | 0.015824416 | 0.088179896 | 0.179456057 | 0.857579618 | 0.926087485 |
| ENSG00000163171 | CDC42EP3 | 18.18226466 | 0.617372059 | 0.433611182 | 1.423791833 | 0.154506741 | 0.297847448 |
| ENSG00000163191 | S100A11 | 8006.103114 | 0.002097769 | 0.043645636 | 0.048063656 | 0.961665511 | 0.982361474 |
| ENSG00000163214 | DHX57 | 912.9754682 | 0.200869622 | 0.069623939 | 2.885065454 | 0.003913323 | 0.012922359 |
| ENSG00000163220 | S100A9 | 5.385605248 | 3.243441684 | 1.045213683 | 3.103137413 | 0.001914807 | 0.006760917 |
| ENSG00000163235 | TGFA | 328.1124653 | 0.600235239 | 0.120321508 | 4.988594723 | 6.08E-07 | 3.62E-06 |
| ENSG00000163249 | CCNYL1 | 291.6188327 | -0.671687661 | 0.119027594 | -5.643125591 | 1.67E-08 | 1.17E-07 |
| ENSG00000163251 | FZD5 | 459.6136419 | 0.525175742 | 0.09206155 | 5.704615487 | 1.17E-08 | 8.31E-08 |
| ENSG00000163257 | DCAF16 | 856.5974696 | -0.090301871 | 0.071510979 | -1.262769331 | 0.20667209 | 0.372519662 |
| ENSG00000163273 | NPPC | 4.604500206 | -0.386792367 | 0.809812398 | -0.477632064 | 0.632912105 | 0.783582692 |
| ENSG00000163281 | GNPDA2 | 311.1822163 | 0.170866571 | 0.108104584 | 1.580567311 | 0.113977005 | 0.235679872 |
| ENSG00000163291 | PAQR3 | 313.2281992 | 0.203683581 | 0.1087773 | 1.872482419 | 0.061139892 | 0.141728893 |
| ENSG00000163293 | NIPAL1 | 391.7101593 | 0.340567625 | 0.098241101 | 3.46665116 | 0.000526985 | 0.002076442 |
| ENSG00000163297 | ANTXR2 | 415.0842311 | -0.267337958 | 0.097045218 | -2.754777243 | 0.005873211 | 0.018573401 |
| ENSG00000163312 | HELQ | 420.7468311 | -0.212916252 | 0.097184135 | -2.190854003 | 0.028462359 | 0.074213062 |
| ENSG00000163319 | MRPS18C | 577.4499571 | 0.022333448 | 0.080637985 | 0.276959406 | 0.781811277 | 0.881574709 |

| | | | | | | | |
|---|---|---|---|---|---|---|---|
| ENSG00000163320 | CGGBP1 | 3945.066612 | -0.101153215 | 0.054094862 | -1.869922784 | 0.061494541 | 0.142464026 |
| ENSG00000163322 | ABRAXAS1 | 573.3346334 | 0.457044711 | 0.099394916 | 4.598270497 | 4.26E-06 | 2.29E-05 |
| ENSG00000163328 | GPR155 | 42.28056911 | 1.098872127 | 0.28729831 | 3.824847162 | 0.000130853 | 0.00057253 |
| ENSG00000163344 | PMVK | 952.04693 | 0.069937623 | 0.078053922 | 0.896016777 | 0.370243803 | 0.562915671 |
| ENSG00000163346 | PBXIP1 | 1416.317054 | 0.589891092 | 0.068455004 | 8.617209242 | 6.86E-18 | 1.00E-16 |
| ENSG00000163347 | CLDN1 | 373.0687341 | 1.434649146 | 0.105408976 | 13.61031296 | 3.48E-42 | 1.55E-40 |
| ENSG00000163348 | PYGO2 | 2891.043482 | 0.328882515 | 0.055607576 | 5.914347266 | 3.33E-09 | 2.53E-08 |
| ENSG00000163349 | HIPK1 | 25517.20284 | 0.085593485 | 0.047963836 | 1.784542098 | 0.074335621 | 0.166708294 |
| ENSG00000163354 | DCST2 | 132.7201046 | 0.376531557 | 0.159439558 | 2.361594334 | 0.018196542 | 0.050341132 |
| ENSG00000163357 | DCST1 | 3.03483496 | 0.927391014 | 1.053372632 | 0.880401659 | 0.378641758 | 0.571322415 |
| ENSG00000163359 | COL6A3 | 5.559886228 | -0.847614751 | 0.766700346 | -1.105535892 | 0.268927451 | 0.450503558 |
| ENSG00000163362 | INAVA | 240.3542108 | 2.819854555 | 0.1631118 | 17.28786367 | 5.81E-67 | 4.86E-65 |
| ENSG00000163364 | LINC01116 | 40.03816105 | -0.741033225 | 0.288384395 | -2.569602366 | 0.010181531 | 0.030215306 |
| ENSG00000163374 | YY1AP1 | 2104.263934 | 0.290213079 | 0.055431869 | 5.235491498 | 1.65E-07 | 1.05E-06 |
| ENSG00000163376 | KBTBD8 | 74.47513265 | -0.030593819 | 0.207055737 | -0.147756443 | 0.882534986 | 0.939121523 |
| ENSG00000163378 | EOGT | 417.3460237 | 0.224349207 | 0.099141417 | 2.262921125 | 0.023640552 | 0.063302704 |
| ENSG00000163382 | NAXE | 2078.305971 | 0.116956892 | 0.060817068 | 1.92309324 | 0.054468341 | 0.128946937 |
| ENSG00000163389 | POGLUT1 | 680.1015401 | -0.094370764 | 0.078745548 | -1.198426652 | 0.230750962 | 0.40417096 |
| ENSG00000163393 | SLC22A15 | 51.26945865 | 0.080328561 | 0.247797169 | 0.324170615 | 0.745808868 | 0.859439164 |
| ENSG00000163395 | IGFN1 | 4.986091729 | 2.648110618 | 0.974378715 | 2.717742678 | 0.006572893 | 0.020522958 |
| ENSG00000163399 | ATP1A1 | 22539.36729 | -0.269282633 | 0.04158289 | -6.475803685 | 9.43E-11 | 8.17E-10 |
| ENSG00000163406 | SLC15A2 | 120.7603429 | -0.138608008 | 0.162601953 | -0.85243754 | 0.393971292 | 0.58690184 |
| ENSG00000163412 | EIF4E3 | 463.2065309 | 0.202346185 | 0.098398247 | 2.056400293 | 0.039743948 | 0.098646064 |
| ENSG00000163428 | LRRC58 | 2783.643325 | 0.025820198 | 0.062093109 | 0.415830325 | 0.677534167 | 0.816641527 |
| ENSG00000163430 | FSTL1 | 39.98526682 | 0.439801993 | 0.279052063 | 1.576057127 | 0.115012638 | 0.237350659 |
| ENSG00000163431 | LMOD1 | 5.086740185 | -0.645608184 | 0.803472942 | -0.803521998 | 0.421673087 | 0.612653152 |
| ENSG00000163435 | ELF3 | 1321.942221 | 0.398133041 | 0.071540439 | 5.565146702 | 2.62E-08 | 1.80E-07 |
| ENSG00000163444 | TMEM183A | 2568.305256 | 0.361802401 | 0.050892791 | 7.109109007 | 1.17E-12 | 1.19E-11 |
| ENSG00000163449 | TMEM169 | 19.79044402 | -0.149346472 | 0.39337024 | -0.379658797 | 0.70419871 | 0.834546593 |
| ENSG00000163462 | TRIM46 | 231.7445406 | -0.141497341 | 0.135475328 | -1.044450996 | 0.29627678 | 0.482506697 |
| ENSG00000163463 | KRTCAP2 | 190.7921333 | -0.135080458 | 0.153689594 | -0.878917393 | 0.379446067 | 0.571911484 |
| ENSG00000163466 | ARPC2 | 6082.049203 | -0.004445274 | 0.042327055 | -0.105022056 | 0.916358308 | 0.95873664 |
| ENSG00000163467 | TSACC | 16.37834741 | -0.087256213 | 0.443339389 | -0.196815838 | 0.843971657 | 0.917708773 |

| | | | | | | | |
|---|---|---|---|---|---|---|---|
| ENSG00000163468 | CCT3 | 14807.44538 | 0.131166323 | 0.043005621 | 3.049980944 | 0.002288559 | 0.00795156 |
| ENSG00000163472 | TMEM79 | 418.4660389 | -0.003469886 | 0.103994699 | -0.033365994 | 0.973382728 | 0.987970845 |
| ENSG00000163479 | SSR2 | 5618.554839 | 0.352220803 | 0.04556043 | 7.730849035 | 1.07E-14 | 1.25E-13 |
| ENSG00000163481 | RNF25 | 891.7671488 | -0.128634707 | 0.072378197 | -1.777257687 | 0.075525852 | 0.168783541 |
| ENSG00000163482 | STK36 | 1285.851423 | -0.075133004 | 0.070459473 | -1.066329352 | 0.286274785 | 0.470021728 |
| ENSG00000163485 | ADORA1 | 12.9383761 | 0.490081209 | 0.48827317 | 1.003702925 | 0.315521827 | 0.504207396 |
| ENSG00000163491 | NEK10 | 13.15195212 | 0.406349813 | 0.49788344 | 0.816154505 | 0.414411774 | 0.606520741 |
| ENSG00000163507 | CIP2A | 962.691468 | -0.148178707 | 0.069522692 | -2.131371833 | 0.033058521 | 0.084431361 |
| ENSG00000163508 | EOMES | 317.6663513 | -0.458367982 | 0.104393775 | -4.390759702 | 1.13E-05 | 5.79E-05 |
| ENSG00000163510 | CWC22 | 691.6242599 | -0.078416875 | 0.080657545 | -0.972219961 | 0.330941132 | 0.521736621 |
| ENSG00000163512 | AZI2 | 1406.136661 | -0.061896105 | 0.059527291 | -1.039793761 | 0.298435728 | 0.485007829 |
| ENSG00000163513 | TGFBR2 | 454.408042 | 1.374479674 | 0.100492617 | 13.67741951 | 1.39E-42 | 6.23E-41 |
| ENSG00000163516 | ANKZF1 | 1383.464155 | 0.155193964 | 0.069141296 | 2.244591476 | 0.02479437 | 0.066007063 |
| ENSG00000163517 | HDAC11 | 781.901121 | 0.088119914 | 0.082465213 | 1.068570744 | 0.285263137 | 0.469166645 |
| ENSG00000163521 | GLB1L | 127.773449 | -0.008475874 | 0.160214821 | -0.052903182 | 0.957809049 | 0.980830956 |
| ENSG00000163527 | STT3B | 5791.253231 | 0.129169593 | 0.050993417 | 2.533064081 | 0.011307028 | 0.033109934 |
| ENSG00000163528 | CHCHD4 | 1038.831623 | 0.008609594 | 0.066915953 | 0.128662806 | 0.897624467 | 0.948077385 |
| ENSG00000163531 | NFASC | 8.406932198 | 1.323907655 | 0.658823959 | 2.009501379 | 0.044483989 | 0.108398417 |
| ENSG00000163535 | SGO2 | 715.8322489 | -0.085647382 | 0.080096685 | -1.069299965 | 0.284934527 | 0.468827861 |
| ENSG00000163536 | SERPINI1 | 22.34899702 | 0.108110716 | 0.368278078 | 0.293557294 | 0.769096213 | 0.873981628 |
| ENSG00000163539 | CLASP2 | 1753.978561 | 0.20704349 | 0.062876616 | 3.292853578 | 0.000991761 | 0.003714019 |
| ENSG00000163541 | SUCLG1 | 2047.16389 | 0.17119765 | 0.057664321 | 2.968866109 | 0.002989008 | 0.010129686 |
| ENSG00000163545 | NUAK2 | 138.3126465 | 1.072045754 | 0.160530572 | 6.678140769 | 2.42E-11 | 2.19E-10 |
| ENSG00000163558 | PRKCI | 2221.232483 | -0.124951985 | 0.055855835 | -2.237044434 | 0.025283436 | 0.067109816 |
| ENSG00000163576 | EFHB | 39.08137789 | 0.125600513 | 0.289503936 | 0.43384734 | 0.664399302 | 0.806248413 |
| ENSG00000163577 | EIF5A2 | 166.3272636 | 0.196760234 | 0.146417408 | 1.343830875 | 0.179003083 | 0.333812913 |
| ENSG00000163584 | RPL22L1 | 897.8846662 | -0.087714144 | 0.076760725 | -1.142695623 | 0.253164988 | 0.431939753 |
| ENSG00000163590 | PPM1L | 816.459319 | 0.147917431 | 0.086769045 | 1.704725815 | 0.088245573 | 0.191410121 |
| ENSG00000163596 | ICA1L | 225.4919852 | 0.392814605 | 0.128936729 | 3.046568727 | 0.002314696 | 0.008038013 |
| ENSG00000163597 | SNHG16 | 3089.100613 | 0.322787837 | 0.05779473 | 5.585073885 | 2.34E-08 | 1.61E-07 |
| ENSG00000163602 | RYBP | 1945.657939 | -0.06396733 | 0.055191561 | -1.159005639 | 0.246453886 | 0.423684964 |
| ENSG00000163605 | PPP4R2 | 2040.976973 | -0.026738308 | 0.057616574 | -0.464073199 | 0.642595303 | 0.790451834 |
| ENSG00000163607 | GTPBP8 | 245.7209966 | 0.145307534 | 0.122506105 | 1.186124843 | 0.235573001 | 0.41056975 |

| | | | | | | | |
|---|---|---|---|---|---|---|---|
| ENSG00000163608 | NEPRO | 1694.695482 | -0.005675577 | 0.066705134 | -0.085084566 | 0.932194161 | 0.967526001 |
| ENSG00000163611 | SPICE1 | 26.09024267 | -0.554026988 | 0.344266045 | -1.609298959 | 0.107550987 | 0.224762652 |
| ENSG00000163612 | FAM86KP | 2.27604896 | 0.627363026 | 1.176591173 | 0.53320392 | 0.593892428 | 0.755791207 |
| ENSG00000163617 | CCDC191 | 205.2220478 | 0.237917982 | 0.13085588 | 1.818168057 | 0.069038451 | 0.156883809 |
| ENSG00000163618 | CADPS | 36.15024393 | 2.856959881 | 0.379419604 | 7.52981621 | 5.08E-14 | 5.66E-13 |
| ENSG00000163624 | CDS1 | 1648.798971 | 0.230836503 | 0.058560711 | 3.941832353 | 8.09E-05 | 0.000366593 |
| ENSG00000163625 | WDFY3 | 2218.742843 | -0.093384898 | 0.066278328 | -1.408980902 | 0.158840815 | 0.304183621 |
| ENSG00000163626 | COX18 | 400.8394625 | -0.042325383 | 0.095564191 | -0.442900032 | 0.657838044 | 0.801849348 |
| ENSG00000163629 | PTPN13 | 2716.972483 | -0.244609751 | 0.061838061 | -3.95565042 | 7.63E-05 | 0.000347408 |
| ENSG00000163630 | SYNPR | 1.700641511 | 3.016953312 | 1.606502639 | 1.877963496 | 0.060386159 | NA |
| ENSG00000163631 | ALB | 10.17661341 | -0.277589988 | 0.591847827 | -0.469022569 | 0.639053504 | 0.787361694 |
| ENSG00000163632 | C3orf49 | 2.638182069 | 0.185059735 | 1.059354087 | 0.174691104 | 0.861322358 | 0.927857943 |
| ENSG00000163634 | THOC7 | 7502.908255 | 0.328018652 | 0.048650704 | 6.742320797 | 1.56E-11 | 1.44E-10 |
| ENSG00000163635 | ATXN7 | 3737.738296 | 0.173308494 | 0.047862892 | 3.620936507 | 0.000293539 | 0.00121024 |
| ENSG00000163636 | PSMD6 | 24961.58421 | 0.130097667 | 0.055951941 | 2.325168095 | 0.020062974 | 0.054823659 |
| ENSG00000163637 | PRICKLE2 | 1349.480705 | -0.532803086 | 0.063202237 | -8.430130166 | 3.45E-17 | 4.80E-16 |
| ENSG00000163638 | ADAMTS9 | 390.9936259 | 2.453569474 | 0.113643072 | 21.59013692 | 2.22E-103 | 3.86E-101 |
| ENSG00000163644 | PPM1K | 288.8359713 | -0.633962305 | 0.113056156 | -5.607499218 | 2.05E-08 | 1.43E-07 |
| ENSG00000163645 | ERICH6 | 12.42124759 | 0.040778133 | 0.510958384 | 0.079807152 | 0.936390636 | 0.969525239 |
| ENSG00000163655 | GMPS | 5233.020269 | 0.244212165 | 0.049153843 | 4.968322924 | 6.75E-07 | 4.00E-06 |
| ENSG00000163659 | TIPARP | 748.9088983 | 0.106980629 | 0.080692582 | 1.325780222 | 0.184912506 | 0.341757403 |
| ENSG00000163660 | CCNL1 | 2537.84824 | -0.171372016 | 0.137471084 | -1.246604094 | 0.2125427 | 0.380501286 |
| ENSG00000163666 | HESX1 | 13.17278854 | 0.037018225 | 0.51981181 | 0.071214667 | 0.943226908 | 0.973002306 |
| ENSG00000163681 | SLMAP | 1298.740278 | 0.098951601 | 0.061246581 | 1.615626523 | 0.106175086 | 0.222479143 |
| ENSG00000163682 | RPL9 | 608.0662894 | 0.22210658 | 0.08022487 | 2.76855021 | 0.005630631 | 0.0178896 |
| ENSG00000163683 | SMIM14 | 3103.686513 | -0.094512684 | 0.0523341 | -1.805948396 | 0.070926401 | 0.160348047 |
| ENSG00000163684 | RPP14 | 446.5802569 | 0.093805747 | 0.090791605 | 1.033198467 | 0.30151103 | 0.488353285 |
| ENSG00000163686 | ABHD6 | 69.57739059 | 0.586236832 | 0.223047786 | 2.628301505 | 0.008581242 | 0.026004451 |
| ENSG00000163689 | CFAP20DC | 108.1686998 | -0.121009869 | 0.171957182 | -0.703720934 | 0.481606581 | 0.665986796 |
| ENSG00000163694 | RBM47 | 1898.873878 | 0.071365802 | 0.055932563 | 1.27592584 | 0.201981736 | 0.365847692 |
| ENSG00000163697 | APBB2 | 6793.205052 | 0.727009544 | 0.046766153 | 15.54563482 | 1.70E-54 | 1.06E-52 |
| ENSG00000163701 | IL17RE | 87.3995534 | 0.161841979 | 0.193850997 | 0.834878238 | 0.403786269 | 0.596617498 |
| ENSG00000163702 | IL17RC | 632.1951105 | -0.010028055 | 0.084731701 | -0.118350685 | 0.905789799 | 0.95262805 |

| | | | | | | | |
|---|---|---|---|---|---|---|---|
| ENSG00000163703 | CRELD1 | 76.46987162 | -0.279533071 | 0.209538861 | -1.334039283 | 0.182190983 | 0.338089398 |
| ENSG00000163704 | PRRT3 | 1643.230217 | 0.00606363 | 0.073411174 | 0.08259819 | 0.934171041 | 0.968450997 |
| ENSG00000163705 | FANCD2OS | 5.813981071 | -1.07884392 | 0.738149786 | -1.461551491 | 0.143864156 | 0.282337559 |
| ENSG00000163710 | PCOLCE2 | 16.98179192 | -0.741926273 | 0.425579308 | -1.74333258 | 0.081275537 | 0.178952746 |
| ENSG00000163714 | U2SURP | 4461.161822 | -0.064334938 | 0.056995841 | -1.128765476 | 0.25899678 | 0.438819022 |
| ENSG00000163719 | MTMR14 | 2679.157014 | -0.064464509 | 0.053687186 | -1.20074294 | 0.229850932 | 0.403056137 |
| ENSG00000163728 | TTC14 | 693.4819609 | -0.041534708 | 0.096781807 | -0.42915822 | 0.667808085 | 0.808785741 |
| ENSG00000163738 | MTHFD2L | 182.021563 | -0.008277112 | 0.133233858 | -0.062124691 | 0.950463534 | 0.977185792 |
| ENSG00000163743 | RCHY1 | 504.1546373 | -0.280366215 | 0.085837461 | -3.266245439 | 0.001089837 | 0.004052662 |
| ENSG00000163749 | CCDC158 | 8.850823839 | -0.084499306 | 0.626437672 | -0.134888608 | 0.892699947 | 0.945104613 |
| ENSG00000163754 | GYG1 | 2536.314543 | 0.037393949 | 0.051922129 | 0.720192914 | 0.471406226 | 0.657149904 |
| ENSG00000163755 | HPS3 | 2415.01486 | -0.08089686 | 0.056596801 | -1.429353925 | 0.152902534 | 0.295699219 |
| ENSG00000163762 | TM4SF18 | 2.610778786 | 0.149843448 | 1.076081552 | 0.139249156 | 0.889253266 | 0.942979241 |
| ENSG00000163781 | TOPBP1 | 3735.3826 | -0.048699513 | 0.059170731 | -0.823033821 | 0.410488761 | 0.6029719 |
| ENSG00000163785 | RYK | 2761.596038 | 0.039555299 | 0.05108338 | 0.774328156 | 0.438736758 | 0.628292729 |
| ENSG00000163788 | SNRK | 443.7755207 | -0.109481904 | 0.093716704 | -1.168221883 | 0.242717276 | 0.419006978 |
| ENSG00000163793 | DNAJC5G | 2.379907968 | -0.711498359 | 1.173626446 | -0.606239201 | 0.544355925 | 0.717823209 |
| ENSG00000163794 | UCN | 21.73296557 | 0.308762911 | 0.402017722 | 0.768033085 | 0.442467529 | 0.631470633 |
| ENSG00000163795 | ZNF513 | 63.36584559 | 0.175247726 | 0.22828718 | 0.767663458 | 0.442687151 | 0.631641123 |
| ENSG00000163798 | SLC4A1AP | 1241.725329 | -0.183377166 | 0.063699614 | -2.878779838 | 0.003992169 | 0.013154318 |
| ENSG00000163806 | SPDYA | 6.376972349 | 0.434839094 | 0.716271779 | 0.607086733 | 0.543793354 | 0.717344424 |
| ENSG00000163807 | KIAA1143 | 1139.475471 | 0.271740008 | 0.06478377 | 4.194569265 | 2.73E-05 | 0.000133126 |
| ENSG00000163808 | KIF15 | 699.5907152 | -0.524721195 | 0.079536771 | -6.597215241 | 4.19E-11 | 3.72E-10 |
| ENSG00000163811 | WDR43 | 2414.213298 | 0.122789872 | 0.06135421 | 2.00132756 | 0.045357102 | 0.110163251 |
| ENSG00000163812 | ZDHHC3 | 1694.011504 | 0.149079861 | 0.0603266 | 2.471212707 | 0.01346557 | 0.038631837 |
| ENSG00000163817 | SLC6A20 | 1.807500844 | 0.076862381 | 1.277770881 | 0.060153492 | 0.952033386 | 0.977646532 |
| ENSG00000163818 | LZTFL1 | 367.717378 | 0.318004574 | 0.097924757 | 3.247437952 | 0.001164491 | 0.00430931 |
| ENSG00000163820 | FYCO1 | 1817.419765 | 0.329458432 | 0.068119477 | 4.836479214 | 1.32E-06 | 7.53E-06 |
| ENSG00000163823 | CCR1 | 42.10233563 | -0.673172215 | 0.284431577 | -2.366728132 | 0.01794611 | 0.049762331 |
| ENSG00000163827 | LRRC2 | 4.452765109 | 0.124565584 | 0.824989092 | 0.150990583 | 0.879983144 | 0.937787482 |
| ENSG00000163832 | ELP6 | 659.4247913 | -0.424507588 | 0.08050967 | -5.272752793 | 1.34E-07 | 8.61E-07 |
| ENSG00000163840 | DTX3L | 451.0315321 | 0.218358392 | 0.093266146 | 2.341239575 | 0.019219829 | 0.052828048 |
| ENSG00000163848 | ZNF148 | 2172.259893 | 0.031050549 | 0.065077845 | 0.477129337 | 0.633270025 | 0.783898898 |

| | | | | | | | |
|---|---|---|---|---|---|---|---|
| ENSG00000163864 | NMNAT3 | 185.9144372 | 1.234048723 | 0.138882355 | 8.885568816 | 6.36E-19 | 9.98E-18 |
| ENSG00000163866 | SMIM12 | 949.9166342 | -0.147234527 | 0.06983237 | -2.108399383 | 0.034996458 | 0.088601561 |
| ENSG00000163867 | ZMYM6 | 149.2808021 | -0.144998242 | 0.151096558 | -0.959639612 | 0.337236626 | 0.528429521 |
| ENSG00000163870 | TPRA1 | 1208.484662 | 0.04782057 | 0.067712158 | 0.70623314 | 0.480043164 | 0.664385532 |
| ENSG00000163872 | YEATS2 | 1338.164964 | -0.008923369 | 0.062574365 | -0.142604229 | 0.886602758 | 0.941561375 |
| ENSG00000163873 | GRIK3 | 3.392342975 | 1.916638514 | 1.074176589 | 1.784286247 | 0.074377164 | 0.166785155 |
| ENSG00000163874 | ZC3H12A | 716.9064263 | 0.002412035 | 0.077258735 | 0.03122022 | 0.975093914 | 0.988713507 |
| ENSG00000163875 | MEAF6 | 2469.872398 | 0.031604309 | 0.052696253 | 0.599744911 | 0.548676253 | 0.721462412 |
| ENSG00000163877 | SNIP1 | 826.1080643 | 0.042819831 | 0.072784817 | 0.588307181 | 0.556326126 | 0.72693858 |
| ENSG00000163879 | DNALI1 | 21.87023219 | -0.090085033 | 0.374497393 | -0.24054916 | 0.809904558 | 0.899408839 |
| ENSG00000163882 | POLR2H | 1673.581543 | -0.31926381 | 0.058173035 | -5.488175237 | 4.06E-08 | 2.74E-07 |
| ENSG00000163884 | KLF15 | 32.68738937 | 0.676806448 | 0.331187479 | 2.043574989 | 0.040995552 | 0.101209424 |
| ENSG00000163888 | CAMK2N2 | 300.6369033 | 0.224258811 | 0.112352787 | 1.996023566 | 0.045931358 | 0.111381115 |
| ENSG00000163898 | LIPH | 372.7035224 | -0.675466309 | 0.106582481 | -6.337498432 | 2.34E-10 | 1.96E-09 |
| ENSG00000163900 | TMEM41A | 1012.518754 | -0.101849522 | 0.066433939 | -1.533094732 | 0.125252515 | 0.253914166 |
| ENSG00000163902 | RPN1 | 8713.736634 | -0.122041674 | 0.042859882 | -2.847457083 | 0.004407004 | 0.014399183 |
| ENSG00000163904 | SENP2 | 2103.649833 | 0.037925074 | 0.056089823 | 0.676148934 | 0.498946091 | 0.680567986 |
| ENSG00000163909 | HEYL | 138.6236256 | 0.796586792 | 0.16134753 | 4.937086993 | 7.93E-07 | 4.66E-06 |
| ENSG00000163913 | IFT122 | 1304.04643 | 0.378812328 | 0.06730569 | 5.628236315 | 1.82E-08 | 1.28E-07 |
| ENSG00000163918 | RFC4 | 593.3758277 | -0.530559365 | 0.082312399 | -6.44567975 | 1.15E-10 | 9.89E-10 |
| ENSG00000163923 | RPL39L | 287.1191506 | -0.12917147 | 0.11935836 | -1.08221552 | 0.279156774 | 0.462305544 |
| ENSG00000163930 | BAP1 | 3793.100826 | 0.077442308 | 0.053000175 | 1.4611708 | 0.143968574 | 0.282478262 |
| ENSG00000163931 | TKT | 15428.29664 | 0.288581221 | 0.042044767 | 6.863665676 | 6.71E-12 | 6.39E-11 |
| ENSG00000163932 | PRKCD | 4715.328635 | 0.224773446 | 0.055637748 | 4.039945062 | 5.35E-05 | 0.000248926 |
| ENSG00000163933 | RFT1 | 750.882645 | 0.053212815 | 0.077945332 | 0.682694057 | 0.494800186 | 0.67711556 |
| ENSG00000163935 | SFMBT1 | 707.5270218 | -0.075424445 | 0.076004118 | -0.992373137 | 0.321015538 | 0.510544679 |
| ENSG00000163938 | GNL3 | 2392.916192 | -0.005139558 | 0.051380905 | -0.100028561 | 0.920321651 | 0.961348694 |
| ENSG00000163939 | PBRM1 | 2057.980158 | -0.086151453 | 0.057281883 | -1.503991279 | 0.132583612 | 0.26556082 |
| ENSG00000163945 | UVSSA | 611.4325956 | -0.115921097 | 0.084082403 | -1.378660613 | 0.167999419 | 0.317992838 |
| ENSG00000163946 | TASOR | 2535.91576 | -0.026164736 | 0.058146136 | -0.449982364 | 0.652723157 | 0.798422458 |
| ENSG00000163947 | ARHGEF3 | 295.7151941 | 0.387204632 | 0.108396722 | 3.572106478 | 0.000354121 | 0.001436016 |
| ENSG00000163950 | SLBP | 1650.290129 | 0.054463177 | 0.05657119 | 0.96273699 | 0.335679529 | 0.526875672 |
| ENSG00000163956 | LRPAP1 | 2670.888579 | -0.038600134 | 0.070114246 | -0.550531964 | 0.581954559 | 0.74680029 |

| | | | | | | | |
|---|---|---|---|---|---|---|---|
| ENSG00000163959 | SLC51A | 5.085519895 | -0.957629923 | 0.830403124 | -1.153210887 | 0.248823836 | 0.42673431 |
| ENSG00000163960 | UBXN7 | 1564.713508 | -0.298971675 | 0.06243201 | -4.78875621 | 1.68E-06 | 9.46E-06 |
| ENSG00000163961 | RNF168 | 1069.262495 | -0.336566157 | 0.068606339 | -4.905758857 | 9.31E-07 | 5.41E-06 |
| ENSG00000163964 | PIGX | 791.9656974 | -0.055006114 | 0.071946152 | -0.764545596 | 0.444542176 | 0.633324898 |
| ENSG00000163975 | MELTF | 208.1104594 | -0.282083151 | 0.136842445 | -2.061371752 | 0.039267588 | 0.097700484 |
| ENSG00000163993 | S100P | 101.2026595 | -0.963462598 | 0.183551694 | -5.24899868 | 1.53E-07 | 9.75E-07 |
| ENSG00000163995 | ABLIM2 | 108.5039224 | -0.171471571 | 0.17475363 | -0.981218935 | 0.326484789 | 0.517056337 |
| ENSG00000164002 | EXO5 | 325.5227699 | -0.187830673 | 0.102935207 | -1.824746636 | 0.068039272 | 0.1550433 |
| ENSG00000164008 | C1orf50 | 186.1475939 | -0.008843261 | 0.133766759 | -0.066109553 | 0.947290605 | 0.975439738 |
| ENSG00000164010 | ERMAP | 1048.001514 | 0.707253347 | 0.066117099 | 10.69698096 | 1.05E-26 | 2.57E-25 |
| ENSG00000164011 | ZNF691 | 281.2178454 | -0.242663793 | 0.112365546 | -2.159592526 | 0.030804228 | 0.079479134 |
| ENSG00000164022 | AIMP1 | 2140.426785 | 0.220766954 | 0.058542716 | 3.771040544 | 0.000162568 | 0.000700341 |
| ENSG00000164023 | SGMS2 | 450.0073883 | 0.289004225 | 0.091741165 | 3.150213149 | 0.001631514 | 0.005849785 |
| ENSG00000164024 | METAP1 | 2408.90884 | 0.196925027 | 0.052779347 | 3.731100111 | 0.000190645 | 0.00081275 |
| ENSG00000164031 | DNAJB14 | 708.6666916 | -0.16124903 | 0.088751649 | -1.816856727 | 0.069239054 | 0.157246203 |
| ENSG00000164032 | H2AZ1 | 7028.770277 | -0.229012295 | 0.053255877 | -4.3002258 | 1.71E-05 | 8.55E-05 |
| ENSG00000164035 | EMCN | 106.4304502 | 0.072722166 | 0.176090446 | 0.412981896 | 0.679619878 | 0.81798632 |
| ENSG00000164037 | SLC9B1 | 11.04497993 | -0.259422294 | 0.566484664 | -0.457951133 | 0.646987549 | 0.793684192 |
| ENSG00000164038 | SLC9B2 | 292.0308117 | 0.958464642 | 0.11267559 | 8.506408879 | 1.79E-17 | 2.55E-16 |
| ENSG00000164039 | BDH2 | 42.31592333 | -0.985590234 | 0.282350724 | -3.490659494 | 0.00048183 | 0.001913639 |
| ENSG00000164040 | PGRMC2 | 1654.408596 | 0.065041012 | 0.060694991 | 1.071604284 | 0.283897808 | 0.467759514 |
| ENSG00000164045 | CDC25A | 547.0583194 | -0.1477837 | 0.082726949 | -1.786403378 | 0.074033969 | 0.166210535 |
| ENSG00000164047 | CAMP | 2.161602431 | -0.365113546 | 1.207728074 | -0.302314365 | 0.762412429 | 0.869703686 |
| ENSG00000164048 | ZNF589 | 289.0921694 | 0.026999012 | 0.11114571 | 0.24291547 | 0.808070881 | 0.898335159 |
| ENSG00000164050 | PLXNB1 | 5387.340981 | 0.252061206 | 0.054597572 | 4.616710938 | 3.90E-06 | 2.11E-05 |
| ENSG00000164051 | CCDC51 | 418.2840688 | -0.307807207 | 0.095499811 | -3.223118477 | 0.001268031 | 0.004654926 |
| ENSG00000164053 | ATRIP | 529.7569651 | -0.283754201 | 0.085907506 | -3.303019882 | 0.000956496 | 0.003592341 |
| ENSG00000164054 | SHISA5 | 4093.449959 | -0.376800528 | 0.054407024 | -6.925586037 | 4.34E-12 | 4.21E-11 |
| ENSG00000164056 | SPRY1 | 153.5619202 | -0.529425833 | 0.157003224 | -3.372069822 | 0.000746055 | 0.002860045 |
| ENSG00000164061 | BSN | 56.42550521 | 0.421420112 | 0.247529145 | 1.702507041 | 0.08866036 | 0.192110008 |
| ENSG00000164062 | APEH | 3348.561762 | 0.233186295 | 0.057559447 | 4.051225403 | 5.10E-05 | 0.000238141 |
| ENSG00000164066 | INTU | 103.0281632 | 0.153110106 | 0.18085831 | 0.846574901 | 0.397232109 | 0.590149898 |
| ENSG00000164068 | RNF123 | 1453.500656 | 0.139903594 | 0.065233225 | 2.144667759 | 0.03197942 | 0.08211416 |

| | | | | | | | |
|---|---|---|---|---|---|---|---|
| ENSG00000164070 | HSPA4L | 1100.770993 | 0.579428525 | 0.069659689 | 8.317988981 | 8.95E-17 | 1.22E-15 |
| ENSG00000164073 | MFSD8 | 664.4616213 | 0.016400788 | 0.078297088 | 0.209468689 | 0.834082376 | 0.912384329 |
| ENSG00000164074 | ABHD18 | 504.8892051 | -0.056479946 | 0.091818301 | -0.615127328 | 0.538470636 | 0.713234967 |
| ENSG00000164077 | MON1A | 142.2089336 | -0.089364656 | 0.151552844 | -0.589660039 | 0.555418591 | 0.726496891 |
| ENSG00000164078 | MST1R | 34.63705457 | 0.118385523 | 0.304391863 | 0.388924729 | 0.697331828 | 0.830053557 |
| ENSG00000164080 | RAD54L2 | 793.1540726 | -0.259203541 | 0.079943506 | -3.242333924 | 0.00118555 | 0.004379472 |
| ENSG00000164081 | TEX264 | 959.0574357 | 0.079856008 | 0.081070657 | 0.985017405 | 0.324615522 | 0.514771193 |
| ENSG00000164082 | GRM2 | 9.216567496 | -0.26385854 | 0.5868849 | -0.449591631 | 0.653004923 | 0.798589026 |
| ENSG00000164086 | DUSP7 | 433.8552277 | 0.196740638 | 0.091772152 | 2.143794543 | 0.032049353 | 0.082266102 |
| ENSG00000164087 | POC1A | 540.410928 | -0.200904 | 0.083077042 | -2.418285414 | 0.01559384 | 0.044014903 |
| ENSG00000164088 | PPM1M | 217.1651387 | 0.046463005 | 0.127481432 | 0.364468806 | 0.715507964 | 0.841687743 |
| ENSG00000164089 | ETNPPL | 10.5429892 | 1.18135968 | 0.566548483 | 2.085187261 | 0.037052306 | 0.093015966 |
| ENSG00000164091 | WDR82 | 4642.000506 | 0.013455548 | 0.045124 | 0.298190493 | 0.765557777 | 0.871774406 |
| ENSG00000164093 | PITX2 | 80.39727156 | -0.123302176 | 0.19838912 | -0.621516824 | 0.534259628 | 0.709706171 |
| ENSG00000164096 | C4orf3 | 1774.148103 | -0.0752514 | 0.062471294 | -1.204575668 | 0.228367155 | 0.401243882 |
| ENSG00000164099 | PRSS12 | 6.933071749 | -0.247193216 | 0.692229051 | -0.357097432 | 0.721018867 | 0.845162383 |
| ENSG00000164100 | NDST3 | 42.99318347 | 0.751748499 | 0.277002016 | 2.713873743 | 0.006650151 | 0.020733142 |
| ENSG00000164104 | HMGB2 | 3496.095193 | -0.408742014 | 0.053517643 | -7.637518982 | 2.21E-14 | 2.54E-13 |
| ENSG00000164105 | SAP30 | 189.8983023 | -0.014680764 | 0.137372931 | -0.106867953 | 0.914893737 | 0.957728499 |
| ENSG00000164109 | MAD2L1 | 1185.365566 | -0.05364289 | 0.074107982 | -0.723847671 | 0.469159259 | 0.655132299 |
| ENSG00000164111 | ANXA5 | 5244.520377 | -0.412165586 | 0.046228062 | -8.915917517 | 4.84E-19 | 7.65E-18 |
| ENSG00000164114 | MAP9 | 782.8784257 | 0.087467454 | 0.076370229 | 1.145308261 | 0.252081485 | 0.430783992 |
| ENSG00000164116 | GUCY1A1 | 435.2285745 | -0.145796198 | 0.094242897 | -1.547025856 | 0.121857011 | 0.2484579 |
| ENSG00000164117 | FBXO8 | 199.8378802 | -0.303355211 | 0.13654787 | -2.22160339 | 0.026310121 | 0.069352729 |
| ENSG00000164118 | CEP44 | 150.3340821 | -0.104493648 | 0.148390814 | -0.704178687 | 0.481321502 | 0.665753106 |
| ENSG00000164120 | HPGD | 21.44330862 | -1.737952007 | 0.406539835 | -4.274985761 | 1.91E-05 | 9.52E-05 |
| ENSG00000164124 | TMEM144 | 115.0664852 | 0.589733192 | 0.169574561 | 3.477722062 | 0.000505694 | 0.001998728 |
| ENSG00000164125 | GASK1B | 334.2517107 | 0.970924573 | 0.106043443 | 9.155913284 | 5.39E-20 | 8.99E-19 |
| ENSG00000164128 | NPY1R | 51.77642197 | 1.606858135 | 0.288203432 | 5.575430265 | 2.47E-08 | 1.70E-07 |
| ENSG00000164134 | NAA15 | 2448.90516 | 0.232861008 | 0.056911481 | 4.091635018 | 4.28E-05 | 0.000202102 |
| ENSG00000164142 | FHIP1A | 555.4406622 | 0.456486579 | 0.087476323 | 5.218401568 | 1.80E-07 | 1.14E-06 |
| ENSG00000164144 | ARFIP1 | 1379.056911 | -0.019003992 | 0.063246006 | -0.300477337 | 0.763813081 | 0.870479014 |
| ENSG00000164151 | ICE1 | 2263.530283 | -0.086861145 | 0.055536718 | -1.564030933 | 0.117810304 | 0.242100545 |

| | | | | | | | |
|---|---|---|---|---|---|---|---|
| ENSG00000164162 | ANAPC10 | 68.32266725 | 0.059485914 | 0.217721222 | 0.273220558 | 0.784683678 | 0.883091372 |
| ENSG00000164163 | ABCE1 | 2370.567314 | 0.06074963 | 0.059133968 | 1.027322063 | 0.304268837 | 0.491613404 |
| ENSG00000164164 | OTUD4 | 1648.383198 | -0.07585922 | 0.062735758 | -1.209186318 | 0.226591273 | 0.398925848 |
| ENSG00000164167 | LSM6 | 276.6533157 | -0.070956499 | 0.112395445 | -0.631311159 | 0.527837092 | 0.703782789 |
| ENSG00000164168 | TMEM184C | 2043.569486 | 0.102386524 | 0.058151556 | 1.760684168 | 0.078291874 | 0.173645517 |
| ENSG00000164169 | PRMT9 | 452.0544852 | -0.190844205 | 0.093022117 | -2.051600326 | 0.040208521 | 0.099544945 |
| ENSG00000164171 | ITGA2 | 577.3164953 | 0.555991213 | 0.083014199 | 6.697543563 | 2.12E-11 | 1.93E-10 |
| ENSG00000164172 | MOCS2 | 882.9947404 | 0.043507463 | 0.071902806 | 0.605087132 | 0.545121104 | 0.718352374 |
| ENSG00000164176 | EDIL3 | 23.50381805 | -0.06360019 | 0.362162838 | -0.175612138 | 0.860598666 | 0.927426758 |
| ENSG00000164180 | TMEM161B | 399.9324351 | 0.017115018 | 0.102548684 | 0.16689652 | 0.867451471 | 0.93096599 |
| ENSG00000164181 | ELOVL7 | 888.6721539 | -0.200038889 | 0.072062122 | -2.77592282 | 0.005504527 | 0.017513711 |
| ENSG00000164182 | NDUFAF2 | 304.708701 | 0.255458775 | 0.107501402 | 2.376329715 | 0.01748583 | 0.048674305 |
| ENSG00000164185 | ZNF474 | 1.809373839 | 0.10605002 | 1.27799841 | 0.08298134 | 0.933866377 | 0.968266427 |
| ENSG00000164187 | LMBRD2 | 975.5412128 | 0.025034747 | 0.072632856 | 0.344675234 | 0.730338539 | 0.850776726 |
| ENSG00000164190 | NIPBL | 3958.789886 | -0.16512766 | 0.054449358 | -3.032683325 | 0.002423898 | 0.008382968 |
| ENSG00000164197 | RNF180 | 51.89896695 | 1.68962999 | 0.270461196 | 6.24721778 | 4.18E-10 | 3.45E-09 |
| ENSG00000164199 | ADGRV1 | 948.8434595 | 0.016651108 | 0.071927554 | 0.231498325 | 0.816927687 | 0.903062617 |
| ENSG00000164209 | SLC25A46 | 1335.587063 | 0.131639364 | 0.073747858 | 1.784992364 | 0.074262556 | 0.166577004 |
| ENSG00000164211 | STARD4 | 1020.325795 | -1.001609895 | 0.088206587 | -11.3552732 | 6.98E-30 | 1.94E-28 |
| ENSG00000164219 | PGGT1B | 877.2942968 | 0.639191557 | 0.074954346 | 8.527745148 | 1.49E-17 | 2.13E-16 |
| ENSG00000164221 | CCDC112 | 262.263761 | 0.403849628 | 0.119318889 | 3.384624437 | 0.000712757 | 0.002742937 |
| ENSG00000164236 | ANKRD33B | 1.979603425 | 0.789143329 | 1.276157834 | 0.618374395 | 0.536328572 | 0.711548056 |
| ENSG00000164237 | CMBL | 991.6110461 | -0.531291916 | 0.072214625 | -7.357123504 | 1.88E-13 | 2.01E-12 |
| ENSG00000164241 | C5orf63 | 117.9740483 | 0.100268624 | 0.173622512 | 0.577509349 | 0.563595425 | 0.732229586 |
| ENSG00000164244 | PRRC1 | 2468.811968 | -0.183542741 | 0.055464307 | -3.309204603 | 0.000935614 | 0.00352141 |
| ENSG00000164251 | F2RL1 | 323.7614828 | -0.14996824 | 0.106600915 | -1.406819443 | 0.159480936 | 0.305148141 |
| ENSG00000164252 | AGGF1 | 447.3590265 | -0.052397913 | 0.096075489 | -0.545382732 | 0.585490314 | 0.749620821 |
| ENSG00000164253 | WDR41 | 938.767862 | -0.022298595 | 0.067516137 | -0.330270596 | 0.741195508 | 0.857309785 |
| ENSG00000164256 | PRDM9 | 3.600865091 | -0.643821068 | 0.933836339 | -0.689436726 | 0.490548478 | 0.673681878 |
| ENSG00000164258 | NDUFS4 | 654.8716786 | 0.134495992 | 0.079167781 | 1.698872826 | 0.089343148 | 0.193260897 |
| ENSG00000164284 | GRPEL2 | 696.6110717 | -0.022266763 | 0.080681511 | -0.275983465 | 0.782560766 | 0.882072719 |
| ENSG00000164287 | CDC20B | 2.684101494 | 4.766467894 | 1.49038605 | 3.198143121 | 0.001383156 | 0.005035228 |
| ENSG00000164291 | ARSK | 103.158967 | 0.483961171 | 0.177734429 | 2.722945547 | 0.006470272 | 0.020235589 |

| | | | | | | | |
|---|---|---|---|---|---|---|---|
| ENSG00000164292 | RHOBTB3 | 2875.714954 | 1.289404509 | 0.05580081 | 23.10727222 | 3.91E-118 | 9.16E-116 |
| ENSG00000164294 | GPX8 | 12.3948043 | 1.054279424 | 0.513475031 | 2.053224325 | 0.040050826 | 0.099208072 |
| ENSG00000164296 | TIGD6 | 232.3040907 | 0.106911844 | 0.125113359 | 0.854519815 | 0.392817035 | 0.585752911 |
| ENSG00000164300 | SERINC5 | 1888.75041 | 0.640180213 | 0.065050361 | 9.841301396 | 7.47E-23 | 1.48E-21 |
| ENSG00000164304 | CAGE1 | 2.485489556 | -0.39496174 | 1.107000924 | -0.356785375 | 0.721252485 | 0.845194729 |
| ENSG00000164305 | CASP3 | 223.5219229 | 0.574408368 | 0.125510517 | 4.576575568 | 4.73E-06 | 2.53E-05 |
| ENSG00000164306 | PRIMPOL | 94.27547854 | -0.101654917 | 0.187543334 | -0.542034283 | 0.587794889 | 0.751198593 |
| ENSG00000164307 | ERAP1 | 709.1371928 | 1.019961833 | 0.077819722 | 13.1067268 | 3.01E-39 | 1.19E-37 |
| ENSG00000164308 | ERAP2 | 1.825900997 | -1.019590615 | 1.330086087 | -0.766559868 | 0.443343244 | 0.632205757 |
| ENSG00000164309 | CMYA5 | 344.2149105 | 0.610404942 | 0.105156645 | 5.804720601 | 6.45E-09 | 4.73E-08 |
| ENSG00000164323 | CFAP97 | 573.2432884 | 0.129192146 | 0.084525112 | 1.528446899 | 0.126401618 | 0.255656244 |
| ENSG00000164327 | RICTOR | 1054.240001 | -0.051287443 | 0.066968096 | -0.765848902 | 0.443766213 | 0.632572978 |
| ENSG00000164329 | TENT2 | 975.0442263 | 0.091280052 | 0.070203463 | 1.300221492 | 0.193525067 | 0.354250321 |
| ENSG00000164331 | ANKRA2 | 319.3674068 | 0.421448815 | 0.10804714 | 3.900601289 | 9.60E-05 | 0.000428747 |
| ENSG00000164332 | UBLCP1 | 582.1179939 | 0.07104601 | 0.087668031 | 0.810398149 | 0.417711382 | 0.609793757 |
| ENSG00000164338 | UTP15 | 515.1648938 | -0.047396705 | 0.093144597 | -0.508850829 | 0.610856788 | 0.768765345 |
| ENSG00000164344 | KLKB1 | 1.837248697 | 2.179346147 | 1.501199556 | 1.451736472 | 0.146574894 | 0.286427486 |
| ENSG00000164346 | NSA2 | 2738.081497 | 0.335782029 | 0.050765897 | 6.614322751 | 3.73E-11 | 3.33E-10 |
| ENSG00000164347 | GFM2 | 1336.039045 | 0.277956108 | 0.067890998 | 4.09415263 | 4.24E-05 | 0.000200124 |
| ENSG00000164362 | TERT | 19.26395978 | -1.230776125 | 0.437231104 | -2.814932686 | 0.004878745 | 0.015767599 |
| ENSG00000164366 | CCDC127 | 683.8685784 | 0.07765321 | 0.076464107 | 1.015551133 | 0.309843185 | 0.497815007 |
| ENSG00000164385 | LINC01600 | 1.56774689 | 1.098391689 | 1.447021356 | 0.759070821 | 0.447810193 | NA |
| ENSG00000164402 | SEPTIN8 | 3588.315639 | -0.144118626 | 0.049438786 | -2.915092342 | 0.003555833 | 0.01185967 |
| ENSG00000164403 | SHROOM1 | 196.5036768 | 0.222426444 | 0.145391394 | 1.529846015 | 0.126054849 | 0.255083188 |
| ENSG00000164404 | GDF9 | 42.50080272 | -0.383130462 | 0.267497716 | -1.43227564 | 0.152064952 | 0.294489413 |
| ENSG00000164405 | UQCRQ | 2104.750044 | 0.034713815 | 0.053311574 | 0.651149697 | 0.514949859 | 0.693942232 |
| ENSG00000164406 | LEAP2 | 26.25820481 | -0.240674051 | 0.409573316 | -0.587621415 | 0.556786434 | 0.727249376 |
| ENSG00000164411 | GJB7 | 1.596187254 | 1.748809174 | 1.530682519 | 1.142502871 | 0.253245053 | NA |
| ENSG00000164414 | SLC35A1 | 537.7254717 | -0.756838446 | 0.085991186 | -8.801349105 | 1.35E-18 | 2.08E-17 |
| ENSG00000164418 | GRIK2 | 67.89884152 | 1.047105235 | 0.226904066 | 4.61474867 | 3.94E-06 | 2.13E-05 |
| ENSG00000164430 | CGAS | 50.71054821 | -1.198253237 | 0.256148418 | -4.677964622 | 2.90E-06 | 1.59E-05 |
| ENSG00000164440 | TXLNB | 9.970824026 | 2.414346846 | 0.671328616 | 3.596371116 | 0.000322687 | 0.001320451 |
| ENSG00000164442 | CITED2 | 2264.948989 | -0.012867425 | 0.055935869 | -0.230038891 | 0.818061549 | 0.90364036 |

| | | | | | | | |
|---|---|---|---|---|---|---|---|
| ENSG00000164451 | CALHM4 | 8.58921238 | -4.731179644 | 1.135431459 | -4.166856228 | 3.09E-05 | 0.000149431 |
| ENSG00000164463 | CREBRF | 363.6737098 | 0.073662159 | 0.101172356 | 0.728085829 | 0.466561048 | 0.652497131 |
| ENSG00000164465 | DCBLD1 | 441.101135 | 0.306301301 | 0.090167623 | 3.397020899 | 0.000681238 | 0.002632681 |
| ENSG00000164466 | SFXN1 | 4289.600142 | -1.010918824 | 0.049105991 | -20.58646628 | 3.63E-94 | 5.24E-92 |
| ENSG00000164494 | PDSS2 | 628.2283938 | -0.026811276 | 0.078487727 | -0.341598326 | 0.732653197 | 0.852241993 |
| ENSG00000164506 | STXBP5 | 1283.284046 | 0.166349477 | 0.064410512 | 2.582644842 | 0.00980462 | 0.029210312 |
| ENSG00000164535 | DAGLB | 927.1248852 | 0.183836555 | 0.072497001 | 2.53578152 | 0.011219671 | 0.032875111 |
| ENSG00000164542 | MATCAP2 | 377.3010644 | 0.77425885 | 0.102150048 | 7.579622947 | 3.47E-14 | 3.91E-13 |
| ENSG00000164543 | STK17A | 324.1987415 | 0.425563261 | 0.110765997 | 3.84200272 | 0.000122034 | 0.0005364 |
| ENSG00000164548 | TRA2A | 2479.448225 | -0.820525776 | 0.058369996 | -14.05732107 | 6.95E-45 | 3.32E-43 |
| ENSG00000164574 | GALNT10 | 4288.386819 | -0.106497591 | 0.060306908 | -1.765926906 | 0.077408112 | 0.172251634 |
| ENSG00000164576 | SAP30L | 1679.118178 | 0.060707864 | 0.056714557 | 1.070410612 | 0.284434525 | 0.468240258 |
| ENSG00000164587 | RPS14 | 10426.49538 | 0.289929 | 0.049066348 | 5.908917407 | 3.44E-09 | 2.61E-08 |
| ENSG00000164597 | COG5 | 943.7009239 | -0.133916884 | 0.07231772 | -1.851785211 | 0.064056672 | 0.14727 |
| ENSG00000164603 | BMT2 | 279.2953877 | 0.186616864 | 0.115943694 | 1.60954734 | 0.107496712 | 0.224690167 |
| ENSG00000164609 | SLU7 | 1959.365357 | 0.164958136 | 0.055927184 | 2.949516193 | 0.003182719 | 0.010719655 |
| ENSG00000164610 | RP9 | 802.2273655 | -0.066037899 | 0.074574876 | -0.885524762 | 0.375873671 | 0.568844307 |
| ENSG00000164611 | PTTG1 | 1056.405839 | -0.197194258 | 0.066270316 | -2.975604602 | 0.002924116 | 0.009930307 |
| ENSG00000164615 | CAMLG | 1216.881806 | 0.215609086 | 0.062951838 | 3.424984747 | 0.000614834 | 0.00239421 |
| ENSG00000164619 | BMPER | 1.667083704 | 0.369517592 | 1.364546011 | 0.270798924 | 0.786545687 | NA |
| ENSG00000164620 | RELL2 | 451.6375281 | 0.346822195 | 0.091350657 | 3.796603162 | 0.000146692 | 0.000636972 |
| ENSG00000164626 | KCNK5 | 49.57894976 | 0.3282037 | 0.289020163 | 1.135573717 | 0.256135014 | 0.435594723 |
| ENSG00000164627 | KIF6 | 15.3294645 | -0.214307748 | 0.478131913 | -0.448218873 | 0.653995243 | 0.799165148 |
| ENSG00000164631 | ZNF12 | 1848.586341 | 0.46310967 | 0.064466623 | 7.183712281 | 6.78E-13 | 7.04E-12 |
| ENSG00000164638 | SLC29A4 | 466.4898093 | 1.679173868 | 0.09819206 | 17.10091286 | 1.46E-65 | 1.19E-63 |
| ENSG00000164647 | STEAP1 | 6.324570387 | 1.400024775 | 0.733202122 | 1.909466343 | 0.05620196 | 0.13230097 |
| ENSG00000164649 | CDCA7L | 2644.274455 | -1.751214699 | 0.060077922 | -29.14905589 | 8.59E-187 | 4.48E-184 |
| ENSG00000164654 | MIOS | 2324.780478 | -1.205074552 | 0.055787992 | -21.60096669 | 1.76E-103 | 3.08E-101 |
| ENSG00000164659 | ELAPOR2 | 1202.324187 | -0.504018058 | 0.07087301 | -7.111565547 | 1.15E-12 | 1.17E-11 |
| ENSG00000164663 | USP49 | 255.6938616 | -0.314932667 | 0.120430483 | -2.615057744 | 0.008921237 | 0.026910346 |
| ENSG00000164669 | INTS4P1 | 8.666036512 | 0.831880251 | 0.666504213 | 1.24812452 | 0.21198546 | 0.379710038 |
| ENSG00000164674 | SYTL3 | 38.60588205 | -1.414764296 | 0.332812438 | -4.250935771 | 2.13E-05 | 0.000105303 |
| ENSG00000164683 | HEY1 | 39.96904736 | -1.862862994 | 0.307453113 | -6.059014906 | 1.37E-09 | 1.08E-08 |

| | | | | | | | |
|---|---|---|---|---|---|---|---|
| ENSG00000164684 | ZNF704 | 1906.295999 | -0.377119616 | 0.064279669 | -5.86685682 | 4.44E-09 | 3.33E-08 |
| ENSG00000164687 | FABP5 | 22.16823289 | 0.196134976 | 0.390467574 | 0.502307973 | 0.615450902 | 0.77213255 |
| ENSG00000164690 | SHH | 11.28573218 | 0.687315589 | 0.534861243 | 1.285035319 | 0.198779942 | 0.361544705 |
| ENSG00000164695 | CHMP4C | 926.559271 | 0.14809398 | 0.067829023 | 2.183342356 | 0.029010608 | 0.075388007 |
| ENSG00000164708 | PGAM2 | 5.132573689 | 0.578748931 | 0.80865714 | 0.715691364 | 0.474181945 | 0.659616385 |
| ENSG00000164713 | BRI3 | 636.4751489 | 0.809790462 | 0.08277965 | 9.782482359 | 1.34E-22 | 2.60E-21 |
| ENSG00000164715 | LMTK2 | 1869.88346 | -0.451459403 | 0.059028415 | -7.648170815 | 2.04E-14 | 2.34E-13 |
| ENSG00000164733 | CTSB | 1961.879741 | 0.825189534 | 0.060866132 | 13.55744997 | 7.16E-42 | 3.15E-40 |
| ENSG00000164741 | DLC1 | 160.5785373 | -0.45438156 | 0.144918175 | -3.135435287 | 0.001715993 | 0.006115407 |
| ENSG00000164742 | ADCY1 | 5245.300642 | 2.295070612 | 0.055369907 | 41.44978273 | 0 | 0 |
| ENSG00000164743 | C8orf48 | 11.78620647 | -1.588803651 | 0.536923481 | -2.959087667 | 0.003085513 | 0.010427471 |
| ENSG00000164749 | HNF4G | 91.38497215 | 0.705959644 | 0.190805957 | 3.699882624 | 0.000215699 | 0.000908916 |
| ENSG00000164751 | PEX2 | 1379.288813 | 0.334579182 | 0.060075571 | 5.569305106 | 2.56E-08 | 1.76E-07 |
| ENSG00000164754 | RAD21 | 14929.23908 | 0.52845646 | 0.045849601 | 11.5258682 | 9.77E-31 | 2.79E-29 |
| ENSG00000164756 | SLC30A8 | 55.6274896 | 5.263752627 | 0.549275029 | 9.583091079 | 9.42E-22 | 1.74E-20 |
| ENSG00000164758 | MED30 | 182.7843935 | 0.182611832 | 0.139862062 | 1.30565666 | 0.191669343 | 0.351667177 |
| ENSG00000164761 | TNFRSF11B | 598.7212979 | -1.836107357 | 0.091640875 | -20.03589955 | 2.68E-89 | 3.64E-87 |
| ENSG00000164776 | PHKG1 | 14.83767296 | 0.194139702 | 0.462209913 | 0.420024963 | 0.674467218 | 0.81390205 |
| ENSG00000164794 | KCNV1 | 27.91246971 | -0.582394443 | 0.363507118 | -1.602154163 | 0.109121523 | 0.227216823 |
| ENSG00000164808 | SPIDR | 2313.744374 | -0.019565193 | 0.056276408 | -0.347662436 | 0.72809371 | 0.849091033 |
| ENSG00000164815 | ORC5 | 689.709945 | -0.417407956 | 0.077718305 | -5.370780521 | 7.84E-08 | 5.12E-07 |
| ENSG00000164818 | DNAAF5 | 2372.730589 | 0.16295131 | 0.055696288 | 2.925712199 | 0.003436687 | 0.011502422 |
| ENSG00000164823 | OSGIN2 | 3054.16471 | 0.317918216 | 0.055405479 | 5.738028487 | 9.58E-09 | 6.91E-08 |
| ENSG00000164828 | SUN1 | 5564.34554 | 0.374567165 | 0.044722726 | 8.375320567 | 5.51E-17 | 7.59E-16 |
| ENSG00000164830 | OXR1 | 1649.063651 | 0.027982543 | 0.058938832 | 0.474772598 | 0.634949066 | 0.784833858 |
| ENSG00000164841 | TMEM74 | 12.61940538 | -0.923852748 | 0.503017735 | -1.836620628 | 0.06626592 | 0.151719101 |
| ENSG00000164845 | FAM86FP | 87.55475697 | -0.020599641 | 0.193330942 | -0.106551184 | 0.915145047 | 0.957860445 |
| ENSG00000164849 | GPR146 | 41.4455417 | 0.877631568 | 0.289853053 | 3.027850005 | 0.002463003 | 0.008500269 |
| ENSG00000164850 | GPER1 | 6277.396932 | 1.430046255 | 0.050391027 | 28.37898595 | 3.67E-177 | 1.69E-174 |
| ENSG00000164855 | TMEM184A | 3633.86452 | 0.146443769 | 0.057571786 | 2.543672502 | 0.010969388 | 0.032228196 |
| ENSG00000164867 | NOS3 | 21.1331949 | 2.686853515 | 0.477830916 | 5.623021499 | 1.88E-08 | 1.31E-07 |
| ENSG00000164877 | MICALL2 | 2276.067787 | 0.523300201 | 0.061448679 | 8.516052914 | 1.65E-17 | 2.35E-16 |
| ENSG00000164880 | INTS1 | 8883.920323 | 0.114042911 | 0.053301356 | 2.139587415 | 0.032388125 | 0.083024201 |

| | | | | | | | |
|---|---|---|---|---|---|---|---|
| ENSG00000164885 | CDK5 | 671.3185241 | 0.404108929 | 0.081151504 | 4.979685029 | 6.37E-07 | 3.78E-06 |
| ENSG00000164889 | SLC4A2 | 7861.973936 | 0.767484674 | 0.05062044 | 15.16155689 | 6.35E-52 | 3.68E-50 |
| ENSG00000164896 | FASTK | 2404.362195 | 0.443108012 | 0.065001539 | 6.816884922 | 9.30E-12 | 8.73E-11 |
| ENSG00000164897 | TMUB1 | 1066.452376 | 0.290088015 | 0.072757608 | 3.98704716 | 6.69E-05 | 0.000307063 |
| ENSG00000164898 | FMC1 | 14.91884308 | -0.336016164 | 0.45083334 | -0.745322349 | 0.456076873 | 0.643397611 |
| ENSG00000164902 | PHAX | 1972.928697 | 0.177135142 | 0.056282709 | 3.147239066 | 0.001648201 | 0.005900396 |
| ENSG00000164904 | ALDH7A1 | 2413.948192 | 0.041502125 | 0.052081014 | 0.79687629 | 0.425522882 | 0.616372729 |
| ENSG00000164916 | FOXK1 | 2218.936595 | 0.248248599 | 0.055309648 | 4.48834171 | 7.18E-06 | 3.77E-05 |
| ENSG00000164919 | COX6C | 3384.409474 | -0.054628355 | 0.054583126 | -1.000828626 | 0.316909668 | 0.505838281 |
| ENSG00000164920 | OSR2 | 634.5387915 | 0.213752268 | 0.081417465 | 2.625385936 | 0.00865508 | 0.026193607 |
| ENSG00000164924 | YWHAZ | 28549.81199 | -0.424089893 | 0.055985179 | -7.575038611 | 3.59E-14 | 4.05E-13 |
| ENSG00000164930 | FZD6 | 2909.065561 | 0.17409841 | 0.051736461 | 3.365100865 | 0.000765157 | 0.002927889 |
| ENSG00000164932 | CTHRC1 | 27.79308066 | -1.80667536 | 0.370441323 | -4.877089155 | 1.08E-06 | 6.21E-06 |
| ENSG00000164933 | SLC25A32 | 39.66599617 | -0.193506912 | 0.282737269 | -0.684405393 | 0.493719211 | 0.676438565 |
| ENSG00000164934 | DCAF13 | 3026.061955 | -0.110940689 | 0.054311783 | -2.042663344 | 0.041085776 | 0.101358648 |
| ENSG00000164938 | TP53INP1 | 4795.049502 | -0.200805715 | 0.051981219 | -3.863043606 | 0.000111983 | 0.000495829 |
| ENSG00000164941 | INTS8 | 1991.177224 | 0.048934578 | 0.057080678 | 0.85728796 | 0.391285761 | 0.584153023 |
| ENSG00000164944 | VIRMA | 5292.440194 | -0.41919582 | 0.050956062 | -8.226613291 | 1.93E-16 | 2.57E-15 |
| ENSG00000164946 | FREM1 | 2.64563662 | 0.419929656 | 1.134273963 | 0.370218898 | 0.711219397 | 0.839311331 |
| ENSG00000164949 | GEM | 57.64486622 | 1.016471735 | 0.248397639 | 4.092115121 | 4.27E-05 | 0.000201725 |
| ENSG00000164951 | PDP1 | 955.0233086 | -0.921684617 | 0.06972472 | -13.21890744 | 6.82E-40 | 2.77E-38 |
| ENSG00000164953 | TMEM67 | 246.2271313 | 0.044311664 | 0.122229161 | 0.362529401 | 0.716956457 | 0.842657092 |
| ENSG00000164961 | WASHC5 | 4561.762485 | -0.380232552 | 0.04716508 | -8.06173866 | 7.52E-16 | 9.61E-15 |
| ENSG00000164967 | RPP25L | 353.604979 | -0.243005198 | 0.102085523 | -2.380408024 | 0.017293478 | 0.048179788 |
| ENSG00000164970 | FAM219A | 635.5274768 | 0.040553008 | 0.081124135 | 0.499888318 | 0.617153718 | 0.773024159 |
| ENSG00000164972 | C9orf24 | 2.737028484 | 1.076789288 | 1.085123095 | 0.992319944 | 0.321041478 | 0.510550537 |
| ENSG00000164975 | SNAPC3 | 507.5444574 | 0.234264013 | 0.09820622 | 2.385429495 | 0.017059192 | 0.047636989 |
| ENSG00000164976 | MYORG | 918.3400203 | -0.291668589 | 0.071261635 | -4.092925839 | 4.26E-05 | 0.000201103 |
| ENSG00000164978 | NUDT2 | 518.1321967 | 0.055326738 | 0.087988477 | 0.628795266 | 0.529483099 | 0.705157753 |
| ENSG00000164983 | TMEM65 | 652.1179386 | -0.118243843 | 0.084003504 | -1.407606074 | 0.159247748 | 0.304758767 |
| ENSG00000164985 | PSIP1 | 2075.955124 | 0.004588953 | 0.060060863 | 0.076405052 | 0.93909685 | 0.971039083 |
| ENSG00000164989 | CCDC171 | 92.30356958 | -0.178960051 | 0.18909482 | -0.946403774 | 0.343942681 | 0.534882049 |
| ENSG00000165006 | UBAP1 | 1407.733734 | 0.088741861 | 0.062330595 | 1.423728777 | 0.154525 | 0.297857617 |

| | | | | | | | |
|---|---|---|---|---|---|---|---|
| ENSG00000165023 | DIRAS2 | 10.86634416 | 0.61856085 | 0.548581065 | 1.127565075 | 0.259503643 | 0.439484833 |
| ENSG00000165025 | SYK | 485.2954213 | 1.13167494 | 0.089654864 | 12.62257157 | 1.59E-36 | 5.67E-35 |
| ENSG00000165028 | NIPSNAP3B | 25.48568008 | 0.702577584 | 0.367992526 | 1.909216995 | 0.056234105 | 0.132363058 |
| ENSG00000165029 | ABCA1 | 619.1855098 | 0.751018531 | 0.084035789 | 8.936889179 | 4.00E-19 | 6.37E-18 |
| ENSG00000165030 | NFIL3 | 368.3160479 | 0.623341858 | 0.099130033 | 6.288123198 | 3.21E-10 | 2.68E-09 |
| ENSG00000165046 | LETM2 | 56.5544624 | -0.51888796 | 0.241142017 | -2.151794059 | 0.031413578 | 0.080815014 |
| ENSG00000165055 | METTL2B | 832.1250876 | 0.200359196 | 0.081697102 | 2.452463919 | 0.014188161 | 0.040472073 |
| ENSG00000165060 | FXN | 141.5662179 | 0.368233128 | 0.16573114 | 2.221870485 | 0.02629206 | 0.069318451 |
| ENSG00000165061 | ZMAT4 | 150.7801615 | 1.261651796 | 0.158342146 | 7.967883631 | 1.61E-15 | 2.01E-14 |
| ENSG00000165072 | MAMDC2 | 2.263223227 | -0.163972514 | 1.135675034 | -0.144383304 | 0.8851978 | 0.940548288 |
| ENSG00000165078 | CPA6 | 21.65087259 | 3.50126997 | 0.563404992 | 6.214481625 | 5.15E-10 | 4.21E-09 |
| ENSG00000165084 | C8orf34 | 62.23061745 | 2.640456282 | 0.285918986 | 9.234980562 | 2.58E-20 | 4.38E-19 |
| ENSG00000165091 | TMC1 | 2.582944812 | 2.603056366 | 1.364850605 | 1.907209738 | 0.056493433 | 0.13278272 |
| ENSG00000165092 | ALDH1A1 | 1.961173528 | -2.356373376 | 1.478168557 | -1.594116831 | 0.11090987 | 0.230271736 |
| ENSG00000165097 | KDM1B | 909.2869912 | -0.092418609 | 0.071552322 | -1.291622779 | 0.196487811 | 0.358158951 |
| ENSG00000165102 | HGSNAT | 835.1765267 | 0.030437817 | 0.073216584 | 0.415722982 | 0.677612722 | 0.816666799 |
| ENSG00000165105 | RASEF | 1883.7527 | 0.058723821 | 0.058547965 | 1.003003619 | 0.315859115 | 0.504653023 |
| ENSG00000165113 | GKAP1 | 168.3069572 | -0.042985167 | 0.142807642 | -0.301000469 | 0.763414137 | 0.870366712 |
| ENSG00000165115 | KIF27 | 331.7940531 | 0.272779815 | 0.108476531 | 2.514643607 | 0.011915276 | 0.034727029 |
| ENSG00000165118 | C9orf64 | 857.1313547 | -0.247713548 | 0.07709573 | -3.213064421 | 0.001313268 | 0.004808679 |
| ENSG00000165119 | HNRNPK | 17668.08347 | -0.135279784 | 0.049660696 | -2.724081493 | 0.006448059 | 0.020179875 |
| ENSG00000165121 | ENSG0000016 | 93.05242016 | -0.063689066 | 0.191898687 | -0.331889014 | 0.739973066 | 0.856567724 |
| ENSG00000165125 | TRPV6 | 37.25500627 | -1.285800231 | 0.297387988 | -4.323645478 | 1.53E-05 | 7.73E-05 |
| ENSG00000165138 | ANKS6 | 2157.496756 | 0.129601088 | 0.065016117 | 1.993368613 | 0.046221098 | 0.111958381 |
| ENSG00000165140 | FBP1 | 8151.691381 | -0.284860822 | 0.046402686 | -6.138886445 | 8.31E-10 | 6.65E-09 |
| ENSG00000165156 | ZHX1 | 2776.817351 | -0.008003751 | 0.056844841 | -0.140799957 | 0.888027977 | 0.942328951 |
| ENSG00000165164 | CFAP47 | 33.69682187 | -0.665780283 | 0.30985032 | -2.148715813 | 0.031656934 | 0.081386314 |
| ENSG00000165168 | CYBB | 13.70483697 | 0.285804114 | 0.481364453 | 0.593737474 | 0.552687715 | 0.7242877 |
| ENSG00000165169 | DYNLT3 | 463.7033911 | 0.284757068 | 0.098573389 | 2.888782354 | 0.003867366 | 0.012789014 |
| ENSG00000165171 | METTL27 | 29.11529474 | 0.468566199 | 0.336313991 | 1.393240281 | 0.163547143 | 0.311169566 |
| ENSG00000165175 | MID1IP1 | 2235.195339 | -0.003642439 | 0.059985769 | -0.060721723 | 0.951580831 | 0.97757565 |
| ENSG00000165182 | CXorf58 | 1.687431957 | -0.231660533 | 1.321653312 | -0.175280863 | 0.860858948 | NA |
| ENSG00000165185 | KIAA1958 | 551.1339167 | 0.125059141 | 0.084303914 | 1.483432191 | 0.13795962 | 0.273605402 |

| | | | | | | | |
|---|---|---|---|---|---|---|---|
| ENSG00000165194 | PCDH19 | 418.5855133 | 0.754816443 | 0.095427407 | 7.909849639 | 2.58E-15 | 3.17E-14 |
| ENSG00000165195 | PIGA | 481.6125703 | 0.468291135 | 0.092051136 | 5.087293365 | 3.63E-07 | 2.22E-06 |
| ENSG00000165209 | STRBP | 2385.641116 | 0.173407751 | 0.054298471 | 3.193602833 | 0.001405093 | 0.005104169 |
| ENSG00000165215 | CLDN3 | 1495.085614 | 0.800667186 | 0.075880296 | 10.55171409 | 4.99E-26 | 1.16E-24 |
| ENSG00000165219 | GAPVD1 | 2692.955499 | 0.162782168 | 0.055065526 | 2.95615386 | 0.003115017 | 0.010520978 |
| ENSG00000165233 | CARD19 | 539.6255786 | 0.389123219 | 0.094276351 | 4.12747433 | 3.67E-05 | 0.000174703 |
| ENSG00000165238 | WNK2 | 462.465523 | 1.445549135 | 0.09789955 | 14.76563611 | 2.44E-49 | 1.33E-47 |
| ENSG00000165240 | ATP7A | 610.785083 | 0.368906129 | 0.084295829 | 4.376327224 | 1.21E-05 | 6.15E-05 |
| ENSG00000165244 | ZNF367 | 666.4568189 | -0.144944921 | 0.089738355 | -1.615194762 | 0.106268525 | 0.222609803 |
| ENSG00000165259 | HDX | 68.75060019 | 0.28863208 | 0.223179839 | 1.293271299 | 0.195917244 | 0.357394409 |
| ENSG00000165264 | NDUFB6 | 527.2238953 | -0.079696112 | 0.085390028 | -0.93331872 | 0.350655432 | 0.541977875 |
| ENSG00000165269 | AQP7 | 1.815159686 | -0.886703578 | 1.378802932 | -0.643096672 | 0.520161374 | 0.697413646 |
| ENSG00000165271 | NOL6 | 1572.604817 | 0.404286954 | 0.062811501 | 6.436511588 | 1.22E-10 | 1.05E-09 |
| ENSG00000165272 | AQP3 | 544.8712567 | -0.391898849 | 0.09729723 | -4.027852083 | 5.63E-05 | 0.000261389 |
| ENSG00000165275 | TRMT10B | 160.8782361 | -0.220843483 | 0.145591391 | -1.516871852 | 0.129299041 | 0.260117514 |
| ENSG00000165280 | VCP | 14669.14884 | -0.320009173 | 0.042000121 | -7.619244146 | 2.55E-14 | 2.91E-13 |
| ENSG00000165282 | PIGO | 1109.707093 | -0.221110068 | 0.07328262 | -3.017223819 | 0.002551014 | 0.008768393 |
| ENSG00000165283 | STOML2 | 2257.694208 | -0.205220864 | 0.057891967 | -3.544893622 | 0.000392772 | 0.001581566 |
| ENSG00000165288 | BRWD3 | 1048.875313 | -0.476468462 | 0.082982718 | -5.741779405 | 9.37E-09 | 6.76E-08 |
| ENSG00000165300 | SLITRK5 | 170.6643763 | 0.150166689 | 0.1447697 | 1.037279828 | 0.299605464 | 0.486305055 |
| ENSG00000165304 | MELK | 921.7409682 | -0.506022421 | 0.074107004 | -6.828267139 | 8.59E-12 | 8.11E-11 |
| ENSG00000165309 | ARMC3 | 1.983903777 | -1.034672018 | 1.330601952 | -0.777596949 | 0.43680666 | 0.626595539 |
| ENSG00000165312 | OTUD1 | 206.4447864 | -0.116910421 | 0.127982766 | -0.913485653 | 0.360987176 | 0.553508844 |
| ENSG00000165322 | ARHGAP12 | 1130.166955 | -0.521583207 | 0.073514422 | -7.094977986 | 1.29E-12 | 1.31E-11 |
| ENSG00000165338 | HECTD2 | 479.7849393 | -0.601760903 | 0.094598159 | -6.361232694 | 2.00E-10 | 1.69E-09 |
| ENSG00000165355 | FBXO33 | 699.2388393 | -0.308842339 | 0.075552838 | -4.087766236 | 4.36E-05 | 0.000205358 |
| ENSG00000165359 | INTS6L | 377.5919168 | -0.414789307 | 0.120510163 | -3.44194462 | 0.000577548 | 0.002258602 |
| ENSG00000165379 | LRFN5 | 5.749811908 | -4.134607596 | 1.200060496 | -3.44533264 | 0.000570357 | 0.002233144 |
| ENSG00000165389 | SPTSSA | 1752.929231 | -0.108622212 | 0.061091148 | -1.778035218 | 0.075398071 | 0.168602564 |
| ENSG00000165392 | WRN | 543.6765752 | 0.046337998 | 0.083881892 | 0.552419563 | 0.580660931 | 0.745737417 |
| ENSG00000165406 | MARCHF8 | 248.0792225 | -0.026499259 | 0.117036695 | -0.22641838 | 0.820876034 | 0.904991377 |
| ENSG00000165410 | CFL2 | 373.0410457 | -0.149503274 | 0.103190069 | -1.448814555 | 0.147389378 | 0.28770527 |
| ENSG00000165416 | SUGT1 | 725.1736784 | -0.204186236 | 0.076422793 | -2.671797628 | 0.007544613 | 0.023203302 |

| | | | | | | | |
|---|---|---|---|---|---|---|---|
| ENSG00000165417 | GTF2A1 | 1375.218736 | -0.895045255 | 0.06763459 | -13.23354307 | 5.62E-40 | 2.30E-38 |
| ENSG00000165424 | ZCCHC24 | 169.9486165 | -0.233516207 | 0.141571008 | -1.649463483 | 0.099052718 | 0.210375832 |
| ENSG00000165434 | PGM2L1 | 379.75292 | 2.551802133 | 0.118260644 | 21.57777975 | 2.90E-103 | 5.01E-101 |
| ENSG00000165449 | SLC16A9 | 97.89902356 | -0.603588018 | 0.186137308 | -3.242703051 | 0.001184015 | 0.004375212 |
| ENSG00000165458 | INPPL1 | 10418.3023 | 0.214683308 | 0.048852583 | 4.394512984 | 1.11E-05 | 5.70E-05 |
| ENSG00000165474 | GJB2 | 57.40737848 | -0.798950224 | 0.236760888 | -3.374502563 | 0.000739492 | 0.00283773 |
| ENSG00000165475 | CRYL1 | 1019.078436 | 0.227822193 | 0.07185725 | 3.170483042 | 0.001521857 | 0.005485764 |
| ENSG00000165476 | REEP3 | 1186.952912 | -0.196359899 | 0.068286179 | -2.875543813 | 0.004033322 | 0.013278473 |
| ENSG00000165480 | SKA3 | 1009.790534 | -0.052748545 | 0.074232603 | -0.71058461 | 0.477341682 | 0.662124694 |
| ENSG00000165487 | MICU2 | 1554.958971 | 0.310441773 | 0.063600993 | 4.881083727 | 1.06E-06 | 6.09E-06 |
| ENSG00000165490 | DDIAS | 584.7521599 | -0.227495187 | 0.084850499 | -2.681129622 | 0.007337409 | 0.02264803 |
| ENSG00000165494 | PCF11 | 4179.50493 | -0.029853677 | 0.04811885 | -0.620415431 | 0.534984316 | 0.710463136 |
| ENSG00000165495 | PKNOX2 | 6.516105234 | 0.173081006 | 0.694560149 | 0.24919513 | 0.803209847 | 0.895230487 |
| ENSG00000165496 | RPL10L | 5.090049226 | -0.451326821 | 0.771242364 | -0.585194541 | 0.558416915 | 0.728502064 |
| ENSG00000165501 | LRR1 | 621.9858921 | -0.878258701 | 0.082171769 | -10.68808315 | 1.16E-26 | 2.82E-25 |
| ENSG00000165502 | RPL36AL | 3417.382123 | -0.62552156 | 0.046514609 | -13.44785171 | 3.17E-41 | 1.36E-39 |
| ENSG00000165506 | DNAAF2 | 613.8890316 | -0.428403599 | 0.079106979 | -5.415496865 | 6.11E-08 | 4.04E-07 |
| ENSG00000165507 | DEPP1 | 12.59992625 | 0.393146814 | 0.506028337 | 0.776926478 | 0.437202148 | 0.62683858 |
| ENSG00000165509 | MAGEC3 | 85.26719906 | 2.033895059 | 0.217252067 | 9.36191351 | 7.83E-21 | 1.38E-19 |
| ENSG00000165511 | ZNF22-AS1 | 9.850441726 | -2.088259159 | 0.649241819 | -3.216458179 | 0.001297834 | 0.004756722 |
| ENSG00000165512 | ZNF22 | 925.4815276 | -0.614986887 | 0.070229931 | -8.756763388 | 2.01E-18 | 3.05E-17 |
| ENSG00000165516 | KLHDC2 | 1786.959651 | -0.773768859 | 0.057656807 | -13.42025159 | 4.60E-41 | 1.97E-39 |
| ENSG00000165521 | EML5 | 69.43773365 | -1.152383837 | 0.220175961 | -5.233922138 | 1.66E-07 | 1.05E-06 |
| ENSG00000165525 | NEMF | 1921.14222 | -0.511580498 | 0.05714253 | -8.952710003 | 3.47E-19 | 5.54E-18 |
| ENSG00000165526 | RPUSD4 | 626.0760993 | 0.330529571 | 0.079106138 | 4.17830497 | 2.94E-05 | 0.000142406 |
| ENSG00000165527 | ARF6 | 3652.023261 | -0.403057608 | 0.04731475 | -8.51864605 | 1.61E-17 | 2.30E-16 |
| ENSG00000165533 | TTC8 | 723.9882456 | -0.682043234 | 0.079093114 | -8.62329478 | 6.51E-18 | 9.52E-17 |
| ENSG00000165548 | TMEM63C | 410.2972925 | 0.670566871 | 0.094029771 | 7.131431539 | 9.93E-13 | 1.02E-11 |
| ENSG00000165555 | NOXRED1 | 32.47736276 | -0.481039714 | 0.316895569 | -1.517975516 | 0.129020571 | 0.259669985 |
| ENSG00000165568 | AKR1E2 | 185.3564121 | -1.506560261 | 0.143137945 | -10.52523327 | 6.61E-26 | 1.53E-24 |
| ENSG00000165572 | KBTBD6 | 889.1642431 | -0.096958277 | 0.071527042 | -1.355547127 | 0.175243304 | 0.328411192 |
| ENSG00000165591 | FAAH2 | 248.2471943 | -0.478436421 | 0.12190453 | -3.924681229 | 8.68E-05 | 0.000391398 |
| ENSG00000165609 | NUDT5 | 1412.592493 | -0.289725291 | 0.060847278 | -4.761516055 | 1.92E-06 | 1.07E-05 |

| | | | | | | | |
|---|---|---|---|---|---|---|---|
| ENSG00000165617 | DACT1 | 118.1878952 | -0.119611167 | 0.167328314 | -0.714829211 | 0.474714583 | 0.659990209 |
| ENSG00000165621 | OXGR1 | 3.647241493 | -0.394370256 | 0.93068399 | -0.423742388 | 0.671753698 | 0.811951622 |
| ENSG00000165626 | BEND7 | 1.74407409 | 0.155833468 | 1.331901209 | 0.117000771 | 0.906859443 | NA |
| ENSG00000165629 | ATP5F1C | 3888.201555 | -0.912416573 | 0.049729466 | -18.34760454 | 3.45E-75 | 3.68E-73 |
| ENSG00000165630 | PRPF18 | 96.5830654 | -0.537374349 | 0.185639786 | -2.894715415 | 0.003795024 | 0.012573347 |
| ENSG00000165632 | TAF3 | 992.980044 | -0.793407546 | 0.068449295 | -11.59117198 | 4.57E-31 | 1.32E-29 |
| ENSG00000165637 | VDAC2 | 961.7937047 | -0.392905077 | 0.070337877 | -5.585967279 | 2.32E-08 | 1.60E-07 |
| ENSG00000165644 | COMTD1 | 172.4570317 | -0.526100183 | 0.145196187 | -3.623374643 | 0.000290784 | 0.001199099 |
| ENSG00000165646 | SLC18A2 | 3.180894164 | -2.081005332 | 1.132211638 | -1.838000302 | 0.066062361 | 0.151304968 |
| ENSG00000165650 | PDZD8 | 1229.528307 | -0.441918978 | 0.067719097 | -6.525765933 | 6.77E-11 | 5.93E-10 |
| ENSG00000165655 | ZNF503 | 150.7609604 | -0.569569112 | 0.158951345 | -3.583292182 | 0.000339291 | 0.001381738 |
| ENSG00000165660 | ABRAXAS2 | 630.1240024 | -0.619314789 | 0.080972642 | -7.648444873 | 2.03E-14 | 2.34E-13 |
| ENSG00000165661 | QSOX2 | 1075.750302 | 0.328503794 | 0.065338553 | 5.027717576 | 4.96E-07 | 2.99E-06 |
| ENSG00000165669 | FAM204A | 951.7835681 | -0.814518247 | 0.070590302 | -11.53867068 | 8.42E-31 | 2.41E-29 |
| ENSG00000165671 | NSD1 | 5643.410806 | -0.514818517 | 0.059364258 | -8.672196627 | 4.24E-18 | 6.30E-17 |
| ENSG00000165672 | PRDX3 | 2518.972856 | -0.464809414 | 0.052897127 | -8.78704459 | 1.54E-18 | 2.34E-17 |
| ENSG00000165675 | ENOX2 | 761.3179797 | 0.08201459 | 0.076974781 | 1.065473513 | 0.286661706 | 0.470488626 |
| ENSG00000165678 | GHITM | 6599.057301 | -0.349341756 | 0.046663284 | -7.486437423 | 7.08E-14 | 7.80E-13 |
| ENSG00000165684 | SNAPC4 | 1272.274623 | 0.155552823 | 0.075027275 | 2.073283665 | 0.038145888 | 0.09540631 |
| ENSG00000165688 | PMPCA | 2874.049372 | 0.232652823 | 0.053246948 | 4.369317504 | 1.25E-05 | 6.35E-05 |
| ENSG00000165689 | ENTR1 | 2309.657131 | 0.131444111 | 0.052735054 | 2.492537706 | 0.012683386 | 0.036649059 |
| ENSG00000165694 | FRMD7 | 3.212295747 | 1.66012671 | 1.210386251 | 1.37156772 | 0.170198048 | 0.320975274 |
| ENSG00000165695 | AK8 | 50.35984287 | 0.084854649 | 0.252536283 | 0.336009733 | 0.736863527 | 0.854938089 |
| ENSG00000165698 | SPACA9 | 146.0713583 | 0.738177942 | 0.154281148 | 4.784628283 | 1.71E-06 | 9.65E-06 |
| ENSG00000165699 | TSC1 | 1951.050508 | 0.090536021 | 0.058931193 | 1.53630049 | 0.124464699 | 0.252562601 |
| ENSG00000165702 | GFI1B | 2.904314246 | -0.151201712 | 1.009254302 | -0.149815276 | 0.880910357 | 0.938252557 |
| ENSG00000165704 | HPRT1 | 1077.474675 | 0.12078552 | 0.068421345 | 1.765319291 | 0.077510119 | 0.172378365 |
| ENSG00000165714 | BORCS5 | 172.4741574 | -0.36769922 | 0.136199197 | -2.699716503 | 0.006939858 | 0.021536844 |
| ENSG00000165716 | DIPK1B | 185.4823815 | 0.77279655 | 0.142462905 | 5.42454577 | 5.81E-08 | 3.85E-07 |
| ENSG00000165724 | ZMYND19 | 1337.756122 | 0.298550784 | 0.060936236 | 4.899396567 | 9.61E-07 | 5.57E-06 |
| ENSG00000165730 | STOX1 | 202.7455701 | -0.24087528 | 0.127712061 | -1.886080915 | 0.059284052 | 0.138124736 |
| ENSG00000165731 | RET | 856.3668482 | -1.623637342 | 0.09228368 | -17.59398126 | 2.74E-69 | 2.48E-67 |
| ENSG00000165732 | DDX21 | 5376.916924 | -0.330071015 | 0.054233826 | -6.086072813 | 1.16E-09 | 9.16E-09 |

| | | | | | | | |
|---|---|---|---|---|---|---|---|
| ENSG00000165733 | BMS1 | 2862.733736 | -0.509205189 | 0.056957773 | -8.9400474 | 3.89E-19 | 6.19E-18 |
| ENSG00000165752 | STK32C | 532.3632836 | -0.335314985 | 0.089235777 | -3.757629476 | 0.000171531 | 0.000736323 |
| ENSG00000165757 | JCAD | 22.6084542 | -0.286119833 | 0.37451764 | -0.763968911 | 0.44488577 | 0.633538988 |
| ENSG00000165775 | FUNDC2 | 2138.902515 | -0.515040301 | 0.055979912 | -9.20044856 | 3.56E-20 | 6.01E-19 |
| ENSG00000165782 | PIP4P1 | 1399.895125 | -0.429907126 | 0.061736927 | -6.963533 | 3.32E-12 | 3.25E-11 |
| ENSG00000165792 | METTL17 | 1260.060017 | -0.511084916 | 0.074965623 | -6.817590488 | 9.26E-12 | 8.69E-11 |
| ENSG00000165794 | SLC39A2 | 2.181001358 | -0.878779449 | 1.183716339 | -0.742390234 | 0.457850935 | 0.645074343 |
| ENSG00000165795 | NDRG2 | 323.1144209 | -0.680583118 | 0.103315882 | -6.587400764 | 4.48E-11 | 3.97E-10 |
| ENSG00000165801 | ARHGEF40 | 121.0436264 | -0.65681544 | 0.162227947 | -4.048719435 | 5.15E-05 | 0.000240558 |
| ENSG00000165802 | NSMF | 1996.041587 | 0.48680071 | 0.054720587 | 8.896116427 | 5.78E-19 | 9.10E-18 |
| ENSG00000165804 | ZNF219 | 1129.869526 | -0.89934036 | 0.073352745 | -12.26048681 | 1.48E-34 | 4.95E-33 |
| ENSG00000165806 | CASP7 | 1419.172247 | -1.038106335 | 0.061596585 | -16.85330995 | 9.92E-64 | 7.71E-62 |
| ENSG00000165807 | PPP1R36 | 12.61030377 | -0.277592039 | 0.498748398 | -0.556577305 | 0.577816263 | 0.743545771 |
| ENSG00000165810 | BTNL9 | 143.9256364 | -0.114550347 | 0.150171919 | -0.762794719 | 0.445585831 | 0.634142252 |
| ENSG00000165813 | CCDC186 | 1012.003531 | -0.499036224 | 0.069718463 | -7.157877621 | 8.19E-13 | 8.44E-12 |
| ENSG00000165816 | VWA2 | 8.832163692 | -0.153892768 | 0.600814732 | -0.256140137 | 0.797842631 | 0.892109083 |
| ENSG00000165819 | METTL3 | 1394.253527 | -0.559125788 | 0.08464723 | -6.60536429 | 3.97E-11 | 3.53E-10 |
| ENSG00000165821 | SALL2 | 311.2804735 | -1.275466804 | 0.109985738 | -11.59665635 | 4.28E-31 | 1.25E-29 |
| ENSG00000165832 | TRUB1 | 777.3729305 | -0.488233608 | 0.074192019 | -6.580675605 | 4.68E-11 | 4.15E-10 |
| ENSG00000165837 | ERICH6B | 1.918539436 | -0.107656989 | 1.276087586 | -0.08436489 | 0.932766321 | 0.96773548 |
| ENSG00000165861 | ZFYVE1 | 1021.090861 | -0.381095766 | 0.078302839 | -4.866946961 | 1.13E-06 | 6.52E-06 |
| ENSG00000165868 | HSPA12A | 285.5714408 | 0.520866871 | 0.116414913 | 4.474228053 | 7.67E-06 | 4.01E-05 |
| ENSG00000165874 | SHLD2P1 | 9.510352365 | -0.248779665 | 0.631463457 | -0.393973178 | 0.693600842 | 0.827329623 |
| ENSG00000165879 | FRAT1 | 528.0632339 | -1.009263365 | 0.089960344 | -11.21898076 | 3.29E-29 | 8.84E-28 |
| ENSG00000165886 | UBTD1 | 403.8780833 | 0.198410249 | 0.122619702 | 1.618094368 | 0.105642262 | 0.22154068 |
| ENSG00000165887 | ANKRD2 | 2.615523549 | 0.532470726 | 1.090487699 | 0.488286779 | 0.625346725 | 0.778628629 |
| ENSG00000165891 | E2F7 | 625.7361429 | -0.431210598 | 0.086149991 | -5.005346999 | 5.58E-07 | 3.34E-06 |
| ENSG00000165895 | ARHGAP42 | 147.4936616 | 0.787628634 | 0.153385747 | 5.134953206 | 2.82E-07 | 1.75E-06 |
| ENSG00000165898 | ISCA2 | 527.9570287 | -0.817981731 | 0.085824586 | -9.530855552 | 1.56E-21 | 2.85E-20 |
| ENSG00000165899 | OTOGL | 14.03610103 | 1.63063082 | 0.523036318 | 3.117624462 | 0.001823149 | 0.006462152 |
| ENSG00000165905 | LARGE2 | 371.8318187 | 1.517730864 | 0.104938344 | 14.46307235 | 2.07E-47 | 1.06E-45 |
| ENSG00000165912 | PACSIN3 | 1599.402429 | -0.473669307 | 0.058438262 | -8.105465375 | 5.25E-16 | 6.79E-15 |
| ENSG00000165914 | TTC7B | 1217.507467 | -0.857846633 | 0.064379077 | -13.32492901 | 1.66E-40 | 6.95E-39 |

| | | | | | | | |
|---|---|---|---|---|---|---|---|
| ENSG00000165915 | SLC39A13 | 1264.653876 | 0.220841506 | 0.074509618 | 2.963932861 | 0.003037345 | 0.010279833 |
| ENSG00000165916 | PSMC3 | 5445.99302 | -0.23363367 | 0.04765937 | -4.902156094 | 9.48E-07 | 5.50E-06 |
| ENSG00000165917 | RAPSN | 7.170278351 | -0.785254916 | 0.663932161 | -1.182733661 | 0.236914719 | 0.412134956 |
| ENSG00000165923 | AGBL2 | 56.64635103 | -0.02942219 | 0.237213819 | -0.124032362 | 0.901289652 | 0.950225965 |
| ENSG00000165929 | TC2N | 1634.834157 | -0.657579009 | 0.068480299 | -9.602455226 | 7.81E-22 | 1.45E-20 |
| ENSG00000165934 | CPSF2 | 3156.545849 | -0.673493122 | 0.05161113 | -13.04937765 | 6.41E-39 | 2.49E-37 |
| ENSG00000165935 | SMCO2 | 4.508586149 | -0.321480192 | 0.827524321 | -0.388484282 | 0.697657683 | 0.830348323 |
| ENSG00000165943 | MOAP1 | 2151.292393 | -0.296635074 | 0.052581054 | -5.641482099 | 1.69E-08 | 1.18E-07 |
| ENSG00000165948 | IFI27L1 | 74.91914034 | -0.879353157 | 0.218896076 | -4.017217535 | 5.89E-05 | 0.000272693 |
| ENSG00000165949 | IFI27 | 7.969322483 | 0.809683697 | 0.638182859 | 1.268733068 | 0.204536284 | 0.369443167 |
| ENSG00000165959 | CLMN | 1231.581173 | -2.007827491 | 0.071649795 | -28.02279479 | 8.57E-173 | 3.71E-170 |
| ENSG00000165973 | NELL1 | 2.329002235 | -2.701658382 | 1.406283933 | -1.921132936 | 0.05471495 | 0.129411378 |
| ENSG00000165983 | PTER | 330.3670694 | -0.215720705 | 0.103387527 | -2.086525427 | 0.036931054 | 0.092792814 |
| ENSG00000165995 | CACNB2 | 55.68342757 | -2.644633016 | 0.295347348 | -8.954314416 | 3.42E-19 | 5.46E-18 |
| ENSG00000165996 | HACD1 | 451.0738179 | -2.508963465 | 0.108720883 | -23.07710722 | 7.86E-118 | 1.82E-115 |
| ENSG00000165997 | ARL5B | 666.9787949 | -0.609640589 | 0.097144243 | -6.275622425 | 3.48E-10 | 2.89E-09 |
| ENSG00000166002 | SMCO4 | 43.69288636 | -0.51293976 | 0.268140542 | -1.912951158 | 0.055754305 | 0.131449514 |
| ENSG00000166004 | CEP295 | 737.591698 | -0.141538837 | 0.082928188 | -1.706763896 | 0.087865945 | 0.190712914 |
| ENSG00000166006 | KCNC2 | 16.23834409 | 1.455060601 | 0.480420567 | 3.028722543 | 0.002455901 | 0.00848214 |
| ENSG00000166012 | TAF1D | 964.8458541 | 0.109744535 | 0.086371611 | 1.270608865 | 0.203867832 | 0.368623635 |
| ENSG00000166016 | ABTB2 | 604.9179555 | 0.608737818 | 0.083685198 | 7.274139668 | 3.49E-13 | 3.66E-12 |
| ENSG00000166024 | R3HCC1L | 909.4290565 | -0.512427568 | 0.08230368 | -6.226058998 | 4.78E-10 | 3.92E-09 |
| ENSG00000166025 | AMOTL1 | 1649.649461 | -3.078697574 | 0.068113853 | -45.19928689 | 0 | 0 |
| ENSG00000166033 | HTRA1 | 390.2379083 | 0.027173283 | 0.097363621 | 0.279090724 | 0.780175202 | 0.880664282 |
| ENSG00000166037 | CEP57 | 959.0043571 | -0.076539149 | 0.075334169 | -1.015995123 | 0.309631709 | 0.497719249 |
| ENSG00000166046 | TCP11L2 | 200.2793057 | 0.527941401 | 0.129006995 | 4.092347078 | 4.27E-05 | 0.000201565 |
| ENSG00000166068 | SPRED1 | 471.0621369 | -0.105905026 | 0.096622502 | -1.09607 | 0.273048142 | 0.455038495 |
| ENSG00000166073 | GPR176 | 287.192147 | -0.123237588 | 0.114444103 | -1.0768365 | 0.281553316 | 0.464998781 |
| ENSG00000166106 | ADAMTS15 | 14.81551804 | 1.901163427 | 0.50958686 | 3.730793658 | 0.000190878 | 0.000813588 |
| ENSG00000166123 | GPT2 | 2161.064299 | -0.078582834 | 0.05321199 | -1.476788098 | 0.139732448 | 0.276023625 |
| ENSG00000166126 | AMN | 16.91534594 | -0.919861682 | 0.447350194 | -2.056245186 | 0.039758889 | 0.098655372 |
| ENSG00000166128 | RAB8B | 201.8739792 | -0.44876577 | 0.131348779 | -3.416596437 | 0.000634092 | 0.002462931 |
| ENSG00000166130 | IKBIP | 170.0181093 | -0.692769163 | 0.142605546 | -4.857939841 | 1.19E-06 | 6.80E-06 |

| | | | | | | | |
|---|---|---|---|---|---|---|---|
| ENSG00000166133 | RPUSD2 | 318.0708914 | 0.077145939 | 0.104382214 | 0.739071688 | 0.459863467 | 0.64675789 |
| ENSG00000166135 | HIF1AN | 2636.424003 | -0.091420451 | 0.053144296 | -1.720230733 | 0.085390509 | 0.186309518 |
| ENSG00000166136 | NDUFB8 | 101.9965016 | -0.222609733 | 0.175876284 | -1.265717742 | 0.205614154 | 0.371050086 |
| ENSG00000166140 | ZFYVE19 | 1241.864236 | -0.358509226 | 0.072609978 | -4.937464985 | 7.91E-07 | 4.65E-06 |
| ENSG00000166143 | PPP1R14D | 11.94937177 | 1.27985903 | 0.532794693 | 2.402161744 | 0.0162985 | 0.045752275 |
| ENSG00000166145 | SPINT1 | 6305.984993 | 0.27830763 | 0.054885996 | 5.070649119 | 3.96E-07 | 2.42E-06 |
| ENSG00000166147 | FBN1 | 64.11711721 | -0.104638144 | 0.231910402 | -0.451200738 | 0.651844878 | 0.797723206 |
| ENSG00000166153 | DEPDC4 | 21.44629311 | -0.367241173 | 0.381257581 | -0.963236384 | 0.33542891 | 0.526604106 |
| ENSG00000166159 | LRTM2 | 2.273722992 | 2.380638116 | 1.395306513 | 1.706175735 | 0.087975365 | 0.190896224 |
| ENSG00000166164 | BRD7 | 1583.912835 | 0.081628531 | 0.057312659 | 1.424267027 | 0.154369189 | 0.297710522 |
| ENSG00000166165 | CKB | 1272.847122 | -0.292682276 | 0.071666573 | -4.083944059 | 4.43E-05 | 0.000208655 |
| ENSG00000166166 | TRMT61A | 1169.359724 | 0.221996312 | 0.073178024 | 3.033647262 | 0.002416168 | 0.008360013 |
| ENSG00000166167 | BTRC | 670.1905164 | 0.117557615 | 0.077434757 | 1.518150503 | 0.128976462 | 0.259627943 |
| ENSG00000166169 | POLL | 699.3327748 | 0.169839769 | 0.079367246 | 2.139922668 | 0.032361017 | 0.082991808 |
| ENSG00000166170 | BAG5 | 2431.21604 | -0.238564562 | 0.052396649 | -4.553049992 | 5.29E-06 | 2.81E-05 |
| ENSG00000166171 | DPCD | 272.5461994 | 0.289543681 | 0.118304882 | 2.44743646 | 0.014387649 | 0.040954544 |
| ENSG00000166173 | LARP6 | 172.9719593 | -0.402266251 | 0.139627844 | -2.880988782 | 0.003964297 | 0.013068111 |
| ENSG00000166181 | API5 | 3739.124469 | 0.042971724 | 0.052029013 | 0.825918504 | 0.408850323 | 0.601219642 |
| ENSG00000166188 | ZNF319 | 299.5151322 | 0.169859726 | 0.111732086 | 1.520241253 | 0.128450353 | 0.258879627 |
| ENSG00000166189 | HPS6 | 608.4873774 | 0.120317478 | 0.093933764 | 1.280875725 | 0.200237321 | 0.36340539 |
| ENSG00000166192 | SENP8 | 175.3058352 | 0.08040734 | 0.135887665 | 0.591719199 | 0.554038638 | 0.725423029 |
| ENSG00000166197 | NOLC1 | 5476.645646 | 0.19013462 | 0.045175613 | 4.208788931 | 2.57E-05 | 0.000125499 |
| ENSG00000166199 | ALKBH3 | 381.1042039 | -0.017768896 | 0.10254968 | -0.1732711 | 0.862438339 | 0.928536889 |
| ENSG00000166200 | COPS2 | 2162.516067 | -0.013502916 | 0.059025625 | -0.228763623 | 0.819052641 | 0.904059452 |
| ENSG00000166224 | SGPL1 | 4507.960071 | -0.491028958 | 0.04761329 | -10.31285496 | 6.16E-25 | 1.37E-23 |
| ENSG00000166225 | FRS2 | 1520.280866 | 0.1134027 | 0.066696214 | 1.700286898 | 0.089076974 | 0.192866991 |
| ENSG00000166226 | CCT2 | 9262.887613 | 0.13135289 | 0.044837721 | 2.929517506 | 0.003394887 | 0.011377459 |
| ENSG00000166228 | PCBD1 | 1026.488954 | -0.256455504 | 0.067654339 | -3.790673398 | 0.000150239 | 0.000651388 |
| ENSG00000166233 | ARIH1 | 2833.787834 | 0.032275018 | 0.049866758 | 0.647225113 | 0.51748626 | 0.695665852 |
| ENSG00000166246 | DNAAF8 | 24.5435712 | -0.347761278 | 0.350001901 | -0.993598255 | 0.320418494 | 0.509913303 |
| ENSG00000166250 | CLMP | 9.526875966 | -1.901554989 | 0.659536571 | -2.883168385 | 0.003936969 | 0.012992962 |
| ENSG00000166260 | COX11 | 1237.7624 | 0.028927053 | 0.06246945 | 0.463059185 | 0.643321945 | 0.790937773 |
| ENSG00000166261 | ZNF202 | 389.8906144 | -0.050530483 | 0.100655455 | -0.502014347 | 0.615657429 | 0.772264951 |

| | | | | | | | |
|---|---|---|---|---|---|---|---|
| ENSG00000166262 | FAM227B | 603.7071215 | -2.296778777 | 0.090686347 | -25.3266213 | 1.63E-141 | 5.33E-139 |
| ENSG00000166263 | STXBP4 | 488.1359118 | -0.292221271 | 0.095373168 | -3.063977824 | 0.002184151 | 0.007616969 |
| ENSG00000166266 | CUL5 | 863.1834455 | -0.124084113 | 0.07271842 | -1.706364259 | 0.087940281 | 0.190856201 |
| ENSG00000166268 | MYRFL | 12.1411553 | 0.535549725 | 0.514297909 | 1.041321995 | 0.297726132 | 0.484282597 |
| ENSG00000166272 | WBP1L | 1706.083359 | -0.038159724 | 0.060623363 | -0.629455752 | 0.529050728 | 0.704950261 |
| ENSG00000166275 | BORCS7 | 229.7031405 | 0.493121132 | 0.123037046 | 4.007907754 | 6.13E-05 | 0.000283038 |
| ENSG00000166278 | C2 | 10.49878847 | -0.411602182 | 0.538239607 | -0.764719237 | 0.444438748 | 0.633256204 |
| ENSG00000166289 | PLEKHF1 | 225.381928 | -1.199230334 | 0.137589724 | -8.71598766 | 2.88E-18 | 4.33E-17 |
| ENSG00000166295 | ANAPC16 | 2157.179261 | -0.676761307 | 0.055648513 | -12.16135471 | 4.99E-34 | 1.63E-32 |
| ENSG00000166311 | SMPD1 | 510.3515239 | -0.024536631 | 0.091210924 | -0.269009784 | 0.787922154 | 0.885283461 |
| ENSG00000166313 | APBB1 | 12.8799858 | -1.782702289 | 0.53862782 | -3.309710754 | 0.000933924 | 0.003515626 |
| ENSG00000166321 | NUDT13 | 49.87275345 | -1.015860391 | 0.257195372 | -3.949761553 | 7.82E-05 | 0.000355432 |
| ENSG00000166323 | C11orf65 | 14.0121054 | -0.312360183 | 0.480180769 | -0.6505054 | 0.515365815 | 0.694378703 |
| ENSG00000166326 | TRIM44 | 4914.095987 | 0.072826492 | 0.046895465 | 1.552953818 | 0.120434168 | 0.246301134 |
| ENSG00000166333 | ILK | 2020.789579 | -0.490092119 | 0.057600727 | -8.508436275 | 1.76E-17 | 2.51E-16 |
| ENSG00000166337 | TAF10 | 96.71507482 | 0.042181884 | 0.179844318 | 0.234546657 | 0.814560616 | 0.901763793 |
| ENSG00000166340 | TPP1 | 920.8505972 | -0.022758176 | 0.071285792 | -0.319252624 | 0.749534955 | 0.861482633 |
| ENSG00000166341 | DCHS1 | 2.816944089 | 0.312709767 | 1.049203849 | 0.298044815 | 0.765668959 | 0.871857735 |
| ENSG00000166343 | MSS51 | 39.00458482 | -1.071363099 | 0.303604806 | -3.5288081 | 0.000417436 | 0.001675293 |
| ENSG00000166347 | CYB5A | 1093.996715 | 0.601525708 | 0.072345931 | 8.314575579 | 9.21E-17 | 1.25E-15 |
| ENSG00000166348 | USP54 | 868.7568328 | -0.750552263 | 0.073281326 | -10.24206714 | 1.28E-24 | 2.80E-23 |
| ENSG00000166349 | RAG1 | 3.207005184 | -0.06899512 | 0.966705246 | -0.071371414 | 0.94310216 | 0.97291737 |
| ENSG00000166352 | IFTAP | 140.2743272 | 0.08194759 | 0.157361289 | 0.520760799 | 0.602533415 | 0.761829824 |
| ENSG00000166359 | WDR88 | 2.985228626 | -0.513932444 | 1.014305292 | -0.506684179 | 0.612376433 | 0.769537956 |
| ENSG00000166377 | ATP9B | 482.0753431 | 0.732922839 | 0.088547524 | 8.277169248 | 1.26E-16 | 1.71E-15 |
| ENSG00000166387 | PPFIBP2 | 802.601907 | -1.433607328 | 0.077220711 | -18.56506249 | 6.16E-77 | 6.73E-75 |
| ENSG00000166398 | GARRE1 | 1283.838947 | 0.237947443 | 0.067630247 | 3.518358338 | 0.000434226 | 0.001738418 |
| ENSG00000166401 | SERPINB8 | 57.96882917 | 0.578663852 | 0.247367201 | 2.339290942 | 0.019320381 | 0.053066694 |
| ENSG00000166402 | TUB | 2.12092563 | -3.556409019 | 1.552552064 | -2.290685834 | 0.021981591 | 0.059380321 |
| ENSG00000166407 | LMO1 | 11.13105962 | 2.602009776 | 0.646659736 | 4.023769583 | 5.73E-05 | 0.000265857 |
| ENSG00000166411 | IDH3A | 2028.825125 | -0.175396173 | 0.056019782 | -3.130968479 | 0.001742309 | 0.006202446 |
| ENSG00000166415 | WDR72 | 2661.761963 | -1.929632228 | 0.055091083 | -35.02621727 | 8.98E-269 | 1.29E-265 |
| ENSG00000166428 | PLD4 | 1.615962539 | -0.76237723 | 1.386111265 | -0.550011567 | 0.58231144 | NA |

| | | | | | | | |
|---|---|---|---|---|---|---|---|
| ENSG00000166432 | ZMAT1 | 516.5736045 | -0.520001467 | 0.114029334 | -4.560242963 | 5.11E-06 | 2.73E-05 |
| ENSG00000166435 | XRRA1 | 288.8344145 | 0.007175275 | 0.111975447 | 0.064079001 | 0.948907322 | 0.976447045 |
| ENSG00000166436 | TRIM66 | 485.3803027 | -0.188406621 | 0.105296373 | -1.789298297 | 0.073566788 | 0.165372084 |
| ENSG00000166439 | RNF169 | 1650.323864 | -0.234819397 | 0.062061195 | -3.783675084 | 0.00015453 | 0.000668598 |
| ENSG00000166441 | RPL27A | 4385.96437 | 0.311485533 | 0.04979859 | 6.254906662 | 3.98E-10 | 3.29E-09 |
| ENSG00000166444 | DENND2B | 141.49622 | 0.72086467 | 0.164052495 | 4.394109759 | 1.11E-05 | 5.70E-05 |
| ENSG00000166446 | CDYL2 | 3440.784921 | -1.001349887 | 0.06042611 | -16.57147693 | 1.12E-61 | 8.37E-60 |
| ENSG00000166448 | TMEM130 | 8.129602884 | -1.314107902 | 0.66685991 | -1.970590647 | 0.048770717 | 0.117262368 |
| ENSG00000166450 | PRTG | 136.5429533 | -0.06546966 | 0.162724586 | -0.402334166 | 0.687438114 | 0.823383061 |
| ENSG00000166451 | CENPN | 1636.982922 | 0.051186765 | 0.064256004 | 0.796606721 | 0.425679473 | 0.61656062 |
| ENSG00000166452 | AKIP1 | 292.4765715 | 0.019332077 | 0.111228007 | 0.173805838 | 0.862018057 | 0.928215087 |
| ENSG00000166454 | ATMIN | 7319.117614 | 0.195714852 | 0.04731625 | 4.136313719 | 3.53E-05 | 0.000168601 |
| ENSG00000166471 | TMEM41B | 619.0438978 | -0.216539877 | 0.084980139 | -2.548123355 | 0.010830417 | 0.031856617 |
| ENSG00000166473 | PKD1L2 | 26.85377869 | 0.704264387 | 0.347907501 | 2.024286298 | 0.042940703 | 0.105196467 |
| ENSG00000166477 | LEO1 | 10469.50584 | -1.637557012 | 0.047065331 | -34.79327491 | 3.07E-265 | 3.92E-262 |
| ENSG00000166478 | ZNF143 | 446.3369078 | -0.099885403 | 0.0908128 | -1.099904445 | 0.271373758 | 0.453376593 |
| ENSG00000166479 | TMX3 | 1037.832622 | 0.627035648 | 0.06914679 | 9.068181612 | 1.21E-19 | 1.98E-18 |
| ENSG00000166483 | WEE1 | 943.13326 | -0.13406924 | 0.075929368 | -1.765709936 | 0.077444525 | 0.172299234 |
| ENSG00000166484 | MAPK7 | 576.9981068 | -0.150331 | 0.084445633 | -1.780210461 | 0.075041524 | 0.167946591 |
| ENSG00000166492 | FAM86GP | 5.951118914 | -0.036912808 | 0.737834324 | -0.050028586 | 0.960099608 | 0.981799414 |
| ENSG00000166503 | HDGFL3 | 4275.839148 | 0.132740942 | 0.050282762 | 2.639889662 | 0.008293302 | 0.025215156 |
| ENSG00000166507 | NDST2 | 7.161413818 | 0.617575167 | 0.653699932 | 0.944738 | 0.344792652 | 0.535659178 |
| ENSG00000166508 | MCM7 | 9902.361114 | -0.436643822 | 0.043445984 | -10.05026889 | 9.16E-24 | 1.89E-22 |
| ENSG00000166509 | CLEC3A | 1246.05795 | 0.70042185 | 0.076961845 | 9.100897372 | 8.96E-20 | 1.47E-18 |
| ENSG00000166510 | CCDC68 | 143.6643675 | -0.183252172 | 0.152140042 | -1.204496659 | 0.228397673 | 0.401243882 |
| ENSG00000166526 | ZNF3 | 1783.900948 | -0.384506267 | 0.059332581 | -6.480524877 | 9.14E-11 | 7.94E-10 |
| ENSG00000166529 | ZSCAN21 | 403.170031 | -0.730855842 | 0.09929885 | -7.360164233 | 1.84E-13 | 1.97E-12 |
| ENSG00000166530 | HSBP1P2 | 13.14962189 | -0.087069637 | 0.539457879 | -0.161402105 | 0.871776709 | 0.933552316 |
| ENSG00000166532 | RIMKLB | 555.072292 | -0.071510276 | 0.107140875 | -0.667441584 | 0.504490111 | 0.685239099 |
| ENSG00000166546 | BEAN1 | 12.34602025 | 0.714634267 | 0.503692308 | 1.418791306 | 0.155959868 | 0.300068722 |
| ENSG00000166548 | TK2 | 172.5201297 | 0.195810316 | 0.14593809 | 1.3417355 | 0.179681772 | 0.334588775 |
| ENSG00000166557 | TMED3 | 4044.967638 | -0.099643506 | 0.058335945 | -1.708097914 | 0.087618176 | 0.190235581 |
| ENSG00000166562 | SEC11C | 666.2663933 | 0.33951629 | 0.080500366 | 4.217574473 | 2.47E-05 | 0.000120942 |

| | | | | | | | |
|---|---|---|---|---|---|---|---|
| ENSG00000166575 | TMEM135 | 2078.634832 | -0.863641583 | 0.063327776 | -13.63764271 | 2.39E-42 | 1.07E-40 |
| ENSG00000166578 | IQCD | 45.19196303 | 0.046176768 | 0.261476338 | 0.176600178 | 0.859822454 | 0.92728725 |
| ENSG00000166579 | NDEL1 | 669.2303991 | 0.0504404 | 0.076707644 | 0.657566806 | 0.51081653 | 0.690585871 |
| ENSG00000166582 | CENPV | 622.9822668 | -0.299582557 | 0.080448131 | -3.723921887 | 0.000196152 | 0.00083359 |
| ENSG00000166592 | RRAD | 1.512389965 | 1.100339576 | 1.640355449 | 0.67079338 | 0.502352165 | NA |
| ENSG00000166595 | CIAO2B | 857.2378915 | 0.075899071 | 0.080010057 | 0.948619129 | 0.342814358 | 0.533778686 |
| ENSG00000166598 | HSP90B1 | 11109.31667 | -0.140187715 | 0.05188198 | -2.702050202 | 0.006891336 | 0.02139784 |
| ENSG00000166619 | BLCAP | 1131.458004 | -0.242128075 | 0.063000501 | -3.843272191 | 0.000121405 | 0.000533836 |
| ENSG00000166669 | ATF7IP2 | 181.4070818 | -0.133153661 | 0.135007492 | -0.986268677 | 0.324001286 | 0.514117002 |
| ENSG00000166670 | MMP10 | 4.93677661 | -2.808731381 | 0.97862477 | -2.870080001 | 0.00410368 | 0.013490744 |
| ENSG00000166676 | TVP23A | 19.93408806 | -0.227128541 | 0.400816044 | -0.566665294 | 0.570941604 | 0.737840144 |
| ENSG00000166681 | BEX3 | 3007.895483 | -0.775669783 | 0.047972472 | -16.16906022 | 8.34E-59 | 5.74E-57 |
| ENSG00000166682 | TMPRSS5 | 5.642375251 | 1.349158765 | 0.787332119 | 1.713582784 | 0.086605378 | 0.188494057 |
| ENSG00000166685 | COG1 | 1419.213439 | 0.126679957 | 0.06605242 | 1.917870034 | 0.055127493 | 0.130212591 |
| ENSG00000166689 | PLEKHA7 | 1954.76178 | 0.041343051 | 0.06256297 | 0.660823025 | 0.508725815 | 0.688880717 |
| ENSG00000166704 | ZNF606 | 539.0785672 | 0.773639391 | 0.087468819 | 8.844744879 | 9.17E-19 | 1.43E-17 |
| ENSG00000166707 | ZCCHC18 | 44.8639046 | 0.580200203 | 0.265525594 | 2.185100857 | 0.028881452 | 0.075109457 |
| ENSG00000166710 | B2M | 2351.260855 | -0.620619091 | 0.053862985 | -11.52218143 | 1.02E-30 | 2.91E-29 |
| ENSG00000166716 | ZNF592 | 4005.642851 | -0.109638899 | 0.050693409 | -2.162784112 | 0.030557788 | 0.078940953 |
| ENSG00000166734 | GOLM2 | 2318.28564 | 0.124085127 | 0.052627034 | 2.357821032 | 0.018382553 | 0.050794479 |
| ENSG00000166747 | AP1G1 | 7913.675057 | 0.201279629 | 0.048639773 | 4.138169615 | 3.50E-05 | 0.000167348 |
| ENSG00000166750 | SLFN5 | 3.564359783 | -1.884192718 | 1.020729186 | -1.84592813 | 0.064902642 | 0.149004967 |
| ENSG00000166762 | CATSPER2 | 80.89675233 | -0.105594866 | 0.215423799 | -0.490172705 | 0.624011694 | 0.777683938 |
| ENSG00000166780 | BMERB1 | 298.6853972 | -0.446147981 | 0.118932 | -3.751286281 | 0.00017593 | 0.000754219 |
| ENSG00000166783 | MARF1 | 2174.99099 | -0.00684738 | 0.057978538 | -0.11810197 | 0.905986862 | 0.95262805 |
| ENSG00000166788 | SAAL1 | 426.7827064 | -0.336113843 | 0.096690452 | -3.476184439 | 0.000508602 | 0.002009531 |
| ENSG00000166794 | PPIB | 6965.980328 | -0.123667117 | 0.041722445 | -2.964042887 | 0.00303626 | 0.010277674 |
| ENSG00000166797 | CIAO2A | 1209.896224 | -0.034185035 | 0.063112533 | -0.541652086 | 0.588058203 | 0.751401597 |
| ENSG00000166801 | FAM111A | 2303.801811 | -0.258438926 | 0.064090258 | -4.032421372 | 5.52E-05 | 0.000256773 |
| ENSG00000166803 | PCLAF | 255.032636 | -0.584067562 | 0.119387529 | -4.892199102 | 9.97E-07 | 5.77E-06 |
| ENSG00000166813 | KIF7 | 363.4832286 | 0.288057027 | 0.100578166 | 2.864011531 | 0.004183128 | 0.013718506 |
| ENSG00000166816 | LDHD | 673.8462202 | 1.366665682 | 0.093941614 | 14.5480328 | 6.01E-48 | 3.10E-46 |
| ENSG00000166819 | PLIN1 | 10.85736536 | -0.645091276 | 0.570789879 | -1.130172941 | 0.258403359 | 0.438055946 |

| | | | | | | | |
|---|---|---|---|---|---|---|---|
| ENSG00000166821 | PEX11A | 296.3546239 | -0.350272355 | 0.109372223 | -3.202571418 | 0.001362065 | 0.004966742 |
| ENSG00000166822 | TMEM170A | 693.9301094 | 0.091351045 | 0.07528837 | 1.213348685 | 0.224996536 | 0.396879704 |
| ENSG00000166823 | MESP1 | 16.29702785 | -4.047635296 | 0.693278238 | -5.838399466 | 5.27E-09 | 3.91E-08 |
| ENSG00000166825 | ANPEP | 3.131800918 | -0.065453835 | 0.986881666 | -0.066323894 | 0.94711996 | 0.975307801 |
| ENSG00000166828 | SCNN1G | 60.49238806 | -0.251913366 | 0.228095072 | -1.104422662 | 0.269409835 | 0.451016256 |
| ENSG00000166831 | RBPMS2 | 53.69501425 | -1.303052903 | 0.250292891 | -5.206112315 | 1.93E-07 | 1.22E-06 |
| ENSG00000166833 | NAV2 | 26.97707264 | -1.149776288 | 0.348942403 | -3.295031724 | 0.000984106 | 0.003687583 |
| ENSG00000166839 | ANKDD1A | 12.64997779 | 0.266310409 | 0.488784444 | 0.544842236 | 0.585862027 | 0.749814489 |
| ENSG00000166840 | GLYATL1 | 350.2120855 | -1.809952466 | 0.107133856 | -16.89430893 | 4.95E-64 | 3.93E-62 |
| ENSG00000166845 | C18orf54 | 274.2625645 | 0.043748316 | 0.122522502 | 0.357063524 | 0.72104425 | 0.845162383 |
| ENSG00000166847 | DCTN5 | 3500.376808 | 0.284396524 | 0.052920447 | 5.374038559 | 7.70E-08 | 5.04E-07 |
| ENSG00000166848 | TERF2IP | 3144.729027 | 0.130643462 | 0.048621726 | 2.686935926 | 0.007211078 | 0.022306114 |
| ENSG00000166851 | PLK1 | 2618.831601 | -0.29200372 | 0.057036894 | -5.119558573 | 3.06E-07 | 1.89E-06 |
| ENSG00000166855 | CLPX | 2419.315186 | 0.048930636 | 0.052476104 | 0.932436517 | 0.35111098 | 0.542458202 |
| ENSG00000166860 | ZBTB39 | 660.3471 | -0.063621041 | 0.080654793 | -0.788806703 | 0.430224991 | 0.620481249 |
| ENSG00000166881 | NEMP1 | 1351.903082 | -0.216929311 | 0.062178566 | -3.488811765 | 0.000485173 | 0.001925582 |
| ENSG00000166886 | NAB2 | 1789.657051 | 0.037873203 | 0.06216074 | 0.609278506 | 0.542339852 | 0.716168328 |
| ENSG00000166887 | VPS39 | 3014.666379 | 0.037384828 | 0.056661814 | 0.659788765 | 0.509389394 | 0.689403699 |
| ENSG00000166888 | STAT6 | 1458.608468 | 0.068404919 | 0.063646425 | 1.074764499 | 0.282480178 | 0.46595925 |
| ENSG00000166889 | PATL1 | 1446.361993 | -0.267574289 | 0.063263603 | -4.229513931 | 2.34E-05 | 0.000114993 |
| ENSG00000166896 | ATP23 | 150.1242062 | -0.075769053 | 0.149660914 | -0.506271487 | 0.612666077 | 0.769733271 |
| ENSG00000166897 | ELFN2 | 33.43562394 | -2.064772996 | 0.34136533 | -6.048572636 | 1.46E-09 | 1.15E-08 |
| ENSG00000166900 | STX3 | 1860.934668 | 0.420518431 | 0.056421598 | 7.453146393 | 9.11E-14 | 9.97E-13 |
| ENSG00000166902 | MRPL16 | 1020.345042 | -0.387018385 | 0.065677 | -5.892753684 | 3.80E-09 | 2.86E-08 |
| ENSG00000166908 | PIP4K2C | 2672.254919 | -0.003303275 | 0.053376506 | -0.061886316 | 0.950653365 | 0.977235535 |
| ENSG00000166912 | MTMR10 | 956.2240067 | -0.053702784 | 0.068492316 | -0.784070202 | 0.432998919 | 0.623082616 |
| ENSG00000166913 | YWHAB | 11836.63939 | -2.76E-05 | 0.04526782 | -0.000608861 | 0.999514199 | 0.999732101 |
| ENSG00000166920 | C15orf48 | 2.578614716 | 0.563297101 | 1.079864085 | 0.521637036 | 0.601923072 | 0.76144895 |
| ENSG00000166922 | SCG5 | 4.049527884 | 0.045047734 | 0.880415396 | 0.051166455 | 0.959192882 | 0.981308688 |
| ENSG00000166924 | NYAP1 | 21.26718019 | -1.094420866 | 0.397449878 | -2.753607251 | 0.005894245 | 0.018634784 |
| ENSG00000166925 | TSC22D4 | 2660.097019 | -0.512154929 | 0.06138347 | -8.343531781 | 7.21E-17 | 9.87E-16 |
| ENSG00000166938 | DIS3L | 1370.979593 | -0.01596304 | 0.062137501 | -0.256898644 | 0.797257017 | 0.89183367 |
| ENSG00000166946 | CCNDBP1 | 1221.60265 | -0.182101971 | 0.062491238 | -2.91404009 | 0.003567841 | 0.011892803 |

| | | | | | | | |
|---|---|---|---|---|---|---|---|
| ENSG00000166949 | SMAD3 | 4202.026007 | 0.180749891 | 0.055513218 | 3.255979328 | 0.00113002 | 0.004191891 |
| ENSG00000166963 | MAP1A | 84.01440524 | 0.229814822 | 0.194052923 | 1.184289411 | 0.236298519 | 0.411406195 |
| ENSG00000166965 | RCCD1 | 795.3487152 | 0.180180223 | 0.074782878 | 2.40937804 | 0.015979736 | 0.044961995 |
| ENSG00000166971 | AKTIP | 1011.940863 | 0.01991846 | 0.069637419 | 0.286030997 | 0.774854381 | 0.877499635 |
| ENSG00000166974 | MAPRE2 | 41.81107674 | 0.040084553 | 0.274828406 | 0.145853021 | 0.884037419 | 0.939910479 |
| ENSG00000166979 | EVA1C | 30.97358134 | -1.114808534 | 0.323898293 | -3.441847515 | 0.000577756 | 0.002259028 |
| ENSG00000166986 | MARS1 | 5151.682591 | -0.068743506 | 0.049531029 | -1.387887707 | 0.165171242 | 0.313687249 |
| ENSG00000166987 | MBD6 | 3281.197311 | 0.031526543 | 0.061061389 | 0.516308967 | 0.605638634 | 0.76434448 |
| ENSG00000166997 | CNPY4 | 318.7356718 | -0.744441399 | 0.115097002 | -6.467947771 | 9.93E-11 | 8.58E-10 |
| ENSG00000167004 | PDIA3 | 7393.826893 | -0.251943444 | 0.042537889 | -5.922800888 | 3.17E-09 | 2.41E-08 |
| ENSG00000167005 | NUDT21 | 3470.117898 | -0.02861227 | 0.052330373 | -0.546762204 | 0.584542113 | 0.748835431 |
| ENSG00000167011 | NAT16 | 1.685419264 | 0.138067345 | 1.419523127 | 0.097263188 | 0.922517389 | NA |
| ENSG00000167034 | NKX3-1 | 115.1084821 | 0.822338352 | 0.170104627 | 4.834309122 | 1.34E-06 | 7.60E-06 |
| ENSG00000167037 | SGSM1 | 49.63356078 | -2.336148902 | 0.288170363 | -8.106832629 | 5.20E-16 | 6.72E-15 |
| ENSG00000167046 | ENSG0000016 | 4.978181302 | 0.767663439 | 0.843893497 | 0.909668627 | 0.362997294 | 0.555760042 |
| ENSG00000167065 | DUSP18 | 135.0240887 | -0.410978839 | 0.155311312 | -2.646161652 | 0.00814109 | 0.024818154 |
| ENSG00000167074 | TEF | 1967.065345 | -0.131553558 | 0.071143175 | -1.84913813 | 0.06443787 | 0.14808703 |
| ENSG00000167077 | MEI1 | 1.993216504 | 1.059017177 | 1.33208262 | 0.795008629 | 0.426608483 | 0.617497513 |
| ENSG00000167080 | B4GALNT2 | 42.05346202 | -5.477861523 | 0.632924079 | -8.65484772 | 4.94E-18 | 7.29E-17 |
| ENSG00000167081 | PBX3 | 885.5598372 | 0.312821773 | 0.0737039 | 4.244304228 | 2.19E-05 | 0.000108151 |
| ENSG00000167085 | PHB1 | 6390.93487 | -0.035275214 | 0.046303395 | -0.761827793 | 0.44616279 | 0.634727095 |
| ENSG00000167088 | SNRPD1 | 1901.295056 | 0.133149009 | 0.055718204 | 2.389685937 | 0.016862786 | 0.047151671 |
| ENSG00000167094 | TTC16 | 4.507154106 | -1.376632554 | 0.861108406 | -1.598675084 | 0.109892817 | 0.228594597 |
| ENSG00000167100 | SAMD14 | 38.90116828 | 0.047499337 | 0.284297421 | 0.167076214 | 0.867310082 | 0.93096599 |
| ENSG00000167103 | PIP5KL1 | 21.20839418 | -0.67042026 | 0.386744464 | -1.733496723 | 0.08300742 | 0.182114595 |
| ENSG00000167105 | TMEM92 | 4.470775929 | 0.138577473 | 0.82974027 | 0.167013074 | 0.867359762 | 0.93096599 |
| ENSG00000167106 | EEIG1 | 3029.001743 | 0.876636264 | 0.052934189 | 16.56087069 | 1.34E-61 | 9.92E-60 |
| ENSG00000167107 | ACSF2 | 1863.31304 | -2.030213698 | 0.072770137 | -27.89899523 | 2.74E-171 | 1.14E-168 |
| ENSG00000167110 | GOLGA2 | 4045.483713 | 0.071657738 | 0.046261397 | 1.548974786 | 0.121387781 | 0.247677284 |
| ENSG00000167112 | TRUB2 | 2761.298144 | 0.169629217 | 0.051686404 | 3.281892372 | 0.00103113 | 0.003849327 |
| ENSG00000167113 | COQ4 | 1671.917705 | 0.075888821 | 0.065067118 | 1.166315988 | 0.243486712 | 0.42006356 |
| ENSG00000167114 | SLC27A4 | 2321.691058 | -0.476095802 | 0.060209208 | -7.907358663 | 2.63E-15 | 3.23E-14 |
| ENSG00000167118 | URM1 | 2097.966064 | 0.226230259 | 0.063199206 | 3.579637656 | 0.000344071 | 0.001398722 |

| | | | | | | | |
|---|---|---|---|---|---|---|---|
| ENSG00000167123 | CERCAM | 4048.810257 | 0.229062336 | 0.056532928 | 4.05183922 | 5.08E-05 | 0.000237614 |
| ENSG00000167130 | DOLPP1 | 1166.46592 | -0.099853982 | 0.067527187 | -1.478722664 | 0.139214452 | 0.275332316 |
| ENSG00000167131 | CCDC103 | 2.987599482 | 0.114416787 | 0.993486528 | 0.115166923 | 0.908312816 | 0.954192242 |
| ENSG00000167136 | ENDOG | 521.0554935 | -0.011710701 | 0.095158626 | -0.12306505 | 0.902055587 | 0.950654947 |
| ENSG00000167157 | PRRX2 | 8.508824978 | -1.419049132 | 0.621875137 | -2.281887549 | 0.02249598 | 0.060591497 |
| ENSG00000167165 | UGT1A6 | 4.829712936 | 2.956568032 | 1.063120605 | 2.781027871 | 0.005418708 | 0.017283812 |
| ENSG00000167173 | C15orf39 | 1527.347443 | 0.069333921 | 0.069826708 | 0.992942719 | 0.32073787 | 0.510279959 |
| ENSG00000167182 | SP2 | 1306.118834 | 0.014478778 | 0.067048063 | 0.215946249 | 0.829029646 | 0.909536221 |
| ENSG00000167183 | PRR15L | 1491.418278 | 0.712046752 | 0.064226168 | 11.08655204 | 1.46E-28 | 3.81E-27 |
| ENSG00000167186 | COQ7 | 600.1922743 | 0.034463738 | 0.084225273 | 0.409185234 | 0.68240373 | 0.819871514 |
| ENSG00000167191 | GPRC5B | 35.52220922 | 0.859048563 | 0.301567574 | 2.848610518 | 0.00439106 | 0.014349133 |
| ENSG00000167193 | CRK | 2524.039912 | 0.096889746 | 0.051614734 | 1.877172237 | 0.060494491 | 0.1404741 |
| ENSG00000167196 | FBXO22 | 1216.309101 | 0.009271858 | 0.062701628 | 0.147872682 | 0.882443248 | 0.939121523 |
| ENSG00000167202 | TBC1D2B | 1539.161021 | -0.173505282 | 0.061061442 | -2.841486804 | 0.004490371 | 0.014640294 |
| ENSG00000167207 | NOD2 | 41.36862313 | 1.249986973 | 0.28526982 | 4.381770825 | 1.18E-05 | 6.01E-05 |
| ENSG00000167216 | KATNAL2 | 168.6254069 | 1.009490341 | 0.144481774 | 6.986973601 | 2.81E-12 | 2.77E-11 |
| ENSG00000167220 | HDHD2 | 158.315832 | 0.694665933 | 0.14634331 | 4.746823983 | 2.07E-06 | 1.15E-05 |
| ENSG00000167232 | ZNF91 | 564.3790078 | 0.165439382 | 0.087400987 | 1.892877734 | 0.058374131 | 0.136434291 |
| ENSG00000167257 | RNF214 | 644.6789896 | -0.230739834 | 0.077695072 | -2.969812967 | 0.002979811 | 0.01010001 |
| ENSG00000167258 | CDK12 | 2380.06202 | -0.290185105 | 0.052193299 | -5.559815298 | 2.70E-08 | 1.85E-07 |
| ENSG00000167261 | DPEP2 | 2.744582476 | -3.941775932 | 1.513666394 | -2.604124626 | 0.009210926 | 0.02768588 |
| ENSG00000167264 | DUS2 | 440.8912272 | -0.20223009 | 0.089983065 | -2.247423892 | 0.024612949 | 0.065614444 |
| ENSG00000167272 | POP5 | 700.4378534 | 0.23260753 | 0.082064536 | 2.834446408 | 0.004590516 | 0.014920152 |
| ENSG00000167280 | ENGASE | 666.2245963 | 0.779796476 | 0.08155906 | 9.561126287 | 1.16E-21 | 2.14E-20 |
| ENSG00000167283 | ATP5MG | 623.3280256 | 0.286998157 | 0.081417101 | 3.525035323 | 0.000423426 | 0.001697553 |
| ENSG00000167291 | TBC1D16 | 7022.059495 | 0.804890471 | 0.051663347 | 15.5795262 | 1.00E-54 | 6.24E-53 |
| ENSG00000167302 | TEPSIN | 944.6491313 | 0.221303495 | 0.082888152 | 2.669905029 | 0.00758727 | 0.023315738 |
| ENSG00000167306 | MYO5B | 3742.756083 | 0.582942657 | 0.054395447 | 10.71675454 | 8.49E-27 | 2.09E-25 |
| ENSG00000167315 | ACAA2 | 1360.702686 | -0.114515341 | 0.062408264 | -1.834938741 | 0.066514767 | 0.152174005 |
| ENSG00000167323 | STIM1 | 726.7740374 | 0.288592648 | 0.076164521 | 3.789069278 | 0.000151213 | 0.000655236 |
| ENSG00000167325 | RRM1 | 3862.616313 | -0.143937215 | 0.048403378 | -2.973701873 | 0.002942308 | 0.009983219 |
| ENSG00000167333 | TRIM68 | 74.84763923 | 0.23246637 | 0.208096325 | 1.117109449 | 0.263947531 | 0.444530971 |
| ENSG00000167363 | FN3K | 413.083974 | -0.143017458 | 0.096148643 | -1.48746205 | 0.136892819 | 0.271889287 |

| | | | | | | | |
|---|---|---|---|---|---|---|---|
| ENSG00000167371 | PRRT2 | 505.2188198 | 0.254007663 | 0.103516581 | 2.453787224 | 0.01413606 | 0.040358583 |
| ENSG00000167377 | ZNF23 | 41.20754132 | 0.872431852 | 0.306905392 | 2.842673589 | 0.004473686 | 0.014590043 |
| ENSG00000167378 | IRGQ | 1058.059794 | 0.02314155 | 0.065077866 | 0.355597857 | 0.722141749 | 0.845601795 |
| ENSG00000167380 | ZNF226 | 668.0531626 | 0.164213934 | 0.077813266 | 2.1103591 | 0.034827437 | 0.088241815 |
| ENSG00000167384 | ZNF180 | 567.8849627 | 0.374639848 | 0.091246798 | 4.105786259 | 4.03E-05 | 0.000190823 |
| ENSG00000167393 | PPP2R3B | 507.0274792 | 0.150179563 | 0.086358509 | 1.739024502 | 0.082030453 | 0.180350642 |
| ENSG00000167393 | PPP2R3B | 507.0274792 | 0.150179563 | 0.086358509 | 1.739024502 | 0.082030453 | 0.180350642 |
| ENSG00000167394 | ZNF668 | 362.0962018 | 0.002149793 | 0.105883051 | 0.020303464 | 0.983801293 | 0.992410257 |
| ENSG00000167395 | ZNF646 | 1315.379693 | 0.165902449 | 0.076363995 | 2.172521868 | 0.029816322 | 0.07727787 |
| ENSG00000167397 | VKORC1 | 180.1556089 | -0.246934985 | 0.135989044 | -1.815844703 | 0.069394198 | 0.15754655 |
| ENSG00000167419 | LPO | 4.393321429 | 0.708231796 | 0.895478148 | 0.79089791 | 0.429003568 | 0.619341841 |
| ENSG00000167434 | CA4 | 373.713464 | 0.783596108 | 0.105527175 | 7.425538542 | 1.12E-13 | 1.22E-12 |
| ENSG00000167447 | SMG8 | 2628.078433 | 0.07944103 | 0.053047693 | 1.497539769 | 0.134252866 | 0.267828571 |
| ENSG00000167460 | TPM4 | 11597.1005 | -0.442582201 | 0.043038315 | -10.28344634 | 8.37E-25 | 1.84E-23 |
| ENSG00000167461 | RAB8A | 2065.997231 | -0.26351302 | 0.057855496 | -4.554675644 | 5.25E-06 | 2.80E-05 |
| ENSG00000167468 | GPX4 | 5220.214338 | 0.33386047 | 0.046906641 | 7.117552339 | 1.10E-12 | 1.12E-11 |
| ENSG00000167470 | MIDN | 2776.611834 | 0.207859687 | 0.060298135 | 3.447199301 | 0.000566431 | 0.002218907 |
| ENSG00000167476 | JSRP1 | 45.06658251 | -0.168581253 | 0.267383603 | -0.630484636 | 0.528377552 | 0.70433965 |
| ENSG00000167483 | NIBAN3 | 5.867351369 | 1.078343907 | 0.749970575 | 1.437848289 | 0.150477105 | 0.292265243 |
| ENSG00000167487 | KLHL26 | 199.5680579 | 0.065317582 | 0.130328133 | 0.501177914 | 0.616245916 | 0.772665136 |
| ENSG00000167491 | GATAD2A | 1282.13986 | 0.067927306 | 0.063694982 | 1.066446741 | 0.286221742 | 0.470021728 |
| ENSG00000167508 | MVD | 5884.627155 | -0.899195256 | 0.052435128 | -17.14871863 | 6.43E-66 | 5.25E-64 |
| ENSG00000167513 | CDT1 | 2084.281536 | 0.356841927 | 0.064650214 | 5.519578404 | 3.40E-08 | 2.31E-07 |
| ENSG00000167515 | TRAPPC2L | 2252.918956 | 0.347385207 | 0.05769068 | 6.02151345 | 1.73E-09 | 1.35E-08 |
| ENSG00000167522 | ANKRD11 | 3995.291341 | 0.551610567 | 0.048898446 | 11.28073813 | 1.63E-29 | 4.46E-28 |
| ENSG00000167523 | SPATA33 | 616.2860381 | 0.51785248 | 0.084626133 | 6.119297414 | 9.40E-10 | 7.49E-09 |
| ENSG00000167524 | RSKR | 208.2515603 | 0.116496906 | 0.132480422 | 0.879351867 | 0.379210522 | 0.571706714 |
| ENSG00000167525 | PROCA1 | 154.4339917 | 0.145898625 | 0.150404286 | 0.970043001 | 0.332025058 | 0.523050051 |
| ENSG00000167526 | RPL13 | 15066.0288 | 0.450857781 | 0.04302376 | 10.47927437 | 1.08E-25 | 2.46E-24 |
| ENSG00000167528 | ZNF641 | 1150.288876 | 0.074608808 | 0.067028673 | 1.113087962 | 0.265670653 | 0.44674423 |
| ENSG00000167535 | CACNB3 | 2804.24027 | 0.147458131 | 0.055650334 | 2.64972589 | 0.00805571 | 0.02459783 |
| ENSG00000167536 | DHRS13 | 778.0546861 | -0.174944844 | 0.075848904 | -2.306491393 | 0.021083191 | 0.057216183 |
| ENSG00000167543 | TP53I13 | 1191.078999 | 0.104773793 | 0.068487914 | 1.529814346 | 0.12606269 | 0.255083188 |

| | | | | | | | |
|---|---|---|---|---|---|---|---|
| ENSG00000167548 | KMT2D | 3739.580182 | -0.172441257 | 0.133034241 | -1.296217091 | 0.194900703 | 0.355975658 |
| ENSG00000167549 | CORO6 | 10.4101477 | 0.584865102 | 0.559350061 | 1.045615515 | 0.295738583 | 0.481835447 |
| ENSG00000167550 | RHEBL1 | 49.60988344 | -0.81287265 | 0.258529259 | -3.144219161 | 0.001665307 | 0.005948637 |
| ENSG00000167552 | TUBA1A | 5249.700402 | -0.366257292 | 0.048728703 | -7.51625367 | 5.64E-14 | 6.27E-13 |
| ENSG00000167553 | TUBA1C | 11236.54777 | -0.268077926 | 0.046253145 | -5.795885411 | 6.80E-09 | 4.98E-08 |
| ENSG00000167554 | ZNF610 | 90.32459674 | 0.788064253 | 0.194174307 | 4.058540314 | 4.94E-05 | 0.000231323 |
| ENSG00000167555 | ZNF528 | 18.39582612 | 2.132704972 | 0.504238143 | 4.229558994 | 2.34E-05 | 0.000114993 |
| ENSG00000167562 | ZNF701 | 293.0590436 | 0.085551968 | 0.110243514 | 0.776027221 | 0.437732913 | 0.627285921 |
| ENSG00000167565 | SERTAD3 | 1065.784543 | -0.129670843 | 0.066272876 | -1.956620132 | 0.050392146 | 0.120642438 |
| ENSG00000167566 | NCKAP5L | 1598.136126 | 0.046709983 | 0.062519935 | 0.747121428 | 0.454990267 | 0.642475746 |
| ENSG00000167578 | RAB4B | 16.50825044 | -0.209943473 | 0.434280453 | -0.483428326 | 0.628791639 | 0.781010353 |
| ENSG00000167588 | GPD1 | 12.01120072 | -0.057273703 | 0.513758018 | -0.11147992 | 0.911235788 | 0.955425246 |
| ENSG00000167595 | PROSER3 | 567.3416752 | -0.157326516 | 0.088866595 | -1.770367327 | 0.076665969 | 0.170931803 |
| ENSG00000167600 | CYP2S1 | 10.67164202 | -0.105574409 | 0.555375602 | -0.190095512 | 0.849234287 | 0.921106747 |
| ENSG00000167601 | AXL | 4.01402135 | -0.672940128 | 0.905252279 | -0.743373029 | 0.457255869 | 0.644513034 |
| ENSG00000167604 | NFKBID | 193.8121946 | -0.137661974 | 0.133468425 | -1.031419784 | 0.302344007 | 0.489261535 |
| ENSG00000167608 | TMC4 | 2556.252424 | -0.122044711 | 0.05695243 | -2.142923673 | 0.032119228 | 0.082436237 |
| ENSG00000167614 | TTYH1 | 7.87392223 | -0.512923885 | 0.625052219 | -0.820609655 | 0.411868647 | 0.604186389 |
| ENSG00000167615 | LENG8 | 3731.149665 | 0.243299093 | 0.153921566 | 1.580669297 | 0.113953672 | 0.235652865 |
| ENSG00000167617 | CDC42EP5 | 3.192317597 | -0.3425096 | 0.966253476 | -0.35447179 | 0.722985342 | 0.845926239 |
| ENSG00000167619 | TMEM145 | 14.30355271 | 0.886085788 | 0.510414897 | 1.736010827 | 0.08256192 | 0.181258536 |
| ENSG00000167625 | ZNF526 | 569.2933695 | -0.01721249 | 0.085246882 | -0.201913421 | 0.83998441 | 0.915757464 |
| ENSG00000167632 | TRAPPC9 | 1559.871638 | 0.086377282 | 0.06056761 | 1.426129937 | 0.153830839 | 0.297018722 |
| ENSG00000167635 | ZNF146 | 3465.495065 | -0.141552204 | 0.053508526 | -2.64541403 | 0.008159102 | 0.024866453 |
| ENSG00000167637 | ZNF283 | 223.1047317 | 0.498844069 | 0.127274044 | 3.919448578 | 8.88E-05 | 0.000399366 |
| ENSG00000167642 | SPINT2 | 5779.023489 | 0.016534449 | 0.049518541 | 0.333904198 | 0.738451852 | 0.855807824 |
| ENSG00000167644 | C19orf33 | 6.535363835 | 0.337202753 | 0.720193328 | 0.468211437 | 0.639633394 | 0.787822285 |
| ENSG00000167645 | YIF1B | 941.036586 | 0.259097156 | 0.074023657 | 3.500193955 | 0.00046492 | 0.001851607 |
| ENSG00000167646 | DNAAF3 | 56.18347443 | -0.761773444 | 0.242012711 | -3.147658811 | 0.001645837 | 0.005893771 |
| ENSG00000167653 | PSCA | 85.84308463 | 0.315797389 | 0.203427627 | 1.552382012 | 0.120570845 | 0.246492753 |
| ENSG00000167656 | LY6D | 2.065100587 | 0.062098665 | 1.211183306 | 0.051271071 | 0.95910952 | 0.981267166 |
| ENSG00000167657 | DAPK3 | 1020.599056 | 0.20563108 | 0.069761689 | 2.947621884 | 0.003202285 | 0.010777643 |
| ENSG00000167658 | EEF2 | 77932.195 | 0.323302297 | 0.042486088 | 7.609603799 | 2.75E-14 | 3.13E-13 |

| | | | | | | | |
|---|---|---|---|---|---|---|---|
| ENSG00000167670 | CHAF1A | 1204.54283 | 0.055938444 | 0.066411433 | 0.842301415 | 0.399619259 | 0.592315423 |
| ENSG00000167671 | UBXN6 | 1966.597397 | 0.32355791 | 0.062101816 | 5.210119955 | 1.89E-07 | 1.19E-06 |
| ENSG00000167674 | HDGFL2 | 2094.820181 | -0.10451378 | 0.066426605 | -1.573372295 | 0.115632638 | 0.238394241 |
| ENSG00000167676 | PLIN4 | 90.89931634 | 0.995571705 | 0.205959859 | 4.833814261 | 1.34E-06 | 7.62E-06 |
| ENSG00000167680 | SEMA6B | 196.204865 | 1.982792731 | 0.149081425 | 13.30006557 | 2.31E-40 | 9.66E-39 |
| ENSG00000167685 | ZNF444 | 1034.553208 | -0.208843117 | 0.07286168 | -2.866295674 | 0.004153062 | 0.013633548 |
| ENSG00000167693 | NXN | 91.37625294 | 0.448432802 | 0.19398528 | 2.311684693 | 0.020795066 | 0.056561395 |
| ENSG00000167695 | TLCD3A | 495.8140842 | 0.119743153 | 0.088019283 | 1.360419552 | 0.173697195 | 0.326206602 |
| ENSG00000167699 | GLOD4 | 1492.875886 | -0.092266733 | 0.059026208 | -1.563148583 | 0.118017651 | 0.242461482 |
| ENSG00000167700 | MFSD3 | 655.2599056 | 0.641360312 | 0.082779695 | 7.747797486 | 9.35E-15 | 1.10E-13 |
| ENSG00000167701 | GPT | 27.74114529 | 0.038991437 | 0.347228512 | 0.112293305 | 0.91059085 | 0.955141934 |
| ENSG00000167702 | KIFC2 | 3781.878983 | 0.365050503 | 0.05239734 | 6.966966385 | 3.24E-12 | 3.18E-11 |
| ENSG00000167703 | SLC43A2 | 2435.986955 | -0.226886566 | 0.061128825 | -3.71161339 | 0.000205942 | 0.000871645 |
| ENSG00000167705 | RILP | 73.08186198 | 0.480914785 | 0.210983126 | 2.279399282 | 0.022643341 | 0.060916881 |
| ENSG00000167711 | SERPINF2 | 3.645639788 | 0.181329994 | 0.983601142 | 0.184353175 | 0.853736401 | 0.923868819 |
| ENSG00000167716 | WDR81 | 1183.786177 | 0.241029302 | 0.070643869 | 3.411892724 | 0.000645135 | 0.002502857 |
| ENSG00000167720 | SRR | 679.5594407 | -0.078487373 | 0.078649877 | -0.997933834 | 0.318311444 | 0.507440204 |
| ENSG00000167721 | TSR1 | 2714.4287 | 0.121175682 | 0.049613328 | 2.442401786 | 0.014589899 | 0.041463365 |
| ENSG00000167723 | TRPV3 | 21.69950889 | 1.560622641 | 0.407886249 | 3.82612222 | 0.000130178 | 0.0005699 |
| ENSG00000167733 | HSD11B1L | 59.49188854 | -0.300611811 | 0.229148419 | -1.311865089 | 0.189565659 | 0.348504264 |
| ENSG00000167740 | CYB5D2 | 422.1828294 | 0.216866617 | 0.093963084 | 2.307998066 | 0.020999244 | 0.057022095 |
| ENSG00000167741 | GGT6 | 151.1327201 | 1.196409354 | 0.153928386 | 7.77250634 | 7.69E-15 | 9.10E-14 |
| ENSG00000167747 | C19orf48P | 3270.502144 | 0.319801726 | 0.051907635 | 6.160976651 | 7.23E-10 | 5.83E-09 |
| ENSG00000167755 | KLK6 | 1.971568224 | 4.336305758 | 1.583783874 | 2.73794034 | 0.006182529 | 0.019471062 |
| ENSG00000167766 | ZNF83 | 1734.580784 | 0.456408016 | 0.079590607 | 5.734445723 | 9.78E-09 | 7.05E-08 |
| ENSG00000167767 | KRT80 | 8667.126578 | 0.21970467 | 0.056604561 | 3.881395144 | 0.000103859 | 0.000461729 |
| ENSG00000167769 | ACER1 | 6.37203208 | -3.694635965 | 1.036349872 | -3.565046964 | 0.000363791 | 0.001472365 |
| ENSG00000167770 | OTUB1 | 3069.756602 | 0.015316046 | 0.058355575 | 0.26246071 | 0.792966273 | 0.888779318 |
| ENSG00000167771 | RCOR2 | 81.92332003 | 1.812121252 | 0.221736826 | 8.172396478 | 3.02E-16 | 3.98E-15 |
| ENSG00000167772 | ANGPTL4 | 77.91384486 | -1.600718663 | 0.217265604 | -7.36756594 | 1.74E-13 | 1.87E-12 |
| ENSG00000167775 | CD320 | 932.2877221 | -0.034097433 | 0.086122815 | -0.395916382 | 0.692166715 | 0.826433973 |
| ENSG00000167778 | SPRYD3 | 2456.194058 | 0.443953102 | 0.058783521 | 7.552339445 | 4.28E-14 | 4.79E-13 |
| ENSG00000167779 | IGFBP6 | 23.98281193 | 0.322316225 | 0.362448583 | 0.889274343 | 0.373855657 | 0.566636335 |

| | | | | | | | |
|---|---|---|---|---|---|---|---|
| ENSG00000167780 | SOAT2 | 1.677402887 | -1.362863379 | 1.489715213 | -0.914848265 | 0.360271293 | NA |
| ENSG00000167785 | ZNF558 | 517.3594575 | -0.137323541 | 0.088947874 | -1.543865346 | 0.122620957 | 0.24952761 |
| ENSG00000167792 | NDUFV1 | 4451.687438 | 0.184102191 | 0.051077544 | 3.604366554 | 0.000312915 | 0.001284587 |
| ENSG00000167797 | CDK2AP2 | 1975.499543 | -0.472573642 | 0.061071492 | -7.738039874 | 1.01E-14 | 1.18E-13 |
| ENSG00000167799 | NUDT8 | 189.019189 | 0.237519585 | 0.140977125 | 1.684809402 | 0.09202535 | 0.198072953 |
| ENSG00000167815 | PRDX2 | 6826.787762 | -0.028469531 | 0.049400159 | -0.576304444 | 0.56440942 | 0.732969584 |
| ENSG00000167840 | ZNF232 | 199.402311 | -0.212873467 | 0.129878327 | -1.639022242 | 0.101208628 | 0.213926558 |
| ENSG00000167842 | MIS12 | 716.3839085 | 0.035500956 | 0.079946967 | 0.444056314 | 0.65700187 | 0.801277136 |
| ENSG00000167861 | HID1 | 8249.803323 | -0.127955639 | 0.051570926 | -2.481158475 | 0.013095614 | 0.037683565 |
| ENSG00000167862 | MRPL58 | 1007.479712 | 0.091371354 | 0.067999355 | 1.343709135 | 0.179042462 | 0.333812913 |
| ENSG00000167863 | ATP5PD | 3024.405709 | 0.101352411 | 0.048543604 | 2.087863328 | 0.036810165 | 0.092529606 |
| ENSG00000167874 | TMEM88 | 21.01312628 | 0.768882342 | 0.402911697 | 1.90831477 | 0.056350545 | 0.132555528 |
| ENSG00000167880 | EVPL | 3335.910227 | 0.699943359 | 0.050642332 | 13.82130969 | 1.90E-43 | 8.65E-42 |
| ENSG00000167881 | SRP68 | 4445.868756 | 0.08098153 | 0.046897229 | 1.726787115 | 0.084205903 | 0.184268189 |
| ENSG00000167889 | MGAT5B | 142.2342383 | 2.001185835 | 0.176805145 | 11.31859502 | 1.06E-29 | 2.92E-28 |
| ENSG00000167895 | TMC8 | 28.89093062 | 1.052801407 | 0.352520062 | 2.98650069 | 0.002821902 | 0.009625938 |
| ENSG00000167900 | TK1 | 3397.394402 | 0.008462239 | 0.055419841 | 0.152693313 | 0.878640134 | 0.93700919 |
| ENSG00000167904 | TMEM68 | 377.3710247 | 0.089634121 | 0.097881403 | 0.915742091 | 0.359802183 | 0.552393393 |
| ENSG00000167916 | KRT24 | 3.35960336 | -1.384275989 | 0.989464274 | -1.399015634 | 0.161808295 | 0.308834536 |
| ENSG00000167920 | KRT10-AS1 | 318.1903448 | -0.66452119 | 0.106495082 | -6.239923739 | 4.38E-10 | 3.61E-09 |
| ENSG00000167925 | GHDC | 26.58671733 | -0.912202523 | 0.350416115 | -2.603197981 | 0.00923586 | 0.027749919 |
| ENSG00000167930 | FAM234A | 3094.143434 | 0.105568872 | 0.055386453 | 1.906041412 | 0.056644833 | 0.133056775 |
| ENSG00000167962 | ZNF598 | 1837.340174 | 0.118670272 | 0.05625461 | 2.109520823 | 0.034899651 | 0.088385735 |
| ENSG00000167964 | RAB26 | 665.4792625 | 0.288724384 | 0.082949397 | 3.480729163 | 0.000500051 | 0.001978811 |
| ENSG00000167965 | MLST8 | 1275.528453 | 0.342930913 | 0.071822852 | 4.774676896 | 1.80E-06 | 1.01E-05 |
| ENSG00000167967 | E4F1 | 1295.374352 | 0.123763544 | 0.065981958 | 1.875717968 | 0.060694017 | 0.140908892 |
| ENSG00000167968 | DNASE1L2 | 92.15582328 | -0.041285648 | 0.186206423 | -0.221719783 | 0.824532029 | 0.90757472 |
| ENSG00000167969 | ECI1 | 1196.979484 | -0.21318187 | 0.066706574 | -3.195814997 | 0.001394365 | 0.005071624 |
| ENSG00000167971 | CASKIN1 | 564.0628141 | 0.207836661 | 0.089186912 | 2.330349344 | 0.019787695 | 0.054207036 |
| ENSG00000167972 | ABCA3 | 5399.341668 | -0.176889694 | 0.053427967 | -3.310807152 | 0.000930273 | 0.003504756 |
| ENSG00000167977 | KCTD5 | 1455.16538 | 0.117752244 | 0.068982897 | 1.706977359 | 0.087826261 | 0.190644817 |
| ENSG00000167978 | SRRM2 | 24975.5249 | 0.092334965 | 0.050962785 | 1.811811588 | 0.070015314 | 0.158585239 |
| ENSG00000167981 | ZNF597 | 401.5109516 | 0.133124257 | 0.096890456 | 1.373966661 | 0.169452033 | 0.320015611 |

| | | | | | | | |
|---|---|---|---|---|---|---|---|
| ENSG00000167985 | SDHAF2 | 147.1387212 | -0.034701129 | 0.152795434 | -0.227108417 | 0.820339439 | 0.904696252 |
| ENSG00000167986 | DDB1 | 14776.05152 | -0.062221679 | 0.043050633 | -1.44531392 | 0.14836973 | 0.289090959 |
| ENSG00000167987 | VPS37C | 1799.400962 | -0.243628071 | 0.063806321 | -3.818243522 | 0.000134405 | 0.000586505 |
| ENSG00000167992 | VWCE | 21.20390768 | 0.082366085 | 0.380759272 | 0.216320629 | 0.82873783 | 0.909446073 |
| ENSG00000167994 | RAB3IL1 | 175.6223843 | 1.012715828 | 0.148372265 | 6.825506311 | 8.76E-12 | 8.25E-11 |
| ENSG00000167995 | BEST1 | 351.9950728 | -0.268964596 | 0.244806273 | -1.098683429 | 0.271906173 | 0.454001573 |
| ENSG00000167996 | FTH1 | 14289.94879 | 0.361526461 | 0.048293905 | 7.485964539 | 7.10E-14 | 7.83E-13 |
| ENSG00000168000 | BSCL2 | 70.64101233 | -0.692348444 | 0.21819933 | -3.173009032 | 0.001508678 | 0.005440824 |
| ENSG00000168002 | POLR2G | 3099.552693 | -0.168900731 | 0.050032315 | -3.375832808 | 0.000735926 | 0.002825464 |
| ENSG00000168003 | SLC3A2 | 6219.696047 | 0.220342368 | 0.046129703 | 4.776583331 | 1.78E-06 | 1.00E-05 |
| ENSG00000168005 | SPINDOC | 1151.138964 | -0.020307793 | 0.074377163 | -0.273038024 | 0.784823987 | 0.883106023 |
| ENSG00000168010 | ATG16L2 | 998.394214 | 0.299209899 | 0.077613273 | 3.85513827 | 0.000115664 | 0.000510748 |
| ENSG00000168014 | C2CD3 | 1252.315294 | 0.014627264 | 0.063491487 | 0.230381501 | 0.817795334 | 0.903530843 |
| ENSG00000168016 | TRANK1 | 514.7773778 | 0.717203196 | 0.102312683 | 7.009914838 | 2.38E-12 | 2.36E-11 |
| ENSG00000168026 | TTC21A | 99.75512203 | 0.223443431 | 0.188237948 | 1.187026492 | 0.235217171 | 0.410208446 |
| ENSG00000168028 | RPSA | 4325.438156 | 0.086090012 | 0.060121839 | 1.431925801 | 0.152165057 | 0.294620731 |
| ENSG00000168032 | ENTPD3 | 184.0272402 | 0.135637714 | 0.147184184 | 0.921550878 | 0.356762891 | 0.548976437 |
| ENSG00000168036 | CTNNB1 | 12604.24386 | -0.382043068 | 0.040936406 | -9.332599108 | 1.03E-20 | 1.80E-19 |
| ENSG00000168038 | ULK4 | 109.2959577 | 0.454400746 | 0.176210543 | 2.578737565 | 0.009916209 | 0.029508086 |
| ENSG00000168040 | FADD | 1150.095972 | 0.150239748 | 0.072954114 | 2.05937321 | 0.0394585 | 0.098079748 |
| ENSG00000168056 | LTBP3 | 3296.628135 | 0.548939283 | 0.057363564 | 9.569476587 | 1.07E-21 | 1.98E-20 |
| ENSG00000168061 | SAC3D1 | 413.3337269 | -0.213538739 | 0.102894312 | -2.075321113 | 0.037956784 | 0.095005851 |
| ENSG00000168062 | BATF2 | 3.939337227 | 0.283871152 | 0.868824335 | 0.326730204 | 0.743871954 | 0.858490754 |
| ENSG00000168066 | SF1 | 9659.294968 | -0.112222738 | 0.044407476 | -2.527113619 | 0.011500429 | 0.033611905 |
| ENSG00000168067 | MAP4K2 | 610.9915735 | 0.238686476 | 0.083682033 | 2.852302533 | 0.004340377 | 0.014197668 |
| ENSG00000168071 | CCDC88B | 95.5523885 | 0.906295305 | 0.189718591 | 4.777050586 | 1.78E-06 | 1.00E-05 |
| ENSG00000168077 | SCARA3 | 512.7864039 | -0.250761365 | 0.095024066 | -2.638924813 | 0.008316942 | 0.025270286 |
| ENSG00000168078 | PBK | 773.7939268 | -0.176222005 | 0.07763589 | -2.269852331 | 0.023216544 | 0.06226181 |
| ENSG00000168090 | COPS6 | 3633.600933 | -0.535611583 | 0.048886589 | -10.95620695 | 6.20E-28 | 1.60E-26 |
| ENSG00000168092 | PAFAH1B2 | 2472.690426 | 0.036607097 | 0.053108595 | 0.689287614 | 0.490642291 | 0.673770377 |
| ENSG00000168096 | ANKS3 | 402.2295535 | -0.178628524 | 0.09676213 | -1.846058202 | 0.064883755 | 0.148977414 |
| ENSG00000168101 | NUDT16L1 | 1061.295165 | 0.074644434 | 0.081881681 | 0.911613358 | 0.361972288 | 0.554611561 |
| ENSG00000168116 | KIAA1586 | 248.6404985 | -0.564226077 | 0.126048077 | -4.476276761 | 7.60E-06 | 3.97E-05 |

| | | | | | | | |
|---|---|---|---|---|---|---|---|
| ENSG00000168118 | RAB4A | 1542.742798 | 0.26766475 | 0.058718923 | 4.558407012 | 5.15E-06 | 2.75E-05 |
| ENSG00000168124 | OR1F1 | 2.388512457 | 0.411423063 | 1.124634269 | 0.365828318 | 0.714493188 | 0.841150775 |
| ENSG00000168135 | KCNJ4 | 56.20177128 | 1.282678621 | 0.247319575 | 5.186320657 | 2.14E-07 | 1.35E-06 |
| ENSG00000168137 | SETD5 | 10686.16831 | -0.150754843 | 0.044435925 | -3.392634305 | 0.00069224 | 0.002671599 |
| ENSG00000168140 | VASN | 307.6156709 | 0.002776374 | 0.123567278 | 0.022468522 | 0.982074222 | 0.991627389 |
| ENSG00000168143 | FAM83B | 1640.842848 | 0.633497573 | 0.066773679 | 9.48723484 | 2.37E-21 | 4.30E-20 |
| ENSG00000168152 | THAP9 | 143.5142102 | -0.22726166 | 0.149680343 | -1.518313327 | 0.12893543 | 0.259568123 |
| ENSG00000168158 | OR2C1 | 1.608454536 | 0.421203291 | 1.453996917 | 0.289686509 | 0.772056079 | NA |
| ENSG00000168159 | RNF187 | 6079.424245 | -0.098946186 | 0.048143303 | -2.055242984 | 0.039855541 | 0.098863118 |
| ENSG00000168172 | HOOK3 | 660.6366024 | -0.200561178 | 0.086319208 | -2.323482598 | 0.020153241 | 0.055044095 |
| ENSG00000168175 | MAPK1IP1L | 2973.159614 | -0.55901686 | 0.052790549 | -10.5893359 | 3.34E-26 | 7.87E-25 |
| ENSG00000168209 | DDIT4 | 3752.544058 | -0.860155278 | 0.048709851 | -17.65875409 | 8.71E-70 | 8.00E-68 |
| ENSG00000168214 | RBPJ | 2154.516863 | 0.04201797 | 0.062105952 | 0.676553023 | 0.498689593 | 0.680381781 |
| ENSG00000168216 | LMBRD1 | 847.5553862 | -0.040274962 | 0.073109 | -0.550889251 | 0.581709597 | 0.746582642 |
| ENSG00000168228 | ZCCHC4 | 199.6829777 | -0.066574394 | 0.127372421 | -0.52267511 | 0.601200362 | 0.761122313 |
| ENSG00000168234 | TTC39C | 585.9482035 | 0.460542975 | 0.084733314 | 5.43520551 | 5.47E-08 | 3.64E-07 |
| ENSG00000168237 | GLYCTK | 108.6103022 | 0.499559878 | 0.188549313 | 2.649491899 | 0.00806129 | 0.02461086 |
| ENSG00000168243 | GNG4 | 90.33947613 | 0.713670281 | 0.194443729 | 3.670317812 | 0.000242249 | 0.001011871 |
| ENSG00000168246 | UBTD2 | 1292.693879 | -0.356946507 | 0.061495817 | -5.804403019 | 6.46E-09 | 4.74E-08 |
| ENSG00000168255 | POLR2J3 | 38.12252065 | -0.401505917 | 0.295607763 | -1.358238742 | 0.174387941 | 0.327289484 |
| ENSG00000168256 | NKIRAS2 | 1890.674003 | 0.16039974 | 0.057781932 | 2.775949767 | 0.005504071 | 0.017513711 |
| ENSG00000168259 | DNAJC7 | 3140.801821 | 0.090735348 | 0.04774395 | 1.900457519 | 0.057373105 | 0.134505776 |
| ENSG00000168264 | IRF2BP2 | 3642.58826 | 0.044691132 | 0.048226041 | 0.926701234 | 0.354081677 | 0.545582595 |
| ENSG00000168268 | NT5DC2 | 4766.288591 | -0.175347178 | 0.047974823 | -3.654983328 | 0.000257199 | 0.001069176 |
| ENSG00000168273 | UQCC5 | 247.2807448 | 0.001702736 | 0.11976298 | 0.014217547 | 0.988656421 | 0.994082067 |
| ENSG00000168275 | COA6 | 309.7456239 | -0.195920142 | 0.104917741 | -1.867369043 | 0.06185007 | 0.143042706 |
| ENSG00000168280 | KIF5C | 2431.807813 | 0.378370981 | 0.055032375 | 6.875425249 | 6.18E-12 | 5.90E-11 |
| ENSG00000168282 | MGAT2 | 786.9125715 | -0.659443399 | 0.074970009 | -8.796096089 | 1.42E-18 | 2.17E-17 |
| ENSG00000168283 | BMI1 | 560.9299571 | -0.168767572 | 0.09818084 | -1.718946111 | 0.085624185 | 0.186748317 |
| ENSG00000168286 | THAP11 | 1060.065612 | -0.317887253 | 0.069758299 | -4.556981119 | 5.19E-06 | 2.77E-05 |
| ENSG00000168288 | MMADHC | 2701.840359 | 0.18154621 | 0.053009125 | 3.424810561 | 0.000615228 | 0.002395339 |
| ENSG00000168291 | PDHB | 1429.586844 | 0.0719343 | 0.060973367 | 1.179765912 | 0.238093331 | 0.413526941 |
| ENSG00000168297 | PXK | 189.7984438 | 0.484610842 | 0.13903297 | 3.485582181 | 0.000491067 | 0.001946283 |

| | | | | | | | |
|---|---|---|---|---|---|---|---|
| ENSG00000168298 | H1-4 | 12.91211529 | 0.241628809 | 0.490969608 | 0.492146165 | 0.622616019 | 0.776597778 |
| ENSG00000168300 | PCMTD1 | 666.6365801 | 0.358170678 | 0.078512358 | 4.56196562 | 5.07E-06 | 2.71E-05 |
| ENSG00000168301 | KCTD6 | 107.1730566 | -0.406071871 | 0.177270291 | -2.29069332 | 0.021981157 | 0.059380321 |
| ENSG00000168303 | MPLKIP | 536.5172818 | 0.625747703 | 0.087632154 | 7.140617634 | 9.29E-13 | 9.55E-12 |
| ENSG00000168306 | ACOX2 | 2.141292486 | 0.007872882 | 1.173590133 | 0.006708374 | 0.994647532 | 0.996733111 |
| ENSG00000168310 | IRF2 | 256.308064 | 0.029318825 | 0.12501194 | 0.234528194 | 0.814574948 | 0.901763793 |
| ENSG00000168329 | CX3CR1 | 7.559159412 | 0.238009902 | 0.642457229 | 0.370468088 | 0.711033749 | 0.839221764 |
| ENSG00000168350 | DEGS2 | 887.1535594 | 0.332057054 | 0.073457474 | 4.520398486 | 6.17E-06 | 3.27E-05 |
| ENSG00000168374 | ARF4 | 2531.717863 | -0.094461965 | 0.052270333 | -1.807181218 | 0.070734028 | 0.160007752 |
| ENSG00000168385 | SEPTIN2 | 8606.756121 | -0.415577833 | 0.047128503 | -8.817972184 | 1.17E-18 | 1.80E-17 |
| ENSG00000168386 | FILIP1L | 84.89633687 | 1.393194863 | 0.204907517 | 6.799139832 | 1.05E-11 | 9.84E-11 |
| ENSG00000168389 | MFSD2A | 102.8375015 | 0.407548256 | 0.179274439 | 2.273320495 | 0.023006873 | 0.061764503 |
| ENSG00000168393 | DTYMK | 1324.392542 | -0.927280736 | 0.06129137 | -15.12905869 | 1.04E-51 | 6.02E-50 |
| ENSG00000168394 | TAP1 | 70.2204777 | 1.073892041 | 0.218399578 | 4.917097603 | 8.78E-07 | 5.13E-06 |
| ENSG00000168395 | ING5 | 1571.933063 | -0.624495499 | 0.062605626 | -9.975069964 | 1.96E-23 | 3.97E-22 |
| ENSG00000168397 | ATG4B | 2150.123819 | -0.52037989 | 0.054512782 | -9.546015941 | 1.35E-21 | 2.47E-20 |
| ENSG00000168398 | BDKRB2 | 79.33211172 | -0.483156915 | 0.216387763 | -2.232829198 | 0.02556021 | 0.067700175 |
| ENSG00000168404 | MLKL | 200.6782396 | 0.599870798 | 0.131032858 | 4.578018111 | 4.69E-06 | 2.51E-05 |
| ENSG00000168405 | CMAHP | 7.064120542 | -0.085717906 | 0.657465086 | -0.130376362 | 0.896268667 | 0.947527685 |
| ENSG00000168411 | RFWD3 | 3405.135014 | 0.488260283 | 0.050514484 | 9.665748283 | 4.22E-22 | 7.91E-21 |
| ENSG00000168421 | RHOH | 93.98932075 | -1.218279446 | 0.193990051 | -6.280113037 | 3.38E-10 | 2.81E-09 |
| ENSG00000168434 | COG7 | 932.011788 | 0.078519148 | 0.074909005 | 1.048193716 | 0.294549365 | 0.48053309 |
| ENSG00000168438 | CDC40 | 963.5554088 | -0.054254392 | 0.074636979 | -0.726910342 | 0.467280881 | 0.653225071 |
| ENSG00000168439 | STIP1 | 8925.746406 | 0.06010543 | 0.041516417 | 1.447750874 | 0.147686736 | 0.288147412 |
| ENSG00000168447 | SCNN1B | 21.72406831 | -0.466507087 | 0.400803887 | -1.16392855 | 0.244452968 | 0.421228261 |
| ENSG00000168453 | HR | 690.7611171 | -0.825713952 | 0.077688375 | -10.62853939 | 2.20E-26 | 5.23E-25 |
| ENSG00000168461 | RAB31 | 494.9894069 | 0.986734966 | 0.089424547 | 11.03427412 | 2.61E-28 | 6.78E-27 |
| ENSG00000168476 | REEP4 | 1206.878697 | -0.3296931 | 0.065207604 | -5.05605296 | 4.28E-07 | 2.60E-06 |
| ENSG00000168477 | TNXB | 32.25374623 | -0.452730739 | 0.321823093 | -1.406768964 | 0.159495909 | 0.305148141 |
| ENSG00000168481 | LGI3 | 3.677007541 | -0.907845273 | 0.96485366 | -0.940914992 | 0.346748424 | 0.537641693 |
| ENSG00000168487 | BMP1 | 550.4787034 | 0.457534005 | 0.087235299 | 5.244826481 | 1.56E-07 | 9.96E-07 |
| ENSG00000168488 | ATXN2L | 6414.884035 | -0.050037407 | 0.06047879 | -0.827354621 | 0.408036093 | 0.600561271 |
| ENSG00000168495 | POLR3D | 1084.583225 | 0.045438336 | 0.067777154 | 0.670407848 | 0.502597833 | 0.683640338 |

| | | | | | | | |
|---|---|---|---|---|---|---|---|
| ENSG00000168496 | FEN1 | 2593.0003 | -0.303905767 | 0.051332996 | -5.920281106 | 3.21E-09 | 2.44E-08 |
| ENSG00000168502 | MTCL1 | 612.1685214 | 0.300821645 | 0.083752472 | 3.59179423 | 0.000328409 | 0.001341711 |
| ENSG00000168505 | GBX2 | 22.00803325 | 0.226305883 | 0.374378068 | 0.604484884 | 0.545521316 | 0.718666765 |
| ENSG00000168517 | HEXIM2 | 119.4550532 | -0.644274449 | 0.186560644 | -3.45343173 | 0.000553502 | 0.002172715 |
| ENSG00000168522 | FNTA | 950.7581754 | 0.141208656 | 0.069755982 | 2.024323256 | 0.042936902 | 0.105196467 |
| ENSG00000168528 | SERINC2 | 1042.735455 | 0.369424769 | 0.073090489 | 5.054348049 | 4.32E-07 | 2.62E-06 |
| ENSG00000168538 | TRAPPC11 | 442.5375167 | 0.0444315 | 0.08984775 | 0.494519894 | 0.62093906 | 0.775560477 |
| ENSG00000168539 | CHRM1 | 449.0860381 | -0.388571068 | 0.093825121 | -4.141439572 | 3.45E-05 | 0.000165324 |
| ENSG00000168542 | COL3A1 | 131.383079 | 0.940404223 | 0.16335428 | 5.756838607 | 8.57E-09 | 6.20E-08 |
| ENSG00000168546 | GFRA2 | 10.59922182 | -1.628458866 | 0.578184107 | -2.816505756 | 0.004854917 | 0.015697224 |
| ENSG00000168556 | ING2 | 143.4840062 | -0.257062166 | 0.1550985 | -1.657412331 | 0.09743614 | 0.207615183 |
| ENSG00000168564 | CDKN2AIP | 200.2719217 | 0.272346593 | 0.132226503 | 2.059697463 | 0.039427472 | 0.098023866 |
| ENSG00000168566 | SNRNP48 | 493.6334324 | -0.814322815 | 0.091343483 | -8.914952496 | 4.88E-19 | 7.70E-18 |
| ENSG00000168569 | TMEM223 | 480.6646929 | -0.271999307 | 0.09436843 | -2.882312524 | 0.00394768 | 0.013018944 |
| ENSG00000168575 | SLC20A2 | 1068.77471 | -0.070198854 | 0.070515671 | -0.995507153 | 0.319489667 | 0.508964789 |
| ENSG00000168589 | DYNLRB2 | 44.44836777 | 0.262017472 | 0.26741001 | 0.979834195 | 0.32716797 | 0.517600364 |
| ENSG00000168591 | TMUB2 | 1112.881415 | 0.068066865 | 0.068608126 | 0.992110833 | 0.321143462 | 0.510676854 |
| ENSG00000168610 | STAT3 | 4256.557539 | 0.093875708 | 0.05247337 | 1.789016181 | 0.07361221 | 0.165425558 |
| ENSG00000168612 | ZSWIM1 | 605.2014965 | -0.089262765 | 0.084008204 | -1.062548188 | 0.287986894 | 0.471988236 |
| ENSG00000168615 | ADAM9 | 2363.271231 | 0.659339181 | 0.058816809 | 11.21004678 | 3.64E-29 | 9.77E-28 |
| ENSG00000168621 | GDNF | 40.4916684 | 1.195609951 | 0.289507407 | 4.129807808 | 3.63E-05 | 0.000173155 |
| ENSG00000168646 | AXIN2 | 102.6957489 | 1.559135026 | 0.193387033 | 8.062252178 | 7.49E-16 | 9.58E-15 |
| ENSG00000168653 | NDUFS5 | 2475.398026 | -0.045661962 | 0.051400353 | -0.888358922 | 0.374347714 | 0.567096122 |
| ENSG00000168661 | ZNF30 | 363.9402478 | -0.960987447 | 0.101210578 | -9.494930918 | 2.20E-21 | 4.00E-20 |
| ENSG00000168672 | LRATD2 | 8158.352036 | -0.9500075 | 0.042471439 | -22.36814948 | 8.04E-111 | 1.60E-108 |
| ENSG00000168675 | LDLRAD4 | 2911.923049 | -0.030441911 | 0.062846291 | -0.484386759 | 0.628111414 | 0.78067269 |
| ENSG00000168676 | KCTD19 | 3.16826424 | 3.948154453 | 1.426132692 | 2.768434154 | 0.005632637 | 0.017891538 |
| ENSG00000168679 | SLC16A4 | 24.35537646 | -0.78193328 | 0.360517045 | -2.168921808 | 0.030088621 | 0.077886816 |
| ENSG00000168701 | TMEM208 | 834.1888589 | 0.562533718 | 0.074125155 | 7.588971869 | 3.22E-14 | 3.65E-13 |
| ENSG00000168702 | LRP1B | 55.77345638 | 2.08705573 | 0.27492468 | 7.591372781 | 3.17E-14 | 3.59E-13 |
| ENSG00000168710 | AHCYL1 | 3709.064025 | -0.247504464 | 0.047892538 | -5.167912838 | 2.37E-07 | 1.48E-06 |
| ENSG00000168724 | DNAJC21 | 2635.862215 | -0.023492978 | 0.053759914 | -0.43699806 | 0.662112755 | 0.80479344 |
| ENSG00000168734 | PKIG | 597.9722896 | 0.705085522 | 0.082124857 | 8.585531213 | 9.04E-18 | 1.31E-16 |

| | | | | | | | |
|---|---|---|---|---|---|---|---|
| ENSG00000168743 | NPNT | 7668.54985 | -0.321226495 | 0.054310861 | -5.91459037 | 3.33E-09 | 2.53E-08 |
| ENSG00000168754 | FAM178B | 9.390047042 | -0.340516792 | 0.591554774 | -0.575630198 | 0.564865165 | 0.733209239 |
| ENSG00000168758 | SEMA4C | 2909.755557 | 0.272475163 | 0.059814978 | 4.555299904 | 5.23E-06 | 2.79E-05 |
| ENSG00000168763 | CNNM3 | 1994.910457 | 0.069761346 | 0.059393087 | 1.17457013 | 0.240166739 | 0.415994035 |
| ENSG00000168765 | GSTM4 | 1054.139724 | -0.288794017 | 0.069746298 | -4.140635765 | 3.46E-05 | 0.0001658 |
| ENSG00000168769 | TET2 | 1378.379171 | 0.09193287 | 0.061648037 | 1.491253807 | 0.135894872 | 0.270492699 |
| ENSG00000168772 | CXXC4 | 46.58536995 | -0.489721673 | 0.260429871 | -1.880435874 | 0.060048698 | 0.139622657 |
| ENSG00000168778 | TCTN2 | 223.2965827 | 0.150367805 | 0.122284674 | 1.229653728 | 0.218826801 | 0.388562198 |
| ENSG00000168779 | SHOX2 | 13.36143765 | -0.565847435 | 0.476800359 | -1.186759666 | 0.235322432 | 0.410329629 |
| ENSG00000168781 | PPIP5K1 | 682.0775799 | 0.009553312 | 0.084725017 | 0.112756684 | 0.910223461 | 0.954995124 |
| ENSG00000168785 | TSPAN5 | 239.5454161 | -0.238433595 | 0.118391313 | -2.013945024 | 0.044015306 | 0.107461804 |
| ENSG00000168792 | ABHD15 | 989.586254 | 0.158076972 | 0.068054877 | 2.322786814 | 0.020190608 | 0.055126463 |
| ENSG00000168795 | ZBTB5 | 947.1357877 | -0.028730401 | 0.069150241 | -0.415477965 | 0.677792043 | 0.816688071 |
| ENSG00000168802 | CHTF8 | 23.51707955 | 0.067610826 | 0.36074703 | 0.18741894 | 0.851332174 | 0.9222591 |
| ENSG00000168803 | ADAL | 354.6928039 | -0.390481538 | 0.101179051 | -3.859312125 | 0.000113707 | 0.000502781 |
| ENSG00000168806 | LCMT2 | 748.4033968 | -0.155193983 | 0.078047658 | -1.988451519 | 0.046761776 | 0.113084033 |
| ENSG00000168807 | SNTB2 | 963.836361 | -0.091557736 | 0.069784873 | -1.311999757 | 0.189520217 | 0.348448647 |
| ENSG00000168811 | IL12A | 5.746781162 | -0.855793953 | 0.761543414 | -1.123762529 | 0.261113782 | 0.441411212 |
| ENSG00000168813 | ZNF507 | 1203.786125 | -0.195765188 | 0.068330674 | -2.864967922 | 0.004170515 | 0.013679098 |
| ENSG00000168818 | STX18 | 754.7984744 | 0.022142592 | 0.074667885 | 0.296547736 | 0.766811817 | 0.872725892 |
| ENSG00000168826 | ZBTB49 | 172.3019489 | -0.097965898 | 0.149595553 | -0.654871726 | 0.512550346 | 0.69212854 |
| ENSG00000168827 | GFM1 | 2237.4881 | -0.073340652 | 0.053400932 | -1.373396493 | 0.169629119 | 0.320165542 |
| ENSG00000168830 | HTR1E | 14.54048522 | 0.757416002 | 0.479259575 | 1.580387833 | 0.114018076 | 0.235743548 |
| ENSG00000168843 | FSTL5 | 538.843411 | 2.046246392 | 0.095532724 | 21.41932426 | 8.83E-102 | 1.47E-99 |
| ENSG00000168852 | TPTE2P5 | 15.46948783 | 0.850717143 | 0.460814941 | 1.846114499 | 0.064875582 | 0.148977414 |
| ENSG00000168872 | DDX19A | 333.5024954 | 0.08730754 | 0.104018927 | 0.839342825 | 0.401276958 | 0.594107762 |
| ENSG00000168874 | ATOH8 | 102.137122 | -1.754572874 | 0.190742639 | -9.198640047 | 3.63E-20 | 6.10E-19 |
| ENSG00000168876 | ANKRD49 | 228.374447 | 0.050621004 | 0.128273314 | 0.394633944 | 0.693113059 | 0.826932295 |
| ENSG00000168883 | USP39 | 2636.52181 | 0.01748036 | 0.052503697 | 0.332935803 | 0.739182743 | 0.856204886 |
| ENSG00000168884 | TNIP2 | 1282.987132 | 0.045553135 | 0.062962424 | 0.723497171 | 0.469374492 | 0.655353064 |
| ENSG00000168887 | C2orf68 | 1769.704176 | 0.11822149 | 0.058985392 | 2.0042503 | 0.045043255 | 0.109516933 |
| ENSG00000168890 | TMEM150A | 594.2363461 | 0.115852992 | 0.08681935 | 1.334414413 | 0.182068079 | 0.337915998 |
| ENSG00000168894 | RNF181 | 1331.539849 | 0.074254441 | 0.066340255 | 1.119296894 | 0.2630135 | 0.443772411 |

| | | | | | | | |
|---|---|---|---|---|---|---|---|
| ENSG00000168899 | VAMP5 | 136.7465876 | 0.070061427 | 0.163258523 | 0.429144068 | 0.667818384 | 0.808785741 |
| ENSG00000168904 | LRRC28 | 443.7756291 | 0.207418509 | 0.097179934 | 2.134375895 | 0.032812023 | 0.08390456 |
| ENSG00000168906 | MAT2A | 8126.542568 | 0.167872065 | 0.06102738 | 2.750766375 | 0.005945603 | 0.018781621 |
| ENSG00000168907 | PLA2G4F | 200.3437871 | 0.902027876 | 0.13152306 | 6.858324887 | 6.97E-12 | 6.62E-11 |
| ENSG00000168913 | ENHO | 39.24914485 | 0.364394183 | 0.29421847 | 1.238515658 | 0.21552492 | 0.384758106 |
| ENSG00000168916 | ZNF608 | 243.3129938 | 0.405329538 | 0.121793618 | 3.328003088 | 0.000874709 | 0.003310179 |
| ENSG00000168918 | INPP5D | 15.23647719 | 0.07990928 | 0.460389125 | 0.173568999 | 0.862204197 | 0.928371945 |
| ENSG00000168924 | LETM1 | 3342.64164 | 0.129001563 | 0.050481057 | 2.555444958 | 0.010605213 | 0.031290494 |
| ENSG00000168936 | TMEM129 | 1442.996249 | 0.343409111 | 0.062127196 | 5.527516648 | 3.25E-08 | 2.21E-07 |
| ENSG00000168938 | PPIC | 1687.714863 | 0.044186115 | 0.056356594 | 0.784045166 | 0.433013609 | 0.623082616 |
| ENSG00000168939 | SPRY3 | 95.17315357 | -0.239085696 | 0.18495712 | -1.292654729 | 0.196130502 | 0.357621311 |
| ENSG00000168939 | SPRY3 | 95.17315357 | -0.239085696 | 0.18495712 | -1.292654729 | 0.196130502 | 0.357621311 |
| ENSG00000168944 | CEP120 | 563.9806201 | -0.157044827 | 0.090220631 | -1.740675336 | 0.081740504 | 0.179816567 |
| ENSG00000168952 | STXBP6 | 3.063640542 | -0.024639248 | 0.98976953 | -0.024893925 | 0.980139573 | 0.991041209 |
| ENSG00000168958 | MFF | 1087.00459 | 0.221107232 | 0.06486134 | 3.408921725 | 0.000652202 | 0.002528564 |
| ENSG00000168961 | LGALS9 | 2.273711851 | 1.670796827 | 1.281865704 | 1.303410195 | 0.192434758 | 0.352620285 |
| ENSG00000168970 | JMJD7-PLA2G | 157.9669922 | -0.358726275 | 0.1535313 | -2.336502561 | 0.019465064 | 0.053431683 |
| ENSG00000168992 | OR7E102P | 12.30616927 | 0.444309877 | 0.533078654 | 0.833479026 | 0.404574622 | 0.597498348 |
| ENSG00000168993 | CPLX1 | 33.60450904 | 0.052508341 | 0.315620407 | 0.166365482 | 0.867869337 | 0.931076558 |
| ENSG00000168994 | PXDC1 | 54.18044383 | -0.257712024 | 0.238478744 | -1.080649873 | 0.279852889 | 0.463190856 |
| ENSG00000169016 | E2F6 | 539.1411574 | 0.047290222 | 0.08734396 | 0.541425214 | 0.588214532 | 0.751525557 |
| ENSG00000169018 | FEM1B | 2705.877669 | -0.121548515 | 0.054441872 | -2.23262925 | 0.025573404 | 0.067721217 |
| ENSG00000169019 | COMMD8 | 268.1379975 | 0.134551001 | 0.112387196 | 1.197209341 | 0.23122497 | 0.404843771 |
| ENSG00000169020 | ATP5ME | 1559.572019 | 0.101112023 | 0.060509681 | 1.671005722 | 0.094720548 | 0.202808416 |
| ENSG00000169021 | UQCRFS1 | 1958.24407 | -0.038938957 | 0.056516147 | -0.688988187 | 0.490830702 | 0.673915141 |
| ENSG00000169026 | SLC49A3 | 9.476866514 | -0.554099441 | 0.596390025 | -0.929089049 | 0.352842939 | 0.544269266 |
| ENSG00000169031 | COL4A3 | 53.18554828 | 0.987886869 | 0.248934037 | 3.968468434 | 7.23E-05 | 0.000330358 |
| ENSG00000169032 | MAP2K1 | 1988.859855 | 0.083430718 | 0.055042236 | 1.515758154 | 0.129580517 | 0.26059236 |
| ENSG00000169045 | HNRNPH1 | 14556.60447 | -0.191630542 | 0.134338214 | -1.426478262 | 0.153730337 | 0.296874647 |
| ENSG00000169047 | IRS1 | 2553.931388 | 1.695828245 | 0.054397089 | 31.17498172 | 2.33E-213 | 1.72E-210 |
| ENSG00000169057 | MECP2 | 4124.175452 | -0.611208127 | 0.05645349 | -10.82675549 | 2.57E-27 | 6.47E-26 |
| ENSG00000169062 | UPF3A | 1012.931758 | 0.046225629 | 0.068813609 | 0.671751262 | 0.501742061 | 0.682833786 |
| ENSG00000169064 | ZBBX | 5.525062142 | -0.286221603 | 0.746608653 | -0.38336229 | 0.70145116 | 0.832949765 |

| | | | | | | | |
|---|---|---|---|---|---|---|---|
| ENSG00000169071 | ROR2 | 10.35413196 | -2.357448667 | 0.638699225 | -3.691015387 | 0.000223361 | 0.000939302 |
| ENSG00000169083 | AR | 7515.934451 | -0.468703327 | 0.055116022 | -8.503939724 | 1.83E-17 | 2.61E-16 |
| ENSG00000169084 | DHRSX | 415.1402703 | -0.00178554 | 0.092341479 | -0.01933627 | 0.98457285 | 0.992970245 |
| ENSG00000169084 | DHRSX | 415.1402703 | -0.00178554 | 0.092341479 | -0.01933627 | 0.98457285 | 0.992970245 |
| ENSG00000169087 | HSPBAP1 | 160.6850579 | 0.03006115 | 0.147058802 | 0.204415846 | 0.838028548 | 0.914820294 |
| ENSG00000169093 | ASMTL | 1010.954245 | 0.208293662 | 0.067584984 | 3.081951762 | 0.002056482 | 0.007207897 |
| ENSG00000169093 | ASMTL | 1010.954245 | 0.208293662 | 0.067584984 | 3.081951762 | 0.002056482 | 0.007207897 |
| ENSG00000169100 | SLC25A6 | 11591.29697 | 0.538557842 | 0.045521096 | 11.83095073 | 2.70E-32 | 8.25E-31 |
| ENSG00000169100 | SLC25A6 | 11591.29697 | 0.538557842 | 0.045521096 | 11.83095073 | 2.70E-32 | 8.25E-31 |
| ENSG00000169105 | CHST14 | 612.4532592 | -0.052288514 | 0.090347441 | -0.578749255 | 0.562758376 | 0.731592267 |
| ENSG00000169116 | PARM1 | 282.8284331 | -0.485707763 | 0.113260519 | -4.288411961 | 1.80E-05 | 8.99E-05 |
| ENSG00000169118 | CSNK1G1 | 1472.543345 | -0.167692081 | 0.065163481 | -2.573405825 | 0.010070305 | 0.029919307 |
| ENSG00000169122 | FAM110B | 490.3665924 | -0.758106692 | 0.090876135 | -8.342197779 | 7.29E-17 | 9.98E-16 |
| ENSG00000169126 | ODAD2 | 18.71626884 | 1.834624448 | 0.456112418 | 4.022307608 | 5.76E-05 | 0.000267351 |
| ENSG00000169129 | AFAP1L2 | 166.1325823 | -1.963074686 | 0.1631897 | -12.0294031 | 2.49E-33 | 7.86E-32 |
| ENSG00000169131 | ZNF354A | 587.6374938 | -0.496109185 | 0.086705105 | -5.721799005 | 1.05E-08 | 7.57E-08 |
| ENSG00000169136 | ATF5 | 2091.107744 | 0.331809148 | 0.058224436 | 5.698795365 | 1.21E-08 | 8.59E-08 |
| ENSG00000169139 | UBE2V2 | 1703.716912 | -0.01270145 | 0.061990256 | -0.204894294 | 0.837654713 | 0.914602528 |
| ENSG00000169155 | ZBTB43 | 547.4025187 | 0.111715579 | 0.084085234 | 1.328599253 | 0.183980219 | 0.340445771 |
| ENSG00000169169 | CPT1C | 19.85781 | 0.209558119 | 0.397066456 | 0.527765859 | 0.597661856 | 0.758106883 |
| ENSG00000169180 | XPO6 | 4342.638666 | 0.129565047 | 0.05125535 | 2.527834588 | 0.011476841 | 0.03355579 |
| ENSG00000169181 | GSG1L | 1.825497472 | 1.175637575 | 1.363099647 | 0.862473684 | 0.388426908 | 0.581170967 |
| ENSG00000169184 | MN1 | 6.992590535 | -0.461902881 | 0.659258942 | -0.700639538 | 0.483527997 | 0.66755731 |
| ENSG00000169188 | APEX2 | 1616.299973 | 0.047821733 | 0.060435652 | 0.791283477 | 0.428778588 | 0.61920798 |
| ENSG00000169189 | NSMCE1 | 775.9956661 | -0.224019466 | 0.074375653 | -3.012000001 | 0.002595326 | 0.008910024 |
| ENSG00000169193 | CCDC126 | 177.1543552 | -1.716768778 | 0.15252561 | -11.25561001 | 2.17E-29 | 5.89E-28 |
| ENSG00000169194 | IL13 | 1.943693365 | 1.367295597 | 1.336921899 | 1.022719127 | 0.306440665 | 0.494094221 |
| ENSG00000169203 | NPIPB12 | 21.29262852 | -0.014045489 | 0.381833739 | -0.036784307 | 0.970656987 | 0.986772291 |
| ENSG00000169213 | RAB3B | 30.15103852 | -0.158801617 | 0.320611821 | -0.495308053 | 0.620382687 | 0.775267138 |
| ENSG00000169217 | CD2BP2 | 3205.815233 | 0.163897352 | 0.051510695 | 3.181812102 | 0.001463567 | 0.005293951 |
| ENSG00000169220 | RGS14 | 56.20840254 | 0.779319681 | 0.241173907 | 3.23135986 | 0.001232027 | 0.004535094 |
| ENSG00000169221 | TBC1D10B | 2690.235795 | -0.036723086 | 0.059604253 | -0.616115195 | 0.537818494 | 0.712782729 |
| ENSG00000169223 | LMAN2 | 3881.66606 | -0.282354406 | 0.05537743 | -5.098727111 | 3.42E-07 | 2.09E-06 |

| | | | | | | | |
|---|---|---|---|---|---|---|---|
| ENSG00000169228 | RAB24 | 78.7249673 | -0.642702954 | 0.22851332 | -2.812540442 | 0.004915184 | 0.015864679 |
| ENSG00000169230 | PRELID1 | 3923.449366 | -0.250058407 | 0.046165605 | -5.416552196 | 6.08E-08 | 4.01E-07 |
| ENSG00000169231 | THBS3 | 1320.768087 | -0.092875925 | 0.071805236 | -1.293442247 | 0.195858146 | 0.357338497 |
| ENSG00000169239 | CA5B | 662.3227692 | -0.390891443 | 0.079603133 | -4.910503267 | 9.08E-07 | 5.29E-06 |
| ENSG00000169241 | SLC50A1 | 2419.759401 | -0.189921781 | 0.051712795 | -3.672626496 | 0.00024007 | 0.001003684 |
| ENSG00000169242 | EFNA1 | 2463.748346 | 0.570789264 | 0.05960062 | 9.576901455 | 1.00E-21 | 1.84E-20 |
| ENSG00000169246 | NPIPB3 | 151.4566208 | -0.133943081 | 0.166400226 | -0.804945308 | 0.420851241 | 0.61204054 |
| ENSG00000169247 | SH3TC2 | 98.74508481 | -1.093055384 | 0.189026914 | -5.782538378 | 7.36E-09 | 5.36E-08 |
| ENSG00000169249 | ZRSR2 | 313.0762836 | 0.069195405 | 0.108875203 | 0.635547882 | 0.525071148 | 0.701269943 |
| ENSG00000169251 | NMD3 | 7556.145746 | -0.399375734 | 0.047883422 | -8.340584547 | 7.39E-17 | 1.01E-15 |
| ENSG00000169253 | RPL36P14 | 14.97246671 | -0.384913887 | 0.459015352 | -0.838564299 | 0.401713852 | 0.594410925 |
| ENSG00000169255 | B3GALNT1 | 1831.609717 | 0.076348435 | 0.055885891 | 1.366148665 | 0.171892306 | 0.323585221 |
| ENSG00000169258 | GPRIN1 | 614.9546898 | -0.119564811 | 0.079568377 | -1.50266746 | 0.132924818 | 0.266091209 |
| ENSG00000169288 | MRPL1 | 448.3984106 | -0.024507651 | 0.089926298 | -0.272530414 | 0.785214209 | 0.883118404 |
| ENSG00000169299 | PGM2 | 1481.843247 | 0.154338186 | 0.060095815 | 2.568201898 | 0.01022276 | 0.030302378 |
| ENSG00000169330 | MINAR1 | 105.3452331 | 0.697862836 | 0.17967786 | 3.883966758 | 0.000102766 | 0.000457402 |
| ENSG00000169347 | GP2 | 50.56983787 | 1.840229355 | 0.28399875 | 6.47970934 | 9.19E-11 | 7.98E-10 |
| ENSG00000169359 | SLC33A1 | 439.8646004 | 0.855860943 | 0.097840419 | 8.747519229 | 2.18E-18 | 3.30E-17 |
| ENSG00000169371 | SNUPN | 184.8232925 | -0.024089866 | 0.136997964 | -0.175841052 | 0.860418817 | 0.927407211 |
| ENSG00000169372 | CRADD | 193.7377971 | 0.175698455 | 0.131770773 | 1.333364372 | 0.182412258 | 0.3382811 |
| ENSG00000169375 | SIN3A | 5465.608412 | -0.11497586 | 0.049503759 | -2.322568243 | 0.020202358 | 0.055151982 |
| ENSG00000169379 | ARL13B | 335.6210526 | -0.068665344 | 0.110885074 | -0.619247853 | 0.535753092 | 0.711031295 |
| ENSG00000169398 | PTK2 | 5569.932304 | 0.174205149 | 0.044002642 | 3.95897021 | 7.53E-05 | 0.000343091 |
| ENSG00000169402 | RSPH10B2 | 6.256986657 | 1.157064551 | 0.717980935 | 1.611553308 | 0.107059178 | 0.223927516 |
| ENSG00000169403 | PTAFR | 16.80525675 | -0.258353331 | 0.429023003 | -0.602189928 | 0.54704772 | 0.719888603 |
| ENSG00000169410 | PTPN9 | 1712.7597 | -0.236758252 | 0.058553651 | -4.043441347 | 5.27E-05 | 0.00024549 |
| ENSG00000169427 | KCNK9 | 7.895697641 | 0.304101587 | 0.624643744 | 0.486840043 | 0.626371691 | 0.779566329 |
| ENSG00000169429 | CXCL8 | 3.864166709 | 1.387856326 | 0.963281367 | 1.440759028 | 0.149652772 | 0.291107826 |
| ENSG00000169439 | SDC2 | 633.8480826 | 0.323834151 | 0.085688959 | 3.779181766 | 0.000157345 | 0.000679368 |
| ENSG00000169446 | MMGT1 | 1663.1215 | -0.154689716 | 0.06603239 | -2.342633919 | 0.019148161 | 0.052649983 |
| ENSG00000169490 | TM2D2 | 682.1838851 | 0.16335361 | 0.077639877 | 2.103991109 | 0.035379223 | 0.089441736 |
| ENSG00000169495 | HTRA4 | 5.046549004 | 1.182138907 | 0.825696787 | 1.431686456 | 0.152233574 | 0.294703645 |
| ENSG00000169499 | PLEKHA2 | 444.7828136 | 0.137096166 | 0.090127832 | 1.521130187 | 0.128227171 | 0.258596158 |

| | | | | | | | |
|---|---|---|---|---|---|---|---|
| ENSG00000169504 | CLIC4 | 3452.170853 | 0.221925093 | 0.062452825 | 3.553483677 | 0.000380165 | 0.001534033 |
| ENSG00000169507 | SLC38A11 | 5.657971686 | 2.425488105 | 0.887142923 | 2.73404436 | 0.006256162 | 0.019667856 |
| ENSG00000169519 | METTL15 | 252.031607 | -0.130505211 | 0.121225209 | -1.076551753 | 0.281680569 | 0.46514197 |
| ENSG00000169564 | PCBP1 | 6508.198392 | -0.130594011 | 0.045471577 | -2.871992113 | 0.004078931 | 0.013415152 |
| ENSG00000169567 | HINT1 | 3025.307161 | -0.075193649 | 0.050966721 | -1.475347988 | 0.140119011 | 0.276644299 |
| ENSG00000169570 | DTWD2 | 464.6232765 | 0.064841644 | 0.090627706 | 0.715472643 | 0.474317039 | 0.6597243 |
| ENSG00000169583 | CLIC3 | 332.1659417 | 1.214680204 | 0.11279673 | 10.7687537 | 4.84E-27 | 1.20E-25 |
| ENSG00000169592 | INO80E | 1427.068347 | -0.314669096 | 0.061587615 | -5.109291793 | 3.23E-07 | 1.99E-06 |
| ENSG00000169594 | BNC1 | 41.76336606 | 0.713282681 | 0.27917457 | 2.554970107 | 0.010619692 | 0.031329183 |
| ENSG00000169598 | DFFB | 163.543266 | -0.045280405 | 0.140978473 | -0.32118666 | 0.748068941 | 0.860436606 |
| ENSG00000169599 | NFU1 | 512.9394699 | -0.122099622 | 0.085511232 | -1.427878181 | 0.153326925 | 0.296241419 |
| ENSG00000169604 | ANTXR1 | 2153.221596 | 0.139024707 | 0.053922622 | 2.578225999 | 0.009930902 | 0.029544145 |
| ENSG00000169607 | CKAP2L | 674.2107558 | -0.166381846 | 0.077462506 | -2.147901669 | 0.031721568 | 0.081525069 |
| ENSG00000169609 | C15orf40 | 839.7678002 | -0.261167295 | 0.072677471 | -3.593511066 | 0.000326252 | 0.001333372 |
| ENSG00000169612 | RAMAC | 318.8440146 | -0.090338159 | 0.105230645 | -0.858477672 | 0.390628754 | 0.58343796 |
| ENSG00000169629 | RGPD8 | 52.29675447 | -0.302124085 | 0.241713363 | -1.249927108 | 0.211326176 | 0.378854523 |
| ENSG00000169635 | HIC2 | 160.7146992 | -1.096874008 | 0.147557391 | -7.433541612 | 1.06E-13 | 1.15E-12 |
| ENSG00000169641 | LUZP1 | 967.4242805 | 0.228950161 | 0.074217252 | 3.084864407 | 0.00203645 | 0.007145328 |
| ENSG00000169660 | HEXD | 796.9447066 | 0.452175918 | 0.078286111 | 5.775940513 | 7.65E-09 | 5.56E-08 |
| ENSG00000169679 | BUB1 | 1437.186507 | -0.358815409 | 0.060307881 | -5.949726693 | 2.69E-09 | 2.05E-08 |
| ENSG00000169682 | SPNS1 | 24.55816397 | -0.088015198 | 0.357358686 | -0.246293714 | 0.805454876 | 0.89673064 |
| ENSG00000169683 | LRRC45 | 1940.139258 | -0.01319826 | 0.056625015 | -0.233081786 | 0.815697898 | 0.902352305 |
| ENSG00000169684 | CHRNA5 | 488.9555093 | -0.303419706 | 0.091058768 | -3.332130587 | 0.000861838 | 0.003266242 |
| ENSG00000169689 | CENPX | 3682.647609 | -0.131691184 | 0.057133312 | -2.304980756 | 0.021167651 | 0.057397863 |
| ENSG00000169692 | AGPAT2 | 1436.481139 | 0.448695389 | 0.073422176 | 6.11116989 | 9.89E-10 | 7.87E-09 |
| ENSG00000169696 | ASPSCR1 | 1532.07428 | 0.054004663 | 0.057846468 | 0.933586171 | 0.350517402 | 0.54191058 |
| ENSG00000169710 | FASN | 109920.8611 | -0.392419409 | 0.039196764 | -10.01152575 | 1.36E-23 | 2.77E-22 |
| ENSG00000169714 | CNBP | 9462.914956 | -0.148058341 | 0.044225333 | -3.347817462 | 0.000814506 | 0.003099664 |
| ENSG00000169718 | DUS1L | 4251.87838 | 0.349227965 | 0.050585044 | 6.903779012 | 5.06E-12 | 4.88E-11 |
| ENSG00000169727 | GPS1 | 4729.429325 | 0.219048951 | 0.05470013 | 4.00454167 | 6.21E-05 | 0.000286636 |
| ENSG00000169733 | RFNG | 1289.71229 | 0.092070896 | 0.071330068 | 1.290772572 | 0.196782551 | 0.358553751 |
| ENSG00000169738 | DCXR | 2573.040762 | 0.219384229 | 0.066991328 | 3.274815335 | 0.001057311 | 0.003939371 |
| ENSG00000169740 | ZNF32 | 755.2516558 | -0.367839015 | 0.076814443 | -4.788669942 | 1.68E-06 | 9.46E-06 |

| | | | | | | | |
|---|---|---|---|---|---|---|---|
| ENSG00000169750 | RAC3 | 835.0957976 | 0.138147258 | 0.079306332 | 1.741944872 | 0.081518091 | 0.179430533 |
| ENSG00000169752 | NRG4 | 2.878810244 | -0.668995226 | 1.067836096 | -0.626496171 | 0.530989547 | 0.706555758 |
| ENSG00000169756 | LIMS1 | 881.7984672 | 0.033902873 | 0.074802314 | 0.453232945 | 0.650381012 | 0.796739315 |
| ENSG00000169758 | TMEM266 | 2.417924428 | 1.791738644 | 1.249620596 | 1.433826115 | 0.151621891 | 0.293976042 |
| ENSG00000169762 | TAPT1 | 1321.756275 | -1.106066665 | 0.068092535 | -16.24358188 | 2.48E-59 | 1.73E-57 |
| ENSG00000169764 | UGP2 | 2057.670016 | -0.07139 | 0.056167589 | -1.2710177 | 0.203722352 | 0.36844771 |
| ENSG00000169783 | LINGO1 | 222.5722014 | 0.299923328 | 0.12833607 | 2.337015047 | 0.019438402 | 0.053377656 |
| ENSG00000169813 | HNRNPF | 9180.800402 | -0.557181639 | 0.042167477 | -13.21353988 | 7.33E-40 | 2.97E-38 |
| ENSG00000169814 | BTD | 927.647219 | 0.204360812 | 0.08002977 | 2.553559912 | 0.010662793 | 0.031436124 |
| ENSG00000169826 | CSGALNACT2 | 349.7060629 | -0.602103794 | 0.106377863 | -5.660047834 | 1.51E-08 | 1.07E-07 |
| ENSG00000169836 | TACR3 | 30.07237829 | -0.667880024 | 0.324204522 | -2.060057705 | 0.039393025 | 0.097948839 |
| ENSG00000169851 | PCDH7 | 1340.881196 | 1.328949349 | 0.068620705 | 19.36659425 | 1.48E-83 | 1.78E-81 |
| ENSG00000169855 | ROBO1 | 2806.101885 | 0.753380578 | 0.053895947 | 13.97842742 | 2.11E-44 | 9.94E-43 |
| ENSG00000169856 | ONECUT1 | 438.4722351 | -3.56639093 | 0.125949343 | -28.31607403 | 2.19E-176 | 9.86E-174 |
| ENSG00000169857 | AVEN | 427.6852087 | -0.03647934 | 0.092399047 | -0.39480212 | 0.69298893 | 0.826907354 |
| ENSG00000169860 | P2RY1 | 23.57573015 | -0.828246773 | 0.374107357 | -2.213928052 | 0.026833729 | 0.070608595 |
| ENSG00000169862 | CTNND2 | 4454.989495 | -0.300435403 | 0.056878409 | -5.282064159 | 1.28E-07 | 8.22E-07 |
| ENSG00000169871 | TRIM56 | 1798.514433 | -0.368528542 | 0.06324717 | -5.826798886 | 5.65E-09 | 4.18E-08 |
| ENSG00000169884 | WNT10B | 19.81703592 | 0.358782577 | 0.403076596 | 0.890110168 | 0.373406733 | 0.566156673 |
| ENSG00000169885 | CALML6 | 20.7183104 | -0.035638482 | 0.393169299 | -0.090644111 | 0.92777538 | 0.96531056 |
| ENSG00000169891 | REPS2 | 454.3430269 | 0.9138272 | 0.093841786 | 9.737956168 | 2.08E-22 | 3.98E-21 |
| ENSG00000169894 | MUC3A | 496.8265004 | -1.033858109 | 0.107348427 | -9.630864055 | 5.92E-22 | 1.10E-20 |
| ENSG00000169895 | SYAP1 | 3623.303982 | 0.179780193 | 0.047997164 | 3.745642022 | 0.000179933 | 0.000769656 |
| ENSG00000169896 | ITGAM | 2.235431564 | 0.441253825 | 1.255300019 | 0.351512641 | 0.725203791 | 0.847271452 |
| ENSG00000169902 | TPST1 | 201.5105198 | 0.642758213 | 0.131077206 | 4.903661244 | 9.41E-07 | 5.46E-06 |
| ENSG00000169905 | TOR1AIP2 | 1499.876526 | -0.196102079 | 0.057904126 | -3.386668464 | 0.000707468 | 0.002723955 |
| ENSG00000169908 | TM4SF1 | 103.2420198 | 2.192466837 | 0.206501641 | 10.61718846 | 2.48E-26 | 5.89E-25 |
| ENSG00000169914 | OTUD3 | 450.0117155 | -0.040520984 | 0.091810406 | -0.441355024 | 0.658955997 | 0.802779253 |
| ENSG00000169918 | OTUD7A | 115.9546582 | -0.097807039 | 0.169845678 | -0.575858274 | 0.564710981 | 0.733133554 |
| ENSG00000169919 | GUSB | 1640.183258 | 0.187488614 | 0.064329505 | 2.914504242 | 0.00356254 | 0.011878584 |
| ENSG00000169925 | BRD3 | 2430.970249 | 0.091855422 | 0.051969525 | 1.767486265 | 0.077146827 | 0.171853585 |
| ENSG00000169926 | KLF13 | 948.5041674 | -0.555506391 | 0.07107862 | -7.815379537 | 5.48E-15 | 6.58E-14 |
| ENSG00000169951 | ZNF764 | 308.2758074 | -0.195682964 | 0.105000003 | -1.863647225 | 0.062371261 | 0.144044773 |

| | | | | | | | |
|---|---|---|---|---|---|---|---|
| ENSG00000169955 | ZNF747 | 57.59504696 | -0.641450941 | 0.235327503 | -2.725779745 | 0.006414979 | 0.020095537 |
| ENSG00000169957 | ZNF768 | 1587.15392 | -0.163267822 | 0.058399224 | -2.795719041 | 0.00517844 | 0.016608159 |
| ENSG00000169962 | TAS1R3 | 8.289629927 | 0.60935196 | 0.663500481 | 0.918389629 | 0.358414919 | 0.550890334 |
| ENSG00000169964 | TMEM42 | 180.4068813 | 0.358325835 | 0.135422141 | 2.645991506 | 0.008145186 | 0.024827341 |
| ENSG00000169967 | MAP3K2 | 2062.652434 | 0.025917729 | 0.063673463 | 0.407041303 | 0.683977653 | 0.820758873 |
| ENSG00000169972 | PUSL1 | 404.2801439 | 0.378263352 | 0.097201918 | 3.891521484 | 9.96E-05 | 0.000444078 |
| ENSG00000169976 | SF3B5 | 2023.135767 | 0.19711041 | 0.061799597 | 3.189509644 | 0.001425144 | 0.005165555 |
| ENSG00000169981 | ZNF35 | 253.1936418 | -0.088761454 | 0.117762296 | -0.753734067 | 0.451008921 | 0.639118987 |
| ENSG00000169989 | TIGD4 | 8.130598304 | -0.490225112 | 0.61678851 | -0.794802601 | 0.426728338 | 0.617533943 |
| ENSG00000169991 | IFFO2 | 336.1995999 | 0.574679387 | 0.103790521 | 5.536915898 | 3.08E-08 | 2.10E-07 |
| ENSG00000169992 | NLGN2 | 2107.637194 | -0.05530528 | 0.060308635 | -0.917037509 | 0.359122985 | 0.551682936 |
| ENSG00000170004 | CHD3 | 11096.40768 | -0.144860917 | 0.050937287 | -2.843907213 | 0.004456402 | 0.014539876 |
| ENSG00000170006 | TMEM154 | 483.1667751 | -0.104594552 | 0.093391744 | -1.119955014 | 0.262732933 | 0.443429479 |
| ENSG00000170011 | MYRIP | 73.16384792 | 1.456708105 | 0.219934103 | 6.623384391 | 3.51E-11 | 3.14E-10 |
| ENSG00000170017 | ALCAM | 6334.425683 | 0.018788002 | 0.047595201 | 0.394745732 | 0.693030549 | 0.826907354 |
| ENSG00000170027 | YWHAG | 6055.884616 | 0.514881629 | 0.042962632 | 11.98440604 | 4.29E-33 | 1.35E-31 |
| ENSG00000170035 | UBE2E3 | 829.141728 | 0.262149992 | 0.072702079 | 3.605811515 | 0.000311179 | 0.001277917 |
| ENSG00000170037 | CNTROB | 1299.29145 | -0.312677713 | 0.063903114 | -4.892996528 | 9.93E-07 | 5.75E-06 |
| ENSG00000170043 | TRAPPC1 | 1040.428184 | -0.191990536 | 0.076438135 | -2.511711418 | 0.012014729 | 0.034973338 |
| ENSG00000170049 | KCNAB3 | 86.64555983 | -0.879680064 | 0.199671954 | -4.405626558 | 1.05E-05 | 5.43E-05 |
| ENSG00000170075 | GPR37L1 | 568.7066525 | 1.956143358 | 0.093453487 | 20.9317321 | 2.75E-97 | 4.21E-95 |
| ENSG00000170085 | SIMC1 | 386.9834433 | -0.320778941 | 0.101515241 | -3.159909157 | 0.001578183 | 0.005674534 |
| ENSG00000170088 | TMEM192 | 1136.332525 | 0.279644585 | 0.06453228 | 4.333406225 | 1.47E-05 | 7.41E-05 |
| ENSG00000170089 | ENSG0000017 | 638.4293726 | -0.463806926 | 0.081038763 | -5.72327255 | 1.04E-08 | 7.50E-08 |
| ENSG00000170100 | ZNF778 | 2975.095219 | 0.812215653 | 0.051243163 | 15.85022493 | 1.40E-56 | 9.13E-55 |
| ENSG00000170113 | NIPA1 | 768.4658866 | -0.205695442 | 0.075740998 | -2.715774133 | 0.006612101 | 0.020634131 |
| ENSG00000170122 | FOXD4 | 11.09413697 | 0.015969457 | 0.51853388 | 0.030797325 | 0.975431173 | 0.988740606 |
| ENSG00000170142 | UBE2E1 | 1891.893625 | 0.006296588 | 0.060849076 | 0.103478773 | 0.917582995 | 0.959405374 |
| ENSG00000170144 | HNRNPA3 | 5467.516027 | 0.174751554 | 0.063939401 | 2.733080878 | 0.006274492 | 0.019711977 |
| ENSG00000170145 | SIK2 | 1059.414777 | 0.027392041 | 0.066801773 | 0.410049609 | 0.681769557 | 0.819321553 |
| ENSG00000170153 | RNF150 | 232.1702636 | 0.844026356 | 0.125889746 | 6.704488508 | 2.02E-11 | 1.84E-10 |
| ENSG00000170161 | FAM88B | 104.9487176 | -0.074490012 | 0.179365515 | -0.415297287 | 0.677924287 | 0.816699042 |
| ENSG00000170175 | CHRNB1 | 60.10737984 | -0.667510021 | 0.239752951 | -2.784157677 | 0.005366694 | 0.017153679 |

| | | | | | | | |
|---|---|---|---|---|---|---|---|
| ENSG00000170178 | HOXD12 | 4.037026567 | -0.727278103 | 0.879535631 | -0.826888732 | 0.408300131 | 0.600718638 |
| ENSG00000170185 | USP38 | 1516.793405 | -0.127408367 | 0.058423508 | -2.180772274 | 0.029200264 | 0.075835397 |
| ENSG00000170190 | SLC16A5 | 60.84974754 | -5.262483157 | 0.495917291 | -10.61161458 | 2.63E-26 | 6.24E-25 |
| ENSG00000170191 | NANP | 459.2440013 | -0.331691147 | 0.097702187 | -3.394920381 | 0.000686486 | 0.002650283 |
| ENSG00000170222 | ADPRM | 92.90387204 | 0.375853554 | 0.185629018 | 2.024756466 | 0.042892377 | 0.105156689 |
| ENSG00000170234 | PWWP2A | 724.7033708 | 0.109564002 | 0.076572817 | 1.430847215 | 0.152474008 | 0.295069491 |
| ENSG00000170236 | USP50 | 97.39196257 | -1.546151975 | 0.201240456 | -7.683107081 | 1.55E-14 | 1.80E-13 |
| ENSG00000170242 | USP47 | 1763.481332 | -0.042764382 | 0.058320656 | -0.733263049 | 0.463398007 | 0.649830069 |
| ENSG00000170248 | PDCD6IP | 4243.42848 | -0.017403247 | 0.047699479 | -0.364851921 | 0.715221945 | 0.841653402 |
| ENSG00000170260 | ZNF212 | 629.1839486 | 0.37337475 | 0.07838912 | 4.763094029 | 1.91E-06 | 1.07E-05 |
| ENSG00000170262 | MRAP | 3.613128694 | 0.061499967 | 0.945995283 | 0.06501086 | 0.948165354 | 0.976023831 |
| ENSG00000170264 | FAM161A | 119.4002439 | 0.310393846 | 0.164900587 | 1.882308918 | 0.059794084 | 0.139157583 |
| ENSG00000170265 | ZNF282 | 1874.956757 | 0.474324811 | 0.063485484 | 7.47138998 | 7.94E-14 | 8.71E-13 |
| ENSG00000170266 | GLB1 | 3081.268127 | 0.065069578 | 0.052740796 | 1.233761774 | 0.217291687 | 0.386858464 |
| ENSG00000170270 | GON7 | 455.7178045 | -0.650470519 | 0.093777819 | -6.936293966 | 4.03E-12 | 3.91E-11 |
| ENSG00000170271 | FAXDC2 | 264.1517344 | -1.75195834 | 0.12537386 | -13.97387254 | 2.25E-44 | 1.06E-42 |
| ENSG00000170275 | CRTAP | 6294.122294 | 0.807267237 | 0.043399716 | 18.6007491 | 3.17E-77 | 3.51E-75 |
| ENSG00000170291 | ELP5 | 614.3132345 | -0.246148506 | 0.085907857 | -2.865261873 | 0.004166645 | 0.013670315 |
| ENSG00000170293 | CMTM8 | 167.2227191 | 0.366118946 | 0.139412012 | 2.626165006 | 0.008635295 | 0.02613935 |
| ENSG00000170296 | GABARAP | 131.872207 | 0.045932368 | 0.177506341 | 0.258764661 | 0.795816826 | 0.890886101 |
| ENSG00000170310 | STX8 | 202.3792391 | 0.204138709 | 0.132413532 | 1.541675581 | 0.123152449 | 0.250409251 |
| ENSG00000170312 | CDK1 | 1402.040791 | -0.853863931 | 0.068422432 | -12.47929809 | 9.68E-36 | 3.36E-34 |
| ENSG00000170315 | UBB | 15971.83827 | -0.093032421 | 0.040303344 | -2.308305274 | 0.020982163 | 0.056989205 |
| ENSG00000170322 | NFRKB | 2140.101412 | 0.226312545 | 0.056826822 | 3.982495197 | 6.82E-05 | 0.000312693 |
| ENSG00000170325 | PRDM10 | 573.0270351 | -0.311626579 | 0.08180339 | -3.809458031 | 0.000139272 | 0.000606587 |
| ENSG00000170340 | B3GNT2 | 428.2421437 | 0.25860288 | 0.093079667 | 2.77829614 | 0.005464479 | 0.017417695 |
| ENSG00000170345 | FOS | 182.5687744 | -0.524548153 | 0.13394847 | -3.916044389 | 9.00E-05 | 0.000404489 |
| ENSG00000170348 | TMED10 | 8620.30834 | -0.577673065 | 0.046532893 | -12.41429508 | 2.19E-35 | 7.53E-34 |
| ENSG00000170356 | OR2A20P | 16.3519918 | 0.047976652 | 0.439742596 | 0.109101671 | 0.91312185 | 0.956615757 |
| ENSG00000170364 | SETMAR | 451.747908 | 0.007436525 | 0.08931811 | 0.083258868 | 0.933645705 | 0.968138388 |
| ENSG00000170365 | SMAD1 | 583.1737471 | 0.244280446 | 0.082774556 | 2.951153807 | 0.003165892 | 0.010673952 |
| ENSG00000170374 | SP7 | 1.977741571 | 1.346690038 | 1.324055861 | 1.017094578 | 0.309108439 | 0.49719167 |
| ENSG00000170379 | TCAF2 | 64.30301712 | -0.973707871 | 0.228183416 | -4.267215764 | 1.98E-05 | 9.83E-05 |

| | | | | | | | |
|---|---|---|---|---|---|---|---|
| ENSG00000170381 | SEMA3E | 190.705883 | 0.421898345 | 0.131150945 | 3.216891398 | 0.001295876 | 0.004751064 |
| ENSG00000170382 | LRRN2 | 181.9018859 | 1.337853775 | 0.158428726 | 8.444515162 | 3.05E-17 | 4.26E-16 |
| ENSG00000170385 | SLC30A1 | 1315.293807 | -0.147159442 | 0.073406786 | -2.004711676 | 0.04499388 | 0.109420078 |
| ENSG00000170390 | DCLK2 | 16.22411726 | 1.635491573 | 0.478057944 | 3.421115776 | 0.000623648 | 0.002424827 |
| ENSG00000170396 | ZNF804A | 32.77305522 | -1.183025605 | 0.326979872 | -3.618038013 | 0.000296845 | 0.001223212 |
| ENSG00000170412 | GPRC5C | 764.8544673 | 0.633742534 | 0.09102633 | 6.962189259 | 3.35E-12 | 3.28E-11 |
| ENSG00000170417 | TMEM182 | 49.65637205 | -0.040611703 | 0.255459896 | -0.158974866 | 0.873688676 | 0.934676032 |
| ENSG00000170419 | VSTM2A | 12.49416718 | 2.158388326 | 0.570342745 | 3.784370615 | 0.000154098 | 0.000667109 |
| ENSG00000170421 | KRT8 | 69936.73398 | 0.120600691 | 0.043037081 | 2.802250683 | 0.005074742 | 0.016327432 |
| ENSG00000170425 | ADORA2B | 79.64680334 | -0.169148667 | 0.202059741 | -0.837122062 | 0.40252396 | 0.595236231 |
| ENSG00000170430 | MGMT | 732.3391388 | -0.595141427 | 0.080333355 | -7.408397517 | 1.28E-13 | 1.38E-12 |
| ENSG00000170439 | METTL7B | 92.4513378 | 0.170972395 | 0.186991833 | 0.914330815 | 0.360543043 | 0.553086626 |
| ENSG00000170442 | KRT86 | 393.5141185 | 1.153532394 | 0.109990581 | 10.4875561 | 9.85E-26 | 2.26E-24 |
| ENSG00000170445 | HARS1 | 2833.905166 | 0.039821353 | 0.052304001 | 0.761344307 | 0.446451443 | 0.634987357 |
| ENSG00000170448 | NFXL1 | 686.0986655 | -0.068292331 | 0.085615837 | -0.797660031 | 0.425067804 | 0.616024728 |
| ENSG00000170456 | DENND5B | 319.7863742 | 0.437055886 | 0.1055497 | 4.140759147 | 3.46E-05 | 0.000165746 |
| ENSG00000170458 | CD14 | 3.113849179 | 0.9281487 | 1.009696642 | 0.919235205 | 0.357972561 | 0.550321039 |
| ENSG00000170464 | DNAJC18 | 204.2376981 | 0.716406775 | 0.139162365 | 5.147992228 | 2.63E-07 | 1.64E-06 |
| ENSG00000170468 | RIOX1 | 707.9521024 | -0.355872787 | 0.082580526 | -4.309403257 | 1.64E-05 | 8.21E-05 |
| ENSG00000170469 | SPATA24 | 32.95793739 | 0.707235046 | 0.315809077 | 2.23943863 | 0.02512739 | 0.06676191 |
| ENSG00000170471 | RALGAPB | 2041.714646 | -0.559187702 | 0.054567148 | -10.24769898 | 1.21E-24 | 2.65E-23 |
| ENSG00000170473 | PYM1 | 1130.288434 | -0.242287515 | 0.069363332 | -3.493020157 | 0.00047759 | 0.00189713 |
| ENSG00000170476 | MZB1 | 17.70564666 | -1.091632976 | 0.460409036 | -2.371006845 | 0.017739703 | 0.049289536 |
| ENSG00000170482 | SLC23A1 | 11.1311669 | -0.629318172 | 0.537523372 | -1.170773597 | 0.241689793 | 0.417816067 |
| ENSG00000170485 | NPAS2 | 222.5832621 | 2.148176254 | 0.141328185 | 15.19991399 | 3.54E-52 | 2.07E-50 |
| ENSG00000170498 | KISS1 | 12.16135376 | 0.662999304 | 0.516024099 | 1.284822366 | 0.198854365 | 0.361612204 |
| ENSG00000170500 | LONRF2 | 2066.954493 | 0.378093661 | 0.05726531 | 6.602490447 | 4.04E-11 | 3.60E-10 |
| ENSG00000170502 | NUDT9 | 803.8224546 | -0.208911356 | 0.071766413 | -2.910990646 | 0.003602848 | 0.011995549 |
| ENSG00000170509 | HSD17B13 | 3.428694867 | 0.209844514 | 0.938749474 | 0.223536226 | 0.823118192 | 0.906409914 |
| ENSG00000170515 | PA2G4 | 4138.575829 | 0.003032437 | 0.046743793 | 0.064873582 | 0.948274655 | 0.976058715 |
| ENSG00000170522 | ELOVL6 | 3354.919627 | -0.602569254 | 0.056392037 | -10.68536073 | 1.19E-26 | 2.90E-25 |
| ENSG00000170523 | KRT83 | 18.2767987 | 0.876477815 | 0.426668236 | 2.054237322 | 0.039952728 | 0.099029236 |
| ENSG00000170525 | PFKFB3 | 1015.304984 | -0.821171913 | 0.068536211 | -11.98157735 | 4.44E-33 | 1.39E-31 |

| | | | | | | | |
|---|---|---|---|---|---|---|---|
| ENSG00000170537 | TMC7 | 150.2323213 | 0.256011962 | 0.147515882 | 1.735487455 | 0.082654502 | 0.181427066 |
| ENSG00000170540 | ARL6IP1 | 4957.393758 | -0.066776273 | 0.051220434 | -1.303703774 | 0.192334602 | 0.352473834 |
| ENSG00000170542 | SERPINB9 | 42.25169042 | -1.617815157 | 0.29835794 | -5.422396852 | 5.88E-08 | 3.89E-07 |
| ENSG00000170545 | SMAGP | 393.269903 | 0.414253072 | 0.101471481 | 4.082458109 | 4.46E-05 | 0.000209864 |
| ENSG00000170558 | CDH2 | 240.6532984 | 1.236683537 | 0.125174876 | 9.879646592 | 5.10E-23 | 1.02E-21 |
| ENSG00000170561 | IRX2 | 392.4257232 | 0.281688593 | 0.09969402 | 2.825531487 | 0.004720225 | 0.015307034 |
| ENSG00000170571 | EMB | 1756.722701 | 0.34812926 | 0.071428979 | 4.873781844 | 1.09E-06 | 6.31E-06 |
| ENSG00000170577 | SIX2 | 583.8181874 | -1.08211532 | 0.085606058 | -12.64063954 | 1.26E-36 | 4.52E-35 |
| ENSG00000170579 | DLGAP1 | 23.81547045 | -1.73877538 | 0.419603201 | -4.143856334 | 3.42E-05 | 0.000163693 |
| ENSG00000170581 | STAT2 | 1648.048153 | -0.082341743 | 0.060775078 | -1.354860335 | 0.175462059 | 0.328705402 |
| ENSG00000170584 | NUDCD2 | 609.1969126 | -0.062986826 | 0.081897391 | -0.769094423 | 0.441837259 | 0.631129174 |
| ENSG00000170604 | IRF2BP1 | 858.6118651 | -0.042946602 | 0.081436405 | -0.527363672 | 0.597941066 | 0.758335253 |
| ENSG00000170606 | HSPA4 | 6752.104075 | 0.138187873 | 0.042627051 | 3.241788255 | 0.001187822 | 0.004386453 |
| ENSG00000170608 | FOXA3 | 2.450453717 | -0.344028496 | 1.102637587 | -0.31200505 | 0.755036685 | 0.864705683 |
| ENSG00000170619 | COMMD5 | 1055.615584 | 0.312547343 | 0.072776814 | 4.294600497 | 1.75E-05 | 8.75E-05 |
| ENSG00000170629 | DPY19L2P2 | 22.46442563 | -3.346256582 | 0.503608446 | -6.644560091 | 3.04E-11 | 2.74E-10 |
| ENSG00000170631 | ZNF16 | 543.7600799 | 0.15973059 | 0.087126834 | 1.833311091 | 0.066756321 | 0.152650518 |
| ENSG00000170632 | ARMC10 | 753.5380679 | -0.259320164 | 0.084958548 | -3.052313984 | 0.002270844 | 0.007900086 |
| ENSG00000170633 | RNF34 | 1018.507318 | -0.265800504 | 0.067408844 | -3.943110227 | 8.04E-05 | 0.000364861 |
| ENSG00000170634 | ACYP2 | 203.6454797 | 0.261235909 | 0.129041982 | 2.024425738 | 0.042926366 | 0.105196467 |
| ENSG00000170638 | TRABD | 802.2460208 | 0.197178384 | 0.073730456 | 2.674313897 | 0.007488232 | 0.023051537 |
| ENSG00000170653 | ATF7 | 1137.626216 | 0.044529809 | 0.065391947 | 0.680967785 | 0.495891875 | 0.677941809 |
| ENSG00000170667 | RASA4B | 34.03360201 | 0.310955703 | 0.325276568 | 0.955973264 | 0.339085742 | 0.530240418 |
| ENSG00000170677 | SOCS6 | 580.5645532 | 0.800015817 | 0.082736626 | 9.669427675 | 4.07E-22 | 7.65E-21 |
| ENSG00000170681 | CAVIN4 | 18.14673375 | 0.275035856 | 0.418099239 | 0.657824339 | 0.510651013 | 0.690540806 |
| ENSG00000170684 | ZNF296 | 83.92053805 | 0.493557123 | 0.199307967 | 2.476354209 | 0.013273182 | 0.038161023 |
| ENSG00000170734 | POLH | 686.5438705 | -0.043637857 | 0.075895097 | -0.574975964 | 0.565307553 | 0.733481521 |
| ENSG00000170745 | KCNS3 | 556.8987601 | 2.452169167 | 0.099219545 | 24.7145779 | 7.46E-135 | 2.28E-132 |
| ENSG00000170759 | KIF5B | 5407.041839 | -0.590334964 | 0.048318078 | -12.21768317 | 2.50E-34 | 8.28E-33 |
| ENSG00000170775 | GPR37 | 732.0571726 | 2.537935765 | 0.100829086 | 25.1706712 | 8.39E-140 | 2.71E-137 |
| ENSG00000170776 | AKAP13 | 2232.740693 | -0.156255893 | 0.074293291 | -2.103230205 | 0.035445651 | 0.089560881 |
| ENSG00000170779 | CDCA4 | 1357.205357 | -0.221522913 | 0.064788119 | -3.419190358 | 0.000628078 | 0.00244081 |
| ENSG00000170786 | SDR16C5 | 7.270794847 | 0.375790414 | 0.672794036 | 0.55855194 | 0.576467548 | 0.74273769 |

| | | | | | | | |
|---|---|---|---|---|---|---|---|
| ENSG00000170791 | CHCHD7 | 348.6667898 | -0.280160527 | 0.100179586 | -2.796582989 | 0.005164615 | 0.016579381 |
| ENSG00000170802 | FOXN2 | 321.3907912 | -0.04016692 | 0.109683439 | -0.366207697 | 0.714210099 | 0.840989046 |
| ENSG00000170819 | BFSP2 | 17.06733029 | -0.270854029 | 0.420089703 | -0.644752841 | 0.519087367 | 0.696692266 |
| ENSG00000170832 | USP32 | 19778.97099 | 0.168328747 | 0.046678553 | 3.606126074 | 0.000310802 | 0.001276598 |
| ENSG00000170835 | CEL | 45.37570393 | 0.418236777 | 0.263690287 | 1.586090946 | 0.11271868 | 0.233477791 |
| ENSG00000170836 | PPM1D | 6847.496 | 0.359572132 | 0.04333471 | 8.2975548 | 1.06E-16 | 1.44E-15 |
| ENSG00000170837 | GPR27 | 315.1080646 | -0.015357522 | 0.108720055 | -0.141257486 | 0.887666534 | 0.9421241 |
| ENSG00000170846 | ENSG0000017 | 1820.857982 | -0.006952284 | 0.05625992 | -0.123574376 | 0.901652282 | 0.950420022 |
| ENSG00000170852 | KBTBD2 | 2749.753787 | -0.132397494 | 0.058975687 | -2.244950438 | 0.024771314 | 0.065953337 |
| ENSG00000170854 | RIOX2 | 953.7338185 | 0.166996977 | 0.068428435 | 2.440461742 | 0.0146685 | 0.041660936 |
| ENSG00000170855 | TRIAP1 | 994.7051864 | -0.009819106 | 0.068135103 | -0.1441123 | 0.885411792 | 0.940732088 |
| ENSG00000170860 | LSM3 | 1815.718321 | -0.012115966 | 0.059796565 | -0.202619766 | 0.839432239 | 0.915367 |
| ENSG00000170871 | KIAA0232 | 2060.042244 | 0.148266535 | 0.057262983 | 2.589221282 | 0.009619326 | 0.028749972 |
| ENSG00000170873 | MTSS1 | 338.9580911 | 0.204027151 | 0.101883177 | 2.002559767 | 0.045224562 | 0.109899518 |
| ENSG00000170876 | TMEM43 | 3893.450871 | 0.198404267 | 0.045667792 | 4.344511921 | 1.40E-05 | 7.06E-05 |
| ENSG00000170881 | RNF139 | 2357.214022 | -0.526585232 | 0.064396731 | -8.177204427 | 2.91E-16 | 3.83E-15 |
| ENSG00000170889 | RPS9 | 9608.994686 | 0.254149601 | 0.043200018 | 5.883090228 | 4.03E-09 | 3.03E-08 |
| ENSG00000170892 | TSEN34 | 2494.953916 | 0.114619126 | 0.055720922 | 2.057021337 | 0.039684174 | 0.098544592 |
| ENSG00000170899 | GSTA4 | 109.4700944 | -0.833257221 | 0.173215902 | -4.810512257 | 1.51E-06 | 8.51E-06 |
| ENSG00000170903 | MSANTD4 | 366.4288403 | 0.085244568 | 0.103670544 | 0.822264114 | 0.410926596 | 0.603383225 |
| ENSG00000170906 | NDUFA3 | 1043.764578 | 0.065109373 | 0.072849633 | 0.89375018 | 0.37145557 | 0.564018453 |
| ENSG00000170909 | OSCAR | 15.74934592 | 0.680595801 | 0.455926525 | 1.492775181 | 0.135496046 | 0.269933069 |
| ENSG00000170915 | PAQR8 | 174.4202206 | 0.006395245 | 0.13831195 | 0.046237833 | 0.963120688 | 0.983178092 |
| ENSG00000170917 | NUDT6 | 209.6233388 | -0.390962112 | 0.135374476 | -2.88800463 | 0.003876942 | 0.012815136 |
| ENSG00000170919 | TPT1-AS1 | 170.9372172 | 0.38850837 | 0.140986507 | 2.755642217 | 0.005857703 | 0.01852947 |
| ENSG00000170921 | TANC2 | 9200.274848 | 0.721849513 | 0.11138699 | 6.480554997 | 9.14E-11 | 7.94E-10 |
| ENSG00000170935 | NCBP2L | 15.25785466 | -1.065509197 | 0.456172713 | -2.33575829 | 0.019503843 | 0.053506117 |
| ENSG00000170946 | DNAJC24 | 245.6449883 | -0.073009718 | 0.126564164 | -0.576859322 | 0.564034492 | 0.73258698 |
| ENSG00000170949 | ZNF160 | 1116.565863 | 0.154603222 | 0.064718719 | 2.388848605 | 0.016901265 | 0.047230482 |
| ENSG00000170950 | PGK2 | 2.232932151 | 0.330255095 | 1.219467839 | 0.270819028 | 0.786530224 | 0.884114237 |
| ENSG00000170965 | PLAC1 | 6.586982861 | 0.916672472 | 0.695159746 | 1.318650105 | 0.187286115 | 0.345138505 |
| ENSG00000171004 | HS6ST2 | 484.5319699 | -2.245131272 | 0.1076053 | -20.86450463 | 1.13E-96 | 1.70E-94 |
| ENSG00000171016 | PYGO1 | 20.96334347 | 0.193433831 | 0.384412936 | 0.503192825 | 0.614828709 | 0.771600111 |

| | | | | | | | |
|---|---|---|---|---|---|---|---|
| ENSG00000171017 | LRRC8E | 543.4341981 | -0.149447089 | 0.085159964 | -1.754898446 | 0.07927668 | 0.175490402 |
| ENSG00000171033 | PKIA | 250.9353662 | -0.33925717 | 0.118522788 | -2.862379251 | 0.004204734 | 0.013787394 |
| ENSG00000171045 | TSNARE1 | 438.8724225 | -0.009943003 | 0.099103527 | -0.100329462 | 0.920082769 | 0.961195531 |
| ENSG00000171055 | FEZ2 | 935.4694588 | 0.380048821 | 0.072304776 | 5.256206318 | 1.47E-07 | 9.40E-07 |
| ENSG00000171056 | SOX7 | 4.730821633 | 0.731703857 | 0.809272179 | 0.904150514 | 0.365915596 | 0.558601436 |
| ENSG00000171067 | C11orf24 | 808.7940271 | -0.057980666 | 0.071788703 | -0.80765725 | 0.41928792 | 0.610870102 |
| ENSG00000171084 | FAM86JP | 237.2421183 | -0.840273877 | 0.12668508 | -6.632776945 | 3.29E-11 | 2.96E-10 |
| ENSG00000171094 | ALK | 1.591464773 | 2.854039207 | 1.638389885 | 1.741978044 | 0.081512286 | NA |
| ENSG00000171097 | KYAT1 | 3.971877657 | -0.653889126 | 0.87272767 | -0.749247616 | 0.453707975 | 0.641379041 |
| ENSG00000171100 | MTM1 | 471.3411686 | -0.633140316 | 0.08883896 | -7.126831671 | 1.03E-12 | 1.05E-11 |
| ENSG00000171103 | TRMT61B | 393.0881396 | 0.167622931 | 0.097902183 | 1.712147019 | 0.086869578 | 0.188961514 |
| ENSG00000171105 | INSR | 1229.019471 | 1.157605745 | 0.068289718 | 16.95139147 | 1.88E-64 | 1.50E-62 |
| ENSG00000171109 | MFN1 | 1428.698301 | -0.144980574 | 0.064061783 | -2.263136728 | 0.023627262 | 0.063281891 |
| ENSG00000171116 | HSFX1 | 10.59997545 | 0.436048875 | 0.565569395 | 0.770990932 | 0.440712306 | 0.630020584 |
| ENSG00000171119 | NRTN | 9.009083563 | -0.730508902 | 0.599176126 | -1.219188935 | 0.222772491 | 0.393868113 |
| ENSG00000171121 | KCNMB3 | 100.8935542 | 0.050539209 | 0.184778866 | 0.273511848 | 0.784459786 | 0.883041734 |
| ENSG00000171124 | FUT3 | 2.598974109 | 0.555458048 | 1.105543761 | 0.502429725 | 0.615365274 | 0.772067347 |
| ENSG00000171126 | KCNG3 | 2.272755063 | -0.904108776 | 1.187641351 | -0.761264144 | 0.446499313 | 0.634997544 |
| ENSG00000171130 | ATP6V0E2 | 2873.068388 | 0.439962178 | 0.060722075 | 7.245506334 | 4.31E-13 | 4.50E-12 |
| ENSG00000171132 | PRKCE | 441.6114846 | 0.340172241 | 0.092721676 | 3.668745603 | 0.000243743 | 0.001017557 |
| ENSG00000171135 | JAGN1 | 1608.397405 | -0.029794489 | 0.061730361 | -0.482655343 | 0.629340476 | 0.7810577 |
| ENSG00000171148 | TADA3 | 5377.958015 | -0.40643969 | 0.05212294 | -7.797712358 | 6.30E-15 | 7.52E-14 |
| ENSG00000171150 | SOCS5 | 491.6677049 | 0.049076553 | 0.087881177 | 0.55844215 | 0.576542497 | 0.74273769 |
| ENSG00000171155 | C1GALT1C1 | 728.4241894 | -0.176734388 | 0.075830638 | -2.330646206 | 0.019772023 | 0.054170573 |
| ENSG00000171159 | BBLN | 1308.970637 | 0.390785867 | 0.085991217 | 4.544485829 | 5.51E-06 | 2.93E-05 |
| ENSG00000171160 | MORN4 | 339.069484 | -0.379962406 | 0.105311403 | -3.607989209 | 0.000308579 | 0.001268604 |
| ENSG00000171161 | ZNF672 | 892.2026592 | -0.50096677 | 0.083775348 | -5.97988289 | 2.23E-09 | 1.72E-08 |
| ENSG00000171163 | ZNF692 | 1095.829296 | -0.216479534 | 0.074103557 | -2.921310956 | 0.003485617 | 0.011647496 |
| ENSG00000171169 | NAIF1 | 271.0352264 | -0.222798657 | 0.112190959 | -1.985887806 | 0.047045783 | 0.113639072 |
| ENSG00000171174 | RBKS | 167.3028473 | -0.149935607 | 0.144877736 | -1.034911311 | 0.300710332 | 0.487482358 |
| ENSG00000171189 | GRIK1 | 4.714027646 | -0.845499562 | 0.837053645 | -1.010090055 | 0.312452146 | 0.500674227 |
| ENSG00000171202 | TMEM126A | 259.2757133 | 0.093587798 | 0.117001635 | 0.79988453 | 0.423777702 | 0.614665 |
| ENSG00000171204 | TMEM126B | 652.6862969 | 0.112488013 | 0.080694272 | 1.394002448 | 0.163316865 | 0.310886142 |

| | | | | | | | |
|---|---|---|---|---|---|---|---|
| ENSG00000171206 | TRIM8 | 1618.396251 | 0.500768357 | 0.058007364 | 8.63284106 | 5.98E-18 | 8.77E-17 |
| ENSG00000171208 | NETO2 | 831.3213162 | 0.188853748 | 0.073242716 | 2.578464573 | 0.009924048 | 0.029527581 |
| ENSG00000171219 | CDC42BPG | 2017.264406 | 0.327079057 | 0.067283791 | 4.86118654 | 1.17E-06 | 6.70E-06 |
| ENSG00000171222 | SCAND1 | 536.4472508 | -0.491012644 | 0.101594121 | -4.833081297 | 1.34E-06 | 7.65E-06 |
| ENSG00000171223 | JUNB | 1095.900883 | 0.112590733 | 0.075469965 | 1.491861472 | 0.135735465 | 0.270316135 |
| ENSG00000171224 | FAM241B | 468.182099 | -0.685563201 | 0.091628359 | -7.481998036 | 7.32E-14 | 8.06E-13 |
| ENSG00000171227 | TMEM37 | 18.30705246 | -0.645430961 | 0.41122176 | -1.569544765 | 0.116521057 | 0.239923979 |
| ENSG00000171236 | LRG1 | 35.235931 | -0.577688886 | 0.303554732 | -1.903079824 | 0.05703013 | 0.133822503 |
| ENSG00000171241 | SHCBP1 | 653.0088242 | -0.18665179 | 0.085920518 | -2.172377379 | 0.02982721 | 0.077294294 |
| ENSG00000171246 | NPTX1 | 14.91354017 | -0.820032515 | 0.457359338 | -1.79297206 | 0.072977392 | 0.164267221 |
| ENSG00000171262 | FAM98B | 1142.711554 | -0.064375996 | 0.065224972 | -0.986983882 | 0.32365054 | 0.513738126 |
| ENSG00000171291 | ZNF439 | 4.410761115 | -0.009297769 | 0.846204725 | -0.010987612 | 0.991233331 | 0.995224641 |
| ENSG00000171295 | ZNF440 | 283.4300432 | -0.089028757 | 0.113776639 | -0.782487143 | 0.433928339 | 0.623821275 |
| ENSG00000171298 | GAA | 5550.713712 | 0.173761364 | 0.054489666 | 3.188886572 | 0.001428219 | 0.005175066 |
| ENSG00000171302 | CANT1 | 3668.276497 | 0.125022063 | 0.054690339 | 2.285999059 | 0.022254314 | 0.0600534 |
| ENSG00000171307 | ZDHHC16 | 2468.335235 | -0.774048054 | 0.058826153 | -13.15823015 | 1.53E-39 | 6.05E-38 |
| ENSG00000171310 | CHST11 | 300.0901579 | 0.880517565 | 0.111773561 | 7.877690907 | 3.33E-15 | 4.06E-14 |
| ENSG00000171311 | EXOSC1 | 1018.344555 | -0.766392627 | 0.065897086 | -11.63014445 | 2.90E-31 | 8.47E-30 |
| ENSG00000171314 | PGAM1 | 1822.442587 | -0.64239805 | 0.06297278 | -10.20120196 | 1.96E-24 | 4.22E-23 |
| ENSG00000171316 | CHD7 | 1795.808352 | 0.314075271 | 0.067186618 | 4.674670041 | 2.94E-06 | 1.62E-05 |
| ENSG00000171320 | ESCO2 | 371.4473241 | -0.070464404 | 0.104631717 | -0.673451661 | 0.500659992 | 0.681732072 |
| ENSG00000171345 | KRT19 | 30798.55746 | -0.045012409 | 0.042179031 | -1.067175039 | 0.285892802 | 0.469663483 |
| ENSG00000171346 | KRT15 | 330.5313418 | 1.364965477 | 0.113692066 | 12.00581119 | 3.31E-33 | 1.04E-31 |
| ENSG00000171357 | LURAP1 | 8.353930577 | 0.374255102 | 0.613192791 | 0.610338392 | 0.54163767 | 0.715650991 |
| ENSG00000171365 | CLCN5 | 353.0542181 | 0.945043921 | 0.107682059 | 8.776243055 | 1.69E-18 | 2.58E-17 |
| ENSG00000171368 | TPPP | 31.57584352 | -0.064558693 | 0.309919877 | -0.208307686 | 0.83498873 | 0.913001315 |
| ENSG00000171385 | KCND3 | 41.55403874 | 1.436892143 | 0.301834719 | 4.760526388 | 1.93E-06 | 1.08E-05 |
| ENSG00000171388 | APLN | 33.88653972 | 0.399173287 | 0.302990978 | 1.317442817 | 0.187690237 | 0.345721377 |
| ENSG00000171401 | KRT13 | 8.1397675 | 0.824303913 | 0.642377507 | 1.283207933 | 0.199419237 | 0.362321979 |
| ENSG00000171408 | PDE7B | 39.56036468 | -1.445644141 | 0.292833476 | -4.936744796 | 7.94E-07 | 4.67E-06 |
| ENSG00000171421 | MRPL36 | 803.6120847 | 0.058362554 | 0.075990557 | 0.768023761 | 0.442473069 | 0.631470633 |
| ENSG00000171425 | ZNF581 | 741.6035999 | -0.002462846 | 0.082658398 | -0.029795466 | 0.976230175 | 0.98920978 |
| ENSG00000171428 | NAT1 | 304.9919921 | 0.468075252 | 0.109181567 | 4.287127062 | 1.81E-05 | 9.04E-05 |

| | | | | | | | |
|---|---|---|---|---|---|---|---|
| ENSG00000171431 | KRT20 | 55.85363228 | 1.47245837 | 0.254672024 | 5.781782975 | 7.39E-09 | 5.38E-08 |
| ENSG00000171433 | GLOD5 | 4.962868806 | 0.274253709 | 0.819370988 | 0.334712497 | 0.737841974 | 0.855509243 |
| ENSG00000171435 | KSR2 | 380.312979 | -0.006853859 | 0.108475959 | -0.063183207 | 0.949620617 | 0.97669911 |
| ENSG00000171443 | ZNF524 | 218.1752324 | 0.556120069 | 0.143225627 | 3.882825155 | 0.00010325 | 0.000459199 |
| ENSG00000171444 | MCC | 1145.434834 | 0.974755404 | 0.070208706 | 13.88368283 | 7.95E-44 | 3.69E-42 |
| ENSG00000171448 | ZBTB26 | 389.7781621 | 0.338918634 | 0.096747618 | 3.503121223 | 0.00045984 | 0.001832013 |
| ENSG00000171450 | CDK5R2 | 4.07305662 | 0.450242703 | 0.908175753 | 0.495766047 | 0.620059482 | 0.775116589 |
| ENSG00000171451 | DSEL | 401.914765 | 0.865732521 | 0.101478078 | 8.531227004 | 1.45E-17 | 2.07E-16 |
| ENSG00000171453 | POLR1C | 827.6527604 | 0.128854717 | 0.073167364 | 1.761095518 | 0.078222238 | 0.173568925 |
| ENSG00000171456 | ASXL1 | 2775.068953 | -0.376661969 | 0.051654059 | -7.292011094 | 3.05E-13 | 3.23E-12 |
| ENSG00000171462 | DLK2 | 16.97607472 | -2.090941907 | 0.485085525 | -4.310460322 | 1.63E-05 | 8.18E-05 |
| ENSG00000171466 | ZNF562 | 1120.381249 | 0.274212329 | 0.064241532 | 4.268458726 | 1.97E-05 | 9.78E-05 |
| ENSG00000171467 | ZNF318 | 1846.937599 | -0.11580576 | 0.058456665 | -1.981053154 | 0.047585313 | 0.114821403 |
| ENSG00000171469 | ZNF561 | 1272.928062 | 0.215576014 | 0.062390762 | 3.455255339 | 0.000549772 | 0.002159919 |
| ENSG00000171475 | WIPF2 | 1643.274199 | -0.05055218 | 0.059789685 | -0.845500025 | 0.397831719 | 0.590849387 |
| ENSG00000171476 | HOPX | 248.2453513 | -2.023969835 | 0.133898048 | -15.11575309 | 1.28E-51 | 7.33E-50 |
| ENSG00000171488 | LRRC8C | 108.147867 | -0.354132303 | 0.170643894 | -2.075270876 | 0.037961437 | 0.095007131 |
| ENSG00000171490 | RSL1D1 | 6869.89913 | 0.317726877 | 0.046026742 | 6.903092953 | 5.09E-12 | 4.90E-11 |
| ENSG00000171492 | LRRC8D | 956.6642981 | 0.325194088 | 0.069235564 | 4.696922614 | 2.64E-06 | 1.46E-05 |
| ENSG00000171497 | PPID | 739.143229 | -0.017356213 | 0.075115759 | -0.23105955 | 0.817268539 | 0.903254867 |
| ENSG00000171502 | COL24A1 | 17.15194199 | -1.230655863 | 0.443202431 | -2.776735364 | 0.005490786 | 0.017484542 |
| ENSG00000171503 | ETFDH | 331.8087814 | -0.051133908 | 0.102736783 | -0.49771763 | 0.61868308 | 0.773944261 |
| ENSG00000171517 | LPAR3 | 2.338019964 | 4.572297092 | 1.538572278 | 2.971779199 | 0.002960795 | 0.010041491 |
| ENSG00000171522 | PTGER4 | 848.4904203 | -2.514074987 | 0.08238944 | -30.51452933 | 1.67E-204 | 1.07E-201 |
| ENSG00000171530 | TBCA | 1714.845792 | 0.079936159 | 0.057957884 | 1.379211137 | 0.167829665 | 0.317736446 |
| ENSG00000171533 | MAP6 | 1.573490427 | 2.873825708 | 1.667853126 | 1.723068814 | 0.084876082 | NA |
| ENSG00000171540 | OTP | 2.799456464 | 0.667111223 | 1.051496405 | 0.634439851 | 0.525793809 | 0.701792633 |
| ENSG00000171552 | BCL2L1 | 3441.04742 | -0.318622324 | 0.057866769 | -5.506136461 | 3.67E-08 | 2.49E-07 |
| ENSG00000171566 | PLRG1 | 1238.958675 | 0.067128736 | 0.063330993 | 1.05996657 | 0.289159808 | 0.473369443 |
| ENSG00000171574 | ZNF584 | 539.852981 | -0.470570385 | 0.086576303 | -5.435325482 | 5.47E-08 | 3.64E-07 |
| ENSG00000171587 | DSCAM | 211.033241 | 0.278429817 | 0.133394018 | 2.087273634 | 0.036863407 | 0.092632988 |
| ENSG00000171603 | CLSTN1 | 6599.485041 | 0.281526476 | 0.052628388 | 5.349327388 | 8.83E-08 | 5.75E-07 |
| ENSG00000171604 | CXXC5 | 2163.18378 | 0.982341988 | 0.060986678 | 16.10748488 | 2.26E-58 | 1.54E-56 |

| | | | | | | | |
|---|---|---|---|---|---|---|---|
| ENSG00000171606 | ZNF274 | 745.4875348 | 0.263275571 | 0.074842964 | 3.517706349 | 0.000435294 | 0.001741782 |
| ENSG00000171608 | PIK3CD | 50.88122453 | -1.010822641 | 0.267507248 | -3.778673841 | 0.000157666 | 0.000680627 |
| ENSG00000171612 | SLC25A33 | 188.2245883 | 0.438450928 | 0.135993827 | 3.22405022 | 0.001263912 | 0.004641292 |
| ENSG00000171617 | ENC1 | 2044.307885 | 0.458892136 | 0.054952372 | 8.350724792 | 6.78E-17 | 9.31E-16 |
| ENSG00000171621 | SPSB1 | 410.0032116 | 1.303859404 | 0.100974954 | 12.91270111 | 3.82E-38 | 1.45E-36 |
| ENSG00000171631 | P2RY6 | 483.0346062 | -3.904643707 | 0.134021514 | -29.13445443 | 1.31E-186 | 6.70E-184 |
| ENSG00000171634 | BPTF | 8454.063091 | -0.001682546 | 0.057680873 | -0.029169917 | 0.976729074 | 0.989278333 |
| ENSG00000171680 | PLEKHG5 | 602.0104776 | -0.018471673 | 0.084220926 | -0.219324035 | 0.82639764 | 0.908407776 |
| ENSG00000171681 | ATF7IP | 1892.475568 | 0.020591636 | 0.055425754 | 0.371517472 | 0.710252145 | 0.83868761 |
| ENSG00000171695 | LKAAEAR1 | 1.657544406 | -2.130539903 | 1.520089619 | -1.401588352 | 0.161038207 | NA |
| ENSG00000171700 | RGS19 | 636.3264256 | -0.206513549 | 0.096568554 | -2.138517567 | 0.03247476 | 0.083203554 |
| ENSG00000171703 | TCEA2 | 2024.104905 | -0.365686048 | 0.060259519 | -6.068519209 | 1.29E-09 | 1.02E-08 |
| ENSG00000171714 | ANO5 | 19.09879466 | -1.70875273 | 0.4305282 | -3.968968185 | 7.22E-05 | 0.000329797 |
| ENSG00000171720 | HDAC3 | 1817.29497 | 0.166507513 | 0.05644649 | 2.949829333 | 0.003179495 | 0.01071037 |
| ENSG00000171723 | GPHN | 1012.991627 | -0.610085282 | 0.06996595 | -8.71974553 | 2.79E-18 | 4.19E-17 |
| ENSG00000171724 | VAT1L | 1009.721763 | 0.080990588 | 0.079378186 | 1.020312904 | 0.307580085 | 0.495493293 |
| ENSG00000171729 | TMEM51 | 162.5917978 | 0.232363764 | 0.156300646 | 1.486646212 | 0.137108276 | 0.272199381 |
| ENSG00000171735 | CAMTA1 | 727.0446996 | -0.111144706 | 0.075611053 | -1.46995317 | 0.141574438 | 0.278965607 |
| ENSG00000171747 | LGALS4 | 5.292377409 | -0.127462326 | 0.811024473 | -0.157162121 | 0.87511708 | 0.935208508 |
| ENSG00000171759 | PAH | 24.46221997 | 3.032105224 | 0.479609762 | 6.322025666 | 2.58E-10 | 2.16E-09 |
| ENSG00000171763 | SPATA5L1 | 275.7996868 | -0.246402402 | 0.112129872 | -2.197473307 | 0.027986661 | 0.07318901 |
| ENSG00000171766 | GATM | 8.423665949 | -0.219714021 | 0.63019009 | -0.348647217 | 0.727354175 | 0.848598564 |
| ENSG00000171777 | RASGRP4 | 92.46858019 | -0.259740345 | 0.194185548 | -1.337588441 | 0.181030634 | 0.336455523 |
| ENSG00000171786 | NHLH1 | 3.063640542 | -0.024639248 | 0.98976953 | -0.024893925 | 0.980139573 | 0.991041209 |
| ENSG00000171790 | SLFNL1 | 7.59792642 | 0.683938314 | 0.645896371 | 1.058897904 | 0.289646279 | 0.473962882 |
| ENSG00000171791 | BCL2 | 3100.21199 | 0.943052366 | 0.054357542 | 17.3490619 | 2.00E-67 | 1.70E-65 |
| ENSG00000171792 | RHNO1 | 1023.12467 | -0.423168972 | 0.065816657 | -6.429511843 | 1.28E-10 | 1.10E-09 |
| ENSG00000171793 | CTPS1 | 2560.187904 | 0.002635646 | 0.058284499 | 0.045220361 | 0.963931666 | 0.983656246 |
| ENSG00000171798 | KNDC1 | 144.3105013 | 2.481696391 | 0.185910189 | 13.34889929 | 1.20E-40 | 5.05E-39 |
| ENSG00000171804 | WDR87 | 3.921362881 | 0.311090381 | 0.881796024 | 0.352791771 | 0.724244553 | 0.846509142 |
| ENSG00000171806 | METTL18 | 157.8318961 | -0.222179497 | 0.146907042 | -1.512381524 | 0.130436832 | 0.262084693 |
| ENSG00000171811 | CFAP46 | 35.11160657 | -0.329048625 | 0.305769241 | -1.07613383 | 0.281867408 | 0.465366562 |
| ENSG00000171813 | PWWP2B | 1256.873532 | -0.220334137 | 0.082543323 | -2.669315075 | 0.007600611 | 0.023353606 |

| | | | | | | | |
|---|---|---|---|---|---|---|---|
| ENSG00000171817 | ZNF540 | 5.593666928 | 0.485130322 | 0.738984521 | 0.656482387 | 0.511513797 | 0.691177858 |
| ENSG00000171823 | FBXL14 | 307.6434999 | 0.144060317 | 0.116758321 | 1.233833407 | 0.217264988 | 0.386840952 |
| ENSG00000171824 | EXOSC10 | 3197.689021 | -0.349959504 | 0.053890961 | -6.493844138 | 8.37E-11 | 7.30E-10 |
| ENSG00000171827 | ZNF570 | 90.77756061 | 0.286013278 | 0.186701594 | 1.531927347 | 0.125540364 | 0.254295449 |
| ENSG00000171843 | MLLT3 | 399.3798175 | 0.554135285 | 0.097705859 | 5.671464226 | 1.42E-08 | 1.00E-07 |
| ENSG00000171848 | RRM2 | 5655.208739 | -0.184655028 | 0.047551334 | -3.883277598 | 0.000103058 | 0.000458434 |
| ENSG00000171853 | TRAPPC12 | 846.0860687 | -0.017154433 | 0.072097391 | -0.237934169 | 0.811932153 | 0.900445907 |
| ENSG00000171858 | RPS21 | 5545.872905 | -0.010673387 | 0.053139784 | -0.200854911 | 0.840812025 | 0.91647005 |
| ENSG00000171861 | MRM3 | 524.6861657 | -0.102057329 | 0.097417399 | -1.047629378 | 0.294809397 | 0.480662112 |
| ENSG00000171862 | PTEN | 2810.812697 | -0.20944846 | 0.054816715 | -3.820886766 | 0.000132973 | 0.000580696 |
| ENSG00000171863 | RPS7 | 4328.772738 | 0.226175328 | 0.051923129 | 4.355964872 | 1.32E-05 | 6.73E-05 |
| ENSG00000171865 | RNASEH1 | 749.1505211 | 0.362755877 | 0.081828508 | 4.433123439 | 9.29E-06 | 4.82E-05 |
| ENSG00000171867 | PRNP | 721.1680825 | 1.247621549 | 0.081417824 | 15.32369069 | 5.31E-53 | 3.16E-51 |
| ENSG00000171877 | FRMD5 | 237.5060529 | 1.093087397 | 0.122424083 | 8.928695844 | 4.31E-19 | 6.84E-18 |
| ENSG00000171903 | CYP4F11 | 62.00677334 | 1.031915793 | 0.232294581 | 4.442272341 | 8.90E-06 | 4.62E-05 |
| ENSG00000171914 | TLN2 | 1162.543505 | -0.273367744 | 0.073485168 | -3.720039726 | 0.000199191 | 0.000845319 |
| ENSG00000171928 | TVP23B | 620.8655016 | 0.145551535 | 0.082429805 | 1.765763434 | 0.077435546 | 0.172295966 |
| ENSG00000171931 | FBXW10 | 1.767207864 | -0.94574637 | 1.337144998 | -0.70728782 | 0.479387633 | NA |
| ENSG00000171940 | ZNF217 | 20914.33006 | 0.466415109 | 0.044021921 | 10.59506494 | 3.14E-26 | 7.41E-25 |
| ENSG00000171943 | SRGAP2C | 363.0526019 | 0.079545061 | 0.10051867 | 0.791346133 | 0.428742035 | 0.61920798 |
| ENSG00000171951 | SCG2 | 6.821557357 | 1.950299101 | 0.773066531 | 2.52280887 | 0.011642166 | 0.034008823 |
| ENSG00000171953 | ATPAF2 | 488.1640661 | 0.144502228 | 0.093479271 | 1.545821076 | 0.122147786 | 0.248855268 |
| ENSG00000171954 | CYP4F22 | 71.79553289 | 1.254843035 | 0.221428993 | 5.667022274 | 1.45E-08 | 1.03E-07 |
| ENSG00000171960 | PPIH | 752.6916358 | -0.094451602 | 0.075206362 | -1.255899098 | 0.209152556 | 0.375927261 |
| ENSG00000171962 | DRC3 | 60.08610329 | 0.171917422 | 0.245519726 | 0.70021837 | 0.483790941 | 0.667759578 |
| ENSG00000171970 | ZNF57 | 250.2037154 | 0.098413784 | 0.118644582 | 0.82948401 | 0.406830581 | 0.599671884 |
| ENSG00000171984 | SHLD1 | 55.895381 | 0.2437004 | 0.251951883 | 0.967249767 | 0.333419189 | 0.52456184 |
| ENSG00000171988 | JMJD1C | 1353.024939 | -0.335757932 | 0.071057023 | -4.725189988 | 2.30E-06 | 1.28E-05 |
| ENSG00000171992 | SYNPO | 152.6968788 | 0.235104882 | 0.159463268 | 1.474351338 | 0.140387019 | 0.277078054 |
| ENSG00000172006 | ZNF554 | 275.2025368 | -0.192498836 | 0.11187762 | -1.720619684 | 0.085319859 | 0.186245998 |
| ENSG00000172007 | RAB33B | 381.2044621 | 0.086458022 | 0.10946666 | 0.78981145 | 0.429637891 | 0.619945479 |
| ENSG00000172009 | THOP1 | 1853.611623 | -0.175447368 | 0.062739872 | -2.796425323 | 0.005167135 | 0.016582831 |
| ENSG00000172031 | EPHX4 | 48.71864657 | 2.398157978 | 0.309070047 | 7.759270109 | 8.54E-15 | 1.01E-13 |

| | | | | | | | |
|---|---|---|---|---|---|---|---|
| ENSG00000172037 | LAMB2 | 5464.082203 | -0.590035287 | 0.058078867 | -10.15920798 | 3.02E-24 | 6.40E-23 |
| ENSG00000172046 | USP19 | 2070.830374 | 0.128047682 | 0.06230517 | 2.05516946 | 0.03986264 | 0.098870035 |
| ENSG00000172053 | QARS1 | 4593.192956 | 0.060117282 | 0.048770488 | 1.232656971 | 0.217703772 | 0.387214171 |
| ENSG00000172057 | ORMDL3 | 1305.340266 | -0.079044095 | 0.064105768 | -1.233026253 | 0.21756597 | 0.387106441 |
| ENSG00000172059 | KLF11 | 557.589831 | 0.593518646 | 0.089891907 | 6.602581545 | 4.04E-11 | 3.60E-10 |
| ENSG00000172061 | LRRC15 | 15.12220121 | -0.470317451 | 0.458793458 | -1.025118041 | 0.305307494 | 0.492805651 |
| ENSG00000172062 | SMN1 | 170.4022412 | -0.045454952 | 0.144732718 | -0.314061345 | 0.753474448 | 0.864088279 |
| ENSG00000172071 | EIF2AK3 | 1095.503677 | -0.098993347 | 0.067447578 | -1.467707961 | 0.142183567 | 0.279805356 |
| ENSG00000172081 | MOB3A | 829.2044545 | -0.057777167 | 0.073221744 | -0.789071173 | 0.430070409 | 0.620374469 |
| ENSG00000172086 | KRCC1 | 872.442607 | 0.089233642 | 0.071262965 | 1.252174143 | 0.210506416 | 0.377812764 |
| ENSG00000172113 | NME6 | 899.734625 | 0.024537321 | 0.069230557 | 0.354429063 | 0.723017357 | 0.845926239 |
| ENSG00000172115 | CYCS | 1203.190702 | -1.040760331 | 0.075020513 | -13.87301002 | 9.23E-44 | 4.27E-42 |
| ENSG00000172159 | FRMD3 | 85.59441021 | 0.681734597 | 0.199798009 | 3.41211908 | 0.000644599 | 0.002501474 |
| ENSG00000172164 | SNTB1 | 17.48193092 | -0.113545633 | 0.433282706 | -0.262059001 | 0.793275955 | 0.889039543 |
| ENSG00000172167 | MTBP | 204.4477205 | -0.189223699 | 0.135938865 | -1.391976449 | 0.16392953 | 0.311793667 |
| ENSG00000172171 | TEFM | 185.1282647 | -0.09480438 | 0.134416106 | -0.705305211 | 0.480620318 | 0.665015749 |
| ENSG00000172172 | MRPL13 | 1004.632105 | -0.238152817 | 0.0660167 | -3.607463225 | 0.000309205 | 0.001270722 |
| ENSG00000172175 | MALT1 | 620.2326945 | 0.677972322 | 0.082666242 | 8.201320215 | 2.38E-16 | 3.15E-15 |
| ENSG00000172183 | ISG20 | 66.92269932 | 0.421611736 | 0.217474737 | 1.938669947 | 0.05254154 | 0.12500549 |
| ENSG00000172186 | HMGN1P35 | 1.616486058 | 0.432069063 | 1.538783999 | 0.280786039 | 0.778874512 | NA |
| ENSG00000172197 | MBOAT1 | 145.6101196 | -0.251496232 | 0.15060963 | -1.669854922 | 0.094948071 | 0.203238663 |
| ENSG00000172201 | ID4 | 44.74433286 | -3.156800328 | 0.353304377 | -8.935072798 | 4.07E-19 | 6.47E-18 |
| ENSG00000172216 | CEBPB | 596.9216856 | -0.114450499 | 0.084372662 | -1.356487942 | 0.174943969 | 0.328015068 |
| ENSG00000172232 | AZU1 | 5.242874382 | 0.912081865 | 0.797590137 | 1.143547071 | 0.252811523 | 0.431655097 |
| ENSG00000172239 | PAIP1 | 1551.338318 | -0.071667352 | 0.05977313 | -1.198989447 | 0.230532049 | 0.403910884 |
| ENSG00000172244 | C5orf34 | 175.2451931 | -0.260179608 | 0.146569756 | -1.775124796 | 0.075877284 | 0.169519369 |
| ENSG00000172250 | SERHL | 19.20384276 | -0.424190181 | 0.411555495 | -1.030699836 | 0.302681602 | 0.489670669 |
| ENSG00000172260 | NEGR1 | 3.559075475 | -2.240497068 | 1.068531462 | -2.09680028 | 0.036011259 | 0.090819951 |
| ENSG00000172262 | ZNF131 | 2092.346814 | 0.191423126 | 0.054263766 | 3.527641765 | 0.000419279 | 0.001681809 |
| ENSG00000172264 | MACROD2 | 269.1172757 | -0.83344822 | 0.116209612 | -7.171938773 | 7.39E-13 | 7.64E-12 |
| ENSG00000172269 | DPAGT1 | 2643.14559 | 0.18143404 | 0.05295133 | 3.426430284 | 0.000611571 | 0.002381907 |
| ENSG00000172270 | BSG | 10448.62402 | 0.055655297 | 0.045204957 | 1.231176864 | 0.218256723 | 0.387918951 |
| ENSG00000172273 | HINFP | 1376.434871 | 0.332444807 | 0.060986976 | 5.451078677 | 5.01E-08 | 3.34E-07 |

| | | | | | | | |
|---|---|---|---|---|---|---|---|
| ENSG00000172292 | CERS6 | 3606.254993 | -0.073342123 | 0.052140562 | -1.406623172 | 0.159539158 | 0.30518915 |
| ENSG00000172296 | SPTLC3 | 7.986048826 | 3.020911889 | 0.818041075 | 3.692860884 | 0.000221745 | 0.000933323 |
| ENSG00000172301 | COPRS | 936.6910566 | 0.401663558 | 0.069181725 | 5.805919978 | 6.40E-09 | 4.70E-08 |
| ENSG00000172315 | TP53RK | 900.8417003 | -0.12879749 | 0.06896445 | -1.867592496 | 0.061818893 | 0.143013857 |
| ENSG00000172318 | B3GALT1 | 8.765074144 | 0.372922203 | 0.606108976 | 0.61527253 | 0.538374756 | 0.713216899 |
| ENSG00000172331 | BPGM | 138.7826605 | -0.245192856 | 0.1527075 | -1.605637293 | 0.108353626 | 0.226089884 |
| ENSG00000172336 | POP7 | 1082.950871 | -0.555979184 | 0.070809597 | -7.851749033 | 4.10E-15 | 4.97E-14 |
| ENSG00000172339 | ALG14 | 91.66615142 | -0.04410929 | 0.185081467 | -0.238323647 | 0.811630082 | 0.900285 |
| ENSG00000172340 | SUCLG2 | 1091.472823 | 0.2766643 | 0.073062899 | 3.786659206 | 0.000152686 | 0.000661122 |
| ENSG00000172345 | STARD5 | 80.43553972 | -0.755298739 | 0.206250489 | -3.662045804 | 0.000250209 | 0.001041895 |
| ENSG00000172346 | CSDC2 | 1.990921381 | -0.188172377 | 1.214238306 | -0.154971537 | 0.876843756 | 0.935974786 |
| ENSG00000172348 | RCAN2 | 39.68615499 | -1.370102636 | 0.297986288 | -4.597871412 | 4.27E-06 | 2.30E-05 |
| ENSG00000172349 | IL16 | 74.80623921 | -0.369061972 | 0.205262432 | -1.798000583 | 0.072176915 | 0.162838163 |
| ENSG00000172350 | ABCG4 | 6.779840898 | -0.265483316 | 0.660448758 | -0.401974132 | 0.687703064 | 0.823383061 |
| ENSG00000172354 | GNB2 | 9012.017055 | -0.450940099 | 0.045711574 | -9.864899782 | 5.91E-23 | 1.17E-21 |
| ENSG00000172361 | CFAP53 | 36.89438433 | 0.753212654 | 0.298147371 | 2.526309898 | 0.011526775 | 0.033684615 |
| ENSG00000172366 | MCRIP2 | 937.6620866 | 0.194782704 | 0.07498181 | 2.597732774 | 0.009384149 | 0.028140181 |
| ENSG00000172375 | C2CD2L | 1610.103234 | 0.37073935 | 0.061477665 | 6.030472178 | 1.63E-09 | 1.28E-08 |
| ENSG00000172379 | ARNT2 | 5695.921423 | -1.390913274 | 0.056056085 | -24.81288644 | 6.51E-136 | 2.02E-133 |
| ENSG00000172380 | GNG12 | 1792.424342 | 0.386501723 | 0.060051716 | 6.436147868 | 1.23E-10 | 1.05E-09 |
| ENSG00000172382 | PRSS27 | 76.38154096 | -0.768359963 | 0.206585712 | -3.719327701 | 0.000199754 | 0.000847172 |
| ENSG00000172399 | MYOZ2 | 2.319062869 | 1.827937735 | 1.315104527 | 1.389956233 | 0.164542168 | 0.312725546 |
| ENSG00000172403 | SYNPO2 | 194.0018911 | 2.629292955 | 0.164558512 | 15.97786053 | 1.82E-57 | 1.23E-55 |
| ENSG00000172409 | CLP1 | 286.7196037 | -0.115636714 | 0.113500509 | -1.018821103 | 0.308287905 | 0.496324271 |
| ENSG00000172425 | TTC36 | 2.029212809 | 1.601778067 | 1.327152999 | 1.206927965 | 0.227459887 | 0.400209373 |
| ENSG00000172426 | RSPH9 | 9.728962857 | -0.803414866 | 0.570481833 | -1.408309293 | 0.159039505 | 0.304487252 |
| ENSG00000172428 | COPS9 | 748.8121644 | -0.263362469 | 0.084185319 | -3.12836576 | 0.001757813 | 0.006251818 |
| ENSG00000172432 | GTPBP2 | 1579.2096 | 0.157263166 | 0.064474237 | 2.439162885 | 0.014721331 | 0.04179029 |
| ENSG00000172456 | FGGY | 227.777217 | -0.209440356 | 0.124196235 | -1.686366394 | 0.09172525 | 0.197519688 |
| ENSG00000172458 | IL17D | 166.1644197 | 0.755974386 | 0.144518003 | 5.231004938 | 1.69E-07 | 1.07E-06 |
| ENSG00000172461 | FUT9 | 560.070638 | 0.04942892 | 0.093687852 | 0.527591553 | 0.597782857 | 0.758176522 |
| ENSG00000172465 | TCEAL1 | 714.5949257 | -0.393997796 | 0.076750775 | -5.133469935 | 2.84E-07 | 1.76E-06 |
| ENSG00000172466 | ZNF24 | 3833.44316 | 0.500439789 | 0.049934575 | 10.02190944 | 1.22E-23 | 2.50E-22 |

| | | | | | | | |
|---|---|---|---|---|---|---|---|
| ENSG00000172469 | MANEA | 270.2157214 | -0.086239346 | 0.115597788 | -0.746029382 | 0.455649666 | 0.642999713 |
| ENSG00000172476 | RAB40A | 6.785547961 | -0.021970548 | 0.712747876 | -0.030825133 | 0.975408997 | 0.988740606 |
| ENSG00000172478 | MAB21L4 | 163.163743 | 0.738476528 | 0.155047697 | 4.762899044 | 1.91E-06 | 1.07E-05 |
| ENSG00000172493 | AFF1 | 1216.772033 | -0.256557407 | 0.065619448 | -3.909776984 | 9.24E-05 | 0.000414074 |
| ENSG00000172500 | FIBP | 3295.785725 | 0.165489468 | 0.055498589 | 2.981868025 | 0.002864955 | 0.009758287 |
| ENSG00000172508 | CARNS1 | 42.22500762 | 0.290179496 | 0.277179442 | 1.046901219 | 0.295145139 | 0.481018631 |
| ENSG00000172530 | BANP | 614.2380699 | 0.235897703 | 0.080257273 | 2.93926886 | 0.003289876 | 0.011054599 |
| ENSG00000172531 | PPP1CA | 8226.596702 | -0.181188728 | 0.043632946 | -4.152566955 | 3.29E-05 | 0.000158211 |
| ENSG00000172534 | HCFC1 | 5946.949659 | -0.203474236 | 0.047328486 | -4.299191749 | 1.71E-05 | 8.59E-05 |
| ENSG00000172551 | MUCL1 | 27.0512623 | -0.149069816 | 0.332152857 | -0.448798836 | 0.653576779 | 0.79899021 |
| ENSG00000172575 | RASGRP1 | 157.4657299 | 2.505702859 | 0.185921238 | 13.47722769 | 2.13E-41 | 9.23E-40 |
| ENSG00000172586 | CHCHD1 | 793.16694 | -0.717373064 | 0.073101923 | -9.813326874 | 9.87E-23 | 1.93E-21 |
| ENSG00000172590 | MRPL52 | 801.647521 | -0.504989078 | 0.073193461 | -6.899374187 | 5.22E-12 | 5.03E-11 |
| ENSG00000172594 | SMPDL3A | 285.8238665 | 0.011247229 | 0.108634621 | 0.10353264 | 0.917540246 | 0.959404405 |
| ENSG00000172602 | RND1 | 168.0354976 | 0.105415914 | 0.138782619 | 0.759575763 | 0.447508211 | 0.635891871 |
| ENSG00000172613 | RAD9A | 1040.568581 | 0.33169071 | 0.077255348 | 4.293433647 | 1.76E-05 | 8.79E-05 |
| ENSG00000172638 | EFEMP2 | 28.66475163 | -0.608576063 | 0.347617932 | -1.750703883 | 0.07999693 | 0.1766416 |
| ENSG00000172650 | AGAP5 | 4.294430126 | -1.024759073 | 0.874544909 | -1.171762664 | 0.241292356 | 0.41728207 |
| ENSG00000172661 | WASHC2C | 1508.631014 | -1.002897218 | 0.06503015 | -15.42203457 | 1.16E-53 | 7.04E-52 |
| ENSG00000172663 | TMEM134 | 922.7208603 | 0.240365849 | 0.073657469 | 3.263292274 | 0.001101259 | 0.00409248 |
| ENSG00000172667 | ZMAT3 | 398.0071802 | 0.156810001 | 0.096213039 | 1.629820682 | 0.103139401 | 0.217404931 |
| ENSG00000172671 | ZFAND4 | 91.45663222 | -0.953766132 | 0.193761713 | -4.922366325 | 8.55E-07 | 5.00E-06 |
| ENSG00000172689 | MS4A10 | 1.63776912 | 0.394057253 | 1.338899878 | 0.294314205 | 0.768517819 | NA |
| ENSG00000172717 | GARIN2 | 20.88698471 | 0.286830148 | 0.388859109 | 0.737619723 | 0.460745567 | 0.647561775 |
| ENSG00000172725 | CORO1B | 6019.084395 | -0.250763859 | 0.059723375 | -4.198755693 | 2.68E-05 | 0.000130772 |
| ENSG00000172728 | FUT10 | 153.7087933 | -0.272212208 | 0.149451241 | -1.82141149 | 0.06854433 | 0.155930873 |
| ENSG00000172731 | LRRC20 | 522.974938 | -0.500015007 | 0.088376396 | -5.657789049 | 1.53E-08 | 1.08E-07 |
| ENSG00000172732 | MUS81 | 1235.162484 | -0.032558707 | 0.070921761 | -0.45907923 | 0.646177271 | 0.793284095 |
| ENSG00000172738 | TMEM217 | 18.05455229 | -0.16348171 | 0.411540556 | -0.397243255 | 0.691188088 | 0.825797152 |
| ENSG00000172748 | ZNF596 | 287.1288756 | 0.161126804 | 0.118188797 | 1.363300149 | 0.172787935 | 0.32489797 |
| ENSG00000172757 | CFL1 | 20339.22633 | -0.243104045 | 0.038988557 | -6.235266514 | 4.51E-10 | 3.71E-09 |
| ENSG00000172765 | TMCC1 | 1668.284138 | 0.336933262 | 0.062485971 | 5.3921425 | 6.96E-08 | 4.57E-07 |
| ENSG00000172766 | NAA16 | 567.6208345 | 0.489637248 | 0.089579891 | 5.465928177 | 4.60E-08 | 3.09E-07 |

| | | | | | | | |
|---|---|---|---|---|---|---|---|
| ENSG00000172771 | EFCAB12 | 7.18506365 | -0.124667255 | 0.647304651 | -0.192594406 | 0.847276626 | 0.920028676 |
| ENSG00000172775 | PSME3IP1 | 1358.963039 | -0.096857667 | 0.061478429 | -1.575474009 | 0.115147073 | 0.237563977 |
| ENSG00000172780 | RAB43 | 31.15889876 | 0.578512574 | 0.33038488 | 1.751026182 | 0.0799414 | 0.176586974 |
| ENSG00000172785 | ZNG1A | 429.2586011 | -0.205071555 | 0.095030209 | -2.15796173 | 0.030930807 | 0.079769839 |
| ENSG00000172789 | HOXC5 | 4.678329014 | -0.20836306 | 0.815084103 | -0.255633817 | 0.798233604 | 0.892259454 |
| ENSG00000172794 | RAB37 | 10.01093176 | 0.450450828 | 0.563978993 | 0.798701429 | 0.424463558 | 0.615226766 |
| ENSG00000172795 | DCP2 | 5697.117966 | 0.601259836 | 0.048624384 | 12.3653974 | 4.02E-35 | 1.38E-33 |
| ENSG00000172799 | ZBTB8OSP2 | 51.98626215 | 0.047675644 | 0.257855952 | 0.184892549 | 0.853313315 | 0.92356723 |
| ENSG00000172803 | SNX32 | 203.0754212 | -0.494458747 | 0.159004944 | -3.109706745 | 0.001872732 | 0.006619486 |
| ENSG00000172809 | RPL38 | 5143.284233 | 0.338392283 | 0.053389432 | 6.338188482 | 2.32E-10 | 1.96E-09 |
| ENSG00000172818 | OVOL1 | 646.9935861 | 0.787036386 | 0.083484791 | 9.427302545 | 4.21E-21 | 7.49E-20 |
| ENSG00000172819 | RARG | 2637.183162 | 0.622737865 | 0.067700749 | 9.198389586 | 3.63E-20 | 6.11E-19 |
| ENSG00000172828 | CES3 | 19.86431371 | 0.300993331 | 0.394999774 | 0.762008869 | 0.446054711 | 0.634689852 |
| ENSG00000172830 | SSH3 | 3427.797599 | 0.045311224 | 0.05574193 | 0.81287504 | 0.416289706 | 0.608341038 |
| ENSG00000172831 | CES2 | 1785.332077 | 0.057642753 | 0.061877287 | 0.931565617 | 0.351561059 | 0.542829016 |
| ENSG00000172840 | PDP2 | 568.0788154 | 0.077405283 | 0.086581975 | 0.894011513 | 0.371315731 | 0.563955434 |
| ENSG00000172845 | SP3 | 3103.696724 | -0.041876302 | 0.048573778 | -0.862117449 | 0.38862289 | 0.581314225 |
| ENSG00000172867 | KRT2 | 1.585245656 | 1.113915549 | 1.452095432 | 0.767109052 | 0.443016681 | NA |
| ENSG00000172869 | DMXL1 | 1238.470062 | -0.139967832 | 0.072265845 | -1.936846261 | 0.052764139 | 0.125496045 |
| ENSG00000172878 | METAP1D | 275.7041407 | 0.196612005 | 0.112899958 | 1.74147101 | 0.081601051 | 0.179561448 |
| ENSG00000172888 | ZNF621 | 1417.724188 | -0.239510349 | 0.063248502 | -3.786814547 | 0.000152591 | 0.000660834 |
| ENSG00000172889 | EGFL7 | 500.2778057 | 0.515323904 | 0.099335969 | 5.187686868 | 2.13E-07 | 1.34E-06 |
| ENSG00000172890 | NADSYN1 | 3126.233552 | 0.143340276 | 0.056229919 | 2.549181604 | 0.010797606 | 0.031780482 |
| ENSG00000172893 | DHCR7 | 14091.11659 | -0.350740078 | 0.044399697 | -7.899605234 | 2.80E-15 | 3.43E-14 |
| ENSG00000172901 | LVRN | 4.776591646 | -0.29176441 | 0.806426705 | -0.361799043 | 0.717502208 | 0.842913738 |
| ENSG00000172915 | NBEA | 1322.554809 | 0.045045082 | 0.076778875 | 0.58668588 | 0.557414692 | 0.727697077 |
| ENSG00000172922 | RNASEH2C | 1187.262358 | 0.280570476 | 0.068641795 | 4.087458319 | 4.36E-05 | 0.000205562 |
| ENSG00000172927 | MYEOV | 14.00375443 | 0.307642495 | 0.475426845 | 0.647086925 | 0.517575687 | 0.695665852 |
| ENSG00000172932 | ANKRD13D | 1483.202815 | 0.007177986 | 0.069662843 | 0.103038943 | 0.917932063 | 0.95968162 |
| ENSG00000172935 | MRGPRF | 7.206365315 | 0.752484213 | 0.672224783 | 1.119393738 | 0.262972201 | 0.443735366 |
| ENSG00000172936 | MYD88 | 1752.269304 | 0.068403086 | 0.057036228 | 1.199291884 | 0.23041447 | 0.403766552 |
| ENSG00000172939 | OXSR1 | 2570.222561 | -0.104836287 | 0.053116111 | -1.97371919 | 0.048413682 | 0.116574983 |
| ENSG00000172940 | SLC22A13 | 4.211281538 | -0.723696545 | 0.894666331 | -0.808901062 | 0.418572058 | 0.61039531 |

| | | | | | | | |
|---|---|---|---|---|---|---|---|
| ENSG00000172943 | PHF8 | 2262.711218 | 0.080494999 | 0.055429109 | 1.452215284 | 0.146441754 | 0.286269607 |
| ENSG00000172954 | LCLAT1 | 757.2416513 | 0.002990484 | 0.074319834 | 0.040238029 | 0.967903359 | 0.985764746 |
| ENSG00000172965 | MIR4435-2HG | 625.7567519 | 1.220003685 | 0.083154477 | 14.6715334 | 9.81E-49 | 5.19E-47 |
| ENSG00000172974 | VDAC2P5 | 1766.834939 | -0.43720786 | 0.066239201 | -6.600439795 | 4.10E-11 | 3.65E-10 |
| ENSG00000172977 | KAT5 | 2539.707797 | -0.074465222 | 0.053342058 | -1.395994553 | 0.162716136 | 0.309999846 |
| ENSG00000172986 | GXYLT2 | 56.75675171 | 0.255687557 | 0.2395173 | 1.067511855 | 0.285740763 | 0.469595837 |
| ENSG00000172987 | HPSE2 | 10.74786331 | -2.274039525 | 0.613672939 | -3.705621317 | 0.000210873 | 0.000890518 |
| ENSG00000172992 | DCAKD | 1273.605352 | 0.001388983 | 0.070966408 | 0.019572394 | 0.984384486 | 0.992823922 |
| ENSG00000172995 | ARPP21 | 8.137690215 | 1.097535846 | 0.634202751 | 1.730575662 | 0.083527476 | 0.183080479 |
| ENSG00000173011 | TADA2B | 1346.329138 | -0.00545167 | 0.06612887 | -0.082440098 | 0.934296751 | 0.968469969 |
| ENSG00000173013 | CCDC96 | 82.34558264 | 0.194376608 | 0.210277398 | 0.92438184 | 0.355287546 | 0.54703306 |
| ENSG00000173020 | GRK2 | 4244.290429 | -0.233090205 | 0.050113957 | -4.651203396 | 3.30E-06 | 1.80E-05 |
| ENSG00000173039 | RELA | 3758.724522 | 0.082046014 | 0.054558974 | 1.503804192 | 0.132631791 | 0.265587753 |
| ENSG00000173041 | ZNF680 | 438.7387918 | 0.427899956 | 0.100660709 | 4.250913388 | 2.13E-05 | 0.000105303 |
| ENSG00000173064 | HECTD4 | 1544.07929 | -0.056968242 | 0.070769118 | -0.804987317 | 0.420826999 | 0.61204054 |
| ENSG00000173065 | FAM222B | 731.5329233 | -0.189234569 | 0.075585963 | -2.503567599 | 0.012294819 | 0.035679083 |
| ENSG00000173083 | HPSE | 29.8395697 | 0.612495009 | 0.329416643 | 1.859332316 | 0.062980049 | 0.145231436 |
| ENSG00000173085 | COQ2 | 639.2429308 | 0.201411698 | 0.078159732 | 2.576924134 | 0.009968383 | 0.029644114 |
| ENSG00000173093 | CCDC63 | 209.8053196 | -0.472899278 | 0.142436275 | -3.320076143 | 0.000899929 | 0.00339937 |
| ENSG00000173110 | HSPA6 | 8.943349118 | 0.213426338 | 0.621103789 | 0.343624273 | 0.731128868 | 0.851119615 |
| ENSG00000173113 | TRMT112 | 2537.032807 | -0.099887974 | 0.052350924 | -1.908046052 | 0.056385264 | 0.132623599 |
| ENSG00000173120 | KDM2A | 7187.906565 | -0.090492443 | 0.048418193 | -1.868976049 | 0.061626147 | 0.142697467 |
| ENSG00000173137 | ADCK5 | 366.6674983 | -0.02511293 | 0.102176749 | -0.245779299 | 0.805853082 | 0.89699984 |
| ENSG00000173141 | MRPL57 | 1629.84115 | 0.04370344 | 0.056892609 | 0.7681743 | 0.442383639 | 0.631470633 |
| ENSG00000173145 | NOC3L | 712.1758626 | -0.34985558 | 0.080929071 | -4.322990164 | 1.54E-05 | 7.74E-05 |
| ENSG00000173153 | ESRRA | 1146.517198 | 0.10635016 | 0.07117579 | 1.494190078 | 0.135125945 | 0.269305262 |
| ENSG00000173156 | RHOD | 1186.934771 | 0.223967776 | 0.066946868 | 3.345455649 | 0.000821475 | 0.003125147 |
| ENSG00000173163 | COMMD1 | 346.1803841 | -0.075629628 | 0.104227282 | -0.725622189 | 0.468070414 | 0.654009947 |
| ENSG00000173166 | RAPH1 | 3371.153013 | -1.116226068 | 0.055235854 | -20.20836068 | 8.26E-91 | 1.14E-88 |
| ENSG00000173171 | MTX1 | 524.8428616 | -0.01613471 | 0.084238124 | -0.191536906 | 0.848104969 | 0.920492429 |
| ENSG00000173175 | ADCY5 | 155.091174 | -1.043629914 | 0.15967744 | -6.535863253 | 6.32E-11 | 5.56E-10 |
| ENSG00000173193 | PARP14 | 216.8408303 | 0.024410679 | 0.126714515 | 0.192643119 | 0.847238473 | 0.920028676 |
| ENSG00000173207 | CKS1B | 662.0592298 | -0.177188556 | 0.077627106 | -2.282560379 | 0.022456277 | 0.060505872 |

| | | | | | | | |
|---|---|---|---|---|---|---|---|
| ENSG00000173209 | AHSA2P | 1241.632578 | 0.141069887 | 0.175218654 | 0.805107697 | 0.420757534 | 0.611996403 |
| ENSG00000173210 | ABLIM3 | 1320.328755 | -0.92985136 | 0.064860041 | -14.3362746 | 1.30E-46 | 6.53E-45 |
| ENSG00000173212 | MAB21L3 | 3.304792235 | -0.152792612 | 1.010476723 | -0.151208443 | 0.87981129 | 0.937691461 |
| ENSG00000173213 | TUBB8B | 5.684824757 | -0.096325607 | 0.728517955 | -0.132221322 | 0.894809236 | 0.946420945 |
| ENSG00000173214 | MFSD4B | 211.6666016 | 0.006622058 | 0.125356755 | 0.052825694 | 0.957870789 | 0.980830956 |
| ENSG00000173218 | VANGL1 | 2787.629724 | -0.050795842 | 0.054115899 | -0.938649143 | 0.347910912 | 0.538897794 |
| ENSG00000173221 | GLRX | 57.0462773 | -0.891089179 | 0.241057437 | -3.696584482 | 0.00021852 | 0.000920125 |
| ENSG00000173226 | IQCB1 | 1057.665445 | 0.301696993 | 0.074848659 | 4.030760195 | 5.56E-05 | 0.000258385 |
| ENSG00000173227 | SYT12 | 1180.076533 | -0.857742822 | 0.080383719 | -10.6706038 | 1.40E-26 | 3.38E-25 |
| ENSG00000173230 | GOLGB1 | 6454.65489 | -0.066593091 | 0.050531395 | -1.317855791 | 0.187551927 | 0.345494356 |
| ENSG00000173253 | DMRT2 | 31.52528676 | 0.19577003 | 0.309800813 | 0.631922262 | 0.527437676 | 0.703507931 |
| ENSG00000173258 | ZNF483 | 127.1723519 | -0.165740315 | 0.166540738 | -0.995193829 | 0.319642003 | 0.509101405 |
| ENSG00000173261 | PLAC8L1 | 7.184099179 | 0.192601465 | 0.661839124 | 0.291009488 | 0.771044063 | 0.87520429 |
| ENSG00000173264 | GPR137 | 1067.305903 | 0.02062994 | 0.072838331 | 0.283229171 | 0.777001168 | 0.878958864 |
| ENSG00000173267 | SNCG | 1087.316235 | -1.620922097 | 0.157498434 | -10.2916712 | 7.68E-25 | 1.70E-23 |
| ENSG00000173269 | MMRN2 | 101.3882972 | -1.919355336 | 0.208622734 | -9.200125524 | 3.58E-20 | 6.02E-19 |
| ENSG00000173272 | MZT2A | 899.9192292 | -0.145404747 | 0.080942685 | -1.796391448 | 0.072432283 | 0.163269782 |
| ENSG00000173273 | TNKS | 1203.953217 | -0.03297746 | 0.069316892 | -0.47574926 | 0.634253022 | 0.784391108 |
| ENSG00000173275 | ZNF449 | 260.9846691 | -0.517335324 | 0.128265817 | -4.033306283 | 5.50E-05 | 0.000255911 |
| ENSG00000173276 | ZBTB21 | 834.4044142 | -0.170279502 | 0.072904576 | -2.335649013 | 0.019509542 | 0.053515352 |
| ENSG00000173281 | PPP1R3B | 563.9644144 | -0.582938212 | 0.090181446 | -6.464059266 | 1.02E-10 | 8.79E-10 |
| ENSG00000173295 | FAM86B3P | 150.1538276 | -0.185519414 | 0.164998147 | -1.124372713 | 0.260854943 | 0.441163976 |
| ENSG00000173320 | STOX2 | 75.42163306 | 0.05139088 | 0.211503178 | 0.242979232 | 0.808021486 | 0.898323781 |
| ENSG00000173327 | MAP3K11 | 4156.270923 | 0.03670643 | 0.05622347 | 0.652866668 | 0.513842239 | 0.692979404 |
| ENSG00000173334 | TRIB1 | 1227.84635 | -0.285578726 | 0.063452972 | -4.500635935 | 6.78E-06 | 3.57E-05 |
| ENSG00000173335 | CST9 | 1.960231664 | 0.747628916 | 1.279772469 | 0.584188935 | 0.559093205 | 0.728807456 |
| ENSG00000173338 | KCNK7 | 4.244240092 | -0.190505875 | 0.84178314 | -0.226312296 | 0.820958536 | 0.904991377 |
| ENSG00000173349 | SFT2D3 | 638.0382273 | 0.202751692 | 0.081335277 | 2.492789102 | 0.01267441 | 0.036627737 |
| ENSG00000173402 | DAG1 | 6568.754768 | 0.177297463 | 0.057992077 | 3.057270447 | 0.002233626 | 0.007782411 |
| ENSG00000173409 | ARV1 | 408.4090575 | 0.152013358 | 0.093622461 | 1.623684711 | 0.104443116 | 0.219588037 |
| ENSG00000173418 | NAA20 | 1884.149281 | -0.362917054 | 0.056967311 | -6.370619394 | 1.88E-10 | 1.59E-09 |
| ENSG00000173436 | MICOS10 | 120.0906071 | 0.249710164 | 0.168730523 | 1.479934751 | 0.138890661 | 0.274857812 |
| ENSG00000173442 | EHBP1L1 | 3117.540009 | -0.145689219 | 0.051548527 | -2.826253785 | 0.004709593 | 0.015276877 |

| | | | | | | | |
|---|---|---|---|---|---|---|---|
| ENSG00000173451 | THAP2 | 85.23347885 | 0.066499783 | 0.198531637 | 0.334958115 | 0.737656682 | 0.855502745 |
| ENSG00000173456 | RNF26 | 1836.065185 | -0.091842893 | 0.058880071 | -1.559829868 | 0.118800091 | 0.243632585 |
| ENSG00000173457 | PPP1R14B | 996.6333173 | 0.293555945 | 0.069170446 | 4.243950441 | 2.20E-05 | 0.000108299 |
| ENSG00000173465 | ZNRD2 | 702.1897521 | 0.071877375 | 0.079445405 | 0.904739236 | 0.36560355 | 0.558348008 |
| ENSG00000173467 | AGR3 | 2.17299958 | 3.391866208 | 1.538269491 | 2.204988286 | 0.027454917 | 0.071995403 |
| ENSG00000173473 | SMARCC1 | 5720.229802 | 0.070439726 | 0.04942387 | 1.425216724 | 0.154094563 | 0.297327722 |
| ENSG00000173480 | ZNF417 | 377.2963275 | 0.362176124 | 0.100732039 | 3.595441215 | 0.000323842 | 0.001324468 |
| ENSG00000173482 | PTPRM | 1.598060248 | 1.787834183 | 1.515571986 | 1.179643197 | 0.238142155 | NA |
| ENSG00000173486 | FKBP2 | 11.42032852 | 0.332887116 | 0.540825683 | 0.615516471 | 0.538213695 | 0.713100506 |
| ENSG00000173511 | VEGFB | 1251.475635 | 0.347895432 | 0.06640539 | 5.238963794 | 1.61E-07 | 1.03E-06 |
| ENSG00000173517 | PEAK1 | 1249.069603 | 0.228936711 | 0.07740183 | 2.957768716 | 0.003098745 | 0.010469104 |
| ENSG00000173531 | MST1 | 197.3119139 | -0.483888678 | 0.129939004 | -3.72396788 | 0.000196116 | 0.00083359 |
| ENSG00000173535 | TNFRSF10C | 9.955903948 | -3.55725844 | 0.796566493 | -4.465739486 | 7.98E-06 | 4.16E-05 |
| ENSG00000173540 | GMPPB | 865.4023641 | -0.10676066 | 0.072242915 | -1.477801111 | 0.139461021 | 0.275629863 |
| ENSG00000173542 | MOB1B | 915.5236141 | -0.199363394 | 0.07203539 | -2.767575678 | 0.005647493 | 0.017923837 |
| ENSG00000173545 | ZNF622 | 1554.463022 | 0.160233431 | 0.061823616 | 2.591783571 | 0.009547982 | 0.02856798 |
| ENSG00000173546 | CSPG4 | 10.01605409 | 0.394141941 | 0.549780695 | 0.716907568 | 0.473431132 | 0.658891596 |
| ENSG00000173548 | SNX33 | 2420.360129 | -0.029438452 | 0.06584418 | -0.447092706 | 0.654808125 | 0.79955812 |
| ENSG00000173559 | NABP1 | 77.8704469 | -0.865621928 | 0.208754045 | -4.146611526 | 3.37E-05 | 0.000161939 |
| ENSG00000173567 | ADGRF3 | 18.59970587 | 0.431496434 | 0.406236459 | 1.062180474 | 0.288153763 | 0.472126799 |
| ENSG00000173575 | CHD2 | 4033.86635 | -0.054145889 | 0.048265953 | -1.121823667 | 0.261937421 | 0.44229273 |
| ENSG00000173581 | CCDC106 | 56.13188887 | -0.111762898 | 0.233916671 | -0.477789368 | 0.632800129 | 0.78358033 |
| ENSG00000173588 | CEP83 | 335.8977756 | -0.347780358 | 0.112415406 | -3.093707258 | 0.001976724 | 0.006959186 |
| ENSG00000173598 | NUDT4 | 1235.513843 | 0.174230651 | 0.066488016 | 2.620482024 | 0.008780556 | 0.026531343 |
| ENSG00000173599 | PC | 1606.233332 | -0.053560463 | 0.068280338 | -0.784420006 | 0.432793703 | 0.622853673 |
| ENSG00000173611 | SCAI | 300.5051072 | 0.064823975 | 0.113724583 | 0.570008464 | 0.568671957 | 0.736073456 |
| ENSG00000173614 | NMNAT1 | 324.1579134 | -0.216570714 | 0.104054617 | -2.081317678 | 0.037404838 | 0.09380857 |
| ENSG00000173621 | LRFN4 | 1967.567944 | 0.198493593 | 0.065259594 | 3.041600166 | 0.002353243 | 0.008160753 |
| ENSG00000173638 | SLC19A1 | 1500.984968 | 0.20449712 | 0.06414237 | 3.188175292 | 0.001431737 | 0.005185357 |
| ENSG00000173653 | RCE1 | 916.8854205 | 0.021045363 | 0.074880805 | 0.281051506 | 0.778670895 | 0.879849785 |
| ENSG00000173660 | UQCRH | 2085.399792 | 0.183806208 | 0.053250235 | 3.451744514 | 0.000556975 | 0.002184867 |
| ENSG00000173674 | EIF1AX | 1169.812057 | 0.229553717 | 0.071850232 | 3.194891783 | 0.001398833 | 0.005083984 |
| ENSG00000173681 | BCLAF3 | 392.3330364 | 0.092340001 | 0.099264613 | 0.930240886 | 0.352246377 | 0.543727512 |

| | | | | | | | |
|---|---|---|---|---|---|---|---|
| ENSG00000173692 | PSMD1 | 3237.56431 | -0.264234555 | 0.050492889 | -5.233104305 | 1.67E-07 | 1.06E-06 |
| ENSG00000173698 | ADGRG2 | 5.500322879 | -1.359981451 | 0.776379644 | -1.751696431 | 0.079826021 | 0.17640005 |
| ENSG00000173706 | HEG1 | 172.7303776 | -1.979182689 | 0.153091493 | -12.92810368 | 3.12E-38 | 1.19E-36 |
| ENSG00000173715 | C11orf80 | 916.328104 | -0.593793764 | 0.072237048 | -8.220072431 | 2.03E-16 | 2.71E-15 |
| ENSG00000173726 | TOMM20 | 6686.254958 | 0.379500572 | 0.055790742 | 6.802214154 | 1.03E-11 | 9.63E-11 |
| ENSG00000173727 | FAUP4 | 37.00740752 | -0.141490591 | 0.286130561 | -0.494496603 | 0.620955505 | 0.775560477 |
| ENSG00000173744 | AGFG1 | 1413.262949 | -0.021371447 | 0.058874714 | -0.362998736 | 0.716605829 | 0.842417634 |
| ENSG00000173757 | STAT5B | 1707.822363 | -0.023603679 | 0.059493779 | -0.396741977 | 0.691557742 | 0.826095324 |
| ENSG00000173786 | CNP | 2468.460939 | 0.094886499 | 0.06397376 | 1.483209659 | 0.138018716 | 0.273627979 |
| ENSG00000173801 | JUP | 19660.92192 | -0.188898512 | 0.048425512 | -3.900805667 | 9.59E-05 | 0.000428468 |
| ENSG00000173805 | HAP1 | 95.40279187 | 0.514879562 | 0.18292259 | 2.814740175 | 0.004881668 | 0.015774824 |
| ENSG00000173809 | TDRD12 | 40.59939191 | 0.694366681 | 0.28282523 | 2.455108697 | 0.014084199 | 0.04022554 |
| ENSG00000173812 | EIF1 | 10229.43468 | -0.014660395 | 0.05203578 | -0.281736817 | 0.778145321 | 0.879573576 |
| ENSG00000173818 | ENDOV | 893.9512627 | 0.134094282 | 0.070162915 | 1.911184595 | 0.055980863 | 0.13191587 |
| ENSG00000173821 | RNF213 | 3438.84186 | 0.020813057 | 0.067447231 | 0.308582821 | 0.757638891 | 0.866391318 |
| ENSG00000173825 | TIGD3 | 51.1472954 | 0.103604848 | 0.249674923 | 0.414958965 | 0.678171943 | 0.81682581 |
| ENSG00000173838 | MARCHF10 | 28.07658556 | 0.868004397 | 0.354273151 | 2.450099297 | 0.014281682 | 0.040683198 |
| ENSG00000173846 | PLK3 | 82.91343125 | 0.308448545 | 0.206534923 | 1.49344499 | 0.135320744 | 0.269607249 |
| ENSG00000173848 | NET1 | 2566.420267 | -1.131940072 | 0.058663903 | -19.29534206 | 5.88E-83 | 7.02E-81 |
| ENSG00000173852 | DPY19L1 | 1617.491001 | -0.128431737 | 0.061576373 | -2.085730783 | 0.037003016 | 0.09293291 |
| ENSG00000173868 | PHOSPHO1 | 3.474554805 | -2.206769771 | 1.055758493 | -2.090222134 | 0.03659785 | 0.092106932 |
| ENSG00000173875 | ZNF791 | 935.0573498 | 0.134228576 | 0.080950055 | 1.658165339 | 0.097284099 | 0.207355897 |
| ENSG00000173889 | PHC3 | 2130.808686 | -0.320018731 | 0.059936078 | -5.339333905 | 9.33E-08 | 6.06E-07 |
| ENSG00000173890 | GPR160 | 1262.837842 | -0.393694517 | 0.072644986 | -5.419431366 | 5.98E-08 | 3.95E-07 |
| ENSG00000173894 | CBX2 | 3785.629565 | 0.240274811 | 0.051366922 | 4.677617438 | 2.90E-06 | 1.59E-05 |
| ENSG00000173898 | SPTBN2 | 3353.786052 | 0.501447028 | 0.056804357 | 8.827615557 | 1.07E-18 | 1.66E-17 |
| ENSG00000173905 | GOLIM4 | 1402.699138 | -0.117716182 | 0.06308095 | -1.866113029 | 0.062025554 | 0.143362482 |
| ENSG00000173914 | RBM4B | 1240.998525 | 0.017093739 | 0.064731504 | 0.264071403 | 0.791724899 | 0.887821733 |
| ENSG00000173915 | ATP5MK | 822.5631184 | 0.061265277 | 0.072032536 | 0.850522287 | 0.395034775 | 0.588013388 |
| ENSG00000173926 | MARCHF3 | 42.39940163 | 0.315991111 | 0.275542456 | 1.146796452 | 0.251465756 | 0.429916861 |
| ENSG00000173928 | SWSAP1 | 116.9351218 | -0.243124246 | 0.172760692 | -1.40728914 | 0.159341668 | 0.304913069 |
| ENSG00000173930 | SLCO4C1 | 478.2473063 | -0.99533631 | 0.091304599 | -10.90127241 | 1.14E-27 | 2.91E-26 |
| ENSG00000173933 | RBM4 | 929.688718 | 0.30852428 | 0.085590685 | 3.604647851 | 0.000312576 | 0.001283427 |

| | | | | | | | |
|---|---|---|---|---|---|---|---|
| ENSG00000173947 | PIFO | 19.36590757 | 1.181397549 | 0.435316644 | 2.713880953 | 0.006650006 | 0.020733142 |
| ENSG00000173950 | XXYLT1 | 848.9338112 | 0.209165345 | 0.070566527 | 2.964087286 | 0.003035822 | 0.010277674 |
| ENSG00000173960 | UBXN2A | 929.8005756 | 0.22692882 | 0.070587354 | 3.21486511 | 0.001305058 | 0.004780907 |
| ENSG00000173991 | TCAP | 28.44681644 | 0.284837045 | 0.341342541 | 0.834461022 | 0.404021243 | 0.596873217 |
| ENSG00000173992 | CCS | 712.7892536 | -0.069013875 | 0.076495028 | -0.902200789 | 0.366950216 | 0.559399478 |
| ENSG00000174004 | NRROS | 3.371505424 | -0.772889642 | 0.955244486 | -0.80910139 | 0.418456829 | 0.61039531 |
| ENSG00000174007 | CEP19 | 42.14972828 | -0.257030493 | 0.281080057 | -0.914438739 | 0.360486354 | 0.553036645 |
| ENSG00000174010 | KLHL15 | 210.3674754 | -0.12172942 | 0.1276578 | -0.953560383 | 0.340306217 | 0.531279749 |
| ENSG00000174013 | FBXO45 | 1231.400978 | -0.214924869 | 0.06448857 | -3.332759127 | 0.000859894 | 0.003259411 |
| ENSG00000174015 | CBY2 | 18.5677671 | 1.047062262 | 0.45365929 | 2.308036637 | 0.020997099 | 0.057022095 |
| ENSG00000174021 | GNG5 | 127.7119555 | -0.083432438 | 0.181848231 | -0.458802579 | 0.646375942 | 0.793412953 |
| ENSG00000174028 | FAM3C2P | 432.0614148 | 0.463136516 | 0.091318775 | 5.071646189 | 3.94E-07 | 2.40E-06 |
| ENSG00000174032 | SLC25A30 | 967.7149626 | 0.887261594 | 0.071462605 | 12.41574651 | 2.15E-35 | 7.41E-34 |
| ENSG00000174080 | CTSF | 837.7798627 | 0.750045227 | 0.074407894 | 10.08018357 | 6.76E-24 | 1.40E-22 |
| ENSG00000174099 | MSRB3 | 28.41046109 | 0.157769364 | 0.330309527 | 0.477640974 | 0.632905762 | 0.783582692 |
| ENSG00000174106 | LEMD3 | 1330.878697 | -0.105041538 | 0.061684583 | -1.702881537 | 0.08859024 | 0.192030625 |
| ENSG00000174109 | UQCC4 | 493.4651807 | 0.111452529 | 0.089895277 | 1.239804062 | 0.215047875 | 0.384026021 |
| ENSG00000174132 | FAM174A | 272.7594836 | 0.094069645 | 0.116392085 | 0.808213416 | 0.418967736 | 0.610656322 |
| ENSG00000174136 | RGMB | 1140.629855 | 0.24629965 | 0.065421852 | 3.76479174 | 0.000166688 | 0.000717415 |
| ENSG00000174137 | FAM53A | 79.94740815 | 0.026923857 | 0.197022823 | 0.136653492 | 0.891304692 | 0.944238 |
| ENSG00000174151 | CYB561D1 | 1158.850459 | 0.554170102 | 0.066739012 | 8.303540714 | 1.01E-16 | 1.37E-15 |
| ENSG00000174165 | ZDHHC24 | 821.6796359 | 0.516145978 | 0.077772186 | 6.636639681 | 3.21E-11 | 2.89E-10 |
| ENSG00000174171 | ENSG0000017 | 4.106264827 | -1.201988233 | 0.920913284 | -1.305213264 | 0.191820239 | 0.351803348 |
| ENSG00000174173 | TRMT10C | 866.160727 | -0.033607762 | 0.073066655 | -0.459960324 | 0.6455447 | 0.792669964 |
| ENSG00000174177 | CTU2 | 715.91394 | 0.263203288 | 0.080401483 | 3.273612355 | 0.001061822 | 0.003954252 |
| ENSG00000174197 | MGA | 2130.747731 | -0.184050471 | 0.072623898 | -2.534296239 | 0.011267343 | 0.033006367 |
| ENSG00000174206 | KICS2 | 596.4203197 | 0.097156029 | 0.083491517 | 1.163663488 | 0.244560412 | 0.421377185 |
| ENSG00000174227 | PIGG | 1822.903482 | 0.015279633 | 0.057176584 | 0.267235844 | 0.78928758 | 0.886041453 |
| ENSG00000174231 | PRPF8 | 12304.33961 | -0.00511871 | 0.044740995 | -0.114407599 | 0.90891469 | 0.954387467 |
| ENSG00000174233 | ADCY6 | 3386.590345 | 0.399248444 | 0.055499874 | 7.193682012 | 6.31E-13 | 6.55E-12 |
| ENSG00000174236 | REP15 | 5.474953469 | 1.320937498 | 0.803546884 | 1.643883542 | 0.10020026 | 0.21230202 |
| ENSG00000174238 | PITPNA | 2749.390531 | 0.213115255 | 0.052672848 | 4.046017351 | 5.21E-05 | 0.000243045 |
| ENSG00000174243 | DDX23 | 5959.618321 | 0.102572833 | 0.048200489 | 2.128045479 | 0.033333313 | 0.085000689 |

| | | | | | | | |
|---|---|---|---|---|---|---|---|
| ENSG00000174276 | ZNHIT2 | 216.4572944 | 0.07619494 | 0.139244879 | 0.547201022 | 0.584240634 | 0.748574628 |
| ENSG00000174282 | ZBTB4 | 2885.080816 | -0.037041707 | 0.05439305 | -0.681000726 | 0.495871031 | 0.677941809 |
| ENSG00000174292 | TNK1 | 476.8091886 | 0.248974109 | 0.091643228 | 2.716775862 | 0.006592123 | 0.0205802 |
| ENSG00000174306 | ZHX3 | 2659.453494 | -0.067672262 | 0.056994647 | -1.187344172 | 0.235091891 | 0.410117476 |
| ENSG00000174307 | PHLDA3 | 2647.624418 | 0.303767184 | 0.062717392 | 4.84342817 | 1.28E-06 | 7.29E-06 |
| ENSG00000174326 | SLC16A11 | 5.35258578 | 1.692647997 | 0.828511948 | 2.04299769 | 0.041052667 | 0.101327717 |
| ENSG00000174327 | SLC16A13 | 254.9905071 | -0.215387477 | 0.121648438 | -1.770573309 | 0.076631684 | 0.17087197 |
| ENSG00000174353 | STAG3L3 | 82.37841636 | 0.730867235 | 0.208307473 | 3.508598255 | 0.000450475 | 0.001796886 |
| ENSG00000174365 | SNHG11 | 229.2438672 | -0.645497461 | 0.129737245 | -4.975421383 | 6.51E-07 | 3.86E-06 |
| ENSG00000174370 | KCNJ5-AS1 | 140.243834 | 0.141882848 | 0.152758103 | 0.928807347 | 0.352988936 | 0.544447401 |
| ENSG00000174371 | EXO1 | 635.0350373 | -0.472276476 | 0.089028809 | -5.304760127 | 1.13E-07 | 7.29E-07 |
| ENSG00000174373 | RALGAPA1 | 899.2005222 | -0.535633382 | 0.072831987 | -7.354370018 | 1.92E-13 | 2.05E-12 |
| ENSG00000174405 | LIG4 | 326.3037483 | 0.071617082 | 0.105197632 | 0.680786061 | 0.496006871 | 0.677981461 |
| ENSG00000174428 | GTF2IRD2B | 260.959173 | 0.279292292 | 0.118743261 | 2.35206857 | 0.018669333 | 0.051487677 |
| ENSG00000174437 | ATP2A2 | 7763.747817 | 0.0817831 | 0.046531976 | 1.757567754 | 0.078821084 | 0.174650407 |
| ENSG00000174442 | ZWILCH | 1595.515251 | -0.424830827 | 0.060441088 | -7.028841493 | 2.08E-12 | 2.08E-11 |
| ENSG00000174444 | RPL4 | 31318.37319 | 0.178173388 | 0.055743345 | 3.196316735 | 0.001391942 | 0.005064418 |
| ENSG00000174446 | SNAPC5 | 799.7863525 | -0.17310701 | 0.073135877 | -2.366923283 | 0.01793665 | 0.049746588 |
| ENSG00000174453 | VWC2L | 3.702047512 | 5.239100785 | 1.406499387 | 3.724922196 | 0.000195375 | 0.000831061 |
| ENSG00000174456 | C12orf76 | 210.0142043 | 0.350458224 | 0.126248212 | 2.775946044 | 0.005504134 | 0.017513711 |
| ENSG00000174460 | ZCCHC12 | 2.03441087 | 0.959527754 | 1.308244409 | 0.73344686 | 0.463285927 | 0.649778624 |
| ENSG00000174469 | CNTNAP2 | 2224.382143 | -1.079569464 | 0.065538002 | -16.47241949 | 5.79E-61 | 4.24E-59 |
| ENSG00000174473 | GALNTL6 | 24.25542959 | -1.137247045 | 0.385423728 | -2.950640977 | 0.003171153 | 0.010685624 |
| ENSG00000174483 | BBS1 | 40.03288664 | 0.18073766 | 0.282188699 | 0.640485111 | 0.521857266 | 0.69885614 |
| ENSG00000174485 | DENND4A | 959.8598058 | 0.039403951 | 0.07224122 | 0.545449693 | 0.585444271 | 0.7496144 |
| ENSG00000174498 | IGDCC3 | 858.0502965 | -0.141034418 | 0.074327021 | -1.897485135 | 0.057763939 | 0.135214771 |
| ENSG00000174501 | ANKRD36C | 60.36210547 | -0.107100989 | 0.226873136 | -0.472074354 | 0.636873717 | 0.786213586 |
| ENSG00000174514 | MFSD4A | 6.521272125 | 0.707093514 | 0.688100735 | 1.027601743 | 0.304137204 | 0.491469954 |
| ENSG00000174516 | PELI3 | 401.617544 | 0.677212107 | 0.101438175 | 6.676106955 | 2.45E-11 | 2.22E-10 |
| ENSG00000174521 | TTC9B | 3.301830217 | 0.011522125 | 1.011621948 | 0.011389754 | 0.990912487 | 0.995176993 |
| ENSG00000174529 | TMEM81 | 87.82408647 | 0.658793883 | 0.193076374 | 3.412089577 | 0.000644669 | 0.002501474 |
| ENSG00000174547 | MRPL11 | 1644.869388 | -0.186827659 | 0.057781774 | -3.233332025 | 0.001223553 | 0.00450839 |
| ENSG00000174564 | IL20RB | 40.6439086 | -0.912195733 | 0.283555097 | -3.216996421 | 0.001295402 | 0.004750084 |

| | | | | | | | |
|---|---|---|---|---|---|---|---|
| ENSG00000174567 | GOLT1A | 482.5980612 | 0.166531561 | 0.090158537 | 1.847096974 | 0.064733087 | 0.148691 |
| ENSG00000174574 | AKIRIN1 | 1478.245251 | 0.05488037 | 0.059332541 | 0.92496241 | 0.354985461 | 0.546644725 |
| ENSG00000174579 | MSL2 | 1282.365071 | -0.227293115 | 0.061557664 | -3.692360934 | 0.000222182 | 0.000934859 |
| ENSG00000174586 | ZNF497 | 128.4645489 | 0.173858098 | 0.16195115 | 1.073521844 | 0.283037043 | 0.466676473 |
| ENSG00000174599 | TRAM1L1 | 109.6881746 | 2.51E-05 | 0.169933323 | 0.000147676 | 0.999882172 | 0.999914377 |
| ENSG00000174606 | ANGEL2 | 530.6851203 | -0.067336686 | 0.092060964 | -0.73143581 | 0.464512999 | 0.650624508 |
| ENSG00000174607 | UGT8 | 117.581745 | -0.055267721 | 0.166141756 | -0.332654009 | 0.73939547 | 0.856263924 |
| ENSG00000174628 | IQCK | 270.5833761 | 0.445556936 | 0.118505461 | 3.759800864 | 0.000170049 | 0.000730919 |
| ENSG00000174640 | SLCO2A1 | 180.311744 | -0.554395656 | 0.141612096 | -3.914889138 | 9.04E-05 | 0.000406032 |
| ENSG00000174652 | ZNF266 | 549.3807568 | 0.138835596 | 0.084572991 | 1.641606782 | 0.100671521 | 0.212985769 |
| ENSG00000174669 | SLC29A2 | 2795.633679 | -0.284637079 | 0.060319299 | -4.718839334 | 2.37E-06 | 1.31E-05 |
| ENSG00000174672 | BRSK2 | 389.2062194 | -0.188079186 | 0.100108541 | -1.878752647 | 0.060278276 | 0.140071276 |
| ENSG00000174680 | GRIK1-AS1 | 3.285538194 | -2.076355762 | 1.111896862 | -1.867399606 | 0.061845805 | 0.143042706 |
| ENSG00000174684 | B4GAT1 | 580.8775904 | -0.214776637 | 0.087030818 | -2.467822798 | 0.013593759 | 0.038946028 |
| ENSG00000174695 | TMEM167A | 2194.91536 | -0.103200433 | 0.058886875 | -1.75252012 | 0.079684413 | 0.176121056 |
| ENSG00000174705 | SH3PXD2B | 613.9024495 | 0.465601836 | 0.082119454 | 5.669811642 | 1.43E-08 | 1.01E-07 |
| ENSG00000174718 | RESF1 | 532.991863 | 0.105753664 | 0.095849932 | 1.103325399 | 0.269885881 | 0.451515614 |
| ENSG00000174720 | LARP7 | 1440.091284 | -0.04669015 | 0.064922874 | -0.719163322 | 0.472040296 | 0.657714066 |
| ENSG00000174721 | FGFBP3 | 119.7070025 | -0.18546 | 0.162665098 | -1.140133944 | 0.254230502 | 0.433255161 |
| ENSG00000174738 | NR1D2 | 2223.419569 | 0.027759466 | 0.054978767 | 0.504912487 | 0.613620303 | 0.770552319 |
| ENSG00000174740 | PABPC5 | 18.66039655 | 0.520207924 | 0.414060704 | 1.256356661 | 0.208986687 | 0.375688003 |
| ENSG00000174744 | BRMS1 | 1919.238817 | 0.089285777 | 0.065399051 | 1.365245753 | 0.172175823 | 0.324012582 |
| ENSG00000174748 | RPL15 | 18732.58632 | 0.303034061 | 0.052208051 | 5.80435493 | 6.46E-09 | 4.74E-08 |
| ENSG00000174749 | FAM241A | 443.4500835 | -0.176169075 | 0.102626652 | -1.716601593 | 0.086051991 | 0.187467491 |
| ENSG00000174775 | HRAS | 1304.349196 | -0.25820458 | 0.071093721 | -3.631890054 | 0.000281353 | 0.001162866 |
| ENSG00000174780 | SRP72 | 5242.599974 | 0.056875034 | 0.050021064 | 1.137021673 | 0.255529226 | 0.43491916 |
| ENSG00000174788 | PCP2 | 15.82971082 | 0.628202512 | 0.440664297 | 1.42558069 | 0.153989413 | 0.297224518 |
| ENSG00000174791 | RIN1 | 148.2796163 | 0.785116456 | 0.153633801 | 5.110310693 | 3.22E-07 | 1.98E-06 |
| ENSG00000174792 | ODAPH | 3.431072778 | -1.079236628 | 0.970316674 | -1.112251965 | 0.266029831 | 0.446987791 |
| ENSG00000174796 | THAP6 | 366.7751256 | 0.103526397 | 0.099032132 | 1.045381886 | 0.295846505 | 0.481960269 |
| ENSG00000174799 | CEP135 | 405.2366782 | 0.001709186 | 0.098471127 | 0.017357225 | 0.986151633 | 0.993350589 |
| ENSG00000174804 | FZD4 | 2671.664791 | -0.80537804 | 0.052735575 | -15.27200646 | 1.18E-52 | 6.93E-51 |
| ENSG00000174808 | BTC | 60.15431742 | -0.723107141 | 0.233820426 | -3.092574729 | 0.001984283 | 0.006983652 |

| | | | | | | | |
|---|---|---|---|---|---|---|---|
| ENSG00000174827 | PDZK1 | 181.1005539 | 1.042066426 | 0.149399134 | 6.975049979 | 3.06E-12 | 3.01E-11 |
| ENSG00000174837 | ADGRE1 | 2.730934548 | 0.489226763 | 1.114544434 | 0.438947743 | 0.660699402 | 0.803842179 |
| ENSG00000174839 | DENND6A | 1203.68412 | -0.024100508 | 0.069668555 | -0.345930929 | 0.729394624 | 0.850046366 |
| ENSG00000174840 | PDE12 | 795.178542 | 0.016731405 | 0.074843742 | 0.223551155 | 0.823106574 | 0.906409914 |
| ENSG00000174842 | GLMN | 156.8591646 | 0.333924248 | 0.145075205 | 2.301732052 | 0.021350287 | 0.057838403 |
| ENSG00000174844 | DNAH12 | 3.337016895 | 0.056958822 | 0.954688418 | 0.059662211 | 0.952424669 | 0.977917014 |
| ENSG00000174851 | YIF1A | 1605.549853 | 0.002237082 | 0.064562008 | 0.034650133 | 0.972358725 | 0.987467756 |
| ENSG00000174871 | CNIH2 | 187.6601795 | 0.344544468 | 0.140060635 | 2.459966479 | 0.013894999 | 0.03973464 |
| ENSG00000174885 | NLRP6 | 1.944119172 | 0.310824265 | 1.225621769 | 0.253605372 | 0.799800446 | 0.893253273 |
| ENSG00000174886 | NDUFA11 | 179.7529313 | 0.222179344 | 0.150036369 | 1.480836583 | 0.138650125 | 0.274591882 |
| ENSG00000174891 | RSRC1 | 957.6553715 | -0.073402795 | 0.06795946 | -1.080096798 | 0.280099078 | 0.463464573 |
| ENSG00000174899 | SLC66A1L | 2.906590766 | -1.513997712 | 1.087504596 | -1.392175921 | 0.163869132 | 0.311704635 |
| ENSG00000174903 | RAB1B | 12835.42019 | -0.1717491 | 0.048190781 | -3.563940986 | 0.000365328 | 0.001478065 |
| ENSG00000174912 | METTL15P1 | 58.98524164 | 0.128220008 | 0.241482328 | 0.530970563 | 0.595439178 | 0.756704466 |
| ENSG00000174915 | PTDSS2 | 849.1869209 | -0.048905224 | 0.070626906 | -0.692444667 | 0.488658124 | 0.671850006 |
| ENSG00000174917 | MICOS13 | 485.3744821 | -0.932538655 | 0.098562723 | -9.461372648 | 3.04E-21 | 5.47E-20 |
| ENSG00000174928 | C3orf33 | 35.8058355 | 0.538223664 | 0.302546942 | 1.778975719 | 0.075243743 | 0.168333476 |
| ENSG00000174938 | SEZ6L2 | 3170.0856 | 0.381345351 | 0.064150501 | 5.944542081 | 2.77E-09 | 2.12E-08 |
| ENSG00000174939 | ASPHD1 | 239.364374 | 1.230418048 | 0.137102152 | 8.974461978 | 2.85E-19 | 4.57E-18 |
| ENSG00000174943 | KCTD13 | 542.247026 | 0.268702995 | 0.083349392 | 3.223814682 | 0.001264952 | 0.004644367 |
| ENSG00000174945 | AMZ1 | 118.5948045 | 2.019795376 | 0.18539278 | 10.89468194 | 1.22E-27 | 3.12E-26 |
| ENSG00000174948 | GPR149 | 3.570588938 | 4.153944894 | 1.398696034 | 2.969869646 | 0.002979261 | 0.010099639 |
| ENSG00000174950 | CD164L2 | 8.68856159 | 1.340767887 | 0.631211636 | 2.124117826 | 0.033660292 | 0.085691611 |
| ENSG00000174951 | FUT1 | 400.1781709 | -0.08457931 | 0.096243392 | -0.87880642 | 0.379506245 | 0.57196065 |
| ENSG00000174953 | DHX36 | 3800.574532 | 0.349730705 | 0.050242366 | 6.960872531 | 3.38E-12 | 3.31E-11 |
| ENSG00000174963 | ZIC4 | 275.8458777 | -0.134826109 | 0.117450123 | -1.14794353 | 0.250991877 | 0.429330673 |
| ENSG00000174977 | PAIP1P2 | 1756.321813 | -0.039133802 | 0.059962935 | -0.652633187 | 0.513992785 | 0.693073347 |
| ENSG00000174989 | FBXW8 | 1336.417603 | 0.184725401 | 0.060563601 | 3.050105979 | 0.002287606 | 0.00794996 |
| ENSG00000174996 | KLC2 | 3220.942255 | -0.328850025 | 0.050449872 | -6.51835201 | 7.11E-11 | 6.22E-10 |
| ENSG00000175003 | SLC22A1 | 2.563949411 | 1.020669569 | 1.171606822 | 0.871170729 | 0.38366094 | 0.576144406 |
| ENSG00000175029 | CTBP2 | 3073.335753 | -0.048886538 | 0.047630647 | -1.026367302 | 0.304718484 | 0.492131937 |
| ENSG00000175048 | ZDHHC14 | 175.4028701 | 0.271816534 | 0.141541423 | 1.920402723 | 0.05480705 | 0.129602487 |
| ENSG00000175054 | ATR | 1329.998573 | 0.001644258 | 0.073171447 | 0.022471305 | 0.982072002 | 0.991627389 |

| | | | | | | | |
|---|---|---|---|---|---|---|---|
| ENSG00000175061 | SNHG29 | 8801.566532 | 0.251143389 | 0.044987374 | 5.582530501 | 2.37E-08 | 1.63E-07 |
| ENSG00000175063 | UBE2C | 2110.134329 | -0.450778863 | 0.053154325 | -8.480567886 | 2.24E-17 | 3.16E-16 |
| ENSG00000175066 | GK5 | 697.969627 | 0.199136477 | 0.075679097 | 2.631327348 | 0.008505207 | 0.025784254 |
| ENSG00000175073 | VCPIP1 | 835.0907136 | 0.176610492 | 0.08006156 | 2.205933682 | 0.027388643 | 0.071854469 |
| ENSG00000175087 | PDIK1L | 496.9637698 | 0.106166908 | 0.090424727 | 1.174091544 | 0.24035836 | 0.416168827 |
| ENSG00000175093 | SPSB4 | 30.59520207 | -0.496501609 | 0.32451482 | -1.52998131 | 0.126021355 | 0.255044542 |
| ENSG00000175104 | TRAF6 | 657.3652065 | -0.04709564 | 0.079958156 | -0.589003578 | 0.555858873 | 0.726680327 |
| ENSG00000175105 | ZNF654 | 672.875503 | -0.221538789 | 0.078423315 | -2.824909761 | 0.004729393 | 0.015325933 |
| ENSG00000175106 | TVP23C | 25.47738632 | 0.153456676 | 0.35148055 | 0.436600763 | 0.662400909 | 0.804963395 |
| ENSG00000175110 | MRPS22 | 1031.306393 | -0.110370966 | 0.069139113 | -1.596360763 | 0.110408271 | 0.229437969 |
| ENSG00000175115 | PACS1 | 7367.888733 | -0.283842315 | 0.048852682 | -5.810168508 | 6.24E-09 | 4.59E-08 |
| ENSG00000175130 | MARCKSL1 | 8794.747652 | 0.258569253 | 0.044196934 | 5.850388872 | 4.90E-09 | 3.66E-08 |
| ENSG00000175137 | SH3BP5L | 1803.266017 | -0.186753258 | 0.060825403 | -3.070316826 | 0.002138318 | 0.007467349 |
| ENSG00000175147 | TMEM51-AS1 | 35.38084558 | 0.841368015 | 0.310846011 | 2.70670359 | 0.00679549 | 0.021131699 |
| ENSG00000175155 | YPEL2 | 1553.978524 | 0.486467103 | 0.061540217 | 7.904864883 | 2.68E-15 | 3.29E-14 |
| ENSG00000175161 | CADM2 | 150.8250694 | -0.21926344 | 0.149810382 | -1.463606444 | 0.143301511 | 0.281425962 |
| ENSG00000175166 | PSMD2 | 9302.499942 | -0.164406887 | 0.043020489 | -3.821595049 | 0.000132591 | 0.00057914 |
| ENSG00000175170 | FAM182B | 33.51390988 | -0.128583196 | 0.309800459 | -0.415051665 | 0.678104082 | 0.81682581 |
| ENSG00000175175 | PPM1E | 380.0249443 | -1.039359791 | 0.106858378 | -9.726516626 | 2.32E-22 | 4.44E-21 |
| ENSG00000175182 | FAM131A | 254.2847671 | 0.509253057 | 0.117598777 | 4.330428176 | 1.49E-05 | 7.50E-05 |
| ENSG00000175183 | CSRP2 | 24.03605587 | -0.153474798 | 0.359810092 | -0.426543895 | 0.66971157 | 0.810251749 |
| ENSG00000175189 | INHBC | 3.519475456 | -0.030034656 | 0.956805379 | -0.031390559 | 0.97495807 | 0.98863534 |
| ENSG00000175193 | PARL | 164.1232591 | -0.138625685 | 0.143679008 | -0.964829076 | 0.33463043 | 0.525746323 |
| ENSG00000175197 | DDIT3 | 363.9931008 | 0.228850137 | 0.102645688 | 2.229515338 | 0.025779636 | 0.068171164 |
| ENSG00000175198 | PCCA | 495.9601262 | -0.340928287 | 0.086651431 | -3.9344796 | 8.34E-05 | 0.000377104 |
| ENSG00000175202 | HIGD2B | 3.240917067 | -0.426516595 | 0.973345128 | -0.438196671 | 0.661243722 | 0.804222818 |
| ENSG00000175203 | DCTN2 | 3827.447755 | -0.068936138 | 0.046833881 | -1.471928794 | 0.141040107 | 0.278127745 |
| ENSG00000175213 | ZNF408 | 694.4532149 | -0.048075603 | 0.082288321 | -0.584233614 | 0.559063149 | 0.728807456 |
| ENSG00000175215 | CTDSP2 | 9800.772991 | -0.036458394 | 0.045571431 | -0.800027417 | 0.423694913 | 0.614617509 |
| ENSG00000175216 | CKAP5 | 10049.51327 | -0.110279698 | 0.045628711 | -2.416892689 | 0.015653631 | 0.044163609 |
| ENSG00000175220 | ARHGAP1 | 3006.154556 | -0.268652226 | 0.060553788 | -4.436588292 | 9.14E-06 | 4.74E-05 |
| ENSG00000175221 | MED16 | 928.2783234 | 0.011006094 | 0.079818231 | 0.137889475 | 0.890327769 | 0.943813263 |
| ENSG00000175224 | ATG13 | 3956.964199 | 0.193414196 | 0.049907245 | 3.875473273 | 0.000106418 | 0.000472281 |

| | | | | | | | |
|---|---|---|---|---|---|---|---|
| ENSG00000175262 | C1orf127 | 7.547973741 | -3.13733779 | 0.845639299 | -3.71001891 | 0.000207244 | 0.000876668 |
| ENSG00000175264 | CHST1 | 11.79286857 | 0.061015598 | 0.516740593 | 0.118077811 | 0.906006005 | 0.95262805 |
| ENSG00000175265 | GOLGA8A | 794.0331061 | 0.214338938 | 0.282296718 | 0.759268263 | 0.447692098 | 0.636060411 |
| ENSG00000175267 | VWA3A | 4.741562944 | 0.637516141 | 0.812012171 | 0.785106633 | 0.432391051 | 0.622430391 |
| ENSG00000175274 | TP53I11 | 3503.71368 | 0.211929426 | 0.049842555 | 4.25197755 | 2.12E-05 | 0.000104849 |
| ENSG00000175279 | CENPS | 21.35278766 | -0.243849771 | 0.380874607 | -0.640236356 | 0.52201895 | 0.699031855 |
| ENSG00000175283 | DOLK | 713.9198258 | 0.051598648 | 0.080338058 | 0.642269049 | 0.520698507 | 0.697950759 |
| ENSG00000175287 | PHYHD1 | 1.955022462 | 0.814733275 | 1.25415114 | 0.649629258 | 0.515931729 | 0.694610826 |
| ENSG00000175302 | ANKRD30BP1 | 16.45278324 | 0.211872511 | 0.439235781 | 0.482366236 | 0.629545802 | 0.781143482 |
| ENSG00000175305 | CCNE2 | 1089.496995 | -0.513111259 | 0.076447082 | -6.711979668 | 1.92E-11 | 1.75E-10 |
| ENSG00000175309 | PHYKPL | 1270.957607 | -0.583057311 | 0.06474033 | -9.006091085 | 2.14E-19 | 3.45E-18 |
| ENSG00000175318 | GRAMD2A | 2.932686559 | -3.059392181 | 1.337593733 | -2.287235731 | 0.022182067 | 0.059879575 |
| ENSG00000175322 | ZNF519 | 235.1639889 | 0.351921453 | 0.121520829 | 2.895976405 | 0.003779808 | 0.012526553 |
| ENSG00000175324 | LSM1 | 761.6078747 | -0.182020688 | 0.074892689 | -2.430419979 | 0.015081336 | 0.042701289 |
| ENSG00000175334 | BANF1 | 4094.413484 | -0.144827867 | 0.048553112 | -2.982875075 | 0.002855545 | 0.009728683 |
| ENSG00000175344 | CHRNA7 | 4.182264203 | -0.741644564 | 0.883979561 | -0.838983837 | 0.40147838 | 0.594226339 |
| ENSG00000175348 | TMEM9B | 758.1241564 | 0.043281658 | 0.074057058 | 0.584436639 | 0.558926582 | 0.728758429 |
| ENSG00000175354 | PTPN2 | 1297.596066 | 0.47226588 | 0.067562966 | 6.990011052 | 2.75E-12 | 2.71E-11 |
| ENSG00000175356 | SCUBE2 | 1522.928924 | 0.200123126 | 0.059545968 | 3.36081743 | 0.000777122 | 0.002969216 |
| ENSG00000175376 | EIF1AD | 1315.143802 | -0.196782919 | 0.064864868 | -3.033736529 | 0.002415453 | 0.0083588 |
| ENSG00000175387 | SMAD2 | 2281.15328 | 0.457258798 | 0.058122322 | 7.867180526 | 3.63E-15 | 4.41E-14 |
| ENSG00000175390 | EIF3F | 2523.359255 | 0.182025068 | 0.050355729 | 3.614783703 | 0.000300598 | 0.001236902 |
| ENSG00000175395 | ZNF25 | 189.6089514 | -0.165820069 | 0.13307712 | -1.246044918 | 0.212747905 | 0.380808126 |
| ENSG00000175414 | ARL10 | 19.67840073 | -0.514716421 | 0.436144576 | -1.180150917 | 0.237940197 | 0.413417262 |
| ENSG00000175416 | CLTB | 1824.441681 | -0.546162268 | 0.06237546 | -8.756043959 | 2.02E-18 | 3.06E-17 |
| ENSG00000175449 | RFESD | 19.13537696 | 0.394718911 | 0.40389995 | 0.977269027 | 0.328435983 | 0.518749755 |
| ENSG00000175455 | CCDC14 | 1554.111713 | -0.252671536 | 0.081442498 | -3.102453174 | 0.001919239 | 0.006772396 |
| ENSG00000175463 | TBC1D10C | 2.184743668 | 1.174859105 | 1.234092801 | 0.952002236 | 0.341095848 | 0.532222742 |
| ENSG00000175467 | SART1 | 3134.259374 | -0.007472963 | 0.054158373 | -0.137983531 | 0.890253433 | 0.943797108 |
| ENSG00000175470 | PPP2R2D | 1009.135552 | -0.42200124 | 0.066063264 | -6.387835113 | 1.68E-10 | 1.43E-09 |
| ENSG00000175471 | MCTP1 | 45.8634233 | 0.707111213 | 0.262618464 | 2.692541881 | 0.007090963 | 0.021960635 |
| ENSG00000175482 | POLD4 | 173.3417007 | -0.259923875 | 0.143293862 | -1.813921905 | 0.06968975 | 0.158035079 |
| ENSG00000175505 | CLCF1 | 65.40609797 | 1.488493731 | 0.233946721 | 6.362533005 | 1.98E-10 | 1.68E-09 |

| | | | | | | | |
|---|---|---|---|---|---|---|---|
| ENSG00000175536 | LIPT2 | 176.1544588 | -0.381768719 | 0.140453496 | -2.718114745 | 0.006565506 | 0.020504719 |
| ENSG00000175544 | CABP4 | 3.657635781 | -0.934372847 | 0.964987541 | -0.968274519 | 0.332907287 | 0.524052325 |
| ENSG00000175548 | ALG10B | 229.8345095 | -0.069358187 | 0.123204531 | -0.562951591 | 0.573467848 | 0.74022645 |
| ENSG00000175550 | DRAP1 | 1916.774944 | 0.189787317 | 0.057514849 | 3.299796839 | 0.000967548 | 0.003631473 |
| ENSG00000175564 | UCP3 | 26.50201564 | 0.220159128 | 0.341530348 | 0.644625373 | 0.519169988 | 0.696723975 |
| ENSG00000175567 | UCP2 | 4612.004989 | -0.321333758 | 0.049523389 | -6.488525175 | 8.67E-11 | 7.55E-10 |
| ENSG00000175573 | C11orf68 | 1016.102923 | 0.252234207 | 0.076542079 | 3.295366576 | 0.000982934 | 0.003684395 |
| ENSG00000175575 | PAAF1 | 1193.863619 | 0.023490463 | 0.06254095 | 0.375601312 | 0.707213315 | 0.836532294 |
| ENSG00000175581 | MRPL48 | 1070.978017 | 0.06651245 | 0.065118703 | 1.021403165 | 0.307063467 | 0.494908728 |
| ENSG00000175582 | RAB6A | 5770.014303 | -0.066228569 | 0.049475741 | -1.338606915 | 0.180698674 | 0.335972408 |
| ENSG00000175591 | P2RY2 | 1118.191725 | 0.392858903 | 0.070627087 | 5.562439541 | 2.66E-08 | 1.83E-07 |
| ENSG00000175592 | FOSL1 | 9.217058303 | 0.595881727 | 0.586257863 | 1.016415751 | 0.309431448 | 0.497467056 |
| ENSG00000175595 | ERCC4 | 512.7757709 | 0.054218852 | 0.085531273 | 0.633906766 | 0.52614167 | 0.702175223 |
| ENSG00000175600 | SUGCT | 3.376683456 | 0.542430644 | 0.971298621 | 0.558459194 | 0.576530862 | 0.74273769 |
| ENSG00000175602 | CCDC85B | 369.0917811 | 0.35874694 | 0.107270846 | 3.344309776 | 0.000824876 | 0.003136524 |
| ENSG00000175606 | TMEM70 | 571.2932159 | 0.44300514 | 0.082067961 | 5.398027825 | 6.74E-08 | 4.43E-07 |
| ENSG00000175611 | ERCC6L2-AS1 | 198.6539597 | 0.387103087 | 0.130102873 | 2.975361569 | 0.002926434 | 0.009935237 |
| ENSG00000175634 | RPS6KB2 | 2438.92224 | -0.011501789 | 0.057737668 | -0.199207722 | 0.842100259 | 0.917009895 |
| ENSG00000175643 | RMI2 | 758.5122661 | -0.48003519 | 0.074836393 | -6.414461889 | 1.41E-10 | 1.21E-09 |
| ENSG00000175662 | TOM1L2 | 3622.696613 | 0.29430975 | 0.069501954 | 4.234553624 | 2.29E-05 | 0.00011271 |
| ENSG00000175691 | ZNF77 | 208.4909727 | -0.07874379 | 0.127711716 | -0.616574518 | 0.537515406 | 0.712545705 |
| ENSG00000175697 | GPR156 | 48.48416153 | 1.094088879 | 0.268521729 | 4.074489178 | 4.61E-05 | 0.00021678 |
| ENSG00000175701 | MTLN | 259.5983078 | 0.068137346 | 0.128530705 | 0.530125046 | 0.596025235 | 0.756993626 |
| ENSG00000175707 | KDF1 | 491.6849251 | 0.17277213 | 0.090796125 | 1.902857971 | 0.05705908 | 0.133865341 |
| ENSG00000175711 | B3GNTL1 | 144.8028802 | 0.135577188 | 0.154537292 | 0.877310493 | 0.380318014 | 0.572774111 |
| ENSG00000175727 | MLXIP | 2293.413381 | -0.513776583 | 0.055135722 | -9.318397711 | 1.18E-20 | 2.04E-19 |
| ENSG00000175730 | BAK1P1 | 137.1336097 | 0.744988154 | 0.162727482 | 4.578133604 | 4.69E-06 | 2.51E-05 |
| ENSG00000175741 | RWDD4P2 | 84.80775485 | 0.182478917 | 0.190753861 | 0.95661978 | 0.338759201 | 0.530045518 |
| ENSG00000175745 | NR2F1 | 282.5261821 | 0.938993204 | 0.120103292 | 7.818213737 | 5.36E-15 | 6.44E-14 |
| ENSG00000175756 | AURKAIP1 | 1413.175542 | 9.34E-05 | 0.07067565 | 0.001320851 | 0.998946114 | 0.999338182 |
| ENSG00000175764 | TTLL11 | 148.6195376 | 0.27688257 | 0.14745961 | 1.877684138 | 0.060424388 | 0.140353935 |
| ENSG00000175768 | TOMM5 | 37.5881016 | -0.012722038 | 0.297742437 | -0.042728334 | 0.965918093 | 0.984632112 |
| ENSG00000175772 | LINC01106 | 53.12441317 | 0.729740896 | 0.252170004 | 2.893844958 | 0.00380556 | 0.012600974 |

| | | | | | | | |
|---|---|---|---|---|---|---|---|
| ENSG00000175773 | ZBTB44-DT | 3.210287939 | 0.676269283 | 1.004551339 | 0.673205297 | 0.500816692 | 0.681904968 |
| ENSG00000175782 | SLC35E3 | 638.3527018 | -0.359931967 | 0.079719691 | -4.514969392 | 6.33E-06 | 3.35E-05 |
| ENSG00000175785 | PRIMA1 | 25.90309156 | -1.962957163 | 0.382794204 | -5.127969918 | 2.93E-07 | 1.81E-06 |
| ENSG00000175787 | ZNF169 | 102.4515169 | 0.207806183 | 0.175499022 | 1.184087411 | 0.236378463 | 0.411514149 |
| ENSG00000175792 | RUVBL1 | 3574.904128 | -0.072671732 | 0.047229799 | -1.538683903 | 0.123881484 | 0.251668548 |
| ENSG00000175793 | SFN | 2172.950455 | -0.75149431 | 0.062129601 | -12.09559216 | 1.11E-33 | 3.56E-32 |
| ENSG00000175806 | MSRA | 290.4049289 | 0.497694734 | 0.114345468 | 4.352553205 | 1.35E-05 | 6.82E-05 |
| ENSG00000175820 | CCDC168 | 3.431506047 | -0.032988719 | 0.923935332 | -0.035704575 | 0.971517922 | 0.987006252 |
| ENSG00000175826 | CTDNEP1 | 983.1607709 | -0.108413515 | 0.071191471 | -1.522844144 | 0.127797703 | 0.25788875 |
| ENSG00000175832 | ETV4 | 136.7910188 | -0.050787743 | 0.159371468 | -0.318675253 | 0.749972783 | 0.861813136 |
| ENSG00000175854 | SWI5 | 1000.918304 | 0.096801851 | 0.069274599 | 1.397364293 | 0.162304051 | 0.309497501 |
| ENSG00000175866 | BAIAP2 | 1505.490925 | 0.58861363 | 0.060028844 | 9.805513366 | 1.07E-22 | 2.08E-21 |
| ENSG00000175874 | CREG2 | 2.957297265 | 0.160483226 | 1.0107469 | 0.158776867 | 0.873844675 | 0.934676032 |
| ENSG00000175877 | TMEM270 | 3.378606223 | 1.116634075 | 0.977258154 | 1.142619348 | 0.253196669 | 0.431939753 |
| ENSG00000175879 | HOXD8 | 14.239276 | 0.190996415 | 0.468000833 | 0.408111272 | 0.683191985 | 0.820223767 |
| ENSG00000175886 | RPL7AP66 | 275.6335582 | 0.10746881 | 0.112607079 | 0.954369927 | 0.339896423 | 0.53082061 |
| ENSG00000175893 | ZDHHC21 | 561.7310032 | 0.072528117 | 0.094418546 | 0.768155404 | 0.442394864 | 0.631470633 |
| ENSG00000175894 | TSPEAR | 10.68257506 | 0.682486927 | 0.572341715 | 1.192446591 | 0.233086181 | 0.407390247 |
| ENSG00000175895 | PLEKHF2 | 3654.024392 | -0.586726163 | 0.054069383 | -10.85135675 | 1.96E-27 | 4.98E-26 |
| ENSG00000175906 | ARL4D | 101.8581349 | -0.615593633 | 0.187520431 | -3.282808332 | 0.001027785 | 0.003839341 |
| ENSG00000175920 | DOK7 | 164.512778 | 0.901905318 | 0.161669951 | 5.578682456 | 2.42E-08 | 1.67E-07 |
| ENSG00000175931 | UBE2O | 2916.77148 | 0.364803308 | 0.053504573 | 6.818170584 | 9.22E-12 | 8.66E-11 |
| ENSG00000175938 | ORAI3 | 205.5301251 | 0.137795348 | 0.157205337 | 0.876530979 | 0.380741443 | 0.573223646 |
| ENSG00000175970 | UNC119B | 3527.303627 | 0.117177275 | 0.046440464 | 2.523171944 | 0.011630152 | 0.033978055 |
| ENSG00000175984 | DENND2C | 129.5409841 | 0.162570901 | 0.157420211 | 1.032719373 | 0.301735245 | 0.488621101 |
| ENSG00000175985 | PLEKHD1 | 190.4704388 | -2.785242468 | 0.167438892 | -16.63438186 | 3.93E-62 | 2.95E-60 |
| ENSG00000176014 | TUBB6 | 1253.488342 | -0.885990629 | 0.062830056 | -14.10138203 | 3.72E-45 | 1.82E-43 |
| ENSG00000176018 | LYSMD3 | 915.4763485 | 0.371855178 | 0.075249021 | 4.941661371 | 7.75E-07 | 4.56E-06 |
| ENSG00000176022 | B3GALT6 | 709.5449281 | 0.262400135 | 0.080690736 | 3.251923931 | 0.001146267 | 0.004250099 |
| ENSG00000176024 | ZNF613 | 61.57085765 | -0.171093122 | 0.222679266 | -0.768338811 | 0.442285923 | 0.631470633 |
| ENSG00000176043 | RPL36P3 | 4.783548336 | -0.523043621 | 0.819983709 | -0.637870747 | 0.523557822 | 0.700001783 |
| ENSG00000176046 | NUPR1 | 1156.882098 | 1.931611082 | 0.070814739 | 27.27696407 | 7.96E-164 | 3.09E-161 |
| ENSG00000176049 | JAKMIP2 | 16.61957525 | 0.53157724 | 0.432123596 | 1.230150923 | 0.218640594 | 0.388355805 |

| | | | | | | | |
|---|---|---|---|---|---|---|---|
| ENSG00000176054 | RPL23P2 | 20.86540119 | -0.027837495 | 0.422764814 | -0.06584629 | 0.947500202 | 0.975544584 |
| ENSG00000176055 | MBLAC2 | 381.5981367 | 0.64653869 | 0.104580711 | 6.182198244 | 6.32E-10 | 5.13E-09 |
| ENSG00000176058 | TPRN | 1240.54606 | 0.158176904 | 0.074632757 | 2.119403181 | 0.034056408 | 0.086517608 |
| ENSG00000176087 | SLC35A4 | 3470.1908 | 0.097801781 | 0.053443556 | 1.830001377 | 0.067249733 | 0.153564491 |
| ENSG00000176092 | CRYBG2 | 87.80446665 | 1.151488983 | 0.205030119 | 5.616194287 | 1.95E-08 | 1.36E-07 |
| ENSG00000176095 | IP6K1 | 1742.66897 | 0.205427061 | 0.057988018 | 3.542577748 | 0.000396237 | 0.00159356 |
| ENSG00000176101 | SSNA1 | 1396.819649 | -0.315098076 | 0.074478975 | -4.230698323 | 2.33E-05 | 0.000114487 |
| ENSG00000176102 | CSTF3 | 1374.627771 | 0.182758352 | 0.060249209 | 3.033373476 | 0.002418361 | 0.008365815 |
| ENSG00000176105 | YES1 | 2479.369591 | 0.333012637 | 0.056020677 | 5.944459358 | 2.77E-09 | 2.12E-08 |
| ENSG00000176108 | CHMP6 | 865.9895524 | 0.312923878 | 0.076991067 | 4.064417982 | 4.82E-05 | 0.000225847 |
| ENSG00000176124 | DLEU1 | 208.5229218 | -0.075497759 | 0.130509293 | -0.578485699 | 0.56293625 | 0.731740569 |
| ENSG00000176125 | UFSP1 | 40.95591847 | -0.57516194 | 0.276497143 | -2.080173173 | 0.037509652 | 0.094040592 |
| ENSG00000176136 | MC5R | 20.39142283 | 1.389241957 | 0.432727558 | 3.210430975 | 0.001325361 | 0.004847541 |
| ENSG00000176142 | TMEM39A | 758.8497112 | -0.278386146 | 0.077306077 | -3.601090073 | 0.000316886 | 0.001299492 |
| ENSG00000176148 | TCP11L1 | 279.5283582 | -0.300094241 | 0.111095755 | -2.701221498 | 0.006908531 | 0.021445427 |
| ENSG00000176153 | GPX2 | 19.86329376 | -0.476275563 | 0.393655543 | -1.209878971 | 0.226325338 | 0.398488237 |
| ENSG00000176155 | CCDC57 | 1361.571248 | -0.07849344 | 0.067564374 | -1.161757814 | 0.245333854 | 0.422329177 |
| ENSG00000176170 | SPHK1 | 481.5622198 | 1.4273792 | 0.097741582 | 14.60360234 | 2.66E-48 | 1.39E-46 |
| ENSG00000176171 | BNIP3 | 899.3149644 | -0.66231457 | 0.071639474 | -9.245106508 | 2.35E-20 | 4.01E-19 |
| ENSG00000176182 | MYPOP | 172.0107013 | -0.132203027 | 0.138055429 | -0.957608311 | 0.338260307 | 0.529559793 |
| ENSG00000176183 | ENSG0000017 | 9.027113613 | -0.191648976 | 0.584875149 | -0.32767502 | 0.743157393 | 0.858367182 |
| ENSG00000176204 | LRRTM4 | 6.440970341 | -5.2193217 | 1.321216198 | -3.950391848 | 7.80E-05 | 0.000354708 |
| ENSG00000176208 | ATAD5 | 283.3157229 | -0.359749633 | 0.112526906 | -3.197009914 | 0.001388602 | 0.005053398 |
| ENSG00000176209 | SMIM19 | 524.5006942 | -0.266056857 | 0.085126987 | -3.125411399 | 0.001775565 | 0.006308111 |
| ENSG00000176222 | ZNF404 | 34.42408399 | 0.557815389 | 0.316453311 | 1.762709918 | 0.077949427 | 0.173120327 |
| ENSG00000176225 | RTTN | 291.4394336 | 0.382950611 | 0.115005367 | 3.329849918 | 0.000868928 | 0.003290394 |
| ENSG00000176236 | RPP38-DT | 2.646628257 | -0.891520396 | 1.111847664 | -0.801836821 | 0.422647356 | 0.613491717 |
| ENSG00000176243 | CDV3P1 | 3.738421603 | -0.73305473 | 0.910524783 | -0.805090365 | 0.420767535 | 0.611996403 |
| ENSG00000176244 | ACBD7 | 563.2656451 | -1.164895769 | 0.086505684 | -13.46611828 | 2.48E-41 | 1.07E-39 |
| ENSG00000176248 | ANAPC2 | 2159.210382 | -0.087686877 | 0.057649514 | -1.521034108 | 0.128251278 | 0.25862204 |
| ENSG00000176261 | ZBTB8OS | 418.7762539 | 0.111133839 | 0.096932048 | 1.146512861 | 0.251583009 | 0.430085269 |
| ENSG00000176268 | CYCSP34 | 9.670459277 | 0.396462446 | 0.568510014 | 0.697371086 | 0.485570592 | 0.669321308 |
| ENSG00000176273 | SLC35G1 | 147.7184299 | -0.02691966 | 0.150261733 | -0.179151801 | 0.857818509 | 0.926188651 |

| | | | | | | | |
|---|---|---|---|---|---|---|---|
| ENSG00000176289 | IDSP1 | 2.215018719 | 1.755017259 | 1.303892059 | 1.345983547 | 0.178307823 | 0.332821925 |
| ENSG00000176293 | ZNF135 | 5.845587099 | 0.124948765 | 0.727048282 | 0.171857589 | 0.863549489 | 0.929172842 |
| ENSG00000176318 | FOXN3P1 | 1.945566359 | 1.397621102 | 1.31365176 | 1.063920549 | 0.287364693 | 0.471279314 |
| ENSG00000176320 | ENSG0000017 | 4.542623214 | -0.548309202 | 0.851689047 | -0.643790364 | 0.519711383 | 0.69707672 |
| ENSG00000176340 | COX8A | 1691.81024 | -0.144482326 | 0.056469684 | -2.558582155 | 0.010509998 | 0.031037508 |
| ENSG00000176343 | RPL37AP8 | 11.60995703 | 0.347161765 | 0.521450041 | 0.665762275 | 0.505563062 | 0.686015208 |
| ENSG00000176371 | ZSCAN2 | 391.9542362 | -0.137939605 | 0.102314159 | -1.348196634 | 0.177595147 | 0.331788637 |
| ENSG00000176383 | B3GNT4 | 68.13798095 | 0.15233675 | 0.223552706 | 0.681435501 | 0.495595966 | 0.677854248 |
| ENSG00000176386 | CDC26 | 115.1254214 | 0.042646689 | 0.166200522 | 0.256597801 | 0.797489273 | 0.891887866 |
| ENSG00000176387 | HSD11B2 | 44.90092077 | 0.720471092 | 0.268509597 | 2.683222873 | 0.007291638 | 0.022537075 |
| ENSG00000176390 | CRLF3 | 234.2985242 | -0.295988253 | 0.118706985 | -2.493435861 | 0.012651344 | 0.036565683 |
| ENSG00000176393 | RNPEP | 3411.003261 | -0.247600553 | 0.048472965 | -5.10801341 | 3.26E-07 | 2.00E-06 |
| ENSG00000176396 | EID2 | 262.7970789 | -0.216210259 | 0.119704549 | -1.806199188 | 0.070887232 | 0.160275291 |
| ENSG00000176399 | DMRTA1 | 571.1914911 | -0.254696446 | 0.08181688 | -3.113006094 | 0.001851922 | 0.006554009 |
| ENSG00000176401 | EID2B | 68.87312052 | -0.048155809 | 0.219858333 | -0.219031084 | 0.826625834 | 0.908528011 |
| ENSG00000176402 | GJC3 | 6.201649028 | -2.026047282 | 0.80492912 | -2.517050546 | 0.011834185 | 0.034516982 |
| ENSG00000176406 | RIMS2 | 401.5620356 | -2.152696572 | 0.108596505 | -19.82288996 | 1.89E-87 | 2.51E-85 |
| ENSG00000176407 | KCMF1 | 1890.29728 | 0.071753351 | 0.05986162 | 1.198653684 | 0.230662635 | 0.404102018 |
| ENSG00000176410 | DNAJC30 | 426.9699143 | 0.574862225 | 0.096467422 | 5.959133257 | 2.54E-09 | 1.94E-08 |
| ENSG00000176422 | SPRYD4 | 519.665083 | 0.12855607 | 0.084495036 | 1.521463 | 0.12814369 | 0.258495976 |
| ENSG00000176428 | VPS37D | 74.09018973 | 0.183221879 | 0.206659808 | 0.8865869 | 0.375301349 | 0.568315595 |
| ENSG00000176438 | SYNE3 | 84.76998472 | -0.491490759 | 0.207052317 | -2.37375155 | 0.017608394 | 0.04897388 |
| ENSG00000176444 | CLK2 | 2056.093247 | -0.189226331 | 0.069229092 | -2.733335461 | 0.006269644 | 0.019704841 |
| ENSG00000176454 | LPCAT4 | 674.3534451 | 0.124565762 | 0.079108717 | 1.57461487 | 0.115345368 | 0.237866123 |
| ENSG00000176463 | SLCO3A1 | 351.3524091 | 0.399101215 | 0.101903662 | 3.916456063 | 8.99E-05 | 0.000403878 |
| ENSG00000176472 | ZNF575 | 24.63139885 | 0.216168713 | 0.358650933 | 0.602727313 | 0.546690109 | 0.719675835 |
| ENSG00000176473 | WDR25 | 290.0931182 | -0.362975519 | 0.118161417 | -3.071861595 | 0.002127283 | 0.00743334 |
| ENSG00000176476 | SGF29 | 550.4832837 | -0.027494621 | 0.08285752 | -0.331830117 | 0.740017542 | 0.856567724 |
| ENSG00000176485 | PLAAT3 | 288.2325977 | -0.599950941 | 0.111167769 | -5.396806504 | 6.78E-08 | 4.46E-07 |
| ENSG00000176490 | DIRAS1 | 58.66314208 | -0.514530047 | 0.229516838 | -2.24179651 | 0.024974527 | 0.066425003 |
| ENSG00000176531 | PHLDB3 | 583.1311008 | 0.030808481 | 0.081330579 | 0.378805632 | 0.704832205 | 0.834995392 |
| ENSG00000176532 | PRR15 | 368.0906096 | -1.693156332 | 0.109038144 | -15.52811029 | 2.24E-54 | 1.38E-52 |
| ENSG00000176533 | GNG7 | 134.4467625 | 1.085837776 | 0.163577928 | 6.638045774 | 3.18E-11 | 2.86E-10 |

| | | | | | | | |
|---|---|---|---|---|---|---|---|
| ENSG00000176542 | USF3 | 1256.910953 | -0.203762966 | 0.165384387 | -1.232056842 | 0.217927851 | 0.387454385 |
| ENSG00000176563 | CNTD1 | 17.69476993 | -0.289499849 | 0.426372865 | -0.678982816 | 0.49714875 | 0.679040564 |
| ENSG00000176593 | ENSG0000017 | 300.6231975 | 1.567395456 | 0.119829457 | 13.08021824 | 4.27E-39 | 1.67E-37 |
| ENSG00000176595 | KBTBD11 | 216.4734936 | 0.693607029 | 0.129001252 | 5.376746499 | 7.58E-08 | 4.97E-07 |
| ENSG00000176597 | B3GNT5 | 146.9846652 | -1.746383432 | 0.176963949 | -9.868583093 | 5.70E-23 | 1.13E-21 |
| ENSG00000176619 | LMNB2 | 4474.096176 | -0.172186613 | 0.049334097 | -3.490215163 | 0.000482632 | 0.001916492 |
| ENSG00000176623 | RMDN1 | 2893.775941 | 0.339026182 | 0.054158629 | 6.259873825 | 3.85E-10 | 3.19E-09 |
| ENSG00000176624 | MEX3C | 1315.747413 | 0.4180822 | 0.064914347 | 6.440520807 | 1.19E-10 | 1.02E-09 |
| ENSG00000176641 | RNF152 | 846.3459197 | 0.298985462 | 0.08015143 | 3.730257374 | 0.000191284 | 0.00081517 |
| ENSG00000176658 | MYO1D | 3426.658879 | 0.413688603 | 0.049475642 | 8.361460076 | 6.19E-17 | 8.51E-16 |
| ENSG00000176681 | LRRC37A | 11.85338708 | -0.043645241 | 0.512412965 | -0.085175911 | 0.932121541 | 0.967506817 |
| ENSG00000176697 | BDNF | 12.49910536 | -0.597254948 | 0.501149412 | -1.191770227 | 0.233351353 | 0.407702936 |
| ENSG00000176700 | SCAND2P | 569.3798663 | -0.484262128 | 0.092486148 | -5.236050359 | 1.64E-07 | 1.04E-06 |
| ENSG00000176714 | CCDC121 | 98.08164913 | 0.113274645 | 0.192887065 | 0.587258896 | 0.557029843 | 0.727416829 |
| ENSG00000176715 | ACSF3 | 1360.11337 | 0.203083499 | 0.061824637 | 3.284831246 | 0.001020435 | 0.003814989 |
| ENSG00000176720 | BOK | 1429.821871 | -0.379385983 | 0.070363197 | -5.39182409 | 6.97E-08 | 4.58E-07 |
| ENSG00000176723 | ZNF843 | 21.9121929 | -0.729705543 | 0.389931275 | -1.871369623 | 0.061293866 | 0.142042761 |
| ENSG00000176731 | RBIS | 780.5849785 | 0.082703044 | 0.078624925 | 1.05186802 | 0.292860106 | 0.478198507 |
| ENSG00000176732 | PFN4 | 23.31938702 | -0.235313723 | 0.376135672 | -0.625608631 | 0.531571676 | 0.707118252 |
| ENSG00000176749 | CDK5R1 | 252.9973059 | 0.684696238 | 0.123632599 | 5.538152918 | 3.06E-08 | 2.09E-07 |
| ENSG00000176753 | PAK6-AS1 | 23.54730849 | 0.007945784 | 0.376098322 | 0.021126879 | 0.983144443 | 0.99214031 |
| ENSG00000176754 | LINC00303 | 3.890586918 | 0.290711402 | 0.886044558 | 0.328100206 | 0.742835898 | 0.858082255 |
| ENSG00000176771 | NCKAP5 | 99.53168579 | -0.071862879 | 0.181323045 | -0.396325127 | 0.691865193 | 0.826372889 |
| ENSG00000176783 | RUFY1 | 1816.155883 | -0.247122829 | 0.057736316 | -4.280197349 | 1.87E-05 | 9.31E-05 |
| ENSG00000176788 | BASP1 | 6434.880178 | -3.00782909 | 0.057570774 | -52.24576408 | 0 | 0 |
| ENSG00000176809 | LRRC37A3 | 460.6999269 | 0.86904276 | 0.094754801 | 9.171490484 | 4.67E-20 | 7.81E-19 |
| ENSG00000176812 | METTL1P1 | 2.051411775 | 0.069304411 | 1.240944525 | 0.055848114 | 0.955462805 | 0.979458252 |
| ENSG00000176826 | FKBP9P1 | 20.54860965 | 0.388349781 | 0.419849405 | 0.924973995 | 0.354979434 | 0.546644725 |
| ENSG00000176834 | VSIG10 | 1945.820441 | 0.115709981 | 0.059093671 | 1.958077389 | 0.050220933 | 0.120294427 |
| ENSG00000176842 | IRX5 | 1379.822121 | 0.407166931 | 0.064230345 | 6.339167775 | 2.31E-10 | 1.94E-09 |
| ENSG00000176845 | METRNL | 1771.06421 | 0.510955411 | 0.067578707 | 7.560893616 | 4.00E-14 | 4.50E-13 |
| ENSG00000176853 | FAM91A1 | 3801.160667 | -0.335278435 | 0.054260974 | -6.178997678 | 6.45E-10 | 5.23E-09 |
| ENSG00000176855 | KRT18P28 | 133.0682835 | 0.212523593 | 0.158983918 | 1.336761574 | 0.181300476 | 0.336654761 |

| | | | | | | | |
|---|---|---|---|---|---|---|---|
| ENSG00000176868 | ENSG0000017 | 41.91281589 | -0.827187138 | 0.288665777 | -2.865553193 | 0.004162813 | 0.013659696 |
| ENSG00000176871 | WSB2 | 2570.468166 | 0.149346595 | 0.055329594 | 2.699217246 | 0.006950279 | 0.021563348 |
| ENSG00000176884 | GRIN1 | 13.45665351 | 0.333417674 | 0.511481107 | 0.651867039 | 0.51448695 | 0.693561182 |
| ENSG00000176890 | TYMS | 2721.252031 | 0.276895992 | 0.057173711 | 4.843064921 | 1.28E-06 | 7.30E-06 |
| ENSG00000176894 | PXMP2 | 86.60622646 | -0.394407593 | 0.196510088 | -2.007060286 | 0.044743244 | 0.10890292 |
| ENSG00000176896 | TCEANC | 112.0786108 | -0.080620883 | 0.170533989 | -0.472755512 | 0.636387617 | 0.78589856 |
| ENSG00000176903 | PNMA1 | 2770.207063 | -0.847508469 | 0.049896448 | -16.98534679 | 1.05E-64 | 8.46E-63 |
| ENSG00000176907 | TCIM | 20.37667502 | -0.03232136 | 0.397608851 | -0.081289336 | 0.935211854 | 0.969069964 |
| ENSG00000176909 | MAMSTR | 11.77996783 | 1.020514794 | 0.544844497 | 1.873038636 | 0.06106305 | 0.141608003 |
| ENSG00000176912 | TYMSOS | 38.63942359 | 0.038799751 | 0.28338651 | 0.136914601 | 0.891098297 | 0.944150152 |
| ENSG00000176915 | ANKLE2 | 2960.413497 | -0.180437481 | 0.056291981 | -3.20538516 | 0.001348819 | 0.004926269 |
| ENSG00000176919 | C8G | 15.98749144 | 1.180756872 | 0.467137983 | 2.52764047 | 0.011483188 | 0.033570068 |
| ENSG00000176927 | EFCAB5 | 8.866744063 | 0.3983212 | 0.598032819 | 0.666052409 | 0.505377602 | 0.68591659 |
| ENSG00000176928 | GCNT4 | 35.97110028 | -0.224200336 | 0.295918687 | -0.757641697 | 0.448665511 | 0.63682631 |
| ENSG00000176933 | TOB2P1 | 64.07584016 | -0.071140495 | 0.219334129 | -0.324347585 | 0.745674898 | 0.859386985 |
| ENSG00000176945 | MUC20 | 31.23996737 | 1.703284108 | 0.349569747 | 4.872515781 | 1.10E-06 | 6.35E-06 |
| ENSG00000176946 | THAP4 | 1390.256763 | -0.420751553 | 0.061546272 | -6.836345054 | 8.12E-12 | 7.68E-11 |
| ENSG00000176953 | NFATC2IP | 1824.81213 | -0.123123983 | 0.069102246 | -1.781765287 | 0.074787515 | 0.167590661 |
| ENSG00000176973 | FAM89B | 170.6181108 | 0.577706103 | 0.144204193 | 4.006167158 | 6.17E-05 | 0.000284844 |
| ENSG00000176974 | SHMT1 | 1941.06416 | -0.279241659 | 0.053713572 | -5.198717022 | 2.01E-07 | 1.27E-06 |
| ENSG00000176978 | DPP7 | 3139.91226 | 0.551165397 | 0.067547334 | 8.159691319 | 3.36E-16 | 4.40E-15 |
| ENSG00000176986 | SEC24C | 4589.787158 | -0.513607156 | 0.047913508 | -10.71946464 | 8.25E-27 | 2.03E-25 |
| ENSG00000176994 | SMCR8 | 2694.431072 | -0.150939555 | 0.057069566 | -2.644834484 | 0.008173088 | 0.024892545 |
| ENSG00000176998 | HCG4 | 11.62183266 | 1.885059388 | 0.576965413 | 3.267196518 | 0.001086183 | 0.004039726 |
| ENSG00000177000 | MTHFR | 1027.560274 | -0.193138122 | 0.066176147 | -2.918545878 | 0.003516681 | 0.011739329 |
| ENSG00000177025 | C19orf18 | 7.034227886 | 2.060401137 | 0.762117419 | 2.703521905 | 0.006860893 | 0.021311967 |
| ENSG00000177030 | DEAF1 | 979.1421614 | 0.102402472 | 0.074287893 | 1.378454389 | 0.168063041 | 0.318022036 |
| ENSG00000177034 | MTX3 | 941.6326001 | 0.191677612 | 0.074100979 | 2.586708228 | 0.00968976 | 0.028924281 |
| ENSG00000177042 | TMEM80 | 594.9520852 | 0.378016689 | 0.084564837 | 4.470140322 | 7.82E-06 | 4.08E-05 |
| ENSG00000177045 | SIX5 | 684.0630587 | 0.289206692 | 0.082679715 | 3.497915913 | 0.000468909 | 0.001864905 |
| ENSG00000177051 | FBXO46 | 367.3210581 | -0.172575996 | 0.100348631 | -1.719764326 | 0.08547529 | 0.186476761 |
| ENSG00000177054 | ZDHHC13 | 593.9278359 | 0.182993094 | 0.088706291 | 2.062909983 | 0.039121181 | 0.097378462 |
| ENSG00000177058 | SLC38A9 | 300.5486459 | -0.274817437 | 0.11016932 | -2.494500628 | 0.01261345 | 0.036465905 |

| | | | | | | | |
|---|---|---|---|---|---|---|---|
| ENSG00000177076 | ACER2 | 199.1957079 | 0.027562127 | 0.134612763 | 0.204751219 | 0.837766501 | 0.91468105 |
| ENSG00000177082 | WDR73 | 2425.840677 | -0.009164955 | 0.054725914 | -0.167470115 | 0.86700016 | 0.930846618 |
| ENSG00000177084 | POLE | 4797.568514 | -0.285085815 | 0.04481317 | -6.361652552 | 2.00E-10 | 1.69E-09 |
| ENSG00000177096 | PHETA2 | 5.030979735 | 0.091980599 | 0.777000984 | 0.118379 | 0.905767365 | 0.95262805 |
| ENSG00000177098 | SCN4B | 2.109117274 | 0.046140211 | 1.176957329 | 0.03920296 | 0.968728574 | 0.985876114 |
| ENSG00000177103 | DSCAML1 | 6.94390787 | -1.137255193 | 0.678967763 | -1.674976715 | 0.0939388 | 0.201397765 |
| ENSG00000177105 | RHOG | 518.8757248 | -0.0669085 | 0.105180215 | -0.636131999 | 0.524690391 | 0.701013253 |
| ENSG00000177106 | EPS8L2 | 4556.402206 | 0.225190918 | 0.053046796 | 4.245137035 | 2.18E-05 | 0.000107797 |
| ENSG00000177108 | ZDHHC22 | 199.3155126 | 4.350043108 | 0.226545759 | 19.2016091 | 3.59E-82 | 4.20E-80 |
| ENSG00000177119 | ANO6 | 676.7739222 | 0.649868509 | 0.079640562 | 8.160019169 | 3.35E-16 | 4.39E-15 |
| ENSG00000177125 | ZBTB34 | 504.1963658 | -0.061096534 | 0.091191282 | -0.669982181 | 0.502869149 | 0.683928279 |
| ENSG00000177133 | PRDM16-DT | 17.59150525 | 3.001225942 | 0.561728449 | 5.342841277 | 9.15E-08 | 5.95E-07 |
| ENSG00000177150 | FAM210A | 780.2592622 | 0.396186742 | 0.076280937 | 5.193784415 | 2.06E-07 | 1.30E-06 |
| ENSG00000177156 | TALDO1 | 4760.589676 | 0.034989857 | 0.049214616 | 0.710964739 | 0.477106087 | 0.661968041 |
| ENSG00000177169 | ULK1 | 2961.42574 | 0.774589756 | 0.057306266 | 13.51666779 | 1.25E-41 | 5.46E-40 |
| ENSG00000177173 | NAP1L4P1 | 14.14932658 | -0.426556754 | 0.50637079 | -0.842380252 | 0.399575143 | 0.592315423 |
| ENSG00000177181 | RIMKLA | 346.45469 | 0.146667733 | 0.102882033 | 1.425591317 | 0.153986344 | 0.297224518 |
| ENSG00000177182 | CLVS1 | 2.463701578 | 1.421869075 | 1.210056685 | 1.175043362 | 0.239977368 | 0.415728804 |
| ENSG00000177189 | RPS6KA3 | 875.7612079 | 0.708106855 | 0.070046283 | 10.10912828 | 5.03E-24 | 1.06E-22 |
| ENSG00000177191 | B3GNT8 | 50.09166016 | 0.954951834 | 0.255324611 | 3.740148003 | 0.000183912 | 0.000785943 |
| ENSG00000177192 | PUS1 | 859.9305859 | 0.013520537 | 0.073819739 | 0.183156123 | 0.854675521 | 0.924486249 |
| ENSG00000177197 | PCNPP5 | 131.519261 | -0.005581661 | 0.15536557 | -0.035925985 | 0.971341376 | 0.986958904 |
| ENSG00000177200 | CHD9 | 869.938358 | 0.078458038 | 0.08851486 | 0.886382671 | 0.375411354 | 0.568360856 |
| ENSG00000177202 | SPACA4 | 4.142151178 | 1.030180176 | 0.939551956 | 1.096458976 | 0.272877967 | 0.454862706 |
| ENSG00000177225 | GATD1 | 1121.200237 | 0.548014568 | 0.070318299 | 7.79334222 | 6.53E-15 | 7.78E-14 |
| ENSG00000177239 | MAN1B1 | 3367.177936 | 0.169862554 | 0.053997165 | 3.145768017 | 0.001656513 | 0.005923724 |
| ENSG00000177283 | FZD8 | 105.4935612 | -1.455285679 | 0.188285574 | -7.729140606 | 1.08E-14 | 1.27E-13 |
| ENSG00000177302 | TOP3A | 1913.79564 | -0.118671859 | 0.058184123 | -2.039591773 | 0.041391002 | 0.101966236 |
| ENSG00000177303 | CASKIN2 | 2846.592744 | 0.296928141 | 0.055594399 | 5.340972247 | 9.24E-08 | 6.01E-07 |
| ENSG00000177311 | ZBTB38 | 621.545859 | 0.005210182 | 0.088398836 | 0.058939488 | 0.953000306 | 0.978201576 |
| ENSG00000177335 | LINC02904 | 19.37198823 | -0.591211163 | 0.400201556 | -1.47728352 | 0.139599654 | 0.275832563 |
| ENSG00000177337 | DLGAP1-AS1 | 70.66684712 | 0.246585636 | 0.220502562 | 1.11828921 | 0.263443494 | 0.44412799 |
| ENSG00000177350 | RPL13AP3 | 15.91243666 | 0.478526064 | 0.474825491 | 1.007793544 | 0.313553586 | 0.501878263 |

| | | | | | | | |
|---|---|---|---|---|---|---|---|
| ENSG00000177352 | CCDC71 | 505.1586242 | 0.061077747 | 0.095633587 | 0.63866419 | 0.523041413 | 0.699706471 |
| ENSG00000177359 | ENSG0000017 | 2.061881796 | -1.192662143 | 1.268424503 | -0.940270501 | 0.347078827 | 0.537972181 |
| ENSG00000177363 | LRRN4CL | 4.467511776 | 1.05888872 | 0.878163608 | 1.20579891 | 0.227895036 | 0.400729122 |
| ENSG00000177369 | FLJ40194 | 8.150362449 | -1.098319703 | 0.641781425 | -1.711361003 | 0.087014491 | 0.189222906 |
| ENSG00000177370 | TIMM22 | 539.6674567 | -0.047785128 | 0.085657577 | -0.55786224 | 0.576938458 | 0.743036617 |
| ENSG00000177374 | HIC1 | 36.63760963 | -0.057520576 | 0.301339551 | -0.190882929 | 0.848617315 | 0.920874229 |
| ENSG00000177380 | PPFIA3 | 969.8025317 | -0.202578538 | 0.069264926 | -2.924691487 | 0.003447979 | 0.011531802 |
| ENSG00000177383 | MAGEF1 | 2091.005638 | -0.141612947 | 0.056209575 | -2.519374096 | 0.011756368 | 0.034311841 |
| ENSG00000177398 | UMODL1 | 33.0201881 | -4.201171052 | 0.506978097 | -8.28669143 | 1.16E-16 | 1.58E-15 |
| ENSG00000177406 | NINJ2-AS1 | 119.1599577 | 0.858105323 | 0.169735692 | 5.055538484 | 4.29E-07 | 2.61E-06 |
| ENSG00000177410 | ZFAS1 | 760.0198183 | 0.507955404 | 0.075602008 | 6.718808333 | 1.83E-11 | 1.67E-10 |
| ENSG00000177425 | PAWR | 3570.797724 | -0.428352095 | 0.054826371 | -7.81288433 | 5.59E-15 | 6.70E-14 |
| ENSG00000177426 | TGIF1 | 1866.076067 | 0.705484108 | 0.056247311 | 12.54253936 | 4.37E-36 | 1.54E-34 |
| ENSG00000177427 | MIEF2 | 408.6761747 | 0.489137385 | 0.096781191 | 5.054054189 | 4.33E-07 | 2.63E-06 |
| ENSG00000177432 | NAP1L5 | 27.40146829 | 0.80055523 | 0.353131988 | 2.267014196 | 0.023389359 | 0.062695943 |
| ENSG00000177447 | CBX3P1 | 11.35865211 | -2.089648366 | 0.584119843 | -3.577430886 | 0.000346988 | 0.001408832 |
| ENSG00000177453 | NIM1K | 10.49278456 | 0.536758307 | 0.539649663 | 0.994642161 | 0.319910334 | 0.509422676 |
| ENSG00000177459 | ERICH5 | 15.18841216 | 1.183520813 | 0.480017852 | 2.465576661 | 0.01367929 | 0.039176394 |
| ENSG00000177463 | NR2C2 | 2673.668082 | -0.228444001 | 0.05027054 | -4.544291757 | 5.51E-06 | 2.93E-05 |
| ENSG00000177465 | ACOT4 | 494.6510116 | -0.72839899 | 0.09570678 | -7.610735492 | 2.73E-14 | 3.11E-13 |
| ENSG00000177469 | CAVIN1 | 137.8611971 | 0.413728132 | 0.158259435 | 2.614239913 | 0.008942622 | 0.026957128 |
| ENSG00000177479 | ARIH2 | 2153.408117 | 0.227936287 | 0.056533525 | 4.0318782 | 5.53E-05 | 0.000257315 |
| ENSG00000177483 | RBM44 | 7.143462079 | 1.077184505 | 0.686815682 | 1.56837494 | 0.116793656 | 0.240413233 |
| ENSG00000177485 | ZBTB33 | 1858.768091 | 0.308443181 | 0.05870671 | 5.253968103 | 1.49E-07 | 9.50E-07 |
| ENSG00000177508 | IRX3 | 1709.687049 | -0.296734758 | 0.059082525 | -5.02237775 | 5.10E-07 | 3.07E-06 |
| ENSG00000177519 | RPRM | 211.773008 | 0.10460747 | 0.127167324 | 0.822597089 | 0.410737155 | 0.603259513 |
| ENSG00000177542 | SLC25A22 | 1073.316633 | 0.284239584 | 0.073404211 | 3.872251738 | 0.000107834 | 0.000478291 |
| ENSG00000177548 | RABEP2 | 891.9541767 | 0.027268453 | 0.075901341 | 0.359261805 | 0.719399244 | 0.844059737 |
| ENSG00000177556 | ATOX1 | 1178.580219 | 0.028431564 | 0.063687269 | 0.44642461 | 0.655290559 | 0.799891737 |
| ENSG00000177565 | TBL1XR1 | 1722.123835 | -0.116263878 | 0.063936234 | -1.818434875 | 0.068997692 | 0.156822259 |
| ENSG00000177570 | SAMD12 | 532.277431 | 0.338875803 | 0.091519298 | 3.702779748 | 0.00021325 | 0.000899422 |
| ENSG00000177576 | C18orf32 | 71.35320524 | 0.982462652 | 0.220827138 | 4.449012296 | 8.63E-06 | 4.48E-05 |
| ENSG00000177595 | PIDD1 | 576.2693137 | -0.08450414 | 0.095610345 | -0.883838878 | 0.376783196 | 0.569544446 |

| | | | | | | | |
|---|---|---|---|---|---|---|---|
| ENSG00000177600 | RPLP2 | 5572.790963 | 0.220432665 | 0.047394437 | 4.651024021 | 3.30E-06 | 1.80E-05 |
| ENSG00000177602 | HASPIN | 253.1880456 | -0.155634832 | 0.121644973 | -1.279418527 | 0.200749713 | 0.364161522 |
| ENSG00000177606 | JUN | 1859.258867 | 0.165516394 | 0.069001698 | 2.398729302 | 0.016452072 | 0.046138207 |
| ENSG00000177613 | CSTF2T | 979.1716887 | -0.915265891 | 0.069929744 | -13.08836333 | 3.84E-39 | 1.50E-37 |
| ENSG00000177614 | PGBD5 | 51.69161243 | -1.030144099 | 0.2498736 | -4.122660809 | 3.75E-05 | 0.000178136 |
| ENSG00000177628 | GBA1 | 1687.708957 | 0.334550713 | 0.06618862 | 5.054504999 | 4.32E-07 | 2.62E-06 |
| ENSG00000177640 | CASC2 | 48.39725089 | -0.627381109 | 0.251717882 | -2.492397856 | 0.012688382 | 0.036658299 |
| ENSG00000177646 | ACAD9 | 2377.400637 | -0.023615945 | 0.05858528 | -0.403103729 | 0.686871918 | 0.823089733 |
| ENSG00000177663 | IL17RA | 1140.715815 | -0.501134204 | 0.066756558 | -7.506890983 | 6.05E-14 | 6.71E-13 |
| ENSG00000177666 | PNPLA2 | 1962.4522 | 0.396113278 | 0.065239917 | 6.071639853 | 1.27E-09 | 9.99E-09 |
| ENSG00000177674 | AGTRAP | 373.6560334 | 0.261322298 | 0.104076859 | 2.510858807 | 0.012043786 | 0.035043689 |
| ENSG00000177675 | CD163L1 | 2.831066171 | 1.637396747 | 1.136887393 | 1.440245319 | 0.149798006 | 0.291340928 |
| ENSG00000177679 | SRRM3 | 522.8666629 | 1.234884428 | 0.102473407 | 12.0507795 | 1.92E-33 | 6.10E-32 |
| ENSG00000177683 | THAP5 | 572.969417 | -0.080313773 | 0.096169744 | -0.835125163 | 0.40364724 | 0.596512767 |
| ENSG00000177685 | CRACR2B | 832.9196557 | 0.057684014 | 0.074948225 | 0.769651512 | 0.44150664 | 0.630841627 |
| ENSG00000177688 | SUMO4 | 2.274126518 | 0.245404882 | 1.139284262 | 0.215402679 | 0.829453382 | 0.909761443 |
| ENSG00000177692 | DNAJC28 | 24.25601769 | -0.780724229 | 0.37267906 | -2.094896961 | 0.036180153 | 0.091195771 |
| ENSG00000177694 | NAALADL2 | 335.8491557 | 2.269878274 | 0.120600162 | 18.8215193 | 5.03E-79 | 5.69E-77 |
| ENSG00000177697 | CD151 | 4118.95793 | 0.261515467 | 0.056492441 | 4.629211662 | 3.67E-06 | 1.99E-05 |
| ENSG00000177700 | POLR2L | 1830.713904 | 0.114667709 | 0.070390429 | 1.629024143 | 0.103307908 | 0.217660122 |
| ENSG00000177706 | FAM20C | 2409.91511 | -0.243772189 | 0.058009562 | -4.202275959 | 2.64E-05 | 0.000128946 |
| ENSG00000177707 | NECTIN3 | 216.970453 | -2.433660565 | 0.147480735 | -16.50154897 | 3.58E-61 | 2.64E-59 |
| ENSG00000177710 | SLC35G5 | 1.653809558 | -1.404769855 | 1.430994578 | -0.981673779 | 0.326260587 | NA |
| ENSG00000177721 | ANXA2R | 90.62510823 | 0.380287325 | 0.185804982 | 2.046701439 | 0.040687408 | 0.10055032 |
| ENSG00000177728 | TMEM94 | 4605.859004 | -0.135382733 | 0.052976818 | -2.555508972 | 0.010603263 | 0.031288764 |
| ENSG00000177731 | FLII | 7640.546543 | 0.190701385 | 0.047574743 | 4.008458563 | 6.11E-05 | 0.000282492 |
| ENSG00000177732 | SOX12 | 2394.289194 | -0.176564125 | 0.054431718 | -3.243772791 | 0.001179578 | 0.004360922 |
| ENSG00000177733 | HNRNPA0 | 5444.844684 | -0.236796858 | 0.046383792 | -5.10516381 | 3.31E-07 | 2.03E-06 |
| ENSG00000177738 | ANXA2R-OT1 | 515.3486314 | -0.559745478 | 0.090261536 | -6.201373286 | 5.60E-10 | 4.56E-09 |
| ENSG00000177770 | CDKN2AIPNLP | 13.25570197 | -0.014707821 | 0.483441523 | -0.030423163 | 0.975729572 | 0.988877242 |
| ENSG00000177788 | RAB4A-AS1 | 784.78878 | 0.117704595 | 0.075957219 | 1.549616963 | 0.121233479 | 0.247480178 |
| ENSG00000177791 | MYOZ1 | 8.581730117 | -1.293631075 | 0.633805669 | -2.041053178 | 0.041245542 | 0.101706194 |
| ENSG00000177803 | YWHAQP4 | 9.369422869 | 0.054717941 | 0.592026203 | 0.092424864 | 0.926360485 | 0.964663921 |

| | | | | | | | |
|---|---|---|---|---|---|---|---|
| ENSG00000177830 | CHID1 | 1091.683958 | 0.42625962 | 0.071449058 | 5.965923623 | 2.43E-09 | 1.87E-08 |
| ENSG00000177839 | PCDHB9 | 21.34846738 | 0.832942729 | 0.388166679 | 2.14583779 | 0.031885922 | 0.081912265 |
| ENSG00000177842 | ZNF620 | 341.9008898 | -0.967947857 | 0.102981311 | -9.399257501 | 5.49E-21 | 9.73E-20 |
| ENSG00000177853 | ZNF518A | 775.4943547 | -0.515334555 | 0.081897552 | -6.292429308 | 3.13E-10 | 2.61E-09 |
| ENSG00000177854 | TMEM187 | 348.2997555 | -0.82499111 | 0.103196137 | -7.994399152 | 1.30E-15 | 1.64E-14 |
| ENSG00000177855 | CACYBPP2 | 716.2733489 | -0.297225941 | 0.075464539 | -3.938617328 | 8.20E-05 | 0.0003711 |
| ENSG00000177868 | SVBP | 407.5734258 | -0.097670537 | 0.095729797 | -1.020273108 | 0.307598953 | 0.495493293 |
| ENSG00000177873 | ZNF619 | 308.3149796 | -0.423253322 | 0.106473376 | -3.975203346 | 7.03E-05 | 0.000321852 |
| ENSG00000177875 | CCDC184 | 12.16640363 | -0.63340705 | 0.516071506 | -1.227362958 | 0.219686202 | 0.389760362 |
| ENSG00000177879 | AP3S1 | 1150.804469 | 0.053941445 | 0.068407993 | 0.788525469 | 0.430389407 | 0.620600477 |
| ENSG00000177885 | GRB2 | 7293.953182 | 0.137604342 | 0.043647231 | 3.152647716 | 0.001617969 | 0.005803943 |
| ENSG00000177888 | ZBTB41 | 1219.395554 | 0.249843344 | 0.074387686 | 3.358665331 | 0.000783199 | 0.002989944 |
| ENSG00000177889 | UBE2N | 2600.691645 | -0.140944312 | 0.052609815 | -2.679049758 | 0.007383142 | 0.022767748 |
| ENSG00000177917 | ARL6IP6 | 349.7153148 | -0.184289396 | 0.104967056 | -1.755687953 | 0.079141705 | 0.175276184 |
| ENSG00000177932 | ZNF354C | 425.8713637 | -0.254159689 | 0.101339029 | -2.50801386 | 0.01214119 | 0.035309192 |
| ENSG00000177943 | MAMDC4 | 620.9023502 | 0.003794191 | 0.096276548 | 0.039409293 | 0.968564071 | 0.985876114 |
| ENSG00000177946 | CENPBD1P | 922.1007358 | 0.463753429 | 0.071114744 | 6.52119943 | 6.97E-11 | 6.11E-10 |
| ENSG00000177951 | BET1L | 1460.97434 | 0.086001542 | 0.063493167 | 1.354500752 | 0.175576674 | 0.328874736 |
| ENSG00000177954 | RPS27 | 556.6229668 | 0.603963382 | 0.088766876 | 6.803927405 | 1.02E-11 | 9.52E-11 |
| ENSG00000177963 | RIC8A | 3146.786857 | -0.018706587 | 0.057326363 | -0.326317357 | 0.744184257 | 0.858515809 |
| ENSG00000177971 | IMP3 | 1670.739315 | -0.070246104 | 0.060799353 | -1.155375839 | 0.247936552 | 0.425500448 |
| ENSG00000177981 | ASB8 | 985.5944257 | -0.094939577 | 0.066665581 | -1.424116862 | 0.154412646 | 0.297737844 |
| ENSG00000177989 | ODF3B | 20.32550504 | -0.23383317 | 0.393016807 | -0.594969899 | 0.551863592 | 0.723703813 |
| ENSG00000178026 | LRRC75B | 270.1871736 | -0.215917259 | 0.113367375 | -1.904580209 | 0.056834661 | 0.133407053 |
| ENSG00000178028 | DMAP1 | 1621.252688 | -0.333670465 | 0.069851754 | -4.776837291 | 1.78E-06 | 1.00E-05 |
| ENSG00000178031 | ADAMTSL1 | 5.590338183 | 1.583136662 | 0.795956027 | 1.988975029 | 0.046703959 | 0.112968033 |
| ENSG00000178035 | IMPDH2 | 5229.915424 | 0.102836344 | 0.047182331 | 2.179552014 | 0.029290685 | 0.076053007 |
| ENSG00000178038 | ALS2CL | 370.1328778 | 0.660517685 | 0.1027117 | 6.430793029 | 1.27E-10 | 1.09E-09 |
| ENSG00000178053 | MLF1 | 270.4869649 | 0.012335585 | 0.117310698 | 0.105153115 | 0.916254313 | 0.958671559 |
| ENSG00000178057 | NDUFAF3 | 627.7583527 | 0.25643782 | 0.084720583 | 3.026865599 | 0.002471038 | 0.008526717 |
| ENSG00000178074 | C2orf69 | 316.0327678 | -0.136089247 | 0.110063379 | -1.236462564 | 0.216286673 | 0.385667802 |
| ENSG00000178075 | GRAMD1C | 45.00066491 | -1.222302436 | 0.271948982 | -4.494601989 | 6.97E-06 | 3.67E-05 |
| ENSG00000178078 | STAP2 | 1662.627023 | -0.127225048 | 0.057572419 | -2.209826332 | 0.027117218 | 0.07123199 |

| | | | | | | | |
|---|---|---|---|---|---|---|---|
| ENSG00000178082 | TWF1P1 | 507.1568094 | 0.089165476 | 0.087770465 | 1.015893858 | 0.309679934 | 0.49772738 |
| ENSG00000178093 | TSSK6 | 13.93550372 | -0.326027697 | 0.46791503 | -0.696766882 | 0.485948695 | 0.669580266 |
| ENSG00000178096 | BOLA1 | 715.049147 | 0.15122511 | 0.076408412 | 1.979168347 | 0.047797055 | 0.115283824 |
| ENSG00000178104 | PDE4DIP | 2940.710677 | 0.32806055 | 0.057674447 | 5.688143795 | 1.28E-08 | 9.12E-08 |
| ENSG00000178105 | DDX10 | 642.2384358 | 0.149028184 | 0.079085922 | 1.884383215 | 0.059513159 | 0.138588149 |
| ENSG00000178125 | PPP1R42 | 1.595689392 | 0.409100308 | 1.472659049 | 0.277797029 | 0.781168172 | NA |
| ENSG00000178127 | NDUFV2 | 31.0091187 | 0.276686551 | 0.329599241 | 0.839463557 | 0.401209231 | 0.594057947 |
| ENSG00000178146 | GK4P | 108.2548759 | 0.218875558 | 0.182025763 | 1.202442743 | 0.229192037 | 0.402238729 |
| ENSG00000178149 | DALRD3 | 729.2152963 | -0.242929541 | 0.087478672 | -2.777014493 | 0.005486073 | 0.017474385 |
| ENSG00000178150 | ZNF114 | 202.7997398 | -0.585876461 | 0.13280588 | -4.411525004 | 1.03E-05 | 5.30E-05 |
| ENSG00000178162 | FAR2P2 | 4.811631163 | -1.290829978 | 0.82439581 | -1.565789105 | 0.117397998 | 0.241447917 |
| ENSG00000178177 | LCORL | 580.7515207 | 0.079848132 | 0.084234767 | 0.947923699 | 0.343168299 | 0.534159855 |
| ENSG00000178184 | PARD6G | 227.5420165 | 1.209425432 | 0.130390453 | 9.275414006 | 1.77E-20 | 3.04E-19 |
| ENSG00000178187 | ZNF454 | 5.341204332 | -0.207845951 | 0.752704172 | -0.276132323 | 0.782446435 | 0.882030527 |
| ENSG00000178188 | SH2B1 | 1532.007975 | 0.233108121 | 0.062709223 | 3.717286064 | 0.000201374 | 0.000853571 |
| ENSG00000178199 | ZC3H12D | 5.417105694 | -0.149418921 | 0.766790381 | -0.1948628 | 0.845500357 | 0.918969875 |
| ENSG00000178201 | VN1R1 | 7.245599813 | 0.650857097 | 0.673194953 | 0.966818147 | 0.333634951 | 0.52468539 |
| ENSG00000178202 | POGLUT3 | 562.8885426 | 0.365389681 | 0.083301733 | 4.386339484 | 1.15E-05 | 5.90E-05 |
| ENSG00000178209 | PLEC | 11134.17865 | 0.298626874 | 0.059067 | 5.055731194 | 4.29E-07 | 2.61E-06 |
| ENSG00000178217 | SH2D4B | 5.449560302 | -0.431718659 | 0.791758494 | -0.545265586 | 0.585570869 | 0.749650179 |
| ENSG00000178226 | PRSS36 | 19.55801586 | 0.331017532 | 0.418393673 | 0.791162853 | 0.428848966 | 0.61920798 |
| ENSG00000178229 | ZNF543 | 182.0558628 | 0.849083157 | 0.144959505 | 5.857381724 | 4.70E-09 | 3.51E-08 |
| ENSG00000178234 | GALNT11 | 1569.838157 | 1.08960563 | 0.059067228 | 18.44687253 | 5.53E-76 | 5.95E-74 |
| ENSG00000178248 | FAM230I | 4.114413016 | -5.617115067 | 1.411565777 | -3.979350562 | 6.91E-05 | 0.000316542 |
| ENSG00000178252 | WDR6 | 10107.91343 | -0.012194072 | 0.051127092 | -0.238505103 | 0.811489358 | 0.900221351 |
| ENSG00000178295 | GEN1 | 793.4718027 | -0.038080196 | 0.160522307 | -0.237226815 | 0.812480835 | 0.900923741 |
| ENSG00000178297 | TMPRSS9 | 2.370526971 | 0.055299415 | 1.143503675 | 0.04835963 | 0.961429632 | 0.982282802 |
| ENSG00000178301 | AQP11 | 106.3251491 | 0.275691458 | 0.178965203 | 1.540475206 | 0.123444562 | 0.250936486 |
| ENSG00000178307 | TMEM11 | 900.3067403 | -0.205783407 | 0.072273881 | -2.84727213 | 0.004409565 | 0.0144055 |
| ENSG00000178338 | ZNF354B | 270.5110256 | -0.426711279 | 0.121677441 | -3.506905434 | 0.00045335 | 0.001808041 |
| ENSG00000178372 | CALML5 | 93.3507921 | -2.16283697 | 0.227545474 | -9.50507575 | 2.00E-21 | 3.63E-20 |
| ENSG00000178381 | ZFAND2A | 1241.423785 | 0.267213482 | 0.063418828 | 4.213472406 | 2.51E-05 | 0.000123108 |
| ENSG00000178385 | PLEKHM3 | 194.7829802 | 0.462362135 | 0.130661942 | 3.538613681 | 0.000402234 | 0.001615981 |

| | | | | | | | |
|---|---|---|---|---|---|---|---|
| ENSG00000178386 | ZNF223 | 36.28612723 | 0.480799054 | 0.302964102 | 1.586983574 | 0.112516365 | 0.233185058 |
| ENSG00000178395 | CCDC185 | 2.778508959 | 0.523376092 | 1.10738845 | 0.472621953 | 0.636482918 | 0.785967494 |
| ENSG00000178401 | DNAJC22 | 1540.211628 | -0.827200631 | 0.062330406 | -13.27122165 | 3.40E-40 | 1.40E-38 |
| ENSG00000178409 | BEND3 | 583.6882853 | 0.171627954 | 0.08299253 | 2.067992798 | 0.0386407 | 0.096392011 |
| ENSG00000178412 | CTDP1-DT | 1.62686583 | -0.181307749 | 1.331067891 | -0.136212247 | 0.891653494 | NA |
| ENSG00000178425 | NT5DC1 | 1036.471924 | 0.21213289 | 0.067195343 | 3.156958215 | 0.001594242 | 0.005726888 |
| ENSG00000178429 | RPS3AP5 | 158.6224517 | 0.179794952 | 0.152623751 | 1.178027346 | 0.238785704 | 0.414322862 |
| ENSG00000178440 | TIMM23B-AG/ | 11.28941012 | 0.061180732 | 0.517400131 | 0.118246456 | 0.905872381 | 0.95262805 |
| ENSG00000178445 | GLDC | 4.435270022 | -0.876030551 | 0.84367178 | -1.038354691 | 0.299104955 | 0.485802016 |
| ENSG00000178449 | COX14 | 645.9599298 | 0.113649348 | 0.078671688 | 1.444602889 | 0.148569461 | 0.289393176 |
| ENSG00000178458 | H3P16 | 41.10157813 | 0.050828652 | 0.301683443 | 0.1684834 | 0.866203002 | 0.930450312 |
| ENSG00000178460 | MCMDC2 | 21.15334324 | 0.851978616 | 0.38927474 | 2.188630624 | 0.028623698 | 0.074574405 |
| ENSG00000178462 | TUBAL3 | 17.45644276 | -1.962840257 | 0.463033507 | -4.239089023 | 2.24E-05 | 0.000110504 |
| ENSG00000178464 | RPL10P16 | 2066.60062 | -0.267891866 | 0.061109134 | -4.38382692 | 1.17E-05 | 5.96E-05 |
| ENSG00000178467 | P4HTM | 1901.604453 | 0.353938229 | 0.057647274 | 6.139721903 | 8.27E-10 | 6.62E-09 |
| ENSG00000178498 | DTX3 | 1577.323727 | 0.095002669 | 0.0674117 | 1.409290502 | 0.158749286 | 0.304083887 |
| ENSG00000178502 | KLHL11 | 357.4662127 | 0.162112833 | 0.107906465 | 1.502345884 | 0.133007804 | 0.266171007 |
| ENSG00000178503 | NECAP1P1 | 2.870277181 | 0.192432556 | 1.05347002 | 0.182665432 | 0.855060541 | 0.924716614 |
| ENSG00000178531 | CTXN1 | 516.4024292 | 1.003575037 | 0.104281888 | 9.6236754 | 6.35E-22 | 1.18E-20 |
| ENSG00000178537 | SLC25A20 | 216.2781905 | -0.142466535 | 0.135186605 | -1.053850968 | 0.291951154 | 0.476936684 |
| ENSG00000178538 | CA8 | 546.9291293 | 0.225738417 | 0.090252548 | 2.501186065 | 0.012377813 | 0.035874545 |
| ENSG00000178556 | CKS1BP6 | 8.249264568 | -0.174461343 | 0.604039037 | -0.288824616 | 0.772715594 | 0.876166615 |
| ENSG00000178567 | EPM2AIP1 | 1710.499688 | 0.037765305 | 0.060024865 | 0.629161014 | 0.52924365 | 0.705079806 |
| ENSG00000178568 | ERBB4 | 436.5960644 | -0.33776807 | 0.099665103 | -3.389030476 | 0.000701402 | 0.002701959 |
| ENSG00000178573 | MAF | 10.13311566 | -1.121819935 | 0.590085947 | -1.901112779 | 0.057287242 | 0.134345669 |
| ENSG00000178585 | CTNNBIP1 | 1042.723964 | 0.32355421 | 0.068661812 | 4.712287706 | 2.45E-06 | 1.36E-05 |
| ENSG00000178605 | GTPBP6 | 1737.217023 | -0.116185162 | 0.061435734 | -1.891165837 | 0.05860221 | 0.136841887 |
| ENSG00000178605 | GTPBP6 | 1737.217023 | -0.116185162 | 0.061435734 | -1.891165837 | 0.05860221 | 0.136841887 |
| ENSG00000178607 | ERN1 | 809.5446357 | 0.215239321 | 0.072224272 | 2.980152179 | 0.002881052 | 0.009798567 |
| ENSG00000178623 | GPR35 | 42.73945784 | 0.349541304 | 0.288111882 | 1.213213775 | 0.225048099 | 0.396909617 |
| ENSG00000178631 | ACTG1P1 | 22.71636056 | 0.03354244 | 0.384217571 | 0.08730064 | 0.930432545 | 0.966409607 |
| ENSG00000178660 | ARMC10P1 | 63.43670838 | -0.040384767 | 0.222150457 | -0.181790157 | 0.855747409 | 0.925175552 |
| ENSG00000178665 | ZNF713 | 130.9258685 | 0.784721942 | 0.162216291 | 4.837503921 | 1.31E-06 | 7.49E-06 |

| | | | | | | | |
|---|---|---|---|---|---|---|---|
| ENSG00000178685 | PARP10 | 48.02350001 | 0.637336256 | 0.29958888 | 2.127369531 | 0.033389392 | 0.085115307 |
| ENSG00000178691 | SUZ12 | 1743.88405 | 0.078726804 | 0.065375497 | 1.204224941 | 0.228502649 | 0.401366827 |
| ENSG00000178694 | NSUN3 | 491.5829703 | 0.101033702 | 0.096414795 | 1.047906626 | 0.294681629 | 0.480591253 |
| ENSG00000178695 | KCTD12 | 393.8811664 | 0.258780209 | 0.100123299 | 2.584615283 | 0.009748771 | 0.029070168 |
| ENSG00000178700 | DHFR2 | 339.6179119 | 0.132535847 | 0.100761205 | 1.315345989 | 0.188393649 | 0.346822109 |
| ENSG00000178715 | ENSG0000017 | 56.68194726 | 0.475034869 | 0.26321146 | 1.804765144 | 0.071111441 | 0.160734698 |
| ENSG00000178718 | RPP25 | 736.9847096 | 0.820696357 | 0.081908654 | 10.01965384 | 1.25E-23 | 2.56E-22 |
| ENSG00000178719 | GRINA | 8760.034502 | 0.119434475 | 0.046244309 | 2.582684813 | 0.009803484 | 0.029210312 |
| ENSG00000178723 | GLULP4 | 9.754485137 | -0.093296932 | 0.615996757 | -0.151456855 | 0.879615343 | 0.937615842 |
| ENSG00000178726 | THBD | 128.1026351 | 3.479678749 | 0.229363045 | 15.17105228 | 5.50E-52 | 3.20E-50 |
| ENSG00000178741 | COX5A | 2445.436258 | 0.12146627 | 0.053226513 | 2.282063276 | 0.022485605 | 0.060570664 |
| ENSG00000178750 | STX19 | 2.931333708 | -1.068764133 | 1.041802952 | -1.025879348 | 0.304948458 | 0.492364698 |
| ENSG00000178752 | ERFE | 41.07973425 | -0.200071167 | 0.282716129 | -0.707675108 | 0.479147039 | 0.663653377 |
| ENSG00000178761 | FAM219B | 1622.184044 | -0.072813463 | 0.057548947 | -1.265244045 | 0.205783858 | 0.371268714 |
| ENSG00000178764 | ZHX2 | 2254.146967 | -0.751754505 | 0.061862767 | -12.15197028 | 5.60E-34 | 1.82E-32 |
| ENSG00000178773 | CPNE7 | 383.5194197 | 1.607086813 | 0.112684087 | 14.26187905 | 3.78E-46 | 1.87E-44 |
| ENSG00000178796 | RIIAD1 | 3.88397284 | -1.121487916 | 0.91771963 | -1.222037624 | 0.221693399 | 0.392592758 |
| ENSG00000178802 | MPI | 1348.144284 | -0.419954559 | 0.070174688 | -5.984416462 | 2.17E-09 | 1.68E-08 |
| ENSG00000178809 | TRIM73 | 5.464525759 | 1.088884255 | 0.788450831 | 1.381042687 | 0.167265836 | 0.316799726 |
| ENSG00000178814 | OPLAH | 504.2411973 | 0.138805362 | 0.090111764 | 1.540368942 | 0.123470448 | 0.250966867 |
| ENSG00000178821 | TMEM52 | 35.02949903 | 0.270036159 | 0.299541054 | 0.90149966 | 0.367322715 | 0.559855363 |
| ENSG00000178826 | TMEM139 | 13.90615844 | -0.808071382 | 0.492097169 | -1.642097195 | 0.100569863 | 0.212888498 |
| ENSG00000178852 | EFCAB13 | 45.04067108 | -0.563789628 | 0.264666778 | -2.13018661 | 0.03315621 | 0.084633744 |
| ENSG00000178878 | APOLD1 | 137.2486736 | 0.036906704 | 0.156319884 | 0.236097313 | 0.813357159 | 0.901448243 |
| ENSG00000178882 | RFLNA | 2.508159217 | -0.359184003 | 1.099224671 | -0.326761228 | 0.743848487 | 0.858490754 |
| ENSG00000178896 | EXOSC4 | 658.1650128 | 0.124326052 | 0.080755236 | 1.539541683 | 0.12367211 | 0.251332229 |
| ENSG00000178904 | DPY19L3 | 1653.493155 | -0.100414495 | 0.060940221 | -1.64775402 | 0.099403158 | 0.210983386 |
| ENSG00000178913 | TAF7 | 4512.963377 | 0.16790591 | 0.051697187 | 3.247873223 | 0.001162711 | 0.004304112 |
| ENSG00000178917 | ZNF852 | 66.90372583 | -0.094297021 | 0.217125694 | -0.434296924 | 0.664072837 | 0.806108101 |
| ENSG00000178921 | PFAS | 1030.397167 | 0.079143983 | 0.06917572 | 1.144100598 | 0.25258192 | 0.431342905 |
| ENSG00000178922 | HYI | 290.46234 | 0.048257 | 0.108912395 | 0.443080883 | 0.657707232 | 0.801775104 |
| ENSG00000178927 | CYBC1 | 2237.045844 | 0.22270697 | 0.05818087 | 3.827838411 | 0.000129274 | 0.000566266 |
| ENSG00000178935 | ZNF552 | 377.0112781 | -0.036246335 | 0.098222385 | -0.369023164 | 0.712110459 | 0.839972059 |

| | | | | | | | |
|---|---|---|---|---|---|---|---|
| ENSG00000178947 | SMIM10L2A | 26.44230796 | 0.337902328 | 0.340336552 | 0.992847596 | 0.320784231 | 0.510318326 |
| ENSG00000178950 | GAK | 3429.062802 | 0.051383376 | 0.055030295 | 0.933728888 | 0.35044376 | 0.541833244 |
| ENSG00000178951 | ZBTB7A | 2087.892956 | 0.252889937 | 0.055886729 | 4.525044524 | 6.04E-06 | 3.20E-05 |
| ENSG00000178952 | TUFM | 5865.845244 | 0.100553629 | 0.04895171 | 2.054139249 | 0.039962216 | 0.099042053 |
| ENSG00000178966 | RMI1 | 571.4354777 | -0.189090873 | 0.084530284 | -2.236960107 | 0.025288947 | 0.067109816 |
| ENSG00000178971 | CTC1 | 594.3204015 | -0.224342924 | 0.089149943 | -2.516467391 | 0.011853787 | 0.03456099 |
| ENSG00000178974 | FBXO34 | 1211.973276 | -0.326373289 | 0.066829637 | -4.883660965 | 1.04E-06 | 6.02E-06 |
| ENSG00000178977 | LINC00324 | 10.74124107 | -0.671766995 | 0.532685966 | -1.261093849 | 0.207275036 | 0.373342442 |
| ENSG00000178980 | SELENOW | 2271.526368 | 0.064848465 | 0.05974814 | 1.085363742 | 0.277760585 | 0.46055857 |
| ENSG00000178982 | EIF3K | 3374.277986 | 0.208329253 | 0.048243481 | 4.318288163 | 1.57E-05 | 7.90E-05 |
| ENSG00000178988 | MRFAP1L1 | 3802.847178 | -0.264064899 | 0.046300602 | -5.703271416 | 1.18E-08 | 8.37E-08 |
| ENSG00000178996 | SNX18 | 512.0762294 | 0.127954701 | 0.085859729 | 1.490276088 | 0.136151657 | 0.270792354 |
| ENSG00000178999 | AURKB | 841.3526951 | -0.333338761 | 0.076037479 | -4.383874428 | 1.17E-05 | 5.96E-05 |
| ENSG00000179008 | C14orf39 | 36.71641395 | -0.344947266 | 0.294132979 | -1.172759568 | 0.240892236 | 0.416810069 |
| ENSG00000179010 | MRFAP1 | 15250.82917 | -0.202213218 | 0.047827089 | -4.228005977 | 2.36E-05 | 0.000115692 |
| ENSG00000179021 | C3orf38 | 690.3906471 | -0.122116022 | 0.077115834 | -1.583540182 | 0.113298396 | 0.234509176 |
| ENSG00000179023 | KLHDC7A | 28.74314354 | -1.1903385 | 0.339789385 | -3.503165641 | 0.000459763 | 0.001832013 |
| ENSG00000179029 | TMEM107 | 89.6046356 | -0.336177997 | 0.191750071 | -1.753209246 | 0.079566097 | 0.175910395 |
| ENSG00000179031 | NSA2P3 | 9.504703165 | 0.148808608 | 0.578582603 | 0.257195095 | 0.79702817 | 0.891799572 |
| ENSG00000179041 | RRS1 | 843.5969108 | 0.417679039 | 0.083547162 | 4.99932049 | 5.75E-07 | 3.43E-06 |
| ENSG00000179044 | EXOC3L1 | 1.65094893 | -1.377286077 | 1.391340877 | -0.989898378 | 0.322223793 | NA |
| ENSG00000179051 | RCC2 | 9451.375355 | 0.052481872 | 0.044250969 | 1.186005032 | 0.235620312 | 0.410570504 |
| ENSG00000179057 | IGSF22 | 1.641503968 | -0.135634195 | 1.35465169 | -0.100124774 | 0.920245267 | NA |
| ENSG00000179058 | C9orf50 | 36.57438885 | -0.089010816 | 0.296230495 | -0.300478234 | 0.763812397 | 0.870479014 |
| ENSG00000179071 | CCDC89 | 10.57704229 | 0.837102936 | 0.57369541 | 1.459141771 | 0.144526091 | 0.283200524 |
| ENSG00000179082 | LINC02913 | 38.33822403 | -0.128295555 | 0.304412565 | -0.421452888 | 0.673424407 | 0.813071362 |
| ENSG00000179085 | DPM3 | 372.2944055 | 0.288142672 | 0.105554769 | 2.729793022 | 0.00633741 | 0.019892879 |
| ENSG00000179088 | C12orf42 | 11.57065201 | 0.261309622 | 0.508346898 | 0.514037999 | 0.607225424 | 0.765665434 |
| ENSG00000179091 | CYC1 | 4720.055793 | 0.104375592 | 0.051741098 | 2.017266664 | 0.043667693 | 0.106783591 |
| ENSG00000179094 | PER1 | 663.3699274 | -0.044619295 | 0.087544757 | -0.509674103 | 0.610279799 | 0.768261612 |
| ENSG00000179097 | HTR1F | 17.0667016 | -0.938393513 | 0.441671887 | -2.124639447 | 0.03361671 | 0.085618665 |
| ENSG00000179101 | H3P47 | 29.81776488 | 0.06512184 | 0.352093132 | 0.18495629 | 0.853263319 | 0.923556693 |
| ENSG00000179104 | TMTC2 | 821.8529158 | 0.079641881 | 0.072629979 | 1.096542805 | 0.272841301 | 0.454862706 |

| | | | | | | | |
|---|---|---|---|---|---|---|---|
| ENSG00000179111 | HES7 | 2.822186713 | 1.45153507 | 1.139936415 | 1.2733474 | 0.202894794 | 0.367095715 |
| ENSG00000179115 | FARSA | 2084.550896 | 0.057542505 | 0.060261018 | 0.954887708 | 0.339634487 | 0.530711987 |
| ENSG00000179119 | SPTY2D1 | 1671.527045 | -0.044460689 | 0.058911403 | -0.754704309 | 0.45042642 | 0.638625693 |
| ENSG00000179131 | RPL31P38 | 2.158791252 | 0.020881113 | 1.161392925 | 0.01797937 | 0.985655311 | 0.993350589 |
| ENSG00000179134 | SAMD4B | 3278.527584 | 0.324440146 | 0.052769411 | 6.148261663 | 7.83E-10 | 6.28E-09 |
| ENSG00000179148 | ALOXE3 | 44.78522275 | 1.732134616 | 0.291340121 | 5.945403644 | 2.76E-09 | 2.11E-08 |
| ENSG00000179151 | EDC3 | 2612.766344 | 0.063131534 | 0.052481068 | 1.202939197 | 0.228999851 | 0.402045919 |
| ENSG00000179152 | TCAIM | 685.2075523 | 0.075550449 | 0.08005611 | 0.943718716 | 0.345313407 | 0.536243449 |
| ENSG00000179157 | RPS2P28 | 6.057423884 | -0.094987285 | 0.717844589 | -0.132322911 | 0.894728887 | 0.946400434 |
| ENSG00000179163 | FUCA1 | 629.7152958 | 0.314574486 | 0.080558317 | 3.904928707 | 9.43E-05 | 0.000421475 |
| ENSG00000179165 | PXT1 | 1.807561433 | 1.117511324 | 1.46193322 | 0.764406547 | 0.444625009 | 0.633374776 |
| ENSG00000179168 | GGN | 13.25117121 | 0.713602805 | 0.495852799 | 1.439142437 | 0.150110171 | 0.291756083 |
| ENSG00000179172 | HNRNPCL1 | 10.56885096 | 0.410106428 | 0.54948933 | 0.746341023 | 0.455461436 | 0.642892281 |
| ENSG00000179178 | TMEM125 | 810.6938222 | -0.220849211 | 0.074595142 | -2.960637994 | 0.003070026 | 0.010379718 |
| ENSG00000179195 | ZNF664 | 8819.501075 | 0.276395104 | 0.045661788 | 6.053094233 | 1.42E-09 | 1.11E-08 |
| ENSG00000179218 | CALR | 12129.71114 | 0.022195984 | 0.040818853 | 0.543767965 | 0.586601156 | 0.750355818 |
| ENSG00000179222 | MAGED1 | 4889.519321 | -0.667505201 | 0.047347322 | -14.09805597 | 3.90E-45 | 1.90E-43 |
| ENSG00000179240 | GVQW3 | 248.5699657 | 0.003471422 | 0.130915501 | 0.026516509 | 0.978845366 | 0.99043965 |
| ENSG00000179241 | LDLRAD3 | 374.0423383 | 1.37152254 | 0.112628033 | 12.17745271 | 4.10E-34 | 1.34E-32 |
| ENSG00000179242 | CDH4 | 13.03512019 | -2.095476662 | 0.548715124 | -3.818878992 | 0.00013406 | 0.000585107 |
| ENSG00000179262 | RAD23A | 3126.223291 | 0.089532865 | 0.051997775 | 1.72185954 | 0.085094966 | 0.185894535 |
| ENSG00000179271 | GADD45GIP1 | 639.4306046 | -0.128424337 | 0.080965334 | -1.586164487 | 0.112702001 | 0.233464323 |
| ENSG00000179277 | MEIS3P1 | 318.6190068 | 0.055988134 | 0.111770713 | 0.500919539 | 0.616427751 | 0.772766413 |
| ENSG00000179292 | TMEM151A | 10.66096425 | -0.467024992 | 0.535301954 | -0.872451498 | 0.382962117 | 0.575434005 |
| ENSG00000179295 | PTPN11 | 7659.03343 | 0.097616365 | 0.043826256 | 2.227348955 | 0.025923961 | 0.06850658 |
| ENSG00000179299 | NSUN7 | 353.623606 | -0.02655839 | 0.101123049 | -0.262634382 | 0.792832397 | 0.88871611 |
| ENSG00000179304 | FAM156B | 5.429082864 | 0.34256192 | 0.816952956 | 0.419316581 | 0.67498478 | 0.814355257 |
| ENSG00000179335 | CLK3 | 2204.286895 | -0.204659091 | 0.054901799 | -3.727730107 | 0.000193212 | 0.000822622 |
| ENSG00000179342 | ENSG0000017 | 22.3242229 | 0.253780998 | 0.380619857 | 0.666757115 | 0.50492729 | 0.685508198 |
| ENSG00000179344 | HLA-DQB1 | 23.59149371 | 2.145872069 | 0.435855236 | 4.923359624 | 8.51E-07 | 4.98E-06 |
| ENSG00000179348 | GATA2 | 931.9941059 | 0.608127622 | 0.071552442 | 8.499047762 | 1.91E-17 | 2.71E-16 |
| ENSG00000179361 | ARID3B | 236.1024326 | -0.451366728 | 0.121536708 | -3.713830445 | 0.000204146 | 0.000864519 |
| ENSG00000179362 | HMGN2P46 | 12.84396911 | 0.112958249 | 0.4991201 | 0.226314767 | 0.820956614 | 0.904991377 |

| | | | | | | | |
|---|---|---|---|---|---|---|---|
| ENSG00000179363 | TMEM31 | 33.2388751 | 0.153278299 | 0.302225572 | 0.507165219 | 0.612038897 | 0.769411385 |
| ENSG00000179364 | PACS2 | 4449.726532 | -0.242421611 | 0.052246157 | -4.639989292 | 3.48E-06 | 1.90E-05 |
| ENSG00000179387 | ELMOD2 | 718.44072 | -0.036806206 | 0.076412525 | -0.481677659 | 0.630034949 | 0.781665859 |
| ENSG00000179388 | EGR3 | 159.6987189 | 2.303020725 | 0.178926159 | 12.87134723 | 6.53E-38 | 2.46E-36 |
| ENSG00000179397 | CATSPERE | 4.342664378 | -0.207990979 | 0.845044382 | -0.246130242 | 0.805581413 | 0.896827993 |
| ENSG00000179399 | GPC5 | 20.9717833 | -0.815374165 | 0.390423462 | -2.088435365 | 0.03675858 | 0.09245059 |
| ENSG00000179403 | VWA1 | 996.8000584 | 0.610537275 | 0.092425924 | 6.605692921 | 3.96E-11 | 3.53E-10 |
| ENSG00000179406 | LINC00174 | 209.4335097 | 0.311895926 | 0.127622811 | 2.44388855 | 0.014529914 | 0.041313366 |
| ENSG00000179409 | GEMIN4 | 1328.372837 | 0.052797745 | 0.063448005 | 0.832141927 | 0.405328838 | 0.598073169 |
| ENSG00000179412 | HNRNPCL4 | 5.886328168 | 0.454026302 | 0.718096861 | 0.632263315 | 0.527214831 | 0.703321018 |
| ENSG00000179431 | FJX1 | 321.2040457 | 0.582983469 | 0.108493058 | 5.373463315 | 7.72E-08 | 5.05E-07 |
| ENSG00000179454 | KLHL28 | 525.2557402 | -0.6449007 | 0.088170042 | -7.314283704 | 2.59E-13 | 2.75E-12 |
| ENSG00000179456 | ZBTB18 | 15.17431604 | 0.777311702 | 0.456096434 | 1.704270509 | 0.088330562 | 0.191522031 |
| ENSG00000179476 | C14orf28 | 183.8732502 | -0.654703271 | 0.139302632 | -4.699862882 | 2.60E-06 | 1.44E-05 |
| ENSG00000179520 | SLC17A8 | 20.71583881 | 6.746813576 | 1.223159307 | 5.515891132 | 3.47E-08 | 2.36E-07 |
| ENSG00000179523 | EIF3J-DT | 223.9473381 | -0.043980429 | 0.124777324 | -0.352471327 | 0.724484818 | 0.846603929 |
| ENSG00000179526 | SHARPIN | 1735.259891 | -0.060093671 | 0.065650078 | -0.915363276 | 0.360000951 | 0.552550636 |
| ENSG00000179528 | LBX2 | 121.4546438 | -0.216236659 | 0.16299687 | -1.326630747 | 0.18463086 | 0.341429417 |
| ENSG00000179532 | DNHD1 | 213.2901019 | -0.150659206 | 0.135912726 | -1.108499627 | 0.2676461 | 0.448914348 |
| ENSG00000179542 | SLITRK4 | 103.2825872 | -0.285642646 | 0.177948014 | -1.605202776 | 0.108449187 | 0.226227548 |
| ENSG00000179546 | HTR1D | 10.81228887 | 1.653237218 | 0.608236718 | 2.718081907 | 0.006566158 | 0.020504719 |
| ENSG00000179562 | GCC1 | 1181.461918 | 0.07379185 | 0.069534192 | 1.061231147 | 0.288584868 | 0.472563128 |
| ENSG00000179564 | LSMEM2 | 4.553737205 | -0.236646 | 0.863513801 | -0.274050049 | 0.78404616 | 0.882792721 |
| ENSG00000179580 | RNF151 | 1.722829335 | 0.079591421 | 1.450918081 | 0.054855902 | 0.956253264 | NA |
| ENSG00000179583 | CIITA | 20.83598482 | 1.061516608 | 0.400670721 | 2.649349088 | 0.008064698 | 0.024614712 |
| ENSG00000179588 | ZFPM1 | 488.1530499 | 0.285224618 | 0.094716642 | 3.0113464 | 0.00260092 | 0.00892522 |
| ENSG00000179598 | PLD6 | 383.5734027 | 0.437698357 | 0.101448974 | 4.314468067 | 1.60E-05 | 8.03E-05 |
| ENSG00000179604 | CDC42EP4 | 2387.614523 | 0.126245627 | 0.060794307 | 2.076602763 | 0.037838238 | 0.094740141 |
| ENSG00000179611 | DGKZP1 | 1573.570236 | -0.019933939 | 0.067554288 | -0.295080287 | 0.767932548 | 0.873221605 |
| ENSG00000179627 | ZBTB42 | 2808.986699 | -0.112563679 | 0.067801649 | -1.660190871 | 0.096876061 | 0.206652114 |
| ENSG00000179630 | LACC1 | 218.2317372 | 0.546686166 | 0.138825968 | 3.937924389 | 8.22E-05 | 0.000372099 |
| ENSG00000179632 | MAF1 | 4648.143384 | -0.080860798 | 0.058177382 | -1.389900937 | 0.164558961 | 0.312731552 |
| ENSG00000179698 | WDR97 | 549.0720629 | 0.254466926 | 0.099322601 | 2.562024381 | 0.010406401 | 0.030775892 |

| | | | | | | | |
|---|---|---|---|---|---|---|---|
| ENSG00000179715 | PCED1B | 121.728998 | 1.484479305 | 0.179888911 | 8.252200204 | 1.56E-16 | 2.10E-15 |
| ENSG00000179743 | SPEN-AS1 | 150.5726303 | 0.380310143 | 0.159154257 | 2.389569407 | 0.016868136 | 0.047160883 |
| ENSG00000179750 | APOBEC3B | 642.2806844 | -0.644063107 | 0.079756107 | -8.075408051 | 6.73E-16 | 8.62E-15 |
| ENSG00000179761 | PIPOX | 18.28323524 | -1.562232231 | 0.443098275 | -3.525701454 | 0.000422363 | 0.001693586 |
| ENSG00000179766 | ATP8B5P | 2.547340891 | -0.16916061 | 1.098296512 | -0.154020894 | 0.877593261 | 0.936395015 |
| ENSG00000179818 | PCBP1-AS1 | 499.0804664 | -0.209072567 | 0.089609465 | -2.333152722 | 0.019640131 | 0.053851666 |
| ENSG00000179820 | MYADM | 827.8388302 | -0.354072587 | 0.074653391 | -4.742886847 | 2.11E-06 | 1.17E-05 |
| ENSG00000179832 | MROH1 | 3218.378498 | 0.269520091 | 0.057497509 | 4.687509009 | 2.77E-06 | 1.52E-05 |
| ENSG00000179833 | SERTAD2 | 571.2595819 | -0.129503855 | 0.084907201 | -1.525239958 | 0.127199258 | 0.256974986 |
| ENSG00000179841 | AKAP5 | 425.7488193 | -0.506489937 | 0.101033953 | -5.013066592 | 5.36E-07 | 3.21E-06 |
| ENSG00000179846 | NKPD1 | 52.90256398 | -0.333857837 | 0.251252022 | -1.328776718 | 0.183921646 | 0.340364839 |
| ENSG00000179859 | RNF227 | 53.68952964 | 0.390315624 | 0.244119549 | 1.598870822 | 0.109849308 | 0.228566271 |
| ENSG00000179862 | CITED4 | 178.9141594 | 0.024491836 | 0.137319535 | 0.178356535 | 0.858442982 | 0.92649707 |
| ENSG00000179869 | ABCA13 | 3.250037696 | 4.012735445 | 1.423394467 | 2.819130985 | 0.004815386 | 0.015582588 |
| ENSG00000179886 | TIGD5 | 632.765795 | 0.006132923 | 0.0885052 | 0.069294493 | 0.944755209 | 0.973711185 |
| ENSG00000179889 | PDXDC1 | 4260.249904 | -0.017228645 | 0.04912222 | -0.350730185 | 0.725790778 | 0.847784533 |
| ENSG00000179899 | PHC1P1 | 484.6461182 | -0.233094563 | 0.088326266 | -2.639017519 | 0.008314668 | 0.025266723 |
| ENSG00000179902 | CFAP276 | 8.468490012 | -1.041341138 | 0.632774089 | -1.64567601 | 0.099830481 | 0.21166383 |
| ENSG00000179912 | R3HDM2 | 1351.521098 | 0.092352422 | 0.062262771 | 1.483268743 | 0.138003024 | 0.273620515 |
| ENSG00000179913 | B3GNT3 | 53.84066405 | 1.645955234 | 0.264350008 | 6.226423995 | 4.77E-10 | 3.91E-09 |
| ENSG00000179918 | SEPHS2 | 3446.308326 | -0.146555438 | 0.047870761 | -3.061481242 | 0.002202448 | 0.007678443 |
| ENSG00000179921 | GPBAR1 | 4.370561843 | 1.29762761 | 0.879511021 | 1.475396646 | 0.140105937 | 0.276642295 |
| ENSG00000179922 | ZNF784 | 241.584229 | 0.078580516 | 0.119588183 | 0.65709265 | 0.511121345 | 0.690810325 |
| ENSG00000179933 | C14orf119 | 1666.259115 | -0.371030261 | 0.055822353 | -6.646625292 | 3.00E-11 | 2.70E-10 |
| ENSG00000179941 | BBS10 | 509.271737 | -0.235129378 | 0.088338886 | -2.661674695 | 0.007775298 | 0.023845634 |
| ENSG00000179943 | FIZ1 | 499.4264602 | -0.142341719 | 0.091373471 | -1.557801374 | 0.11928034 | 0.244355331 |
| ENSG00000179950 | PUF60 | 6070.027898 | -0.06682043 | 0.049714436 | -1.344085054 | 0.178920885 | 0.333721855 |
| ENSG00000179954 | SSC5D | 13.60775429 | -1.106666771 | 0.487156394 | -2.271686843 | 0.023105431 | 0.062007323 |
| ENSG00000179958 | DCTPP1 | 2320.033868 | -0.072432392 | 0.053128453 | -1.363344627 | 0.172773924 | 0.32489797 |
| ENSG00000179965 | ZNF771 | 152.0820104 | -0.08045641 | 0.152257051 | -0.528424856 | 0.597204489 | 0.757729408 |
| ENSG00000179967 | PPP1R14BP3 | 1404.113632 | 0.292396867 | 0.068808225 | 4.249446438 | 2.14E-05 | 0.000105903 |
| ENSG00000179978 | NAIPP2 | 18.26759477 | -0.693290971 | 0.424013179 | -1.635069393 | 0.102034501 | 0.215492636 |
| ENSG00000179981 | TSHZ1 | 1135.918304 | 0.040729067 | 0.070963665 | 0.57394255 | 0.566006678 | 0.73394626 |

| | | | | | | | |
|---|---|---|---|---|---|---|---|
| ENSG00000179988 | PSTK | 83.76433332 | -0.185968472 | 0.192076561 | -0.968199717 | 0.332944636 | 0.524066828 |
| ENSG00000179997 | RPA3P1 | 6.205853944 | -1.225740422 | 0.751577244 | -1.630890813 | 0.102913361 | 0.217028176 |
| ENSG00000180008 | SOCS4 | 1244.445077 | -0.718624297 | 0.066465845 | -10.81193349 | 3.02E-27 | 7.57E-26 |
| ENSG00000180011 | PTGR3 | 1790.393207 | 0.371219009 | 0.057652702 | 6.43888306 | 1.20E-10 | 1.03E-09 |
| ENSG00000180015 | ENSG0000018 | 106.9565501 | 0.05530184 | 0.183670386 | 0.301092852 | 0.763343691 | 0.870333215 |
| ENSG00000180035 | ZNF48 | 814.4763118 | -0.122633065 | 0.078910997 | -1.554068113 | 0.12016817 | 0.245888665 |
| ENSG00000180044 | C3orf80 | 33.87746809 | 0.562128921 | 0.316305712 | 1.777169682 | 0.075540326 | 0.168799443 |
| ENSG00000180066 | LINC02870 | 28.29397335 | -1.92230178 | 0.372146008 | -5.165450489 | 2.40E-07 | 1.50E-06 |
| ENSG00000180071 | ANKRD18A | 184.6458659 | -0.688970085 | 0.154713939 | -4.453186891 | 8.46E-06 | 4.40E-05 |
| ENSG00000180089 | TMEM86B | 46.12698888 | 0.276864353 | 0.268787242 | 1.0300502 | 0.302986441 | 0.490093707 |
| ENSG00000180096 | SEPTIN1 | 52.18379241 | 0.127437899 | 0.244151789 | 0.521961766 | 0.601696952 | 0.761330837 |
| ENSG00000180098 | TRNAU1AP | 384.9517628 | -0.498032733 | 0.096165367 | -5.17891992 | 2.23E-07 | 1.40E-06 |
| ENSG00000180104 | EXOC3 | 3492.40292 | 0.07757156 | 0.050335505 | 1.541090342 | 0.1232948 | 0.250654264 |
| ENSG00000180105 | ACER2P1 | 5.490598945 | 0.06589418 | 0.742051253 | 0.088800039 | 0.929240827 | 0.965981609 |
| ENSG00000180138 | CSNK1A1L | 7.370336849 | 0.911327904 | 0.691474349 | 1.317948967 | 0.187520732 | 0.345464632 |
| ENSG00000180155 | LYNX1 | 11.19202424 | -2.811887481 | 0.660950858 | -4.254306426 | 2.10E-05 | 0.000103831 |
| ENSG00000180172 | RPS12P23 | 8.1561157 | 0.725729307 | 0.648786516 | 1.11859493 | 0.263312987 | 0.44412799 |
| ENSG00000180176 | TH | 5.683317609 | -1.043667248 | 0.76668084 | -1.361279941 | 0.17342524 | 0.325829238 |
| ENSG00000180178 | FAR2P1 | 40.13539097 | 0.111648151 | 0.286180105 | 0.390132468 | 0.696438595 | 0.829463755 |
| ENSG00000180182 | MED14 | 2225.145345 | -0.179084598 | 0.05445018 | -3.288962494 | 0.001005574 | 0.003763725 |
| ENSG00000180185 | FAHD1 | 924.0688768 | -0.05027686 | 0.069977083 | -0.718476082 | 0.47246379 | 0.658168305 |
| ENSG00000180198 | RCC1 | 4503.072115 | 0.013709359 | 0.04601635 | 0.297923644 | 0.765761441 | 0.871876487 |
| ENSG00000180209 | MYL11 | 3.91711598 | 0.530297791 | 0.880236519 | 0.602449205 | 0.546875165 | 0.719836831 |
| ENSG00000180211 | ENSG0000018 | 160.0131715 | 0.340136865 | 0.157723885 | 2.156533646 | 0.031042019 | 0.080011677 |
| ENSG00000180228 | PRKRA | 810.1121394 | 0.127638858 | 0.073070881 | 1.746781415 | 0.080675262 | 0.177899693 |
| ENSG00000180229 | HERC2P3 | 16.65003904 | -0.083445402 | 0.436608192 | -0.191121933 | 0.848430064 | 0.920714588 |
| ENSG00000180233 | ZNRF2 | 83.85927985 | -1.193624881 | 0.203915518 | -5.853526465 | 4.81E-09 | 3.59E-08 |
| ENSG00000180245 | RRH | 4.608797183 | -0.752443399 | 0.876044766 | -0.858909759 | 0.390390305 | 0.583343425 |
| ENSG00000180251 | SLC9A4 | 9.238046471 | 6.558994136 | 1.270099648 | 5.164157119 | 2.42E-07 | 1.51E-06 |
| ENSG00000180257 | ZNF816 | 511.5248587 | 0.307972546 | 0.085953244 | 3.583024114 | 0.000339639 | 0.001382912 |
| ENSG00000180263 | FGD6 | 345.3720523 | 0.547064792 | 0.109089537 | 5.014823679 | 5.31E-07 | 3.19E-06 |
| ENSG00000180269 | GPR139 | 207.657608 | 0.685917722 | 0.128547922 | 5.335891182 | 9.51E-08 | 6.18E-07 |
| ENSG00000180279 | TMEM277P | 7.883729034 | -1.011133554 | 0.636940268 | -1.587485679 | 0.112402688 | 0.233012621 |

| | | | | | | | |
|---|---|---|---|---|---|---|---|
| ENSG00000180304 | OAZ2 | 2195.420469 | -0.072158421 | 0.052623214 | -1.371227949 | 0.170303908 | 0.321144644 |
| ENSG00000180316 | PNPLA1 | 1.621728682 | 2.87567047 | 1.634974346 | 1.758847457 | 0.078603422 | NA |
| ENSG00000180329 | CCDC43 | 1198.040497 | -0.070349877 | 0.063367341 | -1.110191398 | 0.266916559 | 0.448051065 |
| ENSG00000180336 | MEIOC | 25.27078692 | -0.607451967 | 0.364672516 | -1.6657465 | 0.095763916 | 0.204703291 |
| ENSG00000180340 | FZD2 | 476.4406907 | 0.485266985 | 0.099387343 | 4.882583359 | 1.05E-06 | 6.05E-06 |
| ENSG00000180346 | TIGD2 | 474.218541 | -0.193953053 | 0.090927144 | -2.13305998 | 0.032919806 | 0.084105174 |
| ENSG00000180347 | ITPRID1 | 16.90427802 | -0.849874599 | 0.437625894 | -1.942011683 | 0.052135686 | 0.124142894 |
| ENSG00000180353 | HCLS1 | 12.32295507 | 0.474203363 | 0.50423017 | 0.940450197 | 0.346986684 | 0.537873618 |
| ENSG00000180354 | MTURN | 395.5045478 | -0.822196561 | 0.100721695 | -8.163053282 | 3.27E-16 | 4.29E-15 |
| ENSG00000180357 | ZNF609 | 3979.761864 | -0.100249644 | 0.05624333 | -1.782427247 | 0.074679586 | 0.167397859 |
| ENSG00000180370 | PAK2 | 5680.786013 | -0.204055252 | 0.053475749 | -3.81584656 | 0.000135717 | 0.000591778 |
| ENSG00000180376 | CCDC66 | 226.5500818 | -0.140613982 | 0.126176693 | -1.114421204 | 0.265098531 | 0.446099541 |
| ENSG00000180385 | EMC3-AS1 | 562.3794446 | 0.094664674 | 0.082926679 | 1.141546669 | 0.253642503 | 0.43241372 |
| ENSG00000180389 | ATP5F1EP2 | 101.1988654 | -0.282427566 | 0.18119508 | -1.558693349 | 0.119068977 | 0.244031299 |
| ENSG00000180398 | MCFD2 | 3282.339542 | -0.021802325 | 0.049503465 | -0.440420174 | 0.659632816 | 0.803140853 |
| ENSG00000180422 | LINC00304 | 16.75580009 | 1.767756833 | 0.476729626 | 3.708090999 | 0.000208828 | 0.000882717 |
| ENSG00000180423 | HARBI1 | 146.2523084 | -0.140060165 | 0.148902651 | -0.940615655 | 0.346901857 | 0.537843241 |
| ENSG00000180425 | C11orf71 | 228.666321 | 0.258015863 | 0.120428263 | 2.142485972 | 0.032154396 | 0.082498809 |
| ENSG00000180438 | TPRXL | 3.528400682 | 1.688035612 | 1.046196129 | 1.613498238 | 0.106636306 | 0.223196793 |
| ENSG00000180447 | GAS1 | 6.113008163 | -0.616997802 | 0.731632422 | -0.843316648 | 0.399051375 | 0.591894383 |
| ENSG00000180448 | ARHGAP45 | 502.0222213 | 0.002678425 | 0.095231449 | 0.028125428 | 0.977562113 | 0.98951043 |
| ENSG00000180479 | ZNF571 | 80.77266273 | -0.016903156 | 0.2054805 | -0.082261607 | 0.934438684 | 0.968469969 |
| ENSG00000180481 | GLIPR1L2 | 8.57828428 | -0.160764092 | 0.60197011 | -0.267063247 | 0.789420465 | 0.886103903 |
| ENSG00000180488 | MIGA1 | 546.7746879 | 0.130991839 | 0.088454277 | 1.480898876 | 0.138633523 | 0.274591882 |
| ENSG00000180530 | NRIP1 | 1283.981472 | 0.537717354 | 0.073828994 | 7.283281611 | 3.26E-13 | 3.43E-12 |
| ENSG00000180535 | BHLHA15 | 3.079869244 | -1.084865009 | 1.167589434 | -0.929149388 | 0.352811672 | 0.544269266 |
| ENSG00000180537 | RNF182 | 1.899658075 | -2.378180974 | 1.464589106 | -1.623787153 | 0.104421243 | 0.219562173 |
| ENSG00000180539 | LINC02908 | 12.31200947 | 0.688351104 | 0.504178931 | 1.365291291 | 0.172161515 | 0.324012237 |
| ENSG00000180543 | TSPYL5 | 4398.785274 | -0.227146527 | 0.045259659 | -5.018741482 | 5.20E-07 | 3.13E-06 |
| ENSG00000180549 | FUT7 | 4.432899166 | -1.357792313 | 0.856133847 | -1.585957987 | 0.11274884 | 0.233519176 |
| ENSG00000180573 | H2AC6 | 1633.69478 | 0.609554854 | 0.062736593 | 9.716097431 | 2.57E-22 | 4.90E-21 |
| ENSG00000180574 | EIF2S3B | 1268.490528 | 0.10792536 | 0.0612212 | 1.762875605 | 0.077921472 | 0.173091757 |
| ENSG00000180581 | SRP9P1 | 870.4139531 | -0.019574591 | 0.071032388 | -0.275572761 | 0.782876233 | 0.882291026 |

| | | | | | | | |
|---|---|---|---|---|---|---|---|
| ENSG00000180592 | SKIDA1 | 65.79724596 | 1.035906418 | 0.225441928 | 4.595003363 | 4.33E-06 | 2.32E-05 |
| ENSG00000180596 | H2BC4 | 157.5445028 | 0.448017503 | 0.146714014 | 3.053679003 | 0.002260538 | 0.007869005 |
| ENSG00000180611 | MB21D2 | 458.886424 | -0.382378912 | 0.095073367 | -4.021935082 | 5.77E-05 | 0.000267666 |
| ENSG00000180616 | SSTR2 | 3.643159198 | -1.89023316 | 1.028105652 | -1.838559254 | 0.065980039 | 0.151161699 |
| ENSG00000180626 | ZNF594 | 346.619993 | -0.226870135 | 0.107830628 | -2.103948939 | 0.035382901 | 0.089441736 |
| ENSG00000180628 | PCGF5 | 1327.441001 | 0.159417262 | 0.0679022 | 2.347748107 | 0.018887289 | 0.052038725 |
| ENSG00000180667 | YOD1 | 9.844065548 | 0.184085691 | 0.55356462 | 0.332545983 | 0.739477024 | 0.856315141 |
| ENSG00000180673 | EXOC5P1 | 147.5462203 | -0.833065298 | 0.156812816 | -5.312482242 | 1.08E-07 | 6.99E-07 |
| ENSG00000180694 | TMEM64 | 2529.999123 | 2.011313645 | 0.06492819 | 30.97750974 | 1.08E-210 | 7.53E-208 |
| ENSG00000180708 | OR10K2 | 6.669239254 | -0.349067801 | 0.676276323 | -0.5161615 | 0.605741618 | 0.764382678 |
| ENSG00000180712 | LINC02363 | 3.028804989 | 0.386019247 | 1.026547031 | 0.376036592 | 0.706889692 | 0.836412774 |
| ENSG00000180720 | CHRM4 | 2.189073765 | 4.475739586 | 1.535281544 | 2.915256555 | 0.003553963 | 0.011855155 |
| ENSG00000180730 | SHISA2 | 2845.276604 | 0.952791515 | 0.052525111 | 18.13973353 | 1.55E-73 | 1.55E-71 |
| ENSG00000180739 | S1PR5 | 49.67170961 | 0.394973957 | 0.263050224 | 1.501515381 | 0.133222311 | 0.266351736 |
| ENSG00000180747 | SMG1P3 | 62.36362716 | -0.40667778 | 0.228251829 | -1.781706555 | 0.074797097 | 0.167595762 |
| ENSG00000180758 | GPR157 | 594.6841835 | 0.525284964 | 0.086517502 | 6.07143004 | 1.27E-09 | 1.00E-08 |
| ENSG00000180764 | PIPSL | 792.9634267 | 0.084695434 | 0.077009371 | 1.099806854 | 0.271416285 | 0.453414622 |
| ENSG00000180769 | WDFY3-AS2 | 272.3683491 | 0.625634501 | 0.11217775 | 5.577171085 | 2.44E-08 | 1.68E-07 |
| ENSG00000180773 | SLC36A4 | 204.3595224 | 0.071378802 | 0.131632749 | 0.542257171 | 0.587641356 | 0.751132537 |
| ENSG00000180776 | ZDHHC20 | 3411.148759 | -0.017594905 | 0.05138834 | -0.342390999 | 0.732056661 | 0.85176404 |
| ENSG00000180777 | ANKRD30B | 194.6124139 | 0.161948271 | 0.137160957 | 1.180716979 | 0.237715174 | 0.413151458 |
| ENSG00000180787 | ZFP3 | 488.3193793 | -0.025602059 | 0.087989601 | -0.290966874 | 0.771076654 | 0.87520429 |
| ENSG00000180801 | ARSJ | 23.72342099 | 0.073352138 | 0.37010008 | 0.198195413 | 0.842892176 | 0.917308531 |
| ENSG00000180806 | HOXC9 | 260.7773069 | -0.03752509 | 0.119497271 | -0.314024659 | 0.75350231 | 0.864088279 |
| ENSG00000180815 | MAP3K15 | 9.684716971 | -0.052182304 | 0.566169733 | -0.092167244 | 0.926565162 | 0.964663921 |
| ENSG00000180817 | PPA1 | 2731.999405 | -0.48680765 | 0.053807693 | -9.047175654 | 1.47E-19 | 2.40E-18 |
| ENSG00000180818 | HOXC10 | 137.8334727 | -0.177906597 | 0.156976639 | -1.133331675 | 0.257074995 | 0.43670767 |
| ENSG00000180822 | PSMG4 | 304.8354533 | -0.390062409 | 0.108835609 | -3.58395945 | 0.000338425 | 0.0013787 |
| ENSG00000180834 | MAP6D1 | 8.359481763 | -0.239177978 | 0.647342436 | -0.369476748 | 0.711772402 | 0.839747937 |
| ENSG00000180855 | ZNF443 | 219.4954664 | 0.073704749 | 0.124414688 | 0.59241196 | 0.553574759 | 0.725118768 |
| ENSG00000180867 | PDIA3P1 | 2341.807095 | -0.219764496 | 0.052207612 | -4.20943395 | 2.56E-05 | 0.000125168 |
| ENSG00000180875 | GREM2 | 8.053713038 | -6.57126423 | 1.288980298 | -5.098033103 | 3.43E-07 | 2.10E-06 |
| ENSG00000180879 | SSR4 | 2992.034131 | -0.314326614 | 0.05264849 | -5.970287365 | 2.37E-09 | 1.82E-08 |

| | | | | | | | |
|---|---|---|---|---|---|---|---|
| ENSG00000180881 | CAPS2 | 68.84769633 | 1.833844699 | 0.237915189 | 7.707976541 | 1.28E-14 | 1.49E-13 |
| ENSG00000180884 | ZNF792 | 304.4062473 | -0.213399418 | 0.105174771 | -2.028998168 | 0.042458477 | 0.104248899 |
| ENSG00000180891 | CUEDC1 | 1269.649698 | 1.020453789 | 0.070704828 | 14.43258998 | 3.23E-47 | 1.64E-45 |
| ENSG00000180900 | SCRIB | 4973.076439 | 0.050861892 | 0.05974247 | 0.851352338 | 0.394573659 | 0.587570253 |
| ENSG00000180901 | KCTD2 | 2377.158981 | 0.016061347 | 0.063185134 | 0.254195028 | 0.799344892 | 0.892991047 |
| ENSG00000180902 | D2HGDH | 1102.530034 | -0.552765292 | 0.074490808 | -7.420583864 | 1.17E-13 | 1.27E-12 |
| ENSG00000180914 | OXTR | 66.4779608 | -0.104642417 | 0.222059386 | -0.47123618 | 0.637472084 | 0.786553874 |
| ENSG00000180917 | CMTR2 | 1083.379025 | 0.260854053 | 0.074548839 | 3.499102823 | 0.000466827 | 0.001857267 |
| ENSG00000180921 | FAM83H | 7355.910567 | 0.041281337 | 0.046674798 | 0.884445966 | 0.376455519 | 0.569311728 |
| ENSG00000180938 | ZNF572 | 432.1043257 | -1.232569123 | 0.10094021 | -12.21088329 | 2.72E-34 | 8.98E-33 |
| ENSG00000180953 | ST20 | 37.19310442 | -0.219597833 | 0.298462616 | -0.735763277 | 0.461874774 | 0.648552813 |
| ENSG00000180957 | PITPNB | 1838.994726 | -0.67846405 | 0.062364649 | -10.87898455 | 1.45E-27 | 3.69E-26 |
| ENSG00000180979 | LRRC57 | 309.8743679 | 0.10210131 | 0.106975886 | 0.954432948 | 0.339864535 | 0.53082061 |
| ENSG00000180992 | MRPL14 | 986.1327269 | -0.09940571 | 0.079975629 | -1.242950023 | 0.21388625 | 0.38227897 |
| ENSG00000180998 | GPR137C | 197.6546591 | -0.395280135 | 0.135123904 | -2.925316133 | 0.003441065 | 0.011513714 |
| ENSG00000181004 | BBS12 | 55.93848593 | 0.085829151 | 0.242892141 | 0.353363229 | 0.723816149 | 0.846416769 |
| ENSG00000181016 | LSMEM1 | 26.16948424 | -0.680373701 | 0.345893438 | -1.967003786 | 0.049182774 | 0.118129289 |
| ENSG00000181019 | NQO1 | 6460.220869 | -0.120018562 | 0.044517907 | -2.695961463 | 0.007018579 | 0.021751715 |
| ENSG00000181026 | AEN | 1249.020818 | 0.237685709 | 0.063579627 | 3.738394231 | 0.000185199 | 0.000790708 |
| ENSG00000181027 | FKRP | 406.2524975 | 0.182598359 | 0.096003007 | 1.902006673 | 0.057170282 | 0.134098801 |
| ENSG00000181029 | TRAPPC5 | 137.9254038 | 0.526017659 | 0.154425067 | 3.406297102 | 0.000658505 | 0.002551275 |
| ENSG00000181031 | RPH3AL | 163.5094356 | -0.513000403 | 0.147191015 | -3.485269819 | 0.000491641 | 0.00194822 |
| ENSG00000181035 | SLC25A42 | 138.8830833 | -0.265315644 | 0.152682486 | -1.737695341 | 0.08226451 | 0.180747522 |
| ENSG00000181036 | FCRL6 | 2.63346705 | 3.696930873 | 1.479661213 | 2.498498197 | 0.012472078 | 0.036120372 |
| ENSG00000181038 | METTL23 | 410.0240058 | 0.176057658 | 0.105736825 | 1.665055274 | 0.095901729 | 0.204897612 |
| ENSG00000181045 | SLC26A11 | 1424.308069 | 0.149522842 | 0.06374128 | 2.345777224 | 0.018987453 | 0.052264478 |
| ENSG00000181061 | HIGD1A | 355.0474146 | -0.164503151 | 0.104385893 | -1.57591363 | 0.115045709 | 0.237393116 |
| ENSG00000181085 | MAPK15 | 73.56532047 | 0.373511934 | 0.206699977 | 1.807024557 | 0.07075845 | 0.160031436 |
| ENSG00000181090 | EHMT1 | 3252.032859 | -0.033273863 | 0.048953007 | -0.679710304 | 0.496687911 | 0.678619456 |
| ENSG00000181097 | LINC02878 | 43.17199243 | 0.557409487 | 0.275352598 | 2.024348022 | 0.042934356 | 0.105196467 |
| ENSG00000181101 | ENTR1P2 | 26.20371177 | -0.267875443 | 0.350543207 | -0.764172397 | 0.444764513 | 0.633405633 |
| ENSG00000181104 | F2R | 31.06851123 | 1.173751338 | 0.341120846 | 3.440866635 | 0.000579854 | 0.002266847 |
| ENSG00000181135 | ZNF707 | 668.1044249 | 0.107417431 | 0.082097245 | 1.308416998 | 0.19073191 | 0.350311451 |

| | | | | | | | |
|---|---|---|---|---|---|---|---|
| ENSG00000181143 | MUC16 | 4.560977377 | 0.744445519 | 0.884910554 | 0.841266403 | 0.400198708 | 0.592982712 |
| ENSG00000181163 | NPM1 | 11628.9355 | -0.075917373 | 0.060846329 | -1.24769028 | 0.212144502 | 0.379935583 |
| ENSG00000181191 | PJA1 | 537.526636 | -0.333998467 | 0.08513737 | -3.923053626 | 8.74E-05 | 0.000393818 |
| ENSG00000181192 | DHTKD1 | 2645.476671 | -0.950803198 | 0.05113046 | -18.59563159 | 3.49E-77 | 3.84E-75 |
| ENSG00000181218 | H2AC25 | 804.2640301 | 0.522873545 | 0.072131237 | 7.248919691 | 4.20E-13 | 4.39E-12 |
| ENSG00000181220 | ZNF746 | 1000.25403 | 0.581248641 | 0.071840151 | 8.090860487 | 5.92E-16 | 7.62E-15 |
| ENSG00000181222 | POLR2A | 6598.47119 | -0.063864018 | 0.045659836 | -1.398691372 | 0.161905553 | 0.308943053 |
| ENSG00000181227 | DLSTP1 | 322.1508226 | -0.658522741 | 0.10404279 | -6.329345241 | 2.46E-10 | 2.07E-09 |
| ENSG00000181240 | SLC25A41 | 8.47202508 | 0.478965431 | 0.598084827 | 0.800831936 | 0.423228948 | 0.614135749 |
| ENSG00000181260 | MTHFD2P7 | 22.33993523 | 0.229295885 | 0.391547253 | 0.585614847 | 0.558134369 | 0.728263604 |
| ENSG00000181264 | TLCD5 | 239.5387998 | 0.368382054 | 0.127795668 | 2.882586395 | 0.00394425 | 0.013013244 |
| ENSG00000181274 | FRAT2 | 1853.795299 | -1.109350682 | 0.057216437 | -19.38867112 | 9.62E-84 | 1.17E-81 |
| ENSG00000181284 | TMEM102 | 177.1458521 | -0.080230941 | 0.13843955 | -0.57953772 | 0.562226403 | 0.731024977 |
| ENSG00000181291 | TMEM132E | 2.570644107 | 1.84590794 | 1.241522313 | 1.486810121 | 0.137064968 | 0.272182753 |
| ENSG00000181315 | ZNF322 | 294.3942718 | 0.070967624 | 0.113191714 | 0.626968363 | 0.530679973 | 0.706329284 |
| ENSG00000181322 | NME9 | 2.667017394 | 2.086354526 | 1.22143471 | 1.708117927 | 0.087614463 | 0.190235581 |
| ENSG00000181323 | SPEM1 | 1.543663789 | 2.863129113 | 1.648397742 | 1.736916425 | 0.082401924 | NA |
| ENSG00000181333 | HEPHL1 | 1.610825392 | 1.797504541 | 1.514108409 | 1.187170305 | 0.235160451 | NA |
| ENSG00000181350 | LRRC75A | 220.518829 | 0.145949246 | 0.124791272 | 1.16954691 | 0.242183353 | 0.418449689 |
| ENSG00000181355 | OFCC1 | 16.66117511 | -2.11018446 | 0.520452529 | -4.054518601 | 5.02E-05 | 0.000235098 |
| ENSG00000181358 | CTAGE10P | 5.712250321 | 0.259750724 | 0.736304202 | 0.352776371 | 0.7242561 | 0.846509142 |
| ENSG00000181359 | HSP90AA6P | 25.41795488 | -0.208302114 | 0.356097589 | -0.584957946 | 0.558575994 | 0.728591193 |
| ENSG00000181378 | CFAP65 | 4.536369249 | 1.03197166 | 0.88499414 | 1.166077393 | 0.243583156 | 0.420166749 |
| ENSG00000181381 | DDX60L | 19.23104329 | 1.004253597 | 0.413182737 | 2.430531355 | 0.015076701 | 0.042693437 |
| ENSG00000181392 | SYNE4 | 1079.65923 | -0.284190531 | 0.070033426 | -4.057927024 | 4.95E-05 | 0.000231883 |
| ENSG00000181396 | OGFOD3 | 1406.003595 | 0.507562167 | 0.059991054 | 8.460630987 | 2.66E-17 | 3.74E-16 |
| ENSG00000181404 | WASHC1 | 70.64397657 | -0.217856771 | 0.229394515 | -0.949703486 | 0.342262938 | 0.53329254 |
| ENSG00000181409 | AATK | 47.65600612 | 0.466904121 | 0.287660749 | 1.623106809 | 0.104566574 | 0.219807314 |
| ENSG00000181418 | DDN | 12.86319522 | -0.334098034 | 0.487935348 | -0.684717833 | 0.493521993 | 0.676267519 |
| ENSG00000181433 | SAGE1 | 9.863666237 | 3.283471322 | 0.795579679 | 4.127143272 | 3.67E-05 | 0.000174918 |
| ENSG00000181444 | ZNF467 | 1274.773977 | 0.66574361 | 0.072727308 | 9.15397014 | 5.49E-20 | 9.15E-19 |
| ENSG00000181449 | SOX2 | 509.5257536 | 2.07996645 | 0.09917432 | 20.9728329 | 1.16E-97 | 1.79E-95 |
| ENSG00000181450 | ZNF678 | 416.6949029 | -0.292487738 | 0.098374127 | -2.973218117 | 0.00294695 | 0.009996011 |

| | | | | | | | |
|---|---|---|---|---|---|---|---|
| ENSG00000181458 | TMEM45A | 11.7155801 | -1.279425107 | 0.549651648 | -2.32770176 | 0.019927946 | 0.054532634 |
| ENSG00000181467 | RAP2B | 1443.385244 | 0.168226292 | 0.059676112 | 2.818988826 | 0.00481752 | 0.015587292 |
| ENSG00000181472 | ZBTB2 | 3053.643076 | 0.34002914 | 0.050559857 | 6.725278967 | 1.75E-11 | 1.61E-10 |
| ENSG00000181481 | RNF135 | 417.399496 | 0.140571739 | 0.096153204 | 1.461955848 | 0.143753309 | 0.282160624 |
| ENSG00000181511 | GLRX5P1 | 2.211709678 | 1.190755865 | 1.228115415 | 0.969579773 | 0.332255999 | 0.523341982 |
| ENSG00000181513 | ACBD4 | 258.3572714 | 0.114513057 | 0.117320727 | 0.976068421 | 0.329030559 | 0.519478277 |
| ENSG00000181523 | SGSH | 1778.385224 | 0.081886894 | 0.062040918 | 1.319885275 | 0.186873325 | 0.344576326 |
| ENSG00000181524 | RPL24P4 | 1366.041961 | 0.119569124 | 0.074318604 | 1.608872043 | 0.107644324 | 0.224916731 |
| ENSG00000181544 | FANCB | 113.7462272 | -0.095065109 | 0.172309182 | -0.551712378 | 0.581145428 | 0.746192551 |
| ENSG00000181555 | SETD2 | 2630.032606 | 0.008657167 | 0.055991261 | 0.154616392 | 0.877123746 | 0.936150792 |
| ENSG00000181585 | TMIE | 3.576284129 | -0.632930776 | 0.96795996 | -0.653881154 | 0.513188376 | 0.692502432 |
| ENSG00000181588 | MEX3D | 678.2265334 | 0.176497232 | 0.097393424 | 1.812208935 | 0.069953919 | 0.158479187 |
| ENSG00000181610 | MRPS23 | 2834.249952 | 0.105794052 | 0.049668839 | 2.129988435 | 0.033172568 | 0.08465666 |
| ENSG00000181619 | GPR135 | 74.50546368 | -0.234126199 | 0.211097864 | -1.10908843 | 0.267392036 | 0.448648611 |
| ENSG00000181626 | ANKRD62 | 3.146355861 | -1.636839926 | 1.086637057 | -1.506335455 | 0.131981081 | 0.264515638 |
| ENSG00000181634 | TNFSF15 | 6.170890241 | 1.16955334 | 0.732381551 | 1.596918081 | 0.110283969 | 0.229241958 |
| ENSG00000181638 | ZFP41 | 412.5207702 | -0.280470299 | 0.095097862 | -2.949280816 | 0.003185144 | 0.010726249 |
| ENSG00000181649 | PHLDA2 | 538.9434341 | 0.795762933 | 0.095676528 | 8.317222053 | 9.00E-17 | 1.23E-15 |
| ENSG00000181652 | ATG9B | 10.43646424 | 1.877184713 | 0.615792317 | 3.048405544 | 0.002300592 | 0.007990248 |
| ENSG00000181666 | ZNF875 | 1323.311031 | 0.437775562 | 0.06134499 | 7.136288773 | 9.59E-13 | 9.85E-12 |
| ENSG00000181690 | PLAG1 | 3.779999791 | -0.060541693 | 0.909564624 | -0.066561178 | 0.946931053 | 0.975200824 |
| ENSG00000181704 | YIPF6 | 2021.511791 | -0.019540116 | 0.057115406 | -0.342116386 | 0.732263307 | 0.851874855 |
| ENSG00000181722 | ZBTB20 | 13.76664221 | -0.108648803 | 0.476799829 | -0.227870893 | 0.819746612 | 0.904433786 |
| ENSG00000181741 | FDX1P1 | 2.565849572 | 0.556577246 | 1.090618365 | 0.510331811 | 0.60981902 | 0.767828639 |
| ENSG00000181744 | DIPK2A | 436.3253648 | 0.02528465 | 0.096478885 | 0.262074438 | 0.793264054 | 0.889039543 |
| ENSG00000181751 | MACIR | 591.8422266 | -0.111225823 | 0.081610937 | -1.362878885 | 0.172920684 | 0.325067647 |
| ENSG00000181754 | AMIGO1 | 635.5688053 | 0.555296807 | 0.082135396 | 6.760749135 | 1.37E-11 | 1.27E-10 |
| ENSG00000181773 | GPR3 | 10.50921964 | 0.51126991 | 0.551409594 | 0.927205321 | 0.353819941 | 0.545362456 |
| ENSG00000181788 | SIAH2 | 2311.623914 | 0.174855752 | 0.0548436 | 3.188261749 | 0.001431309 | 0.005184625 |
| ENSG00000181789 | COPG1 | 7671.76982 | -0.256384593 | 0.043687497 | -5.868603325 | 4.39E-09 | 3.29E-08 |
| ENSG00000181790 | ADGRB1 | 165.5720558 | 1.009362771 | 0.147133586 | 6.860179239 | 6.88E-12 | 6.53E-11 |
| ENSG00000181798 | LINC00471 | 26.27276192 | 0.038297221 | 0.340298542 | 0.112540067 | 0.910395203 | 0.95506154 |
| ENSG00000181817 | LSM10 | 879.3351935 | -0.35162862 | 0.077404464 | -4.542743462 | 5.55E-06 | 2.95E-05 |

| | | | | | | | |
|---|---|---|---|---|---|---|---|
| ENSG00000181819 | KCTD9P2 | 23.38766224 | -0.725671001 | 0.364877885 | -1.988805108 | 0.046722719 | 0.113001494 |
| ENSG00000181826 | RELL1 | 796.7586212 | -0.3318805 | 0.07374829 | -4.500178899 | 6.79E-06 | 3.58E-05 |
| ENSG00000181827 | RFX7 | 2437.906212 | -0.454310915 | 0.060583342 | -7.498941119 | 6.43E-14 | 7.12E-13 |
| ENSG00000181830 | SLC35C1 | 1514.242418 | 0.558292651 | 0.070268717 | 7.945109441 | 1.94E-15 | 2.40E-14 |
| ENSG00000181852 | RNF41 | 1986.021776 | 0.040493835 | 0.05801745 | 0.697959582 | 0.485202471 | 0.669063758 |
| ENSG00000181856 | SLC2A4 | 16.65754039 | 0.871523662 | 0.467983196 | 1.862296915 | 0.06256125 | 0.144410855 |
| ENSG00000181873 | IBA57 | 705.6384091 | -0.128591445 | 0.076670706 | -1.67719136 | 0.093505068 | 0.200617869 |
| ENSG00000181885 | CLDN7 | 1536.164441 | 0.089556988 | 0.068610061 | 1.305304006 | 0.191789351 | 0.351774822 |
| ENSG00000181894 | ZNF329 | 988.2005421 | 1.082700416 | 0.070575116 | 15.3411072 | 4.06E-53 | 2.43E-51 |
| ENSG00000181896 | ZNF101 | 128.7151292 | 0.19081419 | 0.158374536 | 1.204828725 | 0.228269429 | 0.401172197 |
| ENSG00000181904 | C5orf24 | 1212.415072 | 0.103599581 | 0.070550269 | 1.468450543 | 0.141981881 | 0.279552343 |
| ENSG00000181915 | ADO | 805.0161356 | -0.543445731 | 0.077556489 | -7.007095536 | 2.43E-12 | 2.41E-11 |
| ENSG00000181924 | COA4 | 1841.558368 | 0.148508766 | 0.055389135 | 2.681189462 | 0.007336097 | 0.022647027 |
| ENSG00000181929 | PRKAG1 | 1735.792172 | -0.187490674 | 0.059033123 | -3.17602499 | 0.001493081 | 0.005393052 |
| ENSG00000181938 | GINS3 | 314.572442 | -0.078184171 | 0.10635214 | -0.735144313 | 0.462251609 | 0.64880396 |
| ENSG00000181982 | CCDC149 | 696.6620065 | -0.020065775 | 0.075937515 | -0.264240604 | 0.791594525 | 0.887763734 |
| ENSG00000181991 | MRPS11 | 1544.517382 | 0.089995674 | 0.060878325 | 1.478287609 | 0.139330813 | 0.275514942 |
| ENSG00000182004 | SNRPE | 878.9483466 | 0.281742439 | 0.076876605 | 3.664865768 | 0.000247468 | 0.00103123 |
| ENSG00000182010 | RTKN2 | 321.5238054 | -1.204641191 | 0.110079019 | -10.9434223 | 7.15E-28 | 1.84E-26 |
| ENSG00000182022 | CHST15 | 2188.332114 | -0.440192163 | 0.057959639 | -7.59480508 | 3.08E-14 | 3.49E-13 |
| ENSG00000182040 | USH1G | 22.68792753 | -0.70236447 | 0.378483499 | -1.855733401 | 0.063491567 | 0.146234594 |
| ENSG00000182054 | IDH2 | 8148.898823 | 0.180454201 | 0.046480374 | 3.882374154 | 0.000103442 | 0.000459963 |
| ENSG00000182057 | OGFRP1 | 177.258103 | 0.50524832 | 0.140102194 | 3.60628414 | 0.000310613 | 0.00127605 |
| ENSG00000182087 | TMEM259 | 4650.188813 | 0.163655147 | 0.052149246 | 3.138207324 | 0.001699846 | 0.006061895 |
| ENSG00000182093 | GET1 | 600.4045996 | 0.108952689 | 0.086996491 | 1.252380276 | 0.21043133 | 0.377752149 |
| ENSG00000182095 | TNRC18 | 9570.188555 | 0.310924011 | 0.060189463 | 5.165754857 | 2.39E-07 | 1.49E-06 |
| ENSG00000182103 | FAM181B | 7.678490368 | -5.501352205 | 1.290966427 | -4.261421592 | 2.03E-05 | 0.000100796 |
| ENSG00000182107 | TMEM30B | 2662.954365 | -0.664510988 | 0.051754133 | -12.8397666 | 9.82E-38 | 3.67E-36 |
| ENSG00000182108 | DEXI | 760.2148919 | 0.429669272 | 0.076327789 | 5.629263994 | 1.81E-08 | 1.27E-07 |
| ENSG00000182111 | ZNF716 | 204.5655588 | 3.223009958 | 0.176668592 | 18.24325378 | 2.34E-74 | 2.42E-72 |
| ENSG00000182117 | NOP10 | 1576.557356 | -0.371193773 | 0.060438582 | -6.141669087 | 8.17E-10 | 6.54E-09 |
| ENSG00000182118 | FAM89A | 122.3522806 | -0.200061017 | 0.164416981 | -1.21679048 | 0.223683943 | 0.395080817 |
| ENSG00000182134 | TDRKH | 2149.801525 | 0.160230259 | 0.055210752 | 2.902156769 | 0.00370603 | 0.012299817 |

| | | | | | | | |
|---|---|---|---|---|---|---|---|
| ENSG00000182141 | ZNF708 | 119.130015 | -0.182252599 | 0.186658936 | -0.976393648 | 0.328869428 | 0.519327093 |
| ENSG00000182149 | IST1 | 5864.968877 | 0.407810229 | 0.047257837 | 8.629473044 | 6.16E-18 | 9.03E-17 |
| ENSG00000182150 | ERCC6L2 | 1102.652328 | 0.018286004 | 0.065887368 | 0.277534289 | 0.781369881 | 0.881401824 |
| ENSG00000182154 | MRPL41 | 1564.077144 | 0.033250632 | 0.068030419 | 0.488761241 | 0.625010741 | 0.778425521 |
| ENSG00000182158 | CREB3L2 | 2536.925937 | -0.400543777 | 0.058193803 | -6.882928412 | 5.86E-12 | 5.62E-11 |
| ENSG00000182165 | TP53TG1 | 354.1207769 | -0.672764451 | 0.104466239 | -6.440017927 | 1.19E-10 | 1.03E-09 |
| ENSG00000182168 | UNC5C | 341.5283561 | 1.437663154 | 0.108638848 | 13.23341681 | 5.63E-40 | 2.30E-38 |
| ENSG00000182173 | TSEN54 | 2263.083695 | 0.211204664 | 0.059773494 | 3.53341675 | 0.000410225 | 0.001646932 |
| ENSG00000182175 | RGMA | 4.718504902 | 0.967234218 | 0.822943417 | 1.175335021 | 0.239860709 | 0.415558089 |
| ENSG00000182179 | UBA7 | 6.689578943 | 0.61641222 | 0.681275825 | 0.904790978 | 0.365576132 | 0.558348008 |
| ENSG00000182180 | MRPS16 | 2359.239582 | -0.797103734 | 0.053060261 | -15.02261241 | 5.22E-51 | 2.96E-49 |
| ENSG00000182185 | RAD51B | 202.5393404 | -1.323429116 | 0.137087734 | -9.653884256 | 4.73E-22 | 8.86E-21 |
| ENSG00000182195 | LDOC1 | 10596.44197 | 0.347566025 | 0.046865241 | 7.416285854 | 1.20E-13 | 1.31E-12 |
| ENSG00000182196 | ARL6IP4 | 14.47230408 | 0.283187502 | 0.482310967 | 0.587147135 | 0.557104894 | 0.727416829 |
| ENSG00000182197 | EXT1 | 1097.157142 | 0.660759968 | 0.064845039 | 10.18983066 | 2.20E-24 | 4.73E-23 |
| ENSG00000182199 | SHMT2 | 5939.730464 | 0.043947913 | 0.045653961 | 0.962630877 | 0.335732797 | 0.526875672 |
| ENSG00000182208 | MOB2 | 333.6415324 | 0.249083277 | 0.102797953 | 2.423037324 | 0.015391345 | 0.043530694 |
| ENSG00000182218 | HHIPL1 | 6.264818993 | -1.11528528 | 0.721035972 | -1.546781747 | 0.121915883 | 0.248533756 |
| ENSG00000182220 | ATP6AP2 | 2839.181819 | -0.103739058 | 0.051143023 | -2.028410758 | 0.042518343 | 0.104379301 |
| ENSG00000182224 | CYB5D1 | 116.9863496 | -0.179994071 | 0.168033094 | -1.071182267 | 0.284087483 | 0.467937594 |
| ENSG00000182240 | BACE2 | 706.4447543 | 0.517269168 | 0.082157735 | 6.296049539 | 3.05E-10 | 2.55E-09 |
| ENSG00000182247 | UBE2E2 | 640.1120895 | -0.255211084 | 0.082926333 | -3.077563844 | 0.002087001 | 0.007304823 |
| ENSG00000182253 | SYNM | 306.3080513 | 1.159222088 | 0.115291369 | 10.05471697 | 8.76E-24 | 1.81E-22 |
| ENSG00000182257 | PRR34 | 10.05673199 | -0.419323659 | 0.550406998 | -0.761842891 | 0.446153778 | 0.634727095 |
| ENSG00000182263 | FIGN | 68.13175915 | -1.577474702 | 0.237199689 | -6.650407963 | 2.92E-11 | 2.64E-10 |
| ENSG00000182264 | IZUMO1 | 27.47211765 | -1.298914978 | 0.355909476 | -3.649565592 | 0.000262684 | 0.001090082 |
| ENSG00000182272 | B4GALNT4 | 1725.079646 | 0.461499518 | 0.05932519 | 7.779149462 | 7.30E-15 | 8.66E-14 |
| ENSG00000182287 | AP1S2 | 204.2155529 | -0.030411442 | 0.141753071 | -0.214538152 | 0.830127419 | 0.909932761 |
| ENSG00000182307 | C8orf33 | 3079.201149 | 0.015253033 | 0.047714994 | 0.319669602 | 0.749218806 | 0.861323026 |
| ENSG00000182308 | DCAF4L1 | 7.878711881 | 0.232350584 | 0.643764974 | 0.360924551 | 0.71815585 | 0.843411181 |
| ENSG00000182310 | SPACA6 | 257.3949362 | 0.263690551 | 0.128814559 | 2.047055492 | 0.040652636 | 0.100499776 |
| ENSG00000182318 | ZSCAN22 | 242.8095382 | 0.001132205 | 0.125371326 | 0.009030815 | 0.99279455 | 0.996094602 |
| ENSG00000182324 | KCNJ14 | 49.03836036 | -0.241076781 | 0.2563245 | -0.940514003 | 0.34695397 | 0.537873618 |

| | | | | | | | |
|---|---|---|---|---|---|---|---|
| ENSG00000182325 | FBXL6 | 736.8514674 | 0.124019736 | 0.081912958 | 1.514042951 | 0.130014946 | 0.261291788 |
| ENSG00000182326 | C1S | 8.27160018 | -0.277049887 | 0.617280969 | -0.448822986 | 0.653559356 | 0.79899021 |
| ENSG00000182327 | GLTPD2 | 5.116130503 | -0.795575404 | 0.800104675 | -0.994339152 | 0.320057781 | 0.509515996 |
| ENSG00000182329 | KIAA2012 | 5.44284741 | 0.231797861 | 0.75545339 | 0.306832776 | 0.758970659 | 0.867296921 |
| ENSG00000182359 | KBTBD3 | 69.37935296 | 0.179333207 | 0.218412532 | 0.821075628 | 0.411603191 | 0.603914203 |
| ENSG00000182362 | YBEY | 326.7519053 | -0.233498651 | 0.103222508 | -2.262090454 | 0.023691815 | 0.063410365 |
| ENSG00000182372 | CLN8 | 1210.258539 | 0.382454119 | 0.062178899 | 6.150866697 | 7.71E-10 | 6.19E-09 |
| ENSG00000182378 | PLCXD1 | 497.6463341 | 0.014261622 | 0.087919822 | 0.162211675 | 0.871139166 | 0.933087382 |
| ENSG00000182378 | PLCXD1 | 497.6463341 | 0.014261622 | 0.087919822 | 0.162211675 | 0.871139166 | 0.933087382 |
| ENSG00000182379 | NXPH4 | 216.240667 | 0.932950485 | 0.134886396 | 6.916564721 | 4.63E-12 | 4.47E-11 |
| ENSG00000182383 | RPL27AP5 | 15.18569418 | 0.009662041 | 0.457047588 | 0.021140121 | 0.98313388 | 0.99214031 |
| ENSG00000182389 | CACNB4 | 5.051863275 | 0.174571221 | 0.789378656 | 0.221150167 | 0.82497551 | 0.90772273 |
| ENSG00000182400 | TRAPPC6B | 1468.990315 | -0.270697897 | 0.07279478 | -3.718644354 | 0.000200295 | 0.000849309 |
| ENSG00000182405 | PGBD4 | 150.3429961 | -0.042723122 | 0.151896201 | -0.281265242 | 0.778506967 | 0.879751223 |
| ENSG00000182446 | NPLOC4 | 6168.151252 | -0.130930761 | 0.046005381 | -2.845987995 | 0.004427386 | 0.014455486 |
| ENSG00000182459 | TEX19 | 27.09483388 | 0.576847945 | 0.339422598 | 1.69949776 | 0.089225436 | 0.193070799 |
| ENSG00000182472 | CAPN12 | 52.41308961 | 0.399504633 | 0.245304869 | 1.628604582 | 0.103396754 | 0.217807303 |
| ENSG00000182473 | EXOC7 | 9688.850485 | 0.186177265 | 0.051040126 | 3.647664694 | 0.000264635 | 0.001097779 |
| ENSG00000182481 | KPNA2 | 13093.73585 | -0.099500108 | 0.045395074 | -2.191870151 | 0.028388884 | 0.074055151 |
| ENSG00000182484 | WASH6P | 148.487558 | 0.091057793 | 0.147346309 | 0.617984892 | 0.536585298 | 0.711682859 |
| ENSG00000182484 | WASH6P | 148.487558 | 0.091057793 | 0.147346309 | 0.617984892 | 0.536585298 | 0.711682859 |
| ENSG00000182489 | XKRX | 14.9181178 | 1.350374935 | 0.482079859 | 2.801143651 | 0.005092185 | 0.016372069 |
| ENSG00000182492 | BGN | 42.80845772 | 0.084462466 | 0.27530913 | 0.306791372 | 0.759002176 | 0.867296921 |
| ENSG00000182504 | CEP97 | 319.75742 | -0.167171993 | 0.103611487 | -1.61345037 | 0.106646697 | 0.223198179 |
| ENSG00000182511 | FES | 14.06044225 | -0.029463737 | 0.468907976 | -0.062834797 | 0.949898057 | 0.976896863 |
| ENSG00000182512 | GLRX5 | 1861.443496 | -0.512196842 | 0.055485439 | -9.231193796 | 2.68E-20 | 4.54E-19 |
| ENSG00000182518 | FAM104B | 530.2471761 | 0.312397234 | 0.089544876 | 3.488722633 | 0.000485334 | 0.00192589 |
| ENSG00000182534 | MXRA7 | 150.0458512 | 0.858108642 | 0.148854471 | 5.764748842 | 8.18E-09 | 5.93E-08 |
| ENSG00000182541 | LIMK2 | 1551.855689 | -0.754513257 | 0.065804896 | -11.46591366 | 1.96E-30 | 5.53E-29 |
| ENSG00000182544 | MFSD5 | 1013.367231 | 0.059194357 | 0.070756181 | 0.836596268 | 0.402819544 | 0.595519775 |
| ENSG00000182551 | ADI1 | 2689.643324 | 0.315748662 | 0.049940949 | 6.322440138 | 2.57E-10 | 2.16E-09 |
| ENSG00000182552 | RWDD4 | 124.2830043 | 0.066235809 | 0.164786168 | 0.401950052 | 0.687720785 | 0.823383061 |
| ENSG00000182568 | SATB1 | 985.6117597 | 1.758752531 | 0.075769018 | 23.21202753 | 3.44E-119 | 8.23E-117 |

| | | | | | | | |
|---|---|---|---|---|---|---|---|
| ENSG00000182575 | NXPH3 | 19.98724833 | -0.477380893 | 0.388701033 | -1.228144132 | 0.219392867 | 0.389341291 |
| ENSG00000182578 | CSF1R | 1.939800216 | -1.164807602 | 1.29752799 | -0.89771289 | 0.369338635 | 0.561957173 |
| ENSG00000182580 | EPHB3 | 911.1471636 | 0.790908743 | 0.090503865 | 8.738949896 | 2.35E-18 | 3.55E-17 |
| ENSG00000182583 | VCX | 16.24161938 | 1.230894868 | 0.463856821 | 2.653609504 | 0.007963593 | 0.024363365 |
| ENSG00000182585 | EPGN | 2.556224027 | -1.718302568 | 1.207934158 | -1.422513435 | 0.154877254 | 0.298411239 |
| ENSG00000182586 | LINC00334 | 4.110903639 | 0.20999217 | 0.893967965 | 0.234898988 | 0.814287135 | 0.901759557 |
| ENSG00000182600 | SNORC | 53.26499577 | 1.058087795 | 0.251269668 | 4.210965061 | 2.54E-05 | 0.000124376 |
| ENSG00000182606 | TRAK1 | 1643.389381 | 0.05287757 | 0.059771749 | 0.884658229 | 0.376340991 | 0.569311728 |
| ENSG00000182612 | TSPAN10 | 85.94300902 | 2.868056448 | 0.248360734 | 11.54794639 | 7.56E-31 | 2.17E-29 |
| ENSG00000182621 | PLCB1 | 868.6104022 | -0.197143448 | 0.069876087 | -2.821329246 | 0.004782509 | 0.01548712 |
| ENSG00000182625 | ENSG0000018 | 4.352557754 | -0.418100516 | 0.826253042 | -0.506019941 | 0.612842651 | 0.769891065 |
| ENSG00000182628 | SKA2 | 4976.955195 | -0.07429806 | 0.056971196 | -1.304133762 | 0.192187979 | 0.352308794 |
| ENSG00000182648 | RNF32-DT | 340.0790004 | 0.827063234 | 0.105336197 | 7.85165266 | 4.11E-15 | 4.97E-14 |
| ENSG00000182670 | TTC3 | 12365.60295 | -0.010224196 | 0.043393737 | -0.235614553 | 0.813731779 | 0.901448243 |
| ENSG00000182674 | KCNB2 | 2.163069323 | -3.581545357 | 1.550551429 | -2.309852669 | 0.020896312 | 0.056789646 |
| ENSG00000182676 | PPP1R27 | 2.078890791 | -0.183514376 | 1.343561518 | -0.136587996 | 0.891356465 | 0.944249241 |
| ENSG00000182685 | BRICD5 | 460.3185492 | -0.339416006 | 0.106211331 | -3.195666623 | 0.001395082 | 0.005073428 |
| ENSG00000182687 | GALR2 | 5.332065508 | -1.133669321 | 0.799611394 | -1.417775346 | 0.156256364 | 0.30041242 |
| ENSG00000182700 | IGIP | 84.87619069 | 0.195411819 | 0.192911191 | 1.012962589 | 0.31107803 | 0.499134785 |
| ENSG00000182704 | TSKU | 1638.189899 | -0.177622766 | 0.060464883 | -2.937618605 | 0.003307436 | 0.011107098 |
| ENSG00000182718 | ANXA2 | 10824.22629 | -0.219181928 | 0.042595964 | -5.145603206 | 2.67E-07 | 1.65E-06 |
| ENSG00000182722 | SEPHS1P1 | 3.513355235 | -0.072205325 | 0.926123537 | -0.077965111 | 0.937855806 | 0.970297758 |
| ENSG00000182732 | RGS6 | 661.7681727 | 1.09526847 | 0.089232453 | 12.27432884 | 1.24E-34 | 4.20E-33 |
| ENSG00000182747 | SLC35D3 | 3.311870428 | 0.834805237 | 0.973276006 | 0.857727132 | 0.391043155 | 0.583899955 |
| ENSG00000182749 | PAQR7 | 606.8937215 | 1.016086533 | 0.083733167 | 12.13481546 | 6.91E-34 | 2.24E-32 |
| ENSG00000182759 | MAFA | 41.70497087 | 0.127049659 | 0.271298583 | 0.468301962 | 0.639568665 | 0.787784857 |
| ENSG00000182768 | NGRN | 727.0142476 | 0.281529193 | 0.074895059 | 3.758982191 | 0.000170606 | 0.000732766 |
| ENSG00000182771 | GRID1 | 71.38590279 | 0.331859417 | 0.21365061 | 1.553280926 | 0.120356035 | 0.246185238 |
| ENSG00000182774 | RPS17 | 248.0925211 | 0.173204115 | 0.119643792 | 1.447664879 | 0.147710797 | 0.288161041 |
| ENSG00000182782 | HCAR2 | 131.7141442 | -0.376673042 | 0.170485312 | -2.209416383 | 0.027145693 | 0.071298625 |
| ENSG00000182791 | CCDC87 | 45.29030117 | 0.30696962 | 0.265521703 | 1.156099922 | 0.247640288 | 0.425119225 |
| ENSG00000182795 | C1orf116 | 5.2982076 | 4.74779937 | 1.340237011 | 3.542507281 | 0.000396342 | 0.001593707 |
| ENSG00000182796 | TMEM198B | 661.1329778 | 0.111332621 | 0.100065493 | 1.112597542 | 0.265881317 | 0.446828286 |

| | | | | | | | |
|---|---|---|---|---|---|---|---|
| ENSG00000182809 | CRIP2 | 2510.43042 | -0.400897289 | 0.064773791 | -6.189189804 | 6.05E-10 | 4.91E-09 |
| ENSG00000182810 | DDX28 | 475.2422961 | -0.072072949 | 0.088147307 | -0.817642108 | 0.413561576 | 0.605778849 |
| ENSG00000182814 | FUNDC2P2 | 13.45931584 | -0.674073895 | 0.500653815 | -1.346387214 | 0.178177672 | 0.332660194 |
| ENSG00000182827 | ACBD3 | 1914.460701 | -0.047613299 | 0.06449458 | -0.738252727 | 0.460360887 | 0.647219388 |
| ENSG00000182831 | HAPSTR1 | 1583.652055 | -0.08741691 | 0.060216913 | -1.451700296 | 0.146584957 | 0.286427486 |
| ENSG00000182836 | PLCXD3 | 2.713382331 | 1.527399671 | 1.14531474 | 1.333606927 | 0.182332712 | 0.338215606 |
| ENSG00000182841 | RRP7BP | 153.1519071 | -0.026209722 | 0.148554727 | -0.176431423 | 0.85995502 | 0.927319435 |
| ENSG00000182853 | VMO1 | 38.63424769 | -0.146422742 | 0.2835153 | -0.516454464 | 0.605537035 | 0.764292687 |
| ENSG00000182858 | ALG12 | 822.7155082 | -0.129690892 | 0.075814991 | -1.710623323 | 0.087150671 | 0.189465161 |
| ENSG00000182866 | LCK | 3.036169208 | -3.176036071 | 1.332021619 | -2.384372765 | 0.017108263 | 0.047744958 |
| ENSG00000182871 | COL18A1 | 5132.206943 | -0.045441267 | 0.060078802 | -0.756361065 | 0.449432745 | 0.637718016 |
| ENSG00000182872 | RBM10 | 3657.91481 | -0.058813975 | 0.056729821 | -1.036738238 | 0.299857866 | 0.486576451 |
| ENSG00000182873 | PRKCZ-AS1 | 20.4728075 | 0.862597124 | 0.397221157 | 2.171579006 | 0.029887432 | 0.077418438 |
| ENSG00000182888 | RPS27AP20 | 6.104685647 | 0.070933871 | 0.710794639 | 0.099795169 | 0.920506943 | 0.961454687 |
| ENSG00000182890 | GLUD2 | 973.4735425 | -0.713616681 | 0.068470636 | -10.42222953 | 1.96E-25 | 4.44E-24 |
| ENSG00000182898 | TCHHL1 | 2.699638442 | -0.989331775 | 1.193644023 | -0.828833183 | 0.407198809 | 0.599983343 |
| ENSG00000182899 | RPL35A | 5969.366103 | 0.220012749 | 0.047912537 | 4.591966204 | 4.39E-06 | 2.36E-05 |
| ENSG00000182902 | SLC25A18 | 336.6147536 | -5.536335299 | 0.231325441 | -23.93310164 | 1.39E-126 | 3.70E-124 |
| ENSG00000182903 | ZNF721 | 1200.336007 | -0.034606735 | 0.065291699 | -0.530032699 | 0.59608926 | 0.75703303 |
| ENSG00000182912 | TSPEAR-AS2 | 1.938812583 | -1.110228839 | 1.305446016 | -0.850459403 | 0.395069722 | 0.588013388 |
| ENSG00000182919 | C11orf54 | 206.0567208 | -0.218498245 | 0.128402968 | -1.701660386 | 0.08881905 | 0.19241751 |
| ENSG00000182923 | CEP63 | 1050.950014 | 0.016570591 | 0.068057913 | 0.243477799 | 0.807635281 | 0.897981454 |
| ENSG00000182934 | SRPRA | 3618.772773 | 0.171187035 | 0.050866357 | 3.365427474 | 0.000764252 | 0.002924913 |
| ENSG00000182944 | EWSR1 | 8632.595309 | -0.428496397 | 0.047759618 | -8.971939284 | 2.91E-19 | 4.67E-18 |
| ENSG00000182952 | HMGN4 | 1724.556944 | -0.086769362 | 0.056677859 | -1.530921667 | 0.125788754 | 0.254663667 |
| ENSG00000182957 | SPATA13 | 610.051605 | 0.332440034 | 0.080405184 | 4.13455969 | 3.56E-05 | 0.000169823 |
| ENSG00000182963 | GJC1 | 3.912970145 | 0.388400585 | 0.908418955 | 0.427556672 | 0.668973917 | 0.809848108 |
| ENSG00000182973 | CNOT10 | 1107.27702 | 0.114283185 | 0.068296268 | 1.673344506 | 0.094259496 | 0.202009794 |
| ENSG00000182979 | MTA1 | 4137.897959 | -0.219975844 | 0.05006599 | -4.393718026 | 1.11E-05 | 5.71E-05 |
| ENSG00000182985 | CADM1 | 255.7317004 | 3.11389189 | 0.154030055 | 20.21613183 | 7.06E-91 | 9.82E-89 |
| ENSG00000182986 | ZNF320 | 6.845539392 | 1.80302692 | 0.756653775 | 2.382895558 | 0.017177067 | 0.047902008 |
| ENSG00000182993 | C12orf60 | 6.043312943 | -0.499131984 | 0.772976273 | -0.645727433 | 0.518455891 | 0.696066421 |
| ENSG00000183010 | PYCR1 | 7782.559792 | 0.016814154 | 0.046751791 | 0.359647272 | 0.719110928 | 0.843883997 |

| | | | | | | | |
|---|---|---|---|---|---|---|---|
| ENSG00000183011 | NAA38 | 645.863513 | 0.280191326 | 0.084310152 | 3.323340297 | 0.000889463 | 0.003362605 |
| ENSG00000183018 | SPNS2 | 31.78027729 | 2.041651271 | 0.363514855 | 5.61641771 | 1.95E-08 | 1.36E-07 |
| ENSG00000183019 | MCEMP1 | 6.756210991 | -3.308119129 | 0.928531917 | -3.562741431 | 0.000367002 | 0.001484052 |
| ENSG00000183020 | AP2A2 | 2092.069515 | 0.022773536 | 0.056931487 | 0.400016529 | 0.689144342 | 0.82436022 |
| ENSG00000183022 | TPM3P8 | 45.01865786 | -0.197984691 | 0.265291615 | -0.746290795 | 0.455491771 | 0.64289554 |
| ENSG00000183023 | SLC8A1 | 284.9367895 | 0.32938761 | 0.112151041 | 2.937000009 | 0.003314041 | 0.011124393 |
| ENSG00000183032 | SLC25A21 | 77.96318568 | -0.29364563 | 0.208411608 | -1.408969648 | 0.158844143 | 0.304183621 |
| ENSG00000183036 | PCP4 | 338.1886336 | -0.262465755 | 0.109944772 | -2.387250901 | 0.016974902 | 0.047413156 |
| ENSG00000183044 | ABAT | 2820.135296 | -0.035105761 | 0.059388203 | -0.591123484 | 0.554437686 | 0.72573819 |
| ENSG00000183048 | SLC25A10 | 402.0749111 | 0.891332303 | 0.099294627 | 8.976641881 | 2.79E-19 | 4.48E-18 |
| ENSG00000183049 | CAMK1D | 890.1348077 | -0.559624848 | 0.0748381 | -7.477806757 | 7.56E-14 | 8.30E-13 |
| ENSG00000183055 | FAM133CP | 2.025459359 | -1.20421587 | 1.343133554 | -0.89657195 | 0.369947371 | 0.562601577 |
| ENSG00000183060 | LYSMD4 | 318.2212024 | 0.231818279 | 0.112122751 | 2.067540061 | 0.038683293 | 0.096476924 |
| ENSG00000183066 | WBP2NL | 16.49394072 | -0.504077827 | 0.432863194 | -1.164519953 | 0.244213362 | 0.421001275 |
| ENSG00000183067 | IGSF5 | 72.24515529 | -0.253300389 | 0.210257022 | -1.204717856 | 0.22831224 | 0.401194557 |
| ENSG00000183077 | AFMID | 631.2821417 | 0.238598764 | 0.079990371 | 2.982843555 | 0.002855839 | 0.009728683 |
| ENSG00000183087 | GAS6 | 188.2862953 | 0.713925168 | 0.141713757 | 5.03779721 | 4.71E-07 | 2.85E-06 |
| ENSG00000183091 | NEB | 192.9921006 | 0.572446632 | 0.133918766 | 4.274581139 | 1.91E-05 | 9.53E-05 |
| ENSG00000183092 | BEGAIN | 73.01388714 | -0.626867708 | 0.219143279 | -2.86053814 | 0.004229227 | 0.013863741 |
| ENSG00000183098 | GPC6 | 399.8345193 | 0.912322396 | 0.098761631 | 9.237619778 | 2.52E-20 | 4.29E-19 |
| ENSG00000183111 | ARHGEF37 | 689.1463122 | 0.375393919 | 0.082039139 | 4.575790566 | 4.74E-06 | 2.54E-05 |
| ENSG00000183117 | CSMD1 | 78.31905509 | 3.385545126 | 0.291809097 | 11.60191771 | 4.03E-31 | 1.17E-29 |
| ENSG00000183137 | CEP57L1 | 197.831772 | -0.130608121 | 0.137998784 | -0.946443997 | 0.343922174 | 0.534882049 |
| ENSG00000183145 | RIPPLY3 | 259.3615551 | -0.199322576 | 0.11311131 | -1.762180776 | 0.078038759 | 0.173285174 |
| ENSG00000183150 | GPR19 | 16.24020181 | -1.262576015 | 0.48740969 | -2.590379387 | 0.009587021 | 0.028673567 |
| ENSG00000183153 | GJD3 | 127.1251265 | 0.062033497 | 0.160155157 | 0.387333747 | 0.698509145 | 0.830851384 |
| ENSG00000183154 | ENSG0000018 | 2.486936744 | 0.374932585 | 1.100757716 | 0.340613179 | 0.733394807 | 0.852715503 |
| ENSG00000183155 | RABIF | 1810.627714 | 0.41022469 | 0.056063796 | 7.317105103 | 2.53E-13 | 2.69E-12 |
| ENSG00000183161 | FANCF | 884.9799781 | 0.04131035 | 0.072032907 | 0.57349275 | 0.566311106 | 0.734213676 |
| ENSG00000183171 | ENSG0000018 | 180.0112698 | 0.216138079 | 0.139450029 | 1.549932124 | 0.121157808 | 0.247362062 |
| ENSG00000183172 | SMDT1 | 806.8531505 | -0.172380594 | 0.071219904 | -2.42039913 | 0.01550348 | 0.043808172 |
| ENSG00000183186 | C2CD4C | 55.53041408 | -0.995671987 | 0.241400252 | -4.124568964 | 3.71E-05 | 0.000176776 |
| ENSG00000183196 | CHST6 | 3.088759842 | -0.492267873 | 1.064382325 | -0.462491589 | 0.643728833 | 0.791270263 |

| | | | | | | | |
|---|---|---|---|---|---|---|---|
| ENSG00000183199 | HSP90AB3P | 1213.432039 | -0.043902371 | 0.072018158 | -0.609601418 | 0.542125872 | 0.715968193 |
| ENSG00000183206 | POTEC | 8.247945568 | 3.42865678 | 0.889315316 | 3.855389328 | 0.000115546 | 0.000510323 |
| ENSG00000183207 | RUVBL2 | 3564.732748 | 0.106060382 | 0.052765698 | 2.010025194 | 0.044428522 | 0.108297769 |
| ENSG00000183208 | GDPGP1 | 59.31721871 | 0.203850753 | 0.228792437 | 0.890985538 | 0.372936929 | 0.565706086 |
| ENSG00000183230 | CTNNA3 | 37.97088971 | -0.660343539 | 0.293832808 | -2.24734448 | 0.02461802 | 0.065614444 |
| ENSG00000183239 | RPL29P16 | 2.42579765 | -0.699287743 | 1.142218631 | -0.612218821 | 0.540392991 | 0.714625884 |
| ENSG00000183248 | PRR36 | 1170.253745 | 0.070142791 | 0.074077249 | 0.946887094 | 0.343696315 | 0.534643892 |
| ENSG00000183250 | LINC01547 | 485.8817452 | 0.224258167 | 0.090374122 | 2.481442269 | 0.01308519 | 0.037658295 |
| ENSG00000183255 | PTTG1IP | 6666.218242 | 0.158422448 | 0.044948396 | 3.524540647 | 0.000424218 | 0.001700429 |
| ENSG00000183258 | DDX41 | 3472.173913 | -0.400681885 | 0.050157872 | -7.988414693 | 1.37E-15 | 1.71E-14 |
| ENSG00000183260 | ABHD16B | 5.865923005 | -0.40891906 | 0.743474977 | -0.550010522 | 0.582312157 | 0.746938094 |
| ENSG00000183281 | PLGLB1 | 14.76776782 | 0.025811442 | 0.452193834 | 0.057080483 | 0.954481083 | 0.979088978 |
| ENSG00000183283 | DAZAP2 | 8085.209034 | 0.068812501 | 0.047494047 | 1.448865813 | 0.14737506 | 0.28770527 |
| ENSG00000183291 | SELENOF | 1734.349274 | 0.013315892 | 0.058585407 | 0.227290246 | 0.820198058 | 0.904627311 |
| ENSG00000183298 | RPSAP19 | 71.16397938 | -0.143357533 | 0.225796383 | -0.634897384 | 0.525495343 | 0.701516712 |
| ENSG00000183307 | TMEM121B | 13.25031282 | -0.690604975 | 0.483626174 | -1.427972703 | 0.153299716 | 0.296217611 |
| ENSG00000183308 | ENSG0000018 | 14.88290994 | 0.275032086 | 0.453691141 | 0.606209955 | 0.544375343 | 0.717823209 |
| ENSG00000183309 | ZNF623 | 1957.64191 | -0.031485571 | 0.05817366 | -0.541234153 | 0.5883462 | 0.751651917 |
| ENSG00000183317 | EPHA10 | 371.74269 | 0.306157005 | 0.100318233 | 3.051858039 | 0.002274296 | 0.007908497 |
| ENSG00000183323 | CCDC125 | 992.8642956 | -0.765598823 | 0.072223823 | -10.60036408 | 2.97E-26 | 7.03E-25 |
| ENSG00000183337 | BCOR | 2560.059659 | -0.247567514 | 0.058704593 | -4.217174546 | 2.47E-05 | 0.00012113 |
| ENSG00000183340 | JRKL | 117.4714209 | 0.210975575 | 0.168641442 | 1.251030426 | 0.210923377 | 0.378368961 |
| ENSG00000183346 | CABCOCO1 | 1.578650181 | 1.862448741 | 1.548709583 | 1.202581014 | 0.229138498 | NA |
| ENSG00000183354 | KIAA2026 | 613.2761733 | -0.159394078 | 0.081950095 | -1.94501395 | 0.051773299 | 0.123433744 |
| ENSG00000183386 | FHL3 | 154.1571702 | 0.164261212 | 0.151947308 | 1.081040618 | 0.279679046 | 0.462969932 |
| ENSG00000183396 | TMEM89 | 2.603184537 | 0.164698736 | 1.070809706 | 0.153807661 | 0.877761393 | 0.936395015 |
| ENSG00000183397 | TEKTIP1 | 32.99893015 | -0.004753198 | 0.315461595 | -0.015067439 | 0.987978378 | 0.994030148 |
| ENSG00000183401 | CCDC159 | 566.6762733 | 0.220769203 | 0.096812226 | 2.280385567 | 0.022584831 | 0.060802256 |
| ENSG00000183421 | RIPK4 | 290.0881494 | 0.069729674 | 0.112532066 | 0.619642705 | 0.535493044 | 0.710974097 |
| ENSG00000183423 | LRIT3 | 3.468852153 | -0.039425683 | 0.954554151 | -0.04130272 | 0.967054565 | 0.985295635 |
| ENSG00000183426 | NPIPA1 | 189.6525168 | 0.027286019 | 0.143522089 | 0.190117212 | 0.849217283 | 0.921106747 |
| ENSG00000183431 | SF3A3 | 3578.561969 | -0.186129531 | 0.04801851 | -3.876203833 | 0.000106099 | 0.000471048 |
| ENSG00000183432 | ZBTB8OSP1 | 5.432343085 | 0.10260986 | 0.762841375 | 0.134510088 | 0.892999235 | 0.94527961 |

| | | | | | | | |
|---|---|---|---|---|---|---|---|
| ENSG00000183444 | OR7E38P | 5.396486152 | -0.370731216 | 0.768037003 | -0.482699681 | 0.62930899 | 0.7810577 |
| ENSG00000183458 | PKD1P3 | 166.6753714 | 0.1255848 | 0.149090261 | 0.842340733 | 0.399597257 | 0.592315423 |
| ENSG00000183474 | GTF2H2C | 209.0662085 | -0.532409316 | 0.132223777 | -4.026577728 | 5.66E-05 | 0.000262756 |
| ENSG00000183475 | ASB7 | 1027.800968 | 0.228285861 | 0.067928233 | 3.360691866 | 0.000777475 | 0.002970071 |
| ENSG00000183479 | TREX2 | 49.51986364 | -0.206188205 | 0.251020402 | -0.821400186 | 0.411418358 | 0.603757493 |
| ENSG00000183486 | MX2 | 3.65428884 | 2.508014126 | 1.120757368 | 2.237785088 | 0.025235073 | 0.067001456 |
| ENSG00000183495 | EP400 | 2789.819262 | -0.00685046 | 0.056064047 | -0.122189895 | 0.902748629 | 0.950907167 |
| ENSG00000183496 | MEX3B | 291.7670135 | -0.283566946 | 0.11481254 | -2.469825562 | 0.013517895 | 0.038752876 |
| ENSG00000183506 | PI4KAP2 | 206.4207145 | -0.767916645 | 0.131986755 | -5.818134138 | 5.95E-09 | 4.39E-08 |
| ENSG00000183508 | TENT5C | 108.0143611 | 1.115423584 | 0.18442837 | 6.04800435 | 1.47E-09 | 1.15E-08 |
| ENSG00000183513 | COA5 | 783.8406433 | 0.112448731 | 0.07455272 | 1.508311582 | 0.1314748 | 0.263777499 |
| ENSG00000183520 | UTP11 | 1166.73607 | -0.085469924 | 0.062911199 | -1.358580428 | 0.174279581 | 0.327112877 |
| ENSG00000183527 | PSMG1 | 1181.900893 | 0.47432347 | 0.067849379 | 6.990829903 | 2.73E-12 | 2.70E-11 |
| ENSG00000183530 | PRR14L | 2069.396044 | -0.607556998 | 0.064670415 | -9.394666804 | 5.74E-21 | 1.01E-19 |
| ENSG00000183566 | BPIFA4P | 1.980025553 | 2.308961056 | 1.482043708 | 1.557957463 | 0.119243332 | 0.244323154 |
| ENSG00000183569 | SERHL2 | 25.39193967 | -1.338650144 | 0.378947898 | -3.532544056 | 0.000411582 | 0.001652088 |
| ENSG00000183570 | PCBP3 | 78.24539236 | -0.864543431 | 0.203382604 | -4.250822905 | 2.13E-05 | 0.000105323 |
| ENSG00000183571 | PGPEP1L | 9.815271107 | 0.581324707 | 0.561735243 | 1.034873126 | 0.300728166 | 0.487482358 |
| ENSG00000183576 | SETD3 | 2450.599698 | -0.279143854 | 0.051961728 | -5.372104964 | 7.78E-08 | 5.09E-07 |
| ENSG00000183578 | TNFAIP8L3 | 153.5185589 | -1.542707887 | 0.159068396 | -9.698393412 | 3.06E-22 | 5.81E-21 |
| ENSG00000183579 | ZNRF3 | 909.3687974 | 1.274384719 | 0.071315225 | 17.86974276 | 2.03E-71 | 1.92E-69 |
| ENSG00000183580 | FBXL7 | 7.923991169 | -0.059109099 | 0.615546963 | -0.096026952 | 0.923499167 | 0.963176672 |
| ENSG00000183586 | HMGN3P1 | 13.73933623 | -0.678009521 | 0.4780658 | -1.418234732 | 0.156122245 | 0.300280397 |
| ENSG00000183597 | TANGO2 | 806.330209 | -0.653478491 | 0.077832588 | -8.395949613 | 4.62E-17 | 6.38E-16 |
| ENSG00000183598 | H3C13 | 19.95302185 | 1.020271488 | 0.403204559 | 2.530406629 | 0.01139304 | 0.033336681 |
| ENSG00000183604 | SMG1P5 | 33.53140034 | 0.143804289 | 0.305614041 | 0.470542155 | 0.637967724 | 0.786742264 |
| ENSG00000183605 | SFXN4 | 614.6572831 | -0.26280829 | 0.07953236 | -3.30441961 | 0.000951733 | 0.003577966 |
| ENSG00000183615 | FAM167B | 15.93615582 | 0.617937982 | 0.458799814 | 1.346857526 | 0.178026123 | 0.332431361 |
| ENSG00000183617 | MRPL54 | 555.8249981 | 0.0407334 | 0.09440585 | 0.431471138 | 0.666125834 | 0.807489253 |
| ENSG00000183621 | ZNF438 | 71.8414409 | -0.493308482 | 0.214713888 | -2.297515479 | 0.021589381 | 0.058451599 |
| ENSG00000183624 | HMCES | 3431.853087 | 0.028081351 | 0.048192311 | 0.582693605 | 0.560099579 | 0.729502234 |
| ENSG00000183625 | CCR3 | 9.806579743 | 0.252980463 | 0.564067927 | 0.448492905 | 0.653797505 | 0.799132287 |
| ENSG00000183628 | DGCR6 | 97.88253005 | -0.41876209 | 0.194869908 | -2.148931533 | 0.031639828 | 0.081353031 |

| | | | | | | | |
|---|---|---|---|---|---|---|---|
| ENSG00000183638 | RP1L1 | 2.625254945 | -0.28340329 | 1.146542735 | -0.247180747 | 0.804768345 | 0.896314309 |
| ENSG00000183647 | ZNF530 | 211.8495337 | 0.684725683 | 0.13098187 | 5.227637109 | 1.72E-07 | 1.09E-06 |
| ENSG00000183648 | NDUFB1 | 965.6485559 | -0.499977597 | 0.072772614 | -6.870408696 | 6.40E-12 | 6.10E-11 |
| ENSG00000183654 | MARCHF11 | 3.911552701 | 2.068913267 | 1.040042497 | 1.989258393 | 0.04667269 | 0.112904302 |
| ENSG00000183655 | KLHL25 | 492.5003431 | 0.365565088 | 0.097379215 | 3.754036081 | 0.00017401 | 0.000746407 |
| ENSG00000183665 | TRMT12 | 1024.611521 | -0.347912713 | 0.07230559 | -4.81169868 | 1.50E-06 | 8.46E-06 |
| ENSG00000183666 | GUSBP1 | 375.3855295 | -0.408644129 | 0.099393919 | -4.111359451 | 3.93E-05 | 0.000186583 |
| ENSG00000183668 | PSG9 | 14.98927928 | 1.02486202 | 0.479343999 | 2.138051216 | 0.032512587 | 0.083278108 |
| ENSG00000183682 | BMP8A | 84.1647991 | 0.837704306 | 0.203156929 | 4.123434578 | 3.73E-05 | 0.000177612 |
| ENSG00000183684 | ALYREF | 3654.225119 | -0.146172363 | 0.046906663 | -3.116238814 | 0.001831739 | 0.006489588 |
| ENSG00000183688 | RFLNB | 58.64606161 | 0.257269066 | 0.236040513 | 1.089936057 | 0.275741312 | 0.458019466 |
| ENSG00000183690 | EFHC2 | 6.856816767 | -0.184086549 | 0.688995426 | -0.26718109 | 0.789329735 | 0.886045416 |
| ENSG00000183691 | NOG | 3.748766851 | 1.769255817 | 0.997769121 | 1.773211638 | 0.076193644 | 0.170060537 |
| ENSG00000183696 | UPP1 | 61.31947492 | 0.494836571 | 0.228691921 | 2.163769362 | 0.030482055 | 0.07877192 |
| ENSG00000183718 | TRIM52 | 760.931708 | -0.151342312 | 0.07970314 | -1.898824976 | 0.057587492 | 0.1348738 |
| ENSG00000183722 | LHFPL6 | 104.6714289 | -0.34206758 | 0.177997061 | -1.921759716 | 0.054635999 | 0.12927796 |
| ENSG00000183723 | CMTM4 | 3970.422401 | 0.412522723 | 0.053032953 | 7.778611175 | 7.33E-15 | 8.69E-14 |
| ENSG00000183726 | TMEM50A | 1654.177737 | 0.062886268 | 0.060966305 | 1.031492203 | 0.302310063 | 0.489241117 |
| ENSG00000183734 | ASCL2 | 197.2992246 | -0.393785743 | 0.13726879 | -2.868720142 | 0.004121363 | 0.013539176 |
| ENSG00000183735 | TBK1 | 1505.341749 | 0.057152453 | 0.060678382 | 0.941891524 | 0.34624818 | 0.537083863 |
| ENSG00000183741 | CBX6 | 11331.89187 | -0.149840341 | 0.050501981 | -2.967019065 | 0.003007023 | 0.010186224 |
| ENSG00000183742 | MACC1 | 443.9988115 | -1.27890953 | 0.099589986 | -12.8417482 | 9.57E-38 | 3.58E-36 |
| ENSG00000183751 | TBL3 | 1428.834992 | -0.043431509 | 0.067435251 | -0.644047566 | 0.51954459 | 0.696979701 |
| ENSG00000183760 | ACP7 | 39.94551251 | 0.383469411 | 0.278667833 | 1.376080647 | 0.168796667 | 0.318977299 |
| ENSG00000183762 | KREMEN1 | 517.4237847 | 1.874660386 | 0.098510113 | 19.03013124 | 9.60E-81 | 1.10E-78 |
| ENSG00000183763 | TRAIP | 335.4480937 | -0.243750384 | 0.112612335 | -2.164508752 | 0.030425326 | 0.078643039 |
| ENSG00000183765 | CHEK2 | 607.0410585 | -0.397038442 | 0.081946014 | -4.845122063 | 1.27E-06 | 7.24E-06 |
| ENSG00000183773 | AIFM3 | 140.0611564 | -0.648051113 | 0.159736924 | -4.056990065 | 4.97E-05 | 0.00023272 |
| ENSG00000183778 | B3GALT5 | 1051.651646 | 0.552757354 | 0.068389653 | 8.082470536 | 6.35E-16 | 8.15E-15 |
| ENSG00000183779 | ZNF703 | 1871.305242 | 0.013715347 | 0.060420946 | 0.226996558 | 0.820426419 | 0.90474868 |
| ENSG00000183783 | KCTD8 | 11.49425001 | -2.682230392 | 0.637727261 | -4.205920861 | 2.60E-05 | 0.000127021 |
| ENSG00000183784 | DOCK8-AS1 | 20.43269878 | 0.945425876 | 0.398934563 | 2.369877077 | 0.017794 | 0.049423695 |
| ENSG00000183785 | TUBA8 | 2.931755836 | -0.181666546 | 0.998851792 | -0.181875377 | 0.855680529 | 0.925169031 |

| | | | | | | | |
|---|---|---|---|---|---|---|---|
| ENSG00000183793 | NPIPA5 | 4.058905202 | 0.644642336 | 0.908813146 | 0.709323295 | 0.478123875 | 0.662882047 |
| ENSG00000183798 | EMILIN3 | 10.34234478 | -0.029874224 | 0.569258522 | -0.052479187 | 0.958146879 | 0.980976892 |
| ENSG00000183808 | RBM12B | 1437.366953 | -0.313773779 | 0.06267377 | -5.006460903 | 5.54E-07 | 3.32E-06 |
| ENSG00000183813 | CCR4 | 3.314719915 | 0.515316725 | 0.990841444 | 0.520079906 | 0.603007883 | 0.762148806 |
| ENSG00000183814 | LIN9 | 324.8745366 | -0.104797399 | 0.103646647 | -1.01110265 | 0.311967302 | 0.500246743 |
| ENSG00000183826 | BTBD9 | 1021.869057 | 0.109898144 | 0.073741866 | 1.490308688 | 0.136143089 | 0.270792354 |
| ENSG00000183828 | NUDT14 | 789.702368 | 0.513826012 | 0.07694248 | 6.678053675 | 2.42E-11 | 2.19E-10 |
| ENSG00000183831 | ANKRD45 | 4.00735382 | -1.438034903 | 0.943819938 | -1.523632681 | 0.127600494 | 0.257581428 |
| ENSG00000183840 | GPR39 | 157.9481899 | -0.273242888 | 0.147613552 | -1.851069124 | 0.064159609 | 0.147477098 |
| ENSG00000183844 | FAM3B | 29.79250653 | -1.094775039 | 0.341910961 | -3.201930225 | 0.001365101 | 0.004976229 |
| ENSG00000183850 | ZNF730 | 40.03043012 | 0.152740673 | 0.278152412 | 0.549125826 | 0.582919103 | 0.747298772 |
| ENSG00000183856 | IQGAP3 | 2825.262219 | -0.181087949 | 0.055135553 | -3.284413408 | 0.001021949 | 0.003818783 |
| ENSG00000183862 | CNGA2 | 23.60789388 | 6.937788238 | 1.21556116 | 5.707477719 | 1.15E-08 | 8.18E-08 |
| ENSG00000183864 | TOB2 | 4394.650459 | -0.436065356 | 0.0534532 | -8.157890502 | 3.41E-16 | 4.46E-15 |
| ENSG00000183873 | SCN5A | 9.23677778 | -2.145328784 | 0.649409875 | -3.303505025 | 0.000954843 | 0.003586718 |
| ENSG00000183888 | SRARP | 19.17778922 | -2.387883863 | 0.477875394 | -4.996875533 | 5.83E-07 | 3.47E-06 |
| ENSG00000183889 | NPIPA9 | 23.6829844 | -0.175855776 | 0.379135152 | -0.463834003 | 0.64276668 | 0.790526103 |
| ENSG00000183891 | TTC32 | 88.64120342 | 0.426968045 | 0.192601543 | 2.216846441 | 0.026633587 | 0.070146324 |
| ENSG00000183914 | DNAH2 | 7.940128994 | -0.0573946 | 0.630623625 | -0.091012448 | 0.9274827 | 0.965137362 |
| ENSG00000183929 | DUSP5P1 | 2.426348335 | -0.381149665 | 1.155236198 | -0.329932239 | 0.741451163 | 0.857515143 |
| ENSG00000183935 | HTR7P1 | 534.9480029 | -0.701059331 | 0.08771918 | -7.992087101 | 1.33E-15 | 1.67E-14 |
| ENSG00000183943 | PRKX | 1205.474297 | 0.006520189 | 0.07154634 | 0.091132386 | 0.927387399 | 0.965137362 |
| ENSG00000183955 | KMT5A | 1801.486211 | -0.129585306 | 0.055965404 | -2.315453788 | 0.020588111 | 0.056051657 |
| ENSG00000183960 | KCNH8 | 31.30484739 | 0.993735443 | 0.323517879 | 3.071655405 | 0.002128753 | 0.007436211 |
| ENSG00000183963 | SMTN | 2678.226224 | -0.986093794 | 0.061069464 | -16.1470846 | 1.19E-58 | 8.18E-57 |
| ENSG00000183971 | NPW | 7.18984672 | -0.013456896 | 0.653393915 | -0.020595381 | 0.983568425 | 0.9923499 |
| ENSG00000183977 | PP2D1 | 6.155712476 | -0.543583773 | 0.724871961 | -0.749903158 | 0.453313032 | 0.641003573 |
| ENSG00000183978 | COA3 | 824.7046075 | -0.209375626 | 0.072975961 | -2.869104044 | 0.004116364 | 0.013527443 |
| ENSG00000183979 | NPB | 23.8959729 | 0.175311156 | 0.368116074 | 0.476238797 | 0.633904263 | 0.784236938 |
| ENSG00000184007 | PTP4A2 | 8297.258476 | -0.355701216 | 0.051469864 | -6.910863752 | 4.82E-12 | 4.65E-11 |
| ENSG00000184009 | ACTG1 | 48053.84761 | -0.101079122 | 0.051615376 | -1.958314173 | 0.050193159 | 0.120241339 |
| ENSG00000184012 | TMPRSS2 | 370.7873356 | 0.860551103 | 0.106148521 | 8.107047522 | 5.19E-16 | 6.71E-15 |
| ENSG00000184014 | DENND5A | 1367.316679 | -0.184812204 | 0.060689587 | -3.045204529 | 0.002325221 | 0.008069679 |

| | | | | | | | |
|---|---|---|---|---|---|---|---|
| ENSG00000184047 | DIABLO | 57.58399483 | 0.017488042 | 0.234641516 | 0.074530892 | 0.940587961 | 0.971722037 |
| ENSG00000184056 | VPS33B | 113.2403018 | 0.182344845 | 0.17099513 | 1.066374492 | 0.286254388 | 0.470021728 |
| ENSG00000184058 | TBX1 | 3.308991198 | -1.061942627 | 0.974210331 | -1.090054779 | 0.275689015 | 0.457984358 |
| ENSG00000184060 | ADAP2 | 37.92489482 | -0.406416508 | 0.290996585 | -1.396636691 | 0.162522851 | 0.309734502 |
| ENSG00000184068 | SREBF2-AS1 | 11.84987397 | -1.054889519 | 0.528856486 | -1.99466121 | 0.046079843 | 0.111682155 |
| ENSG00000184076 | UQCR10 | 1653.452307 | -0.770993195 | 0.058729596 | -13.1278478 | 2.28E-39 | 9.00E-38 |
| ENSG00000184083 | FAM120C | 1151.881169 | 0.352717303 | 0.066359947 | 5.315213762 | 1.07E-07 | 6.89E-07 |
| ENSG00000184100 | BRD7P2 | 382.4271188 | 0.161622576 | 0.097670973 | 1.6547657 | 0.097972034 | 0.20852463 |
| ENSG00000184110 | EIF3C | 633.729391 | 0.216297032 | 0.084864129 | 2.548745092 | 0.010811129 | 0.031815823 |
| ENSG00000184117 | NIPSNAP1 | 4050.382263 | -0.506307503 | 0.050985052 | -9.930508746 | 3.07E-23 | 6.18E-22 |
| ENSG00000184154 | LRRC51 | 95.03837202 | -0.497719463 | 0.194314412 | -2.561413011 | 0.010424734 | 0.030820753 |
| ENSG00000184160 | ADRA2C | 860.2883894 | -0.044167265 | 0.081811395 | -0.539866906 | 0.589288822 | 0.752395257 |
| ENSG00000184162 | NR2C2AP | 344.5127062 | -0.136385245 | 0.10254396 | -1.330017347 | 0.183512556 | 0.339717401 |
| ENSG00000184163 | C1QTNF12 | 12.17010793 | -0.454670657 | 0.50348614 | -0.903045031 | 0.366501996 | 0.559103204 |
| ENSG00000184164 | CRELD2 | 930.5016522 | -0.261773286 | 0.07212115 | -3.629632722 | 0.000283825 | 0.001172005 |
| ENSG00000184178 | SCFD2 | 463.8135133 | 0.026847802 | 0.089208747 | 0.300954812 | 0.763448953 | 0.870366712 |
| ENSG00000184182 | UBE2F | 140.2883787 | 0.231794639 | 0.154601916 | 1.499299915 | 0.133795845 | 0.267242201 |
| ENSG00000184188 | ENSG0000018 | 9.378712108 | 0.149118925 | 0.587868107 | 0.253660512 | 0.799757844 | 0.893253273 |
| ENSG00000184194 | GPR173 | 165.9109409 | 0.933094643 | 0.146643868 | 6.362998043 | 1.98E-10 | 1.67E-09 |
| ENSG00000184203 | PPP1R2 | 1062.14157 | -0.441798909 | 0.078089393 | -5.657604624 | 1.54E-08 | 1.08E-07 |
| ENSG00000184205 | TSPYL2 | 3396.971464 | -0.898620671 | 0.068790545 | -13.06314221 | 5.35E-39 | 2.08E-37 |
| ENSG00000184206 | GOLGA6L4 | 3.301897861 | -1.376663033 | 0.994560287 | -1.384192644 | 0.166299475 | 0.315438231 |
| ENSG00000184207 | PGP | 2070.549678 | -0.25813289 | 0.062414756 | -4.13576704 | 3.54E-05 | 0.000168967 |
| ENSG00000184208 | C22orf46P | 1523.479871 | -0.368510656 | 0.057604699 | -6.39723255 | 1.58E-10 | 1.35E-09 |
| ENSG00000184209 | SNRNP35 | 694.5872237 | 0.018520636 | 0.081690136 | 0.226718144 | 0.820642918 | 0.904900429 |
| ENSG00000184216 | IRAK1 | 9965.045001 | -0.239255908 | 0.045586418 | -5.248403387 | 1.53E-07 | 9.77E-07 |
| ENSG00000184220 | CMSS1 | 880.829398 | 0.185749819 | 0.070324637 | 2.641319284 | 0.008258385 | 0.025128977 |
| ENSG00000184226 | PCDH9 | 643.2762997 | -1.418535551 | 0.086890881 | -16.32548246 | 6.50E-60 | 4.63E-58 |
| ENSG00000184227 | ACOT1 | 65.81322717 | -1.332675335 | 0.239828471 | -5.556785359 | 2.75E-08 | 1.88E-07 |
| ENSG00000184232 | OAF | 169.0562286 | 0.475098673 | 0.140621589 | 3.378561411 | 0.000728662 | 0.002798978 |
| ENSG00000184254 | ALDH1A3 | 49.50078122 | 0.508910612 | 0.2638955 | 1.928455058 | 0.053798549 | 0.127678328 |
| ENSG00000184260 | H2AC20 | 61.53256768 | -0.243106719 | 0.253949843 | -0.957302103 | 0.338414794 | 0.529660598 |
| ENSG00000184261 | KCNK12 | 25.11094267 | 0.264601635 | 0.357771744 | 0.739582259 | 0.45955351 | 0.6464805 |

| | | | | | | | |
|---|---|---|---|---|---|---|---|
| ENSG00000184270 | H2AC21 | 12.54384259 | 0.927377407 | 0.519606809 | 1.784767619 | 0.074299018 | 0.166642498 |
| ENSG00000184271 | POU6F1 | 8.439484975 | 1.076041562 | 0.646660623 | 1.663997348 | 0.096112959 | 0.205245836 |
| ENSG00000184277 | TM2D3 | 865.49385 | 0.253841191 | 0.07000787 | 3.625895096 | 0.000287962 | 0.00118789 |
| ENSG00000184281 | TSSC4 | 691.0054469 | 0.020151906 | 0.087921656 | 0.229202983 | 0.818711155 | 0.903986346 |
| ENSG00000184292 | TACSTD2 | 5240.657376 | 0.717296077 | 0.049040072 | 14.62673366 | 1.90E-48 | 9.96E-47 |
| ENSG00000184304 | PRKD1 | 342.4223094 | -0.469350076 | 0.106125957 | -4.422575697 | 9.75E-06 | 5.05E-05 |
| ENSG00000184305 | CCSER1 | 358.9346291 | 0.025767275 | 0.100347095 | 0.256781479 | 0.797347468 | 0.89183367 |
| ENSG00000184307 | ZDHHC23 | 1195.536577 | -0.110975591 | 0.064678133 | -1.715813153 | 0.086196246 | 0.187728272 |
| ENSG00000184319 | RPL23AP82 | 141.6841371 | -0.611201963 | 0.154032532 | -3.968005678 | 7.25E-05 | 0.000330934 |
| ENSG00000184343 | SRPK3 | 13.49735521 | -0.229452441 | 0.472257068 | -0.485863433 | 0.627063996 | 0.780131681 |
| ENSG00000184349 | EFNA5 | 428.8061333 | 0.541832609 | 0.093007728 | 5.825672973 | 5.69E-09 | 4.21E-08 |
| ENSG00000184357 | H1-5 | 8.356074861 | 0.62949538 | 0.610115491 | 1.031764297 | 0.302182548 | 0.489138277 |
| ENSG00000184363 | PKP3 | 2336.761236 | 0.204814687 | 0.062405958 | 3.281973309 | 0.001030834 | 0.003848848 |
| ENSG00000184368 | MAP7D2 | 4.952514995 | 0.0120993 | 0.801916589 | 0.015087978 | 0.987961992 | 0.994030148 |
| ENSG00000184371 | CSF1 | 41.142658 | -0.482031917 | 0.283545771 | -1.700014481 | 0.089128202 | 0.192923282 |
| ENSG00000184374 | COLEC10 | 39.06418362 | -2.953548748 | 0.362644224 | -8.14448033 | 3.81E-16 | 4.97E-15 |
| ENSG00000184378 | ACTRT3 | 118.4620691 | 0.073284101 | 0.166837466 | 0.439254461 | 0.660477167 | 0.803707149 |
| ENSG00000184381 | PLA2G6 | 492.3552116 | -0.343002827 | 0.098000297 | -3.500018245 | 0.000465226 | 0.001852507 |
| ENSG00000184384 | MAML2 | 49.91889041 | 1.688686366 | 0.272578745 | 6.19522393 | 5.82E-10 | 4.74E-09 |
| ENSG00000184402 | SS18L1 | 1162.86471 | -0.104366174 | 0.069456876 | -1.50260393 | 0.132941209 | 0.266091209 |
| ENSG00000184408 | KCND2 | 8.069266011 | 1.635488614 | 0.670605796 | 2.438822662 | 0.014735197 | 0.041824478 |
| ENSG00000184428 | TOP1MT | 1436.933008 | 0.775833644 | 0.060739199 | 12.77319523 | 2.31E-37 | 8.56E-36 |
| ENSG00000184432 | COPB2 | 8716.723789 | -0.114487617 | 0.04427491 | -2.585835102 | 0.009714339 | 0.028990106 |
| ENSG00000184436 | THAP7 | 683.3283631 | -0.675011239 | 0.08690119 | -7.767571852 | 8.00E-15 | 9.46E-14 |
| ENSG00000184441 | ENSG0000018 | 493.1105094 | 0.621442757 | 0.095351041 | 6.51741971 | 7.15E-11 | 6.26E-10 |
| ENSG00000184445 | KNTC1 | 1714.771956 | -0.762339017 | 0.074701766 | -10.20510035 | 1.88E-24 | 4.07E-23 |
| ENSG00000184451 | CCR10 | 2.421139133 | -0.32489314 | 1.165695443 | -0.278711855 | 0.780465964 | 0.880880158 |
| ENSG00000184454 | NCMAP | 43.3700846 | -3.626815168 | 0.385125647 | -9.417225777 | 4.63E-21 | 8.24E-20 |
| ENSG00000184465 | WDR27 | 758.7522899 | 0.108865806 | 0.077109004 | 1.411842967 | 0.157996204 | 0.302919592 |
| ENSG00000184470 | TXNRD2 | 809.5825531 | -1.004439499 | 0.078436818 | -12.8057145 | 1.52E-37 | 5.67E-36 |
| ENSG00000184481 | FOXO4 | 290.5587241 | -0.090168862 | 0.118588522 | -0.760350667 | 0.447045004 | 0.635548611 |
| ENSG00000184486 | POU3F2 | 141.0911792 | 1.875454176 | 0.176798484 | 10.60786345 | 2.74E-26 | 6.49E-25 |
| ENSG00000184489 | PTP4A3 | 367.6440443 | 0.003283286 | 0.100164578 | 0.032778909 | 0.973850897 | 0.988106737 |

| | | | | | | | |
|---|---|---|---|---|---|---|---|
| ENSG00000184492 | FOXD4L1 | 11.53063137 | 0.524377261 | 0.545792561 | 0.960762932 | 0.33667138 | 0.527888882 |
| ENSG00000184497 | TMEM255B | 3.49433009 | -5.358205176 | 1.413630134 | -3.790386924 | 0.000150413 | 0.000652016 |
| ENSG00000184500 | PROS1 | 406.074198 | 0.193351156 | 0.10096587 | 1.915015008 | 0.05549059 | 0.130975834 |
| ENSG00000184508 | HDDC3 | 850.0349146 | 0.049036997 | 0.071207657 | 0.688647802 | 0.491044931 | 0.674019811 |
| ENSG00000184515 | BEX5 | 26.64169988 | -0.87949744 | 0.363631015 | -2.418653538 | 0.01557807 | 0.043982882 |
| ENSG00000184517 | ZFP1 | 1109.971616 | 0.224684875 | 0.063997485 | 3.510839144 | 0.000446695 | 0.001783668 |
| ENSG00000184523 | PTGER4P2 | 5.113688324 | 0.54532541 | 0.788333478 | 0.69174458 | 0.489097746 | 0.672373841 |
| ENSG00000184530 | C6orf58 | 6.524446028 | 0.323121154 | 0.707147055 | 0.456936294 | 0.647716836 | 0.794451384 |
| ENSG00000184545 | DUSP8 | 584.4355917 | 0.688512356 | 0.088089073 | 7.816092603 | 5.45E-15 | 6.55E-14 |
| ENSG00000184557 | SOCS3 | 1967.210823 | -0.655350001 | 0.057867576 | -11.32499479 | 9.87E-30 | 2.72E-28 |
| ENSG00000184560 | SPEM2 | 2.302001376 | 1.841465998 | 1.279732891 | 1.438945589 | 0.150165939 | 0.29183384 |
| ENSG00000184564 | SLITRK6 | 806.3583595 | 1.595122268 | 0.083183349 | 19.17598037 | 5.88E-82 | 6.77E-80 |
| ENSG00000184574 | LPAR5 | 5.274106221 | 2.656355743 | 0.954777828 | 2.782171584 | 0.005399648 | 0.017236506 |
| ENSG00000184575 | XPOT | 9432.170044 | 0.061967889 | 0.061532836 | 1.007070263 | 0.31390101 | 0.502154057 |
| ENSG00000184584 | STING1 | 66.79534985 | 2.205426819 | 0.250825685 | 8.792667367 | 1.46E-18 | 2.23E-17 |
| ENSG00000184588 | PDE4B | 159.2952871 | -2.319915308 | 0.166972983 | -13.89395621 | 6.89E-44 | 3.21E-42 |
| ENSG00000184599 | TAFA3 | 2.129974529 | 0.933482451 | 1.219582539 | 0.765411459 | 0.444026571 | 0.632807922 |
| ENSG00000184602 | SNN | 1032.540829 | 0.143042082 | 0.070984806 | 2.015108447 | 0.043893287 | 0.107209541 |
| ENSG00000184612 | RPL7P26 | 38.83031995 | 0.503712708 | 0.29973433 | 1.680530585 | 0.092854129 | 0.199538521 |
| ENSG00000184613 | NELL2 | 22.49168735 | -0.671029598 | 0.397304839 | -1.688954001 | 0.091228245 | 0.196634027 |
| ENSG00000184619 | KRBA2 | 12.89924803 | -0.684777389 | 0.508343572 | -1.347075928 | 0.17795578 | 0.332327059 |
| ENSG00000184634 | MED12 | 2848.415087 | 0.07365979 | 0.057015576 | 1.29192399 | 0.196383469 | 0.358025652 |
| ENSG00000184635 | ZNF93 | 20.51973889 | 0.654425987 | 0.403621354 | 1.621385933 | 0.104934896 | 0.220359438 |
| ENSG00000184640 | SEPTIN9 | 10651.77309 | -0.410914105 | 0.0478224 | -8.592502806 | 8.51E-18 | 1.24E-16 |
| ENSG00000184661 | CDCA2 | 574.3909527 | -0.36058738 | 0.082272968 | -4.382817192 | 1.17E-05 | 5.98E-05 |
| ENSG00000184669 | OR7E14P | 251.8791828 | -0.163438561 | 0.121198743 | -1.348516971 | 0.177492165 | 0.331650263 |
| ENSG00000184672 | RALYL | 12.72981163 | -1.226801742 | 0.526628834 | -2.329537739 | 0.019830596 | 0.05431159 |
| ENSG00000184675 | AMER1 | 444.8942019 | 0.224053798 | 0.096985471 | 2.310178993 | 0.020878247 | 0.056766862 |
| ENSG00000184677 | ZBTB40 | 1312.630964 | -0.107050068 | 0.064561714 | -1.658104498 | 0.097296376 | 0.207355897 |
| ENSG00000184678 | H2BC21 | 4640.503916 | 0.918856548 | 0.04790633 | 19.18027416 | 5.41E-82 | 6.27E-80 |
| ENSG00000184697 | CLDN6 | 12.61550337 | -0.482247503 | 0.500975397 | -0.962617138 | 0.335739694 | 0.526875672 |
| ENSG00000184702 | SEPTIN5 | 6.842886409 | 0.57222616 | 0.688569182 | 0.831036553 | 0.40595298 | 0.598679911 |
| ENSG00000184708 | EIF4ENIF1 | 1292.173349 | -0.649872757 | 0.066739883 | -9.737397361 | 2.09E-22 | 4.00E-21 |

| | | | | | | | |
|---|---|---|---|---|---|---|---|
| ENSG00000184709 | LRRC26 | 8.189315971 | 0.655770228 | 0.649033309 | 1.010379928 | 0.3123133 | 0.500521665 |
| ENSG00000184716 | SERINC4 | 12.66457872 | -0.717851627 | 0.491798939 | -1.45964452 | 0.144387797 | 0.283010051 |
| ENSG00000184719 | RNLS | 134.2980837 | -0.074188664 | 0.158079308 | -0.469312937 | 0.638845969 | 0.787290808 |
| ENSG00000184730 | APOBR | 36.58935715 | -1.15312256 | 0.316460341 | -3.643813805 | 0.000268628 | 0.001113136 |
| ENSG00000184731 | FAM110C | 486.0023314 | 0.496624505 | 0.088183069 | 5.631744374 | 1.78E-08 | 1.25E-07 |
| ENSG00000184743 | ATL3 | 2062.700902 | -0.019741052 | 0.055299924 | -0.356981547 | 0.72110562 | 0.845162383 |
| ENSG00000184752 | NDUFA12 | 1017.76069 | -0.098785633 | 0.069610709 | -1.419115444 | 0.155865362 | 0.299912046 |
| ENSG00000184774 | MGAT4EP | 9.322186917 | 1.039039714 | 0.59837547 | 1.73643434 | 0.082487065 | 0.181160968 |
| ENSG00000184786 | DYNLT2 | 82.86640845 | 0.547757991 | 0.206448281 | 2.653245595 | 0.007972184 | 0.024377753 |
| ENSG00000184787 | UBE2G2 | 3536.239835 | 0.515004003 | 0.070456328 | 7.309549278 | 2.68E-13 | 2.85E-12 |
| ENSG00000184788 | SATL1 | 7.119326953 | 0.008445442 | 0.670543456 | 0.012594922 | 0.989950972 | 0.994622626 |
| ENSG00000184792 | OSBP2 | 346.0567715 | -0.100672906 | 0.106921644 | -0.94155778 | 0.346419094 | 0.537239623 |
| ENSG00000184809 | B3GALT5-AS1 | 320.6091472 | 1.104073843 | 0.109522017 | 10.08083917 | 6.72E-24 | 1.40E-22 |
| ENSG00000184828 | ZBTB7C | 9.968558323 | 3.351839634 | 0.800200979 | 4.188747227 | 2.80E-05 | 0.000136414 |
| ENSG00000184831 | APOO | 347.3390325 | 0.138675884 | 0.100682902 | 1.377352876 | 0.168403175 | 0.318506788 |
| ENSG00000184838 | PRR16 | 14.79210583 | -1.752025889 | 0.529497739 | -3.308844894 | 0.000936817 | 0.003525236 |
| ENSG00000184840 | TMED9 | 4253.274568 | -0.340245688 | 0.050253407 | -6.770599375 | 1.28E-11 | 1.19E-10 |
| ENSG00000184844 | CYCSP45 | 5.97674464 | -1.31098008 | 0.775038206 | -1.691503812 | 0.09074062 | 0.195785349 |
| ENSG00000184845 | DRD1 | 237.9645502 | -3.697404787 | 0.171015167 | -21.62033253 | 1.16E-103 | 2.05E-101 |
| ENSG00000184857 | TMEM186 | 387.2596532 | 0.099566167 | 0.097642848 | 1.019697483 | 0.307871956 | 0.495863418 |
| ENSG00000184860 | SDR42E1 | 644.7589459 | -0.100218011 | 0.07801529 | -1.284594483 | 0.198934027 | 0.361669566 |
| ENSG00000184863 | RBM33 | 1697.929763 | 0.150207453 | 0.057247981 | 2.623803505 | 0.008695393 | 0.026301731 |
| ENSG00000184867 | ARMCX2 | 181.2693934 | 0.796482821 | 0.141032782 | 5.647501317 | 1.63E-08 | 1.14E-07 |
| ENSG00000184887 | BTBD6 | 2994.389494 | -0.248216354 | 0.052878006 | -4.694132242 | 2.68E-06 | 1.48E-05 |
| ENSG00000184897 | H1-10 | 3721.786222 | 0.13078576 | 0.05721576 | 2.285834532 | 0.022263941 | 0.060065247 |
| ENSG00000184898 | RBM43 | 127.6210059 | 0.365684375 | 0.160518394 | 2.278146238 | 0.022717866 | 0.061076742 |
| ENSG00000184900 | SUMO3 | 6996.287045 | 0.163169772 | 0.044100238 | 3.699974897 | 0.000215621 | 0.000908916 |
| ENSG00000184903 | IMMP2L | 94.54426257 | 0.381401807 | 0.184483402 | 2.067404458 | 0.038696059 | 0.09649827 |
| ENSG00000184905 | TCEAL2 | 3.591333254 | -0.187514041 | 0.921114954 | -0.203572898 | 0.838687274 | 0.915120151 |
| ENSG00000184916 | JAG2 | 1632.881574 | 0.224292532 | 0.058159628 | 3.856498753 | 0.000115023 | 0.000508111 |
| ENSG00000184922 | FMNL1 | 182.457465 | -0.354431246 | 0.135225541 | -2.621037744 | 0.008766256 | 0.026495112 |
| ENSG00000184923 | NUTM2A | 6.937191299 | 0.039751556 | 0.659190745 | 0.060303572 | 0.951913857 | 0.977646532 |
| ENSG00000184924 | PTRHD1 | 495.0178382 | 0.745052408 | 0.095040417 | 7.839321731 | 4.53E-15 | 5.47E-14 |

| | | | | | | | |
|---|---|---|---|---|---|---|---|
| ENSG00000184925 | LCN12 | 48.1748397 | 0.051262209 | 0.25242085 | 0.203082308 | 0.8390707 | 0.915286602 |
| ENSG00000184939 | ZFP90 | 1666.517035 | -0.281214341 | 0.06065682 | -4.636153723 | 3.55E-06 | 1.93E-05 |
| ENSG00000184949 | FAM227A | 90.14371363 | 0.398711477 | 0.199757452 | 1.995977985 | 0.045936319 | 0.111381372 |
| ENSG00000184967 | NOC4L | 576.0556038 | 0.117347208 | 0.083936202 | 1.398052393 | 0.162097334 | 0.309206124 |
| ENSG00000184979 | USP18 | 104.087435 | -0.601342923 | 0.180033815 | -3.340166535 | 0.000837282 | 0.003180007 |
| ENSG00000184983 | NDUFA6 | 1078.095823 | -0.160940636 | 0.065438815 | -2.459406319 | 0.013916701 | 0.03979174 |
| ENSG00000184985 | SORCS2 | 5.285849985 | 1.473968472 | 0.809314396 | 1.821255718 | 0.068567994 | 0.15596924 |
| ENSG00000184986 | TMEM121 | 68.73746683 | -0.93722079 | 0.221421588 | -4.232743508 | 2.31E-05 | 0.000113597 |
| ENSG00000184990 | SIVA1 | 2024.585126 | -0.352824224 | 0.061762014 | -5.712641189 | 1.11E-08 | 7.95E-08 |
| ENSG00000184992 | BRI3BP | 4633.72913 | -0.032187029 | 0.048745125 | -0.660312789 | 0.509053125 | 0.689192557 |
| ENSG00000185000 | DGAT1 | 733.8924054 | 0.337378505 | 0.08081497 | 4.174703089 | 2.98E-05 | 0.000144617 |
| ENSG00000185008 | ROBO2 | 463.0529726 | 2.145526108 | 0.104107024 | 20.60885063 | 2.29E-94 | 3.32E-92 |
| ENSG00000185009 | AP3M1 | 1688.419556 | -0.245607468 | 0.058666025 | -4.186536714 | 2.83E-05 | 0.000137661 |
| ENSG00000185010 | F8 | 133.0072729 | -0.711199191 | 0.158200415 | -4.495558319 | 6.94E-06 | 3.65E-05 |
| ENSG00000185015 | CA13 | 30.5233735 | 0.258157417 | 0.317227377 | 0.813792995 | 0.415763549 | 0.607839896 |
| ENSG00000185019 | UBOX5 | 343.3664995 | 0.158327853 | 0.106135916 | 1.491746233 | 0.135765684 | 0.27034877 |
| ENSG00000185022 | MAFF | 80.95175698 | -0.181673401 | 0.202406831 | -0.897565563 | 0.369417205 | 0.562039441 |
| ENSG00000185024 | BRF1 | 3203.155227 | -0.164396092 | 0.05629319 | -2.920354864 | 0.00349633 | 0.011681592 |
| ENSG00000185033 | SEMA4B | 2357.169056 | 0.6907052 | 0.063958625 | 10.79925037 | 3.47E-27 | 8.67E-26 |
| ENSG00000185038 | MROH2A | 7.728657834 | 1.150891895 | 0.684747702 | 1.680753206 | 0.092810861 | 0.199482916 |
| ENSG00000185043 | CIB1 | 2397.319126 | 0.1226168 | 0.062761417 | 1.953697127 | 0.050737046 | 0.121366824 |
| ENSG00000185046 | ANKS1B | 153.9577051 | 0.86364858 | 0.155744986 | 5.545273741 | 2.93E-08 | 2.01E-07 |
| ENSG00000185049 | NELFA | 1160.564194 | -0.023384428 | 0.064873944 | -0.360459485 | 0.718503549 | 0.843432038 |
| ENSG00000185052 | SLC24A3 | 1924.499724 | -0.177660575 | 0.056548981 | -3.14171134 | 0.001679635 | 0.005994218 |
| ENSG00000185053 | SGCZ | 6.628120318 | -0.047874605 | 0.686090102 | -0.069778888 | 0.944369651 | 0.973580639 |
| ENSG00000185055 | EFCAB10 | 6.473075855 | 0.018951494 | 0.702322554 | 0.026984031 | 0.97847247 | 0.990210919 |
| ENSG00000185065 | UFD1-AS1 | 58.66111752 | -0.545464078 | 0.236576312 | -2.305658046 | 0.021129747 | 0.057309104 |
| ENSG00000185085 | INTS5 | 1316.561806 | -0.058175155 | 0.071865911 | -0.80949583 | 0.41823 | 0.610279003 |
| ENSG00000185088 | RPS27L | 992.3171257 | 0.097513469 | 0.068272125 | 1.428305744 | 0.153203875 | 0.296107246 |
| ENSG00000185090 | MANEAL | 1960.649145 | 0.089042735 | 0.055490398 | 1.604651212 | 0.108570584 | 0.226432911 |
| ENSG00000185100 | ADSS1 | 273.1918665 | 0.150441368 | 0.113281675 | 1.328029156 | 0.184168475 | 0.340711679 |
| ENSG00000185101 | ANO9 | 959.4989273 | 0.075578677 | 0.074502371 | 1.014446594 | 0.310369701 | 0.498346814 |
| ENSG00000185104 | FAF1 | 1998.680193 | -0.167557573 | 0.054519717 | -3.073338999 | 0.002116779 | 0.007401144 |

| | | | | | | | |
|---|---|---|---|---|---|---|---|
| ENSG00000185112 | FAM43A | 55.72600549 | -0.131399067 | 0.239776289 | -0.548006925 | 0.583687148 | 0.748011283 |
| ENSG00000185115 | NSMCE3 | 204.6010891 | -0.134733364 | 0.13033801 | -1.033722735 | 0.301265802 | 0.488132583 |
| ENSG00000185122 | HSF1 | 3576.803457 | 0.051961084 | 0.055144208 | 0.94227636 | 0.346051168 | 0.5369335 |
| ENSG00000185127 | C6orf120 | 1343.624081 | 0.274511411 | 0.063904822 | 4.295629075 | 1.74E-05 | 8.71E-05 |
| ENSG00000185129 | PURA | 1056.334022 | -0.018995371 | 0.076138356 | -0.249484914 | 0.802985709 | 0.895135709 |
| ENSG00000185130 | H2BC13 | 16.05096683 | -0.251314488 | 0.442231187 | -0.568287571 | 0.569839717 | 0.737080657 |
| ENSG00000185133 | INPP5J | 1516.742926 | -0.624644374 | 0.072283134 | -8.641633859 | 5.54E-18 | 8.16E-17 |
| ENSG00000185158 | LRRC37B | 188.9558404 | -0.157947371 | 0.134067369 | -1.178119418 | 0.238749002 | 0.414320455 |
| ENSG00000185163 | DDX51 | 1318.565188 | -0.069387259 | 0.069460234 | -0.99894939 | 0.317819209 | 0.506972578 |
| ENSG00000185164 | NOMO2 | 796.4695847 | 0.195828487 | 0.073800353 | 2.653489842 | 0.007966417 | 0.024363365 |
| ENSG00000185168 | LINC00482 | 35.54639197 | 1.463538101 | 0.324696357 | 4.507405362 | 6.56E-06 | 3.46E-05 |
| ENSG00000185187 | SIGIRR | 1663.918719 | 0.009227565 | 0.063107263 | 0.146220336 | 0.883747452 | 0.93974721 |
| ENSG00000185189 | NRBP2 | 1025.242793 | 0.428441916 | 0.081605173 | 5.250180867 | 1.52E-07 | 9.70E-07 |
| ENSG00000185198 | PRSS57 | 10.01898982 | 0.482256261 | 0.571941387 | 0.843191753 | 0.39912121 | 0.591930681 |
| ENSG00000185201 | IFITM2 | 1230.376861 | -0.410860995 | 0.074916482 | -5.484253716 | 4.15E-08 | 2.80E-07 |
| ENSG00000185203 | WASIR1 | 2.062794324 | -1.808533116 | 1.31749158 | -1.37270943 | 0.169842697 | 0.320412952 |
| ENSG00000185203 | WASIR1 | 2.062794324 | -1.808533116 | 1.31749158 | -1.37270943 | 0.169842697 | 0.320412952 |
| ENSG00000185215 | TNFAIP2 | 55.61178596 | 0.419259815 | 0.240083846 | 1.746305808 | 0.080757827 | 0.17804753 |
| ENSG00000185219 | ZNF445 | 734.361375 | 0.072662701 | 0.079552722 | 0.913390504 | 0.361037198 | 0.553548545 |
| ENSG00000185220 | PGBD2 | 395.1675191 | -0.246382196 | 0.102076903 | -2.413691916 | 0.015791809 | 0.04448233 |
| ENSG00000185236 | RAB11B | 1579.803343 | -0.243284295 | 0.066562863 | -3.654955361 | 0.000257227 | 0.001069176 |
| ENSG00000185238 | PRMT3 | 700.4678255 | 0.083359237 | 0.078005929 | 1.06862694 | 0.285237804 | 0.469158617 |
| ENSG00000185245 | GP1BA | 1.84110464 | -3.306509741 | 1.603389423 | -2.062200045 | 0.039188694 | 0.097527866 |
| ENSG00000185246 | PRPF39 | 1317.088631 | -0.499128382 | 0.0810421 | -6.158877688 | 7.33E-10 | 5.90E-09 |
| ENSG00000185250 | PPIL6 | 50.78664555 | 1.233670815 | 0.263691664 | 4.678459663 | 2.89E-06 | 1.59E-05 |
| ENSG00000185252 | ZNF74 | 529.3264518 | -0.43466877 | 0.088607541 | -4.905550533 | 9.32E-07 | 5.42E-06 |
| ENSG00000185261 | KIAA0825 | 57.451858 | 0.068141075 | 0.233040154 | 0.292400575 | 0.769980369 | 0.874596438 |
| ENSG00000185262 | UBALD2 | 971.6841176 | 0.424915018 | 0.070302766 | 6.04407254 | 1.50E-09 | 1.18E-08 |
| ENSG00000185267 | CDNF | 31.17470871 | -0.868032143 | 0.330936178 | -2.622959353 | 0.008716967 | 0.02636351 |
| ENSG00000185269 | NOTUM | 7.245950107 | -1.386375546 | 0.671811202 | -2.063638626 | 0.039051992 | 0.097237893 |
| ENSG00000185272 | RBM11 | 3.971470128 | 1.196807705 | 0.917818436 | 1.303969998 | 0.192243812 | 0.352382984 |
| ENSG00000185275 | CD24P4 | 135.1940493 | -0.062271473 | 0.164550024 | -0.378434909 | 0.705107539 | 0.834997903 |
| ENSG00000185278 | ZBTB37 | 663.4637526 | -0.477295464 | 0.081750538 | -5.838438231 | 5.27E-09 | 3.91E-08 |

| | | | | | | | |
|---|---|---|---|---|---|---|---|
| ENSG00000185291 | IL3RA | 2.086398166 | -0.360242041 | 1.234729656 | -0.291757826 | 0.770471793 | 0.874956824 |
| ENSG00000185291 | IL3RA | 2.086398166 | -0.360242041 | 1.234729656 | -0.291757826 | 0.770471793 | 0.874956824 |
| ENSG00000185296 | ENSG0000018 | 24.35553645 | -2.793945346 | 0.449532731 | -6.215221163 | 5.13E-10 | 4.19E-09 |
| ENSG00000185298 | CCDC137 | 1991.389197 | 0.246230211 | 0.064370004 | 3.825232182 | 0.000130649 | 0.000571854 |
| ENSG00000185304 | RGPD2 | 12.17733578 | 0.023719971 | 0.533158499 | 0.044489529 | 0.964514198 | 0.983900556 |
| ENSG00000185305 | ARL15 | 543.9748833 | -0.038620769 | 0.090388498 | -0.427275263 | 0.669178848 | 0.809991704 |
| ENSG00000185324 | CDK10 | 3705.26042 | 0.664335896 | 0.062728496 | 10.59065556 | 3.29E-26 | 7.76E-25 |
| ENSG00000185332 | TMEM105 | 90.12404613 | 0.934746493 | 0.194764195 | 4.799375439 | 1.59E-06 | 8.99E-06 |
| ENSG00000185338 | SOCS1 | 64.75617694 | -0.312732874 | 0.223086611 | -1.401845104 | 0.160961506 | 0.307627839 |
| ENSG00000185339 | TCN2 | 16.14773023 | -1.237206143 | 0.463060931 | -2.671799887 | 0.007544563 | 0.023203302 |
| ENSG00000185340 | GAS2L1 | 1314.766454 | -1.019463197 | 0.072090314 | -14.1414725 | 2.11E-45 | 1.03E-43 |
| ENSG00000185344 | ATP6V0A2 | 986.4822639 | -0.05786567 | 0.069135722 | -0.836986568 | 0.402600118 | 0.595272105 |
| ENSG00000185345 | PRKN | 22.00230895 | 0.35692272 | 0.380567306 | 0.937870159 | 0.348311141 | 0.539408463 |
| ENSG00000185347 | TEDC1 | 647.9484799 | -0.775506015 | 0.083948295 | -9.237900683 | 2.51E-20 | 4.28E-19 |
| ENSG00000185352 | HS6ST3 | 658.1390813 | 0.592720619 | 0.085313409 | 6.947566937 | 3.72E-12 | 3.62E-11 |
| ENSG00000185359 | HGS | 4203.70834 | 0.409939119 | 0.058957766 | 6.953097926 | 3.57E-12 | 3.49E-11 |
| ENSG00000185361 | TNFAIP8L1 | 209.6482967 | -0.571806046 | 0.129580989 | -4.412731009 | 1.02E-05 | 5.27E-05 |
| ENSG00000185379 | RAD51D | 424.7121747 | -0.078904122 | 0.093159396 | -0.846979747 | 0.39700641 | 0.589887251 |
| ENSG00000185386 | MAPK11 | 403.0273615 | 0.379493224 | 0.10063189 | 3.771103026 | 0.000162528 | 0.000700341 |
| ENSG00000185404 | SP140L | 236.9176495 | -0.09802562 | 0.121555032 | -0.80642996 | 0.419994978 | 0.611284724 |
| ENSG00000185414 | MRPL30 | 1410.069504 | 0.195908453 | 0.058677406 | 3.338737458 | 0.000841601 | 0.003193767 |
| ENSG00000185418 | TARS3 | 456.5842312 | 0.568586822 | 0.091589074 | 6.208020187 | 5.37E-10 | 4.38E-09 |
| ENSG00000185420 | SMYD3 | 630.106484 | -0.304917661 | 0.086670604 | -3.518120881 | 0.000434614 | 0.001739671 |
| ENSG00000185432 | METTL7A | 666.9941602 | 0.722462691 | 0.080867654 | 8.93388953 | 4.11E-19 | 6.53E-18 |
| ENSG00000185436 | IFNLR1 | 175.5593525 | 0.589165941 | 0.140217166 | 4.201810364 | 2.65E-05 | 0.000129185 |
| ENSG00000185437 | SH3BGR | 6.85068768 | 1.268052255 | 0.725961906 | 1.746720104 | 0.080685901 | 0.177906053 |
| ENSG00000185442 | FAM174B | 1998.85578 | 0.076727802 | 0.056247006 | 1.364122417 | 0.172529041 | 0.324527758 |
| ENSG00000185453 | ZSWIM9 | 257.0910172 | -0.051487699 | 0.117244188 | -0.439149268 | 0.660553382 | 0.803707149 |
| ENSG00000185467 | KPNA7 | 1.857619232 | 3.155978558 | 1.592354839 | 1.981956836 | 0.047484072 | 0.114613279 |
| ENSG00000185475 | TMEM179B | 1321.439966 | -0.082552686 | 0.062296654 | -1.325154417 | 0.185119941 | 0.342113223 |
| ENSG00000185477 | GPRIN3 | 342.0889562 | -3.160311415 | 0.130540152 | -24.209497 | 1.77E-129 | 5.00E-127 |
| ENSG00000185480 | PARPBP | 607.4763109 | -0.123717509 | 0.084064215 | -1.471702428 | 0.141101252 | 0.278200491 |
| ENSG00000185482 | STAC3 | 212.3789767 | 1.084136657 | 0.136271872 | 7.955689171 | 1.78E-15 | 2.21E-14 |

| | | | | | | | |
|---|---|---|---|---|---|---|---|
| ENSG00000185483 | ROR1 | 37.55315536 | -1.486493355 | 0.30163079 | -4.928188378 | 8.30E-07 | 4.86E-06 |
| ENSG00000185485 | SDHAP1 | 403.234928 | -0.195471581 | 0.105899139 | -1.845827854 | 0.064917205 | 0.149009474 |
| ENSG00000185495 | SEPTIN7P13 | 23.22757724 | 0.173347636 | 0.365039202 | 0.474874028 | 0.634876764 | 0.784786775 |
| ENSG00000185499 | MUC1 | 1095.162121 | 0.385571843 | 0.073117903 | 5.273289149 | 1.34E-07 | 8.59E-07 |
| ENSG00000185504 | FAAP100 | 2036.964404 | 0.568094427 | 0.061997379 | 9.163200666 | 5.04E-20 | 8.41E-19 |
| ENSG00000185507 | IRF7 | 193.5085656 | 0.144027207 | 0.138993252 | 1.036217257 | 0.300100797 | 0.48680172 |
| ENSG00000185513 | L3MBTL1 | 41.00836255 | 1.426991302 | 0.300568838 | 4.747635554 | 2.06E-06 | 1.15E-05 |
| ENSG00000185515 | BRCC3 | 1516.631595 | -0.892495596 | 0.062103406 | -14.37112148 | 7.85E-47 | 3.98E-45 |
| ENSG00000185518 | SV2B | 4.61108822 | -1.977057068 | 0.895425685 | -2.207952151 | 0.027247609 | 0.071517179 |
| ENSG00000185519 | FAM131C | 29.30245353 | 1.309948797 | 0.340294156 | 3.849460162 | 0.000118378 | 0.000521547 |
| ENSG00000185522 | LMNTD2 | 157.005387 | 0.096619878 | 0.149885943 | 0.644622676 | 0.519171737 | 0.696723975 |
| ENSG00000185523 | SPATA45 | 2.017439302 | -0.217097162 | 1.223195175 | -0.177483664 | 0.859128496 | 0.926792302 |
| ENSG00000185527 | PDE6G | 5.094793989 | -0.257588092 | 0.788483974 | -0.326687796 | 0.743904032 | 0.858490754 |
| ENSG00000185532 | PRKG1 | 360.8646247 | -0.490054524 | 0.105938557 | -4.62583725 | 3.73E-06 | 2.02E-05 |
| ENSG00000185551 | NR2F2 | 4175.390894 | 0.244261906 | 0.047045916 | 5.191989535 | 2.08E-07 | 1.31E-06 |
| ENSG00000185559 | DLK1 | 1.956714874 | -4.548747859 | 1.61713314 | -2.812846851 | 0.004910503 | 0.01585237 |
| ENSG00000185561 | TLCD2 | 379.5088826 | 0.239680277 | 0.100512456 | 2.384582828 | 0.017098498 | 0.047723513 |
| ENSG00000185565 | LSAMP | 47.37565335 | -0.689024142 | 0.261152096 | -2.638401732 | 0.008329784 | 0.025305952 |
| ENSG00000185567 | AHNAK2 | 437.6237792 | -0.114712809 | 0.103106752 | -1.112563502 | 0.265895943 | 0.446828286 |
| ENSG00000185585 | OLFML2A | 128.7181981 | 1.086157727 | 0.168493172 | 6.446301142 | 1.15E-10 | 9.85E-10 |
| ENSG00000185591 | SP1 | 5232.316916 | -0.117764673 | 0.047661973 | -2.470830864 | 0.013479956 | 0.038663439 |
| ENSG00000185596 | WASH3P | 215.6495919 | -0.357371521 | 0.132236165 | -2.702524845 | 0.006881504 | 0.021373098 |
| ENSG00000185607 | ACTBP7 | 2.319078014 | 0.654777536 | 1.152891117 | 0.567943951 | 0.570073027 | 0.737257596 |
| ENSG00000185608 | MRPL40 | 542.7901314 | -1.061241928 | 0.087889087 | -12.07478606 | 1.44E-33 | 4.57E-32 |
| ENSG00000185614 | INKA1 | 2.013575897 | 1.04012804 | 1.275199251 | 0.81565923 | 0.414695063 | 0.606780504 |
| ENSG00000185615 | PDIA2 | 135.5279293 | 0.303655134 | 0.167265339 | 1.815409784 | 0.069460959 | 0.157626974 |
| ENSG00000185619 | PCGF3 | 2874.283861 | 0.275489427 | 0.050123384 | 5.49622557 | 3.88E-08 | 2.62E-07 |
| ENSG00000185621 | LMLN | 257.3795737 | 0.112866668 | 0.118170057 | 0.955120702 | 0.339516661 | 0.530665194 |
| ENSG00000185624 | P4HB | 28238.30228 | 0.005045377 | 0.03903037 | 0.129267989 | 0.8971456 | 0.947984339 |
| ENSG00000185627 | PSMD13 | 2479.340807 | -0.191611944 | 0.052638661 | -3.640137114 | 0.000272493 | 0.001128745 |
| ENSG00000185630 | PBX1 | 6988.273712 | -0.44286904 | 0.052683145 | -8.40627559 | 4.23E-17 | 5.86E-16 |
| ENSG00000185633 | NDUFA4L2 | 57.50039367 | 0.105712899 | 0.23195314 | 0.455751102 | 0.64856897 | 0.795130996 |
| ENSG00000185634 | SHC4 | 12.12826429 | -1.234488684 | 0.578221341 | -2.134975998 | 0.032762971 | 0.083807153 |

| | | | | | | | |
|---|---|---|---|---|---|---|---|
| ENSG00000185641 | ENSG0000018 | 3229.62149 | 0.387297727 | 0.057586271 | 6.725521948 | 1.75E-11 | 1.60E-10 |
| ENSG00000185650 | ZFP36L1 | 2532.825812 | 0.423423005 | 0.054943703 | 7.706488328 | 1.29E-14 | 1.51E-13 |
| ENSG00000185651 | UBE2L3 | 3030.045827 | -0.557280819 | 0.049877144 | -11.17307 | 5.52E-29 | 1.47E-27 |
| ENSG00000185658 | BRWD1 | 5728.437477 | -0.119922004 | 0.049133247 | -2.440750612 | 0.014656772 | 0.041632783 |
| ENSG00000185664 | PMEL | 95.72929763 | 0.173074849 | 0.180968859 | 0.956379181 | 0.338880699 | 0.530045518 |
| ENSG00000185669 | SNAI3 | 41.4384555 | 0.966960936 | 0.296719033 | 3.258843651 | 0.001118673 | 0.004152485 |
| ENSG00000185670 | ZBTB3 | 239.3034817 | 0.068914576 | 0.13768576 | 0.500520724 | 0.616708468 | 0.772907138 |
| ENSG00000185684 | EP400P1 | 490.7335728 | 0.15796023 | 0.093456588 | 1.69019898 | 0.090989894 | 0.196230907 |
| ENSG00000185686 | PRAME | 44.61371372 | -2.029397116 | 0.294774019 | -6.884586118 | 5.80E-12 | 5.56E-11 |
| ENSG00000185689 | TEX56P | 6.712087098 | -1.267853952 | 0.728961311 | -1.739260964 | 0.08198887 | 0.180276496 |
| ENSG00000185697 | MYBL1 | 365.9795678 | 0.319140823 | 0.114687644 | 2.782695793 | 0.005390933 | 0.017211969 |
| ENSG00000185710 | SMG1P4 | 73.87906275 | 0.048004448 | 0.209149934 | 0.229521698 | 0.818463458 | 0.903931045 |
| ENSG00000185716 | MOSMO | 265.2108249 | 0.161791679 | 0.134018446 | 1.207234401 | 0.227341886 | 0.400063122 |
| ENSG00000185721 | DRG1 | 1626.60684 | -0.611765043 | 0.057849494 | -10.57511487 | 3.89E-26 | 9.12E-25 |
| ENSG00000185722 | ANKFY1 | 2178.258339 | 0.066965406 | 0.053758154 | 1.245679053 | 0.212882247 | 0.380989136 |
| ENSG00000185728 | YTHDF3 | 2387.167146 | 0.4746205 | 0.054857154 | 8.651934446 | 5.06E-18 | 7.47E-17 |
| ENSG00000185730 | ZNF696 | 490.4338545 | -0.094088133 | 0.089526362 | -1.050954504 | 0.293279486 | 0.478751972 |
| ENSG00000185737 | NRG3 | 190.9285761 | -2.421619866 | 0.15389684 | -15.73534497 | 8.66E-56 | 5.55E-54 |
| ENSG00000185745 | IFIT1 | 56.16421585 | 2.443106369 | 0.582215558 | 4.196223091 | 2.71E-05 | 0.000132215 |
| ENSG00000185753 | CXorf38 | 1211.620802 | -0.294230748 | 0.065090797 | -4.520312546 | 6.17E-06 | 3.27E-05 |
| ENSG00000185760 | KCNQ5 | 24.88007865 | -3.761789779 | 0.524781858 | -7.168292356 | 7.59E-13 | 7.84E-12 |
| ENSG00000185761 | ADAMTSL5 | 137.479508 | 0.260284563 | 0.157421159 | 1.653428069 | 0.098243774 | 0.208928542 |
| ENSG00000185787 | MORF4L1 | 3847.853913 | -0.132069221 | 0.055633673 | -2.37390799 | 0.017600936 | 0.04896501 |
| ENSG00000185798 | WDR53 | 305.7051226 | -0.243196534 | 0.11086326 | -2.193662123 | 0.028259709 | 0.073776911 |
| ENSG00000185800 | DMWD | 1014.279305 | 0.032108258 | 0.072378282 | 0.44361729 | 0.657319303 | 0.80147779 |
| ENSG00000185803 | SLC52A2 | 1776.56613 | 0.179716658 | 0.064323681 | 2.793942373 | 0.005206976 | 0.016681159 |
| ENSG00000185808 | PIGP | 426.8441188 | -0.180095858 | 0.093290408 | -1.930486342 | 0.053546606 | 0.127146169 |
| ENSG00000185813 | PCYT2 | 4305.21646 | -0.048540169 | 0.054203172 | -0.895522655 | 0.370507761 | 0.563213043 |
| ENSG00000185818 | NAT8L | 998.4178461 | 1.084987374 | 0.069637262 | 15.5805577 | 9.87E-55 | 6.15E-53 |
| ENSG00000185825 | BCAP31 | 8201.792139 | -0.368112035 | 0.043053291 | -8.550148537 | 1.23E-17 | 1.77E-16 |
| ENSG00000185829 | ARL17A | 44.53775365 | 0.316395227 | 0.276983722 | 1.14228816 | 0.253334261 | 0.431990496 |
| ENSG00000185834 | RPL12P4 | 1596.942383 | 0.221344096 | 0.065763644 | 3.365751703 | 0.000763354 | 0.002922453 |
| ENSG00000185837 | HDHD5-AS1 | 4.387608737 | -1.049942513 | 0.87870218 | -1.194878693 | 0.232134429 | 0.406129026 |

| | | | | | | | |
|---|---|---|---|---|---|---|---|
| ENSG00000185838 | GNB1L | 205.5011726 | -0.540806785 | 0.133166301 | -4.061138458 | 4.88E-05 | 0.000228903 |
| ENSG00000185839 | ENSG0000018 | 599.993142 | 0.09133123 | 0.080064196 | 1.140724997 | 0.253984381 | 0.432900044 |
| ENSG00000185842 | DNAH14 | 207.7184297 | 0.220344557 | 0.13029837 | 1.691076853 | 0.090822126 | 0.195905929 |
| ENSG00000185864 | NPIPB4 | 76.89829648 | 0.155485567 | 0.213098281 | 0.72964252 | 0.465608725 | 0.651602254 |
| ENSG00000185869 | ZNF829 | 66.37229867 | 0.845318537 | 0.230385864 | 3.669142385 | 0.000243366 | 0.001016164 |
| ENSG00000185875 | THNSL1 | 533.0051621 | -0.565845518 | 0.089107859 | -6.350119094 | 2.15E-10 | 1.81E-09 |
| ENSG00000185880 | TRIM69 | 256.3457307 | -0.413141151 | 0.121874457 | -3.389891216 | 0.000699204 | 0.002694847 |
| ENSG00000185883 | ATP6V0C | 5.978093014 | 0.610955849 | 0.741624501 | 0.823807531 | 0.410048928 | 0.602595926 |
| ENSG00000185885 | IFITM1 | 19.89933333 | 1.275790795 | 0.439374706 | 2.903650979 | 0.003688391 | 0.012248362 |
| ENSG00000185896 | LAMP1 | 4132.372095 | 0.146991662 | 0.04637523 | 3.169615821 | 0.001526406 | 0.005501297 |
| ENSG00000185900 | POMK | 100.0833522 | -1.190984865 | 0.183276386 | -6.498299599 | 8.12E-11 | 7.09E-10 |
| ENSG00000185909 | KLHDC8B | 267.3702475 | -0.139648555 | 0.115833173 | -1.205600709 | 0.227971486 | 0.400832826 |
| ENSG00000185917 | SETD4 | 566.6162848 | -0.13925263 | 0.089780668 | -1.551031336 | 0.120894175 | 0.246955687 |
| ENSG00000185920 | PTCH1 | 342.8148402 | 0.077108691 | 0.103378682 | 0.7458858 | 0.455736404 | 0.643082556 |
| ENSG00000185924 | RTN4RL1 | 1215.989696 | 0.002838966 | 0.070932009 | 0.040023758 | 0.968074186 | 0.985820278 |
| ENSG00000185933 | CALHM1 | 2.267014985 | 2.538747147 | 1.457501634 | 1.741848578 | 0.081534944 | 0.179450409 |
| ENSG00000185942 | NKAIN3 | 3.60883202 | -0.179484477 | 0.915652735 | -0.196018065 | 0.844596027 | 0.918248003 |
| ENSG00000185946 | RNPC3 | 240.7704531 | 0.260701632 | 0.122255895 | 2.132425858 | 0.032971854 | 0.084228766 |
| ENSG00000185947 | ZNF267 | 342.924332 | -0.122260263 | 0.101524045 | -1.204249325 | 0.228493227 | 0.401366827 |
| ENSG00000185950 | IRS2 | 369.9464754 | -1.554220188 | 0.110358779 | -14.08333989 | 4.81E-45 | 2.33E-43 |
| ENSG00000185955 | SPACDR | 7.425672594 | -1.121725531 | 0.676498347 | -1.658134918 | 0.097290237 | 0.207355897 |
| ENSG00000185963 | BICD2 | 1536.573228 | -0.054282416 | 0.058392734 | -0.929609082 | 0.352573523 | 0.544055279 |
| ENSG00000185973 | TMLHE | 434.6885784 | -0.743786982 | 0.09304651 | -7.993711733 | 1.31E-15 | 1.65E-14 |
| ENSG00000185986 | SDHAP3 | 582.8621692 | 0.043307702 | 0.093657834 | 0.462403416 | 0.64379205 | 0.791270263 |
| ENSG00000185988 | PLK5 | 23.33775418 | -0.270793702 | 0.364583942 | -0.7427472 | 0.457634748 | 0.644848963 |
| ENSG00000185989 | RASA3 | 305.1621416 | -0.908053851 | 0.108818266 | -8.344682257 | 7.14E-17 | 9.79E-16 |
| ENSG00000186001 | LRCH3 | 1022.344821 | -0.337110926 | 0.07070996 | -4.767516866 | 1.87E-06 | 1.05E-05 |
| ENSG00000186010 | NDUFA13 | 27.02013531 | -0.084786617 | 0.333981559 | -0.253866164 | 0.799598956 | 0.893202184 |
| ENSG00000186017 | ZNF566 | 308.6089342 | 0.300094242 | 0.107272345 | 2.797498673 | 0.005149998 | 0.016541718 |
| ENSG00000186019 | ZNF225-AS1 | 35.62332529 | -0.116245143 | 0.292293588 | -0.397699941 | 0.690851381 | 0.825640142 |
| ENSG00000186020 | ZNF529 | 255.0278364 | 0.68044446 | 0.121990451 | 5.577850209 | 2.44E-08 | 1.68E-07 |
| ENSG00000186026 | ZNF284 | 59.6144876 | -0.059867891 | 0.241720946 | -0.247673577 | 0.804386978 | 0.895992144 |
| ENSG00000186051 | TAL2 | 2.011269634 | -0.690326079 | 1.216777977 | -0.567339393 | 0.570483618 | 0.737372898 |

| | | | | | | | |
|---|---|---|---|---|---|---|---|
| ENSG00000186056 | MATN1-AS1 | 101.4247539 | 0.005044794 | 0.177193424 | 0.028470547 | 0.977286859 | 0.989387067 |
| ENSG00000186063 | AIDA | 364.4131651 | -0.216929872 | 0.102727685 | -2.111698247 | 0.034712339 | 0.087989068 |
| ENSG00000186073 | CDIN1 | 226.1344059 | -0.010914715 | 0.122745421 | -0.088921569 | 0.929144242 | 0.965981609 |
| ENSG00000186076 | ENSG0000018 | 453.6216155 | -0.7614138 | 0.095378594 | -7.983068016 | 1.43E-15 | 1.79E-14 |
| ENSG00000186081 | KRT5 | 1.943655058 | 1.478962616 | 1.451278222 | 1.01907587 | 0.308166949 | 0.49616436 |
| ENSG00000186082 | KRT18P14 | 5.86589672 | -0.165758703 | 0.725948466 | -0.22833398 | 0.81938661 | 0.904358204 |
| ENSG00000186088 | GSAP | 218.8688188 | 2.80303566 | 0.158060377 | 17.73395531 | 2.29E-70 | 2.13E-68 |
| ENSG00000186094 | AGBL4 | 15.25100548 | 0.832852555 | 0.45457488 | 1.832157014 | 0.066928031 | 0.152997413 |
| ENSG00000186104 | CYP2R1 | 259.1607702 | -0.434147291 | 0.120584512 | -3.600356984 | 0.000317781 | 0.001302231 |
| ENSG00000186106 | ANKRD46 | 603.1459779 | -0.205229004 | 0.086153116 | -2.382142562 | 0.017212233 | 0.047988407 |
| ENSG00000186111 | PIP5K1C | 1715.299189 | 0.384568822 | 0.057972321 | 6.633662677 | 3.27E-11 | 2.94E-10 |
| ENSG00000186118 | TEX38 | 7.615797541 | 0.773856502 | 0.647352275 | 1.195417907 | 0.231923793 | 0.40588433 |
| ENSG00000186130 | ZBTB6 | 484.5792745 | 0.143492087 | 0.095514008 | 1.502314577 | 0.133015886 | 0.266171007 |
| ENSG00000186132 | C2orf76 | 190.6106843 | -0.281684705 | 0.133962253 | -2.102716985 | 0.035490516 | 0.089644621 |
| ENSG00000186141 | POLR3C | 3489.371187 | 0.495991332 | 0.052151185 | 9.510643619 | 1.89E-21 | 3.45E-20 |
| ENSG00000186153 | WWOX | 1574.069509 | 0.720981486 | 0.062440198 | 11.54675207 | 7.67E-31 | 2.20E-29 |
| ENSG00000186160 | CYP4Z1 | 4.389462908 | 0.547998977 | 0.849132623 | 0.645363236 | 0.518691822 | 0.696318646 |
| ENSG00000186162 | CIDECP1 | 390.237893 | 0.120326972 | 0.10074039 | 1.194426307 | 0.232311251 | 0.406370643 |
| ENSG00000186166 | CENATAC | 790.2970804 | -0.309062668 | 0.173485669 | -1.781488179 | 0.074832734 | 0.167626494 |
| ENSG00000186174 | BCL9L | 8235.363091 | 0.02722485 | 0.057183167 | 0.476099024 | 0.634003833 | 0.784236938 |
| ENSG00000186184 | POLR1D | 3356.19355 | 0.14363889 | 0.047372416 | 3.032120836 | 0.00242842 | 0.008397339 |
| ENSG00000186185 | KIF18B | 1058.161036 | -0.34379427 | 0.073207107 | -4.696187088 | 2.65E-06 | 1.46E-05 |
| ENSG00000186187 | ZNRF1 | 1895.262148 | 0.743713352 | 0.055578325 | 13.38135591 | 7.77E-41 | 3.29E-39 |
| ENSG00000186193 | SAPCD2 | 4400.519442 | -0.179960872 | 0.050855788 | -3.538650756 | 0.000402178 | 0.001615981 |
| ENSG00000186197 | EDARADD | 33.10320548 | -2.743835336 | 0.37579926 | -7.301332468 | 2.85E-13 | 3.02E-12 |
| ENSG00000186205 | MTARC1 | 564.8903383 | -0.739300977 | 0.083181741 | -8.887779604 | 6.23E-19 | 9.79E-18 |
| ENSG00000186212 | SOWAHB | 948.9416758 | 0.391607649 | 0.081224676 | 4.821289126 | 1.43E-06 | 8.09E-06 |
| ENSG00000186222 | BLOC1S4 | 584.8268536 | 0.014605051 | 0.082863919 | 0.17625344 | 0.860094839 | 0.927319435 |
| ENSG00000186230 | ZNF749 | 271.7570874 | 0.053756315 | 0.116481818 | 0.461499619 | 0.644440195 | 0.791744755 |
| ENSG00000186231 | KLHL32 | 20.22529727 | -0.935227481 | 0.40663294 | -2.299930452 | 0.021452161 | 0.058093798 |
| ENSG00000186260 | MRTFB | 1875.804854 | 0.146081125 | 0.066058133 | 2.211402556 | 0.027007973 | 0.070977535 |
| ENSG00000186272 | ZNF17 | 200.5820253 | 0.21766721 | 0.138615124 | 1.57029914 | 0.116345534 | 0.239670129 |
| ENSG00000186280 | KDM4D | 23.14125527 | -0.825701142 | 0.366258511 | -2.254421718 | 0.02416965 | 0.064561222 |

| | | | | | | | |
|---|---|---|---|---|---|---|---|
| ENSG00000186281 | GPAT2 | 3.281624713 | -0.737191495 | 0.997056348 | -0.739367937 | 0.459683606 | 0.646623861 |
| ENSG00000186283 | TOR3A | 2098.175915 | -0.019166097 | 0.054252547 | -0.353275526 | 0.723881892 | 0.846416769 |
| ENSG00000186298 | PPP1CC | 3200.147395 | 0.03988654 | 0.05379296 | 0.741482526 | 0.458400923 | 0.645532055 |
| ENSG00000186300 | ZNF555 | 237.4385108 | -0.140066282 | 0.131176784 | -1.067767311 | 0.285625487 | 0.469526956 |
| ENSG00000186301 | MST1P2 | 125.1950428 | -0.077538221 | 0.159786192 | -0.485262337 | 0.627490268 | 0.780187429 |
| ENSG00000186306 | OR10T2 | 1.943693365 | 1.367295597 | 1.336921899 | 1.022719127 | 0.306440665 | 0.494094221 |
| ENSG00000186310 | NAP1L3 | 31.66689579 | -0.710080636 | 0.320457091 | -2.215836866 | 0.026702677 | 0.070296007 |
| ENSG00000186312 | CA5BP1 | 287.7358111 | 0.469409518 | 0.115995527 | 4.046789822 | 5.19E-05 | 0.000242351 |
| ENSG00000186314 | PRELID2 | 88.42420902 | -0.416464894 | 0.188835231 | -2.205440644 | 0.027423189 | 0.071928796 |
| ENSG00000186318 | BACE1 | 2490.176025 | 0.383106173 | 0.055666575 | 6.882158117 | 5.90E-12 | 5.64E-11 |
| ENSG00000186326 | RGS9BP | 39.23125582 | -0.424832579 | 0.284944712 | -1.490929859 | 0.135979911 | 0.270615005 |
| ENSG00000186350 | RXRA | 6858.627942 | 0.034586526 | 0.053328714 | 0.648553531 | 0.516627 | 0.695057379 |
| ENSG00000186352 | ANKRD37 | 6.238387318 | -0.782048951 | 0.70207302 | -1.113913979 | 0.265316092 | 0.44634432 |
| ENSG00000186364 | NUDT17 | 308.5349616 | 0.418951981 | 0.11106134 | 3.772257586 | 0.000161777 | 0.000697326 |
| ENSG00000186367 | MINAR2 | 6.938811607 | 0.65515719 | 0.677683743 | 0.966759492 | 0.333664279 | 0.524695542 |
| ENSG00000186376 | ZNF75D | 1637.603669 | -0.073128386 | 0.059002325 | -1.239415319 | 0.215191731 | 0.384253001 |
| ENSG00000186377 | CYP4X1 | 10.01563942 | 0.685635072 | 0.555797151 | 1.233606669 | 0.217349507 | 0.38691722 |
| ENSG00000186395 | KRT10 | 609.8471947 | -0.113692834 | 0.081624986 | -1.392868036 | 0.1636597 | 0.311332077 |
| ENSG00000186399 | GOLGA8R | 2.051476367 | -0.722899703 | 1.224751311 | -0.590242032 | 0.555028396 | 0.726193544 |
| ENSG00000186409 | CCDC30 | 83.45393859 | 0.0411326 | 0.194291437 | 0.211705675 | 0.832336661 | 0.911334752 |
| ENSG00000186416 | NKRF | 1333.808241 | 0.207765765 | 0.065071022 | 3.192907673 | 0.00140848 | 0.005115663 |
| ENSG00000186417 | GLDN | 946.2728377 | -1.249669958 | 0.070708609 | -17.67351918 | 6.71E-70 | 6.20E-68 |
| ENSG00000186432 | KPNA4 | 3443.680821 | -0.112897262 | 0.052132965 | -2.16556382 | 0.030344534 | 0.078478422 |
| ENSG00000186448 | ZNF197 | 506.1122924 | 0.213986358 | 0.08854576 | 2.416675363 | 0.015662979 | 0.044184549 |
| ENSG00000186462 | NAP1L2 | 129.1641336 | -0.259230951 | 0.159088688 | -1.629474443 | 0.103212621 | 0.217491195 |
| ENSG00000186468 | RPS23 | 7897.015546 | 0.304126744 | 0.051165555 | 5.943974278 | 2.78E-09 | 2.13E-08 |
| ENSG00000186470 | BTN3A2 | 1.888351259 | -0.477390658 | 1.309112556 | -0.364667389 | 0.715359704 | 0.841656152 |
| ENSG00000186472 | PCLO | 213.5713249 | -0.222608572 | 0.136841894 | -1.626757469 | 0.103788618 | 0.218432192 |
| ENSG00000186479 | RGS7BP | 8.956221573 | 0.946949681 | 0.604272761 | 1.567089802 | 0.117093703 | 0.240951699 |
| ENSG00000186480 | INSIG1 | 7713.877032 | -0.115747394 | 0.047355071 | -2.444244964 | 0.014515567 | 0.041277687 |
| ENSG00000186481 | ANKRD20A5P | 1180.225201 | 0.235051219 | 0.083077415 | 2.829303459 | 0.004664944 | 0.015142751 |
| ENSG00000186493 | IRX2-DT | 263.7236294 | 0.179670914 | 0.11339291 | 1.584498664 | 0.113080285 | 0.234142228 |
| ENSG00000186496 | ZNF396 | 107.6893567 | 0.250892811 | 0.171141789 | 1.465993854 | 0.142649961 | 0.280389475 |

| | | | | | | | |
|---|---|---|---|---|---|---|---|
| ENSG00000186501 | TMEM222 | 774.4171702 | -0.074170946 | 0.08168978 | -0.907958693 | 0.363900046 | 0.556598684 |
| ENSG00000186510 | CLCNKA | 11.79449071 | 0.386675395 | 0.530770304 | 0.728517387 | 0.466296929 | 0.652247046 |
| ENSG00000186517 | ARHGAP30 | 41.43889749 | 1.120719358 | 0.28751352 | 3.897970977 | 9.70E-05 | 0.000433092 |
| ENSG00000186522 | SEPTIN10 | 876.0316767 | -0.814726657 | 0.073647453 | -11.06252325 | 1.91E-28 | 4.98E-27 |
| ENSG00000186523 | FAM86B1 | 60.32383831 | 0.640032275 | 0.23059717 | 2.775542623 | 0.005510968 | 0.017529339 |
| ENSG00000186532 | SMYD4 | 509.7718337 | -0.09287958 | 0.089017357 | -1.043387304 | 0.296768951 | 0.483068171 |
| ENSG00000186564 | FOXD2 | 18.31421865 | -0.979940803 | 0.437145027 | -2.241683521 | 0.024981834 | 0.066436734 |
| ENSG00000186566 | GPATCH8 | 2577.03519 | -0.094667346 | 0.06342368 | -1.49261831 | 0.135537128 | 0.269991465 |
| ENSG00000186567 | CEACAM19 | 105.4172758 | 0.520599521 | 0.174683643 | 2.980241942 | 0.002880208 | 0.009797149 |
| ENSG00000186575 | NF2 | 1951.524873 | -0.860980773 | 0.066429127 | -12.96089242 | 2.04E-38 | 7.83E-37 |
| ENSG00000186577 | SMIM29 | 652.4690306 | 0.818608852 | 0.084008578 | 9.74434839 | 1.95E-22 | 3.75E-21 |
| ENSG00000186591 | UBE2H | 4091.637828 | 0.139049927 | 0.051917601 | 2.678281049 | 0.00740011 | 0.022804744 |
| ENSG00000186594 | MIR22HG | 161.3636227 | 0.315708861 | 0.15039852 | 2.099148726 | 0.035803792 | 0.090366266 |
| ENSG00000186603 | HPDL | 144.5064068 | 0.515175103 | 0.155646444 | 3.309906035 | 0.000933273 | 0.00351375 |
| ENSG00000186615 | KTN1-AS1 | 218.8588171 | -0.210920012 | 0.122003872 | -1.728797686 | 0.08384531 | 0.183671832 |
| ENSG00000186625 | KATNA1 | 566.2072085 | 0.087219288 | 0.086707326 | 1.005904488 | 0.314461517 | 0.502910429 |
| ENSG00000186628 | FSD2 | 2.788002488 | 0.047942015 | 1.172660901 | 0.040883102 | 0.967389089 | 0.985430982 |
| ENSG00000186635 | ARAP1 | 2361.838671 | -0.063605361 | 0.057071196 | -1.114491473 | 0.265068401 | 0.446091197 |
| ENSG00000186638 | KIF24 | 268.4481452 | -0.096219607 | 0.117201338 | -0.820977036 | 0.411659349 | 0.60391798 |
| ENSG00000186642 | PDE2A | 7.904257869 | -0.154424544 | 0.619058979 | -0.249450454 | 0.803012361 | 0.895135709 |
| ENSG00000186645 | SPDYE17 | 3.309084906 | 1.156951185 | 1.034342482 | 1.118537821 | 0.263337363 | 0.44412799 |
| ENSG00000186648 | CARMIL3 | 150.4255386 | -0.457687591 | 0.163482728 | -2.799608225 | 0.005116466 | 0.016438618 |
| ENSG00000186654 | PRR5 | 225.3945712 | -0.517248416 | 0.129408914 | -3.997007624 | 6.41E-05 | 0.000295079 |
| ENSG00000186660 | ZFP91 | 2045.034063 | -0.02288287 | 0.06097501 | -0.375282758 | 0.707450188 | 0.836584737 |
| ENSG00000186665 | C17orf58 | 1091.208871 | -0.47277049 | 0.063890166 | -7.39973801 | 1.36E-13 | 1.47E-12 |
| ENSG00000186666 | BCDIN3D | 481.4842432 | -0.121000479 | 0.087029943 | -1.390331584 | 0.164428211 | 0.312586655 |
| ENSG00000186676 | EEF1GP1 | 36.62687437 | 0.214957669 | 0.311208214 | 0.690719779 | 0.489741657 | 0.672936425 |
| ENSG00000186687 | LYRM7 | 384.7107431 | 0.161488123 | 0.102261258 | 1.579172079 | 0.114296592 | 0.236170404 |
| ENSG00000186710 | CFAP73 | 27.54898269 | 0.427985578 | 0.350871079 | 1.219780153 | 0.222548227 | 0.393650731 |
| ENSG00000186714 | CCDC73 | 33.38389905 | -0.134006542 | 0.301226409 | -0.444869834 | 0.656413824 | 0.800836735 |
| ENSG00000186715 | MST1L | 30.19434985 | 0.344937421 | 0.327379901 | 1.053630417 | 0.292052157 | 0.476981097 |
| ENSG00000186716 | BCR | 3436.294504 | -0.285051101 | 0.061556277 | -4.630739758 | 3.64E-06 | 1.98E-05 |
| ENSG00000186743 | TPI1P3 | 11.75226169 | 0.235545232 | 0.544866134 | 0.432299271 | 0.665523917 | 0.807143466 |

| | | | | | | | |
|---|---|---|---|---|---|---|---|
| ENSG00000186765 | FSCN2 | 70.76426886 | -0.00206673 | 0.211094473 | -0.009790544 | 0.992188401 | 0.99592006 |
| ENSG00000186767 | SPIN4 | 827.8253892 | -0.07932555 | 0.074268011 | -1.068098486 | 0.285476089 | 0.469386575 |
| ENSG00000186777 | ZNF732 | 78.9970675 | 0.149899621 | 0.211390863 | 0.709111164 | 0.478255495 | 0.66295052 |
| ENSG00000186787 | SPIN2B | 129.9297003 | 0.106054529 | 0.157469298 | 0.673493373 | 0.500633463 | 0.681732072 |
| ENSG00000186792 | HYAL3 | 108.3680793 | 0.25117405 | 0.169545478 | 1.481455319 | 0.138485283 | 0.274434381 |
| ENSG00000186806 | VSIG10L | 63.47266806 | 0.284732182 | 0.229486042 | 1.240738562 | 0.214702344 | 0.383528404 |
| ENSG00000186812 | ZNF397 | 795.2006025 | 0.371580176 | 0.082063894 | 4.527937421 | 5.96E-06 | 3.16E-05 |
| ENSG00000186814 | ZSCAN30 | 189.2396948 | 0.55718144 | 0.140038802 | 3.978764685 | 6.93E-05 | 0.000317259 |
| ENSG00000186815 | TPCN1 | 857.8681896 | 0.077914035 | 0.071621012 | 1.087865593 | 0.276654448 | 0.459189134 |
| ENSG00000186825 | CDRT15P3 | 4.319931078 | 0.00071498 | 0.847791505 | 0.000843344 | 0.999327109 | 0.999588553 |
| ENSG00000186832 | KRT16 | 15.09494296 | 2.797295165 | 0.577595071 | 4.843003874 | 1.28E-06 | 7.30E-06 |
| ENSG00000186834 | HEXIM1 | 1658.441195 | -0.221773032 | 0.061009553 | -3.635054204 | 0.000277922 | 0.001150611 |
| ENSG00000186847 | KRT14 | 10.23790465 | 2.136347664 | 0.635057997 | 3.364019783 | 0.00076816 | 0.002938402 |
| ENSG00000186854 | TRABD2A | 10.32466606 | -0.92815104 | 0.572044539 | -1.62251534 | 0.104693051 | 0.220032854 |
| ENSG00000186862 | PDZD7 | 34.64151017 | 0.259567813 | 0.29799837 | 0.871037694 | 0.383733572 | 0.576215758 |
| ENSG00000186866 | POFUT2 | 1336.182433 | 0.1157967 | 0.066754584 | 1.734662889 | 0.082800533 | 0.181678071 |
| ENSG00000186868 | MAPT | 1697.113835 | 0.229669084 | 0.060170281 | 3.816985395 | 0.000135092 | 0.000589278 |
| ENSG00000186871 | ERCC6L | 531.4856778 | -0.197125268 | 0.086314083 | -2.283813495 | 0.022382494 | 0.06034251 |
| ENSG00000186889 | TMEM17 | 24.90502327 | 1.015935217 | 0.400008582 | 2.539783551 | 0.011092109 | 0.032551232 |
| ENSG00000186891 | TNFRSF18 | 4.159206266 | 5.407279562 | 1.3959439 | 3.873565092 | 0.000107255 | 0.000475904 |
| ENSG00000186897 | C1QL4 | 16.91843395 | 0.417726151 | 0.438879249 | 0.951802008 | 0.341197403 | 0.532308789 |
| ENSG00000186907 | RTN4RL2 | 33.7295988 | 0.420437204 | 0.300011926 | 1.401401636 | 0.161094003 | 0.307829774 |
| ENSG00000186908 | ZDHHC17 | 720.4392868 | -0.171617111 | 0.092023474 | -1.864927532 | 0.062191563 | 0.1437021 |
| ENSG00000186910 | SERPINA11 | 27.94383382 | -2.229229396 | 0.390775812 | -5.704624821 | 1.17E-08 | 8.31E-08 |
| ENSG00000186918 | ZNF395 | 822.1626379 | -0.003884439 | 0.073018693 | -0.053197872 | 0.957574251 | 0.980789986 |
| ENSG00000186919 | ZACN | 111.7247014 | -0.190559413 | 0.187431055 | -1.016690711 | 0.309300586 | 0.497346456 |
| ENSG00000186940 | CHCHD2P9 | 206.1364757 | 0.633046019 | 0.136475325 | 4.638538292 | 3.51E-06 | 1.91E-05 |
| ENSG00000186951 | PPARA | 776.6329899 | -0.080934594 | 0.085216224 | -0.949755698 | 0.342236401 | 0.53329254 |
| ENSG00000186952 | TMEM232 | 15.34400798 | -0.638602969 | 0.451273242 | -1.415113747 | 0.157035148 | 0.301505381 |
| ENSG00000186973 | CFAP144 | 33.79776097 | 1.109888002 | 0.316263528 | 3.509377166 | 0.000449157 | 0.001792567 |
| ENSG00000186976 | EFCAB6 | 9.633542982 | -0.134228072 | 0.558467733 | -0.24035063 | 0.810058448 | 0.899408839 |
| ENSG00000186994 | KANK3 | 16.53910998 | -0.80298307 | 0.436963806 | -1.837642064 | 0.066115166 | 0.151395679 |
| ENSG00000186998 | EMID1 | 506.4879173 | -0.068633681 | 0.086568071 | -0.792829038 | 0.427877436 | 0.618611482 |

| | | | | | | | |
|---|---|---|---|---|---|---|---|
| ENSG00000187013 | LINC02875 | 66.23211543 | 0.299041865 | 0.227284933 | 1.315713548 | 0.188270205 | 0.346650494 |
| ENSG00000187017 | ESPN | 958.6700858 | 3.112043866 | 0.092732223 | 33.55946574 | 6.55E-247 | 7.51E-244 |
| ENSG00000187024 | PTRH1 | 131.9608807 | 0.17734116 | 0.159390065 | 1.112623676 | 0.265870088 | 0.446828286 |
| ENSG00000187045 | TMPRSS6 | 15.78206929 | 2.998822007 | 0.592977308 | 5.057228949 | 4.25E-07 | 2.59E-06 |
| ENSG00000187049 | TMEM216 | 460.0050448 | -0.276924625 | 0.08989381 | -3.08057503 | 0.002066013 | 0.007240198 |
| ENSG00000187051 | RPS19BP1 | 1137.191785 | -0.211616222 | 0.081556324 | -2.594724856 | 0.009466667 | 0.028354268 |
| ENSG00000187066 | TMEM262 | 17.32016484 | -0.66918184 | 0.435539163 | -1.536444704 | 0.12442935 | 0.252513206 |
| ENSG00000187068 | C3orf70 | 163.7803352 | -1.262040711 | 0.145505698 | -8.67347965 | 4.19E-18 | 6.24E-17 |
| ENSG00000187079 | TEAD1 | 2164.752845 | -0.164149905 | 0.063362082 | -2.590664625 | 0.009579079 | 0.028657287 |
| ENSG00000187091 | PLCD1 | 221.8335255 | 0.57029374 | 0.127065184 | 4.488198274 | 7.18E-06 | 3.77E-05 |
| ENSG00000187097 | ENTPD5 | 1881.348865 | -1.123905017 | 0.056904991 | -19.75055253 | 7.93E-87 | 1.03E-84 |
| ENSG00000187098 | MITF | 48.1247259 | -0.845056888 | 0.268369753 | -3.148852944 | 0.001639127 | 0.005873409 |
| ENSG00000187105 | HEATR4 | 11.593273 | -1.000299579 | 0.526791095 | -1.898854384 | 0.057583624 | 0.1348738 |
| ENSG00000187109 | NAP1L1 | 10333.48713 | 0.427597649 | 0.05357909 | 7.980681452 | 1.46E-15 | 1.82E-14 |
| ENSG00000187118 | CMC1 | 446.4558166 | -0.313020875 | 0.09290542 | -3.369242339 | 0.000753751 | 0.002887135 |
| ENSG00000187122 | SLIT1 | 6542.360993 | -2.213003981 | 0.113394879 | -19.51590766 | 8.04E-85 | 9.97E-83 |
| ENSG00000187123 | LYPD6 | 580.4165272 | 0.324280159 | 0.082975847 | 3.908127133 | 9.30E-05 | 0.000416667 |
| ENSG00000187134 | AKR1C1 | 46.33510725 | -0.570919478 | 0.279841935 | -2.040149837 | 0.041335404 | 0.101886148 |
| ENSG00000187140 | FOXD3 | 21.62792267 | -0.075094275 | 0.384052829 | -0.195531108 | 0.844977187 | 0.918575332 |
| ENSG00000187144 | SPATA21 | 25.67463566 | -0.867417641 | 0.417210514 | -2.079088643 | 0.037609204 | 0.094259274 |
| ENSG00000187147 | RNF220 | 2565.500758 | 0.060821658 | 0.051170379 | 1.188610663 | 0.23459291 | 0.409524493 |
| ENSG00000187164 | SHTN1 | 2667.269954 | -0.2529492 | 0.053707354 | -4.70976846 | 2.48E-06 | 1.37E-05 |
| ENSG00000187172 | BAGE2 | 186.8520924 | 0.147047256 | 0.140367849 | 1.04758502 | 0.294829842 | 0.480662112 |
| ENSG00000187186 | ENSG0000018 | 1.919591662 | -1.052099565 | 1.290308603 | -0.815385996 | 0.414851398 | 0.606854423 |
| ENSG00000187187 | ZNF546 | 27.67131107 | -0.782404148 | 0.344625031 | -2.270305627 | 0.023189046 | 0.062195337 |
| ENSG00000187189 | TSPYL4 | 2177.745147 | 0.00744785 | 0.053926728 | 0.138110558 | 0.890153042 | 0.943797108 |
| ENSG00000187193 | MT1X | 48.73687304 | -0.290105755 | 0.249635643 | -1.162116719 | 0.245188057 | 0.422139653 |
| ENSG00000187210 | GCNT1 | 343.0723959 | -1.477692473 | 0.109205657 | -13.53128136 | 1.02E-41 | 4.48E-40 |
| ENSG00000187231 | SESTD1 | 686.7809694 | 0.887571787 | 0.078408069 | 11.3199037 | 1.05E-29 | 2.88E-28 |
| ENSG00000187239 | FNBP1 | 2293.157807 | 0.204729044 | 0.056640434 | 3.614538733 | 0.000300883 | 0.00123785 |
| ENSG00000187240 | DYNC2H1 | 504.0425914 | -0.03284634 | 0.086215321 | -0.380980318 | 0.703217853 | 0.833796972 |
| ENSG00000187243 | MAGED4B | 4.79170809 | -1.102155283 | 0.85925231 | -1.282691092 | 0.199600322 | 0.362564842 |
| ENSG00000187244 | BCAM | 9003.360225 | -0.577146781 | 0.06510645 | -8.864663627 | 7.67E-19 | 1.20E-17 |

| | | | | | | | |
|---|---|---|---|---|---|---|---|
| ENSG00000187257 | RSBN1L | 683.5372584 | 0.61967072 | 0.084937515 | 7.295606856 | 2.97E-13 | 3.15E-12 |
| ENSG00000187260 | WDR86 | 8.087975353 | 0.062818919 | 0.609965033 | 0.102987739 | 0.917972702 | 0.95968162 |
| ENSG00000187266 | EPOR | 173.474181 | 0.429891248 | 0.144516015 | 2.974696253 | 0.002932788 | 0.00995239 |
| ENSG00000187288 | CIDEC | 1.756888682 | 0.616889481 | 1.311517553 | 0.470363114 | 0.638095613 | NA |
| ENSG00000187323 | DCC | 2.368748312 | 3.497103948 | 1.501380475 | 2.329258976 | 0.01984535 | 0.054343789 |
| ENSG00000187325 | TAF9B | 1538.593471 | -0.059535141 | 0.062187235 | -0.957353075 | 0.338389075 | 0.529660598 |
| ENSG00000187372 | PCDHB13 | 44.17782239 | 0.249271023 | 0.274336595 | 0.908632054 | 0.363544381 | 0.556345471 |
| ENSG00000187391 | MAGI2 | 68.23748722 | 0.725784807 | 0.21782054 | 3.332031066 | 0.000862146 | 0.003266871 |
| ENSG00000187398 | LUZP2 | 150.3853927 | -0.45368039 | 0.149211214 | -3.040524757 | 0.002361663 | 0.008188716 |
| ENSG00000187446 | CHP1 | 3671.646539 | 0.000559509 | 0.046948413 | 0.011917518 | 0.990491422 | 0.994958761 |
| ENSG00000187474 | FPR3 | 25.28127469 | -1.450393566 | 0.365965218 | -3.963200588 | 7.40E-05 | 0.000337266 |
| ENSG00000187475 | H1-6 | 3.247147325 | 0.396766094 | 0.955343622 | 0.415312443 | 0.677913193 | 0.816699042 |
| ENSG00000187479 | C11orf96 | 1.963479792 | -0.143004848 | 1.229103583 | -0.116348898 | 0.907376033 | 0.953426448 |
| ENSG00000187486 | KCNJ11 | 351.7648444 | -0.377185034 | 0.10566893 | -3.569498009 | 0.000357666 | 0.001449365 |
| ENSG00000187504 | RPL7P48 | 7.279677355 | 0.068983518 | 0.658883733 | 0.104697558 | 0.9166158 | 0.958874843 |
| ENSG00000187514 | PTMA | 13006.61789 | -0.137885649 | 0.053840414 | -2.56100647 | 0.010436941 | 0.030841612 |
| ENSG00000187522 | HSPA14 | 512.4700855 | -0.404779267 | 0.086013209 | -4.706012835 | 2.53E-06 | 1.40E-05 |
| ENSG00000187531 | SIRT7 | 2851.211974 | 0.198979885 | 0.062196705 | 3.199202993 | 0.001378081 | 0.005019558 |
| ENSG00000187534 | PRR13P5 | 539.3507111 | -0.121057431 | 0.087625483 | -1.381532252 | 0.167115368 | 0.316661723 |
| ENSG00000187535 | IFT140 | 1318.66261 | 0.265263361 | 0.062428876 | 4.249049125 | 2.15E-05 | 0.000106046 |
| ENSG00000187536 | TPM3P7 | 2.02117415 | -0.656059617 | 1.253189205 | -0.523512024 | 0.600617986 | 0.760636889 |
| ENSG00000187537 | POTEG | 2.552651159 | -0.802086753 | 1.117791408 | -0.717563892 | 0.473026228 | 0.658601347 |
| ENSG00000187550 | SBK2 | 56.37799321 | -4.759138097 | 0.450237486 | -10.570284 | 4.09E-26 | 9.59E-25 |
| ENSG00000187554 | TLR5 | 64.33328472 | 0.607478333 | 0.234170967 | 2.594165882 | 0.009482073 | 0.028390523 |
| ENSG00000187555 | USP7 | 5776.161904 | -0.025602973 | 0.049064231 | -0.52182563 | 0.601791743 | 0.761366787 |
| ENSG00000187556 | NANOS3 | 3.790390703 | -0.06491141 | 0.886115205 | -0.073253917 | 0.941604062 | 0.972202592 |
| ENSG00000187566 | NHLRC1 | 148.7522599 | -0.457610979 | 0.155803962 | -2.937094628 | 0.00331303 | 0.011122626 |
| ENSG00000187583 | PLEKHN1 | 214.9818476 | 0.686494053 | 0.141586598 | 4.848580754 | 1.24E-06 | 7.12E-06 |
| ENSG00000187589 | TERF1P2 | 8.943553816 | 1.326734628 | 0.629000002 | 2.109276031 | 0.034920762 | 0.088429439 |
| ENSG00000187595 | ZNF385C | 28.30914723 | -0.44853713 | 0.331480954 | -1.353130924 | 0.176013808 | 0.32937081 |
| ENSG00000187601 | MAGEH1 | 518.5710352 | -0.108970916 | 0.092615743 | -1.176591718 | 0.239358505 | 0.41496865 |
| ENSG00000187605 | TET3 | 3504.257837 | -0.022374037 | 0.064676316 | -0.345938644 | 0.729388826 | 0.850046366 |
| ENSG00000187607 | ZNF286A | 333.7686006 | 0.052815306 | 0.105860383 | 0.498914749 | 0.617839443 | 0.773479417 |

| | | | | | | | |
|---|---|---|---|---|---|---|---|
| ENSG00000187608 | ISG15 | 527.2937495 | -0.737373031 | 0.109569439 | -6.729732659 | 1.70E-11 | 1.56E-10 |
| ENSG00000187609 | EXD3 | 460.1905609 | 0.128004345 | 0.088805026 | 1.441408797 | 0.149469226 | 0.290849441 |
| ENSG00000187624 | C17orf97 | 97.31211693 | 0.061111685 | 0.182174573 | 0.335456719 | 0.737280589 | 0.855251527 |
| ENSG00000187626 | ZKSCAN4 | 288.3288391 | -0.094000625 | 0.11589868 | -0.811058632 | 0.417332001 | 0.609511434 |
| ENSG00000187627 | RGPD1 | 2.15829339 | -0.831511516 | 1.245620389 | -0.667548094 | 0.5044221 | 0.685187291 |
| ENSG00000187630 | DHRS4L2 | 614.1074034 | -0.187402497 | 0.085696358 | -2.186819843 | 0.028755678 | 0.074833268 |
| ENSG00000187634 | SAMD11 | 1988.051474 | 0.184171573 | 0.059210675 | 3.110445427 | 0.001868054 | 0.006607807 |
| ENSG00000187642 | PERM1 | 16.2939731 | 0.853177135 | 0.443520425 | 1.923647902 | 0.054398732 | 0.128827775 |
| ENSG00000187650 | VMAC | 139.4308894 | -0.18583372 | 0.155479971 | -1.195226111 | 0.2319987 | 0.405953483 |
| ENSG00000187653 | TMSB4XP8 | 1926.215383 | -0.213563285 | 0.060258346 | -3.544127909 | 0.000393914 | 0.001584775 |
| ENSG00000187672 | ERC2 | 22.00352656 | -0.59272776 | 0.37840656 | -1.566378128 | 0.11726012 | 0.241229232 |
| ENSG00000187676 | B3GLCT | 424.3190297 | 0.052754018 | 0.091852424 | 0.574334518 | 0.565741455 | 0.733806909 |
| ENSG00000187678 | SPRY4 | 65.47223551 | -0.735500594 | 0.221206591 | -3.324948826 | 0.000884348 | 0.003343817 |
| ENSG00000187686 | KRT18P59 | 12.19980714 | -1.116380348 | 0.523293313 | -2.133374 | 0.032894058 | 0.084067479 |
| ENSG00000187688 | TRPV2 | 2.520924361 | -0.34840478 | 1.143755188 | -0.304614819 | 0.760659532 | 0.86855119 |
| ENSG00000187689 | AMTN | 14.18402161 | 1.769808251 | 0.520927368 | 3.397418451 | 0.000680249 | 0.002629301 |
| ENSG00000187713 | TMEM203 | 1198.748724 | -0.023389874 | 0.065155062 | -0.35898782 | 0.719604201 | 0.844168774 |
| ENSG00000187720 | THSD4 | 630.0131755 | 1.521574513 | 0.086782347 | 17.53322615 | 7.99E-69 | 7.13E-67 |
| ENSG00000187726 | DNAJB13 | 4.621042184 | -1.70653113 | 0.863807323 | -1.975592338 | 0.048200967 | 0.11613593 |
| ENSG00000187730 | GABRD | 16.65465943 | 1.4180913 | 0.455445414 | 3.113636142 | 0.001847972 | 0.006541041 |
| ENSG00000187735 | TCEA1 | 1056.950814 | -0.047438166 | 0.079363672 | -0.597731497 | 0.550019106 | 0.722358636 |
| ENSG00000187736 | NHEJ1 | 6.149717437 | -0.553480299 | 0.747762309 | -0.74018213 | 0.459189489 | 0.646364784 |
| ENSG00000187741 | FANCA | 2883.241873 | 0.145571703 | 0.056784436 | 2.563584548 | 0.010359747 | 0.030652984 |
| ENSG00000187742 | SECISBP2 | 2056.081759 | 0.01127379 | 0.057185824 | 0.197143084 | 0.843715569 | 0.917672106 |
| ENSG00000187764 | SEMA4D | 1184.816906 | 0.617796326 | 0.066337038 | 9.312992295 | 1.24E-20 | 2.15E-19 |
| ENSG00000187773 | DIPK1C | 5.673355962 | 0.177802916 | 0.75575318 | 0.235265853 | 0.814002397 | 0.901565034 |
| ENSG00000187775 | DNAH17 | 10.06767271 | -0.371490499 | 0.562828417 | -0.660042186 | 0.509226758 | 0.689264918 |
| ENSG00000187778 | MCRS1 | 3136.528904 | -0.093069133 | 0.05639691 | -1.650252348 | 0.098891334 | 0.210091425 |
| ENSG00000187790 | FANCM | 456.6416073 | -0.467192355 | 0.091508754 | -5.105438923 | 3.30E-07 | 2.03E-06 |
| ENSG00000187792 | ZNF70 | 181.0233872 | -0.662567281 | 0.139568526 | -4.747254258 | 2.06E-06 | 1.15E-05 |
| ENSG00000187796 | CARD9 | 36.36317198 | 0.723087931 | 0.317383737 | 2.278276565 | 0.022710105 | 0.061068441 |
| ENSG00000187801 | ZFP69B | 154.7385338 | 0.016454864 | 0.145136885 | 0.113374793 | 0.909733423 | 0.954699394 |
| ENSG00000187808 | SOWAHD | 10.59168043 | 0.846299273 | 0.557534034 | 1.517932936 | 0.129031306 | 0.259669985 |

| | | | | | | | |
|---|---|---|---|---|---|---|---|
| ENSG00000187815 | ZFP69 | 175.1285727 | -0.12022913 | 0.138826618 | -0.866038021 | 0.386469312 | 0.578997258 |
| ENSG00000187837 | H1-2 | 1270.996906 | 0.699189368 | 0.148359595 | 4.712801824 | 2.44E-06 | 1.35E-05 |
| ENSG00000187838 | PLSCR3 | 5.588861252 | -0.176747069 | 0.732822858 | -0.241186621 | 0.80941048 | 0.899083692 |
| ENSG00000187840 | EIF4EBP1 | 1274.613636 | -0.460414443 | 0.062672206 | -7.346389617 | 2.04E-13 | 2.18E-12 |
| ENSG00000187848 | P2RX2 | 159.4030012 | 0.683798403 | 0.157262088 | 4.34814526 | 1.37E-05 | 6.95E-05 |
| ENSG00000187860 | CCDC157 | 128.1017119 | -0.508696895 | 0.157867257 | -3.222307809 | 0.001271624 | 0.004665877 |
| ENSG00000187866 | PABIR1 | 251.0032562 | 0.235225423 | 0.11818214 | 1.990363553 | 0.046550902 | 0.112662762 |
| ENSG00000187867 | PALM3 | 501.6016405 | 0.284757266 | 0.096599884 | 2.947801327 | 0.003200427 | 0.010774551 |
| ENSG00000187870 | RNFT1P3 | 6.260329919 | 0.098323924 | 0.739503631 | 0.132959353 | 0.894225527 | 0.946021656 |
| ENSG00000187889 | FYB2 | 19.72716249 | 4.140051333 | 0.676110258 | 6.123337553 | 9.16E-10 | 7.31E-09 |
| ENSG00000187942 | LDLRAD2 | 1.713821321 | 0.689241395 | 1.352019556 | 0.509786557 | 0.610201005 | NA |
| ENSG00000187944 | C2orf66 | 2.250054558 | 1.68392578 | 1.25365967 | 1.343208066 | 0.179204611 | 0.333925251 |
| ENSG00000187950 | OVCH1 | 9.829063342 | 4.140539751 | 0.958573559 | 4.319480454 | 1.56E-05 | 7.86E-05 |
| ENSG00000187951 | ENSG0000018 | 263.9192874 | -0.961195041 | 0.120048556 | -8.00671892 | 1.18E-15 | 1.49E-14 |
| ENSG00000187952 | HS6ST1P1 | 31.93804254 | 0.213193117 | 0.317410088 | 0.671664593 | 0.501797247 | 0.682834788 |
| ENSG00000187953 | PMS2CL | 319.0732033 | 0.091827794 | 0.107529389 | 0.853978577 | 0.393116859 | 0.586009536 |
| ENSG00000187954 | ZFTRAF1 | 2920.629534 | 0.231963736 | 0.059053765 | 3.928009256 | 8.57E-05 | 0.00038663 |
| ENSG00000187955 | COL14A1 | 23.0195237 | 0.113678993 | 0.378623795 | 0.300242602 | 0.763992112 | 0.870544448 |
| ENSG00000187957 | DNER | 4.380888163 | 1.70845038 | 0.919348205 | 1.858327858 | 0.06312247 | 0.145501353 |
| ENSG00000187959 | CPSF4L | 6.035603111 | -0.112785738 | 0.718443027 | -0.156986336 | 0.875255617 | 0.935312985 |
| ENSG00000187961 | KLHL17 | 775.190092 | 0.077432383 | 0.07805689 | 0.991999337 | 0.321197849 | 0.510693003 |
| ENSG00000187988 | KCTD9P3 | 6.760996336 | -0.637166661 | 0.674125464 | -0.945175186 | 0.344569445 | 0.535530019 |
| ENSG00000187994 | RINL | 52.79503051 | 0.150840185 | 0.246434324 | 0.612090812 | 0.540477676 | 0.714696674 |
| ENSG00000187999 | HNRNPA1P61 | 4.894395154 | 0.808009845 | 0.805832575 | 1.002701889 | 0.316004718 | 0.504745038 |
| ENSG00000188001 | TPRG1 | 12.56266822 | 0.520739867 | 0.500192603 | 1.041078704 | 0.297839022 | 0.484397531 |
| ENSG00000188002 | PDCD6P1 | 835.5069314 | 0.024925162 | 0.077002416 | 0.323693246 | 0.746170285 | 0.859596562 |
| ENSG00000188010 | MORN2 | 218.1162567 | 0.189509589 | 0.124712391 | 1.519573055 | 0.128618314 | 0.259126756 |
| ENSG00000188013 | MEIS3P2 | 48.47599368 | -0.182183788 | 0.254425038 | -0.716060767 | 0.473953828 | 0.659459017 |
| ENSG00000188021 | UBQLN2 | 1698.652789 | 0.038206628 | 0.056608106 | 0.674932099 | 0.499718906 | 0.681057017 |
| ENSG00000188026 | RILPL1 | 521.8927123 | 0.226892875 | 0.092140785 | 2.462458682 | 0.013798807 | 0.039498956 |
| ENSG00000188033 | ZNF490 | 124.8507559 | -0.40654156 | 0.162198389 | -2.506446352 | 0.012195156 | 0.035434691 |
| ENSG00000188039 | NWD1 | 9.280077853 | 1.586468594 | 0.622079592 | 2.550266261 | 0.010764068 | 0.031693969 |
| ENSG00000188042 | ARL4C | 353.0027748 | 0.007510921 | 0.100142984 | 0.075001969 | 0.940213145 | 0.971553584 |

| | | | | | | | |
|---|---|---|---|---|---|---|---|
| ENSG00000188050 | RNF133 | 1.903486631 | 1.4147723 | 1.383173341 | 1.022845263 | 0.306381013 | 0.494086498 |
| ENSG00000188051 | TMEM221 | 9.579655526 | 0.259721151 | 0.571619423 | 0.454360263 | 0.649569548 | 0.796021186 |
| ENSG00000188064 | WNT7B | 1417.570966 | 0.53367865 | 0.067113334 | 7.951901925 | 1.84E-15 | 2.28E-14 |
| ENSG00000188070 | ZFTA | 710.8186959 | 0.416378119 | 0.079437575 | 5.241576385 | 1.59E-07 | 1.01E-06 |
| ENSG00000188078 | ENSG0000018 | 4.062194688 | -0.498717229 | 0.910194302 | -0.547923919 | 0.583744144 | 0.748022046 |
| ENSG00000188092 | GPR89B | 385.2090379 | 0.306212978 | 0.09859819 | 3.105665312 | 0.001898515 | 0.00670752 |
| ENSG00000188095 | MESP2 | 6.619460428 | -1.735733898 | 0.747647078 | -2.32159524 | 0.02025474 | 0.055258949 |
| ENSG00000188112 | C6orf132 | 605.9724686 | -0.135825892 | 0.087337023 | -1.5551926 | 0.119900206 | 0.245406025 |
| ENSG00000188130 | MAPK12 | 990.3094997 | 0.030092795 | 0.073758762 | 0.407989426 | 0.683281438 | 0.820223767 |
| ENSG00000188153 | COL4A5 | 1728.031729 | 1.490880764 | 0.068417409 | 21.79095631 | 2.83E-105 | 5.23E-103 |
| ENSG00000188157 | AGRN | 8623.987335 | 0.817145157 | 0.059427052 | 13.75039029 | 5.07E-43 | 2.29E-41 |
| ENSG00000188158 | NHS | 357.8939922 | 0.654424421 | 0.099977925 | 6.545689146 | 5.92E-11 | 5.22E-10 |
| ENSG00000188163 | FAM166A | 52.59397992 | -0.424500112 | 0.266038302 | -1.595635325 | 0.110570236 | 0.229691317 |
| ENSG00000188167 | TMPPE | 53.19286883 | -0.952490216 | 0.24974274 | -3.813885501 | 0.000136799 | 0.000596383 |
| ENSG00000188176 | SMTNL2 | 4.445796954 | -0.041607623 | 0.854873901 | -0.048671064 | 0.961181436 | 0.982116705 |
| ENSG00000188177 | ZC3H6 | 271.6248229 | -0.065821352 | 0.111241209 | -0.59169936 | 0.554051925 | 0.725423029 |
| ENSG00000188185 | LINC00265 | 349.3708813 | 0.770572814 | 0.125030815 | 6.163063192 | 7.14E-10 | 5.76E-09 |
| ENSG00000188186 | LAMTOR4 | 1641.995762 | -0.299240756 | 0.064079514 | -4.669834958 | 3.01E-06 | 1.65E-05 |
| ENSG00000188191 | PRKAR1B | 2353.268501 | 0.074063368 | 0.060141162 | 1.231492137 | 0.218138856 | 0.387794897 |
| ENSG00000188199 | NUTM2B | 32.41420592 | -0.510770375 | 0.321302867 | -1.589685081 | 0.111905808 | 0.232087446 |
| ENSG00000188211 | NCR3LG1 | 759.8524323 | 0.732856802 | 0.084541487 | 8.668605507 | 4.37E-18 | 6.48E-17 |
| ENSG00000188215 | DCUN1D3 | 230.0932823 | 0.028631859 | 0.125631872 | 0.227902825 | 0.819721786 | 0.904433786 |
| ENSG00000188219 | POTEE | 74.35338778 | -0.158079677 | 0.222816873 | -0.709460083 | 0.478039014 | 0.662842676 |
| ENSG00000188227 | ZNF793 | 238.2091117 | -0.243901353 | 0.145105916 | -1.680850514 | 0.092791954 | 0.199460966 |
| ENSG00000188229 | TUBB4B | 11359.17969 | -0.036854942 | 0.04158744 | -0.886203661 | 0.375507791 | 0.56836558 |
| ENSG00000188234 | AGAP4 | 88.63019573 | -0.561464186 | 0.203518722 | -2.758783966 | 0.005801687 | 0.018372544 |
| ENSG00000188243 | COMMD6 | 443.3803825 | 0.06239058 | 0.094861903 | 0.657699015 | 0.510731555 | 0.690568306 |
| ENSG00000188263 | IL17REL | 91.59618407 | -3.148778833 | 0.252991021 | -12.44620785 | 1.47E-35 | 5.07E-34 |
| ENSG00000188266 | HYKK | 47.07819384 | -0.330078435 | 0.255079127 | -1.294023697 | 0.195657235 | 0.357042159 |
| ENSG00000188277 | C15orf62 | 37.47651921 | -0.247747564 | 0.289178088 | -0.85673007 | 0.391594081 | 0.584537234 |
| ENSG00000188283 | ZNF383 | 211.0735659 | 0.236823873 | 0.12902082 | 1.835547723 | 0.066424575 | 0.152013143 |
| ENSG00000188290 | HES4 | 198.1271864 | 0.079404159 | 0.135777297 | 0.584811755 | 0.558674299 | 0.728636559 |
| ENSG00000188295 | ZNF669 | 458.1536463 | -0.031597208 | 0.091985466 | -0.343502177 | 0.731220704 | 0.851136743 |

| | | | | | | | |
|---|---|---|---|---|---|---|---|
| ENSG00000188306 | LRRIQ4 | 4.865974495 | -0.698844189 | 0.796807836 | -0.877054865 | 0.380456838 | 0.57294557 |
| ENSG00000188312 | CENPP | 289.8758198 | 0.171051616 | 0.115950498 | 1.475212431 | 0.14015544 | 0.2766686 |
| ENSG00000188313 | PLSCR1 | 582.1152625 | 0.439700723 | 0.082730219 | 5.314874385 | 1.07E-07 | 6.90E-07 |
| ENSG00000188315 | C3orf62 | 200.2023933 | -0.031156121 | 0.136639302 | -0.228017276 | 0.81963281 | 0.904433786 |
| ENSG00000188316 | ENO4 | 11.28971931 | -0.782946882 | 0.523382186 | -1.4959372 | 0.134670021 | 0.268544009 |
| ENSG00000188321 | ZNF559 | 135.1149559 | 0.149277483 | 0.171655111 | 0.869636106 | 0.3844993 | 0.576987894 |
| ENSG00000188322 | SBK1 | 1851.546329 | -0.454668376 | 0.060511393 | -7.513764851 | 5.75E-14 | 6.38E-13 |
| ENSG00000188338 | SLC38A3 | 9.981308867 | 1.563368207 | 0.60014514 | 2.604983534 | 0.009187868 | 0.02763191 |
| ENSG00000188342 | GTF2F2 | 939.1454107 | 0.308931782 | 0.07103001 | 4.349313509 | 1.37E-05 | 6.92E-05 |
| ENSG00000188343 | CIBAR1 | 343.0992186 | 0.673963624 | 0.10647294 | 6.329905279 | 2.45E-10 | 2.06E-09 |
| ENSG00000188352 | FOCAD | 1429.169204 | 0.127665527 | 0.064988821 | 1.96442286 | 0.049481074 | 0.118746295 |
| ENSG00000188365 | ENSG0000018 | 2.124765328 | 0.994043223 | 1.241110336 | 0.800930581 | 0.423171835 | 0.614118651 |
| ENSG00000188368 | PRR19 | 129.2426626 | 0.038586558 | 0.171024449 | 0.225620125 | 0.821496887 | 0.905363372 |
| ENSG00000188372 | ZP3 | 452.893381 | 1.168252983 | 0.098003085 | 11.92057355 | 9.25E-33 | 2.86E-31 |
| ENSG00000188375 | H3-5 | 138.052086 | 0.183903853 | 0.172983528 | 1.063129277 | 0.287723329 | 0.471631541 |
| ENSG00000188385 | JAKMIP3 | 53.11925372 | 1.494164269 | 0.258986509 | 5.769274532 | 7.96E-09 | 5.78E-08 |
| ENSG00000188419 | CHM | 854.1643217 | -0.039087591 | 0.073004619 | -0.535412576 | 0.592364596 | 0.754736754 |
| ENSG00000188428 | BLOC1S5 | 304.8921499 | -0.695134339 | 0.110336993 | -6.300102264 | 2.97E-10 | 2.49E-09 |
| ENSG00000188451 | SRP72P2 | 20.75561694 | 0.118067471 | 0.390122006 | 0.302642429 | 0.762162376 | 0.869504919 |
| ENSG00000188452 | CERKL | 24.3768183 | 1.522868833 | 0.386236751 | 3.942837727 | 8.05E-05 | 0.000365204 |
| ENSG00000188459 | WASF4P | 233.6299621 | -0.133789073 | 0.121796309 | -1.098465739 | 0.27200117 | 0.454041488 |
| ENSG00000188460 | ACTBP11 | 11.52725582 | -0.30230724 | 0.551376359 | -0.548277479 | 0.583501389 | 0.747919867 |
| ENSG00000188467 | SLC24A5 | 2.882647363 | -1.089595017 | 1.061328574 | -1.026633074 | 0.304593274 | 0.491998993 |
| ENSG00000188483 | IER5L | 509.8792436 | 0.050840344 | 0.102825249 | 0.494434436 | 0.620999399 | 0.775573073 |
| ENSG00000188486 | H2AX | 4277.151287 | 0.256639385 | 0.052362005 | 4.901252067 | 9.52E-07 | 5.52E-06 |
| ENSG00000188488 | SERPINA5 | 4.506218195 | 1.426986697 | 0.90622143 | 1.574655651 | 0.11533595 | 0.237866123 |
| ENSG00000188493 | ACTMAP | 760.6489499 | -0.628902946 | 0.078178621 | -8.044436368 | 8.66E-16 | 1.10E-14 |
| ENSG00000188501 | LCTL | 13.880563 | -0.639841987 | 0.493614442 | -1.296238384 | 0.194893369 | 0.355975658 |
| ENSG00000188505 | NCCRP1 | 3.608850623 | 1.6823083 | 1.021146869 | 1.647469479 | 0.099461585 | 0.211048816 |
| ENSG00000188508 | KRTDAP | 4.357280235 | -0.645490703 | 0.833987311 | -0.773981443 | 0.438941765 | 0.628429389 |
| ENSG00000188512 | ENSG0000018 | 12.55641833 | -1.006579645 | 0.534276664 | -1.884004511 | 0.059564365 | 0.138693315 |
| ENSG00000188522 | FAM83G | 2015.369744 | 0.202834545 | 0.057913752 | 3.502355457 | 0.000461164 | 0.001836968 |
| ENSG00000188529 | SRSF10 | 3089.018087 | -0.110706275 | 0.060161058 | -1.840165041 | 0.065744011 | 0.150651044 |

| | | | | | | | |
|---|---|---|---|---|---|---|---|
| ENSG00000188542 | DUSP28 | 141.1656356 | 0.173951858 | 0.161648839 | 1.076109541 | 0.281878269 | 0.465366562 |
| ENSG00000188549 | CCDC9B | 74.2039904 | 1.285806305 | 0.220686034 | 5.82640542 | 5.66E-09 | 4.19E-08 |
| ENSG00000188554 | NBR1 | 5037.91764 | 0.21788598 | 0.046179753 | 4.718214503 | 2.38E-06 | 1.32E-05 |
| ENSG00000188559 | RALGAPA2 | 1233.802349 | -0.188450291 | 0.075404642 | -2.49918686 | 0.012447866 | 0.036054803 |
| ENSG00000188566 | NDOR1 | 714.8209075 | -0.21181134 | 0.084024263 | -2.52083545 | 0.011707659 | 0.034178377 |
| ENSG00000188581 | KRTAP1-1 | 2.762042715 | 2.845766318 | 1.356905691 | 2.097246947 | 0.035971721 | 0.090730211 |
| ENSG00000188599 | NPIPP1 | 48.03555846 | -0.108976436 | 0.253684524 | -0.429574632 | 0.667505094 | 0.808691146 |
| ENSG00000188603 | CLN3 | 21.54298436 | 0.694445333 | 0.392638386 | 1.768663885 | 0.076949983 | 0.171481698 |
| ENSG00000188610 | FAM72B | 169.2530056 | -0.204992604 | 0.142970745 | -1.43380804 | 0.15162705 | 0.293976042 |
| ENSG00000188611 | ASAH2 | 36.04100414 | -0.162243295 | 0.290204999 | -0.559064439 | 0.576117743 | 0.742521548 |
| ENSG00000188612 | SUMO2 | 2537.757451 | 0.091252325 | 0.053659868 | 1.700569311 | 0.089023891 | 0.192788451 |
| ENSG00000188613 | NANOS1 | 350.7657302 | 0.61480867 | 0.102587356 | 5.993025764 | 2.06E-09 | 1.59E-08 |
| ENSG00000188620 | HMX3 | 3.073537596 | -1.959634142 | 1.13894859 | -1.720564175 | 0.085329939 | 0.186245998 |
| ENSG00000188636 | RTL6 | 2245.040306 | -0.00057272 | 0.059684482 | -0.009595797 | 0.992343779 | 0.99594725 |
| ENSG00000188641 | DPYD | 164.8971022 | 0.70133398 | 0.145691991 | 4.813812858 | 1.48E-06 | 8.38E-06 |
| ENSG00000188643 | S100A16 | 4367.685791 | -1.244321714 | 0.05911622 | -21.04873609 | 2.35E-98 | 3.69E-96 |
| ENSG00000188647 | PTAR1 | 627.3328656 | -0.171693877 | 0.081012943 | -2.11933884 | 0.034061841 | 0.08652183 |
| ENSG00000188649 | CC2D2B | 2.877340775 | -2.395815416 | 1.197696259 | -2.000353093 | 0.04546215 | 0.110395016 |
| ENSG00000188659 | SAXO2 | 37.85064422 | 1.145769708 | 0.296784272 | 3.860614654 | 0.000113102 | 0.000500205 |
| ENSG00000188672 | RHCE | 4.32602123 | -2.183617195 | 0.961471059 | -2.271121084 | 0.023139649 | 0.062078507 |
| ENSG00000188677 | PARVB | 203.3923622 | 0.204518329 | 0.126706454 | 1.614111373 | 0.106503271 | 0.222989159 |
| ENSG00000188681 | TEKT4P2 | 742.0982732 | 0.309133404 | 0.082236823 | 3.759063053 | 0.000170551 | 0.000732666 |
| ENSG00000188687 | SLC4A5 | 15.73727735 | -0.70144412 | 0.473843862 | -1.480327542 | 0.138785857 | 0.27476781 |
| ENSG00000188690 | UROS | 1121.785381 | -0.392951312 | 0.063196314 | -6.21794669 | 5.04E-10 | 4.12E-09 |
| ENSG00000188693 | CYP51A1-AS1 | 7.515975433 | -0.767695309 | 0.643386462 | -1.193210232 | 0.232787048 | 0.40696044 |
| ENSG00000188706 | ZDHHC9 | 3307.21274 | 0.29368485 | 0.049160169 | 5.974040667 | 2.31E-09 | 1.78E-08 |
| ENSG00000188707 | ZBED10P | 684.7735278 | -0.209149023 | 0.078281826 | -2.671744298 | 0.007545812 | 0.023203879 |
| ENSG00000188725 | SMIM15 | 1354.072709 | -0.068637271 | 0.061791043 | -1.110796455 | 0.266655973 | 0.447744695 |
| ENSG00000188732 | FAM221A | 438.532389 | -1.786527103 | 0.120035921 | -14.88327061 | 4.23E-50 | 2.36E-48 |
| ENSG00000188735 | TMEM120B | 616.2271125 | 0.730363157 | 0.093628288 | 7.800667689 | 6.16E-15 | 7.36E-14 |
| ENSG00000188738 | FSIP2 | 9.7616159 | -0.049941386 | 0.585002643 | -0.085369505 | 0.931967636 | 0.967452912 |
| ENSG00000188739 | RBM34 | 15.93968292 | 0.155671089 | 0.462455431 | 0.336618576 | 0.736404451 | 0.854568904 |
| ENSG00000188747 | NOXA1 | 442.6148452 | -0.046336646 | 0.094449546 | -0.490596809 | 0.623711644 | 0.777539482 |

| | | | | | | | |
|---|---|---|---|---|---|---|---|
| ENSG00000188760 | TMEM198 | 163.704003 | 0.033911522 | 0.1468231 | 0.230968572 | 0.817339218 | 0.903254867 |
| ENSG00000188761 | BCL2L15 | 31.79589024 | 0.22346522 | 0.31246638 | 0.715165646 | 0.474506694 | 0.659868063 |
| ENSG00000188763 | FZD9 | 10.2219627 | 1.239659183 | 0.586711351 | 2.112894494 | 0.034609799 | 0.087777644 |
| ENSG00000188765 | TMSB4XP2 | 15.56329964 | -0.610246352 | 0.450861029 | -1.353513193 | 0.175891738 | 0.32919609 |
| ENSG00000188766 | SPRED3 | 106.9842532 | -0.239820377 | 0.178826676 | -1.341077194 | 0.179895391 | 0.334858834 |
| ENSG00000188779 | SKOR1 | 16.45562496 | -0.020783074 | 0.433517077 | -0.047940613 | 0.961763572 | 0.982361474 |
| ENSG00000188782 | CATSPER4 | 2.055806464 | 1.287369868 | 1.361011971 | 0.945891656 | 0.34420385 | 0.535142796 |
| ENSG00000188785 | ZNF548 | 262.9860266 | 0.4546005 | 0.113850939 | 3.992944658 | 6.53E-05 | 0.000299883 |
| ENSG00000188786 | MTF1 | 1104.875717 | -0.114060917 | 0.070743816 | -1.612309368 | 0.106894636 | 0.223635471 |
| ENSG00000188801 | ZNF322P1 | 4.677903207 | 0.190377613 | 0.807522836 | 0.235755083 | 0.813622724 | 0.901448243 |
| ENSG00000188807 | TMEM201 | 614.5235127 | 0.421049529 | 0.084266426 | 4.996646352 | 5.83E-07 | 3.48E-06 |
| ENSG00000188811 | NHLRC3 | 666.812369 | -0.084975837 | 0.091040503 | -0.933384977 | 0.350621234 | 0.541977875 |
| ENSG00000188817 | SNTN | 43.18791743 | 1.493445418 | 0.30349595 | 4.920808399 | 8.62E-07 | 5.03E-06 |
| ENSG00000188818 | ZDHHC11 | 205.8161941 | -0.018842437 | 0.128690957 | -0.14641617 | 0.883592863 | 0.939712848 |
| ENSG00000188822 | CNR2 | 1.782833636 | -0.95890723 | 1.311433181 | -0.731190307 | 0.46466292 | NA |
| ENSG00000188825 | LINC00910 | 93.98641471 | -0.537935681 | 0.184789604 | -2.911071125 | 0.00360192 | 0.011995547 |
| ENSG00000188827 | SLX4 | 686.7847248 | -0.219195885 | 0.076024545 | -2.883225218 | 0.003936259 | 0.012992487 |
| ENSG00000188833 | ENTPD8 | 68.70904779 | 3.69852683 | 0.332017237 | 11.13956269 | 8.05E-29 | 2.13E-27 |
| ENSG00000188846 | RPL14 | 8090.697446 | 0.259816094 | 0.045186456 | 5.749866539 | 8.93E-09 | 6.45E-08 |
| ENSG00000188848 | BEND4 | 2.461187566 | -3.818633536 | 1.526617455 | -2.501368973 | 0.012371421 | 0.035860551 |
| ENSG00000188856 | RPSAP47 | 165.3772041 | -0.133869439 | 0.147492641 | -0.907634698 | 0.364071256 | 0.55674919 |
| ENSG00000188859 | FAM78B | 54.37310649 | 1.821775558 | 0.266675892 | 6.831422022 | 8.41E-12 | 7.94E-11 |
| ENSG00000188868 | ZNF563 | 83.49812207 | -0.147822962 | 0.200033717 | -0.738990225 | 0.459912933 | 0.646782557 |
| ENSG00000188873 | RPL10AP2 | 44.00169141 | 0.222614239 | 0.277461016 | 0.802326189 | 0.422364297 | 0.613322251 |
| ENSG00000188878 | FBF1 | 164.0696838 | -0.072259389 | 0.144957318 | -0.498487345 | 0.618140587 | 0.773612145 |
| ENSG00000188883 | KLRG2 | 68.4287623 | 0.546556621 | 0.216173085 | 2.528328724 | 0.011460699 | 0.033512866 |
| ENSG00000188886 | ASTL | 25.79780319 | -0.932797713 | 0.365743582 | -2.550414439 | 0.010759493 | 0.031684566 |
| ENSG00000188895 | MSL1 | 5275.806287 | 0.016710298 | 0.045544157 | 0.366903225 | 0.713691206 | 0.840800958 |
| ENSG00000188897 | ENSG0000018 | 142.4041097 | -0.732438238 | 0.17291285 | -4.235880895 | 2.28E-05 | 0.00011207 |
| ENSG00000188906 | LRRK2 | 14.15589995 | 0.28925893 | 0.485249114 | 0.596103983 | 0.551105763 | 0.723161782 |
| ENSG00000188910 | GJB3 | 83.04927917 | 2.110108446 | 0.225306904 | 9.365485056 | 7.57E-21 | 1.33E-19 |
| ENSG00000188916 | INSYN2A | 14.96944709 | 0.86114871 | 0.467541632 | 1.841865304 | 0.065494855 | 0.150185123 |
| ENSG00000188917 | TRMT2B | 1897.640695 | -0.004185641 | 0.058860436 | -0.071111274 | 0.943309195 | 0.973031555 |

| | | | | | | | |
|---|---|---|---|---|---|---|---|
| ENSG00000188931 | CFAP126 | 2.714799774 | -0.554151751 | 1.057100723 | -0.524218495 | 0.600126578 | 0.760140451 |
| ENSG00000188938 | FAM120AOS | 3350.046771 | 0.080846657 | 0.049242171 | 1.641817467 | 0.100627838 | 0.21295227 |
| ENSG00000188958 | UTS2B | 2.00180239 | -0.701213744 | 1.276558814 | -0.54929999 | 0.582799594 | 0.747270845 |
| ENSG00000188959 | C9orf152 | 199.428982 | -0.064539332 | 0.129336109 | -0.499004745 | 0.617776041 | 0.773442252 |
| ENSG00000188976 | NOC2L | 4434.388426 | 0.14620259 | 0.049946126 | 2.927205791 | 0.003420225 | 0.011450665 |
| ENSG00000188981 | MSANTD1 | 14.51475833 | 0.368068764 | 0.455924869 | 0.807301354 | 0.419492884 | 0.610929766 |
| ENSG00000188985 | DHFRP1 | 653.1289783 | 0.023695835 | 0.080628507 | 0.293889045 | 0.768842689 | 0.873855116 |
| ENSG00000188986 | NELFB | 2713.265206 | -0.121473684 | 0.061054017 | -1.989610034 | 0.04663391 | 0.112822389 |
| ENSG00000188993 | LRRC66 | 3.987099903 | 0.301904963 | 0.877685844 | 0.343978389 | 0.730862539 | 0.850978918 |
| ENSG00000188994 | ZNF292 | 1106.341624 | 0.00671626 | 0.070148267 | 0.095743781 | 0.923724068 | 0.963279648 |
| ENSG00000188997 | KCTD21 | 669.5343017 | -0.442048064 | 0.079058108 | -5.591432337 | 2.25E-08 | 1.56E-07 |
| ENSG00000189001 | SBSN | 16.35766453 | -0.248523948 | 0.446317235 | -0.556832514 | 0.577641867 | 0.743483389 |
| ENSG00000189007 | ADAT2 | 107.30257 | 0.375239736 | 0.188262749 | 1.99317038 | 0.046242793 | 0.111994265 |
| ENSG00000189014 | SHLD2P3 | 4.190366827 | 0.289187951 | 0.903997761 | 0.319898968 | 0.74904492 | 0.861221455 |
| ENSG00000189042 | ZNF567 | 302.6125312 | 0.695888326 | 0.108289774 | 6.426168411 | 1.31E-10 | 1.12E-09 |
| ENSG00000189043 | NDUFA4 | 4069.207949 | -1.214714374 | 0.052910349 | -22.9579731 | 1.23E-116 | 2.71E-114 |
| ENSG00000189045 | ANKDD1B | 23.69824673 | 0.294086839 | 0.360952368 | 0.814752485 | 0.415214004 | 0.607307399 |
| ENSG00000189046 | ALKBH2 | 649.1445484 | -0.671867427 | 0.079858176 | -8.413257909 | 3.99E-17 | 5.53E-16 |
| ENSG00000189050 | RNFT1 | 345.3351724 | -0.264699659 | 0.121531691 | -2.178029922 | 0.029403809 | 0.076320817 |
| ENSG00000189057 | FAM111B | 1305.378979 | -0.360097742 | 0.064903502 | -5.548202016 | 2.89E-08 | 1.98E-07 |
| ENSG00000189058 | APOD | 197.0019731 | 1.385853508 | 0.153501693 | 9.028262067 | 1.74E-19 | 2.84E-18 |
| ENSG00000189060 | H1-0 | 10330.8749 | -0.271840691 | 0.039573752 | -6.869217007 | 6.46E-12 | 6.15E-11 |
| ENSG00000189067 | LITAF | 4108.312186 | -0.648999461 | 0.0468105 | -13.86439927 | 1.04E-43 | 4.80E-42 |
| ENSG00000189077 | TMEM120A | 596.1541655 | 0.572109223 | 0.082685548 | 6.919095738 | 4.55E-12 | 4.40E-11 |
| ENSG00000189079 | ARID2 | 1711.624224 | -0.107225083 | 0.070129319 | -1.528962286 | 0.126273794 | 0.25542023 |
| ENSG00000189089 | RIMKLBP1 | 5.773566996 | -0.146922743 | 0.745773664 | -0.197007148 | 0.843821944 | 0.917708773 |
| ENSG00000189091 | SF3B3 | 11945.95121 | -0.106544238 | 0.04230685 | -2.518368484 | 0.01178999 | 0.034405595 |
| ENSG00000189108 | IL1RAPL2 | 42.23493166 | -1.740117634 | 0.291357988 | -5.972438398 | 2.34E-09 | 1.80E-08 |
| ENSG00000189114 | BLOC1S3 | 816.2717589 | -0.127050832 | 0.070784288 | -1.794901605 | 0.072669378 | 0.163707703 |
| ENSG00000189120 | SP6 | 126.378948 | 0.319977019 | 0.171262262 | 1.868345164 | 0.061713975 | 0.142833655 |
| ENSG00000189127 | ANKRD34B | 127.1384186 | -0.311274987 | 0.160065033 | -1.944678244 | 0.051813716 | 0.12350443 |
| ENSG00000189136 | UBE2Q2P1 | 105.9252791 | 0.21005043 | 0.194715899 | 1.078753359 | 0.280697692 | 0.464119665 |
| ENSG00000189143 | CLDN4 | 4420.367829 | 0.82717604 | 0.061129228 | 13.53159635 | 1.02E-41 | 4.47E-40 |

| | | | | | | | |
|---|---|---|---|---|---|---|---|
| ENSG00000189144 | ZNF573 | 59.95386404 | -0.001710138 | 0.230658384 | -0.007414161 | 0.99408441 | 0.996533822 |
| ENSG00000189149 | ABCA15P | 6.561355186 | 0.451621702 | 0.705764718 | 0.63990405 | 0.52223498 | 0.69911709 |
| ENSG00000189157 | FAM47E | 30.10302205 | 0.396586288 | 0.326266445 | 1.215528885 | 0.224164439 | 0.395666747 |
| ENSG00000189159 | JPT1 | 4219.351667 | -0.004174302 | 0.045999599 | -0.090746479 | 0.927694038 | 0.965269707 |
| ENSG00000189164 | ZNF527 | 197.2989215 | 0.527291995 | 0.134321185 | 3.925605578 | 8.65E-05 | 0.000389975 |
| ENSG00000189166 | TNRC18P3 | 11.42921992 | -0.416052663 | 0.552018043 | -0.753693957 | 0.451033011 | 0.639118987 |
| ENSG00000189167 | ZAR1L | 4.694968704 | -0.551936023 | 0.832030692 | -0.663360172 | 0.507099908 | 0.687359482 |
| ENSG00000189171 | S100A13 | 818.5004723 | 0.089431179 | 0.072677003 | 1.230529267 | 0.218498974 | 0.38819443 |
| ENSG00000189180 | ZNF33A | 1478.956061 | -0.229529692 | 0.063189499 | -3.632402481 | 0.000280795 | 0.001160827 |
| ENSG00000189184 | PCDH18 | 10.0469638 | -0.316155731 | 0.565196287 | -0.559373333 | 0.575906957 | 0.742333292 |
| ENSG00000189190 | ZNF600 | 508.4269036 | 0.366352647 | 0.088647666 | 4.13268237 | 3.59E-05 | 0.000171145 |
| ENSG00000189195 | BTBD8 | 92.537671 | 0.457228142 | 0.187109576 | 2.443638379 | 0.014539992 | 0.041336897 |
| ENSG00000189212 | DPY19L2P1 | 30.67862972 | -3.951492704 | 0.494019301 | -7.998660566 | 1.26E-15 | 1.58E-14 |
| ENSG00000189221 | MAOA | 899.4782198 | -1.664970415 | 0.079181568 | -21.02724726 | 3.69E-98 | 5.77E-96 |
| ENSG00000189223 | PAX8-AS1 | 1507.974573 | 0.188188435 | 0.061096763 | 3.080170318 | 0.002068823 | 0.00724783 |
| ENSG00000189227 | C15orf61 | 227.6522751 | 0.292887262 | 0.122794971 | 2.385173102 | 0.017071087 | 0.047658601 |
| ENSG00000189241 | TSPYL1 | 7020.094417 | -0.264803005 | 0.044792289 | -5.911798952 | 3.38E-09 | 2.57E-08 |
| ENSG00000189266 | PNRC2 | 2038.034429 | -0.031720862 | 0.060672063 | -0.522824843 | 0.60109615 | 0.761032379 |
| ENSG00000189269 | DRICH1 | 9.975340081 | -1.379759169 | 0.578375832 | -2.38557542 | 0.017052425 | 0.047623891 |
| ENSG00000189283 | FHIT | 97.26610724 | -0.726536202 | 0.181742375 | -3.997615863 | 6.40E-05 | 0.00029444 |
| ENSG00000189298 | ZKSCAN3 | 101.1477749 | -0.913633665 | 0.179492104 | -5.090105055 | 3.58E-07 | 2.19E-06 |
| ENSG00000189306 | RRP7A | 1933.40464 | 0.008924185 | 0.063844364 | 0.139780303 | 0.888833576 | 0.942772834 |
| ENSG00000189308 | LIN54 | 895.6224608 | -0.272166037 | 0.070716558 | -3.848688962 | 0.000118752 | 0.000523073 |
| ENSG00000189319 | FAM53B | 1775.372644 | -0.40725933 | 0.066176492 | -6.15413902 | 7.55E-10 | 6.07E-09 |
| ENSG00000189334 | S100A14 | 2411.15153 | -0.150128666 | 0.054202425 | -2.76977767 | 0.005609457 | 0.017832724 |
| ENSG00000189337 | KAZN | 342.6123201 | -0.815594456 | 0.103215818 | -7.901835893 | 2.75E-15 | 3.37E-14 |
| ENSG00000189339 | SLC35E2B | 2196.455939 | 0.094019078 | 0.057968134 | 1.621909702 | 0.104822684 | 0.220197986 |
| ENSG00000189343 | RPS2P46 | 2140.63941 | 0.138915845 | 0.072074484 | 1.927392857 | 0.053930689 | 0.127897735 |
| ENSG00000189350 | TOGARAM2 | 20.3795269 | 2.875846161 | 0.51263182 | 5.609964205 | 2.02E-08 | 1.41E-07 |
| ENSG00000189362 | NEMP2 | 152.4135086 | -0.732218797 | 0.145991732 | -5.015481265 | 5.29E-07 | 3.18E-06 |
| ENSG00000189366 | ALG1L1P | 277.2972415 | -0.400513693 | 0.113890797 | -3.516646698 | 0.000437035 | 0.001747836 |
| ENSG00000189369 | GSPT2 | 2.216421018 | 0.268153754 | 1.18676648 | 0.225953259 | 0.821237774 | 0.905208271 |
| ENSG00000189376 | C8orf76 | 74.92214371 | -0.065598259 | 0.20308877 | -0.323002888 | 0.746693055 | 0.859607712 |

| | | | | | | | |
|---|---|---|---|---|---|---|---|
| ENSG00000189403 | HMGB1 | 7338.55821 | -0.272968872 | 0.065000914 | -4.19946204 | 2.68E-05 | 0.000130393 |
| ENSG00000189409 | MMP23B | 2.478328576 | 0.992463939 | 1.149240252 | 0.863582647 | 0.3878172 | 0.580492583 |
| ENSG00000189410 | SH2D5 | 47.6929113 | 0.896526649 | 0.260300727 | 3.444195712 | 0.000572761 | 0.002242173 |
| ENSG00000189419 | SPATA41 | 10.91855543 | 0.221787644 | 0.555075811 | 0.399562797 | 0.689478562 | 0.824525492 |
| ENSG00000189423 | USP32P3 | 44.43089767 | 0.049301134 | 0.296448897 | 0.166305676 | 0.8679164 | 0.931076558 |
| ENSG00000189431 | RASSF10 | 16.05008152 | -1.260246264 | 0.472458276 | -2.667423404 | 0.007643531 | 0.023472905 |
| ENSG00000196071 | OR2L13 | 2.762031574 | 2.159874812 | 1.219116175 | 1.771672673 | 0.076448908 | 0.170530723 |
| ENSG00000196072 | BLOC1S2 | 730.4250962 | -0.087563081 | 0.074439939 | -1.176291687 | 0.239478337 | 0.41508372 |
| ENSG00000196074 | SYCP2 | 1840.171851 | -0.50390514 | 0.078781498 | -6.396237099 | 1.59E-10 | 1.35E-09 |
| ENSG00000196081 | ZNF724 | 37.6662727 | -0.088269852 | 0.291490284 | -0.302822621 | 0.762025043 | 0.86943472 |
| ENSG00000196083 | IL1RAP | 145.2738066 | 0.12085695 | 0.147866791 | 0.817336663 | 0.41373606 | 0.605918362 |
| ENSG00000196090 | PTPRT | 7.499954006 | 2.360603842 | 0.763452228 | 3.092012514 | 0.001988045 | 0.006993674 |
| ENSG00000196092 | PAX5 | 35.73631193 | 0.328691939 | 0.294697059 | 1.11535534 | 0.264698179 | 0.445631604 |
| ENSG00000196104 | SPOCK3 | 3.338287179 | -5.288498101 | 1.44247582 | -3.666264647 | 0.000246119 | 0.001025981 |
| ENSG00000196110 | ZNF699 | 91.38985065 | 0.102260855 | 0.188615787 | 0.542164878 | 0.587704928 | 0.751132537 |
| ENSG00000196116 | TDRD7 | 821.4675618 | 0.323400195 | 0.073268452 | 4.413907839 | 1.02E-05 | 5.25E-05 |
| ENSG00000196118 | CFAP119 | 188.5122653 | 0.177020981 | 0.143830634 | 1.230759928 | 0.218412667 | 0.388071147 |
| ENSG00000196119 | OR8A1 | 14.38380843 | 0.406365771 | 0.461304618 | 0.88090549 | 0.378368976 | 0.571100536 |
| ENSG00000196123 | MATCAP1 | 1040.6885 | 0.237040658 | 0.07529433 | 3.148187352 | 0.001642864 | 0.005884961 |
| ENSG00000196126 | HLA-DRB1 | 19.48793606 | 4.391613408 | 0.72130409 | 6.08843547 | 1.14E-09 | 9.03E-09 |
| ENSG00000196132 | MYT1 | 11.87288189 | 0.214573262 | 0.514462873 | 0.417082113 | 0.676618348 | 0.815811253 |
| ENSG00000196136 | SERPINA3 | 3.515759211 | 2.463449194 | 1.139993051 | 2.160933518 | 0.030700475 | 0.079282776 |
| ENSG00000196139 | AKR1C3 | 30.45388606 | 1.921692897 | 0.357500976 | 5.37535007 | 7.64E-08 | 5.00E-07 |
| ENSG00000196141 | SPATS2L | 1129.806626 | -0.506657097 | 0.064386488 | -7.868997219 | 3.57E-15 | 4.35E-14 |
| ENSG00000196150 | ZNF250 | 526.9714294 | 0.193268105 | 0.089330654 | 2.16351383 | 0.030501681 | 0.078813761 |
| ENSG00000196151 | WDSUB1 | 283.7089385 | -0.046063995 | 0.112781102 | -0.408437172 | 0.682952748 | 0.820000839 |
| ENSG00000196152 | ZNF79 | 242.0721997 | -0.200110866 | 0.118548488 | -1.688008589 | 0.09140958 | 0.196980671 |
| ENSG00000196154 | S100A4 | 63.32707682 | 0.117379256 | 0.224742611 | 0.522283049 | 0.60147327 | 0.761283686 |
| ENSG00000196155 | PLEKHG4 | 1125.933009 | 0.625256133 | 0.066917993 | 9.343617555 | 9.31E-21 | 1.62E-19 |
| ENSG00000196159 | FAT4 | 330.8070123 | -0.938295759 | 0.118285917 | -7.932438487 | 2.15E-15 | 2.65E-14 |
| ENSG00000196172 | ZNF681 | 148.0015459 | 0.305773877 | 0.150846775 | 2.027049476 | 0.042657353 | 0.104681181 |
| ENSG00000196177 | ACADSB | 759.778646 | -0.59713617 | 0.075706349 | -7.887530975 | 3.08E-15 | 3.77E-14 |
| ENSG00000196182 | STK40 | 1833.308033 | -0.198388959 | 0.060206463 | -3.295143886 | 0.000983713 | 0.003686714 |

| | | | | | | | |
|---|---|---|---|---|---|---|---|
| ENSG00000196183 | RPS2P4 | 11.50516785 | 0.118829133 | 0.535068455 | 0.222082113 | 0.824249962 | 0.907438409 |
| ENSG00000196187 | TMEM63A | 1719.769657 | 0.134117832 | 0.062821826 | 2.134892304 | 0.032769808 | 0.083815297 |
| ENSG00000196189 | SEMA4A | 956.7324622 | -0.427618888 | 0.070606591 | -6.056359381 | 1.39E-09 | 1.09E-08 |
| ENSG00000196199 | MPHOSPH8 | 2054.193975 | 0.45059135 | 0.057956092 | 7.774702066 | 7.56E-15 | 8.95E-14 |
| ENSG00000196204 | RNF216P1 | 377.17278 | 0.157746846 | 0.098408164 | 1.602985368 | 0.108937885 | 0.226937439 |
| ENSG00000196205 | EEF1A1P5 | 14996.08559 | 0.099191364 | 0.12100548 | 0.819726217 | 0.412372201 | 0.604534362 |
| ENSG00000196208 | GREB1 | 399.1989493 | 0.215452168 | 0.217472616 | 0.990709414 | 0.321827493 | 0.511552293 |
| ENSG00000196214 | ZNF766 | 742.0602246 | 0.026902452 | 0.074679314 | 0.360239681 | 0.718667903 | 0.843510457 |
| ENSG00000196218 | RYR1 | 2.641042696 | 0.167910459 | 1.052988318 | 0.159460894 | 0.873305766 | 0.934604743 |
| ENSG00000196220 | SRGAP3 | 1330.122976 | -0.728989304 | 0.066317492 | -10.9924137 | 4.16E-28 | 1.07E-26 |
| ENSG00000196227 | FAM217B | 1257.397689 | -0.273245739 | 0.065053761 | -4.200306559 | 2.67E-05 | 0.000129963 |
| ENSG00000196230 | TUBB | 25136.26854 | -0.443039866 | 0.043072108 | -10.28600386 | 8.15E-25 | 1.80E-23 |
| ENSG00000196233 | LCOR | 2139.328978 | -1.358462593 | 0.063064897 | -21.54070914 | 6.47E-103 | 1.11E-100 |
| ENSG00000196235 | SUPT5H | 5505.110796 | -0.043500217 | 0.045657104 | -0.952758993 | 0.340712196 | 0.531768796 |
| ENSG00000196236 | XPNPEP3 | 730.7993029 | -0.254478453 | 0.077467268 | -3.28498035 | 0.001019895 | 0.003814214 |
| ENSG00000196247 | ZNF107 | 975.5600313 | 0.61637226 | 0.072171207 | 8.540417808 | 1.34E-17 | 1.92E-16 |
| ENSG00000196262 | PPIA | 10190.32331 | 0.415227012 | 0.062135433 | 6.6826123 | 2.35E-11 | 2.13E-10 |
| ENSG00000196267 | ZNF836 | 308.931369 | 0.293396492 | 0.105879438 | 2.771043165 | 0.005587702 | 0.017766027 |
| ENSG00000196268 | ZNF493 | 119.0047217 | 0.433239857 | 0.167814309 | 2.581662201 | 0.009832577 | 0.02927821 |
| ENSG00000196275 | GTF2IRD2 | 102.7281995 | 0.58665391 | 0.177555888 | 3.304052128 | 0.000952981 | 0.003581485 |
| ENSG00000196284 | SUPT3H | 142.6075514 | -0.933923711 | 0.160858596 | -5.805867604 | 6.40E-09 | 4.70E-08 |
| ENSG00000196290 | NIF3L1 | 968.1649471 | -0.163001843 | 0.066774741 | -2.441070402 | 0.0146438 | 0.041601085 |
| ENSG00000196295 | GARS1-DT | 242.9230318 | -1.279335399 | 0.130857991 | -9.776517192 | 1.42E-22 | 2.75E-21 |
| ENSG00000196296 | ATP2A1 | 39.22221284 | -0.435564732 | 0.293476554 | -1.484155126 | 0.137767772 | 0.273295804 |
| ENSG00000196305 | IARS1 | 6984.832616 | -0.021954337 | 0.056659225 | -0.387480366 | 0.698400617 | 0.830810911 |
| ENSG00000196312 | MFSD14CP | 1006.963531 | 0.422858499 | 0.071900168 | 5.88118928 | 4.07E-09 | 3.06E-08 |
| ENSG00000196313 | POM121 | 2428.734175 | 0.675649843 | 0.058713266 | 11.50761803 | 1.21E-30 | 3.43E-29 |
| ENSG00000196323 | ZBTB44 | 1048.205562 | 0.195922909 | 0.07044533 | 2.781205064 | 0.005415752 | 0.017276782 |
| ENSG00000196335 | STK31 | 74.06971711 | -1.086564142 | 0.216907213 | -5.00934997 | 5.46E-07 | 3.27E-06 |
| ENSG00000196337 | CGB7 | 7.807319298 | -1.80995671 | 0.709305264 | -2.551731675 | 0.010718903 | 0.031577198 |
| ENSG00000196338 | NLGN3 | 30.61519995 | -0.407739852 | 0.313733276 | -1.299638526 | 0.193724889 | 0.354474671 |
| ENSG00000196352 | CD55 | 1793.083344 | 0.337209026 | 0.058713463 | 5.743299868 | 9.28E-09 | 6.70E-08 |
| ENSG00000196353 | CPNE4 | 92.28535906 | 2.866442149 | 0.243069353 | 11.79269257 | 4.26E-32 | 1.28E-30 |

| | | | | | | | |
|---|---|---|---|---|---|---|---|
| ENSG00000196357 | ZNF565 | 110.8378435 | 0.112191723 | 0.16917533 | 0.663168344 | 0.507222744 | 0.687444744 |
| ENSG00000196358 | NTNG2 | 2.395673438 | -1.067559832 | 1.145019752 | -0.932350582 | 0.351155375 | 0.542458202 |
| ENSG00000196363 | WDR5 | 2807.048059 | 0.093314694 | 0.049013697 | 1.903849336 | 0.056929809 | 0.13360305 |
| ENSG00000196365 | LONP1 | 4090.589053 | -0.543898365 | 0.045145473 | -12.04768336 | 1.99E-33 | 6.32E-32 |
| ENSG00000196366 | C9orf163 | 41.28560074 | 0.679132576 | 0.279404909 | 2.430639381 | 0.015072208 | 0.042685981 |
| ENSG00000196367 | TRRAP | 4999.937233 | -0.416037911 | 0.067985704 | -6.119491072 | 9.39E-10 | 7.48E-09 |
| ENSG00000196368 | NUDT11 | 60.34965845 | 0.170305087 | 0.228519707 | 0.745253392 | 0.456118551 | 0.643397611 |
| ENSG00000196369 | SRGAP2B | 475.665127 | 0.147207967 | 0.089569871 | 1.643498708 | 0.100279792 | 0.21243129 |
| ENSG00000196371 | FUT4 | 71.56074689 | -0.454662576 | 0.216439296 | -2.100647078 | 0.035671958 | 0.090062144 |
| ENSG00000196372 | ASB13 | 822.7511388 | -0.61648607 | 0.071802719 | -8.585831875 | 9.02E-18 | 1.31E-16 |
| ENSG00000196376 | SLC35F1 | 35.33072695 | 1.004172567 | 0.306867668 | 3.272330946 | 0.001066646 | 0.00397093 |
| ENSG00000196378 | ZNF34 | 226.8364456 | -0.16700014 | 0.121252685 | -1.377290244 | 0.16842253 | 0.318517135 |
| ENSG00000196381 | ZNF781 | 7.635614037 | 1.590953898 | 0.686752835 | 2.316632444 | 0.020523762 | 0.055896368 |
| ENSG00000196387 | ZNF140 | 585.0054181 | 0.35394585 | 0.080642743 | 4.389060138 | 1.14E-05 | 5.83E-05 |
| ENSG00000196388 | INCA1 | 30.76982094 | 0.220865941 | 0.314656107 | 0.701928029 | 0.482724048 | 0.666848709 |
| ENSG00000196391 | ZNF774 | 192.1207013 | -0.075450624 | 0.133697322 | -0.564339082 | 0.572523389 | 0.73930124 |
| ENSG00000196396 | PTPN1 | 4358.67322 | 0.053277538 | 0.048259986 | 1.103969203 | 0.269606497 | 0.451179826 |
| ENSG00000196405 | EVL | 3736.148745 | 0.605366952 | 0.059499107 | 10.17438717 | 2.58E-24 | 5.51E-23 |
| ENSG00000196408 | NOXO1 | 16.6809889 | -0.12018007 | 0.429411106 | -0.279871825 | 0.779575844 | 0.880351967 |
| ENSG00000196411 | EPHB4 | 4100.198537 | 0.027153845 | 0.05137455 | 0.528546615 | 0.597120002 | 0.75769879 |
| ENSG00000196415 | PRTN3 | 2.024045919 | -0.186164929 | 1.210987483 | -0.153729854 | 0.877822745 | 0.936395015 |
| ENSG00000196417 | ZNF765 | 647.0214207 | 0.281162818 | 0.084395577 | 3.331487604 | 0.000863831 | 0.003272716 |
| ENSG00000196418 | ZNF124 | 297.3466183 | 0.050809308 | 0.110895909 | 0.458171166 | 0.646829474 | 0.793575149 |
| ENSG00000196419 | XRCC6 | 9226.650871 | -0.221641734 | 0.042752528 | -5.184295378 | 2.17E-07 | 1.36E-06 |
| ENSG00000196421 | C20orf204 | 137.7386567 | -0.25902401 | 0.152414275 | -1.699473424 | 0.089230018 | 0.193070799 |
| ENSG00000196422 | PPP1R26 | 1595.439221 | 0.111429134 | 0.061094427 | 1.823883767 | 0.068169646 | 0.155309533 |
| ENSG00000196427 | NBPF4 | 30.38866269 | -0.071057016 | 0.31575725 | -0.225036846 | 0.82195061 | 0.905776385 |
| ENSG00000196428 | TSC22D2 | 1136.693636 | 0.177008178 | 0.063919868 | 2.76922 | 0.005619068 | 0.017858329 |
| ENSG00000196436 | NPIPB15 | 63.28066317 | -0.258096543 | 0.235284368 | -1.096955761 | 0.272660731 | 0.454731892 |
| ENSG00000196437 | ZNF569 | 106.4355577 | 0.166477474 | 0.173787978 | 0.957934349 | 0.338095864 | 0.52937814 |
| ENSG00000196440 | ARMCX4 | 27.76426767 | -0.517984947 | 0.363361432 | -1.425536397 | 0.154002207 | 0.297224518 |
| ENSG00000196449 | YRDC | 689.998439 | 0.179273218 | 0.075288472 | 2.381150967 | 0.017258637 | 0.048100248 |
| ENSG00000196453 | ZNF777 | 944.9155282 | 0.493016339 | 0.071816509 | 6.864944317 | 6.65E-12 | 6.33E-11 |

| | | | | | | | |
|---|---|---|---|---|---|---|---|
| ENSG00000196455 | PIK3R4 | 1492.276756 | -0.170876902 | 0.058972774 | -2.897555801 | 0.003760828 | 0.012467254 |
| ENSG00000196456 | ZNF775 | 56.15326389 | 0.393362087 | 0.242534762 | 1.621879202 | 0.104829216 | 0.220197986 |
| ENSG00000196458 | ZNF605 | 624.0834343 | 0.719496709 | 0.083713345 | 8.594767178 | 8.34E-18 | 1.21E-16 |
| ENSG00000196459 | TRAPPC2 | 350.3389949 | 0.268359388 | 0.103276756 | 2.598449041 | 0.009364594 | 0.028096642 |
| ENSG00000196465 | MYL6B | 1373.421918 | -0.710602627 | 0.064367646 | -11.03974859 | 2.46E-28 | 6.39E-27 |
| ENSG00000196466 | ZNF799 | 152.3243027 | 0.092361393 | 0.158743924 | 0.581826321 | 0.560683673 | 0.729860281 |
| ENSG00000196470 | SIAH1 | 437.8295964 | -0.107440232 | 0.090847182 | -1.18264794 | 0.236948704 | 0.412162821 |
| ENSG00000196476 | C20orf96 | 143.9931456 | 0.289737159 | 0.151589523 | 1.911327069 | 0.055962562 | 0.131886294 |
| ENSG00000196482 | ESRRG | 10.69269741 | 1.151620283 | 0.554778613 | 2.075819538 | 0.037910645 | 0.094900719 |
| ENSG00000196497 | IPO4 | 32.52005866 | -0.274245771 | 0.304980387 | -0.899224287 | 0.368533207 | 0.561103853 |
| ENSG00000196498 | NCOR2 | 6082.425064 | 0.570918365 | 0.049651798 | 11.49844281 | 1.34E-30 | 3.81E-29 |
| ENSG00000196502 | SULT1A1 | 280.0438512 | 0.059328642 | 0.109852532 | 0.540075329 | 0.589145084 | 0.752295476 |
| ENSG00000196503 | ARL9 | 1.859044137 | 3.147274764 | 1.577999992 | 1.99447071 | 0.046100638 | 0.111720751 |
| ENSG00000196504 | PRPF40A | 4095.917046 | 0.118625182 | 0.053920425 | 2.200004581 | 0.02780657 | 0.072775912 |
| ENSG00000196505 | GDAP2 | 633.9296135 | -0.237324856 | 0.079624657 | -2.980544792 | 0.002877362 | 0.009788918 |
| ENSG00000196507 | TCEAL3 | 1281.069208 | -0.180688234 | 0.061169717 | -2.953883768 | 0.003138022 | 0.010588147 |
| ENSG00000196510 | ANAPC7 | 1098.943178 | -0.044898888 | 0.065000577 | -0.690745992 | 0.489725181 | 0.672936425 |
| ENSG00000196511 | TPK1 | 186.5862666 | -0.777919794 | 0.137289653 | -5.666266736 | 1.46E-08 | 1.03E-07 |
| ENSG00000196517 | SLC6A9 | 1236.394578 | 0.352540307 | 0.064633595 | 5.454443682 | 4.91E-08 | 3.28E-07 |
| ENSG00000196526 | AFAP1 | 1114.014889 | -0.391725031 | 0.069927042 | -5.601910497 | 2.12E-08 | 1.47E-07 |
| ENSG00000196531 | NACA | 6133.148727 | 0.220256693 | 0.047634995 | 4.623842046 | 3.77E-06 | 2.04E-05 |
| ENSG00000196535 | MYO18A | 1666.003656 | 0.181521176 | 0.068692803 | 2.642506459 | 0.00822949 | 0.025047697 |
| ENSG00000196542 | SPTSSB | 16718.2314 | -0.712156371 | 0.056886142 | -12.51897825 | 5.88E-36 | 2.06E-34 |
| ENSG00000196544 | BORCS6 | 176.4177092 | -0.275224819 | 0.145635121 | -1.889824491 | 0.058781437 | 0.137170768 |
| ENSG00000196547 | MAN2A2 | 5197.348408 | 0.048845914 | 0.04706087 | 1.03793053 | 0.299302399 | 0.485998083 |
| ENSG00000196549 | MME | 1851.79913 | 1.459159966 | 0.073711837 | 19.79546326 | 3.26E-87 | 4.27E-85 |
| ENSG00000196550 | FAM72A | 206.3706231 | 0.031785913 | 0.128751418 | 0.24687816 | 0.805002519 | 0.896404708 |
| ENSG00000196557 | CACNA1H | 2313.344802 | -0.261172915 | 0.065195919 | -4.005970284 | 6.18E-05 | 0.000285024 |
| ENSG00000196562 | SULF2 | 30344.26605 | 0.282974925 | 0.040473406 | 6.991626137 | 2.72E-12 | 2.68E-11 |
| ENSG00000196569 | LAMA2 | 7.627926552 | -0.943073309 | 0.636608072 | -1.481403316 | 0.138499132 | 0.274438117 |
| ENSG00000196576 | PLXNB2 | 9076.244179 | 0.099421048 | 0.056025871 | 1.77455604 | 0.075971221 | 0.169663144 |
| ENSG00000196584 | XRCC2 | 567.5911566 | 0.483158983 | 0.085403215 | 5.657386387 | 1.54E-08 | 1.08E-07 |
| ENSG00000196586 | MYO6 | 6076.852311 | 0.161532497 | 0.049333584 | 3.274290715 | 0.001059276 | 0.003945411 |

| | | | | | | | |
|---|---|---|---|---|---|---|---|
| ENSG00000196588 | MRTFA | 1007.244751 | -0.001024304 | 0.069653865 | -0.014705632 | 0.988267026 | 0.994030148 |
| ENSG00000196591 | HDAC2 | 2920.617995 | 0.496175939 | 0.057891954 | 8.570723647 | 1.03E-17 | 1.49E-16 |
| ENSG00000196593 | ANKRD20A19I | 4.614966852 | -0.538942431 | 0.854456482 | -0.630742984 | 0.528208589 | 0.704196259 |
| ENSG00000196597 | ZNF782 | 251.5112199 | 0.016864023 | 0.117509739 | 0.143511708 | 0.885886065 | 0.94106165 |
| ENSG00000196604 | POTEF | 31.63762465 | -0.406434869 | 0.318891637 | -1.274523449 | 0.202477968 | 0.366601782 |
| ENSG00000196605 | ZNF846 | 166.7333567 | 0.14312209 | 0.140938977 | 1.015489776 | 0.309872418 | 0.497827107 |
| ENSG00000196611 | MMP1 | 7.719921803 | -1.708386287 | 0.703215117 | -2.429393576 | 0.015124104 | 0.042811816 |
| ENSG00000196620 | UGT2B15 | 7.331156683 | 0.671159424 | 0.648711538 | 1.034603801 | 0.300853978 | 0.487607055 |
| ENSG00000196642 | RABL6 | 4575.916263 | 0.09108819 | 0.048435096 | 1.880623721 | 0.060023123 | 0.139593288 |
| ENSG00000196646 | ZNF136 | 277.5383961 | 0.609310123 | 0.119094249 | 5.116201057 | 3.12E-07 | 1.92E-06 |
| ENSG00000196652 | ZKSCAN5 | 889.2788206 | -0.717879609 | 0.073866455 | -9.71861459 | 2.51E-22 | 4.78E-21 |
| ENSG00000196655 | TRAPPC4 | 1984.687691 | 0.207727846 | 0.053153535 | 3.908072049 | 9.30E-05 | 0.00041668 |
| ENSG00000196656 | ENSG0000019 | 472.2380836 | 0.112909492 | 0.096755786 | 1.166953388 | 0.243229194 | 0.41965085 |
| ENSG00000196659 | TTC30B | 153.147626 | 0.489370996 | 0.147396507 | 3.320099007 | 0.000899855 | 0.00339937 |
| ENSG00000196663 | TECPR2 | 1120.001544 | -0.321628349 | 0.069649735 | -4.617797169 | 3.88E-06 | 2.10E-05 |
| ENSG00000196666 | FAM180B | 2.260502298 | -0.466679878 | 1.266552831 | -0.368464597 | 0.712526839 | 0.840033184 |
| ENSG00000196668 | LINC00173 | 66.26501967 | 1.491221398 | 0.234085095 | 6.370424388 | 1.89E-10 | 1.60E-09 |
| ENSG00000196670 | ZFP62 | 1898.224987 | -0.324471108 | 0.057821723 | -5.611577962 | 2.00E-08 | 1.39E-07 |
| ENSG00000196678 | ERI2 | 795.1095061 | -0.222249857 | 0.087853212 | -2.529786358 | 0.0114132 | 0.033382482 |
| ENSG00000196683 | TOMM7 | 3481.786182 | -1.213615928 | 0.052693615 | -23.03155587 | 2.25E-117 | 5.17E-115 |
| ENSG00000196684 | HSH2D | 217.1006437 | -0.617775625 | 0.128590261 | -4.804217822 | 1.55E-06 | 8.78E-06 |
| ENSG00000196689 | TRPV1 | 99.3658898 | 0.105554952 | 0.178588177 | 0.591052291 | 0.554485385 | 0.72573819 |
| ENSG00000196693 | ZNF33B | 868.5255821 | -0.437955282 | 0.0722895 | -6.058352603 | 1.38E-09 | 1.08E-08 |
| ENSG00000196696 | PDXDC2P-NPI | 106.3227748 | -0.521718522 | 0.191122854 | -2.729754775 | 0.006338145 | 0.019892879 |
| ENSG00000196700 | ZNF512B | 6217.866584 | -0.014218072 | 0.049433977 | -0.287617411 | 0.773639616 | 0.876625971 |
| ENSG00000196704 | AMZ2 | 3218.147403 | -0.058966498 | 0.049375625 | -1.194243056 | 0.232382905 | 0.406408771 |
| ENSG00000196705 | ZNF431 | 258.8544727 | 0.285491159 | 0.119684559 | 2.385363348 | 0.01706226 | 0.047639757 |
| ENSG00000196712 | NF1 | 2770.9422 | -0.463317705 | 0.059323468 | -7.810023981 | 5.72E-15 | 6.85E-14 |
| ENSG00000196715 | VKORC1L1 | 1846.002913 | 0.502342448 | 0.055651872 | 9.026514803 | 1.77E-19 | 2.88E-18 |
| ENSG00000196724 | ZNF418 | 177.3988695 | 0.229319091 | 0.137734607 | 1.664934445 | 0.095925835 | 0.204910947 |
| ENSG00000196730 | DAPK1 | 2.123766554 | 0.546405903 | 1.22717371 | 0.445255548 | 0.656135088 | 0.800666999 |
| ENSG00000196739 | COL27A1 | 302.7243734 | 0.586547846 | 0.113343524 | 5.174956846 | 2.28E-07 | 1.43E-06 |
| ENSG00000196741 | LINC01560 | 63.07407875 | 0.653391542 | 0.227559241 | 2.871303052 | 0.004087834 | 0.013440579 |

| | | | | | | | |
|---|---|---|---|---|---|---|---|
| ENSG00000196743 | GM2A | 543.1486966 | 0.460567049 | 0.091331396 | 5.042811875 | 4.59E-07 | 2.78E-06 |
| ENSG00000196747 | H2AC13 | 153.3670375 | 0.19081746 | 0.147812135 | 1.29094584 | 0.196722458 | 0.358509096 |
| ENSG00000196748 | CLPSL2 | 1.722851617 | 1.255891399 | 1.385167605 | 0.906671073 | 0.364580765 | NA |
| ENSG00000196754 | S100A2 | 6.265033323 | 0.748587512 | 0.743098187 | 1.007387079 | 0.313748798 | 0.501980571 |
| ENSG00000196756 | SNHG17 | 500.3160105 | -0.148347684 | 0.095343874 | -1.555922563 | 0.119726507 | 0.245174947 |
| ENSG00000196757 | ZNF700 | 238.4827744 | -0.1031704 | 0.122809686 | -0.840083573 | 0.40086153 | 0.593736061 |
| ENSG00000196776 | CD47 | 4291.577775 | 0.851539921 | 0.055936489 | 15.22333511 | 2.48E-52 | 1.45E-50 |
| ENSG00000196781 | TLE1 | 2128.666991 | 0.241665247 | 0.053563588 | 4.511744937 | 6.43E-06 | 3.40E-05 |
| ENSG00000196782 | MAML3 | 639.6912323 | -0.282798253 | 0.080325267 | -3.520663698 | 0.000430468 | 0.001723977 |
| ENSG00000196787 | H2AC11 | 487.3968219 | 0.527099682 | 0.093687996 | 5.626117585 | 1.84E-08 | 1.29E-07 |
| ENSG00000196792 | STRN3 | 1305.543935 | -0.33086924 | 0.060895768 | -5.433370062 | 5.53E-08 | 3.67E-07 |
| ENSG00000196793 | ZNF239 | 622.1732448 | -0.650687481 | 0.089057161 | -7.306402671 | 2.74E-13 | 2.91E-12 |
| ENSG00000196810 | CTBP1-DT | 1164.319974 | -0.024791193 | 0.066023661 | -0.375489517 | 0.707296441 | 0.836532294 |
| ENSG00000196812 | ZSCAN16 | 254.684395 | 0.197312601 | 0.114183527 | 1.72803036 | 0.083982782 | 0.183931808 |
| ENSG00000196814 | MVB12B | 458.7995346 | 0.488246193 | 0.092166426 | 5.297440876 | 1.17E-07 | 7.58E-07 |
| ENSG00000196821 | ILRUN | 4435.38224 | -0.054123724 | 0.050218153 | -1.077772079 | 0.281135483 | 0.46454282 |
| ENSG00000196834 | POTEI | 25.28926636 | -0.243607335 | 0.359593437 | -0.677452117 | 0.498119141 | 0.67994219 |
| ENSG00000196839 | ADA | 25.5992585 | -0.584994301 | 0.349689072 | -1.672898433 | 0.094347292 | 0.202162579 |
| ENSG00000196843 | ARID5A | 205.0243772 | 1.285235011 | 0.137368752 | 9.356094386 | 8.27E-21 | 1.45E-19 |
| ENSG00000196844 | PATE2 | 4.179803422 | -0.594626944 | 0.874932655 | -0.679625958 | 0.496741329 | 0.678652021 |
| ENSG00000196850 | PPTC7 | 467.7429469 | 0.061887384 | 0.093478842 | 0.66204697 | 0.50794112 | 0.688052512 |
| ENSG00000196860 | TOMM20L | 3.673547661 | -1.267145321 | 0.945494777 | -1.34019283 | 0.180182662 | 0.335194496 |
| ENSG00000196862 | RGPD4 | 4.334655463 | 0.463184909 | 0.843493798 | 0.549126633 | 0.582918549 | 0.747298772 |
| ENSG00000196865 | NHLRC2 | 839.6974114 | -1.172366327 | 0.074359528 | -15.76618836 | 5.32E-56 | 3.43E-54 |
| ENSG00000196872 | CRACDL | 151.9723348 | 0.984955333 | 0.155283017 | 6.342968797 | 2.25E-10 | 1.90E-09 |
| ENSG00000196873 | ZNG1C | 40.25067679 | 0.105101519 | 0.28347029 | 0.370767316 | 0.710810847 | 0.839088192 |
| ENSG00000196876 | SCN8A | 237.6216874 | -0.572043785 | 0.122496872 | -4.669864437 | 3.01E-06 | 1.65E-05 |
| ENSG00000196878 | LAMB3 | 22.82999498 | 3.455597803 | 0.538082158 | 6.422063524 | 1.34E-10 | 1.15E-09 |
| ENSG00000196890 | H2BC26 | 85.3449092 | 0.190863252 | 0.194903728 | 0.979269378 | 0.327446897 | 0.51777302 |
| ENSG00000196911 | KPNA5 | 222.5561527 | -0.129134534 | 0.133556875 | -0.966887955 | 0.333600048 | 0.524670219 |
| ENSG00000196912 | ANKRD36B | 52.98634484 | -0.494941198 | 0.241749945 | -2.047327032 | 0.040625985 | 0.100459212 |
| ENSG00000196914 | ARHGEF12 | 4605.31473 | -0.026111771 | 0.053964697 | -0.483867652 | 0.628479799 | 0.780792125 |
| ENSG00000196917 | HCAR1 | 624.7445232 | 0.532127094 | 0.088942638 | 5.982812162 | 2.19E-09 | 1.69E-08 |

| | | | | | | | |
|---|---|---|---|---|---|---|---|
| ENSG00000196922 | ZNF252P | 1841.3019 | 0.00017337 | 0.057717745 | 0.003003758 | 0.997603351 | 0.998604568 |
| ENSG00000196923 | PDLIM7 | 1664.624912 | -0.243685584 | 0.065680357 | -3.710174468 | 0.000207116 | 0.000876291 |
| ENSG00000196924 | FLNA | 60513.92807 | -1.038542408 | 0.050497827 | -20.5660811 | 5.53E-94 | 7.92E-92 |
| ENSG00000196932 | TMEM26 | 3.057629174 | 1.659984608 | 1.107695719 | 1.4985926 | 0.133979354 | 0.267469008 |
| ENSG00000196935 | SRGAP1 | 787.121439 | -0.193958268 | 0.073199649 | -2.649715822 | 0.00805595 | 0.02459783 |
| ENSG00000196937 | FAM3C | 291.5813281 | 0.518479723 | 0.127243537 | 4.074703801 | 4.61E-05 | 0.000216625 |
| ENSG00000196943 | NOP9 | 3385.587542 | -0.229268556 | 0.0495881 | -4.623459227 | 3.77E-06 | 2.04E-05 |
| ENSG00000196950 | SLC39A10 | 992.6617043 | 0.262560538 | 0.071875454 | 3.652993095 | 0.000259201 | 0.001076797 |
| ENSG00000196951 | SCOC-AS1 | 8.763630635 | -0.329367174 | 0.590801695 | -0.557491925 | 0.577191375 | 0.743153737 |
| ENSG00000196954 | CASP4 | 56.39888459 | 0.562834261 | 0.24186225 | 2.327086018 | 0.019960688 | 0.054589672 |
| ENSG00000196961 | AP2A1 | 4299.964395 | 0.004347264 | 0.051650942 | 0.084166201 | 0.932924291 | 0.967770789 |
| ENSG00000196967 | ZNF585A | 114.1618519 | 0.249599834 | 0.171717474 | 1.453549419 | 0.146071269 | 0.285642734 |
| ENSG00000196968 | FUT11 | 321.6725314 | -1.187015085 | 0.114428815 | -10.37339314 | 3.28E-25 | 7.36E-24 |
| ENSG00000196972 | SMIM10L2B | 5.595917487 | 0.218581056 | 0.776207022 | 0.281601493 | 0.778249095 | 0.879589804 |
| ENSG00000196975 | ANXA4 | 1356.688231 | 0.036177722 | 0.062421842 | 0.579568329 | 0.562205756 | 0.731024977 |
| ENSG00000196976 | LAGE3 | 1454.616771 | -0.810012426 | 0.066933246 | -12.1017951 | 1.03E-33 | 3.31E-32 |
| ENSG00000196981 | WDR5B | 450.6479392 | 0.103111962 | 0.092817877 | 1.110906271 | 0.266608696 | 0.447698081 |
| ENSG00000196993 | NPIPB9 | 3.442918339 | 1.115803796 | 1.002090893 | 1.113475637 | 0.265504206 | 0.446566011 |
| ENSG00000196998 | WDR45 | 633.4355466 | 0.355335091 | 0.080871696 | 4.393812761 | 1.11E-05 | 5.71E-05 |
| ENSG00000197006 | METTL9 | 3056.263112 | 0.099851513 | 0.04766019 | 2.095071662 | 0.036164623 | 0.091186683 |
| ENSG00000197008 | ZNF138 | 525.8084367 | 0.480165143 | 0.091565563 | 5.243949015 | 1.57E-07 | 1.00E-06 |
| ENSG00000197013 | ZNF429 | 278.5011072 | -0.144565099 | 0.111865085 | -1.292316532 | 0.196247549 | 0.357806293 |
| ENSG00000197019 | SERTAD1 | 307.8187572 | -0.273730612 | 0.108740682 | -2.517278785 | 0.011826521 | 0.034507809 |
| ENSG00000197020 | ZNF100 | 180.5190459 | 0.354645065 | 0.138035088 | 2.56923852 | 0.010192228 | 0.030243139 |
| ENSG00000197021 | EOLA2 | 1724.277969 | 0.100468677 | 0.058367152 | 1.721322237 | 0.085192367 | 0.186071874 |
| ENSG00000197024 | ZNF398 | 1225.05929 | 0.245340186 | 0.068587632 | 3.577032464 | 0.000347517 | 0.001410731 |
| ENSG00000197037 | ZSCAN25 | 822.299978 | -0.345973866 | 0.076128667 | -4.544593781 | 5.50E-06 | 2.93E-05 |
| ENSG00000197043 | ANXA6 | 2574.631202 | -1.91520291 | 0.060169331 | -31.83021779 | 2.47E-222 | 2.27E-219 |
| ENSG00000197045 | GMFB | 2110.845068 | -0.544070652 | 0.072026943 | -7.553710131 | 4.23E-14 | 4.74E-13 |
| ENSG00000197046 | SIGLEC15 | 38.22956464 | 3.087243885 | 0.384578703 | 8.027599717 | 9.94E-16 | 1.26E-14 |
| ENSG00000197050 | ZNF420 | 278.4771089 | 0.351411777 | 0.112361512 | 3.127510229 | 0.001762937 | 0.006266156 |
| ENSG00000197054 | ZNF763 | 11.12049549 | 0.651318259 | 0.534790801 | 1.217893533 | 0.223264434 | 0.394498427 |
| ENSG00000197056 | ZMYM1 | 378.561673 | -0.335107134 | 0.099788453 | -3.358175454 | 0.000784588 | 0.00299475 |

| | | | | | | | |
|---|---|---|---|---|---|---|---|
| ENSG00000197061 | H4C3 | 4.134048554 | -0.025467188 | 0.895456144 | -0.028440464 | 0.977310852 | 0.989387067 |
| ENSG00000197062 | ZSCAN26 | 268.3218653 | -0.075626344 | 0.11454898 | -0.660209671 | 0.509119287 | 0.689241454 |
| ENSG00000197063 | MAFG | 1442.948106 | 0.318105447 | 0.061615352 | 5.162762819 | 2.43E-07 | 1.52E-06 |
| ENSG00000197070 | ARRDC1 | 1463.015219 | 0.063649811 | 0.063908483 | 0.995952457 | 0.319273246 | 0.508689685 |
| ENSG00000197077 | KIAA1671 | 1462.313153 | 0.138443714 | 0.08266518 | 1.67475247 | 0.093982808 | 0.201473284 |
| ENSG00000197081 | IGF2R | 3831.5936 | 0.135312538 | 0.057680758 | 2.345886973 | 0.018981863 | 0.052255362 |
| ENSG00000197093 | GAL3ST4 | 8.927257691 | 0.614850745 | 0.615156609 | 0.999502786 | 0.31755119 | 0.506650737 |
| ENSG00000197102 | DYNC1H1 | 8303.230511 | -0.522583837 | 0.057184173 | -9.13860975 | 6.33E-20 | 1.05E-18 |
| ENSG00000197106 | SLC6A17 | 1.999518408 | -1.658275448 | 1.363020075 | -1.216618506 | 0.223749398 | 0.395135581 |
| ENSG00000197111 | PCBP2 | 5290.794171 | 0.049549517 | 0.06199503 | 0.799249833 | 0.424145562 | 0.615019183 |
| ENSG00000197114 | ZGPAT | 199.5922181 | -0.18947662 | 0.137562451 | -1.377386187 | 0.168392881 | 0.318506788 |
| ENSG00000197119 | SLC25A29 | 3248.339367 | 0.179970233 | 0.057632623 | 3.122714581 | 0.001791914 | 0.006362253 |
| ENSG00000197121 | PGAP1 | 271.8780518 | -0.220163013 | 0.115866645 | -1.90014144 | 0.057414561 | 0.134589211 |
| ENSG00000197122 | SRC | 1146.334261 | 0.387822579 | 0.070952661 | 5.465934238 | 4.60E-08 | 3.09E-07 |
| ENSG00000197124 | ZNF682 | 111.3948088 | 1.039597873 | 0.175242938 | 5.932323939 | 2.99E-09 | 2.28E-08 |
| ENSG00000197128 | ZNF772 | 322.1234498 | 0.39610657 | 0.104379298 | 3.794876724 | 0.000147717 | 0.000640814 |
| ENSG00000197136 | PCNX3 | 6531.73368 | 0.322197895 | 0.050521366 | 6.377458165 | 1.80E-10 | 1.53E-09 |
| ENSG00000197140 | ADAM32 | 62.99859855 | 0.959366721 | 0.240679548 | 3.986074971 | 6.72E-05 | 0.000308139 |
| ENSG00000197147 | LRRC8B | 1037.648488 | 0.066409433 | 0.067488547 | 0.984010413 | 0.325110393 | 0.515289196 |
| ENSG00000197149 | ENSG0000019 | 12.94579387 | -0.104666182 | 0.509060703 | -0.205606485 | 0.837098311 | 0.914148686 |
| ENSG00000197150 | ABCB8 | 740.7909106 | 0.293146671 | 0.074401663 | 3.940055371 | 8.15E-05 | 0.000369174 |
| ENSG00000197153 | H3C12 | 35.67698262 | 0.403113602 | 0.311186651 | 1.295407758 | 0.195179603 | 0.356397365 |
| ENSG00000197157 | SND1 | 6837.581277 | 0.074788542 | 0.046692574 | 1.601722409 | 0.109217008 | 0.227374367 |
| ENSG00000197162 | ZNF785 | 420.7177096 | 0.183393258 | 0.093216841 | 1.967383312 | 0.049139037 | 0.118048958 |
| ENSG00000197165 | SULT1A2 | 7.292024457 | 0.735067917 | 0.672298002 | 1.093366208 | 0.274233044 | 0.456259503 |
| ENSG00000197168 | NEK5 | 53.92470428 | -1.173521424 | 0.246279359 | -4.765001133 | 1.89E-06 | 1.06E-05 |
| ENSG00000197170 | PSMD12 | 4643.552947 | 0.07654398 | 0.052891052 | 1.447200951 | 0.14784065 | 0.288340802 |
| ENSG00000197180 | ATP6AP1-DT | 183.524016 | -1.472654964 | 0.139903707 | -10.52620405 | 6.54E-26 | 1.52E-24 |
| ENSG00000197182 | MIRLET7BHG | 28.16426738 | 0.146986982 | 0.327132214 | 0.449319804 | 0.653200973 | 0.798701046 |
| ENSG00000197183 | NOL4L | 1453.73896 | -0.977592646 | 0.064809727 | -15.08404197 | 2.06E-51 | 1.18E-49 |
| ENSG00000197191 | CYSRT1 | 74.00110801 | 0.681523198 | 0.214777688 | 3.173156413 | 0.001507913 | 0.005439773 |
| ENSG00000197208 | SLC22A4 | 144.0966496 | 0.51005261 | 0.152935311 | 3.335087281 | 0.000852726 | 0.003234382 |
| ENSG00000197217 | ENTPD4 | 980.5287635 | 0.386289885 | 0.067802551 | 5.697276588 | 1.22E-08 | 8.66E-08 |

| | | | | | | | |
|---|---|---|---|---|---|---|---|
| ENSG00000197223 | C1D | 145.869474 | 0.172351079 | 0.15239539 | 1.130946799 | 0.258077484 | 0.437729944 |
| ENSG00000197226 | TBC1D9B | 11230.73497 | -0.111686584 | 0.044758899 | -2.495293354 | 0.012585304 | 0.036393151 |
| ENSG00000197238 | H4C11 | 5.414427076 | -0.95928675 | 0.786228546 | -1.220111829 | 0.222422485 | 0.393488995 |
| ENSG00000197245 | FAM110D | 2.69405656 | -1.731303295 | 1.194291097 | -1.449649335 | 0.147156331 | 0.287348164 |
| ENSG00000197249 | SERPINA1 | 96.79100676 | -0.370088348 | 0.185264364 | -1.997622967 | 0.045757551 | 0.111038277 |
| ENSG00000197256 | KANK2 | 2145.917403 | 0.520562316 | 0.057900354 | 8.990658645 | 2.46E-19 | 3.96E-18 |
| ENSG00000197258 | EIF4BP6 | 3319.767854 | 0.221436245 | 0.048099732 | 4.603689841 | 4.15E-06 | 2.24E-05 |
| ENSG00000197261 | C6orf141 | 468.2778283 | 1.22089261 | 0.203640442 | 5.995334694 | 2.03E-09 | 1.57E-08 |
| ENSG00000197265 | GTF2E2 | 673.1307479 | 0.073187906 | 0.076974603 | 0.95080589 | 0.341702922 | 0.532807574 |
| ENSG00000197275 | RAD54B | 135.5743984 | -0.843825071 | 0.163434604 | -5.163074716 | 2.43E-07 | 1.51E-06 |
| ENSG00000197279 | ZNF165 | 196.9828993 | -0.18801621 | 0.13610048 | -1.381451481 | 0.167140186 | 0.316661723 |
| ENSG00000197283 | SYNGAP1 | 1592.081924 | -0.025519421 | 0.062544143 | -0.408022557 | 0.683257115 | 0.820223767 |
| ENSG00000197291 | RAMP2-AS1 | 7.992505623 | -0.654992106 | 0.627965304 | -1.043038687 | 0.296930376 | 0.483262349 |
| ENSG00000197296 | FITM2 | 1208.52254 | -0.16522432 | 0.064640818 | -2.556036974 | 0.010587187 | 0.031249365 |
| ENSG00000197299 | BLM | 917.1969551 | 0.055703688 | 0.072951639 | 0.763570067 | 0.445123493 | 0.633720159 |
| ENSG00000197302 | KRBOX5 | 749.7103533 | -0.063967212 | 0.080472819 | -0.794892141 | 0.426676246 | 0.617497513 |
| ENSG00000197308 | GATA3-AS1 | 820.9203191 | 0.369273009 | 0.074842014 | 4.934033596 | 8.05E-07 | 4.73E-06 |
| ENSG00000197312 | DDI2 | 1593.961809 | -0.021695538 | 0.064403328 | -0.336869822 | 0.736215036 | 0.854478772 |
| ENSG00000197321 | SVIL | 1946.006072 | -0.346048764 | 0.057367757 | -6.032112508 | 1.62E-09 | 1.26E-08 |
| ENSG00000197323 | TRIM33 | 15404.8475 | 0.386041446 | 0.047381735 | 8.147473901 | 3.72E-16 | 4.85E-15 |
| ENSG00000197324 | LRP10 | 6301.976655 | -0.215646962 | 0.054745013 | -3.939116075 | 8.18E-05 | 0.000370475 |
| ENSG00000197329 | PELI1 | 100.1234084 | 1.050392204 | 0.186768638 | 5.624028819 | 1.87E-08 | 1.30E-07 |
| ENSG00000197332 | ENSG0000019 | 12.22032146 | -0.475581522 | 0.495911018 | -0.95900576 | 0.337555844 | 0.528785427 |
| ENSG00000197345 | MRPL21 | 1183.800185 | 0.111995375 | 0.062476481 | 1.792600551 | 0.073036818 | 0.164349909 |
| ENSG00000197355 | UAP1L1 | 1993.540332 | 0.475806622 | 0.057152503 | 8.325210585 | 8.42E-17 | 1.15E-15 |
| ENSG00000197358 | BNIP3P1 | 175.3756206 | -0.850106828 | 0.138683207 | -6.129846916 | 8.80E-10 | 7.03E-09 |
| ENSG00000197360 | ZNF98 | 4.29917121 | 0.24929599 | 0.847885869 | 0.294020692 | 0.768742092 | 0.873855116 |
| ENSG00000197361 | FBXL22 | 66.22307204 | -0.036036396 | 0.21612536 | -0.166738398 | 0.867575891 | 0.93096599 |
| ENSG00000197362 | ZNF786 | 569.676561 | 0.34942999 | 0.090295273 | 3.86985915 | 0.000108898 | 0.000482823 |
| ENSG00000197363 | ZNF517 | 328.6193614 | 0.030114128 | 0.111799928 | 0.269357313 | 0.787654732 | 0.885118034 |
| ENSG00000197372 | ZNF675 | 279.4853691 | 0.948411758 | 0.116235666 | 8.159386763 | 3.37E-16 | 4.41E-15 |
| ENSG00000197375 | SLC22A5 | 1155.691101 | -0.005103217 | 0.076635146 | -0.066591072 | 0.946907253 | 0.975200824 |
| ENSG00000197380 | DACT3 | 7.371291362 | 0.149221164 | 0.640838306 | 0.232853066 | 0.815875505 | 0.902352305 |

| | | | | | | | |
|---|---|---|---|---|---|---|---|
| ENSG00000197381 | ADARB1 | 1787.892724 | 0.355293398 | 0.056826694 | 6.252227097 | 4.05E-10 | 3.35E-09 |
| ENSG00000197385 | ZNF860 | 295.618554 | -0.2086664 | 0.109861383 | -1.89936076 | 0.057517059 | 0.134788164 |
| ENSG00000197386 | HTT | 5868.521354 | 0.173596594 | 0.055018047 | 3.15526638 | 0.001603516 | 0.005757499 |
| ENSG00000197405 | C5AR1 | 25.62885116 | 0.364833733 | 0.345450714 | 1.056109361 | 0.290918254 | 0.475772777 |
| ENSG00000197406 | DIO3 | 6.296959356 | 0.296547746 | 0.721302863 | 0.411127921 | 0.680978738 | 0.818916557 |
| ENSG00000197415 | VEPH1 | 11.51152671 | -1.172468286 | 0.534733134 | -2.192623219 | 0.028334537 | 0.073930195 |
| ENSG00000197417 | SHPK | 85.78842735 | 0.203365361 | 0.196351542 | 1.035720722 | 0.300332452 | 0.486968225 |
| ENSG00000197429 | IPP | 322.6697538 | -0.136657569 | 0.105194577 | -1.299093294 | 0.193911914 | 0.354618886 |
| ENSG00000197442 | MAP3K5 | 985.0618656 | -0.1032123 | 0.070772813 | -1.458360863 | 0.144741102 | 0.283549179 |
| ENSG00000197444 | OGDHL | 145.6538443 | 0.186264026 | 0.154360881 | 1.206678952 | 0.227555808 | 0.400316736 |
| ENSG00000197448 | GSTK1 | 1510.159141 | -0.179847358 | 0.060774284 | -2.959267427 | 0.003083714 | 0.010422925 |
| ENSG00000197451 | HNRNPAB | 7697.148864 | -0.352566754 | 0.044450051 | -7.931751345 | 2.16E-15 | 2.66E-14 |
| ENSG00000197457 | STMN3 | 466.7010678 | 0.204935538 | 0.093381461 | 2.194606242 | 0.028191855 | 0.073633287 |
| ENSG00000197461 | PDGFA | 616.800936 | 0.538454104 | 0.082983312 | 6.488703478 | 8.66E-11 | 7.54E-10 |
| ENSG00000197467 | COL13A1 | 3.519502622 | -0.36405978 | 0.93724022 | -0.388438067 | 0.697691878 | 0.830348323 |
| ENSG00000197472 | ZNF695 | 124.5232093 | -0.591603794 | 0.168378598 | -3.513533199 | 0.000442189 | 0.001767216 |
| ENSG00000197479 | PCDHB11 | 18.38589397 | -0.135417208 | 0.403696639 | -0.335442992 | 0.737290942 | 0.855251527 |
| ENSG00000197483 | ZNF628 | 221.7247324 | 0.133808333 | 0.124815489 | 1.072049104 | 0.283697977 | 0.467564595 |
| ENSG00000197496 | SLC2A10 | 1232.378358 | -0.602310869 | 0.0656005 | -9.181498126 | 4.25E-20 | 7.13E-19 |
| ENSG00000197498 | RPF2 | 824.8930076 | 0.186888338 | 0.07222663 | 2.587526762 | 0.009666769 | 0.028870677 |
| ENSG00000197530 | MIB2 | 776.7709166 | -0.062693567 | 0.074293861 | -0.843859316 | 0.398748025 | 0.591597444 |
| ENSG00000197535 | MYO5A | 8978.63017 | -1.1372011 | 0.055920313 | -20.33610042 | 6.16E-92 | 8.68E-90 |
| ENSG00000197536 | IRF1-AS1 | 12.9519301 | 0.297637523 | 0.515386996 | 0.577502974 | 0.56359973 | 0.732229586 |
| ENSG00000197548 | ATG7 | 1071.368432 | -0.144637207 | 0.066380037 | -2.178926276 | 0.029337145 | 0.076156401 |
| ENSG00000197555 | SIPA1L1 | 4453.428504 | -0.860381307 | 0.051553948 | -16.68895088 | 1.58E-62 | 1.19E-60 |
| ENSG00000197557 | TTC30A | 157.5165626 | 0.381971502 | 0.159647718 | 2.392589797 | 0.016729934 | 0.046831565 |
| ENSG00000197558 | SSPOP | 473.8000275 | 0.940232472 | 0.097132853 | 9.679860576 | 3.67E-22 | 6.94E-21 |
| ENSG00000197562 | RAB40C | 1443.865266 | 0.140707888 | 0.060690873 | 2.318435727 | 0.020425652 | 0.055655596 |
| ENSG00000197563 | PIGN | 797.395843 | 0.782483957 | 0.078179334 | 10.0088337 | 1.39E-23 | 2.84E-22 |
| ENSG00000197565 | COL4A6 | 50.0705557 | 1.713410687 | 0.275288761 | 6.224048807 | 4.84E-10 | 3.97E-09 |
| ENSG00000197566 | ZNF624 | 278.5295728 | -0.113594357 | 0.112050298 | -1.013780053 | 0.310687716 | 0.498682913 |
| ENSG00000197568 | HHLA3 | 196.2724633 | -0.463671272 | 0.132993152 | -3.486429686 | 0.000489514 | 0.001940797 |
| ENSG00000197575 | RPS17P2 | 1.620270354 | 0.378330089 | 1.353830353 | 0.279451623 | 0.779898259 | NA |

| | | | | | | | |
|---|---|---|---|---|---|---|---|
| ENSG00000197576 | HOXA4 | 5.705934919 | -5.06488525 | 1.330673437 | -3.806257126 | 0.000141086 | 0.000614021 |
| ENSG00000197579 | TOPORS | 777.1892131 | 0.079453207 | 0.076290345 | 1.04145822 | 0.297662934 | 0.484214133 |
| ENSG00000197580 | BCO2 | 8.260608589 | 1.007304205 | 0.626839358 | 1.606957495 | 0.108063693 | 0.225608196 |
| ENSG00000197585 | ENSG0000019 | 1.559289561 | 2.863132081 | 1.645364964 | 1.740119756 | 0.081837992 | NA |
| ENSG00000197586 | ENTPD6 | 2060.012224 | -0.08876176 | 0.056046516 | -1.583715933 | 0.113258377 | 0.234454528 |
| ENSG00000197587 | DMBX1 | 70.37246996 | 0.001224045 | 0.222084364 | 0.005511623 | 0.995602384 | 0.997210788 |
| ENSG00000197594 | ENPP1 | 1064.673817 | -0.540213149 | 0.070302832 | -7.684087984 | 1.54E-14 | 1.79E-13 |
| ENSG00000197599 | CCDC154 | 57.44036515 | 0.985559049 | 0.255194202 | 3.86199624 | 0.000112464 | 0.000497768 |
| ENSG00000197601 | FAR1 | 2494.612422 | -0.132046553 | 0.054962127 | -2.402500777 | 0.0162834 | 0.045715481 |
| ENSG00000197603 | CPLANE1 | 1226.2843 | 1.978608316 | 0.07730483 | 25.59488616 | 1.74E-144 | 5.95E-142 |
| ENSG00000197608 | ZNF841 | 552.0247292 | 0.27128477 | 0.094655152 | 2.866032798 | 0.004156512 | 0.013642922 |
| ENSG00000197619 | ZNF615 | 5.220682521 | -4.022857177 | 1.245565069 | -3.229744698 | 0.001239008 | 0.004558108 |
| ENSG00000197620 | EOLA1 | 1059.762111 | -0.025900735 | 0.068103499 | -0.380314307 | 0.703712116 | 0.834185405 |
| ENSG00000197622 | CDC42SE1 | 10444.65131 | 0.296655298 | 0.04299262 | 6.900144657 | 5.19E-12 | 5.00E-11 |
| ENSG00000197635 | DPP4 | 1.964554299 | -0.020575584 | 1.374942484 | -0.014964687 | 0.988060353 | 0.994030148 |
| ENSG00000197647 | ZNF433 | 310.8574477 | -0.122530461 | 0.109058231 | -1.123532445 | 0.261211429 | 0.441485571 |
| ENSG00000197653 | DNAH10 | 16.52350516 | 0.885432057 | 0.453292404 | 1.953335306 | 0.050779877 | 0.12143129 |
| ENSG00000197670 | ENSG0000019 | 32.36376939 | -0.371531258 | 0.33045897 | -1.124288617 | 0.260890606 | 0.441163976 |
| ENSG00000197694 | SPTAN1 | 9728.814359 | -0.035161616 | 0.046787779 | -0.751512828 | 0.452344084 | 0.640066851 |
| ENSG00000197696 | NMB | 230.6806411 | -0.013514819 | 0.122225142 | -0.110573152 | 0.911954838 | 0.955697231 |
| ENSG00000197702 | PARVA | 2206.007055 | 0.248177594 | 0.055230608 | 4.493479289 | 7.01E-06 | 3.68E-05 |
| ENSG00000197705 | KLHL14 | 4.818354095 | 0.931320104 | 0.843309959 | 1.104362747 | 0.269435814 | 0.4510258 |
| ENSG00000197712 | FAM114A1 | 130.8583226 | -0.093514282 | 0.158644997 | -0.589456228 | 0.555555267 | 0.726586404 |
| ENSG00000197713 | RPE | 1324.077298 | -0.146490653 | 0.063479705 | -2.307676981 | 0.021017109 | 0.057057099 |
| ENSG00000197714 | ZNF460 | 58.11627058 | 0.032924319 | 0.248864826 | 0.132298001 | 0.894748588 | 0.946400434 |
| ENSG00000197724 | PHF2 | 3212.876282 | 0.018943422 | 0.053370367 | 0.354942703 | 0.722632516 | 0.845926239 |
| ENSG00000197728 | RPS26 | 158.2028277 | -0.293852846 | 0.147416557 | -1.993350355 | 0.046223096 | 0.111958381 |
| ENSG00000197734 | C14orf178 | 5.501382567 | -0.644713191 | 0.756525406 | -0.852202961 | 0.394101454 | 0.587019502 |
| ENSG00000197744 | PTMAP2 | 1567.351093 | -0.142073986 | 0.061505782 | -2.309928928 | 0.020892089 | 0.056784897 |
| ENSG00000197746 | PSAP | 12565.86463 | -0.148267707 | 0.047722401 | -3.1068786 | 0.00189074 | 0.006681082 |
| ENSG00000197747 | S100A10 | 7151.157868 | -0.306184549 | 0.046167507 | -6.632035639 | 3.31E-11 | 2.97E-10 |
| ENSG00000197748 | CFAP43 | 3.382371917 | -0.823188336 | 1.004551785 | -0.819458338 | 0.412524963 | 0.604603922 |
| ENSG00000197753 | LHFPL5 | 13.68098949 | -0.431142877 | 0.490623475 | -0.878765284 | 0.379528553 | 0.57196065 |

| | | | | | | | |
|---|---|---|---|---|---|---|---|
| ENSG00000197756 | RPL37A | 8695.211263 | 0.250094205 | 0.043693274 | 5.72386048 | 1.04E-08 | 7.48E-08 |
| ENSG00000197757 | HOXC6 | 243.9450557 | -0.553734104 | 0.120252348 | -4.604767488 | 4.13E-06 | 2.23E-05 |
| ENSG00000197763 | TXNRD3 | 418.2617783 | 0.273841786 | 0.09516274 | 2.877615603 | 0.004006931 | 0.013197271 |
| ENSG00000197766 | CFD | 50.53567072 | 0.114114826 | 0.264426746 | 0.431555537 | 0.66606448 | 0.807457548 |
| ENSG00000197768 | STPG3 | 1.585649181 | -1.333691995 | 1.494096118 | -0.892641363 | 0.372049261 | NA |
| ENSG00000197771 | MCMBP | 2427.556954 | -0.696083126 | 0.052462624 | -13.26817217 | 3.54E-40 | 1.46E-38 |
| ENSG00000197774 | EME2 | 3396.208617 | 0.513275832 | 0.065144715 | 7.879009502 | 3.30E-15 | 4.03E-14 |
| ENSG00000197776 | KLHDC1 | 41.37611914 | -0.975469379 | 0.280439561 | -3.478358667 | 0.000504494 | 0.001994331 |
| ENSG00000197779 | ZNF81 | 272.8111649 | -0.449748662 | 0.117290018 | -3.834500763 | 0.00012582 | 0.000551663 |
| ENSG00000197780 | TAF13 | 595.0886776 | 0.397272325 | 0.08308383 | 4.781584168 | 1.74E-06 | 9.79E-06 |
| ENSG00000197782 | ZNF780A | 437.1780215 | -0.410031278 | 0.098711622 | -4.153829794 | 3.27E-05 | 0.000157406 |
| ENSG00000197785 | ATAD3A | 1117.287205 | 0.672927808 | 0.067323009 | 9.99550994 | 1.59E-23 | 3.24E-22 |
| ENSG00000197798 | FAM118B | 219.1714746 | 0.014667746 | 0.122562133 | 0.119675999 | 0.904739814 | 0.952267525 |
| ENSG00000197808 | ZNF461 | 202.8924819 | 0.801115284 | 0.129929432 | 6.16577223 | 7.01E-10 | 5.66E-09 |
| ENSG00000197813 | ENSG0000019 | 10.01098153 | 0.629657927 | 0.567954491 | 1.108641515 | 0.267584861 | 0.448844451 |
| ENSG00000197816 | CCDC180 | 28.55955878 | 0.248303072 | 0.330153023 | 0.752084805 | 0.452000061 | 0.639856921 |
| ENSG00000197818 | SLC9A8 | 1520.866552 | -0.336130033 | 0.061805848 | -5.438482702 | 5.37E-08 | 3.58E-07 |
| ENSG00000197822 | OCLN | 1055.999079 | 0.118797305 | 0.068594349 | 1.731881796 | 0.08329461 | 0.182639874 |
| ENSG00000197837 | H4C16 | 41.83289301 | -0.049834244 | 0.287305657 | -0.173453753 | 0.862294776 | 0.928425897 |
| ENSG00000197841 | ZNF181 | 351.3270939 | 0.080073826 | 0.100189238 | 0.799225824 | 0.424159481 | 0.615019183 |
| ENSG00000197847 | SLC22A20P | 64.32305033 | 0.043564837 | 0.224953727 | 0.193661324 | 0.846441077 | 0.919600204 |
| ENSG00000197852 | INKA2 | 362.9034952 | -0.62427737 | 0.101535933 | -6.148339302 | 7.83E-10 | 6.28E-09 |
| ENSG00000197857 | ZNF44 | 319.739163 | -0.137176379 | 0.106653501 | -1.2861873 | 0.198377702 | 0.360915575 |
| ENSG00000197858 | GPAA1 | 3264.598915 | 0.204614201 | 0.064250275 | 3.184643189 | 0.001449326 | 0.005245746 |
| ENSG00000197860 | SGTB | 76.80270521 | -0.451734319 | 0.208255815 | -2.169131836 | 0.030072676 | 0.077854327 |
| ENSG00000197863 | ZNF790 | 148.1986898 | 0.233877046 | 0.157206742 | 1.487703666 | 0.13682906 | 0.27180972 |
| ENSG00000197870 | PRB3 | 10.73090023 | -0.749703981 | 0.557426319 | -1.344938255 | 0.178645175 | 0.333316008 |
| ENSG00000197872 | CYRIA | 21.2271731 | 1.645689586 | 0.417715671 | 3.939736289 | 8.16E-05 | 0.000369592 |
| ENSG00000197879 | MYO1C | 6507.028249 | -0.089819782 | 0.051602313 | -1.740615444 | 0.081751009 | 0.179822432 |
| ENSG00000197882 | OR7E13P | 2.187615436 | 1.712361777 | 1.284231145 | 1.333375058 | 0.182408753 | 0.3382811 |
| ENSG00000197885 | NKIRAS1 | 371.37168 | 0.280212134 | 0.103765116 | 2.700446393 | 0.00692465 | 0.021492553 |
| ENSG00000197889 | MEIG1 | 11.63363804 | -0.500481476 | 0.511993619 | -0.977515066 | 0.328314223 | 0.518700294 |
| ENSG00000197892 | KIF13B | 807.2934022 | -0.773328197 | 0.074708384 | -10.3512907 | 4.13E-25 | 9.20E-24 |

| | | | | | | | |
|---|---|---|---|---|---|---|---|
| ENSG00000197894 | ADH5 | 3063.226934 | -0.229014373 | 0.04957824 | -4.619251769 | 3.85E-06 | 2.08E-05 |
| ENSG00000197903 | H2BC12 | 2086.267263 | 0.326862913 | 0.058265793 | 5.609859535 | 2.02E-08 | 1.41E-07 |
| ENSG00000197905 | TEAD4 | 479.2177124 | -0.118211512 | 0.09241652 | -1.279116675 | 0.200855972 | 0.364297597 |
| ENSG00000197912 | SPG7 | 4594.022405 | 0.315133637 | 0.053570329 | 5.882615296 | 4.04E-09 | 3.04E-08 |
| ENSG00000197927 | NBEAP2 | 8.994501453 | -0.085469216 | 0.5913612 | -0.144529631 | 0.88508226 | 0.940548288 |
| ENSG00000197930 | ERO1A | 956.3068681 | -0.679168856 | 0.071640801 | -9.480196255 | 2.54E-21 | 4.59E-20 |
| ENSG00000197933 | ZNF823 | 212.4162537 | -0.022470362 | 0.126551835 | -0.177558562 | 0.85906967 | 0.926792302 |
| ENSG00000197935 | ZNF311 | 143.1276372 | -0.33302607 | 0.154751197 | -2.152009649 | 0.031396594 | 0.080809115 |
| ENSG00000197943 | PLCG2 | 225.2919327 | 0.546655713 | 0.124406325 | 4.394115106 | 1.11E-05 | 5.70E-05 |
| ENSG00000197948 | FCHSD1 | 1174.500657 | 0.337442043 | 0.064529776 | 5.229245513 | 1.70E-07 | 1.08E-06 |
| ENSG00000197956 | S100A6 | 1114.931714 | -1.606102191 | 0.070849092 | -22.66934035 | 8.99E-114 | 1.83E-111 |
| ENSG00000197958 | RPL12 | 5947.272419 | 0.249203435 | 0.051466232 | 4.842076592 | 1.28E-06 | 7.33E-06 |
| ENSG00000197959 | DNM3 | 17.53065674 | 1.269576168 | 0.453316915 | 2.800637096 | 0.005100184 | 0.01639169 |
| ENSG00000197961 | ZNF121 | 1107.246557 | -0.071661426 | 0.071891305 | -0.996802401 | 0.318860433 | 0.508174123 |
| ENSG00000197965 | MPZL1 | 2312.815344 | 0.235351302 | 0.052039927 | 4.522514029 | 6.11E-06 | 3.24E-05 |
| ENSG00000197969 | VPS13A | 967.0501854 | -0.006570535 | 0.072163759 | -0.091050342 | 0.92745259 | 0.965137362 |
| ENSG00000197971 | MBP | 772.5024057 | 0.865541563 | 0.07436501 | 11.639097 | 2.61E-31 | 7.66E-30 |
| ENSG00000197976 | AKAP17A | 1120.178947 | 0.015304173 | 0.065326811 | 0.234270928 | 0.814774654 | 0.901912272 |
| ENSG00000197976 | AKAP17A | 1120.178947 | 0.015304173 | 0.065326811 | 0.234270928 | 0.814774654 | 0.901912272 |
| ENSG00000197977 | ELOVL2 | 97.95270549 | 1.258120553 | 0.212320795 | 5.92556444 | 3.11E-09 | 2.37E-08 |
| ENSG00000197978 | GOLGA6L9 | 126.529245 | 0.047615298 | 0.168955249 | 0.281821951 | 0.778080039 | 0.879572073 |
| ENSG00000197980 | LEKR1 | 44.72579554 | -0.079186561 | 0.269514732 | -0.293811624 | 0.768901852 | 0.873855116 |
| ENSG00000197982 | C1orf122 | 822.154435 | -0.023061193 | 0.082222596 | -0.280472694 | 0.779114872 | 0.880184615 |
| ENSG00000197989 | SNHG12 | 343.4356431 | 0.225183588 | 0.110759294 | 2.033089763 | 0.042043455 | 0.103340497 |
| ENSG00000198000 | NOL8 | 1354.812035 | 0.252428985 | 0.064967342 | 3.88547501 | 0.00010213 | 0.000454817 |
| ENSG00000198001 | IRAK4 | 596.7801315 | 0.011608478 | 0.082143796 | 0.141318991 | 0.887617948 | 0.942116121 |
| ENSG00000198003 | ODAD3 | 20.27979189 | 1.101252118 | 0.405087193 | 2.718555748 | 0.006556761 | 0.020480949 |
| ENSG00000198010 | DLGAP2 | 1.928078734 | 2.125978349 | 1.443583188 | 1.472709273 | 0.140829445 | 0.277807848 |
| ENSG00000198015 | MRPL42 | 1014.387162 | 0.138230382 | 0.072956302 | 1.894701054 | 0.058132018 | 0.135923809 |
| ENSG00000198018 | ENTPD7 | 625.2858591 | -0.053958445 | 0.082546176 | -0.653675883 | 0.513320644 | 0.6925587 |
| ENSG00000198026 | ZNF335 | 1847.850849 | -0.212065688 | 0.064627388 | -3.281359421 | 0.00103308 | 0.003855353 |
| ENSG00000198028 | ZNF560 | 1.568756805 | 2.870142487 | 1.64424151 | 1.745572332 | 0.080885293 | NA |
| ENSG00000198034 | RPS4X | 20030.29661 | 0.241962022 | 0.052264381 | 4.629577868 | 3.66E-06 | 1.99E-05 |

| | | | | | | | |
|---|---|---|---|---|---|---|---|
| ENSG00000198039 | ZNF273 | 155.1448652 | 0.267107401 | 0.149184602 | 1.790448859 | 0.073381782 | 0.16498854 |
| ENSG00000198040 | ZNF84 | 1637.109918 | 0.570855668 | 0.062530768 | 9.12919656 | 6.90E-20 | 1.14E-18 |
| ENSG00000198042 | MAK16 | 895.4676871 | 0.094885119 | 0.071207949 | 1.332507397 | 0.182693512 | 0.33864132 |
| ENSG00000198053 | SIRPA | 130.9362251 | 1.39604445 | 0.165667968 | 8.426761488 | 3.55E-17 | 4.94E-16 |
| ENSG00000198054 | DSCR8 | 3.447979753 | 0.125862744 | 0.972750483 | 0.129388518 | 0.897050232 | 0.947984339 |
| ENSG00000198055 | GRK6 | 2388.780295 | -0.475205673 | 0.058914224 | -8.066060069 | 7.26E-16 | 9.29E-15 |
| ENSG00000198056 | PRIM1 | 387.280318 | -0.502084563 | 0.096565015 | -5.199445816 | 2.00E-07 | 1.26E-06 |
| ENSG00000198060 | MARCHF5 | 1738.40661 | -0.083951143 | 0.063248275 | -1.327327013 | 0.184400532 | 0.34108597 |
| ENSG00000198064 | NPIPB13 | 6.813877963 | -0.413561902 | 0.666437762 | -0.620555926 | 0.534891846 | 0.71038146 |
| ENSG00000198075 | SULT1C4 | 6.152841809 | -5.170102143 | 1.315158126 | -3.931163897 | 8.45E-05 | 0.000382118 |
| ENSG00000198081 | ZBTB14 | 739.2301488 | 0.061352818 | 0.076328399 | 0.803800664 | 0.421512106 | 0.612499166 |
| ENSG00000198087 | CD2AP | 1384.555956 | 0.040382918 | 0.063366402 | 0.637292264 | 0.523934489 | 0.700288829 |
| ENSG00000198088 | NUP62CL | 91.99459122 | -0.747808002 | 0.193607937 | -3.862486282 | 0.000112239 | 0.000496866 |
| ENSG00000198089 | SFI1 | 964.3995208 | -0.552404821 | 0.081248227 | -6.798976926 | 1.05E-11 | 9.85E-11 |
| ENSG00000198105 | ZNF248 | 497.2345116 | -0.406823136 | 0.0882725 | -4.608718867 | 4.05E-06 | 2.19E-05 |
| ENSG00000198108 | CHSY3 | 93.45918831 | 0.768217501 | 0.186938168 | 4.10947379 | 3.97E-05 | 0.000188035 |
| ENSG00000198113 | TOR4A | 79.3113851 | 0.00890421 | 0.198129292 | 0.044941411 | 0.964154009 | 0.983666795 |
| ENSG00000198121 | LPAR1 | 18.60710802 | 0.766701494 | 0.423158769 | 1.811853023 | 0.07000891 | 0.158585239 |
| ENSG00000198125 | MB | 477.0548936 | 1.578144306 | 0.097340567 | 16.2126065 | 4.11E-59 | 2.86E-57 |
| ENSG00000198130 | HIBCH | 521.8953601 | 0.091848549 | 0.086343095 | 1.063762523 | 0.287436293 | 0.471338739 |
| ENSG00000198131 | ZNF544 | 1192.541643 | 0.347398158 | 0.064677327 | 5.371250994 | 7.82E-08 | 5.11E-07 |
| ENSG00000198133 | TMEM229B | 610.1205944 | 0.411253347 | 0.09004069 | 4.567416673 | 4.94E-06 | 2.64E-05 |
| ENSG00000198134 | PTMAP9 | 19.37141231 | -0.426436289 | 0.402534794 | -1.059377465 | 0.289427908 | 0.473706922 |
| ENSG00000198142 | SOWAHC | 652.4962583 | -0.514231296 | 0.082287051 | -6.249237108 | 4.12E-10 | 3.41E-09 |
| ENSG00000198146 | ZNF770 | 1416.920802 | 0.024246226 | 0.066035831 | 0.36716772 | 0.713493916 | 0.840698055 |
| ENSG00000198153 | ZNF849P | 4.222979311 | 0.277545012 | 0.841867524 | 0.329677774 | 0.741643449 | 0.857515143 |
| ENSG00000198155 | ZNF876P | 52.32835729 | 0.615492963 | 0.246124889 | 2.500734344 | 0.012393611 | 0.035915794 |
| ENSG00000198156 | NPIPB6 | 8.347701504 | -0.224467363 | 0.651217852 | -0.34468859 | 0.730328497 | 0.850776726 |
| ENSG00000198157 | HMGN5 | 215.9189511 | -0.777713302 | 0.1272783 | -6.110336951 | 9.94E-10 | 7.90E-09 |
| ENSG00000198160 | MIER1 | 804.4634869 | 0.072798556 | 0.076679566 | 0.949386646 | 0.342424 | 0.533424321 |
| ENSG00000198162 | MAN1A2 | 4595.953542 | -0.062335054 | 0.052240593 | -1.193230218 | 0.232779223 | 0.40696044 |
| ENSG00000198168 | SVIP | 1163.801862 | 0.0817231 | 0.067071912 | 1.218439996 | 0.223056814 | 0.394246345 |
| ENSG00000198169 | ZNF251 | 1074.880795 | -0.095833833 | 0.085312647 | -1.123325039 | 0.261299473 | 0.4415299 |

| | | | | | | | |
|---|---|---|---|---|---|---|---|
| ENSG00000198171 | DDRGK1 | 906.9512534 | 0.030348986 | 0.074503348 | 0.407350631 | 0.683750481 | 0.820572118 |
| ENSG00000198176 | TFDP1 | 3827.672962 | 0.074308501 | 0.048350536 | 1.536870252 | 0.124325085 | 0.252413263 |
| ENSG00000198182 | ZNF607 | 336.3276743 | 0.042114468 | 0.104742622 | 0.402075745 | 0.687628282 | 0.823383061 |
| ENSG00000198189 | HSD17B11 | 83.98888491 | -0.942828477 | 0.203600578 | -4.630775061 | 3.64E-06 | 1.98E-05 |
| ENSG00000198198 | SZT2 | 1616.825959 | -0.161735828 | 0.060489971 | -2.673762717 | 0.00750055 | 0.02308326 |
| ENSG00000198205 | ZXDA | 210.6545919 | 0.124139832 | 0.127188346 | 0.976031504 | 0.329048853 | 0.519478277 |
| ENSG00000198208 | RPS6KL1 | 665.1227148 | 0.178634698 | 0.081350519 | 2.195864283 | 0.028101659 | 0.073439514 |
| ENSG00000198218 | QRICH1 | 2545.159439 | 0.033050001 | 0.050844906 | 0.650015977 | 0.515681901 | 0.6944375 |
| ENSG00000198221 | AFDN-DT | 11.23292634 | -0.874595469 | 0.532983187 | -1.640943822 | 0.100809077 | 0.213237458 |
| ENSG00000198223 | CSF2RA | 2.144114806 | 0.072668849 | 1.192378547 | 0.060944445 | 0.951403453 | 0.977495703 |
| ENSG00000198223 | CSF2RA | 2.144114806 | 0.072668849 | 1.192378547 | 0.060944445 | 0.951403453 | 0.977495703 |
| ENSG00000198225 | FKBP1C | 401.861343 | 0.098370052 | 0.096369558 | 1.020758574 | 0.307368836 | 0.495261719 |
| ENSG00000198231 | DDX42 | 9835.283058 | 0.187726408 | 0.052278699 | 3.59087756 | 0.000329566 | 0.001346199 |
| ENSG00000198242 | RPL23A | 3422.521896 | 0.34468535 | 0.053228279 | 6.475605765 | 9.44E-11 | 8.17E-10 |
| ENSG00000198246 | SLC29A3 | 367.488433 | -0.872304916 | 0.104540397 | -8.344189783 | 7.17E-17 | 9.82E-16 |
| ENSG00000198252 | STYX | 1021.057596 | -0.723173976 | 0.070794339 | -10.2151384 | 1.70E-24 | 3.67E-23 |
| ENSG00000198258 | UBL5 | 1595.243508 | -0.066784825 | 0.062073793 | -1.075894069 | 0.281974635 | 0.465393058 |
| ENSG00000198265 | HELZ | 4306.886731 | 0.062253458 | 0.057646934 | 1.079909262 | 0.280182588 | 0.463535885 |
| ENSG00000198270 | TMEM116 | 387.5539832 | -0.056225147 | 0.101614609 | -0.553317555 | 0.580045982 | 0.745573757 |
| ENSG00000198276 | UCKL1 | 1873.079593 | -0.198002745 | 0.060172003 | -3.290612503 | 0.000999695 | 0.003742331 |
| ENSG00000198298 | ZNF485 | 239.921851 | -0.668239868 | 0.12670101 | -5.2741479 | 1.33E-07 | 8.55E-07 |
| ENSG00000198301 | SDAD1 | 1448.117507 | 0.090432987 | 0.061933203 | 1.460169713 | 0.144243437 | 0.282839939 |
| ENSG00000198315 | ZKSCAN8 | 1187.214442 | -0.076747968 | 0.067994228 | -1.128742395 | 0.259006519 | 0.438819022 |
| ENSG00000198324 | PHETA1 | 266.0292926 | 0.357207294 | 0.118882419 | 3.00471085 | 0.002658334 | 0.009109978 |
| ENSG00000198331 | HYLS1 | 288.6455882 | -0.095969002 | 0.112367881 | -0.854060797 | 0.393071304 | 0.586009536 |
| ENSG00000198342 | ZNF442 | 125.5283419 | -0.175651027 | 0.160793898 | -1.092398591 | 0.274657942 | 0.456767666 |
| ENSG00000198346 | ZNF813 | 482.9819256 | 0.187245531 | 0.089755796 | 2.086166454 | 0.036963548 | 0.092843949 |
| ENSG00000198353 | HOXC4 | 137.9555424 | -0.567272694 | 0.159925184 | -3.547112976 | 0.000389478 | 0.001568577 |
| ENSG00000198354 | DCAF12L2 | 51.02036472 | 0.212218502 | 0.249349926 | 0.851087086 | 0.394720979 | 0.587751477 |
| ENSG00000198355 | PIM3 | 1717.375933 | -0.017438196 | 0.058013167 | -0.300590311 | 0.763726921 | 0.870479014 |
| ENSG00000198356 | GET3 | 2103.692104 | -0.022539287 | 0.060972651 | -0.36966225 | 0.711634163 | 0.839694845 |
| ENSG00000198363 | ASPH | 1323.58791 | 0.679102652 | 0.06642561 | 10.22350648 | 1.56E-24 | 3.38E-23 |
| ENSG00000198369 | SPRED2 | 2182.277716 | -0.405553198 | 0.053903855 | -7.523639943 | 5.33E-14 | 5.93E-13 |

| | | | | | | | |
|---|---|---|---|---|---|---|---|
| ENSG00000198373 | WWP2 | 2502.18685 | 0.078966672 | 0.05791313 | 1.363536584 | 0.172713463 | 0.324837816 |
| ENSG00000198380 | GFPT1 | 2210.902213 | -0.02220367 | 0.059079478 | -0.375827116 | 0.707045427 | 0.836461297 |
| ENSG00000198382 | UVRAG | 913.9812589 | 0.239011871 | 0.070322627 | 3.398790408 | 0.000676846 | 0.002616589 |
| ENSG00000198393 | ZNF26 | 492.1984124 | 0.489341355 | 0.093635046 | 5.226049161 | 1.73E-07 | 1.10E-06 |
| ENSG00000198399 | ITSN2 | 1420.400861 | -0.113077723 | 0.0656983 | -1.721166653 | 0.085220587 | 0.186098074 |
| ENSG00000198406 | BZW1P2 | 2755.451983 | -0.466403104 | 0.058512078 | -7.971056866 | 1.57E-15 | 1.96E-14 |
| ENSG00000198408 | OGA | 4550.990416 | 0.31235168 | 0.054972704 | 5.681941388 | 1.33E-08 | 9.45E-08 |
| ENSG00000198416 | ZNF658B | 19.73932358 | -0.160098275 | 0.39812624 | -0.402129423 | 0.68758878 | 0.823383061 |
| ENSG00000198417 | MT1F | 195.8784802 | -0.692847085 | 0.133298766 | -5.19770067 | 2.02E-07 | 1.27E-06 |
| ENSG00000198420 | TCAF1 | 879.4580229 | 0.022334068 | 0.070848928 | 0.315235081 | 0.752583172 | 0.863428755 |
| ENSG00000198431 | TXNRD1 | 4744.496717 | -0.387021636 | 0.048948472 | -7.906715404 | 2.64E-15 | 3.25E-14 |
| ENSG00000198435 | NRARP | 459.5283113 | 0.26207134 | 0.090082131 | 2.9092489 | 0.003622983 | 0.012052092 |
| ENSG00000198440 | ZNF583 | 25.97241703 | 0.991727842 | 0.355860464 | 2.786844683 | 0.005322398 | 0.017021583 |
| ENSG00000198443 | KRTAP4-1 | 28.29601069 | 1.787106669 | 0.362762983 | 4.926375496 | 8.38E-07 | 4.90E-06 |
| ENSG00000198453 | ZNF568 | 68.98779521 | 0.822190638 | 0.218314518 | 3.766083198 | 0.000165829 | 0.00071385 |
| ENSG00000198455 | ZXDB | 772.0755809 | 0.263976134 | 0.077241952 | 3.41752283 | 0.000631938 | 0.002455396 |
| ENSG00000198464 | ZNF480 | 1099.709789 | 0.215746122 | 0.072523693 | 2.974836385 | 0.002931449 | 0.009949316 |
| ENSG00000198466 | ZNF587 | 1094.04128 | 0.329890799 | 0.070095836 | 4.706282413 | 2.52E-06 | 1.39E-05 |
| ENSG00000198467 | TPM2 | 330.6535935 | -0.010481191 | 0.112190486 | -0.093423171 | 0.925567383 | 0.964412962 |
| ENSG00000198468 | FLVCR1-DT | 31.67505666 | -0.144148998 | 0.318653786 | -0.452368695 | 0.651003395 | 0.797203752 |
| ENSG00000198478 | SH3BGRL2 | 841.7253587 | 0.043766245 | 0.074654709 | 0.586248956 | 0.557708228 | 0.727915708 |
| ENSG00000198482 | ZNF808 | 229.3195069 | -0.023562403 | 0.127938747 | -0.184169407 | 0.853880559 | 0.923919627 |
| ENSG00000198483 | ANKRD35 | 59.4043706 | 0.033120077 | 0.235046494 | 0.140908622 | 0.887942131 | 0.942285816 |
| ENSG00000198492 | YTHDF2 | 3275.485421 | 0.216445297 | 0.048041288 | 4.505401638 | 6.62E-06 | 3.49E-05 |
| ENSG00000198496 | NBR2 | 292.4381121 | 0.105602561 | 0.119362167 | 0.884723889 | 0.376305568 | 0.569310145 |
| ENSG00000198498 | TMA16 | 332.9670827 | 0.0724393 | 0.108673944 | 0.666574689 | 0.505043841 | 0.685625856 |
| ENSG00000198502 | HLA-DRB5 | 1.538504036 | 3.973318516 | 1.68509989 | 2.35791275 | 0.018378012 | NA |
| ENSG00000198513 | ATL1 | 27.93354973 | -1.511969259 | 0.36743464 | -4.114933907 | 3.87E-05 | 0.000183829 |
| ENSG00000198515 | CNGA1 | 57.86362096 | 0.63131488 | 0.239685931 | 2.633925482 | 0.008440401 | 0.025604708 |
| ENSG00000198517 | MAFK | 1551.222657 | 0.331892788 | 0.065990034 | 5.02943806 | 4.92E-07 | 2.96E-06 |
| ENSG00000198520 | ARMH1 | 44.473516 | -1.265372412 | 0.282058359 | -4.486207801 | 7.25E-06 | 3.80E-05 |
| ENSG00000198522 | GPN1 | 1086.148966 | -0.060187182 | 0.065631563 | -0.917046312 | 0.359118373 | 0.551682936 |
| ENSG00000198535 | C2CD4A | 15.58560307 | 1.151575244 | 0.460750247 | 2.499348079 | 0.012442203 | 0.036042953 |

| | | | | | | | |
|---|---|---|---|---|---|---|---|
| ENSG00000198538 | ZNF28 | 1058.998892 | 0.492029614 | 0.067430209 | 7.296872152 | 2.95E-13 | 3.12E-12 |
| ENSG00000198546 | ZNF511 | 184.1464628 | -0.745410412 | 0.136224636 | -5.471920763 | 4.45E-08 | 2.99E-07 |
| ENSG00000198551 | ZNF627 | 746.5995972 | 0.086141307 | 0.077107173 | 1.117163346 | 0.26392449 | 0.444524801 |
| ENSG00000198553 | KCNRG | 2.174896282 | -2.580347041 | 1.431959757 | -1.801968965 | 0.071550288 | 0.161615858 |
| ENSG00000198554 | WDHD1 | 1001.022774 | -0.629690897 | 0.073238317 | -8.597834022 | 8.12E-18 | 1.18E-16 |
| ENSG00000198556 | ZNF789 | 499.1459946 | -0.713635224 | 0.102897004 | -6.93543248 | 4.05E-12 | 3.93E-11 |
| ENSG00000198561 | CTNND1 | 195.8938957 | -0.007842479 | 0.136139994 | -0.057605992 | 0.954062477 | 0.97885385 |
| ENSG00000198563 | DDX39B | 3798.623733 | -0.661847951 | 0.184957177 | -3.578384805 | 0.000345724 | 0.001404198 |
| ENSG00000198569 | SLC34A3 | 29.65249763 | 0.2702429 | 0.324264281 | 0.833403232 | 0.404617353 | 0.597522986 |
| ENSG00000198576 | ARC | 47.20400354 | -0.200699322 | 0.259927871 | -0.772134675 | 0.440034667 | 0.629444114 |
| ENSG00000198580 | ENSG0000019 | 13.43698121 | 1.350581162 | 0.508678967 | 2.655075694 | 0.007929061 | 0.024271773 |
| ENSG00000198585 | NUDT16 | 3104.583133 | 0.159893091 | 0.048495166 | 3.297093365 | 0.00097691 | 0.003664212 |
| ENSG00000198586 | TLK1 | 2185.544994 | -0.085710509 | 0.05458018 | -1.57035959 | 0.116331478 | 0.239662695 |
| ENSG00000198589 | LRBA | 5024.797172 | -0.285547133 | 0.062613176 | -4.560495906 | 5.10E-06 | 2.73E-05 |
| ENSG00000198590 | APRG1 | 21.61852713 | -0.105581142 | 0.388930951 | -0.271465003 | 0.786033413 | 0.883788848 |
| ENSG00000198598 | MMP17 | 203.8511683 | 0.268768002 | 0.137707571 | 1.95173003 | 0.050970265 | 0.121827912 |
| ENSG00000198604 | BAZ1A | 2849.768181 | -0.315570282 | 0.050935755 | -6.195456992 | 5.81E-10 | 4.73E-09 |
| ENSG00000198612 | COPS8 | 886.8266355 | 0.100287495 | 0.070142153 | 1.429774984 | 0.152781611 | 0.29554011 |
| ENSG00000198618 | PPIAP22 | 8531.568343 | 0.261360732 | 0.060976434 | 4.286258086 | 1.82E-05 | 9.07E-05 |
| ENSG00000198624 | CCDC69 | 221.7704721 | 1.87234827 | 0.139343606 | 13.43691561 | 3.67E-41 | 1.57E-39 |
| ENSG00000198625 | MDM4 | 2812.956055 | 0.196562067 | 0.073548379 | 2.672554719 | 0.00752761 | 0.023160323 |
| ENSG00000198633 | ZNF534 | 3.432527102 | 0.813328506 | 0.970708415 | 0.837871078 | 0.402103114 | 0.59476726 |
| ENSG00000198642 | KLHL9 | 942.2446209 | -0.005521817 | 0.069188028 | -0.079808845 | 0.936389289 | 0.969525239 |
| ENSG00000198646 | NCOA6 | 2324.096934 | -0.682010195 | 0.058683765 | -11.62178667 | 3.19E-31 | 9.33E-30 |
| ENSG00000198648 | STK39 | 970.7103235 | 0.030687592 | 0.070042355 | 0.438129073 | 0.661292721 | 0.804222818 |
| ENSG00000198650 | TAT | 2.717690146 | 2.69506828 | 1.340456206 | 2.010560486 | 0.044371901 | 0.108171245 |
| ENSG00000198658 | ABHD17AP1 | 88.82485466 | 0.070555298 | 0.192573177 | 0.366381752 | 0.714080235 | 0.840989046 |
| ENSG00000198663 | C6orf89 | 3458.063741 | 0.02508566 | 0.05096515 | 0.492212032 | 0.622569459 | 0.776597778 |
| ENSG00000198668 | CALM1 | 13328.27386 | -0.703109909 | 0.0430853 | -16.31902092 | 7.23E-60 | 5.13E-58 |
| ENSG00000198673 | TAFA2 | 20.76024299 | -0.093670968 | 0.388235997 | -0.241273268 | 0.809343328 | 0.899052634 |
| ENSG00000198677 | SKIC3 | 2444.6356 | 0.564801063 | 0.06189885 | 9.124580873 | 7.20E-20 | 1.19E-18 |
| ENSG00000198680 | TUSC1 | 254.3251872 | -0.010945622 | 0.119832112 | -0.091341312 | 0.927221393 | 0.965137362 |
| ENSG00000198681 | MAGEA1 | 4.896226939 | -1.10803857 | 0.823213429 | -1.345991856 | 0.178305143 | 0.332821925 |

| | | | | | | | |
|---|---|---|---|---|---|---|---|
| ENSG00000198682 | PAPSS2 | 275.4134074 | 0.729556111 | 0.113629255 | 6.420495447 | 1.36E-10 | 1.16E-09 |
| ENSG00000198689 | SLC9A6 | 734.8477659 | -0.403670752 | 0.075765252 | -5.327914086 | 9.93E-08 | 6.44E-07 |
| ENSG00000198690 | FAN1 | 1162.255636 | -0.41220106 | 0.069537071 | -5.927788656 | 3.07E-09 | 2.34E-08 |
| ENSG00000198691 | ABCA4 | 6.813903621 | -0.39580174 | 0.672181825 | -0.588831363 | 0.555974405 | 0.726719554 |
| ENSG00000198695 | MT-ND6 | 1363.949192 | -0.500996025 | 0.086445495 | -5.795513383 | 6.81E-09 | 4.99E-08 |
| ENSG00000198700 | IPO9 | 3236.809011 | -0.049207379 | 0.048464189 | -1.015334836 | 0.309946244 | 0.497910843 |
| ENSG00000198707 | CEP290 | 584.2770623 | -0.178913827 | 0.084996104 | -2.104965047 | 0.035294351 | 0.089286767 |
| ENSG00000198712 | MT-CO2 | 49781.9563 | -0.052853148 | 0.12358867 | -0.427653666 | 0.668903288 | 0.809829082 |
| ENSG00000198715 | GLMP | 624.3784362 | 0.57813226 | 0.085883075 | 6.731620406 | 1.68E-11 | 1.54E-10 |
| ENSG00000198718 | TOGARAM1 | 1354.644023 | -0.497656946 | 0.067968179 | -7.321910796 | 2.44E-13 | 2.60E-12 |
| ENSG00000198719 | DLL1 | 312.3446644 | 0.143196675 | 0.107117897 | 1.336813724 | 0.181283448 | 0.336654761 |
| ENSG00000198720 | ANKRD13B | 748.343728 | 0.626207761 | 0.076031553 | 8.236156352 | 1.78E-16 | 2.38E-15 |
| ENSG00000198721 | ECI2 | 480.0145033 | -0.904744407 | 0.092626813 | -9.767629644 | 1.55E-22 | 3.00E-21 |
| ENSG00000198722 | UNC13B | 2068.782539 | -0.549315661 | 0.058146273 | -9.447134435 | 3.48E-21 | 6.25E-20 |
| ENSG00000198723 | TEX45 | 22.38641891 | -0.601408082 | 0.372797538 | -1.613229758 | 0.106694601 | 0.223278064 |
| ENSG00000198727 | MT-CYB | 62935.26151 | -0.166669976 | 0.133880303 | -1.244917834 | 0.213161955 | 0.381340969 |
| ENSG00000198728 | LDB1 | 3061.113544 | 0.337284524 | 0.054592194 | 6.178255557 | 6.48E-10 | 5.25E-09 |
| ENSG00000198730 | CTR9 | 1629.166492 | -0.136095828 | 0.058846413 | -2.312729363 | 0.020737524 | 0.056424954 |
| ENSG00000198732 | SMOC1 | 1.785268457 | 0.057034606 | 1.312660724 | 0.043449618 | 0.965343126 | NA |
| ENSG00000198736 | MSRB1 | 1014.138172 | -0.068647323 | 0.072557565 | -0.946108418 | 0.344093291 | 0.535012331 |
| ENSG00000198738 | SMIM11P1 | 1.650085851 | -0.222697489 | 1.478581275 | -0.150615656 | 0.88027891 | NA |
| ENSG00000198739 | LRRTM3 | 2.552085654 | 0.205553961 | 1.080143252 | 0.190302499 | 0.849072095 | 0.921106302 |
| ENSG00000198740 | ZNF652 | 1755.017874 | 0.025999868 | 0.063041793 | 0.412422731 | 0.680029605 | 0.818289926 |
| ENSG00000198742 | SMURF1 | 1790.146232 | -0.222348407 | 0.058811716 | -3.780682177 | 0.000156399 | 0.000675795 |
| ENSG00000198743 | SLC5A3 | 687.844945 | 0.018951503 | 0.088356079 | 0.214490086 | 0.830164898 | 0.909932761 |
| ENSG00000198744 | MTCO3P12 | 267.3185598 | -0.086930061 | 0.139756683 | -0.622010044 | 0.533935263 | 0.709453571 |
| ENSG00000198746 | GPATCH3 | 303.4433987 | -0.269523215 | 0.112612177 | -2.393375405 | 0.01669415 | 0.046737101 |
| ENSG00000198752 | CDC42BPB | 7280.167481 | -0.340207494 | 0.042952209 | -7.920605242 | 2.36E-15 | 2.91E-14 |
| ENSG00000198753 | PLXNB3 | 820.8360535 | -0.289901622 | 0.081138792 | -3.572910259 | 0.000353036 | 0.001431867 |
| ENSG00000198754 | OXCT2 | 10.9106596 | 0.467564703 | 0.538243855 | 0.868685632 | 0.385019103 | 0.577659856 |
| ENSG00000198755 | RPL10A | 11308.7303 | 0.266601639 | 0.046707717 | 5.70787127 | 1.14E-08 | 8.16E-08 |
| ENSG00000198756 | COLGALT2 | 15.03851653 | 0.198673003 | 0.455311345 | 0.436345383 | 0.662586159 | 0.805070259 |
| ENSG00000198763 | MT-ND2 | 36959.08331 | -0.116750244 | 0.122499372 | -0.953068097 | 0.340555569 | 0.531560506 |

| | | | | | | | |
|---|---|---|---|---|---|---|---|
| ENSG00000198780 | FAM169A | 523.0645208 | -0.155482955 | 0.088628807 | -1.754316241 | 0.079376334 | 0.17566015 |
| ENSG00000198783 | ZNF830 | 283.1538771 | -0.026974872 | 0.110536627 | -0.244035599 | 0.807203249 | 0.897631612 |
| ENSG00000198785 | GRIN3A | 2.748335299 | -0.008762444 | 1.052525322 | -0.008325162 | 0.993357558 | 0.996354439 |
| ENSG00000198786 | MT-ND5 | 77581.03397 | 0.051545017 | 0.113905389 | 0.452524828 | 0.650890939 | 0.797170242 |
| ENSG00000198788 | MUC2 | 5.506812878 | -4.101369897 | 1.206169219 | -3.400327113 | 0.000673053 | 0.002602804 |
| ENSG00000198791 | CNOT7 | 2419.195982 | 0.027489547 | 0.056596482 | 0.485711223 | 0.627171926 | 0.780154165 |
| ENSG00000198792 | TMEM184B | 2655.476376 | 0.361329822 | 0.062526928 | 5.778787393 | 7.52E-09 | 5.47E-08 |
| ENSG00000198793 | MTOR | 2993.611682 | -0.073282783 | 0.054236117 | -1.351180498 | 0.17663762 | 0.330349503 |
| ENSG00000198794 | SCAMP5 | 1516.539388 | 0.40991578 | 0.06727456 | 6.093176657 | 1.11E-09 | 8.77E-09 |
| ENSG00000198795 | ZNF521 | 13.73802846 | 1.958606939 | 0.537251028 | 3.645608544 | 0.00026676 | 0.001106195 |
| ENSG00000198797 | BRINP2 | 194.3812184 | -0.114929497 | 0.132489645 | -0.867460221 | 0.385689898 | 0.578320561 |
| ENSG00000198799 | LRIG2 | 594.6320788 | -0.174138991 | 0.085242919 | -2.042855785 | 0.041066716 | 0.101341488 |
| ENSG00000198804 | MT-CO1 | 103619.0377 | -0.201359498 | 0.158063406 | -1.273915971 | 0.202693198 | 0.366817762 |
| ENSG00000198805 | PNP | 1905.215139 | -0.41636695 | 0.059781234 | -6.964843634 | 3.29E-12 | 3.23E-11 |
| ENSG00000198807 | PAX9 | 280.8890298 | -0.773705856 | 0.110387738 | -7.008983728 | 2.40E-12 | 2.38E-11 |
| ENSG00000198814 | GK | 73.54791832 | 0.245623782 | 0.223813889 | 1.097446561 | 0.272446229 | 0.454523114 |
| ENSG00000198815 | FOXJ3 | 2322.686085 | -0.049721967 | 0.052243194 | -0.951740567 | 0.34122857 | 0.532321211 |
| ENSG00000198816 | ZNF358 | 445.9665918 | -0.006980791 | 0.095249377 | -0.073289626 | 0.941575647 | 0.972202592 |
| ENSG00000198818 | SFT2D1 | 539.1467231 | -0.154529288 | 0.083274964 | -1.8556512 | 0.063503291 | 0.146246912 |
| ENSG00000198821 | CD247 | 2.370580422 | -1.068894375 | 1.191584392 | -0.897036234 | 0.369699583 | 0.562357167 |
| ENSG00000198824 | CHAMP1 | 1173.812113 | -0.180227328 | 0.071308885 | -2.527417563 | 0.01149048 | 0.033587105 |
| ENSG00000198825 | INPP5F | 403.1380402 | -0.595909612 | 0.099744569 | -5.97435646 | 2.31E-09 | 1.78E-08 |
| ENSG00000198826 | ARHGAP11A | 2121.519961 | -0.437555407 | 0.061154855 | -7.15487604 | 8.37E-13 | 8.62E-12 |
| ENSG00000198830 | HMGN2 | 4141.837803 | -0.403933203 | 0.117851216 | -3.427484379 | 0.000609201 | 0.002374291 |
| ENSG00000198832 | SELENOM | 493.1649784 | -0.077719959 | 0.097298053 | -0.798782265 | 0.424416676 | 0.615226766 |
| ENSG00000198833 | UBE2J1 | 2779.408045 | 0.216815224 | 0.051274823 | 4.228492852 | 2.35E-05 | 0.000115467 |
| ENSG00000198835 | GJC2 | 16.48583862 | 0.558434187 | 0.445692801 | 1.252957611 | 0.210221134 | 0.377439447 |
| ENSG00000198836 | OPA1 | 2779.174446 | -0.229734162 | 0.058988917 | -3.894530945 | 9.84E-05 | 0.000439028 |
| ENSG00000198837 | DENND4B | 3368.475516 | 0.172004263 | 0.055930612 | 3.075315257 | 0.002102802 | 0.007356759 |
| ENSG00000198839 | ZNF277 | 401.8060977 | 0.16277175 | 0.097886035 | 1.662869995 | 0.096338462 | 0.205658321 |
| ENSG00000198840 | MT-ND3 | 7858.584718 | -0.083911836 | 0.065045146 | -1.290055317 | 0.197031452 | 0.358950248 |
| ENSG00000198841 | KTI12 | 107.3157169 | 0.17277376 | 0.170637226 | 1.01252091 | 0.311289054 | 0.499403517 |
| ENSG00000198843 | SELENOT | 3747.60071 | -0.049090788 | 0.054257346 | -0.904776805 | 0.365583642 | 0.558348008 |

| | | | | | | | |
|---|---|---|---|---|---|---|---|
| ENSG00000198853 | RUSC2 | 166.7920957 | -0.051242939 | 0.141744786 | -0.361515511 | 0.717714112 | 0.843020378 |
| ENSG00000198855 | FICD | 199.2646479 | 0.061053056 | 0.13451809 | 0.453865023 | 0.64992598 | 0.796349063 |
| ENSG00000198856 | OSTC | 1552.478282 | -0.102878934 | 0.065537246 | -1.569778101 | 0.116466744 | 0.239855203 |
| ENSG00000198858 | R3HDM4 | 1654.828961 | 0.30366126 | 0.06042324 | 5.025570618 | 5.02E-07 | 3.02E-06 |
| ENSG00000198860 | TSEN15 | 681.489246 | -0.234034721 | 0.080977072 | -2.890135649 | 0.003850756 | 0.012739594 |
| ENSG00000198862 | LTN1 | 1407.620833 | -0.04604857 | 0.062175403 | -0.740623587 | 0.458921703 | 0.646137686 |
| ENSG00000198863 | RUNDC1 | 1165.899226 | -0.042730035 | 0.070263433 | -0.608140439 | 0.543094334 | 0.71671081 |
| ENSG00000198865 | CCDC152 | 24.54604624 | -0.938894265 | 0.384501976 | -2.441845095 | 0.014612415 | 0.04152221 |
| ENSG00000198868 | MTND4LP30 | 11.68036646 | 0.321558226 | 0.545014272 | 0.589999643 | 0.555190889 | 0.726323296 |
| ENSG00000198870 | STKLD1 | 68.82351749 | 0.034227034 | 0.218273259 | 0.156808187 | 0.87539602 | 0.935410563 |
| ENSG00000198873 | GRK5 | 112.3559253 | -0.461281566 | 0.173987589 | -2.651232589 | 0.008019859 | 0.024497413 |
| ENSG00000198874 | TYW1 | 709.2544694 | 0.51034504 | 0.077998143 | 6.543040927 | 6.03E-11 | 5.30E-10 |
| ENSG00000198876 | DCAF12 | 2666.735409 | 0.276139225 | 0.052776808 | 5.232207814 | 1.67E-07 | 1.06E-06 |
| ENSG00000198879 | SFMBT2 | 186.0944675 | -2.705424981 | 0.165459767 | -16.35095364 | 4.28E-60 | 3.07E-58 |
| ENSG00000198886 | MT-ND4 | 90658.64785 | 0.054432938 | 0.126412313 | 0.430598386 | 0.666760415 | 0.808130391 |
| ENSG00000198887 | SMC5 | 1032.285212 | 0.047153707 | 0.067774597 | 0.695743082 | 0.48658974 | 0.67008339 |
| ENSG00000198888 | MT-ND1 | 34189.77049 | -0.110648567 | 0.131298658 | -0.842724274 | 0.399382668 | 0.592163734 |
| ENSG00000198889 | DCAF12L1 | 2.241570062 | 4.50285244 | 1.588124297 | 2.835327467 | 0.004577873 | 0.01488961 |
| ENSG00000198890 | PRMT6 | 9482.990744 | 0.446967916 | 0.041605212 | 10.74307495 | 6.39E-27 | 1.58E-25 |
| ENSG00000198892 | SHISA4 | 278.9265296 | 0.277899782 | 0.112904139 | 2.461378167 | 0.01384044 | 0.039603303 |
| ENSG00000198894 | CIPC | 1131.362216 | 0.052686836 | 0.065295556 | 0.806897724 | 0.41972541 | 0.611023031 |
| ENSG00000198898 | CAPZA2 | 2200.131939 | 0.00190321 | 0.054264228 | 0.035073007 | 0.972021525 | 0.987343862 |
| ENSG00000198899 | MT-ATP6 | 29194.47072 | -0.15649723 | 0.145159272 | -1.078107024 | 0.280985997 | 0.464439713 |
| ENSG00000198900 | TOP1 | 9042.278728 | -0.114246667 | 0.041436249 | -2.757167182 | 0.005830453 | 0.018448359 |
| ENSG00000198901 | PRC1 | 57.08389281 | 0.094139291 | 0.23718782 | 0.396897662 | 0.691442929 | 0.826001186 |
| ENSG00000198909 | MAP3K3 | 2601.750052 | 0.097937091 | 0.058676952 | 1.669089602 | 0.095099624 | 0.203525083 |
| ENSG00000198910 | L1CAM | 2829.413418 | -1.190610224 | 0.131779082 | -9.034895413 | 1.64E-19 | 2.67E-18 |
| ENSG00000198911 | SREBF2 | 8988.48266 | 0.015850373 | 0.045694843 | 0.346874435 | 0.728685653 | 0.849651768 |
| ENSG00000198912 | C1orf174 | 453.4554241 | -0.048761572 | 0.091651664 | -0.532031496 | 0.594704182 | 0.756386934 |
| ENSG00000198914 | POU3F3 | 4.679671352 | -1.720903543 | 0.899469402 | -1.913243005 | 0.05571695 | 0.131393527 |
| ENSG00000198915 | RASGEF1A | 7.982562799 | -0.637779884 | 0.646735576 | -0.986152468 | 0.3240583 | 0.514171905 |
| ENSG00000198917 | SPOUT1 | 6.954431425 | -0.313770128 | 0.695015256 | -0.451457901 | 0.651659561 | 0.797581514 |
| ENSG00000198918 | RPL39 | 498.7750895 | 0.165907588 | 0.096122322 | 1.726004794 | 0.08434655 | 0.184470384 |

| | | | | | | | |
|---|---|---|---|---|---|---|---|
| ENSG00000198919 | DZIP3 | 1201.542376 | 0.095289766 | 0.076275443 | 1.249285001 | 0.211560852 | 0.379127095 |
| ENSG00000198920 | KIAA0753 | 485.3764188 | -0.019316159 | 0.089238801 | -0.21645471 | 0.828633324 | 0.909383747 |
| ENSG00000198924 | DCLRE1A | 735.4189468 | -1.826782475 | 0.084039318 | -21.73723574 | 9.12E-105 | 1.66E-102 |
| ENSG00000198925 | ATG9A | 1359.461904 | 0.458315019 | 0.067582574 | 6.78155609 | 1.19E-11 | 1.11E-10 |
| ENSG00000198929 | NOS1AP | 161.8766828 | 0.653374051 | 0.145002222 | 4.505958898 | 6.61E-06 | 3.49E-05 |
| ENSG00000198931 | APRT | 3340.424632 | 0.504191811 | 0.059351007 | 8.495084377 | 1.98E-17 | 2.80E-16 |
| ENSG00000198932 | GPRASP1 | 128.9383101 | -0.379031676 | 0.168804518 | -2.245388219 | 0.024743221 | 0.065886186 |
| ENSG00000198933 | TBKBP1 | 446.4241516 | 0.512926519 | 0.096109657 | 5.336888442 | 9.46E-08 | 6.14E-07 |
| ENSG00000198934 | MAGEE1 | 192.2503841 | 0.184130864 | 0.141004054 | 1.305855109 | 0.191601835 | 0.351599559 |
| ENSG00000198937 | CCDC167 | 334.7081495 | 0.102240505 | 0.103568047 | 0.987181935 | 0.323553456 | 0.513619561 |
| ENSG00000198938 | MT-CO3 | 46238.32862 | -0.242826215 | 0.152919336 | -1.587936629 | 0.11230067 | 0.232822174 |
| ENSG00000198939 | ZFP2 | 71.59662459 | -0.371599558 | 0.222975447 | -1.666549227 | 0.095604072 | 0.204433018 |
| ENSG00000198944 | SOWAHA | 41.26290836 | -0.278593079 | 0.279893601 | -0.995353512 | 0.31956436 | 0.509048429 |
| ENSG00000198945 | L3MBTL3 | 41.70211419 | -0.650203768 | 0.282589514 | -2.300877191 | 0.021398574 | 0.057962366 |
| ENSG00000198947 | DMD | 21.68835991 | -0.045515628 | 0.372952746 | -0.122041274 | 0.902866331 | 0.950907167 |
| ENSG00000198948 | MFAP3L | 190.4955655 | 0.246119996 | 0.137167942 | 1.794296785 | 0.072765811 | 0.163876664 |
| ENSG00000198951 | NAGA | 904.2383472 | -0.11007848 | 0.070786622 | -1.555074623 | 0.119928298 | 0.245441624 |
| ENSG00000198952 | SMG5 | 7064.29926 | 0.25857974 | 0.047934577 | 5.39443047 | 6.87E-08 | 4.51E-07 |
| ENSG00000198954 | KIFBP | 1239.734583 | -0.849559084 | 0.064421073 | -13.18759599 | 1.03E-39 | 4.16E-38 |
| ENSG00000198959 | TGM2 | 33.49706775 | -1.716477741 | 0.332959632 | -5.155212756 | 2.53E-07 | 1.58E-06 |
| ENSG00000198960 | ARMCX6 | 567.9824163 | -0.338269832 | 0.08393898 | -4.029949279 | 5.58E-05 | 0.000259143 |
| ENSG00000198961 | PJA2 | 3159.560734 | -0.175480956 | 0.05140007 | -3.414021731 | 0.000640115 | 0.002485483 |
| ENSG00000198964 | SGMS1 | 790.4268782 | -0.638464997 | 0.074110288 | -8.615065667 | 6.99E-18 | 1.02E-16 |
| ENSG00000198976 | MIR429 | 11.80098329 | 0.459123149 | 0.509408371 | 0.901287012 | 0.367435738 | 0.559916024 |
| ENSG00000199023 | MIR339 | 2.738366818 | -0.007914549 | 1.07375212 | -0.007370927 | 0.994118904 | 0.996533822 |
| ENSG00000199436 | SNORD9 | 2.29870748 | 0.225595538 | 1.154337414 | 0.195432926 | 0.845054042 | 0.91861535 |
| ENSG00000199477 | SNORA31 | 2.312049269 | -0.461095167 | 1.195914096 | -0.385558769 | 0.699823474 | 0.831642103 |
| ENSG00000199568 | RNU5A-1 | 2.781946557 | 0.292735595 | 1.067886065 | 0.274126242 | 0.783987607 | 0.882792721 |
| ENSG00000199691 | RN7SKP173 | 5.297613452 | -0.553353166 | 0.783790662 | -0.70599612 | 0.480190551 | 0.66446925 |
| ENSG00000199753 | SNORD104 | 43.98141968 | 0.162693777 | 0.266293044 | 0.610957667 | 0.541227606 | 0.715358452 |
| ENSG00000199804 | RNA5SP383 | 4.10418817 | -0.983251679 | 0.875150588 | -1.123522846 | 0.261215503 | 0.441485571 |
| ENSG00000199961 | SNORD1B | 1.586255571 | 2.883044105 | 1.662018375 | 1.734664399 | 0.082800266 | NA |
| ENSG00000200087 | SNORA73B | 8.867203617 | 0.630250661 | 0.592793516 | 1.063187507 | 0.287696927 | 0.471631541 |

| | | | | | | | |
|---|---|---|---|---|---|---|---|
| ENSG00000200090 | Y_RNA | 8.444783997 | -0.967133587 | 0.65945707 | -1.466560344 | 0.142495694 | 0.280182107 |
| ENSG00000200209 | Y_RNA | 1.80091651 | 1.205022327 | 1.368761886 | 0.880373964 | 0.378656756 | 0.571322415 |
| ENSG00000200403 | RNU6-1099P | 1.894509787 | -0.500349905 | 1.30861583 | -0.382350491 | 0.702201408 | 0.833169561 |
| ENSG00000200534 | SNORA33 | 15.97356924 | 0.276608977 | 0.450002974 | 0.614682553 | 0.538764384 | 0.713418088 |
| ENSG00000200879 | SNORD14E | 4.576161316 | 1.782849963 | 0.904438305 | 1.971223414 | 0.048698327 | 0.117125143 |
| ENSG00000200913 | SNORD46 | 4.432503103 | 0.799341159 | 0.842924448 | 0.948295142 | 0.342979223 | 0.533999144 |
| ENSG00000201183 | RNVU1-3 | 2.769203695 | 1.043366226 | 1.07639919 | 0.969311605 | 0.332389741 | 0.523408886 |
| ENSG00000201208 | Y_RNA | 3.621669219 | 0.871873624 | 0.950944283 | 0.916850377 | 0.35922105 | 0.551759685 |
| ENSG00000201264 | SNORD73B | 1.631610592 | 0.421056821 | 1.408135206 | 0.299017324 | 0.764926827 | NA |
| ENSG00000201302 | SNORA65 | 7.473700556 | 0.403054652 | 0.683838019 | 0.589400767 | 0.555592462 | 0.726586404 |
| ENSG00000201451 | Y_RNA | 1.620281495 | 1.023679263 | 1.387078762 | 0.738010913 | 0.460507818 | NA |
| ENSG00000201457 | SNORA55 | 8.769103615 | 1.109239433 | 0.638816977 | 1.736396296 | 0.082493787 | 0.181160968 |
| ENSG00000201499 | RNU6-312P | 2.072261568 | -1.798781655 | 1.35564673 | -1.326880828 | 0.184548107 | 0.341303901 |
| ENSG00000201555 | Y_RNA | 2.391406507 | 1.826784154 | 1.269281067 | 1.439227451 | 0.15008609 | 0.291756083 |
| ENSG00000201772 | SNORA5C | 22.975921 | 0.734489313 | 0.368441004 | 1.993505891 | 0.046206079 | 0.111940802 |
| ENSG00000201998 | SNORA23 | 2.293487137 | -0.1414521 | 1.151239235 | -0.122869422 | 0.9022105 | 0.950654947 |
| ENSG00000202058 | RN7SKP80 | 3.082077493 | -0.297606642 | 1.009128486 | -0.294914519 | 0.76805918 | 0.87332231 |
| ENSG00000202111 | VTRNA1-2 | 1.815159686 | -0.886703578 | 1.378802932 | -0.643096672 | 0.520161374 | 0.697413646 |
| ENSG00000202314 | SNORD6 | 2.645787459 | 0.549267208 | 1.092421871 | 0.502797704 | 0.615106508 | 0.771827114 |
| ENSG00000202337 | RNU6-8 | 60.70050742 | -0.744890693 | 0.233078808 | -3.195874812 | 0.001394076 | 0.005071376 |
| ENSG00000202343 | ENSG0000020 | 4.183707712 | 0.75483365 | 0.875559872 | 0.862115401 | 0.388624017 | 0.581314225 |
| ENSG00000202474 | RNA5SP283 | 7.955619478 | 1.06010171 | 0.650795372 | 1.628932467 | 0.103327316 | 0.217681021 |
| ENSG00000202515 | VTRNA1-3 | 3.793657434 | 1.162999709 | 0.978196491 | 1.188922389 | 0.234470209 | 0.4093726 |
| ENSG00000203279 | ENSG0000020 | 6.450289428 | 0.70769721 | 0.703183306 | 1.006419242 | 0.314213942 | 0.502584564 |
| ENSG00000203286 | Metazoa_SRP | 2.840089005 | -0.034232071 | 1.016802892 | -0.033666379 | 0.97314319 | 0.987960542 |
| ENSG00000203288 | TDRKH-AS1 | 360.44183 | 0.380189144 | 0.102167196 | 3.72124476 | 0.000198243 | 0.000841699 |
| ENSG00000203326 | ZNF525 | 554.9199481 | 0.657637412 | 0.08497484 | 7.739201558 | 1.00E-14 | 1.17E-13 |
| ENSG00000203387 | SNF8P1 | 6.115665552 | 0.162763328 | 0.720630022 | 0.225862541 | 0.821308333 | 0.905242548 |
| ENSG00000203392 | ENSG0000020 | 56.06985865 | 0.152160149 | 0.246332139 | 0.617703195 | 0.536771006 | 0.711888008 |
| ENSG00000203397 | SHISA5P2 | 1.98770259 | -1.61824391 | 1.478212151 | -1.094730489 | 0.273634727 | 0.455627542 |
| ENSG00000203414 | BTBD7P1 | 44.04757733 | -0.3338618 | 0.275785945 | -1.210583085 | 0.22605523 | 0.398222672 |
| ENSG00000203441 | LINC00449 | 4.459914949 | -0.901046126 | 0.84418312 | -1.067358615 | 0.285809929 | 0.469628233 |
| ENSG00000203469 | ENSG0000020 | 5.369568082 | 0.010604597 | 0.789017371 | 0.013440257 | 0.989276549 | 0.994278654 |

| | | | | | | | |
|---|---|---|---|---|---|---|---|
| ENSG00000203485 | INF2 | 3091.148149 | -0.041007107 | 0.052253638 | -0.784770369 | 0.432588216 | 0.622597006 |
| ENSG00000203497 | PDCD4-AS1 | 130.2576674 | -0.337329879 | 0.156677797 | -2.153016479 | 0.031317383 | 0.080648942 |
| ENSG00000203499 | IQANK1 | 3111.816737 | -0.164698831 | 0.05524013 | -2.981506923 | 0.002868336 | 0.009766902 |
| ENSG00000203546 | ENSG0000020 | 10.48566116 | -0.331356738 | 0.550425968 | -0.602000556 | 0.547173767 | 0.719940707 |
| ENSG00000203585 | LINC02408 | 2.15031532 | -2.561552011 | 1.452230062 | -1.763874801 | 0.07775306 | 0.172818061 |
| ENSG00000203616 | RHOT1P2 | 10.20990344 | -0.808630152 | 0.552868795 | -1.46260769 | 0.143574758 | 0.281938448 |
| ENSG00000203635 | ENSG0000020 | 115.0728595 | -0.350085234 | 0.166916581 | -2.09736643 | 0.03596115 | 0.090713524 |
| ENSG00000203644 | ENSG0000020 | 200.5983098 | -0.197518228 | 0.145884283 | -1.353937686 | 0.175756259 | 0.329028738 |
| ENSG00000203666 | EFCAB2 | 172.71337 | -0.203449299 | 0.138671877 | -1.467127318 | 0.142341424 | 0.280067953 |
| ENSG00000203667 | COX20 | 684.4214067 | -0.226883014 | 0.079266584 | -2.862278198 | 0.004206075 | 0.01378982 |
| ENSG00000203668 | CHML | 1065.287335 | -0.347494113 | 0.069767751 | -4.98072691 | 6.33E-07 | 3.76E-06 |
| ENSG00000203684 | IBA57-DT | 2.889250301 | 0.454964284 | 1.060664351 | 0.428942751 | 0.667964888 | 0.808906431 |
| ENSG00000203685 | STUM | 3.245722744 | 0.708525351 | 0.983199068 | 0.720632651 | 0.471135559 | 0.657018237 |
| ENSG00000203697 | CAPN8 | 112.3726442 | -0.54032121 | 0.17528016 | -3.082614767 | 0.002051906 | 0.007192959 |
| ENSG00000203705 | TATDN3 | 116.3500734 | -0.293394889 | 0.165483364 | -1.772957007 | 0.076235831 | 0.170138143 |
| ENSG00000203706 | SERTAD4-AS1 | 11.98059283 | -0.419439387 | 0.518210169 | -0.809400147 | 0.418285018 | 0.610291228 |
| ENSG00000203709 | MIR29B2CHG | 47.19401289 | -0.024883214 | 0.258278647 | -0.096342512 | 0.923248548 | 0.96304664 |
| ENSG00000203711 | LINC02901 | 33.51153918 | -0.547675421 | 0.305523897 | -1.792578019 | 0.073040424 | 0.164349909 |
| ENSG00000203721 | LINC00862 | 4.758666748 | -0.105268411 | 0.800788977 | -0.131455869 | 0.895414695 | 0.946930348 |
| ENSG00000203722 | RAET1G | 4.992186766 | -0.09463478 | 0.816801314 | -0.115860221 | 0.907763323 | 0.953746021 |
| ENSG00000203724 | C1orf53 | 55.27727672 | 0.732356394 | 0.240676766 | 3.042904415 | 0.002343067 | 0.008129154 |
| ENSG00000203739 | PRDX6-AS1 | 34.88216919 | 0.034682587 | 0.293947552 | 0.117989033 | 0.906076348 | 0.95262805 |
| ENSG00000203740 | NTMT2 | 2.15829339 | -0.831511516 | 1.245620389 | -0.667548094 | 0.5044221 | 0.685187291 |
| ENSG00000203760 | CENPW | 287.0976448 | -0.301551997 | 0.109309076 | -2.7587096 | 0.005803008 | 0.018374189 |
| ENSG00000203761 | MSTO2P | 131.5668675 | -0.100266881 | 0.168123909 | -0.596386804 | 0.550916854 | 0.722999578 |
| ENSG00000203772 | SPRN | 39.65764583 | -0.015631642 | 0.286790583 | -0.054505423 | 0.956532488 | 0.980195505 |
| ENSG00000203778 | FAM229B | 13.36930762 | -0.352530596 | 0.495941741 | -0.710830661 | 0.477189178 | 0.662033248 |
| ENSG00000203780 | FANK1 | 72.28786889 | 0.590202576 | 0.213634966 | 2.762668433 | 0.005733097 | 0.018172897 |
| ENSG00000203791 | EEF1AKMT2 | 106.5122136 | -0.29170152 | 0.172090785 | -1.695044387 | 0.090067001 | 0.19455633 |
| ENSG00000203799 | CCDC162P | 222.9718348 | -0.6608884 | 0.123963145 | -5.331329731 | 9.75E-08 | 6.33E-07 |
| ENSG00000203804 | ADAMTSL4-AS | 11.51026614 | 0.866446407 | 0.536343626 | 1.615468824 | 0.106209206 | 0.222526185 |
| ENSG00000203814 | H2BC18 | 270.4215164 | 0.808406105 | 0.115382629 | 7.006306868 | 2.45E-12 | 2.42E-11 |
| ENSG00000203865 | ATP1A1-AS1 | 377.4911409 | -0.417435341 | 0.103034135 | -4.051427631 | 5.09E-05 | 0.000237984 |

| | | | | | | | |
|---|---|---|---|---|---|---|---|
| ENSG00000203867 | RBM20 | 124.1460815 | 4.59402892 | 0.304266418 | 15.0987051 | 1.65E-51 | 9.47E-50 |
| ENSG00000203872 | C6orf163 | 10.07977095 | -0.065412776 | 0.557177178 | -0.117400315 | 0.906542834 | 0.952856477 |
| ENSG00000203875 | SNHG5 | 1606.449862 | 0.539868215 | 0.067290047 | 8.02300247 | 1.03E-15 | 1.31E-14 |
| ENSG00000203879 | GDI1 | 8339.64688 | -0.446946853 | 0.045540045 | -9.814370083 | 9.76E-23 | 1.91E-21 |
| ENSG00000203880 | PCMTD2 | 2539.898278 | -0.023469919 | 0.055627202 | -0.421914419 | 0.673087485 | 0.81285826 |
| ENSG00000203883 | SOX18 | 13.17954382 | 0.284447475 | 0.482481969 | 0.589550478 | 0.555492061 | 0.726551564 |
| ENSG00000203896 | LIME1 | 174.5684464 | -0.252840788 | 0.14327404 | -1.764735522 | 0.077608226 | 0.172563022 |
| ENSG00000203907 | OOEP | 60.74938728 | 0.07673493 | 0.227613142 | 0.33712873 | 0.73601986 | 0.854386347 |
| ENSG00000203914 | HSP90B3P | 16.83959061 | -0.100998559 | 0.436381188 | -0.231445723 | 0.816968548 | 0.903062617 |
| ENSG00000203930 | LINC00632 | 410.7916108 | 0.869754274 | 0.102625277 | 8.475049226 | 2.35E-17 | 3.31E-16 |
| ENSG00000203943 | SAMD13 | 14.25623154 | 0.512109364 | 0.462804384 | 1.106535249 | 0.268494916 | 0.449943266 |
| ENSG00000203950 | RTL8A | 2915.802496 | -0.202771074 | 0.056582341 | -3.583645892 | 0.000338831 | 0.001380112 |
| ENSG00000203952 | CCDC160 | 2.532249779 | 1.002441112 | 1.169304091 | 0.857297191 | 0.391280661 | 0.584153023 |
| ENSG00000203965 | EFCAB7 | 94.84821581 | -0.042191462 | 0.184811002 | -0.228295184 | 0.819416769 | 0.904358204 |
| ENSG00000203993 | ARRDC1-AS1 | 248.1707202 | 0.038243552 | 0.115331204 | 0.331597616 | 0.74019312 | 0.856646281 |
| ENSG00000203995 | ZYG11A | 452.5883109 | 0.227790577 | 0.090955319 | 2.504422831 | 0.012265136 | 0.035606456 |
| ENSG00000203999 | LINC01270 | 27.64985622 | -0.58525169 | 0.352998048 | -1.657945967 | 0.097328373 | 0.207404819 |
| ENSG00000204019 | CT83 | 264.0065492 | 0.264996604 | 0.11944893 | 2.218492907 | 0.026521243 | 0.069858458 |
| ENSG00000204022 | LIPJ | 3.398388971 | 1.613079349 | 1.039544977 | 1.551716746 | 0.120730014 | 0.246752185 |
| ENSG00000204044 | SLC12A5-AS1 | 119.9112232 | -0.979740041 | 0.18059577 | -5.425044242 | 5.79E-08 | 3.84E-07 |
| ENSG00000204049 | ENSG0000020 | 3.724649122 | 1.069153283 | 0.979115993 | 1.091957736 | 0.274851679 | 0.456957349 |
| ENSG00000204052 | LRRC73 | 76.06781618 | 0.785547283 | 0.213443293 | 3.680355904 | 0.000232909 | 0.00097659 |
| ENSG00000204054 | LINC00963 | 1765.221129 | -0.374579034 | 0.061984292 | -6.043128365 | 1.51E-09 | 1.18E-08 |
| ENSG00000204055 | IER5L-AS1 | 10.86443579 | -0.591376979 | 0.55448199 | -1.066539563 | 0.286179804 | 0.470021728 |
| ENSG00000204060 | FOXO6 | 108.317823 | 0.999496225 | 0.176960093 | 5.648144775 | 1.62E-08 | 1.14E-07 |
| ENSG00000204065 | TCEAL5 | 6.230942482 | 1.208340056 | 0.764689121 | 1.580171631 | 0.114067567 | 0.23582462 |
| ENSG00000204070 | SYS1 | 1020.521025 | -0.178906958 | 0.075394168 | -2.372954854 | 0.017646421 | 0.049073693 |
| ENSG00000204071 | TCEAL6 | 10.99379296 | -0.383524614 | 0.549264281 | -0.698251511 | 0.485019918 | 0.668886808 |
| ENSG00000204072 | ARMCX7P | 2.055730731 | 1.564330164 | 1.349572461 | 1.159130176 | 0.246403127 | 0.423647518 |
| ENSG00000204084 | INPP5B | 820.0302217 | -0.124561272 | 0.075321096 | -1.653736846 | 0.098180993 | 0.208872482 |
| ENSG00000204086 | RPA4 | 2.583796751 | 0.993770445 | 1.128645956 | 0.880497945 | 0.378589618 | 0.571296266 |
| ENSG00000204099 | NEU4 | 3.749174055 | -0.536104432 | 0.893599896 | -0.599937885 | 0.548547633 | 0.721375985 |
| ENSG00000204103 | MAFB | 245.8016162 | 0.011240734 | 0.117663919 | 0.095532547 | 0.92389184 | 0.963279648 |

| | | | | | | | |
|---|---|---|---|---|---|---|---|
| ENSG00000204104 | TRAF3IP1 | 616.1061852 | 0.193237311 | 0.081251621 | 2.378257928 | 0.017394654 | 0.04844282 |
| ENSG00000204110 | LINC02520 | 6.053715729 | 1.942163993 | 0.798637442 | 2.431846907 | 0.015022055 | 0.042564964 |
| ENSG00000204116 | CHIC1 | 228.8648462 | 0.065713604 | 0.121874926 | 0.539188875 | 0.589756536 | 0.752741052 |
| ENSG00000204118 | NAP1L6P | 71.63487537 | 0.204609946 | 0.217621028 | 0.940212202 | 0.347108724 | 0.537982172 |
| ENSG00000204120 | GIGYF2 | 2825.657616 | 0.104364134 | 0.053183072 | 1.962356276 | 0.04972102 | 0.11923481 |
| ENSG00000204128 | C2orf72 | 91.19619282 | -0.141963273 | 0.198608169 | -0.714790704 | 0.47473838 | 0.659990209 |
| ENSG00000204130 | RUFY2 | 452.7302954 | -0.459649317 | 0.092992279 | -4.94287615 | 7.70E-07 | 4.53E-06 |
| ENSG00000204131 | NHSL2 | 59.13318757 | 0.78699812 | 0.242341256 | 3.247478914 | 0.001164323 | 0.00430931 |
| ENSG00000204138 | PHACTR4 | 1897.927347 | -0.325394084 | 0.057892362 | -5.620673788 | 1.90E-08 | 1.33E-07 |
| ENSG00000204147 | ASAH2B | 212.7866223 | -0.513195164 | 0.129558134 | -3.961118846 | 7.46E-05 | 0.000340085 |
| ENSG00000204149 | AGAP6 | 26.448074 | -0.002498301 | 0.342222064 | -0.007300233 | 0.994175308 | 0.996533822 |
| ENSG00000204152 | TIMM23B | 41.09709369 | -0.809082929 | 0.278660358 | -2.903473369 | 0.003690484 | 0.012251765 |
| ENSG00000204160 | ZDHHC18 | 1383.240042 | 0.204488643 | 0.064091009 | 3.190597954 | 0.001419787 | 0.005149394 |
| ENSG00000204172 | AGAP9 | 90.0829559 | -0.966143588 | 0.197006766 | -4.904113738 | 9.39E-07 | 5.45E-06 |
| ENSG00000204173 | LRRC37A5P | 6.716473224 | 0.206009115 | 0.697411843 | 0.295390905 | 0.76769528 | 0.87314707 |
| ENSG00000204177 | BMS1P1 | 66.36626903 | -0.884100224 | 0.223816048 | -3.950119898 | 7.81E-05 | 0.000355041 |
| ENSG00000204178 | MACO1 | 1079.317737 | -0.060069909 | 0.068030939 | -0.882979269 | 0.377247472 | 0.570022519 |
| ENSG00000204186 | ZDBF2 | 129.5373656 | -0.239708947 | 0.162253388 | -1.477374066 | 0.139575394 | 0.275808385 |
| ENSG00000204188 | GGNBP1 | 8.734356936 | 0.612515077 | 0.610667548 | 1.003025425 | 0.315848594 | 0.504653023 |
| ENSG00000204194 | RPL12P1 | 14.41879387 | 0.379883131 | 0.478722533 | 0.793535095 | 0.42746613 | 0.618217808 |
| ENSG00000204209 | DAXX | 1831.358393 | -0.125543218 | 0.055516282 | -2.26137655 | 0.02373595 | 0.063506261 |
| ENSG00000204217 | BMPR2 | 2469.673676 | -0.069368722 | 0.063967188 | -1.084442263 | 0.278168754 | 0.46100211 |
| ENSG00000204219 | TCEA3 | 38.51925443 | 0.084062482 | 0.295526165 | 0.284450218 | 0.77606538 | 0.878333406 |
| ENSG00000204220 | PFDN6 | 630.4959841 | 0.064392062 | 0.078957485 | 0.815528284 | 0.414769981 | 0.606812715 |
| ENSG00000204227 | RING1 | 1383.835998 | 0.016712615 | 0.069059434 | 0.242003356 | 0.808777563 | 0.898836887 |
| ENSG00000204228 | HSD17B8 | 146.4422759 | -0.232911719 | 0.155077613 | -1.501904202 | 0.133121851 | 0.266266917 |
| ENSG00000204231 | RXRB | 1521.759658 | -0.001528061 | 0.06355556 | -0.024042916 | 0.980818376 | 0.991149032 |
| ENSG00000204237 | OXLD1 | 789.3418289 | 0.393530792 | 0.080254445 | 4.903538882 | 9.41E-07 | 5.46E-06 |
| ENSG00000204241 | LINC02731 | 5.674030622 | 0.145216619 | 0.764487173 | 0.189952983 | 0.849345973 | 0.921142049 |
| ENSG00000204248 | COL11A2 | 466.6283203 | 0.61552143 | 0.093769428 | 6.564201598 | 5.23E-11 | 4.62E-10 |
| ENSG00000204253 | HNRNPCP2 | 2339.332814 | -0.040899701 | 0.054694909 | -0.747778946 | 0.454593504 | 0.642180859 |
| ENSG00000204256 | BRD2 | 13161.64925 | -0.086549701 | 0.040824426 | -2.12004697 | 0.034002084 | 0.08640831 |
| ENSG00000204257 | HLA-DMA | 38.11621043 | -0.923595129 | 0.301161759 | -3.06677426 | 0.002163822 | 0.007549517 |

| | | | | | | | |
|---|---|---|---|---|---|---|---|
| ENSG00000204262 | COL5A2 | 98.07518694 | -0.476316881 | 0.19033356 | -2.502537552 | 0.012330654 | 0.035760457 |
| ENSG00000204267 | TAP2 | 17.36009536 | 1.671502955 | 0.46464184 | 3.597400858 | 0.000321413 | 0.00131594 |
| ENSG00000204271 | SPIN3 | 1054.365768 | 0.07249013 | 0.077626931 | 0.933827077 | 0.3503931 | 0.541791434 |
| ENSG00000204272 | NBDY | 1281.509239 | -0.252179958 | 0.060611461 | -4.16059854 | 3.17E-05 | 0.000153391 |
| ENSG00000204291 | COL15A1 | 2.938332709 | -4.071168312 | 1.449692087 | -2.808298638 | 0.004980402 | 0.016053171 |
| ENSG00000204301 | NOTCH4 | 11.59164751 | -0.848479195 | 0.541564051 | -1.566719937 | 0.117180169 | 0.241086374 |
| ENSG00000204304 | PBX2 | 1735.04838 | -0.156597645 | 0.05783054 | -2.707871054 | 0.006771633 | 0.021060365 |
| ENSG00000204305 | AGER | 275.9107657 | -0.363501851 | 0.117439036 | -3.095238722 | 0.001966545 | 0.006925476 |
| ENSG00000204308 | RNF5 | 867.9965636 | -0.346054934 | 0.075173289 | -4.603429463 | 4.16E-06 | 2.24E-05 |
| ENSG00000204310 | AGPAT1 | 1287.518897 | -0.034431667 | 0.066429658 | -0.518317691 | 0.604236632 | 0.763029194 |
| ENSG00000204311 | PJVK | 23.57017598 | 0.136930466 | 0.361960306 | 0.378302438 | 0.705205934 | 0.835050025 |
| ENSG00000204314 | PRRT1 | 145.7395407 | -0.656710124 | 0.163450208 | -4.017799251 | 5.87E-05 | 0.000272131 |
| ENSG00000204315 | FKBPL | 174.1899915 | -0.344972691 | 0.14131038 | -2.441240983 | 0.014636884 | 0.041586589 |
| ENSG00000204316 | MRPL38 | 96.11018854 | 0.446229298 | 0.186307028 | 2.395128633 | 0.016614535 | 0.046531247 |
| ENSG00000204323 | SMIM5 | 79.29045606 | 1.517239942 | 0.223814716 | 6.778999911 | 1.21E-11 | 1.12E-10 |
| ENSG00000204334 | ERICH2 | 46.6359443 | 0.944614876 | 0.271318791 | 3.48156821 | 0.000498487 | 0.001973302 |
| ENSG00000204335 | SP5 | 1.739389916 | 0.602181066 | 1.365643262 | 0.44095049 | 0.65924884 NA | |
| ENSG00000204338 | CYP21A1P | 2.913344542 | 0.148858634 | 1.021542505 | 0.145719471 | 0.88414285 | 0.939910479 |
| ENSG00000204344 | STK19 | 434.5023894 | -0.244346696 | 0.09437172 | -2.589194052 | 0.009620087 | 0.028749972 |
| ENSG00000204348 | DXO | 636.2559426 | -0.063702738 | 0.087746902 | -0.725982756 | 0.467849342 | 0.653820524 |
| ENSG00000204351 | SKIC2 | 1857.066971 | 0.19668636 | 0.064656031 | 3.042041967 | 0.002349791 | 0.008151249 |
| ENSG00000204352 | FAM120A2P | 2.998863987 | 0.333647564 | 1.044286085 | 0.319498238 | 0.749348727 | 0.861330224 |
| ENSG00000204356 | NELFE | 1822.406545 | -0.031009838 | 0.05732844 | -0.54091543 | 0.588565875 | 0.751806959 |
| ENSG00000204366 | ZBTB12 | 512.2216611 | -0.059060521 | 0.089796929 | -0.657712035 | 0.510723187 | 0.690568306 |
| ENSG00000204370 | SDHD | 479.4048347 | -0.108753821 | 0.09347979 | -1.163393941 | 0.244669707 | 0.42153384 |
| ENSG00000204371 | EHMT2 | 3712.021011 | -0.007928669 | 0.051022892 | -0.155394354 | 0.876510435 | 0.935869182 |
| ENSG00000204380 | PKP4-AS1 | 9.907374803 | -0.626928423 | 0.565365055 | -1.108891357 | 0.267477053 | 0.448729238 |
| ENSG00000204385 | SLC44A4 | 2.662254029 | -0.9001964 | 1.106671437 | -0.813426976 | 0.415973298 | 0.608069041 |
| ENSG00000204386 | NEU1 | 3306.692512 | 0.641453757 | 0.050330908 | 12.74472844 | 3.34E-37 | 1.22E-35 |
| ENSG00000204387 | SNHG32 | 253.5309399 | 0.44758419 | 0.115438783 | 3.877242809 | 0.000105647 | 0.000469315 |
| ENSG00000204388 | HSPA1B | 1093.140829 | 0.210912982 | 0.067354161 | 3.131402427 | 0.001739736 | 0.006194248 |
| ENSG00000204389 | HSPA1A | 2578.835673 | 0.146654029 | 0.062867791 | 2.3327371 | 0.019661948 | 0.053901181 |
| ENSG00000204390 | HSPA1L | 29.62856747 | -1.016742487 | 0.336387615 | -3.022532465 | 0.002506692 | 0.008636755 |

| | | | | | | | |
|---|---|---|---|---|---|---|---|
| ENSG00000204392 | LSM2 | 708.9881762 | -0.276723333 | 0.07648825 | -3.617854172 | 0.000297056 | 0.001223861 |
| ENSG00000204394 | VARS1 | 2797.896251 | 0.38069656 | 0.052446983 | 7.258693232 | 3.91E-13 | 4.09E-12 |
| ENSG00000204396 | VWA7 | 32.76180067 | 1.014222409 | 0.329909818 | 3.074241365 | 0.002110386 | 0.007381043 |
| ENSG00000204406 | MBD5 | 875.9084524 | -0.018692084 | 0.072131751 | -0.259138093 | 0.795528694 | 0.890780897 |
| ENSG00000204410 | MSH5 | 44.32412091 | 0.099237799 | 0.27263507 | 0.363994988 | 0.715861752 | 0.841891334 |
| ENSG00000204420 | MPIG6B | 4.549741107 | 3.596071795 | 1.230190218 | 2.92318354 | 0.003464722 | 0.011581047 |
| ENSG00000204421 | LY6G6C | 16.43956596 | 2.393069749 | 0.541729119 | 4.417465599 | 9.99E-06 | 5.17E-05 |
| ENSG00000204427 | ABHD16A | 31.83513049 | -0.121683034 | 0.315541476 | -0.385632454 | 0.699768894 | 0.831642103 |
| ENSG00000204428 | LY6G5C | 131.1087709 | 0.099774438 | 0.159073423 | 0.627222547 | 0.530513365 | 0.70616088 |
| ENSG00000204429 | ENSG0000020 | 3.991811243 | -0.188360029 | 0.877740079 | -0.214596591 | 0.830081852 | 0.909932761 |
| ENSG00000204434 | POTEKP | 189.108915 | 0.647659034 | 0.133244588 | 4.86067798 | 1.17E-06 | 6.72E-06 |
| ENSG00000204435 | CSNK2B | 306.6795349 | 0.041537265 | 0.113930398 | 0.364584573 | 0.715421533 | 0.841656152 |
| ENSG00000204438 | GPANK1 | 353.5088965 | 0.112088318 | 0.102940139 | 1.088868929 | 0.27621169 | 0.458719768 |
| ENSG00000204439 | C6orf47 | 761.5340797 | 0.001247658 | 0.081761531 | 0.015259717 | 0.98782498 | 0.994021364 |
| ENSG00000204444 | APOM | 62.14802958 | -0.217673701 | 0.22801476 | -0.954647413 | 0.339756031 | 0.530782032 |
| ENSG00000204463 | BAG6 | 8417.763524 | -0.031012438 | 0.044258043 | -0.700718699 | 0.483478584 | 0.66755731 |
| ENSG00000204469 | PRRC2A | 16464.31979 | 0.001884918 | 0.048383804 | 0.038957623 | 0.968924175 | 0.985925152 |
| ENSG00000204475 | NCR3 | 1.805552744 | -0.453228033 | 1.341723286 | -0.337795458 | 0.735517333 | 0.854146138 |
| ENSG00000204482 | LST1 | 2.439550427 | -0.733385018 | 1.112851826 | -0.659014076 | 0.50988673 | 0.689870603 |
| ENSG00000204498 | NFKBIL1 | 418.199534 | 0.241790912 | 0.106918683 | 2.261446779 | 0.023731605 | 0.063502043 |
| ENSG00000204514 | ZNF814 | 63.58365356 | 0.624053939 | 0.234483371 | 2.661399557 | 0.007781655 | 0.02386194 |
| ENSG00000204516 | MICB | 730.8247729 | 0.123017503 | 0.07473108 | 1.64613576 | 0.099735811 | 0.211552429 |
| ENSG00000204519 | ZNF551 | 623.745646 | 0.513954404 | 0.085516326 | 6.010015044 | 1.86E-09 | 1.44E-08 |
| ENSG00000204520 | MICA | 1152.347581 | -0.013662048 | 0.062494094 | -0.218613419 | 0.826951198 | 0.908591807 |
| ENSG00000204524 | ZNF805 | 183.8699133 | 0.432075774 | 0.135365644 | 3.191916052 | 0.001413324 | 0.00513082 |
| ENSG00000204525 | HLA-C | 73.96809192 | -0.207106372 | 0.204736777 | -1.011573864 | 0.311741848 | 0.499990071 |
| ENSG00000204531 | POU5F1 | 2.491734958 | -0.280621811 | 1.148898005 | -0.244253023 | 0.807034864 | 0.89748787 |
| ENSG00000204536 | CCHCR1 | 570.7347888 | -0.508413646 | 0.087007586 | -5.843325474 | 5.12E-09 | 3.81E-08 |
| ENSG00000204539 | CDSN | 1.794769448 | 3.039999326 | 1.685919699 | 1.803169705 | 0.071361567 | 0.161220636 |
| ENSG00000204540 | PSORS1C1 | 12.75985168 | 4.120199751 | 0.835572773 | 4.930988518 | 8.18E-07 | 4.79E-06 |
| ENSG00000204560 | DHX16 | 2207.04611 | -0.09055954 | 0.052442539 | -1.726833646 | 0.084197544 | 0.184267474 |
| ENSG00000204564 | C6orf136 | 386.6729414 | -0.090410687 | 0.104456542 | -0.865533984 | 0.386745771 | 0.579318671 |
| ENSG00000204568 | MRPS18B | 1763.846497 | -0.107579916 | 0.062400233 | -1.724030688 | 0.084702303 | 0.185151689 |

| | | | | | | | |
|---|---|---|---|---|---|---|---|
| ENSG00000204569 | PPP1R10 | 4699.579306 | -0.332104727 | 0.047820314 | -6.944846211 | 3.79E-12 | 3.69E-11 |
| ENSG00000204572 | KRTAP5-10 | 15.78597762 | 1.037930729 | 0.485097281 | 2.139634192 | 0.032384341 | 0.083024201 |
| ENSG00000204574 | ABCF1 | 4136.130977 | 0.145978092 | 0.047223943 | 3.091188106 | 0.001993573 | 0.007009896 |
| ENSG00000204576 | PRR3 | 753.0969444 | -0.048367614 | 0.07682078 | -0.629616275 | 0.528945673 | 0.704892166 |
| ENSG00000204580 | DDR1 | 13942.54481 | -0.324962034 | 0.05521197 | -5.885717039 | 3.96E-09 | 2.98E-08 |
| ENSG00000204581 | ACOXL-AS1 | 14.90160667 | 1.341338144 | 0.475973965 | 2.818091416 | 0.004831005 | 0.015628721 |
| ENSG00000204584 | ENSG0000020 | 5.25591787 | 0.119560384 | 0.789981764 | 0.151345751 | 0.879702981 | 0.937619587 |
| ENSG00000204588 | LINC01123 | 49.26753139 | 0.058699955 | 0.25165063 | 0.233259717 | 0.815559737 | 0.902331454 |
| ENSG00000204590 | GNL1 | 2111.876982 | 0.075477923 | 0.056896641 | 1.326579576 | 0.184647796 | 0.341433212 |
| ENSG00000204592 | HLA-E | 2051.399668 | 0.752185695 | 0.064578675 | 11.6475863 | 2.36E-31 | 6.96E-30 |
| ENSG00000204599 | TRIM39 | 347.8431418 | -0.202547273 | 0.104379772 | -1.940483959 | 0.052320902 | 0.124545137 |
| ENSG00000204603 | LINC01257 | 2.065616644 | -1.726589461 | 1.358501375 | -1.270951575 | 0.203745877 | 0.368461207 |
| ENSG00000204604 | ZNF468 | 1294.329458 | 0.553295892 | 0.069472942 | 7.964192579 | 1.66E-15 | 2.07E-14 |
| ENSG00000204611 | ZNF616 | 565.0642305 | -0.319731751 | 0.084104057 | -3.801621045 | 0.000143752 | 0.000625034 |
| ENSG00000204616 | TRIM31 | 1.993356202 | 0.743977245 | 1.279894069 | 0.581280329 | 0.561051536 | 0.730103564 |
| ENSG00000204618 | RNF39 | 14.65715386 | 0.074894681 | 0.49261865 | 0.15203379 | 0.879160285 | 0.937346018 |
| ENSG00000204619 | PPP1R11 | 1515.177357 | 0.140817286 | 0.058352587 | 2.413214097 | 0.015812528 | 0.044529756 |
| ENSG00000204620 | ENSG0000020 | 15.20140337 | -0.546555534 | 0.491914764 | -1.11107772 | 0.266534897 | 0.447606921 |
| ENSG00000204623 | POLR1HASP | 148.1362281 | -0.113209361 | 0.147159911 | -0.769294845 | 0.441718297 | 0.631058764 |
| ENSG00000204624 | DISP3 | 71.28129578 | 1.020153824 | 0.225007691 | 4.533862015 | 5.79E-06 | 3.07E-05 |
| ENSG00000204628 | RACK1 | 38792.70199 | 0.021434929 | 0.055361818 | 0.387178925 | 0.698623752 | 0.830862695 |
| ENSG00000204634 | TBC1D8 | 808.5892463 | -0.242453312 | 0.072288738 | -3.353956892 | 0.000796648 | 0.003036235 |
| ENSG00000204637 | ARPP19P2 | 37.48292718 | -1.648024874 | 0.321027487 | -5.133594296 | 2.84E-07 | 1.76E-06 |
| ENSG00000204642 | HLA-F | 16.85452706 | 0.684821237 | 0.457278422 | 1.497602345 | 0.134236597 | 0.267828571 |
| ENSG00000204650 | LINC02210 | 448.0909567 | 0.017127162 | 0.091389758 | 0.187407895 | 0.851340833 | 0.9222591 |
| ENSG00000204661 | C5orf60 | 3.032053117 | -0.188271684 | 1.005057107 | -0.187324365 | 0.851406321 | 0.922260215 |
| ENSG00000204666 | ZNF473CR | 5.009599286 | -0.458679957 | 0.803671779 | -0.570730452 | 0.568182373 | 0.735799496 |
| ENSG00000204673 | AKT1S1 | 3440.326229 | 0.066947348 | 0.053769201 | 1.245087283 | 0.213099668 | 0.381289009 |
| ENSG00000204677 | FAM153CP | 11.86050166 | -0.639207928 | 0.514625047 | -1.242084761 | 0.21420529 | 0.382729914 |
| ENSG00000204681 | GABBR1 | 225.2634729 | 0.345832715 | 0.130609039 | 2.647846709 | 0.008100625 | 0.024714501 |
| ENSG00000204682 | MIR1915HG | 48.69249576 | 1.65835809 | 0.280401748 | 5.914221646 | 3.33E-09 | 2.53E-08 |
| ENSG00000204685 | STARD7-AS1 | 159.0071973 | 0.168978745 | 0.156416284 | 1.080314276 | 0.280002254 | 0.46340464 |
| ENSG00000204706 | MAMDC2-AS1 | 2.792721291 | 1.25416617 | 1.109174594 | 1.130720246 | 0.258172857 | 0.437826964 |

| | | | | | | | |
|---|---|---|---|---|---|---|---|
| ENSG00000204713 | TRIM27 | 2631.154709 | -0.00259686 | 0.05199311 | -0.049946233 | 0.960165235 | 0.98182181 |
| ENSG00000204740 | MALRD1 | 353.6932889 | -2.257259096 | 0.11481315 | -19.66028371 | 4.72E-86 | 6.05E-84 |
| ENSG00000204745 | ENSG0000020 | 26.81545472 | 0.466627301 | 0.341519041 | 1.3663288 | 0.171835785 | 0.32353192 |
| ENSG00000204758 | RPL26L1-AS1 | 14.25819556 | -0.555658894 | 0.48238233 | -1.151905572 | 0.249359878 | 0.427335695 |
| ENSG00000204764 | RANBP17 | 316.7879408 | -0.525988979 | 0.111104046 | -4.734201836 | 2.20E-06 | 1.22E-05 |
| ENSG00000204778 | ZNG1DP | 12.9986598 | 0.245471116 | 0.490328554 | 0.500625783 | 0.616634513 | 0.772898903 |
| ENSG00000204789 | ZNF204P | 43.67212232 | -0.211592726 | 0.266915403 | -0.79273329 | 0.42793323 | 0.618614173 |
| ENSG00000204792 | LINC01291 | 192.4274835 | -0.125643484 | 0.13070565 | -0.961270484 | 0.336416185 | 0.527540315 |
| ENSG00000204815 | ODAD4 | 79.50958649 | -0.037688434 | 0.202850555 | -0.185794087 | 0.852606242 | 0.923046535 |
| ENSG00000204839 | MROH6 | 569.4787412 | 0.19716138 | 0.102122584 | 1.930634461 | 0.053528273 | 0.127115796 |
| ENSG00000204842 | ATXN2 | 1414.069283 | 0.179066023 | 0.061512382 | 2.911056564 | 0.003602088 | 0.011995547 |
| ENSG00000204843 | DCTN1 | 5387.41783 | -0.428612791 | 0.048679329 | -8.80482135 | 1.31E-18 | 2.02E-17 |
| ENSG00000204852 | TCTN1 | 465.3737736 | 0.607424764 | 0.099974244 | 6.075812522 | 1.23E-09 | 9.74E-09 |
| ENSG00000204856 | FAM216A | 197.5378507 | 0.241761399 | 0.135217239 | 1.787948057 | 0.073784388 | 0.165740532 |
| ENSG00000204859 | ZBTB48 | 435.4819652 | -0.229851594 | 0.095344458 | -2.41074939 | 0.015919783 | 0.044809795 |
| ENSG00000204860 | FAM201A | 229.4778987 | 0.478613555 | 0.125911317 | 3.801195674 | 0.000144 | 0.00062599 |
| ENSG00000204899 | MZT1 | 439.008675 | 0.045167897 | 0.102455251 | 0.440854877 | 0.659318062 | 0.802897431 |
| ENSG00000204904 | LINC01545 | 3.482645814 | 3.130750633 | 1.274958037 | 2.455571511 | 0.014066076 | 0.040183785 |
| ENSG00000204920 | ZNF155 | 356.8121421 | 0.041364118 | 0.098871105 | 0.418364068 | 0.675680953 | 0.814879865 |
| ENSG00000204922 | UQCC3 | 572.3175177 | -0.053392934 | 0.093649189 | -0.570137715 | 0.568584297 | 0.736037682 |
| ENSG00000204923 | FBXO48 | 29.51232636 | -0.326285138 | 0.326686745 | -0.998770666 | 0.317905799 | 0.507067749 |
| ENSG00000204929 | LINC02934 | 2.806033337 | -2.300248743 | 1.204942519 | -1.909011182 | 0.056260649 | 0.132387335 |
| ENSG00000204934 | ATP6V0E2-AS1 | 143.6853027 | 0.466191752 | 0.167392096 | 2.785028459 | 0.005352303 | 0.017112449 |
| ENSG00000204946 | ZNF783 | 1176.872841 | 0.645165404 | 0.068877108 | 9.366906117 | 7.47E-21 | 1.32E-19 |
| ENSG00000204947 | ZNF425 | 362.0018835 | -0.010292498 | 0.101190538 | -0.101714037 | 0.91898366 | 0.960475883 |
| ENSG00000204950 | LRRC10B | 64.82327207 | 1.108004408 | 0.234491989 | 4.725126923 | 2.30E-06 | 1.28E-05 |
| ENSG00000204954 | UQCC6 | 328.6719273 | -0.123165758 | 0.107450874 | -1.1462518 | 0.25169098 | 0.430237786 |
| ENSG00000204956 | PCDHGA1 | 43.26901741 | -0.118904761 | 0.276777583 | -0.429604015 | 0.667483716 | 0.808691146 |
| ENSG00000204959 | ARHGEF34P | 29.86216763 | 0.519674951 | 0.326489474 | 1.591705071 | 0.111450988 | 0.23122779 |
| ENSG00000204962 | PCDHA8 | 2.736455517 | 0.039813085 | 1.05359075 | 0.037787998 | 0.969856714 | 0.986499047 |
| ENSG00000204963 | PCDHA7 | 36.86873619 | 0.011075042 | 0.300167976 | 0.036896149 | 0.97056781 | 0.986772291 |
| ENSG00000204965 | PCDHA5 | 12.40271551 | 0.477894035 | 0.50525549 | 0.945846297 | 0.344226988 | 0.535142796 |
| ENSG00000204977 | TRIM13 | 338.8451507 | 0.086149064 | 0.102086875 | 0.84387992 | 0.39873651 | 0.591597444 |

| | | | | | | | |
|---|---|---|---|---|---|---|---|
| ENSG00000204991 | SPIRE2 | 703.8655693 | 1.028323049 | 0.07742513 | 13.28151539 | 2.96E-40 | 1.23E-38 |
| ENSG00000205002 | AARD | 3594.727576 | 6.667250772 | 0.113345696 | 58.82226693 | 0 | 0 |
| ENSG00000205018 | ENSG0000020 | 26.987229 | -0.71131372 | 0.369062423 | -1.927353415 | 0.053935601 | 0.127897735 |
| ENSG00000205022 | PABPN1L | 5.999715333 | 0.367447523 | 0.707658625 | 0.51924404 | 0.603590572 | 0.762610445 |
| ENSG00000205035 | GRM5P1 | 4.963294938 | 0.090880255 | 0.813434896 | 0.111724067 | 0.911042196 | 0.955316516 |
| ENSG00000205041 | ENSG0000020 | 13.99746468 | -0.944435366 | 0.472746378 | -1.997763304 | 0.045742328 | 0.111016609 |
| ENSG00000205054 | LINC01121 | 3.67776923 | -1.275875249 | 0.941501024 | -1.355150145 | 0.175369725 | 0.328567584 |
| ENSG00000205060 | SLC35B4 | 790.6243138 | -0.544434935 | 0.072036933 | -7.557719475 | 4.10E-14 | 4.60E-13 |
| ENSG00000205078 | SYCE1L | 91.11817452 | -0.273144195 | 0.201220895 | -1.357434547 | 0.174643177 | 0.327661282 |
| ENSG00000205084 | TMEM231 | 507.4878947 | 1.292259038 | 0.094818187 | 13.62880979 | 2.70E-42 | 1.20E-40 |
| ENSG00000205085 | GARIN1A | 30.95941383 | 0.97583517 | 0.326662996 | 2.987284091 | 0.00281468 | 0.009602731 |
| ENSG00000205089 | CCNI2 | 20.07114223 | 0.388290309 | 0.396167489 | 0.980116541 | 0.327028595 | 0.517460158 |
| ENSG00000205090 | TMEM240 | 4.256541122 | 0.290771624 | 0.866107437 | 0.33572235 | 0.73708025 | 0.855093604 |
| ENSG00000205100 | HSP90AA4P | 57.91858823 | 0.198362196 | 0.265619728 | 0.746790147 | 0.455190244 | 0.642658895 |
| ENSG00000205105 | COX17P1 | 177.0199239 | -0.1146321 | 0.138997477 | -0.824706336 | 0.409538334 | 0.602065433 |
| ENSG00000205111 | CDKL4 | 2.8425078 | 1.513147628 | 1.175645785 | 1.287077832 | 0.198067161 | 0.360464949 |
| ENSG00000205133 | TRIQK | 2007.358339 | -0.610355681 | 0.060549012 | -10.08035742 | 6.75E-24 | 1.40E-22 |
| ENSG00000205138 | SDHAF1 | 424.2498334 | 0.10601274 | 0.094349353 | 1.123619147 | 0.26117463 | 0.441481542 |
| ENSG00000205143 | ARID3C | 2.977230851 | 0.803474057 | 1.045240453 | 0.76869782 | 0.442072719 | 0.631335876 |
| ENSG00000205155 | PSENEN | 240.2845705 | -0.307213286 | 0.125285734 | -2.45210109 | 0.014202476 | 0.040502833 |
| ENSG00000205189 | ZBTB10 | 877.0299879 | 0.138836454 | 0.078269997 | 1.773814469 | 0.076093844 | 0.169887375 |
| ENSG00000205208 | C4orf46 | 327.2804007 | -0.216511418 | 0.111072324 | -1.949283217 | 0.051261612 | 0.122395666 |
| ENSG00000205209 | SCGB2B2 | 4.154894668 | 0.170645882 | 0.884584919 | 0.192910685 | 0.847028917 | 0.919933877 |
| ENSG00000205213 | LGR4 | 644.5534926 | -0.143335994 | 0.082402068 | -1.739470832 | 0.081951979 | 0.180229929 |
| ENSG00000205220 | PSMB10 | 64.11741217 | 0.122355551 | 0.227006514 | 0.538995769 | 0.589889774 | 0.752869223 |
| ENSG00000205250 | E2F4 | 2192.68164 | 0.091091023 | 0.05698437 | 1.598526453 | 0.109925863 | 0.228600729 |
| ENSG00000205268 | PDE7A | 1552.34694 | 0.317301761 | 0.06356948 | 4.991416665 | 5.99E-07 | 3.57E-06 |
| ENSG00000205269 | TMEM170B | 370.3726103 | -0.390386206 | 0.103327252 | -3.778153373 | 0.000157996 | 0.000681923 |
| ENSG00000205302 | SNX2 | 2413.096416 | 0.113529059 | 0.052168195 | 2.176212148 | 0.029539401 | 0.076629408 |
| ENSG00000205309 | NT5M | 115.8896676 | -0.429704048 | 0.167295731 | -2.568529672 | 0.010213097 | 0.030285476 |
| ENSG00000205323 | SARNP | 33.60659862 | -0.824912114 | 0.315305856 | -2.616228336 | 0.008890708 | 0.0268359 |
| ENSG00000205336 | ADGRG1 | 4682.193106 | 1.110974356 | 0.053798489 | 20.65066104 | 9.63E-95 | 1.41E-92 |
| ENSG00000205339 | IPO7 | 6641.017671 | 0.126270802 | 0.049788612 | 2.536138241 | 0.011208248 | 0.032845836 |

| | | | | | | | |
|---|---|---|---|---|---|---|---|
| ENSG00000205352 | PRR13 | 1184.841596 | -0.116647981 | 0.066129984 | -1.763919685 | 0.077745502 | 0.172818006 |
| ENSG00000205356 | TECPR1 | 649.82293 | 0.036038406 | 0.079033718 | 0.455987737 | 0.648398797 | 0.795075283 |
| ENSG00000205363 | INSYN1 | 27.96528056 | 0.690934023 | 0.333131754 | 2.074056331 | 0.03807408 | 0.095247479 |
| ENSG00000205413 | SAMD9 | 6.719454252 | 5.095981585 | 1.304067342 | 3.90775953 | 9.32E-05 | 0.000417093 |
| ENSG00000205423 | CNEP1R1 | 157.5779476 | -0.208813139 | 0.145550283 | -1.434646049 | 0.151387985 | 0.293760817 |
| ENSG00000205426 | KRT81 | 81.7117758 | 0.488796677 | 0.199398109 | 2.451360641 | 0.014231728 | 0.040571126 |
| ENSG00000205456 | TP53TG3D | 12.49749593 | 1.534055941 | 0.532084016 | 2.883108485 | 0.003937718 | 0.012993563 |
| ENSG00000205464 | ATP6AP1L | 110.8540391 | -0.502996806 | 0.179552129 | -2.801397065 | 0.005088187 | 0.01636151 |
| ENSG00000205476 | CCDC85C | 3285.93982 | -0.110926458 | 0.058541392 | -1.894838063 | 0.058113859 | 0.135908065 |
| ENSG00000205488 | CALML3-AS1 | 45.30866737 | -1.631086404 | 0.286616524 | -5.690831717 | 1.26E-08 | 8.99E-08 |
| ENSG00000205531 | NAP1L4 | 3818.138073 | -0.069734982 | 0.046284769 | -1.506650761 | 0.131900199 | 0.264399734 |
| ENSG00000205534 | SMG1P2 | 37.34650913 | -0.119979415 | 0.29337256 | -0.408966041 | 0.682564582 | 0.819871514 |
| ENSG00000205542 | TMSB4X | 4213.022874 | -0.226976362 | 0.054121025 | -4.193866678 | 2.74E-05 | 0.000133511 |
| ENSG00000205544 | TMEM256 | 64.55545302 | 0.276483948 | 0.230274295 | 1.200672218 | 0.229878375 | 0.403062894 |
| ENSG00000205559 | CHKB-DT | 68.46763282 | 0.063311926 | 0.217041312 | 0.291704493 | 0.770512574 | 0.874956824 |
| ENSG00000205560 | CPT1B | 23.65375869 | -0.382816767 | 0.363263736 | -1.053825991 | 0.291962592 | 0.476936684 |
| ENSG00000205571 | SMN2 | 123.853282 | 0.035648941 | 0.1679736 | 0.212229427 | 0.83192805 | 0.91112525 |
| ENSG00000205572 | SERF1B | 7.85554068 | 0.537771244 | 0.634864524 | 0.847064569 | 0.396959132 | 0.589858952 |
| ENSG00000205578 | POM121B | 5.736782716 | 0.151979792 | 0.742731054 | 0.204622913 | 0.837866752 | 0.914703435 |
| ENSG00000205579 | DYNLL1P1 | 19.83371063 | -0.602575054 | 0.418470677 | -1.439945704 | 0.149882761 | 0.291456348 |
| ENSG00000205581 | HMGN1 | 11075.34087 | -0.137138233 | 0.125422588 | -1.09340937 | 0.274214101 | 0.456259503 |
| ENSG00000205583 | STAG3L1 | 14.9019125 | 0.997878793 | 0.49535888 | 2.014456252 | 0.043961654 | 0.107342239 |
| ENSG00000205592 | MUC19 | 4.245291216 | 3.434301842 | 1.228420192 | 2.795706114 | 0.005178647 | 0.016608159 |
| ENSG00000205593 | DENND6B | 273.445048 | 0.092749015 | 0.111875214 | 0.829039883 | 0.40708184 | 0.599849661 |
| ENSG00000205609 | EIF3CL | 112.7919708 | 0.450466298 | 0.382455079 | 1.177827991 | 0.238865187 | 0.414396687 |
| ENSG00000205622 | ENSG0000020 | 4.800591551 | -1.810649173 | 0.877239785 | -2.064029932 | 0.039014879 | 0.097186071 |
| ENSG00000205629 | LCMT1 | 1184.695941 | -0.041256478 | 0.062474057 | -0.660377765 | 0.509011438 | 0.689192557 |
| ENSG00000205634 | LINC00898 | 17.78920346 | 0.007750687 | 0.413094443 | 0.018762507 | 0.985030564 | 0.993193404 |
| ENSG00000205639 | MFSD2B | 3.938384222 | 1.438697707 | 0.969021808 | 1.484690741 | 0.137625766 | 0.273132197 |
| ENSG00000205643 | CDPF1 | 120.2443022 | -0.267937627 | 0.164741219 | -1.626415227 | 0.103861354 | 0.218545171 |
| ENSG00000205659 | LIN52 | 492.6362916 | -0.648136116 | 0.091726581 | -7.0659574 | 1.60E-12 | 1.61E-11 |
| ENSG00000205667 | ARSH | 1.971051759 | -1.186476354 | 1.306616451 | -0.908052514 | 0.363850477 | 0.556559975 |
| ENSG00000205683 | DPF3 | 58.95428929 | -2.732792515 | 0.287848712 | -9.493850077 | 2.23E-21 | 4.04E-20 |

| | | | | | | | |
|---|---|---|---|---|---|---|---|
| ENSG00000205702 | CYP2D7 | 74.21760128 | 0.368142926 | 0.215530021 | 1.708081892 | 0.087621148 | 0.190235581 |
| ENSG00000205704 | SMIM45 | 8.018218002 | 0.134325845 | 0.624118893 | 0.21522477 | 0.82959208 | 0.909783073 |
| ENSG00000205707 | ETFRF1 | 110.875066 | 0.295919861 | 0.180258603 | 1.641640714 | 0.100664485 | 0.212985769 |
| ENSG00000205710 | C17orf107 | 13.38077231 | -0.452789098 | 0.486326109 | -0.931040079 | 0.351832833 | 0.543212087 |
| ENSG00000205726 | ITSN1 | 863.3704347 | -0.045096192 | 0.06968157 | -0.64717532 | 0.517518482 | 0.695665852 |
| ENSG00000205730 | ITPRIPL2 | 847.421158 | -0.634286732 | 0.071142461 | -8.915726549 | 4.85E-19 | 7.66E-18 |
| ENSG00000205740 | ENSG0000020 | 17.29087915 | -1.722341793 | 0.451206425 | -3.817192523 | 0.000134979 | 0.000588896 |
| ENSG00000205744 | DENND1C | 11.20188103 | -0.179012577 | 0.527852568 | -0.339133667 | 0.734509037 | 0.853578385 |
| ENSG00000205746 | PKD1P4 | 93.31370762 | 0.153549911 | 0.190640693 | 0.805441427 | 0.420564994 | 0.611818185 |
| ENSG00000205758 | CRYZL1 | 359.4824482 | -0.098656577 | 0.121931279 | -0.809116236 | 0.41844829 | 0.61039531 |
| ENSG00000205763 | RP9P | 248.0723382 | 0.353494783 | 0.1169618 | 3.022309697 | 0.002508538 | 0.008641816 |
| ENSG00000205765 | RIMOC1 | 2226.871635 | -0.137592525 | 0.059740471 | -2.303171084 | 0.021269219 | 0.057635455 |
| ENSG00000205771 | CATSPER2P1 | 20.92159912 | -0.468194403 | 0.411826079 | -1.1368741 | 0.255590921 | 0.434927358 |
| ENSG00000205790 | DPP9-AS1 | 5.252236149 | -0.375562303 | 0.76572918 | -0.490463617 | 0.623805869 | 0.777554154 |
| ENSG00000205791 | LOH12CR2 | 9.127423868 | -0.127212164 | 0.591645249 | -0.215014258 | 0.829756202 | 0.909797661 |
| ENSG00000205794 | ENSG0000020 | 9.023235388 | 0.450869012 | 0.583639699 | 0.772512584 | 0.439810897 | 0.629163256 |
| ENSG00000205795 | CYS1 | 8.463560288 | 0.073968019 | 0.599101561 | 0.123464908 | 0.901738961 | 0.950420022 |
| ENSG00000205808 | PLPP6 | 578.8187562 | 0.174817543 | 0.089750524 | 1.947816398 | 0.051436936 | 0.122708331 |
| ENSG00000205838 | TTC23L | 37.85306666 | -0.572705789 | 0.293753121 | -1.949616018 | 0.051221903 | 0.122335289 |
| ENSG00000205853 | RFPL3S | 3.957250659 | -0.423173983 | 0.861802414 | -0.491033648 | 0.623402649 | 0.777266998 |
| ENSG00000205861 | PCOTH | 13.55480064 | 0.460832122 | 0.50923016 | 0.904958422 | 0.365487415 | 0.558319324 |
| ENSG00000205867 | KRTAP5-2 | 8.965040117 | 0.676759178 | 0.58846937 | 1.150032971 | 0.250130292 | 0.428208126 |
| ENSG00000205869 | KRTAP5-1 | 3.614002915 | -0.163329981 | 0.914327282 | -0.178634045 | 0.85822506 | 0.926435598 |
| ENSG00000205871 | RPS3AP47 | 68.6409372 | 0.007089766 | 0.242902781 | 0.02918767 | 0.976714915 | 0.989278333 |
| ENSG00000205890 | ENSG0000020 | 11.33225859 | -0.599564204 | 0.534169308 | -1.122423536 | 0.261682402 | 0.442014159 |
| ENSG00000205898 | ENSG0000020 | 3.378172954 | 0.060897928 | 0.9542306 | 0.063818879 | 0.949114445 | 0.976558753 |
| ENSG00000205903 | ZNF316 | 2260.10146 | 0.417597608 | 0.054504763 | 7.661671821 | 1.84E-14 | 2.12E-13 |
| ENSG00000205913 | SRRM2-AS1 | 122.8687373 | 0.107625928 | 0.162545898 | 0.662126384 | 0.507890228 | 0.688024202 |
| ENSG00000205918 | PDPK2P | 3.631976936 | -0.446720347 | 0.919781921 | -0.485680721 | 0.627193554 | 0.780154165 |
| ENSG00000205923 | CEMP1 | 20.75564621 | 1.036653647 | 0.401392599 | 2.582642654 | 0.009804682 | 0.029210312 |
| ENSG00000205930 | C21orf62-AS1 | 12.58526237 | -0.45640586 | 0.49665012 | -0.918968589 | 0.358112002 | 0.550498514 |
| ENSG00000205937 | RNPS1 | 4952.376261 | 0.051604288 | 0.049324569 | 1.046218744 | 0.295460051 | 0.481450033 |
| ENSG00000205940 | HSP90AB2P | 60.42729913 | -0.072033253 | 0.249053906 | -0.28922756 | 0.772407243 | 0.876008412 |

| | | | | | | | |
|---|---|---|---|---|---|---|---|
| ENSG00000205955 | HSP90AA5P | 2.413627754 | 1.29338847 | 1.20075268 | 1.077148102 | 0.281414106 | 0.464802333 |
| ENSG00000205959 | ENSG000002C | 1.89594183 | 2.071948341 | 1.469546121 | 1.409923997 | 0.158562126 | 0.303801484 |
| ENSG00000205978 | NYNRIN | 785.02756 | -0.611378914 | 0.077538881 | -7.88480447 | 3.15E-15 | 3.85E-14 |
| ENSG00000205981 | DNAJC19 | 341.5707651 | -0.253545772 | 0.110576881 | -2.292936555 | 0.021851658 | 0.059071062 |
| ENSG00000206028 | ENSG000002C | 1.840964942 | -2.307024396 | 1.500293529 | -1.537715354 | 0.124118228 | 0.252060211 |
| ENSG00000206052 | DOK6 | 29.21690144 | 0.662648538 | 0.328314346 | 2.018335614 | 0.04355632 | 0.106545821 |
| ENSG00000206053 | JPT2 | 7545.081944 | -0.159406273 | 0.043367848 | -3.675678614 | 0.000237218 | 0.000992845 |
| ENSG00000206075 | SERPINB5 | 1.852852187 | 0.813915363 | 1.408980409 | 0.577662654 | 0.563491897 | 0.732172423 |
| ENSG00000206077 | ZDHHC11B | 11.03553041 | 0.283766064 | 0.544512598 | 0.521137739 | 0.602270823 | 0.761636862 |
| ENSG00000206113 | CFAP99 | 9.147117434 | 1.339395855 | 0.639510958 | 2.094406419 | 0.036223791 | 0.091295735 |
| ENSG00000206120 | EGFEM1P | 2.025907772 | -0.652281175 | 1.225911832 | -0.532078375 | 0.594671714 | 0.756386934 |
| ENSG00000206140 | TMEM191C | 12.33685127 | -1.228793736 | 0.514140444 | -2.389996255 | 0.016848545 | 0.047120502 |
| ENSG00000206144 | MAPK6P1 | 5.471208582 | -1.146499341 | 0.774287457 | -1.480715374 | 0.138682435 | 0.274611573 |
| ENSG00000206149 | HERC2P9 | 285.016209 | 0.084668825 | 0.117850643 | 0.718441776 | 0.472484936 | 0.658168305 |
| ENSG00000206172 | HBA1 | 4.281958553 | -0.025783795 | 0.87387463 | -0.029505142 | 0.976461718 | 0.989269612 |
| ENSG00000206187 | LINC02346 | 20.34397651 | -0.764785344 | 0.393341711 | -1.944328105 | 0.051855898 | 0.123579295 |
| ENSG00000206195 | DUXAP8 | 976.4429525 | -0.10998354 | 0.071801984 | -1.53176186 | 0.125581211 | 0.254355729 |
| ENSG00000206199 | ANKUB1 | 1.860887388 | -1.602953731 | 1.358930215 | -1.179570307 | 0.238171158 | 0.413536662 |
| ENSG00000206203 | TSSK2 | 1.612178243 | -0.708192159 | 1.448841194 | -0.488799022 | 0.62498399 | NA |
| ENSG00000206228 | HNRNPA1P4 | 58.46107841 | 0.450300578 | 0.248734819 | 1.810364063 | 0.070239348 | 0.159029871 |
| ENSG00000206262 | FOXL2NB | 4.31744313 | 0.388767897 | 0.900004621 | 0.431962112 | 0.665768951 | 0.807339718 |
| ENSG00000206337 | HCP5 | 3.367280805 | 0.071814406 | 0.945086507 | 0.075987125 | 0.939429342 | 0.971224891 |
| ENSG00000206341 | HLA-H | 7.026657344 | 0.425266043 | 0.655426176 | 0.648838966 | 0.516442468 | 0.69491147 |
| ENSG00000206344 | HCG27 | 17.26324243 | -0.887485412 | 0.479250005 | -1.851821393 | 0.064051475 | 0.14727 |
| ENSG00000206417 | H1-10-AS1 | 276.4959988 | 0.052233995 | 0.136411793 | 0.382914073 | 0.701783476 | 0.833061472 |
| ENSG00000206418 | RAB12 | 1022.04132 | 0.488727693 | 0.070330653 | 6.948999789 | 3.68E-12 | 3.59E-11 |
| ENSG00000206503 | HLA-A | 332.1829085 | -0.069529262 | 0.106439998 | -0.653224948 | 0.513611269 | 0.692787823 |
| ENSG00000206527 | HACD2 | 4896.61131 | -0.280994054 | 0.047548446 | -5.909636989 | 3.43E-09 | 2.60E-08 |
| ENSG00000206530 | CFAP44 | 505.9079708 | 0.270451145 | 0.097939699 | 2.7614047 | 0.005755331 | 0.018235814 |
| ENSG00000206535 | LNP1 | 95.66898993 | -0.05062051 | 0.180492442 | -0.28045778 | 0.779126313 | 0.880184615 |
| ENSG00000206538 | VGLL3 | 34.08492581 | -3.886729586 | 0.459673535 | -8.455413001 | 2.78E-17 | 3.90E-16 |
| ENSG00000206557 | TRIM71 | 9.207045258 | 0.166782905 | 0.610470088 | 0.273204058 | 0.784696361 | 0.883091372 |
| ENSG00000206559 | ZCWPW2 | 15.33416663 | 0.023371514 | 0.444328881 | 0.052599584 | 0.958050948 | 0.980927862 |

| | | | | | | | |
|---|---|---|---|---|---|---|---|
| ENSG00000206560 | ANKRD28 | 2674.388535 | 0.115096165 | 0.053486129 | 2.151888089 | 0.03140617 | 0.080809115 |
| ENSG00000206561 | COLQ | 51.99645906 | 0.674282596 | 0.249173046 | 2.706081599 | 0.006808231 | 0.021168451 |
| ENSG00000206562 | METTL6 | 830.8496486 | -0.261186538 | 0.07188889 | -3.633197541 | 0.000279931 | 0.001157881 |
| ENSG00000206567 | ENSG0000020 | 75.48097122 | 0.079278334 | 0.214991649 | 0.368750761 | 0.712313509 | 0.839972059 |
| ENSG00000206573 | THUMPD3-AS | 1243.077935 | 0.030379049 | 0.085434449 | 0.35558313 | 0.72215278 | 0.845601795 |
| ENSG00000206603 | SNORA22B | 1.991931296 | 0.770966977 | 1.277219554 | 0.603629169 | 0.546090216 | 0.719092449 |
| ENSG00000206625 | RNU6-1 | 8.088111675 | 0.207272424 | 0.620821879 | 0.333867783 | 0.738479332 | 0.855807824 |
| ENSG00000206634 | SNORA22 | 2.753539616 | 1.125329676 | 1.113528211 | 1.010598263 | 0.312208748 | 0.500424027 |
| ENSG00000206754 | SNORD101 | 7.467914709 | 0.639372519 | 0.65127009 | 0.981731741 | 0.326232024 | 0.516727379 |
| ENSG00000206828 | RNVU1-30 | 16.98237045 | 1.141154901 | 0.446761267 | 2.554283429 | 0.01064066 | 0.031382164 |
| ENSG00000206838 | SNORA5A | 5.743136646 | 0.916374115 | 0.755980329 | 1.212166614 | 0.225448609 | 0.39749621 |
| ENSG00000206885 | SNORA75 | 6.651731925 | 0.772215418 | 0.691095253 | 1.117379137 | 0.263832252 | 0.444499989 |
| ENSG00000206897 | SNORA9B | 1.749294432 | 0.6408351 | 1.365612906 | 0.469265556 | 0.638879832 | NA |
| ENSG00000206899 | RNU6-36P | 2.780532792 | 0.676125779 | 1.06015265 | 0.637762665 | 0.523628186 | 0.700001783 |
| ENSG00000206932 | RNU6-4P | 10.0103136 | -0.059599995 | 0.574300683 | -0.103778382 | 0.917345223 | 0.959375401 |
| ENSG00000206965 | RNU6-5P | 2.862321393 | -0.022031526 | 1.037448645 | -0.021236257 | 0.983057192 | 0.992139551 |
| ENSG00000207031 | SNORD59A | 2.503926507 | -0.744648108 | 1.166031395 | -0.638617546 | 0.523071764 | 0.699706471 |
| ENSG00000207041 | RNU6-3P | 5.84326481 | -0.098659697 | 0.730063848 | -0.13513845 | 0.89250241 | 0.944983167 |
| ENSG00000207067 | SNORA72 | 9.192455279 | -1.061303867 | 0.60939914 | -1.741557869 | 0.081585839 | 0.179545198 |
| ENSG00000207088 | SNORA7B | 2.971525297 | -0.224427987 | 1.054124905 | -0.212904548 | 0.831401415 | 0.91071463 |
| ENSG00000207181 | SNORA14B | 10.09108596 | 0.342965173 | 0.576618297 | 0.594787184 | 0.551985735 | 0.723822612 |
| ENSG00000207205 | RNVU1-15 | 4.367234199 | -0.443896132 | 0.835029567 | -0.531593311 | 0.595007699 | 0.756497068 |
| ENSG00000207280 | SNORD20 | 7.03053189 | 0.206088042 | 0.667624498 | 0.308688556 | 0.75755845 | 0.866391318 |
| ENSG00000207357 | RNU6-2 | 14.27696104 | 0.032642597 | 0.465146183 | 0.070177072 | 0.944052723 | 0.973493866 |
| ENSG00000207359 | RNU6-925P | 7.475346979 | 0.44497889 | 0.6348958 | 0.700869167 | 0.483384668 | 0.667507389 |
| ENSG00000207383 | Y_RNA | 2.862306573 | 1.17852777 | 1.082830578 | 1.088376884 | 0.276428762 | 0.458980588 |
| ENSG00000207422 | RNU6-813P | 270.1662723 | -1.466154926 | 0.136746046 | -10.72173545 | 8.05E-27 | 1.98E-25 |
| ENSG00000207425 | Y_RNA | 23.07560043 | 0.390863274 | 0.366175987 | 1.067419185 | 0.285782589 | 0.469616948 |
| ENSG00000207445 | SNORD15B | 9.261705087 | 0.143604287 | 0.587727141 | 0.244338362 | 0.806968775 | 0.89748787 |
| ENSG00000207493 | SNORA46 | 1.919037297 | 0.843706922 | 1.299044355 | 0.649482766 | 0.516026382 | 0.694679946 |
| ENSG00000207507 | RNU6-9 | 6.309004797 | -0.194638399 | 0.707417427 | -0.27513939 | 0.78320915 | 0.882490914 |
| ENSG00000207523 | SNORA66 | 2.862310252 | -0.366286852 | 1.01513456 | -0.36082591 | 0.718229592 | 0.843411181 |
| ENSG00000207524 | RNU6-33P | 7.125401961 | -0.786855378 | 0.660775412 | -1.190806081 | 0.233729721 | 0.4082351 |

| | | | | | | | |
|---|---|---|---|---|---|---|---|
| ENSG00000207547 | MIR25 | 40.51213813 | -0.409878551 | 0.292857802 | -1.399582145 | 0.161638484 | 0.308638823 |
| ENSG00000207554 | MIR647 | 27.84685245 | -0.312676761 | 0.36420479 | -0.858519079 | 0.3906059 | 0.58343796 |
| ENSG00000207561 | MIR635 | 12.19837896 | 0.205487853 | 0.513351783 | 0.400286625 | 0.688945418 | 0.824262433 |
| ENSG00000207563 | MIR23B | 5.777610984 | 0.731649491 | 0.739999976 | 0.98871556 | 0.322802327 | 0.51274653 |
| ENSG00000207586 | MIR135A2 | 2.441434562 | -0.302496206 | 1.168697919 | -0.258831817 | 0.795765008 | 0.890871567 |
| ENSG00000207607 | MIR200A | 6.127513609 | 1.047618776 | 0.785530134 | 1.33364556 | 0.182320044 | 0.338215606 |
| ENSG00000207631 | MIR641 | 2.61041011 | -2.168356779 | 1.281398393 | -1.692180037 | 0.090611652 | 0.195543865 |
| ENSG00000207650 | MIR570 | 24.72161241 | 0.053628139 | 0.353864618 | 0.151549877 | 0.879541969 | 0.937615842 |
| ENSG00000207696 | MIR659 | 3.627363308 | 1.212778082 | 1.003261243 | 1.208835775 | 0.226725944 | 0.399101685 |
| ENSG00000207697 | MIR573 | 4.35493975 | 0.241256383 | 0.857077275 | 0.281487318 | 0.778336654 | 0.879645425 |
| ENSG00000207721 | MIR186 | 1.864192425 | 0.848506213 | 1.314073829 | 0.645706652 | 0.518469352 | 0.696066421 |
| ENSG00000207730 | MIR200B | 3.857981896 | 1.829592472 | 1.004436471 | 1.82151139 | 0.068529157 | 0.155911818 |
| ENSG00000207741 | MIR590 | 8.858746613 | 0.968983232 | 0.606189433 | 1.598482553 | 0.109935625 | 0.228600729 |
| ENSG00000207751 | ENSG0000002C | 12.19085298 | -0.267002266 | 0.510461235 | -0.523060808 | 0.600931938 | 0.760908465 |
| ENSG00000207757 | MIR93 | 8.83602067 | 0.075227394 | 0.659798004 | 0.114015795 | 0.909225272 | 0.954587195 |
| ENSG00000207839 | MIR33B | 4.245151111 | -0.12067153 | 0.89286939 | -0.135150259 | 0.892493074 | 0.944983167 |
| ENSG00000207864 | MIR27B | 11.09800152 | 1.802772827 | 0.597002291 | 3.019708392 | 0.002530182 | 0.00870592 |
| ENSG00000207923 | MIR559 | 2.369667895 | -0.611191366 | 1.196083545 | -0.510993875 | 0.609355346 | 0.767549507 |
| ENSG00000208028 | MIR616 | 3.910968593 | 0.807108555 | 0.906056445 | 0.890792797 | 0.37304034 | 0.565817244 |
| ENSG00000208036 | MIR106B | 3.817790307 | 1.114483996 | 0.921766367 | 1.209074269 | 0.226634314 | 0.398971004 |
| ENSG00000208772 | SNORD94 | 8.45374665 | -0.297996048 | 0.629041459 | -0.473730378 | 0.635692188 | 0.785317815 |
| ENSG00000208892 | SNORA49 | 1.747847245 | -0.431630956 | 1.350443013 | -0.319621748 | 0.749255086 | NA |
| ENSG00000209042 | SNORD12C | 2.184747347 | -0.82669271 | 1.217950887 | -0.678757016 | 0.497291833 | 0.67908058 |
| ENSG00000209482 | SNORD83A | 8.921091398 | 0.296644662 | 0.584489491 | 0.507527794 | 0.61178454 | 0.76921553 |
| ENSG00000210082 | MT-RNR2 | 80440.1996 | 0.076169212 | 0.113156835 | 0.673129569 | 0.500864864 | 0.681930083 |
| ENSG00000210100 | MT-TI | 5.76672677 | -0.304706304 | 0.812749972 | -0.3749078 | 0.707729038 | 0.836785081 |
| ENSG00000210107 | MT-TQ | 49.49039044 | -0.335947516 | 0.251899807 | -1.333655314 | 0.182316846 | 0.338215606 |
| ENSG00000210117 | MT-TW | 8.313640648 | -0.179634011 | 0.617400179 | -0.290952315 | 0.77108779 | 0.87520429 |
| ENSG00000210127 | MT-TA | 4.112717228 | -0.746222929 | 0.952803801 | -0.783186348 | 0.433517691 | 0.623465351 |
| ENSG00000210144 | MT-TY | 12.99237319 | -0.484406081 | 0.496621266 | -0.97540342 | 0.329360188 | 0.519780043 |
| ENSG00000210151 | MT-TS1 | 29.67883929 | -0.427008401 | 0.332298011 | -1.285016421 | 0.198786546 | 0.361544705 |
| ENSG00000210154 | MT-TD | 1.741775288 | 1.243137086 | 1.52793317 | 0.813606976 | 0.41587014 | NA |
| ENSG00000210194 | MT-TE | 11.62316393 | -0.918569676 | 0.524923281 | -1.749912245 | 0.080133457 | 0.176874965 |

| | | | | | | | |
|---|---|---|---|---|---|---|---|
| ENSG00000210195 | MT-TT | 82.6170285 | 0.087509999 | 0.206591138 | 0.423590287 | 0.67186464 | 0.812021148 |
| ENSG00000210196 | MT-TP | 40.21334289 | -0.843419544 | 0.281868971 | -2.992239771 | 0.002769387 | 0.009460868 |
| ENSG00000211445 | GPX3 | 999.1612883 | -0.58800879 | 0.07446349 | -7.896605321 | 2.87E-15 | 3.51E-14 |
| ENSG00000211448 | DIO2 | 10.0842891 | 1.763453916 | 0.60960374 | 2.892787231 | 0.003818399 | 0.012638013 |
| ENSG00000211450 | SELENOH | 100.0028432 | -0.239876123 | 0.176791355 | -1.356831747 | 0.174834678 | 0.327940107 |
| ENSG00000211451 | GNRHR2 | 128.9647675 | 0.314409129 | 0.162510451 | 1.934700981 | 0.053027 | 0.126068958 |
| ENSG00000211452 | DIO1 | 14.11777106 | 0.088815336 | 0.471024933 | 0.188557611 | 0.850439558 | 0.921772038 |
| ENSG00000211454 | AKR7L | 16.79030259 | -0.109343352 | 0.43659641 | -0.250444918 | 0.802243297 | 0.894714402 |
| ENSG00000211455 | STK38L | 365.4476948 | 0.530254283 | 0.106124298 | 4.996539853 | 5.84E-07 | 3.48E-06 |
| ENSG00000211456 | SACM1L | 694.7384537 | -0.05706228 | 0.080360342 | -0.710080109 | 0.477654461 | 0.662398339 |
| ENSG00000211459 | MT-RNR1 | 15275.8617 | 0.103355079 | 0.062361895 | 1.657343475 | 0.097450052 | 0.207625541 |
| ENSG00000211460 | TSN | 3040.957126 | -0.206276687 | 0.051486712 | -4.006406328 | 6.16E-05 | 0.00028467 |
| ENSG00000211584 | SLC48A1 | 604.2779727 | 0.552847257 | 0.089359103 | 6.186804014 | 6.14E-10 | 4.99E-09 |
| ENSG00000211591 | MIR762 | 8.979937996 | -0.437824097 | 0.617511739 | -0.7090134 | 0.478316162 | 0.66295052 |
| ENSG00000211683 | ENSG0000021 | 2.47080197 | -0.750687215 | 1.129297603 | -0.664738164 | 0.50621798 | 0.686625533 |
| ENSG00000211689 | TRGC1 | 296.8844002 | -1.365755063 | 0.117899253 | -11.58408576 | 4.96E-31 | 1.44E-29 |
| ENSG00000211695 | TRGV9 | 33.45464641 | -0.868907859 | 0.315476662 | -2.754269851 | 0.005882325 | 0.018599659 |
| ENSG00000212123 | PRR22 | 88.39188201 | 0.726147779 | 0.196088622 | 3.703161211 | 0.000212929 | 0.000898235 |
| ENSG00000212124 | TAS2R19 | 2.404138229 | 0.399929117 | 1.123335389 | 0.356019334 | 0.721826087 | 0.845596895 |
| ENSG00000212125 | TAS2R15P | 2.185384906 | -0.316925344 | 1.280929176 | -0.24741832 | 0.804584498 | 0.896153058 |
| ENSG00000212127 | TAS2R14 | 5.318378949 | 0.307516111 | 0.825347998 | 0.372589637 | 0.709453887 | 0.838090126 |
| ENSG00000212135 | SNORD67 | 6.715389084 | -0.16502978 | 0.691636931 | -0.23860753 | 0.811409926 | 0.90021491 |
| ENSG00000212195 | U3 | 10.02470078 | -1.125303693 | 0.58098554 | -1.936887609 | 0.052759083 | 0.125496045 |
| ENSG00000212232 | SNORD17 | 17.40781826 | 0.33106525 | 0.424653545 | 0.77961259 | 0.435618943 | 0.625535029 |
| ENSG00000212371 | ENSG0000021 | 3.41449768 | -1.100414596 | 0.973256519 | -1.130652171 | 0.25820152 | 0.437843204 |
| ENSG00000212385 | RNU6-817P | 2.630128265 | -0.559251684 | 1.073344654 | -0.521036446 | 0.602341383 | 0.761646618 |
| ENSG00000212452 | SNORD69 | 3.722795832 | -0.726844992 | 0.904055607 | -0.803982615 | 0.421407015 | 0.612421725 |
| ENSG00000212464 | SNORA12 | 1.961606797 | -0.170338128 | 1.217468439 | -0.13991174 | 0.888729725 | 0.942772712 |
| ENSG00000212493 | SNORD19 | 4.854320657 | -0.428838942 | 0.855217174 | -0.501438646 | 0.616062447 | 0.772581194 |
| ENSG00000212505 | RNA5SP299 | 3.458510363 | -0.074429496 | 0.95702002 | -0.077772142 | 0.938009307 | 0.970412804 |
| ENSG00000212588 | SNORA26 | 7.9711929 | -0.235416153 | 0.61677152 | -0.381691024 | 0.702690559 | 0.833448189 |
| ENSG00000212607 | SNORA3B | 4.318008311 | -0.408281737 | 0.848683759 | -0.481076411 | 0.630462193 | 0.782069037 |
| ENSG00000212664 | RPL17P39 | 202.3144115 | 0.821518644 | 0.131806973 | 6.232740377 | 4.58E-10 | 3.77E-09 |

| | | | | | | | |
|---|---|---|---|---|---|---|---|
| ENSG00000212694 | LINC01089 | 184.7495993 | -0.333969328 | 0.144326057 | -2.313991904 | 0.020668166 | 0.056256257 |
| ENSG00000212695 | ENSG0000021 | 5.625852653 | 0.926510363 | 0.804976114 | 1.150978703 | 0.249740997 | 0.427797079 |
| ENSG00000212710 | CTAGE1 | 34.09944431 | -0.526920069 | 0.328621085 | -1.603427451 | 0.108840314 | 0.226775369 |
| ENSG00000212712 | ENSG0000021 | 1.740361524 | 2.034338219 | 1.512943106 | 1.34462308 | 0.178746986 | NA |
| ENSG00000212719 | LINC02693 | 8.696577968 | 1.674420083 | 0.643908876 | 2.600399135 | 0.009311539 | 0.027955333 |
| ENSG00000212743 | LINC02656 | 1.64341527 | -0.218676861 | 1.429479627 | -0.15297655 | 0.878416768 | NA |
| ENSG00000212747 | RTL8B | 1911.792786 | 0.040091037 | 0.061577807 | 0.651063083 | 0.515005766 | 0.69397511 |
| ENSG00000212766 | EWSAT1 | 34.24226433 | -1.456986147 | 0.316984641 | -4.596393516 | 4.30E-06 | 2.31E-05 |
| ENSG00000212789 | ST13P5 | 187.8104529 | 0.329241316 | 0.133161755 | 2.472491563 | 0.013417488 | 0.03851798 |
| ENSG00000212802 | RPL15P3 | 5009.323 | 0.094701974 | 0.059848062 | 1.582373289 | 0.113564379 | 0.234996108 |
| ENSG00000212829 | RPS26P3 | 5.521699649 | 0.001375837 | 0.735980437 | 0.001869393 | 0.998508441 | 0.999161772 |
| ENSG00000212864 | RNF208 | 676.5167662 | -0.070419799 | 0.09284959 | -0.758428759 | 0.448194347 | 0.63636819 |
| ENSG00000212901 | KRTAP3-1 | 149.1062435 | 1.560253605 | 0.162174653 | 9.620822807 | 6.53E-22 | 1.21E-20 |
| ENSG00000212907 | MT-ND4L | 6936.671047 | 0.075624565 | 0.071241276 | 1.061527372 | 0.2884503 | 0.472443945 |
| ENSG00000212916 | MAP10 | 498.2176627 | 0.062719483 | 0.089206557 | 0.703081533 | 0.48200494 | 0.666287182 |
| ENSG00000212978 | C2orf74-DT | 133.6516192 | -0.550559827 | 0.162646678 | -3.385005062 | 0.000711769 | 0.002739596 |
| ENSG00000212993 | POU5F1B | 35.56433969 | -0.5887383 | 0.293650777 | -2.004892701 | 0.044974519 | 0.109384592 |
| ENSG00000213003 | BTF3P12 | 10.91549431 | 0.654154978 | 0.568486213 | 1.150696293 | 0.249857202 | 0.427932225 |
| ENSG00000213005 | PTTG3P | 7.994428065 | -0.546895522 | 0.610470457 | -0.895859112 | 0.370328014 | 0.562977114 |
| ENSG00000213013 | RPS15AP36 | 7.026659922 | 1.810405698 | 0.757750762 | 2.389183606 | 0.016885861 | 0.047198934 |
| ENSG00000213015 | ZNF580 | 700.7399394 | -0.160791641 | 0.083613108 | -1.923043477 | 0.05447459 | 0.128946937 |
| ENSG00000213018 | PABPN1P1 | 9.895490714 | -0.015896851 | 0.600169533 | -0.026487267 | 0.97886869 | 0.99043965 |
| ENSG00000213020 | ZNF611 | 436.7573079 | -0.044784336 | 0.093552399 | -0.478708582 | 0.632145959 | 0.783268246 |
| ENSG00000213023 | SYT3 | 16.61291397 | 0.615279088 | 0.441157106 | 1.394693816 | 0.16310819 | 0.310540451 |
| ENSG00000213024 | NUP62 | 2363.032083 | -0.157452724 | 0.0519341 | -3.031779186 | 0.00243117 | 0.008404315 |
| ENSG00000213025 | COX20P1 | 121.4668093 | -0.348515913 | 0.160737075 | -2.168235999 | 0.030140735 | 0.077998792 |
| ENSG00000213026 | CFL1P4 | 6.978837131 | -0.22533961 | 0.694531335 | -0.324448443 | 0.74559855 | 0.859379806 |
| ENSG00000213036 | ENSG0000021 | 24.13098577 | -0.107321082 | 0.367836942 | -0.291762653 | 0.770468102 | 0.874956824 |
| ENSG00000213047 | DENND1B | 1133.678861 | 0.350681007 | 0.078092332 | 4.490594651 | 7.10E-06 | 3.73E-05 |
| ENSG00000213051 | RPL5P5 | 26.51650458 | 0.278120663 | 0.379563155 | 0.732738832 | 0.463717735 | 0.650026571 |
| ENSG00000213057 | C1orf220 | 30.66985714 | 0.045699162 | 0.324248445 | 0.140938722 | 0.887918351 | 0.942285816 |
| ENSG00000213058 | ENSG0000021 | 12.21786293 | 0.177842187 | 0.500534253 | 0.355304728 | 0.722361314 | 0.845703316 |
| ENSG00000213060 | ENSG0000021 | 1.845272757 | -0.97000489 | 1.303243237 | -0.744300728 | 0.456694561 | 0.643880107 |

| | | | | | | | |
|---|---|---|---|---|---|---|---|
| ENSG00000213062 | ENSG0000021 | 136.1997129 | -0.009075621 | 0.152938116 | -0.059341786 | 0.952679879 | 0.978056463 |
| ENSG00000213064 | SFT2D2 | 922.4835553 | 0.151605644 | 0.07772532 | 1.95053098 | 0.051112865 | 0.122101234 |
| ENSG00000213066 | CEP43 | 124.3242818 | 0.096784439 | 0.162770278 | 0.594607565 | 0.552105822 | 0.7239387 |
| ENSG00000213070 | HMGB3P6 | 55.06707216 | -0.754256355 | 0.248397791 | -3.036485764 | 0.002393534 | 0.008293446 |
| ENSG00000213071 | LPAL2 | 12.79417514 | -0.327099668 | 0.491595321 | -0.665384015 | 0.505804906 | 0.686293521 |
| ENSG00000213073 | CHP1P2 | 42.09961377 | -0.329339799 | 0.277795932 | -1.185545798 | 0.235801718 | 0.410790661 |
| ENSG00000213078 | ENSG0000021 | 4.056119356 | 0.651782391 | 0.881032324 | 0.739793959 | 0.459425025 | 0.646458326 |
| ENSG00000213079 | SCAF8 | 2846.166752 | 0.053006812 | 0.050868336 | 1.042039429 | 0.297393399 | 0.483809983 |
| ENSG00000213080 | ENSG0000021 | 811.5142943 | -0.370419144 | 0.071721848 | -5.164662539 | 2.41E-07 | 1.50E-06 |
| ENSG00000213082 | PPP1R14BP2 | 9.823795451 | 0.09684983 | 0.611924233 | 0.15827095 | 0.874243298 | 0.934789447 |
| ENSG00000213085 | CFAP45 | 40.86781079 | 0.329275601 | 0.273541195 | 1.203751418 | 0.228685672 | 0.401626804 |
| ENSG00000213087 | ENSG0000021 | 82.37994373 | 0.38123614 | 0.195127547 | 1.953779182 | 0.050727337 | 0.121356254 |
| ENSG00000213089 | PDCL3P5 | 101.2412763 | -0.151749243 | 0.176713449 | -0.858730583 | 0.390489173 | 0.583343425 |
| ENSG00000213090 | SCYL2P1 | 36.13951758 | 0.144445723 | 0.294990616 | 0.489662094 | 0.624373029 | 0.778004121 |
| ENSG00000213096 | ZNF254 | 222.1068492 | -0.096456264 | 0.124088379 | -0.77731907 | 0.436970547 | 0.626676508 |
| ENSG00000213100 | KRT18P68 | 5.480068335 | 0.188823865 | 0.742636034 | 0.254261652 | 0.799293424 | 0.892991047 |
| ENSG00000213104 | NPM1P46 | 39.49679695 | -0.217327453 | 0.281949033 | -0.770804036 | 0.440823094 | 0.630100429 |
| ENSG00000213108 | BTF3L4P3 | 2.254279177 | 0.27440394 | 1.229942316 | 0.223103097 | 0.823455268 | 0.906737572 |
| ENSG00000213110 | RNF11P1 | 2.470813111 | -0.349172566 | 1.123082374 | -0.31090557 | 0.75587241 | 0.865149344 |
| ENSG00000213117 | MEMO1P2 | 4.652388063 | -0.339391486 | 0.820636545 | -0.413571011 | 0.679188307 | 0.81757778 |
| ENSG00000213123 | DYNLT2B | 36.91196784 | -0.168185809 | 0.289791013 | -0.580369305 | 0.5616656 | 0.730585669 |
| ENSG00000213128 | RPL32P31 | 6.646527608 | 0.306429884 | 0.675676954 | 0.453515371 | 0.650177678 | 0.79653276 |
| ENSG00000213130 | EEF1DP5 | 2.315719525 | -0.120886491 | 1.199594681 | -0.10077278 | 0.919730835 | 0.960994095 |
| ENSG00000213131 | YWHAZP4 | 106.5116174 | -0.329644909 | 0.190771555 | -1.727956296 | 0.08399606 | 0.183931808 |
| ENSG00000213137 | ARF1P2 | 4.336434121 | -0.845606391 | 0.85192775 | -0.992579935 | 0.320914707 | 0.510419703 |
| ENSG00000213139 | CRYGS | 23.05885112 | -1.146364109 | 0.401820866 | -2.852923293 | 0.004331908 | 0.014173897 |
| ENSG00000213144 | ENSG0000021 | 353.6970749 | -0.496279191 | 0.103410751 | -4.799106338 | 1.59E-06 | 9.00E-06 |
| ENSG00000213145 | CRIP1 | 14.97451791 | -0.829912359 | 0.470118235 | -1.765326885 | 0.077508844 | 0.172378365 |
| ENSG00000213148 | JPT1P1 | 3.580037579 | 0.909826521 | 0.945125516 | 0.962651527 | 0.33572243 | 0.526875672 |
| ENSG00000213149 | CNN2P9 | 3.274954132 | -0.738749744 | 0.96199331 | -0.767936468 | 0.44252493 | 0.631470633 |
| ENSG00000213152 | RPL7AP60 | 3.473585775 | -0.037423762 | 0.936237671 | -0.039972502 | 0.968115049 | 0.985820278 |
| ENSG00000213160 | KLHL23 | 497.472123 | -0.247750827 | 0.087684635 | -2.825475944 | 0.004721043 | 0.015307524 |
| ENSG00000213169 | RPL8P4 | 23.20801633 | 0.164098183 | 0.361663387 | 0.453731808 | 0.65002187 | 0.79638441 |

| | | | | | | | |
|---|---|---|---|---|---|---|---|
| ENSG00000213171 | LINGO4 | 2.768190101 | 2.142378038 | 1.247434928 | 1.717426689 | 0.085901239 | 0.187227973 |
| ENSG00000213174 | RPL28P1 | 10.74751559 | 0.171660261 | 0.53012311 | 0.323812068 | 0.746080319 | 0.859596562 |
| ENSG00000213176 | RPL13P6 | 5.502411084 | 0.852642438 | 0.75903183 | 1.123328962 | 0.261297808 | 0.4415299 |
| ENSG00000213177 | PRDX2P4 | 7.09987425 | 0.391385177 | 0.688303709 | 0.568622793 | 0.569612153 | 0.736952388 |
| ENSG00000213178 | RPL22P1 | 1454.572299 | 0.242418505 | 0.07380655 | 3.284512064 | 0.001021591 | 0.003818068 |
| ENSG00000213183 | RPS7P8 | 2.376757228 | 1.249710064 | 1.20068124 | 1.040834172 | 0.297952517 | 0.484513415 |
| ENSG00000213184 | ENSG0000021 | 12.13423813 | 0.423472478 | 0.532676845 | 0.79498946 | 0.426619633 | 0.617497513 |
| ENSG00000213185 | FAM24B | 50.74942613 | -0.080236493 | 0.24848506 | -0.322902686 | 0.746768942 | 0.85963488 |
| ENSG00000213186 | TRIM59 | 355.4361934 | -0.156759862 | 0.100395392 | -1.561424861 | 0.11842354 | 0.243077667 |
| ENSG00000213187 | COPS5P1 | 4.145479923 | -0.308204256 | 0.86382176 | -0.356791493 | 0.721247905 | 0.845194729 |
| ENSG00000213189 | BTF3L4P2 | 502.5719888 | -0.027362296 | 0.087942144 | -0.311139745 | 0.755694387 | 0.865149344 |
| ENSG00000213190 | MLLT11 | 185.4391932 | -0.402469863 | 0.133803259 | -3.007922711 | 0.0026304 | 0.009019639 |
| ENSG00000213194 | CDK7P1 | 11.06901583 | 0.174725689 | 0.540456268 | 0.323292928 | 0.746473409 | 0.859604617 |
| ENSG00000213197 | NUDCP1 | 11.94738551 | 0.140008927 | 0.514657043 | 0.272043158 | 0.785588835 | 0.883359045 |
| ENSG00000213199 | ASIC3 | 133.796391 | 0.406371429 | 0.161612209 | 2.514484722 | 0.011920647 | 0.034738267 |
| ENSG00000213204 | ENSG0000021 | 29.38963676 | -1.272108743 | 0.355049816 | -3.582902132 | 0.000339798 | 0.001383312 |
| ENSG00000213205 | STRADBP1 | 2.78339342 | 0.657643374 | 1.084736556 | 0.606270131 | 0.54433539 | 0.717823209 |
| ENSG00000213211 | KRT18P64 | 4.473071051 | 0.692554097 | 0.837186021 | 0.827240398 | 0.408100819 | 0.60057947 |
| ENSG00000213212 | NCLP1 | 7.300337374 | -0.198152297 | 0.726643614 | -0.272695298 | 0.78508745 | 0.883106023 |
| ENSG00000213213 | CCDC183 | 62.02810488 | 0.117849455 | 0.232198509 | 0.507537518 | 0.611777719 | 0.76921553 |
| ENSG00000213214 | ARHGEF35 | 169.6406039 | 0.259588873 | 0.140057192 | 1.853449076 | 0.063818019 | 0.146839052 |
| ENSG00000213216 | ENSG0000021 | 14.49840734 | 0.477682845 | 0.494044009 | 0.966883186 | 0.333602433 | 0.524670219 |
| ENSG00000213221 | DNLZ | 4.706100973 | 0.877685131 | 0.862200649 | 1.017959256 | 0.308697319 | 0.496818242 |
| ENSG00000213222 | TOMM40P4 | 219.2624078 | 0.130419972 | 0.129942689 | 1.003673029 | 0.315536241 | 0.504207396 |
| ENSG00000213225 | NOC2LP1 | 593.843372 | -0.136660557 | 0.0868547 | -1.573438813 | 0.115617246 | 0.238383931 |
| ENSG00000213228 | RPL12P38 | 15.35926601 | 1.250245709 | 0.47264131 | 2.645231557 | 0.008163503 | 0.024873258 |
| ENSG00000213234 | ST13P10 | 3.801233405 | -0.279502118 | 0.906083988 | -0.308472638 | 0.757722717 | 0.866391318 |
| ENSG00000213235 | EEF1A1P16 | 16.3279458 | 0.0797733 | 0.430128973 | 0.185463674 | 0.85286537 | 0.923198544 |
| ENSG00000213236 | YWHAZP2 | 861.5619589 | -0.346328932 | 0.071781082 | -4.824793961 | 1.40E-06 | 7.95E-06 |
| ENSG00000213237 | PTMAP10 | 2.06418806 | 0.544692118 | 1.228991073 | 0.443202664 | 0.657619152 | 0.801710333 |
| ENSG00000213239 | NPM1P32 | 8.692018893 | 0.449528916 | 0.609835118 | 0.737131895 | 0.461042145 | 0.647899216 |
| ENSG00000213246 | SUPT4H1 | 53.78482777 | 0.362392397 | 0.242541352 | 1.494146848 | 0.135137242 | 0.269305262 |
| ENSG00000213250 | RBMS2P1 | 14.47297382 | -1.088981172 | 0.486474885 | -2.23851468 | 0.025187511 | 0.066898403 |

| | | | | | | | |
|---|---|---|---|---|---|---|---|
| ENSG00000213252 | ENSG0000021 | 47.62835793 | 0.458920624 | 0.260531573 | 1.761477957 | 0.078157541 | 0.173489864 |
| ENSG00000213253 | RPL12P42 | 31.44804906 | 0.110833349 | 0.335068563 | 0.330778119 | 0.74081209 | 0.857082375 |
| ENSG00000213260 | YWHAZP5 | 30.70792614 | -0.333830956 | 0.315692098 | -1.057457435 | 0.290302871 | 0.474800218 |
| ENSG00000213261 | EEF1B2P6 | 576.2238785 | 0.203617584 | 0.082289355 | 2.474409764 | 0.013345653 | 0.038345351 |
| ENSG00000213262 | VDAC2P3 | 1.772503313 | 0.044737944 | 1.347441507 | 0.033202142 | 0.97351339 | NA |
| ENSG00000213269 | RPS6P26 | 5.499097957 | -0.641893324 | 0.780961698 | -0.821926767 | 0.411118578 | 0.603581246 |
| ENSG00000213280 | ENSG0000021 | 38.27497495 | 0.521993485 | 0.308207618 | 1.693642383 | 0.090333259 | 0.195018991 |
| ENSG00000213281 | NRAS | 15200.49736 | -0.346708826 | 0.053588978 | -6.469778705 | 9.81E-11 | 8.48E-10 |
| ENSG00000213290 | PGK1P2 | 40.41916899 | -0.231472749 | 0.30844247 | -0.7504568 | 0.452979633 | 0.640611108 |
| ENSG00000213293 | ENSG0000021 | 14.67839108 | -0.215329788 | 0.48490977 | -0.444061558 | 0.656998078 | 0.801277136 |
| ENSG00000213295 | RPL22P10 | 4.157740151 | 0.733621018 | 0.873444691 | 0.83991697 | 0.400954942 | 0.593796409 |
| ENSG00000213296 | ENSG0000021 | 2.011291915 | 0.294180458 | 1.277422973 | 0.23029213 | 0.817864775 | 0.903530843 |
| ENSG00000213300 | HNRNPA3P6 | 3460.126619 | 0.125051189 | 0.054571566 | 2.291508177 | 0.02193404 | 0.059265826 |
| ENSG00000213303 | ENSG0000021 | 14.8477466 | 0.587124525 | 0.469006751 | 1.251846639 | 0.210625752 | 0.377953281 |
| ENSG00000213304 | ENSG0000021 | 7.573096907 | 0.131917099 | 0.667564364 | 0.197609557 | 0.843350557 | 0.917499611 |
| ENSG00000213305 | HNRNPCP6 | 20.73543772 | -0.065071976 | 0.416366999 | -0.156285143 | 0.875808266 | 0.935564696 |
| ENSG00000213307 | RPL18P11 | 2.393800443 | -1.095375338 | 1.202612692 | -0.910829684 | 0.36238512 | 0.555140503 |
| ENSG00000213309 | RPL9P18 | 19.87087836 | 0.166821764 | 0.432623254 | 0.385605172 | 0.699789102 | 0.831642103 |
| ENSG00000213312 | RPL7AP34 | 6.325489969 | 0.190781564 | 0.696671965 | 0.273847052 | 0.784202163 | 0.882881705 |
| ENSG00000213315 | ENSG0000021 | 51.18586965 | 0.547681278 | 0.274867312 | 1.992529683 | 0.046312972 | 0.11214055 |
| ENSG00000213316 | LTC4S | 20.22716775 | 0.106632373 | 0.414058619 | 0.257529654 | 0.796769926 | 0.891676203 |
| ENSG00000213326 | RPS7P11 | 293.4780494 | 0.194893912 | 0.112680495 | 1.729615329 | 0.083699025 | 0.183421441 |
| ENSG00000213331 | ENSG0000021 | 8.25392646 | -0.505142577 | 0.616026761 | -0.820001027 | 0.412215521 | 0.604463313 |
| ENSG00000213332 | SLC25A5P6 | 3.247147325 | 0.396766094 | 0.955343622 | 0.415312443 | 0.677913193 | 0.816699042 |
| ENSG00000213337 | ANKRD39 | 107.0402476 | -0.049897469 | 0.173093564 | -0.288268773 | 0.773141009 | 0.876364005 |
| ENSG00000213339 | QTRT1 | 822.7874299 | 0.236530199 | 0.077909546 | 3.035959128 | 0.002397719 | 0.008304947 |
| ENSG00000213341 | CHUK | 595.0491305 | -0.092497668 | 0.083708348 | -1.104999333 | 0.269159878 | 0.450813539 |
| ENSG00000213347 | MXD3 | 702.6746259 | -0.530678035 | 0.080168207 | -6.619557208 | 3.60E-11 | 3.22E-10 |
| ENSG00000213352 | GAPDHP44 | 7.307525047 | 0.689846351 | 0.663093403 | 1.040345669 | 0.298179334 | 0.484779158 |
| ENSG00000213361 | RPL36P12 | 3.552562567 | 0.087951864 | 0.946991577 | 0.092875023 | 0.926002849 | 0.964567541 |
| ENSG00000213362 | FTH1P12 | 80.03914769 | 0.370832694 | 0.20299788 | 1.826781119 | 0.067732686 | 0.154421369 |
| ENSG00000213363 | RPS3P6 | 50.37691249 | 0.08185435 | 0.257765322 | 0.317553771 | 0.750823445 | 0.862329879 |
| ENSG00000213366 | GSTM2 | 57.79326502 | -0.058970189 | 0.244966125 | -0.240727932 | 0.809765989 | 0.899347944 |

| | | | | | | | |
|---|---|---|---|---|---|---|---|
| ENSG00000213370 | RANP6 | 33.31128317 | -0.181785637 | 0.314025717 | -0.578887739 | 0.562664924 | 0.731512233 |
| ENSG00000213371 | NAP1L1P3 | 93.03718759 | 0.376185368 | 0.195227129 | 1.926911333 | 0.053990681 | 0.127975431 |
| ENSG00000213376 | GAPDHP71 | 18.33452574 | 0.292029874 | 0.42690517 | 0.684062633 | 0.493935616 | 0.676511018 |
| ENSG00000213380 | COG8 | 749.6898497 | 0.111799079 | 0.07452765 | 1.500102025 | 0.133587977 | 0.266919971 |
| ENSG00000213383 | PP1P | 146.1131185 | -0.601639254 | 0.154517316 | -3.893668799 | 9.87E-05 | 0.000440367 |
| ENSG00000213385 | RPL7AP39 | 11.11825459 | -1.077141195 | 0.544594606 | -1.97787709 | 0.047942574 | 0.115610496 |
| ENSG00000213386 | ENSG0000021 | 3.917589726 | 0.589393809 | 0.936560175 | 0.629317608 | 0.529141146 | 0.704988843 |
| ENSG00000213390 | ARHGAP19 | 1024.854827 | -1.128545636 | 0.07053284 | -16.00028642 | 1.27E-57 | 8.58E-56 |
| ENSG00000213394 | RPSAP46 | 1.922323732 | -0.149475534 | 1.2723929 | -0.117475926 | 0.906482921 | 0.952837161 |
| ENSG00000213397 | HAUS7 | 17.82824503 | -0.561254918 | 0.425197293 | -1.319987046 | 0.186839343 | 0.344541361 |
| ENSG00000213398 | LCAT | 4.56798406 | -2.322527408 | 0.965074378 | -2.406578666 | 0.016102736 | 0.045280308 |
| ENSG00000213399 | RPS2P17 | 31.26381953 | 0.052983889 | 0.316132965 | 0.167600013 | 0.866897961 | 0.930846618 |
| ENSG00000213406 | ANXA2P1 | 32.05618956 | -0.043020684 | 0.386741166 | -0.111238958 | 0.91142686 | 0.955494569 |
| ENSG00000213409 | C1QBPP2 | 9.34823036 | -0.363369312 | 0.591568771 | -0.614246947 | 0.539052154 | 0.713716751 |
| ENSG00000213411 | RBM22P2 | 1.968213413 | -0.13620778 | 1.269648646 | -0.1072799 | 0.91456693 | 0.957607799 |
| ENSG00000213412 | HNRNPA1P33 | 12.75412638 | 0.03705572 | 0.524285356 | 0.070678533 | 0.943653606 | 0.973179901 |
| ENSG00000213414 | ENSG0000021 | 5.701399855 | -0.084364159 | 0.753919694 | -0.111900724 | 0.910902123 | 0.955293714 |
| ENSG00000213420 | GPC2 | 28.82651691 | 0.080914552 | 0.329301451 | 0.245715748 | 0.805902281 | 0.897011078 |
| ENSG00000213423 | RBMX2P2 | 1.71381018 | 0.128638839 | 1.33594975 | 0.096290178 | 0.923290112 | NA |
| ENSG00000213430 | HSPD1P1 | 892.8719668 | 0.158740033 | 0.069716847 | 2.27692503 | 0.022790703 | 0.061248679 |
| ENSG00000213432 | RPL17P34 | 5.372850837 | 0.452809893 | 0.774365156 | 0.58474983 | 0.558715943 | 0.728649445 |
| ENSG00000213433 | RPLP1P6 | 4.144929563 | -0.270194683 | 0.881974108 | -0.306352171 | 0.759336521 | 0.867532237 |
| ENSG00000213434 | VTI1BP2 | 5.162818671 | -0.812287695 | 0.798973229 | -1.016664471 | 0.309313073 | 0.497346456 |
| ENSG00000213440 | H2AZP1 | 6.262236335 | 0.586815853 | 0.71400484 | 0.821865371 | 0.411153524 | 0.603581246 |
| ENSG00000213442 | RPL18AP3 | 4484.772657 | 0.328921803 | 0.052380821 | 6.279432023 | 3.40E-10 | 2.82E-09 |
| ENSG00000213445 | SIPA1 | 403.6009996 | 0.370541392 | 0.102923922 | 3.600148379 | 0.000318036 | 0.001303043 |
| ENSG00000213448 | RPS23P2 | 1.661852221 | -0.742944775 | 1.345541199 | -0.552153123 | 0.580843448 | NA |
| ENSG00000213450 | VDAC1P7 | 5.871229594 | 0.059118357 | 0.738132749 | 0.080091768 | 0.93616427 | 0.969509654 |
| ENSG00000213453 | FTH1P3 | 51.63204699 | 0.148003764 | 0.24419025 | 0.60610022 | 0.544448205 | 0.717839549 |
| ENSG00000213462 | ERV3-1 | 497.9531319 | 1.001694705 | 0.093626871 | 10.69879506 | 1.03E-26 | 2.52E-25 |
| ENSG00000213463 | SYNJ2BP | 1844.431705 | -0.393945717 | 0.057149254 | -6.893278372 | 5.45E-12 | 5.24E-11 |
| ENSG00000213465 | ARL2 | 429.3811566 | -0.076081286 | 0.099550491 | -0.764248232 | 0.444719329 | 0.633380605 |
| ENSG00000213467 | HMGB1P37 | 26.85670262 | -0.610043488 | 0.415931634 | -1.466691731 | 0.142459933 | 0.280180973 |

| | | | | | | | |
|---|---|---|---|---|---|---|---|
| ENSG00000213468 | FIRRE | 341.53982 | -0.150133708 | 0.11595788 | -1.294726225 | 0.19541469 | 0.356656296 |
| ENSG00000213478 | CFL1P2 | 12.92128391 | -0.016275349 | 0.497827027 | -0.03269278 | 0.973919582 | 0.988106737 |
| ENSG00000213480 | ENSG0000021 | 7.953994759 | -0.477342893 | 0.67103695 | -0.711351131 | 0.476866674 | 0.661946116 |
| ENSG00000213492 | NT5C3AP1 | 70.18344905 | 0.094546538 | 0.218476441 | 0.432753927 | 0.665193548 | 0.806901013 |
| ENSG00000213493 | ACTN4P1 | 68.84013881 | 0.052172642 | 0.221539829 | 0.235500056 | 0.813820636 | 0.901448243 |
| ENSG00000213500 | LAP3P2 | 28.31338677 | -0.55825348 | 0.354037983 | -1.576818045 | 0.114837398 | 0.237052994 |
| ENSG00000213509 | PPIAP16 | 1.648672411 | 1.013679713 | 1.412022659 | 0.717891959 | 0.472823906 | NA |
| ENSG00000213512 | GBP7 | 4.400584257 | -1.126920207 | 0.867428804 | -1.299150088 | 0.193892427 | 0.354618886 |
| ENSG00000213513 | IMPDH1P5 | 25.19026612 | -1.184981912 | 0.376923088 | -3.14382947 | 0.001667526 | 0.005955635 |
| ENSG00000213514 | ENSG0000021 | 2.444770769 | -0.382436507 | 1.109221874 | -0.344779089 | 0.730260455 | 0.850776726 |
| ENSG00000213516 | RBMXL1 | 1078.615263 | -0.172773388 | 0.064376523 | -2.683794955 | 0.007279173 | 0.022501582 |
| ENSG00000213522 | RAC1P5 | 10.34998848 | 0.040416739 | 0.565921535 | 0.071417566 | 0.94306543 | 0.97291737 |
| ENSG00000213523 | SRA1 | 746.176566 | -0.21152927 | 0.076537804 | -2.76372276 | 0.005714607 | 0.018124303 |
| ENSG00000213526 | SETP8 | 24.68282175 | -0.296692454 | 0.363844522 | -0.815437464 | 0.414821947 | 0.60685004 |
| ENSG00000213530 | MTHFD2P5 | 4.734609305 | 0.310676034 | 0.849379871 | 0.365768067 | 0.71453815 | 0.841150775 |
| ENSG00000213533 | STIMATE | 13.6867226 | 0.274964907 | 0.497035652 | 0.553209625 | 0.580119877 | 0.745585185 |
| ENSG00000213538 | KRT8P41 | 29.8984768 | 0.090479599 | 0.331100846 | 0.273269006 | 0.784646439 | 0.883091372 |
| ENSG00000213539 | YBX1P6 | 17.5997746 | 0.14269435 | 0.455181821 | 0.313488684 | 0.753909416 | 0.86430195 |
| ENSG00000213542 | RPL7AP43 | 12.90024603 | 0.445923727 | 0.498717215 | 0.894141436 | 0.371246222 | 0.563924535 |
| ENSG00000213551 | DNAJC9 | 1585.723711 | -0.808449331 | 0.059952619 | -13.48480432 | 1.92E-41 | 8.37E-40 |
| ENSG00000213553 | RPLP0P6 | 7764.103773 | 0.040336118 | 0.052866082 | 0.76298671 | 0.445471322 | 0.634097661 |
| ENSG00000213560 | ENSG0000021 | 8.587269688 | -0.62217259 | 0.601214561 | -1.034859483 | 0.300734539 | 0.487482358 |
| ENSG00000213563 | C8orf82 | 1070.731544 | -0.038363663 | 0.067971381 | -0.564409056 | 0.572475777 | 0.739291922 |
| ENSG00000213569 | GTF3C6P2 | 117.4615632 | -0.127918821 | 0.168766967 | -0.757961247 | 0.448474183 | 0.636633525 |
| ENSG00000213574 | LDHAP5 | 15.95674714 | -0.07401591 | 0.466872663 | -0.158535541 | 0.874034817 | 0.934773263 |
| ENSG00000213585 | VDAC1 | 6465.372746 | -0.080004829 | 0.045481928 | -1.759046565 | 0.0785696 | 0.174194126 |
| ENSG00000213588 | ZBTB9 | 1319.888768 | 0.610252374 | 0.0654281 | 9.327068561 | 1.09E-20 | 1.89E-19 |
| ENSG00000213590 | ENSG0000021 | 2.370591563 | -0.625078532 | 1.130183235 | -0.553077159 | 0.580210577 | 0.74560514 |
| ENSG00000213593 | TMX2 | 1102.284823 | -0.021697258 | 0.06999861 | -0.309966987 | 0.756586061 | 0.86559694 |
| ENSG00000213598 | RPL31P4 | 173.2529055 | 0.262100564 | 0.137944396 | 1.900045027 | 0.057427211 | 0.134605111 |
| ENSG00000213609 | RPL7AP50 | 200.9821074 | 0.293733929 | 0.132145725 | 2.222803123 | 0.026229078 | 0.069176254 |
| ENSG00000213613 | RPL11P3 | 25.08045446 | 0.285444886 | 0.374115355 | 0.762986287 | 0.445471574 | 0.634097661 |
| ENSG00000213614 | HEXA | 1335.932313 | -0.101868086 | 0.062128035 | -1.639647621 | 0.101078455 | 0.213708734 |

| | | | | | | | |
|---|---|---|---|---|---|---|---|
| ENSG00000213619 | NDUFS3 | 2042.704538 | -0.107471505 | 0.05783775 | -1.858155015 | 0.063147004 | 0.145543281 |
| ENSG00000213621 | RPSAP54 | 29.24094355 | -0.161408913 | 0.322196048 | -0.500964906 | 0.616395822 | 0.772766413 |
| ENSG00000213625 | LEPROT | 1598.736243 | -0.116029378 | 0.057387527 | -2.021857069 | 0.04319112 | 0.105719621 |
| ENSG00000213630 | BOLA3P3 | 2.827889366 | -1.001086077 | 1.10831359 | -0.903251649 | 0.366392352 | 0.559031629 |
| ENSG00000213638 | ADAT3 | 32.7829336 | 0.339063261 | 0.306618373 | 1.105815209 | 0.268806511 | 0.450366737 |
| ENSG00000213639 | PPP1CB | 2747.970804 | 0.0100237 | 0.063238477 | 0.158506341 | 0.874057824 | 0.934773263 |
| ENSG00000213640 | EEF1DP4 | 1.841112102 | 0.548233717 | 1.334901929 | 0.41069213 | 0.681298298 | 0.819086157 |
| ENSG00000213650 | ENSG0000021 | 19.43055085 | 0.339019705 | 0.47485424 | 0.713944779 | 0.475261326 | 0.66046014 |
| ENSG00000213654 | GPSM3 | 36.14623949 | -0.263866191 | 0.293492943 | -0.899054634 | 0.368623561 | 0.561204173 |
| ENSG00000213658 | LAT | 8.910025974 | 0.216563734 | 0.640362486 | 0.33818929 | 0.735220546 | 0.853967465 |
| ENSG00000213661 | ENSG0000021 | 12.27794203 | 0.302590194 | 0.550580598 | 0.54958383 | 0.582604852 | 0.747090374 |
| ENSG00000213667 | PGGT1BP1 | 14.98384415 | 0.452561912 | 0.461395665 | 0.980854278 | 0.326664608 | 0.517185142 |
| ENSG00000213671 | OLA1P2 | 2.916216311 | 0.469162473 | 1.046371197 | 0.448370974 | 0.653885486 | 0.799165148 |
| ENSG00000213672 | NCKIPSD | 909.5076774 | 0.162388468 | 0.069656399 | 2.331278558 | 0.019738676 | 0.05408567 |
| ENSG00000213673 | SLC25A5P3 | 11.85134965 | 0.009511578 | 0.565785552 | 0.016811278 | 0.986587173 | 0.993497695 |
| ENSG00000213676 | ATF6B | 2385.56171 | 0.759024426 | 0.06575154 | 11.5438274 | 7.93E-31 | 2.27E-29 |
| ENSG00000213683 | ENSG0000021 | 172.468615 | 0.143458743 | 0.138228493 | 1.037837714 | 0.299345616 | 0.485998083 |
| ENSG00000213693 | SEC14L1P1 | 29.86427198 | -0.057447283 | 0.325414885 | -0.176535512 | 0.859873252 | 0.92729844 |
| ENSG00000213694 | S1PR3 | 502.89876 | -0.767607395 | 0.089633034 | -8.563889478 | 1.09E-17 | 1.58E-16 |
| ENSG00000213695 | RPS7P14 | 2.491659225 | -0.019391128 | 1.101141894 | -0.017610017 | 0.985949965 | 0.993350589 |
| ENSG00000213698 | CAPZA1P3 | 33.27359293 | 0.266924144 | 0.305366672 | 0.874110268 | 0.382058206 | 0.574414422 |
| ENSG00000213699 | SLC35F6 | 1251.260773 | 0.113810169 | 0.064846686 | 1.75506531 | 0.079248137 | 0.175478018 |
| ENSG00000213700 | RPL17P50 | 9.893636625 | 0.35449569 | 0.552722912 | 0.641362395 | 0.52128726 | 0.698337406 |
| ENSG00000213701 | SETP22 | 3.367642019 | -0.006572328 | 0.959739489 | -0.006848033 | 0.994536103 | 0.996664986 |
| ENSG00000213707 | HMGB1P10 | 52.27523001 | -0.436520932 | 0.247728086 | -1.762097065 | 0.078052899 | 0.173299797 |
| ENSG00000213711 | PHB1P7 | 4.971713635 | -0.411110902 | 0.810427204 | -0.507276779 | 0.611960629 | 0.769394762 |
| ENSG00000213713 | PIGCP1 | 3.755780671 | -0.517141469 | 0.921334739 | -0.561295962 | 0.574595794 | 0.741059623 |
| ENSG00000213714 | FAM209B | 4.038194783 | 1.157115507 | 0.908312523 | 1.273917818 | 0.202692543 | 0.366817762 |
| ENSG00000213719 | CLIC1 | 6595.084989 | -0.000787421 | 0.045100228 | -0.017459366 | 0.986070149 | 0.993350589 |
| ENSG00000213722 | DDAH2 | 1797.742227 | 0.310908069 | 0.066908488 | 4.646765756 | 3.37E-06 | 1.84E-05 |
| ENSG00000213727 | ENSG0000021 | 1.997591962 | -0.190674141 | 1.300333417 | -0.146634808 | 0.883420278 | 0.93961708 |
| ENSG00000213729 | RPS16P2 | 10.8964012 | 0.049307952 | 0.534211744 | 0.092300388 | 0.92645938 | 0.964663921 |
| ENSG00000213730 | POLD2P1 | 5.035720819 | 1.174470244 | 0.824288515 | 1.424829076 | 0.154206615 | 0.297468866 |

| | | | | | | | |
|---|---|---|---|---|---|---|---|
| ENSG00000213735 | ANAPC10P1 | 9.133543136 | -0.043935368 | 0.605220903 | -0.072593937 | 0.942129251 | 0.972530993 |
| ENSG00000213740 | SERBP1P1 | 770.3124879 | -0.053557493 | 0.074912125 | -0.714937572 | 0.47464762 | 0.659944021 |
| ENSG00000213741 | RPS29 | 3088.639205 | -0.207602495 | 0.055199074 | -3.760977832 | 0.00016925 | 0.000727761 |
| ENSG00000213742 | ZNF337-AS1 | 143.603347 | -0.90881709 | 0.160521404 | -5.661656743 | 1.50E-08 | 1.06E-07 |
| ENSG00000213750 | ENSG0000021 | 7.281965016 | -0.310135832 | 0.650751623 | -0.476580957 | 0.633660546 | 0.784143138 |
| ENSG00000213753 | CENPBD2P | 805.374883 | 0.081240529 | 0.079824963 | 1.01773337 | 0.308804684 | 0.496900138 |
| ENSG00000213754 | KPNA4P1 | 1.935672984 | 2.099242558 | 1.448876193 | 1.448876425 | 0.147372096 | 0.28770527 |
| ENSG00000213755 | RPL29P15 | 2.808912567 | 0.308692186 | 1.043315248 | 0.295876234 | 0.767324601 | 0.873006316 |
| ENSG00000213757 | RPS3AP20 | 47.67523107 | 0.252584273 | 0.277662966 | 0.909679374 | 0.362991625 | 0.555760042 |
| ENSG00000213760 | ATP6V1G2 | 27.47272435 | 0.248159667 | 0.337912232 | 0.734390894 | 0.462710535 | 0.649249475 |
| ENSG00000213762 | ZNF134 | 1153.676055 | 0.626158 | 0.067985685 | 9.210144763 | 3.26E-20 | 5.51E-19 |
| ENSG00000213763 | ACTBP2 | 346.2147671 | -0.265847093 | 0.120412779 | -2.207798002 | 0.027258357 | 0.071537206 |
| ENSG00000213771 | KRT8P37 | 49.53881913 | 0.345035007 | 0.264851015 | 1.30275131 | 0.192659679 | 0.352947854 |
| ENSG00000213772 | EIF1P7 | 22.22117007 | 0.04647127 | 0.379872919 | 0.122333726 | 0.902634723 | 0.950907167 |
| ENSG00000213774 | ENSG0000021 | 2.008898778 | -1.773753027 | 1.395836744 | -1.270745332 | 0.203819263 | 0.368564866 |
| ENSG00000213777 | ENSG0000021 | 6.478227045 | 1.06082437 | 0.721799701 | 1.469693557 | 0.141644769 | 0.279056251 |
| ENSG00000213780 | GTF2H4 | 5.813076634 | -0.218042775 | 0.751489576 | -0.290147438 | 0.771703445 | 0.875561078 |
| ENSG00000213781 | PSMC1P2 | 2.588575735 | -1.622295495 | 1.242136144 | -1.306052885 | 0.191534574 | 0.351528276 |
| ENSG00000213782 | DDX47 | 10.63146904 | 0.269963196 | 0.564319745 | 0.478386941 | 0.632374827 | 0.783432036 |
| ENSG00000213786 | NHP2P2 | 2.390950957 | -0.630608153 | 1.118754969 | -0.563669589 | 0.572979018 | 0.739747367 |
| ENSG00000213790 | OLA1P1 | 392.6906151 | 0.412204033 | 0.09944924 | 4.144868626 | 3.40E-05 | 0.000163107 |
| ENSG00000213791 | ENSG0000021 | 3.754978505 | 0.685862154 | 0.909974799 | 0.753715548 | 0.451020044 | 0.639118987 |
| ENSG00000213793 | ZNF888 | 366.5743131 | 1.082567914 | 0.101252641 | 10.69174985 | 1.11E-26 | 2.72E-25 |
| ENSG00000213799 | ZNF845 | 779.1897502 | 0.192805147 | 0.082791366 | 2.328807401 | 0.019869271 | 0.054391537 |
| ENSG00000213801 | ZNF321P | 28.39506795 | 0.885640017 | 0.340882645 | 2.598078929 | 0.009374694 | 0.028119368 |
| ENSG00000213816 | CNN2P4 | 7.736420793 | 0.365501398 | 0.65082701 | 0.561595313 | 0.574391774 | 0.740858848 |
| ENSG00000213820 | RPL13P2 | 8.933596069 | 0.068473777 | 0.617099619 | 0.110960654 | 0.911647548 | 0.955638583 |
| ENSG00000213830 | CFL1P5 | 12.41582501 | -0.9270413 | 0.510824958 | -1.814792495 | 0.069555804 | 0.157770508 |
| ENSG00000213839 | TMX2P1 | 34.25746468 | 0.016156395 | 0.302829335 | 0.053351486 | 0.957451859 | 0.980753638 |
| ENSG00000213842 | SUGT1P2 | 1.590966911 | 1.052117804 | 1.477616389 | 0.712037178 | 0.476441754 | NA |
| ENSG00000213853 | EMP2 | 9154.109544 | 0.308150092 | 0.041196612 | 7.479986258 | 7.43E-14 | 8.17E-13 |
| ENSG00000213856 | VDAC1P2 | 327.875454 | -0.052135915 | 0.111947599 | -0.465717139 | 0.641417983 | 0.789596379 |
| ENSG00000213857 | ACTBP15 | 3.163489734 | -0.066381858 | 0.99936306 | -0.066424166 | 0.947040131 | 0.975269375 |

| | | | | | | | |
|---|---|---|---|---|---|---|---|
| ENSG00000213859 | KCTD11 | 567.7600468 | 0.170259758 | 0.090418397 | 1.883021187 | 0.059697497 | 0.138961705 |
| ENSG00000213860 | RPL21P75 | 12.350671 | 0.405864657 | 0.510120084 | 0.795625716 | 0.426249615 | 0.61707473 |
| ENSG00000213862 | RPL7AP62 | 603.6518855 | 0.154486665 | 0.080391271 | 1.921684564 | 0.054645461 | 0.129287012 |
| ENSG00000213864 | EEF1B2P2 | 27.79361822 | 0.165636258 | 0.351740279 | 0.470905006 | 0.637708573 | 0.78659182 |
| ENSG00000213866 | YBX1P10 | 1580.552915 | 0.116475894 | 0.057812977 | 2.01470155 | 0.04393593 | 0.107302271 |
| ENSG00000213867 | ENSG0000021 | 24.41177131 | 0.187344409 | 0.356560122 | 0.525421653 | 0.599290104 | 0.759416394 |
| ENSG00000213871 | TAF9BP1 | 5.11848959 | -0.297907155 | 0.788892707 | -0.37762696 | 0.705707734 | 0.835385491 |
| ENSG00000213881 | NPM1P6 | 568.974059 | -0.135243676 | 0.09013197 | -1.500507258 | 0.133483055 | 0.266756797 |
| ENSG00000213885 | RPL13AP7 | 179.5255027 | 0.538505024 | 0.140269758 | 3.839067172 | 0.000123503 | 0.000542437 |
| ENSG00000213888 | LINC01521 | 282.2870769 | -0.74551888 | 0.110709079 | -6.73403561 | 1.65E-11 | 1.52E-10 |
| ENSG00000213889 | PPM1N | 66.17648794 | -0.370653109 | 0.22880459 | -1.619954863 | 0.105241973 | 0.220911002 |
| ENSG00000213891 | RPL3P6 | 9.461472316 | 0.168688792 | 0.628348009 | 0.268463956 | 0.788342216 | 0.885492107 |
| ENSG00000213901 | SLC23A3 | 18.10025313 | -0.301823314 | 0.431482024 | -0.699503797 | 0.484237241 | 0.668094203 |
| ENSG00000213903 | LTB4R | 1075.373984 | 0.283006198 | 0.151477596 | 1.868303998 | 0.06171971 | 0.142833655 |
| ENSG00000213904 | LIPE-AS1 | 33.88380361 | -0.304195713 | 0.306103853 | -0.993766362 | 0.320336626 | 0.509889134 |
| ENSG00000213906 | LTB4R2 | 309.1115892 | 0.095285734 | 0.114636106 | 0.831201765 | 0.405859658 | 0.598625381 |
| ENSG00000213918 | DNASE1 | 1457.220561 | -0.217626893 | 0.064368736 | -3.380940889 | 0.000722381 | 0.002777177 |
| ENSG00000213923 | CSNK1E | 1780.369275 | 0.080097291 | 0.066075702 | 1.212204908 | 0.225433953 | 0.39749621 |
| ENSG00000213928 | IRF9 | 48.35568946 | 0.536078554 | 0.25865864 | 2.072532953 | 0.038215766 | 0.095518597 |
| ENSG00000213930 | GALT | 138.6913357 | -0.068747165 | 0.154446285 | -0.445120221 | 0.656232876 | 0.800743733 |
| ENSG00000213935 | RPL22P16 | 31.14410392 | 0.097639275 | 0.326611688 | 0.298946052 | 0.764981208 | 0.871320362 |
| ENSG00000213937 | CLDN9 | 794.211704 | 1.010988417 | 0.086759943 | 11.65270949 | 2.22E-31 | 6.56E-30 |
| ENSG00000213938 | SEPHS1P6 | 17.87565473 | -1.042502973 | 0.422382394 | -2.468149687 | 0.013581351 | 0.038915338 |
| ENSG00000213939 | ENSG0000021 | 46.37670095 | 0.500527732 | 0.308700035 | 1.621404843 | 0.104930843 | 0.220359438 |
| ENSG00000213942 | RPL31P10 | 30.74428451 | 0.421907502 | 0.347444696 | 1.214315563 | 0.224627245 | 0.39631972 |
| ENSG00000213943 | KRT18P17 | 285.8809523 | 0.271854266 | 0.123697804 | 2.197729122 | 0.027968415 | 0.073149634 |
| ENSG00000213946 | DNAJB1P1 | 3.723824349 | 1.624996053 | 1.012700814 | 1.604616122 | 0.108578311 | 0.226432911 |
| ENSG00000213949 | ITGA1 | 3.711531327 | -0.955969322 | 0.955024645 | -1.000989166 | 0.316832046 | 0.505749575 |
| ENSG00000213950 | RPS10P2 | 2.516251328 | 0.359118392 | 1.09861372 | 0.326883221 | 0.743756212 | 0.858490754 |
| ENSG00000213958 | KRT18P29 | 84.18836321 | 0.296574265 | 0.197369743 | 1.502632875 | 0.132933741 | 0.266091209 |
| ENSG00000213959 | ENSG0000021 | 7.257413797 | 0.693337897 | 0.690234654 | 1.004495925 | 0.315139635 | 0.503784196 |
| ENSG00000213962 | API5P2 | 6.225753309 | 0.295023482 | 0.710901923 | 0.414998853 | 0.678142743 | 0.81682581 |
| ENSG00000213964 | CHCHD4P4 | 3.08022678 | 0.308354358 | 0.992657089 | 0.310635326 | 0.756077869 | 0.865184627 |

| | | | | | | | |
|---|---|---|---|---|---|---|---|
| ENSG00000213965 | NUDT19 | 1082.362585 | 0.116604205 | 0.064835459 | 1.798463478 | 0.072103591 | 0.162688737 |
| ENSG00000213967 | ZNF726 | 74.19854244 | 0.083139394 | 0.225564758 | 0.368583259 | 0.712438376 | 0.839972059 |
| ENSG00000213970 | ENSG0000021 | 14.27091809 | -0.122931257 | 0.488600891 | -0.251598512 | 0.801351409 | 0.894398618 |
| ENSG00000213977 | TAX1BP3 | 29.21481419 | 0.235086783 | 0.33296478 | 0.706040993 | 0.480162645 | 0.66446925 |
| ENSG00000213983 | AP1G2 | 4349.180408 | -0.449078242 | 0.067461308 | -6.656826805 | 2.80E-11 | 2.53E-10 |
| ENSG00000213985 | ENSG0000021 | 198.1215729 | -0.011307246 | 0.131520719 | -0.085973113 | 0.931487791 | 0.967155333 |
| ENSG00000213995 | NAXD | 1728.806387 | -0.155777655 | 0.061393641 | -2.537358149 | 0.011169262 | 0.032744137 |
| ENSG00000213997 | PGAM1P7 | 53.93447859 | -0.633083785 | 0.242477949 | -2.610892198 | 0.009030636 | 0.027204567 |
| ENSG00000214003 | ATP5PBP3 | 4.045570142 | -0.712387136 | 0.894547675 | -0.796365757 | 0.425819475 | 0.616646598 |
| ENSG00000214009 | PCNAP3 | 2.171058534 | -0.819625908 | 1.24419989 | -0.658757418 | 0.510051555 | 0.690014901 |
| ENSG00000214012 | KRT18P38 | 158.8264112 | 0.340787621 | 0.145867303 | 2.336285199 | 0.019476382 | 0.05344996 |
| ENSG00000214013 | GANC | 175.2141732 | -0.182630916 | 0.137040099 | -1.33268231 | 0.182636081 | 0.338586689 |
| ENSG00000214016 | RPSAP61 | 5.123618174 | 0.399458003 | 0.795558694 | 0.502110034 | 0.615590122 | 0.772264951 |
| ENSG00000214018 | RRM2P3 | 25.81449253 | -0.127996022 | 0.376748345 | -0.339738776 | 0.734053258 | 0.853308084 |
| ENSG00000214019 | RBM39P1 | 111.5452996 | -0.340708899 | 0.172841118 | -1.971225967 | 0.048698035 | 0.117125143 |
| ENSG00000214021 | TTLL3 | 746.8463605 | 0.125525488 | 0.173656554 | 0.722837611 | 0.469779657 | 0.655719217 |
| ENSG00000214022 | REPIN1 | 5013.891843 | 1.063913791 | 0.049429479 | 21.52387208 | 9.31E-103 | 1.58E-100 |
| ENSG00000214026 | MRPL23 | 640.1357309 | -0.074857736 | 0.085615231 | -0.874350691 | 0.381927302 | 0.574255247 |
| ENSG00000214027 | ARPC3P5 | 6.970436932 | -0.987832093 | 0.684584913 | -1.442965035 | 0.149030321 | 0.290118429 |
| ENSG00000214029 | ZNF891 | 163.290859 | 0.285895845 | 0.149749505 | 1.909160535 | 0.056241386 | 0.132366614 |
| ENSG00000214035 | ENSG0000021 | 2.305803868 | -0.577870003 | 1.16262009 | -0.497041129 | 0.619160048 | 0.77424538 |
| ENSG00000214045 | IMPDH1P2 | 3.988935471 | 0.412122486 | 0.88858465 | 0.46379654 | 0.642793522 | 0.790526103 |
| ENSG00000214046 | SMIM7 | 724.9832259 | 0.127625226 | 0.078490883 | 1.625987903 | 0.103952229 | 0.218716329 |
| ENSG00000214047 | RPS11P7 | 1.74407409 | 0.155833468 | 1.331901209 | 0.117000771 | 0.906859443 | NA |
| ENSG00000214049 | UCA1 | 3.038200504 | -0.529064375 | 1.014991282 | -0.521250166 | 0.602192511 | 0.761606804 |
| ENSG00000214050 | FBXO16 | 41.29416324 | -0.228494555 | 0.278896268 | -0.819281508 | 0.41262582 | 0.604639816 |
| ENSG00000214063 | TSPAN4 | 485.1864614 | -0.226189441 | 0.109837219 | -2.05931508 | 0.039464065 | 0.098082952 |
| ENSG00000214077 | GNAQP1 | 45.52214851 | -0.079274754 | 0.260787207 | -0.303982525 | 0.761141205 | 0.868858441 |
| ENSG00000214078 | CPNE1 | 2008.244447 | -1.029492684 | 0.062705781 | -16.41782732 | 1.43E-60 | 1.04E-58 |
| ENSG00000214087 | ARL16 | 731.3446107 | 0.254732863 | 0.079039598 | 3.222851189 | 0.001269215 | 0.004658526 |
| ENSG00000214089 | RPL9P21 | 2.922822927 | 0.491211384 | 1.056160075 | 0.465091794 | 0.641865722 | 0.789935605 |
| ENSG00000214093 | ENSG0000021 | 1.863759156 | -1.006426433 | 1.283690887 | -0.784009954 | 0.43303427 | 0.623082616 |
| ENSG00000214100 | PLAC9P1 | 2.17863113 | -3.578980725 | 1.569026665 | -2.281019695 | 0.022547281 | 0.060722545 |

| | | | | | | | |
|---|---|---|---|---|---|---|---|
| ENSG00000214106 | PAXIP1-AS2 | 108.984278 | 0.1274755 | 0.17033845 | 0.748365972 | 0.454239442 | 0.641838792 |
| ENSG00000214108 | TPT1P5 | 1.897316963 | 0.844966854 | 1.309465621 | 0.645276089 | 0.518748285 | 0.696318646 |
| ENSG00000214111 | PAFAH1B2P1 | 5.484470161 | 1.514541538 | 0.830787066 | 1.823020121 | 0.068300344 | 0.155499194 |
| ENSG00000214113 | LYRM4 | 292.6498008 | 0.756694655 | 0.11144207 | 6.790026891 | 1.12E-11 | 1.04E-10 |
| ENSG00000214114 | MYCBP | 784.7807728 | -0.157936662 | 0.076417542 | -2.066759243 | 0.038756846 | 0.0966069 |
| ENSG00000214121 | PRDX1P1 | 45.30467347 | 0.099678302 | 0.277269768 | 0.359499354 | 0.719221561 | 0.843935684 |
| ENSG00000214132 | ENSG0000021 | 9.248931853 | -0.266281236 | 0.591471369 | -0.450201397 | 0.652565229 | 0.798306653 |
| ENSG00000214135 | SDHAP4 | 329.1908618 | 0.139778131 | 0.108437625 | 1.289018736 | 0.197391573 | 0.359463578 |
| ENSG00000214144 | ENSG0000021 | 26.27168158 | 0.168999112 | 0.355163309 | 0.475834942 | 0.634191974 | 0.784362944 |
| ENSG00000214146 | LINC02026 | 7.545297479 | -0.061551186 | 0.631155961 | -0.097521358 | 0.922312374 | 0.962245128 |
| ENSG00000214147 | ENAHP1 | 3.254745578 | -0.663397371 | 0.980064289 | -0.676891688 | 0.498474678 | 0.680169469 |
| ENSG00000214160 | ALG3 | 1367.49129 | 0.140964498 | 0.061920916 | 2.276524749 | 0.022814621 | 0.061291417 |
| ENSG00000214174 | AMZ2P1 | 406.1167139 | -0.167563855 | 0.096546521 | -1.73557631 | 0.082638778 | 0.18140991 |
| ENSG00000214176 | PLEKHM1P1 | 1122.487134 | -0.025946752 | 0.070784006 | -0.366562361 | 0.713945488 | 0.840918855 |
| ENSG00000214182 | PTMAP5 | 998.6542045 | -0.123864265 | 0.069171461 | -1.790684522 | 0.073343935 | 0.16491961 |
| ENSG00000214184 | GCC2-AS1 | 7.866746105 | 0.226290142 | 0.640330224 | 0.353396004 | 0.723791581 | 0.846416769 |
| ENSG00000214185 | XPOTP1 | 259.4272588 | 0.030699964 | 0.12550569 | 0.244610134 | 0.806758317 | 0.897397846 |
| ENSG00000214192 | UBE2V1P2 | 214.3443077 | -0.076086669 | 0.130493412 | -0.583069044 | 0.559846822 | 0.729336481 |
| ENSG00000214193 | SH3D21 | 891.2349463 | 0.072419508 | 0.079054072 | 0.91607561 | 0.359627239 | 0.55219872 |
| ENSG00000214194 | SMIM30 | 216.2596653 | -0.414456533 | 0.123563579 | -3.354196583 | 0.000795958 | 0.00303411 |
| ENSG00000214198 | TTC41P | 21.87553588 | -0.389168563 | 0.393075306 | -0.990061081 | 0.322144265 | 0.511949389 |
| ENSG00000214199 | EEF1A1P12 | 74.036243 | 0.223462874 | 0.222832526 | 1.002828801 | 0.31594347 | 0.504717493 |
| ENSG00000214203 | RPS4XP1 | 56.34564063 | 0.442697757 | 0.250624668 | 1.766377429 | 0.077332549 | 0.172166991 |
| ENSG00000214204 | HNRNPA1P43 | 4.669811096 | -0.192752294 | 0.797464401 | -0.241706455 | 0.809007627 | 0.898853827 |
| ENSG00000214207 | KRT18P10 | 144.7049122 | 0.362975316 | 0.14936406 | 2.430138261 | 0.015093064 | 0.042729222 |
| ENSG00000214212 | C19orf38 | 20.75958006 | -2.191195422 | 0.437952122 | -5.003276185 | 5.64E-07 | 3.37E-06 |
| ENSG00000214222 | TUBBP2 | 20.2934039 | -0.666696517 | 0.44216181 | -1.507811172 | 0.131602862 | 0.263919952 |
| ENSG00000214223 | HNRNPA1P10 | 185.704505 | 0.302768005 | 0.157764618 | 1.919112219 | 0.054970133 | 0.129934556 |
| ENSG00000214226 | C17orf67 | 146.3993942 | 0.548657682 | 0.152542433 | 3.596754496 | 0.000322212 | 0.001318742 |
| ENSG00000214243 | ENSG0000021 | 2.339870676 | 2.542786975 | 1.389603563 | 1.829865037 | 0.067270123 | 0.153580475 |
| ENSG00000214244 | SETP21 | 15.92266959 | 0.18380097 | 0.443448555 | 0.414480932 | 0.678521929 | 0.817032861 |
| ENSG00000214248 | ENSG0000021 | 31.30863245 | -0.189335494 | 0.327178931 | -0.578690974 | 0.562797708 | 0.731601939 |
| ENSG00000214249 | CTAGE11P | 5.862569401 | -1.307430146 | 0.748003853 | -1.747892262 | 0.080482685 | 0.177543302 |

| | | | | | | | |
|---|---|---|---|---|---|---|---|
| ENSG00000214253 | FIS1 | 2090.787735 | -0.410376963 | 0.056297225 | -7.289470579 | 3.11E-13 | 3.29E-12 |
| ENSG00000214264 | KCTD9P4 | 38.84093546 | -0.425575537 | 0.286693879 | -1.484424916 | 0.137696229 | 0.273201132 |
| ENSG00000214269 | LGMNP1 | 16.10838179 | -0.226599314 | 0.43524982 | -0.520618972 | 0.60263223 | 0.7618839 |
| ENSG00000214273 | AGGF1P1 | 2.042532318 | -0.223322462 | 1.268783732 | -0.176013024 | 0.86028371 | 0.927382816 |
| ENSG00000214274 | ANG | 5.975650428 | -0.175738831 | 0.718319518 | -0.244652731 | 0.806725331 | 0.897397846 |
| ENSG00000214278 | ENSG0000021 | 2.091629649 | 0.498147685 | 1.187859683 | 0.419365765 | 0.67494884 | 0.814354725 |
| ENSG00000214279 | SCART1 | 67.94028531 | -0.90886345 | 0.239463379 | -3.795417296 | 0.000147395 | 0.000639661 |
| ENSG00000214280 | DTWD1P1 | 2.918503971 | -0.470815818 | 1.045188437 | -0.450460224 | 0.652378629 | 0.798160975 |
| ENSG00000214282 | KRT8P14 | 36.06233437 | 0.268982853 | 0.293688771 | 0.915877214 | 0.359731299 | 0.552321531 |
| ENSG00000214289 | RPL39P5 | 6.897622125 | 0.216093809 | 0.661871781 | 0.326488929 | 0.744054464 | 0.858490754 |
| ENSG00000214293 | APTR | 194.7435882 | 0.771184832 | 0.132318714 | 5.828237049 | 5.60E-09 | 4.15E-08 |
| ENSG00000214297 | ALDOAP2 | 24.79286457 | 0.15545801 | 0.377968892 | 0.411298425 | 0.680853724 | 0.818841625 |
| ENSG00000214300 | SPDYE3 | 3.470680259 | -1.503788253 | 1.001487636 | -1.501554487 | 0.133212205 | 0.266351736 |
| ENSG00000214309 | MBLAC1 | 45.86331848 | -0.529076881 | 0.264229113 | -2.002341355 | 0.045248031 | 0.109944903 |
| ENSG00000214321 | CBX1P4 | 2.250043417 | 1.12751079 | 1.191325791 | 0.946433628 | 0.34392746 | 0.534882049 |
| ENSG00000214322 | CBX1P2 | 20.23055177 | -0.121897026 | 0.391952408 | -0.31099956 | 0.755800956 | 0.865149344 |
| ENSG00000214331 | PDPR2P | 168.1900044 | 0.388859577 | 0.14442294 | 2.692505611 | 0.007091734 | 0.021960635 |
| ENSG00000214336 | FOXI3 | 13.54061198 | -3.12923779 | 0.620642193 | -5.041935317 | 4.61E-07 | 2.79E-06 |
| ENSG00000214353 | VAC14-AS1 | 8.971609527 | -0.71328141 | 0.59195393 | -1.204961018 | 0.228218351 | 0.40111316 |
| ENSG00000214354 | PAFAH1B1P1 | 2.908199933 | -0.356726264 | 1.045173418 | -0.341308206 | 0.732871571 | 0.852366348 |
| ENSG00000214357 | NEURL1B | 914.1299387 | -0.804282834 | 0.071609801 | -11.23146312 | 2.86E-29 | 7.70E-28 |
| ENSG00000214362 | RPS3AP36 | 6.527995833 | 0.159068156 | 0.711377596 | 0.223605799 | 0.823064051 | 0.906409914 |
| ENSG00000214366 | EIF4EP5 | 5.203024777 | -0.504752084 | 0.818289644 | -0.616837922 | 0.537341636 | 0.712397684 |
| ENSG00000214367 | HAUS3 | 582.4575958 | -0.004889527 | 0.085719906 | -0.05704074 | 0.954512742 | 0.979088978 |
| ENSG00000214376 | VSTM5 | 3.459987295 | 4.086609793 | 1.409068394 | 2.900221033 | 0.003728996 | 0.012370668 |
| ENSG00000214380 | NFYBP1 | 8.483700792 | -0.803040312 | 0.603255719 | -1.331177287 | 0.183130683 | 0.339174703 |
| ENSG00000214389 | RPS3AP26 | 1934.933434 | 0.149299727 | 0.071902405 | 2.076421878 | 0.03785495 | 0.094771641 |
| ENSG00000214391 | TUBAP2 | 284.1321802 | -0.259541032 | 0.115120562 | -2.254514985 | 0.024163789 | 0.064556939 |
| ENSG00000214401 | KANSL1-AS1 | 13.11247277 | 0.066512752 | 0.486598889 | 0.136689074 | 0.891276566 | 0.944238 |
| ENSG00000214413 | BBIP1 | 362.1377127 | 0.229895719 | 0.100366633 | 2.290559252 | 0.021988918 | 0.059393121 |
| ENSG00000214417 | KRT18P13 | 13.81641632 | 0.072493021 | 0.50989706 | 0.142171875 | 0.886944247 | 0.941793234 |
| ENSG00000214424 | FAM149B1P1 | 8.326327482 | -1.148888466 | 0.663283032 | -1.732124011 | 0.083251485 | 0.182562763 |
| ENSG00000214425 | LRRC37A4P | 5.015466495 | 0.197685998 | 0.794188172 | 0.248915817 | 0.8034259 | 0.895340789 |

| | | | | | | | |
|---|---|---|---|---|---|---|---|
| ENSG00000214432 | VPS33B-DT | 13.56948084 | -0.753944029 | 0.48051996 | -1.569017091 | 0.116643957 | 0.240133929 |
| ENSG00000214433 | GOLGA2P8 | 4.027291492 | 0.889351899 | 0.886071546 | 1.00370213 | 0.31552221 | 0.504207396 |
| ENSG00000214439 | FAM185BP | 34.73901796 | 1.544155537 | 0.323874499 | 4.767758934 | 1.86E-06 | 1.04E-05 |
| ENSG00000214455 | RCN1P2 | 263.1577257 | 0.10499465 | 0.117304753 | 0.895058792 | 0.370755661 | 0.563440534 |
| ENSG00000214456 | PLIN5 | 127.0811641 | 1.05925346 | 0.1830301 | 5.787318383 | 7.15E-09 | 5.22E-08 |
| ENSG00000214460 | TPT1P6 | 24.67243534 | 0.152817245 | 0.369604087 | 0.413461999 | 0.679268158 | 0.817630998 |
| ENSG00000214465 | SMARCE1P6 | 30.15604816 | 0.206222349 | 0.320607048 | 0.643224627 | 0.520078355 | 0.697413646 |
| ENSG00000214485 | RPL7P1 | 2109.715497 | 0.447276274 | 0.066062408 | 6.770511228 | 1.28E-11 | 1.19E-10 |
| ENSG00000214517 | PPME1 | 1590.582418 | -0.093587583 | 0.058364831 | -1.60349275 | 0.108825909 | 0.226775369 |
| ENSG00000214530 | STARD10 | 6605.476078 | -0.597737284 | 0.045350437 | -13.18040835 | 1.14E-39 | 4.56E-38 |
| ENSG00000214533 | KRT18P33 | 9.539059784 | 0.580642177 | 0.573159891 | 1.013054448 | 0.311034154 | 0.499099294 |
| ENSG00000214534 | ZNF705EP | 3.543619145 | -0.058434907 | 0.931391333 | -0.062739372 | 0.949974045 | 0.976903526 |
| ENSG00000214535 | RPS15AP1 | 154.5641136 | 0.093831141 | 0.184584089 | 0.508338185 | 0.611216195 | 0.76888021 |
| ENSG00000214541 | RPS4XP3 | 65.99912867 | 0.413985055 | 0.227186445 | 1.822226038 | 0.068420695 | 0.155695938 |
| ENSG00000214544 | GTF2IRD2P1 | 11.8719437 | 0.290586474 | 0.507392677 | 0.572705298 | 0.56684425 | 0.734738789 |
| ENSG00000214546 | ENSG0000021 | 2.093491503 | 0.05809022 | 1.178494139 | 0.049291904 | 0.960686672 | 0.981920343 |
| ENSG00000214548 | MEG3 | 7.301520514 | -6.393633861 | 1.315945787 | -4.858584544 | 1.18E-06 | 6.78E-06 |
| ENSG00000214549 | SDHCP2 | 142.8422412 | 0.014738376 | 0.148822141 | 0.099033491 | 0.921111679 | 0.9619656 |
| ENSG00000214552 | COPS8P2 | 111.1923249 | 0.187429412 | 0.170910063 | 1.096655215 | 0.272792141 | 0.454862706 |
| ENSG00000214553 | LRRC37A11P | 5.987851169 | -2.186734924 | 0.801595546 | -2.727977889 | 0.006372388 | 0.019976155 |
| ENSG00000214558 | ZC3H11C | 219.4855057 | 0.085046899 | 0.124940966 | 0.680696665 | 0.496063447 | 0.678008548 |
| ENSG00000214559 | ENSG0000021 | 3.434047445 | -1.421399313 | 1.059169239 | -1.341994518 | 0.179597773 | 0.334547025 |
| ENSG00000214561 | RBBP4P4 | 8.469110918 | -0.444969619 | 0.644088209 | -0.690851987 | 0.489658561 | 0.672936425 |
| ENSG00000214562 | NUTM2D | 45.6475342 | -0.719686031 | 0.265015429 | -2.715638236 | 0.006614815 | 0.020637001 |
| ENSG00000214575 | CPEB1 | 3.937456771 | -1.119892368 | 0.896512642 | -1.249165171 | 0.211604668 | 0.379146378 |
| ENSG00000214578 | HMGN2P15 | 12.09228013 | -0.271922163 | 0.518506718 | -0.524433249 | 0.599977236 | 0.760035219 |
| ENSG00000214584 | PGGT1BP2 | 26.25021939 | 0.42065857 | 0.350376308 | 1.200590796 | 0.229909973 | 0.403062894 |
| ENSG00000214593 | ENSG0000021 | 8.916545088 | 0.353666151 | 0.635030845 | 0.556927515 | 0.577576954 | 0.743451272 |
| ENSG00000214595 | EML6 | 438.6916738 | 0.015534699 | 0.094605697 | 0.164204693 | 0.869570006 | 0.932287011 |
| ENSG00000214612 | RPS19P1 | 66.65121753 | -0.008642557 | 0.255643383 | -0.033807084 | 0.973030987 | 0.987931789 |
| ENSG00000214617 | SLC6A10P | 2.441996388 | 0.402120396 | 1.120992985 | 0.358718031 | 0.719806038 | 0.844319196 |
| ENSG00000214653 | HNRNPA3P3 | 98.57280085 | -0.105275513 | 0.181423989 | -0.580273387 | 0.561730271 | 0.730628362 |
| ENSG00000214654 | B3GALT9 | 104.5193045 | -0.340809446 | 0.178672731 | -1.907450817 | 0.056462234 | 0.132763802 |

| | | | | | | | |
|---|---|---|---|---|---|---|---|
| ENSG00000214655 | ZSWIM8 | 3185.898525 | -0.539238684 | 0.054938898 | -9.815243913 | 9.68E-23 | 1.89E-21 |
| ENSG00000214702 | COMMD4P2 | 2.198500124 | -0.86377862 | 1.181807607 | -0.730896142 | 0.464842593 | 0.650918504 |
| ENSG00000214706 | IFRD2 | 1674.71836 | 0.297695961 | 0.064124211 | 4.642489252 | 3.44E-06 | 1.88E-05 |
| ENSG00000214708 | ENSG0000021 | 16.22612365 | -0.609119745 | 0.442656845 | -1.37605405 | 0.1688049 | 0.318977299 |
| ENSG00000214711 | CAPN14 | 2.589986124 | -1.231298034 | 1.120859251 | -1.098530464 | 0.271972923 | 0.454041488 |
| ENSG00000214717 | ZBED1 | 920.0810371 | 0.189527244 | 0.076102993 | 2.49040462 | 0.012759774 | 0.036823401 |
| ENSG00000214717 | ZBED1 | 920.0810371 | 0.189527244 | 0.076102993 | 2.49040462 | 0.012759774 | 0.036823401 |
| ENSG00000214719 | ENSG0000021 | 8.229764575 | 0.125972864 | 0.617561227 | 0.203984413 | 0.83836568 | 0.91499129 |
| ENSG00000214748 | ENSG0000021 | 2.007900004 | -2.471434637 | 1.4634426 | -1.688781396 | 0.09126133 | 0.196686858 |
| ENSG00000214753 | HNRNPUL2 | 1623.787301 | 0.069040342 | 0.057659444 | 1.197381348 | 0.23115795 | 0.404791097 |
| ENSG00000214756 | CSKMT | 130.4950902 | 0.001123011 | 0.15633091 | 0.007183548 | 0.994268407 | 0.996533822 |
| ENSG00000214760 | RPL21P1 | 1.974383082 | 0.334310122 | 1.241382131 | 0.269304764 | 0.787695166 | 0.885120113 |
| ENSG00000214761 | HNRNPA1P15 | 2.700534316 | 0.033641565 | 1.086920206 | 0.030951274 | 0.975308399 | 0.988740606 |
| ENSG00000214765 | SEPTIN7P2 | 444.6076426 | 0.609024938 | 0.115540755 | 5.271083238 | 1.36E-07 | 8.69E-07 |
| ENSG00000214770 | ENSG0000021 | 12.27976514 | -0.673027048 | 0.514217343 | -1.308837706 | 0.190589332 | 0.35007761 |
| ENSG00000214783 | POLR2J4 | 41.17760285 | 1.918473194 | 0.325522183 | 5.89352522 | 3.78E-09 | 2.85E-08 |
| ENSG00000214784 | RPS3AP21 | 444.8673062 | 0.158114271 | 0.094154017 | 1.679315199 | 0.09309063 | 0.199896945 |
| ENSG00000214796 | TUBA5P | 1325.073511 | 0.531891625 | 0.06157506 | 8.638101582 | 5.72E-18 | 8.40E-17 |
| ENSG00000214807 | NPM1P31 | 2.563554449 | -0.416941877 | 1.11350464 | -0.374441078 | 0.708076186 | 0.837109241 |
| ENSG00000214810 | CYCSP55 | 12.56530955 | -1.004136008 | 0.50301709 | -1.996226425 | 0.045909282 | 0.111351125 |
| ENSG00000214812 | ARL8BP2 | 4.940579508 | -0.964845736 | 0.796135634 | -1.211911255 | 0.225546353 | 0.397545342 |
| ENSG00000214814 | FER1L6 | 2.226397264 | 1.696235323 | 1.296532524 | 1.308285979 | 0.190776329 | 0.350364982 |
| ENSG00000214815 | IMPDH1P3 | 1.753093245 | 0.108957461 | 1.415946333 | 0.076950276 | 0.938663101 | NA |
| ENSG00000214820 | MPRIPP1 | 116.727371 | 0.15286137 | 0.165091313 | 0.925920129 | 0.354487489 | 0.54613452 |
| ENSG00000214821 | HMGB1P4 | 6.332123752 | 0.017626774 | 0.722494878 | 0.024397092 | 0.980535868 | 0.991070363 |
| ENSG00000214826 | DDX12P | 246.0629678 | -0.21439531 | 0.12998728 | -1.649356082 | 0.099074706 | 0.210403052 |
| ENSG00000214827 | MTCP1 | 15.94702588 | -0.552659897 | 0.473381696 | -1.167472045 | 0.243019791 | 0.41941574 |
| ENSG00000214832 | UPF3AP2 | 86.33350711 | -0.010947987 | 0.1935104 | -0.056575705 | 0.954883188 | 0.979301696 |
| ENSG00000214837 | LINC01347 | 17.66818782 | 0.445515834 | 0.41820465 | 1.065305786 | 0.286737575 | 0.470545817 |
| ENSG00000214846 | ENSG0000021 | 9.479560074 | 0.226599564 | 0.603801315 | 0.375288291 | 0.707446073 | 0.836584737 |
| ENSG00000214851 | LINC00612 | 1.92104999 | 0.534520419 | 1.33785612 | 0.399535055 | 0.689498999 | 0.824525492 |
| ENSG00000214857 | SEM1P1 | 5.225170919 | -0.870611382 | 0.795727003 | -1.094108129 | 0.273907558 | 0.455920105 |
| ENSG00000214860 | EVPLL | 13.61327389 | 2.173052503 | 0.548306029 | 3.963211035 | 7.39E-05 | 0.000337266 |

| | | | | | | | |
|---|---|---|---|---|---|---|---|
| ENSG00000214867 | SRSF9P1 | 3.697638224 | -0.27637206 | 0.907967406 | -0.304385442 | 0.760834258 | 0.868680961 |
| ENSG00000214869 | TPM3P4 | 1.783843551 | 0.084799186 | 1.30041342 | 0.065209405 | 0.948007273 | NA |
| ENSG00000214870 | LINC02981 | 4.642819021 | -4.742919338 | 1.357736643 | -3.49325428 | 0.000477172 | 0.001895796 |
| ENSG00000214881 | TMEM14DP | 161.4554595 | -0.778374911 | 0.142531451 | -5.461074784 | 4.73E-08 | 3.17E-07 |
| ENSG00000214889 | RPS9P1 | 7.270074092 | -0.426270604 | 0.650967166 | -0.654826582 | 0.512579414 | 0.69212854 |
| ENSG00000214892 | USP8P1 | 454.778947 | -1.100032438 | 0.098202984 | -11.20161925 | 4.00E-29 | 1.07E-27 |
| ENSG00000214894 | LINC00243 | 101.5237868 | -2.685885107 | 0.219443885 | -12.23950766 | 1.91E-34 | 6.37E-33 |
| ENSG00000214896 | RPSAP55 | 4.652334612 | 0.222042776 | 0.810468833 | 0.27396831 | 0.784108975 | 0.882820117 |
| ENSG00000214900 | LINC01588 | 38.88729509 | -0.577303572 | 0.285003831 | -2.025599336 | 0.04280586 | 0.104999991 |
| ENSG00000214914 | RPL23AP3 | 6.195059588 | 1.172511465 | 0.768310183 | 1.526091272 | 0.126987135 | 0.256659461 |
| ENSG00000214922 | HLA-F-AS1 | 58.97741607 | 0.879599527 | 0.243152427 | 3.617481995 | 0.000297483 | 0.001225401 |
| ENSG00000214925 | RPL36AP52 | 3.010265139 | 1.319229989 | 1.053605927 | 1.252109498 | 0.210529967 | 0.377812764 |
| ENSG00000214941 | ZSWIM7 | 547.5142317 | 0.190058564 | 0.084245429 | 2.256010409 | 0.024069979 | 0.064347433 |
| ENSG00000214944 | ARHGEF28 | 207.5300968 | 1.403388032 | 0.13803978 | 10.16654787 | 2.80E-24 | 5.96E-23 |
| ENSG00000214954 | LRRC69 | 16.80242241 | -0.340521475 | 0.426977559 | -0.797516094 | 0.42515136 | 0.616060327 |
| ENSG00000214955 | ENSG0000021 | 1.905846346 | 2.111551983 | 1.455441627 | 1.450798124 | 0.146836082 | 0.286844905 |
| ENSG00000214960 | CRPPA | 92.6652199 | -1.114080352 | 0.199839511 | -5.574875299 | 2.48E-08 | 1.70E-07 |
| ENSG00000214961 | NT5DC1P2 | 6.273434298 | -0.640634154 | 0.709060047 | -0.903497746 | 0.366261784 | 0.558981127 |
| ENSG00000214973 | CHCHD3P3 | 102.7133914 | -0.224981103 | 0.176641995 | -1.27365581 | 0.202785424 | 0.366926772 |
| ENSG00000214975 | PPIAP29 | 229.3920659 | 0.556814929 | 0.126377859 | 4.405953164 | 1.05E-05 | 5.43E-05 |
| ENSG00000214980 | ENSG0000021 | 2.225952854 | -0.441000938 | 1.166169353 | -0.378162002 | 0.705310251 | 0.835087337 |
| ENSG00000214988 | RPL7AP26 | 5.305060867 | -1.305601901 | 0.795791555 | -1.640633019 | 0.100873617 | 0.213354302 |
| ENSG00000214998 | CCNB2P1 | 6.863513006 | -0.166747533 | 0.679630312 | -0.245350347 | 0.806185172 | 0.897151831 |
| ENSG00000214999 | ENSG0000021 | 17.28402799 | -2.394404474 | 0.495847956 | -4.82890863 | 1.37E-06 | 7.80E-06 |
| ENSG00000215003 | RPL15P20 | 9.472500159 | 0.446601514 | 0.654202891 | 0.682665149 | 0.494818456 | 0.67711556 |
| ENSG00000215004 | MESTP4 | 5.963619807 | -0.193770872 | 0.748321827 | -0.258940558 | 0.795681104 | 0.890864587 |
| ENSG00000215005 | HSPD1P7 | 5.860304022 | -0.438232437 | 0.720214731 | -0.608474693 | 0.542872686 | 0.716541968 |
| ENSG00000215006 | CHCHD2P2 | 44.85812374 | 0.541291952 | 0.289806942 | 1.867767376 | 0.061794503 | 0.142986271 |
| ENSG00000215007 | DNAJA1P3 | 29.28927384 | 0.044735325 | 0.347260347 | 0.128823591 | 0.897497238 | 0.948060798 |
| ENSG00000215012 | RTL10 | 1188.692043 | -0.530503966 | 0.066866626 | -7.933763002 | 2.13E-15 | 2.62E-14 |
| ENSG00000215014 | ENSG0000021 | 8.903765752 | 0.559617185 | 0.591930808 | 0.945409796 | 0.344449703 | 0.535416465 |
| ENSG00000215016 | RPL24P7 | 14.45633312 | 0.154846873 | 0.468312193 | 0.330648818 | 0.740909767 | 0.857108928 |
| ENSG00000215018 | COL28A1 | 581.3861704 | -1.05801281 | 0.099465694 | -10.63696199 | 2.01E-26 | 4.80E-25 |

| | | | | | | | |
|---|---|---|---|---|---|---|---|
| ENSG00000215021 | PHB2 | 4072.289144 | -0.058336777 | 0.046777926 | -1.247100556 | 0.212360629 | 0.380292961 |
| ENSG00000215022 | ENSG0000021 | 7.628726091 | -0.631786953 | 0.661972467 | -0.954400651 | 0.339880877 | 0.53082061 |
| ENSG00000215030 | RPL13P12 | 5768.077607 | 0.348913927 | 0.045926924 | 7.597154277 | 3.03E-14 | 3.43E-13 |
| ENSG00000215032 | GNL3LP1 | 5.54219569 | 0.381072844 | 0.763552759 | 0.499078603 | 0.617724011 | 0.773419321 |
| ENSG00000215035 | FDPSP5 | 18.53611688 | -0.905377168 | 0.432958903 | -2.091138814 | 0.036515622 | 0.091950425 |
| ENSG00000215039 | CD27-AS1 | 15.05234647 | -0.640455821 | 0.496292387 | -1.290480851 | 0.196883756 | 0.358709663 |
| ENSG00000215041 | NEURL4 | 907.3399408 | -0.081221035 | 0.079669651 | -1.019472705 | 0.307978605 | 0.495934609 |
| ENSG00000215045 | GRID2IP | 42.39818256 | 0.478894999 | 0.273039873 | 1.753937964 | 0.079441138 | 0.175733747 |
| ENSG00000215049 | PRDX2P1 | 9.790888972 | -0.733277454 | 0.598522072 | -1.225146889 | 0.220519881 | 0.390755915 |
| ENSG00000215057 | PKMP5 | 16.02060491 | -0.336055742 | 0.450021999 | -0.746754031 | 0.455212048 | 0.642658895 |
| ENSG00000215063 | RPL21P2 | 1.594690618 | -0.129312409 | 1.383274945 | -0.093482796 | 0.925520017 | NA |
| ENSG00000215067 | ALOX12-AS1 | 63.47224159 | 0.047277868 | 0.238372784 | 0.198335846 | 0.842782308 | 0.917308531 |
| ENSG00000215068 | ANXA2R-AS1 | 381.4109665 | -0.172901421 | 0.096806069 | -1.786059728 | 0.074089588 | 0.166302852 |
| ENSG00000215086 | NPM1P24 | 212.2981606 | -0.220257984 | 0.136190784 | -1.617275247 | 0.10581888 | 0.221870497 |
| ENSG00000215089 | KRT18P11 | 278.4521342 | 0.272348662 | 0.110881348 | 2.456217099 | 0.014040831 | 0.040121656 |
| ENSG00000215093 | EEF1A1P29 | 7.141068942 | 0.495588294 | 0.676408835 | 0.732675668 | 0.463756268 | 0.650040865 |
| ENSG00000215097 | ENSG0000021 | 2.753566783 | 0.662454498 | 1.053754614 | 0.628661066 | 0.529570973 | 0.705233839 |
| ENSG00000215102 | TERF1P4 | 23.22995613 | 0.598503213 | 0.386959477 | 1.546681883 | 0.121939974 | 0.248560778 |
| ENSG00000215105 | TTC3P1 | 1674.863009 | 0.15223583 | 0.06307463 | 2.413582626 | 0.015796546 | 0.04449021 |
| ENSG00000215114 | UBXN2B | 815.0931598 | -0.044181094 | 0.078560336 | -0.562384222 | 0.573854266 | 0.740493824 |
| ENSG00000215120 | SOCS6P1 | 2.229311343 | 0.36891481 | 1.249433114 | 0.295265753 | 0.767790876 | 0.87314707 |
| ENSG00000215126 | ZNG1F | 89.04959759 | -0.319822681 | 0.193742304 | -1.650763281 | 0.098786921 | 0.209927926 |
| ENSG00000215146 | ENSG0000021 | 170.5154348 | 0.229882532 | 0.15302912 | 1.502214295 | 0.133041774 | 0.266193092 |
| ENSG00000215154 | ENSG0000021 | 97.7487914 | -0.023931881 | 0.182873358 | -0.13086587 | 0.895881414 | 0.947292908 |
| ENSG00000215158 | GUSBP18 | 118.0984459 | 0.072736252 | 0.167541778 | 0.434138001 | 0.664188232 | 0.806205515 |
| ENSG00000215179 | MAPK6P4 | 38.11391785 | -0.845481866 | 0.310899667 | -2.719468548 | 0.006538691 | 0.020427288 |
| ENSG00000215183 | MSMP | 10.83079306 | -0.014544497 | 0.572142643 | -0.025421103 | 0.979719079 | 0.990906735 |
| ENSG00000215187 | FAM166B | 3.674569042 | -0.187180412 | 0.908326859 | -0.206071647 | 0.836734946 | 0.913925996 |
| ENSG00000215190 | LINC00680 | 248.9014753 | -0.004618671 | 0.11864986 | -0.038926896 | 0.968948673 | 0.985925152 |
| ENSG00000215193 | PEX26 | 2305.88333 | -0.710369137 | 0.052263359 | -13.59210638 | 4.46E-42 | 1.98E-40 |
| ENSG00000215196 | BASP1-AS1 | 121.0375275 | -4.41721655 | 0.280420319 | -15.75212723 | 6.64E-56 | 4.27E-54 |
| ENSG00000215198 | ENSG0000021 | 1.604157862 | -0.114693616 | 1.43790865 | -0.079764188 | 0.936424807 | NA |
| ENSG00000215199 | YWHAZP6 | 105.5315342 | -0.243781061 | 0.192323436 | -1.267557746 | 0.204955929 | 0.370007793 |

| | | | | | | | |
|---|---|---|---|---|---|---|---|
| ENSG00000215208 | KRT18P60 | 2.634458362 | 0.9269706 | 1.095221021 | 0.846377655 | 0.3973421 | 0.590236856 |
| ENSG00000215210 | RBMXP2 | 234.1106755 | -0.156183563 | 0.120462275 | -1.296535062 | 0.194791209 | 0.355943152 |
| ENSG00000215217 | CFAP90 | 37.3586158 | 1.89266915 | 0.322468073 | 5.869322604 | 4.38E-09 | 3.28E-08 |
| ENSG00000215218 | UBE2QL1 | 44.25292459 | 0.896083642 | 0.271787072 | 3.29700613 | 0.000977214 | 0.003664751 |
| ENSG00000215244 | LINC02649 | 6.076679654 | -1.74809459 | 0.761501609 | -2.295588832 | 0.021699402 | 0.058680217 |
| ENSG00000215251 | FASTKD5 | 533.5787073 | -0.121509754 | 0.086456797 | -1.405438999 | 0.159890777 | 0.305734302 |
| ENSG00000215252 | GOLGA8B | 191.4100478 | -0.074785379 | 0.282790341 | -0.264455212 | 0.791429172 | 0.887707081 |
| ENSG00000215256 | DHRS4-AS1 | 1158.729704 | -0.636098389 | 0.064894135 | -9.802093588 | 1.10E-22 | 2.15E-21 |
| ENSG00000215268 | ENSG0000021 | 3.984235272 | 1.20069233 | 0.91268104 | 1.315566204 | 0.188319683 | 0.346713767 |
| ENSG00000215270 | TOMM40P2 | 20.88461473 | 0.207469153 | 0.395426938 | 0.524671267 | 0.599811734 | 0.759867528 |
| ENSG00000215283 | HMGB3P24 | 10.64735271 | -0.668469735 | 0.611153105 | -1.093784405 | 0.274049546 | 0.456119609 |
| ENSG00000215284 | CHMP5P1 | 28.06862537 | 0.291580505 | 0.36853634 | 0.791185219 | 0.428835915 | 0.61920798 |
| ENSG00000215301 | DDX3X | 12698.78415 | -0.028902681 | 0.046798926 | -0.617592823 | 0.536843777 | 0.711908785 |
| ENSG00000215302 | WHAMMP4 | 9.822341755 | -0.706460334 | 0.560188165 | -1.261112566 | 0.207268294 | 0.373342442 |
| ENSG00000215305 | VPS16 | 811.2244487 | 0.03328373 | 0.072676882 | 0.45796861 | 0.646974994 | 0.793684192 |
| ENSG00000215311 | NPM1P12 | 5.893775582 | -0.173334975 | 0.708070857 | -0.244798912 | 0.806612137 | 0.897382528 |
| ENSG00000215319 | EIF5P1 | 58.15678287 | -0.31113455 | 0.233376741 | -1.333185767 | 0.182470849 | 0.338362402 |
| ENSG00000215325 | ASS1P10 | 44.30678744 | 0.872403706 | 0.271467617 | 3.21365662 | 0.001310563 | 0.004799539 |
| ENSG00000215333 | KRT18P23 | 21.78288627 | 0.511490256 | 0.38375273 | 1.33286415 | 0.182576389 | 0.338503384 |
| ENSG00000215347 | SLC25A5P1 | 2.763961071 | -0.017946152 | 1.076123631 | -0.016676664 | 0.986694564 | 0.993497695 |
| ENSG00000215349 | MRPL3P1 | 63.63248846 | 0.166439483 | 0.230542755 | 0.721946276 | 0.470327508 | 0.656115263 |
| ENSG00000215374 | FAM66B | 7.559856508 | 0.267099443 | 0.63905688 | 0.417958793 | 0.675977245 | 0.815166798 |
| ENSG00000215375 | MYL5 | 492.2309905 | -0.138168391 | 0.088487291 | -1.561448979 | 0.118417853 | 0.243077667 |
| ENSG00000215386 | MIR99AHG | 105.5125087 | 0.429115273 | 0.174790941 | 2.455020094 | 0.014087671 | 0.040230447 |
| ENSG00000215388 | ACTG1P3 | 2.672641262 | 0.708075732 | 1.092257022 | 0.648268418 | 0.516811357 | 0.695223862 |
| ENSG00000215414 | PSMA6P1 | 556.8349336 | -0.485016141 | 0.084097731 | -5.767291671 | 8.06E-09 | 5.84E-08 |
| ENSG00000215417 | MIR17HG | 12.96154946 | -0.494532618 | 0.548907079 | -0.9009405 | 0.367619956 | 0.560159535 |
| ENSG00000215421 | ZNF407 | 574.8173304 | 0.26620266 | 0.085597921 | 3.109919684 | 0.001871382 | 0.006615736 |
| ENSG00000215424 | MCM3AP-AS1 | 748.671311 | -0.161582478 | 0.074240486 | -2.176473881 | 0.029519845 | 0.076587337 |
| ENSG00000215440 | NPEPL1 | 768.775912 | 0.100621185 | 0.074320929 | 1.35387416 | 0.175776529 | 0.329034155 |
| ENSG00000215441 | CTAGE12P | 3.730762762 | -0.274434664 | 0.913137765 | -0.300540262 | 0.76376509 | 0.870479014 |
| ENSG00000215450 | ENSG0000021 | 3.570148207 | 0.203702405 | 0.940157765 | 0.216668322 | 0.828466837 | 0.90928207 |
| ENSG00000215464 | ENSG0000021 | 3.429135191 | -0.572419068 | 0.967588395 | -0.591593565 | 0.554122784 | 0.725463798 |

| | | | | | | | |
|---|---|---|---|---|---|---|---|
| ENSG00000215467 | RPL27AP | 10.39058617 | 0.705887028 | 0.591999099 | 1.192378551 | 0.233112847 | 0.407405813 |
| ENSG00000215477 | RCN1P1 | 1.695889286 | -1.407568059 | 1.373896159 | -1.02450833 | 0.305595239 | NA |
| ENSG00000215478 | CES5AP1 | 12.38616944 | -4.665208244 | 0.941845705 | -4.953261685 | 7.30E-07 | 4.31E-06 |
| ENSG00000215481 | BCRP3 | 4.364279236 | -1.926502436 | 0.931596551 | -2.067957888 | 0.038643983 | 0.096392011 |
| ENSG00000215482 | CALM2P3 | 89.98613341 | -0.145293068 | 0.202827803 | -0.716337039 | 0.473783261 | 0.659301669 |
| ENSG00000215483 | LINC00598 | 4.575716906 | 0.664642544 | 0.817648135 | 0.812871106 | 0.416291962 | 0.608341038 |
| ENSG00000215492 | HNRNPA1P7 | 3741.283738 | 0.298029379 | 0.060312014 | 4.941459554 | 7.75E-07 | 4.56E-06 |
| ENSG00000215493 | ENSG0000021 | 34.08476049 | 0.734497184 | 0.321080338 | 2.287580701 | 0.022161951 | 0.059832312 |
| ENSG00000215515 | IFIT1P1 | 2.038571525 | -0.364524378 | 1.335194355 | -0.273012223 | 0.78484382 | 0.883106023 |
| ENSG00000215529 | EFCAB8 | 3.523652136 | -1.513674565 | 0.987572396 | -1.532722636 | 0.125344209 | 0.254055148 |
| ENSG00000215548 | FRG1JP | 12.50471972 | -0.18309162 | 0.518804932 | -0.352910331 | 0.724155665 | 0.846509142 |
| ENSG00000215553 | KRT18P3 | 8.16742652 | -0.039078316 | 0.616896235 | -0.06334666 | 0.949490461 | 0.976618218 |
| ENSG00000215559 | ANKRD20A11I | 44.7832204 | -0.911290781 | 0.273368735 | -3.333558902 | 0.000857425 | 0.003250592 |
| ENSG00000215571 | GRK6P1 | 10.19479004 | -0.707485812 | 0.545809832 | -1.296213023 | 0.194902104 | 0.355975658 |
| ENSG00000215572 | ESRRAP1 | 4.039597082 | 0.368906922 | 0.943913101 | 0.390827209 | 0.695924962 | 0.829110289 |
| ENSG00000215604 | ZNF962P | 16.80476532 | -6.642945204 | 1.230498341 | -5.398581196 | 6.72E-08 | 4.42E-07 |
| ENSG00000215630 | GUSBP9 | 7.743530376 | -1.636796293 | 0.687152361 | -2.381999081 | 0.01721894 | 0.048001275 |
| ENSG00000215644 | GCGR | 2.586657379 | 0.973611079 | 1.104327412 | 0.881632628 | 0.377975505 | 0.570782574 |
| ENSG00000215692 | ENSG0000021 | 2.081242416 | -1.735873687 | 1.369749537 | -1.267292771 | 0.205050624 | 0.370120491 |
| ENSG00000215712 | TMEM242 | 205.5563222 | 0.092386828 | 0.130213962 | 0.709500169 | 0.478014147 | 0.662842676 |
| ENSG00000215717 | TMEM167B | 1136.064395 | 0.221568195 | 0.065235285 | 3.396447122 | 0.000682667 | 0.002636873 |
| ENSG00000215769 | ARHGAP27P1- | 133.3413042 | 0.124742222 | 0.164809723 | 0.756886301 | 0.449117983 | 0.637356203 |
| ENSG00000215784 | FAM72D | 143.8542393 | 0.144211325 | 0.148938034 | 0.968263922 | 0.332912578 | 0.524052325 |
| ENSG00000215788 | TNFRSF25 | 291.9698899 | -0.211188841 | 0.122156311 | -1.728841019 | 0.083837553 | 0.183671832 |
| ENSG00000215790 | SLC35E2A | 512.2148669 | 0.044466287 | 0.094167833 | 0.47220251 | 0.636782248 | 0.786213586 |
| ENSG00000215795 | ENSG0000021 | 3.821574603 | 1.094363442 | 0.925470782 | 1.182493779 | 0.237009833 | 0.412237892 |
| ENSG00000215796 | ENSG0000021 | 1.999453816 | -0.657270364 | 1.258916981 | -0.522091904 | 0.601606343 | 0.761300171 |
| ENSG00000215807 | KRT18P65 | 23.54507745 | 0.334906474 | 0.382855297 | 0.874759934 | 0.381704542 | 0.574033184 |
| ENSG00000215808 | LINC01139 | 24.59423571 | 0.023826898 | 0.353907232 | 0.067325264 | 0.946322765 | 0.974832509 |
| ENSG00000215817 | ZC3H11B | 233.5661819 | 0.073675673 | 0.128536626 | 0.573188167 | 0.566517295 | 0.734439489 |
| ENSG00000215819 | KRT18P12 | 3.186581198 | 0.426123601 | 0.971454819 | 0.438644796 | 0.660918933 | 0.803981351 |
| ENSG00000215833 | QRSL1P1 | 2.476033453 | -0.008632667 | 1.085654502 | -0.007951578 | 0.993655625 | 0.996522691 |
| ENSG00000215835 | ENSG0000021 | 1341.286164 | -0.278988692 | 0.062326963 | -4.476211901 | 7.60E-06 | 3.97E-05 |

| | | | | | | | |
|---|---|---|---|---|---|---|---|
| ENSG00000215840 | ENSG0000021 | 21.79028176 | 0.069912106 | 0.396596449 | 0.176280212 | 0.860073807 | 0.927319435 |
| ENSG00000215841 | ENSG0000021 | 3.032912518 | 0.628782339 | 1.038651719 | 0.605383236 | 0.544924387 | 0.718175654 |
| ENSG00000215845 | TSTD1 | 784.3763874 | 0.609405286 | 0.076557293 | 7.960120604 | 1.72E-15 | 2.13E-14 |
| ENSG00000215861 | ENSG0000021 | 34.189853 | -1.398724866 | 0.332966003 | -4.200803853 | 2.66E-05 | 0.000129733 |
| ENSG00000215867 | KRT18P57 | 7.785890805 | 0.765998256 | 0.654410555 | 1.170516353 | 0.241793237 | 0.417959224 |
| ENSG00000215871 | ENSG0000021 | 2.362080782 | 1.300932701 | 1.207625725 | 1.077264814 | 0.281361977 | 0.464749694 |
| ENSG00000215873 | FEN1P1 | 4.456978588 | -0.72311248 | 0.842216921 | -0.858582227 | 0.390571047 | 0.583427741 |
| ENSG00000215883 | CYB5RL | 246.9585317 | 0.298801077 | 0.116685945 | 2.560728945 | 0.010445282 | 0.030862282 |
| ENSG00000215895 | HSPA5P1 | 17.57791369 | 0.172049165 | 0.463406689 | 0.371270353 | 0.710436177 | 0.838775394 |
| ENSG00000215899 | EEF1A1P46 | 2.362080782 | 1.300932701 | 1.207625725 | 1.077264814 | 0.281361977 | 0.464749694 |
| ENSG00000215900 | SELENOWP1 | 3.952539319 | 0.066300942 | 0.867843274 | 0.076397368 | 0.939102963 | 0.971039083 |
| ENSG00000215908 | CROCCP2 | 465.3169971 | -0.117121701 | 0.09513741 | -1.231079349 | 0.218293188 | 0.387918951 |
| ENSG00000215909 | BRWD1P1 | 4.74251227 | 0.658513695 | 0.837633933 | 0.786159287 | 0.431774174 | 0.621776495 |
| ENSG00000215910 | C1orf167 | 3.701000392 | -0.992341452 | 0.934568444 | -1.061817845 | 0.288318387 | 0.472316334 |
| ENSG00000215912 | TTC34 | 35.91682721 | 0.851786581 | 0.327852756 | 2.598076627 | 0.009374757 | 0.028119368 |
| ENSG00000215915 | ATAD3C | 22.35427605 | 1.331354862 | 0.407415664 | 3.2678048 | 0.001083851 | 0.004031707 |
| ENSG00000216285 | ENSG0000021 | 539.220269 | -0.834176954 | 0.091443395 | -9.122331415 | 7.35E-20 | 1.21E-18 |
| ENSG00000216368 | RPL7AP36 | 4.49111304 | -0.301163613 | 0.846351988 | -0.355837308 | 0.721962407 | 0.845596895 |
| ENSG00000216412 | RPL12P2 | 1.57535228 | 1.860209278 | 1.549136809 | 1.200803743 | 0.22982734 | NA |
| ENSG00000216439 | DUTP5 | 1.581925474 | -0.142326456 | 1.385504367 | -0.102725375 | 0.918180934 | NA |
| ENSG00000216490 | IFI30 | 3.779083888 | 0.708729097 | 0.912698983 | 0.776520091 | 0.437441963 | 0.626920679 |
| ENSG00000216621 | ENSG0000021 | 1.735627902 | 2.029317422 | 1.49455969 | 1.357802861 | 0.174526247 | NA |
| ENSG00000216624 | GAPDHP72 | 25.31434455 | -0.113711089 | 0.366252727 | -0.310471653 | 0.756202313 | 0.865244205 |
| ENSG00000216657 | GLRX3P2 | 136.6886943 | -0.57320667 | 0.162775142 | -3.521463196 | 0.000429172 | 0.001719387 |
| ENSG00000216676 | RPL8P1 | 3.193165532 | -0.154268043 | 0.970987689 | -0.158877445 | 0.873765431 | 0.934676032 |
| ENSG00000216740 | ANXA2P3 | 11.0458604 | 0.037914638 | 0.521737664 | 0.072669927 | 0.94206878 | 0.972530993 |
| ENSG00000216775 | ENSG0000021 | 184.5844533 | -0.26210299 | 0.138061452 | -1.898451639 | 0.057636613 | 0.134971814 |
| ENSG00000216802 | MTCH1P1 | 2.91378149 | -0.143815232 | 1.035343273 | -0.138905845 | 0.889524552 | 0.943223317 |
| ENSG00000216835 | RBMXP1 | 11.47631329 | -0.693920513 | 0.537953008 | -1.289927749 | 0.197075745 | 0.35900243 |
| ENSG00000216854 | RPL17P26 | 3.304343822 | -0.466400928 | 0.955008791 | -0.488373439 | 0.625285352 | 0.778594473 |
| ENSG00000216866 | RPS2P55 | 318.3066458 | 0.17612124 | 0.103921981 | 1.694744825 | 0.090123838 | 0.194637625 |
| ENSG00000216867 | RPL22P12 | 8.301950011 | -0.154423032 | 0.636439897 | -0.242635688 | 0.808287632 | 0.898401971 |
| ENSG00000216895 | RBM33-DT | 34.61769526 | 0.927491357 | 0.314654965 | 2.947645711 | 0.003202038 | 0.010777643 |

| | | | | | | | |
|---|---|---|---|---|---|---|---|
| ENSG00000216906 | ENSG0000021 | 14.58856485 | 0.289165463 | 0.45763727 | 0.631866069 | 0.527474397 | 0.703507931 |
| ENSG00000216915 | GPR89P | 2.982793806 | -1.157432471 | 1.034984223 | -1.11830929 | 0.26343492 | 0.44412799 |
| ENSG00000216937 | CCDC7 | 63.5305529 | -1.118136773 | 0.229788785 | -4.865932753 | 1.14E-06 | 6.55E-06 |
| ENSG00000217026 | RPL10P1 | 24.5773426 | -0.247861399 | 0.361575305 | -0.685504224 | 0.493025796 | 0.675789422 |
| ENSG00000217027 | TPT1P4 | 69.940372 | 0.333703325 | 0.243705236 | 1.369290748 | 0.170908409 | 0.322023475 |
| ENSG00000217060 | ENSG0000021 | 43.83654561 | -0.194957664 | 0.270586702 | -0.720499797 | 0.471217324 | 0.657086221 |
| ENSG00000217094 | PPIAP31 | 664.7724505 | 0.354277116 | 0.077088077 | 4.595744642 | 4.31E-06 | 2.32E-05 |
| ENSG00000217128 | FNIP1 | 1190.811403 | 0.039908626 | 0.065875893 | 0.605815337 | 0.544637384 | 0.717913747 |
| ENSG00000217130 | ENSG0000021 | 151.5764414 | 0.14729653 | 0.150247264 | 0.980360817 | 0.326908044 | 0.517430524 |
| ENSG00000217139 | ENSG0000021 | 4.172020126 | 0.14686084 | 0.90355199 | 0.162537233 | 0.870882811 | 0.932975557 |
| ENSG00000217159 | LARP1P1 | 2.646801378 | -0.032843115 | 1.173240799 | -0.027993499 | 0.977667337 | 0.989573275 |
| ENSG00000217165 | ANKRD18EP | 3.572892845 | -3.381513978 | 1.280661332 | -2.640443569 | 0.008279758 | 0.025187329 |
| ENSG00000217239 | ENSG0000021 | 4.90192849 | 0.229551749 | 0.819353575 | 0.280162016 | 0.779353207 | 0.880273883 |
| ENSG00000217241 | CBX3P9 | 401.2640875 | -1.831329093 | 0.104293936 | -17.55930563 | 5.05E-69 | 4.54E-67 |
| ENSG00000217275 | ENSG0000021 | 43.2370641 | 0.590424986 | 0.267318358 | 2.208695992 | 0.027195793 | 0.071405689 |
| ENSG00000217289 | ENSG0000021 | 106.5142988 | 0.084034094 | 0.176514804 | 0.476073915 | 0.63402172 | 0.784236938 |
| ENSG00000217325 | PRELID1P1 | 61.1506275 | -0.494755339 | 0.228540949 | -2.164843284 | 0.030399689 | 0.078612204 |
| ENSG00000217385 | PSMC1P11 | 5.135532805 | -1.441777252 | 0.832778593 | -1.731285199 | 0.08340091 | 0.182820533 |
| ENSG00000217414 | DDX18P3 | 6.77139471 | 0.148919474 | 0.663585386 | 0.224416446 | 0.822433275 | 0.906177682 |
| ENSG00000217416 | ISCA1P1 | 128.9369621 | 0.259192107 | 0.158138633 | 1.639018262 | 0.101209457 | 0.213926558 |
| ENSG00000217442 | SYCE3 | 11.91488251 | 0.417961449 | 0.507342135 | 0.823825619 | 0.410038648 | 0.602595926 |
| ENSG00000217539 | IQCB2P | 9.4189949 | 0.175149552 | 0.597695577 | 0.293041407 | 0.769490502 | 0.874256457 |
| ENSG00000217555 | CKLF | 15.48754878 | -0.199589784 | 0.446506581 | -0.447003007 | 0.654872889 | 0.799594639 |
| ENSG00000217624 | YWHAZP10 | 430.2734415 | -0.438717644 | 0.094163288 | -4.659115617 | 3.18E-06 | 1.74E-05 |
| ENSG00000217643 | PTGES3P2 | 10.27663004 | 0.161831721 | 0.57444128 | 0.281720216 | 0.778158052 | 0.879573576 |
| ENSG00000217648 | PARLP2 | 52.15881057 | 1.187189648 | 0.26388051 | 4.498966777 | 6.83E-06 | 3.60E-05 |
| ENSG00000217702 | ENSG0000021 | 14.56271645 | -0.381872092 | 0.469971286 | -0.812543453 | 0.416479863 | 0.608557597 |
| ENSG00000217716 | RPS10P3 | 174.861515 | 0.3053113 | 0.142543418 | 2.141882829 | 0.032202912 | 0.082595572 |
| ENSG00000217733 | CCT7P1 | 4.789990345 | 0.66384311 | 0.848054549 | 0.782783503 | 0.433754257 | 0.623610094 |
| ENSG00000217769 | ENSG0000021 | 2.663263943 | -0.186935958 | 1.055076101 | -0.177177701 | 0.85936881 | 0.926885162 |
| ENSG00000217791 | ASS1P9 | 26.54548079 | 0.739513575 | 0.348707457 | 2.120727735 | 0.033944721 | 0.086300776 |
| ENSG00000217801 | ENSG0000021 | 128.9233561 | 0.698334497 | 0.159831769 | 4.369184568 | 1.25E-05 | 6.35E-05 |
| ENSG00000217824 | SNRPEP6 | 1.666574702 | -1.389132866 | 1.396164003 | -0.994963961 | 0.319753793 | NA |

| | | | | | | | |
|---|---|---|---|---|---|---|---|
| ENSG00000217835 | RBMXP5 | 58.75047976 | -0.144054525 | 0.234830976 | -0.613439195 | 0.539585977 | 0.714103677 |
| ENSG00000217889 | KRT18P48 | 60.29957758 | 0.264777432 | 0.225441605 | 1.174483442 | 0.24020144 | 0.416022729 |
| ENSG00000217897 | HSPE1P8 | 1.642491601 | -0.196061353 | 1.341263149 | -0.146176649 | 0.883781939 | NA |
| ENSG00000217930 | PAM16 | 92.06678955 | 0.092695903 | 0.199117919 | 0.465532704 | 0.641550023 | 0.789631802 |
| ENSG00000217950 | NOC2LP2 | 89.11177215 | -0.093551553 | 0.189672956 | -0.493225574 | 0.62185321 | 0.776234318 |
| ENSG00000218014 | KRT19P1 | 312.431967 | 0.108679001 | 0.112345109 | 0.967367442 | 0.33336038 | 0.52456184 |
| ENSG00000218018 | RBM38-AS1 | 11.51538902 | -0.23832562 | 0.51150681 | -0.465928538 | 0.641266653 | 0.789579573 |
| ENSG00000218027 | ENSG0000021 | 3.606336611 | -1.203325176 | 1.002963716 | -1.199769401 | 0.230228911 | 0.403541328 |
| ENSG00000218069 | RSL24D1P1 | 162.1476561 | -0.146975378 | 0.149201068 | -0.985082615 | 0.324583492 | 0.514760705 |
| ENSG00000218073 | ENSG0000021 | 6.886062672 | -1.005927969 | 0.681292316 | -1.476499802 | 0.139809768 | 0.276128807 |
| ENSG00000218175 | ENSG0000021 | 960.8499932 | -0.101743254 | 0.072843594 | -1.396735772 | 0.162493043 | 0.309729157 |
| ENSG00000218180 | SLC25A5P7 | 6.273117019 | 0.26774001 | 0.742513588 | 0.360586007 | 0.718408951 | 0.843432038 |
| ENSG00000218186 | KRT8P43 | 3.896258725 | 0.065068256 | 0.875800863 | 0.07429572 | 0.940775082 | 0.971736581 |
| ENSG00000218208 | RPS27AP11 | 35.58876006 | 0.421208919 | 0.348040048 | 1.210231185 | 0.226190195 | 0.398342014 |
| ENSG00000218226 | TATDN2P2 | 6.39138902 | -2.337399799 | 0.825693193 | -2.830833316 | 0.004642691 | 0.01507905 |
| ENSG00000218261 | RPA3P2 | 2.53079513 | -2.141067884 | 1.243203063 | -1.722218958 | 0.085029862 | 0.185770905 |
| ENSG00000218265 | RPS4XP7 | 3.417817234 | -0.02795702 | 0.964267619 | -0.028993009 | 0.976870166 | 0.989309285 |
| ENSG00000218283 | MORF4L1P1 | 4261.528276 | -0.160821255 | 0.050274303 | -3.198875883 | 0.001379646 | 0.005023662 |
| ENSG00000218305 | CDC14C | 4.587510443 | 0.21129172 | 0.813276713 | 0.259802987 | 0.795015746 | 0.890293445 |
| ENSG00000218336 | TENM3 | 56.78676826 | 2.183941703 | 0.286559031 | 7.621262878 | 2.51E-14 | 2.87E-13 |
| ENSG00000218350 | LYPLA1P3 | 264.6297837 | -0.161327533 | 0.117019581 | -1.378637079 | 0.168006678 | 0.317992838 |
| ENSG00000218357 | LINC01644 | 2.327075789 | -1.043016098 | 1.201465219 | -0.868120094 | 0.385328592 | 0.577892122 |
| ENSG00000218358 | RAET1K | 16.50459138 | -1.168768655 | 0.441438418 | -2.647636921 | 0.008105653 | 0.024723265 |
| ENSG00000218416 | GPC1-AS1 | 43.66702355 | 2.679809512 | 0.335714288 | 7.98241126 | 1.44E-15 | 1.80E-14 |
| ENSG00000218418 | ENSG0000021 | 138.5803452 | -0.512104923 | 0.156650254 | -3.269097307 | 0.001078912 | 0.004015289 |
| ENSG00000218426 | ENSG0000021 | 2299.20296 | 0.248230699 | 0.057732365 | 4.299680064 | 1.71E-05 | 8.57E-05 |
| ENSG00000218502 | H2AZP3 | 66.05970476 | -0.312078679 | 0.256449323 | -1.216921435 | 0.223634109 | 0.395023213 |
| ENSG00000218510 | LINC00339 | 73.45265467 | -0.010334987 | 0.209455387 | -0.049342189 | 0.960646598 | 0.981920343 |
| ENSG00000218537 | MIF-AS1 | 161.0505604 | -0.632086695 | 0.169730569 | -3.724059236 | 0.000196045 | 0.00083359 |
| ENSG00000218549 | OR4K12P | 6.173717667 | -1.577972957 | 0.738863931 | -2.135674636 | 0.032705943 | 0.083689274 |
| ENSG00000218582 | GAPDHP63 | 48.9889735 | -0.195985296 | 0.272807466 | -0.718401509 | 0.472509757 | 0.658168305 |
| ENSG00000218586 | ENSG0000021 | 35.00691342 | -0.326979486 | 0.301090194 | -1.085985171 | 0.277485553 | 0.460302161 |
| ENSG00000218631 | ENSG0000021 | 4.964806393 | -0.032368596 | 0.829846084 | -0.039005542 | 0.96888597 | 0.985925152 |

| | | | | | | | |
|---|---|---|---|---|---|---|---|
| ENSG00000218632 | RPL7P28 | 2.111404935 | -1.308540507 | 1.239700247 | -1.055529763 | 0.291183105 | 0.47613803 |
| ENSG00000218676 | BRD7P4 | 16.66706751 | 0.42207551 | 0.444100532 | 0.950405323 | 0.34190634 | 0.533052293 |
| ENSG00000218690 | H2AC10P | 12.20741878 | -0.667524896 | 0.55346047 | -1.206093175 | 0.227781566 | 0.400621722 |
| ENSG00000218739 | CEBPZOS | 1083.219792 | 0.416335836 | 0.074484905 | 5.589532989 | 2.28E-08 | 1.57E-07 |
| ENSG00000218748 | DBIP1 | 15.0902739 | -0.318439869 | 0.468054957 | -0.680347178 | 0.496284658 | 0.678189674 |
| ENSG00000218772 | FAM8A6P | 2.949669268 | -1.162191497 | 1.039261788 | -1.118285605 | 0.263445033 | 0.44412799 |
| ENSG00000218803 | GSTM2P1 | 39.7484369 | -0.61819464 | 0.290285773 | -2.129607086 | 0.033204066 | 0.084710952 |
| ENSG00000218823 | PAPOLB | 3.167213116 | -0.962351201 | 1.013837108 | -0.949216786 | 0.342510366 | 0.533486406 |
| ENSG00000218857 | ENSG0000021 | 3.706690802 | 0.480588386 | 0.918458036 | 0.523255682 | 0.600796337 | 0.760778758 |
| ENSG00000218890 | NUFIP1P1 | 39.60774035 | 0.459009725 | 0.28954099 | 1.58530136 | 0.11289788 | 0.233785644 |
| ENSG00000218891 | ZNF579 | 289.6980859 | 0.179472142 | 0.111986387 | 1.602624627 | 0.109017553 | 0.227062164 |
| ENSG00000218976 | IMPDH1P9 | 5.324089388 | 0.273372833 | 0.75461481 | 0.362268046 | 0.717151734 | 0.842757071 |
| ENSG00000218980 | FTH1P15 | 2.567722566 | 0.577409842 | 1.091222067 | 0.529140547 | 0.596707956 | 0.75760906 |
| ENSG00000218996 | ARL4AP5 | 5.564419988 | -0.296760721 | 0.783252873 | -0.37888239 | 0.704775202 | 0.834970982 |
| ENSG00000219023 | ENSG0000021 | 270.8138227 | 0.208756117 | 0.114131728 | 1.829080492 | 0.067387551 | 0.15377205 |
| ENSG00000219027 | RPS3AP2 | 2.260788407 | -0.968493398 | 1.18718812 | -0.815787643 | 0.414621602 | 0.606711715 |
| ENSG00000219039 | ENSG0000021 | 2.544144382 | -0.374601685 | 1.125494802 | -0.332832888 | 0.739260431 | 0.856237216 |
| ENSG00000219085 | NPM1P37 | 10.06060253 | -0.240236636 | 0.563681286 | -0.426192321 | 0.669967712 | 0.810390684 |
| ENSG00000219102 | HNRNPA3P12 | 31.27343856 | 0.044194498 | 0.316659749 | 0.139564622 | 0.889003994 | 0.942802073 |
| ENSG00000219133 | RPL21P19 | 13.44360602 | 0.90939464 | 0.496958185 | 1.829921849 | 0.067261626 | 0.153576361 |
| ENSG00000219146 | RPS4XP8 | 6.750047055 | 0.387233623 | 0.688013005 | 0.56282893 | 0.573551379 | 0.740253664 |
| ENSG00000219149 | PPIAP87 | 4.483175855 | 1.003719728 | 0.892090047 | 1.125132749 | 0.260532785 | 0.440851376 |
| ENSG00000219163 | HMGB1P20 | 4.500482897 | -0.954889465 | 0.829065766 | -1.151765643 | 0.24941739 | 0.427402324 |
| ENSG00000219200 | RNASEK | 25.70927536 | -0.047985925 | 0.350394488 | -0.136948287 | 0.891071671 | 0.944150152 |
| ENSG00000219201 | ENSG0000021 | 292.2325647 | -0.387913055 | 0.11677171 | -3.321978029 | 0.000893817 | 0.003378509 |
| ENSG00000219222 | RPL12P47 | 2.096363271 | 0.501588147 | 1.194053593 | 0.42007172 | 0.674433061 | 0.81390205 |
| ENSG00000219294 | PIP5K1P1 | 14.46135913 | 0.595248013 | 0.485912199 | 1.225011462 | 0.220570902 | 0.390816134 |
| ENSG00000219329 | ENSG0000021 | 10.47946858 | -0.456609471 | 0.622922058 | -0.733012204 | 0.463550986 | 0.649941774 |
| ENSG00000219391 | ENSG0000021 | 7.858480719 | -0.098496682 | 0.640459619 | -0.15379062 | 0.87777483 | 0.936395015 |
| ENSG00000219395 | HSPA8P15 | 61.51229548 | 0.019521092 | 0.237033087 | 0.082355979 | 0.934363641 | 0.968469969 |
| ENSG00000219410 | ENSG0000021 | 20.21613171 | -0.915549418 | 0.40949377 | -2.235807931 | 0.025364355 | 0.067259081 |
| ENSG00000219438 | TAFA5 | 56.84004754 | -0.61400325 | 0.238195375 | -2.577729523 | 0.009945181 | 0.029578951 |
| ENSG00000219451 | RPL23P8 | 220.4130248 | 0.266161531 | 0.135938669 | 1.957953047 | 0.050235522 | 0.120305135 |

| | | | | | | | |
|---|---|---|---|---|---|---|---|
| ENSG00000219470 | ENSG0000021 | 9.894040151 | -0.026671793 | 0.552808461 | -0.048247801 | 0.961518755 | 0.982286361 |
| ENSG00000219481 | NBPF1 | 3174.430014 | -0.436145071 | 0.049950357 | -8.73157066 | 2.51E-18 | 3.79E-17 |
| ENSG00000219487 | ENSG0000021 | 19.70125337 | -0.185684782 | 0.400805907 | -0.463278556 | 0.643164715 | 0.790855411 |
| ENSG00000219507 | FTH1P8 | 894.1938387 | 0.37603223 | 0.069226313 | 5.431926273 | 5.57E-08 | 3.70E-07 |
| ENSG00000219545 | UMAD1 | 509.1503745 | -1.129099902 | 0.089907688 | -12.55843555 | 3.57E-36 | 1.26E-34 |
| ENSG00000219553 | ENSG0000021 | 12.31865744 | 0.446434222 | 0.522748607 | 0.854013223 | 0.393097663 | 0.586009536 |
| ENSG00000219565 | ZPR1P1 | 17.59808312 | 0.344355639 | 0.429386491 | 0.80197129 | 0.422569566 | 0.613489389 |
| ENSG00000219626 | FAM228B | 47.49259123 | -0.169745395 | 0.266201095 | -0.637658515 | 0.523695996 | 0.700010847 |
| ENSG00000219642 | BMPR1AP1 | 159.7508244 | -0.2012688 | 0.144090993 | -1.396817361 | 0.1624685 | 0.309715931 |
| ENSG00000219665 | ZNF433-AS1 | 24.25699758 | 0.323096483 | 0.365683861 | 0.88354045 | 0.376944338 | 0.569750485 |
| ENSG00000219682 | ENSG0000021 | 81.1931934 | 0.02624197 | 0.196939705 | 0.133248753 | 0.893996655 | 0.945823146 |
| ENSG00000219699 | ENSG0000021 | 2.663686072 | 0.716149572 | 1.106779066 | 0.647057388 | 0.517594802 | 0.695665852 |
| ENSG00000219703 | RAP1BP3 | 2.421214867 | -0.608317857 | 1.143882643 | -0.531800934 | 0.594863876 | 0.756439984 |
| ENSG00000219747 | RPL32P16 | 65.19437926 | 0.440775131 | 0.225569642 | 1.954053426 | 0.050694899 | 0.1212913 |
| ENSG00000219790 | OSTCP6 | 3.157768154 | -0.328429112 | 0.995352207 | -0.32996271 | 0.741428138 | 0.857515143 |
| ENSG00000219806 | ATP5PBP6 | 9.086533585 | 0.614811377 | 0.593228686 | 1.036381737 | 0.300024087 | 0.486743462 |
| ENSG00000219863 | RPL31P28 | 7.236131363 | 0.491063216 | 0.65847107 | 0.745762781 | 0.455810727 | 0.643117857 |
| ENSG00000219891 | ZSCAN12P1 | 66.80982703 | 0.077817315 | 0.228136042 | 0.341100487 | 0.733027935 | 0.852461767 |
| ENSG00000219928 | ENSG0000021 | 79.14573162 | -0.149312517 | 0.239508042 | -0.623413377 | 0.53301291 | 0.708463324 |
| ENSG00000219932 | RPL12P8 | 9.761193469 | -0.047180609 | 0.593060082 | -0.079554518 | 0.93659157 | 0.96955824 |
| ENSG00000219951 | ENSG0000021 | 8.448861561 | -0.143554876 | 0.597312366 | -0.240334679 | 0.810070813 | 0.899408839 |
| ENSG00000219986 | BTF3P7 | 16.42999618 | 0.572012395 | 0.433913753 | 1.318262884 | 0.187415661 | 0.345298793 |
| ENSG00000220008 | LINGO3 | 5.648858195 | 1.026145056 | 0.764193421 | 1.34278185 | 0.179342622 | 0.334128137 |
| ENSG00000220105 | ENSG0000022 | 1.75215506 | 0.612973503 | 1.318949331 | 0.464743784 | 0.64211495 | NA |
| ENSG00000220157 | HNRNPA1P12 | 12.04446032 | 0.288682037 | 0.522283536 | 0.552730494 | 0.580447969 | 0.745630889 |
| ENSG00000220161 | LINC02076 | 2.775814315 | -0.201831212 | 1.137514135 | -0.177431828 | 0.859169209 | 0.926792302 |
| ENSG00000220201 | ZGLP1 | 43.99750703 | -0.167765541 | 0.271122234 | -0.618781936 | 0.536060024 | 0.711332959 |
| ENSG00000220205 | VAMP2 | 618.406072 | 0.069692773 | 0.086668112 | 0.80413397 | 0.421319608 | 0.61233349 |
| ENSG00000220267 | ACTBP8 | 4.42109081 | -0.067029818 | 0.83907776 | -0.079885109 | 0.936328634 | 0.969525239 |
| ENSG00000220305 | HNRNPH1P1 | 1.865217485 | 0.599355046 | 1.303171281 | 0.459920392 | 0.645573363 | 0.792669964 |
| ENSG00000220323 | H2BC19P | 1875.297873 | 0.420063186 | 0.061648309 | 6.813863912 | 9.50E-12 | 8.91E-11 |
| ENSG00000220343 | PPIAP74 | 2.10953194 | -1.334949828 | 1.26217696 | -1.057656629 | 0.290212015 | 0.474685464 |
| ENSG00000220378 | KRT8P42 | 9.031692717 | 0.246406677 | 0.579611614 | 0.425123775 | 0.670746447 | 0.811118793 |

| | | | | | | | |
|---|---|---|---|---|---|---|---|
| ENSG00000220412 | PTPN11P3 | 63.21879448 | -0.179910303 | 0.221454081 | -0.81240455 | 0.416559536 | 0.608615741 |
| ENSG00000220472 | ENSG0000022 | 596.5301259 | 0.225773517 | 0.099833817 | 2.261493378 | 0.023728722 | 0.063501737 |
| ENSG00000220494 | YAP1P1 | 2.067546549 | 1.529813367 | 1.287353974 | 1.188339337 | 0.234699746 | 0.409617481 |
| ENSG00000220517 | ASS1P1 | 198.3700655 | 0.971898235 | 0.133662464 | 7.271287746 | 3.56E-13 | 3.74E-12 |
| ENSG00000220541 | PPIHP2 | 13.00652501 | 0.330997497 | 0.492510313 | 0.672062063 | 0.501544187 | 0.682733423 |
| ENSG00000220583 | RPL35P2 | 198.6411483 | 0.307495082 | 0.133017211 | 2.311693944 | 0.020794555 | 0.056561395 |
| ENSG00000220585 | DDX18P6 | 3.833446924 | -0.351836518 | 0.903805392 | -0.389283491 | 0.697066447 | 0.829782025 |
| ENSG00000220666 | RCC2P7 | 2.536448742 | -0.156174262 | 1.098943002 | -0.142113159 | 0.886990624 | 0.941798885 |
| ENSG00000220685 | ENSG0000022 | 4.092025736 | 1.56326912 | 0.921846044 | 1.695802818 | 0.089923229 | 0.194295833 |
| ENSG00000220695 | ENSG0000022 | 7.675908118 | 0.044925995 | 0.632546465 | 0.071024024 | 0.943378635 | 0.973031555 |
| ENSG00000220744 | RPL5P18 | 15.44854401 | 0.532242386 | 0.444559102 | 1.197236505 | 0.231214385 | 0.404843771 |
| ENSG00000220749 | RPL21P28 | 65.57343472 | 0.422018309 | 0.226658176 | 1.861915223 | 0.062615041 | 0.14450594 |
| ENSG00000220785 | MTMR9LP | 67.02572542 | 0.174908466 | 0.22557457 | 0.775390889 | 0.438108717 | 0.627656854 |
| ENSG00000220804 | LINC01881 | 75.15272579 | -0.522913246 | 0.206676903 | -2.530100066 | 0.011403 | 0.033360333 |
| ENSG00000220842 | RPL21P16 | 831.2390902 | 0.247626295 | 0.080172015 | 3.088687454 | 0.002010428 | 0.007062668 |
| ENSG00000220848 | RPS18P9 | 149.1408238 | 0.851151812 | 0.154447318 | 5.510952359 | 3.57E-08 | 2.42E-07 |
| ENSG00000220875 | H3C9P | 4.677280164 | -0.224031252 | 0.887659386 | -0.252384254 | 0.80074407 | 0.893871969 |
| ENSG00000220884 | MESTP1 | 118.6107508 | -0.038141628 | 0.17339769 | -0.219966181 | 0.825897493 | 0.90820615 |
| ENSG00000220924 | OSTCP4 | 39.38033622 | -0.223384588 | 0.29728296 | -0.751420758 | 0.452399474 | 0.640066851 |
| ENSG00000220925 | IGBP1-AS2 | 21.54823673 | -0.847166734 | 0.390989829 | -2.166723203 | 0.030255966 | 0.078275839 |
| ENSG00000220937 | HNRNPA1P41 | 6.037562352 | 0.205497315 | 0.699001721 | 0.293986851 | 0.768767951 | 0.873855116 |
| ENSG00000220960 | ENSG0000022 | 19.13526449 | -0.818271691 | 0.429133117 | -1.906801544 | 0.056546291 | 0.132893343 |
| ENSG00000221164 | SNORA11F | 4.973066486 | -1.184476122 | 0.850120542 | -1.393303729 | 0.163527963 | 0.311158882 |
| ENSG00000221184 | ENSG0000022 | 13.25792621 | -1.180774442 | 0.538832265 | -2.191358089 | 0.028425889 | 0.074134823 |
| ENSG00000221496 | U3 | 3.950763712 | 0.800682882 | 0.977543946 | 0.819076099 | 0.412742997 | 0.604772889 |
| ENSG00000221500 | SNORD100 | 7.64055305 | -0.506577379 | 0.657495051 | -0.770465692 | 0.441023699 | 0.63030862 |
| ENSG00000221539 | SNORD99 | 9.554750147 | 0.412961509 | 0.571966658 | 0.722002767 | 0.470292776 | 0.656115263 |
| ENSG00000221676 | RNU6ATAC | 4.552029869 | -0.952557911 | 0.829902717 | -1.147794665 | 0.25105334 | 0.42940379 |
| ENSG00000221716 | SNORA11 | 8.886910852 | -0.580066021 | 0.600219058 | -0.966423864 | 0.333832128 | 0.524818618 |
| ENSG00000221792 | MIR1282 | 101.647636 | -0.560840091 | 0.190904889 | -2.937798472 | 0.003305518 | 0.011102282 |
| ENSG00000221817 | PPP3CB-AS1 | 114.4763283 | -0.649638292 | 0.171708717 | -3.783373975 | 0.000154717 | 0.000669155 |
| ENSG00000221818 | EBF2 | 4.419251563 | 0.813860812 | 0.928998909 | 0.876062183 | 0.38099623 | 0.573306233 |
| ENSG00000221821 | C6orf226 | 97.09035663 | 0.009807617 | 0.187700853 | 0.052251319 | 0.958328442 | 0.981036836 |

| | | | | | | | |
|---|---|---|---|---|---|---|---|
| ENSG00000221823 | PPP3R1 | 1237.025077 | -0.029235729 | 0.071006282 | -0.4117344 | 0.680534108 | 0.818639352 |
| ENSG00000221829 | FANCG | 1218.289143 | -0.465639003 | 0.066901965 | -6.960019778 | 3.40E-12 | 3.33E-11 |
| ENSG00000221838 | AP4M1 | 578.3625875 | -0.226024219 | 0.082504229 | -2.73954706 | 0.006152391 | 0.019384129 |
| ENSG00000221843 | C2orf16 | 2.52277107 | 0.222905701 | 1.114708119 | 0.199967774 | 0.841505785 | 0.916713017 |
| ENSG00000221844 | DPP3P2 | 2.766248731 | -1.032003647 | 1.080767937 | -0.954879962 | 0.339638404 | 0.530711987 |
| ENSG00000221866 | PLXNA4 | 13.94751699 | 1.883823721 | 0.52857521 | 3.563965324 | 0.000365294 | 0.001478065 |
| ENSG00000221869 | CEBPD | 60.8873219 | -0.325256226 | 0.226818896 | -1.433990869 | 0.151574869 | 0.293948007 |
| ENSG00000221879 | MRPS21P3 | 3.725090954 | -0.043977943 | 0.889016281 | -0.049468096 | 0.960546262 | 0.981920343 |
| ENSG00000221883 | ARIH2OS | 23.83182848 | -0.200668589 | 0.370392539 | -0.541772761 | 0.587975058 | 0.751345134 |
| ENSG00000221886 | FAM200C | 115.1134164 | -0.076615718 | 0.164840467 | -0.464787076 | 0.642083944 | 0.790034632 |
| ENSG00000221890 | NPTXR | 3548.041944 | -0.654667802 | 0.067630627 | -9.680049343 | 3.67E-22 | 6.93E-21 |
| ENSG00000221909 | FAM200A | 199.7684867 | -0.280788314 | 0.13150513 | -2.13518905 | 0.032745571 | 0.083771986 |
| ENSG00000221914 | PPP2R2A | 1573.936001 | -0.091459622 | 0.059471675 | -1.537868622 | 0.124080741 | 0.252007329 |
| ENSG00000221916 | C19orf73 | 25.3728681 | 0.096693069 | 0.345255958 | 0.280061987 | 0.779429948 | 0.880308728 |
| ENSG00000221923 | ZNF880 | 35.24845218 | 1.012712463 | 0.315925913 | 3.205537821 | 0.001348103 | 0.004924441 |
| ENSG00000221926 | TRIM16 | 392.7211829 | -0.142621312 | 0.097557186 | -1.461925235 | 0.143761699 | 0.282160624 |
| ENSG00000221930 | DENND10P1 | 324.5901939 | -0.437956304 | 0.104624323 | -4.185989361 | 2.84E-05 | 0.000137964 |
| ENSG00000221944 | TIGD1 | 289.8527197 | 0.354198977 | 0.116652762 | 3.036353117 | 0.002394587 | 0.008295354 |
| ENSG00000221947 | XKR9 | 5.979031199 | 0.454138128 | 0.721009785 | 0.629864028 | 0.52878355 | 0.704798944 |
| ENSG00000221949 | LINC01465 | 11.84723602 | 0.402045285 | 0.518829012 | 0.774909027 | 0.438393417 | 0.627918644 |
| ENSG00000221953 | LINC02897 | 37.30911694 | 0.607316498 | 0.29745422 | 2.041714175 | 0.041179892 | 0.101555227 |
| ENSG00000221955 | SLC12A8 | 158.0988412 | 0.449923775 | 0.147400244 | 3.052395052 | 0.002270231 | 0.007899151 |
| ENSG00000221963 | APOL6 | 2.960225943 | 3.850711021 | 1.452377779 | 2.65131502 | 0.008017902 | 0.024497413 |
| ENSG00000221968 | FADS3 | 623.9824386 | 0.033189428 | 0.07963013 | 0.416794853 | 0.676828468 | 0.815935942 |
| ENSG00000221971 | TTC4P1 | 5.385718881 | 0.126452975 | 0.766169029 | 0.165045793 | 0.868907939 | 0.931785159 |
| ENSG00000221978 | CCNL2 | 5063.535328 | 0.142919171 | 0.113101823 | 1.26363278 | 0.206361862 | 0.372077428 |
| ENSG00000221983 | UBA52 | 6120.768985 | 0.245204392 | 0.044228893 | 5.543986686 | 2.96E-08 | 2.02E-07 |
| ENSG00000221988 | PPT2 | 310.7918534 | -0.34794741 | 0.110251377 | -3.155946149 | 0.001599784 | 0.005745897 |
| ENSG00000221990 | EXOC3-AS1 | 310.4044291 | 0.188347754 | 0.106950934 | 1.761066928 | 0.078227076 | 0.173568925 |
| ENSG00000221994 | ZNF630 | 239.5535154 | -0.275030071 | 0.117390224 | -2.342870308 | 0.019136034 | 0.052622946 |
| ENSG00000222005 | LINC01118 | 1.74407409 | 0.155833468 | 1.331901209 | 0.117000771 | 0.906859443 | NA |
| ENSG00000222009 | BTBD19 | 5.509985629 | 2.406240149 | 0.883530698 | 2.723436951 | 0.006460655 | 0.020213777 |
| ENSG00000222011 | FAM185A | 219.9250789 | -0.367001462 | 0.121985449 | -3.008567552 | 0.002624824 | 0.009001865 |

| | | | | | | | |
|---|---|---|---|---|---|---|---|
| ENSG00000222014 | RAB6C | 83.61625698 | -0.663438319 | 0.204775161 | -3.239837864 | 0.001195977 | 0.004413724 |
| ENSG00000222017 | ENSG0000022 | 30.508168 | 0.121337856 | 0.327450818 | 0.370552917 | 0.710970556 | 0.839190355 |
| ENSG00000222019 | URAHP | 79.79296207 | 0.946721876 | 0.205236415 | 4.61283577 | 3.97E-06 | 2.15E-05 |
| ENSG00000222020 | HDAC4-AS1 | 5.162490251 | 0.115104368 | 0.831770649 | 0.138384743 | 0.889936354 | 0.943616361 |
| ENSG00000222033 | LINC01124 | 2.037712449 | -1.211723128 | 1.300448017 | -0.931773598 | 0.351453542 | 0.542773146 |
| ENSG00000222035 | KIAA2012-AS1 | 29.62611264 | -0.751562123 | 0.32552642 | -2.308759216 | 0.020956946 | 0.056927454 |
| ENSG00000222038 | POTEJ | 1.834953575 | 0.576739108 | 1.287342752 | 0.448007423 | 0.65414784 | 0.799219825 |
| ENSG00000222041 | CYTOR | 340.8282993 | 0.672631715 | 0.105395523 | 6.381976182 | 1.75E-10 | 1.48E-09 |
| ENSG00000222043 | ENSG0000022 | 11.9230667 | 0.41559447 | 0.535211397 | 0.776505269 | 0.437450711 | 0.626920679 |
| ENSG00000222345 | SNORD19C | 6.254688076 | -0.694276881 | 0.741813536 | -0.935918323 | 0.349315243 | 0.540598507 |
| ENSG00000222489 | SNORA79B | 3.9990223 | -1.385550126 | 1.021365651 | -1.356566206 | 0.174919086 | 0.328015068 |
| ENSG00000222612 | RNU2-52P | 40.16111286 | 0.158773092 | 0.312207265 | 0.508550279 | 0.611067488 | 0.768826384 |
| ENSG00000222726 | RNU2-7P | 2.167244902 | 1.16534665 | 1.249844651 | 0.932393197 | 0.351133359 | 0.542458202 |
| ENSG00000222881 | Y_RNA | 2.780494485 | 0.749943137 | 1.139288012 | 0.658255971 | 0.510373664 | 0.690287829 |
| ENSG00000223198 | RNU2-22P | 3.777576111 | -0.277180654 | 0.896540998 | -0.309166736 | 0.757194696 | 0.866120485 |
| ENSG00000223343 | ENSG0000022 | 30.38249702 | -0.211663145 | 0.315219362 | -0.671478882 | 0.501915508 | 0.682955202 |
| ENSG00000223356 | ENSG0000022 | 2.208454088 | -0.454748564 | 1.167665126 | -0.389451183 | 0.696942416 | 0.829719188 |
| ENSG00000223361 | FTH1P10 | 1282.972313 | 0.353505928 | 0.067669668 | 5.223993811 | 1.75E-07 | 1.11E-06 |
| ENSG00000223374 | ENSG0000022 | 4.766150961 | -0.647808961 | 0.827019771 | -0.78330529 | 0.433447858 | 0.623443091 |
| ENSG00000223382 | LINC01778 | 4.972290606 | -0.551883928 | 0.790425245 | -0.698211414 | 0.485044989 | 0.668886808 |
| ENSG00000223393 | ENSG0000022 | 27.16768916 | 0.156324287 | 0.345735927 | 0.452149386 | 0.651161368 | 0.797312078 |
| ENSG00000223396 | RPS10P7 | 238.1163896 | 0.391097579 | 0.126276294 | 3.097157573 | 0.00195386 | 0.00688503 |
| ENSG00000223416 | RPS26P15 | 2.285455615 | -0.147430852 | 1.145124336 | -0.128746589 | 0.89755817 | 0.948060798 |
| ENSG00000223427 | RBM7P1 | 9.567918492 | -0.082360462 | 0.559990768 | -0.147074678 | 0.883073076 | 0.939422017 |
| ENSG00000223442 | TH2LCRR | 4.860750149 | -0.169418421 | 0.806838035 | -0.209978228 | 0.833684666 | 0.912221618 |
| ENSG00000223459 | TCAF1P1 | 111.3219863 | 0.181292585 | 0.175018156 | 1.035850162 | 0.300272052 | 0.486904705 |
| ENSG00000223460 | GAPDHP69 | 2.303942014 | -0.177369624 | 1.180027564 | -0.150309729 | 0.880520257 | 0.938252557 |
| ENSG00000223461 | ENSG0000022 | 1.660050956 | 0.922719835 | 1.525525496 | 0.604853762 | 0.545276168 | NA |
| ENSG00000223467 | CALM1P1 | 12.17859548 | -0.947327586 | 0.524280254 | -1.806910673 | 0.070776208 | 0.160055818 |
| ENSG00000223476 | VN1R42P | 2.928476539 | 2.25721142 | 1.200056527 | 1.880920914 | 0.059982678 | 0.139525718 |
| ENSG00000223478 | ZDHHC12-DT | 89.9141815 | -1.186778674 | 0.195537284 | -6.069321667 | 1.28E-09 | 1.01E-08 |
| ENSG00000223482 | NUTM2A-AS1 | 537.9517461 | -0.367046301 | 0.083268745 | -4.407972061 | 1.04E-05 | 5.38E-05 |
| ENSG00000223496 | EXOSC6 | 1188.6449 | 0.156492056 | 0.062312298 | 2.511415278 | 0.012024815 | 0.034992927 |

| | | | | | | | |
|---|---|---|---|---|---|---|---|
| ENSG00000223500 | RNF14P4 | 2.485063749 | 0.350134025 | 1.146686995 | 0.305344027 | 0.760104147 | 0.868106598 |
| ENSG00000223501 | VPS52 | 2969.193451 | -0.269295172 | 0.050560552 | -5.326191351 | 1.00E-07 | 6.50E-07 |
| ENSG00000223502 | ENSG0000022 | 8.660457582 | -0.04955842 | 0.601888843 | -0.082338159 | 0.934377811 | 0.968469969 |
| ENSG00000223508 | RPL23AP53 | 75.37752487 | 0.257880513 | 0.206861484 | 1.246633778 | 0.21253181 | 0.380501286 |
| ENSG00000223509 | WHAMMP1 | 26.78414354 | -0.016843708 | 0.354376596 | -0.047530531 | 0.962090398 | 0.982564051 |
| ENSG00000223518 | CSNK1A1P1 | 7.246062418 | 0.543046692 | 0.692768331 | 0.78387921 | 0.43311099 | 0.623153921 |
| ENSG00000223523 | ENSG0000022 | 532.8930799 | 0.326177837 | 0.084459463 | 3.861945444 | 0.000112488 | 0.000497775 |
| ENSG00000223528 | ENSG0000022 | 3.321745201 | -1.391586892 | 1.025956357 | -1.356380204 | 0.174978229 | 0.328031993 |
| ENSG00000223529 | EEF1A1P8 | 28.16670037 | 0.223748614 | 0.332061227 | 0.673817344 | 0.500427447 | 0.681707808 |
| ENSG00000223540 | RPS15AP5 | 19.12416833 | 0.159479065 | 0.413647069 | 0.385543805 | 0.699834559 | 0.831642103 |
| ENSG00000223546 | LINC00630 | 176.0960713 | -0.201765917 | 0.145153567 | -1.390016937 | 0.164523734 | 0.312716419 |
| ENSG00000223551 | TMSB4XP4 | 55.73267726 | 0.018793019 | 0.234115491 | 0.080272429 | 0.936020586 | 0.969415691 |
| ENSG00000223553 | SMPD4P1 | 1.986764405 | -1.059793715 | 1.317703304 | -0.804273399 | 0.421239097 | 0.612294062 |
| ENSG00000223559 | ENSG0000022 | 405.4832457 | 0.157634975 | 0.098030116 | 1.608026006 | 0.107829483 | 0.225201051 |
| ENSG00000223566 | TNRC18P2 | 8.283877976 | -0.187760638 | 0.644269801 | -0.291431691 | 0.77072118 | 0.875046557 |
| ENSG00000223571 | DHRSX-IT1 | 2.206964915 | 0.704757595 | 1.198621105 | 0.587973624 | 0.556549997 | 0.727106152 |
| ENSG00000223571 | DHRSX-IT1 | 2.206964915 | 0.704757595 | 1.198621105 | 0.587973624 | 0.556549997 | 0.727106152 |
| ENSG00000223572 | CKMT1A | 446.1607277 | -0.209372145 | 0.092737882 | -2.257676597 | 0.023965829 | 0.06408394 |
| ENSG00000223573 | TINCR | 382.4953661 | 0.828342627 | 0.10892093 | 7.604990408 | 2.85E-14 | 3.24E-13 |
| ENSG00000223575 | RBMX2P3 | 9.519115056 | 0.281862412 | 0.56903284 | 0.495335932 | 0.620363011 | 0.775267138 |
| ENSG00000223599 | ENSG0000022 | 2.37670778 | 0.863661605 | 1.161952618 | 0.743284701 | 0.457309332 | 0.644548788 |
| ENSG00000223612 | ENSG0000022 | 87.22190584 | 0.645294468 | 0.191937569 | 3.362001873 | 0.000773796 | 0.002957494 |
| ENSG00000223635 | NUP133-DT | 1.760763228 | -0.274060877 | 1.355810909 | -0.202137979 | 0.839808858 | NA |
| ENSG00000223642 | ENSG0000022 | 5.679160005 | -1.050402798 | 0.757003484 | -1.387579873 | 0.165265014 | 0.313819511 |
| ENSG00000223652 | PRDM6-AS1 | 141.6980407 | -0.061561534 | 0.15048243 | -0.409094501 | 0.682470311 | 0.819871514 |
| ENSG00000223653 | BCL10-AS1 | 6.009149155 | -0.114240265 | 0.71695313 | -0.15934133 | 0.87339996 | 0.934636147 |
| ENSG00000223657 | KRT8P51 | 5.310837523 | 0.116557333 | 0.786531118 | 0.148191636 | 0.882191533 | 0.939050334 |
| ENSG00000223658 | C1GALT1C1L | 28.84922793 | -0.017589202 | 0.322958913 | -0.054462663 | 0.956566555 | 0.980195505 |
| ENSG00000223668 | EEF1A1P24 | 15.05934647 | -0.238056142 | 0.467761662 | -0.50892615 | 0.61080399 | 0.768741072 |
| ENSG00000223685 | LINC00571 | 18.73677934 | 0.131887925 | 0.408852847 | 0.322580425 | 0.74701302 | 0.859750631 |
| ENSG00000223692 | DIP2A-IT1 | 1.780545975 | 0.6067217 | 1.314677949 | 0.461498347 | 0.644441107 | NA |
| ENSG00000223694 | ADH5P3 | 5.848891729 | -1.113628461 | 0.753545055 | -1.477852523 | 0.139447256 | 0.275629863 |
| ENSG00000223697 | ENSG0000022 | 12.69099006 | -0.230662781 | 0.499529379 | -0.461760189 | 0.644253303 | 0.791557537 |

| | | | | | | | |
|---|---|---|---|---|---|---|---|
| ENSG00000223703 | IGSF3P2 | 10.8675158 | -0.271735183 | 0.60759129 | -0.447233506 | 0.654706472 | 0.799489924 |
| ENSG00000223705 | NSUN5P1 | 522.0345592 | 0.594244465 | 0.10442947 | 5.690390509 | 1.27E-08 | 9.00E-08 |
| ENSG00000223711 | ENSG0000022 | 2.64253187 | -0.848244175 | 1.124535225 | -0.754306452 | 0.450665229 | 0.638834602 |
| ENSG00000223715 | LINC01208 | 11.19514986 | -0.333978824 | 0.543280047 | -0.614745242 | 0.538722977 | 0.713404438 |
| ENSG00000223718 | RPL15P11 | 4.953951042 | 0.199276511 | 0.77775602 | 0.256219824 | 0.797781102 | 0.892083767 |
| ENSG00000223722 | IFITM3P2 | 26.45544343 | -0.399282742 | 0.341863339 | -1.167960106 | 0.242822857 | 0.419138928 |
| ENSG00000223723 | TFDP1P2 | 115.0797869 | 0.09093176 | 0.170039338 | 0.534768964 | 0.592809626 | 0.754957688 |
| ENSG00000223741 | PSMD4P1 | 4.203170278 | 0.245665962 | 0.922751382 | 0.266232017 | 0.790060527 | 0.886605444 |
| ENSG00000223745 | CCDC18-AS1 | 163.8496879 | 0.123260312 | 0.17058014 | 0.722594738 | 0.469928902 | 0.655887625 |
| ENSG00000223749 | MIR503HG | 111.9220017 | 0.774445517 | 0.177452568 | 4.364239564 | 1.28E-05 | 6.49E-05 |
| ENSG00000223751 | ENSG0000022 | 31.82824941 | 0.401299015 | 0.315547346 | 1.271755316 | 0.20346007 | 0.368031383 |
| ENSG00000223764 | LINC02593 | 1169.218472 | 1.277969186 | 0.072953375 | 17.51761567 | 1.05E-68 | 9.28E-67 |
| ENSG00000223768 | LINC00205 | 1894.316903 | -0.033505559 | 0.057474534 | -0.582963564 | 0.559917828 | 0.729374442 |
| ENSG00000223773 | CD99P1 | 46.17347533 | 0.597163967 | 0.268470499 | 2.22431876 | 0.026127005 | 0.068938751 |
| ENSG00000223773 | CD99P1 | 46.17347533 | 0.597163967 | 0.268470499 | 2.22431876 | 0.026127005 | 0.068938751 |
| ENSG00000223774 | ENSG0000022 | 5.006911779 | -0.008230282 | 0.772410577 | -0.01065532 | 0.991498446 | 0.995447256 |
| ENSG00000223776 | LGALS8-AS1 | 21.59106363 | -0.865420839 | 0.388559731 | -2.227253034 | 0.025930367 | 0.06850658 |
| ENSG00000223797 | ENTPD3-AS1 | 97.04284387 | -0.642386073 | 0.185469598 | -3.463565341 | 0.000533067 | 0.002099324 |
| ENSG00000223799 | IL10RB-DT | 3.983191935 | -0.044554559 | 0.90396019 | -0.049288187 | 0.960689633 | 0.981920343 |
| ENSG00000223802 | CERS1 | 57.28472992 | 0.840075043 | 0.240969751 | 3.486226131 | 0.000489887 | 0.001941939 |
| ENSG00000223803 | RPS20P14 | 607.0074371 | 0.17534212 | 0.084589806 | 2.072851666 | 0.038186086 | 0.095486028 |
| ENSG00000223804 | ENSG0000022 | 20.29545983 | -1.124901192 | 0.402773282 | -2.792889307 | 0.005223956 | 0.01672541 |
| ENSG00000223808 | ENSG0000022 | 361.9910389 | 0.779533176 | 0.101010168 | 7.717373359 | 1.19E-14 | 1.39E-13 |
| ENSG00000223813 | PRR15-DT | 37.13230401 | -1.925646546 | 0.32401555 | -5.943068305 | 2.80E-09 | 2.14E-08 |
| ENSG00000223820 | CFL1P1 | 7.73205239 | -0.102376404 | 0.643620948 | -0.159063195 | 0.873619085 | 0.934676032 |
| ENSG00000223822 | EEF1A1P1 | 5.799334369 | 0.232242955 | 0.730088308 | 0.318102553 | 0.750407147 | 0.862139528 |
| ENSG00000223823 | LINC01342 | 2.133222657 | 0.091474321 | 1.233104719 | 0.074182119 | 0.940865473 | 0.971736581 |
| ENSG00000223825 | DAZAP2P1 | 25.63427001 | -0.141726493 | 0.346977442 | -0.40846025 | 0.682935808 | 0.820000839 |
| ENSG00000223828 | BANF1P4 | 6.231388318 | -0.022486204 | 0.688224786 | -0.032672761 | 0.973935546 | 0.988106737 |
| ENSG00000223855 | PDGFA-DT | 192.0121848 | 1.399603822 | 0.144679635 | 9.673813591 | 3.90E-22 | 7.35E-21 |
| ENSG00000223861 | ENSG0000022 | 8.046182786 | 0.249764037 | 0.633107938 | 0.394504669 | 0.693208481 | 0.826947611 |
| ENSG00000223864 | NPM1P19 | 16.76067002 | -0.300713847 | 0.468789947 | -0.641468208 | 0.52121853 | 0.698300881 |
| ENSG00000223873 | SAP18P2 | 8.602563989 | 0.246395046 | 0.592862286 | 0.415602497 | 0.677700899 | 0.816687257 |

| | | | | | | | |
|---|---|---|---|---|---|---|---|
| ENSG00000223878 | PPIAP53 | 1.945981026 | -0.1580355 | 1.219165852 | -0.129625924 | 0.896862392 | 0.947893263 |
| ENSG00000223886 | ENSG0000022 | 6.410998273 | 0.126111423 | 0.698474637 | 0.180552616 | 0.856718751 | 0.925767966 |
| ENSG00000223891 | OSER1-DT | 289.4013294 | -0.505363808 | 0.112884876 | -4.47680705 | 7.58E-06 | 3.96E-05 |
| ENSG00000223901 | ENSG0000022 | 15.70387944 | -1.788660554 | 0.482145218 | -3.709796319 | 0.000207426 | 0.000877278 |
| ENSG00000223908 | ENSG0000022 | 5.468388839 | 0.42202203 | 0.758317167 | 0.556524431 | 0.577852398 | 0.743545771 |
| ENSG00000223916 | RPL18AP9 | 19.84852866 | 1.03014978 | 0.428453551 | 2.404344127 | 0.016201514 | 0.045513437 |
| ENSG00000223935 | LGALSL-DT | 1.878386154 | -1.581977719 | 1.354797855 | -1.167685432 | 0.242933675 | 0.419298661 |
| ENSG00000223940 | KRT8P8 | 6.662529738 | -0.089998068 | 0.674577189 | -0.13341404 | 0.893865942 | 0.945728472 |
| ENSG00000223956 | LINC01767 | 2.656114104 | 1.573963882 | 1.159181143 | 1.357823919 | 0.174519563 | 0.327482927 |
| ENSG00000223959 | AFG3L1P | 1143.557634 | 0.278585758 | 0.065538133 | 4.250742988 | 2.13E-05 | 0.000105337 |
| ENSG00000223960 | CHROMR | 67.67415812 | -1.08522755 | 0.249480106 | -4.349956262 | 1.36E-05 | 6.90E-05 |
| ENSG00000223973 | PAFAH1B1P2 | 4.494444363 | 0.356346066 | 0.848840155 | 0.419803498 | 0.674629009 | 0.814054467 |
| ENSG00000223977 | ENSG0000022 | 1.857174822 | 0.046502754 | 1.271096494 | 0.036584755 | 0.970816099 | 0.98680905 |
| ENSG00000223979 | SMCR2 | 16.30180229 | -0.696441937 | 0.445770086 | -1.562334395 | 0.118209233 | 0.242728378 |
| ENSG00000223984 | HNRNPRP1 | 40.98728144 | -0.252481512 | 0.275559456 | -0.916250583 | 0.359535481 | 0.55209478 |
| ENSG00000223989 | ENSG0000022 | 2.062401939 | 0.747610297 | 1.298607535 | 0.575701493 | 0.564816966 | 0.733188162 |
| ENSG00000224003 | YES1P1 | 5.882958538 | -0.014957768 | 0.719350363 | -0.02079344 | 0.983410431 | 0.9923499 |
| ENSG00000224004 | ATP5F1CP1 | 48.37253102 | -0.832177279 | 0.257300608 | -3.234260829 | 0.00121958 | 0.004495047 |
| ENSG00000224008 | LINC01441 | 42.22359696 | -0.196501231 | 0.274455375 | -0.715967872 | 0.474011188 | 0.659498826 |
| ENSG00000224011 | RPL36AP46 | 5.609299754 | -0.629833424 | 0.776735679 | -0.810872271 | 0.417439026 | 0.609583723 |
| ENSG00000224014 | ENSG0000022 | 15.69108716 | 0.089334393 | 0.449836698 | 0.198592943 | 0.842581175 | 0.917231156 |
| ENSG00000224016 | ENSG0000022 | 8.649018156 | 4.516597905 | 1.129679102 | 3.998124685 | 6.38E-05 | 0.000293867 |
| ENSG00000224017 | LINC01449 | 4.762593319 | 1.699115115 | 0.904371491 | 1.87878005 | 0.060274532 | 0.140071276 |
| ENSG00000224023 | EDRF1-DT | 19.08161812 | -0.683347613 | 0.429299799 | -1.591772499 | 0.111435832 | 0.231217255 |
| ENSG00000224032 | EPB41L4A-AS1 | 975.4381074 | 1.294704347 | 0.074896943 | 17.28647787 | 5.95E-67 | 4.96E-65 |
| ENSG00000224040 | HMGN1P4 | 250.7155479 | -0.276958602 | 0.117565654 | -2.355778179 | 0.018483952 | 0.051019354 |
| ENSG00000224043 | CCNT2-AS1 | 136.6965901 | 0.805981369 | 0.155904952 | 5.169697025 | 2.34E-07 | 1.47E-06 |
| ENSG00000224046 | DMTF1-AS1 | 18.08119056 | -0.841648819 | 0.418164359 | -2.01272251 | 0.04414383 | 0.107741191 |
| ENSG00000224051 | CPTP | 652.9917067 | 0.060470078 | 0.092992807 | 0.650266186 | 0.515520295 | 0.6944375 |
| ENSG00000224059 | HSPA8P16 | 7.303255864 | 0.529937013 | 0.658684199 | 0.804538827 | 0.421085855 | 0.612148892 |
| ENSG00000224061 | SRSF3P4 | 3.440547483 | 0.513815399 | 0.939470286 | 0.54692033 | 0.584433468 | 0.748738063 |
| ENSG00000224063 | CALCRL-AS1 | 62.77442099 | -2.067285498 | 0.254518184 | -8.122348909 | 4.57E-16 | 5.93E-15 |
| ENSG00000224066 | ENSG0000022 | 23.59920557 | -0.044670181 | 0.400974968 | -0.111403916 | 0.911296055 | 0.955444767 |

| | | | | | | | |
|---|---|---|---|---|---|---|---|
| ENSG00000224078 | SNHG14 | 5.318782799 | -0.229829776 | 0.76072142 | -0.302120816 | 0.762559964 | 0.869742236 |
| ENSG00000224080 | UBE2FP1 | 20.40963775 | 0.020910916 | 0.387146928 | 0.054012869 | 0.956924911 | 0.98047512 |
| ENSG00000224081 | SLC44A3-AS1 | 24.82642862 | 0.604157505 | 0.352439557 | 1.714215935 | 0.086489075 | 0.188294523 |
| ENSG00000224086 | PPM1F-AS1 | 112.9215792 | -1.013136622 | 0.180329075 | -5.618265505 | 1.93E-08 | 1.35E-07 |
| ENSG00000224090 | ENSG0000022 | 1.608963538 | 2.866608479 | 1.629665598 | 1.759016379 | 0.078574727 | NA |
| ENSG00000224093 | BCAR3-AS1 | 4.660271798 | -1.009255143 | 0.830845629 | -1.214732446 | 0.224468153 | 0.396099957 |
| ENSG00000224094 | RPS24P8 | 194.9996758 | -0.20892911 | 0.132870557 | -1.57242593 | 0.115851805 | 0.238803163 |
| ENSG00000224097 | ENSG0000022 | 8.088168178 | -0.456141704 | 0.619929748 | -0.735795799 | 0.461854978 | 0.648552813 |
| ENSG00000224114 | ENSG0000022 | 55.63336508 | 0.264983538 | 0.245398133 | 1.079810733 | 0.280226471 | 0.463539661 |
| ENSG00000224116 | INHBA-AS1 | 3.296226054 | -1.069301791 | 0.983043925 | -1.087745689 | 0.276707392 | 0.459243783 |
| ENSG00000224117 | PTPN2P2 | 4.975327812 | 1.845148709 | 0.902285153 | 2.044972925 | 0.040857527 | 0.100933845 |
| ENSG00000224126 | UBE2SP2 | 191.9098442 | 0.0970549 | 0.133374083 | 0.727689354 | 0.466803769 | 0.652756992 |
| ENSG00000224141 | MIR548XHG | 180.3262064 | -0.874717438 | 0.141280181 | -6.191366913 | 5.96E-10 | 4.85E-09 |
| ENSG00000224152 | BAZ2B-AS1 | 88.89115892 | 0.423710504 | 0.198341321 | 2.136269451 | 0.032657457 | 0.083583853 |
| ENSG00000224155 | ENSG0000022 | 89.90335245 | -0.907986259 | 0.193236865 | -4.698825245 | 2.62E-06 | 1.44E-05 |
| ENSG00000224159 | HMGB1P9 | 5.024305068 | -1.397072905 | 0.815095077 | -1.713999931 | 0.086528739 | 0.188362997 |
| ENSG00000224165 | DNAJC27-AS1 | 76.05603904 | -0.381422181 | 0.201632521 | -1.891669947 | 0.05853497 | 0.136726628 |
| ENSG00000224172 | ENSG0000022 | 2.606994491 | 0.179750354 | 1.098239103 | 0.163671421 | 0.869989817 | 0.932552984 |
| ENSG00000224183 | SDHDP6 | 4.8541921 | -0.072235539 | 0.806366032 | -0.089581575 | 0.928619727 | 0.965737636 |
| ENSG00000224186 | PITX1-AS1 | 59.95362377 | -0.03408717 | 0.228050309 | -0.149472151 | 0.881181083 | 0.938366635 |
| ENSG00000224189 | HAGLR | 138.9804531 | -0.494041148 | 0.156605179 | -3.154692266 | 0.001606675 | 0.005767937 |
| ENSG00000224208 | ZCRB1P1 | 5.40016387 | 0.739167549 | 0.777808959 | 0.95032018 | 0.341949588 | 0.533056811 |
| ENSG00000224209 | LINC00466 | 1.838677282 | -0.449060575 | 1.272796557 | -0.352814102 | 0.724227811 | 0.846509142 |
| ENSG00000224216 | ENSG0000022 | 2.244898808 | 0.45127912 | 1.21360656 | 0.371849605 | 0.710004828 | 0.838611408 |
| ENSG00000224217 | PRPF4BP1 | 5.413951496 | -0.931102493 | 0.752223404 | -1.237800484 | 0.215790049 | 0.385021679 |
| ENSG00000224221 | RPS6P8 | 2.370226345 | 0.966151561 | 1.291329416 | 0.748183654 | 0.454349389 | 0.64191507 |
| ENSG00000224236 | MRRFP1 | 7.368985099 | -0.3205003 | 0.639715973 | -0.50100406 | 0.616368266 | 0.772766413 |
| ENSG00000224244 | KRT224P | 5.878742074 | -5.099664069 | 1.333842114 | -3.82328914 | 0.000131683 | 0.000575612 |
| ENSG00000224255 | PDCL3P6 | 2.252331077 | -0.149454172 | 1.143056511 | -0.130749591 | 0.895973401 | 0.947302844 |
| ENSG00000224261 | RPSAP18 | 6.548215853 | 0.401064285 | 0.679724201 | 0.590039732 | 0.555164013 | 0.726323296 |
| ENSG00000224271 | EPIC1 | 162.7113365 | 0.520167761 | 0.149415737 | 3.481345215 | 0.000498902 | 0.001974605 |
| ENSG00000224280 | ENSG0000022 | 21.04064793 | -0.416110332 | 0.410495 | -1.013679416 | 0.310735749 | 0.498725117 |
| ENSG00000224281 | SLC25A5-AS1 | 92.37605116 | 0.167461715 | 0.186315421 | 0.898807592 | 0.368755155 | 0.561255524 |

| | | | | | | | |
|---|---|---|---|---|---|---|---|
| ENSG00000224287 | MSL3P1 | 15.30641648 | 0.555702094 | 0.493676258 | 1.125640709 | 0.26031763 | 0.440584803 |
| ENSG00000224316 | GTF2IP5 | 4.354920927 | 2.005213867 | 0.959357375 | 2.090163603 | 0.036603106 | 0.092110054 |
| ENSG00000224321 | RPL12P14 | 22.88459527 | 0.519643405 | 0.375720627 | 1.383057963 | 0.166647094 | 0.315966967 |
| ENSG00000224324 | THAP5P1 | 7.450329069 | 0.572173849 | 0.653034615 | 0.876176906 | 0.380933869 | 0.57330213 |
| ENSG00000224328 | MDC1-AS1 | 18.44378638 | -0.146348943 | 0.446708248 | -0.327616389 | 0.743201729 | 0.858375172 |
| ENSG00000224334 | ENSG0000022 | 8.074911125 | 0.134595928 | 0.696934067 | 0.19312577 | 0.846860471 | 0.919794479 |
| ENSG00000224356 | ENSG0000022 | 4.837180054 | 0.096133014 | 0.790322977 | 0.12163763 | 0.90318601 | 0.951114904 |
| ENSG00000224358 | TMCO1-AS1 | 13.87835461 | 0.90221401 | 0.488083404 | 1.848483278 | 0.064532461 | 0.148289558 |
| ENSG00000224376 | ENSG0000022 | 21.06783699 | 0.228957257 | 0.388711132 | 0.589016465 | 0.555850228 | 0.726680327 |
| ENSG00000224383 | PRR29 | 11.65473206 | 0.677513041 | 0.525624507 | 1.288967756 | 0.197409296 | 0.359467317 |
| ENSG00000224389 | C4B | 7.233343264 | -0.54571923 | 0.661054446 | -0.825528415 | 0.409071657 | 0.601506558 |
| ENSG00000224401 | RPL7P57 | 29.69413968 | -0.112712124 | 0.322096339 | -0.349932955 | 0.726389014 | 0.847922043 |
| ENSG00000224407 | ENSG0000022 | 8.400095227 | 0.112451057 | 0.602873914 | 0.186525001 | 0.852033075 | 0.922573338 |
| ENSG00000224411 | HSP90AA2P | 1259.034107 | -0.127837131 | 0.06548184 | -1.952253181 | 0.050908153 | 0.121711777 |
| ENSG00000224415 | ENSG0000022 | 70.39518177 | 0.453729165 | 0.245779737 | 1.846080443 | 0.064880526 | 0.148977414 |
| ENSG00000224418 | STK24-AS1 | 11.45213994 | -0.388202838 | 0.528009578 | -0.73521931 | 0.462205941 | 0.648779556 |
| ENSG00000224420 | ADM5 | 152.2892572 | 0.148991246 | 0.146028151 | 1.02029126 | 0.307590347 | 0.495493293 |
| ENSG00000224424 | PRKAR2A-AS1 | 34.39342171 | -0.252182233 | 0.300709257 | -0.838624777 | 0.401679903 | 0.594409558 |
| ENSG00000224425 | SSBP3P2 | 9.706953037 | -0.492623287 | 0.561124612 | -0.877921368 | 0.379986392 | 0.572424996 |
| ENSG00000224426 | SLC31A1P1 | 2.214634898 | 0.432920354 | 1.186271205 | 0.364942141 | 0.715154596 | 0.841653402 |
| ENSG00000224429 | LINC00539 | 3.605112317 | -0.617377745 | 0.954674411 | -0.646689319 | 0.517833037 | 0.69574209 |
| ENSG00000224430 | MKRN5P | 3.013046575 | -1.523245386 | 1.068143632 | -1.426067938 | 0.153848732 | 0.297028272 |
| ENSG00000224437 | PIGUP1 | 27.07173694 | -0.137955288 | 0.351816965 | -0.39212233 | 0.694967829 | 0.828271093 |
| ENSG00000224448 | ENSG0000022 | 2.88210414 | 0.504040261 | 1.024303726 | 0.492080862 | 0.622662181 | 0.776597778 |
| ENSG00000224452 | RSL24D1P6 | 4.539860298 | 0.309600054 | 0.837795268 | 0.369541422 | 0.711724205 | 0.839734262 |
| ENSG00000224463 | RPL39P14 | 2.422635769 | 0.788482048 | 1.181050843 | 0.667610588 | 0.504382197 | 0.685187291 |
| ENSG00000224464 | PGAM1P6 | 4.502442138 | -0.50361156 | 0.851771244 | -0.591252127 | 0.5543515 | 0.725680405 |
| ENSG00000224470 | ATXN1L | 4050.197347 | 0.392633306 | 0.055727483 | 7.045595505 | 1.85E-12 | 1.85E-11 |
| ENSG00000224479 | CDHR18P | 18.42740676 | 0.636712294 | 0.42338668 | 1.503855278 | 0.132618634 | 0.26558459 |
| ENSG00000224481 | ENSG0000022 | 2.299728535 | 1.646575929 | 1.277025755 | 1.289383493 | 0.197264797 | 0.359318282 |
| ENSG00000224505 | HEXIM2-AS1 | 24.98341402 | 0.309121581 | 0.350801589 | 0.881186377 | 0.378216952 | 0.571034413 |
| ENSG00000224509 | MRPS9-AS2 | 1.87696871 | 1.453388234 | 1.353758211 | 1.073595138 | 0.283004178 | 0.466655825 |
| ENSG00000224511 | LINC00365 | 21.65909579 | -1.12256411 | 0.389966303 | -2.878618236 | 0.003994215 | 0.01315917 |

| | | | | | | | |
|---|---|---|---|---|---|---|---|
| ENSG00000224520 | KRT8P45 | 1002.419539 | 0.093374459 | 0.066856078 | 1.396648769 | 0.162519217 | 0.309734502 |
| ENSG00000224531 | SMIM13 | 393.6503082 | -0.634010962 | 0.097849059 | -6.47947942 | 9.20E-11 | 7.99E-10 |
| ENSG00000224533 | TMLHE-AS1 | 1.978145096 | -0.696096698 | 1.258662992 | -0.553044542 | 0.58023291 | 0.74560514 |
| ENSG00000224536 | CSRP1-AS1 | 153.4517324 | 0.481640567 | 0.155858746 | 3.090237653 | 0.001999964 | 0.007029137 |
| ENSG00000224540 | TFDP1P1 | 2.568671892 | 0.615701559 | 1.150860983 | 0.534992122 | 0.592655305 | 0.754844967 |
| ENSG00000224543 | SNRPGP15 | 60.73683506 | -0.453677317 | 0.226491922 | -2.003061798 | 0.045170656 | 0.109780151 |
| ENSG00000224546 | EIF4BP3 | 1412.59628 | 0.185928754 | 0.06236715 | 2.981196894 | 0.002871241 | 0.009773894 |
| ENSG00000224549 | ENSG0000022 | 5.369489297 | 0.643727092 | 0.773111383 | 0.832644695 | 0.405045144 | 0.597811154 |
| ENSG00000224550 | NDUFA4P1 | 912.2616439 | -1.26295865 | 0.074650984 | -16.91817815 | 3.30E-64 | 2.63E-62 |
| ENSG00000224555 | ENSG0000022 | 18.71739016 | -0.322288362 | 0.414205658 | -0.778087784 | 0.436517264 | 0.626326372 |
| ENSG00000224559 | LINC01087 | 57.90253058 | 0.785099101 | 0.240484709 | 3.26465289 | 0.001095983 | 0.004074194 |
| ENSG00000224565 | LINC01754 | 12.01850996 | -1.068398023 | 0.522048726 | -2.046548472 | 0.040702439 | 0.100572376 |
| ENSG00000224566 | CIAO2AP2 | 4.034716625 | -0.5217457 | 0.913802533 | -0.570961101 | 0.568026011 | 0.735690408 |
| ENSG00000224574 | COL18A1-AS2 | 3.990421186 | -3.530344991 | 1.262203714 | -2.796969263 | 0.005158444 | 0.016564209 |
| ENSG00000224577 | LINC01117 | 30.93890569 | -0.320499773 | 0.319634512 | -1.002707034 | 0.316002235 | 0.504745038 |
| ENSG00000224578 | HNRNPA1L3 | 3960.248852 | 0.368758641 | 0.053108252 | 6.943528046 | 3.82E-12 | 3.72E-11 |
| ENSG00000224594 | RPL29P19 | 1.935088876 | -0.13968511 | 1.251272971 | -0.111634402 | 0.911113293 | 0.955340477 |
| ENSG00000224596 | ZMIZ1-AS1 | 4.013599304 | 1.440338941 | 0.978364905 | 1.472189909 | 0.140969602 | 0.278036511 |
| ENSG00000224597 | SVIL-AS1 | 349.1025556 | -1.14901551 | 0.106237823 | -10.8155032 | 2.91E-27 | 7.29E-26 |
| ENSG00000224609 | FGGY-DT | 2.246161409 | -0.556326972 | 1.185309964 | -0.469351468 | 0.638818432 | 0.787290808 |
| ENSG00000224614 | TNK2-AS1 | 14.69952425 | -0.280059867 | 0.450261314 | -0.621994069 | 0.533945768 | 0.709453571 |
| ENSG00000224616 | RTCA-AS1 | 40.69408793 | 0.848518948 | 0.280056494 | 3.029813505 | 0.002447048 | 0.008454108 |
| ENSG00000224621 | ENSG0000022 | 2.546918763 | -1.188373294 | 1.111895099 | -1.068781844 | 0.285167982 | 0.469107664 |
| ENSG00000224628 | ENSG0000022 | 136.3749975 | 0.008351413 | 0.159280396 | 0.052432146 | 0.95818436 | 0.980976892 |
| ENSG00000224631 | RPS27AP16 | 3020.429085 | 0.155004953 | 0.058710636 | 2.640151151 | 0.008286906 | 0.025202389 |
| ENSG00000224635 | ENSG0000022 | 17.31154399 | -0.545901441 | 0.437860705 | -1.24674682 | 0.212490345 | 0.380495553 |
| ENSG00000224645 | ENSG0000022 | 5.091529836 | 0.50042988 | 0.804916428 | 0.621716569 | 0.534128255 | 0.709572745 |
| ENSG00000224646 | RPS28P5 | 15.10092691 | 0.415676359 | 0.505474011 | 0.822349617 | 0.410877945 | 0.603369294 |
| ENSG00000224652 | LINC00885 | 43.83983522 | 1.242710825 | 0.296236543 | 4.194995031 | 2.73E-05 | 0.000132905 |
| ENSG00000224660 | SH3BP5-AS1 | 557.9581422 | -0.030340271 | 0.101401809 | -0.29920838 | 0.764781056 | 0.871149383 |
| ENSG00000224668 | IPO8P1 | 9.334934014 | 0.247615324 | 0.604435449 | 0.409663801 | 0.682052587 | 0.819563478 |
| ENSG00000224669 | ENSG0000022 | 2.004674158 | -0.227531308 | 1.224484325 | -0.185818065 | 0.852587437 | 0.923046535 |
| ENSG00000224680 | PLA2G12AP1 | 52.81050891 | -0.040597201 | 0.249536131 | -0.162690674 | 0.87076199 | 0.932944751 |

| | | | | | | | |
|---|---|---|---|---|---|---|---|
| ENSG00000224687 | RASAL2-AS1 | 4.248924266 | -0.372365458 | 0.843858283 | -0.441265394 | 0.659020876 | 0.802779253 |
| ENSG00000224688 | E2F6P2 | 2.091142929 | 0.105339699 | 1.294135462 | 0.081397738 | 0.935125648 | 0.969062396 |
| ENSG00000224699 | LAMTOR5-AS1 | 81.77483362 | -0.311963487 | 0.197009536 | -1.583494354 | 0.113308832 | 0.234509619 |
| ENSG00000224712 | NPIPA3 | 2.696875202 | 0.130096844 | 1.065633381 | 0.122084055 | 0.90283245 | 0.950907167 |
| ENSG00000224713 | ENSG0000022 | 6.281567211 | -0.732671665 | 0.718680656 | -1.01946763 | 0.307981013 | 0.495934609 |
| ENSG00000224721 | FLVCR2-AS1 | 2.05605915 | -0.825052758 | 1.285182274 | -0.641973341 | 0.520890493 | 0.698050467 |
| ENSG00000224722 | ENSG0000022 | 59.76927121 | -0.235240733 | 0.227389777 | -1.03452642 | 0.300890132 | 0.487631199 |
| ENSG00000224725 | CEP57L1P1 | 14.47138961 | -0.334288738 | 0.470081691 | -0.711129031 | 0.477004281 | 0.661968041 |
| ENSG00000224727 | FCF1P7 | 2.700187921 | -1.19936961 | 1.215870586 | -0.986428674 | 0.3239228 | 0.514063579 |
| ENSG00000224728 | IMPDH1P8 | 21.40312726 | 0.091622811 | 0.378718725 | 0.241928389 | 0.808835652 | 0.898836887 |
| ENSG00000224729 | PCOLCE-AS1 | 23.94595018 | -0.532438387 | 0.363194188 | -1.465988181 | 0.142651507 | 0.280389475 |
| ENSG00000224738 | ENSG0000022 | 458.0882731 | -0.207859203 | 0.091523014 | -2.271114049 | 0.023140075 | 0.062078507 |
| ENSG00000224751 | SHMT1P1 | 2.776748496 | 0.704629547 | 1.080125921 | 0.652358705 | 0.514169797 | 0.693174374 |
| ENSG00000224758 | LINC01167 | 11.10912518 | -0.088277616 | 0.517215366 | -0.170678641 | 0.864476458 | 0.929771201 |
| ENSG00000224763 | FDPSP8 | 1.924823145 | -0.051659411 | 1.326222613 | -0.038952293 | 0.968928425 | 0.985925152 |
| ENSG00000224769 | MUC20P1 | 16.99705578 | 0.714157297 | 0.434382318 | 1.644075433 | 0.100160622 | 0.212237638 |
| ENSG00000224773 | HSPA8P7 | 36.39924246 | 0.358340007 | 0.298821757 | 1.19917643 | 0.23045935 | 0.403814352 |
| ENSG00000224775 | BRAFP1 | 4.843277668 | -0.471708919 | 0.789927154 | -0.597154961 | 0.550403926 | 0.722615811 |
| ENSG00000224781 | EIF4A2P4 | 38.51280173 | 0.495421276 | 0.30449261 | 1.627038754 | 0.103728867 | 0.218346505 |
| ENSG00000224785 | ENSG0000022 | 21.66876977 | 0.285232163 | 0.385639477 | 0.739634244 | 0.459521957 | 0.6464805 |
| ENSG00000224789 | ENSG0000022 | 1.664712848 | -0.769799577 | 1.365581469 | -0.5637156 | 0.572947699 | NA |
| ENSG00000224790 | HLCS-AS1 | 79.45061731 | 0.192097662 | 0.199948257 | 0.960736863 | 0.336684491 | 0.527888882 |
| ENSG00000224791 | KRT18P39 | 38.35139534 | 0.369078447 | 0.311427327 | 1.185119016 | 0.235970393 | 0.411053293 |
| ENSG00000224805 | LINC00853 | 23.16773787 | 0.049233144 | 0.363694231 | 0.135369605 | 0.892319655 | 0.944983167 |
| ENSG00000224806 | ARL5AP4 | 1.719990989 | 1.284343266 | 1.388086563 | 0.925261651 | 0.354829821 | NA |
| ENSG00000224820 | BTF3P4 | 32.05101808 | -0.019888125 | 0.33042006 | -0.06019043 | 0.952003968 | 0.977646532 |
| ENSG00000224829 | TOMM20P1 | 7.131195898 | -0.499564426 | 0.670600419 | -0.74495096 | 0.456301366 | 0.643478245 |
| ENSG00000224831 | TMEM183BP | 2151.32922 | 0.402758352 | 0.0573357 | 7.024564974 | 2.15E-12 | 2.14E-11 |
| ENSG00000224834 | BTF3P9 | 8.077030077 | 0.496051849 | 0.657571655 | 0.75436927 | 0.450627519 | 0.63882062 |
| ENSG00000224837 | GCSHP5 | 652.437664 | 0.288138506 | 0.079317709 | 3.632713406 | 0.000280456 | 0.001159638 |
| ENSG00000224839 | RPL12P17 | 8.516952786 | 0.507228824 | 0.626246727 | 0.809950459 | 0.417968649 | 0.609975241 |
| ENSG00000224842 | LMX1B-DT | 25.05870472 | 1.453287699 | 0.371955734 | 3.90715229 | 9.34E-05 | 0.000417943 |
| ENSG00000224843 | LINC00240 | 17.26269079 | -0.888540295 | 0.43116604 | -2.060784505 | 0.039323603 | 0.097827631 |

| | | | | | | | |
|---|---|---|---|---|---|---|---|
| ENSG00000224848 | ENSG0000022 | 3.702010631 | -0.211793592 | 0.904118974 | -0.234254118 | 0.814787704 | 0.901912272 |
| ENSG00000224855 | OPA1-AS1 | 4.424987638 | 0.023381995 | 0.890466403 | 0.026258143 | 0.97905144 | 0.990493497 |
| ENSG00000224856 | RPL35AP36 | 12.69500744 | 0.193139648 | 0.51779767 | 0.373002158 | 0.709146838 | 0.837943158 |
| ENSG00000224858 | RPL29P11 | 213.4603434 | -0.015445946 | 0.147160547 | -0.104959828 | 0.916407685 | 0.958744575 |
| ENSG00000224861 | YBX1P1 | 1875.279354 | -0.020124179 | 0.061247112 | -0.328573514 | 0.74247807 | 0.857884906 |
| ENSG00000224870 | MRPL20-AS1 | 801.3226372 | 0.832832883 | 0.080973101 | 10.28530302 | 8.21E-25 | 1.81E-23 |
| ENSG00000224877 | NDUFAF8 | 652.2538797 | 0.35946963 | 0.082870412 | 4.337731886 | 1.44E-05 | 7.27E-05 |
| ENSG00000224885 | EIPR1-IT1 | 3.411180076 | 0.854311215 | 1.024061879 | 0.834237885 | 0.404146947 | 0.597020475 |
| ENSG00000224886 | ENSG0000022 | 24.44933536 | -0.571240214 | 0.376863236 | -1.515775909 | 0.129576026 | 0.26059236 |
| ENSG00000224887 | NFYCP2 | 2.748844302 | 1.052626514 | 1.077780218 | 0.976661565 | 0.328736729 | 0.519153282 |
| ENSG00000224888 | ENSG0000022 | 19.70782771 | 0.691089546 | 0.424907426 | 1.626447325 | 0.10385453 | 0.218545171 |
| ENSG00000224891 | ARL14EPP1 | 1.706792576 | -0.77586848 | 1.327128211 | -0.584622099 | 0.558801845 | NA |
| ENSG00000224892 | RPS4XP16 | 28.00753205 | 0.378878397 | 0.353726403 | 1.071105787 | 0.284121867 | 0.46795449 |
| ENSG00000224897 | POT1-AS1 | 28.39751641 | -0.013644202 | 0.330646086 | -0.041265275 | 0.967084416 | 0.985295635 |
| ENSG00000224899 | LINC02830 | 73.39938012 | 0.342217152 | 0.208385806 | 1.642228707 | 0.100542615 | 0.212870106 |
| ENSG00000224903 | RNF32-AS1 | 9.848364799 | 1.183854782 | 0.577213248 | 2.050983384 | 0.040268565 | 0.099657629 |
| ENSG00000224905 | ENSG0000022 | 2.454226872 | -0.778572239 | 1.138638018 | -0.683775024 | 0.49411724 | 0.676647708 |
| ENSG00000224906 | SRP14P1 | 5.984168346 | -0.187725504 | 0.709778285 | -0.26448471 | 0.791406445 | 0.887707081 |
| ENSG00000224908 | TIMM8BP2 | 4.877739913 | -0.731031012 | 0.864891427 | -0.845228649 | 0.397983189 | 0.590997823 |
| ENSG00000224914 | LINC00863 | 150.8757842 | -0.374880759 | 0.14796875 | -2.533513051 | 0.011292553 | 0.03307177 |
| ENSG00000224928 | KRT8P30 | 12.89106048 | -0.260259334 | 0.52455761 | -0.496150144 | 0.619788483 | 0.77486227 |
| ENSG00000224934 | GOT1-DT | 53.79126973 | 0.063317651 | 0.264756895 | 0.239153925 | 0.810986228 | 0.899962465 |
| ENSG00000224936 | SUCLA2P1 | 14.25523582 | 0.250982762 | 0.46292839 | 0.542163254 | 0.587706047 | 0.751132537 |
| ENSG00000224939 | LINC00184 | 4.71562913 | 1.55194204 | 0.864435644 | 1.79532398 | 0.072602097 | 0.163588262 |
| ENSG00000224940 | PRRT4 | 40.73939926 | -0.702013825 | 0.285850271 | -2.455879512 | 0.014054027 | 0.040154363 |
| ENSG00000224945 | ENSG0000022 | 22.19697954 | -0.615017535 | 0.37125537 | -1.656588925 | 0.097602613 | 0.207873358 |
| ENSG00000224950 | ENSG0000022 | 15.24123352 | 0.487726886 | 0.465518946 | 1.04770577 | 0.294774189 | 0.480662112 |
| ENSG00000224953 | SRIP3 | 2.921308096 | 1.374967484 | 1.150269599 | 1.195343671 | 0.231952784 | 0.405904102 |
| ENSG00000224969 | ENSG0000022 | 1.909912665 | -3.441809484 | 1.652719864 | -2.082512324 | 0.037295698 | 0.093575776 |
| ENSG00000224971 | SUMO2P3 | 10.7639933 | -0.085580462 | 0.533376931 | -0.16045025 | 0.872526408 | 0.934137108 |
| ENSG00000224975 | INE1 | 63.30872464 | -0.169616191 | 0.219814423 | -0.771633582 | 0.440331479 | 0.629711622 |
| ENSG00000224976 | PARP4P2 | 5.336922175 | -0.534701438 | 0.799742844 | -0.668591714 | 0.50375596 | 0.684769004 |
| ENSG00000224977 | RC3H1-DT | 2.008409006 | -0.668128227 | 1.216577932 | -0.549186541 | 0.58287744 | 0.747298772 |

| | | | | | | | |
|---|---|---|---|---|---|---|---|
| ENSG00000224984 | ENSG0000022 | 43.75222704 | 1.920670762 | 0.297653806 | 6.452700155 | 1.10E-10 | 9.46E-10 |
| ENSG00000224985 | ENSG0000022 | 18.89548811 | -0.198169351 | 0.405495763 | -0.488708807 | 0.625047868 | 0.778425521 |
| ENSG00000224986 | PPP1R8P1 | 2.145102439 | 0.025433591 | 1.243670202 | 0.020450431 | 0.983684054 | 0.99237927 |
| ENSG00000224988 | ENSG0000022 | 1.666574702 | -1.389132866 | 1.396164003 | -0.994963961 | 0.319753793 | NA |
| ENSG00000224993 | RPL29P12 | 17.38996012 | 0.122197235 | 0.454845301 | 0.268656694 | 0.78819388 | 0.885432696 |
| ENSG00000225008 | TWF1P2 | 9.426588824 | 0.072319734 | 0.570196184 | 0.126833072 | 0.899072519 | 0.949048573 |
| ENSG00000225014 | KCTD9P1 | 3.673042745 | -0.036399546 | 0.958036126 | -0.037993917 | 0.969692532 | 0.986463267 |
| ENSG00000225022 | UBE2D3P1 | 528.9517851 | -0.042165725 | 0.087488628 | -0.481956634 | 0.629836753 | 0.781462227 |
| ENSG00000225026 | PFDN1P1 | 6.954320761 | -0.453161477 | 0.68811791 | -0.658552074 | 0.510183447 | 0.690152629 |
| ENSG00000225031 | EIF4BP7 | 2214.206593 | 0.166972992 | 0.051398182 | 3.248616707 | 0.001159676 | 0.004294265 |
| ENSG00000225032 | ENSG0000022 | 60.02277891 | 0.276141598 | 0.248830107 | 1.109759591 | 0.267102637 | 0.448265015 |
| ENSG00000225039 | LINC01058 | 1.802789504 | 1.236999313 | 1.361512961 | 0.908547585 | 0.363588985 | 0.556345471 |
| ENSG00000225043 | RPL18AP2 | 1.79761861 | 1.198707335 | 1.468378383 | 0.816347713 | 0.414301293 | 0.606477249 |
| ENSG00000225051 | HMGB3P22 | 11.16632168 | -0.339890128 | 0.520399645 | -0.653132897 | 0.513670606 | 0.692827122 |
| ENSG00000225053 | PRPF38AP1 | 3.328012256 | -0.174280086 | 0.970553112 | -0.179567798 | 0.857491887 | 0.926087485 |
| ENSG00000225057 | ENSG0000022 | 4.732773737 | 0.213992309 | 0.859295312 | 0.249032325 | 0.803335778 | 0.89531017 |
| ENSG00000225062 | CATIP-AS1 | 7.352390297 | 0.550098273 | 0.66295216 | 0.829770694 | 0.406668443 | 0.59947141 |
| ENSG00000225067 | RPL23AP2 | 42.2844143 | 0.340458701 | 0.274019154 | 1.24246315 | 0.214065728 | 0.382510345 |
| ENSG00000225071 | RPS26P58 | 5.817723381 | -0.273652544 | 0.716401697 | -0.381981988 | 0.702474725 | 0.83327836 |
| ENSG00000225075 | ENSG0000022 | 2.970036124 | 0.627296856 | 1.046168151 | 0.599613795 | 0.548763651 | 0.721494622 |
| ENSG00000225077 | ICMT-DT | 24.75072956 | -0.975823816 | 0.393763311 | -2.478198927 | 0.01320475 | 0.037978557 |
| ENSG00000225082 | DAP3P1 | 1.991931296 | 0.770966977 | 1.277219554 | 0.603629169 | 0.546090216 | 0.719092449 |
| ENSG00000225083 | GRTP1-AS1 | 4.87787656 | -0.316059101 | 0.793301594 | -0.398409764 | 0.690328162 | 0.825271147 |
| ENSG00000225087 | ENSG0000022 | 5.858385666 | 0.637754617 | 0.724185684 | 0.880650682 | 0.378506918 | 0.57122288 |
| ENSG00000225092 | ENSG0000022 | 1.837764754 | 0.103774166 | 1.306392717 | 0.079435659 | 0.936686107 | 0.96961235 |
| ENSG00000225093 | RPL3P7 | 183.5692191 | 0.192404588 | 0.146599298 | 1.31245232 | 0.189367564 | 0.348223801 |
| ENSG00000225094 | SETP20 | 8.101388872 | 0.214280729 | 0.616189867 | 0.34775114 | 0.728027087 | 0.849088048 |
| ENSG00000225096 | ENSG0000022 | 6.864471523 | -1.599267113 | 0.702451845 | -2.276692879 | 0.022804572 | 0.061278773 |
| ENSG00000225099 | ATP6V1E1P1 | 36.90318112 | -0.579605261 | 0.314327486 | -1.843953476 | 0.065189921 | 0.149570049 |
| ENSG00000225117 | ARSDP1 | 1.624506115 | -0.724127987 | 1.493110262 | -0.48497958 | 0.627690831 | NA |
| ENSG00000225125 | RANP4 | 8.207192302 | 0.462345962 | 0.616608415 | 0.749821039 | 0.453362495 | 0.641034002 |
| ENSG00000225131 | PSME2P2 | 193.9833567 | -0.3684953 | 0.131333718 | -2.805793565 | 0.005019284 | 0.016162601 |
| ENSG00000225137 | DYNC1I2P1 | 530.3980296 | -0.31276081 | 0.092220895 | -3.391431102 | 0.000695286 | 0.002682454 |

| | | | | | | | |
|---|---|---|---|---|---|---|---|
| ENSG00000225138 | SLC9A3-AS1 | 2570.543107 | 0.43886825 | 0.063127893 | 6.952049676 | 3.60E-12 | 3.52E-11 |
| ENSG00000225151 | GOLGA2P7 | 20.53924226 | 0.437264792 | 0.391147635 | 1.117902175 | 0.263608776 | 0.444253991 |
| ENSG00000225155 | TOMM22P5 | 1.867128786 | 0.519931971 | 1.344357185 | 0.386751361 | 0.698940289 | 0.831090713 |
| ENSG00000225159 | NPM1P39 | 770.9082386 | -0.032722304 | 0.077481115 | -0.422326194 | 0.672786942 | 0.812686664 |
| ENSG00000225165 | RTRAFP2 | 2.507661355 | -1.118918466 | 1.20054165 | -0.932011369 | 0.35133065 | 0.542655879 |
| ENSG00000225177 | NHSL1-AS1 | 94.59963229 | 0.48263515 | 0.18530066 | 2.604605672 | 0.009198005 | 0.027657916 |
| ENSG00000225190 | PLEKHM1 | 791.9761783 | 0.147698219 | 0.081494935 | 1.812360732 | 0.069930476 | 0.158455662 |
| ENSG00000225193 | RPS12P26 | 6.924964494 | -0.157029378 | 0.673649972 | -0.23310233 | 0.815681946 | 0.902352305 |
| ENSG00000225200 | ENSG0000022 | 644.3107866 | 0.154960256 | 0.08523685 | 1.817996042 | 0.069064738 | 0.156928 |
| ENSG00000225205 | PDK1-AS1 | 4.123714552 | 1.55967548 | 0.92896884 | 1.678931965 | 0.093165304 | 0.200038569 |
| ENSG00000225210 | DUXAP9 | 94.31009069 | 0.242015961 | 0.181854867 | 1.330819263 | 0.183248488 | 0.339365489 |
| ENSG00000225215 | SMARCE1P1 | 11.10589124 | -0.094048457 | 0.536261502 | -0.175377976 | 0.860782645 | 0.927524107 |
| ENSG00000225218 | ENSG0000022 | 5.643276313 | 0.755452801 | 0.746181446 | 1.012425068 | 0.311334857 | 0.499442072 |
| ENSG00000225224 | RPS27AP12 | 11.41513935 | -0.136195083 | 0.523100272 | -0.260361331 | 0.794585068 | 0.89001146 |
| ENSG00000225235 | INTS6L-AS1 | 3.032954504 | -0.863397265 | 1.017793298 | -0.848303154 | 0.396269162 | 0.58921536 |
| ENSG00000225255 | PSLNR | 8.0904633 | -0.143323981 | 0.61260281 | -0.233959066 | 0.815016757 | 0.902078762 |
| ENSG00000225259 | ST13P6 | 660.1146088 | 0.268361462 | 0.077235236 | 3.474598854 | 0.000511618 | 0.002020131 |
| ENSG00000225264 | ZNRF2P2 | 10.59161563 | -0.626620273 | 0.587139496 | -1.067242585 | 0.285862307 | 0.469663483 |
| ENSG00000225265 | TAF1A-AS1 | 39.8105984 | -0.481556142 | 0.285912492 | -1.684278073 | 0.092127941 | 0.198237967 |
| ENSG00000225282 | ENSG0000022 | 1.910146699 | 0.017617566 | 1.276691871 | 0.013799387 | 0.988990031 | 0.994147115 |
| ENSG00000225285 | LINC01770 | 11.90023981 | 0.767583542 | 0.523625775 | 1.465900989 | 0.142675262 | 0.280412142 |
| ENSG00000225294 | OSTCP2 | 2.578041749 | -0.481236385 | 1.104225499 | -0.435813504 | 0.662972045 | 0.805368498 |
| ENSG00000225308 | ASS1P11 | 16.21977787 | 0.972765573 | 0.450884168 | 2.157462252 | 0.030969665 | 0.079861075 |
| ENSG00000225329 | LHFPL3-AS2 | 6.626183359 | -0.045717455 | 0.680970597 | -0.067135725 | 0.946473653 | 0.974904836 |
| ENSG00000225330 | ENSG0000022 | 7.647384966 | 0.828518746 | 0.643499773 | 1.287519874 | 0.197913147 | 0.360270401 |
| ENSG00000225333 | ENSG0000022 | 2.008909919 | -1.172261444 | 1.343903417 | -0.872281021 | 0.383055089 | 0.575532096 |
| ENSG00000225335 | ENSG0000022 | 40.56028584 | -0.312469072 | 0.276233334 | -1.131178005 | 0.257980178 | 0.437629614 |
| ENSG00000225338 | RPL23AP18 | 23.33156826 | 0.336067462 | 0.362071522 | 0.928179771 | 0.353314326 | 0.544876009 |
| ENSG00000225339 | ENSG0000022 | 223.5828426 | 0.991465618 | 0.125938468 | 7.872619347 | 3.47E-15 | 4.23E-14 |
| ENSG00000225345 | SNX18P3 | 48.59727399 | -0.713558049 | 0.256883121 | -2.777753734 | 0.005473609 | 0.017440874 |
| ENSG00000225347 | SLC25A5P8 | 3.81119851 | 0.20043901 | 0.888261201 | 0.225653231 | 0.821471136 | 0.905363372 |
| ENSG00000225355 | ARL6IP1P2 | 29.35732794 | 0.16197226 | 0.349098483 | 0.463972971 | 0.642667111 | 0.790455392 |
| ENSG00000225356 | ENSG0000022 | 394.6955325 | -0.190399763 | 0.099873 | -1.906418776 | 0.056595895 | 0.132955435 |

| | | | | | | | |
|---|---|---|---|---|---|---|---|
| ENSG00000225357 | RPF2P1 | 15.78572152 | -0.26900526 | 0.444971895 | -0.604544385 | 0.54548177 | 0.718666765 |
| ENSG00000225361 | PPP1R26-AS1 | 150.9227435 | 0.13697004 | 0.151107923 | 0.906438506 | 0.3647038 | 0.557456367 |
| ENSG00000225362 | CT62 | 4.067520507 | 1.447329401 | 0.928409975 | 1.558933488 | 0.119012123 | 0.243958369 |
| ENSG00000225377 | NRSN2-AS1 | 177.409667 | -0.47153934 | 0.140378667 | -3.35905269 | 0.000782102 | 0.002986253 |
| ENSG00000225392 | MRPL36P1 | 3.50251591 | -0.364512895 | 0.981007148 | -0.371570071 | 0.710212976 | 0.83868453 |
| ENSG00000225402 | ENSG0000022 | 6.069239702 | -0.106293264 | 0.701492677 | -0.151524409 | 0.879562057 | 0.937615842 |
| ENSG00000225405 | RPS15AP17 | 9.254930654 | 0.398305793 | 0.592122972 | 0.67267411 | 0.501154642 | 0.682243635 |
| ENSG00000225406 | ELF2P4 | 4.609667097 | 0.343557215 | 0.820594538 | 0.418668659 | 0.675458304 | 0.814669479 |
| ENSG00000225408 | ENSG0000022 | 2.084600905 | -0.718325486 | 1.221447231 | -0.588093753 | 0.556469366 | 0.727042218 |
| ENSG00000225411 | ENSG0000022 | 11.12474492 | -1.814715032 | 0.575951133 | -3.150814241 | 0.00162816 | 0.005839586 |
| ENSG00000225416 | RPL36AP28 | 23.77302783 | -0.21884529 | 0.383800431 | -0.570205952 | 0.56853802 | 0.736037682 |
| ENSG00000225420 | ENSG0000022 | 9.341920994 | -0.385750436 | 0.577049397 | -0.668487721 | 0.503822318 | 0.684818627 |
| ENSG00000225422 | RBMS1P1 | 135.5958004 | 0.479412905 | 0.158793215 | 3.019101943 | 0.002535252 | 0.00872075 |
| ENSG00000225423 | TNPO1P1 | 4.880166797 | 0.855772298 | 0.834157123 | 1.025912594 | 0.304932786 | 0.492364698 |
| ENSG00000225431 | LINC01671 | 92.8259394 | 1.235775159 | 0.199875035 | 6.182738932 | 6.30E-10 | 5.11E-09 |
| ENSG00000225434 | LINC01504 | 6.751973502 | 0.019683482 | 0.664652772 | 0.029614685 | 0.976374354 | 0.989268481 |
| ENSG00000225439 | BOLA3-DT | 13.57305693 | -0.976143723 | 0.513969199 | -1.89922611 | 0.057534753 | 0.134815857 |
| ENSG00000225447 | RPS15AP10 | 2.632661101 | 0.66633964 | 1.210958015 | 0.550258252 | 0.582142254 | 0.746825138 |
| ENSG00000225449 | RAB6C-AS1 | 36.13359507 | -1.155487576 | 0.331577786 | -3.484816006 | 0.000492476 | 0.001950853 |
| ENSG00000225450 | ENSG0000022 | 12.02170386 | 0.184435334 | 0.518655609 | 0.355602698 | 0.722138123 | 0.845601795 |
| ENSG00000225455 | TPI1P4 | 3.251880947 | 0.398984146 | 0.962032449 | 0.414730445 | 0.678339242 | 0.816898641 |
| ENSG00000225462 | FDPSP1 | 17.19400885 | -0.393251372 | 0.435507545 | -0.902972581 | 0.366540447 | 0.559108841 |
| ENSG00000225465 | RFPL1S | 1.913421993 | -1.649766241 | 1.355067714 | -1.21747882 | 0.22342209 | 0.394709492 |
| ENSG00000225470 | JPX | 838.9441587 | 0.235442253 | 0.071520919 | 3.291935525 | 0.000995004 | 0.003725379 |
| ENSG00000225475 | ENSG0000022 | 5.665448112 | 0.435753441 | 0.747829222 | 0.582691112 | 0.560101257 | 0.729502234 |
| ENSG00000225484 | NUTM2B-AS1 | 403.1786753 | -0.491764585 | 0.097208699 | -5.0588537 | 4.22E-07 | 2.57E-06 |
| ENSG00000225486 | ENSG0000022 | 24.46240296 | 0.411660623 | 0.375581884 | 1.096060915 | 0.273052118 | 0.455038495 |
| ENSG00000225489 | ENSG0000022 | 41.47926591 | -1.002496856 | 0.279506242 | -3.586670727 | 0.000334927 | 0.001365663 |
| ENSG00000225496 | RTRAFP1 | 14.08265099 | -0.954389267 | 0.523502232 | -1.823085383 | 0.06829046 | 0.155495108 |
| ENSG00000225506 | CYP4A22-AS1 | 22.88651341 | -0.829229129 | 0.375998363 | -2.205406222 | 0.027425602 | 0.071928796 |
| ENSG00000225511 | LINC00475 | 9.361239627 | -0.00087732 | 0.586659448 | -0.001495451 | 0.998806803 | 0.999329557 |
| ENSG00000225518 | LINC01703 | 45.19106473 | 0.85011255 | 0.270373193 | 3.144219071 | 0.001665307 | 0.005948637 |
| ENSG00000225526 | MKRN2OS | 75.77493996 | -0.224669177 | 0.204585193 | -1.098169292 | 0.272130572 | 0.454173972 |

| | | | | | | | |
|---|---|---|---|---|---|---|---|
| ENSG00000225528 | TMA7B | 1.72715197 | -0.78382461 | 1.367980575 | -0.572979342 | 0.566658681 | NA |
| ENSG00000225530 | SP3P | 3.450025868 | 0.820703501 | 0.991914866 | 0.827393085 | 0.408014299 | 0.600561271 |
| ENSG00000225531 | ENSG0000022 | 48.23220652 | 0.132257531 | 0.253492332 | 0.52174174 | 0.601850159 | 0.761398701 |
| ENSG00000225533 | PAWRP1 | 1.982878718 | -0.691464857 | 1.258040194 | -0.549636538 | 0.582568693 | 0.747090374 |
| ENSG00000225535 | LINC01393 | 4.708581783 | -0.406681 | 0.799939972 | -0.508389397 | 0.611180287 | 0.768877202 |
| ENSG00000225536 | STIP1P3 | 105.7637884 | 0.072569436 | 0.197142742 | 0.36810605 | 0.71279416 | 0.84015416 |
| ENSG00000225544 | ENSG0000022 | 16.6497303 | 0.2450199 | 0.431860664 | 0.567358688 | 0.570470512 | 0.737372898 |
| ENSG00000225555 | ENSG0000022 | 9.134987322 | -1.239165857 | 0.624806348 | -1.983279879 | 0.047336176 | 0.114280348 |
| ENSG00000225556 | C2CD4D | 16.86793962 | 0.798443893 | 0.436876524 | 1.827619128 | 0.067606733 | 0.154195512 |
| ENSG00000225558 | UBE2D3P2 | 189.656444 | -0.072334238 | 0.131293572 | -0.550935108 | 0.58167816 | 0.746582642 |
| ENSG00000225563 | ENSG0000022 | 9.61568863 | 0.472842465 | 0.567016731 | 0.833912723 | 0.404330167 | 0.597252675 |
| ENSG00000225568 | ENSG0000022 | 24.04497134 | 0.039302877 | 0.377855509 | 0.104015625 | 0.917156949 | 0.959222231 |
| ENSG00000225573 | RPL35P5 | 291.9420229 | 0.306494559 | 0.119966712 | 2.554830034 | 0.010623966 | 0.031337763 |
| ENSG00000225578 | NCBP2-AS1 | 8.559326883 | -0.206622442 | 0.665121937 | -0.310653477 | 0.756064069 | 0.865184627 |
| ENSG00000225580 | PA2G4P5 | 430.7246722 | 0.00715521 | 0.092271287 | 0.077545361 | 0.938189707 | 0.9705119 |
| ENSG00000225591 | RPS27P7 | 7.455503642 | 0.003672122 | 0.631392742 | 0.005815907 | 0.995359603 | 0.997076653 |
| ENSG00000225602 | MTOR-AS1 | 1.917164303 | 0.814408333 | 1.265103039 | 0.643748618 | 0.519738458 | 0.69707672 |
| ENSG00000225606 | ENSG0000022 | 3.599346174 | -2.278624353 | 1.050314925 | -2.169467746 | 0.030047191 | 0.077797128 |
| ENSG00000225614 | ZNF469 | 122.5397263 | 1.577771603 | 0.176612163 | 8.933538776 | 4.13E-19 | 6.55E-18 |
| ENSG00000225616 | ENSG0000022 | 66.03807494 | 0.405177484 | 0.216437233 | 1.872032268 | 0.061202139 | 0.141858854 |
| ENSG00000225630 | MTND2P28 | 1078.504456 | -0.021950975 | 0.074333 | -0.295305925 | 0.767760191 | 0.87314707 |
| ENSG00000225639 | PGAM1P2 | 2.318557871 | -1.010776127 | 1.183888136 | -0.853776718 | 0.393228717 | 0.586100108 |
| ENSG00000225642 | SNRPEP5 | 3.974319615 | 0.905740143 | 0.919210901 | 0.985345303 | 0.324454487 | 0.514658134 |
| ENSG00000225643 | ENSG0000022 | 2.622982104 | -0.504189037 | 1.101811839 | -0.457599945 | 0.647239883 | 0.793951282 |
| ENSG00000225648 | SBDSP1 | 586.0818527 | 0.347591608 | 0.084164078 | 4.129928293 | 3.63E-05 | 0.0001731 |
| ENSG00000225663 | MCRIP1 | 2691.788846 | 0.369177558 | 0.063116692 | 5.849127181 | 4.94E-09 | 3.68E-08 |
| ENSG00000225671 | FCF1P6 | 3.832470431 | -0.069310879 | 0.888031697 | -0.078050006 | 0.937788275 | 0.97027165 |
| ENSG00000225674 | IPO7P2 | 289.2477861 | 0.021873752 | 0.114754956 | 0.190612701 | 0.848829039 | 0.920956268 |
| ENSG00000225675 | LRP8-DT | 15.8978344 | 0.545763687 | 0.453999151 | 1.202124908 | 0.229315137 | 0.402362648 |
| ENSG00000225684 | FAM225B | 2.345019289 | 1.316287988 | 1.231624827 | 1.068741032 | 0.285186377 | 0.469107664 |
| ENSG00000225689 | ENSG0000022 | 6.170066014 | -0.562734054 | 0.721351418 | -0.780110831 | 0.435325642 | 0.625250312 |
| ENSG00000225693 | LAGE3P1 | 1.662850995 | -0.20722742 | 1.407831711 | -0.147196159 | 0.882977192 | NA |
| ENSG00000225695 | HNRNPA1P35 | 44.42431154 | 0.252784995 | 0.270402553 | 0.934846927 | 0.349867192 | 0.541124149 |

| | | | | | | | |
|---|---|---|---|---|---|---|---|
| ENSG00000225697 | SLC26A6 | 878.5458184 | 0.240429101 | 0.074043413 | 3.247136942 | 0.001165723 | 0.004313175 |
| ENSG00000225712 | ATP5MC2P1 | 2.722394024 | -0.569093777 | 1.112470044 | -0.511558742 | 0.608959867 | 0.767188235 |
| ENSG00000225715 | PSMA1P1 | 7.957104241 | -0.296560696 | 0.628306658 | -0.471999926 | 0.636926841 | 0.786219422 |
| ENSG00000225721 | ENSG0000022 | 30.27178287 | -0.133446242 | 0.326235054 | -0.409049365 | 0.682503435 | 0.819871514 |
| ENSG00000225733 | FGD5-AS1 | 8671.303459 | -0.136437746 | 0.044224342 | -3.085127791 | 0.002034647 | 0.007140096 |
| ENSG00000225739 | NPM1P18 | 11.98718463 | -0.136829361 | 0.511706663 | -0.267398044 | 0.789162705 | 0.885944625 |
| ENSG00000225742 | LINC02036 | 2.783242906 | 0.866026758 | 1.132634846 | 0.764612497 | 0.444502325 | 0.633307455 |
| ENSG00000225756 | DBH-AS1 | 4.491452601 | -1.176009792 | 0.839830662 | -1.400293946 | 0.161425313 | 0.308334445 |
| ENSG00000225762 | LINC01389 | 65.40324948 | -0.712810967 | 0.224634665 | -3.173201101 | 0.001507681 | 0.005439773 |
| ENSG00000225766 | DHRS4L1 | 4.743629088 | -0.028268114 | 0.812886495 | -0.034774983 | 0.972259169 | 0.987410366 |
| ENSG00000225770 | RPS29P3 | 20.86476914 | -0.302576558 | 0.40262034 | -0.751518313 | 0.452340784 | 0.640066851 |
| ENSG00000225774 | SIRPAP1 | 14.95679047 | 1.147364228 | 0.473617262 | 2.422555767 | 0.01541176 | 0.043577687 |
| ENSG00000225783 | MIAT | 4.029217938 | 0.249631703 | 0.876524768 | 0.28479709 | 0.775799602 | 0.878249253 |
| ENSG00000225787 | LSM3P3 | 19.26690287 | 0.364800103 | 0.420030196 | 0.868509231 | 0.385115621 | 0.577672438 |
| ENSG00000225791 | TRAM2-AS1 | 325.8061815 | -0.009053851 | 0.106484032 | -0.085025437 | 0.932241168 | 0.967526001 |
| ENSG00000225792 | SNX10-AS1 | 2.611219738 | -1.638802028 | 1.186424763 | -1.381294524 | 0.167188421 | 0.316679249 |
| ENSG00000225793 | TMEM30A-DT | 63.50769185 | 0.73612557 | 0.230738499 | 3.190302324 | 0.00142124 | 0.00515385 |
| ENSG00000225798 | ENSG0000022 | 20.05582965 | -0.247946522 | 0.402487686 | -0.616035049 | 0.537871388 | 0.712811649 |
| ENSG00000225806 | ENSG0000022 | 14.06044255 | -0.204799643 | 0.479024138 | -0.427535122 | 0.66898961 | 0.809848108 |
| ENSG00000225808 | DNAJC19P5 | 3.66698193 | 1.029149516 | 0.935134679 | 1.100536146 | 0.27109859 | 0.453048856 |
| ENSG00000225814 | GRPEL2P2 | 6.578446016 | -0.178358001 | 0.684824345 | -0.260443429 | 0.794521747 | 0.890000922 |
| ENSG00000225828 | FAM229A | 147.6464818 | -0.359202958 | 0.158729347 | -2.262990205 | 0.023636293 | 0.063298688 |
| ENSG00000225830 | ERCC6 | 660.3804811 | -0.470680099 | 0.080789272 | -5.82602226 | 5.68E-09 | 4.20E-08 |
| ENSG00000225850 | ENSG0000022 | 7.922009322 | -0.669492969 | 0.624303092 | -1.072384515 | 0.28354736 | 0.467417106 |
| ENSG00000225855 | RUSC1-AS1 | 153.5463319 | -0.07343185 | 0.163932258 | -0.44794021 | 0.654196348 | 0.799227929 |
| ENSG00000225857 | LINC02816 | 2.004101191 | -1.74529785 | 1.312365886 | -1.329886634 | 0.183555626 | 0.339742299 |
| ENSG00000225871 | ENSG0000022 | 97.42201171 | -0.516053599 | 0.182017415 | -2.835188045 | 0.004579872 | 0.014893998 |
| ENSG00000225880 | LINC00115 | 15.81790682 | -0.369354581 | 0.457706213 | -0.806968686 | 0.419684524 | 0.611023031 |
| ENSG00000225882 | LINC01456 | 150.0936929 | -0.909994571 | 0.156501917 | -5.814590588 | 6.08E-09 | 4.48E-08 |
| ENSG00000225891 | DHDDS-AS1 | 9.241800616 | 0.916536245 | 0.593083513 | 1.54537468 | 0.122255661 | 0.248960929 |
| ENSG00000225912 | RPL26P36 | 98.20877272 | 0.391763592 | 0.182055997 | 2.151885119 | 0.031406404 | 0.080809115 |
| ENSG00000225918 | RPL7P59 | 13.73895444 | 0.624475302 | 0.494950477 | 1.261692495 | 0.207059459 | 0.373042015 |
| ENSG00000225921 | NOL7 | 1450.739361 | -0.441821668 | 0.058322174 | -7.575534937 | 3.58E-14 | 4.04E-13 |

| | | | | | | | |
|---|---|---|---|---|---|---|---|
| ENSG00000225934 | FAM133FP | 1.581936615 | 0.458802961 | 1.396486174 | 0.328540998 | 0.742502651 | NA |
| ENSG00000225945 | ZFAND3-DT | 5.37368826 | 0.094755454 | 0.763294333 | 0.124140125 | 0.901204329 | 0.950225965 |
| ENSG00000225950 | NTF4 | 12.90557445 | -0.069869748 | 0.551802145 | -0.126621016 | 0.899240362 | 0.949051063 |
| ENSG00000225951 | ODF2-AS1 | 18.51629718 | -0.383092194 | 0.408982628 | -0.936695515 | 0.348915204 | 0.540161613 |
| ENSG00000225953 | SATB2-AS1 | 162.4131865 | 1.123991951 | 0.146695496 | 7.662075416 | 1.83E-14 | 2.11E-13 |
| ENSG00000225963 | ENSG0000022 | 1.681695557 | 1.26181049 | 1.47785216 | 0.853813746 | 0.393208197 | NA |
| ENSG00000225968 | ELFN1 | 138.1389737 | 0.804936196 | 0.160474868 | 5.015964219 | 5.28E-07 | 3.17E-06 |
| ENSG00000225969 | BICDL3P | 7.697428166 | -0.09454594 | 0.640467946 | -0.147620097 | 0.882642594 | 0.939121523 |
| ENSG00000225972 | MTND1P23 | 71.61374707 | -0.128646407 | 0.231695973 | -0.555237991 | 0.578731886 | 0.744317998 |
| ENSG00000225973 | PIGBOS1 | 240.6173887 | -0.679139108 | 0.122321659 | -5.552075702 | 2.82E-08 | 1.93E-07 |
| ENSG00000225975 | ZNF567-DT | 104.5685442 | 0.631549042 | 0.184459342 | 3.423784534 | 0.000617556 | 0.002403584 |
| ENSG00000225976 | CKS1BP2 | 20.02898555 | 0.002390669 | 0.431572092 | 0.005539444 | 0.995580186 | 0.997210788 |
| ENSG00000225984 | HMGB1P26 | 1.923273058 | -0.098291461 | 1.325745161 | -0.074140539 | 0.940898559 | 0.971736581 |
| ENSG00000226002 | GTF2IP14 | 3.679638546 | -0.034865284 | 0.931682946 | -0.037421833 | 0.970148664 | 0.98657174 |
| ENSG00000226007 | FAM88F | 12.42553255 | -0.182946307 | 0.511553153 | -0.357629126 | 0.720620879 | 0.844878037 |
| ENSG00000226009 | KCNIP2-AS1 | 1.715834013 | 0.350391311 | 1.370189915 | 0.255724631 | 0.798163475 | NA |
| ENSG00000226015 | CCT8P1 | 246.6425933 | 0.06206295 | 0.120101124 | 0.516755777 | 0.605326654 | 0.764069189 |
| ENSG00000226017 | PRICKLE2-AS3 | 21.79303075 | -1.397989954 | 0.398024963 | -3.512317273 | 0.000444217 | 0.001774703 |
| ENSG00000226022 | ENSG0000022 | 2.336550494 | 1.288889438 | 1.253077574 | 1.028579128 | 0.30367749 | 0.49089998 |
| ENSG00000226026 | RWDD3-DT | 5.138294619 | 0.375988395 | 0.782316257 | 0.480609206 | 0.630794272 | 0.782326489 |
| ENSG00000226029 | LINC01772 | 13.29506886 | 0.217386462 | 0.497362525 | 0.437078491 | 0.662054425 | 0.80479344 |
| ENSG00000226031 | FGF13-AS1 | 14.82263508 | 1.141991018 | 0.483019815 | 2.364273642 | 0.018065463 | 0.050014691 |
| ENSG00000226040 | ENSG0000022 | 6.979869327 | 0.313122734 | 0.708275606 | 0.442091653 | 0.658422884 | 0.802336104 |
| ENSG00000226045 | ENSG0000022 | 1.92706882 | 0.853112344 | 1.377618303 | 0.619266122 | 0.535741059 | 0.711031295 |
| ENSG00000226047 | ENSG0000022 | 1.979979784 | -0.554502935 | 1.394218101 | -0.397716063 | 0.690839495 | 0.825640142 |
| ENSG00000226049 | TLK2P1 | 1645.99748 | 0.10320431 | 0.058429844 | 1.766294455 | 0.077346461 | 0.172174899 |
| ENSG00000226051 | ZNF503-AS1 | 2.928951711 | -2.410597498 | 1.195090515 | -2.017083616 | 0.043686788 | 0.106796134 |
| ENSG00000226054 | MEMO1P1 | 335.0574577 | 0.14950797 | 0.105662716 | 1.41495483 | 0.15708174 | 0.30151913 |
| ENSG00000226055 | PAICSP1 | 2.138431858 | 0.02896043 | 1.174032204 | 0.024667492 | 0.980320185 | 0.991070363 |
| ENSG00000226059 | HMGB3P20 | 4.883974174 | -0.896207294 | 0.811420367 | -1.104491987 | 0.269379777 | 0.451016256 |
| ENSG00000226067 | LINC00623 | 95.13805516 | -0.755220307 | 0.184029243 | -4.103805981 | 4.06E-05 | 0.000192346 |
| ENSG00000226068 | HNRNPA3P4 | 6.8483019 | -0.572941882 | 0.682633388 | -0.83931125 | 0.401294672 | 0.594107762 |
| ENSG00000226081 | USP12PX | 25.43901695 | -0.415577123 | 0.371408548 | -1.118921805 | 0.2631735 | 0.443977062 |

| | | | | | | | |
|---|---|---|---|---|---|---|---|
| ENSG00000226082 | RPS10P10 | 3.74783572 | 0.221116936 | 0.933256508 | 0.236930505 | 0.812710704 | 0.90104797 |
| ENSG00000226084 | RPL17P6 | 832.0234121 | 0.63138115 | 0.089108838 | 7.085505372 | 1.39E-12 | 1.40E-11 |
| ENSG00000226085 | UQCRFS1P1 | 82.72528381 | -0.029478584 | 0.210085418 | -0.140317135 | 0.888409427 | 0.94247758 |
| ENSG00000226086 | EIF3LP3 | 14.68917728 | 0.088961698 | 0.469510619 | 0.189477499 | 0.849718589 | 0.921328375 |
| ENSG00000226098 | SEC11B | 23.45233016 | 0.064948733 | 0.36503678 | 0.177923805 | 0.858782815 | 0.926723937 |
| ENSG00000226102 | SEPTIN7P3 | 1.700208242 | 0.369259188 | 1.340940333 | 0.275373317 | 0.783029441 | NA |
| ENSG00000226107 | RPL36AP54 | 5.476884475 | 0.044943062 | 0.766501073 | 0.058634049 | 0.95324359 | 0.978321944 |
| ENSG00000226121 | AHCTF1P1 | 731.6217803 | -0.04734285 | 0.077796431 | -0.608547846 | 0.542824183 | 0.716541968 |
| ENSG00000226124 | FTCDNL1 | 106.3125594 | -0.120935331 | 0.176284678 | -0.686022929 | 0.49269865 | 0.675455731 |
| ENSG00000226126 | PSMC1P12 | 4.928838796 | -0.754429459 | 0.828380308 | -0.910728383 | 0.362438505 | 0.555140503 |
| ENSG00000226131 | PPIAP66 | 3.108678284 | 0.908816032 | 1.017638755 | 0.893063504 | 0.371823166 | 0.564464524 |
| ENSG00000226137 | BAIAP2-DT | 1170.375434 | 0.697100686 | 0.068344456 | 10.19981327 | 1.99E-24 | 4.28E-23 |
| ENSG00000226138 | ENSG0000022 | 3.55965102 | 1.297178124 | 1.043900761 | 1.242625901 | 0.214005721 | 0.382432907 |
| ENSG00000226161 | EIF4A1P5 | 13.74204202 | -0.444749199 | 0.479301431 | -0.927911268 | 0.353453599 | 0.545017514 |
| ENSG00000226167 | AP4B1-AS1 | 1367.827184 | -0.059352178 | 0.061649143 | -0.962741341 | 0.335677345 | 0.526875672 |
| ENSG00000226174 | TEX22 | 64.2528448 | -0.222911567 | 0.226185009 | -0.985527591 | 0.324364986 | 0.514551744 |
| ENSG00000226179 | LINC00685 | 32.89545508 | -0.318988261 | 0.319267255 | -0.999126144 | 0.317733588 | 0.506871245 |
| ENSG00000226179 | LINC00685 | 32.89545508 | -0.318988261 | 0.319267255 | -0.999126144 | 0.317733588 | 0.506871245 |
| ENSG00000226180 | ENSG0000022 | 41.85492722 | 0.39456655 | 0.301678517 | 1.307904034 | 0.190905861 | 0.350574804 |
| ENSG00000226200 | SGMS1-AS1 | 195.8056139 | -0.530792879 | 0.137801657 | -3.85186136 | 0.000117223 | 0.000516937 |
| ENSG00000226210 | WASH8P | 455.8911672 | 0.003950215 | 0.097655041 | 0.040450702 | 0.967733809 | 0.985650827 |
| ENSG00000226221 | RPL26P19 | 1542.55876 | 0.439216399 | 0.063540157 | 6.912422305 | 4.76E-12 | 4.60E-11 |
| ENSG00000226226 | PRELID3BP1 | 2.643405462 | -0.534897259 | 1.135886706 | -0.470907227 | 0.637706987 | 0.78659182 |
| ENSG00000226234 | KRT18P24 | 11.07860858 | 0.27803709 | 0.524233382 | 0.530368915 | 0.595856174 | 0.756993626 |
| ENSG00000226241 | ENSG0000022 | 2.004165156 | -1.780174332 | 1.397061314 | -1.274227776 | 0.202582704 | 0.366704587 |
| ENSG00000226243 | RPL37AP1 | 81.08262908 | 0.300380899 | 0.21192405 | 1.417398818 | 0.156366358 | 0.30055979 |
| ENSG00000226245 | ZNF32-AS1 | 1.775325633 | 0.126973474 | 1.348675534 | 0.094146791 | 0.924992551 | NA |
| ENSG00000226247 | SUPT4H1P1 | 7.389690887 | 0.090289393 | 0.666461863 | 0.135475708 | 0.892235769 | 0.944962537 |
| ENSG00000226259 | GTF2H2B | 20.10154099 | -0.615262569 | 0.403406824 | -1.52516649 | 0.127217577 | 0.256989363 |
| ENSG00000226266 | ENSG0000022 | 15.32628363 | 1.072707452 | 0.509487118 | 2.10546531 | 0.035250824 | 0.08918649 |
| ENSG00000226268 | ENSG0000022 | 133.4604436 | 0.541725431 | 0.157299933 | 3.443901222 | 0.000573385 | 0.002243851 |
| ENSG00000226276 | ENSG0000022 | 7.019892102 | -0.500689542 | 0.685865481 | -0.730011286 | 0.465383286 | 0.651356659 |
| ENSG00000226279 | RPL12P10 | 2.80606308 | 0.689058847 | 1.05262431 | 0.654610425 | 0.51271861 | 0.692235002 |

| | | | | | | | |
|---|---|---|---|---|---|---|---|
| ENSG00000226284 | ARPC3P1 | 94.39078471 | -0.06897961 | 0.180638704 | -0.381865064 | 0.702561455 | 0.833338148 |
| ENSG00000226287 | TMEM191A | 130.8722065 | -0.460112775 | 0.168229736 | -2.735026431 | 0.006237527 | 0.019617338 |
| ENSG00000226299 | RPSAP62 | 2.942131521 | -5.115109565 | 1.467811457 | -3.484854639 | 0.000492404 | 0.001950853 |
| ENSG00000226308 | ENSG0000022 | 4.928835745 | -0.19257273 | 0.784388069 | -0.24550696 | 0.80606392 | 0.897100998 |
| ENSG00000226309 | SMARCE1P2 | 7.4331992 | -0.493688461 | 0.637773251 | -0.774081479 | 0.438882609 | 0.628383913 |
| ENSG00000226310 | ENSG0000022 | 11.49164437 | 0.708766745 | 0.545829415 | 1.298513282 | 0.194111015 | 0.354869835 |
| ENSG00000226312 | CFLAR-AS1 | 3.027330636 | 0.1347691 | 1.01765323 | 0.13243126 | 0.89464319 | 0.946400434 |
| ENSG00000226314 | ZKSCAN8P1 | 3.431992767 | 0.210725454 | 0.950715739 | 0.221649275 | 0.824586921 | 0.907591592 |
| ENSG00000226318 | RPS3AP38 | 1.903815051 | -1.146701785 | 1.356543762 | -0.84531131 | 0.397937048 | 0.590967558 |
| ENSG00000226320 | LINC01811 | 12.50850135 | -1.917227322 | 0.5509815 | -3.479658252 | 0.000502054 | 0.001985709 |
| ENSG00000226327 | RPSAP49 | 2.168197907 | -0.796190258 | 1.252665638 | -0.63559679 | 0.525039263 | 0.701269943 |
| ENSG00000226328 | NUP50-DT | 125.5973352 | 0.455495703 | 0.162301004 | 2.806487286 | 0.005008489 | 0.016130105 |
| ENSG00000226330 | PPP1R15B-AS1 | 7.464921845 | 1.53819951 | 0.700390824 | 2.196201688 | 0.028077511 | 0.073384766 |
| ENSG00000226332 | ENSG0000022 | 119.4893624 | 0.015697814 | 0.166921002 | 0.094043373 | 0.925074702 | 0.964249985 |
| ENSG00000226334 | ENSG0000022 | 4.551525278 | 0.220863394 | 0.856691162 | 0.25780982 | 0.796553685 | 0.891493465 |
| ENSG00000226340 | USP8P2 | 15.94195308 | -1.275131886 | 0.46582117 | -2.737385005 | 0.006192977 | 0.019493261 |
| ENSG00000226342 | NMD3P1 | 30.04025341 | -0.627810091 | 0.338279337 | -1.85589252 | 0.063468879 | 0.146197017 |
| ENSG00000226348 | VN2R10P | 1.926036298 | -1.728799188 | 1.373974075 | -1.258247313 | 0.208302329 | 0.374751426 |
| ENSG00000226352 | PSPC1-AS2 | 3.611692648 | -0.192238161 | 0.942675844 | -0.203928171 | 0.83840963 | 0.91499129 |
| ENSG00000226359 | ACTG1P24 | 2.261350233 | -0.186863497 | 1.160057295 | -0.161081265 | 0.872029396 | 0.933692151 |
| ENSG00000226360 | RPL10AP6 | 1083.372565 | 0.371092827 | 0.065848148 | 5.635584873 | 1.74E-08 | 1.22E-07 |
| ENSG00000226361 | TERF1P5 | 35.53123555 | 0.20775322 | 0.302672699 | 0.686395636 | 0.492463655 | 0.675180269 |
| ENSG00000226363 | HAGLROS | 34.30024688 | 0.029254799 | 0.297509171 | 0.098332428 | 0.921668329 | 0.962142046 |
| ENSG00000226386 | PARD3-DT | 4.835786319 | -0.866271985 | 0.830971689 | -1.042480743 | 0.297188849 | 0.483577128 |
| ENSG00000226389 | MAPK6P6 | 8.062068298 | -1.059040206 | 0.640663271 | -1.653037178 | 0.098323297 | 0.209039522 |
| ENSG00000226396 | ENSG0000022 | 56.99074126 | 0.485239925 | 0.277243937 | 1.750227362 | 0.080079089 | 0.176771969 |
| ENSG00000226415 | TPI1P1 | 1939.955175 | -0.021094005 | 0.057409663 | -0.367429514 | 0.71329866 | 0.84060482 |
| ENSG00000226416 | MRPL23-AS1 | 6.787891717 | -6.307194177 | 1.310550045 | -4.812631307 | 1.49E-06 | 8.43E-06 |
| ENSG00000226419 | SLC16A1-AS1 | 12.76878775 | 0.149692335 | 0.489981691 | 0.305505976 | 0.759980819 | 0.868095404 |
| ENSG00000226421 | SLC25A5P5 | 68.65708997 | 0.804754822 | 0.224761422 | 3.580484652 | 0.000342957 | 0.001394937 |
| ENSG00000226427 | HMGN2P7 | 4.102502812 | -0.654180678 | 0.975435983 | -0.670654651 | 0.502440559 | 0.683507467 |
| ENSG00000226432 | AKTIPP2 | 196.0965323 | 0.021664318 | 0.132671639 | 0.163292759 | 0.870287935 | 0.932790446 |
| ENSG00000226436 | H3P43 | 2.032925376 | 0.103937853 | 1.230325039 | 0.084479995 | 0.932674808 | 0.967687023 |

| | | | | | | | |
|---|---|---|---|---|---|---|---|
| ENSG00000226439 | ENSG0000022 | 6.422911478 | 0.561965466 | 0.717520893 | 0.783204324 | 0.433507136 | 0.623465351 |
| ENSG00000226445 | THBS2-AS1 | 74.9406033 | -0.286048512 | 0.204750023 | -1.397062175 | 0.162394875 | 0.309619218 |
| ENSG00000226450 | CYP2D8P | 22.16664664 | 0.579742717 | 0.415049522 | 1.396803722 | 0.162472603 | 0.309715931 |
| ENSG00000226453 | LINC02542 | 61.01585772 | 0.299070256 | 0.224901855 | 1.329781188 | 0.183590376 | 0.339779203 |
| ENSG00000226457 | RPL22P3 | 7.759000529 | 0.688789932 | 0.631235197 | 1.091177957 | 0.275194585 | 0.457355635 |
| ENSG00000226465 | ENSG0000022 | 43.2912502 | -0.368403984 | 0.271620687 | -1.356317841 | 0.174998061 | 0.328031993 |
| ENSG00000226466 | RPA2P2 | 3.514794333 | -0.597504313 | 1.011449237 | -0.590740782 | 0.554694118 | 0.725963206 |
| ENSG00000226468 | ENSG0000022 | 4.320028465 | 0.685479368 | 0.866043996 | 0.791506403 | 0.428648541 | 0.61920798 |
| ENSG00000226471 | ENSG0000022 | 43.17083271 | -0.114711934 | 0.280687494 | -0.408682026 | 0.682773026 | 0.819956717 |
| ENSG00000226472 | ENSG0000022 | 1.840101863 | -1.004164065 | 1.385303365 | -0.724869433 | 0.468532135 | 0.654495626 |
| ENSG00000226476 | LINC01748 | 68.00136689 | 0.062831372 | 0.222676054 | 0.282164925 | 0.777817052 | 0.879532114 |
| ENSG00000226478 | UPF3AP1 | 115.0478975 | -0.119376297 | 0.167561505 | -0.712432712 | 0.476196865 | 0.661256421 |
| ENSG00000226479 | TMEM185B | 2215.5959 | -0.068092694 | 0.05341242 | -1.274847586 | 0.202363194 | 0.366422896 |
| ENSG00000226499 | ENSG0000022 | 7.871580793 | 0.051198631 | 0.643227922 | 0.079596406 | 0.936558254 | 0.96955824 |
| ENSG00000226508 | LINC01918 | 24.91760016 | 0.857712793 | 0.361197949 | 2.374633617 | 0.017566377 | 0.048880724 |
| ENSG00000226510 | UPK1A-AS1 | 5.959519741 | 0.54649568 | 0.715062395 | 0.764262928 | 0.444710573 | 0.633380605 |
| ENSG00000226519 | LINC00390 | 2.167259722 | -0.3762964 | 1.227531429 | -0.306547263 | 0.759188 | 0.867448957 |
| ENSG00000226521 | ENSG0000022 | 23.40141491 | 0.27327741 | 0.401311619 | 0.680960624 | 0.495896406 | 0.677941809 |
| ENSG00000226525 | RPS7P10 | 232.1187497 | 0.326998835 | 0.133000926 | 2.458620745 | 0.013947187 | 0.039873936 |
| ENSG00000226526 | ENSG0000022 | 6.856344563 | -0.428259283 | 0.675375382 | -0.634105557 | 0.526011936 | 0.702042927 |
| ENSG00000226535 | ENSG0000022 | 19.04232539 | 0.386341134 | 0.437303507 | 0.883462235 | 0.376986578 | 0.569776789 |
| ENSG00000226540 | GAPDHP73 | 2.172057308 | -0.403089296 | 1.329069467 | -0.303286853 | 0.761671265 | 0.869290488 |
| ENSG00000226543 | MYL6P1 | 8.653895122 | -0.935650104 | 0.664676615 | -1.407677181 | 0.159226682 | 0.304743874 |
| ENSG00000226544 | RPL7P22 | 132.4054485 | -0.103993235 | 0.153902125 | -0.675710196 | 0.499224661 | 0.680666492 |
| ENSG00000226549 | SCDP1 | 611.8588929 | -0.71783214 | 0.095541012 | -7.513340352 | 5.76E-14 | 6.40E-13 |
| ENSG00000226553 | IMPDH1P7 | 19.80992768 | -0.043138609 | 0.412855269 | -0.104488454 | 0.916781731 | 0.95891724 |
| ENSG00000226557 | TRAF6P1 | 1.825414276 | -1.562490429 | 1.381113723 | -1.131326409 | 0.257917733 | 0.4375795 |
| ENSG00000226562 | CYP4F26P | 18.40931994 | 2.827468894 | 0.52290275 | 5.40725574 | 6.40E-08 | 4.22E-07 |
| ENSG00000226564 | FTH1P20 | 403.7716079 | 0.389507422 | 0.100936 | 3.858954418 | 0.000113873 | 0.000503421 |
| ENSG00000226578 | ENSG0000022 | 2.365880223 | 0.210942025 | 1.17359074 | 0.179740703 | 0.857356137 | 0.926087485 |
| ENSG00000226581 | LINC02848 | 12.83318763 | 0.197085797 | 0.504936765 | 0.390317779 | 0.696301578 | 0.829343624 |
| ENSG00000226592 | ENSG0000022 | 1.681280891 | 4.103822121 | 1.632553167 | 2.513744853 | 0.011945682 | NA |
| ENSG00000226605 | ENSG0000022 | 1.960657471 | -0.21469238 | 1.26997037 | -0.169053062 | 0.865754905 | 0.9302381 |

| | | | | | | | |
|---|---|---|---|---|---|---|---|
| ENSG00000226608 | FTLP3 | 558.8866096 | 0.581830546 | 0.090314223 | 6.442291419 | 1.18E-10 | 1.01E-09 |
| ENSG00000226609 | SNX30-DT | 4.796060793 | 0.323461855 | 0.8064078 | 0.401114492 | 0.688335832 | 0.82383388 |
| ENSG00000226624 | RPL13AP16 | 3.087821029 | 0.296520922 | 1.008240781 | 0.29409733 | 0.768683531 | 0.873855116 |
| ENSG00000226632 | UBE2V1P1 | 3.148873877 | 0.543432598 | 0.994064813 | 0.546677229 | 0.584600501 | 0.74886841 |
| ENSG00000226647 | ENSG0000022 | 1.723330875 | -1.461043842 | 1.415091926 | -1.032472742 | 0.30185071 | NA |
| ENSG00000226648 | PLCG1-AS1 | 142.1558436 | -0.339237364 | 0.164136295 | -2.066802859 | 0.038752734 | 0.0966069 |
| ENSG00000226650 | KIF4B | 69.00121733 | -0.521330415 | 0.216951844 | -2.402977568 | 0.016262185 | 0.045667098 |
| ENSG00000226659 | TSPAN14-AS1 | 9.328648901 | -2.6991638 | 0.705400493 | -3.826427434 | 0.000130016 | 0.000569303 |
| ENSG00000226666 | HSPA9P1 | 127.3169834 | 0.102550359 | 0.19911497 | 0.515030886 | 0.606531437 | 0.765042677 |
| ENSG00000226677 | IGBP1P1 | 21.0773192 | 0.002962048 | 0.388153049 | 0.007631134 | 0.993911295 | 0.996533822 |
| ENSG00000226688 | ENTPD1-AS1 | 841.5085168 | -0.793173597 | 0.078457454 | -10.10960155 | 5.01E-24 | 1.05E-22 |
| ENSG00000226696 | LENG8-AS1 | 66.01955323 | 0.036472719 | 0.235311611 | 0.154997533 | 0.876823262 | 0.935974786 |
| ENSG00000226701 | RPL15P14 | 14.93960858 | -0.032944177 | 0.457187897 | -0.072058288 | 0.942555521 | 0.97274715 |
| ENSG00000226706 | ENSG0000022 | 12.84190297 | 0.359520388 | 0.517857639 | 0.6942456 | 0.4875282 | 0.670899634 |
| ENSG00000226709 | FGF12-AS3 | 10.95094545 | 0.236643775 | 0.52590595 | 0.449973565 | 0.652729502 | 0.798422458 |
| ENSG00000226711 | FAM66C | 70.81417658 | 0.418998673 | 0.216736347 | 1.933218304 | 0.053209311 | 0.126463074 |
| ENSG00000226715 | LINC01709 | 2.293498278 | 0.282244264 | 1.167251345 | 0.241802475 | 0.808933221 | 0.898853827 |
| ENSG00000226716 | ENSG0000022 | 2.329404659 | 1.88515272 | 1.315260253 | 1.433292549 | 0.15177425 | 0.294112291 |
| ENSG00000226721 | EEF1DP2 | 28.54144778 | 0.178584883 | 0.331840392 | 0.538164996 | 0.590463144 | 0.753103247 |
| ENSG00000226723 | ENSG0000022 | 140.1212582 | 0.099461143 | 0.153899983 | 0.646271304 | 0.518103669 | 0.695932225 |
| ENSG00000226742 | HSBP1L1 | 414.0616008 | 0.691238638 | 0.093896099 | 7.361739645 | 1.82E-13 | 1.95E-12 |
| ENSG00000226745 | OAZ1P1 | 2.855216915 | -0.690517817 | 1.055818061 | -0.654012128 | 0.513103992 | 0.692429295 |
| ENSG00000226752 | CUTALP | 1751.08997 | 0.245475369 | 0.065622772 | 3.740704044 | 0.000183506 | 0.000784352 |
| ENSG00000226754 | MAGOH-DT | 60.88147365 | -0.021309881 | 0.239129383 | -0.089114439 | 0.928990963 | 0.965918174 |
| ENSG00000226756 | ENSG0000022 | 4.503399554 | 0.351917698 | 0.83042993 | 0.423777715 | 0.671727932 | 0.811951622 |
| ENSG00000226763 | SRRM5 | 34.5409948 | -0.240501449 | 0.316595457 | -0.759649083 | 0.447464372 | 0.635888882 |
| ENSG00000226781 | TBCAP1 | 54.86000991 | -0.107525849 | 0.238145591 | -0.451513081 | 0.6516198 | 0.797581514 |
| ENSG00000226783 | TLK1P1 | 11.59637104 | -0.724893984 | 0.536608356 | -1.350880908 | 0.176733585 | 0.330475093 |
| ENSG00000226784 | PGAM4 | 186.7048102 | -0.709741377 | 0.138873841 | -5.110691618 | 3.21E-07 | 1.98E-06 |
| ENSG00000226790 | HNRNPA3P1 | 30.26371219 | -0.004054759 | 0.366457482 | -0.011064746 | 0.99117179 | 0.995222268 |
| ENSG00000226791 | LINC02611 | 1.997996115 | -4.546234175 | 3.445180781 | -1.319592342 | 0.186971161 | 0.344676864 |
| ENSG00000226798 | ENSG0000022 | 1.852855866 | -1.611992293 | 1.360829452 | -1.184565995 | 0.236189089 | 0.411340548 |
| ENSG00000226800 | CACTIN-AS1 | 9.135315389 | 0.216517364 | 0.608873304 | 0.355603313 | 0.722137662 | 0.845601795 |

| | | | | | | | |
|---|---|---|---|---|---|---|---|
| ENSG00000226803 | ZNF451-AS1 | 3.836744824 | -0.350829467 | 0.907654314 | -0.386523219 | 0.69910921 | 0.831105311 |
| ENSG00000226810 | ENSG0000022 | 2.27223492 | -3.673647466 | 1.528065716 | -2.40411615 | 0.016211622 | 0.045536255 |
| ENSG00000226823 | SUGT1P1 | 2.719540858 | 1.509590987 | 1.122103124 | 1.345322863 | 0.178520993 | 0.333156579 |
| ENSG00000226824 | ENSG0000022 | 71.26352711 | 0.025264044 | 0.210208774 | 0.120185488 | 0.904336213 | 0.951932856 |
| ENSG00000226833 | PPP1CB-DT | 14.98625354 | -0.297963072 | 0.45382735 | -0.656556006 | 0.511466444 | 0.691154585 |
| ENSG00000226840 | PAICSP3 | 2.573394373 | 0.236610833 | 1.143796523 | 0.206864445 | 0.836115725 | 0.913443886 |
| ENSG00000226849 | ENSG0000022 | 31.90907663 | -0.508883206 | 0.316452846 | -1.608085413 | 0.107816474 | 0.225194383 |
| ENSG00000226853 | GPR155-DT | 14.4088754 | -0.159253482 | 0.491098817 | -0.324279914 | 0.745726125 | 0.859386985 |
| ENSG00000226855 | RPSAP17 | 7.195729227 | -0.081120979 | 0.665347554 | -0.121922713 | 0.902960228 | 0.950920792 |
| ENSG00000226856 | THORLNC | 7.456405029 | -0.26190974 | 0.634222819 | -0.412961711 | 0.679634667 | 0.81798632 |
| ENSG00000226862 | ENSG0000022 | 5.00090409 | 0.058457519 | 0.815971279 | 0.071641638 | 0.942887103 | 0.972914274 |
| ENSG00000226877 | ENSG0000022 | 1.824050284 | 0.045682524 | 1.275151195 | 0.035825182 | 0.971421754 | 0.986996857 |
| ENSG00000226881 | H3P44 | 2.919940018 | -0.159347496 | 1.012875452 | -0.157321905 | 0.874991157 | 0.935161067 |
| ENSG00000226887 | ERVMER34-1 | 140.8216818 | 0.969189348 | 0.156628862 | 6.187808138 | 6.10E-10 | 4.96E-09 |
| ENSG00000226890 | ENSG0000022 | 3.839638874 | 0.411098475 | 0.895092662 | 0.459280355 | 0.646032853 | 0.793149246 |
| ENSG00000226891 | LINC01359 | 2.689868739 | -0.08474188 | 1.167439009 | -0.072587844 | 0.9421341 | 0.972530993 |
| ENSG00000226900 | ENSG0000022 | 3.717504167 | 1.444126135 | 0.983671817 | 1.4680975 | 0.142077741 | 0.279645077 |
| ENSG00000226913 | BSN-DT | 10.49990273 | -0.842704432 | 0.637495616 | -1.321898396 | 0.186201987 | 0.343559364 |
| ENSG00000226915 | RPL19P4 | 1.813670513 | 0.603613634 | 1.346776553 | 0.448191375 | 0.654015087 | 0.799165148 |
| ENSG00000226925 | IL1R1-AS1 | 2.32952175 | 0.176013198 | 1.208285917 | 0.145671811 | 0.884180475 | 0.939910479 |
| ENSG00000226937 | CEP164P1 | 22.84239862 | -1.423059121 | 0.387319875 | -3.674118512 | 0.000238672 | 0.000998383 |
| ENSG00000226942 | IL9RP3 | 2.838249433 | 1.041601413 | 1.138292091 | 0.915056356 | 0.360162046 | 0.552723931 |
| ENSG00000226944 | RNF207-AS1 | 6.742498878 | 0.797423594 | 0.735660041 | 1.083956652 | 0.278384019 | 0.461225582 |
| ENSG00000226945 | RPL13AP8 | 6.52413533 | 2.389541067 | 0.83107919 | 2.875226687 | 0.004037375 | 0.013288004 |
| ENSG00000226948 | RPS4XP2 | 100.0215574 | 0.246089017 | 0.178828126 | 1.376120312 | 0.168784388 | 0.318977299 |
| ENSG00000226950 | DANCR | 1565.255572 | 0.722177762 | 0.057498175 | 12.56001181 | 3.50E-36 | 1.24E-34 |
| ENSG00000226953 | NCKAP5-AS2 | 1.71609784 | -1.516532392 | 1.475717235 | -1.027657843 | 0.304110805 | NA |
| ENSG00000226964 | RHEBP2 | 16.88678378 | 0.252329141 | 0.442791126 | 0.569860429 | 0.568772366 | 0.736073456 |
| ENSG00000226965 | ENSG0000022 | 2.738868055 | -0.014081737 | 1.088138299 | -0.012941128 | 0.989674762 | 0.994530359 |
| ENSG00000226968 | LINC00423 | 9.230287153 | -0.878596026 | 0.576928801 | -1.522884668 | 0.127787562 | 0.25788875 |
| ENSG00000226970 | NEDD8P1 | 27.91779236 | -0.173717388 | 0.334678751 | -0.519057118 | 0.603720911 | 0.762713828 |
| ENSG00000226971 | ENSG0000022 | 10.50972856 | -0.215239459 | 0.55592716 | -0.387172052 | 0.698628839 | 0.830862695 |
| ENSG00000226972 | RPL12P19 | 3.628776748 | 0.60356272 | 0.985612795 | 0.612373057 | 0.540290964 | 0.714573347 |

| | | | | | | | |
|---|---|---|---|---|---|---|---|
| ENSG00000226976 | COX6A1P2 | 1647.297507 | -0.003886011 | 0.063009808 | -0.061673107 | 0.950823157 | 0.977330339 |
| ENSG00000226979 | LTA | 2.72383007 | -0.219341396 | 1.062147352 | -0.206507502 | 0.836394505 | 0.913666841 |
| ENSG00000226981 | ABHD17AP6 | 2.210240209 | -0.566776568 | 1.254554837 | -0.451775045 | 0.651431049 | 0.797514584 |
| ENSG00000226982 | CENPCP1 | 23.17213506 | 0.5443214 | 0.380190457 | 1.431707163 | 0.152227645 | 0.294703645 |
| ENSG00000226986 | PRELID1P5 | 49.24061111 | -0.359845938 | 0.252654423 | -1.424261382 | 0.154370822 | 0.297710522 |
| ENSG00000227001 | NBPF2P | 42.47505541 | 0.040004398 | 0.278607179 | 0.143587104 | 0.885826525 | 0.941041978 |
| ENSG00000227008 | RPS24P19 | 741.8978515 | -0.120146823 | 0.080415815 | -1.494069589 | 0.135157432 | 0.269305262 |
| ENSG00000227014 | FKBP14-AS1 | 7.66336992 | -2.189994296 | 0.72509487 | -3.020286568 | 0.002525356 | 0.008691924 |
| ENSG00000227018 | IL6STP1 | 3.283003933 | 0.886943866 | 1.036316623 | 0.855861853 | 0.392074197 | 0.584949407 |
| ENSG00000227028 | SLC8A1-AS1 | 1.666563561 | -2.187546828 | 1.57092534 | -1.392521193 | 0.163764629 | NA |
| ENSG00000227034 | ENSG0000022 | 6.247054974 | 1.367042011 | 0.739567788 | 1.848433684 | 0.06453963 | 0.148291176 |
| ENSG00000227036 | LINC00511 | 23.32625343 | 0.116728044 | 0.417995358 | 0.279256795 | 0.780047761 | 0.880624785 |
| ENSG00000227038 | GTF2IP7 | 4.970459597 | -1.602100097 | 0.834844189 | -1.919040844 | 0.054979165 | 0.129942514 |
| ENSG00000227039 | ITGB2-AS1 | 4.912376637 | 1.855418057 | 0.870622452 | 2.131139683 | 0.033077636 | 0.084470776 |
| ENSG00000227040 | ENSG0000022 | 16.85950096 | -0.071402765 | 0.49250105 | -0.144979925 | 0.884726722 | 0.940265509 |
| ENSG00000227045 | ENSG0000022 | 2.760588065 | -0.343169655 | 1.061009395 | -0.323436962 | 0.746364341 | 0.859604617 |
| ENSG00000227050 | ENSG0000022 | 3.091107789 | -0.014492893 | 1.0285425 | -0.014090708 | 0.988757613 | 0.994082067 |
| ENSG00000227051 | C14orf132 | 5131.159308 | -0.566330781 | 0.058377043 | -9.70125845 | 2.98E-22 | 5.66E-21 |
| ENSG00000227057 | WDR46 | 1910.102994 | 0.345911164 | 0.058991951 | 5.86370101 | 4.53E-09 | 3.39E-08 |
| ENSG00000227060 | LINC00629 | 7.026631059 | 0.621280129 | 0.672293936 | 0.924119787 | 0.355423952 | 0.547136322 |
| ENSG00000227063 | RPL41P1 | 483.0402703 | -0.1855814 | 0.196787201 | -0.943056247 | 0.345652134 | 0.536594705 |
| ENSG00000227070 | EPS15-AS1 | 25.93787615 | 0.688251961 | 0.34656793 | 1.985907818 | 0.047043561 | 0.113639072 |
| ENSG00000227073 | SDHDP2 | 15.69792418 | -0.288156886 | 0.450216099 | -0.640041275 | 0.522145766 | 0.699077575 |
| ENSG00000227077 | ENSG0000022 | 21.90279486 | 0.533937413 | 0.383904614 | 1.39080749 | 0.164283811 | 0.312360975 |
| ENSG00000227081 | RPS27P3 | 1973.047697 | 0.277372868 | 0.073327828 | 3.782641291 | 0.000155173 | 0.000671002 |
| ENSG00000227082 | LINC02798 | 34.3442742 | 1.58946776 | 0.33279293 | 4.776146421 | 1.79E-06 | 1.00E-05 |
| ENSG00000227087 | RBMX2P5 | 122.0788464 | 0.10807687 | 0.165648177 | 0.652448288 | 0.514112022 | 0.69315823 |
| ENSG00000227096 | HMGB3P8 | 2.79091594 | -0.347953713 | 1.090271466 | -0.319144107 | 0.749617239 | 0.861534041 |
| ENSG00000227097 | RPS28P7 | 2285.183138 | 0.211903719 | 0.052480453 | 4.037764694 | 5.40E-05 | 0.000251199 |
| ENSG00000227105 | PARP1P1 | 44.44473999 | 0.053268137 | 0.262549538 | 0.202887946 | 0.839222616 | 0.915309377 |
| ENSG00000227110 | LMCD1-AS1 | 21.70868393 | -0.643727072 | 0.377678319 | -1.704432159 | 0.08830038 | 0.191474688 |
| ENSG00000227118 | BTF3P13 | 9.427554175 | -0.145005488 | 0.573767444 | -0.252725194 | 0.800480578 | 0.893708301 |
| ENSG00000227121 | LINC02672 | 93.77766474 | -2.826656079 | 0.229224079 | -12.33140992 | 6.14E-35 | 2.09E-33 |

| | | | | | | | |
|---|---|---|---|---|---|---|---|
| ENSG00000227124 | ZNF717 | 17.75828232 | 2.015146866 | 0.470115606 | 4.286492175 | 1.82E-05 | 9.06E-05 |
| ENSG00000227141 | ENSG0000022 | 10.53572642 | 0.410031318 | 0.543645763 | 0.754225169 | 0.450714027 | 0.638864297 |
| ENSG00000227173 | MYL6P3 | 11.57199189 | -0.104660589 | 0.554622435 | -0.188706013 | 0.850323238 | 0.921722586 |
| ENSG00000227176 | NSA2P5 | 6.096677613 | 0.578183162 | 0.727601187 | 0.794642961 | 0.42682122 | 0.61762108 |
| ENSG00000227183 | HDGFP1 | 5.071615651 | -0.643639298 | 0.81665859 | -0.788137547 | 0.430616255 | 0.620756944 |
| ENSG00000227192 | ENSG0000022 | 2.914814012 | 1.634108882 | 1.114796469 | 1.465836077 | 0.14269295 | 0.28042288 |
| ENSG00000227198 | C6orf47-AS1 | 7.893282849 | -0.629188718 | 0.626353051 | -1.004527266 | 0.315124536 | 0.503784196 |
| ENSG00000227199 | ST7-AS1 | 33.36196178 | -0.055898675 | 0.311895944 | -0.179222193 | 0.857763239 | 0.926188651 |
| ENSG00000227203 | SUB1P1 | 31.32242609 | -0.040606273 | 0.314933923 | -0.12893585 | 0.89740841 | 0.948060798 |
| ENSG00000227209 | WARS2P1 | 15.83183442 | -5.056396123 | 0.916376105 | -5.517817513 | 3.43E-08 | 2.33E-07 |
| ENSG00000227210 | WDR35-DT | 15.9150239 | 0.486143964 | 0.467144654 | 1.040671148 | 0.298028198 | 0.484602131 |
| ENSG00000227212 | PFN1P6 | 1.667072563 | -0.222528607 | 1.337332752 | -0.166397336 | 0.86784427 | NA |
| ENSG00000227214 | HCG15 | 10.08782523 | -0.996092406 | 0.562711813 | -1.770164378 | 0.076699762 | 0.170990528 |
| ENSG00000227218 | ENSG0000022 | 4.125100826 | 1.31011975 | 0.929067236 | 1.410145251 | 0.158496798 | 0.303727053 |
| ENSG00000227227 | ENSG0000022 | 1.967765325 | -0.190545709 | 1.298844863 | -0.146703979 | 0.883365678 | 0.93961708 |
| ENSG00000227232 | WASH7P | 39.90779567 | -0.339209776 | 0.287162498 | -1.181246779 | 0.237504703 | 0.412848214 |
| ENSG00000227237 | ZNF692-DT | 1.623590536 | 1.806608311 | 1.53193139 | 1.17930106 | 0.238278316 | NA |
| ENSG00000227242 | ENSG0000022 | 6.417280473 | 0.351771248 | 0.724815936 | 0.485324936 | 0.62744587 | 0.780187429 |
| ENSG00000227252 | COPS8-DT | 45.98748352 | 0.657194313 | 0.266588499 | 2.465201291 | 0.01369363 | 0.039212567 |
| ENSG00000227253 | ENSG0000022 | 3.806370552 | -1.341391469 | 0.935941956 | -1.43319942 | 0.151800855 | 0.294139 |
| ENSG00000227256 | MIS18A-AS1 | 1.9502777 | 0.289706197 | 1.254933674 | 0.230853792 | 0.81742839 | 0.90328838 |
| ENSG00000227258 | SMIM2-AS1 | 13.18887983 | -1.350351292 | 0.520522128 | -2.594224568 | 0.009480454 | 0.028390523 |
| ENSG00000227268 | KLLN | 99.8716141 | -0.158572744 | 0.182016397 | -0.871200324 | 0.383644783 | 0.576144406 |
| ENSG00000227278 | ENSG0000022 | 3.192732263 | -1.292798499 | 0.997289931 | -1.296311593 | 0.194868156 | 0.355975658 |
| ENSG00000227288 | ARID3BP1 | 19.94687962 | -0.535771559 | 0.39265967 | -1.364468011 | 0.172420315 | 0.324392851 |
| ENSG00000227295 | ELL2P1 | 15.70450363 | -0.206499583 | 0.480966751 | -0.429342741 | 0.667673816 | 0.808785741 |
| ENSG00000227304 | ENSG0000022 | 7.168827485 | -0.436657489 | 0.654422505 | -0.667240942 | 0.504618243 | 0.685322056 |
| ENSG00000227311 | RPL21P20 | 8.728203293 | 0.254027932 | 0.593402332 | 0.428087182 | 0.668587648 | 0.80953239 |
| ENSG00000227337 | RPL23AP43 | 11.37125868 | 0.181245742 | 0.526155182 | 0.344472026 | 0.73049133 | 0.850819131 |
| ENSG00000227339 | THRAP3P1 | 9.687517312 | -0.494538706 | 0.560254094 | -0.882704314 | 0.37739605 | 0.570152414 |
| ENSG00000227344 | HAUS6P1 | 44.546761 | -0.239306108 | 0.277525181 | -0.862286109 | 0.388530095 | 0.581286796 |
| ENSG00000227345 | PARG | 1033.560894 | -0.642164544 | 0.069781595 | -9.202491673 | 3.50E-20 | 5.90E-19 |
| ENSG00000227347 | HNRNPKP2 | 326.5011548 | -0.163577135 | 0.105763158 | -1.546636269 | 0.121950979 | 0.248561124 |

| | | | | | | | |
|---|---|---|---|---|---|---|---|
| ENSG00000227354 | RBM26-AS1 | 35.94437748 | 0.606563052 | 0.297318937 | 2.040109047 | 0.041339466 | 0.101886148 |
| ENSG00000227359 | LINC02936 | 11.63252522 | -0.502523505 | 0.513807692 | -0.978038113 | 0.328055474 | 0.518398607 |
| ENSG00000227361 | RPS24P7 | 6.526985918 | -0.125412489 | 0.682438413 | -0.183771146 | 0.854192993 | 0.924103822 |
| ENSG00000227372 | TP73-AS1 | 1.78383241 | -0.454052336 | 1.323396462 | -0.343096229 | 0.731526068 | NA |
| ENSG00000227373 | RABGAP1L-DT | 3.557794051 | 0.605593298 | 0.929582424 | 0.651468103 | 0.514744361 | 0.693785878 |
| ENSG00000227375 | DLG1-AS1 | 7.062043885 | 0.012345795 | 0.704491407 | 0.017524408 | 0.986018261 | 0.993350589 |
| ENSG00000227376 | FTH1P16 | 110.7570105 | 0.431929747 | 0.173775607 | 2.485560288 | 0.012934768 | 0.03727683 |
| ENSG00000227382 | EIF4A2P2 | 3.747451899 | -0.486387931 | 0.932787017 | -0.521435142 | 0.602063676 | 0.761584821 |
| ENSG00000227388 | ENSG0000022 | 28.90620801 | -0.698375726 | 0.334002837 | -2.090927525 | 0.036534561 | 0.091988018 |
| ENSG00000227398 | KIF9-AS1 | 81.44941208 | 0.471991783 | 0.211073577 | 2.23614812 | 0.02534207 | 0.067215527 |
| ENSG00000227401 | RPL37P1 | 3.75120535 | 0.994356968 | 0.932669637 | 1.066140602 | 0.286360088 | 0.470128134 |
| ENSG00000227403 | LINC01806 | 57.57699296 | 0.385141982 | 0.23327636 | 1.651011625 | 0.098736201 | 0.209839583 |
| ENSG00000227404 | KRT8P20 | 9.729686662 | -0.490198586 | 0.578632733 | -0.84716705 | 0.396902016 | 0.589812286 |
| ENSG00000227407 | ENSG0000022 | 3.440125355 | -0.201058106 | 0.973641554 | -0.206501156 | 0.836399462 | 0.913666841 |
| ENSG00000227409 | ZMYM4-AS1 | 3.178105266 | -0.972255643 | 0.994820075 | -0.977318078 | 0.328411706 | 0.518747128 |
| ENSG00000227431 | CSE1L-DT | 3.353068473 | -0.50811725 | 0.961285763 | -0.528580855 | 0.597096244 | 0.75769879 |
| ENSG00000227440 | ATP5MC1P4 | 1.760114528 | -1.527266478 | 1.410000177 | -1.083167579 | 0.278734047 | NA |
| ENSG00000227449 | FGF7P6 | 4.390276215 | -0.250099476 | 0.842244871 | -0.296943899 | 0.76650934 | 0.872468208 |
| ENSG00000227456 | LINC00310 | 3.199922987 | -0.046576115 | 0.991674394 | -0.046967145 | 0.962539413 | 0.982803851 |
| ENSG00000227462 | PSME2P6 | 3.689649013 | -1.314197852 | 0.972210169 | -1.351763121 | 0.176451105 | 0.330054498 |
| ENSG00000227470 | RPL39P16 | 5.113646338 | 1.501149335 | 0.857871091 | 1.749854205 | 0.080143475 | 0.176880057 |
| ENSG00000227471 | AKR1B15 | 1.589968137 | 0.465938893 | 1.395263328 | 0.333943338 | 0.738422316 | NA |
| ENSG00000227473 | TSSK5P | 3.514780141 | -0.086778263 | 0.932741875 | -0.093035668 | 0.925875225 | 0.964514676 |
| ENSG00000227477 | STK4-DT | 20.98826109 | -0.431747743 | 0.393181446 | -1.098087785 | 0.272166158 | 0.454173972 |
| ENSG00000227482 | ENSG0000022 | 2.423935796 | -0.319372635 | 1.125837575 | -0.283675587 | 0.776659005 | 0.878763805 |
| ENSG00000227486 | ENSG0000022 | 56.05380018 | -0.12240245 | 0.237098003 | -0.516252556 | 0.605678028 | 0.76434448 |
| ENSG00000227492 | CHUK-DT | 2.332235542 | -1.502327223 | 1.219030245 | -1.232395365 | 0.217801432 | 0.387345131 |
| ENSG00000227495 | KIF1C-AS1 | 15.17228217 | -0.152908649 | 0.451450605 | -0.338705158 | 0.734831854 | 0.853737287 |
| ENSG00000227499 | ENSG0000022 | 2.401345245 | -1.506763297 | 1.22471265 | -1.230299449 | 0.218584991 | 0.388287106 |
| ENSG00000227500 | SCAMP4 | 2432.838069 | 0.01329462 | 0.057989995 | 0.22925714 | 0.818669063 | 0.903986346 |
| ENSG00000227513 | ILRUNP1 | 3.67686384 | 0.236011043 | 0.966118149 | 0.244287971 | 0.807007799 | 0.89748787 |
| ENSG00000227525 | RPL7P6 | 12.56274214 | 0.782483819 | 0.549467009 | 1.424077889 | 0.154423926 | 0.297737844 |
| ENSG00000227533 | SLC2A1-DT | 6.887192581 | 0.112906049 | 0.688535701 | 0.163979949 | 0.869746928 | 0.932336193 |

| | | | | | | | |
|---|---|---|---|---|---|---|---|
| ENSG00000227534 | AP1M2P1 | 2.102023937 | -0.370457165 | 1.262369731 | -0.293461698 | 0.769169272 | 0.874004379 |
| ENSG00000227536 | SOCS5P4 | 7.565561738 | 0.541854057 | 0.664073772 | 0.815954612 | 0.414526095 | 0.606649354 |
| ENSG00000227540 | DNAJC9-AS1 | 41.55078319 | -0.897103804 | 0.287930921 | -3.115691082 | 0.001835144 | 0.006499646 |
| ENSG00000227542 | MYO1B-AS1 | 55.08596404 | -0.240150863 | 0.253477184 | -0.947425956 | 0.34342177 | 0.53437586 |
| ENSG00000227543 | SPAG5-AS1 | 70.579492 | -0.317058514 | 0.25009928 | -1.267730613 | 0.204894169 | 0.369954524 |
| ENSG00000227579 | ENSG0000022 | 4.582339548 | 0.198069377 | 0.822097445 | 0.240931751 | 0.809608013 | 0.899259566 |
| ENSG00000227581 | STAU2P1 | 5.350128353 | 0.639232516 | 0.785810365 | 0.813469184 | 0.415949108 | 0.608069041 |
| ENSG00000227582 | ADGRF5P1 | 17.34530435 | 0.334388937 | 0.42214854 | 0.792112031 | 0.428295356 | 0.618922284 |
| ENSG00000227615 | ENSG0000022 | 2217.03078 | 0.394608071 | 0.054961312 | 7.1797426 | 6.98E-13 | 7.24E-12 |
| ENSG00000227617 | CERS6-AS1 | 29.17292399 | -0.367813154 | 0.347607104 | -1.058128989 | 0.28999664 | 0.474427288 |
| ENSG00000227621 | PHB1P11 | 8.983421667 | -0.233783643 | 0.584898014 | -0.399699841 | 0.689377609 | 0.824478513 |
| ENSG00000227624 | SNRPEP3 | 3.772438964 | 0.74244994 | 0.920763879 | 0.806341296 | 0.420046085 | 0.611295895 |
| ENSG00000227627 | ENSG0000022 | 24.55683278 | 0.203047485 | 0.351792131 | 0.577180292 | 0.563817669 | 0.732388297 |
| ENSG00000227630 | LINC01132 | 20.14086539 | -0.387976235 | 0.406028273 | -0.955539948 | 0.339304713 | 0.530474349 |
| ENSG00000227632 | ENSG0000022 | 5.73366194 | -0.037640307 | 0.732698359 | -0.051372174 | 0.959028958 | 0.981228504 |
| ENSG00000227638 | HNRNPA1P14 | 14.15981899 | 0.515754589 | 0.486901967 | 1.059257557 | 0.289482499 | 0.473762468 |
| ENSG00000227653 | ISCA1P6 | 15.43469064 | -0.091868701 | 0.454920026 | -0.201944728 | 0.839959934 | 0.915757464 |
| ENSG00000227666 | CYCSP24 | 3.427835791 | -0.248347109 | 0.95578947 | -0.259834531 | 0.794991414 | 0.890293445 |
| ENSG00000227671 | ZNF731P | 716.9237888 | 0.02178702 | 0.172558171 | 0.126258985 | 0.899526921 | 0.949178821 |
| ENSG00000227682 | ATP5F1AP2 | 6.45794018 | -0.130199483 | 0.693781349 | -0.187666451 | 0.851138131 | 0.9222591 |
| ENSG00000227694 | RPL23AP74 | 34.44944782 | 0.302587998 | 0.321899006 | 0.940009109 | 0.347212888 | 0.538107259 |
| ENSG00000227698 | ENSG0000022 | 4.140245389 | -0.083564883 | 0.849729497 | -0.098342923 | 0.921659996 | 0.962142046 |
| ENSG00000227704 | ENSG0000022 | 19.29657194 | 0.718509968 | 0.405474031 | 1.772024623 | 0.07639047 | 0.170433513 |
| ENSG00000227709 | SNRNP40P1 | 8.751742845 | 0.449799496 | 0.651471702 | 0.690435968 | 0.489920064 | 0.673011336 |
| ENSG00000227719 | ENSG0000022 | 6.393910715 | -1.858821834 | 0.749709899 | -2.479388142 | 0.0131608 | 0.037866394 |
| ENSG00000227725 | POLR2MP1 | 62.91510655 | 0.03698248 | 0.221679661 | 0.166828475 | 0.867505013 | 0.93096599 |
| ENSG00000227726 | ENSG0000022 | 2.581339649 | -0.475274861 | 1.146893754 | -0.414401822 | 0.678579855 | 0.817059725 |
| ENSG00000227729 | RD3L | 111.9200755 | 10.15617312 | 1.186277997 | 8.561376968 | 1.12E-17 | 1.61E-16 |
| ENSG00000227740 | LINC02803 | 9.347047 | -0.9152692 | 0.58309922 | -1.569662878 | 0.116493562 | 0.239888896 |
| ENSG00000227741 | ENSG0000022 | 6.810580063 | -0.414803644 | 0.679990812 | -0.610013601 | 0.541852798 | 0.715752496 |
| ENSG00000227742 | CALR4P | 35.58492691 | 0.360788615 | 0.292836028 | 1.232049955 | 0.217930424 | 0.387454385 |
| ENSG00000227751 | RCC2-AS1 | 17.60489227 | -0.437395462 | 0.450445259 | -0.971029117 | 0.331533779 | 0.522419625 |
| ENSG00000227755 | ENSG0000022 | 10.54681945 | -2.547589914 | 0.640925826 | -3.9748592 | 7.04E-05 | 0.000322189 |

| | | | | | | | |
|---|---|---|---|---|---|---|---|
| ENSG00000227765 | MYL12BP2 | 17.49191617 | 0.261176242 | 0.448678689 | 0.582100841 | 0.560498759 | 0.729849205 |
| ENSG00000227766 | ENSG0000022 | 5.245093364 | -0.699764724 | 0.786708361 | -0.889484285 | 0.373742865 | 0.566553976 |
| ENSG00000227775 | ENSG0000022 | 13.21461051 | -0.135740936 | 0.485588586 | -0.279538976 | 0.779831231 | 0.880534036 |
| ENSG00000227782 | ENSG0000022 | 4.347409466 | 0.234537983 | 0.847784402 | 0.276648146 | 0.782050292 | 0.881757284 |
| ENSG00000227791 | RPS10P9 | 2.693516713 | -0.535711133 | 1.080808715 | -0.495657673 | 0.620135955 | 0.775127707 |
| ENSG00000227799 | CDPF1P1 | 26.93831844 | 0.498237694 | 0.34076626 | 1.462109815 | 0.14371112 | 0.282109625 |
| ENSG00000227811 | INKA2-AS1 | 11.29140021 | -0.25158096 | 0.524863099 | -0.479326821 | 0.631706144 | 0.782934732 |
| ENSG00000227825 | SLC9A7P1 | 12.46882521 | 0.604294275 | 0.507374362 | 1.191022488 | 0.233644757 | 0.408117774 |
| ENSG00000227827 | PKD1P2 | 7.998313124 | -0.425135711 | 0.663603306 | -0.640647368 | 0.521751817 | 0.698796513 |
| ENSG00000227850 | SEPTIN2P1 | 20.26513769 | -0.115136339 | 0.395496661 | -0.29111836 | 0.770960799 | 0.87520429 |
| ENSG00000227854 | ENSG0000022 | 2.112000184 | 0.985882621 | 1.246212625 | 0.79110306 | 0.428883854 | 0.61920798 |
| ENSG00000227855 | DPY19L2P3 | 4.050741037 | -0.693544932 | 0.921926279 | -0.752278081 | 0.451883846 | 0.639731883 |
| ENSG00000227857 | ENSG0000022 | 11.53285098 | -0.331929674 | 0.521535755 | -0.636446631 | 0.524485355 | 0.700820949 |
| ENSG00000227885 | ENSG0000022 | 61.96866519 | -0.010754748 | 0.228619635 | -0.047042101 | 0.962479673 | 0.982786597 |
| ENSG00000227887 | RPS26P13 | 5.232848363 | 0.012867179 | 0.756252379 | 0.017014398 | 0.986425129 | 0.993497695 |
| ENSG00000227888 | FAM66A | 3.720952581 | 1.295357096 | 0.951756692 | 1.361017061 | 0.173508298 | 0.325958591 |
| ENSG00000227890 | PSMA2P3 | 19.16709456 | 0.814021524 | 0.426784522 | 1.907336098 | 0.056477079 | 0.13278272 |
| ENSG00000227896 | WAPL-DT | 29.64731214 | -0.052810295 | 0.324379211 | -0.16280419 | 0.870672609 | 0.932942429 |
| ENSG00000227906 | SNAP25-AS1 | 2.150273334 | 0.054839183 | 1.252289648 | 0.043791133 | 0.965070895 | 0.984162095 |
| ENSG00000227908 | IL6ST-DT | 2.124743046 | 0.034290118 | 1.193526321 | 0.028730089 | 0.977079858 | 0.989371527 |
| ENSG00000227910 | ENSG0000022 | 29.71664652 | 1.130743068 | 0.345117828 | 3.276397149 | 0.001051406 | 0.003918007 |
| ENSG00000227934 | ENSG0000022 | 148.1695062 | 0.037981367 | 0.147533618 | 0.257442119 | 0.796837492 | 0.891680589 |
| ENSG00000227939 | RPL3P2 | 497.5090359 | 0.139028975 | 0.093257019 | 1.490815135 | 0.136010037 | 0.270648295 |
| ENSG00000227942 | FRMD8P1 | 3.722720099 | -0.537021661 | 0.93302166 | -0.575572556 | 0.564904136 | 0.733218336 |
| ENSG00000227945 | PTPRK-AS1 | 11.74944746 | 0.545807622 | 0.509010289 | 1.072291924 | 0.283588933 | 0.467452046 |
| ENSG00000227946 | INO80D-AS1 | 87.79385519 | 0.045451369 | 0.190221472 | 0.238939213 | 0.811152718 | 0.900037232 |
| ENSG00000227953 | LINC01341 | 68.61209593 | 0.163381307 | 0.22109929 | 0.738949941 | 0.459937395 | 0.646782557 |
| ENSG00000227959 | EPHA2-AS1 | 2.622107884 | -0.193354843 | 1.077225864 | -0.179493317 | 0.857550365 | 0.926087485 |
| ENSG00000227963 | RBM15-AS1 | 15.62246723 | -0.029947025 | 0.445128162 | -0.067277309 | 0.946360941 | 0.974832509 |
| ENSG00000227968 | BUB3P1 | 4.037761514 | 0.260485404 | 0.886918896 | 0.293696983 | 0.76898946 | 0.873903607 |
| ENSG00000227972 | THAP12P3 | 3.135637157 | -0.5174022 | 1.041967041 | -0.496562923 | 0.619497305 | 0.774603696 |
| ENSG00000227973 | PIN4P1 | 29.89236316 | 0.252831155 | 0.327557308 | 0.771868463 | 0.440192338 | 0.629591136 |
| ENSG00000227992 | ENSG0000022 | 16.25373105 | 2.530038028 | 0.538704661 | 4.696521508 | 2.65E-06 | 1.46E-05 |

| | | | | | | | |
|---|---|---|---|---|---|---|---|
| ENSG00000228002 | DHX9P1 | 41.94692825 | -0.316253509 | 0.293346663 | -1.078087973 | 0.280994498 | 0.464439713 |
| ENSG00000228007 | RPL36AP7 | 12.17165997 | 0.3719829 | 0.507064343 | 0.733600982 | 0.463191962 | 0.649726282 |
| ENSG00000228010 | ENSG0000022 | 7.102283738 | 0.770360736 | 0.666219522 | 1.156316666 | 0.247551654 | 0.424998873 |
| ENSG00000228013 | IL6R-AS1 | 4.476014874 | 1.992741264 | 0.929197702 | 2.144582643 | 0.031986231 | 0.082122456 |
| ENSG00000228022 | HCG20 | 40.3551207 | -1.297729014 | 0.289242258 | -4.486650817 | 7.24E-06 | 3.79E-05 |
| ENSG00000228034 | HMGN2P21 | 3.043846653 | -0.873652221 | 1.013686859 | -0.861856118 | 0.3887667 | 0.581413918 |
| ENSG00000228043 | SPOPL-DT | 20.87025899 | -0.342488595 | 0.413829376 | -0.827608226 | 0.407892408 | 0.600511443 |
| ENSG00000228049 | POLR2J2 | 8.806257045 | 0.135707049 | 0.590545102 | 0.229799636 | 0.818247468 | 0.903779523 |
| ENSG00000228050 | TOP3BP1 | 3.03439055 | -0.543779783 | 1.020985107 | -0.532603051 | 0.594308389 | 0.756068903 |
| ENSG00000228051 | WBP11P3 | 2.718530944 | 0.711712031 | 1.066779498 | 0.667159457 | 0.504670285 | 0.685322056 |
| ENSG00000228053 | RPL21P87 | 4.014488041 | 0.931898872 | 0.895205954 | 1.040988241 | 0.297881006 | 0.484431467 |
| ENSG00000228054 | RANP5 | 1.849580572 | 0.067212731 | 1.272260561 | 0.052829376 | 0.957867856 | 0.980830956 |
| ENSG00000228057 | SEC63P1 | 156.6803873 | -0.165690733 | 0.147418995 | -1.12394426 | 0.261036673 | 0.441313382 |
| ENSG00000228060 | PABPC4-AS1 | 25.82570357 | 0.130159277 | 0.346383481 | 0.375766411 | 0.707090561 | 0.836461297 |
| ENSG00000228061 | ENSG0000022 | 2.053262488 | -0.833901617 | 1.328310345 | -0.627791254 | 0.530140696 | 0.705746726 |
| ENSG00000228065 | LINC01515 | 117.5035439 | -1.022548389 | 0.168633667 | -6.063726215 | 1.33E-09 | 1.05E-08 |
| ENSG00000228071 | RPL7P47 | 8.982557078 | 0.106325175 | 0.595911654 | 0.178424393 | 0.858389693 | 0.92649707 |
| ENSG00000228084 | ENSG0000022 | 31.94174684 | 0.390745612 | 0.328055189 | 1.191097184 | 0.233615435 | 0.40809763 |
| ENSG00000228089 | PNKDP1 | 4.137881995 | 0.734762401 | 0.898048496 | 0.818176751 | 0.413256269 | 0.605408953 |
| ENSG00000228107 | ENSG0000022 | 4.223416584 | 0.290619071 | 0.861861735 | 0.33719918 | 0.735966755 | 0.854386347 |
| ENSG00000228109 | MELTF-AS1 | 62.42150712 | 0.344210953 | 0.227729316 | 1.511491619 | 0.130663242 | 0.262493631 |
| ENSG00000228110 | ST13P19 | 203.2822549 | 0.211461609 | 0.131302818 | 1.610487974 | 0.107291369 | 0.224322275 |
| ENSG00000228118 | MYL6P5 | 16.73917671 | -0.553196561 | 0.435482924 | -1.270305976 | 0.20397566 | 0.368731414 |
| ENSG00000228121 | PHB1P3 | 12.52656861 | -0.319451065 | 0.496153044 | -0.643855901 | 0.51966888 | 0.69707672 |
| ENSG00000228124 | MDN1-AS1 | 4.304308357 | -0.635605734 | 0.858462381 | -0.740400219 | 0.459057186 | 0.646246487 |
| ENSG00000228126 | FALEC | 17.62547048 | 1.341630597 | 0.443084276 | 3.027935473 | 0.002462307 | 0.008500269 |
| ENSG00000228137 | ENSG0000022 | 50.98625887 | -0.143621478 | 0.246086528 | -0.583621862 | 0.559474748 | 0.72907705 |
| ENSG00000228143 | ENSG0000022 | 5.515967253 | -0.155874116 | 0.737822492 | -0.211262354 | 0.832682557 | 0.911472914 |
| ENSG00000228149 | RPL3P1 | 3.878861025 | -0.30700833 | 0.966121823 | -0.317773932 | 0.750656426 | 0.862253188 |
| ENSG00000228157 | ENSG0000022 | 2.027321212 | -1.773207182 | 1.36717152 | -1.296989555 | 0.194634781 | 0.355713979 |
| ENSG00000228158 | TLE1P1 | 172.833949 | 0.405757933 | 0.147937727 | 2.742761712 | 0.006092488 | 0.019221795 |
| ENSG00000228169 | PPIAP19 | 15.07370258 | 0.310862117 | 0.456475853 | 0.681004516 | 0.495868633 | 0.677941809 |
| ENSG00000228170 | ENSG0000022 | 2.059454438 | 0.541245474 | 1.229335769 | 0.44027473 | 0.659738141 | 0.803183832 |

| | | | | | | | |
|---|---|---|---|---|---|---|---|
| ENSG00000228172 | ENSG0000022 | 64.70479885 | 0.613275458 | 0.228033314 | 2.689411678 | 0.007157809 | 0.022153282 |
| ENSG00000228175 | GEMIN8P4 | 7.732023053 | 0.429658113 | 0.625772456 | 0.686604386 | 0.492332064 | 0.6751209 |
| ENSG00000228187 | ENSG0000022 | 35.64764197 | 0.077690176 | 0.310224219 | 0.250432337 | 0.802253025 | 0.894714402 |
| ENSG00000228192 | ZNF691-DT | 37.00242258 | -1.190792695 | 0.295382325 | -4.031360692 | 5.55E-05 | 0.00025783 |
| ENSG00000228195 | RPL5P27 | 2.602708957 | 0.216748546 | 1.088261846 | 0.19916948 | 0.842130173 | 0.917009895 |
| ENSG00000228196 | PTPN2P1 | 55.36415166 | 0.408474265 | 0.247828542 | 1.64821316 | 0.099308937 | 0.210822415 |
| ENSG00000228201 | ENSG0000022 | 5.302905442 | -2.215095892 | 0.867672076 | -2.552918266 | 0.010682456 | 0.031481956 |
| ENSG00000228204 | ENSG0000022 | 1.697758601 | 1.270342261 | 1.423117953 | 0.892647204 | 0.372046132 | NA |
| ENSG00000228205 | RPS3P3 | 384.7559934 | 0.297626915 | 0.108134497 | 2.752377107 | 0.005916435 | 0.018699781 |
| ENSG00000228212 | OFD1P17 | 5.150615353 | -0.467401855 | 0.766292886 | -0.609951969 | 0.541893625 | 0.715752496 |
| ENSG00000228217 | PNRC2P1 | 90.80870811 | 0.044454229 | 0.192371861 | 0.231084881 | 0.81724886 | 0.903254867 |
| ENSG00000228218 | ATF4P3 | 75.84983286 | -0.051386746 | 0.236472095 | -0.217305749 | 0.827970077 | 0.909090629 |
| ENSG00000228221 | LINC00578 | 4.87267482 | 0.808081138 | 0.833563443 | 0.969429676 | 0.332330852 | 0.523388009 |
| ENSG00000228223 | HCG11 | 406.7211929 | -0.094432057 | 0.103331168 | -0.913877764 | 0.360781079 | 0.553344457 |
| ENSG00000228224 | NACA4P | 82.58815898 | 0.216255231 | 0.195908117 | 1.103860497 | 0.269653657 | 0.451225829 |
| ENSG00000228232 | GAPDHP1 | 885.9069692 | -0.01825899 | 0.07285596 | -0.250617654 | 0.802109731 | 0.894662789 |
| ENSG00000228236 | TXNP5 | 8.039484411 | 0.590638336 | 0.631173487 | 0.935778114 | 0.349387442 | 0.540637296 |
| ENSG00000228237 | EFCAB14-AS1 | 5.573314589 | -0.623170728 | 0.753951454 | -0.826539592 | 0.408498068 | 0.600855712 |
| ENSG00000228242 | XPC-AS1 | 33.32892995 | -0.157089975 | 0.315037575 | -0.498638852 | 0.618033829 | 0.77359751 |
| ENSG00000228247 | UBBP2 | 2.193777643 | -0.404350846 | 1.168536277 | -0.346031916 | 0.729318729 | 0.850044287 |
| ENSG00000228251 | ENSG0000022 | 3.233982659 | 0.552219177 | 1.036459158 | 0.532793958 | 0.594176217 | 0.756026536 |
| ENSG00000228252 | COL6A4P2 | 4.491943001 | -2.265417885 | 0.954355515 | -2.37376727 | 0.017607645 | 0.04897388 |
| ENSG00000228253 | MT-ATP8 | 1249.760643 | -0.110960608 | 0.082334655 | -1.347678058 | 0.177761952 | 0.332019149 |
| ENSG00000228261 | ITPRIP-AS1 | 2.130339747 | -0.808239422 | 1.235564392 | -0.654145933 | 0.513017791 | 0.692381325 |
| ENSG00000228264 | PSMD8P1 | 10.7636797 | 0.04470791 | 0.54938268 | 0.081378449 | 0.935140987 | 0.969062396 |
| ENSG00000228274 | ENSG0000022 | 10.23263706 | -0.805988865 | 0.551684532 | -1.460959694 | 0.144026503 | 0.282535315 |
| ENSG00000228275 | ARMCX3-AS1 | 6.536717314 | -0.454396708 | 0.683740316 | -0.664574982 | 0.506322375 | 0.686670724 |
| ENSG00000228283 | KATNBL1P6 | 64.28182686 | -0.203206879 | 0.227704162 | -0.892416182 | 0.372169901 | 0.564841384 |
| ENSG00000228285 | LYPLA2P1 | 182.1535826 | 0.304448514 | 0.136448822 | 2.231228596 | 0.025665991 | 0.067901952 |
| ENSG00000228288 | PCAT6 | 105.7863634 | 1.298475992 | 0.183943484 | 7.05910296 | 1.68E-12 | 1.69E-11 |
| ENSG00000228300 | FAM174C | 674.5343647 | 0.263529211 | 0.082896699 | 3.179007292 | 0.001477804 | 0.005341235 |
| ENSG00000228302 | ENSG0000022 | 7.018151669 | -1.312197714 | 0.684059883 | -1.918249773 | 0.055079348 | 0.130152476 |
| ENSG00000228305 | PRELID1P6 | 438.8743291 | -0.240963011 | 0.093867949 | -2.567042466 | 0.010257005 | 0.030392107 |

| | | | | | | | |
|---|---|---|---|---|---|---|---|
| ENSG00000228314 | CYP4F29P | 65.84083875 | 0.05230112 | 0.236552561 | 0.221097246 | 0.825016715 | 0.90772273 |
| ENSG00000228315 | GUSBP11 | 228.4054866 | -0.713677378 | 0.12978447 | -5.4989428 | 3.82E-08 | 2.59E-07 |
| ENSG00000228323 | MYADM-AS1 | 9.823830928 | -0.998835878 | 0.574254465 | -1.739361101 | 0.081971266 | 0.180255065 |
| ENSG00000228327 | ENSG0000022 | 107.0817887 | -0.119981005 | 0.176009049 | -0.68167521 | 0.495444346 | 0.677768102 |
| ENSG00000228328 | PGK1P1 | 60.8533927 | -0.199648048 | 0.459644857 | -0.434352838 | 0.66403224 | 0.806101475 |
| ENSG00000228331 | RPL17P43 | 3.273676387 | 0.688848329 | 0.979853455 | 0.703011584 | 0.48204853 | 0.666287182 |
| ENSG00000228335 | ENSG0000022 | 103.2196418 | -0.064725354 | 0.18584877 | -0.348268937 | 0.72763822 | 0.848818753 |
| ENSG00000228339 | AMD1P1 | 1.682200473 | -1.400825913 | 1.417719948 | -0.988083659 | 0.323111678 | NA |
| ENSG00000228343 | ENSG0000022 | 3.138919913 | 0.260634422 | 0.995994413 | 0.261682615 | 0.793566145 | 0.889277874 |
| ENSG00000228348 | EFCAB14P1 | 36.29596239 | -0.221609752 | 0.293596843 | -0.754809722 | 0.450363159 | 0.638625693 |
| ENSG00000228350 | LINC02585 | 2.179150645 | 0.011707396 | 1.188513607 | 0.009850452 | 0.992140603 | 0.995917623 |
| ENSG00000228354 | ENSG0000022 | 1.626854689 | -0.782001371 | 1.398983792 | -0.558978149 | 0.576176633 | NA |
| ENSG00000228360 | ENSG0000022 | 2.371590337 | -0.247102782 | 1.13872437 | -0.216999643 | 0.828208623 | 0.909138327 |
| ENSG00000228363 | CHMP3-AS1 | 46.99158594 | 0.046956758 | 0.275988316 | 0.170140382 | 0.864899736 | 0.929799912 |
| ENSG00000228364 | SAR1AP1 | 3.482593464 | -0.611128744 | 0.955930804 | -0.639302282 | 0.522626304 | 0.699355272 |
| ENSG00000228366 | ENSG0000022 | 4.509535475 | -0.301662184 | 0.817601267 | -0.368960025 | 0.712157522 | 0.839972059 |
| ENSG00000228373 | ENSG0000022 | 3.45706718 | -1.795152931 | 1.007832557 | -1.781201568 | 0.074879528 | 0.167682191 |
| ENSG00000228393 | LINC01004 | 40.76896307 | -0.187269384 | 0.288830468 | -0.648371292 | 0.516744833 | 0.695175139 |
| ENSG00000228395 | ENSG0000022 | 21.53571555 | -0.455417695 | 0.400182874 | -1.13802395 | 0.255110482 | 0.434625655 |
| ENSG00000228398 | HMGN2P25 | 4.307666846 | -0.222995509 | 0.866010201 | -0.257497554 | 0.796794703 | 0.891676203 |
| ENSG00000228399 | ENSG0000022 | 9.073424387 | -1.015491042 | 1.186055536 | -0.856191814 | 0.39189169 | 0.584770973 |
| ENSG00000228404 | ENSG0000022 | 2.015558846 | -4.544790592 | 1.584620477 | -2.868062517 | 0.004129939 | 0.013565407 |
| ENSG00000228407 | ENSG0000022 | 6.56717037 | -0.437222241 | 0.685483983 | -0.637829989 | 0.523584356 | 0.700001783 |
| ENSG00000228409 | CCT6P1 | 161.0253918 | 0.636573338 | 0.14893703 | 4.274110592 | 1.92E-05 | 9.55E-05 |
| ENSG00000228412 | LNC-LBCS | 104.2834049 | -2.525005919 | 0.208824974 | -12.09149397 | 1.17E-33 | 3.74E-32 |
| ENSG00000228414 | REL-DT | 1.982452911 | 0.246392563 | 1.23835083 | 0.198968303 | 0.842287539 | 0.917009895 |
| ENSG00000228420 | LINC01768 | 13.47693553 | -0.548007307 | 0.49245112 | -1.112815638 | 0.265787618 | 0.446828286 |
| ENSG00000228434 | ENSG0000022 | 19.40749927 | 0.137522534 | 0.406351113 | 0.338432774 | 0.73503708 | 0.853889229 |
| ENSG00000228436 | RRAGC-DT | 33.33635296 | 0.429888307 | 0.306280198 | 1.403578517 | 0.160444396 | 0.306716204 |
| ENSG00000228444 | ENSG0000022 | 2.489819653 | 1.227607948 | 1.214585411 | 1.010721796 | 0.312149602 | 0.500364186 |
| ENSG00000228446 | ANKRD49P1 | 2.886382211 | -1.446776456 | 1.066329424 | -1.356781894 | 0.174850522 | 0.327943012 |
| ENSG00000228451 | SDAD1P1 | 31.95787449 | 0.232197848 | 0.32164347 | 0.721910653 | 0.470349411 | 0.656115263 |
| ENSG00000228463 | ENSG0000022 | 35.28631655 | 0.048056676 | 0.315620686 | 0.152260856 | 0.878981197 | 0.937198636 |

| | | | | | | | |
|---|---|---|---|---|---|---|---|
| ENSG00000228474 | OST4 | 2253.980272 | -0.21035805 | 0.053862988 | -3.905428505 | 9.41E-05 | 0.000420718 |
| ENSG00000228477 | ENSG0000022 | 37.93707618 | 0.21421389 | 0.30786318 | 0.695808737 | 0.486548617 | 0.670074159 |
| ENSG00000228486 | C2orf92 | 120.7429252 | 0.041215636 | 0.182122101 | 0.226307714 | 0.820962099 | 0.904991377 |
| ENSG00000228492 | RAB11FIP1P1 | 2.426838734 | -2.108586542 | 1.292812643 | -1.631007056 | 0.102888831 | 0.216996394 |
| ENSG00000228499 | TMSB10P1 | 5.373706455 | -0.879387861 | 0.802756508 | -1.095460269 | 0.273315042 | 0.455220404 |
| ENSG00000228501 | RPL15P18 | 33.6330718 | 0.11000466 | 0.328837187 | 0.334526215 | 0.737982512 | 0.855534609 |
| ENSG00000228502 | EEF1A1P11 | 147.6071272 | 0.293726798 | 0.158055621 | 1.858376162 | 0.063115615 | 0.145500172 |
| ENSG00000228503 | ENSG0000022 | 20.37749302 | 1.019480775 | 0.409253417 | 2.491074562 | 0.012735739 | 0.036763288 |
| ENSG00000228506 | PNISR-AS1 | 85.90465857 | -0.492909405 | 0.202658502 | -2.432216761 | 0.015006723 | 0.042526774 |
| ENSG00000228507 | DAP3P2 | 8.914009202 | 0.305014135 | 0.587910652 | 0.518810356 | 0.603892995 | 0.762847209 |
| ENSG00000228509 | ENSG0000022 | 1.674628506 | -0.130494464 | 1.391585657 | -0.093773936 | 0.925288736 | NA |
| ENSG00000228510 | ENSG0000022 | 7.71649121 | -0.308653124 | 0.631153762 | -0.489029999 | 0.624820458 | 0.778298345 |
| ENSG00000228519 | RANBP1P1 | 372.5526017 | -0.587800186 | 0.098142569 | -5.989248037 | 2.11E-09 | 1.63E-08 |
| ENSG00000228526 | MIR34AHG | 135.3553015 | 0.085095801 | 0.168260884 | 0.505737274 | 0.613041098 | 0.769993582 |
| ENSG00000228532 | SUMO2P21 | 864.6279544 | 0.08530244 | 0.073217073 | 1.165062151 | 0.243993836 | 0.420716972 |
| ENSG00000228544 | CCDC183-AS1 | 231.1027304 | 0.130552979 | 0.126837605 | 1.029292372 | 0.303342307 | 0.490496406 |
| ENSG00000228561 | ENSG0000022 | 1.84447427 | -0.910735499 | 1.401586037 | -0.649789221 | 0.515828382 | 0.694553213 |
| ENSG00000228589 | SPCS2P4 | 832.411265 | 0.000450574 | 0.076288341 | 0.005906195 | 0.995287566 | 0.9970696 |
| ENSG00000228594 | FNDC10 | 1263.222787 | 0.355411231 | 0.073572621 | 4.830753947 | 1.36E-06 | 7.73E-06 |
| ENSG00000228606 | DCAF8-DT | 7.468897457 | 1.091328204 | 0.699869885 | 1.559330137 | 0.118918262 | 0.243853128 |
| ENSG00000228612 | HK2P1 | 126.1909046 | 0.354985592 | 0.165444231 | 2.145651073 | 0.031900827 | 0.08193987 |
| ENSG00000228613 | ENSG0000022 | 88.20048678 | -0.292279016 | 0.194643158 | -1.501614644 | 0.133196659 | 0.266346876 |
| ENSG00000228624 | HDAC2-AS2 | 8.661576978 | 0.54117073 | 0.598852774 | 0.90367909 | 0.36616559 | 0.558908685 |
| ENSG00000228630 | HOTAIR | 366.0587864 | 0.177145482 | 0.102213049 | 1.733100461 | 0.083077814 | 0.182251608 |
| ENSG00000228643 | ENSG0000022 | 4.147315817 | 0.027865101 | 0.961598761 | 0.028977888 | 0.976882226 | 0.989309285 |
| ENSG00000228649 | SNHG26 | 8.174639002 | -2.475955649 | 0.737889064 | -3.355457844 | 0.000792337 | 0.003021815 |
| ENSG00000228656 | MYO5BP3 | 1.724724611 | 1.287950336 | 1.387597097 | 0.92818754 | 0.353310297 | NA |
| ENSG00000228657 | RPL23AP58 | 10.10964185 | 0.561034421 | 0.582408315 | 0.963300842 | 0.335396571 | 0.526589373 |
| ENSG00000228661 | STIM1-AS1 | 2.067037546 | 0.057328307 | 1.265622643 | 0.045296524 | 0.963870958 | 0.983656246 |
| ENSG00000228665 | RPS27P6 | 10.67854558 | 0.114619341 | 0.531861216 | 0.215506109 | 0.82937275 | 0.909755986 |
| ENSG00000228666 | KRT18P1 | 18.73791076 | 0.473607903 | 0.405778915 | 1.167157498 | 0.243146771 | 0.419590311 |
| ENSG00000228672 | PROB1 | 116.1784002 | 0.386499486 | 0.16714404 | 2.312373714 | 0.020757098 | 0.056471516 |
| ENSG00000228677 | TTC3-AS1 | 21.77782738 | -0.463582472 | 0.375937498 | -1.23313709 | 0.217524621 | 0.387066571 |

| | | | | | | | |
|---|---|---|---|---|---|---|---|
| ENSG00000228696 | ARL17B | 4.868846263 | -0.501043471 | 0.781859446 | -0.640835733 | 0.521629414 | 0.698673367 |
| ENSG00000228700 | ENSG0000022 | 1.668436555 | -2.14940984 | 1.520230646 | -1.413870879 | 0.157399818 | NA |
| ENSG00000228701 | TNKS2-DT | 29.11376483 | -1.518646782 | 0.346069955 | -4.388265322 | 1.14E-05 | 5.85E-05 |
| ENSG00000228702 | ENSG0000022 | 3.138780215 | 0.46521155 | 1.006029923 | 0.462423174 | 0.643777884 | 0.791270263 |
| ENSG00000228703 | ENSG0000022 | 5.206853961 | 0.395759358 | 0.777468679 | 0.509035757 | 0.610727161 | 0.768686551 |
| ENSG00000228705 | LINC00659 | 27.61280873 | -1.879551292 | 0.366045466 | -5.134748184 | 2.83E-07 | 1.75E-06 |
| ENSG00000228708 | ENSG0000022 | 2.748399892 | -0.603058322 | 1.068370238 | -0.56446567 | 0.572437258 | 0.739291922 |
| ENSG00000228709 | LINC02575 | 2.384272286 | -2.76445404 | 1.42165588 | -1.944531078 | 0.051831442 | 0.123533846 |
| ENSG00000228716 | DHFR | 1274.025324 | 0.076525054 | 0.065651184 | 1.165630983 | 0.243763675 | 0.42044652 |
| ENSG00000228719 | ENSG0000022 | 3.058409058 | -0.611147885 | 1.004388546 | -0.608477553 | 0.542870789 | 0.716541968 |
| ENSG00000228727 | SAPCD1 | 6.478650275 | -0.560667132 | 0.684226982 | -0.81941687 | 0.412548613 | 0.604603922 |
| ENSG00000228737 | ENSG0000022 | 8.827441211 | -0.045554388 | 0.601601137 | -0.075721912 | 0.939640343 | 0.971233034 |
| ENSG00000228748 | ENSG0000022 | 15.85550218 | -1.852403257 | 0.48587497 | -3.812510153 | 0.000137563 | 0.000599598 |
| ENSG00000228753 | EIF4BP2 | 33.12370582 | 0.075710955 | 0.308965251 | 0.24504683 | 0.806420171 | 0.897326287 |
| ENSG00000228766 | RPL7L1P8 | 134.9385556 | -0.021018529 | 0.155880958 | -0.134837054 | 0.892740709 | 0.945104613 |
| ENSG00000228767 | KRT8P18 | 7.334035506 | -0.044758039 | 0.670930411 | -0.066710404 | 0.946812251 | 0.975200824 |
| ENSG00000228775 | WEE2-AS1 | 143.4185353 | -0.334371985 | 0.151145037 | -2.212259115 | 0.026948766 | 0.070854407 |
| ENSG00000228782 | ENSG0000022 | 159.4858277 | 0.089482907 | 0.156163966 | 0.573006112 | 0.566640556 | 0.734516265 |
| ENSG00000228791 | THRB-AS1 | 11.22655601 | 0.234582942 | 0.521946072 | 0.449439041 | 0.653114973 | 0.798681033 |
| ENSG00000228793 | ENSG0000022 | 4.455607134 | -1.380084798 | 0.859144514 | -1.606347682 | 0.108197539 | 0.225846365 |
| ENSG00000228794 | LINC01128 | 410.4485757 | 0.255714884 | 0.098242869 | 2.602884932 | 0.009244297 | 0.027771632 |
| ENSG00000228797 | FAM207BP | 147.0541525 | 0.085753356 | 0.150822723 | 0.568570532 | 0.569647627 | 0.736956721 |
| ENSG00000228801 | PCMTD1-DT | 195.5113286 | -0.218523276 | 0.132084009 | -1.65442644 | 0.098040898 | 0.208632486 |
| ENSG00000228808 | HMGB3P4 | 1.706792576 | -0.77586848 | 1.327128211 | -0.584622099 | 0.558801845 | NA |
| ENSG00000228812 | LAMA5-AS1 | 3.949158548 | -0.886189656 | 0.893473364 | -0.991847873 | 0.32127174 | 0.510775087 |
| ENSG00000228817 | BACH1-IT2 | 4.559281507 | 0.510349229 | 0.845411536 | 0.603669582 | 0.546063341 | 0.719092449 |
| ENSG00000228818 | ENSG0000022 | 239.4031526 | -0.176479496 | 0.119627066 | -1.47524721 | 0.140146093 | 0.2766686 |
| ENSG00000228819 | AGKP2 | 8.00481379 | 0.151136008 | 0.646864856 | 0.23364387 | 0.815261467 | 0.902135938 |
| ENSG00000228824 | MIR4500HG | 7.702744794 | 2.455155235 | 0.78675214 | 3.120620981 | 0.001804702 | 0.006400719 |
| ENSG00000228825 | LAMTOR3P1 | 1.973000487 | -1.630676076 | 1.364483612 | -1.195086597 | 0.232053198 | 0.406017875 |
| ENSG00000228828 | TLK2P2 | 12.74696235 | 0.522590589 | 0.491198088 | 1.063910063 | 0.287369444 | 0.471279314 |
| ENSG00000228830 | ENSG0000022 | 41.97056742 | -0.215879213 | 0.288524818 | -0.748217136 | 0.454329196 | 0.64191507 |
| ENSG00000228834 | ATP5MFP2 | 39.06038484 | -0.623376103 | 0.310428829 | -2.008112795 | 0.044631307 | 0.10868813 |

| | | | | | | | |
|---|---|---|---|---|---|---|---|
| ENSG00000228836 | CT45A5 | 7.727522039 | 4.342130341 | 1.140271319 | 3.807979967 | 0.000140107 | 0.000609992 |
| ENSG00000228838 | ENSG0000022 | 3.87502879 | -0.839863288 | 0.890300334 | -0.943348279 | 0.345502789 | 0.53639916 |
| ENSG00000228839 | PIK3IP1-DT | 7.953092475 | -1.266403421 | 0.686068021 | -1.845886095 | 0.064908746 | 0.149004967 |
| ENSG00000228843 | ENSG0000022 | 5.13355386 | -0.463455479 | 0.777521738 | -0.596067553 | 0.551130099 | 0.723161782 |
| ENSG00000228844 | ENSG0000022 | 3.198483262 | 1.031513479 | 1.022380974 | 1.008932585 | 0.313006966 | 0.501213166 |
| ENSG00000228847 | ATP5MC2P4 | 23.02783224 | 0.591678423 | 0.377786376 | 1.566171944 | 0.117308369 | 0.24130685 |
| ENSG00000228861 | RPL23AP12 | 5.92408894 | -0.032368191 | 0.705497901 | -0.045879925 | 0.963405955 | 0.983294741 |
| ENSG00000228862 | ENSG0000022 | 3.07551544 | 0.973651611 | 1.036610343 | 0.939264804 | 0.347594804 | 0.538517242 |
| ENSG00000228868 | MYLKP1 | 3.21649103 | 2.277979629 | 1.162296731 | 1.959895067 | 0.050008056 | 0.119847975 |
| ENSG00000228873 | ENSG0000022 | 30.08952358 | -1.526818831 | 0.348141268 | -4.385630118 | 1.16E-05 | 5.91E-05 |
| ENSG00000228878 | SEPTIN7-DT | 48.15916319 | 0.10168654 | 0.255719188 | 0.397649238 | 0.69088876 | 0.825640142 |
| ENSG00000228887 | EEF1DP1 | 122.9272864 | 0.347664296 | 0.192046048 | 1.810317369 | 0.070246585 | 0.15903056 |
| ENSG00000228889 | UBAC2-AS1 | 54.12265872 | 0.337004742 | 0.244991724 | 1.375576027 | 0.168952933 | 0.319204438 |
| ENSG00000228903 | RASA4CP | 68.12010773 | 0.391210884 | 0.220014653 | 1.778112862 | 0.07538532 | 0.168600882 |
| ENSG00000228925 | ENSG0000022 | 7.65671243 | -0.444008277 | 0.653078836 | -0.67986934 | 0.496587197 | 0.678522264 |
| ENSG00000228928 | KPNA2P2 | 7.349084632 | 0.161739295 | 0.660859367 | 0.244740868 | 0.806657082 | 0.897382528 |
| ENSG00000228929 | RPS13P2 | 770.1247311 | 0.284862797 | 0.074052537 | 3.846766236 | 0.000119687 | 0.000526788 |
| ENSG00000228933 | ENSG0000022 | 4.159149032 | 1.298243591 | 0.910807847 | 1.42537594 | 0.154048559 | 0.297288966 |
| ENSG00000228938 | SNRPD2P1 | 7.246348852 | 0.505831294 | 0.712255458 | 0.710182405 | 0.477591031 | 0.662370438 |
| ENSG00000228941 | UBE3AP2 | 3.117113332 | -0.339902927 | 0.977532767 | -0.347715124 | 0.728054137 | 0.849088048 |
| ENSG00000228950 | ENSG0000022 | 3.779487413 | -0.312372281 | 0.893776195 | -0.349497204 | 0.726716069 | 0.848088042 |
| ENSG00000228952 | LINC02041 | 9.534195027 | -0.122598405 | 0.569863958 | -0.215136267 | 0.829661079 | 0.909788127 |
| ENSG00000228956 | SATB1-AS1 | 20.14967165 | 0.869538492 | 0.399064941 | 2.178939825 | 0.029336139 | 0.076156401 |
| ENSG00000228974 | ENSG0000022 | 12.81025646 | 0.085182632 | 0.494402996 | 0.172293924 | 0.863206461 | 0.92901507 |
| ENSG00000228981 | ENSG0000022 | 7.397820424 | 0.273594965 | 0.641422144 | 0.426544308 | 0.669711269 | 0.810251749 |
| ENSG00000228982 | ENSG0000022 | 1.800886766 | -4.389818957 | 1.622484926 | -2.705614633 | 0.006817811 | 0.021192492 |
| ENSG00000228986 | PHF10P1 | 12.73399283 | -0.113890455 | 0.511262902 | -0.222762995 | 0.82371997 | 0.906941978 |
| ENSG00000228998 | PERPP2 | 14.33795445 | 1.462233648 | 0.495883848 | 2.94874224 | 0.0031907 | 0.010743381 |
| ENSG00000229000 | SEPTIN7P8 | 9.589243237 | -0.297823517 | 0.582434987 | -0.511342079 | 0.609111545 | 0.76732668 |
| ENSG00000229017 | LINC01277 | 11.11696839 | 1.399718179 | 0.566592669 | 2.4704135 | 0.013495695 | 0.038698906 |
| ENSG00000229018 | PMS2P7 | 4.523209468 | 0.42473047 | 0.825836885 | 0.5143031 | 0.607040095 | 0.765515902 |
| ENSG00000229021 | ENSG0000022 | 5.563790843 | 0.340616839 | 0.775042451 | 0.439481526 | 0.660312665 | 0.80367002 |
| ENSG00000229022 | RPL6P3 | 17.4304916 | 0.091522709 | 0.418675337 | 0.21860067 | 0.826961131 | 0.908591807 |

| | | | | | | | |
|---|---|---|---|---|---|---|---|
| ENSG00000229023 | RAB1AP1 | 9.707388884 | 0.733402113 | 0.581894442 | 1.260369681 | 0.207536034 | 0.373724516 |
| ENSG00000229028 | KRT223P | 3.374290946 | -1.081395771 | 0.96451119 | -1.121185303 | 0.262208994 | 0.442708054 |
| ENSG00000229036 | VDAC1P8 | 7.513193997 | 0.313206431 | 0.633698419 | 0.494251558 | 0.621128532 | 0.775649892 |
| ENSG00000229043 | ZFAND2A-DT | 561.5477393 | 0.255780585 | 0.093439399 | 2.73739544 | 0.00619278 | 0.019493261 |
| ENSG00000229044 | ENSG0000022 | 12.86698506 | -0.723510627 | 0.532419013 | -1.358912078 | 0.174174451 | 0.326995818 |
| ENSG00000229065 | ENSG0000022 | 3.47458087 | 1.565719527 | 1.019046783 | 1.536455003 | 0.124426826 | 0.252513206 |
| ENSG00000229067 | ENSG0000022 | 2.94983871 | 1.151296719 | 1.078971804 | 1.06703133 | 0.285957689 | 0.46973644 |
| ENSG00000229081 | LINC01165 | 11.66770753 | 2.328379112 | 0.604098144 | 3.854306021 | 0.000116058 | 0.000512291 |
| ENSG00000229083 | PSMA6P2 | 40.34657833 | -0.279909482 | 0.292280964 | -0.957672637 | 0.338227859 | 0.529548668 |
| ENSG00000229091 | HSPA8P8 | 94.13183207 | -0.022420318 | 0.196722178 | -0.113969449 | 0.909262012 | 0.954587195 |
| ENSG00000229097 | CALM2P2 | 270.3853885 | -0.093137187 | 0.121580895 | -0.766051176 | 0.443645853 | 0.632545835 |
| ENSG00000229106 | BTBD6P1 | 11.43536242 | -0.231346729 | 0.558843994 | -0.413973723 | 0.678893351 | 0.817351395 |
| ENSG00000229109 | FRMD3-AS1 | 5.162604292 | -0.047485919 | 0.816293305 | -0.058172618 | 0.953611132 | 0.978522068 |
| ENSG00000229111 | MED4-AS1 | 6.866298853 | -0.170872908 | 0.696900119 | -0.245189954 | 0.806309354 | 0.8972465 |
| ENSG00000229117 | RPL41 | 2188.10029 | -0.011927594 | 0.065865103 | -0.18109125 | 0.856295953 | 0.925565113 |
| ENSG00000229119 | RPLP0P9 | 630.3862616 | 0.189423737 | 0.08150133 | 2.324179701 | 0.020115865 | 0.054961641 |
| ENSG00000229127 | KANSL1L-AS1 | 6.668270224 | 0.607096143 | 0.693520925 | 0.87538259 | 0.381365769 | 0.573598921 |
| ENSG00000229129 | ACTG1P2 | 1.650545405 | 1.0481031 | 1.402489086 | 0.747316404 | 0.454872593 | NA |
| ENSG00000229132 | EIF4A1P10 | 1850.624132 | -0.030553342 | 0.059500431 | -0.513497826 | 0.607603131 | 0.765889105 |
| ENSG00000229133 | RPS7P4 | 62.21463958 | 0.17538593 | 0.222888528 | 0.78687733 | 0.431353679 | 0.621444037 |
| ENSG00000229140 | CCDC26 | 7.208367968 | -1.357148718 | 0.70424797 | -1.927089286 | 0.053968504 | 0.12796253 |
| ENSG00000229152 | ANKRD10-IT1 | 32.29719013 | -0.114076398 | 0.309443893 | -0.3686497 | 0.712388845 | 0.839972059 |
| ENSG00000229161 | TCP1P1 | 34.11069358 | 0.140467425 | 0.339396463 | 0.413874158 | 0.67896627 | 0.817353392 |
| ENSG00000229165 | GDI2P1 | 2.560184819 | -1.62834347 | 1.202355026 | -1.354295058 | 0.175642262 | 0.328902689 |
| ENSG00000229178 | ENSG0000022 | 10.27285156 | -0.527302859 | 0.563002331 | -0.936590898 | 0.348969036 | 0.540208494 |
| ENSG00000229180 | RABGEF1P1 | 340.9264083 | 0.572998995 | 0.112662329 | 5.085985715 | 3.66E-07 | 2.23E-06 |
| ENSG00000229184 | ATP5PDP2 | 11.57838413 | 0.233391327 | 0.569197696 | 0.410035614 | 0.681779822 | 0.819321553 |
| ENSG00000229190 | ENSG0000022 | 3.431920713 | -0.861206277 | 0.943858434 | -0.912431617 | 0.361541552 | 0.554099627 |
| ENSG00000229204 | PTGES3P3 | 172.2988634 | -0.087045375 | 0.147837163 | -0.588792244 | 0.556000649 | 0.726719554 |
| ENSG00000229207 | SERPINH1P1 | 2.904363694 | 0.114337957 | 1.065535782 | 0.1073056 | 0.914546542 | 0.957607799 |
| ENSG00000229211 | KCTD9P6 | 7.371597501 | -0.754911718 | 0.659662645 | -1.144390582 | 0.252461692 | 0.431178901 |
| ENSG00000229212 | ENSG0000022 | 27.24769739 | 0.253716742 | 0.352221131 | 0.720333677 | 0.471319575 | 0.657148912 |
| ENSG00000229214 | LINC00242 | 5.492510246 | 0.040310524 | 0.760982539 | 0.052971681 | 0.957754472 | 0.980830956 |

| | | | | | | | |
|---|---|---|---|---|---|---|---|
| ENSG00000229222 | KRT18P4 | 25.93060601 | 0.317883151 | 0.367568541 | 0.86482687 | 0.387133821 | 0.57972778 |
| ENSG00000229237 | HMGN1P37 | 199.398059 | -0.256857909 | 0.135660809 | -1.89338328 | 0.058306917 | 0.136291082 |
| ENSG00000229241 | PNPT1P1 | 6.14966431 | 0.030044209 | 0.694461866 | 0.043262576 | 0.965492223 | 0.984416712 |
| ENSG00000229251 | HNRNPA1P8 | 19.54166983 | 0.209610421 | 0.429508827 | 0.488023546 | 0.625533163 | 0.778818494 |
| ENSG00000229256 | ST13P13 | 8.206792455 | 0.410385374 | 0.62773201 | 0.653758877 | 0.513267164 | 0.692527276 |
| ENSG00000229257 | ENSG0000022 | 39.9958199 | -0.497258937 | 0.27767725 | -1.790780259 | 0.073328564 | 0.164905503 |
| ENSG00000229267 | SNHG31 | 31.19024495 | 0.685801329 | 0.319460881 | 2.146745875 | 0.031813518 | 0.081743068 |
| ENSG00000229268 | PES1P2 | 3.057037604 | -1.572728253 | 1.050510324 | -1.497108802 | 0.13436495 | 0.26800556 |
| ENSG00000229272 | LINC03036 | 8.424775085 | 0.294093896 | 0.631332649 | 0.46583033 | 0.641336953 | 0.789596379 |
| ENSG00000229273 | ENSG0000022 | 6.722191428 | 0.610889982 | 0.686447936 | 0.889929083 | 0.373503967 | 0.566266671 |
| ENSG00000229298 | TUBB8P1 | 5.393048879 | -0.068472589 | 0.77132444 | -0.088772746 | 0.929262517 | 0.965981609 |
| ENSG00000229299 | ZBTB46-AS2 | 44.84537487 | -0.559901526 | 0.277566123 | -2.017182499 | 0.043676472 | 0.106793675 |
| ENSG00000229312 | ENSG0000022 | 18.36926023 | 0.298979416 | 0.418157866 | 0.714991729 | 0.474614154 | 0.659944021 |
| ENSG00000229316 | HMGB1P45 | 1.630675783 | -0.151402754 | 1.471084925 | -0.102919112 | 0.918027169 | NA |
| ENSG00000229320 | KRT8P12 | 335.4224813 | 0.032049977 | 0.105520397 | 0.303732526 | 0.761331676 | 0.868989385 |
| ENSG00000229321 | ENSG0000022 | 5.434766765 | 0.254479325 | 0.755852425 | 0.336678585 | 0.736359208 | 0.854559632 |
| ENSG00000229324 | NUTF2P7 | 2.350755364 | -0.180975871 | 1.171381283 | -0.154497834 | 0.877217219 | 0.936188091 |
| ENSG00000229334 | TMEM44-AS2 | 20.98551995 | -0.864222005 | 0.400764233 | -2.156434963 | 0.031049717 | 0.080022527 |
| ENSG00000229336 | SREK1IP1P1 | 2.715221903 | 0.342993963 | 1.047905057 | 0.327313968 | 0.743430429 | 0.858490754 |
| ENSG00000229337 | ENSG0000022 | 4.645132747 | -0.810181184 | 0.808688678 | -1.001845588 | 0.316418176 | 0.505194387 |
| ENSG00000229344 | MTCO2P12 | 347.6436096 | 0.120649435 | 0.133080346 | 0.906590939 | 0.364623155 | 0.557370239 |
| ENSG00000229348 | HYI-AS1 | 1.872703206 | -1.644456104 | 1.427484087 | -1.151996102 | 0.249322675 | 0.427335695 |
| ENSG00000229349 | ACTG1P9 | 6.919390074 | 0.384999487 | 0.698754967 | 0.550979249 | 0.5816479 | 0.746582642 |
| ENSG00000229358 | DPY19L1P1 | 68.74968644 | 0.391574724 | 0.21344679 | 1.834530862 | 0.066575231 | 0.152297148 |
| ENSG00000229368 | ENSG0000022 | 9.818465782 | 0.663068393 | 0.572550146 | 1.158096627 | 0.246824606 | 0.42406804 |
| ENSG00000229372 | SZT2-AS1 | 3.316626656 | 1.925902522 | 1.114903087 | 1.727416979 | 0.084092804 | 0.184090936 |
| ENSG00000229388 | TAF12-DT | 11.75665032 | -0.3985027 | 0.542556307 | -0.734490955 | 0.462649571 | 0.649203643 |
| ENSG00000229391 | HLA-DRB6 | 1.698779657 | 4.114809883 | 1.623456526 | 2.534598135 | 0.011257639 | NA |
| ENSG00000229399 | ENSG0000022 | 6.820058145 | 0.088953115 | 0.69559714 | 0.12788022 | 0.898243765 | 0.948554546 |
| ENSG00000229404 | LINC00858 | 109.6052325 | 0.542505801 | 0.177420027 | 3.057748387 | 0.002230067 | 0.00777119 |
| ENSG00000229419 | RALGAPA1P1 | 746.7807794 | -0.537758348 | 0.076217818 | -7.055546388 | 1.72E-12 | 1.73E-11 |
| ENSG00000229422 | ENSG0000022 | 42.31179861 | 0.799096707 | 0.274499973 | 2.9110994 | 0.003601594 | 0.011995547 |
| ENSG00000229425 | ENSG0000022 | 4.307938356 | -4.649955185 | 1.374488566 | -3.383043918 | 0.000716872 | 0.002757384 |

| | | | | | | | |
|---|---|---|---|---|---|---|---|
| ENSG00000229431 | MED8-AS1 | 5.432309338 | -0.606699044 | 0.753879047 | -0.804769739 | 0.420952567 | 0.612069605 |
| ENSG00000229447 | ENSG0000022 | 7.151155142 | -0.221288034 | 0.687489579 | -0.321878383 | 0.74754483 | 0.860186517 |
| ENSG00000229453 | SPINK8 | 3.783845001 | 0.638820683 | 0.962395629 | 0.663781779 | 0.506829989 | 0.687115416 |
| ENSG00000229456 | RLIMP1 | 2.787128268 | 0.333929762 | 1.062223646 | 0.314368601 | 0.753241101 | 0.864010743 |
| ENSG00000229465 | ACTG1P11 | 3.996605454 | 0.258195256 | 0.881807629 | 0.292802248 | 0.76967331 | 0.874359188 |
| ENSG00000229468 | RPL39P18 | 2.138017192 | 1.503460188 | 1.300810904 | 1.155786889 | 0.247768337 | 0.425275395 |
| ENSG00000229473 | RGS17P1 | 4.963467734 | 0.361816499 | 0.830130706 | 0.435854856 | 0.662942041 | 0.805368498 |
| ENSG00000229474 | PATL2 | 3.830604899 | 1.377561176 | 0.938415123 | 1.46796566 | 0.142113551 | 0.279691563 |
| ENSG00000229502 | ENSG0000022 | 2.74037951 | -0.240579142 | 1.057561511 | -0.227484774 | 0.820046807 | 0.904547471 |
| ENSG00000229503 | RPL21P37 | 2.670292688 | 0.738411271 | 1.093714713 | 0.675140657 | 0.499586405 | 0.680997809 |
| ENSG00000229509 | ENSG0000022 | 4.027333803 | -0.014014053 | 0.899385405 | -0.01558181 | 0.987568017 | 0.993927018 |
| ENSG00000229512 | ENSG0000022 | 7.494617673 | 0.642420604 | 0.666014692 | 0.964574223 | 0.334758116 | 0.5258749 |
| ENSG00000229525 | DNPEP-AS1 | 15.6670724 | -0.939943533 | 0.459702869 | -2.04467624 | 0.040886788 | 0.100995252 |
| ENSG00000229534 | HNRNPA1P53 | 7.202072016 | 0.356787331 | 0.653114989 | 0.546285627 | 0.584869615 | 0.749045833 |
| ENSG00000229536 | LINC02572 | 5.680155728 | -0.380155057 | 0.730925398 | -0.520101037 | 0.602993156 | 0.762148806 |
| ENSG00000229539 | PANK2-AS1 | 16.59206152 | -0.305915378 | 0.46115948 | -0.663361356 | 0.50709915 | 0.687359482 |
| ENSG00000229544 | NKX1-2 | 4.130272822 | -2.102165924 | 0.973405405 | -2.1595996 | 0.030803679 | 0.079479134 |
| ENSG00000229558 | SACS-AS1 | 2.033498343 | 1.55035612 | 1.363663634 | 1.136905086 | 0.255577966 | 0.434927358 |
| ENSG00000229562 | ZFYVE9P1 | 4.861252014 | -0.493730284 | 0.795019018 | -0.621029526 | 0.534580197 | 0.710090892 |
| ENSG00000229567 | ENSG0000022 | 16.48401845 | 0.488926513 | 0.43945137 | 1.112583886 | 0.265887184 | 0.446828286 |
| ENSG00000229586 | TNPO1P3 | 9.038571267 | -0.485259226 | 0.613267618 | -0.791268301 | 0.428787442 | 0.61920798 |
| ENSG00000229589 | ACVR2B-AS1 | 66.80730162 | 0.446916752 | 0.216244606 | 2.066718615 | 0.038760676 | 0.0966069 |
| ENSG00000229591 | ENSG0000022 | 9.000764406 | 0.35619474 | 0.660204271 | 0.539522018 | 0.589526709 | 0.752573328 |
| ENSG00000229593 | SUCLA2P3 | 3.425909345 | 0.492170473 | 0.960760667 | 0.512271672 | 0.608460889 | 0.766675425 |
| ENSG00000229598 | PRDX3P1 | 3.423474524 | -0.036399458 | 0.927802379 | -0.039231909 | 0.968705494 | 0.985876114 |
| ENSG00000229604 | MTATP8P2 | 11.05575283 | 0.013289349 | 0.566932511 | 0.023440795 | 0.981298664 | 0.99136792 |
| ENSG00000229605 | RPL21P93 | 5.052736868 | 0.308445453 | 0.785705984 | 0.392571088 | 0.694636297 | 0.828005022 |
| ENSG00000229619 | MBNL1-AS1 | 85.34011431 | -0.064553187 | 0.192123184 | -0.335998946 | 0.736871661 | 0.854938089 |
| ENSG00000229638 | RPL4P4 | 5609.650806 | 0.200604906 | 0.059286331 | 3.383662018 | 0.00071526 | 0.002752107 |
| ENSG00000229642 | ENSG0000022 | 5.014460038 | 1.676417235 | 0.868854337 | 1.929457176 | 0.053674131 | 0.127396231 |
| ENSG00000229644 | NAMPTP1 | 518.8667256 | -0.049385223 | 0.084657428 | -0.583353687 | 0.559655228 | 0.729169701 |
| ENSG00000229656 | ITGB1-DT | 4.50766248 | -0.313952091 | 0.825784234 | -0.380186589 | 0.703806914 | 0.834211563 |
| ENSG00000229659 | RPL26P6 | 20.45765947 | 0.705451769 | 0.405116609 | 1.741354844 | 0.081621398 | 0.179588996 |

| | | | | | | | |
|---|---|---|---|---|---|---|---|
| ENSG00000229661 | BUD31P2 | 2.314355533 | 1.127580367 | 1.218956461 | 0.925037442 | 0.354946431 | 0.546644725 |
| ENSG00000229666 | MAST4-AS1 | 16.26505757 | 1.626543259 | 0.469884375 | 3.461581926 | 0.000537011 | 0.002113403 |
| ENSG00000229670 | PKP4P1 | 32.94488115 | 0.414628955 | 0.315663827 | 1.313514313 | 0.189009701 | 0.347732981 |
| ENSG00000229676 | ZNF492 | 15.98045163 | -0.496459095 | 0.450409276 | -1.102239944 | 0.270357371 | 0.452139541 |
| ENSG00000229689 | ANKRD33BP1 | 169.0895156 | 0.343788732 | 0.140157901 | 2.452867292 | 0.014172261 | 0.040436775 |
| ENSG00000229692 | SOS1-IT1 | 26.83434446 | -0.588780689 | 0.34621057 | -1.700643306 | 0.089009987 | 0.192776539 |
| ENSG00000229700 | RPL29P17 | 9.502204077 | 0.221591299 | 0.561531215 | 0.394619734 | 0.693123547 | 0.826932295 |
| ENSG00000229704 | EIF2S2P2 | 1.546939083 | 0.428893115 | 1.458400979 | 0.294084495 | 0.768693339 | NA |
| ENSG00000229719 | MIR194-2HG | 3.624329235 | 0.267078595 | 0.967497056 | 0.276051067 | 0.782508844 | 0.882057534 |
| ENSG00000229728 | ENSG0000022 | 13.18488383 | 0.729335481 | 0.48540007 | 1.502545067 | 0.132956398 | 0.266098392 |
| ENSG00000229738 | ENSG0000022 | 2.601861022 | -0.004145552 | 1.168581977 | -0.003547507 | 0.997169505 | 0.99830788 |
| ENSG00000229739 | PDC-AS1 | 4.913782615 | 0.175324384 | 0.808611574 | 0.216821512 | 0.828347447 | 0.909208431 |
| ENSG00000229743 | LINC01159 | 11.08749315 | -1.174334232 | 0.541612538 | -2.168218327 | 0.030142079 | 0.077998792 |
| ENSG00000229750 | ENSG0000022 | 2.095918861 | -1.797922202 | 1.295777021 | -1.387524376 | 0.165281924 | 0.313819511 |
| ENSG00000229752 | RPL7P10 | 7.765169872 | 0.690110541 | 0.63301228 | 1.090200875 | 0.275624668 | 0.457910623 |
| ENSG00000229759 | MRPS18AP1 | 4.790716778 | -0.154027817 | 0.815327392 | -0.188915298 | 0.850159203 | 0.921588324 |
| ENSG00000229766 | TMX4-AS1 | 1.858896977 | -0.065960091 | 1.390358008 | -0.047441084 | 0.962161686 | 0.982593112 |
| ENSG00000229780 | UBE2Q1-AS1 | 28.21679855 | -0.04210126 | 0.35343206 | -0.119121224 | 0.905179316 | 0.952558076 |
| ENSG00000229785 | SLC25A38P1 | 25.94429818 | 0.522675019 | 0.357735588 | 1.461065202 | 0.143997549 | 0.282502674 |
| ENSG00000229798 | KRT18P26 | 12.0014041 | 0.381444355 | 0.526352652 | 0.724693519 | 0.468640072 | 0.654566686 |
| ENSG00000229806 | RPS15P5 | 4.12565519 | 0.403407863 | 0.90153443 | 0.447468061 | 0.654537144 | 0.79944 |
| ENSG00000229808 | ENSG0000022 | 2.58378561 | 0.587035381 | 1.107314947 | 0.530143102 | 0.596012717 | 0.756993626 |
| ENSG00000229809 | ZNF688 | 265.6315124 | -0.087453046 | 0.119922944 | -0.729243651 | 0.465852634 | 0.651744797 |
| ENSG00000229814 | RPL35AP21 | 15.08346708 | 0.296583171 | 0.454332706 | 0.652788511 | 0.513892631 | 0.693004348 |
| ENSG00000229816 | DDX50P1 | 22.24650568 | -0.064895941 | 0.378757405 | -0.17133907 | 0.863957161 | 0.929386977 |
| ENSG00000229817 | ENSG0000022 | 1.801331176 | -0.437180065 | 1.289883546 | -0.338929871 | 0.734662561 | 0.853692306 |
| ENSG00000229820 | DNAJC8P4 | 9.238307535 | -0.991660293 | 0.58174581 | -1.704628165 | 0.088263795 | 0.191413448 |
| ENSG00000229833 | PET100 | 52.05448172 | -0.13606856 | 0.2498656 | -0.544567 | 0.586051356 | 0.749932924 |
| ENSG00000229835 | KHSRPP1 | 248.2267868 | 0.055504613 | 0.121015868 | 0.458655661 | 0.646481459 | 0.793463804 |
| ENSG00000229836 | ENSG0000022 | 3.969228457 | -0.123357595 | 0.894149518 | -0.137960814 | 0.890271387 | 0.943797108 |
| ENSG00000229846 | ENSG0000022 | 1.694460701 | 1.267202249 | 1.411518986 | 0.89775785 | 0.369314661 | NA |
| ENSG00000229848 | ENSG0000022 | 45.96436785 | -0.047947049 | 0.26527062 | -0.180747677 | 0.856565634 | 0.925689718 |
| ENSG00000229851 | ARSD-AS1 | 28.01015802 | -0.688317405 | 0.33833334 | -2.034435642 | 0.041907691 | 0.103039918 |

| | | | | | | | |
|---|---|---|---|---|---|---|---|
| ENSG00000229852 | KHDC1-AS1 | 48.35247655 | 0.086457879 | 0.259357012 | 0.333354699 | 0.738866555 | 0.856040341 |
| ENSG00000229867 | STEAP3-AS1 | 5.113183001 | -0.842334942 | 0.781197067 | -1.078261783 | 0.280916947 | 0.464389101 |
| ENSG00000229869 | LARP4B-DT | 5.4655371 | -0.786604633 | 0.80249588 | -0.98019772 | 0.32698853 | 0.517460158 |
| ENSG00000229873 | OGFR-AS1 | 2.879797877 | -0.706025194 | 1.022495808 | -0.690492018 | 0.489884828 | 0.673011336 |
| ENSG00000229880 | IMMTP1 | 33.88584605 | -0.112350949 | 0.301422241 | -0.372736097 | 0.709344868 | 0.838047653 |
| ENSG00000229881 | ENSG0000022 | 2.71151334 | 0.157463571 | 1.075842058 | 0.146363093 | 0.883634761 | 0.939712848 |
| ENSG00000229887 | HNRNPA1P6 | 6.884868122 | 0.356836221 | 0.660690153 | 0.54009617 | 0.589130712 | 0.752295476 |
| ENSG00000229891 | LINC01315 | 6.569513839 | 2.399752697 | 0.81273688 | 2.95268094 | 0.003150274 | 0.010627541 |
| ENSG00000229893 | TAX1BP1-AS1 | 2.090687378 | -3.537843912 | 1.575206746 | -2.245955282 | 0.024706872 | 0.065803856 |
| ENSG00000229894 | GK3 | 8.127182988 | -0.16337897 | 0.615229683 | -0.265557684 | 0.790579879 | 0.887101468 |
| ENSG00000229897 | SEPTIN7P7 | 21.56226451 | 0.25230795 | 0.379543952 | 0.664766092 | 0.506200114 | 0.686625533 |
| ENSG00000229914 | ENSG0000022 | 5.467446567 | -0.738659042 | 0.782183502 | -0.94435518 | 0.344988177 | 0.535890356 |
| ENSG00000229917 | RPL7P46 | 8.260156717 | -0.178547987 | 0.600734756 | -0.29721601 | 0.7663016 | 0.872318316 |
| ENSG00000229927 | RHEBP1 | 4.886864545 | 0.598727409 | 0.838481766 | 0.714061334 | 0.475189254 | 0.66046014 |
| ENSG00000229931 | ATXN1-AS1 | 13.55651806 | 0.556350217 | 0.530856987 | 1.048022783 | 0.29462811 | 0.480572301 |
| ENSG00000229932 | YWHAZP3 | 1672.251889 | -0.36705937 | 0.058322899 | -6.293572159 | 3.10E-10 | 2.59E-09 |
| ENSG00000229939 | ENSG0000022 | 3.437697997 | 0.830624736 | 0.96205279 | 0.863387898 | 0.387924231 | 0.580608198 |
| ENSG00000229944 | EIF4EP2 | 880.1234801 | 0.101850732 | 0.069436223 | 1.466824205 | 0.142423883 | 0.280134089 |
| ENSG00000229950 | TFAP2A-AS1 | 51.3695154 | -1.191076865 | 0.25275866 | -4.712308827 | 2.45E-06 | 1.36E-05 |
| ENSG00000229951 | FOSL2-AS1 | 26.45999908 | 0.612471779 | 0.345628339 | 1.772053126 | 0.076385739 | 0.170433513 |
| ENSG00000229953 | BCAN-AS2 | 7.170749845 | -2.054436674 | 0.747326683 | -2.749047667 | 0.005976869 | 0.018872592 |
| ENSG00000229956 | ZRANB2-DT | 16.23990779 | 0.381624261 | 0.460320841 | 0.82903972 | 0.407081932 | 0.599849661 |
| ENSG00000229965 | RPS3AP7 | 10.93059657 | 0.466615796 | 0.54215196 | 0.860673447 | 0.389417921 | 0.582084259 |
| ENSG00000229970 | ENSG0000022 | 64.38727682 | 0.40255228 | 0.227336425 | 1.770733751 | 0.076604987 | 0.170845655 |
| ENSG00000229980 | TOB1-AS1 | 85.65032502 | -0.218399706 | 0.191166027 | -1.142460875 | 0.2532625 | 0.431958494 |
| ENSG00000229989 | MIR181A1HG | 21.43694922 | -0.344706744 | 0.388366045 | -0.887582085 | 0.374765594 | 0.567579238 |
| ENSG00000229994 | RPL5P4 | 102.1470397 | 0.367916848 | 0.180106595 | 2.042772773 | 0.041074937 | 0.101350872 |
| ENSG00000230002 | ALMS1-IT1 | 3.272293792 | -0.401584323 | 0.974211715 | -0.412214631 | 0.680182114 | 0.818387585 |
| ENSG00000230021 | ENSG0000023 | 25.94790136 | -0.018364327 | 0.347114217 | -0.052905719 | 0.957807027 | 0.980830956 |
| ENSG00000230022 | FNTAP2 | 2.086473899 | -0.691450189 | 1.220670993 | -0.566450905 | 0.571087298 | 0.737986853 |
| ENSG00000230026 | ENSG0000023 | 1.849995239 | -1.586589152 | 1.339496195 | -1.184467084 | 0.236228219 | 0.411377464 |
| ENSG00000230037 | UBBP1 | 9.370560917 | -0.357195109 | 0.579374243 | -0.616518793 | 0.537552172 | 0.712553267 |
| ENSG00000230039 | CCNYL5 | 14.60029707 | -0.829528283 | 0.462309776 | -1.794312659 | 0.072763279 | 0.163876664 |

| | | | | | | | |
|---|---|---|---|---|---|---|---|
| ENSG00000230042 | AK3P3 | 10.29557515 | -2.183438723 | 0.622205111 | -3.509194449 | 0.000449466 | 0.001793487 |
| ENSG00000230043 | TMSB4XP6 | 228.4159158 | -0.196853063 | 0.122494573 | -1.607034978 | 0.108046695 | 0.225608196 |
| ENSG00000230067 | HSPD1P6 | 32.43569166 | 0.054294764 | 0.322456817 | 0.168378404 | 0.866285597 | 0.930495463 |
| ENSG00000230069 | LRRC37A15P | 5.111787839 | 0.584140957 | 0.816109394 | 0.71576306 | 0.474137666 | 0.659616385 |
| ENSG00000230071 | RPL4P6 | 3.605033208 | 0.077601087 | 0.917022419 | 0.08462289 | 0.932561201 | 0.967612915 |
| ENSG00000230074 | ENSG0000023 | 91.13108814 | -0.639107282 | 0.193740151 | -3.298785923 | 0.000971039 | 0.003643979 |
| ENSG00000230076 | RPL10P6 | 257.9801269 | -0.198453974 | 0.115106619 | -1.724088295 | 0.084691905 | 0.185151689 |
| ENSG00000230079 | STK24P1 | 2.425986796 | -0.690676288 | 1.188107247 | -0.581324867 | 0.561021524 | 0.730103564 |
| ENSG00000230082 | PRRT3-AS1 | 7.211992233 | -0.058676339 | 0.65087071 | -0.090150529 | 0.928167597 | 0.965455911 |
| ENSG00000230084 | ZBTB47-AS1 | 3.780010932 | 0.196605844 | 0.902593215 | 0.217823312 | 0.82756678 | 0.90886547 |
| ENSG00000230091 | TMEM254-AS | 241.0905614 | -0.341843145 | 0.124275306 | -2.750692445 | 0.005946945 | 0.018783273 |
| ENSG00000230092 | ENSG0000023 | 5.910321343 | 0.534900103 | 0.730814203 | 0.731923518 | 0.464215251 | 0.650485483 |
| ENSG00000230097 | ME2P1 | 1.551246898 | 1.805313277 | 1.583700403 | 1.139933585 | 0.254313972 | NA |
| ENSG00000230102 | LINC02028 | 12.71739913 | -0.609100611 | 0.499209292 | -1.220130756 | 0.222415311 | 0.393488995 |
| ENSG00000230124 | ACBD6 | 1662.096117 | -0.050501476 | 0.057807783 | -0.873610319 | 0.382330505 | 0.574617273 |
| ENSG00000230126 | FGF12-AS2 | 9.577426945 | 0.894680069 | 0.57667472 | 1.551446662 | 0.12079468 | 0.246818381 |
| ENSG00000230146 | SEPHS1P4 | 34.95300622 | -1.121992836 | 0.312125404 | -3.594686054 | 0.000324783 | 0.001328079 |
| ENSG00000230149 | ENSG0000023 | 12.16681782 | 0.053947159 | 0.49912617 | 0.108083211 | 0.913929685 | 0.957243493 |
| ENSG00000230154 | AIDAP1 | 32.09472474 | -0.207738079 | 0.308305112 | -0.673806796 | 0.500434153 | 0.681707808 |
| ENSG00000230155 | CEP250-AS1 | 1.848133385 | -0.994979136 | 1.285219222 | -0.774170755 | 0.43882982 | 0.628347548 |
| ENSG00000230165 | AURKAP2 | 1.950714648 | -0.153435842 | 1.23040244 | -0.124703786 | 0.90075806 | 0.949953563 |
| ENSG00000230170 | HNRNPA1P28 | 9.933047923 | 0.091488431 | 0.583594963 | 0.156766999 | 0.875428482 | 0.935410563 |
| ENSG00000230183 | CNOT6LP1 | 51.6374814 | -0.069724925 | 0.252380721 | -0.276268822 | 0.782341601 | 0.881955689 |
| ENSG00000230185 | HSDL2-AS1 | 13.50270378 | -0.362371955 | 0.48841445 | -0.741935369 | 0.458126495 | 0.64527349 |
| ENSG00000230191 | IFITM3P4 | 3.397468982 | 0.259786731 | 0.949030529 | 0.273739067 | 0.784285153 | 0.882931806 |
| ENSG00000230199 | ENSG0000023 | 12.35133043 | -0.259248824 | 0.522122594 | -0.496528644 | 0.619521483 | 0.774603696 |
| ENSG00000230202 | RPL29P4 | 3057.774965 | 0.167137668 | 0.050054717 | 3.339099234 | 0.000840505 | 0.003190666 |
| ENSG00000230204 | FTH1P5 | 7.85763887 | 0.053710222 | 0.666476794 | 0.080588286 | 0.935769383 | 0.969261113 |
| ENSG00000230207 | RPL4P5 | 842.3830585 | 0.199908063 | 0.079725043 | 2.507468869 | 0.012159929 | 0.035345765 |
| ENSG00000230212 | CBR1-AS1 | 12.98847926 | 0.199396381 | 0.489160002 | 0.407630182 | 0.683545202 | 0.820454528 |
| ENSG00000230216 | HSPB1P2 | 71.3057558 | 0.936865736 | 0.225644302 | 4.151958314 | 3.30E-05 | 0.000158545 |
| ENSG00000230221 | ENSG0000023 | 13.67661414 | -0.022390155 | 0.506364383 | -0.044217477 | 0.964731051 | 0.984034251 |
| ENSG00000230224 | PHB1P9 | 6.101712488 | -9.12E-05 | 0.714172176 | -0.00012777 | 0.999898054 | 0.999914377 |

| | | | | | | | |
|---|---|---|---|---|---|---|---|
| ENSG00000230228 | H3P1 | 2.775301309 | 0.008121305 | 1.072815436 | 0.007570086 | 0.993960003 | 0.996533822 |
| ENSG00000230267 | HERC2P4 | 3.311372566 | 0.278720179 | 0.98260601 | 0.283654055 | 0.776675508 | 0.878763805 |
| ENSG00000230282 | RPL7P61 | 2.243873748 | 0.666738307 | 1.159129348 | 0.57520613 | 0.565151898 | 0.733481521 |
| ENSG00000230289 | ENSG0000023 | 6.652586915 | -0.069158894 | 0.700865853 | -0.098676364 | 0.921395236 | 0.9619656 |
| ENSG00000230291 | ENSG0000023 | 68.08515525 | 0.179434521 | 0.22215513 | 0.807699202 | 0.419263763 | 0.610870102 |
| ENSG00000230295 | GTF2IP23 | 14.65678582 | 0.400456213 | 0.481819976 | 0.831132442 | 0.405898814 | 0.598644645 |
| ENSG00000230299 | SNRPEP11 | 11.75143441 | -0.41899646 | 0.526435498 | -0.795912247 | 0.426083042 | 0.616893852 |
| ENSG00000230303 | ENSG0000023 | 3.512929429 | 0.466595009 | 0.937065357 | 0.497932194 | 0.618531835 | 0.773839458 |
| ENSG00000230306 | BANF1P2 | 3.272727061 | 0.662659353 | 0.981016729 | 0.675482215 | 0.499369447 | 0.680823435 |
| ENSG00000230311 | TOMM20P4 | 336.6436699 | 0.085915456 | 0.115623949 | 0.743059351 | 0.457445748 | 0.644622241 |
| ENSG00000230314 | ELOVL2-AS1 | 4.980329265 | 0.316768599 | 0.780916797 | 0.405636811 | 0.685009478 | 0.821567283 |
| ENSG00000230316 | FEZF1-AS1 | 122.1375268 | -1.148315957 | 0.169334519 | -6.781345976 | 1.19E-11 | 1.11E-10 |
| ENSG00000230318 | XRCC6P3 | 1.62686583 | -0.181307749 | 1.331067891 | -0.136212247 | 0.891653494 | NA |
| ENSG00000230319 | TTLL1-AS1 | 1.620270354 | 0.378330089 | 1.353830353 | 0.279451623 | 0.779898259 | NA |
| ENSG00000230325 | ENSG0000023 | 4.676911895 | 1.186457099 | 0.837704735 | 1.416318959 | 0.156682139 | 0.301029169 |
| ENSG00000230330 | HMGN2P3 | 695.9464902 | -0.376008434 | 0.081139387 | -4.634104948 | 3.58E-06 | 1.95E-05 |
| ENSG00000230333 | ENSG0000023 | 6.446370214 | -2.085333284 | 0.771025231 | -2.70462392 | 0.006838176 | 0.021247156 |
| ENSG00000230337 | EXOSC10-AS1 | 21.16121122 | -0.720663662 | 0.397584781 | -1.812603742 | 0.06989296 | 0.158386299 |
| ENSG00000230356 | NCAPD2P1 | 3.316095048 | -0.003560336 | 0.994461584 | -0.003580165 | 0.997143448 | 0.99830788 |
| ENSG00000230359 | TPI1P2 | 64.19380117 | -0.493570543 | 0.22195661 | -2.223725361 | 0.026166928 | 0.069022567 |
| ENSG00000230360 | DDX10P2 | 8.226040543 | 0.236505146 | 0.61887771 | 0.382151663 | 0.702348873 | 0.833215242 |
| ENSG00000230362 | ACTG1P23 | 6.230442368 | -0.326378517 | 0.722878709 | -0.451498312 | 0.651630441 | 0.797581514 |
| ENSG00000230364 | RPL4P3 | 55.93176091 | 0.286548998 | 0.260953623 | 1.098084 | 0.272167811 | 0.454173972 |
| ENSG00000230366 | DSCR9 | 4.720781422 | 0.1357269 | 0.797726188 | 0.170142215 | 0.864898295 | 0.929799912 |
| ENSG00000230373 | GOLGA6L5P | 46.11548065 | 0.207252364 | 0.278102313 | 0.74523783 | 0.456127957 | 0.643397611 |
| ENSG00000230383 | RPL6P19 | 189.4183195 | 0.343991711 | 0.142908234 | 2.407081122 | 0.016080597 | 0.0452236 |
| ENSG00000230392 | ENSG0000023 | 1.895934368 | -1.050621469 | 1.321688515 | -0.794908525 | 0.426666715 | 0.617497513 |
| ENSG00000230397 | SPTLC1P1 | 64.93920382 | -0.251073067 | 0.219834508 | -1.142100343 | 0.253412313 | 0.432053397 |
| ENSG00000230406 | RPL23P5 | 2.144114806 | 0.072668849 | 1.192378547 | 0.060944445 | 0.951403453 | 0.977495703 |
| ENSG00000230408 | BZW1-AS1 | 8.478480775 | -0.791649454 | 0.636048113 | -1.24463769 | 0.213264959 | 0.38149549 |
| ENSG00000230409 | TCEA1P2 | 1509.849467 | -0.019073445 | 0.061832651 | -0.308468815 | 0.757725626 | 0.866391318 |
| ENSG00000230415 | LINC01786 | 17.0246448 | -0.701254648 | 0.436465401 | -1.606667208 | 0.108127391 | 0.225720479 |
| ENSG00000230423 | ENSG0000023 | 22.27858045 | -0.073224277 | 0.371251339 | -0.1972364 | 0.843642547 | 0.917646279 |

| | | | | | | | |
|---|---|---|---|---|---|---|---|
| ENSG00000230424 | EMC1-AS1 | 14.76823043 | -0.025358601 | 0.453059959 | -0.055971843 | 0.955364238 | 0.979455025 |
| ENSG00000230438 | SERPINB9P1 | 3.032012232 | -5.144777915 | 1.449665498 | -3.54894141 | 0.000386783 | 0.001557999 |
| ENSG00000230444 | TFAMP1 | 2.346010601 | -0.606926574 | 1.127623326 | -0.538235207 | 0.590414677 | 0.753103247 |
| ENSG00000230445 | LRRC37A6P | 57.51935604 | -1.079655413 | 0.240265045 | -4.493601688 | 7.00E-06 | 3.68E-05 |
| ENSG00000230448 | LINC00276 | 2.279328257 | -2.687858727 | 1.421892245 | -1.890339255 | 0.058712602 | 0.137043863 |
| ENSG00000230450 | NEK2P4 | 8.037399989 | 0.31532322 | 0.633952978 | 0.497392127 | 0.618912556 | 0.774119712 |
| ENSG00000230453 | ANKRD18B | 35.48417331 | 3.023828442 | 0.399564289 | 7.567814562 | 3.80E-14 | 4.27E-13 |
| ENSG00000230454 | ENSG0000023 | 82.16020602 | 0.175772965 | 0.200896213 | 0.874944142 | 0.381604299 | 0.573920057 |
| ENSG00000230457 | PA2G4P4 | 311.9702555 | 0.0615336 | 0.106509225 | 0.577730236 | 0.563446262 | 0.732154597 |
| ENSG00000230487 | PSMG3-AS1 | 954.8324765 | -0.051986631 | 0.073991587 | -0.702601921 | 0.482303865 | 0.666388645 |
| ENSG00000230490 | ENSG0000023 | 70.59854602 | -1.008069719 | 0.230875492 | -4.36629159 | 1.26E-05 | 6.43E-05 |
| ENSG00000230500 | MKX-AS1 | 3.539875734 | 3.14908532 | 1.296438159 | 2.429028564 | 0.01513934 | 0.042849654 |
| ENSG00000230507 | RPL7AP8 | 2.732276259 | -1.33092495 | 1.090020314 | -1.221009308 | 0.222082495 | 0.39313006 |
| ENSG00000230508 | RPL19P21 | 6.770873058 | -0.208394606 | 0.76395971 | -0.272782194 | 0.785020648 | 0.883106023 |
| ENSG00000230510 | PPP5D1P | 14.5677641 | -0.198557914 | 0.459879258 | -0.43176097 | 0.665915149 | 0.807361848 |
| ENSG00000230513 | THAP7-AS1 | 73.86695969 | -0.110739887 | 0.205025894 | -0.540126349 | 0.589109901 | 0.752295476 |
| ENSG00000230524 | COL6A4P1 | 24.0585336 | -0.267230833 | 0.366081656 | -0.729976027 | 0.465404838 | 0.651356659 |
| ENSG00000230530 | LIMD1-AS1 | 28.57754463 | -0.60167618 | 0.329094544 | -1.828277589 | 0.067507901 | 0.154016038 |
| ENSG00000230532 | ENSG0000023 | 55.68575204 | -0.019298413 | 0.24045175 | -0.080258983 | 0.93603128 | 0.969415691 |
| ENSG00000230534 | ENSG0000023 | 2.06647572 | -0.766414491 | 1.301794782 | -0.588736798 | 0.556037849 | 0.72672673 |
| ENSG00000230537 | LINC02937 | 2.862743521 | 0.831401651 | 1.037748857 | 0.801158822 | 0.423039707 | 0.614016506 |
| ENSG00000230547 | HMGB1P11 | 5.385982708 | -0.449537174 | 0.756656187 | -0.59411022 | 0.552438396 | 0.724126438 |
| ENSG00000230550 | ERLNC1 | 2.726086886 | 0.77184952 | 1.124235547 | 0.686554986 | 0.492363203 | 0.675123244 |
| ENSG00000230551 | ENSG0000023 | 316.0842645 | -0.158740048 | 0.130089793 | -1.220234458 | 0.222376008 | 0.39346746 |
| ENSG00000230555 | LINC02916 | 87.28890155 | -0.769287088 | 0.210784081 | -3.649645099 | 0.000262603 | 0.001090082 |
| ENSG00000230561 | CCDC192 | 4.632834294 | 0.557563083 | 0.835994044 | 0.666946238 | 0.504806475 | 0.685384738 |
| ENSG00000230562 | FAM133DP | 184.8291708 | -0.646237104 | 0.137103276 | -4.71350591 | 2.43E-06 | 1.35E-05 |
| ENSG00000230565 | ZNF32-AS2 | 7.485662075 | -0.887051224 | 0.657430863 | -1.349269214 | 0.17725051 | 0.331225701 |
| ENSG00000230568 | SF3A3P1 | 27.89627374 | -0.561842468 | 0.344106267 | -1.632758602 | 0.102519777 | 0.216317484 |
| ENSG00000230580 | RPL19P5 | 16.51480361 | -0.048991358 | 0.434581107 | -0.11273237 | 0.910242737 | 0.954995124 |
| ENSG00000230581 | ACTG1P14 | 4.836587857 | -1.550189602 | 0.849078686 | -1.825731382 | 0.067890734 | 0.154735562 |
| ENSG00000230583 | GTF2IRD1P1 | 15.49862 | 0.36319736 | 0.467599352 | 0.776727681 | 0.437319452 | 0.626920679 |
| ENSG00000230584 | CCT5P2 | 59.18458127 | -0.015721981 | 0.253931772 | -0.061914194 | 0.950631164 | 0.977235535 |

| | | | | | | | |
|---|---|---|---|---|---|---|---|
| ENSG00000230590 | FTX | 775.6207216 | -1.012255966 | 0.089447193 | -11.31679974 | 1.08E-29 | 2.97E-28 |
| ENSG00000230592 | RPSAP8 | 12.817877 | -0.025773763 | 0.489354739 | -0.052668874 | 0.95799574 | 0.980915117 |
| ENSG00000230593 | PPIAP40 | 10.94434597 | 0.634787032 | 0.536246974 | 1.183758722 | 0.236508587 | 0.411646963 |
| ENSG00000230595 | RSL24D1P2 | 22.92429561 | -0.214067818 | 0.368004171 | -0.581699434 | 0.560769153 | 0.729892663 |
| ENSG00000230606 | APPAT | 5.765275023 | 0.068727618 | 0.754624628 | 0.091075238 | 0.927432808 | 0.965137362 |
| ENSG00000230613 | HM13-AS1 | 11.15403579 | -0.750032281 | 0.521140515 | -1.439213148 | 0.150090141 | 0.291756083 |
| ENSG00000230615 | ENSG0000023 | 29.66999813 | -0.204367457 | 0.332798409 | -0.614087844 | 0.53915728 | 0.713814741 |
| ENSG00000230621 | ENSG0000023 | 2.700161636 | -0.583153712 | 1.070315236 | -0.544842951 | 0.585861535 | 0.749814489 |
| ENSG00000230626 | ENSG0000023 | 219.5424697 | -0.089295964 | 0.127254412 | -0.701712124 | 0.482858711 | 0.666954409 |
| ENSG00000230629 | RPS23P8 | 1591.991315 | 0.220774675 | 0.068287173 | 3.233032859 | 0.001224835 | 0.004512238 |
| ENSG00000230648 | ENSG0000023 | 2.776267288 | -1.387540514 | 1.125014654 | -1.23335328 | 0.217443988 | 0.387009471 |
| ENSG00000230650 | ENSG0000023 | 16.70328118 | 1.527382242 | 0.463096796 | 3.2981922 | 0.000973095 | 0.003651096 |
| ENSG00000230658 | KLHL7-DT | 13.91751187 | -1.904592634 | 0.515917097 | -3.691664114 | 0.000222792 | 0.000937081 |
| ENSG00000230662 | TNPO1P2 | 10.14623514 | -0.288068627 | 0.575262852 | -0.500760002 | 0.616540038 | 0.772864944 |
| ENSG00000230665 | THAP12P1 | 7.387693339 | -0.140782764 | 0.655215646 | -0.214864778 | 0.829872747 | 0.909829899 |
| ENSG00000230666 | CEACAM22P | 4.141658829 | -0.555918588 | 0.86592353 | -0.641995013 | 0.520876421 | 0.698050467 |
| ENSG00000230667 | SETSIP | 71.08152646 | -0.161238579 | 0.232477589 | -0.693566117 | 0.487954349 | 0.671365247 |
| ENSG00000230668 | ENSG0000023 | 2.957851629 | -1.087228333 | 1.051303586 | -1.03417162 | 0.301055939 | 0.487865446 |
| ENSG00000230679 | ENO1-AS1 | 5.474091816 | -0.584847731 | 0.758118173 | -0.771446658 | 0.440442229 | 0.62981551 |
| ENSG00000230699 | ENSG0000023 | 124.4680609 | 1.015586651 | 0.182267718 | 5.571950227 | 2.52E-08 | 1.73E-07 |
| ENSG00000230701 | FBXW4P1 | 14.86446081 | -0.384824986 | 0.473224857 | -0.813196898 | 0.416105178 | 0.608184312 |
| ENSG00000230707 | ENSG0000023 | 1.869939966 | 0.057249993 | 1.269699175 | 0.045089415 | 0.964036039 | 0.983666795 |
| ENSG00000230711 | CTAGE13P | 13.21304244 | -0.7436581 | 0.486000534 | -1.530159018 | 0.125977373 | 0.254978024 |
| ENSG00000230715 | ENSG0000023 | 181.9439906 | 0.054717305 | 0.139777716 | 0.391459426 | 0.695457674 | 0.828682684 |
| ENSG00000230716 | KRT8P7 | 33.61463517 | 0.161508429 | 0.316911299 | 0.509632916 | 0.610308659 | 0.768261612 |
| ENSG00000230721 | RPL17P5 | 6.953988965 | 0.493302413 | 0.697250284 | 0.707496898 | 0.479257739 | 0.663653377 |
| ENSG00000230724 | LINC01001 | 10.72628628 | -0.289888363 | 0.540218578 | -0.536613095 | 0.591534897 | 0.753962137 |
| ENSG00000230732 | ENSG0000023 | 3.432515961 | 0.5105495 | 0.946152082 | 0.539606168 | 0.589468662 | 0.752541104 |
| ENSG00000230733 | ENSG0000023 | 23.74464187 | -0.389210304 | 0.374932252 | -1.038081688 | 0.299232026 | 0.485939592 |
| ENSG00000230734 | RPL10P3 | 18.16230004 | -0.521532602 | 0.421222941 | -1.23813912 | 0.215664481 | 0.384917388 |
| ENSG00000230750 | SDAD1P2 | 12.1105143 | -0.256335539 | 0.512346416 | -0.500316838 | 0.616852 | 0.772960336 |
| ENSG00000230753 | ZNF341-AS1 | 2.058071843 | -1.214973561 | 1.241221732 | -0.978852956 | 0.32765264 | 0.51794656 |
| ENSG00000230756 | RHOQP3 | 97.96855291 | 0.154750036 | 0.180427175 | 0.857686964 | 0.39106534 | 0.583899955 |

| | | | | | | | |
|---|---|---|---|---|---|---|---|
| ENSG00000230758 | SNAP23P1 | 3.559233776 | -0.343601949 | 0.915767485 | -0.37520654 | 0.707506867 | 0.836608636 |
| ENSG00000230773 | ENSG0000023 | 4.790868245 | -0.425082897 | 0.815849003 | -0.521031338 | 0.602344941 | 0.761646618 |
| ENSG00000230777 | RPS29P5 | 17.48850566 | -0.355462286 | 0.456075651 | -0.779393254 | 0.435748097 | 0.625614053 |
| ENSG00000230778 | ANKRD63 | 5.77000159 | 1.184087285 | 0.750205011 | 1.578351608 | 0.114484856 | 0.236474212 |
| ENSG00000230793 | SMARCE1P5 | 173.6016746 | 0.062223647 | 0.152510908 | 0.407994734 | 0.683277541 | 0.820223767 |
| ENSG00000230797 | YY2 | 48.28522187 | 0.715911214 | 0.275282912 | 2.600638043 | 0.009305057 | 0.027939531 |
| ENSG00000230798 | FOXD3-AS1 | 68.0063493 | -0.278490947 | 0.221463987 | -1.257499925 | 0.208572666 | 0.375120107 |
| ENSG00000230807 | UBA52P4 | 4.333592096 | 0.615873248 | 0.849940706 | 0.724607309 | 0.468692973 | 0.654600719 |
| ENSG00000230823 | CBX1P1 | 32.0603634 | -0.282547723 | 0.309808195 | -0.912008553 | 0.361764215 | 0.554366807 |
| ENSG00000230832 | SSBL4P | 13.39351039 | -0.131407543 | 0.522571862 | -0.251463106 | 0.801456083 | 0.894398618 |
| ENSG00000230836 | LINC01293 | 431.3383578 | -0.122970793 | 0.094108073 | -1.306697596 | 0.191315435 | 0.351214476 |
| ENSG00000230837 | RPL31P2 | 14.71532509 | 0.548049359 | 0.487505779 | 1.124190487 | 0.260932224 | 0.441169312 |
| ENSG00000230841 | RPL23AP15 | 1.834953575 | 0.576739108 | 1.287342752 | 0.448007423 | 0.65414784 | 0.799219825 |
| ENSG00000230844 | ZNF674-AS1 | 147.8195466 | 0.38719014 | 0.151663899 | 2.552948611 | 0.010681525 | 0.031481956 |
| ENSG00000230847 | OCLNP1 | 9.938619292 | -0.006806635 | 0.548651503 | -0.012406118 | 0.990101604 | 0.994697854 |
| ENSG00000230849 | GOT2P2 | 40.38044087 | 0.015300068 | 0.285084589 | 0.053668521 | 0.957199264 | 0.980624826 |
| ENSG00000230863 | ENSG0000023 | 5.943407249 | 0.409119436 | 0.735764272 | 0.55604689 | 0.5781788 | 0.743882315 |
| ENSG00000230865 | TSEN15P1 | 3.097198348 | 1.084051815 | 1.039918516 | 1.042439189 | 0.297208105 | 0.483577128 |
| ENSG00000230869 | AGAP10P | 1.67839052 | -1.429670759 | 1.407533307 | -1.015727835 | 0.309759009 NA | |
| ENSG00000230870 | FBXW11P1 | 15.73448537 | -0.291506957 | 0.473311719 | -0.615887893 | 0.537968513 | 0.712899179 |
| ENSG00000230873 | STMND1 | 2.500116554 | -0.764991866 | 1.093477582 | -0.699595382 | 0.484180028 | 0.668055445 |
| ENSG00000230882 | ENSG0000023 | 14.04055665 | -1.121092326 | 0.50235778 | -2.23166112 | 0.025637369 | 0.067849705 |
| ENSG00000230896 | ENSG0000023 | 25.73043678 | 0.691595578 | 0.350159527 | 1.975087139 | 0.04825826 | 0.116261761 |
| ENSG00000230897 | RPS18P12 | 333.8358527 | 0.395450019 | 0.107523792 | 3.677790853 | 0.000235263 | 0.00098556 |
| ENSG00000230910 | ADGRB3-DT | 2.972497229 | 0.800587032 | 1.065222801 | 0.751567682 | 0.452311085 | 0.640066851 |
| ENSG00000230911 | PPIHP1 | 75.8816282 | -0.037542475 | 0.211575444 | -0.177442495 | 0.859160831 | 0.926792302 |
| ENSG00000230913 | NPM1P51 | 2.296412357 | -0.908712868 | 1.210330766 | -0.750797132 | 0.452774756 | 0.640439813 |
| ENSG00000230914 | KIF19BP | 2.617392865 | 2.732987555 | 1.354894174 | 2.017122523 | 0.043682729 | 0.106796134 |
| ENSG00000230916 | MTCO1P53 | 6.014914973 | 0.520442638 | 0.722947966 | 0.719889483 | 0.471593044 | 0.657330443 |
| ENSG00000230922 | ENSG0000023 | 4.567633359 | -1.450803321 | 0.858166494 | -1.690584906 | 0.090916109 | 0.196090217 |
| ENSG00000230927 | TMBIM7P | 3.370928779 | -0.288117684 | 0.95496486 | -0.301705012 | 0.762876948 | 0.87001726 |
| ENSG00000230932 | ENSG0000023 | 1.802778363 | 0.62449302 | 1.293457205 | 0.482809185 | 0.629231228 | 0.7810577 |
| ENSG00000230935 | RPS3P1 | 4.927561678 | -0.125419666 | 0.790240058 | -0.158710843 | 0.873896695 | 0.934688091 |

| | | | | | | | |
|---|---|---|---|---|---|---|---|
| ENSG00000230937 | MIR205HG | 42.63596518 | 1.667315641 | 0.297927785 | 5.596375105 | 2.19E-08 | 1.52E-07 |
| ENSG00000230953 | ENSG0000023 | 1.635907267 | 1.003117142 | 1.413630861 | 0.709603313 | 0.477950165 | NA |
| ENSG00000230955 | ENSG0000023 | 7.292759778 | -0.713598152 | 0.646128602 | -1.10442124 | 0.269410451 | 0.451016256 |
| ENSG00000230956 | CAPZA1P1 | 86.57997573 | 0.144205945 | 0.189519005 | 0.760904931 | 0.446713853 | 0.63515655 |
| ENSG00000230965 | SNX18P13 | 10.26039851 | -0.535320063 | 0.554004333 | -0.966274145 | 0.33390702 | 0.524897351 |
| ENSG00000230979 | ENSG0000023 | 96.71164022 | 0.45323523 | 0.201410283 | 2.250308292 | 0.024429382 | 0.065186695 |
| ENSG00000230989 | HSBP1 | 5412.816397 | 0.15762784 | 0.046662397 | 3.378048499 | 0.000730022 | 0.002803265 |
| ENSG00000230992 | FAM201B | 5.751975219 | -0.43832079 | 0.730153628 | -0.600313103 | 0.548297588 | 0.721128035 |
| ENSG00000231006 | RPL7P32 | 8.527747223 | 0.053763697 | 0.597237999 | 0.090020557 | 0.928270879 | 0.965519563 |
| ENSG00000231007 | CDC20P1 | 311.5214503 | -0.346753114 | 0.106821212 | -3.246107288 | 0.001169947 | 0.004328107 |
| ENSG00000231028 | AHI1-DT | 43.50602499 | 0.462537489 | 0.272478604 | 1.697518564 | 0.089598664 | 0.193704021 |
| ENSG00000231035 | RPL7L1P9 | 6.605378603 | -0.616498759 | 0.701773031 | -0.878487391 | 0.379679277 | 0.572075049 |
| ENSG00000231043 | ENSG0000023 | 940.9736808 | -0.090722168 | 0.067843091 | -1.33723518 | 0.181145882 | 0.336558397 |
| ENSG00000231050 | GNB1-DT | 3.758864192 | 0.537725536 | 0.961515819 | 0.559247727 | 0.575992665 | 0.742402052 |
| ENSG00000231064 | THBS3-AS1 | 63.6840024 | 0.77033077 | 0.229150402 | 3.361681949 | 0.000774693 | 0.002960429 |
| ENSG00000231066 | NPM1P9 | 6.33309536 | 0.33901421 | 0.705595965 | 0.480465063 | 0.63089674 | 0.782396541 |
| ENSG00000231073 | ENSG0000023 | 4.15256212 | -0.325835928 | 0.850821806 | -0.382966123 | 0.701744883 | 0.833061472 |
| ENSG00000231074 | HCG18 | 947.3928514 | 0.22995717 | 0.068041181 | 3.379676365 | 0.000725712 | 0.00278905 |
| ENSG00000231086 | TSSK1A | 2.546937366 | 1.405992275 | 1.147272029 | 1.225509068 | 0.220383476 | 0.390635764 |
| ENSG00000231093 | TRMT1P1 | 1.537494121 | 1.868782351 | 1.573810421 | 1.187425325 | 0.235059895 | NA |
| ENSG00000231095 | GYG1P3 | 7.30420519 | 0.542466691 | 0.681139742 | 0.796410278 | 0.425793606 | 0.616646598 |
| ENSG00000231107 | LINC01508 | 25.00025419 | -0.815132007 | 0.373408483 | -2.182949894 | 0.0290395 | 0.075452048 |
| ENSG00000231113 | LINC02976 | 39.31237122 | -0.110829452 | 0.278583204 | -0.397832497 | 0.690753661 | 0.825640142 |
| ENSG00000231119 | ENSG0000023 | 3.063651683 | 0.289938718 | 0.998183146 | 0.290466453 | 0.771459411 | 0.87553945 |
| ENSG00000231120 | BTF3P10 | 94.410436 | 0.336144992 | 0.188272442 | 1.785417924 | 0.074193553 | 0.16647106 |
| ENSG00000231124 | UBE2V1P11 | 1.785290738 | 1.224373604 | 1.344897122 | 0.910384582 | 0.362619726 | NA |
| ENSG00000231125 | ENSG0000023 | 4.856042812 | -0.466756551 | 0.798060395 | -0.584863694 | 0.558639372 | 0.728632431 |
| ENSG00000231128 | ENSG0000023 | 4.966662734 | 0.722912503 | 0.820258189 | 0.881323116 | 0.378142958 | 0.570970942 |
| ENSG00000231131 | LNCAROD | 491.1161197 | -2.10711955 | 0.096464608 | -21.84344696 | 8.97E-106 | 1.69E-103 |
| ENSG00000231133 | HAR1B | 1.807915511 | -1.584310957 | 1.385463917 | -1.143523796 | 0.252821181 | 0.431655097 |
| ENSG00000231154 | MORF4L2-AS1 | 58.08402069 | -0.363634203 | 0.254774608 | -1.427278036 | 0.153499769 | 0.296504269 |
| ENSG00000231160 | KLF3-AS1 | 60.90318374 | 1.099332145 | 0.243506303 | 4.514594202 | 6.34E-06 | 3.35E-05 |
| ENSG00000231162 | COX11P1 | 18.64734304 | 0.373400259 | 0.41783159 | 0.893662109 | 0.371502705 | 0.564052687 |

| | | | | | | | |
|---|---|---|---|---|---|---|---|
| ENSG00000231164 | RPL7P56 | 2.324749821 | 0.258921863 | 1.182906875 | 0.218886092 | 0.82673878 | 0.908541056 |
| ENSG00000231167 | YBX1P2 | 684.4938881 | 0.059660339 | 0.07911134 | 0.754131314 | 0.450770376 | 0.638904691 |
| ENSG00000231169 | EEF1B2P1 | 30.60325715 | 0.444770374 | 0.342763287 | 1.297602136 | 0.194424088 | 0.355385544 |
| ENSG00000231177 | LINC00852 | 18.36719955 | 1.101345387 | 0.42098522 | 2.616114142 | 0.008893682 | 0.026841346 |
| ENSG00000231181 | ENSG0000023 | 11.41002481 | 0.45325443 | 0.536398479 | 0.84499574 | 0.398113218 | 0.591114383 |
| ENSG00000231184 | CCNQP3 | 4.970493345 | -0.707091015 | 0.825937764 | -0.856106896 | 0.391938654 | 0.584785218 |
| ENSG00000231185 | SPRY4-AS1 | 3.021553352 | -1.988462383 | 1.120942913 | -1.773919403 | 0.076076483 | 0.169865146 |
| ENSG00000231187 | SYT15-AS1 | 5.019190202 | -0.172238598 | 0.786034065 | -0.219123579 | 0.826553784 | 0.908492348 |
| ENSG00000231194 | FARP1-AS1 | 1.863250154 | -3.377747778 | 1.629554847 | -2.072803983 | 0.038190526 | 0.09548672 |
| ENSG00000231203 | KRT8P10 | 465.3066801 | 0.253828604 | 0.093147694 | 2.725012212 | 0.006429911 | 0.020134064 |
| ENSG00000231228 | ARPP19P1 | 15.99585072 | -1.430924446 | 0.479397125 | -2.984841523 | 0.002837253 | 0.009672548 |
| ENSG00000231235 | CTNNBL1P1 | 2.745986725 | 0.027330705 | 1.155477419 | 0.023653171 | 0.98112926 | 0.991284088 |
| ENSG00000231240 | KLF2P1 | 12.12263763 | -0.541767077 | 0.551929336 | -0.981587753 | 0.326302984 | 0.51680409 |
| ENSG00000231244 | PSMC1P3 | 9.608128277 | -0.374835867 | 0.596041294 | -0.628875668 | 0.529430457 | 0.705157753 |
| ENSG00000231245 | C1DP1 | 123.3550264 | 0.033287319 | 0.160728306 | 0.207103028 | 0.835929398 | 0.913370821 |
| ENSG00000231249 | ITPR1-DT | 45.62577198 | 0.015942724 | 0.29582621 | 0.053892197 | 0.957021054 | 0.980529834 |
| ENSG00000231255 | ENSG0000023 | 2.06940094 | -1.773651086 | 1.320292048 | -1.343377845 | 0.179149657 | 0.333903242 |
| ENSG00000231290 | APCDD1L-DT | 6.589244457 | -1.349632078 | 0.703018307 | -1.919768041 | 0.054887205 | 0.129778653 |
| ENSG00000231300 | EZH2P1 | 6.600622226 | 0.782364741 | 0.702169418 | 1.114210789 | 0.265188768 | 0.446195565 |
| ENSG00000231302 | RPL36P2 | 3.324703215 | -0.470534022 | 0.975194987 | -0.482502503 | 0.629449021 | 0.781143482 |
| ENSG00000231304 | SGO1-AS1 | 13.08878573 | -0.487623506 | 0.485721545 | -1.003915744 | 0.315419227 | 0.504196019 |
| ENSG00000231305 | IFT122P3 | 50.81019882 | -0.653939147 | 0.256311119 | -2.551349112 | 0.010730678 | 0.031607826 |
| ENSG00000231310 | TBL1XR1-AS1 | 1.64004564 | -2.156794451 | 1.527170034 | -1.412281804 | 0.157867002 | NA |
| ENSG00000231312 | MAP4K3-DT | 92.8760781 | 0.27289046 | 0.18880329 | 1.4453692 | 0.14835421 | 0.289090959 |
| ENSG00000231313 | CLIC1P1 | 36.88226299 | -0.00463157 | 0.308798947 | -0.014998657 | 0.988033252 | 0.994030148 |
| ENSG00000231324 | ENSG0000023 | 17.69632425 | -0.571509335 | 0.431548131 | -1.324323508 | 0.185395626 | 0.342494912 |
| ENSG00000231327 | LINC01816 | 17.14878031 | -1.365081723 | 0.461986034 | -2.954811663 | 0.0031286 | 0.010560635 |
| ENSG00000231329 | ENSG0000023 | 3.873606787 | 1.789866026 | 1.022259158 | 1.750892631 | 0.079964406 | 0.176598898 |
| ENSG00000231333 | RPL34P6 | 8.95750237 | 0.08545899 | 0.578283699 | 0.14778039 | 0.882516086 | 0.939121523 |
| ENSG00000231340 | ACTG1P10 | 4.200697476 | 0.086782598 | 0.877331095 | 0.098916588 | 0.921204499 | 0.9619656 |
| ENSG00000231341 | VDAC1P6 | 80.74159774 | -0.02869068 | 0.201521774 | -0.142370124 | 0.88678766 | 0.941714143 |
| ENSG00000231344 | ENSG0000023 | 6.93552227 | -0.433825257 | 0.681583932 | -0.636495723 | 0.524453368 | 0.700819029 |
| ENSG00000231345 | BEND3P1 | 3.541312882 | -1.114897526 | 0.976092408 | -1.142204895 | 0.253368862 | 0.432011423 |

| | | | | | | | |
|---|---|---|---|---|---|---|---|
| ENSG00000231346 | LINC01160 | 4.191355088 | -0.097563252 | 0.882759713 | -0.110520734 | 0.911996407 | 0.955697231 |
| ENSG00000231351 | WBP1P1 | 14.52745815 | -0.363487837 | 0.481323179 | -0.755184569 | 0.450138246 | 0.638403176 |
| ENSG00000231357 | ENSG0000023 | 17.25886059 | 0.198219687 | 0.430530746 | 0.460407738 | 0.645223581 | 0.792410137 |
| ENSG00000231359 | RPL31P34 | 9.441329966 | 1.466676109 | 0.622565899 | 2.355856802 | 0.01848004 | 0.051014695 |
| ENSG00000231365 | WARS2-AS1 | 458.6781471 | 0.253646532 | 0.097632895 | 2.597961814 | 0.009377892 | 0.028125094 |
| ENSG00000231369 | RPL15P22 | 59.3182486 | 0.274158098 | 0.241080779 | 1.137204302 | 0.255452889 | 0.434821494 |
| ENSG00000231386 | MOB4P1 | 3.102851552 | -1.258144475 | 1.034553364 | -1.216123323 | 0.223937947 | 0.395377242 |
| ENSG00000231397 | ENSG0000023 | 8.992951366 | -0.093696742 | 0.624919797 | -0.149934027 | 0.880816665 | 0.938252557 |
| ENSG00000231407 | GORAB-AS1 | 2.44204984 | -0.677944265 | 1.186797731 | -0.571238255 | 0.56783815 | 0.73560104 |
| ENSG00000231409 | RPL36AP8 | 47.61283242 | 0.188126192 | 0.29033497 | 0.647962567 | 0.517009162 | 0.695286093 |
| ENSG00000231414 | RPL39P15 | 45.33483338 | 0.422065887 | 0.294910434 | 1.431166342 | 0.152382547 | 0.294942257 |
| ENSG00000231416 | ENSG0000023 | 44.33855542 | 0.447575715 | 0.263784333 | 1.696748666 | 0.089744188 | 0.193945517 |
| ENSG00000231419 | LINC00689 | 9.20998467 | -0.199794171 | 0.577037489 | -0.346241232 | 0.72916143 | 0.850013636 |
| ENSG00000231424 | ENSG0000023 | 1.852452341 | 0.589706451 | 1.2975846 | 0.454464742 | 0.649494363 | 0.796021186 |
| ENSG00000231431 | FAR2P4 | 5.289137845 | -1.28573121 | 0.794742408 | -1.617796152 | 0.105706536 | 0.221655204 |
| ENSG00000231434 | HSP90AA3P | 58.96079557 | -0.018600187 | 0.26019078 | -0.071486727 | 0.943010388 | 0.97291737 |
| ENSG00000231437 | LINC01750 | 14.7299343 | 1.781897508 | 0.504234574 | 3.533866178 | 0.000409528 | 0.001644422 |
| ENSG00000231439 | WASIR2 | 2.15634529 | -1.333126833 | 1.225338285 | -1.087966359 | 0.276609959 | 0.459148513 |
| ENSG00000231441 | DST-AS1 | 7.403530539 | 0.128386212 | 0.647704644 | 0.198217216 | 0.842875118 | 0.917308531 |
| ENSG00000231445 | TIMM8AP1 | 7.479232038 | 0.57464076 | 0.71660831 | 0.801889612 | 0.422616816 | 0.613491717 |
| ENSG00000231458 | RNASEH1P2 | 20.93546147 | -0.172492724 | 0.396480027 | -0.43506031 | 0.663518652 | 0.805776489 |
| ENSG00000231466 | ENSG0000023 | 12.23622525 | -0.886323105 | 0.527651192 | -1.679751926 | 0.093005592 | 0.199733036 |
| ENSG00000231468 | PRDX3P2 | 2.889242839 | -1.463045433 | 1.073816184 | -1.362472884 | 0.173048696 | 0.325281636 |
| ENSG00000231482 | ENSG0000023 | 7.757534738 | -0.364496069 | 0.61715162 | -0.590610244 | 0.5547816 | 0.726016596 |
| ENSG00000231500 | RPS18 | 9860.055667 | 0.277690338 | 0.052129727 | 5.326909471 | 9.99E-08 | 6.48E-07 |
| ENSG00000231503 | PTMAP4 | 67.05825724 | -0.079935485 | 0.257047298 | -0.310975783 | 0.755819032 | 0.865149344 |
| ENSG00000231504 | NMD3P2 | 13.59244817 | -0.453495149 | 0.481661206 | -0.941523093 | 0.346436861 | 0.537239623 |
| ENSG00000231528 | FAM225A | 50.87792251 | 0.565306716 | 0.251629465 | 2.246583939 | 0.02466663 | 0.065712749 |
| ENSG00000231530 | ENSG0000023 | 14.87790103 | 0.360070876 | 0.453640956 | 0.793735378 | 0.4273495 | 0.61819886 |
| ENSG00000231531 | HINT1P1 | 12.46281832 | -0.182122021 | 0.513361803 | -0.354763483 | 0.722766788 | 0.845926239 |
| ENSG00000231542 | TAB3-AS1 | 1.838684744 | 3.144231015 | 1.593826446 | 1.972756207 | 0.048523345 | 0.116780838 |
| ENSG00000231544 | RSL24D1P11 | 49.56255704 | -0.480892832 | 0.253322348 | -1.898343493 | 0.057650848 | 0.13499137 |
| ENSG00000231549 | ATP5MKP1 | 10.9211698 | 0.17451354 | 0.578855518 | 0.301480308 | 0.763048264 | 0.870126121 |

| | | | | | | | |
|---|---|---|---|---|---|---|---|
| ENSG00000231551 | PDE4DIPP6 | 12.16295898 | -0.069138883 | 0.516614783 | -0.133830633 | 0.893536503 | 0.945685231 |
| ENSG00000231552 | IGBP1P3 | 2.007534786 | -0.257824251 | 1.322630416 | -0.194932952 | 0.845445437 | 0.918953723 |
| ENSG00000231563 | ENSG0000023 | 8.91445729 | 0.316782057 | 0.588947709 | 0.53787807 | 0.59066123 | 0.753267101 |
| ENSG00000231574 | LINC02015 | 2.184340143 | 4.473434023 | 1.539572941 | 2.905633052 | 0.00366511 | 0.012179867 |
| ENSG00000231579 | RPL7P21 | 97.74922667 | -0.190369847 | 0.185408276 | -1.026760244 | 0.304533374 | 0.491971521 |
| ENSG00000231584 | FAHD2CP | 251.8609567 | -0.060249202 | 0.125952411 | -0.478348935 | 0.632401872 | 0.783432036 |
| ENSG00000231587 | SNORD62B | 15.29375623 | 0.076822411 | 0.457571466 | 0.167891611 | 0.86666855 | 0.930776055 |
| ENSG00000231588 | SHQ1P1 | 7.456430362 | -0.369236447 | 0.655323126 | -0.563441808 | 0.573134075 | 0.73988158 |
| ENSG00000231607 | DLEU2 | 122.5923761 | -0.140152188 | 0.160284063 | -0.87439877 | 0.381901127 | 0.57425353 |
| ENSG00000231609 | EHBP1-AS1 | 6.336887663 | -0.754819525 | 0.763770118 | -0.988281038 | 0.323015029 | 0.512905833 |
| ENSG00000231615 | ENSG0000023 | 2.431081957 | -0.377645812 | 1.13214374 | -0.333567019 | 0.738706309 | 0.855984378 |
| ENSG00000231616 | ENSG0000023 | 1.697321329 | 1.240205373 | 1.473201709 | 0.841843561 | 0.399875525 | NA |
| ENSG00000231651 | DLG3-AS1 | 3.797011836 | -0.266661006 | 0.975780255 | -0.273279772 | 0.784638163 | 0.883091372 |
| ENSG00000231652 | CD109-AS1 | 2.460822348 | -1.179161088 | 1.158648955 | -1.017703493 | 0.308818887 | 0.496900138 |
| ENSG00000231663 | COA6-AS1 | 27.46175011 | 0.006875656 | 0.345691304 | 0.019889582 | 0.984131456 | 0.992635461 |
| ENSG00000231672 | DIRC3 | 12.87488608 | -1.459917883 | 0.528079851 | -2.764577894 | 0.00569965 | 0.018084366 |
| ENSG00000231684 | EIF1P3 | 170.9184214 | -0.056560533 | 0.140467997 | -0.402657788 | 0.687199991 | 0.823290568 |
| ENSG00000231686 | ANKRD11P2 | 89.48061666 | 0.237348264 | 0.192542449 | 1.232706166 | 0.217685411 | 0.387214171 |
| ENSG00000231690 | LINC00574 | 9.848677667 | -0.264491653 | 0.555108614 | -0.476468291 | 0.633740792 | 0.784143138 |
| ENSG00000231691 | ENSG0000023 | 1.810275226 | -1.008155262 | 1.353027664 | -0.745110605 | 0.456204858 | 0.643466518 |
| ENSG00000231700 | ENSG0000023 | 8.467800053 | -0.54505768 | 0.603720385 | -0.902831334 | 0.366615419 | 0.559186017 |
| ENSG00000231702 | ENSG0000023 | 15.00848787 | 0.369670528 | 0.486642029 | 0.759635433 | 0.447472533 | 0.635888882 |
| ENSG00000231703 | ENSG0000023 | 9.813337523 | 0.388662343 | 0.556159403 | 0.698832639 | 0.484656629 | 0.668511999 |
| ENSG00000231705 | ENSG0000023 | 2.050058924 | 2.244656206 | 1.428026993 | 1.571858387 | 0.115983397 | 0.239031455 |
| ENSG00000231707 | PABPC1P1 | 23.98427636 | 0.214521902 | 0.383760365 | 0.55899963 | 0.576161973 | 0.742536835 |
| ENSG00000231713 | ENSG0000023 | 27.84569799 | 0.289348281 | 0.328685518 | 0.880319531 | 0.378686235 | 0.571329315 |
| ENSG00000231721 | LINC-PINT | 11.96173048 | 0.770147208 | 0.517366112 | 1.488592295 | 0.136594762 | 0.271438309 |
| ENSG00000231728 | TMSB15B-AS1 | 26.70046951 | 0.274469569 | 0.353848741 | 0.775669198 | 0.437944331 | 0.627472712 |
| ENSG00000231731 | ENSG0000023 | 10.61015743 | 0.054617026 | 0.575686631 | 0.09487284 | 0.92441583 | 0.963698865 |
| ENSG00000231741 | ENSG0000023 | 2.761537391 | -0.311296711 | 1.056579389 | -0.294626902 | 0.768278909 | 0.873528855 |
| ENSG00000231744 | ENSG0000023 | 9.632611224 | -0.513024663 | 0.603115519 | -0.850624212 | 0.394978135 | 0.588013388 |
| ENSG00000231747 | ENSG0000023 | 19.45683705 | 0.606718731 | 0.400437868 | 1.515138252 | 0.129737396 | 0.260884982 |
| ENSG00000231752 | EMBP1 | 47.32858524 | 0.55406244 | 0.269176417 | 2.058361745 | 0.039555421 | 0.098269814 |

| | | | | | | | |
|---|---|---|---|---|---|---|---|
| ENSG00000231760 | ENSG0000023 | 19.97506506 | -0.299289871 | 0.435094818 | -0.687872756 | 0.491532914 | 0.674428199 |
| ENSG00000231767 | RPS27AP5 | 79.51663525 | 0.114122972 | 0.20065909 | 0.568740605 | 0.569532188 | 0.73689049 |
| ENSG00000231769 | ENSG0000023 | 9.886551378 | 0.649938878 | 0.563740267 | 1.152904832 | 0.248949448 | 0.426853124 |
| ENSG00000231770 | TMEM44-AS1 | 210.1379049 | -0.261068982 | 0.129973442 | -2.008633281 | 0.044576039 | 0.108599654 |
| ENSG00000231776 | LINC01611 | 9.628280653 | 0.23619225 | 0.574311478 | 0.411261588 | 0.680880732 | 0.818841625 |
| ENSG00000231784 | DBIL5P | 18.90832177 | -0.413477667 | 0.416381384 | -0.993026303 | 0.320697136 | 0.510250541 |
| ENSG00000231793 | DOC2GP | 2.943650438 | -1.474546123 | 1.108489003 | -1.330230718 | 0.183442266 | 0.339642097 |
| ENSG00000231799 | PA2G4P6 | 42.40616378 | -0.324983776 | 0.277581799 | -1.170767596 | 0.241692206 | 0.417816067 |
| ENSG00000231806 | PCAT7 | 694.3644174 | 0.119573091 | 0.076381946 | 1.565462758 | 0.117474443 | 0.241540174 |
| ENSG00000231811 | ENSG0000023 | 3.809653936 | -1.081059415 | 0.933446431 | -1.158137605 | 0.246807886 | 0.42406804 |
| ENSG00000231822 | SMC3P1 | 78.62481941 | -0.444210325 | 0.211070118 | -2.104562836 | 0.035329379 | 0.089335975 |
| ENSG00000231829 | ENSG0000023 | 5.839111292 | 1.170922025 | 0.770675375 | 1.519345322 | 0.128675597 | 0.2591512 |
| ENSG00000231831 | MTHFD1P1 | 121.6277191 | -0.982814011 | 0.168317483 | -5.839048898 | 5.25E-09 | 3.90E-08 |
| ENSG00000231840 | TMEM139-AS | 5.597786699 | -1.603915116 | 0.783318478 | -2.047590044 | 0.040600186 | 0.100406238 |
| ENSG00000231841 | FAM192BP | 156.6631006 | -0.118354793 | 0.149167556 | -0.793435225 | 0.427524294 | 0.618217808 |
| ENSG00000231848 | KRTCAP2P1 | 191.4094763 | 0.476906233 | 0.134593065 | 3.543319516 | 0.000395124 | 0.001589363 |
| ENSG00000231851 | MRPS9-AS1 | 14.32798298 | -0.01274225 | 0.458092065 | -0.027815916 | 0.977808972 | 0.989672967 |
| ENSG00000231852 | CYP21A2 | 7.396294453 | 0.468197216 | 0.64723882 | 0.723376289 | 0.469448735 | 0.655385199 |
| ENSG00000231856 | ENSG0000023 | 51.38239562 | -0.236563411 | 0.24351771 | -0.971442327 | 0.331328059 | 0.52224246 |
| ENSG00000231863 | ENSG0000023 | 1.862771523 | -0.948918291 | 1.343384887 | -0.706363678 | 0.479962002 | 0.664313282 |
| ENSG00000231864 | ENSG0000023 | 34.12292867 | -0.625780971 | 0.354486035 | -1.765319108 | 0.07751015 | 0.172378365 |
| ENSG00000231871 | IPO9-AS1 | 11.04061778 | -0.213847495 | 0.52665949 | -0.40604508 | 0.684709478 | 0.821336301 |
| ENSG00000231875 | ENSG0000023 | 1.804689665 | 0.546986814 | 1.353637476 | 0.404086636 | 0.686149012 | 0.822589932 |
| ENSG00000231878 | SNRPFP1 | 43.03214488 | 0.06833429 | 0.273915386 | 0.24947226 | 0.802995496 | 0.895135709 |
| ENSG00000231880 | ENSG0000023 | 17.84864571 | 0.523970533 | 0.429244008 | 1.220682229 | 0.222206359 | 0.393227947 |
| ENSG00000231884 | NDUFB1P1 | 60.80438569 | -0.398353834 | 0.231432423 | -1.721253352 | 0.08520486 | 0.186081445 |
| ENSG00000231887 | PRH1 | 7.576628434 | -0.739362006 | 0.651302072 | -1.135205979 | 0.256289025 | 0.43578394 |
| ENSG00000231889 | TRAF3IP2-AS1 | 41.8493204 | 0.128947325 | 0.274157451 | 0.470340399 | 0.638111839 | 0.786835389 |
| ENSG00000231890 | DARS1-AS1 | 41.9885173 | -0.275470367 | 0.271328467 | -1.015265263 | 0.309979398 | 0.497921249 |
| ENSG00000231892 | ENSG0000023 | 3.407878497 | 2.528235084 | 1.172152359 | 2.156916773 | 0.03101215 | 0.079952654 |
| ENSG00000231908 | IDH1-AS1 | 16.63653148 | -0.645716707 | 0.427031106 | -1.512106959 | 0.130506655 | 0.26220202 |
| ENSG00000231909 | MAP1LC3BP1 | 1.565885037 | 1.85120512 | 1.530583309 | 1.209476877 | 0.226479689 | NA |
| ENSG00000231916 | ENSG0000023 | 4.406803373 | -0.709078509 | 0.835470775 | -0.848717311 | 0.396038611 | 0.589025269 |

| | | | | | | | |
|---|---|---|---|---|---|---|---|
| ENSG00000231920 | NEBL-AS1 | 8.821219599 | -0.007125619 | 0.605238692 | -0.011773237 | 0.990606533 | 0.994991588 |
| ENSG00000231925 | TAPBP | 3613.722807 | 0.111764348 | 0.054683988 | 2.043822166 | 0.040971119 | 0.101170878 |
| ENSG00000231929 | HMGB3P31 | 3.678896561 | -1.583596253 | 1.019811802 | -1.552831856 | 0.12046331 | 0.246338771 |
| ENSG00000231937 | ENSG0000023 | 1.572065846 | 3.996101089 | 1.665395765 | 2.399490364 | 0.016417912 | NA |
| ENSG00000231940 | RPS7P3 | 11.51739219 | 0.404804297 | 0.568128108 | 0.712522918 | 0.476141025 | 0.661218906 |
| ENSG00000231952 | DPY19L1P2 | 12.6644249 | 1.674869514 | 0.579096598 | 2.892210939 | 0.00382541 | 0.012657567 |
| ENSG00000231955 | ENSG0000023 | 2.177726064 | 0.469954656 | 1.196996707 | 0.392611486 | 0.694606455 | 0.828005022 |
| ENSG00000231966 | LINC02818 | 1.7338195 | 0.902586875 | 1.457606262 | 0.619225437 | 0.535767857 | NA |
| ENSG00000231969 | MMADHC-DT | 7.792559437 | -0.506990578 | 0.625575782 | -0.810438307 | 0.417688309 | 0.609793757 |
| ENSG00000231970 | ENSG0000023 | 4.629543774 | -0.557321728 | 0.84936464 | -0.656163092 | 0.511719193 | 0.691292521 |
| ENSG00000231989 | PPP1R2B | 118.4891212 | -0.272863928 | 0.16964765 | -1.608415608 | 0.107744186 | 0.225084384 |
| ENSG00000231991 | ANXA2P2 | 5660.767641 | -0.207475872 | 0.047639589 | -4.355114652 | 1.33E-05 | 6.75E-05 |
| ENSG00000231993 | EP300-AS1 | 4.427807788 | 0.751510056 | 0.871189441 | 0.862625304 | 0.388343513 | 0.581121996 |
| ENSG00000231995 | RPL7AP76 | 4.914765689 | 1.026999354 | 0.816094289 | 1.258432228 | 0.208235483 | 0.374748723 |
| ENSG00000231999 | LRRC8C-DT | 9.610823873 | -0.224077668 | 0.559732647 | -0.400329817 | 0.688913609 | 0.824262433 |
| ENSG00000232000 | CLCN3P1 | 6.03705335 | -0.251827328 | 0.712553323 | -0.353415414 | 0.723777032 | 0.846416769 |
| ENSG00000232004 | CAP1P2 | 33.5211712 | -0.990365483 | 0.345477983 | -2.866652963 | 0.004148376 | 0.013622066 |
| ENSG00000232022 | FAAHP1 | 46.55723888 | -0.175228135 | 0.260369565 | -0.672997766 | 0.500948712 | 0.682003766 |
| ENSG00000232023 | LINC01807 | 28.67999093 | -0.204273745 | 0.339175208 | -0.602266144 | 0.546996993 | 0.719888603 |
| ENSG00000232024 | LSM12P1 | 662.0281796 | -0.006502617 | 0.080639127 | -0.08063848 | 0.935729463 | 0.969261113 |
| ENSG00000232037 | RPL21P29 | 2.127614814 | 0.474088761 | 1.247939366 | 0.379897272 | 0.704021672 | 0.834379891 |
| ENSG00000232040 | SCAND3 | 152.089732 | 0.01437952 | 0.146829093 | 0.097933722 | 0.921984923 | 0.962216621 |
| ENSG00000232041 | PSMD10P3 | 2.004685299 | 0.264596633 | 1.218130787 | 0.217215291 | 0.828040569 | 0.909090629 |
| ENSG00000232043 | ENSG0000023 | 14.16907168 | -0.171162429 | 0.477615349 | -0.358368779 | 0.720067353 | 0.844366665 |
| ENSG00000232044 | SILC1 | 13.08892962 | 1.112654408 | 0.52736504 | 2.109837252 | 0.034872377 | 0.088326412 |
| ENSG00000232054 | NPM1P34 | 1.792938439 | -0.327654201 | 1.373131732 | -0.238618184 | 0.811401663 | 0.90021491 |
| ENSG00000232059 | ENSG0000023 | 2.430105465 | 0.057363019 | 1.112146408 | 0.051578658 | 0.958864425 | 0.981197069 |
| ENSG00000232060 | SLC4A1APP1 | 147.9974882 | -0.243757765 | 0.151347754 | -1.610580658 | 0.107271152 | 0.224322275 |
| ENSG00000232065 | LINC01063 | 8.565323409 | -0.755343738 | 0.630690017 | -1.197646574 | 0.231054637 | 0.404641069 |
| ENSG00000232070 | TMEM253 | 14.01158676 | -0.68051909 | 0.47647162 | -1.428246849 | 0.15322082 | 0.296115048 |
| ENSG00000232077 | LINC01031 | 1.590382803 | -1.329749526 | 1.452601468 | -0.915426258 | 0.359967899 | NA |
| ENSG00000232093 | DCST1-AS1 | 35.04028616 | 1.437623551 | 0.32093071 | 4.479544977 | 7.48E-06 | 3.92E-05 |
| ENSG00000232098 | ZNF584-DT | 514.0016634 | -0.164420827 | 0.095244975 | -1.726293989 | 0.084294536 | 0.184391786 |

| | | | | | | | |
|---|---|---|---|---|---|---|---|
| ENSG00000232104 | RFX3-DT | 25.00199553 | -0.013118585 | 0.355719525 | -0.036879014 | 0.970581473 | 0.986772291 |
| ENSG00000232112 | TMA7 | 1161.635883 | 0.019228527 | 0.063130382 | 0.304584357 | 0.760682736 | 0.86855119 |
| ENSG00000232118 | BACH1-AS1 | 99.50217388 | -1.238536008 | 0.187155282 | -6.617691979 | 3.65E-11 | 3.26E-10 |
| ENSG00000232119 | MCTS1 | 1336.312745 | -0.022465605 | 0.070827524 | -0.317187493 | 0.75110134 | 0.862460033 |
| ENSG00000232124 | ENSG0000023 | 11.4996649 | -0.429942463 | 0.519505661 | -0.827599188 | 0.407897528 | 0.600511443 |
| ENSG00000232133 | IMPDH1P10 | 2.65608436 | -1.29645726 | 1.09834368 | -1.180374853 | 0.237851159 | 0.413356484 |
| ENSG00000232134 | RPS15AP12 | 22.54505719 | 0.036898493 | 0.373750353 | 0.098724971 | 0.921356642 | 0.9619656 |
| ENSG00000232139 | LINC00867 | 4.047560553 | -1.454712107 | 0.932578851 | -1.559881082 | 0.118787986 | 0.243632585 |
| ENSG00000232140 | ENSG0000023 | 22.60275862 | 1.234471402 | 0.406865231 | 3.034103945 | 0.002412513 | 0.008349966 |
| ENSG00000232149 | FERP1 | 11.83529949 | -0.834521446 | 0.554901802 | -1.503908335 | 0.132604971 | 0.265580411 |
| ENSG00000232150 | ST13P4 | 299.8019126 | 0.310838636 | 0.113728686 | 2.73315949 | 0.006272995 | 0.019709971 |
| ENSG00000232160 | RAP2C-AS1 | 157.9917985 | -0.079987815 | 0.148927478 | -0.537092389 | 0.591203798 | 0.753709593 |
| ENSG00000232164 | LINC01873 | 2.394685805 | -1.024750188 | 1.153897573 | -0.888077254 | 0.374499197 | 0.567288139 |
| ENSG00000232176 | ENSG0000023 | 28.17714348 | 0.240150215 | 0.362122103 | 0.663174696 | 0.507218676 | 0.687444744 |
| ENSG00000232177 | MTND4P24 | 16.77766795 | 0.382552316 | 0.428647622 | 0.892463406 | 0.372144599 | 0.564841384 |
| ENSG00000232186 | TERF1P7 | 4.537402871 | -0.740707974 | 0.840796773 | -0.880959582 | 0.378339697 | 0.571100536 |
| ENSG00000232187 | FTH1P7 | 531.4319542 | 0.364172147 | 0.085175031 | 4.275573991 | 1.91E-05 | 9.50E-05 |
| ENSG00000232202 | CHORDC1P1 | 9.447047438 | 0.337889271 | 0.580553716 | 0.582012071 | 0.56055855 | 0.729849205 |
| ENSG00000232203 | SLC25A6P2 | 15.50207393 | 0.023040174 | 0.446910809 | 0.0515543 | 0.958883834 | 0.981197069 |
| ENSG00000232212 | LINC01701 | 8.586565537 | 0.05789804 | 0.605925488 | 0.095553069 | 0.92387554 | 0.963279648 |
| ENSG00000232220 | ENSG0000023 | 3.701361931 | -0.773035425 | 0.92859462 | -0.832478897 | 0.405138685 | 0.597851052 |
| ENSG00000232228 | RPL36AP16 | 15.25378323 | 0.568860164 | 0.452362132 | 1.257532679 | 0.208560814 | 0.375120107 |
| ENSG00000232229 | LINC00865 | 210.4623118 | 0.366884032 | 0.129583753 | 2.831250255 | 0.004636643 | 0.015063672 |
| ENSG00000232233 | LINC02043 | 2.929471854 | -0.722564572 | 1.03004618 | -0.701487551 | 0.482998801 | 0.667074097 |
| ENSG00000232237 | ASCL5 | 5.741658614 | 1.640279756 | 0.803056716 | 2.042545345 | 0.041097466 | 0.101362851 |
| ENSG00000232260 | BTF3L4P1 | 48.98309156 | 0.014784703 | 0.259827693 | 0.056901951 | 0.9546233 | 0.979122709 |
| ENSG00000232267 | ACTR3P2 | 2.652360653 | -0.544347921 | 1.056304295 | -0.515332488 | 0.6063207 | 0.764860977 |
| ENSG00000232284 | GNG12-AS1 | 16.71585635 | -0.368742263 | 0.443386108 | -0.831650466 | 0.405606268 | 0.59836706 |
| ENSG00000232298 | GRHL3-AS1 | 2.040519625 | 0.086118566 | 1.242330822 | 0.069320156 | 0.944734782 | 0.973711185 |
| ENSG00000232303 | DFFBP1 | 9.561509111 | 0.315161115 | 0.593714954 | 0.530829 | 0.595537282 | 0.756709053 |
| ENSG00000232316 | LINC02518 | 9.543712264 | -1.389485871 | 0.621155799 | -2.236936163 | 0.025290513 | 0.067109816 |
| ENSG00000232320 | MXRA7P1 | 20.3659932 | 0.692989157 | 0.398111822 | 1.74068972 | 0.081737981 | 0.179816567 |
| ENSG00000232328 | ENSG0000023 | 2.639645282 | 0.083156696 | 1.189197749 | 0.069926718 | 0.944251987 | 0.973534408 |

| | | | | | | | |
|---|---|---|---|---|---|---|---|
| ENSG00000232332 | ARL5AP5 | 4.897580198 | 0.491585816 | 0.821330607 | 0.598523678 | 0.549490564 | 0.721912465 |
| ENSG00000232333 | RPS27AP2 | 5.477671821 | -0.166543636 | 0.750795759 | -0.221822825 | 0.82445181 | 0.907529967 |
| ENSG00000232335 | ENSG0000023 | 6.306159314 | -0.577086573 | 0.715134048 | -0.806962799 | 0.419687916 | 0.611023031 |
| ENSG00000232341 | RPL4P2 | 28.37483877 | 0.336834657 | 0.341488817 | 0.986370976 | 0.323951102 | 0.51407293 |
| ENSG00000232346 | ENSG0000023 | 602.5318033 | 0.213580468 | 0.083388166 | 2.561280323 | 0.010428717 | 0.030822919 |
| ENSG00000232354 | VIPR1-AS1 | 3.146427916 | -0.350931809 | 0.967146417 | -0.362852825 | 0.716714829 | 0.842466582 |
| ENSG00000232362 | ATP5MGP2 | 12.9980297 | 0.56241546 | 0.48708616 | 1.154652926 | 0.248232584 | 0.425881047 |
| ENSG00000232368 | FTLP2 | 9.487467125 | 1.189670613 | 0.603639003 | 1.970831255 | 0.04874318 | 0.117208444 |
| ENSG00000232380 | ZDHHC20P4 | 147.6297054 | -0.103708648 | 0.148354455 | -0.699059885 | 0.484514607 | 0.668356288 |
| ENSG00000232383 | ENSG0000023 | 2.38901337 | 0.027036775 | 1.203194604 | 0.022470824 | 0.982072385 | 0.991627389 |
| ENSG00000232385 | RPS3AP25 | 12.86141442 | 0.314727218 | 0.517879207 | 0.607723219 | 0.543371061 | 0.716907232 |
| ENSG00000232386 | ENSG0000023 | 34.787376 | -0.588199983 | 0.299106711 | -1.966522186 | 0.049238322 | 0.118213198 |
| ENSG00000232387 | SKA2P1 | 268.3682538 | -0.153179752 | 0.119874728 | -1.277831904 | 0.201308703 | 0.364934116 |
| ENSG00000232388 | SMIM26 | 314.3940702 | -0.284182695 | 0.107780699 | -2.636675187 | 0.008372296 | 0.025416481 |
| ENSG00000232389 | ENSG0000023 | 220.6777776 | -0.172827463 | 0.138225106 | -1.250333377 | 0.211177791 | 0.378677286 |
| ENSG00000232391 | RANP2 | 1.72715197 | -0.78382461 | 1.367980575 | -0.572979342 | 0.566658681 | NA |
| ENSG00000232393 | RPL5P6 | 3.014109941 | -1.91637679 | 1.106778887 | -1.731490195 | 0.083364372 | 0.182757903 |
| ENSG00000232400 | RAD17P1 | 4.594990652 | 0.371428071 | 0.849911379 | 0.437019765 | 0.662097014 | 0.80479344 |
| ENSG00000232412 | STAG2-AS1 | 12.09739131 | -0.391053315 | 0.514598594 | -0.759919128 | 0.447302927 | 0.635757692 |
| ENSG00000232422 | KNOP1P4 | 1.917231947 | -1.624166283 | 1.447672335 | -1.121915674 | 0.261898296 | 0.442281133 |
| ENSG00000232434 | AJM1 | 371.2869917 | 0.424360995 | 0.100610479 | 4.217860815 | 2.47E-05 | 0.000120814 |
| ENSG00000232442 | MHENCR | 250.4794615 | -0.050103203 | 0.118734256 | -0.421977655 | 0.673041328 | 0.81285826 |
| ENSG00000232445 | EMSLR | 184.0414292 | -0.330935088 | 0.134677428 | -2.45724241 | 0.014000818 | 0.040012303 |
| ENSG00000232450 | ENSG0000023 | 45.56490171 | -0.036369606 | 0.26103407 | -0.13932896 | 0.889190206 | 0.94295596 |
| ENSG00000232453 | LINC02777 | 8.389168228 | 1.210236602 | 0.659546489 | 1.834952687 | 0.0665127 | 0.152174005 |
| ENSG00000232454 | ENSG0000023 | 49.75738083 | -0.257774082 | 0.250280006 | -1.029942765 | 0.303036874 | 0.490140724 |
| ENSG00000232456 | FAM131C2P | 1.908209113 | 0.826361067 | 1.355836603 | 0.609484258 | 0.542203504 | 0.716029496 |
| ENSG00000232460 | BMPR1AP2 | 131.7566936 | -0.146846267 | 0.156320034 | -0.939395052 | 0.347527952 | 0.538486405 |
| ENSG00000232467 | TMA16P2 | 22.84697184 | -0.318285706 | 0.382921416 | -0.831203723 | 0.405858552 | 0.598625381 |
| ENSG00000232472 | EEF1B2P3 | 2485.18921 | 0.146810728 | 0.064405074 | 2.279490077 | 0.022637949 | 0.060909518 |
| ENSG00000232479 | KRT8P52 | 5.429437569 | -0.784095139 | 0.751162091 | -1.043842798 | 0.296558127 | 0.482862051 |
| ENSG00000232480 | TGFB2-AS1 | 2.933195561 | -1.444409217 | 1.072994584 | -1.346147724 | 0.17825488 | 0.332777259 |
| ENSG00000232485 | RPL37A-DT | 9.272739237 | -0.785327617 | 0.60160559 | -1.30538617 | 0.191761385 | 0.351761268 |

| | | | | | | | |
|---|---|---|---|---|---|---|---|
| ENSG00000232486 | DEPDC1P2 | 30.00615301 | -0.645492863 | 0.321565889 | -2.007342461 | 0.044713211 | 0.108841369 |
| ENSG00000232499 | ENSG0000023 | 541.6132115 | 0.124235998 | 0.089928901 | 1.381491347 | 0.167127937 | 0.316661723 |
| ENSG00000232504 | ST3GAL5-AS1 | 7.01994893 | -1.180269791 | 0.675681405 | -1.746784479 | 0.08067473 | 0.177899693 |
| ENSG00000232531 | KMT5AP2 | 132.5154408 | -0.097919181 | 0.154146552 | -0.635234325 | 0.5252756 | 0.701327047 |
| ENSG00000232533 | FAM131B-AS2 | 92.96190324 | 0.204637065 | 0.189442999 | 1.080203892 | 0.280051395 | 0.463447298 |
| ENSG00000232536 | ENSG0000023 | 3.413531701 | 0.001485864 | 0.971478036 | 0.001529488 | 0.998779646 | 0.999329557 |
| ENSG00000232545 | ENSG0000023 | 10.7960376 | -2.271231497 | 0.607450272 | -3.738958726 | 0.000184784 | 0.000789375 |
| ENSG00000232546 | ENSG0000023 | 17.05055791 | 0.39857614 | 0.439856171 | 0.906151068 | 0.364855898 | 0.557629459 |
| ENSG00000232549 | SRD5A1P1 | 7.649794047 | -0.323912768 | 0.637611277 | -0.508009785 | 0.611446481 | 0.769085551 |
| ENSG00000232553 | CLK2P1 | 81.54542908 | -0.286699256 | 0.200211824 | -1.431979639 | 0.152149648 | 0.294615762 |
| ENSG00000232573 | RPL3P4 | 5295.578375 | 0.141120605 | 0.050840071 | 2.775775121 | 0.005507028 | 0.017519239 |
| ENSG00000232581 | ENSG0000023 | 11.77668467 | -0.159001891 | 0.51889314 | -0.306425117 | 0.759280987 | 0.867511996 |
| ENSG00000232586 | KIAA1614-AS1 | 66.87636064 | 1.000901425 | 0.221082008 | 4.527285761 | 5.97E-06 | 3.17E-05 |
| ENSG00000232587 | EEF1A1P3 | 6.972227138 | 0.021170053 | 0.656335306 | 0.032254935 | 0.974268747 | 0.988357363 |
| ENSG00000232593 | KANTR | 233.5589966 | -0.008797847 | 0.12887711 | -0.068265399 | 0.945574368 | 0.974261906 |
| ENSG00000232600 | TONSL-AS1 | 28.58449091 | -0.23779137 | 0.3248684 | -0.731962141 | 0.464191676 | 0.650485483 |
| ENSG00000232611 | ENSG0000023 | 19.19070221 | -0.321958425 | 0.417372927 | -0.77139269 | 0.440474206 | 0.62981551 |
| ENSG00000232615 | PPP4R2P5 | 60.6638724 | 0.001168613 | 0.226793101 | 0.005152773 | 0.9958887 | 0.997366925 |
| ENSG00000232618 | HIVEP2-DT | 6.479155599 | 0.534559979 | 0.697954759 | 0.765894884 | 0.44373885 | 0.632572978 |
| ENSG00000232624 | LINC01517 | 14.82624663 | -0.760907588 | 0.473158721 | -1.608144485 | 0.107803538 | 0.225187868 |
| ENSG00000232626 | ENSG0000023 | 8.512268287 | 0.489597959 | 0.627608127 | 0.780101369 | 0.435331211 | 0.625250312 |
| ENSG00000232630 | PRPS1P2 | 4.729822859 | 0.534862175 | 0.801049876 | 0.667701464 | 0.504324175 | 0.685175989 |
| ENSG00000232638 | ENSG0000023 | 232.4174208 | 0.750874612 | 0.124864471 | 6.013516923 | 1.82E-09 | 1.41E-08 |
| ENSG00000232645 | LINC01431 | 15.09248686 | 0.191794702 | 0.45164732 | 0.424655906 | 0.671087532 | 0.81140294 |
| ENSG00000232648 | ING2-DT | 6.772694737 | -0.24454345 | 0.677808525 | -0.360785445 | 0.718259844 | 0.843411181 |
| ENSG00000232653 | GOLGA8N | 4.850407804 | -0.053353621 | 0.839375072 | -0.063563505 | 0.949317792 | 0.976618218 |
| ENSG00000232654 | FAM136BP | 8.048409858 | -0.318084896 | 0.609114307 | -0.522208874 | 0.601524908 | 0.761283686 |
| ENSG00000232656 | IDI2-AS1 | 45.80679663 | -2.465631343 | 0.314358392 | -7.843376872 | 4.39E-15 | 5.30E-14 |
| ENSG00000232665 | PHB1P10 | 4.964771137 | -0.545401668 | 0.816649846 | -0.667852534 | 0.504227729 | 0.685166693 |
| ENSG00000232671 | ZNF687-AS1 | 43.65253911 | 0.163874564 | 0.274951327 | 0.596012993 | 0.551166547 | 0.723161782 |
| ENSG00000232675 | ENSG0000023 | 7.623852772 | -0.528816658 | 0.630716094 | -0.838438503 | 0.401784473 | 0.594410925 |
| ENSG00000232676 | ADH5P2 | 2.753080062 | 0.345594452 | 1.056792171 | 0.327022154 | 0.743651129 | 0.858490754 |
| ENSG00000232677 | LINC00665 | 10.95784563 | 0.614821275 | 0.543056904 | 1.132148898 | 0.25757184 | 0.437260269 |

| | | | | | | | |
|---|---|---|---|---|---|---|---|
| ENSG00000232678 | SPTLC1P4 | 1.917231947 | -1.624166283 | 1.447672335 | -1.121915674 | 0.261898296 | 0.442281133 |
| ENSG00000232682 | ENSG0000023 | 17.63149511 | -1.029664833 | 0.438605224 | -2.347589074 | 0.018895355 | 0.052054694 |
| ENSG00000232690 | HSPE1P9 | 3.168083658 | 0.147399859 | 0.985222093 | 0.149610793 | 0.881071692 | 0.938337262 |
| ENSG00000232699 | BDH2P1 | 12.99134162 | -0.919851944 | 0.491243935 | -1.87249527 | 0.061138116 | 0.141728893 |
| ENSG00000232702 | ENSG0000023 | 110.0808263 | 0.037623858 | 0.181732249 | 0.207029067 | 0.835987158 | 0.913390427 |
| ENSG00000232710 | ENSG0000023 | 5.292423178 | 0.8337501 | 0.767769375 | 1.085938208 | 0.277506331 | 0.460303343 |
| ENSG00000232713 | RPS12P3 | 13.48066953 | 0.45929468 | 0.484533872 | 0.947910368 | 0.343175087 | 0.534159855 |
| ENSG00000232721 | ENSG0000023 | 3.26756017 | -0.404232187 | 0.97429453 | -0.414897318 | 0.678217074 | 0.81683728 |
| ENSG00000232725 | PLXNB3-AS1 | 7.519570583 | -0.781710296 | 0.65047134 | -1.201759782 | 0.229456612 | 0.40258009 |
| ENSG00000232727 | YWHAEP1 | 13.95490051 | -0.198645125 | 0.491680773 | -0.404012392 | 0.686203607 | 0.822612392 |
| ENSG00000232729 | GTF2I-AS1 | 52.46397198 | 0.021457762 | 0.257773185 | 0.0832428 | 0.933658482 | 0.968138388 |
| ENSG00000232732 | ENSG0000023 | 1.762625081 | -0.828837225 | 1.36374174 | -0.607767 | 0.543342019 | NA |
| ENSG00000232742 | RHOQP2 | 116.4447289 | 0.483670869 | 0.17452066 | 2.771424701 | 0.005581158 | 0.01774768 |
| ENSG00000232748 | ENSG0000023 | 5.082530082 | -0.264541305 | 0.805974739 | -0.328225306 | 0.742741316 | 0.858016204 |
| ENSG00000232757 | ETF1P1 | 2.594688901 | 0.978992909 | 1.133519376 | 0.863675496 | 0.387766178 | 0.580492583 |
| ENSG00000232760 | ENSG0000023 | 2.115701609 | -0.847455828 | 1.211305565 | -0.699621839 | 0.4841635 | 0.668055445 |
| ENSG00000232765 | ENSG0000023 | 1.689304951 | -0.200122462 | 1.321338585 | -0.151454339 | 0.879617327 | NA |
| ENSG00000232774 | LINC03033 | 4.687724528 | -0.803851476 | 0.813477384 | -0.988166962 | 0.323070885 | 0.512924501 |
| ENSG00000232788 | ITGA6-AS1 | 1.769155964 | -0.389763134 | 1.33294219 | -0.292408131 | 0.769974592 | NA |
| ENSG00000232794 | HNRNPDP1 | 26.53753511 | 0.374355193 | 0.368123626 | 1.016927922 | 0.309187719 | 0.497224342 |
| ENSG00000232801 | SDCBPP3 | 4.700102172 | -0.432897765 | 0.818632966 | -0.528805683 | 0.596940255 | 0.75769879 |
| ENSG00000232803 | SLCO4A1-AS1 | 2.307665721 | -0.997788215 | 1.170385577 | -0.852529486 | 0.39392028 | 0.58690184 |
| ENSG00000232807 | ENSG0000023 | 7.654991376 | -2.599216696 | 0.761923374 | -3.41138858 | 0.000646329 | 0.002507066 |
| ENSG00000232810 | TNF | 3.614082106 | 2.537730977 | 1.115037484 | 2.275915395 | 0.022851074 | 0.061367787 |
| ENSG00000232815 | DUX4L50 | 43.62097489 | -0.144050015 | 0.272541634 | -0.528543155 | 0.597122402 | 0.75769879 |
| ENSG00000232828 | ENSG0000023 | 7.706131143 | -1.060962893 | 0.649560466 | -1.633355091 | 0.102394336 | 0.216112437 |
| ENSG00000232838 | PET117 | 6.041263453 | -0.56970993 | 0.729216817 | -0.781262742 | 0.434647978 | 0.624660107 |
| ENSG00000232842 | KAT7P1 | 63.05288879 | 0.252831286 | 0.230160982 | 1.098497599 | 0.271987266 | 0.454041488 |
| ENSG00000232848 | RPL7AP18 | 1.59942424 | -0.124247863 | 1.382468455 | -0.089873923 | 0.928387404 | NA |
| ENSG00000232850 | PTGES2-AS1 | 9.238978924 | 0.726961602 | 0.631847093 | 1.150534061 | 0.249923974 | 0.427982678 |
| ENSG00000232855 | ENSG0000023 | 4.667429727 | -1.204360721 | 0.839271806 | -1.435006766 | 0.151285169 | 0.29361098 |
| ENSG00000232858 | RPL34P27 | 6.097765431 | 0.298315763 | 0.810356908 | 0.368128858 | 0.712777154 | 0.84015416 |
| ENSG00000232859 | LYRM9 | 88.10039756 | -0.288631341 | 0.193716765 | -1.489965731 | 0.136233247 | 0.270896354 |

| | | | | | | | |
|---|---|---|---|---|---|---|---|
| ENSG00000232860 | SMG7-AS1 | 9.597380236 | 0.394458853 | 0.571159649 | 0.690628013 | 0.489799339 | 0.672975373 |
| ENSG00000232864 | NUCKS1P1 | 15.57998022 | -0.17979669 | 0.452382134 | -0.397444277 | 0.69103987 | 0.825734692 |
| ENSG00000232871 | SEC1P | 6.874372361 | -0.839629431 | 0.688681138 | -1.21918459 | 0.22277414 | 0.393868113 |
| ENSG00000232872 | CTAGE3P | 20.91474027 | -1.005674034 | 0.400282845 | -2.512408533 | 0.011991018 | 0.034912294 |
| ENSG00000232874 | LINC02924 | 2.459472872 | -0.378324148 | 1.156362299 | -0.327167488 | 0.74354121 | 0.858490754 |
| ENSG00000232878 | DPYD-AS1 | 1.546535558 | 3.978764357 | 1.656828799 | 2.401433606 | 0.016330972 | NA |
| ENSG00000232882 | PHKA1P1 | 85.40038992 | 0.596804727 | 0.197179191 | 3.026712524 | 0.00247229 | 0.00852847 |
| ENSG00000232883 | RPLP0P5 | 9.187898236 | 0.167715331 | 0.579439955 | 0.289443848 | 0.772241744 | 0.875910441 |
| ENSG00000232888 | RPS11P5 | 938.1481059 | 0.089979426 | 0.074495762 | 1.207846245 | 0.22710641 | 0.399679404 |
| ENSG00000232893 | IQCA1-AS1 | 2.746538038 | -0.25706721 | 1.067305715 | -0.24085621 | 0.809666562 | 0.899281056 |
| ENSG00000232901 | CYCSP10 | 1.57002341 | -1.319285218 | 1.509978825 | -0.873711072 | 0.38227562 | NA |
| ENSG00000232903 | LINC01166 | 15.32460458 | 0.381290013 | 0.446653176 | 0.85366014 | 0.393293326 | 0.586120243 |
| ENSG00000232905 | ENSG0000023 | 6.687185806 | 0.037583183 | 0.685414702 | 0.054832765 | 0.956271697 | 0.979980913 |
| ENSG00000232907 | DLGAP4-AS1 | 20.7828519 | -0.636407886 | 0.421818283 | -1.508725229 | 0.131369016 | 0.263588316 |
| ENSG00000232916 | HMGN2P27 | 7.824255389 | 0.047988551 | 0.62749368 | 0.076476548 | 0.939039971 | 0.971039083 |
| ENSG00000232917 | HSPE1P6 | 4.030679318 | 0.277298163 | 0.955102054 | 0.290333543 | 0.77156108 | 0.875561078 |
| ENSG00000232927 | USP12PY | 2.174266185 | -0.118915372 | 1.211205794 | -0.098179328 | 0.921789897 | 0.962181382 |
| ENSG00000232931 | LINC00342 | 186.4860461 | -0.105072409 | 0.14907549 | -0.704826859 | 0.480917992 | 0.665315326 |
| ENSG00000232936 | PANK1-AS1 | 30.09713536 | 0.06911748 | 0.321006268 | 0.215315049 | 0.829521697 | 0.909783073 |
| ENSG00000232938 | RPL23AP87 | 67.5421471 | 0.344122818 | 0.241490898 | 1.424992914 | 0.15415925 | 0.297402505 |
| ENSG00000232940 | HCG25 | 29.22504168 | -0.535119566 | 0.335539303 | -1.594804429 | 0.110755976 | 0.230014675 |
| ENSG00000232951 | IPO7P1 | 33.13266343 | 0.037306106 | 0.315823264 | 0.118123363 | 0.905969912 | 0.95262805 |
| ENSG00000232952 | ENSG0000023 | 245.413427 | 0.085132124 | 0.118046108 | 0.721176876 | 0.470800697 | 0.656664923 |
| ENSG00000232953 | HSPA8P18 | 1.820691795 | -0.945474899 | 1.328282809 | -0.711802406 | 0.476587143 | 0.661638166 |
| ENSG00000232956 | SNHG15 | 1108.364994 | 0.862274632 | 0.066543739 | 12.95801295 | 2.12E-38 | 8.12E-37 |
| ENSG00000232969 | ENSG0000023 | 2.25949952 | 0.653945177 | 1.168142763 | 0.559816143 | 0.575604851 | 0.741985575 |
| ENSG00000232973 | CYP1B1-AS1 | 32.72040488 | 0.441833299 | 0.315956956 | 1.398397128 | 0.161993845 | 0.309060114 |
| ENSG00000232992 | AHCYP4 | 2.334681504 | -0.215654845 | 1.154580551 | -0.186781983 | 0.851831574 | 0.922505405 |
| ENSG00000232995 | ENSG0000023 | 14.05905014 | 0.289628766 | 0.475604218 | 0.608970138 | 0.542544233 | 0.71631532 |
| ENSG00000233006 | MIR3936HG | 20.18119517 | -0.242842572 | 0.388803893 | -0.624588839 | 0.532240947 | 0.707691769 |
| ENSG00000233008 | LINC01725 | 18.1113342 | 0.413017994 | 0.408948341 | 1.00995151 | 0.312518523 | 0.500745611 |
| ENSG00000233012 | HDAC1P2 | 5.926451706 | -0.344647247 | 0.718245812 | -0.479845815 | 0.631337033 | 0.782730991 |
| ENSG00000233014 | TBC1D3P7 | 2.118584518 | 0.053858741 | 1.194322158 | 0.045095656 | 0.964031064 | 0.983666795 |

| | | | | | | | |
|---|---|---|---|---|---|---|---|
| ENSG00000233016 | SNHG7 | 1696.81548 | 0.571683248 | 0.056723235 | 10.0784669 | 6.88E-24 | 1.43E-22 |
| ENSG00000233017 | ENSG0000023 | 16.29960811 | 0.722189512 | 0.439116873 | 1.644640768 | 0.100043914 | 0.212009921 |
| ENSG00000233020 | CHORDC1P5 | 1.691166805 | -0.762702098 | 1.341419856 | -0.568578208 | 0.569642417 | NA |
| ENSG00000233029 | ENSG0000023 | 5.506097586 | -2.04320077 | 0.83036028 | -2.460619589 | 0.013869734 | 0.039667335 |
| ENSG00000233030 | ENSG0000023 | 52.88026585 | 0.222990491 | 0.248735788 | 0.896495402 | 0.369988235 | 0.562609711 |
| ENSG00000233033 | CASK-AS1 | 1.841526769 | -1.031524223 | 1.328372165 | -0.776532549 | 0.43743461 | 0.626920679 |
| ENSG00000233040 | FAM204BP | 4.737771186 | -0.440990346 | 0.809802753 | -0.544565136 | 0.586052639 | 0.749932924 |
| ENSG00000233045 | RPL15P5 | 8.99302493 | 0.163988424 | 0.585024739 | 0.280310238 | 0.779239497 | 0.880232128 |
| ENSG00000233055 | ENSG0000023 | 2.609836515 | -2.909918912 | 1.384735747 | -2.101425429 | 0.035603638 | 0.089910551 |
| ENSG00000233057 | EEF1A1P14 | 15.11560335 | 0.405255809 | 0.489565695 | 0.827786369 | 0.407791496 | 0.600511443 |
| ENSG00000233060 | ENSG0000023 | 2.175276099 | 0.711625928 | 1.241793617 | 0.573062962 | 0.566602064 | 0.734507875 |
| ENSG00000233073 | GRM3-AS1 | 6.309420416 | -0.760826935 | 0.740872669 | -1.026933463 | 0.304451796 | 0.491874372 |
| ENSG00000233080 | LINC01399 | 6.359390712 | -0.133666484 | 0.77700705 | -0.172027376 | 0.863416007 | 0.929172842 |
| ENSG00000233081 | ENSG0000023 | 2.907246928 | 1.15136026 | 1.069989643 | 1.076048041 | 0.281905771 | 0.465366562 |
| ENSG00000233087 | RAB6D | 53.55582038 | -0.133438756 | 0.245375563 | -0.543814365 | 0.586569223 | 0.750355818 |
| ENSG00000233098 | CCDC144NL-A | 11.6112997 | -0.107967091 | 0.517271992 | -0.208724024 | 0.834663685 | 0.912770412 |
| ENSG00000233108 | GLCCI1-DT | 4.876070785 | -2.437490054 | 0.953590523 | -2.556118161 | 0.010584717 | 0.031249365 |
| ENSG00000233111 | RAB1C | 155.7453037 | -0.178749863 | 0.158598188 | -1.127061191 | 0.25971661 | 0.439697346 |
| ENSG00000233122 | CTAGE7P | 15.91273566 | -0.21089753 | 0.44333493 | -0.475707001 | 0.634283133 | 0.784391108 |
| ENSG00000233124 | LINC00456 | 4.726491536 | -0.087730978 | 0.794162392 | -0.110469822 | 0.912036782 | 0.955697231 |
| ENSG00000233131 | RPS15AP14 | 1.783904464 | 1.849077124 | 1.533770867 | 1.205575855 | 0.227981074 | NA |
| ENSG00000233144 | ENSG0000023 | 17.6658485 | -1.42435742 | 0.441781126 | -3.224124655 | 0.001263584 | 0.004640828 |
| ENSG00000233154 | LINC01762 | 21.64694988 | -4.243172376 | 0.629521102 | -6.740317939 | 1.58E-11 | 1.46E-10 |
| ENSG00000233155 | HMGA1P8 | 28.4197682 | -0.152194686 | 0.330051329 | -0.461124293 | 0.644709433 | 0.79186349 |
| ENSG00000233170 | ENSG0000023 | 57.50586775 | -0.032084037 | 0.242435678 | -0.132340412 | 0.894715044 | 0.946400434 |
| ENSG00000233175 | FMNL1-AS1 | 71.93468568 | -0.02844114 | 0.237259728 | -0.11987344 | 0.904583405 | 0.952149367 |
| ENSG00000233178 | ENSG0000023 | 3.171528393 | 1.386568059 | 1.04019929 | 1.332983086 | 0.182537355 | 0.338458367 |
| ENSG00000233184 | DPH5-DT | 104.6648185 | 0.168199942 | 0.184454108 | 0.911879618 | 0.361832092 | 0.554433785 |
| ENSG00000233198 | RNF224 | 44.77501069 | 0.229408429 | 0.26901036 | 0.852786595 | 0.393777659 | 0.586765801 |
| ENSG00000233203 | DHCR24-DT | 5.772873358 | 1.38361222 | 0.765417486 | 1.807656926 | 0.070659912 | 0.159887393 |
| ENSG00000233205 | WBP1P2 | 57.86784882 | -0.136930335 | 0.233611841 | -0.586144669 | 0.557778301 | 0.727915708 |
| ENSG00000233215 | LINC01687 | 1.702077557 | 4.116674767 | 1.63024956 | 2.525180726 | 0.01156388 | NA |
| ENSG00000233217 | MROH3P | 3.229678302 | 4.011128688 | 1.421860975 | 2.821041408 | 0.004786802 | 0.015498835 |

| | | | | | | | |
|---|---|---|---|---|---|---|---|
| ENSG00000233220 | PRDM10-DT | 2.659370795 | -2.273305893 | 1.261569092 | -1.801967017 | 0.071550595 | 0.161615858 |
| ENSG00000233223 | ENSG0000023 | 70.99495467 | 0.067161533 | 0.218144811 | 0.307875912 | 0.758176756 | 0.866725209 |
| ENSG00000233225 | SSBP3P1 | 726.6820312 | -0.114083542 | 0.081794028 | -1.39476616 | 0.163086367 | 0.310540451 |
| ENSG00000233229 | CNOT7P1 | 12.2862841 | -0.461859454 | 0.546241335 | -0.845522709 | 0.397819059 | 0.590849387 |
| ENSG00000233230 | ENSG0000023 | 8.050339762 | 0.445280766 | 0.618778805 | 0.719612181 | 0.47176381 | 0.657466637 |
| ENSG00000233237 | LINC00472 | 108.7743871 | -0.055081059 | 0.1738386 | -0.316851715 | 0.751356121 | 0.862482741 |
| ENSG00000233242 | ENSG0000023 | 2.317619686 | -0.606732626 | 1.13363436 | -0.535210159 | 0.592504543 | 0.754736754 |
| ENSG00000233246 | PHC2-AS1 | 2.563042396 | -0.432148534 | 1.113382829 | -0.388140111 | 0.69791235 | 0.83051669 |
| ENSG00000233247 | UBE2E4P | 109.5046612 | 0.242063302 | 0.191368253 | 1.264908351 | 0.205904183 | 0.371368973 |
| ENSG00000233250 | ENSG0000023 | 10.52975053 | -0.132339095 | 0.556329198 | -0.237879111 | 0.811974857 | 0.900449735 |
| ENSG00000233251 | ENSG0000023 | 437.2662594 | -0.394937009 | 0.093042664 | -4.244687218 | 2.19E-05 | 0.00010799 |
| ENSG00000233254 | RPL21P134 | 1.807500844 | 0.076862381 | 1.277770881 | 0.060153492 | 0.952033386 | 0.977646532 |
| ENSG00000233260 | RPSAP63 | 2.109128415 | 0.509934481 | 1.186689329 | 0.429711862 | 0.667405254 | 0.808691146 |
| ENSG00000233264 | ENSG0000023 | 2.691055607 | -0.655794812 | 1.174918298 | -0.558162055 | 0.576733729 | 0.742898071 |
| ENSG00000233266 | HMGB1P31 | 2.149760956 | -0.392073285 | 1.189965734 | -0.329482837 | 0.741790765 | 0.857590509 |
| ENSG00000233270 | SNRPEP4 | 1275.975064 | 0.225044822 | 0.068587971 | 3.281112108 | 0.001033986 | 0.003858108 |
| ENSG00000233287 | ENSG0000023 | 12.95437014 | 0.001066106 | 0.5141305 | 0.002073609 | 0.998345501 | 0.999129473 |
| ENSG00000233296 | TMEM18-DT | 8.059671311 | 0.110175914 | 0.619230738 | 0.177923845 | 0.858782784 | 0.926723937 |
| ENSG00000233297 | RASA4DP | 4.995348647 | -0.809392038 | 0.803359616 | -1.007508993 | 0.313690238 | 0.501980571 |
| ENSG00000233309 | RPS6P12 | 13.84583381 | -0.060542214 | 0.48158072 | -0.12571561 | 0.899957045 | 0.949471761 |
| ENSG00000233325 | MIPEPP3 | 30.79132023 | -0.153246145 | 0.316610771 | -0.484020632 | 0.628371227 | 0.780741805 |
| ENSG00000233328 | PFN1P1 | 582.5027999 | -0.197421369 | 0.08377878 | -2.356460309 | 0.018450039 | 0.050950271 |
| ENSG00000233347 | ERP29P1 | 38.1070061 | 0.193950093 | 0.317376137 | 0.611104838 | 0.541130177 | 0.715279242 |
| ENSG00000233360 | PDXP-DT | 14.24617978 | -0.557132748 | 0.468914556 | -1.188132766 | 0.234781107 | 0.40969714 |
| ENSG00000233369 | GTF2IP4 | 1168.534449 | 0.420238161 | 0.076584984 | 5.487213543 | 4.08E-08 | 2.75E-07 |
| ENSG00000233370 | ACTR1AP1 | 1.718129136 | 2.01685227 | 1.489572801 | 1.353980328 | 0.175742654 | NA |
| ENSG00000233380 | RPS2P48 | 23.65456473 | -0.033530093 | 0.378213875 | -0.088653789 | 0.929357059 | 0.965993019 |
| ENSG00000233381 | AK4P3 | 11.79537136 | -0.076263832 | 0.525087236 | -0.145240308 | 0.884521142 | 0.94022126 |
| ENSG00000233382 | NKAPP1 | 54.43383159 | 0.064561837 | 0.240037704 | 0.2689654 | 0.787956308 | 0.885283461 |
| ENSG00000233383 | ENSG0000023 | 2.060453212 | 0.994408862 | 1.320111659 | 0.753276328 | 0.451283881 | 0.639355992 |
| ENSG00000233384 | CNIH3-AS2 | 5.706797404 | 2.008676609 | 0.879181941 | 2.284710952 | 0.022329782 | 0.060221632 |
| ENSG00000233396 | LINC01719 | 118.4635871 | 0.298867451 | 0.167294634 | 1.786473622 | 0.074022605 | 0.166201287 |
| ENSG00000233401 | PRKAR1AP1 | 2.031629027 | -0.697942334 | 1.244831395 | -0.560672181 | 0.575021035 | 0.741399648 |

| | | | | | | | |
|---|---|---|---|---|---|---|---|
| ENSG00000233406 | ENSG0000023 | 12.93235624 | -0.314727223 | 0.538347268 | -0.58461748 | 0.558804951 | 0.728707409 |
| ENSG00000233426 | EIF3FP3 | 1042.608703 | 0.171158532 | 0.065567929 | 2.610400166 | 0.009043636 | 0.027240155 |
| ENSG00000233427 | ENSG0000023 | 3.326139262 | -0.191000986 | 0.991077326 | -0.192720569 | 0.847177814 | 0.920028676 |
| ENSG00000233429 | HOTAIRM1 | 4.189421505 | -0.535205627 | 0.842168458 | -0.635508991 | 0.525096505 | 0.701269943 |
| ENSG00000233432 | MTIF2P1 | 2.766343067 | 1.061182117 | 1.073478022 | 0.988545731 | 0.322885449 | 0.512843052 |
| ENSG00000233452 | STXBP5-AS1 | 24.01701704 | 0.110559603 | 0.371529075 | 0.297579949 | 0.76602378 | 0.872131893 |
| ENSG00000233457 | ENSG0000023 | 2.663286225 | 0.559316942 | 1.13616991 | 0.492282833 | 0.622519414 | 0.776597778 |
| ENSG00000233459 | ENSG0000023 | 11.16640392 | 0.126493706 | 0.558767152 | 0.226379997 | 0.820905884 | 0.904991377 |
| ENSG00000233461 | ENSG0000023 | 31.31547696 | 0.60734129 | 0.321397445 | 1.889689227 | 0.058799536 | 0.137176991 |
| ENSG00000233467 | HAX1P1 | 13.6759575 | 0.251804923 | 0.482239271 | 0.522157646 | 0.601560573 | 0.761284247 |
| ENSG00000233476 | EEF1A1P6 | 1299.948328 | 0.140537944 | 0.074431383 | 1.888154412 | 0.059005224 | 0.137572907 |
| ENSG00000233483 | EFCAB15P | 16.56721344 | 0.626255002 | 0.431407468 | 1.451655448 | 0.146597433 | 0.286427486 |
| ENSG00000233487 | RPSAP69 | 17.93685393 | -0.141564624 | 0.410597723 | -0.344776933 | 0.730262075 | 0.850776726 |
| ENSG00000233493 | TMEM238 | 80.57951218 | 0.100747222 | 0.202204963 | 0.498243072 | 0.618312727 | 0.773690709 |
| ENSG00000233503 | HNRNPLP1 | 25.068386 | -0.152816404 | 0.379121084 | -0.403080733 | 0.686888834 | 0.823089733 |
| ENSG00000233509 | ZNF197-AS1 | 2.339437407 | 0.64443214 | 1.168236921 | 0.551627952 | 0.581203282 | 0.746225068 |
| ENSG00000233514 | ENSG0000023 | 74.31957294 | -0.019471391 | 0.224676469 | -0.086664131 | 0.930938488 | 0.96680379 |
| ENSG00000233527 | ZNF529-AS1 | 84.8294982 | 0.226297018 | 0.195217585 | 1.159204064 | 0.246373015 | 0.423641179 |
| ENSG00000233539 | ENSG0000023 | 3.305245209 | -1.096780585 | 1.012752924 | -1.082969556 | 0.278821936 | 0.461817705 |
| ENSG00000233543 | CHTF8P1 | 8.526797269 | 0.221082612 | 0.641690076 | 0.344531762 | 0.730446414 | 0.850819131 |
| ENSG00000233554 | B4GALT1-AS1 | 5.832038612 | -0.259727855 | 0.775659938 | -0.334847583 | 0.737740065 | 0.855509243 |
| ENSG00000233558 | ENSG0000023 | 255.5941931 | 0.319466054 | 0.131064303 | 2.4374757 | 0.014790208 | 0.041970235 |
| ENSG00000233560 | KRT8P39 | 13.67024881 | -0.529175134 | 0.47455259 | -1.115103248 | 0.26480618 | 0.445748002 |
| ENSG00000233585 | ACAA2P1 | 76.6299818 | 0.080071093 | 0.205426038 | 0.389780642 | 0.69669876 | 0.829644391 |
| ENSG00000233586 | ENSG0000023 | 1.84447427 | -0.910735499 | 1.401586037 | -0.649789221 | 0.515828382 | 0.694553213 |
| ENSG00000233588 | CYP51A1P2 | 345.7389809 | -1.069286052 | 0.10387497 | -10.29397216 | 7.50E-25 | 1.66E-23 |
| ENSG00000233593 | LINC02609 | 9.453940488 | 0.314108149 | 0.567781108 | 0.553220501 | 0.58011243 | 0.745585185 |
| ENSG00000233594 | BTF3P5 | 19.19128794 | 0.340145228 | 0.419892913 | 0.810076135 | 0.41789642 | 0.609908631 |
| ENSG00000233597 | RPL19P6 | 35.58443115 | 0.247887238 | 0.324292441 | 0.76439413 | 0.444632406 | 0.633374776 |
| ENSG00000233602 | ERI3-IT1 | 2.0669513 | -0.841211028 | 1.276154605 | -0.659176423 | 0.509782485 | 0.689772927 |
| ENSG00000233611 | GBX2-AS1 | 5.278776352 | -0.01806117 | 0.759037918 | -0.023794819 | 0.981016272 | 0.99123506 |
| ENSG00000233621 | ZC3H12A-DT | 141.0631739 | -0.236742811 | 0.152812016 | -1.549242111 | 0.12132353 | 0.247571378 |
| ENSG00000233622 | CYP2T1P | 155.172976 | 1.070655027 | 0.151834418 | 7.051464628 | 1.77E-12 | 1.78E-11 |

| | | | | | | | |
|---|---|---|---|---|---|---|---|
| ENSG00000233635 | ENSG0000023 | 4.353002164 | 0.590716367 | 0.834078938 | 0.708225972 | 0.47880494 | 0.66343231 |
| ENSG00000233639 | PANTR1 | 1.54822797 | -2.084717743 | 1.649252738 | -1.264037764 | 0.206216473 | NA |
| ENSG00000233654 | NEMP2-DT | 27.06364733 | 0.55522761 | 0.365919515 | 1.517349 | 0.129178593 | 0.259921479 |
| ENSG00000233661 | SPIN4-AS1 | 1.686444324 | -0.172489906 | 1.381534262 | -0.124853875 | 0.900639235 | NA |
| ENSG00000233662 | CALM2P4 | 4.12838726 | 0.793610022 | 0.911911908 | 0.870270489 | 0.384152603 | 0.576618511 |
| ENSG00000233668 | ENSG0000023 | 29.88640492 | 0.291651665 | 0.331764107 | 0.879093484 | 0.37935059 | 0.571805148 |
| ENSG00000233672 | RNASEH2B-AS | 9.490404284 | 0.004133721 | 0.577456452 | 0.007158498 | 0.994288394 | 0.996533822 |
| ENSG00000233673 | ANAPC1P1 | 2.328119451 | 1.636533917 | 1.273490579 | 1.285077365 | 0.19876525 | 0.361544705 |
| ENSG00000233680 | HNRNPA1P27 | 2.129963388 | 0.436223144 | 1.211257676 | 0.360140664 | 0.718741945 | 0.843545493 |
| ENSG00000233695 | GAS6-AS1 | 7.683812509 | -1.353422174 | 0.668501298 | -2.024561774 | 0.042912383 | 0.105183253 |
| ENSG00000233706 | ENSG0000023 | 1.689304951 | -0.200122462 | 1.321338585 | -0.151454339 | 0.879617327 | NA |
| ENSG00000233707 | RPL22P11 | 11.40765514 | 0.089659055 | 0.527801124 | 0.1698728 | 0.865110172 | 0.929929589 |
| ENSG00000233718 | MYCNOS | 1.858113007 | -0.437294889 | 1.361669999 | -0.321146012 | 0.748099743 | 0.860436606 |
| ENSG00000233719 | GOT2P3 | 6.171870412 | 0.259145515 | 0.722768612 | 0.358545613 | 0.71993504 | 0.844366665 |
| ENSG00000233723 | LINC01122 | 24.63484534 | -0.071102453 | 0.359434923 | -0.197817319 | 0.843187995 | 0.917412854 |
| ENSG00000233725 | NRAD1 | 2.575810998 | 4.722233996 | 1.50640517 | 3.134770174 | 0.001719888 | 0.006127385 |
| ENSG00000233728 | ENSG0000023 | 21.77060814 | -0.570930009 | 0.455042255 | -1.254674709 | 0.20959687 | 0.376607833 |
| ENSG00000233741 | RPL27P7 | 2.655221281 | -0.561549262 | 1.082766754 | -0.518624403 | 0.604022688 | 0.762927016 |
| ENSG00000233747 | RPL36AP13 | 3.992715681 | -1.460594865 | 0.967532639 | -1.509607849 | 0.131143517 | 0.263227954 |
| ENSG00000233757 | ZNF892 | 37.28654113 | 0.166512176 | 0.286564127 | 0.581064273 | 0.561197136 | 0.730157957 |
| ENSG00000233762 | RPS15P4 | 1717.988612 | 0.117069651 | 0.057015198 | 2.053306064 | 0.040042903 | 0.099199157 |
| ENSG00000233765 | CDRT15P12 | 4.452693055 | -0.702176471 | 0.823877793 | -0.852282313 | 0.39405742 | 0.586992028 |
| ENSG00000233778 | ENSG0000023 | 3.535139535 | -0.0913863 | 0.925884891 | -0.098701578 | 0.921375216 | 0.9619656 |
| ENSG00000233780 | HNRNPA1P30 | 6.823457111 | -0.115885588 | 0.736356306 | -0.157377056 | 0.874947694 | 0.935158184 |
| ENSG00000233786 | CDC27P1 | 10.33627801 | -0.29458267 | 0.548234418 | -0.537329762 | 0.591039851 | 0.753624266 |
| ENSG00000233791 | BTG2-DT | 4.148702718 | -0.528101662 | 0.8612361 | -0.613190346 | 0.539750488 | 0.714187932 |
| ENSG00000233797 | UFL1-AS1 | 11.34753029 | -0.19264061 | 0.516951537 | -0.372647331 | 0.709410941 | 0.838082552 |
| ENSG00000233799 | ENSG0000023 | 11.69827489 | -0.531511013 | 0.517700119 | -1.0266774 | 0.304572394 | 0.491998993 |
| ENSG00000233806 | LINC01237 | 25.66434961 | 0.329321749 | 0.346767535 | 0.949690255 | 0.342269663 | 0.53329254 |
| ENSG00000233820 | ENSG0000023 | 2.414095872 | -1.069418253 | 1.171200272 | -0.913095974 | 0.361192068 | 0.553748983 |
| ENSG00000233822 | H2BC15 | 115.5832494 | 0.090798497 | 0.168513906 | 0.538819018 | 0.59001174 | 0.752932545 |
| ENSG00000233830 | EIF4HP1 | 1212.880577 | 0.430978811 | 0.06505813 | 6.624518909 | 3.48E-11 | 3.12E-10 |
| ENSG00000233833 | ETF1P3 | 2.591794526 | -0.513526421 | 1.074995648 | -0.477700931 | 0.632863082 | 0.783582692 |

| | | | | | | | |
|---|---|---|---|---|---|---|---|
| ENSG00000233834 | GIRGL | 24.99920304 | -2.447778786 | 0.437947911 | -5.589200739 | 2.28E-08 | 1.58E-07 |
| ENSG00000233837 | EIF3LP2 | 25.27036662 | 0.324684002 | 0.422794364 | 0.7679478 | 0.442518197 | 0.631470633 |
| ENSG00000233846 | SELENOTP1 | 309.1279656 | -0.10648534 | 0.110658248 | -0.962290132 | 0.335903884 | 0.526989133 |
| ENSG00000233849 | ENSG0000023 | 5.336022622 | -0.233276159 | 0.808354181 | -0.288581619 | 0.772901563 | 0.876182887 |
| ENSG00000233859 | ADH5P4 | 161.2964843 | -0.142205462 | 0.159660041 | -0.8906766 | 0.373102692 | 0.565845283 |
| ENSG00000233860 | SHROOM3-AS | 22.37412225 | 0.665727739 | 0.410644549 | 1.621177587 | 0.104979558 | 0.220433049 |
| ENSG00000233868 | LLPHP3 | 188.0595169 | -0.247571209 | 0.139076627 | -1.780106502 | 0.075058533 | 0.16796827 |
| ENSG00000233871 | DLG5-AS1 | 31.08394049 | -0.518605924 | 0.323301118 | -1.604095671 | 0.108692966 | 0.226571253 |
| ENSG00000233873 | RPL7P44 | 22.18305947 | 0.308307618 | 0.426212091 | 0.723366664 | 0.469454646 | 0.655385199 |
| ENSG00000233874 | MFN1P1 | 2.777649883 | -0.022266037 | 1.02508339 | -0.021721195 | 0.982670356 | 0.992011 |
| ENSG00000233885 | YEATS2-AS1 | 26.13114062 | 0.594859836 | 0.349508868 | 1.701987817 | 0.088757652 | 0.192302658 |
| ENSG00000233892 | PAIP1P1 | 8.107528021 | 1.175311146 | 0.669922684 | 1.754398193 | 0.079362301 | 0.175646038 |
| ENSG00000233893 | EZR-AS1 | 8.725366373 | -0.877261736 | 0.600907972 | -1.459893658 | 0.144319303 | 0.28294033 |
| ENSG00000233901 | LINC01503 | 72.63118234 | -0.189888446 | 0.21421762 | -0.886427764 | 0.375387064 | 0.568360856 |
| ENSG00000233912 | ENSG0000023 | 2.171545255 | -0.421992682 | 1.170440083 | -0.360541892 | 0.718441935 | 0.843432038 |
| ENSG00000233913 | RPL10P9 | 2520.275906 | -0.242891149 | 0.055624525 | -4.366619776 | 1.26E-05 | 6.42E-05 |
| ENSG00000233921 | RPS15AP40 | 2.111977902 | 0.024761319 | 1.206380455 | 0.020525299 | 0.983624331 | 0.992362659 |
| ENSG00000233922 | LINC01694 | 7.65313629 | 2.177765349 | 0.733579147 | 2.968684917 | 0.002990771 | 0.010134163 |
| ENSG00000233926 | TLE1-DT | 1.601286094 | -0.691616497 | 1.401609918 | -0.493444351 | 0.621698651 | NA |
| ENSG00000233927 | RPS28 | 716.1123184 | 0.247650427 | 0.077631877 | 3.190061064 | 0.001422427 | 0.005156524 |
| ENSG00000233929 | MT1XP1 | 8.856307381 | -0.165948122 | 0.587683169 | -0.28237685 | 0.777654564 | 0.879394445 |
| ENSG00000233930 | KRTAP5-AS1 | 447.289895 | 0.622530599 | 0.093902195 | 6.629563852 | 3.37E-11 | 3.02E-10 |
| ENSG00000233932 | CTXN2 | 3.978054463 | 0.671059348 | 0.890691421 | 0.753413957 | 0.451201199 | 0.639278333 |
| ENSG00000233937 | CTC-338M12.4 | 271.4221039 | -0.251704947 | 0.114841957 | -2.191750774 | 0.028397507 | 0.074069223 |
| ENSG00000233947 | ASS1P3 | 66.97771921 | 1.156474032 | 0.231323797 | 4.999373369 | 5.75E-07 | 3.43E-06 |
| ENSG00000233951 | RCC2P3 | 1.933215882 | -0.167962526 | 1.232231916 | -0.13630756 | 0.891578147 | 0.944396856 |
| ENSG00000233954 | UQCRHL | 1313.488027 | 0.204442406 | 0.062353319 | 3.278773437 | 0.001042593 | 0.003888957 |
| ENSG00000233966 | UBE2SP1 | 386.9509493 | -0.033704525 | 0.095567342 | -0.35267827 | 0.724329652 | 0.846509568 |
| ENSG00000233967 | ENSG0000023 | 27.64711266 | -0.040189078 | 0.335995649 | -0.1196119 | 0.904790592 | 0.952267525 |
| ENSG00000233971 | RPS20P10 | 51.50816413 | 0.280486413 | 0.286271688 | 0.979790962 | 0.327189315 | 0.517600364 |
| ENSG00000233990 | ENSG0000023 | 4.743997764 | 1.04745529 | 0.831778432 | 1.259296046 | 0.207923421 | 0.374333957 |
| ENSG00000233994 | GDI2P2 | 154.3580383 | -1.130650906 | 0.15271998 | -7.40342492 | 1.33E-13 | 1.44E-12 |
| ENSG00000233997 | LINC01425 | 9.11836651 | 1.552055059 | 0.635454396 | 2.442433428 | 0.01458862 | 0.041463365 |

| | | | | | | | |
|---|---|---|---|---|---|---|---|
| ENSG00000234004 | ENSG0000023 | 29.89665984 | 0.282375825 | 0.32881636 | 0.858764525 | 0.390470443 | 0.583343425 |
| ENSG00000234005 | GAPDHP22 | 3.054172973 | -0.323447351 | 1.073992369 | -0.301163547 | 0.763289786 | 0.87031501 |
| ENSG00000234009 | RPL5P34 | 770.3379918 | 0.312926692 | 0.07838184 | 3.992336651 | 6.54E-05 | 0.000300593 |
| ENSG00000234017 | ENSG0000023 | 2.137519331 | 0.505741819 | 1.201730401 | 0.420844658 | 0.673868519 | 0.813479126 |
| ENSG00000234026 | ENSG0000023 | 4.87365194 | 0.26183066 | 0.805845548 | 0.324914199 | 0.74524601 | 0.859265354 |
| ENSG00000234028 | EIF2AK3-DT | 57.20669278 | -0.051645497 | 0.233641182 | -0.221046207 | 0.825056455 | 0.90772273 |
| ENSG00000234031 | RPS3AP44 | 14.14074849 | 0.255503347 | 0.466243329 | 0.548004296 | 0.583688953 | 0.748011283 |
| ENSG00000234036 | TXNP6 | 149.9038784 | 0.015976976 | 0.154174844 | 0.103628943 | 0.917463818 | 0.959385307 |
| ENSG00000234040 | RPL10P12 | 3.802646845 | -0.801809239 | 0.914689971 | -0.876591265 | 0.380708685 | 0.573211948 |
| ENSG00000234043 | NUDT9P1 | 4.605442478 | 0.974947765 | 0.835454472 | 1.16696696 | 0.243223713 | 0.41965085 |
| ENSG00000234044 | ENSG0000023 | 4.256541122 | 0.290771624 | 0.866107437 | 0.33572235 | 0.73708025 | 0.855093604 |
| ENSG00000234062 | TM9SF5P | 49.54649903 | 0.400765896 | 0.249578931 | 1.605768137 | 0.108324864 | 0.22605043 |
| ENSG00000234070 | SPATA17-AS1 | 3.888227203 | 0.061917684 | 0.872988752 | 0.070926096 | 0.943456574 | 0.973031555 |
| ENSG00000234072 | GTF3C2-AS2 | 427.0345194 | 0.160943945 | 0.095551112 | 1.684375429 | 0.092109136 | 0.198216096 |
| ENSG00000234080 | ENSG0000023 | 1.710086473 | 1.247461975 | 1.407129934 | 0.886529343 | 0.37533235 | NA |
| ENSG00000234093 | RPS15AP11 | 80.71086695 | 0.04979885 | 0.200228945 | 0.248709545 | 0.803585465 | 0.895414541 |
| ENSG00000234106 | SRP14P2 | 58.44042157 | -0.663109757 | 0.236140207 | -2.808118806 | 0.004983184 | 0.016059882 |
| ENSG00000234107 | TPT1P1 | 2.319066873 | 0.217532793 | 1.124952951 | 0.19337057 | 0.846668763 | 0.919716896 |
| ENSG00000234115 | ENSG0000023 | 8.087115952 | -0.240847165 | 0.612520866 | -0.393206466 | 0.694166995 | 0.827678065 |
| ENSG00000234118 | RPL13AP6 | 19.96133996 | 0.361593693 | 0.395010976 | 0.915401636 | 0.35998082 | 0.552550636 |
| ENSG00000234123 | RHBDF1P1 | 2.964465707 | 0.797839629 | 1.032585065 | 0.772662375 | 0.439722219 | 0.629163256 |
| ENSG00000234127 | TRIM26 | 2437.032301 | -0.061535972 | 0.058072435 | -1.059641659 | 0.289307654 | 0.473577678 |
| ENSG00000234129 | HCCS-DT | 17.18077844 | -1.579026718 | 0.477038126 | -3.31006398 | 0.000932746 | 0.003512731 |
| ENSG00000234130 | AP2B1P1 | 144.595057 | -0.060077058 | 0.149713347 | -0.401280574 | 0.688213564 | 0.823819752 |
| ENSG00000234134 | ENSG0000023 | 5.320655793 | -0.218585335 | 0.769429124 | -0.28408768 | 0.776343188 | 0.878561133 |
| ENSG00000234136 | ENSG0000023 | 2.355531297 | -1.988466572 | 1.267055178 | -1.569360677 | 0.116563921 | 0.239990698 |
| ENSG00000234141 | ENSG0000023 | 6.346310019 | -1.638824953 | 0.738442935 | -2.219298034 | 0.026466455 | 0.06972215 |
| ENSG00000234145 | NAP1L4P3 | 144.5066557 | 0.428944933 | 0.154086358 | 2.783795641 | 0.005372687 | 0.017165661 |
| ENSG00000234149 | ENSG0000023 | 3.024880996 | 0.622844094 | 1.005772317 | 0.619269474 | 0.53573885 | 0.711031295 |
| ENSG00000234156 | ENSG0000023 | 8.773844748 | -0.096025466 | 0.609166549 | -0.157634175 | 0.874745072 | 0.935028748 |
| ENSG00000234160 | ENSG0000023 | 62.48195703 | 0.403139221 | 0.228938438 | 1.760906668 | 0.078254201 | 0.173578744 |
| ENSG00000234166 | ARHGEF19-AS | 2.881678333 | 1.205818203 | 1.062498326 | 1.134889509 | 0.256421616 | 0.435903139 |
| ENSG00000234171 | RNASEH1-DT | 164.7854667 | -0.302060653 | 0.139770851 | -2.161113357 | 0.030686584 | 0.079255825 |

| | | | | | | | |
|---|---|---|---|---|---|---|---|
| ENSG00000234176 | HSPA8P1 | 96.51843896 | 0.081132948 | 0.221532105 | 0.36623562 | 0.714189265 | 0.840989046 |
| ENSG00000234203 | ENSG0000023 | 2.06277462 | 1.62522156 | 1.351568643 | 1.202470602 | 0.22918125 | 0.402238729 |
| ENSG00000234213 | FHP1 | 5.373258042 | -1.08936153 | 0.823082843 | -1.323513835 | 0.185664559 | 0.342926326 |
| ENSG00000234222 | LIX1L-AS1 | 70.4032958 | -0.403710189 | 0.221961734 | -1.818827878 | 0.068937694 | 0.156701417 |
| ENSG00000234235 | BOK-AS1 | 6.278265632 | -0.034466697 | 0.703831078 | -0.048970127 | 0.960943103 | 0.982116705 |
| ENSG00000234241 | ENSG0000023 | 349.1870405 | -0.103166211 | 0.10864877 | -0.949538692 | 0.342346703 | 0.533376349 |
| ENSG00000234255 | ENSG0000023 | 51.91342627 | -0.526219703 | 0.245097311 | -2.146982771 | 0.031794653 | 0.081703747 |
| ENSG00000234259 | ENSG0000023 | 1.700622908 | -1.401178414 | 1.401680819 | -0.99964157 | 0.317483998 | NA |
| ENSG00000234261 | ENSG0000023 | 2.842949632 | -0.050462772 | 1.022418421 | -0.049356282 | 0.960635368 | 0.981920343 |
| ENSG00000234264 | DEPDC1-AS1 | 12.48937139 | -1.148131472 | 0.509180666 | -2.254860696 | 0.024142074 | 0.064525129 |
| ENSG00000234268 | RPS27P19 | 35.04493471 | 0.281400032 | 0.300693889 | 0.935835551 | 0.349357864 | 0.540627995 |
| ENSG00000234284 | ZNF879 | 127.8251474 | -0.47897917 | 0.158307122 | -3.025632476 | 0.002481137 | 0.00855513 |
| ENSG00000234286 | ENSG0000023 | 17.22440751 | -1.961666262 | 0.4718858 | -4.157078393 | 3.22E-05 | 0.000155446 |
| ENSG00000234287 | RPS27P4 | 1452.512853 | 0.409616599 | 0.065169263 | 6.285426271 | 3.27E-10 | 2.72E-09 |
| ENSG00000234289 | H2BC12L | 5.821876354 | 0.560923388 | 0.750606892 | 0.747293149 | 0.454886627 | 0.642397145 |
| ENSG00000234297 | PSIP1P1 | 87.88158394 | 0.105734425 | 0.189339166 | 0.558439266 | 0.576544466 | 0.74273769 |
| ENSG00000234311 | LINC03068 | 13.89056005 | -2.105563043 | 0.537402406 | -3.918037992 | 8.93E-05 | 0.000401473 |
| ENSG00000234322 | ST13P18 | 147.0881013 | 0.228732357 | 0.149029828 | 1.534809241 | 0.124830692 | 0.253238068 |
| ENSG00000234327 | ZNF232-AS1 | 32.10680466 | 0.197221568 | 0.323276574 | 0.610070707 | 0.54181497 | 0.715752496 |
| ENSG00000234329 | RPL7AP16 | 8.652139478 | 1.479234024 | 0.647101507 | 2.285938153 | 0.022257877 | 0.060055951 |
| ENSG00000234332 | BCAS2P2 | 267.597997 | 0.266674531 | 0.114593311 | 2.327138717 | 0.019957883 | 0.054588511 |
| ENSG00000234335 | RPS4XP11 | 113.5158289 | 0.359377129 | 0.176836276 | 2.03225909 | 0.042127434 | 0.103535818 |
| ENSG00000234337 | ENSG0000023 | 252.5093288 | 0.003728661 | 0.118857227 | 0.031370927 | 0.974973727 | 0.98863534 |
| ENSG00000234338 | ENSG0000023 | 1.680798174 | 0.435105272 | 1.427057021 | 0.304896907 | 0.760444671 | NA |
| ENSG00000234345 | ELF2P1 | 13.10784172 | -0.143047404 | 0.504071048 | -0.283784209 | 0.776575757 | 0.878737624 |
| ENSG00000234350 | ERICH2-DT | 6.358189003 | 0.099540729 | 0.71290116 | 0.139627672 | 0.888954176 | 0.942802073 |
| ENSG00000234353 | ENSG0000023 | 4.987140172 | -0.124081818 | 0.803786182 | -0.154371674 | 0.877316687 | 0.936250688 |
| ENSG00000234354 | RPS26P47 | 1.862778985 | 2.195528678 | 1.498974871 | 1.46468678 | 0.143006393 | 0.280942593 |
| ENSG00000234356 | ENSG0000023 | 3.580806323 | -1.222123643 | 0.977970231 | -1.249653215 | 0.211426255 | 0.378974706 |
| ENSG00000234369 | TATDN1P1 | 231.2243259 | -0.211814643 | 0.120904131 | -1.751922296 | 0.07978717 | 0.176331183 |
| ENSG00000234371 | RPSAP31 | 2.787117127 | -0.016731231 | 1.034820511 | -0.016168244 | 0.987100169 | 0.99369967 |
| ENSG00000234373 | SNX18P7 | 9.070192301 | 4.015682741 | 0.973575186 | 4.124676552 | 3.71E-05 | 0.00017673 |
| ENSG00000234377 | OBI1-AS1 | 2.734160394 | -0.900839561 | 1.103148291 | -0.816607856 | 0.414152564 | 0.606412192 |

| | | | | | | | |
|---|---|---|---|---|---|---|---|
| ENSG00000234383 | CTBP2P8 | 48.58354077 | 0.173649502 | 0.252072673 | 0.68888666 | 0.490894595 | 0.673915141 |
| ENSG00000234390 | USP27X-DT | 37.588938 | -0.025407764 | 0.294516864 | -0.0862693 | 0.931252343 | 0.967031267 |
| ENSG00000234397 | PPIAP34 | 1.622143349 | 0.410567706 | 1.341968565 | 0.305944354 | 0.759647014 | NA |
| ENSG00000234405 | ENSG0000023 | 8.644026186 | -0.527469715 | 0.604766344 | -0.872187614 | 0.383106036 | 0.575537162 |
| ENSG00000234420 | ZNF37BP | 589.1477852 | -0.245613499 | 0.094492888 | -2.599280269 | 0.009341947 | 0.028035617 |
| ENSG00000234432 | LINC02983 | 1.916677582 | 0.361703143 | 1.283052053 | 0.281908393 | 0.778013754 | 0.879540484 |
| ENSG00000234444 | ZNF736 | 755.4241333 | 0.572260576 | 0.081208803 | 7.046780063 | 1.83E-12 | 1.83E-11 |
| ENSG00000234450 | ENSG0000023 | 1.85759695 | 1.412408392 | 1.372584459 | 1.029013831 | 0.303473176 | 0.490673432 |
| ENSG00000234465 | PINLYP | 28.78915354 | -0.930659515 | 0.343402683 | -2.710111364 | 0.006726062 | 0.020949574 |
| ENSG00000234473 | ZNF101P2 | 1.794789152 | -1.529150084 | 1.465021898 | -1.043772852 | 0.296590495 | 0.482880479 |
| ENSG00000234477 | ENSG0000023 | 39.43667242 | -3.201606719 | 0.380328959 | -8.417993538 | 3.83E-17 | 5.32E-16 |
| ENSG00000234478 | ACBD3-AS1 | 16.71567467 | -0.055070583 | 0.426006183 | -0.129271793 | 0.89714259 | 0.947984339 |
| ENSG00000234488 | RPL27P6 | 6.62256818 | 0.84530932 | 0.714192628 | 1.183587293 | 0.236576473 | 0.411733881 |
| ENSG00000234491 | HNRNPA1P51 | 4.618286406 | 0.228841747 | 0.811738189 | 0.281915709 | 0.778008144 | 0.879540484 |
| ENSG00000234494 | SP2-AS1 | 233.0068071 | -0.418892107 | 0.125108558 | -3.34822903 | 0.000813298 | 0.003095578 |
| ENSG00000234498 | RPL13AP20 | 36.54440533 | 0.159265656 | 0.302575512 | 0.52636664 | 0.59863349 | 0.75887321 |
| ENSG00000234509 | SOD1-DT | 2.890762081 | 0.577798791 | 1.178665895 | 0.490214227 | 0.623982314 | 0.777683938 |
| ENSG00000234515 | PPP1R2P1 | 3.730040008 | -0.41177384 | 0.959360681 | -0.429216923 | 0.667765368 | 0.808785741 |
| ENSG00000234518 | PTGES3P1 | 1010.646538 | -0.139654874 | 0.074394205 | -1.877227859 | 0.060486871 | 0.140470623 |
| ENSG00000234521 | ENSG0000023 | 2.12276778 | 0.146653436 | 1.366593892 | 0.107313107 | 0.914540587 | 0.957607799 |
| ENSG00000234537 | ENSG0000023 | 5.796926715 | -0.279453562 | 0.715955775 | -0.39032238 | 0.696298176 | 0.829343624 |
| ENSG00000234545 | FAM133B | 229.7862603 | -0.516624548 | 0.139504047 | -3.703294337 | 0.000212818 | 0.000897929 |
| ENSG00000234546 | LNCTAM34A | 36.89956218 | 0.206601093 | 0.289955235 | 0.71252755 | 0.476138158 | 0.661218906 |
| ENSG00000234558 | API5P1 | 276.7202119 | 0.084942937 | 0.115004006 | 0.738608509 | 0.460144756 | 0.646955179 |
| ENSG00000234569 | RAD1P1 | 2.912345769 | -0.14535067 | 1.031013453 | -0.140978442 | 0.887886973 | 0.942285816 |
| ENSG00000234585 | CCT6P3 | 60.84285325 | 0.440315788 | 0.236878746 | 1.858823532 | 0.063052156 | 0.145383099 |
| ENSG00000234589 | RPL13AP24 | 68.5459111 | 0.185831857 | 0.240182901 | 0.773709768 | 0.439102442 | 0.628541746 |
| ENSG00000234602 | MCIDAS | 44.00376078 | 2.240471291 | 0.313255336 | 7.15222067 | 8.54E-13 | 8.78E-12 |
| ENSG00000234604 | ENSG0000023 | 2.985737628 | 0.430364214 | 1.003851364 | 0.428713083 | 0.668132038 | 0.80906614 |
| ENSG00000234608 | MAPKAPK5-AS | 336.4814185 | 0.283585392 | 0.104091167 | 2.724394412 | 0.006441952 | 0.020166265 |
| ENSG00000234614 | C2CD4D-AS1 | 67.86802899 | 0.766308414 | 0.219156404 | 3.496627981 | 0.000471178 | 0.001873606 |
| ENSG00000234616 | JRK | 1784.469903 | 0.042128228 | 0.059575686 | 0.707137931 | 0.479480766 | 0.66373081 |
| ENSG00000234617 | SNRK-AS1 | 51.50611506 | -0.08699435 | 0.249411906 | -0.348797902 | 0.727241038 | 0.848528021 |

| | | | | | | | |
|---|---|---|---|---|---|---|---|
| ENSG00000234618 | RPSAP9 | 25.90329093 | -0.246086456 | 0.342324047 | -0.718869906 | 0.47222108 | 0.657886043 |
| ENSG00000234629 | WDR82P1 | 71.70969787 | -0.015896933 | 0.207147966 | -0.076741923 | 0.938828853 | 0.970997921 |
| ENSG00000234636 | MED14OS | 4.590898268 | -0.313020212 | 0.821679415 | -0.380951751 | 0.703239051 | 0.833796972 |
| ENSG00000234645 | YWHAEP5 | 29.31317678 | -0.132758111 | 0.361946598 | -0.366789222 | 0.713776248 | 0.840814785 |
| ENSG00000234648 | ENSG0000023 | 523.0666903 | 0.080756465 | 0.086524816 | 0.933332992 | 0.350648066 | 0.541977875 |
| ENSG00000234664 | HMGN2P5 | 2354.866166 | -0.256703128 | 0.053864223 | -4.765744577 | 1.88E-06 | 1.05E-05 |
| ENSG00000234665 | LERFS | 3.81400969 | -0.016678911 | 0.926443423 | -0.018003162 | 0.985636331 | 0.993350589 |
| ENSG00000234678 | ELF3-AS1 | 137.6700336 | -0.380013359 | 0.178625583 | -2.12742963 | 0.033384402 | 0.085112046 |
| ENSG00000234684 | SDCBP2-AS1 | 66.9810027 | 0.331783768 | 0.21723909 | 1.527274709 | 0.126692716 | 0.25617725 |
| ENSG00000234685 | NUS1P2 | 72.21235878 | -0.020647133 | 0.206262574 | -0.100101207 | 0.920263977 | 0.961332224 |
| ENSG00000234690 | EPCAM-DT | 8.879050198 | -1.240552896 | 0.604669861 | -2.051620193 | 0.040206588 | 0.099544945 |
| ENSG00000234694 | CDC20-DT | 12.59897546 | -0.703581839 | 0.57677298 | -1.219859223 | 0.222518246 | 0.393628051 |
| ENSG00000234695 | TFPI2-DT | 11.69589657 | -1.195939348 | 0.523957747 | -2.282511049 | 0.022459186 | 0.060506603 |
| ENSG00000234699 | ENSG0000023 | 2.39281281 | -1.048213594 | 1.138334326 | -0.920831052 | 0.35713864 | 0.549444062 |
| ENSG00000234702 | VDAC1P3 | 24.04297876 | 0.269408949 | 0.361093282 | 0.746092386 | 0.455611609 | 0.642985561 |
| ENSG00000234705 | HMGA1P4 | 5.123690228 | 1.169418888 | 0.794640851 | 1.471631979 | 0.141120285 | 0.278214106 |
| ENSG00000234709 | UPF3AP3 | 3.286943071 | -0.074153882 | 1.009565781 | -0.073451264 | 0.941447025 | 0.97217172 |
| ENSG00000234718 | ENSG0000023 | 6.510934339 | 0.164384441 | 0.684071157 | 0.240303132 | 0.810095267 | 0.899408839 |
| ENSG00000234719 | NPIPB2 | 96.36777505 | 0.983100465 | 0.186355962 | 5.275390472 | 1.32E-07 | 8.50E-07 |
| ENSG00000234737 | KRT18P15 | 333.2173074 | 0.328455238 | 0.112526254 | 2.918920927 | 0.003512453 | 0.011726921 |
| ENSG00000234740 | ENSG0000023 | 2.998491632 | 0.10260427 | 1.002761368 | 0.102321722 | 0.918501314 | 0.960059241 |
| ENSG00000234741 | GAS5 | 2684.638322 | 0.416935158 | 0.059785546 | 6.973845435 | 3.08E-12 | 3.03E-11 |
| ENSG00000234742 | RPL17P18 | 41.73410244 | 0.613626118 | 0.281780087 | 2.177677366 | 0.029430065 | 0.076367135 |
| ENSG00000234743 | EIF5AP4 | 77.17671473 | 0.042925161 | 0.208332256 | 0.206041836 | 0.836758232 | 0.913925996 |
| ENSG00000234745 | HLA-B | 20.32207933 | 0.091385523 | 0.386186504 | 0.236635724 | 0.812939403 | 0.901214416 |
| ENSG00000234750 | ENSG0000023 | 3.903852975 | 0.055414007 | 0.879405726 | 0.063013015 | 0.94975614 | 0.976794703 |
| ENSG00000234751 | RPL15P16 | 5.528392837 | 0.522953123 | 0.767854808 | 0.6810573 | 0.495835234 | 0.677941809 |
| ENSG00000234752 | LINC02676 | 5.771878588 | 0.566530292 | 0.765988549 | 0.739606739 | 0.459538652 | 0.6464805 |
| ENSG00000234753 | FOXP4-AS1 | 12.27844908 | -0.275489112 | 0.494284536 | -0.557349242 | 0.577288838 | 0.743237509 |
| ENSG00000234764 | E2F6P1 | 71.9446263 | -0.031624393 | 0.218295097 | -0.144869918 | 0.884813578 | 0.940314253 |
| ENSG00000234769 | WASH4P | 11.32822658 | 0.422792378 | 0.525422703 | 0.804670936 | 0.421009596 | 0.612076819 |
| ENSG00000234771 | SLC25A25-AS1 | 548.8198373 | 0.344150774 | 0.190301639 | 1.808448814 | 0.070536677 | 0.159624283 |
| ENSG00000234773 | ENSG0000023 | 8.184571533 | 0.647568865 | 0.623968917 | 1.037822314 | 0.299352786 | 0.485998083 |

| | | | | | | | |
|---|---|---|---|---|---|---|---|
| ENSG00000234776 | FREY1 | 21.06493383 | 1.047101564 | 0.391235572 | 2.676396625 | 0.007441852 | 0.022911835 |
| ENSG00000234782 | TPT1P9 | 311.1864652 | 0.20138343 | 0.130272791 | 1.545859483 | 0.122138508 | 0.248855268 |
| ENSG00000234785 | EEF1GP5 | 46.77672464 | 0.076575203 | 0.299341461 | 0.255812217 | 0.79809584 | 0.892259454 |
| ENSG00000234787 | LINC00458 | 12.33949572 | 2.618101737 | 0.617308368 | 4.24115705 | 2.22E-05 | 0.000109538 |
| ENSG00000234793 | ENSG0000023 | 1.553119892 | 1.841604954 | 1.532135108 | 1.201986003 | 0.229368951 | NA |
| ENSG00000234797 | RPS3AP6 | 6017.642586 | 0.167906499 | 0.061378837 | 2.735576419 | 0.006227113 | 0.01958727 |
| ENSG00000234800 | PCMTD1P3 | 11.83784009 | -0.326532774 | 0.506760721 | -0.644352966 | 0.519346577 | 0.696836315 |
| ENSG00000234801 | MORF4 | 39.71214812 | -0.395886288 | 0.333199615 | -1.18813549 | 0.234780034 | 0.40969714 |
| ENSG00000234807 | LINC01135 | 134.5281156 | -0.874060912 | 0.161080771 | -5.426227504 | 5.76E-08 | 3.82E-07 |
| ENSG00000234814 | SVIL2P | 16.63383211 | -0.653938401 | 0.4733832 | -1.381414466 | 0.167151561 | 0.316661723 |
| ENSG00000234817 | ECI2-DT | 2.205084459 | -1.914544829 | 1.274898792 | -1.501722992 | 0.133168663 | 0.266314109 |
| ENSG00000234818 | ENSG0000023 | 21.92485372 | -0.093286051 | 0.379381578 | -0.245889774 | 0.80576756 | 0.896948167 |
| ENSG00000234819 | ENSG0000023 | 2.412097696 | -1.119560494 | 1.194669431 | -0.937129942 | 0.348691721 | 0.539939594 |
| ENSG00000234825 | XRCC6P2 | 607.3004929 | -0.193177856 | 0.080759804 | -2.392005016 | 0.016756613 | 0.046877648 |
| ENSG00000234828 | IQCM | 2.938332709 | -4.071168312 | 1.449692087 | -2.808298638 | 0.004980402 | 0.016053171 |
| ENSG00000234832 | NXT1-AS1 | 1.680252373 | -2.193606088 | 1.542877962 | -1.421762538 | 0.155095198 | NA |
| ENSG00000234840 | LINC01239 | 132.9386674 | 0.058795705 | 0.154496442 | 0.380563488 | 0.703527179 | 0.834095482 |
| ENSG00000234851 | RPL23AP42 | 4998.949711 | 0.179148143 | 0.059870044 | 2.992283448 | 0.00276899 | 0.009460868 |
| ENSG00000234859 | ENSG0000023 | 1.834369467 | -1.575287751 | 1.363112049 | -1.155655364 | 0.247822153 | 0.425335941 |
| ENSG00000234869 | LINC02925 | 17.99570026 | 1.100193976 | 0.429342996 | 2.562505934 | 0.010391981 | 0.030744396 |
| ENSG00000234899 | SOX9-AS1 | 8.39303725 | 2.328414155 | 0.739111733 | 3.150287095 | 0.001631101 | 0.005849219 |
| ENSG00000234911 | TEX21P | 20.41917069 | -0.692787535 | 0.395047872 | -1.753679958 | 0.079485362 | 0.175799672 |
| ENSG00000234912 | SNHG20 | 206.4453451 | 0.482305722 | 0.13118838 | 3.6764363 | 0.000236515 | 0.000990082 |
| ENSG00000234917 | ENSG0000023 | 4.048145289 | -0.866182323 | 0.915185429 | -0.946455544 | 0.343916287 | 0.534882049 |
| ENSG00000234925 | ATP5PDP4 | 41.59640879 | 0.05003168 | 0.314525426 | 0.159070384 | 0.873613422 | 0.934676032 |
| ENSG00000234928 | LINC01659 | 16.19760141 | -3.808150166 | 0.655736756 | -5.80743741 | 6.34E-09 | 4.66E-08 |
| ENSG00000234936 | ENSG0000023 | 30.73381847 | -0.208025723 | 0.349017476 | -0.596032396 | 0.551153585 | 0.723161782 |
| ENSG00000234945 | GTF3C2-AS1 | 23.65201657 | -0.6388638 | 0.386299597 | -1.653803952 | 0.098167353 | 0.208862833 |
| ENSG00000234949 | RAB17-DT | 47.8015963 | 0.024459987 | 0.257832728 | 0.094867659 | 0.924419945 | 0.963698865 |
| ENSG00000234964 | FABP5P7 | 145.9522818 | 0.234150377 | 0.152790898 | 1.532489045 | 0.125401799 | 0.254126967 |
| ENSG00000234965 | SHISA8 | 10.25675767 | 0.007526371 | 0.568547707 | 0.013237889 | 0.989438002 | 0.994379556 |
| ENSG00000234967 | ENSG0000023 | 102.7713042 | 0.819210384 | 0.178014449 | 4.601931974 | 4.19E-06 | 2.25E-05 |
| ENSG00000234969 | AARSD1P1 | 4.169563024 | -0.764346911 | 0.871759443 | -0.8767865 | 0.380602613 | 0.573127474 |

| | | | | | | | |
|---|---|---|---|---|---|---|---|
| ENSG00000234975 | FTH1P2 | 968.9426732 | 0.263373698 | 0.078274557 | 3.364742123 | 0.000766152 | 0.00293121 |
| ENSG00000234981 | RPL7AP20 | 24.24736588 | 0.057697779 | 0.365719664 | 0.157765044 | 0.874641945 | 0.934962079 |
| ENSG00000234983 | ENSG0000023 | 1.726138051 | 0.115237674 | 1.425781942 | 0.080824193 | 0.935581768 | NA |
| ENSG00000234985 | ENSG0000023 | 5.22002263 | -0.323041289 | 0.774432038 | -0.417133168 | 0.676581006 | 0.815809108 |
| ENSG00000234996 | ACTG1P25 | 164.8572429 | 0.34472286 | 0.142813752 | 2.413793171 | 0.015787421 | 0.044475432 |
| ENSG00000235001 | EIF4A1P2 | 538.1965986 | -0.040778694 | 0.08598623 | -0.474246798 | 0.635323925 | 0.785001122 |
| ENSG00000235010 | JAKMIP3-AS1 | 1.698335247 | 0.335361142 | 1.358694923 | 0.246825933 | 0.80504294 | NA |
| ENSG00000235012 | JCADP1 | 8.704271031 | 0.511268561 | 0.617680069 | 0.827723908 | 0.407826876 | 0.600511443 |
| ENSG00000235016 | SEMA3F-AS1 | 32.13561655 | -0.237875594 | 0.311843298 | -0.762804895 | 0.445579761 | 0.634142252 |
| ENSG00000235023 | ENSG0000023 | 1.575243752 | -0.743410824 | 1.585345544 | -0.468926681 | 0.639122044 | NA |
| ENSG00000235027 | PRADX | 25.31187833 | 0.057446245 | 0.365398567 | 0.157215299 | 0.875075171 | 0.935207288 |
| ENSG00000235028 | HMGN1P30 | 162.7736742 | -0.180160098 | 0.144210376 | -1.249286653 | 0.211560248 | 0.379127095 |
| ENSG00000235029 | MNX1-AS2 | 1.74407409 | 0.155833468 | 1.331901209 | 0.117000771 | 0.906859443 | NA |
| ENSG00000235033 | DAAM2-AS1 | 2.698737055 | -0.21611998 | 1.063110957 | -0.203290144 | 0.838908258 | 0.915274206 |
| ENSG00000235034 | C19orf81 | 23.55718048 | -0.250430139 | 0.366045405 | -0.68415048 | 0.493880148 | 0.676475436 |
| ENSG00000235036 | ENSG0000023 | 93.63227044 | -0.868683935 | 0.187526331 | -4.632330465 | 3.62E-06 | 1.97E-05 |
| ENSG00000235043 | TECRP1 | 855.5950992 | -0.015892648 | 0.08162292 | -0.194708149 | 0.845621432 | 0.919043732 |
| ENSG00000235044 | PPIAP3 | 4.023488593 | -0.66847164 | 0.88319453 | -0.756879281 | 0.44912219 | 0.637356203 |
| ENSG00000235045 | RPL7P8 | 9.797246685 | 0.664021106 | 0.610130643 | 1.088326105 | 0.27645117 | 0.458984573 |
| ENSG00000235049 | LINC00940 | 6.38365102 | 1.053160572 | 0.708404819 | 1.486664889 | 0.13710334 | 0.272199381 |
| ENSG00000235052 | ENSG0000023 | 5.944334921 | -0.153536123 | 0.754167627 | -0.203583551 | 0.838678949 | 0.915120151 |
| ENSG00000235060 | VDAC1P4 | 2.79987113 | -0.356122511 | 1.023211045 | -0.348044045 | 0.727807106 | 0.848972593 |
| ENSG00000235064 | SLC25A5P2 | 31.09138625 | -0.18582363 | 0.320491577 | -0.579808153 | 0.562043999 | 0.730912093 |
| ENSG00000235065 | RPL24P2 | 44.92478336 | 0.304011017 | 0.280200957 | 1.084974943 | 0.277932753 | 0.460677555 |
| ENSG00000235066 | ENSG0000023 | 5.810121042 | -0.926247145 | 0.770225096 | -1.202566821 | 0.229143994 | 0.402216177 |
| ENSG00000235071 | ENSG0000023 | 3.593239996 | 0.929802616 | 0.961483922 | 0.967049573 | 0.333519253 | 0.524611331 |
| ENSG00000235072 | ARNILA | 21.51170175 | 1.692565188 | 0.421150403 | 4.018909106 | 5.85E-05 | 0.000271017 |
| ENSG00000235077 | ENSG0000023 | 2.362529196 | 1.873090063 | 1.277204108 | 1.466554994 | 0.142497151 | 0.280182107 |
| ENSG00000235078 | ENSG0000023 | 8.843937199 | 0.235372192 | 0.615848237 | 0.382191875 | 0.702319048 | 0.833215242 |
| ENSG00000235079 | ZRANB2-AS1 | 7.800115379 | -0.487674462 | 0.621399211 | -0.78480058 | 0.4325705 | 0.622597006 |
| ENSG00000235081 | ENSG0000023 | 8.039876795 | -0.012360478 | 0.620286477 | -0.019927048 | 0.984101568 | 0.992635461 |
| ENSG00000235082 | SUMO1P3 | 413.7939087 | -0.207368724 | 0.092313914 | -2.246343117 | 0.024682039 | 0.065746165 |
| ENSG00000235084 | CHCHD2P6 | 122.9205938 | 0.550729911 | 0.163252119 | 3.373493187 | 0.000742209 | 0.002846726 |

| | | | | | | | |
|---|---|---|---|---|---|---|---|
| ENSG00000235089 | ENSG0000023 | 2.283582621 | -0.170937981 | 1.140701152 | -0.149853431 | 0.880880253 | 0.938252557 |
| ENSG00000235091 | ENSG0000023 | 5.411980013 | 0.911343787 | 0.805756029 | 1.131041846 | 0.258037479 | 0.437694452 |
| ENSG00000235092 | ID2-AS1 | 341.6653626 | -1.36568907 | 0.110524264 | -12.35646379 | 4.49E-35 | 1.54E-33 |
| ENSG00000235095 | ENSG0000023 | 10.28202726 | 1.118573654 | 0.629329983 | 1.777404041 | 0.075501786 | 0.168746198 |
| ENSG00000235098 | ANKRD65 | 6.116670907 | 1.214840032 | 0.756782 | 1.60527078 | 0.108434227 | 0.226216912 |
| ENSG00000235100 | ENSG0000023 | 12.9373067 | -0.774279858 | 0.499549306 | -1.549956827 | 0.121151879 | 0.247362062 |
| ENSG00000235105 | ENSG0000023 | 112.8405701 | -0.140501067 | 0.171131903 | -0.82101037 | 0.411640361 | 0.60391798 |
| ENSG00000235106 | BRD3OS | 1896.142316 | 0.620996695 | 0.058671811 | 10.58424282 | 3.53E-26 | 8.29E-25 |
| ENSG00000235109 | ZSCAN31 | 149.2946822 | -0.797798846 | 0.152919321 | -5.217122618 | 1.82E-07 | 1.15E-06 |
| ENSG00000235121 | ENSG0000023 | 5.082089759 | 0.263805714 | 0.767824455 | 0.343575555 | 0.731165511 | 0.851119615 |
| ENSG00000235123 | DSCAM-AS1 | 7230.48525 | 0.176376245 | 0.051620647 | 3.41677712 | 0.000633671 | 0.002461714 |
| ENSG00000235124 | KRT8P17 | 8.609148323 | 0.025344804 | 0.589254849 | 0.043011617 | 0.965692273 | 0.984576922 |
| ENSG00000235142 | LINC02532 | 1.649087077 | -0.756697124 | 1.347966346 | -0.56136203 | 0.574550763 | NA |
| ENSG00000235151 | ENSG0000023 | 2.792713829 | -0.657806742 | 1.057696651 | -0.62192382 | 0.533991961 | 0.709473856 |
| ENSG00000235159 | LINC02939 | 8.359798568 | 0.404218453 | 0.615377109 | 0.656862998 | 0.511269012 | 0.690928491 |
| ENSG00000235162 | C12orf75 | 510.2936567 | -0.009406416 | 0.100881246 | -0.093242463 | 0.925710941 | 0.964514676 |
| ENSG00000235169 | SMIM1 | 15.62437455 | 0.387673444 | 0.462996464 | 0.837314049 | 0.402416065 | 0.595191781 |
| ENSG00000235173 | HGH1 | 1501.199708 | 0.000663894 | 0.071796242 | 0.009246919 | 0.992622131 | 0.996008734 |
| ENSG00000235174 | RPL39P3 | 4360.270505 | 0.1489409 | 0.067268411 | 2.214128416 | 0.026819946 | 0.070580426 |
| ENSG00000235175 | RPL26P37 | 3.937425926 | 0.317605768 | 0.895857205 | 0.354527224 | 0.722943805 | 0.845926239 |
| ENSG00000235183 | SRP14P3 | 9.991465663 | -0.121325423 | 0.548459869 | -0.221211123 | 0.824928049 | 0.90772273 |
| ENSG00000235189 | ENSG0000023 | 11.61221422 | -0.632267357 | 0.579479316 | -1.091095644 | 0.2752308 | 0.457355635 |
| ENSG00000235191 | NUCB1-AS1 | 24.495116 | -0.23669637 | 0.364129127 | -0.650034157 | 0.515670158 | 0.6944375 |
| ENSG00000235192 | ENSG0000023 | 8.905130372 | 0.071432361 | 0.609182769 | 0.117259326 | 0.906654556 | 0.952929042 |
| ENSG00000235194 | PPP1R3E | 357.3185549 | -0.003904371 | 0.112985419 | -0.034556412 | 0.972433459 | 0.987499936 |
| ENSG00000235205 | TATDN2P3 | 356.8565444 | 0.611218683 | 0.100496554 | 6.081986499 | 1.19E-09 | 9.38E-09 |
| ENSG00000235209 | ENSG0000023 | 16.79977611 | 0.338069994 | 0.449083811 | 0.752799333 | 0.451570507 | 0.639485613 |
| ENSG00000235211 | TMSB10P2 | 1.572895178 | -0.691493359 | 1.414720534 | -0.488784422 | 0.624994328 | NA |
| ENSG00000235214 | FAM83C-AS1 | 1.772081184 | -1.460970921 | 1.377296634 | -1.060752553 | 0.28880237 | NA |
| ENSG00000235217 | TSPY26P | 103.0921147 | 2.494323764 | 0.215947413 | 11.55060731 | 7.33E-31 | 2.11E-29 |
| ENSG00000235225 | RPL31P17 | 5.792161923 | 0.495380585 | 0.745483101 | 0.664509476 | 0.506364286 | 0.686686966 |
| ENSG00000235236 | ENSG0000023 | 22.90649872 | -0.219227226 | 0.365758772 | -0.599376536 | 0.54892182 | 0.721583944 |
| ENSG00000235237 | ENSG0000023 | 3.451948635 | 1.444129394 | 1.013242977 | 1.425254778 | 0.154083567 | 0.297327722 |

| | | | | | | | |
|---|---|---|---|---|---|---|---|
| ENSG00000235238 | SUMO2P1 | 514.1908607 | 0.106667798 | 0.089334352 | 1.194028905 | 0.232466661 | 0.406524258 |
| ENSG00000235244 | DANT2 | 9.803410725 | 0.18007709 | 0.572227265 | 0.314695055 | 0.752993199 | 0.863812772 |
| ENSG00000235246 | ENSG0000023 | 1.60279387 | 1.791269211 | 1.521769907 | 1.177095961 | 0.239157207 | NA |
| ENSG00000235251 | ENSG0000023 | 13.52790092 | 0.117105427 | 0.502784009 | 0.232913984 | 0.815828199 | 0.902352305 |
| ENSG00000235253 | EEF1A1P50 | 1.984751713 | -0.663467049 | 1.24905246 | -0.531176288 | 0.595296623 | 0.756595136 |
| ENSG00000235254 | TMEM185AP1 | 3.529375969 | 0.959034663 | 0.961431915 | 0.997506581 | 0.318518681 | 0.507664736 |
| ENSG00000235256 | FKBP4P7 | 3.240065129 | 0.420245904 | 0.99308068 | 0.42317398 | 0.67216833 | 0.812215855 |
| ENSG00000235257 | ITGA9-AS1 | 3.444783244 | -0.012705813 | 0.994626552 | -0.012774456 | 0.989807736 | 0.994576849 |
| ENSG00000235269 | LINC02331 | 2.320016199 | 0.255518109 | 1.183166547 | 0.215961235 | 0.829017965 | 0.909536221 |
| ENSG00000235272 | RAMACL | 276.4336172 | -0.004236216 | 0.116319514 | -0.03641879 | 0.970948432 | 0.98680905 |
| ENSG00000235275 | KRT18P16 | 49.21427864 | 0.408664414 | 0.261271032 | 1.564139776 | 0.117784747 | 0.242069709 |
| ENSG00000235277 | ENSG0000023 | 99.58549569 | 1.121836163 | 0.190143734 | 5.899937601 | 3.64E-09 | 2.75E-08 |
| ENSG00000235280 | MCF2L-AS1 | 124.891872 | 0.02406853 | 0.159301203 | 0.151088185 | 0.879906153 | 0.937748996 |
| ENSG00000235296 | ENSG0000023 | 5.215933705 | 0.732079446 | 0.784459141 | 0.933228268 | 0.350702122 | 0.542013521 |
| ENSG00000235297 | FAUP1 | 935.2056711 | 0.148451325 | 0.067274834 | 2.206639782 | 0.027339235 | 0.07173305 |
| ENSG00000235298 | HNRNPK-AS1 | 53.24625558 | -0.748676042 | 0.273077077 | -2.741629033 | 0.006113534 | 0.019280242 |
| ENSG00000235313 | HM13-IT1 | 7.868362954 | -0.343173542 | 0.643986228 | -0.532889567 | 0.594110028 | 0.755984249 |
| ENSG00000235314 | LINC00957 | 76.23686222 | 0.40418137 | 0.207618243 | 1.946752679 | 0.051564393 | 0.122974027 |
| ENSG00000235315 | RPL23AP69 | 5.025781999 | 0.496872092 | 0.789007568 | 0.629743125 | 0.528862662 | 0.704863437 |
| ENSG00000235316 | DUSP8P5 | 45.19376587 | -0.542557932 | 0.264425138 | -2.051839456 | 0.040185267 | 0.099508855 |
| ENSG00000235319 | ENSG0000023 | 3.108083036 | -0.627613938 | 0.978428139 | -0.641451234 | 0.521229555 | 0.698300881 |
| ENSG00000235325 | SRSF3P5 | 3.704460272 | -0.646225644 | 0.953954028 | -0.677418015 | 0.498140772 | 0.67994219 |
| ENSG00000235330 | RPL12P25 | 3.185167758 | 1.076196538 | 1.019643918 | 1.055463107 | 0.291213574 | 0.476153912 |
| ENSG00000235351 | ENSG0000023 | 3.455589147 | -0.570890123 | 0.935209618 | -0.610440817 | 0.541569837 | 0.715645856 |
| ENSG00000235354 | RPS29P16 | 14.65949401 | -0.248472449 | 0.471999491 | -0.526425249 | 0.598592777 | 0.75887321 |
| ENSG00000235363 | SNRPGP10 | 335.8405944 | -0.150588246 | 0.104595076 | -1.439725959 | 0.149944946 | 0.291552557 |
| ENSG00000235368 | SAPCD2P2 | 6.405805097 | -0.178837329 | 0.765102175 | -0.233743067 | 0.815184451 | 0.902135938 |
| ENSG00000235371 | ENSG0000023 | 20.86340332 | -0.154255586 | 0.402134372 | -0.383592144 | 0.701280764 | 0.832852595 |
| ENSG00000235374 | SSR4P1 | 93.53454979 | 0.636663147 | 0.190346962 | 3.344750767 | 0.000823565 | 0.00313258 |
| ENSG00000235376 | RPEL1 | 87.07536941 | -0.415079131 | 0.19738045 | -2.102939429 | 0.035471065 | 0.089605354 |
| ENSG00000235381 | ENSG0000023 | 12.42650912 | 0.578172228 | 0.514659744 | 1.123406746 | 0.261264786 | 0.4415299 |
| ENSG00000235400 | ENSG0000023 | 15.22012105 | -0.94431629 | 0.471790267 | -2.001559496 | 0.045332129 | 0.110125917 |
| ENSG00000235408 | SNORA71B | 14.14928956 | -0.393145081 | 0.489682136 | -0.802857717 | 0.422056978 | 0.61309442 |

| | | | | | | | |
|---|---|---|---|---|---|---|---|
| ENSG00000235411 | DRG1P1 | 31.39474295 | -0.696585193 | 0.338418936 | -2.058351702 | 0.039556384 | 0.098269814 |
| ENSG00000235415 | ENSG0000023 | 35.69353833 | 1.171642555 | 0.309606765 | 3.784292486 | 0.000154146 | 0.000667192 |
| ENSG00000235437 | LINC01278 | 1044.410387 | -0.200271346 | 0.080357037 | -2.492268928 | 0.012692989 | 0.036662953 |
| ENSG00000235438 | ESRRAP2 | 2.033502022 | -0.670209133 | 1.252166496 | -0.535239631 | 0.592484165 | 0.754736754 |
| ENSG00000235448 | LURAP1L-AS1 | 1.631173645 | 0.999025143 | 1.414225278 | 0.706411601 | 0.479932209 | NA |
| ENSG00000235449 | COX7CP2 | 9.878158014 | 0.842958315 | 0.570551686 | 1.477444264 | 0.139556588 | 0.27579498 |
| ENSG00000235453 | SMIM27 | 70.46925466 | 0.351615731 | 0.211890442 | 1.659422328 | 0.09703072 | 0.20694354 |
| ENSG00000235454 | HAUS6P3 | 5.534973796 | 0.51240883 | 0.752455748 | 0.680982013 | 0.495882872 | 0.677941809 |
| ENSG00000235472 | EIF4A1P7 | 62.35585432 | -0.010198808 | 0.246925539 | -0.041303171 | 0.967054205 | 0.985295635 |
| ENSG00000235481 | UBE2R2-AS1 | 4.119448723 | -0.091781707 | 0.858053811 | -0.106964978 | 0.914816764 | 0.957703529 |
| ENSG00000235485 | ENSG0000023 | 10.29171001 | -0.215635529 | 0.559756748 | -0.38523078 | 0.700066441 | 0.831719009 |
| ENSG00000235488 | JARID2-AS1 | 1.952161835 | 0.839029448 | 1.262317521 | 0.664673851 | 0.506259123 | 0.686625533 |
| ENSG00000235489 | DBF4P1 | 167.3783478 | -0.823072959 | 0.140765631 | -5.847115895 | 5.00E-09 | 3.73E-08 |
| ENSG00000235491 | LINC01889 | 60.00458986 | -0.926403535 | 0.229806437 | -4.031234051 | 5.55E-05 | 0.000257917 |
| ENSG00000235499 | ENSG0000023 | 76.04271998 | -0.384036221 | 0.210080358 | -1.8280444 | 0.067542888 | 0.154080535 |
| ENSG00000235500 | SNX19P2 | 13.60184009 | 0.430422183 | 0.488614882 | 0.880902729 | 0.37837047 | 0.571100536 |
| ENSG00000235508 | RPS2P7 | 24.92652866 | 0.440474137 | 0.360043229 | 1.223392363 | 0.221181536 | 0.39177704 |
| ENSG00000235512 | TAB3-AS2 | 5.300478084 | -1.27028076 | 0.779526775 | -1.629553726 | 0.103195851 | 0.217491195 |
| ENSG00000235513 | L3MBTL2-AS1 | 7.711222929 | -0.696219488 | 0.626070198 | -1.112047003 | 0.266117941 | 0.447089532 |
| ENSG00000235522 | ENSG0000023 | 13.72170523 | -0.235917857 | 0.473543702 | -0.498196589 | 0.618345487 | 0.773690709 |
| ENSG00000235527 | HIPK1-AS1 | 64.25967203 | 0.471012658 | 0.228616597 | 2.060273241 | 0.039372427 | 0.097908235 |
| ENSG00000235529 | AGAP1-IT1 | 4.981708402 | -0.704784559 | 0.851946197 | -0.827264165 | 0.40808735 | 0.60057947 |
| ENSG00000235530 | TMEM11-DT | 11.56324949 | 1.048191926 | 0.531041847 | 1.97384054 | 0.048399878 | 0.116553978 |
| ENSG00000235545 | DOCK7-DT | 2.083651579 | -0.767560023 | 1.269036801 | -0.604836694 | 0.54528751 | 0.718492252 |
| ENSG00000235546 | ENSG0000023 | 9.089842626 | 0.726701033 | 0.589097919 | 1.233582754 | 0.217358423 | 0.38691722 |
| ENSG00000235552 | RPL6P27 | 5129.30581 | 0.181084576 | 0.061354993 | 2.951423616 | 0.003163128 | 0.010667767 |
| ENSG00000235554 | SRSF6P2 | 198.499629 | -0.274849642 | 0.129435127 | -2.123454804 | 0.033715759 | 0.085794732 |
| ENSG00000235555 | SUMO1P4 | 4.992288157 | -0.195656212 | 0.862555699 | -0.226833134 | 0.820553498 | 0.904845323 |
| ENSG00000235559 | NOP56P1 | 6.628078007 | 0.485030239 | 0.695548895 | 0.697334496 | 0.485593485 | 0.669321308 |
| ENSG00000235560 | ZNF747-DT | 51.09180498 | -0.659683006 | 0.251179455 | -2.626341417 | 0.00863082 | 0.026137427 |
| ENSG00000235568 | NFAM1 | 6.789745415 | -0.255775605 | 0.663480586 | -0.385505787 | 0.69986272 | 0.831642103 |
| ENSG00000235574 | ENSG0000023 | 2.299208392 | -0.181333522 | 1.161785188 | -0.156081799 | 0.875968545 | 0.935616916 |
| ENSG00000235582 | ENSG0000023 | 9.089499607 | 0.190247797 | 0.588603399 | 0.323218992 | 0.746529399 | 0.859604617 |

| | | | | | | | |
|---|---|---|---|---|---|---|---|
| ENSG00000235584 | ENSG0000023 | 39.22754705 | -1.190038719 | 0.310418508 | -3.833659037 | 0.000126251 | 0.000553449 |
| ENSG00000235587 | GAPDHP65 | 349.3793495 | 0.00349992 | 0.105018228 | 0.033326784 | 0.973413995 | 0.987970845 |
| ENSG00000235590 | GNAS-AS1 | 68.19779817 | 0.094349194 | 0.216126438 | 0.436546286 | 0.662440425 | 0.804963395 |
| ENSG00000235597 | LINC01102 | 7.42523227 | -0.725043779 | 0.640069046 | -1.132758697 | 0.257315601 | 0.436954614 |
| ENSG00000235605 | ENSG0000023 | 126.9964994 | 0.08080948 | 0.158228172 | 0.510714866 | 0.609550729 | 0.767711306 |
| ENSG00000235606 | AK4P6 | 1.591829991 | -0.095023634 | 1.440492585 | -0.065966069 | 0.947404839 | NA |
| ENSG00000235609 | ENSG0000023 | 24.88529767 | 0.509903829 | 0.374941718 | 1.359954908 | 0.173844194 | 0.326455943 |
| ENSG00000235613 | NSRP1P1 | 4.237237005 | -0.770576931 | 0.931060427 | -0.827633641 | 0.40787801 | 0.600511443 |
| ENSG00000235618 | FAM21EP | 14.44240234 | 0.219758069 | 0.461517364 | 0.476164249 | 0.633957368 | 0.784236938 |
| ENSG00000235621 | LINC00494 | 27.68670022 | 3.241561446 | 0.466696149 | 6.945764295 | 3.76E-12 | 3.67E-11 |
| ENSG00000235631 | RNF148 | 4.851388382 | 1.259558692 | 0.82514123 | 1.526476495 | 0.12689124 | 0.256488857 |
| ENSG00000235636 | NUS1P1 | 493.6998673 | 0.125226945 | 0.087291965 | 1.434575859 | 0.151407998 | 0.293769423 |
| ENSG00000235643 | LINC01647 | 6.687536507 | -0.369792343 | 0.674823693 | -0.547983638 | 0.583703137 | 0.748011283 |
| ENSG00000235644 | RPL10P5 | 1.972510088 | 0.30688992 | 1.234081108 | 0.24867889 | 0.803609179 | 0.895414541 |
| ENSG00000235651 | G3BP1P1 | 27.48574869 | 0.311128816 | 0.359245748 | 0.866061235 | 0.386456582 | 0.578997258 |
| ENSG00000235652 | EPM2A-DT | 35.22167198 | 0.279410735 | 0.306187244 | 0.91254858 | 0.361480008 | 0.554079339 |
| ENSG00000235655 | H3P6 | 2572.818158 | -0.132587597 | 0.062031807 | -2.137413085 | 0.032564408 | 0.083373607 |
| ENSG00000235660 | LINC00345 | 8.702575838 | -1.729236531 | 0.63691852 | -2.715004317 | 0.006627491 | 0.020673733 |
| ENSG00000235663 | SAPCD1-AS1 | 5.891076934 | 0.049874813 | 0.723932185 | 0.068894316 | 0.945073743 | 0.973900169 |
| ENSG00000235672 | ENSG0000023 | 1.817032681 | -0.852814895 | 1.377472954 | -0.619115528 | 0.535840255 | 0.711105834 |
| ENSG00000235674 | LDHAP2 | 9.089922837 | -0.963119647 | 0.590251061 | -1.631711844 | 0.102740204 | 0.216722784 |
| ENSG00000235677 | NPM1P26 | 4.931171345 | -0.33016348 | 0.8130317 | -0.406089308 | 0.684676981 | 0.821336301 |
| ENSG00000235695 | HIGD2AP2 | 1.633461305 | -0.743396591 | 1.350161272 | -0.550598367 | 0.581909029 | NA |
| ENSG00000235698 | PA2G4P2 | 147.8170561 | 0.089009912 | 0.156799948 | 0.567665445 | 0.570262159 | 0.737339443 |
| ENSG00000235703 | EOLA2-DT | 75.7551047 | 0.825059347 | 0.205777023 | 4.009482377 | 6.09E-05 | 0.000281327 |
| ENSG00000235706 | DICER1-AS1 | 177.7004558 | -0.764706946 | 0.144969073 | -5.274966089 | 1.33E-07 | 8.52E-07 |
| ENSG00000235720 | GABPAP | 119.2051355 | -0.062331812 | 0.164712288 | -0.378428425 | 0.705112355 | 0.834997903 |
| ENSG00000235721 | ENSG0000023 | 12.51154139 | 1.932782427 | 0.570090677 | 3.390307025 | 0.000698144 | 0.002691215 |
| ENSG00000235726 | ENSG0000023 | 9.55278312 | -0.443239704 | 0.570378062 | -0.777098092 | 0.4371009 | 0.626732586 |
| ENSG00000235734 | HMGN1P36 | 459.4823121 | -0.252265617 | 0.095877118 | -2.631134753 | 0.008510029 | 0.025795462 |
| ENSG00000235742 | ENSG0000023 | 1.634460079 | -0.204852114 | 1.412997107 | -0.144977023 | 0.884729013 | NA |
| ENSG00000235748 | SEPTIN14P12 | 2.858446847 | 0.512940368 | 1.063299804 | 0.482404272 | 0.629518787 | 0.781143482 |
| ENSG00000235750 | KIAA0040 | 654.8518138 | 0.328304198 | 0.078957878 | 4.157966341 | 3.21E-05 | 0.000154941 |

| | | | | | | | |
|---|---|---|---|---|---|---|---|
| ENSG00000235760 | MSH2-OT1 | 3.244286697 | 0.410796685 | 0.957282082 | 0.429128146 | 0.66782997 | 0.808785741 |
| ENSG00000235763 | SNRPGP5 | 1.920024931 | 0.789731354 | 1.27525228 | 0.619274607 | 0.53573547 | 0.711031295 |
| ENSG00000235768 | BRD7P5 | 42.89747203 | 0.240760265 | 0.28955418 | 0.831486062 | 0.405699099 | 0.598465521 |
| ENSG00000235770 | LINC00607 | 23.42844356 | -0.215250609 | 0.371094725 | -0.580042223 | 0.561886145 | 0.730748238 |
| ENSG00000235776 | RPL7AP70 | 278.7027747 | 0.128032375 | 0.112518973 | 1.137873656 | 0.255173243 | 0.434668018 |
| ENSG00000235781 | LINC02569 | 7.415377527 | -0.497469767 | 0.665105327 | -0.747956372 | 0.454486474 | 0.642069202 |
| ENSG00000235786 | ZNRF3-IT1 | 7.492341153 | 1.196522416 | 0.679386663 | 1.761180312 | 0.078207889 | 0.173564887 |
| ENSG00000235795 | ENSG0000023 | 1.910146699 | 0.017617566 | 1.276691871 | 0.013799387 | 0.988990031 | 0.994147115 |
| ENSG00000235808 | MYL6P2 | 4.140184475 | -0.739557322 | 0.877118993 | -0.843166467 | 0.39913535 | 0.591930681 |
| ENSG00000235823 | OLMALINC | 1243.329516 | -0.597636707 | 0.066381366 | -9.003079334 | 2.19E-19 | 3.54E-18 |
| ENSG00000235828 | RPL36AP26 | 4.771551561 | 0.946480135 | 0.879497372 | 1.076160276 | 0.281855582 | 0.465366562 |
| ENSG00000235829 | TBCAP2 | 4.709116443 | 0.218239427 | 0.82937143 | 0.263138346 | 0.792443949 | 0.888454337 |
| ENSG00000235830 | SRGAP3-AS4 | 4.105684805 | -0.308573253 | 0.870628857 | -0.354425713 | 0.723019868 | 0.845926239 |
| ENSG00000235831 | BHLHE40-AS1 | 40.76787942 | -0.345479856 | 0.276205538 | -1.250807129 | 0.211004853 | 0.378455929 |
| ENSG00000235837 | ENSG0000023 | 15.19656127 | -2.057488128 | 0.522378614 | -3.938691351 | 8.19E-05 | 0.000371058 |
| ENSG00000235838 | HSP90AB7P | 1.692165579 | -0.227551257 | 1.333515397 | -0.170640143 | 0.864506732 | NA |
| ENSG00000235847 | LDHAP7 | 30.35886618 | -0.545746996 | 0.326381202 | -1.672115283 | 0.094501592 | 0.202396276 |
| ENSG00000235852 | ENSG0000023 | 3.180328659 | -1.016049326 | 1.110432924 | -0.915002882 | 0.360190117 | 0.552730034 |
| ENSG00000235859 | ENSG0000023 | 69.65099724 | -1.387540892 | 0.223671821 | -6.203467601 | 5.52E-10 | 4.51E-09 |
| ENSG00000235863 | B3GALT4 | 25.48319748 | 0.336943742 | 0.361412666 | 0.932296441 | 0.351183347 | 0.542464882 |
| ENSG00000235872 | ENSG0000023 | 2.594719746 | -1.226750928 | 1.110716991 | -1.104467598 | 0.269390352 | 0.451016256 |
| ENSG00000235876 | FEM1AP4 | 2.230184936 | 0.675207096 | 1.273203511 | 0.530321422 | 0.595889097 | 0.756993626 |
| ENSG00000235897 | TM4SF19-AS1 | 13.06901445 | -0.464461399 | 0.512575516 | -0.906132628 | 0.364865658 | 0.557629459 |
| ENSG00000235903 | CPB2-AS1 | 5.393169974 | -1.484510412 | 0.801971853 | -1.851075455 | 0.064158699 | 0.147477098 |
| ENSG00000235912 | ENSG0000023 | 222.0458593 | -0.042544923 | 0.129614786 | -0.328241281 | 0.742729238 | 0.858016204 |
| ENSG00000235916 | MRPS18CP7 | 34.80689806 | -0.22823313 | 0.298783298 | -0.76387513 | 0.444941659 | 0.633579247 |
| ENSG00000235919 | ASH1L-AS1 | 69.82405535 | 0.43918014 | 0.21911972 | 2.004293093 | 0.045038673 | 0.109516933 |
| ENSG00000235920 | THRAP3P3 | 3.937425601 | 0.085863115 | 0.950873946 | 0.090299156 | 0.928049491 | 0.965376834 |
| ENSG00000235939 | ENSG0000023 | 4.604341906 | -1.732971402 | 0.877601157 | -1.974668548 | 0.048305774 | 0.11636401 |
| ENSG00000235944 | ZNF815P | 112.1283803 | -0.381169533 | 0.17062566 | -2.233951989 | 0.025486232 | 0.067519825 |
| ENSG00000235946 | H2BW3P | 7.086310944 | 0.559215254 | 0.665453882 | 0.840351631 | 0.400711261 | 0.593627144 |
| ENSG00000235954 | TTC28-AS1 | 246.2516204 | -0.253124577 | 0.118856501 | -2.129665391 | 0.033199248 | 0.084710952 |
| ENSG00000235957 | COX7CP1 | 121.9819095 | 0.059575935 | 0.165274906 | 0.360465705 | 0.718498899 | 0.843432038 |

| | | | | | | | |
|---|---|---|---|---|---|---|---|
| ENSG00000235961 | PNMA6A | 3.910459591 | 0.070990819 | 0.875916206 | 0.0810475 | 0.935404177 | 0.969116241 |
| ENSG00000235962 | RPL7AP53 | 10.2242626 | -0.248847258 | 0.560933407 | -0.443630661 | 0.657309634 | 0.80147779 |
| ENSG00000235978 | ENSG0000023 | 4.98805575 | -0.914209735 | 0.857327294 | -1.066348571 | 0.286266101 | 0.470021728 |
| ENSG00000235989 | MORC2-AS1 | 6.260182759 | -0.630970069 | 0.697293321 | -0.904884717 | 0.365526465 | 0.558341796 |
| ENSG00000235996 | ENSG0000023 | 2.2864967 | -1.443720365 | 1.262585103 | -1.143463804 | 0.252846075 | 0.431658492 |
| ENSG00000236008 | LINC01814 | 63.8521357 | -0.87019737 | 0.22260014 | -3.909239999 | 9.26E-05 | 0.000414914 |
| ENSG00000236015 | ENSG0000023 | 2.733263892 | -1.373574169 | 1.162898869 | -1.181163905 | 0.237537617 | 0.412874143 |
| ENSG00000236017 | ASMTL-AS1 | 55.01663541 | 0.239006716 | 0.25605522 | 0.933418644 | 0.350603858 | 0.541977875 |
| ENSG00000236017 | ASMTL-AS1 | 55.01663541 | 0.239006716 | 0.25605522 | 0.933418644 | 0.350603858 | 0.541977875 |
| ENSG00000236018 | ENSG0000023 | 8.771138638 | -0.550756962 | 0.58919269 | -0.934765436 | 0.349909196 | 0.54115263 |
| ENSG00000236021 | COP1-DT | 6.697464511 | 0.098520713 | 0.699953745 | 0.140753177 | 0.888064934 | 0.942328951 |
| ENSG00000236035 | NSA2P1 | 14.49305965 | 0.610618325 | 0.494414566 | 1.235033043 | 0.216818207 | 0.386271044 |
| ENSG00000236048 | SSBL5P | 13.44159843 | -0.270180585 | 0.476568118 | -0.566929626 | 0.570761995 | 0.737691147 |
| ENSG00000236051 | MYCBP2-AS1 | 6.031769042 | -0.379368056 | 0.715484059 | -0.530225728 | 0.595955435 | 0.756993626 |
| ENSG00000236056 | GAPDHP14 | 3.174151528 | -5.238515489 | 1.455370958 | -3.599436597 | 0.000318907 | 0.001306148 |
| ENSG00000236058 | RPL17P36 | 88.71856267 | 0.762642601 | 0.190016117 | 4.01356797 | 5.98E-05 | 0.000276723 |
| ENSG00000236060 | HSPB1P1 | 47.38973398 | 1.157175377 | 0.280099132 | 4.131306545 | 3.61E-05 | 0.000172101 |
| ENSG00000236086 | HMGN2P28 | 18.99478171 | -0.466231567 | 0.425416806 | -1.09594064 | 0.273104753 | 0.455038495 |
| ENSG00000236088 | COX10-DT | 322.828734 | -0.050436448 | 0.103000987 | -0.489669561 | 0.624367745 | 0.778004121 |
| ENSG00000236090 | LDHAP3 | 13.21161949 | 0.02195648 | 0.50058775 | 0.0438614 | 0.965014884 | 0.984148726 |
| ENSG00000236104 | ZBTB22 | 987.9756363 | 0.094417142 | 0.066723956 | 1.415041131 | 0.157056437 | 0.30150735 |
| ENSG00000236105 | PRELID3BP10 | 12.40795037 | 0.476765335 | 0.503122652 | 0.947612542 | 0.343326739 | 0.534322618 |
| ENSG00000236111 | RPL6P1 | 2.763471299 | 0.68582553 | 1.063612881 | 0.644807469 | 0.519051961 | 0.696685506 |
| ENSG00000236137 | CD101-AS1 | 29.61593666 | -0.909302243 | 0.331404482 | -2.743783785 | 0.006073553 | 0.01916469 |
| ENSG00000236138 | DUX4L26 | 2.594688901 | 0.978992909 | 1.133519376 | 0.863675496 | 0.387766178 | 0.580492583 |
| ENSG00000236140 | ENSG0000023 | 3.211290717 | -0.102942756 | 1.004170705 | -0.102515196 | 0.918347752 | 0.959952416 |
| ENSG00000236144 | TMEM147-AS | 725.0383869 | 0.849843993 | 0.089762434 | 9.467702241 | 2.86E-21 | 5.16E-20 |
| ENSG00000236148 | RPL23AP37 | 24.60341652 | 0.034660964 | 0.35348254 | 0.09805566 | 0.921888096 | 0.962216621 |
| ENSG00000236152 | MRPS36P1 | 105.4033394 | -0.01567741 | 0.173409116 | -0.090407072 | 0.927963737 | 0.965331908 |
| ENSG00000236182 | ENOPH1P1 | 4.352100777 | 1.088412525 | 0.894359285 | 1.21697459 | 0.223613884 | 0.395017904 |
| ENSG00000236184 | TCEA1P4 | 3.356335204 | 0.854544827 | 0.988038695 | 0.864890041 | 0.387099144 | 0.579713694 |
| ENSG00000236200 | KDM4A-AS1 | 62.01215544 | -0.554846369 | 0.229630879 | -2.416253303 | 0.015681148 | 0.044219489 |
| ENSG00000236204 | LINC01376 | 88.44886449 | -0.522237577 | 0.191493518 | -2.727181481 | 0.00638779 | 0.0200213 |

| | | | | | | | |
|---|---|---|---|---|---|---|---|
| ENSG00000236212 | ENSG0000023 | 18.32312305 | 0.221033241 | 0.409048354 | 0.540359687 | 0.588949003 | 0.752212579 |
| ENSG00000236229 | VEZF1P1 | 19.95377552 | -0.39716416 | 0.430893089 | -0.921723206 | 0.356672973 | 0.548874891 |
| ENSG00000236255 | ENSG0000023 | 29.20778717 | -0.339470215 | 0.354609324 | -0.957307638 | 0.338412001 | 0.529660598 |
| ENSG00000236257 | EI24P2 | 12.69910934 | -0.184208916 | 0.496949383 | -0.370679434 | 0.71087631 | 0.839122288 |
| ENSG00000236263 | ENSG0000023 | 19.5157412 | 0.117041368 | 0.403424355 | 0.290119737 | 0.771724637 | 0.875561078 |
| ENSG00000236264 | RPL26P30 | 163.4662454 | -0.58843763 | 0.141675348 | -4.15342287 | 3.28E-05 | 0.000157653 |
| ENSG00000236266 | ENSG0000023 | 3.008746222 | -0.274473244 | 1.035283281 | -0.265118977 | 0.790917809 | 0.887307049 |
| ENSG00000236281 | NDUFB9P2 | 33.44309432 | -0.236475866 | 0.327323184 | -0.722453763 | 0.470015543 | 0.655940953 |
| ENSG00000236283 | ENSG0000023 | 13.7946442 | 0.876268444 | 0.49008576 | 1.787990011 | 0.07377762 | 0.165740532 |
| ENSG00000236285 | NPM1P8 | 2.763983352 | 0.698006274 | 1.080424214 | 0.646048344 | 0.518248047 | 0.695932225 |
| ENSG00000236287 | ZBED5 | 620.1317849 | -0.188749359 | 0.094492978 | -1.997496129 | 0.045771315 | 0.111051714 |
| ENSG00000236296 | GUSBP5 | 22.81562191 | 0.729945273 | 0.383915344 | 1.901318309 | 0.057260332 | 0.134296291 |
| ENSG00000236305 | SLC12A9-AS1 | 21.38097797 | -0.336442204 | 0.382255491 | -0.880150088 | 0.378778008 | 0.571354252 |
| ENSG00000236307 | EEF1E1P1 | 8.705571058 | 0.198896329 | 0.588090068 | 0.338207258 | 0.735207007 | 0.853967465 |
| ENSG00000236312 | RPL34P34 | 18.68468657 | -0.074739865 | 0.402341047 | -0.185762465 | 0.852631041 | 0.923046535 |
| ENSG00000236319 | ENSG0000023 | 2.540319609 | 0.927779984 | 1.172710089 | 0.791141811 | 0.428861243 | 0.61920798 |
| ENSG00000236325 | RPL32P5 | 2.42823247 | 0.034769451 | 1.09916229 | 0.031632681 | 0.97476498 | 0.988554759 |
| ENSG00000236326 | ENSG0000023 | 1.676490359 | -0.68616016 | 1.452322547 | -0.472457142 | 0.636600526 | NA |
| ENSG00000236330 | RPL5P9 | 16.11666595 | 0.469869536 | 0.441714896 | 1.063739396 | 0.287446773 | 0.471338739 |
| ENSG00000236334 | PPIAL4G | 1.640652029 | 1.779632164 | 1.586688761 | 1.121601292 | 0.262032002 | NA |
| ENSG00000236337 | FMR1-IT1 | 4.292926436 | -0.17951776 | 0.849593655 | -0.211298376 | 0.83265445 | 0.911472914 |
| ENSG00000236348 | PSMC1P10 | 6.761422142 | -0.928154511 | 0.677857988 | -1.369246254 | 0.170922312 | 0.322023475 |
| ENSG00000236349 | SUCLG2P2 | 228.3865632 | 0.387981638 | 0.123721242 | 3.135933906 | 0.001713078 | 0.006106917 |
| ENSG00000236352 | ENSG0000023 | 4.179987684 | 0.167639859 | 0.874052157 | 0.191796173 | 0.847901869 | 0.920424395 |
| ENSG00000236354 | LINC00437 | 1.869777986 | -0.598132821 | 1.317794191 | -0.453889405 | 0.64990843 | 0.796349063 |
| ENSG00000236358 | ENSG0000023 | 5.889091408 | -0.042209356 | 0.720565568 | -0.058578091 | 0.953288161 | 0.978321944 |
| ENSG00000236360 | RRAS2P1 | 103.89781 | 0.042781331 | 0.187837119 | 0.227757597 | 0.819834693 | 0.904443968 |
| ENSG00000236364 | ENSG0000023 | 8.182517631 | -0.395230683 | 0.6110281 | -0.646828981 | 0.517742634 | 0.69574209 |
| ENSG00000236376 | ENSG0000023 | 1.749362076 | -2.192949374 | 1.52060294 | -1.442157789 | 0.149257866 | NA |
| ENSG00000236377 | ENSG0000023 | 1.920996538 | 2.16924081 | 1.508509408 | 1.438002838 | 0.150433249 | 0.292251889 |
| ENSG00000236384 | LINC00479 | 2.196083906 | 1.207146431 | 1.242654909 | 0.97142531 | 0.33133653 | 0.52224246 |
| ENSG00000236397 | DDX11L2 | 40.52049525 | 0.046146529 | 0.27822223 | 0.16586212 | 0.868265459 | 0.931226781 |
| ENSG00000236404 | VLDLR-AS1 | 39.17331366 | 0.881224991 | 0.311811505 | 2.826146496 | 0.004711171 | 0.015279834 |

| | | | | | | | |
|---|---|---|---|---|---|---|---|
| ENSG00000236409 | NRADDP | 26.5374609 | -0.786249411 | 0.351573888 | -2.236370325 | 0.025327523 | 0.067200253 |
| ENSG00000236423 | LINC01134 | 12.4780376 | 1.239758844 | 0.524078862 | 2.365595968 | 0.018001077 | 0.049878574 |
| ENSG00000236432 | MFF-DT | 27.68236491 | 0.546841989 | 0.33708993 | 1.622243622 | 0.104751195 | 0.22009456 |
| ENSG00000236438 | FAM157A | 24.47729801 | -0.413551913 | 0.351495853 | -1.176548486 | 0.239375769 | 0.41496865 |
| ENSG00000236439 | RPS27P8 | 561.0393656 | 0.438796861 | 0.085905311 | 5.107913081 | 3.26E-07 | 2.00E-06 |
| ENSG00000236444 | UBE2L5 | 2.44288175 | 0.469880215 | 1.165892732 | 0.403021823 | 0.68693217 | 0.823098667 |
| ENSG00000236449 | ENSG0000023 | 88.16525758 | 0.145100802 | 0.196050416 | 0.740119837 | 0.459227283 | 0.646378321 |
| ENSG00000236451 | ENSG0000023 | 3.851948467 | -1.04804053 | 0.941237527 | -1.11347083 | 0.26550627 | 0.446566011 |
| ENSG00000236452 | ENSG0000023 | 2.956329336 | -2.417846423 | 1.193200017 | -2.026354667 | 0.042728453 | 0.104844444 |
| ENSG00000236453 | ENSG0000023 | 2.089813785 | -4.581235141 | 1.58606968 | -2.888419846 | 0.003871827 | 0.012800073 |
| ENSG00000236456 | ENSG0000023 | 423.0606659 | 0.199571284 | 0.093543028 | 2.13347042 | 0.032886156 | 0.084056648 |
| ENSG00000236459 | HNRNPA1P22 | 2.474182741 | 0.785354137 | 1.130083167 | 0.694952513 | 0.487085062 | 0.670520307 |
| ENSG00000236471 | LINC02920 | 2.02595722 | -0.256269435 | 1.27033591 | -0.201733599 | 0.840124994 | 0.915861206 |
| ENSG00000236472 | ENSG0000023 | 27.91099402 | -0.477816698 | 0.343933893 | -1.389269008 | 0.164750964 | 0.313044572 |
| ENSG00000236480 | PKMP1 | 45.02989134 | -0.344277883 | 0.303306519 | -1.135082371 | 0.256340807 | 0.435815469 |
| ENSG00000236489 | RPL34P10 | 2.416390995 | 0.014897907 | 1.171584864 | 0.012716029 | 0.989854351 | 0.994580124 |
| ENSG00000236491 | ENSG0000023 | 5.424565301 | -0.18636287 | 0.804035627 | -0.231784344 | 0.816705518 | 0.902945905 |
| ENSG00000236493 | EIF2S2P3 | 60.3360507 | -0.317744036 | 0.230031092 | -1.381309079 | 0.167183948 | 0.316679249 |
| ENSG00000236496 | GPS2P1 | 71.22745283 | -0.131906486 | 0.214204569 | -0.615796791 | 0.538028646 | 0.712906237 |
| ENSG00000236498 | ENSG0000023 | 11.20144927 | -0.606004894 | 0.540835881 | -1.120496836 | 0.2625021 | 0.443137698 |
| ENSG00000236514 | ENSG0000023 | 10.78592874 | -0.294447066 | 0.531640377 | -0.55384632 | 0.579684025 | 0.745192017 |
| ENSG00000236519 | LINC01424 | 8.777838963 | 0.268604343 | 0.593168794 | 0.452829525 | 0.650671501 | 0.797010051 |
| ENSG00000236523 | NPM1P40 | 4.450405394 | -0.075945462 | 0.81586176 | -0.093086189 | 0.925835089 | 0.964514676 |
| ENSG00000236526 | BTBD3-AS1 | 3.862214198 | -1.071411643 | 0.909333398 | -1.178238527 | 0.238701527 | 0.414269408 |
| ENSG00000236527 | ARF4P2 | 2.310032898 | 1.846414466 | 1.288937886 | 1.432508491 | 0.15199835 | 0.294397344 |
| ENSG00000236528 | ENSG0000023 | 4.477830959 | 0.38727126 | 0.863974349 | 0.448243933 | 0.65397716 | 0.799165148 |
| ENSG00000236529 | ENSG0000023 | 6.921525051 | 0.496139081 | 0.728485374 | 0.681055652 | 0.495836277 | 0.677941809 |
| ENSG00000236530 | KPNA2P1 | 8.454419328 | -0.145667733 | 0.653234213 | -0.222994648 | 0.823539672 | 0.906786987 |
| ENSG00000236532 | LINC01695 | 1.588117424 | 1.869930002 | 1.547578847 | 1.208293849 | 0.226934254 | NA |
| ENSG00000236533 | RPS12P4 | 6.887762172 | 0.797675904 | 0.687054739 | 1.161007789 | 0.245638732 | 0.422701956 |
| ENSG00000236534 | H3P36 | 11.39625736 | -0.310615122 | 0.551361762 | -0.563359927 | 0.573189819 | 0.739911904 |
| ENSG00000236539 | HNRNPA1P54 | 3.212122627 | 0.752203601 | 0.990528611 | 0.759396137 | 0.447615624 | 0.636005104 |
| ENSG00000236540 | ENSG0000023 | 2.127030706 | -1.318367133 | 1.234319036 | -1.068092685 | 0.285478705 | 0.469386575 |

| | | | | | | | |
|---|---|---|---|---|---|---|---|
| ENSG00000236552 | RPL13AP5 | 9268.066429 | 0.171123656 | 0.05792734 | 2.954108618 | 0.003135736 | 0.010583167 |
| ENSG00000236565 | HNRNPA3P5 | 1044.398535 | 0.126581077 | 0.072810005 | 1.73851214 | 0.082120613 | 0.180496968 |
| ENSG00000236567 | TCF3P1 | 1.628229822 | -2.107157765 | 1.584148714 | -1.330151485 | 0.183468365 | NA |
| ENSG00000236570 | RAD23BP1 | 28.78442228 | -0.164201743 | 0.333692404 | -0.492075158 | 0.622666213 | 0.776597778 |
| ENSG00000236582 | PRPF38AP2 | 5.208742981 | -0.004110542 | 0.757635525 | -0.005425488 | 0.995671108 | 0.997217798 |
| ENSG00000236591 | UST-AS2 | 2.732690925 | -2.993083211 | 1.343729976 | -2.227443953 | 0.025917618 | 0.068504453 |
| ENSG00000236592 | S100A11P2 | 3.187519383 | 0.153715527 | 1.008911307 | 0.15235782 | 0.878904724 | 0.937160656 |
| ENSG00000236603 | RANP1 | 1001.678691 | -0.104850022 | 0.068734723 | -1.525430208 | 0.127151829 | 0.256901793 |
| ENSG00000236608 | EIF4A1P6 | 7.160972459 | -0.386417852 | 0.67118927 | -0.575721141 | 0.564803683 | 0.733188162 |
| ENSG00000236609 | ZNF853 | 14.97522547 | 0.387541309 | 0.449108051 | 0.862913297 | 0.388185139 | 0.580922898 |
| ENSG00000236617 | CHFR-DT | 22.88303956 | 0.264883851 | 0.367574882 | 0.720625549 | 0.47113993 | 0.657018237 |
| ENSG00000236618 | PITPNA-AS1 | 131.9801488 | 0.051408561 | 0.156043605 | 0.329449971 | 0.741815602 | 0.857590509 |
| ENSG00000236643 | ENSG0000023 | 2.649986746 | -0.20929168 | 1.092644235 | -0.191546043 | 0.848097811 | 0.920492429 |
| ENSG00000236646 | LAMTOR5P1 | 1.957298982 | -1.147622593 | 1.267181449 | -0.905649774 | 0.365121255 | 0.557837983 |
| ENSG00000236651 | DLX2-DT | 3.323704441 | -0.742035203 | 0.966742355 | -0.767562525 | 0.442747133 | 0.631641123 |
| ENSG00000236654 | ENSG0000023 | 3.933600828 | 0.917743926 | 0.903507612 | 1.015756717 | 0.309745252 | 0.49772738 |
| ENSG00000236662 | ENSG0000023 | 14.36404571 | -0.08497035 | 0.473639766 | -0.179398682 | 0.857624666 | 0.926092537 |
| ENSG00000236670 | KRT18P5 | 8.684956401 | -0.297538163 | 0.620390119 | -0.479598487 | 0.631512922 | 0.782886131 |
| ENSG00000236671 | PRKG1-AS1 | 82.73454644 | -1.297494687 | 0.206405726 | -6.286137073 | 3.25E-10 | 2.71E-09 |
| ENSG00000236674 | ENSG0000023 | 1.8524412 | 0.042895994 | 1.259698443 | 0.03405259 | 0.972835214 | 0.987776737 |
| ENSG00000236675 | MTX1P1 | 3.638135463 | -0.457865401 | 0.904093553 | -0.506435866 | 0.612550702 | 0.769672624 |
| ENSG00000236680 | ENSG0000023 | 16.00284462 | -0.624347344 | 0.489005329 | -1.276770022 | 0.201683452 | 0.365422825 |
| ENSG00000236682 | MAP3K2-DT | 46.90864626 | 0.315119327 | 0.258134671 | 1.22075553 | 0.222178596 | 0.393209149 |
| ENSG00000236683 | HMGA1P1 | 36.25191505 | 0.601586498 | 0.298060979 | 2.018333634 | 0.043556526 | 0.106545821 |
| ENSG00000236686 | BZW1P1 | 42.77885053 | -0.364140376 | 0.289180775 | -1.25921364 | 0.207953176 | 0.374358147 |
| ENSG00000236698 | EIF1AXP1 | 711.1718012 | 0.177733271 | 0.084338897 | 2.107370115 | 0.03508551 | 0.088807415 |
| ENSG00000236699 | ARHGEF38 | 52.64589218 | -0.228946332 | 0.248560666 | -0.921088344 | 0.357004305 | 0.549274229 |
| ENSG00000236703 | MYB-AS1 | 2.737039624 | 1.517620384 | 1.12565026 | 1.348216615 | 0.177588722 | 0.331788637 |
| ENSG00000236708 | SDK1-AS1 | 10.86718915 | 2.776604469 | 0.687285938 | 4.0399553 | 5.35E-05 | 0.000248926 |
| ENSG00000236723 | ENSG0000023 | 32.586021 | -0.191538167 | 0.320950627 | -0.5967839 | 0.550651669 | 0.722734253 |
| ENSG00000236735 | RPL31P63 | 8.304606673 | 0.225578767 | 0.608160498 | 0.370919794 | 0.710697272 | 0.839040472 |
| ENSG00000236739 | CLIC4P1 | 124.3232237 | 0.263346736 | 0.16503294 | 1.595722261 | 0.110550816 | 0.229671774 |
| ENSG00000236753 | MKLN1-AS | 43.42769729 | 0.004272438 | 0.274560651 | 0.015560998 | 0.987584621 | 0.993927018 |

| | | | | | | | |
|---|---|---|---|---|---|---|---|
| ENSG00000236754 | ENSG0000023 | 4.342144863 | -1.574583544 | 0.890566126 | -1.768070329 | 0.077049147 | 0.171686006 |
| ENSG00000236756 | DNAJC9-AS1 | 12.57882111 | -0.51218852 | 0.529985428 | -0.966420006 | 0.333834057 | 0.524818618 |
| ENSG00000236762 | RPL19P16 | 39.14975951 | 0.281157654 | 0.313527969 | 0.89675462 | 0.369849868 | 0.562532712 |
| ENSG00000236764 | COX7A2P2 | 8.784461605 | 0.033503878 | 0.587704286 | 0.057008055 | 0.954538779 | 0.979088978 |
| ENSG00000236772 | ENSG0000023 | 2.343986768 | -0.722124699 | 1.229541221 | -0.587312313 | 0.556993973 | 0.727416829 |
| ENSG00000236773 | ENSG0000023 | 179.5550133 | -0.081778663 | 0.136004374 | -0.60129436 | 0.547643946 | 0.720435377 |
| ENSG00000236778 | INTS6-AS1 | 23.78519876 | -0.003198153 | 0.35391286 | -0.009036556 | 0.992789969 | 0.996094602 |
| ENSG00000236779 | ENSG0000023 | 3.180437815 | -0.297283509 | 1.032025713 | -0.288058239 | 0.773302159 | 0.876416754 |
| ENSG00000236780 | LINC01829 | 3.544587074 | 1.997304461 | 1.05809243 | 1.88764649 | 0.059073425 | 0.137689937 |
| ENSG00000236782 | ZNF593OS | 12.51301539 | -0.347155346 | 0.542433342 | -0.639996325 | 0.522174989 | 0.699077575 |
| ENSG00000236790 | LINC00299 | 3.000346023 | -2.419333529 | 1.185917701 | -2.040051789 | 0.041345168 | 0.101886148 |
| ENSG00000236792 | ENSG0000023 | 39.65757076 | -0.136116714 | 0.28213429 | -0.482453637 | 0.629483726 | 0.781143482 |
| ENSG00000236794 | BCRP8 | 2.251407409 | -0.131546651 | 1.191555782 | -0.110399071 | 0.91209289 | 0.955712369 |
| ENSG00000236801 | RPL24P8 | 816.6667486 | 0.218334854 | 0.071077767 | 3.071774224 | 0.002127906 | 0.007434384 |
| ENSG00000236803 | SDAD1P4 | 1.798957269 | 0.049202548 | 1.343499537 | 0.036622676 | 0.970785863 | 0.98680905 |
| ENSG00000236809 | SNX25P1 | 20.90660653 | 0.429210066 | 0.384694902 | 1.115715502 | 0.264543933 | 0.445404611 |
| ENSG00000236810 | ELOA-AS1 | 50.06569026 | -0.171409033 | 0.267003056 | -0.64197405 | 0.520890032 | 0.698050467 |
| ENSG00000236813 | BTF3P8 | 5.324719485 | -0.508455893 | 0.770298687 | -0.660076281 | 0.509204879 | 0.689264918 |
| ENSG00000236814 | ENSG0000023 | 1053.404876 | -0.67992857 | 0.071471537 | -9.513277577 | 1.85E-21 | 3.37E-20 |
| ENSG00000236824 | BCYRN1 | 36.99594681 | 0.001392605 | 0.293665033 | 0.004742156 | 0.996216322 | 0.997651478 |
| ENSG00000236829 | ENSG0000023 | 57.38612318 | 0.653519883 | 0.237023388 | 2.757195765 | 0.005829944 | 0.018448359 |
| ENSG00000236830 | CBR3-AS1 | 410.8680083 | -0.118338207 | 0.094928586 | -1.246602437 | 0.212543308 | 0.380501286 |
| ENSG00000236844 | RPL24P6 | 13.01301166 | 0.152855431 | 0.500093867 | 0.305653481 | 0.759868496 | 0.868010322 |
| ENSG00000236852 | SSBL2P | 15.48090082 | -0.493195324 | 0.467321979 | -1.055365136 | 0.291258362 | 0.476193202 |
| ENSG00000236857 | RAP1BP1 | 2.198511265 | -0.399876453 | 1.20458167 | -0.331962924 | 0.739917256 | 0.856567724 |
| ENSG00000236859 | NIFK-AS1 | 162.1407044 | -0.141989527 | 0.159674073 | -0.88924598 | 0.373870897 | 0.566636335 |
| ENSG00000236871 | LINC00106 | 31.03478901 | 0.285762041 | 0.324690822 | 0.880105076 | 0.37880239 | 0.571354252 |
| ENSG00000236871 | LINC00106 | 31.03478901 | 0.285762041 | 0.324690822 | 0.880105076 | 0.37880239 | 0.571354252 |
| ENSG00000236876 | TMSB4XP1 | 11.60707413 | 0.181826193 | 0.509983953 | 0.356533166 | 0.721441319 | 0.845329648 |
| ENSG00000236882 | LINC01554 | 2.756400244 | 1.105797345 | 1.09332073 | 1.01141167 | 0.311819438 | 0.500045889 |
| ENSG00000236887 | ENSG0000023 | 1.748807712 | 0.160243423 | 1.331297529 | 0.120366349 | 0.904192947 | NA |
| ENSG00000236893 | ASS1P7 | 49.07532345 | 0.716074936 | 0.255120089 | 2.806815167 | 0.005003394 | 0.016116059 |
| ENSG00000236901 | MIR600HG | 221.7019216 | 0.665643858 | 0.129342493 | 5.1463664 | 2.66E-07 | 1.65E-06 |

| | | | | | | | |
|---|---|---|---|---|---|---|---|
| ENSG00000236914 | LINC01852 | 6.553428734 | -0.273953538 | 0.675102508 | -0.405795468 | 0.684892889 | 0.8214704 |
| ENSG00000236915 | CLCA4-AS1 | 2.188102157 | 2.361975915 | 1.417977997 | 1.665735237 | 0.09576616 | 0.204703291 |
| ENSG00000236924 | ENSG0000023 | 4.145874885 | 0.555664239 | 0.907304391 | 0.6124342 | 0.54025052 | 0.714561055 |
| ENSG00000236937 | PTGES3P4 | 41.67014892 | 0.135034088 | 0.278922493 | 0.48412764 | 0.628295286 | 0.780689734 |
| ENSG00000236942 | ENSG0000023 | 7.06687827 | 0.158824947 | 0.664775281 | 0.238915242 | 0.811171307 | 0.900037232 |
| ENSG00000236972 | FABP5P1 | 10.15972792 | -0.546106331 | 0.553529077 | -0.986590144 | 0.323843604 | 0.513973452 |
| ENSG00000236975 | LINC02814 | 15.78061795 | 0.066036053 | 0.455262138 | 0.145050615 | 0.884670909 | 0.940249752 |
| ENSG00000236976 | RPL6P20 | 41.98841919 | 0.072061951 | 0.311189544 | 0.231569321 | 0.816872538 | 0.903062617 |
| ENSG00000236986 | ENSG0000023 | 5.415748757 | -0.781773615 | 0.756108184 | -1.033944125 | 0.301162285 | 0.488003307 |
| ENSG00000236988 | CTDSPL2P1 | 1.93601592 | -1.116745736 | 1.357033349 | -0.822931681 | 0.410546845 | 0.603018609 |
| ENSG00000236991 | EDRF1-AS1 | 2.319481539 | -1.027030132 | 1.149064593 | -0.893796692 | 0.371430679 | 0.564017991 |
| ENSG00000236992 | RPL12P12 | 39.62172708 | 0.605931772 | 0.290447373 | 2.086201593 | 0.036960366 | 0.092843949 |
| ENSG00000236993 | GAPDHP21 | 9.107862334 | 0.020669128 | 0.585157079 | 0.035322358 | 0.971822695 | 0.987229325 |
| ENSG00000237004 | ZNRF2P1 | 85.27509171 | 0.131704687 | 0.19890669 | 0.662143071 | 0.507879534 | 0.688024202 |
| ENSG00000237008 | LAPTM4BP1 | 4.54926295 | 0.663444659 | 0.836140187 | 0.793461035 | 0.427509262 | 0.618217808 |
| ENSG00000237015 | ENSG0000023 | 4.275001863 | 0.245086538 | 0.84101203 | 0.291418588 | 0.7707312 | 0.875046557 |
| ENSG00000237017 | PRPF31-AS1 | 60.00059891 | -0.417958841 | 0.239809113 | -1.742881395 | 0.081354335 | 0.179087269 |
| ENSG00000237026 | ENSG0000023 | 3.006982383 | 0.475109443 | 1.084975788 | 0.43789866 | 0.661459747 | 0.804340663 |
| ENSG00000237036 | ZEB1-AS1 | 74.91273473 | -0.846612766 | 0.213169886 | -3.971540175 | 7.14E-05 | 0.000326451 |
| ENSG00000237037 | NDUFA6-DT | 34.17754216 | -0.43005974 | 0.300716734 | -1.43011576 | 0.152683798 | 0.295400718 |
| ENSG00000237039 | RPS28P4 | 3.555921056 | 0.588837047 | 0.961689332 | 0.612294456 | 0.540342957 | 0.714600913 |
| ENSG00000237054 | PRMT5-AS1 | 63.12470377 | -0.944896767 | 0.259989286 | -3.634368093 | 0.000278663 | 0.001153469 |
| ENSG00000237064 | EIF3IP1 | 6.821509011 | -0.257575392 | 0.677157177 | -0.380377556 | 0.703665172 | 0.834172862 |
| ENSG00000237077 | RPL38P2 | 3.690628082 | 0.734011666 | 0.927809973 | 0.791122846 | 0.428872308 | 0.61920798 |
| ENSG00000237080 | EHMT2-AS1 | 23.27208289 | -0.426775968 | 0.411554488 | -1.036985334 | 0.299742692 | 0.486458957 |
| ENSG00000237082 | COX5BP6 | 2.829185714 | -0.368396242 | 1.021986168 | -0.360470869 | 0.718495038 | 0.843432038 |
| ENSG00000237090 | MKRN8P | 2.546055683 | -0.429327712 | 1.114492395 | -0.385222649 | 0.700072465 | 0.831719009 |
| ENSG00000237094 | ENSG0000023 | 25.81795023 | 0.033877466 | 0.347720867 | 0.097427186 | 0.922387156 | 0.962279383 |
| ENSG00000237101 | ENSG0000023 | 6.721981101 | -1.751582476 | 0.739953518 | -2.367152035 | 0.017925567 | 0.0497234 |
| ENSG00000237118 | CYP2F2P | 3.814534718 | 0.152509304 | 0.895975476 | 0.170215935 | 0.86484032 | 0.929799912 |
| ENSG00000237125 | HAND2-AS1 | 27.81730685 | 0.762312423 | 0.340127029 | 2.241257994 | 0.025009368 | 0.066486836 |
| ENSG00000237126 | ENSG0000023 | 16.04795279 | -0.80152622 | 0.439489346 | -1.823767124 | 0.068187286 | 0.15531887 |
| ENSG00000237149 | ZNF503-AS2 | 59.35109424 | -0.452541723 | 0.23724786 | -1.907463876 | 0.056460545 | 0.132763802 |

| | | | | | | | |
|---|---|---|---|---|---|---|---|
| ENSG00000237161 | ENSG0000023 | 86.40931821 | -0.370346337 | 0.201127811 | -1.841348223 | 0.065570545 | 0.150313611 |
| ENSG00000237169 | RPL12P27 | 2.923291045 | -1.47004277 | 1.070784738 | -1.372864888 | 0.169794354 | 0.320412952 |
| ENSG00000237172 | B3GNT9 | 335.1575293 | 0.416780031 | 0.105303689 | 3.957886322 | 7.56E-05 | 0.000344514 |
| ENSG00000237174 | ENSG0000023 | 36.0474641 | -0.502773107 | 0.300601333 | -1.672557806 | 0.094414379 | 0.202280715 |
| ENSG00000237176 | ENSG0000023 | 25.20780886 | 0.520535437 | 0.361939981 | 1.438181647 | 0.150382521 | 0.292203094 |
| ENSG00000237187 | NR2F1-AS1 | 147.3580488 | 1.674830755 | 0.177902567 | 9.414314736 | 4.76E-21 | 8.45E-20 |
| ENSG00000237188 | ENSG0000023 | 2.034372563 | 1.055920969 | 1.293747227 | 0.816172547 | 0.414401456 | 0.606520741 |
| ENSG00000237190 | CDKN2AIPNL | 723.7609327 | 0.138572206 | 0.078857625 | 1.757245493 | 0.078875974 | 0.174755152 |
| ENSG00000237200 | ZBTB40-IT1 | 3.287779865 | -0.189656851 | 0.958078729 | -0.197955393 | 0.843079964 | 0.917338822 |
| ENSG00000237206 | IMPDH1P4 | 6.424189224 | -0.151131375 | 0.700199698 | -0.215840388 | 0.829112165 | 0.909561143 |
| ENSG00000237208 | ENSG0000023 | 3.806050222 | 0.99149971 | 0.934440989 | 1.061061878 | 0.288661781 | 0.472631666 |
| ENSG00000237211 | SETP4 | 2.029183066 | -2.49640289 | 1.460774576 | -1.708958337 | 0.087458667 | 0.189990703 |
| ENSG00000237214 | ENSG0000023 | 87.90601046 | 0.276316545 | 0.21780203 | 1.268659183 | 0.204562646 | 0.369443167 |
| ENSG00000237224 | ENSG0000023 | 2.573722793 | -1.703189984 | 1.183484442 | -1.439131706 | 0.15011321 | 0.291756083 |
| ENSG00000237232 | ZNF295-AS1 | 4.20626043 | -1.291475893 | 0.908067349 | -1.422224788 | 0.154961005 | 0.29847233 |
| ENSG00000237248 | LINC00987 | 16.35239429 | -0.788928335 | 0.502372217 | -1.570405984 | 0.116320691 | 0.239661995 |
| ENSG00000237263 | MAPK6P3 | 19.90537693 | -1.92282138 | 0.444165903 | -4.329061202 | 1.50E-05 | 7.54E-05 |
| ENSG00000237264 | FTH1P11 | 583.1638245 | 0.259893878 | 0.093931344 | 2.766849338 | 0.005660091 | 0.017961333 |
| ENSG00000237276 | ANO7L1 | 18.42272692 | 0.548043751 | 0.427672063 | 1.281457917 | 0.200032872 | 0.363149264 |
| ENSG00000237285 | HNRNPA1P2 | 3.162000236 | 0.444485897 | 1.01733957 | 0.436910064 | 0.662176573 | 0.804828366 |
| ENSG00000237289 | CKMT1B | 587.1798818 | -0.299092873 | 0.085028449 | -3.517562386 | 0.00043553 | 0.001742424 |
| ENSG00000237294 | EIF4BP9 | 4.428707342 | 0.36575501 | 0.8743158 | 0.418332838 | 0.675703784 | 0.814879865 |
| ENSG00000237296 | SMG1P1 | 25.1044605 | -0.218412815 | 0.348075428 | -0.627487027 | 0.530340037 | 0.705971128 |
| ENSG00000237298 | TTN-AS1 | 269.8945967 | -0.07072298 | 0.115437966 | -0.612649226 | 0.540108302 | 0.714414141 |
| ENSG00000237301 | ENSG0000023 | 10.84289194 | 0.124027169 | 0.547190284 | 0.226661862 | 0.820686685 | 0.904905193 |
| ENSG00000237307 | SRRM1P3 | 9.703168741 | -0.484320943 | 0.566978513 | -0.854213927 | 0.392986468 | 0.585963435 |
| ENSG00000237310 | LINC03011 | 27.04855973 | 0.408748695 | 0.346494829 | 1.17966752 | 0.238132477 | 0.413526941 |
| ENSG00000237330 | RNF223 | 248.9759033 | -0.02565758 | 0.121002535 | -0.212041668 | 0.832074527 | 0.911242166 |
| ENSG00000237343 | ENSG0000023 | 1.911993408 | 0.782448356 | 1.329977509 | 0.588316984 | 0.556319548 | 0.72693858 |
| ENSG00000237350 | CDC42P6 | 705.1823115 | 0.008809853 | 0.075959337 | 0.115981173 | 0.907667463 | 0.953688985 |
| ENSG00000237356 | ENSG0000023 | 33.57822398 | -0.50102478 | 0.303656623 | -1.649971519 | 0.098948761 | 0.210193961 |
| ENSG00000237357 | FAM88E | 1.717545028 | -0.350475688 | 1.367722169 | -0.25624772 | 0.797759563 | NA |
| ENSG00000237361 | TUSC8 | 8.755207609 | 0.237334258 | 0.613068497 | 0.38712519 | 0.69866353 | 0.830862695 |

| | | | | | | | |
|---|---|---|---|---|---|---|---|
| ENSG00000237371 | ENSG0000023 | 5.448018304 | 0.243039326 | 0.743972244 | 0.326677948 | 0.743911481 | 0.858490754 |
| ENSG00000237390 | ENSG0000023 | 2.377771146 | 0.584495631 | 1.173485181 | 0.498085225 | 0.618423974 | 0.773746712 |
| ENSG00000237399 | PITRM1-AS1 | 38.88780823 | -1.465673775 | 0.31418761 | -4.664963637 | 3.09E-06 | 1.69E-05 |
| ENSG00000237406 | NDUFA9P1 | 1.780568257 | 1.978525724 | 1.479315057 | 1.337460681 | 0.181072308 | NA |
| ENSG00000237415 | NRBF2P3 | 2.453314345 | -0.362012656 | 1.113239135 | -0.325188582 | 0.745038349 | 0.859112282 |
| ENSG00000237417 | XRCC6P1 | 14.87926839 | 0.053773382 | 0.475555026 | 0.113074995 | 0.909971099 | 0.95484114 |
| ENSG00000237418 | ENSG0000023 | 2.769173951 | -1.813455292 | 1.147693881 | -1.580086225 | 0.114087121 | 0.235843792 |
| ENSG00000237423 | LINC01522 | 33.834013 | 3.45608788 | 0.442058114 | 7.818175409 | 5.36E-15 | 6.44E-14 |
| ENSG00000237424 | FOXD2-AS1 | 56.56425908 | -0.35075363 | 0.245712691 | -1.427494968 | 0.153437275 | 0.296408512 |
| ENSG00000237433 | RPSAP11 | 7.193898219 | -0.712080101 | 0.670581252 | -1.061884893 | 0.288287944 | 0.472312915 |
| ENSG00000237436 | CAMTA1-DT | 8.955542804 | -0.228993903 | 0.595249007 | -0.384702705 | 0.700457691 | 0.83200432 |
| ENSG00000237437 | ASS1P12 | 1.926194599 | 1.354713547 | 1.349727248 | 1.003694301 | 0.315525985 | 0.504207396 |
| ENSG00000237438 | CECR7 | 2.241951306 | 0.287407274 | 1.205323545 | 0.238448237 | 0.811533459 | 0.900221351 |
| ENSG00000237440 | ZNF737 | 248.2973826 | 1.258388401 | 0.126845315 | 9.920653354 | 3.39E-23 | 6.81E-22 |
| ENSG00000237441 | RGL2 | 6428.45506 | 0.214260688 | 0.053124888 | 4.033150826 | 5.50E-05 | 0.000256029 |
| ENSG00000237447 | CDC27P2 | 1.734169573 | 0.117890299 | 1.332631045 | 0.08846432 | 0.929507642 | NA |
| ENSG00000237451 | CDK2AP2P2 | 3.957683928 | 0.425887396 | 0.882137591 | 0.482790213 | 0.6292447 | 0.7810577 |
| ENSG00000237456 | ID2B | 5.078787772 | -1.188998171 | 0.843368531 | -1.409820414 | 0.158592717 | 0.30383472 |
| ENSG00000237457 | BRINP3-DT | 7.447996741 | -1.58392634 | 0.689870955 | -2.295974819 | 0.021677321 | 0.058655078 |
| ENSG00000237460 | LINC01661 | 83.33648846 | 0.671698367 | 0.206696664 | 3.249681705 | 0.001155342 | 0.00428029 |
| ENSG00000237461 | ENSG0000023 | 5.998825087 | 0.886173409 | 0.727790382 | 1.217621765 | 0.223367739 | 0.394643865 |
| ENSG00000237476 | LINC01637 | 19.70847852 | -0.652128307 | 0.40019948 | -1.629508133 | 0.103205495 | 0.217491195 |
| ENSG00000237481 | ENSG0000023 | 1.791886214 | 0.644603274 | 1.307831445 | 0.492879473 | 0.622097752 | 0.776353086 |
| ENSG00000237489 | C10orf143 | 112.2304099 | -0.641399848 | 0.173688092 | -3.692825691 | 0.000221776 | 0.000933323 |
| ENSG00000237491 | LINC01409 | 17.33034027 | 0.292620467 | 0.43631883 | 0.670657434 | 0.502438785 | 0.683507467 |
| ENSG00000237493 | ENSG0000023 | 661.6887426 | 0.330986995 | 0.080626324 | 4.105197661 | 4.04E-05 | 0.000191271 |
| ENSG00000237499 | WAKMAR2 | 5.818167791 | 0.47695716 | 0.717371941 | 0.664867321 | 0.506135359 | 0.686620055 |
| ENSG00000237505 | PKN2-AS1 | 25.12286269 | 0.513610028 | 0.352682399 | 1.456296173 | 0.145310763 | 0.284495085 |
| ENSG00000237506 | RPSAP15 | 273.5489323 | 0.01411913 | 0.11111699 | 0.127065451 | 0.898888596 | 0.948941758 |
| ENSG00000237514 | PTP4A1P7 | 3.113754843 | -0.920787783 | 1.017951988 | -0.904549324 | 0.365704192 | 0.558395881 |
| ENSG00000237515 | SHISA9 | 370.3133679 | 0.025427459 | 0.098932954 | 0.257017078 | 0.79716559 | 0.89183367 |
| ENSG00000237517 | CELSR1P1 | 35.3689657 | 0.764564326 | 0.307983382 | 2.482485645 | 0.013046932 | 0.037557615 |
| ENSG00000237522 | NONOP2 | 6.684826719 | -0.324491723 | 0.674198899 | -0.481299693 | 0.630303515 | 0.781914484 |

| | | | | | | | |
|---|---|---|---|---|---|---|---|
| ENSG00000237527 | ENSG0000023 | 22.77158468 | 4.345397388 | 0.663442008 | 6.549777274 | 5.76E-11 | 5.08E-10 |
| ENSG00000237529 | ENSG0000023 | 2.687844906 | -0.203245451 | 1.047788923 | -0.193975568 | 0.846195013 | 0.919463508 |
| ENSG00000237540 | RPL36AP36 | 2.966338702 | 0.816504539 | 1.031189302 | 0.791808583 | 0.428472297 | 0.619042354 |
| ENSG00000237550 | ENSG0000023 | 3674.768976 | 0.157807246 | 0.069755668 | 2.262285643 | 0.023679761 | 0.063385498 |
| ENSG00000237574 | LINC01856 | 5.13258961 | -3.949333165 | 1.204377498 | -3.279148916 | 0.001041207 | 0.003884417 |
| ENSG00000237575 | PYY2 | 20.13101172 | 0.171227112 | 0.386038863 | 0.443548897 | 0.657368759 | 0.80147779 |
| ENSG00000237584 | RAC1P4 | 4.242883154 | -1.027604104 | 0.875975701 | -1.173096586 | 0.240757075 | 0.416639055 |
| ENSG00000237588 | ENSG0000023 | 1.655267886 | 0.409629743 | 1.349124378 | 0.303626374 | 0.761412556 | NA |
| ENSG00000237594 | TIAM1-AS1 | 31.37034316 | -0.497347179 | 0.319664171 | -1.555842741 | 0.119745492 | 0.245175196 |
| ENSG00000237596 | PDE7B-AS1 | 1.65431856 | 0.357195993 | 1.380281624 | 0.258784864 | 0.795801237 | NA |
| ENSG00000237605 | DYRK3-AS1 | 4.728394274 | 1.554990103 | 0.860502949 | 1.807071207 | 0.070751177 | 0.160030764 |
| ENSG00000237614 | ENSG0000023 | 17.42972921 | 1.717702859 | 0.460856648 | 3.727195576 | 0.000193622 | 0.000824062 |
| ENSG00000237624 | OXCT2P1 | 1.811725464 | -1.550213987 | 1.494152701 | -1.037520453 | 0.299493369 | 0.48618701 |
| ENSG00000237629 | UQCRHP2 | 4.024412262 | -0.677060808 | 0.861724453 | -0.785704532 | 0.432040608 | 0.62208207 |
| ENSG00000237632 | S100A11P1 | 9.20747003 | -1.1280746 | 0.612162754 | -1.842769086 | 0.065362732 | 0.149912125 |
| ENSG00000237638 | LINC02245 | 25.41696575 | 1.200418397 | 0.37628965 | 3.190144609 | 0.001422016 | 0.005155848 |
| ENSG00000237649 | KIFC1 | 2219.695366 | -0.280859496 | 0.054061395 | -5.19519517 | 2.05E-07 | 1.29E-06 |
| ENSG00000237651 | C2orf74 | 8.707335525 | -0.233109088 | 0.587641205 | -0.396686083 | 0.691598964 | 0.826101549 |
| ENSG00000237661 | TNIP2P1 | 3.607844387 | -0.151942724 | 0.914455994 | -0.166156408 | 0.868033864 | 0.931081965 |
| ENSG00000237668 | RPS15AP38 | 191.7045088 | 0.237253339 | 0.139620784 | 1.699269495 | 0.089268418 | 0.193117455 |
| ENSG00000237669 | ENSG0000023 | 10.68469177 | -0.088622176 | 0.543989863 | -0.162911447 | 0.870588158 | 0.932942429 |
| ENSG00000237676 | RPL30P4 | 37.25428091 | 0.145512796 | 0.31470742 | 0.462374851 | 0.643812531 | 0.791270263 |
| ENSG00000237682 | PRKCIP1 | 140.8140988 | -0.260587024 | 0.151463324 | -1.720462857 | 0.08534834 | 0.186245998 |
| ENSG00000237686 | SCIRT | 2.842527504 | -0.969694478 | 1.104484845 | -0.877960872 | 0.379964953 | 0.572424996 |
| ENSG00000237701 | ATP5PFP1 | 4.775607064 | -0.857447589 | 0.827715842 | -1.035920234 | 0.300239357 | 0.486886106 |
| ENSG00000237705 | ENSG0000023 | 5.10823044 | 0.7564855 | 0.852502217 | 0.887370713 | 0.374879346 | 0.567714035 |
| ENSG00000237708 | SPIN2P1 | 3.634192867 | -2.299918297 | 1.059001969 | -2.171779057 | 0.029872332 | 0.077388063 |
| ENSG00000237719 | ENSG0000023 | 9.161726303 | -0.742805297 | 0.611940185 | -1.213852783 | 0.224803946 | 0.396600978 |
| ENSG00000237729 | RPS27P17 | 31.18940223 | 0.478079398 | 0.325845523 | 1.467196461 | 0.142322619 | 0.280054974 |
| ENSG00000237730 | RPS5P8 | 9.869150324 | 1.069928557 | 0.584140995 | 1.831627239 | 0.067006976 | 0.1531321 |
| ENSG00000237732 | CT75 | 11.94286254 | 1.581765008 | 0.545715622 | 2.898515168 | 0.003749342 | 0.012434567 |
| ENSG00000237737 | DCTN1-AS1 | 3.773758068 | -1.32833184 | 1.02117306 | -1.300790132 | 0.193330301 | 0.353992189 |
| ENSG00000237745 | RPL22P8 | 4.848018752 | 0.618755102 | 0.793703821 | 0.779579342 | 0.435638519 | 0.625535029 |

| | | | | | | | |
|---|---|---|---|---|---|---|---|
| ENSG00000237748 | UQCRBP1 | 46.25096988 | -0.482521393 | 0.291347124 | -1.656173523 | 0.097686683 | 0.207975176 |
| ENSG00000237749 | RPS27P9 | 37.41001927 | 0.557069407 | 0.307193899 | 1.81341299 | 0.069768147 | 0.158181588 |
| ENSG00000237751 | LINC01143 | 8.628019171 | 0.144240632 | 0.602277831 | 0.23949185 | 0.810724216 | 0.899758757 |
| ENSG00000237753 | SLC20A1-DT | 49.883595 | -0.442136678 | 0.256114224 | -1.726326134 | 0.084288756 | 0.184391786 |
| ENSG00000237758 | BANF1P3 | 441.883348 | -0.202491324 | 0.092541946 | -2.188103147 | 0.02866209 | 0.074658792 |
| ENSG00000237765 | FAM200B | 378.7883017 | 0.321958275 | 0.100234309 | 3.212056624 | 0.001317884 | 0.00482404 |
| ENSG00000237766 | GGTA2P | 3.945921562 | -0.194646468 | 0.868892853 | -0.224016652 | 0.822744346 | 0.906215744 |
| ENSG00000237772 | LINC02631 | 6.430287466 | -0.82348481 | 0.727166061 | -1.132457707 | 0.257442056 | 0.437136992 |
| ENSG00000237773 | ENSG0000023 | 11.85243334 | -0.737252228 | 0.529078515 | -1.39346469 | 0.163479316 | 0.311117928 |
| ENSG00000237774 | ENSG0000023 | 6.48527887 | 0.128912869 | 0.705318739 | 0.182772499 | 0.854976527 | 0.924669348 |
| ENSG00000237781 | ADAMTSL4-AS | 6.879561533 | -0.041071064 | 0.654835085 | -0.062719705 | 0.949989706 | 0.976903526 |
| ENSG00000237786 | GFOD1-AS1 | 3.045794753 | -0.541893001 | 1.022127146 | -0.530162028 | 0.595999596 | 0.756993626 |
| ENSG00000237788 | ENSG0000023 | 11.33422985 | -0.314566341 | 0.522886659 | -0.601595653 | 0.547443323 | 0.720254063 |
| ENSG00000237793 | RPL18AP16 | 16.94586562 | 0.211386108 | 0.43277214 | 0.488446663 | 0.625233497 | 0.778594473 |
| ENSG00000237801 | AMD1P2 | 3.275014721 | -0.196364748 | 0.980065118 | -0.200358879 | 0.841199917 | 0.916625749 |
| ENSG00000237801 | AMD1P2 | 3.275014721 | -0.196364748 | 0.980065118 | -0.200358879 | 0.841199917 | 0.916625749 |
| ENSG00000237804 | HNRNPA1P39 | 7.553863336 | 0.202662146 | 0.651347402 | 0.311142941 | 0.755691958 | 0.865149344 |
| ENSG00000237807 | LINC02984 | 292.3243971 | -0.866271825 | 0.116692141 | -7.423566108 | 1.14E-13 | 1.24E-12 |
| ENSG00000237810 | ENSG0000023 | 3.396934322 | -0.585870898 | 0.952954075 | -0.614794473 | 0.53869046 | 0.71340256 |
| ENSG00000237813 | ENSG0000023 | 2.144114806 | 0.072668849 | 1.192378547 | 0.060944445 | 0.951403453 | 0.977495703 |
| ENSG00000237821 | ST13P7 | 174.5186994 | 0.197607777 | 0.139543888 | 1.416097688 | 0.156746905 | 0.301077953 |
| ENSG00000237828 | PA2G4P1 | 69.81510925 | 0.263038001 | 0.210132663 | 1.251771129 | 0.210653273 | 0.3779731 |
| ENSG00000237836 | PHKA2-AS1 | 26.20933139 | -0.320796068 | 0.339950444 | -0.943655388 | 0.345345778 | 0.536243449 |
| ENSG00000237840 | FAM21FP | 12.26890232 | -1.039985111 | 0.504664232 | -2.060746622 | 0.039327219 | 0.097827631 |
| ENSG00000237842 | SMU1P1 | 73.74362312 | -0.170977149 | 0.204427052 | -0.836372425 | 0.402945421 | 0.595590718 |
| ENSG00000237854 | LINC00674 | 598.2889084 | 0.013709452 | 0.085891109 | 0.159614329 | 0.873184891 | 0.934604743 |
| ENSG00000237861 | ENSG0000023 | 2.622107884 | -0.193354843 | 1.077225864 | -0.179493317 | 0.857550365 | 0.926087485 |
| ENSG00000237863 | IRS4-AS1 | 25.61438883 | 1.419617743 | 0.374868457 | 3.786975717 | 0.000152492 | 0.00066053 |
| ENSG00000237864 | LINC00322 | 7.196860865 | -1.048819576 | 0.671731966 | -1.561366184 | 0.118437376 | 0.243084316 |
| ENSG00000237868 | ENSG0000023 | 31.58235226 | -0.864469847 | 0.325100004 | -2.659089009 | 0.007835226 | 0.024010164 |
| ENSG00000237877 | LINC01473 | 7.518364892 | 0.320863212 | 0.629303998 | 0.509869973 | 0.61014256 | 0.76816148 |
| ENSG00000237882 | PPIAP13 | 6.277325938 | 0.320998637 | 0.802450276 | 0.400023088 | 0.689139512 | 0.82436022 |
| ENSG00000237883 | DGUOK-AS1 | 50.41054323 | 0.225505927 | 0.266260212 | 0.846938134 | 0.397029606 | 0.589887251 |

| | | | | | | | |
|---|---|---|---|---|---|---|---|
| ENSG00000237886 | NALT1 | 28.70859799 | 1.446858988 | 0.348600666 | 4.150476835 | 3.32E-05 | 0.000159462 |
| ENSG00000237888 | RPL29P31 | 1.995643862 | -0.679235796 | 1.218121509 | -0.557609229 | 0.577111253 | 0.743117322 |
| ENSG00000237892 | KLF7-IT1 | 1.73318194 | 0.178188755 | 1.356101334 | 0.131397817 | 0.895460616 | NA |
| ENSG00000237897 | MATR3P3 | 10.85921402 | 0.427060782 | 0.529994026 | 0.805784143 | 0.420367323 | 0.611649401 |
| ENSG00000237899 | ENSG0000023 | 6.96284982 | -0.323126995 | 0.668228161 | -0.483557883 | 0.628699671 | 0.780938405 |
| ENSG00000237911 | SRP68P3 | 2.716182369 | 0.744483495 | 1.105337401 | 0.673535062 | 0.50060695 | 0.681732072 |
| ENSG00000237922 | ENSG0000023 | 4.18753737 | -0.790283465 | 0.861635004 | -0.917190529 | 0.359042809 | 0.551633651 |
| ENSG00000237927 | ENSG0000023 | 2.942670267 | 0.470959131 | 1.006347821 | 0.467988425 | 0.639792868 | 0.787976398 |
| ENSG00000237931 | CLIC4P3 | 24.31062793 | 0.150835425 | 0.357291766 | 0.422163171 | 0.672905921 | 0.812787586 |
| ENSG00000237938 | ENSG0000023 | 1.921388923 | -0.631107342 | 1.297759514 | -0.486305309 | 0.626750715 | 0.779953424 |
| ENSG00000237939 | IDI1P1 | 4.145906055 | -0.532964245 | 0.880791372 | -0.605097032 | 0.545114526 | 0.718352374 |
| ENSG00000237940 | LINC01238 | 19.09592347 | -0.918799448 | 0.436689072 | -2.104012916 | 0.03537732 | 0.089441736 |
| ENSG00000237945 | LINC00649 | 170.7321115 | 0.024630426 | 0.141652059 | 0.173879762 | 0.861959958 | 0.928215087 |
| ENSG00000237950 | LINC02918 | 40.64968018 | 0.320011497 | 0.284649326 | 1.124230651 | 0.26091519 | 0.441169312 |
| ENSG00000237972 | TUBG1P | 2.461357008 | 0.036282063 | 1.110048691 | 0.032685109 | 0.973925699 | 0.988106737 |
| ENSG00000237973 | MTCO1P12 | 759.4452519 | -0.078724803 | 0.084864479 | -0.927653178 | 0.353587504 | 0.545083738 |
| ENSG00000237976 | RFX5-AS1 | 32.64995706 | -0.18575014 | 0.311893155 | -0.595556961 | 0.551471236 | 0.72331333 |
| ENSG00000237977 | EIF4HP2 | 6.602551723 | -0.025150801 | 0.690707769 | -0.036413085 | 0.970952981 | 0.98680905 |
| ENSG00000237978 | KCNMB2-AS1 | 142.6192496 | 0.137000112 | 0.153752749 | 0.891041708 | 0.372906795 | 0.565697783 |
| ENSG00000237984 | PTENP1 | 421.6642174 | -0.320350616 | 0.094069245 | -3.405476655 | 0.000660487 | 0.002558521 |
| ENSG00000237991 | RPL35P1 | 41.53282045 | 0.122282336 | 0.282331389 | 0.433116335 | 0.664930256 | 0.806706252 |
| ENSG00000238000 | PSME2P1 | 205.1962691 | -0.111206626 | 0.126994123 | -0.875683245 | 0.381202257 | 0.573432466 |
| ENSG00000238003 | RPL10P4 | 1.666160036 | 0.387960393 | 1.393481429 | 0.278410882 | 0.780696967 | NA |
| ENSG00000238005 | LNCATV | 15.50838002 | 1.107131666 | 0.47467116 | 2.332418229 | 0.0196787 | 0.053940659 |
| ENSG00000238009 | ENSG0000023 | 3.991936425 | -0.152151099 | 0.908206669 | -0.167529158 | 0.866953707 | 0.930846618 |
| ENSG00000238018 | SPTBN1-AS2 | 8.128600431 | -0.83077908 | 0.615095196 | -1.350651226 | 0.176807183 | 0.330558825 |
| ENSG00000238021 | ODAD2P1 | 2.923291045 | -1.47004277 | 1.070784738 | -1.372864888 | 0.169794354 | 0.320412952 |
| ENSG00000238029 | RALBP1P2 | 3.659899733 | 1.055252736 | 1.000120115 | 1.055125999 | 0.291367703 | 0.47633802 |
| ENSG00000238039 | ENSG0000023 | 20.05780044 | 0.463882172 | 0.39336652 | 1.179261957 | 0.238293882 | 0.413718433 |
| ENSG00000238041 | TMEM230P2 | 14.26950874 | 0.515602718 | 0.465883526 | 1.106720218 | 0.268414911 | 0.449874922 |
| ENSG00000238045 | MVP-DT | 386.6643201 | -0.400621515 | 0.271091152 | -1.477811105 | 0.139458345 | 0.275629863 |
| ENSG00000238058 | CAMSAP1-DT | 119.2352188 | 0.161328765 | 0.169414521 | 0.952272357 | 0.340958873 | 0.532081398 |
| ENSG00000238062 | SPATA3-AS1 | 6.910168055 | -0.98935112 | 0.699009183 | -1.41536212 | 0.15696235 | 0.301390835 |

| | | | | | | | |
|---|---|---|---|---|---|---|---|
| ENSG00000238072 | CDC26P1 | 191.7701522 | -0.536812246 | 0.138354957 | -3.879963965 | 0.000104472 | 0.000464275 |
| ENSG00000238082 | NUDCP2 | 34.90278873 | 0.066613807 | 0.295737383 | 0.225246487 | 0.821787527 | 0.905640175 |
| ENSG00000238083 | LRRC37A2 | 68.19549591 | -0.55122 | 0.219873423 | -2.506987849 | 0.01217649 | 0.035384935 |
| ENSG00000238084 | ENSG0000023 | 49.60129306 | 0.169727067 | 0.250038384 | 0.678804049 | 0.497262028 | 0.679080303 |
| ENSG00000238085 | ENSG0000023 | 4.526619575 | 0.336347959 | 0.838327536 | 0.401213063 | 0.688263264 | 0.823819752 |
| ENSG00000238098 | ABCA17P | 9.455820213 | -0.421903465 | 0.584298626 | -0.722068214 | 0.47025254 | 0.656115263 |
| ENSG00000238099 | LINC01625 | 1.621559241 | -2.112222108 | 1.56382318 | -1.350678347 | 0.176798491 | NA |
| ENSG00000238103 | RPL9P7 | 3.930382037 | 0.288107459 | 0.908913539 | 0.316980049 | 0.751258741 | 0.862469998 |
| ENSG00000238105 | GOLGA2P5 | 710.7804151 | -0.164315539 | 0.086644802 | -1.896426965 | 0.057903609 | 0.135500234 |
| ENSG00000238110 | ENSG0000023 | 33.68942659 | 0.133924384 | 0.301927964 | 0.443564028 | 0.657357818 | 0.80147779 |
| ENSG00000238113 | LINC01410 | 28.51545323 | -1.500668855 | 0.345731139 | -4.340566079 | 1.42E-05 | 7.18E-05 |
| ENSG00000238116 | RAD21P1 | 25.03145066 | 0.888976422 | 0.371616716 | 2.392186311 | 0.016748338 | 0.046865928 |
| ENSG00000238117 | ENSG0000023 | 2.719107589 | 0.143202711 | 1.064928664 | 0.134471647 | 0.89302963 | 0.94527961 |
| ENSG00000238123 | MID1IP1-AS1 | 11.02273582 | -0.605684204 | 0.537165205 | -1.127556658 | 0.2595072 | 0.439484833 |
| ENSG00000238132 | GOLM2P1 | 15.12050814 | -0.099223381 | 0.464980325 | -0.213392643 | 0.831020718 | 0.910609757 |
| ENSG00000238133 | MAP3K20-AS1 | 4.530584827 | -1.224236627 | 0.86458676 | -1.41597892 | 0.156781676 | 0.30109432 |
| ENSG00000238141 | BRWD1-AS1 | 4.335088732 | 1.319151447 | 0.874250131 | 1.508894767 | 0.131325677 | 0.263524409 |
| ENSG00000238150 | ENSG0000023 | 17.5793018 | 0.103148381 | 0.431455048 | 0.23907098 | 0.811050544 | 0.8999903 |
| ENSG00000238165 | ENSG0000023 | 29.81835528 | -0.066635351 | 0.323928591 | -0.205710003 | 0.837017445 | 0.914124265 |
| ENSG00000238172 | RPS2P35 | 4.670821011 | 0.205226707 | 0.828678185 | 0.247655497 | 0.804400968 | 0.895992144 |
| ENSG00000238173 | RPL39P6 | 3.060919612 | -0.236014025 | 1.01416949 | -0.23271655 | 0.815981517 | 0.902406402 |
| ENSG00000238184 | CD81-AS1 | 10.88082929 | 0.876169782 | 0.548546373 | 1.597257452 | 0.110208331 | 0.229105493 |
| ENSG00000238193 | HADHBP1 | 7.057113027 | -0.517020489 | 0.688863834 | -0.75054091 | 0.452928995 | 0.640609 |
| ENSG00000238197 | PAXBP1-AS1 | 28.40180758 | 0.180780435 | 0.33212855 | 0.544308628 | 0.586229111 | 0.750100097 |
| ENSG00000238198 | LRIG2-DT | 1.820277129 | 0.638516441 | 1.310597355 | 0.487194972 | 0.626120168 | 0.779457961 |
| ENSG00000238199 | UBE2V2P3 | 2.443371521 | -0.302138248 | 1.230550253 | -0.245531011 | 0.806045299 | 0.897100998 |
| ENSG00000238207 | ENSG0000023 | 2.869381307 | -0.745610315 | 1.064688508 | -0.700308409 | 0.483734722 | 0.667723994 |
| ENSG00000238213 | TARDBPP2 | 5.863462224 | -0.31330987 | 0.728458408 | -0.430099874 | 0.667122992 | 0.808527125 |
| ENSG00000238221 | ENSG0000023 | 6.695753089 | -1.546154721 | 0.7411245 | -2.086228052 | 0.03695797 | 0.092843949 |
| ENSG00000238222 | MKRN4P | 6.247067541 | -0.442083841 | 0.697464537 | -0.633844186 | 0.526182513 | 0.702188881 |
| ENSG00000238227 | TMEM250 | 1323.965541 | 0.115914658 | 0.068978145 | 1.680454858 | 0.09286885 | 0.199547924 |
| ENSG00000238228 | OR7E7P | 6.768948748 | -0.260742732 | 0.662940287 | -0.393312545 | 0.694088653 | 0.827653275 |
| ENSG00000238245 | MYO5BP2 | 2.435389772 | 0.375542867 | 1.187418984 | 0.316268202 | 0.751798944 | 0.862831088 |

| | | | | | | | |
|---|---|---|---|---|---|---|---|
| ENSG00000238247 | RDXP3 | 9.847870941 | 0.608583462 | 0.582176682 | 1.045358705 | 0.295857215 | 0.481960269 |
| ENSG00000238249 | HMGN2P17 | 13.44726215 | -0.619374898 | 0.505528289 | -1.22520324 | 0.220498654 | 0.390748484 |
| ENSG00000238251 | NSA2P7 | 36.99571115 | 0.336710851 | 0.294170115 | 1.144612706 | 0.252369627 | 0.431076637 |
| ENSG00000238258 | ENSG0000023 | 9.482723381 | -1.499216528 | 0.614742944 | -2.438769801 | 0.014737353 | 0.041825421 |
| ENSG00000238260 | ENSG0000023 | 4.068296713 | 0.783682554 | 0.898698216 | 0.872019706 | 0.383197627 | 0.575627796 |
| ENSG00000238278 | ALG1L6P | 1.642491601 | -0.196061353 | 1.341263149 | -0.146176649 | 0.883781939 | NA |
| ENSG00000238280 | ENSG0000023 | 4.959642878 | -1.147824848 | 0.86801093 | -1.322362206 | 0.186047568 | 0.343412553 |
| ENSG00000238286 | SLC35E1P1 | 10.53456629 | 0.065355726 | 0.570061821 | 0.114646734 | 0.908725136 | 0.95431011 |
| ENSG00000238287 | EXO5-DT | 18.43241353 | 0.339453529 | 0.41584463 | 0.816298936 | 0.414329183 | 0.606477249 |
| ENSG00000238317 | SNORD11 | 2.577243262 | -0.421722418 | 1.112613572 | -0.379037636 | 0.704659916 | 0.834877516 |
| ENSG00000238363 | SNORA13 | 5.446540272 | 0.907292479 | 0.765163626 | 1.185749621 | 0.235721192 | 0.410712756 |
| ENSG00000238365 | RNU7-57P | 2.71151334 | 0.157463571 | 1.075842058 | 0.146363093 | 0.883634761 | 0.939712848 |
| ENSG00000238390 | ENSG0000023 | 7.724997988 | -0.446005533 | 0.627331181 | -0.710957062 | 0.477110844 | 0.661968041 |
| ENSG00000238578 | SNORD4A | 13.3300757 | 0.325170171 | 0.505907851 | 0.642745848 | 0.520389026 | 0.69762297 |
| ENSG00000238832 | ENSG0000023 | 2.821688852 | 0.678204067 | 1.061108495 | 0.63914677 | 0.522727456 | 0.699409032 |
| ENSG00000238961 | SNORA47 | 2.285029808 | 0.669675115 | 1.166804726 | 0.573939323 | 0.566008862 | 0.73394626 |
| ENSG00000239122 | RNU2-11P | 1.825497472 | 1.175637575 | 1.363099647 | 0.862473684 | 0.388426908 | 0.581170967 |
| ENSG00000239198 | RPL5P22 | 4.766241293 | 1.293716455 | 0.839982227 | 1.540171224 | 0.123518622 | 0.251042544 |
| ENSG00000239213 | NCK1-DT | 137.9688305 | -0.635263546 | 0.152897768 | -4.154825512 | 3.26E-05 | 0.000156788 |
| ENSG00000239218 | RPS20P22 | 6.017044658 | -0.707577335 | 0.725914811 | -0.974738804 | 0.329689839 | 0.520121375 |
| ENSG00000239223 | RPL34P31 | 9.969176145 | 0.555059525 | 0.59236201 | 0.937027552 | 0.348744386 | 0.539945204 |
| ENSG00000239246 | ENSG0000023 | 139.8411074 | 0.168666284 | 0.151066172 | 1.116505981 | 0.264205612 | 0.444900296 |
| ENSG00000239264 | TXNDC5 | 19.30298264 | -0.815191803 | 0.433760578 | -1.879358903 | 0.060195504 | 0.139921457 |
| ENSG00000239268 | LINC03051 | 5.958913351 | -0.391732671 | 0.722529973 | -0.542168057 | 0.587702738 | 0.751132537 |
| ENSG00000239269 | RPSAP4 | 5.142180306 | 0.271217521 | 0.787377444 | 0.344456808 | 0.730502773 | 0.850819131 |
| ENSG00000239272 | RPL21P10 | 4.556175704 | -0.793284901 | 0.827428556 | -0.958735224 | 0.337692149 | 0.528890414 |
| ENSG00000239282 | CASTOR1 | 6.753702794 | 0.617729435 | 0.711122729 | 0.868667825 | 0.385028845 | 0.577659856 |
| ENSG00000239300 | ENSG0000023 | 2.301982773 | -1.041968462 | 1.18423075 | -0.879869453 | 0.378930034 | 0.571483594 |
| ENSG00000239305 | RNF103 | 750.8700527 | 0.047497818 | 0.077788104 | 0.610605155 | 0.541461009 | 0.715543266 |
| ENSG00000239306 | RBM14 | 1981.315887 | -0.217904002 | 0.062099279 | -3.508961864 | 0.000449859 | 0.001794743 |
| ENSG00000239317 | RPL35AP17 | 2.259564112 | -0.055187265 | 1.181512346 | -0.046709004 | 0.962745154 | 0.982889182 |
| ENSG00000239332 | LINC01119 | 4.027216387 | 0.646715407 | 0.918274779 | 0.704272209 | 0.48126327 | 0.665720776 |
| ENSG00000239335 | LLPH-DT | 26.17981394 | -0.687593641 | 0.341910095 | -2.011036386 | 0.044321614 | 0.108094601 |

| | | | | | | | |
|---|---|---|---|---|---|---|---|
| ENSG00000239344 | ENSG0000023 | 10.10307722 | 0.111833644 | 0.551068699 | 0.202939569 | 0.839182266 | 0.915309377 |
| ENSG00000239345 | HNRNPA1P26 | 49.19589752 | 1.15757145 | 0.263635473 | 4.390803091 | 1.13E-05 | 5.79E-05 |
| ENSG00000239365 | RPS26P49 | 2.203731607 | -0.008378739 | 1.168272797 | -0.007171903 | 0.994277699 | 0.996533822 |
| ENSG00000239374 | ENSG0000023 | 2.185309173 | -0.009257519 | 1.213973199 | -0.007625802 | 0.993915549 | 0.996533822 |
| ENSG00000239382 | ALKBH6 | 86.58847225 | -0.479011409 | 0.202504472 | -2.365436198 | 0.018008846 | 0.049888048 |
| ENSG00000239388 | ASB14 | 11.54744272 | -0.637848392 | 0.519981899 | -1.226674223 | 0.21994506 | 0.39006878 |
| ENSG00000239389 | PCDHA13 | 26.84323667 | -0.057441459 | 0.360210107 | -0.159466539 | 0.873301319 | 0.934604743 |
| ENSG00000239392 | ENSG0000023 | 13.18861978 | -0.309742872 | 0.476391022 | -0.650186208 | 0.515571949 | 0.6944375 |
| ENSG00000239407 | ENSG0000023 | 76.89110531 | -0.031351093 | 0.210459705 | -0.148964827 | 0.881581387 | 0.938595484 |
| ENSG00000239415 | ENSG0000023 | 7.899315953 | 0.550690579 | 0.664248031 | 0.8290436 | 0.407079736 | 0.599849661 |
| ENSG00000239419 | RN7SL535P | 6.006434481 | 0.469643321 | 0.739770665 | 0.634849884 | 0.525526325 | 0.701517248 |
| ENSG00000239443 | PABPC1P10 | 7.323053432 | 0.283169449 | 0.641195936 | 0.441627018 | 0.658759129 | 0.802588264 |
| ENSG00000239454 | LINC02047 | 1.554981746 | 1.086540709 | 1.434918366 | 0.757214302 | 0.448921483 | NA |
| ENSG00000239455 | PTPN11P1 | 30.93718181 | 0.142995626 | 0.314735014 | 0.454336569 | 0.6495866 | 0.796021186 |
| ENSG00000239470 | RPL21P97 | 5.803468739 | 0.005451899 | 0.742191606 | 0.007345676 | 0.994139051 | 0.996533822 |
| ENSG00000239474 | KLHL41 | 3.759723268 | 0.975121016 | 0.930500678 | 1.047953042 | 0.294660242 | 0.480590541 |
| ENSG00000239494 | RN7SL333P | 2.053699435 | -1.290606476 | 1.339324051 | -0.963625252 | 0.335233842 | 0.526441973 |
| ENSG00000239498 | ENSG0000023 | 3.273092279 | -0.45255975 | 0.948671613 | -0.477045738 | 0.633329553 | 0.783930283 |
| ENSG00000239510 | RPL9P5 | 2.536975939 | -1.135616023 | 1.166475606 | -0.973544596 | 0.330282707 | 0.520913393 |
| ENSG00000239521 | CASTOR3P | 978.563244 | -0.030363882 | 0.06912964 | -0.439231017 | 0.660494153 | 0.803707149 |
| ENSG00000239523 | MYLK-AS1 | 107.8986903 | -0.295805037 | 0.171183684 | -1.727997842 | 0.083988611 | 0.183931808 |
| ENSG00000239528 | RPS14P8 | 3.05571041 | -0.192353843 | 1.005229552 | -0.191353151 | 0.848248922 | 0.920605112 |
| ENSG00000239556 | UPK3BL3P | 9.287603727 | 0.590090901 | 0.58016866 | 1.017102338 | 0.309104748 | 0.49719167 |
| ENSG00000239557 | PPP2R5CP | 83.51841313 | -0.116484947 | 0.19951052 | -0.583853661 | 0.559318772 | 0.729021172 |
| ENSG00000239559 | RPL37P2 | 298.0553378 | 0.312604248 | 0.116520783 | 2.682819654 | 0.007300435 | 0.022550472 |
| ENSG00000239569 | KMT2E-AS1 | 47.76903352 | -0.393474136 | 0.261106707 | -1.50694764 | 0.131824079 | 0.264270241 |
| ENSG00000239572 | ENSG0000023 | 42.47956594 | 0.328626109 | 0.280030126 | 1.173538411 | 0.240579963 | 0.416395379 |
| ENSG00000239620 | PRR20G | 3.592686733 | -1.861359459 | 1.007419391 | -1.847651012 | 0.064652845 | 0.148536431 |
| ENSG00000239622 | ENSG0000023 | 13.12425679 | 0.521510129 | 0.49048234 | 1.063259748 | 0.287664174 | 0.471627798 |
| ENSG00000239636 | TFAP2E-AS1 | 2.384204642 | -0.347612473 | 1.135003927 | -0.306265436 | 0.759402553 | 0.867564471 |
| ENSG00000239653 | PSMD6-AS2 | 63.40041571 | -0.404189625 | 0.230370594 | -1.754519178 | 0.079341587 | 0.175617136 |
| ENSG00000239672 | NME1 | 1410.631723 | 0.057023812 | 0.061115212 | 0.933054312 | 0.350791927 | 0.542115791 |
| ENSG00000239704 | CDRT4 | 1.711526198 | -0.770360864 | 1.356369758 | -0.56795786 | 0.570063583 | NA |

| | | | | | | | |
|---|---|---|---|---|---|---|---|
| ENSG00000239718 | HLTF-AS1 | 2.655274732 | -1.727328926 | 1.189159277 | -1.452563134 | 0.146345088 | 0.286129406 |
| ENSG00000239719 | ENSG0000023 | 60.4784322 | 0.425603701 | 0.246868826 | 1.724007474 | 0.084706494 | 0.185151689 |
| ENSG00000239726 | RN7SL688P | 1.746923577 | -0.411672874 | 1.350660269 | -0.304793798 | 0.760523205 | NA |
| ENSG00000239763 | RPL21P118 | 20.88980118 | 0.644072481 | 0.387286635 | 1.663038234 | 0.096304782 | 0.205605557 |
| ENSG00000239779 | WBP1 | 352.8693784 | -0.187203011 | 0.104288958 | -1.79504154 | 0.072647082 | 0.163673547 |
| ENSG00000239789 | MRPS17 | 136.1638827 | 0.92246238 | 0.16732813 | 5.512894797 | 3.53E-08 | 2.39E-07 |
| ENSG00000239791 | ENSG0000023 | 14.39861974 | -0.489001964 | 0.47382989 | -1.032020087 | 0.302062708 | 0.489013304 |
| ENSG00000239801 | DENND6A-AS1 | 12.57963482 | 0.122310338 | 0.495436485 | 0.246873902 | 0.805005815 | 0.896404708 |
| ENSG00000239809 | ENSG0000023 | 2.706818025 | 0.078969214 | 1.105967004 | 0.071402866 | 0.943077129 | 0.97291737 |
| ENSG00000239827 | SUGT1P3 | 16.61532761 | -0.155889894 | 0.424285281 | -0.367417634 | 0.71330752 | 0.84060482 |
| ENSG00000239830 | RPS4XP22 | 5.190331687 | 0.182831726 | 0.783783545 | 0.233268135 | 0.8155532 | 0.902331454 |
| ENSG00000239831 | RNF7P1 | 10.23495447 | -0.354560264 | 0.547023571 | -0.648162681 | 0.516879737 | 0.69523431 |
| ENSG00000239835 | PARLP1 | 1.651958845 | -0.18602231 | 1.409955488 | -0.131934882 | 0.895035798 | NA |
| ENSG00000239857 | GET4 | 232.1883084 | 0.160564924 | 0.136383788 | 1.177302128 | 0.239074938 | 0.414666496 |
| ENSG00000239880 | RALBP1P1 | 2.932830343 | -0.097976738 | 1.044853897 | -0.093770754 | 0.925291264 | 0.964344264 |
| ENSG00000239883 | PARGP1 | 147.6893031 | -0.68133459 | 0.151085029 | -4.509610211 | 6.49E-06 | 3.43E-05 |
| ENSG00000239884 | RN7SL608P | 2.963007379 | -0.198289543 | 1.007804306 | -0.196754014 | 0.844020039 | 0.917708773 |
| ENSG00000239887 | C1orf226 | 533.0195448 | 0.903533587 | 0.092776444 | 9.738825412 | 2.06E-22 | 3.95E-21 |
| ENSG00000239900 | ADSL | 108.9946893 | -0.388044664 | 0.175325902 | -2.213276308 | 0.026878602 | 0.070710451 |
| ENSG00000239908 | RN7SL75P | 1.790876299 | -0.387945755 | 1.343584083 | -0.288739469 | 0.772780756 | 0.876172122 |
| ENSG00000239911 | PRKAG2-AS1 | 16.49801686 | 0.601557697 | 0.437079043 | 1.376313294 | 0.168724659 | 0.318930758 |
| ENSG00000239912 | RPL39P36 | 11.04445931 | 0.406992986 | 0.535392338 | 0.760177083 | 0.447148742 | 0.635656698 |
| ENSG00000239959 | ENPP7P2 | 2.430689572 | 1.798269656 | 1.249166656 | 1.439575454 | 0.149987548 | 0.291610675 |
| ENSG00000239969 | ENSG0000023 | 2.331459337 | -0.510389213 | 1.199340565 | -0.425558201 | 0.670429804 | 0.81090687 |
| ENSG00000239983 | RPL31P31 | 7.018658617 | 1.092203475 | 0.722988131 | 1.510679675 | 0.130870083 | 0.262771091 |
| ENSG00000240003 | RPL7P24 | 4.275051311 | 0.428120815 | 0.90892272 | 0.471020039 | 0.637626425 | 0.78659182 |
| ENSG00000240005 | STIM2-AS1 | 3.927936075 | -0.414062469 | 0.878274529 | -0.471449935 | 0.637319464 | 0.786492469 |
| ENSG00000240023 | RPLP0P3 | 2.978199882 | -1.486794922 | 1.101137361 | -1.350235651 | 0.176940406 | 0.330753986 |
| ENSG00000240024 | LINC00888 | 170.9386357 | -0.289395619 | 0.146672414 | -1.97307462 | 0.048487061 | 0.116727168 |
| ENSG00000240032 | LNCSRLR | 2.209897272 | 2.342454399 | 1.439205961 | 1.627601929 | 0.10360932 | 0.218134893 |
| ENSG00000240036 | ENSG0000024 | 101.1942422 | 0.440597287 | 0.204468748 | 2.154839264 | 0.031174415 | 0.080307812 |
| ENSG00000240038 | AMY2B | 46.9687351 | 0.96230214 | 0.276644902 | 3.478474146 | 0.000504277 | 0.001993815 |
| ENSG00000240048 | DDX50P2 | 13.25523007 | -0.325982765 | 0.487347337 | -0.668892062 | 0.503564335 | 0.684589655 |

| | | | | | | | |
|---|---|---|---|---|---|---|---|
| ENSG00000240052 | ENSG0000024 | 3.349284177 | -0.483846736 | 0.942073933 | -0.513597414 | 0.607533488 | 0.765843401 |
| ENSG00000240053 | LY6G5B | 80.18906694 | -0.12004419 | 0.216419676 | -0.554682421 | 0.579111903 | 0.744665193 |
| ENSG00000240057 | NEPRO-AS1 | 15.4092352 | -0.150821477 | 0.443047527 | -0.34041828 | 0.733541555 | 0.8528429 |
| ENSG00000240086 | ENSG0000024 | 139.4687362 | 0.110591474 | 0.156807729 | 0.705268007 | 0.480643466 | 0.665015749 |
| ENSG00000240087 | RPSAP12 | 62.04324829 | 0.12917781 | 0.22754877 | 0.567692853 | 0.570243546 | 0.737339443 |
| ENSG00000240096 | ENSG0000024 | 2.799882271 | -0.009697078 | 1.024430538 | -0.009465823 | 0.992447479 | 0.995949353 |
| ENSG00000240125 | RPL23AP75 | 2.916238592 | 1.204216866 | 1.108114282 | 1.086726239 | 0.277157814 | 0.45989152 |
| ENSG00000240132 | ETF1P2 | 79.70892155 | 0.226050224 | 0.205517239 | 1.099908819 | 0.271371852 | 0.453376593 |
| ENSG00000240135 | PSMD12P1 | 45.44411613 | -0.085695611 | 0.293793757 | -0.291686289 | 0.770526494 | 0.874956824 |
| ENSG00000240137 | ERICH6-AS1 | 103.9185432 | -0.272910003 | 0.174344071 | -1.56535293 | 0.117500179 | 0.241571438 |
| ENSG00000240138 | EEF1GP4 | 4.950634864 | -0.906892298 | 0.797263958 | -1.137505702 | 0.255326942 | 0.434762737 |
| ENSG00000240184 | PCDHGC3 | 1004.609167 | 0.168370058 | 0.078803808 | 2.13657261 | 0.032632769 | 0.083529985 |
| ENSG00000240204 | SMKR1 | 226.415645 | 0.492052132 | 0.12114912 | 4.06154111 | 4.87E-05 | 0.000228602 |
| ENSG00000240207 | ENSG0000024 | 6.549132059 | -0.410342943 | 0.685703691 | -0.598426038 | 0.549555696 | 0.721915345 |
| ENSG00000240211 | PPP1R35-AS1 | 7.287249951 | -0.418313252 | 0.645819445 | -0.647724771 | 0.51716298 | 0.695452181 |
| ENSG00000240216 | CPHL1P | 15.12117582 | -0.493362257 | 0.450214639 | -1.095837881 | 0.273149727 | 0.455038495 |
| ENSG00000240219 | ENSG0000024 | 16.30324179 | -0.099167331 | 0.433343053 | -0.228842554 | 0.818991291 | 0.904057533 |
| ENSG00000240230 | COX19 | 1168.679542 | 0.146602921 | 0.064757592 | 2.263872338 | 0.023581968 | 0.063167952 |
| ENSG00000240231 | RPS27P29 | 1.769167105 | 0.154205503 | 1.380944434 | 0.111666696 | 0.911087687 | NA |
| ENSG00000240253 | FAR2P3 | 1.905838884 | -1.010907194 | 1.290269406 | -0.783485363 | 0.433342146 | 0.623330124 |
| ENSG00000240254 | B4GALT4-AS1 | 2.154995815 | -0.393989002 | 1.255012158 | -0.313932419 | 0.753572368 | 0.864088279 |
| ENSG00000240286 | MEAF6P1 | 2.719446522 | -0.65741729 | 1.129535348 | -0.582024539 | 0.560550152 | 0.729849205 |
| ENSG00000240288 | GHRLOS | 22.88991736 | -0.403357935 | 0.379020485 | -1.064211437 | 0.287232927 | 0.47125757 |
| ENSG00000240291 | ENSG0000024 | 15.89303232 | -0.628876486 | 0.443114387 | -1.419219293 | 0.155835093 | 0.299878956 |
| ENSG00000240298 | RPL36AP40 | 2.827677938 | -1.848006963 | 1.140792792 | -1.619932187 | 0.105246845 | 0.220911002 |
| ENSG00000240303 | ACAD11 | 33.78386841 | -0.98965819 | 0.30909386 | -3.201804751 | 0.001365695 | 0.004977606 |
| ENSG00000240311 | ENSG0000024 | 7.280973704 | 0.286188357 | 0.647110371 | 0.442255864 | 0.658304065 | 0.802332125 |
| ENSG00000240322 | RN7SL481P | 2.553514238 | -1.631003084 | 1.15345271 | -1.414018165 | 0.157356569 | 0.301920349 |
| ENSG00000240342 | RPS2P5 | 6685.483051 | 0.084265301 | 0.058986273 | 1.428557808 | 0.153131368 | 0.295992045 |
| ENSG00000240344 | PPIL3 | 619.7508977 | -0.17440003 | 0.079755053 | -2.186695687 | 0.028764747 | 0.074848377 |
| ENSG00000240356 | RPL23AP7 | 76.50461156 | -0.371930197 | 0.203542362 | -1.827286439 | 0.067656713 | 0.154294164 |
| ENSG00000240366 | RPL36AP45 | 3.892451823 | -0.643602835 | 0.923777279 | -0.696707799 | 0.485985677 | 0.669580266 |
| ENSG00000240370 | RPL13P5 | 19.66273554 | -0.226454199 | 0.409644368 | -0.552806816 | 0.580395701 | 0.745630889 |

| | | | | | | | |
|---|---|---|---|---|---|---|---|
| ENSG00000240371 | RPS4XP13 | 81.04366969 | 0.174966449 | 0.199765887 | 0.875857492 | 0.381107511 | 0.573376841 |
| ENSG00000240376 | ENSG0000024 | 436.4176999 | 0.22315239 | 0.109451729 | 2.038820136 | 0.041467982 | 0.102122974 |
| ENSG00000240392 | RPL9P3 | 14.83247732 | 0.18608998 | 0.487785086 | 0.381499938 | 0.702832317 | 0.833573227 |
| ENSG00000240395 | RPL5P23 | 70.2582386 | 0.408534538 | 0.242684667 | 1.683396577 | 0.092298346 | 0.198548766 |
| ENSG00000240399 | ENSG0000024 | 6.895817402 | -0.663109387 | 0.677005287 | -0.979474459 | 0.327345602 | 0.517679904 |
| ENSG00000240401 | ENSG0000024 | 13.90500877 | 0.214579816 | 0.474256771 | 0.452454933 | 0.650941281 | 0.797170242 |
| ENSG00000240409 | MTATP8P1 | 2.476971638 | -0.365416322 | 1.148431713 | -0.318187244 | 0.750342908 | 0.862108901 |
| ENSG00000240418 | ENSG0000024 | 9.360949514 | 0.484961286 | 0.602641684 | 0.804725758 | 0.420977952 | 0.612069605 |
| ENSG00000240423 | LINC00636 | 2.92184676 | 1.250568663 | 1.170823377 | 1.068110432 | 0.285470701 | 0.469386575 |
| ENSG00000240429 | LRRFIP1P1 | 121.5572917 | 0.014345886 | 0.171404738 | 0.083695971 | 0.933298161 | 0.967981872 |
| ENSG00000240445 | FOXO3B | 240.8420986 | -0.044725182 | 0.131771665 | -0.339414257 | 0.73429768 | 0.853505714 |
| ENSG00000240449 | REPIN1-AS1 | 31.89136537 | 0.636483231 | 0.319617439 | 1.99139081 | 0.046437939 | 0.112419409 |
| ENSG00000240459 | ENSG0000024 | 1.785279597 | 0.610841382 | 1.295222962 | 0.471610989 | 0.637204481 | NA |
| ENSG00000240463 | RPS19P3 | 14.50730908 | 0.657205906 | 0.488932966 | 1.344163621 | 0.178895483 | 0.333714462 |
| ENSG00000240471 | PHB1P8 | 10.57654811 | 0.238526716 | 0.558405826 | 0.427156567 | 0.669265294 | 0.810007484 |
| ENSG00000240477 | EIF3JP2 | 4.76197884 | -0.474220971 | 0.849310561 | -0.558359913 | 0.57659864 | 0.742765769 |
| ENSG00000240480 | RPL29P2 | 2.444879297 | 1.263536866 | 1.232098165 | 1.025516393 | 0.305119595 | 0.492571675 |
| ENSG00000240489 | SETP14 | 1127.251262 | -0.138493774 | 0.070868484 | -1.954236448 | 0.050673261 | 0.121252175 |
| ENSG00000240498 | CDKN2B-AS1 | 149.9064748 | -0.167303554 | 0.167637704 | -0.998006712 | 0.318276104 | 0.507419128 |
| ENSG00000240509 | RPL34P18 | 394.8289745 | 0.353331378 | 0.102204535 | 3.457100795 | 0.000546021 | 0.002147388 |
| ENSG00000240520 | UOX | 2.219346238 | -0.470383562 | 1.178109937 | -0.399269667 | 0.689694515 | 0.824642561 |
| ENSG00000240540 | RPL3P9 | 36.28463632 | 0.659543578 | 0.334876986 | 1.969510019 | 0.048894553 | 0.117535476 |
| ENSG00000240567 | LINC02067 | 50.31292398 | -0.948272554 | 0.255903817 | -3.705581899 | 0.000210906 | 0.000890518 |
| ENSG00000240571 | ENSG0000024 | 3.69680143 | -0.176823011 | 0.914989843 | -0.193251338 | 0.846762135 | 0.91973122 |
| ENSG00000240579 | ENSG0000024 | 2.528507469 | -0.764258526 | 1.112297231 | -0.687099189 | 0.492020225 | 0.674935356 |
| ENSG00000240591 | ENSG0000024 | 19.20611701 | -1.245566553 | 0.413559552 | -3.011819087 | 0.002596873 | 0.008914001 |
| ENSG00000240601 | RPL9P15 | 2.857606373 | 2.806207911 | 1.319930208 | 2.126027493 | 0.03350097 | 0.085352317 |
| ENSG00000240616 | RPS6P25 | 65.48685745 | 0.35128318 | 0.24497615 | 1.43394849 | 0.151586963 | 0.293948007 |
| ENSG00000240622 | RPL7P15 | 11.8552274 | 0.451459371 | 0.531143868 | 0.849975681 | 0.395338607 | 0.588289287 |
| ENSG00000240634 | ENSG0000024 | 4.399940067 | 1.007483061 | 0.927436574 | 1.086309392 | 0.277342133 | 0.460157474 |
| ENSG00000240652 | ENSG0000024 | 40.37160504 | 0.196404467 | 0.329001617 | 0.596971129 | 0.550526656 | 0.722652867 |
| ENSG00000240666 | MME-AS1 | 6.063092315 | 0.060621709 | 0.724621808 | 0.083659791 | 0.933326927 | 0.967981872 |
| ENSG00000240668 | KRT8P36 | 55.9332791 | 0.339320893 | 0.257015441 | 1.320235434 | 0.186756425 | 0.344471527 |

| | | | | | | | |
|---|---|---|---|---|---|---|---|
| ENSG00000240674 | RPL21P51 | 8.071497643 | 0.407929052 | 0.612450516 | 0.666060427 | 0.505372478 | 0.68591659 |
| ENSG00000240682 | ISY1 | 416.7564965 | -0.174731031 | 0.093594903 | -1.866886188 | 0.061917483 | 0.143155318 |
| ENSG00000240694 | PNMA2 | 18.70613298 | -2.819851788 | 0.516592575 | -5.458560432 | 4.80E-08 | 3.21E-07 |
| ENSG00000240695 | DNAJC8P2 | 42.10597999 | 0.010124168 | 0.280572798 | 0.036083926 | 0.971215439 | 0.986933608 |
| ENSG00000240710 | ENSG0000024 | 3.702809119 | -0.257858048 | 0.936208936 | -0.275427886 | 0.782987522 | 0.882293632 |
| ENSG00000240731 | ENSG0000024 | 52.44175851 | -0.046589352 | 0.261215794 | -0.178355802 | 0.858443558 | 0.92649707 |
| ENSG00000240738 | RPL6P22 | 3.00983187 | 0.125797101 | 0.998763697 | 0.125952817 | 0.899769274 | 0.949347215 |
| ENSG00000240759 | RPS27P21 | 1.934663069 | 0.826583375 | 1.257087817 | 0.657538291 | 0.510834858 | 0.690585871 |
| ENSG00000240764 | PCDHGC5 | 5.499599905 | 1.022573063 | 0.809847237 | 1.262674015 | 0.206706356 | 0.372552153 |
| ENSG00000240771 | ARHGEF25 | 128.528883 | 0.035130831 | 0.161116763 | 0.218045783 | 0.82739344 | 0.908805627 |
| ENSG00000240776 | RPS10P4 | 21.92051388 | 0.689810081 | 0.415124999 | 1.661692459 | 0.096574454 | 0.206142921 |
| ENSG00000240785 | RPL36AP21 | 6.387906484 | -0.261602477 | 0.715190525 | -0.365780121 | 0.714529155 | 0.841150775 |
| ENSG00000240787 | NECTIN2P1 | 23.18104333 | -0.547284862 | 0.364349925 | -1.502085836 | 0.133074942 | 0.266196301 |
| ENSG00000240809 | CAP1P1 | 65.21565935 | -0.491038831 | 0.229944233 | -2.135469219 | 0.032722702 | 0.083722818 |
| ENSG00000240828 | RPL21P4 | 3.176687822 | 0.72146404 | 1.010570025 | 0.713917909 | 0.475277942 | 0.66046014 |
| ENSG00000240849 | PEDS1 | 1992.473742 | -0.192606274 | 0.058936196 | -3.268047267 | 0.001082923 | 0.004028908 |
| ENSG00000240854 | PSMC2P1 | 7.523079911 | -0.509272621 | 0.629646233 | -0.808823422 | 0.418616722 | 0.61039531 |
| ENSG00000240857 | RDH14 | 310.975185 | -0.020073993 | 0.106984356 | -0.187634848 | 0.851162906 | 0.9222591 |
| ENSG00000240859 | LINC03014 | 29.32961036 | -0.233929629 | 0.339705162 | -0.688625477 | 0.491058984 | 0.674019811 |
| ENSG00000240875 | LINC00886 | 480.860368 | 0.495384321 | 0.090826627 | 5.454175043 | 4.92E-08 | 3.29E-07 |
| ENSG00000240877 | RN7SL521P | 5.169990572 | 1.26143026 | 0.852432344 | 1.479801029 | 0.138926354 | 0.274881021 |
| ENSG00000240882 | PTOV1P1 | 1.905359626 | 1.443528994 | 1.348732586 | 1.070285547 | 0.284490798 | 0.468265689 |
| ENSG00000240889 | NDUFB2-AS1 | 62.20499533 | -0.235786179 | 0.22608078 | -1.042928898 | 0.296981225 | 0.483279629 |
| ENSG00000240890 | G2E3P1 | 4.841351222 | 0.05106047 | 0.820032875 | 0.062266369 | 0.95035071 | 0.977185792 |
| ENSG00000240891 | PLCXD2 | 29.83076186 | -2.064456894 | 0.360824632 | -5.721496564 | 1.06E-08 | 7.58E-08 |
| ENSG00000240898 | RPL21P125 | 4.321403598 | 0.257250249 | 0.86839095 | 0.296237828 | 0.767048462 | 0.872778717 |
| ENSG00000240914 | RPL15P2 | 63.21931637 | 0.124143198 | 0.244428966 | 0.507890696 | 0.61153 | 0.76912463 |
| ENSG00000240919 | RPL21P79 | 1.768206638 | -0.440602056 | 1.325513973 | -0.332400914 | 0.739586548 | NA |
| ENSG00000240950 | RPL8P3 | 102.5485816 | 0.066822405 | 0.190766737 | 0.350283314 | 0.726126087 | 0.847853251 |
| ENSG00000240954 | RPL4P1 | 3.597435279 | 1.334677211 | 0.984012809 | 1.356361623 | 0.174984137 | 0.328031993 |
| ENSG00000240972 | MIF | 841.9836148 | -0.052909313 | 0.078151575 | -0.677008917 | 0.498400297 | 0.680139274 |
| ENSG00000240990 | HOXA11-AS | 26.39831976 | -0.677136745 | 0.345883779 | -1.957700206 | 0.050265201 | 0.120363644 |
| ENSG00000240991 | RPL23AP67 | 2.381505994 | 0.200450136 | 1.137274845 | 0.176254787 | 0.86009378 | 0.927319435 |

| | | | | | | | |
|---|---|---|---|---|---|---|---|
| ENSG00000240996 | ENSG0000024 | 12.53685845 | -0.206681718 | 0.493105683 | -0.419142843 | 0.675111741 | 0.814442859 |
| ENSG00000241007 | SEPTIN7P6 | 31.90036607 | 0.052863264 | 0.312719208 | 0.169043867 | 0.865762137 | 0.9302381 |
| ENSG00000241014 | GPR199P | 7.09247315 | -0.060097321 | 0.670628807 | -0.089613391 | 0.928594443 | 0.965737636 |
| ENSG00000241015 | TPM3P9 | 551.4901635 | 0.120162603 | 0.085773703 | 1.400925908 | 0.161236231 | 0.308075889 |
| ENSG00000241030 | RPL29P24 | 1.563450216 | 0.516463784 | 1.482989596 | 0.348258535 | 0.727646031 | NA |
| ENSG00000241057 | ENSG0000024 | 3.069961049 | 0.385025866 | 1.055192671 | 0.364886789 | 0.715195916 | 0.841653402 |
| ENSG00000241058 | NSUN6 | 171.2029357 | -0.599755638 | 0.14481308 | -4.141584697 | 3.45E-05 | 0.000165254 |
| ENSG00000241061 | RPL5P1 | 427.3773064 | 0.296688099 | 0.092067747 | 3.222497659 | 0.001270782 | 0.004663532 |
| ENSG00000241081 | RPL22P2 | 492.998405 | 0.227014906 | 0.095094334 | 2.387260065 | 0.016974479 | 0.047413156 |
| ENSG00000241095 | CYP51A1P1 | 161.7213584 | -1.115984486 | 0.148467721 | -7.516680938 | 5.62E-14 | 6.25E-13 |
| ENSG00000241111 | PRICKLE2-AS1 | 38.92788349 | -0.243991623 | 0.285488622 | -0.8546457 | 0.39274732 | 0.585725102 |
| ENSG00000241120 | HMGN1P8 | 1.990921381 | -0.188172377 | 1.214238306 | -0.154971537 | 0.876843756 | 0.935974786 |
| ENSG00000241127 | YAE1 | 289.2749684 | 0.598437842 | 0.116201256 | 5.150011822 | 2.60E-07 | 1.62E-06 |
| ENSG00000241155 | ARHGAP31-AS | 3.075504299 | 0.629867875 | 1.005864993 | 0.626195244 | 0.531186886 | 0.706729334 |
| ENSG00000241157 | RPL32P32 | 6.683895995 | 0.60271588 | 0.681927136 | 0.883842054 | 0.376781481 | 0.569544446 |
| ENSG00000241170 | RPL31P46 | 27.10635797 | 0.277171387 | 0.348943039 | 0.794316999 | 0.42701091 | 0.61775629 |
| ENSG00000241185 | RPL21P122 | 1.647214082 | -0.790235479 | 1.403277047 | -0.563135755 | 0.573342446 | NA |
| ENSG00000241217 | RN7SL809P | 2.36064874 | -0.573582449 | 1.147525245 | -0.499842989 | 0.617185638 | 0.773024159 |
| ENSG00000241233 | KRTAP5-8 | 4.438510792 | 0.694392198 | 0.855466732 | 0.811711516 | 0.416957185 | 0.609041568 |
| ENSG00000241255 | ENSG0000024 | 69.86784424 | -0.367149563 | 0.230251958 | -1.594555659 | 0.110811635 | 0.230109433 |
| ENSG00000241258 | CRCP | 1618.423648 | 0.40666964 | 0.056191245 | 7.237242072 | 4.58E-13 | 4.78E-12 |
| ENSG00000241269 | ENSG0000024 | 4.760659736 | 1.186154846 | 0.848910155 | 1.397267824 | 0.162333048 | 0.309527065 |
| ENSG00000241281 | RPL31P26 | 6.313331215 | 1.141972002 | 0.750152232 | 1.522320341 | 0.127928834 | 0.258085256 |
| ENSG00000241282 | RPL34P33 | 23.69231878 | 0.144148398 | 0.358607748 | 0.401966769 | 0.687708483 | 0.823383061 |
| ENSG00000241288 | LINC02614 | 66.15245621 | 0.26291619 | 0.222444331 | 1.181941517 | 0.237228907 | 0.412400055 |
| ENSG00000241293 | PPATP1 | 42.61204243 | 0.332581094 | 0.277856922 | 1.196950903 | 0.231325692 | 0.404961185 |
| ENSG00000241313 | WWTR1-AS1 | 11.86423692 | 0.249809329 | 0.50498502 | 0.494686613 | 0.620821353 | 0.775560477 |
| ENSG00000241316 | SUCLG2-DT | 65.58751426 | -0.757159596 | 0.223969646 | -3.380634875 | 0.000723186 | 0.002779806 |
| ENSG00000241318 | WDR82P2 | 14.24470183 | 0.427562123 | 0.466306235 | 0.91691273 | 0.359188373 | 0.551746436 |
| ENSG00000241319 | SETP6 | 5.233858278 | 0.367423304 | 0.78170767 | 0.47002648 | 0.638336099 | 0.786942719 |
| ENSG00000241322 | FBXW10B | 14.98806194 | -0.18583177 | 0.453114434 | -0.410121055 | 0.681717148 | 0.819321553 |
| ENSG00000241343 | RPL36A | 230.0182548 | -0.153983387 | 0.282358365 | -0.545347355 | 0.585514641 | 0.749620821 |
| ENSG00000241352 | ENSG0000024 | 19.73767775 | 0.315586041 | 0.448500945 | 0.703646325 | 0.481653055 | 0.666010915 |

| | | | | | | | |
|---|---|---|---|---|---|---|---|
| ENSG00000241357 | ENSG0000024 | 10.19673814 | -0.609031977 | 0.557622271 | -1.0921945 | 0.27474762 | 0.45683264 |
| ENSG00000241359 | SYNPR-AS1 | 112.135498 | 1.724039269 | 0.195273507 | 8.828843708 | 1.06E-18 | 1.64E-17 |
| ENSG00000241360 | PDXP | 7.802919504 | 0.186974054 | 0.618711131 | 0.302199273 | 0.762500159 | 0.869742236 |
| ENSG00000241361 | SLC25A24P1 | 17.6401815 | -0.170259326 | 0.425101592 | -0.40051444 | 0.68877765 | 0.824190668 |
| ENSG00000241362 | RPL36AP43 | 4.100092107 | -0.713903504 | 0.895781257 | -0.796961868 | 0.425473178 | 0.61633965 |
| ENSG00000241370 | RPP21 | 43.82440522 | -0.132085378 | 0.273469956 | -0.482997766 | 0.629097323 | 0.7810577 |
| ENSG00000241399 | CD302 | 165.6489345 | -0.399978182 | 0.140809547 | -2.840561539 | 0.004503418 | 0.014680747 |
| ENSG00000241404 | EGFL8 | 43.22982235 | -0.325140654 | 0.266636041 | -1.219417496 | 0.222685773 | 0.393802931 |
| ENSG00000241409 | ENSG0000024 | 4.703357762 | 0.322469992 | 0.809590497 | 0.398312471 | 0.690399869 | 0.825313861 |
| ENSG00000241411 | ENSG0000024 | 2.139381184 | 0.0690094 | 1.19272099 | 0.057858796 | 0.953861104 | 0.978691016 |
| ENSG00000241416 | KRT8P13 | 68.63129026 | 0.215849849 | 0.213556925 | 1.010736827 | 0.312142406 | 0.500364186 |
| ENSG00000241429 | EEF1A1P25 | 16.80401643 | -0.012197533 | 0.422835279 | -0.028847009 | 0.976986609 | 0.989320774 |
| ENSG00000241431 | RPL37P6 | 31.62501425 | 0.229750653 | 0.358012106 | 0.641739902 | 0.521042077 | 0.698172035 |
| ENSG00000241458 | RPL7P19 | 6.881520774 | 0.239532035 | 0.664596098 | 0.360417456 | 0.718534975 | 0.843432038 |
| ENSG00000241464 | RPL39P38 | 3.798433366 | 0.195331486 | 0.885440779 | 0.220603671 | 0.825401042 | 0.907872877 |
| ENSG00000241468 | ATP5MF | 1029.824673 | -0.417712649 | 0.065012803 | -6.425082862 | 1.32E-10 | 1.13E-09 |
| ENSG00000241472 | PTPRG-AS1 | 449.7837244 | -0.140394963 | 0.096410674 | -1.456218043 | 0.145332353 | 0.284513072 |
| ENSG00000241478 | HSPA8P9 | 41.72874993 | 0.192875275 | 0.309884933 | 0.622409333 | 0.533672745 | 0.709242542 |
| ENSG00000241484 | ARHGAP8 | 236.5224485 | -0.67653847 | 0.12043023 | -5.617679813 | 1.94E-08 | 1.35E-07 |
| ENSG00000241489 | ENSG0000024 | 32.63749239 | 0.122053027 | 0.316741369 | 0.385339708 | 0.699985746 | 0.831719009 |
| ENSG00000241494 | ENSG0000024 | 70.75280581 | 0.61557458 | 0.214554322 | 2.869084965 | 0.004116612 | 0.013527443 |
| ENSG00000241506 | PSMC1P1 | 3180.600569 | -0.718916674 | 0.050265594 | -14.30236092 | 2.12E-46 | 1.05E-44 |
| ENSG00000241511 | RPS15AP24 | 21.64414336 | 0.096541056 | 0.381120241 | 0.253308655 | 0.800029709 | 0.893422331 |
| ENSG00000241520 | ENSG0000024 | 1.727596379 | 2.025897996 | 1.518434031 | 1.334202182 | 0.182137605 | NA |
| ENSG00000241527 | CA15P1 | 11.37394682 | -0.15296415 | 0.514619559 | -0.297237342 | 0.766285315 | 0.872318316 |
| ENSG00000241532 | AGGF1P3 | 4.655153253 | 1.228308185 | 0.858627157 | 1.430548958 | 0.152559525 | 0.295185181 |
| ENSG00000241537 | U2SURPP1 | 3.536525808 | -0.31563015 | 0.942899832 | -0.334744094 | 0.737818136 | 0.855509243 |
| ENSG00000241547 | ACTG1P20 | 27.93266421 | -0.028965166 | 0.332711812 | -0.087057823 | 0.930625551 | 0.966566312 |
| ENSG00000241549 | GUSBP2 | 42.7594119 | -0.076342093 | 0.272100008 | -0.280566304 | 0.779043064 | 0.880183604 |
| ENSG00000241553 | ARPC4 | 2559.411278 | 0.027601259 | 0.05141052 | 0.536879603 | 0.591350781 | 0.753853121 |
| ENSG00000241556 | RPL29P26 | 57.07122749 | 0.506905583 | 0.244759541 | 2.07103503 | 0.038355523 | 0.095836585 |
| ENSG00000241612 | RPL21P46 | 12.42405139 | 0.388323049 | 0.512160666 | 0.758205531 | 0.448327951 | 0.636466126 |
| ENSG00000241627 | UBQLN4P1 | 490.9189111 | 0.303516744 | 0.089011608 | 3.409855768 | 0.000649972 | 0.002520346 |

| | | | | | | | |
|---|---|---|---|---|---|---|---|
| ENSG00000241634 | RPL13P8 | 26.04390064 | -0.49961523 | 0.36153252 | -1.381937177 | 0.166990991 | 0.316514364 |
| ENSG00000241635 | UGT1A1 | 1.683142745 | 3.005439253 | 1.614819108 | 1.861161562 | 0.062721365 | NA |
| ENSG00000241651 | RPL15P8 | 7.554300609 | 0.209528639 | 0.648655223 | 0.323020045 | 0.746680062 | 0.859607712 |
| ENSG00000241671 | KIAA1328P1 | 1.754028054 | 0.644196651 | 1.343498946 | 0.479491742 | 0.631588841 | NA |
| ENSG00000241680 | RPL31P49 | 41.33769285 | 0.174245977 | 0.279968667 | 0.622376709 | 0.533694192 | 0.709242542 |
| ENSG00000241685 | ARPC1A | 484.6314332 | -0.570719187 | 0.094506412 | -6.038946728 | 1.55E-09 | 1.21E-08 |
| ENSG00000241717 | VWFP1 | 1.619783634 | -0.129718737 | 1.459537969 | -0.088876576 | 0.92918 | NA |
| ENSG00000241720 | ENSG0000024 | 75.2847402 | -1.318078338 | 0.224435596 | -5.872857795 | 4.28E-09 | 3.22E-08 |
| ENSG00000241723 | RPL15P6 | 3.27695168 | -0.200625384 | 0.993655471 | -0.201906385 | 0.839989911 | 0.915757464 |
| ENSG00000241735 | FABP5P3 | 2.671284325 | -0.550969311 | 1.114727704 | -0.494263585 | 0.621120039 | 0.775649892 |
| ENSG00000241741 | RPL7AP30 | 783.6586864 | 0.127909304 | 0.076077829 | 1.681295399 | 0.092705551 | 0.199312591 |
| ENSG00000241746 | RPL13AP18 | 10.230368 | 0.089539673 | 0.552240465 | 0.16213892 | 0.871196458 | 0.933095623 |
| ENSG00000241764 | CBLL1-AS1 | 57.97428753 | -0.267576938 | 0.235654529 | -1.135462743 | 0.256181484 | 0.435641455 |
| ENSG00000241769 | EOLA1-DT | 18.00333632 | 0.039273348 | 0.421990978 | 0.093066795 | 0.925850497 | 0.964514676 |
| ENSG00000241770 | ENSG0000024 | 12.08174992 | 0.735245383 | 0.505110647 | 1.455612523 | 0.145499764 | 0.284743609 |
| ENSG00000241829 | RPL21P54 | 2.549628552 | -1.282147975 | 1.170457583 | -1.095424554 | 0.273330682 | 0.455220404 |
| ENSG00000241837 | ATP5PO | 451.9241542 | -0.011993754 | 0.092232444 | -0.130038345 | 0.896536089 | 0.947723049 |
| ENSG00000241839 | PLEKHO2 | 153.7715696 | -0.052128118 | 0.145445873 | -0.358402178 | 0.720042362 | 0.844366665 |
| ENSG00000241852 | C8orf58 | 117.5217682 | -0.295413133 | 0.165897406 | -1.780697721 | 0.074961846 | 0.167801009 |
| ENSG00000241860 | ENSG0000024 | 1.944119172 | 0.310824265 | 1.225621769 | 0.253605372 | 0.799800446 | 0.893253273 |
| ENSG00000241878 | PISD | 1753.503896 | -1.429929112 | 0.062966283 | -22.70944119 | 3.61E-114 | 7.47E-112 |
| ENSG00000241889 | CTDNEP1P1 | 65.63209092 | -0.036861094 | 0.224639689 | -0.164089856 | 0.869660407 | 0.932287011 |
| ENSG00000241923 | RPL14P3 | 60.73297455 | 0.175870559 | 0.256989466 | 0.684349292 | 0.493754628 | 0.676438565 |
| ENSG00000241932 | RPL35AP34 | 1.721012045 | 4.128993955 | 1.656649798 | 2.492375853 | 0.012689168 | NA |
| ENSG00000241933 | DENND6A-DT | 2.009369473 | -0.124906851 | 1.299356403 | -0.096129785 | 0.923417496 | 0.963135281 |
| ENSG00000241935 | HOGA1 | 15.58094667 | -1.180906852 | 0.478167263 | -2.469652241 | 0.013524446 | 0.03876681 |
| ENSG00000241945 | PWP2 | 22.70900853 | 0.565943528 | 0.373555661 | 1.515017939 | 0.129767861 | 0.260900503 |
| ENSG00000241962 | ENSG0000024 | 7.158926972 | -0.941153546 | 0.663753623 | -1.41792604 | 0.156212359 | 0.300364147 |
| ENSG00000241973 | PI4KA | 5368.721657 | -0.475064352 | 0.049082092 | -9.678975251 | 3.70E-22 | 6.99E-21 |
| ENSG00000241975 | ELOCP19 | 7.975007485 | -0.872296513 | 0.6684462 | -1.304961436 | 0.19190598 | 0.351928829 |
| ENSG00000241990 | PRR34-AS1 | 32.82391787 | -0.50563449 | 0.303854934 | -1.664065427 | 0.096099355 | 0.205243386 |
| ENSG00000242017 | ALG1L15P | 32.6336035 | -0.070182337 | 0.308913641 | -0.227190798 | 0.820275383 | 0.9046691 |
| ENSG00000242021 | ENSG0000024 | 1.61369716 | 2.870099458 | 1.635877519 | 1.754470872 | 0.079349857 | NA |

| | | | | | | | |
|---|---|---|---|---|---|---|---|
| ENSG00000242028 | HYPK | 93.62150652 | -0.132948603 | 0.20637085 | -0.644221811 | 0.519431609 | 0.696909646 |
| ENSG00000242060 | RPS3AP49 | 11.63588308 | -0.214995042 | 0.529587899 | -0.405966682 | 0.684767082 | 0.82136245 |
| ENSG00000242068 | RNF13P1 | 16.88536897 | -0.059649719 | 0.441687493 | -0.135049599 | 0.892572659 | 0.945013929 |
| ENSG00000242071 | RPL7AP6 | 5701.215865 | 0.159821488 | 0.051290135 | 3.116027809 | 0.00183305 | 0.006493231 |
| ENSG00000242083 | RPL7AP31 | 51.6901772 | 0.077657047 | 0.282333919 | 0.275053907 | 0.783274823 | 0.882490914 |
| ENSG00000242085 | RPS20P33 | 5.527334658 | -0.350643317 | 0.754376986 | -0.46481179 | 0.642066244 | 0.790034632 |
| ENSG00000242086 | MUC20-OT1 | 392.4297172 | -0.137703478 | 0.099725314 | -1.380827723 | 0.167331937 | 0.316898757 |
| ENSG00000242087 | RPL36AP41 | 3.650828553 | -1.560378621 | 1.007591233 | -1.548622665 | 0.121472454 | 0.247778375 |
| ENSG00000242088 | RPS27P2 | 7.245097067 | 0.841218224 | 0.676334806 | 1.24378964 | 0.213576994 | 0.381810339 |
| ENSG00000242100 | RPL9P32 | 21.71112375 | -0.222725736 | 0.395913919 | -0.562561014 | 0.573733845 | 0.740405851 |
| ENSG00000242110 | AMACR | 33.78515087 | 0.164276846 | 0.309802975 | 0.530262327 | 0.595930063 | 0.756993626 |
| ENSG00000242114 | MTFP1 | 37.19819812 | -0.578595279 | 0.295297079 | -1.959366755 | 0.050069851 | 0.119971003 |
| ENSG00000242125 | SNHG3 | 1025.980979 | 0.634298386 | 0.072307562 | 8.772227592 | 1.75E-18 | 2.67E-17 |
| ENSG00000242136 | ENSG0000024 | 6.299903399 | -1.059858013 | 0.7394065 | -1.433390176 | 0.151746364 | 0.294083095 |
| ENSG00000242140 | MATR3P2 | 1.568734523 | 1.032751409 | 1.505027717 | 0.686200923 | 0.492586416 | NA |
| ENSG00000242142 | SERBP1P3 | 3.758185748 | -2.351075297 | 1.036201775 | -2.268935794 | 0.023272231 | 0.06239656 |
| ENSG00000242159 | ABCF2P1 | 9.241140775 | 0.549735588 | 0.597245885 | 0.920451028 | 0.357337113 | 0.54960197 |
| ENSG00000242162 | ENSG0000024 | 3.493493076 | 0.89947645 | 0.990714735 | 0.907906604 | 0.363927568 | 0.556603668 |
| ENSG00000242163 | ENSG0000024 | 2.742187912 | 0.359284949 | 1.075139521 | 0.334175186 | 0.738247367 | 0.855798404 |
| ENSG00000242169 | RPL7AP27 | 15.80319628 | 0.510384252 | 0.444930338 | 1.147110477 | 0.251335964 | 0.429791057 |
| ENSG00000242173 | ARHGDIG | 94.18509834 | 0.492512631 | 0.186530048 | 2.640392988 | 0.008280994 | 0.025187749 |
| ENSG00000242193 | CRYZL2P | 31.73654317 | 0.841342794 | 0.320122684 | 2.628188614 | 0.008584091 | 0.026006212 |
| ENSG00000242208 | RPL5P29 | 19.61193368 | 0.425998601 | 0.39844892 | 1.069142316 | 0.285005546 | 0.468877455 |
| ENSG00000242242 | NECTIN3-AS1 | 6.679314314 | -1.371363895 | 0.746826504 | -1.836254991 | 0.066319953 | 0.151819152 |
| ENSG00000242247 | ARFGAP3 | 641.8580648 | -0.088595849 | 0.078313207 | -1.131301503 | 0.257928212 | 0.4375795 |
| ENSG00000242252 | BGLAP | 3.053325038 | -0.531869041 | 1.02043192 | -0.521219525 | 0.602213854 | 0.761606804 |
| ENSG00000242259 | C22orf39 | 682.7137815 | -0.454714188 | 0.0767993 | -5.920811598 | 3.20E-09 | 2.44E-08 |
| ENSG00000242262 | RPL15P7 | 65.29471394 | 0.209034065 | 0.232076641 | 0.900711353 | 0.367741811 | 0.560307993 |
| ENSG00000242265 | PEG10 | 488.6705055 | -7.090523909 | 0.312212653 | -22.71055911 | 3.52E-114 | 7.35E-112 |
| ENSG00000242272 | AK2P2 | 10.14521754 | 0.284076179 | 0.545765965 | 0.52050915 | 0.602708752 | 0.761938652 |
| ENSG00000242276 | RPL5P3 | 13.86877748 | -0.156178201 | 0.538114527 | -0.290232271 | 0.77163855 | 0.875561078 |
| ENSG00000242294 | STAG3L5P | 366.4780471 | -0.438838916 | 0.114276423 | -3.840152722 | 0.000122958 | 0.000540355 |
| ENSG00000242299 | RPS18P5 | 3229.696026 | 0.35908288 | 0.053118412 | 6.760045461 | 1.38E-11 | 1.27E-10 |

| | | | | | | | |
|---|---|---|---|---|---|---|---|
| ENSG00000242314 | RPL12P32 | 12.69215828 | 0.346386718 | 0.503602486 | 0.687817729 | 0.49156757 | 0.67443541 |
| ENSG00000242327 | ENSG0000024 | 29.49009387 | 0.461086966 | 0.347638647 | 1.326339778 | 0.184727176 | 0.341552465 |
| ENSG00000242337 | INHCAP | 5.955405075 | -0.312873081 | 0.719862316 | -0.434629059 | 0.663831699 | 0.805985984 |
| ENSG00000242338 | BMS1P4 | 7.298134638 | -1.217164652 | 0.671920052 | -1.811472434 | 0.070067752 | 0.158672679 |
| ENSG00000242352 | FAM91A3P | 109.4998418 | -0.5529197 | 0.173872535 | -3.180028976 | 0.001472603 | 0.005325795 |
| ENSG00000242358 | RPS21P4 | 44.92634382 | 0.071863156 | 0.299686374 | 0.239794538 | 0.810489541 | 0.899645977 |
| ENSG00000242372 | EIF6 | 1427.352804 | -0.544658957 | 0.066665046 | -8.170082981 | 3.08E-16 | 4.06E-15 |
| ENSG00000242375 | ENSG0000024 | 1.850428508 | 0.365069921 | 1.313160807 | 0.278008542 | 0.781005802 | 0.881298289 |
| ENSG00000242396 | ENSG0000024 | 16.35528512 | 0.090590824 | 0.448245905 | 0.20210073 | 0.839837978 | 0.915722382 |
| ENSG00000242411 | ENSG0000024 | 3.021157288 | 1.305774012 | 1.040933503 | 1.254425963 | 0.209687221 | 0.376711163 |
| ENSG00000242419 | PCDHGC4 | 6.548608238 | -0.325971913 | 0.684588315 | -0.476157576 | 0.633962121 | 0.784236938 |
| ENSG00000242445 | RPL7AP11 | 168.4701333 | 0.168063864 | 0.143792716 | 1.168792609 | 0.242487199 | 0.418753884 |
| ENSG00000242457 | RBBP4P2 | 234.0784271 | 0.275807826 | 0.121118952 | 2.277164899 | 0.02277638 | 0.06121736 |
| ENSG00000242474 | LINC03015 | 362.4108398 | -0.06398539 | 0.114069299 | -0.56093437 | 0.574842279 | 0.741214298 |
| ENSG00000242477 | ENSG0000024 | 82.1406847 | 0.318345205 | 0.216549798 | 1.470078514 | 0.141540491 | 0.278922675 |
| ENSG00000242479 | EIF2AK1P1 | 12.80660135 | -0.155414285 | 0.485575464 | -0.320062063 | 0.748921284 | 0.861122462 |
| ENSG00000242485 | MRPL20 | 1788.944823 | 0.51863891 | 0.056806064 | 9.129992039 | 6.85E-20 | 1.13E-18 |
| ENSG00000242498 | ARPIN | 396.8634611 | 0.942019523 | 0.107767803 | 8.741196303 | 2.31E-18 | 3.48E-17 |
| ENSG00000242516 | LINC00960 | 203.6726567 | 1.168490143 | 0.134335347 | 8.698307407 | 3.37E-18 | 5.04E-17 |
| ENSG00000242537 | MRE11P1 | 1.818404135 | 0.607887589 | 1.332723362 | 0.456124359 | 0.648300555 | 0.794997312 |
| ENSG00000242553 | ENSG0000024 | 8.538696503 | -0.774767401 | 0.615406313 | -1.258952639 | 0.208047439 | 0.37449845 |
| ENSG00000242562 | DCAF13P1 | 41.54404533 | -0.130523261 | 0.273155536 | -0.477834946 | 0.632767686 | 0.78358033 |
| ENSG00000242571 | RPL21P11 | 23.44133616 | 0.254911152 | 0.376114523 | 0.677748761 | 0.497931004 | 0.679791539 |
| ENSG00000242575 | TUBAP13 | 6.314760207 | 2.934612334 | 0.927994907 | 3.16231513 | 0.001565201 | 0.005629619 |
| ENSG00000242578 | ENSG0000024 | 3.02816743 | 0.0013857 | 1.000523709 | 0.001384974 | 0.998894951 | 0.999333951 |
| ENSG00000242588 | ENSG0000024 | 85.17469712 | -0.078206228 | 0.19555629 | -0.399916711 | 0.689217863 | 0.824373418 |
| ENSG00000242612 | DECR2 | 336.6059952 | -0.103367086 | 0.103912034 | -0.994755676 | 0.319855108 | 0.509370093 |
| ENSG00000242615 | RPL17P47 | 12.42459557 | -0.017304873 | 0.49751837 | -0.03478238 | 0.972253271 | 0.987410366 |
| ENSG00000242616 | GNG10 | 3.684025145 | -0.450079571 | 0.905203157 | -0.497213877 | 0.619038236 | 0.77417746 |
| ENSG00000242622 | GSK3B-DT | 123.2291192 | 0.026511919 | 0.16687791 | 0.15887015 | 0.873771179 | 0.934676032 |
| ENSG00000242628 | ENSG0000024 | 2.595092426 | -0.508695739 | 1.125990077 | -0.451776396 | 0.651430076 | 0.797514584 |
| ENSG00000242640 | RPS29P11 | 10.20514948 | -0.013294143 | 0.576036499 | -0.023078646 | 0.981587539 | 0.991616089 |
| ENSG00000242660 | ENSG0000024 | 21.96272585 | 0.41294806 | 0.397173995 | 1.039715756 | 0.298471978 | 0.485007829 |

| | | | | | | | |
|---|---|---|---|---|---|---|---|
| ENSG00000242667 | ENSG0000024 | 6.311325577 | 0.219387554 | 0.767193702 | 0.285961099 | 0.774907916 | 0.877499635 |
| ENSG00000242670 | RPL22P13 | 2.37155203 | -0.166192893 | 1.156704592 | -0.143677905 | 0.885754819 | 0.941041978 |
| ENSG00000242686 | PDE6B-AS1 | 114.9882113 | -0.090163481 | 0.181278643 | -0.497375088 | 0.61892457 | 0.774119712 |
| ENSG00000242689 | CNTF | 9.362190462 | -0.129434212 | 0.561448716 | -0.230536126 | 0.817675195 | 0.90345193 |
| ENSG00000242692 | RPS27AP1 | 36.5987528 | 0.003230258 | 0.292464715 | 0.011044951 | 0.991187583 | 0.995222268 |
| ENSG00000242697 | RPL5P12 | 15.45597845 | 0.254100521 | 0.48317001 | 0.525902924 | 0.598955657 | 0.759076447 |
| ENSG00000242715 | CCDC169 | 11.05770942 | 1.582238787 | 0.568779 | 2.781816464 | 0.00540556 | 0.017249955 |
| ENSG00000242732 | RTL5 | 506.496055 | 0.697539533 | 0.09664474 | 7.217563322 | 5.29E-13 | 5.52E-12 |
| ENSG00000242737 | ENSG0000024 | 4.620140797 | -1.172007295 | 0.872358274 | -1.343493069 | 0.179112369 | 0.333883929 |
| ENSG00000242741 | LINC02005 | 22.42791357 | -0.738878771 | 0.39176952 | -1.886003718 | 0.059294454 | 0.138134941 |
| ENSG00000242748 | RPL23AP81 | 4.39973978 | 0.714792691 | 0.870010718 | 0.821590673 | 0.4113099 | 0.603671145 |
| ENSG00000242756 | RHOT1P3 | 45.09176023 | -0.09519152 | 0.259945615 | -0.366197828 | 0.714217463 | 0.840989046 |
| ENSG00000242759 | LINC00882 | 4.898560694 | -0.40179428 | 0.829863196 | -0.484169297 | 0.628265724 | 0.780689734 |
| ENSG00000242779 | ZNF702P | 91.89902029 | 0.269593538 | 0.192150817 | 1.403030923 | 0.160607618 | 0.307002646 |
| ENSG00000242791 | PFN2-AS1 | 5.076911727 | -0.611604938 | 0.788331031 | -0.775822483 | 0.437853807 | 0.627419972 |
| ENSG00000242798 | ENSG0000024 | 32.18086391 | -1.195414162 | 0.366360812 | -3.262942219 | 0.00110262 | 0.004096874 |
| ENSG00000242802 | AP5Z1 | 2203.083705 | 0.221063573 | 0.056335538 | 3.924051881 | 8.71E-05 | 0.000392346 |
| ENSG00000242808 | SOX2-OT | 3.934987426 | 0.957809541 | 0.950263724 | 1.007940761 | 0.313482902 | 0.501835155 |
| ENSG00000242814 | ENSG0000024 | 38.39464657 | 0.449726009 | 0.354311159 | 1.269296769 | 0.204335238 | 0.369235949 |
| ENSG00000242841 | KRT8P35 | 44.86394275 | 0.35534743 | 0.276225758 | 1.286438428 | 0.198290094 | 0.360813418 |
| ENSG00000242849 | ALDOAP1 | 3.833830745 | 0.324977448 | 0.891421486 | 0.364560932 | 0.715439182 | 0.841656152 |
| ENSG00000242852 | ZNF709 | 6.4518154 | 0.480528575 | 0.693861924 | 0.692542072 | 0.488596974 | 0.671826354 |
| ENSG00000242861 | ENSG0000024 | 68.20253282 | 0.237826285 | 0.217243148 | 1.094746999 | 0.273627492 | 0.455627542 |
| ENSG00000242866 | STRC | 12.94431279 | 0.389621963 | 0.490333178 | 0.794606566 | 0.426842397 | 0.61762108 |
| ENSG00000242882 | RPL5P11 | 2.614513634 | -0.1792192 | 1.057210997 | -0.169520749 | 0.865387053 | 0.930096468 |
| ENSG00000242899 | RPL7P16 | 16.47452372 | 0.504479518 | 0.458836104 | 1.099476511 | 0.271560274 | 0.453577069 |
| ENSG00000242951 | ENSG0000024 | 135.5749805 | -0.017364596 | 0.157710433 | -0.110104297 | 0.91232666 | 0.955898542 |
| ENSG00000242952 | RPSAP3 | 15.82668381 | 0.035337852 | 0.441770366 | 0.079991449 | 0.936244057 | 0.969525239 |
| ENSG00000242960 | FTH1P23 | 25.56890456 | 0.414442829 | 0.364685157 | 1.136440078 | 0.25577243 | 0.435107123 |
| ENSG00000242968 | SULT1D1P2 | 4.514204504 | 0.055323036 | 0.872947948 | 0.063374953 | 0.949467931 | 0.976618218 |
| ENSG00000242970 | ENSG0000024 | 8.709392153 | 0.316984425 | 0.611482017 | 0.518387158 | 0.604188173 | 0.763029194 |
| ENSG00000242992 | FTH1P4 | 14.88736308 | 0.746279051 | 0.48842591 | 1.527926827 | 0.126530707 | 0.255872215 |
| ENSG00000243004 | ENSG0000024 | 2.504827894 | -1.658277551 | 1.203006019 | -1.378444933 | 0.168065958 | 0.318022036 |

| | | | | | | | |
|---|---|---|---|---|---|---|---|
| ENSG00000243007 | RPL12P35 | 3.899628355 | 0.789938965 | 0.89320232 | 0.884389738 | 0.376485861 | 0.569320082 |
| ENSG00000243014 | PTMAP8 | 16.40778372 | -0.612006526 | 0.43095065 | -1.420131344 | 0.155569447 | 0.299468248 |
| ENSG00000243016 | PRR23E2P | 1.999933074 | -4.535837628 | 1.574948645 | -2.879990813 | 0.003976868 | 0.013107664 |
| ENSG00000243023 | UBA52P3 | 17.25265294 | 0.208411296 | 0.424356303 | 0.491123367 | 0.623339195 | 0.777266998 |
| ENSG00000243024 | ENSG0000024 | 1.730883139 | 1.262949612 | 1.404793428 | 0.899028702 | 0.368637373 | NA |
| ENSG00000243069 | ARHGEF26-AS | 53.86579524 | 0.991685469 | 0.259356066 | 3.823644779 | 0.000131493 | 0.000574892 |
| ENSG00000243071 | RPL21P98 | 5.400669992 | 0.155937171 | 0.857151394 | 0.181924887 | 0.855641673 | 0.925169031 |
| ENSG00000243094 | RPL32P2 | 3.448632132 | -0.514663655 | 0.963989072 | -0.533889512 | 0.593417979 | 0.755522723 |
| ENSG00000243101 | RPS3P7 | 3.169244411 | 0.871027691 | 1.041249399 | 0.836521674 | 0.402861489 | 0.595543405 |
| ENSG00000243107 | ENSG0000024 | 3.102448027 | 0.007196774 | 0.998845676 | 0.007205091 | 0.994251219 | 0.996533822 |
| ENSG00000243137 | PSG4 | 4.106186042 | -0.312910072 | 0.892605875 | -0.350557934 | 0.72592002 | 0.847853251 |
| ENSG00000243144 | LINC02932 | 1.73838368 | -4.351118804 | 1.656924291 | -2.626021495 | 0.008638936 | NA |
| ENSG00000243147 | MRPL33 | 898.118038 | -0.252527927 | 0.069912674 | -3.61204791 | 0.000303788 | 0.00124958 |
| ENSG00000243155 | ENSG0000024 | 64.45156203 | 1.459527775 | 0.238199852 | 6.127324445 | 8.94E-10 | 7.13E-09 |
| ENSG00000243156 | MICAL3 | 987.7627238 | -0.449636653 | 0.085552792 | -5.255663104 | 1.47E-07 | 9.42E-07 |
| ENSG00000243160 | RPL7AP32 | 1.872649755 | -0.120111933 | 1.289319499 | -0.093159169 | 0.925777112 | 0.964514676 |
| ENSG00000243176 | ENSG0000024 | 85.69565123 | 0.046895762 | 0.193193035 | 0.242740441 | 0.808206477 | 0.898401971 |
| ENSG00000243181 | RPL21P80 | 7.411588346 | -0.058818021 | 0.657244449 | -0.089491849 | 0.928691031 | 0.965737636 |
| ENSG00000243199 | RPL6P10 | 1722.517048 | 0.202589814 | 0.060077889 | 3.372119356 | 0.000745921 | 0.002860009 |
| ENSG00000243224 | TWF2-DT | 7.841992243 | -1.884803923 | 0.695852486 | -2.708625693 | 0.006756251 | 0.02102393 |
| ENSG00000243232 | PCDHAC2 | 46.7192225 | 1.27590068 | 0.271388763 | 4.701376235 | 2.58E-06 | 1.43E-05 |
| ENSG00000243244 | STON1 | 272.3382191 | 0.468807206 | 0.114222767 | 4.104323666 | 4.05E-05 | 0.000191955 |
| ENSG00000243256 | RPL30P14 | 3.473672021 | 0.512862697 | 0.963698072 | 0.532181927 | 0.594599999 | 0.756355993 |
| ENSG00000243279 | PRAF2 | 204.2958203 | 0.173742805 | 0.136044521 | 1.277102552 | 0.201566044 | 0.365296654 |
| ENSG00000243289 | AGAP13P | 3.05516461 | -1.592688519 | 1.069606984 | -1.489040874 | 0.136476606 | 0.271274009 |
| ENSG00000243302 | ENSG0000024 | 409.6036099 | -0.089408905 | 0.097857048 | -0.913668534 | 0.360891043 | 0.553435425 |
| ENSG00000243312 | RPL6P13 | 2.595540515 | -0.464241122 | 1.146284686 | -0.404996357 | 0.685480189 | 0.821959935 |
| ENSG00000243317 | STMP1 | 1741.168567 | -0.244281268 | 0.060215264 | -4.056799761 | 4.97E-05 | 0.000232862 |
| ENSG00000243323 | PTPRVP | 8.812344878 | 0.962976349 | 0.668652698 | 1.440174177 | 0.149818127 | 0.291355361 |
| ENSG00000243335 | KCTD7 | 166.5889093 | 0.833968527 | 0.141880379 | 5.877969404 | 4.15E-09 | 3.12E-08 |
| ENSG00000243364 | EFNA4 | 376.9360576 | 0.903330408 | 0.100495113 | 8.988799352 | 2.50E-19 | 4.03E-18 |
| ENSG00000243368 | MCCC1-AS1 | 6.946775959 | -0.334896002 | 0.67539366 | -0.495853044 | 0.619998097 | 0.77508209 |
| ENSG00000243388 | RPL3P3 | 14.35313691 | 0.173967473 | 0.45868307 | 0.379275984 | 0.704482931 | 0.834710937 |

| | | | | | | | |
|---|---|---|---|---|---|---|---|
| ENSG00000243403 | ENSG0000024 | 2.714788634 | -0.935030907 | 1.059308809 | -0.882680196 | 0.377409085 | 0.570152414 |
| ENSG00000243404 | RPL35AP32 | 1.645811783 | 1.044097574 | 1.403033707 | 0.744171412 | 0.456772781 | NA |
| ENSG00000243406 | MRPS31P5 | 11.17780224 | -0.542121009 | 0.536876501 | -1.009768557 | 0.312606188 | 0.500805847 |
| ENSG00000243410 | PSMD6-AS1 | 4.89627929 | 1.043830371 | 0.822674853 | 1.26882494 | 0.204503508 | 0.369423614 |
| ENSG00000243422 | RPL23AP49 | 22.14670115 | 0.751486186 | 0.381271654 | 1.970999358 | 0.048723949 | 0.117174483 |
| ENSG00000243429 | OR7E29P | 1.824000836 | -0.383926289 | 1.33913551 | -0.286697116 | 0.774344246 | 0.877192313 |
| ENSG00000243431 | RPL5P30 | 8.356598055 | 0.757049992 | 0.619854317 | 1.221335355 | 0.221959072 | 0.392941898 |
| ENSG00000243445 | ENSG0000024 | 12.34880168 | 0.058936116 | 0.498737063 | 0.118170716 | 0.905932392 | 0.95262805 |
| ENSG00000243449 | C4orf48 | 106.5453814 | 0.431071087 | 0.189221327 | 2.278131609 | 0.022718738 | 0.061076742 |
| ENSG00000243477 | NAA80 | 162.3650707 | 0.375454102 | 0.141734896 | 2.648988438 | 0.008073309 | 0.024637717 |
| ENSG00000243478 | AOX2P | 3.436134819 | -3.33941861 | 1.297683177 | -2.573369733 | 0.010071355 | 0.029919307 |
| ENSG00000243479 | MNX1-AS1 | 73.52685582 | 0.930258593 | 0.223086018 | 4.169954708 | 3.05E-05 | 0.000147507 |
| ENSG00000243491 | ENSG0000024 | 1.798044741 | 0.620694028 | 1.305963879 | 0.475276566 | 0.63458986 | 0.784558953 |
| ENSG00000243495 | GMFBP1 | 6.340260344 | -0.755080276 | 0.72243739 | -1.045184381 | 0.295937761 | 0.482057249 |
| ENSG00000243498 | UBA52P5 | 5.920293503 | -0.182872355 | 0.714417046 | -0.255974233 | 0.797970734 | 0.892208833 |
| ENSG00000243504 | RPS23P1 | 2.877431107 | 1.174469796 | 1.106895641 | 1.061048352 | 0.288667928 | 0.472631666 |
| ENSG00000243508 | DNAJB6P7 | 1.831583945 | -0.96399992 | 1.304933587 | -0.738734852 | 0.460068018 | 0.646886934 |
| ENSG00000243517 | RPL13AP22 | 78.58527186 | 0.380946593 | 0.224258678 | 1.698692763 | 0.089377088 | 0.193297859 |
| ENSG00000243538 | RPS26P28 | 2.539836567 | -1.156076445 | 1.152758491 | -1.002878273 | 0.315919597 | 0.504714503 |
| ENSG00000243547 | HNRNPKP4 | 2125.107937 | -0.143200731 | 0.056374343 | -2.540175594 | 0.011079683 | 0.032518925 |
| ENSG00000243554 | CCZ1P1 | 3.100134709 | 0.893375147 | 1.040274789 | 0.858787655 | 0.39045768 | 0.583343425 |
| ENSG00000243566 | UPK3B | 167.9824625 | 2.18665893 | 0.172266519 | 12.69346446 | 6.43E-37 | 2.34E-35 |
| ENSG00000243592 | RPL17P22 | 4.245219161 | 1.65455714 | 0.933320337 | 1.772764478 | 0.076267742 | 0.170192801 |
| ENSG00000243613 | ENSG0000024 | 4.599377879 | -0.286466072 | 0.85141236 | -0.336459847 | 0.736524126 | 0.854664548 |
| ENSG00000243646 | IL10RB | 12.9291626 | 0.380194233 | 0.4988038 | 0.762211982 | 0.445933496 | 0.634558302 |
| ENSG00000243648 | ENSG0000024 | 1.814168374 | 1.874659881 | 1.515912188 | 1.236654667 | 0.216215316 | 0.385570533 |
| ENSG00000243659 | ENSG0000024 | 2.909036727 | -0.478342772 | 1.018839803 | -0.469497531 | 0.638714049 | 0.787290808 |
| ENSG00000243660 | ZNF487 | 81.83672706 | -0.559958499 | 0.198468804 | -2.821393019 | 0.004781558 | 0.015486227 |
| ENSG00000243663 | RPS4XP14 | 23.18801352 | 0.361766745 | 0.383633155 | 0.943001773 | 0.345679996 | 0.536601645 |
| ENSG00000243667 | DNAAF10 | 117.8241206 | 0.442766597 | 0.172675478 | 2.564154452 | 0.010342751 | 0.030606644 |
| ENSG00000243678 | NME2 | 79.44717979 | -0.124973097 | 0.20889048 | -0.598270907 | 0.549659186 | 0.722005555 |
| ENSG00000243679 | ENSG0000024 | 210.2083371 | -0.046938832 | 0.131824642 | -0.356070242 | 0.721787962 | 0.845596895 |
| ENSG00000243680 | RPL37P23 | 249.0064962 | 0.277087373 | 0.122388926 | 2.263990567 | 0.023574695 | 0.063155845 |

| | | | | | | | |
|---|---|---|---|---|---|---|---|
| ENSG00000243696 | ENSG0000024 | 10.30973301 | 0.853909278 | 0.572812304 | 1.490731383 | 0.136032033 | 0.270648295 |
| ENSG00000243701 | DUBR | 325.8484093 | 0.900256793 | 0.109387243 | 8.229997996 | 1.87E-16 | 2.50E-15 |
| ENSG00000243708 | PLA2G4B | 5.823775634 | 0.336435134 | 0.736485885 | 0.456811381 | 0.647806625 | 0.794519032 |
| ENSG00000243710 | CFAP57 | 19.24744039 | 1.048859985 | 0.412682179 | 2.541568402 | 0.011035635 | 0.032402082 |
| ENSG00000243711 | RPL21P116 | 3.124718722 | -0.041872945 | 1.001331956 | -0.041817246 | 0.966644387 | 0.985153593 |
| ENSG00000243716 | NPIPB5 | 130.8704862 | -0.151610066 | 0.160404149 | -0.945175462 | 0.344569305 | 0.535530019 |
| ENSG00000243725 | TTC4 | 22.12503895 | -0.281948616 | 0.371607269 | -0.758727399 | 0.448015644 | 0.636304641 |
| ENSG00000243738 | RN7SL181P | 2.400851469 | 0.820643446 | 1.149935833 | 0.713642816 | 0.475448075 | 0.660656541 |
| ENSG00000243742 | RPLP0P2 | 48.93228086 | -0.996039454 | 0.26648621 | -3.737677291 | 0.000185728 | 0.000792818 |
| ENSG00000243746 | EEF1A1P10 | 18.5836607 | 0.103878248 | 0.418987054 | 0.247927106 | 0.804190807 | 0.895880917 |
| ENSG00000243753 | HLA-L | 10.46720483 | 0.454226515 | 0.53821636 | 0.843947805 | 0.398698573 | 0.591597444 |
| ENSG00000243759 | ST13P15 | 172.1751926 | 0.255560809 | 0.138492774 | 1.845300669 | 0.064993813 | 0.14915547 |
| ENSG00000243762 | ENSG0000024 | 4.712165792 | -0.643796529 | 0.827054485 | -0.77842094 | 0.436320898 | 0.626240468 |
| ENSG00000243775 | OSTCP1 | 2.01221896 | -0.644878605 | 1.269771337 | -0.50786987 | 0.611544606 | 0.76912463 |
| ENSG00000243779 | RPL36AP49 | 19.71785788 | 0.133870971 | 0.417632026 | 0.320547664 | 0.748553204 | 0.86082868 |
| ENSG00000243789 | JMJD7 | 13.55697201 | -0.242206825 | 0.47203651 | -0.513110363 | 0.607874123 | 0.766146496 |
| ENSG00000243802 | ENSG0000024 | 31.26271722 | 0.134843308 | 0.352837577 | 0.382168218 | 0.702336595 | 0.833215242 |
| ENSG00000243819 | RN7SL832P | 33.07406243 | -0.47413611 | 0.306416699 | -1.547357278 | 0.121777117 | 0.248317072 |
| ENSG00000243824 | RPL12P6 | 13.8840853 | -0.45016945 | 0.482701155 | -0.932604875 | 0.351024015 | 0.542364849 |
| ENSG00000243829 | ENSG0000024 | 90.5632147 | 0.289723982 | 0.234346103 | 1.236308086 | 0.216344067 | 0.385740159 |
| ENSG00000243830 | RPL15P17 | 7.220619004 | 0.599969592 | 0.667908413 | 0.898281232 | 0.369035636 | 0.561572617 |
| ENSG00000243838 | PSMC1P7 | 14.79127245 | 0.029640394 | 0.465311511 | 0.063700109 | 0.949209018 | 0.97658227 |
| ENSG00000243844 | RPL17P33 | 1.804640217 | 0.101857977 | 1.278529811 | 0.07966805 | 0.936501271 | 0.969552268 |
| ENSG00000243859 | RPL5P17 | 33.39327732 | 0.721353442 | 0.325832526 | 2.213877943 | 0.026837176 | 0.070609569 |
| ENSG00000243885 | ENSG0000024 | 3.282133716 | 0.112413985 | 0.947278266 | 0.1186705 | 0.905536409 | 0.95262805 |
| ENSG00000243896 | OR2A7 | 2.584848977 | 0.320203124 | 1.134996912 | 0.282118057 | 0.777852988 | 0.879532114 |
| ENSG00000243904 | RPSAP5 | 3.589421953 | -0.149233013 | 0.939388376 | -0.15886189 | 0.873777687 | 0.934676032 |
| ENSG00000243915 | THAP12P2 | 3.285993117 | 0.364038952 | 1.044487211 | 0.348533662 | 0.727439438 | 0.84863002 |
| ENSG00000243918 | EIF4BP8 | 8.863092327 | -0.155935535 | 0.607321113 | -0.256759615 | 0.797364348 | 0.89183367 |
| ENSG00000243926 | TIPARP-AS1 | 50.03542161 | 0.389680658 | 0.25359307 | 1.536637643 | 0.124382069 | 0.252484264 |
| ENSG00000243927 | MRPS6 | 607.9193876 | 0.017478154 | 0.079909139 | 0.218725344 | 0.826864005 | 0.908572126 |
| ENSG00000243943 | ZNF512 | 402.5593023 | 0.372515002 | 0.094023954 | 3.961915945 | 7.44E-05 | 0.000339019 |
| ENSG00000243960 | ENSG0000024 | 5.497194745 | 0.067073439 | 0.763384478 | 0.087863247 | 0.929985368 | 0.966032619 |

| | | | | | | | |
|---|---|---|---|---|---|---|---|
| ENSG00000243964 | RPL23AP65 | 57.17914891 | 0.41770417 | 0.253705834 | 1.646411367 | 0.099679094 | 0.211451676 |
| ENSG00000243970 | PPIEL | 62.58949458 | 0.518933105 | 0.241294458 | 2.150621729 | 0.031506069 | 0.081033528 |
| ENSG00000243974 | VTI1BP1 | 17.30337874 | -0.553942368 | 0.438903589 | -1.262104895 | 0.206911044 | 0.372833191 |
| ENSG00000243977 | H3P13 | 4.843288809 | -0.267583919 | 0.801032217 | -0.334048886 | 0.73834267 | 0.855807824 |
| ENSG00000243978 | RTL9 | 12.26647129 | -0.370608745 | 0.502126175 | -0.73807892 | 0.460466493 | 0.647248857 |
| ENSG00000243988 | RPS24P17 | 4.775241845 | -0.092412895 | 0.803355984 | -0.115033555 | 0.908418525 | 0.954215897 |
| ENSG00000243989 | ACY1 | 11.50382588 | -0.58724722 | 0.51799735 | -1.13368769 | 0.256925575 | 0.436518493 |
| ENSG00000244005 | NFS1 | 794.9769436 | -0.317038746 | 0.071984181 | -4.404283566 | 1.06E-05 | 5.46E-05 |
| ENSG00000244009 | B3GAT3P1 | 24.43101645 | 0.818708627 | 0.369335026 | 2.216709949 | 0.026642918 | 0.070162846 |
| ENSG00000244026 | FAM86DP | 369.1594791 | 0.035362372 | 0.100973915 | 0.350212949 | 0.72617889 | 0.847853251 |
| ENSG00000244036 | UBE2H-DT | 1.657118599 | -0.747262851 | 1.378533733 | -0.542070777 | 0.587769749 | NA |
| ENSG00000244038 | DDOST | 5027.353303 | 0.055124 | 0.049536347 | 1.112799056 | 0.265794741 | 0.446828286 |
| ENSG00000244040 | IL12A-AS1 | 1.633450164 | -1.398839893 | 1.427059346 | -0.980225452 | 0.326974843 | NA |
| ENSG00000244041 | LINC01011 | 58.62709369 | -0.360355964 | 0.241572545 | -1.491709102 | 0.135775422 | 0.27034877 |
| ENSG00000244045 | TMEM199 | 681.5103549 | 0.079419323 | 0.078687734 | 1.009297358 | 0.312832047 | 0.501037992 |
| ENSG00000244052 | RPL5P24 | 3.013125766 | 1.30274317 | 1.052668359 | 1.23756277 | 0.215878226 | 0.3851191 |
| ENSG00000244055 | ENSG0000024 | 96.95796777 | -0.365490853 | 0.21281191 | -1.717436083 | 0.085899524 | 0.187227973 |
| ENSG00000244073 | RPS4XP6 | 56.47687881 | 0.362404012 | 0.271795993 | 1.33336775 | 0.18241115 | 0.3382811 |
| ENSG00000244076 | ENSG0000024 | 2.633436981 | -0.536005381 | 1.144195445 | -0.468456139 | 0.63945843 | 0.78769137 |
| ENSG00000244086 | ENSG0000024 | 3.180912442 | -0.141290112 | 1.105513885 | -0.127804919 | 0.898303357 | 0.948554546 |
| ENSG00000244097 | RPS4XP17 | 13.48635648 | 0.220035822 | 0.485559351 | 0.453159479 | 0.650433908 | 0.796761567 |
| ENSG00000244119 | PDCL3P4 | 80.55497215 | 0.151156025 | 0.202932898 | 0.744857174 | 0.456358067 | 0.643478245 |
| ENSG00000244128 | LINC01322 | 6.904228741 | 0.225014759 | 0.65533119 | 0.343360368 | 0.731327371 | 0.851217712 |
| ENSG00000244142 | ATP6V0CP2 | 3.871594568 | -1.129004809 | 0.991499852 | -1.138683789 | 0.254835066 | 0.434220933 |
| ENSG00000244151 | ENSG0000024 | 61.12221934 | 0.364071725 | 0.228478006 | 1.593465082 | 0.111055896 | 0.230512328 |
| ENSG00000244153 | WWP1P1 | 181.18573 | 0.409005362 | 0.139128013 | 2.939777202 | 0.003284483 | 0.011039714 |
| ENSG00000244161 | FLNB-AS1 | 13.49103429 | -1.223180835 | 0.502073457 | -2.436258718 | 0.014840066 | 0.042083008 |
| ENSG00000244165 | P2RY11 | 248.9814447 | 0.142298092 | 0.127877051 | 1.11277271 | 0.265806059 | 0.446828286 |
| ENSG00000244171 | PBX2P1 | 188.2631298 | -0.133080283 | 0.133837477 | -0.994342435 | 0.320056183 | 0.509515996 |
| ENSG00000244184 | PELP1-DT | 11.64883054 | -0.816062444 | 0.524018079 | -1.557317346 | 0.119395159 | 0.24454687 |
| ENSG00000244187 | TMEM141 | 361.9153429 | 0.095711099 | 0.099780408 | 0.959217352 | 0.337449261 | 0.528690482 |
| ENSG00000244192 | ENSG0000024 | 13.32227804 | 0.508790605 | 0.48862901 | 1.04126156 | 0.297754172 | 0.484293868 |
| ENSG00000244198 | ARHGEF35-AS | 45.35234109 | 0.512875996 | 0.26658184 | 1.923896976 | 0.054367498 | 0.128788766 |

| | | | | | | | |
|---|---|---|---|---|---|---|---|
| ENSG00000244218 | RN7SL81P | 3.556199703 | -1.24627684 | 1.034664588 | -1.204522562 | 0.228387667 | 0.401243882 |
| ENSG00000244226 | ILF2P1 | 4.746436264 | 0.507356681 | 0.810974231 | 0.625613813 | 0.531568277 | 0.707118252 |
| ENSG00000244229 | RPL26P35 | 2.530380464 | -0.741791384 | 1.118031421 | -0.663479907 | 0.507023244 | 0.687336793 |
| ENSG00000244234 | GMCL2 | 20.97033358 | 0.276694261 | 0.395104643 | 0.700306276 | 0.483736053 | 0.667723994 |
| ENSG00000244242 | IFITM10 | 50.77380952 | 2.079364017 | 0.296230573 | 7.019410569 | 2.23E-12 | 2.22E-11 |
| ENSG00000244245 | ENSG0000024 | 63.75276492 | 0.001598376 | 0.229412771 | 0.006967248 | 0.994440985 | 0.996613202 |
| ENSG00000244257 | PKD1P1 | 5.504326389 | 0.194795169 | 0.783997151 | 0.248464128 | 0.803775321 | 0.89551267 |
| ENSG00000244265 | SIAH2-AS1 | 29.66374091 | 0.318725363 | 0.323689522 | 0.984663828 | 0.324789226 | 0.514904274 |
| ENSG00000244268 | ENSG0000024 | 1.628716542 | -1.405767744 | 1.480865477 | -0.949287944 | 0.342474184 | NA |
| ENSG00000244274 | DBNDD2 | 46.97170778 | -1.060594599 | 0.271408812 | -3.907738261 | 9.32E-05 | 0.000417093 |
| ENSG00000244300 | GATA2-AS1 | 15.6597494 | 0.186572314 | 0.458473705 | 0.406942235 | 0.684050415 | 0.820803249 |
| ENSG00000244306 | DUXAP10 | 393.4964133 | 0.134671573 | 0.098011502 | 1.374038454 | 0.169429745 | 0.320015611 |
| ENSG00000244313 | ENSG0000024 | 4817.623579 | 0.260475641 | 0.048931755 | 5.323243377 | 1.02E-07 | 6.60E-07 |
| ENSG00000244331 | ENSG0000024 | 3.137483866 | -0.035227784 | 0.988109288 | -0.035651708 | 0.971560078 | 0.987006252 |
| ENSG00000244349 | ENSG0000024 | 2.841961999 | -0.015640943 | 1.038134053 | -0.0150664 | 0.987979207 | 0.994030148 |
| ENSG00000244361 | RPL30P7 | 1.858610868 | 0.563327238 | 1.325595561 | 0.424961621 | 0.670864653 | 0.811185291 |
| ENSG00000244363 | RPL7P23 | 40.31444525 | 0.7842589 | 0.297898212 | 2.632640505 | 0.008472397 | 0.025691578 |
| ENSG00000244380 | ENSG0000024 | 15.42447564 | -0.239322944 | 0.469616964 | -0.509613072 | 0.610322564 | 0.768261612 |
| ENSG00000244381 | SDHDP3 | 3.728422277 | 0.75481931 | 0.936947502 | 0.805615372 | 0.420464659 | 0.611711015 |
| ENSG00000244383 | FAM3D-AS1 | 2.679387576 | 0.497263155 | 1.072750249 | 0.46354047 | 0.642977014 | 0.79066699 |
| ENSG00000244398 | RPL36AP37 | 6177.110735 | 0.137978044 | 0.055060889 | 2.505917489 | 0.012213412 | 0.03548324 |
| ENSG00000244405 | ETV5 | 165.9965647 | -1.049033317 | 0.158341778 | -6.625120225 | 3.47E-11 | 3.11E-10 |
| ENSG00000244411 | KRTAP5-7 | 3.671780144 | 0.589627457 | 0.935813636 | 0.630069316 | 0.528649234 | 0.704660862 |
| ENSG00000244459 | ENSG0000024 | 6.433667934 | 0.139190682 | 0.692883569 | 0.200886106 | 0.840787632 | 0.91647005 |
| ENSG00000244462 | RBM12 | 3079.382199 | -0.425819543 | 0.052687738 | -8.081947728 | 6.37E-16 | 8.18E-15 |
| ENSG00000244468 | TM4SF18-AS1 | 60.47912485 | 0.472960828 | 0.233524108 | 2.025319064 | 0.042834612 | 0.105026294 |
| ENSG00000244471 | TPTE2P3 | 2.706757112 | -0.940930372 | 1.084972922 | -0.867238576 | 0.385811304 | 0.578389186 |
| ENSG00000244480 | ENSG0000024 | 1.948408384 | -2.374193142 | 1.455948076 | -1.630685312 | 0.102956737 | 0.217059788 |
| ENSG00000244485 | RPL18P13 | 29.02858825 | 0.061519218 | 0.330741293 | 0.186004045 | 0.852441589 | 0.922972108 |
| ENSG00000244486 | SCARF2 | 38.77175097 | -0.424329336 | 0.27982549 | -1.516407019 | 0.129416465 | 0.260330912 |
| ENSG00000244490 | RWDD4P1 | 7.297651701 | 0.053850046 | 0.654659076 | 0.082256625 | 0.934442646 | 0.968469969 |
| ENSG00000244491 | ENSG0000024 | 3.740290919 | 0.443273493 | 0.90534566 | 0.489617958 | 0.624404267 | 0.778004121 |
| ENSG00000244502 | HDAC11-AS1 | 1.78534419 | -0.311200623 | 1.429980525 | -0.217625777 | 0.827720699 | NA |

| | | | | | | | |
|---|---|---|---|---|---|---|---|
| ENSG00000244509 | APOBEC3C | 5.893904139 | -0.461444145 | 0.736427709 | -0.626598021 | 0.530922766 | 0.706541841 |
| ENSG00000244513 | EOGT-DT | 36.31308081 | 0.139153362 | 0.308133886 | 0.451600322 | 0.651556938 | 0.797581514 |
| ENSG00000244515 | KRT18P34 | 10.37730897 | 0.698315307 | 0.560692047 | 1.245452491 | 0.212965469 | 0.381108344 |
| ENSG00000244527 | RPL21P128 | 14.97023821 | 0.713197357 | 0.481821137 | 1.480211851 | 0.138816719 | 0.274770863 |
| ENSG00000244538 | RPS27P13 | 15.64049635 | 0.528939436 | 0.477684671 | 1.107298326 | 0.268164968 | 0.449653122 |
| ENSG00000244541 | LINC01213 | 17.21920007 | 1.131499978 | 0.444768716 | 2.544018808 | 0.010958518 | 0.032200386 |
| ENSG00000244558 | KCNK15-AS1 | 15.33717399 | 0.048096601 | 0.450392398 | 0.106788217 | 0.914956995 | 0.957751014 |
| ENSG00000244560 | ENSG0000024 | 191.0854214 | 0.893534412 | 0.146341652 | 6.105810608 | 1.02E-09 | 8.12E-09 |
| ENSG00000244582 | RPL21P120 | 15.99152356 | 0.623762747 | 0.451483706 | 1.381584182 | 0.167099414 | 0.316661723 |
| ENSG00000244585 | RPL12P33 | 6.064528362 | 0.212415552 | 0.704457719 | 0.30153059 | 0.763009928 | 0.870125658 |
| ENSG00000244604 | ENSG0000024 | 3.505786644 | -0.025437613 | 0.980500907 | -0.025943488 | 0.979302414 | 0.990703712 |
| ENSG00000244617 | ASPRV1 | 16.08312266 | 0.202010697 | 0.459854815 | 0.439292337 | 0.660449726 | 0.803707149 |
| ENSG00000244625 | MIATNB | 49.23584564 | 0.07092363 | 0.251522159 | 0.281977661 | 0.77796064 | 0.879540484 |
| ENSG00000244627 | TPTEP2 | 26.49406894 | 0.747179099 | 0.34538871 | 2.163299141 | 0.030518179 | 0.078847509 |
| ENSG00000244668 | SNRPCP3 | 5.412978462 | 0.899528029 | 0.773734553 | 1.162579628 | 0.2450001 | 0.421944617 |
| ENSG00000244687 | UBE2V1 | 84.31054675 | -0.513583682 | 0.196094182 | -2.61906639 | 0.008817079 | 0.026631176 |
| ENSG00000244694 | PTCHD4 | 11.75430623 | -1.809703205 | 0.551390701 | -3.282070592 | 0.001030478 | 0.003848773 |
| ENSG00000244701 | ENSG0000024 | 26.1914823 | 0.087558532 | 0.367783286 | 0.238070991 | 0.811826033 | 0.900415279 |
| ENSG00000244705 | RPLP0P10 | 63.17531796 | 0.208512571 | 0.245022517 | 0.850993507 | 0.39477296 | 0.587790725 |
| ENSG00000244716 | RPL17P7 | 1306.255653 | 0.569544208 | 0.072460025 | 7.860116067 | 3.84E-15 | 4.65E-14 |
| ENSG00000244723 | ASLP1 | 3.144472354 | -3.205968207 | 1.352172854 | -2.37097513 | 0.017741225 | 0.049289536 |
| ENSG00000244731 | C4A | 18.46867766 | 0.039179249 | 0.407373504 | 0.096175252 | 0.923381386 | 0.963135281 |
| ENSG00000244733 | ENSG0000024 | 1.75497738 | 0.693558491 | 1.353285476 | 0.512499767 | 0.608301283 | NA |
| ENSG00000244754 | N4BP2L2 | 1987.774033 | 0.13902968 | 0.062322042 | 2.230826792 | 0.025692605 | 0.067956687 |
| ENSG00000244879 | GABPB1-AS1 | 6545.231438 | -0.875749351 | 0.115864549 | -7.558389181 | 4.08E-14 | 4.58E-13 |
| ENSG00000244921 | MTCYBP18 | 35.04632892 | 0.092265015 | 0.318129755 | 0.290023216 | 0.771798476 | 0.875561078 |
| ENSG00000244945 | RUFY1-AS1 | 15.4650491 | -0.567024467 | 0.447120591 | -1.268168988 | 0.204737609 | 0.369700941 |
| ENSG00000244968 | LIFR-AS1 | 155.1982931 | 0.016987998 | 0.147278154 | 0.115346353 | 0.908170599 | 0.954086533 |
| ENSG00000244998 | ENSG0000024 | 3.771809968 | -1.649449817 | 0.979598179 | -1.683802453 | 0.092219853 | 0.198417129 |
| ENSG00000245025 | ENSG0000024 | 40.4971338 | 0.414101488 | 0.279635145 | 1.480863528 | 0.138642944 | 0.274591882 |
| ENSG00000245060 | LINC00847 | 664.0690813 | -0.090173228 | 0.079435741 | -1.135171984 | 0.256303266 | 0.43578394 |
| ENSG00000245080 | MIR3150BHG | 24.52752198 | 0.595943392 | 0.355510597 | 1.676302753 | 0.093678906 | 0.200953254 |
| ENSG00000245105 | A2M-AS1 | 14.22304351 | -0.269143673 | 0.468613286 | -0.57434068 | 0.565737286 | 0.733806909 |

| | | | | | | | |
|---|---|---|---|---|---|---|---|
| ENSG00000245112 | SMARCA5-AS1 | 10.87562368 | -0.482490128 | 0.550889728 | -0.875837947 | 0.381118137 | 0.573376841 |
| ENSG00000245146 | MALINC1 | 64.51819638 | -0.272899272 | 0.241142166 | -1.13169454 | 0.257762878 | 0.437519824 |
| ENSG00000245148 | ARAP1-AS2 | 3.660714874 | -0.0282864 | 0.945447093 | -0.029918544 | 0.976132017 | 0.989157212 |
| ENSG00000245149 | RNF139-DT | 72.73092019 | 0.027860588 | 0.245140703 | 0.113651414 | 0.909514129 | 0.954587195 |
| ENSG00000245156 | ENSG0000024 | 106.1183658 | -0.625286377 | 0.208399398 | -3.000423143 | 0.002696048 | 0.009226814 |
| ENSG00000245205 | EEF1A1P4 | 90.40323464 | 0.226571529 | 0.204519776 | 1.107822108 | 0.26793865 | 0.449306479 |
| ENSG00000245213 | GALNT7-DT | 43.50976684 | -0.362494308 | 0.273918774 | -1.323364235 | 0.18571428 | 0.342962935 |
| ENSG00000245248 | USP2-AS1 | 29.03666139 | 0.778942559 | 0.328815107 | 2.368937865 | 0.01783925 | 0.049531881 |
| ENSG00000245261 | ENSG0000024 | 8.564509221 | -0.378336322 | 0.605009491 | -0.625339482 | 0.531748272 | 0.707271141 |
| ENSG00000245275 | SAP30L-AS1 | 22.78793287 | -0.009140377 | 0.366601035 | -0.024932763 | 0.980108594 | 0.991041209 |
| ENSG00000245281 | ASAH1-AS1 | 13.84671571 | -0.684046621 | 0.477903088 | -1.431350074 | 0.15232991 | 0.294865254 |
| ENSG00000245293 | CYP2U1-AS1 | 6.423744814 | -0.846503625 | 0.697606915 | -1.213439269 | 0.22496192 | 0.396849158 |
| ENSG00000245317 | MRNIP-DT | 56.99533704 | -0.10078548 | 0.237039197 | -0.425184868 | 0.670701914 | 0.811118793 |
| ENSG00000245322 | H2AZ1-DT | 29.70859798 | -0.193803763 | 0.337886097 | -0.573577206 | 0.56625394 | 0.734181054 |
| ENSG00000245330 | DEPTOR-AS1 | 2.905285854 | 0.559451092 | 1.149476691 | 0.486700684 | 0.62647046 | 0.779646957 |
| ENSG00000245384 | CXXC4-AS1 | 4.603075301 | -0.376184086 | 0.807730862 | -0.465729495 | 0.641409138 | 0.789596379 |
| ENSG00000245385 | ENSG0000024 | 13.78210938 | -0.330039105 | 0.48393271 | -0.681993795 | 0.495242875 | 0.677573301 |
| ENSG00000245468 | LINC02447 | 15.01185192 | -0.851079597 | 0.462440146 | -1.840410279 | 0.065708026 | 0.150583755 |
| ENSG00000245498 | MSANTD2-AS1 | 40.82357326 | -0.089735005 | 0.277592814 | -0.323261268 | 0.746497384 | 0.859604617 |
| ENSG00000245522 | LINC02709 | 9.050081729 | -0.072775903 | 0.58294201 | -0.124842441 | 0.900648287 | 0.949925136 |
| ENSG00000245526 | LINC00461 | 58.60114081 | -0.278748307 | 0.244338932 | -1.14082641 | 0.253942168 | 0.432860257 |
| ENSG00000245532 | NEAT1 | 3493.353291 | 0.226555457 | 0.222711199 | 1.017261178 | 0.3090292 | 0.49713393 |
| ENSG00000245534 | RORA-AS1 | 14.12274024 | 0.860415397 | 0.479599462 | 1.794029111 | 0.072808522 | 0.163956759 |
| ENSG00000245552 | LNCRNA-IUR | 6.424469248 | -0.627327939 | 0.707876214 | -0.886211355 | 0.375503646 | 0.56836558 |
| ENSG00000245556 | SCAMP1-AS1 | 291.4933762 | 0.374274692 | 0.109428619 | 3.420263308 | 0.000625606 | 0.002432027 |
| ENSG00000245571 | FAM111A-DT | 217.7213889 | -0.206147396 | 0.123735754 | -1.666029337 | 0.095707571 | 0.204635259 |
| ENSG00000245573 | BDNF-AS | 87.19539907 | 0.018037554 | 0.194975853 | 0.092511733 | 0.92629147 | 0.964662124 |
| ENSG00000245614 | DDX11-AS1 | 32.99621691 | 0.06627627 | 0.316631499 | 0.209316729 | 0.834200993 | 0.912438411 |
| ENSG00000245662 | LINC02211 | 2.18432154 | 0.038983321 | 1.176987431 | 0.033121272 | 0.97357788 | 0.988001378 |
| ENSG00000245680 | ZNF585B | 256.6085821 | 0.245631327 | 0.114659458 | 2.142268337 | 0.032171895 | 0.082534475 |
| ENSG00000245685 | FRG1-DT | 15.14463278 | 0.444456781 | 0.457546209 | 0.971392119 | 0.331353051 | 0.52224246 |
| ENSG00000245694 | CRNDE | 1432.040673 | 0.629920217 | 0.074179204 | 8.491870861 | 2.03E-17 | 2.87E-16 |
| ENSG00000245748 | ENSG0000024 | 33.74893407 | -0.562816685 | 0.3196136 | -1.760928462 | 0.078250512 | 0.173578744 |

| | | | | | | | |
|---|---|---|---|---|---|---|---|
| ENSG00000245750 | DRAIC | 238.5306304 | -2.432723538 | 0.137861325 | -17.64616386 | 1.09E-69 | 9.91E-68 |
| ENSG00000245768 | ENSG0000024 | 7.082305263 | -0.174783238 | 0.669928082 | -0.260898509 | 0.794170775 | 0.88974732 |
| ENSG00000245848 | CEBPA | 676.9477376 | -0.495699037 | 0.080271163 | -6.175306525 | 6.60E-10 | 5.34E-09 |
| ENSG00000245849 | RAD51-AS1 | 265.4948345 | -0.004640334 | 0.120847541 | -0.038398247 | 0.969370159 | 0.986272649 |
| ENSG00000245864 | MEF2C-AS2 | 2.538887241 | -1.194413864 | 1.122394655 | -1.064165673 | 0.287253655 | 0.471257873 |
| ENSG00000245888 | NSMCE1-DT | 23.54074558 | -0.14934367 | 0.39031412 | -0.382624308 | 0.701998344 | 0.833064766 |
| ENSG00000245904 | BTG1-DT | 30.33847625 | -0.445503668 | 0.31838808 | -1.399247321 | 0.161738831 | 0.308753331 |
| ENSG00000245910 | SNHG6 | 746.7412046 | 0.59134825 | 0.086448712 | 6.840451827 | 7.89E-12 | 7.46E-11 |
| ENSG00000245937 | SLC12A2-DT | 690.0448779 | -0.158690582 | 0.07711188 | -2.057926502 | 0.039597189 | 0.098352349 |
| ENSG00000245958 | SEPTIN7P14 | 379.0063906 | 0.32691146 | 0.098487058 | 3.319334202 | 0.000902324 | 0.003407854 |
| ENSG00000245970 | RPL30-AS1 | 182.0544138 | -0.574371996 | 0.275957995 | -2.081374723 | 0.03739962 | 0.09380574 |
| ENSG00000245975 | MINDY2-DT | 18.34083652 | 0.059862017 | 0.412691286 | 0.145052776 | 0.884669203 | 0.940249752 |
| ENSG00000246067 | RAB30-DT | 200.3539796 | -0.287144823 | 0.129901123 | -2.210487605 | 0.02707134 | 0.071127767 |
| ENSG00000246089 | MCPH1-DT | 113.3276176 | 0.425871719 | 0.168324487 | 2.530063971 | 0.011404173 | 0.033360333 |
| ENSG00000246090 | ENSG0000024 | 30.55427669 | -0.164224342 | 0.317094004 | -0.51790428 | 0.604525056 | 0.763309378 |
| ENSG00000246115 | SUPT4H1P2 | 6.885312532 | 1.021978481 | 0.680371456 | 1.502089001 | 0.133074125 | 0.266196301 |
| ENSG00000246145 | RRS1-DT | 1.814107461 | 0.112263246 | 1.320844976 | 0.084993506 | 0.932266553 | 0.967526001 |
| ENSG00000246174 | KCTD21-AS1 | 43.79768776 | -1.200142057 | 0.289196496 | -4.149919076 | 3.33E-05 | 0.000159817 |
| ENSG00000246203 | ENSG0000024 | 4.994134392 | 3.119327155 | 1.066939284 | 2.923621992 | 0.003459846 | 0.01156812 |
| ENSG00000246214 | RETREG1-AS1 | 33.98648387 | 1.779617347 | 0.334787872 | 5.315656563 | 1.06E-07 | 6.88E-07 |
| ENSG00000246223 | LINC01550 | 3.762564191 | 4.229027596 | 1.396690159 | 3.027892457 | 0.002462657 | 0.008500269 |
| ENSG00000246225 | ENSG0000024 | 36.58340973 | 0.533791157 | 0.290376169 | 1.838274669 | 0.066021942 | 0.151227493 |
| ENSG00000246228 | CASC8 | 8.217477181 | -0.245117351 | 0.674590781 | -0.363357102 | 0.716338143 | 0.842275602 |
| ENSG00000246263 | UBR5-DT | 76.47388049 | 0.269062722 | 0.210452575 | 1.278495744 | 0.201074684 | 0.364607799 |
| ENSG00000246273 | SBF2-AS1 | 126.9217403 | -0.458791128 | 0.157619774 | -2.910746015 | 0.00360567 | 0.012003202 |
| ENSG00000246283 | TTBK2-AS1 | 8.868635577 | -0.066033137 | 0.593315979 | -0.111295059 | 0.911382373 | 0.955491597 |
| ENSG00000246308 | ENSG0000024 | 5.408298292 | 0.414901545 | 0.759184975 | 0.546509163 | 0.584715991 | 0.74893271 |
| ENSG00000246323 | FAM13B-AS1 | 4.470218987 | -0.96916063 | 0.872197546 | -1.111171012 | 0.266494746 | 0.447572258 |
| ENSG00000246334 | PRR7-AS1 | 14.59420211 | 0.671762755 | 0.486648334 | 1.380386428 | 0.167467697 | 0.317129682 |
| ENSG00000246339 | EXTL3-AS1 | 22.14441371 | 0.28538026 | 0.377639588 | 0.755694768 | 0.449832222 | 0.638126966 |
| ENSG00000246350 | ZBED9-AS1 | 8.056623064 | -1.013520432 | 0.656774229 | -1.543179356 | 0.122787265 | 0.249799561 |
| ENSG00000246366 | LACTB2-AS1 | 6.138443741 | 0.835302257 | 0.750658159 | 1.112759846 | 0.265811585 | 0.446828286 |
| ENSG00000246375 | PPM1K-DT | 2.741177998 | -0.306573037 | 1.045278513 | -0.293293159 | 0.769298083 | 0.87408113 |

| | | | | | | | |
|---|---|---|---|---|---|---|---|
| ENSG00000246451 | KLC1-AS1 | 4.825190791 | -0.536106493 | 0.826576177 | -0.648586915 | 0.516605415 | 0.695057379 |
| ENSG00000246465 | ENSG0000024 | 76.92301159 | -0.707840892 | 0.209818709 | -3.373583293 | 0.000741966 | 0.002846271 |
| ENSG00000246523 | FZD4-DT | 18.90971644 | -0.389377992 | 0.410158422 | -0.9493356 | 0.342449953 | 0.533428527 |
| ENSG00000246526 | LINC02481 | 12.19974295 | -0.013784204 | 0.516953245 | -0.026664314 | 0.978727476 | 0.990384134 |
| ENSG00000246548 | LINC02288 | 5.45888071 | -1.140999983 | 0.762993735 | -1.495425099 | 0.134803534 | 0.268763521 |
| ENSG00000246560 | UBE2D3-AS1 | 20.5488237 | 0.320301494 | 0.408123247 | 0.784815606 | 0.432561689 | 0.622597006 |
| ENSG00000246575 | ENSG0000024 | 2.90389958 | 0.832980856 | 1.036306334 | 0.8037979 | 0.421513703 | 0.612499166 |
| ENSG00000246582 | TNFRSF10A-DT | 167.6620333 | -0.329329297 | 0.140004501 | -2.3522765 | 0.018658899 | 0.051465089 |
| ENSG00000246596 | ENSG0000024 | 40.27177305 | -0.188551562 | 0.296959448 | -0.63494044 | 0.525467261 | 0.701516712 |
| ENSG00000246640 | PICART1 | 5.51795366 | -0.017813647 | 0.749474709 | -0.023768177 | 0.981037524 | 0.99123506 |
| ENSG00000246662 | CIBAR1-DT | 2.538883562 | 0.54132079 | 1.115718611 | 0.485176805 | 0.627550934 | 0.780187429 |
| ENSG00000246695 | RASSF8-AS1 | 240.6721736 | 0.907497022 | 0.127405594 | 7.122897762 | 1.06E-12 | 1.08E-11 |
| ENSG00000246705 | H2AJ | 1784.402627 | 0.162889998 | 0.057965225 | 2.810133106 | 0.004952101 | 0.015973173 |
| ENSG00000246731 | MGC16275 | 2.543195056 | -0.409137947 | 1.081184563 | -0.378416379 | 0.705121303 | 0.834997903 |
| ENSG00000246740 | PLA2G4E-AS1 | 3.394224533 | -0.500867957 | 0.937602557 | -0.534200716 | 0.593202674 | 0.755354595 |
| ENSG00000246763 | RGMB-AS1 | 45.19300247 | 0.534253577 | 0.261994096 | 2.039181739 | 0.041431893 | 0.10205601 |
| ENSG00000246777 | DYNC1LI2-DT | 5.631739467 | -0.209427172 | 0.753514296 | -0.277933906 | 0.781063096 | 0.881298289 |
| ENSG00000246790 | SORL1-AS1 | 2.311540267 | -1.918551599 | 1.353087846 | -1.417906165 | 0.156218162 | 0.300364147 |
| ENSG00000246792 | ENSG0000024 | 41.57902682 | 2.557089055 | 0.341768033 | 7.48194333 | 7.32E-14 | 8.06E-13 |
| ENSG00000246859 | STARD4-AS1 | 23.65665905 | -0.914972369 | 0.364813317 | -2.508056387 | 0.012139729 | 0.035309192 |
| ENSG00000246863 | GPR176-DT | 4.878848167 | 0.105653661 | 0.818915343 | 0.129016585 | 0.897344525 | 0.948060798 |
| ENSG00000246877 | DNM1P35 | 12.28626139 | -0.967165794 | 0.506415316 | -1.909827297 | 0.056155453 | 0.13220506 |
| ENSG00000246889 | CTTN-DT | 66.58654085 | 0.397234089 | 0.227548029 | 1.745715359 | 0.080860425 | 0.178222343 |
| ENSG00000246922 | UBAP1L | 104.5339267 | 0.310157524 | 0.177304638 | 1.749291659 | 0.080240617 | 0.177060384 |
| ENSG00000246982 | BRPF3-AS1 | 32.20029585 | 0.664310634 | 0.366355455 | 1.813295325 | 0.069786284 | 0.158191438 |
| ENSG00000246985 | SOCS2-AS1 | 76.38447174 | -0.551760586 | 0.202243077 | -2.728205064 | 0.006368001 | 0.01997019 |
| ENSG00000247011 | ENSG0000024 | 10.19284154 | -2.289390827 | 0.65658436 | -3.486819007 | 0.000488802 | 0.001938309 |
| ENSG00000247033 | ENSG0000024 | 47.43087073 | 1.325882076 | 0.270518297 | 4.901265787 | 9.52E-07 | 5.52E-06 |
| ENSG00000247049 | ENSG0000024 | 22.49674526 | -0.521926348 | 0.368065186 | -1.418026934 | 0.156182901 | 0.300364147 |
| ENSG00000247077 | PGAM5 | 2516.762768 | -0.056751216 | 0.052319295 | -1.084709112 | 0.278050511 | 0.460839442 |
| ENSG00000247081 | BAALC-AS1 | 249.1534337 | -0.129840645 | 0.117382256 | -1.106135204 | 0.268668003 | 0.450167555 |
| ENSG00000247092 | SNHG10 | 346.0491904 | -0.413087031 | 0.112732817 | -3.664301494 | 0.000248015 | 0.001033307 |
| ENSG00000247095 | MIR210HG | 68.94934119 | -0.36518346 | 0.223291 | -1.635459823 | 0.101952691 | 0.21535863 |

| | | | | | | | |
|---|---|---|---|---|---|---|---|
| ENSG00000247121 | ENSG0000024 | 38.27235152 | 0.247622284 | 0.290485564 | 0.85244265 | 0.393968457 | 0.58690184 |
| ENSG00000247131 | ENSG0000024 | 2.290664817 | -0.202604451 | 1.139871843 | -0.177743184 | 0.858924669 | 0.926792302 |
| ENSG00000247137 | ANKRD42-DT | 58.96456909 | -0.03337084 | 0.229047024 | -0.145694276 | 0.884162741 | 0.939910479 |
| ENSG00000247151 | CSTF3-DT | 2.46620467 | -0.301898839 | 1.168696923 | -0.258320899 | 0.796159259 | 0.891182463 |
| ENSG00000247157 | LINC01252 | 56.75217271 | -0.156507553 | 0.237111649 | -0.660058473 | 0.509216306 | 0.689264918 |
| ENSG00000247228 | ENSG0000024 | 21.31382389 | -1.14568566 | 0.396457725 | -2.889805367 | 0.003854804 | 0.012751147 |
| ENSG00000247240 | UBL7-DT | 233.3877484 | -0.238415416 | 0.122048972 | -1.953440587 | 0.050767411 | 0.121414137 |
| ENSG00000247271 | ZBED5-AS1 | 99.55101536 | -0.341411307 | 0.177077534 | -1.928032875 | 0.053851037 | 0.127789676 |
| ENSG00000247287 | PRKCH-AS1 | 57.68581436 | -0.31447925 | 0.250384048 | -1.255987565 | 0.209120479 | 0.375899058 |
| ENSG00000247315 | ZCCHC3 | 504.3186679 | -0.043771878 | 0.091311557 | -0.479368439 | 0.631676542 | 0.782934732 |
| ENSG00000247317 | LY6E-DT | 22.41496842 | -0.463543327 | 0.373593676 | -1.240768665 | 0.21469122 | 0.383528404 |
| ENSG00000247363 | NUP107-DT | 11.50431886 | -0.157028853 | 0.529359884 | -0.296639126 | 0.766742037 | 0.87268977 |
| ENSG00000247372 | ENSG0000024 | 1.878819423 | 0.357295728 | 1.274167294 | 0.280415084 | 0.779159066 | 0.880184615 |
| ENSG00000247373 | TMED2-DT | 72.67585833 | 0.114412122 | 0.218502107 | 0.52362022 | 0.600542715 | 0.760625545 |
| ENSG00000247400 | DNAJC3-DT | 105.1407747 | 0.292163879 | 0.173143545 | 1.687408441 | 0.091524842 | 0.197169214 |
| ENSG00000247473 | CARS1-AS1 | 9.721143087 | -0.495498328 | 0.56266107 | -0.88063375 | 0.378516085 | 0.57122288 |
| ENSG00000247498 | GPRC5D-AS1 | 120.24064 | 0.010767893 | 0.167336002 | 0.064348934 | 0.948692391 | 0.97640107 |
| ENSG00000247516 | MIR4458HG | 135.4842733 | 1.199708984 | 0.170925486 | 7.018900516 | 2.24E-12 | 2.22E-11 |
| ENSG00000247556 | OIP5-AS1 | 2705.859752 | -0.161359513 | 0.054183009 | -2.978046381 | 0.002900921 | 0.009861755 |
| ENSG00000247570 | SDCBPP2 | 4.232435414 | 0.052283909 | 0.851304323 | 0.061416238 | 0.951027721 | 0.977409778 |
| ENSG00000247572 | CKMT2-AS1 | 186.7328202 | -0.056904302 | 0.13332016 | -0.426824437 | 0.669507206 | 0.810089947 |
| ENSG00000247595 | MISFA | 35.70966808 | 0.182256852 | 0.307927981 | 0.591881424 | 0.553929994 | 0.725418397 |
| ENSG00000247596 | TWF2 | 381.9167465 | -0.066174117 | 0.102545896 | -0.645312192 | 0.518724894 | 0.696318646 |
| ENSG00000247624 | CPEB2-DT | 2.534172222 | 1.399753695 | 1.147634528 | 1.219685937 | 0.222583955 | 0.393681694 |
| ENSG00000247626 | MARS2 | 350.5164464 | 0.055858244 | 0.102900476 | 0.542837569 | 0.587241644 | 0.750951742 |
| ENSG00000247627 | MTND4P12 | 259.3876687 | 0.293249462 | 0.313160908 | 0.936417844 | 0.349058094 | 0.540273442 |
| ENSG00000247675 | LRP4-AS1 | 4.875922947 | 0.442108741 | 0.842030895 | 0.525050498 | 0.599548087 | 0.759617426 |
| ENSG00000247679 | ENSG0000024 | 130.9129624 | -0.303927014 | 0.160032736 | -1.899155271 | 0.057544064 | 0.134823902 |
| ENSG00000247708 | STX18-AS1 | 128.5798207 | 0.090807648 | 0.156416153 | 0.580551601 | 0.5615427 | 0.730540816 |
| ENSG00000247728 | ARHGAP11B-[ | 7.22652522 | -0.838125035 | 0.697505684 | -1.201603162 | 0.229517316 | 0.402655799 |
| ENSG00000247735 | KCTD13-DT | 37.7432963 | -0.470731576 | 0.289530577 | -1.625844084 | 0.103982828 | 0.218740584 |
| ENSG00000247746 | USP51 | 178.8567828 | -0.456479959 | 0.137192368 | -3.327298496 | 0.000876924 | 0.003317387 |
| ENSG00000247796 | MOCS2-DT | 59.98202828 | -0.226824312 | 0.231685858 | -0.979016648 | 0.327571754 | 0.517919638 |

| | | | | | | | |
|---|---|---|---|---|---|---|---|
| ENSG00000247809 | NR2F2-AS1 | 27.09351783 | 0.765047496 | 0.343922758 | 2.224474764 | 0.026116519 | 0.068919008 |
| ENSG00000247828 | TMEM161B-D | 247.5833779 | 0.187956771 | 0.119180159 | 1.577081063 | 0.114776874 | 0.236949378 |
| ENSG00000247853 | ENSG0000024 | 54.08916712 | -0.198574086 | 0.251874252 | -0.788385808 | 0.43047107 | 0.620640208 |
| ENSG00000247877 | FAM174A-DT | 5.935911813 | -1.12445309 | 0.73415235 | -1.53163453 | 0.125612647 | 0.254396938 |
| ENSG00000247934 | ENSG0000024 | 16.9742497 | -0.18661607 | 0.423049058 | -0.441121582 | 0.65912498 | 0.802846438 |
| ENSG00000247950 | SEC24B-AS1 | 28.8722936 | 0.201477751 | 0.330322735 | 0.609942125 | 0.541900146 | 0.715752496 |
| ENSG00000247970 | ENSG0000024 | 2.596113482 | 0.573155286 | 1.085437461 | 0.528040819 | 0.597471005 | 0.757960331 |
| ENSG00000247982 | LINC00926 | 4.179467541 | -0.747657747 | 0.864936795 | -0.864407377 | 0.387364142 | 0.580034818 |
| ENSG00000247993 | FOXD1-AS1 | 1.88786086 | 1.431331862 | 1.343170634 | 1.065636656 | 0.286587922 | 0.470401183 |
| ENSG00000248008 | NRAV | 405.2999388 | -0.273218558 | 0.097977087 | -2.788596451 | 0.005293698 | 0.016936881 |
| ENSG00000248015 | ENSG0000024 | 126.9354994 | 0.061687723 | 0.159106296 | 0.387713907 | 0.698227763 | 0.830732062 |
| ENSG00000248019 | FAM13A-AS1 | 69.69335568 | -0.500019288 | 0.223071309 | -2.241522186 | 0.02499227 | 0.066451979 |
| ENSG00000248027 | ARHGAP42-AS | 4.89817945 | 0.792245631 | 0.801169651 | 0.988861261 | 0.322731026 | 0.512668773 |
| ENSG00000248049 | UBA6-DT | 255.3088191 | -0.148893186 | 0.115654088 | -1.287400978 | 0.197954564 | 0.360317201 |
| ENSG00000248079 | DPH6-DT | 3.132799692 | 0.207627125 | 1.066834766 | 0.194619759 | 0.845690634 | 0.919043732 |
| ENSG00000248092 | NNT-AS1 | 594.3721352 | 0.113567467 | 0.082018902 | 1.384649937 | 0.166159534 | 0.315198852 |
| ENSG00000248098 | BCKDHA | 8.323444099 | -0.001616256 | 0.611326257 | -0.002643852 | 0.997890514 | 0.998761274 |
| ENSG00000248099 | INSL3 | 14.91278893 | 0.978636383 | 0.471585482 | 2.075204643 | 0.037967572 | 0.095012121 |
| ENSG00000248113 | PTPN11P5 | 50.43079621 | 0.47106181 | 0.277951498 | 1.694762621 | 0.090120461 | 0.194637625 |
| ENSG00000248121 | SMURF2P1 | 3.084459489 | 0.626420164 | 1.056705814 | 0.592804691 | 0.553311868 | 0.72485719 |
| ENSG00000248124 | RRN3P1 | 2.488300736 | -0.736601896 | 1.138924548 | -0.646752147 | 0.517792368 | 0.69574209 |
| ENSG00000248138 | ENSG0000024 | 41.16296962 | -0.02498778 | 0.283044758 | -0.088282082 | 0.92965248 | 0.965993019 |
| ENSG00000248159 | HSPA8P11 | 8.009563112 | -0.227348307 | 0.655795137 | -0.346675805 | 0.728834889 | 0.849739396 |
| ENSG00000248161 | ENSG0000024 | 4.80354619 | -0.681935594 | 0.848526865 | -0.803670009 | 0.421587579 | 0.612567714 |
| ENSG00000248180 | GAPDHP60 | 18.25281744 | 0.141532108 | 0.437521138 | 0.323486331 | 0.746326958 | 0.859604617 |
| ENSG00000248202 | LINC02234 | 3.646178126 | -0.186134812 | 0.895869085 | -0.207770103 | 0.835408477 | 0.91323662 |
| ENSG00000248206 | ENSG0000024 | 2.696875202 | 0.130096844 | 1.065633381 | 0.122084055 | 0.90283245 | 0.950907167 |
| ENSG00000248208 | WDR45P1 | 10.77545016 | 0.63087819 | 0.543866742 | 1.159986705 | 0.246054219 | 0.423251634 |
| ENSG00000248223 | ENSG0000024 | 28.52073716 | -0.48310378 | 0.335311833 | -1.440759712 | 0.149652579 | 0.291107826 |
| ENSG00000248238 | LINC02438 | 3.626712658 | -2.795010465 | 1.171957953 | -2.384906778 | 0.01708345 | 0.047687313 |
| ENSG00000248240 | ENSG0000024 | 5.162492085 | 0.040674786 | 0.788896269 | 0.051559106 | 0.958880005 | 0.981197069 |
| ENSG00000248243 | LINC02014 | 4.128855378 | -0.515935647 | 0.848176362 | -0.608288169 | 0.542996366 | 0.71662275 |
| ENSG00000248265 | FLJ12825 | 3.446204774 | 0.506966701 | 0.953881927 | 0.531477416 | 0.595087987 | 0.756557217 |

| | | | | | | | |
|---|---|---|---|---|---|---|---|
| ENSG00000248268 | ENSG0000024 | 4.493974296 | -0.672529904 | 0.838052942 | -0.802490952 | 0.42226902 | 0.613285933 |
| ENSG00000248275 | TRIM52-AS1 | 332.4944013 | -0.93351816 | 0.108028268 | -8.641424826 | 5.55E-18 | 8.17E-17 |
| ENSG00000248278 | SUMO2P17 | 22.64213473 | -1.273320592 | 0.383625401 | -3.319176963 | 0.000902832 | 0.003408651 |
| ENSG00000248309 | MEF2C-AS1 | 2.34356464 | -2.034399549 | 1.24936248 | -1.628350124 | 0.103450667 | 0.217880858 |
| ENSG00000248323 | LUCAT1 | 2.408894132 | 1.289985882 | 1.177178982 | 1.095828164 | 0.273153981 | 0.455038495 |
| ENSG00000248327 | NOL8P1 | 2.148313768 | -1.339295186 | 1.314971696 | -1.018497348 | 0.30844166 | 0.49649727 |
| ENSG00000248333 | CDK11B | 1070.279129 | -0.202334344 | 0.066340117 | -3.049954575 | 0.00228876 | 0.00795156 |
| ENSG00000248347 | ENSG0000024 | 3.470273055 | 0.920738544 | 0.962090785 | 0.957018359 | 0.338557989 | 0.52977628 |
| ENSG00000248360 | LINC00504 | 55.71007162 | 4.142589441 | 0.402554217 | 10.2907615 | 7.76E-25 | 1.71E-23 |
| ENSG00000248367 | ENSG0000024 | 46.85518916 | -0.196134939 | 0.265857499 | -0.737744618 | 0.460669653 | 0.647494752 |
| ENSG00000248373 | ENSG0000024 | 77.93812831 | 0.454842493 | 0.20089597 | 2.26406977 | 0.023569824 | 0.063150171 |
| ENSG00000248383 | PCDHAC1 | 35.98231888 | 0.300966483 | 0.295200125 | 1.019533723 | 0.307949651 | 0.495934609 |
| ENSG00000248405 | PRR5-ARHGAI | 73.85638278 | -0.61488694 | 0.207428251 | -2.964335555 | 0.003033373 | 0.010270934 |
| ENSG00000248415 | GAPDHP61 | 7.014785497 | -0.388472257 | 0.678467264 | -0.572573325 | 0.566933625 | 0.734771603 |
| ENSG00000248429 | GASK1B-AS1 | 101.1882364 | 0.732799543 | 0.180436618 | 4.061257354 | 4.88E-05 | 0.000228833 |
| ENSG00000248445 | SEMA6A-AS1 | 16.84731445 | 0.387851827 | 0.445324484 | 0.870942067 | 0.383785786 | 0.576218727 |
| ENSG00000248449 | PCDHGB8P | 1.952161835 | 0.839029448 | 1.262317521 | 0.664673851 | 0.506259123 | 0.686625533 |
| ENSG00000248455 | LINC02217 | 1.934644466 | -3.408598557 | 1.571993352 | -2.168328862 | 0.030133674 | 0.077994638 |
| ENSG00000248468 | ENSG0000024 | 11.64700075 | -0.458997523 | 0.554712511 | -0.827451181 | 0.407981381 | 0.600557809 |
| ENSG00000248473 | LINC01962 | 8.013382582 | -1.302801575 | 0.641802273 | -2.029911126 | 0.042365574 | 0.104043065 |
| ENSG00000248476 | BACH1-IT1 | 8.56462672 | 0.53080201 | 0.611077308 | 0.868633155 | 0.385047814 | 0.577659856 |
| ENSG00000248485 | PCP4L1 | 31.63788655 | -0.591626099 | 0.317927249 | -1.860885158 | 0.062760396 | 0.144783134 |
| ENSG00000248487 | ABHD14A | 47.00339906 | -0.271050454 | 0.254547298 | -1.064833359 | 0.286951347 | 0.470862939 |
| ENSG00000248489 | CHD1-DT | 14.07663664 | 0.855247419 | 0.55663111 | 1.536470749 | 0.124422967 | 0.252513206 |
| ENSG00000248494 | LNX1-AS2 | 4.530746807 | -0.942227629 | 0.824625152 | -1.142613254 | 0.2531992 | 0.431939753 |
| ENSG00000248498 | ASNSP1 | 43.75424933 | 0.296987157 | 0.275539973 | 1.077836926 | 0.281106538 | 0.464528452 |
| ENSG00000248503 | C4orf46P2 | 4.692096936 | -0.761831888 | 0.823230227 | -0.925417779 | 0.354748634 | 0.546426756 |
| ENSG00000248508 | SRP14-DT | 133.9638788 | -0.907159249 | 0.160080184 | -5.66690532 | 1.45E-08 | 1.03E-07 |
| ENSG00000248515 | ENSG0000024 | 28.13504567 | -0.144273486 | 0.337865138 | -0.42701501 | 0.669368395 | 0.810007484 |
| ENSG00000248527 | MTATP6P1 | 4486.434181 | -0.100905906 | 0.068723344 | -1.468291566 | 0.142025041 | 0.27960581 |
| ENSG00000248530 | BCL2L12P1 | 11.68426061 | -0.624173095 | 0.550383166 | -1.134070105 | 0.256765142 | 0.436310545 |
| ENSG00000248538 | PPP1R3B-DT | 3.914244212 | 0.286509568 | 0.892790958 | 0.320914505 | 0.748275182 | 0.860552096 |
| ENSG00000248546 | ANP32C | 45.48601525 | -0.324171902 | 0.277007483 | -1.17026406 | 0.24189472 | 0.41807164 |

| | | | | | | | |
|---|---|---|---|---|---|---|---|
| ENSG00000248554 | C5orf34-AS1 | 12.85403261 | -0.760121347 | 0.500305193 | -1.519315326 | 0.128683144 | 0.2591512 |
| ENSG00000248565 | TECRP2 | 2.714290772 | -1.745079583 | 1.169458716 | -1.492211361 | 0.135643745 | 0.270180386 |
| ENSG00000248568 | KRT8P48 | 51.07844854 | 0.212761647 | 0.252359861 | 0.843088303 | 0.39917906 | 0.591930681 |
| ENSG00000248569 | ENSG0000024 | 162.0551467 | -0.080036914 | 0.151954035 | -0.526717924 | 0.598389487 | 0.758736172 |
| ENSG00000248578 | NPM1P21 | 27.99941571 | -0.108211815 | 0.334688294 | -0.323321182 | 0.746452013 | 0.859604617 |
| ENSG00000248583 | ENSG0000024 | 3.689245487 | -0.217167284 | 0.899014048 | -0.241561614 | 0.809119869 | 0.898891462 |
| ENSG00000248587 | GDNF-AS1 | 15.40291064 | 1.027164233 | 0.470457763 | 2.183329331 | 0.029011566 | 0.075388007 |
| ENSG00000248592 | STIMATE-MUS | 5.399324498 | -0.752490741 | 0.7538201 | -0.998236503 | 0.318164689 | 0.50734728 |
| ENSG00000248593 | DSTNP2 | 54.13517958 | 0.165492221 | 0.242421516 | 0.682663089 | 0.494819758 | 0.67711556 |
| ENSG00000248599 | FLJ42969 | 9.42378275 | -1.729570457 | 0.617251321 | -2.802052257 | 0.005077865 | 0.016335187 |
| ENSG00000248610 | HSPA8P4 | 4.036758736 | 0.892492352 | 0.898053312 | 0.993807762 | 0.320316467 | 0.509889134 |
| ENSG00000248625 | ENSG0000024 | 2.344148748 | -0.207983502 | 1.136275842 | -0.183039623 | 0.854766929 | 0.924486249 |
| ENSG00000248626 | GAPDHP40 | 50.29099193 | 0.264397643 | 0.260295415 | 1.015759893 | 0.309743739 | 0.49772738 |
| ENSG00000248636 | ENSG0000024 | 2.020171373 | 0.566712036 | 1.262266749 | 0.448963768 | 0.653457794 | 0.798929904 |
| ENSG00000248641 | HMGA1P2 | 160.3674276 | 0.561201142 | 0.148729495 | 3.773300939 | 0.000161102 | 0.000694545 |
| ENSG00000248643 | RBM14-RBM4 | 19.42492694 | -0.083998252 | 0.422170379 | -0.198967658 | 0.842288044 | 0.917009895 |
| ENSG00000248648 | ENSG0000024 | 2.36206218 | -1.496657299 | 1.220715287 | -1.226049444 | 0.220180069 | 0.390395021 |
| ENSG00000248663 | LINC00992 | 240.7711172 | 0.263988011 | 0.121683065 | 2.169472061 | 0.030046863 | 0.077797128 |
| ENSG00000248664 | ENSG0000024 | 8.073717252 | -0.807586568 | 0.630651898 | -1.280558372 | 0.200348829 | 0.363550241 |
| ENSG00000248668 | OXCT1-AS1 | 10.15762452 | 0.040636148 | 0.564741098 | 0.071955357 | 0.942637435 | 0.972787924 |
| ENSG00000248671 | ALG1L9P | 37.13344179 | -0.040781389 | 0.286804703 | -0.142192192 | 0.886928199 | 0.941793234 |
| ENSG00000248677 | LINC02102 | 3.711933751 | 1.653142286 | 1.011696739 | 1.63402947 | 0.102252663 | 0.215912747 |
| ENSG00000248693 | LINC02100 | 1.538504036 | 3.973318516 | 1.68509989 | 2.35791275 | 0.018378012 | NA |
| ENSG00000248694 | ENSG0000024 | 2.552150246 | -0.420254468 | 1.136943326 | -0.369635371 | 0.711654193 | 0.839694845 |
| ENSG00000248697 | TOX4P1 | 71.39199735 | -0.550714753 | 0.210027074 | -2.622113155 | 0.008738641 | 0.026425577 |
| ENSG00000248712 | CCDC153 | 43.94225392 | -0.207704979 | 0.262048136 | -0.79262147 | 0.427998396 | 0.618669389 |
| ENSG00000248714 | ZNF652-AS1 | 2.789891508 | -0.711078804 | 1.108227617 | -0.641636062 | 0.521109512 | 0.698221612 |
| ENSG00000248724 | NPHP3-AS1 | 2.750706155 | 0.679425134 | 1.057918025 | 0.642228526 | 0.520724814 | 0.697950759 |
| ENSG00000248727 | LINC01948 | 56.62706405 | 1.677924093 | 0.257188786 | 6.524095084 | 6.84E-11 | 6.00E-10 |
| ENSG00000248734 | ENSG0000024 | 3.143653535 | 0.263059708 | 0.995949286 | 0.264129621 | 0.79168004 | 0.887814827 |
| ENSG00000248740 | LINC02428 | 3.129889617 | -0.023422352 | 0.969007911 | -0.024171476 | 0.98071583 | 0.991128294 |
| ENSG00000248746 | ACTN3 | 2.663686072 | 0.716149572 | 1.106779066 | 0.647057388 | 0.517594802 | 0.695665852 |
| ENSG00000248752 | ENSG0000024 | 4.174260172 | -0.808671519 | 0.914576766 | -0.884202999 | 0.376586639 | 0.56939741 |

| | | | | | | | |
|---|---|---|---|---|---|---|---|
| ENSG00000248771 | SMIM31 | 2.013557294 | -2.4870781 | 1.466270752 | -1.696192942 | 0.089849347 | 0.194154485 |
| ENSG00000248773 | ENSG0000024 | 1.698357529 | 1.720835907 | 1.607867936 | 1.070259483 | 0.284502526 | NA |
| ENSG00000248774 | ENSG0000024 | 5.228028495 | -0.344328554 | 0.770991751 | -0.446604718 | 0.655160488 | 0.799808359 |
| ENSG00000248780 | ARL4AP2 | 4.980883629 | -0.381349182 | 0.774638056 | -0.492293374 | 0.622511964 | 0.776597778 |
| ENSG00000248781 | PSMC1P4 | 3.617722618 | 0.247027366 | 0.947054436 | 0.260837558 | 0.79421778 | 0.88974732 |
| ENSG00000248785 | HIGD1AP14 | 7.965189214 | -0.630029273 | 0.619299041 | -1.017326415 | 0.308998175 | 0.497118881 |
| ENSG00000248787 | ENSG0000024 | 5.394571172 | 0.451787425 | 0.753627338 | 0.59948386 | 0.54885027 | 0.72156715 |
| ENSG00000248794 | ENSG0000024 | 9.015154418 | 0.361516342 | 0.58944231 | 0.613319295 | 0.539665238 | 0.714116501 |
| ENSG00000248795 | ENSG0000024 | 30.79618076 | 0.597865341 | 0.336033393 | 1.779184313 | 0.075209549 | 0.168273389 |
| ENSG00000248801 | C8orf34-AS1 | 30.36701059 | 3.883455222 | 0.510529961 | 7.606713641 | 2.81E-14 | 3.20E-13 |
| ENSG00000248817 | PMPCAP1 | 4.839633153 | 1.773557711 | 0.891081572 | 1.990342709 | 0.046553196 | 0.112662762 |
| ENSG00000248827 | ENSG0000024 | 12.55558194 | -0.041544629 | 0.501284835 | -0.082876293 | 0.933949905 | 0.968309266 |
| ENSG00000248830 | ZNF807P | 3.511854596 | 0.117003207 | 0.994071058 | 0.11770105 | 0.906304535 | 0.952693306 |
| ENSG00000248854 | HNRNPH1P3 | 16.35257212 | -0.308717716 | 0.434716258 | -0.710159121 | 0.477605467 | 0.662370438 |
| ENSG00000248858 | FLJ46284 | 13.77006834 | -0.296445972 | 0.469283432 | -0.631699208 | 0.527583446 | 0.703507931 |
| ENSG00000248866 | USP46-DT | 67.53803496 | 0.019840768 | 0.217375476 | 0.091274177 | 0.927274736 | 0.965137362 |
| ENSG00000248873 | SERBP1P6 | 230.3150159 | 0.218595873 | 0.125273595 | 1.744947706 | 0.080993972 | 0.178482394 |
| ENSG00000248890 | HHIP-AS1 | 2.538039306 | -1.547870943 | 1.209865036 | -1.279374887 | 0.200765073 | 0.364161522 |
| ENSG00000248898 | PELO-AS1 | 1.902891383 | -1.124883944 | 1.372262958 | -0.819729147 | 0.41237053 | 0.604534362 |
| ENSG00000248905 | FMN1 | 175.127702 | -0.365375346 | 0.140212108 | -2.605875843 | 0.009163967 | 0.027570556 |
| ENSG00000248919 | ATP5MF-PTCD | 9.503253725 | -0.864899133 | 0.613639441 | -1.409458184 | 0.15869973 | 0.304014347 |
| ENSG00000248923 | MTND5P11 | 215.635776 | 0.18934477 | 0.303869089 | 0.62311297 | 0.533210287 | 0.708681576 |
| ENSG00000248930 | CHP1P1 | 3.023444949 | 0.314317207 | 1.004214254 | 0.312998153 | 0.754282065 | 0.864498202 |
| ENSG00000248932 | COPB2-DT | 150.9218736 | -0.385354029 | 0.147127271 | -2.619188309 | 0.008813928 | 0.026625165 |
| ENSG00000248936 | ENSG0000024 | 5.805266953 | 0.019049271 | 0.757466992 | 0.025148648 | 0.979936397 | 0.991041209 |
| ENSG00000248946 | MTND3P22 | 4.653336288 | 3.568916525 | 1.236747978 | 2.885726591 | 0.003905112 | 0.01289896 |
| ENSG00000248954 | ENSG0000024 | 5.446169018 | -1.772240621 | 0.847031316 | -2.092296456 | 0.036412005 | 0.091729781 |
| ENSG00000248966 | BCLAF1P1 | 32.70130738 | -0.218883348 | 0.327809524 | -0.667715036 | 0.504315511 | 0.685175989 |
| ENSG00000248971 | KRT8P46 | 47.9093869 | -0.047614951 | 0.253410676 | -0.187896389 | 0.850957873 | 0.922214256 |
| ENSG00000248980 | SPCS3-AS1 | 4.070735212 | 0.179259257 | 0.854200836 | 0.209856101 | 0.833779986 | 0.912258208 |
| ENSG00000248988 | PSME2P3 | 6.339257566 | -0.341043546 | 0.702063407 | -0.48577314 | 0.62712802 | 0.780154165 |
| ENSG00000249001 | ENSG0000024 | 6.631241722 | -0.107085674 | 0.730080097 | -0.14667661 | 0.883387281 | 0.93961708 |
| ENSG00000249007 | ENSG0000024 | 3.830149348 | -0.113734655 | 0.925128042 | -0.122939366 | 0.902155113 | 0.950654947 |

| | | | | | | | |
|---|---|---|---|---|---|---|---|
| ENSG00000249014 | HMGN2P4 | 62.93619338 | -0.273769901 | 0.227929358 | -1.201117326 | 0.229705693 | 0.402924652 |
| ENSG00000249026 | CTNNA1P1 | 39.40236095 | -0.11736945 | 0.295396094 | -0.397329052 | 0.691124827 | 0.825793193 |
| ENSG00000249031 | SUMO2P6 | 7.664049989 | 0.519819935 | 0.687697677 | 0.755884384 | 0.449718517 | 0.638044579 |
| ENSG00000249042 | FAM151B-DT | 619.7187415 | -0.181702686 | 0.07879686 | -2.305963541 | 0.02111267 | 0.057282631 |
| ENSG00000249064 | KRT18P25 | 8.359512134 | 0.440379358 | 0.601262234 | 0.732424777 | 0.463909341 | 0.650169158 |
| ENSG00000249069 | LINC01033 | 2.465706809 | -1.095748207 | 1.2123901 | -0.90379178 | 0.366105821 | 0.558854641 |
| ENSG00000249085 | TNPO1-DT | 1.599009574 | 1.842920759 | 1.562540447 | 1.179438755 | 0.238223511 | NA |
| ENSG00000249087 | ZNF436-AS1 | 24.66806389 | 0.118265306 | 0.350607107 | 0.337315768 | 0.735878874 | 0.854386347 |
| ENSG00000249115 | HAUS5 | 1102.664721 | -0.075446305 | 0.064279667 | -1.173719603 | 0.240507356 | 0.416332534 |
| ENSG00000249116 | IRX4-AS1 | 39.60898587 | 0.293877488 | 0.284472308 | 1.033061846 | 0.301574957 | 0.488396013 |
| ENSG00000249119 | MTND6P4 | 31.35225965 | -0.082868427 | 0.363127005 | -0.228207834 | 0.819484671 | 0.904362263 |
| ENSG00000249129 | SUDS3P1 | 345.4190819 | -0.162499318 | 0.103879197 | -1.564310496 | 0.117744668 | 0.242009022 |
| ENSG00000249142 | TMEM183AP2 | 126.4867249 | 0.426184932 | 0.163704495 | 2.603379529 | 0.00923097 | 0.02773886 |
| ENSG00000249158 | PCDHA11 | 218.660368 | -0.041610327 | 0.127621809 | -0.326044015 | 0.744391054 | 0.858668013 |
| ENSG00000249159 | ENSG0000024 | 2.233991514 | 1.674034815 | 1.313558694 | 1.274427113 | 0.202512088 | 0.366634623 |
| ENSG00000249167 | SEMA6A-AS2 | 3.505292868 | 1.608925546 | 1.009673992 | 1.593509943 | 0.11104584 | 0.230512312 |
| ENSG00000249176 | ENSG0000024 | 69.96558667 | -0.152380478 | 0.214906843 | -0.709053633 | 0.478291195 | 0.66295052 |
| ENSG00000249180 | ENSG0000024 | 12.1942922 | 0.410324086 | 0.504525008 | 0.813287904 | 0.416053011 | 0.608146812 |
| ENSG00000249193 | HSPD1P5 | 17.70923031 | -0.131422355 | 0.44011688 | -0.298607849 | 0.765239276 | 0.871498237 |
| ENSG00000249199 | ENSG0000024 | 27.90031294 | -4.636850473 | 0.614030878 | -7.551493973 | 4.30E-14 | 4.82E-13 |
| ENSG00000249207 | ENSG0000024 | 4.632962851 | 0.20467156 | 0.807398841 | 0.253494988 | 0.799885734 | 0.893305035 |
| ENSG00000249209 | ENSG0000024 | 2.947807414 | -0.823102807 | 1.025917445 | -0.802309007 | 0.422374233 | 0.613322251 |
| ENSG00000249210 | GAPDHP38 | 29.52127597 | -0.138530455 | 0.337192246 | -0.410835233 | 0.681193356 | 0.819013756 |
| ENSG00000249215 | NCOA4P4 | 2.233493977 | 1.157729914 | 1.285489727 | 0.900613898 | 0.367793643 | 0.560312536 |
| ENSG00000249222 | ATP5MGL | 13.86568372 | 0.543353195 | 0.473274971 | 1.148070842 | 0.25093932 | 0.429272782 |
| ENSG00000249242 | TMEM150C | 3529.550736 | -1.363481152 | 0.048862201 | -27.90462003 | 2.34E-171 | 9.96E-169 |
| ENSG00000249245 | NCOA4P3 | 9.378723249 | 0.252720442 | 0.592721706 | 0.426372849 | 0.669836182 | 0.810317039 |
| ENSG00000249249 | TICAM2-AS1 | 6.734428966 | -0.751452292 | 0.693215311 | -1.084009945 | 0.278360389 | 0.461219742 |
| ENSG00000249256 | ATP5MGP3 | 3.305331455 | -0.498084723 | 1.018584846 | -0.488996793 | 0.624843967 | 0.778298345 |
| ENSG00000249264 | EEF1A1P9 | 206.3614411 | 0.226861975 | 0.159508218 | 1.42225885 | 0.15495112 | 0.29847233 |
| ENSG00000249266 | MTHFD2P6 | 6.124445789 | -0.143314697 | 0.720833857 | -0.198817933 | 0.842405168 | 0.917083075 |
| ENSG00000249267 | LINC00939 | 44.00077763 | -1.493329655 | 0.290143342 | -5.146868603 | 2.65E-07 | 1.64E-06 |
| ENSG00000249286 | AMD1P3 | 3.685871854 | -0.049873375 | 0.956132629 | -0.052161566 | 0.958399957 | 0.981066266 |

| | | | | | | | |
|---|---|---|---|---|---|---|---|
| ENSG00000249306 | LINC01411 | 6.3198107 | -0.499605056 | 0.687675671 | -0.726512624 | 0.467524571 | 0.653486088 |
| ENSG00000249310 | APOBEC3B-AS | 4.852975317 | -1.863588914 | 0.908522245 | -2.051230914 | 0.040244465 | 0.099623181 |
| ENSG00000249311 | TERF1P3 | 1.993770868 | -0.70767921 | 1.257186113 | -0.562907276 | 0.573498025 | 0.74022645 |
| ENSG00000249326 | CTD-2194D22 | 5.158160783 | -0.964099419 | 0.832619052 | -1.157911793 | 0.246900035 | 0.424165865 |
| ENSG00000249328 | ENSG0000024 | 2.700526854 | -2.25675531 | 1.22462199 | -1.842817889 | 0.065355604 | 0.149910764 |
| ENSG00000249345 | LINC02405 | 5.015514516 | 4.661219094 | 1.379772487 | 3.378251949 | 0.000729482 | 0.002801661 |
| ENSG00000249346 | LINC01016 | 12.10689129 | 2.128205404 | 0.577148786 | 3.687446731 | 0.000226515 | 0.000951348 |
| ENSG00000249348 | UGDH-AS1 | 130.8924584 | -0.043778199 | 0.159970149 | -0.273664802 | 0.78434223 | 0.882952731 |
| ENSG00000249352 | LINC02198 | 3.089502301 | -3.215867007 | 1.343356813 | -2.39390382 | 0.01667012 | 0.046675521 |
| ENSG00000249353 | NPM1P27 | 5530.404638 | -0.098978874 | 0.066722547 | -1.48343968 | 0.137957631 | 0.273605402 |
| ENSG00000249364 | LINC02997 | 45.29848575 | 3.280102444 | 0.365995837 | 8.962130468 | 3.18E-19 | 5.10E-18 |
| ENSG00000249375 | CASC11 | 39.62388172 | 0.191555199 | 0.299574871 | 0.639423456 | 0.522547494 | 0.699290603 |
| ENSG00000249379 | GCLC-AS1 | 6.231528016 | -0.132804717 | 0.711373079 | -0.186687858 | 0.851905376 | 0.922522155 |
| ENSG00000249395 | CASC9 | 124.8478773 | -0.11396469 | 0.160796401 | -0.708751497 | 0.478478701 | 0.663064417 |
| ENSG00000249413 | ENSG0000024 | 8.171697536 | 0.052181354 | 0.605145128 | 0.086229487 | 0.931283991 | 0.967031267 |
| ENSG00000249417 | ENSG0000024 | 2.516665994 | -0.787332763 | 1.10276129 | -0.713964817 | 0.475248936 | 0.66046014 |
| ENSG00000249421 | ADAMTS19-AS | 90.4625445 | -0.080656921 | 0.191529957 | -0.421119089 | 0.673668123 | 0.813322812 |
| ENSG00000249425 | LINC02477 | 1.780568257 | 1.978525724 | 1.479315057 | 1.337460681 | 0.181072308 | NA |
| ENSG00000249437 | NAIP | 19.85950883 | -0.392604859 | 0.39658523 | -0.989963391 | 0.322192013 | 0.511989802 |
| ENSG00000249456 | ENSG0000024 | 1.749859937 | -0.840535976 | 1.335020388 | -0.629605348 | 0.528952824 | NA |
| ENSG00000249459 | ZNF286B | 79.42607547 | 0.017884262 | 0.20215311 | 0.088468893 | 0.929504007 | 0.965993019 |
| ENSG00000249464 | LINC01091 | 8.919300867 | -0.571065447 | 0.597679511 | -0.955471012 | 0.339339558 | 0.530492671 |
| ENSG00000249465 | RBMXP4 | 82.04776326 | -0.09124092 | 0.195216772 | -0.467382585 | 0.64022618 | 0.788383089 |
| ENSG00000249471 | ZNF324B | 245.8070974 | 0.277797728 | 0.126589088 | 2.194483991 | 0.028200634 | 0.073639446 |
| ENSG00000249476 | ENSG0000024 | 4.62583672 | -0.996431696 | 0.844201508 | -1.180324469 | 0.23787119 | 0.41335998 |
| ENSG00000249485 | RBBP4P1 | 531.1180719 | 0.115787006 | 0.087574787 | 1.322150012 | 0.186118203 | 0.343487657 |
| ENSG00000249489 | GAPDHP70 | 9.843389681 | 0.087087774 | 0.561712693 | 0.155039711 | 0.876790011 | 0.935974786 |
| ENSG00000249492 | ENSG0000024 | 10.24153575 | -0.270081685 | 0.551677241 | -0.489564668 | 0.624441983 | 0.778008858 |
| ENSG00000249493 | ANKRD20A18I | 3.170118632 | 0.586274795 | 1.007512999 | 0.581902959 | 0.560632047 | 0.729860281 |
| ENSG00000249494 | DMXL1-DT | 2.040093818 | 1.005382171 | 1.269974124 | 0.791655635 | 0.428561498 | 0.619132235 |
| ENSG00000249506 | ZEB2P1 | 3.178561141 | 1.056482503 | 1.019088574 | 1.036693502 | 0.299878721 | 0.486576451 |
| ENSG00000249532 | MIR302CHG | 25.20488529 | -0.38620909 | 0.34940947 | -1.105319472 | 0.269021183 | 0.450627671 |
| ENSG00000249541 | ENSG0000024 | 1.605593909 | 0.44927417 | 1.406179018 | 0.319499981 | 0.749347405 | NA |

| | | | | | | | |
|---|---|---|---|---|---|---|---|
| ENSG00000249558 | RCC2P4 | 3.453727294 | -0.302954543 | 0.926185832 | -0.32709909 | 0.74359294 | 0.858490754 |
| ENSG00000249565 | SERBP1P5 | 1065.064282 | 0.006463873 | 0.067528752 | 0.09572031 | 0.92374271 | 0.963279648 |
| ENSG00000249568 | ENSG0000024 | 28.04805091 | -0.426419827 | 0.354690567 | -1.202230524 | 0.229274226 | 0.402321636 |
| ENSG00000249572 | TARS1-DT | 45.9119013 | 0.105618264 | 0.269394547 | 0.392057912 | 0.695015425 | 0.828284786 |
| ENSG00000249582 | RPL7L1P4 | 2.652360653 | -0.544347921 | 1.056304295 | -0.515332488 | 0.6063207 | 0.764860977 |
| ENSG00000249592 | PCGF3-AS1 | 313.6062016 | -0.032841995 | 0.105383935 | -0.311641381 | 0.755313082 | 0.864916992 |
| ENSG00000249593 | ENSG0000024 | 14.6260875 | -0.730629243 | 0.4647134 | -1.572214709 | 0.115900765 | 0.23888262 |
| ENSG00000249599 | BMPR1B-DT | 1.583798469 | -0.108565202 | 1.385430509 | -0.07836207 | 0.937540045 | NA |
| ENSG00000249602 | ENSG0000024 | 48.62587953 | 0.312658807 | 0.257106447 | 1.216067549 | 0.223959191 | 0.395384319 |
| ENSG00000249617 | ENSG0000024 | 4.381826348 | 1.434733988 | 0.897035045 | 1.59941799 | 0.109727758 | 0.228334067 |
| ENSG00000249637 | ANKHD1-DT | 30.7757756 | 0.430433672 | 0.325254568 | 1.323374718 | 0.185710795 | 0.342962935 |
| ENSG00000249641 | HOXC13-AS | 313.9683105 | -0.191387581 | 0.109713349 | -1.744432948 | 0.081083624 | 0.178611325 |
| ENSG00000249650 | CEP72-DT | 1.89078608 | 0.512551232 | 1.319667177 | 0.388394317 | 0.697724249 | 0.830348323 |
| ENSG00000249655 | ENSG0000024 | 3.809325516 | 0.18642198 | 0.881689474 | 0.21143723 | 0.832546108 | 0.911410533 |
| ENSG00000249661 | TNRC18P1 | 41.04826454 | 1.042422767 | 0.306376591 | 3.402423025 | 0.000667912 | 0.002585537 |
| ENSG00000249668 | KRT18P56 | 8.049860503 | 1.024780137 | 0.635878984 | 1.611596174 | 0.107049844 | 0.223927516 |
| ENSG00000249673 | NOP14-AS1 | 925.7372932 | 0.302877113 | 0.071409433 | 4.241415987 | 2.22E-05 | 0.000109459 |
| ENSG00000249681 | KRT19P3 | 5.75293231 | 0.062170283 | 0.74326434 | 0.083644916 | 0.933338754 | 0.967981872 |
| ENSG00000249684 | ENSG0000024 | 132.2919551 | -0.885409898 | 0.167187019 | -5.295924895 | 1.18E-07 | 7.64E-07 |
| ENSG00000249685 | ENSG0000024 | 3.513889895 | 0.790598889 | 0.96763344 | 0.817043785 | 0.413903406 | 0.606086055 |
| ENSG00000249693 | THEGL | 1.831067888 | 0.857336032 | 1.392263718 | 0.61578566 | 0.538035993 | 0.712906237 |
| ENSG00000249700 | SRD5A3-AS1 | 4.165349545 | 0.149802522 | 0.851531381 | 0.175921317 | 0.860355758 | 0.927382816 |
| ENSG00000249709 | ZNF564 | 5.362871215 | 0.268613872 | 0.746964186 | 0.359607431 | 0.719140726 | 0.843883997 |
| ENSG00000249715 | FER1L5 | 6.490877301 | -3.234268704 | 0.92542074 | -3.494917028 | 0.000474209 | 0.001884351 |
| ENSG00000249721 | ENSG0000024 | 4.011269332 | 5.354772017 | 1.404626134 | 3.81224006 | 0.000137713 | 0.00060014 |
| ENSG00000249740 | OSMR-DT | 4.819578389 | 0.455036649 | 0.819189587 | 0.555471721 | 0.578572048 | 0.744179578 |
| ENSG00000249741 | ENSG0000024 | 8.309711026 | -0.774048939 | 0.613385658 | -1.261928656 | 0.206974459 | 0.372918167 |
| ENSG00000249773 | ENSG0000024 | 2.668484286 | 0.133551399 | 1.057058028 | 0.126342543 | 0.899460782 | 0.94915269 |
| ENSG00000249774 | ENSG0000024 | 32.44235512 | -1.875803586 | 0.377411204 | -4.970185216 | 6.69E-07 | 3.97E-06 |
| ENSG00000249779 | ENSG0000024 | 5.326550169 | 0.181110156 | 0.780978333 | 0.231901639 | 0.816614411 | 0.902888693 |
| ENSG00000249780 | ENSG0000024 | 3.127504244 | -0.345373174 | 1.009313967 | -0.342186064 | 0.732210872 | 0.851857055 |
| ENSG00000249784 | SCARNA22 | 2.234474148 | -1.38159266 | 1.301334516 | -1.061673723 | 0.288383833 | 0.472368805 |
| ENSG00000249786 | EAF1-AS1 | 28.4229353 | -0.48248036 | 0.354234915 | -1.362035019 | 0.173186834 | 0.32546129 |

| | | | | | | | |
|---|---|---|---|---|---|---|---|
| ENSG00000249797 | LINC02147 | 1.943704506 | 2.104996097 | 1.438085427 | 1.463748995 | 0.143262543 | 0.281373522 |
| ENSG00000249806 | ENSG0000024 | 2.379482161 | 0.046497199 | 1.137077706 | 0.04089184 | 0.967382123 | 0.985430982 |
| ENSG00000249825 | THBS4-AS1 | 3.980896812 | -0.675482307 | 0.901094371 | -0.749624377 | 0.453480965 | 0.641161993 |
| ENSG00000249839 | ENSG0000024 | 7.120698082 | 0.228704483 | 0.653125556 | 0.350169246 | 0.726211686 | 0.847853251 |
| ENSG00000249846 | LINC02021 | 77.84124872 | -1.575145121 | 0.219537949 | -7.174819321 | 7.24E-13 | 7.50E-12 |
| ENSG00000249849 | ENSG0000024 | 3.476427799 | -2.188930572 | 1.053988302 | -2.076807274 | 0.037819351 | 0.094703188 |
| ENSG00000249850 | KRT18P31 | 7.998044869 | -0.120948158 | 0.668838552 | -0.180833113 | 0.856498571 | 0.925660851 |
| ENSG00000249852 | ENSG0000024 | 4.470742506 | -0.517052369 | 0.846446671 | -0.610850496 | 0.54129856 | 0.715369799 |
| ENSG00000249853 | HS3ST5 | 3.045240389 | 0.615168626 | 0.995236095 | 0.618113259 | 0.536500683 | 0.711679378 |
| ENSG00000249855 | EEF1A1P19 | 217.9160264 | 0.026510114 | 0.140603435 | 0.18854528 | 0.850449223 | 0.921772038 |
| ENSG00000249858 | SNX5P1 | 48.78318605 | -0.138166693 | 0.259416072 | -0.532606527 | 0.594305983 | 0.756068903 |
| ENSG00000249859 | PVT1 | 1075.445577 | -0.113206336 | 0.076337661 | -1.482968363 | 0.138082817 | 0.273707752 |
| ENSG00000249863 | ENSG0000024 | 19.2499595 | 0.706788816 | 0.404569293 | 1.747015477 | 0.080634653 | 0.177861437 |
| ENSG00000249884 | RNF103-CHMI | 1.642491601 | -0.196061353 | 1.341263149 | -0.146176649 | 0.883781939 | NA |
| ENSG00000249894 | ENSG0000024 | 1.859055278 | 4.2435703 | 1.584380116 | 2.678378918 | 0.007397947 | 0.022804208 |
| ENSG00000249896 | JAKMIP1-DT | 1.649573797 | -0.243598062 | 1.395433861 | -0.174567974 | 0.861419115 | NA |
| ENSG00000249898 | MCPH1-AS1 | 26.34017874 | 0.763972168 | 0.359315192 | 2.126189443 | 0.033487489 | 0.085327446 |
| ENSG00000249915 | PDCD6 | 493.5895537 | 0.132276301 | 0.100058469 | 1.321990051 | 0.186171464 | 0.343530677 |
| ENSG00000249916 | ENSG0000024 | 25.95010175 | 0.697605432 | 0.35424033 | 1.969299856 | 0.048918668 | 0.117571243 |
| ENSG00000249917 | LINC00536 | 4.381706679 | 0.2572814 | 0.890617596 | 0.288879763 | 0.77267339 | 0.876166615 |
| ENSG00000249920 | HNRNPA1P55 | 4.553985431 | 0.442140633 | 0.836300944 | 0.528686038 | 0.597023264 | 0.75769879 |
| ENSG00000249921 | RACK1P1 | 18.36230251 | -0.163530374 | 0.460298188 | -0.355270513 | 0.722386943 | 0.845703316 |
| ENSG00000249936 | RAC1P2 | 2370.842406 | 0.054522993 | 0.057384782 | 0.950129815 | 0.342046295 | 0.533161787 |
| ENSG00000249937 | LINC02223 | 7.258036807 | -3.889292473 | 0.999470572 | -3.891352665 | 9.97E-05 | 0.000444301 |
| ENSG00000249947 | XBP1P1 | 1.577202993 | 0.45314451 | 1.383006851 | 0.327651674 | 0.743175047 | NA |
| ENSG00000249961 | TERB1 | 2.787178041 | 0.998111106 | 1.105296741 | 0.903025467 | 0.366512379 | 0.559103204 |
| ENSG00000249971 | ENSG0000024 | 6.477888586 | -0.364463069 | 0.722413218 | -0.504507753 | 0.613904615 | 0.770782763 |
| ENSG00000249978 | TRGV7 | 1.849580572 | 0.067212731 | 1.272260561 | 0.052829376 | 0.957867856 | 0.980830956 |
| ENSG00000249986 | YWHAQP6 | 6.969867423 | -0.104051522 | 0.672549223 | -0.154712129 | 0.877048268 | 0.936138436 |
| ENSG00000249992 | TMEM158 | 12.88213382 | 0.475823184 | 0.491917297 | 0.967282889 | 0.333402635 | 0.52456184 |
| ENSG00000249993 | BFSP2-AS1 | 7.742520461 | -1.97827009 | 0.697886258 | -2.834659758 | 0.004587451 | 0.014916532 |
| ENSG00000249996 | PPIC-AS1 | 11.95148557 | -0.226068211 | 0.519714241 | -0.434985599 | 0.66357288 | 0.805799686 |
| ENSG00000250012 | SLC41A3-AS1 | 47.96149487 | 0.598083316 | 0.259539526 | 2.304401663 | 0.021200107 | 0.057465491 |

| | | | | | | | |
|---|---|---|---|---|---|---|---|
| ENSG00000250030 | ENSG0000025 | 20.53773967 | -0.015252805 | 0.384544317 | -0.039664621 | 0.968360508 | 0.985876114 |
| ENSG00000250031 | PTPN11P2 | 10.70308791 | 0.041534339 | 0.530811174 | 0.078246919 | 0.937631641 | 0.970153346 |
| ENSG00000250056 | LINC01018 | 22.98536656 | -0.542313472 | 0.376354043 | -1.440966246 | 0.149594219 | 0.291043286 |
| ENSG00000250067 | YJEFN3 | 98.67470312 | -0.560725906 | 0.205685056 | -2.726138289 | 0.006408014 | 0.020076462 |
| ENSG00000250071 | ARPC1AP3 | 11.05778923 | -0.966320506 | 0.56214551 | -1.718986433 | 0.085616842 | 0.186748317 |
| ENSG00000250072 | SH3TC2-DT | 32.55289902 | -0.037788952 | 0.322408369 | -0.117208347 | 0.906694952 | 0.952929042 |
| ENSG00000250073 | ESAM-AS1 | 2.244353007 | -1.408797911 | 1.252201761 | -1.125056644 | 0.260565032 | 0.44087342 |
| ENSG00000250075 | ENSG0000025 | 4.3818798 | 0.795333359 | 0.841715501 | 0.944895701 | 0.344712126 | 0.535570347 |
| ENSG00000250076 | MAPRE1P2 | 5.259780951 | -0.533040078 | 0.76420217 | -0.697511862 | 0.485482519 | 0.669289001 |
| ENSG00000250081 | ENSG0000025 | 1.787141451 | 0.086889631 | 1.280558901 | 0.067852897 | 0.945902735 | NA |
| ENSG00000250091 | DNAH10OS | 312.7792684 | 0.736384652 | 0.111081051 | 6.629255364 | 3.37E-11 | 3.02E-10 |
| ENSG00000250101 | ENSG0000025 | 6.393525793 | -0.093746525 | 0.683871759 | -0.137082024 | 0.890965961 | 0.944140759 |
| ENSG00000250120 | PCDHA10 | 31.60198432 | 3.166270303 | 0.430784307 | 7.350013101 | 1.98E-13 | 2.12E-12 |
| ENSG00000250131 | AGA-DT | 7.761477335 | 0.342238567 | 0.628900973 | 0.544185145 | 0.586314073 | 0.750100097 |
| ENSG00000250132 | ENSG0000025 | 54.20456975 | 0.819711485 | 0.25583792 | 3.204026544 | 0.0013552 | 0.004945639 |
| ENSG00000250133 | HOXC-AS2 | 2.631990119 | -0.922850894 | 1.094554105 | -0.843129535 | 0.399156002 | 0.591930681 |
| ENSG00000250138 | ENSG0000025 | 10.38427886 | -0.597513056 | 0.545903429 | -1.094539848 | 0.273718281 | 0.455667442 |
| ENSG00000250144 | EEF1A1P35 | 86.43961681 | 0.098627946 | 0.206430187 | 0.477778699 | 0.632807724 | 0.78358033 |
| ENSG00000250148 | KRT8P31 | 2.072291311 | 2.257285181 | 1.43350715 | 1.574659171 | 0.115335137 | 0.237866123 |
| ENSG00000250182 | EEF1A1P13 | 359.5045881 | 0.226981501 | 0.253612942 | 0.894991788 | 0.370791478 | 0.563457638 |
| ENSG00000250186 | ENSG0000025 | 62.86172747 | -0.238166731 | 0.260612449 | -0.913873194 | 0.360783481 | 0.553344457 |
| ENSG00000250202 | ENSG0000025 | 7.263550671 | 0.20296578 | 0.640955692 | 0.316661171 | 0.751500715 | 0.862575122 |
| ENSG00000250215 | CIR1P2 | 25.39034283 | 0.445498718 | 0.356619836 | 1.249225849 | 0.21158248 | 0.379136236 |
| ENSG00000250220 | ANKRD17-DT | 10.20950737 | 0.093156272 | 0.540128572 | 0.172470551 | 0.863067611 | 0.928909215 |
| ENSG00000250221 | KRT8P32 | 80.24940603 | 0.122322594 | 0.21414916 | 0.571202774 | 0.567862198 | 0.73560104 |
| ENSG00000250222 | TRIM7-AS2 | 40.56638147 | -0.16613166 | 0.282205856 | -0.588689626 | 0.556069498 | 0.72672673 |
| ENSG00000250230 | ENSG0000025 | 8.445691794 | -0.829952492 | 0.620813059 | -1.336879888 | 0.181261847 | 0.336654761 |
| ENSG00000250237 | ENSG0000025 | 3.423953783 | -1.43066551 | 0.989730706 | -1.445509877 | 0.148314721 | 0.289044235 |
| ENSG00000250240 | ENSG0000025 | 23.20524173 | 0.816631578 | 0.369758875 | 2.208551662 | 0.027205841 | 0.071423894 |
| ENSG00000250241 | PCDH10-DT | 11.24685554 | 1.2323823 | 0.552103545 | 2.232157918 | 0.025604528 | 0.067778429 |
| ENSG00000250251 | PKD1P6 | 170.0949335 | -0.018659517 | 0.139598877 | -0.133665234 | 0.893667298 | 0.945700277 |
| ENSG00000250254 | PTTG2 | 2.920399572 | 0.527206125 | 1.066975456 | 0.494112702 | 0.621226588 | 0.775691207 |
| ENSG00000250258 | ENSG0000025 | 5.435664148 | 0.501150053 | 0.762029345 | 0.657651908 | 0.510761832 | 0.69056854 |

| | | | | | | | |
|---|---|---|---|---|---|---|---|
| ENSG00000250260 | ENSG0000025 | 1.668082478 | 0.942369214 | 1.425473611 | 0.66109201 | 0.508553309 | NA |
| ENSG00000250261 | CCDC74BP1 | 1.815958173 | -0.952604554 | 1.299783687 | -0.732894683 | 0.463622666 | 0.649941774 |
| ENSG00000250273 | PSMC1P5 | 3.286845056 | -0.475607613 | 0.96087584 | -0.494973016 | 0.620619168 | 0.775393699 |
| ENSG00000250286 | ENSG0000025 | 6.261701676 | 0.111376046 | 0.689611546 | 0.161505484 | 0.871695293 | 0.933508707 |
| ENSG00000250299 | MRPS31P4 | 18.79017518 | 0.23769069 | 0.415125316 | 0.572575752 | 0.566931981 | 0.734771603 |
| ENSG00000250303 | LINC02762 | 5.81995759 | -0.301789248 | 0.733679151 | -0.411336819 | 0.680825575 | 0.818841625 |
| ENSG00000250305 | TRMT9B | 11.23223333 | -0.259178651 | 0.525130916 | -0.493550548 | 0.621623633 | 0.776040526 |
| ENSG00000250312 | ZNF718 | 538.1714663 | 0.143611951 | 0.08522411 | 1.685109432 | 0.09196746 | 0.197966927 |
| ENSG00000250317 | SMIM20 | 398.332239 | -0.15602582 | 0.096205333 | -1.621800112 | 0.104846154 | 0.220213403 |
| ENSG00000250318 | RPL13AP26 | 7.123468785 | 0.553835028 | 0.674356613 | 0.821279153 | 0.41148728 | 0.603820008 |
| ENSG00000250321 | ENSG0000025 | 36.83676868 | 0.399039351 | 0.299206337 | 1.333659425 | 0.182315498 | 0.338215606 |
| ENSG00000250325 | IGBP1P4 | 5.039044679 | 0.686475669 | 0.791047515 | 0.86780586 | 0.385500621 | 0.578074536 |
| ENSG00000250326 | USP38-DT | 16.59026066 | 0.534148731 | 0.435054505 | 1.227774278 | 0.219531714 | 0.389516399 |
| ENSG00000250328 | MGC32805 | 3.072206399 | 0.630307879 | 1.031439595 | 0.611095291 | 0.541136497 | 0.715279242 |
| ENSG00000250332 | PTPN11P4 | 16.64112992 | 0.074996937 | 0.43986241 | 0.170500901 | 0.864616226 | 0.929799912 |
| ENSG00000250334 | LINC00989 | 2.518516707 | -1.662276225 | 1.187897002 | -1.399343733 | 0.161709931 | 0.308723853 |
| ENSG00000250337 | PURPL | 1.647258971 | 2.8958528 | 1.723237053 | 1.680472686 | 0.092865384 | NA |
| ENSG00000250346 | EEF1GP2 | 4.298172436 | 0.044382859 | 0.831922837 | 0.053349731 | 0.957453257 | 0.980753638 |
| ENSG00000250347 | ENSG0000025 | 4.134584095 | 0.733668474 | 0.894481088 | 0.820216865 | 0.412092489 | 0.604360165 |
| ENSG00000250358 | LINC02200 | 1.61927158 | -0.156916222 | 1.393123162 | -0.112636288 | 0.910318915 | NA |
| ENSG00000250363 | KRT18P21 | 6.058917468 | 0.829204182 | 0.750462878 | 1.10492365 | 0.269192674 | 0.450816173 |
| ENSG00000250365 | HEATR5A-DT | 10.19807648 | -0.904207202 | 0.554469013 | -1.630762369 | 0.102940471 | 0.217053521 |
| ENSG00000250397 | ENSG0000025 | 45.73924983 | 0.540907279 | 0.26592202 | 2.034082317 | 0.041943297 | 0.10311641 |
| ENSG00000250403 | ENSG0000025 | 26.21336166 | -0.494708494 | 0.353398125 | -1.39986168 | 0.161554743 | 0.308530287 |
| ENSG00000250412 | KLHL2P1 | 5.988442739 | -0.367583728 | 0.725454879 | -0.506694129 | 0.612369451 | 0.769537956 |
| ENSG00000250415 | ENSG0000025 | 3.964421679 | 2.728345411 | 1.126396895 | 2.422188326 | 0.015427353 | 0.043616401 |
| ENSG00000250420 | AACSP1 | 2.920279579 | -1.517238653 | 1.096655414 | -1.383514487 | 0.166507168 | 0.31572776 |
| ENSG00000250421 | ENSG0000025 | 3.557834935 | 5.180629731 | 1.404696317 | 3.688078106 | 0.000225954 | 0.000949339 |
| ENSG00000250441 | ENSG0000025 | 8.673544736 | -0.941934564 | 0.618002367 | -1.524160125 | 0.127468716 | 0.257368697 |
| ENSG00000250444 | CCT5P1 | 56.2419793 | -0.08547854 | 0.256081789 | -0.333793905 | 0.738535083 | 0.855829198 |
| ENSG00000250451 | HOXC-AS1 | 28.64490529 | 1.063626977 | 0.358911525 | 2.963479582 | 0.003041822 | 0.010291948 |
| ENSG00000250461 | ENSG0000025 | 108.9038939 | -0.630089803 | 0.174927275 | -3.602010047 | 0.000315766 | 0.001295596 |
| ENSG00000250462 | LRRC37BP1 | 111.0614962 | -0.37837939 | 0.178368223 | -2.121338565 | 0.033893321 | 0.086189808 |

| | | | | | | | |
|---|---|---|---|---|---|---|---|
| ENSG00000250471 | GMPSP1 | 222.6571025 | 0.070414421 | 0.121539207 | 0.579355603 | 0.562349255 | 0.731143273 |
| ENSG00000250474 | WBP1LP2 | 6.672517924 | -1.087417817 | 0.722516565 | -1.505042055 | 0.132313265 | 0.265065731 |
| ENSG00000250479 | CHCHD10 | 208.0422218 | -0.169649319 | 0.135288054 | -1.253985947 | 0.209847113 | 0.376968895 |
| ENSG00000250483 | PPM1AP1 | 3.262188988 | -0.745774718 | 0.962335387 | -0.774963416 | 0.438361278 | 0.627911815 |
| ENSG00000250486 | FAM218A | 25.53828159 | 0.031685442 | 0.358086515 | 0.088485438 | 0.929490858 | 0.965993019 |
| ENSG00000250488 | LINC02233 | 15.67655365 | 1.538176626 | 0.49757222 | 3.091363555 | 0.001992395 | 0.007006829 |
| ENSG00000250492 | INTS6P1 | 33.21899471 | -0.360336479 | 0.31008046 | -1.162074126 | 0.245205356 | 0.422139653 |
| ENSG00000250506 | CDK3 | 15.10715922 | -0.283886206 | 0.4438989 | -0.639528969 | 0.522478874 | 0.699239565 |
| ENSG00000250510 | GPR162 | 13.32070813 | 0.212127574 | 0.508917424 | 0.416821205 | 0.676809191 | 0.815935942 |
| ENSG00000250519 | ENSG0000025 | 21.38882422 | 0.254531378 | 0.377898957 | 0.673543478 | 0.500601598 | 0.681732072 |
| ENSG00000250522 | ENSG0000025 | 4.209690973 | 0.035402511 | 0.855984531 | 0.041358821 | 0.967009841 | 0.985295635 |
| ENSG00000250526 | CCT6P2 | 25.64198708 | 0.474829779 | 0.396162554 | 1.198573094 | 0.230693985 | 0.404102018 |
| ENSG00000250536 | ABHD17AP3 | 8.025751338 | -0.207494526 | 0.623080038 | -0.33301424 | 0.739123534 | 0.856204886 |
| ENSG00000250539 | KRT8P33 | 927.5112782 | 0.182505599 | 0.080658166 | 2.262704531 | 0.023653909 | 0.063323688 |
| ENSG00000250562 | RPL38P4 | 1.55598052 | 1.809239485 | 1.583027839 | 1.142898085 | 0.253080908 | NA |
| ENSG00000250565 | ATP6V1E2 | 251.8474116 | 0.317445722 | 0.120368407 | 2.637284402 | 0.008357273 | 0.025382741 |
| ENSG00000250568 | ENSG0000025 | 127.2954672 | 0.23545798 | 0.176092723 | 1.337124987 | 0.181181842 | 0.336597948 |
| ENSG00000250569 | NTAN1P2 | 17.85497163 | -0.437653747 | 0.423482981 | -1.033462424 | 0.301387547 | 0.488230374 |
| ENSG00000250571 | GLI4 | 432.8510288 | -0.002128961 | 0.096364066 | -0.022092895 | 0.982373854 | 0.991820665 |
| ENSG00000250602 | ENSG0000025 | 3.970049226 | 0.322931588 | 0.916058338 | 0.352522951 | 0.72444611 | 0.846601821 |
| ENSG00000250604 | ENSG0000025 | 3.145941195 | -0.621345408 | 0.992071617 | -0.626311041 | 0.531110946 | 0.706669282 |
| ENSG00000250608 | NUDT16-DT | 14.58745377 | -0.698464577 | 0.475622539 | -1.468527077 | 0.141961107 | 0.279535433 |
| ENSG00000250615 | ENSG0000025 | 3.003669663 | 1.724382918 | 1.090894046 | 1.580706141 | 0.113945244 | 0.235652865 |
| ENSG00000250616 | YPEL3-DT | 53.28462421 | -0.214597562 | 0.240682927 | -0.891619382 | 0.372596978 | 0.565302557 |
| ENSG00000250634 | LINC01182 | 2.211295012 | 3.411140973 | 1.52807599 | 2.232311086 | 0.02559441 | 0.067767285 |
| ENSG00000250643 | ENSG0000025 | 2.132888608 | -0.379912879 | 1.375086532 | -0.276282888 | 0.782330798 | 0.881955689 |
| ENSG00000250644 | ENSG0000025 | 2.972519511 | 1.609430631 | 1.120361153 | 1.436528415 | 0.150852037 | 0.292919056 |
| ENSG00000250669 | ENSG0000025 | 1.516271648 | 3.957578687 | 1.695306666 | 2.334432328 | 0.019573096 | NA |
| ENSG00000250685 | ENSG0000025 | 2.540334428 | -0.391339283 | 1.081189882 | -0.36195241 | 0.717387594 | 0.842904702 |
| ENSG00000250697 | ENSG0000025 | 2.404564036 | -0.354753803 | 1.11603426 | -0.317869993 | 0.750583555 | 0.862253188 |
| ENSG00000250708 | LINC02269 | 17.59178773 | 0.983168049 | 0.461078196 | 2.132323883 | 0.03298023 | 0.084240783 |
| ENSG00000250714 | TNFRSF10A-D | 2.002228197 | -1.77463908 | 1.32889372 | -1.335425891 | 0.181736996 | 0.3373834 |
| ENSG00000250722 | SELENOP | 1266.66672 | -0.608404122 | 0.067411224 | -9.025264482 | 1.79E-19 | 2.91E-18 |

| | | | | | | | |
|---|---|---|---|---|---|---|---|
| ENSG00000250731 | TPM3P6 | 54.46946366 | -0.205158887 | 0.270744244 | -0.757758996 | 0.448595274 | 0.636766016 |
| ENSG00000250740 | ENSG0000025 | 13.70175304 | -0.837158892 | 0.479623017 | -1.745451871 | 0.080906243 | 0.178306199 |
| ENSG00000250746 | ENSG0000025 | 9.445919479 | 0.576124634 | 0.588198753 | 0.979472721 | 0.327346461 | 0.517679904 |
| ENSG00000250762 | ENSG0000025 | 6.447947909 | -0.230108079 | 0.725498351 | -0.317172435 | 0.751112765 | 0.862460033 |
| ENSG00000250764 | ENSG0000025 | 4.488975795 | -0.522988203 | 0.865808118 | -0.604046315 | 0.54581285 | 0.718974837 |
| ENSG00000250770 | ENSG0000025 | 4.976198354 | 3.101004408 | 1.054415798 | 2.94096922 | 0.00327187 | 0.010998931 |
| ENSG00000250786 | SNHG18 | 26.27019207 | 2.859404408 | 0.441155266 | 6.481628189 | 9.07E-11 | 7.89E-10 |
| ENSG00000250790 | ENSG0000025 | 38.85204823 | -0.327699387 | 0.286230373 | -1.14487985 | 0.252258932 | 0.430919644 |
| ENSG00000250796 | CIDECP2 | 4.667383738 | 0.543984363 | 0.803200645 | 0.677270824 | 0.498234139 | 0.679962588 |
| ENSG00000250802 | ZBED3-AS1 | 72.71699711 | -0.476101098 | 0.214056277 | -2.224186581 | 0.026135894 | 0.068954272 |
| ENSG00000250820 | ENSG0000025 | 4.717442637 | -1.76692329 | 0.867864649 | -2.035943383 | 0.041756041 | 0.102722099 |
| ENSG00000250826 | HNRNPA3P13 | 2.209403414 | -0.415242172 | 1.185227777 | -0.350347993 | 0.726077552 | 0.847853251 |
| ENSG00000250834 | KRT18P54 | 3.08164022 | -0.314267185 | 1.010958979 | -0.310860472 | 0.755906696 | 0.865149344 |
| ENSG00000250848 | PPP4R2P1 | 39.65287207 | -0.095042644 | 0.281070027 | -0.338145783 | 0.73525333 | 0.853967465 |
| ENSG00000250853 | RNF138P1 | 4.587384937 | -0.019736949 | 0.870838319 | -0.02266431 | 0.981918045 | 0.991627389 |
| ENSG00000250859 | HNRNPKP1 | 142.9341259 | -0.11178849 | 0.15183333 | -0.736257906 | 0.461573758 | 0.648328558 |
| ENSG00000250874 | ENSG0000025 | 17.17319807 | 0.311129704 | 0.432363365 | 0.719602375 | 0.47176985 | 0.657466637 |
| ENSG00000250891 | LINC02208 | 6.183263001 | 2.274998303 | 0.825815683 | 2.754849963 | 0.005871906 | 0.018571834 |
| ENSG00000250893 | ENSG0000025 | 1.707214704 | 0.654300259 | 1.358998446 | 0.481457695 | 0.63019124 | NA |
| ENSG00000250896 | RNPS1P1 | 756.6045727 | 0.051752893 | 0.078112997 | 0.662538824 | 0.507625962 | 0.687838215 |
| ENSG00000250899 | ENSG0000025 | 193.8343897 | 0.007617463 | 0.138721236 | 0.054912015 | 0.95620856 | 0.979978332 |
| ENSG00000250900 | TRIM7-AS1 | 7.928386959 | -1.524929848 | 0.669335914 | -2.278272862 | 0.022710325 | 0.061068441 |
| ENSG00000250902 | SMAD1-AS1 | 5.320815195 | -1.735533905 | 0.817017864 | -2.124230057 | 0.033650911 | 0.085677236 |
| ENSG00000250903 | GMDS-DT | 73.630273 | 0.069795942 | 0.211261123 | 0.330377595 | 0.741114669 | 0.857259505 |
| ENSG00000250906 | ENSG0000025 | 12.59432211 | 0.385191178 | 0.520484394 | 0.740062878 | 0.459261842 | 0.646387304 |
| ENSG00000250909 | ENSG0000025 | 11.60314605 | -0.355587996 | 0.532873056 | -0.667303387 | 0.504578363 | 0.685318392 |
| ENSG00000250910 | MAP9-AS1 | 5.627711048 | -0.814312269 | 0.751519677 | -1.083554155 | 0.278562527 | 0.461488002 |
| ENSG00000250917 | ENSG0000025 | 1.79332226 | 1.229851099 | 1.362363331 | 0.90273356 | 0.366667321 | 0.559190822 |
| ENSG00000250927 | MESTP3 | 12.34415982 | -0.162931684 | 0.502765597 | -0.324070869 | 0.745884381 | 0.859483007 |
| ENSG00000250938 | MAD2L1-DT | 3.361963075 | -1.073047187 | 1.01969704 | -1.052319605 | 0.29265294 | 0.477894251 |
| ENSG00000250942 | ENPP7P11 | 6.523578388 | 1.285769918 | 0.717455309 | 1.792125449 | 0.073112873 | 0.164480662 |
| ENSG00000250950 | ENSG0000025 | 1.701644288 | 0.965163779 | 1.474331801 | 0.65464489 | 0.512696415 | NA |
| ENSG00000250959 | GLUD1P3 | 35.4756866 | -0.261306179 | 0.299069771 | -0.873729826 | 0.382265404 | 0.574617273 |

| | | | | | | | |
|---|---|---|---|---|---|---|---|
| ENSG00000250966 | H3P14 | 12.70644344 | -0.282049408 | 0.496172659 | -0.568450122 | 0.569729365 | 0.737020901 |
| ENSG00000250979 | ENSG0000025 | 13.80529219 | -1.304372191 | 0.491903203 | -2.651684685 | 0.00800913 | 0.024477676 |
| ENSG00000250982 | GAPDHP35 | 22.11478936 | 0.091054992 | 0.389773159 | 0.233610216 | 0.815287596 | 0.902135938 |
| ENSG00000250983 | FAM76AP1 | 6.854457052 | -0.31278417 | 0.673340222 | -0.464526192 | 0.642270798 | 0.790179777 |
| ENSG00000250986 | LINC02600 | 386.4292625 | 0.136759796 | 0.101547927 | 1.346751235 | 0.178060365 | 0.332468238 |
| ENSG00000250988 | SNHG21 | 20.42366212 | 0.001619372 | 0.38962678 | 0.004156213 | 0.996683831 | 0.99798896 |
| ENSG00000250995 | TMEM30BP1 | 21.97318936 | 0.03441124 | 0.38728079 | 0.088853465 | 0.929198366 | 0.965981609 |
| ENSG00000250999 | ENSG0000025 | 18.55062824 | 0.005472529 | 0.404701209 | 0.013522393 | 0.98921102 | 0.994278654 |
| ENSG00000251000 | GGCTP1 | 6.318887032 | -0.494405432 | 0.692514864 | -0.713927539 | 0.475271987 | 0.66046014 |
| ENSG00000251013 | GAPDHP62 | 2.541717023 | 1.004989894 | 1.139076157 | 0.882285076 | 0.377622665 | 0.570362387 |
| ENSG00000251015 | SLC25A30-AS1 | 1.869412768 | 1.37359796 | 1.428291881 | 0.961706762 | 0.336196924 | 0.527274737 |
| ENSG00000251017 | ENSG0000025 | 27.94430168 | -0.290959452 | 0.354721314 | -0.820248009 | 0.412074738 | 0.604360165 |
| ENSG00000251018 | HMMR-AS1 | 4.926426257 | -0.718177406 | 0.868504292 | -0.826912903 | 0.40828643 | 0.600718638 |
| ENSG00000251022 | THAP9-AS1 | 461.7387618 | 0.008717816 | 0.113801335 | 0.07660557 | 0.938937328 | 0.971022554 |
| ENSG00000251023 | ENSG0000025 | 2.099288491 | -0.286683216 | 1.234120315 | -0.232297623 | 0.816306859 | 0.902679168 |
| ENSG00000251026 | NIHCOLE | 24.41926963 | 0.655568151 | 0.380527378 | 1.722788397 | 0.084926798 | 0.185580181 |
| ENSG00000251056 | ANKRD20A17I | 6.413565957 | -0.783900625 | 0.706049369 | -1.110263191 | 0.26688563 | 0.448031931 |
| ENSG00000251062 | ENSG0000025 | 1.724724611 | 1.287950336 | 1.387597097 | 0.92818754 | 0.353310297 | NA |
| ENSG00000251066 | ENSG0000025 | 17.71869193 | -0.448910363 | 0.423663367 | -1.059592116 | 0.289330202 | 0.473580794 |
| ENSG00000251072 | LMNB1-DT | 2.95260195 | 0.094291514 | 1.026346416 | 0.091871042 | 0.9268005 | 0.964865145 |
| ENSG00000251079 | BMS1P2 | 5.208736147 | -1.423807033 | 0.867169663 | -1.64190134 | 0.100610452 | 0.21295227 |
| ENSG00000251081 | ENSG0000025 | 3.13039494 | 2.962732252 | 1.3053264 | 2.26972522 | 0.02322426 | 0.062275223 |
| ENSG00000251095 | ENSG0000025 | 17.47842439 | -0.488300943 | 0.432381212 | -1.129329697 | 0.258758778 | 0.438528839 |
| ENSG00000251102 | CTBP2P4 | 83.98561803 | -0.0298499 | 0.202137895 | -0.147670974 | 0.88260244 | 0.939121523 |
| ENSG00000251107 | PDCD5P2 | 6.289648181 | -0.598563123 | 0.701139289 | -0.85370073 | 0.39327083 | 0.586120243 |
| ENSG00000251118 | H3P24 | 2.415467326 | 0.026722669 | 1.099811357 | 0.024297503 | 0.980615305 | 0.991070363 |
| ENSG00000251129 | LINC02506 | 66.96453669 | 1.153033969 | 0.23553609 | 4.895360075 | 9.81E-07 | 5.68E-06 |
| ENSG00000251131 | ENSG0000025 | 31.08727567 | -0.123148524 | 0.318657847 | -0.386460038 | 0.699155993 | 0.831105311 |
| ENSG00000251136 | RIPK2-DT | 651.0688203 | 0.762743965 | 0.089810258 | 8.492837957 | 2.02E-17 | 2.85E-16 |
| ENSG00000251138 | LINC02882 | 31.94197356 | 0.004520776 | 0.317999793 | 0.014216288 | 0.988657426 | 0.994082067 |
| ENSG00000251141 | MRPS30-DT | 21.432848 | 1.878496311 | 0.425518739 | 4.414603017 | 1.01E-05 | 5.23E-05 |
| ENSG00000251143 | ENSG0000025 | 311.7415102 | -0.481732072 | 0.133869347 | -3.598524098 | 0.000320028 | 0.001310505 |
| ENSG00000251151 | HOXC-AS3 | 79.78652484 | -0.564464414 | 0.212778641 | -2.652824609 | 0.007982134 | 0.024401672 |

| | | | | | | | |
|---|---|---|---|---|---|---|---|
| ENSG00000251155 | SEPTIN14P4 | 3.464702639 | 1.091725912 | 0.998356827 | 1.09352276 | 0.274164342 | 0.456211374 |
| ENSG00000251161 | ENSG0000025 | 26.94982459 | -0.523786834 | 0.35236074 | -1.486507359 | 0.137144972 | 0.272248672 |
| ENSG00000251169 | LINC01843 | 6.607417988 | 0.83010776 | 0.699283902 | 1.18708261 | 0.235195037 | 0.41020103 |
| ENSG00000251172 | TPTE2P7 | 3.78179768 | -0.284101554 | 0.90800327 | -0.312886048 | 0.754367238 | 0.864498202 |
| ENSG00000251187 | ENSG0000025 | 12.63814737 | -0.485088578 | 0.492550153 | -0.984851137 | 0.324697198 | 0.514865122 |
| ENSG00000251188 | ENSG0000025 | 8.347916208 | -0.574417408 | 0.624704146 | -0.919503115 | 0.357832477 | 0.550179425 |
| ENSG00000251192 | ZNF674 | 131.5503214 | -0.135795935 | 0.157591893 | -0.861693658 | 0.388856116 | 0.581509733 |
| ENSG00000251204 | ENSG0000025 | 8.746056764 | -0.436631189 | 0.598613525 | -0.729404149 | 0.46575448 | 0.651726716 |
| ENSG00000251211 | ENSG0000025 | 2.421210863 | 0.810733639 | 1.186290613 | 0.68341908 | 0.494342067 | 0.676865645 |
| ENSG00000251221 | LINC01337 | 2.929855675 | 0.193802902 | 1.035234722 | 0.187206725 | 0.851498552 | 0.922260215 |
| ENSG00000251224 | CNOT10-AS1 | 2.16496092 | 0.462157205 | 1.180188335 | 0.391596147 | 0.695356635 | 0.828627301 |
| ENSG00000251235 | SNRPCP2 | 15.65549666 | 0.476214549 | 0.453683192 | 1.049663195 | 0.293872991 | 0.479545199 |
| ENSG00000251247 | ZNF345 | 110.6024376 | 0.261645287 | 0.172192278 | 1.519494892 | 0.128637973 | 0.259128477 |
| ENSG00000251248 | ENSG0000025 | 7.155771127 | 2.308222736 | 0.784144943 | 2.94361745 | 0.003244007 | 0.01091006 |
| ENSG00000251254 | GTF2F2P1 | 40.40190118 | -0.13136142 | 0.279927987 | -0.46926862 | 0.638877642 | 0.787290808 |
| ENSG00000251259 | ENSG0000025 | 7.335915963 | 0.688669059 | 0.673221723 | 1.022945392 | 0.306333666 | 0.494044875 |
| ENSG00000251260 | WDFY3-AS1 | 2.510070518 | -0.412035243 | 1.09288071 | -0.377017583 | 0.706160538 | 0.835792114 |
| ENSG00000251273 | LINC02228 | 2.522405852 | 1.880490718 | 1.270526311 | 1.480087977 | 0.13884977 | 0.274800597 |
| ENSG00000251279 | SMIM15-AS1 | 4.903851257 | 0.597161655 | 0.791320978 | 0.75463898 | 0.450465629 | 0.638630587 |
| ENSG00000251287 | ALG1L2 | 3.242244261 | -1.738788704 | 1.071072848 | -1.623408443 | 0.104502121 | 0.219691959 |
| ENSG00000251288 | ENSG0000025 | 1.759699861 | 0.140650368 | 1.329269258 | 0.105810292 | 0.915732871 | NA |
| ENSG00000251307 | CCNO-DT | 6.187913752 | 1.377495391 | 0.747045108 | 1.843925322 | 0.065194024 | 0.149570049 |
| ENSG00000251322 | SHANK3 | 173.2914336 | 0.530744178 | 0.141564742 | 3.749126859 | 0.000177451 | 0.000760032 |
| ENSG00000251323 | LINC02728 | 7.518331772 | -0.395947784 | 0.663055281 | -0.597156521 | 0.550402885 | 0.722615811 |
| ENSG00000251333 | RTN3P1 | 400.297879 | -0.097498207 | 0.095352249 | -1.022505584 | 0.306541671 | 0.494206616 |
| ENSG00000251336 | TEMN3-AS1 | 5.094903249 | 5.699401289 | 1.338828199 | 4.257007206 | 2.07E-05 | 0.000102651 |
| ENSG00000251348 | HSPD1P11 | 3.650941492 | 2.074518202 | 1.037381521 | 1.999763983 | 0.045525755 | 0.110537768 |
| ENSG00000251359 | WWC2-AS2 | 3.588483768 | 0.093280941 | 0.907234525 | 0.102818994 | 0.91810663 | 0.959777894 |
| ENSG00000251364 | SYT9-AS1 | 43.36922098 | -0.810754253 | 0.274286387 | -2.955867632 | 0.003117909 | 0.010529196 |
| ENSG00000251365 | LINC02236 | 21.3626567 | 0.240914039 | 0.375825366 | 0.641026553 | 0.521505432 | 0.698548091 |
| ENSG00000251369 | ZNF550 | 226.6971103 | 0.270856286 | 0.137349896 | 1.972016685 | 0.048607702 | 0.116932164 |
| ENSG00000251396 | LINC01301 | 21.59790081 | 0.359830225 | 0.407044668 | 0.884006729 | 0.376692581 | 0.569482523 |
| ENSG00000251400 | ALDH7A1P1 | 410.7145086 | -0.02054564 | 0.095277652 | -0.215639657 | 0.829268641 | 0.9096893 |

| | | | | | | | |
|---|---|---|---|---|---|---|---|
| ENSG00000251410 | ENSG0000025 | 2.95358102 | 3.864897135 | 1.441785344 | 2.680632836 | 0.007348309 | 0.022675574 |
| ENSG00000251411 | ARPC1AP4 | 15.38301838 | -1.025290649 | 0.469164252 | -2.185355438 | 0.028862795 | 0.075069448 |
| ENSG00000251417 | ENSG0000025 | 5.298412264 | -0.405250496 | 0.788050885 | -0.514244072 | 0.607081358 | 0.765525855 |
| ENSG00000251418 | ASS1P8 | 2.078449839 | 2.232466039 | 1.412035029 | 1.581027377 | 0.113871778 | 0.235559179 |
| ENSG00000251429 | AIDAP2 | 9.655138862 | 0.0201144 | 0.583908871 | 0.034447841 | 0.972520034 | 0.987544136 |
| ENSG00000251432 | LINC02615 | 40.29408936 | 0.326008604 | 0.303627973 | 1.073710702 | 0.282952363 | 0.466603926 |
| ENSG00000251441 | RTEL1P1 | 2.809786788 | 0.014252594 | 1.040357104 | 0.013699713 | 0.989069552 | 0.994183487 |
| ENSG00000251453 | HAUS1P1 | 6.379471762 | 0.476856267 | 0.700891486 | 0.68035677 | 0.496278587 | 0.678189674 |
| ENSG00000251455 | ENSG0000025 | 6.701233107 | 0.544480177 | 0.731024012 | 0.744818457 | 0.456381475 | 0.643478245 |
| ENSG00000251463 | FKBP4P1 | 5.303819919 | -0.166727382 | 0.788519011 | -0.211443706 | 0.832541055 | 0.911410533 |
| ENSG00000251467 | CTDNEP1P2 | 6.1893509 | -0.597284883 | 0.706955941 | -0.844868609 | 0.398184203 | 0.591181516 |
| ENSG00000251474 | RPL32P3 | 364.6688163 | 0.140179815 | 0.112200972 | 1.249363639 | 0.211532101 | 0.379127095 |
| ENSG00000251482 | AKIRIN2P1 | 1.633450164 | -1.398839893 | 1.427059346 | -0.980225452 | 0.326974843 | NA |
| ENSG00000251483 | LYPLA1P2 | 2.623099521 | -1.69541142 | 1.189221888 | -1.425647675 | 0.153970067 | 0.297224518 |
| ENSG00000251484 | ENSG0000025 | 2.05329591 | 0.564189793 | 1.251108926 | 0.450951776 | 0.652024306 | 0.797900222 |
| ENSG00000251485 | NRBF2P6 | 9.654323623 | 0.090103914 | 0.576578268 | 0.156273517 | 0.87581743 | 0.935564696 |
| ENSG00000251493 | FOXD1 | 209.1778719 | 0.448929649 | 0.129367267 | 3.470195036 | 0.000520081 | 0.002050997 |
| ENSG00000251495 | PPIAP11 | 67.12976813 | 0.41490896 | 0.23197402 | 1.78860098 | 0.0736791 | 0.165559659 |
| ENSG00000251513 | LIX1-AS1 | 8.14567339 | -0.59020137 | 0.608413277 | -0.970066552 | 0.33201332 | 0.523050051 |
| ENSG00000251532 | ENSG0000025 | 11.26899838 | -0.852245549 | 0.52206189 | -1.632460759 | 0.102582458 | 0.216429835 |
| ENSG00000251548 | ENSG0000025 | 18.68369783 | 0.609047177 | 0.424133981 | 1.435978264 | 0.151008526 | 0.293123675 |
| ENSG00000251550 | ENSG0000025 | 2.408871851 | 0.402756743 | 1.112485989 | 0.362033093 | 0.717327301 | 0.842877031 |
| ENSG00000251553 | DDX18P4 | 3.375323468 | 0.371514506 | 0.979064288 | 0.379458745 | 0.704347234 | 0.834635491 |
| ENSG00000251556 | ENSG0000025 | 8.589624843 | -0.043534607 | 0.628343851 | -0.069284687 | 0.944763015 | 0.973711185 |
| ENSG00000251562 | MALAT1 | 8466.813136 | 0.144562522 | 0.18342971 | 0.788108545 | 0.430633217 | 0.620756944 |
| ENSG00000251571 | DDX3P3 | 1.605643357 | 0.97247207 | 1.493539876 | 0.651118919 | 0.514969724 | NA |
| ENSG00000251574 | ENSG0000025 | 4.9410462 | 0.328848513 | 0.818625654 | 0.401708047 | 0.687898901 | 0.82352577 |
| ENSG00000251580 | LINC02482 | 122.3698789 | -0.445380241 | 0.167405955 | -2.660480276 | 0.00780293 | 0.023917585 |
| ENSG00000251583 | MFFP2 | 8.172714506 | -0.469557564 | 0.627015351 | -0.748877301 | 0.453931163 | 0.641552294 |
| ENSG00000251587 | LDHAP1 | 5.50853098 | 0.696423945 | 0.753019626 | 0.924841692 | 0.35504826 | 0.546704732 |
| ENSG00000251595 | ABCA11P | 9.010959135 | 0.214722233 | 0.608834595 | 0.352677451 | 0.724330266 | 0.846509568 |
| ENSG00000251596 | HADHAP1 | 15.05415587 | 0.180750055 | 0.484576808 | 0.373005996 | 0.709143982 | 0.837943158 |
| ENSG00000251600 | ENSG0000025 | 9.779220942 | -0.419483855 | 0.559809007 | -0.74933388 | 0.453655992 | 0.641369928 |

| | | | | | | | |
|---|---|---|---|---|---|---|---|
| ENSG00000251602 | MTA1-DT | 85.43032194 | -1.350687155 | 0.200197006 | -6.746789973 | 1.51E-11 | 1.39E-10 |
| ENSG00000251611 | FHIP1A-DT | 14.80276135 | 0.256097353 | 0.453219674 | 0.565062305 | 0.572031387 | 0.738873875 |
| ENSG00000251613 | ENSG0000025 | 2.516673456 | 1.391097855 | 1.160990827 | 1.198198833 | 0.23083962 | 0.404295379 |
| ENSG00000251615 | ENSG0000025 | 64.11317306 | 1.273054629 | 0.24252638 | 5.249138779 | 1.53E-07 | 9.75E-07 |
| ENSG00000251620 | STPG2-AS1 | 2.461862006 | 3.596609091 | 1.512631092 | 2.377717283 | 0.017420176 | 0.048503319 |
| ENSG00000251633 | GYG1P1 | 4.856525529 | 0.397710023 | 0.787477408 | 0.505043089 | 0.613528572 | 0.770479304 |
| ENSG00000251634 | NAIPP4 | 9.975927972 | 0.031837963 | 0.562565456 | 0.056594237 | 0.954868425 | 0.979301696 |
| ENSG00000251635 | LINC02278 | 1.899178817 | 0.346735834 | 1.271339334 | 0.27273272 | 0.785058681 | 0.883106023 |
| ENSG00000251661 | ENSG0000025 | 20.0529315 | -0.942648686 | 0.408129764 | -2.309678857 | 0.02090594 | 0.056809082 |
| ENSG00000251664 | PCDHA12 | 104.8240474 | 0.039005513 | 0.173938283 | 0.224249154 | 0.822563438 | 0.90619104 |
| ENSG00000251666 | ZNF346-IT1 | 2.834330323 | 0.142768665 | 1.087407035 | 0.131292754 | 0.895543724 | 0.947023142 |
| ENSG00000251667 | BRCC3P1 | 52.48153223 | -0.948096708 | 0.245838258 | -3.856587317 | 0.000114981 | 0.000508088 |
| ENSG00000251669 | FAM86EP | 186.4903905 | 0.041741347 | 0.132716665 | 0.314514738 | 0.753130124 | 0.863926645 |
| ENSG00000251675 | ENSG0000025 | 1.790002078 | 0.061687621 | 1.292214865 | 0.047737897 | 0.961925131 | NA |
| ENSG00000251689 | ENSG0000025 | 4.280150151 | -0.41185053 | 0.848371139 | -0.485460327 | 0.627349848 | 0.780187429 |
| ENSG00000251692 | PTX4 | 7.30405803 | -0.083301908 | 0.656120162 | -0.12696136 | 0.898970982 | 0.948985059 |
| ENSG00000251739 | RNU6-1053P | 2.834507448 | -0.259823673 | 1.131632506 | -0.229600751 | 0.818402024 | 0.903906713 |
| ENSG00000251819 | RNU6-322P | 1.729021285 | 1.992576219 | 1.520764604 | 1.310246315 | 0.190112522 | NA |
| ENSG00000251867 | ENSG0000025 | 54.28979556 | 0.969909567 | 0.247708087 | 3.915534528 | 9.02E-05 | 0.000405106 |
| ENSG00000251920 | RNA5SP216 | 2.059869104 | -0.80400519 | 1.250240985 | -0.643080174 | 0.520172078 | 0.697413646 |
| ENSG00000252147 | RNY3P16 | 4.419183595 | 1.718497451 | 0.910517912 | 1.887384563 | 0.05910862 | 0.13774398 |
| ENSG00000252206 | RNU7-40P | 2.184170701 | -0.084071273 | 1.240355435 | -0.067779986 | 0.945960776 | 0.974551588 |
| ENSG00000252311 | RNU1-103P | 2.727139112 | 0.146694901 | 1.085497355 | 0.135140727 | 0.89250061 | 0.944983167 |
| ENSG00000252464 | RN7SKP70 | 2.172035654 | -2.556518901 | 1.445132395 | -1.769055147 | 0.076884672 | 0.17136945 |
| ENSG00000252473 | ENSG0000025 | 2.093970761 | -2.549541921 | 1.481916447 | -1.720435674 | 0.085353277 | 0.186245998 |
| ENSG00000252498 | RNU6-1016P | 8.051775809 | 0.565512861 | 0.622769793 | 0.90806084 | 0.363846079 | 0.556559975 |
| ENSG00000252516 | RNA5SP82 | 6.049932534 | -0.387118289 | 0.716706151 | -0.540135295 | 0.589103731 | 0.752295476 |
| ENSG00000252577 | SCARNA20 | 5.320225982 | 0.746139398 | 0.797838912 | 0.935200561 | 0.349684949 | 0.540951699 |
| ENSG00000252618 | RNA5SP441 | 2.772079467 | 0.120535229 | 1.089710759 | 0.110612131 | 0.911923928 | 0.955697231 |
| ENSG00000252690 | ENSG0000025 | 38.18961704 | -0.058505252 | 0.324203893 | -0.180458205 | 0.856792864 | 0.925804441 |
| ENSG00000252755 | RNU6-703P | 3.205991265 | 0.39380466 | 0.998179154 | 0.394523025 | 0.693194931 | 0.826947611 |
| ENSG00000252835 | SCARNA21 | 2.154494578 | -0.386843812 | 1.236227273 | -0.312922891 | 0.754339246 | 0.864498202 |
| ENSG00000252916 | RNU6-762P | 2.381057581 | -0.242246345 | 1.167330986 | -0.207521558 | 0.835602558 | 0.913361733 |

| | | | | | | | |
|---|---|---|---|---|---|---|---|
| ENSG00000252933 | RNU6-1223P | 3.607494313 | 0.193773617 | 0.967293001 | 0.200325668 | 0.841225889 | 0.916625749 |
| ENSG00000252965 | Y_RNA | 4.55116311 | 0.410001624 | 0.848602296 | 0.483149322 | 0.628989716 | 0.7810577 |
| ENSG00000253047 | ENSG0000025 | 4.262376417 | 0.07958116 | 0.878974182 | 0.090538677 | 0.92785916 | 0.965331908 |
| ENSG00000253102 | ENSG0000025 | 23.40046583 | -0.133860049 | 0.40604428 | -0.3296686 | 0.741650382 | 0.857515143 |
| ENSG00000253106 | ENSG0000025 | 89.74224311 | -0.182091414 | 0.195714571 | -0.930392734 | 0.352167779 | 0.543656047 |
| ENSG00000253125 | ENSG0000025 | 45.20946003 | 0.129092489 | 0.273469421 | 0.472054566 | 0.636887841 | 0.786213586 |
| ENSG00000253133 | GPAT4-AS1 | 12.24716975 | -0.052052435 | 0.507812745 | -0.102503206 | 0.918357268 | 0.959952416 |
| ENSG00000253144 | ENSG0000025 | 322.7531929 | 0.409008565 | 0.121645103 | 3.36231017 | 0.000772932 | 0.002954686 |
| ENSG00000253146 | CIR1P1 | 39.15086169 | 0.050898523 | 0.280023918 | 0.181764914 | 0.85576722 | 0.925175552 |
| ENSG00000253154 | ENSG0000025 | 8.987958735 | 3.509864958 | 0.87513703 | 4.010646147 | 6.06E-05 | 0.000280057 |
| ENSG00000253159 | PCDHGA12 | 3.752117878 | 0.698883974 | 0.903977759 | 0.773120762 | 0.439450915 | 0.628922826 |
| ENSG00000253161 | LINC01605 | 2.553543982 | 1.432104616 | 1.178482505 | 1.215210755 | 0.224285719 | 0.395808478 |
| ENSG00000253167 | WASHC5-AS1 | 12.36709331 | -0.402838812 | 0.510930147 | -0.788442049 | 0.430438183 | 0.620631799 |
| ENSG00000253172 | ENSG0000025 | 1.987186533 | 0.250640223 | 1.231157018 | 0.203581037 | 0.838680913 | 0.915120151 |
| ENSG00000253173 | H2AZP2 | 23.00232903 | 0.084674854 | 0.377089345 | 0.224548518 | 0.822330519 | 0.906107983 |
| ENSG00000253174 | ENSG0000025 | 3.798938282 | -0.981654738 | 0.96434734 | -1.017947266 | 0.308703018 | 0.496818242 |
| ENSG00000253180 | ENSG0000025 | 5.957130282 | -0.820720295 | 0.729941151 | -1.124365017 | 0.260858206 | 0.441163976 |
| ENSG00000253194 | ENSG0000025 | 22.93827276 | 0.898456956 | 0.374782446 | 2.397275986 | 0.016517477 | 0.046287678 |
| ENSG00000253196 | ENSG0000025 | 1.64585009 | 0.915049512 | 1.48261406 | 0.617186587 | 0.537111662 | NA |
| ENSG00000253198 | ENSG0000025 | 1.801395768 | -1.479286619 | 1.380726279 | -1.071382968 | 0.283997268 | 0.467822586 |
| ENSG00000253200 | ENSG0000025 | 114.2220415 | 0.020650776 | 0.170085864 | 0.12141383 | 0.903363263 | 0.951170572 |
| ENSG00000253203 | GUSBP3 | 5.073474129 | -0.463819583 | 0.780319272 | -0.59439719 | 0.552246488 | 0.723957622 |
| ENSG00000253204 | ENSG0000025 | 65.10227876 | -0.012465788 | 0.218710255 | -0.056996814 | 0.954547733 | 0.979088978 |
| ENSG00000253210 | DENND3-AS1 | 74.20304639 | 0.080640947 | 0.210289547 | 0.383475777 | 0.701367027 | 0.832911922 |
| ENSG00000253213 | USP12P1 | 4.937412743 | 0.41199194 | 0.79681672 | 0.517047308 | 0.605123135 | 0.763896364 |
| ENSG00000253219 | KRT18P41 | 5.95331665 | -0.098018941 | 0.720969756 | -0.135954304 | 0.891857405 | 0.944605424 |
| ENSG00000253223 | ENSG0000025 | 82.2664224 | 0.433199952 | 0.210877208 | 2.054275833 | 0.039949002 | 0.099029236 |
| ENSG00000253227 | ENSG0000025 | 7.257395602 | 1.492235512 | 0.697355657 | 2.139848581 | 0.032367006 | 0.082997889 |
| ENSG00000253228 | NRBF2P4 | 4.25887823 | -0.17333605 | 0.830234396 | -0.208779655 | 0.834620255 | 0.912770412 |
| ENSG00000253250 | C8orf88 | 5.454752377 | 3.15892619 | 1.040585541 | 3.035719856 | 0.002399622 | 0.008309031 |
| ENSG00000253251 | SHLD3 | 45.20429821 | -0.895086622 | 0.266677505 | -3.356438412 | 0.000789533 | 0.003012122 |
| ENSG00000253263 | ENSG0000025 | 2.053850274 | -1.200827114 | 1.310463421 | -0.916337758 | 0.35948977 | 0.55209478 |
| ENSG00000253266 | ENSG0000025 | 24.30380868 | 0.01638508 | 0.354773391 | 0.046184635 | 0.963163089 | 0.983178092 |

| | | | | | | | |
|---|---|---|---|---|---|---|---|
| ENSG00000253276 | CCDC71L | 235.8550486 | 0.44067187 | 0.121335142 | 3.631856873 | 0.000281389 | 0.001162866 |
| ENSG00000253293 | HOXA10 | 144.5321939 | -0.895540142 | 0.164543461 | -5.442575088 | 5.25E-08 | 3.50E-07 |
| ENSG00000253304 | TMEM200B | 12.24456143 | 0.808036968 | 0.520390211 | 1.552752051 | 0.120482382 | 0.246348342 |
| ENSG00000253305 | PCDHGB6 | 17.23719927 | 1.896497096 | 0.474175122 | 3.999571059 | 6.35E-05 | 0.000292253 |
| ENSG00000253307 | ENSG0000025 | 1.74077619 | 0.159059439 | 1.393976458 | 0.114104825 | 0.909154698 | NA |
| ENSG00000253313 | C1orf210 | 382.0094683 | 0.107715648 | 0.099210061 | 1.085733114 | 0.277597086 | 0.460354019 |
| ENSG00000253320 | MAILR | 191.4117507 | 0.195973207 | 0.140077178 | 1.399037374 | 0.161801776 | 0.308834536 |
| ENSG00000253327 | RAD21-AS1 | 10.1104232 | 1.429079637 | 0.595951275 | 2.397980669 | 0.016485735 | 0.046210015 |
| ENSG00000253330 | ENSG0000025 | 3.070333404 | 0.611360987 | 1.017123639 | 0.601068507 | 0.547794358 | 0.720591925 |
| ENSG00000253336 | ENSG0000025 | 37.29367131 | -0.187482728 | 0.287650058 | -0.651773648 | 0.514547204 | 0.693601649 |
| ENSG00000253341 | PCBP2P2 | 3621.713753 | 0.039258666 | 0.055466762 | 0.707787239 | 0.479077392 | 0.663653377 |
| ENSG00000253352 | TUG1 | 10018.29287 | -0.642242113 | 0.048744146 | -13.17577925 | 1.21E-39 | 4.83E-38 |
| ENSG00000253355 | ENSG0000025 | 2.044880892 | -0.263948043 | 1.239302802 | -0.212981075 | 0.831341724 | 0.910700499 |
| ENSG00000253356 | ENSG0000025 | 16.33656548 | -0.840910656 | 0.448980739 | -1.872932586 | 0.061077695 | 0.141627648 |
| ENSG00000253358 | ENSG0000025 | 2.027769626 | -1.14718549 | 1.285920958 | -0.892111978 | 0.372332915 | 0.564976656 |
| ENSG00000253361 | ENSG0000025 | 52.87256912 | -0.503014655 | 0.241963664 | -2.078885094 | 0.037627913 | 0.094285563 |
| ENSG00000253368 | TRNP1 | 91.25988095 | 0.61925355 | 0.18693026 | 3.312751776 | 0.000923829 | 0.003482768 |
| ENSG00000253369 | ENSG0000025 | 5.798739121 | -0.577097128 | 0.730234406 | -0.790290246 | 0.429358282 | 0.619714368 |
| ENSG00000253384 | ABRAXAS1P2 | 6.870233662 | -0.278364537 | 0.671055085 | -0.414816224 | 0.678276442 | 0.816865895 |
| ENSG00000253392 | ENSG0000025 | 16.3312994 | 0.382450949 | 0.449729892 | 0.850401444 | 0.395101934 | 0.588013388 |
| ENSG00000253394 | LINC00534 | 28.67683619 | 0.071283469 | 0.325657152 | 0.218891152 | 0.826734839 | 0.908541056 |
| ENSG00000253395 | LINC03044 | 44.48504333 | 0.42072063 | 0.269556988 | 1.560785469 | 0.118574377 | 0.243300198 |
| ENSG00000253404 | CTNNA1-AS1 | 62.06844088 | -0.113551035 | 0.234195899 | -0.484854924 | 0.627779259 | 0.780428993 |
| ENSG00000253406 | ENSG0000025 | 2.725107816 | -2.273298728 | 1.21454913 | -1.87172233 | 0.061245029 | 0.141943924 |
| ENSG00000253414 | LINC01605 | 2.624553845 | 0.897661907 | 1.143222127 | 0.785203405 | 0.432334319 | 0.622408697 |
| ENSG00000253431 | SRPK2P | 62.20250306 | -0.230294941 | 0.23198636 | -0.992708973 | 0.320851801 | 0.510355035 |
| ENSG00000253433 | NCRNA00250 | 3.832014881 | -1.632210669 | 0.951510637 | -1.715388778 | 0.086273972 | 0.18786188 |
| ENSG00000253438 | PCAT1 | 167.443119 | -0.925360037 | 0.148402769 | -6.235463432 | 4.50E-10 | 3.71E-09 |
| ENSG00000253439 | CDC42P5 | 3.581888292 | 0.348655462 | 0.910040139 | 0.383120972 | 0.701630071 | 0.833050046 |
| ENSG00000253445 | ENSG0000025 | 3.582318103 | -1.162898264 | 0.988954136 | -1.175886951 | 0.239640054 | 0.415332641 |
| ENSG00000253476 | ENSG0000025 | 2.984252134 | -0.151659628 | 1.016861118 | -0.149144879 | 0.881439314 | 0.938554487 |
| ENSG00000253477 | ENSG0000025 | 8.902738336 | -2.083109934 | 0.662680266 | -3.143461547 | 0.001669624 | 0.005962199 |
| ENSG00000253485 | PCDHGA5 | 17.07502837 | 1.709803849 | 0.465020696 | 3.676833879 | 0.000236147 | 0.000988722 |

| | | | | | | | |
|---|---|---|---|---|---|---|---|
| ENSG00000253506 | NACA2 | 222.458944 | 0.219216634 | 0.127973216 | 1.712988392 | 0.086714675 | 0.188678236 |
| ENSG00000253516 | HMGB1P41 | 2.009856194 | 0.292536884 | 1.277518287 | 0.228988412 | 0.818877922 | 0.904040596 |
| ENSG00000253517 | XRCC6P4 | 4.317069822 | 0.358182702 | 0.885743309 | 0.404386574 | 0.685928472 | 0.822454482 |
| ENSG00000253537 | PCDHGA7 | 7.598236017 | 0.933859319 | 0.69100269 | 1.351455402 | 0.176549597 | 0.330211802 |
| ENSG00000253540 | FAM86HP | 218.4890254 | -0.399391253 | 0.136820593 | -2.919087285 | 0.003510579 | 0.011724077 |
| ENSG00000253541 | SEPTIN10P1 | 17.87254847 | -1.181795988 | 0.446575084 | -2.646354514 | 0.00813645 | 0.024807304 |
| ENSG00000253549 | CA3-AS1 | 7.420078552 | 1.919970995 | 0.751145649 | 2.556056867 | 0.010586582 | 0.031249365 |
| ENSG00000253553 | ENSG0000025 | 112.803729 | 0.853608236 | 0.176164545 | 4.845516659 | 1.26E-06 | 7.22E-06 |
| ENSG00000253554 | LINC01414 | 19.75282067 | -0.595158627 | 0.396898914 | -1.499521934 | 0.133738283 | 0.267173753 |
| ENSG00000253555 | TARDBPP4 | 1.75215506 | 0.612973503 | 1.318949331 | 0.464743784 | 0.64211495 | NA |
| ENSG00000253559 | OSGEPL1-AS1 | 5.340616546 | -0.098171037 | 0.759509297 | -0.129255873 | 0.897155187 | 0.947984339 |
| ENSG00000253570 | RNF5P1 | 188.6159843 | -0.349136612 | 0.140512962 | -2.484728868 | 0.012965014 | 0.037354612 |
| ENSG00000253582 | OXR1-AS1 | 7.406347231 | -0.619703842 | 0.656677788 | -0.943695452 | 0.345325299 | 0.536243449 |
| ENSG00000253596 | ENSG0000025 | 6.42526263 | 2.137941472 | 0.803809229 | 2.659762285 | 0.007819582 | 0.023965425 |
| ENSG00000253598 | SLC10A5 | 10.0732275 | 0.95463785 | 0.59138429 | 1.614242832 | 0.106474765 | 0.222980747 |
| ENSG00000253606 | AFG3L2P1 | 16.2774533 | 0.437620451 | 0.448005709 | 0.976818916 | 0.32865881 | 0.519065962 |
| ENSG00000253618 | GRPEL2-AS1 | 1.929442726 | 0.372494187 | 1.289363857 | 0.288897649 | 0.772659703 | 0.876166615 |
| ENSG00000253626 | EIF5AL1 | 1447.317367 | 0.033868898 | 0.062940367 | 0.538110913 | 0.590500479 | 0.753103963 |
| ENSG00000253636 | ENSG0000025 | 7.620260656 | 0.089926155 | 0.659558612 | 0.136342932 | 0.891550186 | 0.944396856 |
| ENSG00000253641 | LINC03022 | 2.266592857 | 1.09924887 | 1.216986184 | 0.903255012 | 0.366390567 | 0.559031629 |
| ENSG00000253653 | ENSG0000025 | 3.354537942 | 0.669203704 | 1.021653543 | 0.655020196 | 0.512454751 | 0.69208242 |
| ENSG00000253669 | GASAL1 | 90.85901605 | -0.309866202 | 0.191975809 | -1.61408983 | 0.106507943 | 0.222989159 |
| ENSG00000253671 | ENSG0000025 | 22.29252353 | 0.858438681 | 0.388166441 | 2.211522148 | 0.0269997 | 0.070972054 |
| ENSG00000253676 | TAGLN2P1 | 278.0739844 | 0.573961973 | 0.113996752 | 5.034897607 | 4.78E-07 | 2.89E-06 |
| ENSG00000253677 | UBE2HP1 | 84.51426902 | 0.356883579 | 0.195042744 | 1.829771113 | 0.067284172 | 0.153588121 |
| ENSG00000253683 | ENSG0000025 | 612.0680559 | 0.017234377 | 0.083114839 | 0.207356192 | 0.835731693 | 0.913370821 |
| ENSG00000253696 | KBTBD11-OT1 | 1.577202993 | 0.45314451 | 1.383006851 | 0.327651674 | 0.743175047 | NA |
| ENSG00000253704 | VIRMA-DT | 17.4776453 | -0.713924088 | 0.419611157 | -1.701394437 | 0.088868945 | 0.192507422 |
| ENSG00000253710 | ALG11 | 267.7340874 | -0.347738329 | 0.114158012 | -3.046114097 | 0.002318198 | 0.008048959 |
| ENSG00000253716 | MINCR | 166.4870683 | 0.387616808 | 0.146037088 | 2.654235398 | 0.007948835 | 0.024325812 |
| ENSG00000253719 | ATXN7L3B | 8295.488523 | 0.278120792 | 0.041777927 | 6.657122883 | 2.79E-11 | 2.52E-10 |
| ENSG00000253721 | SUMO2P20 | 18.31141568 | -0.13453059 | 0.425287202 | -0.316328799 | 0.751752954 | 0.862821472 |
| ENSG00000253722 | ENSG0000025 | 5.317023845 | -0.397483808 | 0.770166159 | -0.516101367 | 0.605783613 | 0.764393624 |

| | | | | | | | |
|---|---|---|---|---|---|---|---|
| ENSG00000253729 | PRKDC | 7499.903756 | 0.04577645 | 0.064614568 | 0.708454008 | 0.478663364 | 0.663276205 |
| ENSG00000253731 | PCDHGA6 | 25.29921885 | 1.349827256 | 0.368497699 | 3.663054778 | 0.000249225 | 0.001037986 |
| ENSG00000253738 | OTUD6B-AS1 | 789.7178025 | -0.042962294 | 0.077532696 | -0.554118401 | 0.579497818 | 0.745038089 |
| ENSG00000253741 | LNCOC1 | 104.6059839 | -0.078153943 | 0.176936692 | -0.441705687 | 0.658702194 | 0.802561522 |
| ENSG00000253744 | ENSG0000025 | 1.883617637 | -0.48226713 | 1.309952956 | -0.368156068 | 0.712756866 | 0.84015416 |
| ENSG00000253764 | KBTBD11-AS1 | 2.829769822 | 0.999903725 | 1.099965496 | 0.909031901 | 0.363333287 | 0.556139695 |
| ENSG00000253767 | PCDHGA8 | 15.39612524 | 2.285314425 | 0.553014311 | 4.132468868 | 3.59E-05 | 0.000171268 |
| ENSG00000253768 | ENSG0000025 | 18.05687475 | -3.411936739 | 0.583303542 | -5.849333142 | 4.94E-09 | 3.68E-08 |
| ENSG00000253773 | CFAP418-AS1 | 3.514199084 | -2.247118338 | 1.078612939 | -2.083340795 | 0.037220171 | 0.093406708 |
| ENSG00000253785 | ENSG0000025 | 632.3234127 | -0.571734721 | 0.082258623 | -6.950453346 | 3.64E-12 | 3.55E-11 |
| ENSG00000253802 | SIRLNT | 7.466477561 | 2.884585579 | 0.834416074 | 3.45701104 | 0.000546202 | 0.002147735 |
| ENSG00000253806 | ENSG0000025 | 1.6405803 | -0.105105336 | 1.481112996 | -0.070963753 | 0.943426604 | NA |
| ENSG00000253816 | GUSBP14 | 11.8662452 | -0.478699294 | 0.532057339 | -0.899713733 | 0.368272613 | 0.560818757 |
| ENSG00000253833 | DSTNP3 | 52.57760485 | -0.365406094 | 0.265201288 | -1.377844341 | 0.168251353 | 0.318324621 |
| ENSG00000253846 | PCDHGA10 | 8.408696119 | 2.809693694 | 0.783133571 | 3.587757948 | 0.000333534 | 0.001360708 |
| ENSG00000253848 | RBM12B-DT | 2.13750819 | 0.043911656 | 1.204334728 | 0.036461338 | 0.970914506 | 0.98680905 |
| ENSG00000253869 | PIGFP1 | 2.21936852 | 0.434510924 | 1.204035484 | 0.360878836 | 0.718190025 | 0.843411181 |
| ENSG00000253873 | PCDHGA11 | 5.347403745 | 1.430888003 | 0.810263119 | 1.76595475 | 0.077403441 | 0.172251634 |
| ENSG00000253875 | ENSG0000025 | 4.039039259 | -0.91449595 | 0.873541377 | -1.046883381 | 0.295153368 | 0.481018631 |
| ENSG00000253882 | ENSG0000025 | 8.888279728 | -1.29003017 | 0.618565432 | -2.085519338 | 0.037022185 | 0.092960696 |
| ENSG00000253899 | ENSG0000025 | 8.533980856 | 0.155800522 | 0.63472913 | 0.245459858 | 0.806100386 | 0.897100998 |
| ENSG00000253910 | PCDHGB2 | 46.98157273 | 1.969313197 | 0.317927296 | 6.194224971 | 5.86E-10 | 4.76E-09 |
| ENSG00000253915 | MAPRE1P1 | 4.987053298 | -0.193355953 | 0.773324057 | -0.250032249 | 0.80256241 | 0.894943694 |
| ENSG00000253919 | THAP12P7 | 214.8309932 | -0.087053074 | 0.124796903 | -0.697557964 | 0.485453678 | 0.669289001 |
| ENSG00000253931 | LINC02990 | 27.55675211 | 0.027919348 | 0.352195709 | 0.079272256 | 0.936816074 | 0.969659375 |
| ENSG00000253943 | KRT18P37 | 2.648625805 | -0.216146248 | 1.093013025 | -0.197752674 | 0.843238576 | 0.917424374 |
| ENSG00000253944 | MTUS1-DT | 2.069419543 | 1.557315763 | 1.306668627 | 1.1918215 | 0.233331243 | 0.407702936 |
| ENSG00000253945 | ENSG0000025 | 15.20229339 | 0.118101856 | 0.44849311 | 0.263330369 | 0.792295954 | 0.888331828 |
| ENSG00000253948 | VPS13B-DT | 51.84480514 | 0.478778689 | 0.259203179 | 1.847117351 | 0.064730134 | 0.148691 |
| ENSG00000253953 | PCDHGB4 | 2.736090298 | 1.485412148 | 1.143875887 | 1.298578076 | 0.194088766 | 0.354857439 |
| ENSG00000253954 | HMGN1P38 | 2293.230855 | -0.231092721 | 0.057513593 | -4.018053985 | 5.87E-05 | 0.000271892 |
| ENSG00000253955 | LINC02995 | 3.273722376 | -2.051161983 | 1.095067168 | -1.873092394 | 0.061055627 | 0.141605105 |
| ENSG00000253958 | CLDN23 | 90.84339593 | 1.868007261 | 0.208170727 | 8.973438705 | 2.87E-19 | 4.61E-18 |

| | | | | | | | |
|---|---|---|---|---|---|---|---|
| ENSG00000253966 | ENSG0000025 | 3.148862736 | 0.220980694 | 1.002079094 | 0.220522208 | 0.825464479 | 0.907872877 |
| ENSG00000253971 | CDC42P3 | 4.997060714 | -0.552914007 | 0.808063277 | -0.684245928 | 0.493819885 | 0.676438565 |
| ENSG00000253972 | MAL2-AS1 | 32.12038581 | -0.84208212 | 0.329894124 | -2.55258296 | 0.010692744 | 0.031504181 |
| ENSG00000253973 | NRIP3-DT | 8.996638275 | -0.422125077 | 0.582451161 | -0.724739007 | 0.46861216 | 0.654566686 |
| ENSG00000253978 | TENM2-AS1 | 1.989997713 | -0.171538587 | 1.245174133 | -0.13776273 | 0.890427941 | 0.943868964 |
| ENSG00000253982 | CLN8-AS1 | 171.4751128 | 0.440854037 | 0.141957914 | 3.10552632 | 0.001899407 | 0.00670964 |
| ENSG00000253991 | ENSG0000025 | 2.698694744 | 1.130369504 | 1.124241202 | 1.005451056 | 0.314679706 | 0.503189213 |
| ENSG00000254003 | SRI-AS1 | 2.26265026 | -1.430290143 | 1.232497915 | -1.16048078 | 0.245853114 | 0.422937416 |
| ENSG00000254004 | ZNF260 | 738.3378685 | 0.339430202 | 0.077211142 | 4.396129813 | 1.10E-05 | 5.66E-05 |
| ENSG00000254012 | RPL10P8 | 1.86334449 | 0.568388885 | 1.311825372 | 0.433280906 | 0.664810708 | 0.806662311 |
| ENSG00000254013 | MAP2K1P1 | 87.69630342 | 0.286390066 | 0.201309939 | 1.422632523 | 0.154842711 | 0.298394808 |
| ENSG00000254023 | PKMP4 | 4.820569701 | -0.448747125 | 0.826065691 | -0.543234188 | 0.58696857 | 0.750686235 |
| ENSG00000254024 | ENSG0000025 | 251.4906931 | 0.025105895 | 0.121037015 | 0.207423282 | 0.835679302 | 0.913370821 |
| ENSG00000254027 | LNMICC | 5.140081367 | 0.02569995 | 0.794977208 | 0.032327908 | 0.974210553 | 0.988342035 |
| ENSG00000254028 | ENSG0000025 | 5.279088746 | -0.499425714 | 0.815145962 | -0.612682559 | 0.540086257 | 0.714414141 |
| ENSG00000254035 | ENSG0000025 | 1.802778363 | 0.62449302 | 1.293457205 | 0.482809185 | 0.629231228 | 0.7810577 |
| ENSG00000254039 | ENSG0000025 | 1.806513211 | 0.133445923 | 1.310991999 | 0.101790036 | 0.918923335 | 0.960456593 |
| ENSG00000254042 | SLIT3-AS2 | 5.084935787 | -1.869019209 | 0.871365022 | -2.144932562 | 0.031958239 | 0.082068959 |
| ENSG00000254065 | ENSG0000025 | 5.077291544 | 0.568686291 | 0.798500397 | 0.712192871 | 0.476345351 | 0.661422574 |
| ENSG00000254087 | LYN | 114.8527687 | 4.139295814 | 0.281455036 | 14.70677476 | 5.83E-49 | 3.13E-47 |
| ENSG00000254093 | PINX1 | 155.3708145 | -0.2094521 | 0.159422871 | -1.313814628 | 0.188908594 | 0.347574843 |
| ENSG00000254099 | ENSG0000025 | 1.809351557 | -0.987428665 | 1.352534318 | -0.730058123 | 0.465354658 | 0.651356659 |
| ENSG00000254101 | LINC02055 | 189.4404716 | 0.804112712 | 0.140074984 | 5.740587577 | 9.43E-09 | 6.81E-08 |
| ENSG00000254109 | RBPMS-AS1 | 26.00823786 | 0.095191174 | 0.342509271 | 0.277922911 | 0.781071536 | 0.881298289 |
| ENSG00000254112 | ENSG0000025 | 2.379123998 | -1.043927479 | 1.180280401 | -0.884474128 | 0.376440322 | 0.569311728 |
| ENSG00000254122 | PCDHGB7 | 17.26838765 | 1.131993699 | 0.441348592 | 2.564851727 | 0.01032199 | 0.030557034 |
| ENSG00000254153 | LINC02950 | 11.58110689 | 0.253841847 | 0.515427326 | 0.492488144 | 0.622374302 | 0.77657147 |
| ENSG00000254164 | ENSG0000025 | 2.748316696 | -2.999691341 | 1.347891943 | -2.225468708 | 0.02604979 | 0.068774562 |
| ENSG00000254165 | ENSG0000025 | 2.368653976 | 0.033961423 | 1.175580887 | 0.028889057 | 0.976953073 | 0.989320774 |
| ENSG00000254170 | ENSG0000025 | 2.315343166 | 1.080555251 | 1.204997671 | 0.896728082 | 0.369864032 | 0.562532712 |
| ENSG00000254180 | ENSG0000025 | 13.31352266 | 1.566133664 | 0.520557854 | 3.008567924 | 0.002624821 | 0.009001865 |
| ENSG00000254183 | DNAJC8P3 | 2.588021371 | -0.103568338 | 1.14475421 | -0.090472118 | 0.927912049 | 0.965331908 |
| ENSG00000254185 | MAPK6P5 | 9.786732321 | -0.81287162 | 0.582201644 | -1.396202894 | 0.162653405 | 0.309906073 |

| | | | | | | | |
|---|---|---|---|---|---|---|---|
| ENSG00000254186 | ENSG0000025 | 1.594690618 | -0.129312409 | 1.383274945 | -0.093482796 | 0.925520017 | NA |
| ENSG00000254192 | ENSG0000025 | 3.004928806 | -2.471449361 | 1.185916611 | -2.083999278 | 0.037160233 | 0.093276698 |
| ENSG00000254200 | RPL7AP33 | 1.58624443 | 1.83270152 | 1.556746969 | 1.177263587 | 0.239090316 | NA |
| ENSG00000254206 | NPIPB11 | 27.22882929 | 0.62480298 | 0.347227546 | 1.799404996 | 0.07195464 | 0.162400574 |
| ENSG00000254207 | ENSG0000025 | 6.293816298 | -0.18258243 | 0.684322517 | -0.266807573 | 0.789617323 | 0.886238142 |
| ENSG00000254211 | LINC01485 | 5.965105301 | 0.082468625 | 0.717586101 | 0.114925059 | 0.908504522 | 0.95425968 |
| ENSG00000254213 | PRXL2AP2 | 6.708457727 | -0.114237641 | 0.675621685 | -0.169085219 | 0.865729611 | 0.9302381 |
| ENSG00000254221 | PCDHGB1 | 120.6228362 | 0.329820364 | 0.166508124 | 1.980806433 | 0.047612985 | 0.114876092 |
| ENSG00000254224 | UQCRB-AS1 | 11.72738415 | -1.21755467 | 0.54367927 | -2.239472308 | 0.025125201 | 0.06676191 |
| ENSG00000254226 | LINC01933 | 3.583761287 | 0.36351179 | 0.91406004 | 0.397689182 | 0.690859313 | 0.825640142 |
| ENSG00000254230 | ENSG0000025 | 9.022173448 | -0.768137919 | 0.612760147 | -1.253570297 | 0.209998233 | 0.377122238 |
| ENSG00000254231 | WWP1-AS1 | 146.1841468 | 0.336740029 | 0.151264984 | 2.226159816 | 0.026003479 | 0.068691824 |
| ENSG00000254236 | ENSG0000025 | 1.55685474 | 1.127914975 | 1.490390404 | 0.756791625 | 0.449174711 | NA |
| ENSG00000254238 | ENSG0000025 | 4.927580281 | 1.227970943 | 0.834345466 | 1.471777571 | 0.141080952 | 0.278184377 |
| ENSG00000254244 | PAICSP4 | 169.9915884 | -0.060612379 | 0.141079326 | -0.429633322 | 0.667462394 | 0.808691146 |
| ENSG00000254245 | PCDHGA3 | 9.772419226 | -1.142615605 | 0.585798202 | -1.950527673 | 0.051113258 | 0.122101234 |
| ENSG00000254248 | ENSG0000025 | 4.360622698 | 0.263623731 | 0.857186416 | 0.30754539 | 0.75842828 | 0.866883146 |
| ENSG00000254251 | ENSG0000025 | 239.9972965 | 6.220678651 | 0.355194805 | 17.51342804 | 1.13E-68 | 9.91E-67 |
| ENSG00000254270 | ERHP1 | 28.70033374 | -0.881622365 | 0.350840801 | -2.512884372 | 0.011974858 | 0.034874093 |
| ENSG00000254271 | ANK3-DT | 5.896520219 | -1.94542554 | 0.816256077 | -2.383351982 | 0.017155782 | 0.047853683 |
| ENSG00000254272 | BTF3P3 | 6.577021435 | -0.031965554 | 0.677057696 | -0.047212451 | 0.962343904 | 0.98273545 |
| ENSG00000254275 | LINC00824 | 12.30319575 | -2.007932492 | 0.557674391 | -3.600546348 | 0.000317549 | 0.001301515 |
| ENSG00000254285 | KRT8P3 | 4183.823814 | 0.135513557 | 0.045924935 | 2.950762089 | 0.00316991 | 0.010684357 |
| ENSG00000254290 | ENSG0000025 | 9.732018991 | 1.08413044 | 0.608549149 | 1.781500217 | 0.074830769 | 0.167626494 |
| ENSG00000254294 | IMPDH1P6 | 8.179895923 | -0.173632636 | 0.630370504 | -0.27544537 | 0.782974092 | 0.882293632 |
| ENSG00000254305 | MRPL9P1 | 1.729435951 | 0.113453128 | 1.33326253 | 0.085094364 | 0.932186371 | NA |
| ENSG00000254307 | ENSG0000025 | 5.36998826 | -1.513348582 | 0.814823427 | -1.857271811 | 0.063272493 | 0.145803213 |
| ENSG00000254331 | CKS1BP7 | 53.31987804 | 0.049797946 | 0.245218689 | 0.203075655 | 0.8390759 | 0.915286602 |
| ENSG00000254332 | TPM4P3 | 445.5926734 | -0.507940968 | 0.093613897 | -5.42591415 | 5.77E-08 | 3.82E-07 |
| ENSG00000254337 | ENSG0000025 | 233.9093703 | 3.635509556 | 0.186403685 | 19.50342109 | 1.03E-84 | 1.27E-82 |
| ENSG00000254338 | MAFA-AS1 | 12.79576265 | -0.356717742 | 0.491474261 | -0.725811645 | 0.467954247 | 0.65388746 |
| ENSG00000254340 | ENSG0000025 | 1.699688098 | -2.170934484 | 1.554118695 | -1.396891042 | 0.162446339 | NA |
| ENSG00000254343 | ENSG0000025 | 22.36991634 | 1.280885164 | 0.389936531 | 3.284855515 | 0.001020347 | 0.003814989 |

| | | | | | | | |
|---|---|---|---|---|---|---|---|
| ENSG00000254349 | MIR2052HG | 14.51352036 | 2.696283793 | 0.574826249 | 4.69060659 | 2.72E-06 | 1.50E-05 |
| ENSG00000254363 | ENSG0000025 | 13.86185517 | 0.087055831 | 0.46338105 | 0.187870934 | 0.850977828 | 0.922214256 |
| ENSG00000254373 | ENSG0000025 | 78.30659062 | -0.163075201 | 0.212679839 | -0.766763796 | 0.443221966 | 0.632072106 |
| ENSG00000254387 | MYL12AP1 | 14.43960873 | 0.030276825 | 0.497122616 | 0.060904139 | 0.951435553 | 0.977495703 |
| ENSG00000254389 | RHPN1-AS1 | 249.1406073 | 0.20977836 | 0.1196452 | 1.753337035 | 0.079544172 | 0.175895827 |
| ENSG00000254400 | ENSG0000025 | 4.931224168 | -0.538232438 | 0.897607641 | -0.599629965 | 0.548752873 | 0.721494622 |
| ENSG00000254402 | LRRC24 | 174.9502459 | -0.135709245 | 0.137685393 | -0.985647371 | 0.324306184 | 0.514494043 |
| ENSG00000254406 | ENSG0000025 | 3.367691467 | 0.221711362 | 1.02681991 | 0.215920396 | 0.829049798 | 0.909536221 |
| ENSG00000254409 | ENSG0000025 | 4.071149878 | -0.506708056 | 0.862935791 | -0.587190915 | 0.557075493 | 0.727416829 |
| ENSG00000254413 | CHKB-CPT1B | 13.17943865 | -0.397259912 | 0.480562745 | -0.826655657 | 0.408432262 | 0.600797441 |
| ENSG00000254416 | LINC02732 | 5.739303634 | 5.869478583 | 1.355944097 | 4.328702484 | 1.50E-05 | 7.56E-05 |
| ENSG00000254418 | SPON1-AS1 | 1.945566359 | 1.397621102 | 1.31365176 | 1.063920549 | 0.287364693 | 0.471279314 |
| ENSG00000254419 | ENSG0000025 | 35.76308021 | 0.337088592 | 0.310459633 | 1.085772695 | 0.277579569 | 0.460354019 |
| ENSG00000254420 | ENSG0000025 | 2.26178718 | -0.567247742 | 1.172821931 | -0.483660586 | 0.62862677 | 0.780890133 |
| ENSG00000254422 | ENSG0000025 | 6.581797367 | -0.577735738 | 0.691583474 | -0.835381064 | 0.403503188 | 0.596376724 |
| ENSG00000254427 | LINC02696 | 3.394224533 | -0.500867957 | 0.937602557 | -0.534200716 | 0.593202674 | 0.755354595 |
| ENSG00000254429 | ENSG0000025 | 6.066888077 | 0.361884066 | 0.708725947 | 0.51061213 | 0.60962268 | 0.767759774 |
| ENSG00000254433 | ENSG0000025 | 5.473940977 | -0.628131308 | 0.746960242 | -0.840916655 | 0.400394627 | 0.593234693 |
| ENSG00000254449 | SF3A3P2 | 27.14451473 | -0.130962442 | 0.360317534 | -0.363463973 | 0.716258321 | 0.842224916 |
| ENSG00000254452 | ENSG0000025 | 54.17513733 | -0.667420151 | 0.248976742 | -2.680652603 | 0.007347875 | 0.022675574 |
| ENSG00000254454 | RCC2P6 | 25.34276889 | 0.000978598 | 0.36692389 | 0.002667033 | 0.997872018 | 0.998761274 |
| ENSG00000254455 | HIGD1AP10 | 1.647236364 | 0.403086531 | 1.40484624 | 0.286925729 | 0.77416919 | NA |
| ENSG00000254459 | ENSG0000025 | 3.14830091 | -0.334101147 | 0.970969075 | -0.34409041 | 0.730778295 | 0.850978918 |
| ENSG00000254461 | ENSG0000025 | 7.928193182 | -0.778965207 | 0.644805503 | -1.208062281 | 0.227023307 | 0.399571949 |
| ENSG00000254469 | XNDC1N | 51.24980645 | 0.054361117 | 0.252877187 | 0.214970429 | 0.829790374 | 0.909797661 |
| ENSG00000254470 | AP5B1 | 1684.413836 | 0.278925753 | 0.063419034 | 4.398139426 | 1.09E-05 | 5.61E-05 |
| ENSG00000254473 | UBQLN1-AS1 | 53.03980171 | -0.208607346 | 0.239552561 | -0.87082077 | 0.383852023 | 0.576280458 |
| ENSG00000254477 | ENSG0000025 | 3.312447074 | 0.369950215 | 0.960167174 | 0.385297712 | 0.700016857 | 0.831719009 |
| ENSG00000254479 | SLC25A1P1 | 2.428293059 | 0.779422405 | 1.156865897 | 0.673736175 | 0.500479058 | 0.681728496 |
| ENSG00000254480 | LINC02749 | 6.370008197 | -0.200006465 | 0.687360218 | -0.290977656 | 0.771068408 | 0.87520429 |
| ENSG00000254481 | PTP4A2P2 | 18.34766209 | -0.774330958 | 0.427286851 | -1.812204041 | 0.069954675 | 0.158479187 |
| ENSG00000254485 | ENSG0000025 | 7.830910898 | 0.247654856 | 0.653865237 | 0.378755195 | 0.704869662 | 0.834996645 |
| ENSG00000254501 | ENSG0000025 | 10.66959786 | 0.612455698 | 0.562896516 | 1.088043148 | 0.27657606 | 0.459125466 |

| | | | | | | | |
|---|---|---|---|---|---|---|---|
| ENSG00000254502 | FNTAP1 | 3.211139878 | -0.187229046 | 0.997143291 | -0.187765438 | 0.851060531 | 0.9222591 |
| ENSG00000254505 | CHMP4A | 26.30263165 | -0.623923344 | 0.359675464 | -1.734684198 | 0.082796757 | 0.181678071 |
| ENSG00000254506 | ENSG0000025 | 3.893993346 | 1.384305147 | 0.938784877 | 1.474571204 | 0.140327861 | 0.276985127 |
| ENSG00000254528 | FXYD6-AS1 | 4.084029063 | -0.720593029 | 0.889133435 | -0.810444193 | 0.417684927 | 0.609793757 |
| ENSG00000254531 | FLJ20021 | 340.6251417 | -0.058880728 | 0.106518102 | -0.552776735 | 0.580416301 | 0.745630889 |
| ENSG00000254535 | PABPC4L | 34.38327434 | -1.827008503 | 0.339361783 | -5.38366013 | 7.30E-08 | 4.78E-07 |
| ENSG00000254536 | ENSG0000025 | 27.08691394 | -0.773687118 | 0.354512607 | -2.182396625 | 0.029080273 | 0.07554943 |
| ENSG00000254543 | HSPD1P2 | 6.630068418 | 0.09707532 | 0.726040947 | 0.133705021 | 0.893635834 | 0.945700277 |
| ENSG00000254548 | ENSG0000025 | 41.14275488 | 0.506248126 | 0.278566221 | 1.817334939 | 0.069165843 | 0.157095489 |
| ENSG00000254559 | ENSG0000025 | 5.720236955 | -0.577115433 | 0.736297379 | -0.783807535 | 0.433153052 | 0.623170067 |
| ENSG00000254571 | ENSG0000025 | 2.808479298 | -0.960030491 | 1.091353379 | -0.879669692 | 0.379038273 | 0.571559717 |
| ENSG00000254572 | GLULP3 | 3.086023768 | 0.085274303 | 1.011376021 | 0.084315132 | 0.932805882 | 0.96773548 |
| ENSG00000254574 | ENSG0000025 | 1.926198278 | -1.017606687 | 1.303509629 | -0.780666798 | 0.434998493 | 0.624968087 |
| ENSG00000254577 | ENSG0000025 | 1.771515679 | 0.101620474 | 1.295291607 | 0.078453742 | 0.937467125 | NA |
| ENSG00000254578 | ENSG0000025 | 6.70177112 | 0.205247956 | 0.698323727 | 0.293915196 | 0.768822706 | 0.873855116 |
| ENSG00000254589 | EIF4A2P3 | 7.668620007 | -0.833675085 | 0.646865253 | -1.288792496 | 0.197470235 | 0.359549742 |
| ENSG00000254595 | ENSG0000025 | 3.266515406 | 1.931355414 | 1.080077015 | 1.788164536 | 0.073749466 | 0.165701543 |
| ENSG00000254598 | CSNK2A3 | 816.9242333 | 0.119343326 | 0.072515935 | 1.645753122 | 0.099814597 | 0.21166083 |
| ENSG00000254604 | ENSG0000025 | 2.989528979 | -1.902268886 | 1.141273348 | -1.666795154 | 0.095555144 | 0.204354679 |
| ENSG00000254612 | DNAJB6P1 | 939.964289 | 0.532959278 | 0.071059684 | 7.500163962 | 6.37E-14 | 7.06E-13 |
| ENSG00000254614 | CAPN1-AS1 | 90.41706134 | 0.05583995 | 0.188823789 | 0.295725187 | 0.76743996 | 0.873024431 |
| ENSG00000254618 | TMED10P1 | 10.31721802 | -0.399152947 | 0.564282596 | -0.707363561 | 0.479340575 | 0.663653377 |
| ENSG00000254632 | ENSG0000025 | 3.796624964 | -0.24385756 | 0.91007317 | -0.267953795 | 0.788734885 | 0.885767771 |
| ENSG00000254634 | SMG1P6 | 5.371238619 | 0.368004458 | 0.791494166 | 0.464949046 | 0.641967946 | 0.790019027 |
| ENSG00000254635 | WAC-AS1 | 1020.236684 | -0.55229909 | 0.068066063 | -8.114162411 | 4.89E-16 | 6.34E-15 |
| ENSG00000254648 | ENSG0000025 | 2.764960473 | -0.262486054 | 1.103094315 | -0.237954316 | 0.811916527 | 0.900445907 |
| ENSG00000254662 | ENSG0000025 | 58.38343973 | -0.400359835 | 0.236463387 | -1.69311554 | 0.090433477 | 0.195196082 |
| ENSG00000254676 | ENSG0000025 | 9.359668767 | -0.35366373 | 0.579418396 | -0.610377117 | 0.541612023 | 0.715650991 |
| ENSG00000254677 | OSBPL9P2 | 2.491161363 | -0.754912554 | 1.166969159 | -0.646900175 | 0.517696552 | 0.695721084 |
| ENSG00000254681 | PKD1P5 | 29.10148015 | 0.131867495 | 0.326133758 | 0.404335618 | 0.685965937 | 0.822456415 |
| ENSG00000254682 | ENSG0000025 | 291.0851236 | -0.086336348 | 0.110833045 | -0.778976597 | 0.435993501 | 0.625888042 |
| ENSG00000254685 | FPGT | 135.3119601 | 0.299677144 | 0.168239483 | 1.781253359 | 0.07487107 | 0.16767962 |
| ENSG00000254689 | LINC02235 | 4.684347844 | -0.069583237 | 0.810498986 | -0.085852343 | 0.931583796 | 0.967211237 |

| | | | | | | | |
|---|---|---|---|---|---|---|---|
| ENSG00000254694 | ENSG0000025 | 4.143758396 | -0.664315798 | 0.931885087 | -0.712873086 | 0.475924295 | 0.661056589 |
| ENSG00000254701 | ENSG0000025 | 5.669564204 | -0.724575583 | 0.75033247 | -0.965672701 | 0.334207984 | 0.525154541 |
| ENSG00000254702 | ENSG0000025 | 22.45096451 | 0.151533348 | 0.374987045 | 0.404102889 | 0.686137061 | 0.822589932 |
| ENSG00000254708 | MMADHCP2 | 6.01825335 | -0.007331622 | 0.776022102 | -0.009447697 | 0.99246194 | 0.995949353 |
| ENSG00000254718 | ENSG0000025 | 14.68352212 | -0.099607881 | 0.453228949 | -0.219773871 | 0.82604727 | 0.908261176 |
| ENSG00000254719 | ENSG0000025 | 94.061854 | 0.295580417 | 0.183189263 | 1.613524791 | 0.106630542 | 0.223196793 |
| ENSG00000254721 | ENSG0000025 | 34.05876909 | 0.050488801 | 0.313097796 | 0.161255689 | 0.871892021 | 0.933588638 |
| ENSG00000254726 | MEX3A | 1029.463198 | 0.529185316 | 0.066624235 | 7.942835187 | 1.98E-15 | 2.44E-14 |
| ENSG00000254739 | ENSG0000025 | 29.63261728 | 0.558261239 | 0.331109531 | 1.686031919 | 0.091789652 | 0.197621268 |
| ENSG00000254759 | NAP1L1P1 | 64.41942863 | 0.339427096 | 0.235294815 | 1.442560885 | 0.149144209 | 0.290315498 |
| ENSG00000254760 | SIGLEC10-AS2 | 1.686444324 | -0.172489906 | 1.381534262 | -0.124853875 | 0.900639235 | NA |
| ENSG00000254762 | KLC2-AS2 | 2.122862589 | -3.559993158 | 1.559221448 | -2.28318637 | 0.022419393 | 0.060413587 |
| ENSG00000254780 | ENSG0000025 | 8.978796573 | 0.188075257 | 0.578012215 | 0.325382842 | 0.74489134 | 0.858985941 |
| ENSG00000254783 | ENSG0000025 | 16.26554112 | -1.228886126 | 0.470321453 | -2.612864282 | 0.008978695 | 0.027058757 |
| ENSG00000254786 | ENSG0000025 | 3.242406241 | -1.305654504 | 1.000668268 | -1.30478256 | 0.191966899 | 0.351987904 |
| ENSG00000254812 | ENSG0000025 | 14.28754405 | -0.37202364 | 0.477738203 | -0.778718632 | 0.436145478 | 0.626067034 |
| ENSG00000254815 | LMNTD2-AS1 | 264.8320501 | 0.053541758 | 0.115862961 | 0.462112804 | 0.644000429 | 0.791331611 |
| ENSG00000254827 | SLC22A18AS | 18.56334394 | -0.035965507 | 0.409085326 | -0.087916885 | 0.929942736 | 0.966032078 |
| ENSG00000254831 | KRT18P58 | 6.686684893 | 0.176499965 | 0.667955832 | 0.264238977 | 0.791595779 | 0.887763734 |
| ENSG00000254835 | RNF185-AS1 | 5.032837502 | -1.626560814 | 0.860705934 | -1.889798537 | 0.058784909 | 0.137170768 |
| ENSG00000254837 | LIPT2-AS1 | 140.7147642 | 0.268424043 | 0.151598983 | 1.770619026 | 0.076624076 | 0.170871615 |
| ENSG00000254846 | ENSG0000025 | 3.039105569 | -4.107098679 | 1.44646644 | -2.839401291 | 0.004519828 | 0.014723778 |
| ENSG00000254851 | ENSG0000025 | 11.68176971 | 0.02041018 | 0.513260082 | 0.039765765 | 0.96827987 | 0.985876114 |
| ENSG00000254854 | NECTIN1-DT | 14.65279745 | -0.319980048 | 0.463962161 | -0.689668413 | 0.490402734 | 0.673562371 |
| ENSG00000254855 | KLC2-AS1 | 13.76146009 | -0.972367856 | 0.48497009 | -2.005005825 | 0.044962424 | 0.109366771 |
| ENSG00000254858 | MPV17L2 | 344.2840944 | -0.164385698 | 0.106264129 | -1.546953806 | 0.121874385 | 0.24847124 |
| ENSG00000254860 | TMEM9B-AS1 | 35.9204879 | 0.159865277 | 0.296742717 | 0.538733615 | 0.590070675 | 0.752932545 |
| ENSG00000254864 | ENSG0000025 | 10.31699616 | 0.165020009 | 0.579589122 | 0.284718955 | 0.775859468 | 0.878273683 |
| ENSG00000254872 | LINC02688 | 11.25348187 | 0.609978264 | 0.530805778 | 1.149155283 | 0.250491956 | 0.428699304 |
| ENSG00000254873 | ENSG0000025 | 29.59108169 | -0.858919776 | 0.326925046 | -2.627268195 | 0.008607346 | 0.026073224 |
| ENSG00000254876 | SUGT1P4-STR | 14.23945254 | -0.863999301 | 0.483629579 | -1.786489781 | 0.07401999 | 0.166201287 |
| ENSG00000254877 | ENSG0000025 | 2.622959823 | -1.301836521 | 1.11723501 | -1.165230689 | 0.243925626 | 0.420662597 |
| ENSG00000254893 | RAP1BL | 925.7543718 | 0.080358162 | 0.068107717 | 1.179868665 | 0.238052454 | 0.413525228 |

| | | | | | | | |
|---|---|---|---|---|---|---|---|
| ENSG00000254898 | ENSG0000025 | 5.166191677 | -0.626732262 | 0.796650129 | -0.786709547 | 0.431451914 | 0.621507498 |
| ENSG00000254901 | BORCS8 | 106.1879411 | 0.006648138 | 0.175403529 | 0.037901965 | 0.969765846 | 0.986494103 |
| ENSG00000254905 | ENSG0000025 | 2.111902796 | -0.398598719 | 1.245039125 | -0.320149553 | 0.748854963 | 0.861089366 |
| ENSG00000254907 | ENSG0000025 | 12.94231571 | -0.619281109 | 0.491963315 | -1.258795298 | 0.208104279 | 0.374550959 |
| ENSG00000254910 | ENSG0000025 | 8.782147879 | -0.991398836 | 0.604617218 | -1.639713206 | 0.101064812 | 0.213699584 |
| ENSG00000254963 | ENSG0000025 | 2.499215167 | 0.044709981 | 1.192498752 | 0.037492686 | 0.970092172 | 0.986563569 |
| ENSG00000254964 | ENSG0000025 | 5.290994187 | -0.501172489 | 0.812760371 | -0.616630076 | 0.537478752 | 0.71253829 |
| ENSG00000254966 | IGSF22-AS1 | 44.86938273 | 0.145294765 | 0.271446546 | 0.535261055 | 0.592469352 | 0.754736754 |
| ENSG00000254973 | ENSG0000025 | 65.9833437 | 0.379619086 | 0.221081944 | 1.717096748 | 0.085961496 | 0.187323713 |
| ENSG00000254974 | ENSG0000025 | 2.357761827 | 0.048047458 | 1.164103779 | 0.041274205 | 0.967077297 | 0.985295635 |
| ENSG00000254978 | ALG1L9P | 2.721817378 | 0.012278067 | 1.043358897 | 0.011767827 | 0.990610849 | 0.994991588 |
| ENSG00000254985 | RSF1-IT2 | 1.659990367 | -0.178741223 | 1.338528944 | -0.133535568 | 0.893769837 | NA |
| ENSG00000254986 | DPP3 | 3506.319191 | 0.072826548 | 0.058119198 | 1.253054937 | 0.210185714 | 0.377429376 |
| ENSG00000254990 | ENSG0000025 | 1.77347492 | 1.266404298 | 1.448260657 | 0.874431196 | 0.381883475 | NA |
| ENSG00000254995 | STX16-NPEPL1 | 105.246433 | 0.640361219 | 0.180383195 | 3.550004865 | 0.000385224 | 0.001552537 |
| ENSG00000254996 | ANKHD1-EIF4I | 13.14739567 | -0.867819167 | 0.484189049 | -1.792314735 | 0.073082564 | 0.164428602 |
| ENSG00000254997 | KRTAP5-9 | 14.3347257 | 1.004311192 | 0.476852064 | 2.106127388 | 0.035193288 | 0.089050743 |
| ENSG00000254999 | BRK1 | 6064.486551 | -0.016364471 | 0.048223674 | -0.339345177 | 0.734349713 | 0.853522948 |
| ENSG00000255002 | LINC02324 | 3.590782569 | -0.394981968 | 0.984387911 | -0.40124626 | 0.688238825 | 0.823819752 |
| ENSG00000255007 | LINC02489 | 2.001315669 | -1.155429439 | 1.273521739 | -0.907271076 | 0.364263465 | 0.556894628 |
| ENSG00000255014 | ARL6IP1P3 | 7.254884965 | -0.197381938 | 0.697142051 | -0.283130156 | 0.777077066 | 0.87898243 |
| ENSG00000255020 | TDH-AS1 | 23.81516918 | 0.265893133 | 0.359030057 | 0.740587391 | 0.458943656 | 0.646137686 |
| ENSG00000255031 | ENSG0000025 | 57.15963276 | 0.444478368 | 0.244000684 | 1.821627553 | 0.068511517 | 0.155887146 |
| ENSG00000255035 | SDHCP4 | 79.77751912 | 0.095116779 | 0.20429807 | 0.465578449 | 0.641517272 | 0.789631802 |
| ENSG00000255038 | ENSG0000025 | 12.62620105 | -1.184832301 | 0.525487947 | -2.254727836 | 0.024150417 | 0.064539908 |
| ENSG00000255040 | MORF4L1P3 | 13.41212838 | 0.075599911 | 0.507584553 | 0.148940527 | 0.881600562 | 0.938595484 |
| ENSG00000255042 | SEPTIN7P11 | 2.621195356 | 0.189243222 | 1.11995399 | 0.168974104 | 0.865817011 | 0.930253488 |
| ENSG00000255045 | ENSG0000025 | 2.534586888 | 0.19583038 | 1.097776743 | 0.178388166 | 0.858418142 | 0.92649707 |
| ENSG00000255046 | ENSG0000025 | 15.81689245 | -0.033865167 | 0.448273347 | -0.075545797 | 0.939780461 | 0.971233034 |
| ENSG00000255050 | ENSG0000025 | 10.15079196 | -0.08345461 | 0.539831119 | -0.154593923 | 0.877141461 | 0.936150792 |
| ENSG00000255052 | FAM66D | 35.2363709 | 0.500003022 | 0.336001909 | 1.488095776 | 0.136725637 | 0.271651318 |
| ENSG00000255059 | PPIAP43 | 7.805083035 | 0.148353167 | 0.650763217 | 0.227967967 | 0.819671144 | 0.904433786 |
| ENSG00000255062 | TIRAP-AS1 | 8.225666679 | 0.31135822 | 0.642421314 | 0.484663589 | 0.627914999 | 0.780547313 |

| | | | | | | | |
|---|---|---|---|---|---|---|---|
| ENSG00000255063 | RBMXP3 | 6.846693785 | -0.684269456 | 0.726385682 | -0.942019471 | 0.346182671 | 0.537054878 |
| ENSG00000255070 | OSBPL9P3 | 2.919514211 | 0.470738348 | 1.05259179 | 0.447218335 | 0.654717424 | 0.799489924 |
| ENSG00000255100 | TSKU-AS1 | 6.994006553 | 0.457706375 | 0.67220667 | 0.680901269 | 0.495933965 | 0.677952754 |
| ENSG00000255107 | ENSG0000025 | 1.689719617 | -2.1781547 | 1.538800562 | -1.41548863 | 0.15692528 | NA |
| ENSG00000255108 | ENSG0000025 | 10.26709515 | 0.581271872 | 0.559701261 | 1.038539509 | 0.299018951 | 0.485696718 |
| ENSG00000255112 | CHMP1B | 2060.62999 | 0.443885002 | 0.054073677 | 8.208892576 | 2.23E-16 | 2.96E-15 |
| ENSG00000255114 | ENSG0000025 | 17.71246024 | 0.058246687 | 0.433830144 | 0.134261502 | 0.893195794 | 0.945411873 |
| ENSG00000255121 | CENATAC-DT | 176.1308008 | -0.240967179 | 0.137295131 | -1.755103602 | 0.079241588 | 0.175478018 |
| ENSG00000255128 | HSPD1P3 | 3.682585419 | 0.457148406 | 0.930422455 | 0.491334236 | 0.623190069 | 0.777166632 |
| ENSG00000255129 | TTC12-DT | 3.444735305 | -0.616826764 | 0.974884638 | -0.632717698 | 0.52691801 | 0.703006814 |
| ENSG00000255135 | EMSY-DT | 201.3276281 | 0.101470951 | 0.127934917 | 0.793145087 | 0.427693297 | 0.618384232 |
| ENSG00000255136 | TPBGL-AS1 | 1.945193679 | 0.464570662 | 1.39361806 | 0.333355799 | 0.738865724 | 0.856040341 |
| ENSG00000255138 | GLTPP1 | 19.8505046 | 0.135718511 | 0.401779613 | 0.337793422 | 0.735518867 | 0.854146138 |
| ENSG00000255139 | ENSG0000025 | 17.98217811 | 0.000410563 | 0.410696329 | 0.000999675 | 0.999202375 | 0.999528515 |
| ENSG00000255142 | ENSG0000025 | 11.3390841 | 0.056137319 | 0.518108575 | 0.108350493 | 0.913717671 | 0.95710883 |
| ENSG00000255145 | STX17-DT | 10.13902817 | 0.83959787 | 0.566936748 | 1.480937462 | 0.138623239 | 0.274591882 |
| ENSG00000255150 | EID3 | 2.020171373 | 0.566712036 | 1.262266749 | 0.448963768 | 0.653457794 | 0.798929904 |
| ENSG00000255152 | MSH5-SAPCD1 | 7.281512042 | 0.104434706 | 0.678787468 | 0.153854794 | 0.877724229 | 0.936395015 |
| ENSG00000255153 | TOLLIP-DT | 47.18637786 | -0.129754239 | 0.260189972 | -0.49869039 | 0.617997516 | 0.77359751 |
| ENSG00000255154 | HTD2 | 2.640609427 | -1.211797595 | 1.123322779 | -1.078761705 | 0.280693971 | 0.464119665 |
| ENSG00000255158 | ENSG0000025 | 1.889350033 | -0.002057426 | 1.298540199 | -0.001584415 | 0.99873582 | 0.999329557 |
| ENSG00000255164 | ENSG0000025 | 2.423000987 | -0.704354474 | 1.179734447 | -0.597044933 | 0.550477382 | 0.722629536 |
| ENSG00000255173 | ENSG0000025 | 13.59796576 | -0.400936314 | 0.471367329 | -0.850581468 | 0.395001887 | 0.588013388 |
| ENSG00000255176 | ENSG0000025 | 5.299200338 | -0.328211075 | 0.773491164 | -0.424324272 | 0.67132934 | 0.811609753 |
| ENSG00000255182 | ENSG0000025 | 297.0684303 | 0.451846011 | 0.121929003 | 3.705812389 | 0.000210714 | 0.000890036 |
| ENSG00000255197 | SLC39A13-AS1 | 2.234417645 | 1.141646591 | 1.19358702 | 0.956483752 | 0.338827889 | 0.530045518 |
| ENSG00000255198 | SNHG9 | 229.2250829 | 0.28690915 | 0.127428036 | 2.251538662 | 0.024351441 | 0.065028856 |
| ENSG00000255200 | PGAM1P8 | 47.98796571 | 0.481882568 | 0.270166686 | 1.783649109 | 0.0744807 | 0.167001004 |
| ENSG00000255210 | ENSG0000025 | 2.314784716 | -0.517041416 | 1.33423674 | -0.387518497 | 0.698372394 | 0.830810911 |
| ENSG00000255213 | NPM1P35 | 1.988539384 | -1.769599379 | 1.368426436 | -1.29316369 | 0.195954451 | 0.357413939 |
| ENSG00000255220 | DDX18P5 | 9.762009662 | -0.187176078 | 0.588655652 | -0.317972107 | 0.750506095 | 0.862210027 |
| ENSG00000255222 | SETP17 | 1.859025535 | -1.011494621 | 1.284502557 | -0.787460185 | 0.431012527 | 0.621162456 |
| ENSG00000255224 | ENSG0000025 | 41.81024115 | 0.076630942 | 0.27601877 | 0.27762946 | 0.781296815 | 0.881384261 |

| | | | | | | | |
|---|---|---|---|---|---|---|---|
| ENSG00000255234 | CCDC90B-AS1 | 3.249619571 | 0.877291086 | 1.054501594 | 0.831948563 | 0.405437979 | 0.59815726 |
| ENSG00000255236 | CHKA-DT | 21.18741883 | 0.16883607 | 0.38197532 | 0.442007798 | 0.658483563 | 0.802337756 |
| ENSG00000255240 | ENSG0000025 | 46.42528014 | -1.068827613 | 0.268049794 | -3.987421874 | 6.68E-05 | 0.00030664 |
| ENSG00000255252 | ENSG0000025 | 7.218215354 | -0.286566996 | 0.650045707 | -0.440841302 | 0.65932789 | 0.802897431 |
| ENSG00000255262 | ELOBP2 | 2.549725939 | -0.129828813 | 1.146451798 | -0.113244022 | 0.909837095 | 0.9547442 |
| ENSG00000255276 | RRM1-AS1 | 10.9026477 | -1.135497434 | 0.543103473 | -2.090757083 | 0.036549845 | 0.092016401 |
| ENSG00000255277 | ABCC6P2 | 1.977157463 | -0.643536307 | 1.29064517 | -0.498615981 | 0.618049945 | 0.77359751 |
| ENSG00000255282 | WTAPP1 | 7.425319144 | -0.685893422 | 0.67448439 | -1.01691519 | 0.309193777 | 0.497224342 |
| ENSG00000255284 | GATD1-DT | 93.05400885 | -0.167871922 | 0.187772831 | -0.894016034 | 0.371313312 | 0.563955434 |
| ENSG00000255291 | HMGB1P40 | 2.209403414 | -0.415242172 | 1.185227777 | -0.350347993 | 0.726077552 | 0.847853251 |
| ENSG00000255298 | OR8G5 | 3.613607953 | -1.232789311 | 0.960468344 | -1.283529352 | 0.199306683 | 0.36217485 |
| ENSG00000255302 | EID1 | 2877.759608 | -0.457914301 | 0.058555841 | -7.820130232 | 5.28E-15 | 6.35E-14 |
| ENSG00000255306 | ENSG0000025 | 11.06350806 | -0.863897841 | 0.548522782 | -1.574953437 | 0.115267193 | 0.237769032 |
| ENSG00000255308 | CSRP3-AS1 | 5.301019881 | 0.173321782 | 0.786208935 | 0.220452572 | 0.825518706 | 0.907876653 |
| ENSG00000255320 | ENSG0000025 | 10.58362316 | -0.179102993 | 0.544785632 | -0.328758658 | 0.742338114 | 0.85777393 |
| ENSG00000255326 | ENSG0000025 | 2.745976212 | -0.926893904 | 1.093122606 | -0.847932244 | 0.396475706 | 0.589369667 |
| ENSG00000255337 | TMEM123-DT | 4.805516896 | 0.121208664 | 0.809562667 | 0.149721163 | 0.88098461 | 0.93828808 |
| ENSG00000255340 | ENSG0000025 | 3.472138587 | -0.587413815 | 0.927098052 | -0.633604842 | 0.52633874 | 0.702315652 |
| ENSG00000255341 | OR8Q1P | 1.915728256 | 0.315830352 | 1.259486753 | 0.250761154 | 0.801998777 | 0.894633256 |
| ENSG00000255345 | ENSG0000025 | 6.470143498 | -0.397242249 | 0.678964062 | -0.585071098 | 0.558499911 | 0.728533377 |
| ENSG00000255346 | NOX5 | 206.5311717 | -0.36331032 | 0.135256583 | -2.686082337 | 0.007229527 | 0.022357152 |
| ENSG00000255351 | ENSG0000025 | 17.59683346 | -0.992465664 | 0.431498455 | -2.300044536 | 0.021445697 | 0.058083151 |
| ENSG00000255353 | ASS1P13 | 19.96558638 | 0.899019324 | 0.402515946 | 2.233499897 | 0.025515997 | 0.067590875 |
| ENSG00000255361 | ENSG0000025 | 2.773450596 | 0.705021828 | 1.157926637 | 0.608865714 | 0.542613452 | 0.716364675 |
| ENSG00000255363 | LINC02757 | 2.374401517 | -0.607119153 | 1.158013048 | -0.524276608 | 0.600086164 | 0.760131232 |
| ENSG00000255367 | ENSG0000025 | 3.912321445 | -0.161619482 | 0.871554992 | -0.185438077 | 0.852885445 | 0.923198544 |
| ENSG00000255387 | RDXP1 | 2.632566765 | -1.684698744 | 1.157745729 | -1.45515436 | 0.145626533 | 0.28494308 |
| ENSG00000255389 | ENSG0000025 | 22.05054084 | -0.460666538 | 0.384341591 | -1.19858623 | 0.230688875 | 0.404102018 |
| ENSG00000255397 | NASPP1 | 77.69999589 | -0.162550772 | 0.213903054 | -0.759927309 | 0.447298037 | 0.635757692 |
| ENSG00000255398 | HCAR3 | 41.22447343 | -0.397088441 | 0.288653961 | -1.375655611 | 0.168928281 | 0.31918415 |
| ENSG00000255404 | ENSG0000025 | 6.053157278 | 1.140113376 | 0.752504981 | 1.515090803 | 0.12974941 | 0.260886272 |
| ENSG00000255418 | LINC02718 | 14.17688111 | 0.406997709 | 0.485663945 | 0.838023315 | 0.402017609 | 0.594679131 |
| ENSG00000255422 | ENSG0000025 | 4.141609381 | -0.752684566 | 0.859105979 | -0.876125396 | 0.380961868 | 0.57330213 |

| | | | | | | | |
|---|---|---|---|---|---|---|---|
| ENSG00000255423 | EBLN2 | 42.27365378 | -0.308420563 | 0.322258464 | -0.957059621 | 0.338537163 | 0.52977628 |
| ENSG00000255425 | OR8G3P | 2.151611668 | -1.33789285 | 1.226053388 | -1.09121908 | 0.275176494 | 0.457355635 |
| ENSG00000255435 | TTC36-AS1 | 75.04157009 | 0.624549755 | 0.211355299 | 2.954975624 | 0.003126938 | 0.010556578 |
| ENSG00000255438 | ENSG0000025 | 99.38767235 | -0.042780258 | 0.196475316 | -0.217738587 | 0.827632797 | 0.908894459 |
| ENSG00000255441 | SIGLEC10-AS1 | 3.897759039 | -0.344220913 | 0.881636899 | -0.390433878 | 0.696215741 | 0.829343624 |
| ENSG00000255443 | CD44-AS1 | 3.683524232 | -0.192854731 | 0.93107228 | -0.207131858 | 0.835906882 | 0.913370821 |
| ENSG00000255449 | ENSG0000025 | 2.571597112 | -0.022819686 | 1.107423985 | -0.020606097 | 0.983559877 | 0.9923499 |
| ENSG00000255455 | ENSG0000025 | 54.61754804 | 0.587301545 | 0.251807135 | 2.332346718 | 0.019682459 | 0.053944516 |
| ENSG00000255458 | SMIM14-DT | 13.84561933 | -0.010122053 | 0.474052582 | -0.021352173 | 0.982964726 | 0.992139551 |
| ENSG00000255467 | ENSG0000025 | 4.129303792 | -0.263451405 | 0.891073896 | -0.295656068 | 0.76749275 | 0.873024431 |
| ENSG00000255468 | B4GAT1-DT | 81.84719728 | 0.208347606 | 0.200820563 | 1.037481433 | 0.299511544 | 0.48618701 |
| ENSG00000255471 | PRSS23-AS1 | 54.2024089 | 0.323988189 | 0.244647353 | 1.324306944 | 0.185401125 | 0.342494912 |
| ENSG00000255487 | ENSG0000025 | 1.859895751 | 0.376386828 | 1.347069582 | 0.279411571 | 0.779928992 | 0.880534036 |
| ENSG00000255491 | ENSG0000025 | 9.323596679 | 1.32178427 | 0.61211484 | 2.15937302 | 0.030821239 | 0.079505142 |
| ENSG00000255495 | ENSG0000025 | 2.617818672 | 1.558501061 | 1.184134217 | 1.316152374 | 0.188122905 | 0.346434891 |
| ENSG00000255498 | ENSG0000025 | 9.221433761 | 0.324522901 | 0.571162607 | 0.568179528 | 0.569913071 | 0.737133843 |
| ENSG00000255500 | ENSG0000025 | 16.08809452 | -1.510957061 | 0.470140297 | -3.213842909 | 0.001309713 | 0.004797193 |
| ENSG00000255503 | LINC02951 | 3.587888519 | -1.246801186 | 0.951045402 | -1.310979669 | 0.189864633 | 0.348997971 |
| ENSG00000255507 | UVRAG-DT | 3.561517758 | 0.08162109 | 0.913842928 | 0.089316323 | 0.928830522 | 0.965838909 |
| ENSG00000255508 | ENSG0000025 | 6.269646324 | 0.054438882 | 0.710462942 | 0.076624521 | 0.938922252 | 0.971022554 |
| ENSG00000255517 | DPP3-DT | 154.4201214 | 0.267715702 | 0.148628311 | 1.801242987 | 0.07166459 | 0.161825543 |
| ENSG00000255521 | CD44-DT | 1.857228273 | -1.533555829 | 1.43035974 | -1.072146947 | 0.283654035 | 0.467525762 |
| ENSG00000255524 | NPIPB8 | 1.587145817 | 0.378432341 | 1.414968914 | 0.267449226 | 0.789123302 | NA |
| ENSG00000255529 | POLR2M | 411.2081723 | 0.00921969 | 0.097995226 | 0.094083055 | 0.925043181 | 0.964249985 |
| ENSG00000255539 | ENSG0000025 | 5.180313758 | 0.032788547 | 0.765291433 | 0.042844523 | 0.965825472 | 0.984625204 |
| ENSG00000255541 | CKS1BP4 | 2.239140126 | 0.663725904 | 1.15937113 | 0.572487866 | 0.566991504 | 0.734805102 |
| ENSG00000255547 | RPA2P3 | 3.737892897 | 0.262524209 | 1.008426136 | 0.260330628 | 0.794608749 | 0.89001146 |
| ENSG00000255559 | ZNF252P-AS1 | 12.08687225 | 0.685358464 | 0.521642516 | 1.313847017 | 0.188897692 | 0.347574843 |
| ENSG00000255561 | FDXACB1 | 23.16911115 | -0.320015514 | 0.381653382 | -0.838497782 | 0.401751193 | 0.594410925 |
| ENSG00000255568 | BRWD1-AS2 | 51.076876 | 0.342820058 | 0.250589986 | 1.368051708 | 0.171295891 | 0.322647816 |
| ENSG00000255571 | MIR9-3HG | 427.9655693 | 0.103678977 | 0.093138879 | 1.113165397 | 0.265637401 | 0.446724037 |
| ENSG00000255581 | ENSG0000025 | 1.961629079 | 0.848462051 | 1.321977243 | 0.641812902 | 0.520994672 | 0.698149294 |
| ENSG00000255585 | ENSG0000025 | 12.15297008 | -0.267881534 | 0.529468489 | -0.505944243 | 0.612895792 | 0.769891065 |

| | | | | | | | |
|---|---|---|---|---|---|---|---|
| ENSG00000255587 | RAB44 | 5.928848627 | 2.235390014 | 0.833490817 | 2.68196118 | 0.007319195 | 0.022603977 |
| ENSG00000255606 | LINC02952 | 3.040488164 | -1.553873469 | 1.047394582 | -1.483560729 | 0.137925494 | 0.273585027 |
| ENSG00000255621 | ENSG0000025 | 4.303008958 | -0.377532287 | 0.866255037 | -0.435821174 | 0.66296648 | 0.805368498 |
| ENSG00000255622 | PCDHB17P | 2.683537091 | -0.919067189 | 1.070099702 | -0.858861269 | 0.39041706 | 0.583343425 |
| ENSG00000255624 | C10orf88B | 70.58972741 | -0.308265713 | 0.213041296 | -1.446976335 | 0.147903552 | 0.28837223 |
| ENSG00000255642 | PABPC1P4 | 281.6382779 | 0.142121548 | 0.133648795 | 1.063395656 | 0.287602562 | 0.471560487 |
| ENSG00000255647 | ENSG0000025 | 4.54535866 | -0.581229459 | 0.83420276 | -0.696748424 | 0.485960248 | 0.669580266 |
| ENSG00000255652 | LINC02963 | 11.01876326 | -0.573714791 | 0.530613666 | -1.081228825 | 0.279595339 | 0.462933542 |
| ENSG00000255655 | ENSG0000025 | 2.153635502 | -1.217401094 | 1.275411091 | -0.954516628 | 0.339822196 | 0.5308131 |
| ENSG00000255680 | ENSG0000025 | 2.467941342 | -0.732037094 | 1.126019274 | -0.650110625 | 0.515620767 | 0.6944375 |
| ENSG00000255690 | TRIL | 1.639227448 | 2.888369599 | 1.643961335 | 1.756957136 | 0.078925116 | NA |
| ENSG00000255693 | LINC02389 | 4.378213223 | -0.380483279 | 0.869808332 | -0.437433472 | 0.661797014 | 0.804622827 |
| ENSG00000255703 | ENSG0000025 | 2.043455986 | -0.238815257 | 1.239373053 | -0.192690374 | 0.847201463 | 0.920028676 |
| ENSG00000255710 | ENSG0000025 | 2.886476548 | 0.62777551 | 1.093710122 | 0.573987109 | 0.565976525 | 0.73394626 |
| ENSG00000255717 | SNHG1 | 2172.603268 | 0.15586733 | 0.125939201 | 1.237639498 | 0.215849762 | 0.385098269 |
| ENSG00000255725 | TDGP1 | 174.342673 | -0.049496935 | 0.138037906 | -0.358574949 | 0.71991309 | 0.844366665 |
| ENSG00000255734 | HNRNPABP1 | 9.470398643 | -0.570410967 | 0.588463739 | -0.969322202 | 0.332384455 | 0.523408886 |
| ENSG00000255737 | AGAP2-AS1 | 505.554244 | 0.174784154 | 0.098095368 | 1.781777844 | 0.074785467 | 0.167590661 |
| ENSG00000255769 | GOLGA2P10 | 303.1972256 | -0.05128043 | 0.12237932 | -0.419028559 | 0.675195261 | 0.81448069 |
| ENSG00000255774 | LINC02747 | 38.19774703 | 0.8736895 | 0.291523558 | 2.996977354 | 0.00272671 | 0.009326185 |
| ENSG00000255794 | RMST | 32.70971709 | 0.024229092 | 0.335197246 | 0.072283089 | 0.942376622 | 0.972650037 |
| ENSG00000255807 | PTP4A1P2 | 3.597491782 | -0.199324233 | 0.942711092 | -0.211437242 | 0.832546099 | 0.911410533 |
| ENSG00000255811 | ENSG0000025 | 2.877785184 | -0.461905545 | 1.027021856 | -0.4497524 | 0.652888983 | 0.798532378 |
| ENSG00000255815 | KRT8P11 | 25.36807051 | 0.099119856 | 0.346384171 | 0.286155848 | 0.774758758 | 0.877499635 |
| ENSG00000255836 | ENSG0000025 | 49.67324234 | 0.755789032 | 0.258968378 | 2.918460694 | 0.003517642 | 0.011740828 |
| ENSG00000255847 | ENSG0000025 | 35.37064969 | 0.0895779 | 0.304891379 | 0.293802667 | 0.768908697 | 0.873855116 |
| ENSG00000255850 | RXYLT1-AS1 | 7.356849001 | -1.281414356 | 0.687296335 | -1.864427745 | 0.06226166 | 0.14382061 |
| ENSG00000255856 | ENSG0000025 | 46.94237953 | 0.834385079 | 0.271289676 | 3.075624156 | 0.002100625 | 0.007350263 |
| ENSG00000255857 | PXN-AS1 | 220.3110038 | -0.378364953 | 0.127126768 | -2.976280754 | 0.002917676 | 0.009911372 |
| ENSG00000255867 | DENND5B-AS1 | 1.900626004 | 1.440269632 | 1.349102652 | 1.067576014 | 0.285711808 | 0.469595837 |
| ENSG00000255874 | PRECSIT | 11.84132236 | 1.177952485 | 0.553920127 | 2.126574622 | 0.033455443 | 0.085274207 |
| ENSG00000255875 | PABPN1P2 | 7.734367216 | -0.612112758 | 0.643436574 | -0.951317943 | 0.341443001 | 0.532510874 |
| ENSG00000255883 | FUNDC2P1 | 229.2973303 | -0.538019253 | 0.125450435 | -4.288699798 | 1.80E-05 | 8.98E-05 |

| | | | | | | | |
|---|---|---|---|---|---|---|---|
| ENSG00000255893 | ENSG0000025 | 2.294410805 | -0.154217161 | 1.187502249 | -0.129866837 | 0.896671782 | 0.947822813 |
| ENSG00000255909 | PDCD5P1 | 18.28806452 | -0.499961788 | 0.407962965 | -1.22550778 | 0.220383961 | 0.390635764 |
| ENSG00000255949 | RPS6KB2-AS1 | 17.38976945 | 0.392331628 | 0.431635214 | 0.908942587 | 0.363380432 | 0.556174748 |
| ENSG00000255959 | ENSG0000025 | 14.5556123 | -0.447028371 | 0.473494043 | -0.944105587 | 0.345115695 | 0.536037809 |
| ENSG00000255964 | HSPD1P12 | 19.09923024 | 0.555943017 | 0.410509968 | 1.354274098 | 0.175648947 | 0.328902689 |
| ENSG00000255966 | GAPDH-DT | 4.353330583 | -0.481442938 | 0.865907498 | -0.555998117 | 0.578212142 | 0.743883491 |
| ENSG00000255967 | HADHAP2 | 70.70696892 | 0.230805184 | 0.218859723 | 1.054580447 | 0.29161725 | 0.476576172 |
| ENSG00000255973 | ENSG0000025 | 1.872223948 | 0.853262458 | 1.30009931 | 0.6563056 | 0.511627515 | 0.691250085 |
| ENSG00000255977 | ENSG0000025 | 2.750197153 | -0.33746247 | 1.051107368 | -0.321054233 | 0.748169292 | 0.860473456 |
| ENSG00000255983 | ENSG0000025 | 3.158332558 | 5.001425094 | 1.455849544 | 3.435399705 | 0.00059168 | 0.002309864 |
| ENSG00000255992 | PUS1-AS1 | 12.64152814 | -0.188921912 | 0.490587429 | -0.385093259 | 0.700168322 | 0.831778571 |
| ENSG00000255993 | PSMC1P9 | 9.992471492 | -0.649431395 | 0.592649655 | -1.09580996 | 0.273161949 | 0.455038495 |
| ENSG00000256007 | ARAP1-AS1 | 5.262566473 | -0.718847689 | 0.762710378 | -0.942491029 | 0.345941302 | 0.536831407 |
| ENSG00000256013 | ENSG0000025 | 35.57755063 | -0.160891604 | 0.307734605 | -0.522825844 | 0.601095453 | 0.761032379 |
| ENSG00000256029 | ENSG0000025 | 11.67908629 | 0.854823624 | 0.537491098 | 1.590395871 | 0.1117456 | 0.231797095 |
| ENSG00000256034 | ENSG0000025 | 16.79242646 | -0.122352128 | 0.455263829 | -0.268749943 | 0.788122116 | 0.885426384 |
| ENSG00000256035 | ENSG0000025 | 1.655682552 | -1.369571315 | 1.421012076 | -0.963799912 | 0.335146251 | NA |
| ENSG00000256037 | MRPL40P1 | 1.69547462 | 0.364893817 | 1.34152901 | 0.271998454 | 0.785623208 | NA |
| ENSG00000256043 | CTSO | 198.4908564 | 0.648570435 | 0.129697401 | 5.000643267 | 5.71E-07 | 3.41E-06 |
| ENSG00000256053 | COA8 | 395.2835908 | -0.509583917 | 0.098058461 | -5.196735837 | 2.03E-07 | 1.28E-06 |
| ENSG00000256056 | C12orf71BP | 2.682703755 | 3.692436112 | 1.475119405 | 2.50314388 | 0.012309549 | 0.03571731 |
| ENSG00000256060 | TRAPPC2B | 111.4390941 | 0.500252085 | 0.178902074 | 2.796234126 | 0.005170193 | 0.016590325 |
| ENSG00000256061 | DNAAF4 | 5.504155114 | -0.942300007 | 0.813459785 | -1.158385484 | 0.24670676 | 0.423960827 |
| ENSG00000256069 | A2MP1 | 2.547333429 | -2.88105015 | 1.363104563 | -2.113594384 | 0.034549926 | 0.08765679 |
| ENSG00000256073 | URB1-AS1 | 178.052122 | -0.031304706 | 0.13678587 | -0.228859209 | 0.818978345 | 0.904057533 |
| ENSG00000256075 | ENSG0000025 | 6.812489856 | -0.244320652 | 0.681580897 | -0.358461708 | 0.719997819 | 0.844366665 |
| ENSG00000256081 | ENSG0000025 | 2.249556696 | 0.711769947 | 1.235310773 | 0.57618695 | 0.564488825 | 0.732969584 |
| ENSG00000256087 | ZNF432 | 228.4492951 | 0.00092804 | 0.12744612 | 0.007281821 | 0.994189998 | 0.996533822 |
| ENSG00000256091 | MTRF1LP1 | 2.069823068 | -0.391891697 | 1.236966361 | -0.316816778 | 0.751382632 | 0.862482741 |
| ENSG00000256092 | SBNO1-AS1 | 36.13407843 | 0.20673634 | 0.297509867 | 0.69488902 | 0.487124855 | 0.670520307 |
| ENSG00000256103 | ATP5MFP5 | 4.581343825 | -0.588867623 | 0.858428158 | -0.685983582 | 0.492723461 | 0.675455731 |
| ENSG00000256115 | LINC02443 | 3.450481744 | 4.102784931 | 1.416816798 | 2.895776601 | 0.003782215 | 0.01253272 |
| ENSG00000256116 | ENSG0000025 | 31.43764437 | 0.525835623 | 0.32708415 | 1.607646296 | 0.107912667 | 0.225354262 |

| | | | | | | | |
|---|---|---|---|---|---|---|---|
| ENSG00000256124 | LINC01152 | 8.750913838 | 1.257574598 | 0.621712407 | 2.022759371 | 0.043097962 | 0.105502855 |
| ENSG00000256148 | ENSG0000025 | 35.53187448 | -0.164152058 | 0.314912795 | -0.521261953 | 0.602184302 | 0.761606804 |
| ENSG00000256152 | ENSG0000025 | 18.78325464 | 0.262545111 | 0.400931386 | 0.65483801 | 0.512572055 | 0.69212854 |
| ENSG00000256167 | ATF4P4 | 73.70276078 | 0.370074082 | 0.214014639 | 1.729199852 | 0.083773333 | 0.183566745 |
| ENSG00000256196 | ENSG0000025 | 7.852477271 | 0.657750861 | 0.685900372 | 0.958959767 | 0.337579014 | 0.528785427 |
| ENSG00000256210 | TMEM167AP2 | 195.7606402 | -0.375152326 | 0.129199723 | -2.903661996 | 0.003688261 | 0.012248362 |
| ENSG00000256211 | SKP1P2 | 61.06395018 | 0.163005626 | 0.250713651 | 0.650166535 | 0.515584655 | 0.6944375 |
| ENSG00000256223 | ZNF10 | 99.21814673 | 0.158332603 | 0.183889997 | 0.861018029 | 0.389228113 | 0.581914294 |
| ENSG00000256229 | ZNF486 | 4.587536508 | 4.540742102 | 1.364263843 | 3.328345998 | 0.000873633 | 0.003307664 |
| ENSG00000256235 | SMIM3 | 178.4540716 | 0.283989276 | 0.137153766 | 2.070590432 | 0.038397087 | 0.095902352 |
| ENSG00000256238 | SUPT16HP1 | 331.5808367 | -0.426347727 | 0.108894684 | -3.915229924 | 9.03E-05 | 0.000405539 |
| ENSG00000256249 | ENSG0000025 | 2.215411103 | -0.537324756 | 1.257038295 | -0.427452973 | 0.669049431 | 0.809877787 |
| ENSG00000256263 | DDX11L8 | 2.570436359 | -0.848714416 | 1.160323025 | -0.731446672 | 0.464506366 | 0.650624508 |
| ENSG00000256269 | HMBS | 1696.933332 | -0.033348251 | 0.065858835 | -0.506359568 | 0.612604252 | 0.769697752 |
| ENSG00000256280 | ATP5MGP1 | 2.381057581 | -0.242246345 | 1.167330986 | -0.207521558 | 0.835602558 | 0.913361733 |
| ENSG00000256283 | METTL8P1 | 9.391444132 | -0.417610467 | 0.581946575 | -0.717609631 | 0.472998018 | 0.658601347 |
| ENSG00000256293 | ATP6V1E1P3 | 41.13986478 | -0.353768742 | 0.273111744 | -1.295325998 | 0.195207795 | 0.356420472 |
| ENSG00000256294 | ZNF225 | 204.4332861 | 0.017170574 | 0.135508115 | 0.126712511 | 0.899167943 | 0.949051063 |
| ENSG00000256338 | RPL41P2 | 93.04636862 | 0.120629978 | 0.204361386 | 0.590277744 | 0.555004458 | 0.726193544 |
| ENSG00000256340 | ABCC6P1 | 2.70760905 | -2.289973974 | 1.217670964 | -1.880618033 | 0.060023897 | 0.139593288 |
| ENSG00000256341 | ENSG0000025 | 5.320705241 | -0.075590968 | 0.756101043 | -0.099974691 | 0.920364419 | 0.961349593 |
| ENSG00000256356 | HSPA8P5 | 38.97973034 | 0.15329993 | 0.315731328 | 0.485539181 | 0.627293927 | 0.780163875 |
| ENSG00000256357 | ENSG0000025 | 9.000088115 | -2.101161373 | 0.654398094 | -3.210830521 | 0.00132352 | 0.004843123 |
| ENSG00000256364 | ENSG0000025 | 16.88021617 | -0.547732891 | 0.45808136 | -1.195710936 | 0.231809382 | 0.405746011 |
| ENSG00000256385 | UBE2NP1 | 4.786570943 | -0.280340224 | 0.807144525 | -0.347323453 | 0.728348333 | 0.849344793 |
| ENSG00000256393 | RPL41P5 | 32.89594422 | -0.329929414 | 0.303997276 | -1.085303849 | 0.277787102 | 0.460569248 |
| ENSG00000256417 | ENSG0000025 | 25.19125162 | 3.196434578 | 0.479953197 | 6.659888079 | 2.74E-11 | 2.48E-10 |
| ENSG00000256433 | ENSG0000025 | 18.69059529 | 0.465184014 | 0.432457558 | 1.075675532 | 0.282072394 | 0.465502266 |
| ENSG00000256436 | TAS2R31 | 1.839637749 | 0.137164289 | 1.352076967 | 0.101447101 | 0.919195548 | 0.96065357 |
| ENSG00000256443 | ENSG0000025 | 53.4551714 | -0.278231129 | 0.242229279 | -1.148627161 | 0.250709752 | 0.428944042 |
| ENSG00000256448 | ENSG0000025 | 5.50563693 | 0.162201122 | 0.734561484 | 0.22081354 | 0.82523762 | 0.907785125 |
| ENSG00000256453 | DND1 | 49.46583541 | 0.206476639 | 0.264414511 | 0.780882404 | 0.434871662 | 0.624864136 |
| ENSG00000256458 | ENSG0000025 | 1.635907267 | 1.003117142 | 1.413630861 | 0.709603313 | 0.477950165 | NA |

| | | | | | | | |
|---|---|---|---|---|---|---|---|
| ENSG00000256464 | YWHABP2 | 11.41092187 | 0.488485055 | 0.534039924 | 0.914697635 | 0.360350387 | 0.552902004 |
| ENSG00000256525 | POLG2 | 826.3806458 | -0.167404358 | 0.07633535 | -2.193012261 | 0.028306496 | 0.073873835 |
| ENSG00000256537 | SMIM10L1 | 1007.126153 | -0.294323438 | 0.068520029 | -4.295436561 | 1.74E-05 | 8.72E-05 |
| ENSG00000256542 | ENSG0000025 | 5.876250751 | -2.717198917 | 0.875740919 | -3.102742897 | 0.001917361 | 0.006768893 |
| ENSG00000256546 | LINC02985 | 1.66412874 | -4.308825331 | 1.658871332 | -2.597443966 | 0.009392044 | NA |
| ENSG00000256569 | ENSG0000025 | 14.36505049 | -0.625266609 | 0.478989187 | -1.305387734 | 0.191760853 | 0.351761268 |
| ENSG00000256576 | LINC02361 | 9.434157741 | 0.154013027 | 0.56751612 | 0.271380885 | 0.786098102 | 0.883798765 |
| ENSG00000256591 | ENSG0000025 | 8.749618915 | -0.324691401 | 0.602173127 | -0.53919942 | 0.589749261 | 0.752741052 |
| ENSG00000256594 | ENSG0000025 | 36.33787213 | -0.071623967 | 0.28844037 | -0.24831464 | 0.803890972 | 0.895598024 |
| ENSG00000256596 | ENSG0000025 | 1.866076561 | 1.484430614 | 1.430239007 | 1.037889896 | 0.299321318 | 0.485998083 |
| ENSG00000256612 | CYP2B7P | 9.137352616 | 1.261430982 | 0.621283919 | 2.030361552 | 0.042319802 | 0.103952913 |
| ENSG00000256616 | ADGRA3P1 | 61.89655303 | 0.227693235 | 0.229814609 | 0.990769197 | 0.321798294 | 0.511541325 |
| ENSG00000256625 | ENSG0000025 | 1.880681277 | -0.113295683 | 1.288350507 | -0.087938556 | 0.929925512 | 0.966032078 |
| ENSG00000256628 | ZBTB11-AS1 | 208.2991163 | 0.201418003 | 0.13000534 | 1.549305615 | 0.121308271 | 0.247571378 |
| ENSG00000256637 | LINC01965 | 2.529985502 | -2.835988454 | 1.38438401 | -2.048556206 | 0.040505531 | 0.100182951 |
| ENSG00000256650 | RERG-IT1 | 2.52428285 | 0.362203537 | 1.121415952 | 0.32298768 | 0.746704572 | 0.859607712 |
| ENSG00000256663 | ENSG0000025 | 554.7550853 | -0.056607921 | 0.082309322 | -0.687746167 | 0.491612642 | 0.674456911 |
| ENSG00000256667 | KLRA1P | 20.12449503 | 0.050372216 | 0.387325533 | 0.130051371 | 0.896525784 | 0.947723049 |
| ENSG00000256673 | ENSG0000025 | 39.86738985 | -0.333049258 | 0.281094115 | -1.184831844 | 0.236083939 | 0.411214535 |
| ENSG00000256686 | ENSG0000025 | 2.249094091 | 1.087010498 | 1.241335844 | 0.875678007 | 0.381205105 | 0.573432466 |
| ENSG00000256690 | STX5-DT | 26.95397802 | -0.192128973 | 0.36483269 | -0.526622142 | 0.598456013 | 0.758778585 |
| ENSG00000256704 | ENTR1P1 | 3.757367557 | -0.200920412 | 0.91991109 | -0.218412859 | 0.827107447 | 0.908665525 |
| ENSG00000256737 | RBBP4P5 | 50.75290917 | -0.012755847 | 0.252208093 | -0.050576675 | 0.95966285 | 0.981621201 |
| ENSG00000256742 | KDM2B-DT | 4.526103518 | 1.116711541 | 0.849904557 | 1.313925819 | 0.188871169 | 0.347567446 |
| ENSG00000256745 | ARPC3P3 | 679.0980298 | -0.150533513 | 0.076944555 | -1.956389418 | 0.050419298 | 0.120694844 |
| ENSG00000256747 | ENSG0000025 | 1.841040048 | -1.57310965 | 1.391203781 | -1.130754295 | 0.258158522 | 0.437826964 |
| ENSG00000256751 | PLBD1-AS1 | 78.05536054 | 2.266420496 | 0.242131526 | 9.360286668 | 7.95E-21 | 1.40E-19 |
| ENSG00000256771 | ZNF253 | 255.2835251 | 0.669071298 | 0.121958056 | 5.486077084 | 4.11E-08 | 2.77E-07 |
| ENSG00000256789 | ENSG0000025 | 4.180380068 | -1.007127974 | 0.857680323 | -1.174246333 | 0.240296372 | 0.416155729 |
| ENSG00000256799 | ENSG0000025 | 8.021838485 | 0.009932753 | 0.623114601 | 0.015940492 | 0.987281866 | 0.9938236 |
| ENSG00000256802 | LCIIAR | 4.896693956 | 0.420300158 | 0.79171915 | 0.530870269 | 0.595508682 | 0.756709053 |
| ENSG00000256806 | C17orf100 | 60.14233529 | 0.106464692 | 0.231083369 | 0.460719835 | 0.644999622 | 0.7921775 |
| ENSG00000256813 | CCDC86-AS1 | 1.72471347 | 0.667180768 | 1.335622373 | 0.499527997 | 0.617407468 | NA |

| | | | | | | | |
|---|---|---|---|---|---|---|---|
| ENSG00000256843 | ENSG0000025 | 6.976011434 | 0.009981631 | 0.673132265 | 0.014828633 | 0.988168896 | 0.994030148 |
| ENSG00000256849 | TCP1P3 | 22.9011875 | -0.124226672 | 0.371338245 | -0.334537779 | 0.737973788 | 0.855534609 |
| ENSG00000256894 | ENSG0000025 | 7.07719786 | 1.461757115 | 0.707502202 | 2.066081364 | 0.038820798 | 0.096735726 |
| ENSG00000256913 | ENSG0000025 | 11.42359756 | -0.27361872 | 0.559359407 | -0.48916442 | 0.624725297 | 0.778277306 |
| ENSG00000256928 | ENSG0000025 | 3.097213493 | 0.310242018 | 1.075260011 | 0.28852744 | 0.772943029 | 0.876182887 |
| ENSG00000256940 | PPP1R14B-AS1 | 42.34254123 | 0.991838314 | 0.277762471 | 3.570814694 | 0.000355873 | 0.001442608 |
| ENSG00000256943 | GALNT9-AS1 | 2.095195056 | -4.64953546 | 1.618303356 | -2.873092638 | 0.004064749 | 0.01337426 |
| ENSG00000256955 | ENSG0000025 | 7.912175351 | -0.050526074 | 0.622963407 | -0.081106006 | 0.935357649 | 0.969111805 |
| ENSG00000256967 | ENSG0000025 | 2.512478173 | 0.80644573 | 1.142070435 | 0.70612609 | 0.480109728 | 0.664437571 |
| ENSG00000256968 | SNRPEP2 | 674.4478506 | 0.213054225 | 0.083036992 | 2.565774839 | 0.010294563 | 0.030487642 |
| ENSG00000256973 | ENSG0000025 | 2.399992394 | 0.166228765 | 1.16362967 | 0.142853667 | 0.886405752 | 0.941439323 |
| ENSG00000256982 | ENSG0000025 | 17.41251744 | 1.611783936 | 0.468521883 | 3.440146544 | 0.000581399 | 0.002272112 |
| ENSG00000257000 | ENSG0000025 | 2.827741903 | -1.871780457 | 1.221453545 | -1.532420504 | 0.125418701 | 0.254138769 |
| ENSG00000257002 | ENSG0000025 | 15.22265604 | -0.42986582 | 0.467450936 | -0.919595592 | 0.357784131 | 0.550141965 |
| ENSG00000257027 | ENSG0000025 | 93.43002531 | -0.0552003 | 0.186336832 | -0.296239338 | 0.767047309 | 0.872778717 |
| ENSG00000257038 | ARHGEF17-AS | 7.21717059 | 0.678836929 | 0.654863541 | 1.03660822 | 0.299918481 | 0.486606546 |
| ENSG00000257043 | SRSF3P1 | 39.39834855 | -0.255197925 | 0.289954967 | -0.880129517 | 0.378789151 | 0.571354252 |
| ENSG00000257052 | PRPF19-DT | 5.261277586 | -0.023583678 | 0.764371633 | -0.03085368 | 0.97538623 | 0.988740606 |
| ENSG00000257058 | ENSG0000025 | 2.037658998 | 0.107734039 | 1.22991698 | 0.087594562 | 0.930198925 | 0.966210701 |
| ENSG00000257074 | RPL29P33 | 6.75712179 | -0.395288708 | 0.663427495 | -0.595828045 | 0.551290106 | 0.723235044 |
| ENSG00000257084 | MIR200CHG | 71.44783865 | 0.890600696 | 0.216526357 | 4.113128333 | 3.90E-05 | 0.000185235 |
| ENSG00000257086 | ENSG0000025 | 58.18872678 | -0.152706115 | 0.235358963 | -0.648822177 | 0.516453321 | 0.69491147 |
| ENSG00000257093 | DENND11 | 2303.563311 | -0.300582918 | 0.062092131 | -4.840918041 | 1.29E-06 | 7.37E-06 |
| ENSG00000257103 | LSM14A | 4952.294246 | -0.063715884 | 0.047823783 | -1.332305386 | 0.182759858 | 0.33865195 |
| ENSG00000257108 | NHLRC4 | 1.712973386 | 0.379786346 | 1.359035515 | 0.279452849 | 0.779897318 | NA |
| ENSG00000257122 | RRN3P3 | 133.7255748 | -0.246825164 | 0.157383667 | -1.568302284 | 0.116810603 | 0.240413233 |
| ENSG00000257135 | ODC1-DT | 8.754253724 | 0.700776065 | 0.592113301 | 1.183516845 | 0.236604375 | 0.411744504 |
| ENSG00000257137 | LINC02874 | 5.11462651 | 0.368722645 | 0.781742516 | 0.471667636 | 0.637164041 | 0.786385275 |
| ENSG00000257150 | PGAM1P5 | 1.910550225 | -2.396038691 | 1.485513619 | -1.6129362 | 0.10675837 | 0.223386529 |
| ENSG00000257159 | ENSG0000025 | 3.183849127 | -0.072038962 | 0.995925088 | -0.072333715 | 0.942336334 | 0.972650037 |
| ENSG00000257167 | TMPO-AS1 | 287.7152468 | -0.144663405 | 0.108772135 | -1.329967509 | 0.183528977 | 0.339720384 |
| ENSG00000257175 | ENSG0000025 | 11.6136696 | 0.103330724 | 0.510963874 | 0.202227064 | 0.839739216 | 0.91565822 |
| ENSG00000257176 | ENSG0000025 | 22.15011622 | 0.552127025 | 0.391933775 | 1.4087253 | 0.158916411 | 0.304276977 |

| | | | | | | | |
|---|---|---|---|---|---|---|---|
| ENSG00000257179 | HMGN2P6 | 1.655256745 | -0.183602495 | 1.339256921 | -0.13709281 | 0.890957436 | NA |
| ENSG00000257181 | ENSG0000025 | 13.16634651 | -0.964665832 | 0.496269033 | -1.943836445 | 0.051915177 | 0.123694866 |
| ENSG00000257185 | LINC02293 | 28.80426892 | 0.300421858 | 0.33012015 | 0.910037929 | 0.362802507 | 0.555549664 |
| ENSG00000257191 | ENSG0000025 | 37.46769154 | 0.132726323 | 0.299673847 | 0.44290259 | 0.657836194 | 0.801849348 |
| ENSG00000257194 | ENSG0000025 | 7.933141134 | 0.676173091 | 0.659903397 | 1.02465466 | 0.305526163 | 0.49301985 |
| ENSG00000257195 | HNRNPA1P50 | 9.810502229 | 0.310032635 | 0.570172775 | 0.543752085 | 0.586612085 | 0.750355818 |
| ENSG00000257199 | ENSG0000025 | 17.31462872 | -0.459740679 | 0.429720286 | -1.069860312 | 0.28468219 | 0.468513482 |
| ENSG00000257202 | ENSG0000025 | 6.402952152 | -1.471060525 | 0.720187693 | -2.042607141 | 0.041091344 | 0.101358648 |
| ENSG00000257210 | NACAP8 | 3.295897634 | 0.387417467 | 0.978068553 | 0.396104614 | 0.692027855 | 0.826376122 |
| ENSG00000257218 | GATC | 1811.466487 | 0.189904979 | 0.055493644 | 3.422103215 | 0.000621387 | 0.002417267 |
| ENSG00000257225 | ENSG0000025 | 5.536944502 | 1.288975711 | 0.791565242 | 1.628388468 | 0.103442541 | 0.217880858 |
| ENSG00000257242 | LINC01619 | 6.430190503 | -0.644652268 | 0.725445438 | -0.888629569 | 0.374202194 | 0.566950554 |
| ENSG00000257243 | PARK7P1 | 3.096278359 | -0.289207008 | 1.00886759 | -0.28666498 | 0.774368855 | 0.877192313 |
| ENSG00000257261 | SLC38A4-AS1 | 170.1841967 | -1.04347472 | 0.146054913 | -7.144400011 | 9.04E-13 | 9.29E-12 |
| ENSG00000257267 | ZNF271P | 1022.86203 | 0.603593841 | 0.066469251 | 9.080798034 | 1.08E-19 | 1.77E-18 |
| ENSG00000257268 | ENSG0000025 | 11.07881839 | -0.370075686 | 0.59176266 | -0.625378571 | 0.531722623 | 0.707271141 |
| ENSG00000257270 | ENSG0000025 | 5.611508631 | -0.509296782 | 0.736091134 | -0.691893651 | 0.489004118 | 0.672285417 |
| ENSG00000257279 | ENSG0000025 | 6.410696546 | 3.470753315 | 1.002146384 | 3.463319701 | 0.000533554 | 0.00210088 |
| ENSG00000257285 | PRMT5-DT | 4.582723369 | 0.767495496 | 0.828148029 | 0.926761243 | 0.354050512 | 0.545582595 |
| ENSG00000257286 | ENSG0000025 | 2.820678937 | 0.002326915 | 1.043501316 | 0.002229911 | 0.99822079 | 0.999048249 |
| ENSG00000257303 | CNPY2-AS1 | 80.45008212 | -0.160133693 | 0.200397343 | -0.799080923 | 0.424243491 | 0.615065066 |
| ENSG00000257307 | ENSG0000025 | 36.16931973 | -0.537095933 | 0.296044447 | -1.814240867 | 0.069640651 | 0.157939351 |
| ENSG00000257315 | ZBED6 | 38.26293436 | -0.151507133 | 0.293958529 | -0.515403085 | 0.606271377 | 0.764860977 |
| ENSG00000257316 | ASS1P14 | 18.85909246 | 0.505916679 | 0.415928653 | 1.216354476 | 0.223849918 | 0.395252241 |
| ENSG00000257331 | RACGAP1P1 | 11.91908076 | -0.574479338 | 0.532888471 | -1.078047977 | 0.281012346 | 0.464439713 |
| ENSG00000257335 | MGAM | 2.253789406 | 1.178950239 | 1.239406743 | 0.951221418 | 0.341491988 | 0.532551068 |
| ENSG00000257337 | TNS2-AS1 | 2189.185773 | 0.427074857 | 0.053790315 | 7.939623639 | 2.03E-15 | 2.51E-14 |
| ENSG00000257342 | ENSG0000025 | 23.57625199 | -0.773414455 | 0.37610636 | -2.056371649 | 0.039746707 | 0.098646064 |
| ENSG00000257350 | ENSG0000025 | 4.279226483 | -0.404163764 | 0.832783168 | -0.485316922 | 0.627451554 | 0.780187429 |
| ENSG00000257354 | MIRLET7IHG | 97.89442559 | 0.408698368 | 0.203266303 | 2.010654807 | 0.044361931 | 0.108158432 |
| ENSG00000257359 | ENSG0000025 | 4.012999969 | -1.759010448 | 0.944171325 | -1.863020409 | 0.062459395 | 0.144221456 |
| ENSG00000257365 | FNTB | 270.6407358 | -0.753447522 | 0.127082818 | -5.928791421 | 3.05E-09 | 2.32E-08 |
| ENSG00000257376 | ENSG0000025 | 10.45068072 | 0.38350342 | 0.585314787 | 0.655208835 | 0.512333307 | 0.691999885 |

| | | | | | | | |
|---|---|---|---|---|---|---|---|
| ENSG00000257386 | ENSG0000025 | 31.36562984 | -0.382175149 | 0.323300101 | -1.182106497 | 0.237163447 | 0.41238001 |
| ENSG00000257390 | ENSG0000025 | 9.985825026 | -0.488421384 | 0.555333115 | -0.87951064 | 0.379124466 | 0.571614542 |
| ENSG00000257398 | ENSG0000025 | 5.214076957 | -2.88955452 | 0.984356327 | -2.935476149 | 0.003330362 | 0.011171006 |
| ENSG00000257433 | RPAP3-DT | 14.97843985 | 0.01694629 | 0.453806282 | 0.037342563 | 0.970211868 | 0.98657174 |
| ENSG00000257435 | ENSG0000025 | 3.072184117 | -0.009232406 | 0.989239812 | -0.009332829 | 0.992553588 | 0.995983523 |
| ENSG00000257453 | PHLDA1-AS1 | 8.098516327 | 2.051152798 | 0.719183547 | 2.852057455 | 0.004343725 | 0.014206594 |
| ENSG00000257489 | ENSG0000025 | 37.80754608 | -0.327988213 | 0.285463393 | -1.148967683 | 0.250569307 | 0.428735728 |
| ENSG00000257497 | GLIPR1-AS1 | 49.38757809 | 0.094129277 | 0.259792092 | 0.362325412 | 0.71710887 | 0.842749871 |
| ENSG00000257511 | ENSG0000025 | 517.0969317 | 0.335554169 | 0.090172408 | 3.721251076 | 0.000198238 | 0.000841699 |
| ENSG00000257512 | ENSG0000025 | 35.62531387 | -0.178331083 | 0.293441994 | -0.607721754 | 0.543372033 | 0.716907232 |
| ENSG00000257526 | ENSG0000025 | 7.571167328 | -0.548160763 | 0.642982533 | -0.852528234 | 0.393920976 | 0.58690184 |
| ENSG00000257545 | ENSG0000025 | 2.334194783 | -0.57692458 | 1.186528044 | -0.486229198 | 0.626804672 | 0.779978257 |
| ENSG00000257550 | ENSG0000025 | 8.244885023 | -0.736992033 | 0.620982518 | -1.186816071 | 0.235300178 | 0.410322013 |
| ENSG00000257553 | ENSG0000025 | 52.55733841 | -0.38847961 | 0.308040831 | -1.261130247 | 0.207261925 | 0.373342442 |
| ENSG00000257557 | PPP1R12A-AS | 55.08000393 | -0.21259537 | 0.244307684 | -0.870195185 | 0.384193747 | 0.576642538 |
| ENSG00000257564 | ENSG0000025 | 30.30279457 | 0.676042923 | 0.334195359 | 2.022897397 | 0.043083727 | 0.105490522 |
| ENSG00000257576 | HSPD1P4 | 192.5301558 | 0.0768113 | 0.14951464 | 0.513737653 | 0.607435423 | 0.765803946 |
| ENSG00000257588 | AQP5-AS1 | 8.960498218 | 1.836928785 | 0.645381653 | 2.84626744 | 0.004423503 | 0.014446918 |
| ENSG00000257591 | ZNF625 | 4.737493167 | 0.605465435 | 0.9013695 | 0.671717242 | 0.501763723 | 0.682833786 |
| ENSG00000257594 | GALNT4 | 7.072638784 | 0.03324927 | 0.68046535 | 0.048862547 | 0.961028836 | 0.982116705 |
| ENSG00000257596 | SCAT2 | 1.581914333 | -0.745280894 | 1.472234292 | -0.506224381 | 0.612699141 | NA |
| ENSG00000257599 | OVCH1-AS1 | 25.65114988 | -0.826008854 | 0.398183985 | -2.074440172 | 0.03803845 | 0.095179105 |
| ENSG00000257605 | MYG1-AS1 | 21.71846459 | -0.108174427 | 0.423150343 | -0.255640645 | 0.798228331 | 0.892259454 |
| ENSG00000257607 | ENSG0000025 | 64.71031407 | 0.995482034 | 0.226968301 | 4.385995895 | 1.15E-05 | 5.91E-05 |
| ENSG00000257611 | ENSG0000025 | 3.412062232 | -1.152206057 | 1.002378455 | -1.14947209 | 0.250361369 | 0.428507782 |
| ENSG00000257616 | ENSG0000025 | 11.36313683 | -0.614563554 | 0.532551041 | -1.153999348 | 0.248500436 | 0.426244954 |
| ENSG00000257621 | PSMA3-AS1 | 1487.096315 | -0.08615904 | 0.080136404 | -1.075154812 | 0.282305423 | 0.465749817 |
| ENSG00000257653 | ENSG0000025 | 10.4441584 | -0.395514425 | 0.562684336 | -0.702906408 | 0.482114077 | 0.666287182 |
| ENSG00000257654 | ENSG0000025 | 1.743124764 | 0.107280438 | 1.352916742 | 0.079295669 | 0.936797451 | NA |
| ENSG00000257657 | ENSG0000025 | 9.219884775 | -1.102452058 | 0.60223005 | -1.830616154 | 0.067157856 | 0.153400499 |
| ENSG00000257660 | ADCY6-DT | 5.516005885 | -0.025640944 | 0.805257422 | -0.031841921 | 0.974598115 | 0.988514222 |
| ENSG00000257663 | ENSG0000025 | 18.34422067 | 0.161666065 | 0.413589504 | 0.390885319 | 0.695882006 | 0.829102172 |
| ENSG00000257671 | KRT7-AS | 33.36817377 | 0.540723992 | 0.317994308 | 1.700420351 | 0.089051887 | 0.192830874 |

| | | | | | | | |
|---|---|---|---|---|---|---|---|
| ENSG00000257674 | ENSG0000025 | 2.625405784 | -0.191489085 | 1.056684331 | -0.181216925 | 0.85619731 | 0.925509674 |
| ENSG00000257675 | BTBD10P1 | 9.323456573 | 0.094664967 | 0.593770256 | 0.159430294 | 0.873329873 | 0.934604743 |
| ENSG00000257681 | ENSG0000025 | 6.448054894 | 0.603166643 | 0.713249462 | 0.845660145 | 0.397742363 | 0.590793175 |
| ENSG00000257696 | UHRF1BP1L-D | 3.713845052 | 1.612876886 | 1.011753748 | 1.59413977 | 0.110904733 | 0.230271736 |
| ENSG00000257698 | GIHCG | 47.58702041 | -0.967598119 | 0.260105564 | -3.720020844 | 0.000199206 | 0.000845319 |
| ENSG00000257702 | LBX2-AS1 | 157.6597723 | -0.344735454 | 0.147564245 | -2.336171975 | 0.01948228 | 0.05345975 |
| ENSG00000257704 | INAFM1 | 230.7916043 | 0.554866963 | 0.121496438 | 4.56694018 | 4.95E-06 | 2.65E-05 |
| ENSG00000257711 | ENSG0000025 | 4.688644518 | 0.114536814 | 0.799635988 | 0.143236192 | 0.886103648 | 0.941249198 |
| ENSG00000257715 | ENSG0000025 | 1.821604323 | -1.590359982 | 1.395470574 | -1.139658558 | 0.254428579 | 0.433560511 |
| ENSG00000257727 | CNPY2 | 184.8979967 | 0.118811452 | 0.140994235 | 0.842668866 | 0.399413664 | 0.592163734 |
| ENSG00000257752 | ENSG0000025 | 2.951267294 | -0.397606584 | 1.039616423 | -0.382455081 | 0.702123841 | 0.833120618 |
| ENSG00000257758 | KRT18P20 | 29.17600697 | 0.170583771 | 0.33998012 | 0.501746313 | 0.615845982 | 0.772407485 |
| ENSG00000257771 | LINC02395 | 2.252342218 | 0.277662885 | 1.177332042 | 0.235840761 | 0.813556237 | 0.901448243 |
| ENSG00000257773 | ST13P3 | 59.68769629 | 0.132473177 | 0.22619583 | 0.585657024 | 0.55810602 | 0.728263604 |
| ENSG00000257790 | EIF4A1P4 | 145.9615007 | -0.029921963 | 0.149002686 | -0.200814924 | 0.840843293 | 0.91647005 |
| ENSG00000257800 | FNBP1P1 | 110.7838412 | -0.257825663 | 0.171684163 | -1.501744011 | 0.133163233 | 0.266314109 |
| ENSG00000257808 | ENSG0000025 | 3.672684907 | -0.470731342 | 0.950027806 | -0.495492173 | 0.620252746 | 0.775231447 |
| ENSG00000257809 | MYL6B-AS1 | 8.009258725 | -0.772477446 | 0.6822509 | -1.132248334 | 0.257530045 | 0.437221671 |
| ENSG00000257815 | PRANCR | 41.56851556 | 0.385475919 | 0.275349242 | 1.399952715 | 0.161527479 | 0.308503902 |
| ENSG00000257817 | TBX3-AS1 | 5.906103453 | -0.178176267 | 0.710232496 | -0.250870339 | 0.801914357 | 0.894633256 |
| ENSG00000257818 | C1GALT1P1 | 2.578668167 | -0.440943645 | 1.102035149 | -0.400117588 | 0.689069911 | 0.82436022 |
| ENSG00000257829 | ENSG0000025 | 1.986104564 | -3.488779922 | 1.583145563 | -2.203701291 | 0.027545358 | 0.072199557 |
| ENSG00000257839 | PHLDA1-DT | 5.79139318 | 2.10428417 | 0.855018855 | 2.461096803 | 0.013851299 | 0.039629433 |
| ENSG00000257842 | NOVA1-DT | 90.75752839 | 0.34431479 | 0.192880773 | 1.785117222 | 0.074242305 | 0.166547866 |
| ENSG00000257846 | ENSG0000025 | 31.33015673 | -0.379416713 | 0.325327466 | -1.16626093 | 0.243508964 | 0.42007036 |
| ENSG00000257851 | HNRNPA3P10 | 243.6066175 | 0.069053204 | 0.125448187 | 0.55045199 | 0.582009398 | 0.74680029 |
| ENSG00000257859 | CASC18 | 6.661670662 | -0.325208424 | 0.678461914 | -0.479331878 | 0.631702547 | 0.782934732 |
| ENSG00000257869 | ENSG0000025 | 6.888527237 | 0.3128125 | 0.672227556 | 0.465337217 | 0.641689987 | 0.789761698 |
| ENSG00000257883 | ENSG0000025 | 6.656435803 | -0.324046091 | 0.676281614 | -0.479158511 | 0.631825867 | 0.782956212 |
| ENSG00000257884 | ENSG0000025 | 12.68630494 | -0.117883082 | 0.556338256 | -0.211891022 | 0.832192055 | 0.911283868 |
| ENSG00000257894 | ENSG0000025 | 10.17439344 | -2.307288003 | 0.628409454 | -3.671631589 | 0.000241007 | 0.001006866 |
| ENSG00000257900 | ENSG0000025 | 1.88217045 | -1.629296492 | 1.377439575 | -1.182844259 | 0.236870876 | 0.412089936 |
| ENSG00000257907 | EEF1A1P17 | 4.019636329 | 0.208414522 | 0.891957191 | 0.233659782 | 0.815249113 | 0.902135938 |

| | | | | | | | |
|---|---|---|---|---|---|---|---|
| ENSG00000257913 | DDN-AS1 | 5.369436947 | -1.310755257 | 0.781110977 | -1.67806534 | 0.093334344 | 0.200326521 |
| ENSG00000257918 | ENSG0000025 | 2.528854492 | -0.138969327 | 1.136507362 | -0.122277543 | 0.902679217 | 0.950907167 |
| ENSG00000257923 | CUX1 | 3917.793604 | -0.508040462 | 0.067951195 | -7.476549385 | 7.63E-14 | 8.37E-13 |
| ENSG00000257941 | ENSG0000025 | 26.03713488 | -0.152917821 | 0.339393522 | -0.45056199 | 0.652305266 | 0.798158887 |
| ENSG00000257950 | P2RX5-TAX1BI | 10.65460849 | -0.825968394 | 0.554192226 | -1.490400542 | 0.13611895 | 0.270774255 |
| ENSG00000257954 | ENSG0000025 | 5.720768564 | 0.410767861 | 0.742585118 | 0.553159296 | 0.580154336 | 0.745587702 |
| ENSG00000257966 | OLA1P3 | 12.49026054 | -0.313407049 | 0.491064892 | -0.638219213 | 0.523330992 | 0.699904915 |
| ENSG00000257976 | ENSG0000025 | 1.98282927 | -1.118516025 | 1.303407354 | -0.858147702 | 0.39081091 | 0.583644831 |
| ENSG00000258011 | HMGA1P3 | 94.7984805 | 0.551681901 | 0.184168231 | 2.995532394 | 0.002739662 | 0.009369089 |
| ENSG00000258012 | ENSG0000025 | 8.301862664 | 1.21773414 | 0.645222583 | 1.887308617 | 0.059118828 | 0.137753775 |
| ENSG00000258016 | HIGD1AP1 | 44.31164699 | -0.0344661 | 0.270897765 | -0.127229178 | 0.898759013 | 0.948848624 |
| ENSG00000258017 | TUBA1B-AS1 | 275.2955648 | -0.881342891 | 0.247599114 | -3.55955591 | 0.000371482 | 0.001500582 |
| ENSG00000258021 | ENSG0000025 | 1.769155964 | -0.389763134 | 1.33294219 | -0.292408131 | 0.769974592 | NA |
| ENSG00000258033 | SNX5P2 | 15.67112721 | -0.609037239 | 0.445892282 | -1.365884235 | 0.171975302 | 0.323714895 |
| ENSG00000258034 | ENSG0000025 | 3.302557702 | -0.385370667 | 0.973668036 | -0.395792665 | 0.692257988 | 0.826457434 |
| ENSG00000258048 | PPP1R12A-AS1 | 10.25317404 | -1.340458244 | 0.571284893 | -2.346391896 | 0.018956164 | 0.05219088 |
| ENSG00000258056 | CD63-AS1 | 141.7765969 | 0.407891287 | 0.158516122 | 2.573184876 | 0.010076736 | 0.029931417 |
| ENSG00000258057 | BCDIN3D-AS1 | 22.58020762 | -0.541186828 | 0.370509925 | -1.460654064 | 0.144110401 | 0.282651556 |
| ENSG00000258072 | ENSG0000025 | 42.05636128 | -0.137778398 | 0.27176254 | -0.506980829 | 0.61216827 | 0.769498044 |
| ENSG00000258090 | ENSG0000025 | 8.708766059 | 0.403449584 | 0.624332604 | 0.646209379 | 0.518143766 | 0.695932225 |
| ENSG00000258092 | ENSG0000025 | 48.36031011 | -0.244551368 | 0.255914981 | -0.955596137 | 0.339276314 | 0.530466101 |
| ENSG00000258099 | ATXN2-AS | 3.641945417 | -0.44613968 | 0.936070526 | -0.476609045 | 0.633640541 | 0.784143138 |
| ENSG00000258101 | ENSG0000025 | 26.00853926 | -0.501252245 | 0.348859083 | -1.436833005 | 0.15076545 | 0.292775706 |
| ENSG00000258102 | MAP1LC3B2 | 592.9149412 | 0.451409557 | 0.08165509 | 5.528247655 | 3.23E-08 | 2.20E-07 |
| ENSG00000258111 | ENSG0000025 | 2.217484384 | -0.050543784 | 1.216491863 | -0.041548806 | 0.966858385 | 0.985284143 |
| ENSG00000258128 | MKRN9P | 16.84045715 | 0.30249021 | 0.430345009 | 0.702901633 | 0.482117054 | 0.666287182 |
| ENSG00000258136 | PRDM4-AS1 | 4.607311386 | -0.566290033 | 0.85683346 | -0.660910269 | 0.50866986 | 0.688876946 |
| ENSG00000258153 | HSPE1P4 | 46.54082804 | 0.082469136 | 0.261426377 | 0.31545836 | 0.752413662 | 0.863320632 |
| ENSG00000258168 | ENSG0000025 | 3.898776416 | 1.671145168 | 0.995911195 | 1.678006208 | 0.093345887 | 0.200332552 |
| ENSG00000258181 | ENSG0000025 | 4.980969958 | 2.602579821 | 1.022145944 | 2.546191996 | 0.010890527 | 0.032012903 |
| ENSG00000258199 | ENSG0000025 | 18.13415138 | -0.303340157 | 0.427995102 | -0.708746795 | 0.47848162 | 0.663064417 |
| ENSG00000258203 | ENSG0000025 | 2.00602701 | -2.505344755 | 1.513432701 | -1.655405459 | 0.097842279 | 0.208267782 |
| ENSG00000258204 | ENSG0000025 | 38.4625322 | -0.524338643 | 0.281198949 | -1.864653637 | 0.062229969 | 0.143776362 |

| | | | | | | | |
|---|---|---|---|---|---|---|---|
| ENSG00000258216 | ENSG0000025 | 10.17320525 | 0.768840472 | 0.56768805 | 1.354336192 | 0.175629145 | 0.328902689 |
| ENSG00000258232 | ENSG0000025 | 108.5836512 | -0.676888487 | 0.196424627 | -3.446046951 | 0.000568852 | 0.00222763 |
| ENSG00000258244 | OSTF1P1 | 16.21924431 | -0.019392819 | 0.431513153 | -0.044941431 | 0.964153993 | 0.983666795 |
| ENSG00000258245 | RPL10P13 | 7.908014371 | 0.175516878 | 0.637013106 | 0.275531032 | 0.782908288 | 0.882291026 |
| ENSG00000258279 | LINC00592 | 10.75827204 | -0.076482938 | 0.540275345 | -0.141562887 | 0.887425283 | 0.941998797 |
| ENSG00000258282 | BTF3P2 | 6.936447888 | 1.03045935 | 0.739782454 | 1.392922129 | 0.16364334 | 0.311326772 |
| ENSG00000258289 | CHURC1 | 2172.232955 | -0.258134641 | 0.061293727 | -4.211436494 | 2.54E-05 | 0.00012417 |
| ENSG00000258290 | BRWD1P2 | 27.74572809 | -0.230341197 | 0.337532313 | -0.682427099 | 0.494968925 | 0.677239198 |
| ENSG00000258300 | NUTF2P2 | 10.28480156 | -0.142738846 | 0.589014259 | -0.242335129 | 0.808520496 | 0.898617256 |
| ENSG00000258301 | VASH1-AS1 | 219.6014834 | -0.321806261 | 0.125412974 | -2.565972643 | 0.010288694 | 0.030474196 |
| ENSG00000258302 | ENSG0000025 | 7.588831857 | 0.902206908 | 0.659587777 | 1.367834487 | 0.17136389 | 0.322749396 |
| ENSG00000258303 | ENSG0000025 | 1.669955472 | 0.97792033 | 1.427162631 | 0.685219966 | 0.493205126 | NA |
| ENSG00000258311 | ENSG0000025 | 16.19210535 | -0.018501413 | 0.468353342 | -0.039503109 | 0.968489275 | 0.985876114 |
| ENSG00000258314 | ENSG0000025 | 1.758252674 | -0.932470499 | 1.358617834 | -0.686337597 | 0.492500246 | NA |
| ENSG00000258315 | C17orf49 | 15.76214537 | 0.895170277 | 0.458541622 | 1.952211606 | 0.050913087 | 0.121711777 |
| ENSG00000258317 | ENSG0000025 | 13.74905874 | -0.349265862 | 0.493917613 | -0.707133848 | 0.479483303 | 0.66373081 |
| ENSG00000258334 | TROAP-AS1 | 40.75640961 | 0.256725896 | 0.277298105 | 0.925811938 | 0.354543722 | 0.546184473 |
| ENSG00000258345 | ENSG0000025 | 1.682698335 | -0.237280604 | 1.334966927 | -0.177742683 | 0.858925063 | NA |
| ENSG00000258354 | ENSG0000025 | 5.705300768 | 1.323645922 | 0.779717796 | 1.697596142 | 0.089584011 | 0.193690596 |
| ENSG00000258357 | NSA2P2 | 19.71459027 | -0.106104193 | 0.406178417 | -0.261225581 | 0.793918552 | 0.889585873 |
| ENSG00000258359 | PCNPP1 | 15.69257603 | 0.081711532 | 0.44066095 | 0.185429483 | 0.852892186 | 0.923198544 |
| ENSG00000258365 | ENSG0000025 | 6.354389562 | 0.487544088 | 0.709695835 | 0.686976116 | 0.492097779 | 0.675001378 |
| ENSG00000258366 | RTEL1 | 56.33955599 | -0.154312839 | 0.237677931 | -0.649251859 | 0.516175597 | 0.694735284 |
| ENSG00000258373 | SLC25A3P2 | 6.065888675 | 0.461553073 | 0.732012312 | 0.63052638 | 0.528350249 | 0.70433965 |
| ENSG00000258376 | PAPLN-AS1 | 121.2888126 | -0.613671812 | 0.163853532 | -3.745246172 | 0.000180217 | 0.000770583 |
| ENSG00000258377 | ENSG0000025 | 22.93676668 | -1.063484865 | 0.38020204 | -2.79715718 | 0.005155445 | 0.016556895 |
| ENSG00000258378 | ENSG0000025 | 4.076810545 | -0.975307897 | 0.910213776 | -1.071515202 | 0.283937839 | 0.467791872 |
| ENSG00000258384 | RCCD1-AS1 | 27.87942679 | 0.051346005 | 0.347855248 | 0.147607388 | 0.882652624 | 0.939121523 |
| ENSG00000258387 | ENSG0000025 | 6.117398848 | 0.279562113 | 0.721006899 | 0.387738472 | 0.698209582 | 0.830732062 |
| ENSG00000258388 | PPT2-EGFL8 | 18.64825226 | -0.797227823 | 0.435439798 | -1.830856588 | 0.067121951 | 0.153349026 |
| ENSG00000258401 | ATP5MC2P2 | 2.699087129 | -0.653218973 | 1.112651642 | -0.587083098 | 0.557147899 | 0.727431576 |
| ENSG00000258413 | FBXO34-AS1 | 2.739380737 | -0.570548536 | 1.068285768 | -0.534078571 | 0.593287175 | 0.755398113 |
| ENSG00000258424 | ENSG0000025 | 28.86026044 | -2.021328131 | 0.368798681 | -5.480844256 | 4.23E-08 | 2.85E-07 |

| | | | | | | | |
|---|---|---|---|---|---|---|---|
| ENSG00000258427 | RBM8B | 1262.839841 | 0.146911686 | 0.062321813 | 2.357307636 | 0.01840799 | 0.050852515 |
| ENSG00000258429 | PDF | 13.52256058 | -0.357579325 | 0.515054302 | -0.694255584 | 0.48752194 | 0.670899634 |
| ENSG00000258430 | ENSG0000025 | 66.76775511 | -1.145976086 | 0.249318981 | -4.59642535 | 4.30E-06 | 2.31E-05 |
| ENSG00000258441 | LINC00641 | 282.5811482 | 0.182241886 | 0.117130194 | 1.555891615 | 0.119733868 | 0.245174947 |
| ENSG00000258445 | MAD2L1P1 | 33.99531332 | -0.107392646 | 0.302714089 | -0.354765932 | 0.722764953 | 0.845926239 |
| ENSG00000258446 | ENSG0000025 | 4.78284756 | 0.296008828 | 0.848101701 | 0.349025156 | 0.727070424 | 0.848415257 |
| ENSG00000258450 | ENSG0000025 | 5.485334667 | -1.13907757 | 0.779573171 | -1.461155429 | 0.143972792 | 0.282478262 |
| ENSG00000258454 | ENSG0000025 | 2.477386305 | -1.62101374 | 1.196819389 | -1.354434726 | 0.175597725 | 0.328887313 |
| ENSG00000258457 | AJUBA-DT | 50.12609669 | -1.045067487 | 0.252412079 | -4.140322812 | 3.47E-05 | 0.000165957 |
| ENSG00000258458 | OXA1L-DT | 10.68033939 | -0.634280244 | 0.545524703 | -1.162697566 | 0.24495223 | 0.421893847 |
| ENSG00000258469 | CHMP4BP1 | 8.672895408 | -0.989527796 | 0.625836203 | -1.581129042 | 0.113848536 | 0.235541613 |
| ENSG00000258471 | ENSG0000025 | 6.300540331 | -0.604273178 | 0.71192012 | -0.84879351 | 0.395996202 | 0.589000381 |
| ENSG00000258477 | PPIAP6 | 3.326172685 | 0.711716272 | 1.021155562 | 0.69697145 | 0.485820661 | 0.669513755 |
| ENSG00000258479 | LINC00640 | 1.63776912 | 0.394057253 | 1.338899878 | 0.294314205 | 0.768517819 | NA |
| ENSG00000258484 | SPESP1 | 144.1969749 | 0.046182023 | 0.153512114 | 0.300836345 | 0.763539292 | 0.870426445 |
| ENSG00000258498 | DIO3OS | 9.18303087 | 0.981196222 | 0.633543506 | 1.548743239 | 0.121443455 | 0.247768842 |
| ENSG00000258499 | LINC02287 | 4.75248248 | -3.205001352 | 1.069376657 | -2.997074353 | 0.002725842 | 0.009324608 |
| ENSG00000258500 | ENSG0000025 | 3.668410514 | -0.175972326 | 0.904648337 | -0.194520145 | 0.845768623 | 0.919043732 |
| ENSG00000258501 | EIF3LP1 | 22.7073601 | 0.450865414 | 0.382475522 | 1.178808544 | 0.238474421 | 0.413937895 |
| ENSG00000258504 | ENSG0000025 | 32.39625352 | 0.199677537 | 0.32029895 | 0.623409903 | 0.533015192 | 0.708463324 |
| ENSG00000258509 | ENSG0000025 | 3.709977561 | 0.218622308 | 0.927968946 | 0.235592267 | 0.813749074 | 0.901448243 |
| ENSG00000258525 | ENSG0000025 | 6.415309767 | -0.246418807 | 0.693190277 | -0.355485088 | 0.722226214 | 0.845625137 |
| ENSG00000258526 | ENSG0000025 | 14.57433046 | -0.887588103 | 0.465084058 | -1.908446631 | 0.056333515 | 0.132529056 |
| ENSG00000258527 | ASB9P1 | 1.713388052 | -1.385273719 | 1.398479787 | -0.99055684 | 0.321902021 | NA |
| ENSG00000258539 | ENSG0000025 | 6.118733097 | -1.414883084 | 0.752397718 | -1.880498903 | 0.060040116 | 0.139616853 |
| ENSG00000258548 | LINC00645 | 10.98602453 | 6.807107835 | 1.25590925 | 5.420063462 | 5.96E-08 | 3.94E-07 |
| ENSG00000258551 | CRAT37 | 12.24063618 | -2.329467725 | 0.592579372 | -3.931064484 | 8.46E-05 | 0.000382189 |
| ENSG00000258554 | ENSG0000025 | 9.035348373 | -0.731410838 | 0.599692326 | -1.219643484 | 0.222600054 | 0.393681694 |
| ENSG00000258559 | ENSG0000025 | 20.28569564 | -1.269445779 | 0.412349432 | -3.078568028 | 0.00207998 | 0.007282471 |
| ENSG00000258561 | ENSG0000025 | 13.26798102 | -0.190463775 | 0.473579559 | -0.402179045 | 0.687552263 | 0.823383061 |
| ENSG00000258568 | RHOQP1 | 26.21718979 | -0.405153967 | 0.34047806 | -1.189956168 | 0.23406362 | 0.408756048 |
| ENSG00000258591 | PPIAP4 | 3.115262619 | 0.265838404 | 0.981837756 | 0.270755939 | 0.78657875 | 0.88412546 |
| ENSG00000258592 | LINC03059 | 2.909949255 | -0.829777924 | 1.063885166 | -0.779950648 | 0.435419925 | 0.625299429 |

| | | | | | | | |
|---|---|---|---|---|---|---|---|
| ENSG00000258593 | ENSG0000025 | 7.189774665 | -0.527446503 | 0.670691836 | -0.786421535 | 0.431620571 | 0.621633344 |
| ENSG00000258605 | DYNLL1P2 | 4.414921142 | -0.276164708 | 0.834152379 | -0.331072254 | 0.740589909 | 0.856911758 |
| ENSG00000258608 | DNAJC19P9 | 180.3647876 | -0.128103579 | 0.136964579 | -0.935304434 | 0.349631431 | 0.540914772 |
| ENSG00000258613 | RPL21P7 | 1.895934368 | -1.050621469 | 1.321688515 | -0.794908525 | 0.426666715 | 0.617497513 |
| ENSG00000258616 | LINC02303 | 8.000765226 | -0.893436733 | 0.667585614 | -1.338310343 | 0.180795292 | 0.336121526 |
| ENSG00000258630 | LINC02292 | 1.861313195 | -3.375155688 | 1.588877805 | -2.124238678 | 0.033650191 | 0.085677236 |
| ENSG00000258633 | RRAGAP1-AS1 | 1.710139924 | -0.299574315 | 1.400199607 | -0.213951149 | 0.830585152 | NA |
| ENSG00000258634 | ENSG0000025 | 401.292545 | -0.108516101 | 0.09839308 | -1.102883467 | 0.270077775 | 0.451737816 |
| ENSG00000258640 | RPL21P5 | 1.695026206 | -0.263289883 | 1.445185184 | -0.182184184 | 0.855438185 | NA |
| ENSG00000258643 | BCL2L2-PABPN | 7.257950186 | -0.904745479 | 0.698175703 | -1.295870759 | 0.195020015 | 0.356162658 |
| ENSG00000258645 | HSPE1P2 | 225.1575967 | 0.022007807 | 0.125293354 | 0.175650237 | 0.860568733 | 0.927426758 |
| ENSG00000258646 | ENSG0000025 | 13.41203868 | -0.712525479 | 0.478572322 | -1.488856429 | 0.136525179 | 0.271347047 |
| ENSG00000258647 | LINC00930 | 6.361418632 | 1.04857754 | 0.7098513 | 1.477179149 | 0.139627621 | 0.275864063 |
| ENSG00000258648 | UBE2CP1 | 5.036568671 | -0.813351029 | 0.874444383 | -0.930134661 | 0.352301367 | 0.543727512 |
| ENSG00000258651 | SEC23A-AS1 | 3.647230352 | -0.676024777 | 0.945916077 | -0.714677331 | 0.474808448 | 0.660047619 |
| ENSG00000258655 | ARHGAP5-AS1 | 90.60635301 | -0.726096299 | 0.197282165 | -3.680496407 | 0.00023278 | 0.000976231 |
| ENSG00000258666 | ENSG0000025 | 10.32510606 | -1.324686231 | 0.58757637 | -2.25449201 | 0.024165233 | 0.064556939 |
| ENSG00000258674 | ENSG0000025 | 9.525383841 | -0.828202303 | 0.57304829 | -1.445257436 | 0.148385589 | 0.289090959 |
| ENSG00000258682 | ENSG0000025 | 15.92056825 | -0.273280027 | 0.446531804 | -0.612005741 | 0.540533959 | 0.7147299 |
| ENSG00000258701 | LINC00638 | 246.9047368 | -0.19966364 | 0.129182931 | -1.545588394 | 0.122204006 | 0.248944134 |
| ENSG00000258704 | SRP54-AS1 | 11.79375473 | -0.800894409 | 0.521192004 | -1.536659049 | 0.124376824 | 0.252484264 |
| ENSG00000258708 | SLC25A21-AS1 | 117.7618763 | -0.101975637 | 0.16831972 | -0.605844861 | 0.544617777 | 0.717913747 |
| ENSG00000258725 | PRC1-AS1 | 115.7334735 | -0.207277901 | 0.20550691 | -1.008617673 | 0.313158028 | 0.501385062 |
| ENSG00000258727 | AP1G2-AS1 | 397.0007684 | -0.332916333 | 0.100035718 | -3.327974631 | 0.000874798 | 0.003310179 |
| ENSG00000258733 | LINC02328 | 14.71883211 | -2.089402838 | 0.525626116 | -3.975074246 | 7.04E-05 | 0.000321962 |
| ENSG00000258738 | BAZ1A-AS1 | 93.59786429 | -0.548274902 | 0.193340103 | -2.835805376 | 0.004571029 | 0.014869457 |
| ENSG00000258741 | H2AZ2P1 | 336.5089144 | 0.135922701 | 0.112670969 | 1.206368431 | 0.227675463 | 0.400465812 |
| ENSG00000258744 | ENSG0000025 | 123.5124521 | -0.524084362 | 0.165429613 | -3.168020231 | 0.001534808 | 0.005528974 |
| ENSG00000258745 | ENSG0000025 | 1.977647235 | -1.753521252 | 1.353969577 | -1.295096494 | 0.195286944 | 0.356488274 |
| ENSG00000258748 | KIF26A-DT | 12.86137527 | -1.557255519 | 0.514619317 | -3.026033939 | 0.002477845 | 0.008545307 |
| ENSG00000258749 | ENSG0000025 | 15.66464059 | -0.241857696 | 0.454852767 | -0.531727438 | 0.594914786 | 0.756462789 |
| ENSG00000258751 | ENSG0000025 | 4.772487796 | -1.497892272 | 0.873335817 | -1.715138944 | 0.086319756 | 0.187943735 |
| ENSG00000258759 | ENSG0000025 | 22.44470443 | -0.762398554 | 0.374187786 | -2.037475789 | 0.041602386 | 0.102420984 |

| | | | | | | | |
|---|---|---|---|---|---|---|---|
| ENSG00000258760 | ENSG0000025 | 3.30417438 | -3.315002834 | 1.308905867 | -2.532651826 | 0.011320333 | 0.033144665 |
| ENSG00000258768 | ENSG0000025 | 9.520193242 | -0.135961309 | 0.567613184 | -0.239531627 | 0.810693376 | 0.899758757 |
| ENSG00000258769 | RAP1AP | 4.977020224 | 0.123120772 | 0.791264861 | 0.155599948 | 0.876348366 | 0.93573969 |
| ENSG00000258777 | HIF1A-AS1 | 11.3515595 | -0.813160171 | 0.545254264 | -1.491341242 | 0.135871926 | 0.270470495 |
| ENSG00000258778 | EIF2S2P1 | 1.902902524 | -0.595373999 | 1.287137426 | -0.462556668 | 0.643682175 | 0.791270263 |
| ENSG00000258788 | CKS1BP1 | 3.160629106 | -0.051983252 | 1.02857427 | -0.050539133 | 0.959692766 | 0.981621201 |
| ENSG00000258794 | DUX4L27 | 16.34904019 | -0.610756554 | 0.444216089 | -1.374908674 | 0.16915976 | 0.319568878 |
| ENSG00000258798 | CCDC88C-DT | 5.02806598 | 0.795698557 | 0.833091488 | 0.955115457 | 0.339519313 | 0.530665194 |
| ENSG00000258800 | ENSG0000025 | 4.730207781 | -1.760996477 | 0.857453373 | -2.053751881 | 0.039999712 | 0.099113567 |
| ENSG00000258811 | ENSG0000025 | 11.85899797 | -0.342553468 | 0.503428524 | -0.68044112 | 0.496225192 | 0.678189211 |
| ENSG00000258813 | RBM25-AS1 | 17.6000174 | -0.315814875 | 0.416415676 | -0.75841255 | 0.448204047 | 0.63636819 |
| ENSG00000258815 | LINC02820 | 43.59411771 | -0.11434732 | 0.265394854 | -0.430857337 | 0.666572106 | 0.807944843 |
| ENSG00000258824 | ENSG0000025 | 39.74974115 | -1.608552394 | 0.303433048 | -5.301177329 | 1.15E-07 | 7.42E-07 |
| ENSG00000258828 | KRT8P2 | 27.45808612 | -0.1345226 | 0.3449178 | -0.390013504 | 0.696526561 | 0.829482392 |
| ENSG00000258839 | MC1R | 283.3995901 | 0.788697734 | 0.115262691 | 6.842610802 | 7.78E-12 | 7.36E-11 |
| ENSG00000258846 | EEF1A1P33 | 1.544078456 | 0.456952752 | 1.418201732 | 0.322205749 | 0.747296829 | NA |
| ENSG00000258875 | ENSG0000025 | 1.972498947 | -0.186428566 | 1.246972178 | -0.149504993 | 0.88115517 | 0.938366635 |
| ENSG00000258876 | TGFB3-AS1 | 2.410726242 | -2.754843885 | 1.374794827 | -2.003821829 | 0.04508915 | 0.109605287 |
| ENSG00000258884 | LINC02321 | 72.89884218 | -1.743230559 | 0.226141911 | -7.708569163 | 1.27E-14 | 1.48E-13 |
| ENSG00000258890 | CEP95 | 2160.712977 | 0.189125148 | 0.066313899 | 2.851968447 | 0.004344941 | 0.014208547 |
| ENSG00000258891 | ENSG0000025 | 7.839190695 | -1.37094892 | 0.690280934 | -1.986073862 | 0.047025123 | 0.11362506 |
| ENSG00000258895 | ENSG0000025 | 104.4910325 | 0.183035177 | 0.177159895 | 1.033163725 | 0.301527286 | 0.488353285 |
| ENSG00000258904 | ENSG0000025 | 3.376311101 | 0.341222283 | 0.965797128 | 0.353306376 | 0.723858766 | 0.846416769 |
| ENSG00000258908 | ENSG0000025 | 19.27223266 | -0.763222548 | 0.431950274 | -1.766922246 | 0.077241251 | 0.171997116 |
| ENSG00000258920 | FOXN3-AS1 | 46.69079807 | -0.332408827 | 0.262758494 | -1.265073575 | 0.205844955 | 0.37129134 |
| ENSG00000258925 | NPM1P5 | 6.867358215 | 0.209104893 | 0.669159049 | 0.312489077 | 0.754668863 | 0.864498202 |
| ENSG00000258927 | ENSG0000025 | 2.29828881 | 4.553893652 | 1.523748212 | 2.988612958 | 0.002802469 | 0.009565338 |
| ENSG00000258938 | ENSG0000025 | 45.93186173 | -0.305202842 | 0.270512017 | -1.128241344 | 0.259218008 | 0.439112472 |
| ENSG00000258940 | MIA2-AS1 | 36.80173647 | -0.606559142 | 0.306066114 | -1.981791236 | 0.047502611 | 0.114633904 |
| ENSG00000258944 | ENSG0000025 | 10.82408568 | -0.115125302 | 0.546767322 | -0.210556296 | 0.833233519 | 0.911858454 |
| ENSG00000258947 | TUBB3 | 44.06334382 | 1.72075626 | 0.300842314 | 5.719794656 | 1.07E-08 | 7.65E-08 |
| ENSG00000258957 | GALNT16-AS1 | 7.412564838 | -0.206049658 | 0.660619822 | -0.311903535 | 0.755113836 | 0.864731998 |
| ENSG00000258959 | ENSG0000025 | 15.69662775 | -1.10561232 | 0.45802632 | -2.413861978 | 0.01578444 | 0.044472496 |

| | | | | | | | |
|---|---|---|---|---|---|---|---|
| ENSG00000258969 | LINC02307 | 54.21284915 | 0.561326853 | 0.241011372 | 2.329047171 | 0.019856567 | 0.054363247 |
| ENSG00000258982 | ENSG0000025 | 18.0642202 | -0.91257085 | 0.430621682 | -2.11919392 | 0.034074082 | 0.086543339 |
| ENSG00000258988 | SF3B4P1 | 7.946250723 | -0.200215717 | 0.612313609 | -0.326982308 | 0.743681267 | 0.858490754 |
| ENSG00000258995 | PARP1P2 | 64.79492763 | 0.151362788 | 0.218888933 | 0.691504984 | 0.48924825 | 0.672489708 |
| ENSG00000259001 | ENSG0000025 | 1.730875677 | -2.151299952 | 1.519851615 | -1.415467096 | 0.156931589 | NA |
| ENSG00000259006 | ENSG0000025 | 5.857955855 | 1.667241477 | 0.781239603 | 2.134097491 | 0.032834801 | 0.083953449 |
| ENSG00000259010 | NUP50P1 | 34.04390758 | -0.051741101 | 0.309995339 | -0.166909286 | 0.867441427 | 0.93096599 |
| ENSG00000259015 | ENSG0000025 | 58.82563921 | -1.929835081 | 0.26329777 | -7.329477494 | 2.31E-13 | 2.47E-12 |
| ENSG00000259020 | ENSG0000025 | 12.39816206 | 0.169611641 | 0.528083869 | 0.321183152 | 0.748071599 | 0.860436606 |
| ENSG00000259022 | DNAJC8P1 | 17.71692809 | -0.323797479 | 0.409917298 | -0.789909284 | 0.429580749 | 0.619902019 |
| ENSG00000259030 | FPGT-TNNI3K | 4.867070656 | 0.076614469 | 0.870508931 | 0.088011124 | 0.929867835 | 0.966032078 |
| ENSG00000259031 | ENSG0000025 | 2.60976141 | -3.888014368 | 1.480507234 | -2.62613669 | 0.008636013 | 0.02613935 |
| ENSG00000259032 | ENSAP2 | 1659.739471 | 0.175077905 | 0.059468322 | 2.944053211 | 0.003239443 | 0.010896308 |
| ENSG00000259033 | ENSG0000025 | 12.77581689 | -1.449399355 | 0.542648144 | -2.670974498 | 0.007563139 | 0.023254043 |
| ENSG00000259038 | ENSG0000025 | 4.364327582 | 1.429932524 | 0.900829283 | 1.58735129 | 0.112433105 | 0.233054615 |
| ENSG00000259045 | MTCO1P2 | 15.80082164 | -0.212955227 | 0.452657943 | -0.470455076 | 0.638029923 | 0.786776672 |
| ENSG00000259049 | ENSG0000025 | 30.28422507 | -1.431390466 | 0.341375817 | -4.193004873 | 2.75E-05 | 0.000133963 |
| ENSG00000259051 | HNRNPUP1 | 371.9080769 | -0.110738109 | 0.101489106 | -1.091132971 | 0.275214377 | 0.457355635 |
| ENSG00000259052 | ENSG0000025 | 6.004946817 | 0.682469396 | 0.74456332 | 0.916603568 | 0.359350414 | 0.551921431 |
| ENSG00000259065 | MIDEAS-AS1 | 60.70649337 | -0.66362357 | 0.234481528 | -2.830174199 | 0.004652267 | 0.015105874 |
| ENSG00000259066 | ENSG0000025 | 4.888727026 | 0.094082985 | 0.82130515 | 0.11455302 | 0.908799419 | 0.95431011 |
| ENSG00000259070 | LINC00639 | 14.85372461 | -1.58204654 | 0.489933039 | -3.229107682 | 0.001241771 | 0.004567297 |
| ENSG00000259071 | ENSG0000025 | 1.673242232 | 0.340872787 | 1.398653074 | 0.243715038 | 0.807451526 | NA |
| ENSG00000259078 | PTBP1P | 12.60794639 | -1.347770404 | 0.57936744 | -2.326279163 | 0.020003663 | 0.054681128 |
| ENSG00000259079 | ENSG0000025 | 24.17519606 | -0.401980585 | 0.366413241 | -1.097068938 | 0.272611257 | 0.454715498 |
| ENSG00000259081 | ENSG0000025 | 8.578160608 | -0.326103486 | 0.601336057 | -0.54229824 | 0.587613068 | 0.751132537 |
| ENSG00000259083 | PNN-AS1 | 22.83511585 | -0.283215185 | 0.376633291 | -0.751965351 | 0.452071896 | 0.639919122 |
| ENSG00000259086 | KIAA1143P1 | 76.72184481 | -0.04719745 | 0.200359028 | -0.235564379 | 0.813770717 | 0.901448243 |
| ENSG00000259088 | ENSG0000025 | 11.44587689 | -1.056433898 | 0.537044422 | -1.967125724 | 0.049168718 | 0.118107895 |
| ENSG00000259096 | SHLD2P2 | 21.39034719 | -0.396561872 | 0.378338839 | -1.048165906 | 0.294562176 | 0.48053309 |
| ENSG00000259100 | COILP1 | 4.572773083 | -0.3450023 | 0.829103769 | -0.416114741 | 0.677326044 | 0.816449974 |
| ENSG00000259102 | SMARCE1P3 | 4.827067161 | -0.906556541 | 0.880700891 | -1.029358038 | 0.30331146 | 0.490496406 |
| ENSG00000259113 | ENSG0000025 | 3.73688817 | -1.963935821 | 1.009047837 | -1.946325782 | 0.051615618 | 0.123083398 |

| | | | | | | | |
|---|---|---|---|---|---|---|---|
| ENSG00000259118 | ENSG0000025 | 22.11404535 | -1.111337554 | 0.399815665 | -2.779624843 | 0.005442173 | 0.017353829 |
| ENSG00000259137 | ENSG0000025 | 37.73935923 | 0.051393821 | 0.315867299 | 0.162707 | 0.870749135 | 0.932944751 |
| ENSG00000259144 | RANBP20P | 9.442780101 | -0.200734506 | 0.594578191 | -0.337608256 | 0.73565842 | 0.854264953 |
| ENSG00000259146 | SIPA1L1-AS1 | 129.5321171 | -0.810520142 | 0.177167205 | -4.57488811 | 4.76E-06 | 2.55E-05 |
| ENSG00000259151 | CAP2P1 | 2.708182017 | -0.960449776 | 1.095928943 | -0.876379606 | 0.380823701 | 0.573272243 |
| ENSG00000259153 | MAP3K9-DT | 244.1108859 | -0.818148717 | 0.120692005 | -6.778814551 | 1.21E-11 | 1.12E-10 |
| ENSG00000259158 | ADAM20P1 | 10.2536578 | -0.634958413 | 0.58934395 | -1.077398712 | 0.281302179 | 0.464717845 |
| ENSG00000259165 | DDX18P1 | 5.741979571 | -0.235479907 | 0.742213293 | -0.317267165 | 0.75104089 | 0.862460033 |
| ENSG00000259172 | SNRPA1-DT | 27.72678012 | 0.150134175 | 0.351610022 | 0.426990604 | 0.669386171 | 0.810007484 |
| ENSG00000259175 | ENSG0000025 | 4.011171863 | -0.411787825 | 0.892671983 | -0.461298028 | 0.6445848 | 0.791795197 |
| ENSG00000259179 | UBE2CP4 | 7.999716051 | -1.021948731 | 0.632872467 | -1.614778308 | 0.106358712 | 0.222778384 |
| ENSG00000259185 | ENSG0000025 | 39.33283266 | -1.505099451 | 0.30019118 | -5.013803041 | 5.34E-07 | 3.20E-06 |
| ENSG00000259188 | ENSG0000025 | 5.903194479 | -3.549570012 | 1.024694602 | -3.464027239 | 0.000532153 | 0.002096082 |
| ENSG00000259192 | ENSG0000025 | 2.41131035 | -0.637796986 | 1.144954622 | -0.557050012 | 0.57749326 | 0.743451272 |
| ENSG00000259194 | ENSG0000025 | 2.742176772 | 0.006335667 | 1.052940714 | 0.006017117 | 0.995199064 | 0.997068024 |
| ENSG00000259198 | ENSG0000025 | 9.383276613 | -0.434187851 | 0.604708684 | -0.718011602 | 0.472750133 | 0.658343231 |
| ENSG00000259201 | ENSG0000025 | 32.38329121 | -2.028823683 | 0.35378074 | -5.734692299 | 9.77E-09 | 7.04E-08 |
| ENSG00000259205 | PRKXP1 | 13.49308027 | 0.743690483 | 0.503203208 | 1.477912842 | 0.139431109 | 0.275629863 |
| ENSG00000259209 | ENSG0000025 | 6.664048655 | 0.611027179 | 0.684424466 | 0.892760573 | 0.371985404 | 0.56467345 |
| ENSG00000259211 | ENSG0000025 | 8.968921717 | -0.17722988 | 0.598267507 | -0.296238519 | 0.767047934 | 0.872778717 |
| ENSG00000259215 | LINC02896 | 3.505251984 | -0.876606222 | 0.931106883 | -0.941466805 | 0.346465693 | 0.537239623 |
| ENSG00000259216 | ENSG0000025 | 3.411636425 | -0.581799877 | 0.97373456 | -0.597493301 | 0.550178078 | 0.722460026 |
| ENSG00000259234 | ANKRD34C-AS | 2.256141031 | -0.126742706 | 1.20442134 | -0.105231202 | 0.916192352 | 0.958650454 |
| ENSG00000259238 | ENSG0000025 | 4.380440295 | -1.554803488 | 0.929927631 | -1.671961813 | 0.094531852 | 0.202442186 |
| ENSG00000259240 | MIR4713HG | 290.5775354 | -1.092626911 | 0.117814227 | -9.274150792 | 1.79E-20 | 3.07E-19 |
| ENSG00000259244 | ENSG0000025 | 2.284042175 | 0.717155888 | 1.205077334 | 0.595111922 | 0.55176866 | 0.723620687 |
| ENSG00000259248 | USP3-AS1 | 92.00644757 | -0.250877503 | 0.18763524 | -1.337048966 | 0.181206654 | 0.336616782 |
| ENSG00000259251 | VPS13C-DT | 2.159661793 | 1.757398904 | 1.371166466 | 1.281681653 | 0.199954343 | 0.363064163 |
| ENSG00000259271 | ANKRD62P1 | 1.778536962 | -1.525784435 | 1.426301884 | -1.069748594 | 0.284732487 | NA |
| ENSG00000259275 | ENSG0000025 | 64.76639409 | 1.369086499 | 0.243722811 | 5.617391719 | 1.94E-08 | 1.35E-07 |
| ENSG00000259286 | ENSG0000025 | 2.665570207 | 1.132689925 | 1.115205113 | 1.015678562 | 0.30978248 | 0.497752336 |
| ENSG00000259287 | ENSG0000025 | 7.974595952 | 0.474130455 | 0.646617358 | 0.733247336 | 0.463407589 | 0.649830069 |
| ENSG00000259291 | ZNF710-AS1 | 299.2360311 | 0.514488692 | 0.112199252 | 4.585491276 | 4.53E-06 | 2.43E-05 |

| | | | | | | | |
|---|---|---|---|---|---|---|---|
| ENSG00000259295 | CSPG4P12 | 2.347295809 | -0.30924097 | 1.233952087 | -0.250610192 | 0.802115501 | 0.894662789 |
| ENSG00000259299 | ENSG0000025 | 13.24163529 | 0.281990515 | 0.539644819 | 0.52254836 | 0.601288585 | 0.761191994 |
| ENSG00000259305 | ZHX1-C8orf76 | 3.024060227 | 0.046517484 | 1.05633844 | 0.044036534 | 0.964875282 | 0.984122576 |
| ENSG00000259306 | ENSG0000025 | 110.0575171 | -0.259500417 | 0.169494325 | -1.531027172 | 0.125762677 | 0.254641967 |
| ENSG00000259314 | ENSG0000025 | 4.504812994 | -0.086216843 | 0.81607567 | -0.105648099 | 0.915861561 | 0.958435482 |
| ENSG00000259315 | ACTG1P17 | 9.17901515 | -1.121735425 | 0.600377406 | -1.868383811 | 0.061708592 | 0.142833655 |
| ENSG00000259318 | HMGN1P1 | 13.96933759 | -1.184722969 | 0.499917704 | -2.369835993 | 0.017795977 | 0.049423695 |
| ENSG00000259319 | JDP2-AS1 | 97.3878586 | -0.918577163 | 0.182632353 | -5.029651909 | 4.91E-07 | 2.96E-06 |
| ENSG00000259321 | ENSG0000025 | 14.30657309 | -0.909034838 | 0.479317833 | -1.896517876 | 0.057891598 | 0.135499772 |
| ENSG00000259322 | ENSG0000025 | 7.426679458 | -0.459536979 | 0.6379351 | -0.720350674 | 0.471309112 | 0.657148912 |
| ENSG00000259327 | ENSG0000025 | 21.50370926 | -1.433782196 | 0.415624809 | -3.449703106 | 0.000561203 | 0.002199933 |
| ENSG00000259330 | INAFM2 | 436.128793 | 0.198616942 | 0.093209639 | 2.13086269 | 0.033100456 | 0.084510235 |
| ENSG00000259332 | ST20-MTHFS | 6.368360723 | -0.665160799 | 0.705559173 | -0.942742756 | 0.345812498 | 0.536750313 |
| ENSG00000259343 | TMC3-AS1 | 11.02027063 | -1.358078104 | 0.544896722 | -2.492358733 | 0.01268978 | 0.036658299 |
| ENSG00000259345 | ENSG0000025 | 50.1129255 | -0.447179058 | 0.249037269 | -1.795631071 | 0.072553211 | 0.163510233 |
| ENSG00000259347 | ENSG0000025 | 10.05954726 | 0.197563365 | 0.54460451 | 0.362764835 | 0.716780563 | 0.842493524 |
| ENSG00000259349 | APPBP2-DT | 121.3843867 | 0.641253013 | 0.164290419 | 3.903167433 | 9.49E-05 | 0.000424473 |
| ENSG00000259352 | ENSG0000025 | 6.071516222 | -0.725462822 | 0.715027899 | -1.014593729 | 0.31029953 | 0.498303901 |
| ENSG00000259357 | ENSG0000025 | 168.7464619 | -0.082782817 | 0.165749114 | -0.499446514 | 0.617464858 | 0.773221455 |
| ENSG00000259358 | DEPDC1P1 | 36.94776996 | -0.319837533 | 0.301045039 | -1.062424193 | 0.288043155 | 0.472013 |
| ENSG00000259362 | ENSG0000025 | 13.41590963 | 0.271309207 | 0.482155319 | 0.562700848 | 0.573638607 | 0.740324593 |
| ENSG00000259366 | ENSG0000025 | 24.84945464 | -0.179385647 | 0.362371038 | -0.49503307 | 0.620576777 | 0.775382967 |
| ENSG00000259370 | ENSG0000025 | 3.657621264 | -0.685040399 | 0.978794043 | -0.699882068 | 0.484000957 | 0.667928891 |
| ENSG00000259375 | ENSG0000025 | 6.355860458 | -0.778253325 | 0.703140169 | -1.106825295 | 0.26836947 | 0.449874922 |
| ENSG00000259376 | ENSG0000025 | 1.804640217 | 0.101857977 | 1.278529811 | 0.07966805 | 0.936501271 | 0.969552268 |
| ENSG00000259377 | ENSG0000025 | 1.870787901 | 0.352889401 | 1.296149852 | 0.272259724 | 0.785422322 | 0.883215101 |
| ENSG00000259378 | DCAF13P3 | 25.63908085 | -1.654803598 | 0.38734727 | -4.272144731 | 1.94E-05 | 9.63E-05 |
| ENSG00000259388 | ENSG0000025 | 10.81981434 | -0.268307029 | 0.53125118 | -0.505047403 | 0.613525542 | 0.770479304 |
| ENSG00000259394 | ENSG0000025 | 9.229420395 | -0.198085081 | 0.578529367 | -0.342394168 | 0.732054277 | 0.85176404 |
| ENSG00000259397 | ENSG0000025 | 21.73009687 | -0.188432459 | 0.398589482 | -0.472748198 | 0.636392836 | 0.78589856 |
| ENSG00000259404 | EFL1P1 | 3.767585349 | -0.802099849 | 0.909316099 | -0.882091332 | 0.377727419 | 0.570483047 |
| ENSG00000259405 | ISCA1P4 | 7.651301823 | 0.156840575 | 0.636134316 | 0.246552609 | 0.805254485 | 0.896594568 |
| ENSG00000259407 | ENSG0000025 | 9.4200468 | -0.103302149 | 0.568869393 | -0.181592032 | 0.855902902 | 0.925278632 |

| | | | | | | | |
|---|---|---|---|---|---|---|---|
| ENSG00000259408 | ENSG0000025 | 13.35730186 | 0.41053242 | 0.492923486 | 0.832852221 | 0.404928079 | 0.597789441 |
| ENSG00000259409 | BMF-AS1 | 11.07365607 | -0.383547987 | 0.526010434 | -0.729164219 | 0.465901216 | 0.651756481 |
| ENSG00000259411 | HNRNPA1P45 | 2.549797993 | 1.386498235 | 1.14559249 | 1.210289214 | 0.226167935 | 0.398342014 |
| ENSG00000259416 | LINC02883 | 3.480332089 | -0.222642269 | 0.967957406 | -0.230012465 | 0.818082083 | 0.90364036 |
| ENSG00000259419 | HNRNPCP3 | 11.60033487 | -0.286030264 | 0.518761015 | -0.551371934 | 0.581378736 | 0.746408563 |
| ENSG00000259422 | ENSG0000025 | 6.872521648 | -0.550977087 | 0.665182788 | -0.828309297 | 0.407495359 | 0.600343172 |
| ENSG00000259424 | IRAIN | 6.011225013 | 1.641806722 | 0.780563999 | 2.103359524 | 0.035434354 | 0.089542198 |
| ENSG00000259426 | KIF23-AS1 | 54.8824205 | -0.843446969 | 0.242883479 | -3.472640345 | 0.000515365 | 0.002033801 |
| ENSG00000259429 | UBE2Q2P2 | 82.43510004 | 0.046242901 | 0.211335754 | 0.218812481 | 0.826796124 | 0.908541056 |
| ENSG00000259431 | THTPA | 231.6271228 | -0.087786424 | 0.131403148 | -0.668069417 | 0.504089283 | 0.685059724 |
| ENSG00000259438 | MAPK6-DT | 46.11657227 | -0.515184634 | 0.267352334 | -1.926987609 | 0.053981174 | 0.127971962 |
| ENSG00000259439 | LINC01833 | 2.801744125 | -0.337330801 | 1.023098259 | -0.329714959 | 0.74161535 | 0.857515143 |
| ENSG00000259448 | LINC02352 | 1.745563263 | -1.432052126 | 1.409291954 | -1.016150076 | 0.309557926 | NA |
| ENSG00000259456 | ADNP-AS1 | 49.07624785 | -0.325938169 | 0.259609892 | -1.2554921 | 0.209300174 | 0.376133647 |
| ENSG00000259457 | ENSG0000025 | 15.42028597 | -3.978757501 | 0.715292784 | -5.562418059 | 2.66E-08 | 1.83E-07 |
| ENSG00000259459 | LINC02568 | 11.32246749 | -0.540885771 | 0.52568587 | -1.028914419 | 0.303519892 | 0.490679798 |
| ENSG00000259460 | ENSG0000025 | 2.392409285 | 0.617399241 | 1.167888911 | 0.52864552 | 0.597051376 | 0.75769879 |
| ENSG00000259470 | KRT8P9 | 44.19055129 | -0.131386325 | 0.279208917 | -0.470566363 | 0.637950433 | 0.786742264 |
| ENSG00000259479 | SORD2P | 95.62852004 | -0.619999165 | 0.18934938 | -3.274365969 | 0.001058994 | 0.003945001 |
| ENSG00000259485 | LINC02253 | 112.2684532 | -0.04232235 | 0.173788953 | -0.243527275 | 0.807596959 | 0.897981454 |
| ENSG00000259488 | DUT-AS1 | 28.10377963 | -0.372844496 | 0.333587094 | -1.117682619 | 0.263702569 | 0.444379411 |
| ENSG00000259493 | TFDP1P3 | 7.681031073 | -0.218950631 | 0.639804186 | -0.342215065 | 0.732189049 | 0.851857055 |
| ENSG00000259494 | MRPL46 | 95.88638935 | 0.440588571 | 0.181786585 | 2.423658323 | 0.015365054 | 0.043461694 |
| ENSG00000259495 | ARNT2-DT | 149.7503658 | -1.240855506 | 0.152536502 | -8.134810278 | 4.13E-16 | 5.37E-15 |
| ENSG00000259498 | TPM1-AS | 4.957240852 | -0.550751461 | 0.834066744 | -0.660320609 | 0.509048107 | 0.689192557 |
| ENSG00000259500 | KRT8P24 | 5.772274024 | -0.752181379 | 0.75276942 | -0.999218829 | 0.317688696 | 0.506834877 |
| ENSG00000259502 | ENSG0000025 | 3.990854455 | -1.708524344 | 0.930934481 | -1.835278828 | 0.066464386 | 0.15208908 |
| ENSG00000259515 | ENSG0000025 | 16.38911428 | -0.961630465 | 0.46271402 | -2.078239307 | 0.037687323 | 0.094413804 |
| ENSG00000259516 | ANP32AP1 | 53.01786058 | -0.321384843 | 0.258399407 | -1.243752243 | 0.213590762 | 0.381810339 |
| ENSG00000259520 | SLC28A2-AS1 | 2.062379658 | -0.247498213 | 1.218774282 | -0.203071411 | 0.839079217 | 0.915286602 |
| ENSG00000259522 | ENSG0000025 | 1.82499961 | 0.092123193 | 1.288105551 | 0.071518358 | 0.942985215 | 0.97291737 |
| ENSG00000259523 | ENSG0000025 | 11.49396225 | 0.114737266 | 0.512109396 | 0.224048352 | 0.82271968 | 0.906215744 |
| ENSG00000259527 | LINC00052 | 912.1518045 | 3.241982325 | 0.267941149 | 12.09960598 | 1.06E-33 | 3.40E-32 |

| | | | | | | | |
|---|---|---|---|---|---|---|---|
| ENSG00000259529 | ENSG0000025 | 32.74690072 | -0.112746021 | 0.307609352 | -0.366523385 | 0.713974566 | 0.840918855 |
| ENSG00000259531 | ENSG0000025 | 200.8490663 | -1.401687576 | 0.147819019 | -9.482457594 | 2.48E-21 | 4.49E-20 |
| ENSG00000259536 | ENSG0000025 | 6.897972827 | -0.18169623 | 0.666721846 | -0.272521788 | 0.785220841 | 0.883118404 |
| ENSG00000259539 | ENSG0000025 | 8.685250297 | 0.286264826 | 0.60310541 | 0.474651397 | 0.635035466 | 0.784856074 |
| ENSG00000259544 | ENSG0000025 | 6.658275375 | -0.765836636 | 0.673353919 | -1.137346371 | 0.255393517 | 0.434784955 |
| ENSG00000259548 | ENSG0000025 | 2.053356824 | 2.245987028 | 1.434567378 | 1.565619756 | 0.117437662 | 0.241507841 |
| ENSG00000259556 | ENSG0000025 | 49.38953304 | -1.422998509 | 0.263165465 | -5.407238796 | 6.40E-08 | 4.22E-07 |
| ENSG00000259557 | HMGN1P26 | 1.549810851 | 1.052832841 | 1.523859874 | 0.690898723 | 0.489629189 | NA |
| ENSG00000259562 | ENSG0000025 | 38.82865593 | -0.077116607 | 0.280916321 | -0.274518072 | 0.783686517 | 0.882691054 |
| ENSG00000259564 | ENSG0000025 | 1.802205396 | -0.90239634 | 1.377622289 | -0.65503901 | 0.512442638 | 0.69208242 |
| ENSG00000259570 | ENSG0000025 | 6.774658863 | -0.419240008 | 0.687421113 | -0.609873628 | 0.541945523 | 0.715771219 |
| ENSG00000259577 | CERNA1 | 135.4250295 | -1.032833582 | 0.175280508 | -5.892461137 | 3.80E-09 | 2.87E-08 |
| ENSG00000259581 | TYRO3P | 8.475668642 | -0.60531025 | 0.657481097 | -0.920650424 | 0.357232967 | 0.549552325 |
| ENSG00000259583 | ALDH1A3-AS1 | 7.779956597 | -0.23334413 | 0.660692988 | -0.353180878 | 0.723952842 | 0.846456585 |
| ENSG00000259585 | RBM17P4 | 16.33632378 | -1.148606146 | 0.494753622 | -2.321571982 | 0.020255994 | 0.055258949 |
| ENSG00000259592 | PRELID1P4 | 3.04572711 | 0.914777239 | 1.028782436 | 0.889184347 | 0.373904014 | 0.566648482 |
| ENSG00000259595 | CTDSPL2-DT | 11.27591308 | -0.535030159 | 0.527405825 | -1.014456294 | 0.310365074 | 0.498346814 |
| ENSG00000259601 | ENSG0000025 | 2.109117274 | 0.046140211 | 1.176957329 | 0.03920296 | 0.968728574 | 0.985876114 |
| ENSG00000259605 | ENSG0000025 | 42.22405494 | 0.411175724 | 0.27426853 | 1.499172083 | 0.133828995 | 0.267285143 |
| ENSG00000259612 | EEF1A1P22 | 56.62693211 | 0.08425564 | 0.249091445 | 0.338251842 | 0.735173411 | 0.853967465 |
| ENSG00000259619 | ENSG0000025 | 2.190457461 | -1.37421407 | 1.260354801 | -1.090339061 | 0.275563815 | 0.457842682 |
| ENSG00000259620 | ENSG0000025 | 2.025915234 | 2.302487206 | 1.440392705 | 1.598513515 | 0.10992874 | 0.228600729 |
| ENSG00000259627 | ENSG0000025 | 8.203404547 | -0.576974505 | 0.645515409 | -0.89381988 | 0.371418271 | 0.564017991 |
| ENSG00000259635 | ENSG0000025 | 1.891595708 | 0.89641411 | 1.313060386 | 0.682690697 | 0.494802309 | 0.67711556 |
| ENSG00000259642 | ST20-AS1 | 106.3343897 | -0.161064849 | 0.176578842 | -0.912141269 | 0.361694356 | 0.554296782 |
| ENSG00000259648 | ENSG0000025 | 2.572000637 | -1.657249818 | 1.154032417 | -1.436051357 | 0.150987728 | 0.293108105 |
| ENSG00000259657 | PIGHP1 | 17.37754012 | -1.065549015 | 0.459458892 | -2.319138957 | 0.020387503 | 0.055558247 |
| ENSG00000259659 | ENSG0000025 | 3.424466161 | -1.099688055 | 1.002914729 | -1.096492078 | 0.272863488 | 0.454862706 |
| ENSG00000259661 | ENSG0000025 | 25.25032968 | -0.572892295 | 0.376876115 | -1.520107728 | 0.128483903 | 0.258909059 |
| ENSG00000259663 | ISL1-DT | 4.819974452 | -1.521556183 | 0.863580296 | -1.761916281 | 0.078083443 | 0.173350836 |
| ENSG00000259668 | ENSG0000025 | 34.40693782 | -1.618750368 | 0.324182101 | -4.993336655 | 5.93E-07 | 3.53E-06 |
| ENSG00000259673 | IQCH-AS1 | 489.4577544 | 0.004395153 | 0.088132769 | 0.049869683 | 0.960226237 | 0.981840422 |
| ENSG00000259678 | ENSG0000025 | 15.96555802 | -2.14781183 | 0.493259638 | -4.354323087 | 1.33E-05 | 6.77E-05 |

| | | | | | | | |
|---|---|---|---|---|---|---|---|
| ENSG00000259684 | ENSG0000025 | 4.639593176 | -2.306197865 | 0.939891602 | -2.453684936 | 0.014140081 | 0.04036504 |
| ENSG00000259685 | IDH2-DT | 2.777227755 | -0.942917683 | 1.065958255 | -0.884572804 | 0.37638708 | 0.569311728 |
| ENSG00000259687 | LINC01220 | 3.39980609 | -0.313752203 | 0.957289855 | -0.327750474 | 0.743100337 | 0.858344498 |
| ENSG00000259692 | ENSG0000025 | 29.46447538 | 0.05784661 | 0.319508429 | 0.181048775 | 0.856329292 | 0.925565113 |
| ENSG00000259696 | ENSG0000025 | 7.491793519 | 0.671252633 | 0.654061886 | 1.026283058 | 0.304758181 | 0.492161398 |
| ENSG00000259699 | HMGB1P8 | 5.830914332 | -0.591406903 | 0.717595594 | -0.824150689 | 0.409853942 | 0.602386559 |
| ENSG00000259702 | ENSG0000025 | 3.365321564 | -0.51812672 | 1.049856935 | -0.493521262 | 0.62164432 | 0.776040526 |
| ENSG00000259705 | FBN1-DT | 1.805638991 | 0.597554646 | 1.334120112 | 0.447901685 | 0.654224152 | 0.799227929 |
| ENSG00000259706 | HSP90B2P | 57.38639218 | -0.259685519 | 0.244683677 | -1.06131117 | 0.288548511 | 0.472537325 |
| ENSG00000259707 | ENSG0000025 | 13.49223008 | 0.140000345 | 0.491484239 | 0.284852155 | 0.775757413 | 0.878244833 |
| ENSG00000259709 | ENSG0000025 | 9.044485248 | -1.398890418 | 0.602644436 | -2.321253353 | 0.020273174 | 0.055292666 |
| ENSG00000259711 | ENSG0000025 | 22.68869896 | -0.039062911 | 0.370014928 | -0.105571176 | 0.915922595 | 0.95845563 |
| ENSG00000259715 | ENSG0000025 | 2.222583225 | -1.894832733 | 1.29338434 | -1.465019077 | 0.142915713 | 0.280788494 |
| ENSG00000259723 | ENSG0000025 | 57.18167617 | 0.28826555 | 0.236098313 | 1.220955568 | 0.222102843 | 0.393135743 |
| ENSG00000259727 | ENSG0000025 | 3.853817458 | -0.101580895 | 0.908747312 | -0.111781233 | 0.910996868 | 0.955316516 |
| ENSG00000259728 | LINC00933 | 40.11847825 | 0.637356518 | 0.283079521 | 2.251510518 | 0.024353221 | 0.065028856 |
| ENSG00000259736 | CRTC3-AS1 | 18.88510095 | 0.12534523 | 0.400648736 | 0.312855673 | 0.754390316 | 0.864498202 |
| ENSG00000259746 | HSPE1P3 | 97.07897093 | 0.121365691 | 0.189719414 | 0.6397115 | 0.522360178 | 0.69916229 |
| ENSG00000259750 | NSA2P4 | 8.464517076 | 0.717325318 | 0.613179356 | 1.169845839 | 0.242063013 | 0.418299474 |
| ENSG00000259751 | TUBAP11 | 2.537021708 | 0.926769309 | 1.145072852 | 0.809354014 | 0.418311546 | 0.610291228 |
| ENSG00000259762 | ENSG0000025 | 2.243387028 | 0.293598647 | 1.264774772 | 0.232135123 | 0.816433066 | 0.902775211 |
| ENSG00000259767 | ENSG0000025 | 1.842973956 | 0.031070192 | 1.323311825 | 0.023479116 | 0.981268097 | 0.99136792 |
| ENSG00000259768 | ZFHX3-AS1 | 310.0975925 | -0.18174253 | 0.107461123 | -1.691239814 | 0.09079101 | 0.195857228 |
| ENSG00000259773 | ENSG0000025 | 7.794918744 | -2.190381725 | 0.710369805 | -3.083438666 | 0.002046233 | 0.007176362 |
| ENSG00000259775 | EIF5-DT | 10.69568281 | 0.056122806 | 0.536839594 | 0.104542971 | 0.916738469 | 0.95891724 |
| ENSG00000259780 | ENSG0000025 | 18.96146718 | 0.299101449 | 0.407758261 | 0.733526399 | 0.463237433 | 0.649750334 |
| ENSG00000259781 | HMGB1P6 | 1251.058892 | -0.420034998 | 0.069715428 | -6.024993491 | 1.69E-09 | 1.32E-08 |
| ENSG00000259790 | ANP32BP1 | 166.2961299 | -0.147276699 | 0.15023201 | -0.980328354 | 0.326924063 | 0.517430524 |
| ENSG00000259793 | ENSG0000025 | 6.627144927 | -2.110179086 | 0.758566132 | -2.781799764 | 0.005405838 | 0.017249955 |
| ENSG00000259802 | MARCHF6-DT | 66.24603497 | 0.063022967 | 0.218342376 | 0.288642854 | 0.772854698 | 0.876182887 |
| ENSG00000259803 | SLC22A31 | 9.831294417 | 2.060531955 | 0.653289608 | 3.154086535 | 0.001610013 | 0.005779018 |
| ENSG00000259804 | CTCF-DT | 3.252415606 | 1.401816901 | 1.073153262 | 1.306259741 | 0.191464243 | 0.351431408 |
| ENSG00000259810 | ENSG0000025 | 1.958806758 | 0.772519696 | 1.309942776 | 0.589735453 | 0.555368023 | 0.72647217 |

| | | | | | | | |
|---|---|---|---|---|---|---|---|
| ENSG00000259813 | ENSG0000025 | 2.898133437 | -0.80686286 | 1.041405988 | -0.774782236 | 0.438468348 | 0.627947553 |
| ENSG00000259818 | CZIB-DT | 2.636309075 | 0.165007099 | 1.059106355 | 0.155798422 | 0.876191914 | 0.935703297 |
| ENSG00000259826 | CDK13-DT | 22.28008161 | 0.89392677 | 0.381919945 | 2.340612953 | 0.019252114 | 0.052899516 |
| ENSG00000259838 | ELOCP2 | 88.06176617 | 0.421461147 | 0.190456218 | 2.21290306 | 0.02690433 | 0.070761904 |
| ENSG00000259843 | HEATR3-AS1 | 1.774387448 | 0.630560456 | 1.297553213 | 0.485961154 | 0.626994709 | NA |
| ENSG00000259845 | HERC2P10 | 6.977146448 | -1.289258804 | 0.699217949 | -1.84385828 | 0.065203797 | 0.149577509 |
| ENSG00000259856 | RAB43P1 | 85.96802183 | 0.230703712 | 0.199590998 | 1.155882354 | 0.247729282 | 0.425240177 |
| ENSG00000259865 | ENSG0000025 | 29.2214866 | -0.144350488 | 0.344413223 | -0.419120053 | 0.675128396 | 0.814442859 |
| ENSG00000259867 | DYNLRB2-AS1 | 7.400655312 | -1.237647393 | 0.682662005 | -1.812972428 | 0.069836074 | 0.158288661 |
| ENSG00000259868 | ENSG0000025 | 7.476642022 | -2.30230034 | 0.810085768 | -2.842045165 | 0.004482514 | 0.014616755 |
| ENSG00000259877 | ZNF778-DT | 217.6822801 | 0.524409509 | 0.12605237 | 4.1602511 | 3.18E-05 | 0.00015356 |
| ENSG00000259881 | ENSG0000025 | 10.61469975 | -0.358532755 | 0.551560503 | -0.65003341 | 0.515670641 | 0.6944375 |
| ENSG00000259882 | ENSG0000025 | 2.908633202 | 0.835291151 | 1.03624667 | 0.806073664 | 0.420200376 | 0.611481643 |
| ENSG00000259884 | NR4A1AS | 3.564955031 | -0.37221719 | 0.955303928 | -0.38963222 | 0.696808524 | 0.829719188 |
| ENSG00000259891 | ENSG0000025 | 97.98996217 | 0.412774179 | 0.18100567 | 2.280448882 | 0.022581079 | 0.060799291 |
| ENSG00000259895 | TEDC2-AS1 | 3.49151781 | 0.960055738 | 0.970810056 | 0.988922326 | 0.322701146 | 0.512656808 |
| ENSG00000259905 | PWRN1 | 3.530868925 | 5.17255197 | 1.408126445 | 3.673357593 | 0.000239384 | 0.00100118 |
| ENSG00000259916 | AQP7B | 4.251875226 | 1.9555449 | 0.962676473 | 2.031362514 | 0.042218234 | 0.103736749 |
| ENSG00000259917 | HNRNPLP2 | 1335.557658 | -0.087034676 | 0.060997218 | -1.426863035 | 0.153619378 | 0.296685344 |
| ENSG00000259918 | NDUFA5P11 | 72.21262972 | 0.10678781 | 0.214273724 | 0.498370997 | 0.618222575 | 0.773663514 |
| ENSG00000259920 | ENSG0000025 | 3.166225483 | -0.927438363 | 1.04488716 | -0.887596669 | 0.374757746 | 0.567579238 |
| ENSG00000259924 | ENSG0000025 | 21.12800712 | -0.42004977 | 0.405970199 | -1.034681291 | 0.300817776 | 0.487582829 |
| ENSG00000259933 | PKD1-AS1 | 3.021636547 | -0.223137176 | 1.0109239 | -0.220725987 | 0.825305796 | 0.907816588 |
| ENSG00000259935 | ENSG0000025 | 3.150151623 | -0.956864701 | 1.002031411 | -0.954924856 | 0.339615699 | 0.530711987 |
| ENSG00000259937 | ENSG0000025 | 2.87836183 | -1.054874945 | 1.077672997 | -0.978845112 | 0.327656516 | 0.51794656 |
| ENSG00000259940 | ENSG0000025 | 14.6160971 | 0.998340696 | 0.493330237 | 2.023676275 | 0.043003471 | 0.105316497 |
| ENSG00000259943 | ZMPSTE24-DT | 227.3932134 | -0.075526024 | 0.126802216 | -0.595620695 | 0.551428648 | 0.723298828 |
| ENSG00000259945 | ENSG0000025 | 5.218658638 | 0.193732751 | 0.754672788 | 0.256710927 | 0.797401936 | 0.89183367 |
| ENSG00000259948 | ENSG0000025 | 27.07970878 | -0.195683233 | 0.39547448 | -0.494806222 | 0.620736912 | 0.775498571 |
| ENSG00000259952 | ENSG0000025 | 141.4658258 | -0.526561875 | 0.17960671 | -2.931749452 | 0.003370586 | 0.011300969 |
| ENSG00000259953 | LINC02977 | 42.77060054 | 0.066538717 | 0.28077061 | 0.236986046 | 0.812667616 | 0.901043746 |
| ENSG00000259956 | RBM15B | 3657.098899 | -0.021350424 | 0.053465632 | -0.399329876 | 0.689650156 | 0.824632504 |
| ENSG00000259959 | ENSG0000025 | 72.98744294 | 0.35154355 | 0.207612917 | 1.693264344 | 0.090405162 | 0.195153328 |

| | | | | | | | |
|---|---|---|---|---|---|---|---|
| ENSG00000259972 | ENSG0000025 | 296.9435939 | 0.287685477 | 0.11446112 | 2.513390373 | 0.011957694 | 0.034837375 |
| ENSG00000259977 | ENSG0000025 | 2.306309191 | 3.488682502 | 1.514535676 | 2.303466704 | 0.021252598 | 0.05760097 |
| ENSG00000259985 | ENSG0000025 | 60.24514442 | 0.494096468 | 0.231025058 | 2.138713744 | 0.032458859 | 0.083177639 |
| ENSG00000259993 | ENSG0000025 | 5.982679173 | 0.236863763 | 0.718457322 | 0.329683831 | 0.741638872 | 0.857515143 |
| ENSG00000259994 | ENSG0000025 | 13.23210118 | -0.415395557 | 0.504658468 | -0.823122137 | 0.410438541 | 0.602953271 |
| ENSG00000260000 | ENSG0000026 | 71.52318368 | 0.172691082 | 0.211668441 | 0.815856538 | 0.414582192 | 0.606692746 |
| ENSG00000260001 | TGFBR3L | 52.34825915 | -0.169525291 | 0.247746137 | -0.684270168 | 0.493804581 | 0.676438565 |
| ENSG00000260005 | ENSG0000026 | 13.36254928 | -0.396072762 | 0.494817569 | -0.800441996 | 0.423454758 | 0.614346834 |
| ENSG00000260007 | ENSG0000026 | 53.57751391 | 0.383589133 | 0.239317722 | 1.602844658 | 0.108968955 | 0.226981551 |
| ENSG00000260011 | ENSG0000026 | 4.929825955 | 1.616411934 | 0.851746565 | 1.897761612 | 0.057727492 | 0.135143246 |
| ENSG00000260017 | ENSG0000026 | 2.70777103 | -1.678566373 | 1.182715941 | -1.419247271 | 0.155826939 | 0.299878956 |
| ENSG00000260018 | MBTPS1-DT | 81.85578932 | 0.67897813 | 0.201393152 | 3.371406253 | 0.000747855 | 0.002866465 |
| ENSG00000260025 | CRIM1-DT | 40.34119887 | 0.833905278 | 0.28763693 | 2.899159288 | 0.003741648 | 0.012410844 |
| ENSG00000260027 | HOXB7 | 93.05808167 | -0.073344798 | 0.189316418 | -0.387419108 | 0.69844596 | 0.830819306 |
| ENSG00000260032 | NORAD | 7200.49983 | -0.869015566 | 0.047839353 | -18.16528671 | 9.72E-74 | 9.82E-72 |
| ENSG00000260036 | ENSG0000026 | 5.447004386 | 0.513887337 | 0.760835856 | 0.675424709 | 0.499405972 | 0.680832768 |
| ENSG00000260046 | ENSG0000026 | 2.386643142 | -1.514757064 | 1.176682336 | -1.287311808 | 0.19798563 | 0.360345156 |
| ENSG00000260047 | BCAP31P2 | 2.270817477 | -0.179658018 | 1.159395468 | -0.154958358 | 0.876854146 | 0.935974786 |
| ENSG00000260051 | ENSG0000026 | 2.03967169 | -0.198542567 | 1.221045337 | -0.162600488 | 0.870833003 | 0.932975557 |
| ENSG00000260053 | ABCB10P4 | 2.900496831 | -2.389664685 | 1.216676216 | -1.964092545 | 0.049519361 | 0.118813318 |
| ENSG00000260060 | ENSG0000026 | 3.73461165 | -0.744654025 | 0.907662906 | -0.820408127 | 0.411983484 | 0.604316205 |
| ENSG00000260063 | ENSG0000026 | 31.61964215 | 0.192956495 | 0.311179417 | 0.620081165 | 0.535204352 | 0.710673063 |
| ENSG00000260077 | ENSG0000026 | 33.48159343 | 0.737782478 | 0.312698224 | 2.359407316 | 0.018304153 | 0.050596127 |
| ENSG00000260078 | MPHOSPH10F | 6.133989091 | -0.094515145 | 0.707620165 | -0.133567626 | 0.893744486 | 0.945700277 |
| ENSG00000260081 | MRFAP1P1 | 66.0005058 | -0.250831999 | 0.223321424 | -1.123188245 | 0.261357554 | 0.441595513 |
| ENSG00000260082 | ENSG0000026 | 1.982464052 | 0.766258742 | 1.248729025 | 0.613630921 | 0.539459246 | 0.714049685 |
| ENSG00000260083 | MIR762HG | 118.1128932 | 0.282564595 | 0.168907777 | 1.672892747 | 0.094348412 | 0.202162579 |
| ENSG00000260086 | ENSG0000026 | 4.130313706 | 0.181612222 | 0.860490057 | 0.211056735 | 0.832843 | 0.91160504 |
| ENSG00000260088 | DDX59-AS1 | 4.186119926 | 0.48736902 | 0.858260486 | 0.567856761 | 0.570132234 | 0.737292601 |
| ENSG00000260095 | ENSG0000026 | 8.906570979 | -0.281360442 | 0.595996691 | -0.472083899 | 0.636866904 | 0.786213586 |
| ENSG00000260101 | AFTPH-DT | 7.438867956 | -0.240696027 | 0.643795063 | -0.37387057 | 0.708500613 | 0.837453382 |
| ENSG00000260103 | ENSG0000026 | 22.52062953 | -0.031131906 | 0.367782557 | -0.084647587 | 0.932541566 | 0.967612915 |
| ENSG00000260107 | ENSG0000026 | 9.199491486 | -0.169893394 | 0.58305787 | -0.291383417 | 0.770758096 | 0.875046557 |

| | | | | | | | |
|---|---|---|---|---|---|---|---|
| ENSG00000260111 | ENSG0000026 | 3.976207754 | 0.310749812 | 0.877821371 | 0.354001192 | 0.72333799 | 0.846082679 |
| ENSG00000260118 | ENSG0000026 | 1.74699931 | -0.815427651 | 1.365710507 | -0.597072108 | 0.550459239 | NA |
| ENSG00000260121 | ENSG0000026 | 66.47389652 | 0.098407595 | 0.236055457 | 0.416883372 | 0.676763717 | 0.815935942 |
| ENSG00000260136 | ENSG0000026 | 101.7655133 | -0.248603546 | 0.185481511 | -1.34031443 | 0.180143142 | 0.335184011 |
| ENSG00000260144 | ENSG0000026 | 4.910424451 | 0.004716852 | 0.793655855 | 0.005943196 | 0.995258044 | 0.9970696 |
| ENSG00000260160 | ENSG0000026 | 11.65053645 | 0.479055482 | 0.512197306 | 0.935294809 | 0.34963639 | 0.540914772 |
| ENSG00000260163 | ENSG0000026 | 1.975381856 | 0.801823111 | 1.310697802 | 0.611752846 | 0.540701292 | 0.714868747 |
| ENSG00000260166 | ENSG0000026 | 1.611251199 | 0.43414744 | 1.404493436 | 0.309113186 | 0.757235429 | NA |
| ENSG00000260167 | ENSG0000026 | 10.70049554 | -0.176635221 | 0.559929201 | -0.31545992 | 0.752412478 | 0.863320632 |
| ENSG00000260174 | SYNGR2P1 | 16.9382599 | 0.102564997 | 0.43731599 | 0.234532921 | 0.814571278 | 0.901763793 |
| ENSG00000260179 | ENSG0000026 | 4.967757271 | -0.341506058 | 0.794255938 | -0.429969789 | 0.667217618 | 0.808556374 |
| ENSG00000260185 | ENSG0000026 | 2.318482765 | -1.467386807 | 1.225714747 | -1.197168273 | 0.231240973 | 0.404843771 |
| ENSG00000260190 | ENSG0000026 | 5.456879159 | -0.800207998 | 0.752126369 | -1.063927595 | 0.287361501 | 0.471279314 |
| ENSG00000260192 | LINC02240 | 3.558743377 | 0.629898343 | 0.962570454 | 0.654391936 | 0.512859328 | 0.692343494 |
| ENSG00000260193 | LINC02846 | 22.77637425 | 0.304669565 | 0.371839414 | 0.819357911 | 0.412582241 | 0.604614578 |
| ENSG00000260196 | ENSG0000026 | 136.6165028 | 0.916814543 | 0.158488904 | 5.784723865 | 7.26E-09 | 5.30E-08 |
| ENSG00000260206 | ENSG0000026 | 4.22335567 | -0.331090941 | 0.90124828 | -0.367369291 | 0.713343575 | 0.84060482 |
| ENSG00000260211 | ENSG0000026 | 8.398354248 | 0.963723892 | 0.647572164 | 1.488210806 | 0.136695308 | 0.271614584 |
| ENSG00000260213 | CENPN-AS1 | 7.544819322 | -1.205323684 | 0.688382799 | -1.750949743 | 0.079954567 | 0.176598898 |
| ENSG00000260219 | CD2BP2-DT | 68.6902003 | -0.196934115 | 0.210810664 | -0.9341753 | 0.350213474 | 0.541550192 |
| ENSG00000260228 | CDH13-AS2 | 178.5816071 | -1.210998031 | 0.148169907 | -8.173036313 | 3.01E-16 | 3.96E-15 |
| ENSG00000260231 | KDM7A-DT | 100.1611457 | 0.5089625 | 0.186742821 | 2.725472902 | 0.006420945 | 0.020111479 |
| ENSG00000260233 | ZNRD2-DT | 62.45037601 | 0.255642118 | 0.23208347 | 1.101509372 | 0.270675028 | 0.452506848 |
| ENSG00000260236 | PTPN23-DT | 25.37104518 | 0.066384159 | 0.347884051 | 0.190822657 | 0.848664537 | 0.920881911 |
| ENSG00000260239 | LINC02533 | 21.42531773 | -3.678160941 | 0.55410913 | -6.637972097 | 3.18E-11 | 2.86E-10 |
| ENSG00000260240 | APOOP5 | 2.474160459 | -0.031568166 | 1.119220611 | -0.02820549 | 0.977498259 | 0.989489456 |
| ENSG00000260244 | ENSG0000026 | 81.27229124 | 0.502599692 | 0.203146901 | 2.474070192 | 0.013358345 | 0.038377011 |
| ENSG00000260252 | ENSG0000026 | 2.085399392 | -0.775560635 | 1.294702355 | -0.599026202 | 0.549155412 | 0.721774261 |
| ENSG00000260257 | ENSG0000026 | 19.45163295 | -0.213172119 | 0.395924112 | -0.538416612 | 0.59028946 | 0.753103247 |
| ENSG00000260259 | LINC02166 | 21.77862823 | 0.504491836 | 0.380485774 | 1.325915109 | 0.184867818 | 0.341752453 |
| ENSG00000260260 | SNHG19 | 456.8851632 | 0.497276115 | 0.09242747 | 5.380176617 | 7.44E-08 | 4.87E-07 |
| ENSG00000260261 | ENSG0000026 | 209.0602813 | 0.213122608 | 0.155162058 | 1.373548472 | 0.169581903 | 0.320165542 |
| ENSG00000260267 | ENSG0000026 | 328.1478408 | -0.005236587 | 0.10991424 | -0.047642476 | 0.96200118 | 0.982516676 |

| | | | | | | | |
|---|---|---|---|---|---|---|---|
| ENSG00000260273 | ENSG0000026 | 5.472135626 | -1.55894502 | 0.802585924 | -1.942402643 | 0.052088376 | 0.124055994 |
| ENSG00000260274 | ENSG0000026 | 135.4274078 | 0.252883997 | 0.157195579 | 1.60872207 | 0.107677128 | 0.224964782 |
| ENSG00000260279 | ENSG0000026 | 80.61180474 | 0.34570439 | 0.197448246 | 1.750860785 | 0.079969893 | 0.176598898 |
| ENSG00000260281 | ITFG1-AS1 | 3.691107341 | -0.470130169 | 0.903606855 | -0.520281765 | 0.602867204 | 0.762096966 |
| ENSG00000260284 | TPSP2 | 2.062824068 | 2.25244712 | 1.417280386 | 1.589274178 | 0.111998506 | 0.232258699 |
| ENSG00000260285 | TRMT61A-DT | 187.0363851 | -0.362012742 | 0.132841771 | -2.725142388 | 0.006427376 | 0.020128875 |
| ENSG00000260288 | ENSG0000026 | 5.580947774 | -0.978670541 | 0.780522882 | -1.253865278 | 0.209890977 | 0.377018168 |
| ENSG00000260290 | ENSG0000026 | 986.1244188 | 0.147666886 | 0.069295672 | 2.130968388 | 0.033091747 | 0.084497403 |
| ENSG00000260293 | ENSG0000026 | 29.31502397 | 0.271546978 | 0.329313323 | 0.82458546 | 0.409606979 | 0.602084447 |
| ENSG00000260296 | ENSG0000026 | 13.0353015 | -0.294552611 | 0.50407564 | -0.584342086 | 0.558990182 | 0.728799931 |
| ENSG00000260298 | ACTG1P16 | 2.641992022 | 0.201531051 | 1.107939291 | 0.181897197 | 0.855663405 | 0.925169031 |
| ENSG00000260304 | ENSG0000026 | 48.98018681 | -0.594473062 | 0.255984321 | -2.322302627 | 0.020216646 | 0.05518442 |
| ENSG00000260317 | LINC02986 | 8.218737238 | 1.798714791 | 0.696710688 | 2.581724123 | 0.009830814 | 0.029276758 |
| ENSG00000260318 | COX6CP1 | 10.78772938 | -0.399141045 | 0.552016258 | -0.723060307 | 0.469642834 | 0.655568128 |
| ENSG00000260326 | PHB1P21 | 9.186865087 | -0.180714195 | 0.606552283 | -0.297936716 | 0.765751464 | 0.871876487 |
| ENSG00000260329 | TMEM263-DT | 4.361022545 | 0.364561069 | 0.831586832 | 0.438392065 | 0.661102097 | 0.804118882 |
| ENSG00000260331 | ENSG0000026 | 2.13279685 | 0.998918002 | 1.233661134 | 0.809718305 | 0.418102095 | 0.610131174 |
| ENSG00000260337 | ENSG0000026 | 10.78678782 | -0.147409403 | 0.530676977 | -0.277776141 | 0.781184207 | 0.881300566 |
| ENSG00000260339 | HEXA-AS1 | 32.45184046 | 0.354873901 | 0.314445558 | 1.128570249 | 0.259079167 | 0.438909689 |
| ENSG00000260350 | ENSG0000026 | 16.57939785 | 0.285094268 | 0.435847881 | 0.654114154 | 0.513038264 | 0.692381325 |
| ENSG00000260352 | SMG1-DT | 5.615254619 | -0.488392104 | 0.75363945 | -0.64804477 | 0.516955994 | 0.69525535 |
| ENSG00000260360 | ENSG0000026 | 3.582375013 | 0.592166762 | 0.928472699 | 0.637785863 | 0.523613084 | 0.700001783 |
| ENSG00000260361 | ENSG0000026 | 4.778938386 | -0.228880662 | 0.839241534 | -0.27272323 | 0.785065976 | 0.883106023 |
| ENSG00000260367 | ENSG0000026 | 7.70781845 | 0.098069065 | 0.643295081 | 0.152448025 | 0.878833581 | 0.937128357 |
| ENSG00000260368 | ENSG0000026 | 15.75117436 | 0.424421728 | 0.440997531 | 0.962412934 | 0.335842219 | 0.526943222 |
| ENSG00000260369 | ENSG0000026 | 3.372406811 | -1.436787756 | 1.013537057 | -1.417597656 | 0.156308265 | 0.300487019 |
| ENSG00000260382 | TMEM183AP3 | 16.31988831 | 0.220079106 | 0.433426306 | 0.507765919 | 0.611617514 | 0.769174155 |
| ENSG00000260388 | LINC00562 | 2.252915185 | 1.65962296 | 1.276354196 | 1.300284016 | 0.193503644 | 0.354239374 |
| ENSG00000260389 | WBP11P1 | 10.9351299 | 0.363196542 | 0.546798964 | 0.664223171 | 0.506547485 | 0.686854198 |
| ENSG00000260390 | ENSG0000026 | 1.979592284 | 0.269955721 | 1.239108728 | 0.217862819 | 0.827535997 | 0.90886547 |
| ENSG00000260394 | STUB1-DT | 3.081712275 | 0.896713279 | 1.01786222 | 0.880977073 | 0.378330229 | 0.571100536 |
| ENSG00000260398 | ENSG0000026 | 2.56919692 | 0.885986604 | 1.176495384 | 0.753072741 | 0.451406204 | 0.639390671 |
| ENSG00000260400 | ENSG0000026 | 8.73148259 | -0.453528176 | 0.590494452 | -0.768048158 | 0.442458575 | 0.631470633 |

| | | | | | | | |
|---|---|---|---|---|---|---|---|
| ENSG00000260401 | ENSG0000026 | 1015.474431 | 0.492382054 | 0.078290456 | 6.289170828 | 3.19E-10 | 2.66E-09 |
| ENSG00000260404 | ENSG0000026 | 253.9120722 | 0.037426048 | 0.118634236 | 0.315474265 | 0.752401588 | 0.863320632 |
| ENSG00000260410 | ENSG0000026 | 2.632607649 | 1.877295242 | 1.234682932 | 1.520467477 | 0.128393527 | 0.258840628 |
| ENSG00000260418 | ENSG0000026 | 30.50598877 | 0.232170039 | 0.315100264 | 0.736813217 | 0.461235945 | 0.648052463 |
| ENSG00000260425 | ENSG0000026 | 1.622143349 | 0.410567706 | 1.341968565 | 0.305944354 | 0.759647014 | NA |
| ENSG00000260428 | SCX | 105.7029692 | 0.936143654 | 0.184011889 | 5.087408519 | 3.63E-07 | 2.22E-06 |
| ENSG00000260430 | ENSG0000026 | 58.3090894 | -1.231091958 | 0.248742082 | -4.94927094 | 7.45E-07 | 4.39E-06 |
| ENSG00000260441 | ENSG0000026 | 1.623994061 | -0.753330117 | 1.398720324 | -0.538585237 | 0.590173077 | NA |
| ENSG00000260442 | ATP2A1-AS1 | 24.07643583 | 0.507861118 | 0.359829468 | 1.411393907 | 0.158128498 | 0.303097239 |
| ENSG00000260443 | ENSG0000026 | 14.20119895 | -0.323348161 | 0.466182479 | -0.693608566 | 0.48792772 | 0.671365247 |
| ENSG00000260448 | LCMT1-AS1 | 4.630996149 | -1.462549568 | 0.863100539 | -1.694529782 | 0.090164657 | 0.19470745 |
| ENSG00000260456 | C16orf95 | 15.82160206 | 0.749245297 | 0.473905322 | 1.581002076 | 0.113877563 | 0.235559179 |
| ENSG00000260459 | FTLP14 | 22.30643498 | 0.43129029 | 0.369530361 | 1.167130868 | 0.243157524 | 0.419590311 |
| ENSG00000260464 | ENSG0000026 | 11.88003213 | 0.820616553 | 0.520454345 | 1.576731101 | 0.114857411 | 0.237072972 |
| ENSG00000260465 | ENSG0000026 | 13.36887725 | 0.08942374 | 0.526853841 | 0.16973159 | 0.86522123 | 0.929961814 |
| ENSG00000260466 | ENSG0000026 | 10.70129403 | -0.192355923 | 0.542638917 | -0.354482357 | 0.722977423 | 0.845926239 |
| ENSG00000260468 | LINC01290 | 10.65016839 | 1.029122327 | 0.553369477 | 1.859738148 | 0.062922582 | 0.145113505 |
| ENSG00000260475 | SLC25A28-DT | 4.959460869 | -0.227522615 | 0.795544283 | -0.285996166 | 0.774881058 | 0.877499635 |
| ENSG00000260488 | ENSG0000026 | 2.993312173 | 3.879638258 | 1.446940372 | 2.681270309 | 0.007334325 | 0.022644604 |
| ENSG00000260490 | MYL12BP1 | 43.04570153 | -0.313163249 | 0.283102479 | -1.106183352 | 0.268647167 | 0.450165523 |
| ENSG00000260493 | E2F5-DT | 2.155504817 | 0.948336149 | 1.235611134 | 0.767503725 | 0.442782079 | 0.631641123 |
| ENSG00000260494 | ENSG0000026 | 5.141019552 | -0.144445241 | 0.77025693 | -0.187528648 | 0.851246164 | 0.9222591 |
| ENSG00000260495 | ENSG0000026 | 10.76470517 | 1.108588607 | 0.554883727 | 1.997875506 | 0.045730159 | 0.110998819 |
| ENSG00000260498 | ENSG0000026 | 3.385578058 | 0.017934581 | 0.966985471 | 0.018546898 | 0.985202564 | 0.993255991 |
| ENSG00000260500 | ENSG0000026 | 4.882531072 | 0.186645385 | 0.814450144 | 0.229167355 | 0.818738845 | 0.903986346 |
| ENSG00000260509 | ENSG0000026 | 2.293487137 | -0.1414521 | 1.151239235 | -0.122869422 | 0.9022105 | 0.950654947 |
| ENSG00000260518 | BMS1P8 | 13.34880853 | -0.296638134 | 0.477933739 | -0.620667908 | 0.534818149 | 0.710342986 |
| ENSG00000260526 | AP1AR-DT | 13.86043059 | 0.157378134 | 0.469446109 | 0.335242174 | 0.73744241 | 0.855304663 |
| ENSG00000260528 | FAM157C | 8.732655112 | -0.278341184 | 0.610623483 | -0.455831116 | 0.648511426 | 0.795128387 |
| ENSG00000260545 | ENSG0000026 | 4.312189041 | -0.68332052 | 0.867720157 | -0.787489508 | 0.430995368 | 0.621162456 |
| ENSG00000260552 | COSMOC | 26.93972482 | 0.656283233 | 0.341198334 | 1.923465524 | 0.054421612 | 0.128863727 |
| ENSG00000260563 | ENSG0000026 | 163.0558906 | 0.558180982 | 0.151979404 | 3.672740951 | 0.000239963 | 0.001003417 |
| ENSG00000260565 | ERVK13-1 | 423.3785013 | 0.388264987 | 0.103949434 | 3.735133229 | 0.000187616 | 0.00080028 |

| | | | | | | | |
|---|---|---|---|---|---|---|---|
| ENSG00000260570 | ENSG0000026 | 6.489915105 | -0.304618678 | 0.686107962 | -0.443980678 | 0.657056554 | 0.801279961 |
| ENSG00000260572 | PPM1L-DT | 15.0188512 | -0.077855997 | 0.457394923 | -0.170216137 | 0.864840161 | 0.929799912 |
| ENSG00000260577 | CDH3-AS1 | 15.15951302 | 0.063861417 | 0.471984113 | 0.135304168 | 0.89237139 | 0.944983167 |
| ENSG00000260578 | ENSG0000026 | 4.368277861 | 0.919471614 | 0.884353238 | 1.039710801 | 0.298474281 | 0.485007829 |
| ENSG00000260588 | ENSG0000026 | 10.76977882 | -0.327893538 | 0.538440251 | -0.608969218 | 0.542544843 | 0.71631532 |
| ENSG00000260589 | STAM-DT | 13.66183347 | -0.733746152 | 0.475913437 | -1.541763889 | 0.12313098 | 0.250393034 |
| ENSG00000260593 | ENSG0000026 | 7.185496291 | 0.558624742 | 0.700824931 | 0.79709599 | 0.425395285 | 0.61626573 |
| ENSG00000260597 | ENSG0000026 | 44.88934607 | -0.812489531 | 0.267978721 | -3.031918082 | 0.002430051 | 0.008401715 |
| ENSG00000260604 | ENSG0000026 | 15.10436664 | 0.182938554 | 0.446136244 | 0.410050867 | 0.681768634 | 0.819321553 |
| ENSG00000260608 | ENSG0000026 | 5.315669485 | 0.555790388 | 0.782079194 | 0.710657427 | 0.477296547 | 0.662124694 |
| ENSG00000260612 | ENSG0000026 | 2.813710782 | -0.266763006 | 1.068942738 | -0.249557807 | 0.802929331 | 0.895135709 |
| ENSG00000260615 | RPL23AP97 | 3.2963123 | -0.470216343 | 0.970364779 | -0.484576886 | 0.627976513 | 0.780547313 |
| ENSG00000260617 | ENSG0000026 | 4.810748379 | 0.509783659 | 0.802544786 | 0.635208985 | 0.525292124 | 0.701327047 |
| ENSG00000260618 | ENSG0000026 | 19.70927895 | -2.503147321 | 0.490199307 | -5.106386907 | 3.28E-07 | 2.02E-06 |
| ENSG00000260619 | ENSG0000026 | 6.156901024 | -0.662287838 | 0.747955669 | -0.885464026 | 0.375906414 | 0.568856331 |
| ENSG00000260625 | RUSF1-DT | 6.013433482 | -0.325116199 | 0.723077834 | -0.44962822 | 0.652978537 | 0.798589026 |
| ENSG00000260630 | SNAI3-AS1 | 239.4636392 | 0.031212345 | 0.121459459 | 0.256977477 | 0.79719616 | 0.89183367 |
| ENSG00000260633 | ENSG0000026 | 17.92377151 | 0.175003335 | 0.416423252 | 0.420253514 | 0.674300265 | 0.813829014 |
| ENSG00000260634 | ENSG0000026 | 2.366434587 | -1.429670442 | 1.2227257 | -1.169248706 | 0.242303443 | 0.418557303 |
| ENSG00000260641 | TSPAN5-DT | 2.681664096 | -0.939452565 | 1.070226094 | -0.877807568 | 0.380048157 | 0.572480446 |
| ENSG00000260643 | ENSG0000026 | 25.44176939 | -0.086401594 | 0.374311569 | -0.230828009 | 0.817448421 | 0.90328838 |
| ENSG00000260645 | ENSG0000026 | 5.811583781 | 0.997325586 | 0.789040011 | 1.2639734 | 0.206239575 | 0.371886169 |
| ENSG00000260646 | ENSG0000026 | 3.204208521 | -0.079814776 | 1.027680855 | -0.077664944 | 0.938094581 | 0.970457257 |
| ENSG00000260647 | ENSG0000026 | 5.540656743 | 0.529224787 | 0.752782826 | 0.703024523 | 0.482040467 | 0.666287182 |
| ENSG00000260653 | MIR3147HG | 111.9523179 | 1.363565777 | 0.182064254 | 7.489475523 | 6.91E-14 | 7.63E-13 |
| ENSG00000260658 | ENSG0000026 | 9.20021875 | 1.148173634 | 0.607581073 | 1.889745557 | 0.058791998 | 0.137173356 |
| ENSG00000260659 | ENSG0000026 | 8.159079552 | 0.620185451 | 0.617142507 | 1.0049307 | 0.314930221 | 0.503554699 |
| ENSG00000260668 | ENSG0000026 | 20.27258593 | 0.467652288 | 0.396612026 | 1.179117769 | 0.238351284 | 0.413786776 |
| ENSG00000260669 | ENSG0000026 | 7.081524971 | -0.899171107 | 0.669109563 | -1.343832396 | 0.179002591 | 0.333812913 |
| ENSG00000260698 | ENSG0000026 | 5.854017262 | 0.013192529 | 0.737641631 | 0.017884741 | 0.985730802 | 0.993350589 |
| ENSG00000260708 | TBC1D22A-DT | 129.0551697 | 0.645364493 | 0.159537035 | 4.045233098 | 5.23E-05 | 0.000243719 |
| ENSG00000260710 | ENSG0000026 | 3.5691766 | -0.378293034 | 0.928376157 | -0.407478188 | 0.683656811 | 0.820502629 |
| ENSG00000260711 | ENSG0000026 | 29.15104183 | -0.595283377 | 0.352364631 | -1.689395939 | 0.091143579 | 0.196506926 |

| | | | | | | | |
|---|---|---|---|---|---|---|---|
| ENSG00000260727 | SLC7A5P1 | 33.82997118 | 0.446810277 | 0.304312625 | 1.46826073 | 0.142033414 | 0.27960581 |
| ENSG00000260729 | ENSG0000026 | 4.837666775 | 0.269143269 | 0.837277011 | 0.321450686 | 0.747868878 | 0.860419264 |
| ENSG00000260731 | KRT8P22 | 9.824082513 | -0.081073591 | 0.550745286 | -0.147207054 | 0.882968593 | 0.939354428 |
| ENSG00000260735 | ENSG0000026 | 12.6414944 | -0.495402907 | 0.489460995 | -1.012139704 | 0.311471261 | 0.499625951 |
| ENSG00000260740 | ENSG0000026 | 1.878415898 | 3.168128947 | 1.577616779 | 2.008173968 | 0.044624808 | 0.108683841 |
| ENSG00000260742 | ITPRID2-DT | 7.513615245 | 0.613492444 | 0.662012965 | 0.926707597 | 0.354078372 | 0.545582595 |
| ENSG00000260747 | ENSG0000026 | 268.0318161 | 0.171283889 | 0.11172082 | 1.533142072 | 0.125240853 | 0.253912963 |
| ENSG00000260751 | ENSG0000026 | 24.34885844 | 0.179790495 | 0.353526343 | 0.508563218 | 0.611058417 | 0.768826384 |
| ENSG00000260773 | ENSG0000026 | 3.86449818 | -0.630907475 | 0.910870743 | -0.692642155 | 0.488534148 | 0.671800345 |
| ENSG00000260774 | ENSG0000026 | 12.87003587 | 0.05277062 | 0.496967792 | 0.10618519 | 0.915435421 | 0.958033237 |
| ENSG00000260777 | ENSG0000026 | 5.785081153 | -1.666850417 | 0.779448594 | -2.138499486 | 0.032476226 | 0.083203554 |
| ENSG00000260778 | MIR3677HG | 31.54091962 | 0.843133179 | 0.324893335 | 2.595107653 | 0.00945613 | 0.028326405 |
| ENSG00000260782 | ENSG0000026 | 1.613674879 | 0.982889098 | 1.437624012 | 0.683689956 | 0.494170967 | NA |
| ENSG00000260793 | ENSG0000026 | 295.5188366 | 0.625086746 | 0.120965682 | 5.167471747 | 2.37E-07 | 1.48E-06 |
| ENSG00000260796 | ENSG0000026 | 2.12998567 | 1.49941487 | 1.294637023 | 1.158173947 | 0.246793058 | 0.42406804 |
| ENSG00000260798 | ENSG0000026 | 90.29732894 | -0.448372807 | 0.22785776 | -1.967775011 | 0.049093931 | 0.11795295 |
| ENSG00000260804 | LINC01963 | 169.8386697 | 1.176809245 | 0.154038057 | 7.639730504 | 2.18E-14 | 2.50E-13 |
| ENSG00000260805 | ENSG0000026 | 25.44462094 | -0.80161243 | 0.352374321 | -2.274888897 | 0.022912596 | 0.061518603 |
| ENSG00000260806 | PAPOLA-DT | 51.37971222 | 0.00982654 | 0.245811094 | 0.039975983 | 0.968112274 | 0.985820278 |
| ENSG00000260807 | CEROX1 | 12.87134163 | 1.446164913 | 0.522361316 | 2.768514566 | 0.005631247 | 0.0178896 |
| ENSG00000260808 | ENSG0000026 | 9.363049455 | -1.118500703 | 0.616134564 | -1.815351336 | 0.069469935 | 0.157626974 |
| ENSG00000260809 | VPS35P1 | 3.697789063 | -0.204970484 | 0.933978389 | -0.219459557 | 0.82629208 | 0.908364356 |
| ENSG00000260810 | ENSG0000026 | 1.698335247 | 0.335361142 | 1.358694923 | 0.246825933 | 0.80504294 | NA |
| ENSG00000260815 | PPIAP47 | 2.741297991 | 2.080571203 | 1.286658393 | 1.61703465 | 0.105870802 | 0.221938791 |
| ENSG00000260816 | ENSG0000026 | 2.522420996 | 0.756466284 | 1.132097364 | 0.668198962 | 0.504006598 | 0.684987935 |
| ENSG00000260830 | ENSG0000026 | 10.87765065 | 0.25945946 | 0.560338239 | 0.463040788 | 0.643335131 | 0.790937773 |
| ENSG00000260852 | FBXL19-AS1 | 288.4480231 | 0.719725902 | 0.111157802 | 6.474812259 | 9.49E-11 | 8.21E-10 |
| ENSG00000260853 | NFATC2IP-AS1 | 11.74800019 | -0.514223182 | 0.526017339 | -0.977578387 | 0.328282891 | 0.518686516 |
| ENSG00000260855 | ENSG0000026 | 42.02503348 | 0.510915598 | 0.276680668 | 1.846589435 | 0.064806667 | 0.148845107 |
| ENSG00000260860 | ENSG0000026 | 4.510971521 | -0.095400807 | 0.816078615 | -0.116901491 | 0.906938117 | 0.95304868 |
| ENSG00000260871 | ENSG0000026 | 1.54507723 | 1.047955924 | 1.511351448 | 0.693389963 | 0.48806486 | NA |
| ENSG00000260874 | ENSG0000026 | 28.19133294 | 0.269873031 | 0.338835462 | 0.796472215 | 0.425757619 | 0.616634875 |
| ENSG00000260879 | ENSG0000026 | 7.470168947 | -0.141789014 | 0.632936696 | -0.224017686 | 0.822743541 | 0.906215744 |

| | | | | | | | |
|---|---|---|---|---|---|---|---|
| ENSG00000260880 | HCCAT5 | 12.95990203 | -2.204269706 | 0.560591131 | -3.932045272 | 8.42E-05 | 0.000380869 |
| ENSG00000260886 | TAT-AS1 | 6.700472519 | -1.219833622 | 0.728036441 | -1.675511765 | 0.093833865 | 0.2012104 |
| ENSG00000260892 | ENSG0000026 | 7.508355526 | -0.779458762 | 0.643274211 | -1.211705288 | 0.225625214 | 0.397621757 |
| ENSG00000260894 | ENSG0000026 | 7.206191787 | -1.210310791 | 0.6769369 | -1.787922613 | 0.073788494 | 0.165740532 |
| ENSG00000260895 | ENSG0000026 | 2.190540656 | 0.876988166 | 1.247921372 | 0.702759152 | 0.482205857 | 0.666327572 |
| ENSG00000260903 | XKR7 | 7.942051436 | 0.048092268 | 0.626435196 | 0.076771338 | 0.938805452 | 0.970997921 |
| ENSG00000260911 | ENSG0000026 | 5.541633863 | 0.055974642 | 0.809588056 | 0.069139658 | 0.944878454 | 0.973786411 |
| ENSG00000260912 | ENSG0000026 | 99.66069218 | 0.091174105 | 0.187914374 | 0.485189628 | 0.627541839 | 0.780187429 |
| ENSG00000260916 | CCPG1 | 116.8256424 | -0.272461007 | 0.163763767 | -1.66374414 | 0.096163571 | 0.205323187 |
| ENSG00000260917 | ENSG0000026 | 167.0551581 | -0.379287276 | 0.140650355 | -2.696667749 | 0.007003711 | 0.021708571 |
| ENSG00000260919 | ENSG0000026 | 4.362937525 | -0.65152212 | 0.843285446 | -0.772599745 | 0.439759296 | 0.629163256 |
| ENSG00000260920 | ENSG0000026 | 114.759322 | 0.254450398 | 0.184896889 | 1.37617458 | 0.16876759 | 0.318977299 |
| ENSG00000260923 | LINC02193 | 4.093941041 | 0.609919087 | 0.888208495 | 0.686684591 | 0.492281509 | 0.6751209 |
| ENSG00000260924 | LINC01311 | 28.76455664 | -0.46673598 | 0.326231097 | -1.430691265 | 0.152518718 | 0.295131117 |
| ENSG00000260927 | ENSG0000026 | 19.52585752 | -0.512015663 | 0.393096598 | -1.302518682 | 0.192739136 | 0.353065222 |
| ENSG00000260928 | SPCS2P1 | 4.916544347 | -0.150296522 | 0.801239277 | -0.187580073 | 0.851205848 | 0.9222591 |
| ENSG00000260934 | ENSG0000026 | 7.153868609 | -0.892184447 | 0.663201929 | -1.345268171 | 0.178538648 | 0.333156579 |
| ENSG00000260942 | CAPN10-DT | 84.72365462 | 0.201005997 | 0.21067675 | 0.954096724 | 0.340034684 | 0.53096424 |
| ENSG00000260948 | ENSG0000026 | 12.25968629 | 0.891238872 | 0.512750624 | 1.738152681 | 0.082183914 | 0.180601494 |
| ENSG00000260949 | ENSG0000026 | 3.509631853 | 0.742659539 | 1.007758083 | 0.73694228 | 0.461157451 | 0.648021558 |
| ENSG00000260954 | ENSG0000026 | 3.288375114 | 1.34454466 | 1.042933446 | 1.289195073 | 0.197330277 | 0.359409017 |
| ENSG00000260955 | ENSG0000026 | 3.217908474 | 0.230031066 | 0.974827395 | 0.235971073 | 0.813455117 | 0.901448243 |
| ENSG00000260958 | ENSG0000026 | 5.957722259 | 4.019560245 | 1.192643993 | 3.370293455 | 0.000750882 | 0.002877586 |
| ENSG00000260971 | ENSG0000026 | 3.013186273 | -1.902196628 | 1.178779666 | -1.613699899 | 0.106592536 | 0.2231459 |
| ENSG00000260988 | ENSG0000026 | 3.365400348 | -1.708388243 | 1.025964582 | -1.665153235 | 0.095882188 | 0.204894029 |
| ENSG00000260992 | DOCK9-DT | 24.53622121 | 1.515003048 | 0.389088418 | 3.893724353 | 9.87E-05 | 0.000440367 |
| ENSG00000261000 | ENSG0000026 | 2.532321509 | 2.605892711 | 1.355092351 | 1.923036986 | 0.054475405 | 0.128946937 |
| ENSG00000261002 | VPS39-DT | 4.41831448 | -2.24893686 | 0.979580895 | -2.295815356 | 0.021686441 | 0.05865922 |
| ENSG00000261003 | ENSG0000026 | 6.526999636 | 1.420371508 | 0.776119833 | 1.830093045 | 0.067236027 | 0.153548477 |
| ENSG00000261008 | LINC01572 | 81.32490192 | -0.003091143 | 0.195654999 | -0.015798946 | 0.98739479 | 0.993893658 |
| ENSG00000261019 | ENSG0000026 | 1.780557116 | 1.220857595 | 1.345317003 | 0.907486928 | 0.364149359 | NA |
| ENSG00000261026 | ENSG0000026 | 2.904974087 | 0.982148434 | 1.113908354 | 0.88171386 | 0.377931564 | 0.570753791 |
| ENSG00000261033 | SPECC1-DT | 5.504227169 | -0.257944043 | 0.750773472 | -0.343571068 | 0.731168886 | 0.851119615 |

| | | | | | | | |
|---|---|---|---|---|---|---|---|
| ENSG00000261040 | WFDC21P | 2.056217451 | 2.217760325 | 1.42994546 | 1.550940498 | 0.120915944 | 0.246956175 |
| ENSG00000261051 | ENSG0000026 | 2.424497622 | 0.391419134 | 1.128283605 | 0.346915556 | 0.72865476 | 0.849651768 |
| ENSG00000261052 | SULT1A3 | 2.90726921 | 2.095769065 | 1.183129066 | 1.771378225 | 0.076497827 | 0.170623253 |
| ENSG00000261054 | SYNM-AS2 | 10.67996561 | 0.404554537 | 0.543095759 | 0.744904615 | 0.456329385 | 0.643478245 |
| ENSG00000261056 | ENSG0000026 | 2.269404036 | 0.682238255 | 1.15780371 | 0.589252089 | 0.55569218 | 0.726640663 |
| ENSG00000261061 | ENSG0000026 | 341.2053461 | 0.424433223 | 0.101626855 | 4.176388435 | 2.96E-05 | 0.00014358 |
| ENSG00000261063 | ENSG0000026 | 15.29122332 | -0.815994108 | 0.464411615 | -1.757049309 | 0.078909405 | 0.174812337 |
| ENSG00000261064 | LINC02256 | 1.660477088 | 0.33177744 | 1.365948179 | 0.242891674 | 0.808089316 | NA |
| ENSG00000261065 | ENSG0000026 | 13.7530088 | 0.421039912 | 0.47603627 | 0.884470236 | 0.376442423 | 0.569311728 |
| ENSG00000261067 | ENSG0000026 | 13.2591086 | -0.142978448 | 0.527667401 | -0.270963201 | 0.786419335 | 0.884032907 |
| ENSG00000261071 | ENSG0000026 | 38.65572039 | -0.248136847 | 0.286498916 | -0.866100473 | 0.386435066 | 0.578997258 |
| ENSG00000261079 | ENSG0000026 | 12.39046715 | 1.26189346 | 0.524423148 | 2.406250497 | 0.016117209 | 0.045315453 |
| ENSG00000261087 | ZNNT1 | 55.80219941 | 0.294522877 | 0.240247902 | 1.225912379 | 0.22023165 | 0.390456299 |
| ENSG00000261093 | ENSG0000026 | 12.00879493 | -0.342992748 | 0.549537485 | -0.624148047 | 0.532530362 | 0.707982991 |
| ENSG00000261094 | ENSG0000026 | 45.80964539 | 0.362766331 | 0.25981845 | 1.396230064 | 0.162645226 | 0.309906073 |
| ENSG00000261096 | ENSG0000026 | 4.910489044 | -0.324748378 | 0.823329877 | -0.394432885 | 0.693261469 | 0.826967818 |
| ENSG00000261098 | ENSG0000026 | 6.459297118 | 0.388685372 | 0.692138383 | 0.561571763 | 0.574407824 | 0.740858848 |
| ENSG00000261101 | ENSG0000026 | 8.027112279 | -0.204992779 | 0.618382716 | -0.331498235 | 0.740268174 | 0.856669118 |
| ENSG00000261104 | ENSG0000026 | 27.08725838 | -1.050176598 | 0.361649656 | -2.903850675 | 0.003686039 | 0.012244098 |
| ENSG00000261105 | LMO7-AS1 | 2.303995466 | -1.422783884 | 1.223431107 | -1.16294565 | 0.244851554 | 0.421752114 |
| ENSG00000261114 | ENSG0000026 | 1.980018091 | -0.668069855 | 1.237703363 | -0.539765727 | 0.589358606 | 0.752442477 |
| ENSG00000261115 | TMEM178B | 1672.167008 | -0.058944679 | 0.067912881 | -0.867945493 | 0.385424173 | 0.577997681 |
| ENSG00000261118 | ENSG0000026 | 12.61512213 | -0.025216971 | 0.503517076 | -0.050081661 | 0.960057314 | 0.981799414 |
| ENSG00000261123 | ENSG0000026 | 16.7292754 | -0.146397957 | 0.430480883 | -0.340080043 | 0.73379625 | 0.853084627 |
| ENSG00000261126 | RBFADN | 31.99055102 | 0.557923561 | 0.317245023 | 1.758651898 | 0.078636652 | 0.174275413 |
| ENSG00000261135 | ENSG0000026 | 2.025044692 | 0.250939158 | 1.275141366 | 0.196793207 | 0.843989368 | 0.917708773 |
| ENSG00000261136 | ENSG0000026 | 21.97546629 | 0.381894143 | 0.376333347 | 1.014776253 | 0.310212496 | 0.49819901 |
| ENSG00000261150 | EPPK1 | 1387.27635 | -0.149857224 | 0.080502355 | -1.861525954 | 0.062669939 | 0.144603541 |
| ENSG00000261159 | ENSG0000026 | 20.42920301 | -1.039830381 | 0.408153131 | -2.547647689 | 0.010845194 | 0.031891904 |
| ENSG00000261167 | ENSG0000026 | 86.74122145 | 0.490076118 | 0.199795006 | 2.452894727 | 0.01417118 | 0.040436775 |
| ENSG00000261168 | ENSG0000026 | 4.385610644 | 1.416129635 | 0.892210953 | 1.58721391 | 0.112464206 | 0.233098019 |
| ENSG00000261173 | DNAJA2-DT | 2.97485662 | 0.79038113 | 1.0737287 | 0.736108786 | 0.461664496 | 0.648393573 |
| ENSG00000261183 | SPINT1-AS1 | 434.2488791 | 0.04027178 | 0.095194729 | 0.423046327 | 0.672261461 | 0.812222991 |

| | | | | | | | |
|---|---|---|---|---|---|---|---|
| ENSG00000261186 | LINC01238 | 7.51835743 | -0.378714543 | 0.629799007 | -0.60132604 | 0.547622849 | 0.720435377 |
| ENSG00000261187 | ENSG0000026 | 3.549777673 | -0.070185051 | 0.925876966 | -0.075803863 | 0.939575142 | 0.971233034 |
| ENSG00000261188 | TFIP11-DT | 171.4910836 | -0.25103535 | 0.136401869 | -1.840409898 | 0.065708081 | 0.150583755 |
| ENSG00000261189 | DSP-AS1 | 57.14802233 | -0.194672981 | 0.244213908 | -0.797141253 | 0.425369 | 0.61626573 |
| ENSG00000261202 | TNRC6B-DT | 2.552236493 | 0.335730418 | 1.135446916 | 0.295681298 | 0.767473481 | 0.873024431 |
| ENSG00000261204 | ENSG0000026 | 2.089767796 | 0.971597713 | 1.225600128 | 0.792752621 | 0.427921966 | 0.618614173 |
| ENSG00000261211 | ENSG0000026 | 55.9652164 | -0.715619668 | 0.241912256 | -2.95817864 | 0.003094627 | 0.010456731 |
| ENSG00000261215 | ENSG0000026 | 1.784330271 | 0.557395398 | 1.395192732 | 0.399511397 | 0.689516428 | NA |
| ENSG00000261217 | BCAP31P1 | 7.070526243 | -0.010243135 | 0.696423774 | -0.014708192 | 0.988264984 | 0.994030148 |
| ENSG00000261220 | ST3GAL1-DT | 52.70688111 | 0.133683084 | 0.24529876 | 0.544980673 | 0.58576681 | 0.749814489 |
| ENSG00000261221 | ZNF865 | 435.3747008 | -0.205579654 | 0.092997501 | -2.210593311 | 0.027064012 | 0.07111666 |
| ENSG00000261226 | ENSG0000026 | 17.25488074 | 0.661034543 | 0.452834977 | 1.45976918 | 0.144353521 | 0.282983232 |
| ENSG00000261235 | HSD17B2-AS1 | 55.79832237 | -0.780613513 | 0.237649638 | -3.284724182 | 0.001020823 | 0.003815817 |
| ENSG00000261236 | BOP1 | 1494.565566 | 0.091300231 | 0.063690657 | 1.433494885 | 0.15171646 | 0.294083095 |
| ENSG00000261240 | ENSG0000026 | 1.731320087 | 0.705128679 | 1.363913601 | 0.516989257 | 0.605163658 | NA |
| ENSG00000261242 | ENSG0000026 | 4.38529847 | -1.077431903 | 0.891876725 | -1.208050254 | 0.227027933 | 0.399571949 |
| ENSG00000261243 | ENSG0000026 | 2.655221281 | -0.561549262 | 1.082766754 | -0.518624403 | 0.604022688 | 0.762927016 |
| ENSG00000261266 | ENSG0000026 | 8.393390596 | -0.052085908 | 0.618537861 | -0.084208117 | 0.932890965 | 0.967770789 |
| ENSG00000261270 | ENSG0000026 | 6.687547973 | -0.088043733 | 0.690995928 | -0.127415704 | 0.898611388 | 0.948793422 |
| ENSG00000261278 | ENSG0000026 | 3.830555451 | 1.11941053 | 0.930452675 | 1.20308164 | 0.22894473 | 0.402020215 |
| ENSG00000261286 | ATP2C2-AS1 | 45.63063621 | 0.499808938 | 0.273552377 | 1.827105083 | 0.067683971 | 0.154340984 |
| ENSG00000261292 | ENSG0000026 | 2.640100832 | 2.68435684 | 1.361225371 | 1.972014993 | 0.048607895 | 0.116932164 |
| ENSG00000261303 | GOLGA6GP | 1.773943038 | -2.217187289 | 1.490150379 | -1.487894993 | 0.136778588 | NA |
| ENSG00000261305 | ENSG0000026 | 21.16671614 | -0.839277241 | 0.404627518 | -2.074197141 | 0.038061006 | 0.095225158 |
| ENSG00000261308 | FIGNL2 | 5.257772262 | -1.077610397 | 0.802217526 | -1.343289522 | 0.179178244 | 0.333903242 |
| ENSG00000261310 | ENSG0000026 | 1.769653826 | 0.626738848 | 1.29803591 | 0.482836294 | 0.629211979 | NA |
| ENSG00000261312 | ENSG0000026 | 6.586357869 | -0.920959859 | 0.720301983 | -1.278574655 | 0.20104688 | 0.3645862 |
| ENSG00000261315 | LARP4P | 5.014867242 | -0.065943207 | 0.796827502 | -0.082757193 | 0.934044608 | 0.968363687 |
| ENSG00000261318 | DIS3L-AS1 | 1.874662447 | -0.469857961 | 1.268666352 | -0.370355815 | 0.711117391 | 0.839248612 |
| ENSG00000261324 | ENSG0000026 | 77.3032371 | -0.193198897 | 0.20663081 | -0.934995593 | 0.349790571 | 0.541042121 |
| ENSG00000261326 | LINC01355 | 39.38132294 | 1.036358523 | 0.307023676 | 3.375500346 | 0.000736816 | 0.002828407 |
| ENSG00000261329 | ENSG0000026 | 4.266472804 | 0.032524232 | 0.87147751 | 0.037320793 | 0.970229226 | 0.98657174 |
| ENSG00000261335 | ENSG0000026 | 12.02345796 | -1.439267061 | 0.525187755 | -2.740480995 | 0.006134933 | 0.019334436 |

| | | | | | | | |
|---|---|---|---|---|---|---|---|
| ENSG00000261336 | EIF4BP5 | 11.99104098 | 0.160181348 | 0.501283206 | 0.319542618 | 0.749315079 | 0.861330224 |
| ENSG00000261338 | ENSG0000026 | 26.24833165 | 0.517268978 | 0.365030343 | 1.41705748 | 0.156466123 | 0.300714884 |
| ENSG00000261342 | ENSG0000026 | 40.85446293 | 0.501484456 | 0.297930972 | 1.683223641 | 0.092331806 | 0.198602121 |
| ENSG00000261349 | ENSG0000026 | 5.946872013 | 0.102338565 | 0.728645885 | 0.140450344 | 0.888304183 | 0.942452038 |
| ENSG00000261351 | ENSG0000026 | 52.00770284 | -0.031472781 | 0.243183342 | -0.129419972 | 0.897025345 | 0.947984339 |
| ENSG00000261359 | PYCARD-AS1 | 27.68706128 | 0.662531642 | 0.338771904 | 1.955686507 | 0.050502095 | 0.120867821 |
| ENSG00000261360 | OTULIN-DT | 24.79991921 | 0.960483065 | 0.384678202 | 2.496848174 | 0.012530259 | 0.03625683 |
| ENSG00000261373 | VPS9D1-AS1 | 1443.296789 | 0.927673217 | 0.06199432 | 14.96384223 | 1.27E-50 | 7.10E-49 |
| ENSG00000261377 | PDCD6IPP2 | 3.184700741 | -1.297429811 | 1.020136107 | -1.271820302 | 0.203436974 | 0.368018625 |
| ENSG00000261379 | ENSG0000026 | 24.90941784 | 2.208226901 | 0.412453651 | 5.353878905 | 8.61E-08 | 5.61E-07 |
| ENSG00000261385 | ENSG0000026 | 10.3504792 | -0.203338737 | 0.564808192 | -0.360013789 | 0.718836822 | 0.843613685 |
| ENSG00000261386 | ENSG0000026 | 2.516262469 | 0.772908767 | 1.155019542 | 0.669173758 | 0.503384644 | 0.684426489 |
| ENSG00000261393 | ENSG0000026 | 2.440997614 | 0.043886294 | 1.115560398 | 0.039340132 | 0.96861921 | 0.985876114 |
| ENSG00000261395 | HSPE1P5 | 2.075144802 | -0.239797056 | 1.257781942 | -0.190650738 | 0.848799236 | 0.920956268 |
| ENSG00000261396 | ENSG0000026 | 12.95352036 | 0.251889593 | 0.509331153 | 0.494549747 | 0.620917982 | 0.775560477 |
| ENSG00000261404 | PSMD7-DT | 4.194151448 | 0.490028257 | 0.852072873 | 0.575101347 | 0.565222757 | 0.733481521 |
| ENSG00000261408 | TEN1-CDK3 | 20.71767316 | -0.045257626 | 0.386726523 | -0.117027468 | 0.906838287 | 0.952992363 |
| ENSG00000261416 | CORO1A-AS1 | 1.59747614 | -0.717702222 | 1.437820242 | -0.49915991 | 0.617666735 | NA |
| ENSG00000261420 | MPC1-DT | 6.067807659 | -0.833427727 | 0.711665243 | -1.171095169 | 0.241560526 | 0.417682856 |
| ENSG00000261423 | TMEM202-AS | 34.59260644 | 0.011238721 | 0.298888781 | 0.037601681 | 0.970005268 | 0.986563569 |
| ENSG00000261424 | ATP5PBP7 | 75.94423488 | 0.250900976 | 0.204832292 | 1.224909284 | 0.220609402 | 0.390854162 |
| ENSG00000261437 | LINC02894 | 120.8364537 | -0.415786047 | 0.16606201 | -2.503799917 | 0.01228675 | 0.035664689 |
| ENSG00000261441 | POLG-DT | 9.598121896 | 0.83714853 | 0.588275834 | 1.423054428 | 0.154720378 | 0.298184111 |
| ENSG00000261455 | LINC01003 | 209.6892427 | 0.734842882 | 0.130596888 | 5.626802396 | 1.84E-08 | 1.28E-07 |
| ENSG00000261456 | TUBB8 | 25.96579023 | -0.62524756 | 0.344334419 | -1.815814876 | 0.069398775 | 0.15754655 |
| ENSG00000261457 | ENSG0000026 | 7.337167423 | 0.235966124 | 0.67358492 | 0.350313846 | 0.726103175 | 0.847853251 |
| ENSG00000261460 | ENSG0000026 | 29.84727104 | -0.878931012 | 0.325573363 | -2.699640423 | 0.006941446 | 0.021538856 |
| ENSG00000261461 | UBE2MP1 | 435.4004901 | -0.115736899 | 0.092668549 | -1.248933982 | 0.211689221 | 0.379268255 |
| ENSG00000261465 | COQ7-DT | 3.2589888 | 0.412964131 | 1.002093715 | 0.412101308 | 0.68026517 | 0.818419487 |
| ENSG00000261468 | ENSG0000026 | 1.697343935 | 4.113616272 | 1.659210903 | 2.479260631 | 0.013165506 | NA |
| ENSG00000261474 | ENSG0000026 | 59.71000689 | -0.030268461 | 0.236860712 | -0.12779013 | 0.898315062 | 0.948554546 |
| ENSG00000261476 | ENSG0000026 | 2.987580879 | -2.430054259 | 1.173405398 | -2.070941777 | 0.038364237 | 0.095847921 |
| ENSG00000261485 | PAN3-AS1 | 26.23674191 | 0.374584571 | 0.339952787 | 1.101872334 | 0.270517178 | 0.452340845 |

| | | | | | | | |
|---|---|---|---|---|---|---|---|
| ENSG00000261488 | TBILA | 12.22362193 | 0.213403146 | 0.507708459 | 0.420326158 | 0.674247203 | 0.813807789 |
| ENSG00000261499 | ENSG0000026 | 11.56106399 | 0.018752734 | 0.557063377 | 0.033663556 | 0.97314544 | 0.987960542 |
| ENSG00000261502 | ENSG0000026 | 5.590315901 | 1.145373493 | 0.759019557 | 1.509017102 | 0.131294412 | 0.263484719 |
| ENSG00000261504 | LINC01686 | 4.473128182 | -0.871329796 | 0.855538165 | -1.018458125 | 0.308460291 | 0.49649727 |
| ENSG00000261505 | ENSG0000026 | 73.90726687 | -0.452978825 | 0.246072527 | -1.840834615 | 0.065645799 | 0.150471085 |
| ENSG00000261512 | ENSG0000026 | 44.98390123 | 0.127032195 | 0.282402095 | 0.449827382 | 0.652834913 | 0.798508819 |
| ENSG00000261513 | ENSG0000026 | 4.795111467 | 0.305777183 | 0.788318806 | 0.387885181 | 0.698101005 | 0.830667413 |
| ENSG00000261526 | ENSG0000026 | 129.1538456 | 0.55385062 | 0.161220485 | 3.435361331 | 0.000591764 | 0.002309864 |
| ENSG00000261532 | ENSG0000026 | 11.44805246 | -0.309738598 | 0.539479299 | -0.57414362 | 0.565870618 | 0.733932944 |
| ENSG00000261542 | ENSG0000026 | 5.473138486 | 0.021803482 | 0.790507912 | 0.027581612 | 0.977995848 | 0.98981844 |
| ENSG00000261546 | ENSG0000026 | 1.749859937 | -0.840535976 | 1.335020388 | -0.629605348 | 0.528952824 | NA |
| ENSG00000261549 | ENSG0000026 | 22.75375212 | -0.104468833 | 0.390048143 | -0.267835741 | 0.788825758 | 0.885806405 |
| ENSG00000261556 | SMG1P7 | 169.8459864 | 0.414175855 | 0.142418473 | 2.90816104 | 0.003635611 | 0.012090593 |
| ENSG00000261557 | EEF1A1P38 | 33.48191886 | -0.005331461 | 0.306854755 | -0.017374542 | 0.986137819 | 0.993350589 |
| ENSG00000261559 | FSCN1P1 | 7.073549258 | 2.020087054 | 0.766904969 | 2.63407741 | 0.008436625 | 0.025600024 |
| ENSG00000261560 | ENSG0000026 | 14.07040703 | 0.035248555 | 0.465440589 | 0.075731588 | 0.939632644 | 0.971233034 |
| ENSG00000261572 | ENSG0000026 | 36.33082552 | 0.025515254 | 0.296199438 | 0.086142142 | 0.931353424 | 0.967059591 |
| ENSG00000261575 | ENSG0000026 | 8.446056963 | 0.559932048 | 0.620617282 | 0.902217945 | 0.366941104 | 0.559399478 |
| ENSG00000261578 | ENSG0000026 | 338.8442255 | 0.759942383 | 0.116305855 | 6.533999339 | 6.40E-11 | 5.62E-10 |
| ENSG00000261582 | ENSG0000026 | 6.519795194 | -0.712060205 | 0.679699547 | -1.04761024 | 0.294818218 | 0.480662112 |
| ENSG00000261587 | TMEM249 | 3.867042156 | 0.43476618 | 0.904185363 | 0.480837446 | 0.630632035 | 0.78221733 |
| ENSG00000261594 | TPBGL | 12.59018003 | 0.855247234 | 0.50746605 | 1.685328967 | 0.09192512 | 0.197894356 |
| ENSG00000261595 | PPM1J-DT | 2.621735529 | -0.493193393 | 1.144200483 | -0.431037567 | 0.666441055 | 0.807828682 |
| ENSG00000261596 | ENSG0000026 | 6.567041813 | -0.179178823 | 0.66999145 | -0.267434492 | 0.789134645 | 0.885944625 |
| ENSG00000261600 | ENSG0000026 | 6.14347236 | -0.440914144 | 0.703786916 | -0.626488124 | 0.530994824 | 0.706555758 |
| ENSG00000261602 | ENSG0000026 | 9.554595525 | -0.626280324 | 0.58240774 | -1.075329671 | 0.282227157 | 0.465675823 |
| ENSG00000261604 | ENSG0000026 | 40.54279074 | -0.99313799 | 0.298244052 | -3.329950697 | 0.000868614 | 0.003289747 |
| ENSG00000261606 | ENSG0000026 | 7.172297404 | 1.838619205 | 0.733800845 | 2.505610641 | 0.012224015 | 0.035509548 |
| ENSG00000261609 | GAN | 542.117757 | -0.14035375 | 0.087476522 | -1.604473377 | 0.108609748 | 0.226438937 |
| ENSG00000261612 | SUB1P3 | 24.22290569 | 0.171875988 | 0.366316831 | 0.469200359 | 0.638926429 | 0.787290808 |
| ENSG00000261614 | YBX3P1 | 90.24044811 | 0.19972227 | 0.189905194 | 1.051694617 | 0.292939682 | 0.478260359 |
| ENSG00000261616 | TTC23-AS1 | 7.608652911 | 1.085152474 | 0.675869814 | 1.605564343 | 0.108369665 | 0.226102785 |
| ENSG00000261634 | PAQR5-DT | 46.83761044 | -2.853372328 | 0.323790832 | -8.812393816 | 1.23E-18 | 1.89E-17 |

| | | | | | | | |
|---|---|---|---|---|---|---|---|
| ENSG00000261635 | ENSG0000026 | 2.017865109 | -1.184607296 | 1.244050994 | -0.952217635 | 0.340986619 | 0.5320885 |
| ENSG00000261641 | ENSG0000026 | 14.57520295 | 0.054520028 | 0.455950156 | 0.119574534 | 0.904820194 | 0.952267525 |
| ENSG00000261647 | IMPDH1P11 | 4.394124801 | -0.076953342 | 0.865988866 | -0.088861814 | 0.929191731 | 0.965981609 |
| ENSG00000261652 | PIERCE2 | 28.64342949 | -1.059272624 | 0.336358496 | -3.149237007 | 0.001636974 | 0.005867474 |
| ENSG00000261654 | ENSG0000026 | 18.39467904 | 0.423056159 | 0.412996727 | 1.024357172 | 0.305666603 | 0.493177091 |
| ENSG00000261655 | ENSG0000026 | 2.174861433 | 2.423645731 | 1.414803994 | 1.713061132 | 0.086701294 | 0.188667015 |
| ENSG00000261659 | ENSG0000026 | 76.58290641 | 0.511397373 | 0.205557442 | 2.487856283 | 0.012851566 | 0.03705567 |
| ENSG00000261663 | ECI1-AS1 | 20.74650145 | 0.111528238 | 0.396008174 | 0.281631151 | 0.778226351 | 0.879589804 |
| ENSG00000261664 | TTC39A-AS1 | 64.07382123 | -0.596069091 | 0.224326815 | -2.657145961 | 0.007880532 | 0.024136101 |
| ENSG00000261684 | ENSG0000026 | 13.07821361 | -0.957506545 | 0.504197504 | -1.899070377 | 0.057555223 | 0.134836277 |
| ENSG00000261691 | ENSG0000026 | 10.3064202 | 0.1424781 | 0.542851901 | 0.262462193 | 0.79296513 | 0.888779318 |
| ENSG00000261693 | ENSG0000026 | 17.09854394 | 0.400910243 | 0.423683397 | 0.946249596 | 0.344021295 | 0.534968039 |
| ENSG00000261707 | ENSG0000026 | 5.270077852 | -1.516324887 | 0.817540055 | -1.85474079 | 0.063633251 | 0.146472685 |
| ENSG00000261713 | SSTR5-AS1 | 13.98257143 | 2.638369307 | 0.588103329 | 4.486234264 | 7.25E-06 | 3.80E-05 |
| ENSG00000261716 | H2BC20P | 4006.779731 | 0.574360161 | 0.052886972 | 10.8601446 | 1.78E-27 | 4.53E-26 |
| ENSG00000261731 | ENSG0000026 | 1.718608394 | -0.817057398 | 1.38416372 | -0.590289563 | 0.554996535 | NA |
| ENSG00000261732 | ENSG0000026 | 4.334493483 | 0.155504909 | 0.832364989 | 0.186822981 | 0.851799428 | 0.922505405 |
| ENSG00000261744 | ZFPM1-AS1 | 14.33236425 | -0.700991563 | 0.490945395 | -1.427840182 | 0.153337864 | 0.296241419 |
| ENSG00000261754 | ENSG0000026 | 147.6729655 | -1.338070551 | 0.154298151 | -8.671980486 | 4.25E-18 | 6.31E-17 |
| ENSG00000261758 | STAG1-DT | 3.069004261 | -1.535714529 | 1.084452018 | -1.416120312 | 0.156740282 | 0.301077953 |
| ENSG00000261759 | ENSG0000026 | 2.277326706 | -1.482601514 | 1.259298352 | -1.177323477 | 0.23906642 | 0.414666496 |
| ENSG00000261762 | ENSG0000026 | 7.469226676 | -1.01047759 | 0.658683243 | -1.534087289 | 0.12500818 | 0.253553284 |
| ENSG00000261764 | KRT18P18 | 11.42586096 | 0.39434215 | 0.542940356 | 0.726308417 | 0.467649721 | 0.65362119 |
| ENSG00000261766 | ENSG0000026 | 4.601700168 | 0.004334698 | 0.803631967 | 0.005393885 | 0.995696324 | 0.997217798 |
| ENSG00000261770 | ENSG0000026 | 5.644290556 | 0.660559262 | 0.787584868 | 0.838715025 | 0.401629245 | 0.594372936 |
| ENSG00000261774 | ENSG0000026 | 2.794202374 | -1.047256793 | 1.234645705 | -0.848224546 | 0.396312929 | 0.589242245 |
| ENSG00000261775 | ENSG0000026 | 5.119386092 | -0.076790543 | 0.786443935 | -0.097642743 | 0.922215983 | 0.962243482 |
| ENSG00000261777 | DDX19A-DT | 37.28381201 | -0.036423726 | 0.288109193 | -0.126423337 | 0.89939683 | 0.949128866 |
| ENSG00000261779 | ENSG0000026 | 2.275039045 | -0.192045313 | 1.158779471 | -0.165730683 | 0.868368899 | 0.931294181 |
| ENSG00000261783 | ENSG0000026 | 8.895492734 | -0.596359401 | 0.595037773 | -1.002221083 | 0.316236828 | 0.505010291 |
| ENSG00000261787 | TCF24 | 53.98653618 | 0.549902076 | 0.246362511 | 2.232085044 | 0.025609343 | 0.067783353 |
| ENSG00000261790 | ENSG0000026 | 28.07298382 | 0.282601492 | 0.352987766 | 0.800598547 | 0.423364092 | 0.614254144 |
| ENSG00000261794 | GOLGA8H | 2.573862491 | -2.185789397 | 1.216339588 | -1.797022327 | 0.072332075 | 0.163108012 |

| | | | | | | | |
|---|---|---|---|---|---|---|---|
| ENSG00000261798 | ENSG0000026 | 11.50250583 | 0.119440345 | 0.520522798 | 0.229462275 | 0.818509638 | 0.903938528 |
| ENSG00000261799 | ENSG0000026 | 61.24532001 | 0.750030815 | 0.234769452 | 3.194754725 | 0.001399497 | 0.005085453 |
| ENSG00000261801 | LOXL1-AS1 | 329.1813479 | 0.134615511 | 0.115721202 | 1.163274385 | 0.244718195 | 0.421585716 |
| ENSG00000261804 | ENSG0000026 | 115.2623177 | -0.664990932 | 0.168386435 | -3.949195377 | 7.84E-05 | 0.000356062 |
| ENSG00000261812 | TUBB8P7 | 2.077865731 | 0.071052742 | 1.19822842 | 0.059298161 | 0.952714626 | 0.978056463 |
| ENSG00000261819 | ENSG0000026 | 19.39143487 | 0.125476264 | 0.408732389 | 0.306988795 | 0.758851901 | 0.867237712 |
| ENSG00000261822 | ENSG0000026 | 1.786719323 | -1.41111177 | 1.436324084 | -0.98244664 | 0.325879859 | NA |
| ENSG00000261824 | LINC00662 | 374.8812795 | 0.617472852 | 0.104809395 | 5.891388382 | 3.83E-09 | 2.88E-08 |
| ENSG00000261840 | ENSG0000026 | 34.87732161 | 0.354747409 | 0.298969853 | 1.186565821 | 0.235398924 | 0.410400617 |
| ENSG00000261845 | ENSG0000026 | 15.18208258 | -0.101247279 | 0.490886617 | -0.206253899 | 0.836592587 | 0.913834291 |
| ENSG00000261873 | SMIM36 | 2.90146476 | 0.195895171 | 1.033076875 | 0.189623033 | 0.849604537 | 0.921263493 |
| ENSG00000261879 | ZNF594-DT | 38.76914037 | 0.225764112 | 0.2938174 | 0.768382374 | 0.442260049 | 0.631470633 |
| ENSG00000261884 | ENSG0000026 | 18.91758641 | -0.238734031 | 0.407322275 | -0.586106004 | 0.557804282 | 0.727915708 |
| ENSG00000261886 | GOSR2-DT | 2.132390747 | -1.301348857 | 1.34886707 | -0.96477176 | 0.334659143 | 0.525755427 |
| ENSG00000261888 | ENSG0000026 | 294.827594 | 1.389887311 | 0.120782618 | 11.50734542 | 1.21E-30 | 3.44E-29 |
| ENSG00000261889 | ENSG0000026 | 21.02854288 | 0.861175068 | 0.405236974 | 2.125114741 | 0.03357704 | 0.085527126 |
| ENSG00000261898 | ENSG0000026 | 3.072237569 | -0.873130203 | 1.012896481 | -0.862013266 | 0.388680218 | 0.58132248 |
| ENSG00000261934 | PCDHGA9 | 26.47846676 | 1.475116299 | 0.373197345 | 3.952644141 | 7.73E-05 | 0.000351594 |
| ENSG00000261938 | ENSG0000026 | 2.440488612 | -1.11266991 | 1.246776627 | -0.892437255 | 0.37215861 | 0.564841384 |
| ENSG00000261971 | MMP25-AS1 | 663.7716876 | 0.135683041 | 0.087318613 | 1.55388452 | 0.120211965 | 0.245934404 |
| ENSG00000262001 | DLGAP1-AS2 | 49.34689629 | 0.838712326 | 0.262792068 | 3.191543541 | 0.001415148 | 0.005135816 |
| ENSG00000262049 | ENSG0000026 | 341.023146 | 0.460577462 | 0.10807015 | 4.261837906 | 2.03E-05 | 0.000100631 |
| ENSG00000262050 | ENSG0000026 | 7.847855773 | -0.733006422 | 0.632145494 | -1.159553344 | 0.246230706 | 0.423428216 |
| ENSG00000262075 | DKFZP434A06 | 2.018897631 | 1.341851115 | 1.349296945 | 0.994481697 | 0.319988412 | 0.509511638 |
| ENSG00000262089 | ENSG0000026 | 17.21518474 | 0.004295364 | 0.455503537 | 0.009429924 | 0.992476121 | 0.995949353 |
| ENSG00000262094 | ENSG0000026 | 18.78383344 | 0.206118794 | 0.436255297 | 0.472472874 | 0.636589299 | 0.786056546 |
| ENSG00000262112 | ENSG0000026 | 10.39230487 | -0.144740368 | 0.557671424 | -0.259544172 | 0.795215405 | 0.890473562 |
| ENSG00000262115 | ENSG0000026 | 5.428600147 | -0.419882871 | 0.792601834 | -0.529752585 | 0.596283483 | 0.757195854 |
| ENSG00000262136 | NQO1-DT | 51.56163808 | -0.357255245 | 0.252384401 | -1.415520308 | 0.156915998 | 0.301327055 |
| ENSG00000262152 | GREP1 | 29.83126465 | -0.73068502 | 0.329795104 | -2.215572675 | 0.026720782 | 0.070335598 |
| ENSG00000262155 | LINC02175 | 36.39043943 | -0.039598503 | 0.296509707 | -0.133548757 | 0.893759407 | 0.945700277 |
| ENSG00000262160 | ENSG0000026 | 1404.619256 | 0.145766267 | 0.063072219 | 2.311100962 | 0.020827279 | 0.056641519 |
| ENSG00000262165 | C17orf114 | 10.07852005 | 0.182218181 | 0.56488389 | 0.322576345 | 0.74701611 | 0.859750631 |

| | | | | | | | |
|---|---|---|---|---|---|---|---|
| ENSG00000262172 | ENSG0000026 | 7.838045593 | -0.015376949 | 0.619868588 | -0.024806789 | 0.980209076 | 0.991052765 |
| ENSG00000262185 | LINC02861 | 8.600943681 | -0.23367365 | 0.599894386 | -0.389524649 | 0.69688808 | 0.829719188 |
| ENSG00000262198 | ENSG0000026 | 11.93099242 | -0.217670489 | 0.523931956 | -0.415455647 | 0.677808378 | 0.816688071 |
| ENSG00000262209 | PCDHGB3 | 15.35496003 | 1.082746444 | 0.466470276 | 2.321147775 | 0.02027887 | 0.055295052 |
| ENSG00000262211 | ENSG0000026 | 6.388434007 | -0.490958951 | 0.724685813 | -0.677478353 | 0.4981025 | 0.67994219 |
| ENSG00000262223 | LINC03048 | 59.81440641 | -0.155812153 | 0.243267286 | -0.640497766 | 0.521849041 | 0.69885614 |
| ENSG00000262227 | ENSG0000026 | 21.57493227 | 0.120669511 | 0.373116606 | 0.32340965 | 0.746385022 | 0.859604617 |
| ENSG00000262246 | CORO7 | 151.3848834 | 0.106168601 | 0.150998896 | 0.703108459 | 0.481988161 | 0.666287182 |
| ENSG00000262248 | ENSG0000026 | 3.077408138 | -1.189338807 | 1.055510128 | -1.126790521 | 0.259831059 | 0.439858645 |
| ENSG00000262294 | RPH3AL-AS2 | 2.217062256 | -1.282774106 | 1.329020614 | -0.965202565 | 0.334443362 | 0.525488406 |
| ENSG00000262313 | ENSG0000026 | 2.380857294 | -0.670930359 | 1.182244847 | -0.56750542 | 0.570370845 | 0.737351771 |
| ENSG00000262333 | HNRNPA1P16 | 47.71364355 | -0.052449069 | 0.256834423 | -0.204213548 | 0.838186624 | 0.91492202 |
| ENSG00000262358 | ENSG0000026 | 2.341276979 | -0.610742234 | 1.138459543 | -0.536463713 | 0.591638108 | 0.7540099 |
| ENSG00000262362 | ENSG0000026 | 8.587767549 | -0.408186627 | 0.590175384 | -0.691636144 | 0.489165858 | 0.67242718 |
| ENSG00000262402 | MCUR1P1 | 3.422532253 | -2.166611458 | 1.090406422 | -1.986976063 | 0.04692505 | 0.113419096 |
| ENSG00000262406 | MMP12 | 4.756808898 | -0.630569226 | 0.809940794 | -0.778537432 | 0.436252248 | 0.626181115 |
| ENSG00000262410 | ENSG0000026 | 4.506663707 | -0.510513242 | 0.810788598 | -0.629650248 | 0.52892344 | 0.704892166 |
| ENSG00000262412 | ENSG0000026 | 10.65148524 | -0.412556388 | 0.557503198 | -0.740007215 | 0.459295617 | 0.646395181 |
| ENSG00000262413 | ENSG0000026 | 104.2263641 | -0.234376255 | 0.211331011 | -1.109048092 | 0.267409436 | 0.448648611 |
| ENSG00000262420 | TXNDC11-AS1 | 2.362560041 | -0.630854643 | 1.130821594 | -0.55787283 | 0.576931226 | 0.743036617 |
| ENSG00000262429 | ENSG0000026 | 6.90994591 | -0.833221295 | 0.729302578 | -1.142490538 | 0.253250177 | 0.431958494 |
| ENSG00000262454 | MIR193BHG | 107.924956 | 0.483674117 | 0.186232085 | 2.597157823 | 0.009399872 | 0.028179962 |
| ENSG00000262456 | ENSG0000026 | 5.698008571 | -1.234489542 | 0.794426624 | -1.553937777 | 0.12019926 | 0.245930344 |
| ENSG00000262468 | LINC01569 | 238.3784824 | -0.256844653 | 0.134250025 | -1.913181421 | 0.055724831 | 0.131393527 |
| ENSG00000262470 | TVP23CP2 | 2.373452191 | -0.644283597 | 1.130358996 | -0.569981395 | 0.568690317 | 0.736073456 |
| ENSG00000262482 | ENSG0000026 | 3.306227957 | -0.153924788 | 0.968837157 | -0.15887581 | 0.873766719 | 0.934676032 |
| ENSG00000262484 | CCER2 | 21.47046577 | 0.59617436 | 0.379700783 | 1.570116226 | 0.116388074 | 0.239714709 |
| ENSG00000262497 | FAM187B2P | 2.631059396 | 1.510050649 | 1.209388103 | 1.248607163 | 0.211808791 | 0.379423213 |
| ENSG00000262500 | MAPK8IP1P1 | 3.809747319 | 0.576592467 | 0.962882606 | 0.598819071 | 0.549293544 | 0.721777632 |
| ENSG00000262519 | TXNP4 | 49.92747653 | 0.166107228 | 0.252320152 | 0.658319308 | 0.510332974 | 0.690273492 |
| ENSG00000262521 | ENSG0000026 | 3.760209989 | 1.242477478 | 0.988196192 | 1.257318625 | 0.208638282 | 0.375149882 |
| ENSG00000262528 | ENSG0000026 | 44.53038584 | -3.194782445 | 0.368348912 | -8.673250668 | 4.20E-18 | 6.24E-17 |
| ENSG00000262533 | ENSG0000026 | 19.78482125 | -0.646121213 | 0.391471529 | -1.650493495 | 0.098842043 | 0.210025608 |

| | | | | | | | |
|---|---|---|---|---|---|---|---|
| ENSG00000262576 | PCDHGA4 | 41.22977842 | 0.720115415 | 0.281063064 | 2.562113302 | 0.010403737 | 0.030775207 |
| ENSG00000262580 | ENSG0000026 | 201.4480158 | 0.032602528 | 0.136890454 | 0.238165093 | 0.81175305 | 0.900377863 |
| ENSG00000262583 | TMEM231P1 | 16.45680054 | 0.549608624 | 0.450307637 | 1.220518106 | 0.22226853 | 0.393307627 |
| ENSG00000262585 | LINC01979 | 35.24459575 | 0.894368503 | 0.323807383 | 2.762038641 | 0.005744168 | 0.018205473 |
| ENSG00000262587 | ENSG0000026 | 33.38807015 | 0.215514666 | 0.318350323 | 0.676973291 | 0.4984229 | 0.680139274 |
| ENSG00000262619 | LINC00621 | 18.72613763 | -0.47900278 | 0.41828985 | -1.145145597 | 0.25214885 | 0.430795756 |
| ENSG00000262624 | ENSG0000026 | 2.700534316 | 0.033641565 | 1.086920206 | 0.030951274 | 0.975308399 | 0.988740606 |
| ENSG00000262652 | ENSG0000026 | 3.156483271 | -0.247506996 | 1.032248297 | -0.239774671 | 0.810504944 | 0.899645977 |
| ENSG00000262662 | ENSG0000026 | 1.888286667 | 0.367016558 | 1.294321605 | 0.283559014 | 0.776748351 | 0.878802878 |
| ENSG00000262663 | ENSG0000026 | 5.635713234 | -0.048566511 | 0.745635379 | -0.065134397 | 0.948066994 | 0.975976343 |
| ENSG00000262678 | CAMTA2-AS1 | 3.054685351 | -0.018125728 | 0.995481174 | -0.018208007 | 0.985472915 | 0.993350589 |
| ENSG00000262691 | ENSG0000026 | 23.35377255 | -1.080612657 | 0.378043521 | -2.85843454 | 0.004257369 | 0.013946031 |
| ENSG00000262693 | ZFP3-DT | 2.875086537 | -0.014792232 | 1.041308302 | -0.01420543 | 0.988666088 | 0.994082067 |
| ENSG00000262712 | ENSG0000026 | 34.67772799 | 0.297453287 | 0.305330021 | 0.974202558 | 0.329955975 | 0.520505437 |
| ENSG00000262714 | ENSG0000026 | 60.65868192 | -0.397936541 | 0.2326384 | -1.710536788 | 0.087166657 | 0.189481958 |
| ENSG00000262728 | ARHGAP11A-[ | 5.479684514 | -0.284113554 | 0.736155351 | -0.385942387 | 0.699539337 | 0.831516265 |
| ENSG00000262732 | ENSG0000026 | 6.288273048 | -0.311577711 | 0.70641219 | -0.441070688 | 0.659161823 | 0.802846438 |
| ENSG00000262766 | ENSG0000026 | 30.15344712 | -0.080777351 | 0.322594506 | -0.250399027 | 0.802278782 | 0.894714402 |
| ENSG00000262768 | ENSG0000026 | 5.588830407 | 0.80086177 | 0.764142535 | 1.048052861 | 0.294614253 | 0.480572301 |
| ENSG00000262772 | LINC01977 | 196.0849152 | 0.202941662 | 0.130466143 | 1.555512088 | 0.119824158 | 0.245272257 |
| ENSG00000262791 | ENSG0000026 | 3.500125977 | 0.514766772 | 0.948714196 | 0.542594149 | 0.587409269 | 0.751053167 |
| ENSG00000262801 | ENSG0000026 | 10.78771494 | 0.650553425 | 0.565128028 | 1.151161141 | 0.249665948 | 0.427721961 |
| ENSG00000262814 | MRPL12 | 129.685864 | 0.37201588 | 0.16539648 | 2.249236979 | 0.024497423 | 0.065345451 |
| ENSG00000262831 | ENSG0000026 | 128.4322347 | -0.231015714 | 0.199565103 | -1.157595746 | 0.247029046 | 0.424292177 |
| ENSG00000262877 | ENSG0000026 | 319.3467636 | 0.822863709 | 0.114209833 | 7.204841222 | 5.81E-13 | 6.04E-12 |
| ENSG00000262879 | ENSG0000026 | 1265.575521 | 0.421391633 | 0.067175993 | 6.272949841 | 3.54E-10 | 2.94E-09 |
| ENSG00000262884 | ENSG0000026 | 11.08804498 | 0.786213602 | 0.53893187 | 1.458836721 | 0.144610053 | 0.283316647 |
| ENSG00000262898 | ENSG0000026 | 2.290231548 | -1.964118696 | 1.281931211 | -1.532156078 | 0.125483924 | 0.25422602 |
| ENSG00000262902 | MTCO1P40 | 117.3159573 | -0.070084057 | 0.193151674 | -0.362844677 | 0.716720916 | 0.842466582 |
| ENSG00000262903 | CTNS-AS1 | 35.14507206 | -0.132324337 | 0.294923575 | -0.448673312 | 0.65366734 | 0.799053343 |
| ENSG00000262904 | TMPOP2 | 33.05675827 | -0.03978067 | 0.309282865 | -0.128622288 | 0.89765653 | 0.948077385 |
| ENSG00000262919 | CCNQ | 654.0244046 | -0.708959784 | 0.083291743 | -8.511765516 | 1.71E-17 | 2.44E-16 |
| ENSG00000262943 | ALOX12P2 | 165.5942048 | 0.226204886 | 0.1460716 | 1.548589087 | 0.121480531 | 0.247778375 |

| | | | | | | | |
|---|---|---|---|---|---|---|---|
| ENSG00000262953 | EIF4A1P9 | 7.247502144 | 0.081184374 | 0.668292539 | 0.121480294 | 0.903310622 | 0.951158803 |
| ENSG00000262967 | ENSG0000026 | 8.965393787 | -1.139100584 | 0.63307641 | -1.799309791 | 0.07196969 | 0.162418563 |
| ENSG00000262979 | ENSG0000026 | 4.72277897 | 0.511091184 | 0.811890904 | 0.629507218 | 0.529017045 | 0.704946326 |
| ENSG00000263001 | GTF2I | 3653.497869 | 0.353062699 | 0.046322245 | 7.621882238 | 2.50E-14 | 2.86E-13 |
| ENSG00000263002 | ZNF234 | 337.1356123 | 0.157752876 | 0.100831483 | 1.564520049 | 0.117695488 | 0.241929615 |
| ENSG00000263004 | AKAP1-DT | 122.4891666 | -0.056948328 | 0.163200499 | -0.348947023 | 0.727129082 | 0.848440547 |
| ENSG00000263006 | ROCK1P1 | 26.44796825 | 0.835906165 | 0.361342366 | 2.313335616 | 0.020704194 | 0.05634095 |
| ENSG00000263011 | ENSG0000026 | 4.057872681 | -0.52099939 | 0.878113548 | -0.593316651 | 0.552969257 | 0.72450299 |
| ENSG00000263017 | ENSG0000026 | 2.267108914 | -0.432856721 | 1.192338584 | -0.363031715 | 0.716581193 | 0.842417634 |
| ENSG00000263033 | ENSG0000026 | 2.97109949 | 0.402113632 | 1.042681769 | 0.385653269 | 0.699753477 | 0.831642103 |
| ENSG00000263050 | DPH1-AS1 | 2.094866636 | -0.765731215 | 1.293420786 | -0.592020186 | 0.553837071 | 0.725418397 |
| ENSG00000263063 | ENSG0000026 | 1.789429111 | -1.547507352 | 1.367836901 | -1.131353709 | 0.257906247 | NA |
| ENSG00000263069 | RNF213-AS1 | 88.97814871 | 0.349038036 | 0.193118049 | 1.807381749 | 0.070702778 | 0.159968608 |
| ENSG00000263072 | ZNF213-AS1 | 518.9126818 | 0.173203254 | 0.089746029 | 1.929926653 | 0.053615926 | 0.12728442 |
| ENSG00000263080 | ENSG0000026 | 4.874431824 | -1.106725869 | 0.818604148 | -1.351967092 | 0.176385843 | 0.329959328 |
| ENSG00000263105 | ENSG0000026 | 4.366321672 | -0.210810468 | 0.843808879 | -0.24983201 | 0.802717264 | 0.895072865 |
| ENSG00000263126 | ENSG0000026 | 15.47497936 | 0.126711827 | 0.443772015 | 0.285533613 | 0.775235356 | 0.877740441 |
| ENSG00000263142 | LRRC37A17P | 2.979131012 | 0.407357873 | 1.029618005 | 0.395639811 | 0.692370764 | 0.826464009 |
| ENSG00000263155 | MYZAP | 2.661323306 | 1.524100026 | 1.171624572 | 1.300843344 | 0.193312082 | 0.353992189 |
| ENSG00000263165 | ENSG0000026 | 3.940633576 | 0.685185055 | 0.910779757 | 0.752305976 | 0.451867075 | 0.639731883 |
| ENSG00000263171 | ENSG0000026 | 2.687822624 | -0.957355936 | 1.129754849 | -0.847401485 | 0.396771377 | 0.5896749 |
| ENSG00000263179 | HNRNPCP4 | 19.90346123 | -0.436708689 | 0.400815084 | -1.089551532 | 0.275910743 | 0.458253146 |
| ENSG00000263218 | ENSG0000026 | 5.56627879 | 0.037631159 | 0.746716458 | 0.050395512 | 0.959807212 | 0.981674504 |
| ENSG00000263219 | RYKP1 | 9.839794206 | 0.013946676 | 0.580206922 | 0.024037417 | 0.980822763 | 0.991149032 |
| ENSG00000263232 | ATP5F1AP3 | 21.65701335 | 0.47569708 | 0.403808875 | 1.178025323 | 0.238786511 | 0.414322862 |
| ENSG00000263235 | ENSG0000026 | 42.30337975 | -0.166010592 | 0.270627704 | -0.613427929 | 0.539593423 | 0.714103677 |
| ENSG00000263266 | RPS7P1 | 4342.999137 | 0.184538971 | 0.048794891 | 3.781932194 | 0.000155616 | 0.000672664 |
| ENSG00000263272 | ENSG0000026 | 5.956780208 | -0.617120432 | 0.716466931 | -0.861338333 | 0.389051729 | 0.581688501 |
| ENSG00000263276 | ENSG0000026 | 2.108129641 | 0.095085449 | 1.214629842 | 0.078283479 | 0.937602559 | 0.970153346 |
| ENSG00000263280 | ENSG0000026 | 4.537456323 | -1.40866404 | 0.874940194 | -1.610011804 | 0.107395279 | 0.224519066 |
| ENSG00000263293 | EFCAB13-DT | 2.325247682 | 1.110972332 | 1.19448286 | 0.930086457 | 0.352326322 | 0.543727512 |
| ENSG00000263311 | ENSG0000026 | 1.615061152 | 0.462591465 | 1.461815123 | 0.316450047 | 0.751660935 | NA |
| ENSG00000263325 | ENSG0000026 | 3.988495148 | 1.143694553 | 0.962205178 | 1.188618165 | 0.234589956 | 0.409524493 |

| | | | | | | | |
|---|---|---|---|---|---|---|---|
| ENSG00000263327 | TAPT1-AS1 | 79.49762869 | 0.157973627 | 0.200188881 | 0.789122881 | 0.43004019 | 0.620369888 |
| ENSG00000263335 | ENSG0000026 | 45.21454627 | -0.279506184 | 0.262348637 | -1.065399798 | 0.286695048 | 0.470509687 |
| ENSG00000263345 | SGSM2-AS1 | 36.47025278 | -0.284756003 | 0.294840018 | -0.965798349 | 0.334145095 | 0.525127662 |
| ENSG00000263366 | ABHD17AP5 | 1.85472886 | -1.580021182 | 1.365093087 | -1.157445742 | 0.247090295 | 0.424302071 |
| ENSG00000263369 | ENSG0000026 | 2.247608596 | 0.273042662 | 1.159350002 | 0.235513573 | 0.813810145 | 0.901448243 |
| ENSG00000263388 | ENSG0000026 | 3.997494191 | -0.162157557 | 0.920886996 | -0.176088443 | 0.86022446 | 0.927382816 |
| ENSG00000263412 | NFE2L1-DT | 31.01876795 | -0.199031479 | 0.334783479 | -0.594508069 | 0.552172347 | 0.723944914 |
| ENSG00000263424 | ENSG0000026 | 2.573394373 | 0.236610833 | 1.143796523 | 0.206864445 | 0.836115725 | 0.913443886 |
| ENSG00000263432 | RN7SL689P | 16.3279462 | 0.790267745 | 0.439274712 | 1.799028545 | 0.072014165 | 0.162502946 |
| ENSG00000263465 | SRSF8 | 883.4627362 | -0.294769397 | 0.070517065 | -4.180114388 | 2.91E-05 | 0.000141397 |
| ENSG00000263503 | MAPK8IP1P2 | 12.03976791 | 1.086158189 | 0.540121547 | 2.010951414 | 0.044330589 | 0.108095189 |
| ENSG00000263513 | FAM72C | 2.391334778 | 0.473720737 | 1.173394489 | 0.403718222 | 0.686419939 | 0.822742745 |
| ENSG00000263528 | IKBKE | 529.9181859 | 0.872899534 | 0.088926363 | 9.815981482 | 9.61E-23 | 1.88E-21 |
| ENSG00000263531 | ENSG0000026 | 1.872223948 | 0.853262458 | 1.30009931 | 0.6563056 | 0.511627515 | 0.691250085 |
| ENSG00000263535 | AK4P1 | 212.3656485 | 0.006948894 | 0.12387773 | 0.056094782 | 0.955266301 | 0.979455025 |
| ENSG00000263563 | UBBP4 | 92.80898071 | -0.157356222 | 0.383873704 | -0.409916649 | 0.681867092 | 0.819383504 |
| ENSG00000263585 | ENSG0000026 | 43.29343893 | -0.28758475 | 0.294657891 | -0.975995412 | 0.329066738 | 0.519478277 |
| ENSG00000263606 | CHORDC1P4 | 50.261698 | 0.126113563 | 0.251367835 | 0.50170923 | 0.615872071 | 0.772407485 |
| ENSG00000263618 | ENSG0000026 | 22.78772862 | 1.284353913 | 0.39173751 | 3.278608454 | 0.001043203 | 0.003890598 |
| ENSG00000263627 | PPP4R1-AS1 | 3.8906092 | 0.813224263 | 0.932340246 | 0.872239793 | 0.383077575 | 0.575532096 |
| ENSG00000263639 | MSMB | 20.50897421 | -1.420479005 | 0.415325657 | -3.420157125 | 0.00062585 | 0.002432564 |
| ENSG00000263675 | MIR5581 | 12.43641507 | -0.370076415 | 0.503800422 | -0.734569482 | 0.46260173 | 0.64917622 |
| ENSG00000263718 | SEPTIN9-DT | 3.222046847 | -1.302431323 | 0.999494058 | -1.30309061 | 0.192543829 | 0.352791968 |
| ENSG00000263731 | ENSG0000026 | 64.6125095 | 0.510988166 | 0.226192495 | 2.259085411 | 0.023878072 | 0.063871616 |
| ENSG00000263740 | RN7SL4P | 1.538492895 | 2.847924683 | 1.672351832 | 1.702945892 | 0.088578195 | NA |
| ENSG00000263753 | LINC00667 | 2.346564965 | -2.719648465 | 1.480809725 | -1.836595491 | 0.066269633 | 0.151719101 |
| ENSG00000263756 | ENSG0000026 | 37.4780524 | -0.686424016 | 0.318193898 | -2.157250718 | 0.030986135 | 0.079894563 |
| ENSG00000263766 | KPNB1-DT | 41.68536172 | -0.121141611 | 0.277753098 | -0.436148551 | 0.662728954 | 0.805201112 |
| ENSG00000263781 | ENSG0000026 | 21.033157 | -0.127042223 | 0.416643987 | -0.304917932 | 0.760428658 | 0.868347524 |
| ENSG00000263786 | ENSG0000026 | 14.52660716 | -0.217362871 | 0.461214311 | -0.471283881 | 0.637438025 | 0.786553874 |
| ENSG00000263818 | RDM1P5 | 31.24018143 | -0.61561702 | 0.329670046 | -1.867373233 | 0.061849485 | 0.143042706 |
| ENSG00000263821 | ENSG0000026 | 1.570618658 | 1.854896672 | 1.53002746 | 1.212329007 | 0.225386464 | NA |
| ENSG00000263823 | ENSG0000026 | 11.98439647 | 1.169643135 | 0.52648948 | 2.221588804 | 0.026311107 | 0.069352729 |

| | | | | | | | |
|---|---|---|---|---|---|---|---|
| ENSG00000263826 | ENSG0000026 | 16.53595372 | -0.229970487 | 0.454081031 | -0.506452531 | 0.612539005 | 0.769672624 |
| ENSG00000263829 | SINHCAFP1 | 2.072219582 | 0.551950995 | 1.248487378 | 0.442095775 | 0.658419902 | 0.802336104 |
| ENSG00000263834 | MIR4635 | 2.61883259 | 0.938859204 | 1.096580036 | 0.856170251 | 0.391903615 | 0.584770973 |
| ENSG00000263843 | MIF4GD-DT | 236.1997051 | 0.547153893 | 0.131351251 | 4.165578098 | 3.11E-05 | 0.000150208 |
| ENSG00000263846 | CIAPIN1P | 35.22697833 | -0.21113982 | 0.308089949 | -0.685318753 | 0.4931428 | 0.675909413 |
| ENSG00000263874 | LASP1NB | 61.80296145 | 0.166856262 | 0.237508693 | 0.70252697 | 0.482350589 | 0.666413063 |
| ENSG00000263884 | THOC1-DT | 8.385920646 | -0.054721927 | 0.673683988 | -0.081227887 | 0.935260722 | 0.969069964 |
| ENSG00000263887 | USP32P4 | 7.096897005 | -0.651823564 | 0.714075691 | -0.91282139 | 0.361336487 | 0.553896359 |
| ENSG00000263916 | ENSG0000026 | 6.804523251 | -0.350968832 | 0.68658856 | -0.511177803 | 0.60922656 | 0.767429427 |
| ENSG00000263934 | SNORD3A | 13.42211889 | 0.092974646 | 0.516505723 | 0.180007001 | 0.857147072 | 0.926012707 |
| ENSG00000263946 | ENSG0000026 | 3.606023743 | 4.169761327 | 1.422647901 | 2.930986173 | 0.003378878 | 0.011327118 |
| ENSG00000263956 | NBPF11 | 1092.631687 | 0.332042447 | 0.071835519 | 4.622259986 | 3.80E-06 | 2.06E-05 |
| ENSG00000263968 | RN7SL381P | 27.48037183 | 0.628998269 | 0.342270939 | 1.837720345 | 0.066103624 | 0.151384361 |
| ENSG00000263986 | ENSG0000026 | 14.20131115 | -0.357001951 | 0.461942953 | -0.772826923 | 0.439624818 | 0.629093214 |
| ENSG00000264017 | RN7SL336P | 2.861368388 | 1.579179295 | 1.140465 | 1.384680193 | 0.166150278 | 0.315198852 |
| ENSG00000264019 | ENSG0000026 | 16.37981342 | -0.378314565 | 0.434352739 | -0.870984642 | 0.383762539 | 0.576218727 |
| ENSG00000264044 | ENSG0000026 | 42.43420118 | -0.58837475 | 0.290103675 | -2.028153383 | 0.042544596 | 0.104426817 |
| ENSG00000264057 | ENSG0000026 | 8.392911965 | 0.286803251 | 0.608001939 | 0.471714369 | 0.637130679 | 0.786385275 |
| ENSG00000264058 | ENSG0000026 | 10.50107613 | -0.681485972 | 0.547901845 | -1.243810326 | 0.213569379 | 0.381810339 |
| ENSG00000264080 | ENSG0000026 | 2.515327659 | 0.369214613 | 1.145360906 | 0.32235657 | 0.747182582 | 0.859899083 |
| ENSG00000264107 | MIR4733HG | 9.900628164 | -0.651452257 | 0.561289201 | -1.160635651 | 0.245790101 | 0.422860725 |
| ENSG00000264112 | ENSG0000026 | 327.1004281 | 0.01670467 | 0.285080241 | 0.058596378 | 0.953273595 | 0.978321944 |
| ENSG00000264148 | ENSG0000026 | 8.832503252 | -0.575397793 | 0.601104711 | -0.957233877 | 0.338449221 | 0.529678342 |
| ENSG00000264176 | MAGOH2P | 6.610248872 | -0.304738816 | 0.688590742 | -0.442554332 | 0.658088123 | 0.802111553 |
| ENSG00000264179 | ENSG0000026 | 1.666585843 | -0.73595902 | 1.376671531 | -0.534593041 | 0.592931297 | NA |
| ENSG00000264187 | ENSG0000026 | 6.685012186 | 0.332717191 | 0.725931978 | 0.458331084 | 0.646714595 | 0.793519085 |
| ENSG00000264201 | MIR4701 | 4.592198295 | -0.921604599 | 0.868250437 | -1.061450198 | 0.288485354 | 0.472467625 |
| ENSG00000264204 | AGAP7P | 53.43191356 | -1.323850642 | 0.261126242 | -5.069772501 | 3.98E-07 | 2.43E-06 |
| ENSG00000264215 | ENSG0000026 | 83.62852656 | -0.62943854 | 0.247006767 | -2.548264351 | 0.01082604 | 0.031847827 |
| ENSG00000264229 | RNU4ATAC | 4.891587654 | 0.08514626 | 0.801230807 | 0.106269329 | 0.915368665 | 0.958033237 |
| ENSG00000264235 | MYL12-AS1 | 105.6526987 | -0.397425635 | 0.176848383 | -2.247267561 | 0.024622933 | 0.0656192 |
| ENSG00000264247 | ZNF407-AS1 | 336.4777497 | 0.403561504 | 0.102050929 | 3.954510833 | 7.67E-05 | 0.000348929 |
| ENSG00000264269 | DYM-AS1 | 7.257513646 | 0.853465784 | 0.702693019 | 1.2145642 | 0.22453235 | 0.396182764 |

| | | | | | | | |
|---|---|---|---|---|---|---|---|
| ENSG00000264278 | ZNF236-DT | 41.75181786 | 1.515508897 | 0.297147147 | 5.100196694 | 3.39E-07 | 2.08E-06 |
| ENSG00000264281 | ENSG0000026 | 24.47142859 | 0.050815612 | 0.368980747 | 0.137718872 | 0.890462604 | 0.943868964 |
| ENSG00000264290 | ENSG0000026 | 20.05604517 | 0.208051757 | 0.391208009 | 0.531818757 | 0.59485153 | 0.756439984 |
| ENSG00000264301 | LINC01444 | 26.69042835 | 2.983514813 | 0.451707696 | 6.604967857 | 3.98E-11 | 3.54E-10 |
| ENSG00000264311 | MIX23P1 | 8.041738649 | -0.125877353 | 0.615886993 | -0.204383847 | 0.838053552 | 0.914820294 |
| ENSG00000264324 | ENSG0000026 | 25.44801883 | -0.210237667 | 0.368296719 | -0.570837742 | 0.568109636 | 0.735755381 |
| ENSG00000264343 | NOTCH2NLA | 13.64370088 | 1.070372942 | 0.510302694 | 2.097525556 | 0.035947077 | 0.090687997 |
| ENSG00000264349 | MIR4258 | 4.101074856 | 0.043859378 | 0.900156609 | 0.048724164 | 0.961139119 | 0.982116705 |
| ENSG00000264350 | SNRPGP2 | 267.8501017 | -0.135818844 | 0.112110555 | -1.211472411 | 0.225714402 | 0.397748378 |
| ENSG00000264364 | DYNLL2 | 11369.9758 | 0.339264828 | 0.040242583 | 8.430493407 | 3.44E-17 | 4.79E-16 |
| ENSG00000264425 | MIR4653 | 11.37145519 | 0.366168307 | 0.544965033 | 0.671911562 | 0.501639999 | 0.682818106 |
| ENSG00000264443 | NCMAP-DT | 2.113406486 | -2.527904078 | 1.432254966 | -1.764981891 | 0.077566809 | 0.172487651 |
| ENSG00000264456 | ENSG0000026 | 16.33090892 | -0.7474289 | 0.44907947 | -1.664357756 | 0.096040957 | 0.205137761 |
| ENSG00000264491 | ENSG0000026 | 1.711959467 | 1.2792687 | 1.396189426 | 0.916257261 | 0.359531978 | NA |
| ENSG00000264522 | OTUD7B | 2625.760636 | 0.245530823 | 0.053378702 | 4.599790099 | 4.23E-06 | 2.28E-05 |
| ENSG00000264538 | SUZ12P1 | 206.5934505 | 0.283640702 | 0.129249634 | 2.194518411 | 0.028198162 | 0.073639446 |
| ENSG00000264548 | ENSG0000026 | 35.47702954 | 0.46819533 | 0.299075269 | 1.56547658 | 0.117471205 | 0.241540174 |
| ENSG00000264558 | ENSG0000026 | 61.93886193 | -0.605306441 | 0.238477685 | -2.53820998 | 0.011142111 | 0.032681246 |
| ENSG00000264569 | DCXR-DT | 7.633214538 | -1.491187197 | 0.667214982 | -2.234942616 | 0.025421116 | 0.067378441 |
| ENSG00000264570 | SNX19P3 | 12.49768392 | 0.09875008 | 0.499843334 | 0.197562062 | 0.84338772 | 0.917499611 |
| ENSG00000264577 | ENSG0000026 | 129.8568504 | -0.487693747 | 0.19531183 | -2.497000544 | 0.012524876 | 0.036245826 |
| ENSG00000264578 | ENSG0000026 | 12.31247045 | -0.108135536 | 0.526455432 | -0.20540302 | 0.837257261 | 0.914212079 |
| ENSG00000264589 | MAPT-AS1 | 6.593216797 | 0.67311724 | 0.692768114 | 0.971634269 | 0.331232528 | 0.522160128 |
| ENSG00000264635 | ENSG0000026 | 7.535856521 | -0.250926596 | 0.68831134 | -0.364553918 | 0.715444419 | 0.841656152 |
| ENSG00000264644 | KRT18P8 | 40.54189479 | 0.168565249 | 0.293627878 | 0.574077811 | 0.565915148 | 0.73394626 |
| ENSG00000264655 | PPIAP54 | 24.61292528 | 0.308059225 | 0.371989621 | 0.8281393 | 0.407591615 | 0.600369311 |
| ENSG00000264666 | ENSG0000026 | 5.604072357 | 0.323332467 | 0.735932469 | 0.439350729 | 0.660407422 | 0.803707149 |
| ENSG00000264672 | SEPTIN4-AS1 | 30.77561664 | -1.991817885 | 0.347272203 | -5.735609907 | 9.72E-09 | 7.00E-08 |
| ENSG00000264701 | ENSG0000026 | 19.42698435 | 0.012468659 | 0.400170699 | 0.03115835 | 0.975143256 | 0.988719836 |
| ENSG00000264736 | BDP1P | 1.740339242 | 0.651239229 | 1.34606348 | 0.48381019 | 0.628520582 | NA |
| ENSG00000264769 | ENSG0000026 | 4.140660055 | -0.774133319 | 0.861237212 | -0.898861902 | 0.368726223 | 0.561248726 |
| ENSG00000264772 | ENSG0000026 | 2.564002863 | -0.007247278 | 1.10797696 | -0.006541 | 0.994781074 | 0.99681023 |
| ENSG00000264808 | ENSG0000026 | 2.527143477 | 0.344424781 | 1.103984424 | 0.31198337 | 0.755053162 | 0.864705683 |

| | | | | | | | |
|---|---|---|---|---|---|---|---|
| ENSG00000264812 | ENSG0000026 | 2.444367244 | 1.251561286 | 1.181897283 | 1.058942519 | 0.289625959 | 0.473962882 |
| ENSG00000264859 | DSG2-AS1 | 2.323789354 | -0.200483297 | 1.120658207 | -0.178897808 | 0.858017944 | 0.926299197 |
| ENSG00000264920 | SP2-DT | 103.8309292 | -0.499055719 | 0.179319988 | -2.783045693 | 0.005385122 | 0.017198204 |
| ENSG00000264943 | SH3GL1P2 | 2.30961077 | 0.648326724 | 1.17047914 | 0.553898572 | 0.579648263 | 0.745187802 |
| ENSG00000264964 | TWSG1-DT | 35.10853089 | 0.673325938 | 0.303918967 | 2.215478504 | 0.026727238 | 0.070344521 |
| ENSG00000264982 | ENSG0000026 | 3.256625709 | 0.714033197 | 1.014156004 | 0.70406643 | 0.481391405 | 0.665769508 |
| ENSG00000265008 | ENSG0000026 | 2.385802669 | 0.586284385 | 1.199655901 | 0.488710458 | 0.625046699 | 0.778425521 |
| ENSG00000265010 | ENSG0000026 | 2.979142153 | 0.758745816 | 1.030505236 | 0.73628526 | 0.461557114 | 0.648328558 |
| ENSG00000265015 | ENSG0000026 | 2.414479693 | 0.07000132 | 1.135214618 | 0.061663512 | 0.950830798 | 0.977330339 |
| ENSG00000265018 | AGAP12P | 1.714820095 | 1.251769384 | 1.418572854 | 0.882414591 | 0.377552648 | NA |
| ENSG00000265055 | ENSG0000026 | 8.108749412 | -0.28141955 | 0.613765043 | -0.458513487 | 0.646583575 | 0.793491418 |
| ENSG00000265073 | ENSG0000026 | 4.050406664 | -1.214386663 | 0.921914836 | -1.317243866 | 0.187756895 | 0.345816392 |
| ENSG00000265091 | ENSG0000026 | 1.780557116 | 1.220857595 | 1.345317003 | 0.907486928 | 0.364149359 | NA |
| ENSG00000265095 | FTLP12 | 3.434280428 | -0.591889898 | 0.95680372 | -0.618611618 | 0.536172246 | 0.711422947 |
| ENSG00000265100 | ENSG0000026 | 6.748163548 | 0.013803265 | 0.6680808 | 0.020661071 | 0.983516023 | 0.9923499 |
| ENSG00000265112 | MIR3153 | 3.636635149 | -0.017737159 | 0.975202587 | -0.018188179 | 0.985488733 | 0.993350589 |
| ENSG00000265134 | MIR3190 | 1.592244657 | -2.091222545 | 1.590879634 | -1.314507082 | 0.188675616 | NA |
| ENSG00000265137 | MIR3192 | 3.050592967 | -1.142425015 | 1.062628609 | -1.07509341 | 0.28233291 | 0.465749817 |
| ENSG00000265148 | TSPOAP1-AS1 | 63.49396082 | -0.0955832 | 0.249601248 | -0.382943596 | 0.701761586 | 0.833061472 |
| ENSG00000265158 | LRRC37A7P | 10.89506717 | -0.220410825 | 0.547291383 | -0.402730304 | 0.687146638 | 0.823269645 |
| ENSG00000265168 | ENSG0000026 | 2.554972567 | -0.360247667 | 1.135925815 | -0.317140136 | 0.751137271 | 0.862460033 |
| ENSG00000265182 | SRP72P1 | 8.378489258 | 0.128851943 | 0.614564238 | 0.209663914 | 0.833929992 | 0.912272485 |
| ENSG00000265185 | SNORD3B-1 | 1.756942133 | -0.907154092 | 1.517621569 | -0.597747232 | 0.550008605 | NA |
| ENSG00000265218 | ENSG0000026 | 2.477895307 | -0.379959467 | 1.084603016 | -0.350321234 | 0.726097631 | 0.847853251 |
| ENSG00000265241 | RBM8A | 7254.038656 | 0.40875183 | 0.042976691 | 9.511012182 | 1.89E-21 | 3.44E-20 |
| ENSG00000265246 | ENSG0000026 | 57.46942107 | -0.623198903 | 0.26405643 | -2.360097433 | 0.018270136 | 0.05052519 |
| ENSG00000265254 | ENSG0000026 | 4.093081965 | 0.218753265 | 0.931605611 | 0.234813168 | 0.814353747 | 0.901759557 |
| ENSG00000265273 | PGDP1 | 38.4221766 | -0.460149174 | 0.318888766 | -1.442977061 | 0.149026933 | 0.290118429 |
| ENSG00000265282 | ENSG0000026 | 15.9027503 | -1.648259168 | 0.480276943 | -3.431893185 | 0.000599384 | 0.002338014 |
| ENSG00000265296 | FEM1AP2 | 170.305544 | 1.663175 | 0.151725377 | 10.96174573 | 5.84E-28 | 1.50E-26 |
| ENSG00000265298 | ENSG0000026 | 38.68072724 | -0.02872063 | 0.296665259 | -0.09681157 | 0.922876035 | 0.962730484 |
| ENSG00000265321 | MIR4263 | 5.84185593 | -0.458475224 | 0.72070322 | -0.636149821 | 0.524678775 | 0.701013253 |
| ENSG00000265342 | ENSG0000026 | 2.188094695 | -0.448474454 | 1.180334848 | -0.379955277 | 0.703978614 | 0.834371966 |

| | | | | | | | |
|---|---|---|---|---|---|---|---|
| ENSG00000265349 | ENSG0000026 | 7.465872273 | -0.259600813 | 0.629570737 | -0.412345742 | 0.680086026 | 0.818314893 |
| ENSG00000265354 | TIMM23 | 1448.678133 | -0.595371484 | 0.06006796 | -9.911631514 | 3.71E-23 | 7.42E-22 |
| ENSG00000265359 | ENSG0000026 | 3.476842465 | -4.323817543 | 1.410859736 | -3.064668607 | 0.002179113 | 0.007600556 |
| ENSG00000265366 | GLUD1P2 | 10.07497274 | 0.090230748 | 0.569733301 | 0.158373661 | 0.874162367 | 0.934789447 |
| ENSG00000265386 | RN7SL219P | 6.043086999 | -0.692950922 | 0.726368111 | -0.953994141 | 0.340086608 | 0.531009174 |
| ENSG00000265393 | ENSG0000026 | 52.43267238 | 0.492728602 | 0.263625852 | 1.869045082 | 0.061616543 | 0.142690336 |
| ENSG00000265401 | ENSG0000026 | 428.6277924 | -0.450391264 | 0.257301284 | -1.750443129 | 0.080041879 | 0.17670944 |
| ENSG00000265415 | PRR11-AS1 | 12.93691737 | -0.611980262 | 0.494719773 | -1.23702406 | 0.216078151 | 0.385415813 |
| ENSG00000265417 | ENSG0000026 | 7.659271804 | 0.005639345 | 0.671323429 | 0.008400339 | 0.993297578 | 0.996354439 |
| ENSG00000265437 | ENSG0000026 | 5.023901542 | -0.564390797 | 0.783628053 | -0.720227913 | 0.471384681 | 0.657149904 |
| ENSG00000265445 | ENSG0000026 | 2.186232841 | -0.022119468 | 1.180759034 | -0.018733262 | 0.985053894 | 0.993193404 |
| ENSG00000265451 | CLASP1-AS1 | 4.518490665 | -0.305980472 | 0.8147572 | -0.375548043 | 0.707252923 | 0.836532294 |
| ENSG00000265458 | NARF-AS2 | 86.12988088 | 0.150421474 | 0.192975535 | 0.779484685 | 0.435694256 | 0.625575905 |
| ENSG00000265474 | ENSG0000026 | 5.06639972 | 0.766511074 | 0.825913923 | 0.928076224 | 0.353368032 | 0.5449222 |
| ENSG00000265478 | ENSG0000026 | 5.224726916 | 0.460369153 | 0.779610309 | 0.590511885 | 0.55484752 | 0.726039707 |
| ENSG00000265480 | KRT18P55 | 10.45869915 | -0.6684502 | 0.54246123 | -1.232254331 | 0.217854094 | 0.387378723 |
| ENSG00000265489 | ARHGAP44-AS | 1.804651358 | 0.654850386 | 1.313406118 | 0.498589414 | 0.618068664 | 0.77359751 |
| ENSG00000265490 | ENSG0000026 | 2.382869987 | -0.993392657 | 1.195943061 | -0.830635412 | 0.406179623 | 0.598866287 |
| ENSG00000265491 | RNF115 | 3492.739508 | 0.242686536 | 0.047326099 | 5.127964111 | 2.93E-07 | 1.81E-06 |
| ENSG00000265496 | ENSG0000026 | 14.23389658 | 0.702770915 | 0.485975628 | 1.446103208 | 0.148148256 | 0.288793423 |
| ENSG00000265519 | ENSG0000026 | 3.244802754 | -0.623663952 | 1.016867014 | -0.61331909 | 0.539665373 | 0.714116501 |
| ENSG00000265539 | MIR3164 | 2.038733505 | 0.244330419 | 1.273409598 | 0.191871036 | 0.847843226 | 0.920424395 |
| ENSG00000265545 | ELOCP27 | 6.848807224 | 0.446328255 | 0.677311605 | 0.658970335 | 0.509914818 | 0.689870603 |
| ENSG00000265566 | RN7SL605P | 2.861906726 | 1.036520588 | 1.094441272 | 0.947077394 | 0.343599343 | 0.534565542 |
| ENSG00000265574 | WDR45BP1 | 102.942343 | 0.495255555 | 0.180681956 | 2.741034943 | 0.006124599 | 0.019304522 |
| ENSG00000265590 | CFAP298-TCP1 | 13.59783658 | -0.162584086 | 0.495007319 | -0.328447843 | 0.742573074 | 0.857909488 |
| ENSG00000265618 | ENSG0000026 | 7.150853464 | -0.293500464 | 0.660794881 | -0.444162739 | 0.656924929 | 0.801277136 |
| ENSG00000265626 | ENSG0000026 | 1.553119892 | 1.841604954 | 1.532135108 | 1.201986003 | 0.229368951 | NA |
| ENSG00000265646 | TUFMP1 | 10.92678478 | 0.500184972 | 0.58747899 | 0.851409123 | 0.394542126 | 0.587570253 |
| ENSG00000265660 | MIR4664 | 91.11277098 | -0.337024898 | 0.184854772 | -1.823187439 | 0.068275007 | 0.155487805 |
| ENSG00000265664 | ENSG0000026 | 15.02815841 | 0.496154894 | 0.462863131 | 1.071925719 | 0.283753397 | 0.467622337 |
| ENSG00000265678 | ENSG0000026 | 6.589767976 | -1.010841596 | 0.701273671 | -1.441436686 | 0.149461352 | 0.290849441 |
| ENSG00000265681 | RPL17 | 181.6033467 | 0.584389897 | 0.141004198 | 4.144485789 | 3.41E-05 | 0.000163312 |

| | | | | | | | |
|---|---|---|---|---|---|---|---|
| ENSG00000265683 | SYPL1P2 | 15.63951486 | 0.140772993 | 0.448296035 | 0.31401793 | 0.753507421 | 0.864088279 |
| ENSG00000265688 | MILIP | 161.5108269 | 0.254194592 | 0.152008114 | 1.672243576 | 0.094476301 | 0.202361003 |
| ENSG00000265692 | LINC01970 | 9.080037551 | -0.428211001 | 0.584231453 | -0.73294753 | 0.463590432 | 0.649941774 |
| ENSG00000265702 | ENSG0000026 | 22.94888404 | 0.011579922 | 0.367346402 | 0.031523168 | 0.974852316 | 0.988599625 |
| ENSG00000265737 | ENSG0000026 | 2.345577332 | -2.660165758 | 1.419363537 | -1.874196208 | 0.060903387 | 0.141280583 |
| ENSG00000265749 | ENSG0000026 | 1.814510986 | -2.319031189 | 1.519206524 | -1.52647527 | 0.126891545 | 0.256488857 |
| ENSG00000265750 | ENSG0000026 | 4.722266917 | 0.504013094 | 0.810663408 | 0.621729178 | 0.534119962 | 0.709572745 |
| ENSG00000265766 | CXADRP3 | 5.155431069 | 5.711876839 | 1.34290411 | 4.253376542 | 2.11E-05 | 0.00010424 |
| ENSG00000265778 | ZNF516-AS1 | 36.90312772 | 1.124350782 | 0.298997445 | 3.760402638 | 0.00016964 | 0.0007293 |
| ENSG00000265784 | ENSG0000026 | 14.06089739 | -0.459497959 | 0.474216018 | -0.968963387 | 0.332563457 | 0.523610549 |
| ENSG00000265786 | LINC01906 | 2.423559437 | 0.776975184 | 1.149704176 | 0.675804438 | 0.499164817 | 0.68062535 |
| ENSG00000265787 | CYP4F35P | 223.3045059 | 0.668021781 | 0.137463286 | 4.859637798 | 1.18E-06 | 6.75E-06 |
| ENSG00000265790 | RNASEH1P1 | 2.721372968 | -1.776802894 | 1.130506559 | -1.571687383 | 0.11602307 | 0.239091737 |
| ENSG00000265800 | ENSG0000026 | 7.836005734 | 0.212144467 | 0.636521546 | 0.333287173 | 0.738917521 | 0.856056155 |
| ENSG00000265806 | MIR4292 | 51.23958159 | 0.011119827 | 0.247445952 | 0.044938408 | 0.964156403 | 0.983666795 |
| ENSG00000265808 | SEC22B | 3369.295441 | 0.001542458 | 0.048340131 | 0.031908441 | 0.974545067 | 0.988514222 |
| ENSG00000265817 | FSBP | 84.77330449 | -0.425816806 | 0.207618883 | -2.050954137 | 0.040271413 | 0.099657629 |
| ENSG00000265840 | ENSG0000026 | 5.804863901 | -1.302771109 | 0.761136474 | -1.711613034 | 0.086968005 | 0.189157678 |
| ENSG00000265907 | ENSG0000026 | 3.941217684 | 1.817125487 | 0.967350897 | 1.87845537 | 0.060318897 | 0.140137279 |
| ENSG00000265917 | MIR3685 | 2.344148748 | -0.207983502 | 1.136275842 | -0.183039623 | 0.854766929 | 0.924486249 |
| ENSG00000265936 | ENSG0000026 | 6.528206533 | 0.211111508 | 0.702102431 | 0.300684769 | 0.763654884 | 0.870479014 |
| ENSG00000265939 | UBE2CP2 | 3.082061467 | 0.46493957 | 1.995981064 | 0.232937866 | 0.815809654 | 0.902352305 |
| ENSG00000265943 | ENSG0000026 | 6.499555469 | 0.039371242 | 0.703569778 | 0.055959256 | 0.955374265 | 0.979455025 |
| ENSG00000265962 | GACAT2 | 7.091863385 | -0.60901719 | 0.684035344 | -0.890330003 | 0.373288715 | 0.56605256 |
| ENSG00000265967 | ENSG0000026 | 3.353920412 | -1.415972519 | 1.009739259 | -1.40231501 | 0.1608212 | 0.307385297 |
| ENSG00000265972 | TXNIP | 4952.29935 | -0.916042665 | 0.062686896 | -14.61298488 | 2.32E-48 | 1.21E-46 |
| ENSG00000265975 | ENSG0000026 | 12.38589293 | 0.63872752 | 0.516198653 | 1.237367661 | 0.215950619 | 0.38521829 |
| ENSG00000265982 | ENSG0000026 | 14.97199922 | -0.125903229 | 0.448304115 | -0.280843349 | 0.778830553 | 0.879986844 |
| ENSG00000266010 | GATA6-AS1 | 4.346448999 | -0.013722647 | 0.849746373 | -0.016149109 | 0.987115435 | 0.99369967 |
| ENSG00000266017 | MIR4477B | 9.728779174 | 0.286402371 | 0.609159428 | 0.470159958 | 0.63824074 | 0.786909737 |
| ENSG00000266028 | SRGAP2 | 4882.727289 | 0.314073791 | 0.050123849 | 6.26595512 | 3.71E-10 | 3.07E-09 |
| ENSG00000266036 | SLC9A3R1-AS1 | 23.1336043 | -0.02010884 | 0.366351004 | -0.054889546 | 0.95622646 | 0.979978332 |
| ENSG00000266053 | NDUFV2-AS1 | 27.06767807 | 0.498674381 | 0.345989043 | 1.441301078 | 0.149499643 | 0.290883953 |

| | | | | | | | |
|---|---|---|---|---|---|---|---|
| ENSG00000266066 | POLRMTP1 | 152.1450395 | 0.104928588 | 0.148225703 | 0.707897384 | 0.479008984 | 0.663653377 |
| ENSG00000266074 | BAHCC1 | 1567.672239 | 0.11987311 | 0.060461082 | 1.982649109 | 0.047406638 | 0.114438417 |
| ENSG00000266094 | RASSF5 | 350.7830392 | 0.079630218 | 0.101077134 | 0.787816339 | 0.430804143 | 0.620964312 |
| ENSG00000266104 | MIR4326 | 2.272095222 | -2.718712127 | 1.422622148 | -1.91105708 | 0.055997246 | 0.131935597 |
| ENSG00000266124 | MIR5587 | 7.425300646 | 0.361450744 | 0.656831463 | 0.550294504 | 0.582117394 | 0.746825138 |
| ENSG00000266145 | RHOT1P1 | 252.3064459 | -0.067931014 | 0.128857447 | -0.527179572 | 0.598068893 | 0.758455438 |
| ENSG00000266171 | ENSG0000026 | 1.602733281 | 0.483810775 | 1.457201143 | 0.332013722 | 0.739878897 | NA |
| ENSG00000266173 | STRADA | 177.7119747 | 0.051066111 | 0.136948484 | 0.372885548 | 0.709233628 | 0.838002546 |
| ENSG00000266184 | ENSG0000026 | 4.934922218 | -0.959187047 | 0.810147112 | -1.183966509 | 0.236426321 | 0.411560661 |
| ENSG00000266208 | GJD3-AS1 | 1393.802765 | 0.428690413 | 0.070087403 | 6.116511593 | 9.56E-10 | 7.62E-09 |
| ENSG00000266235 | MIR3176 | 10.52761979 | 0.322133459 | 0.544431957 | 0.591687271 | 0.554060022 | 0.725423029 |
| ENSG00000266236 | NARF-IT1 | 7.302844877 | 0.33307751 | 0.652396372 | 0.510544699 | 0.609669907 | 0.767761904 |
| ENSG00000266242 | GRAMD4P7 | 13.60795951 | 1.256606072 | 0.503190478 | 2.497277128 | 0.012515111 | 0.036226706 |
| ENSG00000266256 | LINC00683 | 5.944059407 | 2.893354594 | 0.939794966 | 3.078708335 | 0.002079001 | 0.007281265 |
| ENSG00000266258 | LINC01909 | 6.812154381 | 1.46783899 | 0.730825551 | 2.008466983 | 0.044593691 | 0.108631122 |
| ENSG00000266261 | ENSG0000026 | 3.660854572 | -0.216334651 | 0.904260243 | -0.23923937 | 0.810919976 | 0.899932478 |
| ENSG00000266282 | UBL5P2 | 29.41887824 | -0.135093709 | 0.335658038 | -0.402474226 | 0.687335053 | 0.823367217 |
| ENSG00000266283 | ENSG0000026 | 1.753104386 | 0.662444994 | 1.358488295 | 0.487633936 | 0.625809154 | NA |
| ENSG00000266296 | ARIH2P1 | 39.07007296 | -0.199781077 | 0.280796067 | -0.711481038 | 0.476786197 | 0.661874455 |
| ENSG00000266306 | ENSG0000026 | 5.79283328 | -1.309476357 | 0.781284632 | -1.676055439 | 0.093727334 | 0.20103834 |
| ENSG00000266338 | NBPF15 | 2389.803064 | 0.179520263 | 0.06020837 | 2.981649584 | 0.002866999 | 0.009763802 |
| ENSG00000266341 | ENSG0000026 | 20.85486342 | -0.36285966 | 0.387963765 | -0.935292655 | 0.349637499 | 0.540914772 |
| ENSG00000266365 | ENSG0000026 | 23.70906942 | 0.18751745 | 0.43080926 | 0.435267919 | 0.663367968 | 0.805636147 |
| ENSG00000266373 | ENSG0000026 | 23.16719553 | 0.250157605 | 0.363374518 | 0.688429136 | 0.491182581 | 0.674149121 |
| ENSG00000266389 | PIK3R5-DT | 9.028904144 | 0.66601731 | 0.585621093 | 1.137283676 | 0.255419716 | 0.434797291 |
| ENSG00000266402 | SNHG25 | 84.71065577 | 0.602058973 | 0.206506155 | 2.91545292 | 0.003551727 | 0.01184942 |
| ENSG00000266405 | CBX3P2 | 21.64244084 | 0.882754419 | 0.395578248 | 2.2315545 | 0.025644422 | 0.067860541 |
| ENSG00000266412 | NCOA4 | 4580.356181 | -0.230867635 | 0.048107023 | -4.799042274 | 1.59E-06 | 9.00E-06 |
| ENSG00000266437 | MIR4746 | 3.550138888 | -0.139241154 | 0.933070187 | -0.149229025 | 0.881372918 | 0.938527352 |
| ENSG00000266469 | ENSG0000026 | 41.95099965 | 0.247015624 | 0.27019072 | 0.914226901 | 0.360597632 | 0.553133378 |
| ENSG00000266472 | MRPS21 | 4059.020766 | 0.337311493 | 0.045639577 | 7.390767289 | 1.46E-13 | 1.58E-12 |
| ENSG00000266473 | HELZ-AS1 | 8.762732707 | 0.891867984 | 0.639833148 | 1.393907125 | 0.163345653 | 0.31091514 |
| ENSG00000266479 | ENSG0000026 | 2.911969409 | 0.771503046 | 1.055610131 | 0.730859835 | 0.464864772 | 0.650918504 |

| | | | | | | | |
|---|---|---|---|---|---|---|---|
| ENSG00000266490 | ENSG0000026 | 25.8606257 | 1.106923183 | 0.372985654 | 2.967736617 | 0.003000013 | 0.010163978 |
| ENSG00000266495 | ENSG0000026 | 10.64729514 | -1.114863759 | 0.570886621 | -1.952863699 | 0.050835748 | 0.121552227 |
| ENSG00000266498 | ENSG0000026 | 2.788545712 | -1.776176462 | 1.139713893 | -1.558440652 | 0.119128826 | 0.244110342 |
| ENSG00000266501 | ATG12P1 | 25.8909238 | -0.213671229 | 0.345592226 | -0.618275565 | 0.536393706 | 0.711593316 |
| ENSG00000266522 | ENSG0000026 | 3.744526679 | -0.037957642 | 0.93542462 | -0.04057798 | 0.967632339 | 0.985635001 |
| ENSG00000266554 | LINC01443 | 10.51261566 | 2.333638318 | 0.636407198 | 3.666894917 | 0.000245514 | 0.001023829 |
| ENSG00000266563 | EIF1P5 | 7.096705232 | 0.289619361 | 0.68816038 | 0.420860267 | 0.673857121 | 0.813479126 |
| ENSG00000266589 | MIR4512 | 2.431104239 | 0.41623172 | 1.111519629 | 0.374470868 | 0.708054027 | 0.837109241 |
| ENSG00000266602 | ENSG0000026 | 94.53623993 | -0.078511978 | 0.187168422 | -0.41947235 | 0.674870958 | 0.814303586 |
| ENSG00000266613 | COP1P1 | 1.934651928 | 0.303699216 | 1.244894335 | 0.243955818 | 0.807265038 | 0.897656809 |
| ENSG00000266642 | ENSG0000026 | 15.0717544 | -0.448500305 | 0.487475131 | -0.92004756 | 0.357547906 | 0.54981559 |
| ENSG00000266643 | MIR3677 | 2.707251294 | 1.447961233 | 1.166467165 | 1.241321897 | 0.214486859 | 0.383203158 |
| ENSG00000266648 | SETP3 | 27.68746495 | 0.075954425 | 0.354064776 | 0.21452127 | 0.830140582 | 0.909932761 |
| ENSG00000266677 | ENSG0000026 | 17.92017979 | -0.777715638 | 0.432661826 | -1.797513881 | 0.072254076 | 0.162980188 |
| ENSG00000266680 | ENSG0000026 | 47.82210156 | 0.262718037 | 0.256417967 | 1.024569535 | 0.305566345 | 0.493050008 |
| ENSG00000266707 | PSMD7P1 | 7.316933211 | 0.263844842 | 0.641434632 | 0.411335511 | 0.680826534 | 0.818841625 |
| ENSG00000266708 | ENSG0000026 | 10.60884994 | -0.190115194 | 0.552109477 | -0.34434329 | 0.730588132 | 0.850832153 |
| ENSG00000266709 | ENSG0000026 | 2.468439204 | 0.00638237 | 1.102610322 | 0.005788418 | 0.995381536 | 0.997076653 |
| ENSG00000266714 | MYO15B | 2019.257101 | -0.383001503 | 0.068281337 | -5.609168174 | 2.03E-08 | 1.41E-07 |
| ENSG00000266751 | MIR3661 | 20.8760213 | -0.37046705 | 0.397151486 | -0.932810435 | 0.350917853 | 0.542237339 |
| ENSG00000266770 | RN7SL619P | 1.682200473 | -1.400825913 | 1.417719948 | -0.988083659 | 0.323111678 | NA |
| ENSG00000266777 | SH3GL1P1 | 14.87879527 | -0.064019929 | 0.44854312 | -0.142728594 | 0.886504533 | 0.941500648 |
| ENSG00000266783 | ENSG0000026 | 3.35500606 | -1.032020827 | 1.05296688 | -0.980107586 | 0.327033015 | 0.517460158 |
| ENSG00000266821 | ENSG0000026 | 2.235706532 | -0.512397721 | 1.253189029 | -0.408875045 | 0.682631363 | 0.819872433 |
| ENSG00000266844 | ENSG0000026 | 2.302427183 | 0.869650637 | 1.218845285 | 0.713503713 | 0.475534116 | 0.660696059 |
| ENSG00000266850 | ENSG0000026 | 2.307590616 | -1.453319739 | 1.229591978 | -1.181952847 | 0.237224411 | 0.412400055 |
| ENSG00000266852 | MIR4482 | 2.64204147 | 0.505057906 | 1.123949536 | 0.449359949 | 0.653172019 | 0.798701046 |
| ENSG00000266865 | ENSG0000026 | 60.10831809 | -0.4432543 | 0.244174792 | -1.815315562 | 0.069475429 | 0.157626974 |
| ENSG00000266869 | ENSG0000026 | 156.4410382 | -0.130199935 | 0.167451542 | -0.777537986 | 0.436841433 | 0.626595539 |
| ENSG00000266876 | ENSG0000026 | 48.08418509 | -0.139592019 | 0.254279705 | -0.54897035 | 0.583025798 | 0.747393786 |
| ENSG00000266896 | ENSG0000026 | 15.31636195 | -0.39205237 | 0.486879897 | -0.805234253 | 0.420684514 | 0.61195325 |
| ENSG00000266897 | ENSG0000026 | 2.853408513 | -1.385817121 | 1.112005223 | -1.246232565 | 0.212679027 | 0.380714544 |
| ENSG00000266903 | CEACAM16-AS | 54.78782078 | -0.048631352 | 0.245616731 | -0.197996902 | 0.843047488 | 0.917338822 |

| | | | | | | | |
|---|---|---|---|---|---|---|---|
| ENSG00000266904 | LINC00663 | 17.38447368 | 0.236840846 | 0.418678123 | 0.565687177 | 0.571606454 | 0.738449741 |
| ENSG00000266907 | ENSG0000026 | 3.914280685 | 0.334272525 | 0.95467897 | 0.350141289 | 0.726232665 | 0.847853251 |
| ENSG00000266910 | ENSG0000026 | 6.183813907 | -0.057860252 | 0.731819559 | -0.07906355 | 0.936982076 | 0.96978744 |
| ENSG00000266916 | ZNF793-AS1 | 34.42314909 | 0.763893339 | 0.305358569 | 2.501627325 | 0.012362398 | 0.035838925 |
| ENSG00000266920 | ACTBP9 | 12.60303147 | -0.22502613 | 0.521328931 | -0.431639444 | 0.666003486 | 0.807426275 |
| ENSG00000266921 | ZNF230-DT | 30.97186412 | -0.098328583 | 0.316586709 | -0.310589739 | 0.756112529 | 0.865184627 |
| ENSG00000266933 | MADCAM1-AS | 6.369925409 | 0.867206739 | 0.709726497 | 1.221888633 | 0.221749745 | 0.392631919 |
| ENSG00000266934 | ENSG0000026 | 136.720872 | -0.793262822 | 0.161856687 | -4.90101976 | 9.53E-07 | 5.53E-06 |
| ENSG00000266947 | ENSG0000026 | 2.1048206 | -0.364328784 | 1.194006687 | -0.305131276 | 0.760266171 | 0.868248418 |
| ENSG00000266954 | ENSG0000026 | 3.508123752 | -0.597947436 | 0.925502711 | -0.646078536 | 0.518228495 | 0.695932225 |
| ENSG00000266962 | HSD17B1-AS1 | 551.5040997 | -0.053494249 | 0.095772415 | -0.558555911 | 0.576464836 | 0.74273769 |
| ENSG00000266967 | AARSD1 | 27.71702971 | -0.405730068 | 0.331675065 | -1.223275764 | 0.221225558 | 0.391824761 |
| ENSG00000266969 | CEP192-DT | 2.012333 | -1.148702715 | 1.379166721 | -0.832896195 | 0.404903276 | 0.597789441 |
| ENSG00000266970 | SOCS3-DT | 119.3162311 | 0.006175016 | 0.167059746 | 0.03696292 | 0.970514571 | 0.986772291 |
| ENSG00000266975 | FARSA-AS1 | 3.924521084 | 0.478660975 | 0.921471597 | 0.519452772 | 0.603445039 | 0.762492237 |
| ENSG00000266978 | LIMASI | 15.0713011 | -0.307198031 | 0.476666967 | -0.644470987 | 0.519270066 | 0.696787882 |
| ENSG00000266980 | ENSG0000026 | 4.561082772 | -0.097394658 | 0.847824203 | -0.114876006 | 0.908543404 | 0.95425968 |
| ENSG00000266983 | RANBP3-DT | 6.534440794 | 0.113163056 | 0.67852829 | 0.166777211 | 0.86754535 | 0.93096599 |
| ENSG00000266984 | POLR3GP2 | 1.623153263 | 1.769280212 | 1.538533801 | 1.149978123 | 0.250152882 | NA |
| ENSG00000266992 | DHX40P1 | 2.199009126 | 0.44419526 | 1.212653227 | 0.366300316 | 0.714140994 | 0.840989046 |
| ENSG00000266993 | ENSG0000026 | 18.42589614 | 1.664008597 | 0.448264786 | 3.7121109 | 0.000205538 | 0.000870094 |
| ENSG00000266994 | ENSG0000026 | 4.087349245 | -0.226744226 | 0.886419586 | -0.255797852 | 0.798106933 | 0.892259454 |
| ENSG00000267002 | ENSG0000026 | 495.2259745 | 0.275798219 | 0.101930327 | 2.705752326 | 0.006814985 | 0.021186578 |
| ENSG00000267009 | ENSG0000026 | 36.62761751 | -0.207979167 | 0.29748976 | -0.699113699 | 0.484480978 | 0.66835009 |
| ENSG00000267011 | ENSG0000026 | 3.582815336 | -0.166185628 | 0.962405715 | -0.172677308 | 0.862905082 | 0.92885434 |
| ENSG00000267023 | LRRC37A16P | 77.52015978 | -0.755005563 | 0.22160877 | -3.406929985 | 0.00065698 | 0.002545797 |
| ENSG00000267024 | ENSG0000026 | 1.92998595 | -2.376123454 | 1.554781071 | -1.528268834 | 0.126445805 | 0.255723068 |
| ENSG00000267026 | ENSG0000026 | 5.938745274 | -0.912266452 | 0.778048739 | -1.172505534 | 0.240994152 | 0.416892077 |
| ENSG00000267030 | ENSG0000026 | 10.74372534 | -0.427008894 | 0.547269634 | -0.780253219 | 0.435241843 | 0.625239393 |
| ENSG00000267040 | ATP8B1-AS1 | 22.72818802 | 0.720488126 | 0.376583809 | 1.913221194 | 0.055719741 | 0.131393527 |
| ENSG00000267041 | ZNF850 | 224.0127679 | 0.00711855 | 0.127033559 | 0.056036765 | 0.955312519 | 0.979455025 |
| ENSG00000267042 | ENSG0000026 | 2.495808739 | -1.621962933 | 1.162838928 | -1.394830268 | 0.163067029 | 0.310539403 |
| ENSG00000267049 | ENSG0000026 | 2.591847978 | -1.678899033 | 1.179331374 | -1.42360245 | 0.154561586 | 0.297903108 |

| | | | | | | | |
|---|---|---|---|---|---|---|---|
| ENSG00000267056 | ENSG0000026 | 10.65123664 | 0.64646361 | 0.549804423 | 1.175806492 | 0.239672212 | 0.415342373 |
| ENSG00000267058 | ENSG0000026 | 11.56356276 | 0.810199812 | 0.532879593 | 1.520418164 | 0.128405912 | 0.258842849 |
| ENSG00000267060 | PTGES3L | 9.256880808 | -0.96850739 | 0.585742838 | -1.6534686 | 0.098235531 | 0.208928542 |
| ENSG00000267062 | ENSG0000026 | 48.46917818 | -0.583209077 | 0.260064522 | -2.242555321 | 0.024925504 | 0.066317683 |
| ENSG00000267064 | UXT-AS1 | 42.40575395 | 0.368878917 | 0.278805942 | 1.323066913 | 0.185813127 | 0.343090239 |
| ENSG00000267072 | ENSG0000026 | 1.807960399 | 1.275337238 | 1.471666043 | 0.866594187 | 0.3861644 | 0.578767237 |
| ENSG00000267076 | MIX23P3 | 29.02295538 | 0.03316752 | 0.327965208 | 0.101131216 | 0.919446298 | 0.960822968 |
| ENSG00000267077 | ENSG0000026 | 6.088517534 | 0.848448004 | 0.753931533 | 1.125364793 | 0.260434484 | 0.44075006 |
| ENSG00000267078 | ENSG0000026 | 3.721721325 | -0.776947464 | 0.922965632 | -0.841794576 | 0.399902948 | 0.592697612 |
| ENSG00000267080 | ASB16-AS1 | 394.2257317 | 0.210923386 | 0.101014236 | 2.08805605 | 0.036792779 | 0.092516316 |
| ENSG00000267082 | ENSG0000026 | 4.234331015 | 0.75942841 | 0.874434205 | 0.868479762 | 0.385131747 | 0.577672438 |
| ENSG00000267091 | CTBP2P7 | 1.865639613 | 2.162514524 | 1.459033572 | 1.482155425 | 0.138298944 | 0.274088793 |
| ENSG00000267092 | ENSG0000026 | 2.819740752 | 0.320635342 | 1.072698756 | 0.298905298 | 0.765012304 | 0.871320362 |
| ENSG00000267096 | ENSG0000026 | 7.693400154 | 1.009501677 | 0.672713827 | 1.500640594 | 0.133448546 | 0.266734307 |
| ENSG00000267100 | ILF3-DT | 641.8689128 | 0.105114581 | 0.078241896 | 1.343456459 | 0.179124216 | 0.333883929 |
| ENSG00000267105 | ENSG0000026 | 13.52245969 | -0.08965366 | 0.486642745 | -0.184228905 | 0.853833884 | 0.923912703 |
| ENSG00000267106 | ZNF561-AS1 | 186.3053554 | -0.426222487 | 0.136149458 | -3.130548532 | 0.001744802 | 0.006210091 |
| ENSG00000267108 | ENSG0000026 | 3.985226909 | 0.288133418 | 0.881173687 | 0.326988223 | 0.743676793 | 0.858490754 |
| ENSG00000267112 | ENSG0000026 | 9.104712221 | 0.359769114 | 0.58715651 | 0.612731202 | 0.540054088 | 0.714414141 |
| ENSG00000267113 | ENSG0000026 | 3.753990872 | 0.713057245 | 0.914139517 | 0.780031092 | 0.435372575 | 0.625270573 |
| ENSG00000267114 | ENSG0000026 | 4.201760842 | -0.08431111 | 0.858628285 | -0.098192793 | 0.921779205 | 0.962181382 |
| ENSG00000267116 | PMM2P2 | 2.384739302 | 0.871064817 | 1.192221927 | 0.730623047 | 0.465009431 | 0.65100803 |
| ENSG00000267119 | RPL10P15 | 33.73132727 | -0.130285848 | 0.312740315 | -0.41659435 | 0.676975142 | 0.816069877 |
| ENSG00000267121 | FMNL1-DT | 10.18126016 | -0.30632865 | 0.556341557 | -0.550612562 | 0.581899296 | 0.746784327 |
| ENSG00000267122 | ENSG0000026 | 2.663263943 | -0.186935958 | 1.055076101 | -0.177177701 | 0.85936881 | 0.926885162 |
| ENSG00000267125 | ENSG0000026 | 1.734667435 | 1.216856015 | 1.400548303 | 0.86884259 | 0.384933234 | NA |
| ENSG00000267127 | ENSG0000026 | 18.27073072 | 0.659355045 | 0.412246548 | 1.599419202 | 0.109727489 | 0.228334067 |
| ENSG00000267128 | RNF157-AS1 | 9.103094084 | -0.427929686 | 0.583897896 | -0.732884447 | 0.46362891 | 0.649941774 |
| ENSG00000267130 | ENSG0000026 | 2.787117127 | -0.016731231 | 1.034820511 | -0.016168244 | 0.987100169 | 0.99369967 |
| ENSG00000267131 | ENSG0000026 | 379.4866765 | 0.568868471 | 0.101418965 | 5.609093632 | 2.03E-08 | 1.41E-07 |
| ENSG00000267132 | HMGB3P27 | 2.189458687 | -1.905974375 | 1.271771158 | -1.49867715 | 0.133957407 | 0.267469008 |
| ENSG00000267135 | ENSG0000026 | 4.052301637 | -0.054596366 | 0.912862206 | -0.059807894 | 0.952308638 | 0.977841644 |
| ENSG00000267138 | ENSG0000026 | 2.458112559 | -1.0756603 | 1.141069424 | -0.94267735 | 0.345845962 | 0.536750313 |

| | | | | | | | |
|---|---|---|---|---|---|---|---|
| ENSG00000267139 | ENSG0000026 | 11.84418183 | -0.27227442 | 0.515295514 | -0.528385001 | 0.597232146 | 0.757729408 |
| ENSG00000267152 | ENSG0000026 | 12.94041821 | 0.958211469 | 0.50750793 | 1.888071914 | 0.059016297 | 0.137584741 |
| ENSG00000267156 | TPMTP1 | 18.15362901 | -1.179714642 | 0.41924376 | -2.81391103 | 0.004894277 | 0.015802212 |
| ENSG00000267160 | ENSG0000026 | 4.644314555 | 0.644993003 | 0.823790758 | 0.782957319 | 0.433652176 | 0.623549332 |
| ENSG00000267165 | CHMP1B-AS1 | 7.673827375 | -0.801847742 | 0.639643855 | -1.253584687 | 0.209993 | 0.377122238 |
| ENSG00000267169 | ADGRL1-AS1 | 42.91017814 | -1.097564566 | 0.279765484 | -3.923159327 | 8.74E-05 | 0.000393725 |
| ENSG00000267174 | ENSG0000026 | 14.43490323 | -0.09439521 | 0.462859943 | -0.203939034 | 0.838401141 | 0.91499129 |
| ENSG00000267178 | PHF5AP1 | 21.57730988 | 0.038778507 | 0.396215426 | 0.097872281 | 0.922033711 | 0.962216621 |
| ENSG00000267184 | ENSG0000026 | 2.512433284 | -1.20146926 | 1.189937784 | -1.009690823 | 0.312643441 | 0.500805847 |
| ENSG00000267185 | PTP4A2P1 | 88.34307602 | -0.680173441 | 0.210556858 | -3.230355207 | 0.001236365 | 0.004549601 |
| ENSG00000267197 | ENSG0000026 | 14.68300823 | -0.077676237 | 0.454663576 | -0.170843324 | 0.864346962 | 0.929719103 |
| ENSG00000267199 | ENSG0000026 | 22.99540536 | -0.23034466 | 0.385530683 | -0.59747426 | 0.550190787 | 0.722460026 |
| ENSG00000267201 | LINC01775 | 4.068392151 | -0.974209493 | 0.924181459 | -1.054132263 | 0.291822367 | 0.476877411 |
| ENSG00000267207 | BCAS3-AS1 | 6.164166126 | -0.140082642 | 0.729674811 | -0.19197955 | 0.847758224 | 0.920377435 |
| ENSG00000267213 | DPY19L3-DT | 17.47293837 | -0.135939671 | 0.418900155 | -0.324515687 | 0.745547648 | 0.859379806 |
| ENSG00000267216 | ZNF8-ERVK3-1 | 10.13314341 | 0.142109856 | 0.571111584 | 0.248830281 | 0.803492066 | 0.895371029 |
| ENSG00000267218 | ENSG0000026 | 1.703057728 | -0.249509062 | 1.364551596 | -0.182850588 | 0.854915254 | NA |
| ENSG00000267219 | ENSG0000026 | 7.076821093 | 0.141650707 | 0.653899941 | 0.216624438 | 0.828501039 | 0.90928207 |
| ENSG00000267221 | C17orf113 | 86.6324071 | -0.533717985 | 0.190486789 | -2.801863514 | 0.005080836 | 0.016342414 |
| ENSG00000267226 | MALT1-AS1 | 2.78190825 | 0.364474914 | 1.121387541 | 0.325021369 | 0.745164899 | 0.859215018 |
| ENSG00000267239 | ENSG0000026 | 5.460757488 | 0.673993559 | 0.775982889 | 0.868567553 | 0.385083708 | 0.577672438 |
| ENSG00000267244 | ENSG0000026 | 4.028093658 | -0.234921137 | 0.910171919 | -0.258106334 | 0.796324844 | 0.891280831 |
| ENSG00000267248 | ENSG0000026 | 105.0697318 | 0.344548198 | 0.179744346 | 1.916879199 | 0.05525328 | 0.130496268 |
| ENSG00000267249 | ENSG0000026 | 14.36110046 | 0.413397726 | 0.45985081 | 0.898982272 | 0.368662104 | 0.561204714 |
| ENSG00000267253 | WHSC1L2P | 8.00396336 | 0.335250883 | 0.622948183 | 0.538168169 | 0.590460953 | 0.753103247 |
| ENSG00000267254 | ZNF790-AS1 | 175.8029063 | 0.750059413 | 0.140139191 | 5.352245923 | 8.69E-08 | 5.66E-07 |
| ENSG00000267255 | ENSG0000026 | 7.116793793 | -0.553775484 | 0.668541354 | -0.828333925 | 0.407481415 | 0.600343172 |
| ENSG00000267259 | ERVE-1 | 8.357364849 | -1.40153695 | 0.64080225 | -2.187159846 | 0.028730857 | 0.074785642 |
| ENSG00000267264 | ENSG0000026 | 2.049561062 | 1.012528725 | 1.281254984 | 0.790263248 | 0.429374046 | 0.619714368 |
| ENSG00000267265 | GP6-AS1 | 8.967185045 | -0.056836326 | 0.595874821 | -0.095382997 | 0.924010621 | 0.963359707 |
| ENSG00000267270 | PARD6G-AS1 | 5.25181292 | 1.733683518 | 0.836457549 | 2.072649737 | 0.038204889 | 0.095501814 |
| ENSG00000267274 | ENSG0000026 | 3.77848496 | 0.575255918 | 0.914467726 | 0.62906093 | 0.529309167 | 0.70508985 |
| ENSG00000267277 | ENSG0000026 | 12.49775957 | -0.803483575 | 0.522099329 | -1.538947727 | 0.123817058 | 0.25160451 |

| | | | | | | | |
|---|---|---|---|---|---|---|---|
| ENSG00000267278 | MAP3K14-AS1 | 340.6626002 | -0.010064127 | 0.101915153 | -0.098750055 | 0.921336725 | 0.9619656 |
| ENSG00000267280 | TBX2-AS1 | 1115.172167 | 0.25283926 | 0.066828485 | 3.783405513 | 0.000154697 | 0.000669155 |
| ENSG00000267281 | ATF7-NPFF | 4.022146883 | 0.521228501 | 0.886288433 | 0.588102565 | 0.556463451 | 0.727042218 |
| ENSG00000267282 | ENSG0000026 | 15.37943239 | -0.111982136 | 0.506628618 | -0.221033973 | 0.825065981 | 0.90772273 |
| ENSG00000267283 | ENSG0000026 | 43.97089563 | 0.133237905 | 0.272453711 | 0.489029509 | 0.624820806 | 0.778298345 |
| ENSG00000267288 | ENSG0000026 | 32.28851486 | -0.303951785 | 0.317764508 | -0.956531573 | 0.338803741 | 0.530045518 |
| ENSG00000267293 | ENSG0000026 | 23.09786247 | 0.252358174 | 0.363408013 | 0.694421051 | 0.487418196 | 0.670828738 |
| ENSG00000267296 | CEBPA-DT | 81.7869749 | 0.145409485 | 0.201288614 | 0.722392996 | 0.470052892 | 0.655940953 |
| ENSG00000267302 | RNFT1-DT | 2.26655455 | 1.187641642 | 1.253735107 | 0.947282752 | 0.343494717 | 0.534439014 |
| ENSG00000267309 | ZNF566-AS1 | 59.37048762 | 0.416514524 | 0.229057955 | 1.818380523 | 0.069005993 | 0.156825588 |
| ENSG00000267312 | ENSG0000026 | 78.25277809 | 0.050448836 | 0.19813474 | 0.254618834 | 0.799017511 | 0.892813527 |
| ENSG00000267315 | UFM1P2 | 2.112437456 | 1.011301118 | 1.247614365 | 0.810587908 | 0.417602363 | 0.609751016 |
| ENSG00000267317 | ENSG0000026 | 319.5406306 | 0.012130242 | 0.111716549 | 0.10858053 | 0.913535205 | 0.956961395 |
| ENSG00000267319 | SELENOKP1 | 7.553369886 | 1.059497098 | 0.654141853 | 1.619674834 | 0.105302145 | 0.220948615 |
| ENSG00000267321 | SNHG30 | 157.5761885 | 0.377132823 | 0.143177186 | 2.634028751 | 0.008437834 | 0.025600307 |
| ENSG00000267322 | SNHG22 | 14.91907528 | 0.016988199 | 0.475786513 | 0.035705507 | 0.971517179 | 0.987006252 |
| ENSG00000267324 | ENSG0000026 | 2.11013833 | 1.526719412 | 1.336554741 | 1.142279748 | 0.253337757 | 0.431990496 |
| ENSG00000267336 | EIF4A2P1 | 4.774709763 | 1.079505241 | 0.839856022 | 1.285345599 | 0.198671542 | 0.361421505 |
| ENSG00000267340 | ENSG0000026 | 5.128488442 | 0.84601799 | 0.793273472 | 1.066489703 | 0.286202331 | 0.470021728 |
| ENSG00000267341 | ENSG0000026 | 5.101020568 | 1.15295463 | 0.819796533 | 1.406391201 | 0.159607992 | 0.305295366 |
| ENSG00000267342 | ENSG0000026 | 3.562550279 | 1.556124247 | 1.02048527 | 1.524886534 | 0.127287401 | 0.257107773 |
| ENSG00000267344 | ENSG0000026 | 2.205091921 | 0.681412493 | 1.229728536 | 0.554116191 | 0.57949933 | 0.745038089 |
| ENSG00000267348 | GEMIN7-AS1 | 60.56578638 | -0.248846564 | 0.230171251 | -1.0811366 | 0.279636355 | 0.462933542 |
| ENSG00000267352 | SH3GL1P3 | 3.123697666 | -0.961087638 | 1.008881573 | -0.952626813 | 0.340779187 | 0.531837169 |
| ENSG00000267370 | ENSG0000026 | 9.788730023 | 0.361281584 | 0.60154882 | 0.600585643 | 0.548116003 | 0.720932349 |
| ENSG00000267374 | MIR924HG | 114.7442178 | 0.375852861 | 0.169311021 | 2.219896008 | 0.026425828 | 0.069639114 |
| ENSG00000267383 | ENSG0000026 | 18.17136008 | -0.072812092 | 0.421876642 | -0.172590953 | 0.862972963 | 0.92885434 |
| ENSG00000267385 | ENSG0000026 | 7.482982438 | 1.88063558 | 0.708583856 | 2.654076244 | 0.007952586 | 0.024334042 |
| ENSG00000267387 | ENSG0000026 | 3.021594561 | 1.326054777 | 1.094937314 | 1.21107826 | 0.225865413 | 0.397953346 |
| ENSG00000267390 | KDSR-DT | 11.61664349 | 1.27820851 | 0.540280578 | 2.365823541 | 0.017990017 | 0.049859971 |
| ENSG00000267394 | ATXN7L3-AS1 | 16.4936647 | -0.52623077 | 0.430064065 | -1.223610185 | 0.221099316 | 0.391661645 |
| ENSG00000267395 | DM1-AS | 36.3517387 | -0.145412994 | 0.300069608 | -0.484597542 | 0.627961857 | 0.780547313 |
| ENSG00000267397 | ENSG0000026 | 3.459965013 | 2.444639826 | 1.139952797 | 2.144509696 | 0.031992069 | 0.082128253 |

| | | | | | | | |
|---|---|---|---|---|---|---|---|
| ENSG00000267414 | SETBP1-DT | 22.37881872 | 1.521580088 | 0.409632511 | 3.714500309 | 0.000203606 | 0.000862392 |
| ENSG00000267416 | HEATR6-DT | 22.76301633 | -0.137359792 | 0.371489014 | -0.369754653 | 0.711565306 | 0.839676344 |
| ENSG00000267418 | ELOCP29 | 34.62078028 | 0.452804447 | 0.321882053 | 1.406740275 | 0.159504419 | 0.305148141 |
| ENSG00000267419 | ZNF56P | 35.61119463 | 0.954719295 | 0.30772181 | 3.102540232 | 0.001918675 | 0.006771446 |
| ENSG00000267422 | ENSG0000026 | 569.880919 | 0.481733464 | 0.08620376 | 5.588311494 | 2.29E-08 | 1.58E-07 |
| ENSG00000267426 | ENSG0000026 | 2.172992442 | 0.464750168 | 1.243134425 | 0.37385351 | 0.708513306 | 0.837453382 |
| ENSG00000267432 | DNAH17-AS1 | 25.00633462 | 0.434776538 | 0.370605002 | 1.173153454 | 0.240734274 | 0.416631025 |
| ENSG00000267436 | ENSG0000026 | 19.30900358 | 0.046697495 | 0.418002818 | 0.111715741 | 0.911048798 | 0.955316516 |
| ENSG00000267439 | ENSG0000026 | 6.981494095 | -0.121674446 | 0.69283913 | -0.175617169 | 0.860594713 | 0.927426758 |
| ENSG00000267449 | ENSG0000026 | 2.590881998 | -0.122408727 | 1.14771728 | -0.106654077 | 0.915063416 | 0.957818706 |
| ENSG00000267458 | ENSG0000026 | 27.88118147 | -1.075347927 | 0.35057307 | -3.067400262 | 0.002159295 | 0.007536015 |
| ENSG00000267469 | ENSG0000026 | 30.57327401 | -0.593014845 | 0.324433936 | -1.827844686 | 0.067572865 | 0.15413359 |
| ENSG00000267470 | ZNF571-AS1 | 21.45266407 | 1.30391822 | 0.407542018 | 3.199469407 | 0.001376808 | 0.005015718 |
| ENSG00000267472 | ARHGAP27P2 | 8.915811243 | -1.654280901 | 0.638112267 | -2.592460584 | 0.00952921 | 0.028515533 |
| ENSG00000267475 | NUDT19-DT | 102.8276086 | 0.26456456 | 0.17642955 | 1.499547895 | 0.133731553 | 0.267173753 |
| ENSG00000267484 | ENSG0000026 | 9.280025723 | -1.085671453 | 0.592637937 | -1.831930399 | 0.066961791 | 0.153044085 |
| ENSG00000267493 | CIRBP-AS1 | 44.92345162 | 0.739986171 | 0.265792613 | 2.784073507 | 0.005368087 | 0.017155741 |
| ENSG00000267500 | ZNF887P | 24.22762039 | -0.117769634 | 0.376972383 | -0.312409183 | 0.754729572 | 0.864498202 |
| ENSG00000267508 | ZNF285 | 43.61122678 | 0.502231032 | 0.27100482 | 1.853218076 | 0.063851108 | 0.146885721 |
| ENSG00000267512 | ENSG0000026 | 4.593099054 | -1.006559126 | 0.843072552 | -1.193917563 | 0.232510217 | 0.406552301 |
| ENSG00000267519 | MIR23AHG | 56.4025622 | 0.42012089 | 0.238333904 | 1.762740772 | 0.077944221 | 0.173120327 |
| ENSG00000267526 | ENSG0000026 | 4.282546665 | 0.050351948 | 0.831864657 | 0.060529014 | 0.951734308 | 0.977646532 |
| ENSG00000267530 | LINC01836 | 31.01920295 | -0.201407175 | 0.326441433 | -0.616977976 | 0.537249252 | 0.712316371 |
| ENSG00000267532 | MIR497HG | 25.04211025 | -0.413070017 | 0.356449898 | -1.158844537 | 0.24651956 | 0.423702607 |
| ENSG00000267533 | ENSG0000026 | 153.3742626 | 0.086223077 | 0.151184816 | 0.570315721 | 0.56846358 | 0.736037682 |
| ENSG00000267534 | S1PR2 | 58.54506158 | 0.280315576 | 0.231431147 | 1.211226663 | 0.225808547 | 0.397883713 |
| ENSG00000267541 | MTCO2P2 | 42.6235158 | 0.032349306 | 0.282381568 | 0.114558843 | 0.908794803 | 0.95431011 |
| ENSG00000267544 | HIKESHIP2 | 45.04277253 | -0.088384434 | 0.264060092 | -0.334713335 | 0.737841341 | 0.855509243 |
| ENSG00000267546 | ENSG0000026 | 9.789110942 | -0.324570532 | 0.564712478 | -0.574753604 | 0.565457949 | 0.733605098 |
| ENSG00000267549 | ZSCAN5A-AS1 | 3.524247385 | -0.081225054 | 0.91697011 | -0.088579827 | 0.929415841 | 0.965993019 |
| ENSG00000267573 | KRT8P5 | 7.72977587 | 0.382753682 | 0.657464535 | 0.582166279 | 0.560454685 | 0.729849205 |
| ENSG00000267575 | LINC02987 | 254.5790552 | 0.729251311 | 0.116762868 | 6.245575552 | 4.22E-10 | 3.48E-09 |
| ENSG00000267576 | ENSG0000026 | 16.67846037 | -0.660187256 | 0.430402987 | -1.533881677 | 0.125058765 | 0.253611038 |

| | | | | | | | |
|---|---|---|---|---|---|---|---|
| ENSG00000267580 | ENSG0000026 | 4.836991382 | -2.807361953 | 1.004793753 | -2.79396836 | 0.005206557 | 0.016681159 |
| ENSG00000267586 | LINC00907 | 2.229197303 | 0.71911862 | 1.197346522 | 0.600593568 | 0.548110723 | 0.720932349 |
| ENSG00000267589 | ENSG0000026 | 1.950725788 | 0.345609646 | 1.299462247 | 0.265963592 | 0.790267249 | 0.886794047 |
| ENSG00000267595 | BRCA1P1 | 4.872720809 | -0.853474346 | 0.823719337 | -1.036122751 | 0.300144879 | 0.48680172 |
| ENSG00000267598 | ENSG0000026 | 22.20688689 | -0.148369337 | 0.399492089 | -0.371394932 | 0.7103434 | 0.838742158 |
| ENSG00000267601 | ENSG0000026 | 1.60279387 | 1.791269211 | 1.521769907 | 1.177095961 | 0.239157207 | NA |
| ENSG00000267607 | ICAM4-AS1 | 2.804615893 | -0.006576386 | 1.04398677 | -0.0062993 | 0.994973919 | 0.996913245 |
| ENSG00000267620 | ENSG0000026 | 3.88299572 | -0.397733026 | 0.902440592 | -0.440730425 | 0.659408168 | 0.802952565 |
| ENSG00000267632 | ENSG0000026 | 12.96751032 | -1.130949 | 0.540078325 | -2.094046265 | 0.036255859 | 0.091366517 |
| ENSG00000267633 | ENSG0000026 | 10.78636193 | -0.986168095 | 0.588425584 | -1.675943605 | 0.093749239 | 0.201066525 |
| ENSG00000267649 | ENSG0000026 | 6.002070637 | -0.830353593 | 0.722286483 | -1.149618071 | 0.250301212 | 0.42846165 |
| ENSG00000267655 | ENSG0000026 | 13.04253616 | 0.408720278 | 0.506295505 | 0.807276134 | 0.419507411 | 0.610929766 |
| ENSG00000267656 | ENSG0000026 | 4.799833948 | 0.101692526 | 0.784408419 | 0.129642319 | 0.89684942 | 0.947893263 |
| ENSG00000267658 | RAB5C-AS1 | 3.198374734 | -0.194237239 | 1.027300707 | -0.189075348 | 0.850033762 | 0.921503648 |
| ENSG00000267666 | ENSG0000026 | 17.58251118 | -0.622071275 | 0.418405153 | -1.486767718 | 0.13707617 | 0.272182753 |
| ENSG00000267676 | THA1P | 2.839090231 | -0.344415424 | 1.05748053 | -0.325694341 | 0.744655626 | 0.858800466 |
| ENSG00000267679 | EIF5AP2 | 1.671806185 | -0.21712742 | 1.323974678 | -0.163996656 | 0.869733776 | NA |
| ENSG00000267680 | ZNF224 | 749.9224187 | 0.135608202 | 0.08837307 | 1.534496911 | 0.124907453 | 0.253371382 |
| ENSG00000267681 | ENSG0000026 | 9.278973602 | 0.639269838 | 0.587415087 | 1.088276163 | 0.27647321 | 0.458987946 |
| ENSG00000267682 | ENSG0000026 | 5.761955469 | -0.544332295 | 0.732530652 | -0.743084666 | 0.457430423 | 0.644622241 |
| ENSG00000267688 | ENSG0000026 | 10.37950998 | -0.854300647 | 0.559953827 | -1.525662665 | 0.127093897 | 0.256851744 |
| ENSG00000267698 | ENSG0000026 | 15.61242735 | -0.145290118 | 0.443968446 | -0.327253252 | 0.743476347 | 0.858490754 |
| ENSG00000267707 | ENSG0000026 | 5.812597375 | 0.559310657 | 0.737228372 | 0.758666755 | 0.448051929 | 0.636309741 |
| ENSG00000267724 | ENSG0000026 | 3.25993475 | 1.047846237 | 1.014093737 | 1.033283413 | 0.301471287 | 0.488331545 |
| ENSG00000267727 | ENSG0000026 | 5.009198811 | -0.25039805 | 0.819925254 | -0.305391313 | 0.760068138 | 0.868106598 |
| ENSG00000267729 | ENSG0000026 | 6.417704331 | -1.644050159 | 0.736377286 | -2.232619326 | 0.025574059 | 0.067721217 |
| ENSG00000267731 | ENSG0000026 | 2.818839365 | 1.086165771 | 1.114287055 | 0.97476298 | 0.329677844 | 0.520121375 |
| ENSG00000267733 | ENSG0000026 | 20.07406018 | 1.336553152 | 0.412861773 | 3.237289669 | 0.001206709 | 0.004451897 |
| ENSG00000267735 | ENSG0000026 | 6.106577245 | 1.146623545 | 0.748452852 | 1.531991684 | 0.125524487 | 0.254285741 |
| ENSG00000267736 | HMGB2P1 | 3.014927031 | 0.340899282 | 1.035952806 | 0.329068352 | 0.742104025 | 0.857711927 |
| ENSG00000267737 | ENSG0000026 | 2.274613238 | 0.625330865 | 1.190665142 | 0.525194568 | 0.599447941 | 0.759532492 |
| ENSG00000267741 | UBE2L4 | 8.861577172 | 0.016762373 | 0.585767484 | 0.028616086 | 0.977170782 | 0.989387067 |
| ENSG00000267742 | SINHCAFP2 | 49.46641978 | -0.50391406 | 0.249255182 | -2.02167937 | 0.043209486 | 0.105742012 |

| | | | | | | | |
|---|---|---|---|---|---|---|---|
| ENSG00000267750 | RUNDC3A-AS1 | 826.5894704 | 0.654114678 | 0.082967206 | 7.884014816 | 3.17E-15 | 3.87E-14 |
| ENSG00000267751 | BSG-AS1 | 194.4171072 | -0.187412611 | 0.138421383 | -1.353928181 | 0.175759292 | 0.329028738 |
| ENSG00000267757 | EML2-AS1 | 27.18776837 | 0.450120741 | 0.34718989 | 1.296468456 | 0.194814141 | 0.355956702 |
| ENSG00000267767 | LINC01801 | 125.054109 | -1.111176032 | 0.166194368 | -6.686002929 | 2.29E-11 | 2.09E-10 |
| ENSG00000267769 | ENSG0000026 | 20.94945966 | 0.534929005 | 0.395213773 | 1.353518128 | 0.175890163 | 0.32919609 |
| ENSG00000267778 | CBARP-DT | 8.609491342 | 0.587531121 | 0.612729756 | 0.958874797 | 0.337621823 | 0.528816374 |
| ENSG00000267786 | ENSG0000026 | 3.083001161 | -0.309561394 | 0.991007003 | -0.312370541 | 0.754758936 | 0.864498202 |
| ENSG00000267787 | ENSG0000026 | 19.24308197 | -0.164986486 | 0.396792629 | -0.415800279 | 0.677556154 | 0.816641527 |
| ENSG00000267791 | ENSG0000026 | 2.264659274 | 0.24045733 | 1.206871698 | 0.199240177 | 0.842074873 | 0.917009895 |
| ENSG00000267795 | SMIM22 | 10.93761112 | 0.873613856 | 0.539318457 | 1.619847875 | 0.105264959 | 0.220911002 |
| ENSG00000267796 | LIN37 | 80.51948283 | -0.25433339 | 0.218486101 | -1.164071256 | 0.244395136 | 0.421187321 |
| ENSG00000267809 | NDUFV2P1 | 543.780611 | 0.219253559 | 0.090946348 | 2.410801142 | 0.015917525 | 0.044808935 |
| ENSG00000267811 | ENSG0000026 | 8.340098388 | 0.990902801 | 0.624707331 | 1.586187248 | 0.11269684 | 0.233464323 |
| ENSG00000267815 | ENSG0000026 | 4.456989729 | -0.499524748 | 0.821918702 | -0.60775445 | 0.543350344 | 0.716907232 |
| ENSG00000267834 | ENSG0000026 | 15.98943922 | 1.182642014 | 0.457963753 | 2.58239218 | 0.009811802 | 0.029227727 |
| ENSG00000267838 | ENSG0000026 | 3.645236262 | 1.297608057 | 0.971225017 | 1.336052957 | 0.181531965 | 0.337057326 |
| ENSG00000267855 | NDUFA7 | 7.815384022 | 0.457717922 | 0.655215896 | 0.698575729 | 0.484817216 | 0.668693298 |
| ENSG00000267858 | MZF1-AS1 | 163.2670322 | -0.432844911 | 0.148105427 | -2.922545915 | 0.003471824 | 0.011603096 |
| ENSG00000267868 | ENSG0000026 | 124.1616875 | 0.056267931 | 0.160715016 | 0.350109978 | 0.726256163 | 0.847853251 |
| ENSG00000267871 | ZNF460-AS1 | 17.44372159 | 0.687855667 | 0.425754055 | 1.615617419 | 0.106177055 | 0.222479143 |
| ENSG00000267882 | ENSG0000026 | 2.949262064 | 1.749858552 | 1.116696371 | 1.566995826 | 0.117115667 | 0.240975281 |
| ENSG00000267886 | ENSG0000026 | 45.76086147 | -0.232123885 | 0.266239463 | -0.871861301 | 0.383284047 | 0.575691485 |
| ENSG00000267892 | ENSG0000026 | 26.79445434 | 1.1192027 | 0.361290532 | 3.097791387 | 0.001949686 | 0.006872434 |
| ENSG00000267898 | ENSG0000026 | 2.636734881 | -0.535767932 | 1.056906902 | -0.506920648 | 0.612210497 | 0.769498044 |
| ENSG00000267904 | ENSG0000026 | 23.04142904 | 0.481041985 | 0.381635333 | 1.260475497 | 0.207497882 | 0.373685147 |
| ENSG00000267909 | CCDC177 | 21.87782741 | -3.576011984 | 0.545753423 | -6.552431616 | 5.66E-11 | 5.00E-10 |
| ENSG00000267920 | SNX6P1 | 8.061491653 | -0.836825376 | 0.6358746 | -1.31602265 | 0.188166441 | 0.346487249 |
| ENSG00000267938 | EIF1P6 | 5.891932331 | 1.068961385 | 0.736892933 | 1.450633241 | 0.146882014 | 0.286910201 |
| ENSG00000267939 | ENSG0000026 | 13.60040316 | 0.347027121 | 0.474569579 | 0.731246029 | 0.46462889 | 0.650747099 |
| ENSG00000267940 | ENSG0000026 | 12.33075591 | -0.347060829 | 0.518540938 | -0.669302659 | 0.503302431 | 0.68435527 |
| ENSG00000267980 | ENSG0000026 | 104.5684469 | 0.234135865 | 0.173761182 | 1.347457828 | 0.177832827 | 0.332124485 |
| ENSG00000267984 | ENSG0000026 | 13.03542941 | -0.569959755 | 0.495308901 | -1.150715754 | 0.249849193 | 0.427932225 |
| ENSG00000267992 | ENSG0000026 | 4.191705487 | -0.16727025 | 0.844615932 | -0.198042973 | 0.843011442 | 0.917338822 |

| | | | | | | | |
|---|---|---|---|---|---|---|---|
| ENSG00000268001 | CARD8-AS1 | 34.62269996 | -0.154771582 | 0.3079697 | -0.502554577 | 0.615277471 | 0.771999409 |
| ENSG00000268006 | PTOV1-AS1 | 140.9431839 | 0.092504719 | 0.152623612 | 0.606097037 | 0.544450318 | 0.717839549 |
| ENSG00000268015 | ENSG0000026 | 18.62947953 | -0.028614099 | 0.429879122 | -0.066563129 | 0.9469295 | 0.975200824 |
| ENSG00000268030 | ENSG0000026 | 2.809889687 | -0.65582808 | 1.136255714 | -0.577183527 | 0.563815484 | 0.732388297 |
| ENSG00000268034 | TPM3P5 | 248.6605978 | -0.104212202 | 0.118933762 | -0.876220517 | 0.380910165 | 0.57330213 |
| ENSG00000268041 | ERFL | 22.39491677 | 0.474972007 | 0.379971236 | 1.250020954 | 0.211291893 | 0.378848255 |
| ENSG00000268043 | NBPF12 | 961.7836754 | 0.220116528 | 0.070573626 | 3.118962998 | 0.001814888 | 0.006433862 |
| ENSG00000268047 | ENSG0000026 | 15.23520618 | -0.077039127 | 0.448591182 | -0.171735714 | 0.863645307 | 0.929172842 |
| ENSG00000268049 | ENSG0000026 | 87.16111221 | 0.265041419 | 0.192086107 | 1.379805251 | 0.167646616 | 0.317416091 |
| ENSG00000268056 | ENSG0000026 | 5.404004993 | -0.088512472 | 0.749999087 | -0.118016773 | 0.906054368 | 0.95262805 |
| ENSG00000268061 | NAPA-AS1 | 63.56149687 | -0.848260362 | 0.23737838 | -3.573452491 | 0.000352305 | 0.001429157 |
| ENSG00000268069 | ENSG0000026 | 13.12845608 | 0.367305131 | 0.479777491 | 0.765573911 | 0.443929873 | 0.632753994 |
| ENSG00000268089 | GABRQ | 214.370169 | 3.872836608 | 0.197391434 | 19.62008442 | 1.04E-85 | 1.32E-83 |
| ENSG00000268093 | ENSG0000026 | 9.268171378 | 0.384568372 | 0.570993977 | 0.67350688 | 0.500624874 | 0.681732072 |
| ENSG00000268104 | SLC6A14 | 1.628738824 | -0.148917016 | 1.343621332 | -0.110832578 | 0.911749111 | NA |
| ENSG00000268112 | ENSG0000026 | 2.217890487 | 2.446812831 | 1.418818695 | 1.724542283 | 0.084609993 | 0.18501127 |
| ENSG00000268120 | ENSG0000026 | 1.925206966 | 1.412840647 | 1.362763536 | 1.036746735 | 0.299853905 | 0.486576451 |
| ENSG00000268129 | ENSG0000026 | 18.94069239 | -0.198179212 | 0.408113609 | -0.485598147 | 0.627252111 | 0.780154165 |
| ENSG00000268140 | PCGF7P | 40.61261721 | 0.064946139 | 0.278875209 | 0.232886027 | 0.815849909 | 0.902352305 |
| ENSG00000268144 | NIFKP6 | 3.537968992 | 1.23657962 | 1.012401502 | 1.221432029 | 0.221922487 | 0.392907451 |
| ENSG00000268149 | ENSG0000026 | 4.352983561 | -0.85906566 | 0.843295112 | -1.018701103 | 0.308344889 | 0.496381175 |
| ENSG00000268186 | ZNF114-AS1 | 6.662575728 | -1.356606817 | 0.701174305 | -1.93476402 | 0.05301926 | 0.126063622 |
| ENSG00000268189 | ENSG0000026 | 13.52699421 | -0.330359808 | 0.476709829 | -0.692999782 | 0.488309688 | 0.67162154 |
| ENSG00000268199 | ENSG0000026 | 3.074045971 | -0.303350809 | 0.98771232 | -0.307124659 | 0.758748489 | 0.867190412 |
| ENSG00000268201 | ENSG0000026 | 8.125476081 | 1.349387094 | 0.676388373 | 1.994988601 | 0.046044123 | 0.111607374 |
| ENSG00000268204 | ENSG0000026 | 13.83213143 | -0.041738129 | 0.479955019 | -0.086962585 | 0.930701253 | 0.966601174 |
| ENSG00000268205 | ENSG0000026 | 183.9640054 | 0.5215024 | 0.142981846 | 3.647332968 | 0.000264976 | 0.001098998 |
| ENSG00000268218 | ENSG0000026 | 20.08712768 | 0.446457369 | 0.398691192 | 1.119807454 | 0.262795822 | 0.443502991 |
| ENSG00000268222 | EEF1A1P7 | 23.50300549 | 0.455761082 | 0.370906293 | 1.228776892 | 0.219155468 | 0.388999259 |
| ENSG00000268230 | ENSG0000026 | 20.23548568 | -0.069377701 | 0.383524653 | -0.180895024 | 0.856449975 | 0.92565194 |
| ENSG00000268234 | FKBP4P6 | 18.18583223 | 0.045348031 | 0.420989519 | 0.107717719 | 0.914219614 | 0.957459731 |
| ENSG00000268266 | ENSG0000026 | 2.494413902 | 2.003374685 | 1.294187687 | 1.547978477 | 0.121627479 | 0.248056042 |
| ENSG00000268279 | ENSG0000026 | 24.474077 | 0.23097932 | 0.373448461 | 0.618503875 | 0.536243244 | 0.711475999 |

| | | | | | | | |
|---|---|---|---|---|---|---|---|
| ENSG00000268282 | ARPC1AP2 | 11.59055953 | -0.576973446 | 0.510324834 | -1.130600371 | 0.258223331 | 0.437847824 |
| ENSG00000268288 | ENSG0000026 | 15.19757837 | 0.53220246 | 0.455488568 | 1.168421113 | 0.242636943 | 0.418944447 |
| ENSG00000268292 | ENSG0000026 | 7.770684805 | -1.849396515 | 0.678295991 | -2.726533163 | 0.006400352 | 0.020055194 |
| ENSG00000268295 | POLR3GP1 | 3.786606407 | -0.044268045 | 0.895446266 | -0.049436853 | 0.96057116 | 0.981920343 |
| ENSG00000268297 | CLEC4GP1 | 1.740180941 | -3.297814754 | 1.691588077 | -1.949537715 | 0.051231243 | NA |
| ENSG00000268307 | LINC02560 | 1.634919633 | 1.068886703 | 1.439211085 | 0.742689321 | 0.457669797 | NA |
| ENSG00000268309 | ENSG0000026 | 8.937225764 | -0.674757749 | 0.606390664 | -1.11274429 | 0.265818268 | 0.446828286 |
| ENSG00000268333 | ENSG0000026 | 13.9890557 | 0.175332332 | 0.46425055 | 0.377667473 | 0.705677634 | 0.835385491 |
| ENSG00000268350 | FAM156A | 10.97011739 | -0.88736223 | 0.538353796 | -1.648288238 | 0.099293537 | 0.210809231 |
| ENSG00000268362 | ENSG0000026 | 124.9347644 | 0.009498438 | 0.164074585 | 0.057890978 | 0.953835469 | 0.978691016 |
| ENSG00000268364 | SMC5-DT | 12.50640204 | -0.66671902 | 0.504120891 | -1.32253797 | 0.185989074 | 0.343332209 |
| ENSG00000268379 | ENSG0000026 | 32.97314509 | 0.063252356 | 0.318934831 | 0.198323762 | 0.842791761 | 0.917308531 |
| ENSG00000268401 | ENSG0000026 | 2.656668468 | 0.15761204 | 1.067917851 | 0.147588169 | 0.882667793 | 0.939121523 |
| ENSG00000268403 | ENSG0000026 | 39.42120494 | -0.220693877 | 0.280600103 | -0.786506756 | 0.431570663 | 0.62160049 |
| ENSG00000268412 | TRMT112P6 | 247.0209521 | -0.063740178 | 0.120976765 | -0.5268795 | 0.598277271 | 0.758658892 |
| ENSG00000268416 | ENSG0000026 | 12.09192881 | 5.959092954 | 1.250343539 | 4.765964529 | 1.88E-06 | 1.05E-05 |
| ENSG00000268471 | MIR4453HG | 188.6209765 | 0.48859927 | 0.139092207 | 3.512772439 | 0.000443457 | 0.001771975 |
| ENSG00000268509 | ENSG0000026 | 2.214646039 | 0.916812203 | 1.267525927 | 0.723308442 | 0.469490407 | 0.655395238 |
| ENSG00000268516 | ZNF8-DT | 84.93510037 | -0.324945369 | 0.205709833 | -1.579629732 | 0.114191686 | 0.236017414 |
| ENSG00000268520 | ENSG0000026 | 2.208454088 | -0.454748564 | 1.167665126 | -0.389451183 | 0.696942416 | 0.829719188 |
| ENSG00000268521 | VN1R83P | 7.626691117 | -0.803555787 | 0.647969644 | -1.24011332 | 0.214933483 | 0.383881509 |
| ENSG00000268530 | ENSG0000026 | 5.501479954 | -0.118231909 | 0.737638027 | -0.160284455 | 0.872657004 | 0.934146135 |
| ENSG00000268543 | ENSG0000026 | 37.81835719 | -0.336118409 | 0.291284033 | -1.153919786 | 0.248533057 | 0.426269033 |
| ENSG00000268565 | ENSG0000026 | 2.827745989 | 0.815466195 | 1.071345438 | 0.761160842 | 0.446561004 | 0.635017941 |
| ENSG00000268573 | ENSG0000026 | 24.60571307 | -0.343074043 | 0.35727479 | -0.960252591 | 0.336928104 | 0.528123295 |
| ENSG00000268575 | ENSG0000026 | 152.6118563 | 0.320279249 | 0.1596069 | 2.006675454 | 0.044784231 | 0.108979555 |
| ENSG00000268583 | ENSG0000026 | 7.655364463 | -0.369266337 | 0.642190568 | -0.575010528 | 0.565284177 | 0.733481521 |
| ENSG00000268584 | ENSG0000026 | 25.81895401 | -0.683197796 | 0.35465579 | -1.926368654 | 0.054058359 | 0.128109375 |
| ENSG00000268592 | RAET1E-AS1 | 2.567274478 | 0.537005312 | 1.120577054 | 0.479222121 | 0.631780619 | 0.782942434 |
| ENSG00000268603 | ENSG0000026 | 5.616487427 | 0.565836301 | 0.783256882 | 0.722414719 | 0.47003954 | 0.655940953 |
| ENSG00000268615 | ENSG0000026 | 4.031090933 | 0.262575826 | 0.886561276 | 0.296173353 | 0.767097697 | 0.872791448 |
| ENSG00000268628 | ENSG0000026 | 15.88563395 | -0.379556605 | 0.461550658 | -0.822350913 | 0.410877208 | 0.603369294 |
| ENSG00000268649 | ENSG0000026 | 19.35806011 | 0.240156509 | 0.411125107 | 0.584144593 | 0.559123035 | 0.728807456 |

| | | | | | | | |
|---|---|---|---|---|---|---|---|
| ENSG00000268655 | ENSG0000026 | 7.644668041 | 0.303082235 | 0.64379418 | 0.470775046 | 0.637801386 | 0.786664003 |
| ENSG00000268660 | LETM1P2 | 5.847492888 | 0.907572196 | 0.791389222 | 1.14680889 | 0.251460614 | 0.429916861 |
| ENSG00000268670 | ENSG0000026 | 22.51140232 | -0.198573207 | 0.369720433 | -0.537090162 | 0.591205336 | 0.753709593 |
| ENSG00000268707 | ENSG0000026 | 2.480616236 | -0.113847396 | 1.154042907 | -0.098650922 | 0.921415438 | 0.9619656 |
| ENSG00000268723 | ENSG0000026 | 4.525629364 | -1.668015477 | 0.882432044 | -1.89024808 | 0.058724789 | 0.137058364 |
| ENSG00000268736 | MTCO3P39 | 13.73243175 | 2.178988007 | 0.549532764 | 3.965164864 | 7.33E-05 | 0.000334701 |
| ENSG00000268743 | ENSG0000026 | 4.124272595 | -0.457742171 | 0.87688768 | -0.522007757 | 0.601664929 | 0.761330837 |
| ENSG00000268744 | ENSG0000026 | 1.864192425 | 0.848506213 | 1.314073829 | 0.645706652 | 0.518469352 | 0.696066421 |
| ENSG00000268745 | ENSG0000026 | 2.30631287 | 0.647039482 | 1.153511119 | 0.560930425 | 0.574844968 | 0.741214298 |
| ENSG00000268747 | ENSG0000026 | 4.239525073 | 1.335123895 | 0.893430575 | 1.494379007 | 0.135076585 | 0.269237716 |
| ENSG00000268751 | SCGB1B2P | 4.82957066 | 0.596513425 | 0.796535189 | 0.748885213 | 0.453926393 | 0.641552294 |
| ENSG00000268785 | RPL7P50 | 5.77422621 | 0.645975864 | 0.737320113 | 0.876113173 | 0.380968512 | 0.57330213 |
| ENSG00000268798 | ENSG0000026 | 11.93517731 | 0.548024901 | 0.521874647 | 1.050108305 | 0.293668321 | 0.479245307 |
| ENSG00000268833 | ENSG0000026 | 1.882105858 | -0.610543382 | 1.364864732 | -0.447328858 | 0.654637634 | 0.799489924 |
| ENSG00000268836 | ENSG0000026 | 2.415467326 | 0.026722669 | 1.099811357 | 0.024297503 | 0.980615305 | 0.991070363 |
| ENSG00000268854 | ENSG0000026 | 16.88677785 | 0.063793055 | 0.432082248 | 0.147641001 | 0.882626096 | 0.939121523 |
| ENSG00000268858 | ENSG0000026 | 549.3089324 | 0.064967627 | 0.086812065 | 0.748370944 | 0.454236444 | 0.641838792 |
| ENSG00000268864 | ENSG0000026 | 20.01410621 | -1.997037335 | 0.43927246 | -4.546238421 | 5.46E-06 | 2.90E-05 |
| ENSG00000268869 | ESPNP | 18.96022459 | -0.052954409 | 0.4270127 | -0.124011322 | 0.901306311 | 0.950225965 |
| ENSG00000268889 | LIM2-AS1 | 1.564426708 | -0.158490845 | 1.445524719 | -0.109642432 | 0.912692958 | NA |
| ENSG00000268895 | A1BG-AS1 | 473.0906635 | -0.274108901 | 0.090293601 | -3.035751137 | 0.002399373 | 0.008309031 |
| ENSG00000268903 | ENSG0000026 | 31.59924362 | -0.288048558 | 0.318497865 | -0.904397139 | 0.365784855 | 0.558469938 |
| ENSG00000268941 | LINC01711 | 2.308717947 | -1.997542043 | 1.311470693 | -1.523131285 | 0.127725863 | 0.257789134 |
| ENSG00000268942 | CKS1BP3 | 131.4840991 | -0.202335126 | 0.155661926 | -1.299836965 | 0.193656853 | 0.354418971 |
| ENSG00000268945 | ENSG0000026 | 4.02304827 | 0.013502917 | 0.871951534 | 0.015485858 | 0.987644567 | 0.993927018 |
| ENSG00000268947 | ENSG0000026 | 15.51912093 | 0.294657417 | 0.452247179 | 0.651540642 | 0.514697551 | 0.693763548 |
| ENSG00000268964 | ERVV-2 | 41.60186156 | 0.78012208 | 0.281821344 | 2.768144064 | 0.005637653 | 0.017900036 |
| ENSG00000268970 | ENSG0000026 | 2.408882992 | 0.82451939 | 1.1257985 | 0.732386293 | 0.463932822 | 0.650169158 |
| ENSG00000268983 | KXD1-AS1 | 14.47198004 | -0.610090326 | 0.493996538 | -1.235009315 | 0.216827037 | 0.386271044 |
| ENSG00000268996 | MAN1B1-DT | 99.56528218 | 0.120848941 | 0.177087055 | 0.682426737 | 0.494969154 | 0.677239198 |
| ENSG00000269001 | ZNF818P | 665.539279 | 0.158339103 | 0.082882085 | 1.910414082 | 0.056079919 | 0.132054336 |
| ENSG00000269012 | KRT18P40 | 4.771008662 | 2.533604261 | 0.981411782 | 2.581591447 | 0.009834593 | 0.029280411 |
| ENSG00000269019 | HOMER3-AS1 | 2.171567537 | 0.490100044 | 1.198115593 | 0.409059065 | 0.682496316 | 0.819871514 |

| | | | | | | | |
|---|---|---|---|---|---|---|---|
| ENSG00000269038 | SF1-DT | 11.74488838 | -0.271798445 | 0.524929155 | -0.517781195 | 0.60461094 | 0.763375803 |
| ENSG00000269044 | ENSG0000026 | 53.1124595 | 0.665957396 | 0.244440248 | 2.724417941 | 0.006441494 | 0.020166265 |
| ENSG00000269054 | ZNF497-AS1 | 24.54314547 | 0.151026153 | 0.361506684 | 0.417768633 | 0.676116287 | 0.815291612 |
| ENSG00000269068 | ENSG0000026 | 7.130572855 | -0.778337997 | 0.661148154 | -1.177252017 | 0.239094933 | 0.414669823 |
| ENSG00000269069 | ENSG0000026 | 68.19325111 | 0.687285781 | 0.218358464 | 3.147511514 | 0.001646666 | 0.005895821 |
| ENSG00000269091 | ENSG0000026 | 14.46025151 | 0.198559704 | 0.465676281 | 0.426389988 | 0.669823696 | 0.810317039 |
| ENSG00000269106 | ENSG0000026 | 15.96179236 | 0.173038638 | 0.451710751 | 0.383073986 | 0.701664908 | 0.833050046 |
| ENSG00000269113 | TRABD2B | 3.551624382 | 0.336205532 | 0.925648106 | 0.363210954 | 0.716447305 | 0.84236078 |
| ENSG00000269119 | HNRNPA1P52 | 2.720521029 | -0.590848744 | 1.112397875 | -0.531148753 | 0.595315702 | 0.756595136 |
| ENSG00000269125 | ENSG0000026 | 3.943497882 | -0.403451389 | 0.872939731 | -0.462175537 | 0.643955444 | 0.791318722 |
| ENSG00000269139 | ENSG0000026 | 8.767998262 | 1.833235981 | 0.681785423 | 2.688875296 | 0.00716932 | 0.022185916 |
| ENSG00000269145 | MAST3-AS1 | 1.644353455 | -0.762112684 | 1.348890524 | -0.564992244 | 0.57207904 | NA |
| ENSG00000269148 | ENSG0000026 | 3.36963243 | -0.78957427 | 0.969979908 | -0.814010954 | 0.415638676 | 0.607773535 |
| ENSG00000269151 | ENSG0000026 | 2.526581651 | -0.358758012 | 1.163114278 | -0.308446056 | 0.757742941 | 0.866391318 |
| ENSG00000269153 | LYPLA2P2 | 46.47551067 | 0.250929556 | 0.260388676 | 0.963673073 | 0.335209858 | 0.526440347 |
| ENSG00000269176 | ENSG0000026 | 98.59770818 | 0.354731907 | 0.180626499 | 1.963897372 | 0.049541995 | 0.118855193 |
| ENSG00000269177 | ENSG0000026 | 10.0445887 | -0.006829661 | 0.60198358 | -0.011345261 | 0.990947985 | 0.995176993 |
| ENSG00000269190 | FBXO17 | 444.0313174 | 0.276323552 | 0.09049103 | 3.053601589 | 0.002261122 | 0.007869842 |
| ENSG00000269194 | ENSG0000026 | 4.971849654 | 0.293872146 | 0.782050022 | 0.375771546 | 0.707086743 | 0.836461297 |
| ENSG00000269226 | TMSB15C | 4.352154228 | 0.468866607 | 0.84855507 | 0.552547057 | 0.580573603 | 0.74566701 |
| ENSG00000269235 | ZNF350-AS1 | 3.990428648 | -1.130471404 | 0.908512111 | -1.24431077 | 0.213385209 | 0.381561828 |
| ENSG00000269243 | ENSG0000026 | 33.33475523 | -0.377302425 | 0.322607739 | -1.169539287 | 0.242186422 | 0.418449689 |
| ENSG00000269246 | ENSG0000026 | 8.675834346 | -0.075135911 | 0.605740293 | -0.12403981 | 0.901283755 | 0.950225965 |
| ENSG00000269275 | ENSG0000026 | 34.79225877 | 0.579205428 | 0.303085485 | 1.911029913 | 0.056000737 | 0.131935597 |
| ENSG00000269289 | ENSG0000026 | 1.718140277 | 3.028410241 | 1.611424033 | 1.879337889 | 0.060198371 | NA |
| ENSG00000269292 | ENSG0000026 | 21.72211797 | -1.049068855 | 0.394601815 | -2.658550504 | 0.007847759 | 0.024045357 |
| ENSG00000269293 | ZSCAN16-AS1 | 44.94297065 | -0.847163762 | 0.264376761 | -3.204380588 | 0.001353534 | 0.004941329 |
| ENSG00000269304 | FKBP1AP1 | 3.179947415 | 0.78517383 | 1.029161446 | 0.76292581 | 0.445507642 | 0.634109655 |
| ENSG00000269313 | MAGIX | 72.01188416 | -0.341725304 | 0.210977279 | -1.619725615 | 0.105291231 | 0.220945924 |
| ENSG00000269318 | ENSG0000026 | 18.6526469 | 0.729803679 | 0.42476185 | 1.718147895 | 0.085769643 | 0.187012223 |
| ENSG00000269335 | IKBKG | 346.3918344 | -0.471524056 | 0.104817039 | -4.498543961 | 6.84E-06 | 3.60E-05 |
| ENSG00000269343 | ZNF587B | 506.7273593 | -0.025057933 | 0.098353062 | -0.254775317 | 0.798896641 | 0.892721951 |
| ENSG00000269352 | PTOV1-AS2 | 297.2857528 | 0.330436886 | 0.121808529 | 2.71275655 | 0.006672611 | 0.020794687 |

| | | | | | | | |
|---|---|---|---|---|---|---|---|
| ENSG00000269356 | ENSG0000026 | 12.29689012 | 0.23013628 | 0.495414006 | 0.464533253 | 0.642265741 | 0.790179777 |
| ENSG00000269371 | ENSG0000026 | 1.709513506 | -0.357075762 | 1.383835368 | -0.258033412 | 0.796381121 | NA |
| ENSG00000269374 | ENSG0000026 | 157.1933584 | 0.302261768 | 0.144427859 | 2.092821776 | 0.036365067 | 0.091621597 |
| ENSG00000269376 | ENSG0000026 | 6.820676303 | 0.879837263 | 0.714781264 | 1.230918194 | 0.218353462 | 0.387996005 |
| ENSG00000269378 | ITGB1P1 | 622.2558624 | -0.291330271 | 0.080924179 | -3.600039852 | 0.000318168 | 0.001303354 |
| ENSG00000269386 | RAB11B-AS1 | 117.8484231 | 0.08609139 | 0.167605128 | 0.513656065 | 0.607492474 | 0.765833783 |
| ENSG00000269388 | ENSG0000026 | 5.166558404 | -1.713351884 | 0.845961418 | -2.025330999 | 0.042833388 | 0.105026294 |
| ENSG00000269392 | ENSG0000026 | 6.230400685 | -0.005871836 | 0.700993641 | -0.008376447 | 0.99331664 | 0.996354439 |
| ENSG00000269397 | ENSG0000026 | 5.036685289 | 0.692303164 | 0.801671619 | 0.863574496 | 0.38782168 | 0.580492583 |
| ENSG00000269399 | ENSG0000026 | 21.84404224 | -0.125787105 | 0.378677782 | -0.332174504 | 0.739757494 | 0.856527588 |
| ENSG00000269421 | ENSG0000026 | 1.810821026 | 1.242762911 | 1.371626282 | 0.906050669 | 0.364909034 | 0.5576586 |
| ENSG00000269425 | ENSG0000026 | 7.842549592 | 0.372058104 | 0.640324718 | 0.581045982 | 0.561209463 | 0.730157957 |
| ENSG00000269439 | PGLS-DT | 16.3110018 | 0.566145796 | 0.435018882 | 1.301428097 | 0.193111962 | 0.353691688 |
| ENSG00000269444 | ENSG0000026 | 1.881691191 | 0.860359105 | 1.299248341 | 0.662197578 | 0.507844606 | 0.688024202 |
| ENSG00000269463 | ENSG0000026 | 1.585256797 | 1.910474675 | 1.601811074 | 1.192696633 | 0.232988205 | NA |
| ENSG00000269473 | ZNF132-DT | 56.79488679 | -0.257891593 | 0.234121093 | -1.101530792 | 0.270665711 | 0.452506848 |
| ENSG00000269480 | ENSG0000026 | 1.989474194 | -1.186941868 | 1.29224596 | -0.918510799 | 0.358351508 | 0.550829777 |
| ENSG00000269486 | ERVK9-11 | 23.75048424 | 0.719806108 | 0.384408785 | 1.872501712 | 0.061137225 | 0.141728893 |
| ENSG00000269487 | ENSG0000026 | 18.43380676 | -0.19985177 | 0.435547689 | -0.458851637 | 0.646340711 | 0.793412953 |
| ENSG00000269489 | ENSG0000026 | 19.5851451 | 0.246409347 | 0.405411 | 0.607801336 | 0.543319243 | 0.716907232 |
| ENSG00000269516 | CYP4F23P | 14.95910842 | 0.611221988 | 0.478687248 | 1.276871257 | 0.201647703 | 0.365400947 |
| ENSG00000269534 | ENSG0000026 | 7.569342273 | -0.279166111 | 0.653793174 | -0.426994534 | 0.669383309 | 0.810007484 |
| ENSG00000269546 | CLIC4P2 | 1.865228626 | 1.199780914 | 1.441256411 | 0.832454867 | 0.405152243 | 0.597851052 |
| ENSG00000269556 | TMEM185A | 854.6718313 | -0.202004251 | 0.07177798 | -2.814292775 | 0.004888468 | 0.015790125 |
| ENSG00000269559 | ENSG0000026 | 3.816396246 | -0.33307101 | 0.932815298 | -0.357059978 | 0.721046905 | 0.845162383 |
| ENSG00000269570 | ENSG0000026 | 72.02967165 | -0.442794218 | 0.240732254 | -1.8393639 | 0.06586168 | 0.150905607 |
| ENSG00000269583 | ENSG0000026 | 14.93729973 | -0.837858339 | 0.463619543 | -1.807211001 | 0.070729386 | 0.160007752 |
| ENSG00000269586 | CT45A10 | 43.39459417 | 5.329366128 | 0.631844947 | 8.434610659 | 3.32E-17 | 4.63E-16 |
| ENSG00000269604 | ENSG0000026 | 27.73409581 | -0.166932384 | 0.364986626 | -0.457365756 | 0.647408173 | 0.794115255 |
| ENSG00000269608 | OSTCP3 | 5.863036092 | -0.158191797 | 0.720935978 | -0.219425583 | 0.826318544 | 0.908364356 |
| ENSG00000269609 | C10orf95-AS1 | 526.3864205 | -0.115332362 | 0.08741798 | -1.319320833 | 0.187061876 | 0.344757728 |
| ENSG00000269621 | ENSG0000026 | 16.404475 | -0.616268242 | 0.444992292 | -1.38489644 | 0.166084136 | 0.315116179 |
| ENSG00000269637 | RPL12P41 | 3.992248191 | -0.407816583 | 0.870314635 | -0.46858523 | 0.639366137 | 0.787619975 |

| | | | | | | | |
|---|---|---|---|---|---|---|---|
| ENSG00000269656 | NOP53-AS1 | 3.103360555 | -0.315422316 | 0.972186048 | -0.324446454 | 0.745600055 | 0.859379806 |
| ENSG00000269680 | ENSG0000026 | 6.814374316 | 0.042585143 | 0.685667724 | 0.062107551 | 0.950477183 | 0.977185792 |
| ENSG00000269688 | ENSG0000026 | 34.05188811 | -0.032065826 | 0.298908516 | -0.107276389 | 0.914569715 | 0.957607799 |
| ENSG00000269692 | FKBP4P2 | 3.413105894 | 0.542420219 | 0.965496922 | 0.561804193 | 0.574249435 | 0.740779511 |
| ENSG00000269707 | ENSG0000026 | 2.571446273 | -0.134706608 | 1.080204258 | -0.124704755 | 0.900757293 | 0.949953563 |
| ENSG00000269713 | NBPF9 | 2341.720054 | 0.41649851 | 0.058462463 | 7.124203901 | 1.05E-12 | 1.07E-11 |
| ENSG00000269737 | ENSG0000026 | 1.693612766 | 0.966489127 | 1.401491964 | 0.689614462 | 0.49043667 | NA |
| ENSG00000269743 | SLC25A53 | 158.4873523 | 0.198641754 | 0.143559636 | 1.383688059 | 0.166453991 | 0.315675557 |
| ENSG00000269765 | ENSG0000026 | 10.40769365 | -0.052317433 | 0.582430078 | -0.089826118 | 0.928425393 | 0.965636494 |
| ENSG00000269800 | PLEKHA3P1 | 12.71919697 | 0.527925907 | 0.502333112 | 1.050947855 | 0.29328254 | 0.478751972 |
| ENSG00000269807 | ENSG0000026 | 7.492766553 | -0.624559968 | 0.637794174 | -0.979250037 | 0.327456451 | 0.51777302 |
| ENSG00000269821 | KCNQ1OT1 | 79.03548272 | 0.009551871 | 0.204768133 | 0.046647256 | 0.962794368 | 0.982889182 |
| ENSG00000269825 | ENSG0000026 | 188.6925993 | -0.381461611 | 0.131822802 | -2.893745273 | 0.003806768 | 0.012603156 |
| ENSG00000269837 | IPO5P1 | 116.2820762 | 0.528579605 | 0.175178104 | 3.017383988 | 0.002549666 | 0.008766388 |
| ENSG00000269855 | RNF225 | 1.798033601 | 0.068148431 | 1.344046298 | 0.050703931 | 0.959561444 | 0.981621201 |
| ENSG00000269858 | EGLN2 | 332.6489578 | -0.062606875 | 0.114828866 | -0.545218965 | 0.58560293 | 0.749650179 |
| ENSG00000269867 | ENSG0000026 | 1.9077082 | 1.400868179 | 1.368756496 | 1.023460479 | 0.306090177 | 0.493756322 |
| ENSG00000269883 | ENSG0000026 | 2.415074942 | 2.516138519 | 1.385909133 | 1.815514783 | 0.069444836 | 0.157619921 |
| ENSG00000269886 | ENSG0000026 | 5.431797284 | -0.61348875 | 0.765110906 | -0.801829832 | 0.422651399 | 0.613491717 |
| ENSG00000269887 | ENSG0000026 | 12.02779143 | -1.241886392 | 0.527120446 | -2.355982207 | 0.018473803 | 0.051009753 |
| ENSG00000269888 | ENSG0000026 | 98.5778549 | -0.303341403 | 0.391387687 | -0.775040741 | 0.438315586 | 0.627885571 |
| ENSG00000269889 | ENSG0000026 | 7.56429288 | 0.298705742 | 0.634645257 | 0.470665682 | 0.637879495 | 0.786718044 |
| ENSG00000269891 | ARHGAP19-SL | 13.80887042 | -1.865088348 | 0.531944486 | -3.506171033 | 0.000454603 | 0.001812723 |
| ENSG00000269892 | ENSG0000026 | 2.592865354 | 1.329208793 | 1.171865991 | 1.13426689 | 0.256682613 | 0.436202619 |
| ENSG00000269893 | SNHG8 | 1096.434357 | 0.769190124 | 0.065954547 | 11.66242752 | 1.98E-31 | 5.87E-30 |
| ENSG00000269896 | ENSG0000026 | 79.07597691 | -0.284806527 | 0.200496644 | -1.420505208 | 0.155460653 | 0.299309055 |
| ENSG00000269903 | ENSG0000026 | 3.662248308 | 1.027086191 | 0.930144891 | 1.104221721 | 0.26949697 | 0.451059856 |
| ENSG00000269904 | MAP2K4P1 | 53.88227104 | 0.042588099 | 0.237249639 | 0.17950754 | 0.857539197 | 0.926087485 |
| ENSG00000269906 | ENSG0000026 | 16.73214459 | -0.597723445 | 0.426820647 | -1.400408927 | 0.161390898 | 0.30829438 |
| ENSG00000269907 | PTMAP11 | 1.995643862 | -0.679235796 | 1.218121509 | -0.557609229 | 0.577111253 | 0.743117322 |
| ENSG00000269910 | ENSG0000026 | 9.874288573 | -0.647364496 | 0.560945207 | -1.154060127 | 0.248475519 | 0.426234084 |
| ENSG00000269915 | ENSG0000026 | 8.639823221 | 0.196829035 | 0.597006349 | 0.329693369 | 0.741631664 | 0.857515143 |
| ENSG00000269921 | ENSG0000026 | 6.05361393 | -0.102215934 | 0.699589034 | -0.146108543 | 0.883835703 | 0.939797489 |

| | | | | | | | |
|---|---|---|---|---|---|---|---|
| ENSG00000269925 | ENSG0000026 | 8.90970169 | -0.172216944 | 0.621193755 | -0.277235473 | 0.781599303 | 0.881489169 |
| ENSG00000269929 | MIRLET7A1HG | 89.68818749 | -0.001853404 | 0.212544511 | -0.008720073 | 0.993042477 | 0.996294293 |
| ENSG00000269934 | ENSG0000026 | 2.014495479 | -3.497564138 | 1.583118249 | -2.209287992 | 0.027154616 | 0.071305734 |
| ENSG00000269935 | ENSG0000026 | 11.06294829 | 1.215735932 | 0.553683949 | 2.195721826 | 0.02811186 | 0.073449438 |
| ENSG00000269937 | ENSG0000026 | 29.14844776 | -0.444976678 | 0.322797976 | -1.378498972 | 0.168049285 | 0.318022036 |
| ENSG00000269938 | ENSG0000026 | 1.6728161 | 0.9432117 | 1.450788884 | 0.650137115 | 0.515603657 | NA |
| ENSG00000269939 | PCF11-AS1 | 15.81239028 | -0.256927277 | 0.45417248 | -0.565704194 | 0.571594884 | 0.738449741 |
| ENSG00000269940 | ENSG0000026 | 118.344559 | 0.00926056 | 0.166097389 | 0.055753797 | 0.955537943 | 0.979491506 |
| ENSG00000269947 | ENSG0000026 | 4.432898065 | 1.744656094 | 0.933685661 | 1.868569013 | 0.0616828 | 0.142814235 |
| ENSG00000269949 | ENSG0000026 | 9.191614586 | 0.92048588 | 0.595606088 | 1.545460831 | 0.122234836 | 0.248960929 |
| ENSG00000269951 | ENSG0000026 | 9.122565065 | -0.144445284 | 0.574545308 | -0.251407995 | 0.801498687 | 0.894398618 |
| ENSG00000269958 | ENSG0000026 | 126.4033058 | 0.129535394 | 0.158359002 | 0.81798567 | 0.41336537 | 0.605530114 |
| ENSG00000269959 | SPACA6-AS1 | 21.44587526 | 0.293685118 | 0.435413636 | 0.674496833 | 0.499995499 | 0.681231617 |
| ENSG00000269961 | ERBIN-DT | 12.52477513 | 0.384837032 | 0.499234157 | 0.770854771 | 0.440793018 | 0.6300967 |
| ENSG00000269968 | ENSG0000026 | 560.8671731 | -0.341732621 | 0.210323038 | -1.624798801 | 0.104205437 | 0.219168673 |
| ENSG00000269972 | ENSG0000026 | 14.9742803 | -0.651803992 | 0.471423892 | -1.382628253 | 0.166778881 | 0.316138451 |
| ENSG00000269973 | ENSG0000026 | 120.7661379 | 0.799806398 | 0.164276018 | 4.868674128 | 1.12E-06 | 6.47E-06 |
| ENSG00000269978 | ENSG0000026 | 17.27521743 | -0.305633521 | 0.421354074 | -0.725360309 | 0.468231016 | 0.654194501 |
| ENSG00000269981 | ENSG0000026 | 72.2162273 | -0.276965355 | 0.206660823 | -1.340192841 | 0.180182659 | 0.335194496 |
| ENSG00000269982 | ENSG0000026 | 9.593939504 | -0.343553187 | 0.595102795 | -0.577300577 | 0.563736424 | 0.732365702 |
| ENSG00000269983 | ENSG0000026 | 3.527820253 | -0.676348493 | 0.964571316 | -0.701190759 | 0.483183976 | 0.66728315 |
| ENSG00000269984 | ENSG0000026 | 20.27021673 | -0.870493887 | 0.418525988 | -2.079904023 | 0.037534337 | 0.094092197 |
| ENSG00000269987 | ENSG0000026 | 27.66919344 | -1.531286103 | 0.368228863 | -4.158517318 | 3.20E-05 | 0.000154633 |
| ENSG00000269993 | ENSG0000026 | 12.54760207 | 4.525371175 | 0.930158943 | 4.865159023 | 1.14E-06 | 6.58E-06 |
| ENSG00000269994 | LINC02893 | 30.53359102 | 1.155763121 | 0.330301176 | 3.499119001 | 0.000466798 | 0.001857267 |
| ENSG00000269997 | ENSG0000026 | 3.32984099 | -2.610761716 | 1.180311045 | -2.211926869 | 0.026971718 | 0.070906626 |
| ENSG00000270001 | ENSG0000027 | 7.477795517 | 2.337680499 | 0.760225558 | 3.074982779 | 0.002105147 | 0.007363842 |
| ENSG00000270006 | C16orf95-DT | 18.81861324 | 0.918533008 | 0.432109366 | 2.125695668 | 0.033528608 | 0.085413244 |
| ENSG00000270012 | ENSG0000027 | 76.0447463 | 0.128374252 | 0.203206879 | 0.631741666 | 0.527555697 | 0.703507931 |
| ENSG00000270017 | ENSG0000027 | 2.112863263 | 0.098829681 | 1.214269004 | 0.081390269 | 0.935131588 | 0.969062396 |
| ENSG00000270019 | ENSG0000027 | 1.88217045 | -1.629296492 | 1.377439575 | -1.182844259 | 0.236870876 | 0.412089936 |
| ENSG00000270021 | ENSG0000027 | 29.86329474 | -0.187962282 | 0.340120402 | -0.55263454 | 0.580513686 | 0.745653906 |
| ENSG00000270035 | ENSG0000027 | 8.413815209 | -0.430802901 | 0.62609676 | -0.688077194 | 0.491404171 | 0.674332218 |

| | | | | | | | |
|---|---|---|---|---|---|---|---|
| ENSG00000270038 | ENSG0000027 | 1.741808711 | 4.141557461 | 1.63351485 | 2.535365664 | 0.011233 | NA |
| ENSG00000270039 | ENSG0000027 | 29.13058636 | -0.034929719 | 0.321518212 | -0.108639939 | 0.913488081 | 0.95695573 |
| ENSG00000270049 | ATP6V0D1-DT | 25.59130436 | -0.120814476 | 0.35045893 | -0.344732194 | 0.730295713 | 0.850776726 |
| ENSG00000270050 | ENSG0000027 | 7.840397218 | -0.37887521 | 0.640573469 | -0.591462539 | 0.554210548 | 0.725537292 |
| ENSG00000270055 | ENSG0000027 | 18.45328982 | -0.638213759 | 0.424050516 | -1.505041819 | 0.132313326 | 0.265065731 |
| ENSG00000270059 | ENSG0000027 | 1.848133385 | -0.994979136 | 1.285219222 | -0.774170755 | 0.43882982 | 0.628347548 |
| ENSG00000270060 | ENSG0000027 | 39.88448094 | -0.11997461 | 0.303183042 | -0.395716757 | 0.692313992 | 0.826457434 |
| ENSG00000270061 | ENSG0000027 | 2.259864738 | -0.947645077 | 1.230132895 | -0.770359919 | 0.441086422 | 0.630358991 |
| ENSG00000270062 | ENSG0000027 | 8.793009184 | -0.340968041 | 0.595933977 | -0.572157411 | 0.567215337 | 0.735053657 |
| ENSG00000270069 | MIR222HG | 1.688306177 | -0.730940994 | 1.45074301 | -0.50383906 | 0.614374477 | NA |
| ENSG00000270074 | ENSG0000027 | 4.287269146 | -0.174316864 | 0.827791934 | -0.21058053 | 0.833214607 | 0.911858454 |
| ENSG00000270083 | ENSG0000027 | 5.145957465 | -0.603213181 | 0.773168342 | -0.780183496 | 0.435282875 | 0.625250312 |
| ENSG00000270084 | GAS5-AS1 | 24.40615696 | -0.017844616 | 0.35236064 | -0.050643047 | 0.95960996 | 0.981621201 |
| ENSG00000270091 | ENSG0000027 | 19.059262 | 0.431106612 | 0.402664212 | 1.07063553 | 0.284333341 | 0.468122053 |
| ENSG00000270094 | ENSG0000027 | 5.358115312 | -0.092970763 | 0.749063274 | -0.124116034 | 0.901223403 | 0.950225965 |
| ENSG00000270096 | ENSG0000027 | 5.289125278 | 0.833088532 | 0.769078407 | 1.083229648 | 0.278706503 | 0.461693182 |
| ENSG00000270100 | ENSG0000027 | 2.225952854 | -0.441000938 | 1.166169353 | -0.378162002 | 0.705310251 | 0.835087337 |
| ENSG00000270104 | ENSG0000027 | 1.620270354 | 0.378330089 | 1.353830353 | 0.279451623 | 0.779898259 | NA |
| ENSG00000270108 | ENSG0000027 | 3.635274836 | -0.445589576 | 0.898949959 | -0.495677842 | 0.620121723 | 0.775127707 |
| ENSG00000270115 | ENSG0000027 | 9.717185974 | -0.996376277 | 0.568918218 | -1.751352383 | 0.07988523 | 0.176513887 |
| ENSG00000270116 | ENSG0000027 | 6.274870345 | -0.490492693 | 0.690234418 | -0.710617552 | 0.477321263 | 0.662124694 |
| ENSG00000270127 | ENSG0000027 | 27.62209381 | 1.273759699 | 0.360144759 | 3.536799207 | 0.000405007 | 0.001626838 |
| ENSG00000270130 | ENSG0000027 | 1.671806185 | -0.21712742 | 1.323974678 | -0.163996656 | 0.869733776 | NA |
| ENSG00000270133 | ENSG0000027 | 2.744517884 | -2.314791593 | 1.21768962 | -1.900970128 | 0.057305925 | 0.134375747 |
| ENSG00000270135 | ENSG0000027 | 3.284071303 | -0.364323683 | 0.998702175 | -0.364797126 | 0.715262851 | 0.841656152 |
| ENSG00000270147 | ENSG0000027 | 4.642784173 | -0.791475565 | 0.822987358 | -0.961710477 | 0.336195058 | 0.527274737 |
| ENSG00000270149 | ENSG0000027 | 6.748642179 | -0.413081379 | 0.689949014 | -0.598712905 | 0.54936435 | 0.721787983 |
| ENSG00000270157 | ENSG0000027 | 57.17999364 | -0.154541821 | 0.230348238 | -0.670905157 | 0.502280951 | 0.683411923 |
| ENSG00000270165 | ENSG0000027 | 3.881707461 | 0.144772031 | 0.889280974 | 0.162796726 | 0.870678486 | 0.932942429 |
| ENSG00000270170 | NCBP2AS2 | 725.7476339 | 0.02067089 | 0.075292811 | 0.274540023 | 0.783669651 | 0.882691054 |
| ENSG00000270171 | ENSG0000027 | 5.234752528 | -0.86251082 | 0.774530458 | -1.113591869 | 0.265454316 | 0.446544109 |
| ENSG00000270175 | ENSG0000027 | 20.83357698 | -0.272874463 | 0.381447023 | -0.715366607 | 0.474382541 | 0.659775402 |
| ENSG00000270177 | PPP2CA-DT | 27.88191266 | -0.195646555 | 0.355819321 | -0.549848037 | 0.582423607 | 0.747039282 |

| | | | | | | | |
|---|---|---|---|---|---|---|---|
| ENSG00000270179 | ENSG0000027 | 7.698513593 | 1.973792401 | 0.708517396 | 2.78580655 | 0.005339473 | 0.017073809 |
| ENSG00000270194 | GOLGA4-AS1 | 63.27404479 | -0.08116747 | 0.22436038 | -0.361772742 | 0.717521863 | 0.842913738 |
| ENSG00000270231 | NBPF8 | 611.4848477 | -0.24241287 | 0.090488142 | -2.67894626 | 0.007385425 | 0.022771726 |
| ENSG00000270236 | ENSG0000027 | 2.911908496 | -0.162944922 | 1.029515935 | -0.158273336 | 0.874241418 | 0.934789447 |
| ENSG00000270244 | ENSG0000027 | 1.857163681 | -0.486239503 | 1.270990025 | -0.382567521 | 0.702040456 | 0.833064766 |
| ENSG00000270248 | ENSG0000027 | 6.157111724 | -0.573254363 | 0.711488911 | -0.805710889 | 0.42040957 | 0.61166966 |
| ENSG00000270264 | NDUFB8P2 | 4.356854428 | -0.214953597 | 0.850065664 | -0.252867051 | 0.800370952 | 0.893665701 |
| ENSG00000270265 | ENSG0000027 | 11.48654458 | 1.102278935 | 0.536010625 | 2.056449784 | 0.039739182 | 0.098646064 |
| ENSG00000270276 | H4C15 | 15.23384381 | 0.699609172 | 0.515752971 | 1.356481129 | 0.174946136 | 0.328015068 |
| ENSG00000270277 | ENSG0000027 | 14.7104729 | -0.652413185 | 0.459118515 | -1.42101258 | 0.155313102 | 0.299050073 |
| ENSG00000270294 | XIAPP1 | 30.60844905 | -0.364297033 | 0.337260347 | -1.080165621 | 0.280068434 | 0.463447298 |
| ENSG00000270307 | MTATP6P2 | 14.31204849 | -0.130326121 | 0.467802205 | -0.278592362 | 0.780557675 | 0.880940326 |
| ENSG00000270333 | ENSG0000027 | 2.064602726 | -0.798328129 | 1.263144404 | -0.632016519 | 0.527376083 | 0.703495223 |
| ENSG00000270344 | POC1B-AS1 | 99.68803074 | 0.194197025 | 0.176384789 | 1.100985101 | 0.270903141 | 0.452821194 |
| ENSG00000270361 | ENSG0000027 | 2.564839657 | -0.15873539 | 1.076061305 | -0.147515192 | 0.88272539 | 0.939139247 |
| ENSG00000270362 | HMGN3-AS1 | 38.2400876 | 0.249378516 | 0.28310303 | 0.880875473 | 0.378385224 | 0.571100536 |
| ENSG00000270372 | ENSG0000027 | 34.76784189 | -0.211267087 | 0.298656816 | -0.70739081 | 0.479323647 | 0.663653377 |
| ENSG00000270393 | ENSG0000027 | 21.29705254 | -0.10482893 | 0.377812672 | -0.277462716 | 0.781424831 | 0.881401824 |
| ENSG00000270419 | CAHM | 8.269221168 | 2.05682714 | 0.71692876 | 2.868942152 | 0.004118471 | 0.013531614 |
| ENSG00000270424 | ENSG0000027 | 1.594690618 | -0.129312409 | 1.383274945 | -0.093482796 | 0.925520017 | NA |
| ENSG00000270426 | ENSG0000027 | 2.329959023 | 0.201897433 | 1.15178225 | 0.175291322 | 0.86085073 | 0.927524107 |
| ENSG00000270441 | LAMB2P1 | 79.21251035 | -0.351010031 | 0.205358312 | -1.709256509 | 0.087403446 | 0.18990671 |
| ENSG00000270442 | ENSG0000027 | 1.8925722 | 0.316109972 | 1.314027844 | 0.240565657 | 0.80989177 | 0.899408839 |
| ENSG00000270460 | ENSG0000027 | 7.095643374 | 0.204409982 | 0.664096016 | 0.307801849 | 0.758233114 | 0.866725443 |
| ENSG00000270482 | ENSG0000027 | 5.584085644 | 0.616687381 | 0.760443006 | 0.810958055 | 0.417389759 | 0.609556982 |
| ENSG00000270503 | YTHDF2P1 | 31.11077044 | 0.218079689 | 0.331243095 | 0.658367503 | 0.510302012 | 0.690272313 |
| ENSG00000270504 | ENSG0000027 | 4.706104977 | 0.160785769 | 0.8140915 | 0.197503314 | 0.843433688 | 0.917506108 |
| ENSG00000270513 | ENSG0000027 | 1.905846346 | 2.111551983 | 1.455441627 | 1.450798124 | 0.146836082 | 0.286844905 |
| ENSG00000270521 | ENSG0000027 | 2.108619413 | -0.807701794 | 1.265896373 | -0.638047324 | 0.523442874 | 0.699998807 |
| ENSG00000270532 | PEBP1P2 | 123.5593951 | 0.199563173 | 0.169598146 | 1.176682514 | 0.23932225 | 0.41496865 |
| ENSG00000270533 | ENSG0000027 | 5.968531434 | -0.347088062 | 0.70437781 | -0.492758371 | 0.622183328 | 0.776417644 |
| ENSG00000270540 | ENSG0000027 | 5.418258209 | 2.056670849 | 0.865555236 | 2.376128945 | 0.017495348 | 0.048694889 |
| ENSG00000270542 | ENSG0000027 | 2.444261767 | -1.629081243 | 1.224732292 | -1.330152927 | 0.18346789 | 0.339662125 |

| | | | | | | | |
|---|---|---|---|---|---|---|---|
| ENSG00000270553 | ENSG0000027 | 5.710457317 | -0.138894871 | 0.858642826 | -0.16176094 | 0.871494113 | 0.933336833 |
| ENSG00000270557 | ENSG0000027 | 4.488772556 | 0.562490842 | 0.831886351 | 0.676163085 | 0.498937107 | 0.680567986 |
| ENSG00000270558 | ENSG0000027 | 24.31325567 | 0.779633518 | 0.368828058 | 2.11381293 | 0.034531248 | 0.087626862 |
| ENSG00000270562 | FOXP1-DT | 15.25792561 | 0.039974727 | 0.453952941 | 0.088059188 | 0.929829634 | 0.966032078 |
| ENSG00000270571 | ENSG0000027 | 80.72630233 | 1.191482093 | 0.205382386 | 5.801286647 | 6.58E-09 | 4.83E-08 |
| ENSG00000270574 | ENSG0000027 | 20.77503983 | 0.211683432 | 0.422231322 | 0.501344692 | 0.616128557 | 0.772602443 |
| ENSG00000270580 | PKD1P6-NPIPI | 16.90030941 | -0.113227021 | 0.44273063 | -0.25574698 | 0.798146217 | 0.892259454 |
| ENSG00000270589 | ENSG0000027 | 4.818256383 | 0.100350411 | 0.802582347 | 0.125034411 | 0.900496308 | 0.949808511 |
| ENSG00000270598 | ENSG0000027 | 1.726575324 | 0.140043282 | 1.334208871 | 0.104963537 | 0.916404742 | NA |
| ENSG00000270604 | HCG17 | 13.0099718 | -0.41134624 | 0.503337073 | -0.817238114 | 0.413792365 | 0.605962136 |
| ENSG00000270605 | ENSG0000027 | 43.60337744 | -0.649325468 | 0.275176064 | -2.359672784 | 0.018291061 | 0.050566032 |
| ENSG00000270612 | ENSG0000027 | 2.439561568 | -0.330151131 | 1.113901931 | -0.29639156 | 0.76693107 | 0.872778717 |
| ENSG00000270624 | ENSG0000027 | 11.96877397 | -0.183517833 | 0.501618802 | -0.365851185 | 0.714476123 | 0.841150775 |
| ENSG00000270629 | NBPF14 | 351.6564676 | 0.448282427 | 0.101650111 | 4.410053476 | 1.03E-05 | 5.33E-05 |
| ENSG00000270638 | ENSG0000027 | 9.754726633 | 0.964301845 | 0.592304933 | 1.628049661 | 0.103514356 | 0.217994981 |
| ENSG00000270647 | TAF15 | 4221.708819 | 0.045837591 | 0.050172227 | 0.913604859 | 0.360924512 | 0.553449753 |
| ENSG00000270659 | ENSG0000027 | 13.5592471 | 0.346303613 | 0.475993117 | 0.72753912 | 0.46689576 | 0.652845833 |
| ENSG00000270665 | HNRNPA1P67 | 1.770603152 | 0.677504785 | 1.364081296 | 0.496674786 | 0.619418406 | NA |
| ENSG00000270670 | MARK2P16 | 2.445720095 | -0.346857759 | 1.113535414 | -0.311492346 | 0.75542636 | 0.865003529 |
| ENSG00000270673 | YTHDF3-DT | 34.01266519 | 0.447431211 | 0.308822887 | 1.448827888 | 0.147385654 | 0.28770527 |
| ENSG00000270681 | ENSG0000027 | 20.27635509 | 0.581902616 | 0.402154217 | 1.446963855 | 0.147907047 | 0.28837223 |
| ENSG00000270706 | PRMT1P1 | 15.61116011 | 0.307354138 | 0.447039066 | 0.687533063 | 0.491746873 | 0.674600722 |
| ENSG00000270714 | MICOS10P2 | 7.621500519 | 0.051382299 | 0.63270975 | 0.081209906 | 0.935275022 | 0.969069964 |
| ENSG00000270720 | ENSG0000027 | 4.08011621 | -0.272637279 | 0.87261903 | -0.312435633 | 0.754709474 | 0.864498202 |
| ENSG00000270722 | RNVU1-31 | 2.002762857 | -0.158901281 | 1.295017196 | -0.122702062 | 0.902343031 | 0.950707258 |
| ENSG00000270761 | CD2AP-DT | 15.39187871 | 0.3494733 | 0.458759809 | 0.76177837 | 0.446192292 | 0.634729702 |
| ENSG00000270765 | GAS2L2 | 6.635031646 | 0.721959838 | 0.712457899 | 1.013336844 | 0.310899295 | 0.498917791 |
| ENSG00000270799 | ENSG0000027 | 2.29023901 | 0.6118666 | 1.189040848 | 0.514588377 | 0.606840689 | 0.765306509 |
| ENSG00000270804 | UBE2CP5 | 93.52571243 | 0.73489734 | 0.187961998 | 3.909818727 | 9.24E-05 | 0.000414074 |
| ENSG00000270806 | C17orf50 | 3.018278058 | -0.826236966 | 1.003038718 | -0.823733871 | 0.410090789 | 0.60261884 |
| ENSG00000270813 | NANOGNBP3 | 5.810645271 | 0.99185351 | 0.759589137 | 1.305776322 | 0.191628634 | 0.351620611 |
| ENSG00000270820 | USP34-DT | 48.15042742 | -0.064349574 | 0.263266932 | -0.244427107 | 0.80690005 | 0.897468468 |
| ENSG00000270823 | ENSG0000027 | 4.975162374 | -0.363194399 | 0.774539119 | -0.468916791 | 0.639129113 | 0.78741256 |

| | | | | | | | |
|---|---|---|---|---|---|---|---|
| ENSG00000270858 | VDAC1P5 | 4.222945889 | -0.406594696 | 0.872984498 | -0.465752481 | 0.641392682 | 0.789596379 |
| ENSG00000270863 | DDX55P1 | 1.845261617 | -1.592107463 | 1.36742216 | -1.164313048 | 0.244297171 | 0.42105012 |
| ENSG00000270872 | SRGAP2D | 2.055297462 | -0.212232073 | 1.267494833 | -0.16744216 | 0.867022154 | 0.930846618 |
| ENSG00000270876 | ZNF30-AS1 | 2.101096893 | 0.505463772 | 1.216107453 | 0.415640716 | 0.677672928 | 0.816687257 |
| ENSG00000270878 | GCATP1 | 7.982111335 | -0.419205946 | 0.617052254 | -0.679368633 | 0.49690432 | 0.678793848 |
| ENSG00000270882 | H4C14 | 22.78415471 | 0.903868729 | 0.403891608 | 2.237899254 | 0.025227626 | 0.066989435 |
| ENSG00000270885 | RASL10B | 133.2136012 | 1.555942954 | 0.166746355 | 9.331196182 | 1.05E-20 | 1.82E-19 |
| ENSG00000270889 | PPP1R14BP4 | 3.54733517 | 1.693030133 | 1.025343116 | 1.651183986 | 0.098701012 | 0.209784232 |
| ENSG00000270906 | MTND4P35 | 5.927469083 | 0.866231131 | 0.766318361 | 1.130380238 | 0.258316037 | 0.437940278 |
| ENSG00000270917 | ENSG0000027 | 2.346519603 | 0.611098245 | 1.176360202 | 0.51948225 | 0.603424487 | 0.762492237 |
| ENSG00000270919 | ENSG0000027 | 5.896643264 | -1.286268599 | 0.780670175 | -1.64764665 | 0.099425202 | 0.211010651 |
| ENSG00000270923 | TAS2R6P | 1.7167354 | -0.851051082 | 1.38554587 | -0.614235227 | 0.539059898 | NA |
| ENSG00000270945 | HSPE1P7 | 6.771262067 | -0.655476618 | 0.680323424 | -0.96347795 | 0.335307724 | 0.526485914 |
| ENSG00000270953 | ENSG0000027 | 4.277278383 | -0.631750556 | 0.851556089 | -0.741877798 | 0.458161379 | 0.64527349 |
| ENSG00000270956 | ENSG0000027 | 11.69321285 | -0.125433336 | 0.50819145 | -0.246822994 | 0.805045214 | 0.896405068 |
| ENSG00000270959 | LPP-AS2 | 74.88119752 | 0.429233452 | 0.212912431 | 2.016009352 | 0.043798998 | 0.107036222 |
| ENSG00000270964 | MAP2K5-DT | 35.76173742 | -0.323613615 | 0.294359956 | -1.099380564 | 0.271602105 | 0.453578063 |
| ENSG00000270966 | ENSG0000027 | 6.145242455 | -0.138381495 | 0.713795052 | -0.193867265 | 0.846279816 | 0.919479255 |
| ENSG00000270976 | ENSG0000027 | 1.904786659 | -0.052629619 | 1.375310949 | -0.038267433 | 0.969474457 | 0.9863289 |
| ENSG00000270977 | ENSG0000027 | 58.7459859 | -0.954281313 | 0.24015514 | -3.97360354 | 7.08E-05 | 0.000323764 |
| ENSG00000270978 | ENSG0000027 | 2.074092576 | 0.580260446 | 1.261576254 | 0.45994877 | 0.645552993 | 0.792669964 |
| ENSG00000270983 | ENSG0000027 | 22.94448784 | -0.02807811 | 0.371903929 | -0.075498287 | 0.939818261 | 0.971233034 |
| ENSG00000270987 | ENSG0000027 | 109.0335656 | -0.345010225 | 0.180145204 | -1.915178519 | 0.055469741 | 0.130940097 |
| ENSG00000270996 | ENSG0000027 | 6.793167868 | -0.006764386 | 0.682930355 | -0.009904943 | 0.992097128 | 0.995917623 |
| ENSG00000271009 | ENSG0000027 | 36.84253045 | 0.294023665 | 0.299292899 | 0.982394389 | 0.32590559 | 0.51635294 |
| ENSG00000271011 | ENSG0000027 | 10.13533154 | -0.136328614 | 0.567195462 | -0.240355615 | 0.810054584 | 0.899408839 |
| ENSG00000271013 | LRRC37A9P | 41.10509129 | -0.961413962 | 0.286800883 | -3.352200142 | 0.00080172 | 0.003054553 |
| ENSG00000271029 | ENSG0000027 | 4.735468381 | 0.636093294 | 0.867497758 | 0.733250648 | 0.463405569 | 0.649830069 |
| ENSG00000271034 | RNPC3P1 | 2.694375789 | 0.048971272 | 1.082732783 | 0.045229324 | 0.963924521 | 0.983656246 |
| ENSG00000271047 | ENSG0000027 | 3.918396777 | -1.420645407 | 0.953746871 | -1.489541356 | 0.136344872 | 0.271074819 |
| ENSG00000271072 | PHB1P20 | 3.539801102 | -1.176659747 | 1.000084463 | -1.176560371 | 0.239371023 | 0.41496865 |
| ENSG00000271086 | NAMA | 5.508648396 | 0.210016674 | 0.779342344 | 0.269479357 | 0.787560825 | 0.885055863 |
| ENSG00000271092 | TLCD4-RWDD: | 1.788053979 | -0.472919793 | 1.408050746 | -0.335868429 | 0.736970085 | NA |

| | | | | | | | |
|---|---|---|---|---|---|---|---|
| ENSG00000271105 | SCML2P2 | 4.465225217 | -1.334228487 | 0.908534396 | -1.46855033 | 0.141954796 | 0.279535433 |
| ENSG00000271109 | LINC03049 | 73.6309213 | -1.105948133 | 0.220275473 | -5.020750233 | 5.15E-07 | 3.09E-06 |
| ENSG00000271119 | LINC03067 | 4.618105823 | -1.480045623 | 0.852127825 | -1.736882166 | 0.082407972 | 0.181007169 |
| ENSG00000271122 | HERPUD2-AS1 | 1018.703714 | 0.691850881 | 0.071993031 | 9.609970161 | 7.26E-22 | 1.35E-20 |
| ENSG00000271127 | ENSG0000027 | 5.481045455 | -0.281423889 | 0.764569801 | -0.368081355 | 0.712812573 | 0.84015416 |
| ENSG00000271133 | ITGB8-AS1 | 21.58839144 | -1.331046067 | 0.405993257 | -3.278493039 | 0.001043629 | 0.003891557 |
| ENSG00000271141 | ENSG0000027 | 6.411518741 | 0.87617884 | 0.71430413 | 1.226618751 | 0.219965918 | 0.390075616 |
| ENSG00000271147 | ARMCX5-GPR/ | 177.4570837 | 0.075909962 | 0.135059487 | 0.56204835 | 0.574083077 | 0.740648209 |
| ENSG00000271151 | ENSG0000027 | 7.305193126 | 0.193280492 | 0.644507119 | 0.299888839 | 0.764261948 | 0.870644539 |
| ENSG00000271153 | RPL23AP88 | 2.670259265 | -0.333585452 | 1.185584722 | -0.281367874 | 0.778428256 | 0.879705611 |
| ENSG00000271155 | ENSG0000027 | 5.464228184 | 0.963340127 | 0.78838444 | 1.221916718 | 0.221739123 | 0.392631919 |
| ENSG00000271180 | COX5AP1 | 6.790803269 | 0.324575548 | 0.702610611 | 0.461956514 | 0.644112506 | 0.791426935 |
| ENSG00000271198 | VDAC3P1 | 13.51290124 | 0.061868862 | 0.492712992 | 0.125567751 | 0.900074092 | 0.949537809 |
| ENSG00000271200 | PATJ-DT | 3.133688429 | -0.317392852 | 1.043541185 | -0.304149809 | 0.761013761 | 0.868799083 |
| ENSG00000271207 | MTCO1P22 | 4.585738514 | -0.111326072 | 0.844574062 | -0.131813274 | 0.895131987 | 0.946675017 |
| ENSG00000271218 | ENSG0000027 | 13.0471921 | -0.30090026 | 0.483583356 | -0.62223039 | 0.533790385 | 0.709329285 |
| ENSG00000271234 | ENSG0000027 | 1.765880671 | 1.290379979 | 1.41294954 | 0.913252698 | 0.361109654 | NA |
| ENSG00000271259 | ENSG0000027 | 1.968688993 | -0.207344955 | 1.247158327 | -0.166253916 | 0.867957131 | 0.931076558 |
| ENSG00000271265 | ENSG0000027 | 2.176290018 | 0.033369751 | 1.171003224 | 0.02849672 | 0.977265984 | 0.989387067 |
| ENSG00000271266 | LAPTM4BP2 | 58.59978668 | 0.414102353 | 0.237322848 | 1.744890371 | 0.081003954 | 0.178487244 |
| ENSG00000271268 | ENSG0000027 | 87.09612818 | -0.280220109 | 0.193497307 | -1.448186096 | 0.147565012 | 0.287999096 |
| ENSG00000271270 | TMCC1-DT | 65.2859576 | 0.641284287 | 0.230975226 | 2.776420218 | 0.005496112 | 0.017499072 |
| ENSG00000271303 | SRXN1 | 2.064176919 | 0.078396755 | 1.22672503 | 0.063907357 | 0.949043994 | 0.976543878 |
| ENSG00000271318 | MOB4P2 | 4.04419501 | -0.279957353 | 0.857320395 | -0.326549275 | 0.744008815 | 0.858490754 |
| ENSG00000271327 | ENSG0000027 | 6.02908642 | -0.525302339 | 0.706163615 | -0.74388191 | 0.45694792 | 0.64419772 |
| ENSG00000271335 | CCNY-AS1 | 21.30746543 | 0.100635386 | 0.376196997 | 0.267507148 | 0.789078711 | 0.885944625 |
| ENSG00000271344 | ENSG0000027 | 8.290526104 | 0.638903465 | 0.632826589 | 1.00960275 | 0.312685652 | 0.500838489 |
| ENSG00000271361 | HTATSF1P2 | 54.34746309 | -4.824643054 | 0.461905805 | -10.44507992 | 1.54E-25 | 3.52E-24 |
| ENSG00000271367 | ENSG0000027 | 1.66944647 | -0.763557367 | 1.412985827 | -0.540385722 | 0.588931052 | NA |
| ENSG00000271380 | ENSG0000027 | 56.70008016 | 0.197598801 | 0.234074483 | 0.84417062 | 0.39857407 | 0.591568916 |
| ENSG00000271383 | NBPF19 | 3.956421327 | 1.063628396 | 0.918097284 | 1.158513825 | 0.246654413 | 0.423902625 |
| ENSG00000271384 | ENSG0000027 | 3.407751366 | -0.869185234 | 1.012940048 | -0.858081617 | 0.390847397 | 0.583650521 |
| ENSG00000271392 | ENSG0000027 | 1.711544801 | 4.12264468 | 1.617648685 | 2.548541421 | 0.010817444 | NA |

| | | | | | | | |
|---|---|---|---|---|---|---|---|
| ENSG00000271425 | NBPF10 | 241.3089401 | 0.242676175 | 0.119245417 | 2.035098559 | 0.041840957 | 0.102897894 |
| ENSG00000271426 | ENSG0000027 | 16.94114777 | -0.017247181 | 0.429006555 | -0.040202606 | 0.9679316 | 0.985764746 |
| ENSG00000271427 | ENSG0000027 | 19.22669492 | 0.496489385 | 0.403223696 | 1.231300119 | 0.218210637 | 0.387892446 |
| ENSG00000271447 | MMP28 | 3.736043693 | 0.41802754 | 0.932553261 | 0.448261303 | 0.653964625 | 0.799165148 |
| ENSG00000271452 | ENSG0000027 | 16.26258296 | 0.102776285 | 0.435941123 | 0.23575726 | 0.813621035 | 0.901448243 |
| ENSG00000271474 | UNC5C-AS1 | 39.36080569 | 1.658521719 | 0.304108708 | 5.453713338 | 4.93E-08 | 3.30E-07 |
| ENSG00000271503 | CCL5 | 10.75711695 | 4.301187046 | 0.95459009 | 4.505794783 | 6.61E-06 | 3.49E-05 |
| ENSG00000271525 | ARF4P1 | 12.53087988 | 0.511067165 | 0.513522275 | 0.995219079 | 0.319629725 | 0.509101405 |
| ENSG00000271533 | ENSG0000027 | 27.23534962 | -0.809750618 | 0.368781136 | -2.195748475 | 0.028109951 | 0.073449438 |
| ENSG00000271538 | LINC02427 | 2.648539559 | -0.945626987 | 1.119660347 | -0.844565934 | 0.398353236 | 0.591380497 |
| ENSG00000271547 | ENSG0000027 | 3.23719336 | 0.13260054 | 0.96662767 | 0.137178507 | 0.890889699 | 0.944103557 |
| ENSG00000271551 | ENSG0000027 | 8.517488954 | 0.710061602 | 0.610864508 | 1.162388047 | 0.245077876 | 0.422015199 |
| ENSG00000271553 | ENSG0000027 | 3.74077764 | 0.678747495 | 0.930646007 | 0.729329401 | 0.465800192 | 0.651744797 |
| ENSG00000271576 | PRKACB-DT | 9.723502802 | -0.402109612 | 0.56297269 | -0.714261313 | 0.47506561 | 0.660365091 |
| ENSG00000271590 | ENSG0000027 | 2.66043306 | 3.70816524 | 1.473149157 | 2.517168898 | 0.01183021 | 0.034514183 |
| ENSG00000271601 | LIX1L | 66.8243848 | -0.79317443 | 0.216828807 | -3.658067583 | 0.000254124 | 0.001056854 |
| ENSG00000271602 | KRT8P49 | 57.4894384 | 0.122869647 | 0.256296634 | 0.479404061 | 0.631651205 | 0.782934732 |
| ENSG00000271614 | ATP2B1-AS1 | 86.69403621 | 0.656007771 | 0.202917092 | 3.232885731 | 0.001225466 | 0.004513838 |
| ENSG00000271643 | PDCD6IP-DT | 270.3087221 | 0.395108419 | 0.112933669 | 3.498588345 | 0.000467728 | 0.001860531 |
| ENSG00000271646 | IRF2-DT | 16.96196576 | 0.134998883 | 0.4353123 | 0.310119614 | 0.756469997 | 0.865507319 |
| ENSG00000271647 | KRT8P47 | 13.72394027 | 0.897958945 | 0.506399428 | 1.773222668 | 0.076191817 | 0.170060537 |
| ENSG00000271659 | ENSG0000027 | 6.222903822 | 0.463039561 | 0.723330951 | 0.640148967 | 0.522075757 | 0.699067118 |
| ENSG00000271676 | ENSG0000027 | 10.4077274 | 0.319380437 | 0.562717503 | 0.56756798 | 0.570328354 | 0.737339443 |
| ENSG00000271687 | MTND5P10 | 2.323789354 | -0.200483297 | 1.120658207 | -0.178897808 | 0.858017944 | 0.926299197 |
| ENSG00000271698 | ENSG0000027 | 2.078291538 | -0.80483686 | 1.251987421 | -0.642847402 | 0.520323122 | 0.697575385 |
| ENSG00000271707 | ATP1B3P1 | 12.14182187 | 0.214201063 | 0.504060232 | 0.424951323 | 0.67087216 | 0.811185291 |
| ENSG00000271711 | SAP30P1 | 2.159778885 | -0.033790723 | 1.285075644 | -0.026294734 | 0.979022255 | 0.990493497 |
| ENSG00000271714 | FAM169A-AS1 | 4.59652071 | 1.775593213 | 0.899321528 | 1.974369742 | 0.048339716 | 0.116433545 |
| ENSG00000271730 | ENSG0000027 | 2.040075215 | -2.512083815 | 1.440525225 | -1.743866592 | 0.081182355 | 0.178794473 |
| ENSG00000271737 | ENSG0000027 | 19.18165277 | -0.06776507 | 0.399118909 | -0.16978667 | 0.865177911 | 0.929958826 |
| ENSG00000271743 | ENSG0000027 | 3.288363973 | 0.983554275 | 1.003152311 | 0.980463548 | 0.326857354 | 0.517416244 |
| ENSG00000271746 | ENSG0000027 | 5.329772963 | 0.011109091 | 0.785131733 | 0.014149334 | 0.988710842 | 0.994082067 |
| ENSG00000271749 | ENSG0000027 | 5.724650247 | 0.931015788 | 0.748106213 | 1.244496799 | 0.213316777 | 0.381522035 |

| | | | | | | | |
|---|---|---|---|---|---|---|---|
| ENSG00000271751 | ENSG0000027 | 12.94796963 | -0.477698486 | 0.510269402 | -0.936169177 | 0.34918609 | 0.54043509 |
| ENSG00000271752 | ENSG0000027 | 1.750730154 | 0.643315143 | 1.392316017 | 0.462046787 | 0.644047769 | NA |
| ENSG00000271754 | ENSG0000027 | 24.52933406 | 0.484921599 | 0.356101997 | 1.361749173 | 0.173277057 | 0.325604168 |
| ENSG00000271780 | ENSG0000027 | 55.52662677 | 0.212516701 | 0.23679373 | 0.897476048 | 0.369464949 | 0.562074801 |
| ENSG00000271781 | ENSG0000027 | 88.64372416 | -0.462302895 | 0.199139478 | -2.321502996 | 0.020259713 | 0.055262522 |
| ENSG00000271784 | TP53RK-DT | 7.18647709 | -0.397325094 | 0.643797375 | -0.617158612 | 0.537130112 | 0.71219957 |
| ENSG00000271788 | ENSG0000027 | 9.60615084 | -0.044045241 | 0.571421532 | -0.077080122 | 0.938559806 | 0.970807194 |
| ENSG00000271795 | ENSG0000027 | 2.099672312 | 1.002412102 | 1.226433181 | 0.817339352 | 0.413734524 | 0.605918362 |
| ENSG00000271797 | ENSG0000027 | 7.85071335 | -0.379157564 | 0.629668405 | -0.602154342 | 0.547071405 | 0.719888603 |
| ENSG00000271806 | ENSG0000027 | 4.38566001 | -0.881198081 | 0.864022273 | -1.019878896 | 0.3077859 | 0.495759622 |
| ENSG00000271816 | BMS1P4 | 16.21203962 | -0.976897731 | 0.467280637 | -2.090601779 | 0.036563776 | 0.092041372 |
| ENSG00000271819 | RNU6-94P | 5.58482232 | -1.330792419 | 0.810272369 | -1.642401333 | 0.100506859 | 0.212814043 |
| ENSG00000271826 | PLS3-AS1 | 4.839867561 | -0.37165907 | 0.816489905 | -0.455191262 | 0.648971646 | 0.795495779 |
| ENSG00000271828 | ENSG0000027 | 6.914521639 | 0.027978068 | 0.672862971 | 0.041580632 | 0.966833013 | 0.985284143 |
| ENSG00000271830 | ENSG0000027 | 3.293007263 | -2.09467567 | 1.075807591 | -1.947072774 | 0.051526011 | 0.122895267 |
| ENSG00000271833 | ENSG0000027 | 3.988008427 | 0.90225269 | 0.902094187 | 1.000175706 | 0.317225484 | 0.506271922 |
| ENSG00000271843 | ENSG0000027 | 59.51287814 | 0.862194541 | 0.240921523 | 3.578736058 | 0.00034526 | 0.00140281 |
| ENSG00000271850 | LINC02343 | 31.29049001 | -0.39407601 | 0.313263959 | -1.257967916 | 0.20840336 | 0.374874388 |
| ENSG00000271851 | ENSG0000027 | 16.43581946 | 0.448168385 | 0.432528278 | 1.036159731 | 0.30012763 | 0.48680172 |
| ENSG00000271855 | ENSG0000027 | 18.47687484 | -0.122048429 | 0.404119081 | -0.302011053 | 0.762643637 | 0.869794424 |
| ENSG00000271857 | ENSG0000027 | 6.032839871 | 0.367148077 | 0.70144588 | 0.523416114 | 0.600684713 | 0.760679399 |
| ENSG00000271858 | ENSG0000027 | 19.82606018 | -0.45316827 | 0.390368127 | -1.160874155 | 0.245693081 | 0.422753252 |
| ENSG00000271860 | ENSG0000027 | 9.250747244 | 1.919037771 | 0.645971357 | 2.970778427 | 0.00297046 | 0.01007278 |
| ENSG00000271862 | ENSG0000027 | 15.06321091 | 0.243531213 | 0.451093531 | 0.539868555 | 0.589287685 | 0.752395257 |
| ENSG00000271869 | DCTN6-DT | 15.14149669 | 0.362385286 | 0.477608246 | 0.758750063 | 0.448002083 | 0.636304641 |
| ENSG00000271871 | ENSG0000027 | 6.736825562 | 1.157443635 | 0.713449995 | 1.622319214 | 0.104735017 | 0.220080726 |
| ENSG00000271882 | ENSG0000027 | 6.248962817 | -0.116762526 | 0.710566336 | -0.164323189 | 0.869476728 | 0.932264367 |
| ENSG00000271886 | MIR98 | 2.605046391 | -0.1851757 | 1.084373073 | -0.170767519 | 0.86440657 | 0.929739624 |
| ENSG00000271888 | JARID2-DT | 13.31388368 | -0.144871115 | 0.50509503 | -0.286819522 | 0.774250514 | 0.877144886 |
| ENSG00000271913 | TAGAP-AS1 | 10.26541498 | 0.17204149 | 0.552843375 | 0.311193907 | 0.755653215 | 0.865149344 |
| ENSG00000271916 | ENSG0000027 | 2.542196282 | -0.768853432 | 1.100953082 | -0.698352586 | 0.484956721 | 0.668845498 |
| ENSG00000271918 | ENSG0000027 | 40.72173982 | 0.556711928 | 0.276067417 | 2.016579628 | 0.043739401 | 0.106901966 |
| ENSG00000271919 | ENSG0000027 | 4.969554532 | -0.199177647 | 0.775522245 | -0.256830347 | 0.797309742 | 0.89183367 |

| | | | | | | | |
|---|---|---|---|---|---|---|---|
| ENSG00000271926 | ENSG0000027 | 6.139762371 | 1.508251283 | 0.765416429 | 1.970497661 | 0.048781363 | 0.117275672 |
| ENSG00000271930 | ENSG0000027 | 3.353927874 | 0.230621934 | 0.978960477 | 0.235578391 | 0.813759843 | 0.901448243 |
| ENSG00000271931 | PNRC1-DT | 1.572065846 | 3.996101089 | 1.665395765 | 2.399490364 | 0.016417912 | NA |
| ENSG00000271933 | ENSG0000027 | 4.66776866 | -1.758730898 | 0.869431801 | -2.022850896 | 0.043088522 | 0.105491004 |
| ENSG00000271936 | ENSG0000027 | 62.80749967 | -0.370662006 | 0.225236536 | -1.645656665 | 0.099834466 | 0.21166383 |
| ENSG00000271943 | TPM4P2 | 308.5744952 | -0.287990209 | 0.106967649 | -2.692311301 | 0.007095868 | 0.021970469 |
| ENSG00000271959 | ENSG0000027 | 2.248023263 | -0.978726475 | 1.199995651 | -0.815608352 | 0.414724171 | 0.606784393 |
| ENSG00000271964 | ENSG0000027 | 5.250385436 | -0.017105918 | 0.764769477 | -0.022367418 | 0.982154871 | 0.991665173 |
| ENSG00000271966 | ARFGEF1-DT | 35.83287705 | 1.006628557 | 0.307735729 | 3.271081194 | 0.001071371 | 0.003987873 |
| ENSG00000271967 | ENSG0000027 | 1.779109929 | 0.080307438 | 1.281525375 | 0.062665508 | 0.950032864 | NA |
| ENSG00000271969 | ENSG0000027 | 2.757867136 | -0.595316099 | 1.052174845 | -0.565795791 | 0.571532608 | 0.738437512 |
| ENSG00000271971 | ENSG0000027 | 23.39628124 | -0.641950144 | 0.371105828 | -1.729830404 | 0.083660581 | 0.183354707 |
| ENSG00000271976 | ENSG0000027 | 78.93325268 | 0.225510893 | 0.208973881 | 1.07913435 | 0.28052784 | 0.463939778 |
| ENSG00000271978 | ENSG0000027 | 4.56579757 | 1.453155255 | 0.900289391 | 1.614097945 | 0.106506183 | 0.222989159 |
| ENSG00000271980 | ENSG0000027 | 67.18259967 | -0.433908402 | 0.240396 | -1.804973471 | 0.071078833 | 0.160676827 |
| ENSG00000271983 | ENSG0000027 | 4.637964304 | -1.240744446 | 0.837825939 | -1.480909565 | 0.138630674 | 0.274591882 |
| ENSG00000271984 | ENSG0000027 | 9.472893497 | -0.125696499 | 0.584761732 | -0.21495336 | 0.829803682 | 0.909797661 |
| ENSG00000271989 | ENSG0000027 | 1.603159088 | -0.648827872 | 1.465997172 | -0.442584668 | 0.658066176 | NA |
| ENSG00000271991 | TTC32-DT | 2.448667597 | -0.229555643 | 1.147029209 | -0.200130599 | 0.841378443 | 0.916700769 |
| ENSG00000271993 | ENSG0000027 | 2.422127394 | -1.062466745 | 1.159545158 | -0.916278885 | 0.35952064 | 0.55209478 |
| ENSG00000272004 | ENSG0000027 | 17.95217172 | 0.449589651 | 0.430991547 | 1.0431519 | 0.296877947 | 0.483211303 |
| ENSG00000272008 | ENSG0000027 | 4.926148238 | 0.278614541 | 0.806909429 | 0.345286015 | 0.72987936 | 0.850481663 |
| ENSG00000272009 | ENSG0000027 | 13.51040646 | -0.312347981 | 0.472116675 | -0.661590657 | 0.508233597 | 0.688367404 |
| ENSG00000272010 | ENSG0000027 | 3.051421198 | 1.318517031 | 1.043485109 | 1.263570529 | 0.206384217 | 0.372088489 |
| ENSG00000272023 | AFF4-DT | 1.774387448 | 0.630560456 | 1.297553213 | 0.485961154 | 0.626994709 | NA |
| ENSG00000272024 | ENSG0000027 | 3.967212086 | 1.160458368 | 0.914789725 | 1.268552035 | 0.204600881 | 0.36948313 |
| ENSG00000272030 | ENSG0000027 | 28.69330839 | 0.13462279 | 0.327690542 | 0.410822935 | 0.681202374 | 0.819013756 |
| ENSG00000272031 | ANKRD34A | 445.2869348 | -0.828330601 | 0.101395774 | -8.169281319 | 3.10E-16 | 4.08E-15 |
| ENSG00000272034 | SNORD14A | 8.625079027 | -0.911939997 | 0.64229182 | -1.419821908 | 0.155659535 | 0.299616524 |
| ENSG00000272037 | ENSG0000027 | 11.94746925 | -0.475739293 | 0.521981178 | -0.91141082 | 0.362078955 | 0.554737943 |
| ENSG00000272043 | ENSG0000027 | 3.155872878 | -0.987035488 | 1.021754805 | -0.966019913 | 0.334034217 | 0.524989376 |
| ENSG00000272046 | ENSG0000027 | 7.047525519 | 4.271048748 | 1.163043574 | 3.672303295 | 0.000240374 | 0.001004405 |
| ENSG00000272047 | GTF2H5 | 417.6708456 | -0.062656771 | 0.096188775 | -0.651393787 | 0.51479232 | 0.693809754 |

| | | | | | | | |
|---|---|---|---|---|---|---|---|
| ENSG00000272049 | ENSG0000027 | 2.335137054 | 2.539885987 | 1.383236999 | 1.836190031 | 0.066329557 | 0.151825986 |
| ENSG00000272054 | ENSG0000027 | 30.77330358 | -0.034549566 | 0.320730734 | -0.107721406 | 0.914216689 | 0.957459731 |
| ENSG00000272055 | RNU6-6P | 5.850866197 | 0.64177717 | 0.747260808 | 0.858839596 | 0.390429019 | 0.583343425 |
| ENSG00000272056 | ENSG0000027 | 3.086822256 | 0.015219384 | 0.995136797 | 0.015293761 | 0.98779782 | 0.994021364 |
| ENSG00000272068 | BCAN-AS1 | 26.21865695 | -1.849010966 | 0.376314951 | -4.913466657 | 8.95E-07 | 5.22E-06 |
| ENSG00000272070 | ENSG0000027 | 56.83260666 | 0.138294836 | 0.235270353 | 0.58781242 | 0.556658207 | 0.7271782 |
| ENSG00000272077 | ENSG0000027 | 12.50918797 | 0.17724309 | 0.511156615 | 0.346749089 | 0.728779828 | 0.849718386 |
| ENSG00000272078 | ENSG0000027 | 14.18941592 | -0.006104296 | 0.474542015 | -0.012863551 | 0.989736654 | 0.994548988 |
| ENSG00000272079 | ENSG0000027 | 279.8070292 | 0.229380598 | 0.122380012 | 1.87433057 | 0.060884877 | 0.141266215 |
| ENSG00000272081 | FCHO2-DT | 8.612313988 | 0.715655841 | 0.612307512 | 1.168785009 | 0.242490262 | 0.418753884 |
| ENSG00000272084 | ENSG0000027 | 4.463679541 | 0.565735876 | 0.836273828 | 0.676495972 | 0.498725802 | 0.680390717 |
| ENSG00000272086 | GOLPH3-DT | 89.6736286 | 0.043901551 | 0.18788695 | 0.233659392 | 0.815249416 | 0.902135938 |
| ENSG00000272088 | ENSG0000027 | 4.604127526 | -0.766849485 | 0.825029713 | -0.929481051 | 0.352639841 | 0.544055279 |
| ENSG00000272092 | ENSG0000027 | 6.65636007 | -0.227439079 | 0.680952936 | -0.334001172 | 0.738378674 | 0.855807824 |
| ENSG00000272097 | TMEM14B-DT | 5.146786797 | -1.915855093 | 0.845304136 | -2.266468376 | 0.023422721 | 0.062778038 |
| ENSG00000272100 | ENSG0000027 | 4.664052415 | -0.080155394 | 0.819666831 | -0.097790213 | 0.922098879 | 0.962216621 |
| ENSG00000272103 | ENSG0000027 | 6.469703174 | 0.020565547 | 0.70447791 | 0.029192607 | 0.976710978 | 0.989278333 |
| ENSG00000272106 | ENSG0000027 | 213.2192551 | 0.644330799 | 0.129660313 | 4.969375612 | 6.72E-07 | 3.98E-06 |
| ENSG00000272108 | ENSG0000027 | 13.82837255 | 0.671621214 | 0.483871687 | 1.388015111 | 0.165132444 | 0.313666886 |
| ENSG00000272109 | ENSG0000027 | 1.923322831 | 0.790231413 | 1.344395058 | 0.587797023 | 0.556668542 | 0.7271782 |
| ENSG00000272112 | ENSG0000027 | 8.240901575 | 0.751928828 | 0.623842853 | 1.205317694 | 0.228080682 | 0.400963356 |
| ENSG00000272114 | ENSG0000027 | 10.93473156 | 0.204094835 | 0.527848571 | 0.386654139 | 0.699012272 | 0.831105086 |
| ENSG00000272115 | ENSG0000027 | 2.652793922 | 0.729684338 | 1.101001926 | 0.662745742 | 0.507493409 | 0.687730316 |
| ENSG00000272129 | ENSG0000027 | 17.25736123 | 0.255398206 | 0.422533222 | 0.604445267 | 0.545547648 | 0.718666765 |
| ENSG00000272138 | LINC01607 | 6.654868947 | -1.410883044 | 0.729658288 | -1.933621624 | 0.053159667 | 0.126358176 |
| ENSG00000272140 | ENSG0000027 | 24.6854064 | -1.145049815 | 0.414877561 | -2.759970463 | 0.005780659 | 0.018308479 |
| ENSG00000272141 | ENSG0000027 | 86.53905018 | 0.363267363 | 0.194989099 | 1.863013703 | 0.062460338 | 0.144221456 |
| ENSG00000272142 | LYRM4-AS1 | 199.9100987 | 0.041992586 | 0.13137715 | 0.319633863 | 0.749245901 | 0.861323026 |
| ENSG00000272144 | ENSG0000027 | 20.42119831 | -0.194847958 | 0.399411659 | -0.487837433 | 0.625664994 | 0.778940354 |
| ENSG00000272145 | NFYC-AS1 | 59.03343958 | 0.35644057 | 0.240438668 | 1.482459426 | 0.138218092 | 0.273952223 |
| ENSG00000272146 | ARF4-AS1 | 17.64326903 | -0.14113755 | 0.413721755 | -0.341141235 | 0.73299726 | 0.852461767 |
| ENSG00000272148 | ENSG0000027 | 4.674469613 | -0.386544258 | 0.82209241 | -0.470195629 | 0.638215256 | 0.786909737 |
| ENSG00000272149 | ENSG0000027 | 8.315904926 | 0.780698813 | 0.650029489 | 1.20102061 | 0.229743207 | 0.402959642 |

| | | | | | | | |
|---|---|---|---|---|---|---|---|
| ENSG00000272150 | NBPF25P | 103.5194674 | -0.226026483 | 0.187557139 | -1.205107328 | 0.228161871 | 0.40104462 |
| ENSG00000272153 | ENSG0000027 | 8.263526347 | 0.153460992 | 0.601626343 | 0.255076915 | 0.798663693 | 0.892592085 |
| ENSG00000272154 | ENSG0000027 | 7.802994609 | 0.303457606 | 0.627342264 | 0.483719371 | 0.628585044 | 0.780880586 |
| ENSG00000272156 | ENSG0000027 | 3.279261947 | -0.168648031 | 0.950717919 | -0.177390189 | 0.859201913 | 0.926792302 |
| ENSG00000272158 | ENSG0000027 | 1.988147 | 0.811221781 | 1.309604553 | 0.619440257 | 0.535626368 | 0.711031295 |
| ENSG00000272159 | ENSG0000027 | 2.012727962 | 0.786913497 | 1.253011094 | 0.628017981 | 0.529992161 | 0.705630887 |
| ENSG00000272160 | RNU4-5P | 2.191818402 | -1.367357728 | 1.272328844 | -1.074688933 | 0.28251402 | 0.465981566 |
| ENSG00000272168 | CASC15 | 8.798421332 | 2.323778637 | 0.722163218 | 3.21780254 | 0.001291767 | 0.00473827 |
| ENSG00000272172 | ENSG0000027 | 21.71343561 | 0.198250646 | 0.395988657 | 0.500647286 | 0.616619377 | 0.772898903 |
| ENSG00000272173 | ENSG0000027 | 52.45604935 | -0.592597802 | 0.254772926 | -2.32598421 | 0.020019393 | 0.054717607 |
| ENSG00000272175 | ENSG0000027 | 1.950299981 | 1.400799832 | 1.31333013 | 1.066601458 | 0.286151842 | 0.470021713 |
| ENSG00000272181 | ENSG0000027 | 4.751728661 | 1.639558783 | 0.940179185 | 1.74387905 | 0.081180182 | 0.178794473 |
| ENSG00000272182 | ENSG0000027 | 11.32279102 | -0.686054538 | 0.522140986 | -1.313925847 | 0.18887116 | 0.347567446 |
| ENSG00000272183 | ENSG0000027 | 3.51472669 | 0.640404363 | 0.932663783 | 0.68664011 | 0.492309546 | 0.6751209 |
| ENSG00000272186 | VPS11-DT | 12.74986161 | 0.578336429 | 0.512827304 | 1.127741101 | 0.259429274 | 0.439437906 |
| ENSG00000272189 | MAP7-AS1 | 165.2030028 | -0.49217471 | 0.140640048 | -3.499534558 | 0.000466071 | 0.001855227 |
| ENSG00000272192 | PDE7A-DT | 4.344982107 | 1.065138509 | 0.862814424 | 1.234493165 | 0.217019192 | 0.386508148 |
| ENSG00000272195 | ENSG0000027 | 7.665799636 | 0.446331681 | 0.645611941 | 0.691331204 | 0.489357427 | 0.672569611 |
| ENSG00000272205 | POU2F1-DT | 3.723282552 | -0.492248312 | 0.907117044 | -0.542651376 | 0.587369859 | 0.751053167 |
| ENSG00000272209 | ENSG0000027 | 9.614294894 | 0.007461562 | 0.55608347 | 0.01341806 | 0.989294258 | 0.994278654 |
| ENSG00000272217 | ENSG0000027 | 5.789332466 | -0.27306466 | 0.713486375 | -0.382718815 | 0.701928263 | 0.833064766 |
| ENSG00000272221 | ENSG0000027 | 46.97542378 | -0.504155181 | 0.261566719 | -1.927443915 | 0.053924331 | 0.127897452 |
| ENSG00000272223 | ENSG0000027 | 8.151545011 | 0.837411486 | 0.679365392 | 1.232637835 | 0.217710915 | 0.387214171 |
| ENSG00000272226 | ENSG0000027 | 1.594690618 | -0.129312409 | 1.383274945 | -0.093482796 | 0.925520017 | NA |
| ENSG00000272247 | ENSG0000027 | 3.074068252 | 0.324226727 | 1.018881063 | 0.318218425 | 0.750319257 | 0.862108901 |
| ENSG00000272256 | ENSG0000027 | 2.521399616 | -0.784034172 | 1.119113115 | -0.700585277 | 0.483561869 | 0.667563897 |
| ENSG00000272263 | ENSG0000027 | 10.18122131 | 0.916239739 | 0.564042431 | 1.624416335 | 0.104286983 | 0.219299972 |
| ENSG00000272264 | ENSG0000027 | 4.995398095 | -0.648339911 | 0.838513996 | -0.77320106 | 0.439403399 | 0.628894058 |
| ENSG00000272269 | NUP153-AS1 | 20.65853274 | -0.537778887 | 0.400743435 | -1.341953079 | 0.17961121 | 0.334547025 |
| ENSG00000272273 | IER3-AS1 | 29.68376506 | -1.178956691 | 0.339268271 | -3.474998377 | 0.000510856 | 0.002017742 |
| ENSG00000272274 | LINC00551 | 3.440547483 | 0.513815399 | 0.939470286 | 0.54692033 | 0.584433468 | 0.748738063 |
| ENSG00000272275 | ENSG0000027 | 5.477845267 | 0.414997181 | 0.753880105 | 0.550481673 | 0.581989044 | 0.74680029 |
| ENSG00000272277 | ENSG0000027 | 28.90099113 | -0.417396815 | 0.326895905 | -1.276849322 | 0.201655449 | 0.365400947 |

| | | | | | | | |
|---|---|---|---|---|---|---|---|
| ENSG00000272281 | TPTE2P2 | 5.206361612 | -0.862803061 | 0.773609121 | -1.115295876 | 0.264723652 | 0.445641783 |
| ENSG00000272288 | ILRUN-AS1 | 47.83492754 | 0.101883858 | 0.254505954 | 0.400320135 | 0.688920739 | 0.824262433 |
| ENSG00000272305 | ENSG0000027 | 6.293386813 | 0.8043151 | 0.728725861 | 1.10372795 | 0.269711168 | 0.451289146 |
| ENSG00000272316 | ENSG0000027 | 24.10118591 | 0.750404563 | 0.362044176 | 2.072687844 | 0.03820134 | 0.095501814 |
| ENSG00000272320 | ENSG0000027 | 5.149262502 | 0.257450802 | 0.767772571 | 0.335321698 | 0.737382428 | 0.855304663 |
| ENSG00000272323 | TTC23L-AS1 | 81.00368953 | -0.282794521 | 0.19692204 | -1.436073487 | 0.150981431 | 0.293108105 |
| ENSG00000272324 | DAP-DT | 10.70545101 | -0.045632473 | 0.535000354 | -0.085294285 | 0.932027435 | 0.967452912 |
| ENSG00000272325 | NUDT3 | 1743.483569 | 0.669895073 | 0.060795979 | 11.01873976 | 3.10E-28 | 8.05E-27 |
| ENSG00000272328 | ENSG0000027 | 7.123046657 | 0.217292898 | 0.649210094 | 0.334703512 | 0.737848752 | 0.855509243 |
| ENSG00000272333 | KMT2B | 3293.645438 | 0.100481808 | 0.058774147 | 1.709625969 | 0.08733506 | 0.1897761 |
| ENSG00000272335 | ENSG0000027 | 139.2521112 | 0.59081642 | 0.163733363 | 3.608405817 | 0.000308084 | 0.001266796 |
| ENSG00000272338 | ENSG0000027 | 3.492554891 | 1.193187416 | 1.005640528 | 1.186494958 | 0.235426892 | 0.410418185 |
| ENSG00000272341 | ENSG0000027 | 34.82513577 | -0.210316574 | 0.296519206 | -0.709284826 | 0.478147743 | 0.662882047 |
| ENSG00000272342 | LINC01115 | 18.71557698 | 0.495747625 | 0.404594352 | 1.225295465 | 0.220463916 | 0.390717108 |
| ENSG00000272345 | ENSG0000027 | 25.58705441 | 0.018446275 | 0.355354751 | 0.051909466 | 0.958600832 | 0.981140547 |
| ENSG00000272356 | ENSG0000027 | 10.53953291 | -0.401960779 | 0.558603056 | -0.719582134 | 0.471782315 | 0.657466637 |
| ENSG00000272361 | ENSG0000027 | 3.494827952 | -3.379434271 | 1.298765163 | -2.602036432 | 0.009267201 | 0.027836792 |
| ENSG00000272368 | ENSG0000027 | 16.02167126 | 0.871664994 | 0.444513367 | 1.960942143 | 0.049885772 | 0.119592394 |
| ENSG00000272369 | ENSG0000027 | 15.72515131 | -0.728690875 | 0.443628367 | -1.642570512 | 0.100471826 | 0.212764456 |
| ENSG00000272374 | SCUBE3-AS1 | 6.788136247 | -0.717099156 | 0.695429869 | -1.031159558 | 0.302466002 | 0.489389906 |
| ENSG00000272375 | ENSG0000027 | 8.323364665 | 0.179920806 | 0.643656005 | 0.279529445 | 0.779838545 | 0.880534036 |
| ENSG00000272382 | ENSG0000027 | 4.520341378 | -0.736152058 | 0.821615955 | -0.895980724 | 0.370263059 | 0.562915671 |
| ENSG00000272384 | ENSG0000027 | 56.45930214 | -0.042058944 | 0.245036243 | -0.171643767 | 0.863717597 | 0.929172842 |
| ENSG00000272391 | POM121C | 2542.184724 | 0.678519839 | 0.056797445 | 11.94630912 | 6.79E-33 | 2.10E-31 |
| ENSG00000272398 | CD24 | 13704.53056 | -0.118153902 | 0.054897625 | -2.1522589 | 0.031376969 | 0.08078425 |
| ENSG00000272402 | ENSG0000027 | 2.733784035 | 0.100130974 | 1.069880621 | 0.093590792 | 0.925434225 | 0.964370625 |
| ENSG00000272405 | ENSG0000027 | 53.4658431 | -1.825945492 | 0.270358791 | -6.753786274 | 1.44E-11 | 1.33E-10 |
| ENSG00000272416 | ENSG0000027 | 2.98283469 | 1.786869021 | 1.145369551 | 1.560080779 | 0.118740793 | 0.243587668 |
| ENSG00000272419 | LINC01145 | 309.9950567 | 0.178634065 | 0.107173271 | 1.66677814 | 0.095558529 | 0.204354679 |
| ENSG00000272420 | ENSG0000027 | 5.11420806 | -0.97490593 | 0.786830291 | -1.239029484 | 0.215334579 | 0.384448225 |
| ENSG00000272425 | ENSG0000027 | 5.643340906 | 0.472889573 | 0.75202661 | 0.628820266 | 0.52946673 | 0.705157753 |
| ENSG00000272434 | ENSG0000027 | 25.6088163 | 0.719361951 | 0.371431677 | 1.936727521 | 0.05277866 | 0.125517568 |
| ENSG00000272438 | ENSG0000027 | 26.1546475 | 0.753440686 | 0.353658683 | 2.130417612 | 0.033137151 | 0.084594508 |

| | | | | | | | |
|---|---|---|---|---|---|---|---|
| ENSG00000272444 | ENSG0000027 | 12.15030636 | -0.03200998 | 0.499051906 | -0.064141584 | 0.948857491 | 0.976445122 |
| ENSG00000272447 | ENSG0000027 | 74.99865819 | -0.080173169 | 0.202415886 | -0.396081409 | 0.692044973 | 0.826376122 |
| ENSG00000272449 | ENSG0000027 | 55.33968719 | 0.843533822 | 0.252999399 | 3.334133702 | 0.000855655 | 0.003244954 |
| ENSG00000272455 | MRPL20-DT | 235.9141366 | 0.60557602 | 0.12306008 | 4.920978586 | 8.61E-07 | 5.03E-06 |
| ENSG00000272459 | FAM193B-DT | 3.60504067 | 1.671650297 | 0.997858421 | 1.675237951 | 0.093887554 | 0.20130671 |
| ENSG00000272462 | LINC02980 | 4.64695229 | -0.226851723 | 0.827678981 | -0.274081773 | 0.784021781 | 0.882792721 |
| ENSG00000272468 | ENSG0000027 | 42.26610033 | 0.997348805 | 0.276948195 | 3.601210701 | 0.000316739 | 0.001299122 |
| ENSG00000272473 | ENSG0000027 | 5.635215372 | -0.369629366 | 0.732198047 | -0.504821568 | 0.613684166 | 0.770590331 |
| ENSG00000272476 | ENSG0000027 | 31.0001951 | 0.266234519 | 0.321912866 | 0.827039076 | 0.408214913 | 0.600670309 |
| ENSG00000272498 | ENSG0000027 | 7.791145996 | -0.255580631 | 0.619022827 | -0.412877554 | 0.679696327 | 0.818017614 |
| ENSG00000272501 | ENSG0000027 | 62.10699419 | -0.395208203 | 0.230366064 | -1.715566066 | 0.086241494 | 0.187808987 |
| ENSG00000272502 | ENSG0000027 | 2.771065549 | 0.671626498 | 1.043578241 | 0.643580396 | 0.519847566 | 0.697182295 |
| ENSG00000272505 | ENSG0000027 | 6.387973979 | 0.983356334 | 0.725815656 | 1.354829322 | 0.175471942 | 0.328705402 |
| ENSG00000272509 | ENSG0000027 | 2.71894561 | -0.320019342 | 1.052347338 | -0.304100491 | 0.761051333 | 0.868799083 |
| ENSG00000272510 | ENSG0000027 | 2.828759907 | 0.289646511 | 1.036601432 | 0.279419362 | 0.779923014 | 0.880534036 |
| ENSG00000272512 | ENSG0000027 | 147.5822607 | 0.390442069 | 0.161710576 | 2.414449812 | 0.015758994 | 0.04441171 |
| ENSG00000272514 | CFAP206 | 4.404017851 | -0.489003923 | 0.836491437 | -0.584589275 | 0.558823921 | 0.728707409 |
| ENSG00000272518 | ENSG0000027 | 10.17679036 | -0.614997853 | 0.551031029 | -1.116085702 | 0.264385452 | 0.445170454 |
| ENSG00000272523 | LINC01023 | 19.50738204 | 0.202467224 | 0.394996692 | 0.512579542 | 0.608245466 | 0.766456043 |
| ENSG00000272525 | BTF3-DT | 21.95417168 | -0.177877535 | 0.390915778 | -0.455027771 | 0.649089259 | 0.795581728 |
| ENSG00000272529 | ENSG0000027 | 3.925595591 | 0.559707137 | 0.897127737 | 0.623887897 | 0.53270121 | 0.708128049 |
| ENSG00000272533 | SNORA28 | 2.852903597 | 0.250848504 | 1.067206251 | 0.235051569 | 0.814168708 | 0.9017057 |
| ENSG00000272537 | ENSG0000027 | 3.111829025 | -0.590012288 | 1.015293231 | -0.581125009 | 0.561156205 | 0.730157957 |
| ENSG00000272540 | ENSG0000027 | 133.1094306 | -0.912830881 | 0.197929524 | -4.61189854 | 3.99E-06 | 2.16E-05 |
| ENSG00000272556 | GTF2IP13 | 187.3091627 | 0.578779372 | 0.144108973 | 4.016261874 | 5.91E-05 | 0.000273697 |
| ENSG00000272562 | H3-3A-DT | 2.663328536 | -0.831376994 | 1.106304067 | -0.751490498 | 0.452357517 | 0.640066851 |
| ENSG00000272564 | ENSG0000027 | 2.772399797 | -5.028779404 | 1.479264684 | -3.399512919 | 0.00067506 | 0.002610125 |
| ENSG00000272572 | ENSG0000027 | 11.28134117 | 0.096360015 | 0.522399521 | 0.184456553 | 0.853655308 | 0.923850203 |
| ENSG00000272574 | ENSG0000027 | 2.596589062 | 0.521531177 | 1.091073897 | 0.477998034 | 0.632651605 | 0.783513891 |
| ENSG00000272576 | ENSG0000027 | 1.600347909 | -0.142989839 | 1.419366328 | -0.100742026 | 0.919755249 | NA |
| ENSG00000272578 | ENSG0000027 | 19.3195171 | -0.387666413 | 0.429180032 | -0.90327225 | 0.366381421 | 0.559031629 |
| ENSG00000272582 | ENSG0000027 | 2.179648507 | 0.895979525 | 1.249702209 | 0.716954422 | 0.47340222 | 0.658891332 |
| ENSG00000272588 | ENSG0000027 | 3.812630553 | 1.407559661 | 0.945164615 | 1.489221706 | 0.136428997 | 0.271202876 |

| | | | | | | | |
|---|---|---|---|---|---|---|---|
| ENSG00000272599 | ENSG0000027 | 29.04020738 | -0.48410571 | 0.337792295 | -1.433146097 | 0.15181609 | 0.294143675 |
| ENSG00000272600 | ENSG0000027 | 3.09633181 | -1.194473925 | 1.018691563 | -1.172557001 | 0.240973501 | 0.416887791 |
| ENSG00000272602 | ZNF595 | 1319.749994 | -0.41386572 | 0.060952477 | -6.789973757 | 1.12E-11 | 1.04E-10 |
| ENSG00000272604 | EFCAB10-AS1 | 37.55231933 | 0.637719572 | 0.290612801 | 2.194396012 | 0.028206952 | 0.073647563 |
| ENSG00000272606 | PPP4R3B-DT | 23.48846937 | 0.36932756 | 0.366606155 | 1.00742324 | 0.313731428 | 0.501980571 |
| ENSG00000272610 | MAGI1-IT1 | 2.334714926 | 1.115750953 | 1.2110989 | 0.921271543 | 0.356908674 | 0.549163927 |
| ENSG00000272620 | AFAP1-AS1 | 16.52439273 | -0.648607357 | 0.432591895 | -1.499351617 | 0.133782438 | 0.267238691 |
| ENSG00000272622 | ENSG0000027 | 237.8367061 | 2.371906281 | 0.14525707 | 16.32902473 | 6.14E-60 | 4.38E-58 |
| ENSG00000272630 | ENSG0000027 | 4.689658436 | -0.204037256 | 0.800656535 | -0.254837433 | 0.798848662 | 0.892721951 |
| ENSG00000272631 | ENSG0000027 | 105.3107215 | 0.406747019 | 0.185903862 | 2.187942814 | 0.028673768 | 0.074670932 |
| ENSG00000272638 | ENSG0000027 | 2.743594215 | -2.299213796 | 1.199512791 | -1.916789727 | 0.055264651 | 0.130509685 |
| ENSG00000272645 | GTF2IP20 | 285.8532093 | -0.131567117 | 0.128565579 | -1.023346357 | 0.306144113 | 0.493773883 |
| ENSG00000272654 | JTB-DT | 44.99438505 | 0.192361903 | 0.284837539 | 0.675339015 | 0.499460402 | 0.680866509 |
| ENSG00000272655 | ENSG0000027 | 90.7036565 | 1.139446905 | 0.203799632 | 5.591015527 | 2.26E-08 | 1.56E-07 |
| ENSG00000272661 | ENSG0000027 | 38.51548073 | 0.426277584 | 0.285875205 | 1.491131711 | 0.135926918 | 0.270533013 |
| ENSG00000272662 | TIMMDC1-DT | 3.763010027 | 0.688848204 | 0.908525502 | 0.758204588 | 0.448328515 | 0.636466126 |
| ENSG00000272663 | PPP1R21-DT | 9.212860442 | -0.467507059 | 0.574403901 | -0.81389952 | 0.415702516 | 0.607789402 |
| ENSG00000272667 | ENSG0000027 | 37.84128199 | 0.201192477 | 0.2876851 | 0.69934966 | 0.484333539 | 0.668186877 |
| ENSG00000272668 | ENSG0000027 | 65.788564 | 1.048171641 | 0.228254308 | 4.592122042 | 4.39E-06 | 2.36E-05 |
| ENSG00000272669 | ENSG0000027 | 37.76353793 | 0.068865082 | 0.292479903 | 0.235452357 | 0.813857653 | 0.901448243 |
| ENSG00000272674 | PCDHB16 | 55.68661112 | 0.473434542 | 0.253784361 | 1.865499277 | 0.062111453 | 0.143531454 |
| ENSG00000272677 | HNRNPD-DT | 4.296310583 | 0.260282143 | 0.837052015 | 0.310950979 | 0.755837889 | 0.865149344 |
| ENSG00000272678 | ENSG0000027 | 1.79332226 | 1.229851099 | 1.362363331 | 0.90273356 | 0.366667321 | 0.559190822 |
| ENSG00000272686 | WASL-DT | 188.0217763 | 0.660511242 | 0.139867905 | 4.722393179 | 2.33E-06 | 1.29E-05 |
| ENSG00000272688 | ENSG0000027 | 74.97898508 | -0.076632282 | 0.203757221 | -0.376096031 | 0.706845505 | 0.836412774 |
| ENSG00000272690 | LINC02018 | 15.06945147 | -0.535927098 | 0.468701965 | -1.143428314 | 0.252860802 | 0.431658492 |
| ENSG00000272692 | ENSG0000027 | 19.55075508 | 0.055571434 | 0.414089518 | 0.134201499 | 0.893243241 | 0.945418471 |
| ENSG00000272693 | LINC03006 | 41.03174971 | 0.591801739 | 0.281727187 | 2.100619915 | 0.035674344 | 0.090062144 |
| ENSG00000272695 | GAS6-DT | 3.211101246 | -0.403036557 | 1.052729796 | -0.382849007 | 0.701831722 | 0.833064766 |
| ENSG00000272696 | ENSG0000027 | 3.825863815 | -0.085091109 | 0.879249305 | -0.096776999 | 0.922903491 | 0.962730484 |
| ENSG00000272701 | MESTIT1 | 3.037197726 | 0.287563379 | 0.999139607 | 0.28781101 | 0.773491409 | 0.876544629 |
| ENSG00000272702 | ENSG0000027 | 2.669293914 | 0.408605267 | 1.093509768 | 0.37366403 | 0.70865429 | 0.837576867 |
| ENSG00000272711 | HK2-DT | 216.1000354 | 0.352804806 | 0.131153642 | 2.690011499 | 0.007144956 | 0.022116487 |

| | | | | | | | |
|---|---|---|---|---|---|---|---|
| ENSG00000272715 | ENSG0000027 | 6.222588377 | 1.44881621 | 0.770495094 | 1.880370455 | 0.060057607 | 0.13962922 |
| ENSG00000272716 | SLC30A6-DT | 9.02141096 | 0.514584337 | 0.58534512 | 0.879112713 | 0.379340165 | 0.571805148 |
| ENSG00000272717 | ENSG0000027 | 5.930669596 | 0.364267139 | 0.728805726 | 0.499813773 | 0.617206212 | 0.773024159 |
| ENSG00000272720 | ENSG0000027 | 58.44538519 | -0.011922144 | 0.254420062 | -0.04686008 | 0.962624745 | 0.982847234 |
| ENSG00000272721 | EIF2B5-DT | 9.252448219 | 0.079120537 | 0.573332435 | 0.138001152 | 0.890239507 | 0.943797108 |
| ENSG00000272732 | ENSG0000027 | 125.5572679 | -0.715987398 | 0.163558519 | -4.377561019 | 1.20E-05 | 6.12E-05 |
| ENSG00000272733 | ENSG0000027 | 3.603525431 | -0.952514461 | 0.93073162 | -1.023403998 | 0.30611687 | 0.49376466 |
| ENSG00000272734 | ADIRF-AS1 | 126.2272827 | -0.602451081 | 0.169233453 | -3.559881745 | 0.000371022 | 0.001499046 |
| ENSG00000272735 | ENSG0000027 | 6.412978171 | -0.675719422 | 0.706886526 | -0.955909326 | 0.339118047 | 0.530254788 |
| ENSG00000272742 | ENSG0000027 | 31.68941629 | -0.689119485 | 0.322925529 | -2.133988872 | 0.032843692 | 0.083966822 |
| ENSG00000272744 | ENSG0000027 | 2.075992737 | 0.045667699 | 1.19804432 | 0.038118539 | 0.96959317 | 0.98640593 |
| ENSG00000272745 | ENSG0000027 | 3.925576988 | -1.085486454 | 0.916491879 | -1.184392877 | 0.236257578 | 0.411397362 |
| ENSG00000272750 | ENSG0000027 | 18.53378338 | 0.195996251 | 0.413460252 | 0.47403892 | 0.635472154 | 0.785141983 |
| ENSG00000272752 | STAG3L5P-PVI | 15.32352261 | 0.021898871 | 0.449066047 | 0.048765368 | 0.961106282 | 0.982116705 |
| ENSG00000272755 | ENSG0000027 | 1.643949929 | 1.786979254 | 1.512638712 | 1.181365543 | 0.23745754 | NA |
| ENSG00000272758 | WDR5B-DT | 87.79799151 | 0.142448576 | 0.188603538 | 0.755280506 | 0.450080693 | 0.638403176 |
| ENSG00000272764 | ENSG0000027 | 34.3367421 | -1.15513415 | 0.313086798 | -3.689501299 | 0.000224694 | 0.00094439 |
| ENSG00000272767 | JMJD1C-AS1 | 45.57720025 | 0.0536416 | 0.258965018 | 0.207138404 | 0.835901771 | 0.913370821 |
| ENSG00000272768 | ENSG0000027 | 7.898486946 | 1.486518472 | 0.663395321 | 2.240773225 | 0.025040768 | 0.066547176 |
| ENSG00000272769 | ENSG0000027 | 6.551144752 | -0.517402539 | 0.704249526 | -0.734686386 | 0.462530513 | 0.64911599 |
| ENSG00000272770 | ENSG0000027 | 2.078431236 | -1.22028515 | 1.239392858 | -0.984583009 | 0.324828939 | 0.514931647 |
| ENSG00000272777 | ENSG0000027 | 4.711951738 | 0.381354655 | 0.852693381 | 0.447235388 | 0.654705113 | 0.799489924 |
| ENSG00000272779 | BMS1P20 | 4.025848308 | -0.440360416 | 0.866557639 | -0.508172101 | 0.611332655 | 0.768984542 |
| ENSG00000272787 | ENSG0000027 | 11.89656955 | 0.710945149 | 0.52741069 | 1.347991542 | 0.177661103 | 0.331884829 |
| ENSG00000272791 | ENSG0000027 | 26.00173004 | -0.377527488 | 0.343706155 | -1.098401881 | 0.272029041 | 0.454041488 |
| ENSG00000272799 | ENSG0000027 | 2.105841981 | 1.552311685 | 1.306200937 | 1.188417219 | 0.234669076 | 0.409617481 |
| ENSG00000272800 | ENSG0000027 | 7.670639836 | -0.336963112 | 0.633384571 | -0.532003978 | 0.59472324 | 0.756386934 |
| ENSG00000272807 | ENSG0000027 | 4.614476452 | 0.220608046 | 0.802723347 | 0.274824504 | 0.783451071 | 0.882599203 |
| ENSG00000272808 | GCAWKR | 195.2225417 | -0.662625485 | 0.134292471 | -4.93419683 | 8.05E-07 | 4.72E-06 |
| ENSG00000272812 | ENSG0000027 | 14.1621444 | 0.35508821 | 0.467451586 | 0.759625641 | 0.447478388 | 0.635888882 |
| ENSG00000272817 | ENSG0000027 | 34.53954165 | -0.815234802 | 0.315302801 | -2.58556156 | 0.009722051 | 0.029009346 |
| ENSG00000272829 | ENSG0000027 | 9.066061578 | -0.574102478 | 0.602267262 | -0.953235406 | 0.340470811 | 0.531464372 |
| ENSG00000272831 | ENSG0000027 | 135.9449821 | 0.18924411 | 0.154238917 | 1.226954351 | 0.219839749 | 0.389942305 |

| | | | | | | | |
|---|---|---|---|---|---|---|---|
| ENSG00000272836 | ENSG0000027 | 9.794797673 | 0.658658355 | 0.561973886 | 1.172044417 | 0.241179224 | 0.417149317 |
| ENSG00000272841 | MAP3K4-AS1 | 34.72053385 | -0.820712657 | 0.309290052 | -2.653537194 | 0.007965299 | 0.024363365 |
| ENSG00000272842 | ENSG0000027 | 2.535121548 | 1.434523766 | 1.180180764 | 1.215511903 | 0.224170912 | 0.395666747 |
| ENSG00000272843 | ENSG0000027 | 1.553545699 | 0.468176289 | 1.43939328 | 0.325259466 | 0.744984706 | NA |
| ENSG00000272844 | ENSG0000027 | 8.545216978 | 0.656384951 | 0.622790381 | 1.053942019 | 0.291909464 | 0.476936684 |
| ENSG00000272848 | ENSG0000027 | 28.1920467 | 0.024736407 | 0.33111566 | 0.074706243 | 0.940448441 | 0.971621655 |
| ENSG00000272851 | ENSG0000027 | 2.093480362 | -0.397487651 | 1.206990124 | -0.329321378 | 0.741912787 | 0.857590509 |
| ENSG00000272853 | SUV39H2-DT | 23.26125651 | -0.598229565 | 0.361854921 | -1.653230428 | 0.098283976 | 0.208975291 |
| ENSG00000272858 | ENSG0000027 | 57.37620374 | 1.394603865 | 0.249640993 | 5.58643775 | 2.32E-08 | 1.60E-07 |
| ENSG00000272861 | ENSG0000027 | 3.835360803 | 2.118639606 | 1.038519727 | 2.040057162 | 0.041344633 | 0.101886148 |
| ENSG00000272870 | SAP30-DT | 8.212921239 | -0.478377973 | 0.615232105 | -0.777556907 | 0.436830274 | 0.626595539 |
| ENSG00000272871 | ENSG0000027 | 1.870787901 | 0.352889401 | 1.296149852 | 0.272259724 | 0.785422322 | 0.883215101 |
| ENSG00000272872 | ENSG0000027 | 55.6687898 | 0.489706508 | 0.235786939 | 2.076902612 | 0.037810549 | 0.094691483 |
| ENSG00000272885 | ENSG0000027 | 6.211172081 | 1.332348052 | 0.771839355 | 1.726198649 | 0.084311681 | 0.184411705 |
| ENSG00000272886 | DCP1A | 683.2084727 | -0.109799988 | 0.07563258 | -1.451755149 | 0.146569699 | 0.286427486 |
| ENSG00000272888 | CHASERR | 688.9303824 | -0.26006254 | 0.082996571 | -3.133413067 | 0.001727861 | 0.006152923 |
| ENSG00000272892 | ENSG0000027 | 8.288442784 | -1.22141874 | 0.625818827 | -1.951712999 | 0.050972288 | 0.121827912 |
| ENSG00000272894 | MIOS-DT | 277.7415769 | -1.292287081 | 0.120950769 | -10.68440567 | 1.20E-26 | 2.93E-25 |
| ENSG00000272899 | ATP6V1FNB | 68.16051251 | 0.63321089 | 0.225141109 | 2.812506754 | 0.004915699 | 0.015864679 |
| ENSG00000272902 | TBC1D8-AS1 | 44.6918681 | -0.25409441 | 0.269145594 | -0.944077911 | 0.345129836 | 0.536037809 |
| ENSG00000272905 | ENSG0000027 | 3.652366073 | 1.672927853 | 1.008690209 | 1.658515012 | 0.09721356 | 0.207256419 |
| ENSG00000272906 | ENSG0000027 | 20.0842544 | 0.46429795 | 0.395850801 | 1.172911481 | 0.240831305 | 0.416736074 |
| ENSG00000272908 | ENSG0000027 | 1.659990367 | -0.178741223 | 1.338528944 | -0.133535568 | 0.893769837 | NA |
| ENSG00000272909 | ENSG0000027 | 45.55374728 | -1.094889007 | 0.269868885 | -4.05711465 | 4.97E-05 | 0.000232644 |
| ENSG00000272910 | ENSG0000027 | 13.8228853 | -0.848804353 | 0.473047066 | -1.794333827 | 0.072759902 | 0.163876664 |
| ENSG00000272913 | LINC03052 | 286.6023122 | 0.406957861 | 0.118230339 | 3.442076425 | 0.000577267 | 0.002257887 |
| ENSG00000272914 | ENSG0000027 | 2.667553156 | -2.202000884 | 1.241169191 | -1.77413434 | 0.076040932 | 0.169802295 |
| ENSG00000272916 | ENSG0000027 | 9.738524762 | -0.120669106 | 0.582367154 | -0.20720452 | 0.835850138 | 0.913370821 |
| ENSG00000272918 | ENSG0000027 | 9.21595773 | -0.683089215 | 0.581121556 | -1.175467006 | 0.23980793 | 0.41549803 |
| ENSG00000272927 | ENSG0000027 | 34.28567301 | -0.045935224 | 0.298365701 | -0.153956116 | 0.877644338 | 0.936395015 |
| ENSG00000272931 | LRRC8D-DT | 33.7088927 | 0.404512009 | 0.305324909 | 1.324857546 | 0.185218404 | 0.342212483 |
| ENSG00000272933 | TRIM8-DT | 4.947788835 | 1.110363338 | 0.805914017 | 1.377768985 | 0.168274625 | 0.318342396 |
| ENSG00000272936 | ENSG0000027 | 2.596589062 | 0.521531177 | 1.091073897 | 0.477998034 | 0.632651605 | 0.783513891 |

| | | | | | | | |
|---|---|---|---|---|---|---|---|
| ENSG00000272940 | ENSG0000027 | 4.234297268 | -0.162392779 | 0.84495061 | -0.192192037 | 0.847591783 | 0.92032733 |
| ENSG00000272941 | ENSG0000027 | 25.85777289 | -0.564322909 | 0.357346112 | -1.579205396 | 0.114288953 | 0.236170404 |
| ENSG00000272944 | ENSG0000027 | 28.2667428 | 0.860956869 | 0.347803865 | 2.475409146 | 0.013308362 | 0.038252577 |
| ENSG00000272945 | ENSG0000027 | 3.121922687 | -0.553593008 | 0.997826645 | -0.554798783 | 0.5790323 | 0.744646314 |
| ENSG00000272948 | ENSG0000027 | 8.232414502 | 0.07870833 | 0.611174042 | 0.128782187 | 0.897530001 | 0.948060798 |
| ENSG00000272953 | ENSG0000027 | 13.22038228 | 0.656637442 | 0.501311322 | 1.309839641 | 0.19025009 | 0.349534617 |
| ENSG00000272954 | ENSG0000027 | 20.2981334 | -0.481713585 | 0.428453018 | -1.124309001 | 0.260881961 | 0.441163976 |
| ENSG00000272966 | ENSG0000027 | 2.653830122 | 0.956711252 | 1.129023919 | 0.847379081 | 0.39678386 | 0.5896749 |
| ENSG00000272969 | ENSG0000027 | 2.803594837 | -1.039119736 | 1.056014443 | -0.984001443 | 0.325114804 | 0.515289196 |
| ENSG00000272970 | ENSG0000027 | 29.19037257 | -0.703444224 | 0.333251822 | -2.110848844 | 0.034785306 | 0.088144806 |
| ENSG00000272973 | ENSG0000027 | 7.730579869 | -0.244710172 | 0.621297899 | -0.393869305 | 0.693677533 | 0.827371387 |
| ENSG00000272977 | ENSG0000027 | 41.4783632 | -0.303358613 | 0.272810614 | -1.111975112 | 0.266148852 | 0.447089532 |
| ENSG00000272982 | ENSG0000027 | 2.701110962 | -0.549692298 | 1.098593848 | -0.50035989 | 0.61682169 | 0.772960336 |
| ENSG00000272983 | ENSG0000027 | 5.418266773 | 0.414156875 | 0.755497029 | 0.548191269 | 0.583560577 | 0.747953941 |
| ENSG00000272986 | ENSG0000027 | 13.01337109 | -0.453367048 | 0.503587463 | -0.900274692 | 0.367974086 | 0.560512984 |
| ENSG00000272989 | LINC02012 | 15.71955705 | 1.533801209 | 0.481413001 | 3.186040276 | 0.001442345 | 0.005222127 |
| ENSG00000272990 | ENSG0000027 | 2.222601827 | 1.173315269 | 1.230765208 | 0.953321773 | 0.340427063 | 0.531432244 |
| ENSG00000272991 | ENSG0000027 | 1.598474914 | -0.178247836 | 1.39679048 | -0.127612436 | 0.898455689 | NA |
| ENSG00000272994 | C2orf49-DT | 107.0209466 | 0.431863414 | 0.19410027 | 2.224950096 | 0.026084588 | 0.068858511 |
| ENSG00000273001 | PFKP-DT | 8.291273668 | -2.508966255 | 0.713205862 | -3.517871052 | 0.000435024 | 0.001741006 |
| ENSG00000273002 | ARHGEF2-AS2 | 7.294802214 | 0.19547572 | 0.644893636 | 0.303113117 | 0.761803658 | 0.869297509 |
| ENSG00000273004 | ENSG0000027 | 6.641755679 | 0.33292436 | 0.674394587 | 0.493664045 | 0.621543462 | 0.775999075 |
| ENSG00000273007 | ENSG0000027 | 8.709443655 | -1.193625898 | 0.63111899 | -1.891285028 | 0.058586306 | 0.136818677 |
| ENSG00000273008 | CSGALNACT2- | 10.85831815 | 0.180935723 | 0.537255876 | 0.336777561 | 0.73628459 | 0.854516265 |
| ENSG00000273010 | ENSG0000027 | 9.215272981 | -0.486624231 | 0.605926696 | -0.80310743 | 0.421912644 | 0.612923567 |
| ENSG00000273013 | ENSG0000027 | 3.17253485 | -0.87419378 | 1.02017503 | -0.856905682 | 0.391497013 | 0.584430368 |
| ENSG00000273014 | ENSG0000027 | 39.48306719 | 0.144598694 | 0.276899744 | 0.52220595 | 0.601526944 | 0.761283686 |
| ENSG00000273015 | LINC00938 | 613.7474313 | 0.259249891 | 0.080931429 | 3.203327752 | 0.001358493 | 0.004956079 |
| ENSG00000273018 | FAM106A | 16.78430942 | -0.140899616 | 0.431147893 | -0.326801124 | 0.743818309 | 0.858490754 |
| ENSG00000273026 | ENSG0000027 | 22.00113683 | 0.320408255 | 0.376349606 | 0.851358019 | 0.394570505 | 0.587570253 |
| ENSG00000273027 | ENSG0000027 | 42.11068413 | 0.393889975 | 0.275590133 | 1.42926008 | 0.152929495 | 0.295726428 |
| ENSG00000273032 | DGCR5 | 4.385640305 | 0.612689987 | 0.90239062 | 0.67896316 | 0.497161205 | 0.679040564 |
| ENSG00000273033 | LINC02035 | 141.2508479 | 0.764748812 | 0.15791532 | 4.842777851 | 1.28E-06 | 7.31E-06 |

| | | | | | | | |
|---|---|---|---|---|---|---|---|
| ENSG00000273045 | C2orf15 | 173.5018396 | 0.051681572 | 0.136565009 | 0.378439343 | 0.705104246 | 0.834997903 |
| ENSG00000273056 | ENSG0000027 | 2.854247885 | 1.678206941 | 1.139129358 | 1.473236493 | 0.140687278 | 0.277622896 |
| ENSG00000273058 | ENSG0000027 | 13.64627097 | 1.076394657 | 0.486738633 | 2.211442826 | 0.027005187 | 0.070977535 |
| ENSG00000273059 | ENSG0000027 | 4.00047544 | 0.893439471 | 2.039026549 | 0.438169612 | 0.661263336 | 0.804222818 |
| ENSG00000273061 | CDC37L1-DT | 16.64693724 | -0.641629518 | 0.447179909 | -1.434835297 | 0.151334037 | 0.293680975 |
| ENSG00000273062 | RALGPS2-AS1 | 4.170444707 | 0.303393284 | 0.857975189 | 0.353615452 | 0.723627093 | 0.846377683 |
| ENSG00000273063 | ENSG0000027 | 1.631599452 | -0.176308599 | 1.343051472 | -0.13127464 | 0.895558052 | NA |
| ENSG00000273064 | ENSG0000027 | 5.074488047 | -0.782414118 | 0.773919905 | -1.010975572 | 0.312028122 | 0.500309297 |
| ENSG00000273066 | ENSG0000027 | 60.58849305 | 0.049720839 | 0.225483353 | 0.220507803 | 0.825475696 | 0.907872877 |
| ENSG00000273073 | ENSG0000027 | 9.313010667 | 0.526872301 | 0.584390392 | 0.901575913 | 0.367282191 | 0.559830795 |
| ENSG00000273076 | ENSG0000027 | 2.193777643 | -0.404350846 | 1.168536277 | -0.346031916 | 0.729318729 | 0.850044287 |
| ENSG00000273080 | ENSG0000027 | 34.29585434 | -0.219547376 | 0.310773553 | -0.706454504 | 0.479905536 | 0.664275206 |
| ENSG00000273090 | ENSG0000027 | 2.620683303 | 0.174962026 | 1.059953967 | 0.165065684 | 0.868892283 | 0.931785159 |
| ENSG00000273091 | ENSG0000027 | 9.14527561 | 0.778466506 | 0.597790445 | 1.302239794 | 0.192834426 | 0.353211572 |
| ENSG00000273108 | ENSG0000027 | 1.774376307 | 0.076095828 | 1.282089647 | 0.05935297 | 0.952670972 | NA |
| ENSG00000273113 | ENSG0000027 | 27.97126003 | 0.240751968 | 0.352657427 | 0.682679422 | 0.494809436 | 0.67711556 |
| ENSG00000273117 | INSIG1-DT | 112.2117523 | 0.198598316 | 0.171337731 | 1.159104389 | 0.246413637 | 0.423647518 |
| ENSG00000273132 | ENSG0000027 | 4.354784826 | -1.575773594 | 0.929556785 | -1.695188094 | 0.090039745 | 0.194529266 |
| ENSG00000273133 | ENSG0000027 | 1.818818801 | -0.977290136 | 1.30713695 | -0.747657034 | 0.454667053 | 0.642205665 |
| ENSG00000273136 | NBPF26 | 386.9587584 | -0.187888095 | 0.096683915 | -1.94332319 | 0.051977121 | 0.123829594 |
| ENSG00000273137 | SELENOO-AS1 | 16.11935056 | -0.252000975 | 0.437356051 | -0.576191811 | 0.564485539 | 0.732969584 |
| ENSG00000273139 | ENSG0000027 | 3.465109843 | -1.420831055 | 0.981912142 | -1.447004262 | 0.14789573 | 0.28837223 |
| ENSG00000273141 | ENSG0000027 | 2.384377763 | 0.611902882 | 1.175091126 | 0.520728026 | 0.602556248 | 0.761829824 |
| ENSG00000273142 | LINC02604 | 156.2493823 | 0.544795769 | 0.157823763 | 3.451924858 | 0.000556603 | 0.002184511 |
| ENSG00000273145 | ENSG0000027 | 40.64406971 | -0.318401742 | 0.289194159 | -1.100996449 | 0.270898203 | 0.452821194 |
| ENSG00000273148 | LCDR | 46.10587049 | 0.229491375 | 0.265726823 | 0.863636468 | 0.387787624 | 0.580492583 |
| ENSG00000273149 | ENSG0000027 | 872.4374459 | -0.206230028 | 0.192234622 | -1.072803775 | 0.283359167 | 0.467174018 |
| ENSG00000273151 | ENSG0000027 | 85.25462665 | 0.374660188 | 0.199494167 | 1.878050839 | 0.060374211 | 0.140251584 |
| ENSG00000273154 | ENSG0000027 | 2.541781616 | 0.372734284 | 1.125063588 | 0.331300638 | 0.740417409 | 0.856755378 |
| ENSG00000273156 | ENSG0000027 | 5.39979119 | 0.438163667 | 0.777376014 | 0.563644439 | 0.572996137 | 0.739747367 |
| ENSG00000273162 | ENSG0000027 | 31.25610666 | -0.01593275 | 0.317159608 | -0.050235749 | 0.959934524 | 0.981760945 |
| ENSG00000273174 | ENSG0000027 | 8.461302696 | 1.498899649 | 0.677484554 | 2.212448446 | 0.026935695 | 0.070832741 |
| ENSG00000273179 | ENSG0000027 | 2.446643764 | -0.360402719 | 1.113340073 | -0.323713058 | 0.746155284 | 0.859596562 |

| | | | | | | | |
|---|---|---|---|---|---|---|---|
| ENSG00000273181 | ENSG0000027 | 1.568734523 | 1.032751409 | 1.505027717 | 0.686200923 | 0.492586416 | NA |
| ENSG00000273183 | ENSG0000027 | 2.22782217 | 1.671629035 | 1.258547221 | 1.328221148 | 0.18410506 | 0.34062183 |
| ENSG00000273186 | ENSG0000027 | 27.66358719 | 0.679547213 | 0.343701171 | 1.977145469 | 0.048025189 | 0.115748881 |
| ENSG00000273188 | ENSG0000027 | 5.043383664 | 0.15100587 | 0.787640163 | 0.191719363 | 0.847962038 | 0.920424395 |
| ENSG00000273192 | ENSG0000027 | 21.10967179 | 0.028449237 | 0.407794042 | 0.06976374 | 0.944381708 | 0.973580639 |
| ENSG00000273199 | ENSG0000027 | 2.533685501 | 1.00142956 | 1.145984751 | 0.873859412 | 0.382194821 | 0.574582161 |
| ENSG00000273204 | ENSG0000027 | 11.41793971 | -0.21179922 | 0.555751587 | -0.381104121 | 0.703125989 | 0.833796972 |
| ENSG00000273209 | ENSG0000027 | 3.34979318 | 0.361066364 | 0.96244558 | 0.375155096 | 0.707545124 | 0.836610749 |
| ENSG00000273210 | ENSG0000027 | 14.7199546 | 0.844437006 | 0.469855569 | 1.797226769 | 0.072299626 | 0.163066899 |
| ENSG00000273211 | ENSG0000027 | 16.39655264 | -0.475803416 | 0.440653365 | -1.07976803 | 0.280245491 | 0.463539661 |
| ENSG00000273212 | ENSG0000027 | 28.77157016 | -0.664741678 | 0.328959716 | -2.020738847 | 0.043306806 | 0.105968868 |
| ENSG00000273216 | ENSG0000027 | 7.491041022 | -0.348907935 | 0.651977197 | -0.535153585 | 0.592543659 | 0.754744671 |
| ENSG00000273221 | ENSG0000027 | 20.80973874 | 0.561944306 | 0.40023564 | 1.404033651 | 0.160308829 | 0.306482585 |
| ENSG00000273226 | ENSG0000027 | 4.424271294 | 0.523822207 | 0.87809168 | 0.596546146 | 0.550810436 | 0.722901276 |
| ENSG00000273230 | ENSG0000027 | 435.3206037 | 0.423887462 | 0.091536796 | 4.63078763 | 3.64E-06 | 1.98E-05 |
| ENSG00000273243 | ENSG0000027 | 3.822505326 | -0.55168732 | 0.881753073 | -0.625670992 | 0.531530764 | 0.707118252 |
| ENSG00000273247 | RAB33B-AS1 | 135.5883537 | 0.113181417 | 0.157836667 | 0.71707937 | 0.473325124 | 0.658863978 |
| ENSG00000273249 | WDR5-DT | 4.897665563 | 0.862748558 | 0.854080998 | 1.010148405 | 0.312424194 | 0.500664409 |
| ENSG00000273253 | TRABD-AS1 | 32.36291644 | 0.422547007 | 0.319393216 | 1.32296801 | 0.185846016 | 0.34312335 |
| ENSG00000273254 | ENSG0000027 | 8.303635066 | -0.017870924 | 0.607225956 | -0.029430435 | 0.9765213 | 0.989278333 |
| ENSG00000273262 | ENSG0000027 | 4.425760468 | -0.050151401 | 0.830183515 | -0.060410018 | 0.95182908 | 0.977646532 |
| ENSG00000273264 | ENSG0000027 | 8.850752296 | 3.196933396 | 0.809425754 | 3.949631426 | 7.83E-05 | 0.000355555 |
| ENSG00000273265 | CNNM3-DT | 20.37429015 | 0.079000463 | 0.39497032 | 0.200016202 | 0.84146791 | 0.916713017 |
| ENSG00000273270 | LINC03072 | 548.9589037 | 0.163004104 | 0.084814707 | 1.921884892 | 0.054620243 | 0.129254011 |
| ENSG00000273271 | ENSG0000027 | 19.31905045 | 1.038820881 | 0.431893238 | 2.405272389 | 0.016160415 | 0.045420231 |
| ENSG00000273274 | ZBTB8B | 8.53884434 | -2.7876847 | 0.748806569 | -3.722836865 | 0.000196997 | 0.000836872 |
| ENSG00000273275 | ENSG0000027 | 8.073632058 | -0.747959092 | 0.654303037 | -1.143138652 | 0.252981029 | 0.431735218 |
| ENSG00000273284 | ENSG0000027 | 4.572894586 | 0.63418424 | 0.825585977 | 0.768162563 | 0.442390612 | 0.631470633 |
| ENSG00000273287 | ENSG0000027 | 2.024460585 | -1.748898714 | 1.32382735 | -1.32109275 | 0.18647044 | 0.343999349 |
| ENSG00000273293 | ENSG0000027 | 2.150778333 | 4.457188784 | 1.546323993 | 2.882441716 | 0.003946061 | 0.013015478 |
| ENSG00000273294 | C1QTNF3-AM/ | 1.658038588 | 2.017942729 | 1.612601251 | 1.251358777 | 0.21080361 | NA |
| ENSG00000273295 | ENSG0000027 | 21.99229227 | -0.813851186 | 0.38546406 | -2.111354264 | 0.034741873 | 0.088044473 |
| ENSG00000273297 | LINC03060 | 3.208791304 | -0.161943987 | 0.970650586 | -0.166840663 | 0.867495423 | 0.93096599 |

| | | | | | | | |
|---|---|---|---|---|---|---|---|
| ENSG00000273300 | ENSG0000027 | 20.26527942 | -1.01294145 | 0.427123208 | -2.371543928 | 0.017713941 | 0.049243554 |
| ENSG00000273305 | ENSG0000027 | 19.59664001 | -0.028825122 | 0.395938962 | -0.072801934 | 0.941963732 | 0.972486409 |
| ENSG00000273306 | ENSG0000027 | 16.5054493 | 0.685992782 | 0.444029245 | 1.544927028 | 0.122363915 | 0.249137145 |
| ENSG00000273308 | ENSG0000027 | 2.629215738 | -0.175395741 | 1.101823407 | -0.159186799 | 0.873521704 | 0.934676032 |
| ENSG00000273311 | DGCR11 | 47.59130925 | -0.507493052 | 0.256290919 | -1.98014449 | 0.047687295 | 0.11504328 |
| ENSG00000273314 | ENSG0000027 | 100.6547918 | 0.15167991 | 0.184560864 | 0.821842219 | 0.411166702 | 0.603581246 |
| ENSG00000273320 | ENSG0000027 | 14.02025779 | 0.114845025 | 0.46197388 | 0.248596359 | 0.803673025 | 0.895442187 |
| ENSG00000273321 | ENSG0000027 | 2.763041488 | 3.76835036 | 1.46625347 | 2.570053839 | 0.010168271 | 0.03017986 |
| ENSG00000273328 | ENSG0000027 | 6.535180201 | -0.370264761 | 0.743058518 | -0.498298252 | 0.61827384 | 0.773685462 |
| ENSG00000273329 | ENSG0000027 | 122.6990746 | 0.169792621 | 0.176828476 | 0.960210848 | 0.336949108 | 0.528123295 |
| ENSG00000273335 | ENSG0000027 | 8.397272906 | 0.09575801 | 0.59641927 | 0.160554856 | 0.872444013 | 0.934092489 |
| ENSG00000273340 | MICE | 13.07512299 | 0.396233354 | 0.480343738 | 0.824895429 | 0.409430962 | 0.601996299 |
| ENSG00000273343 | ENSG0000027 | 20.24193359 | -0.924809197 | 0.403449658 | -2.292254261 | 0.021890976 | 0.059156435 |
| ENSG00000273344 | PAXIP1-DT | 230.5593637 | 0.896731521 | 0.125790342 | 7.128778797 | 1.01E-12 | 1.04E-11 |
| ENSG00000273345 | ENSG0000027 | 3.230085507 | 0.119886799 | 0.989442492 | 0.12116601 | 0.903559546 | 0.951246259 |
| ENSG00000273355 | ENSG0000027 | 20.02619515 | 0.756939569 | 0.406319554 | 1.862916912 | 0.062473957 | 0.144238383 |
| ENSG00000273356 | LINC02019 | 2.672602955 | 0.787786236 | 1.160320684 | 0.678938372 | 0.497176911 | 0.679040564 |
| ENSG00000273361 | ENSG0000027 | 9.914668427 | -0.427192251 | 0.568319531 | -0.751676175 | 0.452245822 | 0.640066851 |
| ENSG00000273367 | ENSG0000027 | 2.242449168 | 1.146731839 | 1.220781075 | 0.939342739 | 0.347554802 | 0.538491636 |
| ENSG00000273373 | SLC16A4-AS1 | 31.6355385 | 0.329547401 | 0.344611329 | 0.956287193 | 0.338927158 | 0.5300647 |
| ENSG00000273374 | ENSG0000027 | 2.323374688 | 1.084499438 | 1.211374576 | 0.895263496 | 0.370646249 | 0.5633489 |
| ENSG00000273381 | ENSG0000027 | 15.29218156 | 0.456869027 | 0.453628288 | 1.00714404 | 0.31386556 | 0.502132363 |
| ENSG00000273382 | TMEM167B-D | 142.5443338 | 0.171715088 | 0.151778561 | 1.131352725 | 0.257906661 | 0.4375795 |
| ENSG00000273387 | ENSG0000027 | 15.20952096 | -0.5598633 | 0.471547651 | -1.187288918 | 0.235113678 | 0.410121494 |
| ENSG00000273391 | ENSG0000027 | 23.33740563 | 0.030741696 | 0.367809388 | 0.083580509 | 0.933389964 | 0.967991219 |
| ENSG00000273394 | ENSG0000027 | 8.090896569 | 0.273527312 | 0.620181226 | 0.441044167 | 0.659181023 | 0.802846438 |
| ENSG00000273398 | ENSG0000027 | 14.26780067 | 0.175813408 | 0.474063278 | 0.370864853 | 0.710738194 | 0.839045604 |
| ENSG00000273402 | ENSG0000027 | 3.248583371 | 0.693972768 | 0.97331843 | 0.712996637 | 0.475847838 | 0.661019853 |
| ENSG00000273403 | ENSG0000027 | 2.825450866 | -0.061398653 | 1.038383686 | -0.059129061 | 0.952849311 | 0.978090353 |
| ENSG00000273419 | ENSG0000027 | 31.89982576 | 0.850569194 | 0.347003682 | 2.45118204 | 0.014238792 | 0.040586219 |
| ENSG00000273424 | ENSG0000027 | 2.303429961 | -0.193620817 | 1.138992095 | -0.169993117 | 0.865015549 | 0.929871448 |
| ENSG00000273428 | ENSG0000027 | 5.171001032 | -0.766480062 | 0.817917635 | -0.93711155 | 0.348701181 | 0.539939594 |
| ENSG00000273432 | ENSG0000027 | 2.628299834 | 0.943135436 | 1.129754448 | 0.834814537 | 0.40382214 | 0.596617498 |

| | | | | | | | |
|---|---|---|---|---|---|---|---|
| ENSG00000273437 | ENSG0000027 | 7.126792645 | 0.232497959 | 0.653452872 | 0.355799123 | 0.721991006 | 0.845596895 |
| ENSG00000273443 | ENSG0000027 | 3.11049069 | 0.325686068 | 1.02095547 | 0.319001247 | 0.749725567 | 0.861615376 |
| ENSG00000273445 | ENSG0000027 | 3.456068406 | -2.18692545 | 1.072120916 | -2.039812318 | 0.041369023 | 0.101923035 |
| ENSG00000273447 | ENSG0000027 | 4.857524303 | 0.58316573 | 0.811980996 | 0.718201205 | 0.472633235 | 0.658220399 |
| ENSG00000273448 | ENSG0000027 | 4.23251861 | 1.228619045 | 0.883410686 | 1.390767697 | 0.164295882 | 0.312360975 |
| ENSG00000273449 | ENSG0000027 | 5.379424659 | -0.366801786 | 0.767069407 | -0.478185915 | 0.632517886 | 0.783513891 |
| ENSG00000273451 | ENSG0000027 | 10.01935677 | -1.381078277 | 0.571283038 | -2.417502682 | 0.015627419 | 0.044100503 |
| ENSG00000273454 | ENSG0000027 | 2.219357379 | -0.020240193 | 1.167006685 | -0.017343682 | 0.986162437 | 0.993350589 |
| ENSG00000273455 | ENSG0000027 | 2.096363271 | 0.501588147 | 1.194053593 | 0.42007172 | 0.674433061 | 0.81390205 |
| ENSG00000273456 | ENSG0000027 | 33.48249949 | -0.254484554 | 0.33036632 | -0.770310224 | 0.441115893 | 0.630361838 |
| ENSG00000273464 | ENSG0000027 | 23.60089181 | -1.871773403 | 0.408662795 | -4.580239321 | 4.64E-06 | 2.49E-05 |
| ENSG00000273472 | ENSG0000027 | 35.11504742 | 0.462665677 | 0.315589603 | 1.466035868 | 0.142638516 | 0.280389475 |
| ENSG00000273473 | ENSG0000027 | 20.56450649 | -0.689865764 | 0.386355875 | -1.785570786 | 0.074168781 | 0.166448036 |
| ENSG00000273483 | ENSG0000027 | 3.457150375 | -0.262231544 | 0.97701042 | -0.268401993 | 0.788389906 | 0.885492107 |
| ENSG00000273485 | ENSG0000027 | 4.333641544 | 0.810839031 | 0.849440962 | 0.954556075 | 0.339802239 | 0.5308131 |
| ENSG00000273486 | ENSG0000027 | 49.64548919 | -0.156154554 | 0.257745928 | -0.605846831 | 0.544616468 | 0.717913747 |
| ENSG00000273487 | ENSG0000027 | 2.109117274 | 0.046140211 | 1.176957329 | 0.03920296 | 0.968728574 | 0.985876114 |
| ENSG00000273489 | ENSG0000027 | 3.117113332 | -0.339902927 | 0.977532767 | -0.347715124 | 0.728054137 | 0.849088048 |
| ENSG00000273492 | APP-DT | 3.043281148 | -0.017927992 | 1.026569701 | -0.017463979 | 0.986066469 | 0.993350589 |
| ENSG00000273507 | ENSG0000027 | 3.075553747 | 0.91156509 | 1.010038045 | 0.902505697 | 0.366788296 | 0.559338132 |
| ENSG00000273509 | CNTNAP3P1 | 2.768204921 | 0.688344041 | 1.053448329 | 0.653419842 | 0.51348565 | 0.692692264 |
| ENSG00000273523 | ENSG0000027 | 2.248972589 | -0.92929003 | 1.262549433 | -0.736042491 | 0.46170484 | 0.648393573 |
| ENSG00000273542 | H4C12 | 26.36289154 | -0.486186836 | 0.339273879 | -1.433021717 | 0.151851631 | 0.294187688 |
| ENSG00000273559 | CWC25 | 721.6930998 | -0.12080945 | 0.078764167 | -1.533812314 | 0.125075833 | 0.25362323 |
| ENSG00000273565 | ENSG0000027 | 2.550277252 | -0.441048165 | 1.114125894 | -0.395869235 | 0.692201498 | 0.826433973 |
| ENSG00000273568 | ENSG0000027 | 26.44900935 | -0.423190064 | 0.367941238 | -1.150156656 | 0.250079355 | 0.428152877 |
| ENSG00000273599 | ENSG0000027 | 17.61069477 | -0.761373896 | 0.424903298 | -1.791875702 | 0.073152879 | 0.164554525 |
| ENSG00000273604 | EPOP | 631.6336577 | 0.174131807 | 0.088049306 | 1.977662461 | 0.047966798 | 0.115653993 |
| ENSG00000273611 | ZNHIT3 | 988.5857757 | 0.00647039 | 0.066181701 | 0.097767052 | 0.922117271 | 0.962216621 |
| ENSG00000273619 | ENSG0000027 | 72.30848373 | 0.31249443 | 0.216407912 | 1.444006492 | 0.148737149 | 0.289621441 |
| ENSG00000273654 | ENSG0000027 | 467.1816915 | -0.141605142 | 0.089700645 | -1.578641302 | 0.114418355 | 0.236358134 |
| ENSG00000273669 | ENSG0000027 | 3.165336443 | 0.384437936 | 1.062730543 | 0.361745448 | 0.717542262 | 0.842913738 |
| ENSG00000273674 | ENSG0000027 | 39.83886792 | -1.244510677 | 0.299178388 | -4.159761283 | 3.19E-05 | 0.000153858 |

| | | | | | | | |
|---|---|---|---|---|---|---|---|
| ENSG00000273679 | ENSG0000027 | 12.6037188 | -0.007268551 | 0.493458152 | -0.014729822 | 0.988247728 | 0.994030148 |
| ENSG00000273680 | ENSG0000027 | 8.437992899 | 0.221403363 | 0.666567136 | 0.332154633 | 0.739772498 | 0.856527588 |
| ENSG00000273682 | BLVRBP1 | 3.479721696 | -0.890045565 | 0.944882465 | -0.941964316 | 0.346210909 | 0.537062366 |
| ENSG00000273693 | CDRT15P10 | 2.10437619 | -3.542992138 | 1.543212761 | -2.295854615 | 0.021684195 | 0.05865922 |
| ENSG00000273702 | ENSG0000027 | 190.5980753 | 0.581319505 | 0.132320795 | 4.393258851 | 1.12E-05 | 5.72E-05 |
| ENSG00000273703 | H2BC14 | 2.075506016 | -0.344208483 | 1.236595929 | -0.278351622 | 0.780742452 | 0.881105517 |
| ENSG00000273706 | LHX1 | 356.5103921 | 0.786977051 | 0.102132763 | 7.705431912 | 1.30E-14 | 1.52E-13 |
| ENSG00000273711 | ENSG0000027 | 43.07812885 | -0.566908524 | 0.270196945 | -2.098130771 | 0.035893595 | 0.09056303 |
| ENSG00000273716 | ENSG0000027 | 39.4179893 | -0.002489545 | 0.292810606 | -0.008502237 | 0.993216278 | 0.996343424 |
| ENSG00000273723 | SUGT1-DT | 7.087037376 | 0.010158025 | 0.701903448 | 0.014472112 | 0.988453329 | 0.994043197 |
| ENSG00000273724 | ENSG0000027 | 1.841134384 | 1.899291355 | 1.488657726 | 1.275841533 | 0.202011543 | 0.365872792 |
| ENSG00000273729 | ENSG0000027 | 51.94404399 | -0.068990083 | 0.243934178 | -0.282822536 | 0.777312878 | 0.879138067 |
| ENSG00000273733 | ENSG0000027 | 6.296186526 | 0.352228486 | 0.715062002 | 0.492584537 | 0.622306176 | 0.776528703 |
| ENSG00000273747 | ENSG0000027 | 15.7364567 | -0.148306661 | 0.445684758 | -0.332761348 | 0.739314437 | 0.856254116 |
| ENSG00000273749 | CYFIP1 | 3957.281436 | -0.186723589 | 0.050500772 | -3.697440284 | 0.000217784 | 0.000917366 |
| ENSG00000273759 | ENSG0000027 | 10.66809439 | -0.278059562 | 0.532380086 | -0.522295198 | 0.601464812 | 0.761283686 |
| ENSG00000273763 | ENSG0000027 | 35.50756822 | -0.173844509 | 0.296262055 | -0.586793031 | 0.557342718 | 0.727644525 |
| ENSG00000273776 | MIR6126 | 1.999868482 | -2.480987223 | 1.494381494 | -1.660210082 | 0.096872197 | 0.206652114 |
| ENSG00000273783 | GTF2A1-AS1 | 71.05872086 | -0.07467932 | 0.210541046 | -0.354701954 | 0.722812887 | 0.845926239 |
| ENSG00000273796 | ENSG0000027 | 46.1208234 | -0.564288557 | 0.265808231 | -2.122916037 | 0.033760888 | 0.085890516 |
| ENSG00000273802 | H2BC8 | 583.4552583 | 0.633938371 | 0.088980739 | 7.124444862 | 1.05E-12 | 1.07E-11 |
| ENSG00000273820 | USP27X | 252.2894348 | -0.005001126 | 0.115425705 | -0.043327663 | 0.96544034 | 0.984407565 |
| ENSG00000273821 | ENSG0000027 | 12.82285016 | -0.003584035 | 0.503307183 | -0.007120968 | 0.994318337 | 0.996533822 |
| ENSG00000273828 | ENSG0000027 | 12.97657462 | -0.581436653 | 0.492801934 | -1.179858708 | 0.238056415 | 0.413525228 |
| ENSG00000273841 | TAF9 | 1020.440279 | -0.104193802 | 0.069087942 | -1.508132953 | 0.131520503 | 0.263846116 |
| ENSG00000273853 | ENSG0000027 | 5.553444968 | -0.987024523 | 0.78437993 | -1.258350049 | 0.208265188 | 0.374751426 |
| ENSG00000273870 | ENSG0000027 | 2.214125895 | -0.8738449 | 1.202697972 | -0.726570528 | 0.467489088 | 0.653476308 |
| ENSG00000273891 | ENSG0000027 | 5.325943779 | -0.900417004 | 0.763303592 | -1.179631557 | 0.238146786 | 0.413526941 |
| ENSG00000273893 | ENSG0000027 | 48.23211503 | 0.139786223 | 0.282342087 | 0.49509524 | 0.620532894 | 0.775370368 |
| ENSG00000273899 | NOL12 | 70.82966936 | 0.26978601 | 0.2198688 | 1.227031803 | 0.219810638 | 0.38993566 |
| ENSG00000273901 | ENSG0000027 | 22.57677237 | 0.436495625 | 0.371905544 | 1.173673349 | 0.240525889 | 0.4163332 |
| ENSG00000273951 | ENSG0000027 | 3.933694757 | -0.561838005 | 0.912948808 | -0.615410196 | 0.53828386 | 0.713152281 |
| ENSG00000273973 | ENSG0000027 | 2.337459564 | -4.764628732 | 1.537853789 | -3.098232593 | 0.001946786 | 0.00686397 |

| | | | | | | | |
|---|---|---|---|---|---|---|---|
| ENSG00000273980 | ENSG0000027 | 1.65947431 | 2.020880847 | 1.63070601 | 1.239267431 | 0.215246476 | NA |
| ENSG00000273982 | ENSG0000027 | 73.8628386 | 0.379957984 | 0.207677654 | 1.829556415 | 0.067316296 | 0.153624735 |
| ENSG00000273983 | H3C8 | 138.3003291 | 0.243571816 | 0.155667358 | 1.564694227 | 0.117654623 | 0.241867286 |
| ENSG00000273998 | ENSG0000027 | 1.754582418 | -1.485188992 | 1.41148961 | -1.052213903 | 0.292701422 | NA |
| ENSG00000274001 | ENSG0000027 | 2.263223227 | -0.163972514 | 1.135675034 | -0.144383304 | 0.8851978 | 0.940548288 |
| ENSG00000274002 | ENSG0000027 | 5.199430255 | -0.80571997 | 0.792946861 | -1.016108404 | 0.309577768 | 0.497667413 |
| ENSG00000274010 | ZFP91P1 | 2.361568729 | 1.28805935 | 1.221092323 | 1.054841903 | 0.291497636 | 0.476478957 |
| ENSG00000274015 | ENSG0000027 | 23.8877691 | -0.525030807 | 0.36076959 | -1.455307822 | 0.145584062 | 0.284884278 |
| ENSG00000274020 | LINC01138 | 19.23976695 | 0.604581175 | 0.407649244 | 1.483091615 | 0.138050072 | 0.273666492 |
| ENSG00000274051 | ENSG0000027 | 1.895903523 | 2.182112985 | 1.467318698 | 1.487143173 | 0.136977001 | 0.272032933 |
| ENSG00000274070 | CASTOR2 | 1479.942163 | 0.352152509 | 0.061144011 | 5.759394958 | 8.44E-09 | 6.11E-08 |
| ENSG00000274080 | ENSG0000027 | 4.100034977 | 0.991347428 | 0.89999114 | 1.101507985 | 0.270675631 | 0.452506848 |
| ENSG00000274099 | ABCB10P1 | 1.801779589 | 0.131220223 | 1.344485691 | 0.097598824 | 0.922250858 | 0.962243482 |
| ENSG00000274104 | ENSG0000027 | 30.72611723 | 0.224095229 | 0.320178164 | 0.699907909 | 0.483984818 | 0.667928891 |
| ENSG00000274105 | ENSG0000027 | 1.735194633 | -0.185381042 | 1.381527764 | -0.134185535 | 0.893255864 | NA |
| ENSG00000274114 | ALOX15P1 | 2.336976626 | 0.842026563 | 1.190334619 | 0.707386435 | 0.479326364 | 0.663653377 |
| ENSG00000274173 | LINC02967 | 613.4101495 | -0.537530823 | 0.086757427 | -6.195790293 | 5.80E-10 | 4.72E-09 |
| ENSG00000274177 | ENSG0000027 | 14.48734607 | 0.088613602 | 0.466719631 | 0.189864741 | 0.849415122 | 0.921173493 |
| ENSG00000274180 | NATD1 | 554.8456978 | 1.083838331 | 0.094961327 | 11.41347082 | 3.58E-30 | 1.00E-28 |
| ENSG00000274191 | ENSG0000027 | 2.452662594 | -0.078925654 | 1.27058434 | -0.062117603 | 0.950469179 | 0.977185792 |
| ENSG00000274210 | RNVU1-27 | 5.453509529 | -1.361128915 | 0.776000614 | -1.754030719 | 0.079425244 | 0.175733747 |
| ENSG00000274211 | SOCS7 | 1448.184931 | 0.07039969 | 0.065480701 | 1.075121191 | 0.282320473 | 0.465749817 |
| ENSG00000274213 | ENSG0000027 | 30.07256932 | 0.421070927 | 0.32129922 | 1.310525829 | 0.190018012 | 0.349195962 |
| ENSG00000274214 | ENSG0000027 | 149.9722029 | -0.070588394 | 0.14814808 | -0.476471877 | 0.633738238 | 0.784143138 |
| ENSG00000274220 | ENSG0000027 | 56.58981199 | 0.093945823 | 0.235158421 | 0.399500143 | 0.689524718 | 0.824525492 |
| ENSG00000274225 | HEPFAL | 3.383332383 | -0.491586051 | 0.942954507 | -0.52132531 | 0.602140173 | 0.761606804 |
| ENSG00000274227 | ENSG0000027 | 10.50225213 | 0.630098202 | 0.558870782 | 1.127448818 | 0.259552769 | 0.439484833 |
| ENSG00000274253 | ENSG0000027 | 12.88743662 | -0.048758257 | 0.481456445 | -0.101272415 | 0.919334213 | 0.96075472 |
| ENSG00000274259 | SYNGAP1-AS1 | 18.41197969 | 0.201914538 | 0.437763565 | 0.461241078 | 0.644625653 | 0.791802982 |
| ENSG00000274265 | ENSG0000027 | 100.2235697 | 0.127466567 | 0.18171091 | 0.701479987 | 0.48300352 | 0.667074097 |
| ENSG00000274270 | ENSG0000027 | 15.86799295 | 0.761649884 | 0.464869501 | 1.63841655 | 0.101334829 | 0.214152094 |
| ENSG00000274272 | ENSG0000027 | 8.976908979 | -0.912954816 | 0.589527336 | -1.548621685 | 0.12147269 | 0.247778375 |
| ENSG00000274276 | ENSG0000027 | 33.89383575 | 0.088723496 | 0.312015798 | 0.284355782 | 0.776137742 | 0.878371968 |

| | | | | | | | |
|---|---|---|---|---|---|---|---|
| ENSG00000274286 | ADRA2B | 22.58041058 | 0.50285099 | 0.377924492 | 1.330559413 | 0.183334025 | 0.339496491 |
| ENSG00000274290 | H2BC6 | 181.6448719 | 0.709393697 | 0.143348035 | 4.948750769 | 7.47E-07 | 4.40E-06 |
| ENSG00000274292 | ENSG0000027 | 20.62843347 | 0.432313196 | 0.386585074 | 1.118287344 | 0.263444291 | 0.44412799 |
| ENSG00000274315 | ENSG0000027 | 3.959112513 | -0.659733114 | 0.867929747 | -0.760122714 | 0.447181237 | 0.635663501 |
| ENSG00000274317 | LINC02334 | 4.835690881 | 0.626586223 | 0.811777981 | 0.771868956 | 0.440192045 | 0.629591136 |
| ENSG00000274340 | ENSG0000027 | 2.45795058 | -1.633778446 | 1.205031833 | -1.35579692 | 0.175163792 | 0.328289002 |
| ENSG00000274349 | ZNF658 | 29.53682472 | 0.732070722 | 0.345098945 | 2.121335728 | 0.03389356 | 0.086189808 |
| ENSG00000274363 | ENSG0000027 | 3.317120107 | -0.164585086 | 0.958086625 | -0.171785183 | 0.863606415 | 0.929172842 |
| ENSG00000274367 | ENSG0000027 | 14.0136817 | -0.345331196 | 0.480804026 | -0.718236905 | 0.472611226 | 0.658220399 |
| ENSG00000274370 | ENSG0000027 | 53.66353604 | 1.423800125 | 0.290205716 | 4.906175341 | 9.29E-07 | 5.40E-06 |
| ENSG00000274372 | LINC02804 | 12.20704621 | 0.330988496 | 0.506263536 | 0.653786956 | 0.513249072 | 0.692527276 |
| ENSG00000274378 | ENSG0000027 | 1.600871428 | 1.093244567 | 1.447434516 | 0.755298119 | 0.450070127 | NA |
| ENSG00000274383 | ENSG0000027 | 33.39348439 | 0.736205036 | 0.319178904 | 2.306559194 | 0.021079407 | 0.057212682 |
| ENSG00000274386 | TMEM269 | 21.77158118 | -0.006247102 | 0.378778813 | -0.016492745 | 0.98684129 | 0.993598104 |
| ENSG00000274391 | TPTE | 2.841950858 | -0.360925165 | 1.021524003 | -0.353320298 | 0.72384833 | 0.846416769 |
| ENSG00000274414 | ENSG0000027 | 5.868840542 | 0.622860779 | 0.724308769 | 0.8599382 | 0.38982311 | 0.582613991 |
| ENSG00000274422 | ENSG0000027 | 5.094443915 | -0.017991584 | 0.783976207 | -0.022949146 | 0.981690838 | 0.991627389 |
| ENSG00000274423 | SEC22B2P | 12.41408846 | 0.012208833 | 0.573799395 | 0.02127718 | 0.983024547 | 0.992139551 |
| ENSG00000274425 | ENSG0000027 | 146.8580599 | -0.107902434 | 0.148409009 | -0.727061211 | 0.467188459 | 0.653190484 |
| ENSG00000274427 | ENSG0000027 | 3.232459738 | 0.130230536 | 0.966660723 | 0.134722072 | 0.892831621 | 0.94515724 |
| ENSG00000274444 | ENSG0000027 | 2.107734679 | -1.833309591 | 1.320898761 | -1.387925892 | 0.165159613 | 0.313687249 |
| ENSG00000274447 | ENSG0000027 | 7.83736347 | -0.6550375 | 0.644873118 | -1.015761832 | 0.309742816 | 0.49772738 |
| ENSG00000274460 | ENSG0000027 | 18.64625539 | 0.608935347 | 0.419369712 | 1.452025099 | 0.146494626 | 0.286348563 |
| ENSG00000274471 | ENSG0000027 | 24.16993466 | -0.418577986 | 0.355999563 | -1.175782302 | 0.23968188 | 0.415342373 |
| ENSG00000274487 | NPEPPSP1 | 24.67760278 | -0.192935232 | 0.359593315 | -0.536537314 | 0.591587254 | 0.753986978 |
| ENSG00000274494 | MIR6832 | 2.375325185 | -0.621040969 | 1.119564262 | -0.554716679 | 0.579088466 | 0.744665193 |
| ENSG00000274499 | ENSG0000027 | 6.056493161 | 0.934656836 | 0.730068289 | 1.280232068 | 0.200463531 | 0.363672076 |
| ENSG00000274512 | TBC1D3L | 67.21109549 | 0.006101856 | 0.226556448 | 0.026933047 | 0.978513135 | 0.990210919 |
| ENSG00000274515 | ENSG0000027 | 4.81545972 | 0.09606183 | 0.790336836 | 0.121545429 | 0.903259035 | 0.951148141 |
| ENSG00000274523 | RCC1L | 1830.872051 | 0.423719367 | 0.055678486 | 7.610109378 | 2.74E-14 | 3.12E-13 |
| ENSG00000274525 | ENSG0000027 | 3.42018809 | 0.51997543 | 0.938369606 | 0.554126462 | 0.579492301 | 0.745038089 |
| ENSG00000274528 | ENSG0000027 | 105.197118 | -0.521473294 | 0.180810566 | -2.884086401 | 0.00392551 | 0.012958873 |
| ENSG00000274549 | ENSG0000027 | 5.54975509 | 2.376940265 | 0.885924859 | 2.683004367 | 0.007296404 | 0.02254284 |

| | | | | | | | |
|---|---|---|---|---|---|---|---|
| ENSG00000274554 | ENSG0000027 | 2.500549823 | 0.593966517 | 1.123071844 | 0.528876688 | 0.596890994 | 0.75769879 |
| ENSG00000274559 | ENSG0000027 | 4.854226403 | 0.581020113 | 0.850803582 | 0.68290746 | 0.49466532 | 0.67711556 |
| ENSG00000274561 | ENSG0000027 | 76.76408895 | 0.427465621 | 0.214057656 | 1.996964885 | 0.045828998 | 0.111168151 |
| ENSG00000274565 | ENSG0000027 | 67.75491727 | -1.80077697 | 0.233391038 | -7.715707434 | 1.20E-14 | 1.40E-13 |
| ENSG00000274600 | RIMBP3B | 6.524980688 | 0.778490919 | 0.693310652 | 1.122860172 | 0.261496885 | 0.441733324 |
| ENSG00000274602 | PI4KAP1 | 274.7580504 | -1.250923797 | 0.132682307 | -9.427962383 | 4.18E-21 | 7.46E-20 |
| ENSG00000274605 | PCCA-DT | 100.7446751 | -0.420877528 | 0.176520541 | -2.38429775 | 0.017111751 | 0.047748883 |
| ENSG00000274629 | ENSG0000027 | 8.278518563 | -0.391556084 | 0.612910193 | -0.638847401 | 0.522922208 | 0.699628803 |
| ENSG00000274630 | ENSG0000027 | 57.2063186 | -0.783768595 | 0.243049196 | -3.224732309 | 0.001260905 | 0.004631731 |
| ENSG00000274641 | H2BC17 | 26.18799017 | 0.837830438 | 0.348954131 | 2.40097584 | 0.016351416 | 0.045875895 |
| ENSG00000274642 | ENSG0000027 | 1.65094893 | -1.377286077 | 1.391340877 | -0.989898378 | 0.322223793 | NA |
| ENSG00000274653 | ENSG0000027 | 22.90001223 | -0.179401856 | 0.365732018 | -0.490528165 | 0.623760205 | 0.777539482 |
| ENSG00000274667 | ENSG0000027 | 14.17076114 | -0.280627733 | 0.474123545 | -0.591887358 | 0.55392602 | 0.725418397 |
| ENSG00000274677 | ENSG0000027 | 8.956300357 | 0.546454758 | 0.597059659 | 0.915243142 | 0.360064001 | 0.552610435 |
| ENSG00000274678 | ENSG0000027 | 1.737056486 | -0.738920852 | 1.426516378 | -0.517989743 | 0.604465426 | NA |
| ENSG00000274695 | ENSG0000027 | 8.054128062 | 0.013558204 | 0.629846056 | 0.021526219 | 0.982825888 | 0.992098874 |
| ENSG00000274698 | ENSG0000027 | 7.083402052 | 0.132810927 | 0.677453873 | 0.196044236 | 0.844575543 | 0.918248003 |
| ENSG00000274712 | ENSG0000027 | 247.3019192 | 0.232112305 | 0.120150018 | 1.93185411 | 0.053377515 | 0.126819494 |
| ENSG00000274750 | H3C6 | 412.0400155 | 0.06156035 | 0.094787551 | 0.649456068 | 0.516043633 | 0.694679946 |
| ENSG00000274751 | ENSG0000027 | 7.929185226 | 0.221191003 | 0.624248089 | 0.354331887 | 0.723090174 | 0.8459455 |
| ENSG00000274767 | ENSG0000027 | 3.65640338 | 0.61965228 | 1.044411004 | 0.593303094 | 0.552978329 | 0.72450299 |
| ENSG00000274798 | ENSG0000027 | 2.927982681 | 0.17405305 | 1.021530227 | 0.17038463 | 0.864707659 | 0.929799912 |
| ENSG00000274818 | ENSG0000027 | 9.372250377 | -0.528478029 | 0.59624253 | -0.886347422 | 0.375430342 | 0.568360856 |
| ENSG00000274825 | ENSG0000027 | 2.073282948 | 0.218386851 | 1.262044488 | 0.173042118 | 0.862618322 | 0.928643498 |
| ENSG00000274828 | ENSG0000027 | 191.8263376 | 1.090543127 | 0.139806865 | 7.80035463 | 6.17E-15 | 7.37E-14 |
| ENSG00000274840 | BALR6 | 18.02158375 | 0.803515237 | 0.442072302 | 1.817610453 | 0.069123693 | 0.157046401 |
| ENSG00000274849 | ENSG0000027 | 11.8296681 | 0.892289711 | 0.528613525 | 1.687981236 | 0.091414831 | 0.196980671 |
| ENSG00000274859 | ENSG0000027 | 9.977728911 | -0.325488004 | 0.552449369 | -0.589172551 | 0.555745528 | 0.726640663 |
| ENSG00000274898 | ENSG0000027 | 2.245302333 | -1.363524227 | 1.202071407 | -1.13431217 | 0.256663626 | 0.436202619 |
| ENSG00000274902 | ENSG0000027 | 3.007363627 | -1.559713264 | 1.060436728 | -1.470821618 | 0.141339364 | 0.278622069 |
| ENSG00000274904 | ENSG0000027 | 6.751016714 | -0.789099091 | 0.673530678 | -1.171585968 | 0.241363324 | 0.417373335 |
| ENSG00000274918 | ENSG0000027 | 2.76777511 | 3.770247271 | 1.46762133 | 2.568950991 | 0.010200688 | 0.030260415 |
| ENSG00000274922 | ENSG0000027 | 62.96735154 | -0.548430124 | 0.222238732 | -2.467752214 | 0.01359644 | 0.038948843 |

| | | | | | | | |
|---|---|---|---|---|---|---|---|
| ENSG00000274925 | ZKSCAN2-DT | 74.1717057 | 0.179757827 | 0.213976533 | 0.84008197 | 0.400862428 | 0.593736061 |
| ENSG00000274928 | KRT89P | 51.53246123 | -0.939795777 | 0.260835255 | -3.603024361 | 0.000314536 | 0.001290779 |
| ENSG00000274943 | ENSG0000027 | 5.231985284 | 0.356713767 | 0.783583643 | 0.45523381 | 0.648941039 | 0.795495779 |
| ENSG00000274964 | ENSG0000027 | 10.26759669 | 0.341950187 | 0.547101108 | 0.625021924 | 0.531956669 | 0.707466287 |
| ENSG00000274985 | PTCHD3P1 | 2.609772551 | -2.879994535 | 1.356448476 | -2.123187563 | 0.033738138 | 0.085842156 |
| ENSG00000274993 | ENSG0000027 | 4.90131096 | -1.149686778 | 0.822272339 | -1.398182481 | 0.162058276 | 0.309157327 |
| ENSG00000274997 | H2AC12 | 2.466079489 | -0.353171183 | 1.11248215 | -0.317462337 | 0.750892813 | 0.862351997 |
| ENSG00000275011 | ALOXE3P1 | 17.45161605 | -2.928739613 | 0.545296659 | -5.370910612 | 7.83E-08 | 5.12E-07 |
| ENSG00000275023 | MLLT6 | 5073.068444 | 0.372531155 | 0.04942216 | 7.537735204 | 4.78E-14 | 5.34E-13 |
| ENSG00000275029 | HMGB1P24 | 7.277634919 | -0.845266449 | 0.659172593 | -1.282314311 | 0.19973241 | 0.362776047 |
| ENSG00000275052 | PPP4R3B | 3317.10115 | -0.021725004 | 0.048133384 | -0.451350031 | 0.651737292 | 0.797634095 |
| ENSG00000275055 | ENSG0000027 | 8.031827402 | 0.748370949 | 0.642732978 | 1.164357478 | 0.244279172 | 0.42105012 |
| ENSG00000275056 | ENSG0000027 | 3.170533298 | -0.321516146 | 0.990960821 | -0.324448898 | 0.745598205 | 0.859379806 |
| ENSG00000275066 | SYNRG | 1112.071174 | -0.166037677 | 0.070187638 | -2.365625648 | 0.017999634 | 0.049878574 |
| ENSG00000275074 | NUDT18 | 221.973476 | 0.285098264 | 0.129748861 | 2.197308411 | 0.027998427 | 0.073211435 |
| ENSG00000275084 | SNORD91B | 1.705345389 | -2.189480563 | 1.529881763 | -1.431143646 | 0.152389051 | NA |
| ENSG00000275091 | ENSG0000027 | 3.73749088 | 0.96702538 | 0.930119306 | 1.03967886 | 0.298489125 | 0.485007829 |
| ENSG00000275092 | ENSG0000027 | 4.254607214 | -0.342762201 | 0.905794146 | -0.378410704 | 0.705125518 | 0.834997903 |
| ENSG00000275111 | ZNF2 | 305.6839347 | -0.147374501 | 0.114949081 | -1.282085076 | 0.199812804 | 0.362864607 |
| ENSG00000275120 | ZSCAN2-AS1 | 10.40568529 | -0.224466365 | 0.548918462 | -0.408924787 | 0.682594858 | 0.819871514 |
| ENSG00000275131 | PDE4DIPP2 | 218.392561 | -0.22011233 | 0.127027602 | -1.732791349 | 0.083132761 | 0.182337273 |
| ENSG00000275160 | RPL7AP45 | 66.8351867 | -0.769059473 | 0.216595357 | -3.550673859 | 0.000384246 | 0.001549054 |
| ENSG00000275180 | ENSG0000027 | 22.38544204 | 0.086945827 | 0.377496656 | 0.230322113 | 0.817841479 | 0.903530843 |
| ENSG00000275183 | LENG9 | 214.2078357 | 0.417623333 | 0.128968284 | 3.238186326 | 0.001202922 | 0.004438642 |
| ENSG00000275185 | ENSG0000027 | 7.047055045 | 0.395509401 | 0.663905312 | 0.595731641 | 0.551354517 | 0.723242945 |
| ENSG00000275197 | ENSG0000027 | 3.195453193 | -1.024312152 | 1.022804621 | -1.001473919 | 0.316597743 | 0.505410732 |
| ENSG00000275198 | LINC02960 | 34.89774017 | -1.23113287 | 0.313473561 | -3.927389816 | 8.59E-05 | 0.000387475 |
| ENSG00000275202 | ENSG0000027 | 17.71763858 | 0.167821991 | 0.446941002 | 0.375490256 | 0.707295892 | 0.836532294 |
| ENSG00000275212 | LINC02826 | 10.68459548 | -1.719746159 | 0.575664592 | -2.987409998 | 0.002813521 | 0.009600205 |
| ENSG00000275221 | H2AC15 | 33.15556648 | 0.45263845 | 0.30454728 | 1.486266597 | 0.137208618 | 0.27235145 |
| ENSG00000275223 | ENSG0000027 | 33.87610587 | -0.547436139 | 0.311251801 | -1.758820795 | 0.078607952 | 0.174262313 |
| ENSG00000275227 | ENSG0000027 | 2.847622665 | -0.677563825 | 1.048654747 | -0.646126695 | 0.518197309 | 0.695932225 |
| ENSG00000275234 | ENSG0000027 | 261.5806715 | 0.197375713 | 0.116443608 | 1.695032615 | 0.090069234 | 0.19455633 |

| | | | | | | | |
|---|---|---|---|---|---|---|---|
| ENSG00000275236 | ENSG0000027 | 7.156608142 | 0.096017749 | 0.645442697 | 0.148762623 | 0.881740945 | 0.938683755 |
| ENSG00000275263 | ENSG0000027 | 3.349356232 | 0.634984343 | 0.998646401 | 0.635845023 | 0.524877439 | 0.701099828 |
| ENSG00000275291 | RNVU1-26 | 1.856160903 | 0.847272517 | 1.394273117 | 0.607680451 | 0.543399431 | 0.716907232 |
| ENSG00000275318 | ENSG0000027 | 6.050297752 | -1.01362452 | 0.719911333 | -1.407985224 | 0.159135445 | 0.304620087 |
| ENSG00000275322 | ENSG0000027 | 52.72175018 | -0.27096283 | 0.245651644 | -1.103036907 | 0.270011138 | 0.45165929 |
| ENSG00000275329 | ENSG0000027 | 45.67213557 | -0.168241161 | 0.279425844 | -0.60209592 | 0.54711029 | 0.719898477 |
| ENSG00000275332 | ENSG0000027 | 1.583798469 | -0.108565202 | 1.385430509 | -0.07836207 | 0.937540045 | NA |
| ENSG00000275340 | FGD5P1 | 2.438175294 | -0.013317067 | 1.114895319 | -0.011944679 | 0.990469751 | 0.994958761 |
| ENSG00000275342 | PRAG1 | 676.761633 | 0.316045246 | 0.076397339 | 4.136861972 | 3.52E-05 | 0.000168234 |
| ENSG00000275367 | ENSG0000027 | 7.120691353 | 1.279174737 | 0.710022804 | 1.801596696 | 0.071608881 | 0.161715667 |
| ENSG00000275383 | ENSG0000027 | 2.30631287 | 0.647039482 | 1.153511119 | 0.560930425 | 0.574844968 | 0.741214298 |
| ENSG00000275389 | ENSG0000027 | 7.781832169 | 0.893181756 | 0.667414817 | 1.338270793 | 0.180808179 | 0.336121526 |
| ENSG00000275393 | ENSG0000027 | 9.468751527 | 2.303784085 | 0.666601196 | 3.45601553 | 0.000548224 | 0.002154575 |
| ENSG00000275395 | FCGBP | 29.08316989 | -0.779634392 | 0.333391788 | -2.338493087 | 0.019361684 | 0.053173354 |
| ENSG00000275401 | ENSG0000027 | 4.496344524 | 0.105432415 | 0.827886321 | 0.127351319 | 0.898662344 | 0.948793422 |
| ENSG00000275437 | ENSG0000027 | 119.1763576 | 0.194023085 | 0.168630348 | 1.150582249 | 0.249904139 | 0.427980661 |
| ENSG00000275438 | ENSG0000027 | 5.882944346 | 0.286565358 | 0.735609386 | 0.389561856 | 0.696860563 | 0.829719188 |
| ENSG00000275441 | ENSG0000027 | 17.93823543 | 0.769174324 | 0.420693164 | 1.828349947 | 0.067497048 | 0.154006593 |
| ENSG00000275450 | PTGER4P2-CD | 3.381331157 | 0.262211942 | 1.043129792 | 0.251370389 | 0.801527759 | 0.894398618 |
| ENSG00000275454 | ENSG0000027 | 17.86495808 | -0.001555915 | 0.413185809 | -0.003765655 | 0.996995449 | 0.998257414 |
| ENSG00000275457 | ENSG0000027 | 18.91525939 | -0.216034892 | 0.403094327 | -0.535941285 | 0.59199913 | 0.754408176 |
| ENSG00000275464 | ENSG0000027 | 336.9943126 | 0.308268434 | 0.102051538 | 3.020713258 | 0.002521801 | 0.008682291 |
| ENSG00000275468 | ENSG0000027 | 4.58000244 | 0.642936914 | 0.831295268 | 0.773415824 | 0.439276328 | 0.628751417 |
| ENSG00000275479 | ENSG0000027 | 5.413970099 | 0.248839447 | 0.738341514 | 0.337024862 | 0.736098158 | 0.854386347 |
| ENSG00000275481 | ENSG0000027 | 16.44045238 | -1.025269649 | 0.44150784 | -2.322200321 | 0.020222152 | 0.05518681 |
| ENSG00000275484 | ENSG0000027 | 6.838062234 | 1.174012656 | 0.722631253 | 1.624635871 | 0.104240169 | 0.219221624 |
| ENSG00000275485 | ENSG0000027 | 2.592340327 | 1.01707453 | 1.199348714 | 0.848022362 | 0.396425517 | 0.589333248 |
| ENSG00000275491 | LINC01730 | 7.68052575 | -1.203358821 | 0.646455499 | -1.861472016 | 0.062677549 | 0.144606555 |
| ENSG00000275494 | ENSG0000027 | 29.48356534 | -0.46650386 | 0.343761126 | -1.357058214 | 0.174762714 | 0.327858739 |
| ENSG00000275527 | ENSG0000027 | 15.3564149 | 0.68365585 | 0.463114698 | 1.476212812 | 0.139886771 | 0.276257105 |
| ENSG00000275532 | ENSG0000027 | 15.76722531 | 0.398512958 | 0.459936723 | 0.866451706 | 0.386242499 | 0.578846471 |
| ENSG00000275549 | STPG3-AS1 | 35.42563279 | -0.222534503 | 0.305608517 | -0.728168524 | 0.466510431 | 0.65246612 |
| ENSG00000275552 | ENSG0000027 | 1.887438732 | 0.071700586 | 1.317906191 | 0.054404924 | 0.956612556 | 0.980198858 |

| | | | | | | | |
|---|---|---|---|---|---|---|---|
| ENSG00000275576 | ENSG0000027 | 2.723743824 | -0.943507163 | 1.094519972 | -0.862028274 | 0.388671959 | 0.58132248 |
| ENSG00000275580 | ENSG0000027 | 4.564229508 | -0.356928196 | 0.821189072 | -0.434648009 | 0.663817941 | 0.805985984 |
| ENSG00000275601 | ENSG0000027 | 8.523964436 | -0.086309427 | 0.59764427 | -0.144416054 | 0.88517194 | 0.940548288 |
| ENSG00000275632 | ENSG0000027 | 9.0747935 | 0.560737177 | 0.584557631 | 0.959250461 | 0.337432585 | 0.528690482 |
| ENSG00000275638 | ENSG0000027 | 2.222612968 | 1.732788018 | 1.293382116 | 1.339734017 | 0.180331834 | 0.335426277 |
| ENSG00000275645 | ENSG0000027 | 43.37481446 | 0.188707649 | 0.276761554 | 0.681841991 | 0.495338869 | 0.677664221 |
| ENSG00000275665 | ENSG0000027 | 13.37889781 | -0.35576635 | 0.52527176 | -0.677299594 | 0.498215889 | 0.679962588 |
| ENSG00000275700 | AATF | 1958.831813 | 0.186108774 | 0.053352391 | 3.488293035 | 0.000486115 | 0.001928654 |
| ENSG00000275703 | ENSG0000027 | 1.80938498 | 0.660784466 | 1.357202456 | 0.486872436 | 0.626348734 | 0.779566329 |
| ENSG00000275709 | ENSG0000027 | 3.752121557 | -0.465097178 | 0.929771868 | -0.5002272 | 0.616915108 | 0.772972842 |
| ENSG00000275710 | ENSG0000027 | 2.717155811 | 1.474255492 | 1.171666825 | 1.258254874 | 0.208299595 | 0.374751426 |
| ENSG00000275713 | H2BC9 | 146.1817612 | 0.413714387 | 0.154196352 | 2.683036151 | 0.00729571 | 0.02254284 |
| ENSG00000275714 | H3C1 | 1.697343935 | 4.113616272 | 1.659210903 | 2.479260631 | 0.013165506 | NA |
| ENSG00000275718 | CCL15 | 2.960165354 | 2.248482318 | 1.186307503 | 1.895362131 | 0.058044442 | 0.135774396 |
| ENSG00000275719 | ENSG0000027 | 102.3783996 | -0.697103417 | 0.190447586 | -3.660342631 | 0.000251878 | 0.001048084 |
| ENSG00000275720 | ENSG0000027 | 1.601297235 | -0.08435445 | 1.438760594 | -0.058629942 | 0.953246861 | NA |
| ENSG00000275734 | ENSG0000027 | 66.23345904 | 1.094436182 | 0.240333013 | 4.553832078 | 5.27E-06 | 2.81E-05 |
| ENSG00000275741 | ENSG0000027 | 2.193755361 | -1.371515362 | 1.259872951 | -1.088614023 | 0.27632413 | 0.458873284 |
| ENSG00000275759 | ENSG0000027 | 2.0169303 | -1.763950346 | 1.393730581 | -1.265632232 | 0.205644781 | 0.371076164 |
| ENSG00000275763 | ZNF516-DT | 6.013448081 | 1.333453339 | 0.764381832 | 1.744485914 | 0.081074396 | 0.178608147 |
| ENSG00000275764 | ENSG0000027 | 65.45093674 | 0.118721964 | 0.218980843 | 0.54215685 | 0.587710458 | 0.751132537 |
| ENSG00000275765 | ENSG0000027 | 8.205583135 | 0.081669623 | 0.656355421 | 0.12442896 | 0.900975645 | 0.95013935 |
| ENSG00000275793 | RIMBP3 | 36.75547185 | -0.75675262 | 0.299251313 | -2.528819713 | 0.01144468 | 0.03347029 |
| ENSG00000275807 | ENSG0000027 | 16.21170547 | 0.436631945 | 0.438903519 | 0.994824436 | 0.319821659 | 0.509352185 |
| ENSG00000275812 | ENSG0000027 | 6.141922273 | -0.451992122 | 0.718166529 | -0.629369517 | 0.52910717 | 0.704984519 |
| ENSG00000275835 | TUBGCP5 | 604.0227389 | -0.043943818 | 0.083499474 | -0.526276586 | 0.598696049 | 0.75887321 |
| ENSG00000275846 | ENSG0000027 | 1.534196546 | 2.855510509 | 1.744847389 | 1.636538833 | 0.101726865 | NA |
| ENSG00000275854 | ENSG0000027 | 8.741033029 | 0.077792017 | 0.582398229 | 0.133571864 | 0.893741134 | 0.945700277 |
| ENSG00000275857 | ENSG0000027 | 42.28261229 | -0.409091978 | 0.30172528 | -1.355842565 | 0.175149265 | 0.328288597 |
| ENSG00000275880 | NAXD-AS1 | 23.73701709 | 0.213140869 | 0.372829479 | 0.571684594 | 0.567535674 | 0.73534416 |
| ENSG00000275882 | IKBKGP1 | 5.108102356 | -1.195768969 | 0.817811416 | -1.462157347 | 0.143698097 | 0.282108202 |
| ENSG00000275888 | ENSG0000027 | 8.613881945 | 0.026241131 | 0.589413371 | 0.04452076 | 0.964489304 | 0.983900556 |
| ENSG00000275894 | ENSG0000027 | 3.678353338 | -0.208085491 | 0.89539648 | -0.232394806 | 0.816231383 | 0.902639217 |

| | | | | | | | |
|---|---|---|---|---|---|---|---|
| ENSG00000275897 | ENSG0000027 | 8.342405973 | -1.937209623 | 0.673697906 | -2.875487078 | 0.004034047 | 0.013278954 |
| ENSG00000275902 | ENSG0000027 | 77.3552336 | 0.564103605 | 0.208562133 | 2.704726874 | 0.006836057 | 0.02124345 |
| ENSG00000275910 | ENSG0000027 | 4.649413396 | -0.146551849 | 0.838450999 | -0.174788806 | 0.861245584 | 0.927818808 |
| ENSG00000275936 | ENSG0000027 | 4.41387075 | 0.02068697 | 0.93750614 | 0.022065956 | 0.982395343 | 0.991820665 |
| ENSG00000275944 | ENSG0000027 | 14.42376829 | 5.310954094 | 1.0861676 | 4.889626699 | 1.01E-06 | 5.84E-06 |
| ENSG00000275945 | EIF3FP1 | 1.662839854 | -0.806208662 | 1.470741636 | -0.54816471 | 0.583578811 | NA |
| ENSG00000275963 | ENSG0000027 | 2.221652501 | 1.134401266 | 1.200453816 | 0.944977016 | 0.34467061 | 0.535542116 |
| ENSG00000275964 | ENSG0000027 | 98.35019816 | 0.573717474 | 0.185711677 | 3.08929134 | 0.002006346 | 0.007049406 |
| ENSG00000275993 | ENSG0000027 | 257.9233933 | -0.372330839 | 0.116516305 | -3.195525632 | 0.001395764 | 0.005075103 |
| ENSG00000276007 | ENSG0000027 | 78.76578783 | -0.120698067 | 0.207652696 | -0.581249699 | 0.561072176 | 0.730103564 |
| ENSG00000276023 | DUSP14 | 498.2796784 | 0.025872441 | 0.088308837 | 0.292976805 | 0.769539881 | 0.874269259 |
| ENSG00000276043 | UHRF1 | 930.5483685 | -0.059240493 | 0.072785394 | -0.813906332 | 0.415698613 | 0.607789402 |
| ENSG00000276045 | ORAI1 | 731.4814754 | 0.112486183 | 0.081575157 | 1.378926953 | 0.167917277 | 0.31787608 |
| ENSG00000276058 | STMN1P1 | 1.911107166 | 0.602350298 | 1.423486209 | 0.423151481 | 0.672184743 | 0.812215855 |
| ENSG00000276071 | ENSG0000027 | 4.619179229 | 1.504514123 | 0.876058359 | 1.717367464 | 0.085912052 | 0.187233753 |
| ENSG00000276075 | ENSG0000027 | 13.49350662 | -0.28699451 | 0.483918252 | -0.593064033 | 0.5531383 | 0.724671193 |
| ENSG00000276101 | ENSG0000027 | 16.10113757 | 0.658225667 | 0.447865278 | 1.469695688 | 0.141644191 | 0.279056251 |
| ENSG00000276103 | U4 | 3.64089014 | 0.774524518 | 0.957597137 | 0.808820837 | 0.418618209 | 0.61039531 |
| ENSG00000276107 | THBS1-IT1 | 29.27989823 | 1.321483981 | 0.345568421 | 3.824087798 | 0.000131257 | 0.000574078 |
| ENSG00000276110 | ENSG0000027 | 2.218282872 | -0.13434663 | 1.18218428 | -0.113642714 | 0.909521026 | 0.954587195 |
| ENSG00000276116 | FUT8-AS1 | 399.9699301 | -0.197477483 | 0.098133636 | -2.012332268 | 0.044184923 | 0.107818544 |
| ENSG00000276131 | ENSG0000027 | 19.29061145 | 0.385334792 | 0.41496376 | 0.928598662 | 0.353097115 | 0.544577641 |
| ENSG00000276136 | ENSG0000027 | 2.366867856 | 0.169512079 | 1.131329628 | 0.149834385 | 0.88089528 | 0.938252557 |
| ENSG00000276141 | WHAMMP3 | 5.661371754 | -0.51029902 | 0.757308651 | -0.673832287 | 0.500417945 | 0.681707808 |
| ENSG00000276148 | ENSG0000027 | 4.639912404 | -1.007214005 | 0.823008388 | -1.223819854 | 0.221020193 | 0.39155172 |
| ENSG00000276168 | RN7SL1 | 233.1916328 | -0.374235502 | 0.13154491 | -2.844925761 | 0.004442177 | 0.014495527 |
| ENSG00000276180 | H4C9 | 103.4596025 | 0.202649621 | 0.185621105 | 1.091738033 | 0.274948263 | 0.4570848 |
| ENSG00000276216 | ENSG0000027 | 7.066803164 | 0.030425087 | 0.651565608 | 0.046695355 | 0.962756033 | 0.982889182 |
| ENSG00000276223 | ENSG0000027 | 24.14077838 | -0.187364283 | 0.351021728 | -0.533768334 | 0.593501825 | 0.755562373 |
| ENSG00000276234 | TADA2A | 460.3385875 | -0.02908583 | 0.093553501 | -0.310900494 | 0.755876269 | 0.865149344 |
| ENSG00000276248 | ENSG0000027 | 5.046670099 | -0.228103152 | 0.777364513 | -0.293431393 | 0.769192433 | 0.874004379 |
| ENSG00000276250 | ENSG0000027 | 3.155480493 | 0.563708729 | 0.985801471 | 0.571827844 | 0.567438612 | 0.735301461 |
| ENSG00000276255 | LINC02809 | 3.22074682 | -0.405902501 | 1.009911762 | -0.401918778 | 0.687743803 | 0.823383061 |

| | | | | | | | |
|---|---|---|---|---|---|---|---|
| ENSG00000276261 | ENSG0000027 | 1.626462304 | 2.879132443 | 1.634463915 | 1.761514841 | 0.078151303 | NA |
| ENSG00000276272 | ENSG0000027 | 5.985626674 | 0.296204473 | 0.720710008 | 0.410989815 | 0.681080003 | 0.818990574 |
| ENSG00000276278 | ENSG0000027 | 12.85772219 | -0.08344072 | 0.493547934 | -0.169063053 | 0.865747046 | 0.9302381 |
| ENSG00000276290 | RBBP4P3 | 2.880656953 | -0.145691633 | 1.030604499 | -0.141365221 | 0.887581428 | 0.942116121 |
| ENSG00000276291 | FRG1HP | 139.2050424 | 0.4025961 | 0.157690352 | 2.553080097 | 0.010677494 | 0.031475419 |
| ENSG00000276292 | ENSG0000027 | 3.825365954 | -0.563367637 | 0.884580072 | -0.636875795 | 0.52420575 | 0.700569759 |
| ENSG00000276293 | PIP4K2B | 4665.930194 | -0.08588049 | 0.046420317 | -1.850062532 | 0.064304537 | 0.14779542 |
| ENSG00000276317 | ENSG0000027 | 2.450868383 | -1.600770478 | 1.185172935 | -1.350664051 | 0.176803073 | 0.330558825 |
| ENSG00000276334 | ENSG0000027 | 6.318162852 | 1.437773503 | 0.808260384 | 1.778849405 | 0.075264455 | 0.168363393 |
| ENSG00000276337 | ENSG0000027 | 2.366818408 | -0.169158791 | 1.182484547 | -0.143053701 | 0.88624777 | 0.9413587 |
| ENSG00000276376 | U7 | 3.353144207 | -0.729553601 | 1.047034423 | -0.696780912 | 0.485939913 | 0.669580266 |
| ENSG00000276380 | UBE2NL | 11.62157962 | -0.272808128 | 0.528419655 | -0.516271726 | 0.605664641 | 0.76434448 |
| ENSG00000276384 | ENSG0000027 | 3.456509137 | 0.647837512 | 0.966105942 | 0.670565705 | 0.502497236 | 0.683544035 |
| ENSG00000276390 | ENSG0000027 | 2.865604148 | 0.815270614 | 1.049833698 | 0.776571199 | 0.437411799 | 0.626920679 |
| ENSG00000276404 | MIR6835 | 3.363424454 | -1.028518987 | 1.030497714 | -0.998079834 | 0.318240648 | 0.507399162 |
| ENSG00000276408 | ENSG0000027 | 6.796871546 | 0.530709913 | 0.672174595 | 0.78954176 | 0.429795432 | 0.620133796 |
| ENSG00000276434 | ENSG0000027 | 3.697608888 | 0.834478311 | 1.044697682 | 0.798774923 | 0.424420934 | 0.615226766 |
| ENSG00000276445 | ENSG0000027 | 3.260824996 | 0.092108982 | 0.974911507 | 0.094479326 | 0.924728404 | 0.963976624 |
| ENSG00000276449 | ENSG0000027 | 14.26725633 | 0.921153307 | 0.49320542 | 1.86768691 | 0.061805724 | 0.142997813 |
| ENSG00000276488 | ENSG0000027 | 3.976174331 | -0.427732198 | 0.892548571 | -0.479225683 | 0.631778085 | 0.782942434 |
| ENSG00000276505 | ENSG0000027 | 2.522835662 | -0.403071774 | 1.08215387 | -0.372471776 | 0.709541622 | 0.838150608 |
| ENSG00000276517 | ENSG0000027 | 10.06246133 | -0.058808793 | 0.546855592 | -0.107539895 | 0.914360677 | 0.95756375 |
| ENSG00000276524 | LINC03065 | 43.08974769 | -1.611538411 | 0.293589635 | -5.489084829 | 4.04E-08 | 2.73E-07 |
| ENSG00000276529 | ENSG0000027 | 82.18859557 | 0.639245307 | 0.201318456 | 3.1752941 | 0.001496847 | 0.005404954 |
| ENSG00000276531 | SNX19P4 | 2.449059981 | -2.808069636 | 1.39493976 | -2.013040073 | 0.044110414 | 0.107682547 |
| ENSG00000276533 | ENSG0000027 | 2.310609544 | 1.077660409 | 1.205211985 | 0.894166688 | 0.371232713 | 0.563924535 |
| ENSG00000276535 | ENSG0000027 | 6.593156209 | 0.390867106 | 0.676438689 | 0.577830796 | 0.563378362 | 0.732154597 |
| ENSG00000276545 | PCDHGB9P | 2.078027711 | 0.731013358 | 1.296369219 | 0.563892869 | 0.572827044 | 0.739651716 |
| ENSG00000276547 | PCDHGB5 | 538.9473386 | 0.885183649 | 0.08979167 | 9.858193461 | 6.32E-23 | 1.25E-21 |
| ENSG00000276550 | HERC2P2 | 1413.755787 | 0.236140907 | 0.137203888 | 1.72109486 | 0.085233612 | 0.1861088 |
| ENSG00000276564 | ENSG0000027 | 9.365194873 | 0.556928958 | 0.590916233 | 0.942483769 | 0.345945017 | 0.536831407 |
| ENSG00000276570 | ENSG0000027 | 71.78287203 | -0.123877086 | 0.210565601 | -0.588306377 | 0.556326666 | 0.72693858 |
| ENSG00000276571 | ENSG0000027 | 3.52899961 | 1.951209043 | 1.042811204 | 1.871104793 | 0.061330557 | 0.142113433 |

| | | | | | | | |
|---|---|---|---|---|---|---|---|
| ENSG00000276600 | RAB7B | 24.39973605 | 3.694404633 | 0.541756251 | 6.819311498 | 9.15E-12 | 8.60E-11 |
| ENSG00000276603 | ENSG0000027 | 6.571417899 | -0.825133 | 0.715933918 | -1.152526761 | 0.24910468 | 0.427057869 |
| ENSG00000276644 | DACH1 | 44.90875538 | -1.645975796 | 0.285345289 | -5.768365068 | 8.00E-09 | 5.81E-08 |
| ENSG00000276651 | ENSG0000027 | 2.758250957 | 0.366375676 | 1.050934818 | 0.348618839 | 0.727375483 | 0.848598564 |
| ENSG00000276667 | ENSG0000027 | 2.006547153 | -0.200189136 | 1.201422595 | -0.166626745 | 0.867663749 | 0.930973171 |
| ENSG00000276672 | ENSG0000027 | 54.53981803 | -0.264644135 | 0.241564317 | -1.095543159 | 0.273278748 | 0.455200006 |
| ENSG00000276698 | ENSG0000027 | 25.52897223 | 0.087176802 | 0.342998062 | 0.254161209 | 0.799371017 | 0.892991047 |
| ENSG00000276710 | CSPG4P10 | 7.396788228 | -0.238842755 | 0.638455547 | -0.374094573 | 0.708333956 | 0.83737083 |
| ENSG00000276712 | MIR7111 | 5.640981515 | 0.478062527 | 0.750780175 | 0.636754329 | 0.524284879 | 0.700634692 |
| ENSG00000276727 | ENSG0000027 | 6.780661667 | -0.008874995 | 0.674207124 | -0.013163604 | 0.989497267 | 0.994395554 |
| ENSG00000276728 | ENSG0000027 | 46.13260304 | 0.180019517 | 0.267937685 | 0.671870838 | 0.501665927 | 0.682818106 |
| ENSG00000276744 | ENSG0000027 | 11.25158618 | -0.594085343 | 0.538818379 | -1.102570674 | 0.270213651 | 0.451932135 |
| ENSG00000276755 | ENSG0000027 | 2.520920357 | 0.995214092 | 1.154218228 | 0.862240838 | 0.388555001 | 0.581286796 |
| ENSG00000276757 | RN7SL192P | 3.523821578 | 0.454443337 | 0.951770932 | 0.477471334 | 0.633026529 | 0.78368206 |
| ENSG00000276791 | ENSG0000027 | 79.01008188 | -0.583196901 | 0.207241897 | -2.814087827 | 0.004891586 | 0.015797971 |
| ENSG00000276805 | ENSG0000027 | 231.1330834 | -0.979583019 | 0.122812935 | -7.976220273 | 1.51E-15 | 1.88E-14 |
| ENSG00000276809 | ENSG0000027 | 1.971560762 | 0.26437471 | 1.233052422 | 0.214406707 | 0.830229911 | 0.909960543 |
| ENSG00000276832 | CDRT15P6 | 5.331337242 | -0.30790912 | 0.781905915 | -0.393793056 | 0.693733831 | 0.827371387 |
| ENSG00000276846 | ENSG0000027 | 60.95463842 | 0.445553113 | 0.231620846 | 1.923631314 | 0.054400813 | 0.128827775 |
| ENSG00000276851 | ENSG0000027 | 1.620281495 | 1.023679263 | 1.387078762 | 0.738010913 | 0.460507818 | NA |
| ENSG00000276853 | ENSG0000027 | 3.809751323 | -0.299036453 | 0.887206397 | -0.337053987 | 0.736076203 | 0.854386347 |
| ENSG00000276855 | ENSG0000027 | 132.6734818 | 0.067768662 | 0.155624463 | 0.435462785 | 0.663226546 | 0.805549688 |
| ENSG00000276863 | ENSG0000027 | 4.141255304 | 0.368525866 | 0.853327984 | 0.431868957 | 0.665836658 | 0.807352024 |
| ENSG00000276900 | ENSG0000027 | 38.54807848 | 0.796465584 | 0.294042554 | 2.708674551 | 0.006755257 | 0.021023686 |
| ENSG00000276903 | H2AC16 | 2.319055732 | -0.204315523 | 1.137920153 | -0.179551722 | 0.857504509 | 0.926087485 |
| ENSG00000276931 | ENSG0000027 | 8.606348285 | 0.237333227 | 0.593734806 | 0.399729348 | 0.689355874 | 0.824478513 |
| ENSG00000276934 | ENSG0000027 | 1.660001508 | 0.41487127 | 1.395322035 | 0.297330121 | 0.766214487 | NA |
| ENSG00000276945 | ENSG0000027 | 3.196101893 | -0.367169458 | 0.971561095 | -0.377917004 | 0.70549225 | 0.835259714 |
| ENSG00000276952 | ENSG0000027 | 12.49231784 | -1.340975967 | 0.520575161 | -2.57595073 | 0.009996489 | 0.029716141 |
| ENSG00000276966 | H4C5 | 35.45754579 | 0.113725574 | 0.293027648 | 0.388105271 | 0.697938132 | 0.83051669 |
| ENSG00000276997 | ENSG0000027 | 7.304522144 | -0.358987937 | 0.644599256 | -0.556916463 | 0.577584506 | 0.743451272 |
| ENSG00000277007 | ENSG0000027 | 2.278932194 | 2.308717744 | 1.413703339 | 1.633099166 | 0.102448142 | 0.216206106 |
| ENSG00000277010 | ENSG0000027 | 7.304661842 | -0.458209904 | 0.645069932 | -0.710325938 | 0.477502039 | 0.662307079 |

| | | | | | | | |
|---|---|---|---|---|---|---|---|
| ENSG00000277013 | SUNO1 | 54.12962255 | -1.855594025 | 0.267210982 | -6.944303007 | 3.80E-12 | 3.70E-11 |
| ENSG00000277020 | ENSG0000027 | 2.550277252 | -0.441048165 | 1.114125894 | -0.395869235 | 0.692201498 | 0.826433973 |
| ENSG00000277022 | ENSG0000027 | 283.5594208 | -2.308638829 | 0.13243264 | -17.43255161 | 4.67E-68 | 4.01E-66 |
| ENSG00000277047 | ENSG0000027 | 9.705969187 | 0.502406327 | 0.561827066 | 0.894236603 | 0.371195312 | 0.563924535 |
| ENSG00000277053 | GTF2IP1 | 246.2450982 | 0.325155116 | 0.119911214 | 2.711632255 | 0.006695283 | 0.020856842 |
| ENSG00000277072 | STAG3L2 | 138.9658866 | 0.711097733 | 0.161205964 | 4.411113067 | 1.03E-05 | 5.31E-05 |
| ENSG00000277075 | H2AC8 | 135.6953759 | 0.279465173 | 0.157041455 | 1.779563065 | 0.075147495 | 0.16815095 |
| ENSG00000277103 | ENSG0000027 | 3.067515088 | -0.540365787 | 1.004100094 | -0.538159283 | 0.590467088 | 0.753103247 |
| ENSG00000277112 | ANKRD20A21I | 15.74991699 | -0.765656936 | 0.455218812 | -1.681953634 | 0.092577831 | 0.199075314 |
| ENSG00000277117 | ENSG0000027 | 93.66565148 | -0.004197582 | 0.18355223 | -0.022868598 | 0.981755089 | 0.991627389 |
| ENSG00000277142 | LINC00235 | 94.28971914 | 0.88729749 | 0.191942787 | 4.622718598 | 3.79E-06 | 2.05E-05 |
| ENSG00000277147 | LINC00869 | 71.90038302 | 0.074569233 | 0.209100707 | 0.356618752 | 0.721377237 | 0.845297738 |
| ENSG00000277149 | TYW1B | 43.26808336 | 0.218841695 | 0.269169113 | 0.813026771 | 0.416202709 | 0.608288109 |
| ENSG00000277150 | F8A3 | 5.115203155 | 0.071258639 | 0.772978301 | 0.092187115 | 0.926549375 | 0.964663921 |
| ENSG00000277151 | ENSG0000027 | 2.489224404 | -0.752332237 | 1.103771896 | -0.681601189 | 0.495491163 | 0.677781102 |
| ENSG00000277152 | ENSG0000027 | 1.7129471 | 1.219302036 | 1.532299966 | 0.795733253 | 0.426187095 | NA |
| ENSG00000277156 | TOMM40P1 | 18.57377511 | 0.487079868 | 0.414838355 | 1.174143765 | 0.240337446 | 0.416168827 |
| ENSG00000277157 | H4C4 | 10.60034835 | -0.623240159 | 0.547934079 | -1.137436387 | 0.255355903 | 0.434762737 |
| ENSG00000277159 | ENSG0000027 | 8.10873081 | -1.084172966 | 0.627876749 | -1.726728961 | 0.084216352 | 0.184273475 |
| ENSG00000277161 | PIGW | 464.2853791 | -0.080292861 | 0.092709107 | -0.866073076 | 0.386450089 | 0.578997258 |
| ENSG00000277170 | ENSG0000027 | 2.897718771 | 0.157473921 | 1.012013026 | 0.155604639 | 0.876344668 | 0.93573969 |
| ENSG00000277182 | ENSG0000027 | 37.20449759 | 0.127795037 | 0.295329895 | 0.43271961 | 0.665218482 | 0.806901013 |
| ENSG00000277186 | ENSG0000027 | 4.213512068 | 0.274048536 | 0.841728439 | 0.325578325 | 0.744743413 | 0.858858531 |
| ENSG00000277200 | CCDC92B | 21.28762666 | -0.244094017 | 0.38271915 | -0.637788876 | 0.523611122 | 0.700001783 |
| ENSG00000277215 | SPANXA2-OT1 | 94.04505279 | 0.623540446 | 0.194147127 | 3.211690318 | 0.001319565 | 0.004829424 |
| ENSG00000277218 | ENSG0000027 | 4.530464377 | -1.880037738 | 0.992148211 | -1.89491622 | 0.058103502 | 0.135898688 |
| ENSG00000277224 | H2BC7 | 53.75685062 | 0.17638885 | 0.240270868 | 0.734124997 | 0.46287256 | 0.649357665 |
| ENSG00000277232 | GTSE1-DT | 26.66843681 | 0.211156561 | 0.342639108 | 0.616265206 | 0.537719499 | 0.712733882 |
| ENSG00000277258 | PCGF2 | 2215.35421 | 0.068635415 | 0.05556826 | 1.235155023 | 0.216772815 | 0.386234437 |
| ENSG00000277268 | LHX1-DT | 118.4965812 | 1.011518202 | 0.175283423 | 5.770757914 | 7.89E-09 | 5.73E-08 |
| ENSG00000277283 | RAB35-AS1 | 156.8748166 | 0.392288399 | 0.143275858 | 2.737993722 | 0.006181526 | 0.019471062 |
| ENSG00000277287 | ENSG0000027 | 18.89805014 | 1.008475868 | 0.413265911 | 2.440259022 | 0.014676734 | 0.041679164 |
| ENSG00000277297 | ATP5F1AP10 | 1.586233289 | 1.047844246 | 1.478325821 | 0.70880467 | 0.478445699 | NA |

| | | | | | | | |
|---|---|---|---|---|---|---|---|
| ENSG00000277301 | ENSG0000027 | 53.11858976 | -3.047787354 | 0.313109859 | -9.733923313 | 2.16E-22 | 4.13E-21 |
| ENSG00000277310 | ENSG0000027 | 2.673651177 | 1.508568113 | 1.141133434 | 1.321990985 | 0.186171153 | 0.343530677 |
| ENSG00000277324 | ENSG0000027 | 11.94727988 | 0.596274214 | 0.531529249 | 1.121808846 | 0.261943724 | 0.44229273 |
| ENSG00000277363 | SRCIN1 | 779.1529453 | 0.413141282 | 0.075456491 | 5.475225197 | 4.37E-08 | 2.94E-07 |
| ENSG00000277368 | ENSG0000027 | 13.62803196 | 0.141357397 | 0.508701516 | 0.277878859 | 0.781105353 | 0.881298289 |
| ENSG00000277369 | ENSG0000027 | 8.390028731 | -0.04566009 | 0.603118396 | -0.075706678 | 0.939652463 | 0.971233034 |
| ENSG00000277383 | ENSG0000027 | 14.74392076 | 0.12033056 | 0.456374067 | 0.263666517 | 0.792036899 | 0.888128187 |
| ENSG00000277386 | ENSG0000027 | 2.160630824 | -1.381196173 | 1.302741388 | -1.060222838 | 0.289043233 | 0.473212373 |
| ENSG00000277399 | GPR179 | 16.76001569 | -0.774706683 | 0.448521568 | -1.727245105 | 0.084123653 | 0.184140898 |
| ENSG00000277406 | SEC22B4P | 17.56338654 | 0.275522638 | 0.41611037 | 0.662138358 | 0.507882555 | 0.688024202 |
| ENSG00000277423 | ENSG0000027 | 1.886900393 | 0.792001734 | 1.333677838 | 0.593847863 | 0.552613873 | 0.724232303 |
| ENSG00000277443 | MARCKS | 3713.267259 | -1.0403391 | 0.072078346 | -14.43344858 | 3.19E-47 | 1.62E-45 |
| ENSG00000277447 | ENSG0000027 | 3.047678888 | -0.196713985 | 1.010648226 | -0.1946414 | 0.845673691 | 0.919043732 |
| ENSG00000277449 | CEBPB-AS1 | 80.54381263 | -0.323153397 | 0.207261468 | -1.559158101 | 0.118958965 | 0.243879598 |
| ENSG00000277459 | ENSG0000027 | 4.940737809 | 0.17416407 | 0.786708432 | 0.22138325 | 0.824794033 | 0.907715593 |
| ENSG00000277462 | ZNF670 | 221.2466172 | -0.389282402 | 0.123272343 | -3.157905435 | 0.001589071 | 0.005710996 |
| ENSG00000277463 | ENSG0000027 | 6.919432385 | -0.131124933 | 0.669562909 | -0.195836614 | 0.844738051 | 0.918358888 |
| ENSG00000277476 | ENSG0000027 | 323.0591916 | 0.859873299 | 0.109586591 | 7.846519312 | 4.28E-15 | 5.17E-14 |
| ENSG00000277496 | SLCO4A1-AS2 | 8.415148031 | -0.211672831 | 0.616591921 | -0.34329485 | 0.731376655 | 0.851231886 |
| ENSG00000277504 | ENSG0000027 | 12.92768314 | 1.731586894 | 0.534131867 | 3.241871529 | 0.001187475 | 0.004385878 |
| ENSG00000277534 | ENSG0000027 | 108.3104928 | -0.129227988 | 0.181131405 | -0.713448825 | 0.47556807 | 0.660703217 |
| ENSG00000277548 | ENSG0000027 | 1.911955101 | 0.883518893 | 1.309646559 | 0.674623918 | 0.499914733 | 0.681205745 |
| ENSG00000277559 | ENSG0000027 | 42.39925045 | -0.23693292 | 0.279932536 | -0.846392935 | 0.397333579 | 0.590236856 |
| ENSG00000277566 | ENSG0000027 | 1.747998084 | -0.287001989 | 1.357763713 | -0.21137845 | 0.832591972 | NA |
| ENSG00000277589 | ENSG0000027 | 1.650534264 | 0.405263448 | 1.349737096 | 0.30025362 | 0.763983708 | NA |
| ENSG00000277597 | ENSG0000027 | 5.084750099 | 0.041039544 | 0.784677508 | 0.05230116 | 0.958288729 | 0.981036836 |
| ENSG00000277619 | ENSG0000027 | 1.866653531 | 1.166829377 | 1.439049055 | 0.810833636 | 0.417461216 | 0.609583723 |
| ENSG00000277639 | CHD9NB | 10.89458534 | -0.589673061 | 0.529044319 | -1.114600498 | 0.265021657 | 0.446045253 |
| ENSG00000277654 | ENSG0000027 | 93.75703335 | 0.099189622 | 0.186720914 | 0.531218597 | 0.595267308 | 0.756595136 |
| ENSG00000277662 | ENSG0000027 | 3.816396571 | -0.090294374 | 0.899526593 | -0.100379883 | 0.92004274 | 0.961195531 |
| ENSG00000277687 | ENSG0000027 | 6.002632464 | -0.528110995 | 0.733434462 | -0.72005206 | 0.471492943 | 0.657230851 |
| ENSG00000277688 | ENSG0000027 | 1.758324728 | 1.20370047 | 1.376632891 | 0.874380147 | 0.381911265 | NA |
| ENSG00000277693 | ENSG0000027 | 18.59281901 | -0.719806518 | 0.445106836 | -1.617154489 | 0.105844937 | 0.22190485 |

| | | | | | | | |
|---|---|---|---|---|---|---|---|
| ENSG00000277701 | ENSG0000027 | 2.029201669 | 1.023383912 | 1.25985259 | 0.812304487 | 0.416616937 | 0.60862208 |
| ENSG00000277715 | ENSG0000027 | 9.887983421 | 1.114838209 | 0.574282764 | 1.941270538 | 0.052225471 | 0.124343779 |
| ENSG00000277728 | ENSG0000027 | 7.29613904 | -0.017676132 | 0.687986475 | -0.025692557 | 0.97950256 | 0.990775111 |
| ENSG00000277741 | GOLGA6L17P | 3.267571311 | -0.097844683 | 0.97496204 | -0.100357429 | 0.920060566 | 0.961195531 |
| ENSG00000277744 | ENSG0000027 | 13.48587895 | -0.503712292 | 0.472002251 | -1.067181969 | 0.285889673 | 0.469663483 |
| ENSG00000277767 | ENSG0000027 | 4.04796328 | 0.229376972 | 0.943800108 | 0.243035543 | 0.807977863 | 0.898318819 |
| ENSG00000277775 | H3C7 | 9.321467671 | 0.228851811 | 0.579680237 | 0.394789742 | 0.692998066 | 0.826907354 |
| ENSG00000277778 | PGM5P2 | 2.356902751 | -0.621003301 | 1.159383434 | -0.535632373 | 0.59221265 | 0.754616353 |
| ENSG00000277782 | ENSG0000027 | 44.35514006 | -0.149005008 | 0.274411993 | -0.542997433 | 0.58713157 | 0.750852838 |
| ENSG00000277791 | PSMB3 | 1841.068425 | -0.004705573 | 0.057154675 | -0.082330495 | 0.934383905 | 0.968469969 |
| ENSG00000277801 | ENSG0000027 | 1.793736926 | -0.417427598 | 1.278370191 | -0.326531079 | 0.744022579 | 0.858490754 |
| ENSG00000277840 | ENSG0000027 | 2.788391194 | -0.128292745 | 1.107810188 | -0.115807515 | 0.907805094 | 0.953746227 |
| ENSG00000277855 | ENSG0000027 | 7.228138148 | 0.463284343 | 0.657957342 | 0.704125197 | 0.48135481 | 0.665759034 |
| ENSG00000277873 | ENSG0000027 | 6.481962692 | -0.694312177 | 0.725958062 | -0.956408108 | 0.33886609 | 0.530045518 |
| ENSG00000277879 | ENSG0000027 | 23.40740127 | 0.148372117 | 0.370412518 | 0.400559132 | 0.68874474 | 0.824190668 |
| ENSG00000277883 | NLRP3P1 | 4.490951061 | -0.571011672 | 0.858758751 | -0.664926758 | 0.50609734 | 0.686620055 |
| ENSG00000277893 | SRD5A2 | 1.897765376 | 1.469798134 | 1.376303228 | 1.067931909 | 0.285551228 | 0.469472168 |
| ENSG00000277895 | ENSG0000027 | 1.875958796 | 0.385734039 | 1.310389738 | 0.294365888 | 0.76847833 | 0.873712291 |
| ENSG00000277914 | DDX6P2 | 7.451131235 | -0.031677882 | 0.641366171 | -0.049391257 | 0.960607495 | 0.981920343 |
| ENSG00000277938 | ENSG0000027 | 15.40338244 | 0.353945348 | 0.47601394 | 0.743560886 | 0.457142174 | 0.644392372 |
| ENSG00000277952 | ENSG0000027 | 20.2732863 | -0.289803881 | 0.396094355 | -0.731653651 | 0.464379993 | 0.650604562 |
| ENSG00000277957 | SENP3-EIF4A1 | 4.331772228 | -0.20040377 | 0.853864156 | -0.23470217 | 0.814439903 | 0.901763793 |
| ENSG00000277959 | ENSG0000027 | 7.149496202 | -0.917828585 | 0.6716964 | -1.366433683 | 0.171802883 | 0.323496522 |
| ENSG00000277969 | ENSG0000027 | 45.64286016 | -0.009220828 | 0.270484262 | -0.034090073 | 0.972805325 | 0.987776737 |
| ENSG00000277972 | CISD3 | 634.8833688 | 0.308935275 | 0.080606702 | 3.832625182 | 0.000126783 | 0.000555569 |
| ENSG00000277978 | ENSG0000027 | 6.366638567 | -0.638394019 | 0.703019904 | -0.90807389 | 0.363839184 | 0.556559975 |
| ENSG00000277998 | ENSG0000027 | 1052.193607 | 0.040925569 | 0.072358579 | 0.565593875 | 0.571669892 | 0.738490107 |
| ENSG00000278000 | ENSG0000027 | 9.021934887 | 2.424894541 | 0.70214056 | 3.453574225 | 0.00055321 | 0.002171939 |
| ENSG00000278002 | ENSG0000027 | 2.90667764 | -1.423958099 | 1.132501225 | -1.257356785 | 0.20862447 | 0.375149882 |
| ENSG00000278009 | ENSG0000027 | 1.820265988 | 0.087867883 | 1.288639263 | 0.068186564 | 0.945637123 | 0.974261906 |
| ENSG00000278017 | ENSG0000027 | 3.063640542 | -0.024639248 | 0.98976953 | -0.024893925 | 0.980139573 | 0.991041209 |
| ENSG00000278023 | RDM1 | 43.64814765 | -0.490063464 | 0.275673607 | -1.777694535 | 0.075454038 | 0.168688785 |
| ENSG00000278053 | DDX52 | 765.3251563 | 0.105252223 | 0.07857929 | 1.339439726 | 0.180427564 | 0.33557713 |

| | | | | | | | |
|---|---|---|---|---|---|---|---|
| ENSG00000278068 | ENSG0000027 | 1.979094422 | -0.651007072 | 1.237637511 | -0.526007871 | 0.598882738 | 0.759025967 |
| ENSG00000278075 | ENSG0000027 | 2.07226903 | 0.959939174 | 1.244456914 | 0.771371964 | 0.440486488 | 0.62981551 |
| ENSG00000278090 | LUNAR1 | 3.353995518 | -1.098280886 | 0.988319106 | -1.111261413 | 0.266455844 | 0.447539688 |
| ENSG00000278098 | MIR7973-1 | 2.053360503 | -0.205392508 | 1.259924832 | -0.163019652 | 0.870502961 | 0.932928658 |
| ENSG00000278107 | ENSG0000027 | 22.90417919 | 1.212241618 | 0.381628931 | 3.176492977 | 0.001490674 | 0.005385206 |
| ENSG00000278112 | ENSG0000027 | 10.00535509 | 0.07767483 | 0.556418828 | 0.139597775 | 0.888977799 | 0.942802073 |
| ENSG00000278129 | ZNF8 | 400.1522066 | 0.079521167 | 0.0964274 | 0.824673964 | 0.409556717 | 0.602065433 |
| ENSG00000278133 | ENSG0000027 | 145.0876503 | 0.324442063 | 0.161425159 | 2.00986057 | 0.044445948 | 0.108317226 |
| ENSG00000278156 | TSC22D1-AS1 | 15.9670822 | 0.370685422 | 0.452124094 | 0.8198754 | 0.412287142 | 0.604529692 |
| ENSG00000278175 | GLIDR | 157.6812968 | -0.158984306 | 0.144993337 | -1.096493881 | 0.2728627 | 0.454862706 |
| ENSG00000278177 | ENSG0000027 | 9.985314597 | 0.403466453 | 0.555728993 | 0.726012963 | 0.467830824 | 0.653820524 |
| ENSG00000278192 | ENSG0000027 | 5.737886967 | 1.599337774 | 0.790310375 | 2.02368313 | 0.043002765 | 0.105316497 |
| ENSG00000278206 | ENSG0000027 | 10.60351098 | -0.493566396 | 0.554224027 | -0.89055395 | 0.373168513 | 0.565907695 |
| ENSG00000278212 | ENSG0000027 | 113.2701837 | 0.555995974 | 0.17175476 | 3.237150298 | 0.001207298 | 0.004453356 |
| ENSG00000278231 | ENSG0000027 | 110.5345457 | -1.76243649 | 0.182859276 | -9.638212107 | 5.51E-22 | 1.03E-20 |
| ENSG00000278238 | CKAP2-DT | 26.53981214 | -0.21158852 | 0.357149961 | -0.592436072 | 0.553558617 | 0.725118768 |
| ENSG00000278259 | MYO19 | 2156.153652 | -0.029744239 | 0.061621925 | -0.482689218 | 0.62931642 | 0.7810577 |
| ENSG00000278266 | ENSG0000027 | 24.8480373 | 3.865836066 | 0.561359624 | 6.886558814 | 5.72E-12 | 5.48E-11 |
| ENSG00000278267 | MIR6859-1 | 5.572361585 | 0.138017669 | 0.747824179 | 0.184558982 | 0.853574961 | 0.923806832 |
| ENSG00000278267 | MIR6859-1 | 5.572361585 | 0.138017669 | 0.747824179 | 0.184558982 | 0.853574961 | 0.923806832 |
| ENSG00000278275 | ENSG0000027 | 2.127170404 | -1.819947704 | 1.366915049 | -1.331427074 | 0.183048526 | 0.339074971 |
| ENSG00000278291 | ENSG0000027 | 584.3214125 | 0.684808261 | 0.084708235 | 8.084317442 | 6.25E-16 | 8.03E-15 |
| ENSG00000278311 | GGNBP2 | 1652.661776 | 0.051737096 | 0.062472149 | 0.828162584 | 0.40757843 | 0.600369311 |
| ENSG00000278318 | ZNF229 | 333.430202 | 1.050815743 | 0.106665274 | 9.851526196 | 6.75E-23 | 1.34E-21 |
| ENSG00000278338 | VWA8-AS1 | 3.317617968 | 0.387778958 | 0.953713767 | 0.406598889 | 0.684302613 | 0.821019976 |
| ENSG00000278341 | ENSG0000027 | 51.78236315 | 0.653004722 | 0.264312903 | 2.470574519 | 0.013489621 | 0.038686324 |
| ENSG00000278344 | ENSG0000027 | 2.075155943 | 0.24837232 | 1.241026966 | 0.200134507 | 0.841375387 | 0.916700769 |
| ENSG00000278356 | ENSG0000027 | 8.61282972 | 0.219681124 | 0.596413618 | 0.368336868 | 0.712622065 | 0.840102275 |
| ENSG00000278376 | ENSG0000027 | 35.77272811 | 0.21215181 | 0.301585702 | 0.70345447 | 0.481772571 | 0.666136024 |
| ENSG00000278383 | ENSG0000027 | 14.53277793 | -1.073344494 | 0.471463932 | -2.276620588 | 0.022808892 | 0.061283203 |
| ENSG00000278385 | ENSG0000027 | 80.22890279 | -0.137016192 | 0.198848299 | -0.689048851 | 0.490792526 | 0.673915141 |
| ENSG00000278390 | ENSG0000027 | 10.86911964 | -0.753453935 | 0.533704449 | -1.411743776 | 0.158025419 | 0.30294021 |
| ENSG00000278396 | ENSG0000027 | 10.57489349 | -0.452051562 | 0.532290097 | -0.849257886 | 0.395737809 | 0.588730567 |

| | | | | | | | |
|---|---|---|---|---|---|---|---|
| ENSG00000278416 | PMS2P2 | 3.284915559 | 1.153005766 | 1.014325512 | 1.136721646 | 0.255654667 | 0.43496672 |
| ENSG00000278463 | H2AC4 | 1.814085179 | -0.982529954 | 1.307978996 | -0.751181753 | 0.45254328 | 0.640191345 |
| ENSG00000278467 | ENSG0000027 | 3.308161865 | 0.693653437 | 0.989993806 | 0.700664421 | 0.483512465 | 0.66755731 |
| ENSG00000278472 | ENSG0000027 | 1.934165208 | -0.120232542 | 1.281752193 | -0.093803266 | 0.925265437 | 0.964344264 |
| ENSG00000278484 | ENSG0000027 | 4.151989153 | -1.007644661 | 0.868739503 | -1.159892761 | 0.246092471 | 0.423284457 |
| ENSG00000278493 | ENSG0000027 | 2.491659225 | -0.019391128 | 1.101141894 | -0.017610017 | 0.985949965 | 0.993350589 |
| ENSG00000278514 | ENSG0000027 | 2.153642964 | 1.485762386 | 1.297529834 | 1.145069923 | 0.252180194 | 0.430817221 |
| ENSG00000278535 | DHRS11 | 265.7176359 | 0.390213407 | 0.117628831 | 3.317327949 | 0.000908829 | 0.003429035 |
| ENSG00000278540 | ACACA | 8459.994483 | -0.338167528 | 0.05291133 | -6.391212023 | 1.65E-10 | 1.40E-09 |
| ENSG00000278558 | TMEM191B | 32.86293002 | -1.037049057 | 0.316463665 | -3.276992499 | 0.001049192 | 0.003911026 |
| ENSG00000278570 | NR2E3 | 3.264231425 | 1.3807506 | 1.021936931 | 1.351111363 | 0.176659762 | 0.33036398 |
| ENSG00000278571 | MIR7161 | 38.629795 | -3.170081517 | 0.385653813 | -8.220018603 | 2.03E-16 | 2.71E-15 |
| ENSG00000278572 | ENSG0000027 | 1.635884985 | -0.239594016 | 1.481304153 | -0.161745321 | 0.871506413 | NA |
| ENSG00000278576 | ENSG0000027 | 9.508458042 | -0.530449245 | 0.570942811 | -0.929075968 | 0.352849718 | 0.544269266 |
| ENSG00000278581 | MIR6080 | 4.593630338 | 0.041740569 | 0.810211932 | 0.051518088 | 0.958912689 | 0.981197069 |
| ENSG00000278588 | H2BC10 | 3.457063501 | -0.356673256 | 0.965856157 | -0.369281962 | 0.71191757 | 0.839858692 |
| ENSG00000278600 | ENSG0000027 | 147.943448 | -0.108662869 | 0.150496884 | -0.722027365 | 0.470277653 | 0.656115263 |
| ENSG00000278601 | ENSG0000027 | 2.312399343 | -0.991317918 | 1.206061041 | -0.821946721 | 0.411107221 | 0.603581246 |
| ENSG00000278615 | C11orf98 | 2.096853671 | -1.225301771 | 1.349813298 | -0.907756482 | 0.364006895 | 0.556687878 |
| ENSG00000278619 | MRM1 | 163.584883 | -0.195706867 | 0.145844576 | -1.3418865 | 0.1796328 | 0.334560073 |
| ENSG00000278621 | THBS1-AS1 | 12.18321947 | -0.1974594 | 0.502806716 | -0.392714326 | 0.694530489 | 0.827977606 |
| ENSG00000278627 | ENSG0000027 | 7.354686521 | -0.854797547 | 0.659209888 | -1.296700131 | 0.194734384 | 0.355867663 |
| ENSG00000278654 | ENSG0000027 | 3.265060757 | -0.455985683 | 0.96383097 | -0.473097148 | 0.636143871 | 0.785675698 |
| ENSG00000278662 | GOLGA6L10 | 66.69909846 | -0.048223155 | 0.231972692 | -0.207882894 | 0.835320406 | 0.913215357 |
| ENSG00000278677 | H2AC17 | 7.701720836 | 0.447825928 | 0.629038632 | 0.711921184 | 0.476513583 | 0.661576082 |
| ENSG00000278690 | ENSG0000027 | 3.528546637 | 4.10845318 | 1.419627986 | 2.894035072 | 0.003803257 | 0.012595165 |
| ENSG00000278703 | ENSG0000027 | 2.649123666 | 0.47752491 | 1.100561638 | 0.433892018 | 0.664366856 | 0.806248413 |
| ENSG00000278709 | NKILA | 168.3196252 | 0.324845722 | 0.141322995 | 2.298604858 | 0.021527387 | 0.058290636 |
| ENSG00000278713 | ENSG0000027 | 4.868382149 | -0.077897894 | 0.833122472 | -0.093501131 | 0.925505451 | 0.964392235 |
| ENSG00000278716 | ENSG0000027 | 2.133211516 | -0.365783691 | 1.184762799 | -0.30874002 | 0.757519298 | 0.866391318 |
| ENSG00000278727 | ENSG0000027 | 4.998400998 | 0.891680953 | 0.795426575 | 1.121009759 | 0.262283708 | 0.44280161 |
| ENSG00000278730 | ENSG0000027 | 5104.445343 | -0.023910826 | 0.04652758 | -0.513906494 | 0.607317367 | 0.765739276 |
| ENSG00000278740 | ENSG0000027 | 2.211103288 | -1.43329045 | 1.311707681 | -1.092690445 | 0.274529736 | 0.45662066 |

| | | | | | | | |
|---|---|---|---|---|---|---|---|
| ENSG00000278743 | ENSG0000027 | 3.301543783 | 0.081100062 | 0.956971564 | 0.084746574 | 0.932462868 | 0.967598417 |
| ENSG00000278765 | ENSG0000027 | 2.600835963 | 0.195473684 | 1.077568874 | 0.181402497 | 0.856051659 | 0.925395837 |
| ENSG00000278768 | BACE1-AS | 200.5771039 | -0.002914884 | 0.128898851 | -0.022613728 | 0.981958394 | 0.991627389 |
| ENSG00000278769 | ENSG0000027 | 3.474512494 | -0.849355547 | 0.993164108 | -0.855201614 | 0.392439543 | 0.585380266 |
| ENSG00000278771 | RN7SL3 | 10.71395778 | -0.142833635 | 0.526412178 | -0.271334216 | 0.786133993 | 0.883798765 |
| ENSG00000278784 | ENSG0000027 | 74.43868359 | -0.531633198 | 0.219036043 | -2.427149392 | 0.015217988 | 0.043066944 |
| ENSG00000278791 | ENSG0000027 | 4.6665172 | -0.958052411 | 0.839401047 | -1.141352414 | 0.253723299 | 0.432519319 |
| ENSG00000278811 | LINC00624 | 3.679162966 | 0.001850379 | 0.919151938 | 0.002013137 | 0.99839375 | 0.999134172 |
| ENSG00000278828 | H3C10 | 645.6440426 | 0.081548442 | 0.085921818 | 0.949100511 | 0.342569494 | 0.533542277 |
| ENSG00000278831 | ENSG0000027 | 10.06998753 | -0.774594936 | 0.549710396 | -1.409096393 | 0.158806667 | 0.304168402 |
| ENSG00000278834 | ENSG0000027 | 398.2782955 | -0.061451499 | 0.098033656 | -0.626840843 | 0.530763568 | 0.706370961 |
| ENSG00000278845 | MRPL45 | 1245.981066 | 0.04948096 | 0.061659087 | 0.802492582 | 0.422268078 | 0.613285933 |
| ENSG00000278861 | ENSG0000027 | 4.80607935 | 0.171551834 | 0.827792257 | 0.207240201 | 0.835822273 | 0.913370821 |
| ENSG00000278864 | ENSG0000027 | 105.8356579 | -0.721848723 | 0.183707912 | -3.929328436 | 8.52E-05 | 0.000384591 |
| ENSG00000278867 | ENSG0000027 | 4.808951118 | 0.372630872 | 0.840176988 | 0.443514733 | 0.657393465 | 0.80147779 |
| ENSG00000278869 | ENSG0000027 | 16.25009223 | 0.015169352 | 0.453743808 | 0.033431536 | 0.973330462 | 0.987970845 |
| ENSG00000278876 | ENSG0000027 | 7.454930675 | -0.3613208 | 0.637994175 | -0.566338712 | 0.571163549 | 0.738043813 |
| ENSG00000278899 | ENSG0000027 | 13.63947888 | 0.716009488 | 0.49177195 | 1.455978705 | 0.145398506 | 0.284594005 |
| ENSG00000278901 | ENSG0000027 | 2.18432154 | 0.038983321 | 1.176987431 | 0.033121272 | 0.97357788 | 0.988001378 |
| ENSG00000278915 | ENSG0000027 | 19.13405953 | 0.267036001 | 0.411571755 | 0.648820036 | 0.516454705 | 0.69491147 |
| ENSG00000278916 | CEP83-DT | 18.95984934 | -0.204327505 | 0.401790332 | -0.508542613 | 0.611072862 | 0.768826384 |
| ENSG00000278917 | ENSG0000027 | 4.82967573 | -0.402822705 | 0.839164364 | -0.480028374 | 0.631207217 | 0.782654644 |
| ENSG00000278918 | ENSG0000027 | 13.7273662 | -0.553025887 | 0.474757831 | -1.164858905 | 0.24407611 | 0.420827206 |
| ENSG00000278921 | EPB41L4A-DT | 49.36910567 | 0.536832298 | 0.255166311 | 2.103852566 | 0.03539131 | 0.089453134 |
| ENSG00000278922 | ENSG0000027 | 8.254560016 | 0.037272221 | 0.609485398 | 0.061153592 | 0.951236889 | 0.97748496 |
| ENSG00000278931 | ENSG0000027 | 34.14904031 | -0.737210814 | 0.312375179 | -2.360017264 | 0.018274085 | 0.05052519 |
| ENSG00000278932 | ENSG0000027 | 93.31441269 | 0.08637203 | 0.186148152 | 0.46399617 | 0.64265049 | 0.790455392 |
| ENSG00000278943 | ENSG0000027 | 5.807787695 | 0.490471544 | 0.747757242 | 0.655923496 | 0.51187335 | 0.691460054 |
| ENSG00000278946 | ENSG0000027 | 2.380005355 | 0.472568188 | 1.234626092 | 0.382762191 | 0.701896098 | 0.833064766 |
| ENSG00000278948 | ENSG0000027 | 162.9450092 | 0.541337009 | 0.146715168 | 3.689713983 | 0.000224506 | 0.000943774 |
| ENSG00000278949 | ENSG0000027 | 13.29600294 | -0.652521772 | 0.49751888 | -1.311551779 | 0.189671412 | 0.348670743 |
| ENSG00000278952 | ENSG0000027 | 96.30993223 | -0.425776816 | 0.179466353 | -2.372460401 | 0.017670057 | 0.049133468 |
| ENSG00000278962 | ENSG0000027 | 2.806546122 | 3.767422442 | 1.476716379 | 2.551215992 | 0.010734778 | 0.031615842 |

| | | | | | | | |
|---|---|---|---|---|---|---|---|
| ENSG00000278970 | HEIH | 1007.516952 | -0.181202657 | 0.074104222 | -2.445240676 | 0.014475551 | 0.041179208 |
| ENSG00000278974 | ENSG0000027 | 1.843410904 | -0.444446244 | 1.272165488 | -0.349361972 | 0.726817579 | 0.848163356 |
| ENSG00000278977 | ENSG0000027 | 2.899165958 | 0.831046387 | 1.046253157 | 0.794307172 | 0.42701663 | 0.61775629 |
| ENSG00000278985 | ENSG0000027 | 2.668506568 | 0.909654836 | 1.133421173 | 0.802574416 | 0.42222076 | 0.613285933 |
| ENSG00000278986 | ENSG0000027 | 3.585208474 | 0.926316528 | 0.942037808 | 0.983311413 | 0.325454195 | 0.515744628 |
| ENSG00000278987 | ENSG0000027 | 10.33294989 | 0.08548324 | 0.548345142 | 0.155893128 | 0.876117262 | 0.935667132 |
| ENSG00000278991 | ENSG0000027 | 4.735574959 | -0.655545717 | 0.883103287 | -0.742320549 | 0.457893144 | 0.645094192 |
| ENSG00000278993 | ENSG0000027 | 7.319710968 | -0.244432219 | 0.653654102 | -0.37394735 | 0.708443487 | 0.837453382 |
| ENSG00000278995 | ENSG0000027 | 24.74115061 | -0.183902478 | 0.347896048 | -0.528613299 | 0.597073733 | 0.75769879 |
| ENSG00000278997 | ENSG0000027 | 24.53368448 | 0.318334923 | 0.363345849 | 0.876120983 | 0.380964267 | 0.57330213 |
| ENSG00000279009 | ENSG0000027 | 6.390800859 | 0.331263729 | 0.69254103 | 0.478330835 | 0.632414753 | 0.783432036 |
| ENSG00000279013 | ENSG0000027 | 4.816805516 | 0.592683891 | 0.805884459 | 0.735445242 | 0.462068377 | 0.648626151 |
| ENSG00000279022 | ENSG0000027 | 4.787931884 | -0.277074017 | 0.786468004 | -0.352301703 | 0.724612011 | 0.846666303 |
| ENSG00000279024 | ENSG0000027 | 7.942573006 | -1.00161101 | 0.640892739 | -1.562837195 | 0.118090893 | 0.242546789 |
| ENSG00000279026 | ENSG0000027 | 11.84164291 | -0.702844218 | 0.515693093 | -1.362911833 | 0.172910299 | 0.325067647 |
| ENSG00000279031 | ENSG0000027 | 4.297219431 | 1.050403053 | 0.871465046 | 1.205330102 | 0.228075893 | 0.400963356 |
| ENSG00000279033 | ENSG0000027 | 4.775241845 | -0.092412895 | 0.803355984 | -0.115033555 | 0.908418525 | 0.954215897 |
| ENSG00000279035 | ENSG0000027 | 11.13610313 | -0.931749671 | 0.548716756 | -1.698052157 | 0.089497918 | 0.193522692 |
| ENSG00000279041 | ENSG0000027 | 42.28545382 | -1.339593323 | 0.295954513 | -4.526348699 | 6.00E-06 | 3.18E-05 |
| ENSG00000279042 | ENSG0000027 | 44.65397587 | 3.258460127 | 0.370624381 | 8.791812672 | 1.47E-18 | 2.25E-17 |
| ENSG00000279047 | PCDHB1-AS1 | 16.89630863 | -0.17598478 | 0.44869157 | -0.392217709 | 0.694897361 | 0.828271093 |
| ENSG00000279048 | ENSG0000027 | 20.98214404 | -1.424398262 | 0.412375266 | -3.454131176 | 0.000552069 | 0.002167829 |
| ENSG00000279058 | AGAP14P | 6.756179926 | 0.363050907 | 0.680193465 | 0.533746537 | 0.593516907 | 0.755562373 |
| ENSG00000279059 | ENSG0000027 | 150.9359765 | 0.168874974 | 0.150007872 | 1.125774079 | 0.260261158 | 0.440521728 |
| ENSG00000279066 | HEXD-IT1 | 6.791289989 | 0.451421462 | 0.679852036 | 0.663999574 | 0.506690583 | 0.686971025 |
| ENSG00000279069 | ENSG0000027 | 128.9813562 | 0.313645893 | 0.15717368 | 1.995536989 | 0.045984344 | 0.111474253 |
| ENSG00000279072 | ENSG0000027 | 1.606080629 | 1.009392252 | 1.467013413 | 0.688059321 | 0.491415426 | NA |
| ENSG00000279075 | ENSG0000027 | 3.226000585 | 0.815958032 | 1.06633057 | 0.765201762 | 0.44415141 | 0.632925417 |
| ENSG00000279080 | ENSG0000027 | 2.339426266 | 0.209013345 | 1.133896492 | 0.184331944 | 0.853753056 | 0.923868819 |
| ENSG00000279081 | ENSG0000027 | 10.76352148 | 0.497499438 | 0.569560906 | 0.873478908 | 0.382402098 | 0.57466781 |
| ENSG00000279086 | ENSG0000027 | 7.186944106 | 0.622584455 | 0.65881028 | 0.945013266 | 0.344652103 | 0.535542116 |
| ENSG00000279088 | ENSG0000027 | 209.9489375 | -0.387494878 | 0.128263331 | -3.021088528 | 0.002518677 | 0.008672839 |
| ENSG00000279089 | ENSG0000027 | 30.33491052 | -0.231520145 | 0.32371472 | -0.715198078 | 0.474486657 | 0.659868063 |

| | | | | | | | |
|---|---|---|---|---|---|---|---|
| ENSG00000279108 | ENSG0000027 | 24.93206816 | 1.183470923 | 0.380020728 | 3.114227294 | 0.001844274 | 0.006529964 |
| ENSG00000279114 | ENSG0000027 | 1.721043215 | 0.265562207 | 1.403792233 | 0.189174866 | 0.849955766 | NA |
| ENSG00000279118 | ENSG0000027 | 11.05205246 | 1.700436236 | 0.584776387 | 2.907840113 | 0.003639344 | 0.012101253 |
| ENSG00000279119 | ENSG0000027 | 9.979171214 | 0.173463881 | 0.561859788 | 0.308731617 | 0.757525691 | 0.866391318 |
| ENSG00000279124 | IGSF3P1 | 15.10235596 | -0.601237481 | 0.53706396 | -1.119489532 | 0.262931354 | 0.443699078 |
| ENSG00000279130 | ENSG0000027 | 4.469382193 | -0.896275717 | 0.836244384 | -1.071786829 | 0.28381579 | 0.467676593 |
| ENSG00000279133 | ENSG0000027 | 22.64393408 | 1.028229712 | 0.380949993 | 2.699119911 | 0.006952312 | 0.02156674 |
| ENSG00000279138 | ENSG0000027 | 2.730930869 | 2.639144505 | 1.344308486 | 1.963198575 | 0.049623105 | 0.119037334 |
| ENSG00000279140 | ENSG0000027 | 10.95708883 | 0.450067604 | 0.533188823 | 0.844105473 | 0.39861047 | 0.591584665 |
| ENSG00000279145 | ENSG0000027 | 7.065506815 | -0.197969256 | 0.663723678 | -0.298270595 | 0.765496645 | 0.871748066 |
| ENSG00000279148 | ENSG0000027 | 16.58043188 | 0.496513598 | 0.43058186 | 1.153122423 | 0.248860139 | 0.42673431 |
| ENSG00000279152 | ENSG0000027 | 2.256141031 | -0.126742706 | 1.20442134 | -0.105231202 | 0.916192352 | 0.958650454 |
| ENSG00000279159 | ENSG0000027 | 44.10123313 | -1.496448217 | 0.300803263 | -4.974840367 | 6.53E-07 | 3.87E-06 |
| ENSG00000279168 | ENSG0000027 | 4.106698723 | -0.696835551 | 0.884096835 | -0.788189171 | 0.430586062 | 0.620756944 |
| ENSG00000279170 | TSTD3 | 121.0025661 | -0.149499314 | 0.164815595 | -0.907070194 | 0.364369679 | 0.557019888 |
| ENSG00000279172 | ENSG0000027 | 2.353658302 | -2.020129003 | 1.308981761 | -1.543282774 | 0.122762181 | 0.249770682 |
| ENSG00000279179 | ENSG0000027 | 5.674878883 | 0.407908472 | 0.748788476 | 0.544757946 | 0.585920006 | 0.749846858 |
| ENSG00000279187 | ENSG0000027 | 1.857163681 | -0.486239503 | 1.270990025 | -0.382567521 | 0.702040456 | 0.833064766 |
| ENSG00000279196 | ENSG0000027 | 24.17798776 | -0.714519807 | 0.370664374 | -1.927673272 | 0.053895779 | 0.12785617 |
| ENSG00000279198 | ENSG0000027 | 42.07382681 | 0.017216909 | 0.280378463 | 0.061405962 | 0.951035904 | 0.977409778 |
| ENSG00000279199 | ENSG0000027 | 6.223507634 | 0.022949537 | 0.720007451 | 0.031874027 | 0.974572511 | 0.988514222 |
| ENSG00000279203 | ENSG0000027 | 4.015582577 | -0.414075916 | 0.952979661 | -0.434506562 | 0.663920631 | 0.806043413 |
| ENSG00000279205 | ENSG0000027 | 4.165782814 | 0.992145162 | 0.87537134 | 1.133399184 | 0.257046657 | 0.436691869 |
| ENSG00000279207 | ENSG0000027 | 1290.026384 | 0.930778814 | 0.069248176 | 13.44120335 | 3.47E-41 | 1.49E-39 |
| ENSG00000279208 | ENSG0000027 | 19.7619256 | 0.065513613 | 0.394075367 | 0.166246404 | 0.867963043 | 0.931076558 |
| ENSG00000279227 | ENSG0000027 | 22.94462111 | 0.140594106 | 0.363733154 | 0.386530906 | 0.699103518 | 0.831105311 |
| ENSG00000279232 | ENSG0000027 | 15.00245155 | -0.223354082 | 0.468765198 | -0.47647326 | 0.633737253 | 0.784143138 |
| ENSG00000279233 | ENSG0000027 | 80.68153501 | -0.437101845 | 0.222266065 | -1.966570319 | 0.049232768 | 0.118212236 |
| ENSG00000279246 | ENSG0000027 | 7.243506723 | -0.52250264 | 0.696382292 | -0.75031006 | 0.453067985 | 0.640696559 |
| ENSG00000279253 | ENSG0000027 | 25.57976524 | -0.769621317 | 0.350984173 | -2.192752198 | 0.028325238 | 0.073914339 |
| ENSG00000279255 | ENSG0000027 | 26.8220465 | 0.466498641 | 0.346112022 | 1.347825593 | 0.177714484 | 0.331957517 |
| ENSG00000279259 | ENSG0000027 | 22.59181555 | 0.21316534 | 0.38053265 | 0.560176217 | 0.575359247 | 0.74175234 |
| ENSG00000279265 | ENSG0000027 | 9.689665225 | 0.910112287 | 0.638051299 | 1.426393596 | 0.153754761 | 0.29689682 |

| | | | | | | | |
|---|---|---|---|---|---|---|---|
| ENSG00000279267 | ENSG0000027 | 7.624119176 | 0.168839736 | 0.660527752 | 0.255613387 | 0.79824938 | 0.892259454 |
| ENSG00000279274 | ENSG0000027 | 13.8670897 | 0.019858271 | 0.472372124 | 0.042039465 | 0.966467238 | 0.98505693 |
| ENSG00000279277 | ENSG0000027 | 35.66729349 | 1.655323495 | 0.321242888 | 5.15287204 | 2.57E-07 | 1.60E-06 |
| ENSG00000279283 | ENSG0000027 | 10.50087667 | 0.717356063 | 0.549888887 | 1.304547299 | 0.192047043 | 0.352078572 |
| ENSG00000279306 | ENSG0000027 | 2.782444419 | 0.993201852 | 1.122774121 | 0.884596316 | 0.376374394 | 0.569311728 |
| ENSG00000279307 | NDUFAF2P1 | 119.8549773 | 0.123068445 | 0.164591096 | 0.747722372 | 0.454627634 | 0.642189527 |
| ENSG00000279320 | ENSG0000027 | 3.180935048 | 0.750847317 | 1.020841109 | 0.735518301 | 0.462023899 | 0.648626151 |
| ENSG00000279322 | ENSG0000027 | 1.800321261 | -1.563766638 | 1.385158284 | -1.128944364 | 0.258921304 | 0.438739453 |
| ENSG00000279328 | ENSG0000027 | 3.255167706 | 0.099625295 | 0.963747003 | 0.103372872 | 0.917667042 | 0.959449521 |
| ENSG00000279329 | ENSG0000027 | 41.58964123 | 0.249945628 | 0.28016403 | 0.89214032 | 0.372317726 | 0.564976656 |
| ENSG00000279330 | ENSG0000027 | 3.397894788 | -0.274539697 | 0.950636299 | -0.288795723 | 0.772737705 | 0.876166615 |
| ENSG00000279331 | RBM12B-AS1 | 2.552651159 | -0.802086753 | 1.117791408 | -0.717563892 | 0.473026228 | 0.658601347 |
| ENSG00000279333 | ENSG0000027 | 2.915545957 | -1.520678424 | 1.10677351 | -1.373974358 | 0.169449643 | 0.320015611 |
| ENSG00000279342 | ENSG0000027 | 10.68415711 | -0.36260204 | 0.533663333 | -0.679458411 | 0.496847451 | 0.678756582 |
| ENSG00000279344 | ENSG0000027 | 28.43094119 | -0.36496826 | 0.35578093 | -1.025822997 | 0.304975024 | 0.492372936 |
| ENSG00000279347 | ENSG0000027 | 11.19163821 | -0.127438731 | 0.540246965 | -0.235889768 | 0.813518208 | 0.901448243 |
| ENSG00000279348 | ENSG0000027 | 68.63571098 | 0.986296744 | 0.220434925 | 4.474321596 | 7.67E-06 | 4.01E-05 |
| ENSG00000279356 | ENSG0000027 | 5.124150884 | -0.998892045 | 0.781925358 | -1.277477492 | 0.201433722 | 0.36508569 |
| ENSG00000279364 | ENSG0000027 | 3.601641296 | -1.268122399 | 0.960999419 | -1.319587062 | 0.186972925 | 0.344676864 |
| ENSG00000279368 | ENSG0000027 | 1.771006677 | -1.542858292 | 1.450774457 | -1.063472192 | 0.28756787 | NA |
| ENSG00000279369 | ENSG0000027 | 3.415081788 | 0.03761544 | 0.953436431 | 0.039452488 | 0.968529633 | 0.985876114 |
| ENSG00000279372 | ENSG0000027 | 4.068862218 | 0.166394415 | 0.860315757 | 0.193410866 | 0.846637206 | 0.919716896 |
| ENSG00000279382 | ENSG0000027 | 11.83971904 | 0.230838632 | 0.517933565 | 0.445691585 | 0.655820044 | 0.80041029 |
| ENSG00000279392 | ENSG0000027 | 166.7570288 | -0.562273107 | 0.143650974 | -3.914161472 | 9.07E-05 | 0.000407099 |
| ENSG00000279399 | ENSG0000027 | 5.39080658 | -0.739364131 | 0.759900734 | -0.972974625 | 0.330565914 | 0.521252548 |
| ENSG00000279406 | ENSG0000027 | 14.63992337 | -1.469631523 | 0.485167487 | -3.029122032 | 0.002452656 | 0.008472207 |
| ENSG00000279407 | ENSG0000027 | 37.70868705 | -0.896693213 | 0.305756581 | -2.932702903 | 0.003360253 | 0.01126962 |
| ENSG00000279413 | ENSG0000027 | 2.770578828 | 0.35599242 | 1.072884348 | 0.331808756 | 0.740033672 | 0.856567724 |
| ENSG00000279414 | CDRT15P9 | 3.456483072 | -4.31880582 | 1.415207369 | -3.05171236 | 0.0022754 | 0.007911137 |
| ENSG00000279415 | ENSG0000027 | 6.453568401 | -0.406892711 | 0.683475961 | -0.595328489 | 0.551623916 | 0.723472222 |
| ENSG00000279419 | ENSG0000027 | 8.493864862 | 3.925017277 | 0.980424523 | 4.003385456 | 6.24E-05 | 0.000287983 |
| ENSG00000279425 | ENSG0000027 | 3.27598265 | 1.934948755 | 1.08007377 | 1.791496848 | 0.0732136 | 0.164674966 |
| ENSG00000279428 | ENSG0000027 | 9.854369178 | -1.038125861 | 0.578949169 | -1.79312091 | 0.072953593 | 0.164235076 |

| | | | | | | | |
|---|---|---|---|---|---|---|---|
| ENSG00000279434 | ENSG0000027 | 7.54514664 | -0.098280303 | 0.642262094 | -0.153022113 | 0.878380837 | 0.936776215 |
| ENSG00000279441 | ENSG0000027 | 1.734234166 | -0.827543744 | 1.336874528 | -0.619013772 | 0.535907287 | NA |
| ENSG00000279443 | ENSG0000027 | 84.21162458 | 0.617072558 | 0.204658341 | 3.015135143 | 0.002568648 | 0.008822396 |
| ENSG00000279453 | ENSG0000027 | 82.76772544 | 0.617789153 | 0.199591676 | 3.095265123 | 0.00196637 | 0.006925476 |
| ENSG00000279457 | WASH9P | 127.7597112 | -0.687199148 | 0.164628037 | -4.174253439 | 2.99E-05 | 0.000144842 |
| ENSG00000279464 | ENSG0000027 | 4.516590505 | -0.058495013 | 0.842355104 | -0.069442226 | 0.944637619 | 0.973711185 |
| ENSG00000279483 | ENSG0000027 | 91.50131948 | 0.086813098 | 0.18724185 | 0.46364153 | 0.642904595 | 0.790620317 |
| ENSG00000279488 | ENSG0000027 | 5.344788342 | -0.41829238 | 0.755726581 | -0.553496979 | 0.579923148 | 0.745457639 |
| ENSG00000279491 | ENSG0000027 | 5.531069506 | -0.506408884 | 0.755500286 | -0.670296086 | 0.502669062 | 0.683696685 |
| ENSG00000279495 | ENSG0000027 | 38.59627619 | -0.760338335 | 0.284548052 | -2.67209116 | 0.007538017 | 0.023189232 |
| ENSG00000279500 | ENSG0000027 | 24.56538186 | 0.017029508 | 0.355909021 | 0.047847923 | 0.961837443 | 0.982393185 |
| ENSG00000279504 | ENSG0000027 | 9.750740101 | -0.172436968 | 0.557595745 | -0.309250869 | 0.757130701 | 0.866096054 |
| ENSG00000279511 | ENSG0000027 | 28.77676403 | 0.773277766 | 0.340299718 | 2.272343244 | 0.023065787 | 0.061915417 |
| ENSG00000279513 | ENSG0000027 | 2.296977862 | -2.594900064 | 1.517344669 | -1.710158619 | 0.087236546 | 0.189601698 |
| ENSG00000279518 | ENSG0000027 | 10.11109823 | -0.132055129 | 0.55788726 | -0.236705762 | 0.812885065 | 0.901197728 |
| ENSG00000279520 | ENSG0000027 | 49.04559983 | 0.124079959 | 0.263454374 | 0.470973234 | 0.637659849 | 0.78659182 |
| ENSG00000279529 | ENSG0000027 | 35.28171646 | 0.205992576 | 0.294246094 | 0.700069026 | 0.483884198 | 0.667848115 |
| ENSG00000279536 | ENSG0000027 | 2.21920654 | -0.149394226 | 1.179648469 | -0.126643004 | 0.899222959 | 0.949051063 |
| ENSG00000279539 | ENSG0000027 | 7.598253331 | 0.214485102 | 0.653029175 | 0.328446432 | 0.742574141 | 0.857909488 |
| ENSG00000279549 | ENSG0000027 | 7.006046366 | -0.834672377 | 0.692103041 | -1.205994379 | 0.227819658 | 0.400658 |
| ENSG00000279555 | ENSG0000027 | 7.36084137 | -0.258683701 | 0.651694624 | -0.396940057 | 0.691411665 | 0.826001186 |
| ENSG00000279561 | ENSG0000027 | 40.85280864 | 0.44516801 | 0.282488868 | 1.575878059 | 0.115053908 | 0.237393116 |
| ENSG00000279569 | ENSG0000027 | 120.6104791 | -0.413732215 | 0.361136493 | -1.145639455 | 0.251944365 | 0.430636065 |
| ENSG00000279571 | ENSG0000027 | 4.469777155 | -0.057290485 | 0.824397821 | -0.069493737 | 0.944596618 | 0.973711185 |
| ENSG00000279579 | LINC01666 | 4.187495687 | -0.294932032 | 0.930282962 | -0.317034755 | 0.75121723 | 0.862469998 |
| ENSG00000279583 | ENSG0000027 | 3.158107412 | -0.709896872 | 1.109153033 | -0.6400351 | 0.52214978 | 0.699077575 |
| ENSG00000279592 | ENSG0000027 | 6.968303144 | 0.129909857 | 0.685160136 | 0.189605101 | 0.84961859 | 0.921263493 |
| ENSG00000279594 | ENSG0000027 | 3.559678186 | 0.916647485 | 0.937053126 | 0.978223603 | 0.327963745 | 0.518289358 |
| ENSG00000279602 | ENSG0000027 | 6.932055254 | -1.309360309 | 0.68915351 | -1.899954495 | 0.057439092 | 0.134619203 |
| ENSG00000279605 | ENSG0000027 | 148.4318602 | -0.104207794 | 0.165794122 | -0.628537325 | 0.529652003 | 0.705300804 |
| ENSG00000279611 | ENSG0000027 | 50.50948346 | -0.494166476 | 0.266134738 | -1.856828164 | 0.063335605 | 0.145933988 |
| ENSG00000279613 | ENSG0000027 | 5.003176931 | 0.166810646 | 0.77854998 | 0.214258108 | 0.830345784 | 0.910033374 |
| ENSG00000279631 | ENSG0000027 | 12.80932228 | -0.105380629 | 0.480855106 | -0.219152563 | 0.826531207 | 0.908492348 |

| | | | | | | | |
|---|---|---|---|---|---|---|---|
| ENSG00000279637 | ENSG0000027 | 1.583798469 | -0.108565202 | 1.385430509 | -0.07836207 | 0.937540045 | NA |
| ENSG00000279641 | ENSG0000027 | 29.37946832 | 0.071114982 | 0.327410695 | 0.217204212 | 0.828049202 | 0.909090629 |
| ENSG00000279649 | ENSG0000027 | 6.201177859 | -0.446773745 | 0.699770214 | -0.638457791 | 0.523175722 | 0.699804727 |
| ENSG00000279653 | ENSG0000027 | 2.5015526 | -0.384368934 | 1.140048331 | -0.337151438 | 0.736002743 | 0.854386347 |
| ENSG00000279656 | ENSG0000027 | 7.004428635 | -0.190576595 | 0.656561123 | -0.290264819 | 0.771613651 | 0.875561078 |
| ENSG00000279659 | ENSG0000027 | 1.995643862 | -0.679235796 | 1.218121509 | -0.557609229 | 0.577111253 | 0.743117322 |
| ENSG00000279660 | ENSG0000027 | 34.92005379 | -0.044675316 | 0.306704235 | -0.145662533 | 0.8841878 | 0.939910479 |
| ENSG00000279662 | ENSG0000027 | 47.26291475 | 0.723606465 | 0.280832715 | 2.576645906 | 0.009976409 | 0.029664138 |
| ENSG00000279667 | ENSG0000027 | 5.951521966 | 1.263328531 | 0.76602065 | 1.649209497 | 0.099104723 | 0.210447315 |
| ENSG00000279672 | ENSG0000027 | 30.13467439 | 0.390739356 | 0.323872635 | 1.206459928 | 0.227640201 | 0.400434492 |
| ENSG00000279673 | ENSG0000027 | 7.350977331 | -0.712834002 | 0.648356458 | -1.099447677 | 0.271572844 | 0.453577069 |
| ENSG00000279675 | ENSG0000027 | 4.267998859 | 2.773261407 | 1.094518321 | 2.5337734 | 0.011284168 | 0.033051431 |
| ENSG00000279689 | ENSG0000027 | 66.99180033 | -1.101469067 | 0.259395957 | -4.246284638 | 2.17E-05 | 0.000107293 |
| ENSG00000279691 | ENSG0000027 | 2.035797144 | 0.550370682 | 1.239437577 | 0.44404873 | 0.657007353 | 0.801277136 |
| ENSG00000279692 | ENSG0000027 | 18.0927909 | 0.082407208 | 0.411092444 | 0.200459068 | 0.841121567 | 0.916599152 |
| ENSG00000279693 | ENSG0000027 | 5.438969406 | -1.119191625 | 0.781255356 | -1.43255546 | 0.151984918 | 0.294396185 |
| ENSG00000279696 | ENSG0000027 | 26.90768185 | 0.436443831 | 0.337600061 | 1.292783625 | 0.196085905 | 0.357611444 |
| ENSG00000279699 | ENSG0000027 | 3.392830797 | -2.168230589 | 1.101020913 | -1.969291013 | 0.048919683 | 0.117571243 |
| ENSG00000279700 | ENSG0000027 | 6.672605205 | 0.717751118 | 0.707684327 | 1.014224974 | 0.310475414 | 0.498458778 |
| ENSG00000279712 | ENSG0000027 | 4.573599365 | -0.994750601 | 0.889468319 | -1.11836541 | 0.263410961 | 0.44412799 |
| ENSG00000279716 | ENSG0000027 | 38.22230535 | 0.09862809 | 0.286394873 | 0.344377987 | 0.730562042 | 0.850832153 |
| ENSG00000279718 | SNX18P12 | 17.14620144 | -0.121439921 | 0.425200602 | -0.285606182 | 0.775179768 | 0.877720823 |
| ENSG00000279720 | SOWAHCP2 | 43.81721557 | 0.340831493 | 0.275547165 | 1.236926146 | 0.216114502 | 0.385420717 |
| ENSG00000279722 | ENSG0000027 | 16.61133335 | 0.574656114 | 0.442340034 | 1.299127527 | 0.193900168 | 0.354618886 |
| ENSG00000279726 | ENSG0000027 | 65.1948945 | 0.662398332 | 0.225474206 | 2.937800928 | 0.003305492 | 0.011102282 |
| ENSG00000279730 | KMT5AP1 | 289.4623354 | -0.087393988 | 0.112184168 | -0.779022476 | 0.435966475 | 0.625888042 |
| ENSG00000279738 | ENSG0000027 | 25.60710917 | 0.228434408 | 0.349071851 | 0.654405124 | 0.512850834 | 0.692343494 |
| ENSG00000279741 | ENSG0000027 | 31.24067722 | -0.360311956 | 0.317193853 | -1.135936125 | 0.255983298 | 0.43543355 |
| ENSG00000279742 | ENSG0000027 | 17.26646358 | 0.523851604 | 0.435477576 | 1.202935886 | 0.229001132 | 0.402045919 |
| ENSG00000279744 | ENSG0000027 | 23.47990902 | 0.08497295 | 0.367532373 | 0.231198546 | 0.817160559 | 0.90323134 |
| ENSG00000279748 | ENSG0000027 | 22.93701734 | 0.49004884 | 0.371080221 | 1.320600809 | 0.186634503 | 0.344274324 |
| ENSG00000279753 | ENSG0000027 | 15.91629297 | -0.9202885 | 0.47231586 | -1.948459869 | 0.051359962 | 0.122572861 |
| ENSG00000279759 | ENSG0000027 | 90.27270498 | -0.863964369 | 0.198472564 | -4.353067 | 1.34E-05 | 6.81E-05 |

| | | | | | | | |
|---|---|---|---|---|---|---|---|
| ENSG00000279762 | ENSG0000027 | 10.81604824 | -0.743890541 | 0.579904998 | -1.282780014 | 0.199569158 | 0.36253694 |
| ENSG00000279766 | ENSG0000027 | 2.193838232 | 0.412638376 | 1.243464846 | 0.33184563 | 0.740005827 | 0.856567724 |
| ENSG00000279773 | ENSG0000027 | 8.916225909 | -2.107057127 | 0.6868755 | -3.067596859 | 0.002157875 | 0.007532206 |
| ENSG00000279789 | ENSG0000027 | 63.35718131 | 0.11914834 | 0.220175062 | 0.541152748 | 0.588402304 | 0.751676167 |
| ENSG00000279791 | ENSG0000027 | 1.851868233 | -1.55220412 | 1.38805473 | -1.118258586 | 0.263456569 | 0.44412799 |
| ENSG00000279792 | ENSG0000027 | 16.24509975 | 0.659137467 | 0.467417049 | 1.410169929 | 0.158489513 | 0.303727053 |
| ENSG00000279794 | ENSG0000027 | 23.94447641 | -0.445231339 | 0.369577005 | -1.204705197 | 0.228317129 | 0.401194557 |
| ENSG00000279799 | ENSG0000027 | 11.8607316 | -0.200726979 | 0.514651844 | -0.390024794 | 0.696518213 | 0.829482392 |
| ENSG00000279800 | BCLAF1P2 | 751.4852144 | 0.148303546 | 0.075431797 | 1.966061415 | 0.049291517 | 0.118316145 |
| ENSG00000279803 | ENSG0000027 | 10.40483398 | -0.105936547 | 0.54458522 | -0.194527033 | 0.84576323 | 0.919043732 |
| ENSG00000279817 | ENSG0000027 | 2.313341614 | 1.876215284 | 1.312156539 | 1.429871535 | 0.152753893 | 0.295511411 |
| ENSG00000279821 | ENSG0000027 | 27.34336171 | -0.211498271 | 0.332543916 | -0.636001023 | 0.524775755 | 0.7010535 |
| ENSG00000279827 | ENSG0000027 | 44.28632754 | -0.556675093 | 0.262346398 | -2.121908653 | 0.03384541 | 0.086095997 |
| ENSG00000279833 | ENSG0000027 | 4.174855421 | 0.332500302 | 0.87635974 | 0.379410745 | 0.704382873 | 0.834635491 |
| ENSG00000279837 | ENSG0000027 | 4.254295447 | -0.108538197 | 0.852299377 | -0.127347502 | 0.898665365 | 0.948793422 |
| ENSG00000279838 | ENSG0000027 | 11.46942055 | 0.051935359 | 0.532254553 | 0.097576166 | 0.922268851 | 0.962243482 |
| ENSG00000279845 | ENSG0000027 | 1.721469022 | -0.844095034 | 1.370608898 | -0.615854045 | 0.537990854 | NA |
| ENSG00000279853 | ENSG0000027 | 1.582423336 | 1.024123817 | 1.44628124 | 0.708108346 | 0.478877977 | NA |
| ENSG00000279859 | ENSG0000027 | 1.588106284 | 1.081034691 | 1.427911794 | 0.757073859 | 0.449005615 | NA |
| ENSG00000279861 | ENSG0000027 | 40.10535002 | 0.00500573 | 0.28000923 | 0.017877017 | 0.985736964 | 0.993350589 |
| ENSG00000279865 | ENSG0000027 | 20.68849276 | 0.146960133 | 0.389578868 | 0.377228195 | 0.706004029 | 0.835693108 |
| ENSG00000279873 | LINC01126 | 32.71685529 | -0.238574399 | 0.305028462 | -0.782138157 | 0.434133385 | 0.624076942 |
| ENSG00000279878 | ENSG0000027 | 40.56047336 | -0.537346612 | 0.28045327 | -1.91599339 | 0.055365937 | 0.13070851 |
| ENSG00000279879 | ENSG0000027 | 6.642660744 | -0.610121304 | 0.676174551 | -0.90231332 | 0.366890452 | 0.559399478 |
| ENSG00000279880 | ENSG0000027 | 3.111878472 | -0.337400285 | 1.033435657 | -0.326484076 | 0.744058136 | 0.858490754 |
| ENSG00000279884 | ENSG0000027 | 2.271891984 | -0.079468933 | 1.252687499 | -0.063438753 | 0.949417129 | 0.976618218 |
| ENSG00000279900 | ENSG0000027 | 4.000712982 | 0.441311059 | 0.87593946 | 0.503814566 | 0.614391691 | 0.771190919 |
| ENSG00000279901 | ENSG0000027 | 2.826400192 | -0.030785264 | 1.043022553 | -0.029515434 | 0.97645351 | 0.989269612 |
| ENSG00000279908 | ENSG0000027 | 19.57820298 | -0.519809604 | 0.394209415 | -1.31861286 | 0.187298572 | 0.345138505 |
| ENSG00000279923 | ENSG0000027 | 1.756888682 | 0.616889481 | 1.311517553 | 0.470363114 | 0.638095613 | NA |
| ENSG00000279930 | ENSG0000027 | 9.000479399 | -0.944782365 | 0.611610956 | -1.544744016 | 0.122408194 | 0.249183065 |
| ENSG00000279943 | FLJ38576 | 2.390027288 | -0.617661094 | 1.209993728 | -0.51046636 | 0.609724776 | 0.767761904 |
| ENSG00000279945 | ENSG0000027 | 13.63754986 | 0.276971502 | 0.479353356 | 0.577802363 | 0.56339756 | 0.732154597 |

| | | | | | | | |
|---|---|---|---|---|---|---|---|
| ENSG00000279953 | ENSG0000027 | 7.405908856 | 1.144046477 | 0.66225329 | 1.727505916 | 0.084076844 | 0.184073563 |
| ENSG00000279954 | ENSG0000027 | 5.129701596 | 0.2135981 | 0.769836503 | 0.277459043 | 0.78142765 | 0.881401824 |
| ENSG00000279968 | CCDC28A-AS1 | 11.84544949 | -0.538036158 | 0.508412601 | -1.058266764 | 0.289933841 | 0.474365759 |
| ENSG00000279970 | ENSG0000027 | 51.60177308 | 1.384431848 | 0.266249358 | 5.199756569 | 2.00E-07 | 1.26E-06 |
| ENSG00000279973 | ENSG0000027 | 3.018108616 | -5.162954739 | 1.468290842 | -3.516302488 | 0.000437602 | 0.001749798 |
| ENSG00000279982 | ENSG0000027 | 7.960354202 | -0.559017667 | 0.623519956 | -0.896551364 | 0.36995836 | 0.562601577 |
| ENSG00000280007 | ENSG0000028 | 2.466566209 | -0.015633847 | 1.11324853 | -0.014043447 | 0.988795319 | 0.994082067 |
| ENSG00000280010 | ENSG0000028 | 5.51182888 | 0.69692484 | 0.744157545 | 0.936528621 | 0.349001083 | 0.54022165 |
| ENSG00000280022 | ENSG0000028 | 1.751073091 | -3.308970536 | 1.680760902 | -1.968733645 | 0.048983686 | NA |
| ENSG00000280023 | ENSG0000028 | 3.131239092 | -0.623754222 | 1.03528275 | -0.602496489 | 0.546843699 | 0.719836717 |
| ENSG00000280027 | ENSG0000028 | 22.78074167 | -0.152000141 | 0.383722629 | -0.396119826 | 0.692016633 | 0.826376122 |
| ENSG00000280029 | ENSG0000028 | 4.019275115 | 0.280453413 | 0.868659888 | 0.322857561 | 0.746803118 | 0.85963488 |
| ENSG00000280032 | ENSG0000028 | 2.292545273 | 2.548822547 | 1.427184893 | 1.785909141 | 0.074113971 | 0.166341306 |
| ENSG00000280033 | ENSG0000028 | 9.923641169 | -0.051047159 | 0.577886435 | -0.08833424 | 0.929611026 | 0.965993019 |
| ENSG00000280046 | ENSG0000028 | 21.76621 | -0.057487636 | 0.383695943 | -0.14982602 | 0.88090188 | 0.938252557 |
| ENSG00000280047 | ENSG0000028 | 8.518246232 | -0.395758312 | 0.604873212 | -0.654283086 | 0.512929441 | 0.692356657 |
| ENSG00000280053 | ENSG0000028 | 2.379146279 | -0.182540052 | 1.189763221 | -0.153425529 | 0.878062714 | 0.936524022 |
| ENSG00000280057 | ENSG0000028 | 4.2347234 | -0.376638487 | 0.87166974 | -0.432088519 | 0.665677079 | 0.80728654 |
| ENSG00000280063 | ENSG0000028 | 81.05145989 | 0.168980049 | 0.217536625 | 0.77678896 | 0.437283291 | 0.626915736 |
| ENSG00000280064 | ENSG0000028 | 4.351061118 | -1.071201416 | 0.867081152 | -1.235410797 | 0.216677657 | 0.386094877 |
| ENSG00000280067 | ENSG0000028 | 3.522761891 | -0.575265462 | 0.942604393 | -0.610293635 | 0.541667313 | 0.715650991 |
| ENSG00000280068 | ENSG0000028 | 7.166464312 | -2.039799477 | 0.729665491 | -2.795526855 | 0.00518152 | 0.016615049 |
| ENSG00000280071 | ENSG0000028 | 102.6476701 | 0.208420049 | 0.200657239 | 1.038686917 | 0.298950368 | 0.485654092 |
| ENSG00000280073 | ENSG0000028 | 4.525214698 | -0.916760714 | 0.84486913 | -1.085091976 | 0.277880921 | 0.460658211 |
| ENSG00000280077 | ENSG0000028 | 26.92437687 | -0.294411705 | 0.347006369 | -0.848433144 | 0.396196791 | 0.589145937 |
| ENSG00000280088 | ENSG0000028 | 62.71619552 | 0.022390502 | 0.222082311 | 0.100820734 | 0.919692767 | 0.960994095 |
| ENSG00000280099 | ENSG0000028 | 73.70605875 | 0.359842549 | 0.213022012 | 1.689227068 | 0.091175924 | 0.196539719 |
| ENSG00000280106 | ENSG0000028 | 9.100742458 | -0.116313399 | 0.583573187 | -0.199312445 | 0.842018345 | 0.917009895 |
| ENSG00000280114 | ENSG0000028 | 13.62225911 | -0.225396234 | 0.498105797 | -0.452506747 | 0.650903962 | 0.797170242 |
| ENSG00000280120 | ENSG0000028 | 44.49584884 | 0.634948252 | 0.27227768 | 2.33198789 | 0.019701328 | 0.053989783 |
| ENSG00000280128 | ENSG0000028 | 418.5611438 | 0.670295664 | 0.10350687 | 6.475856777 | 9.43E-11 | 8.17E-10 |
| ENSG00000280129 | ENSG0000028 | 7.211069193 | -0.266806974 | 0.649876911 | -0.410550012 | 0.681402524 | 0.819131835 |
| ENSG00000280132 | ENSG0000028 | 13.42015891 | -0.719169652 | 0.514447578 | -1.397945451 | 0.162129449 | 0.309215959 |

| | | | | | | | |
|---|---|---|---|---|---|---|---|
| ENSG00000280138 | ENSG0000028 | 13.16377794 | -0.519766307 | 0.494744598 | -1.050575002 | 0.293453827 | 0.478997495 |
| ENSG00000280143 | ENSG0000028 | 2.741264244 | 0.376084314 | 1.148605263 | 0.327426946 | 0.743344988 | 0.858490754 |
| ENSG00000280145 | ENSG0000028 | 2.304504468 | -0.093470489 | 1.229036059 | -0.076051869 | 0.939377832 | 0.971215408 |
| ENSG00000280149 | ENSG0000028 | 15.43444972 | 0.137563657 | 0.445746427 | 0.308614156 | 0.757615051 | 0.866391318 |
| ENSG00000280152 | ENSG0000028 | 47.50331812 | 0.677279913 | 0.264015219 | 2.565306329 | 0.010308475 | 0.030524902 |
| ENSG00000280157 | ENSG0000028 | 1.655279027 | 1.052578984 | 1.448291482 | 0.726772888 | 0.467365094 | NA |
| ENSG00000280160 | ENSG0000028 | 3.147942747 | -1.212652847 | 1.058893501 | -1.145207564 | 0.252123186 | 0.430783992 |
| ENSG00000280161 | ENSG0000028 | 48.23662681 | -0.421959061 | 0.258861058 | -1.630060018 | 0.103088813 | 0.217318266 |
| ENSG00000280169 | ENSG0000028 | 1.738541981 | 0.28560923 | 1.379114414 | 0.20709611 | 0.8359348 | NA |
| ENSG00000280172 | ENSG0000028 | 34.18285224 | 0.988102808 | 0.314822465 | 3.138603243 | 0.001697551 | 0.00605533 |
| ENSG00000280173 | ENSG0000028 | 69.11513222 | -0.143356811 | 0.214916646 | -0.667034471 | 0.504750115 | 0.68534878 |
| ENSG00000280184 | ENSG0000028 | 3.050403822 | -1.138209639 | 1.085104409 | -1.048940203 | 0.294205641 | 0.480053873 |
| ENSG00000280186 | ENSG0000028 | 14.88841281 | 0.773377811 | 0.477579766 | 1.619368882 | 0.105367918 | 0.221046186 |
| ENSG00000280187 | ENSG0000028 | 148.8629909 | -0.089382379 | 0.148125858 | -0.60342185 | 0.54622809 | 0.719191436 |
| ENSG00000280194 | ENSG0000028 | 23.74523039 | 0.104956365 | 0.382279743 | 0.274553823 | 0.783659048 | 0.882691054 |
| ENSG00000280195 | ENSG0000028 | 112.8340974 | -0.680399369 | 0.175515623 | -3.876574379 | 0.000105937 | 0.000470514 |
| ENSG00000280198 | ENSG0000028 | 4.318607563 | -0.10508015 | 0.870525951 | -0.120708808 | 0.903921682 | 0.951583837 |
| ENSG00000280202 | ENSG0000028 | 45.46918967 | 0.497882516 | 0.263119252 | 1.892231423 | 0.058460153 | 0.136579684 |
| ENSG00000280206 | ENSG0000028 | 180.6121072 | -0.168017889 | 0.138631036 | -1.211978886 | 0.225520463 | 0.397545342 |
| ENSG00000280207 | ENSG0000028 | 2.073719896 | -0.214073405 | 1.257928224 | -0.170179348 | 0.864869092 | 0.929799912 |
| ENSG00000280211 | ENSG0000028 | 36.26783756 | -0.101513124 | 0.290040019 | -0.349996957 | 0.726340981 | 0.84790912 |
| ENSG00000280213 | UCKL1-AS1 | 15.42718727 | -0.284307413 | 0.45004462 | -0.631731611 | 0.527562269 | 0.703507931 |
| ENSG00000280214 | ENSG0000028 | 3.54775384 | -0.16513164 | 0.928319285 | -0.177882376 | 0.858815352 | 0.926723937 |
| ENSG00000280216 | ENSG0000028 | 8.958912759 | 0.175206611 | 0.586245256 | 0.298862309 | 0.765045106 | 0.871320362 |
| ENSG00000280219 | ENSG0000028 | 22.45629776 | -0.419564041 | 0.381361057 | -1.100175368 | 0.271255721 | 0.45324541 |
| ENSG00000280225 | ENSG0000028 | 2.603752619 | 3.633072548 | 1.510551772 | 2.405129447 | 0.016166738 | 0.045432436 |
| ENSG00000280231 | ENSG0000028 | 3.729259071 | 0.624820961 | 0.952192039 | 0.656192171 | 0.511700485 | 0.691292521 |
| ENSG00000280233 | ENSG0000028 | 1.749806486 | 0.660527506 | 1.366554381 | 0.483352522 | 0.628845453 | NA |
| ENSG00000280239 | ENSG0000028 | 48.64547952 | -0.271843988 | 0.266706402 | -1.019263075 | 0.308078089 | 0.496056107 |
| ENSG00000280247 | ENSG0000028 | 4.302917199 | 0.274940047 | 0.853228063 | 0.32223512 | 0.747274581 | 0.859961818 |
| ENSG00000280254 | ENSG0000028 | 3.733699122 | -0.463645582 | 0.90345546 | -0.513191411 | 0.607817434 | 0.766117139 |
| ENSG00000280273 | ENSG0000028 | 5.277766437 | -0.367282617 | 0.748737754 | -0.490535724 | 0.623754857 | 0.777539482 |
| ENSG00000280278 | FLJ30679 | 2.519526621 | -0.807113766 | 1.135077243 | -0.711065058 | 0.477043922 | 0.661968041 |

| | | | | | | | |
|---|---|---|---|---|---|---|---|
| ENSG00000280279 | LINC02887 | 10.43182715 | -0.214028551 | 0.541855032 | -0.394992274 | 0.69284859 | 0.826827386 |
| ENSG00000280286 | FRG1KP | 3.984653942 | -0.39798019 | 0.879539427 | -0.452487039 | 0.650918157 | 0.797170242 |
| ENSG00000280287 | ENSG0000028 | 85.88437644 | -0.476165979 | 0.191678833 | -2.484186551 | 0.012984776 | 0.03739746 |
| ENSG00000280300 | ENSG0000028 | 2.823931949 | -1.89326246 | 1.200883046 | -1.576558572 | 0.114897132 | 0.237133621 |
| ENSG00000280303 | ERICD | 65.78608322 | 0.775027967 | 0.21980417 | 3.525993009 | 0.000421898 | 0.001692018 |
| ENSG00000280310 | ENSG0000028 | 3.907720466 | 1.323971641 | 0.977870831 | 1.353933054 | 0.175757737 | 0.329028738 |
| ENSG00000280311 | ENSG0000028 | 5.576661938 | -0.47661322 | 0.750151905 | -0.635355609 | 0.525196513 | 0.701327047 |
| ENSG00000280320 | ENSG0000028 | 3.980346127 | -0.880058007 | 0.897059237 | -0.981047818 | 0.326569161 | 0.517145554 |
| ENSG00000280327 | ENSG0000028 | 7.566029228 | 0.03225892 | 0.663646798 | 0.048608568 | 0.961231242 | 0.98212385 |
| ENSG00000280332 | ENSG0000028 | 29.70033001 | -0.319125753 | 0.336583974 | -0.94813116 | 0.343062687 | 0.534092843 |
| ENSG00000280339 | ENSG0000028 | 52.03133407 | 0.989185374 | 0.256817799 | 3.851701004 | 0.0001173 | 0.000517176 |
| ENSG00000280347 | ENSG0000028 | 47.59968787 | 0.299946798 | 0.27440672 | 1.093073805 | 0.274361396 | 0.456373753 |
| ENSG00000280351 | ENSG0000028 | 140.6960626 | 0.39550111 | 0.155366717 | 2.545597388 | 0.010909093 | 0.032063369 |
| ENSG00000280355 | ENSG0000028 | 63.71881273 | -0.59435938 | 0.222025759 | -2.676983891 | 0.007428821 | 0.022887073 |
| ENSG00000280367 | ENSG0000028 | 3.803166988 | 0.197417167 | 0.897027473 | 0.220079288 | 0.825809405 | 0.908152817 |
| ENSG00000280378 | ENSG0000028 | 5.117087291 | 0.274219932 | 0.764127281 | 0.358866826 | 0.719694718 | 0.844231787 |
| ENSG00000280383 | ENSG0000028 | 4.219286774 | -0.358742943 | 0.877774443 | -0.408696044 | 0.682762737 | 0.819956717 |
| ENSG00000280387 | ENSG0000028 | 10.58739522 | 0.954837602 | 0.556290155 | 1.716438072 | 0.086081893 | 0.187514826 |
| ENSG00000280388 | ENSG0000028 | 15.91046723 | 0.256639673 | 0.437858285 | 0.586124968 | 0.557791539 | 0.727915708 |
| ENSG00000280399 | ENSG0000028 | 3.954047095 | 1.052080001 | 0.948530205 | 1.109168685 | 0.267357419 | 0.448626961 |
| ENSG00000280401 | ENSG0000028 | 4.987976966 | -0.199717915 | 0.771001295 | -0.259037068 | 0.79560664 | 0.890824694 |
| ENSG00000280407 | ENSG0000028 | 32.86295057 | -0.512137272 | 0.320917287 | -1.595854424 | 0.110521299 | 0.229652047 |
| ENSG00000280416 | ENSG0000028 | 6.037984073 | -1.346706523 | 0.760057201 | -1.771848909 | 0.076419641 | 0.170482016 |
| ENSG00000280417 | ENSG0000028 | 25.08202345 | 0.446839858 | 0.359085808 | 1.24438184 | 0.213359063 | 0.381544817 |
| ENSG00000280420 | ENSG0000028 | 3.933193844 | -0.316071741 | 0.925452363 | -0.341532156 | 0.732703001 | 0.852256711 |
| ENSG00000280422 | ENSG0000028 | 1.991909014 | -0.240703035 | 1.266553015 | -0.190045764 | 0.84927327 | 0.921106747 |
| ENSG00000280424 | ENSG0000028 | 3.666043744 | 1.320748619 | 0.996813033 | 1.324971258 | 0.185180684 | 0.34217035 |
| ENSG00000280426 | ENSG0000028 | 2.155919483 | -0.411132664 | 1.171629803 | -0.350906628 | 0.725658399 | 0.847716233 |
| ENSG00000280433 | ENSG0000028 | 46.58341366 | 0.063028487 | 0.268637853 | 0.234622509 | 0.814501738 | 0.901763793 |
| ENSG00000280434 | ENSG0000028 | 2.774886643 | 1.075726981 | 1.135268971 | 0.947552526 | 0.343357304 | 0.534333937 |
| ENSG00000280486 | ENSG0000028 | 6.6649757 | 0.323286561 | 0.681069019 | 0.474675182 | 0.63501851 | 0.784856074 |
| ENSG00000280537 | ENSG0000028 | 12.87415164 | -0.538566319 | 0.515012469 | -1.045734524 | 0.295683619 | 0.481780113 |
| ENSG00000280587 | LINC01348 | 22.86186417 | -0.551756428 | 0.364430132 | -1.514025265 | 0.130019432 | 0.261291788 |

| | | | | | | | |
|---|---|---|---|---|---|---|---|
| ENSG00000280594 | BTG3-AS1 | 3.284493431 | 0.388067739 | 0.968129076 | 0.400842975 | 0.688535736 | 0.823987156 |
| ENSG00000280604 | ENSG0000028 | 10.82836968 | -0.316972255 | 0.533695049 | -0.593920172 | 0.552565506 | 0.724232303 |
| ENSG00000280620 | SCAANT1 | 24.47514046 | 0.940438782 | 0.364121074 | 2.582763949 | 0.009801236 | 0.029210312 |
| ENSG00000280623 | PCAT14 | 18.17551647 | -5.266840742 | 0.904912841 | -5.820274067 | 5.88E-09 | 4.34E-08 |
| ENSG00000280634 | THRIL | 36.48702108 | -0.802977214 | 0.302909311 | -2.65088323 | 0.008028159 | 0.024519501 |
| ENSG00000280670 | CCDC163 | 106.7924704 | -0.508367437 | 0.177966807 | -2.856529509 | 0.004283002 | 0.014021987 |
| ENSG00000280721 | LINC01943 | 7.334470284 | 0.240926537 | 0.666635579 | 0.361406658 | 0.717795472 | 0.843038507 |
| ENSG00000280734 | LINC01232 | 125.4490365 | -0.515571068 | 0.163827909 | -3.147028307 | 0.00164939 | 0.00590373 |
| ENSG00000280739 | EIF1B-AS1 | 35.25175795 | -0.331160918 | 0.292642371 | -1.131623271 | 0.257792852 | 0.437525331 |
| ENSG00000280758 | ENSG0000028 | 29.05547242 | 0.272787299 | 0.322768393 | 0.845148734 | 0.398027801 | 0.591025812 |
| ENSG00000280776 | LINC01202 | 31.01343245 | 0.322117044 | 0.319615512 | 1.007826691 | 0.31353767 | 0.501878263 |
| ENSG00000280789 | PAGR1 | 1183.829028 | -0.376127238 | 0.063646721 | -5.909609028 | 3.43E-09 | 2.60E-08 |
| ENSG00000280798 | LINC00294 | 135.6064789 | -0.026273922 | 0.156678425 | -0.167693299 | 0.866824568 | 0.930846618 |
| ENSG00000280828 | ENSG0000028 | 8.695154488 | 0.187528005 | 0.585228492 | 0.320435535 | 0.748638191 | 0.860883258 |
| ENSG00000280832 | GSEC | 56.28599334 | -0.496825468 | 0.235067837 | -2.113540819 | 0.034554505 | 0.08765679 |
| ENSG00000280836 | ENSG0000028 | 6.383685868 | -0.356070583 | 0.684718928 | -0.520024449 | 0.603046535 | 0.762155667 |
| ENSG00000280852 | ENSG0000028 | 49.18216561 | 0.259125656 | 0.252860541 | 1.024776958 | 0.305468441 | 0.492996063 |
| ENSG00000280870 | MIR325HG | 4.327016325 | -0.658963778 | 0.857574985 | -0.768403685 | 0.442247392 | 0.631470633 |
| ENSG00000280924 | LINC00628 | 18.4333086 | 1.074930077 | 0.422863936 | 2.542023535 | 0.011021275 | 0.032364062 |
| ENSG00000280927 | CTBP1-AS | 32.57864118 | -0.781367271 | 0.33033728 | -2.365362064 | 0.018012452 | 0.049892012 |
| ENSG00000281005 | LINC00921 | 26.39321126 | 0.226124056 | 0.338354145 | 0.668305853 | 0.503938378 | 0.684935797 |
| ENSG00000281021 | ENSG0000028 | 6.314752662 | -0.291967824 | 0.743007507 | -0.392954071 | 0.694353405 | 0.827839878 |
| ENSG00000281026 | N4BP2L2-IT2 | 22.11494574 | 0.554663905 | 0.458041771 | 1.210946118 | 0.225916056 | 0.398012006 |
| ENSG00000281091 | ENSG0000028 | 9.460434166 | -0.950766149 | 0.573284984 | -1.658452907 | 0.097226085 | 0.20726386 |
| ENSG00000281100 | ENSG0000028 | 2.981437275 | 1.715370075 | 1.099699067 | 1.559854078 | 0.118794369 | 0.243632585 |
| ENSG00000281103 | TRG-AS1 | 3.834810916 | -1.09927175 | 0.924564019 | -1.188962287 | 0.234454508 | 0.4093726 |
| ENSG00000281133 | ENSG0000028 | 3.792840668 | -0.223626137 | 0.933640343 | -0.239520645 | 0.81070189 | 0.899758757 |
| ENSG00000281183 | NPTN-IT1 | 74.62572322 | -0.648623548 | 0.21783979 | -2.977525588 | 0.002905854 | 0.009874135 |
| ENSG00000281189 | GHET1 | 23.52524707 | 0.078298452 | 0.361134835 | 0.216812239 | 0.828354673 | 0.909208431 |
| ENSG00000281195 | ENSG0000028 | 6.986975859 | -1.439948508 | 0.719814011 | -2.000445235 | 0.045452208 | 0.110382559 |
| ENSG00000281207 | SLFNL1-AS1 | 12.53424756 | -0.057202087 | 0.50443514 | -0.113398298 | 0.909714789 | 0.954699394 |
| ENSG00000281332 | LINC00997 | 118.3974964 | 1.026579588 | 0.174172738 | 5.894031406 | 3.77E-09 | 2.84E-08 |
| ENSG00000281333 | ENSG0000028 | 7.938825915 | 0.641549807 | 0.642362393 | 0.998735004 | 0.317923079 | 0.507067749 |

| | | | | | | | |
|---|---|---|---|---|---|---|---|
| ENSG00000281344 | HELLPAR | 13.24675823 | -0.123840926 | 0.492894744 | -0.251252274 | 0.801619071 | 0.894413497 |
| ENSG00000281357 | ARRDC3-AS1 | 25.80942189 | 0.345726755 | 0.349842075 | 0.988236637 | 0.323036769 | 0.512905833 |
| ENSG00000281358 | RASSF1-AS1 | 3.38900419 | -0.762851269 | 0.968770687 | -0.787442559 | 0.431022841 | 0.621162456 |
| ENSG00000281371 | INE2 | 2.430659829 | -1.520361246 | 1.183765859 | -1.284342875 | 0.199022011 | 0.361800851 |
| ENSG00000281376 | ABALON | 42.71831885 | -0.585604013 | 0.28927023 | -2.024418528 | 0.042927107 | 0.105196467 |
| ENSG00000281398 | SNHG4 | 242.89987 | 1.356293435 | 0.130975334 | 10.3553348 | 3.96E-25 | 8.83E-24 |
| ENSG00000281404 | LINC01176 | 10.17059463 | -2.153647266 | 0.619046232 | -3.478976456 | 0.000503333 | 0.001990425 |
| ENSG00000281450 | PANDAR | 2.562107262 | -1.221078047 | 1.225474824 | -0.996412185 | 0.319049915 | 0.508405464 |
| ENSG00000281468 | ENSG0000028 | 9.028195582 | 0.428056723 | 0.600485177 | 0.71285144 | 0.47593769 | 0.661056589 |
| ENSG00000281490 | CICP14 | 10.45590552 | -0.216565476 | 0.586745652 | -0.369096005 | 0.712056166 | 0.839953127 |
| ENSG00000281501 | SEPSECS-AS1 | 70.25256081 | -0.622849316 | 0.22108901 | -2.81718805 | 0.004844615 | 0.015669759 |
| ENSG00000281530 | ENSG0000028 | 7.124494945 | -1.374392866 | 0.702594171 | -1.956168899 | 0.050445261 | 0.120744396 |
| ENSG00000281538 | ENSG0000028 | 17.92937605 | -0.426610543 | 0.422216613 | -1.010406814 | 0.312300425 | 0.500521665 |
| ENSG00000281571 | ENSG0000028 | 10.35130005 | 0.99727529 | 0.560079193 | 1.780596929 | 0.074978322 | 0.167821515 |
| ENSG00000281641 | SAMD12-AS1 | 8.189698167 | -0.132388529 | 0.652741462 | -0.202819243 | 0.839276317 | 0.915309377 |
| ENSG00000281649 | EBLN3P | 1739.306276 | 0.124414449 | 0.063924295 | 1.946278015 | 0.051621353 | 0.123084278 |
| ENSG00000281691 | RBM5-AS1 | 7.596324936 | -0.776336258 | 0.646649474 | -1.200551905 | 0.229925067 | 0.403062894 |
| ENSG00000281692 | PACRG-AS1 | 2.586706827 | 1.3504197 | 1.153415491 | 1.170800731 | 0.241678884 | 0.417816067 |
| ENSG00000281706 | LINC01012 | 80.94509 | -0.231362811 | 0.198051547 | -1.168194922 | 0.242728148 | 0.419006978 |
| ENSG00000281731 | METTL14-DT | 7.503783884 | -0.613422517 | 0.641537041 | -0.956176305 | 0.338983169 | 0.530116156 |
| ENSG00000281832 | LINC00602 | 3.469323321 | -3.352217188 | 1.287090307 | -2.604492607 | 0.009201041 | 0.027663417 |
| ENSG00000281849 | ENSG0000028 | 4.054333235 | 0.756378077 | 0.936510003 | 0.807656164 | 0.419288545 | 0.610870102 |
| ENSG00000281849 | ENSG0000028 | 4.054333235 | 0.756378077 | 0.936510003 | 0.807656164 | 0.419288545 | 0.610870102 |
| ENSG00000281903 | LINC02246 | 2.987960696 | 0.02973691 | 1.052981699 | 0.028240672 | 0.977470199 | 0.989489456 |
| ENSG00000281912 | LINC01144 | 42.91296855 | -0.279275909 | 0.271323794 | -1.029308579 | 0.303334693 | 0.490496406 |
| ENSG00000282021 | ENSG0000028 | 7.608779938 | 0.573324518 | 0.656249172 | 0.873638463 | 0.382315173 | 0.574617273 |
| ENSG00000282048 | ENSG0000028 | 6.172816687 | 0.736774183 | 0.721040578 | 1.021820694 | 0.306865773 | 0.494659605 |
| ENSG00000282100 | HSP90AB4P | 26.96808543 | -0.388253362 | 0.388408582 | -0.999600371 | 0.317503944 | 0.506610591 |
| ENSG00000282121 | ENSG0000028 | 1.935077735 | -0.639606682 | 1.242261044 | -0.514873009 | 0.606641763 | 0.7650978 |
| ENSG00000282164 | PEG13 | 43.38863744 | 0.202186159 | 0.265552267 | 0.761379901 | 0.446430189 | 0.634987357 |
| ENSG00000282206 | ENSG0000028 | 4.705295349 | -9.36E-05 | 0.798749656 | -0.000117233 | 0.999906461 | 0.999914377 |
| ENSG00000282221 | ENSG0000028 | 48.36368874 | 1.248508264 | 0.271018607 | 4.606725257 | 4.09E-06 | 2.21E-05 |
| ENSG00000282381 | ENSG0000028 | 3.972419454 | 1.222306639 | 0.959197768 | 1.274300963 | 0.202556775 | 0.366686587 |

| | | | | | | | |
|---|---|---|---|---|---|---|---|
| ENSG00000282386 | ENSG0000028 | 30.81807264 | -0.120889169 | 0.318465988 | -0.379598367 | 0.704243573 | 0.834556647 |
| ENSG00000282393 | ENSG0000028 | 12.53291331 | 0.751236648 | 0.531483011 | 1.413472554 | 0.157516826 | 0.302177286 |
| ENSG00000282508 | LINC01002 | 11.30059584 | 0.416063852 | 0.522669012 | 0.796036961 | 0.426010553 | 0.616845426 |
| ENSG00000282572 | FAM157D | 5.392600466 | -0.257807614 | 0.756406016 | -0.340832315 | 0.733229821 | 0.852610101 |
| ENSG00000282728 | PGR-AS1 | 6.32836247 | 3.011864856 | 0.941155006 | 3.200179395 | 0.001373421 | 0.005004173 |
| ENSG00000282772 | ENSG0000028 | 11.09376366 | 0.003426298 | 0.546876907 | 0.006265209 | 0.995001119 | 0.996913245 |
| ENSG00000282787 | ENSG0000028 | 4.360057193 | 0.857730395 | 0.883227398 | 0.971132007 | 0.331482547 | 0.522374777 |
| ENSG00000282826 | FRG1CP | 15.12905819 | -0.452165945 | 0.475057424 | -0.951813239 | 0.341191707 | 0.532308789 |
| ENSG00000282840 | ENSG0000028 | 1.877436828 | -1.635441169 | 1.378634815 | -1.186275837 | 0.235513386 | 0.410506577 |
| ENSG00000282851 | BISPR | 3.724103321 | -0.016118206 | 0.910108267 | -0.017710208 | 0.985870037 | 0.993350589 |
| ENSG00000282855 | ENSG0000028 | 7.375964396 | -0.086114311 | 0.651770382 | -0.132123694 | 0.894886455 | 0.94645898 |
| ENSG00000282870 | FRG1DP | 34.02271474 | -0.089033981 | 0.306784256 | -0.290216916 | 0.771650296 | 0.875561078 |
| ENSG00000282876 | ENSG0000028 | 5.524978947 | -1.173597021 | 0.75613545 | -1.552098927 | 0.120638555 | 0.2466092 |
| ENSG00000282885 | ENSG0000028 | 5.227681472 | -0.650704453 | 0.765697594 | -0.849819117 | 0.39542566 | 0.588342498 |
| ENSG00000282886 | PTMAP12 | 57.0725437 | -0.030105167 | 0.513411414 | -0.058637511 | 0.953240833 | 0.978321944 |
| ENSG00000282914 | ENSG0000028 | 2.932776892 | 0.800621037 | 1.046524602 | 0.765028395 | 0.444254636 | 0.633033192 |
| ENSG00000282917 | ENSG0000028 | 3.554887654 | -0.82656679 | 1.022992433 | -0.807989153 | 0.419096826 | 0.610728065 |
| ENSG00000282933 | RHOXF1P3 | 11.33126633 | -0.151452512 | 0.534562583 | -0.283320452 | 0.776931201 | 0.878923057 |
| ENSG00000282936 | ENSG0000028 | 53.81981273 | -0.322869741 | 0.249609958 | -1.293497037 | 0.195839208 | 0.357338497 |
| ENSG00000282950 | ENSG0000028 | 14.49587891 | 0.821923513 | 0.469953639 | 1.748945949 | 0.080300363 | 0.177159665 |
| ENSG00000282951 | ENSG0000028 | 1.857163681 | -0.486239503 | 1.270990025 | -0.382567521 | 0.702040456 | 0.833064766 |
| ENSG00000282961 | PRNCR1 | 9.252444133 | -0.688003345 | 0.593279657 | -1.159661109 | 0.24618681 | 0.423384469 |
| ENSG00000282977 | PCBP2-OT1 | 3.670694496 | 0.228430949 | 0.905001925 | 0.252409352 | 0.800724672 | 0.893871969 |
| ENSG00000282978 | BTF3P16 | 91.09743185 | 0.251725942 | 0.193087343 | 1.303689502 | 0.19233947 | 0.352473834 |
| ENSG00000282988 | ENSG0000028 | 58.32085742 | 0.481919733 | 0.237484045 | 2.029272038 | 0.04243059 | 0.104191579 |
| ENSG00000282989 | PDCL3P2 | 2.37626337 | -1.023343419 | 1.189492098 | -0.860319645 | 0.389612867 | 0.58233771 |
| ENSG00000282995 | FRG1EP | 29.79594216 | -0.271881574 | 0.318162749 | -0.854536161 | 0.392807983 | 0.585752911 |
| ENSG00000282996 | MPP7-DT | 2.357826419 | -0.634587879 | 1.13119467 | -0.560989099 | 0.574804968 | 0.741214298 |
| ENSG00000283000 | LINC02635 | 1.735183492 | -0.775030806 | 1.380455131 | -0.561431363 | 0.574503509 | NA |
| ENSG00000283005 | CDC27P3 | 5.061254785 | 0.293946516 | 0.773088891 | 0.380223438 | 0.703779562 | 0.834211563 |
| ENSG00000283016 | ATP5MC1P2 | 7.72907923 | -0.15665666 | 0.634026913 | -0.247082035 | 0.804844737 | 0.896355873 |
| ENSG00000283020 | DUX4L37 | 4.431779297 | 0.096373463 | 0.87023507 | 0.110744173 | 0.911819217 | 0.955697231 |
| ENSG00000283023 | FRG1GP | 46.59740037 | 0.099066211 | 0.261593425 | 0.378702987 | 0.704908435 | 0.834997903 |

| | | | | | | | |
|---|---|---|---|---|---|---|---|
| ENSG00000283031 | ENSG0000028 | 1.780184436 | 0.224065658 | 1.459715518 | 0.153499538 | 0.878004355 | NA |
| ENSG00000283041 | ENSG0000028 | 12069.58987 | 0.138249669 | 0.046935951 | 2.94549628 | 0.00322437 | 0.010850382 |
| ENSG00000283047 | FRG1FP | 9.716383808 | -0.509177358 | 0.564372825 | -0.902200346 | 0.366950451 | 0.559399478 |
| ENSG00000283050 | GTF2IP12 | 244.0269318 | -0.36469538 | 0.120862586 | -3.017438174 | 0.00254921 | 0.008766135 |
| ENSG00000283064 | ENSG0000028 | 8.024337898 | 0.037009304 | 0.614390285 | 0.06023745 | 0.951966519 | 0.977646532 |
| ENSG00000283075 | ENSG0000028 | 35.36118406 | 0.443888477 | 0.296603319 | 1.496572856 | 0.134504437 | 0.268237138 |
| ENSG00000283078 | ENSG0000028 | 42.99851027 | -0.219764985 | 0.266964898 | -0.823198054 | 0.410395375 | 0.602950551 |
| ENSG00000283098 | ENSG0000028 | 1.65094893 | -1.377286077 | 1.391340877 | -0.989898378 | 0.322223793 | NA |
| ENSG00000283103 | ENSG0000028 | 336.3164903 | -0.044234721 | 0.107641042 | -0.410946602 | 0.68111169 | 0.818990574 |
| ENSG00000283117 | MGC4859 | 6.612521713 | -0.221157112 | 0.705188127 | -0.313614344 | 0.753813964 | 0.864272694 |
| ENSG00000283141 | LINC02666 | 2.987610623 | 0.447983114 | 1.013111657 | 0.442185332 | 0.6583551 | 0.802336104 |
| ENSG00000283154 | IQCJ-SCHIP1 | 22.79979609 | 0.601236978 | 0.389052889 | 1.54538623 | 0.122252869 | 0.248960929 |
| ENSG00000283156 | ENSG0000028 | 1.837825343 | 1.150319651 | 1.391416029 | 0.826725887 | 0.408392445 | 0.600793959 |
| ENSG00000283162 | ENSG0000028 | 4.995875845 | -1.354306649 | 0.847532387 | -1.597940881 | 0.110056135 | 0.228830575 |
| ENSG00000283196 | ENSG0000028 | 100.8712822 | 0.424941398 | 0.193238434 | 2.199052172 | 0.027874213 | 0.072928198 |
| ENSG00000283199 | C13orf46 | 3.055548431 | -0.595208672 | 1.017088078 | -0.585208583 | 0.558407475 | 0.728502064 |
| ENSG00000283208 | ENSG0000028 | 3.578092855 | 0.100025257 | 0.956973871 | 0.104522454 | 0.91675475 | 0.95891724 |
| ENSG00000283209 | ENSG0000028 | 1.513825686 | 1.098306035 | 1.590190743 | 0.690675657 | 0.489769391 | NA |
| ENSG00000283235 | ENSG0000028 | 4.899585753 | -0.520738252 | 0.821797305 | -0.633657775 | 0.526304187 | 0.702310398 |
| ENSG00000283240 | ENSG0000028 | 2.094057008 | -1.231364519 | 1.255933006 | -0.980438059 | 0.326869931 | 0.517416244 |
| ENSG00000283294 | ENSG0000028 | 208.2432888 | 0.145765014 | 0.131316777 | 1.110025828 | 0.266987897 | 0.448105235 |
| ENSG00000283303 | ENSG0000028 | 1.822127842 | -0.416462505 | 1.317762697 | -0.316037558 | 0.751974001 | 0.862988823 |
| ENSG00000283312 | ENSG0000028 | 5.934988227 | 0.694805947 | 0.740338547 | 0.938497596 | 0.347988752 | 0.538977059 |
| ENSG00000283317 | ENSG0000028 | 4.626743735 | -0.167973187 | 0.806595214 | -0.20824967 | 0.835034028 | 0.913001315 |
| ENSG00000283341 | ENSG0000028 | 11.85447641 | -0.30644086 | 0.537206388 | -0.570434132 | 0.568383285 | 0.73598513 |
| ENSG00000283342 | NBPF13P | 10.73475923 | -1.978260762 | 0.588090538 | -3.363871093 | 0.000768574 | 0.002939496 |
| ENSG00000283375 | ENSG0000028 | 3.737328493 | -3.475367583 | 1.273360288 | -2.729288495 | 0.006347115 | 0.019915582 |
| ENSG00000283390 | PET100P1 | 2.949800403 | 1.219608284 | 1.098448561 | 1.110300771 | 0.266869441 | 0.448031931 |
| ENSG00000283413 | ENSG0000028 | 7.902007684 | -2.045953936 | 0.695575021 | -2.941385002 | 0.003267481 | 0.010985787 |
| ENSG00000283429 | MIR1244-3 | 2.428809116 | -0.624980274 | 1.179972561 | -0.529656616 | 0.596350033 | 0.757238445 |
| ENSG00000283445 | ENSG0000028 | 3.500590091 | -0.078678133 | 0.938389471 | -0.083843794 | 0.933180628 | 0.967949159 |
| ENSG00000283480 | ENSG0000028 | 3.532693573 | -0.901327333 | 0.932506931 | -0.966563682 | 0.333762198 | 0.524777574 |
| ENSG00000283491 | ENSG0000028 | 15.07594815 | -0.495507154 | 0.469688367 | -1.054970037 | 0.291439029 | 0.476420674 |

| | | | | | | | |
|---|---|---|---|---|---|---|---|
| ENSG00000283498 | MIR1244-2 | 34.44824591 | -0.259010447 | 0.323676239 | -0.800214585 | 0.423586481 | 0.614499075 |
| ENSG00000283511 | ENSG0000028 | 2.814105743 | 1.0837984 | 1.098425432 | 0.986683637 | 0.323797754 | 0.51393624 |
| ENSG00000283526 | PRRT1B | 441.7922696 | -0.831350172 | 0.109950943 | -7.561100907 | 4.00E-14 | 4.49E-13 |
| ENSG00000283529 | ENSG0000028 | 1.811235692 | -0.400157943 | 1.288300991 | -0.310609047 | 0.756097849 | 0.865184627 |
| ENSG00000283537 | ENSG0000028 | 4.023846757 | -0.038674481 | 0.863669142 | -0.044779278 | 0.964283242 | 0.983752449 |
| ENSG00000283538 | ENSG0000028 | 52.93868845 | 0.767948042 | 0.248361361 | 3.092059244 | 0.001987732 | 0.006993645 |
| ENSG00000283566 | ENSG0000028 | 2.033502022 | -0.670209133 | 1.252166496 | -0.535239631 | 0.592484165 | 0.754736754 |
| ENSG00000283627 | ENSG0000028 | 14.10918918 | 0.099291008 | 0.46656711 | 0.212811847 | 0.831473723 | 0.91071463 |
| ENSG00000283635 | ENSG0000028 | 1.724735752 | 2.061724389 | 1.553030597 | 1.327549111 | 0.184327106 NA | |
| ENSG00000283667 | ENSG0000028 | 14.02590652 | 0.649330539 | 0.477483283 | 1.359902141 | 0.173860893 | 0.326460579 |
| ENSG00000283674 | ENSG0000028 | 14.74355065 | 0.581327367 | 0.466943849 | 1.244962041 | 0.213145704 | 0.381340969 |
| ENSG00000283689 | ENSG0000028 | 19.54055147 | 0.45208329 | 0.39566537 | 1.142589986 | 0.253208865 | 0.431939753 |
| ENSG00000283692 | ENSG0000028 | 1.818732555 | -2.340369997 | 1.540811092 | -1.518920787 | 0.128782436 | 0.259305633 |
| ENSG00000283696 | ENSG0000028 | 7.152987334 | 0.106824209 | 0.667733937 | 0.1599802 | 0.872896672 | 0.93435909 |
| ENSG00000283709 | FAM238C | 9.74558803 | -0.887018007 | 0.568192984 | -1.561121013 | 0.118495201 | 0.24317769 |
| ENSG00000283765 | ENSG0000028 | 2.366867856 | 0.169512079 | 1.131329628 | 0.149834385 | 0.88089528 | 0.938252557 |
| ENSG00000283828 | ENSG0000028 | 26.22211859 | 0.246146157 | 0.338579233 | 0.726997208 | 0.467227666 | 0.653190484 |
| ENSG00000283839 | ENSG0000028 | 3.333571532 | -0.564030748 | 1.032230856 | -0.54641919 | 0.584777822 | 0.748970088 |
| ENSG00000283897 | ENSG0000028 | 31.06349761 | 0.129116951 | 0.313020259 | 0.412487524 | 0.679982123 | 0.818289926 |
| ENSG00000283930 | PLD5P1 | 23.78672506 | -1.796081927 | 0.432040734 | -4.157205062 | 3.22E-05 | 0.000155392 |
| ENSG00000283959 | ENSG0000028 | 27.82249992 | 0.329153536 | 0.335109367 | 0.982227202 | 0.325987929 | 0.516447728 |
| ENSG00000283971 | MIR4442 | 8.942717396 | -0.323684461 | 0.596879966 | -0.542294061 | 0.587615946 | 0.751132537 |
| ENSG00000283982 | ENSG0000028 | 5.703307698 | -1.834441051 | 0.804588976 | -2.279972887 | 0.022609297 | 0.060853838 |
| ENSG00000283991 | ENSG0000028 | 1.910093248 | 1.448392118 | 1.368032395 | 1.058741097 | 0.289717706 | 0.474045947 |
| ENSG00000284024 | MSANTD7 | 758.6599712 | -0.747381557 | 0.074205556 | -10.0717736 | 7.36E-24 | 1.53E-22 |
| ENSG00000284052 | ENSG0000028 | 2.4963066 | -0.782547745 | 1.124827345 | -0.695704766 | 0.48661374 | 0.67008339 |
| ENSG00000284128 | ENSG0000028 | 1.838677282 | -0.449060575 | 1.272796557 | -0.352814102 | 0.724227811 | 0.846509142 |
| ENSG00000284194 | SCO2 | 329.1119661 | -0.007312769 | 0.116760342 | -0.062630591 | 0.950060669 | 0.976932707 |
| ENSG00000284308 | C2orf81 | 66.71943913 | -0.403935342 | 0.214717625 | -1.881239798 | 0.059939306 | 0.139453112 |
| ENSG00000284343 | MIR631 | 10.99578411 | 0.493724826 | 0.549354662 | 0.898736026 | 0.368793283 | 0.561276316 |
| ENSG00000284395 | PERCC1 | 6.725331501 | -2.177823094 | 0.766721156 | -2.84043694 | 0.004505178 | 0.014684396 |
| ENSG00000284428 | ENSG0000028 | 37.05717224 | 0.405437248 | 0.297022217 | 1.365006471 | 0.172251016 | 0.324127497 |
| ENSG00000284473 | MIR4720 | 15.11098777 | 0.130009574 | 0.449175852 | 0.289440256 | 0.772244493 | 0.875910441 |

| | | | | | | | |
|---|---|---|---|---|---|---|---|
| ENSG00000284512 | ENSG0000028 | 2.158780111 | -0.431888908 | 1.17158427 | -0.368636657 | 0.712398569 | 0.839972059 |
| ENSG00000284523 | ENSG0000028 | 13.72999079 | -0.534035798 | 0.488315336 | -1.093628971 | 0.274117738 | 0.456166913 |
| ENSG00000284526 | ENSG0000028 | 58.96212871 | -0.545967171 | 0.232586849 | -2.347369053 | 0.018906518 | 0.05207294 |
| ENSG00000284602 | ENSG0000028 | 5.609202367 | -1.187703061 | 0.773327001 | -1.535835498 | 0.124578731 | 0.252749278 |
| ENSG00000284606 | ENSG0000028 | 4.098248856 | 1.119488848 | 0.918122012 | 1.219324701 | 0.222720977 | 0.393834827 |
| ENSG00000284618 | ENSG0000028 | 8.86876381 | -0.359773841 | 0.618509111 | -0.581679128 | 0.560782833 | 0.729892663 |
| ENSG00000284624 | ENSG0000028 | 2.373312493 | -0.332855993 | 1.154918966 | -0.288207228 | 0.773188117 | 0.876374099 |
| ENSG00000284630 | ENSG0000028 | 7.937788509 | 0.427658358 | 0.631334568 | 0.677387838 | 0.498159913 | 0.67994219 |
| ENSG00000284633 | ENSG0000028 | 25.49629547 | 0.185964989 | 0.344857824 | 0.539251182 | 0.589713549 | 0.752741052 |
| ENSG00000284634 | ENSG0000028 | 12.89084368 | 0.131757024 | 0.501297451 | 0.262832025 | 0.79268005 | 0.888675611 |
| ENSG00000284642 | ENSG0000028 | 2.588530373 | 0.996372952 | 1.128517407 | 0.882904372 | 0.377287941 | 0.570044482 |
| ENSG00000284648 | ENSG0000028 | 1.944108031 | -0.184539859 | 1.230989479 | -0.149911809 | 0.880834195 | 0.938252557 |
| ENSG00000284664 | ENSG0000028 | 2.279328257 | -2.687858727 | 1.421892245 | -1.890339255 | 0.058712602 | 0.137043863 |
| ENSG00000284669 | ENSG0000028 | 3.68113886 | -0.473301318 | 0.975054009 | -0.48541036 | 0.627385285 | 0.780187429 |
| ENSG00000284686 | ENSG0000028 | 4.083979615 | -0.92793662 | 0.873569236 | -1.062235919 | 0.288128597 | 0.472119287 |
| ENSG00000284691 | ENSG0000028 | 62.17671868 | 1.404072814 | 0.24216077 | 5.798101862 | 6.71E-09 | 4.92E-08 |
| ENSG00000284693 | LINC02606 | 8.809894506 | -0.282378991 | 0.587885882 | -0.480329601 | 0.630993044 | 0.78247367 |
| ENSG00000284707 | ENSG0000028 | 21.32925414 | 0.477255975 | 0.384728153 | 1.240501822 | 0.214789841 | 0.383654827 |
| ENSG00000284713 | SMIM38 | 10.23463502 | 0.492663775 | 0.561274814 | 0.877758564 | 0.380074755 | 0.572482921 |
| ENSG00000284719 | ENSG0000028 | 16.21217616 | 0.628169297 | 0.446688081 | 1.406281752 | 0.159640477 | 0.305306589 |
| ENSG00000284738 | ENSG0000028 | 13.22310831 | -0.51943534 | 0.48105576 | -1.079781978 | 0.280239278 | 0.463539661 |
| ENSG00000284740 | ENSG0000028 | 8.04872571 | 0.431140834 | 0.637788892 | 0.675993012 | 0.499045082 | 0.680567986 |
| ENSG00000284747 | ENSG0000028 | 10.15954045 | -0.466499019 | 0.580804003 | -0.803195255 | 0.421861888 | 0.612888645 |
| ENSG00000284753 | EEF1AKMT4 | 99.4531502 | -0.100803875 | 0.182332676 | -0.552856886 | 0.580361412 | 0.745630889 |
| ENSG00000284770 | TBCE | 33.31841448 | -0.535190977 | 0.316715547 | -1.689815933 | 0.091063175 | 0.196370486 |
| ENSG00000284773 | ENSG0000028 | 12.54162287 | 0.589667578 | 0.52408591 | 1.125135338 | 0.260531689 | 0.440851376 |
| ENSG00000284820 | ENSG0000028 | 2.615376714 | -0.871690523 | 1.157750177 | -0.752917633 | 0.451499412 | 0.639463882 |
| ENSG00000284830 | ENSG0000028 | 4.685927267 | -0.980791683 | 0.827816041 | -1.184794248 | 0.236098808 | 0.411214535 |
| ENSG00000284862 | CCDC39 | 2.852356287 | -0.674304821 | 1.048459226 | -0.643138812 | 0.520134033 | 0.697413646 |
| ENSG00000284874 | ENSG0000028 | 11.41999848 | -0.040008722 | 0.518868122 | -0.07710769 | 0.938537876 | 0.970807194 |
| ENSG00000284879 | NCAL1 | 3.66662039 | 0.859609781 | 1.009382425 | 0.851619524 | 0.3944253 | 0.58746373 |
| ENSG00000284882 | ENSG0000028 | 3.635490267 | -0.872520071 | 1.008100797 | -0.86550876 | 0.38675961 | 0.579318671 |
| ENSG00000284883 | KRT88P | 10.01270825 | 0.18744365 | 0.569846721 | 0.328936964 | 0.742203334 | 0.85777393 |

| | | | | | | | |
|---|---|---|---|---|---|---|---|
| ENSG00000284946 | ENSG0000028 | 165.4993442 | -0.088199575 | 0.153566639 | -0.574340728 | 0.565737254 | 0.733806909 |
| ENSG00000284948 | ENSG0000028 | 3.442372539 | -0.080290759 | 1.016691131 | -0.078972617 | 0.937054404 | 0.969818544 |
| ENSG00000284954 | ENSG0000028 | 19.82230596 | -0.037210455 | 0.397631899 | -0.093580156 | 0.925442674 | 0.964370625 |
| ENSG00000284959 | ENSG0000028 | 7.448452291 | -0.733504136 | 0.654709277 | -1.120350912 | 0.262564254 | 0.443210007 |
| ENSG00000284966 | ENSG0000028 | 36.6985265 | -0.239649432 | 0.312028203 | -0.768037726 | 0.442464772 | 0.631470633 |
| ENSG00000284968 | ENSG0000028 | 23.77436885 | -1.000845343 | 0.36741384 | -2.724027332 | 0.006449117 | 0.020180431 |
| ENSG00000284976 | ENSG0000028 | 602.1955553 | -0.07845189 | 0.087368492 | -0.897942594 | 0.369216155 | 0.561808079 |
| ENSG00000284985 | ENSG0000028 | 8.950929176 | 0.406230999 | 0.58054235 | 0.699743953 | 0.484087222 | 0.668007753 |
| ENSG00000285018 | ENSG0000028 | 35.33620942 | -1.156285842 | 0.303450451 | -3.810460126 | 0.000138708 | 0.000604248 |
| ENSG00000285041 | ENSG0000028 | 6.710395314 | -0.34586531 | 0.665827678 | -0.519451686 | 0.603445796 | 0.762492237 |
| ENSG00000285043 | ENSG0000028 | 45.7958173 | -0.262525537 | 0.277098664 | -0.947408163 | 0.343430833 | 0.53437586 |
| ENSG00000285053 | ENSG0000028 | 224.4821163 | 0.294418099 | 0.125646233 | 2.343230602 | 0.019117563 | 0.052584053 |
| ENSG00000285077 | ARHGAP11B | 21.51400147 | -0.262022746 | 0.378361968 | -0.69251872 | 0.488611634 | 0.671826354 |
| ENSG00000285091 | ENSG0000028 | 9.503557556 | -0.328576619 | 0.575148896 | -0.571289663 | 0.567803308 | 0.73560104 |
| ENSG00000285094 | LINC01488 | 4.060831021 | 0.391466998 | 0.925335004 | 0.423054349 | 0.672255608 | 0.812222991 |
| ENSG00000285095 | ENSG0000028 | 3.209726113 | 0.126101943 | 0.997040654 | 0.12647623 | 0.899354963 | 0.949128346 |
| ENSG00000285130 | ENSG0000028 | 3.020151052 | -0.806670434 | 1.030343926 | -0.782913757 | 0.433677759 | 0.623549332 |
| ENSG00000285177 | ENSG0000028 | 1.895519702 | 0.516559192 | 1.319196087 | 0.391571198 | 0.695375073 | 0.828627301 |
| ENSG00000285184 | ENSG0000028 | 233.9079764 | 0.428992797 | 0.120845585 | 3.549925268 | 0.000385341 | 0.001552733 |
| ENSG00000285190 | ENSG0000028 | 4.717027971 | -1.035665739 | 0.809711217 | -1.279055689 | 0.200877446 | 0.364307741 |
| ENSG00000285205 | ENSG0000028 | 6.289659322 | -0.438439567 | 0.701967409 | -0.624586785 | 0.532242295 | 0.707691769 |
| ENSG00000285210 | LINC02795 | 2.299219533 | 0.241096473 | 1.173057044 | 0.205528345 | 0.837159355 | 0.914148686 |
| ENSG00000285212 | ENSG0000028 | 7.287558666 | -0.177443904 | 0.669678452 | -0.264968812 | 0.791033488 | 0.887393428 |
| ENSG00000285219 | HULC | 42.17228425 | -0.387937346 | 0.271493348 | -1.428901845 | 0.153032447 | 0.295872867 |
| ENSG00000285230 | RALY-AS1 | 163.2646823 | -0.508560142 | 0.151689938 | -3.352629385 | 0.000800478 | 0.003050327 |
| ENSG00000285257 | ENSG0000028 | 13.54302199 | -0.518663563 | 0.479681455 | -1.081266656 | 0.279578515 | 0.462933542 |
| ENSG00000285278 | TFAP2A-AS2 | 25.18271308 | -0.782969374 | 0.355241454 | -2.204048447 | 0.027520937 | 0.072143789 |
| ENSG00000285329 | ENSG0000028 | 26.76314033 | -0.211083102 | 0.344535615 | -0.612659745 | 0.540101345 | 0.714414141 |
| ENSG00000285331 | TCF12-DT | 19.7005786 | -0.89089715 | 0.409545705 | -2.175330226 | 0.029605379 | 0.076765841 |
| ENSG00000285407 | ENSG0000028 | 22.47029571 | -0.164730657 | 0.36756618 | -0.44816598 | 0.654033413 | 0.799165148 |
| ENSG00000285410 | GABPB1-IT1 | 4053.013903 | -0.817240855 | 0.048784918 | -16.75191624 | 5.48E-63 | 4.22E-61 |
| ENSG00000285427 | SOD2-OT1 | 19.49467535 | -1.607927621 | 0.424921551 | -3.784057595 | 0.000154292 | 0.000667697 |
| ENSG00000285437 | POLR2J3 | 36.14658628 | -0.477487977 | 0.295309458 | -1.616907162 | 0.105898322 | 0.221976198 |

| | | | | | | | |
|---|---|---|---|---|---|---|---|
| ENSG00000285444 | ENSG0000028 | 4.835883706 | -0.244614106 | 0.815032296 | -0.300128115 | 0.764079435 | 0.870566317 |
| ENSG00000285458 | ENSG0000028 | 2.363996088 | -0.230595515 | 1.134773525 | -0.203208403 | 0.838972146 | 0.915286602 |
| ENSG00000285467 | DHRS4L1 | 5.363869989 | 0.434111869 | 0.761358755 | 0.570180439 | 0.568555322 | 0.736037682 |
| ENSG00000285486 | ENSG0000028 | 2.675580674 | -0.567964195 | 1.128815285 | -0.503150695 | 0.614858327 | 0.771600111 |
| ENSG00000285509 | TBCEL-TECTA | 3.310008574 | 1.145933808 | 0.990118445 | 1.157370427 | 0.247121051 | 0.424323122 |
| ENSG00000285518 | ENSG0000028 | 4.588930937 | -1.704431865 | 0.894901567 | -1.904602615 | 0.056831746 | 0.133407053 |
| ENSG00000285531 | ENSG0000028 | 45.98509603 | -0.04280206 | 0.274428022 | -0.155968256 | 0.876058043 | 0.935647446 |
| ENSG00000285533 | RELA-DT | 188.3601392 | -0.003288603 | 0.133855339 | -0.024568336 | 0.980399276 | 0.991070363 |
| ENSG00000285568 | ENSG0000028 | 3.679262187 | 0.680947143 | 0.979256427 | 0.695371636 | 0.486822431 | 0.670250064 |
| ENSG00000285570 | ENSG0000028 | 2.233493977 | 1.157729914 | 1.285489727 | 0.900613898 | 0.367793643 | 0.560312536 |
| ENSG00000285571 | ENSG0000028 | 12.2156364 | -0.434984491 | 0.516914446 | -0.841501905 | 0.400066818 | 0.592902178 |
| ENSG00000285577 | ENSG0000028 | 4.284932037 | 0.290544623 | 0.898740155 | 0.323279895 | 0.746483278 | 0.859604617 |
| ENSG00000285581 | ENSG0000028 | 1.7167354 | -0.851051082 | 1.38554587 | -0.614235227 | 0.539059898 | NA |
| ENSG00000285587 | ENSG0000028 | 8.662612957 | 2.592052521 | 0.727942438 | 3.560793253 | 0.000369736 | 0.001494581 |
| ENSG00000285588 | ENSG0000028 | 3.108497702 | -1.648070915 | 1.074438291 | -1.533890711 | 0.125056542 | 0.253611038 |
| ENSG00000285589 | ENSG0000028 | 102.6999209 | -0.362649559 | 0.184252479 | -1.96822078 | 0.04904264 | 0.117842062 |
| ENSG00000285596 | ENSG0000028 | 10.16027781 | 0.529188035 | 0.548905304 | 0.964078926 | 0.335006357 | 0.526221015 |
| ENSG00000285597 | ENSG0000028 | 2.513880472 | -0.395949963 | 1.109766084 | -0.356786866 | 0.721251369 | 0.845194729 |
| ENSG00000285599 | ENSG0000028 | 7.042305397 | 0.712189427 | 0.665786726 | 1.069696044 | 0.284756148 | 0.46860158 |
| ENSG00000285600 | ENSG0000028 | 38.32627055 | -0.345450387 | 0.315592224 | -1.094609946 | 0.273687556 | 0.455667442 |
| ENSG00000285601 | ENSG0000028 | 2.192828317 | -0.444536318 | 1.179916681 | -0.376752296 | 0.706357695 | 0.835939204 |
| ENSG00000285608 | ENSG0000028 | 13.78141706 | -1.78110457 | 0.507518846 | -3.509435329 | 0.000449059 | 0.001792567 |
| ENSG00000285612 | ENSG0000028 | 31.68647662 | -1.404838192 | 0.326966019 | -4.296587752 | 1.73E-05 | 8.68E-05 |
| ENSG00000285637 | ENSG0000028 | 15.68158885 | 1.321600192 | 0.467067507 | 2.82956997 | 0.004661061 | 0.015132286 |
| ENSG00000285641 | ENSG0000028 | 26.46529715 | 0.407433392 | 0.349707031 | 1.165070632 | 0.243990403 | 0.420716972 |
| ENSG00000285644 | ENSG0000028 | 20.44739852 | -0.118564803 | 0.386087817 | -0.307092837 | 0.75877271 | 0.867190412 |
| ENSG00000285646 | ENSG0000028 | 3.134623239 | -0.02078562 | 0.974534851 | -0.02132876 | 0.982983402 | 0.992139551 |
| ENSG00000285649 | ENSG0000028 | 6.670644456 | 0.612195453 | 0.680688771 | 0.899376455 | 0.368452176 | 0.561017717 |
| ENSG00000285650 | ENSG0000028 | 1.947831738 | -1.156358359 | 1.30918591 | -0.883265203 | 0.377093 | 0.569900086 |
| ENSG00000285665 | TOP1P1 | 323.7206615 | -0.145028717 | 0.104576345 | -1.386821435 | 0.165496219 | 0.314174385 |
| ENSG00000285667 | ENSG0000028 | 19.00453333 | 0.233026519 | 0.400519609 | 0.581810514 | 0.560694322 | 0.729860281 |
| ENSG00000285669 | ENSG0000028 | 40.27212239 | 0.380862247 | 0.285289697 | 1.335001756 | 0.181875772 | 0.337613708 |
| ENSG00000285676 | ENSG0000028 | 3.256524236 | -3.233718969 | 1.311042109 | -2.466525634 | 0.013643096 | 0.039077616 |

| | | | | | | | |
|---|---|---|---|---|---|---|---|
| ENSG00000285677 | ENSG0000028 | 17.38961053 | 0.161342623 | 0.447888102 | 0.36022976 | 0.718675322 | 0.843510457 |
| ENSG00000285679 | ENSG0000028 | 20.22794393 | -0.163941641 | 0.388573855 | -0.421906 | 0.67309363 | 0.81285826 |
| ENSG00000285693 | ENSG0000028 | 89.03829835 | 0.213544926 | 0.191702836 | 1.113937234 | 0.265306115 | 0.44634432 |
| ENSG00000285696 | ENSG0000028 | 6.23507114 | -1.813387757 | 0.765584583 | -2.368631496 | 0.017854032 | 0.049566925 |
| ENSG00000285706 | ENSG0000028 | 2.936045048 | -1.906791654 | 1.10616712 | -1.723782619 | 0.084747094 | 0.185205137 |
| ENSG00000285710 | ENSG0000028 | 12.8412262 | -0.281541634 | 0.493333011 | -0.570692875 | 0.568207849 | 0.735799496 |
| ENSG00000285711 | LINC02835 | 2.235405278 | 1.092276556 | 1.255659188 | 0.86988298 | 0.384364357 | 0.576860867 |
| ENSG00000285712 | ENSG0000028 | 2.3105798 | -2.705213617 | 1.430938806 | -1.890516636 | 0.058688898 | 0.137016417 |
| ENSG00000285725 | ENSG0000028 | 18.87531408 | -0.71212604 | 0.404076556 | -1.76235426 | 0.078009462 | 0.173236888 |
| ENSG00000285728 | ENSG0000028 | 21.03425206 | -0.60595471 | 0.391997209 | -1.545813837 | 0.122149534 | 0.248855268 |
| ENSG00000285730 | ENSG0000028 | 2.730831649 | 1.17911083 | 1.216042011 | 0.969630012 | 0.332230948 | 0.523338456 |
| ENSG00000285737 | LINC02680 | 2.603610344 | -0.538322323 | 1.078619513 | -0.499084539 | 0.617719829 | 0.773419321 |
| ENSG00000285744 | ENSG0000028 | 1.778111155 | -0.403384177 | 1.299758677 | -0.310353133 | 0.75629243 | NA |
| ENSG00000285752 | CDC42-AS1 | 4.85841304 | 0.233693382 | 0.826883622 | 0.282619435 | 0.77746858 | 0.87922747 |
| ENSG00000285755 | ENSG0000028 | 1.639630974 | -0.169059384 | 1.329067842 | -0.12720147 | 0.898780942 | NA |
| ENSG00000285756 | ENSG0000028 | 829.199479 | 0.162724219 | 0.072606032 | 2.241194216 | 0.025013497 | 0.066490108 |
| ENSG00000285761 | ENSG0000028 | 1.987630943 | 3.267641982 | 1.565059536 | 2.087870721 | 0.036809498 | 0.092529606 |
| ENSG00000285793 | ANAPC1P2 | 3.207411286 | 1.458119296 | 1.096090115 | 1.330291438 | 0.183422267 | 0.339632481 |
| ENSG00000285796 | ENSG0000028 | 290.4295608 | 0.31439473 | 0.136172604 | 2.308795756 | 0.020954918 | 0.056927454 |
| ENSG00000285799 | ENSG0000028 | 3.293907223 | 1.202993733 | 1.044519711 | 1.151719512 | 0.249436351 | 0.427402891 |
| ENSG00000285816 | ENSG0000028 | 5.811545149 | 0.853636922 | 0.753434689 | 1.13299392 | 0.257216806 | 0.436883869 |
| ENSG00000285825 | CCDC15-DT | 21.75275269 | -0.931504981 | 0.390986248 | -2.382449476 | 0.017197892 | 0.047954253 |
| ENSG00000285830 | ENSG0000028 | 20.50859817 | -1.213352535 | 0.405445516 | -2.992640163 | 0.002765756 | 0.009452689 |
| ENSG00000285851 | ENSG0000028 | 3.256629713 | -0.334680291 | 1.002417461 | -0.333873165 | 0.73847527 | 0.855807824 |
| ENSG00000285852 | ENSG0000028 | 4.987694536 | -0.880625203 | 0.843490551 | -1.044024976 | 0.296473834 | 0.482781276 |
| ENSG00000285853 | ENSG0000028 | 1.55685474 | 1.127914975 | 1.490390404 | 0.756791625 | 0.449174711 | NA |
| ENSG00000285865 | ENSG0000028 | 18.92282422 | -0.551289178 | 0.41236889 | -1.336883532 | 0.181260657 | 0.336654761 |
| ENSG00000285867 | ENSG0000028 | 18.20484088 | -1.387889662 | 0.441325239 | -3.144822776 | 0.001661875 | 0.005940076 |
| ENSG00000285868 | ENSG0000028 | 2.886777173 | -0.086221248 | 1.096467346 | -0.078635491 | 0.937322557 | 0.970052308 |
| ENSG00000285871 | ENSG0000028 | 1.754593559 | -0.835159534 | 1.334149448 | -0.625986493 | 0.531323801 | NA |
| ENSG00000285872 | ENSG0000028 | 55.6169967 | -0.272191707 | 0.237161191 | -1.14770762 | 0.251089284 | 0.42943325 |
| ENSG00000285873 | ENSG0000028 | 5.361020503 | 0.631608751 | 0.7670833 | 0.823390043 | 0.410286224 | 0.602867408 |
| ENSG00000285877 | ENSG0000028 | 75.31850743 | 0.604549424 | 0.213751357 | 2.828283438 | 0.004679835 | 0.01518679 |

| | | | | | | | |
|---|---|---|---|---|---|---|---|
| ENSG00000285884 | ENSG0000028 | 8.142839929 | -0.731573827 | 0.618791379 | -1.182262475 | 0.23710157 | 0.412334926 |
| ENSG00000285886 | ENSG0000028 | 3.72425416 | 0.063943904 | 0.912345615 | 0.07008737 | 0.94412412 | 0.973493866 |
| ENSG00000285897 | ENSG0000028 | 8.463977857 | 0.861551169 | 0.618984301 | 1.391878869 | 0.163959082 | 0.311824021 |
| ENSG00000285907 | ENSG0000028 | 32.65200465 | -0.97413176 | 0.311874934 | -3.12346923 | 0.001787326 | 0.006346942 |
| ENSG00000285908 | ENSG0000028 | 3.124722726 | -1.192840491 | 1.054404046 | -1.131293545 | 0.25793156 | 0.4375795 |
| ENSG00000285923 | ENSG0000028 | 10.42595506 | -0.231354846 | 0.548933259 | -0.421462613 | 0.673417306 | 0.813071362 |
| ENSG00000285926 | ENSG0000028 | 3.287775779 | -3.243611956 | 1.323688753 | -2.450434023 | 0.014268411 | 0.040655489 |
| ENSG00000285933 | ENSG0000028 | 2.789906025 | -1.033821417 | 1.089423878 | -0.948961591 | 0.342640147 | 0.533616088 |
| ENSG00000285943 | ENSG0000028 | 11.04315918 | -0.612820873 | 0.530845791 | -1.154423532 | 0.248326572 | 0.426010437 |
| ENSG00000285948 | ENSG0000028 | 13.02893122 | -0.515846017 | 0.494107511 | -1.043995499 | 0.296487472 | 0.482781276 |
| ENSG00000285955 | ENSG0000028 | 3.782947293 | 0.022826209 | 0.92240932 | 0.02474629 | 0.980257332 | 0.991057876 |
| ENSG00000285967 | NIPBL-DT | 682.2585755 | 0.503407603 | 0.084594955 | 5.950799321 | 2.67E-09 | 2.04E-08 |
| ENSG00000285969 | ENSG0000028 | 21.19789165 | -3.512727712 | 0.539203657 | -6.514658543 | 7.29E-11 | 6.37E-10 |
| ENSG00000285970 | ENSG0000028 | 1.666160036 | 0.387960393 | 1.393481429 | 0.278410882 | 0.780696967 | NA |
| ENSG00000285976 | ENSG0000028 | 15.25423244 | -0.035342975 | 0.492904825 | -0.071703447 | 0.942837913 | 0.972907279 |
| ENSG00000285999 | ENSG0000028 | 19.54867031 | -0.272515598 | 0.420020498 | -0.648814998 | 0.516457962 | 0.69491147 |
| ENSG00000286010 | ENSG0000028 | 5.941119588 | 0.895894811 | 0.733829612 | 1.22084854 | 0.222143371 | 0.393177142 |
| ENSG00000286011 | ENSG0000028 | 9.627230157 | -1.088860143 | 0.590024609 | -1.845448691 | 0.064972296 | 0.149121007 |
| ENSG00000286017 | ENSG0000028 | 7.462209479 | 0.350370473 | 0.655173824 | 0.534774834 | 0.592805567 | 0.754957688 |
| ENSG00000286018 | ENSG0000028 | 26.0857482 | -0.712041044 | 0.381635035 | -1.865764354 | 0.062074342 | 0.143460145 |
| ENSG00000286032 | ENSG0000028 | 9.253011367 | -3.798973136 | 0.873280344 | -4.350233191 | 1.36E-05 | 6.89E-05 |
| ENSG00000286033 | ENSG0000028 | 19.28073557 | 1.53302539 | 0.441725604 | 3.470537765 | 0.000519417 | 0.002049086 |
| ENSG00000286035 | ENSG0000028 | 2.649500026 | -0.527544556 | 1.056250683 | -0.499450144 | 0.617462301 | 0.773221455 |
| ENSG00000286037 | ENSG0000028 | 79.53769008 | -0.643209917 | 0.234577005 | -2.741999018 | 0.006106652 | 0.019263835 |
| ENSG00000286043 | ENSG0000028 | 2.634873028 | -0.185437792 | 1.056298078 | -0.175554415 | 0.860644017 | 0.927432063 |
| ENSG00000286045 | ENSG0000028 | 4.18282603 | -0.313366347 | 0.839281579 | -0.373374508 | 0.708869729 | 0.837745175 |
| ENSG00000286046 | ENSG0000028 | 26.05833875 | -0.463102391 | 0.339653885 | -1.36345383 | 0.172739526 | 0.324860201 |
| ENSG00000286058 | ENSG0000028 | 113.8637176 | -0.015235542 | 0.176446452 | -0.086346548 | 0.931190936 | 0.967022186 |
| ENSG00000286061 | RCAN3AS | 2.7965015 | -1.409002423 | 1.106144637 | -1.273795827 | 0.202735784 | 0.366865891 |
| ENSG00000286067 | ENSG0000028 | 24.80731773 | -0.319873885 | 0.427894127 | -0.747553809 | 0.454729335 | 0.642254091 |
| ENSG00000286078 | ENSG0000028 | 12.01460457 | -0.436632418 | 0.49990423 | -0.873432134 | 0.382427582 | 0.574668462 |
| ENSG00000286080 | ENSG0000028 | 1.667083704 | 0.369517592 | 1.364546011 | 0.270798924 | 0.786545687 | NA |
| ENSG00000286081 | ENSG0000028 | 4.893362633 | -0.17701213 | 0.775932164 | -0.228128358 | 0.819546455 | 0.904386938 |

| | | | | | | | |
|---|---|---|---|---|---|---|---|
| ENSG00000286084 | ENSG0000028 | 2.381920661 | -1.035266287 | 1.144751562 | -0.904358921 | 0.365805113 | 0.558469938 |
| ENSG00000286086 | ENSG0000028 | 9.85404517 | 0.28327268 | 0.562265515 | 0.503805892 | 0.614397787 | 0.771190919 |
| ENSG00000286089 | ENSG0000028 | 1.718543802 | 0.133076002 | 1.335309503 | 0.099659294 | 0.920614818 | NA |
| ENSG00000286104 | ENSG0000028 | 1.952211283 | 1.32598483 | 1.356676633 | 0.977377215 | 0.328382439 | 0.518747128 |
| ENSG00000286106 | NOTCH2NLR | 1.948841653 | -0.180454934 | 1.230494059 | -0.146652422 | 0.883406374 | 0.93961708 |
| ENSG00000286112 | ENSG0000028 | 5.451760615 | 1.085621831 | 0.785022089 | 1.382918832 | 0.166689755 | 0.315995619 |
| ENSG00000286115 | ENSG0000028 | 7.602038729 | -1.615872621 | 0.674868754 | -2.39435092 | 0.01664981 | 0.046624347 |
| ENSG00000286118 | ENSG0000028 | 9.005356805 | -0.21099346 | 0.581090542 | -0.363099112 | 0.716530848 | 0.842415828 |
| ENSG00000286128 | ENSG0000028 | 9.390916852 | -1.360201375 | 0.603034679 | -2.255593954 | 0.024096072 | 0.064409683 |
| ENSG00000286129 | ENSG0000028 | 14.3270871 | -0.197766248 | 0.457865011 | -0.431931341 | 0.665791315 | 0.807339718 |
| ENSG00000286136 | ENSG0000028 | 5.393184573 | 0.426614147 | 0.755875932 | 0.564397051 | 0.572483945 | 0.739291922 |
| ENSG00000286138 | ENSG0000028 | 21.73163134 | 0.438846198 | 0.38096642 | 1.151928819 | 0.249350324 | 0.427335695 |
| ENSG00000286145 | ENSG0000028 | 3.103360555 | -0.315422316 | 0.972186048 | -0.324446454 | 0.745600055 | 0.859379806 |
| ENSG00000286150 | ENSG0000028 | 58.32457218 | -0.193058223 | 0.238711075 | -0.808752688 | 0.418657415 | 0.610397935 |
| ENSG00000286156 | BMS1P23 | 94.89543119 | -0.226813888 | 0.181330706 | -1.250830005 | 0.210996505 | 0.378455929 |
| ENSG00000286158 | ENSG0000028 | 1.789515358 | -0.393354404 | 1.413045649 | -0.278373459 | 0.780725692 | NA |
| ENSG00000286159 | ENSG0000028 | 127.2437425 | -1.779106016 | 0.195242971 | -9.112266672 | 8.07E-20 | 1.33E-18 |
| ENSG00000286161 | ENSG0000028 | 6.052124757 | 0.314075971 | 0.724613782 | 0.433439136 | 0.664695774 | 0.806565515 |
| ENSG00000286162 | ENSG0000028 | 25.02845927 | 0.019457554 | 0.351428538 | 0.055367029 | 0.955846062 | 0.979763569 |
| ENSG00000286176 | ENSG0000028 | 45.72014219 | -0.300735818 | 0.260405527 | -1.154874945 | 0.248141641 | 0.425788709 |
| ENSG00000286177 | ENSG0000028 | 55.00842146 | 0.039971249 | 0.246586025 | 0.162098598 | 0.87122821 | 0.933095623 |
| ENSG00000286181 | ENSG0000028 | 14.75842367 | -0.810896089 | 0.473964655 | -1.710878818 | 0.087103486 | 0.189380529 |
| ENSG00000286191 | ENSG0000028 | 5.76195179 | 0.206706539 | 0.74301172 | 0.278200912 | 0.780858134 | 0.88119272 |
| ENSG00000286193 | CPLANE1-AS1 | 31.08847194 | 2.395164312 | 0.376734624 | 6.357696258 | 2.05E-10 | 1.73E-09 |
| ENSG00000286195 | ENSG0000028 | 2.620016 | -4.968569129 | 1.522865983 | -3.262643717 | 0.001103782 | 0.004099863 |
| ENSG00000286205 | ENSG0000028 | 2.136072143 | -0.38669853 | 1.191339795 | -0.324591297 | 0.745490415 | 0.859379806 |
| ENSG00000286206 | ENSG0000028 | 6.494694716 | -1.671947327 | 0.730060796 | -2.290148077 | 0.022012734 | 0.05945045 |
| ENSG00000286207 | ENSG0000028 | 14.77924441 | 0.337236038 | 0.485556584 | 0.694534991 | 0.487346765 | 0.670770672 |
| ENSG00000286208 | ENSG0000028 | 102.3118704 | 0.38825931 | 0.184973225 | 2.099002766 | 0.035816657 | 0.090388791 |
| ENSG00000286214 | COPG2IT1 | 21.82972333 | 0.063026273 | 0.373686262 | 0.16866093 | 0.866063352 | 0.930387435 |
| ENSG00000286219 | NOTCH2NLC | 55.66869735 | 0.797485519 | 0.249289526 | 3.199033396 | 0.001378892 | 0.005021715 |
| ENSG00000286223 | ENSG0000028 | 58.91962397 | -0.624770654 | 0.231767286 | -2.695680938 | 0.007024492 | 0.021764159 |
| ENSG00000286230 | ENSG0000028 | 17.30507523 | -0.572065597 | 0.417639173 | -1.369760391 | 0.17076171 | 0.321866195 |

| | | | | | | | |
|---|---|---|---|---|---|---|---|
| ENSG00000286237 | ARMCX5-GPRA | 2.738890337 | 0.703097207 | 1.055494961 | 0.666130331 | 0.505327799 | 0.68591659 |
| ENSG00000286258 | ENSG0000028 | 17.08698628 | -2.915284468 | 0.537290913 | -5.425895726 | 5.77E-08 | 3.82E-07 |
| ENSG00000286262 | ENSG0000028 | 1.583798469 | -0.108565202 | 1.385430509 | -0.07836207 | 0.937540045 | NA |
| ENSG00000286281 | ENSG0000028 | 2.347299488 | -2.700676842 | 1.416407855 | -1.906708462 | 0.056558351 | 0.132895996 |
| ENSG00000286293 | ENSG0000028 | 10.56836118 | 0.59685888 | 0.541027888 | 1.103194296 | 0.269942798 | 0.451577903 |
| ENSG00000286299 | ENSG0000028 | 2.364440498 | 1.812404758 | 1.256733406 | 1.442155313 | 0.149258564 | 0.290488797 |
| ENSG00000286305 | ENSG0000028 | 6.645396191 | -0.630413219 | 0.702530318 | -0.897346639 | 0.369533977 | 0.562142535 |
| ENSG00000286311 | ENSG0000028 | 27.00541089 | -1.145919429 | 0.358613214 | -3.195418863 | 0.00139628 | 0.005076176 |
| ENSG00000286312 | ENSG0000028 | 2.212113203 | -0.543498878 | 1.203158851 | -0.451726617 | 0.65146594 | 0.797514738 |
| ENSG00000286315 | ENSG0000028 | 3.679781922 | -1.532010728 | 0.967301677 | -1.583798276 | 0.113239632 | 0.234451007 |
| ENSG00000286319 | ENSG0000028 | 2.556374866 | -1.649515315 | 1.15058905 | -1.433626815 | 0.151678787 | 0.294051498 |
| ENSG00000286327 | ENSG0000028 | 2.253363599 | 2.39739404 | 1.406377967 | 1.704658418 | 0.088258149 | 0.191413448 |
| ENSG00000286340 | ENSG0000028 | 12.64335114 | 0.700851324 | 0.503657464 | 1.391523752 | 0.164066662 | 0.312002755 |
| ENSG00000286342 | ENSG0000028 | 12.92468971 | -3.681255599 | 0.720823716 | -5.107012323 | 3.27E-07 | 2.01E-06 |
| ENSG00000286351 | ENSG0000028 | 2.27267933 | -0.581980616 | 1.154033746 | -0.504301211 | 0.614049726 | 0.770880574 |
| ENSG00000286360 | ENSG0000028 | 24.84197894 | -0.817779393 | 0.373888771 | -2.187226408 | 0.028726 | 0.074785642 |
| ENSG00000286362 | ENSG0000028 | 3.41644578 | -0.789319988 | 0.948029735 | -0.832589906 | 0.405076054 | 0.597815534 |
| ENSG00000286366 | ENSG0000028 | 23.44670908 | 0.149524737 | 0.377998688 | 0.39556946 | 0.692422671 | 0.826482963 |
| ENSG00000286369 | ENSG0000028 | 14.63673616 | -1.11090414 | 0.474438289 | -2.341514516 | 0.019205679 | 0.05279548 |
| ENSG00000286373 | ENSG0000028 | 9.12565835 | 0.004756596 | 0.592751768 | 0.008024601 | 0.993597363 | 0.996507827 |
| ENSG00000286379 | ENSG0000028 | 18.73535995 | -0.112321681 | 0.402923281 | -0.278766918 | 0.780423704 | 0.8808758 |
| ENSG00000286380 | ENSG0000028 | 2.996037107 | 1.821264796 | 1.200295005 | 1.517347643 | 0.129178935 | 0.259921479 |
| ENSG00000286381 | NHIP | 6.901451359 | -1.341513604 | 0.738166757 | -1.817358466 | 0.069162243 | 0.157095489 |
| ENSG00000286388 | ENSG0000028 | 419.3338494 | 0.001783095 | 0.095677385 | 0.018636538 | 0.985131055 | 0.993227548 |
| ENSG00000286389 | ENSG0000028 | 18.82777901 | 2.032897116 | 0.459115928 | 4.427851426 | 9.52E-06 | 4.93E-05 |
| ENSG00000286406 | ENSG0000028 | 11.19521413 | -0.56700438 | 0.52251087 | -1.08515327 | 0.277853777 | 0.460646502 |
| ENSG00000286408 | ENSG0000028 | 32.2232191 | 0.642705319 | 0.312544376 | 2.056365013 | 0.039747346 | 0.098646064 |
| ENSG00000286416 | ENSG0000028 | 18.67163116 | 0.830970865 | 0.423981164 | 1.959924016 | 0.050004672 | 0.119847975 |
| ENSG00000286417 | ENSG0000028 | 2.105819374 | 0.044830015 | 1.218646191 | 0.036786735 | 0.97065505 | 0.986772291 |
| ENSG00000286424 | ENSG0000028 | 10.20588543 | -1.218926611 | 0.590012819 | -2.065932422 | 0.038834861 | 0.09676026 |
| ENSG00000286427 | ENSG0000028 | 4.325621163 | 1.059542091 | 0.881865732 | 1.201477791 | 0.229565916 | 0.402710264 |
| ENSG00000286430 | ENSG0000028 | 4.292397729 | 0.665772922 | 0.904478303 | 0.736085011 | 0.461678964 | 0.648393573 |
| ENSG00000286431 | ENSG0000028 | 12.39140861 | -0.129493373 | 0.504894492 | -0.256476106 | 0.797583228 | 0.891941177 |

| | | | | | | | |
|---|---|---|---|---|---|---|---|
| ENSG00000286433 | ENSG0000028 | 3.806389155 | 0.352560039 | 0.924942035 | 0.381169874 | 0.703077202 | 0.833796972 |
| ENSG00000286435 | ENSG0000028 | 10.67150122 | 1.215813074 | 0.56738791 | 2.142825135 | 0.032127142 | 0.082447326 |
| ENSG00000286437 | ENSG0000028 | 1.979094422 | -0.651007072 | 1.237637511 | -0.526007871 | 0.598882738 | 0.759025967 |
| ENSG00000286443 | ENSG0000028 | 13.48069008 | -1.138869924 | 0.504143237 | -2.259020532 | 0.023882108 | 0.063874962 |
| ENSG00000286444 | ENSG0000028 | 2.713352587 | -1.325331966 | 1.132375696 | -1.170399516 | 0.241840231 | 0.418008958 |
| ENSG00000286445 | ENSG0000028 | 8.350885363 | -0.122393973 | 0.598122476 | -0.204630286 | 0.837860991 | 0.914703435 |
| ENSG00000286447 | ENSG0000028 | 5.25371308 | 1.455847828 | 0.803092312 | 1.812802595 | 0.069862274 | 0.1583324 |
| ENSG00000286451 | ENSG0000028 | 324.0388085 | -0.305100055 | 0.107221872 | -2.845502042 | 0.004434148 | 0.014473441 |
| ENSG00000286472 | ENSG0000028 | 63.42290904 | -0.724654559 | 0.221884252 | -3.265912528 | 0.001091119 | 0.004056772 |
| ENSG00000286475 | ENSG0000028 | 1.922896699 | 1.351880925 | 1.362692808 | 0.992065796 | 0.32116543 | 0.510676854 |
| ENSG00000286482 | ENSG0000028 | 18.61317317 | 0.411668976 | 0.429563196 | 0.958343219 | 0.337889717 | 0.529131362 |
| ENSG00000286485 | ENSG0000028 | 3.177117633 | -0.94008123 | 0.995305152 | -0.944515587 | 0.344906241 | 0.535799361 |
| ENSG00000286502 | ENSG0000028 | 2.774845758 | -2.312890863 | 1.222500695 | -1.891934191 | 0.05849975 | 0.136658275 |
| ENSG00000286503 | ENSG0000028 | 1.641089302 | 1.819993712 | 1.540972633 | 1.181068159 | 0.237575648 | NA |
| ENSG00000286511 | ENSG0000028 | 9.668684298 | 0.53890515 | 0.587108895 | 0.917896415 | 0.358673099 | 0.551213297 |
| ENSG00000286519 | ENSG0000028 | 2.014495479 | -3.497564138 | 1.583118249 | -2.209287992 | 0.027154616 | 0.071305734 |
| ENSG00000286522 | H3C2 | 23.17811271 | -1.360653472 | 0.378936447 | -3.59071681 | 0.00032977 | 0.00134679 |
| ENSG00000286529 | ENSG0000028 | 12.10905246 | 0.049131957 | 0.541302354 | 0.090766198 | 0.927678369 | 0.965269707 |
| ENSG00000286532 | PARTICL | 75.81511758 | 0.24168302 | 0.210621693 | 1.147474487 | 0.251185572 | 0.429565902 |
| ENSG00000286535 | ENSG0000028 | 16.43856715 | -0.279162003 | 0.426771326 | -0.65412549 | 0.513030961 | 0.692381325 |
| ENSG00000286536 | ENSG0000028 | 23.33751309 | -0.218164093 | 0.365531593 | -0.596840593 | 0.550613814 | 0.722725919 |
| ENSG00000286538 | ENSG0000028 | 8.552030935 | 0.070660259 | 0.621015287 | 0.113781836 | 0.909410738 | 0.954587195 |
| ENSG00000286540 | ENSG0000028 | 3.22164678 | 4.008402534 | 1.431686661 | 2.799776406 | 0.005113801 | 0.016432358 |
| ENSG00000286543 | ENSG0000028 | 3.825882418 | 1.719134488 | 0.972418858 | 1.767895053 | 0.07707845 | 0.17173462 |
| ENSG00000286548 | ENSG0000028 | 40.07066193 | 0.088003381 | 0.27760826 | 0.317005629 | 0.751239331 | 0.862469998 |
| ENSG00000286555 | ENSG0000028 | 6.608823966 | -0.296914308 | 0.687638487 | -0.431788379 | 0.665895226 | 0.807361848 |
| ENSG00000286563 | ENSG0000028 | 5.386136125 | 1.231479098 | 0.795723077 | 1.547622702 | 0.121713162 | 0.248208725 |
| ENSG00000286570 | ENSG0000028 | 6.198367005 | -0.170134309 | 0.78730334 | -0.216097532 | 0.828911724 | 0.909515231 |
| ENSG00000286572 | ENSG0000028 | 10.17504828 | 1.396978579 | 0.593838101 | 2.352456969 | 0.018649847 | 0.051446308 |
| ENSG00000286579 | ENSG0000028 | 3.365354359 | 0.825774332 | 0.968031431 | 0.853044958 | 0.393634373 | 0.586590393 |
| ENSG00000286580 | ENSG0000028 | 1.76533487 | -0.97698965 | 1.35883682 | -0.71898968 | 0.472147278 | NA |
| ENSG00000286585 | ENSG0000028 | 37.48943412 | -0.500155783 | 0.291144812 | -1.71789351 | 0.085816041 | 0.187088431 |
| ENSG00000286591 | ENSG0000028 | 2.236207444 | -0.941598185 | 1.264012034 | -0.744928181 | 0.456315138 | 0.643478245 |

| | | | | | | | |
|---|---|---|---|---|---|---|---|
| ENSG00000286593 | ENSG0000028 | 206.9644271 | 1.45427967 | 0.135383794 | 10.74190364 | 6.47E-27 | 1.60E-25 |
| ENSG00000286595 | ENSG0000028 | 5.906118598 | -0.591562851 | 0.730126234 | -0.810219964 | 0.417813765 | 0.609904408 |
| ENSG00000286596 | ENSG0000028 | 13.73569029 | 0.926369064 | 0.49760114 | 1.861669897 | 0.062649634 | 0.144571231 |
| ENSG00000286599 | ENSG0000028 | 10.06285189 | -0.496101273 | 0.563878177 | -0.879802222 | 0.378966461 | 0.571488996 |
| ENSG00000286602 | ENSG0000028 | 3.357387429 | 0.348766873 | 0.964429574 | 0.361630213 | 0.717628384 | 0.84297174 |
| ENSG00000286607 | ENSG0000028 | 3.664751399 | -0.111553702 | 0.946458458 | -0.11786434 | 0.906175148 | 0.952644604 |
| ENSG00000286613 | ENSG0000028 | 4.063159566 | 1.928117231 | 0.977377853 | 1.972744957 | 0.048524627 | 0.116780838 |
| ENSG00000286614 | ENSG0000028 | 10.39714885 | 0.183233259 | 0.536048315 | 0.341822284 | 0.732484639 | 0.852089128 |
| ENSG00000286619 | ENSG0000028 | 483.0868219 | 0.832022264 | 0.089788146 | 9.26650451 | 1.92E-20 | 3.30E-19 |
| ENSG00000286623 | ENSG0000028 | 2.973412808 | -0.489814832 | 1.068290541 | -0.458503387 | 0.64659083 | 0.793491418 |
| ENSG00000286626 | ENSG0000028 | 3.059483566 | -0.543976114 | 0.98979527 | -0.549584475 | 0.58260441 | 0.747090374 |
| ENSG00000286634 | ENSG0000028 | 4.437229263 | -0.416080947 | 0.854104349 | -0.487154699 | 0.626148706 | 0.779457961 |
| ENSG00000286650 | ENSG0000028 | 3.686320267 | 0.221269663 | 0.904795463 | 0.244552135 | 0.806803229 | 0.89740429 |
| ENSG00000286659 | ENSG0000028 | 2.469437978 | 0.36416624 | 1.107729886 | 0.328750036 | 0.742344631 | 0.85777393 |
| ENSG00000286667 | ENSG0000028 | 14.23967464 | 0.142875651 | 0.459660398 | 0.310828715 | 0.755930839 | 0.865149344 |
| ENSG00000286669 | ENSG0000028 | 12.40937958 | 0.008473426 | 0.499755929 | 0.016955128 | 0.986472413 | 0.993497695 |
| ENSG00000286670 | ENSG0000028 | 1.924174445 | -1.153596103 | 1.311120978 | -0.879854813 | 0.378937966 | 0.571483594 |
| ENSG00000286681 | ENSG0000028 | 52.55601914 | 0.40648677 | 0.245065881 | 1.658683647 | 0.097179556 | 0.207203181 |
| ENSG00000286682 | ENSG0000028 | 2.439464181 | -1.603771357 | 1.25256229 | -1.280392496 | 0.200407132 | 0.363598513 |
| ENSG00000286683 | ENSG0000028 | 5.79445101 | -2.416581343 | 0.852527735 | -2.834607301 | 0.004588205 | 0.014916867 |
| ENSG00000286688 | ENSG0000028 | 5.435693484 | -0.25692739 | 0.75254745 | -0.341410219 | 0.732794783 | 0.852320251 |
| ENSG00000286689 | ENSG0000028 | 56.54580449 | -0.063600769 | 0.24038667 | -0.264576937 | 0.791335389 | 0.887688696 |
| ENSG00000286691 | ENSG0000028 | 1.56444899 | 1.096666791 | 1.455008077 | 0.753718696 | 0.451018153 | NA |
| ENSG00000286705 | ENSG0000028 | 54.95033436 | 0.466877612 | 0.239691452 | 1.947827545 | 0.051435602 | 0.122708331 |
| ENSG00000286707 | ENSG0000028 | 5.362012139 | -0.018364734 | 0.749787946 | -0.024493237 | 0.980459178 | 0.991070363 |
| ENSG00000286718 | ENSG0000028 | 9.004003953 | 0.20348846 | 0.590415112 | 0.344653204 | 0.730355103 | 0.850776726 |
| ENSG00000286721 | ENSG0000028 | 2.597975335 | 0.212741694 | 1.071797535 | 0.198490561 | 0.842661269 | 0.917274818 |
| ENSG00000286724 | ENSG0000028 | 30.46371703 | 0.784927824 | 0.337261762 | 2.327354925 | 0.019946383 | 0.054570068 |
| ENSG00000286729 | ENSG0000028 | 13.49192127 | -0.219632043 | 0.49355271 | -0.445002202 | 0.656318163 | 0.800805205 |
| ENSG00000286732 | ENSG0000028 | 24.64660087 | -0.244199177 | 0.350777913 | -0.696164634 | 0.486325733 | 0.669947245 |
| ENSG00000286734 | ENSG0000028 | 2.144153113 | -0.017052442 | 1.230630824 | -0.013856668 | 0.988944333 | 0.994147115 |
| ENSG00000286753 | ENSG0000028 | 7.090359392 | 0.226174144 | 0.691487102 | 0.327083677 | 0.743604597 | 0.858490754 |
| ENSG00000286760 | ENSG0000028 | 46.35069434 | -0.257208321 | 0.259509243 | -0.991133565 | 0.321620365 | 0.511293914 |

| | | | | | | | |
|---|---|---|---|---|---|---|---|
| ENSG00000286786 | ENSG0000028 | 2.746982448 | 1.471513787 | 1.121422082 | 1.312185493 | 0.189457555 | 0.34836136 |
| ENSG00000286787 | ENSG0000028 | 7.957451263 | -0.104470544 | 0.624208158 | -0.167364912 | 0.867082932 | 0.930846618 |
| ENSG00000286790 | ENSG0000028 | 4.253635606 | -0.837315404 | 0.857929832 | -0.975971894 | 0.329078392 | 0.519478277 |
| ENSG00000286791 | ENSG0000028 | 10.0077243 | 4.79920674 | 1.12270971 | 4.274663966 | 1.91E-05 | 9.53E-05 |
| ENSG00000286808 | ENSG0000028 | 7.370827248 | 0.428968548 | 0.66251158 | 0.647488377 | 0.517315915 | 0.695617062 |
| ENSG00000286813 | ENSG0000028 | 2.960572558 | -0.815652464 | 1.010421186 | -0.807240066 | 0.419528187 | 0.610929766 |
| ENSG00000286817 | ENSG0000028 | 3.91337367 | -0.607332615 | 0.886616693 | -0.685000204 | 0.493343792 | 0.676104104 |
| ENSG00000286825 | ENSG0000028 | 3.911888176 | -1.082803779 | 0.907986307 | -1.192533158 | 0.233052257 | 0.407361992 |
| ENSG00000286828 | ENSG0000028 | 9.582482323 | -1.30284715 | 0.592327945 | -2.199536864 | 0.027839771 | 0.072846395 |
| ENSG00000286833 | ENSG0000028 | 8.987729481 | -0.029794668 | 0.579333718 | -0.051429197 | 0.95898352 | 0.981225778 |
| ENSG00000286834 | ENSG0000028 | 8.977515148 | 1.247418863 | 0.621013151 | 2.008683489 | 0.044570711 | 0.108598206 |
| ENSG00000286835 | ENSG0000028 | 15.16076009 | 0.278733353 | 0.459305939 | 0.606857716 | 0.543945342 | 0.717456866 |
| ENSG00000286836 | ENSG0000028 | 101.6613985 | -1.450170094 | 0.185304876 | -7.825860436 | 5.04E-15 | 6.08E-14 |
| ENSG00000286837 | ENSG0000028 | 3.679800525 | 0.315023256 | 0.911976639 | 0.345429085 | 0.729771815 | 0.850415471 |
| ENSG00000286847 | ENSG0000028 | 2.04531784 | -0.703363984 | 1.236574047 | -0.568800539 | 0.569491509 | 0.736879419 |
| ENSG00000286850 | ENSG0000028 | 2.596016095 | -0.523245834 | 1.10983674 | -0.471461986 | 0.63731086 | 0.786492469 |
| ENSG00000286856 | ENSG0000028 | 2.257543958 | -2.659716224 | 1.455913528 | -1.826836672 | 0.06772433 | 0.154417666 |
| ENSG00000286861 | ENSG0000028 | 1.649087077 | -0.756697124 | 1.347966346 | -0.56136203 | 0.574550763 | NA |
| ENSG00000286872 | ENSG0000028 | 10.46477747 | 0.792707437 | 0.547628438 | 1.447527889 | 0.147749131 | 0.288211316 |
| ENSG00000286873 | ENSG0000028 | 30.868121 | 0.193318848 | 0.322536563 | 0.599370334 | 0.548925955 | 0.721583944 |
| ENSG00000286874 | ENSG0000028 | 12.1493459 | -0.118959504 | 0.506498468 | -0.234866464 | 0.81431238 | 0.901759557 |
| ENSG00000286875 | ENSG0000028 | 6.546737821 | 0.959845229 | 0.719261111 | 1.334487871 | 0.182044019 | 0.337898681 |
| ENSG00000286878 | ENSG0000028 | 2.151611668 | -1.33789285 | 1.226053388 | -1.09121908 | 0.275176494 | 0.457355635 |
| ENSG00000286883 | ENSG0000028 | 23.49413624 | -0.967396689 | 0.372060519 | -2.600105737 | 0.009319504 | 0.027975585 |
| ENSG00000286885 | ENSG0000028 | 3.663206825 | -1.556673258 | 1.01164414 | -1.53875577 | 0.123863931 | 0.251655176 |
| ENSG00000286898 | ENSG0000028 | 1.760763228 | -0.274060877 | 1.355810909 | -0.202137979 | 0.839808858 | NA |
| ENSG00000286900 | ENSG0000028 | 3.96392933 | 0.525705016 | 0.892287461 | 0.589165531 | 0.555750237 | 0.726640663 |
| ENSG00000286905 | ENSG0000028 | 3.361099995 | -0.503161251 | 0.943326065 | -0.533390595 | 0.593763226 | 0.755710631 |
| ENSG00000286909 | ENSG0000028 | 46.54438303 | -0.061443992 | 0.266276281 | -0.23075278 | 0.817506868 | 0.903309453 |
| ENSG00000286912 | ENSG0000028 | 40.14749739 | 1.592093601 | 0.301191556 | 5.285983524 | 1.25E-07 | 8.05E-07 |
| ENSG00000286915 | ENSG0000028 | 2.994740758 | 1.190865665 | 1.085344174 | 1.097223991 | 0.272543488 | 0.454635516 |
| ENSG00000286918 | ENSG0000028 | 2.191339143 | 0.718242663 | 1.200615438 | 0.598228743 | 0.549687316 | 0.722005555 |
| ENSG00000286931 | ENSG0000028 | 83.08193243 | -0.162982753 | 0.209439668 | -0.778184739 | 0.436460112 | 0.626296602 |

| | | | | | | | |
|---|---|---|---|---|---|---|---|
| ENSG00000286935 | ENSG0000028 | 3.430077462 | 1.266812098 | 0.994348305 | 1.274012428 | 0.202659012 | 0.366813771 |
| ENSG00000286938 | ENSG0000028 | 3.249073771 | -0.383280352 | 0.989167839 | -0.387477572 | 0.698402686 | 0.830810911 |
| ENSG00000286957 | ENSG0000028 | 1.879384928 | -1.018334743 | 1.300694543 | -0.782916134 | 0.433676363 | 0.623549332 |
| ENSG00000286964 | ENSG0000028 | 67.29236704 | 0.211116365 | 0.233775106 | 0.903074619 | 0.366486293 | 0.559103204 |
| ENSG00000286977 | ENSG0000028 | 32.25171864 | 0.038056644 | 0.320628664 | 0.118693828 | 0.905517927 | 0.95262805 |
| ENSG00000286984 | ENSG0000028 | 3.871204017 | 0.016203028 | 0.963272514 | 0.016820814 | 0.986579565 | 0.993497695 |
| ENSG00000286989 | ENSG0000028 | 33.88390426 | 0.858738 | 0.308595276 | 2.782732161 | 0.005390328 | 0.017211969 |
| ENSG00000286999 | ENSG0000028 | 2.857056487 | -1.883745348 | 1.168914865 | -1.611533401 | 0.107063513 | 0.223927516 |
| ENSG00000287001 | ENSG0000028 | 4.68160721 | 1.228859318 | 0.850459208 | 1.444936225 | 0.1484758 | 0.289235296 |
| ENSG00000287009 | ENSG0000028 | 9.746033646 | -0.29165347 | 0.593719676 | -0.491230932 | 0.623263124 | 0.777215485 |
| ENSG00000287016 | ENSG0000028 | 22.08481809 | -1.281348819 | 0.389702415 | -3.288018675 | 0.001008951 | 0.003775134 |
| ENSG00000287024 | ENSG0000028 | 28.77003672 | -0.802344485 | 0.330684633 | -2.426313182 | 0.015253101 | 0.043155666 |
| ENSG00000287032 | ENSG0000028 | 4.136966092 | 1.543340912 | 0.919819563 | 1.677873546 | 0.093371788 | 0.200369393 |
| ENSG00000287042 | ENSG0000028 | 4.725089237 | 0.533369909 | 0.800949635 | 0.665921908 | 0.505461017 | 0.685970999 |
| ENSG00000287045 | ENSG0000028 | 2.108269339 | -0.240547996 | 1.311359144 | -0.183434109 | 0.854457415 | 0.924325808 |
| ENSG00000287051 | ENSG0000028 | 6.316870881 | -2.022449524 | 0.793758919 | -2.547939275 | 0.010836133 | 0.031869345 |
| ENSG00000287054 | ENSG0000028 | 2.440613793 | -1.089959284 | 1.142760308 | -0.953795189 | 0.340187325 | 0.531130282 |
| ENSG00000287061 | ENSG0000028 | 1.663714074 | -1.359697858 | 1.418949179 | -0.958242817 | 0.337940331 | NA |
| ENSG00000287064 | ENSG0000028 | 59.72556161 | -0.357825662 | 0.234680199 | -1.52473734 | 0.127324624 | 0.257151431 |
| ENSG00000287071 | CDRT15P4 | 14.4343878 | 0.028940035 | 0.48164596 | 0.060085701 | 0.952087378 | 0.977658212 |
| ENSG00000287074 | ENSG0000028 | 2.796999362 | -0.692628225 | 1.046519762 | -0.661839604 | 0.508074022 | 0.688191902 |
| ENSG00000287077 | ENSG0000028 | 4.339227105 | 0.156998241 | 0.82759723 | 0.189703682 | 0.849541336 | 0.921263493 |
| ENSG00000287078 | ENSG0000028 | 3.526109239 | -0.345115466 | 0.916001044 | -0.37676318 | 0.706349606 | 0.835939204 |
| ENSG00000287086 | ENSG0000028 | 39.25842799 | -0.218611974 | 0.31109425 | -0.70271943 | 0.482230617 | 0.666327572 |
| ENSG00000287089 | ENSG0000028 | 2.883042325 | 0.200666922 | 1.06481283 | 0.188452765 | 0.85052174 | 0.921807083 |
| ENSG00000287090 | ENSG0000028 | 2.346508463 | 0.177289741 | 1.139139725 | 0.155634763 | 0.876320922 | 0.93573969 |
| ENSG00000287091 | ENSG0000028 | 5.053663912 | -0.052269938 | 0.85421362 | -0.06119071 | 0.951207328 | 0.97748496 |
| ENSG00000287104 | ENSG0000028 | 30.25268375 | -0.062158874 | 0.323736832 | -0.192004331 | 0.847738813 | 0.920377435 |
| ENSG00000287108 | ENSG0000028 | 2.586627635 | -2.173380863 | 1.231645876 | -1.764615061 | 0.077628482 | 0.172591334 |
| ENSG00000287110 | ACAD9-DT | 40.09405092 | -0.678753119 | 0.28166396 | -2.409797542 | 0.015961375 | 0.044915842 |
| ENSG00000287120 | ENSG0000028 | 4.303822369 | 2.413078475 | 1.022160549 | 2.360762678 | 0.018237397 | 0.050448076 |
| ENSG00000287125 | ENSG0000028 | 5.19985496 | 1.411673018 | 0.831020379 | 1.698722503 | 0.089371481 | 0.193297859 |
| ENSG00000287126 | ENSG0000028 | 130.0970121 | -0.421814326 | 0.159502169 | -2.644567963 | 0.008179528 | 0.02490885 |

| | | | | | | | |
|---|---|---|---|---|---|---|---|
| ENSG00000287133 | ENSG0000028 | 5.453762166 | 0.779837541 | 0.76894637 | 1.014163759 | 0.310504618 | 0.498458778 |
| ENSG00000287146 | ENSG0000028 | 1.70865443 | -1.391575502 | 1.39968774 | -0.994204252 | 0.320123439 | NA |
| ENSG00000287151 | C2orf27A | 176.6218553 | 0.537421865 | 0.145367861 | 3.696978545 | 0.000218181 | 0.000918867 |
| ENSG00000287165 | ENSG0000028 | 2.104405934 | 1.005507078 | 1.226158438 | 0.820046616 | 0.412189533 | 0.604463313 |
| ENSG00000287166 | ENSG0000028 | 1.657190328 | 0.964437114 | 1.454778711 | 0.662944203 | 0.50736629 | NA |
| ENSG00000287168 | ENSG0000028 | 99.31765402 | 0.158695457 | 0.189107123 | 0.839182863 | 0.401366702 | 0.594137714 |
| ENSG00000287170 | ENSG0000028 | 8.854983393 | 1.098354139 | 0.689693779 | 1.592524352 | 0.111266936 | 0.230929477 |
| ENSG00000287182 | ENSG0000028 | 3.382397574 | -0.782421366 | 0.969662195 | -0.80690097 | 0.41972354 | 0.611023031 |
| ENSG00000287185 | ENSG0000028 | 7.56792593 | -1.326759937 | 0.657697613 | -2.017279538 | 0.04366635 | 0.106783591 |
| ENSG00000287190 | ENSG0000028 | 1.791441804 | -2.191481627 | 1.51307606 | -1.44836184 | 0.147515881 | 0.287927705 |
| ENSG00000287192 | ENSG0000028 | 1.763585548 | -0.18657349 | 2.570883722 | -0.072571734 | 0.94214692 | NA |
| ENSG00000287193 | ENSG0000028 | 4.426353464 | -0.701848011 | 0.931364899 | -0.753569317 | 0.451107874 | 0.639185586 |
| ENSG00000287202 | ENSG0000028 | 12.66306664 | -0.531645668 | 0.523381842 | -1.015789287 | 0.309729738 | 0.49772738 |
| ENSG00000287205 | ENSG0000028 | 1.979116704 | 0.340241291 | 1.282238083 | 0.265349544 | 0.790740201 | 0.887194572 |
| ENSG00000287216 | ENSG0000028 | 2.123744272 | -0.37164909 | 1.224176532 | -0.303591092 | 0.761439439 | 0.869069144 |
| ENSG00000287218 | ENSG0000028 | 1.615061152 | 0.462591465 | 1.461815123 | 0.316450047 | 0.751660935 | NA |
| ENSG00000287221 | ENSG0000028 | 9.919989516 | 0.496333769 | 0.57963712 | 0.856283616 | 0.391840921 | 0.584770973 |
| ENSG00000287222 | ENSG0000028 | 290.5499101 | 0.004023859 | 0.109493538 | 0.036749741 | 0.970684547 | 0.986772291 |
| ENSG00000287234 | ENSG0000028 | 10.19505645 | -0.174579049 | 0.576077185 | -0.303048019 | 0.761853268 | 0.869297509 |
| ENSG00000287236 | ENSG0000028 | 7.654558514 | -1.080378328 | 0.65856004 | -1.640516067 | 0.10089791 | 0.213373899 |
| ENSG00000287242 | ENSG0000028 | 13.33512002 | -0.483327321 | 0.483465726 | -0.999713724 | 0.317449069 | 0.506558266 |
| ENSG00000287249 | ENSG0000028 | 4.214424595 | 0.037213444 | 0.84573692 | 0.044001205 | 0.964903443 | 0.984122576 |
| ENSG00000287250 | ENSG0000028 | 2.032503248 | -1.141957968 | 1.285190976 | -0.888551188 | 0.374244334 | 0.566976953 |
| ENSG00000287261 | HPS1-AS1 | 5.141054401 | -2.186510421 | 0.872886244 | -2.504920241 | 0.012247901 | 0.035560922 |
| ENSG00000287262 | ENSG0000028 | 5.709398257 | -1.062702791 | 0.739057606 | -1.437916046 | 0.150457876 | 0.29225264 |
| ENSG00000287263 | ENSG0000028 | 23.86880318 | 0.874279839 | 0.373030157 | 2.343724287 | 0.019092279 | 0.05252781 |
| ENSG00000287265 | ENSG0000028 | 6.955620788 | -0.849173527 | 0.685994251 | -1.237872658 | 0.215763282 | 0.385021679 |
| ENSG00000287275 | ENSG0000028 | 23.51178491 | 0.515102158 | 0.384678901 | 1.339044479 | 0.180556193 | 0.335761923 |
| ENSG00000287286 | ENSG0000028 | 13.57309827 | -0.613869546 | 0.483709939 | -1.269086072 | 0.204410367 | 0.369342614 |
| ENSG00000287292 | ENSG0000028 | 7.202965941 | -0.656825101 | 0.647720473 | -1.014056415 | 0.310555834 | 0.498506111 |
| ENSG00000287296 | ENSG0000028 | 2.750634101 | -0.653200156 | 1.068717838 | -0.611199826 | 0.541067299 | 0.715270203 |
| ENSG00000287299 | ENSG0000028 | 17.64794654 | -1.110923032 | 0.432211544 | -2.570322441 | 0.01016039 | 0.030164273 |
| ENSG00000287306 | ENSG0000028 | 4.046547263 | -1.462399683 | 0.916024079 | -1.596464238 | 0.110385184 | 0.229410774 |

| | | | | | | | |
|---|---|---|---|---|---|---|---|
| ENSG00000287315 | ENSG0000028 | 253.6575603 | 1.140702968 | 0.127994362 | 8.912134468 | 5.01E-19 | 7.90E-18 |
| ENSG00000287337 | ENSG0000028 | 3.478741525 | 0.6557073 | 0.945797742 | 0.693284908 | 0.488130772 | 0.671528246 |
| ENSG00000287338 | ENSG0000028 | 4.54548274 | 2.933352363 | 1.072706471 | 2.734534042 | 0.006246864 | 0.01964401 |
| ENSG00000287352 | ENSG0000028 | 81.42288814 | 0.074706899 | 0.200118343 | 0.373313598 | 0.708915057 | 0.837755584 |
| ENSG00000287354 | ENSG0000028 | 16.22513322 | 4.448582717 | 0.802525637 | 5.543228168 | 2.97E-08 | 2.03E-07 |
| ENSG00000287356 | ENSG0000028 | 50.74907934 | -0.153725353 | 0.251646267 | -0.610878732 | 0.541279866 | 0.715369799 |
| ENSG00000287358 | ENSG0000028 | 10.47089243 | -2.705836074 | 0.657041959 | -4.118208949 | 3.82E-05 | 0.000181424 |
| ENSG00000287362 | ENSG0000028 | 2.307251055 | 0.246066174 | 1.161064704 | 0.211931491 | 0.832160483 | 0.911283868 |
| ENSG00000287377 | ENSG0000028 | 11.29961379 | -2.370362026 | 0.617989779 | -3.835600698 | 0.000125258 | 0.000549515 |
| ENSG00000287385 | ENSG0000028 | 2.698748196 | 0.150897301 | 1.056049613 | 0.142888458 | 0.886378274 | 0.941439323 |
| ENSG00000287387 | ENSG0000028 | 35.52291546 | -1.092430832 | 0.317844978 | -3.436992582 | 0.000588212 | 0.00229717 |
| ENSG00000287390 | ENSG0000028 | 12.35730846 | -0.024754893 | 0.496992342 | -0.049809405 | 0.960274272 | 0.981845775 |
| ENSG00000287391 | ENSG0000028 | 3.81743623 | 3.213752828 | 1.252983392 | 2.564880626 | 0.010321131 | 0.030557034 |
| ENSG00000287394 | MAGEA3-DT | 22.90657254 | 3.766197665 | 0.576803888 | 6.529424891 | 6.60E-11 | 5.79E-10 |
| ENSG00000287401 | ENSG0000028 | 26.76121025 | 1.349701804 | 0.359438958 | 3.755023695 | 0.000173325 | 0.000743748 |
| ENSG00000287402 | ENSG0000028 | 2.376184261 | 0.045906915 | 1.16023029 | 0.039567071 | 0.96843828 | 0.985876114 |
| ENSG00000287407 | ENSG0000028 | 12.3098243 | -0.163010726 | 0.516271698 | -0.315746005 | 0.752195305 | 0.863199614 |
| ENSG00000287425 | ENSG0000028 | 3.002264787 | -0.256117073 | 1.059869094 | -0.241649723 | 0.80905159 | 0.898859138 |
| ENSG00000287426 | ENSG0000028 | 2.778129142 | -1.823475161 | 1.168477152 | -1.560556967 | 0.118628319 | 0.24338911 |
| ENSG00000287430 | ENSG0000028 | 125.6610358 | 0.152892778 | 0.162032842 | 0.943591288 | 0.345378546 | 0.536243449 |
| ENSG00000287431 | RENO1 | 169.3340943 | 0.432647773 | 0.148194038 | 2.919468149 | 0.003506292 | 0.011711465 |
| ENSG00000287437 | ENSG0000028 | 1.600336768 | -0.748139244 | 1.410662385 | -0.530346065 | 0.595872014 | NA |
| ENSG00000287443 | ENSG0000028 | 10.42797867 | 1.077587113 | 0.560788243 | 1.92155796 | 0.054661403 | 0.129311393 |
| ENSG00000287460 | ENSG0000028 | 44.26439917 | 0.101308776 | 0.269425514 | 0.376017751 | 0.7069037 | 0.836412774 |
| ENSG00000287500 | ENSG0000028 | 4.771009764 | -0.624390816 | 0.799809024 | -0.780674882 | 0.434993737 | 0.624968087 |
| ENSG00000287502 | ENSG0000028 | 2.864605374 | 0.495278286 | 1.024625828 | 0.483374782 | 0.628829651 | 0.781015278 |
| ENSG00000287507 | ENSG0000028 | 5.637129251 | 1.141768856 | 0.762114634 | 1.498158945 | 0.134091959 | 0.267600657 |
| ENSG00000287513 | ENSG0000028 | 1.794302431 | -2.221297794 | 1.505125691 | -1.475822124 | 0.13999165 | 0.276440427 |
| ENSG00000287515 | ENSG0000028 | 4.876877461 | -0.694467157 | 0.826759989 | -0.839986412 | 0.400916005 | 0.593777078 |
| ENSG00000287519 | ENSG0000028 | 673.6562831 | 9.090655392 | 0.549493989 | 16.54368486 | 1.78E-61 | 1.32E-59 |
| ENSG00000287527 | ENSG0000028 | 2.793275655 | -0.031375532 | 1.030359928 | -0.030451041 | 0.975707339 | 0.988877242 |
| ENSG00000287529 | ENSG0000028 | 1.791886214 | 0.644603274 | 1.307831445 | 0.492879473 | 0.622097752 | 0.776353086 |
| ENSG00000287539 | ENSG0000028 | 25.46383141 | 0.249557314 | 0.359098276 | 0.694955477 | 0.487083204 | 0.670520307 |

| | | | | | | | |
|---|---|---|---|---|---|---|---|
| ENSG00000287541 | ENSG0000028 | 2.581453062 | -0.208072935 | 1.172743614 | -0.177424061 | 0.859175309 | 0.926792302 |
| ENSG00000287548 | ENSG0000028 | 3.783199979 | -1.084994144 | 0.917678476 | -1.182324934 | 0.237076796 | 0.4123231 |
| ENSG00000287553 | ENSG0000028 | 16.22729844 | 0.158950716 | 0.434126744 | 0.366138963 | 0.714261385 | 0.840997596 |
| ENSG00000287562 | ENSG0000028 | 48.36979476 | 0.243734908 | 0.27655977 | 0.88131006 | 0.378150023 | 0.570970942 |
| ENSG00000287566 | ENSG0000028 | 27.10957997 | -1.20921017 | 0.352049142 | -3.434776644 | 0.000593042 | 0.002314459 |
| ENSG00000287569 | ENSG0000028 | 9.631706786 | -0.027290938 | 0.565432161 | -0.048265627 | 0.961504549 | 0.982286361 |
| ENSG00000287574 | ENSG0000028 | 28.15656305 | 0.611893566 | 0.348387916 | 1.756357032 | 0.079027465 | 0.175040074 |
| ENSG00000287576 | ENSG0000028 | 47.42842536 | -0.625188747 | 0.254631743 | -2.455266339 | 0.014078024 | 0.040212909 |
| ENSG00000287578 | ENSG0000028 | 5.752840876 | 0.70544289 | 0.774586033 | 0.910735361 | 0.362434828 | 0.555140503 |
| ENSG00000287581 | ENSG0000028 | 2.027769626 | -1.14718549 | 1.285920958 | -0.892111978 | 0.372332915 | 0.564976656 |
| ENSG00000287584 | ENSG0000028 | 9.816128674 | 0.879193857 | 0.581824441 | 1.511098185 | 0.130763437 | 0.262625919 |
| ENSG00000287588 | ENSG0000028 | 3.482665519 | 0.452061968 | 0.953102067 | 0.474305936 | 0.635281759 | 0.784991303 |
| ENSG00000287595 | ENSG0000028 | 10.10879565 | -0.936300494 | 0.583879264 | -1.603585795 | 0.108805384 | 0.226775369 |
| ENSG00000287602 | ENSG0000028 | 46.00901455 | -0.36606205 | 0.25944231 | -1.410957412 | 0.158257172 | 0.303318533 |
| ENSG00000287603 | ENSG0000028 | 9.517475771 | 0.719842268 | 0.578069276 | 1.245252599 | 0.213038913 | 0.381210037 |
| ENSG00000287616 | ENSG0000028 | 4.426661854 | -0.496096336 | 0.844210831 | -0.587645074 | 0.55677055 | 0.727249376 |
| ENSG00000287617 | ENSG0000028 | 4.563825983 | 0.473623278 | 0.850690299 | 0.55675171 | 0.577697082 | 0.743512739 |
| ENSG00000287619 | ENSG0000028 | 8.995263142 | -0.228047748 | 0.575715551 | -0.396111842 | 0.692022522 | 0.826376122 |
| ENSG00000287620 | ENSG0000028 | 30.85422194 | 0.055318724 | 0.318159867 | 0.173870842 | 0.861966968 | 0.928215087 |
| ENSG00000287624 | ENSG0000028 | 10.76465282 | 0.114572145 | 0.535814606 | 0.213827961 | 0.830681219 | 0.910324711 |
| ENSG00000287625 | ENSG0000028 | 4.237670902 | -0.31209335 | 0.857419887 | -0.363991265 | 0.715864532 | 0.841891334 |
| ENSG00000287629 | ENSG0000028 | 2.346010601 | -0.606926574 | 1.127623326 | -0.538235207 | 0.590414677 | 0.753103247 |
| ENSG00000287630 | ENSG0000028 | 13.68526419 | -0.698364231 | 0.473774217 | -1.474044399 | 0.140469637 | 0.277217264 |
| ENSG00000287637 | ENSG0000028 | 9.558398204 | 1.453814717 | 0.613060878 | 2.37140351 | 0.017720673 | 0.049244366 |
| ENSG00000287642 | ENSG0000028 | 17.54814091 | 0.674156664 | 0.431508128 | 1.562326688 | 0.118211048 | 0.242728378 |
| ENSG00000287643 | ENSG0000028 | 4.271982307 | -0.663838425 | 0.908411677 | -0.730768265 | 0.464920711 | 0.650957099 |
| ENSG00000287655 | ENSG0000028 | 1.631610592 | 0.421056821 | 1.408135206 | 0.299017324 | 0.764926827 | NA |
| ENSG00000287670 | ENSG0000028 | 9.679267176 | 0.430431146 | 0.629739197 | 0.683506994 | 0.494286532 | 0.676830004 |
| ENSG00000287672 | ENSG0000028 | 12.84093677 | -1.307078166 | 0.51404289 | -2.542741457 | 0.010998658 | 0.032305917 |
| ENSG00000287676 | ENSG0000028 | 3.687820582 | -0.204248482 | 0.910812938 | -0.224248552 | 0.822563907 | 0.90619104 |
| ENSG00000287683 | ENSG0000028 | 3.186642111 | 1.365160193 | 1.050474052 | 1.29956584 | 0.193749814 | 0.354483437 |
| ENSG00000287688 | ENSG0000028 | 8.634509577 | 0.588254992 | 0.624342075 | 0.942199822 | 0.346090345 | 0.536947959 |
| ENSG00000287697 | ENSG0000028 | 7.506967826 | 0.916088387 | 0.668668169 | 1.370019434 | 0.170680835 | 0.321753501 |

| | | | | | | | |
|---|---|---|---|---|---|---|---|
| ENSG00000287698 | ENSG0000028 | 3.38418778 | 1.268340238 | 1.027038449 | 1.234949129 | 0.216849437 | 0.386280952 |
| ENSG00000287699 | ENSG0000028 | 3.006097022 | 0.422249636 | 1.016133454 | 0.415545453 | 0.677742649 | 0.816688071 |
| ENSG00000287707 | ENSG0000028 | 18.07541227 | -0.267250297 | 0.42116236 | -0.634554087 | 0.52571928 | 0.701733986 |
| ENSG00000287710 | ENSG0000028 | 24.9550045 | 0.05939995 | 0.349138706 | 0.170132812 | 0.864905689 | 0.929799912 |
| ENSG00000287721 | ENSG0000028 | 7.15892955 | 0.248496598 | 0.673951282 | 0.368715966 | 0.712339447 | 0.839972059 |
| ENSG00000287724 | ENSG0000028 | 76.65269185 | -0.882851009 | 0.207763042 | -4.249316917 | 2.14E-05 | 0.000105942 |
| ENSG00000287725 | ENSG0000028 | 25.303622 | 0.531051254 | 0.355942468 | 1.491958113 | 0.135710127 | 0.270289139 |
| ENSG00000287729 | ENSG0000028 | 13.37291832 | 0.30658959 | 0.482065165 | 0.635992003 | 0.524781634 | 0.7010535 |
| ENSG00000287731 | ENSG0000028 | 2.428308204 | -0.22420229 | 1.185840048 | -0.189066216 | 0.85004092 | 0.921503648 |
| ENSG00000287733 | ENSG0000028 | 4.742174438 | -1.736707384 | 0.86513357 | -2.00744422 | 0.044702384 | 0.108832103 |
| ENSG00000287737 | ENSG0000028 | 37.57816775 | 0.095819691 | 0.296969071 | 0.322658823 | 0.74695364 | 0.859750631 |
| ENSG00000287742 | ENSG0000028 | 9.002507318 | -0.097009069 | 0.577020932 | -0.168120537 | 0.866488453 | 0.930626205 |
| ENSG00000287750 | LINC03026 | 37.37865902 | 0.088660381 | 0.305710175 | 0.290014492 | 0.771805151 | 0.875561078 |
| ENSG00000287757 | ENSG0000028 | 10.77500317 | -0.56819236 | 0.530715571 | -1.070615582 | 0.284342314 | 0.468122053 |
| ENSG00000287759 | ENSG0000028 | 3.353463435 | 0.562406497 | 0.963974748 | 0.583424512 | 0.559607561 | 0.729149008 |
| ENSG00000287760 | ENSG0000028 | 13.05742466 | 0.449411918 | 0.505694381 | 0.888702613 | 0.374162926 | 0.566928503 |
| ENSG00000287763 | ENSG0000028 | 85.50805447 | 0.150032624 | 0.196732579 | 0.762622157 | 0.445688767 | 0.634249399 |
| ENSG00000287765 | ENSG0000028 | 19.93164472 | -0.712918473 | 0.40414254 | -1.764027299 | 0.077727383 | 0.172794473 |
| ENSG00000287766 | ENSG0000028 | 1.830231094 | 1.1792681 | 1.362663722 | 0.865413881 | 0.386811665 | 0.579358814 |
| ENSG00000287769 | ENSG0000028 | 84.58602181 | 0.719242948 | 0.196200754 | 3.66585212 | 0.000246516 | 0.00102745 |
| ENSG00000287780 | ENSG0000028 | 6.887182068 | -0.251531951 | 0.676274591 | -0.371937604 | 0.709939306 | 0.838577194 |
| ENSG00000287782 | ENSG0000028 | 3.917481198 | -0.39167055 | 0.899181982 | -0.435585408 | 0.66313756 | 0.805526906 |
| ENSG00000287784 | ENSG0000028 | 14.42631871 | -4.490192739 | 0.819594225 | -5.478555856 | 4.29E-08 | 2.89E-07 |
| ENSG00000287787 | ENSG0000028 | 5.597375491 | 0.570690855 | 0.757839225 | 0.753050035 | 0.451419848 | 0.639390671 |
| ENSG00000287799 | ENSG0000028 | 1.839231646 | -3.317845796 | 1.622948688 | -2.044331913 | 0.040920769 | 0.101057426 |
| ENSG00000287808 | DHX9-AS1 | 27.87988443 | -0.23453648 | 0.336977228 | -0.696000976 | 0.486428219 | 0.669988792 |
| ENSG00000287811 | ENSG0000028 | 4.888229165 | -0.280382474 | 0.801402374 | -0.349864791 | 0.726440172 | 0.847938612 |
| ENSG00000287817 | ENSG0000028 | 21.03707526 | -0.584553415 | 0.386838679 | -1.511103842 | 0.130761996 | 0.262625919 |
| ENSG00000287824 | ENSG0000028 | 3.266478201 | -2.597231361 | 1.145440289 | -2.267452426 | 0.023362602 | 0.06263154 |
| ENSG00000287825 | ENSG0000028 | 23.2877254 | -0.141298158 | 0.365169885 | -0.386938145 | 0.698802001 | 0.830984288 |
| ENSG00000287831 | ENSG0000028 | 2.456661693 | -0.046191003 | 1.168481212 | -0.039530805 | 0.968467194 | 0.985876114 |
| ENSG00000287837 | ENSG0000028 | 95.6877313 | -0.445997736 | 0.182837432 | -2.439313061 | 0.014715214 | 0.041778095 |
| ENSG00000287851 | ENSG0000028 | 4.900489393 | 0.61167846 | 0.852725672 | 0.717321502 | 0.473175743 | 0.658734955 |

| | | | | | | | |
|---|---|---|---|---|---|---|---|
| ENSG00000287853 | ENSG0000028 | 5.08724205 | -0.985448208 | 0.786598428 | -1.252797073 | 0.210279567 | 0.377509256 |
| ENSG00000287859 | ENSG0000028 | 3.655559124 | -0.704299519 | 0.943841676 | -0.74620515 | 0.455543498 | 0.642928992 |
| ENSG00000287865 | ENSG0000028 | 7.264960653 | -1.351849251 | 0.688851612 | -1.962468007 | 0.049708022 | 0.119228568 |
| ENSG00000287875 | ENSG0000028 | 6.610564724 | 0.615129781 | 0.68965588 | 0.891937268 | 0.372426558 | 0.56508137 |
| ENSG00000287878 | ENSG0000028 | 1.576204219 | -0.075657275 | 1.468866049 | -0.051507266 | 0.958921312 | NA |
| ENSG00000287886 | ENSG0000028 | 301.6346659 | 0.308253818 | 0.110739412 | 2.783596313 | 0.00537599 | 0.01717143 |
| ENSG00000287887 | ENSG0000028 | 1.923741176 | -4.49612107 | 1.585164663 | -2.836374779 | 0.004562887 | 0.01484718 |
| ENSG00000287898 | ENSG0000028 | 2.335605172 | -0.229620912 | 1.147620608 | -0.20008434 | 0.84141462 | 0.916700769 |
| ENSG00000287900 | ENSG0000028 | 15.66039399 | -0.348385144 | 0.451486136 | -0.771640847 | 0.440327174 | 0.629711622 |
| ENSG00000287906 | ENSG0000028 | 63.01451318 | -0.38955118 | 0.23675368 | -1.645385955 | 0.099890244 | 0.211762517 |
| ENSG00000287914 | ENSG0000028 | 4.931624318 | -0.960447684 | 0.804163286 | -1.19434411 | 0.23234339 | 0.406370643 |
| ENSG00000287916 | ENSG0000028 | 2.6745819 | -0.911298896 | 1.080154453 | -0.843674619 | 0.398851255 | 0.591712331 |
| ENSG00000287917 | ENSG0000028 | 4.333099747 | -0.890070527 | 0.877637287 | -1.014166718 | 0.310503207 | 0.498458778 |
| ENSG00000287918 | ENSG0000028 | 35.93789112 | -0.0268653 | 0.290197521 | -0.092575912 | 0.926240481 | 0.964662124 |
| ENSG00000287920 | ENSG0000028 | 2.197934619 | 0.301634159 | 1.189144828 | 0.253656369 | 0.799761044 | 0.893253273 |
| ENSG00000287925 | ENSG0000028 | 2.320065647 | 0.607953957 | 1.185430459 | 0.512855016 | 0.608052742 | 0.766329519 |
| ENSG00000287934 | ENSG0000028 | 13.38886354 | -0.339930049 | 0.497605333 | -0.683131844 | 0.494523535 | 0.677033295 |
| ENSG00000287935 | ENSG0000028 | 6.186361716 | -0.311999057 | 0.702154703 | -0.444345179 | 0.656793042 | 0.801214166 |
| ENSG00000287952 | ENSG0000028 | 7.704541207 | -0.829556543 | 0.649852402 | -1.276530703 | 0.20176798 | 0.365547107 |
| ENSG00000287957 | ENSG0000028 | 1.717695867 | -0.202720972 | 1.329719653 | -0.152453919 | 0.878828933 | NA |
| ENSG00000287966 | LARRPM | 41.97896964 | 0.216488809 | 0.27081878 | 0.799386249 | 0.424066481 | 0.615019183 |
| ENSG00000287967 | ENSG0000028 | 29.36375932 | -0.980108453 | 0.359952606 | -2.722881949 | 0.006471518 | 0.020236726 |
| ENSG00000287978 | ENSG0000028 | 31.5567965 | -0.080459813 | 0.31780187 | -0.253176021 | 0.800132196 | 0.893449794 |
| ENSG00000287979 | ENSG0000028 | 2.552715751 | -1.59682602 | 1.192577963 | -1.33896992 | 0.180580466 | 0.335779839 |
| ENSG00000287985 | ENSG0000028 | 55.34187294 | 1.601621487 | 0.256294101 | 6.249154707 | 4.13E-10 | 3.41E-09 |
| ENSG00000288004 | ENSG0000028 | 32.99976623 | -0.287001461 | 0.316283699 | -0.907417809 | 0.364185895 | 0.556813145 |
| ENSG00000288007 | ENSG0000028 | 2.477473179 | -1.517131164 | 1.201350859 | -1.262854355 | 0.206641526 | 0.37249384 |
| ENSG00000288009 | ENSG0000028 | 12.61703038 | 0.18318293 | 0.496757794 | 0.368757032 | 0.712308834 | 0.839972059 |
| ENSG00000288012 | ENSG0000028 | 35.18636475 | 0.903335785 | 0.3075806 | 2.93690755 | 0.003315029 | 0.011126082 |
| ENSG00000288021 | ENSG0000028 | 2.636734881 | -0.535767932 | 1.056906902 | -0.506920648 | 0.612210497 | 0.769498044 |
| ENSG00000288029 | ENSG0000028 | 4.31998758 | -1.322804694 | 0.894461729 | -1.478883502 | 0.139171453 | 0.275271007 |
| ENSG00000288031 | ENSG0000028 | 7.091247804 | -0.131305338 | 0.683955123 | -0.191979464 | 0.847758291 | 0.920377435 |
| ENSG00000288033 | ENSG0000028 | 3.038222785 | 0.123408787 | 1.002059651 | 0.123155131 | 0.901984256 | 0.950634881 |

| | | | | | | | |
|---|---|---|---|---|---|---|---|
| ENSG00000288040 | ENSG0000028 | 3.651827652 | -0.982722429 | 1.000912817 | -0.981826202 | 0.326185478 | 0.516713236 |
| ENSG00000288044 | ENSG0000028 | 116.5137227 | -0.295779119 | 0.167464297 | -1.766221965 | 0.077358617 | 0.172174899 |
| ENSG00000288045 | ENSG0000028 | 9.069088491 | -1.148365293 | 0.60832857 | -1.887738551 | 0.059061058 | 0.137675102 |
| ENSG00000288046 | ENSG0000028 | 1.982452911 | 0.246392563 | 1.23835083 | 0.198968303 | 0.842287539 | 0.917009895 |
| ENSG00000288056 | ENSG0000028 | 2.882583399 | -1.066101654 | 1.066347022 | -0.999769899 | 0.317421876 | 0.506550111 |
| ENSG00000288060 | ENSG0000028 | 2.341306723 | 3.505311818 | 1.502410055 | 2.333125904 | 0.019641538 | 0.053851666 |
| ENSG00000288061 | ENSG0000028 | 6.436155881 | -0.119509008 | 0.683271487 | -0.174907062 | 0.861152661 | 0.927805845 |
| ENSG00000288062 | ENSG0000028 | 2.348946962 | -0.950285109 | 1.193995176 | -0.795886892 | 0.426097781 | 0.616893852 |
| ENSG00000288068 | ENSG0000028 | 7.855918768 | -1.831898597 | 0.695008244 | -2.635794056 | 0.008394066 | 0.025474252 |
| ENSG00000288075 | ENSG0000028 | 56.75207108 | -1.219469513 | 0.245448832 | -4.96832478 | 6.75E-07 | 4.00E-06 |
| ENSG00000288086 | ENSG0000028 | 12.45862753 | 1.249077857 | 0.517218641 | 2.414990022 | 0.015735641 | 0.044356795 |
| ENSG00000288093 | ENSG0000028 | 11.93172654 | -1.240800095 | 0.51916896 | -2.389973576 | 0.016849585 | 0.047120502 |
| ENSG00000288095 | ENSG0000028 | 17.99043588 | -0.069386716 | 0.432849056 | -0.160302338 | 0.872642917 | 0.934146135 |
| ENSG00000288098 | ENSG0000028 | 17.14784828 | 0.140370424 | 0.422311746 | 0.332385793 | 0.739597965 | 0.856411959 |
| ENSG00000288103 | ENSG0000028 | 2.454238013 | -0.375574123 | 1.108653332 | -0.338766062 | 0.73478597 | 0.853727216 |
| ENSG00000288109 | ENSG0000028 | 2.125240907 | 0.929380172 | 1.230518612 | 0.755275185 | 0.450083885 | 0.638403176 |
| ENSG00000288234 | ENSG0000028 | 9.145663991 | 0.602622518 | 0.579170194 | 1.040492975 | 0.298110926 | 0.484702293 |
| ENSG00000288262 | ENSG0000028 | 3.718668599 | 0.849964564 | 0.993188021 | 0.855794217 | 0.392111615 | 0.584967187 |
| ENSG00000288398 | ENSG0000028 | 280.69386 | -0.267304839 | 0.112140481 | -2.383660535 | 0.017141406 | 0.04782 |
| ENSG00000288526 | ENSG0000028 | 2.256216764 | -0.417751617 | 1.207142537 | -0.34606652 | 0.729292724 | 0.850044287 |
| ENSG00000288533 | ENSG0000028 | 9.096935963 | 0.833471701 | 0.597373031 | 1.395228203 | 0.162947035 | 0.310336652 |
| ENSG00000288534 | ENSG0000028 | 1.660902895 | -0.796706106 | 1.379146574 | -0.577680517 | 0.563479835 | NA |
| ENSG00000288538 | ENSG0000028 | 44.53455165 | -0.591543643 | 0.273865133 | -2.159981579 | 0.030774096 | 0.07942819 |
| ENSG00000288548 | ENSG0000028 | 5.795939082 | -0.262488671 | 0.715241635 | -0.366992997 | 0.713624242 | 0.840800958 |
| ENSG00000288552 | ENSG0000028 | 9.529129202 | -0.638942882 | 0.56932644 | -1.122278604 | 0.261744001 | 0.442085656 |
| ENSG00000288559 | ENSG0000028 | 104.7923181 | -0.006008822 | 0.179054873 | -0.033558548 | 0.973229177 | 0.987970845 |
| ENSG00000288560 | ENSG0000028 | 7.398257697 | 0.280676452 | 0.646792073 | 0.433951595 | 0.664323592 | 0.806241836 |
| ENSG00000288568 | ZCCHC14-DT | 14.6068792 | 0.801065647 | 0.483666519 | 1.656235475 | 0.097674142 | 0.207967775 |
| ENSG00000288573 | ENSG0000028 | 13.41732556 | -0.024042287 | 0.484191388 | -0.049654511 | 0.960397707 | 0.981920343 |
| ENSG00000288579 | ENSG0000028 | 2.878319844 | 0.535951344 | 1.073335705 | 0.499332447 | 0.6175452 | 0.773279853 |
| ENSG00000288585 | ENSG0000028 | 35.9495754 | -0.53428899 | 0.291004069 | -1.836018963 | 0.066354852 | 0.151868733 |
| ENSG00000288586 | ENSG0000028 | 3.660865713 | 0.05082884 | 0.905772657 | 0.056116553 | 0.955248957 | 0.979455025 |
| ENSG00000288596 | C8orf44 | 61.67018541 | 0.778666486 | 0.232388138 | 3.35071528 | 0.000806031 | 0.003069958 |

| | | | | | | | |
|---|---|---|---|---|---|---|---|
| ENSG00000288597 | ENSG0000028 | 22.17340034 | 0.3151987 | 0.373487375 | 0.843934015 | 0.398706279 | 0.591597444 |
| ENSG00000288604 | ENSG0000028 | 7.500805212 | -0.039396885 | 0.635996388 | -0.06194514 | 0.95060652 | 0.977235535 |
