## Supplementary Table 5 for "GLYATL1 is associated with metabolic and epigenetic changes and with endocrine resistance in luminal breast cancer"

| Sample | Peak Area (y7 - 817.4666+) | Peak Area (y6 - 718.3981+) | Peak Area (y5 - 617.3505+) |
|---|---|---|---|
| parental_PRM_r1 | 12,738 | 0 | 8,290 |
| parental_PRM_r2 | 0 | 0 | 0 |
| parental_PRM_r3 | 0 | 3,887 | 727 |
| parental_PRM_r4 | 1,028 | 0 | 0 |
| LTED_PRM_r1 | 1,911,542 | 1,174,859 | 163,990 |
| LTED_PRM_r2 | 951,609 | 545,892 | 74,999 |
| LTED_PRM_r3 | 1,049,634 | 560,465 | 82,652 |
| LTED_PRM_r4 | 1,621,848 | 838,739 | 107,674 |
| KO1_PRM_r1 | 18,390 | 0 | 0 |
| KO1_PRM_r2 | 17,272 | 0 | 0 |
| KO1_PRM_r3 | 22,530 | 0 | 0 |
| KO1_PRM_r4 | 41,603 | 16,645 | 1,808 |
| KO2_PRM_r1 | 344,786 | 167,110 | 20,608 |
| KO2_PRM_r3 | 320,796 | 174,550 | 24,950 |
| KO2_PRM_r4 | 393,369 | 205,861 | 29,102 |
| KO2_PRM_r5 | 336,717 | 157,694 | 21,243 |
