## Supplementary Table 6 for "GLYATL1 is associated with metabolic and epigenetic changes and with endocrine resistance in luminal breast cancer"

| Protein | proteinGroup | GeneSymbol | Description | parental_r1 | parental_r2 |
|---|---|---|---|---|---|
| CEACAM5;CEACAM5;CEAC/ | A0A024R0K5;A0A087WYX0;F | CEACAM5;CEACAM5;( | Carcinoembryonic ant | NA | NA |
| RGS6 | A0A087WTW9;A0A0A0MRA9 | RGS6 | Regulator of G-protei | NA | NA |
| PDLIM3 | A0A087WYF8;A0A2U3TZH4;( | PDLIM3 | PDZ and LIM domain | 10802.3457 | 10154.27539 |
| MAPK8IP3 | A0A087WYG2;E9PFH7;Q9UP | MAPK8IP3 | C-Jun-amino-terminal | NA | NA |
| OAS2 | A0A087X0V5;P29728 | OAS2 | 2'-5'-oligoadenylate sy | 7983.85791 | 5961.518066 |
| KRT81;KRT86;KRT83;KRT85 | A0A087X106;O43790;P7838 | KRT81;KRT86;KRT83;K | Keratin, type II cuticul | NA | NA |
| CD99 | A0A096LP69;P14209 | CD99 | CD99 antigen | NA | NA |
| NDRG2 | A0A0A0MS87;G3V4S9;G3V5 | NDRG2 | Protein NDRG2;Protei | NA | NA |
| AKR1C3 | A0A0A0MSS8;P42330 | AKR1C3 | Aldo-keto reductase f | NA | NA |
| ZG16B | A0A0C4DGN4;I3L1H9;I3L3X0 | ZG16B | Zymogen granule prot | NA | NA |
| TAP1 | A0A140T9T7;Q03518 | TAP1 | Antigen peptide trans | 1868.425171 | 2075.86377 |
| ZNF385B | A0A2U3TZT0;C9JKK0;Q569K4 | ZNF385B | Zinc finger protein 38 | 6026.759277 | 2780.671387 |
| GUCY1B1 | A0A494C1B9;E9PCN2;Q0215 | GUCY1B1 | Guanylate cyclase solu | NA | NA |
| DNALI1 | A0A499FIY3;O14645 | DNALI1 | Axonemal dynein light | 20878.35742 | 18412.52734 |
| AMD1 | A0A5F9ZHD5;P17707;Q5VXN | AMD1 | S-adenosylmethionine | 20536.99414 | 19124.88086 |
| BNIPL | A2A492;F8W685;Q7Z465 | BNIPL | Bcl-2/adenovirus E1B | NA | NA |
| MROH6 | A6NGR9 | MROH6 | Maestro heat-like rep | 1669.080566 | 1063.829468 |
| TRAF3 | A6NHG8;H0YL25;H0YL26;Q1 | TRAF3 | TNF receptor-associat | NA | NA |
| MDM2 | A7UKX8;E7EPE2;F5GWH7;J3 | MDM2 | E3 ubiquitin-protein li | 911.1810913 | 1407.638794 |
| NPY1R | B4DKL9;P25929 | NPY1R | Neuropeptide Y recep | 1 | 2.63851738 |
| CSAD | C9JTU8;E9PFW9;F8VV11;F8V | CSAD | Cysteine sulfinic acid | NA | NA |
| BBS1 | E9PMB7;Q8NFJ9 | BBS1 | Bardet-Biedl syndrom | NA | NA |
| TIMP1 | H0Y789;P01033;Q5H9A7 | TIMP1 | Metalloproteinase inh | 8121.140625 | NA |
| VAV3 | H0YDG2;Q9UKW4 | VAV3 | Guanine nucleotide e | NA | NA |
| CDKN2A;CDKN2A;CDKN2A; | J3QRG6;K7ENC6;K7ES20;P42 | CDKN2A;CDKN2A;CDK | Cyclin-dependent kin | NA | NA |
| SOX1;SOX14;SOX3;SOX2;SC | O00570;O95416;P41225;P48 | SOX1;SOX14;SOX3;SO | Transcription factor S | NA | NA |
| TSPAN1 | O60635 | TSPAN1 | Tetraspanin-1 | NA | 1231.616699 |
| ENTPD3 | O75355 | ENTPD3 | Ectonucleoside tripho | NA | NA |
| CYBA | P13498 | CYBA | Cytochrome b-245 ligl | 19207.75391 | 32056.05664 |
| HPGD | P15428 | HPGD | 15-hydroxyprostaglan | 4933.90625 | 21899.20313 |
| CPE | P16870 | CPE | Carboxypeptidase E | NA | NA |
| SPRR1B | P22528 | SPRR1B | Cornifin-B | 5588.347168 | 7385.514648 |

| | | | | | |
|---|---|---|---|---|---|
| CD82 | P27701 | CD82 | CD82 antigen | 13091.84375 | 13328.26465 |
| RNASE4 | P34096 | RNASE4 | Ribonuclease 4 | 1144.686768 | NA |
| FMO5 | P49326 | FMO5 | Dimethylaniline monc | NA | NA |
| PCDH1 | Q08174 | PCDH1 | Protocadherin-1 | NA | NA |
| NEXN | Q0ZGT2 | NEXN | Nexilin | NA | NA |
| IL24 | Q13007 | IL24 | Interleukin-24 | 8006.725586 | 8086.186523 |
| KRT33B;KRT31 | Q14525;Q15323 | KRT33B;KRT31 | Keratin, type I cuticula | NA | NA |
| MYLK | Q15746 | MYLK | Myosin light chain kin | NA | NA |
| DOK7 | Q18PE1 | DOK7 | Protein Dok-7 | 1887.827515 | 2929.999756 |
| ANKRD22 | Q5VYY1 | ANKRD22 | Ankyrin repeat domai | NA | NA |
| SBSN | Q6UWP8 | SBSN | Suprabasin | 345.7049255 | 441.422699 |
| PHLDA1 | Q8WV24;R4GND3 | PHLDA1 | Pleckstrin homology-l | NA | NA |
| FRMD6 | Q96NE9 | FRMD6 | FERM domain-contair | NA | NA |
| SPANXD;SPANXB1;SPANXA | Q9BXN6;Q9NS25;Q9NS26;Q9 | SPANXD;SPANXB1;SPA | Sperm protein associa | 0.506998122 | 2.74260211 |
| PLEKHA4 | Q9H4M7 | PLEKHA4 | Pleckstrin homology d | 3080.536133 | 3820.230469 |
| CLIP2 | Q9UDT6 | CLIP2 | CAP-Gly domain-conta | 719.411438 | 388.4516907 |
| C2CD2 | Q9Y426 | C2CD2 | C2 domain-containing | NA | NA |
| IRS2 | Q9Y4H2 | IRS2 | Insulin receptor subst | NA | NA |
| CXCR4 | P61073 | CXCR4 | C-X-C chemokine rece | NA | 114.5247269 |
| FAM20C | Q8IXL6 | FAM20C | Extracellular serine/th | 263.5974121 | NA |
| BASP1 | P80723 | BASP1 | Brain acid soluble pro | 38462.83594 | 48943.85156 |
| TYMP | A0A494C0L6;C9JGI3;P19971 | TYMP | Thymidine phosphory | 383.2762756 | 293.2657471 |
| GNB5 | H0YNW7;Q96F32 | GNB5 | Guanine nucleotide-b | 35987.71362 | 29195.25781 |
| AGR2 | O95994 | AGR2 | Anterior gradient prot | 58370.89453 | 22890.84766 |
| APOD | C9JF17;P05090 | APOD | Apolipoprotein D (Fra | 1043.610352 | 4084.465332 |
| TNFAIP8L3 | A0A1B0GTK8;Q5GJ75 | TNFAIP8L3 | Tumor necrosis factor | NA | 317.1912537 |
| SPPL2A | Q8TCT8 | SPPL2A | Signal peptide peptida | 29390.24414 | 23974.46484 |
| DTWD1 | Q8N5C7 | DTWD1 | DTW domain-containi | 3533.873047 | 3908.327393 |
| ACSL4 | O60488 | ACSL4 | Long-chain-fatty-acid- | 858.9390869 | 1413.859009 |
| SRGAP3 | O43295 | SRGAP3 | SLIT-ROBO Rho GTPas | 338.8704529 | NA |
| USP8 | P40818 | USP8 | Ubiquitin carboxyl-ter | 44611.05859 | 45398.48047 |
| LXN | Q9BS40 | LXN | Latexin | 17449.98633 | 16102.87012 |
| TMOD2 | Q9NZR1 | TMOD2 | Tropomodulin-2 | 2780.48584 | 5317.661133 |

| | | | | | |
|---|---|---|---|---|---|
| SH3BGRL2 | Q9UJC5 | SH3BGRL2 | SH3 domain-binding g | 9202.578125 | 9275.832031 |
| ARPP19 | H3BMD8 | ARPP19 | cAMP-regulated phos | 214913.2941 | 199650.1094 |
| MYO5A | Q9Y4I1 | MYO5A | Unconventional myos | 65196.10547 | 63749.73828 |
| CLU | P10909 | CLU | Clusterin | 9651.375 | 5212.882324 |
| SEPTIN10 | B5ME97;E7EW69;Q9P0V9 | SEPTIN10 | Septin 10, isoform CR | 2616.991943 | 2187.975098 |
| SLC27A2 | O14975 | SLC27A2 | Very long-chain acyl-C | 22393.76367 | 22637.69922 |
| ST3GAL1 | Q11201 | ST3GAL1 | CMP-N-acetylneurami | 2397.354736 | 2826.053467 |
| NAPRT | Q6XQN6 | NAPRT | Nicotinate phosphorib | 21373.8125 | 14331.65234 |
| SNCG | O76070 | SNCG | Gamma-synuclein | 4830.34668 | 4141.875 |
| RFTN1 | Q14699 | RFTN1 | Raftlin | NA | 916.3873901 |
| TMEM181 | Q9P2C4 | TMEM181 | Transmembrane prote | 1645.650024 | 3674.899902 |
| TGFB1I1 | O43294 | TGFB1I1 | Transforming growth | 2179.354004 | 1549.091187 |
| GLYATL1 | E9PP99;Q969I3 | GLYATL1 | Glycine N-acyltransfer | NA | NA |
| CYP1A1 | E7EMT5;P04798 | CYP1A1 | Cytochrome P450 1A1 | 4949.07959 | 2741.512939 |
| FUT8 | Q9BYC5 | FUT8 | Alpha-(1,6)-fucosyltra | 15305.13965 | 19154.33203 |
| TRPM7 | H0YLN8;Q96QT4 | TRPM7 | Transient receptor po | 3270.064453 | 3428.447266 |
| ESR1 | P03372 | ESR1 | Estrogen receptor | 8111.900879 | 10369.94629 |
| NUPR1 | O60356 | NUPR1 | Nuclear protein 1 | 127.5089264 | 137.836731 |
| TRIQK | Q629K1 | TRIQK | Triple QxxK/R motif-c | 32077.30859 | 15079.99902 |
| SOWAHC | Q53LP3 | SOWAHC | Ankyrin repeat domai | NA | 2126.651855 |
| ITPR1 | A0A3B3ITU8;A0A3B3IU04;A0 | ITPR1 | Inositol 1,4,5-trisphos | 21589.47266 | 21128.97852 |
| MBNL2 | O95205;Q5VZF2 | MBNL2 | Muscleblind-like 2 (Dr | 1457.565063 | NA |
| BCAS1 | G3XAF7 | BCAS1 | Breast carcinoma amp | 9122.325195 | 10890.17987 |
| DMXL2 | Q8TDJ6 | DMXL2 | DmX-like protein 2 | 58103.86328 | 56103.02344 |
| CYSRT1 | A8MQ03;B8A4K4 | CYSRT1 | Cysteine-rich tail prot | 653.1707153 | 236.4849396 |
| SLC2A6 | Q9UGQ3 | SLC2A6 | Solute carrier family 2 | 1293.485107 | NA |
| SLC34A2 | O95436 | SLC34A2 | Sodium-dependent ph | 81282.90625 | 80138.10938 |
| PREX1 | Q8TCU6 | PREX1 | Phosphatidylinositol 3 | 225715.4375 | 175631.8906 |
| CDK6 | Q00534 | CDK6 | Cyclin-dependent kina | 2146.835693 | NA |
| MYO5C | Q9NQX4 | MYO5C | Unconventional myos | 47813.90625 | 42885.31641 |
| NET1 | Q5SQI6 | NET1 | Neuroepithelial cell-tr | 428407.625 | 172342.8906 |
| DCLK1 | O15075 | DCLK1 | Serine/threonine-prot | 13255.73535 | 11096.22852 |
| ARFGEF3 | Q5TH69 | ARFGEF3 | Brefeldin A-inhibited g | 10974.9668 | 10831.39844 |

| | | | | | |
|---|---|---|---|---|---|
| LMCD1 | Q9NZU5 | LMCD1 | LIM and cysteine-rich | 16435.21289 | 13585.28711 |
| TFPI | P10646 | TFPI | Tissue factor pathway | 546.4213257 | 1077.404297 |
| FARP1 | C9JME2;Q9Y4F1 | FARP1 | FERM, ARHGEF and pl | 2170.04541 | 1907.249023 |
| ELOVL6 | D6RBP4;D6RCU0;D6RHI2;Q9I | ELOVL6 | Elongation of very lon | NA | NA |
| LYSMD2 | Q8IV50 | LYSMD2 | LysM and putative pe | 39894.51953 | 35310.82813 |
| ANKMY2 | Q8IV38 | ANKMY2 | Ankyrin repeat and M | 4929.334961 | 4795.010742 |
| FAM221A | A4D161;B8ZZQ8 | FAM221A | Protein FAM221A | 4962.405762 | 3131.289307 |
| CAB39L | Q5TAW7;Q9H9S4 | CAB39L | Calcium-binding prote | 10946.6875 | 15371.75879 |
| AR | P10275 | AR | Androgen receptor | 3088.41748 | 2788.254639 |
| TMEM135 | E9PQL0;Q86UB9 | TMEM135 | Transmembrane prote | 2225.298096 | 2513.493408 |
| GDAP1 | Q8TB36 | GDAP1 | Ganglioside-induced c | 21150.52344 | 22701.48438 |
| ANTXR2 | J3KPY9;P58335 | ANTXR2 | Anthrax toxin recepto | 583.3782349 | 660.0952759 |
| BOK | Q9UMX3 | BOK | Bcl-2-related ovarian l | 624.5980835 | 1523.240967 |
| MYO1B | E9PDF6;O43795 | MYO1B | Unconventional myos | 49286.09766 | 57603.96484 |
| RAB27A | P51159 | RAB27A | Ras-related protein Ra | 30269.26367 | 38988.74219 |
| ACKR3 | P25106 | ACKR3 | Atypical chemokine re | 556.6116943 | 1633.057007 |
| EFEMP1 | A0A0U1RQV3;Q12805 | EFEMP1 | EGF-containing fibulin | 5429.975586 | 1324.092041 |
| CDCA7L | Q96GN5 | CDCA7L | Cell division cycle-ass | 1420.192383 | 918.12323 |
| STC2 | H0YB13;O76061 | STC2 | Stanniocalcin-2 (Fragr | 818.5586548 | NA |
| FLNA | P21333 | FLNA | Filamin-A | 386068.0645 | 392618.9863 |
| AP4E1 | Q9UPM8 | AP4E1 | AP-4 complex subunit | 18092.74414 | 19152.02539 |
| EPB41L2 | E9PHY5 | EPB41L2 | Band 4.1-like protein | 15788.41309 | 14794.72461 |
| NPNT | D6RH31;H0YAK9;Q6UXI9 | NPNT | Nephronectin (Fragme | 718.5405884 | 1854.232178 |
| VWDE | E5RG96;J3KQJ9;Q8N2E2 | VWDE | von Willebrand factor | 132.3049316 | 321.3639526 |
| ATP2A3 | Q93084 | ATP2A3 | Sarcoplasmic/endopla | 23139.375 | 27183.63867 |
| DGLUCY | H0YB09;H0YB11;H0YB62;Q7Z | DGLUCY | D-glutamate cyclase, r | NA | 714.3413086 |
| AKAP5 | P24588 | AKAP5 | A-kinase anchor prote | 795.8441772 | 1311.228027 |
| ABHD17C | Q6PCB6 | ABHD17C | Alpha/beta hydrolase | 3199.823242 | 2765.414307 |
| ASB9 | Q96DX5 | ASB9 | Ankyrin repeat and SC | 2324.706299 | 582.144043 |
| ACSL3 | O95573 | ACSL3 | Long-chain-fatty-acid- | 72797.80469 | 92930.15625 |
| CYP1B1 | Q16678 | CYP1B1 | Cytochrome P450 1B1 | 41170.11328 | 22485.6875 |
| KIF5A | Q12840 | KIF5A | Kinesin heavy chain is | 2058.50415 | 16827.6875 |
| FOLH1;FOLH1B | Q04609;Q9HBA9 | FOLH1;FOLH1B | Glutamate carboxype | NA | 141.2749329 |

| | | | | | |
|---|---|---|---|---|---|
| BAMBI | Q13145 | BAMBI | BMP and activin mem | NA | 494.1544189 |
| PRR15 | Q8IV56 | PRR15 | Proline-rich protein 15 | 5189.682129 | 8976.806641 |
| DKK1 | O94907 | DKK1 | Dickkopf-related prot | 7552.331543 | 6456.3125 |
| TAGLN | Q01995 | TAGLN | Transgelin | 47919.55078 | 57721.41016 |
| L1CAM | P32004 | L1CAM | Neural cell adhesion r | 10063.9502 | 6120.036621 |
| TTC9 | Q92623 | TTC9 | Tetratricopeptide rep | 4612.228516 | 7412.09668 |
| CXADR | P78310 | CXADR | Coxsackievirus and ad | 3195.263672 | 1710.400879 |
| SHANK2 | C9JFP8 | SHANK2 | SH3 and multiple anky | 9124.138184 | 7594.643066 |
| ICAM1 | P05362 | ICAM1 | Intercellular adhesion | 3291.769287 | 2986.483643 |
| PCDH7 | O60245 | PCDH7 | Protocadherin-7 | 1072.985718 | 1500.007568 |
| CCDC126 | C9J0J0;Q96EE4 | CCDC126 | Coiled-coil domain-co | 5075.934082 | 2840.349609 |
| PRKD1 | F8WBA3;Q15139 | PRKD1 | Serine/threonine-prot | 624.4707031 | NA |
| OBSL1 | A6NN50;O75147 | OBSL1 | Obscurin-like protein | 2769.66748 | 1403.902344 |
| DPYSL5 | Q9BPU6 | DPYSL5 | Dihydropyrimidinase- | 5623.144043 | 2910.00293 |
| MAPK12 | B5MDL5;P53778 | MAPK12 | Mitogen-activated pro | NA | 1137.028198 |
| TESK2 | Q96S53 | TESK2 | Dual specificity testis- | 1171.891602 | 933.3110962 |
| RBMS1 | E7ETU5;F6Y5H0;P29558 | RBMS1 | RNA-binding motif, sir | 290.4062195 | 7888.355957 |
| CMC4 | P56277 | CMC4 | Cx9C motif-containing | 5051.23584 | 5917.180176 |
| THBS1 | P07996 | THBS1 | Thrombospondin-1 | 23671.51563 | 20460.46094 |
| MAP9 | A2VCS9;E7ETZ8;Q49MG5 | MAP9 | MAP9 protein (Fragm | 1261.765869 | 394.9317627 |
| TMOD3 | Q9NYL9 | TMOD3 | Tropomodulin-3 | 143049.2344 | 139628.0781 |
| PKN3 | Q6P5Z2 | PKN3 | Serine/threonine-prot | 422.7148743 | 343.4072266 |
| ARNT2 | A0A087WVE9;Q9HBZ2 | ARNT2 | Aryl hydrocarbon rece | 1602.755859 | 1298.440186 |
| MME | P08473 | MME | Neprilysin | 1937.6875 | 1399.675293 |
| BCAM | P50895 | BCAM | Basal cell adhesion mo | 126163.9766 | 91092.57813 |
| SLIT1 | O75093 | SLIT1 | Slit homolog 1 protein | 7324.297852 | 5326.486328 |
| NUDT7 | P0C024 | NUDT7 | Peroxisomal coenzym | 2738.375732 | 2585.611816 |
| EEF1A2 | A0A2U3TZH3;Q05639 | EEF1A2 | Elongation factor 1-alp | 887202.1875 | 612320.25 |
| TNS3 | Q68CZ2 | TNS3 | Tensin-3 | 5967.396484 | 5659.0625 |
| SELENOM | A0A087WWF1;Q8WWX9 | SELENOM | Selenoprotein M | NA | 676.0651245 |
| NPTXR | A0A1X7SBT7;O95502 | NPTXR | Neuronal pentraxin re | 540.0657349 | 500.2240906 |
| AP4B1 | Q9Y6B7 | AP4B1 | AP-4 complex subunit | 25094.59766 | 24235.93945 |
| SELENBP1 | Q13228 | SELENBP1 | Methanethiol oxidase | 233530.1875 | 161053.8594 |

| | | | | | |
|---|---|---|---|---|---|
| SORBS3 | O60504 | SORBS3 | Vinexin | 13759.79004 | 11652.78711 |
| EFNB2 | P52799 | EFNB2 | Ephrin-B2 | 2034.334717 | 2162.002197 |
| FKBP10 | Q96AY3 | FKBP10 | Peptidyl-prolyl cis-trar | 91557.875 | 67671.01563 |
| UPK2 | O00526 | UPK2 | Uroplakin-2 | 1069.328491 | 1486.286133 |
| C16orf46 | Q6P387 | C16orf46 | Uncharacterized prote | 347697.875 | 229442.9219 |
| PPP1R35 | Q8TAP8 | PPP1R35 | Protein phosphatase 1 | 4986.915527 | 9984.048828 |
| EMD | P50402 | EMD | Emerin | 43124.67969 | 94003.4375 |
| MOCOS | Q96EN8 | MOCOS | Molybdenum cofactoı | NA | 1534.369995 |
| MICAL1 | Q8TDZ2 | MICAL1 | [F-actin]-monooxygen | 1971.049316 | 2881.136963 |
| KCNAB2 | A0A590UJB6;Q13303 | KCNAB2 | Voltage-gated potassi | 8953.066406 | 10814.40234 |
| TAPT1 | Q6NXT6 | TAPT1 | Transmembrane ante | 82278.28906 | 105783.0234 |
| ABCG1 | C9JKD9;E9PGV9;P45844 | ABCG1 | ATP-binding cassette s | 6095.637695 | 5205.303223 |
| PAK1 | Q13153 | PAK1 | Serine/threonine-prot | 45827.06641 | 65908.99219 |
| VGF | O15240 | VGF | Neurosecretory prote | 279.7450867 | NA |
| TPK1 | F5GZG6;F8VRJ6;F8WCM7;Q9 | TPK1 | Thiamin pyrophospho | 297.0066528 | 1122.878418 |
| IDNK | Q5T6J7;Q5T6J8 | IDNK | Probable gluconokina | 2551.429932 | 2426.615967 |
| RAB39B | Q96DA2 | RAB39B | Ras-related protein Ra | 9802.255859 | 6027.092285 |
| SMIM20 | Q8N5G0 | SMIM20 | Small integral membra | 24212.94922 | 17204.95313 |
| ABCC5 | O15440 | ABCC5 | Multidrug resistance-a | 9735.283203 | 10814.37402 |
| BBC3 | Q9BXH1 | BBC3 | Bcl-2-binding compon | 3023.862549 | 616.2875366 |
| SEPSECS | Q9HD40 | SEPSECS | O-phosphoseryl-tRNA | 5166.118164 | 4958.449707 |
| DPYSL2 | A0A1C7CYX9;Q16555 | DPYSL2 | Dihydropyrimidinase- | 117474.8906 | 128946.8203 |
| PDP1 | Q9P0J1 | PDP1 | [Pyruvate dehydroger | 13162.06543 | 14028.05762 |
| GRTP1 | A0A087X0Y1;Q5TC63 | GRTP1 | Growth hormone-regu | 2650.714355 | 2436.968018 |
| SIPA1L2 | Q9P2F8 | SIPA1L2 | Signal-induced prolife | 3156.466309 | 1415.117676 |
| TCF12 | Q99081 | TCF12 | Transcription factor 1 | 2124.725586 | 387.4258728 |
| ENPP1 | P22413 | ENPP1 | Ectonucleotide pyropl | 5790.14209 | 6214.160156 |
| ACSF2 | Q96CM8 | ACSF2 | Medium-chain acyl-Co | 19968.58984 | 18052.24219 |
| MAP1B | P46821 | MAP1B | Microtubule-associate | 2368.144531 | 1361.375244 |
| RAPH1 | Q70E73 | RAPH1 | Ras-associated and pl | 25256.33398 | 31731.24805 |
| S100P | P25815 | S100P | Protein S100-P | NA | 2242.266602 |
| PCTP | I3L2M9;I3L3H0;Q9UKL6 | PCTP | Phosphatidylcholine t | 1085.862305 | 893.512085 |
| PDE8A | O60658 | PDE8A | High affinity cAMP-sp | 486.086853 | 918.1456909 |

| | | | | | |
|---|---|---|---|---|---|
| SCD | O00767 | SCD | Acyl-CoA desaturase | 198557.7813 | 280630.3125 |
| HMGB3 | O15347 | HMGB3 | High mobility group p | 245675.5313 | 262161.875 |
| EML2 | O95834 | EML2 | Echinoderm microtub | 18562.14844 | 23939.37695 |
| TMC4 | A0A087WVI4;A0A0G2JMM6; | TMC4 | Transmembrane chan | NA | NA |
| SZT2 | Q5T011 | SZT2 | KICSTOR complex prot | 1293.672974 | 719.380188 |
| PLXNA3 | P51805 | PLXNA3 | Plexin-A3 | 3514.554932 | 4376.293945 |
| IRAK1 | D3YTB5;P51617 | IRAK1 | Interleukin-1 receptor | 12635.25195 | 15765.46191 |
| CYP2J2 | A0A3B3IT99;P51589 | CYP2J2 | Cytochrome P450 2J2 | 3343.645996 | 2794.371338 |
| MARCKS | P29966 | MARCKS | Myristoylated alanine | 290690.4688 | 197498.0156 |
| UGDH | O60701 | UGDH | UDP-glucose 6-dehydr | 261233.3125 | 239121.4844 |
| PIGBOS1 | A0A0B4J2F0 | PIGBOS1 | Protein PIGBOS1 | 7746.620117 | 12299.37598 |
| SLC43A2 | Q8N370 | SLC43A2 | Large neutral amino a | 834.4818115 | 594.3898926 |
| METTL7A | H0YI09;Q9H8H3 | METTL7A | Methyltransferase-like | 12982.52539 | 13566.62305 |
| FAR1 | Q8WVX9 | FAR1 | Fatty acyl-CoA reducta | 13762.52734 | 23162.11719 |
| FAM89A | Q96GI7 | FAM89A | Protein FAM89A | 1904.372681 | 1567.742676 |
| FAM107B | A0A1C7CYX8;C9J6N5;C9JQ40 | FAM107B | Protein FAM107B (Fra | 11253.84277 | 8137.372559 |
| JPH1 | Q9HDC5 | JPH1 | Junctophilin-1 | 4197.788086 | 7210.166992 |
| CENPV | Q7Z7K6 | CENPV | Centromere protein V | 3964.634521 | 3701.654053 |
| MT1F | P04733 | MT1F | Metallothionein-1F | 789.6029053 | NA |
| VIL1 | P09327 | VIL1 | Villin-1 | 2682.443848 | 1406.651245 |
| ICMT | O60725 | ICMT | Protein-S-isoprenylcys | 12204.66211 | 8733.322266 |
| CA5B | Q9Y2D0 | CA5B | Carbonic anhydrase 5 | 10497.49219 | 12643.12012 |
| ACSS2 | Q9NR19 | ACSS2 | Acetyl-coenzyme A sy | 34252.73047 | 37343.55078 |
| MTMR10 | Q9NXD2 | MTMR10 | Myotubularin-related | 1013.12738 | 2194.962646 |
| MBOAT2 | Q6ZWT7 | MBOAT2 | Lysophospholipid acyl | 2243.078369 | 1527.010742 |
| HACD1 | B0YJ81 | HACD1 | Very-long-chain (3R)-3 | 1020.637024 | 1318.50415 |
| RPS6KA5 | O75582 | RPS6KA5 | Ribosomal protein S6 | 9752.457031 | 10806.73535 |
| VAT1L | Q9HCJ6 | VAT1L | Synaptic vesicle meml | 4089.960938 | 3391.698242 |
| LMBRD1 | A0A3B3IRS7;A0A3B3IT31;A0A | LMBRD1 | Probable lysosomal co | 7116.964355 | 8956.806641 |
| CLMN | Q96JQ2 | CLMN | Calmin | 3801.174072 | 6941.92334 |
| GDI1 | P31150 | GDI1 | Rab GDP dissociation | 175049.3125 | 210502.7031 |
| CBX6 | B0QXZ6;O95503 | CBX6 | Chromobox protein ho | 5892.187012 | 8388.170898 |
| LETMD1 | F8VP71;F8VVQ3;F8VZA2;Q6P | LETMD1 | LETM1 domain-contai | 189.6425323 | 2750.338623 |

| | | | | | |
|---|---|---|---|---|---|
| HIBADH | P31937 | HIBADH | 3-hydroxyisobutyrate | 17623.58203 | 20781.90234 |
| IGSF3 | O75054 | IGSF3 | Immunoglobulin supe | 35279.17969 | 41112.50781 |
| PLEKHF2 | Q9H8W4 | PLEKHF2 | Pleckstrin homology d | 90991.39063 | 86473.75 |
| PXMP2 | F5H4N4;Q9NR77 | PXMP2 | Peroxisomal membrar | 19419.92383 | 19052.41016 |
| DHCR24 | A0A0A0MTI1;Q15392 | DHCR24 | Delta(24)-sterol reduc | 253572.3438 | 283903.9688 |
| DOP1B | Q9Y3R5 | DOP1B | Protein dopey-2 | 15245.54785 | 19482.88867 |
| SMAD1 | Q15797 | SMAD1 | Mothers against deca | 481.0163879 | NA |
| ANKIB1 | Q9P2G1 | ANKIB1 | Ankyrin repeat and IB | NA | 1832.111816 |
| GALNT12 | Q8IXK2 | GALNT12 | Polypeptide N-acetylg | 3289.328857 | 2218.647217 |
| C8orf37 | Q96NL8 | C8orf37 | Protein C8orf37 | 2390.790771 | NA |
| SLC9A1 | P19634 | SLC9A1 | Sodium/hydrogen exc | 19631.5332 | 20271.13086 |
| ARSD | P51689 | ARSD | Arylsulfatase D | 4625.015137 | 5330.425293 |
| HEBP1 | Q9NRV9 | HEBP1 | Heme-binding protein | 265129.1563 | 259851.5781 |
| FBP2 | O00757 | FBP2 | Fructose-1,6-bisphosp | 1216986.5 | 589882.375 |
| GLCCI1 | Q86VQ1 | GLCCI1 | Glucocorticoid-induce | 4539.581543 | 6489.452148 |
| FBXW8 | Q8N3Y1 | FBXW8 | F-box/WD repeat-con | 373.826355 | 4654.272461 |
| FAM50B | Q9Y247 | FAM50B | Protein FAM50B | 77586.125 | 81178.71094 |
| PRRT3 | Q5FWE3 | PRRT3 | Proline-rich transmem | 2385.224609 | 2918.369141 |
| SLC25A18 | Q9H1K4 | SLC25A18 | Mitochondrial glutam | 9933.087891 | 5978.266113 |
| RWDD2A | Q9UIY3 | RWDD2A | RWD domain-containi | 470.9031067 | 785.5628052 |
| ICA1 | A0A3B3IST8;E9PDL4;Q05084 | ICA1 | Islet cell autoantigen 1 | 10291.41992 | 11095.66406 |
| PYGL | P06737 | PYGL | Glycogen phosphoryla | 11098.71094 | 8638.763672 |
| MECP2 | B5MCB4 | MECP2 | Methyl-CpG-binding p | 12308.72021 | 15908.78809 |
| TAX1BP1 | B8ZZD4;Q86VP1 | TAX1BP1 | Tax1-binding protein 1 | 2065.525391 | 1320.959351 |
| SPTSSB | Q8NFR3 | SPTSSB | Serine palmitoyltransf | 25214.77344 | 13991.75781 |
| SNX14 | D6RJG9;Q9Y5W7 | SNX14 | Sorting nexin-14 (Frag | 4280.753418 | 5414.316895 |
| PLGRKT | Q9HBL7 | PLGRKT | Plasminogen receptor | 21559.58594 | 17057.5 |
| AGAP1 | A0A087X1U1;E7EUN2;F5GXN | AGAP1 | Arf-GAP with GTPase, | 2773.186279 | 4120.585938 |
| NBEA | A0A0D9SF28;F5GXV7;Q5T32 | NBEA | Neurobeachin | 4092.225342 | 5013.37207 |
| CRYBG1 | A0A0J9YWL0 | CRYBG1 | Beta/gamma crystallir | 22342.55664 | 22154.85156 |
| AP1G2 | O75843 | AP1G2 | AP-1 complex subunit | 25908.91016 | 43831.77344 |
| FAM241A | Q8N8J7 | FAM241A | Uncharacterized prote | 10420.00586 | 11505.02246 |
| DIP2C | E7EPU2;Q9Y2E4 | DIP2C | Disco-interacting prot | 16370.32617 | 20684.35352 |

| | | | | | |
|---|---|---|---|---|---|
| MIOS | Q9NXC5 | MIOS | GATOR complex prote | 21211.79883 | 20457.61914 |
| APOBEC3B | B0QYD3;Q9UH17 | APOBEC3B | DNA dC->dU-editing e | 9737.919922 | 6018.750977 |
| LIMA1 | Q9UHB6 | LIMA1 | LIM domain and actin | 101637.1875 | 82087.98438 |
| UTRN | P46939 | UTRN | Utrophin | 5387.911621 | 4419.249512 |
| DHRS1 | Q96LJ7 | DHRS1 | Dehydrogenase/reduc | 14268.91211 | 12723.31348 |
| TMLHE | Q9NVH6 | TMLHE | Trimethyllysine dioxyg | 13640.15039 | 15163.48242 |
| FUNDC2 | Q9BWH2 | FUNDC2 | FUN14 domain-contai | 37133.61328 | 46560.29688 |
| LEO1 | Q8WVC0 | LEO1 | RNA polymerase-asso | 59428.53125 | 52785.55078 |
| ACACA | Q13085 | ACACA | Acetyl-CoA carboxylas | 82908.41406 | 97145.50781 |
| PLEKHA7 | E9PKC0;Q6IQ23 | PLEKHA7 | Pleckstrin homology d | 5953.287598 | 9162.787109 |
| OXCT1 | P55809 | OXCT1 | Succinyl-CoA:3-ketoac | 171699.625 | 182251.0781 |
| SIPA1L1 | O43166 | SIPA1L1 | Signal-induced prolife | 5327.299316 | 4680.319336 |
| ASAP1 | A0A0A0MRE5;Q9ULH1 | ASAP1 | Arf-GAP with SH3 dom | 16713.54102 | 16740.02344 |
| NAXD | Q8IW45 | NAXD | ATP-dependent (S)-NA | 15034.05273 | 17821.01367 |
| ACYP1 | P07311 | ACYP1 | Acylphosphatase-1 | 34369.44922 | 32795.96094 |
| DIP2A | A0A494C143;Q14689 | DIP2A | Disco-interacting prot | 14931.4082 | 20085.64844 |
| LSS | P48449 | LSS | Lanosterol synthase | 216596.4688 | 253262.7656 |
| AP4S1 | Q9Y587 | AP4S1 | AP-4 complex subunit | 31906.42578 | 35911.09766 |
| CUL7 | Q14999 | CUL7 | Cullin-7 | 9699.94043 | 12994.72852 |
| HIPK1 | J3KP92;Q5SQL3;Q86Z02 | HIPK1 | Homeodomain-intera | 1935.027832 | NA |
| SYNJ2 | A0A1W2PR85;O15056 | SYNJ2 | Synaptojanin-2 | 2087.076172 | 2507.653076 |
| FAM50A | Q14320 | FAM50A | Protein FAM50A | 108950.7266 | 117199.1406 |
| ST8SIA4 | Q92187 | ST8SIA4 | CMP-N-acetylneurami | 13405.29492 | 11812.20996 |
| NR3C1 | P04150 | NR3C1 | Glucocorticoid recept | 17429.29297 | 14213.21973 |
| NEDD4 | A0A3B3IUC1 | NEDD4 | E3 ubiquitin-protein li | 70929.21875 | 78928.11328 |
| MICA | Q29983 | MICA | MHC class I polypepti | 7133.341797 | 5346.984375 |
| ALDH6A1 | Q02252 | ALDH6A1 | Methylmalonate-semi | 92210.75781 | 88450.14844 |
| PAAT | Q9H8K7 | PAAT | ATPase PAAT | 4128.613281 | 4209.088379 |
| DCAKD | K7ESP4;Q8WVC6 | DCAKD | Dephospho-CoA kinas | 8890.532227 | 9866.963867 |
| JMJD7 | P0C870 | JMJD7 | Bifunctional peptidase | 1374.401855 | 1117.983765 |
| BICDL2 | A0A2R8Y2X6;A1A5D9 | BICDL2 | BICD family-like cargo | NA | 3922.616455 |
| RASEF | Q8IZ41 | RASEF | Ras and EF-hand dom | 2482.544678 | 2212.310791 |
| SRR | Q9GZT4 | SRR | Serine racemase | 17660.71289 | 22529.53516 |

| | | | | | |
|---|---|---|---|---|---|
| ENTPD5 | G3V4I0;O75356 | ENTPD5 | Ectonucleoside tripho | 4021.626221 | 3821.010986 |
| TAP2 | A0A087WYD6;A0A0G2JLV0;C | TAP2 | Antigen peptide trans | 17107.06641 | 11769.72852 |
| SLC25A40 | Q8TBP6 | SLC25A40 | Solute carrier family 2 | 23166.6543 | 25113.23047 |
| MAFG | O15525 | MAFG | Transcription factor M | 4863.126465 | 314.924408 |
| DEPTOR | Q8TB45 | DEPTOR | DEP domain-containin | 928.9362793 | 965.3428955 |
| GRHL3 | Q8TE85 | GRHL3 | Grainyhead-like prote | 1288.904541 | 2733.691162 |
| MID1IP1 | Q9NPA3 | MID1IP1 | Mid1-interacting prot | 3207.335205 | 2280.305908 |
| GCHFR | P30047 | GCHFR | GTP cyclohydrolase 1 | 17611.69531 | 18386.50781 |
| FASN | P49327 | FASN | Fatty acid synthase | 1990926.125 | 2716709.25 |
| IDI1 | Q13907 | IDI1 | Isopentenyl-diphosph | 258956.2969 | 334411.2813 |
| PGPEP1 | Q9NXJ5;S4R2Y9;U3KQ24;U3K | PGPEP1 | Pyroglutamyl-peptida | 3443.255127 | 5121.311035 |
| MRFAP1 | Q9Y605 | MRFAP1 | MORF4 family-associa | 20229.88086 | 13806.9209 |
| GPRC5C | A0A0C4DFY5;A8MXZ4;Q9NQ | GPRC5C | G-protein-coupled rec | 5003.390137 | 1900.544922 |
| CAMK2D | D6R938;E9PBG7;E9PF82;Q13 | CAMK2D | Calcium/calmodulin-d | 26346.0293 | 24586.15234 |
| TMEM184C | Q9NVA4 | TMEM184C | Transmembrane prote | 10510.5 | 4310.314941 |
| BCAP31 | P51572 | BCAP31 | B-cell receptor-associa | 250411.0781 | 261942.1719 |
| MPP6 | Q9NZW5 | MPP6 | MAGUK p55 subfamily | 48975.78516 | 50007.15625 |
| FBP1 | P09467 | FBP1 | Fructose-1,6-bisphosp | 1265451.25 | 942361.375 |
| FBXO4 | Q9UKT5 | FBXO4 | F-box only protein 4 | 24876.81055 | 22618.20508 |
| ATN1 | P54259 | ATN1 | Atrophin-1 | 14427.08789 | 17902.91797 |
| GGT7 | Q9UJ14 | GGT7 | Glutathione hydrolase | 1731.342285 | 2830.189941 |
| STBD1 | O95210 | STBD1 | Starch-binding domain | 1675.856689 | 3964.279785 |
| ENO2 | P09104 | ENO2 | Gamma-enolase | 66403.99219 | 53898.40234 |
| ERBB2 | P04626 | ERBB2 | Receptor tyrosine-pro | 31401.38281 | 30351.46875 |
| ABLIM3 | A0A0C4DGA7;O94929 | ABLIM3 | Actin-binding LIM pro | 194.9687653 | 6577.476563 |
| SEMA3C | Q99985 | SEMA3C | Semaphorin-3C | 6169.817871 | 7096.731934 |
| IGFBP2 | P18065 | IGFBP2 | Insulin-like growth fac | 21717.36133 | 16852.80273 |
| ENO3 | P13929 | ENO3 | Beta-enolase | 104218.2344 | 223912.6094 |
| GCNT1 | Q02742 | GCNT1 | Beta-1,3-galactosyl-O- | 960.9059448 | NA |
| PAFAH2 | Q99487 | PAFAH2 | Platelet-activating fac | 3481.235596 | 4395.049316 |
| SHPRH | A0A0D9SFM0;Q149N8 | SHPRH | E3 ubiquitin-protein li | 103586.9922 | 106644.1719 |
| ZNF740 | Q8NDX6 | ZNF740 | Zinc finger protein 74( | 4920.447754 | 3206.85498 |
| HMGN5 | P82970;Q5JSK6;Q5JSL0 | HMGN5 | High mobility group n | 7204.237793 | 6362.497559 |

| | | | | | |
|---|---|---|---|---|---|
| PKP4 | A0A0D9SF60;Q99569 | PKP4 | Plakophilin-4 | 6895.959961 | 9032.756836 |
| ALDH4A1 | P30038 | ALDH4A1 | Delta-1-pyrroline-5-ca | 86735.0625 | 83266.85938 |
| PRKACB | P22694 | PRKACB | cAMP-dependent prot | 12008.12305 | 13594.49316 |
| DHCR7 | Q9UBM7 | DHCR7 | 7-dehydrocholesterol | 401936.6563 | 494360.9688 |
| NEDD4L | Q96PU5 | NEDD4L | E3 ubiquitin-protein li | 34375.69922 | 37031.41016 |
| GBP2 | P32456 | GBP2 | Guanylate-binding pro | 26623.06055 | 23553.62109 |
| NSDHL | Q15738 | NSDHL | Sterol-4-alpha-carbox | 140900.3906 | 178036.3438 |
| AKT1 | P31749 | AKT1 | RAC-alpha serine/thre | 59138.13281 | 71977.10938 |
| OCIAD2 | Q56VL3 | OCIAD2 | OCIA domain-containi | 24207.35938 | 26674.25195 |
| LAMB2 | P55268 | LAMB2 | Laminin subunit beta- | 3468.361084 | 4387.602051 |
| RIT1 | A0A494C0S1;Q92963 | RIT1 | GTP-binding protein R | 792.0847168 | 1030.178467 |
| RAB20 | Q9NX57 | RAB20 | Ras-related protein Ra | 2974.19751 | 2969.385254 |
| KRT20 | P35900 | KRT20 | Keratin, type I cytoske | NA | 1307.162476 |
| GNG12 | Q9UBI6 | GNG12 | Guanine nucleotide-b | 46247.72656 | 37108.91406 |
| F5 | A0A0A0MRJ7;P12259 | F5 | Coagulation factor V | 40162.53906 | 25643.03125 |
| SH3KBP1 | Q96B97 | SH3KBP1 | SH3 domain-containir | 8450.556641 | 7415.134277 |
| SPATA20 | A0A0U1RRL8;Q8TB22 | SPATA20 | Spermatogenesis-asso | 65628.79688 | 61255.8125 |
| TMEM128 | Q5BJH2 | TMEM128 | Transmembrane prote | 22166.6582 | 21142.67383 |
| MFSD10 | Q14728 | MFSD10 | Major facilitator supe | 35427.42188 | 28725.12891 |
| RETREG1 | Q9H6L5 | RETREG1 | Reticulophagy regulat | 2100.494629 | 3106.128906 |
| PICK1 | Q9NRD5 | PICK1 | PRKCA-binding protei | 25527.17188 | 24458.87891 |
| PPP1R3D | O95685 | PPP1R3D | Protein phosphatase 1 | 39045.6875 | 41790.08203 |
| MB21D2 | Q8IYB1 | MB21D2 | Protein MB21D2 | 8078.590332 | 7947.345703 |
| PCMTD2 | A0A0G2JM70;Q9NV79 | PCMTD2 | Protein-L-isoaspartate | 6208.737793 | 6470.010742 |
| TSPO | B1AH87;P30536 | TSPO | Putative peripheral be | 22233.58594 | 15181.54004 |
| TXNDC16 | Q9P2K2 | TXNDC16 | Thioredoxin domain-c | 4427.859375 | 4671.612793 |
| EXOG | Q9Y2C4 | EXOG | Nuclease EXOG, mitoc | 4833.381836 | 4309.952637 |
| HMGCS1 | Q01581 | HMGCS1 | Hydroxymethylglutary | 193346.3594 | 232002.2344 |
| TC2N | Q8N9U0 | TC2N | Tandem C2 domains n | 4856.081543 | 4808.695313 |
| PI4K2B | G5E9Z4;Q8TCG2 | PI4K2B | Phosphatidylinositol 4 | 6255.760742 | 5293.43457 |
| MID2 | A6PVI4;Q9UJV3 | MID2 | Probable E3 ubiquitin- | 1051.656616 | 992.4345703 |
| WDR24 | Q96S15 | WDR24 | GATOR complex prote | 3225.797363 | 3013.002441 |
| MAOA | P21397 | MAOA | Amine oxidase [flavin- | 40241.88672 | 49806.69141 |

| | | | | | |
|---|---|---|---|---|---|
| ORMDL2 | Q53FV1 | ORMDL2 | ORM1-like protein 2 | 25801.24219 | 20177.65039 |
| SMURF1 | Q9HCE7 | SMURF1 | E3 ubiquitin-protein li | 37833.3125 | 39265.57031 |
| CNPY4 | A0A0J9YYJ0;Q8N129 | CNPY4 | Protein canopy homol | 8765.006836 | 7023.366211 |
| COMMD9 | E9PJ95;Q9P000 | COMMD9 | COMM domain-conta | 5975.266602 | 19554.11914 |
| PLPP2 | K7EMK9;O43688 | PLPP2 | Phospholipid phospha | 6949.124512 | 3866.88916 |
| TSPYL2 | Q9H2G4 | TSPYL2 | Testis-specific Y-encoc | 28334.36719 | 19458.33984 |
| PGM1 | P36871 | PGM1 | Phosphoglucomutase- | 75264.89844 | 59684.73828 |
| NDRG4 | H3BQJ4 | NDRG4 | Protein NDRG4 | 4326.088867 | 4849.214844 |
| ERG28 | Q9UKR5 | ERG28 | Ergosterol biosyntheti | 80208.08594 | 104910.1563 |
| BBS2 | H3BRL0;Q9BXC9 | BBS2 | Bardet-Biedl syndrom | 1142.653687 | 2407.496094 |
| AMMECR1 | Q9Y4X0 | AMMECR1 | AMME syndrome canc | 10221.59473 | 14744.24805 |
| PLEKHS1 | A0A0C4DFS9 | PLEKHS1 | Pleckstrin homology d | 7907.942383 | 8521.927734 |
| DPY19L1 | A0A1B0GW05 | DPY19L1 | Probable C-mannosylt | 21002.17773 | 20455.21289 |
| TMSB4X | P62328 | TMSB4X | Thymosin beta-4 | 11185.91113 | 12895.92285 |
| TPM4 | A0A2R8Y5V9;P67936 | TPM4 | Tropomyosin alpha-4 | 2384.953369 | 1271.987671 |
| RAB27B | O00194 | RAB27B | Ras-related protein Ra | 16899.14453 | 15770.21582 |
| AKAP12 | Q02952 | AKAP12 | A-kinase anchor prote | 857.5661011 | 3361.508545 |
| TMEM106B | Q9NUM4 | TMEM106B | Transmembrane prote | 32944.06641 | 24163.60547 |
| CTNND2 | E7EPC8;Q9UQB3 | CTNND2 | Catenin delta-2 | 10140.34039 | 11100.8808 |
| TRIOBP | Q9H2D6 | TRIOBP | TRIO and F-actin-bind | 12747.97461 | 11866.14258 |
| SGSM3 | Q96HU1 | SGSM3 | Small G protein signal | 3143.61499 | 6212.60791 |
| CSRP1 | P21291 | CSRP1 | Cysteine and glycine-r | 664348.6875 | 498164.2813 |
| SQOR | Q9Y6N5 | SQOR | Sulfide:quinone oxido | 64982.16406 | 52188.12891 |
| PEX1 | A0A0C4DG33;O43933 | PEX1 | Peroxisome biogenesi | 5973.037598 | 7567.443848 |
| CORO2A | Q92828 | CORO2A | Coronin-2A | 2988.638672 | 1261.942993 |
| NAT14 | Q8WUY8 | NAT14 | N-acetyltransferase 14 | 10919.99121 | 12370.50391 |
| PRKCD | Q05655 | PRKCD | Protein kinase C delta | 48456.37109 | 56843.84375 |
| ZNF22 | P17026 | ZNF22 | Zinc finger protein 22 | 4055.389404 | 3554.639648 |
| ADAL | Q6DHV7 | ADAL | Adenosine deaminase | 14680.3457 | 30599.02148 |
| TCEAL1 | Q15170 | TCEAL1 | Transcription elongati | 14557.75391 | 14765.69629 |
| MROH1 | E9PHY8;Q8NDA8 | MROH1 | Maestro heat-like rep | 31412.50391 | 42690.50391 |
| RAB28 | H0Y927;H0Y9S6;P51157 | RAB28 | Ras-related protein Ra | NA | NA |
| EOLA1;EOLA2;CXorf40A;CX | D6RA30;Q5HY62;Q8TE69;Q9 | EOLA1;EOLA2;CXorf40 | Endothelium and lymp | 2960.411621 | 3342.79834 |

| | | | | | |
|---|---|---|---|---|---|
| TWISTNB | Q3B726 | TWISTNB | DNA-directed RNA po | 23351.37891 | 14780.43359 |
| USP30 | B3KUS5;Q70CQ3 | USP30 | Ubiquitin carboxyl-ter | 2631.272461 | 2519.526367 |
| MTM1 | Q13496 | MTM1 | Myotubularin | 14500.5918 | 14575.70703 |
| MAP2 | P11137 | MAP2 | Microtubule-associate | 21367.2207 | 7815.70752 |
| ATP2B1 | P20020 | ATP2B1 | Plasma membrane cal | 47097.1875 | 45758.04297 |
| RTL8C | A6ZKI3 | RTL8C | Retrotransposon Gag- | 23717.20703 | 25915.24023 |
| PANK3 | Q9H999 | PANK3 | Pantothenate kinase 3 | 12235.40332 | 14290.38281 |
| GSTT1 | P30711 | GSTT1 | Glutathione S-transfer | 26231.68555 | 25494.13281 |
| RAB24 | Q969Q5 | RAB24 | Ras-related protein Ra | 16515.75 | 17948.23828 |
| EPPK1 | P58107 | EPPK1 | Epiplakin | 321213.5938 | 160193.3281 |
| CERS4 | H0YKR4;H0YN04;H0YNR6;Q9 | CERS4 | Ceramide synthase 4 ( | 5482.254883 | 5969.211426 |
| CUL9 | E9PEZ1;Q8IWT3 | CUL9 | Cullin-9 | 1308.246948 | 5433.734863 |
| ANXA2 | P07355 | ANXA2 | Annexin A2 | 1341855 | 777954.5 |
| PTPRJ | A0A087WTK0;A0A087WVC6; | PTPRJ | Receptor-type tyrosin | 6868.774414 | 7519.061523 |
| FAM102B | Q5T8I3 | FAM102B | Protein FAM102B | 5480.104492 | 10861.64453 |
| TM7SF2 | O76062 | TM7SF2 | Delta(14)-sterol reduc | 33379.5625 | 39403 |
| LAGE3 | Q14657 | LAGE3 | EKC/KEOPS complex s | 25287.54688 | 35817.44141 |
| MTHFR | P42898;Q5SNW7 | MTHFR | Methylenetetrahydro | 3659.803223 | 2347.775879 |
| MBOAT7 | Q96N66 | MBOAT7 | Lysophospholipid acyl | 99513.46094 | 116208.6875 |
| VAMP7 | P51809 | VAMP7 | Vesicle-associated me | 47497.29688 | 55035.52734 |
| PCIF1 | Q9H4Z3 | PCIF1 | mRNA (2'-O-methylad | 28815.74219 | 29197.94336 |
| NR2F2 | P24468 | NR2F2 | COUP transcription fa | 18632.54297 | 20490.56641 |
| GABPB1 | H9ZYI9;Q06547 | GABPB1 | GA binding protein tra | NA | 1604.287964 |
| LRATD1 | Q96KN4 | LRATD1 | Protein LRATD1 | NA | 4180.748535 |
| GALNT10 | Q86SR1 | GALNT10 | Polypeptide N-acetylg | 6845.0625 | 6879.692871 |
| IQSEC2 | A0A1W2PR28;A0A1W2PRJ5;( | IQSEC2 | IQ motif and SEC7 dor | 5847.26123 | 3271.314697 |
| BNIP2 | H7C096;J3KN59;Q12982 | BNIP2 | BCL2/adenovirus E1B | 259.9554443 | 13980.63379 |
| GALM | Q96C23 | GALM | Aldose 1-epimerase | 56664.78906 | 58326.66797 |
| MAPKAP1 | Q9BPZ7 | MAPKAP1 | Target of rapamycin c | 3548.156982 | 8267.696289 |
| SH3BGRL | O75368 | SH3BGRL | SH3 domain-binding g | 29326.54492 | 39132.24219 |
| GCH1 | P30793 | GCH1 | GTP cyclohydrolase 1 | 8547.291992 | 7542.352539 |
| KIF13B | Q9NQT8 | KIF13B | Kinesin-like protein KI | 17838.03906 | 18085.64844 |
| DECR1 | Q16698 | DECR1 | 2,4-dienoyl-CoA reduc | 295358.9688 | 316593.0625 |

| | | | | | |
|---|---|---|---|---|---|
| CCNE2 | O96020 | CCNE2 | G1/S-specific cyclin-E2 | 3528.400146 | 3825.081543 |
| SLC38A1 | F8VX04;Q9H2H9 | SLC38A1 | Sodium-coupled neutr | 33985.01172 | 32654.66211 |
| ARHGAP10 | A1A4S6 | ARHGAP10 | Rho GTPase-activating | 14110.63965 | 14756.37793 |
| HMGCL | P35914 | HMGCL | Hydroxymethylglutary | 33492.21875 | 41250.25391 |
| MTMR1 | F8WA39;Q13613 | MTMR1 | Myotubularin-related | 19574.88086 | 23553.52539 |
| ACLY | P53396 | ACLY | ATP-citrate synthase | 539815.0625 | 605549.125 |
| ROGDI | K7EPS3;Q9GZN7 | ROGDI | Protein rogdi homolog | 9401.373047 | 10182.4502 |
| KIF5C | O60282 | KIF5C | Kinesin heavy chain is | 190804.6719 | 105777.1094 |
| INPP4B | O15327 | INPP4B | Inositol polyphosphat | 100925.1719 | 100827.3594 |
| UNC13B | A0A1B0GUS7;A0A1B0GVW8; | UNC13B | Protein unc-13 homol | 3558.814697 | 5155.463379 |
| GPC4 | O75487 | GPC4 | Glypican-4 | 8575.293945 | 13581.99512 |
| CETN2 | P41208 | CETN2 | Centrin-2 | 58407.45313 | 79677.05469 |
| CAPN5 | A0A3B3IRX8;E7EV01;O15484 | CAPN5 | Calpain-5 (Fragment); | 1725.563843 | 1386.449585 |
| ANTXR1 | Q9H6X2 | ANTXR1 | Anthrax toxin recepto | 1788.609375 | 3417.14917 |
| SMARCA2 | P51531 | SMARCA2 | Probable global transc | 10364.76465 | 11823.71094 |
| XIAP | P98170 | XIAP | E3 ubiquitin-protein li | 17679.06445 | 18040.69922 |
| PPFIBP2 | Q8ND30 | PPFIBP2 | Liprin-beta-2 | 7444.018066 | 5649.780762 |
| SMPD1 | E9PQT3;G3V1E1;P17405 | SMPD1 | Sphingomyelin phospl | 5443.901855 | 3359.709961 |
| ACADS | P16219 | ACADS | Short-chain specific ac | 11289.44727 | 11245.71289 |
| FADS1 | O60427 | FADS1 | Acyl-CoA (8-3)-desatu | 9105.398438 | 15173.71484 |
| SUOX | P51687 | SUOX | Sulfite oxidase, mitocl | 4741.183105 | 6378.311523 |
| ABCC3 | O15438 | ABCC3 | Canalicular multispeci | 8501.036133 | 10193.0625 |
| CARD14 | A0A494C1N2;Q9BXL6 | CARD14 | Caspase recruitment d | 438.9330444 | 526.9387207 |
| NAA40 | Q86UY6 | NAA40 | N-alpha-acetyltransfe | 6552.027832 | 7689.521484 |
| NUP42 | C9JYA1;O15504 | NUP42 | Nucleoporin NUP42 (F | 5368.271973 | 5754.319336 |
| PPP1R2;PPP1R2;PPP1R2;PP | E7EMN6;E7EUI7;P41236;Q6N | PPP1R2;PPP1R2;PPP1 | Protein phosphatase i | 1447.750732 | 1717.803101 |
| SMIM8 | Q96KF7 | SMIM8 | Small integral membra | 1195.352905 | 3105.580566 |
| SLC12A2 | P55011 | SLC12A2 | Solute carrier family 1 | 81286.50781 | 81839.83594 |
| CNN2 | B4DDF4;B4DUT8;Q99439 | CNN2 | Calponin;Calponin;Cal | 431437.5938 | 294814.375 |
| MGST1 | P10620 | MGST1 | Microsomal glutathior | 161174.5938 | 184040.9688 |
| APOM | O95445;Q5SRP5 | APOM | Apolipoprotein M | 8381.958984 | 8166.54541 |
| MINDY1 | Q8N5J2 | MINDY1 | Ubiquitin carboxyl-ter | 11187.93262 | 14435.0625 |
| CRYL1 | Q9Y2S2 | CRYL1 | Lambda-crystallin hon | 32861.35547 | 34255.44531 |

| | | | | | |
|---|---|---|---|---|---|
| MCTP1 | D6RA42;E5RJR1;H0YA70;Q6D | MCTP1 | Multiple C2 and transı | 5830.307129 | 7781.090332 |
| ARHGAP5 | A0A0A0MSK6;Q13017 | ARHGAP5 | Rho GTPase-activating | 13023.84277 | 15086.78809 |
| PPP3CA | Q08209 | PPP3CA | Serine/threonine-prot | 163887.9531 | 192572.6563 |
| CAMK1D | Q8IU85 | CAMK1D | Calcium/calmodulin-d | 7683.859375 | 7460.455566 |
| FLOT1 | O75955 | FLOT1 | Flotillin-1 | 36655.125 | 24647.50977 |
| ARHGAP18 | Q8N392 | ARHGAP18 | Rho GTPase-activating | 2626.593262 | 3513.774414 |
| LMF1 | H3BVI4;Q96S06 | LMF1 | Lipase maturation fac | 6528.937012 | 11148.45215 |
| NOL4L | A0A087WTU5;A0A087X0N3;( | NOL4L | Nucleolar protein 4-lik | 6977.012207 | 6693.012207 |
| AIFM2 | Q9BRQ8 | AIFM2 | Ferroptosis suppresso | 26196.46484 | 23029.48633 |
| PRKAR2B | P31323 | PRKAR2B | cAMP-dependent prot | 9406.078125 | 8570.392578 |
| FAM114A1 | Q8IWE2 | FAM114A1 | Protein NOXP20 | 1649.597412 | 1505.791382 |
| ABHD4 | F5H6T0;Q8TB40 | ABHD4 | (Lyso)-N-acylphosphat | 5314.738281 | 4718.312012 |
| SYK | P43405 | SYK | Tyrosine-protein kinas | 1389.321289 | 2997.087891 |
| DMAC1 | Q96GE9 | DMAC1 | Distal membrane-arm | 9244.796875 | 14894.11328 |
| KLHDC2 | Q9Y2U9 | KLHDC2 | Kelch domain-contain | 9806.301758 | 9763.335938 |
| ING1 | A0A0C4DFW2;Q9UK53 | ING1 | Inhibitor of growth pr | 4928.94873 | 5047.368652 |
| ADAP1 | O75689 | ADAP1 | Arf-GAP with dual PH | 6628.97998 | 8143.278809 |
| SIVA1 | B4DTY2;G3V3U1;G3V439;O1 | SIVA1 | Apoptosis regulatory ı | 6676.02002 | 6703.904297 |
| MSI1 | O43347 | MSI1 | RNA-binding protein M | 17204.22656 | 16018.27246 |
| VMA21 | Q3ZAQ7 | VMA21 | Vacuolar ATPase asseı | 24369.36328 | 36193.53125 |
| FBXW5 | Q969U6 | FBXW5 | F-box/WD repeat-con | 5453.508789 | 3183.587646 |
| STK19 | A0A140T8X5;A0A140TA37;B7 | STK19 | STK19;Serine/threonii | 4111.648926 | 2521.113525 |
| HYI | A0A0A0MTR5;E7EWH8;F6UJ' | HYI | Putative hydroxypyruv | 8144.07666 | 1568.2854 |
| NCAM2 | H9KV31;O15394 | NCAM2 | Neural cell adhesion ı | 22471.9043 | 31071.26367 |
| FCSK | Q8N0W3 | FCSK | L-fucose kinase | 10994.37793 | 12500.44043 |
| SLC38A2 | Q96QD8 | SLC38A2 | Sodium-coupled neutı | 30406.81836 | 41666.26172 |
| DAB2IP | A0A3B3ITG3;H0Y3A3;Q5VW( | DAB2IP | Disabled homolog 2-ir | 4145.713867 | 915.6465454 |
| CPN1 | B1AP58;P15169 | CPN1 | Carboxypeptidase N c | 1009.450378 | NA |
| P2RX4 | Q99571 | P2RX4 | P2X purinoceptor 4 | 10664.64551 | 12398.85449 |
| ACYP2 | P14621 | ACYP2 | Acylphosphatase-2 | 24872.95313 | 26420.82617 |
| SQLE | Q14534 | SQLE | Squalene monooxygeı | 29739.14648 | 32237.26758 |
| SLC22A18 | A0A024RCG2;A0A0J9YWM3; | SLC22A18 | Solute carrier family 2 | 24605.72656 | 23448.28906 |
| TMUB2 | E7ESS3;K7EKW3;Q71RG4 | TMUB2 | Transmembrane and ı | 6273.176758 | 9763.141602 |

| | | | | | |
|---|---|---|---|---|---|
| NDRG1 | Q92597 | NDRG1 | Protein NDRG1 | 22980.93945 | 16265.69043 |
| MPI | P34949 | MPI | Mannose-6-phosphate | 47235.62891 | 59870.92188 |
| TBC1D2 | Q9BYX2 | TBC1D2 | TBC1 domain family m | 2752.49585 | 3438.56665 |
| KIAA1324L | A8MWY0;C9JA41;H7C2N5 | KIAA1324L | UPF0577 protein KIAA | 7004.77002 | 4806.773438 |
| WDR37 | A0A3B3IT47;Q9Y2I8 | WDR37 | WD repeat-containing | 24856.36328 | 26696.63477 |
| SLC9A7 | A0A087WXD1;Q96T83 | SLC9A7 | Sodium/hydrogen exc | 4114.553711 | 5384.041992 |
| MYL6 | F8W1R7;G3V1V0;P60660 | MYL6 | Myosin light polypept | 35517.59766 | 38423.34375 |
| CRBN | J3QT87;Q96SW2 | CRBN | Protein cereblon (Frag | 29281.84375 | 29217.91406 |
| NRP1 | E9PEP6;O14786;Q5JWQ6 | NRP1 | Neuropilin;Neuropilin | 3810.570801 | 4550.055176 |
| BZW2 | E7ETZ4;Q9Y6E2 | BZW2 | Basic leucine zipper ar | 462653.3125 | 537462 |
| TSPAN13 | O95857 | TSPAN13 | Tetraspanin-13 | 113521.5781 | 92939.78906 |
| CYP51A1 | A0A0C4DFL7;Q16850 | CYP51A1 | Lanosterol 14-alpha d | 96384.22656 | 110929.7813 |
| GPRC5A | A0A3B3ITN8;Q8NFJ5 | GPRC5A | Retinoic acid-induced | 15691.91797 | 13789.91699 |
| IRAK4 | Q9NWZ3 | IRAK4 | Interleukin-1 receptor | 10074.61133 | 14787.1377 |
| PACS1 | Q6VY07 | PACS1 | Phosphofurin acidic cl | 22269.57227 | 26794.21484 |
| ABCG2 | Q9UNQ0 | ABCG2 | Broad substrate speci | 12421.02051 | 4297.635254 |
| DNHD1 | Q96M86 | DNHD1 | Dynein heavy chain do | 44435.82422 | 43459.60547 |
| PRKAA1 | Q13131 | PRKAA1 | 5'-AMP-activated prot | 89766.69531 | 91293.60156 |
| SLC12A9 | Q9BXP2 | SLC12A9 | Solute carrier family 1 | 32513.24023 | 31188.38672 |
| ASNS | P08243 | ASNS | Asparagine synthetase | 648110.8125 | 475282.875 |
| ITGB4 | P16144 | ITGB4 | Integrin beta-4 | 26023.14063 | 23080.42188 |
| NRAS | P01111 | NRAS | GTPase NRas | 695679.8125 | 804101.25 |
| PBLD | P30039 | PBLD | Phenazine biosynthes | 22605.72266 | 14250.04199 |
| RMDN1 | Q96DB5 | RMDN1 | Regulator of microtub | 238082.8438 | 241851.8125 |
| FKBP8 | M0R2K9 | FKBP8 | Peptidylprolyl isomera | 86737.33984 | 93795.66797 |
| GNPNAT1 | Q96EK6 | GNPNAT1 | Glucosamine 6-phosp | 337747.5 | 303003.5313 |
| S100A10 | P60903 | S100A10 | Protein S100-A10 | 288857.375 | 100630.2422 |
| C11orf52 | Q96A22 | C11orf52 | Uncharacterized prote | 1746.097046 | 2066.880127 |
| CPNE2 | Q96FN4 | CPNE2 | Copine-2 | 5415.776855 | 2391.76709 |
| SDCBP | G5EA09 | SDCBP | Syndecan binding prot | 18427.43652 | 23222.67676 |
| LMF2 | Q9BU23 | LMF2 | Lipase maturation fact | 14371.11328 | 17804.08203 |
| MANBA | A0A2R8YEC9;O00462 | MANBA | Beta-mannosidase | 15297.78711 | 13128.21289 |
| ARVCF | C9JJX6;E9PDC3;O00192 | ARVCF | Armadillo repeat prot | 12417.72852 | 16099.78223 |

| | | | | | |
|---|---|---|---|---|---|
| HEATR5A | F5H619;Q86XA9 | HEATR5A | HEAT repeat-containi | 5755.342285 | 7339 |
| YBEY | P58557 | YBEY | Endoribonuclease Ybe | 7984.893555 | 9035.466797 |
| CRABP2 | P29373 | CRABP2 | Cellular retinoic acid-b | 797214.125 | 1213582.375 |
| ATP11B | Q9Y2G3 | ATP11B | Probable phospholipi | 3378.737061 | 2823.486084 |
| MVK | F5H8H2;Q03426 | MVK | Mevalonate kinase | 128039.3281 | 139498.125 |
| WDR1 | O75083 | WDR1 | WD repeat-containing | 312014.3438 | 296285.8438 |
| STON1-GTF2A1L;STON1-GT | A0A0A6YYG5;Q53S48;Q9Y6C | STON1-GTF2A1L;STON | Protein STON1-GTF2A | 2469.165039 | 1163.225586 |
| SYTL1 | Q8IYJ3 | SYTL1 | Synaptotagmin-like pr | 17600.0332 | 19861.26563 |
| TSC22D4 | Q9Y3Q8 | TSC22D4 | TSC22 domain family | 33139.625 | 40057.89063 |
| MAP1S | Q66K74 | MAP1S | Microtubule-associate | 17554.52344 | 18950.88867 |
| SVIL | O95425 | SVIL | Supervillin | 22846.10938 | 22885.02344 |
| LRCH4 | O75427 | LRCH4 | Leucine-rich repeat ar | 13611.89941 | 14915.52051 |
| NR1H2 | M0R0K3;M0R2F9;P55055 | NR1H2 | Oxysterols receptor L | 1654.19519 | 2892.428223 |
| FKBP2 | A0A494C1N0;P26885 | FKBP2 | Peptidylprolyl isomera | 115050.7266 | 131667.625 |
| ATM | Q13315 | ATM | Serine-protein kinase | 2131.775879 | 2804.635742 |
| GSTCD | Q8NEC7 | GSTCD | Glutathione S-transfe | 9401.705078 | 13699.8877 |
| MON1B | Q7L1V2 | MON1B | Vacuolar fusion prote | 21105.12695 | 23903.63672 |
| MYO6 | A0A0A0MRM8 | MYO6 | Unconventional myos | 60262.76172 | 62163.24219 |
| MVD | P53602 | MVD | Diphosphomevalonat | 286671.5313 | 353326.5 |
| KIF1A | A0A3B3IT28;A0A3B3ITE5;A0 | KIF1A | Kinesin-like protein KI | 28389.54492 | 24241.04492 |
| KBTBD2 | Q8IY47 | KBTBD2 | Kelch repeat and BTB | 10946.37012 | 13947.9668 |
| MIS18BP1 | Q6P0N0 | MIS18BP1 | Mis18-binding protei | 1521.065063 | 2974.605469 |
| ZNF185 | H0Y704 | ZNF185 | Zinc finger protein 18 | 14320.71899 | 26789.62915 |
| DALRD3 | C9JA38;Q5D0E6 | DALRD3 | DALR anticodon-bindi | 3068.925781 | 1971.861694 |
| MAPK6 | Q16659 | MAPK6 | Mitogen-activated pro | 21949.48828 | 24783.64063 |
| TRAPPC6B | Q86SZ2 | TRAPPC6B | Trafficking protein par | 22696.03125 | 29051.51367 |
| ZFYVE1 | G3V5N8;Q9HBF4 | ZFYVE1 | Zinc finger FYVE doma | 7073.690918 | 8357.782227 |
| FBXL16 | Q8N461 | FBXL16 | F-box/LRR-repeat pro | 329.2080078 | 491.4266968 |
| PEX7 | O00628 | PEX7 | Peroxisomal targeting | 4291.720215 | 4544.101563 |
| ZNF362 | Q5T0B9 | ZNF362 | Zinc finger protein 36 | 2065.57251 | 5943.865234 |
| PTGFRN | Q9P2B2 | PTGFRN | Prostaglandin F2 rece | 40550.0625 | 38590.75 |
| NSD2 | O96028 | NSD2 | Histone-lysine N-meth | 8672.618164 | 8464.008789 |
| NNT | Q13423 | NNT | NAD(P) transhydroger | 144151.7031 | 156339.6563 |

| | | | | | |
|---|---|---|---|---|---|
| HSPA1L | P34931;Q53FA3 | HSPA1L | Heat shock 70 kDa pro | 7688.812988 | 11048.31738 |
| H6PD | O95479 | H6PD | GDH/6PGL endoplasm | 8005.917969 | 10446.75586 |
| MAGED2 | Q9UNF1 | MAGED2 | Melanoma-associated | 120028.7109 | 144243.3594 |
| NMNAT1 | Q9HAN9 | NMNAT1 | Nicotinamide/nicotini | 22121.76367 | 17967.74219 |
| STXBP5 | Q5T5C0 | STXBP5 | Syntaxin-binding prot | 9123.299805 | 9784.092773 |
| ENO1 | P06733 | ENO1 | Alpha-enolase | 5360372.5 | 5393364.5 |
| LRRC58 | Q96CX6 | LRRC58 | Leucine-rich repeat-co | 4735.924805 | 3762.506592 |
| PRKAB1 | Q9Y478 | PRKAB1 | 5'-AMP-activated prot | 17595.58008 | 16234.68262 |
| PTGR2 | Q8N8N7 | PTGR2 | Prostaglandin reducta | 63346.83203 | 61956.25391 |
| STAG2 | Q8N3U4 | STAG2 | Cohesin subunit SA-2 | 51066.57813 | 48907.27344 |
| SEC22B | O75396 | SEC22B | Vesicle-trafficking pro | 359985.2188 | 403854 |
| FBH1 | Q8NFZ0 | FBH1 | F-box DNA helicase 1 | 1793.243408 | 2381.184814 |
| JOSD2 | M0QZL8;Q8TAC2 | JOSD2 | Josephin-2 (Fragment | 4450.291016 | 5239.080078 |
| PIP | P12273 | PIP | Prolactin-inducible pr | 5559.683105 | 6176.166016 |
| ZKSCAN3 | Q9BRR0 | ZKSCAN3 | Zinc finger protein wit | 978.8312988 | 860.4105835 |
| TUBA4A | P68366 | TUBA4A | Tubulin alpha-4A chai | 1534957 | 1642932.375 |
| NOVA1 | P51513 | NOVA1 | RNA-binding protein N | 5373.334473 | 3050.14209 |
| PLOD3 | O60568 | PLOD3 | Multifunctional proco | 95491.41406 | 102890.8828 |
| EPM2AIP1 | Q7L775 | EPM2AIP1 | EPM2A-interacting pr | 46717.54688 | 41309.54297 |
| IFT80 | Q9P2H3 | IFT80 | Intraflagellar transpor | NA | 1439.618042 |
| RALB | P11234 | RALB | Ras-related protein Ra | 377832.2188 | 400793.6875 |
| TSPAN31 | F8VS78;F8VWE0;Q12999 | TSPAN31 | Tetraspanin-31 | 46242.98828 | 44858.96875 |
| SDR39U1 | Q9NRG7 | SDR39U1 | Epimerase family prot | 13203.15332 | 14334.52148 |
| PRKAG1 | P54619 | PRKAG1 | 5'-AMP-activated prot | 54935.10938 | 57551.10547 |
| ATP11C | A0A067XG54;Q8NB49 | ATP11C | Phospholipid-transpor | 10169.23242 | 13270.76563 |
| PPCDC | Q96CD2 | PPCDC | Phosphopantothenoy | 13336.58594 | 12718.82617 |
| ORMDL1 | Q9P0S3 | ORMDL1 | ORM1-like protein 1 | 45146.15625 | 39852.20313 |
| NUDT16 | Q96DE0 | NUDT16 | U8 snoRNA-decapping | 53771.58984 | 51515.88672 |
| MAGI1 | Q96QZ7 | MAGI1 | Membrane-associated | 6321.280762 | 6064.423828 |
| TMEM165 | Q9HC07 | TMEM165 | Transmembrane prote | 50275.25 | 57999.27344 |
| MOSPD1 | Q9UJG1 | MOSPD1 | Motile sperm domain- | 3824.394043 | 3426.122559 |
| LRBA | A0A494C1L5 | LRBA | Lipopolysaccharide-re | 43557.98047 | 51216.92578 |
| MYL6B | P14649 | MYL6B | Myosin light chain 6B | 25220.39648 | 23241.37305 |

| | | | | | |
|---|---|---|---|---|---|
| CDIPT | B3KY94;O14735 | CDIPT | CDP-diacylglycerol--in | 83906.9375 | 105565.2344 |
| MAP3K4 | F5H4R1;F5H538;J3KNB8;Q9Y | MAP3K4 | Mitogen-activated pro | 6037.631348 | 4680.119629 |
| ARFGEF1 | Q9Y6D6 | ARFGEF1 | Brefeldin A-inhibited g | 25031.53516 | 29602.5 |
| PODXL | O00592 | PODXL | Podocalyxin | 1025.466064 | 1290.55957 |
| VPS50 | Q96JG6 | VPS50 | Syndetin | 16269.51465 | 18941.89063 |
| KIAA0513 | O60268 | KIAA0513 | Uncharacterized prote | 10076.08496 | 7764.23877 |
| SLC44A2 | Q8IWA5 | SLC44A2 | Choline transporter-lil | 95920.04688 | 134077.7344 |
| ADGRG6 | Q86SQ4 | ADGRG6 | Adhesion G-protein co | 5967.59082 | 5717.163574 |
| CUL4B | K4DI93 | CUL4B | Cullin 4B, isoform CRA | 70464.33594 | 71503.27344 |
| CXorf38 | Q8TB03 | CXorf38 | Uncharacterized prote | 49656.60156 | 48495.20703 |
| DIP2B | Q9P265 | DIP2B | Disco-interacting prot | 46696.96875 | 49039.94922 |
| LRMDA | A0A087WWI0 | LRMDA | Leucine-rich melanocy | 1749.228394 | NA |
| CD63 | F8VV56;F8VWK8;F8W022;P0 | CD63 | CD63 antigen;Tetrasp | 18737.68945 | 19856.61914 |
| RABEP2 | Q9H5N1 | RABEP2 | Rab GTPase-binding e | 3630.742432 | 4731.704102 |
| LPCAT3 | Q6P1A2 | LPCAT3 | Lysophospholipid acyl | 54157.52734 | 29905.5 |
| MACF1 | A0A0A6YYJ5 | MACF1 | Microtubule-actin cro | 23712.54663 | 29446.15527 |
| ZNF385A | Q96PM9 | ZNF385A | Zinc finger protein 38 | 5936.573242 | 6804.571289 |
| APMAP | Q9HDC9 | APMAP | Adipocyte plasma me | 409228.375 | 432489.875 |
| KIFC3 | B7Z896;F5H3M2;H3BTL6;H3E | KIFC3 | Kinesin-like protein;Ki | 1791.225464 | NA |
| SREBF1 | P36956 | SREBF1 | Sterol regulatory elem | 1098.654785 | 3863.859619 |
| COL9A2 | Q14055 | COL9A2 | Collagen alpha-2(IX) c | 4736.19873 | 4238.578125 |
| BMP2K | H0Y9P1;Q9NSY1 | BMP2K | BMP-2-inducible prot | 4468.657227 | 13352.80078 |
| MPV17 | B5MC53;B5MCF8;P39210 | MPV17 | MpV17 transgene, mu | 8267.90918 | 8692.94043 |
| NEO1 | Q92859 | NEO1 | Neogenin | 18506.38867 | 15351.37988 |
| TRIM32 | Q13049 | TRIM32 | E3 ubiquitin-protein li | 7218.456055 | 11343.49121 |
| PLPP6 | Q8IY26 | PLPP6 | Phospholipid phospha | 11896.27637 | 18807.9707 |
| SIDT2 | C9JBG5;E9PMC3;F5H8L4;Q8N | SIDT2 | SID1 transmembrane | 4744.571289 | 3817.480225 |
| LACTB | P83111 | LACTB | Serine beta-lactamase | 16016.99023 | 18699.91797 |
| FAM120C | Q9NX05 | FAM120C | Constitutive coactivat | 68333.03125 | 70340.98438 |
| SLC35A1 | P78382 | SLC35A1 | CMP-sialic acid transp | 12232.07324 | 14507.44434 |
| QDPR | P09417 | QDPR | Dihydropteridine redu | 71527.50781 | 65454.62891 |
| UBAC2 | Q8NBM4 | UBAC2 | Ubiquitin-associated c | 29249.33203 | 35471.07422 |
| RIPOR1 | A0A0A0MTL6;Q6ZS17 | RIPOR1 | Rho family-interacting | 6619.115723 | 5101.957031 |

| | | | | | |
|---|---|---|---|---|---|
| SLC2A10 | O95528 | SLC2A10 | Solute carrier family 2 | 8559.480469 | 10446.21387 |
| DOCK5 | Q9H7D0 | DOCK5 | Dedicator of cytokines | 6006.584473 | 5531.638184 |
| FANCD2 | Q9BXW9 | FANCD2 | Fanconi anemia group | 29431.70117 | 44872.01953 |
| WDR44 | Q5JSH3 | WDR44 | WD repeat-containing | 51138.79297 | 53968.88672 |
| IDH1 | O75874 | IDH1 | Isocitrate dehydrogen | 491550.4063 | 595298.4375 |
| ACVR1 | Q04771 | ACVR1 | Activin receptor type- | 1899.795288 | 1576.765259 |
| ATP6AP2 | A0A1B0GVW0;O75787 | ATP6AP2 | Renin receptor | 52409.55469 | 56871.19531 |
| RAB4B | P61018 | RAB4B | Ras-related protein Ra | 21178.08398 | 24648.17383 |
| FKBP3 | Q00688 | FKBP3 | Peptidyl-prolyl cis-trar | 874877.0625 | 861774.5 |
| RAF1 | P04049 | RAF1 | RAF proto-oncogene s | 43682.99219 | 54723.29688 |
| SIRT5 | Q9NXA8 | SIRT5 | NAD-dependent prote | 14360.99609 | 16719.50391 |
| UBP1 | Q9NZI7 | UBP1 | Upstream-binding pro | 11761.65039 | 11311.12109 |
| CTBP1 | Q13363 | CTBP1 | C-terminal-binding pr | 281686.3438 | 290144.3125 |
| F8A1 | P23610 | F8A1 | 40-kDa huntingtin-ass | 14623.20801 | 18253.57813 |
| OPTC | Q5T2G3;Q9UBM4 | OPTC | Opticin (Fragment);Op | 63278.00391 | 75175.69531 |
| ZFHX3 | Q15911 | ZFHX3 | Zinc finger homeobox | 5703.331055 | 5947.433594 |
| HYDIN | Q4G0P3 | HYDIN | Hydrocephalus-induci | 732029.75 | 765125.75 |
| C9orf64 | Q5T6V5 | C9orf64 | Queuosine salvage pr | 98249.03906 | 96625.48438 |
| PGM2L1 | Q6PCE3 | PGM2L1 | Glucose 1,6-bisphospl | 1235.662964 | 2603.05249 |
| RARS2 | Q5T160 | RARS2 | Probable arginine--tRI | 11446.21484 | 13712.24805 |
| ZNF512 | G3XAG1;Q96ME7 | ZNF512 | Zinc finger protein 51 | 10053.27832 | 11484.28516 |
| GGCT | O75223 | GGCT | Gamma-glutamylcyclo | 249513.5625 | 250944.1406 |
| ORMDL3 | Q8N138 | ORMDL3 | ORM1-like protein 3 | 13460.29785 | 11101.87793 |
| PHLDB3 | Q6NSJ2 | PHLDB3 | Pleckstrin homology-l | 6844.054199 | 5418.23877 |
| AMBP | P02760 | AMBP | Protein AMBP | NA | 4419.626953 |
| FERMT2 | Q96AC1 | FERMT2 | Fermitin family homol | 69959.29688 | 58670.87891 |
| MAZ | I3L0M3;I3L4Y2 | MAZ | MYC-associated zinc f | 40459.57324 | 40597.35693 |
| LHPP | Q9H008 | LHPP | Phospholysine phosph | 20932.17383 | 30764.01172 |
| NMRAL1 | Q9HBL8 | NMRAL1 | NmrA-like family dom | 52007.19141 | 47336.24609 |
| TJAP1 | Q5JTD0 | TJAP1 | Tight junction-associa | 17627.25195 | 19449.82227 |
| ZNF865 | P0CJ78 | ZNF865 | Zinc finger protein 86 | 2382.128418 | 2319.44458 |
| ABCD4 | O14678 | ABCD4 | ATP-binding cassette s | 125926.2344 | 135171.1094 |
| HRAS | P01112 | HRAS | GTPase HRas | 271327.125 | 306696.5938 |

| | | | | | |
|---|---|---|---|---|---|
| ARHGAP27 | A0A0G2JNY3;A0A0G2JRG8;E! | ARHGAP27 | Rho GTPase-activating | 5591.724609 | 5181.675781 |
| KIF13A | A0A1B0GUA8;Q9H1H9 | KIF13A | Kinesin-like protein KI | 2737.950439 | 2285.816162 |
| ARHGAP35 | Q9NRY4 | ARHGAP35 | Rho GTPase-activating | 42527.125 | 42597.56641 |
| ACOX3 | O15254 | ACOX3 | Peroxisomal acyl-coer | 28128.18555 | 40883.12109 |
| XPNPEP1 | Q9NQW7 | XPNPEP1 | Xaa-Pro aminopeptida | 83493.5625 | 79062.4375 |
| KHNYN | O15037 | KHNYN | Protein KHNYN | 3951.677734 | 4879.71875 |
| OXNAD1 | C9JLB7;Q96HP4;U3KQU8 | OXNAD1 | Oxidoreductase NAD- | 15670.82715 | 421.9755249 |
| C11orf54 | Q9H0W9 | C11orf54 | Ester hydrolase C11or | 18688.92188 | 18537.91602 |
| KNTC1 | P50748 | KNTC1 | Kinetochore-associate | 20075.8457 | 26768.87305 |
| ACOT8 | O14734 | ACOT8 | Acyl-coenzyme A thio | 33979.96484 | 40930.0625 |
| CEP170B | J3KQR7;Q9Y4F5 | CEP170B | Centrosomal protein | 19116.39648 | 17497.76172 |
| MID1 | O15344 | MID1 | E3 ubiquitin-protein li | NA | 1213.06665 |
| STX18 | D6RF48;Q9P2W9 | STX18 | Syntaxin-18 | 18836.33398 | 19181.13867 |
| RAB11A | H3BSC1;P62491 | RAB11A | Ras-related protein Ra | 15320.43555 | 22255.98438 |
| PFN2 | C9J0J7;G5E9Q6 | PFN2 | Profilin | 927180.6719 | 1086394.094 |
| DYNC2H1 | Q8NCM8 | DYNC2H1 | Cytoplasmic dynein 2 | 3772.05249 | 4990.605469 |
| MAN2A2 | P49641 | MAN2A2 | Alpha-mannosidase 2 | 14828.16699 | 13263.70117 |
| CHN1 | P15882 | CHN1 | N-chimaerin | 3237.804199 | 3261.627197 |
| ZNF131 | P52739 | ZNF131 | Zinc finger protein 13 | 5015.02832 | 4929.937012 |
| CBR1 | P16152 | CBR1 | Carbonyl reductase [N | 480254.0625 | 493958.25 |
| EHD4 | Q9H223 | EHD4 | EH domain-containing | 143959.5938 | 153321.6563 |
| NRCAM | C9JYY6;Q92823 | NRCAM | Neuronal cell adhesio | 12623.04199 | 10873.98535 |
| SH3YL1 | H7C499;Q96HL8 | SH3YL1 | SH3 domain-containir | 2728.088135 | 3971.829102 |
| SLC39A6 | Q13433 | SLC39A6 | Zinc transporter ZIP6 | 10948.99805 | 10394.41113 |
| RNF168 | Q8IYW5 | RNF168 | E3 ubiquitin-protein li | 3564.13501 | 2922.690918 |
| LASP1 | Q14847 | LASP1 | LIM and SH3 domain | 412382.0625 | 451509.5313 |
| HSD17B12 | Q53GQ0 | HSD17B12 | Very-long-chain 3-oxo | 75203.91406 | 74633.79688 |
| TAOK3 | Q9H2K8 | TAOK3 | Serine/threonine-prot | 35330.02734 | 44980.49219 |
| TTC39B | Q5VTQ0 | TTC39B | Tetratricopeptide rep | 2854.930664 | 5104.866699 |
| NMD3 | C9JA08;Q96D46 | NMD3 | 60S ribosomal export | 170881.8594 | 143969.1563 |
| ARMT1 | Q9H993 | ARMT1 | Damage-control phos | 1148479.625 | 1354454.5 |
| PHLDA3 | Q9Y5J5 | PHLDA3 | Pleckstrin homology-l | 7243.974121 | 5828.101563 |
| TANGO2 | C9J695;C9JDT9;Q6ICL3 | TANGO2 | Transport and Golgi o | 6507.394531 | 6145.591309 |

| | | | | | |
|---|---|---|---|---|---|
| CDR2L | Q86X02 | CDR2L | Cerebellar degenerati | 5692.338867 | 4916.528809 |
| GARS1 | P41250 | GARS1 | Glycine--tRNA ligase | 158015.9531 | 169296.7344 |
| C1orf43 | A0A1W2PQP6;Q9BWL3;X6R6 | C1orf43 | Uncharacterized prote | 7264.908203 | NA |
| MAN2B2 | H0YA68;Q9Y2E5 | MAN2B2 | Alpha-mannosidase (F | 8538.160156 | 9368.959961 |
| DGUOK | Q16854 | DGUOK | Deoxyguanosine kinas | 5341.366699 | 4712.670898 |
| PARVA | Q9NVD7 | PARVA | Alpha-parvin | 29088.30664 | 29166.35352 |
| FAM169A | D6RB01;Q9Y6X4 | FAM169A | Soluble lamin-associat | 15465.66992 | 14170.98047 |
| GLUD1 | P00367 | GLUD1 | Glutamate dehydroge | 517334.0625 | 519413.8125 |
| G6PD | E7EUI8 | G6PD | Glucose-6-phosphate | 478663.4688 | 694481.1484 |
| BET1 | H7C1N3;O15155 | BET1 | BET1 homolog (Fragm | 13641.69629 | 11960.99219 |
| SERPINB1 | P30740 | SERPINB1 | Leukocyte elastase inh | 62487.32422 | 67271.60938 |
| ACAT2 | Q9BWD1 | ACAT2 | Acetyl-CoA acetyltran | 191408.4063 | 228610.4844 |
| STXBP1 | A0A0D9SG72;P61764 | STXBP1 | Syntaxin-binding prot | 25740.5332 | 29883.8457 |
| VPS37C | A5D8V6;F5H4Q5 | VPS37C | Vacuolar protein sorti | 19620.86133 | 9234.946289 |
| UPRT | E9PSD7;Q96BW1 | UPRT | Uracil phosphoribosyl | 8395.049805 | 8271.21582 |
| PHLDA2 | Q53GA4 | PHLDA2 | Pleckstrin homology-l | 23493.59375 | 11393.29004 |
| NAT1 | P18440 | NAT1 | Arylamine N-acetyltra | 15963.31152 | 18695.28125 |
| AFAP1 | Q8N556 | AFAP1 | Actin filament-associa | 2475.951904 | 3026.936035 |
| CBX5 | P45973 | CBX5 | Chromobox protein ho | 53275.64844 | 52788.37891 |
| PLEKHG2 | H7BXC7;Q9H7P9 | PLEKHG2 | Pleckstrin homology d | 3769.963867 | 4495.40625 |
| FAM172A | Q8WUF8 | FAM172A | Cotranscriptional regu | 10644.45898 | 12723.74414 |
| CDK5RAP2 | Q96SN8 | CDK5RAP2 | CDK5 regulatory subu | 2532.847168 | 2140.432373 |
| HTT | P42858 | HTT | Huntingtin | 28931.58594 | 37635.03906 |
| PHF14 | A0A0U1RRH6;B4DG57;O9488 | PHF14 | PHD finger protein 14 | 3937.779053 | 13134.38477 |
| APOOL | A0A087WYF7;Q6UXV4 | APOOL | MICOS complex subur | 18182.03125 | 22748.92969 |
| TLE5 | Q08117 | TLE5 | TLE family member 5 | 6307.069336 | 9427.97168 |
| AKAP9 | A0A0A0MRF6;Q99996 | AKAP9 | A-kinase anchor prote | 15824.2334 | 19076.80859 |
| PEX26 | E9PRC5;Q7Z412 | PEX26 | Peroxisome assembly | 694.137207 | 1022.750916 |
| GLTP | Q9NZD2 | GLTP | Glycolipid transfer pro | 51935.96484 | 60239.01172 |
| THUMPD3 | Q9BV44 | THUMPD3 | THUMP domain-conta | 97438.63281 | 98037.41406 |
| PGM3 | H0Y987;O95394 | PGM3 | Phosphoacetylglucosa | 106155.1641 | 113610.3047 |
| ZMYND11 | B0QZE2;B7Z2J6;E7ENI9;Q153 | ZMYND11 | Zinc finger MYND dom | 10061.33789 | 3569.962646 |
| RBKS | Q9H477 | RBKS | Ribokinase | 20340.18164 | 23327.91211 |

| | | | | | |
|---|---|---|---|---|---|
| CCDC167 | Q9P0B6 | CCDC167 | Coiled-coil domain-co | 8607.74707 | 9345.981445 |
| RAPGEF1 | Q13905 | RAPGEF1 | Rap guanine nucleotic | 4055.13623 | 6392.612793 |
| SLC30A9 | Q6PML9 | SLC30A9 | Zinc transporter 9 | 12115.74414 | 12796.38574 |
| PHKB | Q93100 | PHKB | Phosphorylase b kinas | 14437.65723 | 13379.3291 |
| INPP5A | Q14642 | INPP5A | Inositol polyphosphat | 4608.239258 | 4852.845215 |
| RNF41 | Q9H4P4 | RNF41 | E3 ubiquitin-protein li | 4238.489746 | 7055.759277 |
| TCEA2 | Q15560 | TCEA2 | Transcription elongati | 86511.72656 | 80904.48438 |
| CD44 | H0Y2P0;H0YD13;P16070 | CD44 | CD44 antigen (Fragme | 22765.03125 | 18502.23633 |
| RABL2A;RABL2B | Q9UBK7;Q9UNT1 | RABL2A;RABL2B | Rab-like protein 2A;Ra | 5331.615234 | 8941.821289 |
| VPS52 | Q8N1B4 | VPS52 | Vacuolar protein sorti | 21732.09766 | 25874.40625 |
| GAMT | Q14353 | GAMT | Guanidinoacetate N-n | 12854.86621 | 13850.02148 |
| STARD3 | J3QLM1;J3QRM3;Q14849 | STARD3 | StAR-related lipid trar | 3143.838135 | 4460.525879 |
| ABCA3 | H0Y3H2;Q99758 | ABCA3 | ATP-binding cassette : | 14615.9209 | 11465.68066 |
| SETD5 | E7EWN3;H0Y3R4;Q9C0A6 | SETD5 | Histone-lysine N-meth | 1929.796875 | 1596.626587 |
| PSMC3IP | K7EK12;K7ELD8;K7EQS1;Q9P | PSMC3IP | Homologous-pairing p | 6261.262207 | 9429.286133 |
| XPOT | O43592 | XPOT | Exportin-T | 71836.40625 | 98135.48438 |
| INPP5K | Q9BT40 | INPP5K | Inositol polyphosphat | 9350.548828 | 12446.89746 |
| SLC25A13 | Q9UJS0 | SLC25A13 | Calcium-binding mitoc | 127569.6094 | 113945.125 |
| RPA2 | P15927 | RPA2 | Replication protein A | 200304.8906 | 208619.1406 |
| RMND1 | Q9NWS8 | RMND1 | Required for meiotic r | 283635.875 | 313249 |
| IRF5 | A0A087WWA0;C9J7M2;C9JB | IRF5 | Interferon regulatory | 5605.477539 | 4076.33252 |
| GNAS | Q5JWF2 | GNAS | Guanine nucleotide-b | 259728 | 314582.5 |
| RIDA | P52758 | RIDA | 2-iminobutanoate/2-i | 155631 | 168446.4531 |
| KPRP | Q5T749 | KPRP | Keratinocyte proline-r | 277.9387512 | 450.3046265 |
| CDK2 | G3V5T9;P24941 | CDK2 | Cyclin-dependent kina | 114100.4609 | 136745.6406 |
| B4GALT4 | C9J3R8;E7ETS9;O60513 | B4GALT4 | Beta-1,4-galactosyltra | 4324.081543 | 4925.668457 |
| AUH | Q13825 | AUH | Methylglutaconyl-CoA | 5266.694824 | 10667.73535 |
| PRDX2 | P32119 | PRDX2 | Peroxiredoxin-2 | 1688337.375 | 1770708.375 |
| ERBB3 | P21860 | ERBB3 | Receptor tyrosine-pro | 9524.478516 | 7652.786133 |
| OPLAH | O14841 | OPLAH | 5-oxoprolinase | 16179.54492 | 18366.02734 |
| RETSAT | Q6NUM9 | RETSAT | All-trans-retinol 13,14 | 25570.97852 | 28201.19336 |
| PIGG | Q5H8A4 | PIGG | GPI ethanolamine phc | 12299.73828 | 19294.72266 |
| UAP1L1 | Q3KQV9 | UAP1L1 | UDP-N-acetylhexosam | 93385.57031 | 84165.36719 |

| | | | | | |
|---|---|---|---|---|---|
| SPATS2L | Q9NUQ6 | SPATS2L | SPATS2-like protein | 9350.875 | 17948.99414 |
| SLC2A8 | A0A087WT42;Q5VVW5;Q9N | SLC2A8 | Solute carrier family 2 | 4764.715332 | 2988.626221 |
| PXMP4 | Q9Y6I8 | PXMP4 | Peroxisomal membrar | 20086.19336 | 17878.82617 |
| CERS5 | F8VXY1;H0YI10;Q49AQ3;Q8N | CERS5 | Ceramide synthase 5; | 17555.62695 | 24408.76563 |
| GRB7 | Q14451 | GRB7 | Growth factor recepto | 1687.029907 | 3827.683105 |
| B3GALT6 | Q96L58 | B3GALT6 | Beta-1,3-galactosyltra | 8619.544922 | 8149.477539 |
| SLC44A1 | Q8WWI5 | SLC44A1 | Choline transporter-lil | 19030.08984 | 28867.63867 |
| ATP2C2 | O75185 | ATP2C2 | Calcium-transporting | 14131.19141 | 12617.22363 |
| MPP7 | Q5T2T1 | MPP7 | MAGUK p55 subfamily | 15327.38184 | 18386.40039 |
| UBE2Q2 | E9PHD0;Q8WVN8 | UBE2Q2 | Ubiquitin-conjugating | 4013.604492 | 3871.157227 |
| NT5C | Q8TCD5 | NT5C | 5'(3')-deoxyribonucle | 33588.99219 | 27476.86719 |
| POFUT2 | Q9Y2G5 | POFUT2 | GDP-fucose protein O | 2994.694336 | 3531.388184 |
| RBPJ | Q06330 | RBPJ | Recombining binding | 12168.57422 | 14307.75586 |
| DOLPP1 | Q86YN1 | DOLPP1 | Dolichyldiphosphatase | 17076.95703 | 18407.11914 |
| FAM210B | Q96KR6 | FAM210B | Protein FAM210B, mit | 2612.385986 | 2351.844727 |
| KIF22 | Q14807 | KIF22 | Kinesin-like protein KI | 63189.19922 | 59991.82813 |
| B3GLCT | Q6Y288 | B3GLCT | Beta-1,3-glucosyltrans | 36409.90234 | 21127.71875 |
| CRACR2B | Q8N4Y2 | CRACR2B | EF-hand calcium-bindi | 2773.164307 | 1307.443237 |
| SERPINB6 | A0A024QZX5;A0A087X1N8;P | SERPINB6 | Serpin B6 | 266530.9375 | 344649.75 |
| KLC4 | Q9NSK0 | KLC4 | Kinesin light chain 4 | 18473.27344 | 20665.56641 |
| SRCIN1 | A0A087X165;Q9C0H9 | SRCIN1 | SRC kinase-signaling ir | 2374.838623 | 2674.168701 |
| ZNF516 | Q92618 | ZNF516 | Zinc finger protein 51 | 1089.037231 | 1582.796509 |
| MAP2K5 | A6NK28;Q13163 | MAP2K5 | Dual-specificity mitog | 69605.83594 | 54724.58203 |
| OXSR1 | O95747 | OXSR1 | Serine/threonine-prot | 104748.2656 | 113326.5156 |
| CNKSR1 | Q969H4 | CNKSR1 | Connector enhancer c | 6392.969727 | 8208.960938 |
| DCAF5 | Q96JK2 | DCAF5 | DDB1- and CUL4-asso | 30308.04102 | 35250.25781 |
| LMBRD2 | Q68DH5 | LMBRD2 | G-protein coupled rec | 6136.157227 | 7313.212402 |
| MYH9 | P35579 | MYH9 | Myosin-9 | 496854.6563 | 443023.25 |
| RAB5A | P20339 | RAB5A | Ras-related protein Ra | 103596.9766 | 113019.2656 |
| SLC7A11 | Q9UPY5 | SLC7A11 | Cystine/glutamate tra | 2031.767456 | 1190.926392 |
| PGD | P52209 | PGD | 6-phosphogluconate c | 965117.375 | 1068316.25 |
| H1-4 | P10412 | H1-4 | Histone H1.4 | 1957405.25 | 509377.375 |
| ZDHHC4 | Q9NPG8 | ZDHHC4 | Probable palmitoyltra | 10852.03516 | 7770.082031 |

| | | | | | |
|---|---|---|---|---|---|
| TUSC3 | A0A0A0MTC2;D6RA37;D6RD | TUSC3 | Tumor suppressor car | 16652.25977 | 13881.76172 |
| ZKSCAN1 | E9PC66;P17029 | ZKSCAN1 | Zinc finger protein wit | 4936.617676 | 5765.042969 |
| PIP4K2B | P78356 | PIP4K2B | Phosphatidylinositol 5 | 18528.63477 | 21950.97461 |
| APOB | P04114 | APOB | Apolipoprotein B-100 | 32410.22852 | 34508.27734 |
| PHF1 | A0A087WVG4;A0A140TA01;/ | PHF1 | PHD finger protein 1;F | 3327.826416 | 2594.315674 |
| PHYKPL | Q8IUZ5 | PHYKPL | 5-phosphohydroxy-L-l | 46780.96484 | 25120.5918 |
| POGLUT1 | Q8NBL1 | POGLUT1 | Protein O-glucosyltrar | 36346.82813 | 34220.375 |
| SERPINC1 | P01008 | SERPINC1 | Antithrombin-III | 56639.63672 | 50367.20313 |
| CAMK1 | Q14012 | CAMK1 | Calcium/calmodulin-d | 6994.403809 | 6790.64502 |
| ANXA3 | P12429 | ANXA3 | Annexin A3 | 195681.5625 | 174584.4531 |
| LPP | Q93052 | LPP | Lipoma-preferred par | 86164.0625 | 67213.96094 |
| ZYG11B | Q9C0D3 | ZYG11B | Protein zyg-11 homolo | 11125.24609 | 13205.14453 |
| DTX3 | A0A0C4DGI8;F8VUZ4;Q8N9I | DTX3 | Probable E3 ubiquitin- | 3119.656982 | 3191.218262 |
| SYNGR3 | O43761 | SYNGR3 | Synaptogyrin-3 | 4662.773438 | 4908.604492 |
| ZNF689 | Q96CS4 | ZNF689 | Zinc finger protein 68 | 5786.762207 | 5686.839355 |
| CASTOR2 | A6NHX0 | CASTOR2 | Cytosolic arginine sen | 10126.5918 | 14408.19434 |
| NECAP2 | Q9NVZ3 | NECAP2 | Adaptin ear-binding c | 48359.80469 | 52202.07031 |
| RICTOR | Q6R327 | RICTOR | Rapamycin-insensitive | 6918.228516 | 6625.026367 |
| GAS2L1 | A0A5E8;Q99501 | GAS2L1 | GAS2-like protein 1 | 6335.510742 | 4025.203125 |
| KCTD21 | Q4G0X4 | KCTD21 | BTB/POZ domain-cont | 12187.93652 | 16668.01953 |
| RAP2C | Q9Y3L5 | RAP2C | Ras-related protein Ra | 38412.00781 | 47945.96875 |
| ST6GALNAC2 | Q9UJ37 | ST6GALNAC2 | Alpha-N-acetylgalacto | 1818.065674 | NA |
| HGFAC | D6RAR4;Q04756 | HGFAC | Hepatocyte growth fa | 30966.70703 | 34601.43359 |
| SGSH | P51688 | SGSH | N-sulphoglucosamine | 21073.91992 | 21445.19727 |
| PRKCA | P17252 | PRKCA | Protein kinase C alpha | 4009.927979 | 4959.566406 |
| TARBP1 | Q13395 | TARBP1 | Probable methyltrans | 9640.810547 | 11601.14746 |
| DDX19B | Q9UMR2 | DDX19B | ATP-dependent RNA h | 19488.01367 | 33115.87891 |
| CT83 | Q5H943 | CT83 | Kita-kyushu lung canc | 23350.41797 | 23469.5 |
| CES2 | O00748 | CES2 | Cocaine esterase | 1629.836548 | 2560.121338 |
| TBC1D1 | Q86TI0 | TBC1D1 | TBC1 domain family m | 14906.48242 | 17683.08984 |
| SLC35E1 | Q96K37 | SLC35E1 | Solute carrier family 3 | 11195.74902 | 10935.98242 |
| LGALS3 | P17931 | LGALS3 | Galectin-3 | 676568.4375 | 610298.8125 |
| CIB1 | Q99828 | CIB1 | Calcium and integrin-l | 1493.715454 | 2420.755615 |

| | | | | | |
|---|---|---|---|---|---|
| AGTPBP1 | J3KNS1;Q9UPW5 | AGTPBP1 | Cytosolic carboxypept | 1245.700317 | 1455.199585 |
| C10orf67 | Q8IYJ2 | C10orf67 | Uncharacterized prote | 328003.5 | 297419.6875 |
| LLGL2 | J3QRV5;Q6P1M3 | LLGL2 | LLGL scribble cell pola | 91824.4375 | 109228.5938 |
| RBL2 | F5H837;Q08999 | RBL2 | Retinoblastoma-like p | 984.6947632 | 3380.583252 |
| APAF1 | C9JLV4;O14727 | APAF1 | Apoptotic protease-ac | 6683.333496 | 8084.59668 |
| PRKDC | P78527 | PRKDC | DNA-dependent prote | 308089.5625 | 455468.3125 |
| DHRSX | C9JB06;Q8N5I4 | DHRSX | Dehydrogenase/reduc | 5603.146973 | 5113.854492 |
| FAM98C | Q17RN3 | FAM98C | Protein FAM98C | 16328.93848 | 16435.2207 |
| ZBTB14 | O43829 | ZBTB14 | Zinc finger and BTB dc | 1193.368286 | 807.6885376 |
| VPS13C | Q709C8 | VPS13C | Vacuolar protein sorti | 19654.69141 | 27808.68164 |
| C12orf10 | Q9HB07 | C12orf10 | UPF0160 protein MYG | 246430.9531 | 227509.7031 |
| HSPA2 | P54652 | HSPA2 | Heat shock-related 70 | 64513.38672 | 67468.52344 |
| H1-2 | P16403 | H1-2 | Histone H1.2 | 9664545 | 3409885 |
| SEC61A1 | B4DR61;P61619 | SEC61A1 | Protein transport prot | 161924.5 | 171175.125 |
| HSD3B7 | Q9H2F3 | HSD3B7 | 3 beta-hydroxysteroid | 7671.34082 | 9144.84082 |
| SLC9A6 | A0A0D9SGH0 | SLC9A6 | Sodium/hydrogen exc | 11795.23633 | 11858.81348 |
| OARD1 | C9JNE2;Q9Y530 | OARD1 | ADP-ribose glycohydr | 45782.23828 | 49462.33203 |
| TFAP2A;TFAP2E;TFAP2B | P05549;Q6VUC0;Q92481 | TFAP2A;TFAP2E;TFAP2 | Transcription factor A | 5932.290039 | 3482.144531 |
| ULK3 | Q6PHR2 | ULK3 | Serine/threonine-prot | 26015.14844 | 28576.04688 |
| WDR59 | Q6PJI9 | WDR59 | GATOR complex prote | 11523.84766 | 13022.77539 |
| TGM2 | P21980 | TGM2 | Protein-glutamine gar | 1939.909546 | 2479.902588 |
| SVIP | Q8NHG7 | SVIP | Small VCP/p97-interac | 26105.5957 | 26430.78516 |
| CARMIL1 | Q5VZK9 | CARMIL1 | F-actin-uncapping pro | 47949.99609 | 55027.875 |
| SRPK2 | P78362 | SRPK2 | SRSF protein kinase 2 | 44859.93359 | 35882.61719 |
| RAB32 | Q13637 | RAB32 | Ras-related protein Ra | 63577.89453 | 49199.50781 |
| USP11 | G5E9A6 | USP11 | Ubiquitin carboxyl-ter | 48552.40234 | 60877.30859 |
| HIGD1A | Q9Y241 | HIGD1A | HIG1 domain family m | 120202.5859 | 110582.5391 |
| PITPNB | P48739 | PITPNB | Phosphatidylinositol t | 93697.57031 | 90880.02344 |
| RAB5B | P61020 | RAB5B | Ras-related protein Ra | 85000.82031 | 88629.21094 |
| YWHAZ | P63104 | YWHAZ | 14-3-3 protein zeta/de | 7648116.5 | 7831599.5 |
| GATAD1 | Q8WUU5 | GATAD1 | GATA zinc finger doma | 5206.621094 | 4440.738281 |
| NR2C2 | P49116 | NR2C2 | Nuclear receptor subf | 6748.194336 | 6850.369141 |
| NDOR1 | Q9UHB4 | NDOR1 | NADPH-dependent dif | 7455.079102 | 10905.36719 |

| | | | | | |
|---|---|---|---|---|---|
| SMG9 | Q9H0W8 | SMG9 | Protein SMG9 | 29240.08789 | 30326.58594 |
| TRADD | Q15628 | TRADD | Tumor necrosis factor | 34655.53906 | 43550.53516 |
| LPIN3 | Q9BQK8 | LPIN3 | Phosphatidate phospl | 9113.094727 | 10203.85938 |
| ARPC1A | Q92747 | ARPC1A | Actin-related protein : | 46180.88672 | 44832.19922 |
| RHOC | P08134;Q5JR08 | RHOC | Rho-related GTP-bind | 818800.8125 | 898337.4375 |
| COX14 | Q96I36 | COX14 | Cytochrome c oxidase | 2987.685547 | 3205.388428 |
| HTATSF1 | O43719 | HTATSF1 | HIV Tat-specific factor | 146953.1094 | 148349.9219 |
| FAM241B | Q96D05 | FAM241B | Protein FAM241B | 81766.33594 | 61961.23828 |
| ZFP62 | Q8NB50 | ZFP62 | Zinc finger protein 62 | 4912.986328 | 4660.52832 |
| DDAH2 | O95865 | DDAH2 | N(G),N(G)-dimethylar; | 29082.61523 | 35295.01172 |
| ZBTB2 | Q8N680 | ZBTB2 | Zinc finger and BTB dc | 177422.9375 | 125755.0234 |
| BSPRY | Q5W0U4 | BSPRY | B box and SPRY doma | 6962.314453 | 9509.366211 |
| TMEM97 | Q5BJF2 | TMEM97 | Sigma intracellular rec | 54318.4375 | 73077.84375 |
| P3H4 | K7ERA3;Q92791 | P3H4 | Endoplasmic reticulur | 30480.58594 | 21113.4707 |
| ADGRG4 | Q8IZF6 | ADGRG4 | Adhesion G-protein cc | 117484.0234 | 189288.25 |
| PALM | O75781 | PALM | Paralemmin-1 | 19381.16797 | 17664.36914 |
| MTMR12 | Q9C0I1 | MTMR12 | Myotubularin-related | 34657.84375 | 34921.77734 |
| ATP6AP1 | Q15904 | ATP6AP1 | V-type proton ATPase | 71754.97656 | 77523.40625 |
| BLOC1S4 | Q9NUP1 | BLOC1S4 | Biogenesis of lysosom | 29272.52344 | 24821.125 |
| ATP9A | O75110 | ATP9A | Probable phospholipic | 44636.49609 | 33742.36328 |
| PIK3CA | A0A2R8Y2F6;P42336 | PIK3CA | Phosphatidylinositol 4 | 2524.613037 | 5185.483398 |
| ROCK1 | Q13464 | ROCK1 | Rho-associated protei | 29593.88867 | 32654.89844 |
| SMYD3 | Q9H7B4 | SMYD3 | Histone-lysine N-meth | 27767.08398 | 26710.73047 |
| TMEM38B | Q9NVV0 | TMEM38B | Trimeric intracellular ( | 590.7979736 | 14099.90527 |
| STARD3NL | C9JKL2;C9JPX5;O95772 | STARD3NL | STARD3 N-terminal-lik | 16045.20605 | 16752.57227 |
| ;EIF4E3 | A0A3B3IRU0;Q8N5X7 | ;EIF4E3 | Uncharacterized prote | 4474.683105 | 6167.004883 |
| MICOS13 | A0A140TA86;Q5XKP0 | MICOS13 | MICOS complex subur | 26934.06836 | 28576.75195 |
| PARP16 | Q8N5Y8 | PARP16 | Protein mono-ADP-rib | 6007.405273 | 6744.559082 |
| DYRK1A | A0A2R8Y6L5;Q13627 | DYRK1A | Dual-specificity tyrosi | 13103.76465 | 13372.08301 |
| GTF2A2 | P52657 | GTF2A2 | Transcription initiatior | 16400.36523 | 28094.00195 |
| NUDT14 | O95848 | NUDT14 | Uridine diphosphate g | 16080.10352 | 18979.23438 |
| PHGDH | A0A286YF22;O43175 | PHGDH | D-3-phosphoglycerate | 1305443.875 | 1236366.5 |
| ANXA11 | P50995 | ANXA11 | Annexin A11 | 137775.0469 | 119012.8906 |

| | | | | | |
|---|---|---|---|---|---|
| VPS37B | Q9H9H4 | VPS37B | Vacuolar protein sorti | 11216.60547 | 10177.7832 |
| AACS | Q86V21 | AACS | Acetoacetyl-CoA syntl | 146029.6875 | 157780.4531 |
| MAN1A2 | O60476 | MAN1A2 | Mannosyl-oligosaccha | 24988.22656 | 28571.78125 |
| VPS51 | Q9UID3 | VPS51 | Vacuolar protein sorti | 28774.04883 | 33641.69141 |
| RALGAPA1 | A0A1B0GUI1 | RALGAPA1 | Ral GTPase-activating | 19504.52539 | 18263.86914 |
| DENND2D | Q9H6A0 | DENND2D | DENN domain-contair | 18706.03711 | 18438.00977 |
| PPP1R7 | Q15435 | PPP1R7 | Protein phosphatase : | 254652.1563 | 270873.5938 |
| SDSL | Q96GA7 | SDSL | Serine dehydratase-lik | 66651.375 | 63401.38672 |
| TMEM230 | A0A087WTT2;Q5JWB9;Q96A | TMEM230 | Transmembrane prote | 17440.36914 | 21521.9082 |
| NUBPL | Q8TB37 | NUBPL | Iron-sulfur protein NU | 15253.14746 | 16330.08496 |
| TSPAN3 | O60637 | TSPAN3 | Tetraspanin-3 | 2191.233887 | NA |
| DYNLT1 | P63172 | DYNLT1 | Dynein light chain Tct | 40092.63672 | 46245.30859 |
| ENDOG | Q14249 | ENDOG | Endonuclease G, mito | 31965.70898 | 34158.49219 |
| STMN1 | P16949 | STMN1 | Stathmin | 1024702 | 1234861 |
| NF1 | J3KSB5;P21359 | NF1 | Neurofibromin (Fragm | 9875.15918 | 11525.4668 |
| FANCI | Q9NVI1 | FANCI | Fanconi anemia group | 59429.67578 | 76253.10156 |
| CST3 | P01034 | CST3 | Cystatin-C | 42485.79688 | 42941.12891 |
| PRUNE1 | Q86TP1 | PRUNE1 | Exopolyphosphatase F | 33541.78516 | 41961.45313 |
| LBR | Q14739 | LBR | Delta(14)-sterol reduc | 76493.28906 | 86763.77344 |
| RAB33B | A0A494C0Z5;Q9H082 | RAB33B | Ras-related protein Ra | 21842.77734 | 24178.18555 |
| CMTM6 | Q9NX76 | CMTM6 | CKLF-like MARVEL trai | 18234.83789 | 17542.24219 |
| CTNND1 | C9JZR2;O60716 | CTNND1 | Catenin delta-1 | 108435.4922 | 115721.4766 |
| RP2 | O75695 | RP2 | Protein XRP2 | 25578.48242 | 28454.08008 |
| ZER1 | Q7Z7L7 | ZER1 | Protein zer-1 homolog | 8246.719727 | 10851.70215 |
| ADAM10 | O14672 | ADAM10 | Disintegrin and metall | 72728.14844 | 66196.24219 |
| TALDO1 | P37837 | TALDO1 | Transaldolase | 1088456.125 | 1369627.625 |
| DIAPH2 | A6NML8;C9J6U3;K4DI95;O6( | DIAPH2 | Diaphanous homolog | 14703.2207 | 19983.93945 |
| VCPKMT | Q9H867 | VCPKMT | Protein-lysine methylt | 7578.150391 | 8261.642578 |
| SCAND1 | P57086 | SCAND1 | SCAN domain-contain | 373.8937683 | NA |
| XPC | Q01831 | XPC | DNA repair protein co | 33713.60938 | 41751.24609 |
| PACS2 | F8VW41;Q86VP3 | PACS2 | Phosphofurin acidic cl | 11376.17969 | 11932.1416 |
| NVL | O15381 | NVL | Nuclear valosin-conta | 53958.21484 | 44634.67969 |
| IGFBP4 | P22692 | IGFBP4 | Insulin-like growth fac | 14256.55762 | 20444.17188 |

| | | | | | |
|---|---|---|---|---|---|
| PARP1 | P09874 | PARP1 | Poly [ADP-ribose] poly | 216559.875 | 377279.25 |
| TIMMDC1 | Q9NPL8 | TIMMDC1 | Complex I assembly fa | 22626.45313 | 29130.05273 |
| MMUT | P22033 | MMUT | Methylmalonyl-CoA m | 47275.11719 | 56191.94141 |
| NFIB | A0A0A0MRX8;A0A1B0GU97; | NFIB | Nuclear factor 1;Nucle | 9016.268555 | 13994.72754 |
| H1FX | Q92522 | H1FX | Histone H1x | 732473.6875 | 689562.75 |
| TACC1 | O75410 | TACC1 | Transforming acidic co | 27147.58984 | 23986.94336 |
| GRN | P28799 | GRN | Progranulin | 91708.23438 | 95454.07813 |
| TTLL5 | G3V2J9;Q6EMB2 | TTLL5 | Tubulin polyglutamyla | 180654.8594 | 237628.9219 |
| PBXIP1 | Q96AQ6 | PBXIP1 | Pre-B-cell leukemia tra | 98533.57031 | 150016.2656 |
| RPL36AL | Q969Q0 | RPL36AL | 60S ribosomal protein | 59708.29297 | 38766.25 |
| PTPRA | P18433 | PTPRA | Receptor-type tyrosin | 3303.217773 | 5171.645508 |
| ABCD3 | A0A3B3ITW3;P28288 | ABCD3 | ATP-binding cassette s | 60107.52344 | 127437.0156 |
| CAPN1 | P07384 | CAPN1 | Calpain-1 catalytic sub | 500438.2813 | 659543.875 |
| ANP32A | P39687 | ANP32A | Acidic leucine-rich nuc | 344047.9375 | 479795.5938 |
| HACD3 | Q9P035 | HACD3 | Very-long-chain (3R)-3 | 293350.0938 | 310093.875 |
| GORASP1 | Q9BQQ3 | GORASP1 | Golgi reassembly-stac | 8968.255859 | 8043.521484 |
| GUK1 | B1ANH0 | GUK1 | Guanylate kinase (Frag | 71874.78271 | 67792.33398 |
| PHF6 | A0A0D9SGE8;Q8IWS0 | PHF6 | PHD finger protein 6 | 35681.55859 | 29926.11328 |
| POGLUT3 | Q7Z4H8 | POGLUT3 | Protein O-glucosyltran | 25570.67969 | 19482.84766 |
| MKNK1 | A0A499FIS5;A0A499FJN1;A0A | MKNK1 | MAP kinase-interactin | 2638.155762 | 4236.002441 |
| INPP1 | P49441 | INPP1 | Inositol polyphosphat | 2977.223389 | 2566.042236 |
| PARP4 | Q9UKK3 | PARP4 | Protein mono-ADP-rib | 12290.65527 | 9189.052734 |
| SNX1 | Q13596 | SNX1 | Sorting nexin-1 | 204393.5313 | 254829.3906 |
| CGN | A2A3M4 | CGN | Cingulin (Fragment) | 108579.4595 | 108597.2837 |
| WDR45 | A0A0U1RQR1;A0A0U1RQS7;A | WDR45 | WD repeat domain ph | 8082.810547 | 6996.226563 |
| PIP4K2C | Q8TBX8 | PIP4K2C | Phosphatidylinositol 5 | 317005.7188 | 359113.7188 |
| MAP3K6 | O95382 | MAP3K6 | Mitogen-activated pro | 4385.651855 | 4151.75293 |
| LAP3 | P28838 | LAP3 | Cytosol aminopeptida | 639998 | 634270.8125 |
| TBC1D30 | Q9Y2I9 | TBC1D30 | TBC1 domain family m | 12250.68555 | 10024.51953 |
| KRAS | P01116 | KRAS | GTPase KRas | 9095.604492 | 16883.83008 |
| RPS6KA1 | E9PGT3;Q15418 | RPS6KA1 | Ribosomal protein S6 | 37359.30469 | 32544.15039 |
| RAB8B | Q92930 | RAB8B | Ras-related protein Ra | 186290.7188 | 203078.1719 |
| MAP3K11 | Q16584 | MAP3K11 | Mitogen-activated pro | 3575.046143 | 2760.681641 |

| | | | | | |
|---|---|---|---|---|---|
| FIG4 | Q92562 | FIG4 | Polyphosphoinositide | 5334.54541 | 6098.396973 |
| CATSPERG | B8ZZI7;Q6ZRH7;X1WI24 | CATSPERG | Cation channel sperm | 43326.45703 | 68158.04688 |
| DAPK3 | O43293 | DAPK3 | Death-associated prot | 3106.781738 | 3909.781982 |
| IRF2BP2 | Q7Z5L9 | IRF2BP2 | Interferon regulatory | 81726.90625 | 75073.38281 |
| CHML | P26374 | CHML | Rab proteins geranylg | 175.0019531 | 513.4577637 |
| PVR | A0A0C4DG49;P15151 | PVR | Poliovirus receptor | 24372.6543 | 34962.29297 |
| SLBP | F8W8D3;Q14493 | SLBP | Histone RNA hairpin-b | 10225.08496 | 7951.939941 |
| EXOC1 | Q9NV70 | EXOC1 | Exocyst complex comp | 30881.01172 | 32385.90039 |
| NECTIN2 | Q92692 | NECTIN2 | Nectin-2 | 72100.24219 | 86248.71094 |
| NUMA1 | A0A087WY61 | NUMA1 | Nuclear mitotic appar | 360484.0117 | 312952.0078 |
| COA3 | Q9Y2R0 | COA3 | Cytochrome c oxidase | 91479.57031 | 95162.10156 |
| DAP | P51397 | DAP | Death-associated prot | 23457.47656 | 25740.66016 |
| ERGIC1 | Q969X5 | ERGIC1 | Endoplasmic reticulum | 276486.875 | 271545.9375 |
| CTSZ | Q9UBR2 | CTSZ | Cathepsin Z | 22192.32227 | 14030.46875 |
| ITIH1 | P19827 | ITIH1 | Inter-alpha-trypsin inh | 6001.364258 | 4803.871582 |
| MYO9B | M0R0P8;M0R300;Q13459 | MYO9B | Unconventional myos | 13180.1709 | 16797.10547 |
| WDTC1 | Q8N5D0 | WDTC1 | WD and tetratricopep | 2732.667236 | 4140.222656 |
| TRIM4 | Q9C037 | TRIM4 | E3 ubiquitin-protein li | 11568.20313 | 12118.02637 |
| GSS | P48637 | GSS | Glutathione synthetas | 226031.3438 | 227195.2969 |
| NBEAL1 | Q6ZS30 | NBEAL1 | Neurobeachin-like pro | 3456.838867 | 5502.712891 |
| RPA3 | P35244 | RPA3 | Replication protein A | 230238.3906 | 277889.5938 |
| SCFD1 | Q8WVM8 | SCFD1 | Sec1 family domain-co | 214903.0156 | 228911.7656 |
| PRDX1 | Q06830 | PRDX1 | Peroxiredoxin-1 | 10688065 | 11837479 |
| ALDH16A1 | Q8IZ83 | ALDH16A1 | Aldehyde dehydrogen | 33088.46094 | 34043.42188 |
| EZR | E7EQR4;P15311 | EZR | Ezrin | 402101.3125 | 407698.0313 |
| CDS1 | Q92903 | CDS1 | Phosphatidate cytidyl | 14251.56641 | 8321.058594 |
| CLYBL | Q5JVB9;Q5JVC0;Q8N0X4 | CLYBL | Citramalyl-CoA lyase, | 2126.635742 | 2751.285156 |
| MICALL1 | Q8N3F8 | MICALL1 | MICAL-like protein 1 | 6862.895508 | 9164.375 |
| HDAC10 | C9J8B8;Q08AP5;Q969S8 | HDAC10 | Polyamine deacetylas | 5634.143066 | 5448.011719 |
| TDP1 | Q9NUW8 | TDP1 | Tyrosyl-DNA phospho | 23145.02344 | 23252.36914 |
| PRKAB2 | O43741 | PRKAB2 | 5'-AMP-activated prot | 12896.01465 | 12560.82324 |
| RAB6D | Q53S08 | RAB6D | Ras-related protein Ra | 100356.6172 | 109659.7188 |
| BIRC2 | A0A087X2B2;E9PIW1;E9PMH | BIRC2 | Baculoviral IAP repeat | NA | 4899.320313 |

| | | | | | |
|---|---|---|---|---|---|
| CAPN7 | Q9Y6W3 | CAPN7 | Calpain-7 | 33520.05469 | 33560.51563 |
| BNIP3L | H0YBC7;O60238 | BNIP3L | BCL2/adenovirus E1B | 2199.646973 | 4416.90332 |
| IGFBP3 | A6XND0;A6XND1;P17936 | IGFBP3 | Insulin-like growth fac | 4470.151855 | 7353.620117 |
| MTARC1 | Q5VT66 | MTARC1 | Mitochondrial amidox | 29437.2168 | 40176.69531 |
| NUB1 | H3BM14;H3BM74;Q9Y5A7 | NUB1 | NEDD8 ultimate buste | 16748.75 | 18557.57031 |
| ASF1A | Q9Y294 | ASF1A | Histone chaperone AS | 49534.1875 | 59078.86328 |
| EXOC6B | A0A0U1RRB6;Q9Y2D4 | EXOC6B | Exocyst complex comp | 29764.41406 | 33010.64453 |
| MICALL2 | Q8IY33 | MICALL2 | MICAL-like protein 2 | 21567.17188 | 20535.86523 |
| PPP1R37 | O75864 | PPP1R37 | Protein phosphatase 1 | 5766.569336 | 8643.019531 |
| ADD1 | E7ENY0;E7EV99;P35611 | ADD1 | Alpha-adducin | 22530.41406 | 29374.24609 |
| ATL3 | F5H6I7 | ATL3 | Atlastin-3 | 96107.8772 | 107532.3684 |
| SPAG1 | Q07617 | SPAG1 | Sperm-associated ant | 20984.62305 | 20111.2168 |
| ARSA | A0A0C4DFZ2;P15289 | ARSA | Arylsulfatase A | 1986.017456 | 2656.058594 |
| CDH3 | P22223 | CDH3 | Cadherin-3 | 1760.223633 | 2417.616943 |
| AP4M1 | C9JC87;C9JWL4;O00189 | AP4M1 | AP-4 complex subunit | 20925.97852 | 20171.25391 |
| ECHDC1 | Q9NTX5 | ECHDC1 | Ethylmalonyl-CoA dec | 58608.5625 | 54247.70313 |
| MOSPD2 | Q8NHP6 | MOSPD2 | Motile sperm domain- | 21527.9082 | 24926.01172 |
| CRELD1 | F8WBY3;Q96HD1 | CRELD1 | Protein disulfide isom | 3594.12085 | 3430.894043 |
| NADK2 | Q4G0N4 | NADK2 | NAD kinase 2, mitocho | 42997.95703 | 41830.16406 |
| GOLPH3 | Q9H4A6 | GOLPH3 | Golgi phosphoprotein | 59442.86719 | 71336.67188 |
| CTDSP1 | Q9GZU7 | CTDSP1 | Carboxy-terminal dom | 13469.01367 | 16470.48242 |
| SLC9A8 | Q9Y2E8 | SLC9A8 | Sodium/hydrogen exc | 4874.358398 | 7144.211426 |
| HP1BP3 | Q5SSJ5 | HP1BP3 | Heterochromatin prot | 344065.6563 | 342591.0938 |
| HINT3 | Q9NQE9 | HINT3 | Histidine triad nucleot | 24197.7832 | 25085.20313 |
| EBP | Q15125 | EBP | 3-beta-hydroxysteroid | 147055.7813 | 175784.9219 |
| EXOC5 | O00471 | EXOC5 | Exocyst complex comp | 39563.65234 | 47294.85938 |
| TEAD3 | A0A1X7SBS4;Q99594 | TEAD3 | Transcriptional enhan | 4311.086426 | 1706.088989 |
| ZC3HAV1L | Q96H79 | ZC3HAV1L | Zinc finger CCCH-type | 24867.65039 | 28746.64648 |
| PHYH | B1ALH6;C9IYS5;O14832 | PHYH | Phytanoyl-CoA dioxyg | 3733.930908 | 6711.456543 |
| DYNC1LI1 | Q9Y6G9 | DYNC1LI1 | Cytoplasmic dynein 1 | 139113.7188 | 134165.4063 |
| RBBP9 | O75884 | RBBP9 | Serine hydrolase RBBP | 59797.65625 | 65595.5 |
| ARRB1 | H0YET9 | ARRB1 | Beta-arrestin-1 (Fragn | 29899.24609 | 32775.99194 |
| LAMC1 | P11047 | LAMC1 | Laminin subunit gamn | 5153.643555 | 4810.220215 |

| | | | | | |
|---|---|---|---|---|---|
| FOXP1 | A0A0B4J2F3;A0A3B3IRS5;A0/ | FOXP1 | Forkhead box protein | 6776.958984 | 7853.98584 |
| STX12 | Q86Y82 | STX12 | Syntaxin-12 | 75580.74219 | 75680.46094 |
| RABL3 | Q5HYI8 | RABL3 | Rab-like protein 3 | 97125.71875 | 95679.96875 |
| GBE1 | Q04446 | GBE1 | 1,4-alpha-glucan-bran | 61891.76172 | 61368.77734 |
| PCK2 | Q16822 | PCK2 | Phosphoenolpyruvate | 255515.6719 | 258719 |
| MSRA | Q9UJ68 | MSRA | Mitochondrial peptide | 24509.12891 | 24804.71094 |
| ATG2B | Q96BY7 | ATG2B | Autophagy-related pr | 14136.06641 | 13241.95117 |
| ZBTB33 | Q86T24 | ZBTB33 | Transcriptional regula | 4141.850098 | 3006.887207 |
| RAB11B | Q15907 | RAB11B | Ras-related protein Ra | 828097.8125 | 978191 |
| NKAP | A0A494C050;Q8N5F7 | NKAP | NF-kappa-B-activating | 26146.57031 | 18995.8457 |
| SCAMP4 | Q969E2 | SCAMP4 | Secretory carrier-asso | 110937.5234 | 120046.6172 |
| TMEM254 | A0A087WZM8;Q8TBM7 | TMEM254 | Transmembrane prote | 14760.93066 | 10169.07227 |
| CPEB2 | A0A5K1VW61;A0A5K1VW71; | CPEB2 | Cytoplasmic polyaden | 4255.518555 | 4663.227539 |
| PRKCZ | Q05513 | PRKCZ | Protein kinase C zeta t | 32967.11719 | 34954.125 |
| UNG | P13051 | UNG | Uracil-DNA glycosylase | 61459.14063 | 74099.40625 |
| LPGAT1 | Q92604 | LPGAT1 | Acyl-CoA:lysophospha | 2290.269043 | 2587.147705 |
| EEFSEC | P57772 | EEFSEC | Selenocysteine-specifi | 51222.83203 | 56717.17578 |
| TMPPE | Q6ZT21 | TMPPE | Transmembrane prote | 6078.474121 | 4986.349609 |
| UFL1 | O94874 | UFL1 | E3 UFM1-protein ligas | 127100.7266 | 123458.0859 |
| SLC6A9 | B7Z589;P48067 | SLC6A9 | Sodium- and chloride- | 6023.347168 | 5696.32959 |
| MTERF1 | B4DPR9;Q99551 | MTERF1 | Transcription termina | 2564.031006 | 1914.701416 |
| ITIH4 | B7ZKJ8;Q14624 | ITIH4 | ITIH4 protein;Inter-alp | 30960.27539 | 23369.00977 |
| CYSTM1 | Q9H1C7 | CYSTM1 | Cysteine-rich and tran | 21471.52148 | 22235.56055 |
| RSPRY1 | H3BPL2;H3BQQ7;H3BSC7;Q9 | RSPRY1 | RING finger and SPRY | 1968.31311 | 1354.125732 |
| RBCK1 | Q9BYM8 | RBCK1 | RanBP-type and C3HC | 12774.58887 | 12388.83984 |
| TF | P02787 | TF | Serotransferrin | 40405.82813 | 40639.41797 |
| RPS6KA6 | Q9UK32 | RPS6KA6 | Ribosomal protein S6 | 838.9633789 | 900.5911865 |
| BRD3 | Q15059 | BRD3 | Bromodomain-contair | 14552.57031 | 7394.727051 |
| RANBP6 | O60518 | RANBP6 | Ran-binding protein 6 | 4550.037109 | 5006.016113 |
| RIPK2 | O43353 | RIPK2 | Receptor-interacting s | 10539.08496 | 18486.34766 |
| MAP2K1 | Q02750 | MAP2K1 | Dual specificity mitog | 142682.7188 | 156087.5156 |
| ANAPC11 | Q9NYG5 | ANAPC11 | Anaphase-promoting | 7595.131836 | 10396.63379 |
| FYTTD1 | Q96QD9 | FYTTD1 | UAP56-interacting fac | 60367.30469 | 49259.4375 |

| | | | | | |
|---|---|---|---|---|---|
| CMC1 | Q7Z7K0 | CMC1 | COX assembly mitoch | 119672.3672 | 103753.7266 |
| TBC1D5 | Q92609 | TBC1D5 | TBC1 domain family m | 80258.66406 | 88060.875 |
| VASP | P50552 | VASP | Vasodilator-stimulate | 150322.8125 | 142399.4219 |
| CLEC3B | E9PHK0;P05452 | CLEC3B | Tetranectin | 12742.83105 | 9385.69043 |
| KRT6A | P02538 | KRT6A | Keratin, type II cytosk | 7391.560547 | 17136.11523 |
| ACTR1B | P42025 | ACTR1B | Beta-centractin | 46976.60547 | 45666.98828 |
| TRPM4 | Q8TD43 | TRPM4 | Transient receptor po | 23522.49219 | 23151.5918 |
| WDR48 | Q8TAF3 | WDR48 | WD repeat-containing | 19873.47656 | 24303.33203 |
| SERGEF | G3V1B4;Q9UGK8 | SERGEF | Secretion regulating g | 2146.576904 | 2278.880127 |
| SBF1 | O95248 | SBF1 | Myotubularin-related | 8404.161133 | 9252.010742 |
| ACADSB | P45954 | ACADSB | Short/branched chain | 43851.69922 | 42468.12109 |
| SSBP2 | A0A087X159;P81877 | SSBP2 | Single-stranded DNA-l | 3099.621094 | 2825.290527 |
| FAM111A | Q96PZ2 | FAM111A | Protein FAM111A | 6887.978516 | 5456.715332 |
| B3GALNT1 | A0A494C0Y5;O75752 | B3GALNT1 | Hexosyltransferase (Fr | 19228.84961 | 15793.14258 |
| C2CD2L | O14523 | C2CD2L | Phospholipid transfer | 9715.118164 | 9501.604492 |
| ILK | A0A0A0MTH3;Q13418 | ILK | Integrin-linked protei | 80773.17188 | 89434.41406 |
| BCOR | H7C231;Q6W2J9 | BCOR | BCL-6 corepressor (Fra | 4475.244141 | 4028.862061 |
| TBCEL | Q5QJ74 | TBCEL | Tubulin-specific chape | 14182.25781 | 9303.304688 |
| ASAH1 | A0A1B0GTM3;A0A1B0GUA4; | ASAH1 | Acid ceramidase;Acid | 58177.73828 | 63628.83594 |
| ATP1B3 | P54709 | ATP1B3 | Sodium/potassium-tra | 161383.0625 | 172240.6563 |
| UAP1 | Q16222 | UAP1 | UDP-N-acetylhexosam | 83048.25 | 68642.33594 |
| SLC33A1 | O00400 | SLC33A1 | Acetyl-coenzyme A tra | 22230.55664 | 25520.85742 |
| FAM3A | D3DWX8;P98173;Q5HY75 | FAM3A | Family with sequence | 7102.397461 | 9681.798828 |
| FBXW9 | Q5XUX1 | FBXW9 | F-box/WD repeat-con | 4059.355469 | 5133.495117 |
| F2 | E9PIT3;P00734 | F2 | Prothrombin | 7119.277344 | 10762.09082 |
| EOGT | Q5NDL2 | EOGT | EGF domain-specific C | 6498.695313 | 7039.713867 |
| ZNF154 | Q13106 | ZNF154 | Zinc finger protein 15 | 1203.17334 | 1413.198853 |
| LONP1 | K7EJE8;K7EKE6 | LONP1 | Lon protease homolog | 270453.5313 | 292557.2188 |
| CHPF | Q8IZ52 | CHPF | Chondroitin sulfate sy | 6056.764648 | 6248.550781 |
| TSG101 | F5H442;Q99816 | TSG101 | Tumor susceptibility g | 99684.32813 | 90163.78906 |
| EMC6 | Q9BV81 | EMC6 | ER membrane protein | 103776.6016 | 110428.0938 |
| RPL26L1 | Q9UNX3 | RPL26L1 | 60S ribosomal protein | 56547.40625 | 54631.40625 |
| FN3K | Q9H479 | FN3K | Fructosamine-3-kinas | 20443.82227 | 20757.60547 |

| | | | | | |
|---|---|---|---|---|---|
| STAU2 | E7EPX0;E7EVJ4;Q9NUL3 | STAU2 | Double-stranded RNA | 56587.32813 | 55389.92578 |
| RALBP1 | Q15311 | RALBP1 | RalA-binding protein 1 | 11881.2334 | 13141.4209 |
| VTN | P04004 | VTN | Vitronectin | 174961.2188 | 69305.5 |
| HDAC8;HDAC8;;HDAC8;HDA | A0A3B3IRI9;A0A3B3IRP8;A0A | HDAC8;HDAC8;;HDAC | Histone deacetylase 8 | 2626.367188 | 4039.03833 |
| DNMT3B | Q9UBC3 | DNMT3B | DNA (cytosine-5)-met | 3641.092773 | 4623.270996 |
| TPM1 | B7Z596;F5H7S3;H0YKX5;H7B | TPM1 | Tropomyosin alpha-1 | 7598.16748 | 8621.71875 |
| SERPINA7 | P05543 | SERPINA7 | Thyroxine-binding glo | 24510.18555 | 16031.43652 |
| PTPN9 | A0A0A0MR09;P43378 | PTPN9 | Tyrosine-protein phos | 14326.50195 | 14527.46484 |
| CCDC90B | Q9GZT6 | CCDC90B | Coiled-coil domain-co | 4768.666016 | 6193.214844 |
| UBE4B | O95155 | UBE4B | Ubiquitin conjugation | 23247.12891 | 28777.97266 |
| GSR | P00390 | GSR | Glutathione reductase | 444235.7813 | 488483.0625 |
| CYBC1 | Q9BQA9 | CYBC1 | Cytochrome b-245 cha | 50329.55859 | 56452.32031 |
| CENPM | B1AHQ6;B1AHQ7;B1AHQ8;Q | CENPM | Centromere protein M | 2392.660645 | 1983.175537 |
| CCDC28A | B4DUJ5;Q8IWP9 | CCDC28A | Coiled-coil domain-co | 1408.055908 | 2291.136475 |
| DSTN | P60981 | DSTN | Destrin | 671577.125 | 840192.5 |
| SPIN1 | Q9Y657 | SPIN1 | Spindlin-1 | 10350.27344 | 9472.990234 |
| SNAP23 | O00161 | SNAP23 | Synaptosomal-associa | 20013.30664 | 9016.993164 |
| NAGK | Q9UJ70 | NAGK | N-acetyl-D-glucosamir | 157931.4531 | 191244.4219 |
| SIGIRR | Q6IA17 | SIGIRR | Single Ig IL-1-related r | 67103.35156 | 72004.72656 |
| C8orf82 | Q6P1X6 | C8orf82 | UPF0598 protein C8or | 44560.30469 | 50627.28125 |
| ATP1A1 | P05023 | ATP1A1 | Sodium/potassium-tra | 659601.75 | 618656.9375 |
| RAB6A | P20340 | RAB6A | Ras-related protein Ra | 506720.4063 | 583984.8125 |
| MYO9A | B2RTY4;H3BMM1;H3BRD5 | MYO9A | Unconventional myos | 1789.401733 | 1091.51416 |
| RPS6KA4 | A0A0A6YYC0;E9PJN1;O7567 | RPS6KA4 | Ribosomal protein S6 | 14391.74316 | 16858.86328 |
| WWP1 | Q9H0M0 | WWP1 | NEDD4-like E3 ubiquit | 8531.269531 | 10120.00488 |
| NSFL1C | Q9UNZ2 | NSFL1C | NSFL1 cofactor p47 | 94945.09375 | 84678.875 |
| TMEM30A | Q9NV96 | TMEM30A | Cell cycle control prot | 29952.55273 | 37510.89844 |
| TMUB1 | A0A590UJA9;C9JCW7;C9JE12 | TMUB1 | Transmembrane and u | 10432.5332 | 15686.69043 |
| DEGS1 | O15121 | DEGS1 | Sphingolipid delta(4)- | 26767.2207 | 17010.90039 |
| C4A | A0A0G2JPR0;A0A140TA32;A0 | C4A | Complement C4-A | 18878.72656 | 16120.20996 |
| DTYMK | P23919 | DTYMK | Thymidylate kinase | 187940.0313 | 210269.7031 |
| POMGNT1 | Q8WZA1 | POMGNT1 | Protein O-linked-mann | 4507.168457 | 5639.28418 |
| C2orf76 | Q3KRA6 | C2orf76 | UPF0538 protein C2or | 32938.11719 | 34511.76563 |

| | | | | | |
|---|---|---|---|---|---|
| TMEM223 | A0PJW6 | TMEM223 | Transmembrane prote | 28136.45898 | 27638.67578 |
| NEDD8-MDP1 | E9PL57 | NEDD8-MDP1 | NEDD8-MDP1 readthr | 20927.03906 | 26562.79297 |
| THEM6 | Q8WUY1 | THEM6 | Protein THEM6 | 347421.75 | 402496.0938 |
| PARL | A0A1W2PP11;Q9H300 | PARL | Presenilins-associated | 20858.29688 | 21747.72656 |
| CYB5R3 | P00387 | CYB5R3 | NADH-cytochrome b5 | 306866.2188 | 283138.8438 |
| INPP4A | Q96PE3 | INPP4A | Inositol polyphosphat | 17671.00781 | 19590.65234 |
| LRATD2 | Q96KN1 | LRATD2 | Protein LRATD2 | 58432.78906 | 52653.50391 |
| NRDC | O43847 | NRDC | Nardilysin | 210503.5938 | 253441.7188 |
| TMEM41B | Q5BJD5 | TMEM41B | Transmembrane prote | 11713.54883 | 14257.77441 |
| CDC45 | O75419 | CDC45 | Cell division control pr | 8592.411133 | 15334.05078 |
| GGPS1 | O95749 | GGPS1 | Geranylgeranyl pyrop | 54758.78516 | 60803.98047 |
| COL12A1 | D6RGG3;Q99715 | COL12A1 | Collagen alpha-1(XII) c | 4674.424316 | 4543.837402 |
| NEB | A0A087X1N7;P20929 | NEB | Nebulin | 21498.08008 | 26063.77539 |
| GDPGP1 | Q6ZNW5 | GDPGP1 | GDP-D-glucose phosp | 12720.49219 | 15516.75977 |
| ITIH3 | A0A087WW43;E7ET33;Q060 | ITIH3 | Inter-alpha-trypsin inh | 35847.72656 | 30211.73047 |
| SMAP2 | Q8WU79 | SMAP2 | Stromal membrane-as | 71500.85938 | 67589.39063 |
| TRMT12 | Q53H54 | TRMT12 | tRNA wybutosine-synt | 11378.8584 | 10172.11133 |
| SUMF2 | J3KQJ1;Q8NBJ7 | SUMF2 | Inactive C-alpha-form | 32019.43555 | 30814.67773 |
| RFC1 | P35251 | RFC1 | Replication factor C su | 55186.91797 | 59071.91406 |
| UBE2E2 | Q96LR5 | UBE2E2 | Ubiquitin-conjugating | 24366.92188 | 21920.60938 |
| DTD2 | Q96FN9 | DTD2 | D-aminoacyl-tRNA dea | 45111.11328 | 51747.91406 |
| MGMT | B4DEE8;P16455 | MGMT | Methylated-DNA--pro | 75242.17188 | 82839.85156 |
| EHD3 | Q9NZN3 | EHD3 | EH domain-containing | 10159.56934 | 9819.426758 |
| ILVBL | A1L0T0 | ILVBL | Acetolactate synthase | 48279.57813 | 50513.62891 |
| REEP5 | Q00765 | REEP5 | Receptor expression-e | 296919.1563 | 332477.125 |
| ZWILCH | Q9H900 | ZWILCH | Protein zwilch homolo | 19568.77344 | 23384.56445 |
| TLE3 | H0YKH0 | TLE3 | Transducin-like enhan | 119766.0742 | 145570.7656 |
| PYCR1 | P32322 | PYCR1 | Pyrroline-5-carboxylat | 364411.5313 | 413367.625 |
| SRBD1 | Q8N5C6 | SRBD1 | S1 RNA-binding doma | 41721.94141 | 35543.8125 |
| USP24 | Q9UPU5 | USP24 | Ubiquitin carboxyl-ter | 27791.41211 | 29494.58984 |
| PNPLA2 | Q96AD5 | PNPLA2 | Patatin-like phospholi | 10806.6748 | 12258.1543 |
| DNTTIP1 | Q9H147 | DNTTIP1 | Deoxynucleotidyltrans | 51113.47266 | 50882.9375 |
| SLC16A3 | O15427 | SLC16A3 | Monocarboxylate trar | 9223.301758 | 5985.582031 |

| | | | | | |
|---|---|---|---|---|---|
| YIF1B | Q5BJH7 | YIF1B | Protein YIF1B | 14363.62207 | 14004.30469 |
| RAP2A | P10114 | RAP2A | Ras-related protein Ra | 151217.2813 | 167286.3125 |
| SLC27A4 | Q6P1M0 | SLC27A4 | Long-chain fatty acid t | 131890.2031 | 131976.9531 |
| DHRS7B | A0A0C4DGQ8;Q6IAN0 | DHRS7B | Dehydrogenase/reduc | 61676.25781 | 61913.27734 |
| SAP30L | Q9HAJ7 | SAP30L | Histone deacetylase c | 14579.64746 | 17013.52148 |
| MON2 | Q7Z3U7 | MON2 | Protein MON2 homolo | 127077.1328 | 146060.1563 |
| TLN2 | Q9Y4G6 | TLN2 | Talin-2 | 110591.75 | 109126.7578 |
| PMS1 | B7ZAA0;P54277 | PMS1 | PMS1 protein homolo | 6795.073242 | 7568.850586 |
| TSPAN15 | H7C285;O95858 | TSPAN15 | Tetraspanin-15 (Fragn | 33981.76172 | 28992.49023 |
| RFT1 | B5MDE0;Q96AA3 | RFT1 | Protein RFT1 homolog | 24541.92578 | 26757.00195 |
| PCYT1A | C9JEJ2;P49585 | PCYT1A | Choline-phosphate cy | 144256.9219 | 128916.4297 |
| CCDC88C | Q9P219 | CCDC88C | Protein Daple | 13531.92188 | 12957.77148 |
| FN3KRP | Q9HA64 | FN3KRP | Ketosamine-3-kinase | 60560.90234 | 63419.02344 |
| SNX9 | Q9Y5X1 | SNX9 | Sorting nexin-9 | 21744.23438 | 21572.60938 |
| IPO13 | O94829 | IPO13 | Importin-13 | 20046.36328 | 23733.33789 |
| TRNAU1AP | Q9NX07 | TRNAU1AP | tRNA selenocysteine 1 | 4505.061035 | 10304.37695 |
| ACAD8 | Q9UKU7 | ACAD8 | Isobutyryl-CoA dehydı | 36423.12891 | 36260.9375 |
| NKIRAS2 | K7ERG2;Q9NYR9 | NKIRAS2 | NF-kappa-B inhibitor-i | 22770.61523 | 22866.04102 |
| C3 | P01024 | C3 | Complement C3 | 76565.04688 | 69041.82813 |
| PTPN12 | Q05209 | PTPN12 | Tyrosine-protein phos | 8037.928711 | 9306.869141 |
| BCAP29 | Q9UHQ4 | BCAP29 | B-cell receptor-associ | 11042.20508 | 10803.85156 |
| CHTF8 | J3KSJ7;P0CG13 | CHTF8 | Chromosome transmi | 9335.339844 | 11448.41406 |
| HOOK3 | H0YDM4 | HOOK3 | Protein Hook homolog | 24853.96484 | 27230.60229 |
| PLG | P00747 | PLG | Plasminogen | 19743.36719 | 20198.20313 |
| SCP2 | P22307 | SCP2 | Non-specific lipid-tran | 419365.1563 | 421843.125 |
| FOXO3 | O43524 | FOXO3 | Forkhead box protein | 8764.666992 | 11336.54199 |
| SH3BP4 | Q9P0V3 | SH3BP4 | SH3 domain-binding p | 12868.89941 | 10138.44141 |
| ARL6IP4 | Q66PJ3 | ARL6IP4 | ADP-ribosylation facto | 39611.24609 | 50658.35547 |
| KRIT1 | A0A0C4DG23;C9J718;C9JF32 | KRIT1 | Krev interaction trapp | 5474.42041 | 6381.050781 |
| FAM162A | F8W7Q4;Q96A26 | FAM162A | Protein FAM162A | 111336.8438 | 132327.9063 |
| FAM160B1 | Q5W0V3 | FAM160B1 | Protein FAM160B1 | 20819.76563 | 19910.46289 |
| LYL1 | K7ER61 | LYL1 | Protein lyl-1 (Fragmer | 2878724.5 | 3284710.5 |
| ARHGEF1 | M0QZR4;Q92888 | ARHGEF1 | Rho guanine nucleotic | 112164.4141 | 120367.4141 |

| | | | | | |
|---|---|---|---|---|---|
| SIRT3 | E9PM75;Q9NTG7 | SIRT3 | NAD-dependent prote | 14666.04004 | 11938.45117 |
| PIP5K1C | O60331 | PIP5K1C | Phosphatidylinositol 4 | 5661.221191 | 6245.126953 |
| RDH11 | Q8TC12 | RDH11 | Retinol dehydrogenas | 112098.3281 | 141057.4844 |
| CYP20A1 | Q6UW02 | CYP20A1 | Cytochrome P450 20A | 16685.72852 | 16657.29297 |
| DYNC1H1 | Q14204 | DYNC1H1 | Cytoplasmic dynein 1 | 402794.4688 | 451655.0938 |
| SLX4IP | Q5VYV7 | SLX4IP | Protein SLX4IP | 894.6401367 | 1156.190063 |
| ARGLU1 | Q9NWB6 | ARGLU1 | Arginine and glutamat | 84034.95313 | 60861.15625 |
| ALB | A0A0C4DGB6;H0YA55;P0276 | ALB | Serum albumin;Serum | 14487474 | 12187360 |
| POLE2 | P56282 | POLE2 | DNA polymerase epsil | 13210.14063 | 17717.37109 |
| AARS1 | P49588 | AARS1 | Alanine--tRNA ligase, | 205471.1875 | 225591.1563 |
| COASY | Q13057 | COASY | Bifunctional coenzym | 967473.0625 | 1072681.75 |
| TTPAL | Q9BTX7 | TTPAL | Alpha-tocopherol trar | 13225.63281 | 13658.82324 |
| CHMP4C | Q96CF2 | CHMP4C | Charged multivesicula | 10498.40625 | 15628.23242 |
| EBAG9 | E9PN10;E9PND3;O00559 | EBAG9 | Receptor-binding canc | 48187.25781 | 59074.79688 |
| SIRT2 | Q8IXJ6 | SIRT2 | NAD-dependent prote | 27202.0918 | 29363.48828 |
| STK32C | Q86UX6 | STK32C | Serine/threonine-prot | 15283.37598 | 17603.90234 |
| BDH1 | Q02338 | BDH1 | D-beta-hydroxybutyra | 210322.2188 | 211865.9688 |
| ECH1 | Q13011 | ECH1 | Delta(3,5)-Delta(2,4)-c | 210200.4219 | 234321.8281 |
| SLC25A5 | P05141 | SLC25A5 | ADP/ATP translocase | 4354488 | 5048970 |
| ARHGEF6 | Q15052 | ARHGEF6 | Rho guanine nucleotic | NA | 2333.350098 |
| MICOS10;MICOS10;MICOS1 | A0A087WU07;Q5TGZ0;R4GN | MICOS10;MICOS10;M | MICOS complex subur | 21808.60742 | 25601.96289 |
| TMEM41A | Q96HV5 | TMEM41A | Transmembrane prote | 23681.59375 | 22282.18359 |
| HIST2H2AC;HIST2H2AA3 | Q16777;Q6FI13 | HIST2H2AC;HIST2H2A | Histone H2A type 2-C; | 16115.87012 | 711.5388794 |
| GRAMD1A | M0QZ12;Q96CP6 | GRAMD1A | Protein Aster-A | 5038.179199 | 5363.061523 |
| SDK1 | A0A087WTQ6;F8W6X9;Q7Z5 | SDK1 | Protein sidekick-1 | 28354.28711 | 26248.91602 |
| AASS | Q9UDR5 | AASS | Alpha-aminoadipic ser | 7982.560059 | 11743.50879 |
| EIF2AK1 | Q9BQI3 | EIF2AK1 | Eukaryotic translation | 8421.449219 | 6758.501953 |
| AGPAT2 | O15120 | AGPAT2 | 1-acyl-sn-glycerol-3-pl | 46652.95313 | 35887.54297 |
| KEAP1 | Q14145 | KEAP1 | Kelch-like ECH-associa | 28261.84375 | 25422.43164 |
| AREL1 | G3V506;O15033 | AREL1 | Apoptosis-resistant E3 | 3360.355957 | 2518.737061 |
| USP25 | Q9UHP3 | USP25 | Ubiquitin carboxyl-ter | 5998.43457 | 7341.274902 |
| PI4K2A | Q9BTU6 | PI4K2A | Phosphatidylinositol 4 | 45602.36719 | 41245.76172 |
| CCDC93 | F8W9X7;Q567U6 | CCDC93 | Coiled-coil domain-co | 40447.44922 | 42037.71094 |

| | | | | | |
|---|---|---|---|---|---|
| RBSN | Q9H1K0 | RBSN | Rabenosyn-5 | 10281.07227 | 12594.75781 |
| PEX11G | Q96HA9 | PEX11G | Peroxisomal membrar | 9180.829102 | 6978.046387 |
| TUBGCP5 | A0A087X1Z1;A0A0G2JPW5;A | TUBGCP5 | Gamma-tubulin comp | 1881.228882 | 2231.506104 |
| SLC11A2 | A0A2R8Y5Q7;A0A2R8YD71;F | SLC11A2 | Natural resistance-ass | 28507.08789 | 32449.62695 |
| B3GAT3 | O94766 | B3GAT3 | Galactosylgalactosylxy | 42881 | 45918.11719 |
| MGAT2 | Q10469 | MGAT2 | Alpha-1,6-mannosyl-g | 17681.87109 | 19841.09375 |
| SEC22A | C9JNZ0;C9JRY4;Q96IW7 | SEC22A | Vesicle-trafficking pro | 46294.39844 | 43184.46875 |
| TBC1D15 | C9JA93 | TBC1D15 | TBC1 domain family m | 74796.6416 | 81853.7085 |
| CDS2 | O95674 | CDS2 | Phosphatidate cytidyl | 13349.69629 | 14499.4541 |
| RAB4A | P20338 | RAB4A | Ras-related protein Ra | 32923.96094 | 35678.16406 |
| HLCS | P50747 | HLCS | Biotin--protein ligase | 4609.131348 | 5570.547363 |
| VPS28 | Q9UK41 | VPS28 | Vacuolar protein sorti | 93749.77344 | 84239.20313 |
| ADD3 | Q9UEY8 | ADD3 | Gamma-adducin | 10973.39355 | 13554.27344 |
| TUBA1C | Q9BQE3 | TUBA1C | Tubulin alpha-1C chai | 265780.7188 | 291790.2188 |
| SLC25A17 | O43808 | SLC25A17 | Peroxisomal membrar | 1978.167847 | 3293.781494 |
| GSTZ1 | A0A0C4DFM0;G3V5T0 | GSTZ1 | Glutathione transfera | 161090.1563 | 146245.9688 |
| OTUD6B | A0A087X0W9;Q8N6M0 | OTUD6B | Deubiquitinase OTUD | 53393.6875 | 59393.56641 |
| ERCC6 | D6R9X7;P0DP91;Q03468 | ERCC6 | Chimeric ERCC6-PGBD | 9026.446289 | 6810.062012 |
| MCF2L | O15068 | MCF2L | Guanine nucleotide ex | NA | 2521.501953 |
| PPM1L | Q5SGD2 | PPM1L | Protein phosphatase 1 | 13379.40234 | 10675.57617 |
| SYTL2 | A0A0U1RQH1 | SYTL2 | Synaptotagmin-like pr | 81121.71924 | 62750.18042 |
| WLS | E9PJW4;E9PKJ0;E9PM62;H0Y | WLS | Protein wntless homo | 3580.993896 | 3915.852051 |
| NCEH1 | A0A0A0MTJ9;A0A0R4J2G3;Q | NCEH1 | Neutral cholesterol es | 32876.10547 | 27851.50586 |
| GLCE | H0YNP1;O94923 | GLCE | D-glucuronyl C5-epim | 26054.40625 | 24000.62891 |
| CACFD1 | Q9UGQ2 | CACFD1 | Calcium channel flowe | 3727.540283 | 4778.667969 |
| BRWD1 | Q9NSI6 | BRWD1 | Bromodomain and WI | 1301.159668 | 5752.245117 |
| ANO10 | Q9NW15 | ANO10 | Anoctamin-10 | 20827.01758 | 18856.00195 |
| SYVN1 | Q86TM6 | SYVN1 | E3 ubiquitin-protein li | 28262.0332 | 36308.14063 |
| NFKB2 | Q00653 | NFKB2 | Nuclear factor NF-kap | 90092.21875 | 78437.88281 |
| PDS5B | A9IYQ1 | PDS5B | Sister chromatid cohe | 90500.95728 | 87838.41162 |
| PANK4 | Q9NVE7 | PANK4 | 4'-phosphopantetheir | 24658.69336 | 26572.32031 |
| VIPAS39 | G3V4K3;Q9H9C1 | VIPAS39 | Spermatogenesis-defe | 37907.20313 | 44795.04688 |
| UNC119B | A6NIH7 | UNC119B | Protein unc-119 homo | 68828.67188 | 78038.84375 |

| | | | | | |
|---|---|---|---|---|---|
| SLC35A4 | L0R6Q1 | SLC35A4 | SLC35A4 upstream op | 93675.95313 | 98057.85156 |
| SNX33 | Q8WV41 | SNX33 | Sorting nexin-33 | 10078.93164 | 11454.85938 |
| ANK3 | Q12955 | ANK3 | Ankyrin-3 | 72940.99219 | 69194.07031 |
| RHOA | P61586 | RHOA | Transforming protein | 305544.1563 | 311761.8125 |
| EIF1B | O60739 | EIF1B | Eukaryotic translation | 20459.14844 | 20289.95508 |
| ARHGAP32 | A7KAX9;G3V174 | ARHGAP32 | Rho GTPase-activating | 6602.555176 | 7355.380859 |
| OGFRL1 | A0A3B3IRI0;Q5TC84 | OGFRL1 | Opioid growth factor r | 1491.021118 | 1269.735229 |
| ZFYVE26 | G3V2D8;Q68DK2 | ZFYVE26 | Zinc finger FYVE doma | 2078.175049 | 2852.432617 |
| FMR1 | G3V0J0 | FMR1 | Fragile X mental retar | 644589.6484 | 674578.9648 |
| C19orf54 | Q5BKX5;Q6ZWG5 | C19orf54 | UPF0692 protein C19o | 5822.734375 | 6459.504395 |
| TJP2 | A0A3B3IRI3 | TJP2 | Tight junction protein | 17094.02148 | 19550.57617 |
| FOXA1 | P55317 | FOXA1 | Hepatocyte nuclear fa | 19546.8418 | 20828.06836 |
| STK16 | B8ZZI5;B8ZZN3;O75716 | STK16 | Serine/threonine-prot | 2596.728027 | 2303.448975 |
| MGST2 | Q99735 | MGST2 | Microsomal glutathion | 234894.1563 | 250754.9688 |
| STX5 | Q13190 | STX5 | Syntaxin-5 | 65626.39844 | 68946.32813 |
| TMEM50B | C9K0I4;F8WBF4;F8WDH1;P5( | TMEM50B | Transmembrane prote | 40353.28125 | 40437.28516 |
| C1QTNF3-AMACR;C1QTNF3 | E9PGA6;Q9BXJ4 | C1QTNF3-AMACR;C1C | C1QTNF3-AMACR rea | 8408.083008 | 7058.79248 |
| TNS2 | F8VV64;Q63HR2 | TNS2 | Tensin-2 | 3506.438965 | 3972.48877 |
| PTBP2 | Q9UKA9 | PTBP2 | Polypyrimidine tract-b | 9143.879883 | 3888.223633 |
| EXOC3 | O60645 | EXOC3 | Exocyst complex comp | 77045.42969 | 89532.98438 |
| ARG2 | P78540 | ARG2 | Arginase-2, mitochond | 2692.875732 | 5553.756348 |
| UBE2J1 | Q9Y385 | UBE2J1 | Ubiquitin-conjugating | 12403.66211 | 11825.1748 |
| CZIB | Q9NWV4 | CZIB | CXXC motif containing | 203173.6094 | 166515.5469 |
| BUD31 | P41223 | BUD31 | Protein BUD31 homol | 108139.1172 | 82181.89063 |
| CBSL;CBS | P0DN79;P35520 | CBSL;CBS | Cystathionine beta-sy | 276822.5938 | 308017.1875 |
| ACAA1 | P09110 | ACAA1 | 3-ketoacyl-CoA thiolas | 115620.6719 | 119948.9609 |
| LARP7 | Q4G0J3 | LARP7 | La-related protein 7 | 51859.69141 | 46376.84766 |
| ;;CFB;CFB | B4E1Z4;E7ETN3;H7C5H1;P00 | ;;CFB;CFB | cDNA FLJ55673, highly | 7056.910156 | 6431.375 |
| AIFM1 | O95831 | AIFM1 | Apoptosis-inducing fa | 426374.5 | 436954.25 |
| POLM | Q9NP87 | POLM | DNA-directed DNA/RN | 6879.933594 | 6076.737305 |
| ARHGAP25 | B7Z8K7;H7C4P1;P42331 | ARHGAP25 | Rho GTPase-activating | 14672.76172 | 14913.36914 |
| RRM2B | Q7LG56 | RRM2B | Ribonucleoside-dipho | 38610.84375 | 38299.32422 |
| LAMTOR4 | Q0VGL1 | LAMTOR4 | Ragulator complex pro | 83495.32813 | 80356.6875 |

| | | | | | |
|---|---|---|---|---|---|
| ZC2HC1A | Q96GY0 | ZC2HC1A | Zinc finger C2HC doma | 13282.44727 | 13106.89063 |
| CNOT6L | H0Y9C1;H0Y9Z5;Q96LI5 | CNOT6L | CCR4-NOT transcriptio | 7155.188965 | 4212.512207 |
| EXOC8 | Q8IYI6 | EXOC8 | Exocyst complex comp | 18756.67188 | 21840.63086 |
| ERLIN2 | E5RHW4;O94905 | ERLIN2 | Erlin-2 (Fragment);Erli | 63753.67969 | 88716.83594 |
| ATP1B1 | P05026 | ATP1B1 | Sodium/potassium-tra | 322913.1875 | 241991.9844 |
| SMARCAD1 | Q9H4L7 | SMARCAD1 | SWI/SNF-related matr | 14382.83105 | 20093.18164 |
| OVOL2 | Q9BRP0 | OVOL2 | Transcription factor O | 753.2295532 | 1685.402832 |
| ITIH2 | P19823;Q5T985 | ITIH2 | Inter-alpha-trypsin inh | 145526.6094 | 123400.4141 |
| PPM1H | Q9ULR3 | PPM1H | Protein phosphatase 1 | 28524.92969 | 34445.95703 |
| MSH3 | P20585 | MSH3 | DNA mismatch repair | 19498.29883 | 23353.11523 |
| NPEPL1 | Q8NDH3 | NPEPL1 | Probable aminopeptic | 106444.3828 | 115004.9141 |
| PDCD6IP | Q8WUM4 | PDCD6IP | Programmed cell deat | 406351 | 440241.6563 |
| NXT2 | Q9NPJ8 | NXT2 | NTF2-related export p | 23308.5332 | 15321.17188 |
| TBC1D22A | Q8WUA7 | TBC1D22A | TBC1 domain family m | 10189.55273 | 14857.83789 |
| AGFG2 | O95081 | AGFG2 | Arf-GAP domain and F | 11292.34766 | 12801.58496 |
| TDRD7 | Q8NHU6 | TDRD7 | Tudor domain-contair | 9887.713867 | 11534.21875 |
| SYS1;SYS1-DBNDD2;SYS1-D | E7ETM6;F2Z2A3;H3BUS1;Q5( | SYS1;SYS1-DBNDD2;SY | Protein SYS1 homolog | NA | 13141.12793 |
| EXOC7 | A0A0A0MSB8;Q9UPT5 | EXOC7 | Exocyst complex comp | 74681.00781 | 82042.41406 |
| PLPBP | O94903 | PLPBP | Pyridoxal phosphate h | 164399.1875 | 160927.0156 |
| GRK3 | P35626 | GRK3 | Beta-adrenergic recep | 1999.2323 | 2904.953125 |
| MEPCE | Q7L2J0 | MEPCE | 7SK snRNA methylpho | 85237.80469 | 88001.39844 |
| ARHGEF16 | Q5VV41 | ARHGEF16 | Rho guanine nucleotic | 77400.74219 | 72994.35938 |
| HSD17B7 | P56937 | HSD17B7 | 3-keto-steroid reducta | 11040.47559 | 19836.66992 |
| COL1A1 | P02452 | COL1A1 | Collagen alpha-1(I) ch | 9172.007813 | 6568.441406 |
| ADCK1 | Q86TW2 | ADCK1 | AarF domain-containi | 11606.0625 | 9568.117188 |
| TMEM184A | Q6ZMB5 | TMEM184A | Transmembrane prote | 18997.16211 | 3083.124023 |
| THBS4 | E7ES19;P35443 | THBS4 | Thrombospondin-4 | 4819.321777 | 3783.12793 |
| PYGB | P11216 | PYGB | Glycogen phosphoryla | 280105.75 | 290822.5625 |
| GPR89A;GPR89B | B7ZAQ6;P0CG08 | GPR89A;GPR89B | Golgi pH regulator A;G | 70201.49219 | 59165.87109 |
| FAM118A | B0QY29;Q9NWS6 | FAM118A | Protein FAM118A | 9013.12207 | 6662.483887 |
| HMCN1 | Q96RW7 | HMCN1 | Hemicentin-1 | 189667.5781 | 198170.2031 |
| SAMD12 | B8ZZB7;F8VYB8;H0YEJ0;Q8N | SAMD12 | Sterile alpha motif do | 2391.674805 | 6863.741699 |
| ;LRRC57 | A0A3B3ISV5;Q8N9N7 | ;LRRC57 | Uncharacterized prote | 43899.67188 | 50249.37891 |

| | | | | | |
|---|---|---|---|---|---|
| CRLF3 | Q8IUI8 | CRLF3 | Cytokine receptor-like | 3459.533203 | 5804.786133 |
| EXOC2 | Q96KP1 | EXOC2 | Exocyst complex comp | 119173.9219 | 140084.9219 |
| CD55 | B1AP13;H3BLV0;H7BY55;P08 | CD55 | Complement decay-ac | 37720.82031 | 29982.92969 |
| CXorf56 | Q9H5V9 | CXorf56 | UPF0428 protein CXor | 32408.98047 | 23509.08008 |
| SLC29A2 | G5E943;Q14542 | SLC29A2 | Equilibrative nucleosi | 10549.35449 | 15147.34863 |
| CDK5RAP3 | Q96JB5 | CDK5RAP3 | CDK5 regulatory subu | 43783.03125 | 58460.30469 |
| KRT80 | Q6KB66 | KRT80 | Keratin, type II cytosk | 71821.91406 | 43217.82422 |
| DLST | P36957 | DLST | Dihydrolipoyllysine-re | 575101.1875 | 606316.125 |
| ASF1B | Q9NVP2 | ASF1B | Histone chaperone AS | 127030.8281 | 136282.1094 |
| DOCK6 | Q96HP0 | DOCK6 | Dedicator of cytokines | 7188.851563 | 8179.655273 |
| ATG7 | O95352 | ATG7 | Ubiquitin-like modifie | 78011.96875 | 94744.5 |
| LRPAP1 | P30533 | LRPAP1 | Alpha-2-macroglobuli | 88244.25 | 102415.3594 |
| NCKIPSD | Q9NZQ3 | NCKIPSD | NCK-interacting prote | 31228.5332 | 36740.64844 |
| AK1 | P00568;Q5T9B7 | AK1 | Adenylate kinase isoe | 295754 | 279458.1563 |
| CTNNB1 | A0A2R8Y543;A0A2R8Y5A3;A( | CTNNB1 | Catenin beta-1;Cateni | 147269.3594 | 193874.7031 |
| ITPR3 | Q14573 | ITPR3 | Inositol 1,4,5-trisphos | 111106.3047 | 135340.8125 |
| KALRN | H7BXZ5;O60229 | KALRN | Kalirin (Fragment);Kal | 71966.09375 | 97472.52344 |
| HDAC6 | Q9UBN7 | HDAC6 | Histone deacetylase 6 | 16691.08398 | 25199.17383 |
| SEC23A | F5H365;Q15436 | SEC23A | Protein transport prot | 125332.2031 | 131771.75 |
| ZHX1 | Q9UKY1 | ZHX1 | Zinc fingers and home | 29782.08398 | 23723.16211 |
| GGACT | Q9BVM4 | GGACT | Gamma-glutamylamir | 13798.61816 | 15219.90527 |
| RNASEL | Q05823 | RNASEL | 2-5A-dependent ribor | 6998.068848 | 10037.68555 |
| MIDEAS | A0A1C7CYX1 | MIDEAS | Mitotic deacetylase-as | 46936.875 | 46484.64844 |
| ATE1 | F5GXE4 | ATE1 | Arginyl-tRNA--protein | 30085.08594 | 31190.25977 |
| SFT2D3 | Q587I9 | SFT2D3 | Vesicle transport prot | 32017.92188 | 37403.48047 |
| PASK | Q96RG2 | PASK | PAS domain-containin | 6944.453125 | 6922.143066 |
| ARAF | P10398 | ARAF | Serine/threonine-prot | 48083.11328 | 53580.8125 |
| ATG2A | Q2TAZ0 | ATG2A | Autophagy-related pr | 12062.62793 | 17286.88672 |
| NCKAP5L | H7C1V4;Q9HCH0 | NCKAP5L | Nck-associated protei | 5363.814941 | 5500.208984 |
| ETV3 | P41162 | ETV3 | ETS translocation vari | 2152.887207 | 2307.794678 |
| PGM2 | Q96G03 | PGM2 | Phosphoglucomutase- | 189190.5781 | 235037.1094 |
| ZDHHC12 | Q5T266;Q5T269;Q96GR4;X6I | ZDHHC12 | Palmitoyltransferase ( | 5209.494629 | 5418.699707 |
| MIA2 | G3V599;Q96PC5 | MIA2 | Melanoma inhibitory a | 34166.32422 | 32870.92578 |

| | | | | | |
|---|---|---|---|---|---|
| NGLY1 | Q96IV0 | NGLY1 | Peptide-N(4)-(N-acety | 14911.65625 | 13354.38574 |
| TOB2 | Q14106 | TOB2 | Protein Tob2 | 3080.850586 | 1941.778687 |
| CHP1 | Q99653 | CHP1 | Calcineurin B homolog | 19112.67969 | 20872.68945 |
| GNA11 | P29992 | GNA11 | Guanine nucleotide-b | 154266.375 | 184702.9688 |
| EXOC4 | Q96A65 | EXOC4 | Exocyst complex comp | 56717.375 | 72157.69531 |
| TRAPPC1 | Q9Y5R8 | TRAPPC1 | Trafficking protein par | 78163.80469 | 75008.67188 |
| ABCB10 | Q9NRK6 | ABCB10 | ATP-binding cassette s | 40253.22266 | 37983.63672 |
| EMC9 | H0YNH6;Q9Y3B6 | EMC9 | ER membrane protein | 17064.71094 | 18950.19922 |
| PYCR2 | Q96C36 | PYCR2 | Pyrroline-5-carboxylat | 47662.35547 | 49468.57422 |
| CHKB | Q9Y259 | CHKB | Choline/ethanolamine | 7034.667969 | 7674.36377 |
| RAB9B | Q9NP90 | RAB9B | Ras-related protein Ra | 6487.698242 | 6027.824219 |
| NIPSNAP1 | Q9BPW8 | NIPSNAP1 | Protein NipSnap homo | 339671.2813 | 348754.7188 |
| BGN | P21810 | BGN | Biglycan | 29053.06445 | 17966.98828 |
| ANXA5 | D6RBL5 | ANXA5 | Annexin | 2515003.453 | 2625775.938 |
| PSPH | C9JBI3;P78330 | PSPH | Phosphoserine phospl | 118024.3125 | 123759.4844 |
| RGL2 | O15211 | RGL2 | Ral guanine nucleotide | 12183.00195 | 15375.5293 |
| EXOC6 | Q8TAG9 | EXOC6 | Exocyst complex comp | 23203.36719 | 27426.31836 |
| NUDT9 | Q9BW91 | NUDT9 | ADP-ribose pyrophosp | 48787.75781 | 45513.33594 |
| TMCO1 | Q9UM00 | TMCO1 | Calcium load-activated | 217956.5 | 257898.7656 |
| CD276 | A0A0C4DGH0;Q5ZPR3 | CD276 | CD276 antigen | 53812.40234 | 85552.89063 |
| FBXO17 | M0QYI3;M0QYV7;Q96EF6 | FBXO17 | F-box only protein 17 | 10122.58789 | 10666.62402 |
| CNIH4 | A6NLH6;Q9P003 | CNIH4 | Protein cornichon hor | 248968.25 | 226133.0938 |
| FBXL8 | Q96CD0 | FBXL8 | F-box/LRR-repeat prot | 3156.59082 | 3757.037842 |
| PSAP | C9JIZ6;P07602;Q5BJH1 | PSAP | Prosaposin;Prosaposin | 60112.17188 | 62619.05859 |
| PRMT6 | Q96LA8 | PRMT6 | Protein arginine N-me | 159661.4531 | 147195.2969 |
| A2M | P01023 | A2M | Alpha-2-macroglobuli | 492663.375 | 444444.3438 |
| PDCL | Q13371 | PDCL | Phosducin-like proteir | 21927.57617 | 27717.43945 |
| MB | B0QYF8;P02144 | MB | Myoglobin (Fragment | 15998.81152 | 15773.60254 |
| SETD1B | A0A0A0MQV9;Q9UPS6 | SETD1B | Histone-lysine N-meth | 4896.953125 | 4928.826172 |
| GLO1 | Q04760 | GLO1 | Lactoylglutathione lya | 891259.125 | 1094463.375 |
| PACSIN2 | Q9UNF0 | PACSIN2 | Protein kinase C and c | 68153.17969 | 72287.99219 |
| NANP | Q8TBE9 | NANP | N-acylneuraminate-9- | 8614.321289 | 8495.991211 |
| APPL2 | Q8NEU8 | APPL2 | DCC-interacting prote | 15254.67676 | 17744.13281 |

| | | | | | |
|---|---|---|---|---|---|
| ZBTB17 | Q13105 | ZBTB17 | Zinc finger and BTB dc | 194.9720306 | 425.5244751 |
| DPY19L4 | E5RGB7;E5RGH1;H0YBV9;Q7 | DPY19L4 | Probable C-mannosylt | 20219.48828 | 17458.80469 |
| TBK1 | Q9UHD2 | TBK1 | Serine/threonine-prot | 37125.61328 | 42980.32031 |
| COPG1 | Q9Y678 | COPG1 | Coatomer subunit gar | 359749.3438 | 403103.0938 |
| WBP2 | K7EMC9;Q969T9 | WBP2 | WW domain-binding p | 100333.0078 | 122453.9375 |
| GC | D6RF35;P02774 | GC | Vitamin D-binding pro | 189931.4063 | 151836.7969 |
| TUFT1 | Q9NNX1 | TUFT1 | Tuftelin | 8464.141602 | 8949.032227 |
| LRRFIP2 | Q9Y608 | LRRFIP2 | Leucine-rich repeat fli | 47890.48438 | 56906.09375 |
| DTD1 | Q8TEA8 | DTD1 | D-aminoacyl-tRNA dea | 170345.8906 | 182202.25 |
| FBLN1 | P23142 | FBLN1 | Fibulin-1 | 45345.62109 | 37323.60156 |
| AMACR | D6RB81;Q9UHK6 | AMACR | Alpha-methylacyl-CoA | 3157.539795 | 5073.922363 |
| UCK1 | Q9HA47 | UCK1 | Uridine-cytidine kinas | 63600.96875 | 53125.07422 |
| ARHGDIA | P52565 | ARHGDIA | Rho GDP-dissociation | 1545980.875 | 1615056.375 |
| FOXRED1 | Q96CU9 | FOXRED1 | FAD-dependent oxido | 24896.75977 | 25310.72852 |
| LRRC45 | Q96CN5 | LRRC45 | Leucine-rich repeat-cc | 2404.740479 | 1292.247192 |
| SLC25A12 | O75746 | SLC25A12 | Calcium-binding mitoc | 18936.50977 | 18120.50391 |
| ATP6V0C | P27449 | ATP6V0C | V-type proton ATPase | 129183.5078 | 96705.32813 |
| SLC6A6 | A0A087WY96;A0A087WYN0; | SLC6A6 | Transporter;Transport | 5591.285156 | 8679.46875 |
| NFKBIA | P25963 | NFKBIA | NF-kappa-B inhibitor a | 17084.25977 | 20508.38281 |
| PHF2 | O75151 | PHF2 | Lysine-specific demetl | 37751.49219 | 37140.10938 |
| PPP1R12C | K7EL81 | PPP1R12C | Protein phosphatase 1 | 23508.9624 | 24938.41406 |
| WIPI1 | G5EA37;K7ELY0;Q5MNZ9 | WIPI1 | WD repeat domain ph | 7274.066406 | 12187.00391 |
| PSMG3 | Q9BT73 | PSMG3 | Proteasome assembly | 46910.83984 | 42413.76953 |
| DPAGT1 | A0A1W2PPC6;Q9H3H5 | DPAGT1 | UDP-N-acetylglucosar | 12056.14453 | 12662.12695 |
| GAB2 | Q9UQC2 | GAB2 | GRB2-associated-bind | 9215.625977 | 8362.189453 |
| NQO1 | P15559 | NQO1 | NAD(P)H dehydrogena | 336870.7813 | 396050.375 |
| EI24 | E9PK61;E9PL33;E9PM05;E9P: | EI24 | Etoposide-induced pro | 22604.03516 | 30249.95898 |
| ASCC3 | Q8N3C0 | ASCC3 | Activating signal coint | 38714.22266 | 45355.60938 |
| TMEM205 | K7ELQ9;Q6UW68 | TMEM205 | Transmembrane prote | 257115.9688 | 251574.4844 |
| CBR3 | O75828 | CBR3 | Carbonyl reductase [N | 151991.7188 | 131874.0469 |
| LTF | E7EQB2;E7ER44;P02788 | LTF | Lactotransferrin (Frag | 1468134.625 | 1004043.063 |
| RFC3 | P40938 | RFC3 | Replication factor C su | 67163.14063 | 73602.07813 |
| NPDC1 | Q5SPY9;Q9NQX5 | NPDC1 | Neural proliferation d | 9477.223633 | 7119.393066 |

| | | | | | |
|---|---|---|---|---|---|
| UROS | P10746 | UROS | Uroporphyrinogen-III | 4813.077148 | 7834.660645 |
| BCAS3 | Q9H6U6 | BCAS3 | Breast carcinoma-amp | 34918.48047 | 42448.37109 |
| PIP4P1 | Q86T03 | PIP4P1 | Type 1 phosphatidylin | 57174.37109 | 59611.39844 |
| SNX21 | Q5JZH3;Q969T3 | SNX21 | Sorting nexin-21 | 4617.036621 | 5962.912109 |
| GRHL2 | Q6ISB3 | GRHL2 | Grainyhead-like prote | 57036.70703 | 61470.48438 |
| SRP14 | P37108 | SRP14 | Signal recognition par | 669946.0625 | 674074.1875 |
| BRMS1L | Q5PSV4 | BRMS1L | Breast cancer metasta | 20344.42969 | 15359.65137 |
| HIRIP3 | Q9BW71 | HIRIP3 | HIRA-interacting prote | 18103.6582 | 16002.8125 |
| SCRN3 | Q0VDG4 | SCRN3 | Secernin-3 | 19515.34766 | 20890.19727 |
| ATP6V1C1 | P21283 | ATP6V1C1 | V-type proton ATPase | 124472.5547 | 120240.9063 |
| GMPPA | Q96IJ6 | GMPPA | Mannose-1-phosphate | 75789.77344 | 87550.63281 |
| EIF4A2 | Q14240 | EIF4A2 | Eukaryotic initiation fa | 85379.89063 | 87586.69531 |
| ROMO1 | P60602 | ROMO1 | Reactive oxygen speci | 74197.64063 | 73791.10156 |
| ACAP2 | Q15057 | ACAP2 | Arf-GAP with coiled-co | 40378.1875 | 45139.16406 |
| ALKBH4 | Q9NXW9 | ALKBH4 | Alpha-ketoglutarate-d | 26804.76953 | 25867.80078 |
| RER1 | A0A0A0MR06;O15258;Q5T0 | RER1 | Protein RER1 (Fragme | 68094.92969 | 69000.90625 |
| RAB6B | Q9NRW1 | RAB6B | Ras-related protein Ra | 10416.72363 | 16305.85059 |
| AP1M2 | Q9Y6Q5 | AP1M2 | AP-1 complex subunit | 300863.4688 | 327076.625 |
| DST | E9PHM6;F6QMI7;Q03001 | DST | Dystonin | 3342.948242 | 4599.703613 |
| CCDC97 | Q96F63 | CCDC97 | Coiled-coil domain-co | 22999.4668 | 19703.80664 |
| AP1M1 | Q9BXS5 | AP1M1 | AP-1 complex subunit | 94674.88281 | 95689.08594 |
| ANKRD28 | O15084 | ANKRD28 | Serine/threonine-prot | 67594.57031 | 83233.78906 |
| AP1B1 | Q10567 | AP1B1 | AP-1 complex subunit | 380963.5 | 406307.0938 |
| GNA12 | Q03113 | GNA12 | Guanine nucleotide-b | 10905.29883 | 16759.2793 |
| ARHGEF7 | B1ALK7 | ARHGEF7 | Rho guanine nucleotic | 54027.46094 | 54611.51563 |
| RECQL4 | O94761 | RECQL4 | ATP-dependent DNA h | 27792.21094 | 33304.85156 |
| THYN1 | Q9P016 | THYN1 | Thymocyte nuclear pr | 73373.60938 | 77358.39063 |
| TOP2B | Q02880 | TOP2B | DNA topoisomerase 2 | 43187.75 | 36717.31641 |
| RAB3IP | H0YI02;J3KPG1;Q96QF0 | RAB3IP | Rab-3A-interacting pr | 5227.124023 | 4877.260254 |
| PPP1R8 | Q12972 | PPP1R8 | Nuclear inhibitor of pr | 95197.85156 | 79874.44531 |
| PPIL3 | Q9H2H8 | PPIL3 | Peptidyl-prolyl cis-trar | 83138.30469 | 83864.46875 |
| GEMIN2 | A0A3B3IU42 | GEMIN2 | Gem-associated prote | 68316.71094 | 59876.41016 |
| PPP1R21 | Q6ZMI0 | PPP1R21 | Protein phosphatase 1 | 10573.00879 | 12439.11133 |

| | | | | | |
|---|---|---|---|---|---|
| DDR1 | A0A0A0MSX3;A0A0G2JNZ7;( | DDR1 | Epithelial discoidin do | 14003.45313 | 14225.84473 |
| PRMT9 | Q6P2P2 | PRMT9 | Protein arginine N-me | 6427.71582 | 6846.883301 |
| STK17B | O94768;U3KQF8 | STK17B | Serine/threonine-prot | 524.6982422 | 2858.303223 |
| DHRS2 | Q13268 | DHRS2 | Dehydrogenase/reduc | 49804.33203 | 82604.07813 |
| SLC37A4 | O43826;U3KPU7;U3KQS2 | SLC37A4 | Glucose-6-phosphate | 20580.23047 | 26655.07813 |
| KRT19 | K7EMS3 | KRT19 | Keratin, type I cytoske | 24863987.64 | 29134771.52 |
| DCTN2 | Q13561 | DCTN2 | Dynactin subunit 2 | 76887.70313 | 68712.29688 |
| PPM1A | P35813 | PPM1A | Protein phosphatase 1 | 116644.5469 | 111349.4766 |
| ALDH7A1 | A0A1B0GTG2;A0A1B0GW77; | ALDH7A1 | Alpha-aminoadipic sei | 288793.0625 | 301899.625 |
| CASC4 | Q6P4E1 | CASC4 | Protein CASC4 | 43635.14063 | 47295.84766 |
| GPD1L | Q8N335 | GPD1L | Glycerol-3-phosphate | 291448.0938 | 424438.0313 |
| MSI2 | Q96DH6 | MSI2 | RNA-binding protein M | 232852.6719 | 288241.25 |
| SLC25A28 | Q96A46 | SLC25A28 | Mitoferrin-2 | 7144.40625 | 8238.724609 |
| HSPA1B;HSPA1A;HSPA1B | A0A0G2JIW1;P0DMV8;P0DM | HSPA1B;HSPA1A;HSPA | Heat shock 70 kDa pro | 7894271 | 7194001 |
| RMI2 | Q96E14 | RMI2 | RecQ-mediated genor | 46979.94141 | 47471.31641 |
| LAMTOR3 | Q9UHA4 | LAMTOR3 | Ragulator complex pro | 59211.73438 | 55879.41016 |
| BRCC3 | P46736 | BRCC3 | Lys-63-specific deubiq | 36482.59375 | 38209.87109 |
| CDC42EP2 | O14613 | CDC42EP2 | Cdc42 effector proteir | 17668.32227 | 13825.99609 |
| CCDC17 | F2Z395;Q96LX7 | CCDC17 | Coiled-coil domain-co | 38025.55469 | 44774.66406 |
| LRSAM1 | Q6UWE0 | LRSAM1 | E3 ubiquitin-protein li | 56077.34375 | 64878.35547 |
| RCCD1 | A6NED2 | RCCD1 | RCC1 domain-contain | 7656.725098 | 10679.4209 |
| SHROOM2 | Q13796 | SHROOM2 | Protein Shroom2 | 18723.79688 | 16707.54492 |
| SGO1 | Q5FBB7 | SGO1 | Shugoshin 1 | 3912.250488 | 3673.454834 |
| D2HGDH | Q8N465 | D2HGDH | D-2-hydroxyglutarate | 38754.6875 | 46531.74219 |
| TUBGCP6 | Q96RT7 | TUBGCP6 | Gamma-tubulin comp | 5852.915527 | 4748.864258 |
| OSBPL8 | Q9BZF1 | OSBPL8 | Oxysterol-binding pro | 90129.58594 | 85440.55469 |
| RAB31 | Q13636 | RAB31 | Ras-related protein Ra | 868.7218018 | 831.4468994 |
| NEK7 | Q8TDX7 | NEK7 | Serine/threonine-prot | 32002.62305 | 34170.01953 |
| ELMOD2 | D6RBS5;Q8IZ81 | ELMOD2 | ELMO domain-contair | 34387.625 | 38559.1875 |
| THAP11 | Q96EK4 | THAP11 | THAP domain-contain | 42862.71875 | 35854.08203 |
| TAB2 | A0A1B0GV57;Q9NYJ8 | TAB2 | TGF-beta-activated kii | 13292.43262 | 14557.41113 |
| KANK2 | Q63ZY3 | KANK2 | KN motif and ankyrin | 6252.878418 | 4887.072754 |
| AP5M1 | E7EQ45;Q9H0R1 | AP5M1 | AP-5 complex subunit | 10676.08105 | 11633.23145 |

| | | | | | |
|---|---|---|---|---|---|
| TUBGCP3 | Q96CW5 | TUBGCP3 | Gamma-tubulin comp | 16332.60059 | 18564.40234 |
| CHAF1B | Q13112 | CHAF1B | Chromatin assembly f | 15027.47754 | 18128.02344 |
| TFB1M | Q8WVM0 | TFB1M | Dimethyladenosine tr | 12341.125 | 14216.41895 |
| CHEK1 | E7EPP6;O14757 | CHEK1 | Serine/threonine-prot | 16650.35156 | 14182.65918 |
| DPM1 | H0Y368;O60762;Q5QPK2 | DPM1 | Dolichol-phosphate m | 310365.25 | 309104.2188 |
| RFC4 | P35249 | RFC4 | Replication factor C su | 108481.2969 | 111847.7188 |
| ZXDC;ZXDA;ZXDB;ZXDC | H0YAA9;P98168;P98169;Q2C | ZXDC;ZXDA;ZXDB;ZXD | Zinc finger protein ZXI | 3128.985107 | 2741.244385 |
| ABRAXAS1 | D6REL5;Q6UWZ7 | ABRAXAS1 | BRCA1-A complex sub | 8571.09668 | 6828.52832 |
| POLD4 | Q9HCU8 | POLD4 | DNA polymerase delta | 14818.16406 | 15811.4043 |
| TJP2;;TJP2;TJP2 | A0A1B0GTW1;A0A2R8YDH4; | TJP2;;TJP2;TJP2 | Tight junction protein | 92545.72656 | 96800.34375 |
| PZP | P20742 | PZP | Pregnancy zone prote | 172142.0156 | 151246.4063 |
| SLC39A9 | Q9NUM3 | SLC39A9 | Zinc transporter ZIP9 | 2925.271484 | 4335.817383 |
| IQGAP3 | Q86VI3 | IQGAP3 | Ras GTPase-activating | 31747.79492 | 38617.73438 |
| MFSD8 | A0A286YEV4;A0A286YEW8;A | MFSD8 | Major facilitator super | 4528.137695 | 5676.411621 |
| GLE1 | Q53GS7 | GLE1 | Nucleoporin GLE1 | 21218.68945 | 21709.0293 |
| MEX3D | Q86XN8;S4R446 | MEX3D | RNA-binding protein M | 10520.92871 | 15229.98828 |
| ZNF652 | Q9Y2D9 | ZNF652 | Zinc finger protein 652 | 3601.45874 | 4206.455078 |
| UBA5 | Q9GZZ9 | UBA5 | Ubiquitin-like modifie | 124038.5469 | 145684.3125 |
| TRAPPC4 | Q9Y296 | TRAPPC4 | Trafficking protein par | 78876.1875 | 88831.64844 |
| H1-0 | P07305 | H1-0 | Histone H1.0 | 771264.125 | 71986.35156 |
| PPP1R11 | O60927 | PPP1R11 | E3 ubiquitin-protein li | 27316.23633 | 24672.22656 |
| MAL2 | Q969L2 | MAL2 | Protein MAL2 | 320784.3125 | 265438.7188 |
| LSR | Q86X29;S4R3V8 | LSR | Lipolysis-stimulated li | 105688.2422 | 108667.8828 |
| VPS39 | Q96JC1 | VPS39 | Vam6/Vps39-like prot | 31849.16211 | 40224.13672 |
| DNM1 | A0A0D9SFE4;A0A0U1RQP1;C | DNM1 | Dynamin-1 | 40730.29297 | 41589.71875 |
| ITPRID2 | E7EUL7;E9PHV5;P28290 | ITPRID2 | Protein ITPRID2 | 5313.967285 | 2623.90332 |
| TOR1B | O14657 | TOR1B | Torsin-1B | 22070.19141 | 21425.4707 |
| TXNRD2 | A0A182DWF2;A0A182DWF3; | TXNRD2 | Thioredoxin reductase | 30357.58594 | 38188.35547 |
| ZDHHC21 | Q8IVQ6 | ZDHHC21 | Palmitoyltransferase Z | 4916.745117 | 7615.040039 |
| BLOC1S6 | H3BST1;Q9UL45 | BLOC1S6 | Biogenesis of lysosom | 13977.56055 | 14375.2666 |
| THOC2 | Q8NI27 | THOC2 | THO complex subunit | 184116.0156 | 149309.6875 |
| SMS | P52788 | SMS | Spermine synthase | 295834.7188 | 303638.8125 |
| CRY2 | A0A0D2X7Z3;Q49AN0 | CRY2 | Cryptochrome-2 | 9394.491211 | 7537.493652 |

| | | | | | |
|---|---|---|---|---|---|
| RALGAPB | Q86X10 | RALGAPB | Ral GTPase-activating | 21091.49023 | 24507.92773 |
| ZNF768 | H3BS42;Q9H5H4 | ZNF768 | Zinc finger protein 768 | 10626.23242 | 9326.449219 |
| LYPLA1 | O75608 | LYPLA1 | Acyl-protein thioester | 305667.5 | 411280.25 |
| TADA2B | Q86TJ2 | TADA2B | Transcriptional adapte | 26039.49219 | 28390.27148 |
| BRIP1 | Q9BX63 | BRIP1 | Fanconi anemia group | 128906.9375 | 126205.3906 |
| TTC9C | Q8N5M4 | TTC9C | Tetratricopeptide repe | 42680.25 | 49513.29688 |
| NSF | I3L0N3;P46459 | NSF | Vesicle-fusing ATPase | 192841.8281 | 236157.9375 |
| ZFPL1 | O95159 | ZFPL1 | Zinc finger protein-like | 64259.88281 | 64513.0625 |
| COMMD7 | A0A2R8Y455;A0A2R8Y600;A( | COMMD7 | COMM domain-conta | 17851.00586 | 22931.36914 |
| RAB13 | P51153 | RAB13 | Ras-related protein Ra | 179855.5 | 199936.5 |
| TPRG1L | Q5T0D9 | TPRG1L | Tumor protein p63-re | 25317.05664 | 25967.47266 |
| MAP3K20 | Q9NYL2 | MAP3K20 | Mitogen-activated pro | 10503.19824 | 9727.324219 |
| SEPTIN8 | A6NFQ9;A6NMH6;F8W8I8;Q | SEPTIN8 | Septin-8 | 83887.08594 | 94832.13281 |
| DNAJC21 | Q5F1R6 | DNAJC21 | DnaJ homolog subfam | 27309.53906 | 26490.82031 |
| LIMS1 | C9JF46 | LIMS1 | LIM and senescent cel | 129793.3203 | 138420.5352 |
| MTARC2 | Q969Z3 | MTARC2 | Mitochondrial amidox | 22852.19922 | 18691.30859 |
| CCDC186 | Q7Z3E2 | CCDC186 | Coiled-coil domain-co | 8691.348633 | 10215.11523 |
| STRN3 | Q13033 | STRN3 | Striatin-3 | 38818.16016 | 39051.49219 |
| POLR2K | P53803 | POLR2K | DNA-directed RNA po | 68062.82813 | 55948.44922 |
| VPS8 | C9JJN9;Q8N3P4 | VPS8 | Vacuolar protein sorti | 13886.2998 | 17882.53906 |
| NEMP1 | O14524 | NEMP1 | Nuclear envelope inte | 19067.79688 | 21514.09375 |
| MORC3 | Q14149 | MORC3 | MORC family CW-type | 18328.39453 | 18031.58203 |
| HES1 | Q14469 | HES1 | Transcription factor H | 4328.097656 | 5832.132324 |
| NCK1 | P16333 | NCK1 | Cytoplasmic protein N | 66833.5 | 64882.14453 |
| USP9Y | O00507 | USP9Y | Probable ubiquitin car | 70647.61719 | 90518.28125 |
| PALM3 | A0A590UJ36 | PALM3 | Paralemmin-3 | 7663.678711 | 7620.85498 |
| PPA2 | A0A0C4DGB9 | PPA2 | Inorganic pyrophosph | 210592.5723 | 242443.0703 |
| DR1 | Q01658 | DR1 | Protein Dr1 | 8042.508789 | 6273.52002 |
| FAM120B | A0A0D9SEJ5;F5GY05;Q96EK7 | FAM120B | Constitutive coactivat | 37482.57813 | 39141.62891 |
| NUP210 | Q8TEM1 | NUP210 | Nuclear pore membra | 176311.1094 | 182585.9531 |
| SESN2 | P58004 | SESN2 | Sestrin-2 | 18998.46094 | 15694.56543 |
| HPRT1 | P00492 | HPRT1 | Hypoxanthine-guanine | 697694.625 | 706797.625 |
| HDDC3 | Q8N4P3 | HDDC3 | Guanosine-3',5'-bis(di | 24004.75781 | 22773.24023 |

| | | | | | |
|---|---|---|---|---|---|
| WDR11 | Q9BZH6 | WDR11 | WD repeat-containing | 55660.23047 | 54299.90234 |
| PDPR | Q8NCN5 | PDPR | Pyruvate dehydrogen | 53396.82813 | 53193.82813 |
| UBXN6 | Q9BZV1 | UBXN6 | UBX domain-containir | 18517.69531 | 19544.80664 |
| GPR107 | G5E994;Q5VW38 | GPR107 | G protein-coupled rec | 22383.76953 | 22761.39844 |
| PAWR | Q96IZ0 | PAWR | PRKC apoptosis WT1 r | 160778.7656 | 164669.375 |
| GNB2 | P62879 | GNB2 | Guanine nucleotide-b | 779518.875 | 881377.6875 |
| KIAA1522 | Q9P206 | KIAA1522 | Uncharacterized prote | 144221.4219 | 146725.9219 |
| HSPA6 | P17066 | HSPA6 | Heat shock 70 kDa pro | 2627529 | 2351150 |
| ALDOA | P04075 | ALDOA | Fructose-bisphosphat | 4850906 | 5406292.5 |
| METTL6 | B4DDX3;M0R0W0;Q8TCB7 | METTL6 | cDNA FLJ57901, highly | NA | 3080.05835 |
| LRRC41 | A0A0B4J2G4;Q15345 | LRRC41 | Leucine-rich repeat-cc | 20448.24414 | 17778.24219 |
| CARHSP1 | Q9Y2V2 | CARHSP1 | Calcium-regulated hea | 120190.4922 | 106777.9844 |
| DENR | O43583 | DENR | Density-regulated pro | 409559.8438 | 406323.5 |
| TTC5 | Q8N0Z6 | TTC5 | Tetratricopeptide rep | 37731.70703 | 41414.00391 |
| COPB1 | P53618 | COPB1 | Coatomer subunit bet | 241875.7344 | 299583 |
| MYO5B | A0A0A0MR36 | MYO5B | Unconventional myos | 19150.43115 | 14335.11108 |
| C5orf51 | A6NDU8 | C5orf51 | UPF0600 protein C5or | 40834.01172 | 54040.72656 |
| CCNDBP1 | O95273 | CCNDBP1 | Cyclin-D1-binding pro | 7753.69873 | 13440.93164 |
| PURA | Q00577 | PURA | Transcriptional activat | 206441.9844 | 205299.6406 |
| ALAD | P13716 | ALAD | Delta-aminolevulinic a | 50347.42578 | 57334.06641 |
| GLB1 | P16278 | GLB1 | Beta-galactosidase | 123429.0938 | 161531.9063 |
| LPCAT1 | Q8NF37 | LPCAT1 | Lysophosphatidylcholi | 187756.25 | 203822.3594 |
| CLEC3A | J3KNC9;O75596 | CLEC3A | C-type lectin domain f | NA | NA |
| TRIP4 | Q15650 | TRIP4 | Activating signal coint | 41241.96094 | 41138.79688 |
| IARS2 | Q9NSE4 | IARS2 | Isoleucine--tRNA ligas | 150526.3125 | 175000.4219 |
| TEPSIN | A0A1B0GV70 | TEPSIN | AP-4 complex accesso | 7993.506348 | 10484.24219 |
| STAT2 | A0A494C164;P52630 | STAT2 | Signal transducer and | 21981.90625 | 32168.25977 |
| TOR1A | O14656 | TOR1A | Torsin-1A | 86451.57031 | 91066.49219 |
| ARHGAP1 | Q07960 | ARHGAP1 | Rho GTPase-activating | 292807.0625 | 289306.7188 |
| PANK1 | Q8TE04 | PANK1 | Pantothenate kinase 1 | 8548.222656 | 3046.495361 |
| RIC8A | Q9NPQ8 | RIC8A | Synembryn-A | 76609.4375 | 84944.64844 |
| PRPS1 | B1ALA9;P60891 | PRPS1 | Ribose-phosphate pyr | 57308.05859 | 61981.41016 |
| POGLUT2 | Q6UW63 | POGLUT2 | Protein O-glucosyltrar | 1136.386353 | 2492.613281 |

| | | | | | |
|---|---|---|---|---|---|
| ITPKC | Q96DU7 | ITPKC | Inositol-trisphosphate | 8155.645508 | 10456.06348 |
| GLT8D1 | Q68CQ7 | GLT8D1 | Glycosyltransferase 8 | 18939.5625 | 21004.73242 |
| SFXN1 | Q9H9B4 | SFXN1 | Sideroflexin-1 | 489860.0938 | 488906.125 |
| AKT2 | P31751 | AKT2 | RAC-beta serine/threo | 49704.71875 | 56252.47266 |
| HPS5 | Q9UPZ3 | HPS5 | Hermansky-Pudlak syr | 3039.603271 | 3691.324463 |
| WDFY1 | Q8IWB7 | WDFY1 | WD repeat and FYVE c | 29325.60352 | 33062.49219 |
| VNN1 | O95497 | VNN1 | Pantetheinase | 7117.233887 | 7836.712891 |
| NCL | P19338 | NCL | Nucleolin | 5276229 | 3675851 |
| LIN7A | O14910 | LIN7A | Protein lin-7 homolog | 89683.14063 | 109729.3594 |
| ITGB5 | P18084 | ITGB5 | Integrin beta-5 | 29080.11523 | 33465.96094 |
| ARHGEF11 | O15085 | ARHGEF11 | Rho guanine nucleotic | 8125.947266 | 8822.337891 |
| FAM120A | A0A494C0Y9 | FAM120A | Constitutive coactivat | 108757.2432 | 125104.1113 |
| GRAMD4 | Q6IC98 | GRAMD4 | GRAM domain-contair | 8018.13623 | 8058.852539 |
| PSMC6 | A0A087X2I1;P62333 | PSMC6 | 26S proteasome regul | 504336.4375 | 509715.2813 |
| TSC1 | A0A2R8Y5N2;A0A2R8Y5S3;A( | TSC1 | Hamartin | 1832.29187 | 3935.621338 |
| TUBB | P07437;Q5JP53 | TUBB | Tubulin beta chain | 7767259 | 8391516 |
| TBX2 | Q13207 | TBX2 | T-box transcription fa | 32734.00195 | 35252.66797 |
| RAB9A | P51151 | RAB9A | Ras-related protein Ra | 64421.89063 | 63101.31641 |
| COQ8B | Q96D53 | COQ8B | Atypical kinase COQ8I | 15364.4375 | 15972.29785 |
| PPP2CB | P62714 | PPP2CB | Serine/threonine-prot | 21071.90039 | 25148.4668 |
| WRB-SH3BGR;GET1;WRB | A0A3B3ISE9;B7Z1T1;O00258 | WRB-SH3BGR;GET1;W | Uncharacterized prote | 43435.79297 | 40093.40234 |
| POLI | J3KQ09;J3KSW2;Q9UNA4;X6I | POLI | DNA polymerase iota; | 5325.039063 | 5274.427246 |
| SEL1L | Q9UBV2 | SEL1L | Protein sel-1 homolog | 71336.41406 | 79570.35156 |
| SFN | P31947 | SFN | 14-3-3 protein sigma | 55234.25 | 84917.36719 |
| RRM1 | P23921 | RRM1 | Ribonucleoside-dipho | 176124.6875 | 224951.1094 |
| EEF2K | O00418 | EEF2K | Eukaryotic elongation | 33506.53125 | 35738.02344 |
| HERC2 | O95714 | HERC2 | E3 ubiquitin-protein li | 11972.88672 | 14315.46191 |
| MACO1 | Q8N5G2 | MACO1 | Macoilin | 28307.14258 | 26928.30273 |
| XYLB | A0A3B3IRM4;O75191 | XYLB | Xylulose kinase | 13696.66309 | 11397.47656 |
| RNF113A | O15541 | RNF113A | E3 ubiquitin-protein li | 46338.04688 | 47221.66406 |
| STK3 | Q13188 | STK3 | Serine/threonine-prot | 60951.29688 | 70306.53125 |
| PFKL | P17858 | PFKL | ATP-dependent 6-pho | 437351.5625 | 523931.875 |
| MTCH1 | Q9NZJ7 | MTCH1 | Mitochondrial carrier | 83836.5625 | 66181.97656 |

| | | | | | |
|---|---|---|---|---|---|
| CGGBP1 | Q9UFW8 | CGGBP1 | CGG triplet repeat-bir | 93641.8125 | 93311.19531 |
| AIF1L | Q9BQI0 | AIF1L | Allograft inflammator | 94547.48438 | 62289.44531 |
| GOLGA5 | Q8TBA6 | GOLGA5 | Golgin subfamily A me | 79257.35156 | 79832.65625 |
| IQGAP1 | P46940 | IQGAP1 | Ras GTPase-activating | 429086.3125 | 368773.125 |
| MCTS1 | Q9ULC4 | MCTS1 | Malignant T-cell-ampl | 353358.5 | 379359.4688 |
| CCDC115 | Q96NT0 | CCDC115 | Coiled-coil domain-co | 24942.62109 | 23841.64648 |
| DDT | P30046 | DDT | D-dopachrome decarb | 748406.125 | 946511.75 |
| RACGAP1 | Q9H0H5 | RACGAP1 | Rac GTPase-activating | 93445.32031 | 80886.54688 |
| RAB3GAP2 | Q9H2M9 | RAB3GAP2 | Rab3 GTPase-activatir | 78079.875 | 86950.5 |
| BROX | Q5VW32 | BROX | BRO1 domain-contain | 58890.11328 | 76978.19531 |
| CSTB | P04080 | CSTB | Cystatin-B | 387114.125 | 498960.125 |
| BPNT1 | O95861 | BPNT1 | 3'(2'),5'-bisphosphate | 112980.4531 | 132690.1563 |
| SPG11 | Q96JI7 | SPG11 | Spatacsin | 8618.021484 | 10782.34668 |
| SLC25A1 | B4DP62 | SLC25A1 | Solute carrier family 2 | 488937.832 | 541418.8438 |
| ATP2C1 | B4E2Q0;H0Y9V7;P98194 | ATP2C1 | Calcium-transporting | 27441.95508 | 30615.22656 |
| NIF3L1 | Q9GZT8 | NIF3L1 | NIF3-like protein 1 | 130017.2656 | 136942.3125 |
| SP3 | A0A494C070;H0Y6K5;H0Y7L | SP3 | Transcription factor S | 13776.74805 | 10662.2373 |
| SLC16A7 | O60669 | SLC16A7 | Monocarboxylate trar | 1902.306763 | 2845.172607 |
| RRAGC | Q9HB90 | RRAGC | Ras-related GTP-bindi | 75654.08594 | 75106.59375 |
| BZW1 | Q7L1Q6 | BZW1 | Basic leucine zipper ar | 286701.2188 | 333178.5625 |
| TMED8 | Q6PL24 | TMED8 | Protein TMED8 | 18847.65039 | 28728.48828 |
| NIBAN2 | Q96TA1 | NIBAN2 | Protein Niban 2 | 73392.38281 | 74586.57031 |
| LANCL1 | O43813 | LANCL1 | Glutathione S-transfer | 123046.8516 | 148106.4375 |
| MIGA1 | A0A2R8YE40;A0A2R8YF99;A( | MIGA1 | Mitoguardin 1 | 2832.998779 | 2886.364014 |
| SLC25A22 | Q9H936 | SLC25A22 | Mitochondrial glutam | 99718.78906 | 98120.82031 |
| CAP1 | Q01518 | CAP1 | Adenylyl cyclase-asso | 605123 | 637059.1875 |
| BCKDK | H3BS02;H3BTL2;O14874 | BCKDK | Protein-serine/threon | 13092.40234 | 15110.8916 |
| LRRC40 | Q9H9A6 | LRRC40 | Leucine-rich repeat-co | 50958.60547 | 54640.99609 |
| UBL4A | P11441 | UBL4A | Ubiquitin-like protein | 130206.6797 | 123184.875 |
| SHISA2 | Q6UWI4 | SHISA2 | Protein shisa-2 homol | 13767.89746 | 9231.335938 |
| PPP1R18 | A0A0G2JHC2;Q6NYC8 | PPP1R18 | PPP1R18;Phostensin | 10083.10742 | 9885.873047 |
| PRC1 | O43663 | PRC1 | Protein regulator of c | 48370.57422 | 47608.95703 |
| FAM174B | Q3ZCQ3 | FAM174B | Membrane protein FA | 16146.38086 | 7076.916992 |

| | | | | | |
|---|---|---|---|---|---|
| ARL8B | Q9NVJ2 | ARL8B | ADP-ribosylation facto | 186468.8906 | 199312.7656 |
| FARSB | Q9NSD9 | FARSB | Phenylalanine--tRNA l | 484450.0625 | 456072.375 |
| CAV1 | Q03135 | CAV1 | Caveolin-1 | 2419.857178 | 2432.374756 |
| RGPD2;RGPD1;RGPD1;RGPI | A0A1W2PNU4;A0A286YES2;I | RGPD2;RGPD1;RGPD1 | RANBP2-like and GRIP | NA | 189.6631927 |
| FAM83G | A6ND36 | FAM83G | Protein FAM83G | 13033.18945 | 12773.89746 |
| FNBP1 | Q96RU3 | FNBP1 | Formin-binding protei | 62796.79688 | 57783.05078 |
| PCBD2 | H0YA52;Q9H0N5 | PCBD2 | Pterin-4-alpha-carbino | 39129.60156 | 33354.80078 |
| EFL1 | Q7Z2Z2 | EFL1 | Elongation factor-like | 62979.85547 | 63713.49219 |
| SPECC1 | Q5M775 | SPECC1 | Cytospin-B | 19812.44531 | 23777.92188 |
| ITGA3 | P26006 | ITGA3 | Integrin alpha-3 | 68801.91406 | 80363.9375 |
| NRM | Q8IXM6 | NRM | Nurim | 4553.936035 | 10435.32715 |
| GMPPB | Q9Y5P6 | GMPPB | Mannose-1-phosphate | 120995.9844 | 143766.5781 |
| WFS1 | O76024 | WFS1 | Wolframin | 40518.41406 | 50217.88281 |
| DERL1 | B4E1G1;Q9BUN8 | DERL1 | Derlin;Derlin-1 | 427168.8125 | 397249.75 |
| HNRNPD | D6RBQ9 | HNRNPD | Heterogeneous nucle | 1305946.9 | 1052026.178 |
| AKAP13 | Q12802 | AKAP13 | A-kinase anchor prote | 21839.32617 | 30860.12695 |
| PKM | H3BR70 | PKM | Pyruvate kinase | 11642780.72 | 13618029.51 |
| TKFC | Q3LXA3 | TKFC | Triokinase/FMN cycla | 194554.7656 | 184733.7188 |
| SMTN | P53814 | SMTN | Smoothelin | 26954.89648 | 17002.75195 |
| SNX6 | Q9UNH7 | SNX6 | Sorting nexin-6 | 107993.2969 | 122026.4063 |
| RFC5 | P40937 | RFC5 | Replication factor C su | 95313.35156 | 99862.05469 |
| TMEM43 | Q9BTV4 | TMEM43 | Transmembrane prote | 313072.8125 | 305928.0313 |
| CEP85L;STMP1;CEP85L | A2A3P3;E0CX11;Q5SZL2 | CEP85L;STMP1;CEP85 | Centrosomal protein c | 3937.7146 | 4088.906494 |
| DAAM1 | Q9Y4D1 | DAAM1 | Disheveled-associated | 11791.25488 | 10784.18945 |
| EML3 | B7WPE2;Q32P44 | EML3 | Echinoderm microtub | 55653.80859 | 58617.26953 |
| MTRES1 | A0A0D9SEI0;Q9P0P8 | MTRES1 | Mitochondrial transcr | 5604.934082 | 8466.449219 |
| USP9X | Q93008 | USP9X | Probable ubiquitin car | 98193.09375 | 126343.6563 |
| TUBA1A | Q71U36 | TUBA1A | Tubulin alpha-1A chai | 11193358 | 15982017 |
| NEU1 | Q99519 | NEU1 | Sialidase-1 | 50661.5 | 44545.22656 |
| PLEKHG3 | A1L390 | PLEKHG3 | Pleckstrin homology d | 22941.79102 | 23627.54102 |
| ARF6 | P62330 | ARF6 | ADP-ribosylation facto | 311151.4375 | 309707.8438 |
| ARFIP1 | P53367 | ARFIP1 | Arfaptin-1 | 137683.6094 | 148286.8594 |
| CLASP2 | E7ERI8;E7EW49 | CLASP2 | CLIP-associating prote | 42883.94434 | 43076.02759 |

| | | | | | |
|---|---|---|---|---|---|
| MMGT1 | Q8N4V1 | MMGT1 | Membrane magnesiur | 42840.79688 | 52082.21094 |
| EYA3 | Q99504 | EYA3 | Eyes absent homolog | 13648.36719 | 17946.50195 |
| ALMS1 | A0A087WTU9;A0A087WV20; | ALMS1 | Alstrom syndrome prc | 266081.375 | 270690.7188 |
| SIPA1L3 | O60292 | SIPA1L3 | Signal-induced prolife | 6458.415039 | 5609.366211 |
| DIABLO | Q9NR28 | DIABLO | Diablo homolog, mito | 211099.5156 | 231049.4375 |
| CHD6 | Q8TD26 | CHD6 | Chromodomain-helica | 8490.092773 | 12164.86035 |
| DGAT1 | O75907 | DGAT1 | Diacylglycerol O-acyltı | 6496.182129 | 6029.316406 |
| ACOT2;ACOT1 | A0A087X0W7;Q86TX2 | ACOT2;ACOT1 | Acyl-coenzyme A thio | 67763.64063 | 70280.42969 |
| TMEM199 | Q8N511 | TMEM199 | Transmembrane prote | 29192.63672 | 31742.61719 |
| SEPTIN2 | Q15019 | SEPTIN2 | Septin-2 | 526864.5625 | 585746.8125 |
| HCFC1 | P51610 | HCFC1 | Host cell factor 1 | 169669.6719 | 184805.7188 |
| WDR7 | A2RRE0;Q9Y4E6 | WDR7 | WD repeat-containing | 10021.19727 | 11435.21289 |
| VPS11 | A0A087WXL6 | VPS11 | Vacuolar protein sorti | 91146.28906 | 101159.6563 |
| ZNF579 | Q8NAF0 | ZNF579 | Zinc finger protein 57! | 25263.85742 | 27431.84375 |
| AKR1C1 | Q04828 | AKR1C1 | Aldo-keto reductase fa | 2462.060059 | 2935.453857 |
| HERPUD1 | H3BP08;H3BTA8;H3BTT7;Q1! | HERPUD1 | Homocysteine-respon | 901.1585083 | 1243.880005 |
| AKTIP | Q9H8T0 | AKTIP | AKT-interacting protei | 26062.93555 | 21956.13867 |
| TCF4 | H3BPJ7;P15884 | TCF4 | Transcription factor 4 | 4024948.25 | 4408425.5 |
| LRR1 | Q96L50 | LRR1 | Leucine-rich repeat pr | 6769.244629 | 4398.599609 |
| ZC3H7B | Q9UGR2 | ZC3H7B | Zinc finger CCCH dom | 23937.2168 | 16892.69922 |
| ACO2 | A2A274;Q99798 | ACO2 | Aconitate hydratase, ı | 396170.5313 | 459768.2813 |
| SEPTIN7 | Q16181 | SEPTIN7 | Septin-7 | 329681.8125 | 329373.3438 |
| FAM89B | Q8N5H3 | FAM89B | Leucine repeat adapte | 1035.574707 | 1514.827393 |
| TMEM179B | Q7Z7N9 | TMEM179B | Transmembrane prote | 38818.52734 | 36168.875 |
| AHNAK2 | Q8IVF2 | AHNAK2 | Protein AHNAK2 | 6085246.5 | 5416163 |
| EHBP1L1 | Q8N3D4 | EHBP1L1 | EH domain-binding pr | 21158.49609 | 25996.15234 |
| PGRMC2 | O15173 | PGRMC2 | Membrane-associatec | 381884.0313 | 354384.9063 |
| BLOC1S2 | Q6QNY1 | BLOC1S2 | Biogenesis of lysosom | 1250522 | 906718.6875 |
| AFP | J3KMX3;P02771 | AFP | Alpha-fetoprotein | 299457.5625 | 238400.75 |
| AGO1 | Q9UL18 | AGO1 | Protein argonaute-1 | 17991.9668 | 15174.11328 |
| VTA1 | A0A087WY55;Q9NP79 | VTA1 | Chromosome 6 open ı | 259684.7031 | 274663.2813 |
| UBE4A | Q14139 | UBE4A | Ubiquitin conjugation | 107670.3516 | 117604.6563 |
| C16orf70 | Q9BSU1 | C16orf70 | UPF0183 protein C16c | 10004.95801 | 10164.26855 |

| | | | | | |
|---|---|---|---|---|---|
| EPB41L4B | Q9H329 | EPB41L4B | Band 4.1-like protein 4 | 4645.95459 | 4376.587891 |
| GAK | O14976 | GAK | Cyclin-G-associated ki | 83457.30469 | 92468.75781 |
| PGRMC1 | O00264 | PGRMC1 | Membrane-associated | 38435.22266 | 69405.70313 |
| MTHFS | P49914 | MTHFS | 5-formyltetrahydrofol | 50555.55078 | 49818.89453 |
| RELL1 | Q8IUW5 | RELL1 | RELT-like protein 1 | 13156.73926 | 17803.02148 |
| CC2D1B | H7C1U3 | CC2D1B | Coiled-coil and C2 dor | 131880.8438 | 138729.6563 |
| ARHGDIB | H0YGX7;P52566 | ARHGDIB | Rho GDP-dissociation | 20326.47852 | 16460.94922 |
| TRAM1 | G3XAN4;Q15629 | TRAM1 | Translocating chain-as | 24467.43945 | 24937.76172 |
| PRORP | O15091 | PRORP | Mitochondrial ribonuc | 36521.41016 | 47850.21875 |
| ZNF384 | Q8TF68 | ZNF384 | Zinc finger protein 384 | 7727.57373 | 12731.15234 |
| TMX1 | Q9H3N1 | TMX1 | Thioredoxin-related tr | 516078.7813 | 557961.6875 |
| ENPP4 | Q9Y6X5 | ENPP4 | Bis(5'-adenosyl)-triph | 15661.60156 | 18844.19727 |
| PIGQ | B8ZZ31;E7ERP4;Q9BRB3 | PIGQ | Phosphatidylinositol N | 13054.61426 | 17224.93018 |
| PCID2 | Q5JVF3 | PCID2 | PCI domain-containing | 31509.42578 | 39650.22656 |
| ADCK5 | Q3MIX3 | ADCK5 | Uncharacterized aarF | 4247.681641 | 7274.253418 |
| RAP2B | P61225 | RAP2B | Ras-related protein Ra | 117860.2344 | 113803.9297 |
| VAV2 | P52735 | VAV2 | Guanine nucleotide ex | 172146.8438 | 182357.9375 |
| ASL | P04424 | ASL | Argininosuccinate lyas | 29963.83203 | 34626.95313 |
| FN1 | P02751 | FN1 | Fibronectin | 8533.206055 | 8348.232422 |
| PFKP | Q01813 | PFKP | ATP-dependent 6-pho | 1036822.313 | 1222613.75 |
| ASCC2 | Q9H1I8 | ASCC2 | Activating signal coint | 56004.49219 | 51885.56641 |
| MRI1 | Q9BV20 | MRI1 | Methylthioribose-1-pl | 32611.67969 | 31832.54883 |
| SERPINF2 | A0A0G2JPA8;A0A0J9YY65;C9 | SERPINF2 | Alpha-2-antiplasmin;A | 12180.44238 | 22654.33984 |
| AMY1A;AMY2A;AMY2B | P04745;P04746;P19961 | AMY1A;AMY2A;AMY2 | Alpha-amylase 1;Panc | 3178.820801 | 1787.165649 |
| PSMG4 | Q5JS54 | PSMG4 | Proteasome assembly | 36318.73438 | 36282.96484 |
| IRF3 | Q14653 | IRF3 | Interferon regulatory | 88430.59375 | 112687.6563 |
| ATOX1 | O00244 | ATOX1 | Copper transport prot | 24300.23438 | 24221.96484 |
| L1RE1 | Q9UN81 | L1RE1 | LINE-1 retrotransposa | 234202.1406 | 188879.2813 |
| BABAM2 | Q9NXR7 | BABAM2 | BRISC and BRCA1-A co | 72157.30469 | 85110.17969 |
| GATD3B;GATD3A | A0A0B4J2D5;P0DPI2 | GATD3B;GATD3A | Glutamine amidotrans | 30719.54688 | 73271.15625 |
| CHD3 | Q12873 | CHD3 | Chromodomain-helica | 39382.14844 | 43692.53906 |
| GNB1 | P62873 | GNB1 | Guanine nucleotide-b | 305118.1563 | 350257.5938 |
| STEAP3 | B8ZZX6;Q658P3 | STEAP3 | Metalloreductase STE | 19695.08984 | 29125.89844 |

| | | | | | |
|---|---|---|---|---|---|
| TMEM245 | Q9H330 | TMEM245 | Transmembrane prote | 19274.64844 | 21758.00195 |
| VRK3 | M0QYA8;M0R073;Q8IV63 | VRK3 | Inactive serine/threor | 13606.31836 | 16031.65039 |
| SGF29 | Q96ES7 | SGF29 | SAGA-associated facto | 35927.72266 | 32378.29492 |
| PRDX5 | P30044 | PRDX5 | Peroxiredoxin-5, mito | 1256085 | 1184680.5 |
| FBXW2 | Q9UKT8 | FBXW2 | F-box/WD repeat-con | 4646.672363 | 3420.665039 |
| APEX1 | P27695 | APEX1 | DNA-(apurinic or apyr | 975265.875 | 986021.0625 |
| C5 | P01031 | C5 | Complement C5 | 11180.80078 | 13206.46289 |
| MDK | E9PLM6;E9PPJ5;P21741 | MDK | Midkine;Midkine (Frag | 3079.321533 | 7325.077148 |
| VAPB | O95292 | VAPB | Vesicle-associated me | 377199.8125 | 423194.5 |
| FAM219B | H3BRU1;Q5XKK7 | FAM219B | Protein FAM219B (Fra | 11206.6084 | 12940.91797 |
| PHKA2 | P46019 | PHKA2 | Phosphorylase b kinas | 20983.60742 | 25301.59961 |
| RAB3GAP1 | Q15042 | RAB3GAP1 | Rab3 GTPase-activatir | 48853.50781 | 54103.03125 |
| STK39 | Q9UEW8 | STK39 | STE20/SPS1-related pr | 27902.63281 | 34832.64844 |
| DHFR | P00374 | DHFR | Dihydrofolate reducta | 117276.0938 | 139123.4531 |
| DDB2 | Q92466 | DDB2 | DNA damage-binding | 104956.5078 | 98961.00781 |
| FBXO2 | A0A286YF37;Q9UK22 | FBXO2 | F-box only protein 2 (I | 1357.126831 | 2984.764893 |
| PGAM1 | P18669 | PGAM1 | Phosphoglycerate mu | 2546987.75 | 2794315 |
| SH3GLB2 | B7ZC38;Q9NR46 | SH3GLB2 | Endophilin-B2 | 70712.78125 | 87128.39063 |
| SLC25A20 | C9JPE1;O43772 | SLC25A20 | Mitochondrial carnitir | 17369.82031 | 16649.73633 |
| ATP6V1H | Q9UI12 | ATP6V1H | V-type proton ATPase | 56813.54688 | 53881.95313 |
| RAB2A | P61019 | RAB2A | Ras-related protein Ra | 249074.0625 | 272144.9375 |
| VRK2 | Q86Y07 | VRK2 | Serine/threonine-prot | 8403.712891 | 7503.339355 |
| C17orf75 | J9JIC5;Q9HAS0 | C17orf75 | Protein Njmu-R1 | 29338.66406 | 30816.91016 |
| LTN1 | O94822 | LTN1 | E3 ubiquitin-protein li | 23416.9668 | 28173.06445 |
| CASP7 | P55210 | CASP7 | Caspase-7 | 147382.5 | 134308.2969 |
| RRAGA | Q7L523 | RRAGA | Ras-related GTP-bindi | 62020.375 | 63326.14844 |
| DYNC1I2 | E7EQL5 | DYNC1I2 | Cytoplasmic dynein 1 | 480681.4844 | 520225.4688 |
| SERPINA10 | G3V2W1;Q9UK55 | SERPINA10 | Protein Z-dependent p | 4976.933105 | 3637.333984 |
| SSRP1 | Q08945 | SSRP1 | FACT complex subunit | 162551.625 | 451871.3125 |
| CCNA2 | P20248 | CCNA2 | Cyclin-A2 | 2847.825439 | 2897.344482 |
| TBCK | Q8TEA7 | TBCK | TBC domain-containin | 4456.375977 | 4777.636719 |
| SEC61B | P60468 | SEC61B | Protein transport prot | 526177.3125 | 457767.7813 |
| SENP1 | Q9P0U3 | SENP1 | Sentrin-specific protea | 1303.795532 | 1351.338745 |

| | | | | | |
|---|---|---|---|---|---|
| NAPA | P54920 | NAPA | Alpha-soluble NSF att | 534391.1875 | 557678.625 |
| PKNOX1 | P55347 | PKNOX1 | Homeobox protein PK | 26814.75391 | 18714.25195 |
| FAM114A2 | Q9NRY5 | FAM114A2 | Protein FAM114A2 | 38555.28516 | 35309.9375 |
| FBXO6 | Q9NRD1 | FBXO6 | F-box only protein 6 | 5187.397949 | 5580.103516 |
| UBE2E1 | C9J2P0;P51965 | UBE2E1 | Ubiquitin-conjugating | 84322.36719 | 117922 |
| RBM23 | Q86U06 | RBM23 | Probable RNA-binding | 31392.20313 | 25597.92969 |
| DHX15 | O43143 | DHX15 | Pre-mRNA-splicing fac | 404547.0938 | 377320.8125 |
| OPA1 | A0A2R8YE54 | OPA1 | Dynamin-like 120 kDa | 69546.78638 | 77046.42139 |
| DHRS13 | Q6UX07 | DHRS13 | Dehydrogenase/reduc | 33759.94141 | 34774.11719 |
| NCS1 | P62166 | NCS1 | Neuronal calcium sen | 2328.715332 | 2484.682129 |
| MCM5 | P33992 | MCM5 | DNA replication licens | 385757.2813 | 381006.7813 |
| ATG16L1 | E7EVC7;Q676U5 | ATG16L1 | Autophagy-related pr | 34379.68359 | 39139.09766 |
| WDR62 | O43379 | WDR62 | WD repeat-containing | 828.9279785 | 899.203064 |
| GOLGA1 | Q92805 | GOLGA1 | Golgin subfamily A me | 26350.6875 | 25702.83398 |
| MTHFD1 | P11586 | MTHFD1 | C-1-tetrahydrofolate s | 1157220.875 | 1267751.625 |
| EIF5 | P55010 | EIF5 | Eukaryotic translation | 797116.4375 | 815958.0625 |
| C11orf68 | Q9H3H3 | C11orf68 | UPF0696 protein C11c | 77704.94531 | 86018.85156 |
| PCYT2 | Q99447 | PCYT2 | Ethanolamine-phosph | 125460.8281 | 150322.6563 |
| CFL1 | P23528 | CFL1 | Cofilin-1 | 8458571 | 8108584 |
| TMEM109 | Q9BVC6 | TMEM109 | Transmembrane prot | 276849.5313 | 246826.6563 |
| ;TATDN2 | H7C1W4;Q93075 | ;TATDN2 | Uncharacterized prot | 1562.807617 | 1505.766479 |
| CDC42 | P60953 | CDC42 | Cell division control p | 1096089.875 | 1239507.5 |
| STT3B | Q8TCJ2 | STT3B | Dolichyl-diphosphooli | 186551.625 | 219624.7344 |
| CBARP | A0A140LJL2;Q8N350 | CBARP | Voltage-dependent ca | 5392.114258 | 3888.003662 |
| A1BG | M0R009;P04217 | A1BG | Alpha-1B-glycoprotei | 277175.8125 | 230092.125 |
| N4BP1 | O75113 | N4BP1 | NEDD4-binding protei | 1430.234619 | 1729.836914 |
| TRAPPC10 | P48553 | TRAPPC10 | Trafficking protein pa | 18000.82422 | 18004.16016 |
| DPP7 | Q9UHL4 | DPP7 | Dipeptidyl peptidase 2 | 42176.79688 | 53530.80078 |
| MAPK13 | O15264 | MAPK13 | Mitogen-activated pr | 35968.53516 | 35687.53906 |
| MAGT1 | A0A087WU53;Q9H0U3 | MAGT1 | Magnesium transport | 82288.4375 | 85518.625 |
| ELOA | A0A024RAC6 | ELOA | Elongin-A | 43055.96094 | 42343.33594 |
| PPP6R2 | O75170 | PPP6R2 | Serine/threonine-prot | 6159.464355 | 7140.646484 |
| AP5B1 | Q2VPB7 | AP5B1 | AP-5 complex subunit | 9334.505859 | 10335.36523 |

| | | | | | |
|---|---|---|---|---|---|
| LZIC | Q8WZA0 | LZIC | Protein LZIC | 51053.23438 | 52840.34766 |
| STAT5B | P51692 | STAT5B | Signal transducer and | 31459.32617 | 38687.29688 |
| GON4L;GON4L;YY1AP1 | A0A087WUE6;Q3T8J9;Q9H8( | GON4L;GON4L;YY1AP | GON-4-like protein;G( | 6230.872559 | 6564.813477 |
| TAF9B | Q9HBM6 | TAF9B | Transcription initiatio | 22143.94727 | 15243.94141 |
| ATP6V0A1 | Q93050 | ATP6V0A1 | V-type proton ATPase | 113802.8359 | 110297.6797 |
| WDR76 | A0A0C4DFX7;Q9H967 | WDR76 | WD repeat-containing | 11190.64648 | 8841.439453 |
| MCM7 | P33993 | MCM7 | DNA replication licens | 375753.4375 | 395408.9688 |
| BRK1 | Q8WUW1 | BRK1 | Protein BRICK1 | 11905.23145 | 17378.95117 |
| NRDE2 | Q9H7Z3 | NRDE2 | Nuclear exosome regu | 19164.89453 | 18559.3125 |
| RNMT | O43148 | RNMT | mRNA cap guanine-N7 | 79026.32031 | 68379.97656 |
| ZFP36L2 | P47974 | ZFP36L2 | mRNA decay activator | 11484.28223 | 9826.418945 |
| TBCB | Q99426 | TBCB | Tubulin-folding cofact | 352786.5625 | 413759.5313 |
| TUBB4A | P04350 | TUBB4A | Tubulin beta-4A chain | 236268.0469 | 261913.2813 |
| UGCG | Q16739 | UGCG | Ceramide glucosyltrar | 6861.151367 | 7066.571777 |
| CHAMP1 | Q96JM3 | CHAMP1 | Chromosome alignme | 47049.00781 | 42233.08594 |
| FARSA | Q9Y285 | FARSA | Phenylalanine--tRNA l | 353901.1563 | 328359.4063 |
| SMIM12 | E5RH51;Q96EX1 | SMIM12 | Small integral membra | 20977.0957 | 24860.55664 |
| DBN1 | Q16643 | DBN1 | Drebrin | 22462.54297 | 8955.803711 |
| SRP9 | P49458 | SRP9 | Signal recognition par | 850820.75 | 837172.375 |
| TRPT1 | F5H6B6;F5H8A0;Q86TN4 | TRPT1 | tRNA 2'-phosphotrans | 12611.50586 | 19258.28516 |
| CNPY3 | Q9BT09 | CNPY3 | Protein canopy homol | 110097.2813 | 114246.9531 |
| OFD1 | A6NF31;O75665 | OFD1 | Oral-facial-digital sync | 797.1486816 | 1388.20752 |
| FIZ1 | Q96SL8 | FIZ1 | Flt3-interacting zinc fi | 8045.499512 | 9313.764648 |
| SCCPDH | Q8NBX0 | SCCPDH | Saccharopine dehydro | 281267.875 | 297134.9688 |
| RAC1 | P63000 | RAC1 | Ras-related C3 botulir | 1113314.875 | 1231018.25 |
| NFKB1 | P19838 | NFKB1 | Nuclear factor NF-kap | 49135.14063 | 48614.28125 |
| NINJ1 | Q92982 | NINJ1 | Ninjurin-1 | 4178.213379 | 5192.720703 |
| FGGY | A0A0U1RQJ1;A0A0U1RRB9;C | FGGY | FGGY carbohydrate ki | 2152.324463 | 2127.828369 |
| SRGAP2 | O75044 | SRGAP2 | SLIT-ROBO Rho GTPas | 48736.53906 | 53041.16406 |
| VPS33B | Q9H267 | VPS33B | Vacuolar protein sorti | 66374.52344 | 71495.78125 |
| DSCC1 | Q9BVC3 | DSCC1 | Sister chromatid cohe | 34846.37109 | 33809.40625 |
| HNRNPUL1 | Q9BUJ2 | HNRNPUL1 | Heterogeneous nucle | 290096.0625 | 312260.25 |
| PTDSS1 | P48651 | PTDSS1 | Phosphatidylserine sy | 65086.9375 | 66818.34375 |

| | | | | | |
|---|---|---|---|---|---|
| ARIH2 | O95376 | ARIH2 | E3 ubiquitin-protein li | 23913.14258 | 25302.27344 |
| CBX4 | O00257 | CBX4 | E3 SUMO-protein liga: | 34213.70313 | 29826.42773 |
| OPTN | Q96CV9 | OPTN | Optineurin | 4672.640625 | 13153.84277 |
| ZDHHC13 | Q8IUH4 | ZDHHC13 | Palmitoyltransferase Z | 28448.08398 | 26938.08594 |
| DAD1 | F5GXX5;P61803 | DAD1 | Dolichyl-diphosphooli | 214746.875 | 274597.4375 |
| PLCB4 | Q15147 | PLCB4 | 1-phosphatidylinosito | 19455.14063 | 21802.28906 |
| L2HGDH | Q9H9P8 | L2HGDH | L-2-hydroxyglutarate ( | 94371.57031 | 108646.125 |
| NCK2 | O43639 | NCK2 | Cytoplasmic protein N | 16827.56445 | 18558.55273 |
| HAGH | Q16775 | HAGH | Hydroxyacylglutathior | 172257.2031 | 196692.4375 |
| ABHD5 | Q8WTS1 | ABHD5 | 1-acylglycerol-3-phosp | 11914.75488 | 10845.13574 |
| HID1 | Q8IV36 | HID1 | Protein HID1 | 157431.5469 | 181045.5 |
| ARV1 | A0A0A0MRI7;H7C484;Q9H2( | ARV1 | Protein ARV1;Protein | 15648.89648 | 15485.33398 |
| SHARPIN | Q9H0F6 | SHARPIN | Sharpin | 2215.509766 | 3737.774414 |
| OSBPL10 | Q9BXB5 | OSBPL10 | Oxysterol-binding pro | 41324.49609 | 55214.32422 |
| FEN1 | P39748 | FEN1 | Flap endonuclease 1 | 751848.375 | 776278.875 |
| TASOR | Q9UK61 | TASOR | Protein TASOR | 25703.3457 | 26658.91992 |
| KATNAL1 | Q9BW62 | KATNAL1 | Katanin p60 ATPase-c | 7731.616211 | 6399.166504 |
| ZFYVE16 | Q7Z3T8 | ZFYVE16 | Zinc finger FYVE doma | 208705.5313 | 250029.9219 |
| ME1 | P48163 | ME1 | NADP-dependent mal | 160054.375 | 168153.0313 |
| ABCF3 | Q9NUQ8 | ABCF3 | ATP-binding cassette : | 118277.6563 | 118912.2344 |
| DIS3L2 | Q8IYB7 | DIS3L2 | DIS3-like exonuclease | 12942.7168 | 14853.65234 |
| IRX5 | P78411 | IRX5 | Iroquois-class homeo( | 14857.83301 | 18601.74609 |
| ADSS2 | P30520 | ADSS2 | Adenylosuccinate syn | 208090.7031 | 221170.4844 |
| DDX19A | Q9NUU7 | DDX19A | ATP-dependent RNA h | 202143.2813 | 221227.3906 |
| FXYD3 | Q14802;S4R445 | FXYD3 | FXYD domain-containi | 57298.97656 | 85399.89844 |
| VPS53 | Q5VIR6 | VPS53 | Vacuolar protein sorti | 41431.82422 | 45789.50781 |
| WIZ | M0QXA7 | WIZ | Protein Wiz | 29634.7832 | 26253.60156 |
| COX7B | A0A3B3IRN8;A0A3B3ITX0;P2 | COX7B | Cytochrome c oxidase | 24863.42383 | 43042.46484 |
| MORF4L2 | Q15014 | MORF4L2 | Mortality factor 4-like | 87235.51563 | 89148.48438 |
| ATP6V1D | Q9Y5K8 | ATP6V1D | V-type proton ATPase | 41885.56641 | 41007.53906 |
| CHMP1B | Q7LBR1 | CHMP1B | Charged multivesicula | 36879.08203 | 53817.80469 |
| TPD52 | P55327 | TPD52 | Tumor protein D52 | 132528.6875 | 120933.6641 |
| GPAT4 | Q86UL3 | GPAT4 | Glycerol-3-phosphate | 11949.09961 | 11604.36328 |

| | | | | | |
|---|---|---|---|---|---|
| CASZ1 | Q86V15 | CASZ1 | Zinc finger protein cas | 10454.84766 | 13931.66797 |
| LPCAT4 | A0A0C4DGT4;Q643R3 | LPCAT4 | Lysophospholipid acyl | 10966.03809 | 9770.212891 |
| ASAP2 | O43150 | ASAP2 | Arf-GAP with SH3 dom | 17655.7207 | 18967.51953 |
| MTR | Q99707 | MTR | Methionine synthase | 24444.80664 | 24878.3125 |
| PLAA | Q9Y263 | PLAA | Phospholipase A-2-act | 68138.86719 | 71903.57813 |
| UEVLD | Q8IX04 | UEVLD | Ubiquitin-conjugating | 47152.83984 | 61594.43359 |
| ZZZ3 | Q8IYH5 | ZZZ3 | ZZ-type zinc finger-cor | 6716.571289 | 6188.460938 |
| ANP32B | Q92688 | ANP32B | Acidic leucine-rich nuc | 1245925 | 1365767.375 |
| CLIC1 | O00299 | CLIC1 | Chloride intracellular | 1713928.875 | 1756892.25 |
| DUT | H0YNJ9 | DUT | Deoxyuridine 5'-tripho | 708190.0781 | 801787.6641 |
| ARHGAP8 | P85298 | ARHGAP8 | Rho GTPase-activating | 23667.17969 | 26652.83594 |
| CCDC25 | Q86WR0 | CCDC25 | Coiled-coil domain-co | 58515.16016 | 52258.13281 |
| HSD17B4 | P51659 | HSD17B4 | Peroxisomal multifunc | 440055.625 | 521174.2188 |
| GOLGA3 | Q08378 | GOLGA3 | Golgin subfamily A me | 66682.14063 | 64568.08203 |
| PSMD3 | O43242 | PSMD3 | 26S proteasome non-A | 187235.0313 | 204630.0469 |
| CARS2 | Q9HA77 | CARS2 | Probable cysteine--tRI | 31418.61133 | 36298.75391 |
| RABL6 | Q3YEC7 | RABL6 | Rab-like protein 6 | 93256.94531 | 106696.3125 |
| SPATS2 | Q86XZ4 | SPATS2 | Spermatogenesis-asso | 25854.66797 | 29891.44727 |
| NF2 | P35240 | NF2 | Merlin | 21957.30273 | 20940.43945 |
| GRIPAP1 | Q4V328 | GRIPAP1 | GRIP1-associated prot | 80895.51563 | 81584.67188 |
| COL6A1 | A0A087X0S5;P12109 | COL6A1 | Collagen alpha-1(VI) c | 26226.40039 | 23766.95508 |
| ZW10 | O43264 | ZW10 | Centromere/kinetoch | 97965.80469 | 110495.6406 |
| PACSIN3 | Q9UKS6 | PACSIN3 | Protein kinase C and c | 82324.16406 | 88137.21875 |
| PEX3 | P56589 | PEX3 | Peroxisomal biogenes | 13898.5625 | 17422.66602 |
| PIK3R1 | P27986 | PIK3R1 | Phosphatidylinositol 3 | 21286.57422 | 26762.38086 |
| RAB18 | Q9NP72 | RAB18 | Ras-related protein Ra | 301235.4375 | 341496.875 |
| HAT1 | O14929 | HAT1 | Histone acetyltransfer | 333198.9375 | 400962.8125 |
| ARRDC1 | Q8N5I2 | ARRDC1 | Arrestin domain-conta | 34387.92969 | 39240.48828 |
| GYS1 | P13807 | GYS1 | Glycogen [starch] synt | 120038.5859 | 127194.2734 |
| ZNF219 | Q9P2Y4 | ZNF219 | Zinc finger protein 219 | 1212.975464 | 1416.321289 |
| CRIP2 | P52943 | CRIP2 | Cysteine-rich protein 2 | 357205.75 | 337593.9375 |
| TOM1L2 | Q6ZVM7 | TOM1L2 | TOM1-like protein 2 | 20240.35938 | 19685.17188 |
| CMTR1 | Q8N1G2 | CMTR1 | Cap-specific mRNA (nu | 92641.70313 | 76294.28906 |

| | | | | | |
|---|---|---|---|---|---|
| CADPS2 | A0A087X1P3;C9IYE1;F8W8P5 | CADPS2 | Calcium-dependent se | 3071.747314 | 2583.15625 |
| NUMB | P49757 | NUMB | Protein numb homolo | 52695.79297 | 53583.05078 |
| NAP1L4 | Q99733 | NAP1L4 | Nucleosome assembly | 251102.9375 | 224658.6875 |
| LDOC1 | O95751 | LDOC1 | Protein LDOC1 | 10136.06543 | 9294.740234 |
| RPTOR | Q8N122 | RPTOR | Regulatory-associated | 40261.80859 | 46170.61328 |
| PAN2 | Q504Q3 | PAN2 | PAN2-PAN3 deadenyla | 10824.96973 | 11373.00586 |
| IMPA1 | P29218 | IMPA1 | Inositol monophospha | 53810.9375 | 53330.12891 |
| RFC2 | P35250 | RFC2 | Replication factor C su | 142172.8125 | 158615.9219 |
| ZYX | Q15942 | ZYX | Zyxin | 261156.9531 | 202561.8594 |
| DMAC2L | A0A5K1VW60;Q99766 | DMAC2L | ATP synthase subunit | 30993.60352 | 31704.41602 |
| NIPSNAP2 | O75323 | NIPSNAP2 | Protein NipSnap homo | 64861.53906 | 68612.96094 |
| SMARCD1 | Q96GM5 | SMARCD1 | SWI/SNF-related matr | 29411.12109 | 30752.41602 |
| CIRBP | Q14011 | CIRBP | Cold-inducible RNA-bi | 513576.5938 | 431859.5 |
| GPSM1 | A0A0A0MRC4;A0A0A0MSK4; | GPSM1 | G-protein-signaling m | 3976.759277 | 6799.090332 |
| MVB12A | Q96EY5 | MVB12A | Multivesicular body su | 41017.53516 | 36304.85156 |
| SUPT16H | Q9Y5B9 | SUPT16H | FACT complex subunit | 114674.0625 | 320499.125 |
| SLC25A10 | F6RGN5 | SLC25A10 | Mitochondrial dicarbo | 41266.26953 | 20692.38867 |
| GPT2 | Q8TD30 | GPT2 | Alanine aminotransfer | 28225.85547 | 31373.14648 |
| STARD4 | Q96DR4 | STARD4 | StAR-related lipid tran | 9917.712891 | 9375.204102 |
| TTF2 | Q9UNY4 | TTF2 | Transcription termina | 27853.16602 | 23372.72461 |
| AZI2 | C9JVK8;Q9H6S1 | AZI2 | 5-azacytidine-induced | 18311.39063 | 17721.3418 |
| SENP8 | Q96LD8 | SENP8 | Sentrin-specific prote | 31137.60547 | 25802.8457 |
| LRRC1 | Q9BTT6 | LRRC1 | Leucine-rich repeat-co | 33174.67578 | 36372.26172 |
| ABHD14B | Q96IU4 | ABHD14B | Protein ABHD14B | 128993.625 | 123282.9688 |
| RNF2 | Q99496 | RNF2 | E3 ubiquitin-protein li | 70126.24219 | 69253.9375 |
| MCRIP1 | C9JLW8;I3L2L5;I3L4Q0 | MCRIP1 | Mapk-regulated corep | 42371.64453 | 36832.00391 |
| GOLGA7 | Q7Z5G4 | GOLGA7 | Golgin subfamily A me | 23441.27734 | 22537.41797 |
| RAB3D | O95716 | RAB3D | Ras-related protein Ra | 198990.0313 | 213514.4219 |
| SKOR2 | A0A087X0E6;Q2VWA4 | SKOR2 | SKI family transcriptio | 34584.20703 | 22434.16211 |
| PTK2 | Q05397 | PTK2 | Focal adhesion kinase | 45236.11328 | 62912.39063 |
| SLC4A7 | E9PFN4 | SLC4A7 | Anion exchange prote | 31716.02832 | 29859.65039 |
| UHRF1 | A0A087WTW0;A0A087WVR3 | UHRF1 | E3 ubiquitin-protein li | 87695.15625 | 99058.21094 |
| JTB | O76095 | JTB | Protein JTB | 79999.94531 | 90040.88281 |

| | | | | | |
|---|---|---|---|---|---|
| HECTD1 | Q9ULT8 | HECTD1 | E3 ubiquitin-protein li | 53350.63281 | 46380.08203 |
| ISOC2 | Q96AB3 | ISOC2 | Isochorismatase doma | 85404.57813 | 74321.46094 |
| SLC41A3 | D6R9S7;D6RDB0;D6RDM3;D6 | SLC41A3 | Solute carrier family 4 | 5675.775391 | 4686.016113 |
| UBTD1 | Q9HAC8 | UBTD1 | Ubiquitin domain-con | 10575.66309 | 14434.04785 |
| GMFB | P60983 | GMFB | Glia maturation factor | 168262.1875 | 164545.875 |
| UPF3B | Q9BZI7 | UPF3B | Regulator of nonsense | 38940.83203 | 37764.44141 |
| GSK3B | P49841 | GSK3B | Glycogen synthase kir | 126736.25 | 139798.4375 |
| NUDT4B | A0A024RBG1 | NUDT4B | Diphosphoinositol pol | 12684.7041 | 13739.47559 |
| RWDD2B | P57060 | RWDD2B | RWD domain-containi | 7132.906738 | 6941.084961 |
| KIF21A | A0A1B0GV47;Q7Z4S6 | KIF21A | Kinesin-like protein KI | 17875.83594 | 16860.05664 |
| BCL2L1 | Q07817;Q5QP56;Q5TE64;Q9 | BCL2L1 | Bcl-2-like protein 1;Bc | 52258.84766 | 41041.11719 |
| DCTN1 | E7EX90 | DCTN1 | Dynactin subunit 1 | 131253.4063 | 139713.9688 |
| ATG4A | A0A0A0MT61;G5E979;H7BYN | ATG4A | Cysteine protease (Fra | 3761.655273 | 5033.71582 |
| CAPNS1 | A0A0C4DGQ5;P04632 | CAPNS1 | Calpain small subunit | 190892.0938 | 293729.5938 |
| KIDINS220 | Q9ULH0 | KIDINS220 | Kinase D-interacting s | 339996.1563 | 320966.4063 |
| PLD3 | Q8IV08 | PLD3 | 5'-3' exonuclease PLD | 99378.30469 | 69777.72656 |
| PIK3C2A | O00443 | PIK3C2A | Phosphatidylinositol 4 | 129301.6172 | 144938.4688 |
| RHOF | Q9HBH0 | RHOF | Rho-related GTP-bind | 10705.35449 | 13705.67676 |
| NLRX1 | Q86UT6 | NLRX1 | NLR family member X | 9024.763672 | 7512.033203 |
| MIA3 | Q5JRA6 | MIA3 | Transport and Golgi o | 36265.05469 | 41432.72656 |
| RHOG | P84095 | RHOG | Rho-related GTP-bind | 27425.37695 | 27590.67969 |
| ALKBH2 | F5H5X2;Q6NS38;S4R3T5 | ALKBH2 | DNA oxidative demeth | 29010.10156 | 40162.14063 |
| STX4 | Q12846 | STX4 | Syntaxin-4 | 58236.69531 | 58030.78906 |
| FBXW11 | Q9UKB1 | FBXW11 | F-box/WD repeat-con | 8151.226563 | 6766.852539 |
| CRTC3 | Q6UUV7 | CRTC3 | CREB-regulated transc | 6893.827148 | 5936.819336 |
| PACSIN1 | F6U236;Q9BY11 | PACSIN1 | Protein kinase C and c | 24712.44336 | 20357.26172 |
| TRIP11 | Q15643 | TRIP11 | Thyroid receptor-inter | 46668.31641 | 55345.14844 |
| TECR | Q9NZ01 | TECR | Very-long-chain enoyl | 91944.88281 | 123062.0703 |
| IPO9 | Q96P70 | IPO9 | Importin-9 | 87958.54688 | 106295.3438 |
| PTPN6 | P29350 | PTPN6 | Tyrosine-protein phos | 29911.875 | 33748.98047 |
| PPP1R9A | Q9ULJ8 | PPP1R9A | Neurabin-1 | 5014.740234 | 6651.913086 |
| FAM49B | Q9NUQ9 | FAM49B | Protein FAM49B | 112579.2031 | 105218.9766 |
| REPS1 | Q96D71 | REPS1 | RalBP1-associated Eps | 334093.7188 | 320886.9375 |

| | | | | | |
|---|---|---|---|---|---|
| PAFAH1B3 | Q15102 | PAFAH1B3 | Platelet-activating fac | 629578.125 | 671652.9375 |
| RASSF8 | Q8NHQ8 | RASSF8 | Ras association domai | 5824.77002 | 4761.969238 |
| ORC3 | Q9UBD5 | ORC3 | Origin recognition cor | 27429.86133 | 33866.17188 |
| SLC25A3 | F8VVM2 | SLC25A3 | Phosphate carrier pro | 1257317.016 | 1438252.125 |
| RNF31 | Q96EP0 | RNF31 | E3 ubiquitin-protein li | 32660.95898 | 30414.17383 |
| NFAT5 | O94916 | NFAT5 | Nuclear factor of activ | 1031.60437 | 2958.904297 |
| TADA3 | O75528 | TADA3 | Transcriptional adapte | 24394.97852 | 23787.23828 |
| FAM53C | D6RE00;Q9NYF3 | FAM53C | Protein FAM53C | 141609.5469 | 115990.3359 |
| LIN37 | Q96GY3 | LIN37 | Protein lin-37 homolo | 11330.30762 | 5237.910156 |
| CLIC4 | Q9Y696 | CLIC4 | Chloride intracellular | 112393.2578 | 127782.0234 |
| MAP2K2 | P36507 | MAP2K2 | Dual specificity mitog | 201223.625 | 240943.9375 |
| ASPSCR1 | Q9BZE9 | ASPSCR1 | Tether containing UB | 131944.0625 | 131530.0781 |
| ZNF3 | P17036 | ZNF3 | Zinc finger protein 3 | 2802.731934 | 2791.875977 |
| MTOR | P42345 | MTOR | Serine/threonine-prot | 46670.45313 | 55568.54297 |
| MGAT5 | Q09328 | MGAT5 | Alpha-1,6-mannosylgl | 13092.1709 | 13181.5791 |
| MAPK9 | P45984 | MAPK9 | Mitogen-activated pro | 66719.14063 | 82574.53906 |
| UGP2 | A0A087WYS1;E7EUC7;Q1685 | UGP2 | UTP--glucose-1-phosp | 266984.0488 | 377712.5391 |
| SMC3 | Q9UQE7 | SMC3 | Structural maintenanc | 146095.5781 | 131992.4688 |
| LRRC20 | Q8TCA0 | LRRC20 | Leucine-rich repeat-co | 25125.39844 | 27980.48633 |
| COMMD2 | F8WBW1;Q86X83 | COMMD2 | COMM domain-conta | 77812.82031 | 91005.04688 |
| NIT1 | Q86X76 | NIT1 | Deaminated glutathio | 19878.76563 | 19905.80859 |
| CAPZA2 | P47755 | CAPZA2 | F-actin-capping protei | 708280.375 | 705987.625 |
| RAB21 | Q9UL25 | RAB21 | Ras-related protein Ra | 271036.2188 | 272969.875 |
| NME7 | Q9Y5B8 | NME7 | Nucleoside diphospha | 11536.45313 | 13402.02051 |
| RNF213 | A0A0A0MTC1;A0A0A0MTR7; | RNF213 | E3 ubiquitin-protein li | 41040.375 | 61269.59375 |
| MED20 | Q9H944 | MED20 | Mediator of RNA poly | 25686.23242 | 26973.48242 |
| SCRN1 | Q12765 | SCRN1 | Secernin-1 | 29010.24805 | 58446.60547 |
| PNPLA8 | A0A0C4DG51;Q9NP80 | PNPLA8 | Calcium-independent | 3358.917969 | 4271.671875 |
| AP3M2 | P53677 | AP3M2 | AP-3 complex subunit | 34372.46875 | 40611.40234 |
| GNS | P15586 | GNS | N-acetylglucosamine- | 165478.0156 | 152418.3438 |
| FAM8A1 | Q9UBU6 | FAM8A1 | Protein FAM8A1 | 3797.425049 | 4500.312012 |
| ZNF691 | A0A0A0MRR1;Q5VV50;Q5VV | ZNF691 | Zinc finger protein 69 | 3099.766602 | 3572.62915 |
| OCRL | Q01968 | OCRL | Inositol polyphosphat | 154369.5625 | 169847.0781 |

| | | | | | |
|---|---|---|---|---|---|
| GDAP2 | Q9NXN4 | GDAP2 | Ganglioside-induced d | 19979.10352 | 17341.62305 |
| ATPAF1 | I3L448;Q5TC12 | ATPAF1 | ATP synthase mitocho | 62323.63281 | 70808.49219 |
| MACC1 | Q6ZN28 | MACC1 | Metastasis-associated | 19493.98633 | 15115.61426 |
| C7orf50 | C9JQV0;Q9BRJ6 | C7orf50 | Uncharacterized prote | 572738.9375 | 639129.4375 |
| CDYL | Q9Y232 | CDYL | Chromodomain Y-like | 14828.30273 | 12324.22656 |
| AGPAT1 | Q99943 | AGPAT1 | 1-acyl-sn-glycerol-3-pl | 1271.519287 | 1806.107666 |
| CASP6 | P55212 | CASP6 | Caspase-6 | 72946.21875 | 90371.70313 |
| NEDD8 | Q15843 | NEDD8 | NEDD8 | 159506.9531 | 201438.2656 |
| TBC1D10B | Q4KMP7 | TBC1D10B | TBC1 domain family m | 44972.59375 | 49643.78906 |
| HTATIP2 | Q9BUP3 | HTATIP2 | Oxidoreductase HTAT | 163464.6563 | 184423.2656 |
| MARF1 | Q9Y4F3 | MARF1 | Meiosis regulator and | 9901.550781 | 12218.43262 |
| MTMR14 | Q8NCE2 | MTMR14 | Myotubularin-related | 34227.86328 | 37844.55078 |
| SMPD4 | F2Z2I0;F2Z2W5;F8WF03 | SMPD4 | Sphingomyelin phospl | 75764.57227 | 74955.31885 |
| YWHAB | P31946 | YWHAB | 14-3-3 protein beta/a | 1672895 | 1649872.875 |
| NUP188 | Q5SRE5 | NUP188 | Nucleoporin NUP188 | 43291.28125 | 49785.87109 |
| TWF2 | Q6IBS0 | TWF2 | Twinfilin-2 | 100965.8281 | 111360.7266 |
| UBR7 | Q8N806 | UBR7 | Putative E3 ubiquitin- | 148096.8438 | 142874.0313 |
| TMEM59 | Q5T6Z8;Q9BXS4 | TMEM59 | Transmembrane prote | 9229.0625 | 10603.69336 |
| TLK1 | Q9UKI8 | TLK1 | Serine/threonine-prot | 4402.598145 | 4955.133301 |
| TAOK2 | Q9UL54 | TAOK2 | Serine/threonine-prot | 24326.24023 | 22537.86328 |
| GPHN | F5H039;Q9NQX3 | GPHN | Molybdopterin molyb | 87598.98438 | 118350.3828 |
| MTDH | Q86UE4 | MTDH | Protein LYRIC | 413313.7813 | 373243.25 |
| PAIP1 | D6RJF2 | PAIP1 | Polyadenylate-binding | 216433.7695 | 262531.3711 |
| MAP11 | Q8WVR3 | MAP11 | Microtubule-associate | 1873.369385 | 2028.712891 |
| AVL9 | Q8NBF6 | AVL9 | Late secretory pathwa | 63394.46484 | 84685.0625 |
| HOOK2 | Q96ED9 | HOOK2 | Protein Hook homolog | 35581.41406 | 34498.57422 |
| FCHSD1 | Q86WN1 | FCHSD1 | F-BAR and double SH3 | 2407.316162 | 5514.90918 |
| TRIM24 | O15164 | TRIM24 | Transcription interme | 55518.58984 | 52519.44141 |
| RHOT1 | Q8IXI2 | RHOT1 | Mitochondrial Rho GT | 40817.30078 | 38521.76563 |
| USP47 | Q96K76 | USP47 | Ubiquitin carboxyl-ter | 41499.98438 | 44499.00391 |
| GCLC | P48506 | GCLC | Glutamate--cysteine li | 100699.1875 | 101183.1563 |
| DCAF11 | Q8TEB1 | DCAF11 | DDB1- and CUL4-asso | 47326.27734 | 55200.21484 |
| COPS7A | Q9UBW8 | COPS7A | COP9 signalosome co | 133390.0625 | 127693.6094 |

| | | | | | |
|---|---|---|---|---|---|
| COBLL1 | A0A0X1KG75;A0A590PXD8;C | COBLL1 | Cordon-bleu protein-l | 7946.030273 | 7832.387695 |
| ARID4A | P29374 | ARID4A | AT-rich interactive do | 8803.145508 | 10985.86914 |
| ORC4 | O43929 | ORC4 | Origin recognition cor | 28461.51563 | 30403.20508 |
| CHMP3 | Q9Y3E7 | CHMP3 | Charged multivesicula | 139567.6094 | 178632.3906 |
| ATP7A | Q04656 | ATP7A | Copper-transporting A | 8184.498535 | 11276.28711 |
| KPNA4 | O00629 | KPNA4 | Importin subunit alph | 282895.5 | 305241.75 |
| COPB2 | P35606 | COPB2 | Coatomer subunit bet | 654477.4375 | 756729.5 |
| RAB8A | P61006 | RAB8A | Ras-related protein Ra | 79054.0625 | 84584.3125 |
| F11R | Q9Y624 | F11R | Junctional adhesion m | 226231.7344 | 262671 |
| DHX40 | Q8IX18 | DHX40 | Probable ATP-depend | 26936.16992 | 33394.88672 |
| PKLR | P30613 | PKLR | Pyruvate kinase PKLR | 50542.94531 | 50148.60938 |
| | A0A096LPI6 | | Utp12 domain-contair | 1348320.816 | 1502291.271 |
| SMC1A | Q14683 | SMC1A | Structural maintenanc | 215221 | 191588.0313 |
| SERPINF1 | P36955 | SERPINF1 | Pigment epithelium-d | 20138.0293 | 16018.78809 |
| BAG5 | Q9UL15 | BAG5 | BAG family molecular | 76148.94531 | 68646.88281 |
| RPN2 | P04844 | RPN2 | Dolichyl-diphosphooli | 305004.9063 | 349374.0625 |
| SMG8 | Q8ND04 | SMG8 | Protein SMG8 | 56710.62109 | 55361.88281 |
| TKT | P29401 | TKT | Transketolase | 2618980.25 | 3014439.25 |
| RETREG2 | Q8NC44 | RETREG2 | Reticulophagy regulat | 14484.5332 | 15829.80176 |
| BRF1 | Q92994 | BRF1 | Transcription factor II | 9115.305664 | 9002.958008 |
| DYNLL2 | Q96FJ2 | DYNLL2 | Dynein light chain 2, c | 123816.4844 | 95352.99219 |
| ZNF732 | B4DXR9 | ZNF732 | Zinc finger protein 73 | 2666.998535 | 3271.993896 |
| PSMC3 | E9PN50 | PSMC3 | 26S proteasome regul | 920414.7891 | 857073.4297 |
| PPP4R3A | Q6IN85 | PPP4R3A | Serine/threonine-prot | 97536.03125 | 95538.73438 |
| KIAA0319L | E7EN73;Q8IZA0 | KIAA0319L | Dyslexia-associated pr | 23527.52148 | 27539.55664 |
| VHL | P40337 | VHL | von Hippel-Lindau dis | 14675.78223 | 12076.43164 |
| FIBP | O43427 | FIBP | Acidic fibroblast grow | 27562.84766 | 31479.2832 |
| PDE4DIP | E9PQG4 | PDE4DIP | Myomegalin | 16276.93652 | 19170.87109 |
| KIAA1324 | Q6UXG2 | KIAA1324 | UPF0577 protein KIAA | 74778.14063 | 60768.13672 |
| PDE6D | O43924 | PDE6D | Retinal rod rhodopsin | 53930.20703 | 66532.5625 |
| NIPSNAP3A | Q9UFN0 | NIPSNAP3A | Protein NipSnap hom | 47517.14453 | 48439.15234 |
| VAC14 | Q08AM6 | VAC14 | Protein VAC14 homol | 48582.46484 | 52748.62891 |
| CCDC149 | A0A0U1RQD2;Q6ZUS6 | CCDC149 | Coiled-coil domain-co | 2480.53833 | 2336.902832 |

| | | | | | |
|---|---|---|---|---|---|
| CLDN3 | O15551 | CLDN3 | Claudin-3 | 66918.10156 | 71511.57813 |
| FDPS | P14324 | FDPS | Farnesyl pyrophospha | 795149.5 | 963723.25 |
| SYNGR1 | B5MCD7;O43759 | SYNGR1 | Synaptogyrin-1 | 45185.11719 | 43808.48438 |
| SEC63 | Q9UGP8 | SEC63 | Translocation protein | 94061.29688 | 99120.64844 |
| RXYLT1 | G3V1K2;Q9Y2B1 | RXYLT1 | Ribitol-5-phosphate x | 4508.400879 | 12404.95703 |
| CIZ1 | F5H2X7;H0Y5D5;Q5SYW2;Q9 | CIZ1 | Cip1-interacting zinc f | 31034.6582 | 30085.15039 |
| UBA1 | P22314 | UBA1 | Ubiquitin-like modifie | 1019741.688 | 1165790.625 |
| CBX8 | Q9HC52 | CBX8 | Chromobox protein h | 51533.17188 | 44207.44531 |
| PLCH1 | A0A2U3TZV8;Q4KWH8 | PLCH1 | Phosphoinositide pho | 50016.11719 | 50309.84375 |
| PIR | O00625 | PIR | Pirin | 129364.4922 | 161372.4531 |
| TRIM41 | Q8WV44 | TRIM41 | E3 ubiquitin-protein li | 5773.594727 | 5879.713379 |
| NUDT4 | Q9NZJ9 | NUDT4 | Diphosphoinositol pol | 37087.48438 | 34571.03516 |
| ESRP1 | Q6NXG1 | ESRP1 | Epithelial splicing regu | 126764.625 | 132483.7969 |
| NT5DC2 | Q9H857 | NT5DC2 | 5'-nucleotidase doma | 46524.10938 | 63665.26172 |
| AP2A2 | O94973 | AP2A2 | AP-2 complex subunit | 214181.0313 | 231621.3594 |
| DCK | D6RFG8;P27707 | DCK | Deoxycytidine kinase | 76227.98438 | 71027.51563 |
| UNC45A | Q9H3U1 | UNC45A | Protein unc-45 homol | 241348.4844 | 262887.7188 |
| DHRS4 | Q9BTZ2 | DHRS4 | Dehydrogenase/reduc | 162699.0156 | 180813.0781 |
| SLC27A1 | Q6PCB7 | SLC27A1 | Long-chain fatty acid t | 10449.91211 | 9848.773438 |
| MCM3 | P25205 | MCM3 | DNA replication licens | 789028.75 | 766911.8125 |
| RNF170 | E9PP55;Q96K19 | RNF170 | E3 ubiquitin-protein li | 4003.960205 | 5074.344238 |
| UBXN4 | Q92575 | UBXN4 | UBX domain-containir | 76261.35938 | 85316.375 |
| GNAI2 | P04899 | GNAI2 | Guanine nucleotide-b | 204437.7344 | 222235.2031 |
| PHKG2 | J3KNN3;P15735 | PHKG2 | Phosphorylase b kinas | 29845.23828 | 35354.67969 |
| MGME1 | Q9BQP7 | MGME1 | Mitochondrial genom | 49737.35938 | 59158.63281 |
| GNPDA2 | Q8TDQ7 | GNPDA2 | Glucosamine-6-phosp | 60150.59766 | 65142.25391 |
| ARF1 | P84077 | ARF1 | ADP-ribosylation facto | 1150566.375 | 1134525.375 |
| DPM3 | Q9P2X0 | DPM3 | Dolichol-phosphate m | 142320.5156 | 155483.5938 |
| SLC39A11 | J3KRX2;J3QLB2;J3QS49;Q8N1 | SLC39A11 | Zinc transporter ZIP11 | 15633.15137 | 20199.0293 |
| NUSAP1 | Q9BXS6 | NUSAP1 | Nucleolar and spindle | 68295.07813 | 67307.45313 |
| FMC1 | Q96HJ9 | FMC1 | Protein FMC1 homolo | 45220.27344 | 54341.54688 |
| DKC1 | O60832 | DKC1 | H/ACA ribonucleoprot | 189717.5469 | 86136.60156 |
| MSL1 | J3KSZ8;Q68DK7 | MSL1 | Male-specific lethal 1 | 16654.88867 | 20074.81641 |

| | | | | | |
|---|---|---|---|---|---|
| ALDH18A1 | P54886 | ALDH18A1 | Delta-1-pyrroline-5-ca | 607268.875 | 677916.375 |
| AFDN | A8MQ02 | AFDN | Afadin | 102808.5558 | 101514.751 |
| ZC3H12A | A0A1W2PQC8;Q5D1E8 | ZC3H12A | Endoribonuclease ZC3 | 94648.13281 | 93571.24219 |
| NOTCH2 | Q04721 | NOTCH2 | Neurogenic locus notc | 36736.18359 | 47914.80469 |
| RBM12B | Q8IXT5 | RBM12B | RNA-binding protein 1 | 61097.59766 | 47315.04688 |
| SLC5A3 | P53794 | SLC5A3 | Sodium/myo-inositol c | 242006.9063 | 326508.9688 |
| ARFRP1 | Q13795 | ARFRP1 | ADP-ribosylation facto | 18802.23828 | 20909.42773 |
| TYK2 | E9PM19;P29597 | TYK2 | Tyrosine-protein kinas | 6077.768066 | 7142.967285 |
| AKR1A1 | P14550 | AKR1A1 | Aldo-keto reductase fa | 265206.0625 | 288276.5313 |
| C22orf39 | A0A0C4DH14;Q6P5X5 | C22orf39 | UPF0545 protein C22o | 2552.939941 | 1679.272949 |
| RAB10 | P61026 | RAB10 | Ras-related protein Ra | 578765.6875 | 599141.875 |
| SCYL1 | E9PS17;Q96KG9 | SCYL1 | N-terminal kinase-like | 152274.3125 | 172286.7813 |
| STK10 | O94804 | STK10 | Serine/threonine-prot | 50221.25391 | 49500.13672 |
| DCAF6 | Q58WW2 | DCAF6 | DDB1- and CUL4-asso | 3339.509277 | 5205.169922 |
| RAB22A | Q9UL26 | RAB22A | Ras-related protein Ra | 125857.8359 | 123615.4766 |
| MPV17L2 | Q567V2 | MPV17L2 | Mpv17-like protein 2 | 15429.69434 | 9947.581055 |
| MCCC1 | Q96RQ3 | MCCC1 | Methylcrotonoyl-CoA | 81483.03125 | 89030.82031 |
| CACNB3 | F8VNV8;P54284 | CACNB3 | Voltage-dependent L- | 6046.810547 | 6943.978027 |
| IKBKB | A0A499FJS7;O14920 | IKBKB | Inhibitor of nuclear fa | 12911.39258 | 15992.79883 |
| TRA2A | Q13595 | TRA2A | Transformer-2 protein | 219634.4844 | 116938.3828 |
| APOH | J3KS17;P02749 | APOH | Beta-2-glycoprotein 1 | 16410.74805 | 15201.36914 |
| PLXNA1 | Q9UIW2 | PLXNA1 | Plexin-A1 | 13080.44727 | 13646.71582 |
| HACD2 | Q6Y1H2 | HACD2 | Very-long-chain (3R)-3 | 8014.262207 | 8702.867188 |
| RNF214 | Q8ND24 | RNF214 | RING finger protein 21 | 40838.1875 | 40362.64453 |
| PPP2R5E | Q16537 | PPP2R5E | Serine/threonine-prot | 88952.25781 | 95687.28125 |
| IVNS1ABP | Q9Y6Y0 | IVNS1ABP | Influenza virus NS1A-b | 60842.01563 | 55795.53125 |
| PLEKHG4 | Q58EX7 | PLEKHG4 | Puratrophin-1 | 4440.786621 | 7512.817871 |
| ATIC | P31939 | ATIC | Bifunctional purine bi | 965188.625 | 993331.0625 |
| MTX2 | O75431 | MTX2 | Metaxin-2 | 81065.70313 | 49236.91406 |
| ORC5 | O43913 | ORC5 | Origin recognition cor | 58204.00781 | 63463.75781 |
| PDK3 | Q15120 | PDK3 | [Pyruvate dehydroger | 33615.53906 | 41835.36328 |
| AAMDC | Q9H7C9 | AAMDC | Mth938 domain-conta | 23531.81055 | 25539.29102 |
| TINF2 | B4DFJ1;H0YKA6;H0YLC9;Q9B | TINF2 | TERF1-interacting nuc | 5606.608887 | 5784.520996 |

| | | | | | |
|---|---|---|---|---|---|
| OTULIN | Q96BN8 | OTULIN | Ubiquitin thioesterase | 71661.40625 | 77206.95313 |
| CYHR1 | E9PJD7 | CYHR1 | Cysteine and histidine | 26892.30664 | 34112.26172 |
| HEBP2 | Q5THN1;Q9Y5Z4 | HEBP2 | Heme-binding protein | 2362.80249 | 4836.562012 |
| HNRNPH2 | P55795 | HNRNPH2 | Heterogeneous nuclea | 590726.4375 | 484101.2813 |
| ZC3H7A | Q8IWR0 | ZC3H7A | Zinc finger CCCH doma | 27692.87305 | 21631.25977 |
| WIPI2 | Q9Y4P8 | WIPI2 | WD repeat domain ph | 35724.64063 | 40984.44531 |
| KDM2B | A0A0C4DGG3;Q8NHM5 | KDM2B | Lysine-specific demetl | 13028.3916 | 13431.95508 |
| FLII | Q13045 | FLII | Protein flightless-1 ho | 139907.8438 | 136144.8438 |
| GFAP | K7EPI4 | GFAP | Glial fibrillary acidic pr | 72718.78125 | 123522.5313 |
| KDELR1 | P24390 | KDELR1 | ER lumen protein-reta | 29864.37305 | 36064.56641 |
| PRPSAP1 | Q14558 | PRPSAP1 | Phosphoribosyl pyrop | 56677.47656 | 61701.53906 |
| MGAT4A | Q9UM21 | MGAT4A | Alpha-1,3-mannosyl-g | 8568.3125 | 15135.19629 |
| TOR2A | Q5JU69 | TOR2A | Torsin-2A | 12074.66797 | 12090.75488 |
| CIP2A | A0A087X211;Q8TCG1 | CIP2A | Protein CIP2A | 53224.28906 | 60022.29688 |
| CHM | P24386 | CHM | Rab proteins geranylg | 20792.90234 | 23397.83203 |
| XRN1 | Q8IZH2 | XRN1 | 5'-3' exoribonuclease | 25855.89258 | 26462.57617 |
| TCEA1 | A0A1W2PPZ5;P23193 | TCEA1 | Transcription elongati | 283034.7813 | 249901.7969 |
| ABHD16A | A0A0G2JJD3;O95870 | ABHD16A | HLA-B associated tran | 45398.71094 | 48431.16797 |
| TIMM22 | Q9Y584 | TIMM22 | Mitochondrial import | 93537.01563 | 74602.35156 |
| ALDH9A1 | P49189 | ALDH9A1 | 4-trimethylaminobuty | 247546.3906 | 248945.7344 |
| CYTH2 | A0A0D9SFG6;Q99418 | CYTH2 | Cytohesin-2 | 13732.11035 | 11298.1416 |
| KLC2 | Q9H0B6 | KLC2 | Kinesin light chain 2 | 57923.125 | 56354.98438 |
| GNG5 | P63218 | GNG5 | Guanine nucleotide-b | 97778.80469 | 137953.7344 |
| ASCC1 | A0A5F9ZH79;A0A5F9ZHP1 | ASCC1 | Activating signal coint | 35208.28125 | 38672.5625 |
| CENPQ | Q7L2Z9 | CENPQ | Centromere protein C | 12256.44336 | 11028.93262 |
| MORF4L1 | B3KTM8 | MORF4L1 | Mortality factor 4-like | 81173.02344 | 75658.11719 |
| OS9 | B4E321 | OS9 | Protein OS-9 | 87302.72168 | 93987.49414 |
| RAB2B | Q8WUD1 | RAB2B | Ras-related protein Ra | 12559.56836 | 12458.78125 |
| DCP2 | Q8IU60 | DCP2 | m7GpppN-mRNA hydr | 21679.32813 | 23393.57617 |
| E2F3 | O00716 | E2F3 | Transcription factor E2 | 1799.55835 | 2712.899658 |
| LIN9 | C9J5J4;H0Y322;H7C4T7;Q5TK | LIN9 | Protein lin-9 homolog | 17584.61914 | 14393.50098 |
| TBC1D24 | Q9ULP9 | TBC1D24 | TBC1 domain family m | 40617.91797 | 48571.69922 |
| CAPZB | B1AK87;B1AK88 | CAPZB | F-actin-capping protei | 1121286.75 | 1153212.758 |

| | | | | | |
|---|---|---|---|---|---|
| JUP | P14923 | JUP | Junction plakoglobin | 199615.9063 | 182092.3125 |
| FREM2 | Q5SZK8 | FREM2 | FRAS1-related extrace | 16997.95898 | 34049.46094 |
| NEBL | O76041 | NEBL | Nebulette | 49199.5 | 55550.53906 |
| PAK2 | Q13177 | PAK2 | Serine/threonine-prot | 381039.7813 | 382275.8125 |
| ARL6IP5 | O75915 | ARL6IP5 | PRA1 family protein 3 | 335618.2813 | 344167.4688 |
| TRMT2A | F2Z2W7;Q8IZ69 | TRMT2A | tRNA (uracil-5-)-meth | 25584.65625 | 31543.03125 |
| NSD3 | Q9BZ95 | NSD3 | Histone-lysine N-meth | 11295.51172 | 10133.05762 |
| PKN1 | Q16512 | PKN1 | Serine/threonine-prot | 52671.52734 | 55611.76563 |
| TBC1D17 | Q9HA65 | TBC1D17 | TBC1 domain family m | 28304.80078 | 31571.95703 |
| CERS2 | Q5SZE1;Q96G23 | CERS2 | Ceramide synthase 2 ( | 285766.9063 | 248080.4375 |
| PGP | A6NDG6 | PGP | Glycerol-3-phosphate | 415526.7188 | 423016.0313 |
| C3orf38 | Q5JPI3 | C3orf38 | Uncharacterized prote | 10660.89453 | 12234.42871 |
| BAD | Q92934 | BAD | Bcl2-associated agoni | 49013.96094 | 47090.10938 |
| TTC33 | Q6PID6 | TTC33 | Tetratricopeptide rep | 8095.367676 | 8757.703125 |
| ZNF335 | Q9H4Z2 | ZNF335 | Zinc finger protein 33 | 3444.52417 | 3849.888672 |
| KATNA1 | O75449 | KATNA1 | Katanin p60 ATPase-c | 22519.41016 | 24154.09375 |
| AIDA | Q96BJ3 | AIDA | Axin interactor, dorsa | 39205.47656 | 43419.37109 |
| MISP3 | A0A044PY82;Q96FF7 | MISP3 | Uncharacterized prote | 9129.003906 | 11059.75098 |
| TIMM29 | Q9BSF4 | TIMM29 | Mitochondrial import | 64807.40625 | 71167.10938 |
| STK4 | Q13043 | STK4 | Serine/threonine-prot | 29841.57813 | 32768.70703 |
| ERLEC1 | Q96DZ1 | ERLEC1 | Endoplasmic reticulur | 56740.72656 | 55494.64453 |
| PCNA | P12004 | PCNA | Proliferating cell nucle | 692841.6875 | 753415.0625 |
| RNPEP | Q9H4A4 | RNPEP | Aminopeptidase B | 381504.3438 | 478202.2813 |
| CPT2 | P23786 | CPT2 | Carnitine O-palmitoylt | 63616.35156 | 69511.14844 |
| ACOT9 | Q9Y305 | ACOT9 | Acyl-coenzyme A thio | 97554.89063 | 93117.63281 |
| SNX15 | Q9NRS6 | SNX15 | Sorting nexin-15 | 21938.62109 | 21902.3457 |
| CDK19 | Q9BWU1 | CDK19 | Cyclin-dependent kina | 25327.91602 | 29404.10352 |
| POLE | Q07864 | POLE | DNA polymerase epsil | 26619.30664 | 29459.72266 |
| MAP7 | Q14244 | MAP7 | Ensconsin | 145077.5625 | 136407.8438 |
| SOX13 | Q9UN79 | SOX13 | Transcription factor S | 614.918457 | 2417.666504 |
| PITPNC1 | A0A0C4DGP0;E7EU35 | PITPNC1 | Cytoplasmic phosphat | 35991.05078 | 35192.77344 |
| TP53I3 | Q53FA7 | TP53I3 | Quinone oxidoreducta | 138048.1094 | 141835.2031 |
| PPARD | Q03181 | PPARD | Peroxisome proliferat | 55475.24609 | 25910.08594 |

| | | | | | |
|---|---|---|---|---|---|
| CAPZA1 | P52907 | CAPZA1 | F-actin-capping protei | 655656.3125 | 648983.1875 |
| ASMTL | O95671 | ASMTL | Probable bifunctional | 66589.51563 | 75549.70313 |
| LIN54 | Q6MZP7 | LIN54 | Protein lin-54 homolo | 16130.70996 | 13796.05078 |
| GDI2 | P50395 | GDI2 | Rab GDP dissociation | 1085838.875 | 1169329.375 |
| CHMP4A | Q9BY43 | CHMP4A | Charged multivesicula | 80386.53906 | 106301.9141 |
| EFCAB14 | A0A5F9ZHQ6;O75071 | EFCAB14 | EF-hand calcium-bindi NA | | 4261.464355 |
| APOA1 | F8W696;P02647 | APOA1 | Apolipoprotein A-I | 32773.26563 | 21354.30664 |
| ZFYVE27 | Q5T4F4 | ZFYVE27 | Protrudin | 7982.546387 | 7499.009766 |
| EIF4G3 | A0A0A0MSA7;A0A0U1RQK7; | EIF4G3 | Eukaryotic translation | 32580.28125 | 40455.53516 |
| FHIT | P49789 | FHIT | Bis(5'-adenosyl)-triph | 7868.555176 | 9494.680664 |
| HBS1L | Q9Y450 | HBS1L | HBS1-like protein | 40983.83594 | 48279.32031 |
| ABR | A0A0G2JQ41;A0A1C7CYZ0;Q | ABR | Active breakpoint clus | 59345.75659 | 74293.59912 |
| ELF2 | D6R9B1 | ELF2 | ETS-related transcript | 6563.377441 | 5170.750488 |
| DTNBP1 | Q96EV8 | DTNBP1 | Dysbindin | 28265.40234 | 26213.60742 |
| SNTB2 | Q13425 | SNTB2 | Beta-2-syntrophin | 65370.60938 | 63606.33984 |
| MESD | Q14696 | MESD | LRP chaperone MESD | 1076869.625 | 1091201.5 |
| SLC25A25 | Q6KCM7 | SLC25A25 | Calcium-binding mito | 6319.769043 | 9735.990234 |
| FBXO21 | O94952;Q4G104 | FBXO21 | F-box only protein 21 | 21038.47461 | 22707.52148 |
| ESYT1 | Q9BSJ8 | ESYT1 | Extended synaptotagr | 418999.6563 | 441385.25 |
| ELOVL7 | A1L3X0;D6RBM2;D6RBR5;D6 | ELOVL7 | Elongation of very lon | 12333.23828 | 7128.112305 |
| NPLOC4 | Q8TAT6 | NPLOC4 | Nuclear protein locali | 206700.3438 | 201318.5156 |
| PDAP1 | Q13442 | PDAP1 | 28 kDa heat- and acid | 429402.6563 | 466740.25 |
| DICER1 | Q9UPY3 | DICER1 | Endoribonuclease Dic | 105069.2188 | 96470.38281 |
| B4GALT7 | Q9UBV7 | B4GALT7 | Beta-1,4-galactosyltra | 16364.29102 | 12528.08887 |
| CRYZ | Q08257 | CRYZ | Quinone oxidoreducta | 180654.25 | 178357.7969 |
| CSK | P41240 | CSK | Tyrosine-protein kinas | 224930.5469 | 230297.8281 |
| TEDC1 | Q86SX3 | TEDC1 | Tubulin epsilon and d | 33168.00781 | 35595.35938 |
| ABRAXAS2 | Q15018 | ABRAXAS2 | BRISC complex subuni | 53384.82031 | 66917.08594 |
| HACL1 | Q9UJ83 | HACL1 | 2-hydroxyacyl-CoA lya | 23019.20313 | 22258.28711 |
| HMGN1 | P05114 | HMGN1 | Non-histone chromos | 442088.75 | 172129.625 |
| EIF4EBP1 | Q13541 | EIF4EBP1 | Eukaryotic translation | 138807.2656 | 150595.8438 |
| LIN7C | Q9NUP9 | LIN7C | Protein lin-7 homolog | 131229.7813 | 111780.5156 |
| SH3GLB1 | A0A087WW40;Q9Y371 | SH3GLB1 | Endophilin-B1 | 20516.71875 | 26052.05273 |

| | | | | | |
|---|---|---|---|---|---|
| ITFG1 | Q8TB96 | ITFG1 | T-cell immunomodula | 14681.95898 | 14637.95508 |
| RPA1 | P27694 | RPA1 | Replication protein A | 158241.1563 | 165686.375 |
| MPDZ | F5H1U9;O75970 | MPDZ | Multiple PDZ domain | 1824.920776 | 2275.231445 |
| ACAD9 | Q9H845 | ACAD9 | Complex I assembly fa | 156526.3594 | 198828.7969 |
| KIN | O60870 | KIN | DNA/RNA-binding pro | 59110.90234 | 43590.35547 |
| MGRN1 | K7EPJ5;O60291 | MGRN1 | E3 ubiquitin-protein li | 10538.5166 | 10415.43457 |
| CDC42SE2 | Q9NRR3 | CDC42SE2 | CDC42 small effector | 32097.28711 | 30707.57422 |
| SLC25A46 | Q96AG3 | SLC25A46 | Solute carrier family 2 | 9765.895508 | 19027.51172 |
| SLC25A44 | E9PGQ0;Q96H78 | SLC25A44 | Solute carrier family 2 | 7270.146484 | 7735.806152 |
| MTFP1 | Q9UDX5 | MTFP1 | Mitochondrial fission | 59067.73047 | 73018.02344 |
| TUBB4B | P68371 | TUBB4B | Tubulin beta-4B chain | 11647086 | 13419420 |
| PEX11B | O96011 | PEX11B | Peroxisomal membrar | 61250.30469 | 64445.16406 |
| NCOA3 | Q9Y6Q9 | NCOA3 | Nuclear receptor coac | 51644.73047 | 59275.63672 |
| COG4 | J3KNI1;Q9H9E3 | COG4 | Conserved oligomeric | 51725.91406 | 63799.57813 |
| SSH3 | Q8TE77 | SSH3 | Protein phosphatase S | 123888.4922 | 130999.9766 |
| AP1S1 | A0A2R8YGH5;P61966 | AP1S1 | AP complex subunit si | 324649.3125 | 346365.6563 |
| ARPC4-TTLL3;ARPC4 | F8WCF6;P59998 | ARPC4-TTLL3;ARPC4 | Actin-related protein | 750967.5625 | 853149.75 |
| WDR34 | Q96EX3 | WDR34 | WD repeat-containing | 24614.21094 | 22622.82031 |
| CAMLG | P49069 | CAMLG | Calcium signal-modula | 33163.98828 | 34870.84766 |
| ABHD12 | Q8N2K0 | ABHD12 | Lysophosphatidylserir | 65150.42188 | 81740.14063 |
| PRDM2 | Q13029 | PRDM2 | PR domain zinc finger | 3934.398682 | 3457.580811 |
| TBC1D7 | Q5SZL3;Q5SZL4;Q5SZL5;Q5SZ | TBC1D7 | TBC1 domain family m | 12560.61426 | 10563.01074 |
| SCAMP2 | O15127 | SCAMP2 | Secretory carrier-asso | 73631.75781 | 78246.78125 |
| SNX7 | Q9UNH6 | SNX7 | Sorting nexin-7 | 19237.03711 | 16930.34375 |
| VPS4A | Q9UN37 | VPS4A | Vacuolar protein sorti | 283815.2813 | 291866.4688 |
| QPCTL | Q9NXS2 | QPCTL | Glutaminyl-peptide cy | 40564.41016 | 36385.26953 |
| SSBP3 | Q9BWW4 | SSBP3 | Single-stranded DNA-I | 19949.125 | 25174.9707 |
| DDRGK1 | Q96HY6 | DDRGK1 | DDRGK domain-conta | 158675.6406 | 131083.3594 |
| ARIH1 | Q9Y4X5 | ARIH1 | E3 ubiquitin-protein li | 58966.81641 | 64970.48047 |
| TTC17 | Q96AE7 | TTC17 | Tetratricopeptide rep | 3536.329834 | 1744.981323 |
| NCOA5 | Q9HCD5 | NCOA5 | Nuclear receptor coac | 152435.7031 | 112019.2422 |
| DDOST | A0A0C4DGS1;P39656 | DDOST | Dolichyl-diphosphooli | 816368.75 | 910955.1875 |
| CDC42BPB | Q9Y5S2 | CDC42BPB | Serine/threonine-prot | 55240.45313 | 61286.71484 |

| | | | | | |
|---|---|---|---|---|---|
| PITRM1 | Q5JRX3 | PITRM1 | Presequence protease | 183747.0469 | 213548.9844 |
| OGDH | Q02218 | OGDH | 2-oxoglutarate dehydı | 322783.875 | 378060.75 |
| METTL9 | Q8TD49;Q9H1A3 | METTL9 | Methyltransferase-like | 13862.79492 | 15757.27246 |
| SURF1 | Q15526 | SURF1 | Surfeit locus protein 1 | 97093.14844 | 99371.72656 |
| KYNU | Q16719 | KYNU | Kynureninase | 436571.9688 | 418307.25 |
| ARHGEF4;;ARHGEF9;ARHGI | A0A087X0P1;A0A0A0MRY4;/ | ARHGEF4;;ARHGEF9;/ | Rho guanine nucleotic | 2508.356445 | 2528.432129 |
| TMEM177 | Q53S58 | TMEM177 | Transmembrane prote | 12916.35938 | NA |
| LAMTOR2 | Q9Y2Q5 | LAMTOR2 | Ragulator complex pro | 101775.1016 | 100022.3203 |
| RNF7 | Q9UBF6 | RNF7 | RING-box protein 2 | 41289.75 | 42317.39063 |
| PNMA1 | Q8ND90 | PNMA1 | Paraneoplastic antiger | 5806.552246 | 9184.414063 |
| PGS1 | Q32NB8 | PGS1 | CDP-diacylglycerol--gl | 11095.6582 | 12787.98926 |
| HPX | P02790;Q9BS19 | HPX | Hemopexin;Epididymi | 18104.05273 | NA |
| ATG5 | Q9H1Y0 | ATG5 | Autophagy protein 5 | 43673.61328 | 40760.36719 |
| TRAPPC9 | H0YBR0;Q96Q05 | TRAPPC9 | Trafficking protein par | 25499.45703 | 25408.36719 |
| UBE2K | P61086 | UBE2K | Ubiquitin-conjugating | 445781.3438 | 475063.0625 |
| MYO19 | Q96H55 | MYO19 | Unconventional myos | 9505.648438 | 7000.373047 |
| HSD17B10 | Q99714 | HSD17B10 | 3-hydroxyacyl-CoA de | 722351.625 | 803131.5 |
| RHOB | P62745 | RHOB | Rho-related GTP-bind | 86790.19531 | 91402.82031 |
| SLC35A2 | P78381 | SLC35A2 | UDP-galactose translo | 12960.0166 | 20657.19336 |
| FAM234A | Q9H0X4 | FAM234A | Protein FAM234A | 25668.33203 | 29017.66602 |
| DCUN1D1 | Q96GG9 | DCUN1D1 | DCN1-like protein 1 | 83952.75781 | 86846.38281 |
| KLHDC3 | E7ERR0;Q9BQ90 | KLHDC3 | Kelch domain-contain | 5103.956543 | 6598.822754 |
| NOP10 | Q9NPE3 | NOP10 | H/ACA ribonucleoprot | 77351.69531 | 34832.90625 |
| IKBKG | A0A087X1B1;Q9Y6K9 | IKBKG | NF-kappa-B essential | 65434.53516 | 71276.28906 |
| MTHFD1L | A0A087WVM4;B7ZM99 | MTHFD1L | Monofunctional C1-te | 156080.8125 | 173091.4844 |
| LRRC8D | Q7L1W4 | LRRC8D | Volume-regulated ani | 2549.930908 | 3486.361084 |
| HMBS | P08397 | HMBS | Porphobilinogen dean | 125588.2422 | 136428.5781 |
| RILPL1 | Q5EBL4 | RILPL1 | RILP-like protein 1 | 19295.35352 | 21703.51953 |
| TONSL | Q96HA7 | TONSL | Tonsoku-like protein | 16658.13672 | 21016.01172 |
| NFRKB | Q6P4R8 | NFRKB | Nuclear factor related | 9504.042969 | 9780.032227 |
| NISCH | Q9Y2I1 | NISCH | Nischarin | 117317.0859 | 110516.4766 |
| XRCC3 | O43542 | XRCC3 | DNA repair protein XR | 41789.01563 | 48109.59375 |
| TAF5L | O75529 | TAF5L | TAF5-like RNA polyme | 9350.795898 | 9203.541992 |

| | | | | | |
|---|---|---|---|---|---|
| IFT27 | Q9BW83 | IFT27 | Intraflagellar transpor | 54245.59766 | 56937.19531 |
| DOCK9 | A0A088AWN3;A0A0A0MSY4; | DOCK9 | Dedicator of cytokines | 5091.229492 | 5140.946777 |
| ZCCHC3 | Q9NUD5 | ZCCHC3 | Zinc finger CCHC doma | 14716.24805 | 10481.17773 |
| DENND10P1 | Q6NSW5 | DENND10P1 | Putative DENN domai | 54304.15234 | 50930.14844 |
| ARF4 | P18085 | ARF4 | ADP-ribosylation facto | 745152.5 | 738011.0625 |
| ERI3 | H0Y4B0;O43414 | ERI3 | ERI1 exoribonuclease | 66091.23438 | 73948.29688 |
| TUBB1 | Q9H4B7 | TUBB1 | Tubulin beta-1 chain | 301170.0625 | 313451.9063 |
| ARCN1 | P48444 | ARCN1 | Coatomer subunit del | 606317.1875 | 652527.875 |
| STXBP3 | O00186 | STXBP3 | Syntaxin-binding prot | 121675.5625 | 138165.3281 |
| PRSS8 | Q16651 | PRSS8 | Prostasin | 10603.97949 | 16859.35742 |
| SEC61G | P60059 | SEC61G | Protein transport prot | 201455.8125 | 234663.5781 |
| COG2 | Q14746 | COG2 | Conserved oligomeric | 53387.09375 | 61456.88281 |
| ARHGEF12 | E9PMR6;Q9NZN5 | ARHGEF12 | Rho guanine nucleotic | 13660.10254 | 15883.98145 |
| TAF6L | Q9Y6J9 | TAF6L | TAF6-like RNA polyme | 20592.38086 | 19516.48438 |
| MFGE8 | Q08431 | MFGE8 | Lactadherin | 52660.53906 | 67351.22656 |
| ZCCHC10 | B3KVL5;Q8TBK6 | ZCCHC10 | Zinc finger CCHC doma | 3322.861084 | 33096.97266 |
| KRT8 | P05787 | KRT8 | Keratin, type II cytosk | 56924452 | 69068848 |
| PGLS | O95336 | PGLS | 6-phosphogluconolact | 268671.3125 | 287082.5625 |
| PIP5K1A | A6PW57 | PIP5K1A | Phosphatidylinositol 4 | 22976.68555 | 21929.49023 |
| TYW5 | A2RUC4 | TYW5 | tRNA wybutosine-synt | 10579.84961 | 18295.61719 |
| HDAC1 | Q13547 | HDAC1 | Histone deacetylase 1 | 223175.3125 | 233086.5469 |
| ARFIP2 | A0A087X1E4;P53365 | ARFIP2 | Arfaptin-2 | 49278.28125 | 49972.69531 |
| C6orf89 | Q6UWU4 | C6orf89 | Bombesin receptor-ac | 14618.79004 | 16973.66992 |
| SMAP1 | Q8IYB5 | SMAP1 | Stromal membrane-as | 35439.55469 | 35852.13281 |
| UBE2J2 | A6NGS0;B1AMF0;D6RD90;Q8 | UBE2J2 | Ubiquitin-conjugating | 5335.176758 | 7591.424805 |
| SCRIB | A0A0G2JNZ2;A0A0G2JPP5;Q1 | SCRIB | Protein scribble homo | 200418.1172 | 244311.7031 |
| SERPINH1 | P50454 | SERPINH1 | Serpin H1 | 953248 | 894595.5 |
| UBE2F | C9J9P8;Q969M7 | UBE2F | NEDD8-conjugating er | 18810.60156 | 20192.15039 |
| STT3A | P46977 | STT3A | Dolichyl-diphosphooli | 423272.4375 | 478184.5938 |
| RBFOX2 | A0A0G2JRA5;B0QYV1;O4325 | RBFOX2 | RNA-binding protein f | 14723.72656 | 14544.24609 |
| OAT | P04181 | OAT | Ornithine aminotrans | 456268.6875 | 596144.5625 |
| CRADD | P78560 | CRADD | Death domain-contair | 18970.08398 | 16856.55469 |
| UACA | Q9BZF9 | UACA | Uveal autoantigen wit | 19334.82617 | 20109.80469 |

| | | | | | |
|---|---|---|---|---|---|
| PIH1D3 | Q9NQM4 | PIH1D3 | Protein PIH1D3 | 52895.70703 | 32990.625 |
| SYNJ2BP-COX16 | A0A087WYV9;A0A087X1F5 | SYNJ2BP-COX16 | SYNJ2BP-COX16 readt | 106797.7266 | 114916.2969 |
| PSMC2 | P35998 | PSMC2 | 26S proteasome regul | 481803.8438 | 586768.5 |
| SAPCD2 | Q86UD0 | SAPCD2 | Suppressor APC doma | 37158.60156 | 30059.25781 |
| TWF1 | Q12792 | TWF1 | Twinfilin-1 | 129788.4922 | 141196.7188 |
| PSMD14 | O00487 | PSMD14 | 26S proteasome non-/ | 466570.4375 | 551392.6875 |
| AIP | O00170 | AIP | AH receptor-interactir | 132619.7813 | 147025.375 |
| PTPN13 | Q12923 | PTPN13 | Tyrosine-protein phos | 4237.188477 | 5345.573242 |
| N6AMT1 | Q9Y5N5 | N6AMT1 | Methyltransferase N6 | 19841.14648 | 20149.68945 |
| CCDC134 | Q9H6E4 | CCDC134 | Coiled-coil domain-co | 15238.93066 | 15792.37207 |
| PKN2 | Q16513 | PKN2 | Serine/threonine-prot | 618110.125 | 686814.25 |
| PTPN11 | Q06124 | PTPN11 | Tyrosine-protein phos | 155307.9531 | 160378.2656 |
| BLVRA | P53004 | BLVRA | Biliverdin reductase A | 1067534.25 | 1129181.125 |
| CSNK1G3 | Q9Y6M4 | CSNK1G3 | Casein kinase I isoforr | 7018.56543 | 9008.699219 |
| TEX264 | Q9Y6I9 | TEX264 | Testis-expressed prote | 78722.15625 | 78374.5 |
| PSMD7 | P51665 | PSMD7 | 26S proteasome non-/ | 400218.1563 | 460726.2813 |
| EFR3A | A0A1B0GUZ7;Q14156 | EFR3A | Protein EFR3 homolog | 6013.241699 | 11111.46582 |
| NFYA | P23511 | NFYA | Nuclear transcription | 35491.64063 | 25998.88477 |
| COL3A1 | P02461 | COL3A1 | Collagen alpha-1(III) c | 7412.203613 | 8557.223633 |
| TTC26 | A0AVF1;B7Z6R6 | TTC26 | Intraflagellar transpor | 5619.585449 | 6935.938477 |
| NHLRC3 | Q5JS37 | NHLRC3 | NHL repeat-containing | 12098.28906 | 11710.31641 |
| PSMC1 | P62191 | PSMC1 | 26S proteasome regul | 834910.6875 | 806565 |
| PEF1 | Q9UBV8 | PEF1 | Peflin | 154669.5625 | 177572.125 |
| BRPF1 | A0A5F9ZH11;P55201 | BRPF1 | Peregrin (Fragment);P | 10085.88477 | 9034.178711 |
| NRBP1 | Q9UHY1 | NRBP1 | Nuclear receptor-bind | 115243.8672 | 120939.4063 |
| ALDH1L2 | A0A494C1M4;Q3SY69 | ALDH1L2 | 10-formyltetrahydrofo | 20685.61133 | 23812.55469 |
| NMT1 | P30419 | NMT1 | Glycylpeptide N-tetra | 196331.5625 | 207016.1406 |
| FRYL | O94915 | FRYL | Protein furry homolog | 68076.78125 | 79794.6875 |
| ARID3A | Q99856 | ARID3A | AT-rich interactive dor | 27208.65234 | 26453.07031 |
| TGFBRAP1 | Q8WUH2 | TGFBRAP1 | Transforming growth | 21120.5293 | 23963.32617 |
| YIF1A | A6NGW1;O95070 | YIF1A | Protein YIF1A | 35039.53516 | 41550.39844 |
| SPATA5 | Q8NB90 | SPATA5 | ATPase family protein | 48321.9375 | 54115.76953 |
| EMC2 | Q15006 | EMC2 | ER membrane protein | 172226.2969 | 173852.2656 |

| | | | | | |
|---|---|---|---|---|---|
| PSPC1 | Q8WXF1 | PSPC1 | Paraspeckle compone | 287988.625 | 212204.125 |
| SCLY | A0A0A0MQU4;Q96I15 | SCLY | Selenocysteine lyase | 17760.12891 | 23249.23438 |
| MAPRE2 | K7EL66;K7ERD8;Q15555 | MAPRE2 | Microtubule-associate | 12177.88867 | 11821.9541 |
| PDS5A | Q29RF7 | PDS5A | Sister chromatid cohe | 152979.1875 | 166156.2031 |
| TMEM132A | Q24JP5 | TMEM132A | Transmembrane prote | 4792.569824 | 4895.146484 |
| COQ8A | Q8NI60 | COQ8A | Atypical kinase COQ8/ | 48523.90625 | 49998.67188 |
| TTC1 | Q99614 | TTC1 | Tetratricopeptide rep | 66904.21875 | 56105.14453 |
| ARL1 | P40616 | ARL1 | ADP-ribosylation facto | 182361.5313 | 216564.5156 |
| HIST2H2AB | Q8IUE6 | HIST2H2AB | Histone H2A type 2-B | 262972.4688 | 65309.98047 |
| COG3 | Q96JB2 | COG3 | Conserved oligomeric | 58593.4375 | 77799.44531 |
| CENPB | P07199 | CENPB | Major centromere aut | 32775.66406 | 31856.87305 |
| FAHD1 | Q6P587 | FAHD1 | Acylpyruvase FAHD1, | 173662.8125 | 194280.9063 |
| PSMD10 | B1AJY7;O75832 | PSMD10 | 26S proteasome non-/ | 134429.0156 | 142798.8438 |
| UBA6 | A0AVT1 | UBA6 | Ubiquitin-like modifie | 157447.9688 | 193282.4375 |
| SECISBP2 | Q5HYY5;Q96T21 | SECISBP2 | Selenocysteine inserti | 8541.992188 | 7372.987305 |
| UBFD1 | H3BRL3;O14562 | UBFD1 | Ubiquitin domain-con | 174797.25 | 203796.625 |
| KDM4A | O75164 | KDM4A | Lysine-specific demetl | 18343.44922 | 17200.5957 |
| PSMF1 | Q5QPM7;Q92530 | PSMF1 | Proteasome inhibitor | 82248.52344 | 87038.125 |
| SFXN5 | Q8TD22 | SFXN5 | Sideroflexin-5 | 11168.9707 | 16528.55664 |
| PHC3 | Q8NDX5 | PHC3 | Polyhomeotic-like pro | 31549.60352 | 37251.94531 |
| PAXX | Q9BUH6 | PAXX | Protein PAXX | 105568.3125 | 157259.4844 |
| FAM133B | G3XAI9;Q5BKY9 | FAM133B | Protein FAM133B | 37166.78516 | 32091.32617 |
| POLR1B | Q9H9Y6 | POLR1B | DNA-directed RNA po | 77872.39844 | 54570.20703 |
| VAMP5 | O95183 | VAMP5 | Vesicle-associated me | 12515.34766 | 11249.84473 |
| OGFOD2 | F5H145;Q6N063 | OGFOD2 | 2-oxoglutarate and irc | 4548.419922 | 5219.038086 |
| KANSL1 | A0A0G2JNB1;A0A0G2JNT7;A | KANSL1 | KAT8 regulatory NSL c | 2743.904297 | 4874.911133 |
| RELA | A0A087X0W8;E9PKH5;Q042( | RELA | Transcription factor p | 91988.27344 | 103846.1563 |
| RP9 | Q8TA86 | RP9 | Retinitis pigmentosa 9 | 27295.9707 | 25949.7207 |
| DPH7 | Q9BTV6 | DPH7 | Diphthine methyltrans | 7321.372559 | 8818.667969 |
| VTI1B | Q9UEU0 | VTI1B | Vesicle transport thro | 67197.0625 | 77551.96094 |
| CERCAM | A0A087WYE2;Q5T4B2 | CERCAM | Inactive glycosyltransf | 6711.04834 | 9032.283203 |
| CPNE1 | F2Z2V0;Q99829 | CPNE1 | Copine-1 (Fragment);( | 249422.125 | 301371.8125 |
| EEF1AKMT1 | Q8WVE0 | EEF1AKMT1 | EEF1A lysine methyltr | 12906.80176 | 17382.72266 |

| | | | | | |
|---|---|---|---|---|---|
| LETM1 | O95202 | LETM1 | Mitochondrial proton/ | 138498.7188 | 157795.0625 |
| RAB35 | Q15286 | RAB35 | Ras-related protein Ra | 78885.20313 | 78353.53125 |
| KDF1 | Q8NAX2 | KDF1 | Keratinocyte different | 13613.33984 | 10221.04492 |
| IDH3G | P51553 | IDH3G | Isocitrate dehydrogen | 123573.875 | 141780.1563 |
| NDUFAF6 | A0A075B6P0;Q330K2 | NDUFAF6 | NADH dehydrogenase | 13098.56543 | 14076.26172 |
| IST1 | H3BUI0 | IST1 | IST1 homolog (Fragme | 109374.1719 | 124128.3633 |
| HEATR5B | Q9P2D3 | HEATR5B | HEAT repeat-containi | 49487.75391 | 60486.63672 |
| EHMT1 | Q9H9B1 | EHMT1 | Histone-lysine N-meth | 34732.98438 | 35281.72266 |
| ZDHHC7 | Q9NXF8 | ZDHHC7 | Palmitoyltransferase Z | 5515.112305 | 5275.887695 |
| YIPF6 | Q96EC8 | YIPF6 | Protein YIPF6 | 68617 | 77518.19531 |
| ADPRHL2 | Q9NX46 | ADPRHL2 | ADP-ribose glycohydr | 143436.0469 | 141385 |
| BCAT2 | O15382 | BCAT2 | Branched-chain-amin | 114639.5234 | 122554.4844 |
| C18orf32 | Q8TCD1 | C18orf32 | UPF0729 protein C18 | 11326.12305 | 14175.40234 |
| PPIL1 | Q9Y3C6 | PPIL1 | Peptidyl-prolyl cis-tra | 150026.3438 | 141949.7344 |
| DENND4C | Q5VZ89 | DENND4C | DENN domain-contain | 16077.39063 | 19944.01563 |
| PLEKHH3 | Q7Z736;X6R3U4 | PLEKHH3 | Pleckstrin homology d | 15001.68164 | 13788.67188 |
| SLC25A32 | Q9H2D1 | SLC25A32 | Mitochondrial folate t | 18585.69531 | 16064.62402 |
| PIGK | Q92643 | PIGK | GPI-anchor transamid | 80702.82813 | 84656.95313 |
| HDDC2 | Q7Z4H3 | HDDC2 | HD domain-containing | 223529.5313 | 247380.6094 |
| GIT1 | Q9Y2X7 | GIT1 | ARF GTPase-activating | 77066.75781 | 81216.51563 |
| CEBPZOS | A8MTT3 | CEBPZOS | Protein CEBPZOS | 6531.390137 | 6545.401367 |
| CHTF18 | E7EXA6;Q8WVB6 | CHTF18 | Chromosome transmi | 23730.25586 | 24551.15234 |
| LNPK | Q9C0E8 | LNPK | Endoplasmic reticulum | 68145.21094 | 69911.21094 |
| PAPOLA | P51003 | PAPOLA | Poly(A) polymerase al | 95316.07031 | 110997.8438 |
| TFG | Q92734 | TFG | Protein TFG | 307280.5625 | 325476.5 |
| CCS | J3KNF4;O14618 | CCS | Copper chaperone for | 8241.584961 | 20268.13086 |
| SFR1 | Q86XK3 | SFR1 | Swi5-dependent reco | 2207.58667 | 2567.927246 |
| HYOU1 | A0A494C039;Q9Y4L1 | HYOU1 | Hypoxia up-regulated | 868945.875 | 935372.875 |
| PPHLN1 | F8W0Q9 | PPHLN1 | Periphilin-1 | 27381.12939 | 29537.32471 |
| DDX41 | Q9UJV9 | DDX41 | Probable ATP-depend | 102649.8594 | 88023.00781 |
| MAP4K5 | Q9Y4K4 | MAP4K5 | Mitogen-activated pro | 18960.32422 | 27131.79883 |
| HLTF | A0A0C4DGA6;Q14527 | HLTF | Helicase-like transcrip | 40884.3125 | 43017.17969 |
| CDK1 | P06493 | CDK1 | Cyclin-dependent kina | 218496.25 | 250215.6563 |

| | | | | | |
|---|---|---|---|---|---|
| HNRNPU | Q00839 | HNRNPU | Heterogeneous nucle | 833141.9375 | 739869.9375 |
| LATS1 | O95835 | LATS1 | Serine/threonine-prot | 10863.14648 | 25448.07617 |
| YIPF5 | Q969M3 | YIPF5 | Protein YIPF5 | 20865.30078 | 28710.11914 |
| CISD3 | P0C7P0 | CISD3 | CDGSH iron-sulfur dor | 6949.450684 | 11081.83594 |
| SH2D4A | Q9H788 | SH2D4A | SH2 domain-containir | 10766.37109 | 9803.398438 |
| PSMD2 | Q13200 | PSMD2 | 26S proteasome non- | 534077.75 | 563382.5 |
| PRKACA | P17612 | PRKACA | cAMP-dependent pro | 146171.625 | 168795.125 |
| PYM1 | Q9BRP8 | PYM1 | Partner of Y14 and ma | 150317.7344 | 134618.9531 |
| PTOV1 | M0QY25;Q86YD1 | PTOV1 | Prostate tumor-overe | 12779.98438 | 13749.39551 |
| OSBPL5 | Q9H0X9 | OSBPL5 | Oxysterol-binding pro | 12768.62695 | 10771.50781 |
| AFM | P43652 | AFM | Afamin | 28774.8125 | 23022.20313 |
| UBL7 | Q96S82 | UBL7 | Ubiquitin-like protein | 6876.004883 | 10744.87891 |
| PPCS | Q9HAB8 | PPCS | Phosphopantothenate | 145046.5313 | 169178.7969 |
| PRKRA | O75569 | PRKRA | Interferon-inducible d | 85107.91406 | 80058.32031 |
| PANK2 | Q9BZ23 | PANK2 | Pantothenate kinase 2 | 4902.983398 | 5918.433105 |
| OSTC | Q9NRP0 | OSTC | Oligosaccharyltransfer | 217291.9688 | 211304.2344 |
| LMO7 | E9PMS6;F8WD26;J3KP06;Q8 | LMO7 | LIM domain only prot | 9035.999023 | 7286.399902 |
| TTC4 | O95801 | TTC4 | Tetratricopeptide rep | 102138.8906 | 108141.4531 |
| ACTR3 | P61158 | ACTR3 | Actin-related protein | 634747.25 | 700968.9375 |
| QSOX1 | O00391 | QSOX1 | Sulfhydryl oxidase 1 | 17409.33398 | 15576.98438 |
| PTDSS2 | Q9BVG9 | PTDSS2 | Phosphatidylserine sy | 7496.334473 | 7840.428223 |
| OSBPL11 | Q9BXB4 | OSBPL11 | Oxysterol-binding pro | 46948.25781 | 50693.01172 |
| NBEAL2 | A0A494C1V1;Q6ZNJ1 | NBEAL2 | Neurobeachin-like pro | 20083.07227 | 30642.40039 |
| RASA1 | E9PGC0;P20936 | RASA1 | Ras GTPase-activating | 48450.375 | 52397.98047 |
| HK1 | P19367 | HK1 | Hexokinase-1 | 906073.5625 | 386595.0625 |
| ABL1 | P00519 | ABL1 | Tyrosine-protein kina | 7414.395508 | 7719.489746 |
| AP2B1 | P63010 | AP2B1 | AP-2 complex subunit | 369652.25 | 411875.6563 |
| GEMIN7 | Q9H840 | GEMIN7 | Gem-associated prote | 19722.75195 | 30041.04102 |
| TARBP2 | Q15633 | TARBP2 | RISC-loading complex | 18107.20508 | 13166.91992 |
| CXXC1 | Q9P0U4 | CXXC1 | CXXC-type zinc finger | 21055.88477 | 20785.52344 |
| TP53I11 | E9PIK4;E9PIN5;E9PKN9;E9PK | TP53I11 | Tumor protein p53-in | 24350.03125 | 35179.95703 |
| SLC12A7 | A0A0G2JNW7;Q9Y666 | SLC12A7 | Solute carrier family 1 | 46545.625 | 41406.56641 |
| CACTIN | Q8WUQ7 | CACTIN | Cactin | 48115.46094 | 40103.89844 |

| | | | | | |
|---|---|---|---|---|---|
| SRP54 | P61011 | SRP54 | Signal recognition par | 264099.2188 | 295164.125 |
| GALNT14 | J3KQQ5;Q96FL9 | GALNT14 | Polypeptide N-acetylg | 6793.918945 | 7886.862793 |
| ACBD7 | Q8N6N7 | ACBD7 | Acyl-CoA-binding dom | 5686.443848 | 4032.232178 |
| DCTN3 | O75935 | DCTN3 | Dynactin subunit 3 | 147692.8438 | 163796.0781 |
| UBA3 | Q8TBC4 | UBA3 | NEDD8-activating enz | 49458.00391 | 53307.96484 |
| FSD1 | M0R366;Q9BTV5 | FSD1 | Fibronectin type III an | 5350.249023 | 7918.057617 |
| WDR54 | Q9H977 | WDR54 | WD repeat-containing | 11667.78906 | 12776.39844 |
| CYCS | C9JFR7;P99999 | CYCS | Cytochrome c (Fragm | 725781.3125 | 625137.625 |
| PLEKHA3 | Q9HB20 | PLEKHA3 | Pleckstrin homology d | 9226.612305 | 10010.23633 |
| ORC6 | H3BT22;Q9Y5N6 | ORC6 | Origin recognition cor | 8724.385742 | 8418.5625 |
| DBR1 | Q9UK59 | DBR1 | Lariat debranching en | 75193.58594 | 76267.64063 |
| POLR1A | O95602 | POLR1A | DNA-directed RNA po | 77613.1875 | 55355.5625 |
| MAPK3 | P27361 | MAPK3 | Mitogen-activated pro | 187152.4688 | 206833.8281 |
| PPP1CA | P62136 | PPP1CA | Serine/threonine-prot | 734342.3125 | 773735.625 |
| ATG12 | O94817 | ATG12 | Ubiquitin-like protein | 26964.05469 | 20381.17578 |
| PSMB7 | Q99436 | PSMB7 | Proteasome subunit b | 475597 | 522506.125 |
| NTPCR | Q9BSD7 | NTPCR | Cancer-related nucleo | 123138.625 | 130006.5313 |
| EGLN1 | Q9GZT9 | EGLN1 | Egl nine homolog 1 | 24184.41992 | 28937.91797 |
| OSBPL9 | Q96SU4 | OSBPL9 | Oxysterol-binding pro | 78141.29688 | 86176.71875 |
| CLK2 | B1AVT0;P49760 | CLK2 | Dual-specificity protei | 7509.786621 | 7329.182129 |
| EHHADH | Q08426 | EHHADH | Peroxisomal bifunctio | 12028.37402 | 10032.52246 |
| DERPC | P0CG12 | DERPC | Decreased expression | 7854.294434 | 12038.6416 |
| HYPK | Q9NX55 | HYPK | Huntingtin-interacting | 42618.82031 | 15025.91699 |
| CCZ1B;CCZ1 | P86790;P86791 | CCZ1B;CCZ1 | Vacuolar fusion protei | 101850.6719 | 102028.7422 |
| KCTD18 | Q6PI47 | KCTD18 | BTB/POZ domain-cont | 8287.246094 | 7658.088379 |
| USP14 | P54578 | USP14 | Ubiquitin carboxyl-ter | 312945.0625 | 323465.8125 |
| CNNM3 | Q8NE01 | CNNM3 | Metal transporter CNI | 35800.82422 | 39718.92969 |
| IMPA2 | O14732 | IMPA2 | Inositol monophospha | 24101.33203 | 33936.82422 |
| BCR | P11274 | BCR | Breakpoint cluster reg | 28549.97461 | 29389.21875 |
| HNRNPA2B1 | P22626 | HNRNPA2B1 | Heterogeneous nucle | 5420360 | 2649236.5 |
| ACBD3 | Q9H3P7 | ACBD3 | Golgi resident protein | 159741.4844 | 173821.6875 |
| SLTM | Q9NWH9 | SLTM | SAFB-like transcriptio | 142742.125 | 113892.9531 |
| CTPS2 | Q9NRF8 | CTPS2 | CTP synthase 2 | 183024.7031 | 208154.4063 |

| | | | | | |
|---|---|---|---|---|---|
| IGBP1 | P78318 | IGBP1 | Immunoglobulin-bind | 132115.1875 | 129577.5 |
| FLG2 | Q5D862 | FLG2 | Filaggrin-2 | 1906.921021 | 1227.654907 |
| RNF13 | O43567 | RNF13 | E3 ubiquitin-protein li | 19742.25195 | 14904.31738 |
| MND1 | A0A087WTC6;D6R9E3;D6RC1 | MND1 | Meiotic nuclear divisi | 12585.72656 | 12334.19336 |
| PPFIBP1 | Q86W92 | PPFIBP1 | Liprin-beta-1 | 12009.33789 | 12053.32129 |
| AP2M1 | A0A087WY71;Q96CW1 | AP2M1 | AP-2 complex subunit | 461284.1563 | 518693.8438 |
| DOCK7 | Q96N67 | DOCK7 | Dedicator of cytokines | 42196.04688 | 41961.17188 |
| LGALS8 | F6V2D4 | LGALS8 | Galectin | 26732.75 | 22467.83984 |
| KAT8 | Q9H7Z6 | KAT8 | Histone acetyltransfer | 30027.66797 | 34080.79688 |
| MIF4GD | A9UHW6;J3QRZ6 | MIF4GD | MIF4G domain-contai | 19348.82227 | 22136.41211 |
| PURB | Q96QR8 | PURB | Transcriptional activat | 185053.4375 | 180880.2344 |
| SMARCA1 | B7ZLQ5;P28370 | SMARCA1 | Probable global transc | 273712.9375 | 310617.5625 |
| GATA3 | P23771 | GATA3 | Trans-acting T-cell-spe | 133542.5625 | 142600.5781 |
| EP400P1 | Q6ZTU2 | EP400P1 | Putative EP400-like pr | 877.999939 | 1656.748657 |
| COG1 | Q8WTW3 | COG1 | Conserved oligomeric | 44157.37109 | 52661.74609 |
| CRYZL1 | O95825 | CRYZL1 | Quinone oxidoreducta | 21546.68555 | 23318.22852 |
| ACP6 | Q9NPH0 | ACP6 | Lysophosphatidic acid | 14040.73145 | 14453.1875 |
| TSPAN14 | Q8NG11 | TSPAN14 | Tetraspanin-14 | 21155.58984 | 20879.19727 |
| ITGAV | P06756 | ITGAV | Integrin alpha-V | 44289.51953 | 47893.69141 |
| RAB43 | Q86YS6 | RAB43 | Ras-related protein Ra | 399437.5625 | 546315.5 |
| ZFYVE19 | H3BRF9;Q96K21 | ZFYVE19 | Abscission/NoCut che | 49385.82422 | 59465.52734 |
| GTF2H3 | F5GWD3;F5GX35;F5GXG6;F5 | GTF2H3 | General transcription | 13248.96875 | 9172.946289 |
| PLEC | H0YDN1 | PLEC | Plectin | 621373.0313 | 431072.2617 |
| H1-3 | P16402 | H1-3 | Histone H1.3 | 448011.8125 | 168201.9531 |
| RTN3 | B7Z4M1 | RTN3 | Reticulon | 310672.8828 | 368329.4531 |
| MMADHC | F8WEC0;Q9H3L0 | MMADHC | Methylmalonic acidur | 13009.46484 | 16026.01074 |
| MMAB | Q96EY8 | MMAB | Corrinoid adenosyltra | 184354.5313 | 192226.6875 |
| GNPTG | Q9UJJ9 | GNPTG | N-acetylglucosamine- | 24042.19922 | 26493.19922 |
| CDC7 | O00311 | CDC7 | Cell division cycle 7-re | 14777.92676 | 12869.10059 |
| OSCP1 | C9JF58;Q8WVF1 | OSCP1 | Protein OSCP1 | 4218.841064 | 4705.510986 |
| ELMO2 | Q5JVZ5;Q96JJ3 | ELMO2 | Engulfment and cell m | 11806.96191 | 10377.28516 |
| IQSEC1 | A0A087WWK8;A0A2R8Y7T6; | IQSEC1 | IQ motif and SEC7 dor | 27594.6875 | 24612.10547 |
| PDCD10 | Q9BUL8 | PDCD10 | Programmed cell deat | 165886 | 198263.375 |

| | | | | | |
|---|---|---|---|---|---|
| MOB4 | Q9Y3A3 | MOB4 | MOB-like protein pho | 117307.6641 | 117935.9766 |
| HSPB11 | Q9Y547 | HSPB11 | Intraflagellar transpor | 33448.57813 | 41425.31641 |
| PSMD8 | P48556;R4GMR5 | PSMD8 | 26S proteasome non-/ | 297004.9688 | 339021.0625 |
| RAI14 | Q9P0K7 | RAI14 | Ankycorbin | 25679.15625 | 20548.82617 |
| PDPK1 | O15530 | PDPK1 | 3-phosphoinositide-de | 47878.69531 | 48345.16016 |
| RETREG3 | Q86VR2 | RETREG3 | Reticulophagy regulat | 79334.875 | 86401.95313 |
| IER5 | Q5VY09 | IER5 | Immediate early respo | 1522401.875 | 1506127.625 |
| STK38 | Q15208 | STK38 | Serine/threonine-prot | 35057.875 | 36159.74219 |
| STRADA | A0A1W2PPJ9;Q7RTN6 | STRADA | STE20-related kinase a | 7487.692871 | 8150.202637 |
| TERF2 | Q15554 | TERF2 | Telomeric repeat-binc | 50914.35938 | 47899.41797 |
| WARS1 | P23381 | WARS1 | Tryptophan--tRNA liga | 384551.75 | 428240.2188 |
| HADH | E9PF18;Q16836 | HADH | Hydroxyacyl-coenzym | 552927.6875 | 605969.6875 |
| WDR83OS | A0A0A0MQS5;Q9Y284 | WDR83OS | PTD008 protein, isofo | 6428.131348 | 6010.296875 |
| MPHOSPH8 | Q99549 | MPHOSPH8 | M-phase phosphopro | 22782.04883 | 21403.74219 |
| HERC1 | Q15751 | HERC1 | Probable E3 ubiquitin- | 118854.5391 | 144002.0469 |
| DUSP3 | P51452 | DUSP3 | Dual specificity protei | 55023.16797 | 63275.45313 |
| PIP4K2A | P48426 | PIP4K2A | Phosphatidylinositol 5 | 45197.22656 | 58017.69531 |
| PSMD11 | O00231 | PSMD11 | 26S proteasome non-/ | 542028.375 | 635519.8125 |
| COPA | P53621 | COPA | Coatomer subunit alp | 695842.75 | 750571.5 |
| RAB7A | P51149 | RAB7A | Ras-related protein Ra | 1407695.75 | 1360371.75 |
| TSPYL1 | Q9H0U9 | TSPYL1 | Testis-specific Y-encoc | 12031.64941 | 11856.61719 |
| TRAF2 | Q12933 | TRAF2 | TNF receptor-associat | 40736.08984 | 43473.38672 |
| HIP1R | O75146 | HIP1R | Huntingtin-interacting | 67863.02344 | 74220.1875 |
| HSF1 | E9PMQ6;Q00613 | HSF1 | Heat shock factor pro | 41911.35547 | 37249.33984 |
| KRT15 | P19012 | KRT15 | Keratin, type I cytoske | 1082.084961 | 3617.314209 |
| UBR1 | Q8IWV7 | UBR1 | E3 ubiquitin-protein li | 68989.375 | 80092.42188 |
| NCKAP1 | Q9Y2A7 | NCKAP1 | Nck-associated protei | 110370.7969 | 124806.9375 |
| SLC25A24 | Q6NUK1 | SLC25A24 | Calcium-binding mitoc | 605039.125 | 632547.75 |
| CDH1 | A0A087WXI5;P12830 | CDH1 | Cadherin-1 | 112396.4688 | 191770.0313 |
| LCMT1 | Q9UIC8 | LCMT1 | Leucine carboxyl meth | 156976.3906 | 156980.1719 |
| FAM111B | Q6SJ93 | FAM111B | Protein FAM111B | 7001.125977 | 8388.807617 |
| PNPLA4 | P41247 | PNPLA4 | Patatin-like phospholi | 6600.16748 | 5772.163086 |
| UCKL1 | Q9NWZ5 | UCKL1 | Uridine-cytidine kinas | 32529.86914 | 33286.42188 |

| | | | | | |
|---|---|---|---|---|---|
| KYAT3 | Q6YP21 | KYAT3 | Kynurenine--oxogluta | 56189.49219 | 53791.84766 |
| G6PC3 | K7ESE6;Q9BUM1 | G6PC3 | Glucose-6-phosphatas | 46387.09375 | 43229.07031 |
| TSFM | P43897 | TSFM | Elongation factor Ts, r | 212065.1094 | 221264.8906 |
| C15orf40 | Q8WUR7 | C15orf40 | UPF0235 protein C15c | 63594.34375 | 67574.3125 |
| RINT1 | Q6NUQ1 | RINT1 | RAD50-interacting prc | 9282.151367 | 10074.76758 |
| DUSP12 | Q9UNI6 | DUSP12 | Dual specificity protei | 42345.18359 | 35403.71094 |
| REEP4 | E5RGS2;Q9H6H4 | REEP4 | Receptor expression-e | 25015.80469 | 31329.81641 |
| RMI1 | A0A0A0MSU3;Q9H9A7 | RMI1 | RecQ-mediated genor | 2843.560791 | 4619.574707 |
| IDH2 | P48735 | IDH2 | Isocitrate dehydrogen | 636631.0625 | 785879.4375 |
| YLPM1 | P49750 | YLPM1 | YLP motif-containing p | 99439.02344 | 96022.66406 |
| HAUS3 | Q68CZ6 | HAUS3 | HAUS augmin-like con | 5511.783691 | 6120.84668 |
| ERBIN | Q96RT1 | ERBIN | Erbin | 27662.99414 | 41127.10938 |
| IVD | A0A0A0MT83;P26440 | IVD | Isovaleryl-CoA dehydr | 210974.375 | 233495.3281 |
| AP3B1 | O00203 | AP3B1 | AP-3 complex subunit | 122775.9609 | 154663.5313 |
| RAD23A | A0A494C0B4;P54725 | RAD23A | UV excision repair prc | 22519.95313 | 42640.41016 |
| FLOT2 | Q14254 | FLOT2 | Flotillin-2 | 45332.4375 | 48816.05078 |
| PSMA6 | P60900 | PSMA6 | Proteasome subunit a | 699053.9375 | 721091.3125 |
| TADA1 | Q96BN2 | TADA1 | Transcriptional adapte | 11232.40332 | 12884.14941 |
| PRAF2 | A6NP52;O60831 | PRAF2 | PRA1 family protein;P | 102475.0703 | 114378.9375 |
| MTMR9 | Q96QG7 | MTMR9 | Myotubularin-related | 13556.23633 | 14604.57617 |
| DCTPP1 | Q9H773 | DCTPP1 | dCTP pyrophosphatas | 182883.7188 | 196402.1563 |
| TMEM167A | Q8TBQ9 | TMEM167A | Protein kish-A | 130328.2656 | 155132.8125 |
| BCKDHB | P21953 | BCKDHB | 2-oxoisovalerate dehy | 23827.21094 | 26457.32031 |
| SRXN1 | Q9BYN0 | SRXN1 | Sulfiredoxin-1 | 68448.76563 | 58888.60938 |
| COPS6 | E7EM64;Q7L5N1 | COPS6 | COP9 signalosome coi | 147416.5938 | 158090.4688 |
| RLF | Q13129 | RLF | Zinc finger protein Rlf | 4538.22168 | 6032.664551 |
| PTPMT1 | Q8WUK0 | PTPMT1 | Phosphatidylglyceropl | 36417.60156 | 36582.04297 |
| SLC30A7 | Q8NEW0 | SLC30A7 | Zinc transporter 7 | 43894.39453 | 42466.73438 |
| PSMA1 | P25786 | PSMA1 | Proteasome subunit a | 879683.0625 | 985241.6875 |
| C1orf198 | Q9H425 | C1orf198 | Uncharacterized prote | 27645.42578 | 26166.45117 |
| TPP2 | P29144 | TPP2 | Tripeptidyl-peptidase | 102653.875 | 118768.7188 |
| RBM47 | A0AV96 | RBM47 | RNA-binding protein 4 | 71530.32031 | 74303.26563 |
| OXR1 | C9JY63;E5RFD1 | OXR1 | Oxidation resistance p | 56912.80469 | 50082.98633 |

| | | | | | |
|---|---|---|---|---|---|
| CEPT1 | A0A0C4DGS9;Q9Y6K0 | CEPT1 | Choline/ethanolamine | 97019.41406 | 120163.3047 |
| GLYR1 | Q49A26 | GLYR1 | Putative oxidoreducta | 57853.72266 | 58903.87109 |
| SAR1B | Q9Y6B6 | SAR1B | GTP-binding protein S | 675506.625 | 782671.9375 |
| CASP2 | P42575 | CASP2 | Caspase-2 | 18592.7793 | 23289.01367 |
| GOLPH3L | Q9H4A5 | GOLPH3L | Golgi phosphoprotein | 67557.66406 | 82542.08594 |
| SYNE2 | Q8WXH0 | SYNE2 | Nesprin-2 | 41023.23438 | 49057.73438 |
| UBTF | P17480 | UBTF | Nucleolar transcriptio | 181112.7656 | 201598.1094 |
| ZFYVE21 | Q9BQ24 | ZFYVE21 | Zinc finger FYVE doma | 34532.08594 | 42693.71484 |
| EIF2S2 | P20042 | EIF2S2 | Eukaryotic translation | 689563.875 | 688895.3125 |
| IGHMBP2 | P38935 | IGHMBP2 | DNA-binding protein S | 10569.05957 | 8689.150391 |
| SOAT1 | B1APM4;P35610 | SOAT1 | Sterol O-acyltransfera | 18976.14648 | 15076.14551 |
| SLC35B2 | Q8TB61 | SLC35B2 | Adenosine 3'-phospho | 135824.5 | 153329.3438 |
| TM9SF3 | Q9HD45 | TM9SF3 | Transmembrane 9 sup | 418594.5938 | 458545.4688 |
| COG6 | Q9Y2V7 | COG6 | Conserved oligomeric | 56805.96094 | 63183.57813 |
| ZDHHC5 | Q9C0B5 | ZDHHC5 | Palmitoyltransferase Z | 17296.28711 | 14698.9375 |
| REEP6 | Q96HR9 | REEP6 | Receptor expression-e | 55974.55469 | 63224.625 |
| TBC1D13 | Q9NVG8 | TBC1D13 | TBC1 domain family m | 11102.83887 | 13303.21582 |
| NDUFA4 | O00483 | NDUFA4 | Cytochrome c oxidase | 637956.0625 | 617101.875 |
| GNAI3 | P08754 | GNAI3 | Guanine nucleotide-b | 320665.4375 | 309879.8125 |
| DYNLRB1 | Q9NP97 | DYNLRB1 | Dynein light chain roa | 45667.72266 | 59610.34375 |
| FAM174A | Q8TBP5 | FAM174A | Membrane protein FA | 15791.63672 | 14356.75488 |
| PABPC1 | P11940 | PABPC1 | Polyadenylate-binding | 2573658.5 | 2336769 |
| TBCC | Q15814 | TBCC | Tubulin-specific chape | 106197.4063 | 101807.1016 |
| RUFY2 | H0YD93;Q8WXA3 | RUFY2 | RUN and FYVE domain | 16863.64453 | 16089.11523 |
| IMPACT | A0A3B3ITH3;Q9P2X3 | IMPACT | Protein IMPACT | 21578.3418 | 23391.52539 |
| AGMAT | Q9BSE5 | AGMAT | Agmatinase, mitochon | 41833.52734 | 32993.50781 |
| VRK1 | Q99986 | VRK1 | Serine/threonine-prot | 127384.0078 | 107779.4375 |
| PPP1CB | P62140 | PPP1CB | Serine/threonine-prot | 320355.5 | 310352.2188 |
| RAP1B | P61224 | RAP1B | Ras-related protein Ra | 454410.8125 | 447688.2188 |
| TOMM34 | Q15785 | TOMM34 | Mitochondrial import | 358875.4688 | 320477.25 |
| DEK | A0A494C028;H0Y993;P35659 | DEK | Protein DEK;DEK onco | 383268.9688 | 370665.0938 |
| THOP1 | P52888 | THOP1 | Thimet oligopeptidase | 456856.5938 | 551067.1875 |
| RAP1GDS1 | D6REZ0 | RAP1GDS1 | Rap1 GTPase-GDP diss | 112578.6143 | 137935.9648 |

| | | | | | |
|---|---|---|---|---|---|
| SLC25A4 | P12235 | SLC25A4 | ADP/ATP translocase | 2567443 | 3038774.75 |
| SEMA4B | Q9NPR2 | SEMA4B | Semaphorin-4B | 117817.0234 | 168198.6406 |
| DDHD2 | H0YF17;O94830 | DDHD2 | Phospholipase DDHD2 | 4253.534668 | 5357.72168 |
| ANXA7 | P20073 | ANXA7 | Annexin A7 | 252975.2188 | 264142.5625 |
| ACTG1 | P63261 | ACTG1 | Actin, cytoplasmic 2 | 5130344 | 3802825.75 |
| RXRB | P28702 | RXRB | Retinoic acid receptor | 28799.41016 | 32696.24219 |
| DRG2 | P55039 | DRG2 | Developmentally-regu | 96266.67969 | 105498.0391 |
| PSMD5 | Q16401 | PSMD5 | 26S proteasome non-A | 269088.5625 | 286962.9375 |
| OSBP | P22059 | OSBP | Oxysterol-binding pro | 102265.2188 | 109505.1953 |
| SLC30A5 | H0Y957;Q8TAD4;Q9BY48 | SLC30A5 | Zinc transporter 5 (Fra | 6222.552246 | 5886.231934 |
| TMEM87A | Q8NBN3 | TMEM87A | Transmembrane prote | 67282.46094 | 75118.39063 |
| CNOT1 | A5YKK6 | CNOT1 | CCR4-NOT transcriptio | 167524.0469 | 178892.1563 |
| PC | P11498 | PC | Pyruvate carboxylase, | 135996.4375 | 153488.7813 |
| HNRNPM | M0QZM1 | HNRNPM | Heterogeneous nuclea | 2480994.641 | 2205140.234 |
| SCRN2 | J3QL71;Q96FV2 | SCRN2 | Secernin-2 | 29903.3418 | 30701.47461 |
| SPRYD4 | Q8WW59 | SPRYD4 | SPRY domain-containi | 72581.17969 | 82370.75781 |
| NPTN | Q9Y639 | NPTN | Neuroplastin | 132114.1875 | 125769.3047 |
| CHD2 | O14647 | CHD2 | Chromodomain-helica | 31544.68359 | 37472.00781 |
| ZMYND8 | A0A087WVZ6 | ZMYND8 | Protein kinase C-bindi | 57677.7793 | 62616.66992 |
| IER3IP1 | A0A1W2PNR9;Q9Y5U9 | IER3IP1 | Immediate early respo | 211676.25 | 214973.0313 |
| PSMA3 | P25788 | PSMA3 | Proteasome subunit a | 615307.3125 | 694796.625 |
| VDAC3 | Q9Y277 | VDAC3 | Voltage-dependent an | 371168.125 | 132149.5469 |
| PTPA | Q15257 | PTPA | Serine/threonine-prot | 526185.125 | 591180 |
| NR2F6 | P10588 | NR2F6 | Nuclear receptor subf | 12221.54102 | 13799.63574 |
| PCP4 | P48539 | PCP4 | Calmodulin regulator | 17336.54297 | 22427.85938 |
| PIK3CB | P42338 | PIK3CB | Phosphatidylinositol 4 | 20193.54492 | 22597.24414 |
| LRRFIP1 | Q32MZ4 | LRRFIP1 | Leucine-rich repeat fli | 47334.91406 | 59617.28906 |
| ARPC2 | O15144 | ARPC2 | Actin-related protein 2 | 349526.6875 | 392594.9063 |
| GOLGA4 | Q13439 | GOLGA4 | Golgin subfamily A me | 41008.16797 | 39371.02344 |
| NPC2 | E7EMS2;G3V2V8;G3V3D1;G3 | NPC2 | NPC intracellular chol | 102378.1563 | 119407.0313 |
| UBE2A | A0A0D9SG71;P49459 | UBE2A | Ubiquitin-conjugating | 187088.8281 | 180065.5938 |
| CARS1 | B4DKY1 | CARS1 | Cysteinyl-tRNA synthe | 278472.291 | 295059.79 |
| FAM76B | F5GX09;Q5HYJ3 | FAM76B | Protein FAM76B | 5502.476563 | 4713.928223 |

| | | | | | |
|---|---|---|---|---|---|
| EPHB4 | P54760 | EPHB4 | Ephrin type-B recepto | 37584.79297 | 36909.69531 |
| FUCA1 | P04066 | FUCA1 | Tissue alpha-L-fucosid | 49861.00391 | 57213.01953 |
| RAB14 | P61106 | RAB14 | Ras-related protein Ra | 347057.5625 | 353475.9375 |
| SAR1A | Q9NR31 | SAR1A | GTP-binding protein S | 412581.7813 | 436884.6563 |
| DGCR6L | B5MCQ4;Q9BY27 | DGCR6L | Protein DGCR6L | 326546.4375 | 289062.1563 |
| RAD50 | Q92878 | RAD50 | DNA repair protein RA | 104472.7578 | 114403.0547 |
| SCPEP1 | Q9HB40 | SCPEP1 | Retinoid-inducible ser | 28647.9668 | 34405.59375 |
| SRM | P19623 | SRM | Spermidine synthase | 768281.3125 | 890451.5 |
| DNAJC3 | Q13217 | DNAJC3 | DnaJ homolog subfam | 68160.35156 | 77961.6875 |
| SETD1A | O15047 | SETD1A | Histone-lysine N-meth | 15896.66797 | 14410.99219 |
| MYL12A;MYL12B;MYL12A | J3QRS3;O14950;P19105 | MYL12A;MYL12B;MYL | Myosin regulatory ligh | 1430971.75 | 1240569.875 |
| CWC25 | Q9NXE8 | CWC25 | Pre-mRNA-splicing fac | 20596.73047 | 15539.14551 |
| TSC22D2 | O75157 | TSC22D2 | TSC22 domain family p | 10557.41992 | 11700.48242 |
| STYX | Q8WUJ0 | STYX | Serine/threonine/tyro | 12662.75781 | 13763.21484 |
| ATRN | O75882 | ATRN | Attractin | 8553.868164 | 9823.478516 |
| UQCR10 | Q9UDW1 | UQCR10 | Cytochrome b-c1 com | 518274.0938 | 494970.625 |
| LDB1 | Q86U70 | LDB1 | LIM domain-binding p | 20871.39453 | 22828.85156 |
| KIF15 | Q9NS87 | KIF15 | Kinesin-like protein KI | 16190.28516 | 19967.42578 |
| PSMB2 | P49721 | PSMB2 | Proteasome subunit b | 680871.9375 | 734194.6875 |
| MADD | A0A0A0MRB5 | MADD | MAP kinase-activating | 44516.75781 | 51080.03125 |
| PBX1 | H0YLF5;P40424 | PBX1 | Pre-B-cell leukemia tra | 17599.61719 | 20560.78125 |
| KAT5 | Q92993 | KAT5 | Histone acetyltransfer | 6287.885254 | 5216.046387 |
| DLG1 | Q12959 | DLG1 | Disks large homolog 1 | 92185.35156 | 111333.6797 |
| DMXL1 | F5H269;Q9Y485 | DMXL1 | DmX-like protein 1 | 1831.888428 | 2184.697998 |
| NAA35 | Q5VZE5 | NAA35 | N-alpha-acetyltransfe | 50405.32813 | 45825.375 |
| SKP1 | E5RJR5;P63208 | SKP1 | S-phase kinase-associa | 48005.71094 | 56988.97266 |
| CDC42BPA | A0A0A0MRJ0;Q5VT25 | CDC42BPA | Non-specific serine/th | 85652.07813 | 91496.85156 |
| NAAA | B4DVL2;Q02083;R4GNC0 | NAAA | N-acylethanolamine-h | 23134.62891 | 23529.56836 |
| TSNAX | Q99598 | TSNAX | Translin-associated pr | 90238.48438 | 97925.38281 |
| MOV10 | Q9HCE1 | MOV10 | Helicase MOV-10 | 144537.2344 | 145275.8438 |
| HAUS5 | O94927 | HAUS5 | HAUS augmin-like con | 21339.93945 | 24217.06055 |
| NUBP2 | Q9Y5Y2 | NUBP2 | Cytosolic Fe-S cluster | 169261.9531 | 190890.2969 |
| MED18 | Q9BUE0 | MED18 | Mediator of RNA poly | 20314.1543 | 24128.85938 |

| | | | | | |
|---|---|---|---|---|---|
| TEX30 | A0A0C4DFW4;Q5JUR7 | TEX30 | Testis-expressed prot | 8368.40625 | 10184.80176 |
| TBL1X | A0A2R8YFW3 | TBL1X | F-box-like/WD repeat- | 19410.87305 | 23047.77539 |
| FAM98B | Q52LJ0 | FAM98B | Protein FAM98B | 165064.375 | 177169.875 |
| PSMC4 | P43686 | PSMC4 | 26S proteasome regul | 431501.0625 | 492937.9688 |
| IFT140 | Q96RY7 | IFT140 | Intraflagellar transpor | 2861.776611 | 6562.98877 |
| PCBP2 | F8VRH0 | PCBP2 | Poly(rC)-binding prote | 3178697.758 | 3292026.074 |
| NUF2 | Q9BZD4 | NUF2 | Kinetochore protein N | 18787.77344 | 21391.00977 |
| TAB1 | Q15750 | TAB1 | TGF-beta-activated ki | 38597.37891 | 39903.69922 |
| TMPO | H0YJH7 | TMPO | Lamina-associated po | 2001298.95 | 1883320.265 |
| TOP2A | P11388 | TOP2A | DNA topoisomerase 2 | 33463.16016 | 43722.4375 |
| BRD1 | O95696 | BRD1 | Bromodomain-contai | 12830.76953 | 9824.172852 |
| CNOT7 | Q9UIV1 | CNOT7 | CCR4-NOT transcriptio | 41371.33203 | 39163.88672 |
| CDC34 | P49427 | CDC34 | Ubiquitin-conjugating | 48493.21484 | 62053.06641 |
| SEPTIN9 | Q9UHD8 | SEPTIN9 | Septin-9 | 621300.25 | 658530.6875 |
| ACTBL2 | Q562R1 | ACTBL2 | Beta-actin-like protei | 7395166.5 | 6568148 |
| RCC2 | Q9P258 | RCC2 | Protein RCC2 | 653517.0625 | 658434.375 |
| KDM5A | F5GZ99 | KDM5A | Lysine-specific demet | 60581.78906 | 56926.94531 |
| ELAVL1 | Q15717 | ELAVL1 | ELAV-like protein 1 | 536564.75 | 460041.8125 |
| GAB1 | Q13480 | GAB1 | GRB2-associated-bind | 5528.061035 | 7066.05957 |
| ZNF592 | Q92610 | ZNF592 | Zinc finger protein 59 | 15276.83203 | 15781.67676 |
| NDC80 | O14777 | NDC80 | Kinetochore protein N | 65119.53125 | 80899.30469 |
| TRAPPC8 | Q9Y2L5 | TRAPPC8 | Trafficking protein pa | 11614.33105 | 12405.16602 |
| CDK16 | A0A087WZU2;E5RGN0;Q005 | CDK16 | Cyclin-dependent kin | 13757.01465 | 17072.99023 |
| PRR36 | Q9H6K5 | PRR36 | Proline-rich protein 3 | 26373.41016 | 31196.86328 |
| GCOM1;POLR2M;GCOM2 | H8Y6P7;P0CAP2;Q9BZD3 | GCOM1;POLR2M;GCC | GRINL1A combined pr | 6710.029785 | 7494.023438 |
| MED13L | A0A3B3IRX3;Q71F56 | MED13L | Mediator of RNA poly | 2811.708984 | 3148.46167 |
| OCIAD1 | Q9NX40 | OCIAD1 | OCIA domain-containi | 170277.9375 | 156004.3906 |
| MMACHC | A0A0C4DGU2;Q9Y4U1 | MMACHC | Methylmalonic acidur | 12139.82324 | 16347.24609 |
| CAMK2G | Q5SWX3 | CAMK2G | Calcium/calmodulin-d | 60379.00781 | 47710.16797 |
| PSMD13 | Q9UNM6 | PSMD13 | 26S proteasome non- | 285724.8125 | 333688.8125 |
| CYFIP1 | Q7L576 | CYFIP1 | Cytoplasmic FMR1-int | 234026.6719 | 254614.5 |
| EPRS1 | P07814 | EPRS1 | Bifunctional glutamat | 368659.125 | 361384.3438 |
| GPAA1 | E9PPZ9;O43292 | GPAA1 | Glycosylphosphatidyli | 108544.5625 | 111196.5938 |

| | | | | | |
|---|---|---|---|---|---|
| SEC13 | P55735 | SEC13 | Protein SEC13 homolo | 321817.375 | 338821.4063 |
| PBRM1 | Q86U86 | PBRM1 | Protein polybromo-1 | 66662.61719 | 62591.13672 |
| HDHD3 | Q9BSH5 | HDHD3 | Haloacid dehalogenas | 31778.5918 | 37492.3125 |
| TUBB8 | Q3ZCM7 | TUBB8 | Tubulin beta-8 chain | 25753.82813 | 29468.4043 |
| ATG14 | Q6ZNE5 | ATG14 | Beclin 1-associated au | 6652.393066 | 4926.430176 |
| MTG1 | Q9BT17 | MTG1 | Mitochondrial ribosor | 75991.35938 | 96877.03125 |
| FLYWCH1 | Q4VC44 | FLYWCH1 | FLYWCH-type zinc fing | 4804.241211 | 3500.500488 |
| ADH5 | P11766 | ADH5 | Alcohol dehydrogenas | 582907.875 | 620260.375 |
| FAIM | Q9NVQ4 | FAIM | Fas apoptotic inhibito | 25828.14063 | 27503.67578 |
| ARHGAP17 | Q68EM7 | ARHGAP17 | Rho GTPase-activating | 52429 | 50978.88672 |
| RRAS | P10301 | RRAS | Ras-related protein R- | 6533.420898 | 7665.39209 |
| EHMT2 | A0A0G2JIS2 | EHMT2 | Histone-lysine N-meth | 43998.81641 | 46421.96875 |
| COPS4 | Q9BT78 | COPS4 | COP9 signalosome coi | 149047.9219 | 165236.4375 |
| TARS1 | P26639 | TARS1 | Threonine--tRNA ligas | 686163.4375 | 775778.75 |
| NAE1 | Q13564 | NAE1 | NEDD8-activating enz | 58805.05469 | 68279.14844 |
| NHLRC2 | Q8NBF2 | NHLRC2 | NHL repeat-containing | 109158.9844 | 124403.5469 |
| ERF | P50548 | ERF | ETS domain-containin | 8535.606445 | 11547.62695 |
| LRRC8A | Q8IWT6 | LRRC8A | Volume-regulated ani | 26435.3418 | 24092.22656 |
| AP2S1 | M0QYZ2;M0R0N4;P53680 | AP2S1 | AP complex subunit si | 229494.5156 | 247906.875 |
| PSEN1 | P49768 | PSEN1 | Presenilin-1 | 30426.77148 | 26839.70313 |
| RPRD1B | Q9NQG5 | RPRD1B | Regulation of nuclear | 210056.6563 | 171617.7656 |
| ZNF629 | Q9UEG4 | ZNF629 | Zinc finger protein 62 | 17111.33203 | 15682.91602 |
| KIAA1191 | A0A087WT18;Q96A73 | KIAA1191 | Putative monooxygen | 24776.30664 | 25680.71289 |
| SUMO3 | A8MU27;A8MUA9;P55854 | SUMO3 | Small ubiquitin-relate | 828238.5625 | 912344.3125 |
| POFUT1 | Q9H488 | POFUT1 | GDP-fucose protein O | 90210.97656 | 90834.25 |
| NHP2 | Q9NX24 | NHP2 | H/ACA ribonucleoprot | 135668.4219 | 69153.45313 |
| ACTB | P60709 | ACTB | Actin, cytoplasmic 1 | 40769376 | 31259846 |
| DNAJA3 | Q96EY1 | DNAJA3 | DnaJ homolog subfam | 197977.3438 | 182468.1406 |
| MYO10 | A0A0A0MQX1;D6RGD1;Q9HI | MYO10 | Unconventional myos | 544.7402954 | NA |
| TRIM26 | Q12899 | TRIM26 | Tripartite motif-conta | 69896.16406 | 75268.80469 |
| TES | Q9UGI8 | TES | Testin | 142756.0313 | 152901.7031 |
| MPC2 | O95563;Q5R3B4 | MPC2 | Mitochondrial pyruva | 83163.04688 | 87293.71875 |
| MIF | P14174 | MIF | Macrophage migratio | 4763816 | 5487902 |

| | | | | | |
|---|---|---|---|---|---|
| RPRD1A | A0A0C4DGQ6;Q96P16 | RPRD1A | Regulation of nuclear | 39481.77344 | 40111.61719 |
| ZNF687 | Q8N1G0 | ZNF687 | Zinc finger protein 68 | 31065.55859 | 39332.75 |
| ACADM | P11310;Q5T4U5 | ACADM | Medium-chain specifi | 253915.6094 | 288917.7813 |
| ATP5F1C | P36542 | ATP5F1C | ATP synthase subunit | 789949.0625 | 895630.4375 |
| DERA | Q9Y315 | DERA | Deoxyribose-phospha | 44327.75781 | 43164.81641 |
| TRRAP | F2Z2U4 | TRRAP | Transformation/trans | 44919.04102 | 49154.1709 |
| ARPC3 | O15145 | ARPC3 | Actin-related protein | 443615.0938 | 474986.4063 |
| FAF1 | Q9UNN5 | FAF1 | FAS-associated factor | 110107.3984 | 118192.1719 |
| KIFC1 | Q9BW19 | KIFC1 | Kinesin-like protein KI | 83029.79688 | 67871.65625 |
| TTYH3 | Q9C0H2 | TTYH3 | Protein tweety homol | 3262.747559 | 4191.17334 |
| SCO2 | O43819 | SCO2 | Protein SCO2 homolo | 39556.39063 | 39845.97266 |
| GALC | P54803 | GALC | Galactocerebrosidase | 28677.73242 | 29671.39453 |
| VPS45 | Q9NRW7 | VPS45 | Vacuolar protein sorti | 117473.75 | 122534.4375 |
| PELO | Q9BRX2 | PELO | Protein pelota homolo | 53493.67969 | 59594.20313 |
| DBI | P07108 | DBI | Acyl-CoA-binding prot | 41743.14844 | 74715.64844 |
| RBM34 | P42696 | RBM34 | RNA-binding protein 3 | 103621.3125 | 58646.47266 |
| RBM45 | Q8IUH3 | RBM45 | RNA-binding protein 4 | 24940.43164 | 18259.0625 |
| CTCF | P49711 | CTCF | Transcriptional repres | 58297.08984 | 47756.50781 |
| PSMC5 | P62195 | PSMC5 | 26S proteasome regul | 650994.25 | 680514.6875 |
| ZFAND3 | Q9H8U3 | ZFAND3 | AN1-type zinc finger p | 17679.53906 | 26055.24805 |
| ARMC7 | Q9H6L4 | ARMC7 | Armadillo repeat-cont | 2117.409668 | 2157.116699 |
| ARHGAP11A | Q6P4F7 | ARHGAP11A | Rho GTPase-activating | 4057.959717 | 3574.81665 |
| ELP1 | O95163 | ELP1 | Elongator complex pr | 83821.95313 | 89056.78125 |
| NME6 | A0A0C4DG91;A0A2R8Y547;A | NME6 | Nucleoside diphospha | 19289.70703 | 20675.4375 |
| BNIP1 | Q12981 | BNIP1 | Vesicle transport prot | 75920.90625 | 73140.65625 |
| STX17 | P56962 | STX17 | Syntaxin-17 | 40559.03906 | 35714.78906 |
| DDB1 | Q16531 | DDB1 | DNA damage-binding | 620709.375 | 657352.75 |
| BCAS2 | O75934 | BCAS2 | Pre-mRNA-splicing fac | 32859.44922 | 36948.14844 |
| AP3D1 | O14617 | AP3D1 | AP-3 complex subunit | 81171.0625 | 93109.72656 |
| CTNNA1 | P35221 | CTNNA1 | Catenin alpha-1 | 519494.3125 | 523802.875 |
| AGO2 | Q9UKV8 | AGO2 | Protein argonaute-2 | 69562.35156 | 62562.52734 |
| CIC | I3L2J0 | CIC | Protein capicua homo | 28441.93945 | 31953.2832 |
| WWP2 | O00308 | WWP2 | NEDD4-like E3 ubiquit | 32885.04688 | 28674.85742 |

| | | | | | |
|---|---|---|---|---|---|
| OPA3 | Q9H6K4 | OPA3 | Optic atrophy 3 prote | 35256.08984 | 38496.37891 |
| GINS1 | Q14691 | GINS1 | DNA replication comp | 48401.23047 | 46158.69531 |
| H2AC4;HIST3H2A;HIST1H2/ | P04908;Q7L7L0;Q93077 | H2AC4;HIST3H2A;HIST | Histone H2A type 1-B/ | 13088.36621 | 1019.906982 |
| SETD7 | Q8WTS6 | SETD7 | Histone-lysine N-meth | 41048.53516 | 45583.68359 |
| NUDT6 | P53370 | NUDT6 | Nucleoside diphospha | 18535.62695 | 19444.33398 |
| PSMB1 | P20618 | PSMB1 | Proteasome subunit b | 594873.125 | 650031.3125 |
| HM13 | A0A3B3IUB5;Q8TCT9 | HM13 | Minor histocompatibi | 522309.8125 | 545265.3125 |
| ALG2 | Q9H553 | ALG2 | Alpha-1,3/1,6-mannos | 37085.27344 | 37507.10547 |
| GCLM | P48507 | GCLM | Glutamate--cysteine li | 42506.03516 | 39581.85156 |
| COPZ1 | F8VVA7;P61923 | COPZ1 | Coatomer subunit zet | 380380.25 | 455999.875 |
| PLEKHA5 | Q9HAU0 | PLEKHA5 | Pleckstrin homology d | 9130.666992 | 9222.428711 |
| CSTF3 | Q12996 | CSTF3 | Cleavage stimulation f | 464897.9063 | 474518.0938 |
| EIF2B2 | P49770 | EIF2B2 | Translation initiation f | 115049.9297 | 113798.9453 |
| GAR1 | Q9NY12 | GAR1 | H/ACA ribonucleoprot | 209123.75 | 92451.11719 |
| PFKM | A0A2R8Y891;P08237 | PFKM | ATP-dependent 6-pho | 123673.8438 | 137164.4219 |
| CENPO | Q9BU64 | CENPO | Centromere protein C | 2136.447021 | 3638.702881 |
| PGK1 | P00558 | PGK1 | Phosphoglycerate kin | 4764985 | 4964065 |
| CAMK2B | Q13554 | CAMK2B | Calcium/calmodulin-d | 103530.6641 | 109460.75 |
| KHDRBS1 | Q07666 | KHDRBS1 | KH domain-containing | 1112478.25 | 847837.4375 |
| FUS | P35637 | FUS | RNA-binding protein F | 1839575 | 1097938.875 |
| ZNF706 | E5RHR6;E5RHV4;H0YAP1;Q9 | ZNF706 | Zinc finger protein 70 | 461562.875 | 434759.5313 |
| PSIP1 | O75475 | PSIP1 | PC4 and SFRS1-intera | 89909.48438 | 69289.10156 |
| DXO | F8WC68;O77932 | DXO | Decapping and exorib | 6236.562988 | 7751.898438 |
| DENND10 | Q8TCE6 | DENND10 | DENN domain-contair | 69846.86719 | 70517.77344 |
| FAHD2A | Q96GK7 | FAHD2A | Fumarylacetoacetate | 72552.32813 | 69142.36719 |
| IFT81 | F8W1J4;Q8WYA0 | IFT81 | Intraflagellar transpor | 14559.74121 | 603.6351929 |
| MTA1 | E7ESY4;Q13330 | MTA1 | Metastasis-associated | 113142.3984 | 128174.3359 |
| PRPF19 | Q9UMS4 | PRPF19 | Pre-mRNA-processing | 336290.8438 | 354023.3438 |
| RAD17 | A0A0G2JNH5;A0A0G2JP78;A | RAD17 | Cell cycle checkpoint p | 11728.22266 | 12139.03613 |
| ATAD1 | Q8NBU5 | ATAD1 | ATPase family AAA do | 92383.96094 | 71414.85938 |
| IDUA | P35475 | IDUA | Alpha-L-iduronidase | 11956.18066 | 20452.39453 |
| UBA2 | Q9UBT2 | UBA2 | SUMO-activating enzy | 310510.4688 | 335333 |
| DNAJB14 | Q8TBM8 | DNAJB14 | DnaJ homolog subfam | 11667.0293 | 12186.19824 |

| | | | | | |
|---|---|---|---|---|---|
| ATP5MPL | A0A0A0MTQ5;G3V556;P5631 | ATP5MPL | ATP synthase subunit | 249536.8594 | 246041.75 |
| NCBP2AS2 | Q69YL0 | NCBP2AS2 | Protein NCBP2AS2 | 35679.53516 | 44673.84766 |
| ARHGEF17 | Q96PE2 | ARHGEF17 | Rho guanine nucleotid | 7956.762207 | 8692.405273 |
| PI4KA | P42356 | PI4KA | Phosphatidylinositol 4 | 38772.29688 | 49628.75781 |
| THEM4 | Q5T1C6 | THEM4 | Acyl-coenzyme A thio | 13748.29199 | 13556.37695 |
| MANBAL | Q9NQG1 | MANBAL | Protein MANBAL | 2125.052979 | 4055.818359 |
| COPS5 | Q92905 | COPS5 | COP9 signalosome co | 107382.9141 | 113170.7969 |
| BLOC1S5 | Q8TDH9 | BLOC1S5 | Biogenesis of lysosom | 40578.30078 | 41205.58203 |
| WAPL | Q7Z5K2 | WAPL | Wings apart-like prote | 39984.86328 | 48283.80469 |
| FLCN | Q8NFG4 | FLCN | Folliculin | 17537.375 | 18393 |
| RBMX2 | A0A1W2PP04;Q9Y388 | RBMX2 | RNA-binding motif pro | 20515.36328 | 13290.92383 |
| MLH1 | P40692 | MLH1 | DNA mismatch repair | 43380.0625 | 47012.60938 |
| PMVK | Q15126 | PMVK | Phosphomevalonate k | 137311.2344 | 143915.75 |
| GSTK1 | Q9Y2Q3 | GSTK1 | Glutathione S-transfe | 206523.0625 | 199971.9063 |
| APOO | H7C1U8;Q9BUR5 | APOO | MICOS complex subur | 117376.5391 | 110744.8203 |
| TBC1D23 | Q9NUY8 | TBC1D23 | TBC1 domain family m | 52550.9375 | 50406.5 |
| PSMA2 | A0A024RA52;P25787 | PSMA2 | Proteasome subunit a | 564274.8125 | 622538.75 |
| NT5C3B | Q969T7 | NT5C3B | 7-methylguanosine ph | 26414.57617 | 25349.19141 |
| NFIC | A0A286YEX4;P08651 | NFIC | Nuclear factor 1;Nucle | 25589.88477 | 36887.74609 |
| PHC2 | A0A0A0MSI2;B3KPJ4;Q8IXK0 | PHC2 | Polyhomeotic-like pro | 32554.07422 | 34538.46875 |
| CISD2 | Q8N5K1 | CISD2 | CDGSH iron-sulfur do | 319628.9063 | 352784.6563 |
| RAC3 | P60763 | RAC3 | Ras-related C3 botulir | 31802.74805 | 29870.61523 |
| PIGT | A0A1W2PNP0;A0A1W2PP57; | PIGT | GPI transamidase com | 60776.78906 | 67220.94531 |
| PSMD1 | A0A087WW66 | PSMD1 | 26S proteasome non-/ | 465399.7461 | 475598.043 |
| CLN3;CLN3;CLN3;;CLN3 | A0A0D9SFH9;A0A1B0GV71;A | CLN3;CLN3;CLN3;;CLN | Battenin | 13550.8916 | 12836.45215 |
| UFD1 | Q92890 | UFD1 | Ubiquitin recognition | 554324.6875 | 557614 |
| SEPTIN11 | D6RER5;D6RGI3;Q9NVA2 | SEPTIN11 | Septin-11;Septin 11, is | 376358.2188 | 391469.125 |
| SAC3D1 | A6NKF1;F8WC89 | SAC3D1 | SAC3 domain-containi | 10766.16895 | 11366.08008 |
| GET4 | Q7L5D6 | GET4 | Golgi to ER traffic prot | 206156.3281 | 221083.8125 |
| RPP21 | A0A096LP39;A0A0G2JJ26;A0 | RPP21 | Ribonuclease P protei | 32358.95313 | 35225.60156 |
| USP15 | Q9Y4E8 | USP15 | Ubiquitin carboxyl-ter | 104676.7578 | 111165.2813 |
| ZMYM4 | Q5VZL5 | ZMYM4 | Zinc finger MYM-type | 11425.6543 | 13845.96191 |
| TRIM11 | Q96F44;R4GMV1;R4GNB9 | TRIM11 | E3 ubiquitin-protein li | 19440.59766 | 21971.75977 |

| | | | | | |
|---|---|---|---|---|---|
| SRPRA | P08240 | SRPRA | Signal recognition par | 87975.54688 | 94553.15625 |
| COPS2 | P61201 | COPS2 | COP9 signalosome coı | 200641.4375 | 214754.625 |
| MLEC | Q14165 | MLEC | Malectin | 150414.2969 | 166163.0156 |
| MEN1 | A0A5F9ZHS3;E7EN32 | MEN1 | Menin | 24389.70508 | 26362.12109 |
| MINPP1 | Q9UNW1 | MINPP1 | Multiple inositol polyı | 4805374.5 | 3660163.75 |
| RMC1 | K7ENL9;Q96DM3 | RMC1 | Regulator of MON1-C | 100472.0078 | 112362.5938 |
| NAMPT | P43490 | NAMPT | Nicotinamide phosphı | 431795.875 | 420436.875 |
| PTER | Q96BW5 | PTER | Phosphotriesterase-re | 11317.25098 | 8661.080078 |
| STUB1 | Q9UNE7 | STUB1 | E3 ubiquitin-protein li | 554096.0625 | 572229.0625 |
| AMZ2 | Q86W34 | AMZ2 | Archaemetzincin-2 | 6743.172852 | 5647.003418 |
| RNF4 | D6RA71;H0YAI5;P78317 | RNF4 | E3 ubiquitin-protein li | 16920.09961 | 12640.47754 |
| UBE2V2 | Q15819 | UBE2V2 | Ubiquitin-conjugating | 199455.8281 | 195744.6563 |
| ZMYM3 | A6NHB5;Q14202 | ZMYM3 | Zinc finger MYM-type | 47950.875 | 50872.78516 |
| WDR74 | E9PS41;Q6RFH5 | WDR74 | WD repeat-containing | 19172.54883 | 9778.851563 |
| EIF2B1 | Q14232 | EIF2B1 | Translation initiation f | 71470.4375 | 81762.5 |
| FAM210A | K7EK00;Q96ND0 | FAM210A | Protein FAM210A (Fra | 49655.01953 | 40998.51172 |
| MAP2K6 | P52564 | MAP2K6 | Dual specificity mitog | 14149.49707 | 16841.76367 |
| MAP2K7 | O14733 | MAP2K7 | Dual specificity mitog | 23873.42969 | 28163.39844 |
| PMM2 | O15305 | PMM2 | Phosphomannomutas | 280212.4688 | 260792.7188 |
| MCM4 | P33991 | MCM4 | DNA replication licens | 379189.1875 | 393561.9063 |
| CYFIP2 | E7EVJ5 | CYFIP2 | Cytoplasmic FMR1-int | 16925.70117 | 16344.69238 |
| P4HB | P07237 | P4HB | Protein disulfide-isom | 713260.125 | 1071584.875 |
| INPPL1 | O15357 | INPPL1 | Phosphatidylinositol 3 | 76858.625 | 84452.77344 |
| YEATS4 | O95619 | YEATS4 | YEATS domain-contair | 20916.78516 | 18447.52539 |
| SCML2 | Q9UQR0 | SCML2 | Sex comb on midleg-li | 24644.5 | 28241.71289 |
| RBM15B | Q8NDT2 | RBM15B | Putative RNA-binding | 19136.3418 | 17392.26758 |
| AKR7A2 | O43488 | AKR7A2 | Aflatoxin B1 aldehyde | 253124.0938 | 274348.9375 |
| VEZF1 | J3QSH4;Q14119 | VEZF1 | Vascular endothelial z | 22123.44141 | 20902.41406 |
| SLC25A39 | B4DFG5;K7EMR4;K7EMW3;C | SLC25A39 | Solute carrier family 2 | 2933.322021 | 3089.76416 |
| CYB5R1 | Q9UHQ9 | CYB5R1 | NADH-cytochrome b5 | 74107.26563 | 80642.02344 |
| PTPN2 | P17706 | PTPN2 | Tyrosine-protein phos | 68562.71875 | 68775.4375 |
| WDR73 | Q6P4I2 | WDR73 | WD repeat-containing | 54067.91797 | 51899.26953 |
| RABGAP1 | Q9Y3P9 | RABGAP1 | Rab GTPase-activating | 76574.25 | 89187.64063 |

| | | | | | |
|---|---|---|---|---|---|
| CERT1 | A0A2R8Y7Q9;Q9Y5P4 | CERT1 | Ceramide transporter | 60510.83203 | 66834.84375 |
| ADARB1 | P78563 | ADARB1 | Double-stranded RNA | 17596.33594 | 14461.92188 |
| TSPYL5 | Q86VY4 | TSPYL5 | Testis-specific Y-encod | 132643.2813 | 150807.125 |
| CPEB4 | A0A590UJI5;B7ZLQ8;E5RFP2; | CPEB4 | Cytoplasmic polyaden | 4036.23877 | 2758.100098 |
| NEMF | O60524 | NEMF | Nuclear export media | 93849.39844 | 105252.4141 |
| FUCA2 | Q9BTY2 | FUCA2 | Plasma alpha-L-fucosi | 35097.38281 | 38335.57422 |
| SRP19 | P09132 | SRP19 | Signal recognition par | 126852.1016 | 130836.3672 |
| MREG | Q8N565 | MREG | Melanoregulin | 31424.00977 | 31247.88672 |
| CNBP | P62633 | CNBP | Cellular nucleic acid-b | 598241.0625 | 527913.625 |
| YARS1 | P54577 | YARS1 | Tyrosine--tRNA ligase, | 317152.5 | 338632.375 |
| MYD88 | A0A0A0MS70;H0Y4G9;Q9983 | MYD88 | Myeloid differentiatio | 57592.16016 | 65897.89844 |
| AP5S1 | Q9NUS5 | AP5S1 | AP-5 complex subunit | 11088.68262 | 9402.603516 |
| STAP2 | Q9UGK3 | STAP2 | Signal-transducing ada | 74141.1875 | 80748.51563 |
| USP46 | H7BZK6;P62068 | USP46 | Ubiquitin carboxyl-ter | 12143.04199 | 15082.84961 |
| MANEA | Q5SRI9;X6R7A2 | MANEA | Glycoprotein endo-alp | 9023.449219 | 17237.66406 |
| C7orf25 | Q9BPX7 | C7orf25 | UPF0415 protein C7or | 3775.463135 | 2644.068359 |
| ATP5F1A | P25705 | ATP5F1A | ATP synthase subunit | 2398156.75 | 2743549.5 |
| ASPH | Q12797 | ASPH | Aspartyl/asparaginyl b | 18235.17188 | 19977.23242 |
| ARID2 | Q68CP9 | ARID2 | AT-rich interactive dor | 25921.57227 | 30516.95898 |
| GIT2 | F8VXI9;F8WAK2;Q14161 | GIT2 | ARF GTPase-activating | 14683.74121 | 14282.4082 |
| CUL5 | Q93034 | CUL5 | Cullin-5 | 98377.24219 | 101087.9922 |
| TCEAL3 | Q969E4 | TCEAL3 | Transcription elongati | 228830.3594 | 252381.3281 |
| MAX | P61244 | MAX | Protein max | 32941.28906 | 28431.09766 |
| ARPC5 | O15511 | ARPC5 | Actin-related protein 2 | 303005.25 | 287934.6563 |
| RRBP1 | Q9P2E9 | RRBP1 | Ribosome-binding pro | 155371.4844 | 109632.9453 |
| NUP54 | Q7Z3B4 | NUP54 | Nucleoporin p54 | 127657.3828 | 125816.5313 |
| LIPA | A0A0A0MT32;P38571 | LIPA | Lysosomal acid lipase/ | 39981.16797 | 35111.64453 |
| GANAB | E9PKU7 | GANAB | Neutral alpha-glucosic | 658479.4883 | 723989.0469 |
| FAAH | O00519 | FAAH | Fatty-acid amide hydr | 35768.99609 | 42197.16797 |
| MED23 | Q5JWT2;Q9ULK4 | MED23 | Mediator of RNA poly | 32268.58789 | 36259.60938 |
| ZADH2 | Q8N4Q0 | ZADH2 | Prostaglandin reducta | 35059.64453 | 37559.67578 |
| EIPR1 | A8MUM1;Q53HC9 | EIPR1 | EARP and GARP comp | 27858.38281 | 31791.85156 |
| DNAJC13 | O75165 | DNAJC13 | DnaJ homolog subfam | 108436 | 111908.9844 |

| | | | | | |
|---|---|---|---|---|---|
| TMPRSS9 | A0A3B3IU58;Q7Z410 | TMPRSS9 | Transmembrane prote | 114853.7578 | 87142.625 |
| NELFA | A0A0C4DFX9;Q9H3P2 | NELFA | Negative elongation fa | 129295.8984 | 124830.0078 |
| ELP2 | Q6IA86 | ELP2 | Elongator complex pro | 44935.54297 | 51472.52734 |
| RAB1B | Q9H0U4 | RAB1B | Ras-related protein Ra | 2223739.25 | 2025815 |
| GNAI1 | P63096 | GNAI1 | Guanine nucleotide-b | 285884.5 | 348737.7188 |
| KATNB1 | Q9BVA0 | KATNB1 | Katanin p80 WD40 rep | 122078.8047 | 142204.5469 |
| ATXN2 | A0A2R8Y5A6;A0A2R8YDM9 | ATXN2 | Ataxin-2 | 50626.60156 | 56294.73438 |
| TUT7 | Q5VYS8 | TUT7 | Terminal uridylyltrans | 25930.50195 | 25131.38672 |
| LSM6 | P62312 | LSM6 | U6 snRNA-associated | 206267.9531 | 171914.75 |
| GOLGB1 | Q14789 | GOLGB1 | Golgin subfamily B me | 55696.10547 | 51993.42969 |
| CNOT9 | Q92600 | CNOT9 | CCR4-NOT transcriptio | 76098.07031 | 75125.21875 |
| FNIP1 | A0A087WX11;J3KNG8;Q8TF4 | FNIP1 | Folliculin-interacting p | 2740.057129 | 4002.919189 |
| UBE2D3 | P61077 | UBE2D3 | Ubiquitin-conjugating | 23890.29883 | 26551.77344 |
| BTF3L4 | Q96K17 | BTF3L4 | Transcription factor B | 146074.8281 | 115975.8516 |
| PPP2R2A | P63151 | PPP2R2A | Serine/threonine-prot | 205044.4375 | 211971.5625 |
| ENOPH1 | Q9UHY7 | ENOPH1 | Enolase-phosphatase | 246735.625 | 249283.7188 |
| DPP3 | Q9NY33 | DPP3 | Dipeptidyl peptidase 3 | 404896.625 | 504712.3125 |
| CRTC1 | M0QX46;Q6UUV9 | CRTC1 | CREB-regulated transc | 11459.07324 | 12237.28418 |
| SERINC1 | Q9NRX5 | SERINC1 | Serine incorporator 1 | 12236.77539 | 12542.89551 |
| RAB23 | Q9ULC3 | RAB23 | Ras-related protein Ra | 28996.08789 | 26513.24219 |
| RPS27L | H0YMV8;Q71UM5 | RPS27L | 40S ribosomal protein | 828932 | 838240.4375 |
| ACTR8 | Q9H981 | ACTR8 | Actin-related protein 8 | 11165.3584 | 10651.22266 |
| TBP | P20226;Q32MN6 | TBP | TATA-box-binding pro | 34260.71484 | 28245.17383 |
| CNOT11 | Q9UKZ1 | CNOT11 | CCR4-NOT transcriptio | 29867.63086 | 32207.36133 |
| PPP4C | P60510 | PPP4C | Serine/threonine-prot | 170317.6719 | 180951.1563 |
| ZNF316 | A6NFI3 | ZNF316 | Zinc finger protein 31 | 25094.05664 | 24678.34766 |
| DCLRE1B | A0A3B3ITQ0;Q9H816 | DCLRE1B | 5' exonuclease Apollo | 4576.022949 | 5677.51709 |
| COG7 | P83436 | COG7 | Conserved oligomeric | 47748.94531 | 57391.93359 |
| MYEF2 | A0A0A0MQW0 | MYEF2 | Myelin expression fac | 289733.9941 | 253804.0986 |
| HNRNPLL | B7WPG3;C9IYN3;Q8WVV9;V9 | HNRNPLL | Heterogeneous nuclea | 82941.42188 | 69634.46875 |
| SMG1 | J3KRA9;Q96Q15 | SMG1 | Serine/threonine-prot | 22035.38477 | 24702.35352 |
| TMEM258 | F5GWH5;P61165 | TMEM258 | Transmembrane prote | 187088.25 | 228232.5938 |
| SEC16A | O15027 | SEC16A | Protein transport prot | 158646.6406 | 186212.6406 |

| | | | | | |
|---|---|---|---|---|---|
| KRT18 | P05783 | KRT18 | Keratin, type I cytoske | 11713992 | 15376857 |
| AHSG | P02765 | AHSG | Alpha-2-HS-glycoprote | 409084.3438 | 330628.4688 |
| QKI | Q96PU8 | QKI | Protein quaking | 32430.23047 | 26973.91406 |
| STK24 | B4DR80 | STK24 | Serine/threonine-prot | 115403.9688 | 119369.8516 |
| PCMT1 | A0A0A0MRJ6;H7BY58;P2206 | PCMT1 | Protein-L-isoaspartate | 500486.75 | 509103.9688 |
| STK26 | B4E0Y9;Q8NBY1;Q9P289 | STK26 | Serine/threonine-prot | 122614.0391 | 125438.9453 |
| CISD1 | Q9NZ45 | CISD1 | CDGSH iron-sulfur dor | 88594.16406 | 88120.17188 |
| ORAI1 | Q96D31 | ORAI1 | Calcium release-activa | 6102.899902 | 4621.455566 |
| PIN1 | Q13526 | PIN1 | Peptidyl-prolyl cis-trar | 287965.625 | 307118.0625 |
| EPB41L5 | Q9HCM4 | EPB41L5 | Band 4.1-like protein 5 | 50141.51563 | 58960.57031 |
| ABRACL | Q9P1F3 | ABRACL | Costars family protein | 148777.7813 | 165099.875 |
| MAN1B1 | Q9UKM7 | MAN1B1 | Endoplasmic reticulur | 86113.9375 | 79710.64844 |
| AAK1 | Q2M2I8 | AAK1 | AP2-associated protei | 30287.40234 | 38902.78906 |
| NASP | P49321 | NASP | Nuclear autoantigenic | 6016.182129 | 10526.84473 |
| CACNG6 | Q9BXT2 | CACNG6 | Voltage-dependent ca | 421100.2813 | 451832.75 |
| ARHGAP12 | Q8IWW6 | ARHGAP12 | Rho GTPase-activating | 40910.54688 | 43541.69141 |
| SLC30A6 | B5MCR8;Q6NXT4 | SLC30A6 | Solute carrier family 3 | 12426.51563 | 8891.347656 |
| CCDC174 | A0A0B4J1R8;Q6PII3 | CCDC174 | Coiled-coil domain-co | 3172.166016 | 4152.200684 |
| VBP1 | P61758 | VBP1 | Prefoldin subunit 3 | 444010.1563 | 474828.3125 |
| MMTAG2 | Q9BU76 | MMTAG2 | Multiple myeloma tur | 97164.0625 | 70441.85156 |
| RAD54B | E5RHN9;E5RI14;E5RJF7;Q9Y6 | RAD54B | DNA repair and recom | 6478.948242 | 6084.561523 |
| RPS6KB2 | Q9UBS0 | RPS6KB2 | Ribosomal protein S6 | 48459.77734 | 48665.83203 |
| PAF1 | Q8N7H5 | PAF1 | RNA polymerase II-ass | 105405.875 | 95659.71094 |
| STIM2 | A0A1W2PRA1;A0A1X7SBY3;I | STIM2 | Stromal interaction m | 7936.10791 | 7410.625 |
| CLPTM1L | Q96KA5 | CLPTM1L | Cleft lip and palate tra | 74359.5625 | 70086.01563 |
| SURF4 | O15260;Q5T8U5 | SURF4 | Surfeit locus protein 4 | 681772.125 | 706553.625 |
| TBPL1 | P62380 | TBPL1 | TATA box-binding pro | 8938.805664 | 10180.46582 |
| CCM2 | A0A0A0MT72;C9JUH3;Q9BS( | CCM2 | Cerebral cavernous m | 6885.838867 | 8316.717773 |
| MKRN2 | C9J494;Q9H000 | MKRN2 | Probable E3 ubiquitin- | 43677.13281 | 39382.76953 |
| BRI3BP | A0A5F9ZHY7;Q8WY22 | BRI3BP | BRI3-binding protein | 208036.9531 | 204467.2188 |
| ATP6V0D1 | F5GYQ1;P61421 | ATP6V0D1 | V-type proton ATPase | 200712.3281 | 191657.4375 |
| RAMAC | Q9BTL3 | RAMAC | RNA guanine-N7 meth | 68723.34375 | 59850.10938 |
| LEMD2 | Q8NC56 | LEMD2 | LEM domain-containi | 138447.9063 | 133431.5 |

| | | | | | |
|---|---|---|---|---|---|
| GABARAP | H6UMI1;O95166 | GABARAP | GABARAP-a;Gamma-a | 46645.78906 | 51079.89844 |
| HNRNPUL2 | Q1KMD3 | HNRNPUL2 | Heterogeneous nucle | 1019284.438 | 787498.6875 |
| RMDN3 | Q96TC7 | RMDN3 | Regulator of microtub | 103044.8438 | 109103.8906 |
| ECT2 | Q9H8V3 | ECT2 | Protein ECT2 | 57348.88281 | 48884.35547 |
| CLK3 | P49761 | CLK3 | Dual specificity protei | 35243.76953 | 35045.36719 |
| RHBDD2 | Q6NTF9 | RHBDD2 | Rhomboid domain-co | 16463.73047 | 16494.15234 |
| COQ6 | Q9Y2Z9 | COQ6 | Ubiquinone biosynthe | 51335.20703 | 49257.125 |
| PRPS2 | P11908 | PRPS2 | Ribose-phosphate pyr | 224910.4219 | 260884.4844 |
| RALGPS2 | Q86X27 | RALGPS2 | Ras-specific guanine n | 11625.60449 | 13173.27539 |
| MCUR1 | Q96AQ8 | MCUR1 | Mitochondrial calcium | 8993.78418 | 9771.398438 |
| MCUB | Q9NWR8 | MCUB | Calcium uniporter reg | 4501.228027 | 6608.276367 |
| ABCB7 | A0A590UJS8;A0A5F9ZA98;O7 | ABCB7 | ATP-binding cassette s | 37651.58984 | 41954.88672 |
| ATG101 | F8VQD9;Q9BSB4 | ATG101 | Autophagy-related pro | 24529.51953 | 27712.50391 |
| CUL4A | Q13619 | CUL4A | Cullin-4A | 50418.23438 | 59096.80859 |
| PDCD6 | O75340 | PDCD6 | Programmed cell deat | 559474.9375 | 370584.75 |
| MRE11 | P49959 | MRE11 | Double-strand break r | 103363.9844 | 112043.6328 |
| XRCC6 | P12956 | XRCC6 | X-ray repair cross-com | 841484 | 984334.4375 |
| LYRM2 | Q9NU23 | LYRM2 | LYR motif-containing p | 31276.46484 | 32306.14258 |
| SELENOT | A0A087WVA1;P62341 | SELENOT | Selenoprotein T;Thior | 52938.88281 | 62312.32813 |
| PIGU | Q9H490 | PIGU | Phosphatidylinositol g | 35846.46094 | 41058.37109 |
| PLCG1 | P19174 | PLCG1 | 1-phosphatidylinosito | 154243.5938 | 168038.5313 |
| FAM20B | O75063 | FAM20B | Glycosaminoglycan xy | 23927.17969 | 23711.31641 |
| CD2BP2 | O95400 | CD2BP2 | CD2 antigen cytoplasr | 52491.89453 | 39906.08984 |
| PSMB5 | P28074 | PSMB5 | Proteasome subunit b | 432428.25 | 448400.8438 |
| KLHL36 | Q8N4N3 | KLHL36 | Kelch-like protein 36 | 27875.37109 | 16173.48926 |
| FBXL15 | A0A0C4DFV0;Q9H469 | FBXL15 | F-box/LRR-repeat pro | 16489.16406 | 15103.24219 |
| CREB1 | P16220 | CREB1 | Cyclic AMP-responsive | 35667.41406 | 25827.61328 |
| ATP5PB | P24539 | ATP5PB | ATP synthase F(0) con | 391967.7813 | 479118.5 |
| C19orf25 | Q9UFG5 | C19orf25 | UPF0449 protein C19c | 39765.50781 | 42418.67188 |
| FBXO3 | E9PJM3;Q9UK99 | FBXO3 | F-box only protein 3 | 16890.08203 | 16612.13086 |
| SSR4 | P51571 | SSR4 | Translocon-associated | 410036.2813 | 436735.5313 |
| MFN2 | O95140 | MFN2 | Mitofusin-2 | 19156.3418 | 17747.2793 |
| PPIA | P62937 | PPIA | Peptidyl-prolyl cis-trar | 16819106 | 15819427 |

| | | | | | |
|---|---|---|---|---|---|
| THADA | Q6YHU6 | THADA | Thyroid adenoma-ass | 20517.05273 | 26938.44531 |
| ABCC4 | O15439 | ABCC4 | Multidrug resistance-a | 4039.013916 | 3452.505127 |
| SKP2 | A0A0A0MTL5;Q13309 | SKP2 | S-phase kinase-associ | 30827.89648 | 40024.44531 |
| YIPF3 | D6RFI3;E7EQR8;Q9GZM5 | YIPF3 | Protein YIPF3 (Fragme | 77785.04688 | 86787.57031 |
| IRF2BPL | Q9H1B7 | IRF2BPL | Probable E3 ubiquitin- | 69326.35156 | 68131.27344 |
| ACSS3 | A0A0B4J1R2;Q9H6R3 | ACSS3 | Acyl-CoA synthetase s | 64007.625 | 77883.35938 |
| PPP4R1 | Q8TF05 | PPP4R1 | Serine/threonine-prot | 54019.99609 | 64096.97656 |
| ATP6V0A2 | Q9Y487 | ATP6V0A2 | V-type proton ATPase | 30208.17383 | 30508.70508 |
| U2AF2 | P26368 | U2AF2 | Splicing factor U2AF 6 | 880737.375 | 680627 |
| WDHD1 | O75717 | WDHD1 | WD repeat and HMG- | 82092.51563 | 87160.95313 |
| SAE1 | Q9UBE0 | SAE1 | SUMO-activating enzy | 396389.7188 | 447071.6563 |
| SMARCAL1 | H7BYI2;Q9NZC9 | SMARCAL1 | SWI/SNF-related matr | 12497.00098 | 12468.43945 |
| ALCAM | Q13740 | ALCAM | CD166 antigen | 234784.6719 | 239734.2969 |
| TRAPPC12 | Q8WVT3 | TRAPPC12 | Trafficking protein par | 27241.64844 | 31832.74414 |
| PPIL4 | Q8WUA2 | PPIL4 | Peptidyl-prolyl cis-trar | 67107.47656 | 63815.00781 |
| PGAM5 | Q96HS1 | PGAM5 | Serine/threonine-prot | 380843.625 | 366051.9688 |
| TPMT | P51580 | TPMT | Thiopurine S-methyltr | 43766.62109 | 43563.63281 |
| AGPAT3 | Q9NRZ7 | AGPAT3 | 1-acyl-sn-glycerol-3-pl | 23113.31836 | 23823.17578 |
| SHTN1 | A0MZ66 | SHTN1 | Shootin-1 | 161658.2344 | 191885.3125 |
| ATP5PO | P48047 | ATP5PO | ATP synthase subunit | 638518 | 757577.625 |
| MT-ATP6 | P00846 | MT-ATP6 | ATP synthase subunit | 86073.53125 | 108991.1172 |
| ELP3 | Q9H9T3 | ELP3 | Elongator complex pro | 52686.34375 | 52089.26953 |
| MAML1 | Q92585 | MAML1 | Mastermind-like prote | 36673.72656 | 48486.16406 |
| FAM91A1 | Q658Y4 | FAM91A1 | Protein FAM91A1 | 95590.4375 | 95572.15625 |
| TDRKH | F6TB26;Q5SZR4;Q9Y2W6 | TDRKH | Tudor and KH domain | 45312.04297 | 54168.97656 |
| CAB39 | Q9Y376 | CAB39 | Calcium-binding prote | 63575.875 | 66302.94531 |
| KIF14 | Q15058 | KIF14 | Kinesin-like protein KI | 6935.804688 | 9628.969727 |
| NFATC2IP | Q8NCF5 | NFATC2IP | NFATC2-interacting pr | 73102.95313 | 68261.57813 |
| CNOT2 | Q9NZN8 | CNOT2 | CCR4-NOT transcriptio | 73634.14063 | 78403.85938 |
| ANKRD27 | Q96NW4 | ANKRD27 | Ankyrin repeat domai | 17495.64063 | 19146.91602 |
| KBTBD4 | A0A075B6T4;Q9NVX7 | KBTBD4 | Kelch repeat and BTB | 6127.949707 | 6700.640137 |
| RTN4 | F8W914 | RTN4 | Reticulon | 771698.4609 | 830284.6641 |
| THOC5 | Q13769 | THOC5 | THO complex subunit | 72687.875 | 61874.75781 |

| | | | | | |
|---|---|---|---|---|---|
| TRAF7 | Q6Q0C0 | TRAF7 | E3 ubiquitin-protein li | 8669.808594 | 7616.525879 |
| ZMPSTE24 | O75844 | ZMPSTE24 | CAAX prenyl protease | 186231.9219 | 210663.0938 |
| AP5Z1 | O43299 | AP5Z1 | AP-5 complex subunit | 28276.35742 | 26516.58203 |
| SPICE1 | Q8N0Z3 | SPICE1 | Spindle and centriole- | 16448.88672 | 20986.37109 |
| NDUFA11 | Q86Y39 | NDUFA11 | NADH dehydrogenase | 55805.73438 | 66313.29688 |
| MFSD1 | A0A494C105;H7C593;H7C5U | MFSD1 | Major facilitator supe | 19683.11328 | 14447.44727 |
| ELOF1 | A0A024R7E8;K7EMV4;K7ENC | ELOF1 | Transcription elongati | 95876.99219 | 64918.42969 |
| TTC37 | Q6PGP7 | TTC37 | Tetratricopeptide rep | 26844.12695 | 29318.12109 |
| UBE2L3 | P68036 | UBE2L3 | Ubiquitin-conjugating | 1129193.5 | 1021323.188 |
| HAUS2 | Q9NVX0 | HAUS2 | HAUS augmin-like con | 2145.634277 | 8887.503906 |
| PARS2 | Q7L3T8 | PARS2 | Probable proline--tRN | 25415.41797 | 25672.0293 |
| PCYOX1L | Q8NBM8 | PCYOX1L | Prenylcysteine oxidas | 45845.67969 | 52202.48047 |
| MINDY3 | Q9H8M7 | MINDY3 | Ubiquitin carboxyl-ter | 39406.50781 | 40515.40234 |
| NKX2-8;NKX2-2 | O15522;O95096 | NKX2-8;NKX2-2 | Homeobox protein Nk | 59839.41016 | 58198.46484 |
| EIF2S1 | P05198 | EIF2S1 | Eukaryotic translation | 1149944.125 | 1180536 |
| DDX39A | O00148 | DDX39A | ATP-dependent RNA h | 482894.0625 | 483047.3125 |
| BRD2 | A0A0G2JK44;A0A140T9E9;H( | BRD2 | Bromodomain-contair | 46265.22266 | 37729.77344 |
| LEMD3 | Q9Y2U8 | LEMD3 | Inner nuclear membra | 52089.66406 | 49998.15625 |
| PFKFB3 | A0A1W2PR17;F2Z2I2;Q1687! | PFKFB3 | 6-phosphofructo-2-kir | 285.0731201 | 527.5877686 |
| CLN6 | A0A0S2Z5D0;A0A1B0GTU6;H | CLN6 | Ceroid-lipofuscinosis r | 43730.00391 | 46239.85547 |
| OXSM | Q9NWU1 | OXSM | 3-oxoacyl-[acyl-carrier | 38499.65625 | 40387.52734 |
| CYB5R4 | Q7L1T6 | CYB5R4 | Cytochrome b5 reduc | 50912.49219 | 62233.77344 |
| EMC8 | O43402 | EMC8 | ER membrane protein | 125305.3984 | 121766.3984 |
| KCTD10 | Q9H3F6 | KCTD10 | BTB/POZ domain-cont | 9438.355469 | 9531.335938 |
| PSMD9 | J3KN29;O00233 | PSMD9 | 26S proteasome non-A | 555566.75 | 541396.0625 |
| ABCD1 | P33897 | ABCD1 | ATP-binding cassette s | 24488.41797 | 31210.95117 |
| STX7 | O15400 | STX7 | Syntaxin-7 | 202568.9688 | 175006.3125 |
| SEC14L1 | Q92503 | SEC14L1 | SEC14-like protein 1 | 37145.21484 | 39605.89063 |
| DNAJB11 | Q9UBS4 | DNAJB11 | DnaJ homolog subfam | 249006.3281 | 256531.9688 |
| CPSF6 | F8WJN3;Q16630 | CPSF6 | Cleavage and polyade | 476185.3125 | 425168.5 |
| KIFAP3 | Q92845 | KIFAP3 | Kinesin-associated pro | 15393.41504 | 16080.1543 |
| SVBP | Q8N300 | SVBP | Small vasohibin-bindir | 5017.887695 | 3987.006104 |
| BANP | Q8N9N5 | BANP | Protein BANP | 9765.499023 | 6083.250977 |

| | | | | | |
|---|---|---|---|---|---|
| SUDS3 | Q9H7L9 | SUDS3 | Sin3 histone deacetyla | 27836.11328 | 32668.86328 |
| SIX4 | G3V2N2;Q9UIU6 | SIX4 | Homeobox protein SIX | 24034.76367 | 26763.02148 |
| PSME3IP1 | Q9GZU8 | PSME3IP1 | PSME3-interacting pro | 37675.1875 | 36832.41016 |
| SEH1L | Q96EE3 | SEH1L | Nucleoporin SEH1 | 106697.8516 | 119815.7734 |
| SIPA1 | F6RY50;Q96FS4 | SIPA1 | Signal-induced prolife | 84999.83594 | 67701.02344 |
| CMAS | Q8NFW8 | CMAS | N-acylneuraminate cy | 67842.08594 | 60073.94531 |
| MCEE | F5GZ54;Q96PE7 | MCEE | Methylmalonyl-CoA e | 8249.242188 | 7518.978516 |
| DNPEP | E5RIA4;E7ETB3;Q9ULA0 | DNPEP | Aspartyl aminopeptida | 466010.8125 | 506726.9375 |
| TNPO2 | O14787 | TNPO2 | Transportin-2 | 109836.5703 | 139083.7656 |
| PPOX | A0A494C146;P50336 | PPOX | Protoporphyrinogen c | 10504.63379 | 13676.36914 |
| MTCH2 | E9PIE4 | MTCH2 | Mitochondrial carrier | 433800.4063 | 294361.7891 |
| PPP2CA | P67775 | PPP2CA | Serine/threonine-prot | 934398.25 | 964279.75 |
| MAP3K7 | O43318 | MAP3K7 | Mitogen-activated pro | 17803.46094 | 18164.23242 |
| TMEM33 | P57088 | TMEM33 | Transmembrane prote | 73937.1875 | 99155.24219 |
| DDX17 | A0A1W2PQ51;A0A1X7SBZ2;A | DDX17 | Probable ATP-depend | 585811.1875 | 533022.625 |
| CHMP4B | Q9H444 | CHMP4B | Charged multivesicula | 417596.5938 | 407909.3438 |
| SPINDOC | Q9BUA3 | SPINDOC | Spindlin interactor and | 8356.058594 | 5447.447754 |
| CTDSPL2 | Q05D32 | CTDSPL2 | CTD small phosphatas | 1712069.875 | 748462.25 |
| PTPN23 | Q9H3S7 | PTPN23 | Tyrosine-protein phos | 72759.48438 | 79956.98438 |
| BTD | A0A5F9ZH57 | BTD | Biotinidase (Fragment | 16980.07031 | 16514.45313 |
| SRGAP1 | G5EA48;Q7Z6B7 | SRGAP1 | SLIT-ROBO Rho GTPas | 5898.027832 | 8874.296875 |
| DCAF8 | Q5TAQ9 | DCAF8 | DDB1- and CUL4-asso | 37025.34375 | 45817.32422 |
| MMS19 | Q96T76 | MMS19 | MMS19 nucleotide ex | 134759.875 | 158387.4219 |
| YWHAH | Q04917 | YWHAH | 14-3-3 protein eta | 340974.3438 | 366154.4688 |
| GPATCH8 | Q9UKJ3 | GPATCH8 | G patch domain-conta | 77270.63281 | 73098.3125 |
| CTSA | P10619;X6R5C5;X6R8A1 | CTSA | Lysosomal protective | 90141.39063 | 112153.5625 |
| TTN | A0A0A0MTS7;Q8WZ42 | TTN | Titin | 226295.2344 | 195075.4063 |
| C19orf53 | Q9UNZ5 | C19orf53 | Leydig cell tumor 10 k | 26385.74219 | 35753.83594 |
| CLASP1 | Q7Z460 | CLASP1 | CLIP-associating prote | 38725.375 | 43368.79297 |
| PPP2R5C | H0YJ75;Q13362 | PPP2R5C | Serine/threonine-prot | 170732.9688 | 214592.5 |
| SRPRB | Q9Y5M8 | SRPRB | Signal recognition par | 162076.7969 | 145864.4219 |
| TSN | E9PGT1;Q15631 | TSN | Translin | 315171.625 | 342431.5938 |
| CDYL2 | A0A0B4J291;Q8N8U2 | CDYL2 | Chromodomain Y-like | 16902.75391 | 11075.29688 |

| | | | | | |
|---|---|---|---|---|---|
| HARS2 | A0A2R8Y5P7;P49590 | HARS2 | Probable histidine--tR | 76248.50781 | 87275.53906 |
| POT1 | A0A590UJF2;A0A590UJF7;A0 | POT1 | Protection of telomere | 1573.116577 | 5124.967285 |
| PSMA8 | Q8TAA3 | PSMA8 | Proteasome subunit a | 625545.1875 | 646035.6875 |
| UBIAD1 | Q9Y5Z9 | UBIAD1 | UbiA prenyltransferas | 20389.02148 | 28699.86914 |
| NCAPD3 | P42695 | NCAPD3 | Condensin-2 complex | 44949.80469 | 48275.87891 |
| PDXK | O00764 | PDXK | Pyridoxal kinase | 341231.875 | 407242.75 |
| RO60 | P10155 | RO60 | 60 kDa SS-A/Ro ribonu | 160949.4219 | 180375.3906 |
| CTR9 | Q6PD62 | CTR9 | RNA polymerase-asso | 77116.32031 | 81384.42969 |
| COX17 | C9J8T6;Q14061 | COX17 | Cytochrome c oxidase | 12085.75977 | 14619.95703 |
| FCHSD2 | E7ENZ2;E9PG19;O94868 | FCHSD2 | F-BAR and double SH3 | 8215.560547 | 7774.456543 |
| OLFML3 | B4DNG0;Q9NRN5 | OLFML3 | Olfactomedin-like pro | 3782.171631 | 3081.713867 |
| TUT1 | F5H0R1;Q9H6E5 | TUT1 | Speckle targeted PIP5 | 13898.77637 | 11746.42871 |
| JUND | P17535 | JUND | Transcription factor ju | 15422.5293 | 13140.01855 |
| ASB6 | F6TX30;Q9NWX5 | ASB6 | Ankyrin repeat and SC | 19188.54688 | 17629.83594 |
| DMAP1 | Q9NPF5 | DMAP1 | DNA methyltransferas | 41664.28125 | 32385.37305 |
| MBNL1 | A0A0A0MQX8;C9JP00;Q9NR! | MBNL1 | Muscleblind-like prote | 288178.375 | 245476.5469 |
| TRIM27 | P14373 | TRIM27 | Zinc finger protein RFI | 32128.72656 | 22465.10352 |
| RBM25 | P49756 | RBM25 | RNA-binding protein 2 | 273987.5313 | 255427.1875 |
| HERPUD2 | Q9BSE4 | HERPUD2 | Homocysteine-respon | 7382.284668 | 7834.492676 |
| PEBP1 | P30086 | PEBP1 | Phosphatidylethanola | 2541628.25 | 2526142.25 |
| CCDC47 | Q96A33 | CCDC47 | Coiled-coil domain-co | 288003 | 286242.5938 |
| ANKLE2 | Q86XL3 | ANKLE2 | Ankyrin repeat and LE | 20663.68945 | 19243.9082 |
| ZC3HAV1 | C9J6P4 | ZC3HAV1 | Zinc finger CCCH-type | 160566.2207 | 165681.2832 |
| TMEM65 | Q6PI78 | TMEM65 | Transmembrane prote | 30750.72852 | 28826.55469 |
| MAPK1 | P28482 | MAPK1 | Mitogen-activated pro | 275365.8438 | 309672.0938 |
| CDK17 | F5H6Z0;Q00537 | CDK17 | Cyclin-dependent kina | 11065.50586 | 11406.49316 |
| RAB25 | P57735 | RAB25 | Ras-related protein Ra | 260890.5 | 280185.25 |
| CAND1 | Q86VP6 | CAND1 | Cullin-associated NED | 691864.4375 | 830635.9375 |
| RBX1 | P62877 | RBX1 | E3 ubiquitin-protein li | 78964.41406 | 86725.36719 |
| PGK2 | P07205 | PGK2 | Phosphoglycerate kina | 45804.11328 | 47802.72656 |
| DHX36 | Q9H2U1 | DHX36 | ATP-dependent DNA/ | 207475.0938 | 218241.9063 |
| METTL26 | Q96S19 | METTL26 | Methyltransferase-like | 65441.09766 | 50879 |
| FBXO38 | Q6PIJ6 | FBXO38 | F-box only protein 38 | 5286.127441 | 4595.899902 |

| | | | | | |
|---|---|---|---|---|---|
| SLC25A21 | Q9BQT8 | SLC25A21 | Mitochondrial 2-oxod | 17348.86523 | 17444.08594 |
| KRTCAP2 | Q8N6L1 | KRTCAP2 | Keratinocyte-associat | 15165.50879 | 17828.65234 |
| NKRF | O15226 | NKRF | NF-kappa-B-repressin | 88368.19531 | 89140.15625 |
| DIDO1 | Q9BTC0 | DIDO1 | Death-inducer oblitera | 75133.60938 | 76602.86719 |
| ASH2L | F5H8F7;Q9UBL3 | ASH2L | Set1/Ash2 histone me | 49298.59766 | 51660.60547 |
| COPS3 | Q9UNS2 | COPS3 | COP9 signalosome co | 182188.5313 | 185396.7344 |
| PHF20L1 | A0A0A0MQS0;F8W9L8 | PHF20L1 | PHD finger protein 20 | 8363.663086 | 6854.945313 |
| PPP5C | H0YDU8;P53041 | PPP5C | Serine/threonine-prot | 157055.875 | 161888.0156 |
| PPP2R5D | E9PFR3;Q14738 | PPP2R5D | Serine/threonine-prot | 287390.2188 | 327704.7813 |
| TBC1D9 | Q6ZT07 | TBC1D9 | TBC1 domain family m | 12315.78223 | 12482.35254 |
| ZNF512B | Q96KM6 | ZNF512B | Zinc finger protein 51 | 47367.46875 | 47056.44141 |
| SDC1 | P18827 | SDC1 | Syndecan-1 | 5339.30127 | 6009.741211 |
| PSMA7 | O14818 | PSMA7 | Proteasome subunit a | 1551799.25 | 1648227.5 |
| DYNLL1 | P63167 | DYNLL1 | Dynein light chain 1, c | 548877.0625 | 606625.375 |
| PARP2 | G3V167;Q9UGN5 | PARP2 | Poly [ADP-ribose] poly | 2670.825195 | 13301.27539 |
| C12orf4 | Q9NQ89 | C12orf4 | Protein C12orf4 | 3359.025391 | 4437.620605 |
| ARL2 | P36404 | ARL2 | ADP-ribosylation facto | 260482.6563 | 304374.0938 |
| COPS8 | E9PGT6;Q99627 | COPS8 | COP9 signalosome co | 184251.5 | 192440.9375 |
| ANKRD16 | Q6P6B7 | ANKRD16 | Ankyrin repeat domai | 9746.799805 | 10456.45996 |
| PPP1R12A | O14974 | PPP1R12A | Protein phosphatase 1 | 170642.4063 | 167354.875 |
| TIMM10 | P62072 | TIMM10 | Mitochondrial import | 4793.628418 | 5847.577637 |
| PES1 | B5MCF9;O00541 | PES1 | Pescadillo homolog | 175965.7031 | 105439.9375 |
| ATP5F1B | P06576 | ATP5F1B | ATP synthase subunit | 1152675.5 | 1458851.75 |
| HDHD2 | Q9H0R4 | HDHD2 | Haloacid dehalogenas | 105347.9453 | 114969.5 |
| SDF2L1 | Q9HCN8 | SDF2L1 | Stromal cell-derived fa | 227878.5156 | 235240.4219 |
| SCYL3 | Q8IZE3;X6RHX1 | SCYL3 | Protein-associating wi | 10391.36719 | 11738.97363 |
| TARDBP | Q13148 | TARDBP | TAR DNA-binding prot | 324061.875 | 299262.625 |
| SLMAP | A0A590UJK3;Q14BN4 | SLMAP | Sarcolemmal membra | 17112.91797 | 16575.3125 |
| POGZ | Q7Z3K3 | POGZ | Pogo transposable ele | 128207.9375 | 133123.4531 |
| IP6K1 | C9JNA8;Q92551 | IP6K1 | Kinase;Inositol hexakis | 5184.153809 | 7147.519531 |
| UIMC1 | Q96RL1 | UIMC1 | BRCA1-A complex sub | 43373.08594 | 43029.49219 |
| ATP5ME | P56385 | ATP5ME | ATP synthase subunit | 647757.5 | 840856.9375 |
| CLTA | F8WF69;P09496 | CLTA | Clathrin light chain;Cla | 42336.55078 | 21640.70703 |

| | | | | | |
|---|---|---|---|---|---|
| ARHGAP19 | Q14CB8 | ARHGAP19 | Rho GTPase-activating | 39239.32422 | 25835.32617 |
| PTBP1 | P26599 | PTBP1 | Polypyrimidine tract-b | 1785640 | 1722119.25 |
| FBXL17 | H7C835;Q9UF56 | FBXL17 | F-box and leucine-rich | 8411.87207 | 7819.010742 |
| PIK3R2 | O00459 | PIK3R2 | Phosphatidylinositol 3 | 15192.45313 | 15451.76953 |
| RING1 | Q06587 | RING1 | E3 ubiquitin-protein li | 57106.28906 | 55533.58984 |
| THOC1 | A0A087WWS1;Q96FV9 | THOC1 | THO complex subunit | 141377.9844 | 116079.2031 |
| RAD21 | O60216 | RAD21 | Double-strand-break r | 124942.5469 | 126392.1875 |
| MANEAL | E9PMU5;Q5VSG8 | MANEAL | Glycoprotein endo-alp | 13623.73145 | 12635.38379 |
| DNAJC6 | O75061 | DNAJC6 | Putative tyrosine-prot | 14043.62402 | 11231.15918 |
| GTPBP6 | O43824 | GTPBP6 | Putative GTP-binding | 24073.58203 | 26361.49805 |
| APIP | Q96GX9 | APIP | Methylthioribulose-1- | 99461.3125 | 99162.42188 |
| PARK7 | Q99497 | PARK7 | Protein/nucleic acid d | 1424395.25 | 1535818.75 |
| ZNF326 | Q5BKZ1 | ZNF326 | DBIRD complex subun | 65174.89453 | 68460.53125 |
| LSM1 | O15116 | LSM1 | U6 snRNA-associated | 23411.54688 | 29822.79297 |
| TRAF6 | Q9Y4K3 | TRAF6 | TNF receptor-associat | 22292.52734 | 20101.80859 |
| FUT11 | Q495W5 | FUT11 | Alpha-(1,3)-fucosyltra | 3817.391602 | 4162.068848 |
| EMC3 | Q9P0I2 | EMC3 | ER membrane protein | 52177.11719 | 54411.96875 |
| DHX34 | Q14147 | DHX34 | Probable ATP-depend | 10603.87012 | 9080.945313 |
| GYG1 | P46976 | GYG1 | Glycogenin-1 | 402660.375 | 413985.7188 |
| KIF23 | H7BYN4;Q02241 | KIF23 | Kinesin-like protein;Ki | 83202.54688 | 77120.55469 |
| SMAD3 | P84022 | SMAD3 | Mothers against deca | 29747.04688 | 28511.3125 |
| GOSR1 | E9PCW1;O95249 | GOSR1 | Golgi SNAP receptor c | 44168.70313 | 46137.67969 |
| WASF2 | Q9Y6W5 | WASF2 | Wiskott-Aldrich syndr | 77852.21875 | 83555.17188 |
| PCCA | P05165 | PCCA | Propionyl-CoA carbox | 34253.15625 | 40293.08594 |
| KIF4A | O95239 | KIF4A | Chromosome-associat | 80316.03125 | 96327.20313 |
| PMM1 | Q92871 | PMM1 | Phosphomannomutas | 7784.586426 | 9505.99707 |
| NUDT3 | O95989 | NUDT3 | Diphosphoinositol pol | 45024.17188 | 47039.73438 |
| EIF2AK3 | Q9NZJ5 | EIF2AK3 | Eukaryotic translation | 9915.188477 | 9056.896484 |
| SUPT7L | O94864 | SUPT7L | STAGA complex 65 su | 7950.192871 | 12197.44629 |
| EPC2 | Q52LR7 | EPC2 | Enhancer of polycomb | 9565.0625 | 7143.609863 |
| TUBGCP4 | Q9UGJ1 | TUBGCP4 | Gamma-tubulin comp | 8211.907227 | 9259.801758 |
| MAGED1 | Q9Y5V3 | MAGED1 | Melanoma-associated | 19642.06445 | 23825.66211 |
| CLPTM1 | O96005 | CLPTM1 | Cleft lip and palate tra | 255886.3281 | 310124.6563 |

| | | | | | |
|---|---|---|---|---|---|
| TMEM209 | Q96SK2 | TMEM209 | Transmembrane prote | 24803.55273 | 24824.95898 |
| UFC1 | Q9Y3C8 | UFC1 | Ubiquitin-fold modifie | 144240.1094 | 158894.7969 |
| MEF2D | Q14814 | MEF2D | Myocyte-specific enha | NA | 3542.578857 |
| ZMYM1 | Q5SVZ6 | ZMYM1 | Zinc finger MYM-type | 8767.049805 | 8459.344727 |
| SEC24D | O94855 | SEC24D | Protein transport prot | 75729.76563 | 96177.24219 |
| PTTG1IP | P53801 | PTTG1IP | Pituitary tumor-transf | 30705.75 | 20809.35938 |
| NABP2 | C9JMP5;C9JT95;Q9BQ15 | NABP2 | SOSS complex subunit | 1129.929199 | 25178.37305 |
| THUMPD1 | Q9NXG2 | THUMPD1 | THUMP domain-conta | 153070.4844 | 175687.7188 |
| NUDCD3 | Q8IVD9 | NUDCD3 | NudC domain-contain | 92719.73438 | 106154.5156 |
| TMEM63A | O94886 | TMEM63A | CSC1-like protein 1 | 54711.22266 | 49317.82813 |
| AP2A1 | O95782 | AP2A1 | AP-2 complex subunit | 266874.5625 | 321142.4375 |
| MPZL1 | O95297 | MPZL1 | Myelin protein zero-li | 20708.33008 | 17142.94141 |
| HNRNPK | P61978 | HNRNPK | Heterogeneous nuclea | 3584435.789 | 2969046.137 |
| ETFB | P38117 | ETFB | Electron transfer flavo | 649620.6875 | 661110.3125 |
| INO80 | Q9ULG1 | INO80 | Chromatin-remodeling | 9772.402344 | 11406.76074 |
| SPATA33 | A0A087WYH0;J3KRC8;Q96N( | SPATA33 | Spermatogenesis-asso | 25361.34375 | 30101.01953 |
| DPP8 | Q6V1X1 | DPP8 | Dipeptidyl peptidase 8 | 15007.24121 | 17422.58789 |
| GNAQ | P50148 | GNAQ | Guanine nucleotide-b | 94752.70313 | 100199.2188 |
| PANX1 | Q96RD7 | PANX1 | Pannexin-1 | 5382.677246 | 5141.122559 |
| EAF1 | Q96JC9 | EAF1 | ELL-associated factor | 15345.88867 | 13636.90527 |
| RNGTT | O60942 | RNGTT | mRNA-capping enzym | 59626.02734 | 53465.20703 |
| TSTA3 | Q13630 | TSTA3 | GDP-L-fucose synthas | 276003.5 | 331201.3125 |
| DDI2 | Q5TDH0 | DDI2 | Protein DDI1 homolog | 56036.74609 | 69716.0625 |
| SIN3B | O75182 | SIN3B | Paired amphipathic he | 1908.028076 | 2003.767456 |
| TASOR2 | A0A2R8YH03;Q5VWN6 | TASOR2 | Protein TASOR 2 | 7433.608887 | 5786.286621 |
| ARL8A | Q96BM9 | ARL8A | ADP-ribosylation facto | 273327.875 | 312218.2188 |
| SKIV2L | Q15477 | SKIV2L | Helicase SKI2W | 33434.76563 | 39535.69141 |
| GALNT2 | Q10471 | GALNT2 | Polypeptide N-acetylg | 84091.26563 | 79889.625 |
| ABI1 | A0A0A0MRT6;B6VEX4;Q8IZP | ABI1 | Abl interactor 1 | 144525.6563 | 141343.1406 |
| DTL | F5GZ90;Q9NZJ0 | DTL | Denticleless protein h | 18082.61133 | 19514.74805 |
| PITHD1 | Q9GZP4 | PITHD1 | PITH domain-containi | 88075.95313 | 91018.17188 |
| ZCCHC17 | A0A087WXF8;A0A087WXU5; | ZCCHC17 | Nucleolar protein of 4 | 17322.68945 | 18997.88672 |
| POMGNT2 | Q8NAT1 | POMGNT2 | Protein O-linked-mann | 2055.476807 | 1459.151123 |

| | | | | | |
|---|---|---|---|---|---|
| STX16 | Q96NX8 | STX16 | Syntaxin-16 | 9134.427734 | 9019.822266 |
| LEPROT | A0A087X0N2;A0A3B3ISI6;O1 | LEPROT | Leptin receptor gene- | 3593.734863 | 3542.925537 |
| PRKRIP1 | Q9H875 | PRKRIP1 | PRKR-interacting prot | 52924.45313 | 55316.51953 |
| STXBP2 | Q15833 | STXBP2 | Syntaxin-binding prot | 106810.2578 | 130894.6484 |
| MICAL3 | Q7RTP6 | MICAL3 | [F-actin]-monooxygen | 18972.7793 | 17764.40625 |
| NQO2 | P16083;Q5TD07 | NQO2 | Ribosyldihydronicotin | 7881.282715 | 8847.333984 |
| MSL3 | A0A3B3IT59;A0A3B3ITL5;A0A | MSL3 | Male-specific lethal 3 | 6232.211426 | 6941.195313 |
| SPATA5L1 | Q9BVQ7 | SPATA5L1 | Spermatogenesis-asso | 35529.56641 | 36046.5625 |
| MLST8 | A0A0A0MR05;Q9BVC4 | MLST8 | Target of rapamycin c | 22638.30664 | 26884.94727 |
| TCEAL4 | Q96EI5 | TCEAL4 | Transcription elongati | 356400.8125 | 386276.2188 |
| HMOX2 | P30519 | HMOX2 | Heme oxygenase 2 | 308089.5 | 312492.25 |
| SLC27A3 | Q5K4L6;X6R3N0 | SLC27A3 | Solute carrier family 2 | 67077.27344 | 73402.22656 |
| PSMD12 | O00232 | PSMD12 | 26S proteasome non-A | 477499.9375 | 507850.8125 |
| ELOVL1 | Q9BW60 | ELOVL1 | Elongation of very lon | 27080.42383 | 25358.86328 |
| MCM2 | P49736 | MCM2 | DNA replication licens | 437144.625 | 472968.7188 |
| BAK1 | Q16611 | BAK1 | Bcl-2 homologous ant | 29129.55078 | 25464.25977 |
| FKBP15 | Q5T1M5 | FKBP15 | FK506-binding proteir | 43007.62891 | 43283.26172 |
| CRTAP | C9JP16;O75718 | CRTAP | Cartilage-associated p | 28133.36328 | 19837.05664 |
| CNOT6 | Q9ULM6 | CNOT6 | CCR4-NOT transcriptic | 45067.73438 | 51482.52734 |
| CCDC22 | O60826 | CCDC22 | Coiled-coil domain-co | 82434.94531 | 88289.08594 |
| MCL1 | Q07820 | MCL1 | Induced myeloid leuke | 40620.37891 | 42126.30078 |
| TOM1 | O60784 | TOM1 | Target of Myb protein | 47899.83203 | 88664.625 |
| PRXL2A | Q9BRX8 | PRXL2A | Peroxiredoxin-like 2A | 342955.0625 | 447349.1563 |
| RBM39 | Q14498 | RBM39 | RNA-binding protein 3 | 427158.9688 | 418976.0313 |
| CMTM4 | J3QRP2;Q8IZR5 | CMTM4 | CKLF-like MARVEL trar | 36119.25391 | 42228.36328 |
| BSCL2 | H7C2D5;J3KQ12;Q96G97 | BSCL2 | Seipin (Fragment);HC( | 11919.99219 | 11690.44043 |
| AK3 | Q9UIJ7 | AK3 | GTP:AMP phosphotra | 168556 | 160651.4219 |
| DFFB | B4DZS0;O76075 | DFFB | DNA fragmentation fa | 8893.396484 | 9625.956055 |
| PPIB | P23284 | PPIB | Peptidyl-prolyl cis-trar | 2425947.5 | 2450665.5 |
| MARK3 | P27448 | MARK3 | MAP/microtubule affi | 53144.12109 | 51787.57422 |
| PSMB3 | P49720 | PSMB3 | Proteasome subunit b | 507144.2188 | 553565.5625 |
| RAD18 | Q9NS91 | RAD18 | E3 ubiquitin-protein li | 40234.19141 | 28917.12695 |
| FOXP4 | B7ZBM3;Q8IVH2 | FOXP4 | Forkhead box protein | 11906.83496 | 11656.12891 |

| | | | | | |
|---|---|---|---|---|---|
| SP1 | P08047 | SP1 | Transcription factor Sp | 21325.58398 | 23722.86719 |
| EPHB3 | P54753 | EPHB3 | Ephrin type-B recepto | 25850.73438 | 17970.46484 |
| WDR5 | P61964 | WDR5 | WD repeat-containing | 186509.0781 | 177430.75 |
| TRMT1L | Q7Z2T5 | TRMT1L | TRMT1-like protein | 34213.35547 | 32996.64063 |
| PIK3C2B | F5GWN5;O00750 | PIK3C2B | Phosphatidylinositol 4 | 17168.45703 | 21217.44141 |
| BCLAF1 | E9PK91 | BCLAF1 | Bcl-2-associated trans | 484638.1602 | 437595.6484 |
| CCNL1 | Q9UK58 | CCNL1 | Cyclin-L1 | 20921.42969 | 17554.10742 |
| ESRP2 | Q9H6T0 | ESRP2 | Epithelial splicing regu | 78815.99219 | 82597 |
| SLC25A19 | Q9HC21 | SLC25A19 | Mitochondrial thiamir | 43385.74219 | 38402.90625 |
| GABPA | Q06546 | GABPA | GA-binding protein alp | 48523.07031 | 50620.72656 |
| PSMA4 | P25789 | PSMA4 | Proteasome subunit a | 1481676 | 1664949.125 |
| PYCR3 | A0A0A0MQS1;Q53H96 | PYCR3 | Pyrroline-5-carboxylat | 141031.4844 | 180086.2656 |
| LUC7L | Q9NQ29 | LUC7L | Putative RNA-binding | 169554.4844 | 173248.5469 |
| WDR45B | Q5MNZ6 | WDR45B | WD repeat domain ph | 25644.25195 | 28073.5 |
| NUP133 | Q8WUM0 | NUP133 | Nuclear pore complex | 73606.07031 | 71296.39063 |
| PRRC1 | Q96M27 | PRRC1 | Protein PRRC1 | 160309.9688 | 160522.8594 |
| ESPL1 | H3BRX7;Q14674 | ESPL1 | Separin (Fragment);Se | 5564.775879 | 7191.65918 |
| SERPINB8 | P50452 | SERPINB8 | Serpin B8 | 68434.35156 | 59252.66797 |
| ALG5 | Q9Y673 | ALG5 | Dolichyl-phosphate be | 179504.4063 | 183344.625 |
| IRF6 | O14896 | IRF6 | Interferon regulatory | 19897.00195 | 16958.29883 |
| WRN | Q14191 | WRN | Werner syndrome ATI | 7176.754883 | 7040.993164 |
| ATPAF2 | Q8N5M1 | ATPAF2 | ATP synthase mitocho | 37701.24219 | 44144.72656 |
| XRCC5 | P13010 | XRCC5 | X-ray repair cross-con | 1702875.375 | 2095297.125 |
| PHIP | Q8WWQ0 | PHIP | PH-interacting proteir | 5572.63916 | 1111.316772 |
| VAPA | Q9P0L0 | VAPA | Vesicle-associated me | 597135.25 | 565913.375 |
| PPIP5K1 | B7WPL9;F8W9A8;Q6PFW1 | PPIP5K1 | Inositol hexakisphospl | 7839.798828 | 8270.767578 |
| COG5 | Q9UP83 | COG5 | Conserved oligomeric | 62503.55371 | 60996.7207 |
| ABI2 | A0A0C4DG21;E7EP65;E7EW7 | ABI2 | Abl interactor 2;Abl in | 68380.5625 | 77908.88281 |
| R3HDM1 | Q15032 | R3HDM1 | R3H domain-containir | 5232.744141 | 7366.455566 |
| CAAP1 | Q9H8G2 | CAAP1 | Caspase activity and a | 15812.52148 | 18071.36523 |
| ECI1 | P42126 | ECI1 | Enoyl-CoA delta isome | 314138.9688 | 374318.1875 |
| SFT2D1 | Q8WV19 | SFT2D1 | Vesicle transport prot | 31330.90039 | 17700.91406 |
| WDR61 | Q9GZS3 | WDR61 | WD repeat-containing | 89918.24219 | 99400.99219 |

| | | | | | |
|---|---|---|---|---|---|
| ACSS1 | Q9NUB1 | ACSS1 | Acetyl-coenzyme A sy | 10589.05664 | 11069.58398 |
| MAPK14 | Q16539 | MAPK14 | Mitogen-activated prc | 93621.29688 | 98728.57031 |
| MYO15A | A0A087WYA1;Q9UKN7 | MYO15A | Unconventional myos | 116842.7813 | 133516.0625 |
| HIRA | P54198 | HIRA | Protein HIRA | 15281.44629 | 18494.04492 |
| ARFGAP2 | E9PN48 | ARFGAP2 | ADP-ribosylation facto | 445711.7344 | 430738.6875 |
| IARS1 | P41252 | IARS1 | Isoleucine--tRNA ligas | 426382.2188 | 495371.5625 |
| TM9SF4 | A0A0C4DFM1;Q92544 | TM9SF4 | Transmembrane 9 sup | 161979.6563 | 189864.375 |
| ATP5MG | O75964 | ATP5MG | ATP synthase subunit | 716902.0625 | 769496.125 |
| FRS2 | Q8WU20 | FRS2 | Fibroblast growth fact | 466.3434448 | 591.6417236 |
| CANT1 | Q8WVQ1 | CANT1 | Soluble calcium-activa | 173994.4375 | 179860.3906 |
| LGMN | Q99538 | LGMN | Legumain | 21964.5 | 20197.32031 |
| PAFAH1B2 | P68402 | PAFAH1B2 | Platelet-activating fac | 528045.625 | 542464.9375 |
| TBC1D9B | Q66K14 | TBC1D9B | TBC1 domain family m | 88162.71094 | 99055.89844 |
| TUBG1 | P23258 | TUBG1 | Tubulin gamma-1 chai | 191352.0938 | 204382.2656 |
| POTEF | A5A3E0 | POTEF | POTE ankyrin domain | 1089702.375 | 1180950.75 |
| WDR91 | A4D1P6;C9J1X0 | WDR91 | WD repeat-containing | 18986.67188 | 17354.18945 |
| EIF4E | P06730 | EIF4E | Eukaryotic translation | 429730.9688 | 428286.9063 |
| LZTFL1 | Q9NQ48 | LZTFL1 | Leucine zipper transcr | 191930.6875 | 195192.5781 |
| AAR2 | A2A2Q9;Q9Y312 | AAR2 | Protein AAR2 homolog | 53380.19141 | 53803.35938 |
| ZNHIT2 | Q9UHR6 | ZNHIT2 | Zinc finger HIT domair | 45850.69922 | 36933.74219 |
| ETFA | P13804 | ETFA | Electron transfer flavo | 713480.0625 | 717136.6875 |
| DNMT3A | Q9Y6K1 | DNMT3A | DNA (cytosine-5)-met | 27730.26172 | 31722.71484 |
| WDR82 | Q6UXN9 | WDR82 | WD repeat-containing | 98810.17969 | 94338.55469 |
| GOT1 | P17174 | GOT1 | Aspartate aminotrans | 743974.75 | 802816.875 |
| TBCE | Q15813 | TBCE | Tubulin-specific chape | 125244.6797 | 124266.1953 |
| EIF4G1 | E7EUU4 | EIF4G1 | Eukaryotic translation | 780294.5234 | 822689.2422 |
| SPOUT1 | Q5T280 | SPOUT1 | Putative methyltransf | 107143.6563 | 103309.8906 |
| FANCA | O15360 | FANCA | Fanconi anemia group | 9825.791992 | 13179.00391 |
| UBE2M | P61081 | UBE2M | NEDD8-conjugating er | 658624.3125 | 653913 |
| AHSA1 | O95433 | AHSA1 | Activator of 90 kDa he | 1018232.625 | 1095045.875 |
| POP1 | Q99575 | POP1 | Ribonucleases P/MRP | 78013.66406 | 92856.94531 |
| DLGAP4 | Q9Y2H0 | DLGAP4 | Disks large-associated | 45098.50391 | 49107.33203 |
| HSP90AA4P | Q58FG1 | HSP90AA4P | Putative heat shock pr | 151613.9844 | 111946.3906 |

| | | | | | |
|---|---|---|---|---|---|
| NUDCD1 | Q96RS6 | NUDCD1 | NudC domain-contain | 172570.5 | 187559.9688 |
| ORC1 | Q13415 | ORC1 | Origin recognition cor | 13612.69727 | 13676.69824 |
| BIVM-ERCC5;ERCC5;ERCC5; | A0A1W2PS85;A0A494C0S2;P | BIVM-ERCC5;ERCC5;E | BIVM-ERCC5 readthro | 1744.630859 | 2043.333374 |
| THOC6 | Q86W42 | THOC6 | THO complex subunit | 90701.49219 | 80357.21875 |
| SMIM4 | Q8WVI0 | SMIM4 | Small integral membra | 35168.94141 | 46937.76172 |
| PAAF1 | Q9BRP4 | PAAF1 | Proteasomal ATPase-a | 136640.6563 | 151197.7188 |
| NAA30 | Q147X3 | NAA30 | N-alpha-acetyltransfe | 51417.59375 | 59589.12891 |
| DHPS | P49366 | DHPS | Deoxyhypusine syntha | 56421.64453 | 67666.48438 |
| DYNC1LI2 | O43237 | DYNC1LI2 | Cytoplasmic dynein 1 | 186266.25 | 210766.4531 |
| DDX39B | Q13838 | DDX39B | Spliceosome RNA heli | 2872799.5 | 2804849.5 |
| SETX | Q7Z333 | SETX | Probable helicase sen | 14996.00488 | 17313.66992 |
| PHACTR2 | J3KP75 | PHACTR2 | Phosphatase and actir | 12677.57031 | 14454.64844 |
| HDAC7 | J3KPH8;Q8WUI4 | HDAC7 | Histone deacetylase;H | 6305.308105 | 4070.553955 |
| AKAP17A | Q02040 | AKAP17A | A-kinase anchor prote | 37990.17969 | 34848.20313 |
| GOT2 | P00505 | GOT2 | Aspartate aminotrans | 633060.25 | 731967.4375 |
| ROBO1 | A0A087WTM1;A0A087WZ85 | ROBO1 | Roundabout homolog | 764842.5625 | 948333.875 |
| SCAP | A0A0A0MTG6;Q12770 | SCAP | Sterol regulatory elem | 15689.63379 | 13275.44531 |
| RSF1 | Q96T23 | RSF1 | Remodeling and spaci | 3502.884277 | 13237.00684 |
| NAA25 | Q14CX7 | NAA25 | N-alpha-acetyltransfe | 78005.47656 | 86393.63281 |
| CNOT10 | Q9H9A5 | CNOT10 | CCR4-NOT transcriptic | 37278.36328 | 40746.14844 |
| CENPT | H3BMP6;H3BN05;H3BQ71;H | CENPT | Centromere protein T | 7671.415039 | 3352.068848 |
| BAIAP2L1 | Q9UHR4 | BAIAP2L1 | Brain-specific angioge | 15935.58789 | 14444.50195 |
| SUB1 | P53999 | SUB1 | Activated RNA polyme | 2158573.5 | 2204032.25 |
| ATG13 | O75143 | ATG13 | Autophagy-related pr | 9838.118164 | 11319.81836 |
| ZNF48 | A0A087WVT1;Q96MX3 | ZNF48 | Zinc finger protein 48 | 9456.810547 | 9567.269531 |
| OSBPL2 | Q9H1P3 | OSBPL2 | Oxysterol-binding pro | 64611.34766 | 69222.78125 |
| NCR3LG1 | Q68D85 | NCR3LG1 | Natural cytotoxicity tr | NA | 11760.89551 |
| SNAP47 | A0A087X0B7;Q5SQN1 | SNAP47 | Synaptosomal-associa | 9318.558594 | 10256.88867 |
| STRN | O43815 | STRN | Striatin | 53027.31641 | 52207.76953 |
| LUZP1 | Q86V48 | LUZP1 | Leucine zipper proteir | 11842.70703 | 11692.86133 |
| DNAJB2 | P25686 | DNAJB2 | DnaJ homolog subfam | 23375.4375 | 18357.48047 |
| ACAD11 | Q709F0 | ACAD11 | Acyl-CoA dehydrogena | 7759.090332 | 12900.08691 |
| PTPRK | E9PGC5;Q5TG12 | PTPRK | Receptor-type tyrosin | 21985.56445 | 25032.3357 |

| | | | | | |
|---|---|---|---|---|---|
| YPEL5 | P62699 | YPEL5 | Protein yippee-like 5 | 10734.48438 | 12936.44824 |
| CHSY1 | Q86X52 | CHSY1 | Chondroitin sulfate sy | 5408.776367 | 4335.294434 |
| DHRS7 | Q9Y394 | DHRS7 | Dehydrogenase/reduc | 59527.22656 | 60008.72656 |
| SLC39A7 | Q92504 | SLC39A7 | Zinc transporter SLC3! | 19080.5332 | 23155.38086 |
| TARS3 | A2RTX5 | TARS3 | Threonine--tRNA ligas | 26644.46484 | 27365.87305 |
| KIF3B | O15066 | KIF3B | Kinesin-like protein KI | 22174.24219 | 23079.07227 |
| U2AF1L5;U2AF1 | P0DN76;Q01081 | U2AF1L5;U2AF1 | Splicing factor U2AF 3 | 548351.9375 | 427936.875 |
| PPP1R13L | Q8WUF5 | PPP1R13L | RelA-associated inhibi | 95823.78906 | 98264.39063 |
| PIGS | Q96S52 | PIGS | GPI transamidase com | 71590.17969 | 77869.35156 |
| TIPRL | O75663 | TIPRL | TIP41-like protein | 147126.0156 | 149085.8906 |
| ATAD3A | Q9NVI7 | ATAD3A | ATPase family AAA do | 219755.0469 | 229381.3906 |
| CDC25B | B4DIG0;G8JLH2;P30305 | CDC25B | M-phase inducer phos | 3649.250488 | 2614.128418 |
| CCDC6 | Q16204 | CCDC6 | Coiled-coil domain-co | 305501.3438 | 275035.5 |
| PPT1 | P50897 | PPT1 | Palmitoyl-protein thio | 228714.9531 | 238497.5938 |
| SLC25A6 | P12236 | SLC25A6 | ADP/ATP translocase : | 624281.3125 | 603013.8125 |
| ZBTB11 | O95625 | ZBTB11 | Zinc finger and BTB dc | 5312.218262 | 6214.460449 |
| PAPSS1 | O43252 | PAPSS1 | Bifunctional 3'-phospl | 357681.875 | 330967.2813 |
| SSR3 | C9JA28;Q9UNL2 | SSR3 | Translocon-associatec | 83950.71875 | 87088.99219 |
| NT5C2 | P49902 | NT5C2 | Cytosolic purine 5'-nu | 30697.76758 | 36878.66406 |
| JADE3 | Q92613 | JADE3 | Protein Jade-3 | 2722.986084 | 3710.2146 |
| NONO | Q15233 | NONO | Non-POU domain-con | 1543432.5 | 1246536.125 |
| HMGB1 | P09429 | HMGB1 | High mobility group p | 6960128.5 | 7097147 |
| FBXO7 | Q9Y3I1 | FBXO7 | F-box only protein 7 | 69812.05469 | 63682.53125 |
| TOLLIP | Q9H0E2 | TOLLIP | Toll-interacting protei | 40638.55859 | 44529.72266 |
| TRMT112 | Q9UI30 | TRMT112 | Multifunctional methy | 150847.2344 | 159461.3125 |
| RAB5C | P51148 | RAB5C | Ras-related protein Ra | 768645.75 | 866360.375 |
| TMA7 | Q9Y2S6 | TMA7 | Translation machinery | 216812.2813 | 251963.4688 |
| NCBP2 | P52298 | NCBP2 | Nuclear cap-binding p | 65088.90625 | 68789.42969 |
| APPL1 | Q9UKG1 | APPL1 | DCC-interacting prote | 50624.97656 | 55455.95703 |
| STAM2 | O75886 | STAM2 | Signal transducing ada | 45692.32813 | 54098.17188 |
| GOLGA2 | A0A2Q2TH77;Q08379 | GOLGA2 | Golgin subfamily A me | 135069.8438 | 130892.125 |
| VPS18 | Q9P253 | VPS18 | Vacuolar protein sorti | 62547.98438 | 70163.33594 |
| HUWE1 | A0A087X1S3 | HUWE1 | E3 ubiquitin-protein li | 430988.459 | 571232.7109 |

| | | | | | |
|---|---|---|---|---|---|
| PLS1 | Q14651 | PLS1 | Plastin-1 | 165867.2031 | 164900.7813 |
| VWA1 | Q6PCB0 | VWA1 | von Willebrand factor | 1378.098511 | 1835.529663 |
| ROCK2 | O75116 | ROCK2 | Rho-associated protei | 107288.0313 | 102832.3359 |
| COPE | O14579 | COPE | Coatomer subunit eps | 156127.9844 | 170430.7813 |
| IMUP | Q9GZP8 | IMUP | Immortalization up-re | 45853.88281 | 27918.15234 |
| GLMN | Q92990 | GLMN | Glomulin | 21487.18555 | 22988.25977 |
| APH1A | Q96BI3 | APH1A | Gamma-secretase sub | 11082.01367 | 13169.03418 |
| TK1 | P04183 | TK1 | Thymidine kinase, cyt | 225673.8281 | 231655.7188 |
| NIT2 | Q9NQR4 | NIT2 | Omega-amidase NIT2 | 190135.5781 | 214938.9063 |
| MCM6 | Q14566 | MCM6 | DNA replication licens | 442876.2188 | 491961.375 |
| EMC7 | Q9NPA0 | EMC7 | ER membrane protein | 156623.4844 | 162790.0313 |
| CD151 | E9PMR4;K4DIA7;P48509 | CD151 | Tetraspanin;Tetraspar | 138988.0938 | 156800.2031 |
| ARPC5L | Q9BPX5 | ARPC5L | Actin-related protein | 232484.0781 | 212208.6719 |
| PPIP5K2 | A0A087WZV0;O43314 | PPIP5K2 | Inositol hexakisphospl | 33160.76172 | 36568.02344 |
| BRD7 | Q9NPI1 | BRD7 | Bromodomain-contair | 22924.75 | 23126.6543 |
| CRIPT | Q9P021 | CRIPT | Cysteine-rich PDZ-bin | 19419.0918 | 22193.74023 |
| RFX7 | H0YLX2;Q2KHR2 | RFX7 | DNA-binding protein F | 46758.61328 | 88325.74219 |
| RHEB | Q15382 | RHEB | GTP-binding protein R | 107386.3516 | 122112.5781 |
| ZNF317 | Q96PQ6 | ZNF317 | Zinc finger protein 31 | 931.9171753 | 1228.96814 |
| MISP | Q8IVT2 | MISP | Mitotic interactor and | 67867.23438 | 48310.69141 |
| WDR89 | Q96FK6 | WDR89 | WD repeat-containing | 16383.61719 | 13666.21191 |
| EPB41L1 | A0A0C4DH22;A0A1B0GTW6; | EPB41L1 | Band 4.1-like protein | 30775.33105 | 33975.09082 |
| GXYLT1 | Q4G148 | GXYLT1 | Glucoside xylosyltrans | 17505.55859 | 16644.12891 |
| RECQL5 | O94762 | RECQL5 | ATP-dependent DNA h | 19689.24609 | 19705.1875 |
| ERCC4 | Q92889 | ERCC4 | DNA repair endonucle | 18099.39648 | 18081.43359 |
| SRP68 | Q9UHB9 | SRP68 | Signal recognition par | 295776.7188 | 324450.0313 |
| HOOK1 | Q9UJC3 | HOOK1 | Protein Hook homolog | 99109.8125 | 89751.67969 |
| PTGES3L-AARSD1;AARSD1 | C9J5N1;Q9BTE6 | PTGES3L-AARSD1;AAF | PTGES3L-AARSD1 reac | 51868.20703 | 47729.41016 |
| TRAFD1 | O14545 | TRAFD1 | TRAF-type zinc finger | 41159.96484 | 46316.58984 |
| TRNT1 | Q96Q11 | TRNT1 | CCA tRNA nucleotidylt | 138655.7656 | 152371.8125 |
| AK2 | F8W1A4;P54819 | AK2 | Adenylate kinase 2, m | 505762 | 522592.5625 |
| RPN1 | P04843 | RPN1 | Dolichyl-diphosphooli | 906318.5 | 1054828.875 |
| UBXN7 | O94888 | UBXN7 | UBX domain-containir | 17486.14063 | 17643.25391 |

| | | | | | |
|---|---|---|---|---|---|
| NCLN | Q969V3 | NCLN | Nicalin | 107841.8281 | 106748.375 |
| EPN1 | Q9Y6I3 | EPN1 | Epsin-1 | 140752 | 161990.5625 |
| USF1 | B1AQP1;E9PME6;P22415 | USF1 | Upstream stimulatory | 46722.56641 | 50356.60547 |
| BAG6 | A0A024RCR6;A0A0G2JK23;P4 | BAG6 | BAG6;Large proline-ri | 207706.2969 | 261697.7813 |
| CDC73 | Q6P1J9 | CDC73 | Parafibromin | 197060.8281 | 178534.8438 |
| CTTNBP2NL | Q9P2B4 | CTTNBP2NL | CTTNBP2 N-terminal-l | 18522.41211 | 21166.00977 |
| EIF2AK4 | H0YME5;Q9P2K8 | EIF2AK4 | eIF-2-alpha kinase GC | 5525.90625 | 6524.094727 |
| ACY1 | A0A1B0GU86;Q03154 | ACY1 | Aminoacylase-1 | 66028.64063 | 67270.46875 |
| VTI1A | Q96AJ9 | VTI1A | Vesicle transport thro | 12057.2002 | 16907.48633 |
| SLC1A4 | P43007 | SLC1A4 | Neutral amino acid tra | 79161.9375 | 69864.3125 |
| MED16 | Q9Y2X0 | MED16 | Mediator of RNA poly | 21931.42188 | 25372.88672 |
| USP38 | Q8NB14 | USP38 | Ubiquitin carboxyl-ter | 7370.959961 | 7386.125977 |
| MSX2 | D6RIS4;P35548 | MSX2 | Homeobox protein M: | 12900.70508 | 12339.68848 |
| SLC25A36 | Q96CQ1 | SLC25A36 | Solute carrier family 2 | 13178.53516 | 12900.95703 |
| FBXO9 | Q9UK97 | FBXO9 | F-box only protein 9 | 16250.59473 | 17576.91406 |
| RNF145 | Q96MT1 | RNF145 | RING finger protein 14 | 10513.10352 | 12197.75488 |
| SUCLG2 | Q96I99 | SUCLG2 | Succinate--CoA ligase | 197991.7969 | 217068.0156 |
| GPS1 | C9JFE4;Q13098 | GPS1 | COP9 signalosome coı | 214749.9375 | 245586.4531 |
| NUDT2 | P50583 | NUDT2 | Bis(5'-nucleosyl)-tetra | 69437.88281 | 78586.59375 |
| NELFB | A0A5H1ZRP4 | NELFB | Negative elongation fa | 141124.9219 | 145133.4063 |
| PPP2R1A | P30153 | PPP2R1A | Serine/threonine-prot | 1033145.375 | 1410829.75 |
| FZD6 | O60353 | FZD6 | Frizzled-6 | 18089.63086 | 20915.01563 |
| PROSER2 | D3DRR9;Q86WR7 | PROSER2 | Chromosome 10 oper | 102334.8594 | 123168.8359 |
| RABGEF1 | E9PFK9;Q9UJ41 | RABGEF1 | Rab5 GDP/GTP exchar | 30008.01172 | 29043.9082 |
| ARMC10 | Q8N2F6 | ARMC10 | Armadillo repeat-cont | 40612.46094 | 47787.84375 |
| RUSC1 | A0A087WYU8 | RUSC1 | RUN and SH3 domain- | 30408.27783 | 30670.97656 |
| TPM3 | A0A087WWU8 | TPM3 | Tropomyosin alpha-3 | 44731.39258 | 62270.85156 |
| UQCC1 | B1AKV3;Q9NVA1 | UQCC1 | Ubiquinol-cytochrome | 51563.65234 | 63474.11328 |
| PCLAF | Q15004 | PCLAF | PCNA-associated facto | 154727.5313 | 130666.7813 |
| MTFR1L | Q9H019 | MTFR1L | Mitochondrial fission | 14788.55566 | 13518.75684 |
| SNRPA | P09012 | SNRPA | U1 small nuclear ribor | 380998.9688 | 264692.5313 |
| RABAC1 | M0R3D4;Q9UI14 | RABAC1 | PRA1 family protein;P | 55856.53516 | 55848.96875 |
| DNAJA2 | O60884 | DNAJA2 | DnaJ homolog subfam | 317175.6563 | 347122.6563 |

| | | | | | |
|---|---|---|---|---|---|
| MGAT4B | E5RFS3;H0YB63;H0YB84;H0Y | MGAT4B | Alpha-1,3-mannosyl-g | 5775.984375 | 5740.817383 |
| SMARCE1 | A0A2R8Y855;B4DGM3;Q969( | SMARCE1 | SWI/SNF-related matr | 78717 | 81509.14063 |
| UBQLN1 | Q9UMX0 | UBQLN1 | Ubiquilin-1 | 32902.29297 | 32862.25391 |
| RPS11 | P62280 | RPS11 | 40S ribosomal protein | 1027472.25 | 1095504.375 |
| GGCX | P38435 | GGCX | Vitamin K-dependent | 21516.97656 | 29670.99609 |
| PHKA1 | A6NMN0;P46020 | PHKA1 | Phosphorylase b kinas | 9624.431641 | 11774.0625 |
| LEPROTL1 | A0A3B3IS19;C9JVM4;E5RHU8 | LEPROTL1 | Leptin receptor overla | 2417.575928 | 3042.401611 |
| TBL1XR1 | Q9BZK7 | TBL1XR1 | F-box-like/WD repeat- | 93844.63281 | 104611.0469 |
| C19orf47 | J3QS80;Q8N9M1 | C19orf47 | Uncharacterized prote | 18098.23242 | 12966.75488 |
| LSM14A | Q8ND56 | LSM14A | Protein LSM14 homol | 112181.0156 | 123553.9063 |
| EMC4 | H0YNK8;Q5J8M3 | EMC4 | ER membrane protein | 158558.3438 | 144335.7188 |
| HNRNPH1 | G8JLB6;P31943 | HNRNPH1 | Heterogeneous nucle | 1119681 | 782386.1875 |
| KRI1 | A0A494C108;Q8N9T8 | KRI1 | Protein KRI1 homolog | 101635.2266 | 82614.76563 |
| VPS41 | P49754 | VPS41 | Vacuolar protein sorti | 22836.56445 | 30255.43164 |
| DNAJC5 | Q9H3Z4 | DNAJC5 | DnaJ homolog subfam | 46935.87891 | 51165.78125 |
| PPP4R2 | Q9NY27 | PPP4R2 | Serine/threonine-prot | 115988.3047 | 128579.8281 |
| PRELID3B | Q9Y3B1 | PRELID3B | PRELI domain contain | 11816.92676 | 9118.157227 |
| RTF2 | Q9BY42 | RTF2 | Replication terminatic | 216061.0156 | 229888.0781 |
| STAG1 | Q8WVM7 | STAG1 | Cohesin subunit SA-1 | 91389.04688 | 84046.28125 |
| MCM3AP | O60318 | MCM3AP | Germinal-center asso | 20154.58203 | 20521.16016 |
| FUBP1 | C9JSZ1 | FUBP1 | Far upstream element | 1105436.469 | 1158851.75 |
| ZNF574 | A0A0C4DFM2;Q6ZN55 | ZNF574 | HCG1643764, isoform | 8849.367188 | 8009.767578 |
| ;AHR;AHR | A0A1B0GVI7;A0A2R8Y7G1;P3 | ;AHR;AHR | Uncharacterized prote | 12531.59473 | 11205.48145 |
| PRSS1 | E7EQ64;P07477 | PRSS1 | Trypsin-1 | 308255.875 | 226913.5625 |
| SECISBP2L | J3KPI1;Q93073 | SECISBP2L | Selenocysteine inserti | 23845.18359 | 23601.85547 |
| DOCK1 | A0A096LNH6;Q14185 | DOCK1 | Dedicator of cytokine | 87780.30469 | 83729.875 |
| FBXO45 | P0C2W1 | FBXO45 | F-box/SPRY domain-co | 7053.737305 | 7689.381836 |
| HS2ST1 | Q7LGA3 | HS2ST1 | Heparan sulfate 2-O-s | 39708.99609 | 44932.09375 |
| AMFR | Q9UKV5 | AMFR | E3 ubiquitin-protein li | 28607.71094 | 36844.54688 |
| CNDP2 | Q96KP4 | CNDP2 | Cytosolic non-specific | 536487.5 | 621159.25 |
| MLLT1 | Q03111 | MLLT1 | Protein ENL | 15780.60156 | 16511.18555 |
| ETHE1 | O95571 | ETHE1 | Persulfide dioxygenas | 95482.5 | 109429.5156 |
| DNAJC10 | Q8IXB1 | DNAJC10 | DnaJ homolog subfam | 48814.57031 | 54269.03516 |

| | | | | | |
|---|---|---|---|---|---|
| AKAP10 | O43572 | AKAP10 | A-kinase anchor prote | 20935.78125 | 18081.32227 |
| CIT | O14578 | CIT | Citron Rho-interacting | 16863.08398 | 20895.21094 |
| PRPF31 | Q8WWY3 | PRPF31 | U4/U6 small nuclear r | 179505.4844 | 177810.3281 |
| SRP72 | O76094 | SRP72 | Signal recognition par | 216409.5781 | 228661.7031 |
| ZNF581 | Q9P0T4 | ZNF581 | Zinc finger protein 58 | 7325.917969 | 5265.26416 |
| SREK1 | A0A087WY03;E5RFV3 | SREK1 | Splicing regulatory glu | 92178.44531 | 78468.78125 |
| UBR4 | Q5T4S7 | UBR4 | E3 ubiquitin-protein li | 128032.4453 | 129227.3203 |
| SSX2IP | Q9Y2D8;S4R403 | SSX2IP | Afadin- and alpha-acti | 4878.479492 | 5516.72998 |
| LENG8 | A0A087WUE4;A0A0G2JNN3; | LENG8 | Leukocyte receptor clu | 9208.820313 | 8875.352539 |
| VAMP3 | Q15836 | VAMP3 | Vesicle-associated me | 149315.5781 | 156693.8125 |
| PRKAR2A | P13861 | PRKAR2A | cAMP-dependent pro | 123905.9688 | 115846.6563 |
| SHMT1 | P34896 | SHMT1 | Serine hydroxymethyl | 205115.0313 | 220033.4219 |
| PLP2 | Q04941 | PLP2 | Proteolipid protein 2 | 373033.5938 | 377955.875 |
| RRAS2 | P62070 | RRAS2 | Ras-related protein R- | 1451466 | 1480762.375 |
| XRCC1 | F5H8D7 | XRCC1 | DNA repair protein XR | 179740.5645 | 173526.1221 |
| MAGOH | P61326 | MAGOH | Protein mago nashi ho | 129908.5781 | 94405.40625 |
| SYNGR2 | O43760 | SYNGR2 | Synaptogyrin-2 | 342498.625 | 370801.7813 |
| POLB | P06746 | POLB | DNA polymerase beta | 13777.67871 | 13489.78613 |
| EML1 | A0A3B3IU69;F8W717;O0042 | EML1 | Echinoderm microtub | 5020.193848 | 3980.345459 |
| ATXN7L3B | Q96GX2 | ATXN7L3B | Ataxin-7-like protein 3 | 49514.52734 | 61480.83203 |
| ILKAP | Q9H0C8 | ILKAP | Integrin-linked kinase- | 63842.61719 | 69452.04688 |
| MYO1D | O94832 | MYO1D | Unconventional myos | 57663.44922 | 68078.69531 |
| BAX | Q07812 | BAX | Apoptosis regulator B. | 322818.5 | 289675.8125 |
| KLHL22 | Q53GT1 | KLHL22 | Kelch-like protein 22 | 15267.76953 | 15181.17285 |
| CYC1 | P08574 | CYC1 | Cytochrome c1, heme | 971323 | 863870.4375 |
| GEMIN8 | Q9NWZ8 | GEMIN8 | Gem-associated prote | 27360.33789 | 32520.49023 |
| ARAP1 | Q96P48 | ARAP1 | Arf-GAP with Rho-GAI | 60623.23828 | 55957.25391 |
| NUDT21 | O43809 | NUDT21 | Cleavage and polyade | 610991.25 | 586925.125 |
| SNRNP70 | P08621 | SNRNP70 | U1 small nuclear ribor | 855578.875 | 559152.5625 |
| CCAR1 | Q8IX12 | CCAR1 | Cell division cycle and | 188365.4219 | 162406.7813 |
| ZNF672 | Q499Z4 | ZNF672 | Zinc finger protein 67. | 4816.820801 | 4506.998535 |
| NAGLU | P54802 | NAGLU | Alpha-N-acetylglucosa | 61899.125 | 65762.54688 |
| HELLS | A0A0B4J1V9;Q9NRZ9 | HELLS | Helicase, lymphoid-sp | 99939.78906 | 109218.7109 |

| | | | | | |
|---|---|---|---|---|---|
| DDX59 | Q5T1V6 | DDX59 | Probable ATP-depend | 40953.96484 | 39595.35547 |
| THOC3 | Q96J01 | THOC3 | THO complex subunit | 215231.125 | 202375.6094 |
| SNAPC4 | Q5SXM2 | SNAPC4 | snRNA-activating prot | 13194.94629 | 13199.25586 |
| PIAS3 | E9PHH8;Q9Y6X2 | PIAS3 | E3 SUMO-protein liga | 12064.51074 | 12541.61621 |
| PRPF4 | O43172 | PRPF4 | U4/U6 small nuclear r | 174180.5625 | 163404.9219 |
| POP7 | C9JYM0;O75817 | POP7 | Ribonuclease P protei | 102124.9922 | 104567.8125 |
| STRIP1 | Q5VSL9 | STRIP1 | Striatin-interacting pr | 44893.53125 | 49309.48047 |
| ZCRB1 | G3V1V1;Q8TBF4 | ZCRB1 | Zinc finger CCHC-type | 9172.717773 | 8047.410156 |
| UBE2I | P63279 | UBE2I | SUMO-conjugating en | 836905.75 | 864754.375 |
| URM1 | Q9BTM9 | URM1 | Ubiquitin-related moc | 47801.19531 | 58394.67578 |
| MATR3 | A0A0R4J2E8;P43243 | MATR3 | Matrin-3 | 549543 | 506609 |
| FLYWCH2 | Q96CP2 | FLYWCH2 | FLYWCH family memb | 290564.6563 | 295407.4063 |
| CHTOP | Q9Y3Y2;X6R700 | CHTOP | Chromatin target of P | 125829.8672 | 105702.9922 |
| PATZ1 | Q9HBE1 | PATZ1 | POZ-, AT hook-, and zi | 36966.87109 | 39999.34375 |
| EMB | D6RDX7;Q6PCB8 | EMB | Embigin | 6849.950195 | 6092.088867 |
| MFSD6 | Q6ZSS7 | MFSD6 | Major facilitator supe | 8088.187988 | 5187.01416 |
| CTNNAL1 | Q9UBT7 | CTNNAL1 | Alpha-catulin | 4334.546875 | 5769.350586 |
| ACBD6 | A0A2R8Y544;Q9BR61 | ACBD6 | Acyl-CoA-binding dom | 6316.032715 | 10227.04199 |
| PPM1B | O75688 | PPM1B | Protein phosphatase 1 | 109903.2969 | 112941.5391 |
| RPL24 | C9JNW5;C9JXB8;P83731 | RPL24 | 60S ribosomal protein | 1069133.375 | 953545.625 |
| MIEF1 | L0R8F8 | MIEF1 | MIEF1 upstream open | 45164.58594 | 38581.55078 |
| COMMD5 | Q9GZQ3 | COMMD5 | COMM domain-conta | 43389.33984 | 48762.96484 |
| TRAPPC2;TRAPPC2B | P0DI81;P0DI82 | TRAPPC2;TRAPPC2B | Trafficking protein pa | 18256.60156 | 24076.86328 |
| CENPE | A0A087X0P0;Q02224 | CENPE | Centromere-associate | 4782.291016 | 5267.208008 |
| FBXO5 | Q9UKT4 | FBXO5 | F-box only protein 5 | 3713.868164 | 4898.164551 |
| UBL5 | Q9BZL1 | UBL5 | Ubiquitin-like protein | 181404.75 | 170179.8281 |
| SHQ1 | Q6PI26 | SHQ1 | Protein SHQ1 homolo | 7902.255371 | 7787.612793 |
| VPS26C | A8MTY9;O14972 | VPS26C | Vacuolar protein sorti | 3222.590576 | 4199.781738 |
| ARFGEF2 | Q9Y6D5 | ARFGEF2 | Brefeldin A-inhibited g | 80838.75781 | 75860.875 |
| IGF2R | P11717 | IGF2R | Cation-independent m | 168607.0156 | 144017.0156 |
| MYO18A | A0A0D9SFK2;Q92614 | MYO18A | Unconventional myos | 30326.57813 | 31131.88672 |
| HMGN2 | P05204 | HMGN2 | Non-histone chromos | 39400.41797 | 9323.139648 |
| SETMAR | Q53H47 | SETMAR | Histone-lysine N-meth | 11190.39063 | 11117.8457 |

| | | | | | |
|---|---|---|---|---|---|
| DLG3 | Q5JUW8;Q92796 | DLG3 | Discs, large homolog 3 | 29761.33789 | 27678.67969 |
| MPLKIP | Q8TAP9 | MPLKIP | M-phase-specific PLK1 | 14281.41602 | 18878.13672 |
| PRPF39 | Q86UA1 | PRPF39 | Pre-mRNA-processing | 45844.31641 | 46938.07422 |
| NME3 | Q13232 | NME3 | Nucleoside diphospha | 124485.3125 | 94186.59375 |
| AKR1C2 | B4DK69;P52895 | AKR1C2 | Aldo-keto reductase fa | 302.822998 | 807.3808594 |
| LMNB2 | Q03252 | LMNB2 | Lamin-B2 | 283393.4063 | 265462.8125 |
| NT5C3A | Q9H0P0;X6RM59 | NT5C3A | Cytosolic 5'-nucleotida | 72643.625 | 70483.80469 |
| SH3BGRL3 | Q5T123;Q9H299 | SH3BGRL3 | SH3 domain-binding g | 183108 | 220384.6719 |
| PRDM10 | E9PLV1;Q9NQV6 | PRDM10 | PR domain zinc finger | 11135.91504 | 7105.346191 |
| TTI1 | O43156 | TTI1 | TELO2-interacting pro | 14772.16211 | 16549.29492 |
| FNBP1L | Q5T0N5;S4R347 | FNBP1L | Formin-binding protei | 28263.76953 | 30020.84766 |
| ERCC1 | P07992 | ERCC1 | DNA excision repair pr | 9978.126953 | 10732.16992 |
| CCNT2 | O60583 | CCNT2 | Cyclin-T2 | 8124.310547 | 8591.774414 |
| KAT6A | A0A3B3IS53;A0A3F2YNX6;A5 | KAT6A | Histone acetyltransfer | 5751.464844 | 5533.875488 |
| DCAF7 | P61962 | DCAF7 | DDB1- and CUL4-asso | 390884.75 | 455008.9688 |
| MAF1 | E9PR76;E9PSH4;Q9H063 | MAF1 | Repressor of RNA poly | 14338.2373 | 16575.41992 |
| GSK3A | P49840 | GSK3A | Glycogen synthase kir | 91429.96875 | 95027.57813 |
| RAB29 | O14966 | RAB29 | Ras-related protein Ra | 23930.23828 | 29405.80859 |
| APLP2 | Q06481 | APLP2 | Amyloid-like protein 2 | 27301.45508 | 28069.10938 |
| ITPR2 | Q14571 | ITPR2 | Inositol 1,4,5-trisphos | 2211.546875 | 3255.44043 |
| SYMPK | Q92797 | SYMPK | Symplekin | 90432.38281 | 106288.5547 |
| VARS2 | A0A0G2JL52;B4E0K6 | VARS2 | VARS2;Valine--tRNA li | 18059.70313 | 18529.45703 |
| MTX1 | Q13505 | MTX1 | Metaxin-1 | 108410.5 | 96587.89844 |
| VCP | P55072 | VCP | Transitional endoplası | 1685802.25 | 1833833.125 |
| HPS3 | Q969F9 | HPS3 | Hermansky-Pudlak syı | 15225.51074 | 15968.56055 |
| CEP78 | A0A2R8Y432;A0A2R8Y4C1;A( | CEP78 | Centrosomal protein o | 14882.19043 | 12194.80176 |
| STAT6 | A0A1W2PNW1;A0A494C0T4; | STAT6 | Signal transducer and | 47417.55859 | 56031.28906 |
| RBM17 | Q96I25 | RBM17 | Splicing factor 45 | 175123.8281 | 165188.5625 |
| SIN3A | Q96ST3 | SIN3A | Paired amphipathic he | 88509.10938 | 97850.67969 |
| CHST14 | H0YN65;Q8NCH0 | CHST14 | Carbohydrate sulfotra | 19447.20508 | 16182.30469 |
| TOPBP1 | A0A2R8YD63;Q92547 | TOPBP1 | DNA topoisomerase 2 | 35831.51172 | 45662.03516 |
| GNA13 | Q14344 | GNA13 | Guanine nucleotide-b | 205765.3594 | 236213.25 |
| MTPAP | Q9NVV4 | MTPAP | Poly(A) RNA polymera | 31264.6875 | 37338.31641 |

| | | | | | |
|---|---|---|---|---|---|
| EEF1AKNMT | Q8N6R0 | EEF1AKNMT | eEF1A lysine and N-te | 102713.8672 | 101085.7031 |
| TCP1 | P17987 | TCP1 | T-complex protein 1 s | 1074750.875 | 1206452.25 |
| TP53BP2 | Q13625 | TP53BP2 | Apoptosis-stimulating | 16334.45508 | 17030.65039 |
| HERC4 | Q5GLZ8 | HERC4 | Probable E3 ubiquitin- | 40770.73047 | 44864.86328 |
| CWF19L2 | Q2TBE0 | CWF19L2 | CWF19-like protein 2 | 9429.714844 | 9142.146484 |
| ECHS1 | P30084 | ECHS1 | Enoyl-CoA hydratase, | 616380.5625 | 660214 |
| ADSS1 | Q8N142 | ADSS1 | Adenylosuccinate syn | 16176.88281 | 19964.58789 |
| CNP | P09543 | CNP | 2',3'-cyclic-nucleotide | 163388.3281 | 175934.8594 |
| PSMD6 | Q15008 | PSMD6 | 26S proteasome non-A | 484018.8125 | 443281.4688 |
| RLIM | Q9NVW2 | RLIM | E3 ubiquitin-protein li | 2851.958008 | 2179.747559 |
| CLTB | P09497 | CLTB | Clathrin light chain B | 126331.3203 | 103319.3672 |
| PTPN1 | P18031 | PTPN1 | Tyrosine-protein phos | 445822.7188 | 481043.9063 |
| STN1 | Q9H668 | STN1 | CST complex subunit S | 5509.852051 | 5764.344238 |
| RPS27 | P42677 | RPS27 | 40S ribosomal protein | 360296.4375 | 344742.6563 |
| ARHGAP39 | Q9C0H5 | ARHGAP39 | Rho GTPase-activating | 6359.597656 | 5558.724609 |
| TMEM9 | B1ALM5;Q9P0T7 | TMEM9 | Transmembrane prote | 43509.41016 | 50416.58203 |
| DCPS | Q96C86 | DCPS | m7GpppX diphosphat | 114717.3672 | 126674.6875 |
| KIF20B | Q96Q89 | KIF20B | Kinesin-like protein KI | 13118.23438 | 13584.73926 |
| VCPIP1 | Q96JH7 | VCPIP1 | Deubiquitinating prot | 98984.55469 | 109095.0547 |
| CLEC16A | Q2KHT3 | CLEC16A | Protein CLEC16A | 9634.429688 | 11380.01758 |
| LAMA5 | O15230 | LAMA5 | Laminin subunit alpha | 15988.22559 | 16118.04688 |
| GZF1 | Q9H116 | GZF1 | GDNF-inducible zinc fi | 6406.461914 | 5878.209961 |
| CASK | A0A2R8YE77;A0A2U3TZM1;C | CASK | Calcium/calmodulin-d | 311751.625 | 312395.5625 |
| PDE5A | G5E9C5;O76074 | PDE5A | Phosphodiesterase;cG | 1897.658569 | 2531.007813 |
| MIEN1 | Q9BRT3 | MIEN1 | Migration and invasio | 18076.25 | 25972.7832 |
| HSDL1 | Q3SXM5 | HSDL1 | Inactive hydroxystero | 118417.25 | 135393.3438 |
| HEXIM1 | O94992 | HEXIM1 | Protein HEXIM1 | 85651.29688 | 94446.71875 |
| AJUBA | Q96IF1 | AJUBA | LIM domain-containin | 17763.4043 | 19665.2207 |
| SEPTIN5 | E7EQM7;E7EX32;Q99719 | SEPTIN5 | Septin-5 | 5565.308594 | 7164.5625 |
| ANAPC4 | Q9UJX5 | ANAPC4 | Anaphase-promoting | 45327.66797 | 51118.51563 |
| SNX16 | A0A0C4DGW2;E5RGQ6;E5RH | SNX16 | Sorting nexin-16;Sorti | 1013.828186 | 1369.22522 |
| PPP1CC | F8VYE8;P36873 | PPP1CC | Serine/threonine-prot | 789764.0625 | 820058.5625 |
| CPSF4 | O95639 | CPSF4 | Cleavage and polyade | 77621.67969 | 71703.25781 |

| | | | | | |
|---|---|---|---|---|---|
| PPP1R13B | A0A3B3ISJ2;Q96KQ4 | PPP1R13B | Apoptosis-stimulating | 26932.12695 | 21958.86914 |
| ATP5F1E;ATP5F1EP2 | P56381;Q5VTU8 | ATP5F1E;ATP5F1EP2 | ATP synthase subunit | 117518.1484 | 133280.5625 |
| MARCHF5 | Q9NX47 | MARCHF5 | E3 ubiquitin-protein li | 88763.28906 | 94544.03906 |
| RMND5B | F5H6G4;Q96G75 | RMND5B | E3 ubiquitin-protein tr | 4784.253906 | 4239.348633 |
| CCDC85B | Q15834 | CCDC85B | Coiled-coil domain-co | 19091.01758 | 26541.17383 |
| INTS12 | Q96CB8 | INTS12 | Integrator complex su | 21100.32813 | 21704.99414 |
| DCUN1D4 | Q92564 | DCUN1D4 | DCN1-like protein 4 | 12966.7041 | 10890.56152 |
| RPL10 | F8W7C6 | RPL10 | 60S ribosomal protein | 1077882.625 | 980319.2969 |
| C6orf47 | O95873 | C6orf47 | Uncharacterized prote | 16912.91016 | 20973.43945 |
| HNRNPA3 | P51991 | HNRNPA3 | Heterogeneous nuclea | 1104447.625 | 571930.5 |
| SASS6 | Q6UVJ0 | SASS6 | Spindle assembly abn | 21037.25 | 13917.52539 |
| OGT | O15294 | OGT | UDP-N-acetylglucosar | 124244.7656 | 135189.0781 |
| MBD4 | O95243 | MBD4 | Methyl-CpG-binding d | 14066.53711 | 12713.85645 |
| C17orf49;BAP18 | F8W038;Q8IXM2 | C17orf49;BAP18 | Chromosome 17 oper | 63759.90234 | 54370.86719 |
| GORASP2 | Q9H8Y8 | GORASP2 | Golgi reassembly-stac | 59450.77344 | 71261.57813 |
| PABPN1 | Q86U42 | PABPN1 | Polyadenylate-binding | 654967 | 599951.6875 |
| RNF123 | C9J266;Q5XPI4 | RNF123 | E3 ubiquitin-protein li | 3213.127686 | 4339.459961 |
| BAG1 | J3QTA2;Q99933 | BAG1 | BAG family molecular | 50315.02344 | 44564.09766 |
| EPS8L2 | Q9H6S3 | EPS8L2 | Epidermal growth fact | 296393.7188 | 318001.125 |
| WDR70 | Q9NW82 | WDR70 | WD repeat-containing | 55491.67969 | 45131.58594 |
| TFPT | A0A0G2JP55;A8MTQ3;P0C1Z | TFPT | TCF3 fusion partner | 9961.827148 | 6897.817871 |
| GRK2 | P25098 | GRK2 | Beta-adrenergic recep | 106692.7578 | 125482.875 |
| RPS9 | P46781 | RPS9 | 40S ribosomal protein | 2632020 | 2460294.5 |
| TMEM231 | H3BTN6;Q9H6L2 | TMEM231 | Transmembrane prote | 7261.972656 | 8431.424805 |
| MKKS | Q9HB66 | MKKS | Alternative protein M | 183316.5781 | 185573.8594 |
| KAT7 | O95251 | KAT7 | Histone acetyltransfer | 39436.23438 | 43391.75781 |
| ARL6IP1 | H3BPJ2;Q15041 | ARL6IP1 | ADP-ribosylation facto | 102755.3594 | 107249.4141 |
| KMT2D | O14686 | KMT2D | Histone-lysine N-meth | 13302.83398 | 13624.86426 |
| BUB3 | O43684 | BUB3 | Mitotic checkpoint pro | 456065.5938 | 421232.125 |
| DMAC2 | Q9NW81 | DMAC2 | Distal membrane-arm | 14802.56836 | 16694.80859 |
| RNF14 | Q9UBS8 | RNF14 | E3 ubiquitin-protein li | 36062.24609 | 36418.44141 |
| USF2 | M0QXT0;Q15853 | USF2 | Upstream stimulatory | 43059.83203 | 33589.43359 |
| CETN3 | E5RJF8;E5RK82;O15182 | CETN3 | Centrin-3 | 4563.165527 | 2276.232178 |

| | | | | | |
|---|---|---|---|---|---|
| NUP205 | Q92621 | NUP205 | Nuclear pore complex | 82833.14063 | 93749.46875 |
| SYNCRIP | B7Z645 | SYNCRIP | Heterogeneous nucle | 2243398.352 | 1942324.961 |
| PLD2 | O14939 | PLD2 | Phospholipase D2 | 4554.959961 | 7794.541992 |
| STAMBP | O95630 | STAMBP | STAM-binding protein | 106044.1641 | 102084.3984 |
| KIF1B | A0A087WWA3;O60333;Q4R | KIF1B | Kinesin-like protein KI | 54123.28516 | 60859.46094 |
| COMMD3 | Q9UBI1 | COMMD3 | COMM domain-conta | 17852.01758 | 29774.71094 |
| GCN1 | Q92616 | GCN1 | eIF-2-alpha kinase act | 386212.6875 | 453595.0313 |
| ACTR2 | P61160 | ACTR2 | Actin-related protein | 847011.625 | 821549.8125 |
| SUPT20H | F6S7C4;Q8NEM7 | SUPT20H | Transcription factor S | 10464.11621 | 9460.782227 |
| SHOC2 | Q9UQ13 | SHOC2 | Leucine-rich repeat pr | 30118.99805 | 31495.57031 |
| FAF2 | Q96CS3 | FAF2 | FAS-associated factor | 278115.0938 | 265808.7188 |
| MTX3 | Q5HYI7 | MTX3 | Metaxin-3 | 20234.68945 | 13503.94727 |
| PRXL2B | A0A0A0MT35;A0A2P0CTB1;J | PRXL2B | Prostamide/prostagla | 21848.375 | 27032.21094 |
| BCL10 | A0A087WWW9;O95999 | BCL10 | B-cell lymphoma/leuk | 76342.6875 | 77594.54688 |
| SACM1L | A0A5F9ZHN7;Q9NTJ5 | SACM1L | Phosphatidylinositide | 88901.91406 | 94611.21875 |
| HSPA5 | P11021 | HSPA5 | Endoplasmic reticulur | 1694154.375 | 1756862 |
| SF3B4 | Q15427 | SF3B4 | Splicing factor 3B sub | 327591.2188 | 255916.2344 |
| SRRT | Q9BXP5 | SRRT | Serrate RNA effector r | 317668.25 | 333080.4688 |
| BICD2 | Q8TD16 | BICD2 | Protein bicaudal D ho | 22596.96289 | 26359.63672 |
| SMARCB1 | A0A2R8YEE8 | SMARCB1 | SWI/SNF-related matr | 308583.3203 | 317532.0117 |
| EPS15 | P42566 | EPS15 | Epidermal growth fac | 80732.28125 | 92862.53906 |
| CHFR | F5GWH4;Q96EP1 | CHFR | E3 ubiquitin-protein li | 6853.57666 | 5770.953125 |
| TUSC1 | Q2TAM9 | TUSC1 | Tumor suppressor car | 4009.092773 | 3925.811035 |
| RFX1 | P22670 | RFX1 | MHC class II regulator | 20060.78125 | 19585.08203 |
| DNAJC8 | O75937 | DNAJC8 | DnaJ homolog subfam | 301177.6875 | 308777.375 |
| LIG1 | P18858 | LIG1 | DNA ligase 1 | 173838.6719 | 214871.9531 |
| QARS1 | P47897 | QARS1 | Glutamine--tRNA ligas | 265968.5 | 305342.625 |
| CHPF2 | G5E9W2;Q9P2E5 | CHPF2 | Hexosyltransferase;Cl | 2133.227295 | 4059.62085 |
| TUBGCP2 | Q9BSJ2 | TUBGCP2 | Gamma-tubulin comp | 79259.88281 | 88360.52344 |
| GUF1 | Q8N442 | GUF1 | Translation factor GU | 13476.54492 | 15438.91309 |
| PRPSAP2 | O60256 | PRPSAP2 | Phosphoribosyl pyrop | 118696.8906 | 160378.5 |
| AARS2 | Q5JTZ9 | AARS2 | Alanine--tRNA ligase, | 46905.17578 | 51846.67969 |
| ATRX | P46100 | ATRX | Transcriptional regula | 40556.97656 | 46636.47656 |

| | | | | | |
|---|---|---|---|---|---|
| CORO7-PAM16 | A0A0A6YYL4 | CORO7-PAM16 | Coronin | 124928.2031 | 134811.125 |
| LMNB1 | P20700 | LMNB1 | Lamin-B1 | 1844185.625 | 1552671 |
| CPNE3 | O75131 | CPNE3 | Copine-3 | 1119245.25 | 1010365.063 |
| BAG2 | O95816 | BAG2 | BAG family molecular | 54015.91016 | 59703.83984 |
| COX20 | Q5RI15 | COX20 | Cytochrome c oxidase | 92610.9375 | 98554 |
| TRPS1 | Q9UHF7 | TRPS1 | Zinc finger transcriptio | 70631.44531 | 70589.64063 |
| PSMB6 | P28072 | PSMB6 | Proteasome subunit b | 796510.25 | 847469.25 |
| LCMT2 | O60294 | LCMT2 | tRNA wybutosine-synt | 11049.3291 | 10557.84277 |
| MAD2L1 | Q13257 | MAD2L1 | Mitotic spindle assem | 178759.7969 | 216615.0781 |
| QRICH1 | Q2TAL8 | QRICH1 | Glutamine-rich protei | 49046.28906 | 58435.94141 |
| AP3M1 | Q9Y2T2 | AP3M1 | AP-3 complex subunit | 95638.97656 | 115271.2422 |
| CREBBP | Q92793 | CREBBP | CREB-binding protein | 54782.82813 | 54543.33203 |
| PRADC1 | Q9BSG0 | PRADC1 | Protease-associated d | 5704.967285 | 9286.368164 |
| YTHDC1 | J3QR07;Q96MU7 | YTHDC1 | YTH domain-containir | 41926.14453 | 30838.0918 |
| MBTD1 | Q05BQ5 | MBTD1 | MBT domain-containi | 6394.807617 | 7539.320801 |
| CCDC154 | A0A590PWR5;A6NI56;B7ZBA | CCDC154 | Coiled-coil domain-co | 416908.3438 | 231793.9219 |
| NUP93 | Q8N1F7 | NUP93 | Nuclear pore complex | 193436.75 | 207978.2188 |
| DNAJB1 | P25685 | DNAJB1 | DnaJ homolog subfam | 286603.875 | 291866.0313 |
| SNRPB | P14678 | SNRPB | Small nuclear ribonuc | 934292.8125 | 787148.5625 |
| FIS1 | Q9Y3D6 | FIS1 | Mitochondrial fission | 275158.125 | 293584.875 |
| ERO1A | Q96HE7 | ERO1A | ERO1-like protein alph | 221565.5625 | 213275.3438 |
| CHMP2A | M0R1T5;O43633 | CHMP2A | Charged multivesicula | 106951.6172 | 97633.28125 |
| ACTN4 | K7EJH8 | ACTN4 | Alpha-actinin-4 (Fragr | 564633.1372 | 295700.2419 |
| SRSF11 | Q05519;Q5T760 | SRSF11 | Serine/arginine-rich sp | 133233.3594 | 130962.3906 |
| DDX23 | Q9BUQ8 | DDX23 | Probable ATP-depend | 223910.9688 | 206799.125 |
| CPSF7 | Q8N684 | CPSF7 | Cleavage and polyade | 194100.9531 | 178922.9531 |
| PPFIA1 | A0A2R8Y7R9;Q13136 | PPFIA1 | Liprin-alpha-1 (Fragme | 52316.10547 | 55591.83984 |
| RPL22 | P35268 | RPL22 | 60S ribosomal protein | 2302442.75 | 1734722.5 |
| AFF1 | F5GXF9;P51825 | AFF1 | AF4/FMR2 family mer | 24660.17969 | 25965.04297 |
| ITSN2 | A0A087WVF7;Q9NZM3 | ITSN2 | Intersectin-2 | 31024.94336 | 27505.5625 |
| DNAJC30 | Q96LL9 | DNAJC30 | DnaJ homolog subfam | 11953.6416 | 9264.549805 |
| FBXO28 | Q9NVF7 | FBXO28 | F-box only protein 28 | 15167.07129 | 14832.83594 |
| TIMELESS | Q9UNS1 | TIMELESS | Protein timeless homo | 35187.92188 | 39778.26953 |

| | | | | | |
|---|---|---|---|---|---|
| LSM4 | Q9Y4Z0;V9GZ56 | LSM4 | U6 snRNA-associated | 446546.6875 | 373551.0313 |
| TRAF4 | F6SA91;Q9BUZ4 | TRAF4 | TNF receptor-associat | 10302.58008 | 6105.178711 |
| SARNP | P82979 | SARNP | SAP domain-containin | 552471.125 | 550043.8125 |
| RPS6 | P62753 | RPS6 | 40S ribosomal protein | 1214362.5 | 1139206.5 |
| PCGF5 | Q86SE9 | PCGF5 | Polycomb group RING | 10071.44824 | 10333.11523 |
| CSNK2A2 | P19784 | CSNK2A2 | Casein kinase II subun | 184055.6875 | 197788.9375 |
| ATP6V1E1 | P36543 | ATP6V1E1 | V-type proton ATPase | 128220.6484 | 115282.2734 |
| PTBP3 | O95758 | PTBP3 | Polypyrimidine tract-b | 62093.125 | 61496.74609 |
| EFNB1 | P98172 | EFNB1 | Ephrin-B1 | 3894.491699 | 4552.490723 |
| UXS1 | Q8NBZ7 | UXS1 | UDP-glucuronic acid d | 20981.93945 | 17795.16602 |
| ATXN3 | A0A087X1A8;A0A0A0MS38;A | ATXN3 | Ataxin-3;Ataxin-3;Ata | 47170.47656 | 61246.59766 |
| S100A13 | Q99584 | S100A13 | Protein S100-A13 | 1822555.75 | 1910126.125 |
| HIST3H2BB | Q8N257 | HIST3H2BB | Histone H2B type 3-B | 18268.54102 | 1730.612793 |
| REEP3 | Q6NUK4 | REEP3 | Receptor expression-e | 22546.7832 | 20341.17383 |
| EXD2 | Q9NVH0 | EXD2 | Exonuclease 3'-5' dom | 55970.91406 | 57588.90625 |
| COQ3 | Q9NZJ6 | COQ3 | Ubiquinone biosynthe | 40876.16406 | 37464.17969 |
| ADAM9 | A0AVL1;Q13443 | ADAM9 | ADAM9 protein;Disint | 36466.05078 | 44404.875 |
| PRPF3 | O43395 | PRPF3 | U4/U6 small nuclear r | 110401.6953 | 93933.17188 |
| UPK3BL1;UPK3BL2 | B0FP48;E5RIL1 | UPK3BL1;UPK3BL2 | Uroplakin-3b-like prot | 8065.134766 | 7621.396973 |
| CCT6A | P40227 | CCT6A | T-complex protein 1 s | 1355402.875 | 1469933.25 |
| SPC25 | Q9HBM1 | SPC25 | Kinetochore protein S | 36478.19922 | 48085.33594 |
| SDHAF4 | Q5VUM1 | SDHAF4 | Succinate dehydroger | 17684.66992 | 13310.28906 |
| NELFCD | H0UI80;Q8IXH7;X6RLT1 | NELFCD | Negative elongation fa | 192862.0938 | 228259.625 |
| CHCHD6 | J3QTA6;Q9BRQ6 | CHCHD6 | MICOS complex subur | 24640.36133 | 16402.91406 |
| PAGR1 | Q9BTK6 | PAGR1 | PAXIP1-associated glu | 373.4301147 | 719.081543 |
| KANSL3 | A0A590UJ53;C9J5A4;Q9P2N6 | KANSL3 | KAT8 regulatory NSL c | 11749.30078 | 10123.69238 |
| MTERF3 | Q96E29 | MTERF3 | Transcription termina | 13706.65137 | 18151.32227 |
| BABAM1 | Q9NWV8 | BABAM1 | BRISC and BRCA1-A cc | 15887.81934 | 20154.16602 |
| CCNC | E5RFK5;E5RFX8;E5RI39;G5E9 | CCNC | Cyclin-C;Cyclin-C (Frag | 14231.02246 | 13468.09863 |
| RUFY1 | Q96T51 | RUFY1 | RUN and FYVE domair | 149183.5625 | 166384.6875 |
| GOLT1B | F5H6U7;Q9Y3E0 | GOLT1B | Vesicle transport prot | 55698.21094 | 75615.6875 |
| CUL3 | Q13618 | CUL3 | Cullin-3 | 122156.0391 | 127495.0547 |
| EHBP1 | Q8NDI1 | EHBP1 | EH domain-binding pr | 11801.54004 | 11689.63379 |

| | | | | | |
|---|---|---|---|---|---|
| MITD1 | H7C3Q6;Q8WV92 | MITD1 | MIT domain-containir | 7540.748535 | 6286.22168 |
| TRIR | Q9BQ61 | TRIR | Telomerase RNA comp | 84048.22656 | 77098.52344 |
| CPOX | P36551 | CPOX | Oxygen-dependent cc | 131907.1875 | 125230.1719 |
| NDUFAF3 | Q9BU61 | NDUFAF3 | NADH dehydrogenase | 35301.83594 | 43902.90234 |
| P3H1 | Q32P28 | P3H1 | Prolyl 3-hydroxylase 1 | 39931.68359 | 33900.24219 |
| CBWD2;CBWD1 | Q8IUF1;Q9BRT8 | CBWD2;CBWD1 | COBW domain-contai | 27388.12695 | 37288.55469 |
| CSNK2A1 | A0A2R8YD58;E7EU96;P6840( | CSNK2A1 | Casein kinase II subun | 258092.0313 | 291233 |
| AGRN | O00468 | AGRN | Agrin | 20932.93555 | 30414.85547 |
| YY1 | P25490 | YY1 | Transcriptional repres | 136097.7656 | 152603.0313 |
| CKS1B | P61024 | CKS1B | Cyclin-dependent kina | 173045.0469 | 187575.25 |
| TRIM28 | Q13263 | TRIM28 | Transcription interme | 1694371.125 | 1944529.625 |
| UROD | P06132 | UROD | Uroporphyrinogen de | 113945.2344 | 120580.9375 |
| PDZD11 | Q5EBL8 | PDZD11 | PDZ domain-containir | 55302.86719 | 62563.48438 |
| SF3A2 | Q15428 | SF3A2 | Splicing factor 3A subu | 183960.9063 | 122980.8594 |
| BCL7C | I3L1Q2;Q8WUZ0 | BCL7C | B-cell CLL/lymphoma | 46671.83984 | 42952.14063 |
| AP1G1 | O43747 | AP1G1 | AP-1 complex subunit | 265480.0313 | 296405.1875 |
| GATD1 | Q8NB37 | GATD1 | Glutamine amidotrans | 58008.43359 | 71410.6875 |
| RFXAP | O00287 | RFXAP | Regulatory factor X-as | 15775.50684 | 15771.64453 |
| RAN | P62826 | RAN | GTP-binding nuclear p | 4481440 | 4938730 |
| EXOSC7 | Q15024 | EXOSC7 | Exosome complex cor | 94802.40625 | 92108.64063 |
| USP4 | Q13107 | USP4 | Ubiquitin carboxyl-ter | 70278.875 | 68722.98438 |
| TIMM10B | Q9Y5J6 | TIMM10B | Mitochondrial import | 10047.94531 | 14143.38965 |
| CAMKMT | B5MC16;B5MC79;Q7Z624 | CAMKMT | Calmodulin-lysine N-n | 7127.409668 | 9288.476563 |
| COMMD3-BMI1 | R4GMX3 | COMMD3-BMI1 | COMMD3-BMI1 readt | 27368.35938 | 41210.21875 |
| LARP4B | H0Y4V9 | LARP4B | La-related protein 4B | 212726.9023 | 195386.459 |
| NANS | Q9NR45 | NANS | Sialic acid synthase | 495993.2813 | 550760.6875 |
| WWC3 | Q9ULE0 | WWC3 | Protein WWC3 | 170791.2188 | 189069.4531 |
| ANXA4 | P09525 | ANXA4 | Annexin A4 | 240751.1094 | 239249.3281 |
| H1-5 | P16401 | H1-5 | Histone H1.5 | 5660141 | 711740.9375 |
| TRIM33 | Q9UPN9 | TRIM33 | E3 ubiquitin-protein li | 282985.1563 | 289097.6875 |
| JAK1 | A0A5F9ZH07;A0A5F9ZHN8;P. | JAK1 | Tyrosine-protein kinas | 11283.66602 | 15208.61621 |
| NDUFB6 | O95139 | NDUFB6 | NADH dehydrogenase | 117529.8984 | 121872.6953 |
| RBM15 | A0A087WWP4;A0A499FJY3;( | RBM15 | RNA-binding protein 1 | 90088.5625 | 85826.07031 |

| | | | | | |
|---|---|---|---|---|---|
| CAMKK2 | Q96RR4 | CAMKK2 | Calcium/calmodulin-d | 16756.25977 | 16314.2168 |
| WDR13 | Q9H1Z4 | WDR13 | WD repeat-containing | 28477.9707 | 28249.96094 |
| ANAPC5 | Q9UJX4 | ANAPC5 | Anaphase-promoting | 24309.30859 | 27036.17969 |
| MUS81 | E9PL60;E9PRI1;H0YDE3;H0YD | MUS81 | Crossover junction en | 3911.36084 | 5687.082031 |
| USP19 | A0A0A0MR08 | USP19 | Ubiquitin carboxyl-ter | 44951.58105 | 44943.7041 |
| PDLIM7 | Q9NR12 | PDLIM7 | PDZ and LIM domain p | 244371.0313 | 223282.125 |
| FOXJ3 | Q9UPW0 | FOXJ3 | Forkhead box protein | 40119.08203 | 86439.84375 |
| GPATCH2 | Q9NW75 | GPATCH2 | G patch domain-conta | 16251.69336 | 13130.39941 |
| DNM2 | P50570 | DNM2 | Dynamin-2 | 486448.3438 | 499133.9375 |
| ZNF451 | D6RIA1;E9PH99;Q9Y4E5 | ZNF451 | E3 SUMO-protein liga: NA | | 2235.89917 |
| CIAO1 | O76071 | CIAO1 | Probable cytosolic iro | 139300 | 150076.1094 |
| NFX1 | Q12986 | NFX1 | Transcriptional repres | 20526.32227 | 21947.96289 |
| PPP1R9B | Q96SB3 | PPP1R9B | Neurabin-2 | 69287.92969 | 71012.375 |
| NOL12; | Q9UGY1;V9GYY5 | NOL12; | Nucleolar protein 12;U | 20978.74805 | 14124.06738 |
| RPL14 | P50914 | RPL14 | 60S ribosomal protein | 2792409 | 2283106.75 |
| GMEB1 | Q9Y692 | GMEB1 | Glucocorticoid modula | 16820.85547 | 15341.54199 |
| RPL9 | P32969 | RPL9 | 60S ribosomal protein | 963329.25 | 1103446.75 |
| RAD1 | O60671 | RAD1 | Cell cycle checkpoint p | 52087.97266 | 47912.54688 |
| RPS8 | P62241 | RPS8 | 40S ribosomal protein | 2021136.875 | 1899712.75 |
| BAZ1A | Q9NRL2 | BAZ1A | Bromodomain adjacer | 236003.7188 | 273551.8438 |
| C9orf78 | Q9NZ63 | C9orf78 | Telomere length and s | 20045.02148 | 27542.23047 |
| TRA2B | P62995 | TRA2B | Transformer-2 protein | 296413.375 | 169128.1719 |
| SLC66A2 | Q8N2U9 | SLC66A2 | Solute carrier family 6 | 637.2432251 | 664.9790039 |
| MED27 | A0A494C0K7;A0A494C0P0;Q | MED27 | Mediator of RNA poly | 14290.76855 | 14154.65234 |
| RPS27A | P62979 | RPS27A | Ubiquitin-40S ribosom | 7061801 | 6283174.5 |
| TUBD1 | Q9UJT1 | TUBD1 | Tubulin delta chain | 6684.932129 | 7558.95166 |
| NBN | O60934 | NBN | Nibrin | 45082.06641 | 41293.16016 |
| TELO2 | Q9Y4R8 | TELO2 | Telomere length regu | 62488.11328 | 70876.42188 |
| TRMT6 | Q9UJA5 | TRMT6 | tRNA (adenine(58)-N( | 39075.18359 | 37845.30469 |
| SRSF8 | Q9BRL6 | SRSF8 | Serine/arginine-rich sp | 12329.05566 | 8390.639648 |
| GSPT1 | P15170 | GSPT1 | Eukaryotic peptide ch | 495565.125 | 530423.625 |
| RGPD8 | O14715 | RGPD8 | RANBP2-like and GRIP | 19832.30273 | 21074.37305 |
| RPS3 | P23396 | RPS3 | 40S ribosomal protein | 5618744.5 | 4406407.5 |

| | | | | | |
|---|---|---|---|---|---|
| RTCB | Q9Y3I0 | RTCB | RNA-splicing ligase Rt | 575911.5625 | 598442.5 |
| RECQL | P46063 | RECQL | ATP-dependent DNA h | 108993.5547 | 126511.8672 |
| FAM32A | K7EIY1;Q9Y421 | FAM32A | Protein FAM32A | 24700.60742 | 21231.66211 |
| HECTD4 | F8VWT9;J3KPF0;Q9Y4D8 | HECTD4 | Probable E3 ubiquitin- | 8728.297852 | 11444.12207 |
| HSP90AB3P | Q58FF7 | HSP90AB3P | Putative heat shock pr | 287731.1563 | 311625.8125 |
| G3BP2 | Q9UN86 | G3BP2 | Ras GTPase-activating | 177513.0313 | 202547.3438 |
| YEATS2 | Q9ULM3 | YEATS2 | YEATS domain-contair | 6546.852051 | 3479.926025 |
| ZFAND2B | B4DEN4;B8ZZ56;E7EV05;E9P | ZFAND2B | AN1-type zinc finger p | 20314.74414 | 23247.02344 |
| MAPT | A0A0G2JPD5;A0A0G2JQJ7;P1 | MAPT | Microtubule-associate | 228034.2266 | 252242.0078 |
| RPL23 | P62829 | RPL23 | 60S ribosomal protein | 1995831.75 | 1709786.875 |
| RPS15A | P62244 | RPS15A | 40S ribosomal protein | 1301329.875 | 1158407 |
| CORO1B | Q9BR76 | CORO1B | Coronin-1B | 1106642.625 | 1108047.125 |
| NUP107 | P57740 | NUP107 | Nuclear pore complex | 120996.3516 | 115310.4688 |
| DLGAP5 | Q15398 | DLGAP5 | Disks large-associated | 38280.52734 | 39444.51172 |
| E4F1 | H3BUJ7;Q66K89 | E4F1 | Transcription factor E4 | 365.5020142 | 3247.134033 |
| SETD2 | Q9BYW2 | SETD2 | Histone-lysine N-meth | 6313.865234 | 4497.702148 |
| RPS13 | P62277 | RPS13 | 40S ribosomal protein | 2364120.5 | 1860700.875 |
| YME1L1 | Q96TA2 | YME1L1 | ATP-dependent zinc m | 140083.4688 | 150397.8281 |
| TOP3B | H0Y6W0;O95985 | TOP3B | DNA topoisomerase (I | 9116.148438 | 12550.65918 |
| CHMP6 | I3L4A1;Q96FZ7 | CHMP6 | Charged multivesicula | 51537.17578 | 57154.78125 |
| GTF2H1 | P32780 | GTF2H1 | General transcription | 16897.68164 | 15372.19922 |
| GMEB2 | Q5JTV1;Q9UKD1 | GMEB2 | Glucocorticoid modula | 34558.71875 | 32334.07031 |
| SMARCC2 | Q8TAQ2 | SMARCC2 | SWI/SNF complex sub | 96655.6875 | 103374.6875 |
| NIPBL | A0A590UJS4;Q6KC79 | NIPBL | Nipped-B protein;Nipp | 19940.94922 | 20214.09961 |
| FASTKD5 | Q7L8L6 | FASTKD5 | FAST kinase domain-c | 8760.868164 | 9223.507813 |
| CLP1 | Q92989 | CLP1 | Polyribonucleotide 5'- | 44037.53516 | 50424.11719 |
| SH3PXD2B | A0A1B0GUF2;A1X283;G3V14 | SH3PXD2B | SH3 and PX domain-co | 11376.92383 | 10926.19727 |
| RPS20 | P60866 | RPS20 | 40S ribosomal protein | 1628855.375 | 1406326 |
| TRMT61A | Q96FX7 | TRMT61A | tRNA (adenine(58)-N( | 56687.25391 | 57238.75781 |
| TMED7 | Q9Y3B3 | TMED7 | Transmembrane emp | 188834.1875 | 203904.8906 |
| ;BCKDHA | F5H5P2;P12694 | ;BCKDHA | 2-oxoisovalerate dehy | 67138.86719 | 72869.29688 |
| WDR12 | Q9GZL7 | WDR12 | Ribosome biogenesis | 90545.34375 | 76427.25 |
| EP300 | Q09472 | EP300 | Histone acetyltransfer | 17153.02148 | 17465.52344 |

| | | | | | |
|---|---|---|---|---|---|
| CD58 | B1AMW1;P19256 | CD58 | CD58 antigen, (Lymph | 18091.44531 | 15920.77148 |
| CBFB | J3KTD8 | CBFB | Core-binding factor su | 160971.9922 | 153339.668 |
| LSM2 | Q9Y333 | LSM2 | U6 snRNA-associated | 217643.3281 | 211713.4375 |
| THAP4 | Q8WY91 | THAP4 | THAP domain-contain | 1851.339111 | 2078.39502 |
| KIF5B | P33176 | KIF5B | Kinesin-1 heavy chain | 289812.4063 | 297835.5313 |
| UBE3A | Q05086 | UBE3A | Ubiquitin-protein ligas | 45052.31641 | 46799.61719 |
| VPS26B | Q4G0F5 | VPS26B | Vacuolar protein sorti | 74465.60156 | 82093.80469 |
| STX16-NPEPL1;STX16 | H3BU86;O14662 | STX16-NPEPL1;STX16 | STX16-NPEPL1 readth | 129517.2656 | 115979.0859 |
| PHF5A | Q7RTV0 | PHF5A | PHD finger-like domai | 162858.9531 | 113335.8281 |
| IMPAD1 | Q9NX62 | IMPAD1 | Golgi-resident adenos | 69724.64844 | 72203.75781 |
| SLC25A15 | Q9Y619 | SLC25A15 | Mitochondrial ornithi | 82879.10938 | 103580.4922 |
| ZNF207 | J3QRS9;O43670 | ZNF207 | BUB3-interacting and | 239884.5781 | 206940.6094 |
| TMEM147 | A8MU21;Q9BVK8 | TMEM147 | Transmembrane prot | 9092.59668 | 9973.615234 |
| RAVER1 | A0A087WZ13;E9PAU2 | RAVER1 | Ribonucleoprotein PT | 184907.1563 | 200342.3594 |
| KLHL12 | J3QSX3;Q53G59 | KLHL12 | Kelch-like protein 12 ( | 10042.88867 | 11633.40625 |
| ARL2BP | H3BU49;Q9Y2Y0 | ARL2BP | ADP-ribosylation facto | 5924.759766 | 11608.33008 |
| ERH | P84090 | ERH | Enhancer of rudiment | 46577.95313 | 82670.375 |
| SLC35B1 | D6RHG0;P78383 | SLC35B1 | Solute carrier family 3 | 29054.06445 | 32176.32422 |
| TUBAL3 | A6NHL2 | TUBAL3 | Tubulin alpha chain-lil | 40633.55859 | 69945.42969 |
| REXO2 | F5GYG5;H0YGR4;Q9Y3B8 | REXO2 | Oligoribonuclease, mi | 75639.70313 | 71602.59375 |
| MIER3 | Q7Z3K6 | MIER3 | Mesoderm induction | 2418.302002 | 2197.732422 |
| CSNK2B | P67870;Q5SRQ6 | CSNK2B | Casein kinase II subun | 224569.9063 | 222640.0469 |
| SPECC1L | A0A494C1J1;Q69YQ0 | SPECC1L | Cytospin-A | 43057.73438 | 38887.11719 |
| CABLES1 | Q8TDN4 | CABLES1 | CDK5 and ABL1 enzym | 250044.6406 | 140013.875 |
| RPL35A | P18077 | RPL35A | 60S ribosomal protein | 883926.9375 | 734257.5625 |
| AFTPH | A0A0A0MSN6;B8ZZI9;Q6ULP | AFTPH | Aftiphilin;Aftiphilin (Fr | 5458.975586 | 5142.341797 |
| NRF1 | Q16656 | NRF1 | Nuclear respiratory fa | 60187.37891 | 47901.02344 |
| PDIA3 | H7BZJ3 | PDIA3 | Protein disulfide-isom | 4586303.625 | 4072068 |
| MTIF3 | Q9H2K0 | MTIF3 | Translation initiation f | 40354.19531 | 40723.41797 |
| KTN1 | Q86UP2 | KTN1 | Kinectin | 314551.7813 | 308443.9063 |
| SPNS1 | H3BMF4;Q9H2V7 | SPNS1 | Protein spinster homo | 38111.89453 | 42416.64453 |
| NDUFB3 | O43676 | NDUFB3 | NADH dehydrogenase | 142914.875 | 162248.8125 |
| ERCC3 | P19447 | ERCC3 | General transcription | 33439.65625 | 32037.4375 |

| | | | | | |
|---|---|---|---|---|---|
| FTO | A0A1B0GTC3;A0A1B0GTC5;A | FTO | Alpha-ketoglutarate-d | 59972.94531 | 60890.14063 |
| ITPKB | P27987 | ITPKB | Inositol-trisphosphate | NA | 1901.564697 |
| DNAJB12 | Q9NXW2 | DNAJB12 | DnaJ homolog subfam | 81774.95313 | 85591.17188 |
| ID1 | P41134 | ID1 | DNA-binding protein i | 5290.113281 | 3316.574463 |
| PSMB4 | P28070 | PSMB4 | Proteasome subunit b | 543179.8125 | 592622.4375 |
| KDM5C | P41229 | KDM5C | Lysine-specific demetl | 15259.43457 | 19415.28125 |
| GALNT7 | Q86SF2 | GALNT7 | N-acetylgalactosamin | 78750.66406 | 76620.82813 |
| TMEM189 | A5PLL7 | TMEM189 | Plasmanylethanolami | 28061.60938 | 22622.98633 |
| RPS10-NUDT3 | A0A1W2PQS6 | RPS10-NUDT3 | RPS10-NUDT3 readthr | 2655523.75 | 2128771.75 |
| WASL | O00401 | WASL | Neural Wiskott-Aldricl | 50418.41016 | 49689.38281 |
| ALDH5A1 | P51649 | ALDH5A1 | Succinate-semialdehy | 73138.0625 | 84384.10938 |
| MOB2 | Q70IA6 | MOB2 | MOB kinase activator | 25026.03906 | 25715.82813 |
| SENP7 | J3QT09;Q9BQF6 | SENP7 | Sentrin-specific prote | 17114.6875 | 12522.95898 |
| MAPRE1 | Q15691 | MAPRE1 | Microtubule-associate | 746218.8125 | 784150.3125 |
| ZFAND1 | Q8TCF1 | ZFAND1 | AN1-type zinc finger p | 45112.39063 | 34444.99609 |
| TRIO | O75962 | TRIO | Triple functional dom | 19692.80664 | 20807.38086 |
| CCNY | Q8ND76 | CCNY | Cyclin-Y | 23412.95898 | 24307.58594 |
| RTRAF | Q9Y224 | RTRAF | RNA transcription, tra | 544480.75 | 517889.2813 |
| NAA20 | P61599 | NAA20 | N-alpha-acetyltransfe | 41954.86719 | 46150.84766 |
| YIPF4 | Q9BSR8 | YIPF4 | Protein YIPF4 | 83091.54688 | 97270.07813 |
| ING5 | A0A1B0GW41;E9PEN0;Q8W | ING5 | Inhibitor of growth pr | 16898.51953 | 15038.97266 |
| PRDX6 | P30041 | PRDX6 | Peroxiredoxin-6 | 3891866.75 | 4337075 |
| HAPLN1 | P10915 | HAPLN1 | Hyaluronan and prote | 556938.375 | 564098.0625 |
| NUBP1 | P53384 | NUBP1 | Cytosolic Fe-S cluster | 126566.4609 | 129837.5781 |
| RNF10 | H0YG28;Q8N5U6 | RNF10 | RING finger protein 1( | 2086.618652 | 2130.293701 |
| GNPDA1 | P46926 | GNPDA1 | Glucosamine-6-phosp | 207672.5938 | 226465.8438 |
| SWAP70 | Q9UH65 | SWAP70 | Switch-associated pro | 31164.91602 | 29623.67383 |
| FKBP1A | P62942 | FKBP1A | Peptidyl-prolyl cis-tra | 1173760.375 | 1190973.75 |
| UHRF1BP1 | H7C1J4;Q6BDS2 | UHRF1BP1 | Chromosome 6 open | 15959.37695 | 17485.14258 |
| NDUFB5 | H0Y886;O43674 | NDUFB5 | NADH dehydrogenase | 129531.3203 | 140306.5625 |
| NDUFAF4 | Q9P032 | NDUFAF4 | NADH dehydrogenase | 51939.55469 | 52552.88281 |
| RPS24 | A0A2R8Y849;A0A2R8YD14;E | RPS24 | 40S ribosomal protein | 1134257.625 | 959719.875 |
| ATAD3B | Q5T9A4 | ATAD3B | ATPase family AAA do | 40716.28516 | 46834.69922 |

| | | | | | |
|---|---|---|---|---|---|
| CSDE1 | O75534 | CSDE1 | Cold shock domain-co | 675084.375 | 642992.0625 |
| PPP6C | O00743 | PPP6C | Serine/threonine-prot | 182643.9063 | 192307.4375 |
| FH | P07954 | FH | Fumarate hydratase, r | 278163.1875 | 292382.8438 |
| BAHD1 | Q8TBE0 | BAHD1 | Bromo adjacent homo | 4750.536133 | 5904.57666 |
| KDELR3 | O43731 | KDELR3 | ER lumen protein-reta | 16665.375 | 19782.97852 |
| CDK11B | A0A0D9SEI3;A0A0D9SEN2;A( | CDK11B | Cyclin-dependent kina | 202674.4688 | 188367.4531 |
| SAMM50 | Q9Y512 | SAMM50 | Sorting and assembly | 254178.9219 | 131897.4688 |
| MTO1 | H7C2S9 | MTO1 | Protein MTO1 homolo | 21432.71094 | 30554.43262 |
| IFT74 | Q96LB3 | IFT74 | Intraflagellar transpor | 12546.58008 | 13735.51172 |
| SRSF9 | Q13242 | SRSF9 | Serine/arginine-rich sp | 548315 | 261518.4063 |
| ALS2 | Q96Q42 | ALS2 | Alsin | 8201.158203 | 4534.390137 |
| LRRC47 | Q8N1G4 | LRRC47 | Leucine-rich repeat-co | 308166.2188 | 323630.9375 |
| TBC1D2B | Q9UPU7 | TBC1D2B | TBC1 domain family m | 12607.02441 | 14252.36426 |
| C5orf24 | Q7Z6I8 | C5orf24 | UPF0461 protein C5or | 8237.270508 | 19599.77734 |
| ZNF668 | Q96K58 | ZNF668 | Zinc finger protein 668 | 23306.71484 | 27794.58984 |
| PON2 | A0A0J9YXF2 | PON2 | Paraoxonase 2, isofor | 58635.73047 | 44598.82031 |
| MOCS3 | O95396 | MOCS3 | Adenylyltransferase a | 158472.8438 | 180498.3594 |
| HINT2 | Q9BX68 | HINT2 | Histidine triad nucleot | 38519.11719 | 49817.29297 |
| RC3H1 | Q5TC82 | RC3H1 | Roquin-1 | 9999.202148 | 11180.68848 |
| VPS16 | Q9H269 | VPS16 | Vacuolar protein sorti | 66338.32031 | 70804.84375 |
| IDE | A0A3B3ISG5;P14735 | IDE | Insulin-degrading enzy | 483692.0625 | 561572.125 |
| BRD4 | O60885 | BRD4 | Bromodomain-contair | 52338.07813 | 35009.26953 |
| CTSL | P07711 | CTSL | Cathepsin L1 | 85345.85938 | 58773.86719 |
| IDH3B | A0A087WZN1;O43837 | IDH3B | Isocitrate dehydrogen | 178093.4375 | 185406.1406 |
| PAFAH1B1 | P43034 | PAFAH1B1 | Platelet-activating fac | 123706.3125 | 126316.2266 |
| ACP2 | A0A5F9ZHR7;P11117 | ACP2 | Lysosomal acid phosp | 71423.9375 | 60084.53906 |
| GAN | Q9H2C0 | GAN | Gigaxonin | 93942.58594 | 107418.2734 |
| LYPLA2 | O95372 | LYPLA2 | Acyl-protein thioester | 71491.75781 | 90696.38281 |
| ERI1 | Q8IV48 | ERI1 | 3'-5' exoribonuclease | 25880.57422 | 28760.43555 |
| MED14 | O60244 | MED14 | Mediator of RNA poly | 126725.1563 | 138079.4063 |
| TMEM222 | H0Y861;H0Y870;H0Y8D0;Q5S | TMEM222 | Chromosome 1 open r | 14145.50977 | 14891.2998 |
| EXOSC2 | Q13868 | EXOSC2 | Exosome complex cor | 91485.84375 | 88334.8125 |
| GALNT3 | Q14435 | GALNT3 | Polypeptide N-acetylg | 44066.04297 | 51096.58984 |

| | | | | | |
|---|---|---|---|---|---|
| RBM4B | Q9BQ04 | RBM4B | RNA-binding protein 4 | 27392.30859 | 20867.78906 |
| HIF1AN | Q9NWT6 | HIF1AN | Hypoxia-inducible fact | 48713.05469 | 49908.93359 |
| COPS7B | J3KQ34;Q9H9Q2 | COPS7B | COP9 signalosome cor | 51182.50391 | 59430.51953 |
| RPL3 | P39023 | RPL3 | 60S ribosomal protein | 1558402.625 | 1326522.625 |
| MED12 | Q93074 | MED12 | Mediator of RNA poly | 41324.36719 | 44267.18359 |
| ATAD2 | Q6PL18 | ATAD2 | ATPase family AAA do | 56518.61719 | 54255.49219 |
| RPS4X | P62701 | RPS4X | 40S ribosomal protein | 1936176 | 1601603.625 |
| CFAP20 | Q9Y6A4 | CFAP20 | Cilia- and flagella-asso | 73871.875 | 74164.46094 |
| LLPH | Q9BRT6 | LLPH | Protein LLP homolog | 50851.94531 | 33011.75781 |
| PPIAL4C | A0A0B4J2A2 | PPIAL4C | Peptidyl-prolyl cis-tra | 526344.375 | 47356.88672 |
| SARAF | H0YAN9;H0YAZ1;Q96BY9 | SARAF | Store-operated calciu | 7801.53418 | 6329.621094 |
| HNRNPH3 | P31942 | HNRNPH3 | Heterogeneous nucle | 540239.1875 | 282394.625 |
| VAT1 | Q99536 | VAT1 | Synaptic vesicle meml | 734856.8125 | 837530.5 |
| C2CD5 | Q86YS7 | C2CD5 | C2 domain-containing | 20139.5332 | 20682.33594 |
| CHKA | P35790 | CHKA | Choline kinase alpha | 41450.08594 | 48787.59375 |
| TSC2 | A0A2R8Y5F1;A0A2R8Y7C8;A( | TSC2 | Tuberin;Tuberin;Tube | 103279.7813 | 97976.44531 |
| FYCO1 | Q9BQS8 | FYCO1 | FYVE and coiled-coil d | 27330.2168 | 27207.00781 |
| CLCC1 | Q96S66 | CLCC1 | Chloride channel CLIC | 87175.24219 | 98863.14844 |
| TMEM102 | Q8N9M5 | TMEM102 | Transmembrane prote | 7130.240723 | 9704.600586 |
| NUP58 | H7BYF2;Q5JRG1;Q9BVL2 | NUP58 | Nucleoporin p58/p45 | 100370.0625 | 83245.71094 |
| DNLZ | Q5SXM8 | DNLZ | DNL-type zinc finger p | 30121.21875 | 31274.29102 |
| CHDH | Q8NE62 | CHDH | Choline dehydrogenas | 47276.74219 | 41271.62891 |
| SGTA | K7EMD6;O43765 | SGTA | Small glutamine-rich t | 7210.294922 | 12301.34473 |
| UBE2N | P61088 | UBE2N | Ubiquitin-conjugating | 2147369.25 | 2273169.25 |
| SNRPD3 | P62318 | SNRPD3 | Small nuclear ribonuc | 1449369.625 | 1076911.25 |
| TEFM | Q96QE5 | TEFM | Transcription elongati | 14020.1709 | 12828.85254 |
| SLC19A1 | P41440 | SLC19A1 | Reduced folate transp | 48626.23828 | 43151.48828 |
| ATP5MF-PTCD1 | G3V325 | ATP5MF-PTCD1 | ATP5MF-PTCD1 readt | 436537.25 | 512909.5 |
| EXOSC4 | Q9NPD3 | EXOSC4 | Exosome complex cor | 94630.55469 | 85349.25781 |
| MYCBP2 | A0A499FJI4;O75592 | MYCBP2 | E3 ubiquitin-protein li | 6680.84668 | 8259.616211 |
| MAN2C1 | Q9NTJ4 | MAN2C1 | Alpha-mannosidase 2( | 34310.32813 | 45219.11719 |
| API5 | Q9BZZ5 | API5 | Apoptosis inhibitor 5 | 483679.1875 | 536641.75 |
| STRN4 | Q9NRL3 | STRN4 | Striatin-4 | 88702.625 | 97006.30469 |

| | | | | | |
|---|---|---|---|---|---|
| EXOSC6 | Q5RKV6 | EXOSC6 | Exosome complex cor | 127898.0938 | 111871.0391 |
| PA2G4 | Q9UQ80 | PA2G4 | Proliferation-associate | 637639.75 | 600026.9375 |
| ZNF281 | Q9Y2X9 | ZNF281 | Zinc finger protein 28 | 9286.042969 | 8923.573242 |
| EMC1 | Q8N766 | EMC1 | ER membrane protein | 224780.1406 | 226227.1563 |
| ACTN1 | P12814 | ACTN1 | Alpha-actinin-1 | 1362156.5 | 1020067 |
| PREP | P48147 | PREP | Prolyl endopeptidase | 212721.8906 | 237801.3906 |
| CCAR2 | Q8N163 | CCAR2 | Cell cycle and apoptos | 354093.0625 | 358275.0625 |
| RAPGEF6;RAPGEF6;;RAPGE | A0A5F9ZHG7;B7Z7Y1;E9PCH | RAPGEF6;RAPGEF6;;R | Rap guanine nucleotic | 14775.07617 | 13464.55176 |
| GMDS | O60547 | GMDS | GDP-mannose 4,6 deh | 84947.85938 | 121100.625 |
| RUNDC1 | Q96C34 | RUNDC1 | RUN domain-containi | 19551.76172 | 18328.65039 |
| RPL34 | P49207 | RPL34 | 60S ribosomal protein | 841863.875 | 686420.8125 |
| PET100 | A0A0A0MSK2;P0DJ07 | PET100 | Protein PET100 homo | 22314.56055 | 29205.06836 |
| BTAF1 | A0A0A0MTH9;O14981 | BTAF1 | TATA-binding protein- | 54128.56641 | 59126.47266 |
| C1D | J3KPZ4;Q13901 | C1D | Nuclear nucleic acid-b | 22735.39648 | 22071.09766 |
| PPIG | Q13427 | PPIG | Peptidyl-prolyl cis-trar | 147365.1094 | 112413.6094 |
| SNRPD2 | P62316 | SNRPD2 | Small nuclear ribonuc | 1423971.375 | 1003489.25 |
| PRPF6 | O94906 | PRPF6 | Pre-mRNA-processing | 363687.5938 | 320963.1875 |
| POLDIP3 | Q9BY77 | POLDIP3 | Polymerase delta-inte | 93390.20313 | 90788.08594 |
| DDX1 | Q92499 | DDX1 | ATP-dependent RNA h | 583125.625 | 618963.1875 |
| RAB12 | A0A3B3ITT1 | RAB12 | Ras-related protein Ra | 191476.0313 | 214870.0469 |
| NEXMIF | A0A2R8YEQ5;Q5QGS0 | NEXMIF | Neurite extension anc | NA | NA |
| GATB | O75879 | GATB | Glutamyl-tRNA(Gln) a | 34876.64063 | 34387.42969 |
| CALM2;CALM1;CALM1;CAL | F8WBR5;G3V479;M0QZ52;P( | CALM2;CALM1;CALM | Calmodulin-2;Calmod | 3232.863281 | 8517.868164 |
| TMEM189-UBE2V1 | I3L0A0 | TMEM189-UBE2V1 | HCG2044781 | 2271224.25 | 2436660.75 |
| UBN2 | H7C4P8;Q6ZU65 | UBN2 | Ubinuclein-2 (Fragme | 26892.5625 | 24680.10352 |
| POLE3 | Q9NRF9 | POLE3 | DNA polymerase epsil | 130196.8438 | 107604.7578 |
| ZDHHC20 | B4DRN8;Q5W0Z9 | ZDHHC20 | Palmitoyltransferase; | 29842.11914 | 29238.26172 |
| LRCH1 | Q9Y2L9 | LRCH1 | Leucine-rich repeat ar | 34581.33594 | 28913.6875 |
| STIP1 | P31948 | STIP1 | Stress-induced-phosp | 3198069 | 3111412.75 |
| MAP7D3 | Q8IWC1 | MAP7D3 | MAP7 domain-contair | 53837.49609 | 48056.54297 |
| DAZAP1 | K7EK33;K7EQ55;Q96EP5 | DAZAP1 | DAZ-associated protei | 288634.5625 | 238754.4531 |
| KPNA6 | O60684 | KPNA6 | Importin subunit alph | 349199.4375 | 385335.4063 |
| CDC37 | Q16543 | CDC37 | Hsp90 co-chaperone ( | 1436316.75 | 1434764.625 |

| | | | | | |
|---|---|---|---|---|---|
| RTF1 | Q92541 | RTF1 | RNA polymerase-asso | 122737.8359 | 112876.7109 |
| NSMCE3 | Q96MG7 | NSMCE3 | Non-structural mainte | 10891.76465 | 9411.485352 |
| CCT6B | Q92526 | CCT6B | T-complex protein 1 s | 2002967 | 1853310.625 |
| ERCC6L | Q2NKX8 | ERCC6L | DNA excision repair pı | 32046.01563 | 36255.58594 |
| SNRNP48 | Q6IEG0 | SNRNP48 | U11/U12 small nuclea | 4638.245605 | 4980.841797 |
| PUM1 | H0YEH2;Q14671 | PUM1 | Pumilio homolog 1 (Fr | 68203.53125 | 83937.76563 |
| PPFIA3 | O75145 | PPFIA3 | Liprin-alpha-3 | 135411.1719 | 110191.2578 |
| NUP88 | Q99567 | NUP88 | Nuclear pore complex | 118356.3828 | 116252.4453 |
| TACC3 | A0A494BZT8;Q9Y6A5 | TACC3 | Transforming acidic cc | 8363893 | 8056065.5 |
| ADNP | Q9H2P0 | ADNP | Activity-dependent ne | 108140.1875 | 111745.0625 |
| MAIP1 | Q8WWC4 | MAIP1 | m-AAA protease-inter | 208085.4844 | 218339.625 |
| PAK4 | O96013 | PAK4 | Serine/threonine-prot | 172737.5781 | 197893.7344 |
| ORC2 | Q13416 | ORC2 | Origin recognition cor | 57764.10547 | 59800.53125 |
| SCAF8 | A0A0A0MT33;Q9UPN6 | SCAF8 | SR-related and CTD-as | 69662.54688 | 74402.45313 |
| TPRN | Q4KMQ1 | TPRN | Taperin | 24990.9707 | 21016.71875 |
| PEPD | P12955 | PEPD | Xaa-Pro dipeptidase | 203013.7813 | 242464.3906 |
| RPL18A | Q02543 | RPL18A | 60S ribosomal protein | 1487839.625 | 1333651.375 |
| RABGAP1L | Q5R372 | RABGAP1L | Rab GTPase-activating | 18747.26953 | 19216.92773 |
| ENY2 | Q9NPA8 | ENY2 | Transcription and mRI | 286910.5313 | 298510.2813 |
| TRPC4AP | Q8TEL6 | TRPC4AP | Short transient recept | 12055.6123 | 11904.02051 |
| GRK6 | F8W9W2;P43250 | GRK6 | G protein-coupled rec | 102247.6328 | 108698.0234 |
| NDUFAF5 | B3KR61;Q5TEU4 | NDUFAF5 | Arginine-hydroxylase | 14549.36328 | 17103.70898 |
| RPL11 | P62913 | RPL11 | 60S ribosomal protein | 850894.3125 | 724523.875 |
| CKS2 | P33552 | CKS2 | Cyclin-dependent kina | 20208.32031 | 22047.08203 |
| MCCC2 | Q9HCC0 | MCCC2 | Methylcrotonoyl-CoA | 446563.9688 | 466270.5625 |
| CAD | F8VPD4;P27708 | CAD | CAD protein | 588366.1875 | 682766 |
| CTPS1 | P17812 | CTPS1 | CTP synthase 1 | 461576.7188 | 497774.125 |
| ;ALG9 | A0A087WZ62;Q9H6U8 | ;ALG9 | Mannosyltransferase; | 11087.94336 | 12997.9541 |
| WWC1 | H0YBE8;H3BLZ3;Q8IX03 | WWC1 | Protein KIBRA (Fragm | 4786.622559 | 6017.614258 |
| RASA2 | A0A0A0MSJ9;Q15283 | RASA2 | Ras GTPase-activating | 3772.170898 | 2893.30127 |
| GRPEL1 | Q9HAV7 | GRPEL1 | GrpE protein homolog | 162076.1094 | 178493.7031 |
| RSL24D1 | Q9UHA3 | RSL24D1 | Probable ribosome bi | 36865.64063 | 19280.25781 |
| RIOX1 | Q9H6W3 | RIOX1 | Ribosomal oxygenase | 48571.45313 | 56396.61328 |

| | | | | | |
|---|---|---|---|---|---|
| ALG13 | A0A087WTT9;A0A087WX43;. | ALG13 | Putative bifunctional U | 42377.78223 | 53841.69531 |
| HMCES | Q96FZ2 | HMCES | Abasic site processing | 119933.0938 | 119385.0469 |
| UBR5 | O95071 | UBR5 | E3 ubiquitin-protein li | 70283.07813 | 73677.88281 |
| USO1 | O60763 | USO1 | General vesicular tran | 204586.2813 | 216505.2813 |
| MED4 | Q9NPJ6 | MED4 | Mediator of RNA poly | 5052.982422 | 8289.608398 |
| TAMM41 | Q96BW9 | TAMM41 | Phosphatidate cytidyl | 89059.78906 | 100779.4375 |
| SNRPD1 | P62314 | SNRPD1 | Small nuclear ribonuc | 632183.125 | 508749.3438 |
| ELOC | E5RHG8;Q15369 | ELOC | Elongin-C (Fragment); | 109606.2109 | 162142.875 |
| DNAJC19 | Q96DA6 | DNAJC19 | Mitochondrial import | 86816.51563 | 87643.71094 |
| EMSY | Q7Z589 | EMSY | BRCA2-interacting tra | 25210.25 | 25987.85352 |
| PDSS2 | Q86YH6 | PDSS2 | Decaprenyl-diphospha | 21349.31445 | 29342.88086 |
| ACTA2;ACTG2 | P62736;P63267 | ACTA2;ACTG2 | Actin, aortic smooth n | 7321230.5 | 4800644.5 |
| SEC24B | O95487 | SEC24B | Protein transport prot | 70100.33594 | 74859.375 |
| PPM1F | P49593 | PPM1F | Protein phosphatase 1 | 28363.48633 | 34380.6875 |
| PPP1R16A | A0A087WWZ0;Q96I34 | PPP1R16A | Protein phosphatase 1 | 9647.610352 | 6509.717773 |
| VNN2 | B4DQB2;E9PL76;E9PQ75;E9P | VNN2 | Vascular non-inflamm | NA | 3960.637207 |
| COG8 | B4DYU2;Q96MW5 | COG8 | Conserved oligomeric | 43352.53125 | 50376.61719 |
| ZC3H4 | M0QY97;Q9UPT8 | ZC3H4 | Zinc finger CCCH dom | 74490.54688 | 72698.28906 |
| PTEN | P60484 | PTEN | Phosphatidylinositol 3 | 32961.74219 | 36339.73438 |
| HAUS8 | Q9BT25 | HAUS8 | HAUS augmin-like con | 5516.541016 | 6656.638672 |
| CC2D1A | Q6P1N0 | CC2D1A | Coiled-coil and C2 dor | 116123.0781 | 121531.5625 |
| SEC31A | D6REX3;O94979 | SEC31A | Protein transport prot | 346942.625 | 359188.0625 |
| ITCH | A0A590UJQ1;A0A590UJW8;( | ITCH | E3 ubiquitin-protein li | 53529.42578 | 51517.22656 |
| RB1 | P06400 | RB1 | Retinoblastoma-assoc | 53465.32031 | 59889.51563 |
| HSP90AB1 | P08238 | HSP90AB1 | Heat shock protein HS | 9970059 | 11595688 |
| RPS10P5 | Q9NQ39 | RPS10P5 | Putative 40S ribosoma | 725457.6875 | 550793.875 |
| RNF34 | H7BYJ1;Q969K3 | RNF34 | E3 ubiquitin-protein li | 6035.675781 | 6610.481934 |
| PPP6R1 | Q9UPN7 | PPP6R1 | Serine/threonine-prot | 27373.76367 | 40783.30859 |
| RBM10 | P98175 | RBM10 | RNA-binding protein 1 | 123292.4219 | 115192.0156 |
| TXNDC5 | Q8NBS9 | TXNDC5 | Thioredoxin domain-c | 683674.25 | 727040.6875 |
| LHFPL4 | Q7Z7J7 | LHFPL4 | LHFPL tetraspan subfa | 64750.99219 | 66927.97656 |
| CHMP7 | Q8WUX9 | CHMP7 | Charged multivesicula | 42844.75781 | 46807.19922 |
| GMPR2 | A0A087WWM4;F8WAN9;H0Y | GMPR2 | GMP reductase;GMP r | 120224.5469 | 133821.625 |

| | | | | | |
|---|---|---|---|---|---|
| CLDND1 | A0A0R4J2F2;D6R9S8;D6RA76 | CLDND1 | Claudin domain-conta | 12749.89648 | 9427.875 |
| KAT2A | Q92830 | KAT2A | Histone acetyltransfer | 12918.44629 | 13722.94141 |
| TOMM20 | Q15388 | TOMM20 | Mitochondrial import | 165082.4063 | 107871.1641 |
| CMPK1 | P30085 | CMPK1 | UMP-CMP kinase | 541979.875 | 543996.125 |
| USP16 | Q9Y5T5 | USP16 | Ubiquitin carboxyl-ter | 23091.06641 | 22226.21094 |
| RPS18 | P62269 | RPS18 | 40S ribosomal protein | 5108934 | 4304848 |
| FEZ2 | F8W6C0;G3V0F5;Q9UHY8 | FEZ2 | Fasciculation and elor | 7779.012695 | 7673.668945 |
| BRAP | Q7Z569 | BRAP | BRCA1-associated pro | 22070.28516 | 22253.12695 |
| YTHDC2 | Q9H6S0 | YTHDC2 | 3'-5' RNA helicase YTH | 44365.55078 | 46477.89844 |
| SIK3 | Q9Y2K2 | SIK3 | Serine/threonine-prot | 7478.878418 | 8210.392578 |
| SLC4A11 | A0A2R8Y6X5 | SLC4A11 | Sodium bicarbonate tr | 184203.5 | 152438.0938 |
| CPSF1 | Q10570 | CPSF1 | Cleavage and polyade | 102918.9453 | 102246.5703 |
| LDLR | J3KMZ9;P01130 | LDLR | Low-density lipoprote | 90777 | 79534.29688 |
| HSPA13 | P48723 | HSPA13 | Heat shock 70 kDa pro | 31340.06836 | 28067.17578 |
| RPL12 | P30050 | RPL12 | 60S ribosomal protein | 2518571.25 | 1952902.75 |
| MAD2L1BP | Q15013 | MAD2L1BP | MAD2L1-binding prote | 12871.23828 | 12214.02832 |
| SFXN2 | Q96NB2 | SFXN2 | Sideroflexin-2 | 28877.55078 | 44504.48438 |
| ERCC2 | P18074 | ERCC2 | General transcription | 51234.25 | 55484.42969 |
| RSAD1 | Q9HA92 | RSAD1 | Radical S-adenosyl me | 3866.656982 | 2797.620605 |
| ASNA1 | A0A087WXS7;O43681 | ASNA1 | ATPase ASNA1 | 647746.5 | 697600.3125 |
| RPUSD3 | Q6P087 | RPUSD3 | Mitochondrial mRNA | 12055.74219 | 16182.87207 |
| SLC31A1 | O15431 | SLC31A1 | High affinity copper u | 2658.797607 | 2503.494385 |
| C1orf112 | Q9NSG2 | C1orf112 | Uncharacterized prote | 6106.40918 | 7128.835938 |
| PATL1 | Q86TB9 | PATL1 | Protein PAT1 homolog | 25506.69141 | 27346.42188 |
| NUDT18 | Q6ZVK8 | NUDT18 | 8-oxo-dGDP phosphat | 4761.053223 | 4152.814941 |
| KIF1C | O43896 | KIF1C | Kinesin-like protein KI | 36113.87109 | 39945.16016 |
| KIF2C | Q99661 | KIF2C | Kinesin-like protein KI | 88801.22656 | 95777.14844 |
| DNMT1 | P26358 | DNMT1 | DNA (cytosine-5)-met | 182090.4219 | 208297.7344 |
| MALSU1 | Q96EH3 | MALSU1 | Mitochondrial assemb | 17506.74219 | 20893.60547 |
| SPTLC2 | O15270 | SPTLC2 | Serine palmitoyltransf | 113774.2891 | 121143.4922 |
| XPO1 | O14980 | XPO1 | Exportin-1 | 259633.4531 | 296336.25 |
| BAP1 | F8W6N3;Q92560 | BAP1 | Ubiquitin carboxyl-ter | 12039.45898 | 11859.50879 |
| RPS14 | A0A2R8Y811;P62263 | RPS14 | 40S ribosomal protein | 2390928 | 2086586.875 |

| | | | | | |
|---|---|---|---|---|---|
| TTL | Q8NG68 | TTL | Tubulin--tyrosine ligas | 20094.08008 | 19588.09766 |
| MPP5 | Q8N3R9 | MPP5 | MAGUK p55 subfamily | 43669.1875 | 44597.78516 |
| PPIC | P45877 | PPIC | Peptidyl-prolyl cis-trar | 139078.1875 | 162954.5938 |
| XXYLT1 | Q8NBI6 | XXYLT1 | Xyloside xylosyltransfe | 18922.68945 | 22291.4707 |
| ELF1 | A0A0U1RRL5;P32519 | ELF1 | ETS-related transcript | 22363.82227 | 46324.15625 |
| EIF2B4 | A0A087WTA5 | EIF2B4 | Translation initiation f | 163373.3564 | 167655.6426 |
| AGK | Q53H12 | AGK | Acylglycerol kinase, m | 174518 | 173337.7188 |
| RPS7 | P62081 | RPS7 | 40S ribosomal protein | 2065547.5 | 1693451 |
| CTBP2 | P56545 | CTBP2 | C-terminal-binding pro | 117601.6406 | 127440.2109 |
| SMC5 | Q8IY18 | SMC5 | Structural maintenanc | 14935.0127 | 12392.55566 |
| HDAC3 | O15379 | HDAC3 | Histone deacetylase 3 | 15076.26953 | 16053.13867 |
| COMMD10 | Q9Y6G5 | COMMD10 | COMM domain-conta | 73261.44531 | 73439.70313 |
| COMMD4 | Q9H0A8 | COMMD4 | COMM domain-conta | 33939.58594 | 37346.3125 |
| RPS3A | P61247 | RPS3A | 40S ribosomal protein | 2913362.5 | 2380496.5 |
| FUBP3 | Q96I24 | FUBP3 | Far upstream element | 301164.3125 | 247925.7969 |
| ADCY3 | A0A0A0MSC1;O60266;U3KQ | ADCY3 | Adenylate cyclase typ | 3123.986816 | 1785.403931 |
| GCDH | Q92947 | GCDH | Glutaryl-CoA dehydro | 92735.91406 | 108455.0234 |
| MARS2 | Q96GW9 | MARS2 | Methionine--tRNA liga | 21756.39063 | 23115.95508 |
| RBPMS | B4E3T4;Q93062 | RBPMS | HCG2043421, isoform | 32278.37305 | 45544.85938 |
| RPL28 | P46779 | RPL28 | 60S ribosomal protein | 1428005.25 | 1215759.125 |
| NCAPH2 | A0A0A6YYG7;Q6IBW4 | NCAPH2 | Condensin-2 complex | 7005.162598 | 8143.773926 |
| NSUN4 | Q96CB9 | NSUN4 | 5-methylcytosine rRN | 72133.26563 | 76815.07031 |
| TRIM56 | Q9BRZ2 | TRIM56 | E3 ubiquitin-protein li | 31561.61133 | 30150.39258 |
| RFFL | C9JE27;Q8WZ73 | RFFL | E3 ubiquitin-protein li | NA | 4170.609863 |
| PTP4A2 | Q12974 | PTP4A2 | Protein tyrosine phos | 17277.375 | 20728.11328 |
| ZFP64 | Q9NTW7 | ZFP64 | Zinc finger protein 64 | 6434.590332 | 6380.878906 |
| SNX12 | A0A087X0R6;Q9UMY4 | SNX12 | Sorting nexin-12 | 321049.5625 | 322607.5625 |
| PRKCSH | K7ELL7;P14314 | PRKCSH | Glucosidase 2 subunit | 986147.75 | 874940.9375 |
| RPL15 | P61313 | RPL15 | 60S ribosomal protein | 2059065.25 | 1846501.875 |
| CCT3 | P49368 | CCT3 | T-complex protein 1 s | 2119216.75 | 2150087.25 |
| TOE1 | Q96GM8 | TOE1 | Target of EGR1 protei | 34743.26563 | 35598.02344 |
| CA8 | P35219 | CA8 | Carbonic anhydrase-re | 66110.85938 | 84566.39844 |
| HNRNPA1 | F8W6I7;P09651 | HNRNPA1 | Heterogeneous nuclea | 4205025.5 | 2686139.5 |

| | | | | | |
|---|---|---|---|---|---|
| RPL26 | P61254 | RPL26 | 60S ribosomal protein | 2722141.75 | 2242738.25 |
| SAFB2 | Q14151 | SAFB2 | Scaffold attachment fa | 230488.8438 | 174519.6719 |
| ZHX3 | Q9H4I2 | ZHX3 | Zinc fingers and home | 37740.53125 | 39817.125 |
| GINS3 | Q9BRX5 | GINS3 | DNA replication comp | 4767.727051 | 5281.736816 |
| RAB17 | Q9H0T7 | RAB17 | Ras-related protein Ra | 3428814 | 3631186.75 |
| RPLP0 | P05388 | RPLP0 | 60S acidic ribosomal p | 1979526.875 | 1544665.375 |
| SPAG9 | A0A087X2D8 | SPAG9 | C-Jun-amino-terminal | 133660.915 | 144874.25 |
| FAM83B | Q5T0W9 | FAM83B | Protein FAM83B | 71573.60156 | 70202.55469 |
| SYPL1 | C9JYN0 | SYPL1 | Synaptophysin-like pr | 408680.291 | 402581.376 |
| SRRM1 | A9Z1X7;Q8IYB3 | SRRM1 | Serine/arginine repeti | 161529.5625 | 137750.1094 |
| GPR180 | Q86V85 | GPR180 | Integral membrane pr | 7693.352051 | 7695.847656 |
| EXOSC10 | Q01780 | EXOSC10 | Exosome component | 96716.8125 | 87395.54688 |
| MAPKAPK2 | P49137 | MAPKAPK2 | MAP kinase-activated | 57929.64453 | 59239.40625 |
| WASH3P;WASH2P | C4AMC7;Q6VEQ5 | WASH3P;WASH2P | Putative WAS protein | 85614.85938 | 84370.82031 |
| RXRA | A0A5F9ZHH6;P19793 | RXRA | Retinoic acid receptor | 32752.24609 | 33050.19141 |
| CSNK1D | H7BYT1;P48730 | CSNK1D | Casein kinase I isoform | 80447.92188 | 85974.23438 |
| SMARCA5 | O60264 | SMARCA5 | SWI/SNF-related matr | 171822.6719 | 195984.3281 |
| SEPHS2 | Q99611 | SEPHS2 | Selenide, water dikina | 42813.03906 | 51979.26953 |
| SPTBN1 | Q01082 | SPTBN1 | Spectrin beta chain, n | 245510.125 | 185201.375 |
| PLIN5 | Q00G26 | PLIN5 | Perilipin-5 | 1128.986084 | 1691.307495 |
| NSMCE1 | Q8WV22 | NSMCE1 | Non-structural mainte | 18544.50586 | 13122.79492 |
| ULK1 | O75385 | ULK1 | Serine/threonine-prot | 8086.75293 | 11843.99316 |
| SMN2;SMN1;SMN1 | A0A1W2PRV5;E7EQZ4;Q166 | SMN2;SMN1;SMN1 | Survival motor neuron | 97272.26563 | 93425.67188 |
| SEZ6L2 | A0A087WYL5;H3BN26;Q6UX | SEZ6L2 | Seizure 6-like protein | 15291.16699 | 12366.41309 |
| NPC1 | O15118 | NPC1 | NPC intracellular chol | 47607.85156 | 44184.26172 |
| RSRC2 | Q7L4I2 | RSRC2 | Arginine/serine-rich c | 94406.86719 | 82902.54688 |
| CENPS | Q8N2Z9 | CENPS | Centromere protein S | 113693.2109 | 84038.54688 |
| EDC4 | Q6P2E9 | EDC4 | Enhancer of mRNA-de | 170166.4531 | 188949.2188 |
| XRCC4 | Q13426 | XRCC4 | DNA repair protein XR | 38676.82422 | 39097.20703 |
| FAAP100 | Q0VG06 | FAAP100 | Fanconi anemia core c | 4789.467285 | 6081.526367 |
| SEC23IP | Q9Y6Y8 | SEC23IP | SEC23-interacting pro | 176050.4844 | 176290.2969 |
| LYAR | Q9NX58 | LYAR | Cell growth-regulating | 73870.46875 | 75599.89063 |
| TNRC6B | H0Y720;Q9UPQ9 | TNRC6B | Trinucleotide repeat-c | 9735.286133 | 10484.7207 |

| | | | | | |
|---|---|---|---|---|---|
| LONP2 | Q86WA8 | LONP2 | Lon protease homolog | 24878.71094 | 27034.79883 |
| CLINT1 | Q14677 | CLINT1 | Clathrin interactor 1 | 197032.8906 | 190559.7813 |
| SRSF4 | Q08170 | SRSF4 | Serine/arginine-rich sp | 1184970.125 | 987105.3125 |
| SPTLC1 | O15269 | SPTLC1 | Serine palmitoyltransf | 157160.7188 | 182073.8281 |
| RNH1 | P13489 | RNH1 | Ribonuclease inhibitor | 694095.75 | 774880.8125 |
| EPN3 | Q9H201 | EPN3 | Epsin-3 | 48547.43359 | 51606.26563 |
| RAD51C | O43502 | RAD51C | DNA repair protein RA | 41389.91016 | 43117.60547 |
| NDUFB10 | O96000 | NDUFB10 | NADH dehydrogenase | 363879.0313 | 372861.6563 |
| C12orf73 | Q69YU5 | C12orf73 | Uncharacterized prote | 41063.74609 | 57238.73828 |
| RPL27A | E9PLL6;P46776 | RPL27A | 60S ribosomal protein | 5100190 | 4353945.5 |
| TBCD | J3KR97;Q9BTW9 | TBCD | Tubulin-specific chape | 212233.3125 | 262634.7188 |
| EXOSC1 | Q9Y3B2 | EXOSC1 | Exosome complex cor | 75883.76563 | 73694.46875 |
| NME2P1 | O60361 | NME2P1 | Putative nucleoside di | 55379.03906 | 58816.47266 |
| ANKRD13D | Q6ZTN6 | ANKRD13D | Ankyrin repeat domai | 6851.586914 | 7623.721191 |
| ATG4B | Q9Y4P1 | ATG4B | Cysteine protease ATG | 78493.875 | 87425.45313 |
| SH3GL1 | Q99961 | SH3GL1 | Endophilin-A2 | 293801.3125 | 278545.0938 |
| GIPC1 | O14908 | GIPC1 | PDZ domain-containir | 113558.8047 | 116764.1094 |
| ARHGEF18 | A0A087WZG4;A0A3B3IPE9;A | ARHGEF18 | Rho guanine nucleotic | 3355.683838 | 5170.017578 |
| MYH10 | P35580 | MYH10 | Myosin-10 | 790585.5 | 829695.8125 |
| STK25 | O00506 | STK25 | Serine/threonine-prot | 111807.1016 | 125365.9531 |
| EIF2B5 | A0A3B3IUB1;Q13144 | EIF2B5 | Translation initiation f | 165416.7188 | 181243.625 |
| ATXN1L | P0C7T5 | ATXN1L | Ataxin-1-like | 3525.143799 | 7794.694824 |
| POLE4 | Q9NR33 | POLE4 | DNA polymerase epsil | 39952.17578 | 35525.84766 |
| LUC7L3 | O95232 | LUC7L3 | Luc7-like protein 3 | 154998.5156 | 111471.0781 |
| KIF3A | A0A087X011;E9PES4;J3KPF9; | KIF3A | Kinesin-like protein;Ki | 26715.44531 | 26126.92969 |
| BET1L | Q9NYM9 | BET1L | BET1-like protein | 19984.375 | 21945.70508 |
| YKT6 | O15498 | YKT6 | Synaptobrevin homol | 155722.4688 | 171959.4531 |
| NUP214 | A0A494C1F2;P35658 | NUP214 | Nuclear pore complex | 149868.7031 | 143668.4531 |
| CENPJ | F6VUX8;Q9HC77 | CENPJ | Centromere protein J | 2506.723877 | 2933.314209 |
| EEF1AKMT2 | A0A494BZY7;G3V238;Q5JPI9 | EEF1AKMT2 | EEF1A lysine methyltr | 51320.67578 | 57168.46875 |
| SLC7A6OS | Q96CW6 | SLC7A6OS | Probable RNA polyme | 32067.5293 | 30109.08398 |
| LRCH3 | Q96II8 | LRCH3 | DISP complex protein | 6074.150391 | 5370.731445 |
| EDF1 | O60869 | EDF1 | Endothelial differentia | 690943.125 | 667333.4375 |

| | | | | | |
|---|---|---|---|---|---|
| RPP25L | Q8N5L8 | RPP25L | Ribonuclease P protei | 73814.85938 | 70229.09375 |
| RPL30 | P62888 | RPL30 | 60S ribosomal protein | 1614575.75 | 1380710.375 |
| MIS12 | Q9H081 | MIS12 | Protein MIS12 homolo | 13352.30371 | 14893.29395 |
| EPHX1 | P07099 | EPHX1 | Epoxide hydrolase 1 | 115459.0938 | 119480.2969 |
| COX4I1 | P13073 | COX4I1 | Cytochrome c oxidase | 1282901.625 | 1334085.25 |
| CIAO3 | Q9H6Q4 | CIAO3 | Cytosolic iron-sulfur a | 37482.3125 | 39733.13281 |
| RPS23 | P62266 | RPS23 | 40S ribosomal protein | 1591763.375 | 1588399.75 |
| CKAP5 | Q14008 | CKAP5 | Cytoskeleton-associat | 280218.1563 | 248119.5625 |
| BRD9 | Q9H8M2 | BRD9 | Bromodomain-contair | 8609.142578 | 6439.286133 |
| EIF2B3 | Q9NR50 | EIF2B3 | Translation initiation f | 164789.0625 | 166603.1719 |
| CFDP1 | Q9UEE9 | CFDP1 | Craniofacial developm | 93850.82031 | 98372.52344 |
| POP4 | O95707 | POP4 | Ribonuclease P protei | 32522.60742 | 37744.55469 |
| ATAD2B | Q9ULI0 | ATAD2B | ATPase family AAA do | 5524.294922 | 7617.327637 |
| RBM12 | Q9NTZ6 | RBM12 | RNA-binding protein 1 | 170083.625 | 166294.2031 |
| FARP2 | O94887 | FARP2 | FERM, ARHGEF and pl | 19189.54688 | 19281.87305 |
| MED6 | A0A087WYL7;G3V4A5;O7558 | MED6 | Mediator of RNA poly | 20836.94531 | 18521.10352 |
| PPIH | O43447 | PPIH | Peptidyl-prolyl cis-trar | 356541.375 | 358844.875 |
| ZBTB1 | Q9Y2K1 | ZBTB1 | Zinc finger and BTB dc | 6867.65918 | 5389.563965 |
| PTGES2 | Q9H7Z7 | PTGES2 | Prostaglandin E synth | 191143.5156 | 194877.2344 |
| LACTB2 | Q53H82 | LACTB2 | Endoribonuclease LAC | 75961.53906 | 79987.33594 |
| RPL37A | C9J4Z3;P61513 | RPL37A | 60S ribosomal protein | 396773.4063 | 306109.7813 |
| UCHL5 | Q5LJA5 | UCHL5 | Ubiquitin carboxyl-ter | 343627.4414 | 341583.2988 |
| CTU1 | Q7Z7A3 | CTU1 | Cytoplasmic tRNA 2-tl | 29946.5957 | 30332.41992 |
| POLR2H | C9JLU1;P52434 | POLR2H | DNA-directed RNA po | 101087.6094 | 101981.8984 |
| DGKQ | H0YA20;P52824 | DGKQ | Diacylglycerol kinase ( | 7650.487305 | 8777.84082 |
| RPSA | A0A0C4DG17;C9J9K3;P08865 | RPSA | 40S ribosomal protein | 3106341.75 | 2301087 |
| RYBP | Q8N488 | RYBP | RING1 and YY1-bindin | 18782.7832 | 15316.31934 |
| ADAT1 | Q9BUB4 | ADAT1 | tRNA-specific adenosi | 14374.13477 | 13310.62305 |
| PSMG2 | Q969U7 | PSMG2 | Proteasome assembly | 165812.1719 | 180652.1406 |
| RPL6 | Q02878 | RPL6 | 60S ribosomal protein | 4743207 | 3704990.25 |
| ARID1A | O14497 | ARID1A | AT-rich interactive do | 56424.02734 | 63818.63281 |
| H3-3A;H3-3A;H3-3B;H3-3B; | A0A590UJJ6;B4DEB1;K7EK07 | H3-3A;H3-3A;H3-3B;H | Histone H3;Histone H3 | 3241510 | 195436.6094 |
| PRPF38A | Q8NAV1 | PRPF38A | Pre-mRNA-splicing fac | 81142.4375 | 53136.84375 |

| | | | | | |
|---|---|---|---|---|---|
| DNAJC9 | Q8WXX5 | DNAJC9 | DnaJ homolog subfam | 392118.25 | 365420.5313 |
| ;RPS17 | H3BNC9;P08708 | ;RPS17 | Uncharacterized prote | 1155495.375 | 961671.9375 |
| TRIM37 | O94972 | TRIM37 | E3 ubiquitin-protein li | 27726.26953 | 32965.98828 |
| MACROD1 | Q9BQ69 | MACROD1 | ADP-ribose glycohydr | 307204.7188 | 314656.0313 |
| TOP3A | Q13472 | TOP3A | DNA topoisomerase 3 | 12020.83594 | 13383.12012 |
| RPS29 | P62273 | RPS29 | 40S ribosomal protein | 930098.5625 | 782816.6875 |
| RPL13A | P40429 | RPL13A | 60S ribosomal protein | 2279338.75 | 1977402.375 |
| CSNK1E | P49674 | CSNK1E | Casein kinase I isoforr | 9611.239258 | 6685.312988 |
| SLC25A11 | I3L1P8;Q02978 | SLC25A11 | Mitochondrial 2-oxogl | 284528.5 | 285240.5 |
| HCCS | P53701 | HCCS | Cytochrome c-type he | 206691.6719 | 175876.7344 |
| USP7 | Q93009 | USP7 | Ubiquitin carboxyl-ter | 245980.2813 | 248970 |
| STARD10 | F5H1Z6;Q9Y365 | STARD10 | START domain-contai | 1427219.625 | 1406262.375 |
| ACOT7 | O00154 | ACOT7 | Cytosolic acyl coenzyr | 639227.0625 | 695567.5625 |
| CBX2 | Q14781 | CBX2 | Chromobox protein h | 29039.5957 | 30846.02148 |
| ACAD10 | Q6JQN1 | ACAD10 | Acyl-CoA dehydrogen | 27429.93945 | 28644.70898 |
| MGST3 | O14880;Q5VV89 | MGST3 | Microsomal glutathio | 39662.9375 | 19754.78906 |
| HMGN4 | O00479 | HMGN4 | High mobility group n | 15194.88965 | 1877.737549 |
| WDR83 | Q9BRX9 | WDR83 | WD repeat domain-cc | 10250.03516 | 9047.710938 |
| KPNA1 | P52294 | KPNA1 | Importin subunit alph | 197705.0313 | 202034.2969 |
| GDPD3 | Q7L5L3 | GDPD3 | Lysophospholipase D | 29637.33984 | 26802.16016 |
| ZNF554 | K7EQV8 | ZNF554 | Zinc finger protein 55 | 35399.16016 | 45797.46484 |
| RPS16 | P62249 | RPS16 | 40S ribosomal protein | 3730297.25 | 3116778.75 |
| TTC21A | A0A0B4J1Y2;Q8NDW8 | TTC21A | Tetratricopeptide rep | 76573.8125 | 54218.21484 |
| UGGT1 | Q9NYU2 | UGGT1 | UDP-glucose:glycopro | 139023.2031 | 156284.0781 |
| SHMT2 | P34897 | SHMT2 | Serine hydroxymethyl | 904377.4375 | 907689.875 |
| NCAPG | Q9BPX3 | NCAPG | Condensin complex su | 91284.01563 | 105210.8359 |
| ANAPC2 | Q9UJX6 | ANAPC2 | Anaphase-promoting | 28738.72266 | 33220.14453 |
| GALNT6 | Q8NCL4 | GALNT6 | Polypeptide N-acetylg | 57072.10547 | 66710.98438 |
| IMPDH1 | P20839 | IMPDH1 | Inosine-5'-monophos | 266181.0938 | 293819.875 |
| RPL21 | P46778 | RPL21 | 60S ribosomal protein | 835014.4375 | 733485.8125 |
| NUDT1 | P36639 | NUDT1 | 7,8-dihydro-8-oxogua | 90304.96875 | 106504.2344 |
| EIF6 | P56537 | EIF6 | Eukaryotic translation | 573530.8125 | 616686.9375 |
| METTL14 | Q9HCE5 | METTL14 | N6-adenosine-methyl | 24460.63281 | 25138.64844 |

| | | | | | |
|---|---|---|---|---|---|
| PBK | Q96KB5 | PBK | Lymphokine-activatec | 100929.1172 | 138496.8125 |
| WRNIP1 | Q96S55 | WRNIP1 | ATPase WRNIP1 | 71922.78906 | 72340.21094 |
| KNOP1 | Q1ED39 | KNOP1 | Lysine-rich nucleolar p | 41708.89453 | 27373.97852 |
| PRPF40A | H0YG38 | PRPF40A | Pre-mRNA-processing | 203672.1367 | 210074.9961 |
| ZNF143 | E7EN86;P52747 | ZNF143 | Zinc finger protein 143 | 42562.23438 | 52000.48438 |
| USP34 | Q70CQ2 | USP34 | Ubiquitin carboxyl-ter | 14507.6416 | 18753.06641 |
| DNAJC11 | Q9NVH1 | DNAJC11 | DnaJ homolog subfam | 63516.72266 | 39813.94922 |
| HNRNPA1L2 | Q32P51 | HNRNPA1L2 | Heterogeneous nucle | 2683.10376 | 1912.697144 |
| DIAPH3 | Q9NSV4 | DIAPH3 | Protein diaphanous ho | 16498.08203 | 18781.81055 |
| HMGN3 | Q15651 | HMGN3 | High mobility group n | 381594 | 116118.3281 |
| TMEM167B | A0A494C0V7;Q9NRX6 | TMEM167B | Protein kish-B | 28777.36719 | 20443.27148 |
| PFDN5 | Q99471 | PFDN5 | Prefoldin subunit 5 | 52349.88672 | 72849.94531 |
| KIFBP | A0A1B0GUA3;Q96EK5 | KIFBP | KIF-binding protein | 169034.9375 | 181553.2656 |
| TAF6 | J3KR72;P49848 | TAF6 | Transcription initiatio | 46859.27734 | 49247.14453 |
| RPL10A | P62906 | RPL10A | 60S ribosomal protein | 1533693.125 | 1205726.5 |
| GSTM4 | A0A0A0MR85;Q03013 | GSTM4 | Glutathione S-transfe | 33556.75781 | 31750.72266 |
| DDX3X | A0A0D9SF53;A0A0D9SFB3;A0 | DDX3X | ATP-dependent RNA h | 1716163.25 | 1756449.75 |
| UFM1 | P61960 | UFM1 | Ubiquitin-fold modifie | 182017.2344 | 227692.3594 |
| PEX14 | O75381 | PEX14 | Peroxisomal membrar | 44802.27344 | 48914.67578 |
| B3GNT2 | Q9NY97 | B3GNT2 | N-acetyllactosaminide | 3598.883057 | 3219.897705 |
| ;SRCAP | A0A0C4DFX4;Q6ZRS2 | ;SRCAP | Uncharacterized prote | 21747.10938 | 14297.65137 |
| HGS | O14964 | HGS | Hepatocyte growth fa | 204065.7188 | 204738.2188 |
| SGPL1 | O95470 | SGPL1 | Sphingosine-1-phosph | 198384.4219 | 188802.4219 |
| GMNN | E2QRF9;O75496 | GMNN | Geminin (Fragment);G | 50390.54297 | 46709.73438 |
| ARMC1 | Q9NVT9 | ARMC1 | Armadillo repeat-cont | 62503.81641 | 65698.76563 |
| ABHD10 | Q9NUJ1 | ABHD10 | Mycophenolic acid acy | 106764.375 | 115564.4063 |
| MAPRE3 | C9JB30;Q9UPY8 | MAPRE3 | Microtubule-associate | 42056.98828 | 42824.58203 |
| KCT2 | Q8NC54 | KCT2 | Keratinocyte-associat | 31142.81641 | 28200.62305 |
| PLEKHB2 | A0A0A0MSE9;A0A0A0MSI4;B | PLEKHB2 | Pleckstrin homology d | 6031.518555 | 5160.54541 |
| NUP98 | P52948 | NUP98 | Nuclear pore complex | 104100.7969 | 111898.2734 |
| PEA15 | Q15121 | PEA15 | Astrocytic phosphopro | 3238.905518 | 4679.63916 |
| STAT3 | P40763 | STAT3 | Signal transducer and | 94735.24219 | 105751.6563 |
| CEP131 | Q9UPN4 | CEP131 | Centrosomal protein c | 36074.14063 | 41129.1875 |

| | | | | | |
|---|---|---|---|---|---|
| RPL8 | P62917 | RPL8 | 60S ribosomal protein | 901212.25 | 806632.6875 |
| CDKAL1 | Q5VV42 | CDKAL1 | Threonylcarbamoylad | 23283.64453 | 28973.45898 |
| EXOSC8 | Q96B26 | EXOSC8 | Exosome complex cor | 108762.0781 | 90243.65625 |
| GDE1 | Q9NZC3 | GDE1 | Glycerophosphodieste | 39004.875 | 37897.34766 |
| AGFG1 | B8ZZY2;P52594 | AGFG1 | Arf-GAP domain and F | 46674.68359 | 73776.72656 |
| DNAJA4 | Q8WW22 | DNAJA4 | DnaJ homolog subfam | 56422.32422 | 59811.02734 |
| SLC35F2 | Q8IXU6 | SLC35F2 | Solute carrier family 3 | 5923.946289 | 6725.283203 |
| MECR | Q9BV79 | MECR | Enoyl-[acyl-carrier-pro | 23999.83203 | 29095.17383 |
| ADPGK | Q9BRR6 | ADPGK | ADP-dependent gluco | 96806.19531 | 98159.20313 |
| INTS11 | Q5TA45 | INTS11 | Integrator complex su | 122395.8047 | 130359.4453 |
| MAD1L1 | Q9Y6D9 | MAD1L1 | Mitotic spindle assem | 71869.78125 | 65139.35156 |
| UNK | Q9C0B0 | UNK | RING finger protein ur | 21846.90039 | 16468.65234 |
| ARF5 | P84085 | ARF5 | ADP-ribosylation facto | 372152 | 397886.9375 |
| TERF2IP | Q9NYB0 | TERF2IP | Telomeric repeat-binc | 75411.03906 | 67009.99219 |
| TARS2 | Q9BW92 | TARS2 | Threonine--tRNA ligas | 143381.6875 | 154337.9844 |
| VEZT | F8W8C2 | VEZT | Vezatin (Fragment) | 5722.654297 | 6213.438477 |
| FAM186B | Q8IYM0 | FAM186B | Protein FAM186B | 52417.04297 | 47094.53516 |
| LIG3 | P49916 | LIG3 | DNA ligase 3 | 114967.0391 | 121111.1953 |
| LPIN1 | Q14693 | LPIN1 | Phosphatidate phospl | 13665.14648 | 12450.67969 |
| GCAT | O75600 | GCAT | 2-amino-3-ketobutyra | 15586.19727 | 21085.45313 |
| PTPN18 | E7EMB8;Q99952 | PTPN18 | Tyrosine-protein phos | 4500.230957 | 5085.044434 |
| TRIP13 | Q15645 | TRIP13 | Pachytene checkpoint | 211822.2813 | 240362.2656 |
| ATF7 | P17544 | ATF7 | Cyclic AMP-dependen | 4536.396484 | 4641.853027 |
| HSPA8 | A8K7Q2 | HSPA8 | Heat shock cognate 7 | 16260903.13 | 16969967.88 |
| DRAP1 | C9JCC6;Q14919 | DRAP1 | Dr1-associated corepr | 142109.3125 | 136495.5469 |
| CDC23 | Q9UJX2 | CDC23 | Cell division cycle prot | 70233.64063 | 77641.69531 |
| TOR1AIP2 | Q8NFQ8 | TOR1AIP2 | Torsin-1A-interacting | 115143.3867 | 131614.7422 |
| C20orf27 | Q9GZN8 | C20orf27 | UPF0687 protein C20c | 236205.2031 | 277411.5313 |
| CNOT3 | O75175 | CNOT3 | CCR4-NOT transcriptic | 97057.67969 | 93460.25781 |
| DNAJC25-GNG10;GNG10 | A0A024R161;P50151 | DNAJC25-GNG10;GNG | Guanine nucleotide-b | 31555.17969 | 38466.40234 |
| RPS26 | P62854 | RPS26 | 40S ribosomal protein | 925849.125 | 814612.1875 |
| RPS2 | P15880 | RPS2 | 40S ribosomal protein | 2359707.5 | 1949292.125 |
| GTF3A | A0A1X7SBT5;H7BZH7;Q9266 | GTF3A | Transcription factor II | 6364.281738 | 7205.367188 |

| | | | | | |
|---|---|---|---|---|---|
| HMGXB4 | Q9UGU5 | HMGXB4 | HMG domain-contain | 6674.166016 | 4180.202637 |
| USP48 | Q86UV5 | USP48 | Ubiquitin carboxyl-ter | 44851.77734 | 46723.15625 |
| CCDC50 | Q8IVM0 | CCDC50 | Coiled-coil domain-co | 56201.37109 | 61307.04297 |
| TMBIM6 | F8VPI1;F8VQQ5;F8VQW0;F8V | TMBIM6 | Bax inhibitor 1 (Fragm | 39886.98828 | 37061.39063 |
| ZNF654 | A0A1B0GWA0 | ZNF654 | Zinc finger protein 65 | 4800.615234 | 6435.099121 |
| VPS35 | Q96QK1 | VPS35 | Vacuolar protein sorti | 380310.7813 | 424669.375 |
| ANLN | Q9NQW6 | ANLN | Anillin | 154251.4063 | 107265.5 |
| PRMT7 | Q9NVM4 | PRMT7 | Protein arginine N-me | 26125.94336 | 33612.12891 |
| TTI2 | E5RIH5;Q6NXR4 | TTI2 | TELO2-interacting pro | 29239.29297 | 37017.76953 |
| XPO7 | Q9UIA9 | XPO7 | Exportin-7 | 108814.1406 | 114548.5859 |
| HIBCH | Q6NVY1 | HIBCH | 3-hydroxyisobutyryl-C | 123992.9453 | 124151.3906 |
| ATP5IF1 | Q9UII2 | ATP5IF1 | ATPase inhibitor, mitc | 15541.36719 | 23626.48438 |
| HIKESHI | Q53FT3 | HIKESHI | Protein Hikeshi | 57420.19922 | 70330.02344 |
| LSM7 | Q9UK45 | LSM7 | U6 snRNA-associated | 205698.6719 | 173168.6094 |
| NELFE | P18615 | NELFE | Negative elongation fa | 124608.4141 | 106700.5156 |
| LRWD1 | Q9UFC0 | LRWD1 | Leucine-rich repeat ar | 73376.21094 | 75646.39844 |
| PLIN3 | O60664 | PLIN3 | Perilipin-3 | 352315.8125 | 318034.1563 |
| AGPAT5 | Q9NUQ2 | AGPAT5 | 1-acyl-sn-glycerol-3-pl | 32024.75781 | 22396.16211 |
| AP3S1 | Q92572 | AP3S1 | AP-3 complex subunit | 74841.23438 | 107524.3438 |
| MRRF | Q96E11 | MRRF | Ribosome-recycling fa | 122362.9922 | 132123 |
| RPL5 | P46777 | RPL5 | 60S ribosomal protein | 1008508.313 | 804303 |
| DHODH | Q02127 | DHODH | Dihydroorotate dehyc | 69752.22656 | 64331.62109 |
| SMURF2 | Q9HAU4 | SMURF2 | E3 ubiquitin-protein li | 4247.580566 | 4893.575195 |
| KIF16B | A0A1B0GTU3;Q96L93 | KIF16B | Kinesin-like protein KI | 98201.75781 | 106196.1328 |
| DNAJC7 | Q99615 | DNAJC7 | DnaJ homolog subfam | 260737.0625 | 278389.75 |
| TMED10 | P49755 | TMED10 | Transmembrane emp | 491924.4063 | 502734.7813 |
| SENP6 | Q9GZR1 | SENP6 | Sentrin-specific protea | 2459.776367 | 1677.287231 |
| SDF2 | Q99470 | SDF2 | Stromal cell-derived fa | 37672.86719 | 38584.17969 |
| DARS1 | P14868 | DARS1 | Aspartate--tRNA ligas | 996710.75 | 1019119.125 |
| TXNIP | Q9H3M7 | TXNIP | Thioredoxin-interactir | 44372.31641 | 41548.46484 |
| CSNK1A1 | A0A590UJ43;P48729;U3KPX3 | CSNK1A1 | Casein kinase I isoforr | 238195.9688 | 239946.3281 |
| TRAPPC11 | Q7Z392 | TRAPPC11 | Trafficking protein par | 12130.86426 | 11376.11914 |
| SP2 | Q02086 | SP2 | Transcription factor S | 7722.987793 | 5902.211914 |

| | | | | | |
|---|---|---|---|---|---|
| RPL7L1 | Q6DKI1 | RPL7L1 | 60S ribosomal protein | 100840.4375 | 44369.53516 |
| OTUD7B | Q6GQQ9 | OTUD7B | OTU domain-containi | 44769.13672 | 55792.86719 |
| FSCN1 | Q16658 | FSCN1 | Fascin | 64766.24219 | 60404.44141 |
| PPP3R1 | D3YTA9;F6U1T9;P63098 | PPP3R1 | Calcineurin subunit B | 2366.98291 | 2611.428223 |
| ARPC1B | O15143 | ARPC1B | Actin-related protein | 308224.75 | 326277.1875 |
| PFN1 | P07737 | PFN1 | Profilin-1 | 13057397 | 13794073 |
| RFX5 | F8W689;P48382 | RFX5 | DNA-binding protein | 27451.16211 | 26840.35352 |
| MRPS36 | P82909 | MRPS36 | 28S ribosomal protein | 118232.6641 | 128718.9688 |
| SDHAF1 | A6NFY7 | SDHAF1 | Succinate dehydroger | 6635.875488 | 6375.891113 |
| UBB;UBC;UBC;UBC;UBC;UB | B4DV12;F5GXK7;F5GYU3;F5 | UBB;UBC;UBC;UBC;U | Polyubiquitin-B;Polyu | 28749.42383 | 23010.27148 |
| NDUFB4 | O95168 | NDUFB4 | NADH dehydrogenase | 135336.8281 | 136136.5469 |
| SEPHS1 | P49903 | SEPHS1 | Selenide, water dikina | 320600.25 | 346949.125 |
| BANF1 | O75531 | BANF1 | Barrier-to-autointegra | 39809.02734 | 100507.8047 |
| EIF2A | Q9BY44 | EIF2A | Eukaryotic translation | 323088.7188 | 299189.1875 |
| LENG1 | Q96BZ8 | LENG1 | Leukocyte receptor cl | 24715.69727 | 22374.59961 |
| CENPU | Q71F23 | CENPU | Centromere protein U | 8224.117188 | 7856.643555 |
| MAN1A1 | P33908 | MAN1A1 | Mannosyl-oligosaccha | 16662.37305 | 19618.66992 |
| LRMP | A0A1B0GU45 | LRMP | Lymphoid-restricted n | 133105.8125 | 148336.4531 |
| MYO1C | F5H6E2;O00159 | MYO1C | Unconventional myos | 303424.8438 | 285699.5938 |
| RANBP3 | B7Z7F3 | RANBP3 | Ran-binding protein 3 | 70895.27344 | 82568.59375 |
| SMC4 | Q9NTJ3 | SMC4 | Structural maintenanc | 135604.8594 | 151974.0625 |
| NDUFB1 | O75438 | NDUFB1 | NADH dehydrogenase | 42784.16406 | 48601.39844 |
| EXOSC9 | Q06265 | EXOSC9 | Exosome complex cor | 103708.4063 | 102652.2734 |
| POLD1 | M0R2B7;P28340 | POLD1 | DNA polymerase;DNA | 125064.0078 | 143296.5781 |
| UBQLN2 | Q9UHD9 | UBQLN2 | Ubiquilin-2 | 67007.1875 | 68485.34375 |
| CASKIN2 | Q8WXE0 | CASKIN2 | Caskin-2 | 56472.38281 | 45562.41406 |
| PPP2R2D | Q66LE6 | PPP2R2D | Serine/threonine-prot | 41218.23047 | 37628.9375 |
| PXN | A0A1B0GTU4;F5GZ78;P4902 | PXN | Paxillin | 100247.7578 | 108707.0703 |
| SSU72 | Q9NP77 | SSU72 | RNA polymerase II sub | 100461.1719 | 91296.21875 |
| RBM42 | K7EP90;Q9BTD8 | RBM42 | RNA-binding protein 4 | 166293.7656 | 155641.2344 |
| KIAA2013 | Q8IYS2 | KIAA2013 | Uncharacterized prote | 38880.92969 | 44636.91797 |
| VWA8 | A3KMH1 | VWA8 | von Willebrand factor | 86750.24219 | 95036.94531 |
| RPL7A | P62424 | RPL7A | 60S ribosomal protein | 2929283.25 | 2362739 |

| | | | | | |
|---|---|---|---|---|---|
| RPL4 | P36578 | RPL4 | 60S ribosomal protein | 3903241.5 | 3177608 |
| NMT2 | O60551;Q5VUC6 | NMT2 | Glycylpeptide N-tetra | 9967.737305 | 11562.16309 |
| MFF | Q9GZY8 | MFF | Mitochondrial fission | 39291.92578 | 40723.10156 |
| SSB | P05455 | SSB | Lupus La protein | 576345.875 | 635381.75 |
| ABCB6 | Q9NP58 | ABCB6 | ATP-binding cassette | 44048.17969 | 42347.30078 |
| PPAT | Q06203 | PPAT | Amidophosphoribosyl | 389472.6563 | 454176 |
| EHD1 | Q9H4M9 | EHD1 | EH domain-containing | 164358.125 | 170703.9844 |
| DHX9 | Q08211 | DHX9 | ATP-dependent RNA h | 818644.9375 | 568059.875 |
| ATF1 | P18846 | ATF1 | Cyclic AMP-dependen | 11871.14746 | NA |
| VPS33A | A0A2R8YF87;Q96AX1 | VPS33A | Vacuolar protein sorti | 146315.6094 | 166153.0625 |
| TOMM70 | O94826 | TOMM70 | Mitochondrial import | 290571.5625 | 293990.8125 |
| SOD1 | P00441 | SOD1 | Superoxide dismutase | 116951.3516 | 142133.9219 |
| RCC1 | P18754 | RCC1 | Regulator of chromos | 807997.4375 | 432305.4375 |
| RALA | P11233 | RALA | Ras-related protein Ra | 156931.3281 | 153970.7813 |
| RBMX | P38159 | RBMX | RNA-binding motif pro | 966070.4375 | 399690.75 |
| NDUFS7 | F5GXJ1;F5H5N1;O75251 | NDUFS7 | NADH dehydrogenase | 138783.2031 | 157020.7344 |
| BLOC1S1 | G8JLQ3;P78537 | BLOC1S1 | Biogenesis of lysosom | 43948.74609 | 39343.65234 |
| SPINT1 | H3BR01 | SPINT1 | Kunitz-type protease i | 152567.4922 | 126129.5918 |
| PATJ | A0A2R8Y549;Q8NI35 | PATJ | InaD-like protein | 24750.97266 | 29009.06641 |
| ATP5MD | Q96IX5 | ATP5MD | ATP synthase membra | 252484.8281 | 262128.2813 |
| UHRF2 | Q96PU4 | UHRF2 | E3 ubiquitin-protein li | 67922.71094 | 78779.36719 |
| ABCF1 | Q8NE71 | ABCF1 | ATP-binding cassette | 385626.3438 | 324732.625 |
| TNRC6A | Q8NDV7 | TNRC6A | Trinucleotide repeat-c | 1324.308105 | 2525.222168 |
| EPC1 | A0A590UJN0;Q9H2F5 | EPC1 | Enhancer of polycomb | 16796.38477 | 9113.310547 |
| GTF2H4 | Q92759 | GTF2H4 | General transcription | 25057.35742 | 23925.90625 |
| PGGT1B | P53609 | PGGT1B | Geranylgeranyl transfe | 19343.47656 | 18838.06836 |
| SAP130 | A0A2R8YDB8;H7BXF5;Q9H0E | SAP130 | Histone deacetylase c | 50034.78516 | 49917.62109 |
| DPF2 | Q92785 | DPF2 | Zinc finger protein ubi | 52598.99609 | 51326.41016 |
| NAPG | Q99747 | NAPG | Gamma-soluble NSF a | 210859.1875 | 269767.1875 |
| SETD3 | Q86TU7 | SETD3 | Actin-histidine N-meth | 117450.8359 | 116852.9453 |
| POU2F1 | A0A0C4DG88;P14859 | POU2F1 | POU domain protein ( | 23241.95703 | 34437.67969 |
| RPS15 | P62841 | RPS15 | 40S ribosomal protein | 1054550.75 | 907620.625 |
| ZZEF1 | O43149 | ZZEF1 | Zinc finger ZZ-type an | 21484.27148 | 25215.08594 |

| | | | | | |
|---|---|---|---|---|---|
| ERP29 | P30040 | ERP29 | Endoplasmic reticulur | 2241613.5 | 2243613.25 |
| RPS25 | P62851 | RPS25 | 40S ribosomal protein | 3842346 | 2972496.75 |
| WASHC2C | Q9Y4E1 | WASHC2C | WASH complex subun | 5896.567871 | 6173.361328 |
| LRRC8E | Q6NSJ5 | LRRC8E | Volume-regulated ani | 16941.44141 | 16977.98047 |
| TFIP11 | Q9UBB9 | TFIP11 | Tuftelin-interacting pr | 59127.21875 | 56629.23828 |
| RDH14 | Q9HBH5 | RDH14 | Retinol dehydrogenas | 67963.71094 | 70887.89063 |
| ABCC1 | I3L4X2;P33527 | ABCC1 | Multidrug resistance-a | 33124.98047 | 29100.78711 |
| RPL36 | Q9Y3U8 | RPL36 | 60S ribosomal protein | 2609097.5 | 2000863.75 |
| HUS1 | O60921 | HUS1 | Checkpoint protein HU | 12759.26074 | 13565.38379 |
| WASHC5 | Q12768 | WASHC5 | WASH complex subun | 97367.55469 | 101420.3203 |
| EXOSC5 | Q9NQT4 | EXOSC5 | Exosome complex cor | 86057.625 | 88892.45313 |
| LUC7L2 | Q9Y383 | LUC7L2 | Putative RNA-binding | 384266.9375 | 358828.5 |
| CPSF2 | Q9P2I0 | CPSF2 | Cleavage and polyade | 111375.5313 | 112304.6875 |
| ENSA | O43768 | ENSA | Alpha-endosulfine | 981243.1152 | 1007800.578 |
| TPRKB | Q9Y3C4 | TPRKB | EKC/KEOPS complex s | 75844.14844 | 78936.23438 |
| MED24 | A0A0B4J1W0;O75448 | MED24 | Mediator of RNA poly | 35680.41406 | 37133.44531 |
| DHTKD1 | Q96HY7 | DHTKD1 | Probable 2-oxoglutara | 144726.9688 | 166963.9375 |
| EXOSC3 | Q9NQT5 | EXOSC3 | Exosome complex cor | 98888.72656 | 87844.98438 |
| XPO6 | Q96QU8 | XPO6 | Exportin-6 | 20693.29688 | 24493.39453 |
| RPL7 | P18124 | RPL7 | 60S ribosomal protein | 2566875 | 2022821.375 |
| ARMCX3 | Q9UH62 | ARMCX3 | Armadillo repeat-cont | 60135.89844 | 78446.10156 |
| ALKBH5 | Q6P6C2 | ALKBH5 | RNA demethylase ALK | 92488.82031 | 90816.57813 |
| SNIP1 | Q8TAD8 | SNIP1 | Smad nuclear-interact | 47920.45703 | 31338.84766 |
| NOTCH3 | Q9UM47 | NOTCH3 | Neurogenic locus notc | 10866.625 | 12760.97754 |
| COQ5 | Q5HYK3 | COQ5 | 2-methoxy-6-polyprer | 42070.19531 | 37792.86328 |
| TRUB1 | Q8WWH5 | TRUB1 | Probable tRNA pseudo | 63616.30469 | 68343.44531 |
| MED1 | Q15648 | MED1 | Mediator of RNA poly | 46976.30078 | 46339.04688 |
| EEF2 | P13639 | EEF2 | Elongation factor 2 | 7813211.5 | 8061347 |
| TRIM65 | Q6PJ69 | TRIM65 | Tripartite motif-conta | 30720.84961 | 39069.38672 |
| INTS1 | Q8N201 | INTS1 | Integrator complex su | 64505.92578 | 66352.36719 |
| STK11 | K7EP59;Q15831 | STK11 | Serine/threonine-prot | 51781.08984 | 52854.39063 |
| KDM6A | A0A087X0R0;F5H5V6;F5H6S1 | KDM6A | Lysine-specific demetl | 8006.891113 | 10328.05957 |
| COX7A2 | D6RGV5;D6RIE3;H0UI06;P14 | COX7A2 | Cytochrome c oxidase | 412880.3438 | 389280.9688 |

| | | | | | |
|---|---|---|---|---|---|
| ATP6V1G1 | O75348 | ATP6V1G1 | V-type proton ATPase | 271486.2813 | 257586.9688 |
| TMEM256-PLSCR3;TMEM2! | K7ENI6;K7ERE1;Q8N2U0 | TMEM256-PLSCR3;TM | TMEM256-PLSCR3 rea | 30770.14844 | 34262.04688 |
| MTMR6 | Q9Y217 | MTMR6 | Myotubularin-related | 52249.08984 | 53383.50781 |
| ARMC9 | A0A2Q3DP09;Q7Z3E5 | ARMC9 | LisH domain-containir | 25598.41016 | 26487.46875 |
| PITPNA | Q00169 | PITPNA | Phosphatidylinositol t | 121564.2969 | 130608.5469 |
| RACK1 | P63244 | RACK1 | Receptor of activated | 3345373.25 | 2821361.25 |
| VPS29 | Q9UBQ0 | VPS29 | Vacuolar protein sorti | 126654.6016 | 145578.875 |
| CHID1 | Q9BWS9 | CHID1 | Chitinase domain-con | 67028.35156 | 84743.0625 |
| COQ10B;COQ10B;COQ10A; | B8ZZV9;B8ZZX2;H0YHF1;Q96 | COQ10B;COQ10B;COC | Coenzyme Q-binding p | 2537.222168 | 4962.891113 |
| KMT2A | Q03164 | KMT2A | Histone-lysine N-meth | 19524.86523 | 21031.76563 |
| ST13 | P50502 | ST13 | Hsc70-interacting pro | 1365561.5 | 1410215.25 |
| CCSMST1 | Q4G0I0 | CCSMST1 | Protein CCSMST1 | 19053.14844 | 18700.52734 |
| ;IL10RB;IL10RB | H0Y3Z8;H7C0Z5;Q08334 | ;IL10RB;IL10RB | Uncharacterized prote | 12259.56738 | 10989.98828 |
| ;VKORC1;;VKORC1 | E9PLN8;F8W9H0;I3L3B4;Q9B | ;VKORC1;;VKORC1 | VKc domain-containin | 87122.49219 | 103468.8516 |
| PFN3 | P60673 | PFN3 | Profilin-3 | 102564.9766 | 82369.82031 |
| NTHL1 | A0A494BZZ6;P78549 | NTHL1 | Endonuclease III-like p | 53227.96875 | 49640.86328 |
| RPL27 | P61353 | RPL27 | 60S ribosomal protein | 3028147.75 | 2306045.5 |
| FAM83H | A0A494C1T9;Q6ZRV2 | FAM83H | Protein FAM83H | 91055.65625 | 92030.32813 |
| FAM185A | Q8N0U4 | FAM185A | Protein FAM185A | 3617.848145 | 2842.655273 |
| ALYREF | E9PB61 | ALYREF | THO complex subunit | 1649001.377 | 1515279.965 |
| COX16 | Q9P0S2 | COX16 | Cytochrome c oxidase | 64277.38672 | 67779.625 |
| MLYCD | O95822 | MLYCD | Malonyl-CoA decarbo: | 47277.61328 | 55615.25 |
| STAM | Q92783 | STAM | Signal transducing ada | 59803.5 | 49566.50391 |
| VIRMA | Q69YN4 | VIRMA | Protein virilizer homo | 70938.47656 | 62299.21484 |
| ECPAS | Q5VYK3 | ECPAS | Proteasome adapter a | 179320.2813 | 182509.4844 |
| DCXR | Q7Z4W1 | DCXR | L-xylulose reductase | 635279.4375 | 796126.8125 |
| SPG21 | Q9NZD8 | SPG21 | Maspardin | 69920.74219 | 76090.75781 |
| LARP4 | Q71RC2 | LARP4 | La-related protein 4 | 135719.2188 | 144347.25 |
| NFIA | A0A590UK09;A0A590UK73;B | NFIA | Nuclear factor 1;Nucle | 33053.39453 | 52096.48828 |
| RAD51 | Q06609 | RAD51 | DNA repair protein RA | 33095.49609 | 36273.05469 |
| CIAPIN1 | Q6FI81 | CIAPIN1 | Anamorsin | 119437.5547 | 116725.4063 |
| HPS6 | Q86YV9 | HPS6 | Hermansky-Pudlak syr | 29804.04297 | 31492.55469 |
| ZNF639 | J3KR59;J3KR68;Q9UID6 | ZNF639 | Zinc finger protein 63! | 4667.046875 | 3326.503662 |

| | | | | | |
|---|---|---|---|---|---|
| UBLCP1 | Q8WVY7 | UBLCP1 | Ubiquitin-like domain- | 77363.88281 | 76739.85938 |
| TLN1 | Q9Y490 | TLN1 | Talin-1 | 720546.6875 | 685095.9375 |
| FBRSL1 | A0A3B3IRR3;Q9HCM7 | FBRSL1 | Fibrosin-1-like protein | 11250.13086 | 10637.63281 |
| NCOA7 | Q8NI08 | NCOA7 | Nuclear receptor coac | 2251.649902 | 3228.497314 |
| MAGOHB | Q96A72 | MAGOHB | Protein mago nashi ho | 403918.7813 | 395446.5 |
| ARMC8 | Q8IUR7 | ARMC8 | Armadillo repeat-cont | 207201.2031 | 226042.625 |
| SMC2 | O95347 | SMC2 | Structural maintenanc | 208925.8906 | 230532.8438 |
| FOXK2 | Q01167 | FOXK2 | Forkhead box protein | 72300.50781 | 79911.09375 |
| GLRX3 | O76003 | GLRX3 | Glutaredoxin-3 | 912736.4375 | 895091 |
| PPIL2 | Q13356 | PPIL2 | RING-type E3 ubiquiti | 34463.78516 | 32974.62891 |
| MAT2B | Q9NZL9 | MAT2B | Methionine adenosylt | 233086.125 | 244410.2656 |
| TAF15 | Q92804 | TAF15 | TATA-binding protein- | 2161402.75 | 2104213.25 |
| DLAT | P10515 | DLAT | Dihydrolipoyllysine-re | 236874.2969 | 241130.7969 |
| TLK2 | J3KRK0 | TLK2 | Serine/threonine-prot | 86494.62891 | 93088.14453 |
| SFPQ | P23246 | SFPQ | Splicing factor, proline | 1944563.375 | 1553947.625 |
| CS | B4DJV2;O75390 | CS | Citrate synthase;Citra | 2249387.5 | 2451122.5 |
| STARD5 | Q9NSY2 | STARD5 | StAR-related lipid tran | 8384.561523 | 7866.871094 |
| PDHX | O00330 | PDHX | Pyruvate dehydrogena | 69797.375 | 66661.69531 |
| HDAC4 | P56524 | HDAC4 | Histone deacetylase 4 | 5498.510254 | 5738.17334 |
| SRSF5 | Q13243 | SRSF5 | Serine/arginine-rich sp | 246544.5313 | 205166.9219 |
| SLC25A29 | Q8N8R3 | SLC25A29 | Mitochondrial basic ar | 29979.55273 | 44552.87891 |
| PLCB3 | Q01970 | PLCB3 | 1-phosphatidylinosito | 64286.62109 | 65531.61719 |
| TP53RK | Q96S44 | TP53RK | EKC/KEOPS complex s | 74262.5 | 71338.89844 |
| CRKL | P46109 | CRKL | Crk-like protein | 421794.875 | 413777.75 |
| SLC4A1AP | A0A087X0M4 | SLC4A1AP | Kanadaptin | 122707.3594 | 112783.6953 |
| TNPO3 | Q9Y5L0 | TNPO3 | Transportin-3 | 93336.80469 | 103687.9141 |
| HNRNPL | A0A3B3ITJ4 | HNRNPL | Heterogeneous nuclea | 1069428.285 | 575015.3223 |
| EAPP | A0A087X060;G3V2W9;Q56P( | EAPP | E2F-associated phospl | 19896.55273 | 19819.67188 |
| MFN1 | Q8IWA4 | MFN1 | Mitofusin-1 | 87538.39844 | 65792.55469 |
| PRR12 | Q9ULL5 | PRR12 | Proline-rich protein 12 | 11547.1084 | 13502.37793 |
| CABIN1 | A0A087WWW8;Q9Y6J0 | CABIN1 | Calcineurin-binding pr | 5125.643555 | 5376.386719 |
| TRMT5 | Q32P41 | TRMT5 | tRNA (guanine(37)-N1 | 54053.67969 | 53460.77344 |
| ILF3 | Q12906 | ILF3 | Interleukin enhancer- | 1013000.563 | 631929.0625 |

| | | | | | |
|---|---|---|---|---|---|
| SCARB2 | A0A1W2PRS1;A0A1W2PS43; | SCARB2 | Lysosome membrane | 144938.6406 | 119329.7344 |
| PPP2R5A | Q15172 | PPP2R5A | Serine/threonine-prot | 31445.2207 | 37914.57031 |
| GTPBP10 | A4D1E9 | GTPBP10 | GTP-binding protein 1 | 96038.02344 | 97225.98438 |
| DNAJC1 | Q96KC8 | DNAJC1 | DnaJ homolog subfam | 37001.16797 | 36542.12891 |
| ZFP91-CNTF;ZFP91 | A0A0A6YYC7;Q96JP5 | ZFP91-CNTF;ZFP91 | HCG2042749, isoform | 40394.92578 | 38106.71875 |
| PRKAR1A | P10644 | PRKAR1A | cAMP-dependent prot | 460417.4375 | 437903.6875 |
| CFL2 | Q9Y281 | CFL2 | Cofilin-2 | 63748.85156 | 51978.76172 |
| RPL32 | D3YTB1;F8W727;P62910 | RPL32 | 60S ribosomal protein | 938600.8125 | 795226.125 |
| CDK4 | P11802 | CDK4 | Cyclin-dependent kina | 94818.35938 | 100337.5234 |
| SAFB | Q15424 | SAFB | Scaffold attachment fa | 350691.3438 | 208942.2969 |
| RPL29 | A0A3B3ITT5;P47914 | RPL29 | 60S ribosomal protein | 1849778.625 | 1431241.125 |
| OSGEP | Q9NPF4 | OSGEP | Probable tRNA N6-ade | 93789.02344 | 107172.1953 |
| NSL1 | Q96IY1 | NSL1 | Kinetochore-associate | 9897.952148 | 12139.17578 |
| AGL | P35573 | AGL | Glycogen debranching | 100693.3594 | 118803.4688 |
| METAP1 | P53582 | METAP1 | Methionine aminopep | 223796.3281 | 199376.6875 |
| C2orf49 | Q9BVC5 | C2orf49 | Ashwin | 48984.91797 | 51099.6875 |
| TLE1 | Q04724 | TLE1 | Transducin-like enhan | 36187.37891 | 29610.31055 |
| ATP5MF | P56134 | ATP5MF | ATP synthase subunit | 127036.6172 | 131155.1719 |
| SSR1 | A0A3B3IRT8;P43307 | SSR1 | Translocon-associated | 410842.5 | 350257.4688 |
| PIN4 | Q9Y237 | PIN4 | Peptidyl-prolyl cis-tran | 200037.0156 | 221680.4531 |
| GRHL1 | Q9NZI5 | GRHL1 | Grainyhead-like prote | 12372.99609 | 16782.47266 |
| IDH3A | P50213 | IDH3A | Isocitrate dehydrogen | 590640.8125 | 600789.4375 |
| GTF3C5 | Q9Y5Q8 | GTF3C5 | General transcription | 87221.76563 | 90576.14844 |
| TPD52L2 | O43399 | TPD52L2 | Tumor protein D54 | 1208844.875 | 1158947.5 |
| TPR | P12270 | TPR | Nucleoprotein TPR | 298771.3438 | 326718.7188 |
| KRT13 | K7ERE3;P13646 | KRT13 | Keratin, type I cytoske | 16410.97852 | 16780.9707 |
| NECAP1 | A0A1W2PR09;A0A1W2PRL5; | NECAP1 | Adaptin ear-binding co | 51790.73438 | 53927.12891 |
| WASHC4 | A0A087X256;Q2M389 | WASHC4 | WASH complex subun | 207649.4844 | 209103.6719 |
| SON | H7C1M2 | SON | Protein SON (Fragmer | 659204.9961 | 620534.0225 |
| THRAP3 | Q9Y2W1 | THRAP3 | Thyroid hormone rece | 409363.9688 | 400082.7188 |
| APC | P25054 | APC | Adenomatous polypos | 10635.55664 | 12049.76074 |
| ZNF148 | Q9UQR1 | ZNF148 | Zinc finger protein 14 | 24207.07227 | 29274.40234 |
| SMAD5 | Q99717 | SMAD5 | Mothers against deca | 28689.27344 | 28505.46875 |

| | | | | | |
|---|---|---|---|---|---|
| PNPO | Q9NVS9 | PNPO | Pyridoxine-5'-phospha | 347786.9688 | 353894.8438 |
| CBX1 | J3KS05;P83916 | CBX1 | Chromobox protein h | 123497.75 | 106268.2266 |
| ELP6 | C9IYN7;C9J2D6;C9J9N8;C9JA | ELP6 | Elongator complex pr | 17848.16016 | 20088.84766 |
| TMED2 | Q15363 | TMED2 | Transmembrane emp | 169668.9219 | 177542.0469 |
| PTK7 | Q13308 | PTK7 | Inactive tyrosine-prot | 21235.21484 | 25663.49414 |
| BAZ2A | A0A0C4DGI9;F8VU39;J3KPG5 | BAZ2A | Bromodomain adjacer | 9130.867188 | 12948.18262 |
| CHD8 | Q9HCK8 | CHD8 | Chromodomain-helica | 102766.2188 | 117206.6719 |
| TUBB2A | Q13885 | TUBB2A | Tubulin beta-2A chain | 2226273.5 | 2392692.25 |
| BUB1B | O60566 | BUB1B | Mitotic checkpoint se | 35357.5 | 35053.84375 |
| KANSL2 | F8VX10;F8VXI8;H0YIQ8;Q9H | KANSL2 | KAT8 regulatory NSL c | 4978.512695 | 5097.741211 |
| FABP6 | P51161 | FABP6 | Gastrotropin | 2737.707764 | 5504.351563 |
| AGTRAP | Q6RW13 | AGTRAP | Type-1 angiotensin II | 35694.19922 | 39490.10156 |
| EED | A0A5F9ZI63 | EED | Polycomb protein EED | 157806.2188 | 139883.8438 |
| CDC16 | Q13042 | CDC16 | Cell division cycle pro | 31738.86914 | 35312.27734 |
| EIF5B | A0A087WUT6;O60841 | EIF5B | Eukaryotic translation | 308012.4375 | 320608.625 |
| MAP4 | E7EVA0 | MAP4 | Microtubule-associate | 207151.1875 | 229856.8281 |
| SERPINB9 | P50453 | SERPINB9 | Serpin B9 | 5831.526855 | 9374.763672 |
| STRBP | Q96SI9 | STRBP | Spermatid perinuclea | 102811.9844 | 113450.2109 |
| CDC26 | Q8NHZ8 | CDC26 | Anaphase-promoting | 25949.15234 | 24713.00977 |
| KIAA1143 | Q96AT1 | KIAA1143 | Uncharacterized prote | 52558.87109 | 44924.54297 |
| SYNJ1 | A0A0D9SGJ6;C9JFZ1;J3KPK1; | SYNJ1 | Synaptojanin-1 | 9925.666992 | 10034.65039 |
| RALY | Q9UKM9 | RALY | RNA-binding protein R | 229920.5625 | 111628.0078 |
| TATDN1 | E5RG17;Q6P1N9 | TATDN1 | Putative deoxyribonu | 313821.0625 | 321669.9063 |
| GPRASP2 | Q96D09 | GPRASP2 | G-protein coupled rec | 13176.33789 | 12222.42383 |
| ZCCHC8 | Q6NZY4 | ZCCHC8 | Zinc finger CCHC dom | 60959.30859 | 61030.68359 |
| OTUB1 | J3KR44;Q96FW1 | OTUB1 | Ubiquitin thioesterase | 574572.625 | 626426.5 |
| MIB1 | Q86YT6 | MIB1 | E3 ubiquitin-protein li | 11138.32324 | 11613.46484 |
| ATXN7 | A0A2R8YDD6;O15265 | ATXN7 | Ataxin 7, isoform CRA | 13726.65137 | 12303.12207 |
| CUEDC2 | Q9H467 | CUEDC2 | CUE domain-containir | 30016.30664 | 39683.36328 |
| CDK13 | A0A2R8Y4Z0;Q14004 | CDK13 | Cyclin-dependent kina | 42288.8125 | 42927.17578 |
| TXNRD1 | A0A087WSW9;A0A087WSY9 | TXNRD1 | Thioredoxin reductase | 257833.9375 | 249211.8438 |
| LSM12 | K7ELG9;Q3MHD2 | LSM12 | Protein LSM12 homol | 200560.4219 | 194633.2188 |
| HECTD3 | Q5T447 | HECTD3 | E3 ubiquitin-protein li | 47016.91016 | 50148.15625 |

| | | | | | |
|---|---|---|---|---|---|
| TOMM40 | O96008 | TOMM40 | Mitochondrial import | 888243 | 566049.625 |
| TRAPPC3 | O43617 | TRAPPC3 | Trafficking protein par | 250283.4844 | 274247.4063 |
| CCNK | O75909 | CCNK | Cyclin-K | 104456.7031 | 112394.9063 |
| RPL38 | P63173 | RPL38 | 60S ribosomal protein | 1099699.75 | 869397.1875 |
| CROT | C9J3D7;Q9UKG9 | CROT | Peroxisomal carnitine | 1858.619507 | 1589.877319 |
| HNRNPDL | A0A087WU03 | HNRNPDL | Heterogeneous nuclea | 512176.1094 | 248530.4297 |
| DERL2 | I3L1T3;Q9GZP9 | DERL2 | Derlin (Fragment);Der | 13676.4082 | 17218.50977 |
| EIF4E2 | O60573 | EIF4E2 | Eukaryotic translation | 69471.38281 | 67757.91406 |
| CBX3 | Q13185 | CBX3 | Chromobox protein ho | 718490.3125 | 617356.5 |
| GTF2B | Q00403 | GTF2B | Transcription initiatio | 52477.72266 | 49015.05469 |
| PXDN | Q92626 | PXDN | Peroxidasin homolog | 12596.9209 | 13152.87109 |
| RPP30 | P78346 | RPP30 | Ribonuclease P protei | 87983.11719 | 104070.0547 |
| MARS1 | P56192 | MARS1 | Methionine--tRNA liga | 406038.5313 | 426969.7813 |
| U2SURP | O15042 | U2SURP | U2 snRNP-associated | 220593.8438 | 196007.1875 |
| CFAP298 | P57076 | CFAP298 | Cilia- and flagella-assc | 64174.20313 | 64565.98438 |
| PCCB | E9PDR0;P05166 | PCCB | Propionyl-CoA carbox | 41617.19922 | 49104.71875 |
| LIMD2 | J3QQM5;Q9BT23 | LIMD2 | LIM domain-containin | 9223.835938 | 12164.65723 |
| HABP4 | Q5JVS0 | HABP4 | Intracellular hyaluron | 26863.54102 | 25860.4043 |
| PSMD4 | P55036 | PSMD4 | 26S proteasome non-A | 627923.7852 | 693250.0977 |
| CACYBP | Q9HB71 | CACYBP | Calcyclin-binding prot | 654547.5 | 666597.0625 |
| TEAD1 | H0YE88;H0YEJ9;P28347 | TEAD1 | Transcriptional enhan | 20889.16016 | 26999.58398 |
| FXR1 | P51114 | FXR1 | Fragile X mental retar | 225284.7031 | 223767.0781 |
| BCCIP | Q9P287 | BCCIP | BRCA2 and CDKN1A-ir | 142959.8125 | 129819.5 |
| PCF11 | O94913 | PCF11 | Pre-mRNA cleavage co | 48312.14063 | 45584.55078 |
| RPP25 | Q9BUL9 | RPP25 | Ribonuclease P protei | 41352.4375 | 43650.79297 |
| HEATR6 | Q6AI08 | HEATR6 | HEAT repeat-containir | 354926.625 | 409740.4375 |
| ZNF414 | M0R0Q5;M0R398 | ZNF414 | Zinc finger protein 41 | 3810.15918 | 3663.740234 |
| HNRNPA0 | Q13151 | HNRNPA0 | Heterogeneous nuclea | 470581.4688 | 357651.2188 |
| IBTK | E7EPI0;Q9P2D0 | IBTK | Inhibitor of Bruton tyr | 5997.91748 | 6043.963867 |
| VPS26A | O75436 | VPS26A | Vacuolar protein sorti | 115990.125 | 122644.7188 |
| CDCA5 | B5MBX0;Q96FF9 | CDCA5 | Sororin | 45692.89453 | 31285.04492 |
| IMMT | C9J406 | IMMT | MICOS complex subur | 604764.5234 | 353507.207 |
| TRMT11 | Q7Z4G4 | TRMT11 | tRNA (guanine(10)-N2 | 20100.66211 | 19586.06836 |

| | | | | | |
|---|---|---|---|---|---|
| PNKP | M0QYH2;Q96T60 | PNKP | Bifunctional polynucle | 54341.66016 | 53341.4375 |
| SCUBE2;SCUBE1;SCUBE2 | A0A3B3ISZ7;Q8IWY4;Q9NQ3 | SCUBE2;SCUBE1;SCUE | Signal peptide, CUB ar | 2028.508423 | 2470.502197 |
| CANX | P27824 | CANX | Calnexin | 4107134.25 | 4408108 |
| RPL19 | J3KTE4;J3QR09;P84098 | RPL19 | Ribosomal protein L19 | 1497782.125 | 1144921.875 |
| NDUFA10 | A0A087WXC5;E7ESZ7;O9529 | NDUFA10 | NADH dehydrogenase | 214323.125 | 232507.2344 |
| DSN1 | Q9H410 | DSN1 | Kinetochore-associate | 20255.45508 | 20979.56836 |
| BOD1L1 | Q8NFC6 | BOD1L1 | Biorientation of chron | 50171.04297 | 57257.5625 |
| NCBP1 | Q09161 | NCBP1 | Nuclear cap-binding p | 231007.375 | 238440.1406 |
| DIS3L | Q8TF46 | DIS3L | DIS3-like exonuclease | 4271.323242 | 3243.757568 |
| PCM1 | A0A5H1ZRS1 | PCM1 | Pericentriolar materia | 114959.9063 | 161503.2813 |
| RNF40 | O75150 | RNF40 | E3 ubiquitin-protein li | 89406.44531 | 84487.16406 |
| SRSF6;SRSF6;SRSF6;;SRSF6 | A0A590UJK4;A0A590UJP7;A0 | SRSF6;SRSF6;SRSF6;;S | Serine/arginine-rich-s | 942387.6875 | 703134.9375 |
| RTCA | O00442 | RTCA | RNA 3'-terminal phos | 43106.54688 | 45514.13281 |
| CWC27 | Q6UX04 | CWC27 | Spliceosome-associate | 79162.85938 | 71856.71094 |
| HSP90AA1 | P07900 | HSP90AA1 | Heat shock protein HS | 14768103 | 15061243 |
| C6orf136 | A0A140T9G8;H7C4U4;H7C5B | C6orf136 | Uncharacterized prote | 4400.148438 | 4348.022461 |
| CD2AP | Q9Y5K6 | CD2AP | CD2-associated protei | 101808.2031 | 85373.83594 |
| WARS2 | Q9UGM6 | WARS2 | Tryptophan--tRNA liga | 32653.60938 | 28351.9707 |
| ZNFX1 | Q9P2E3 | ZNFX1 | NFX1-type zinc finger- | 16403.01758 | 16816.23242 |
| RPL18 | J3QQ67;Q07020 | RPL18 | 60S ribosomal protein | 3441580.75 | 2800209.5 |
| KTI12 | Q96EK9 | KTI12 | Protein KTI12 homolo | 25831.76367 | 23897.79688 |
| UBE2D2 | P62837 | UBE2D2 | Ubiquitin-conjugating | 598175.9375 | 615913.3125 |
| RPP38 | P78345 | RPP38 | Ribonuclease P protei | 57681.49609 | 59530.73047 |
| NUP43 | Q8NFH3 | NUP43 | Nucleoporin Nup43 | 56695.20703 | 76807.125 |
| PMS2P1 | A4D2B8 | PMS2P1 | Putative postmeiotic s | 184802.5781 | 190908.125 |
| MARK2 | E9PC69 | MARK2 | Non-specific serine/th | 72644.60156 | 70405.92188 |
| NAXE | Q8NCW5 | NAXE | NAD(P)H-hydrate epir | 644668.0625 | 704757.1875 |
| ZNRD2 | H0YEB6;H0YEK1;O60232 | ZNRD2 | Protein ZNRD2 (Fragm | 13004.53516 | 13942.89844 |
| NDUFB11 | Q9NX14 | NDUFB11 | NADH dehydrogenase | 329629.2813 | 323473.125 |
| TMEM131 | Q92545 | TMEM131 | Transmembrane prote | 9140.229492 | 9777.084961 |
| CIAO2A | Q9H5X1 | CIAO2A | Cytosolic iron-sulfur a | 66183.33594 | 71824.20313 |
| CNOT4 | O95628 | CNOT4 | CCR4-NOT transcriptic | 13852.51172 | 16619.71289 |
| HPF1 | Q9NWY4 | HPF1 | Histone PARylation fa | 43500.98828 | 34666.25781 |

| | | | | | |
|---|---|---|---|---|---|
| RUVBL2 | Q9Y230 | RUVBL2 | RuvB-like 2 | 1051089.5 | 1159217.125 |
| SEC11A | H0YNG3;P67812 | SEC11A | Signal peptidase comp | 225989.2344 | 195542.1094 |
| INTS14 | B4DJL6;B4DVT3;Q96SY0 | INTS14 | Integrator complex su | 40597.68359 | 41070.79688 |
| SRI | C9J0K6;P30626 | SRI | Sorcin | 60192.25 | 102959.5313 |
| RNF20 | Q5VTR2 | RNF20 | E3 ubiquitin-protein li | 840360.9375 | 836318.375 |
| CDAN1 | Q8IWY9 | CDAN1 | Codanin-1 | 11120.13672 | 12709.18848 |
| GFPT1 | Q06210 | GFPT1 | Glutamine--fructose-6 | 363549.3438 | 365988.6875 |
| ERP44 | Q9BS26 | ERP44 | Endoplasmic reticulum | 322843.25 | 307736.875 |
| NUP85 | Q9BW27 | NUP85 | Nuclear pore complex | 114470.3047 | 124516.7969 |
| RANBP2 | P49792 | RANBP2 | E3 SUMO-protein ligas | 153308.8125 | 144391.0938 |
| PDE12 | Q6L8Q7 | PDE12 | 2',5'-phosphodiestera | 110247.0625 | 119229.7109 |
| MOB1B | Q7L9L4 | MOB1B | MOB kinase activator | 84690.71094 | 94864.08594 |
| GTSE1 | Q9NYZ3 | GTSE1 | G2 and S phase-expre | 7852.336914 | 7137.788574 |
| RRP9 | O43818 | RRP9 | U3 small nucleolar RN | 65062.51953 | 38127.55859 |
| GTPBP4 | Q9BZE4 | GTPBP4 | Nucleolar GTP-binding | 276415.4375 | 230595.3125 |
| HMGCR | P04035 | HMGCR | 3-hydroxy-3-methylglu | 7932.247559 | 6785.146484 |
| NFYC | Q13952 | NFYC | Nuclear transcription | 98992.46875 | 96289.79688 |
| TMEM70 | Q9BUB7 | TMEM70 | Transmembrane prote | 69887.69531 | 60245.23828 |
| MT-CO1 | P00395 | MT-CO1 | Cytochrome c oxidase | 44974.22266 | 55535.24219 |
| LAMTOR1 | Q6IAA8 | LAMTOR1 | Ragulator complex pro | 110788.2891 | 100664.9609 |
| TANC1 | Q9C0D5 | TANC1 | Protein TANC1 | 13508.13867 | 17955.17383 |
| LTBP3 | E9PJR2;E9PQR0;E9PR84;H0Y | LTBP3 | Latent-transforming g | 8059.177734 | 5748.349609 |
| MED13 | Q9UHV7 | MED13 | Mediator of RNA poly | 20525.63477 | 22157.55273 |
| MTFR2 | Q6P444 | MTFR2 | Mitochondrial fission | 771.5126953 | 725.5744629 |
| RPS5 | M0R0F0;P46782 | RPS5 | 40S ribosomal protein | 1743036.75 | 1503166.5 |
| KLC1 | G3V3H3;G3V5R9 | KLC1 | Kinesin light chain 1 | 251205.4946 | 246643.4004 |
| WDR26 | A0A499FIZ0 | WDR26 | WD repeat-containing | 88890.89844 | 105399.0469 |
| COA5 | B9A057;Q86WW8 | COA5 | Cytochrome c oxidase | 7356.593262 | 7423.797852 |
| XPO5 | Q9HAV4 | XPO5 | Exportin-5 | 161456.5 | 180331.5938 |
| ACTR10 | Q9NZ32 | ACTR10 | Actin-related protein | 55308.20313 | 60923.9375 |
| MNAT1 | P51948 | MNAT1 | CDK-activating kinase | 38921.09375 | 36501.25 |
| PUF60 | A0A0J9YXX5;Q9UHX1 | PUF60 | Poly(U)-binding-splicir | 932732.2422 | 902995.5859 |
| POR | P16435 | POR | NADPH--cytochrome P | 331883.8438 | 348670.2188 |

| | | | | | |
|---|---|---|---|---|---|
| KHSRP | Q92945 | KHSRP | Far upstream element | 1480766.125 | 1372072 |
| CHERP | Q8IWX8 | CHERP | Calcium homeostasis | 133851.2344 | 128591.9688 |
| SF3B3 | Q15393 | SF3B3 | Splicing factor 3B subu | 783989.6875 | 697358.75 |
| IRF2BP1 | Q8IU81 | IRF2BP1 | Interferon regulatory | 86805.57031 | 76043.21875 |
| CWF19L1 | Q69YN2 | CWF19L1 | CWF19-like protein 1 | 123240.0156 | 138898.1563 |
| MED19 | A0A494C0G4;A0A494C0Y4;A | MED19 | Mediator of RNA poly | 21422.90234 | 23746.69922 |
| PTGES3 | Q15185 | PTGES3 | Prostaglandin E synth | 969560.3125 | 1378414.25 |
| PHACTR4 | Q8IZ21 | PHACTR4 | Phosphatase and actir | 39176.82813 | 45050.46094 |
| DDX6 | P26196 | DDX6 | Probable ATP-depend | 675522 | 744236.9375 |
| CCDC85C | A6NKD9 | CCDC85C | Coiled-coil domain-co | 34544.96875 | 33974.05078 |
| MT-ATP8 | P03928 | MT-ATP8 | ATP synthase protein | 90781.59375 | 125894.9688 |
| RPL17;RPL17-C18orf32;RPL | A0A087WXM6;A0A0A0MRF8 | RPL17;RPL17-C18orf3 | 60S ribosomal protein | 1417387.375 | 1183893.375 |
| MTIF2 | P46199 | MTIF2 | Translation initiation f | 148087.9844 | 138256.4844 |
| INPP5F | A0A3B3IRJ9;A0A3B3ISI3;A0A | INPP5F | Phosphatidylinositide | 235977.7969 | 337259.5 |
| UMPS | P11172 | UMPS | Uridine 5'-monophosp | 340615.5 | 361275.7188 |
| SLC39A1 | Q5T4K2;Q5T4K3;Q5T4K4;Q9I | SLC39A1 | Zinc transporter ZIP1 ( | 11589.12793 | 14182.68262 |
| GPR108 | M0R1G1;Q9NPR9 | GPR108 | Protein GPR108 (Fragi | 20187.0293 | 21138.20313 |
| ZNF787 | Q6DD87 | ZNF787 | Zinc finger protein 787 | 32555.55078 | 30848.85742 |
| KRR1 | Q13601 | KRR1 | KRR1 small subunit pr | 156487.125 | 79118.625 |
| PTMA | B8ZZQ6;P06454 | PTMA | Prothymosin alpha | 105145.7813 | 235505.8906 |
| EIF4ENIF1 | Q9NRA8 | EIF4ENIF1 | Eukaryotic translation | 51108.48047 | 50665.10938 |
| ATP6V1B2 | P21281 | ATP6V1B2 | V-type proton ATPase | 220222.6875 | 202337.6406 |
| TAF1 | P21675 | TAF1 | Transcription initiatio | 24876.32227 | 22203.4668 |
| TSPAN6 | A0A087WYV6;A0A087WZU5; | TSPAN6 | Tetraspanin-6 | 31293.17578 | 28249.11914 |
| CDK9 | P50750 | CDK9 | Cyclin-dependent kin | 156305.6406 | 152663.9688 |
| ELP4 | A0A1W2PNY5;A0A1W2PPP6; | ELP4 | Elongator complex pr | 16552.63672 | 15974.11328 |
| TOB1 | P50616 | TOB1 | Protein Tob1 | 22714.3457 | 23176.09766 |
| UQCRH | P07919 | UQCRH | Cytochrome b-c1 com | 3748.544189 | 5249.96875 |
| TOMM22 | Q9NS69 | TOMM22 | Mitochondrial import | 477950.2813 | 347829.625 |
| EIF2S3 | P41091 | EIF2S3 | Eukaryotic translation | 561610.875 | 558123.75 |
| SAMD1 | Q6SPF0 | SAMD1 | Atherin | 59615.10547 | 65532.54688 |
| MT-ND5 | P03915 | MT-ND5 | NADH-ubiquinone oxi | 26491.20117 | 37095.10547 |
| SRSF2 | J3KP15;Q01130 | SRSF2 | Serine/arginine-rich-s | 1575209.25 | 1366033.875 |

| | | | | | |
|---|---|---|---|---|---|
| SUCLG1 | P53597 | SUCLG1 | Succinate--CoA ligase | 317403.125 | 357947.625 |
| OIP5 | O43482 | OIP5 | Protein Mis18-beta | 14820.57617 | 15457.25391 |
| CSTF1 | Q05048 | CSTF1 | Cleavage stimulation f | 160358.5625 | 155378.2188 |
| BARD1 | A0A087X2H0;C9IYG1;F6MDI1 | BARD1 | BRCA1-associated RIN | 7660.338379 | 4993.021973 |
| CAMSAP1 | Q5T5Y3 | CAMSAP1 | Calmodulin-regulated | 16348.7666 | 17849.21484 |
| AGAP3 | Q96P47 | AGAP3 | Arf-GAP with GTPase, | 23396.87109 | 25234.03516 |
| PIK3R4 | Q99570 | PIK3R4 | Phosphoinositide 3-ki | 35449.12109 | 43462.98828 |
| ATG3 | Q9NT62 | ATG3 | Ubiquitin-like-conjuga | 171533.6563 | 203587.9219 |
| UBE2H | C9JZY6;P62256 | UBE2H | Ubiquitin-conjugating | 41035.32813 | 52771.75781 |
| ALKBH1 | G3V4M4;H0YJF3;Q13686 | ALKBH1 | Nucleic acid dioxygena | 9686.869141 | 12673.69824 |
| RPL13 | P26373 | RPL13 | 60S ribosomal protein | 3142967.75 | 2342811.5 |
| ESD | P10768 | ESD | S-formylglutathione h | 1144311.625 | 1159767.125 |
| EDC3 | Q96F86 | EDC3 | Enhancer of mRNA-de | 124339.4063 | 124129.5469 |
| RBM3 | P98179 | RBM3 | RNA-binding protein 3 | 615902.6875 | 390645.9063 |
| TMED3 | Q9Y3Q3 | TMED3 | Transmembrane emp2 | 70782.96094 | 78286.46875 |
| INTS5 | Q6P9B9 | INTS5 | Integrator complex su | 50314.3125 | 49582.79688 |
| BMPR2 | A0A1W2PPB5;Q13873 | BMPR2 | Serine/threonine-prot | NA | 2961.900635 |
| FNTB | P49356 | FNTB | Protein farnesyltransf | 22264.73438 | 24230.86328 |
| PROSER1 | Q5JS36;Q86XN7 | PROSER1 | Proline and serine-rich | 20013.09766 | 21308.79688 |
| TMED1 | Q13445 | TMED1 | Transmembrane emp2 | 97705.94531 | 94106.50781 |
| FKBP9 | O95302 | FKBP9 | Peptidyl-prolyl cis-tran | 67701.25781 | 90275.23438 |
| ZRANB2 | O95218 | ZRANB2 | Zinc finger Ran-bindin | 132562.7188 | 123427.4297 |
| RBBP6 | Q7Z6E9 | RBBP6 | E3 ubiquitin-protein li | 27989.55273 | 24352.74023 |
| DDX46 | A0A0C4DG89 | DDX46 | Probable ATP-depend | 605007.1719 | 602919.5469 |
| DOK1 | Q99704 | DOK1 | Docking protein 1 | NA | 9335.494141 |
| VPS4B | O75351 | VPS4B | Vacuolar protein sorti | 510205.6875 | 521040.2188 |
| ILF2 | B4DY09;Q12905 | ILF2 | Interleukin enhancer- | 838375 | 620492.1875 |
| KDM1B | H0Y6H0;Q8NB78 | KDM1B | Lysine-specific histone | 14754.23926 | 17194.98242 |
| NXF1 | Q9UBU9 | NXF1 | Nuclear RNA export fa | 192919.125 | 194697.5625 |
| CHRAC1 | Q9NRG0 | CHRAC1 | Chromatin accessibilit | 67439.60156 | 58477.19922 |
| NSMCE4A | Q9NXX6 | NSMCE4A | Non-structural mainte | 11917.4248 | 10721.39746 |
| RAE1 | P78406 | RAE1 | mRNA export factor | 245170.1563 | 238070.4844 |
| INTS4 | Q96HW7 | INTS4 | Integrator complex su | 57942.14063 | 63267.38672 |

| | | | | | |
|---|---|---|---|---|---|
| BOP1 | Q14137 | BOP1 | Ribosome biogenesis | 121577.1328 | 101462.4453 |
| JPT2 | H3BMT0 | JPT2 | Jupiter microtubule-as | 1201430.395 | 1449659.133 |
| NDC1 | Q9BTX1 | NDC1 | Nucleoporin NDC1 | 33823.32031 | 33208.04688 |
| DDX24 | G3V529;Q9GZR7 | DDX24 | ATP-dependent RNA h | 138182.3438 | 112354.4297 |
| MT-ND1 | P03886 | MT-ND1 | NADH-ubiquinone oxi | 73132.82031 | 76268.94531 |
| RCOR3 | Q9P2K3 | RCOR3 | REST corepressor 3 | 92677.17969 | 98141.46875 |
| EP400 | Q96L91 | EP400 | E1A-binding protein p | 36314.52734 | 29537.2793 |
| ATP5PD | O75947 | ATP5PD | ATP synthase subunit | 225246.2344 | 235629.1094 |
| EPG5 | Q9HCE0 | EPG5 | Ectopic P granules prc | 4061.826172 | 5744.760742 |
| MED25 | M0QZQ2;Q71SY5 | MED25 | Mediator of RNA poly | 15928.16016 | 17315.66211 |
| BID | P55957 | BID | BH3-interacting doma | 50461.16797 | 53569.20313 |
| GTPBP1 | O00178 | GTPBP1 | GTP-binding protein 1 | 65049.23438 | 66776.54688 |
| NOL7 | Q9UMY1 | NOL7 | Nucleolar protein 7 | 32398.82813 | 17843.28906 |
| PREPL | Q4J6C6 | PREPL | Prolyl endopeptidase- | 79251.69531 | 83249.0625 |
| MTHFSD | A0A087WX22 | MTHFSD | Methenyltetrahydrofc | 32865.3291 | 29153.87207 |
| NDUFB7 | P17568 | NDUFB7 | NADH dehydrogenase | 78373.10938 | 86326.75781 |
| BTN2A1 | Q7KYR7 | BTN2A1 | Butyrophilin subfamily | 10139.9082 | 11470.61328 |
| INTS8 | Q75QN2 | INTS8 | Integrator complex su | 33875.72656 | 35244.86719 |
| HK2 | P52789 | HK2 | Hexokinase-2 | 71173.02344 | 42754.17578 |
| TMEM168 | H7C268;Q9H0V1 | TMEM168 | Transmembrane prote | 8112.216797 | 12914.31738 |
| YTHDF2 | Q9Y5A9 | YTHDF2 | YTH domain-containir | 141927.2188 | 136948.8594 |
| PABPC4 | B1ANR0;Q13310 | PABPC4 | Polyadenylate-binding | 331688.1875 | 340436.7188 |
| NDUFC2 | O95298 | NDUFC2 | NADH dehydrogenase | 147114.6719 | 151460.1563 |
| NDE1 | Q9NXR1 | NDE1 | Nuclear distribution p | 25691.75195 | 34047.74219 |
| PPP1R14B | Q96C90 | PPP1R14B | Protein phosphatase 1 | 240968.5938 | 244988.7031 |
| RPL23A | P62750 | RPL23A | 60S ribosomal protein | 2901736.5 | 2407970.25 |
| EIF4G2 | H0Y3P2 | EIF4G2 | Eukaryotic translation | 423405.666 | 479399.6055 |
| TRIP10 | Q15642 | TRIP10 | Cdc42-interacting pro | 32061.51172 | 32328.62695 |
| PCBP1 | Q15365 | PCBP1 | Poly(rC)-binding prote | 2234545 | 2198004.25 |
| BOLA2 | Q9H3K6 | BOLA2 | BolA-like protein 2 | 265769.5313 | 381725.7188 |
| LLGL1 | Q15334 | LLGL1 | Lethal(2) giant larvae | 84514.66406 | 95343.00781 |
| PMS2 | P54278 | PMS2 | Mismatch repair endc | 43199.86328 | 47531.12109 |
| PWP1 | Q13610 | PWP1 | Periodic tryptophan p | 118073.8516 | 114559.8359 |

| | | | | | |
|---|---|---|---|---|---|
| SUMO1 | B8ZZN6;P63165 | SUMO1 | Small ubiquitin-relate | 40401.15234 | 49419.03516 |
| MMP21 | Q8N119 | MMP21 | Matrix metalloprotein | 23633.25195 | 31341.69727 |
| ASXL2 | E7EWD6;Q76L83 | ASXL2 | Putative Polycomb gro | 18526.5332 | 17782.38477 |
| IPO7 | O95373 | IPO7 | Importin-7 | 406778.4063 | 459878.4063 |
| USP32 | Q8NFA0 | USP32 | Ubiquitin carboxyl-ter | 149187.5156 | 174905.2969 |
| KCTD9 | Q7L273 | KCTD9 | BTB/POZ domain-cont | 14901.17773 | 13421.99902 |
| LRP8 | A0A590UJ39;A0A590UJ57;A0 | LRP8 | Low-density lipoprote | 9503.850586 | 6745.842773 |
| TMCC1 | O94876 | TMCC1 | Transmembrane and | 7257.033691 | 6675.880859 |
| PPP6R3 | E9PKF6;H7BXH2;Q5H9R7 | PPP6R3 | Serine/threonine-prot | 185108.6875 | 196803.0156 |
| BLMH | Q13867 | BLMH | Bleomycin hydrolase | 264109.5625 | 283379.0625 |
| ACOT13 | Q9NPJ3 | ACOT13 | Acyl-coenzyme A thio | 111340.6484 | 136031.5313 |
| MED7 | O43513 | MED7 | Mediator of RNA poly | 2793.253174 | 2853.197754 |
| RPL31 | P62899 | RPL31 | 60S ribosomal protein | 1960852.375 | 1556186.125 |
| EIF3I | Q13347 | EIF3I | Eukaryotic translation | 614223.0625 | 647350.375 |
| VCL | P18206 | VCL | Vinculin | 784305.1875 | 663934.3125 |
| CMC2 | Q9NRP2 | CMC2 | COX assembly mitoch | 191203 | 212102.4844 |
| EIF4A3 | P38919 | EIF4A3 | Eukaryotic initiation f | 1068553.375 | 914878.3125 |
| AFG3L2 | Q9Y4W6 | AFG3L2 | AFG3-like protein 2 | 204283.4844 | 209757.5938 |
| SAMD4B | A0A087WZT0;M0QZ22;Q5PR | SAMD4B | Protein Smaug homol | 19436.66797 | 20691.48438 |
| CALR | P27797 | CALR | Calreticulin | 1277844.625 | 1640971.375 |
| MRTFB | Q9ULH7 | MRTFB | Myocardin-related tra | 34714.67578 | 56733.96484 |
| DUS2 | Q9NX74 | DUS2 | tRNA-dihydrouridine( | 73022.85156 | 77013.35156 |
| DCTN5 | Q9BTE1 | DCTN5 | Dynactin subunit 5 | 31928.36914 | 33723.54297 |
| SEC24A | O95486 | SEC24A | Protein transport prot | 85426.53125 | 94259.83594 |
| AK4 | P27144 | AK4 | Adenylate kinase 4, m | 272356.25 | 298187.8125 |
| TIAL1 | Q01085 | TIAL1 | Nucleolysin TIAR | 210728.7656 | 176790.8438 |
| RNF126 | K7EIY6;Q9BV68 | RNF126 | E3 ubiquitin-protein li | 54581.125 | 47474.1875 |
| SNRPGP15;SNRPG | A8MWD9;P62308 | SNRPGP15;SNRPG | Putative small nuclear | 438358.0625 | 321152.875 |
| NEPRO | Q6NW34 | NEPRO | Nucleolus and neural | 18479.70898 | 19303.44922 |
| GOLIM4 | O00461 | GOLIM4 | Golgi integral membra | 18806.83984 | 22102.38086 |
| NUP160 | Q12769 | NUP160 | Nuclear pore complex | 121042.1328 | 126197.8828 |
| LARS2 | Q15031 | LARS2 | Probable leucine--tRN | 37853.46484 | 41796.84375 |
| SH3BP5 | O60239 | SH3BP5 | SH3 domain-binding p | 6155.575195 | 6008.618164 |

| | | | | | |
|---|---|---|---|---|---|
| BCS1L | Q9Y276 | BCS1L | Mitochondrial chaper | 34160.11719 | 37556.96875 |
| NACA | E9PAV3;F8VZJ2;F8W0W4;H0 | NACA | Nascent polypeptide-a | 466320.5313 | 403103.375 |
| BTBD1 | Q9H0C5 | BTBD1 | BTB/POZ domain-cont | 11771.07031 | 8165.714844 |
| PEAK1 | H0YN99;Q9H792 | PEAK1 | Inactive tyrosine-prot | 2776.911133 | 3962.54126 |
| TMEM160 | Q9NX00 | TMEM160 | Transmembrane prot | 54166.11719 | 56659.21484 |
| NDUFV1 | G3V0I5;P49821 | NDUFV1 | NADH dehydrogenase | 302772.5313 | 337895.375 |
| ARHGEF26 | Q96DR7 | ARHGEF26 | Rho guanine nucleotic | 10611.20898 | 8220.146484 |
| DAPK1 | P53355 | DAPK1 | Death-associated prot | 58780.25 | 61387.74609 |
| CPSF3 | Q9UKF6 | CPSF3 | Cleavage and polyade | 178178.6094 | 191410.1094 |
| TNK1 | Q13470 | TNK1 | Non-receptor tyrosine | 12614.43457 | 12650.18457 |
| PIAS1 | O75925 | PIAS1 | E3 SUMO-protein liga | 14427.89453 | 16225.05566 |
| EIF4B | E7EX17;P23588 | EIF4B | Eukaryotic translation | 1276696.125 | 1169926.75 |
| SAP18 | O00422 | SAP18 | Histone deacetylase c | 425852 | 374020.3438 |
| AKAP8L | Q9ULX6 | AKAP8L | A-kinase anchor prote | 81149.65625 | 82811.39063 |
| EFTUD2 | Q15029 | EFTUD2 | 116 kDa U5 small nucl | 2072669.625 | 1899181 |
| RNF5 | A0A140TA09;Q99942 | RNF5 | E3 ubiquitin-protein li | 5906.33252 | 9903.646484 |
| MYADM | A0A494C017;A0A499FIY8;C9 | MYADM | Myeloid-associated di | 66032.36719 | 56341.75 |
| RUVBL1 | Q9Y265 | RUVBL1 | RuvB-like 1 | 998324.5625 | 1025583.063 |
| TTC12 | J3KR69;Q9H892 | TTC12 | Tetratricopeptide rep | 2087.248535 | 3008.797119 |
| WASHC3 | F5GWI9;F5GXZ7;F5GZ97;Q9Y | WASHC3 | WASH complex subun | 10082.03418 | 15014.79102 |
| TYMS | P04818 | TYMS | Thymidylate synthase | 122421.7891 | 115974.5781 |
| MRPS21 | P82921 | MRPS21 | 28S ribosomal protein | 104714.4063 | 129975.5391 |
| GK | P32189 | GK | Glycerol kinase | 16366.0127 | 8882.912109 |
| IBA57 | Q5T440 | IBA57 | Putative transferase C | 36104.97656 | 38158.69141 |
| SLK | Q9H2G2 | SLK | STE20-like serine/thre | 76473.78125 | 78310.96875 |
| CDC40 | O60508 | CDC40 | Pre-mRNA-processing | 52816.69141 | 39041.72266 |
| ADO | Q96SZ5 | ADO | 2-aminoethanethiol d | 47347.29297 | 54693.39453 |
| YTHDF3 | A0A087X0Q1;Q7Z739 | YTHDF3 | YTH domain-containir | 37328.06641 | 39198.33984 |
| MDH1 | P40925 | MDH1 | Malate dehydrogenas | 2724338.5 | 3027909.5 |
| RCOR1 | Q9UKL0 | RCOR1 | REST corepressor 1 | 80773.95313 | 79616.60156 |
| SARS1 | P49591 | SARS1 | Serine--tRNA ligase, cy | 1582590.375 | 1364066 |
| DCP1B | Q8IZD4 | DCP1B | mRNA-decapping enzy | 8122.533691 | 7707.302246 |
| TVP23C;TVP23B;TVP23C;TV | C9JYS5;J3KS67;J3QL47;J3QL6 | TVP23C;TVP23B;TVP2 | Golgi apparatus meml | 15217.82129 | 16516.15625 |

| | | | | | |
|---|---|---|---|---|---|
| ZNF439 | A0A0C4DG37;Q8NDP4 | ZNF439 | Zinc finger protein 43! | 28632.71094 | 25747.93359 |
| HSPD1 | P10809 | HSPD1 | 60 kDa heat shock pro | 4501242 | 4849136.5 |
| SLC7A1 | P30825 | SLC7A1 | High affinity cationic a | 314217.5313 | 328975.4688 |
| XRN2 | Q9H0D6 | XRN2 | 5'-3' exoribonuclease | 263560.7188 | 263083.7813 |
| DHX38 | Q92620 | DHX38 | Pre-mRNA-splicing fac | 74939.60938 | 71123.71094 |
| SHKBP1 | M0R2P6;Q8TBC3 | SHKBP1 | SH3KBP1 binding prot | 13868.19629 | 16568.875 |
| ZNF217 | O75362 | ZNF217 | Zinc finger protein 21 | 60480.35156 | 52597.73828 |
| PDLIM5 | Q96HC4 | PDLIM5 | PDZ and LIM domain p | 226271.875 | 245172.0938 |
| POGK | Q9P215 | POGK | Pogo transposable ele | 224617.75 | 186318.8594 |
| MED30 | Q96HR3 | MED30 | Mediator of RNA poly | 43413.48828 | 43301.55078 |
| PUS10 | Q3MIT2 | PUS10 | Putative tRNA pseudo | 3363.067627 | 2018.933228 |
| TBC1D4 | O60343 | TBC1D4 | TBC1 domain family m | 57146.51172 | 60238.40234 |
| NDUFS2 | O75306 | NDUFS2 | NADH dehydrogenase | 254909.3906 | 301486.3438 |
| NUP155 | E9PF10;O75694 | NUP155 | Nuclear pore complex | 741995.625 | 796888.0625 |
| DYM | Q7RTS9 | DYM | Dymeclin | 23819.41016 | 30650.23633 |
| AKAP8 | O43823 | AKAP8 | A-kinase anchor prote | 47378.52734 | 47707.24609 |
| HAUS4 | Q9H6D7 | HAUS4 | HAUS augmin-like con | 35916.42969 | 38666.67578 |
| PNP | P00491 | PNP | Purine nucleoside pho | 466233.5625 | 558945.1875 |
| KDELR2 | P33947 | KDELR2 | ER lumen protein-reta | 9996.797852 | 13733.90332 |
| RPL36A | J3KQN4;P83881 | RPL36A | 60S ribosomal protein | 1177321.625 | 1214324.875 |
| URGCP | Q8TCY9 | URGCP | Up-regulator of cell pr | 11430.50977 | 12193.18164 |
| RPL35 | P42766 | RPL35 | 60S ribosomal protein | 1313732.625 | 1135206.375 |
| MAP2K3 | P46734 | MAP2K3 | Dual specificity mitoge | 92985.60156 | 73798.28906 |
| SNX3 | O60493 | SNX3 | Sorting nexin-3 | 489612.2813 | 507718.8438 |
| ITPA | Q9BY32 | ITPA | Inosine triphosphate p | 97883.10938 | 114921.7734 |
| BOD1 | Q96IK1 | BOD1 | Biorientation of chron | 52366.62109 | 51930.60547 |
| RAB3A | P20336 | RAB3A | Ras-related protein Ra | 15055.65723 | 14809.74219 |
| PM20D2 | Q8IYS1 | PM20D2 | Peptidase M20 domai | 42143.73047 | 38145.99609 |
| RHOT2 | Q8IXI1 | RHOT2 | Mitochondrial Rho GT | 142132.0313 | 149685.4063 |
| PKP2 | Q99959 | PKP2 | Plakophilin-2 | 33629.64063 | 37210.01563 |
| HDHD5 | Q9BXW7 | HDHD5 | Haloacid dehalogenas | 155919.6719 | 163884.1875 |
| KCMF1 | Q9P0J7 | KCMF1 | E3 ubiquitin-protein li | 13905.57813 | 13915.29395 |
| INTS10 | Q9NVR2 | INTS10 | Integrator complex su | 43898.65625 | 47198.47266 |

| | | | | | |
|---|---|---|---|---|---|
| BCL9L | A0A087WZX0;Q86UU0 | BCL9L | B-cell CLL/lymphoma 9 | 28860.64063 | 32613.63086 |
| FTH1 | P02794 | FTH1 | Ferritin heavy chain | 7014.016113 | 10035.26953 |
| ANKS1A | A0A3B3ISP1;Q92625 | ANKS1A | Ankyrin repeat and SA | 28843.60156 | 28778.58594 |
| UPF2 | Q9HAU5 | UPF2 | Regulator of nonsense | 81670.67969 | 85903.23438 |
| ETFDH | Q16134 | ETFDH | Electron transfer flavo | 30597.89063 | 30703.87109 |
| TIA1 | F8W8I6;P31483 | TIA1 | Nucleolysin TIA-1 isofo | 290609.9688 | 253102.6875 |
| CAT | P04040 | CAT | Catalase | 162936.0938 | 171157.1563 |
| LARS1 | Q9P2J5 | LARS1 | Leucine--tRNA ligase, | 409886.5313 | 470531.7813 |
| MACIR | Q96GV9 | MACIR | Macrophage immuno | 8360.939453 | 8989.889648 |
| UBE2Q1 | Q7Z7E8 | UBE2Q1 | Ubiquitin-conjugating | 111354.7578 | 126728.4531 |
| CLSPN | E7ESG2;Q9HAW4 | CLSPN | Claspin | 6206.072266 | 4338.489746 |
| ENOSF1 | Q7L5Y1 | ENOSF1 | Mitochondrial enolase | 20290.76758 | 25776.86328 |
| ZSWIM8 | A7E2V4;S4R393;S4R3H3;S4R4 | ZSWIM8 | Zinc finger SWIM dom | 19052.32227 | 21884.72461 |
| PCNP | Q8WW12 | PCNP | PEST proteolytic signa | 262918.5625 | 279714.2813 |
| TIMM44 | O43615 | TIMM44 | Mitochondrial import | 170233.5781 | 169867.7656 |
| ETNK1 | Q9HBU6 | ETNK1 | Ethanolamine kinase 1 | 179926 | 113756.8359 |
| MT-CYB | P00156 | MT-CYB | Cytochrome b | 35927.59766 | 39564.90625 |
| SCAMP3 | O14828 | SCAMP3 | Secretory carrier-asso | 249679.9688 | 282965.8125 |
| KDM1A | O60341 | KDM1A | Lysine-specific histone | 148032.1875 | 152844.7813 |
| DBT | P11182 | DBT | Lipoamide acyltransfe | 54125.75391 | 67827 |
| TYSND1 | Q2T9J0 | TYSND1 | Peroxisomal leader pe | 37911.30469 | 39306.00781 |
| ATR | A0A590UJ01;Q13535 | ATR | Serine/threonine-prot | 7632.549805 | 9671.322266 |
| MAN2A1 | Q16706 | MAN2A1 | Alpha-mannosidase 2 | 73120.39063 | 81962.53125 |
| DYNLT3 | A6NGJ0;P51808 | DYNLT3 | Dynein light chain Tct | 18102.41016 | 17285.60352 |
| MCAT | Q8IVS2 | MCAT | Malonyl-CoA-acyl carr | 45864.04688 | 43879.28516 |
| B4GAT1 | O43505 | B4GAT1 | Beta-1,4-glucuronyltra | 93682.26563 | 81176.46875 |
| RREB1 | Q92766 | RREB1 | Ras-responsive eleme | 38717.65625 | 42062.14844 |
| SF3B1 | O75533 | SF3B1 | Splicing factor 3B subu | 983099.75 | 746070.9375 |
| BCL7B | F2Z3H6 | BCL7B | B-cell CLL/lymphoma 7 | 18529.38281 | 16024.95605 |
| FAU | P62861 | FAU | 40S ribosomal protein | 1509214.625 | 1334199.25 |
| RNF25 | Q96BH1 | RNF25 | E3 ubiquitin-protein li | 40230.60938 | 46379.83984 |
| DCTN4 | Q9UJW0 | DCTN4 | Dynactin subunit 4 | 93087.42969 | 102800.1406 |
| DAGLB | Q8NCG7 | DAGLB | Sn1-specific diacylglyc | 62352.64453 | 64575.6875 |

| | | | | | |
|---|---|---|---|---|---|
| DEGS2 | Q6QHC5 | DEGS2 | Sphingolipid delta(4)-( | 6360.956543 | 3116.756592 |
| CAMSAP2 | Q08AD1 | CAMSAP2 | Calmodulin-regulated | 31919.65234 | 27497.0332 |
| DROSHA | Q9NRR4 | DROSHA | Ribonuclease 3 | 21232.67773 | 20882.36328 |
| GSPT2 | Q8IYD1 | GSPT2 | Eukaryotic peptide ch | 588010.75 | 831885.8125 |
| EDRF1 | Q3B7T1 | EDRF1 | Erythroid differentiati | 12363.90625 | 20294.375 |
| SNAP29 | O95721 | SNAP29 | Synaptosomal-associa | 160392.2031 | 169459.7969 |
| UTP23 | Q9BRU9 | UTP23 | rRNA-processing prot( | 24690.64648 | 14523.65918 |
| CSE1L | P55060 | CSE1L | Exportin-2 | 2010105.25 | 2300886.25 |
| TTF1 | Q15361 | TTF1 | Transcription termina | 2145.495117 | 1371.583496 |
| SPTY2D1 | Q68D10 | SPTY2D1 | Protein SPT2 homolog | 46534.57813 | 41786.60938 |
| PLXDC2 | Q6UX71 | PLXDC2 | Plexin domain-contair | 9258.733398 | 10093.65137 |
| RAB1A | P62820 | RAB1A | Ras-related protein Ra | 465284.9063 | 489975.7813 |
| CCT8 | P50990 | CCT8 | T-complex protein 1 s | 1988707.75 | 2187082.25 |
| SDE2 | Q6IQ49 | SDE2 | Replication stress resp | 23282.41602 | 18920.93945 |
| WDR81 | A0A0J9YXM6;Q562E7 | WDR81 | WD repeat-containing | 57817.37109 | 49567.21094 |
| RANBP9 | Q96S59 | RANBP9 | Ran-binding protein 9 | 77649.25781 | 98083.96094 |
| RBM8A | Q9Y5S9 | RBM8A | RNA-binding protein 8 | 682213.25 | 602834.8125 |
| COMMD1 | Q8N668 | COMMD1 | COMM domain-conta | 88078.95313 | 94766.60938 |
| CRAT | P43155 | CRAT | Carnitine O-acetyltrar | 986.1401367 | NA |
| NEDD1 | Q8NHV4 | NEDD1 | Protein NEDD1 | 27419.05078 | 29787.42188 |
| MRPS15 | P82914 | MRPS15 | 28S ribosomal protein | 137843.8906 | 154269.5625 |
| PFDN2 | Q9UHV9 | PFDN2 | Prefoldin subunit 2 | 404021.25 | 407085.1563 |
| RNPC3 | Q96LT9;X6RLP6 | RNPC3 | RNA-binding region-co | 12780.96387 | 10677.0791 |
| TDP2 | O95551 | TDP2 | Tyrosyl-DNA phospho | 16216.78418 | 11953.00977 |
| RBM7;RBM7;;RBM7;RBM7 | F5GXV8;F5GY08;I3L521;J3KP | RBM7;RBM7;;RBM7;R | RNA-binding protein 7 | 12614.81641 | 11726.06055 |
| USP10 | Q14694 | USP10 | Ubiquitin carboxyl-ter | 296570.9688 | 304674.8438 |
| TTK | P33981 | TTK | Dual specificity protei | 17616.21484 | 22743.60156 |
| CD46 | P15529 | CD46 | Membrane cofactor p | 41900.00488 | 50493.1084 |
| CCNH | A0A2R8YEM2;P51946 | CCNH | Cyclin-H | 33001.38281 | 32129.46484 |
| POLD3 | Q15054 | POLD3 | DNA polymerase delta | 71423.51563 | 60235.75 |
| SRSF10 | O75494;Q5JRI1 | SRSF10 | Serine/arginine-rich sp | 181200.4688 | 93823.60156 |
| MAP2K4 | P45985 | MAP2K4 | Dual specificity mitoge | 44143.98438 | 42255.09375 |
| MCU | Q8NE86 | MCU | Calcium uniporter pro | 27728.44336 | 29793.52344 |

| | | | | | |
|---|---|---|---|---|---|
| MSMO1 | Q15800 | MSMO1 | Methylsterol monoox | 55326.93359 | 70806.59375 |
| SNRNP40 | Q96DI7 | SNRNP40 | U5 small nuclear ribor | 193837.7969 | 171288.1719 |
| SEC24C | P53992 | SEC24C | Protein transport prot | 298985.5313 | 322122.6563 |
| APEH | C9JIF9;P13798 | APEH | Acylamino-acid-releas | 382222.9063 | 447842.4688 |
| TOX4 | O94842 | TOX4 | TOX high mobility gro | 32341.33984 | 58814.78906 |
| NEURL4 | I3L100;I3L2W2;I3L2Z9;Q96JN | NEURL4 | Neuralized-like protei | 5906.23877 | 6351.595703 |
| NSRP1 | A0A024QZ33;Q9H0G5 | NSRP1 | Coiled-coil domain co | 4590.609863 | 3986.54126 |
| FIP1L1 | Q6UN15 | FIP1L1 | Pre-mRNA 3'-end-pro | 197067.8125 | 193036.0313 |
| TOM1L1 | I3L4I4;J3KQU4;O75674 | TOM1L1 | TOM1-like protein 1 | 61531.28906 | 78497.48438 |
| NDUFA12 | Q9UI09 | NDUFA12 | NADH dehydrogenase | 185676.625 | 195726.2188 |
| MBD1 | Q9UIS9 | MBD1 | Methyl-CpG-binding d | 8583.124023 | 13284.87695 |
| RABIF | P47224 | RABIF | Guanine nucleotide ex | 571968.3125 | 591181.625 |
| STAU1 | O95793 | STAU1 | Double-stranded RNA | 347189.4375 | 299254.4688 |
| OLA1 | J3KQ32;Q9NTK5 | OLA1 | Obg-like ATPase 1 | 785774.1875 | 818666.3125 |
| SMAD2 | Q15796 | SMAD2 | Mothers against deca | 111617.8047 | 110976.8047 |
| PYROXD2 | Q8N2H3 | PYROXD2 | Pyridine nucleotide-di | 21399.24414 | 38488.5 |
| OGA | O60502 | OGA | Protein O-GlcNAcase | 165849.9688 | 183364.6406 |
| EIF3J | O75822 | EIF3J | Eukaryotic translation | 603937.125 | 625898.4375 |
| CELF1 | G5EA30 | CELF1 | CUG triplet repeat, RN | 281950.9375 | 303043.125 |
| NOMO2 | Q5JPE7 | NOMO2 | Nodal modulator 2 | 228562.1875 | 231518.2656 |
| PFDN6 | O15212 | PFDN6 | Prefoldin subunit 6 | 221600.0625 | 226669.2969 |
| SCAF1 | Q9H7N4 | SCAF1 | Splicing factor, argini | 30876.29492 | 33700.39844 |
| IRGQ | Q8WZA9 | IRGQ | Immunity-related GTP | 98458.02344 | 102964.9922 |
| EIF3G | K7ER90 | EIF3G | Eukaryotic translation | 495325.3672 | 478568.75 |
| CNOT8 | E5RIL2;Q9UFF9 | CNOT8 | CCR4-NOT transcriptic | 13808.62402 | 15311.47168 |
| SMARCD2 | B9EGA3;J3KMX2;Q92925 | SMARCD2 | SMARCD2 protein;SW | 59321.89453 | 61464.31641 |
| MRPS11 | P82912 | MRPS11 | 28S ribosomal protein | 12524.76563 | 15679.91309 |
| CCDC9 | A0A2R8Y4Z8 | CCDC9 | Coiled-coil domain-co | 108164.3984 | 116227.0195 |
| ELOVL5 | A0A0A0MTI6;Q9NYP7 | ELOVL5 | Elongation of very lon | 84978.39844 | 114637.0391 |
| R3HCC1L | A0A384DVK4;Q7Z5L2 | R3HCC1L | Coiled-coil domain-co | 4098.061523 | 8981.598633 |
| KHDC4 | Q7Z7F0 | KHDC4 | KH homology domain- | 14869.17969 | 16086.33984 |
| RRM2 | P31350 | RRM2 | Ribonucleoside-dipho | 536791.4375 | 435091.2813 |
| TUFM | P49411 | TUFM | Elongation factor Tu, r | 1093199.25 | 1287949.75 |

| | | | | | |
|---|---|---|---|---|---|
| RBBP7 | Q16576 | RBBP7 | Histone-binding prote | 506219.1875 | 780176.75 |
| PISD | Q9UG56 | PISD | Phosphatidylserine de | 158399.0313 | 154240.0156 |
| CCDC51 | Q96ER9 | CCDC51 | Mitochondrial potassi | 30520.27148 | 32710.40625 |
| KLHDC10 | Q6PID8 | KLHDC10 | Kelch domain-contain | 18304.36523 | 10636.12695 |
| ANAPC1 | Q9H1A4 | ANAPC1 | Anaphase-promoting | 55112.83984 | 63317.39844 |
| GPATCH11 | Q8N954 | GPATCH11 | G patch domain-conta | 7751.189941 | 7668.187012 |
| APPBP2 | Q92624 | APPBP2 | Amyloid protein-bindi | 130670.5313 | 157296.4063 |
| CKMT1A | P12532 | CKMT1A | Creatine kinase U-typ | 981979.9375 | 873314.6875 |
| UBE2Z | Q9H832 | UBE2Z | Ubiquitin-conjugating | 366970.6563 | 357448.9375 |
| RNF114 | Q9Y508 | RNF114 | E3 ubiquitin-protein li | 378447.1875 | 387124.5313 |
| PAICS | P22234 | PAICS | Multifunctional protei | 1523522.75 | 1786553.25 |
| TIMM23;TIMM23B | O14925;Q5SRD1 | TIMM23;TIMM23B | Mitochondrial import | 111744.2734 | 122791.5313 |
| AAAS | Q9NRG9 | AAAS | Aladin | 118559.2031 | 113628.1172 |
| PRKD2 | Q9BZL6 | PRKD2 | Serine/threonine-prot | 50770.17969 | 48644.79688 |
| ICE2 | H0YK97;H0YNU9;Q659A1 | ICE2 | Little elongation comp | 2460.428223 | 7126.885742 |
| CHD1L | Q86WJ1 | CHD1L | Chromodomain-helica | 118745.4922 | 134946.2813 |
| PHF10 | Q8WUB8 | PHF10 | PHD finger protein 10 | 11286.47949 | 9612.856445 |
| ACSF3 | Q4G176 | ACSF3 | Malonate--CoA ligase | 78073.28125 | 84479.65625 |
| THG1L | Q9NWX6 | THG1L | Probable tRNA(His) gu | 30284.42188 | 35370.76563 |
| QSER1 | A0A3B3IRW6;Q2KHR3 | QSER1 | Glutamine and serine- | 8888.165039 | 11193.88574 |
| UBE2D1;UBE2D4;UBE2D1;U | A0A087WW00;C9J9H9;P516 | UBE2D1;UBE2D4;UBE | Ubiquitin-conjugating | 73054.98438 | 72795.01563 |
| ADK | P55263 | ADK | Adenosine kinase | 474300.6875 | 550750.9375 |
| POLR2L | P62875 | POLR2L | DNA-directed RNA po | 52510.82031 | 39750.82422 |
| TMED4 | Q7Z7H5 | TMED4 | Transmembrane emp | 191375.6563 | 187573.6094 |
| TIMM9 | Q9Y5J7 | TIMM9 | Mitochondrial import | 29008.3457 | 38455.40625 |
| NDUFB9 | Q9Y6M9 | NDUFB9 | NADH dehydrogenase | 195695.0625 | 201995.2031 |
| STX3 | A0A0C4DGE4;A0A0J9YW33;A | STX3 | Syntaxin-3 (Fragment) | 37504.74219 | 32552.41406 |
| APBB2 | G5E9Y1;H0YAJ5;Q92870 | APBB2 | Amyloid beta (A4) pro | 31247.03125 | 38333.22656 |
| NCDN | C9J5H8 | NCDN | Neurochondrin (Fragn | 155644.7891 | 160349.4941 |
| NDUFS5 | O43920 | NDUFS5 | NADH dehydrogenase | 196825.5469 | 197358.2188 |
| NCAPD2 | Q15021 | NCAPD2 | Condensin complex su | 118247.2734 | 125909.1406 |
| SYNRG | Q9UMZ2 | SYNRG | Synergin gamma | 16660.80273 | 25363.35547 |
| FCHO2 | H7BXJ4;Q0JRZ9 | FCHO2 | F-BAR domain only pr | 20051.11914 | 18154.92969 |

| | | | | | |
|---|---|---|---|---|---|
| NDUFS1 | P28331 | NDUFS1 | NADH-ubiquinone oxi | 268628.4688 | 302811.6875 |
| ITSN1 | Q15811 | ITSN1 | Intersectin-1 | 9284.43457 | 8040.224121 |
| SNF8 | Q96H20 | SNF8 | Vacuolar-sorting prote | 98698.84375 | 106806.1641 |
| GPATCH1 | Q9BRR8 | GPATCH1 | G patch domain-conta | 82736.17188 | 77450.98438 |
| REXO1 | Q8N1G1 | REXO1 | RNA exonuclease 1 hc | 1266.426514 | 1239.232666 |
| FADD | Q13158 | FADD | FAS-associated death | 171674.7188 | 156888.2031 |
| SUZ12 | J3QQW9;Q15022 | SUZ12 | Polycomb protein SUZ | 29363.38086 | 29021.23633 |
| ANKZF1 | Q9H8Y5 | ANKZF1 | Ankyrin repeat and zir | 42126.08594 | 45229.83594 |
| PDHA1 | P08559 | PDHA1 | Pyruvate dehydrogena | 607298.25 | 649137.6875 |
| GTF3C4 | Q9UKN8 | GTF3C4 | General transcription | 80677.13281 | 79640.86719 |
| WDR33 | Q9C0J8 | WDR33 | pre-mRNA 3' end proc | 75827.8125 | 79504.55469 |
| IFT43 | G3V4X2;Q96FT9 | IFT43 | Intraflagellar transpor | 973.8365479 | 11554.60352 |
| MRTO4 | Q9UKD2 | MRTO4 | mRNA turnover prote | 238003.6875 | 173801.8438 |
| BOLA3 | Q53S33 | BOLA3 | BolA-like protein 3 | 63188.08984 | 64233.02734 |
| CDC37L1 | B1AL69;Q7L3B6 | CDC37L1 | Hsp90 co-chaperone C | 4241.638672 | 6563.338867 |
| DDX31 | F8WAJ0 | DDX31 | RNA helicase | 208557.0781 | 219771.8906 |
| NEK3 | A0A087WY58;P51956 | NEK3 | Serine/threonine-prot | 4326.303223 | 6594.023438 |
| AASDHPPT | Q9NRN7 | AASDHPPT | L-aminoadipate-semia | 112398.5 | 117744.7422 |
| RBBP4 | Q09028 | RBBP4 | Histone-binding prote | 127369 | 189386.0781 |
| BLM | A0A3B3IT82;H0YNU5;P54132 | BLM | Bloom syndrome prot | 628636.3125 | 652788.9375 |
| RPS19 | P39019 | RPS19 | 40S ribosomal protein | 1387160.375 | 1066446.5 |
| EML4 | Q9HC35 | EML4 | Echinoderm microtub | 76367.32813 | 86372.24219 |
| EIF1 | K7EM18;P41567 | EIF1 | Eukaryotic translation | 392689.5625 | 362964.2813 |
| BEND3 | Q5T5X7 | BEND3 | BEN domain-containir | 11262.07715 | 15346.75879 |
| TMED9 | Q9BVK6 | TMED9 | Transmembrane emp | 860826.4375 | 853918.875 |
| SUMF1 | Q8NBK3 | SUMF1 | Formylglycine-genera | 15960.53027 | 16652.73828 |
| GLOD4 | F6TLX2 | GLOD4 | Glyoxalase domain-co | 818907.1055 | 780209.6406 |
| SNRNP27 | Q8WVK2 | SNRNP27 | U4/U6.U5 small nucle | 111677.3984 | 102633.4063 |
| RANGAP1 | P46060 | RANGAP1 | Ran GTPase-activating | 288111.0625 | 328714.5938 |
| HSPE1 | P61604 | HSPE1 | 10 kDa heat shock pro | 6338761 | 6379283.5 |
| RIPK1 | Q13546 | RIPK1 | Receptor-interacting s | 41264.49219 | 44380.70313 |
| PRIM1 | A0A5F9ZHB6;P49642 | PRIM1 | DNA primase;DNA pri | 83750.49219 | 97601.9375 |
| GNB1L | Q9BYB4 | GNB1L | Guanine nucleotide-b | 46025.85938 | 56931.27344 |

| | | | | | |
|---|---|---|---|---|---|
| ANGEL2 | Q5VTE6 | ANGEL2 | Protein angel homolog | 16081.7627 | 16033.30762 |
| FCF1 | G3V1S4;G3V2M5;G3V5S9;Q9 | FCF1 | Chromosome 14 oper | 29479.66797 | 9125.498047 |
| NOSIP | Q9Y314 | NOSIP | Nitric oxide synthase- | 79466.46875 | 77678.26563 |
| NDUFA6 | A0A2Y9D025;P56556 | NDUFA6 | NADH dehydrogenase | 129349.7969 | 150887.9219 |
| TRIM16 | O95361 | TRIM16 | Tripartite motif-conta | 111956.2266 | 98558.21094 |
| PARN | A0A494C0W0;O95453 | PARN | Poly(A)-specific ribonu | 68863.82813 | 46347.59766 |
| EIF3H | B3KS98;O15372 | EIF3H | Eukaryotic translation | 455309.5313 | 453658.4063 |
| TAOK1 | Q7L7X3 | TAOK1 | Serine/threonine-prot | 36321.36328 | 35862.39844 |
| DDHD1 | Q8NEL9 | DDHD1 | Phospholipase DDHD1 | 30287.19141 | 37607.36328 |
| RRP8 | O43159 | RRP8 | Ribosomal RNA-proce | 51727.53516 | 32969.79297 |
| AAK1; | A0A096LP25;Q6ZSR9 | AAK1; | AP2-associated protei | 18471.17969 | 26699.36523 |
| ZNF114 | Q8NC26 | ZNF114 | Zinc finger protein 114 | 214.2544861 | 297.864624 |
| PNISR | Q8TF01 | PNISR | Arginine/serine-rich p | 10592.43555 | 9881.99707 |
| LSG1 | Q9H089 | LSG1 | Large subunit GTPase | 115366.1563 | 111824.5234 |
| RBBP5 | Q15291 | RBBP5 | Retinoblastoma-bindi | 482145.625 | 550302 |
| TOMM6 | Q96B49 | TOMM6 | Mitochondrial import | 141491.6719 | 71025.14063 |
| TIMM17B | O60830 | TIMM17B | Mitochondrial import | 66568.75781 | 66649.38281 |
| CDCA8 | Q53HL2 | CDCA8 | Borealin | 36734.70313 | 30797.3125 |
| METTL15 | A6NJ78 | METTL15 | Probable methyltrans | 15351.96973 | 18669.26953 |
| LYSMD3 | Q7Z3D4 | LYSMD3 | LysM and putative pe | 9680.239258 | 11931.95703 |
| EIF1AX | P47813 | EIF1AX | Eukaryotic translation | 438452.5313 | 467473.25 |
| NUDCD2 | Q8WVJ2 | NUDCD2 | NudC domain-contain | 127063.8594 | 151649.7969 |
| PARD3 | Q5VWU8 | PARD3 | Partitioning defective | 57511.63672 | 64475.63086 |
| DHX8 | Q14562 | DHX8 | ATP-dependent RNA h | 247259.3438 | 214536.4375 |
| TRIP12 | G5E9G6 | TRIP12 | E3 ubiquitin-protein li | 61903.39648 | 63579.97949 |
| POP5 | Q969H6 | POP5 | Ribonuclease P/MRP p | 7601.208984 | 9392.371094 |
| PDCD4 | Q53EL6 | PDCD4 | Programmed cell deat | 112212.1641 | 210071.875 |
| ALG1 | Q9BT22 | ALG1 | Chitobiosyldiphospho | 24140.7832 | 24020.54297 |
| VKORC1L1 | A0A3B3ISV4 | VKORC1L1 | Vitamin K epoxide red | 99251.23438 | 110762.5547 |
| TAF1C | Q15572 | TAF1C | TATA box-binding pro | 59849.44141 | 58516.0625 |
| ABCC10 | Q5T3U5 | ABCC10 | Multidrug resistance-a | 13379.59961 | 15545.41309 |
| ATP6V1A | P38606 | ATP6V1A | V-type proton ATPase | 255754.4375 | 263312.9063 |
| RTN4IP1 | Q8WWV3 | RTN4IP1 | Reticulon-4-interactin | 32334.22656 | 29497.76172 |

| | | | | | |
|---|---|---|---|---|---|
| GFM1 | Q96RP9 | GFM1 | Elongation factor G, m | 232180.6406 | 248959.3438 |
| CDC5L | Q99459 | CDC5L | Cell division cycle 5-lik | 232906.1094 | 202106.9375 |
| MRM1 | Q6IN84 | MRM1 | rRNA methyltransfera | 26828.08203 | 31769.94531 |
| GRB2 | P62993 | GRB2 | Growth factor recepto | 518805.3438 | 551980.1875 |
| ELOB | Q15370 | ELOB | Elongin-B | 123272.3906 | 198674.6094 |
| SF3A3 | Q12874 | SF3A3 | Splicing factor 3A subu | 580767.9375 | 345006.2813 |
| SOS1 | G5E9C8;Q07889 | SOS1 | Son of sevenless hom | 15087.91992 | 16419.71484 |
| CDK11A | Q5QPR3;Q5QPR4;Q9UQ88 | CDK11A | Cyclin-dependent kina | 19649.69141 | 30034.94922 |
| SQSTM1 | Q13501 | SQSTM1 | Sequestosome-1 | 228952.6719 | 199741.1719 |
| WNK2 | A6PVV2;F8W9F9;H0Y7T5;Q9 | WNK2 | Serine/threonine-prot | 1242.179565 | 1496.713867 |
| SMARCC1 | Q92922 | SMARCC1 | SWI/SNF complex sub | 65494.25781 | 66881.74219 |
| NAP1L1 | F5H4R6;F8VY35;H0YIV4;P552 | NAP1L1 | Nucleosome assembly | 564022.4297 | 611704.6953 |
| ATP5PF | P18859 | ATP5PF | ATP synthase-coupling | 16842.51563 | 18358.00195 |
| NDUFA3 | O95167;Q6FGG4 | NDUFA3 | NADH dehydrogenase | 57598 | 55703.30078 |
| PLOD1 | Q02809 | PLOD1 | Procollagen-lysine,2-c | 76272.99219 | 75684.85156 |
| PDHB | P11177 | PDHB | Pyruvate dehydrogena | 263033.0313 | 280761.5625 |
| ADRM1 | Q16186 | ADRM1 | Proteasomal ubiquitin | 717740.4375 | 721280.125 |
| SRSF3 | P84103 | SRSF3 | Serine/arginine-rich s | 3718110.75 | 2540374.25 |
| NDUFA13 | Q9P0J0 | NDUFA13 | NADH dehydrogenase | 202007.1719 | 209100.8594 |
| EIF3F | O00303 | EIF3F | Eukaryotic translation | 447856.1563 | 455965.0625 |
| POLD2 | P49005 | POLD2 | DNA polymerase delta | 78289.39063 | 90440.3125 |
| ZNF598 | Q86UK7 | ZNF598 | E3 ubiquitin-protein li | 69403.82031 | 70231.65625 |
| SCAF11 | F8VXG7;Q99590 | SCAF11 | Protein SCAF11 | 45117.12109 | 48695.84375 |
| RNASEH2A | O75792 | RNASEH2A | Ribonuclease H2 subu | 99933.97656 | 94063.3125 |
| PUDP | Q08623 | PUDP | Pseudouridine-5'-pho | 28263.05859 | 34017.74219 |
| SMC6 | Q96SB8 | SMC6 | Structural maintenanc | 13998.02734 | 11840.53613 |
| TPX2 | Q9ULW0 | TPX2 | Targeting protein for X | 119968.2891 | 86222.25781 |
| PAXBP1 | Q9Y5B6 | PAXBP1 | PAX3- and PAX7-bindi | 343292.4063 | 313739.6563 |
| HNRNPR | O43390 | HNRNPR | Heterogeneous nuclea | 518949.875 | 287777.625 |
| EZH2 | Q15910 | EZH2 | Histone-lysine N-meth | 31773.66602 | 29180.77148 |
| CLPP | Q16740 | CLPP | ATP-dependent Clp pr | 96515.04688 | 89300.84375 |
| CEP104 | O60308 | CEP104 | Centrosomal protein c | 94100.27344 | 78017.13281 |
| RABGGTA | Q92696 | RABGGTA | Geranylgeranyl transf | 137017.6094 | 145747.2188 |

| | | | | | |
|---|---|---|---|---|---|
| RNPS1 | H3BMM9;H3BTC0;H3BV80;Q | RNPS1 | RNA-binding protein w | 539787.3125 | 421800.4375 |
| SMARCA4 | A0A2R8Y4P4;A0A2R8Y7S2 | SMARCA4 | Transcription activato | 214807.1406 | 216318.1563 |
| NDUFA1 | O15239 | NDUFA1 | NADH dehydrogenase | 39044.85938 | 54425.21484 |
| RPP14 | O95059 | RPP14 | Ribonuclease P protei | 19392.3418 | 19572.53516 |
| SF3A1 | Q15459 | SF3A1 | Splicing factor 3A subu | 418941.0625 | 315245.875 |
| RDH13 | Q8NBN7 | RDH13 | Retinol dehydrogenas | 64199.55859 | 70521.34375 |
| GSTM3 | P21266 | GSTM3 | Glutathione S-transfer | 1645766.25 | 1829573.625 |
| CYB5A | P00167 | CYB5A | Cytochrome b5 | 38889.76953 | 64779.99609 |
| PPP1R10 | Q96QC0 | PPP1R10 | Serine/threonine-prot | 141058.1719 | 135089.625 |
| DIPK2A | Q8NDZ4 | DIPK2A | Divergent protein kina | 15447.48828 | 13435.69727 |
| CDT1 | Q9H211 | CDT1 | DNA replication factor | 7799.672363 | 9944.126953 |
| SDF4 | A0A5F9UP49;Q9BRK5 | SDF4 | 45 kDa calcium-bindin | 55924.27344 | 49116.49219 |
| STX10 | K7EIY4;K7ELY2;K7EPP9;K7EQ | STX10 | Syntaxin-10 (Fragmen | 8998.637695 | 8575.46875 |
| CTU2 | Q2VPK5 | CTU2 | Cytoplasmic tRNA 2-th | 72360.40625 | 76761.57031 |
| CAMSAP3 | Q9P1Y5 | CAMSAP3 | Calmodulin-regulated | 55944.61328 | 58062.39844 |
| JUNB | P17275 | JUNB | Transcription factor ju | 10312.98926 | 5306.225586 |
| CHMP5 | Q9NZZ3 | CHMP5 | Charged multivesicula | 141514.5469 | 144522.9063 |
| GCC1 | Q96CN9 | GCC1 | GRIP and coiled-coil d | 15593.87793 | 14908.41504 |
| TNFRSF12A | Q9NP84 | TNFRSF12A | Tumor necrosis factor | 13409.0459 | 9381.584961 |
| NDFIP2 | A0A0C4DGW0;A0A3B3IRK3;E | NDFIP2 | NEDD4 family-interact | 5672.424316 | 5648.262695 |
| METTL3 | Q86U44 | METTL3 | N6-adenosine-methylt | 42423.36328 | 43770.57031 |
| PIAS4 | Q8N2W9 | PIAS4 | E3 SUMO-protein ligas | 23815.38672 | 26819.70898 |
| ALG3 | H7C0X4;Q92685 | ALG3 | Dol-P-Man:Man(5)Glc | 18563.33008 | 20195.64648 |
| SF3B6 | Q9Y3B4 | SF3B6 | Splicing factor 3B subu | 421111.1563 | 328362.6563 |
| GPN3 | Q9UHW5 | GPN3 | GPN-loop GTPase 3 | 20844.25 | 26263.63086 |
| NDUFAF7 | Q7L592 | NDUFAF7 | Protein arginine meth | 15391.17773 | 18778.40625 |
| SF1 | H7BZT1 | SF1 | Splicing factor 1 (Frag | 616586.625 | 598978.1797 |
| EIF3D | O15371 | EIF3D | Eukaryotic translation | 391608.8125 | 382601.25 |
| SIAH2 | O43255 | SIAH2 | E3 ubiquitin-protein li | 2076.986328 | 2996.874023 |
| SNAPC3 | Q5T282;Q92966 | SNAPC3 | snRNA-activating prot | 5277.345703 | 6332.138184 |
| RPL22L1 | C9JYQ9;H0Y8C2;Q6P5R6 | RPL22L1 | 60S ribosomal protein | 41992.29688 | 49857.39063 |
| GPI | A0A2U3TZU2;P06744 | GPI | Glucose-6-phosphate | 2515438 | 3022118.5 |
| DHX30 | H7BXY3;Q7L2E3 | DHX30 | ATP-dependent RNA h | 126188.7188 | 129758.5703 |

| | | | | | |
|---|---|---|---|---|---|
| PYCARD | Q9ULZ3 | PYCARD | Apoptosis-associated | 583674.6875 | 487317.5 |
| GLRX5 | Q86SX6 | GLRX5 | Glutaredoxin-related p | 19607.3125 | 21097.79883 |
| GTF3C3 | Q9Y5Q9 | GTF3C3 | General transcription | 78019.51563 | 81460.39063 |
| GALK2 | Q01415 | GALK2 | N-acetylgalactosamin | 67742.27344 | 74583.11719 |
| HDGFL2 | Q7Z4V5 | HDGFL2 | Hepatoma-derived gr | 91609.9375 | 107441.2344 |
| KIF11 | P52732 | KIF11 | Kinesin-like protein KI | 196494.9844 | 191152.9219 |
| EIF3L | B0QY89;Q9Y262 | EIF3L | Eukaryotic translation | 257764.7969 | 265415.3438 |
| CLTC | A0A087WVQ6;Q00610 | CLTC | Clathrin heavy chain;C | 3989024.5 | 4304853.5 |
| TFDP1 | Q14186 | TFDP1 | Transcription factor D | 12710.2168 | 12991.99805 |
| GTF2H2C | Q6P1K8 | GTF2H2C | General transcription | 41030.1875 | 36592.57422 |
| ACTL6A | O96019 | ACTL6A | Actin-like protein 6A | 280342.9375 | 286268.9375 |
| SNAPC1 | Q16533 | SNAPC1 | snRNA-activating prot | 3854.4646 | 2980.813965 |
| ACAT1 | A0A5F9ZHJ0 | ACAT1 | Acetyl-CoA acetyltran | 368643.9268 | 376672.6338 |
| KYAT1 | A0A494C066;B7Z4W5;Q1677 | KYAT1 | Kynurenine--oxogluta | 5886.312988 | 5578.726563 |
| PCNT | O95613 | PCNT | Pericentrin | 14642.4707 | 13983.53027 |
| EIF3M | Q7L2H7 | EIF3M | Eukaryotic translation | 207628.8906 | 207404.6406 |
| CBLL1 | Q75N03 | CBLL1 | E3 ubiquitin-protein li | 13390.12695 | 11456.86621 |
| TMEM126A | Q9H061 | TMEM126A | Transmembrane prote | 66919.07031 | 58719.58594 |
| YWHAQ | P27348 | YWHAQ | 14-3-3 protein theta | 1202638.5 | 1388494.5 |
| DDX42 | Q86XP3 | DDX42 | ATP-dependent RNA h | 478726.7188 | 439170.2813 |
| NCSTN | Q92542 | NCSTN | Nicastrin | 135696.8281 | 118473.4063 |
| PCBD1 | P61457 | PCBD1 | Pterin-4-alpha-carbin | 125731.3594 | 138305.9063 |
| LSM14B | Q5TBP9 | LSM14B | Protein LSM14 homol | 114915.6758 | 133554.3203 |
| TMEM50A | O95807 | TMEM50A | Transmembrane prote | 94194.03125 | 74145.92969 |
| MLPH | H7C2D8 | MLPH | Melanophilin (Fragme | 132555.1028 | 136776.6394 |
| SERBP1 | Q8NC51 | SERBP1 | Plasminogen activator | 1892088.5 | 1488845.5 |
| CEP85 | H7BZW2;Q6P2H3 | CEP85 | Centrosomal protein c | 58991.8125 | 77951.33594 |
| NARS1 | K7ENF0 | NARS1 | Asparagine--tRNA liga | 1323689.438 | 1193620.219 |
| INTS13 | Q9NVM9 | INTS13 | Integrator complex su | 98724.28125 | 108820.3047 |
| ZMIZ2 | A0A087X127;E7EWM3;Q8NF | ZMIZ2 | Zinc finger MIZ domai | 40340.97656 | 48470.3125 |
| DTWD2 | Q8NBA8 | DTWD2 | DTW domain-containi | 39100.01953 | 44663.8125 |
| NUP35 | Q8NFH5 | NUP35 | Nucleoporin NUP35 | 58866.36328 | 61345.60156 |
| NUDC | Q9Y266 | NUDC | Nuclear migration pro | 646763.125 | 688238.6875 |

| | | | | | |
|---|---|---|---|---|---|
| SNX27 | Q96L92 | SNX27 | Sorting nexin-27 | 166016.75 | 179841.6563 |
| ARID1B | A0A3F2YNW7;Q8NFD5 | ARID1B | AT-rich interactive do | 11961.04297 | 15695.56641 |
| UQCC2 | Q9BRT2;V5IRT4 | UQCC2 | Ubiquinol-cytochrome | 112154.2656 | 108270.0391 |
| CSTF2T | Q9H0L4 | CSTF2T | Cleavage stimulation f | 23556.9043 | 28949.72266 |
| DTNA | Q9Y4J8 | DTNA | Dystrobrevin alpha | 34964.12891 | 30879.26563 |
| DCTN6 | O00399 | DCTN6 | Dynactin subunit 6 | 33522.34766 | 36325.41406 |
| RBM22 | Q9NW64 | RBM22 | Pre-mRNA-splicing fac | 736460.6875 | 541028.75 |
| NDUFS3 | O75489 | NDUFS3 | NADH dehydrogenase | 256381.5469 | 273094.25 |
| CENPC | Q03188 | CENPC | Centromere protein C | 236321.0625 | 103736.5625 |
| SMCHD1 | A6NHR9 | SMCHD1 | Structural maintenanc | 133703.2656 | 120693.5859 |
| NT5DC1 | Q5TFE4 | NT5DC1 | 5'-nucleotidase doma | 180763.2344 | 217143.2344 |
| SHC1 | P29353 | SHC1 | SHC-transforming pro | 37745.57422 | 42432.85547 |
| KIAA1671 | Q9BY89 | KIAA1671 | Uncharacterized prote | 7806.92041 | 8332.075195 |
| YJU2 | Q9BW85 | YJU2 | Splicing factor YJU2 | 48997.98438 | 45822.18359 |
| PPP3CB | P16298 | PPP3CB | Serine/threonine-prot | 320995.8125 | 414337.7188 |
| MTMR4 | J3QR65;Q9NYA4 | MTMR4 | Myotubularin-related | 21446.11719 | 24020.67773 |
| POMK | Q9H5K3 | POMK | Protein O-mannose ki | 9828.466797 | 10275.60742 |
| CHMP1A | Q9HD42 | CHMP1A | Charged multivesicula | 216753.0313 | 254057.5625 |
| MAPKAPK3 | Q16644 | MAPKAPK3 | MAP kinase-activated | 74820.10156 | 76422.59375 |
| MEAF6 | Q9HAF1 | MEAF6 | Chromatin modificatio | 67257.5 | 55243.97656 |
| COA7 | Q96BR5 | COA7 | Cytochrome c oxidase | 125981.9531 | 120337.4141 |
| HJURP | Q8NCD3 | HJURP | Holliday junction reco | 9774.347656 | 9295.310547 |
| ZNF830 | Q96NB3 | ZNF830 | Zinc finger protein 830 | 13239.83887 | 10893.39453 |
| PRMT3 | O60678 | PRMT3 | Protein arginine N-me | 108409.3516 | 118530.0469 |
| NFKBIE | O00221;Q7LC14 | NFKBIE | NF-kappa-B inhibitor e | 4992.26709 | 9046.338867 |
| SH3BP5L | Q7L8J4 | SH3BP5L | SH3 domain-binding p | 9588.956055 | 11345.00977 |
| TOP1 | P11387 | TOP1 | DNA topoisomerase 1 | 1236989.875 | 548583.375 |
| CDK7 | P50613 | CDK7 | Cyclin-dependent kina | 71830.40625 | 75722.84375 |
| POLA2 | Q14181 | POLA2 | DNA polymerase alph | 65863.83594 | 72253.59375 |
| PDXP | Q96GD0 | PDXP | Pyridoxal phosphate p | 58123.80859 | 58475.90234 |
| INTS9 | Q9NV88 | INTS9 | Integrator complex su | 32615.27148 | 38070.80859 |
| LRP10 | Q7Z4F1 | LRP10 | Low-density lipoprote | 20273.29102 | 17793.0332 |
| PLK4 | J3KR82;J3KR84;O00444 | PLK4 | Serine/threonine-prot | 38581.80859 | 38930.14453 |

| | | | | | |
|---|---|---|---|---|---|
| ZHX2 | Q9Y6X8 | ZHX2 | Zinc fingers and home | 51557.70313 | 72010.73438 |
| TEX10 | Q9NXF1 | TEX10 | Testis-expressed prot | 62483.17969 | 55017.39063 |
| EVL | Q9UI08 | EVL | Ena/VASP-like protein | 36573.49219 | 46551.80859 |
| CLCN7 | P51798 | CLCN7 | H(+)/Cl(-) exchange tr | 27961.33789 | 30016.80273 |
| FBXL18 | Q96ME1 | FBXL18 | F-box/LRR-repeat prot | 71315.70313 | 72229.55469 |
| SELENOI | A0A087WW58;Q9C0D9 | SELENOI | Ethanolaminephospho | 127516.7266 | 117760.0781 |
| THNSL1 | Q8IYQ7 | THNSL1 | Threonine synthase-lil | 68728.64844 | 67201.44531 |
| PPWD1 | Q96BP3 | PPWD1 | Peptidylprolyl isomera | 167855.4688 | 133938.5313 |
| MGA | H3BU53 | MGA | MAX gene-associated | 15841.88428 | 14142.4292 |
| ACP1 | D3YTI2 | ACP1 | Low molecular weight | 707058.4453 | 695167.1641 |
| HMG20B | Q9P0W2 | HMG20B | SWI/SNF-related matr | 25094.70703 | 27091.68359 |
| AQR | O60306 | AQR | RNA helicase aquarius | 100502.5625 | 89733.9375 |
| KDM3B | Q7LBC6 | KDM3B | Lysine-specific demetl | 256589.0313 | 320950.7813 |
| EIF3A | Q14152 | EIF3A | Eukaryotic translation | 605846.0625 | 603798.1875 |
| TMX4 | Q9H1E5 | TMX4 | Thioredoxin-related tr | 71140.84375 | 64499.40625 |
| RARS1 | P54136 | RARS1 | Arginine--tRNA ligase, | 677675.5 | 705483.125 |
| ENAH | A0A0U1RRM6 | ENAH | Protein enabled homo | 137103.6875 | 155019.8125 |
| FNTA | P49354 | FNTA | Protein farnesyltransf | 56382.38672 | 58673.95313 |
| FANCG | O15287 | FANCG | Fanconi anemia group | 3603.122559 | 4520.627441 |
| ACADVL | P49748 | ACADVL | Very long-chain specif | 145536.9531 | 183319.5469 |
| NFXL1 | Q6ZNB6 | NFXL1 | NF-X1-type zinc finger | 31273.16211 | 33620.03125 |
| FARS2 | A0A3B3ITR6;O95363 | FARS2 | Phenylalanine--tRNA l | 27715.50977 | 29542.39648 |
| ERMP1 | Q7Z2K6 | ERMP1 | Endoplasmic reticulum | 70192.38281 | 86447.67969 |
| PDF | Q9HBH1 | PDF | Peptide deformylase, | 55355.14844 | 50716.69531 |
| HSCB | Q8IWL3 | HSCB | Iron-sulfur cluster co-c | 83391.5625 | 76412.60156 |
| SFSWAP | Q12872 | SFSWAP | Splicing factor, suppre | 31143.19727 | 30016.92969 |
| PRMT5 | O14744 | PRMT5 | Protein arginine N-me | 457535.1875 | 510037.5313 |
| ABL2 | P42684 | ABL2 | Tyrosine-protein kinas | 9227 | 9469.503906 |
| MYO1E | Q12965 | MYO1E | Unconventional myos | 28479.29883 | 19051.3418 |
| PAK1IP1 | Q9NWT1 | PAK1IP1 | p21-activated protein | 80842.04688 | 37845.00781 |
| FHOD1 | Q9Y613 | FHOD1 | FH1/FH2 domain-cont | 70350.72656 | 73784.0625 |
| GGA2 | Q9UJY4 | GGA2 | ADP-ribosylation facto | 26589.16797 | 31867.89258 |
| CCT5 | P48643 | CCT5 | T-complex protein 1 s | 1140879.5 | 1335163.25 |

| | | | | | |
|---|---|---|---|---|---|
| SNX4 | O95219 | SNX4 | Sorting nexin-4 | 97193.60156 | 111293.3047 |
| MARCKSL1 | P49006 | MARCKSL1 | MARCKS-related prote | 938189.1875 | 905874.75 |
| XAB2 | Q9HCS7 | XAB2 | Pre-mRNA-splicing fac | 99958.74219 | 93180.42969 |
| TSEN2 | C9J7Z4;Q8NCE0 | TSEN2 | tRNA-splicing endonu | 14055.69141 | 14746.12695 |
| USP3 | Q9Y6I4 | USP3 | Ubiquitin carboxyl-ter | 34468.78906 | 34174.08594 |
| SPR | P35270 | SPR | Sepiapterin reductase | 393123.7813 | 401050.0625 |
| ACTR5 | Q9H9F9 | ACTR5 | Actin-related protein ! | 16552.13477 | 17550.89258 |
| PHF3 | Q92576 | PHF3 | PHD finger protein 3 | 25429.55664 | 25489.11914 |
| SND1 | Q7KZF4 | SND1 | Staphylococcal nuclea | 982378.1875 | 1026278.438 |
| ZC3H13 | Q5T200 | ZC3H13 | Zinc finger CCCH dom | 296948.875 | 220658.3125 |
| AIMP1 | Q12904 | AIMP1 | Aminoacyl tRNA synth | 588443.6875 | 555107 |
| NDUFB8 | O95169 | NDUFB8 | NADH dehydrogenase | 128310.25 | 144326.1563 |
| SAMD11 | A0A087WYW1;A0A087X1J5;I | SAMD11 | Sterile alpha motif do | 5719.441406 | 6353.891602 |
| LAS1L | Q9Y4W2 | LAS1L | Ribosomal biogenesis | 150673.8594 | 144347.8281 |
| RSU1 | Q15404 | RSU1 | Ras suppressor protei | 326709.5313 | 301109.9688 |
| RPS21 | P63220;Q8WVC2 | RPS21 | 40S ribosomal protein | 3396804 | 2639006.75 |
| RBM41 | Q96IZ5 | RBM41 | RNA-binding protein 4 | 8844.21582 | 8081.472656 |
| DHX35 | Q5THR1;Q9H5Z1 | DHX35 | Probable ATP-depend | 23040.69531 | 19733.42578 |
| MTFR1 | Q15390 | MTFR1 | Mitochondrial fission | 16803.0957 | 15207.6748 |
| BECN1 | Q14457 | BECN1 | Beclin-1 | 28438.58203 | 30404.80664 |
| MAN2B1 | O00754 | MAN2B1 | Lysosomal alpha-man | 84778.79688 | 92436.80469 |
| PSMG1 | O95456 | PSMG1 | Proteasome assembly | 315885.4688 | 368456.4375 |
| VMP1 | Q96GC9 | VMP1 | Vacuole membrane pr | 100829.6328 | 108325.9141 |
| COMT | P21964 | COMT | Catechol O-methyltra | 631655.5 | 773245.5625 |
| DHX16 | O60231 | DHX16 | Pre-mRNA-splicing fac | 504364.2188 | 510726.9375 |
| TACO1 | Q9BSH4 | TACO1 | Translational activato | 210296.1406 | 212890.1875 |
| CCNB2 | O95067 | CCNB2 | G2/mitotic-specific cy | 29340.63086 | 31471.30664 |
| DDX11L8;DDX12P;DDX11 | A8MPP1;Q92771;Q96FC9 | DDX11L8;DDX12P;DD) | Putative ATP-depende | 5317.508301 | 4934.468262 |
| RANBP1 | P43487 | RANBP1 | Ran-specific GTPase-a | 2193581.5 | 2142322.25 |
| TMED5 | Q9Y3A6 | TMED5 | Transmembrane emp | 165385.5313 | 195177.75 |
| MED31 | Q9Y3C7 | MED31 | Mediator of RNA poly | 22850.69531 | 24234.88281 |
| CCT7 | Q99832 | CCT7 | T-complex protein 1 s | 832943.8125 | 917521.5625 |
| SSBP1 | Q04837 | SSBP1 | Single-stranded DNA-l | 808452 | 721519.25 |

| | | | | | |
|---|---|---|---|---|---|
| TAGLN2 | P37802 | TAGLN2 | Transgelin-2 | 1690334.5 | 1549620.625 |
| ARFGAP3 | Q9NP61 | ARFGAP3 | ADP-ribosylation facto | 33061.625 | 37185.53125 |
| SCYL2 | A0A0U1RQQ9;Q6P3W7 | SCYL2 | SCY1-like protein 2 | 80105.01563 | 89399.49219 |
| SMYD2 | Q9NRG4 | SMYD2 | N-lysine methyltransfe | 10243.14453 | 9364.536133 |
| SLC38A10 | Q9HBR0 | SLC38A10 | Putative sodium-coup | 3180.676758 | 6897.687988 |
| DLD | P09622 | DLD | Dihydrolipoyl dehydro | 479499.625 | 544166.4375 |
| AIMP2 | Q13155 | AIMP2 | Aminoacyl tRNA synth | 763011.4375 | 850701.0625 |
| SPAST | A0A2R8YCL5;A0A2U3TZR0;Q | SPAST | Spastin (Fragment);Sp | 15269.96484 | 17004.83789 |
| NIFK | Q9BYG3 | NIFK | MKI67 FHA domain-in | 57875.71484 | 25423.33008 |
| GFER | P55789 | GFER | FAD-linked sulfhydryl | 64550.14063 | 59444.61328 |
| INTS3 | Q68E01 | INTS3 | Integrator complex su | 94976.6875 | 92648.6875 |
| TRMU | O75648 | TRMU | Mitochondrial tRNA-s | 13754.12402 | 15346.75586 |
| IREB2 | P48200 | IREB2 | Iron-responsive eleme | 76428.65625 | 100720.75 |
| NSUN2 | Q08J23 | NSUN2 | RNA cytosine C(5)-me | 491349.5938 | 489823.5 |
| PTPRF | P10586 | PTPRF | Receptor-type tyrosin | 94631.14063 | 139626.2813 |
| NAGA | P17050 | NAGA | Alpha-N-acetylgalacto | 21785.48438 | 24967.91016 |
| INTS7 | Q9NVH2 | INTS7 | Integrator complex su | 41790.76172 | 45713.35938 |
| WTAP | Q15007 | WTAP | Pre-mRNA-splicing reg | 71294.53125 | 48736.83203 |
| VPS72 | Q15906 | VPS72 | Vacuolar protein sorti | 47047.9375 | 43344.04297 |
| ACTR6 | F8W057;Q9GZN1 | ACTR6 | Actin-related protein | 16659.18945 | 15829.50488 |
| IQCB1 | Q15051 | IQCB1 | IQ calmodulin-binding | 5334.527832 | 4302.095703 |
| EIF3C | Q99613 | EIF3C | Eukaryotic translation | 768801.125 | 759713.5 |
| TFDP3 | Q5H9I0 | TFDP3 | Transcription factor D | 13312.67969 | 10312.20996 |
| KCNK1 | O00180 | KCNK1 | Potassium channel su | 5544.100586 | 8829.734375 |
| SBNO1 | A3KN83 | SBNO1 | Protein strawberry no | 59468.03125 | 63842.25391 |
| MRGBP | Q9NV56 | MRGBP | MRG/MORF4L-binding | 205342.2344 | 171774.2656 |
| CSNK1G1 | A0A0U1RQT7;A0A0U1RQY7;( | CSNK1G1 | Casein kinase I isoforr | 53376.54297 | 55736.57813 |
| BAZ1B | Q9UIG0 | BAZ1B | Tyrosine-protein kinas | 418682.9375 | 419920.0313 |
| HSPA9 | P38646 | HSPA9 | Stress-70 protein, mit | 3656422.5 | 3388252 |
| MCMBP | Q9BTE3 | MCMBP | Mini-chromosome ma | 103747.5156 | 121197.1953 |
| ZNF609 | O15014 | ZNF609 | Zinc finger protein 60 | 31708.33984 | 29704.67773 |
| RAD9A | Q99638 | RAD9A | Cell cycle checkpoint | 16713.53125 | 15407.52148 |
| COP1 | A0A3B3ISQ9;Q8NHY2 | COP1 | E3 ubiquitin-protein li | 6652.891602 | 6587.640137 |

| | | | | | |
|---|---|---|---|---|---|
| ATXN2L | Q8WWM7 | ATXN2L | Ataxin-2-like protein | 324039.9063 | 323619.6563 |
| EXT2 | Q93063 | EXT2 | Exostosin-2 | 17309.01172 | 17838.49805 |
| PRR15L | Q9BU68 | PRR15L | Proline-rich protein 15 | 18329.91602 | 23025.56836 |
| ZBTB7B | O15156 | ZBTB7B | Zinc finger and BTB dc | 16660.60938 | 16773.2832 |
| hCG_2039566;;H2AC11;H2 | A0A0U1RR32;A0A0U1RRH7;F | hCG_2039566;;H2AC1 | Histone H2A;Histone I | 10929695 | 780632.125 |
| SNAPIN | O95295 | SNAPIN | SNARE-associated pro | 123702.2813 | 121803.7344 |
| TSEN54 | Q7Z6J9 | TSEN54 | tRNA-splicing endonu | 54675.28125 | 54944.60938 |
| C7orf26 | H7BZK9;H7C0Q5;Q96N11 | C7orf26 | Uncharacterized prote | 6636.945801 | 6531.068359 |
| VDAC1 | P21796 | VDAC1 | Voltage-dependent ar | 1327419.5 | 537675.5 |
| SAP30BP | J3QQJ0;Q9UHR5 | SAP30BP | SAP30-binding protein | 271854.5938 | 274827.4375 |
| LAMP1 | P11279 | LAMP1 | Lysosome-associated | 556933.6875 | 500537.25 |
| TSEN34 | A0A590UJW4;E7EQB3;Q9BS\ | TSEN34 | tRNA-splicing endonu | 71808.42969 | 75145.3125 |
| FAM234B | A2RU67 | FAM234B | Protein FAM234B | 17065.61914 | 17910.03125 |
| KPNB1 | Q14974 | KPNB1 | Importin subunit beta | 1597861.625 | 1741670.125 |
| ELL | P55199 | ELL | RNA polymerase II elo | 9025.348633 | 8261.40625 |
| PPP4R3B | Q5MIZ7 | PPP4R3B | Serine/threonine-prot | 22660.39648 | 31700.5332 |
| ARL3 | P36405 | ARL3 | ADP-ribosylation facto | 393963.1563 | 438539.0938 |
| XPO4 | F2Z2X4;Q9C0E2 | XPO4 | Exportin-4 | 67554.85156 | 67307.75 |
| HDLBP | A0A024R4E5 | HDLBP | High density lipoprote | 279315.6563 | 301788.625 |
| ENGASE | Q8NFI3 | ENGASE | Cytosolic endo-beta-N | 11225.60059 | 11410.08789 |
| MANF | A8K878;P55145 | MANF | Mesencephalic astroc | 303550.9063 | 322959.2188 |
| RPLP1 | P05386 | RPLP1 | 60S acidic ribosomal p | 72714.10156 | 92305.71094 |
| PARG | Q86W56 | PARG | Poly(ADP-ribose) glyco | 56802.46484 | 53104.37891 |
| TRAPPC5 | Q8IUR0 | TRAPPC5 | Trafficking protein par | 72520.16406 | 72367.65625 |
| ALDH3A2 | A0A5F9ZHN9;P51648 | ALDH3A2 | Aldehyde dehydrogen | 167606.0938 | 177937.125 |
| SUMO2 | P61956 | SUMO2 | Small ubiquitin-relate | 76759.78125 | 78797.78906 |
| ZNF24 | P17028 | ZNF24 | Zinc finger protein 24 | 22111.60352 | 22358.3457 |
| FIGNL1 | Q6PIW4 | FIGNL1 | Fidgetin-like protein 1 | 12889.56152 | 15320.42285 |
| POLR1C | O15160 | POLR1C | DNA-directed RNA po | 232830.4375 | 202014.6563 |
| CWC15 | Q9P013 | CWC15 | Spliceosome-associate | 140321.1719 | 113507.8125 |
| GLUL | P15104 | GLUL | Glutamine synthetase | 50870.41016 | 57637.98047 |
| TTC19 | Q6DKK2 | TTC19 | Tetratricopeptide rep | 80256.96875 | 82647.89063 |
| COX6B1 | P14854 | COX6B1 | Cytochrome c oxidase | 930831.375 | 1169216.25 |

| | | | | | |
|---|---|---|---|---|---|
| H4C1 | P62805 | H4C1 | Histone H4 | 14205993 | 696213.5 |
| PUS7L | F8VWC0;F8VZA0;Q9H0K6 | PUS7L | Pseudouridylate synth | 1019.735901 | 889.84375 |
| PLRG1 | O43660 | PLRG1 | Pleiotropic regulator 1 | 81913.89063 | 80697.85938 |
| RNF169 | Q8NCN4 | RNF169 | E3 ubiquitin-protein li | 21016.11719 | 23042.23438 |
| TMEM87B | A0A494BZZ8;H7C0B3;H7C1G | TMEM87B | Transmembrane prote | 5474.962891 | 5443.053223 |
| ZNF638 | Q14966 | ZNF638 | Zinc finger protein 638 | 102707.4609 | 107433.5 |
| IPO8 | O15397 | IPO8 | Importin-8 | 80455.40625 | 85356.13281 |
| PEX16 | Q9Y5Y5 | PEX16 | Peroxisomal membrar | 16638.2168 | 16417.69922 |
| SPCS2 | A0A087WUC6;Q15005 | SPCS2 | Signal peptidase comp | 499478.1875 | 548063.4375 |
| SKA2 | Q8WVK7 | SKA2 | Spindle and kinetocho | 7965.911621 | 10138.53516 |
| CIAO2B | Q9Y3D0 | CIAO2B | Cytosolic iron-sulfur a | 48012.88672 | 62220.01563 |
| AAGAB | Q6PD74 | AAGAB | Alpha- and gamma-ad | 82226.45313 | 123126.9141 |
| ATXN10 | Q9UBB4 | ATXN10 | Ataxin-10 | 190957.6094 | 194136.9375 |
| KDM2A | I3VM54;Q9Y2K7 | KDM2A | Lysine-specific demetl | 96613.54688 | 79580.35156 |
| RNF167 | I3L0L6;Q9H6Y7 | RNF167 | E3 ubiquitin-protein li | 12659.25586 | 11190.79199 |
| RPS12 | P25398 | RPS12 | 40S ribosomal protein | 2591340.25 | 2061875.125 |
| GPKOW | Q92917 | GPKOW | G-patch domain and K | 144140.7344 | 155132.1406 |
| CEP55 | Q53EZ4 | CEP55 | Centrosomal protein d | 24413.58203 | 35987.70703 |
| YBX1 | P67809 | YBX1 | Y-box-binding protein | 2044557 | 2006099 |
| STARD7 | Q9NQZ5 | STARD7 | StAR-related lipid trar | 19903.47656 | 18642.49805 |
| YES1 | J3QRU1;P07947 | YES1 | Tyrosine-protein kinas | 102639.8203 | 96949.875 |
| PUM2 | A0A0A0MR59;A0A0C4DG68 | PUM2 | Pumilio homolog 2;Pu | 76695.59375 | 87482.75391 |
| PI4KB | A0A0B4J1S8;Q9UBF8 | PI4KB | Phosphatidylinositol 4 | 26974.03711 | 30980.3125 |
| GNL3 | Q9BVP2 | GNL3 | Guanine nucleotide-b | 171814.625 | 86675.38281 |
| RPS28 | P62857 | RPS28 | 40S ribosomal protein | 2639776 | 1857272.625 |
| NUP37 | Q8NFH4 | NUP37 | Nucleoporin Nup37 | 97840.83594 | 121588.4688 |
| WIPF2 | Q8TF74 | WIPF2 | WAS/WASL-interactin | 81948.71094 | 81750.96094 |
| ARHGEF5;;ARHGEF5 | A0A2R8YF49;A0A2R8YFR7;Q | ARHGEF5;;ARHGEF5 | Rho guanine nucleotic | 11415.60547 | 11184.87207 |
| STK38L | Q9Y2H1 | STK38L | Serine/threonine-prot | 33892.95703 | 24393.05273 |
| HIP1 | O00291 | HIP1 | Huntingtin-interacting | 16561.07031 | 18423.06445 |
| PICALM | H0YEF7;H0YEH1 | PICALM | Phosphatidylinositol-b | 175469.623 | 204526.8164 |
| SLC35F6 | Q8N357 | SLC35F6 | Solute carrier family 3 | 23255.10352 | 24972.76172 |
| ATP2A2 | P16615 | ATP2A2 | Sarcoplasmic/endopla | 587070 | 632267.125 |

| | | | | | |
|---|---|---|---|---|---|
| NDUFA9 | Q16795 | NDUFA9 | NADH dehydrogenase | 303712.8438 | 338527.5938 |
| EIF4A1 | P60842 | EIF4A1 | Eukaryotic initiation fa | 3994650.75 | 4218925.5 |
| NSMAF | Q92636 | NSMAF | Protein FAN | 7448.063477 | 6632.080078 |
| BCAR1 | P56945 | BCAR1 | Breast cancer anti-est | 38745.34766 | 39157.07422 |
| SKA1 | Q96BD8 | SKA1 | Spindle and kinetocho | 12537.12305 | 19958.7832 |
| TGS1 | Q96RS0 | TGS1 | Trimethylguanosine sy | 26071.67188 | 24093.5625 |
| CLK4 | A0A087WX74;B7ZL31;Q9HAZ | CLK4 | Dual-specificity protei | 15250.84473 | 9603.474609 |
| OGFOD3 | Q6PK18 | OGFOD3 | 2-oxoglutarate and irc | 34531.875 | 44158.35938 |
| MT-CO2 | P00403 | MT-CO2 | Cytochrome c oxidase | 1440324.875 | 1445974.75 |
| TM9SF2 | Q99805 | TM9SF2 | Transmembrane 9 sup | 180517.4375 | 216057.9219 |
| COQ9 | O75208 | COQ9 | Ubiquinone biosynthe | 27238.21094 | 32301.28711 |
| PMPCB | O75439 | PMPCB | Mitochondrial-process | 414452.7188 | 401716.5938 |
| GTF2E2 | P29084 | GTF2E2 | Transcription initiatior | 38707.64844 | 39346.28125 |
| NOL9 | Q5SY16 | NOL9 | Polynucleotide 5'-hyd | 100960.4609 | 100173.6094 |
| ANKRD54 | Q6NXT1 | ANKRD54 | Ankyrin repeat domai | 31688.34375 | 31492.31445 |
| EFHD2 | Q96C19 | EFHD2 | EF-hand domain-conta | 106523.1484 | 111336.4297 |
| MPST | P25325 | MPST | 3-mercaptopyruvate s | 175729.0156 | 220563.1875 |
| NACC1 | Q96RE7 | NACC1 | Nucleus accumbens-a | 105647.2813 | 103661.1016 |
| CAPN15 | O75808 | CAPN15 | Calpain-15 | 4567.692383 | 4926.671387 |
| BRIX1 | Q8TDN6 | BRIX1 | Ribosome biogenesis | 265990.0625 | 201343.5625 |
| NPEPPS | E9PLK3;P55786 | NPEPPS | Aminopeptidase;Puro | 339865.2813 | 366598.3438 |
| MTERF4 | Q7Z6M4 | MTERF4 | Transcription termina | 8089.42041 | 8110.541016 |
| ZFR | Q96KR1 | ZFR | Zinc finger RNA-bindir | 145615.1875 | 93434.28125 |
| INTS6 | Q9UL03 | INTS6 | Integrator complex su | 43578.27344 | 42454.10547 |
| AHCYL2 | Q96HN2 | AHCYL2 | Adenosylhomocystein | 345835.4063 | 417672.0938 |
| GRWD1 | Q9BQ67 | GRWD1 | Glutamate-rich WD re | 268284.6563 | 273011.4063 |
| CINP | H0YN33;Q9BW66 | CINP | Cyclin-dependent kina | 5695.668457 | 5253.486816 |
| RBBP8NL | Q8NC74 | RBBP8NL | RBBP8 N-terminal-like | 23790.74023 | 22661.71875 |
| MRPL48 | Q96GC5 | MRPL48 | 39S ribosomal protein | 79360.38281 | 89822.33594 |
| SET | Q01105 | SET | Protein SET | 908637.875 | 1043055.313 |
| SBDS | Q9Y3A5 | SBDS | Ribosome maturation | 180355.4531 | 173240.6094 |
| SNRNP200 | O75643 | SNRNP200 | U5 small nuclear ribor | 481817.5313 | 388067.9688 |
| ZNF318 | Q5VUA4 | ZNF318 | Zinc finger protein 31ε | 15542.67285 | 17674.19922 |

| | | | | | |
|---|---|---|---|---|---|
| TMX2 | Q9Y320 | TMX2 | Thioredoxin-related tr | 118907.7891 | 119677.6016 |
| IPO5 | O00410 | IPO5 | Importin-5 | 300826.375 | 399075.3125 |
| TROAP | F8W130;F8W1U0;Q12815 | TROAP | Tastin | 8983.916016 | 6678.864746 |
| CHCHD1 | Q96BP2 | CHCHD1 | Coiled-coil-helix-coilec | 105629.0703 | 102292.1016 |
| ZNF444 | Q8N0Y2 | ZNF444 | Zinc finger protein 44 | 3584.378906 | 3757.150146 |
| SUPT5H | O00267 | SUPT5H | Transcription elongati | 104626.2109 | 85302.90625 |
| EEF1E1 | O43324 | EEF1E1 | Eukaryotic translation | 418016.9063 | 416399.625 |
| MTPN | C9JL85;P58546 | MTPN | Myotrophin | 27981.93164 | 58657.80078 |
| GMPS | P49915 | GMPS | GMP synthase [glutan | 690299.9375 | 761704.125 |
| GTF3C1 | Q12789 | GTF3C1 | General transcription | 272955.4063 | 282082.5938 |
| TMEM11 | P17152 | TMEM11 | Transmembrane prote | 72950.71875 | 65757.70313 |
| TAF5 | Q15542 | TAF5 | Transcription initiatior | 25506.98438 | 26947.95703 |
| SNRPC | A0A0A0MRR7;P09234 | SNRPC | U1 small nuclear ribor | 711042.875 | 468415.4063 |
| TRIM3 | O75382 | TRIM3 | Tripartite motif-conta | 41224.96094 | 39305.80078 |
| BORCS5 | G3V1P3;Q969J3 | BORCS5 | BLOC-1-related compl | 21303.38867 | 20884.11914 |
| PHF12 | C9J9G2;Q96QT6 | PHF12 | PHD finger protein 12 | 5895.379883 | 7661.615234 |
| ACIN1 | G3V3B0 | ACIN1 | Apoptotic chromatin c | 294659.1973 | 259855.2207 |
| IL17RA | Q96F46 | IL17RA | Interleukin-17 recepto | 11674.35352 | 4929.77002 |
| TMEM201 | Q5SNT2 | TMEM201 | Transmembrane prote | 34900.62891 | 30626.21484 |
| SCAI | Q8N9R8 | SCAI | Protein SCAI | 3832.406006 | 4436.726563 |
| CMBL | Q96DG6 | CMBL | Carboxymethylenebut | 337298.375 | 299466.1563 |
| ZMAT2 | Q96NC0 | ZMAT2 | Zinc finger matrin-typ | 28282.58594 | 25658.79492 |
| TAGLN3 | C9J5W6;Q9UI15 | TAGLN3 | Transgelin;Transgelin- | 31085.03906 | 31299.68164 |
| DNAJC17 | Q9NVM6 | DNAJC17 | DnaJ homolog subfam | 135004.9688 | 120894.9844 |
| MPG | P29372 | MPG | DNA-3-methyladenine | 155517.7031 | 154115.3594 |
| ANKRD24 | M0QZ18;Q8TF21 | ANKRD24 | Ankyrin repeat domai | 162461.125 | 147513.8906 |
| GOPC | Q9HD26 | GOPC | Golgi-associated PDZ a | 54587.37109 | 62593.3125 |
| CLPX | O76031 | CLPX | ATP-dependent Clp pr | 199071.2344 | 194315.2188 |
| TNPO1 | Q92973 | TNPO1 | Transportin-1 | 351489.25 | 413814.8438 |
| ZC3H11B | A0A1B0GTU1;A0A494C0C4 | ZC3H11B | Zinc finger CCCH doma | 6669.537109 | 5671.056641 |
| HDGF | P51858 | HDGF | Hepatoma-derived gro | 915240.0625 | 889061.875 |
| APTX | Q7Z2E3 | APTX | Aprataxin | 27583.55664 | 32471.78125 |
| RBM33 | H0Y3K4 | RBM33 | RNA-binding protein 3 | 56072.43945 | 53400.71777 |

| | | | | | |
|---|---|---|---|---|---|
| DSP | P15924 | DSP | Desmoplakin | 399915.4688 | 379436.625 |
| LGALSL | Q3ZCW2 | LGALSL | Galectin-related prote | 7134.125488 | 8915.673828 |
| DIAPH1 | A0A0G2JH68;O60610 | DIAPH1 | Protein diaphanous hc | 330481.6875 | 375432.7813 |
| POLR2E | A0A087WVZ9;P19388 | POLR2E | DNA-directed RNA po | 186050.0469 | 145436.1406 |
| SLC26A6 | C9J6Y3;C9JKR6;F8W9I8;G3XA | SLC26A6 | Solute carrier family 2 | 12778.41504 | 14112.48047 |
| PREB | Q9HCU5 | PREB | Prolactin regulatory e | 157633.2188 | 165479.625 |
| GAPVD1 | F8W9S7 | GAPVD1 | GTPase-activating pro | 173244.4766 | 186763.8301 |
| ISG15 | P05161 | ISG15 | Ubiquitin-like protein | 432820.9375 | 438744.9063 |
| EPB41 | A0A2R8Y783 | EPB41 | Protein 4.1 (Fragment | 48535.41797 | 59553.55273 |
| DUS1L | Q6P1R4 | DUS1L | tRNA-dihydrouridine( | 42693.28516 | 44320.76172 |
| EWSR1 | B0QYK0;Q01844 | EWSR1 | RNA-binding protein E | 1749177.361 | 1428985.164 |
| TXNL1 | O43396 | TXNL1 | Thioredoxin-like prote | 964287.25 | 801746.75 |
| ANP32E | Q9BTT0 | ANP32E | Acidic leucine-rich nuc | 198484.7969 | 308151.8125 |
| YTHDF1 | Q9BYJ9 | YTHDF1 | YTH domain-containir | 60051.76563 | 58256.92969 |
| TSEN15 | Q8WW01 | TSEN15 | tRNA-splicing endonu | 59352.08594 | 53744.90625 |
| CDCA3 | F5GX58;F8WDL1;J3KMY0;Q9 | CDCA3 | Cell division cycle-asso | 25093.43359 | 25302.46094 |
| SORT1 | Q99523 | SORT1 | Sortilin | 97723.66406 | 105879.7578 |
| DBNL | B4DDD6 | DBNL | Drebrin-like protein | 219330.666 | 255600.1563 |
| SDHAF2 | F5GYJ5;Q9NX18 | SDHAF2 | Succinate dehydroger | 12757.69238 | 19492.60938 |
| HARS1 | P12081 | HARS1 | Histidine--tRNA ligase | 776923.6875 | 833753.25 |
| ITM2B | Q9Y287 | ITM2B | Integral membrane pr | 9733.493164 | 11490.26563 |
| JAGN1 | Q8N5M9 | JAGN1 | Protein jagunal homol | 37204.82813 | 35290.90625 |
| INCENP | Q9NQS7 | INCENP | Inner centromere pro | 35683.83203 | 29676.05273 |
| TAF9 | Q16594 | TAF9 | Transcription initiatior | 110016.7969 | 112142.9766 |
| SCAF4 | O95104 | SCAF4 | SR-related and CTD-as | 129651.7344 | 117383.5781 |
| SRSF7 | A0A0B4J1Z1;C9JAB2;Q16629 | SRSF7 | Serine/arginine-rich-s | 2678241.25 | 1810184.25 |
| NCAPH | Q15003 | NCAPH | Condensin complex su | 113403.75 | 129876.2891 |
| RAP1A | P62834 | RAP1A | Ras-related protein Ra | 62704.6875 | 71190.03125 |
| PPM1D | O15297 | PPM1D | Protein phosphatase 1 | 61034.15234 | 56944.19141 |
| TXLNA | P40222 | TXLNA | Alpha-taxilin | 167751.8125 | 154074.6094 |
| EIF3E | P60228 | EIF3E | Eukaryotic translation | 595460.875 | 571732.25 |
| COLGALT1 | Q8NBJ5 | COLGALT1 | Procollagen galactosy | 115088.3984 | 121483.9063 |
| PPIE | Q9UNP9 | PPIE | Peptidyl-prolyl cis-trar | 136581.6719 | 128360.9297 |

| | | | | | |
|---|---|---|---|---|---|
| EPS8 | A0A2R8Y4W2;A0A2R8YE63;C | EPS8 | Epidermal growth fact | 55136.14063 | 62392.91406 |
| CDC42SE1 | Q9NRR8 | CDC42SE1 | CDC42 small effector | 139590.9844 | 146854.625 |
| GPALPP1 | A0A0A0MRI1;Q8IXQ4 | GPALPP1 | GPALPP motifs-contai | 52538.54688 | 53760.48047 |
| MDC1 | Q14676 | MDC1 | Mediator of DNA dam | 62891.12109 | 56884.75391 |
| ST14 | Q9Y5Y6 | ST14 | Suppressor of tumorig | 52357.9375 | 59965.86328 |
| ITGAE | P38570 | ITGAE | Integrin alpha-E | 156468.9688 | 152953.9531 |
| URB2 | Q14146 | URB2 | Unhealthy ribosome b | 40545.13281 | 40435.65234 |
| ANKRD52 | Q8NB46 | ANKRD52 | Serine/threonine-prot | 26104.21289 | 35236.24609 |
| GORAB | Q5T7V8 | GORAB | RAB6-interacting golg | 5588.789551 | 7072.989258 |
| PDIA5 | Q14554 | PDIA5 | Protein disulfide-isom | 104253.4219 | 109893.0469 |
| NTAN1 | Q96AB6 | NTAN1 | Protein N-terminal asp | 15211.52441 | 17360.47656 |
| C11orf58;SMAP | E9PM92;O00193 | C11orf58;SMAP | Chromosome 11 oper | 17024.69922 | 17193.94531 |
| VPS35L | C9J7I2;E7EWW0;F5H7K1;H3E | VPS35L | VPS35 endosomal pro | 420.8912354 | 548.2581177 |
| CCP110 | O43303 | CCP110 | Centriolar coiled-coil | 6524.928223 | 5738.299316 |
| MRPS26 | Q9BYN8 | MRPS26 | 28S ribosomal protein | 104607.2813 | 111275.6094 |
| EMC10 | M0R2A0;Q5UCC4 | EMC10 | ER membrane protein | 35957.99609 | 30130.85547 |
| PDCD2 | F5H4V9;Q16342 | PDCD2 | Programmed cell deat | 74423.05469 | 60339.01563 |
| FAM3C | Q92520 | FAM3C | Protein FAM3C | 132368.875 | 129238.0625 |
| FRA10AC1 | Q70Z53 | FRA10AC1 | Protein FRA10AC1 | 19318.07813 | 19119.49609 |
| RSL1D1 | O76021 | RSL1D1 | Ribosomal L1 domain- | 644557.6875 | 354733.375 |
| NDUFA7 | O95182 | NDUFA7 | NADH dehydrogenase | 49590.34766 | 59896.44531 |
| SART1 | O43290 | SART1 | U4/U6.U5 tri-snRNP-a | 226560.0938 | 197155.0313 |
| PSME3 | P61289 | PSME3 | Proteasome activator | 384744.8438 | 468725.5938 |
| FAH | P16930 | FAH | Fumarylacetoacetase | 187382.2188 | 223531.5156 |
| CLCN5 | P51795 | CLCN5 | H(+)/Cl(-) exchange tr | 38617.02734 | 41739.53516 |
| KIF2A | D6R9M0 | KIF2A | Kinesin-like protein KI | 185617.4512 | 192211.9219 |
| IWS1 | Q96ST2 | IWS1 | Protein IWS1 homolog | 101223.6875 | 91342.03906 |
| POLR2B | C9J2Y9;P30876 | POLR2B | DNA-directed RNA po | 128036.0313 | 84738.58594 |
| MT-ND2 | P03891 | MT-ND2 | NADH-ubiquinone oxi | 69713.40625 | 71868 |
| HSPA14 | Q0VDF9 | HSPA14 | Heat shock 70 kDa pro | 151026.0156 | 163172.9375 |
| EEF1A1 | P68104 | EEF1A1 | Elongation factor 1-al | 43630392 | 49987868 |
| RRN3 | F5H148;H3BMT1;Q9NYV6 | RRN3 | RNA polymerase I-spe | 12840.43359 | 13345.42578 |
| USP13 | Q92995 | USP13 | Ubiquitin carboxyl-ter | 47532.71094 | 52830.35156 |

| | | | | | |
|---|---|---|---|---|---|
| VPS25 | Q9BRG1 | VPS25 | Vacuolar protein-sorti | 195843.1094 | 203100.5625 |
| ANKHD1 | Q8IWZ3 | ANKHD1 | Ankyrin repeat and KH | 136924.6094 | 152913.4063 |
| MSH2 | P43246 | MSH2 | DNA mismatch repair | 288587.3438 | 300446.6875 |
| KCTD5 | Q9NXV2 | KCTD5 | BTB/POZ domain-cont | 21435.48828 | 20165.04688 |
| HGH1 | Q9BTY7 | HGH1 | Protein HGH1 homolo | 27851.18945 | 56923.92578 |
| NOP56 | O00567 | NOP56 | Nucleolar protein 56 | 307892.25 | 71517.55469 |
| RPF2 | Q9H7B2 | RPF2 | Ribosome production | 173527.2344 | 78934.96875 |
| SMNDC1 | O75940 | SMNDC1 | Survival of motor neu | 151259.4844 | 111211.2188 |
| HMG20A | Q9NP66 | HMG20A | High mobility group p | 6064.303711 | 11796.48242 |
| UBXN1 | Q04323 | UBXN1 | UBX domain-containir | 135632.875 | 147115.2969 |
| EDEM3 | Q9BZQ6 | EDEM3 | ER degradation-enhar | 31218.99414 | 28043.40625 |
| MOB1A | Q9H8S9 | MOB1A | MOB kinase activator | 173050.875 | 181634.9219 |
| HAUS6 | Q5VY60;Q7Z4H7 | HAUS6 | HAUS augmin-like con | 3616.628174 | 5438.924316 |
| YIPF1 | Q9Y548 | YIPF1 | Protein YIPF1 | 33439.08984 | 31654.66992 |
| C6orf120 | Q7Z4R8 | C6orf120 | UPF0669 protein C6or | 9025.616211 | 9326.661133 |
| MED15 | Q96RN5 | MED15 | Mediator of RNA poly | 50404.08594 | 50206.16016 |
| TTC39C | Q8N584 | TTC39C | Tetratricopeptide rep | 9593.81543 | 11681.87402 |
| ARHGEF2 | A0A5F9ZI21;Q92974;V9GYM | ARHGEF2 | Rho guanine nucleotic | 57876.62891 | 64395.31641 |
| POLR3F | Q9H1D9 | POLR3F | DNA-directed RNA po | 33405.97656 | 35415.65625 |
| SPINT2 | O43291 | SPINT2 | Kunitz-type protease i | 223191.6094 | 282095.6563 |
| FTSJ1 | Q9UET6 | FTSJ1 | Putative tRNA (cytidin | 25182.37695 | 27075.74805 |
| CCND3 | P30281 | CCND3 | G1/S-specific cyclin-D | 15331.34668 | 17738.92773 |
| MRPS17;hCG_1984214;MR | E9PE17;I3L0E3;Q9Y2R5 | MRPS17;hCG_198421 | 28S ribosomal protein | 133645.4688 | 148439.3125 |
| NTMT1 | Q9BV86 | NTMT1 | N-terminal Xaa-Pro-Ly | 255772.8906 | 204555.3594 |
| RPAIN | Q86UA6 | RPAIN | RPA-interacting prote | 24907.85352 | 20158.40625 |
| HSPBP1 | Q9NZL4 | HSPBP1 | Hsp70-binding proteir | 218335.8281 | 226444.2344 |
| BIRC5 | A0A0B4J1S3;H3BLT4;K7EMW | BIRC5 | Baculoviral IAP repeat | 14308.68945 | 14576.27637 |
| TMF1 | P82094 | TMF1 | TATA element modula | 48650.15625 | 44470.68359 |
| SURF2 | Q15527 | SURF2 | Surfeit locus protein 2 | 78535.50781 | 66908.5 |
| TUBB6 | Q9BUF5 | TUBB6 | Tubulin beta-6 chain | 1602543.5 | 1905886 |
| TXNL4A | P83876 | TXNL4A | Thioredoxin-like prote | 56205.70313 | 55208.89844 |
| MAEA | E7ESC7;Q7L5Y9 | MAEA | E3 ubiquitin-protein tr | 29255.64063 | 30745.23242 |
| CHUK | O15111 | CHUK | Inhibitor of nuclear fa | 21073.32031 | 19475.55859 |

| | | | | | |
|---|---|---|---|---|---|
| NAA50 | E7EQ69;Q9GZZ1 | NAA50 | N-alpha-acetyltransfe | 187578 | 193350.4375 |
| KDM3A | Q9Y4C1 | KDM3A | Lysine-specific demetl | 7053.297363 | 6338.372559 |
| TOR1AIP1 | J3KN66 | TOR1AIP1 | Torsin-1A-interacting | 260851.7031 | 280118.1289 |
| CCNL2 | Q96S94 | CCNL2 | Cyclin-L2 | 2769.450195 | 2298.972412 |
| NMRK1 | Q9NWW6 | NMRK1 | Nicotinamide riboside | 12976.96191 | 24930.70508 |
| CYB5B | O43169 | CYB5B | Cytochrome b5 type E | 45011.57813 | 112997.1875 |
| USP39 | Q53GS9 | USP39 | U4/U6.U5 tri-snRNP-a | 175083.8438 | 179474.3906 |
| EBNA1BP2 | H7C2Q8;Q99848 | EBNA1BP2 | EBNA1 binding protei | 207535.4219 | 146458.3594 |
| SLC30A1 | Q9Y6M5 | SLC30A1 | Zinc transporter 1 | 84176.8125 | 74305.04688 |
| ZC3H11A | O75152 | ZC3H11A | Zinc finger CCCH dom | 159166.3906 | 153216.0781 |
| SLC4A2 | P04920 | SLC4A2 | Anion exchange prote | 53521.69531 | 42022.69141 |
| MRPL18 | Q9H0U6 | MRPL18 | 39S ribosomal protein | 130093.625 | 132634.1406 |
| PLS3 | P13797 | PLS3 | Plastin-3 | 181084.2813 | 187265.6094 |
| PIK3C3 | A0A1W2PS05;Q8NEB9 | PIK3C3 | Phosphatidylinositol 3 | 29410.75977 | 32049.00781 |
| RBM6 | P78332 | RBM6 | RNA-binding protein 6 | 66424.07813 | 68819.78906 |
| SNX2 | O60749 | SNX2 | Sorting nexin-2 | 202745.0313 | 215150.6563 |
| MDN1 | Q9NU22 | MDN1 | Midasin | 64582.06641 | 67632.14844 |
| BRCA1 | A0A2R8Y7V5;B7ZA85;E7EUM | BRCA1 | Breast cancer type 1 s | 2646.857178 | 1823.097046 |
| RPAP2 | Q8IXW5 | RPAP2 | Putative RNA polymer | 7363.257324 | 13456.55273 |
| FAM98A | Q8NCA5 | FAM98A | Protein FAM98A | 107579.875 | 120129.1953 |
| NAA10 | P41227 | NAA10 | N-alpha-acetyltransfe | 181707.8438 | 177686.375 |
| PLK1 | P53350 | PLK1 | Serine/threonine-prot | 57851.45313 | 55865.99609 |
| PRPF8 | Q6P2Q9 | PRPF8 | Pre-mRNA-processing | 552467.125 | 470760.75 |
| GFM2 | Q969S9 | GFM2 | Ribosome-releasing fa | 37220.70703 | 41762.17969 |
| SLC39A10 | Q9ULF5 | SLC39A10 | Zinc transporter ZIP1C | 22290.35156 | 19638.67383 |
| COL18A1 | H7BXV5;P39060 | COL18A1 | Collagen alpha-1(XVIII | 14330.88965 | 9435.024414 |
| LARP1B | A0A3B3ISF0;Q659C4 | LARP1B | La-related protein 1B | 19638.06836 | 17682.22656 |
| ZBTB40 | B1AKC8;Q9NUA8 | ZBTB40 | Zinc finger and BTB dc | 5210.382813 | 6394.316406 |
| ZNF800 | A0A087WXB4;C9JHX9;C9JLY3 | ZNF800 | Zinc finger protein 80( | 31559.49219 | 28444.66602 |
| ZBTB7A | O95365 | ZBTB7A | Zinc finger and BTB dc | 102387 | 105061.8984 |
| HELZ | J3QS41 | HELZ | Probable helicase witl | 41793.27734 | 43265.36133 |
| GAPDH | P04406 | GAPDH | Glyceraldehyde-3-phc | 21844280 | 21054910 |
| NOMO1 | A0A087X117;A0A0G2JN29;Q | NOMO1 | Nodal modulator 1 | 11658.30371 | 10599.58887 |

| | | | | | |
|---|---|---|---|---|---|
| WDR6 | Q9NNW5 | WDR6 | WD repeat-containing | 75703.57813 | 71927.66406 |
| RANBP10 | A0A0D9SEU5;Q6VN20;R4GM | RANBP10 | Ran-binding protein 1 | 24612.64453 | 27572.02734 |
| H2BC11;HIST1H2BO;HIST1H | P06899;P23527;P33778;Q16 | H2BC11;HIST1H2BO;H | Histone H2B type 1-J;I | 11599807 | 832643.4375 |
| SART3 | A0A499FI31 | SART3 | Squamous cell carcinc | 491753.6563 | 501059.9063 |
| MIS18A | Q9NYP9 | MIS18A | Protein Mis18-alpha | 25182.65625 | 13978.61328 |
| CTNNBL1 | A0A087WUB9;Q8WYA6 | CTNNBL1 | Beta-catenin-like prot | 371197 | 384291.375 |
| GIGYF2 | Q6Y7W6 | GIGYF2 | GRB10-interacting GY | 129999.5781 | 113396.7344 |
| NDUFS8 | O00217 | NDUFS8 | NADH dehydrogenase | 232102.5313 | 286203.375 |
| MIPEP | Q99797 | MIPEP | Mitochondrial interm | 58660.74609 | 81297.01563 |
| NDUFAF8 | A1L188 | NDUFAF8 | NADH dehydrogenase | 13273.2832 | 11143.90332 |
| RELCH | Q9P260 | RELCH | RAB11-binding protei | 54206.72266 | 60576.16016 |
| SIRT7 | I3L2A4;Q9NRC8 | SIRT7 | NAD-dependent prote | 9602.226563 | 10136.97754 |
| TXNL4B | Q9NX01 | TXNL4B | Thioredoxin-like prote | 21233.14453 | 21155.36328 |
| SMG5 | Q9UPR3 | SMG5 | Protein SMG5 | 22061.71484 | 22335.98438 |
| SNW1 | Q13573 | SNW1 | SNW domain-containi | 215226.8438 | 160092.8438 |
| PUS7 | Q96PZ0 | PUS7 | Pseudouridylate synth | 106212.9141 | 107166.4766 |
| MRPS12 | O15235 | MRPS12 | 28S ribosomal protein | 81241.40625 | 100801.0781 |
| ICOSLG | K4DIA0;O75144 | ICOSLG | ICOS ligand | 9184.332031 | 9257.583008 |
| ANKRD50 | Q9ULJ7 | ANKRD50 | Ankyrin repeat domai | 23294.68164 | 36707.22266 |
| ARL14EP | Q8N8R7 | ARL14EP | ARL14 effector protei | 20344.62109 | 19643.96289 |
| CPTP | Q5TA50 | CPTP | Ceramide-1-phosphat | 17129.11328 | 17163.51563 |
| FAM193B | Q96PV7 | FAM193B | Protein FAM193B | 5013.779297 | 4645.618652 |
| SNU13 | P55769 | SNU13 | NHP2-like protein 1 | 758792.0625 | 334273.9688 |
| CHD1 | O14646 | CHD1 | Chromodomain-helica | 5585.133301 | 22720.21484 |
| MRPL13 | Q9BYD1 | MRPL13 | 39S ribosomal protein | 108122.1563 | 120657.4063 |
| TYW3 | Q6IPR3 | TYW3 | tRNA wybutosine-syn | NA | 7749.519043 |
| UBE2C | O00762 | UBE2C | Ubiquitin-conjugating | 239426.6719 | 210327.375 |
| TIMM13 | Q9Y5L4 | TIMM13 | Mitochondrial import | 347056.3125 | 329477.9688 |
| SULF2 | Q8IWU5 | SULF2 | Extracellular sulfatase | 16385.50195 | 24066.14453 |
| AMBRA1 | A0A075B6T1;Q9C0C7 | AMBRA1 | Activating molecule ir | 9595.171875 | 9114.819336 |
| ASNSD1 | C9IYZ1;Q9NWL6 | ASNSD1 | Asparagine synthetase | 3615.311523 | 3691.138428 |
| SPCS3 | P61009 | SPCS3 | Signal peptidase comp | 273873.6875 | 278373.7188 |
| NUTF2 | P61970 | NUTF2 | Nuclear transport fact | 146826.7656 | 312133.2188 |

| | | | | | |
|---|---|---|---|---|---|
| AP1S3 | B8ZZZ1;F8W8K9;G5E966;Q96 | AP1S3 | AP complex subunit si | 14850.08984 | 17521.28906 |
| RNF115 | Q9Y4L5 | RNF115 | E3 ubiquitin-protein li | 13008.44141 | 11796.66602 |
| SRFBP1 | Q8NEF9 | SRFBP1 | Serum response facto | 28398.63281 | 21271.40039 |
| SPIRE1 | Q08AE8 | SPIRE1 | Protein spire homolog | 10777.16699 | 9575.049805 |
| ARNT | P27540 | ARNT | Aryl hydrocarbon rece | 18734.73828 | 24990.20703 |
| GCC2 | Q8IWJ2 | GCC2 | GRIP and coiled-coil d | 16162.27344 | 17326.47852 |
| YWHAE | P62258 | YWHAE | 14-3-3 protein epsilor | 2244603.5 | 2329126 |
| POLR2G | P62487 | POLR2G | DNA-directed RNA po | 75636.19531 | 51326.59375 |
| INTS2 | J3KMZ7 | INTS2 | Integrator complex su | 37214.52734 | 38258.58594 |
| DGKE | I3L112;P52429 | DGKE | Diacylglycerol kinase ( | 28280.14258 | 28427.58008 |
| FPGT | O14772 | FPGT | Fucose-1-phosphate g | 3823.154053 | 5343.879883 |
| CDK12 | Q9NYV4 | CDK12 | Cyclin-dependent kina | 14872.43652 | 15382.87988 |
| ELMO3 | F8W9E7;Q96BJ8 | ELMO3 | Engulfment and cell m | 55992.62891 | 59851.78906 |
| CDC42EP4 | Q9H3Q1 | CDC42EP4 | Cdc42 effector proteir | 107831.3906 | 111382.2891 |
| RPL39;RPL39P5 | P62891;Q59GN2 | RPL39;RPL39P5 | 60S ribosomal protein | 260796.7969 | 246527.7031 |
| LNPEP | Q9UIQ6 | LNPEP | Leucyl-cystinyl aminop | 144202.0938 | 141722.1719 |
| H3-2 | Q5TEC6 | H3-2 | Histone H3 | 266500.9688 | 5211.216309 |
| CASC3 | O15234 | CASC3 | Protein CASC3 | 25310.00977 | 23695.1582 |
| BPTF | A0A2R8Y7Q1;E9PE19 | BPTF | Nucleosome-remodel | 245320.2368 | 251119.4043 |
| TPI1 | P60174 | TPI1 | Triosephosphate isom | 7751553.5 | 7296411.5 |
| ADI1 | Q9BV57 | ADI1 | 1,2-dihydroxy-3-keto- | 122363.0078 | 201495.8438 |
| TRIM16L | Q309B1 | TRIM16L | Tripartite motif-conta | 13425.16895 | 11373.76367 |
| ADSL | A0A1B0GTJ7;A0A1B0GWJ0;P | ADSL | Adenylosuccinate lyas | 196479.2656 | 218205.2969 |
| GOSR2;GOSR2;GOSR2;GOS | A0A1W2PPG1;A0A1W2PPG5 | GOSR2;GOSR2;GOSR2 | Golgi SNAP receptor c | 75609.84375 | 89398.77344 |
| C11orf98 | E9PRG8 | C11orf98 | Uncharacterized prote | 297006.4063 | 332151.3125 |
| PRMT1 | E9PKG1;Q99873 | PRMT1 | Protein arginine N-me | 1039466.688 | 1261043.75 |
| NUP50 | Q9UKX7 | NUP50 | Nuclear pore complex | 166986.2656 | 158253.9531 |
| PHAX | Q9H814 | PHAX | Phosphorylated adapt | 137012.2656 | 137845.8438 |
| SCFD2 | Q8WU76 | SCFD2 | Sec1 family domain-co | 28593.54492 | 33820.48047 |
| FLAD1 | Q8NFF5 | FLAD1 | FAD synthase | 127462.2969 | 136420.6406 |
| STK11IP | Q8N1F8 | STK11IP | Serine/threonine-prot | 41225.20313 | 46019.63672 |
| MIGA2 | Q7L4E1 | MIGA2 | Mitoguardin 2 | 15717.69043 | 13433.21973 |
| POLR2M | Q6EEV4 | POLR2M | DNA-directed RNA po | 6599.111328 | 8636.042969 |

| | | | | | |
|---|---|---|---|---|---|
| TSR3 | Q9UJK0 | TSR3 | Ribosome biogenesis | 13644.30957 | 19605.28711 |
| MICU1 | A0A0U1RRK1;A0A286YF11;F( | MICU1 | Calcium uptake protei | 18387.27539 | 23234.29688 |
| DAG1 | Q14118 | DAG1 | Dystroglycan | 120515.1953 | 149362.375 |
| ANKRD13A | Q8IZ07 | ANKRD13A | Ankyrin repeat domai | 6552.161621 | 7924.187012 |
| BIRC6 | A0A3B3IUB9 | BIRC6 | Baculoviral IAP repeat | 66716.32422 | 74291.80176 |
| POLG2 | A0A5F9ZH93;A0A5F9ZHX9;Q | POLG2 | DNA polymerase subu | 32866.11328 | 52660.17578 |
| E2F4 | Q16254 | E2F4 | Transcription factor E | 16134.5752 | 7523.8125 |
| MRPL46 | Q9H2W6 | MRPL46 | 39S ribosomal protein | 171522.1875 | 183284.1094 |
| MRPS33 | Q9Y291 | MRPS33 | 28S ribosomal protein | 83835.5625 | 91091.60938 |
| TRIM25 | Q14258 | TRIM25 | E3 ubiquitin/ISG15 lig | 422046.8438 | 454231.3438 |
| TAF10 | Q12962 | TAF10 | Transcription initiatio | 52388.52734 | 53421.51563 |
| ZMIZ1 | F6WR09;Q9ULJ6 | ZMIZ1 | Zinc finger MIZ domai | 8069.698242 | 11325.9541 |
| CCT2 | P78371 | CCT2 | T-complex protein 1 s | 1466582 | 1594740.125 |
| SARS2 | M0QWZ7;Q9NP81 | SARS2 | Serine--tRNA ligase, m | 86928.07031 | 84808.02344 |
| RDX | A0A2R8Y5S7;P35241 | RDX | Radixin | 1891294.75 | 1996414.75 |
| ACTR1A | P61163 | ACTR1A | Alpha-centractin | 280259.8125 | 328212.4063 |
| PTP4A1 | Q93096 | PTP4A1 | Protein tyrosine phos | 74115.25 | 73312.5 |
| FEM1B | Q9UK73 | FEM1B | Protein fem-1 homolc | 11510.6748 | 11635.16992 |
| TBC1D10A | Q9BXI6 | TBC1D10A | TBC1 domain family m | 98075.13281 | 98164.875 |
| CARM1 | Q86X55 | CARM1 | Histone-arginine meth | 185804.0781 | 218091.8125 |
| WDYHV1 | Q96HA8 | WDYHV1 | Protein N-terminal glu | 24130.69922 | 23615.24414 |
| GBF1 | Q92538 | GBF1 | Golgi-specific brefeldi | 70107.15625 | 88721.79688 |
| MRPL55 | Q7Z7F7 | MRPL55 | 39S ribosomal protein | 123710.1563 | 129515.2031 |
| GEMIN4 | I3L2C7;P57678 | GEMIN4 | Gem-associated prote | 79601.29688 | 113816.625 |
| RPP40 | O75818 | RPP40 | Ribonuclease P protei | 77587.53125 | 78394.07813 |
| KDSR | K7ERC8;Q06136 | KDSR | 3-ketodihydrosphingo | 26847.92188 | 28082.23828 |
| LPCAT2 | Q7L5N7 | LPCAT2 | Lysophosphatidylcholi | 7192.456543 | 7815.324707 |
| AURKA | O14965 | AURKA | Aurora kinase A | 60683.14453 | 43777.19531 |
| KMT2B | Q9UMN6 | KMT2B | Histone-lysine N-meth | 10202.71875 | 9115.063477 |
| PRDX4 | Q13162 | PRDX4 | Peroxiredoxin-4 | 7105735.5 | 7780168 |
| SLC7A6 | Q92536 | SLC7A6 | Y+L amino acid transp | 20240.05273 | 20042.08789 |
| SPCS1 | A0A5F9YFS9 | SPCS1 | Signal peptidase comp | 141895.0469 | 167566.9375 |
| RBL1 | P28749 | RBL1 | Retinoblastoma-like p | 8436.595703 | 7293.195801 |

| | | | | | |
|---|---|---|---|---|---|
| REXO4 | Q9GZR2 | REXO4 | RNA exonuclease 4 | 30368.0332 | 25862.82031 |
| SPEN | Q96T58 | SPEN | Msx2-interacting prot | 25475.86523 | 11664.52344 |
| HSP90AB4P | Q58FF6 | HSP90AB4P | Putative heat shock pr | 46635.24609 | 52693.84766 |
| GTF2F2 | P13984 | GTF2F2 | General transcription | 88946.33594 | 86602.42969 |
| CHD7 | Q9P2D1 | CHD7 | Chromodomain-helica | 2176.47998 | 3145.196045 |
| GAA | P10253 | GAA | Lysosomal alpha-gluco | 324815.3438 | 343198.25 |
| APP | E9PG40;P05067 | APP | Amyloid-beta precurs | 57724.41406 | 50267.89453 |
| TMEM126B | E9PJQ6;E9PKZ7;E9PKZ9;Q8IU | TMEM126B | Complex I assembly fa | 19529.82617 | 17991.67773 |
| SF3B2 | Q13435 | SF3B2 | Splicing factor 3B subu | 567564.6875 | 438325.1875 |
| TACSTD2 | P09758 | TACSTD2 | Tumor-associated calc | 201977.7031 | 238852.6875 |
| MRPL1 | Q9BYD6 | MRPL1 | 39S ribosomal protein | 137619.3281 | 136734.1875 |
| AFF4 | Q9UHB7 | AFF4 | AF4/FMR2 family mer | 70016.70313 | 67438.75 |
| NPM1 | P06748 | NPM1 | Nucleophosmin | 2988224.25 | 2009476.5 |
| MAT2A | P31153 | MAT2A | S-adenosylmethionine | 877585.5 | 804561 |
| WBP4 | O75554 | WBP4 | WW domain-binding p | 58202.55078 | 54295.12109 |
| CDK5 | Q00535 | CDK5 | Cyclin-dependent-like | 112794.8672 | 120553.4219 |
| ZNRF2 | Q8NHG8 | ZNRF2 | E3 ubiquitin-protein li | 18593.97852 | 21368.21875 |
| OSBPL1A | Q9BXW6 | OSBPL1A | Oxysterol-binding pro | 17536.05273 | 17615.19141 |
| TCF3 | P15923;X6REB3 | TCF3 | Transcription factor E | 12404.21777 | 11129.10742 |
| HENMT1 | A0A0A0MRN1;Q5T8I9 | HENMT1 | Small RNA 2'-O-methy | 9232.445313 | 7634.959473 |
| ARL15 | A0A087WUW9;Q9NXU5 | ARL15 | ADP-ribosylation facto | 11804.0293 | 15295.72754 |
| RPAP1 | Q9BWH6 | RPAP1 | RNA polymerase II-ass | 30834.17188 | 29504.86914 |
| COQ7 | H3BP28;Q99807 | COQ7 | 5-demethoxyubiquinc | 19476.87305 | 19791.43359 |
| MRPS24 | Q96EL2 | MRPS24 | 28S ribosomal protein | 88750.82813 | 107049.4531 |
| CRK | P46108 | CRK | Adapter molecule crk | 165599.4375 | 176522.5625 |
| MT-CO3 | P00414 | MT-CO3 | Cytochrome c oxidase | 20661.27734 | 25413 |
| RBM26 | Q5T8P6 | RBM26 | RNA-binding protein 2 | 56407.50781 | 54156.98438 |
| MGAT1 | P26572 | MGAT1 | Alpha-1,3-mannosyl-g | 74354.99219 | 87069.77344 |
| CHCHD3 | C9JRZ6;Q9NX63 | CHCHD3 | MICOS complex subur | 558077.25 | 320462.0313 |
| CAST | A0A0C4DGB5 | CAST | Calpastatin | 107330.5889 | 154771.4482 |
| RPS6KB1 | P23443 | RPS6KB1 | Ribosomal protein S6 | 810374.5 | 951394.0625 |
| SNRPB2 | P08579 | SNRPB2 | U2 small nuclear ribor | 204311.4531 | 155638.0156 |
| SNRPE | P62304 | SNRPE | Small nuclear ribonuc | 670952.6875 | 505077.0625 |

| | | | | | |
|---|---|---|---|---|---|
| RNASEH2B | A0A2R8YCX2;A0A2R8YGP2;Q | RNASEH2B | Ribonuclease H2 subu | 60789.51563 | 65240.78125 |
| SAP30 | O75446 | SAP30 | Histone deacetylase c | 25262.23242 | 26365.84766 |
| CCNT1 | O60563 | CCNT1 | Cyclin-T1 | 42732.28906 | 43105.90625 |
| TBRG4 | Q969Z0 | TBRG4 | FAST kinase domain-c | 187334.9688 | 221632.25 |
| CHD4 | A0A2R8Y521;A0A2R8Y5J0;A0 | CHD4 | Chromodomain-helica | 425690.2813 | 429941 |
| UPF1 | Q92900 | UPF1 | Regulator of nonsense | 387146.3125 | 341450.4375 |
| GPN1 | Q9HCN4 | GPN1 | GPN-loop GTPase 1 | 73837.71875 | 86203.82031 |
| NDUFS4 | O43181 | NDUFS4 | NADH dehydrogenase | 126720.0859 | 142480.0313 |
| PRELID1 | D6RC47;D6REW1;Q9Y255 | PRELID1 | PRELI domain-contain | 22971.81641 | 20336.2832 |
| RCHY1 | Q96PM5 | RCHY1 | RING finger and CHY z | 12135.0752 | 10228.67578 |
| KNL1 | E9PPJ1;Q8NG31 | KNL1 | Kinetochore scaffold 1 | 8366.974609 | 7780.843262 |
| ACOX1 | Q15067 | ACOX1 | Peroxisomal acyl-coer | 117980.6719 | 133403.3125 |
| FBXO22 | Q8NEZ5 | FBXO22 | F-box only protein 22 | 147801.9375 | 167786.3594 |
| SORD | Q00796 | SORD | Sorbitol dehydrogena | 916539.625 | 1063502.75 |
| MRPL14 | Q6P1L8 | MRPL14 | 39S ribosomal protein | 73224.875 | 83347.41406 |
| MRPL4 | K7ES61;Q9BYD3 | MRPL4 | 39S ribosomal protein | 140816.2031 | 150135.5625 |
| VPS37A | Q8NEZ2 | VPS37A | Vacuolar protein sorti | 8234.654297 | 10697.03027 |
| PRIM2 | A0A5F9ZHU7;P49643 | PRIM2 | DNA primase large sul | 85157.55469 | 109863.3047 |
| ZNF346 | Q9UL40 | ZNF346 | Zinc finger protein 34 | 19878.32227 | 19091.83594 |
| PRKCI | P41743 | PRKCI | Protein kinase C iota t | 208357.4844 | 229688.25 |
| XYLT2 | Q9H1B5 | XYLT2 | Xylosyltransferase 2 | 45448.03906 | 48902.79297 |
| FAM118B | E9PMJ2;G3V179;J3KP39;Q9B | FAM118B | Protein FAM118B (Fra | 6984.14209 | 10393.89551 |
| CNPY2 | F8VXJ7;Q9Y2B0 | CNPY2 | Protein canopy homol | 16310.38086 | 21731.78711 |
| DCP1A | Q9NPI6 | DCP1A | mRNA-decapping enzy | 15695.8457 | 16952.79492 |
| C16orf58 | H3BSM7;Q96GQ5 | C16orf58 | RUS1 family protein C | 40515.56641 | 44472.66797 |
| POLG | P54098 | POLG | DNA polymerase subu | 39067.14063 | 42885.6875 |
| SRA1 | Q9HD15 | SRA1 | Steroid receptor RNA | 96131.52344 | 104161.5938 |
| MRPS5 | P82675 | MRPS5 | 28S ribosomal protein | 132017.3594 | 145053.9844 |
| ZNF8 | P17098 | ZNF8 | Zinc finger protein 8 | 152.6986237 | 476.8701477 |
| PRPF4B | Q13523 | PRPF4B | Serine/threonine-prot | 192979.4688 | 131516.3438 |
| ZC3HC1 | C9J0I9;Q86WB0 | ZC3HC1 | Nuclear-interacting pa | 45341.375 | 46104.56641 |
| DDX51 | Q8N8A6 | DDX51 | ATP-dependent RNA h | 58372.82813 | 53551.19531 |
| C5orf22 | Q49AR2 | C5orf22 | UPF0489 protein C5or | 17703.81836 | 29096.18945 |

| | | | | | |
|---|---|---|---|---|---|
| PMPCA | Q10713 | PMPCA | Mitochondrial-proces | 151651.6875 | 161561.7344 |
| SCAMP1 | O15126 | SCAMP1 | Secretory carrier-asso | 190557.4063 | 214364.0938 |
| TIMM50 | Q3ZCQ8 | TIMM50 | Mitochondrial import | 369739.1563 | 398958.625 |
| YWHAG | P61981 | YWHAG | 14-3-3 protein gamma | 1162802.875 | 1253046.625 |
| MYDGF | Q969H8 | MYDGF | Myeloid-derived grow | 184271.4688 | 196647.5313 |
| MFSD3 | Q96ES6 | MFSD3 | Major facilitator supe | 14685.0957 | 14948.61133 |
| TP53BP1 | Q12888 | TP53BP1 | TP53-binding protein | 38467.48438 | 34712.21094 |
| DUSP11 | O75319 | DUSP11 | RNA/RNP complex-1-i | 19975.47852 | 20702.79688 |
| GGA3 | Q9NZ52 | GGA3 | ADP-ribosylation facto | 78600.92188 | 69723.17969 |
| GTPBP3 | Q969Y2 | GTPBP3 | tRNA modification GT | 15300.52832 | 16695.85352 |
| TMEM192 | Q8IY95 | TMEM192 | Transmembrane prote | 110904.1406 | 93872.58594 |
| MT-ND3 | P03897 | MT-ND3 | NADH-ubiquinone oxi | 50475.62891 | 69805.86719 |
| HAS1 | G3V1S7;M0QXH4;M0R2V0;Q | HAS1 | Hyaluronan synthase | 3120840.75 | 3629670.5 |
| DDX20 | Q9UHI6 | DDX20 | Probable ATP-depend | 170325.0313 | 237769.1563 |
| RCN2 | Q14257 | RCN2 | Reticulocalbin-2 | 2833.722168 | 4745.839355 |
| MPHOSPH6 | H3BNT4;Q99547 | MPHOSPH6 | M-phase phosphopro | 54440.27344 | 41997.31641 |
| TPP1 | A0A2R8YGD1;O14773 | TPP1 | Tripeptidyl-peptidase | 20105.62109 | 27827.79102 |
| WDR77 | Q9BQA1 | WDR77 | Methylosome protein | 468311.8438 | 500848.9063 |
| RNF216 | Q9NWF9 | RNF216 | E3 ubiquitin-protein li | 2676.113037 | 2691.721924 |
| DIS3 | Q9Y2L1 | DIS3 | Exosome complex exo | 157570.6875 | 152613.2656 |
| PDXDC1 | Q6P996 | PDXDC1 | Pyridoxal-dependent | 154190.6406 | 188660.625 |
| SRSF1 | J3KTL2;Q07955 | SRSF1 | Serine/arginine-rich-s | 4215129.5 | 2338289.25 |
| NDUFA8 | P51970 | NDUFA8 | NADH dehydrogenase | 313951.4375 | 296044.9063 |
| LCLAT1 | Q6UWP7 | LCLAT1 | Lysocardiolipin acyltra | 63572.95703 | 59462.68359 |
| MRPS18B | Q9Y676 | MRPS18B | 28S ribosomal protein | 122784.2031 | 122921.7422 |
| HMGB2 | P26583 | HMGB2 | High mobility group p | 2121284 | 2016362.625 |
| DNAJC2 | Q99543 | DNAJC2 | DnaJ homolog subfam | 124514.5313 | 123693.6563 |
| BUD13 | Q9BRD0 | BUD13 | BUD13 homolog | 29115.01172 | 19654.96094 |
| NBAS | A2RRP1 | NBAS | Neuroblastoma-ampli | 24990.40039 | 24656.48633 |
| SLAIN2 | Q9P270 | SLAIN2 | SLAIN motif-containin | 55028.10938 | 55811.14844 |
| DDX55 | F5H5U2;Q8NHQ9 | DDX55 | RNA helicase;ATP-dep | 75500.89844 | 76435.85156 |
| ISY1 | Q9ULR0 | ISY1 | Pre-mRNA-splicing fac | 96484.03906 | 87642.49219 |
| VPS13A | Q96RL7 | VPS13A | Vacuolar protein sorti | 24379.57422 | 28512.57617 |

| | | | | | |
|---|---|---|---|---|---|
| MRPL45 | A0A087X2D5;Q9BRJ2 | MRPL45 | 39S ribosomal protein | 171904.2656 | 193189.6406 |
| LRIF1 | Q5T3J3 | LRIF1 | Ligand-dependent nuc | 5580.103027 | 4127.777344 |
| MRPL40 | Q9NQ50 | MRPL40 | 39S ribosomal protein | 154345.875 | 143572.2188 |
| DRG1 | Q9Y295 | DRG1 | Developmentally-regu | 252289.1875 | 273598.2813 |
| RPLP2 | P05387 | RPLP2 | 60S acidic ribosomal p | 23339.95117 | 30655.04102 |
| HSPA4L | O95757 | HSPA4L | Heat shock 70 kDa pro | 351019.375 | 395685.8438 |
| MRPL39 | Q9NYK5 | MRPL39 | 39S ribosomal protein | 141111.0156 | 155305.5469 |
| NDRG3 | Q9UGV2 | NDRG3 | Protein NDRG3 | 116293.75 | 143626.6719 |
| NUDT19 | A8MXV4 | NUDT19 | Nucleoside diphospha | 16112.91211 | 18254.50391 |
| PDLIM2 | A0A087WVI1;Q96JY6 | PDLIM2 | PDZ and LIM domain p | 15876.25293 | 12249.8457 |
| MRPL28 | A2IDC6;Q13084 | MRPL28 | 39S ribosomal protein | 88398.00781 | 104403.1094 |
| MRPL16 | Q9NX20 | MRPL16 | 39S ribosomal protein | 124948.4766 | 142632.3281 |
| AMOTL2 | Q9Y2J4 | AMOTL2 | Angiomotin-like prote | 56567.5625 | 51195.26563 |
| SETDB1 | Q15047 | SETDB1 | Histone-lysine N-meth | 45875.40625 | 47715.9375 |
| GPSM2 | P81274 | GPSM2 | G-protein-signaling m | 10030.54492 | 9446.844727 |
| DCD | P81605 | DCD | Dermcidin | 3822.293945 | 7738.114746 |
| NME1-NME2 | Q32Q12 | NME1-NME2 | Nucleoside diphospha | 18655164 | 19486252 |
| SMG7 | E9PD50;Q92540 | SMG7 | Protein SMG7 | 24639.65039 | 24719.7832 |
| EPHA1 | P21709 | EPHA1 | Ephrin type-A recepto | 5588.322266 | 7800.29834 |
| CIR1 | Q86X95 | CIR1 | Corepressor interactir | 9777.427734 | 12579.48145 |
| TRABD | J3KPT4;Q9H4I3 | TRABD | TraB domain-containi | 13994.66602 | 14260.625 |
| NDUFA5 | Q16718 | NDUFA5 | NADH dehydrogenase | 102124.4766 | 117160.9219 |
| SPTAN1 | A0A0D9SGF6 | SPTAN1 | Spectrin alpha chain, r | 1004353.665 | 633162.6914 |
| ZFAND6 | Q6FIF0 | ZFAND6 | AN1-type zinc finger p | 58111.75781 | 74155.32031 |
| MRPS9 | P82933 | MRPS9 | 28S ribosomal protein | 257119.5781 | 290995.5 |
| LMAN2 | Q12907 | LMAN2 | Vesicular integral-mer | 1154221.625 | 1353857.625 |
| CENPH | B3KVZ3;Q9H3R5 | CENPH | Centromere protein H | 13264.98828 | 14442.31934 |
| RPUSD2 | Q8IZ73 | RPUSD2 | RNA pseudouridylate | 20475.33203 | 21117.51563 |
| FRMPD1 | Q5SYB0 | FRMPD1 | FERM and PDZ domai | 82924.78906 | 89141.97656 |
| SLC1A5 | Q15758 | SLC1A5 | Neutral amino acid tra | 905922 | 965075.125 |
| AURKB | Q96GD4 | AURKB | Aurora kinase B | 45232.25 | 40748.76563 |
| AGGF1 | Q8N302 | AGGF1 | Angiogenic factor with | 19478.12891 | 19252.66992 |
| SAAL1 | Q96ER3 | SAAL1 | Protein SAAL1 | 94125.83594 | 105907.7344 |

| | | | | | |
|---|---|---|---|---|---|
| PPME1 | Q9Y570 | PPME1 | Protein phosphatase r | 501387.1563 | 505250.8125 |
| GUSB | P08236 | GUSB | Beta-glucuronidase | 78858.55469 | 90508.15625 |
| GART | P22102 | GART | Trifunctional purine b | 613295.4375 | 723125.375 |
| SYF2 | O95926 | SYF2 | Pre-mRNA-splicing fac | 32435.16016 | 24770.84375 |
| CENPA | F8WD88;P49450 | CENPA | Histone H3-like centro | 2148.566162 | 600.6226196 |
| MRPS25 | P82663 | MRPS25 | 28S ribosomal protein | 113448.4766 | 127537.4297 |
| POLDIP2 | Q9Y2S7 | POLDIP2 | Polymerase delta-inte | 248580.6094 | 279558.25 |
| POLR3C | Q9BUI4 | POLR3C | DNA-directed RNA po | 1922922.5 | 1769514.75 |
| CHEK2 | O96017 | CHEK2 | Serine/threonine-prot | 18125.51172 | 23217.77344 |
| NDFIP1 | Q9BT67 | NDFIP1 | NEDD4 family-interact | 25444.46484 | 14515.74609 |
| HS1BP3 | Q53T59 | HS1BP3 | HCLS1-binding proteir | 72225.64063 | 69393.46875 |
| VANGL1 | Q8TAA9 | VANGL1 | Vang-like protein 1 | 64331.13672 | 69945.19531 |
| UQCR11 | O14957 | UQCR11 | Cytochrome b-c1 com | 22176.37695 | 25053.76563 |
| LMNA | P02545 | LMNA | Prelamin-A/C | 2575035.469 | 2172639.844 |
| POLR3H | Q9Y535 | POLR3H | DNA-directed RNA po | 48554.33984 | 34452.27344 |
| MRPL44 | Q9H9J2 | MRPL44 | 39S ribosomal protein | 115555.1016 | 126627.0156 |
| CYB561 | F5H757;J3KSL5;J3QRH5;J3QS | CYB561 | Cytochrome b561;Cyt | 45636.49609 | 40020.24219 |
| SENP3 | Q9H4L4 | SENP3 | Sentrin-specific protea | 55695.48828 | 64978.92578 |
| PPA1 | Q15181 | PPA1 | Inorganic pyrophosph | 1952925 | 1990066.75 |
| WRAP53 | E9PMG4;Q9BUR4 | WRAP53 | Telomerase Cajal body | 15297.52344 | 15581.72852 |
| SZRD1 | Q7Z422 | SZRD1 | SUZ domain-containin | 72930.85938 | 65836.15625 |
| KRT6C | P48668 | KRT6C | Keratin, type II cytosk | 14836495 | 22475294 |
| KPNA3 | O00505 | KPNA3 | Importin subunit alph | 168965.1094 | 184304.5781 |
| C12orf43;CUSTOS | F5H7W8;Q96C57 | C12orf43;CUSTOS | Chromosome 12 oper | 75027.08594 | 77197.42188 |
| RAB11FIP1 | Q6WKZ4 | RAB11FIP1 | Rab11 family-interacti | 45466.24219 | 46378.64063 |
| PYGO2 | Q5T171;Q9BRQ0 | PYGO2 | Pygopus homolog 2 | 21923.5918 | 24260.52734 |
| PSMA5 | P28066 | PSMA5 | Proteasome subunit a | 345742.375 | 541114.75 |
| NLE1 | Q9NVX2 | NLE1 | Notchless protein hon | 71638.375 | 50411.34766 |
| ASPHD1 | Q5U4P2 | ASPHD1 | Aspartate beta-hydro: | 5577.560547 | 4751.129395 |
| MRPS30 | Q9NP92 | MRPS30 | 39S ribosomal protein | 62233.73047 | 66817.64844 |
| INIP | Q9NRY2 | INIP | SOSS complex subunit | 22655.61523 | 22622.1875 |
| DLG5 | Q8TDM6 | DLG5 | Disks large homolog 5 | 59897.58594 | 69785.0625 |
| TRMT10C | Q7L0Y3 | TRMT10C | tRNA methyltransfera | 255409.5625 | 253514.3906 |

| | | | | | |
|---|---|---|---|---|---|
| ZNF460 | Q14592 | ZNF460 | Zinc finger protein 46( | 3213.133057 | 3777.322754 |
| CDC42BPG | Q6DT37 | CDC42BPG | Serine/threonine-prot | 38977.13281 | 43627.47266 |
| NUDT12 | Q9BQG2 | NUDT12 | Peroxisomal NADH py | 26162.05664 | 24863.38477 |
| PHPT1 | Q9NRX4 | PHPT1 | 14 kDa phosphohistid | 37273.50391 | 49233.33984 |
| MRPS22 | G5E9V5;G5E9W7;P82650 | MRPS22 | 28S ribosomal protein | 219114.8906 | 220218.0625 |
| SMPD2 | O60906 | SMPD2 | Sphingomyelin phospl | 5176.523926 | 5458.626465 |
| MAFK | O60675 | MAFK | Transcription factor M | 6974.408691 | 7347.88623 |
| GTF2I | P78347 | GTF2I | General transcription | 494655.7188 | 517775.2813 |
| NAT10 | Q9H0A0 | NAT10 | RNA cytidine acetyltra | 1014323.188 | 812629 |
| RHPN2 | Q8IUC4 | RHPN2 | Rhophilin-2 | 41931.22656 | 30407.69727 |
| AXIN1 | O15169 | AXIN1 | Axin-1 | 7930.687012 | 7040.580566 |
| POLR3D | P05423 | POLR3D | DNA-directed RNA po | 128213.125 | 130669.2578 |
| ZNF883;ZNF614 | P0CG24;Q8N883 | ZNF883;ZNF614 | Zinc finger protein 88: | 922.4788818 | 1149.039063 |
| DDX5 | P17844 | DDX5 | Probable ATP-depend | 3046984 | 2869598.5 |
| TMEM63B | Q5T3F8 | TMEM63B | CSC1-like protein 2 | 22523.71875 | 29417.13477 |
| DOLK | Q9UPQ8 | DOLK | Dolichol kinase | 10301.27637 | 11250.91113 |
| LARP1 | Q6PKG0 | LARP1 | La-related protein 1 | 385971.7813 | 329386.8438 |
| C8orf33 | A0A3B3IT54;Q9H7E9 | C8orf33 | UPF0488 protein C8or | 130333.8438 | 128100.5469 |
| GBA | P04062 | GBA | Lysosomal acid glucos | 174082.8438 | 166448.1094 |
| COX15 | Q7KZN9 | COX15 | Cytochrome c oxidase | 31947.98438 | 31017.63281 |
| KMT2C | Q8NEZ4 | KMT2C | Histone-lysine N-meth | 10534.83887 | 10499.59375 |
| PRR14 | Q9BWN1 | PRR14 | Proline-rich protein 14 | 13197.9375 | 10225.81934 |
| IMPDH2 | P12268 | IMPDH2 | Inosine-5'-monophosp | 976584.4375 | 1118263.625 |
| MRPL30 | Q8TCC3 | MRPL30 | 39S ribosomal protein | 30462.91602 | 44121.24609 |
| CCT4 | P50991 | CCT4 | T-complex protein 1 s | 1383439 | 1451745.25 |
| JMJD6 | B2WTI3;B2WTI4;K7EJU9;Q6N | JMJD6 | Bifunctional arginine d | 122350.7266 | 122968.6016 |
| BPHL | Q86WA6 | BPHL | Valacyclovir hydrolase | 59728.53906 | 68022.23438 |
| MRPL37 | Q9BZE1 | MRPL37 | 39S ribosomal protein | 186406.3125 | 215951.4063 |
| NDUFV2 | E7EPT4;P19404 | NDUFV2 | NADH dehydrogenase | 144256.4531 | 154661.1563 |
| EPHA2 | P29317 | EPHA2 | Ephrin type-A recepto | 6738.11377 | 3148.253418 |
| DCUN1D5 | Q9BTE7 | DCUN1D5 | DCN1-like protein 5 | 140925.3906 | 149947.5781 |
| TXN | P10599 | TXN | Thioredoxin | 540477.125 | 696031 |
| UBR3 | Q6ZT12 | UBR3 | E3 ubiquitin-protein li | 18455.82617 | 18591.90234 |

| | | | | | |
|---|---|---|---|---|---|
| SUPT4H1 | P63272 | SUPT4H1 | Transcription elongati | 138223.4219 | 109139.5625 |
| NFKBIB | G5E9C2;I3L4X3;Q15653 | NFKBIB | NF-kappa-B inhibitor b | 33219.85938 | 42756.875 |
| KCTD15 | Q96SI1 | KCTD15 | BTB/POZ domain-cont | 24538.48242 | 27908.41602 |
| NAA16 | Q6N069 | NAA16 | N-alpha-acetyltransfe | 16085.28711 | 19126.43555 |
| SOD2 | P04179 | SOD2 | Superoxide dismutase | 150558.7344 | 171503.3125 |
| PRCC | Q92733 | PRCC | Proline-rich protein PI | 60069.16797 | 57719.87891 |
| NDUFA2 | O43678 | NDUFA2 | NADH dehydrogenase | 166544.1406 | 183909.4219 |
| ERC1 | G8JLD3;Q8IUD2;X6RLX0 | ERC1 | ELKS/Rab6-interacting | 36108.125 | 38666.21484 |
| RSBN1 | A0A087WWP8;Q5VWQ0 | RSBN1 | Lysine-specific demetl | 88628.14063 | 85446.69531 |
| TTC13 | Q8NBP0 | TTC13 | Tetratricopeptide rep | 12240.50391 | 16663.29102 |
| YAP1 | P46937 | YAP1 | Transcriptional coacti | 43869.55469 | 38897.80469 |
| MRPL27 | H7C5U8;Q9P0M9 | MRPL27 | 39S ribosomal protein | 56295.09766 | 60897.60547 |
| POLR2D | E9PHV4;O15514 | POLR2D | DNA-directed RNA po | 10208.93457 | 5223.989258 |
| TCERG1 | O14776 | TCERG1 | Transcription elongati | 272801.625 | 267513.5625 |
| DPH2 | Q9BQC3 | DPH2 | 2-(3-amino-3-carboxy | 57654.48047 | 55649.81641 |
| TM2D3 | B3KT51;E7EPS7;H0YNF6;H0Y | TM2D3 | TM2 domain containi | 4359.90918 | 4212.644043 |
| PROCR | Q9UNN8 | PROCR | Endothelial protein C | 4763.967285 | 3897.201904 |
| LAMP2 | P13473 | LAMP2 | Lysosome-associated | 676973.5 | 759345.875 |
| MAU2 | Q9Y6X3 | MAU2 | MAU2 chromatid coh | 17962.69336 | 15664.35938 |
| MT-ND4 | P03905 | MT-ND4 | NADH-ubiquinone oxi | 41067.48828 | 52366.59766 |
| NDST1 | E7EVJ3;P52848 | NDST1 | Bifunctional heparan s | 8739.052734 | 12072.59473 |
| DHRS11 | Q6UWP2 | DHRS11 | Dehydrogenase/reduc | 12544.44141 | 18498.30078 |
| USP22 | Q9UPT9 | USP22 | Ubiquitin carboxyl-ter | 30022.50977 | 30896.25977 |
| ANKHD1-EIF4EBP3 | H7C0V5 | ANKHD1-EIF4EBP3 | ANKHD1-EIF4EBP3 rea | 3649.738525 | 5689.013184 |
| ZNF277 | C9JH62;Q9NRM2 | ZNF277 | Zinc finger protein 27 | 7061.172363 | 7751.452148 |
| ETF1 | P62495 | ETF1 | Eukaryotic peptide ch | 727565.75 | 795439.9375 |
| PRRC2A | P48634 | PRRC2A | Protein PRRC2A | 126044.0938 | 127079.625 |
| MTREX | P42285 | MTREX | Exosome RNA helicase | 1533128.625 | 1425996.625 |
| L3HYPDH | Q96EM0 | L3HYPDH | Trans-3-hydroxy-L-pro | 9285.6875 | 8915.043945 |
| MT-ND6 | P03923 | MT-ND6 | NADH-ubiquinone oxi | 23195.61719 | 21808.13867 |
| ZGPAT | Q8N5A5 | ZGPAT | Zinc finger CCCH-type | 32063.03125 | 28240.73438 |
| NUP153 | P49790 | NUP153 | Nuclear pore complex | 159379.7813 | 160119.1094 |
| PRKAR1B | P31321 | PRKAR1B | cAMP-dependent pro | 112315.9844 | 96276.71875 |

| | | | | | |
|---|---|---|---|---|---|
| COX6C | P09669 | COX6C | Cytochrome c oxidase | 649454.875 | 754195.5 |
| PRCP | P42785 | PRCP | Lysosomal Pro-X carbo | 52859.35938 | 45336.58203 |
| SFT2D2 | O95562;Q5TIH2 | SFT2D2 | Vesicle transport prot | 30191.94336 | 26903.15234 |
| GTF3C2 | Q8WUA4 | GTF3C2 | General transcription | 92101.96094 | 89608.70313 |
| USE1 | Q9NZ43 | USE1 | Vesicle transport prot | 22626.37891 | 27517.23633 |
| COX5B | P10606 | COX5B | Cytochrome c oxidase | 112937.6641 | 122812.9219 |
| POC5 | D6RBC1;D6RDG4;Q8NA72 | POC5 | Centrosomal protein P | 16932.57422 | 13797.7207 |
| UBE2G1 | P62253 | UBE2G1 | Ubiquitin-conjugating | 50677.76563 | 48721.70703 |
| AHCTF1 | Q8WYP5 | AHCTF1 | Protein ELYS | 66084.48438 | 68859.47656 |
| COA6 | Q5JTJ3 | COA6 | Cytochrome c oxidase | 223439.5313 | 209334.0781 |
| MRPL15 | Q9P015 | MRPL15 | 39S ribosomal protein | 167935.1563 | 186684.8281 |
| MSH6 | P52701 | MSH6 | DNA mismatch repair | 247223.1094 | 264186.2188 |
| DDA1 | Q9BW61 | DDA1 | DET1- and DDB1-asso | 35899.30469 | 39699.53125 |
| GLB1L2 | Q8IW92 | GLB1L2 | Beta-galactosidase-1-l | 8928.277344 | 10983.35742 |
| ASH1L | Q9NR48 | ASH1L | Histone-lysine N-meth | 12698.53223 | 8573.608398 |
| MSRB2 | Q9Y3D2 | MSRB2 | Methionine-R-sulfoxid | 82450.36719 | 84568.92188 |
| THOC7 | Q6I9Y2 | THOC7 | THO complex subunit | 177728.2188 | 165192.4844 |
| MRPL3 | E7ETU7;H0Y9G6;P09001 | MRPL3 | 39S ribosomal protein | 122254.1328 | 134579.125 |
| CCDC124 | Q96CT7 | CCDC124 | Coiled-coil domain-co | 185484.3906 | 178436.6875 |
| NOC3L | Q8WTT2 | NOC3L | Nucleolar complex pro | 73404.79688 | 66139.61719 |
| GLRX2 | Q9NS18 | GLRX2 | Glutaredoxin-2, mitoc | 16581.63867 | 13308.17188 |
| C15orf39 | H3BMJ2;Q6ZRI6 | C15orf39 | Uncharacterized prote | 10699.30566 | 9315.709961 |
| TTLL12 | Q14166 | TTLL12 | Tubulin--tyrosine ligas | 380148.4063 | 419503.1875 |
| AVEN | Q9NQS1 | AVEN | Cell death regulator A | 16750.80469 | 26518.58789 |
| LYRM7 | Q5U5X0 | LYRM7 | Complex III assembly f | 66111.05469 | 64949.63281 |
| MAP7D1 | Q3KQU3 | MAP7D1 | MAP7 domain-contain | 66790.72656 | 60920.39063 |
| GNL1 | P36915 | GNL1 | Guanine nucleotide-b | 107896.4297 | 126470.9219 |
| MRPL19 | P49406 | MRPL19 | 39S ribosomal protein | 209931.2813 | 233271.6094 |
| GPD2 | P43304 | GPD2 | Glycerol-3-phosphate | 273036.6563 | 280736.4375 |
| ITFG2 | Q969R8 | ITFG2 | KICSTOR complex prot | 8120.764648 | 7452.288574 |
| PFDN4 | E9PQY2;Q9NQP4 | PFDN4 | Prefoldin subunit 4 | 10779.98633 | 11668.62988 |
| RRS1 | Q15050 | RRS1 | Ribosome biogenesis | 262138.3906 | 113018.8438 |
| EIF3K | K7ERF1;Q9UBQ5 | EIF3K | Eukaryotic translation | 331361.8438 | 339625.7188 |

| | | | | | |
|---|---|---|---|---|---|
| HSP90B1 | P14625 | HSP90B1 | Endoplasmin | 5487788 | 5683442 |
| DEDD2 | M0QXK7;M0QYK9;Q8WXF8 | DEDD2 | DNA-binding death ef | 230593.4219 | 243745.75 |
| DDX18 | Q9NVP1 | DDX18 | ATP-dependent RNA h | 271542.7188 | 180191.1094 |
| VDAC2 | P45880 | VDAC2 | Voltage-dependent ar | 2059675 | 854900.5625 |
| FOXK1 | P85037 | FOXK1 | Forkhead box protein | 118783.2813 | 130223.1875 |
| SNX11 | J3KTN6;J3QLV8;J3QRB9;Q9Y5 | SNX11 | Sorting nexin-11 (Frag | 16230.11621 | 15680.10742 |
| ACBD5 | B7Z2R7 | ACBD5 | Acyl-CoA-binding dom | 53973.17578 | 54744.91797 |
| UQCC3 | Q6UW78 | UQCC3 | Ubiquinol-cytochrome | 83507.84375 | 87657.875 |
| VAMP4 | O75379 | VAMP4 | Vesicle-associated me | 9951.662109 | 10958.08691 |
| MSTO1 | Q9BUK6 | MSTO1 | Protein misato homol | 54765.21094 | 60704.62109 |
| PHF8 | H0Y3N9;Q9UPP1 | PHF8 | Histone lysine demeth | 14425.33398 | 15367.09961 |
| CFAP47 | Q6ZTR5 | CFAP47 | Cilia- and flagella-assc | 97123.82031 | 97713 |
| MRPS6 | P82932 | MRPS6 | 28S ribosomal protein | 327384.6875 | 341724.0625 |
| MRPS35 | P82673 | MRPS35 | 28S ribosomal protein | 133008.1563 | 143541.1406 |
| COIL | P38432 | COIL | Coilin | 125674.5703 | 79856.84375 |
| RPAP3 | Q9H6T3 | RPAP3 | RNA polymerase II-ass | 92300.07813 | 93487.5625 |
| MRPS28 | Q9Y2Q9 | MRPS28 | 28S ribosomal protein | 268418.0625 | 306610.6875 |
| WRAP73 | A0A0A0MRV3;Q5T0D5;Q9P2 | WRAP73 | WD repeat domain 8, | 2286.021484 | 2209.967041 |
| PTCD3 | Q96EY7 | PTCD3 | Pentatricopeptide rep | 175530.0313 | 174814.0781 |
| PELP1 | Q8IZL8 | PELP1 | Proline-, glutamic acic | 229499.4844 | 219648.6406 |
| PDK2 | Q15119 | PDK2 | [Pyruvate dehydroger | 21343.96289 | 27682.46484 |
| MRPL43 | B1AL05;H0Y6Y8;Q8N983 | MRPL43 | 39S ribosomal protein | 147575.125 | 158296.5938 |
| SKA3 | Q8IX90 | SKA3 | Spindle and kinetochc | 26609.16406 | 40859.13281 |
| THAP12 | O43422 | THAP12 | 52 kDa repressor of th | 12615.05371 | 12323.14648 |
| MICU2 | Q8IYU8 | MICU2 | Calcium uptake protei | 28348.58008 | 30764.26563 |
| MRPL38 | Q96DV4 | MRPL38 | 39S ribosomal protein | 171110.6406 | 186811.7188 |
| TXNDC12 | O95881 | TXNDC12 | Thioredoxin domain-c | 28379.19336 | 35550.77344 |
| GRSF1 | Q12849 | GRSF1 | G-rich sequence facto | 192531.5938 | 198849.375 |
| SLC39A14 | E5RIP4;Q15043 | SLC39A14 | Zinc transporter ZIP14 | 51278.23047 | 52370.12109 |
| TXNDC9 | O14530 | TXNDC9 | Thioredoxin domain-c | 91706.20313 | 102623.6094 |
| HAX1 | O00165 | HAX1 | HCLS1-associated pro | 27572.93359 | 46539.17578 |
| MPRIP | Q6WCQ1 | MPRIP | Myosin phosphatase R | 57359.35547 | 54207.41016 |
| MFAP1 | P55081 | MFAP1 | Microfibrillar-associat | 57823.72656 | 39862.86719 |

| | | | | | |
|---|---|---|---|---|---|
| ST3GAL4 | E9PMC7;Q11206 | ST3GAL4 | CMP-N-acetylneurami | 2279.048828 | 2487.757813 |
| MRPS31 | Q92665 | MRPS31 | 28S ribosomal protein | 347388.25 | 349499.875 |
| TMEM186 | Q96B77 | TMEM186 | Transmembrane prote | 5774.108398 | 6983.70166 |
| TTC27 | Q6P3X3 | TTC27 | Tetratricopeptide rep | 216809.1406 | 223881.5156 |
| HIGD2A | Q9BW72 | HIGD2A | HIG1 domain family m | 6576.455078 | 9590.696289 |
| TAF12 | Q16514 | TAF12 | Transcription initiatio | 32178.43555 | 28239.77539 |
| COX11 | Q9Y6N1 | COX11 | Cytochrome c oxidase | 32911.84375 | 30735.69922 |
| TEX2 | A0A087WVA8;Q8IWB9 | TEX2 | Testis-expressed prot | 66085.95313 | 72371.39844 |
| PTGR1 | F2Z3J9;Q14914 | PTGR1 | Prostaglandin reducta | 93867.38281 | 88346.13281 |
| NGRN | Q9NPE2 | NGRN | Neugrin | 10575.35254 | 10357.89355 |
| NUDT8 | Q8WV74 | NUDT8 | Nucleoside diphospha | 43544.13672 | 41420.30469 |
| PRPF38B | Q5VTL8 | PRPF38B | Pre-mRNA-splicing fac | 131752.4844 | 104220.0703 |
| TFAP4 | Q01664 | TFAP4 | Transcription factor A | 14640.72168 | 12374.9043 |
| ESS2 | Q96DF8 | ESS2 | Splicing factor ESS-2 h | 20368.2832 | 19680.96289 |
| MRPL22 | E7ESL0;J3KQY1;Q9NWU5 | MRPL22 | 39S ribosomal protein | 133870.6563 | 146051.0156 |
| MOGS | Q13724 | MOGS | Mannosyl-oligosaccha | 190348.875 | 200311.6094 |
| FBXO30 | Q8TB52 | FBXO30 | F-box only protein 30 | 14313.7666 | 15324.48828 |
| HADHA | H0YFD6;P40939 | HADHA | Trifunctional enzyme | 549073.1875 | 583235.125 |
| DGKZ | Q13574 | DGKZ | Diacylglycerol kinase | 23346.74414 | 26206.72266 |
| NOP14 | P78316 | NOP14 | Nucleolar protein 14 | 99002.57813 | 40930.5625 |
| PIH1D1 | Q9NWS0 | PIH1D1 | PIH1 domain-containi | 52825.93359 | 51748.03516 |
| DSTYK | Q6XUX3 | DSTYK | Dual serine/threonine | 4708.000488 | 7002.366211 |
| EIF3B | P55884 | EIF3B | Eukaryotic translation | 1069163.25 | 1088083.5 |
| HSD17B8 | Q92506 | HSD17B8 | Estradiol 17-beta-deh | 50097.21484 | 46470.55859 |
| PRDX3 | P30048 | PRDX3 | Thioredoxin-depende | 1060365.5 | 1176943 |
| ZNF280C | Q8ND82 | ZNF280C | Zinc finger protein 28 | 3769.591797 | NA |
| AKAP11 | Q9UKA4 | AKAP11 | A-kinase anchor prote | 21522.99414 | 20898.27344 |
| DPH1 | I3L1H5;Q9BZG8 | DPH1 | 2-(3-amino-3-carboxy | 67910.34375 | 66926.28906 |
| OVCA2 | Q8WZ82 | OVCA2 | Esterase OVCA2 | 148282.9844 | 166837.7188 |
| NDUFS6 | O75380 | NDUFS6 | NADH dehydrogenase | 520731.0313 | 544596.4375 |
| FKBP5 | Q13451 | FKBP5 | Peptidyl-prolyl cis-tra | 125128.5859 | 128257.6172 |
| HNRNPC | G3V251;G3V3K6;G3V5X6 | HNRNPC | Heterogeneous nucle | 4177827.516 | 1818268.656 |
| MRPL10 | Q7Z7H8 | MRPL10 | 39S ribosomal protein | 91819.17969 | 95689.01563 |

| | | | | | |
|---|---|---|---|---|---|
| ISG20L2 | Q9H9L3 | ISG20L2 | Interferon-stimulated | 30624.86133 | 9767.96582 |
| HNRNPF | P52597 | HNRNPF | Heterogeneous nuclea | 1126666.375 | 842100.6875 |
| TST | Q16762 | TST | Thiosulfate sulfurtrans | 452805.7188 | 765619.8125 |
| SEC62 | F8WF48;Q99442 | SEC62 | Translocation protein | 79023.65625 | 98488.13281 |
| RPRD2 | Q5VT52 | RPRD2 | Regulation of nuclear | 96462.80469 | 104399.1797 |
| POLR1D | P0DPB5 | POLR1D | Protein POLR1D, isofo | 54401.94141 | 64672.37109 |
| FLNB | O75369 | FLNB | Filamin-B | 598066.5 | 687611.5625 |
| RIF1 | Q5UIP0 | RIF1 | Telomere-associated | 101771.7891 | 108220.5625 |
| DIMT1 | Q9UNQ2 | DIMT1 | Probable dimethylade | 90501.96875 | 89821.5625 |
| PRRC2C | E7EPN9 | PRRC2C | Protein PRRC2C | 495194.7813 | 491038.4063 |
| BLVRB | P30043 | BLVRB | Flavin reductase (NAD | 968120.4375 | 1107017.125 |
| CWC22 | Q9HCG8 | CWC22 | Pre-mRNA-splicing fac | 39817.95313 | 34283.94922 |
| MRPL33 | O75394 | MRPL33 | 39S ribosomal protein | 57016.5625 | 66741.89063 |
| QRSL1 | Q9H0R6 | QRSL1 | Glutamyl-tRNA(Gln) a | 33030.94922 | 33755.3125 |
| MRPS2 | Q9Y399 | MRPS2 | 28S ribosomal protein | 229723.9688 | 256340.8281 |
| UVRAG | E9PK00;E9PR71;Q9P2Y5 | UVRAG | UV radiation resistanc | NA | 17991.19336 |
| MRPS18A | Q5QPA5;Q9NVS2 | MRPS18A | 39S ribosomal protein | 156867.6563 | 181035.6719 |
| TUSC2 | O75896 | TUSC2 | Tumor suppressor car | 71794.02344 | 77470.79688 |
| SLU7 | O95391 | SLU7 | Pre-mRNA-splicing fac | 18095.10352 | 13872.57813 |
| LGALS1 | P09382 | LGALS1 | Galectin-1 | 42229.89063 | 48322.74609 |
| ETFRF1 | A0A0C4DGJ8;Q6IPR1 | ETFRF1 | Electron transfer flavo | 8028.669434 | 10148.82129 |
| NOC4L | Q9BVI4 | NOC4L | Nucleolar complex pro | 126768.4922 | 59103.67578 |
| HASPIN | Q8TF76 | HASPIN | Serine/threonine-prot | 14561.42285 | 12482.93262 |
| MAP1LC3B2;MAP1LC3B;MA | A6NCE7;H3BTL1;Q9GZQ8 | MAP1LC3B2;MAP1LC3 | Microtubule-associate | 165104.3906 | 185868.9844 |
| RBM5 | P52756 | RBM5 | RNA-binding protein 5 | 310854.9375 | 227374.7188 |
| PDIA4 | A0A499FI48 | PDIA4 | Protein disulfide-isom | 1421021.831 | 1566116.403 |
| ADPRM | Q3LIE5;V9GYR6 | ADPRM | Manganese-dependen | 3064.480713 | 3041.739014 |
| MAP4K3 | F5H5A3;Q8IVH8 | MAP4K3 | Mitogen-activated pro | 17227 | 19526.14648 |
| EPCAM | B5MCA4;P16422 | EPCAM | Epithelial cell adhesio | 598263.125 | 748130 |
| DDX27 | B7Z6D5;Q96GQ7 | DDX27 | Probable ATP-depend | 379305.9688 | 274046.2188 |
| MRPS10 | P82664 | MRPS10 | 28S ribosomal protein | 87027.26563 | 87815.72656 |
| OGFR | Q9NZT2 | OGFR | Opioid growth factor r | 324791.0313 | 306610.7813 |
| ADAM17 | P78536 | ADAM17 | Disintegrin and metall | 2384.449951 | 2664.925781 |

| | | | | | |
|---|---|---|---|---|---|
| AAMP | C9JEH3;C9JG97;Q13685 | AAMP | Angio-associated migr | 134835.0313 | 158233.8906 |
| INPP5B | P32019 | INPP5B | Type II inositol 1,4,5-t | 46828.52344 | 77794.73438 |
| FAM136A | Q96C01 | FAM136A | Protein FAM136A | 119440.2578 | 131726.4219 |
| STXBP4 | B5MCT9;E7EPP7;Q6ZWJ1 | STXBP4 | Syntaxin-binding prote | 9900.095703 | 9610.5625 |
| GPRIN1 | Q7Z2K8 | GPRIN1 | G protein-regulated ir | 8846.460938 | 10495.9873 |
| SMU1 | Q2TAY7 | SMU1 | WD40 repeat-contain | 272449.0938 | 180232.6719 |
| TSSC4 | C9JJV1;Q9Y5U2 | TSSC4 | Protein TSSC4 (Fragm | 45094.88672 | 40799.89063 |
| ;MYCBP | A0A087WV05;Q99417 | ;MYCBP | Uncharacterized prote | 78324.07813 | 80979.02344 |
| DPP9 | Q86TI2 | DPP9 | Dipeptidyl peptidase 9 | 89186.39063 | 119815.7734 |
| MRPL57 | Q9BQC6 | MRPL57 | Ribosomal protein 63, | 47036.39844 | 54545.44922 |
| LDHA | P00338 | LDHA | L-lactate dehydrogena | 4011560 | 4096107.25 |
| METTL5 | Q9NRN9 | METTL5 | Methyltransferase-like | 25754.01758 | 26034.33984 |
| MRPL9 | Q9BYD2 | MRPL9 | 39S ribosomal protein | 198067.4219 | 202236.0625 |
| MBLAC2 | Q68D91 | MBLAC2 | Metallo-beta-lactama | 37528.53125 | 33649.57813 |
| TDG | B4E127;G8JL98;Q13569 | TDG | G/T mismatch-specific | 30416.55859 | 30576.62305 |
| COX6A1 | P12074 | COX6A1 | Cytochrome c oxidase | 44856.875 | 49310.94141 |
| PPL | K7EKI8;O60437 | PPL | Periplakin | 59887.72656 | 40911.71875 |
| PET117 | Q6UWS5 | PET117 | Protein PET117 homo | 5801.476563 | 8025.23584 |
| MRPL23 | Q16540 | MRPL23 | 39S ribosomal protein | 198941.375 | 209817.3125 |
| PLA2G15 | Q8NCC3 | PLA2G15 | Group XV phospholipa | 31394.95703 | 28061.80469 |
| MRPL58 | J3KS15;Q14197 | MRPL58 | Peptidyl-tRNA hydrola | 183223.0156 | 193227.2813 |
| STIM1 | Q13586 | STIM1 | Stromal interaction m | 23671.06445 | 23973.55469 |
| TBL2 | E9PF19;Q9Y4P3 | TBL2 | Transducin beta-like p | 69222.92188 | 70781.59375 |
| MRPS16 | Q9Y3D3 | MRPS16 | 28S ribosomal protein | 167943.7969 | 237467.5625 |
| TRMT61B | Q9BVS5 | TRMT61B | tRNA (adenine(58)-N( | 16103.81445 | 16303.24219 |
| CDC42EP1 | Q00587 | CDC42EP1 | Cdc42 effector proteir | 21063.05859 | 20242.32422 |
| POLR2C | P19387 | POLR2C | DNA-directed RNA po | 80154.8125 | 58002.9375 |
| HLA-E | P13747 | HLA-E | HLA class I histocompa | 9143.924805 | 6288.858887 |
| GSE1 | Q14687 | GSE1 | Genetic suppressor ele | 39201.12109 | 38141.00781 |
| ZNHIT3 | Q15649 | ZNHIT3 | Zinc finger HIT domair | 28672.04102 | 28419.68164 |
| TNRC18 | H9KVB4;O15417 | TNRC18 | Trinucleotide repeat-c | 4104.147949 | 4408.397949 |
| ING3 | E7ET07;Q9NXR8 | ING3 | Inhibitor of growth pr | 18166.45508 | 13139.16309 |
| RABGGTB | P53611 | RABGGTB | Geranylgeranyl transfe | 56634.77344 | 68292.32031 |

| | | | | | |
|---|---|---|---|---|---|
| CHMP2B | Q9UQN3 | CHMP2B | Charged multivesicula | 87527.41406 | 84060.15625 |
| CMIP | A0A087WY48;Q8IY22 | CMIP | C-Maf-inducing protei | 11432.45703 | 13558.57813 |
| MRPS27 | Q92552 | MRPS27 | 28S ribosomal protein | 220163.8281 | 248910.4219 |
| DCAF13 | A0A087WT20;Q9NV06 | DCAF13 | DDB1- and CUL4-asso | 80767.57813 | 41203.92188 |
| METTL2B | Q6P1Q9 | METTL2B | tRNA N(3)-methylcytic | 86427.69531 | 100278.6953 |
| DARS2 | Q6PI48 | DARS2 | Aspartate--tRNA ligase | 435187.5313 | 494416.9375 |
| CAPRIN1 | Q14444 | CAPRIN1 | Caprin-1 | 1332386.375 | 1301133.125 |
| ARID4B | Q4LE39 | ARID4B | AT-rich interactive do | 8786.400391 | 9465.28125 |
| TJP1 | G5E9E7 | TJP1 | Tight junction protein | 60599.57813 | 55983.03516 |
| NPRL3 | B7Z6Q0;Q12980 | NPRL3 | GATOR complex prote | 11849.97461 | 12229.09863 |
| MRPL20 | Q9BYC9 | MRPL20 | 39S ribosomal protein | 79933.17188 | 95608.30469 |
| CLN5 | A0A024R644;A0A1B0GWI2;C | CLN5 | Ceroid-lipofuscinosis r | 31248.53906 | 24804.125 |
| ADAR | A0A3B3IRQ9 | ADAR | Double-stranded RNA | 490098.3984 | 487872.7813 |
| GTF2E1 | P29083 | GTF2E1 | General transcription | 131563.2031 | 166080.8906 |
| CPS1 | P31327 | CPS1 | Carbamoyl-phosphate | 1827.190186 | 2773.157959 |
| LRRC59 | Q96AG4 | LRRC59 | Leucine-rich repeat-co | 2493200.75 | 2453792.5 |
| ANKRD17 | O75179 | ANKRD17 | Ankyrin repeat domai | 41944.6875 | 44199.30469 |
| CERS6 | Q6ZMG9 | CERS6 | Ceramide synthase 6 | 42015.91797 | 49591.01172 |
| IPO11 | Q9UI26 | IPO11 | Importin-11 | 135337.7031 | 173803.1406 |
| ACAA2 | P42765 | ACAA2 | 3-ketoacyl-CoA thiolas | 393843.25 | 369768.75 |
| HSPB1 | P04792 | HSPB1 | Heat shock protein be | 14839563 | 14378580 |
| H2AZ1;H2AFV | P0C0S5;Q71UI9 | H2AZ1;H2AFV | Histone H2A.Z;Histone | 1157009.125 | 132666.3906 |
| VPS36 | Q86VN1 | VPS36 | Vacuolar protein-sorti | 87320.57031 | 95812.17969 |
| RSRC1 | Q96IZ7 | RSRC1 | Serine/Arginine-relate | 30396.9375 | 29960.52344 |
| UBA52 | M0R2S1 | UBA52 | Ubiquitin-60S ribosom | 39273.26953 | 39174.52734 |
| TIMM21 | Q9BVV7 | TIMM21 | Mitochondrial import | 38747.86328 | 30934.15234 |
| MRPS34 | C9JJ19;P82930 | MRPS34 | 28S ribosomal protein | 391071.5938 | 413804.125 |
| TRUB2 | O95900 | TRUB2 | Mitochondrial mRNA | 26469.53516 | 31179.25 |
| NRBF2 | Q96F24 | NRBF2 | Nuclear receptor-bind | 13070.38672 | 14511.34082 |
| ENTPD2 | Q9Y5L3 | ENTPD2 | Ectonucleoside tripho | 17555.39648 | 17181.30664 |
| NOP9 | Q86U38 | NOP9 | Nucleolar protein 9 | 109065.6328 | 92631.83594 |
| HADHB | P55084 | HADHB | Trifunctional enzyme | 1077852.375 | 1163807.5 |
| MRPL21 | Q7Z2W9 | MRPL21 | 39S ribosomal protein | 204861.625 | 226707.0938 |

| | | | | | |
|---|---|---|---|---|---|
| POC1B-GALNT4;GALNT4 | F8VUJ3;Q8N4A0 | POC1B-GALNT4;GALN | Polypeptide N-acetylg | 4031.170654 | 3280.90918 |
| CLPB | A0A2R8Y602 | CLPB | Caseinolytic peptidase | 347014.707 | 356720.5898 |
| SNRPF | P62306 | SNRPF | Small nuclear ribonuc | 189029.0938 | 180414.8125 |
| NOP58 | Q9Y2X3 | NOP58 | Nucleolar protein 58 | 376648.4063 | 150084.1094 |
| PSME4 | Q14997 | PSME4 | Proteasome activator | 87675.16406 | 101703.3828 |
| TANC2 | J3KTM5;J3QQY7 | TANC2 | Protein TANC2 (Fragm | 57230.98633 | 61551.11279 |
| PQBP1 | O60828 | PQBP1 | Polyglutamine-binding | 529244.1875 | 443594.875 |
| CCDC12 | J3KR35;Q8WUD4 | CCDC12 | Coiled-coil domain co | 79643.71094 | 78176.03906 |
| SUCLA2 | A0A2R8YDQ9;Q9P2R7 | SUCLA2 | Succinate--CoA ligase | 231199.625 | 233779.2813 |
| RPS19BP1 | Q86WX3 | RPS19BP1 | Active regulator of SIF | 59320.83594 | 55782.70313 |
| ZC3H14 | G3V5I6 | ZC3H14 | Zinc finger CCCH dom | 187039.8984 | 166293.8984 |
| HNRNPAB | D6R9P3;D6RBZ0;D6RD18 | HNRNPAB | Heterogeneous nucle | 1287666.625 | 828720.5625 |
| GALNT1 | Q10472 | GALNT1 | Polypeptide N-acetylg | 16775.8125 | 16490.03125 |
| MEMO1 | Q9Y316 | MEMO1 | Protein MEMO1 | 108032.6016 | 115937.0781 |
| NSMCE2 | A0A087WTZ8 | NSMCE2 | E3 SUMO-protein liga | 72409.30469 | 70808.08203 |
| BRD8 | B5MCW3 | BRD8 | Bromodomain-contair | 105103.8398 | 80348.28125 |
| DAP3 | P51398 | DAP3 | 28S ribosomal protein | 142496.7813 | 163095.6875 |
| BRMS1 | E9PJF5;G5E9I4;H0YCF7;Q9H( | BRMS1 | Breast cancer metasta | 15857.19531 | 14603.87988 |
| CSTF2 | E7EWR4;P33240 | CSTF2 | Cleavage stimulation f | 102674.7266 | 114147.9297 |
| GLA | P06280 | GLA | Alpha-galactosidase A | 75645.84375 | 103408.3125 |
| MDH2 | P40926 | MDH2 | Malate dehydrogenas | 3986461.25 | 4170142.25 |
| SCO1 | J3QL56;O75880 | SCO1 | Protein SCO1 homolog | 144843.0156 | 153365.9844 |
| MORC2 | Q9Y6X9 | MORC2 | ATPase MORC2 | 98155.08594 | 103343.0078 |
| SMG6 | Q86US8 | SMG6 | Telomerase-binding p | 14109.85645 | 14359.10254 |
| GGA1 | Q9UJY5 | GGA1 | ADP-ribosylation facto | 68779.33594 | 69029.89063 |
| ADNP2 | Q6IQ32 | ADNP2 | Activity-dependent ne | 6786.924805 | 7048.086426 |
| RRP1B | Q14684 | RRP1B | Ribosomal RNA proce | 155516.4531 | 144491.9531 |
| AHCY | P23526 | AHCY | Adenosylhomocystein | 3756613 | 4331542 |
| ZNF622 | Q969S3 | ZNF622 | Zinc finger protein 62 | 63105.55859 | 61942.50781 |
| RRP12 | Q5JTH9 | RRP12 | RRP12-like protein | 426541.8125 | 355681.375 |
| DNASE2 | O00115 | DNASE2 | Deoxyribonuclease-2- | 141746.7188 | 158995.6094 |
| CREG1 | A0A3B3IRL2;O75629 | CREG1 | Cellular repressor of E | 113885.9375 | 115575.625 |
| TMEM68 | E5RGT2;Q96MH6 | TMEM68 | Transmembrane prote | 21852.51172 | 19233.1582 |

| | | | | | |
|---|---|---|---|---|---|
| MRPL41 | Q8IXM3 | MRPL41 | 39S ribosomal protein | 220518.6094 | 209481.1094 |
| FXR2 | P51116 | FXR2 | Fragile X mental retar | 89198.32031 | 90000.42969 |
| CKAP2 | Q8WWK9 | CKAP2 | Cytoskeleton-associat | 51062.20313 | 41603.02344 |
| TAF3 | Q5VWG9 | TAF3 | Transcription initiatio | 14103.4541 | 11270.05957 |
| KIF20A | O95235 | KIF20A | Kinesin-like protein KI | 29322.12695 | 30017.24219 |
| MALT1 | A0A3B3IS32;Q9UDY8 | MALT1 | Mucosa-associated lyı | 4230.193359 | 4133.812988 |
| CRNKL1 | Q5JY65;Q9BZJ0 | CRNKL1 | Crooked neck-like pro | 108857.2656 | 89416.70313 |
| PLCG2 | P16885 | PLCG2 | 1-phosphatidylinosito | 11704.67578 | 12734.91504 |
| DDX28 | Q9NUL7 | DDX28 | Probable ATP-depend | 51136.71484 | 57656.21094 |
| PAXIP1 | Q6ZW49 | PAXIP1 | PAX-interacting prote | 15449.43262 | 14257.18555 |
| SUPT6H | Q7KZ85 | SUPT6H | Transcription elongati | 138946.9375 | 120780.8047 |
| SNX8 | Q9Y5X2 | SNX8 | Sorting nexin-8 | 32935.24609 | 33393.85547 |
| SPDEF | O95238 | SPDEF | SAM pointed domain- | 32569.61719 | 26903.0625 |
| CCDC117 | Q8IWD4 | CCDC117 | Coiled-coil domain-co | 54294.38672 | 77294.28125 |
| UBR2 | Q8IWV8 | UBR2 | E3 ubiquitin-protein li | 91334.71094 | 90936.38281 |
| MRPL2 | Q5T653 | MRPL2 | 39S ribosomal protein | 73560.25 | 85808.78906 |
| MRPS14 | O60783 | MRPS14 | 28S ribosomal protein | 49488.48047 | 56404.20703 |
| ZNHIT6 | Q9NWK9 | ZNHIT6 | Box C/D snoRNA prote | 13135.8291 | 12074.68555 |
| FTL | P02792 | FTL | Ferritin light chain | 2341.805176 | 1570.329224 |
| MFAP3 | P55082 | MFAP3 | Microfibril-associated | 7940.516113 | 9843.092773 |
| FBL | P22087 | FBL | rRNA 2'-O-methyltran | 590353.375 | 262316.8438 |
| GRHPR | Q9UBQ7 | GRHPR | Glyoxylate reductase/ | 346778.2813 | 391256.4375 |
| TMEM214 | Q6NUQ4 | TMEM214 | Transmembrane prote | 94210.30469 | 99342.84375 |
| DNM1L | O00429 | DNM1L | Dynamin-1-like protei | 229003.8125 | 245999.25 |
| TBL3 | Q12788 | TBL3 | Transducin beta-like p | 64779.77734 | 31576.48242 |
| KDM5B | A0A3B3IS40;A0A3B3ISE6;A0A | KDM5B | Lysine-specific demetl | 12677.72168 | 12294.53418 |
| C1orf174 | Q8IYL3 | C1orf174 | UPF0688 protein C1or | 7086.076172 | 7876.294922 |
| PTRH2 | J3KQ48;Q9Y3E5 | PTRH2 | Peptidyl-tRNA hydrola | 315636 | 322762.1875 |
| ATG9A | Q7Z3C6 | ATG9A | Autophagy-related pro | 33262.53516 | 43541.05859 |
| CHORDC1 | Q9UHD1 | CHORDC1 | Cysteine and histidine | 180339.2656 | 183624.7188 |
| EPS15L1 | M0QY01 | EPS15L1 | Epidermal growth fact | 233889.6787 | 256999.9531 |
| PEX13 | Q92968 | PEX13 | Peroxisomal membrar | 14553.46777 | 18889.47852 |
| ARFGAP1 | Q8N6T3 | ARFGAP1 | ADP-ribosylation facto | 221526.625 | 236071.4844 |

| | | | | | |
|---|---|---|---|---|---|
| POLR3B | Q9NW08 | POLR3B | DNA-directed RNA po | 28337.03125 | 28580.44922 |
| GATAD2B | A0A0U1RRM1;Q8WXI9 | GATAD2B | Transcriptional repres | 131731.0625 | 122296.6641 |
| ATP5F1D | P30049 | ATP5F1D | ATP synthase subunit | 9085.741211 | 13359.00879 |
| OXA1L | A0A087X0L7;J3KNA0;Q1507( | OXA1L | Mitochondrial inner m | 75717.79688 | 84100.35156 |
| CHCHD7 | E5RFN2;E5RHM5;E5RII1;E5RJ | CHCHD7 | Coiled-coil-helix-coile | 24904.98438 | 25616.06445 |
| RBM28 | Q9NW13 | RBM28 | RNA-binding protein 2 | 441108.4375 | 302873.5625 |
| NSA2 | O95478 | NSA2 | Ribosome biogenesis | 178495.6719 | 87087.5625 |
| LRPPRC | P42704 | LRPPRC | Leucine-rich PPR moti | 1522610.125 | 1726735.25 |
| LYPLAL1 | Q5VWZ2 | LYPLAL1 | Lysophospholipase-lik | 83492.36719 | 86276.07031 |
| STRAP | Q9Y3F4 | STRAP | Serine-threonine kina | 241676.2969 | 249540.8438 |
| ZNF746 | A0A2R8YDQ5;Q6NUN9 | ZNF746 | Zinc finger protein 74( | 1305.484253 | 1059.643555 |
| RIOK3 | B4E1Q4;O14730 | RIOK3 | Serine/threonine-prot | 1018.075989 | 1515.685181 |
| RPF1 | Q9H9Y2 | RPF1 | Ribosome production | 18515.51367 | 10908.7666 |
| POLR1E | Q9GZS1 | POLR1E | DNA-directed RNA po | 113513.7109 | 89927.33594 |
| TAF4 | O00268 | TAF4 | Transcription initiatio | 43415.14453 | 39326.17969 |
| EIF2D | P41214 | EIF2D | Eukaryotic translation | 123647.9375 | 123639.2969 |
| GSTO1 | P78417 | GSTO1 | Glutathione S-transfe | 1260505.75 | 1309186.125 |
| MNT | Q99583;V9GY02 | MNT | Max-binding protein M | 10143.86426 | 11678.20898 |
| STX6 | O43752 | STX6 | Syntaxin-6 | 41220.60156 | 65907.45313 |
| USP42 | C9JPC9;Q9H9J4 | USP42 | Ubiquitin carboxyl-ter | 11547.88867 | 11254.38867 |
| BCL2L13 | Q9BXK5 | BCL2L13 | Bcl-2-like protein 13 | 21802.24805 | 29295.7168 |
| TMEM259 | A0A0A0MTU3;Q4ZIN3;U3KPY | TMEM259 | Membralin | 20604.75 | 25452.48633 |
| SIRT6 | Q8N6T7 | SIRT6 | NAD-dependent prote | 10185.53516 | 7959.781738 |
| AHNAK | Q09666 | AHNAK | Neuroblast differentia | 3237010.25 | 2105211.5 |
| DHX33 | I3L1L6;Q9H6R0 | DHX33 | ATP-dependent RNA h | 29926.75586 | 23986.55469 |
| METTL2A | Q96IZ6 | METTL2A | tRNA N(3)-methylcytid | 55332.19531 | 60680.96875 |
| RPL37 | D6RG19;P61927 | RPL37 | Ribosomal protein L37 | 23032.14648 | 23910.79688 |
| SPG7 | A0A2R8Y4Y7;A0A2R8Y726;A( | SPG7 | Paraplegin;Paraplegin | 32687.94727 | 36743.51563 |
| DVL2 | O14641 | DVL2 | Segment polarity prot | 43647.88281 | 57517.55859 |
| MZT2A | Q6P582 | MZT2A | Mitotic-spindle organi | 65504.5 | 81813.25 |
| PRNP | A2A2V1;P04156 | PRNP | Major prion protein (F | 14753.20605 | 13388.80566 |
| COA1 | Q9GZY4 | COA1 | Cytochrome c oxidase | 12485.14648 | 14128.12109 |
| SNRPA1 | P09661 | SNRPA1 | U2 small nuclear ribor | 486546.9063 | 368778.8438 |

| | | | | | |
|---|---|---|---|---|---|
| TRIT1 | Q9H3H1;S4R2Z0 | TRIT1 | tRNA dimethylallyltra | 2652.489746 | 3362.904541 |
| MRPL49 | Q13405 | MRPL49 | 39S ribosomal protein | 128145.3203 | 130576.0156 |
| ANKRD40 | Q6AI12 | ANKRD40 | Ankyrin repeat domai | 10160.0498 | 13827.94434 |
| GHITM | Q9H3K2 | GHITM | Growth hormone-indu | 139290.0625 | 143783.5781 |
| MAK16 | Q9BXY0 | MAK16 | Protein MAK16 homo | 76059.86719 | 43323.70313 |
| NUCKS1 | Q9H1E3 | NUCKS1 | Nuclear ubiquitous ca | 374187.375 | 902374.375 |
| INF2 | Q27J81 | INF2 | Inverted formin-2 | 165901.6094 | 183562.7969 |
| POLA1 | A6NMQ1;P09884 | POLA1 | DNA polymerase;DNA | 296161.3125 | 371908.9063 |
| CUL2 | A0A0A0MTN0;Q13617 | CUL2 | Cullin-2 | 127382.2813 | 130123.8594 |
| DHX32 | Q7L7V1 | DHX32 | Putative pre-mRNA-sp | 19038.54688 | 22312.09766 |
| PDCD5 | O14737 | PDCD5 | Programmed cell deat | 120457.0625 | 124120.2813 |
| LOC102724159;PWP2 | A0A0B4J2E5;Q15269 | LOC102724159;PWP2 | Uncharacterized prote | 63203.53906 | 25180.83594 |
| CPPED1 | Q9BRF8 | CPPED1 | Serine/threonine-prot | 22720.74414 | 25440.6543 |
| LANCL2 | Q9NS86 | LANCL2 | LanC-like protein 2 | 77719.75781 | 81924.85938 |
| ZWINT | O95229 | ZWINT | ZW10 interactor | 58746.23438 | 56559.20313 |
| SINHCAF | F5GZ82;F5H6U2;Q9NP50 | SINHCAF | SIN3-HDAC complex-a | 4014.392334 | 3308.335693 |
| MASTL | Q96GX5 | MASTL | Serine/threonine-prot | 19186.2793 | 24072.92383 |
| NME1 | P15531 | NME1 | Nucleoside diphospha | 2684564.75 | 2848799 |
| RBM4 | Q9BWF3 | RBM4 | RNA-binding protein 4 | 709414.9375 | 446332.8125 |
| PSME2 | A0A087X1Z3;Q9UL46 | PSME2 | Proteasome activator | 487991.9063 | 605845.6875 |
| MRPL35 | Q9NZE8 | MRPL35 | 39S ribosomal protein | 56516.375 | 67484.69531 |
| TRAPPC6A | O75865 | TRAPPC6A | Trafficking protein par | 38146.25 | 37493.6875 |
| DDX54 | Q8TDD1 | DDX54 | ATP-dependent RNA h | 225239.4375 | 198979.8594 |
| MRPL47 | Q9HD33 | MRPL47 | 39S ribosomal protein | 233640.4688 | 250464.3594 |
| PKP1 | Q13835 | PKP1 | Plakophilin-1 | 6237.635254 | 2875.693604 |
| CCDC59 | Q9P031 | CCDC59 | Thyroid transcription | 67939.85938 | 43477.82813 |
| TYW1 | A0A087WUV9;Q9NV66 | TYW1 | S-adenosyl-L-methion | 8819.558594 | 14089.61133 |
| MKI67 | P46013 | MKI67 | Proliferation marker p | 272745.7188 | 190709.2031 |
| CEP170 | Q5SW79 | CEP170 | Centrosomal protein | 11103.42578 | 11872.66016 |
| RFK | Q969G6 | RFK | Riboflavin kinase | 12612.24219 | 19750.10352 |
| RMND5A | Q9H871 | RMND5A | E3 ubiquitin-protein tr | 16543.79492 | 17177.92578 |
| DCAF1 | Q9Y4B6 | DCAF1 | DDB1- and CUL4-asso | 55476.46094 | 64907.19531 |
| PML | P29590 | PML | Protein PML | 94156.41406 | 87121.9375 |

| | | | | | |
|---|---|---|---|---|---|
| C18orf21 | Q32NC0 | C18orf21 | UPF0711 protein C18c | 1266.46167 | 1802.656128 |
| TMEM30B | Q3MIR4 | TMEM30B | Cell cycle control prot | 18692.16602 | 18642.41016 |
| CDC20 | Q12834 | CDC20 | Cell division cycle prot | 435295.4375 | 316682.5 |
| PDCL3 | Q9H2J4 | PDCL3 | Phosducin-like proteir | 75397.32031 | 79665.67188 |
| FRMD8 | Q9BZ67 | FRMD8 | FERM domain-contair | 13754.27539 | 17262.29492 |
| YRDC | Q86U90 | YRDC | YrdC domain-containi | 96549.76563 | 98465.53125 |
| MRPL17 | Q9NRX2 | MRPL17 | 39S ribosomal protein | 278904.9375 | 297375.6875 |
| MLF2 | Q15773 | MLF2 | Myeloid leukemia fact | 75493.5 | 75158.78906 |
| NUDT15 | Q9NV35 | NUDT15 | Nucleotide triphospha | 20198.08789 | 20564.41797 |
| QTRT2 | Q9H974 | QTRT2 | Queuine tRNA-ribosyl | 55292.55078 | 65650.45313 |
| MRPS23 | Q9Y3D9 | MRPS23 | 28S ribosomal protein | 391935.25 | 397853.6875 |
| TMEM134 | Q9H6X4 | TMEM134 | Transmembrane prote | 7771.486328 | 7657.625488 |
| WDR36 | A0A0A0MTB8;Q8NI36 | WDR36 | WD repeat-containing | 67547.9375 | 30835.25195 |
| SDHA | P31040 | SDHA | Succinate dehydroger | 411890.5938 | 587538.75 |
| URB1 | O60287 | URB1 | Nucleolar pre-ribosom | 26981.98242 | 23528.01953 |
| CKAP4 | Q07065 | CKAP4 | Cytoskeleton-associat | 136516.4063 | 116897.8438 |
| SIRT1 | Q96EB6 | SIRT1 | NAD-dependent prote | 27362.25781 | 27656.13867 |
| MRPL51 | Q4U2R6 | MRPL51 | 39S ribosomal protein | 51913.22266 | 54252.71484 |
| UBXN8 | A0A087WT23;A0A087WV67; | UBXN8 | UBX domain-containir | 3540.907715 | 2506.80127 |
| NSD1 | Q96L73 | NSD1 | Histone-lysine N-meth | 10387.73242 | 9387.915039 |
| VARS1 | P26640 | VARS1 | Valine--tRNA ligase | 427574.9063 | 483038.375 |
| GTF2IRD1 | E9PFE2;Q9UHL9 | GTF2IRD1 | General transcription | 1710.458862 | 3512.978271 |
| EIF5A2 | C9J7B5;F8WCJ1;Q9GZV4 | EIF5A2 | Eukaryotic translation | 59806.23438 | 35728.46094 |
| ADAT2 | Q7Z6V5 | ADAT2 | tRNA-specific adenosi | 11794.09668 | 10938.14551 |
| MYH14 | M0QY43 | MYH14 | Myosin-14 (Fragment) | 615406.2383 | 619004.5078 |
| HOMER3 | Q9NSC5 | HOMER3 | Homer protein homol | 7928.57959 | 10245.66699 |
| FER | P16591;W0S4B9 | FER | Tyrosine-protein kinas | 3464.428223 | 3127.953369 |
| DSG2 | Q14126 | DSG2 | Desmoglein-2 | 48816.10156 | 31236.08594 |
| DDX50 | Q9BQ39 | DDX50 | ATP-dependent RNA h | 195951.7969 | 184490.7188 |
| RCE1 | E9PI08;E9PKA7;Q9Y256 | RCE1 | CAAX prenyl protease | 40928.31641 | 39915.25781 |
| LAD1 | O00515 | LAD1 | Ladinin-1 | 413776.5 | 400436.1563 |
| NUFIP1 | Q9UHK0 | NUFIP1 | Nuclear fragile X ment | 11485.82129 | 12893.19727 |
| CBR4 | Q8N4T8 | CBR4 | Carbonyl reductase fa | 21546.86133 | 19989.46289 |

| | | | | | |
|---|---|---|---|---|---|
| NAA15 | Q9BXJ9 | NAA15 | N-alpha-acetyltransfe | 352393.1563 | 350367 |
| PNN | Q9H307 | PNN | Pinin | 324935.3125 | 300599.375 |
| ERGIC2 | A0A087WU02;Q96RQ1 | ERGIC2 | Endoplasmic reticulur | 53301.41406 | 55591.69531 |
| CMSS1 | Q9BQ75 | CMSS1 | Protein CMSS1 | 105289.4531 | 96625.89844 |
| SUGP1 | Q8IWZ8 | SUGP1 | SURP and G-patch dor | 43885.02344 | 41270.14063 |
| NECTIN4 | Q96NY8 | NECTIN4 | Nectin-4 | 26569.17383 | 33499.60547 |
| WBP11 | Q9Y2W2 | WBP11 | WW domain-binding p | 207714.7031 | 181933.4844 |
| UTP11 | Q9Y3A2 | UTP11 | Probable U3 small nuc | 30135.64648 | 12514.6543 |
| H2BC12;HIST1H2BD;H2BC4 | O60814;P58876;P62807;Q5C | H2BC12;HIST1H2BD;H | Histone H2B type 1-K; | 1112342.375 | 63452.80859 |
| ZMYM2 | Q9UBW7 | ZMYM2 | Zinc finger MYM-type | 53765.94531 | 56908.83984 |
| LRRC42 | Q9Y546 | LRRC42 | Leucine-rich repeat-cc | 11451.64355 | 12098.27441 |
| PCYOX1 | Q9UHG3 | PCYOX1 | Prenylcysteine oxidas | 257040.7344 | 298184.3125 |
| TOMM5 | Q8N4H5 | TOMM5 | Mitochondrial import | 41904.54297 | 34097.73047 |
| ERGIC3 | H0Y5K5;H0Y621;Q9Y282 | ERGIC3 | Endoplasmic reticulur | 71578.57031 | 67936.77344 |
| TSTD1 | Q8NFU3 | TSTD1 | Thiosulfate:glutathion | 26565.19922 | 33899.63672 |
| GINS2 | Q9Y248 | GINS2 | DNA replication comp | 44281.07813 | 58100.49609 |
| KLHDC4 | Q8TBB5 | KLHDC4 | Kelch domain-contain | 53038.31641 | 63255.60156 |
| PUM3 | Q15397 | PUM3 | Pumilio homolog 3 | 91846.32813 | 33271.62109 |
| BOLA1 | Q9Y3E2 | BOLA1 | BolA-like protein 1 | 9759.913086 | 13774.83887 |
| PHF23 | Q9BUL5 | PHF23 | PHD finger protein 23 | 65898.21875 | 54240.8125 |
| FKBP4 | Q02790 | FKBP4 | Peptidyl-prolyl cis-trar | 1795650.5 | 2172228 |
| IPO4 | Q8TEX9 | IPO4 | Importin-4 | 377073.2813 | 493199.2813 |
| MRPL24 | Q96A35 | MRPL24 | 39S ribosomal proteir | 208576.7813 | 221753 |
| PNPLA6 | A0A384DVU0;Q8IY17 | PNPLA6 | Neuropathy target est | 13679.86133 | 16823.4707 |
| CDKN1B | E7ES52;P46527 | CDKN1B | Cyclin-dependent kina | 16095.45508 | 21422.36133 |
| TAF8 | A0A0A0MRR3;Q7Z7C8 | TAF8 | Transcription initiatio | 28268.92578 | 28525.93164 |
| BCL7A | Q4VC05 | BCL7A | B-cell CLL/lymphoma | 2482.847168 | 2716.081055 |
| GEN1 | Q17RS7 | GEN1 | Flap endonuclease GE | 6922.693359 | 11897.58691 |
| UBE2B | H0YA80;P63146 | UBE2B | Ubiquitin-conjugating | 95375.74219 | 95017.45313 |
| TMTC3 | Q6ZXV5 | TMTC3 | Protein O-mannosyl-tr | 37977.23438 | 44475.3125 |
| NCAPG2 | Q86XI2 | NCAPG2 | Condensin-2 complex | 32713.44141 | 37258.14063 |
| MRPS18C | Q9Y3D5 | MRPS18C | 28S ribosomal proteir | 68233.125 | 65660.88281 |
| SHFL | Q9NUL5 | SHFL | Shiftless antiviral inhil | 8121.820313 | 6073.627441 |

| | | | | | |
|---|---|---|---|---|---|
| SMYD5 | Q6GMV2 | SMYD5 | SET and MYND domai | 86152.50781 | 94849.67188 |
| EARS2 | Q5JPH6 | EARS2 | Probable glutamate--t | 39554.73438 | 45414.64453 |
| SPC24 | Q8NBT2 | SPC24 | Kinetochore protein S | 2536.300049 | 6397.563965 |
| RBM14 | Q96PK6 | RBM14 | RNA-binding protein 1 | 728977.875 | 712381.1875 |
| WDR3 | Q9UNX4 | WDR3 | WD repeat-containing | 172014.9375 | 81469.79688 |
| HEXA | H3BP20;P06865 | HEXA | Beta-hexosaminidase; | 261636.3594 | 254717.2344 |
| OAS1 | F8VXY3;P00973 | OAS1 | 2'-5'-oligoadenylate sy | 229814.5625 | 291688.1875 |
| LYRM4 | C9JRX8;C9JY28;F5H189;H7C4 | LYRM4 | LYR motif-containing p | 14434.83398 | 15837.38867 |
| DDX52 | Q9Y2R4 | DDX52 | Probable ATP-depend | 70052.375 | 57355.22266 |
| MEST | C9JW74;Q5EB52 | MEST | Mesoderm-specific tra | 14046.25195 | 17102.91211 |
| COTL1 | Q14019 | COTL1 | Coactosin-like protein | 210358.5469 | 250204.5625 |
| BUD23 | O43709 | BUD23 | Probable 18S rRNA (gu | 37647.60156 | 40804.22656 |
| GTF2F1 | P35269 | GTF2F1 | General transcription | 214490.3281 | 194600 |
| RNF149 | F8WCD0;Q8NC42 | RNF149 | E3 ubiquitin-protein li | 9907.06543 | 8919.21582 |
| GNL2 | Q13823 | GNL2 | Nucleolar GTP-binding | 113794.8516 | 89913.53906 |
| G3BP1 | Q13283 | G3BP1 | Ras GTPase-activating | 709454.0625 | 823085.25 |
| ZC3H15 | Q8WU90 | ZC3H15 | Zinc finger CCCH doma | 196614.1875 | 208466.4531 |
| C12orf29 | Q8N999 | C12orf29 | Uncharacterized prote | 20692.79102 | 19458.75977 |
| LENG9 | A0A087WVD1 | LENG9 | Leukocyte receptor clu | 14614.82227 | 13408.31543 |
| CDK8 | P49336 | CDK8 | Cyclin-dependent kina | 6412.211426 | 8053.312988 |
| RFXANK | A0A024R7P0;F5GY33;H0YGQ | RFXANK | DNA-binding protein R | 3507.001953 | 6896.244629 |
| ADGRG1 | Q9Y653 | ADGRG1 | Adhesion G-protein co | 15019.71191 | 16623.71484 |
| BORCS6 | Q96GS4 | BORCS6 | BLOC-1-related compl | 18900.71875 | 16785.58789 |
| BCL2 | P10415 | BCL2 | Apoptosis regulator Bc | 43704.12109 | 65529.62891 |
| MRPS7 | J3QLS3;Q9Y2R9 | MRPS7 | 28S ribosomal protein | 268910.25 | 312647.5625 |
| SLC29A1 | Q99808 | SLC29A1 | Equilibrative nucleosid | 40108.16797 | 48969 |
| ACO1 | P21399 | ACO1 | Cytoplasmic aconitate | 91302.89063 | 104603.6484 |
| ERCC8 | A0A2R8YD24;G3XAG7;Q1321 | ERCC8 | DNA excision repair pr | 5471.82666 | 4817.415039 |
| PLEKHA6 | A0A1B0GUN5;Q5VTI5;Q9Y2H | PLEKHA6 | Pleckstrin homology d | 106396.9629 | 128492.0195 |
| DNAAF5 | Q86Y56 | DNAAF5 | Dynein assembly facto | 289811.625 | 336376.5 |
| RCC1L | Q96I51 | RCC1L | RCC1-like G exchangir | 25750.03711 | 31405.55273 |
| UQCRC1 | P31930 | UQCRC1 | Cytochrome b-c1 com | 695170.1875 | 721670.1875 |
| RIOX2 | Q8IUF8 | RIOX2 | Ribosomal oxygenase | 28481.96289 | 23401.66992 |

| | | | | | |
|---|---|---|---|---|---|
| CNNM4 | Q6P4Q7 | CNNM4 | Metal transporter CNI | 50070.17578 | 62727.97266 |
| SELENOH | A0A087WYP2;Q8IZQ5 | SELENOH | Selenoprotein H | 2617.66333 | 3526.390625 |
| SLC22A5 | O76082 | SLC22A5 | Solute carrier family 2 | 17355.28906 | 17768.16406 |
| LSM8 | C9JIZ0;C9JNV3;F2Z2Y6;O957 | LSM8 | U6 snRNA-associated | 12462.21484 | 23398.875 |
| FAM117A | Q9C073 | FAM117A | Protein FAM117A | 6317.779785 | 12502.78516 |
| ESRRB;ESRRB;ESRRA;ESRRB | A0A2R8Y491;E7EWD9;H0YG | ESRRB;ESRRB;ESRRA;E | Steroid hormone rece | 18302.61133 | 16341.76758 |
| METAP2 | P50579 | METAP2 | Methionine aminopep | 82700.17188 | 84911.07813 |
| PTMS | P20962 | PTMS | Parathymosin | 33851.78516 | 57456.07031 |
| MRPL11 | Q9Y3B7 | MRPL11 | 39S ribosomal protein | 174433.5781 | 188188.2813 |
| BBX | C9J8D0;C9JA69;Q8WY36 | BBX | HMG box transcriptio | 10604.66876 | 20986.9292 |
| METTL1 | Q9UBP6 | METTL1 | tRNA (guanine-N(7)-)- | 121382.3594 | 147173.3125 |
| MRPL32 | Q9BYC8 | MRPL32 | 39S ribosomal protein | 38672.80469 | 36698.24609 |
| RPIA | P49247 | RPIA | Ribose-5-phosphate is | 111844.1016 | 112485.6641 |
| KLF16 | Q9BXK1 | KLF16 | Krueppel-like factor 1 | 13488.43164 | 9909.837891 |
| ESYT2 | H7BXI1 | ESYT2 | Extended synaptotagr | 84661.14063 | 89621.13281 |
| METTL16 | Q86W50 | METTL16 | RNA N6-adenosine-m | 31470.39258 | 32007.27734 |
| CDCA2 | E9PEI0;Q69YH5 | CDCA2 | Cell division cycle-ass | 13372.34473 | 8642.351563 |
| SUN2 | Q9UH99 | SUN2 | SUN domain-containi | 15664.0332 | 15988.87012 |
| CALCOCO2 | Q13137 | CALCOCO2 | Calcium-binding and c | 25370.08008 | 24250.74023 |
| TAF7 | Q15545 | TAF7 | Transcription initiatio | 86752.4375 | 99260.10156 |
| DNPH1 | H0Y8X4;O43598 | DNPH1 | 2'-deoxynucleoside 5' | 495207 | 485192.3438 |
| HSPA4 | P34932 | HSPA4 | Heat shock 70 kDa pro | 1162623.125 | 1201507.875 |
| SERF2 | P84101 | SERF2 | Small EDRK-rich facto | 71473.00781 | 78477.78906 |
| MTA2 | O94776 | MTA2 | Metastasis-associated | 388786.75 | 374774.3438 |
| ISCA1 | Q9BUE6 | ISCA1 | Iron-sulfur cluster ass | 19203.32813 | 21408.45313 |
| WASHC2A | Q641Q2 | WASHC2A | WASH complex subun | 60553.875 | 70741.49219 |
| SIAE | Q9HAT2 | SIAE | Sialate O-acetylestera | 13089.48438 | 16216.15332 |
| PEG10 | A0A087WUL4;A0A087WX23; | PEG10 | Retrotransposon-deri | 20909.01758 | 20422.51953 |
| GSN | A0A0A0MS51;A0A0A0MT01; | GSN | Gelsolin | 76993.68652 | 94966.57422 |
| B4GALT3 | O60512 | B4GALT3 | Beta-1,4-galactosyltra | 4262.1875 | 5453.75 |
| NARS2 | Q96I59 | NARS2 | Probable asparagine-- | 101692.9375 | 118430.1016 |
| EEA1 | Q15075 | EEA1 | Early endosome antig | 93904.4375 | 85555.52344 |
| MRPL53 | Q96EL3 | MRPL53 | 39S ribosomal protein | 144739.8281 | 169009.5781 |

| | | | | | |
|---|---|---|---|---|---|
| PBDC1 | Q9BVG4 | PBDC1 | Protein PBDC1 | 35393.66016 | 54088.25 |
| PAN3 | A0A0C4DFZ9;Q58A45 | PAN3 | PAN2-PAN3 deadenyla | 15987.86621 | 17018.94531 |
| CNEP1R1 | H3BUT5;Q8N9A8 | CNEP1R1 | Nuclear envelope pho | 3674.272217 | 3960.151123 |
| REL | Q04864 | REL | Proto-oncogene c-Rel | 21955.33008 | 23157.35742 |
| AKT1S1 | H9KV91;M0R2V8;Q96B36 | AKT1S1 | Proline-rich AKT1 subs | 10143.02051 | 12766.19141 |
| SPDL1 | Q96EA4 | SPDL1 | Protein Spindly | 75244.75781 | 78704.08594 |
| CD9 | A0A2R8Y478;A6NNI4;G8JLH6 | CD9 | Tetraspanin;Tetraspar | 330613.7813 | 276481.2188 |
| EPHA4 | E9PG71;P54764 | EPHA4 | Ephrin type-A recepto | 21256.56836 | 18615.4082 |
| ATL2 | Q8NHH9 | ATL2 | Atlastin-2 | 106817.9063 | 106422.5 |
| PCDH11X;PCDH11Y | Q9BZA7;Q9BZA8 | PCDH11X;PCDH11Y | Protocadherin-11 X-lir | 23179.92773 | 24524.18359 |
| ALDH1B1 | P30837 | ALDH1B1 | Aldehyde dehydrogen | 162910.0313 | 157260.0938 |
| SDHD | O14521 | SDHD | Succinate dehydroger | 37372.78516 | 43440.91406 |
| AHCYL1 | O43865 | AHCYL1 | S-adenosylhomocyste | 228734.3906 | 265695.25 |
| COX7C | D6R9Z7;P15954 | COX7C | Cytochrome c oxidase | 134252.5781 | 167351.1406 |
| IL1R1 | B8ZZ73;B8ZZW4;B9A040;C9J | IL1R1 | Interleukin-1 receptor | 1586.335693 | NA |
| DFFA | O00273 | DFFA | DNA fragmentation fa | 6497.342773 | 6646.721191 |
| EIF4H | Q15056 | EIF4H | Eukaryotic translation | 1575522.125 | 1456897.75 |
| DDAH1 | O94760 | DDAH1 | N(G),N(G)-dimethylarg | 130041.7734 | 134322.2656 |
| USP5 | P45974 | USP5 | Ubiquitin carboxyl-ter | 940663.625 | 997448.9375 |
| SULT1A1 | P50225 | SULT1A1 | Sulfotransferase 1A1 | 23309.07031 | 24500.33984 |
| ATRIP | A0A0U1RQJ8;Q8WXE1 | ATRIP | ATR-interacting prote | 142455.7813 | 45201.07031 |
| BTF3 | D6RDG3 | BTF3 | Transcription factor B | 545785.5938 | 454039.9219 |
| CUL1 | Q13616 | CUL1 | Cullin-1 | 130557.1641 | 143366.2344 |
| IK | Q13123 | IK | Protein Red | 336324.125 | 204201.9063 |
| S100A11 | P31949 | S100A11 | Protein S100-A11 | 783686.875 | 979732.375 |
| ARHGEF10 | H0YAN8;O15013 | ARHGEF10 | Rho guanine nucleotic | 13885.99316 | 19334.50391 |
| UCHL3 | P15374 | UCHL3 | Ubiquitin carboxyl-ter | 33120.24219 | 45713.10156 |
| CRTC2 | Q53ET0 | CRTC2 | CREB-regulated transc | 22141.5918 | 24570.00977 |
| WWOX | Q9NZC7 | WWOX | WW domain-containir | 13337.86133 | 16707.75586 |
| NKTR | P30414 | NKTR | NK-tumor recognition | 99345.02344 | 165152.1719 |
| POMT1 | Q9Y6A1 | POMT1 | Protein O-mannosyl-tr | 12712.0332 | 14560.32617 |
| TRAPPC2L | A0A0B4J294;Q9UL33 | TRAPPC2L | Trafficking protein par | 106172.4141 | 116134.0156 |
| ILRUN | Q9H6K1 | ILRUN | Protein ILRUN | 23228.7168 | 27933.53516 |

| | | | | | |
|---|---|---|---|---|---|
| CCDC137 | I3L0U5;Q6PK04 | CCDC137 | Coiled-coil domain-co | 195161.6563 | 164313.125 |
| CLASRP | A0A0A0MQS2;F5H0Q6;Q8N2 | CLASRP | CLK4-associating serin | 14631.47266 | 10122.48926 |
| P4HA1 | P13674 | P4HA1 | Prolyl 4-hydroxylase s | 160114.0781 | 190890.1094 |
| PHB2 | Q99623 | PHB2 | Prohibitin-2 | 806165.125 | 820860.125 |
| PINX1 | Q96BK5 | PINX1 | PIN2/TERF1-interactir | 26811.32813 | 25782.27734 |
| UQCRC2 | P22695 | UQCRC2 | Cytochrome b-c1 com | 360395 | 386404.9063 |
| RABEPK | Q7Z6M1 | RABEPK | Rab9 effector protein | 29637.64648 | 24952.39258 |
| MAGI3 | Q5TCQ9 | MAGI3 | Membrane-associatec | 24327.95898 | 31607.97461 |
| NECTIN1 | Q15223 | NECTIN1 | Nectin-1 | 23856.84375 | 27195.58594 |
| PKMYT1 | A6NHV6;B4DZM6;Q99640 | PKMYT1 | Membrane-associatec | 13642.06641 | 15054.91699 |
| ARMH3 | Q5T2E6 | ARMH3 | Armadillo-like helical | 31970.89648 | 36157.46094 |
| NOC2L | Q9Y3T9 | NOC2L | Nucleolar complex pro | 195010.9844 | 155466.0781 |
| PARVB | A0A087WZB5;Q9HBI1 | PARVB | Beta-parvin | 8071.862793 | 9653.546875 |
| CEBPB | P17676 | CEBPB | CCAAT/enhancer-binc | 21977.06641 | 20768.76953 |
| TFAM | Q00059 | TFAM | Transcription factor A | 449033.3438 | 477915.4375 |
| WDR4 | P57081 | WDR4 | tRNA (guanine-N(7)-)- | 101625.3906 | 117808.5703 |
| MED8 | Q96G25 | MED8 | Mediator of RNA poly | 28482.24414 | 32250.8125 |
| SNX5 | Q9Y5X3 | SNX5 | Sorting nexin-5 | 243874.0938 | 302826 |
| PARD6B | Q9BYG5 | PARD6B | Partitioning defective | 555177.4375 | 606520.1875 |
| SUN1 | E9PHI4 | SUN1 | SUN domain-containir | 89101.71094 | 61859.29688 |
| EEF1B2 | P24534 | EEF1B2 | Elongation factor 1-be | 102436.1719 | 185657.7813 |
| PDSS1 | Q5T2R2 | PDSS1 | Decaprenyl-diphospha | 8674.789063 | 9051.588867 |
| TMSB10 | P63313 | TMSB10 | Thymosin beta-10 | 75485.9375 | 90912.58594 |
| ZNF511-PRAP1;ZNF511 | H7BY64;Q8NB15 | ZNF511-PRAP1;ZNF51 | ZNF511-PRAP1 readth | 10390.86816 | 8601.832031 |
| CLIP1 | J3KP58 | CLIP1 | CAP-Gly domain-conta | 68239.66602 | 66782.72363 |
| UCK2 | A0A2R8Y653;Q9BZX2 | UCK2 | Uridine-cytidine kinas | 116968.5938 | 107804.1484 |
| KCTD1 | A0A2U3U043;J3KSG1;J3QRK1 | KCTD1 | BTB/POZ domain-cont | 27152.99023 | 29536.17188 |
| PPT2;PPT2;PPT2;PPT2;PPT2 | A0A024RCQ8;A0A0G2JH76;A | PPT2;PPT2;PPT2;PPT2 | HCG1999928, isoform | 22461.42383 | 24768.77344 |
| FPGS | Q05932 | FPGS | Folylpolyglutamate sy | 8845.311523 | 11087.50781 |
| REPIN1 | Q9BWE0 | REPIN1 | Replication initiator 1 | 31362.63086 | 33737.73828 |
| BYSL | Q13895 | BYSL | Bystin | 205897.375 | 129692.0703 |
| HMGA1 | P17096 | HMGA1 | High mobility group p | 35426.27734 | 36459.89844 |
| CENPF | P49454 | CENPF | Centromere protein F | 67115.11719 | 48296.03906 |

| | | | | | |
|---|---|---|---|---|---|
| SLIRP | Q9GZT3 | SLIRP | SRA stem-loop-interac | 849747.5625 | 914103.375 |
| METTL17 | Q9H7H0 | METTL17 | Methyltransferase-like | 13283.95117 | 16923.38086 |
| YAF2 | Q8IY57 | YAF2 | YY1-associated factor | 4603.963867 | 6359.970215 |
| EPM2A | B3EWF7 | EPM2A | Laforin, isoform 9 | 65731.15625 | 63328.41016 |
| POLR3A | O14802 | POLR3A | DNA-directed RNA po | 50170.65234 | 49876 |
| CARD19 | Q96LW7 | CARD19 | Caspase recruitment c | 56982.19922 | 25834.9375 |
| COMMD8 | Q9NX08 | COMMD8 | COMM domain-conta | 2555.077881 | 6624.030273 |
| PSME1 | Q06323 | PSME1 | Proteasome activator | 1922407.5 | 2397338 |
| VGLL4 | A0A075B6E4;A0A0A6YYI5;E7 | VGLL4 | Transcription cofactor | 2405.579102 | 4273.087891 |
| HELZ2 | Q9BYK8 | HELZ2 | Helicase with zinc fing | 33046.59766 | 27489.07617 |
| MACROH2A1 | O75367 | MACROH2A1 | Core histone macro-H | 295361.375 | 16792.68945 |
| SRRM2 | Q9UQ35 | SRRM2 | Serine/arginine repeti | 382814.5938 | 381378.75 |
| HEATR3 | Q7Z4Q2 | HEATR3 | HEAT repeat-containin | 32985.22266 | 34470.80859 |
| SELENOS | Q9BQE4 | SELENOS | Selenoprotein S | 35228.91797 | 34012.96484 |
| FOXJ2 | Q9P0K8 | FOXJ2 | Forkhead box protein | 7811.554199 | 8371.818359 |
| WNK1 | F5GWT4;Q9H4A3 | WNK1 | Serine/threonine-prot | 84462.71094 | 89193.76563 |
| IAH1 | Q2TAA2 | IAH1 | Isoamyl acetate-hydro | 218524.2656 | 182526.7813 |
| HECW2 | A0A2R8Y6F3;A0A2R8YE75;Q9 | HECW2 | E3 ubiquitin-protein li | 120929.1641 | 127617.2188 |
| ALG8 | H0YDD3;H0YDQ1;Q9BVK2 | ALG8 | Alpha-1,3-glucosyltran | 17132.4043 | 30094.90039 |
| OSTF1 | Q92882 | OSTF1 | Osteoclast-stimulating | 147615.7656 | 151624.8281 |
| C2orf69 | Q8N8R5 | C2orf69 | UPF0565 protein C2or | 9811.850586 | 10246.48633 |
| LMAN2L | Q9H0V9 | LMAN2L | VIP36-like protein | 105716.75 | 110216 |
| LZTS2 | Q9BRK4 | LZTS2 | Leucine zipper putativ | 2038.789795 | 2032.504395 |
| FGFR1OP | A0A087WV25;O95684 | FGFR1OP | FGFR1 oncogene partn | 16493.97461 | 19599.25 |
| ANKFY1 | Q9P2R3 | ANKFY1 | Rabankyrin-5 | 80054.24219 | 93937.1875 |
| DECR2 | Q4VXZ8;Q9NUI1 | DECR2 | Peroxisomal 2,4-dienc | 78903.32031 | 87709.69531 |
| NGDN | Q8NEJ9 | NGDN | Neuroguidin | 89250.71875 | 61976.53516 |
| CCDC43 | Q96MW1 | CCDC43 | Coiled-coil domain-co | 163985.4063 | 179667.2188 |
| ALDOC | P09972 | ALDOC | Fructose-bisphosphat | 1974444.75 | 3221445.25 |
| HSPH1 | Q92598 | HSPH1 | Heat shock protein 10 | 1478945.625 | 1646419.375 |
| DNA2 | F8VR31;P51530 | DNA2 | DNA replication ATP-c | 4919.533691 | 5271.560059 |
| DHX29 | A0A087WYN9;Q7Z478 | DHX29 | ATP-dependent RNA h | 77393.82031 | 77854.90625 |
| EIF5A | P63241 | EIF5A | Eukaryotic translation | 6248498.5 | 5881968 |

| | | | | | |
|---|---|---|---|---|---|
| LSM3 | P62310 | LSM3 | U6 snRNA-associated | 16074.87305 | 38068.61328 |
| NSUN5 | Q96P11 | NSUN5 | Probable 28S rRNA (cy | 55212.58984 | 44490.59766 |
| LTA4H | P09960 | LTA4H | Leukotriene A-4 hydro | 811683.875 | 841390.875 |
| PALLD | Q8WX93 | PALLD | Palladin | 44425.96875 | 41714.83984 |
| PHTF2 | Q8N3S3 | PHTF2 | Putative homeodoma | 3111.398438 | 3577.252686 |
| PPP1R15B | Q5SWA1 | PPP1R15B | Protein phosphatase 1 | 4128.347168 | 3758.934082 |
| HINT1 | P49773 | HINT1 | Histidine triad nucleot | 175458.2656 | 263355.3125 |
| LIMD1 | C9JRJ5;Q9UGP4 | LIMD1 | LIM domain-containin | 26177.30859 | 26671.79883 |
| NUDT16L1 | Q9BRJ7 | NUDT16L1 | Tudor-interacting repa | 184029.4063 | 175638.0156 |
| CTDP1 | A0A0J9YWB6;Q9Y5B0 | CTDP1 | RNA polymerase II sub | 50543.66016 | 56440.29688 |
| MRPL54 | Q6P161 | MRPL54 | 39S ribosomal protein | 129586.4531 | 128656.5938 |
| B2M | P61769 | B2M | Beta-2-microglobulin | 48206.54688 | 60297.82031 |
| CBL | A0A0U1RQX8;A0A1B0GW38; | CBL | E3 ubiquitin-protein li | 65891.44531 | 77106.59375 |
| XPNPEP3 | Q9NQH7 | XPNPEP3 | Xaa-Pro aminopeptida | 45914.75 | 55162.66406 |
| PHLDB1 | Q86UU1 | PHLDB1 | Pleckstrin homology-l | 7986.913574 | 14113.93945 |
| NCOR1 | O75376 | NCOR1 | Nuclear receptor core | 31101.95508 | 30611.75195 |
| MBD3 | O95983 | MBD3 | Methyl-CpG-binding d | 124694.7578 | 129255.0625 |
| MRPL34 | M0R226;Q9BQ48 | MRPL34 | 39S ribosomal protein | 96146.09375 | 105342.75 |
| POLR3E | A0A0C4DH01;Q9NVU0 | POLR3E | DNA-directed RNA po | 17009.96289 | 17274.91016 |
| FAM207A | Q9NSI2 | FAM207A | Protein FAM207A | 149855.1719 | 120521.9844 |
| PLCB1 | A0A087WT80;A0A087WW73 | PLCB1 | 1-phosphatidylinosito | 34757.54688 | 21382.85547 |
| RAVER2 | Q9HCJ3 | RAVER2 | Ribonucleoprotein PT | 13567.6084 | 11189.8291 |
| NUDT5 | A6NFX8;Q9UKK9 | NUDT5 | ADP-sugar pyrophosp | 520873.1563 | 589286.5 |
| PHB | P35232 | PHB | Prohibitin | 2031704.75 | 2069171 |
| UPF3A | Q9H1J1 | UPF3A | Regulator of nonsense | 6199.8125 | 6247.907715 |
| WAC | A0A0A0MRT2 | WAC | WW domain-containin | 48913.24023 | 47588.3623 |
| TIPIN | Q9BVW5 | TIPIN | TIMELESS-interacting | 23107.1875 | 25522.60938 |
| ALG6 | Q9Y672;S4R350 | ALG6 | Dolichyl pyrophospha | 15745.31934 | 15151.57715 |
| RBM27 | Q9P2N5 | RBM27 | RNA-binding protein 2 | 94743.5 | 89591.85938 |
| RDH10 | Q8IZV5 | RDH10 | Retinol dehydrogenas | 5392.349609 | 5221.047852 |
| SEC23B | A0A2R8YFH5;Q15437 | SEC23B | Protein transport prot | 414943.0938 | 456947.5938 |
| WDR18 | Q9BV38;U3KQC1 | WDR18 | WD repeat-containing | 219419.7656 | 236722.3281 |
| EEF1D | E9PL71 | EEF1D | Elongation factor 1-de | 745509.6685 | 1183359.74 |

| | | | | | |
|---|---|---|---|---|---|
| UBE2S | Q16763 | UBE2S | Ubiquitin-conjugating | 142384.0625 | 120979.2422 |
| DGCR8 | Q8WYQ5 | DGCR8 | Microprocessor comp | 14481.07422 | 11971.12891 |
| GFRA1 | P56159 | GFRA1 | GDNF family receptor | 34200.56641 | 45062.55859 |
| KBTBD7 | Q8WVZ9 | KBTBD7 | Kelch repeat and BTB | 18689.04492 | 19855.44727 |
| M6PR | P20645 | M6PR | Cation-dependent ma | 205140.4375 | 180891.875 |
| POLRMT | O00411 | POLRMT | DNA-directed RNA po | 12767.03613 | 16516.47266 |
| POM121C | A8CG34 | POM121C | Nuclear envelope por | 43745.26953 | 38714.74219 |
| ERLIN1 | O75477 | ERLIN1 | Erlin-1 | 36453.89844 | 42124.53516 |
| LMAN1 | P49257 | LMAN1 | Protein ERGIC-53 | 281787.5625 | 326309.0938 |
| CDK18 | Q07002 | CDK18 | Cyclin-dependent kina | 14926.41699 | 17391.29102 |
| MTA3 | E7EQY4;E7EV10;Q9BTC8 | MTA3 | Metastasis-associated | 93842.83594 | 89757.27344 |
| GLUD2 | P49448 | GLUD2 | Glutamate dehydroge | 8835.170898 | 7311.711914 |
| MRPL52 | Q86TS9 | MRPL52 | 39S ribosomal protein | 119941.6406 | 113683.9922 |
| UBTD2 | Q8WUN7 | UBTD2 | Ubiquitin domain-con | 9119.336914 | 9581.44043 |
| CD81 | A6NMH8;E9PIF1;E9PJK1;E9PI | CD81 | Tetraspanin;Tetraspar | 232692.0625 | 224682.9375 |
| SUGT1 | Q9Y2Z0 | SUGT1 | Protein SGT1 homolog | 390605.6563 | 424661.125 |
| NOA1 | Q8NC60 | NOA1 | Nitric oxide-associated | 114084.8125 | 120140.1641 |
| GNL3L | Q9NVN8 | GNL3L | Guanine nucleotide-b | 70763.54688 | 73409.92969 |
| MRPL50 | Q8N5N7 | MRPL50 | 39S ribosomal protein | 18029.24609 | 21590.17969 |
| PDIA6 | Q15084 | PDIA6 | Protein disulfide-isom | 2950282 | 3073537.5 |
| PTPN14 | Q15678 | PTPN14 | Tyrosine-protein phos | 4737.393555 | 4352.755859 |
| ABT1 | Q9ULW3 | ABT1 | Activator of basal trar | 29116.75391 | 12166.99219 |
| SDC4 | P31431 | SDC4 | Syndecan-4 | 27425.00781 | 85475.75 |
| METTL8 | B3KW44;E7ETE0 | METTL8 | Methyltransferase-like | 5330.852539 | 4757.966797 |
| PAPSS2 | O95340 | PAPSS2 | Bifunctional 3'-phosph | 80698.1875 | 81650.40625 |
| PRR11 | Q96HE9 | PRR11 | Proline-rich protein 11 | 29430.81445 | 32478.10742 |
| ISOC1 | Q96CN7 | ISOC1 | Isochorismatase doma | 1118834 | 1509062.125 |
| HSDL2 | Q6YN16 | HSDL2 | Hydroxysteroid dehyd | 120212.8672 | 146966.4375 |
| DNAJB6 | O75190 | DNAJB6 | DnaJ homolog subfam | 44872.70313 | 49440.80078 |
| UTP3 | Q9NQZ2 | UTP3 | Something about siler | 84782 | 54544.94531 |
| RBM19 | Q9Y4C8 | RBM19 | Probable RNA-binding | 54564.87109 | 21319.92969 |
| TJP3 | O95049 | TJP3 | Tight junction protein | 61606.33984 | 66685.77344 |
| AGA | P20933 | AGA | N(4)-(beta-N-acetylglu | 66406.57813 | 70234.17188 |

| | | | | | |
|---|---|---|---|---|---|
| PKP3 | Q9Y446 | PKP3 | Plakophilin-3 | 373547.1563 | 394909.4063 |
| FNDC3B | Q53EP0 | FNDC3B | Fibronectin type III do | 10472.74121 | 7599.212891 |
| SURF6 | O75683 | SURF6 | Surfeit locus protein 6 | 89627.67969 | 180931.3281 |
| HDAC2 | Q92769 | HDAC2 | Histone deacetylase 2 | 740550.4375 | 781498 |
| PLXNB2 | O15031 | PLXNB2 | Plexin-B2 | 25555.18555 | 25495.86328 |
| CCDC58 | Q4VC31 | CCDC58 | Coiled-coil domain-co | 226038.6563 | 217277.1094 |
| UBE2G2 | P60604 | UBE2G2 | Ubiquitin-conjugating | 14538.36328 | 21278.29297 |
| UBE2R2 | Q712K3 | UBE2R2 | Ubiquitin-conjugating | 33415.43359 | 54152.28906 |
| KARS1 | Q15046 | KARS1 | Lysine--tRNA ligase | 786822.375 | 851724.3125 |
| AP1S2 | A0A5F9ZHW1;A0A5F9ZHX2;A | AP1S2 | AP complex subunit si | 24409.57617 | 28417.30078 |
| SLAIN1 | A0A0A0MSS3;A0A0C4DFN6;( | SLAIN1 | SLAIN motif-containin NA | | 10510.31934 |
| KRT3 | P12035 | KRT3 | Keratin, type II cytosk | 21008.26758 | 27904.64844 |
| MTG2 | Q5JXJ0;Q9H4K7 | MTG2 | Mitochondrial ribosor | 7116.766602 | 7713.527344 |
| GRPEL2 | Q8TAA5 | GRPEL2 | GrpE protein homolog | 12361.14258 | 13553.5127 |
| FOPNL | I3L269;I3NI25;Q96NB1 | FOPNL | LisH domain-containir | 6658.712891 | 7151.507324 |
| FOCAD | Q5VW36 | FOCAD | Focadhesin | 36196.71484 | 39757.61719 |
| GABARAPL2 | P60520 | GABARAPL2 | Gamma-aminobutyric | 183225.125 | 197242.5 |
| NAF1 | Q96HR8 | NAF1 | H/ACA ribonucleoprot | 32735.73438 | 37025.78906 |
| WDR92 | Q96MX6 | WDR92 | WD repeat-containing | 39358.78125 | 39515.14453 |
| NDUFAB1 | H3BNK3;O14561 | NDUFAB1 | Acyl carrier protein (F | 8233.678711 | 24687.45508 |
| CD3EAP | O15446 | CD3EAP | DNA-directed RNA po | 155642.5469 | 131024.7969 |
| TRAP1 | Q12931 | TRAP1 | Heat shock protein 75 | 1113809.5 | 1278612.5 |
| SDAD1 | Q9NVU7 | SDAD1 | Protein SDA1 homolo | 50191.61719 | 50120.35938 |
| SF3B5 | Q9BWJ5 | SF3B5 | Splicing factor 3B sub | 11672.67969 | 22185.18359 |
| FDFT1 | P37268 | FDFT1 | Squalene synthase | 343363.8438 | 837846.6875 |
| GPATCH4 | Q5T3I0 | GPATCH4 | G patch domain-conta | 61296.83203 | 51187.96484 |
| GALE | Q14376 | GALE | UDP-glucose 4-epimer | 180586.8906 | 220404.7969 |
| SIL1 | D6REA1;Q9H173 | SIL1 | Nucleotide exchange f | 35069.67578 | 36311.92188 |
| UBE3C | Q15386 | UBE3C | Ubiquitin-protein ligas | 44909.40234 | 48370.11719 |
| TRMT1 | Q9NXH9 | TRMT1 | tRNA (guanine(26)-N( | 32007.63672 | 36286.16016 |
| IMP3 | Q9NV31 | IMP3 | U3 small nucleolar rib | 83407.39844 | 39090.44531 |
| TSR1 | Q2NL82 | TSR1 | Pre-rRNA-processing p | 172455.8125 | 165596.8594 |
| CTTN | Q14247 | CTTN | Src substrate cortactir | 1042046.625 | 1158826.25 |

| | | | | | |
|---|---|---|---|---|---|
| MKLN1 | Q9UL63 | MKLN1 | Muskelin | 24365.46875 | 22729.01563 |
| TACC2 | E7EMZ9;E9PBC6 | TACC2 | Transforming acidic co | 915112.625 | 921293.75 |
| AMPD2 | A0A5F9UK94 | AMPD2 | AMP deaminase | 137329.0938 | 149774.4688 |
| SNX17 | Q15036 | SNX17 | Sorting nexin-17 | 101356.1094 | 99932.80469 |
| EPN2 | E9PBC1;F6PQP6;O95208 | EPN2 | Epsin-2;Epsin-2 (Fragr | 44903.97266 | 50320.42969 |
| NLN | Q9BYT8 | NLN | Neurolysin, mitochond | 216793.5469 | 239384.0156 |
| JMY | Q8N9B5 | JMY | Junction-mediating ar | 11493.06348 | 12406.04297 |
| DPH5 | Q9H2P9 | DPH5 | Diphthine methyl este | 43492.92188 | 38007.83203 |
| AHDC1 | Q5TGY3 | AHDC1 | AT-hook DNA-binding | 4280.349121 | 4261.063477 |
| SFXN4 | Q6P4A7 | SFXN4 | Sideroflexin-4 | 47164.53906 | 47244.86719 |
| C1orf131 | Q8NDD1 | C1orf131 | Uncharacterized prote | 27065.05664 | 13373.40918 |
| OGFOD1 | Q8N543 | OGFOD1 | Prolyl 3-hydroxylase C | 74339.14063 | 81698.80469 |
| AK6 | Q9Y3D8 | AK6 | Adenylate kinase isoe | 103482.7656 | 99665.78125 |
| UBAC1 | Q9BSL1 | UBAC1 | Ubiquitin-associated d | 3541.536865 | 8001.038574 |
| UBL3 | O95164 | UBL3 | Ubiquitin-like protein | 90624.83594 | 117402.5703 |
| LCP1 | P13796 | LCP1 | Plastin-2 | 2461.234863 | 1994.363037 |
| POLR2A | A0A590UJN4;A0A590UKB6;P | POLR2A | DNA-directed RNA po | 145750.6719 | 73776.03906 |
| CDC27 | I3L328 | CDC27 | Cell division cycle prot | 98309.52344 | 103285.5508 |
| SRPK1 | H3BLV9;Q96SB4 | SRPK1 | SRSF protein kinase 1 | 64946.57422 | 65297.92578 |
| INSR | P06213 | INSR | Insulin receptor | 4846.644043 | 8148.838867 |
| TNIP1 | A0A0A0MRZ4;Q15025 | TNIP1 | TNFAIP3-interacting p | 38170.71094 | 39505.17969 |
| PUS1 | G8JLB3;Q9Y606 | PUS1 | tRNA pseudouridine s | 155988.4063 | 173619.5156 |
| CHAC2 | Q8WUX2 | CHAC2 | Glutathione-specific g | 23568.59961 | 24116.82813 |
| SELENOF | A0A087X1G7;O60613 | SELENOF | Selenoprotein F | 9382.613281 | 14016.90332 |
| TCF25 | Q9BQ70 | TCF25 | Transcription factor 25 | 119966.1797 | 145545.4063 |
| USP36 | Q9P275 | USP36 | Ubiquitin carboxyl-ter | 21691.43555 | 14212.75098 |
| DSG1 | Q02413 | DSG1 | Desmoglein-1 | 3101.287109 | 3715.665771 |
| CDV3 | Q9UKY7 | CDV3 | Protein CDV3 homolog | 1843096.25 | 2083995.5 |
| EVPL | Q92817 | EVPL | Envoplakin | 60634.49219 | 41549.8125 |
| PHLPP2 | H3BMS5;Q6ZVD8 | PHLPP2 | PH domain leucine-ric | 9343.330078 | 8120.461426 |
| C1orf122 | Q6ZSJ8 | C1orf122 | Uncharacterized prote | 32609.05859 | 33163.59375 |
| NCBP3 | Q53F19 | NCBP3 | Nuclear cap-binding p | 102104.5391 | 90162.01563 |
| RNF138 | Q8WVD3 | RNF138 | E3 ubiquitin-protein li | 10539.28516 | 10863.34082 |

| | | | | | |
|---|---|---|---|---|---|
| GADD45GIP1 | Q8TAE8 | GADD45GIP1 | Growth arrest and DN | 131900.8125 | 123023.9141 |
| TCTN3 | A0A0C4DFN5;Q6NUS6 | TCTN3 | Chromosome 10 oper | 25120.00391 | 22009.27539 |
| EPHX2 | E5RFU2;H0YAW7;P34913 | EPHX2 | Bifunctional epoxide h | 9208.871094 | 11005.78125 |
| NOL8 | Q76FK4 | NOL8 | Nucleolar protein 8 | 38026.32813 | 29340.44727 |
| UQCRQ | O14949 | UQCRQ | Cytochrome b-c1 com | 148564.9688 | 167249.0313 |
| LDAH | B5MDU6;Q9H6V9 | LDAH | Lipid droplet-associat | 36176.8125 | 39932.08984 |
| CDK5RAP1 | Q96SZ6 | CDK5RAP1 | CDK5 regulatory subu | 9492.527344 | 7880.600098 |
| TOR3A | Q9H497 | TOR3A | Torsin-3A | 49858.48047 | 49656.53125 |
| CEP97 | E9PG22;Q8IW35 | CEP97 | Centrosomal protein c | 24213.19531 | 30885.73047 |
| DGKH | A0A0D9SFR6;Q86XP1 | DGKH | Diacylglycerol kinase;I | 16060.48535 | 18151.60352 |
| NDUFV3 | P56181 | NDUFV3 | NADH dehydrogenase | 71646.05469 | 84355.375 |
| SHROOM3 | A0A2R8Y5P9;Q8TF72 | SHROOM3 | Protein Shroom3 | 134405.1406 | 125286.4688 |
| AMDHD2 | Q9Y303 | AMDHD2 | N-acetylglucosamine-( | 35748.76563 | 35815.59766 |
| UQCRFS1 | P47985 | UQCRFS1 | Cytochrome b-c1 com | 663183.625 | 704232.3125 |
| PIGO | Q8TEQ8 | PIGO | GPI ethanolamine pho | 33219.76953 | 34907.30469 |
| MAPK8 | P45983 | MAPK8 | Mitogen-activated pro | 21353.33984 | 26326.52734 |
| ATP8B1 | A0A2R8Y5C5;O43520 | ATP8B1 | Phospholipid-transpor | 15807.82617 | 8161.358398 |
| SDHC | Q99643 | SDHC | Succinate dehydroger | 20257.05273 | 26691.86523 |
| CFAP36 | Q96G28 | CFAP36 | Cilia- and flagella-asso | 28078.10352 | 36153.50391 |
| UBE2O | Q9C0C9 | UBE2O | (E3-independent) E2 u | 241350.5781 | 245782.7188 |
| QTRT1 | Q9BXR0 | QTRT1 | Queuine tRNA-ribosyl | 90979.41406 | 106097.4141 |
| GALK1 | P51570 | GALK1 | Galactokinase | 565123.0625 | 618231.375 |
| FAM83D | A0A087WXK8;Q9H4H8 | FAM83D | Chromosome 20 oper | 34994.25391 | 31999.82813 |
| UNC93B1 | Q9H1C4 | UNC93B1 | Protein unc-93 homol | 22226.62891 | 23828.54297 |
| WDR47 | A0A0A0MRE6;O94967 | WDR47 | WD repeat-containing | 13083.5752 | 12292.72363 |
| WDR55 | Q9H6Y2 | WDR55 | WD repeat-containing | 76890.26563 | 73346.60156 |
| TIMM17A | Q99595 | TIMM17A | Mitochondrial import | 44855.28125 | 39984.48828 |
| NOB1 | Q9ULX3 | NOB1 | RNA-binding protein N | 55670.33203 | 69416.75781 |
| CCNB1 | P14635 | CCNB1 | G2/mitotic-specific cy | 149300.4844 | 131277.5469 |
| ZC3H18 | E7ERS3;Q86VM9 | ZC3H18 | Zinc finger CCCH dom | 72795.21094 | 87794.61719 |
| ABCF2 | Q9UG63 | ABCF2 | ATP-binding cassette s | 181913.75 | 198846.9375 |
| GLG1 | Q92896 | GLG1 | Golgi apparatus prote | 272097.5938 | 324939.8438 |
| RBIS | A0A087WWE7;Q8N0T1 | RBIS | Ribosomal biogenesis | 17624.67188 | 17239.45898 |

| | | | | | |
|---|---|---|---|---|---|
| TNFRSF10A | F8U8C0;O00220 | TNFRSF10A | Tumor necrosis factor | 28305.85156 | 17324.67969 |
| MCM10 | Q5T670;Q7L590 | MCM10 | MCM10 minichromos | 74751.46875 | 66610.24219 |
| ZNRF1 | H3BRB6;Q8ND25 | ZNRF1 | E3 ubiquitin-protein li | 65850.5 | 62135.51953 |
| PDCD2L | Q9BRP1 | PDCD2L | Programmed cell deat | 144801.5 | 140326.0625 |
| OTUD4 | Q01804 | OTUD4 | OTU domain-containi | 27801.97852 | 30032.08984 |
| BRAT1 | Q6PJG6 | BRAT1 | BRCA1-associated ATI | 81224.57813 | 100122.9297 |
| ABCE1 | H0Y990 | ABCE1 | ATP-binding cassette : | 504605.375 | 505879.7822 |
| FTSJ3 | Q8IY81 | FTSJ3 | pre-rRNA 2'-O-ribose | 545874.6875 | 460657.2813 |
| CRCP | O75575 | CRCP | DNA-directed RNA po | 29722.69141 | 28377.82227 |
| MEAK7 | Q6P9B6 | MEAK7 | MTOR-associated pro | 79726.76563 | 98357.07031 |
| ANAPC7 | Q9UJX3 | ANAPC7 | Anaphase-promoting | 69773.46875 | 70393.02344 |
| UTP6 | Q9NYH9 | UTP6 | U3 small nucleolar RN | 30834.03711 | 7749.350098 |
| COBL | H7C1N2;J3KR05;O75128 | COBL | Protein cordon-bleu (I | 6466.289551 | 5248.885254 |
| TNFRSF10B | O14763 | TNFRSF10B | Tumor necrosis factor | 6420.765137 | 5899.309082 |
| CPT1A | P50416 | CPT1A | Carnitine O-palmitoylt | 454902.2813 | 433651.0313 |
| DHX37 | Q8IY37 | DHX37 | Probable ATP-depend | 76109.64844 | 71417.82813 |
| RARA | A8MUP8;P10276 | RARA | Retinoic acid receptor | 13900.56445 | 11740.93164 |
| NADSYN1 | Q6IA69 | NADSYN1 | Glutamine-dependent | 51145.57813 | 75902.02344 |
| NHEJ1 | C9JWV4;Q9H9Q4 | NHEJ1 | Non-homologous end | 15520.33203 | 17473.97852 |
| NFS1 | Q9Y697 | NFS1 | Cysteine desulfurase, | 41273.22656 | 46138.50781 |
| ECE1 | A0A3B3ISF9;P42892 | ECE1 | Endothelin-converting | 27296.55859 | 26582.94141 |
| RRP7A | Q9Y3A4 | RRP7A | Ribosomal RNA-proce | 42730.11328 | 24703.80273 |
| CAP2 | P40123 | CAP2 | Adenylyl cyclase-asso | 642266.6875 | 698594.8125 |
| COMTD1 | Q86VU5 | COMTD1 | Catechol O-methyltra | 41925.19922 | 47986.91797 |
| C15orf38-AP3S2 | A0A0A6YYH1 | C15orf38-AP3S2 | Protein C15orf38-AP3 | 21704.65625 | 27354.62695 |
| HOMER1 | Q86YM7 | HOMER1 | Homer protein homol | 36575.07422 | 35718.34766 |
| FNBP4 | Q8N3X1 | FNBP4 | Formin-binding protei | 38850.76172 | 46021.81641 |
| CNST | Q6PJW8 | CNST | Consortin | 3984.10083 | 6895.207031 |
| C18orf25 | Q96B23 | C18orf25 | Uncharacterized prote | 17282.28711 | 18018.77539 |
| PPM1G | O15355 | PPM1G | Protein phosphatase : | 142374.4375 | 230365.6406 |
| ABCA2 | A0A087WXK5;J3QSS3;Q9BZC | ABCA2 | ATP-binding cassette : | 2066.83374 | 4656.19873 |
| CDKN1A | P38936 | CDKN1A | Cyclin-dependent kina | 40092.33984 | 31232.19727 |
| MTHFD2 | P13995 | MTHFD2 | Bifunctional methylen | 443068.5 | 559722.25 |

| | | | | | |
|---|---|---|---|---|---|
| METTL7B | A0A087WZT2;Q6UX53 | METTL7B | Methyltransferase-like | 29206.56445 | 27222.96875 |
| TXNDC15 | H0Y928;H0Y997;Q96J42 | TXNDC15 | Thioredoxin domain-c | 6438.696777 | 15138.11816 |
| MYORG | Q6NSJ0 | MYORG | Myogenesis-regulating | 13989.51465 | 13087.88477 |
| RCL1 | Q9Y2P8 | RCL1 | RNA 3'-terminal phos | 107924.9375 | 65634.36719 |
| DCDC2 | Q9UHG0 | DCDC2 | Doublecortin domain- | 6398.304199 | 9976.060547 |
| CCDC86 | Q9H6F5 | CCDC86 | Coiled-coil domain-co | 99404.69531 | 68340.47656 |
| PROM2 | Q8N271 | PROM2 | Prominin-2 | 1384.656128 | 2893.961426 |
| PRPF18 | Q99633 | PRPF18 | Pre-mRNA-splicing fac | 27777.34375 | 27180.89453 |
| ISYNA1 | Q9NPH2 | ISYNA1 | Inositol-3-phosphate s | 219924.5938 | 322876.7813 |
| DHX57 | Q6P158 | DHX57 | Putative ATP-depende | 43771.25781 | 44453.78516 |
| ZNF692 | H0YEY1;Q9BU19 | ZNF692 | Zinc finger protein 69 | 1000.328003 | 834.2283936 |
| AUP1 | Q9Y679 | AUP1 | Ancient ubiquitous pr | 84313.91406 | 98670.67969 |
| KDM4B | A0A0C4DFL8;F5GX28 | KDM4B | Lysine-specific demet | 8603.847656 | 13394.63965 |
| HSPA12A | O43301 | HSPA12A | Heat shock 70 kDa pro | 7767.084473 | 8981.899414 |
| SS18L1 | O75177 | SS18L1 | Calcium-responsive tr | 1757.94397 | 4068.16748 |
| AKAP1 | Q92667 | AKAP1 | A-kinase anchor prote | 106019.9375 | 112131.5234 |
| ADCY9 | O60503 | ADCY9 | Adenylate cyclase typ | 5594.510742 | 7341.850586 |
| PLBD2 | Q8NHP8 | PLBD2 | Putative phospholipas | 116408.6719 | 128947.6875 |
| MTFMT | Q96DP5 | MTFMT | Methionyl-tRNA form | 6506.692383 | 7479.459961 |
| FBXO44 | R4GMZ8;R4GNH4 | FBXO44 | F-box only protein 44 | 8685.271484 | 8972.261719 |
| EEF1G | P26641 | EEF1G | Elongation factor 1-ga | 5958146 | 6047349.5 |
| CCND1 | P24385 | CCND1 | G1/S-specific cyclin-D | 29811.37695 | 28993.2207 |
| MTSS2 | A0A087WXN9;Q765P7 | MTSS2 | Protein MTSS 2 | 26587.25 | 24180.66406 |
| RGS10 | O43665 | RGS10 | Regulator of G-protei | 55303.05078 | 64372.92969 |
| MAP4K4 | A0A0D9SEY1;E7EN19;E7ENQ | MAP4K4 | Mitogen-activated pro | 22697.67773 | 25982.66016 |
| DUSP23 | Q9BVJ7 | DUSP23 | Dual specificity protei | 147564.875 | 165813.2656 |
| NOP16 | Q9Y3C1 | NOP16 | Nucleolar protein 16 | 127068.5469 | 88095 |
| CEP41 | Q9BYV8 | CEP41 | Centrosomal protein | 27808.4707 | 32428.0918 |
| ELAC2 | Q9BQ52 | ELAC2 | Zinc phosphodiestera | 242743.7813 | 273823.625 |
| DNAJA1 | P31689 | DNAJA1 | DnaJ homolog subfam | 516896.75 | 524370.1875 |
| BARGIN | Q6ZT62 | BARGIN | Bargin | 73024.14844 | 73764.42188 |
| IFIT5 | Q13325 | IFIT5 | Interferon-induced pr | 34573.41016 | 31543.7207 |
| FDXR | P22570 | FDXR | NADPH:adrenodoxin | 242021.2813 | 278068.4375 |

| | | | | | |
|---|---|---|---|---|---|
| BAIAP2 | I3L4C2 | BAIAP2 | Brain-specific angioge | 355215.2344 | 365098.3906 |
| FRMD8P1 | Q9BZ68 | FRMD8P1 | Putative FERM domai | 6648.145996 | NA |
| JPT1 | J3KT51;Q9UK76 | JPT1 | Jupiter microtubule-a | 318915.4688 | 428281.75 |
| IMP4 | H0Y714;Q96G21 | IMP4 | U3 small nucleolar rib | 98318.57813 | 53435.76172 |
| PNKD | Q8N490 | PNKD | Probable hydrolase PN | 25692.36914 | 28514.89453 |
| SELENOO | A0A087X0Q9;Q9BVL4 | SELENOO | Protein adenylyltransf | 16043.77441 | 18899.85938 |
| SOGA1 | O94964 | SOGA1 | Protein SOGA1 | 34866.86719 | 39524.19141 |
| TAF11 | Q15544 | TAF11 | Transcription initiatio | 5028.562988 | 4051.748779 |
| MRPL42 | J3KPP0;Q9Y6G3;S4R2Z7;S4R3 | MRPL42 | 39S ribosomal protein | 23550.0332 | 31513.18164 |
| FECH | P22830 | FECH | Ferrochelatase, mitoc | 78734.17188 | 79662.59375 |
| KRT72 | Q14CN4 | KRT72 | Keratin, type II cytosk | 4107.166504 | NA |
| GCA | H7BXD5;P28676 | GCA | Grancalcin | 229055.1719 | 262704.9063 |
| PFAS | O15067 | PFAS | Phosphoribosylformyl | 323352.0625 | 344923.0625 |
| TFRC | P02786 | TFRC | Transferrin receptor p | 1018337.563 | 1083919.625 |
| FAM117B | Q6P1L5 | FAM117B | Protein FAM117B | 3080.748047 | 4048.585938 |
| SLC9A3R2 | Q15599 | SLC9A3R2 | Na(+)/H(+) exchange r | 201569.4688 | 184546.5313 |
| ZC3H8 | A0A0C4DGZ1;Q8N5P1 | ZC3H8 | Zinc finger CCCH dom | 38065.61328 | 33569.46484 |
| HTRA2 | O43464 | HTRA2 | Serine protease HTRA | 157307.7031 | 151766.5938 |
| LPAR2 | Q9HBW0 | LPAR2 | Lysophosphatidic acid | 6624.810547 | 6611.195801 |
| DAXX | Q9UER7 | DAXX | Death domain-associa | 52302.87109 | 51594.71484 |
| KHK | C9JDL1;P50053 | KHK | Ketohexokinase (Frag | 18091.49609 | 16788.31836 |
| PPP1R12B | O60237 | PPP1R12B | Protein phosphatase 1 | 4756.22168 | 4552.617676 |
| PPP2R1B | P30154 | PPP2R1B | Serine/threonine-prot | 23985.70898 | 31637.57617 |
| CALD1 | E7EX44;E9PGZ1 | CALD1 | Caldesmon | 51411.85938 | 33179.75 |
| NOL6 | Q9H6R4 | NOL6 | Nucleolar protein 6 | 151088.7344 | 89467.27344 |
| POLR3K | Q9Y2Y1 | POLR3K | DNA-directed RNA po | 56112.80078 | 44592.91797 |
| PPIF | P30405 | PPIF | Peptidyl-prolyl cis-tra | 420196.0625 | 370994.875 |
| OBI1 | Q5W0B1 | OBI1 | ORC ubiquitin ligase 1 | 17452.53125 | 20413.89258 |
| NXT1 | Q9UKK6 | NXT1 | NTF2-related export p | 20635.23047 | 28635.875 |
| NOP53 | Q9NZM5 | NOP53 | Ribosome biogenesis | 127546.625 | 70405.28125 |
| LDHD | Q86WU2 | LDHD | Probable D-lactate de | 21988.86328 | 20469.34961 |
| GNE | Q9Y223 | GNE | Bifunctional UDP-N-ac | 5944.81543 | 4941.45459 |
| ECD | O95905 | ECD | Protein ecdysoneless | 72072.375 | 72524.14063 |

| | | | | | |
|---|---|---|---|---|---|
| SLC7A2 | A0A1W2PR06;P52569 | SLC7A2 | Cationic amino acid tr | 8761.234375 | 14499.55371 |
| PDLIM1 | O00151 | PDLIM1 | PDZ and LIM domain p | 142375.5781 | 187053.5625 |
| ODR4 | Q5SWX8 | ODR4 | Protein odr-4 homolog | 38539.42188 | 41215.37891 |
| ZNF593 | O00488 | ZNF593 | Zinc finger protein 593 | 89180.9375 | 63191.67969 |
| MON1A | A0A0A0MSQ9;Q86VX9 | MON1A | Vacuolar fusion protei | 1073.192749 | 1830.377075 |
| ISCU | Q9H1K1 | ISCU | Iron-sulfur cluster ass | 79307.89844 | 99180.72656 |
| SPAG7 | O75391 | SPAG7 | Sperm-associated ant | 246161.2031 | 228257.3125 |
| DDX10 | Q13206 | DDX10 | Probable ATP-depend | 49393.77734 | 15220.82715 |
| ATP2B4 | P23634 | ATP2B4 | Plasma membrane cal | 24686.48633 | 23766.29688 |
| PYROXD1 | B4DEW4;Q8WU10 | PYROXD1 | Pyridine nucleotide-di | 11608.30176 | 10656.52637 |
| TMX3 | Q96JJ7 | TMX3 | Protein disulfide-isom | 84754.52344 | 88421.25 |
| SLCO4C1 | Q6ZQN7 | SLCO4C1 | Solute carrier organic | 10190.48926 | 10220.56543 |
| ;PSMB10;PSMB10 | J3QL48;J3QQN1;P40306 | ;PSMB10;PSMB10 | Uncharacterized prote | 24980.93359 | 24967.875 |
| SPRYD7 | Q5W111 | SPRYD7 | SPRY domain-containi | 70067.86719 | 74245.4375 |
| RBFA | Q8N0V3 | RBFA | Putative ribosome-bir | 17238.79102 | 27383.66211 |
| PNO1 | Q9NRX1 | PNO1 | RNA-binding protein P | 79673.69531 | 67128.48438 |
| C9orf40 | Q8IXQ3 | C9orf40 | Uncharacterized prote | 10537.78809 | 6814.066406 |
| MAGED4 | Q96JG8 | MAGED4 | Melanoma-associated | 10284.1123 | 10640.02148 |
| UTP18 | Q9Y5J1 | UTP18 | U3 small nucleolar RN | 95963.63281 | 60185.79688 |
| RRP36 | Q96EU6 | RRP36 | Ribosomal RNA proce | 55991.67969 | 37580.375 |
| EPS8L1 | Q8TE68 | EPS8L1 | Epidermal growth fact | 76057.34375 | 73530.41406 |
| TBCA | E5RJD8;O75347 | TBCA | Tubulin-specific chape | 2039656.5 | 1961003.75 |
| SORL1 | Q92673 | SORL1 | Sortilin-related recept | 13862.25781 | 12826.52637 |
| CACUL1 | Q86Y37 | CACUL1 | CDK2-associated and c | 11965.9209 | 19925.64844 |
| EEF1AKMT4 | P0DPD7 | EEF1AKMT4 | EEF1A lysine methyltr | 65979.24219 | 68346.48438 |
| MAVS | Q7Z434 | MAVS | Mitochondrial antivira | 258954.2031 | 313879.1563 |
| RERG | F5H252;Q96A58 | RERG | Ras-related and estrog | 10679.53223 | 11768.18945 |
| CTBS | Q01459 | CTBS | Di-N-acetylchitobiase | 13458.2959 | 10088.03711 |
| SNX18 | Q96RF0 | SNX18 | Sorting nexin-18 | 8941.916016 | 10515.96387 |
| BCL2L11 | O43521 | BCL2L11 | Bcl-2-like protein 11 | 30721.41602 | 34217.69531 |
| CHCHD2 | Q9Y6H1 | CHCHD2 | Coiled-coil-helix-coile | 151077.5938 | 205946.0469 |
| ERAP1 | Q9NZ08 | ERAP1 | Endoplasmic reticulum | 47392.44922 | 50324.00391 |
| KBTBD6 | Q86V97 | KBTBD6 | Kelch repeat and BTB | 34974.66016 | 34922.05469 |

| | | | | | |
|---|---|---|---|---|---|
| RBBP8 | I6L8A6;Q99708 | RBBP8 | DNA endonuclease RE | 17287.64844 | 32295.64648 |
| DDX56 | Q9NY93 | DDX56 | Probable ATP-depend | 233218.875 | 128475.6719 |
| NUMBL | A0A0C4DGH3;Q9Y6R0 | NUMBL | Numb-like protein | 33638.37891 | 30591.83398 |
| ARMCX1 | Q9P291 | ARMCX1 | Armadillo repeat-cont | 10346.8291 | 12142.86035 |
| ALG11 | Q2TAA5 | ALG11 | GDP-Man:Man(3)GlcN | 18573.22656 | 17751.0293 |
| IFRD2 | A0A0R4J2F6;Q12894 | IFRD2 | Interferon-related dev | 30619.0957 | 33334.74609 |
| SDHB | P21912 | SDHB | Succinate dehydroger | 183955.2188 | 185753.5 |
| MDM4 | A0A087WZ58;O15151;Q5T0Y | MDM4 | Protein Mdm4;Protei | 4129.550293 | 5911.237793 |
| SH3BP1 | Q9Y3L3 | SH3BP1 | SH3 domain-binding p | 38448.33984 | 35290.23828 |
| KIAA0100 | Q08E86;Q14667 | KIAA0100 | KIAA0100 protein;Pro | 22109.25781 | 25788.6582 |
| CUTC | Q9NTM9 | CUTC | Copper homeostasis p | 76405.57813 | 67896.05469 |
| ZCCHC7 | Q8N3Z6 | ZCCHC7 | Zinc finger CCHC dom | 670.4041748 | 918.3079834 |
| PTPRS | Q13332 | PTPRS | Receptor-type tyrosin | 2563.329834 | 3410.72583 |
| WDR41 | Q9HAD4 | WDR41 | WD repeat-containing | 21715.55273 | 26385.61914 |
| HMMR | O75330 | HMMR | Hyaluronan mediated | 34759.87109 | 29482.57617 |
| RPUSD1 | H3BQN8;Q9UJJ7 | RPUSD1 | RNA pseudouridylate | 6320.600098 | 6269.423828 |
| TAX1BP3 | O14907 | TAX1BP3 | Tax1-binding protein 3 | 341281.3438 | 311226.625 |
| OCLN | A0A0G2JMZ8;Q16625 | OCLN | Occludin | 220501.5469 | 213855.7969 |
| BMS1 | Q14692 | BMS1 | Ribosome biogenesis | 137737 | 88464.71094 |
| TCAF1 | Q9Y4C2 | TCAF1 | TRPM8 channel-assoc | 141046.7969 | 161843.8594 |
| UBE2T | Q9NPD8 | UBE2T | Ubiquitin-conjugating | 420910.5 | 366423.6875 |
| TRMT10A | Q8TBZ6 | TRMT10A | tRNA methyltransfera | 7412.024902 | 8607.769531 |
| SLC25A30 | Q5SVS4 | SLC25A30 | Kidney mitochondrial | 15880.97656 | 18854.23633 |
| RAD23B | P54727 | RAD23B | UV excision repair pro | 60012.80859 | 80222.73438 |
| SAMHD1 | Q9Y3Z3 | SAMHD1 | Deoxynucleoside triph | 137059.3281 | 120630.0859 |
| ARF3 | P61204 | ARF3 | ADP-ribosylation facto | 7889.118652 | 11663.4043 |
| GLS | B8ZZC5 | GLS | Glutaminase kidney is | 129037.1641 | 157175.1836 |
| SNX30 | Q5VWJ9 | SNX30 | Sorting nexin-30 | 27024.16016 | 27727.06836 |
| UBQLN4 | Q9NRR5 | UBQLN4 | Ubiquilin-4 | 171215.0313 | 157617.9063 |
| CDK2AP1 | O14519 | CDK2AP1 | Cyclin-dependent kina | 51927.78906 | 51374.81641 |
| OTUD5 | A0A087WXV6;H7BZQ3;Q96G | OTUD5 | OTU domain-containi | 24504.86523 | 23486.23828 |
| GALNS | P34059 | GALNS | N-acetylgalactosamin | 46532.83203 | 46526.4375 |
| MNX1 | P50219 | MNX1 | Motor neuron and pa | 10643.84668 | 10484.83105 |

| | | | | | |
|---|---|---|---|---|---|
| NAB2 | Q15742 | NAB2 | NGFI-A-binding protei | 98831.42188 | 94720.10156 |
| ERAL1 | O75616 | ERAL1 | GTPase Era, mitochon | 88643.61719 | 102449.4375 |
| CTSD | A0A1B0GVD5;A0A1B0GW44; | CTSD | Cathepsin D | 1537987.625 | 2036656.75 |
| PHRF1 | A0A0J9YWD5;A0A0J9YYB5;B | PHRF1 | PHD and RING finger ( | 15411.94336 | 19652.84375 |
| TTC38 | Q5R3I4 | TTC38 | Tetratricopeptide rep | 15315.20996 | 20373.01172 |
| COX19 | Q49B96 | COX19 | Cytochrome c oxidase | 104930.5625 | 95336.82813 |
| ABCA12 | Q86UK0 | ABCA12 | ATP-binding cassette s | 8811.771484 | 4931.601563 |
| GALT | P07902 | GALT | Galactose-1-phosphat | 9445.569336 | 11404.33691 |
| HOMER2 | Q9NSB8 | HOMER2 | Homer protein homol | 11478.12402 | 12122.64648 |
| PRICKLE3 | H0Y413;O43900 | PRICKLE3 | Prickle planar cell pola | 9036.267578 | 10124.46777 |
| PTRH1 | Q86Y79 | PTRH1 | Probable peptidyl-tRN | 35573.08984 | 35797.72266 |
| MYOF | Q9NZM1 | MYOF | Myoferlin | 246086.0625 | 358803.375 |
| BNIP3 | A0A0J9YW18;F6RP06;Q1298 | BNIP3 | BCL2/adenovirus E1B | 4092.756104 | 4891.433105 |
| ARRB2 | P32121 | ARRB2 | Beta-arrestin-2 | 28470.92578 | 34324.11328 |
| MRM3 | Q9HC36 | MRM3 | rRNA methyltransfera | 338303.7188 | 385985 |
| TMEM120A | A0A024R4K9;A0A087WZA9;A | TMEM120A | Transmembrane prote | 22504.57031 | 26008.57813 |
| RBM38 | F6VZ39;Q9H0Z9 | RBM38 | RNA-binding protein 3 | 101277.3359 | 90885.02344 |
| ZBTB8OS | A0A087X0V4;A8K0B5;D3DPC | ZBTB8OS | Protein archease (Frag | 30606.76367 | 42540.30469 |
| UTP14A | Q9BVJ6 | UTP14A | U3 small nucleolar RN | 140065.2188 | 91986.33594 |
| PNPT1 | Q8TCS8 | PNPT1 | Polyribonucleotide nu | 328044.2188 | 326048.6563 |
| TCOF1 | A0A3B3IS06 | TCOF1 | Treacle protein (Fragn | 254927.0254 | 272810.6367 |
| SUPV3L1 | Q8IYB8 | SUPV3L1 | ATP-dependent RNA h | 162068.2344 | 151991.5469 |
| NDUFAF1 | H0YL22;Q9Y375 | NDUFAF1 | Complex I intermediat | 24343.16016 | 25202.19531 |
| DPCD | Q9BVM2 | DPCD | Protein DPCD | 33997.74609 | 33182.19922 |
| NUFIP2 | Q7Z417 | NUFIP2 | Nuclear fragile X men | 98824.17969 | 111708.4531 |
| RRP1 | P56182 | RRP1 | Ribosomal RNA proce | 259317.4219 | 196423.8438 |
| KPNA2 | P52292 | KPNA2 | Importin subunit alph | 1553155.875 | 1590705.875 |
| RNASET2 | A0A087WZM2;D6RHI9;O005 | RNASET2 | Ribonuclease T2;Ribo | 17189.26367 | 20993.0918 |
| EMG1 | Q92979 | EMG1 | Ribosomal RNA small | 334876.3125 | 175115.4375 |
| HPCAL1 | P37235 | HPCAL1 | Hippocalcin-like prote | 134269.5625 | 165551.5625 |
| HBB | P68871 | HBB | Hemoglobin subunit b | 101113.2031 | 126701.5703 |
| CYTH1 | Q15438 | CYTH1 | Cytohesin-1 | 44948.98047 | 50443.54688 |
| BSG | P35613 | BSG | Basigin | 481325.2813 | 525126.8125 |

| | | | | | |
|---|---|---|---|---|---|
| TENT4B | H3BQM0;Q8NDF8 | TENT4B | Terminal nucleotidyltr | 16917.94336 | 16898.79297 |
| CORO1C | Q9ULV4 | CORO1C | Coronin-1C | 329293.4375 | 234217.7188 |
| CD59;;CD59 | A0A2U3TZL5;E9PNW4;P1398 | CD59;;CD59 | CD59 glycoprotein (Fr | 6559.234863 | 9620.099609 |
| NHS | A0A087WU78;A0A087WWC4 | NHS | Nance-Horan syndrom | 3627.30542 | 3737.902832 |
| ABCB8 | Q9NUT2 | ABCB8 | Mitochondrial potassi | 11207.32422 | 13192.1748 |
| CLUH | A0A494C0R8 | CLUH | Clustered mitochondr | 414562.7168 | 410732.8281 |
| TGFB1 | A0A499FJK2;P01137 | TGFB1 | Transforming growth | 7471.258789 | 8080.591309 |
| CFAP45 | Q9UL16 | CFAP45 | Cilia- and flagella-assc | 67564.77344 | 59516.44141 |
| ARL6IP6 | Q8N6S5 | ARL6IP6 | ADP-ribosylation facto | 24772.62695 | 28746.07617 |
| ADAM15 | Q13444 | ADAM15 | Disintegrin and metall | 54701.24219 | 64014.24609 |
| APRT | P07741 | APRT | Adenine phosphoribo | 2198463.5 | 2512541.5 |
| KCTD12 | Q96CX2 | KCTD12 | BTB/POZ domain-cont | 74712.02344 | 62964.9375 |
| UBAP2 | Q5T6F2 | UBAP2 | Ubiquitin-associated p | 73949.82813 | 84524.54688 |
| UBAP2L | Q14157 | UBAP2L | Ubiquitin-associated p | 214831.6289 | 324882.4658 |
| MYBBP1A | Q9BQG0 | MYBBP1A | Myb-binding protein 1 | 293821.7813 | 293362.6563 |
| MBD2 | Q9UBB5 | MBD2 | Methyl-CpG-binding d | 68253.80469 | 84813.24219 |
| TAF13 | Q15543 | TAF13 | Transcription initiation | 4.243143082 | NA |
| CEP350 | Q5VT06 | CEP350 | Centrosome-associate | 3722.411621 | 3408.215088 |
| RIOK1 | Q9BRS2 | RIOK1 | Serine/threonine-prot | 32273.53516 | 32221.13867 |
| PRRC2B | H0Y412 | PRRC2B | Protein PRRC2B (Fragi | 104206.9219 | 113417.7627 |
| PDCD11 | A0A3B3IUD7;Q14690 | PDCD11 | Programmed cell deat | 103996.1719 | 45295.14453 |
| MRS2 | Q9HD23 | MRS2 | Magnesium transport | 29187.66602 | 28184.35156 |
| DNTTIP2 | Q5QJE6 | DNTTIP2 | Deoxynucleotidyltrans | 71431.10156 | 19594.70313 |
| CD47 | H0YC40 | CD47 | Leukocyte surface ant | 344530.4766 | 392223.3047 |
| UQCRB | P14927 | UQCRB | Cytochrome b-c1 com | 438905.7813 | 442258.1875 |
| FHL2 | Q14192 | FHL2 | Four and a half LIM dc | 10483.68262 | 5157.865723 |
| DDX47 | Q9H0S4 | DDX47 | Probable ATP-depend | 148343.8906 | 119498.8906 |
| MRNIP | Q6NTE8 | MRNIP | MRN complex-interac | 22720.21094 | 21010.29492 |
| PARP12 | Q9H0J9 | PARP12 | Protein mono-ADP-rib | 21675.42578 | 20031.69336 |
| UTP15 | Q8TED0 | UTP15 | U3 small nucleolar RN | 48447.76172 | 25444.07227 |
| P4HTM | Q9NXG6 | P4HTM | Transmembrane proly | 37020.59375 | 32014.94141 |
| ENDOD1 | O94919 | ENDOD1 | Endonuclease domain | 32915.66797 | 37340.34766 |
| IFIT3 | O14879 | IFIT3 | Interferon-induced pr | 8846.05957 | 8033.356445 |

| | | | | | |
|---|---|---|---|---|---|
| MTRR | Q9UBK8 | MTRR | Methionine synthase | 19619.13086 | 19456.55859 |
| COPG2 | Q9UBF2 | COPG2 | Coatomer subunit gar | 79918.04688 | 92886.34375 |
| MVP | Q14764 | MVP | Major vault protein | 7108.719727 | 10580.72168 |
| HEATR1 | Q9H583 | HEATR1 | HEAT repeat-containi | 109091.2031 | 64266.22266 |
| FOSL2 | C9JCN8;P15408 | FOSL2 | Fos-related antigen 2 | 14107.7832 | 12487.81152 |
| SUGP2 | M0R2Z9;Q8IX01 | SUGP2 | SURP and G-patch dor | 22558.65039 | 26790.11719 |
| SMAD4 | K7EIU8;Q13485 | SMAD4 | Mothers against deca | 16109.88574 | 18580.53516 |
| CRELD2 | Q6UXH1 | CRELD2 | Protein disulfide isom | 82744.89063 | 87192.53906 |
| NOL10 | Q9BSC4 | NOL10 | Nucleolar protein 10 | 139094.4219 | 56064.5625 |
| UTP4 | Q969X6 | UTP4 | U3 small nucleolar RN | 50535.51172 | 25174.0332 |
| RASAL2 | Q9UJF2 | RASAL2 | Ras GTPase-activating | 4641.624512 | 4912.570801 |
| TXLNG | Q9NUQ3 | TXLNG | Gamma-taxilin | 123419.8594 | 147078.4844 |
| CPD | O75976 | CPD | Carboxypeptidase D | 39723.51563 | 43275.01953 |
| CTSC | P53634 | CTSC | Dipeptidyl peptidase 1 | 36820.10938 | 42173.21875 |
| STOML2 | Q9UJZ1 | STOML2 | Stomatin-like protein | 423692.1563 | 436085.0938 |
| RIOK2 | Q9BVS4 | RIOK2 | Serine/threonine-prot | 23587.37695 | 24865.9668 |
| IKBIP | Q70UQ0 | IKBIP | Inhibitor of nuclear fa | 21420.47266 | 17005.68555 |
| CAPN2 | P17655 | CAPN2 | Calpain-2 catalytic sub | 68790.59375 | 54719.20703 |
| CUEDC1 | Q9NWM3 | CUEDC1 | CUE domain-containir | 29304.77734 | 32859.41016 |
| TPT1 | A0A0B4J2C3;P13693;Q5W0H | TPT1 | Translationally-contro | 610169.0625 | 569361.0625 |
| CDC123 | O75794 | CDC123 | Cell division cycle prot | 71846.83594 | 85425.17188 |
| DCTD | P32321 | DCTD | Deoxycytidylate deam | 68084.45313 | 56735.38281 |
| PDRG1 | Q9NUG6 | PDRG1 | p53 and DNA damage | 46531.70703 | 50020.67188 |
| HEXD | J3KT84;J3QKL0;K7EIV9;Q8W\ | HEXD | Hexosaminidase D (Fr | 7345.505859 | 9502.18457 |
| ITPK1 | Q13572 | ITPK1 | Inositol-tetrakisphosp | 325986 | 486164.4063 |
| PIMREG | I3L156;I3L2R2;I3L419;Q9BSJ6 | PIMREG | Protein PIMREG;Prote | 7206.318848 | 5679.626465 |
| S100A16 | Q96FQ6 | S100A16 | Protein S100-A16 | 473288.6875 | 405080.1563 |
| TSPAN4 | A8MVV6;E9PN41;J3KQ42;O1 | TSPAN4 | Tetraspanin;Tetraspar | 12227.67578 | 19019.80273 |
| MZT2B | B8ZZ87;Q6NZ67 | MZT2B | Mitotic-spindle organi | 350089 | 258647.2031 |
| NAGPA | K7EL75;K7EPH3;K7ERI4;Q9UI | NAGPA | N-acetylglucosamine- | 8862.489258 | 6846.097168 |
| ESF1 | Q9H501 | ESF1 | ESF1 homolog | 2330517 | 1056414.75 |
| RTKN | Q9BST9 | RTKN | Rhotekin | 97512.89844 | 111859.2656 |
| QSOX2 | Q6ZRP7 | QSOX2 | Sulfhydryl oxidase 2 | 90945.42188 | 92225.0625 |

| | | | | | |
|---|---|---|---|---|---|
| KRT5 | P13647 | KRT5 | Keratin, type II cytosk | 15849.14648 | 17437.01758 |
| STAT1 | P42224 | STAT1 | Signal transducer and | 360985.6563 | 362938.4688 |
| PACC1 | Q9H813 | PACC1 | Proton-activated chlo | 9347.788086 | 8739.149414 |
| CEBPZ | Q03701 | CEBPZ | CCAAT/enhancer-binc | 211029.4375 | 127503.3906 |
| CD320 | Q9NPF0 | CD320 | CD320 antigen | 2881.704346 | 6662.749512 |
| PTRHD1 | Q6GMV3 | PTRHD1 | Putative peptidyl-tRN | 43193.57422 | 47919.16797 |
| SIGMAR1 | Q99720 | SIGMAR1 | Sigma non-opioid intr | 37641.90234 | 40033.42188 |
| PFKFB2 | O60825 | PFKFB2 | 6-phosphofructo-2-kir | 106386.0469 | 99437.84375 |
| CT45A10;CT45A8;CT45A9;( | P0DMU9;P0DMV1;P0DMV2;( | CT45A10;CT45A8;CT4 | Cancer/testis antigen | 2717.010254 | 1477.370117 |
| DNAJC15 | Q9Y5T4 | DNAJC15 | DnaJ homolog subfam | 47880.40234 | 51228.64063 |
| DDX49 | Q9Y6V7 | DDX49 | Probable ATP-depend | 61176.70313 | 51554.07422 |
| ZNF703 | Q9H7S9 | ZNF703 | Zinc finger protein 70 | 16664.20313 | 24252.53711 |
| SP110 | G5E9C0;Q9HB58 | SP110 | SP110 nuclear body p | 9384.703125 | 7763.680664 |
| PPAN-P2RY11;PPAN-P2RY1 | A0A0A6YYI3;A0A0B4J1V8;Q9 | PPAN-P2RY11;PPAN-F | Protein PPAN-P2RY11 | 108376.4844 | 66060.50781 |
| DDX21 | Q9NR30 | DDX21 | Nucleolar RNA helicas | 662110.125 | 410334.75 |
| CBWD6 | Q4V339 | CBWD6 | COBW domain-contai | 10400.02344 | 10247.0918 |
| GPBP1 | D4PHA4;Q86WP2 | GPBP1 | GC-rich promoter binc | 6360.141113 | 6913.040527 |
| KRT27 | Q7Z3Y8 | KRT27 | Keratin, type I cytoske | 10776.6377 | 5001.292969 |
| SYAP1 | Q96A49 | SYAP1 | Synapse-associated pr | 29485.44922 | 53125.91016 |
| SCARB1 | B7ZKQ9 | SCARB1 | SCARB1 protein | 155404.875 | 158198.5313 |
| IGF1R | C9J5X1;P08069 | IGF1R | Tyrosine-protein kinas | 32111.19336 | 39989.56641 |
| FA2H | J3QS89;Q7L5A8 | FA2H | Fatty acid 2-hydroxyla | 5929.70166 | 7363.170898 |
| TDRD1 | H9KV62;H9KV63;Q9BXT4 | TDRD1 | Tudor domain-contair | 4043.004395 | 3213.209229 |
| MT2A | P02795 | MT2A | Metallothionein-2 | 223704.6719 | 215416.1094 |
| MINK1 | Q8N4C8 | MINK1 | Misshapen-like kinase | 2587.114258 | 2867.38501 |
| KRT14 | P02533 | KRT14 | Keratin, type I cytoske | 32296.83398 | 58145.10156 |
| ZNF106 | H3BSS6;Q9H2Y7 | ZNF106 | Zinc finger protein 10 | 412781.2813 | 322533.5625 |
| MPHOSPH10 | O00566 | MPHOSPH10 | U3 small nucleolar rib | 65452.26953 | 33271.96484 |
| YBX2 | Q9Y2T7 | YBX2 | Y-box-binding protein | 47812.32031 | 44579.28125 |
| NR2C2AP | Q86WQ0 | NR2C2AP | Nuclear receptor 2C2- | 135340.2656 | 139692.1406 |
| DUS3L | Q96G46 | DUS3L | tRNA-dihydrouridine( | 39389.07422 | 34204.61719 |
| TFB2M | Q9H5Q4 | TFB2M | Dimethyladenosine tr | 185119.5938 | 196497.5625 |
| WDR46 | O15213 | WDR46 | WD repeat-containing | 54945.05859 | 22782.16602 |

| | | | | | |
|---|---|---|---|---|---|
| GATAD2A | Q86YP4 | GATAD2A | Transcriptional repres | 77433.3125 | 83437.89844 |
| FASTKD2 | Q9NYY8 | FASTKD2 | FAST kinase domain-c | 38203.47266 | 49791.30859 |
| NARF | E7EP87;Q9UHQ1 | NARF | Nuclear prelamin A re | 17404.83398 | 19883.20898 |
| GULP1 | Q9UBP9 | GULP1 | PTB domain-containin | 31128.60742 | 30921.05078 |
| PPID | Q08752 | PPID | Peptidyl-prolyl cis-trai | 523067.1875 | 536358.5 |
| ZFC3H1 | O60293 | ZFC3H1 | Zinc finger C3H1 doma | 6603.657227 | 6879.798828 |
| KRT10 | P13645 | KRT10 | Keratin, type I cytoske | 341504.7813 | 290366.0313 |
| CRACR2A | Q9BSW2 | CRACR2A | EF-hand calcium-bindi | 3543.995117 | 2369.254639 |
| KCTD3 | Q9Y597 | KCTD3 | BTB/POZ domain-cont | 110887.4375 | 113021.2344 |
| ANXA9 | O76027 | ANXA9 | Annexin A9 | 57066.10938 | 60248.66016 |
| CTIF | O43310 | CTIF | CBP80/20-dependent | 1561.553467 | 2412.718018 |
| KNSTRN | Q9Y448;V9GY01 | KNSTRN | Small kinetochore-ass | 72638.00781 | 95848.99219 |
| ECSIT | Q9BQ95 | ECSIT | Evolutionarily conserv | 34793.17578 | 34984.20313 |
| TRIM21 | P19474 | TRIM21 | E3 ubiquitin-protein li | 9454.044922 | 10282.08789 |
| CEMIP2 | F5H6B2;Q9UHN6 | CEMIP2 | Cell surface hyaluroni | 13309.29688 | 11249.49707 |
| YARS2 | Q9Y2Z4 | YARS2 | Tyrosine--tRNA ligase, | 97967.36719 | 100820.0156 |
| SLC5A6 | Q9Y289 | SLC5A6 | Sodium-dependent m | 88069.41406 | 101383.5391 |
| ABHD11 | Q8NFV4 | ABHD11 | Protein ABHD11 | 391263.3125 | 428986.25 |
| NOL11 | Q9H8H0 | NOL11 | Nucleolar protein 11 | 95860.32813 | 71317.875 |
| TUBB3 | Q13509 | TUBB3 | Tubulin beta-3 chain | 7292719 | 7493061 |
| USP28 | B4E3L3;G3V1N5 | USP28 | Ubiquitin carboxyl-ter | 28409.4126 | 30283.06006 |
| SLC2A1 | P11166 | SLC2A1 | Solute carrier family 2 | 335244.875 | 325869.5 |
| IRX2 | Q9BZI1 | IRX2 | Iroquois-class homeo | 26385.52148 | 24554.68164 |
| SULT2B1 | O00204 | SULT2B1 | Sulfotransferase 2B1 | 542201.5 | 615888.5625 |
| PGM5 | Q15124 | PGM5 | Phosphoglucomutase- | 9771.055664 | 5175.584473 |
| NDUFAF2 | Q8N183 | NDUFAF2 | NADH dehydrogenase | 358610.5 | 385937.4688 |
| NIP7 | Q9Y221 | NIP7 | 60S ribosome subunit | 85011.33594 | 55855.40234 |
| IDS | B3KWA1;P22304 | IDS | Iduronate 2-sulfatase | 4507.878906 | 4148.643066 |
| NMI | Q13287 | NMI | N-myc-interactor | 14384.07324 | 13298.16016 |
| MCRIP2 | Q9BUT9 | MCRIP2 | MAPK regulated corep | 40950.52734 | 39420.75 |
| ITGB1 | P05556 | ITGB1 | Integrin beta-1 | 680922.75 | 668267.9375 |
| WWTR1 | C9J038;C9JR84;Q9GZV5 | WWTR1 | WW domain-containi | 5664.133789 | 2901.218262 |
| WDR75 | Q8IWA0 | WDR75 | WD repeat-containing | 45772.24609 | 24524.13281 |

| | | | | | |
|---|---|---|---|---|---|
| OSBPL6 | Q9BZF3 | OSBPL6 | Oxysterol-binding pro | 7121.978027 | 7484.241699 |
| CD97 | P48960 | CD97 | CD97 antigen | 7641.880859 | 5699.750488 |
| ING2 | Q9H160 | ING2 | Inhibitor of growth pr | 43126.65234 | 39177.39063 |
| ATF7IP | Q6VMQ6 | ATF7IP | Activating transcriptic | 54053.05859 | 56775.69141 |
| ME2 | A0A1W2PPH1;A0A1W2PQH3 | ME2 | Malic enzyme;Malic e | 99761.46094 | 108058.2344 |
| TNKS1BP1 | Q9C0C2 | TNKS1BP1 | 182 kDa tankyrase-1-k | 129639.1641 | 165913.6563 |
| SNX24 | A8MXB7;Q9Y343 | SNX24 | Sorting nexin-24 | 49841.74609 | 60968.34766 |
| TXNDC17 | Q9BRA2 | TXNDC17 | Thioredoxin domain-c | 320425.7813 | 424655.75 |
| YBX3 | P16989 | YBX3 | Y-box-binding protein | 313352.5625 | 286038.7813 |
| PSRC1 | Q6PGN9 | PSRC1 | Proline/serine-rich coi | 9574.731445 | 12572.02539 |
| CUX1 | A0A2R8Y852 | CUX1 | Homeobox protein cu | 211336.7988 | 211323.188 |
| AATF | Q9NY61 | AATF | Protein AATF | 72511.96094 | 32437.47852 |
| CLDN7 | F5H496;I3L3X1;K7EP40;O954 | CLDN7 | Claudin;Claudin (Fragr | 38160.65625 | 53380.23828 |
| ZFAND5 | O76080 | ZFAND5 | AN1-type zinc finger p | 12078.33887 | 11527.11914 |
| HEXB | P07686 | HEXB | Beta-hexosaminidase | 313949.9063 | 340626.6875 |
| TCIRG1 | Q13488 | TCIRG1 | V-type proton ATPase | 35381.11328 | 26381.44336 |
| HYAL2 | Q12891 | HYAL2 | Hyaluronidase-2 | 12761.86133 | 10498.47168 |
| URI1 | O94763 | URI1 | Unconventional prefo | 38921.57813 | 49172.79688 |
| HBA1 | P69905 | HBA1 | Hemoglobin subunit a | 72008.71094 | 34450.72266 |
| RNASEH1 | O60930 | RNASEH1 | Ribonuclease H1 | 18230.97852 | 19673.49609 |
| CBFA2T3 | O75081 | CBFA2T3 | Protein CBFA2T3 | 4858.210449 | 5248.530762 |
| RBP4 | P02753;Q5VY30 | RBP4 | Retinol-binding protei | 15930.89648 | 11211.70996 |
| CASP8 | C9JB29 | CASP8 | Caspase-8 (Fragment) | 81455.90918 | 80644.45313 |
| DIEXF | Q68CQ4 | DIEXF | Digestive organ expar | 133263.1719 | 117248.8125 |
| RHBDF2 | Q6PJF5 | RHBDF2 | Inactive rhomboid prc | 4341.618652 | 4545.775391 |
| LTV1 | Q96GA3 | LTV1 | Protein LTV1 homolog | 51956.19922 | 56984.88281 |
| MPDU1 | I3L295;J3KT75;J3QRD5;J3QS | MPDU1 | Mannose-P-dolichol u | 220246.9375 | 262447.1563 |
| DES | P17661 | DES | Desmin | 87405.11719 | 113711.9609 |
| FAS | P25445 | FAS | Tumor necrosis factor | 4516.015625 | 2852.169678 |
| VAMP8 | Q9BV40 | VAMP8 | Vesicle-associated me | 131141.875 | 172379.8438 |
| CARNMT1 | Q8N4J0 | CARNMT1 | Carnosine N-methyltra | 15747.85156 | 15212.03613 |
| MEX3A | A1L020 | MEX3A | RNA-binding protein M | 13267.2373 | 15015.93457 |
| ASDURF | L0R819 | ASDURF | ASNSD1 upstream ope | 33987.74219 | 33064.17969 |

| | | | | | |
|---|---|---|---|---|---|
| SLC19A2 | A0A2R8Y5B5;O60779 | SLC19A2 | Solute carrier family 1 | 11112.32813 | 14274.46582 |
| CALB2 | H3BN14;P22676 | CALB2 | Calretinin (Fragment); | 5293.747559 | 15123.81348 |
| LONRF2 | Q1L5Z9 | LONRF2 | LON peptidase N-term | 24611.75781 | 29356.82422 |
| C1QBP | Q07021 | C1QBP | Complement compon | 39992.8125 | 94159.69531 |
| GGH | Q92820 | GGH | Gamma-glutamyl hydr | 80788.10156 | 90142.21094 |
| TAPBP | A0A0A0MSV9;A0A0G2JH37;A | TAPBP | Tapasin | 61218.82031 | 65446.60547 |
| KRT4 | P19013 | KRT4 | Keratin, type II cytosk | 19673.01758 | NA |
| N4BP3 | O15049 | N4BP3 | NEDD4-binding protei | 80287.84375 | 89597.91406 |
| ANKRD39 | Q53RE8 | ANKRD39 | Ankyrin repeat domai | 10672.86914 | 16153.64648 |
| ST7 | B7Z4L1;B7Z4U3;C9JU30;E7EN | ST7 | Suppression of tumori | 381.8377075 | 482.1340942 |
| VASN | Q6EMK4 | VASN | Vasorin | 21925.08984 | 20124.33008 |
| OGDHL | Q9ULD0 | OGDHL | 2-oxoglutarate dehydr | 13867.42871 | 10349.70215 |
| ARSB | A0A2U3U034;P15848 | ARSB | Arylsulfatase B | 9225.138672 | 10796.3252 |
| NCOR2 | C9J0Q5;C9JE98;Q9Y618 | NCOR2 | Nuclear receptor core | 27339.69922 | 31023.17383 |
| ZNF330 | Q9Y3S2 | ZNF330 | Zinc finger protein 33( | 14640.59961 | 14478.36914 |
| LAMB1 | G3XAI2;P07942 | LAMB1 | Laminin subunit beta- | 332708.125 | 337314.1563 |
| ABLIM1 | A0A0A0MRL6;A0A3B3IS55;F8 | ABLIM1 | Actin-binding LIM pro | 8324.813477 | 9454.30957 |
| ANGEL1 | Q9UNK9 | ANGEL1 | Protein angel homolo | 18991.17773 | 14803.51465 |
| FABP5 | I6L8B7 | FABP5 | Fatty acid-binding pro | 1489749.008 | 1400855.344 |
| HRNR | Q86YZ3 | HRNR | Hornerin | 1354.181396 | 5740.552246 |
| SPAG5 | Q96R06 | SPAG5 | Sperm-associated ant | 13619.82227 | 28646.89258 |
| UBE2L6 | O14933 | UBE2L6 | Ubiquitin/ISG15-conju | 47557.57422 | 46037.24219 |
| TEAD4 | A0A0A0MRF3;Q15561;Q53G | TEAD4 | Transcriptional enhan | 2929.51416 | 3857.678955 |
| NOM1 | Q5C9Z4 | NOM1 | Nucleolar MIF4G dom | 35283.83984 | 36466.94531 |
| KRT78 | Q8N1N4 | KRT78 | Keratin, type II cytosk | 36004.76172 | 46856.44922 |
| RAI1 | A0A1W2PQJ7;Q7Z5J4 | RAI1 | Retinoic acid-induced | 16038.46387 | 14920.77246 |
| TGOLN2 | A0A5F9UY30;A0A5H1ZRP2;O | TGOLN2 | Trans-Golgi network i | 84549.46094 | 79772.32813 |
| TMA16 | H0Y9X1;Q96EY4 | TMA16 | Translation machinery | 106963.3281 | 111728.6328 |
| SLC9A3R1 | O14745 | SLC9A3R1 | Na(+)/H(+) exchange r | 2945708.5 | 4319983.5 |
| NOP2 | P46087 | NOP2 | Probable 28S rRNA (cy | 215881.2813 | 121026.7188 |
| TFAP2C | Q92754 | TFAP2C | Transcription factor A | 37303.24219 | 43136.8125 |
| WDR43 | Q15061 | WDR43 | WD repeat-containing | 91040.21875 | 49656.01172 |
| CAPG | P40121 | CAPG | Macrophage-capping | 65788.96875 | 38594.59375 |

| | | | | | |
|---|---|---|---|---|---|
| RRP15 | Q9Y3B9 | RRP15 | RRP15-like protein | 344851.75 | 226703.9688 |
| SPTBN2 | O15020 | SPTBN2 | Spectrin beta chain, n | 152153.5156 | 87907.57813 |
| STARD13 | Q9Y3M8 | STARD13 | StAR-related lipid trar | 9105.885742 | 10957.81152 |
| NRIP1 | P48552 | NRIP1 | Nuclear receptor-inte | 1087.310059 | 1433.574951 |
| APOL2 | J3KQL8 | APOL2 | Apolipoprotein L2 | 47845.52344 | 63416.0625 |
| CHURC1 | A0A0C4DGJ7;Q8WUH1 | CHURC1 | Protein Churchill | 11858.54688 | 13901.09277 |
| ASRGL1 | Q7L266 | ASRGL1 | Isoaspartyl peptidase/ | 31832.9375 | 23171.125 |
| CTSH | A0A087X0D5;P09668 | CTSH | Pro-cathepsin H | 37116.17188 | 48553.01172 |
| TMEM238 | C9JI98 | TMEM238 | Transmembrane prote | 14997.61035 | 6867.03418 |
| ARMCX2 | Q7L311 | ARMCX2 | Armadillo repeat-cont | 11907.04785 | 13405.07715 |
| KRT2 | P35908 | KRT2 | Keratin, type II cytosk | 114562.5 | 129278.6641 |
| TPD52L1 | Q16890 | TPD52L1 | Tumor protein D53 | 865584 | 1267212.375 |
| KRT1 | P04264 | KRT1 | Keratin, type II cytosk | 268714.75 | 430393.2813 |
| CDKN2AIP | Q9NXV6 | CDKN2AIP | CDKN2A-interacting p | 50905.92188 | 51005.36719 |
| TIGAR | Q9NQ88 | TIGAR | Fructose-2,6-bisphosp | 264248 | 278686.1563 |
| GEMIN5 | Q8TEQ6 | GEMIN5 | Gem-associated prote | 181256.9375 | 189013.0156 |
| GPX4 | A0A087WT12;A0A087X2I2;P3 | GPX4 | Glutathione peroxidas | 126370.0313 | 187879.1563 |
| KRT6B | P04259 | KRT6B | Keratin, type II cytosk | 230921.2344 | 412084.4375 |
| F10 | P00742 | F10 | Coagulation factor X | 17978.58203 | 3109.254883 |
| FERMT1 | Q9BQL6 | FERMT1 | Fermitin family homol | 11085.44141 | 7724.072754 |
| NOS1AP | O75052 | NOS1AP | Carboxyl-terminal PDZ | 14936.84668 | 21528.77344 |
| RPS6KA3 | P51812 | RPS6KA3 | Ribosomal protein S6 | 213695.9531 | 271569.2188 |
| UNC13D | Q70J99 | UNC13D | Protein unc-13 homol | 39956.53125 | 40329.8125 |
| UTP20 | O75691 | UTP20 | Small subunit process | 152274.9531 | 121767.8984 |
| NOLC1 | A0A0A0MRM9 | NOLC1 | Nucleolar and coiled-b | 369827.0313 | 176198.125 |
| HPDL | Q96IR7 | HPDL | 4-hydroxyphenylpyruv | 221959.9063 | 252722.3125 |
| IRF2 | D6R9N5;D6RB08;K4DIA5;P14 | IRF2 | Interferon regulatory | 11139.16895 | 18349.57422 |
| P4HA2 | O15460 | P4HA2 | Prolyl 4-hydroxylase s | 136157.3438 | 149023.3125 |
| GMIP | Q9P107 | GMIP | GEM-interacting prote | 10454.6084 | 10943.16699 |
| KRT9 | P35527 | KRT9 | Keratin, type I cytoske | 48499.86328 | 115117.7109 |
| POM121 | Q96HA1 | POM121 | Nuclear envelope pore | 20434.17969 | 16679.34375 |
| MRC2 | Q9UBG0 | MRC2 | C-type mannose recep | 9474.623047 | 11519.77246 |
| C6orf132 | Q5T0Z8 | C6orf132 | Uncharacterized prote | 33710.1875 | 36602.98828 |

| | | | | | |
|---|---|---|---|---|---|
| PIEZO1 | Q92508 | PIEZO1 | Piezo-type mechanose | 35814.65234 | 49296.85547 |
| ABAT | H3BRN4;P80404 | ABAT | 4-aminobutyrate amir | 418852.25 | 556206.75 |
| SLC16A1 | P53985;Q5T8R3;Q5T8R5 | SLC16A1 | Monocarboxylate trar | 71485.0625 | 75543.32813 |
| CTSB | P07858 | CTSB | Cathepsin B | 70427.91406 | 113320.7656 |
| GOLM1 | Q8NBJ4 | GOLM1 | Golgi membrane prot | 34369.42188 | 28298.54688 |
| S100A6 | P06703;R4GN98 | S100A6 | Protein S100-A6;Prote | 717746.25 | 561982.625 |
| TPBG | Q13641 | TPBG | Trophoblast glycoprot | 95489.39063 | 104560.5703 |
| STX8 | Q9UNK0 | STX8 | Syntaxin-8 | 20748.06641 | 27208.52734 |
| RWDD4 | K4DI92;Q6NW29 | RWDD4 | RWD domain containi | 111553.8672 | 106267.2422 |
| SCD5 | Q86SK9 | SCD5 | Stearoyl-CoA desatura | 19581.94531 | 28346.77148 |
| PARP10 | E9PK67;E9PNI7;Q53GL7 | PARP10 | Poly [ADP-ribose] poly | 33510.25 | 46108.17969 |
| ATP13A1 | Q9HD20 | ATP13A1 | Manganese-transport | 71542.46875 | 78101.28125 |
| SRC | P12931 | SRC | Proto-oncogene tyros | 189003.4063 | 204489.1094 |
| IFI30 | P13284 | IFI30 | Gamma-interferon-in | 59066.29688 | 72480.89063 |
| RESF1 | J3KPI3;Q9HCM1 | RESF1 | Retroelement silencin | 812.824646 | 648.8781128 |
| SCIN | Q9Y6U3 | SCIN | Adseverin | 172668.5156 | 143762.8438 |
| ARMC6 | Q6NXE6 | ARMC6 | Armadillo repeat-cont | 39791.01953 | 41231.32031 |
| UFSP2 | Q9NUQ7 | UFSP2 | Ufm1-specific proteas | 39327.07031 | 42553.30078 |
| COX7A2L | O14548 | COX7A2L | Cytochrome c oxidase | 156363.5313 | 148314.5313 |
| MORC4 | Q8TE76 | MORC4 | MORC family CW-type | 11903.7373 | 10464.42578 |
| PPM1E | Q8WY54 | PPM1E | Protein phosphatase 1 | 76390.6875 | 93730.02344 |
| KLC3 | Q6P597 | KLC3 | Kinesin light chain 3 | 10164.4043 | 11868.49121 |
| CD109 | Q6YHK3 | CD109 | CD109 antigen | 33709.07422 | 21011.80859 |
| ANXA6 | E5RK69 | ANXA6 | Annexin | 2906584.656 | 1155726.578 |
| COA8 | A0A0U1RQS9;H7BZ67;Q96IL( | COA8 | Cytochrome c oxidase | 2602.530518 | 2040.257813 |
| FHL3 | Q13643 | FHL3 | Four and a half LIM do | 57899.6875 | 59345.42969 |
| MT1X | P80297 | MT1X | Metallothionein-1X | 55446.35938 | 30821.32031 |
| EIF2AK2 | P19525 | EIF2AK2 | Interferon-induced, d | 237152.8906 | 175103.3125 |
| ECI2 | A0A0C4DGA2;O75521 | ECI2 | Enoyl-CoA delta isome | 121864.3047 | 128414.6719 |
| UNC119 | Q13432 | UNC119 | Protein unc-119 homo | 141218.9688 | 216964.6875 |
| CORO1A | P31146 | CORO1A | Coronin-1A | 138408.2031 | 145252.1563 |
| KIAA1217 | Q5T5P2 | KIAA1217 | Sickle tail protein hom | 15914.0791 | 18755.15234 |
| UXT | Q9UBK9 | UXT | Protein UXT | 38836.17969 | 34891.33984 |

| | | | | | |
|---|---|---|---|---|---|
| IGSF8 | Q969P0 | IGSF8 | Immunoglobulin supe | 29743.5918 | 28509.88281 |
| KRT16 | P08779 | KRT16 | Keratin, type I cytoske | 3213785 | 4942529 |
| MLKL | Q8NB16 | MLKL | Mixed lineage kinase | 190621.4375 | 192048.375 |
| NOTCH1 | P46531 | NOTCH1 | Neurogenic locus notc | 14779.28125 | 18339.94141 |
| POLR3G | D6R921;D6R9U7;D6RIT0;D6R | POLR3G | DNA-directed RNA po | 16511.4707 | 14338.1875 |
| ISG20 | Q96AZ6 | ISG20 | Interferon-stimulated | 17659.02148 | 12026.9707 |
| BAG3 | C9JFK9 | BAG3 | BAG family molecular | 347294.4023 | 468817.8672 |
| SEC11C | A0A0A0MR04;B4DI03;K7EJQ | SEC11C | Signal peptidase comp | 15673.5918 | 15188.96191 |
| ARHGEF10L | Q5VXI4;Q9HCE6 | ARHGEF10L | Rho guanine nucleotic | 15425.16016 | 19542.39648 |
| SULT1A4;SULT1A3;SULT1A | A0A0A6YYL2;P0DMM9;P0DM | SULT1A4;SULT1A3;SU | Sulfotransferase;Sulfo | 28173.75195 | 21591.83594 |
| SLC3A2 | F5GZS6;P08195 | SLC3A2 | 4F2 cell-surface antige | 8786059.34 | 9062316.605 |
| SPART | Q8N0X7 | SPART | Spartin | 79074.97656 | 87192.84375 |
| S100A9 | P06702 | S100A9 | Protein S100-A9 | 8037.64209 | 17730.16797 |
| SEMA4C | Q9C0C4 | SEMA4C | Semaphorin-4C | 100705.5156 | 147508.6094 |
| RIN1 | E9PNR2;Q13671 | RIN1 | Ras and Rab interacto | 35689.80859 | 30651.88477 |
| AREG | P15514 | AREG | Amphiregulin | 7489.313965 | 8272.74707 |
| FRG1 | Q14331 | FRG1 | Protein FRG1 | 108791.4531 | 105059.1563 |
| SYNPO2 | H0Y9Y3;Q9UMS6 | SYNPO2 | Synaptopodin-2 (Fragi | 20984.62305 | 43542.74219 |
| SLC7A5 | Q01650 | SLC7A5 | Large neutral amino a | 1922592.375 | 2474031.25 |
| C1orf226 | A1L170 | C1orf226 | Uncharacterized prote | 25160.98438 | 28406.70898 |
| PSTPIP2 | Q9H939 | PSTPIP2 | Proline-serine-threoni | 4327.108887 | 3889.233643 |
| TTC39A | E9PQX3;F8W961 | TTC39A | Tetratricopeptide repe | 224594.4585 | 283668.2388 |
| ACSL1 | P33121 | ACSL1 | Long-chain-fatty-acid- | 96299.125 | 105844.6406 |
| IFIH1 | Q9BYX4 | IFIH1 | Interferon-induced he | 6018.468262 | 6197.787109 |
| B4GALT1 | P15291 | B4GALT1 | Beta-1,4-galactosyltra | 58539.09375 | 60544.26953 |
| RCN1 | Q15293 | RCN1 | Reticulocalbin-1 | 219748.625 | 226168.2969 |
| NEIL2 | Q969S2 | NEIL2 | Endonuclease 8-like 2 | 17358.24609 | 28830.25977 |
| APOE | P02649 | APOE | Apolipoprotein E | 427017.2188 | 415289.0938 |
| ASS1 | P00966 | ASS1 | Argininosuccinate syn | 2070571.75 | 2345388.75 |
| CHCHD10 | B5MBW9;Q8WYQ3 | CHCHD10 | Coiled-coil-helix-coile | 26711.99023 | 31127.4707 |
| PARP9 | Q8IXQ6 | PARP9 | Protein mono-ADP-rib | 17839.2168 | 15386.33594 |
| DTX3L | Q8TDB6 | DTX3L | E3 ubiquitin-protein li | 16825.5293 | 16651.63672 |
| OAS3 | Q9Y6K5 | OAS3 | 2'-5'-oligoadenylate sy | 129019.0703 | 125806.8359 |

| | | | | | |
|---|---|---|---|---|---|
| SYT12 | Q8IV01 | SYT12 | Synaptotagmin-12 | 8735.417969 | 9624.907227 |
| STOM | P27105 | STOM | Erythrocyte band 7 int | 236961.6094 | 205439.6406 |
| PARP14 | Q460N5 | PARP14 | Protein mono-ADP-rib | 35504.08594 | 40756.65234 |
| CA12 | O43570 | CA12 | Carbonic anhydrase 1 | 159886.1719 | 232596.9219 |
| KSR1 | A0A2R8Y5H9;Q8IVT5 | KSR1 | Kinase suppressor of R | 1152.88562 | 1312.090576 |
| TCEA3 | O75764;S4R3M9 | TCEA3 | Transcription elongati | 14271.07129 | 12353.88672 |
| ZCCHC14;ZCCHC2;ZCCHC2;Z | A0A590UJW6;K7EKB8;K7ESN | ZCCHC14;ZCCHC2;ZCC | Zinc finger CCHC doma | 2628.549316 | 5093.605957 |
| TMEM164 | Q5U3C3;Q9H617 | TMEM164 | Transmembrane prote | 14118.26367 | 20465.91016 |
| HLA-A | A0A140T913;A0A140T9I0;A0 | HLA-A | HLA class I histocompa | 154826.375 | 153994.9531 |
| NT5DC3 | Q86UY8 | NT5DC3 | 5'-nucleotidase doma | 27924.40039 | 32508.91602 |
| NPM3 | O75607 | NPM3 | Nucleoplasmin-3 | 129385.375 | 153208.0938 |
| PGR | P06401 | PGR | Progesterone recepto | 9247.214844 | 16463.88867 |
| IGFBP5 | P24593 | IGFBP5 | Insulin-like growth fac | 7279.224121 | 2616.614502 |
| DNASE1L2 | Q92874 | DNASE1L2 | Deoxyribonuclease-1- | 30787.46094 | 27404.84375 |
| ADA | F5GWI4;P00813 | ADA | Adenosine deaminase | 6361.224609 | 10257.84082 |
| IRS1 | P35568 | IRS1 | Insulin receptor subst | 66510.24219 | 81518.70313 |
| PLOD2 | O00469 | PLOD2 | Procollagen-lysine,2-c | 84789.45313 | 119575.1797 |
| VAMP2;VAMP2;;VAMP2 | F8WCA0;J3QRU4;L7N2F9;P63 | VAMP2;VAMP2;;VAM | Vesicle-associated me | 53789.94531 | 51235.32031 |
| QPRT | Q15274 | QPRT | Nicotinate-nucleotide | 49589.24219 | 60272.20313 |
| PLBD1 | Q6P4A8 | PLBD1 | Phospholipase B-like 1 | 7147.272461 | 6968.096191 |
| SYTL5 | Q8TDW5 | SYTL5 | Synaptotagmin-like pr | 5970.790527 | 6039.160156 |
| PRTFDC1 | Q9NRG1 | PRTFDC1 | Phosphoribosyltransfe | 18203.54297 | 17733.21875 |
| EFHD1 | Q9BUP0 | EFHD1 | EF-hand domain-conta | 1493511.625 | 1137451.25 |
| S100A14 | Q9HCY8 | S100A14 | Protein S100-A14 | 8150.79834 | 8659.930664 |
| KLF13 | A0A087X1K6;Q9Y2Y9 | KLF13 | Krueppel-like factor 1 | 6175.574219 | 9233.751953 |
| SFXN3 | A0A1P0AYU5;Q9BWM7 | SFXN3 | Sidoreflexin;Sideroflex | 144645.9531 | 150514.8906 |
| GPC6 | Q9Y625 | GPC6 | Glypican-6 | 10594.80762 | 33274.08984 |
| KRT17 | Q04695 | KRT17 | Keratin, type I cytoske | 87660576 | 63794800 |
| SMPDL3B | Q92485 | SMPDL3B | Acid sphingomyelinas | 167070.1719 | 132883.8906 |
| CDC42EP5 | Q6NZY7 | CDC42EP5 | Cdc42 effector protein | 34522.02344 | 26226.10938 |
| FGFR4 | J3KPQ0;P22455 | FGFR4 | Fibroblast growth fact | 29329.5625 | 29435.11523 |
| INAFM1 | C9JVW0 | INAFM1 | Putative transmembra | 5303.597168 | 9336.170898 |
| CXXC5 | Q7LFL8 | CXXC5 | CXXC-type zinc finger | 78696.85156 | 94575.79688 |

| | | | | | |
|---|---|---|---|---|---|
| HLA-C | A0A0G2JH50 | HLA-C | HLA class I histocompa | 116161.1875 | 126021.2266 |
| S100A7 | P31151 | S100A7 | Protein S100-A7 | 69722.35156 | 519101.75 |
| RHOD | O00212 | RHOD | Rho-related GTP-bind | 112131.0391 | 111159.9375 |
| DMD | A0A075B6G3;A0A087WTU7;A | DMD | Dystrophin | 2964226.5 | 2923260.75 |
| PLCD3 | Q8N3E9 | PLCD3 | 1-phosphatidylinosito | 65997.65625 | 91715.80469 |
| STMN3 | A0A0J9YW36;Q9NZ72 | STMN3 | Stathmin-3 (Fragment | 164130.2969 | 201525.2656 |
| ANXA1 | P04083 | ANXA1 | Annexin A1 | 10035.36523 | 10391.80859 |
| TENM3 | Q9P273 | TENM3 | Teneurin-3 | 18786.06055 | 27395.63672 |
| NUDT10;NUDT11 | Q8NFP7;Q96G61 | NUDT10;NUDT11 | Diphosphoinositol pol | 30618.67773 | 24499.04297 |
| MOCS2 | O96007 | MOCS2 | Molybdopterin syntha | 312387.0469 | 523790.7578 |
| SYNE1 | A0A0C4DG40;E7ENN3;Q8NF | SYNE1 | Nesprin-1 | 3080886.25 | 2379160.5 |
| HNMT | P50135 | HNMT | Histamine N-methyltr | 240710.9219 | 190347.5938 |
| IFITM3 | Q01628 | IFITM3 | Interferon-induced tra | 99854 | 160672.3906 |
| FKBP11 | F8VU90;Q9NYL4 | FKBP11 | Peptidylprolyl isomera | 160373.1094 | 150778.3906 |
| GNB4 | Q9HAV0 | GNB4 | Guanine nucleotide-b | 105605.6484 | 156354.3438 |
| ITGA2 | E7EMF1 | ITGA2 | Integrin alpha-2 | 765430.7969 | 925149.7969 |
| DDX58 | O95786 | DDX58 | Antiviral innate immu | 41510.36328 | 36864.35156 |
| MUC1 | A0A087WWM1 | MUC1 | Mucin-1 | 75150.9375 | 85893.47656 |
| RAPGEFL1 | A0A087WVW6;Q9UHV5 | RAPGEFL1 | Rap guanine nucleotic | 12622.71973 | 25341.57813 |
| ATP6V0A4 | Q9HBG4 | ATP6V0A4 | V-type proton ATPase | 4680.077148 | 5463.648926 |
| CELSR2 | Q9HCU4 | CELSR2 | Cadherin EGF LAG sev | 28255.51758 | 34198.23438 |
| LGALS3BP | Q08380 | LGALS3BP | Galectin-3-binding pro | 405009.4063 | 518186.6875 |
| FHL1 | A0A494C0D6;Q13642;Q5JXI8 | FHL1 | Four and a half LIM do | 11496.04297 | 8985.78125 |
| S100A4 | P26447 | S100A4 | Protein S100-A4 | 109871.1094 | 84803.75 |
| RNF223 | E7ERA6 | RNF223 | RING finger protein 22 | 12428.06738 | 13875.41699 |
| POF1B | Q8WVV4 | POF1B | Protein POF1B | 61621.4375 | 73772.07031 |
| DDX60 | Q8IY21 | DDX60 | Probable ATP-depend | 15715.6709 | 12591.9541 |
| SPHK1 | Q9NYA1 | SPHK1 | Sphingosine kinase 1 | 24454.83008 | 19961.70313 |
| IRF9 | H0YNP4;Q00978 | IRF9 | Interferon regulatory | 9327.032227 | 9482.438477 |
| HSPB8 | Q9UJY1 | HSPB8 | Heat shock protein be | 245112.0781 | 239633.5 |
| CA2 | P00918 | CA2 | Carbonic anhydrase 2 | 218310.6563 | 263187.9063 |
| PADI2 | Q9Y2J8 | PADI2 | Protein-arginine deim | 83449.9375 | 114577.875 |
| ESPN | B1AK53 | ESPN | Espin | 51858.32813 | 80014.48438 |

| | | | | | |
|---|---|---|---|---|---|
| SUSD2 | Q9UGT4 | SUSD2 | Sushi domain-contain | 82008.26563 | 71194.05469 |
| IFI35 | P80217 | IFI35 | Interferon-induced 35 | 57747.12891 | 49380.08984 |
| GREB1 | Q4ZG55 | GREB1 | Protein GREB1 | 252642.7656 | 379112.375 |
| SLC26A2 | P50443 | SLC26A2 | Sulfate transporter | 175190.7813 | 166182.0156 |
| CLIC3 | O95833 | CLIC3 | Chloride intracellular | 209706.7188 | 298553.5625 |
| MAOB | P27338 | MAOB | Amine oxidase [flavin- | 35601.32031 | 59292.83984 |
| IFIT1 | P09914 | IFIT1 | Interferon-induced pr | 3502.716797 | 3365.447021 |
| S100A8 | P05109 | S100A8 | Protein S100-A8 | 72350.70313 | 124588.3203 |
| EGR3 | Q06889 | EGR3 | Early growth response | 1986.485596 | 3002.179932 |
| TRIP6 | Q15654 | TRIP6 | Thyroid receptor-inter | 195165.875 | 204931.4219 |
| GAL3ST2 | A0A0G2JPZ6;A0A0J9YWU2;Q | GAL3ST2 | Galactose-3-O-sulfotra | 38294.92578 | 40087.00391 |
| FABP3 | P05413;S4R371 | FABP3 | Fatty acid-binding pro | 25697.65234 | 28833.18945 |
| PDZK1 | Q5T2W1 | PDZK1 | Na(+)/H(+) exchange r | 870177.875 | 1240092.625 |

| parental_r3 | LTED_r1 | LTED_r2 | LTED_r3 | KO1_r1 | KO1_r2 | KO1_r3 |
|---|---|---|---|---|---|---|
| NA | 715.2779541 | NA | 2131.451416 | 24468.95508 | 10535.25977 | 17656.5957 |
| NA | NA | NA | 2225.947754 | 6557.917969 | 5875.529785 | 4963.834961 |
| 2714.862793 | NA | NA | NA | 155.3779144 | NA | NA |
| NA | NA | 394.4131165 | 222.7645416 | 411.7068481 | 1293.189453 | 970.4672852 |
| NA | NA | NA | NA | 11668.87988 | NA | 1727.049194 |
| NA | NA | NA | NA | 741.3161621 | 97.15795898 | 99.78037262 |
| NA | 11.15878963 | NA | NA | 2.345895052 | NA | NA |
| NA | 1866.786133 | 4372.04248 | NA | 2085.071777 | NA | 2582.868164 |
| NA | NA | NA | NA | 84805.70313 | 33837.45703 | 46286.83594 |
| NA | NA | NA | NA | NA | NA | NA |
| 1056.633789 | NA | NA | NA | 1919.086548 | 242.7609406 | 567.7671509 |
| 2576.413086 | NA | NA | NA | NA | NA | NA |
| NA | 1339.813965 | 960.0352783 | 844.3025513 | 1291.776978 | 402.1470947 | 249.2745819 |
| 15230.38281 | NA | NA | NA | NA | NA | NA |
| NA | NA | NA | NA | NA | NA | NA |
| NA | NA | NA | NA | NA | 374.5022278 | NA |
| 935.8546143 | NA | NA | NA | NA | NA | NA |
| NA | 260.6226196 | 363.289032 | 284.2433472 | NA | 249.589386 | 294.8302307 |
| 544.8516235 | NA | NA | NA | 687.6620483 | NA | 780.6821899 |
| 1.705773592 | NA | NA | NA | NA | NA | NA |
| NA | 945.3991089 | 1362.589844 | 1796.764038 | 4110.882813 | 2545.80249 | 3745.355713 |
| NA | 1687.473511 | 1539.040405 | NA | 3642.172119 | 2646.80249 | 4112.509277 |
| 4781.602051 | NA | NA | NA | 4086.495117 | NA | NA |
| NA | NA | 196.0921631 | NA | 796.3924561 | 858.843689 | 822.3571167 |
| NA | 896.6276245 | 1396.71582 | 1587.131836 | 1110.517212 | 1045.462891 | 740.2753296 |
| NA | NA | NA | NA | 3695.706543 | 2070.577148 | 2873.452148 |
| NA | NA | NA | NA | 13792.37988 | 2222.862793 | NA |
| NA | NA | NA | 546.3461914 | 2155.428955 | NA | NA |
| 20176.625 | NA | NA | NA | 2156.088623 | 4430.724609 | 1573.391357 |
| 4412.559082 | NA | NA | NA | NA | NA | NA |
| NA | 3058.24292 | 5836.8125 | 6515.361328 | 8445.167969 | 4166.568848 | 6024.314453 |
| 5763.611328 | NA | NA | NA | 282.0969238 | NA | 190.7924957 |

| | | | | | | |
|---|---|---|---|---|---|---|
| 11525.75781 | NA | NA | NA | 5187.726563 | 3782.268555 | 3216.606201 |
| 1567.442871 | NA | NA | NA | NA | NA | NA |
| NA | NA | NA | NA | 1347.486572 | NA | 1335.199951 |
| NA | 12053.88965 | 6320.994141 | 4648.571777 | 14790.5918 | 12767.0752 | 13648.65234 |
| 1448.520264 | NA | NA | NA | 4948.63623 | 9484.671875 | 4350.614746 |
| 6250.883301 | NA | NA | NA | NA | NA | NA |
| NA | NA | NA | NA | NA | NA | NA |
| NA | NA | NA | NA | NA | NA | NA |
| 3244.950684 | NA | NA | NA | 2014.986206 | NA | NA |
| NA | 417.7215881 | NA | NA | 3225.90625 | 1445.741333 | 1330.180908 |
| 380.138031 | NA | NA | NA | 5329.817383 | NA | 289.367981 |
| NA | 864.7929688 | NA | NA | 6155.878418 | 3110.160156 | 2888.812256 |
| NA | 24461.53516 | 9150.996094 | 7620.470215 | 7748.456543 | 10848.25684 | 6594.462891 |
| 1 | NA | NA | NA | NA | NA | NA |
| 2046.769897 | NA | NA | NA | 975.4564209 | NA | NA |
| 681.4842529 | NA | NA | NA | NA | 92.3058548 | NA |
| NA | 4675.815918 | 4141.710938 | 2311.22583 | 4740.254883 | 4767.438965 | 3523.712158 |
| NA | 1481.701294 | 5404.183594 | 7410.492676 | 2502.900635 | 1068.240601 | 1724.182983 |
| NA | 4977.371094 | 4803.494629 | 6492.72168 | 3903.567383 | 4453.361816 | 4213.466797 |
| NA | 15093.9082 | 8074.121094 | 12785.21777 | 11155.79883 | 6949.558594 | 10123.83691 |
| 61494.32422 | 826720.4375 | 1678597.5 | 2307713.5 | 1003747.125 | 280084.9063 | 372802.375 |
| 166.0476532 | 11371.54395 | 6715.794922 | 7130.97998 | 4807.145996 | 4180.65625 | 4284.98584 |
| 107930.9766 | 1740234.346 | 1438060.232 | 1410527.412 | 867886.459 | 797605.1484 | 783914.5732 |
| 23710.5 | 624099.4375 | 664174.9375 | 1376723.375 | 563718 | 287217.7188 | 522916.6875 |
| NA | 9943.206055 | 104559.9922 | NA | 9164.583008 | 6086.808105 | 9206.331055 |
| NA | 7900.935547 | 7211.125977 | 6059.781738 | 2442.390625 | 1914.177734 | 2115.35376 |
| 25440.16992 | 640446.25 | 428667.4375 | 531286.8125 | 168003.6875 | 189796.1563 | 183342.1406 |
| 4123.918945 | 65042.07813 | 79935.8125 | 82448.10938 | 29781.22656 | 26572.10547 | 29843.25977 |
| 982.7218628 | 13452.91992 | 22131.47656 | 23164.08203 | 7823.346191 | 6933.542969 | 7216.706055 |
| NA | 7280.230957 | 4784.262695 | 4970.564941 | 2724.189453 | 5231.59375 | 22092.92969 |
| 49505.68359 | 694419.5 | 702656.125 | 755325.9375 | 264833.6875 | 252374.3125 | 254014.6875 |
| 18958.91602 | 306660.0938 | 216171.6719 | 273257.9063 | 674175.9375 | 425422.6563 | 446806.2188 |
| 3865.851807 | 77458.17188 | 50541.42578 | 50585.60547 | 21647.6543 | 26926.74219 | 23870.74219 |

| | | | | | | |
|---|---|---|---|---|---|---|
| 1259.854248 | 85685.67188 | 105654.9844 | 96741.03906 | 115428.9063 | 105562.7969 | 98643.95313 |
| 213821.1244 | 2502114.621 | 2917628.465 | 3271449.701 | 1120865.604 | 1148380.092 | 1213974.909 |
| 63480.86328 | 1008990.938 | 850862.9375 | 783862.875 | 347470.2188 | 389713.9375 | 306808.5938 |
| 8185.786621 | 116694.8438 | 91003.39063 | 98833.13281 | 80546.09375 | 53333.14844 | 60978.85547 |
| 1692.214355 | 31395.00586 | 25421.00195 | 27881.55664 | 16009.85059 | 19030.95898 | 18030.28516 |
| 15175.82129 | 290997.2188 | 234769.9688 | 252690.7969 | 130700.9688 | 138409.625 | 135547.3594 |
| 2574.575928 | 37605.39063 | 35297.37891 | 27470.95117 | 27792.76172 | 35000.17578 | 25400.12891 |
| 16542.3418 | 216627.0938 | 227290.6719 | 224586.3281 | 462924.625 | 425850.6563 | 429250.3438 |
| 2898.205811 | 60658.64844 | 53717.51953 | 35129.98828 | 17111.0957 | 21145.58398 | 14281.21582 |
| 916.0923462 | 7901.996094 | 9810.700195 | 16597.74805 | 11020.15137 | 3525.441162 | 5858.739746 |
| 2844.743896 | 36817.76953 | 33774.98828 | 30488.69922 | 6123.439453 | 8295.003906 | 10289.875 |
| 1557.942261 | 24584.37891 | 19413.46484 | 21416.92773 | 6156.600586 | 7124.279297 | 7421.09668 |
| 1013.963074 | 16509.09961 | 10442.17383 | 9757.046875 | NA | NA | NA |
| 2324.414795 | 71244.16406 | 43752.66797 | 3012.12915 | 2345.70874 | 7643.433594 | 3061.374512 |
| 15219.37402 | 218411.75 | 182950.4063 | 171849.5313 | 129509.9766 | 144997 | 128137.5156 |
| 3893.696777 | 44925.51563 | 35825.0625 | 34178.32422 | 14763.46484 | 15568.20703 | 14278.6582 |
| 8172.246582 | 81285.82031 | 97265.24219 | 103341.7578 | 110211.5078 | 133246.5781 | 126516.9219 |
| 180.1045685 | 702.6072998 | 1240.190186 | 2695.34668 | 10081.2373 | 8552.822266 | 17409.45898 |
| 60330.26953 | 315988.125 | 353800.1563 | 419386.5938 | 268559.875 | 317949.5625 | 302432.5313 |
| NA | 25648.13281 | 19787.89648 | 15445.89355 | 12952.56445 | 14623.75391 | 12920.89063 |
| 18828.60938 | 186110.0313 | 201889.875 | 189924.5313 | 41858.71094 | 50230.69922 | 41055.38281 |
| NA | 14726.2334 | 14636.87695 | 10944.27734 | 12021.63672 | 12002.6543 | 10711.08691 |
| 8881.674835 | 120397.6206 | 54810.28125 | 85643.40771 | 275403.8311 | 123231.8706 | 151057.4199 |
| 53293.375 | 539062.3125 | 518844.0313 | 444527.8125 | 176200.9063 | 199445.8594 | 184530.7188 |
| NA | 3982.094482 | NA | NA | 4226.702637 | 3468.252441 | 3838.235352 |
| NA | 13283.97656 | 8432.405273 | 12559.32031 | 11849.84375 | 12641.16016 | 12415.47363 |
| 87918.28906 | 642603.6875 | 613875.6875 | 844737.1875 | 291869.1563 | 265242.75 | 236020.0313 |
| 149092.1719 | 1650777.875 | 1212250.25 | 1766285.125 | 710098.9375 | 517167.8125 | 508807.0313 |
| NA | 21807.93555 | 15970.08594 | 16049.84961 | 9193.47168 | 5852.983398 | 5340.931152 |
| 41887.00391 | 349239.7813 | 365809.25 | 372915.6563 | 135568.5156 | 183314.4688 | 115143.9531 |
| 312155.5938 | 2576875.25 | 2573453.5 | 2218528.25 | 2984133.25 | 2688658.25 | 3155178.25 |
| 17650.40234 | 123700.6406 | 127794.4141 | 84940.92969 | 38902.16797 | 79514.36719 | 66335.4375 |
| 8583.007813 | 80446.07031 | 87220.01563 | 73169.21094 | 70030.5 | 66123.92188 | 64623.52734 |

| | | | | | | |
|---|---|---|---|---|---|---|
| 13806.52734 | 149017.9531 | 102493.875 | 84078.57031 | 30565.77734 | 25696.89063 | 21522.3125 |
| 335.8495178 | 6728.491211 | 3447.505615 | 4690.819824 | 1716.224731 | 1893.133789 | 1713.265869 |
| 3188.131836 | 18558.87109 | 18503.89258 | 17967.39063 | 36004.77344 | 22168.98438 | 23042.15039 |
| 715.1704712 | NA | 5669.349121 | 4981.496582 | 2520.801514 | 5230.690918 | 4616.559082 |
| 33159.19531 | 270077.9375 | 282139.5 | 244577.625 | 115673.8438 | 126118.3281 | 145083.9219 |
| 6152.916016 | 34601.70313 | 45885.55469 | 36227.82422 | 10374.48438 | 9434.135742 | 9704.608398 |
| 3106.110352 | 23639.36914 | 31345.50586 | 26917.49805 | 6015.029297 | 7441.175781 | 6335.925781 |
| 12437.91797 | 107553.6016 | 84161.73438 | 85451.5625 | 62266.375 | 62909.43359 | 49082.41797 |
| 2970.946777 | 22879.77344 | 18325.71875 | 21779.71094 | 23878.47852 | 20703.03125 | 20586.35547 |
| 2281.230469 | 14182.26855 | 17104.23438 | 18510.60938 | 26693.55664 | 22344.75391 | 24746.46484 |
| 23866.41602 | 125745.6016 | 173103.8438 | 171357.75 | 236369.0938 | 325229 | 323167.8438 |
| 796.4755249 | 4861.103027 | 4887.949707 | 4157.610352 | 4848.89209 | 4592.708984 | 4441.229004 |
| NA | 8013.540039 | 5285.549316 | 8414.688477 | 3674.812012 | 2497.642334 | 1792.743408 |
| 55945.16797 | 368268.0313 | 340217.75 | 383945.3438 | 264490.4688 | 204270.5469 | 199333.7344 |
| 27723.9707 | 208252.0625 | 213178.1875 | 224186.6875 | 123143.0234 | 113570.9844 | 105856.6641 |
| 1555.075806 | 14747.75098 | 5086.324219 | 5052.975586 | 16295.69531 | 23655.16602 | 16850.70703 |
| 4712.020508 | 35986.91406 | 22578.59375 | 17479.56836 | 7313.229492 | 12867.87598 | 9977.157227 |
| 1787.674316 | 9633.383789 | 9731.271484 | 7914.132324 | 2331.061279 | 1771.989624 | 2649.052246 |
| NA | 5459.614258 | 3999.092285 | 6533.8125 | 7984.608887 | 8176.725098 | 6736.698242 |
| 418646.7148 | 2858785.172 | 2344287.188 | 2552429.922 | 911719.6602 | 921082.8711 | 832937.0977 |
| 20036.77148 | 132101.5938 | 120864.6641 | 115649.7266 | 47434.68359 | 55906.01172 | 55906.28516 |
| 19370.62305 | 136931.3594 | 100895.1172 | 81807.54688 | 73148.125 | 100601.9844 | 87159.82813 |
| 1783.857666 | 12272.84473 | 7031.970215 | 8521.241211 | 7239.458984 | 8284.550781 | 9652.458984 |
| NA | 1093.068237 | 1523.718262 | 1685.125122 | 1373.238403 | 1272.022339 | 1034.782837 |
| 16118.28516 | 157311.9219 | 131274.0156 | 131175.1563 | 263467.5 | 172273.2031 | 172137.3906 |
| 275.1817322 | 3517.146973 | 2847.133789 | 3008.811035 | 6889.141113 | 4210.744629 | 4817.749512 |
| 377.4507446 | 3367.284912 | 6316.227051 | 5967.614258 | 5524.304199 | 6740.751953 | 4688.195801 |
| 2763.083008 | 17113.83789 | 18591.71484 | 19003.22266 | 16195.94043 | 9895.5 | 8953.294922 |
| 277.8351135 | 6675.463379 | 7113.152344 | 6099.331543 | 28154.22461 | 51801.47266 | 41337.625 |
| 77508.29688 | 447413.25 | 524018.2188 | 545505.25 | 136853.4844 | 161888.7344 | 174543.2969 |
| 19741.30469 | 225308.9375 | 281312.0938 | 10122.88867 | 5000.421387 | 62223.36328 | 10792.44141 |
| 23559.80469 | 89645.03125 | 90900.72656 | 82305.05469 | 102738.4297 | 124874.625 | 114189.625 |
| NA | 906.8364868 | 370.1672974 | 1325.386597 | 218.0867767 | NA | 130.1285248 |

| | | | | | | |
|---|---|---|---|---|---|---|
| NA | 1884.815674 | 3443.84375 | 3722.89917 | 6332.565918 | 4607.749023 | 5105.63916 |
| 8014.893066 | 37112.92578 | 34617.82813 | 63617.48828 | 27016.89844 | 25211.11133 | 33291.73438 |
| 3936.484863 | 40009.79688 | 35852.32813 | 32862.0625 | 26890.16602 | 19574.63477 | 20902.75391 |
| 27094.73828 | 361679.5313 | 201349.875 | 238198.8906 | 122509.7188 | 164085.5625 | 127653.9688 |
| 8477.256836 | 70286.375 | 42326.03906 | 35305.05469 | 8328.671875 | 6079.172363 | 4437.949707 |
| 3360.942139 | 32292.60156 | 30070.90234 | 29869.20703 | 31305.38477 | 25342.55664 | 15424.09668 |
| 992.9935303 | 11611.50684 | 11600.53613 | 11964.6377 | 18727.69531 | 17415.70508 | 19112.31055 |
| 4853.063965 | 47551.26758 | 37681.75391 | 42882.83301 | 34390.09863 | 36972.17969 | 32810.02344 |
| 2320.092773 | 15587.86719 | 18357.38867 | 17042.62891 | 21665.50391 | 16165.02344 | 20009.06055 |
| NA | 9081.892578 | 6864.020996 | 6865.79541 | 16406.10742 | 25274.67383 | 20155.47266 |
| 4192.824707 | 19785.48633 | 25447.24414 | 25719.62695 | 7581.475098 | 7844.582031 | 9852.834961 |
| NA | 6839.452637 | 2870.765137 | 1256.357666 | 2990.943604 | NA | 1763.680908 |
| 3157.773193 | 15763.90137 | 14464.10645 | 12534.80762 | 8576.960938 | 8854.383789 | 9017.767578 |
| 7606.529785 | 33547.36719 | 31486.82227 | 28912.04883 | 9771.350586 | 11824.08105 | 8233.920898 |
| 2595.256592 | 12112.91406 | 11845.38086 | 8449.871094 | 12735.31348 | 15275.68164 | 12393.16406 |
| NA | 6744.622559 | 5545.693848 | 5809.42627 | 13668.68359 | 15895.85547 | 15065.31738 |
| 10662.91602 | 44533.96875 | 31819.01953 | 31018.57227 | 46774.10547 | 39321.66016 | 34199.07031 |
| 6795.658203 | 26005.92578 | 31679.6543 | 41815.70703 | 17730.27539 | 13047.95996 | 17500.5 |
| 33597.66406 | 136742.3281 | 122769.6797 | 164751.0938 | 134222.4844 | 143836.7031 | 147947.1094 |
| 1065.713379 | 8258.231445 | 4526.86377 | 1898.846802 | 5704.966309 | 4810.632813 | 8453.540039 |
| 168180.5 | 941985.8125 | 737420.125 | 746527.4375 | 457927 | 471241.375 | 466367.1563 |
| 237.0960693 | 2037.432617 | 1797.332886 | 1521.907593 | 628.401062 | 311.3312683 | 541.5952148 |
| 738.7749023 | 8217.174805 | 5969.089355 | 4874.205078 | 3319.736816 | 2702.993164 | 2655.528564 |
| NA | 11289.15332 | 5857.084473 | 9034.0625 | 15804.17578 | 38467.44531 | 27925.06055 |
| 171977.0938 | 799459 | 602010.875 | 598373.625 | 425007.3438 | 429131.125 | 378487.0313 |
| 9026.807617 | 41640.68359 | 27065.99219 | 40684.28516 | 14409.43262 | 11267.97168 | 14206.47266 |
| 2169.285645 | 8963.446289 | 14456.05664 | 13668.13477 | 5166.232422 | 5556.568848 | 5896.04834 |
| 763335.8125 | 3905963.5 | 3556533.5 | 3664502.5 | 3037700.5 | 3191055.25 | 3402929.75 |
| 6254.69873 | 43999.53516 | 23134.58203 | 19579.0918 | 30115.16797 | 26131.08203 | 21418.33203 |
| NA | NA | NA | 3258.443848 | 6307.370117 | 4630.390137 | 6928.873535 |
| 305.4093628 | 3206.82959 | 1799.888428 | 1298.866577 | 551.1906128 | 1948.339355 | 1326.814453 |
| 22947.70313 | 114172.9609 | 117198.5547 | 106293.9844 | 51065.69531 | 62635.58984 | 56817.38672 |
| 206627.2813 | 960218.3125 | 792046.3125 | 1053766 | 2205150.25 | 2174724.25 | 2231454.5 |

| | | | | | | |
|---|---|---|---|---|---|---|
| 13708.14355 | 69738.75781 | 57715.55469 | 53759.08203 | 33766.46094 | 33598.21875 | 32091.19141 |
| 1858.01123 | 9293.94043 | 10079.42773 | 8473.816406 | 12805.34277 | 14629.70508 | 12929.58691 |
| 91158.74219 | 353999.5625 | 420742.8125 | 376829.9375 | 403738.1563 | 419529.7188 | 406720.6563 |
| 2044.738159 | 8664.625 | 5665.299805 | 6579.524902 | 28194.92969 | 13447.10742 | 13466.64258 |
| 300540.0625 | 1399602.5 | 1124901.5 | 1453609.125 | 3174462 | 2900157 | 2998764 |
| 2539.340576 | 15046.23828 | 26853.00391 | 37109.91797 | 18308.37305 | 17185.66406 | 12179.83594 |
| 75639.6875 | 662065.1875 | 140250.8438 | 154110.4063 | 148242.7813 | 116452.3047 | 163562.4375 |
| 1247.477539 | 5470.157227 | 6064.021484 | 7217.083984 | 5932.567383 | 7373.006836 | 6464.29834 |
| 1815.42395 | 12113.32813 | 10765.29883 | 7061.087891 | 7498.653809 | 11017.5791 | 8376.146484 |
| 4971.109375 | 35935.32813 | 43206.69141 | 31328.54492 | 44542.31641 | 44414.57422 | 43802.56641 |
| 81241.71875 | 419157.125 | 371535.0938 | 411875.3125 | 345120.9688 | 353784.2188 | 344521.9375 |
| 4054.281738 | 27434.96484 | 21792.18359 | 19246.69336 | 19057.40039 | 24566.75391 | 24659.45117 |
| 43802.18359 | 223151.5781 | 247802.1406 | 221756.4531 | 191290.8438 | 209554.125 | 220151.3594 |
| NA | 1580.779053 | 1371.067749 | 782.6569824 | 678.7915649 | 904.538208 | 983.9694214 |
| 1393.650146 | 2815.244873 | 3593.737061 | 6105.485352 | 5219.137695 | 2429.616211 | 3822.889893 |
| 3477.267822 | 10058.51367 | 13345.45898 | 14027.25391 | 10107.6748 | 6266.686035 | 9321.842773 |
| 7230.362793 | 33779.55469 | 33241.58984 | 34903.64844 | 21172.50391 | 24941.23633 | 25206.05078 |
| 13119.88672 | 73252.0625 | 84868.35156 | 82082.20313 | 105437.4844 | 105671.4688 | 120291.4766 |
| 10163.14746 | 48646.28906 | 48588.45313 | 36851.21094 | 16662.33203 | 22265.33203 | 19243.46094 |
| 1159.251343 | 6932.947754 | 5967.146973 | 8021.936523 | 24495.75391 | 23941.57031 | 30310.0918 |
| 6287.462891 | 24203.81641 | 24097.125 | 22810.89453 | 27693.5 | 27467.5625 | 29947.21484 |
| 122088.0469 | 497141.625 | 511470.75 | 586259.5 | 316706.25 | 311992.625 | 326779.7813 |
| 13182.75195 | 61812.4375 | 51141.69531 | 59470.40234 | 58851.69531 | 46006.64844 | 50849.125 |
| 2693.697266 | 10059.61816 | 11749.35352 | 11317.50977 | 6696.004883 | 9511.984375 | 8444.734375 |
| 1405.455322 | 11453.23828 | 8174.89502 | 5706.037109 | 3199.482422 | 4113.413574 | 4127.083496 |
| 2279.347168 | 10424.44922 | 5048.954102 | 4739.225586 | 5725.065918 | 4464.217285 | 4635.759277 |
| 5715.872559 | 24455.89063 | 23564.46094 | 26148.00195 | 25181.47656 | 27940.1582 | 25771.09375 |
| 20310.4375 | 69257.64063 | 76677.79688 | 97571.66406 | 42655.40625 | 27889.20508 | 32932.23438 |
| 2203.228027 | 12496.77148 | 8359.849609 | 3755.509277 | 8776.982422 | 2903.88208 | 4354.632324 |
| 28695.55859 | 97337.38281 | 114285.9609 | 143378.6406 | 88746 | 94531.4375 | 102930.8203 |
| 1974.516602 | 1326.089966 | 8518.985352 | 16291.63379 | 87491.66406 | 12501.85254 | 39447.06641 |
| 1511.250366 | 2372.296387 | 5986.289551 | 6055.563965 | 13281.35742 | 9488.399414 | 9992.332031 |
| 607.2293701 | 2646.14917 | 2748.715576 | 2911.032471 | 3323.043701 | 2682.498779 | 2720.79248 |

| | | | | | | |
|---|---|---|---|---|---|---|
| 199080.3281 | 784617.5 | 972589.75 | 1033406.938 | 806409.8125 | 952815.9375 | 941658 |
| 293460.5938 | 1007497.625 | 1106839 | 1166139.75 | 588329.9375 | 546377.0625 | 590140.3125 |
| 18681.91797 | 85802.64063 | 78989.39844 | 85444.5625 | 92752.19531 | 94497.38281 | 95058.1875 |
| 4256.525879 | 26532.32031 | 14414.68652 | 11233.63281 | 20960.04102 | 20144.08008 | 18757.26367 |
| 1472.676514 | 6303.236328 | 1477.351807 | 6304.170898 | 11953.35156 | 4692.723633 | 9019.120117 |
| 4328.964355 | 16372.93457 | 16474.16016 | 16305.21484 | 14669.5293 | 16899.74023 | 14455.58496 |
| 11558.95801 | 58706.47656 | 52694.36719 | 48715.75391 | 42804.9375 | 37390.86719 | 40754.91016 |
| 1880.36731 | 10659.50488 | 8044.996582 | 13239.6084 | 9751.56543 | 6658.078125 | 4396.834473 |
| 285997.2813 | 838906.6875 | 1017646.875 | 1213799.625 | 491438.9063 | 449557.5 | 368343 |
| 254888.3594 | 783871.8125 | 977778.625 | 1232197.5 | 1091316.75 | 819767.375 | 985230.9375 |
| 9380.808594 | 37184.32422 | 37460.45703 | 41939.08984 | 33510.63672 | 29267.79492 | 35167.97266 |
| 547.8795776 | 2080.797852 | 2824.981445 | 2885.939453 | 4186.132813 | 3454.62085 | 2993.374756 |
| 10930.54688 | 51078.10938 | 44943.92969 | 51673.45313 | 65728.90625 | 94311.11719 | 89425.59375 |
| 20097.7207 | 48163.91797 | 80902.92969 | 95425.35156 | 83937.47656 | 65836.89063 | 78728.08594 |
| 2016.371338 | 6619.686523 | 6797.264648 | 8091.214844 | 5294.020996 | 4839.414551 | 6213.785156 |
| 12041.50098 | 33511.875 | 40036.68359 | 49346.39453 | 37227.03125 | 32273.23438 | 32988.4375 |
| 6286.058594 | 14776.82227 | 26368.25586 | 27716.6543 | 57303.34375 | 93232.96875 | 83026.85156 |
| 4260.574707 | 11613.73242 | 17587.66797 | 17108.9668 | 12906.81445 | 11934.23828 | 13705.99023 |
| 885.7057495 | 3491.009766 | 2969.516357 | 3131.183105 | 1857.967163 | 3462.287842 | 2820.525391 |
| 1072.236938 | 9324.704102 | 5669.308105 | 4621.808594 | NA | 2605.036865 | 1485.708496 |
| 2276.319336 | 35171.38281 | 22610.64258 | 29992.22656 | 30905.48438 | 26620.3457 | 26737.66797 |
| 15529.2627 | 38453.50391 | 51045.55469 | 56251.64063 | 81104.45313 | 62453.23438 | 74900.03906 |
| 26723.18555 | 89739.61719 | 122990.0078 | 154721.75 | 120791.7188 | 89018.78906 | 111760.1641 |
| 1700.581299 | 6029.257813 | 6389.859375 | 5910.643066 | 4093.03125 | 3873.425293 | 3912.12915 |
| 1697.711304 | 6553.003418 | 6617.708984 | 7223.240234 | 9047.102539 | 7591.608398 | 7158.248047 |
| 1473.00061 | 3976.694092 | 5056.149902 | 5073.710938 | 2276.943115 | 1629.026611 | 1545.373047 |
| 10879.92188 | 43544.97266 | 33976.89453 | 38784.76563 | 32667.80469 | 28159.04297 | 32051.68359 |
| 4464.576172 | 10615.24023 | 12821.46094 | 20522.12109 | 57208.13672 | 31175.21094 | 31265.75391 |
| 9612.22168 | 31489.74414 | 29653.51172 | 32718.43555 | NA | 26512.79102 | 11664.99707 |
| 4779.786621 | 32371.20898 | 10088.41602 | 14202.19238 | 20790.14063 | 11927.97559 | 14701.79004 |
| 212831.9688 | 770352.8125 | 732363.375 | 677105.375 | 498731.8125 | 593464.625 | 582992.9375 |
| 5795.915527 | 32558.98242 | 22343.73047 | 18173.45898 | 10150.16113 | 15822.56152 | 12857.50781 |
| 3247.427002 | 7810.333008 | 6760.445313 | 7835.211426 | 7249.819824 | 7929.297363 | 7651.884766 |

| | | | | | | |
|---|---|---|---|---|---|---|
| 19313.13086 | 66731.25781 | 68175.40625 | 73392.19531 | 37094.72266 | 29898.54688 | 33210.94141 |
| 38658.91406 | 104913.1953 | 130889 | 178417.0469 | 144491.2188 | 120856.1094 | 124424.6016 |
| 80649.95313 | 301133.625 | 300056.4063 | 326014.25 | 264484.9688 | 242760.3594 | 260599.5 |
| 18392.38281 | 77844.38281 | 58832.59766 | 66575.75781 | 44978.61328 | 47119.10938 | 44680.48047 |
| 210740.4688 | 987704.375 | 783405.6875 | 902532.375 | 984277.4375 | 894611.875 | 860925.4375 |
| 11829.16797 | 63710.90625 | 52249.03516 | 50185.65625 | 42510.52734 | 42313.125 | 39866.13281 |
| NA | NA | 1716.193726 | NA | NA | NA | NA |
| 570.5210571 | 9820.87207 | 2170.381836 | 850.6744385 | 4119.791504 | 2754.808105 | 3475.421875 |
| 3474.231689 | 11164.18555 | 9797.65918 | 10892.36621 | 8225.539063 | 6759.402344 | 7522.215332 |
| NA | 8468.53125 | NA | NA | NA | NA | NA |
| 16435.42383 | 85157.60156 | 57354.16016 | 54079.07031 | 69207.46875 | 87257.39844 | 78983.59375 |
| 6262.974609 | 22228.1582 | 15400.4541 | 18752.64844 | 28011.34766 | 20432.32617 | 24160.76953 |
| 263030.7188 | 963706.9375 | 890400.5 | 882814.625 | 449293.6563 | 552414.75 | 504906.6563 |
| 661670.5 | 3246522.5 | 2491535.75 | 2831576 | 2196248.25 | 2335631 | 2423265.25 |
| 9093.264648 | 59197.96875 | 5254.78125 | 5237.415527 | 20046.43359 | 20466.80859 | 22522.16406 |
| 2814.930664 | 11470.27441 | 9464.228516 | 6146.143555 | 7168.12793 | 10280.41016 | 11255.69531 |
| 76126.92188 | 297534.2813 | 248755.3906 | 263855.0313 | 142024.0625 | 192827.5156 | 190171.75 |
| 1052.276611 | 8410.78125 | 6442.224609 | 6962.011719 | 12332.3125 | 11043.11328 | 10496.61426 |
| 16896.69727 | 43034.96094 | 34535.23047 | 34840.83203 | NA | 1092.65271 | 1076.463623 |
| 437.2405701 | 2359.478516 | 1665.028198 | 1753.291016 | 2734.54126 | 2161.666992 | 2219.407715 |
| 9840.098633 | 36594.875 | 35903.88281 | 34021.61328 | 29795.97461 | 31581.45313 | 30744.48828 |
| 15722.68652 | 32267.83008 | 47784.67969 | 40585.70703 | 13006.25684 | 12272.42969 | 12449.18652 |
| 10930.30908 | 39553.12695 | 41413.38184 | 51924.49023 | 60120.5293 | 17179.56201 | 58199.11914 |
| 1697.313232 | 5923.797363 | 5461.01416 | 5862.448242 | 2100.336182 | 2881.072266 | 2405.282471 |
| 11963.97852 | 60748.09375 | 51575.78125 | 61217.01172 | 45798.24609 | 36709.82813 | 36492.91406 |
| 4698.791992 | 16810.19336 | 16524.375 | 15295.39063 | 7238.471191 | 7997.710449 | 6732.848633 |
| 25316.28711 | 77683.98438 | 68666.86719 | 69099.35156 | 51395.40625 | 42146.66406 | 41316.30078 |
| 5451.641602 | 16131.70703 | 11245.37012 | 14106.41406 | 16985.41016 | 16208.13965 | 13571.72852 |
| 4596.744629 | 17029.5918 | 15498.88867 | 13303.4834 | 14973.30566 | 18194.48633 | 14907.22949 |
| 16038.55078 | 65427.13281 | 73221.5625 | 62746.15234 | 23112.37891 | 41556.54297 | 27788.55078 |
| 32950.22266 | 119384.7891 | 104934.8594 | 117216.2266 | 21166.27148 | 22407.26367 | 20045.73828 |
| 9170.433594 | 33487.08984 | 35761.61328 | 34017.27344 | 21017.87891 | 22973.51563 | 21507.9043 |
| 18854.42969 | 73063.60156 | 58930.68359 | 53399.10938 | 24953.26758 | 29916.26953 | 27474.76953 |

| | | | | | | |
|---|---|---|---|---|---|---|
| 19963.23047 | 69891.88281 | 66494.92188 | 67872.90625 | 51414.26563 | 46176.37109 | 48304.62109 |
| 6001.258301 | 28249.73242 | 26869.49609 | 16760.89453 | 4655.550293 | 10105.04102 | 11092.27734 |
| 62643.67969 | 310384.9375 | 236555.4531 | 265122.625 | 339459.1875 | 285670 | 254514.3906 |
| 4756.033203 | 16201.4834 | 13521.22754 | 18025.78516 | 21169.08008 | 25737.54688 | 22829.80859 |
| 12949.79688 | 45594.40234 | 43802.72656 | 41556.07813 | 47517.125 | 49595.98047 | 44643.79688 |
| 16152.51172 | 47218.85547 | 43821.37891 | 55784.31641 | 54572.59375 | 48090.67578 | 50415.25391 |
| 39906.40234 | 150694.2188 | 128336.7656 | 123970.9375 | 94050.19531 | 98407.13281 | 78470.97656 |
| 52938.96094 | 167168 | 187591.7188 | 181561.0156 | 68232.48438 | 76385.21094 | 75622.52344 |
| 88310.54688 | 257221.3438 | 283614.9688 | 329640.9063 | 325737.625 | 371121.6563 | 368514.375 |
| 5163.368652 | 24926.8418 | 20459.93164 | 20320.56055 | 22991.88086 | 20130.45898 | 20011.42578 |
| 168093.9063 | 552640.0625 | 539461.4375 | 576181 | 701235.9375 | 741416.3125 | 727567.125 |
| 5113.315918 | 14603.35547 | 15560.57715 | 18036.72656 | 8295.691406 | 8024.126953 | 7878.799805 |
| 16436.31055 | 55742.53906 | 50262.75391 | 52190.33984 | 56485.42188 | 44249.62109 | 48422.92188 |
| 15850.64746 | 51440.41406 | 44864.76563 | 58127.62891 | 86354.17969 | 59903.21484 | 69811.29688 |
| 34467.54688 | 98260.88281 | 116646.4219 | 107056.7969 | 75366.11719 | 90284.57813 | 92464.64844 |
| 17456.5625 | 60069.17969 | 51767.49609 | 54348.09766 | 38085.18359 | 41354.12109 | 40906.84766 |
| 206813.3125 | 632819.4375 | 656382.3125 | 842786.25 | 876694.4375 | 700310.625 | 829191.375 |
| 34619.88281 | 111482.4766 | 103117.4766 | 107863.1484 | 54220.95703 | 72369.49219 | 60951.66406 |
| 12406.72754 | 38694.36328 | 34612.55859 | 37177.14063 | 42465.12109 | 44419.98438 | 45847.78516 |
| NA | 6810.577637 | 6349.804688 | 4971.436035 | 3550.50293 | 4076.570801 | 3242.958984 |
| 2427.994873 | 8859.348633 | 6813.621582 | 6254.891113 | 5014.610352 | 5336.014648 | 5473.899902 |
| 110805.1563 | 366675.5938 | 335687.375 | 344338.2188 | 232648.8438 | 241708.0469 | 238783.8281 |
| 18099.34766 | 44302.02734 | 49147.30859 | 41041.88672 | 65424.53516 | 80970.84375 | 73244.21094 |
| 14636.83887 | 66249.00781 | 35878.03125 | 41380.79688 | 41504.69922 | 29671.91211 | 33802.76953 |
| 59437.82227 | 224455.9141 | 203107.2188 | 218601.8359 | 133572.6836 | 141590.6367 | 132736.625 |
| 2873.683594 | 12997.85742 | 17236.97266 | 17024.96484 | 19513.04883 | 16696.69336 | 18027.29297 |
| 92362.50781 | 253147.5156 | 271594.625 | 311611.6875 | 283250.6875 | 221107.4375 | 240883.3281 |
| 3420.71582 | 15715.10742 | 2256.28418 | 18039.7207 | 21403.5 | 1492.563599 | 4111.569336 |
| 10914.92676 | 31850.41211 | 27708.25586 | 31235.96094 | 28850.35742 | 28515.57031 | 29480.4375 |
| 2486.503662 | 5050.726563 | 3552.397949 | 6614.158203 | 17734.9668 | 14270.93066 | 15824.2666 |
| 3036.954346 | 11793.15137 | 11275.7334 | 8719.637695 | 7760.040527 | 10107.69336 | NA |
| 1686.041992 | 6731.95166 | 6137.70459 | 6501.322754 | 6379.483398 | 6284.566895 | 6098.717285 |
| 18922.86719 | 57795.32422 | 57923.41406 | 63562.80859 | 64790.92188 | 64226.28906 | 62564.02734 |

| | | | | | | |
|---|---|---|---|---|---|---|
| 5712.323242 | 13340.16895 | 13437.5791 | 14320.52051 | 7498.833984 | 6236.551758 | 6349.388672 |
| 5907.984375 | NA | 45002.4375 | 25298.04102 | 24239.33789 | 15977.54883 | 29371.15039 |
| 21043.75977 | 73242.47656 | 66822.38281 | 69547.25781 | 39223.37109 | 39128.90234 | 41891.67969 |
| 4151.624023 | 14888.42285 | 9979.148438 | 3322.849854 | 12385.31641 | 11278.90137 | 11721.86719 |
| 182.388031 | 2410.34668 | 1180.104736 | 2671.478516 | 9595.418945 | 9213.479492 | 9338.779297 |
| 4222.625488 | 11302.44043 | 3183.150146 | 10367.49316 | 9595.326172 | 7794.685059 | 4421.999512 |
| 2879.839355 | 6164.308105 | 9549.578125 | 9491.799805 | 15283.73242 | 14549.96973 | 23149.97461 |
| 22837.63477 | 59222.58594 | 55382.19141 | 62495.05078 | 30389.18164 | 22719.54297 | 24193.35352 |
| 1898276.125 | 6469991 | 6178341.5 | 7161669.5 | 7465147 | 7502095.5 | 7577360 |
| 267109.1563 | 639263 | 900096.0625 | 1037861.188 | 398361.5625 | 336804.0313 | 397218 |
| 4219.894531 | 11608.31152 | 10894.91211 | 15754.74023 | 29512.05469 | 19669.60938 | 28910.35938 |
| 15138.07617 | 9368.179688 | 71086.64844 | 65761.69531 | 12580.10449 | 17382.00977 | 15936.41016 |
| 3814.301758 | 12877.17578 | 11106.5957 | 7847.947754 | 11963.93457 | 15867.86328 | 11945.52148 |
| 17640.57813 | 63611.23828 | 60168.01953 | 79813.48438 | 108662.6094 | 84107.17969 | 89810.3125 |
| 15831.86426 | 34606.96094 | 16958.60352 | 39374.85156 | 67325.20313 | 206110.0469 | 35719.91016 |
| 284900.8125 | 857065.125 | 709645.6875 | 794463.375 | 595515.25 | 600592.8125 | 578033.8125 |
| 57209.88672 | 162520.0625 | 161042.6406 | 137038.9219 | 58249.33594 | 50198.12891 | 54705.41797 |
| 910407.3125 | 3554671.75 | 2639447.25 | 2996599.25 | 2834444.5 | 3084028 | 3026951 |
| 23468.02148 | 69277.17188 | 68380.4375 | 70830.03125 | 78699.99219 | 71476.88281 | 73406.07031 |
| 13171.37695 | 47596.15625 | 40836.65234 | 45053.75391 | 76602.1875 | 63691.33203 | 73185.74219 |
| 1711.071655 | 6324.944824 | 6490.952637 | 5557.637695 | 3556.179688 | 4004.239014 | 3773.579346 |
| 4059.218018 | 17286.1582 | 6216.004395 | 4899.00293 | 13771.65332 | 16122.81445 | 14629.9248 |
| 37155.05859 | 157877.8906 | 156366.7344 | 146591.7031 | 94813.39844 | 116751.3203 | 124844.2656 |
| 32188.14648 | 97216.53906 | 88344.70313 | 88960.9375 | 96535.96094 | 90482.13281 | 85150.96875 |
| 5665.750488 | 13571.58496 | 10350.86426 | 12385.06641 | 29770.60352 | 11356.03125 | 22307.19727 |
| 4456.441406 | 11667.01367 | 16379.32324 | 23612.49805 | 64318.25 | 55419.28125 | 73375.67188 |
| 15159.97461 | 48347.04688 | 54043.47656 | 53773.21094 | 93046.22656 | 59839.26953 | 73603.64844 |
| 51723.63672 | 544873.375 | 248635.9844 | 304787.6875 | 1930759.125 | 569569.4375 | 940936.5625 |
| 1250.311523 | 3634.88501 | 2743.698486 | NA | 2197.87915 | 2377.956787 | 9602.017578 |
| 3829.858398 | 11512.91016 | 10856.62305 | 11267.01074 | 11637.21973 | 11525.59277 | 10296.14648 |
| 86362.25 | 270975.25 | 260388.0469 | 320196 | 289894.2188 | 292931.5625 | 329028.0625 |
| 3735.354004 | 14370.07324 | 10219.4502 | 9469.698242 | 14568.33105 | 9122.507813 | 14312.58105 |
| 6137.50293 | 14786.27441 | 22396.59961 | 19293.60938 | 19064.33398 | 17309.90234 | 20694.98828 |

| | | | | | | |
|---|---|---|---|---|---|---|
| 7759.445313 | 32609.16211 | 16049.41309 | 19154.88672 | 29094.29492 | 23687.13672 | 27500.19531 |
| 85625.03125 | 246201.5156 | 237262.0938 | 246831.1563 | 307322.25 | 245327.9844 | 262465.2188 |
| 10254.10742 | 34430.91797 | 33812.16797 | 34129.05469 | 34979.5 | 38179.1875 | 33381.51953 |
| 370170.5 | 1051817.875 | 1124381 | 1438262.875 | 1372933 | 1272024.625 | 1419542.625 |
| 25344.4668 | 95537.375 | 95724.42188 | 84598.39063 | 85158.82031 | 77125.83594 | 76893.02344 |
| 21802.1875 | 74997.59375 | 71857.4375 | 57702.36719 | 18747.14648 | 22019.88672 | 17466.94531 |
| 159852.7813 | 469928.6563 | 435972.5938 | 453053.5 | 368064.5313 | 314254.2813 | 345536.2813 |
| 65080.54688 | 192067.8594 | 182220.1094 | 182371.4531 | 151187.9531 | 153637.0625 | 147630.6094 |
| 24439.5918 | 69211.05469 | 65663.35938 | 77412.25781 | 109969.3359 | 89213.8125 | 93341.34375 |
| 5582.424316 | 13484.87012 | 12760.0293 | 11544.38672 | 12654.45508 | 11793.43555 | 13170.37988 |
| 1140.296875 | 1762.652588 | 3287.766113 | 3249.02124 | 4453.808105 | 3907.443359 | 4164.92041 |
| 4460.703125 | 13276.20605 | 6140.668945 | 9704.546875 | 10489.19043 | 10643.62598 | 8969.256836 |
| NA | 3903.677734 | 2944.195801 | 4107.397949 | 23327.70508 | 21559.65234 | 21244.16602 |
| 37724.375 | 119021.0547 | 116090.7266 | 102276.2891 | 143799.5313 | 144912.8281 | 141763.75 |
| 29684.55664 | 98389.125 | 82356.72656 | 85189.45313 | 78347.01563 | 110388.0938 | 86453.32813 |
| 6364.384766 | 20269.98242 | 20895.20508 | 20130.9668 | 17865.67969 | 22407.60547 | 22626.55859 |
| 57710.30859 | 181184.8125 | 161206.3125 | 165884.2969 | 154081.9844 | 143172.375 | 140341.0781 |
| 25664.77539 | 66787.14844 | 63790.18359 | 59220.08984 | 51152.99609 | 42471.72656 | 44877.17578 |
| 42662.00391 | 99947.26563 | 88163.25 | 105636.9688 | 109587.2969 | 118143.8359 | 144441.6094 |
| 3358.681885 | 15848.34375 | 2643.94873 | 4981.930664 | 16723.4668 | 29175.98047 | 22517.92383 |
| 25053.81836 | 65453.14844 | 70050.40625 | 70042.46094 | 52615.57031 | 62281.62891 | 55443.51953 |
| 43709.72266 | 115439.2813 | 123443.5703 | 102149.1328 | 88168.60938 | 110362.0781 | 104782.8828 |
| 8171.946289 | 26205.10742 | 22866.86914 | 17153.49023 | 11927.92773 | 12681.69238 | 9969.271484 |
| 11032.66797 | 24753.66602 | 22694.93555 | 17421.72656 | 27322.63672 | 21958.41406 | 23618.72852 |
| 39925.89453 | 96017.65625 | 56820.46484 | 58151.08203 | 103010.2109 | 116237.375 | 107136.8906 |
| 4235.824707 | 11348.6748 | 11148.69043 | 13876.97559 | 12555.05273 | 12797.875 | 12573.82129 |
| 4183.236816 | 11816.96094 | 10987.56543 | 13545.16992 | 17486.88477 | 19180.19336 | 16432.68164 |
| 180832.7031 | 392699.3438 | 627845 | 631714.375 | 395776.9375 | 354936.375 | 401017.6563 |
| 5250.359863 | 12824.12793 | 13875.18555 | 13918.4043 | 26748.88477 | 14286.3916 | 17986.76172 |
| 6035.28125 | 16539.86133 | 15871.56836 | 15377.84961 | 15033.49023 | 12044.875 | 12821.2334 |
| NA | 2217.688232 | 2668.465332 | 3438.791016 | 3638.114746 | 2704.341797 | 2590.217285 |
| 3062.679932 | 7810.359863 | 8838.786133 | 8600.875977 | 8373.871094 | 8790.616211 | 8291.609375 |
| 49009.12891 | 97483.47656 | 86400.57813 | 193360.6875 | 468778.1875 | 135292.5938 | 197017.9063 |

| | | | | | | |
|---|---|---|---|---|---|---|
| 20622.66992 | 66792.63281 | 52916.09375 | 60735.48047 | 49424.37891 | 48490.87891 | 47530.30078 |
| 33199.21875 | 101314.8203 | 101386.0938 | 96027.79688 | 73657.02344 | 75521.64063 | 66893.35156 |
| 9961.224609 | 18285.88086 | 27605.89844 | 23755.5 | 22170.4707 | 15173.43164 | 18508.76563 |
| 13133.46875 | 32321.99609 | 40568.95703 | 31132.50391 | 34482.12109 | 13968.86426 | 40151.52734 |
| 5102.040039 | 15275.32324 | 16149.7998 | 11398.06445 | 13868.06543 | 19410.78516 | 16469.76367 |
| 31635.26172 | 72725.27344 | 57664.04688 | 82679.10156 | 73134.69531 | 93357 | 90793.57813 |
| 60170.64453 | 170613.0156 | 174881.7969 | 177099.5313 | 192935.375 | 213695.625 | 216109.4375 |
| 5013.583984 | 13141.42578 | 14706.44238 | 10128.0957 | 4804.506836 | 5117.682129 | 4071.75293 |
| 44018.82813 | 149764.4219 | 221584.6563 | 241565.4063 | 234612.3594 | 133528.9844 | 132248.5 |
| 2315.150146 | 5241.402832 | 4616.551758 | 5809.85791 | 12417.37891 | 10510.8877 | 12541.91602 |
| 11163.67285 | 32400.49609 | 34276.75781 | 29761.52148 | 25510.75586 | 27983.32617 | 24150.18164 |
| 5836.836914 | 18204.27393 | 18014.47021 | 23171.48438 | 15567.28345 | 10498.29688 | 8535.395508 |
| 19977.52344 | 59634.90234 | 48199.51172 | 56024.38672 | 56565.74609 | 51965.24609 | 49333.86719 |
| 9350.421875 | 34905.59766 | 27481.83789 | 26659.21289 | 16305.33691 | 20464.41797 | 25604.19141 |
| 1342.173828 | 7005.932129 | 3423.481201 | 2860.023682 | 1926.453491 | 5078.663574 | 2605.227295 |
| 13412.56836 | 39146.68359 | 36392 | 46871.86719 | 143009.6875 | 82144.23438 | 92833.92188 |
| 2141.439209 | 10740.69824 | 2177.678955 | 3974.049072 | 4664.898438 | 10401.5791 | 6612.585938 |
| 27067.15039 | 81545.82813 | 66745.24219 | 75175.80469 | 45528.35938 | 44602.89063 | 41586.6875 |
| 8619.633484 | 30455.14185 | 22934.81689 | 25644.57471 | 39160.79297 | 31450.57056 | 33160.91626 |
| 9496.491211 | 25731.11719 | 32471.61914 | 31601.82031 | 26696.45703 | 23986.66406 | 22838.34961 |
| 2939.491943 | 12521.33496 | 9170.435547 | 10668.52734 | 12165.3623 | 10249.87598 | 12318.48926 |
| 497621.7188 | 1368921.125 | 1436424.5 | 1544215 | 1959372.375 | 1834165.75 | 1814915.125 |
| 35298.53906 | 140327.7031 | 128410.0391 | 129135.9922 | 211347.1563 | 161756.9219 | 169033.5156 |
| 5769.39502 | 17142.53125 | 18392.47656 | 14412.62891 | 15051.94043 | 13676.08203 | 12937.87012 |
| 2827.476318 | 10016.68164 | 4827.967773 | 3407.870605 | 5324.762207 | 5857.39502 | 3744.420654 |
| 9428.575195 | 30016.26953 | 26583.56055 | 27762.98438 | 37495.22656 | 28185.59766 | 32997.375 |
| 43805.4375 | 144904.75 | 124021.6875 | 115489.7813 | 144464.3125 | 149624.1094 | 134612.2344 |
| 3347.115723 | 8859.524414 | 8639.160156 | 10749.01367 | 5384.217285 | 3787.14502 | 4624.708984 |
| 38379.60938 | 73093.53906 | 56215.82422 | 86270.05469 | 142992.4063 | 134818.8906 | 131183.5781 |
| 15441.28125 | 22346.0293 | 45055.47656 | 47645.34766 | 27217.69922 | 27519.53906 | 28548.07227 |
| 26620.51563 | 98849.71875 | 72908.1875 | 87056.90625 | 75039.38281 | 71555.53125 | 74718.44531 |
| 5051.085449 | 10613.25684 | 13881.38672 | 14422.35352 | 14251.85156 | 11156.65527 | 15369.08789 |
| 2316.223877 | 6722.847168 | 6721.829102 | 8587.061523 | 8393.805664 | 6971.246094 | 8032.593262 |

| | | | | | | |
|---|---|---|---|---|---|---|
| 20825.79297 | 49064.53125 | 51203.36719 | 50420.41406 | 15871.22656 | 16626.66797 | 14019.45996 |
| 2193.955322 | 10424.5332 | 4217.299805 | 4121.30957 | 6396.129395 | 6037.080566 | 4598.058105 |
| 14835.64746 | 35979.06641 | 38529.79688 | 37645.39453 | 26403.41797 | 25493.65625 | 25636.55078 |
| 9388.291992 | 37431.25 | 35408.88281 | 25402.73633 | 44691.25 | 40049.07813 | 37842.37891 |
| 48659.12891 | 126630.4844 | 109592.3906 | 123051.8984 | 174483.8438 | 166951.1719 | 154384.1719 |
| 25965.60352 | 63722.14453 | 66347.86719 | 61784.60547 | 54674.71484 | 61530.33594 | 64294.32422 |
| 16279.27246 | 33361.29688 | 37640.72266 | 37164.44922 | 18560.88281 | 27341.75781 | 23354.89453 |
| 23062.84375 | 67552.15625 | 54868.08984 | 66480.13281 | 24894.30078 | 18872.0625 | 21096.02344 |
| 16558.13477 | 48172.83984 | 41911.33203 | 38780.93359 | 29358.44336 | 31410.52539 | 28960.7832 |
| 226439.625 | 783843 | 536961.0625 | 465528 | 279704.8125 | 444110.3125 | 314289.0625 |
| 8861.732422 | 17084.17773 | 15984.87207 | 17930.01172 | 14568.62109 | 24980.17969 | 16842.1543 |
| 3643.664795 | 8185.61377 | 8372.055664 | 9506.086914 | 13940.69824 | 16596.10547 | 14014.2998 |
| 1072864.625 | 5152104 | 1888425.25 | 960864.75 | 1644906.875 | 511643.6875 | 776609.625 |
| 4173.075684 | 15936.62207 | 15094.10059 | 15181.44141 | 8457.833008 | 7110.20459 | 6262.576172 |
| 6445.190918 | 20772.62109 | 18747.39063 | 17182.3125 | 10586.58008 | 12215.66309 | 12229.11523 |
| 31295.24023 | 90594.57813 | 71747.20313 | 96354.25 | 122825.5547 | 115139.5313 | 111801.5859 |
| 19669.88867 | 65896.01563 | 62691.32813 | 72098.15625 | 52141.44531 | 42728.26563 | 41017.25 |
| 3820.632568 | 6129.514648 | 8218.436523 | 10017.92676 | 10615.47363 | 11755.52441 | 15589.87598 |
| 86719.54688 | 321136.3438 | 212439.9531 | 215435.25 | 231730.3125 | 238405.6563 | 214980.1406 |
| 53901.23438 | 121742.5469 | 123348.5781 | 142221.9531 | 89492.86719 | 101930.5313 | 90186.96094 |
| 30326.71875 | 75778.0625 | 70999.10156 | 71336.07031 | 52992.34766 | 47633.71094 | 47820.73828 |
| 18054.46289 | 43641.71875 | 42267.77734 | 54770.49219 | 81412.55469 | 58159.64063 | 64428.07813 |
| 440.4890442 | 1814.62793 | 2397.009766 | 3328.986572 | 834.0512695 | 1791.285522 | 1091.932373 |
| 4050.610352 | 15395.3291 | 11865.03027 | 3067.466797 | 9091.40332 | 22393.12305 | 16751.00586 |
| 5124.853027 | 18701.80273 | 13641.30664 | 13936.74023 | 18803.90234 | 15652.06934 | 13634.10059 |
| 2141.845703 | 11266.97168 | 7869.024414 | 8467.323242 | 5019.757324 | 7919.499023 | 6669.890137 |
| 15483.72168 | 22816.71289 | 24621.44727 | 25393.34961 | 20291.13281 | 20784.2793 | 21769.08203 |
| 63963.23047 | 152626.3125 | 145059.9219 | 140115.75 | 194962.1563 | 224017.2188 | 211741.4375 |
| 6551.150879 | 15771.67969 | 14363.04297 | 14756.44727 | 14365.94043 | 12882.02344 | 14837.71582 |
| 25411.82617 | 82985.10156 | 63933.01563 | 82332.67969 | 208143.6719 | 195760.6719 | 188018.3906 |
| 7783.515137 | 21308.55273 | 19970.05469 | 16835.62109 | 8821.663086 | 7506.757813 | 8039.844238 |
| 13533.9209 | 35168.83984 | 40080.01953 | 44756.49609 | 43996.83594 | 31436.23242 | 34437.80859 |
| 297914.9375 | 721462 | 691061.25 | 794989.4375 | 1372602.625 | 1121574.625 | 1228998.25 |

| | | | | | | |
|---|---|---|---|---|---|---|
| 3492.983887 | 10732.29492 | 8383.082031 | 7193.265137 | 3400.822021 | 5361.196777 | 3813.800293 |
| 32483.33789 | 95167.03125 | 74454.32813 | 70636.82031 | 46199.18359 | 70485.04688 | 50109.10938 |
| 11504.34863 | 34508.76172 | 28369.27148 | 34639.51563 | 36077.28125 | 21485.75391 | 29259.30859 |
| 34847.90234 | 82970.98438 | 85934 | 94988.29688 | 94642.17969 | 90841 | 93272.47656 |
| 22651.96289 | 54753.51172 | 50028.87109 | 53592.85938 | 40479.09766 | 38601.77734 | 37856.14844 |
| 481608.5313 | 1112607 | 1260200.875 | 1543473.25 | 1483846.875 | 1354563.75 | 1553714.75 |
| 8847.797852 | 22915.08008 | 22057.16797 | 23422.85156 | 18380.74414 | 19739.75195 | 18397.375 |
| 126984.7422 | 331229.625 | 368999.6563 | 316373.7813 | 208530.7656 | 232468.2813 | 231636.4844 |
| 82838.75 | 254926.6406 | 200500.7813 | 227328.2656 | 314396.75 | 315141.3438 | 290914.3125 |
| 5785.060059 | 8391.535156 | 9609.401367 | 16739.36523 | 18285.00586 | 14437.0293 | 16693.68555 |
| 8582.275391 | 14902.14355 | 30647.36914 | 28014.61719 | 16205.32422 | 11886.72949 | 19753.36914 |
| 71017.13281 | 145999.3906 | 176682.2344 | 176934.6563 | 106382.1484 | 107165.5781 | 124079.7422 |
| 1094.926636 | 4020.314453 | 3241.141846 | 2778.883789 | 17090.39063 | 6172.177246 | 7141.051758 |
| 4017.589844 | 8768.476563 | 6259.612793 | 6963.624023 | 12041.50879 | 10933.48145 | 11257.33691 |
| 9957.316406 | 25845.08789 | 24671.53711 | 26070.20703 | 22466.34375 | 23100.92578 | 23936.19531 |
| 17326.01758 | 46821.59375 | 41138.48828 | 38204.46094 | 42011.55859 | 41182.875 | 42635.24219 |
| 6032.435059 | 21600.61719 | 11174.49023 | 12661.41895 | 18648.125 | 10486.93945 | 11949.60645 |
| 4591.79834 | 13098.21582 | 10186.91797 | 8527.636719 | 14894.75391 | 16708.22852 | 11972.31836 |
| 11507.66797 | 26439.64844 | 25871.50391 | 28076.07617 | 61514.01953 | 43223.54297 | 48071.07031 |
| 16132.09277 | 30120.62695 | 31158.94531 | 34124.74219 | 17317.08594 | 20945.14258 | 17411.21875 |
| 5696.450684 | 12316.73535 | 13341.55566 | 13999.14258 | 19689.5332 | 14575.06836 | 15625.45313 |
| 4321.430664 | 26000.94727 | 15483.23145 | 12734.25195 | 7298.616211 | 9481.652344 | 8770.517578 |
| 435.5675354 | 1455.18457 | 988.4813843 | 855.6450806 | 1979.866333 | 1857.434448 | 1616.341064 |
| 6542.63916 | 15565.14063 | 18408.36719 | 14933.32031 | 12416.80859 | 8231.476563 | 14559.00586 |
| 7122.237793 | 15288.53906 | 14851.93066 | 12783.12109 | 7351.956055 | 10014.40918 | 7170.367676 |
| 1404.751099 | 990.2688599 | 6328.021484 | 3413.206787 | 2402.568115 | 4577.893066 | 3255.014648 |
| 1643.345947 | 3305.21167 | 4917.061035 | 5716.932129 | 4834.510254 | 5812.982422 | 5108.265625 |
| 91171.97656 | 235147.5781 | 166034.8125 | 194647.0469 | 216159.4063 | 234210.6875 | 209370.4063 |
| 354228.2188 | 1003817.375 | 796677.75 | 730638.875 | 612260.125 | 651383.625 | 596770.9375 |
| 141183.0781 | 398627.0313 | 373058.0938 | 366572.9375 | 254002.5781 | 363984.5625 | 271300.3125 |
| 20125.20313 | 33780.45703 | 15428.79004 | 36200.78125 | 31417.25195 | 42072.48047 | 28979.75 |
| 11202.69238 | 30863.92383 | 29083.63281 | 25765.05078 | 44659.55859 | 40915.41406 | 44483.625 |
| 22340.52344 | 67830.03906 | 65774.57813 | 74182.97656 | 112980.6953 | 94754.50781 | 102030.6328 |

| | | | | | | | |
|---|---|---|---|---|---|---|---|
| | 5218.034668 | 19270.60742 | 13698.51953 | 10711.70117 | 8429.594727 | 13146.87598 | 8324.188477 |
| | 14520.12207 | 33167.96875 | 30990.62109 | 34464.03125 | 30176.11914 | 27051.57227 | 26686.23828 |
| | 183002.7969 | 409667.8438 | 382279.7813 | 454415.3125 | 463108 | 380524.7813 | 430677.5625 |
| | 6014.48877 | 16011.56934 | 15833.12207 | 17028.7793 | 12628.96387 | 10368.69727 | 10200.13184 |
| | 34000.13281 | 73609.22656 | 95946.32031 | 49838.40625 | 55595.10547 | 100330.6094 | 57405.53906 |
| | 1883.970093 | 8770.991211 | 5272.862305 | 4387.102051 | 6314.547852 | 5351.303223 | 4740.974609 |
| | 5265.182129 | 18844.02344 | 15470.05566 | 18299.0293 | 25759.30273 | 32883.4375 | 24282.14453 |
| | 8095.623535 | 16717.32813 | 17636.52148 | 15557.94043 | 12290.4668 | 10071.78906 | 10010.83984 |
| | 23197.26758 | 51681.17969 | 64224.44141 | 50046.26953 | 23488.99219 | 28540.62695 | 25523.4668 |
| | 10012.6875 | 22009.91211 | 20803.06055 | 21272.99219 | 9680.244141 | 12254.64551 | 12334.78027 |
| | 1098.635864 | 1575.433594 | 3964.161377 | 4188.52832 | 6459.181641 | 3097.835449 | 3324.563232 |
| | 4212.194824 | 13618.60547 | 11000.66895 | 7942.84082 | 8537.152344 | 8608.46875 | 7781.291504 |
| NA | | 6043.470215 | 4492.308594 | 4482.47998 | 10970.38379 | 10137.70703 | 8545.664063 |
| | 15326.55859 | 31185.82227 | 28446.40234 | 30375.76563 | 41336.98828 | 61954.33984 | 42084.87891 |
| | 12120.57715 | 24326.62109 | 25458.3125 | 22458.41016 | 17983.54492 | 18097.48242 | 16901.64648 |
| | 5517.859375 | 11231.32715 | 11166.82813 | 12796.60547 | 11760.13379 | 11664.36426 | 11634.37695 |
| | 8050.249023 | 21372.99609 | 15161.01074 | 15301.62305 | 22046.52734 | 23085.34961 | 22750.62891 |
| | 6612.772949 | 16419.70508 | 14391.33398 | 14596.05664 | 13420.95703 | 11234.01758 | 11436.45508 |
| | 18721.57422 | 40557.40234 | 35182.14844 | 42047.46484 | 11694.27832 | 10231.78223 | 13570.46777 |
| | 43801.49609 | 89395.21875 | 67789.66406 | 79465.3125 | 62311.73438 | 75338.15625 | 73490.375 |
| | 6171.928223 | 13283.31152 | 7354.808105 | 12932.87207 | 8018.142578 | 11349.30566 | 8932.152344 |
| | 3490.939941 | 7051.050781 | 8271.945313 | 7622.489746 | 4982.871094 | 5330.771484 | 5313.389648 |
| NA | | 9631.015625 | 11777.78516 | 11580.34668 | 13963.85547 | 13097.69531 | 16529.16406 |
| | 35295.10156 | 51761.52344 | 59590.15625 | 89389.09375 | 178727.0938 | 195334.1406 | 197840.1563 |
| | 10475.06152 | 26515.86914 | 24464.37891 | 25770.7793 | 45026.35547 | 42066.73438 | 43219.28516 |
| | 34140.97266 | 78328.96094 | 77078.03125 | 84523.51563 | 92631.73438 | 71772.78906 | 77719.07031 |
| | 4019.676514 | 10100.55566 | 5492.984863 | 4900.825684 | 10324.43164 | 10073.86523 | 9448.600586 |
| NA | | 2202.323486 | 1136.68396 | 3484.288818 | 1783.095337 | 4694.37207 | 2246.27002 |
| | 9724.863281 | 27366.95508 | 21134.62109 | 25324.15625 | 43895.01172 | 35145.91406 | 31651.97656 |
| | 19654.43164 | 48973.10547 | 50175.95703 | 60553.03906 | 76176.45313 | 78305.41406 | 79955.79688 |
| | 29400.35938 | 70543.14844 | 62943.54297 | 72002.86719 | 43879.96875 | 38587.89844 | 39190.89844 |
| | 24256.72266 | 62892.82031 | 53041.65234 | 46619.10156 | 46649.96875 | 52396.72656 | 47242.76172 |
| | 7358.248535 | 17707.66602 | 18660.57422 | 16193.15137 | 12358.47754 | 11177.68848 | 11591.93848 |

| | | | | | | |
|---|---|---|---|---|---|---|
| 20516.7168 | 41984.11719 | 51949.89844 | 40192.12109 | 51210.89844 | 35550.51953 | 40096.76172 |
| 47574.125 | 105255.8984 | 114719.7891 | 127164.2734 | 98058.00781 | 83430.14844 | 92798.35156 |
| 2809.491211 | 7566.913086 | 5843.241699 | 6786.491211 | 8434.90918 | 6064.700684 | 5363.057129 |
| 4240.491699 | 14535.63379 | 10443.71289 | 11027.20703 | 8330.870117 | 13052.75293 | 11066.58789 |
| 23934.27734 | 59444.68359 | 52955.05859 | 56849.88281 | 29092.22852 | 26177.33789 | 26978.80469 |
| 3255.577637 | 9720.102539 | 8838.170898 | 10013.06152 | 13357.04785 | 11933.96777 | 9798.063477 |
| 25143.37891 | 135765.9688 | 33050.03125 | 52827.19531 | 48064.37109 | 103514.3516 | 60842.08203 |
| 25666.38086 | 72204.5625 | 56234.72266 | 59671.75 | 65597.07031 | 57856.49219 | 62785.19141 |
| 3060.356934 | 8408.020508 | 7113.030273 | 9967.46875 | 6493.213379 | 4542.10498 | 3865.44165 |
| 551927.75 | 1100631.875 | 1195940.75 | 1165916.125 | 376803.9375 | 397823.3125 | 389754.1563 |
| 102410.75 | 217577.6406 | 234198.5938 | 236200.7969 | 74314.26563 | 69853.66406 | 63554.91016 |
| 111053.1094 | 251487.125 | 220731.7813 | 236400.5781 | 128496.75 | 95845.75781 | 116187.1641 |
| 7011.210938 | 42035.43359 | 22571.68164 | 16611.94141 | 27744.39844 | 29618.99414 | 18186.97461 |
| 15408.34375 | 27642.90234 | 27360.29883 | 34512.57422 | 39865.44922 | 39067.91406 | 44847.58984 |
| 23171.57617 | 62436.66406 | 44526.16797 | 53594.04297 | 54916.83594 | 46112.34375 | 46904.62891 |
| 910.0238037 | 15313.29395 | 13845.72559 | 9947.526367 | 6128.643066 | 7825.443359 | 2892.37085 |
| 42229.82031 | 106461.8203 | 86425.74219 | 95716.73438 | 90812.14063 | 100387.1875 | 102840.9766 |
| 90254.92188 | 205092.3438 | 196427.1719 | 200038.4531 | 199530 | 190466.5469 | 188275.4375 |
| 20565.99609 | 60856.73438 | 63890.45703 | 61893.89453 | 47205.57031 | 49435.17578 | 47304.41797 |
| 721202 | 797258.3125 | 1540067.625 | 1741099.375 | 944337 | 1012327.75 | 1204224 |
| 21477.76172 | 73747.54688 | 43500.69141 | 38666.46875 | 36194.23047 | 37265.29297 | 33758.50781 |
| 911247.875 | 1717909.625 | 1824319.875 | 1777627.5 | 1353320.375 | 1359316.25 | 1397102.125 |
| 11430.66992 | 43135.44922 | 23700.75586 | 39704.07422 | 27077.57617 | 24570.57227 | 27166.59375 |
| 245016.8281 | 540753.6875 | 523313.9375 | 533566.25 | 673615.4375 | 641795.125 | 642166 |
| 104413.7305 | 214085.4688 | 213308.5234 | 199573.1094 | 160850.9063 | 188918.625 | 183080.5938 |
| 314105.2188 | 627318.25 | 738701.1875 | 733589.4375 | 491463.4375 | 453083.875 | 493062.0938 |
| 164776.9688 | 877452.875 | 237127.5469 | 102137.5391 | 118703.2578 | 63964.05469 | 62857.54297 |
| 1899.456177 | 3288.796143 | 4772.261719 | 4443.919434 | 3991.453613 | 3564.129639 | 4127.072754 |
| 3895.951416 | 14303.64941 | 8172.538086 | 3116.100098 | 2658.929688 | 6421.068848 | 2705.740723 |
| 21092.69873 | 63135.52539 | 35916.88672 | 38083.73926 | 76655.79688 | 63210.81055 | 65273.75 |
| 12942.92383 | 35152.52734 | 30936.70508 | 32447.80469 | 29523.00977 | 29021.66992 | 26502.57813 |
| 9132.273438 | 25407.53711 | 25992.32617 | 30507.77734 | 43027.54297 | 43785.51953 | 41556.23828 |
| 12836.04785 | 34518.33203 | 26927.08594 | 28707.91992 | 16637.46289 | 15464.71582 | 16272.04102 |

| | | | | | | |
|---|---|---|---|---|---|---|
| 5634.619629 | 11974.67383 | 14098.56641 | 14678.79883 | 11887.5625 | 12629.59082 | 10850.90234 |
| 10344.42871 | 25167.07227 | 18168.17188 | 16036.79102 | 15634.23145 | 12998.32129 | 17040.76563 |
| 2077726.375 | 3584685.5 | 2799618.25 | 2484062 | 3047812.25 | 3605288.25 | 3130054.75 |
| 2633.836914 | 6792.01416 | 6372.209961 | 6000.976074 | 7319.13916 | 6294.537598 | 6589.27832 |
| 121084.8438 | 244135.3906 | 271467.2813 | 326836.5313 | 242218.9844 | 228079.0469 | 235014.8594 |
| 302479.1875 | 689569.125 | 642230 | 642058.375 | 590440.25 | 651527.625 | 583759.1875 |
| NA | 5266.991699 | 3110.946045 | 3421.068115 | 3878.150635 | 3461.560303 | 2726.764648 |
| 21884.46484 | 36528.40234 | 43669.98438 | 48154.66797 | 69845.32813 | 51030.46875 | 59249.9375 |
| 34996.91797 | 100161.7344 | 72585.22656 | 60140.05078 | 82062.91406 | 80284.45313 | 87310.96875 |
| 17098.49414 | 39282.625 | 37381.44141 | 38563.22656 | 48829.90625 | 47024.56641 | 45247.45703 |
| 23707.2168 | 56896.87109 | 43474.66406 | 48835.94141 | 53624.71484 | 44596.17188 | 49813.375 |
| 15814.64258 | 30272.1543 | 34137.19531 | 30805.52344 | 23207.12891 | 26355.51563 | 26826.60547 |
| 2711.0979 | 5726.457031 | 4351.193848 | 5504.470215 | 6284.668457 | 8643.453125 | 7768.13916 |
| 118523.1406 | 282617.4375 | 218462.7344 | 282210.9375 | 301947.3438 | 277711.625 | 299089.9375 |
| 2097.868652 | 4600.547852 | 4189.182129 | 6266.448242 | 3988.927002 | 3006.269287 | 3716.650879 |
| 9367.682617 | 22481.94727 | 22906.1875 | 24071.37695 | 23653.82422 | 23109.05273 | 24325.41211 |
| 22209.61328 | 44731.09766 | 50128.81641 | 48767.85156 | 34046.74219 | 34586.10938 | 34339.84766 |
| 65637.89844 | 134950.2656 | 126280.1265 | 139847.6411 | 236070.7617 | 221889.4141 | 186490.5957 |
| 260883.2656 | 494035.9375 | 625767.9375 | 798645.625 | 685931.6875 | 559009.25 | 690112.3125 |
| 30069.56445 | 62098.33203 | 55459.44141 | 58540.07031 | 34247.89453 | 34692.01563 | 34728.62109 |
| 13899.36914 | 28282.55273 | 26757.78516 | 27403.24805 | 28624.33789 | 26852.76172 | 25923.37109 |
| 3053.978516 | 6190.786621 | 4578.273926 | 5275.179688 | 3667.412598 | 2168.270996 | 2006.010376 |
| 24121.36194 | 91334.57227 | 23103.86914 | 24173.35913 | 124573.0811 | 54162.54199 | 67790.63965 |
| 3265.937744 | 5816.870117 | 6104.481934 | 5699.463379 | 5735.831543 | 4636.120117 | 4765.626953 |
| 2613.369629 | 30138.4375 | 32978.07031 | 41450.98438 | 19168.81836 | 16742.04883 | 21680.54297 |
| 27465.34961 | 60444.88672 | 57694.78516 | 49497.62891 | 53395.14453 | 58772.05859 | 55215.83594 |
| 7059.043945 | 15624.26758 | 14506.78711 | 17387.57031 | 15595.36133 | 14091.85352 | 14782.52051 |
| 657.2623291 | 1146.850098 | 1108.267944 | 865.7731934 | 1246.52002 | 1534.977173 | 1184.739136 |
| 4121.51416 | 8075.513672 | 10288.11914 | 8974.645508 | 11843.95605 | 13717.62891 | 13399.97266 |
| 3172.068115 | 7691.02832 | 5980.371094 | 9904.001953 | 5217.710938 | NA | 5720.38916 |
| 40824.85547 | 77522.09375 | 82239.89844 | 92737.47656 | 138684.9688 | 144345.7031 | 148661.5469 |
| 10623.4502 | 21129.89453 | 18746.64844 | 18492.33008 | 10890.79492 | 10154.10156 | 9578.060547 |
| 147400.7969 | 307809.25 | 285676.4688 | 346226 | 384095.4688 | 343131.875 | 348085.8438 |

| | | | | | | |
|---|---|---|---|---|---|---|
| 10444.51855 | 18613.91211 | 23258.68359 | 19348.31641 | 14529.00879 | 15967.13867 | 15076.32715 |
| 9901.416992 | 19300.60156 | 18161.52734 | 21996.30078 | 23344.40039 | 16638.46484 | 17846.69336 |
| 138967.2031 | 249841.1094 | 293759.7813 | 300580.625 | 235199.7344 | 229004.2031 | 262952.1875 |
| 18793.64258 | 34337.91016 | 44155.22656 | 44474.75 | 30457.98438 | 29869.08594 | 26696.91797 |
| 8418.920898 | 19708.17188 | 18752.24414 | 18549.53516 | 24983.13281 | 21977.77148 | 22743.11133 |
| 5156742 | 13868761 | 10125387 | 9184307 | 7648462 | 7910456 | 8040302 |
| 7367.548828 | 27239.69727 | 4024.695557 | 1803.459229 | 16918.48242 | 13858.05566 | 6736.811523 |
| 17210.63672 | 34236.62109 | 36740.05859 | 35387.33984 | 33662.29688 | 34490.77734 | 29589.51758 |
| 54805.88281 | 121641.8359 | 124562.1172 | 128719.7734 | 113193.1484 | 93786.71094 | 100630.9922 |
| 56156.95313 | 99450.49219 | 112063.1328 | 112813.2734 | 93786.10938 | 107705.3359 | 102489.8203 |
| 348014.2188 | 786272.5 | 715462.125 | 805752 | 1001224.813 | 838247.625 | 892683.875 |
| 2748.662842 | 5000.818359 | 4906.515625 | 4441.26416 | 1957.671265 | 2337.283447 | 1847.648315 |
| 3077.169434 | 8521.3125 | 8982.787109 | 8939.40625 | 7445.327148 | 7842.975098 | 8823.262695 |
| 7471.491699 | 8632.442383 | 20622.58594 | 10521.15723 | 10546.63184 | 11644.68359 | 7888.165039 |
| 911.7093506 | 1881.404175 | 2184.737793 | 1625.333252 | 1529.478149 | 1596.394653 | 2381.080078 |
| 2384081.5 | 4551310 | 3797019.5 | 3120389.25 | 5920195 | 6765214.5 | 6652645.5 |
| 4170.277344 | 8992.439453 | 8989.451172 | 7973.412109 | 7093.672852 | 9356.56543 | 8801.617188 |
| 97078.24219 | 203976.5 | 221140.9531 | 183223.2813 | 96576.94531 | 103454.7578 | 100914.75 |
| 49047.49219 | 92533.25 | 93434.82031 | 95997.24219 | 114597.0859 | 118099.1875 | 111854.5859 |
| 1408.067993 | 2445.705322 | 3038.444336 | 3294.47583 | 3021.035645 | 3404.935059 | 3799.338135 |
| 379956.2188 | 851755.6875 | 749974.8125 | 777163.5625 | 762224.125 | 746551.1875 | 741202.8125 |
| 43994.74219 | 90694.51563 | 80050.625 | 106550.3906 | 59987.34375 | 52458.79688 | 53223.41406 |
| 11494.87793 | 25963.13281 | 24598.04492 | 29472.74414 | 26382.84375 | 20935.50781 | 21454.97852 |
| 54645.75391 | 110869.2734 | 112182.6875 | 119514.2188 | 115147.3047 | 113690.8359 | 110948.7422 |
| 12493.21582 | 27348.27539 | 23824.84961 | 22467.83789 | 15687.69141 | 14559.42383 | 14053.68164 |
| 12788.63867 | 17849.5332 | 30739.41602 | 31007.64453 | 19004.02539 | 16050.41406 | 17708.41992 |
| 41108.41016 | 90972.07813 | 86023.30469 | 81393.875 | 69407.17969 | 65725.91406 | 59959.53125 |
| 54769.85547 | 106852.3047 | 112562.8281 | 108129.5469 | 94150.67969 | 95754.57813 | 93622.67969 |
| 5097.300293 | 12106.33398 | 11482.9873 | 12183.85645 | 12678.87598 | 10668.48438 | 11665.18457 |
| 51894.60156 | 114586.5 | 100541.6875 | 112574.8203 | 111284.9219 | 104517.9375 | 109944.6641 |
| 3474.640137 | 7491.861328 | 7481.845703 | 6945.206543 | 4843.732422 | 5728.838379 | 5834.178223 |
| 42958.74609 | 100266.9844 | 87961.34375 | 93233.10156 | 82987.21875 | 93887.46875 | 91376.07031 |
| 19896.42969 | 34632.27734 | 51433.24219 | 53547.42188 | 42332.41406 | 37809.09375 | 44424.55859 |

| | | | | | | |
|---|---|---|---|---|---|---|
| 76832.70313 | 195195.3281 | 164466.2656 | 183297.4531 | 225107.625 | 270834.4688 | 230191.1406 |
| 1078.25 | 9271.636719 | 7718.383301 | 7035.390625 | 5148.989746 | 5786.841309 | 6156.266113 |
| 21886.46094 | 53298.44531 | 50469.67188 | 51858.10938 | 76288.64063 | 75899.28906 | 73808.42188 |
| 674.2705688 | 3088.969238 | 1325.410278 | 1664.438232 | 869.5014648 | 1245.829224 | 1907.421021 |
| 16352.81934 | 34719.12109 | 35377.08594 | 34422.71875 | 26237.06055 | 27482.39648 | 27101.87695 |
| 8099.156738 | 23679.7832 | 14183.58789 | 14713.51953 | 25899.82031 | 22261.52148 | 20426.01563 |
| 109567.4531 | 252999.625 | 214589.9063 | 219614.8281 | 265477.3438 | 259544.8906 | 237650.875 |
| 6517.564453 | 11853.21289 | 11017.84082 | 13961.09668 | 13090.21582 | 9963.623047 | 11535.79395 |
| 68400.61719 | 139907.2969 | 140357.5781 | 145300.2188 | 138836.9531 | 141245.75 | 142352.8438 |
| 51776.48047 | 98180.05469 | 104781.5078 | 100226.6797 | 79297.47656 | 89855.63281 | 83985.04688 |
| 45630.33203 | 95326.60156 | 99784.77344 | 90540.60938 | 90349.97656 | 100434.8594 | 98039.57031 |
| NA | 3533.630371 | NA | NA | 7756.603027 | 5697.180176 | 5560.616211 |
| 18175.50586 | 40827.27734 | 31954.79492 | 41776.11719 | 50835.17578 | 50532.27344 | 50470.08984 |
| 2450.650879 | 11206.46289 | 3284.41626 | 7324.130859 | 9694.479492 | 9757.552734 | 9247.291016 |
| 51164.42188 | 96679.5625 | 92928.64844 | 83107.75781 | 103694.6875 | 116077.4453 | 94516.35156 |
| 31775.98193 | 51176.375 | 55938.51855 | 64048.75879 | 54227.06445 | 55503.99414 | 58152.49121 |
| 7370.762695 | 13017.28418 | 10421.08789 | 17082.61523 | 22562.57422 | 15336.96387 | 17474.32617 |
| 386009.75 | 853592.875 | 778498.5 | 838143.9375 | 903258.25 | 811513.75 | 803296.125 |
| NA | 3504.471191 | 5742.854004 | 1561.923828 | NA | NA | NA |
| 3568.215576 | 9426.723633 | 4869.751465 | 2854.929932 | 13689.54688 | 12332.85254 | 13851.17578 |
| 2349.22583 | 6544.518066 | 6849.414551 | 9321.123047 | 4725.569336 | 1799.582031 | 4250.37793 |
| 11953.61621 | 19889.9375 | NA | NA | 11589.83301 | 11053.30371 | 11320.73145 |
| 9890.44043 | 20793.42578 | 13352.91016 | 19585.81445 | 20231.89648 | 26624.74414 | 23552.68359 |
| 23618.46094 | 37670.27344 | 35600.42578 | 41739.24609 | 24758.18359 | 24205.13281 | 25446.72656 |
| 10093.81445 | 16098.41406 | 20381.68945 | 20821.73242 | 20657.60352 | 18975.02344 | 17654.95703 |
| 16636.77734 | 37168.55469 | 29646.94727 | 27796.79492 | 55770.05469 | 43555.28516 | 44581.74609 |
| 2023.471436 | 8547.772461 | 4894.850586 | 7705.196777 | 16853.04883 | 14213.88184 | 11675.54688 |
| 16673.94531 | 36607.99609 | 31194.35352 | 34767.08984 | 49411.44922 | 70730.99219 | 52263.51563 |
| 78567.96094 | 139314.3906 | 144900.5 | 149272.5469 | 218932 | 221592.6094 | 225344.5781 |
| 12470.31348 | 24903.70703 | 27939.74414 | 25362.84766 | 22212.5918 | 27805.75195 | 20924.6875 |
| 61583.53906 | 116424.3984 | 122847.8672 | 156517.7813 | 155678.9531 | 104322.6172 | 119126.875 |
| 32331.91602 | 67567.85156 | 58553.12891 | 66986.80469 | 64470.26953 | 60406.50391 | 68670.47656 |
| 5289.700195 | 13272.64941 | 10336.99805 | 10226.45313 | 8218.263672 | 9616.874023 | 9730.00293 |

| | | | | | | |
|---|---|---|---|---|---|---|
| 7520.486328 | 22176.32031 | 15026.53027 | 15529.26855 | 11358.45996 | 11914.83984 | 10590.54297 |
| 4605.974609 | 10565.62695 | 10471.54785 | 11012.66699 | 7273.503906 | 8292.154297 | 7431.523438 |
| 36078.78516 | 68798.55469 | 77295.28906 | 72974.11719 | 39885.42969 | 49952.72656 | 36976.35938 |
| 51627.69531 | 103896.5156 | 104268.0156 | 102853.5547 | 94024.36719 | 91152.92969 | 97342.82031 |
| 487459.3125 | 992982 | 997084.875 | 1133553 | 1450967.375 | 1222541.5 | 1335909.625 |
| 2285.458008 | 4626.840332 | 2941.740479 | 3855.952148 | 4546.237305 | 3853.218018 | 4408.843262 |
| 59238.73047 | 121295.6875 | 104786.1875 | 107666.0859 | 84796.21875 | 84539.71094 | 88629.125 |
| 21841.36914 | 40794.42578 | 41746.87109 | 51428.05078 | 47418.0625 | 48343.3125 | 49558.12109 |
| 913402.0625 | 1564292.875 | 1786984.125 | 1881262.75 | 1121867.375 | 1078738.375 | 1169857.375 |
| 47981.53516 | 102979.1875 | 94854.83594 | 91045.53906 | 82565.47656 | 81606.64844 | 80607.75781 |
| 13768.08984 | 24724.28516 | 29277.98438 | 34464.96484 | 23934.60352 | 18251.83008 | 18330.40234 |
| 14432.79688 | 23936.13281 | 23520.88477 | 26502.65625 | 26713.36523 | 20396.86523 | 25828.45117 |
| 287014.4063 | 566772.25 | 565103.625 | 560056.125 | 533708.8125 | 544133.4375 | 524597.625 |
| 16004.87695 | 32267.24023 | 31632.4707 | 32157.8457 | 30144.16211 | 32078.19922 | 28928.67383 |
| 59554.42578 | 108962.9531 | 129622.4766 | 150348.6094 | 128398.6641 | 126134.6094 | 147442.2188 |
| 5701.592285 | 11136.84961 | 11116.76172 | 11823.44043 | 11553.23633 | 9867.621094 | 9173.737305 |
| 773784 | 1363885.75 | 1433483.375 | 1660536 | 2409972.25 | 1938558.75 | 1662157.75 |
| 78751.23438 | 168464.5469 | 182318 | 186072.9844 | 164420.3594 | 166324.9531 | 175569.4063 |
| 1049.817139 | 3279.134766 | 2230.680176 | 4080.954834 | 16885.9375 | 18808.625 | 17177.89844 |
| 13711.72656 | 23849.30078 | 25637.99414 | 26702.64844 | 23604.52734 | 22337.29883 | 24128.08594 |
| 9575.992188 | 21842.21094 | 18367.54102 | 20751.70508 | 32428.83203 | 26091.05469 | 28349.84961 |
| 253263.5625 | 469294.625 | 490265.7813 | 515584.4688 | 235907.4375 | 224898.75 | 232406.6719 |
| 12366.06055 | 27646.25 | 20192.30859 | 24426.83008 | 19324.24023 | 20393.12109 | 15075.24707 |
| 5999.635254 | 10785.31348 | 12945.53125 | 11979.13867 | 11035.47266 | 12824.70801 | 11219.38281 |
| 35010.54688 | 35171.59766 | NA | 41894.33594 | 21114.50586 | 50984.24219 | 31392.03711 |
| 68201.90625 | 129388.6406 | 129411.9844 | 125856.6797 | 55038.65625 | 64917.64844 | 61594.08594 |
| 44253.58594 | 72797.18848 | 81789.76758 | 90074.55469 | 59310.29102 | 64717.51367 | 59004.69629 |
| 21220.71484 | 45272.78125 | 43679.88281 | 53393.82813 | 41835.87891 | 34799.79297 | 38986.69141 |
| 49131.30859 | 99472.85938 | 103975.7813 | 86016.28125 | 100472.0703 | 92972.1875 | 94194.84375 |
| 17985.76563 | 34500.20313 | 38206.35938 | 34582.98438 | 41583.80078 | 39265.41406 | 40395.13281 |
| 2595.521973 | 4726.306641 | 4381.438965 | 5107.313965 | 5392.805664 | 5429.582031 | 4914.987793 |
| 157493.1406 | 273900.375 | 261902.2656 | 279313.6875 | 130529.9531 | 123866.2344 | 136506.9844 |
| 337480.9063 | 577340.375 | 608720.1875 | 596382.375 | 470420.125 | 476081.4375 | 482191.1875 |

| | | | | | | |
|---|---|---|---|---|---|---|
| 3111.138428 | 9671.3125 | 8096.598633 | 9260.161133 | 14188.26855 | 12984.2627 | 11608.92773 |
| 2406.673096 | 5180.038086 | 4706.114746 | 4575.923828 | 4437.464844 | 4821.258789 | 4690.679199 |
| 37965.16016 | 89975.25781 | 72898.14063 | 76692.86719 | 89045.00781 | 80684.57031 | 76369.97656 |
| 30716.58789 | 66014.71875 | 63676.34766 | 64236.40234 | 119690.4453 | 119429.8906 | 116126.2578 |
| 75261.99219 | 154780.0313 | 154461.4375 | 153061.5313 | 136983.6094 | 125176.3672 | 132792.8281 |
| 4918.660156 | 12565.66211 | 7728.725586 | 6431.416504 | 7549.772949 | 9099.734375 | 8078.888184 |
| 9133.443359 | 12836.07031 | 20084.88867 | 16104.57422 | 23538.24414 | 27445.97656 | 21979.07227 |
| 17693.39258 | 33160.15234 | 33449.03125 | 40061.97656 | 49051.05469 | 43764.83594 | 46816.35547 |
| 26772.05273 | 46698.96484 | 47493.15234 | 48308.51563 | 23678.20898 | 29535.06836 | 27723.49609 |
| 32725.19336 | 74121.15625 | 64030.69922 | 70040.89063 | 78859.91406 | 81991.57031 | 86066.22656 |
| 18569.45313 | 53913.00391 | 29182.43359 | 23602.04883 | 27939.86328 | 27135.62891 | 31087.25977 |
| NA | NA | 1578.005371 | 3112.775391 | 9877.396484 | 2043.707886 | 3773.298828 |
| 20135.91992 | 36203.51953 | 38504.88672 | 37631.28906 | 29884.52148 | 27442.33594 | 26867.17969 |
| 12872.47656 | 49211.58203 | 26787.47266 | 21059.87305 | 25133.29492 | 35051.42578 | 28315.98633 |
| 944474.5 | 1800753.813 | 2004424.25 | 1881213.75 | 1877369.5 | 1684170.75 | 1711661.063 |
| 3926.001953 | 8244.295898 | 8042.599121 | 8047.42627 | 8451.990234 | 9482.566406 | 8816.833984 |
| 13462.01074 | 29789.19141 | 24403.25586 | 25338.78906 | 31515.65039 | 25906.13086 | 27779.00195 |
| 1815.727661 | 6323.19873 | 4865.151367 | 4704.718262 | 4786.174805 | 6883.111328 | 5393.052246 |
| 4042.608643 | 11761.37793 | 7877.89502 | 7077.875977 | 8306.606445 | 8853.149414 | 9748.041992 |
| 449146.8438 | 937380.625 | 907996.3125 | 869975.375 | 1059424.375 | 838111.125 | 903804.875 |
| 153633.7031 | 296859.1875 | 293206.4688 | 268624.5313 | 183444.3594 | 144218.625 | 143922.0156 |
| 13081.9209 | 25240.67578 | 21514.44531 | 22893.79883 | 25356.10352 | 32410.92188 | 26652.32422 |
| 2936.826904 | 5581.023438 | 6640.216309 | 6125.720215 | 9683.162109 | 7470.616211 | 8122.777344 |
| 11580.37988 | 19265.81055 | 19750.58398 | 23650.60352 | 20764.20313 | 20940.23242 | 22803.45508 |
| 3009.010986 | 6041.673828 | 6014.703613 | 6010.51123 | 4138.659668 | 3579.137695 | 4155.980957 |
| 453168.6875 | 878861.125 | 738840.1875 | 886713.0625 | 1273208.875 | 987800 | 1144764.875 |
| 73598.80469 | 123313.3672 | 142187.5469 | 158926.75 | 144402.6719 | 146033.3906 | 147939.8125 |
| 34602.80078 | 85441.17188 | 66009.72656 | 66828.10156 | 75877.96875 | 62978.94531 | 71175.70313 |
| 3341.594727 | 7964.039551 | 6941.465332 | 6544.501953 | 5663.125 | 9278.15918 | 8790.441406 |
| 157342.1875 | 282543.0625 | 300413.8125 | 312707.1875 | 263213.25 | 214461.7813 | 222508.3906 |
| 1480046.75 | 2345909.75 | 2823174.5 | 2385238.5 | 2280612.5 | 3197595 | 2946220.5 |
| 9402.02832 | 14386.3623 | 13871.50879 | 14350.53906 | 17500.94727 | 17191.78125 | 17128.14063 |
| 6252.212402 | 11831.03711 | 10878.59375 | 13122.57617 | 10693.22461 | 8772.885742 | 10592.80273 |

| | | | | | | | |
|---|---|---|---|---|---|---|---|
| | 4806.162109 | 10227.74512 | 9256.90918 | 9685.091797 | 9392.510742 | 10803.8125 | 10259.08008 |
| | 172926.5781 | 262539.125 | 325459.5625 | 358277.0938 | 133714.7188 | 141634.9375 | 157351.6875 |
| NA | | 15074.32324 | 12180.64941 | 13952.81738 | 25342.73828 | 16576.60352 | 19711.48828 |
| | 9253.647461 | 18213.45703 | 16173.2334 | 16896.47266 | 31610.58789 | 29949.67773 | 29216.4082 |
| | 5314.023926 | 9145.735352 | 9029.810547 | 10837.42383 | 13523.38281 | 12183.83789 | 15885.59961 |
| | 28084.27734 | 59482.22656 | 50889.03906 | 52608.57422 | 45561.20703 | 45474.625 | 48775.21875 |
| | 11517.83984 | 29317.15234 | 21739.96094 | 26622.0332 | 34171.25 | 35973.46094 | 36560.97656 |
| | 472115.0625 | 961834 | 845638.4375 | 1040141.188 | 1202023.125 | 770929.6875 | 900751 |
| | 625884.4023 | 1165719.953 | 1177977.188 | 1048293.336 | 855783.4219 | 897720.7031 | 873463.5313 |
| | 14585.58691 | 21004.51367 | 29202.56055 | 25535.58594 | 18814.75391 | 16359.93848 | 20146.375 |
| | 46982.74219 | 96153.92188 | 108260.9531 | 128541.8359 | 118762.0625 | 95115.67188 | 101931.9375 |
| | 195092.6719 | 337860 | 392784.8125 | 428128.25 | 300639.8125 | 285612.9063 | 317194.3125 |
| | 23486.66406 | 46818.46875 | 52346.98047 | 49854.64844 | 56836.00391 | 60848.8125 | 60520.25781 |
| | 19779.29297 | 38281.95313 | 30639.69727 | 22659.24414 | 23496.91211 | 19574.0293 | 22296.5293 |
| | 10885.80859 | 14565.30859 | 17747.87891 | 19531.24805 | 16706.25977 | 21080.51953 | 17216.82617 |
| | 23241.69727 | 46459.20703 | 35960.78906 | 26941.59375 | 25969.19922 | 29446.1543 | 28770.07227 |
| | 14300.89453 | 34415.02344 | 27370.05859 | 30320.94531 | 43177.77734 | 36296.03906 | 39207.52344 |
| | 2940.020752 | 6753.405273 | 4817.977051 | 4311.708496 | 3970.925537 | 2332.208008 | 1915.38147 |
| | 59091.3125 | 99360.15625 | 121417.6328 | 89730.5625 | 104948.4688 | 116263.4844 | 150939.0781 |
| | 5545.275879 | 7593.605469 | 9135.321289 | 9231.438477 | 4095.014648 | 5972.416992 | 4615.641113 |
| | 12171.02148 | 23379.6543 | 22759.61914 | 20645.81836 | 33715.12109 | 30838.48047 | 30375.13477 |
| | 2771.322021 | 5161.338867 | 4737.828125 | 4088.695313 | 4253.815918 | 3842.628174 | 3361.148926 |
| | 33455.96875 | 62718.85156 | 63179.12109 | 62006.61328 | 61256.53516 | 66189.09375 | 64962.49609 |
| | 5237.379395 | 8414.191406 | 20254.35547 | 13242.21094 | 8201.051758 | 5669.513672 | 17440.44531 |
| | 21730.62109 | 51580.98828 | 30250.43359 | 35848.60156 | 71986.78906 | 62790.02734 | 69024.58594 |
| | 8357.619141 | 14440.72168 | 16143.72559 | 14585.51367 | 15605.52734 | 15383.11426 | 14285.13086 |
| | 16864 | 44139.57422 | 28476.50977 | 24435.15234 | 26199.87305 | 26324.01367 | 26005.67578 |
| | 607.4297485 | 1766.729736 | 1143.938477 | 1445.978149 | 1506.430786 | 1093.827637 | 1268.431396 |
| | 52708.48047 | 101919.7734 | 104724.0938 | 102379.8125 | 84495.80469 | 113852.0313 | 82099.61719 |
| | 98370.76563 | 183265.4375 | 197018.3438 | 170374.5 | 147082.1094 | 155781.5 | 149340.4219 |
| | 96828.16406 | 177414.2344 | 188732.8594 | 226396.0938 | 239392.8906 | 186311.75 | 197501.3281 |
| | 7330.442871 | 19419.13867 | 11662.91113 | 8132.039063 | 2598.614258 | 3291.582031 | 2239.559082 |
| | 18937.89063 | 39972.41797 | 37221.53906 | 39588.28125 | 35549.12109 | 33401.05469 | 33210.55859 |

| | | | | | | |
|---|---|---|---|---|---|---|
| 7998.468262 | 15755.96289 | 24979.875 | 7667.900879 | 17097.11914 | 23598.03711 | 25082.03125 |
| 5248.372559 | 10419.00195 | 9750.545898 | 9087.667969 | 21668.03711 | 23782.0625 | 19417.4043 |
| 12671.32617 | 24819.24414 | 23571.09766 | 21600.97461 | 25925.4375 | 25296.33984 | 25743.61328 |
| 13706.7959 | 20217.51953 | 27435.87305 | 29665.55469 | 34054.44141 | 35466.38281 | 31386.26563 |
| 3671.447266 | 8503.701172 | 7786.6875 | 8158.540039 | 5446.755371 | 5023.605957 | 4175.868164 |
| 6441.140137 | 9014.624023 | 12651.43848 | 11345.25391 | 8880.338867 | 8768.509766 | 9343.913086 |
| 90344.82813 | 142746.7813 | 166833.75 | 169679.6719 | 77434.21875 | 79037.5625 | 79174.71875 |
| 13857.64258 | 33998.25 | 35194.67188 | 33281.17969 | 78749.66406 | 78065.29688 | 84127.46094 |
| 7768.938477 | 13302.50977 | 13217.60254 | 14446.80957 | 15138.69922 | 13776.06738 | 14508.57324 |
| 25762.01172 | 46053.17188 | 45807.89453 | 44496.73047 | 36597.28516 | 36411.64063 | 37778.14453 |
| 9704.006836 | 24555.75977 | 20472.53906 | 22468.43164 | 30885.99609 | 33331.42188 | 32819.82031 |
| 3906.88501 | 7716.033691 | 5445.53125 | 8150.337891 | 7458.317871 | 6567.383301 | 8077.010742 |
| 12466.81055 | 26502.21484 | 20981.03711 | 23848.11328 | 22749.71875 | 27949.94727 | 23173.60352 |
| 1894.445679 | 2613.433105 | 3742.282715 | 3674.746094 | 2670.451416 | 3067.856201 | 2772.439697 |
| 9002.072266 | 13828.90625 | 16216.34961 | 15599.89746 | 8087.138184 | 7620.907227 | 10555.37988 |
| 84592.71094 | 132601.4375 | 159496.3281 | 178400.625 | 155376.0156 | 165670.0938 | 191801.0938 |
| 8146.681152 | 19583.85742 | 17858.89063 | 17892.1543 | 17153.59961 | 15660.19238 | 16137.45801 |
| 126521.7969 | 222190.5625 | 238675.8438 | 218819.0625 | 168167.625 | 170074.9375 | 161436.8125 |
| 208977.0156 | 370125.0625 | 391374.3438 | 378858.5625 | 275347.4375 | 288809.1875 | 271207.875 |
| 302490.375 | 553505.375 | 553102.25 | 551674.5 | 609504.6875 | 717800.875 | 684754.75 |
| 6402.471191 | 10427.25879 | 11546.44043 | 7648.297852 | 7859.944824 | 8581.214844 | 8534.816406 |
| 282123.6563 | 475186.375 | 554009.125 | 547634.875 | 427343.5313 | 453498.875 | 438233.875 |
| 165446.3281 | 222305.2031 | 324679.25 | 353883.4063 | 204129.2656 | 181176.5 | 197029.6719 |
| 399.3089294 | NA | 690.8720093 | NA | NA | NA | 4331.372559 |
| 130135.9922 | 248694.9844 | 223414.9063 | 228010.1406 | 141576.0469 | 153908.7188 | 137747.8906 |
| 2741.155762 | 7203.16748 | 8423.519531 | 6383.737305 | 10355.45508 | 10266.51953 | 9428.62207 |
| 8712.169922 | 16526.29297 | 15058.58398 | 13622.91211 | 26294.76758 | 29645.02148 | 25284.10547 |
| 1506510.5 | 3087826.25 | 2869526.25 | 3146049.5 | 3255311 | 3249770 | 3384548 |
| 9793.299805 | 19050.73633 | 15342.12988 | 14898.62988 | 20583.125 | 17168.75391 | 16050.89063 |
| 16924.34375 | 35871.48047 | 23537.24023 | 34654.80859 | 57569.19141 | 48742.4375 | 50071.49219 |
| 24053.64258 | 45781.10156 | 42616.57422 | 53550.70703 | 53351.83203 | 51664.92969 | 53691.41797 |
| 15384.55859 | 29706.8125 | 27284.0293 | 28674.53711 | 40697.82422 | 39427.07813 | 39026.86328 |
| 86103.41406 | 175242.0781 | 136472.7344 | 168843.8125 | 241488.1094 | 233566.7188 | 248693.0313 |

| | | | | | | |
|---|---|---|---|---|---|---|
| 13374.66113 | 26450.30078 | 22247.27734 | 25420.08203 | 29569.85742 | 20130.51367 | 22809.61133 |
| 3478.703857 | 7197.160645 | 6257.080078 | 7012.111328 | 8414.12793 | 9155.731445 | 9113.919922 |
| 16045.09668 | 38030.41797 | 32233.94727 | 28123.59766 | 13205.9502 | 15882.34473 | 11106.10742 |
| 24246.95898 | 40025.8125 | 40751.64063 | 39783.91016 | 28606.27539 | 33376.71484 | 31624.51563 |
| 1527.85437 | 2611.327148 | 4417.049316 | 5781.255371 | 10932.89453 | 9265.097656 | 7186.705566 |
| 10556.63477 | 19465.4668 | 15348.63867 | 14817.44141 | 16348.95117 | 20124.81641 | 17983.62109 |
| 24598.02148 | 45183.25781 | 39910.16406 | 46552.65234 | 42497.00781 | 43467.28125 | 44327.19141 |
| 7615.815918 | 24926.88867 | 20933.53125 | 16536.92773 | 20733.48242 | 26679.52148 | 24561.67969 |
| 15653.8877 | 29401.93555 | 31571.22266 | 28608.30664 | 22572.74805 | 23289.13477 | 23131.11133 |
| 3514.444824 | 8149.71582 | 6217.901855 | 6294.255371 | 6483.455566 | 5926.094727 | 6342.811523 |
| 27544.79688 | 50567.23047 | 52870.21484 | 57104.26563 | 67017.73438 | 61723.97266 | 67857.82031 |
| 4356.414063 | 7078.617676 | 5681.908203 | 6944.611816 | 9330.331055 | 10849.44238 | 9304.348633 |
| 12745.80078 | 24782.33984 | 22930.8418 | 23216.13477 | 23311.10938 | 15469.42383 | 25960.9668 |
| 16461.74219 | 32867.98828 | 30000.49414 | 30920.01563 | 26139.45703 | 31262.7832 | 29021.76172 |
| 2427.876953 | 4996.049805 | 4073.588623 | 4276.366699 | 9035.289063 | 9505.674805 | 6583.241699 |
| 77424.85156 | 147669.9688 | 112269.5313 | 102224.9219 | 45555.70313 | 48377.79297 | 44207.33984 |
| 22763.25977 | 52866.61719 | 47716.03516 | 44369.42188 | 44403.16406 | 42210.06641 | 47950.4375 |
| 1777.925781 | 4440.009277 | 3264.351807 | 2869.008789 | 3251.45459 | 5400.573242 | 3796.579834 |
| 217281.3906 | 442920.5 | 485571.0938 | 564786.5625 | 381170.0625 | 349022.5313 | 360041.125 |
| 17451.48633 | 31234.85938 | 32864.20313 | 37894.29688 | 40412.9375 | 45876.67188 | 49141.78125 |
| 4016.744629 | 7179.037598 | 4327.597168 | 4830.266113 | 7513.537598 | 8584.676758 | 9169.924805 |
| 1765.544067 | 2665.35791 | NA | NA | 2104.871826 | 3311.0354 | 2877.332031 |
| 81758.80469 | 136478.125 | 84058.27344 | 150765.9063 | 172446.7969 | 196717.125 | 162880.125 |
| 113454.7891 | 208622.7031 | 192906.3281 | 195654 | 211614.0156 | 202534.3594 | 208546.9844 |
| 6362.38623 | 13731.18555 | 12842.12891 | 11169.59082 | 19461.12695 | 12836.0791 | 15888.93164 |
| 30762.5293 | 62726.50781 | 58741.3125 | 51787.32031 | 50685.22266 | 40808.03906 | 47175.13281 |
| 6784.281738 | 15489.13184 | 11340.65234 | 9559.480469 | 10519.7002 | 13883.68945 | 13427.79785 |
| 446985.25 | 979667.6875 | 770444.375 | 743447.125 | 703701.75 | 713742.4375 | 639989.75 |
| 106684.2656 | 204222.75 | 187372.5938 | 189522.0469 | 195553.4688 | 185568.7031 | 198090.4531 |
| 2757.196045 | 1940.871216 | 3348.426025 | 5459.153809 | 2864.374756 | 6814.858887 | 6229.539551 |
| 938255.375 | 1634442.125 | 1827864.625 | 1878113 | 1580353.875 | 1481350 | 1530399.5 |
| 1757808.375 | 2778662.75 | 2249149.25 | 2558312.75 | 2381700.25 | 2452127.75 | 1506787 |
| 10966.79004 | 23397.1543 | 18837.98828 | 10894.84961 | 11583.11426 | 9320.926758 | 9326.367188 |

| | | | | | | |
|---|---|---|---|---|---|---|
| 13601.65918 | 30441.79492 | 26403.20508 | 22392.66992 | 20870.72461 | 26944.39063 | 20176.27734 |
| 5273.570313 | 8385.771484 | 10916.88672 | 9354.299805 | 9544.274414 | 10156.08105 | 10016.48145 |
| 21776.2832 | 37784.79688 | 34622.59375 | 39252.23828 | 45980.73828 | 38926.71484 | 42215.27344 |
| 60325.29297 | 78858.52344 | 61178.57813 | 88118.1875 | 62321.34766 | 72036.625 | 59900.26953 |
| 2700.800049 | 4598.224121 | 5842.697266 | 5018.788574 | 3925.031738 | 3667.064941 | 3580.914063 |
| 39281.36719 | 59891.00391 | 63664.08203 | 75765.96875 | 68437.32031 | 46793.64453 | 56450.33203 |
| 33072.51563 | 64695.75781 | 59328.69531 | 61727.83984 | 68062.79688 | 59538.28906 | 63736.70703 |
| 131586.0938 | 158082.8906 | 83751.54688 | 185737.1875 | 134174.3438 | 221365.7813 | 155804.0313 |
| 7912.158691 | 14519.61328 | 12054.94629 | 12293.04492 | 6608.910156 | 10483.06055 | 9288.526367 |
| 149334.2344 | 277848.9375 | 260012.0156 | 392857.125 | 668221.0625 | 308930.2813 | 339763.4375 |
| 74076.64063 | 123067.9219 | 144727.5625 | 138892.2344 | 140292.5 | 147929.0625 | 138879.3906 |
| 13401.84473 | 23331.97461 | 21981.33008 | 22108.80273 | 25049.80664 | 23893.66797 | 24283.71289 |
| 5169.957031 | 5969.205566 | 7136.619629 | 7385.455566 | 7046.194336 | 5914.550293 | 5917.226563 |
| 4739.521973 | 9963.375977 | 6242.105957 | 9332.981445 | 12498.23438 | 15982.68848 | 10612.43652 |
| 3728.144531 | 9731.15625 | 9004.848633 | 8390.341797 | 6103.06543 | 4559.217773 | 6362.112305 |
| 12595.75293 | 22025.59961 | 22845.78711 | 21356.35156 | 18470.28516 | 23388.50977 | 19356.57617 |
| 47844.95703 | 96591.29688 | 82959.51563 | 85123.41406 | 73895.53906 | 77168.83594 | 72291.01563 |
| 6733.612305 | 11957.15137 | 12629.05273 | 11571.10547 | 12116.88477 | 9464.741211 | 11850.31055 |
| 5422.371094 | 11305.78809 | 9063.105469 | 7772.260254 | 2998.552734 | 4205.623047 | 2944.589111 |
| 15481.16602 | 24710.98047 | 25654.37891 | 28677.96484 | 24344.58789 | 24476.875 | 24974.65625 |
| 45952.01563 | 65920.21094 | 91277.78906 | 78457.51563 | 62937.87109 | 89602.07031 | 82531.95313 |
| NA | 4091.407227 | 2405.965576 | 3212.256592 | 3590.123047 | 2388.922119 | 1867.059937 |
| 72737.03125 | 76514.71875 | 77431.74219 | 92237.67188 | 65849.63281 | 73132.07813 | 60143.46875 |
| 17430.36719 | 34199.52344 | 34025.29688 | 38481.79297 | 94625.69531 | 61785.19922 | 70515.69531 |
| 3380.131104 | 8699.605469 | 7078.403809 | 6196.190918 | 9785.480469 | 11974.40918 | 7553.964355 |
| 11088.8457 | 18787.04297 | 17573.95117 | 21142.08984 | 20414.24023 | 21263.8125 | 19054.38867 |
| 23841.17578 | 64441.13281 | 36223.02344 | 35222.27344 | 58179.97656 | 37499.33594 | 47337.85938 |
| 29140.18359 | 44295.55469 | 44369.82813 | 46322.24219 | 38282.47656 | 35295.26953 | 35764.27344 |
| 2396.952148 | 4841.83252 | 3657.267822 | 3202.672363 | 7712.329102 | 5451.567383 | 8143.781738 |
| 12280.54102 | 27428.75 | 24719.66797 | 27506.55273 | 50276.88672 | 44773.55469 | 43883.73828 |
| 10852.95996 | 25988.01953 | 16834.36523 | 15710.44727 | 15572.65332 | 18426.28906 | 15448.02246 |
| 654033 | 1102647.625 | 1071700.125 | 1268048.125 | 2575209.5 | 2097377.75 | 1933555.125 |
| 1526.831055 | 1866.170654 | 3433.840332 | 4349.216797 | 9452.500977 | 8938.40332 | 7718.483398 |

| | | | | | | |
|---|---|---|---|---|---|---|
| 1414.646973 | 2423.799072 | 2323.885986 | 2548.749023 | 1527.839478 | 1790.897217 | 1451.596313 |
| 318709.375 | 599115.0625 | 550870.75 | 522809.3125 | 467275.1563 | 541985.125 | 434531.8438 |
| 82035.11719 | 169523.0156 | 163059.1875 | 168879.7813 | 230310.5469 | 225867 | 222461.3594 |
| 2622.20752 | 6236.501465 | 2757.200439 | 3374.113037 | 19954.99023 | 6832.931641 | 10102.40527 |
| 8371.375977 | 13522.38281 | 12061.55176 | 15357.66504 | 11518.75098 | 10151.33594 | 11100.58789 |
| 288280.0313 | 629055.5625 | 575500.9375 | 656035.9375 | 608149 | 529819.0625 | 649823.5 |
| 4314.899414 | 9818.234375 | 8392.802734 | 8377.750977 | 10559.02344 | 13205.5957 | 11809.78418 |
| 17857.77148 | 31771.16602 | 27404.9043 | 30315.70898 | 16238.29297 | 13092.64746 | 15698.2207 |
| NA | 1803.483765 | 1989.239136 | 1510.214478 | 1552.888062 | 1417.786987 | 1621.888916 |
| 17872.70703 | 38471.35547 | 36546.34375 | 40380.64063 | 49620.58203 | 49991.23828 | 50602.49219 |
| 225193.6719 | 412259.1563 | 412437.3438 | 410077.5313 | 443129.5 | 403731.4375 | 430917.75 |
| 75454.01563 | 140143.1875 | 125360 | 100813.125 | 47224.14063 | 45081.99219 | 47115.72266 |
| 8675168 | 17063880 | 9939820 | 11398833 | 7929145.5 | 9931827 | 5299077.5 |
| 150614.4375 | 261836.3906 | 288904.9688 | 303099.6563 | 256962.5156 | 261781.8125 | 268229.2813 |
| 7841.529297 | 16969.04102 | 13331.29688 | 13222.76074 | 18083.23633 | 14547.80957 | 16535.91211 |
| 10073.23535 | 19990.98828 | 19491.91797 | 19961.39063 | 17987.7168 | 17625.26953 | 17711.04883 |
| 51099.41016 | 70913.54688 | 93092.04688 | 93652.85938 | 49592.20313 | 44777.14453 | 40776.50391 |
| 3760.04126 | 8909.235352 | 7204.991699 | 7077.008301 | 2681.131348 | NA | 2231.785645 |
| 23991.21094 | 45571.64063 | 46913.01563 | 45681.86719 | 42006.94141 | 47443.81641 | 43213.65625 |
| 13162.31055 | 21655.25781 | 21975.50391 | 22613.97656 | 25680.76172 | 23902.93555 | 23495.63672 |
| 3839.510742 | 9409.995117 | 2988.873535 | 2110.359863 | 8068.185059 | 2433.023926 | 3493.583984 |
| 22921.48633 | 49471.34766 | 29458.03125 | 53479.90234 | 44923.00391 | 47868.375 | 52416.10547 |
| 52533.42969 | 86010.22656 | 94942.09375 | 91721.3125 | 66296.61719 | 67136.67188 | 70599.03125 |
| 46301.75391 | 67379.89063 | 84941.5 | 70334.17969 | 46002.96875 | 47495.46094 | 45422.64844 |
| 52675.88281 | 91748.27344 | 92501.89844 | 105647.5 | 109432.6406 | 87851.70313 | 90265.99219 |
| 55038.35547 | 100322.0234 | 93268.38281 | 94525.94531 | 51633.35547 | 57860.59766 | 56875.50391 |
| 66608.32813 | 139547.2188 | 184128.8281 | 197287.1875 | 462107.75 | 489410.3438 | 365380.9063 |
| 91706.96875 | 150581.9219 | 163788.7031 | 168290.4844 | 111704.5938 | 105669.0078 | 109773.8828 |
| 81058.875 | 157632.5313 | 135966.0313 | 151215.25 | 191320.3594 | 173144.9844 | 177883 |
| 8275412.5 | 12514594 | 14178046 | 14746919 | 13150227 | 12234785 | 12703224 |
| 4453.238281 | 7573.71875 | 8857.050781 | 8152.729492 | 4049.181885 | 4189.915527 | 4909.490234 |
| 6721.992676 | 11564.51074 | 11848.13379 | 12005.56445 | 11161.72461 | 11452.7373 | 11444.86035 |
| 8730.748047 | 15382.10254 | 16727.97266 | 15092.19238 | 9396.155273 | 11079.98438 | 10882.29004 |

| | | | | | | |
|---|---|---|---|---|---|---|
| 27709.91797 | 50368.95703 | 49350.75391 | 52290.37891 | 44195.03125 | 40654.58594 | 46024.66016 |
| 37189.26563 | 66837.60938 | 67364.77344 | 66651.30469 | 59455.02344 | 59486.15234 | 60517.12891 |
| 13629.81934 | 20377.00977 | 18271.0625 | 18676.46094 | 17326.70703 | 13602.9082 | 14634.39551 |
| 50298.5 | 81286.66406 | 80901.76563 | 83508.97656 | 71060.28906 | 71904.16406 | 69664.55469 |
| 751556.3125 | 1563194.875 | 1337616.875 | 1390927 | 1341654.375 | 1420514.125 | 1371795.375 |
| 3301.569336 | 5319.397461 | 5324.275879 | 5851.366699 | 4833.314941 | 20042.85938 | 5671.875488 |
| 145561.3438 | 261782.5313 | 256105.0313 | 247239.3438 | 240054.8125 | 233918.5 | 239999.125 |
| 62471.15625 | 106556.0313 | 128259.3438 | 122657.0078 | 58793.81641 | 77118.82813 | 69332.85156 |
| 4790.255859 | 7664.57373 | 8370.027344 | 8859.016602 | 5866.626953 | 4772.191406 | 5049.820801 |
| 31110.6582 | 41829.33984 | 57458.47656 | 66165.22656 | 110503.5 | 85055.15625 | 115346.7109 |
| 246500.25 | 286084.6563 | 421650.6875 | 244530.6406 | 182789.7031 | 288271.5938 | 294776.375 |
| 6833.684082 | 12979.20703 | 13551.47949 | 13803.92676 | 14837.36328 | 13486.4082 | 13561.46094 |
| 56653.94922 | 106832.0156 | 103785.6563 | 107903.6094 | 82380.02344 | 71796.58594 | 76015.39063 |
| 33323.63281 | 47047.14063 | 52146.46875 | 47577.92969 | 74730.1875 | 52137.94922 | 25677.04297 |
| 118187.5156 | 243770.8906 | 234961.4063 | 255550.25 | 202054.5 | 169988.3906 | 222811.1563 |
| 18841.07422 | 35771.08984 | 32517.31836 | 28229.60547 | 50284.34766 | 45450.48828 | 47369.29297 |
| 37930.0625 | 65338.13672 | 57957.85156 | 62338.375 | 52927.625 | 49749.43359 | 53016.34375 |
| 84832.17969 | 158333.7969 | 126302.8984 | 119440.8594 | 104271.5313 | 129106.1797 | 111939.3047 |
| 30251.1582 | 36528.97266 | 51175.67188 | 57850.85938 | 46571.75 | 41118.83984 | 51939.5 |
| 35953.0625 | 77951.78125 | 62031.55859 | 57201.72266 | 71916.05469 | 77106.03125 | 71867.26563 |
| 5439.018555 | 6728.281738 | 7498 | 8430.213867 | 8989.620117 | 10205.93066 | 7964.793457 |
| 29630.01563 | 52976.81641 | 51404.17969 | 53919.34375 | 55449.65625 | 51741.26563 | 53269.95703 |
| 33371.32813 | 53634.52734 | 48081.75391 | 49615.01563 | 30826.53125 | 37055.36719 | 33889.32813 |
| 11102.58594 | 14324.68164 | 14887.85352 | 15212.9873 | 10683.20703 | 16897.64453 | 11968.39844 |
| 15316.81738 | 26584.05469 | 25817.43164 | 30463.875 | 43905.97656 | 37843.48047 | 38397.84766 |
| 5870.742188 | 8960.682617 | 8926.698242 | 10545.52637 | 12046.79297 | 11657.31738 | 11134.07129 |
| 29070.89844 | 56537.26563 | 41090.64063 | 47923.13281 | 54033.55078 | 54842.20703 | 51975.01563 |
| 7037.006836 | 11517.26855 | 12466.01563 | 10066.17578 | 14226.41016 | 11984.01074 | 14907.14063 |
| 13521.46094 | 25095.69922 | 22857.65234 | 20825.18164 | 20150.45117 | 18572.38281 | 19510.01172 |
| 33901.46484 | 42245.23047 | 45574.71484 | 46923.02734 | 24982.79102 | 33591.82031 | 30419.25977 |
| 14038.07324 | 32386.56641 | 23514.49219 | 28430.98438 | 40913.6875 | 36086.24609 | 39320.98438 |
| 1348558.5 | 1959167.125 | 2168586.25 | 2545346.5 | 2484233.5 | 3292267 | 3496742.5 |
| 129879.6328 | 285004.4063 | 257598.8281 | 120572.3359 | 81762.54688 | 27533.69531 | 20141.78125 |

| | | | | | | |
|---|---|---|---|---|---|---|
| 11708.04883 | 16963.78711 | 20643.10547 | 19166.86719 | 21313.76953 | 21151.36328 | 22730.60156 |
| 125781.4219 | 202837.5469 | 265212 | 268725.3125 | 258944.4688 | 262960.4063 | 256599.5156 |
| 24036.10156 | 47589.76563 | 41401.01953 | 44071.41406 | 59680.03125 | 60677.65625 | 59876.05078 |
| 31409.95313 | 54674.13672 | 51146.64453 | 55037.55859 | 44394.73828 | 46721.80469 | 45327.37109 |
| 18289.26953 | 35107.95313 | 31919.36133 | 29070.83008 | 24324.8418 | 23323.46875 | 23618.04688 |
| 16624.37695 | 33233.50391 | 27397.62109 | 31537.55664 | 40156.86328 | 37205.29297 | 37825.46875 |
| 255866.4844 | 391060.5938 | 438294.4375 | 509692.8125 | 335319.9375 | 310698.9063 | 357583.875 |
| 54016.70313 | 100089.6641 | 107020.6172 | 108057.875 | 104581.0391 | 107246.3438 | 111192.6484 |
| 24489.9707 | 35321.0625 | 37141.01563 | 36170.30859 | 21313.33594 | 22440.62109 | 23454.82422 |
| 16843.25586 | 28087.29102 | 27736.31836 | 26973.90039 | 25794.91406 | 21484.60547 | 22861.01953 |
| 2410.106445 | NA | NA | 3928.848633 | 3679.75708 | 5655.90918 | 4635.768066 |
| 38964.98438 | 65000.37109 | 74110.80469 | 74764.63281 | 56386.96875 | 60747.60156 | 58432.27344 |
| 30524.21289 | 76486.30469 | 40368.89844 | 48037.00391 | 46603.12891 | 37748.03516 | 40702.875 |
| 1351922 | 1821154.875 | 2194450.5 | 2143293 | 1473943.625 | 1432757.5 | 1666117.75 |
| 8329.996094 | 16983.06445 | 17679.39258 | 16034.58984 | 11599.92773 | 10334.33789 | 11933.88672 |
| 69014.125 | 111470.0625 | 120794.9219 | 116609.9531 | 77895.17188 | 90645.29688 | 78519.60938 |
| 37914.14844 | 60242.05469 | 66161.70313 | 83722.46875 | 83935.83594 | 73115.01563 | 74684.28125 |
| 40289.90234 | 64849.3125 | 63568.23438 | 68798.50781 | 99600.47656 | 93056.19531 | 98955.26563 |
| 88729.71875 | 150876.1875 | 140537.9844 | 137390.1719 | 84472.88281 | 74176.57813 | 82876.4375 |
| 25174.22266 | 43654.24219 | 38191.81641 | 39253.28516 | 40015.74609 | 45507.23438 | 40614.53516 |
| 21650.52344 | 37614.55469 | 31562.75977 | 28421.73828 | 37271.78906 | 47705.19531 | 42908.32422 |
| 106896.7188 | 216370.5313 | 177837.0781 | 168365.8125 | 170139.2969 | 150849.9688 | 157256.2188 |
| 27985.40039 | 46593.19922 | 49157.57813 | 43538.64844 | 34672.90234 | 43840.41016 | 43292 |
| 8825.783203 | 16630.26367 | 16635.58008 | 14138.67676 | 20914.61133 | 20422.96094 | 20264.63672 |
| 66024.44531 | 114738.4844 | 113504.6797 | 119535.0078 | 127867.9453 | 101027.5 | 99396.71875 |
| 1169428.875 | 2053202.25 | 2072278.75 | 2026602.875 | 2270981.25 | 2254332 | 2139831.5 |
| 15380.04199 | 23899.18164 | 27454.6543 | 33544.55859 | 39151.89844 | 39422.51953 | 42254.15234 |
| 7886.501953 | 10846.22461 | 14802.79688 | 14574.74902 | 7867.919434 | 7674.967285 | 7593.276855 |
| 68.95477295 | 671.8074341 | NA | 78.88459778 | 221.5680237 | NA | 161.776886 |
| 31552.80078 | 56884.41797 | 62746.51563 | 61767.52344 | 55854.29688 | 41434.83594 | 61467.07813 |
| 9241.986328 | 20718.83203 | 16730.65039 | 17718.06445 | 14253.74805 | 14510.2666 | 11473.16699 |
| 42656.93359 | 99364.08594 | 76869.3125 | 63156.73438 | 26631.99219 | 30073.32813 | 27679.92773 |
| 15752.88281 | 22193.03711 | 27131.61328 | 36165.82813 | 46863.22266 | 25730.62695 | 31029.0625 |

| | | | | | | |
|---|---|---|---|---|---|---|
| 229586.8125 | 460881.6563 | 444880.5625 | 488611.375 | 386646.9688 | 350728.9375 | 412173.6875 |
| 29508.16211 | 50539.69531 | 42616.55859 | 44309.05469 | 51290.10547 | 53481.59766 | 52913.09375 |
| 53010.45313 | 89898.53125 | 93400.99219 | 81382.27344 | 68134.25 | 68461.86719 | 65790.5 |
| 10141.68555 | 21521.01367 | 18017.16211 | 16519.93555 | 35992.35938 | 25463.75 | 37770.47266 |
| 842381.75 | 1281256.25 | 1205062.375 | 1341991.25 | 1172045.5 | 1182317.625 | 922167.5625 |
| 24746.45703 | 80534.34375 | 23850.7832 | 23716.66992 | 63933.12109 | 57872.34766 | 60591.14063 |
| 80269.04688 | 140283.3906 | 145276.9219 | 165836.3438 | 206813.2344 | 187038.6406 | 194507.4688 |
| 147180.4219 | 346775.4375 | 265991.6875 | 340915.125 | 292732.9688 | 352219.6875 | 285752.3125 |
| 108087.8594 | 174760.5156 | 193468.8594 | 233056.6406 | 456885.0313 | 354729.0625 | 390214.4063 |
| 66837.10156 | 98163.38281 | 83596.77344 | 96827.57813 | 65804.80469 | 61690.125 | 81778.59375 |
| 3510.464355 | 6868.580078 | 6071.133789 | 7247.838867 | 7166.273438 | 7951.974121 | 8158.268555 |
| 67973.77344 | 147106.0156 | 134155.5938 | 149109.8281 | 104525.0703 | 84174.22656 | 125547.3203 |
| 504497.7188 | 943278.0625 | 876841.75 | 983134.75 | 1233275.5 | 1016367 | 1040605.125 |
| 320981.7188 | 1005780.5 | 414692 | 507121.0938 | 661936.5 | 829474.5625 | 669288.8125 |
| 290814.3438 | 528780.5 | 458176.2813 | 518676.6875 | 446240.5313 | 398347.0938 | 388858.5 |
| 8216.439453 | 12142.65332 | 14354.29102 | 15961.51367 | 19057.70703 | 21588.21875 | 22122.9707 |
| 62283.29785 | 112164.5322 | 111930.5996 | 115782.085 | 153679.9023 | 147666.5596 | 143216.5586 |
| 39109.68359 | 58731.47266 | 59414.41406 | 58085.08594 | 53632.56641 | 52414.68359 | 49601.90625 |
| 23869.49023 | 34395.73438 | 41110.91016 | 40356.89844 | 47005.57813 | 46320.86719 | 45226.44141 |
| 3306.949951 | 6381.850098 | 6235.185547 | 4489.682617 | 5418.175293 | 4986.571289 | 5077.791016 |
| 1834.736328 | 4715.564453 | 4083.324951 | 3596.85376 | 5970.711426 | 6953.471191 | 5962.27002 |
| 8476.542969 | 19963.54102 | 16117.39453 | 14200.00977 | 13002.64453 | 14817.39063 | 12224.12793 |
| 202140.5625 | 400216.7813 | 340112.5625 | 368122.6563 | 291152.6563 | 282948.375 | 292584.8125 |
| 98348.85303 | 202583.7109 | 187794.9941 | 138303.0034 | 164180.5869 | 252288.3936 | 227509.5166 |
| 7079.121094 | 11023.3125 | 13876.79004 | 12223.30273 | 9622.012695 | 10757.16895 | 8589.411133 |
| 296042.7188 | 557303.8125 | 531469.1875 | 539957 | 590571.875 | 593500.3125 | 563388.8125 |
| 5090.960449 | 8203.579102 | 7937.934082 | 6669.654785 | 9505.03418 | 9374.348633 | 8535.211914 |
| 559923.125 | 1029809.063 | 996254.6875 | 1043158.063 | 1326171.75 | 992283.75 | 1134371.875 |
| 7612.882813 | 18300.65039 | 16875.06055 | 14834.79785 | 18169.11914 | 17297.70313 | 14618.47559 |
| 11599.93359 | 22169.89844 | 19762.45117 | 20925.35156 | 20691.83984 | 33136.76563 | 24772.37891 |
| 29896.74219 | 60407.21484 | 52189.66406 | 54196.1875 | 50035.92969 | 51391.87891 | 49530.83594 |
| 174821.1406 | 323618.5313 | 317829.8125 | 301326 | 202510.2813 | 198013.5938 | 177178.7656 |
| 3270.515869 | 7155.564941 | 6740.697754 | 2136.966309 | 5654.808105 | 6324.280273 | 6597.405273 |

| | | | | | | |
|---|---|---|---|---|---|---|
| 4281.097168 | 8702.260742 | 7730.878906 | 9783.376953 | 11387.00391 | 8570.516602 | 9669.365234 |
| 54631.00391 | 92337 | 102076.3438 | 82717.61719 | 111093.3828 | 43963.48047 | 74375.48438 |
| 3858.088135 | 7149.818848 | 4803.141602 | 6178.936523 | 7697.668457 | 5549.334961 | 6846.905762 |
| 78314.58594 | 138055.0625 | 125430.7734 | 127960.7109 | 145445.8906 | 139096.0938 | 136414.2188 |
| 438.2516785 | 491.5086975 | 622.4011841 | 760.5886841 | 498.2127991 | 486.1552734 | NA |
| 23686.07031 | 47440.32031 | 43792.50391 | 46782.23438 | 46163.88281 | 46527.52344 | 47545.97266 |
| 11176.17383 | 17682.41406 | 15123.41211 | 15983.26758 | 11242.28809 | 12246.66699 | 11668.88965 |
| 31067.89648 | 49478.39453 | 52389.78516 | 54863.07422 | 50475.75391 | 47850.71094 | 48556.36719 |
| 81251.67969 | 121144.3594 | 131269.5 | 145412.8594 | 149536.9219 | 150294.3281 | 149061.6875 |
| 291998.6914 | 486023.5586 | 543331.8672 | 572251.2031 | 400672.6641 | 398750.4063 | 405910.0625 |
| 86970.98438 | 151912.875 | 147619.6094 | 154026.5938 | 153065.3438 | 143936.8438 | 157146.5313 |
| 22646.85352 | 31739.49805 | 41770.22266 | 45528.21484 | 71638.20313 | 80177.65625 | 88444.19531 |
| 229530.8906 | 424130.4688 | 395386.4688 | 467794.0313 | 355376.6563 | 294320.9375 | 316962.2813 |
| 19166.3125 | 36431.94531 | 31172.10742 | 23998.13281 | 17274.7832 | 22298.01172 | 17315.25 |
| 12476.82715 | 15115.88672 | 8229.924805 | 15155.06055 | 13925.51465 | 18948.8418 | 12181.93555 |
| 15726.64453 | 27248.27539 | 24806.14453 | 23449.75586 | 23262.0625 | 24994.76367 | 25122.27734 |
| 2755.554443 | 4800.897461 | 5777.214355 | 5323.751953 | 9627.899414 | 8435.991211 | 7308.773438 |
| 9586.99707 | 18286.45117 | 18954.70117 | 17704.20508 | 19419.19531 | 19467.15234 | 16158.85938 |
| 219605.4844 | 325970.0313 | 396675.0313 | 387990.0313 | 208344.5469 | 194893.0781 | 204754.2813 |
| 3417.525146 | 6489.875 | 7557.44873 | 6382.077637 | 7221.260742 | 6679.018066 | 7192.854004 |
| 239007.4688 | 437499.6563 | 387213.5 | 408378.125 | 331687.3125 | 367479.25 | 330609.8125 |
| 217943.2656 | 366931.6563 | 345286.9063 | 379937.375 | 338702.5625 | 288297.5 | 302848.7188 |
| 10893224 | 17315832 | 19170682 | 18651352 | 19719554 | 19749674 | 19993504 |
| 25041.40625 | 43539.01953 | 52918.51953 | 55561.67969 | 50005.30859 | 44921.27734 | 43576.47266 |
| 348912.3438 | 642454.25 | 635906.9375 | 632671 | 673209.375 | 599128.5 | 588616.5 |
| 7442.907715 | 18035.68164 | 18188.51953 | 13192.52832 | 26809.91211 | 26203.05469 | 28861.37305 |
| NA | 3461.970459 | 3518.37915 | 5064.993164 | 9006.370117 | 5846.500488 | 8251.260742 |
| 7943.092285 | 14122.25586 | 12708.94434 | 12629.14551 | 11202.47461 | 11863.33594 | 12837.83301 |
| 6417.337402 | 10740.05762 | 8224.167969 | 9824.360352 | 13797.05859 | 13185.05469 | 11573.65625 |
| 22685.61719 | 37557.19141 | 40371.81641 | 35700.75391 | 16396.48047 | 16919.41211 | 15963.1748 |
| 13454.7041 | 22936.58398 | 19063.75781 | 21947.08008 | 31628.75586 | 28262.82031 | 33505.65234 |
| 113913.9375 | 159324.5 | 176882.7344 | 196080.9844 | 160622.375 | 162120.6563 | 176317.0469 |
| 6253.109863 | 9631.880859 | NA | 8690.354492 | 10453.71973 | 10128.86523 | 12229.75 |

| | | | | | | |
|---|---|---|---|---|---|---|
| 33500.67969 | 55384.63281 | 54924.13672 | 54930.32031 | 86741.05469 | 95145.65625 | 86855.6875 |
| 2219.976318 | 6059.308594 | 3686.622803 | 4766.130371 | 7617.027832 | 6008.999512 | 7231.284668 |
| 4239.900879 | 12647.42676 | 6879.461914 | 6851.889648 | 181415.3281 | 58031.65625 | 95873.00781 |
| 34264.25781 | 62794.14453 | 43152.45313 | 64618.5 | 36242.99609 | 48832.55469 | 36586.30859 |
| 17354.51563 | 27239.23242 | 29250.69531 | 29932.3125 | 34575.03906 | 29824.10938 | 30444.85547 |
| 58357.44922 | 74555.9375 | 104150.1172 | 95147.89844 | 79511.07031 | 97202.85156 | 100655.4844 |
| 33302.48047 | 52941.55469 | 48367.16797 | 56197.17969 | 48237.29688 | 54322.42188 | 48784.92188 |
| 17400.10547 | 34923.62891 | 31966.2168 | 30555.17773 | 36301.48047 | 40440.83984 | 38694.20313 |
| 7310.131836 | 16017.62793 | 9760.399414 | 9737.599609 | 8807.188477 | 10820.75 | 9347.380859 |
| 25374.35156 | 43903.06641 | 41908.62891 | 40407.60156 | 37309.14063 | 35231.19531 | 36042.42969 |
| 98475.52368 | 179596.1748 | 151766.2324 | 161977.5527 | 157363.0635 | 177309.9863 | 155674.9561 |
| 18840.27344 | 36357.03906 | 28177.39648 | 33338.21094 | 28977.41211 | 34871.96875 | 32009.5625 |
| NA | 4061.359131 | 2196.164551 | 5108.666992 | 17495.26563 | 12982.90625 | 16600.13086 |
| 1425.692139 | 3790.508789 | 2208.654053 | 3141.481445 | 3028.976318 | 3189.839355 | 2162.880127 |
| 19866.42773 | 32829.38281 | 35486.75781 | 31113.83008 | 33832.37891 | 32081.67383 | 34158.70703 |
| 55435.73438 | 82663.5625 | 97437.96094 | 94188.70313 | 111302.0703 | 99358.25 | 105287.1641 |
| 22744.3418 | 35275.64063 | 39207.1875 | 38215.21875 | 38252.73828 | 43591.94531 | 42312.10156 |
| 3399.579346 | 6308.835938 | 5833.362793 | 4824.36377 | 4744.869629 | 3483.47876 | 4604.268555 |
| 39864.12891 | 66499.54688 | 65414.53125 | 70976.48438 | 91328.40625 | 74312.03906 | 87090.875 |
| 61036.42578 | 106576.3125 | 105044.5313 | 100442.2344 | 116911.3438 | 124316.4922 | 111465.5156 |
| 17666.22656 | 27186.40039 | 23750.38281 | 26483.9375 | 22444.49805 | 21579.01172 | 20044.51563 |
| 6514.547852 | 13007.37891 | 8187.201172 | 8940.767578 | 11144.16699 | 10189.7998 | 10514.96484 |
| 339534.9375 | 610910.1875 | 489351.9063 | 566750.25 | 660106.5625 | 521092 | 519302.5938 |
| 29513.24023 | 39018.64453 | 46404.90625 | 42527.96875 | 65431.70313 | 55725.54297 | 61110.33203 |
| 128633.4063 | 255198.9375 | 220227.7969 | 257295.5469 | 228833.5938 | 229881.2031 | 222485.125 |
| 42949.37109 | 71776.05469 | 68740.6875 | 70040.84375 | 52423.02344 | 56130.98438 | 54510.35547 |
| 3829.121582 | 6419.65918 | 3885.127441 | 5666.412109 | 9685.692383 | 11830.57324 | 4992.92041 |
| 27668.76563 | 43651.32031 | 44202.52344 | 43970.30859 | 48269.94531 | 51237.85156 | 50731.00391 |
| 4101.781738 | 10006.64551 | 6899.289063 | 6683.12207 | NA | 7946.016113 | 6120.748535 |
| 144666.5469 | 229884.1563 | 225088.2969 | 222372.2813 | 182377.2656 | 197269.0781 | 199392.1406 |
| 60535.00391 | 89320.08594 | 104624.5703 | 107339.2578 | 71897.9375 | 59856.46875 | 59154.67578 |
| 23668.22369 | 46841.05054 | 46111.59961 | 46890.23193 | 105299.1162 | 77976.28711 | 74644.22461 |
| 5331.699707 | 7351.12793 | 8303.995117 | 9112.032227 | 10075.5791 | 8364.125 | 7523.135742 |

| | | | | | | |
|---|---|---|---|---|---|---|
| 7737.566406 | 12921.99414 | 10261.56836 | 13028.89258 | 19799.625 | 14847.94336 | 16272.85547 |
| 75429.92188 | 140302.4844 | 119498.0547 | 107094.0859 | 180569.3281 | 162326.6406 | 183719.5469 |
| 92933.70313 | 155455.4375 | 145871.7969 | 161017.7344 | 155322.75 | 158600.8594 | 151440.1875 |
| 50680.625 | 93380.54688 | 85039.82031 | 102890.1797 | 125051.8359 | 112819.8594 | 119085.5313 |
| 252189.4531 | 334401.9688 | 381528.4688 | 522184.7813 | 500571.5625 | 467431.4688 | 583139.5 |
| 23038.63477 | 39981.60547 | 36233.32422 | 40569.90625 | 50771.30859 | 51699.96094 | 52614.35938 |
| 11866.54688 | 20124.73242 | 22095.94141 | 21073.16211 | 15665.15918 | 13037.7168 | 15642.4375 |
| 5171.101563 | 5976.624023 | 5881.990234 | 7991.121582 | 8846.489258 | 8037.078125 | 7041.956543 |
| 822045.3125 | 1451974.25 | 1370010.125 | 1410563.5 | 1205280.125 | 1243526.75 | 1137142.75 |
| 25964.08594 | 36156.82422 | 39840.32422 | 38486.92188 | 35453.22266 | 37779.13672 | 38409.21875 |
| 84649.28125 | 193381.75 | 161901.3281 | 152709.3594 | 186726.6875 | 207847.1875 | 181932.5625 |
| 11795.49512 | 17985.72266 | 18778.79102 | 22266.09766 | 26022.66406 | 22902.30664 | 22169.03711 |
| 3217.036865 | 9409.961914 | 5793.650879 | 4297.144043 | 11116.63672 | 11394.30957 | 11963.23242 |
| 32404.30469 | 59343.82422 | 48952.55078 | 52865.47266 | 50442.85938 | 49399.88672 | 48372.55859 |
| 65737.35156 | 118328.9531 | 99236.75781 | 105622.5625 | 60014.36719 | 50163.33594 | 47347.70313 |
| 3929.384277 | 2046.44397 | 4061.92041 | 8028.715332 | 3220.935303 | 10536.41211 | 10934.37012 |
| 49436.87891 | 86711.52344 | 88879.8125 | 76962.6875 | 61830.46094 | 61790.71094 | 60263.69531 |
| 4569.484375 | 9329.996094 | 8175.02002 | 7563.804199 | 6417.220703 | 7448.404785 | 6692.658691 |
| 123323.9375 | 189168.0156 | 206428.875 | 203777.7969 | 232614.2344 | 210595.2656 | 221018.5938 |
| 4698.290527 | 7506.144043 | 8403.24707 | 10405.86621 | 17814.17383 | 19983.87109 | 23333.20508 |
| 2373.310791 | 4062.948242 | 4307.274902 | 2608.214355 | 2986.269775 | 3893.898193 | 8098.704102 |
| 50251.89453 | 61147.55078 | 33562.93359 | 72807.82813 | 52587.80469 | 78792.53906 | 55565.11719 |
| 19788.6582 | 37504.28516 | 34676.51172 | 29476.23242 | 28257.20313 | 33680.85547 | 28131.42969 |
| 1955.720947 | 3604.445068 | 2972.789063 | 1871.54541 | 3743.643311 | 2520.964844 | 2313.579102 |
| 9157.96582 | 21543.66211 | 17153.17188 | 16240.28027 | 21469.11133 | 20449.96484 | 22134.53906 |
| 79088.86719 | 88211.20313 | 50676.75391 | 117420.6953 | 87714.28906 | 119183.0703 | 83889.64844 |
| 1076.325439 | 1524.680786 | 1633.431763 | 1347.447998 | NA | 1122.060669 | 914.6777344 |
| 11909.05566 | 21996.07031 | 17087.39063 | 15082.19434 | 14480.43066 | 14167.90625 | 11852.21875 |
| 5258.12207 | 7673.233887 | 7898.959473 | 8113.55127 | 8841.355469 | 7705.956543 | 8546.508789 |
| 13732.50684 | 34520.69922 | 15125.66113 | 18701.47656 | 46208.34375 | 38251.18359 | 40954.80859 |
| 137579.9688 | 242958.9219 | 230128.2813 | 224197.5313 | 251068.2969 | 241881.8906 | 235217.4531 |
| 5919.056152 | 11500.73047 | 15083.46973 | 11621.63574 | 11638.1748 | 13750.03418 | 11851.43945 |
| 58121.41797 | 103335.9688 | 83504.625 | 81163.19531 | 57315.66797 | 54658.98438 | 58859.375 |

| | | | | | | |
|---|---|---|---|---|---|---|
| 116879.9063 | 168800.625 | 185802.7969 | 189057.9844 | 221612.6094 | 208360.4844 | 227548.5469 |
| 75708.19531 | 151250.2813 | 111908.8047 | 126687.3359 | 148140.3281 | 127410.2266 | 136289.8906 |
| 132678.4688 | 221370.7969 | 220478.625 | 237726.9063 | 200366.4375 | 198526.8125 | 204150.8906 |
| 20299.9082 | 27902.80078 | 14795.22656 | 25076.42188 | 17679.72461 | 26119.98828 | 19976.35547 |
| 33166.81641 | 13709.96387 | 31213.80469 | 47202.26953 | 101801.8984 | 50051.94531 | 44846.25 |
| 40688.43359 | 74066.625 | 64147.45703 | 74567.88281 | 90100.47656 | 94090.98438 | 89540.125 |
| 26657.1543 | 51467.64844 | 35309.07813 | 30197.57813 | 26783.04102 | 36018.81641 | 26088.56641 |
| 21840.24219 | 32771.53125 | 37484.89063 | 35046.83984 | 37484.20313 | 36039.69141 | 35683.28516 |
| 2292.186035 | 2754.214355 | 3551.052002 | 4404.235352 | 5346.740234 | 4959.652832 | 4913.150391 |
| 8445.006836 | 14080.21777 | 13610.84961 | 13910.30371 | 14857.24219 | 14470.04004 | 14199.52539 |
| 39776.78906 | 70442.25 | 64395.44531 | 66110.20313 | 50011.94141 | 42073.07813 | 44939.0625 |
| 2499.858398 | 4339.316406 | 4732.137207 | 4353.416992 | 5719.529297 | 7448.927734 | 6005.330566 |
| 9536.68457 | 11998.78516 | 11825.23145 | 11026.76953 | 9590.44043 | 10365.89551 | 8375.614258 |
| 14401.96582 | 23146.38477 | 30132.56836 | 25436.0957 | 40688.30469 | 41204.37109 | 36814.63672 |
| 8320.301758 | 15605.01367 | 13946.99902 | 14298.95117 | 23959.89844 | 24752.24609 | 25876.39063 |
| 79437.53906 | 143557.125 | 126651.2813 | 127327.3203 | 98836.08594 | 99580.33594 | 95805.60938 |
| 4045.760254 | 6734.466797 | 6286.348633 | 6957.510254 | 4854.994141 | 5232.30957 | 4297.652832 |
| 11892.46875 | 21256.31445 | 17632.13281 | 17423.32031 | 32743.96484 | 27934.26367 | 30646.25 |
| 47429.78906 | 82917.89063 | 85039.14844 | 100993.2578 | 246755.1875 | 180061.2969 | 176519.3438 |
| 164934.7031 | 295225.4688 | 251172.9375 | 245775.0156 | 206020.5 | 259087.9375 | 224626.2031 |
| 68162.375 | 107470.1484 | 117981.7891 | 123637.6563 | 103288.9219 | 101241.7891 | 109083.3359 |
| 25847.56836 | 35763.68359 | 40841.49219 | 40247.15625 | 52796.87109 | 68425.13281 | 63504.01953 |
| 8320.516602 | 16833.21094 | 11847.72656 | 11172.19336 | 8238.772461 | 10747.72363 | 8349.387695 |
| 5201.175781 | 8953.728516 | 6183.873535 | 7702.315918 | 16139.06348 | 12590.09375 | 15332.93359 |
| 16752.38477 | 27288.85156 | 5911.494141 | 21708.65234 | 21014.58203 | 28349.39844 | 16475.27734 |
| 2679.72583 | 9166.050781 | 8806.186523 | 7708.010742 | 11378.08984 | 10786.89648 | 11048.16406 |
| 1030.936646 | 1936.12793 | 1774.947876 | 2063.600342 | 2718.178711 | 2640.750732 | 2319.835938 |
| 257187.8438 | 440649.5313 | 417012 | 440796.3125 | 300534.7188 | 324185.7813 | 346398.4688 |
| 5328.277344 | 11059.65332 | 8493.22168 | 8360.132813 | 11261.09082 | 9560.003906 | 10737.45801 |
| 95148.01563 | 165927.2656 | 150682.7188 | 134480.375 | 129129.2109 | 129235.9063 | 133301.1719 |
| 125992.0859 | 182090.6875 | 194317.75 | 161980.125 | 202364.375 | 192291.1719 | 214347.2344 |
| 60424.69922 | 96395.10156 | 100452.8828 | 74718.32813 | 72149.71875 | 80457.07031 | 56512.20313 |
| 19540.33789 | 39156.60156 | 28597.68945 | 28358.71289 | 26480.23047 | 23286.4707 | 24851.28906 |

| | | | | | | |
|---|---|---|---|---|---|---|
| 52295.79688 | 85215.4375 | 92134.85938 | 82577.71875 | 108170.8125 | 109443.2422 | 108601.5781 |
| 12228.59375 | 18471.4375 | 21570.39844 | 18861.94336 | 29697.39453 | 30204.27148 | 30720.81445 |
| 94732.90625 | 234288.3125 | 134477.8906 | 167100.375 | 256437.6406 | 130004.4219 | 227062.3906 |
| 3000.376465 | 4472.074707 | 5202.53418 | 5601.281738 | 10738.69141 | 10359.24023 | 10688.65332 |
| 5426.512207 | 10028.38184 | 7250.930664 | 4347.956543 | 477.6147461 | 1943.914673 | 1907.0802 |
| 10502.13672 | 15443.56543 | 11896.0918 | 14869.74121 | 12150.30762 | 29829.28125 | 17828.35156 |
| 37245.17969 | 43323.0625 | 26494.57031 | 52973.59766 | 43192.19141 | 59109.28516 | 44634.08203 |
| 15590.56641 | 24669.6582 | 21893.64648 | 23590.88086 | 15693.70508 | 15390.12988 | 14967.94043 |
| 3298.854492 | NA | 6054.264648 | 8950.084961 | 24975.8418 | 24283.82422 | 27207.83789 |
| 23814.79297 | 37741.21094 | 42565.46875 | 39376.29688 | 37978.31641 | 39525.16406 | 40280.98828 |
| 406229 | 661730.625 | 722226.5 | 728932.75 | 843485.625 | 774802 | 801068.8125 |
| 48786.67188 | 81280.51563 | 77389.16406 | 86795.66406 | 111722.6484 | 118919.3438 | 98674.94531 |
| 2125.858398 | 4568.821289 | 3137.329102 | 2550.105225 | 2429.010742 | 2911.754639 | 1505.28125 |
| 1671.779419 | 3290.927979 | 2482.198975 | 2691.816406 | 4578.83252 | 5010.446777 | 5738.436035 |
| 675290.5625 | 1178736.75 | 1079427.625 | 1185304.25 | 1076967 | 1074953.5 | 926248.875 |
| 13143.47168 | 18758.04492 | 16833.82031 | 16295.0791 | 12077.34668 | 15277.17383 | 11286.95313 |
| 21664.36328 | 41822.60547 | 20542.93164 | 17396.60742 | 17789.20117 | 16110.46191 | 12080.87402 |
| 165003.2031 | 282499.125 | 255649.9375 | 270724.4688 | 255982.8281 | 220523.2969 | 228468.7969 |
| 59504.02734 | 121691.9609 | 91818.78125 | 98774.8125 | 96504.52344 | 100924.6484 | 97874.07813 |
| 42727.89453 | 76285.23438 | 65602.4375 | 74820 | 87133.8125 | 83863.72656 | 80695.52344 |
| 613000.5 | 1137615.875 | 873283.6875 | 958360.375 | 1254919.75 | 1142470.875 | 1123762.875 |
| 552054.4375 | 846424.5625 | 859122.75 | 872485.0625 | 648324.75 | 710375.125 | 727693.1875 |
| 1386.848267 | 2257.070313 | 2421.731445 | 2018.197144 | 2615.926758 | 2269.229736 | 2184.598145 |
| 15324.02539 | 29572.19141 | 21215.61914 | 22292.39063 | 17647.49219 | 20693.61133 | 15617.97949 |
| 8924.204102 | 17421.85938 | 13434.03516 | 12407.42773 | 25085.9082 | 22248.00977 | 21358.9375 |
| 89059.83594 | 104062.1094 | 163458.7969 | 153944.4219 | 133992.7656 | 117869.8594 | 134759.6094 |
| 32349.39063 | 54736.03125 | 48672.41016 | 53133.79688 | 92502.55469 | 72975.44531 | 80736.6875 |
| 9793.02832 | 13189.73926 | 21892.25781 | 21211.31445 | 25738.69531 | 23405.28125 | 30884.41797 |
| 25130.12891 | 44645.39063 | 33671.66406 | 29669.55078 | 13039.80273 | 18841.58594 | 15461.70898 |
| 29903.40625 | 40597.53516 | 19100.92969 | 41929.49219 | 32683.04688 | 40083.89453 | 32447.45117 |
| 198349.7813 | 323048.5625 | 310997.9063 | 299795.9063 | 137352.5156 | 143756.6094 | 139620.4688 |
| 4968.265137 | 8906.12207 | 7650.442871 | 7088.569824 | 10584.91211 | 10098.25781 | 8267.933594 |
| 33852.57813 | 51788.07031 | 50943.45703 | 55649.54688 | 40164.61719 | 44474.60938 | 39789.05859 |

| | | | | | | |
|---|---|---|---|---|---|---|
| 31438.99023 | 43152.95313 | 49375.46484 | 43790.71875 | 27944.78125 | 31536.85156 | 30138.17188 |
| 24439.95117 | 25601.49219 | 40808.32031 | 45991.13281 | 34686.23828 | 27662.26367 | 35549.21875 |
| 320257.1875 | 562784.4375 | 516371.75 | 593105.1875 | 636274.25 | 540146.8125 | 560847.5625 |
| 19672.08203 | 30688.43164 | 33550.86719 | 33063.36719 | 31596.375 | 34237.29688 | 32202.62109 |
| 265657.4063 | 477498.75 | 453937.4375 | 405021.4688 | 366423.5625 | 366685.9375 | 342605.9375 |
| 16831.08203 | 29805.63672 | 27003.29102 | 27641.0625 | 36420.15234 | 32333.97656 | 36444.01953 |
| 51399.78516 | 89504.40625 | 80276.03125 | 83893.79688 | 67918.10938 | 64665.52734 | 64578.96875 |
| 228278.7188 | 328679.6875 | 379310.0625 | 372697.3125 | 343558.9375 | 365750.5938 | 352491.9063 |
| 11031.18652 | 19511.41406 | 18121.32227 | 20114.01172 | 21059.94922 | 19109.45898 | 18421.41016 |
| 10441.70996 | 18381.04883 | 18190.32422 | 17036.78711 | 5271.157715 | 6292.71582 | 5898.923828 |
| 51813.23828 | 72251.77344 | 82618.80469 | 105927.2031 | 91039.98438 | 76841.08594 | 83148.46094 |
| 5867.392578 | 6554.326172 | 6216.820801 | 10728.58691 | 58821.89844 | 42435.92969 | 54649.23828 |
| 22793.875 | 42907.67578 | 33255.1875 | 33408.57422 | 32007.95117 | 35687.72656 | 37094.28516 |
| 13212.31934 | 19784.39453 | 24142.49023 | 20614.11328 | 32358.7793 | 25824.16016 | 29894.23438 |
| 68675.52344 | 80562.16406 | 39154.75 | 89926.96875 | 67815.33594 | 99898.67188 | 69297.78906 |
| 69223.17188 | 108919.4531 | 111634.2422 | 103566.6016 | 80508.39844 | 80721.13281 | 81239.42188 |
| 11816.1875 | 14415.40039 | 18858.80859 | 18636.85938 | 9829.117188 | 6062.67041 | 11034.25977 |
| 36392.85547 | 51905.40625 | 53211.5625 | 49156.82813 | 43604.03906 | 44375.60938 | 45255.14453 |
| 62691.0625 | 89778.36719 | 92685.76563 | 92401.20313 | 69793.71094 | 67338.16406 | 70667.59375 |
| 25772.79102 | 43633.00781 | 31045.55273 | 37239.50391 | 32477.74609 | 27242.63672 | 27260.9375 |
| 46439.73828 | 76936.38281 | 70223.28906 | 75336.67188 | 51437.22266 | 38282.64063 | 43592.88281 |
| 82677.17188 | 130268.9688 | 122302.25 | 121180.375 | 61328.50391 | 57924.99609 | 57096.82813 |
| 7380.214355 | 17629.46289 | 11030.10742 | 13810.9834 | 19266.74609 | 16626.60156 | 15650.87988 |
| 48596.99219 | 92725.58594 | 68351.92969 | 67686.76563 | 96288.10156 | 75703.35156 | 75327.23438 |
| 278953.5625 | 517829.1875 | 419986.25 | 471994.8125 | 858102.8125 | 960034.5 | 892743.125 |
| 23326.32031 | 33805.13672 | 34572.01563 | 34487.87891 | 17208.43945 | 20697.64063 | 18854.06641 |
| 128738.1426 | 227344.2383 | 191069.7832 | 193171.4434 | 171568.5664 | 190192.7383 | 184497.1094 |
| 390903.3125 | 565986.5625 | 578674.4375 | 669072.8125 | 964227.9375 | 940478.5 | 954936.5 |
| 34011.28906 | 61245.8125 | 56528.41406 | 54883.99219 | 41863.87109 | 40557.56641 | 44773.01172 |
| 26475.44727 | 43203.85547 | 42869.44141 | 43871.36719 | 40918.04688 | 40922.40625 | 40957.04297 |
| 12587.28125 | 14081.4668 | 20752.36133 | 20473.9082 | 34667.08594 | 40618.38672 | 39961.91797 |
| 51269.83203 | 79303.98438 | 80055.07813 | 78328.22656 | 49179.51563 | 48269.18359 | 53064.64453 |
| 7596.544434 | 13770.03516 | 11972.2627 | 9620.591797 | 6351.804199 | 10122.94043 | 9154.769531 |

| | | | | | | |
|---|---|---|---|---|---|---|
| 16893.19531 | 28096.01758 | 19958.33203 | 22128.79102 | 27802.08594 | 30454.63477 | 26197.58398 |
| 149957.6094 | 216837.5156 | 267847 | 241495.3438 | 242676.4219 | 308373.4688 | 283016.625 |
| 117970.5781 | 203450.2031 | 191174.5781 | 196788.6094 | 144501.4375 | 156611.2969 | 151629.3438 |
| 51497.10938 | 92997.47656 | 84914.5 | 93124.08594 | 87224.63281 | 60722.61719 | 69214.63281 |
| 14823.32031 | 22303.71289 | 24404.67969 | 25059.04102 | 25862.32813 | 24622.0332 | 25023.17969 |
| 132079.3594 | 215751.1094 | 205062.3438 | 205634.4844 | 204188.125 | 209581.4375 | 216129.0469 |
| 113690.3828 | 176889.5781 | 164940.875 | 173529.2031 | 149881.5625 | 149835.75 | 139129.75 |
| 3937.996094 | 10228.32813 | 7250.42627 | 10810.79199 | 6675.585449 | 5091.552246 | 4539.344238 |
| 26330.52539 | 43726.26563 | 52354.36719 | 41943.76172 | 52096.80078 | 31380.02539 | 30618.76953 |
| 23809.67578 | 40970.86719 | 37825.27344 | 37250.82031 | 35923.07031 | 33617.71875 | 33551.92969 |
| 138363.25 | 200289.5469 | 232472.2188 | 202995.5313 | 174430.4531 | 169191.0156 | 184313.4688 |
| 11475.31445 | 19831.44336 | 19456.56641 | 19320.17578 | 14308.15039 | 14734.68555 | 14095.78418 |
| 59158.04297 | 97776.70313 | 90454.19531 | 94479.28906 | 112556.6406 | 126506.3906 | 125150.875 |
| 22611.74805 | 33340.625 | 33678.85938 | 34734.42578 | 30426.95313 | 30415.22852 | 28155.93945 |
| 17743.97461 | 30982.14258 | 32337.02344 | 31634.41602 | 28338.44531 | 26970.17383 | 30749.10742 |
| 9772.523438 | 13784.48926 | 11294.2334 | 12838.07715 | 11616.71094 | 13770.53516 | 13327.32715 |
| 32047.9707 | 53737.74219 | 52514.19141 | 55281.55078 | 59561.83203 | 43781.23047 | 48542.75 |
| 23412.48242 | 38342.76953 | 34902.44141 | 33222.24219 | 30601.6543 | 31452.39453 | 28297.01758 |
| 160293.375 | 173444.3906 | 87969.91406 | 210226.7656 | 153421.75 | 215634.7969 | 155349.9688 |
| 8255.236328 | 13078.77832 | 13826.59082 | 12563.60938 | 16498.33008 | 17160.75781 | 16684.11719 |
| 11474.87402 | 14964.85938 | 18454.51172 | 17947.91016 | 15379.38965 | 15165.69629 | 14301.37891 |
| 10735.51074 | 14191.34668 | 16519.51758 | 17866.33203 | 19728.35156 | 22861.24609 | 16966.99414 |
| 22566.52148 | 36256.69678 | 35943.28125 | 42782.44604 | 48445.78809 | 41750.30713 | 46633.80811 |
| 42185.30859 | 45855.63672 | 27147.30273 | 53492.27734 | 43900.69141 | 57093.69141 | 41726.20313 |
| 410153.1875 | 574307.8125 | 691543.0625 | 661428.75 | 629288.6875 | 580354 | 607724.25 |
| 8719.458008 | 11155.96582 | 17121.70898 | 16099.12012 | 20890.36133 | 21557.21094 | 22539.17383 |
| 10391.47363 | 16509.14453 | 18002.74805 | 16913.66016 | 28860.61328 | 19784.54688 | 19129.70703 |
| 43034.72266 | 82363.27344 | 63266.49609 | 59615.13672 | 54281.42578 | 55977 | 56920.58203 |
| 6737.883301 | 10662.34863 | 9750.827148 | 8176.619141 | 8672.03125 | 9689.933594 | 8365.213867 |
| 114259.9609 | 159462 | 188734.1563 | 201707.6406 | 185052.4531 | 160587.9844 | 171322.5313 |
| 20326.06641 | 30592.20313 | 30008.97461 | 33104.96875 | 20249.36133 | 21873.00977 | 21120.59961 |
| 2782567 | 4063447.25 | 4019759.5 | 5644129.5 | 6021790 | 4442987 | 4806580.5 |
| 116182.7578 | 177264.875 | 175466.75 | 182204.0313 | 144012.0625 | 145814.1875 | 147491.3594 |

| | | | | | | |
|---|---|---|---|---|---|---|
| 11919.81641 | 19728.42969 | 19768.40625 | 19589.76953 | 30850.46289 | 26320.81836 | 28792.80859 |
| 4727.959961 | 9084.450195 | 8325.782227 | 8098.462402 | 8183.285645 | 9653.850586 | 9483.689453 |
| 118421.9063 | 194664.7188 | 194568.6719 | 180502.7188 | 110207.5 | 112933.1641 | 117074.9688 |
| 14602.10645 | 26602.50391 | 22522.87695 | 24379.5625 | 22088.87891 | 24062.81055 | 23008.9707 |
| 410204.5938 | 647069.875 | 638639.125 | 651304.8125 | 491117.75 | 506093.75 | 516593.4688 |
| 755.3782959 | 1805.412476 | 1114.976563 | 1377.28418 | 1357.454224 | 1000.947449 | 964.4389648 |
| 73520.03906 | 122083.5703 | 107076.5078 | 105313.5625 | 99301.9375 | 92578.42188 | 97653.9375 |
| 29909446 | 31125108 | 16466913 | 39050588 | 29720126 | 42283340 | 29022736 |
| 16346.73047 | 24874.10547 | 24944.91016 | 22566.16211 | 8029.157715 | 7876.976563 | 7595.459473 |
| 208123.7031 | 269084.0938 | 320783 | 388246.0938 | 343156.8125 | 309783.7188 | 380806.0313 |
| 953111.6875 | 1400512.5 | 1623395.5 | 1556429.875 | 1258409.25 | 1346434.75 | 1383537 |
| 15362.14941 | 23544.89258 | 18901.32422 | 22187 | 17061.58398 | 15447.90918 | 16317.87695 |
| 11512.94336 | 13851.39648 | 17710.48828 | 26003.21875 | 31923.85938 | 31544.68164 | 32114.64453 |
| 42853.01563 | 165845.8125 | 31390.35156 | 32304.75977 | 125115.8906 | 113084.6641 | 137605.0469 |
| 19898.26758 | 37257.87109 | 39834.28125 | 39821.66797 | 37839.41016 | 34350.89453 | 33997.99609 |
| 19356.77539 | 28466.83203 | 24753.07227 | 26591.83008 | 12750.89355 | 17707.97461 | 15094.38672 |
| 191375.7656 | 317994.9063 | 331999.2188 | 287185.5 | 455769.8125 | 464207.4063 | 448365.4063 |
| 218009.7344 | 303946.1875 | 345796.9375 | 361736.875 | 309284.9688 | 261204.4688 | 263644.9063 |
| 4568268.5 | 7448111 | 6822087.5 | 7053475 | 6715088 | 6069439 | 6448563 |
| 1391.084839 | 3387.097656 NA | | 2297.100586 | 2440.709229 | 4823.626953 | 4696.285645 |
| 22285.21094 | 34272.61328 | 35066.65625 | 37024.40234 | 49660.63281 | 50551.36328 | 45575.33203 |
| 20187.08594 | 37891.09375 | 32142.37891 | 30892.86719 | 35237.60547 | 35079.85938 | 32814.5625 |
| 968.355957 | 12529.54688 | 14093.69629 | 519.3699951 | 6919.251465 | 3717.331787 | 2229.776123 |
| 4313.13623 | 8892.048828 | 8074.908203 | 5475.047363 | 13891.10645 | 16050.53418 | 12952.2334 |
| 13509.22559 | 39823.43359 | 31006.18164 | 33009.81641 | 52395.67969 | 66393.95313 | 56151.63281 |
| 8320.170898 | 8665.836914 | 10261.13965 | 23817.54492 | 31971.13867 | 9302.243164 | 10796.86621 |
| 8609.924805 | 9501.113281 | 16773.20508 | 9982.663086 | 17036.25586 | 13970.44629 | 6042.889648 |
| 45979.86719 | 75867.89844 | 56874.09375 | 63091.79688 | 63487.78125 | 72046.53125 | 67410.625 |
| 28629.07422 | 43540.35547 | 46171.95703 | 35707.21094 | 29204.53516 | 31046.93164 | 26280.98633 |
| 3481.789307 | 5172.093262 | 4666.000977 | 4422.024902 | 2697.79126 | 2400.415039 | 2417.369873 |
| 6726.211426 | 13760.50781 | 7986.83252 | 8818.666016 | 16975.11328 | 15677.46777 | 14679.5166 |
| 40663.26563 | 67451.3125 | 62933.23047 | 63809.65234 | 47481.35938 | 47223.08594 | 42814.375 |
| 38083.70313 | 63546.54688 | 56488.28125 | 63579.65234 | 60075.98438 | 62655.13281 | 60891.37891 |

| | | | | | | | |
|---|---|---|---|---|---|---|---|
| | 11925.34961 | 18058.12695 | 17009.92383 | 17929.25586 | 19400.17969 | 17054.58789 | 18156.7832 |
| | 7009.033203 | 12410.97168 | 10016.37988 | 12844.49316 | 6146.228027 | 9164.360352 | 7101.069824 |
| | 2221.672852 | 3933.132568 | 2721.314941 | 2987.442383 | 2752.106445 | 2671.456543 | 2640.118896 |
| | 27502.70508 | 44988.58594 | 45177.10938 | 44445.48047 | 43201.58203 | 51248.78906 | 47904.69141 |
| | 39297.81641 | 73748.45313 | 61700.00781 | 59443.38672 | 104570.4766 | 100671.4375 | 100760.1797 |
| | 19298.33984 | 26971.53125 | 27665.51367 | 31745.40234 | 27122.16211 | 24697.20508 | 27258.08398 |
| | 43830.46875 | 67476.13281 | 68606.8125 | 66484.77344 | 52831.30078 | 52545.66797 | 49869.08594 |
| | 83436.37549 | 118999.5151 | 117835.3555 | 127981.8047 | 111476.6982 | 115617.8047 | 110284.6724 |
| | 10540.40039 | 21668.76758 | 18428.29492 | 18227.20703 | 26097.16992 | 17576.56445 | 17912.14258 |
| | 33626.55859 | 54970.21484 | 51133.10938 | 49135.50781 | 52286.86328 | 51874.77344 | 49683.96875 |
| | 5055.834961 | 8229.261719 | 7865.088867 | 7025.869141 | 5739.257324 | 7861.512695 | 7132.570801 |
| | 91935.25 | 136808.0938 | 145225.3125 | 127522.2344 | 117290.0156 | 125106.1406 | 125654.8125 |
| | 9370.283203 | 14278.1123 | 17561.12305 | 19591.79102 | 10956.10156 | 9062.474609 | 9076.96582 |
| | 255908.0625 | 403429.3125 | 370182.875 | 460011.9375 | 377932.875 | 408030.1563 | 399687.0938 |
| | 2419.259033 | 3563.359375 | 3718.341553 | 4369.60791 | 3751.168457 | 4030.762939 | 4542.799316 |
| | 138475.6094 | 223998.5 | 218891.3594 | 232267.7813 | 168506 | 168286.4375 | 162870.8125 |
| | 55869.26953 | 81065.57813 | 89310.25781 | 84918.98438 | 86937.75 | 110626.0234 | 114393.25 |
| | 7439.065918 | 6705.574219 | 24299.89453 | 4217.117188 | 9856.124023 | 8029.931152 | 9154.65918 |
| NA | | 3995.071777 | 3203.601074 | 4240.767578 | 1567.95874 | 2600.963135 | 2632.897217 |
| | 8433.662109 | 17406.83594 | 16470.66211 | 15246.74609 | 24368.04688 | 17225.47266 | 16883.59766 |
| | 75734.83643 | 140203.8252 | 98067.07568 | 93772.33008 | 175197.0762 | 146944.2246 | 134746.6025 |
| | 3409.791016 | 7825.675293 | 3678.278076 | 4980.810059 | 8433.358398 | 8858.782227 | 7248.831543 |
| | 26364.46484 | 46793.10156 | 44926.75 | 39823.94531 | 43298.70313 | 47148.46484 | 43056.58203 |
| | 22067.14648 | 38535.55859 | 32997.51563 | 37351.85156 | 33064.55469 | 37838.24219 | 33356.83594 |
| | 2886.796387 | 7208.651367 | 3486.005615 | 6505.123535 | 11873.85938 | 8520.699219 | 6943.721191 |
| | 4810.86377 | 4660.82959 | 8854.915039 | 4393.116211 | 15270.16992 | 9001.855469 | 14326.51758 |
| | 17203.63672 | 27869.38672 | 28789.57031 | 29209.80078 | 33238.86328 | 33143.46094 | 31943.92578 |
| | 29371.19141 | 48077.54688 | 45769.20703 | 47912.08594 | 55527.55469 | 54628.12109 | 56764.57031 |
| | 69131.24219 | 123345.7734 | 124268.8672 | 110749.2344 | 140376.9844 | 115071.0313 | 115840.3438 |
| | 90807.27539 | 130111.3086 | 127198.5947 | 148334.835 | 120426.1367 | 94800.75781 | 98069.25439 |
| | 25167.45898 | 38462.36719 | 38136.58594 | 38526.75781 | 44442.625 | 45510.60547 | 45333.67578 |
| | 42948.80078 | 70702.35938 | 59276.05859 | 59345.34766 | 53936.43359 | 46024.19531 | 46825.44531 |
| | 82315.25781 | 110780.6328 | 110104.5078 | 124091.1016 | 115090.9844 | 104147.4688 | 108440.1172 |

| | | | | | | |
|---|---|---|---|---|---|---|
| 104325.6875 | 142047.7031 | 145162.6719 | 158401.0625 | 199299.2188 | 180935 | 186948.6563 |
| 11061.02344 | 14400.89844 | 17397.7832 | 17251.6875 | 19528.77148 | 18111.38867 | 19412.41602 |
| 59592.13672 | 111679.25 | 93964.85938 | 97779.03125 | 56507.91797 | 52590.55469 | 55146.82422 |
| 272703.0313 | 467217.5313 | 381954.3438 | 489484.375 | 458196.875 | 374723.625 | 372271.5313 |
| 19969.7832 | 27121.78906 | 30252.2793 | 33943.90625 | 24026.17773 | 23706.52344 | 24635.5625 |
| 6447.756348 | 9737.69043 | 10106.38281 | 10817.89355 | 8055.207031 | 9872.530273 | 9896.560547 |
| 853.4979858 | NA | 1810.175049 | NA | 1467.349121 | 1586.654419 | NA |
| 2851.59082 | 4451.547363 | 3732.900879 | 3507.805176 | 3328.842529 | 3294.96167 | 3589.165039 |
| 660187.1016 | 936042.9297 | 1015652.453 | 1020792.137 | 956794.7656 | 933779.6094 | 967938.2461 |
| 5852.362305 | 7585.992188 | 9924.958984 | 9708.083008 | 7238.216797 | 7223.116699 | 6762.815918 |
| 14653.2207 | 23011.71484 | 34140.94141 | 19830.30273 | 87605.32031 | 21047.71094 | 40028.55859 |
| 20252.26563 | 34778.86328 | 27021.74609 | 29142.85156 | 32420.17383 | 26937.79297 | 29407.53711 |
| 1687.810791 | 3146.988525 | 3253.735107 | 3480.465088 | 4100.168945 | 3455.63623 | 3294.631836 |
| 192973.25 | 394492.875 | 292248.125 | 330940.9688 | 286098.2188 | 349414.375 | 287023.25 |
| 66194.97656 | 99554.65625 | 99228.94531 | 102074.9688 | 102957.4531 | 89177.95313 | 94987.88281 |
| 40896.27344 | 72243.65625 | 52524.01563 | 57557.46875 | 59391.58984 | 65767.96875 | 55452.80469 |
| 5927.839355 | 13422.01367 | 8167.509277 | 10464.00781 | 6133.286621 | 7697.595215 | 7887.452148 |
| 3997.354004 | 7405.993164 | 3628.782959 | 6158.181641 | 5577.605957 | 5903.331055 | 6474.117676 |
| 6480.42041 | 8564.167969 | 15335.29102 | 5330.923828 | 8698.59375 | 9448.678711 | 6251.912109 |
| 76640.84375 | 118259.2578 | 122364.6406 | 123575.3047 | 111421.1719 | 108757.8203 | 108216.5547 |
| 6006.137207 | 6197.519043 | 9490.927734 | 5649.674316 | 1860.964233 | 2650.462646 | 1601.134277 |
| 9722.795898 | 15694.73633 | 17017.46289 | 18116.29102 | 16098.78027 | 15726.89941 | 15188.53711 |
| 174361.0469 | 237413.375 | 297313.4375 | 279702.2813 | 246927.3438 | 241494.3438 | 248006.5469 |
| 97370.22656 | 133285.4063 | 156934.5781 | 140376.4531 | 91346.63281 | 99605.69531 | 98364.52344 |
| 330125.5313 | 439195.4375 | 468643.5625 | 461279.4688 | 487953.5625 | 525459.375 | 545571.75 |
| 107048.0703 | 180195.25 | 165191.0625 | 167081.1406 | 202671.3438 | 181655.5781 | 192071.0313 |
| 48786.76563 | 71540.78125 | 72876.76563 | 75381.23438 | 79241.60156 | 72109.25781 | 74726.15625 |
| 9524.536133 | 14563.96777 | 4798.70166 | 15032.38086 | 12062.1416 | 18958.56641 | 14414.05762 |
| 416657.9375 | 591001.9375 | 635417.4375 | 686390.3125 | 783473.875 | 786262.6875 | 792065.375 |
| 5165.98584 | 11744.47754 | 8114.770508 | 7220.75 | 10880.43359 | 7497.128906 | 9211.141602 |
| 19379.16797 | 17187.58789 | 27839.57813 | 28086.48242 | 42597.13672 | 44993.52734 | 50702.32813 |
| 36824.53516 | 52585.11719 | 56142.12109 | 61005.33984 | 58660.89063 | 61331.51953 | 61996.78906 |
| 75151.03906 | 110723.6328 | 125556.6016 | 120378.6406 | 119054.2109 | 128164.4531 | 117420.5781 |

| | | | | | | |
|---|---|---|---|---|---|---|
| 15405.1123 | 18170.12109 | 21837.83594 | 22298.49805 | 29176.65625 | 23935.85547 | 28850.67188 |
| 5508.883301 | 8293.494141 | 8981.241211 | 7882.210449 | 12255.44043 | 13220.63867 | 9109.828125 |
| 18172.67383 | 30314.9375 | 27938.14844 | 29342.12109 | 25358.39453 | 24540.17969 | 23847.91211 |
| 76061.92969 | 111965.1406 | 113658.4609 | 114994.6719 | 117056.0938 | 105607.7109 | 116185.3281 |
| 256202.125 | 476695.8125 | 353872.5 | 392801.4688 | 738507.6875 | 540084.875 | 627166.75 |
| 12421.16797 | 25328.27734 | 22110.85352 | 22418.23633 | 16998.5 | 13878.75195 | 18763.95313 |
| 695.2211304 | 1726.13916 | 1104.562012 | 1834.755005 | 1266.94458 | 1071.671143 | 1281.513306 |
| 282891.4063 | 298569.5 | 161649.125 | 361127.5938 | 272649.7813 | 398453.3125 | 275228.2813 |
| 19896.41992 | 41684.72266 | 37784.24219 | 43746.60938 | 51651.30469 | 50454.92969 | 51881.40625 |
| 21233.91797 | 32071.81641 | 32575.76758 | 30638.19727 | 27123.70508 | 30979.78125 | 29194.02734 |
| 89139.24219 | 167023.625 | 138725.3906 | 155982.0469 | 256783.0938 | 205779.2188 | 214281.9219 |
| 424621.9063 | 657510.875 | 598103.125 | 633865.3125 | 645681.5 | 671497.375 | 674979.4375 |
| 16786.78711 | 26203.11133 | 28000.69141 | 28131.98828 | 26855.37109 | 29521.82031 | 27230.53906 |
| 12373.96289 | 21789.71094 | 14791.22168 | 19014.25 | 22330.50977 | 21823.40234 | 19907.20898 |
| 13201.22754 | 20122.33594 | 18391.99805 | 16890.5957 | 13035.37207 | 14433.85156 | 14010.8623 |
| 7082.727051 | 15073.05176 | 13421.02832 | 13849.80469 | 16677.96484 | 16825.53125 | 15296.46094 |
| 9159.566406 | 17578.55664 | 17633.8125 | 14451.05273 | 11639.2334 | 17073.19336 | 12674.06152 |
| 81412.60938 | 120639.5234 | 112683.2578 | 120079.4766 | 96326.59375 | 99867.60938 | 100158.3516 |
| 160390.6719 | 217627.6094 | 240235.1094 | 262699.75 | 278462.625 | 234936.6563 | 280203.5313 |
| 2444.543945 | 3557.734375 | 3527.058838 | 3817.067627 | 1625.348267 | 1819.484497 | 1844.295166 |
| 80370.47656 | 110611.1719 | 134291.3438 | 131081.4531 | 92225.91406 | 83766.85938 | 85796.45313 |
| 72544.89063 | 117408.4766 | 103525.8359 | 109543.1406 | 98248.44531 | 99770.53125 | 94995.32813 |
| 14475.49902 | 24498.05078 | 19049.10352 | 23681.68945 | 24284.02148 | 23223.88477 | 24356.78516 |
| 16923.11523 | 16981.48242 | 10138.11035 | 21294.88672 | 17066.58008 | 24819.35938 | 17624.70313 |
| 8340.951172 | 16278.74219 | 13123.89258 | 14313.45703 | 9844.805664 | 10011.44824 | 9354.004883 |
| 7879.946777 | 15746.97852 | 19220.69141 | 9395.787109 | 21992.97461 | 16628.24609 | 14916.11133 |
| 9205.915039 | 10501.09473 | 4249.063477 | 11616.61035 | 9982.603516 | 14457.48828 | 7203.168457 |
| 272009.9063 | 439613.4688 | 388857.75 | 418265.5625 | 341790.75 | 367119.9375 | 358939 |
| 57791.20313 | 97072.64844 | 89291.99219 | 90386.96875 | 96827.80469 | 99001.29688 | 93061.09375 |
| 9805.522461 | 12541.64258 | 13049.59766 | 12080.26465 | 9595.797852 | 9748.783203 | 12607.1582 |
| 358392.375 | 394163.1875 | 225542.1719 | 483079.4375 | 412307.0313 | 645078.9375 | 473186.1875 |
| 4361.148926 | 6137.001465 | 6838.166504 | 7147.275391 | 15462.51172 | 10955.94238 | 15054.56641 |
| 46830.85938 | 76024.53125 | 64927.50391 | 67250.82031 | 60153.36328 | 63604.71484 | 62153.62109 |

| | | | | | | |
|---|---|---|---|---|---|---|
| 6155.141602 | 7002.819336 | 8147.44043 | 7596.072754 | 4141.5625 | 4666.243164 | 5155.322266 |
| 129771.75 | 172853.4688 | 204150.2969 | 196759.6719 | 151495.4063 | 160388.7344 | 164834.8906 |
| 34799.73047 | 47893.77344 | 59508.46875 | 43715.66797 | 62221.44531 | 60949.67578 | 54403.82031 |
| 28872.7832 | 55655.38281 | 35861.40234 | 33483.48828 | 29916.55859 | 29365.55273 | 29268.13867 |
| 9448.851563 | 17380.51367 | 13416.54883 | 20997.87109 | 21368.04102 | 20707.42969 | 22439.35352 |
| 56397.30078 | 84492.82031 | 75644.53906 | 73642.10156 | 112454.6641 | 104087.1641 | 111643.4219 |
| 35352.48438 | 97413.14844 | 55751.8125 | 68426.88281 | 92629.03125 | 64376.56641 | 61262.82031 |
| 581826.6875 | 854020.375 | 836284.375 | 906791.375 | 787375.1875 | 648051.5 | 710677.1875 |
| 121837.8594 | 187743.7969 | 182033.6406 | 197099.1719 | 114888.9219 | 114865.1953 | 116416.4688 |
| 8130.908691 | 13225.91504 | 10802.59473 | 10555.41211 | 19218.69727 | 19645.18555 | 17827.82813 |
| 83042.35938 | 127094.3828 | 121278.5313 | 128027.3203 | 122020.9688 | 115772.3047 | 116533.5703 |
| 87923.17969 | 127589.9922 | 138140.9219 | 144171.0313 | 143464.8281 | 126684.7188 | 137134.8281 |
| 31087.76172 | 52181.58984 | 44697.3125 | 48860.67969 | 51393.06641 | 53450.22656 | 51145.24609 |
| 269091.8438 | 418607 | 419011.1563 | 404485.125 | 399997.25 | 402723.625 | 406001.9063 |
| 155257.3125 | 237180.1875 | 232904.4688 | 260120.1719 | 236095.9688 | 220927.1875 | 222699.4375 |
| 124943.9063 | 181054.7031 | 179715.7031 | 185416.5625 | 196406.625 | 239015.4844 | 204434.3281 |
| 46761.30469 | 125230.9688 | 101355.0547 | 91346.8125 | 145337.5625 | 237728.6563 | 179720.6563 |
| 21515.90234 | 25384.86328 | 34096.23828 | 33708.89453 | 31038.59375 | 33176.96484 | 33108.97656 |
| 124912.8594 | 186578.6094 | 185418.4844 | 189387.9219 | 211018.1875 | 179166.9844 | 185869.3906 |
| 25429.04297 | 40252.83984 | 39037.98828 | 36622.625 | 34823.78906 | 34016.89453 | 35651.76953 |
| 9899.463867 | 19461.91797 | 17595.84766 | 20089.15039 | 30720.47266 | 27088.40234 | 28993.05859 |
| 7538.692383 | 12380.13672 | 11455.9209 | 12247.95508 | 20014.94727 | 16013.28613 | 16875 |
| 41935.64063 | 79253.89063 | 63090.98828 | 56376.28125 | 44764.80859 | 40216.65234 | 42772.46094 |
| 30030.26367 | 44792.29688 | 44535.01563 | 44720.26563 | 28372.20703 | 26251.99414 | 28509.31836 |
| 39548.46094 | 53675.44531 | 54337.82031 | 51963.85938 | 63310.84375 | 63227.16406 | 61911.1875 |
| 7162.958984 | 10491.71777 | 10914.48047 | 9466.219727 | 5211.51416 | 7209.326172 | 9045.850586 |
| 50778.31641 | 73162.51563 | 74490.76563 | 76137.14844 | 83217.46875 | 73388.58594 | 78692.71094 |
| 13602.50684 | 21370.51953 | 22103.91797 | 19568.44727 | 13883.98047 | 16621.19531 | 14537.7627 |
| 5781.269531 | 7475.942871 | 9081.423828 | 7867.275879 | 7498.208984 | 6740.606445 | 8460.700195 |
| 2287.425537 | 5482.277832 | 2553.104004 | 1861.991089 | 4136.959961 | 4526.869141 | 4606.540039 |
| 205777.4063 | 303164.375 | 299695.3125 | 320628.4375 | 553512.4375 | 420325.2188 | 427078.1875 |
| 6329.939453 | 10885.03516 | 9128.985352 | 4842.151855 | 7035.430664 | 8662.477539 | 6227.703613 |
| 32298.55078 | 47753.91406 | 47860.86328 | 49968.03516 | 53701.37891 | 44638.56641 | 51355.85938 |

| | | | | | | |
|---|---|---|---|---|---|---|
| 12417.6416 | 16622.61914 | 21167.8457 | 21827.7207 | 36822.89453 | 32799.64063 | 33743.98047 |
| 3114.932373 | 4070.804443 | 4343.88623 | 3507.496094 | 7492.285645 | 6053.766113 | 5748.844238 |
| 15373.21582 | 24457.93164 | 25061.58984 | 31572.22266 | 42785.78906 | 45032.40234 | 48173.8125 |
| 167700.8594 | 236289.7344 | 261263.7656 | 244330.9844 | 253943.625 | 279776.6563 | 266023.6875 |
| 63304.1875 | 90361.36719 | 95013.38281 | 95929 | 85805.07031 | 87204.54688 | 87548.32813 |
| 77032.53125 | 118532.3359 | 115932.7344 | 102492.7734 | 89968.22656 | 90603.20313 | 80725.49219 |
| 40061.76953 | 62095.32422 | 55659.66797 | 55382.73828 | 47126.15625 | 52530.94141 | 54873.53516 |
| 11019.87988 | 25458.93555 | 23777.7832 | 19575.51367 | 11925.41699 | 9519.893555 | 11492.13281 |
| 44033.57813 | 76678.03125 | 67559.49219 | 62261.34766 | 67696.78125 | 72209.76563 | 72172.80469 |
| 6539.963867 | 11246.48145 | 8797.587891 | 11038.04395 | 16355.61426 | 12668.09961 | 11759.48828 |
| 6062.746582 | 8138.028809 | 9148.688477 | 9876.491211 | 20617.85742 | 18070.97656 | 18068.83203 |
| 346782.4375 | 513450.9063 | 485991.75 | 513982.875 | 524673.5 | 426710.4063 | 461906.875 |
| 33535.8125 | 34481.68359 | NA | 43996.01563 | 37717.74609 | 43627.55469 | 36244.07813 |
| 2137132.988 | 3235077.656 | 3317909.266 | 4081908.75 | 4757020.688 | 3492486.484 | 3865547.734 |
| 119928.6797 | 113005.4766 | 195482.4844 | 220001.9531 | 306880.1875 | 377994.6563 | 471476.6563 |
| 13550.84863 | 19685.45508 | 17470.0957 | 22898.06055 | 24631.08398 | 26024.2793 | 25601.12305 |
| 25477.18359 | 36660.91406 | 38301.37891 | 36201.68359 | 23668.24805 | 26336.375 | 22305.77539 |
| 43461.54688 | 65593.3125 | 64488.96094 | 71059.11719 | 62179.86328 | 52087.25391 | 58438.82031 |
| 226885.5781 | 351471.2188 | 333520.125 | 341036.3125 | 191497.5156 | 200486.0781 | 208842.4063 |
| 62869.25781 | 112122.0703 | 78145.34375 | 104962.7109 | 91642.34375 | 113770.8359 | 103698.5391 |
| 10919.05273 | 15448.69629 | 16590.07422 | 14247.60742 | 4749.67334 | 4483.061035 | 6987.401367 |
| 200475.5781 | 346526.125 | 303847.25 | 335755.9688 | 257542 | 265021.4063 | 274194.3125 |
| 2374.181641 | 3881.234863 | NA | 5154.872559 | 14327.53711 | 10773.01758 | 13242.7998 |
| 59579.42188 | 88332.89844 | 77828.88281 | 99863.82813 | 125422.7656 | 122689.3984 | 120598.6953 |
| 153256.8438 | 230933.6094 | 215816.7813 | 224602 | 343879.7188 | 358568.0313 | 349798.875 |
| 949544.5625 | 987888.6875 | 556911.4375 | 1207327.375 | 863342.5625 | 1258804.25 | 848120.4375 |
| 25322.24805 | 38977.29297 | 34916.07031 | 35453.32813 | 32928.78906 | 38425.98438 | 34522.97656 |
| 9506.373047 | 15403.0459 | 17472.9375 | 27330.98047 | 101385.0156 | 108936.4375 | 108519.2734 |
| 3865.003418 | 6830.235352 | 6727.054688 | 6404.845703 | 9161.871094 | 8077.662109 | 7517.56543 |
| 991376.25 | 1131176.875 | 1376165 | 1832907 | 1459924.125 | 1232991.125 | 1353393.25 |
| 73301.75781 | 105446.4453 | 96399.39844 | 109527.6172 | 90767.85156 | 85689.01563 | 97696.96094 |
| 6456.480957 | 11648.85449 | 11387.90234 | 11294.41699 | 9148.922852 | 8494.90918 | 8746.833984 |
| 16113.05078 | 17964.49023 | 24639.11133 | 28928.77734 | 36482.92578 | 24615.88867 | 26518.1875 |

| | | | | | | |
|---|---|---|---|---|---|---|
| 507.7922363 | 759.1032104 | 394.296814 | 489.744751 | 482.1105347 | 470.2208252 | 450.9933167 |
| 23562.92188 | 29725.05469 | 27801.1543 | 31615.10742 | 22703.25781 | 24358.76172 | 20517.51367 |
| 39258.92188 | 62199.26563 | 57713.57422 | 53799.98828 | 60932.42969 | 60306.625 | 57958.99219 |
| 367021.3125 | 558257.375 | 543578.3125 | 542443.4375 | 608243.5625 | 575014.9375 | 596534.6875 |
| 101097.8672 | 143293.9219 | 166050.8438 | 161934.1094 | 189040.1563 | 217671.2344 | 216020.2344 |
| 377750.3438 | 366536.0938 | 215579.9688 | 464611.9688 | 345450.0313 | 502608.4688 | 365353.1563 |
| 8424.835938 | 20776.8125 | 8865.922852 | 7914.706543 | 23980.89648 | 23848.06836 | 25788.7168 |
| 53992.77344 | 65453.71094 | 86090.92969 | 79144.82031 | 78165.17969 | 78304.00781 | 84617.39844 |
| 169770.1406 | 252564.0469 | 263919.5625 | 242138.0469 | 182439.7656 | 194291.8906 | 189178.8125 |
| 82560.03906 | 93033.76563 | 45471.38281 | 101451.8438 | 83871.41406 | 100754.7109 | 75984.4375 |
| 3605.566162 | 6231.90918 | 4912.113281 | 6035.492676 | 13864.71973 | 8801.891602 | 10886.89746 |
| 52318.28125 | 91144.80469 | 78360.21094 | 75664.39063 | 52196.59766 | 54685.55859 | 50635.83594 |
| 1571673.625 | 2427182 | 2214371 | 2222412.75 | 2736055.75 | 2570378.75 | 2657087 |
| 24883.7168 | 37452.87109 | 36875.25391 | 34545.01172 | 28569.84961 | 28755.60938 | 27087.80273 |
| 1854.815918 | 3209.828369 | 2373.550537 | 2461.90332 | 2202.526123 | 2420.237549 | 1840.089844 |
| 19260.61328 | 27522.17188 | 26930.29102 | 27148.53906 | 30448.13477 | 27676.64844 | 27505.99023 |
| 112355.9219 | 237884.5 | 123224.4141 | 128721.9453 | 129028.3984 | 181641.4844 | 103045.6641 |
| 5922.13623 | 9030.998047 | 9764.722656 | 10440.14258 | 7779.236816 | 8253.571289 | 6036.612305 |
| 16667.9668 | 28944.43359 | 24843.94922 | 24766.26563 | 26144.78125 | 28377.74414 | 29443.62109 |
| 39520.20313 | 55989.05469 | 53058.35938 | 56471.74609 | 49503 | 49395.40234 | 50338.71484 |
| 20596.73193 | 31544.17188 | 34007.26758 | 34299.57715 | 37959.66211 | 36365.50195 | 34170.57422 |
| 7688.966309 | 12912.53711 | 12114.08105 | 14224.59082 | 25751.22266 | 20080.45313 | 22185.41992 |
| 37239.3125 | 53404.86328 | 64993.86328 | 64516.23828 | 42519.11328 | 41001.87109 | 43092.53125 |
| 10683.45898 | 18648.48242 | 15337.79199 | 17167.83594 | 26792.96484 | 34512.70703 | 28692.82227 |
| 6507.686035 | 10795.03613 | 11794.96582 | 12202.69629 | 14026.41406 | 10988.25781 | 15664.33789 |
| 371219.8125 | 512649.125 | 548497.0625 | 533806.125 | 633388 | 613840.75 | 607562.875 |
| 30592.83008 | 46147.33594 | 37468 | 36907.48828 | 35016.66406 | 38252.38672 | 32135.40039 |
| 41379.74219 | 58523.80469 | 64311.87891 | 58339.43359 | 49120.79688 | 53162.125 | 50435.36719 |
| 231415.1406 | 390837.5938 | 313895.0625 | 364052.8125 | 463557.5625 | 433110.3438 | 409758.5938 |
| 110841.6953 | 175923 | 212917.7188 | 181010.9531 | 215632.1406 | 195837.0469 | 189191.0625 |
| 2360874 | 2396455.5 | 1496276.625 | 3081402.75 | 2688656.25 | 3133652.25 | 2473457.5 |
| 74354.125 | 102103.6016 | 105218.7188 | 103013.8672 | 69456.21094 | 74637.46875 | 63243.22266 |
| 8824.332031 | 14484.24707 | 10650.73242 | 11530.68262 | 20234.125 | 14927.87207 | 19590.63867 |

| | | | | | | |
|---|---|---|---|---|---|---|
| 4468.910645 | 7959.13916 | 8370.833008 | 8354.341797 | 10409.97461 | 8104.621094 | 8882.000977 |
| 38397.03906 | 56084.66406 | 54792.04297 | 55805.49219 | 64376.20313 | 53727.24219 | 58942.17578 |
| 52560.48438 | 73499.5 | 80588.80469 | 89679.17969 | 81421.60156 | 73488.85938 | 76201.1875 |
| 6700.961914 | 9013.004883 | 7716.40332 | 8137.879395 | 14650.55664 | 13549.1582 | 14930.01758 |
| 63855.33203 | 99731.54688 | 78562.17188 | 83889.125 | 79615.27344 | 54830.32813 | 74141.79688 |
| 719160.5 | 983379.75 | 928160.125 | 1054562.5 | 926884.625 | 904466.0625 | 987220.1875 |
| 18836.83008 | 23400.0625 | 27422.14844 | 27580.80273 | 19855.04883 | 19413.45508 | 19892.55078 |
| 17746.68555 | 23065.54688 | 25713.4375 | 25756.15039 | 23050.28125 | 19623.31055 | 21701.94922 |
| 21101.33789 | 31182.55859 | 29129.98828 | 28076.22461 | 42874.46875 | 34567.80078 | 38178.82031 |
| 115942.0547 | 175185.2031 | 175182.2969 | 167886.1094 | 138357.7344 | 150609.0156 | 136022.0781 |
| 77404.88281 | 109960.8125 | 114111.7422 | 121853.625 | 126624.2578 | 120953.0078 | 121234.7109 |
| 85245.875 | 116628.4063 | 124428.1172 | 129938.5703 | 185918.25 | 204647.3125 | 189655.8906 |
| 78490.40625 | 94961.25 | 124135.75 | 106233.2188 | 84433.47656 | 108339.8984 | 104484.3984 |
| 45744.37891 | 61095.14063 | 68231.02344 | 59150.35938 | 57214.83984 | 51860.85547 | 57432.08203 |
| 25398.37891 | 32091.0332 | 44875.94922 | 35130.66016 | 23673.41211 | 21359.9082 | 24894.71289 |
| 67457.0625 | 104261.8125 | 95632.9375 | 93760.72656 | 108229.0313 | 109800.1406 | 100533.2734 |
| 21195.44336 | 24661.93164 | 25087.87305 | 19026.98047 | 9771.674805 | 21440.875 | 22421.12305 |
| 279780.4688 | 441332.5625 | 422719.5313 | 438136.875 | 340572.125 | 371331.5 | 344524.4063 |
| 2548.224365 | 5589.712402 | 5086.120117 | 4364.195313 | 5438.73584 | 5555.856934 | 4936.807129 |
| 29543.89063 | 18092.80078 | 42865.66797 | 42582.09375 | 33333.5625 | 36241.19141 | 36470.18359 |
| 105813.5938 | 141045.7188 | 143694.0469 | 139707.2656 | 103660.3047 | 107099.0938 | 104122.5781 |
| 72001.28125 | 103362.2891 | 105546.7813 | 110351.7969 | 122843.1563 | 115238.6797 | 116599.4609 |
| 370440.5 | 537770.625 | 546082.5 | 574443.9375 | 400116.75 | 407709.4375 | 397679.4688 |
| 16394.89648 | 22553.69141 | 21603.00586 | 18945.96875 | 16593.25 | 15699.56055 | 17695.86719 |
| 51315.58984 | 78216.75781 | 74916.54688 | 75882.00781 | 77983.89063 | 80096.20313 | 79008.53125 |
| 26970.6582 | 44248.99609 | 38808.71094 | 43013.99219 | 28163.22852 | 27568.23828 | 24310.45313 |
| 67624.17969 | 96133.67188 | 104583.2734 | 111794.6875 | 83815.36719 | 70960 | 73059.46094 |
| 47429.0625 | 17736.72461 | 141176.4844 | 23282.72656 | 27969.63867 | 61347.12891 | 74061.35938 |
| 3401.401855 | 7139.127441 | 4887.506836 | 7297.675293 | 8539.077148 | 7041.912109 | 8204.094727 |
| 91883.13281 | 133507.0625 | 124031.7656 | 124385.7813 | 110120.4766 | 106563.3281 | 102831.1875 |
| 86740.75 | 112133.2891 | 124314.25 | 126567.4297 | 104053.9141 | 90603.71094 | 101231.7969 |
| 73174.53906 | 79813.42969 | 105961.7031 | 102160.3438 | 76420.09375 | 76920.32031 | 84400.16406 |
| 9276.845703 | 16157.88867 | 14064.29004 | 15920.89063 | 15713.99512 | 14743.25 | 15278.30566 |

| | | | | | | |
|---|---|---|---|---|---|---|
| 13423.30762 | 16824.02539 | 22342.88086 | 20342.42578 | 29796.39844 | 29530.36328 | 28365.55664 |
| 6820.109375 | 9983.183594 | 9673.854492 | 9043.179688 | 9302.750977 | 9398.120117 | 9878.72168 |
| 1734.334106 | 2872.020264 | NA | 1998.44812 | 1776.128052 | 1814.445435 | 942.4735718 |
| 28604.49023 | 121611.6875 | 38917.04297 | 69335.76563 | 205255.6875 | 159261.3125 | 185154.9844 |
| 19552.03125 | 31661.44141 | 31040.84375 | 32632.77539 | 45141.10156 | 42394.42578 | 41781.26563 |
| 10748513.12 | 45285514.61 | 17842372.68 | 29291429.65 | 48303761.15 | 31636967.6 | 35077699.63 |
| 64026.70703 | 60469.76172 | 122434.3125 | 116299.8984 | 56543.14063 | 61177.08984 | 59900.78125 |
| 110443.25 | 145167.0313 | 171869.6563 | 165833.4688 | 153578.1406 | 123577.1641 | 137622.9063 |
| 285640.5625 | 393471 | 395087.8125 | 461714.375 | 534690.5 | 370110.75 | 398159.2813 |
| 38259.37891 | 59721.49609 | 60387.08203 | 64181.76172 | 82019.42188 | 80752.59375 | 84569.10156 |
| 332996.2813 | 494819.1563 | 469316.9063 | 532015.625 | 501328.0938 | 456639.6875 | 499235.6563 |
| 258899.3594 | 312417.6875 | 385683.75 | 413966.7813 | 508878.75 | 479038.5938 | 502725.8125 |
| 8008.852539 | 14724.05957 | 11903.09277 | 6717.340332 | 6203.29248 | 9381.109375 | 7611.333496 |
| 6943815.5 | 10320681 | 10557139 | 10528009 | 12956382 | 11980482 | 13209512 |
| 49789.10547 | 71749.33594 | 67842.32813 | 65911.30469 | 35879.90625 | 40137.26953 | 37789.85547 |
| 47990.5625 | 76944.66406 | 70494.6875 | 84891.17188 | 93374.44531 | 94077.76563 | 86517.58594 |
| 33284.64063 | 50843.55859 | 50775.72266 | 52193.89844 | 38470.17578 | 35626.92969 | 36898.74219 |
| 16792.80469 | 16913.36719 | 26280.09375 | 25587.58594 | 10449.19531 | 18910.65234 | 16774.98828 |
| 69036.6875 | 73534.75781 | 70451.78906 | 72262.35938 | 18293.72852 | 58820.21875 | 17062.63281 |
| 49417.74609 | 79200.46875 | 80187.71094 | 83170.49219 | 63232.10547 | 68935.90625 | 67686.63281 |
| 9604.666992 | 13704.15332 | 13244.23828 | 12824.94922 | 13197.61719 | 13647.26953 | 12400.20508 |
| 16704.49219 | 23464.19336 | 24690.13086 | 26042.77734 | 31783.60547 | 29819.55078 | 33176.46094 |
| 5630.204102 | 4919.227539 | 8467.953125 | 5420.975586 | 59763.37891 | 14988.7666 | 11994.28027 |
| 36781.5 | 69639.85938 | 49337.92578 | 54736.61328 | 44590.08594 | 37858.05469 | 37816.33594 |
| 5067.179688 | 6947.183594 | 7120.870605 | 8223.760742 | 3811.747803 | 7599.847168 | 5901.834473 |
| 95124.73438 | 129773.0391 | 125309.6328 | 129673 | 137502.5156 | 139316.7344 | 144233.5156 |
| NA | 1256.638672 | 1250.474365 | 1117.643433 | NA | 1042.124512 | 999.6122437 |
| 31282.67188 | 47032.26172 | 44351.31641 | 47117.64453 | 55232.71875 | 48809.71094 | 51253.84766 |
| 31977.26953 | 50908.24219 | 49258.25391 | 48903.44141 | 50653.42188 | 49646.64063 | 50517.54688 |
| 42503.24219 | 51809.16016 | 58307.96094 | 62060.73438 | 40566.75781 | 40684 | 39814.96094 |
| 16102.48145 | 19844.45117 | 20984.75586 | 21587.82422 | 25609.44922 | 25172.42383 | 25308.76563 |
| 6321.35791 | 10106.67969 | 7673.755371 | 7013.771973 | 6760.195313 | 8574.77832 | 7508.967773 |
| 10319.94727 | 15654.76953 | 15586.86719 | 15089.99121 | 13940.99512 | 17049.74609 | 14020.01758 |

| | | | | | | |
|---|---|---|---|---|---|---|
| 17993.61523 | 25943.87695 | 25326.27734 | 23824.45898 | 20761.88281 | 24182.50586 | 23218.58984 |
| 14428.53516 | 23704.3457 | 24114.3418 | 19740.71875 | 15479.31934 | 11945.68945 | 17440.47656 |
| 14145.96875 | 19531.31055 | 20304.06641 | 17952.46094 | 17705.80273 | 18645.0957 | 19210.54688 |
| 15768.66309 | 26317.02344 | 19370.74805 | 20463.92383 | 11332.05664 | 10796.31348 | 10541.89258 |
| 295557.9688 | 449672.8438 | 397372.7813 | 451781.75 | 335663.4063 | 322687.75 | 325649.25 |
| 116286.2813 | 157489.7188 | 166096.8438 | 154206.25 | 107155.5781 | 108921.2578 | 104113.3438 |
| 2756.970703 | 3923.387451 | 4436.213379 | 3880.043701 | 3314.243408 | 3349.192627 | 3640.07666 |
| 7575.895996 | 10237.41992 | 10892.22852 | 11460.5459 | 10154.95215 | 9437.422852 | 10296.83691 |
| 7466.826172 | 14711.88477 | 19861.41406 | 19433.2793 | 30109.19141 | 26979.23047 | 26499.24805 |
| 81565.46094 | 139505.7344 | 126991.2422 | 117547.9219 | 111763.9922 | 115015.6406 | 108391.7813 |
| 305505.9688 | 329787.1563 | 176998.3594 | 384563.4688 | 316211.0938 | 414518.9688 | 280812.8125 |
| 4245.985352 | 6090.089844 | 5021.198242 | 5194.164063 | 3726.676758 | 4849.038086 | 5088.03125 |
| 32163.5293 | 48716.51172 | 47227.17578 | 49323.69922 | 30675.38867 | 35821.73828 | 30437.55664 |
| 5430.487793 | 8484.511719 | 6175.606445 | 7488.387695 | 7520.78125 | 9911.585938 | 7148.250488 |
| 20552.02539 | 30434.49609 | 28686.85938 | 30740.16602 | 25015.70898 | 28765.53516 | 25786.56641 |
| 17325.16992 | 20584.75977 | 18547.55859 | 21840.5332 | 16427.23438 | 21461.19531 | 19763.19727 |
| 2818.938232 | 4165.529785 | 5259.54248 | 5602.925293 | 9089.928711 | 6054.056641 | 7345.655762 |
| 113961.7266 | 184981.1094 | 177386.8125 | 180054.1563 | 164236.4063 | 170588.75 | 175282.1719 |
| 93868.28906 | 128397.9766 | 124675.1016 | 116623.3359 | 106303.0781 | 113192.3281 | 102000.2969 |
| 678089.3125 | 1428776.5 | 340906.0938 | 380462.875 | 582325.5 | 855453.1875 | 121088.6875 |
| 30180.98828 | 23993.73828 | 45231.85547 | 46894.92969 | 70966.78906 | 61807.53125 | 69515.65625 |
| 281134.75 | 477228.6563 | 370018.5938 | 378446.0625 | 419087.3438 | 391558.1563 | 332995.6563 |
| 98781.95313 | 149839.8281 | 146375.4688 | 145812.6719 | 136180.8125 | 135336.5781 | 141929.6406 |
| 32960.9375 | 50135.43359 | 46709.73438 | 51413.47266 | 66630.07031 | 68563.90625 | 62997.65234 |
| 50786.75781 | 68355.6875 | 64288.72266 | 55146.95313 | 41415.43359 | 53519.30859 | 50828.85938 |
| 3074.165771 | 6655.171387 | 4589.480469 | 4287.963867 | 4089.6875 | 8469.205078 | 4409.196289 |
| 22285.50586 | 37115.99219 | 27910.73047 | 27743.22266 | 37684.62109 | 42829.65234 | 36076.82031 |
| 28315.53516 | 46043.59375 | 42079.31641 | 48409.73047 | 38101.76172 | 27631.75781 | 31262.08984 |
| 7720.209473 | 4757.366211 | 10349.71875 | 13429.8916 | 3488.343262 | 16882.20508 | 5227.144043 |
| 12923.03906 | 18203.79492 | 20940.13477 | 19013.86133 | 17901.85742 | 21898.39258 | 17915.19141 |
| 157084.1719 | 256255.8438 | 224652.25 | 210133.5 | 198501.0938 | 204463.0156 | 171623.2813 |
| 305442.3438 | 431400.4063 | 429038.0625 | 414348.8438 | 396759.5938 | 319400.9063 | 362778.0938 |
| 2731.908691 | 11255.00391 | 5256.79248 | 11189.37891 | 21243.02734 | 16439.97461 | 18175.49219 |

| | | | | | | |
|---|---|---|---|---|---|---|
| 20556.15625 | 33027.04688 | 28600.38086 | 31502.33008 | 26220.82617 | 24823.88086 | 24174.9043 |
| 12418.85156 | 15094.2793 | 14035.60742 | 16414.4668 | 11297.95215 | 13315.08789 | 16490.25 |
| 336010.1875 | 463112.0313 | 483793.1563 | 534442 | 597217.25 | 554009.6875 | 550851.75 |
| 27240.14844 | 45152.47266 | 32981.64063 | 36733.97656 | 43013.50781 | 37133.21875 | 43946.5625 |
| 149224.7188 | 237981.375 | 164557.4219 | 166044.1719 | 135982.2813 | 126861.2734 | 120249.2813 |
| 43480.53125 | 55203.65234 | 64954.51563 | 70604.89063 | 71898.89063 | 66495.47656 | 73401.82031 |
| 190358 | 301893.9688 | 282762.0625 | 286081.2813 | 379143.4063 | 345118.4063 | 341761.375 |
| 56069.38281 | 82355.29688 | 87165.96875 | 90261.88281 | 88210.39844 | 86788.47656 | 84941.19531 |
| 18049.0625 | 25993.84961 | 28378.11133 | 28275.44336 | 23230.56445 | 25462.67578 | 23430.42969 |
| 195978.8906 | 298673.9063 | 280008.375 | 230151.8281 | 203748.375 | 218871.5938 | 211969.0625 |
| 23155.38281 | 36095.10547 | 33950.79688 | 34495.08594 | 27779.4707 | 38786.28516 | 28544.60547 |
| 13252.78516 | 15062.38086 | 16688.89648 | 15266.21875 | 8352.585938 | 10719.83301 | 8149.787109 |
| 83778.96875 | 126115.0313 | 113790.6172 | 128570.375 | 204605.4688 | 139156.9375 | 154179.4375 |
| 24618.40625 | 36377.98047 | 37327.77734 | 36311.27344 | 32643.89844 | 30920.41211 | 35078.73438 |
| 120204.5039 | 192948.4102 | 181344.2891 | 170611.707 | 137757.0234 | 128160.0156 | 140321.4961 |
| 18580.05859 | 32835.95313 | 24295.49219 | 27179.31641 | 29662.35156 | 25102.50391 | 24739.4043 |
| 8433.603516 | 14377.44238 | 11728.71484 | 12232.03613 | 10913.99707 | 10020.35742 | 9056.803711 |
| 41624.35547 | 53543.55078 | 59216.23828 | 54676.67969 | 44980.52734 | 48067.28125 | 51918.08594 |
| 81401.39063 | 87403.28906 | 115395.7656 | 84999.5 | 53209.13672 | 83347.79688 | 57882.57813 |
| 12519.50098 | 20580.75195 | 19492 | 21969.98242 | 24967.33203 | 23401.41406 | 22721.51758 |
| 18525.16406 | 25180.74414 | 29454.14258 | 28129.94727 | 11564.90039 | 11243.8252 | 12473.79883 |
| 17981.59961 | 30349.81641 | 23910.47852 | 21817.38867 | 21211.58398 | 21950.44141 | 19870.16406 |
| 6925.82959 | 7255.919434 | 7598.299805 | 9063.833984 | 7618.594238 | 10486.79395 | 8624.454102 |
| 67093.42188 | 86693.53906 | 95194.96094 | 96369.53125 | 94699.96094 | 89811.80469 | 95319.86719 |
| 84704.05469 | 109522.3984 | 115542.9844 | 119042.7813 | 113501.7813 | 124587.8359 | 122020.9375 |
| 6879.855957 | 13024.62012 | 9472.837891 | 8517.056641 | 16923.5 | 18592.91602 | 19810.0625 |
| 231277.8887 | 293795.5156 | 309384.5195 | 354367.6055 | 538275.1797 | 510686.9766 | 535296.6797 |
| 3946.405273 | 7051.365234 | 12164.56836 | 6326.6875 | 20378.12891 | 8128.461426 | 12875.69141 |
| 35401.14063 | 54637.76953 | 51177.35938 | 50816.41016 | 61826.42578 | 58873.42188 | 59996.47656 |
| 152940.1875 | 251443 | 232568.2188 | 231271.2813 | 227365.3281 | 291193.6563 | 253467.1563 |
| 17226.08984 | 21185.00195 | 23622.68359 | 27674.00781 | 35683.42578 | 32991.80078 | 35733.47266 |
| 730534.375 | 971205.875 | 969523.9375 | 1039245.75 | 1143837.5 | 1099030.75 | 1131858 |
| 19257.61133 | 29307.36719 | 28644.36328 | 34198.23047 | 45491.87891 | 40228.21484 | 44518.78906 |

| | | | | | | |
|---|---|---|---|---|---|---|
| 55815.07422 | 70845.07031 | 80184.77344 | 80262.47656 | 55436.57031 | 54974.41016 | 53403.01172 |
| 52733.42188 | 73739.16406 | 72623.40625 | 75885.40625 | 82348.71094 | 68406.13281 | 69522.76563 |
| 20412.05273 | 30162.9668 | 24489.37109 | 26905.41602 | 42210.0625 | 40629.17188 | 40310.25391 |
| 18808.69141 | 30139.96484 | 28539.66602 | 30466.66797 | 42895.80859 | 45650.47656 | 41939.23828 |
| 195086.25 | 252978.0469 | 234361.0156 | 238091.9531 | 261528.9688 | 235989.6094 | 269061.875 |
| 810527.4375 | 1109153.5 | 1185158 | 1149416.75 | 988054.8125 | 999475.9375 | 965939.9375 |
| 136898.2344 | 216496.6719 | 184300.3281 | 195325.75 | 252867.2969 | 252886.5 | 247890.7813 |
| 2277074.25 | 3256486.5 | 3500884.25 | 3351249.5 | 4291327 | 3940632.25 | 4345037.5 |
| 5099249 | 7151094.5 | 7140783 | 7098920 | 7696767.5 | 7479690 | 7643923 |
| 4747.305176 | 4774.937988 | NA | 6126.978516 | 2134.445068 | 1056.172974 | NA |
| 19091.54883 | 26467.39453 | 28753.93555 | 24596.55859 | 26919.36133 | 24498 | 24880.67188 |
| 109685.2422 | 89542.80469 | 182193.9063 | 196649.3438 | 171412.0625 | 161641.5313 | 163388.6563 |
| 407682.25 | 581634.5625 | 532541.25 | 588127.25 | 702197.875 | 687730.375 | 757165.25 |
| 41037.21094 | 51795.24219 | 59343.87891 | 56057.85938 | 34052.48828 | 33431.17578 | 33874.21484 |
| 272196.0313 | 395220.5938 | 367956.4375 | 368727.1875 | 393312.5938 | 373334.0625 | 403297.4063 |
| 14614.104 | 27314.79834 | 21712.76758 | 17883.1604 | 33233.86035 | 50367.85547 | 26970.14014 |
| 57069.97266 | 53731.70703 | 79230.67969 | 78268.03125 | 92689.88281 | 81256.17969 | 90043.80469 |
| 10244.95117 | 9995.741211 | 17986.23828 | 15710.63379 | 15842.3418 | 15515.78809 | 19108.36914 |
| 190914.5469 | 257276.6563 | 292671.9375 | 287328.1875 | 303741 | 321372.0313 | 316071.1563 |
| 49067.67969 | 79520.15625 | 70849.07031 | 67394.72656 | 134310.0313 | 109596.0781 | 112821.1016 |
| 127669.2422 | 217682.0781 | 159780.9219 | 195672.1563 | 263680.9688 | 245923.125 | 231634.3438 |
| 182108.0156 | 286388.4375 | 243205.7969 | 266903.3125 | 269055.6563 | 245216.3438 | 235791.0938 |
| 9703.62207 | 18773.42578 | 8170.399902 | NA | 24135.10547 | 27086.39648 | 37788.19922 |
| 39072.52734 | 54142.85938 | 60330.74219 | 54008.30469 | 46379.33203 | 45756.67188 | 47087.90625 |
| 158638.1406 | 236867.9844 | 214673.5938 | 219882.125 | 284832.9375 | 231075.9844 | 258368.5781 |
| 8472.358398 | 16991.13672 | 10992.79297 | 9389.328125 | 12202.31152 | 12038.77148 | 12323.98242 |
| 22424.67773 | 36128.67969 | 33262.20313 | 36778.46094 | 41454.71094 | 42092.44141 | 37596.12891 |
| 93156.67188 | 127850.75 | 125133.5547 | 122233.4844 | 122450.0547 | 139436.4531 | 126558.2891 |
| 281284.0625 | 408809.2813 | 394420.3125 | 393211.625 | 300354.625 | 269723.125 | 264615 |
| 4802.611328 | 8289.759766 | 9714.674805 | 4717.049805 | 8558.764648 | 14063.93555 | 6517.188477 |
| 82629.48438 | 123179.1641 | 106357.875 | 108744.8828 | 124195.1406 | 124610.2656 | 119864.3594 |
| 67347.55469 | 66343.01563 | 95047.23438 | 97167.1875 | 116120.5938 | 84388.26563 | 91717.30469 |
| 2762.484375 | 2430.555664 | 3335.078613 | 3086.633301 | 2150.090576 | 3371.066895 | 1605.33252 |

| | | | | | | |
|---|---|---|---|---|---|---|
| 12671.91113 | 13372.70898 | 15170.83203 | 14783.5166 | 18320.03711 | 14901.59766 | 16810.68945 |
| 21897.00586 | 30705.10938 | 29391.45313 | 25539.39648 | 27853.35547 | 25872.21484 | 25747.85156 |
| 418993.1875 | 691182.875 | 623477.5625 | 620755.3125 | 448370.125 | 389726.0313 | 397523.6563 |
| 55864.46875 | 77606.51563 | 71382.51563 | 74994.30469 | 86197.125 | 85655.77344 | 84710.03906 |
| 3787.529541 | 5033.833008 | 5138.188477 | 4384.960938 | 3685.044922 | 3825.533447 | 2924.868896 |
| 29817.46875 | 40231.21094 | 42264.71484 | 45082.37109 | 45780.35547 | 45185.09766 | 45282.27344 |
| 14937.81641 | 14450.79785 | 9152.949219 | 17746.76758 | 17006.75 | 26941.30078 | 14890.74219 |
| 5063404 | 5827371 | 6670313 | 6875231.5 | 6331549.5 | 6712496.5 | 6046445 |
| 94326.57031 | 150654.5313 | 138063.7969 | 117199.4844 | 91065.03125 | 98224.70313 | 95490 |
| 16561.55664 | 39864.83594 | 36025.91016 | 33353.32813 | 39158.13281 | 37841.39453 | 35070.52344 |
| 8950.80957 | 14402.05273 | 10837.58984 | 10524.98926 | 15502.69824 | 13685.90137 | 14512.61035 |
| 114829.6543 | 173109.6953 | 154498.252 | 153874.0078 | 173017.8008 | 167432 | 165528.2891 |
| 8609.407227 | 14149.2666 | 9770.910156 | 10165.0332 | 8784.345703 | 8632.798828 | 8470.177734 |
| 496097.3125 | 650705.8125 | 762910.375 | 671278.125 | 453141.125 | 443662.4063 | 477148.8125 |
| 3655.510498 | 4936.941406 | 3809.483643 | 4258.066895 | 5682.637207 | 5858.453613 | 5573.471191 |
| 7959349.5 | 10010167 | 11061863 | 12210481 | 8419278 | 7308757.5 | 8107437 |
| 38017.11328 | 59119.17188 | 43588.86719 | 43548.62891 | 42776.65234 | 37531.82422 | 38342.25781 |
| 68952.13281 | 97754.85156 | 87770.13281 | 85397.09375 | 111135.1953 | 104147.2891 | 107124.4922 |
| 19330.92969 | 30519.24609 | 22336.80273 | 16968.74609 | 31496.63477 | 26303.55859 | 33120.44141 |
| 24080.41602 | 31487.36914 | 31745.73438 | 33626.71484 | 36031.23828 | 31952.66211 | 36622.58203 |
| 40427.54297 | 60157.46094 | 57024.10938 | 53557.19531 | 54962.66797 | 59309.41797 | 54796.21484 |
| 5193.243164 | 7972.575684 | 6827.666016 | 6950.931641 | 11117.97754 | 9733.552734 | 10433.33301 |
| 69773.14063 | 99957.17969 | 97748.0625 | 106211.1797 | 119821.9688 | 112307.7656 | 113726.1094 |
| 62870.74609 | 101703.1563 | 81051.625 | 96775.48438 | 82353.59375 | 99088.32031 | 78591.75 |
| 194405.4375 | 257426 | 283404.8438 | 278978.2188 | 178677.5781 | 218616.6094 | 199333.0625 |
| 36401.96094 | 51226.39844 | 47053.23438 | 47128.75781 | 50928.21875 | 50165.63672 | 52480.00781 |
| 11658.77148 | 17696.80664 | 18792.46484 | 15737.23535 | 12310.77441 | 13056.54688 | 12739.9873 |
| 22184.70508 | 34354.51953 | 38796.5 | 33401.34375 | 22992.07422 | 23165.86133 | 25376.80859 |
| 15753.98145 | 18974.29688 | 17575.49609 | 19668.76953 | 28023.87109 | 24209.96094 | 26584.47656 |
| 49930.33203 | 66926.48438 | 65722.78125 | 64798.49609 | 92310.75 | 82241.13281 | 95865.75781 |
| 68872.3125 | 104106.3359 | 84566.25 | 86464.95313 | 89989.42188 | 97607.70313 | 103013.1563 |
| 442602.0313 | 684601.6875 | 608714.5 | 636359 | 620115.75 | 602403.625 | 615081 |
| 77083.27344 | 110485.5703 | 103390.9766 | 98205.82031 | 123006.3438 | 122120.9063 | 108078.8516 |

| | | | | | | |
|---|---|---|---|---|---|---|
| 105221.6797 | 122015.0313 | 134976.7188 | 144411.3281 | 120395.5391 | 117249.5938 | 128509.6406 |
| 98898.80469 | 153897.0313 | 93375.69531 | 103984.8047 | 121330.8438 | 198937.0313 | 113704.3281 |
| 79028.90625 | 108713.5313 | 105477.5469 | 112767.0938 | 102453.8047 | 87243.28125 | 101534.1953 |
| 413854.4688 | 688197.9375 | 473313.5625 | 501682.5313 | 591108 | 686009.75 | 550715.75 |
| 354387.9688 | 479632.9375 | 490074.9688 | 522404.5 | 555890 | 548603.6875 | 585959.5 |
| 27019.17969 | 36212.03906 | 35535.42188 | 32278.33203 | 32700.56836 | 30088.77539 | 32985.45313 |
| 739965 | 896552.0625 | 1140243.125 | 1304608.875 | 1231514.5 | 1002094 | 1301186.5 |
| 97182.42188 | 124994.7578 | 137644.7969 | 109908.8359 | 59583.15625 | 69102.13281 | 62247.76563 |
| 81474.8125 | 118461.9063 | 108679.6719 | 111066.4453 | 110170.2344 | 105120.3828 | 103714.875 |
| 66231.34375 | 92859.78906 | 86473.86719 | 97826.85938 | 100108.2422 | 97222.01563 | 93340.76563 |
| 322389.4688 | 689058.5625 | 451855 | 516182.5313 | 664873.0625 | 655867.6875 | 602863.875 |
| 118107.0234 | 151601.8438 | 173258.1719 | 173810.9063 | 156522.3281 | 136083.6563 | 149410.6406 |
| 9989.097656 | 12094.69043 | 13525.41309 | 14665.95801 | 14678.45117 | 11614.00879 | 11899.65039 |
| 434250.0039 | 690389.2813 | 594972.0898 | 721530.8906 | 694218.8203 | 565961.9453 | 604619.1133 |
| 30071.70313 | 41512.04688 | 40637 | 38589.78125 | 53339.75781 | 54584.28125 | 52779.19531 |
| 135698.1719 | 174954.6719 | 189914.5 | 186725.1719 | 176799.2031 | 176444.9375 | 181535.2188 |
| 13108.33691 | 15416.54199 | 17796.54297 | 18197.94727 | 13240.17676 | 12898.74023 | 11762.86621 |
| 3768.388672 | 6478.536133 | 1294.066162 | NA | 4036.45874 | 3871.150879 | 3136.580078 |
| 62594.73438 | 92246.21094 | 91323.04688 | 108396.9609 | 126787.9141 | 136333.375 | 124724.9844 |
| 333838.75 | 446137.9688 | 415031.7188 | 443208.625 | 311742.6875 | 360360.625 | 322849.3125 |
| 21147.52344 | 26111.83008 | 32500.63281 | 35363.87109 | 29278.5918 | 30579.81836 | 32923.92578 |
| 75235.26563 | 105587.5625 | 101899.9375 | 97732.33594 | 178133.7031 | 160825.8594 | 158006.7344 |
| 132845.75 | 179094.2031 | 183510.3594 | 189646.1719 | 253385.8281 | 250370.3906 | 240903.7813 |
| 2895.462646 | 3547.258545 | 4652.695801 | 3573.302979 | 5512.488281 | 5912.017578 | 5426.671875 |
| 96739.89063 | 131864.4219 | 132515.3438 | 138097.8594 | 136327.0781 | 133767.1094 | 134802.6406 |
| 594411.125 | 859841.75 | 804966.125 | 841764.125 | 787678.4375 | 752315.25 | 737101.6875 |
| 14598.27246 | 20962.81055 | 18464.06641 | 18957.94531 | 22298.43164 | 21578.60742 | 20678.625 |
| 57172.37109 | 71574.125 | 76931 | 73519.21094 | 48518.60547 | 47528.03125 | 47140.03516 |
| 135368.5938 | 140804.8281 | 195651.9688 | 193752.4219 | 198711.4844 | 215695.125 | 213929.8906 |
| 12920.92188 | 18903.1875 | 15140.23438 | 14943.96191 | 23592.99023 | 21292.90039 | 17925.46094 |
| 12103.10254 | 17994.58203 | 12761.36133 | 12965.99805 | 17393.53711 | 20353.75781 | 20438.44141 |
| 57217.82422 | 75496.65625 | 68347.07813 | 64993.90234 | 48355.90234 | 48080.51563 | 46843.40625 |
| 5414.223145 | 10725.25293 | 15447.01172 | 12863.07129 | 28490.41406 | 20966.80664 | 26031.05859 |

| | | | | | | |
|---|---|---|---|---|---|---|
| 179252.5781 | 275816.5 | 233333.0156 | 260955 | 271937.3125 | 306593.625 | 263863.8125 |
| 500486.7188 | 659889.1875 | 660264.0625 | 643517.625 | 277408.1875 | 287096.0938 | 285333.125 |
| NA | 1293.619995 | NA | 5317.534668 | 25859.61133 | 3512.485352 | 8939.462891 |
| 153.5724182 | 233.7580414 | NA | NA | NA | NA | 158.6143646 |
| 12773.61523 | 17830.81445 | 18313.875 | 16402.23828 | 15322.73828 | 16710.6543 | 13594.6084 |
| 57356.25781 | 77189.71875 | 83250.21875 | 81845.17969 | 77410.42969 | 86047.90625 | 92771.375 |
| 32593.43555 | 50070.78516 | 44081.67188 | 48908.33594 | 53611.95703 | 48285.98438 | 53718.73047 |
| 55322.33203 | 77854.16406 | 80705.38281 | 89221.89844 | 75438.1875 | 75867.92188 | 77485.40625 |
| 20029.85156 | 39954.48828 | 22983.91797 | 23650.02344 | 43891.45313 | 42913.85156 | 37965.01563 |
| 59920.44141 | 96222.90625 | 89628.55469 | 98706.51563 | 47637.48047 | 51541.07813 | 47788.39453 |
| 12542.26465 | 16920.98242 | 12462.47168 | 8083.956543 | 24226.35742 | 14661.71387 | 5727.387695 |
| 130462.0391 | 180696.8281 | 174827.8594 | 182188.9688 | 202418.3438 | 192495.625 | 200487.5156 |
| 40758.92188 | 64897.76563 | 60893.25781 | 53071.57031 | 69539.375 | 84372.03125 | 71675.82031 |
| 394597.5625 | 566640.6875 | 537087.75 | 554030.75 | 331827.875 | 394749.9063 | 363498.6875 |
| 1325626.185 | 1836563.587 | 1624329.31 | 1548263.353 | 2039875.62 | 1742831.728 | 1921278.529 |
| 26223.5293 | 56070.375 | 27912.00977 | 23340.26563 | 36061.91016 | 32645.38281 | 33834.38672 |
| 12309981.07 | 16491206.44 | 16583480.66 | 18005208.11 | 17842844.61 | 16807843.94 | 17488009.44 |
| 186846.0156 | 262771.2813 | 247517.6719 | 259113.625 | 268843.625 | 227085 | 234912.3125 |
| 24365.91797 | 33017.37109 | 31776.69336 | 28037.39258 | 6971.651367 | 9139.847656 | 8383.119141 |
| 118310.7422 | 154991.8125 | 156837.9688 | 161413.4219 | 158545.3125 | 147166.7031 | 151822.2969 |
| 104404.9141 | 137369.7813 | 136718.7344 | 132871.8438 | 90051.17188 | 91578.95313 | 86387.80469 |
| 328187.125 | 494702.9688 | 386862.75 | 405092.75 | 464103.8438 | 461436.7188 | 446023.125 |
| 4236.9375 | 3755.768066 | 5993.584961 | 6907.245117 | 8046.423828 | 5753.54541 | 6958.168945 |
| 8887.947266 | 16506.67969 | 13367.55078 | 12858.50098 | 9135.21582 | 9175.824219 | 9093.063477 |
| 51440.74609 | 78951.32031 | 72557.77344 | 73410.21094 | 72714.1875 | 67539.10156 | 67811.10938 |
| 8859.771484 | 16121.48535 | 3838.896973 | 11161.64844 | 20153.08008 | 27869.90625 | 19269.73047 |
| 109630.6328 | 148842.5313 | 150442.0469 | 154135.1563 | 148825.0625 | 154929.0781 | 159355.1563 |
| 12805430 | 19477096 | 16419156 | 18346544 | 14922079 | 14519693 | 14934090 |
| 39282.94922 | 60401.25781 | 59822.79297 | 62165.125 | 93341.03125 | 97153.5625 | 86972.49219 |
| 24511.46289 | 33418.22266 | 30397.04688 | 32547.5918 | 18974.59375 | 21138.22852 | 19154.09961 |
| 303393.375 | 412284.1875 | 420454.7813 | 419947.5313 | 346604.7813 | 325668.0625 | 298055.1563 |
| 131570.1875 | 217792.125 | 164477.0938 | 183641.4375 | 190784.6094 | 202484.5 | 196696.4844 |
| 41205.96716 | 64981.51855 | 51436.43286 | 55877.65723 | 62811.28418 | 66753.85742 | 66953.80859 |

| | | | | | | |
|---|---|---|---|---|---|---|
| 52158.76172 | 62105.48828 | 72125.76563 | 64817.04688 | 61143.74609 | 61131.92578 | 71692.98438 |
| 16283.75293 | 22240.61523 | 21170.70703 | 21377.68164 | 26801.86523 | 28651.89453 | 25559.30273 |
| 284557.375 | 372395.875 | 369527.0313 | 369346.125 | 318361.2813 | 337288.8125 | 400142.25 |
| 4875.733398 | 7622.941895 | 8007.425781 | 7293.596191 | 5968.090332 | 7453.728027 | 6853.596191 |
| 215176.5 | 284694.6875 | 292496.5313 | 312093.25 | 451895.0938 | 437920.6875 | 422103.0625 |
| 10387.58887 | 14787.68555 | 13854.58203 | 13314.88672 | 15638.90137 | 12899.18359 | 16362.05859 |
| 6346.312012 | 9177.155273 | 6722.353516 | 9595.064453 | 10473.6709 | 11168.73145 | 10304.14258 |
| 65436.37891 | 89032.57813 | 86677.70313 | 99137.57813 | 53400.40234 | 40448.97266 | 44652.51172 |
| 30325.9043 | 41208.39063 | 41839.27734 | 40220.13672 | 37401.70313 | 42336.00781 | 40644.00781 |
| 515023.9688 | 737388.3125 | 676503.4375 | 783568 | 833970.4375 | 724457.6875 | 762969.125 |
| 192408.4531 | 252506.2344 | 241518.9688 | 244257.2656 | 211921.0781 | 209195.3906 | 214344.1875 |
| 8696.076172 | 14452.66895 | 13540.19043 | 12704.45313 | 16802.41797 | 17027.85156 | 16260.36621 |
| 89273.14063 | 129765.0547 | 119003.5313 | 131194.0781 | 168146.5313 | 160080.5625 | 160388.7813 |
| 24103.21289 | 29810.28711 | 36540.52734 | 37280.67578 | 28780.02539 | 28194.86914 | 27598.94336 |
| 5169.483887 | 3236.175049 | 4132.700684 | 6887.376953 | 27217.11328 | 24171.35742 | 26833.02734 |
| 834.6799316 | 1133.900391 | 1338.235229 | 1546.328735 | 2052.839355 | 1707.258423 | 1924.934082 |
| 20903.42578 | 29758.51758 | 33108.05469 | 30070.36719 | 33901.78125 | 28392.73633 | 39114.76563 |
| 3956660.5 | 5626035 | 5451893.5 | 5628192 | 6027230.5 | 5623505.5 | 5960432 |
| 4476.114746 | 9156.939453 | 6298.69043 | 5637.167969 | 2227.630615 | 4658.654297 | 1931.062744 |
| 21194.26172 | 26898.39063 | 29664.95898 | 27052.28516 | 25018.10352 | 24265.63281 | 21303.36133 |
| 419086.5938 | 607037.9375 | 553612.875 | 557426.0625 | 584772.5625 | 603969 | 584371.5 |
| 321256.5625 | 439005.5938 | 417858.125 | 463974.8438 | 521425.9063 | 450368.1563 | 473402.8438 |
| 1227.426514 | 649.2988281 | 2124.772217 | 2314.841309 | 3040.35083 | 1747.46814 | 1727.595825 |
| 38789.04297 | 47445.60156 | 51040.69141 | 54767.51953 | 58363.80078 | 50238.61328 | 51506.25781 |
| 2746860 | 9049713 | 4432199 | 5705352.5 | 10932473 | 8319493 | 7902590 |
| 22948.19922 | 40050.19922 | 26730.41016 | 27611.29297 | 27271.50977 | 27261.64258 | 25117.43164 |
| 369398.6875 | 388545.875 | 481814.8438 | 618343.5 | 813775.75 | 630990.5625 | 751615.625 |
| 785286.125 | 1103491.375 | 1359925.75 | 1498076.875 | 1429372 | 1375715.125 | 1337681.75 |
| 605617.125 | 560797.9375 | 298057.5 | 679866.9375 | 512678.0625 | 743054.6875 | 540984 |
| 13971.77051 | 21809.46289 | 21599.36133 | 20019.58203 | 31159.45898 | 25270.11719 | 22859.37695 |
| 271845.5938 | 349310 | 385460.2188 | 350004.75 | 463529.2188 | 477769.25 | 480341.375 |
| 115415.8672 | 150107.25 | 157336.4375 | 150736.6563 | 152821.0313 | 147240.0313 | 149050.0156 |
| 9066.233398 | 13494.79492 | 12155.1543 | 13661.72461 | 12811.43066 | 11901.88184 | 12792.71875 |

| | | | | | | |
|---|---|---|---|---|---|---|
| 4851.757813 | 6536.078613 | 7502.601563 | 4613.534668 | 5979.20752 | 8696.152344 | 8211.634766 |
| 84097.33594 | 120149.75 | 115131.1094 | 114264.3359 | 118676.3594 | 128981.6875 | 124637.3594 |
| 45792.79297 | 47054.49609 | 77126.75 | 82340.73438 | 83610.375 | 90913.1875 | 100164.9922 |
| 48137.86719 | 67561.82031 | 66903.3125 | 65158.73438 | 61283.23828 | 54669.25391 | 54692.65234 |
| 15086.54004 | 26482.98438 | 18235.36914 | 17165.95898 | 10209.7627 | 16889.1582 | 9640.200195 |
| 119569.2109 | 162511.9688 | 192736.4375 | 169028.2813 | 164462.25 | 150707.7969 | 145895.0313 |
| 18465.89844 | 23982.65625 | 25465.32422 | 24768.70703 | 30518.24414 | 26240.0957 | 28413.63672 |
| 23280.61328 | 31467.79492 | 30736.92773 | 35339.3125 | 44495.24609 | 52007.70313 | 37090.93359 |
| 40687.44531 | 56604.55859 | 60405.87109 | 50814.74219 | 33275.42969 | 37101.39453 | 34639.60547 |
| 6414.012207 | 12019.54297 | 9506.598633 | 14519.76563 | 13777.38672 | 4419.129395 | 15648.42188 |
| 516748.125 | 691201.0625 | 695633 | 746722.5625 | 468628 | 405963.7813 | 458573.9063 |
| 10744.67285 | 21836.25 | 18609.50586 | 20241.16016 | 29188.75977 | 30245.09375 | 37431.92188 |
| 14333.24072 | 20696.39551 | 19785.3335 | 19318.86328 | 25581.21875 | 25211.08447 | 23984.75488 |
| 36027.84375 | 49970.62109 | 46331.77734 | 47349.46875 | 35804.74609 | 38276.13672 | 36671.53516 |
| 4184.00293 | 9171.384766 | 4234.683105 | 7634.706055 | 8218.307617 | 8841.676758 | 7042.608398 |
| 132203.0781 | 162320.0469 | 167217.3906 | 157863.0625 | 191862.3125 | 189218.1406 | 185967.8594 |
| 165717.9688 | 258818.3438 | 215185.375 | 222592.7813 | 269916.0938 | 244271.0313 | 250956.0781 |
| 32638.08789 | 48364.28516 | 40588.0625 | 41174.99219 | 79115.85938 | 68914.24219 | 67862.25 |
| 12409.30664 | 15050.44727 | 8186.130859 | 15921.19727 | 17382.69922 | 17815.00195 | 14679.79199 |
| 1153166 | 1551469.875 | 1534260.875 | 1476450.75 | 559104.6875 | 598850.5625 | 614324.875 |
| 54424.05078 | 69561.97656 | 76054.42969 | 71333.49219 | 52214.30859 | 59265.13672 | 56532.98047 |
| 38076.76953 | 45244.70703 | 47730.5625 | 44011.58594 | 68334.71875 | 74296.92969 | 71591.51563 |
| 16342.91504 | 28130.38477 | 15231.00586 | 24992.94727 | 29699.48242 | 43629.77344 | 38344.54688 |
| 5069.330566 | 4443.896484 | 3184.952148 | 5772.633301 | 5284.092773 | 9083.138672 | 5368.158203 |
| 33439.16406 | 37755.02734 | 51700.1875 | 52017.41016 | 28630.83594 | 38797.42578 | 32692.13281 |
| 98613.17969 | 133692.6719 | 133894.6563 | 132216.4688 | 117883.7656 | 123619.6563 | 124071.8828 |
| 26236.25 | 36146.80469 | 32808.77734 | 30668.27148 | 47784.27734 | 40580.50391 | 51604.72266 |
| 235219.4219 | 301113.1875 | 317301.0313 | 258396.4063 | 282466.0313 | 325873.5313 | 303470.7813 |
| 77825.02344 | 106239.0859 | 104446.6172 | 102430.3516 | 79593.70313 | 80913.95313 | 81690.77344 |
| 27695.45313 | 70486.85938 | 51134.76563 | 53747.01953 | 98284.91406 | 86236.98438 | 72080.99219 |
| 33633.62891 | 49712.11328 | 53483.15234 | 52155.51563 | 48255.79297 | 50776.30078 | 50065.78516 |
| 329967 | 452011.5938 | 445120.8125 | 414359.5938 | 367587.3438 | 415179.3438 | 420064.5938 |
| 27436.33398 | 31109.27148 | 33830.54297 | 36549.875 | 37736.10156 | 35051.51953 | 33761.37109 |

| | | | | | | |
|---|---|---|---|---|---|---|
| 20775.02539 | 30165.62891 | 26535.55859 | 25542.25781 | 44038.73828 | 40961.89063 | 37604.42188 |
| 16669.88477 | 21169.11328 | 24100.65234 | 16338.84766 | 7715.886719 | 13634.375 | 10366.12402 |
| 34969.37109 | 45780.20703 | 47306.40625 | 44259.23047 | 40181.95313 | 34493.01563 | 37231.51172 |
| 1178758.75 | 1485148.25 | 1576659.625 | 1750788.625 | 1695829.875 | 1463900.625 | 1593860.875 |
| 4608.957031 | 4674.945313 | 5834.740234 | 6339.373535 | 6798.916992 | 7015.447754 | 6713.104004 |
| 953582.4375 | 1220994 | 1294654 | 1356652.125 | 1062743 | 880326.25 | 988403.1875 |
| 24835.60938 | 28375.24609 | 13586.40039 | 23384.27539 | 18441.30859 | 23736.88672 | 21055.41211 |
| 7027.724121 | 5649.646484 | 9150.091797 | 8339.166992 | 15623.7832 | 13446.03516 | 16409.36133 |
| 388624.5313 | 691758.5 | 442691.6563 | 443708.6563 | 668898.75 | 615884.6875 | 638804.8125 |
| 7846.85791 | 15611.73047 | 14297.32129 | 12553.67969 | 15763.26855 | 14130.4248 | 14526.55762 |
| 21536.25195 | 28164.46289 | 32387.70508 | 29458.72656 | 28930.86719 | 27471.36719 | 27847.30664 |
| 51594.04688 | 67306.875 | 68212.58594 | 69568.16406 | 65760.09375 | 65033.96484 | 64362.55859 |
| 30667.11719 | 47464.01563 | 41237.98047 | 35215.05078 | 44249.77344 | 43220.09375 | 45197.81641 |
| 149558.5313 | 185248.6563 | 175761.5781 | 177523.3125 | 131042.3281 | 135018.3438 | 125773.4609 |
| 94063.53125 | 132648.6563 | 130246.7266 | 132302.7656 | 71966.60156 | 68009.875 | 69048.15625 |
| 4067.807373 | 3731.601807 | 3007.943604 | 4413.355469 | 8846.953125 | 13089.02148 | 11945.69727 |
| 2613758.75 | 3507521 | 3355991.5 | 3682894.25 | 1938205 | 1947673.25 | 2007517.125 |
| 80130.51563 | 122150.0469 | 97941.39844 | 95349.79688 | 128923.4453 | 136408.0781 | 137599.9531 |
| 14793.2041 | 24216.9707 | 20186.85352 | 20297.45117 | 27057.1875 | 23470.27539 | 22394.03516 |
| 50751.65625 | 72316.32813 | 73997.78906 | 67651.75 | 61420.32813 | 65176.85156 | 60663.42969 |
| 252455.6875 | 338431.1875 | 332367.375 | 354467.1563 | 466038.125 | 464858.0313 | 479784.8125 |
| 8814.375977 | 13318.46191 | 9827.179688 | 9611.663086 | 12804.2998 | 11770.5752 | 11180.65527 |
| 29863.36914 | 39644.60938 | 42225.36719 | 37397.19531 | 35798.76563 | 38789.3125 | 41682.14453 |
| 23306.70508 | 32831.78125 | 33874.67188 | 32478.8125 | 27465.55664 | 29518.4043 | 29840.08984 |
| 122909.0391 | 184788.4375 | 164889.1719 | 186128.8906 | 138249.7344 | 90361.58594 | 93984.46875 |
| 56823.57813 | 78549.24219 | 77316.22656 | 85353.53906 | 97736.34375 | 106521.2578 | 96276.82031 |
| 464650.6875 | 608033.1875 | 621250.4375 | 711149.9375 | 583427.9219 | 601517.1563 | 617627.25 |
| 8330.512695 | 8656.241211 | 4229.168457 | 9545.084961 | 8361.357422 | 10200.68945 | 7844.495605 |
| 207127.7031 | 230868.8281 | 402360.5625 | 453864 | 339072.1875 | 81239.75 | 393394.25 |
| 3565.184814 | 5177.933105 | 3973.020264 | 3167.759766 | 2092.385742 | 2075.965332 | 2166.102295 |
| 3849.877686 | 5856.927246 | 6470.601074 | 4981.246582 | 5599.929688 | 3895.924805 | 5075.662598 |
| 505777.8125 | 565832.5 | 715318.125 | 689356.125 | 686516.125 | 645515.625 | 780840.4375 |
| 1557.983765 | 1998.829468 | 1832.808716 | 1740.751465 | 2797.5625 | 3078.811523 | 2033.407837 |

| | | | | | | |
|---|---|---|---|---|---|---|
| 529440.9375 | 697322.125 | 718357.8125 | 727963.8125 | 746210.1875 | 817861.625 | 752910.0625 |
| 27603.83203 | 28945.49023 | 29036.55078 | 38682.49609 | 37133.60938 | 35539.0625 | 37894.29688 |
| 32762.45508 | 45752.66016 | 46155.91016 | 49023.62109 | 50148.80859 | 44085.90234 | 47543 |
| 4883.239746 | 8187.238281 | 5928.845215 | 6563.397461 | 8284.112305 | 10160.97949 | 8895.663086 |
| 106820.4141 | 149002.9063 | 121082.3594 | 138280.2969 | 158820.9531 | 160355.9375 | 126647.3359 |
| 33859.20313 | 43333.51172 | 37218.80078 | 39485.89063 | 18219.31836 | 21117.52539 | 18755.15625 |
| 378340.9375 | 513923.2188 | 501077.0313 | 517777.0938 | 475671.8125 | 485305.25 | 457901.1875 |
| 78267.47729 | 101442.7109 | 98531.61572 | 97069.45557 | 110303.1338 | 96475.17188 | 104082.4619 |
| 33586.08984 | 49363.91406 | 41682.13281 | 43835.25 | 56550.81641 | 57621.98047 | 55704.82031 |
| 1995.678589 | 2974.541748 | 3041.853271 | 2974.717529 | 3628.829102 | 6225.337402 | 4636.602051 |
| 390237.125 | 536125.25 | 505045.4375 | 486593.4688 | 366650.875 | 403918.2813 | 340799.0313 |
| 32536.90234 | 44426.13281 | 48217.19922 | 47392.79297 | 42668.16406 | 42106.02734 | 43801.62109 |
| 697.435791 | 1193.544189 | 950.8776245 | 1057.007935 | 271.8546448 | NA | 260.8873901 |
| 24580.46875 | 36811.88672 | 31758.53711 | 32546.74023 | 39291.48828 | 37955.02734 | 38416.10156 |
| 1226781.25 | 1595977.125 | 1589712.25 | 1632108.375 | 751057.8125 | 715413.25 | 728867.1875 |
| 821636.75 | 1007346.813 | 1064044.75 | 1140055.25 | 752410.1875 | 840174.5625 | 902526.8125 |
| 74336.47656 | 95849.11719 | 105914.8594 | 112232.5703 | 133531.7969 | 132936 | 133800.25 |
| 117996.8438 | 158607.4219 | 175411.5938 | 185365.6719 | 205883.7344 | 193161.5938 | 205960.25 |
| 8286288.5 | 10778172 | 10947391 | 11036458 | 9810620 | 10213050 | 9556390 |
| 229228.8438 | 311167.9688 | 331718.75 | 349442.6875 | 376711.5313 | 279838.1875 | 309125.875 |
| 1761.63269 | 1874.229614 | 1951.325928 | 2537.192139 | 2843.641846 | 2591.649902 | 3114.929443 |
| 1107734.5 | 1485621.25 | 1557365.5 | 1492365.625 | 1573914.75 | 1537513.375 | 1465858.75 |
| 207283.5625 | 302573.8438 | 254471.375 | 250726.8125 | 299549.0625 | 288546.7188 | 269526.3438 |
| 5721.085938 | 7830.467773 | 5414.367676 | 6504.302246 | 6093.474121 | 6686.599121 | 7292.19043 |
| 568930.125 | 442983.2188 | 287507.7813 | 686256.75 | 513578.4063 | 753223.75 | 574994.6875 |
| 1227.586304 | 2171.027832 | 1773.731201 | 1829.54248 | 3665.212402 | 1934.818237 | 2857.130859 |
| 16426.63672 | 23867.59766 | 23420.66992 | 21712.4707 | 24850.33398 | 23335.13867 | 23911.22656 |
| 44649.47266 | 61478.84766 | 51059.94922 | 72135.59375 | 163044.4375 | 147480.7813 | 138363.1719 |
| 33186.73828 | 49412.05469 | 44414.03125 | 44087.33984 | 28075.86133 | 28721.38281 | 26666.63867 |
| 93773.42969 | 123229.5234 | 109470.5547 | 111365.0625 | 133262.2344 | 124982.2109 | 124162.2266 |
| 42822.69141 | 50706.05469 | 59345.43359 | 58584.42578 | 49756.60156 | 53089.55469 | 55271.64844 |
| 8267.889648 | 10595.24414 | 9007.81543 | 8759.788086 | 13429.25879 | 13117.84961 | 12632.62207 |
| 8297.787109 | 12136.80469 | 12195.00488 | 12444.82617 | 12786.60449 | 12437.14258 | 11769.05957 |

| | | | | | | |
|---|---|---|---|---|---|---|
| 58111.74609 | 56253 | 79567.59375 | 77137.22656 | 81166.28125 | 72128.21094 | 80334.3125 |
| 36530.23828 | 45374.61719 | 48498.80469 | 46336.98828 | 44257.07422 | 40538.53906 | 44259.64844 |
| 7462.213867 | 12178.20313 | 8162.787598 | 6281.499512 | 13115.66797 | 10754.18164 | 12203.57715 |
| 21020.52734 | 28362.99023 | 23254.49805 | 25140.8457 | 31044.65039 | 23532.32227 | 24344.48438 |
| 105686.0781 | 153592.6719 | 140314.375 | 139325.125 | 138871.2969 | 152165.3438 | 131760.625 |
| 9025.354492 | 14621.95898 | 12143.0459 | 11401.38574 | 7504.536621 | 7298.364258 | 7302.973145 |
| 402199.4375 | 555849.3125 | 498611.5313 | 486525.0313 | 336488.75 | 361201.9063 | 306032.5625 |
| 12503.6748 | 18612.33984 | 19542.57813 | 16716.25195 | 22318.17578 | 22225.7168 | 21651.91406 |
| 19356.97461 | 25972.42578 | 24584.10938 | 24368.19922 | 17896.52344 | 14176.32324 | 15688.20801 |
| 71114.29688 | 93138.21875 | 97293.42188 | 96338.85156 | 143139.5938 | 122476.1641 | 130399.8047 |
| 9303.210938 | 13714.47852 | 11870.375 | 14589.64453 | 14277.64063 | 10148.0293 | 10175.65918 |
| 399079.7188 | 532466.4375 | 513159.0313 | 482959.7813 | 414417.5313 | 464877.125 | 456399.1563 |
| 213903.7031 | 280934.9375 | 323044.5313 | 329629.9688 | 245540.1094 | 278520.8125 | 316139.6875 |
| 7021.082031 | 8748.960938 | 9520.206055 | 9194.058594 | 18021.06055 | 18181.86914 | 18836.2832 |
| 40128.25391 | 53138.00391 | 56671.60156 | 59789.48438 | 61389.12891 | 60233.53125 | 61770.59375 |
| 358701.7188 | 450778.3125 | 460873.9688 | 451609.75 | 204650.875 | 224913.6094 | 217012.2344 |
| 21835.77734 | 28753.23438 | 29382.6543 | 30489.95508 | 34611.69141 | 33583.3125 | 34630.34766 |
| 32428.22656 | 33776.64453 | 28469.36328 | 21348.73047 | 10624.59668 | 26965.00391 | 13980.78516 |
| 828715.8125 | 992289.1875 | 1123640.875 | 1179111 | 1062330 | 1110492.125 | 1083995.375 |
| 16668.16602 | 14364.9248 | 21472.11914 | 27684.49414 | 30018.84375 | 42619.51563 | 35045.21875 |
| 104777.9063 | 129636 | 141382.5156 | 159559.0781 | 140533.1406 | 125955.7188 | 134675.5625 |
| 1433.205566 | 2809.206299 | 925.2848511 | 999.369873 | 984.1487427 | 1242.081787 | 1627.062622 |
| 9670.297852 | 9784.629883 | 12986.39551 | 12583.63574 | 9403.69043 | 10448.39355 | 8303.981445 |
| 227179.7188 | 475710.8125 | 268352.4063 | 309083.2188 | 506850.0625 | 370081.8438 | 360582.75 |
| 1050459.625 | 1533486.75 | 1471837.125 | 1430921.375 | 1706539.125 | 1488702.875 | 1458541.25 |
| 44628.07422 | 63824.38281 | 56152.46875 | 66041.10156 | 70334.54688 | 60675.98828 | 65046.28906 |
| 3911.05835 | 5467.725586 | 5677.935059 | 6202.492676 | 11994.21484 | 11471.71875 | 9145.095703 |
| 1583.944946 | 2668.795166 | 2167.70752 | 2821.58667 | 1036.112427 | 1520.840576 | 1487.828491 |
| 53716.50781 | 66823.0625 | 66521.25 | 69678.17969 | 75637.71094 | 86156.21094 | 82944.77344 |
| 67316.8125 | 89584.67969 | 91273.86719 | 87045.67969 | 76411.58594 | 70903.72656 | 76294.60156 |
| 33587.01953 | 44485.19141 | 43252.88672 | 45740.39453 | 22952.28125 | 26777.03125 | 22820.47266 |
| 285016.2188 | 405739.0938 | 378398.0625 | 374178.7188 | 403181.0938 | 332779.7813 | 380405.9688 |
| 68927.94531 | 96085.92188 | 87291.39844 | 78773.85938 | 69456.14063 | 74445.64063 | 62658.16797 |

| | | | | | | |
|---|---|---|---|---|---|---|
| 23749.99609 | 31386.00781 | 32703.57813 | 31093.02148 | 37197.54688 | 40404.1875 | 39071.96484 |
| 30474.42188 | 38779.0625 | 42521.47656 | 41963.48047 | 37105.82813 | 42940.07031 | 39576.38672 |
| 5408.870117 | 12693.88379 | 8722.202148 | 8885.755859 | 12665.93555 | 10336.53516 | 6768.239746 |
| 29194.20117 | 38244.16016 | 36603.03906 | 35438.28516 | 39363.60938 | 44638.27734 | 42956.34766 |
| 244571.9688 | 371860.5938 | 292386.6875 | 292458.9063 | 301979.125 | 325659.5938 | 274648.625 |
| 13468.0293 | 25641.50195 | 25700 | 19986.52734 | 11657.51367 | 7368.128906 | 17539.39063 |
| 105226.6719 | 129450.8281 | 133009.6875 | 139082.6563 | 112708.3438 | 102253.5703 | 107178.5 |
| 19881.10352 | 24373.59961 | 22852.25977 | 24735.66211 | 22667.6543 | 19498.50391 | 21053.31641 |
| 151426.4531 | 230696.6094 | 214587.2969 | 231859.25 | 254947.6719 | 271232.0625 | 260557 |
| 12097.36621 | 14910.05957 | 16314.75879 | 14132.21582 | 11604.85547 | 15526.45605 | 11156.90137 |
| 130114.7031 | 231740.4688 | 191882.1563 | 185859.2656 | 221294.8906 | 216221.5781 | 205047.3125 |
| 13813.62598 | 23665.73047 | 17872.11914 | 16915.16016 | 22115.42969 | 17439.01758 | 18792.91797 |
| 2690.946045 | 4682.143066 | 2972.686035 | 3586.051025 | 4590.002441 | 5714.310547 | 5304.049805 |
| 46178.86719 | 63226.19141 | 60984.12891 | 61312.54688 | 75732.34375 | 79627.375 | 74144.9375 |
| 766976.3125 | 1070552.125 | 962013.125 | 950597.625 | 655761.1875 | 630242.125 | 608971.5 |
| 27277.47852 | 36655.94531 | 31991.26758 | 34857.66406 | 36193.9375 | 36257.24609 | 36805.56641 |
| 10437.19922 | 7594.517578 | 13683.63672 | 10650.67188 | 7831.78418 | 10818.77441 | 10640.06055 |
| 260880.8281 | 447363.7813 | 230281.6563 | 257406.3906 | 341221.5 | 312208.9688 | 335796.125 |
| 156205.75 | 160902.75 | 262834 | 205614.875 | 169800.9375 | 229076.375 | 222120.8281 |
| 111791.7266 | 152178.1406 | 163836.7344 | 137312.8906 | 128387.125 | 137416.1406 | 129480.9297 |
| 12917.10938 | 18030.64844 | 17659.5957 | 17176.40625 | 21531.18164 | 22936.53516 | 22272.28516 |
| 12506.01367 | 21406.19727 | 18371.05859 | 19881.5957 | 13516.11035 | 19610.95117 | 15958.36621 |
| 206153 | 299145.875 | 264341.4688 | 261040.8906 | 211292.8125 | 188883.6094 | 187803.4063 |
| 222604.5156 | 301390.5625 | 265833.7188 | 270334.9063 | 273597.375 | 262179.2188 | 264383.2813 |
| 56828.69922 | 94872.14063 | 80661.75 | 83118.98438 | 169598.7031 | 110729.5 | 135497.7969 |
| 42494.94922 | 58703.9375 | 54274.66016 | 55168.5625 | 51512.72656 | 47870.94922 | 47200.3125 |
| 28843.95508 | 34116.08203 | 39616.87891 | 36092.14844 | 31332.44531 | 31988.54297 | 34081.66406 |
| 39612.75391 | 51516.72656 | 44976.28516 | 42864.00781 | 55047.29688 | 80233.94531 | 52383.75781 |
| 92152.86719 | 105114.6094 | 120205.3594 | 122719.1016 | 90488.58594 | 76930.33594 | 85489.71875 |
| 39576.03906 | 56134.29297 | 51509.15234 | 51059.30859 | 44682.85938 | 46604.25 | 41365.21094 |
| 46672.34375 | 56794.26953 | 65266.95313 | 55922.85156 | 77163.49219 | 84211.52344 | 71099.82813 |
| 104135.7656 | 110988.2578 | 175322.0313 | 176999.1719 | 160386.5313 | 150605.0313 | 170103.125 |
| 11412.63574 | 17179.13281 | 14718.55469 | 13397.97852 | 16828.52148 | 14614.4043 | 14610.3418 |

| | | | | | | |
|---|---|---|---|---|---|---|
| 8151.663086 | 17042.8125 | 11962.92285 | 13141.74316 | 13124.26855 | 7031.577148 | 14407.75781 |
| 11953.72363 | 14106.74121 | 14842.61816 | 13387.62305 | 12599.56348 | 9119.361328 | 12403.68359 |
| 17225.50781 | 24208.98828 | 22679.68359 | 22843.50195 | 24558.13086 | 25591.67969 | 23316.04688 |
| 31659.31055 | 34479.20313 | 36654.14844 | 33726.75391 | 31161.95898 | 31375.10547 | 32205.08398 |
| 70259.54688 | 88940.57813 | 94109.19531 | 89256.85938 | 86670.67969 | 89785.3125 | 87492.79688 |
| 50579.51563 | 73665.5625 | 67509.91406 | 65122.33594 | 62968.58594 | 68369.64844 | 63306.01172 |
| 8589.505859 | 10772.3623 | 7092.023926 | 9966.613281 | 11194.87695 | 13553.0957 | 12614.44531 |
| 1241956 | 1900412.125 | 1456949.75 | 1631866.375 | 1451336.625 | 1422905.875 | 1612542.375 |
| 1741172.25 | 1907900.25 | 2416433.75 | 2422216.5 | 2831001 | 2687593.75 | 2666393.75 |
| 830951.4336 | 799546.4766 | 1066550.008 | 1163636.773 | 667458.5586 | 662904.8164 | 694514.8008 |
| 24031.35156 | 36762.47656 | 33764.21484 | 25695.36719 | 14419.6748 | 20135.20313 | 16069.22949 |
| 62955.9375 | 68435.67188 | 76479.24219 | 79905.0625 | 65484.95703 | 67606.96094 | 64502.94531 |
| 459047.375 | 565185.75 | 626040.5 | 646463.8125 | 695241.125 | 652912.4375 | 673747.3125 |
| 64613.39453 | 78891.88281 | 89942.28906 | 84529.71875 | 77370.65625 | 77580.96094 | 81655.25781 |
| 178643.0781 | 242580.5781 | 252265.2344 | 243097.3281 | 209806.9375 | 215689.5469 | 217924.75 |
| 31406.28516 | 43376.19531 | 43038.1875 | 41800.30469 | 40437.72656 | 38083.28125 | 40240.22266 |
| 97016.26563 | 133221.5313 | 123061.4141 | 127832.7109 | 133698.9688 | 140718.4688 | 138464.6094 |
| 26848.19141 | 33084.03906 | 37059.75391 | 36683.46094 | 24371.49609 | 25185.03125 | 22533.11133 |
| 22499.12305 | 30825.87695 | 27677.26563 | 26069.10547 | 19258.33789 | 19478.35742 | 19500.91406 |
| 74858.375 | 92837.69531 | 107422.9297 | 106650.6719 | 110721.3359 | 116724.5156 | 126108.2813 |
| 44412.14063 | 44343.27344 | 26797.76367 | 50931.86719 | 36300.88672 | 42926.14063 | 40022.5625 |
| 107196.7578 | 131683.9844 | 142129.5156 | 134271.8125 | 100118.4219 | 104086.0156 | 104736.9453 |
| 90801.59375 | 108947.9141 | 115573.5 | 113210.1016 | 65293.94531 | 71522.88281 | 73022.22656 |
| 15676.53125 | 20865.71094 | 20512.87695 | 19343.49414 | 25846.95117 | 25624.30078 | 28843.39453 |
| 22773.73047 | 27528.1582 | 29891.04492 | 34042.35156 | 51063.50391 | 44509.48438 | 51877.44531 |
| 298722.7188 | 407376.8438 | 391524.1563 | 416839 | 241518.2969 | 280918.2188 | 263132.9688 |
| 363133 | 475238.6875 | 468178.4688 | 472488.125 | 360066.2188 | 368565.2813 | 345898.7188 |
| 31364.34961 | 43989.75 | 44818.99609 | 46653.08594 | 45380.34375 | 41815.64453 | 43298.01172 |
| 110181.5547 | 153167.8125 | 154228.7188 | 153412.6094 | 126109.7578 | 144236.2188 | 146586.4219 |
| 1351.674194 | NA | 1337.79834 | 2082.88916 | 1132.399658 | 921.7765503 | NA |
| 343173.4688 | 459426.4063 | 409934.9375 | 468323 | 309938.75 | 358838.6875 | 352249.8438 |
| 16567.86328 | 20622.50391 | 26568.55469 | 25609.33008 | 39117.5625 | 33017.01172 | 33102.55859 |
| 78550.25 | 96486.25781 | 110715.3594 | 111595.2422 | 105571.6016 | 95171.49219 | 93869.42188 |

| | | | | | | |
|---|---|---|---|---|---|---|
| 3810.45874 | 5400.447266 | 3685.856445 | 3104.814697 | 6241.466797 | 4910.729492 | 5691.191895 |
| 57725.53516 | 70716.96875 | 70135.21094 | 70364.70313 | 50944.12891 | 43732.72656 | 47387.43359 |
| 227958.2031 | 250388.6406 | 332376.6875 | 323231.0938 | 279628.0938 | 268017.1875 | 264590.625 |
| 11240.29199 | 12355.23926 | 12969.30566 | 14159.83496 | 42362.07031 | 31517.64063 | 25371.45117 |
| 41314.69141 | 53286.01172 | 54171.04297 | 56974.24609 | 66938.91406 | 66382.96094 | 69502.28125 |
| 11400.97852 | 14211.29004 | 14829.49512 | 14196.94141 | 11764.80762 | 11911.7041 | 12414.05273 |
| 55104.51953 | 59669.95703 | 73948.78125 | 75038.21094 | 76430.39063 | 73099.44531 | 77539.03906 |
| 165928.7969 | 190340.6406 | 210682.6563 | 198995.3906 | 143484.5625 | 149942.6406 | 139780.1563 |
| 218731.1406 | 311697.4063 | 282126.1875 | 283433.9375 | 354173.8125 | 312743.4375 | 305145.625 |
| 32215.30664 | 41232.48438 | 40533.78516 | 40235.75391 | 44530.98047 | 35496.12891 | 36115.06641 |
| 61296.42578 | 88077.99219 | 76041.78906 | 86209.60938 | 152189.2344 | 141457 | 151807.4531 |
| 31114.47461 | 40966.85547 | 40471.96094 | 35853.60938 | 30995.00781 | 29804.42188 | 30835.33984 |
| 614903.5 | 675741.3125 | 658475.75 | 670038.5 | 602670 | 471948 | 667225.1875 |
| 7506.125488 | 15184.02637 | 4668.618164 | 3630.387207 | 8632.117188 | 8463.017578 | 8501.192383 |
| 39651.25 | 52659.28906 | 49119.37891 | 48411.76172 | 54397.46875 | 51636.60156 | 50846.57422 |
| 152307.0938 | 163184.5938 | 278543.1875 | 312547 | 228131.2656 | 56791.68359 | 265280.375 |
| 29911.19531 | 53374.8125 | 40560.19531 | 23978.29297 | 55011.66016 | 80253.22656 | 71123.74219 |
| 31860.39063 | 28880.3418 | 45075.90234 | 43428.85938 | 40256.20313 | 39205.83984 | 46099.98438 |
| 10971.58105 | 10473.34277 | 13148.04199 | 15220.64746 | 12380.80371 | 10517.93848 | 12975.94531 |
| 27634.19922 | 37692.92188 | 31995.72266 | 31517.21484 | 26789.26953 | 24555 | 25819.60742 |
| 19106.43945 | 24603.82031 | 23457.75391 | 22688.73633 | 24200.07813 | 22065.64453 | 25454.86719 |
| 22363.00195 | 31781.43359 | 38350.13672 | 31586.17383 | 53338.47656 | 39798.44922 | 41683.03906 |
| 35582.89063 | 51978.5625 | 45502.57031 | 37353.14844 | 50903.24609 | 51901.80859 | 48626.72656 |
| 98489.19531 | 142500.75 | 143650.7344 | 163702.1563 | 228343.6563 | 203282.4531 | 189483.0781 |
| 70965.60938 | 92059.15625 | 88007.60156 | 89607.91406 | 72991.42188 | 76423.58594 | 74685.60156 |
| 50868.17578 | 54635.00391 | 58755.57031 | 53356.58594 | 80763.05469 | 75484.08594 | 89079.98438 |
| 23397.54297 | 34458.38672 | 28549.33984 | 25920.89258 | 26087.83594 | 30935.49219 | 25005.49023 |
| 186814.8438 | 245625.4219 | 271581.3438 | 250722.3906 | 270542.4375 | 277190.9063 | 251024.3906 |
| 58197.32422 | 58348.67578 | 26187.64063 | 63084.66797 | 52333.99609 | 62431.76953 | 52619.60547 |
| 64195.71094 | 99639.39844 | 62868.99609 | 58233.46875 | 95433.88281 | 98302.04688 | 103540.5625 |
| 33256.2666 | 43219.33984 | 42122.0625 | 36119.82129 | 46927.84766 | 59662.58594 | 53640.39453 |
| 96525.22656 | 134566.1406 | 116853.2656 | 111323.875 | 65755.78125 | 69214.66406 | 63730.96875 |
| 82442.375 | 115546.125 | 98620.14844 | 109128.2578 | 135434.9375 | 143167.9688 | 143964.0938 |

| | | | | | | |
|---|---|---|---|---|---|---|
| 54445.81641 | 71079.25781 | 68424.52344 | 57912.96484 | 44068.96875 | 42736.89844 | 40888.91797 |
| 72574.96875 | 92925.50781 | 91411.46094 | 113101.4297 | 145568.9688 | 117718.3125 | 131729.2969 |
| 3897.26123 | 6332.29541 | 6090.13623 | 5831.007324 | 11330.4668 | 7849.927734 | 7821.311523 |
| 16358.52441 | 17693.52148 | 17717.60938 | 17538.83398 | 9809.133789 | 16210.75 | 17910.80078 |
| 154392.1719 | 196809 | 216342.3594 | 210445.7188 | 184180.6719 | 157426.7344 | 180117.8281 |
| 43081.07422 | 46977.59375 | 52336.62109 | 53914.12109 | 37308.32031 | 36090.63672 | 39422.14844 |
| 131401.3281 | 161467.5938 | 169888.0625 | 177561.4531 | 158328.6094 | 156207.6406 | 160277.7188 |
| 15521.0293 | 21425.25781 | 12086.50098 | 20127.19727 | 57922.26953 | 24243.01563 | 30456.58789 |
| 6522.779297 | 8096.463867 | 9227.262695 | 9013.191406 | 6081.338379 | 6727.206543 | 6081.007324 |
| 16345.62891 | 23202.39648 | 21727.61328 | 20376.74609 | 17411.64063 | 20603.00977 | 18735.67969 |
| 44950.01953 | 59437.71875 | 62565.94531 | 54674.42578 | 63191.76953 | 50102.54688 | 55630.97656 |
| 135585.2344 | 169757.7031 | 176918.9063 | 172787.0313 | 123244.5938 | 127189.3594 | 126175.2734 |
| 3549.72168 | 5095.91748 | 5435.154785 | 5241.869629 | 4456.317871 | 6216.898926 | 5148.569824 |
| 223837.6719 | 189482.7344 | 330676.1563 | 384993.9688 | 602636.625 | 409986.8125 | 507843.5313 |
| 345834.2813 | 492275.7188 | 411699.9063 | 381983.75 | 294762.9063 | 351423.0938 | 282656 |
| 82511.98438 | 132050.0938 | 97637.89844 | 91556.90625 | 93228.60156 | 100813.8125 | 88865.17188 |
| 133917.0156 | 174499.8438 | 176513.9844 | 169807.75 | 187990.3125 | 184261.4531 | 186292.0938 |
| 13907.00195 | 17776.68945 | 16918.61133 | 14189.25879 | 13774.01367 | 19970.45313 | 17387.78125 |
| 7136.919922 | 10932.77637 | 8577.708008 | 10680.64258 | 10492.62891 | 8250.548828 | 8353.451172 |
| 40139.98828 | 59240.84375 | 43299.94141 | 47730.46484 | 72370.64063 | 55179.71094 | 68686.10156 |
| 23571.59375 | 34590.58984 | 32145.16797 | 33479.06641 | 35252.67969 | 31384.45508 | 33110.26563 |
| 47439.4375 | 44685.86328 | 53111.76953 | 50883.23438 | 34716.66797 | 37314.15234 | 35778.78516 |
| 48671.21484 | 68975.40625 | 68398.50781 | 72784.05469 | 79414.0625 | 77656.25 | 85915.5 |
| 9008.393555 | 10799.42676 | 10738.53906 | 8945.904297 | 7470.968262 | 8219.568359 | 7023.349121 |
| 7413.943359 | 11384.75879 | 6209.147949 | 8198.552734 | 17517.8457 | 13236.38184 | 13930.93945 |
| 21455.11719 | 23404.05664 | 30051.01367 | 31272.89258 | 50150.71484 | 49864.875 | 50732.14063 |
| 49944.58984 | 85608.96875 | 56253.33203 | 51610.01172 | 76702.21875 | 59881.24609 | 74391.61719 |
| 97394.07813 | 147514.7031 | 118616.4609 | 131572.5625 | 150001.3906 | 143508.2656 | 135098.6563 |
| 98341.5 | 125484.1328 | 126843.4297 | 120158.9688 | 93366.05469 | 106835.2656 | 96935.08594 |
| 29750.46289 | 41980.53516 | 38054.0625 | 38878.04297 | 40469.73047 | 39419.67188 | 39132.55469 |
| 4904.056152 | 5794.069336 | 9950.74707 | 5347.479004 | 24124.18555 | 20360.11133 | 7454.233887 |
| 103289.2734 | 132669.5156 | 140013.6719 | 135955.0938 | 121017.2344 | 123877.6094 | 116577.8359 |
| 336963.5313 | 479168.1563 | 387534.2813 | 395453.7813 | 568063.75 | 608158.1875 | 628637.6875 |

| | | | | | | |
|---|---|---|---|---|---|---|
| 680398.3125 | 656896 | 926937.4375 | 937188.0625 | 951816.5 | 838013.875 | 1006164.875 |
| 6225.709961 | 7985.057617 | 6851.375 | 6551.337891 | 5335.45459 | 9223.041016 | 8205.276367 |
| 29619.02344 | 36199.75391 | 41013.54688 | 38438.45703 | 28453.89063 | 33425.92578 | 30287.33594 |
| 1411604.492 | 1878037.063 | 1673645.219 | 1670819.016 | 1642056.328 | 1723388.891 | 1636368.25 |
| 28018.38477 | 41233.47656 | 36222.26563 | 38372.92578 | 44411.43359 | 39473.53125 | 41885.01172 |
| 802.4807129 | 4040.195557 | 880.1341553 | 1173.769409 | 3587.869385 | 508.9846802 | 3007.686523 |
| 23531.86328 | 29663.53125 | 31105.97852 | 30392.62305 | 27458.27344 | 27643.37305 | 31429.39258 |
| 295347.8438 | 259632.6563 | 132175.6719 | 311017.5 | 213965.9375 | 392948.8438 | 231493.5313 |
| 10890.79492 | 11153.06934 | 13057.67773 | 10690.61719 | 9939.503906 | 9355.777344 | 10972.27832 |
| 138584.5 | 129302.0859 | 159443.0313 | 192588.3438 | 238913.4219 | 250272.9063 | 264267.375 |
| 209402.4063 | 291393.7188 | 262536.1875 | 274081.0625 | 289535.375 | 294717.1875 | 288601.8125 |
| 111072.9297 | 149848.375 | 161558.875 | 164433.375 | 202107.9219 | 186724.4219 | 190441.7188 |
| 2803.369629 | 2837.100586 | 3976.402344 | 3853.073486 | 3163.438232 | 2961.576904 | 2696.099121 |
| 48330.75781 | 63163.28906 | 62655.45703 | 65383.72266 | 63971.68359 | 68397.4375 | 67673.38281 |
| 14498.89551 | 18294.64258 | 16635.91992 | 16844.22656 | 15561.33984 | 16143.79395 | 17629.0293 |
| 73408.40625 | 88514.88281 | 98437.92188 | 95839.38281 | 76740.57813 | 90527.39844 | 83043.42969 |
| 293452.0859 | 442543.0938 | 374367.2969 | 373860.4453 | 640128.9375 | 407928.1563 | 429018.4375 |
| 145968.7969 | 175196.6094 | 179893.2344 | 182865.9844 | 133632 | 134740.3906 | 135400.4375 |
| 25530.23438 | 24150.44531 | 34043.76172 | 41547.38672 | 21071.04492 | 22249.00195 | 22412.64258 |
| 81762.05469 | 95935.48438 | 110516.5 | 111340.4453 | 69836.02344 | 101171.4219 | 77332.73438 |
| 22043.85547 | 23452.30664 | 26862.92188 | 28092.48438 | 45542.32422 | 46661.78125 | 45377.3125 |
| 739170.8125 | 888659.375 | 934273.1875 | 907205.4375 | 906849.125 | 867597 | 832404.125 |
| 264464.7188 | 320576.2188 | 359205.8125 | 345008.5625 | 310382.9688 | 331644.2813 | 322751.625 |
| 13149.37207 | 15077.30176 | 15819 | 17377.33203 | 12088.85449 | 11556.29102 | 11234.13184 |
| 39697.87109 | 62657.14844 | 60948.59375 | 56369.10547 | 53324.31641 | 48666.39844 | 50731.37891 |
| 27049.41016 | 32314.09375 | 32585.54297 | 36118.6875 | 31671.99609 | 32819.97266 | 31686.08398 |
| 38104.62109 | 50010.13672 | 50183.0625 | 58800.26172 | 31521.59961 | 32267.52148 | 40706.82031 |
| 2889.186523 | 5890.756348 | 6483.219727 | 940.5458374 | 8848.772461 | 6451.745117 | 9194.682617 |
| 35346.01953 | 51218.65625 | 49062.87891 | 39276.85547 | 76320.75 | 74579.96094 | 79426.98438 |
| 144125.8438 | 178594.0938 | 191108.4531 | 214648.1719 | 345194.6563 | 287134.875 | 307897.8125 |
| 2969.501221 | 4887.665039 | 4597.416504 | 4762.66748 | 5680.14502 | 5030.229492 | 5754.916016 |
| 3968.944092 | 3224.961426 | 4870.706055 | 5359.69873 | 3807.688721 | 3256.090332 | 3296.603516 |
| 165733.6406 | 216866.4531 | 198685.8594 | 203626.7969 | 269984.2188 | 287705.1875 | 272897.4063 |

| | | | | | | |
|---|---|---|---|---|---|---|
| 17184.96875 | 23828.22656 | 23010.20117 | 22037.85156 | 27082.17188 | 24437.76172 | 26033.73047 |
| 69092.9375 | 79278.72656 | 92082.67188 | 84156.98438 | 69045.30469 | 77029.98438 | 69392.35938 |
| 18868.93945 | 16088.64941 | 24952.67773 | 26529.72266 | 20808.09766 | 8083.835938 | 9881.577148 |
| 569222.375 | 799788.5 | 790645.125 | 659894.25 | 662032.25 | 834424 | 779798 |
| 17510.20117 | 17767.18555 | 19699.34961 | 18959.53711 | 16871.90625 | 18367.39453 | 17106.9707 |
| 2234.241211 | 1844.401978 | 2490.781494 | 2371.327148 | 2011.007568 | 1855.989136 | 2321.304443 |
| 82196.875 | 101807.5703 | 102465.3359 | 105685.3359 | 84338.30469 | 94942.07813 | 90729.4375 |
| 178225.5156 | 255316.4063 | 237738.5625 | 187441.2188 | 175772.75 | 187571.5313 | 160597.0938 |
| 42918.0625 | 61953.1875 | 55999.90234 | 55601.59375 | 56165.94922 | 53523.33984 | 56361.98438 |
| 153297.9688 | 211836.0781 | 223860.7813 | 196587.1875 | 177094.3125 | 184323.1406 | 180260.4063 |
| 7877.765625 | 13130.10254 | 12554.97559 | 12138.125 | 15871.71289 | 15371.24512 | 15289.74609 |
| 33929.76172 | 52800.71875 | 38431.58594 | 42417.50781 | 51433.28516 | 48675.27734 | 47378.19922 |
| 82108.23584 | 104984.6963 | 93978.67773 | 94574.4541 | 61856.09863 | 61997.74512 | 56184.88623 |
| 1611755.875 | 1998308.5 | 2047836.375 | 2173056.5 | 2192268.25 | 2209097.75 | 2364733.75 |
| 39263.19141 | 60360.16016 | 50496.14453 | 55916.29297 | 43936.03125 | 42183.10156 | 45107.44531 |
| 107984.0703 | 135739.3125 | 133700.6406 | 134122.1406 | 116145.5938 | 121645.8984 | 122134.5 |
| 153199.6875 | 174792.5938 | 202879.9063 | 181644.2344 | 126087.7969 | 113675.5313 | 115201.0625 |
| 10321.53418 | 11587.74805 | 14069.18848 | 12311.94531 | 16687.88672 | 14983.81055 | 16768.76367 |
| 5337.177246 | 5813.475586 | 6336.208496 | 6351.265625 | 5577.908203 | 5499.424316 | 5158.697266 |
| 24002.41406 | 33420.64844 | 27427.20117 | 28355.58789 | 22396.09961 | 20410.29297 | 21339.4707 |
| 111826.7188 | 152855.3281 | 127633.0938 | 119444.7969 | 111561.0625 | 97137.67969 | 116234.5547 |
| 406812.8125 | 480801.125 | 503827.5938 | 516873.9688 | 452523.9688 | 450270.1875 | 464769.9688 |
| 261698.3398 | 247846.9648 | 342758.7344 | 341223.8438 | 376265.1328 | 396581.8984 | 419619.1875 |
| 1964.904175 | 3336.429443 | 1910.276611 | 2134.267334 | 2200.143311 | 2793.843018 | 2088.929688 |
| 75400.04688 | 93422.4375 | 96990.0625 | 90656.75781 | 75302.51563 | 91689.98438 | 87703.73438 |
| 31779.58594 | 42732.27344 | 41403.89844 | 43947.96875 | 36713.83203 | 38771.46484 | 40416.01563 |
| 5615.297852 | 6183.506836 | 5031.870605 | 5798.547363 | 10458.0957 | 6821.953125 | 9854.654297 |
| 57539.91797 | 83281.22656 | 65441.86328 | 59339.85938 | 64361.33984 | 75073.42188 | 70634.4375 |
| 35956.67969 | 54654.69531 | 45351.30469 | 44787.61328 | 63251.22266 | 55053.19141 | 56240.33984 |
| 45441.42969 | 55987.82813 | 55980.45703 | 53098.53906 | 58659.89844 | 61521.41406 | 59645.59766 |
| 99676.57813 | 117355.4219 | 130974.6328 | 130303.0703 | 157213.6406 | 149936.7344 | 153886.4844 |
| 49519.72266 | 57217.67969 | 64565.07422 | 69087.75 | 56927.95703 | 58944.32031 | 60384.13672 |
| 132754.8281 | 169968.1094 | 165521.8438 | 158803.4219 | 160913.125 | 155971.8594 | 153185.1406 |

| | | | | | | |
|---|---|---|---|---|---|---|
| 6709.295898 | 10721.94629 | 7961.598145 | 9539.142578 | 23947.00586 | 18363.29492 | 20061.19336 |
| 10030.76367 | 15123.65039 | 11009.33398 | 11281.66797 | 10518.16895 | 9669.148438 | 9544.929688 |
| 30011.08984 | 35263.81641 | 37779.34766 | 38466.51172 | 31285.73242 | 33914.99219 | 31655.21289 |
| 163254.4531 | 195552.7344 | 197041.3594 | 211459.8438 | 198727.125 | 180694 | 192000.5938 |
| 7373.785645 | 13354.29688 | 10486.51953 | 9825.957031 | 19722.5293 | 17164.05078 | 20159.26953 |
| 279763.0938 | 344028.375 | 373742.3438 | 370705.625 | 336687.4375 | 341650.6563 | 325065.2813 |
| 681600.375 | 842535.375 | 880546.6875 | 899890.125 | 958181.5625 | 952774.875 | 958188.75 |
| 85200.51563 | 112365.3359 | 103276.2344 | 96211.38281 | 67247.59375 | 72832.75781 | 69989.66406 |
| 219426.5625 | 273700.6875 | 277896.3125 | 335749.4063 | 336357.3125 | 281794.375 | 318532.7188 |
| 33499.87891 | 40251.01172 | 39671.375 | 37603.82031 | 42291.8125 | 42545.75 | 41975.31641 |
| 47582.06641 | 59079.76172 | 62725.71094 | 63905.58203 | 66225.58594 | 48883.39844 | 58926.13281 |
| 1448363.399 | 1781139.281 | 1827658.953 | 1775310.244 | 1882781.244 | 2245660.649 | 1935348.232 |
| 214519.2031 | 251986.0313 | 260203.8594 | 265534.625 | 190076.7813 | 196406.0781 | 194958.2344 |
| 40930.39844 | 36073.66797 | 18121.05469 | 42216.36328 | 36511.56641 | 52140.21484 | 34261.11719 |
| 71442.08594 | 88526.89844 | 89840.14844 | 92076.24219 | 66315.07031 | 64162.79297 | 63264.03516 |
| 310119.7188 | 415428.4063 | 384574.1563 | 406111.4688 | 401013.0625 | 394745.7813 | 372491.7813 |
| 59000.71094 | 66355.60938 | 74120.95313 | 73388.27344 | 64145.29297 | 64989.89063 | 67014.50781 |
| 2893775.25 | 3502347.75 | 3562766.75 | 3593822 | 4656342.5 | 4719185 | 4726718.5 |
| 16373.9082 | 19898.67383 | 18827.67578 | 19607.50781 | 22992.86133 | 24719.73828 | 21782.98438 |
| 9183.47168 | 11775.17383 | 10993.20898 | 11339.21387 | 9097.266602 | 8947.798828 | 8007.371582 |
| 107548.5156 | 141291.6563 | 130741.7422 | 135992.1875 | 178255.4063 | 145052.5625 | 147159.0625 |
| 3518.358643 | 4553.962402 | 3719.999023 | 3535.540039 | 4740.165039 | 5620.996094 | 5015.479004 |
| 905475.6172 | 1045968.781 | 1218716.602 | 1084621.082 | 761479.1289 | 742946.8242 | 779181.9219 |
| 98465.91406 | 119410.625 | 124962.1328 | 119535.7031 | 95454.5625 | 93678.02344 | 96024.02344 |
| 26494.10156 | 31464.87695 | 33603.08203 | 31720.73633 | 31011.00195 | 33879.16797 | 30161.32813 |
| 18306.10938 | 15278.76172 | 19452.7793 | 21473.34766 | 18996.38867 | 19017.0332 | 20532.55469 |
| 29688.96875 | 38103.64063 | 36818.70703 | 35739.69531 | 29100.41016 | 27442.71484 | 26033.24609 |
| 18956.16406 | 35067.78125 | 18049.22656 | 14722.7959 | 25332.62109 | 21914.28906 | 25779.80469 |
| 52468.84766 | 55774.125 | 79424.875 | 99245.0625 | 153092.0156 | 117977.3281 | 126541.0234 |
| 61854.89844 | 73050.33594 | 78142.35156 | 76105.89844 | 70531.41406 | 72857.19531 | 74339.45313 |
| 41487.28906 | 55825.34766 | 53358 | 62169.49219 | 89908.26563 | 79207.04688 | 83686.03125 |
| 50120.23047 | 62241.15234 | 63159.63281 | 63401.42188 | 74744.46875 | 65961.99219 | 71937.89063 |
| 1868.027344 | 3245.336426 | 2542.121338 | 2546.456543 | 4230.453613 | 3143.450928 | 3155.980225 |

| | | | | | | |
|---|---|---|---|---|---|---|
| 73916.77344 | 127233.4375 | 82818.75 | 54632.83594 | 146195.5938 | 168728.2188 | 126849.1094 |
| 721757.75 | 841708.3125 | 1015830.188 | 1233407.5 | 1174761 | 959281.25 | 1070820.875 |
| 50180.60156 | 61311.70703 | 56215.76953 | 55869.70703 | 50311.12109 | 53845.42578 | 52659.92188 |
| 104618.0859 | 113705.0859 | 129530.7578 | 127713.6563 | 92702.16406 | 102029.8438 | 98489.67969 |
| 6453.328125 | 9516.964844 | 8198.962891 | 11384.62207 | 6816.971191 | 9560.964844 | 8318.598633 |
| 30672.45117 | 41825.74609 | 39699.90625 | 32787.35156 | 32886.78516 | 34330.05859 | 32458.04883 |
| 1037952.813 | 1340483.25 | 1309776.25 | 1363188.5 | 1341745.625 | 1249393.375 | 1313180.125 |
| 46809.17969 | 57865.74609 | 60209.49219 | 59335.20313 | 61639.77734 | 57804.99609 | 60387.03906 |
| 47553.36719 | 65517.50781 | 67850.96094 | 50552.55078 | 65229.93359 | 97199.125 | 68105.59375 |
| 121881.4922 | 139649.2813 | 205729.3906 | 167674.7344 | 119278.5156 | 121460.7109 | 121990.7422 |
| 4969.067871 | 6925.034668 | 7205.041016 | 6529.737305 | 7274.925781 | 6983.042969 | 7643.759277 |
| 38500.11719 | 47598.125 | 39557.80859 | 49714.96484 | 153289.75 | 68254.46875 | 80769.66406 |
| 123403.7422 | 158884.6406 | 161339.8438 | 155209.3125 | 130768.6328 | 134653.7188 | 135020.25 |
| 53529.32813 | 76832.50781 | 65242.00781 | 61267.77734 | 56428.16797 | 58537.51172 | 56701.94141 |
| 217397.0938 | 279723.4063 | 275821.625 | 268157.9063 | 264745.8438 | 280229.375 | 268697.625 |
| 74529.65625 | 83758.17188 | 101392.5391 | 90206.85938 | 71390.22656 | 72630.17188 | 69533.99219 |
| 245465.4531 | 323456.3438 | 313956.4688 | 293213.1563 | 363927.1563 | 401684.6563 | 388613.625 |
| 138391.3594 | 193152.7969 | 193043.8125 | 211829.4688 | 170670.3906 | 181719.1719 | 182556 |
| 9345.875977 | 14378.00098 | 11689.43359 | 10718.87012 | 21588.95117 | 18751.79883 | 18727.2832 |
| 826567 | 1036668.5 | 985452.125 | 932243.125 | 683526.0625 | 762808.5 | 652608.9375 |
| 3561.852295 | 6148.593262 | 4422.085449 | 5100.736816 | 9653.311523 | 6240.980469 | 6225.848633 |
| 76520.21875 | 103216.5625 | 99339.53906 | 92595.1875 | 103518.1328 | 102241.5469 | 104014.6172 |
| 212139.625 | 245516.0781 | 287225.2813 | 259081.4531 | 236339.4063 | 250523.9063 | 255868.2188 |
| 32752.14453 | 41729.32813 | 39555.38672 | 40096.16016 | 42599.89063 | 45863.76953 | 41472.78125 |
| 52490.86719 | 70183.96094 | 63865.25391 | 65937.91406 | 35307.99219 | 35982.90234 | 37529.3125 |
| 59781.59766 | 74347.55469 | 75475.07813 | 79499.17969 | 100323.5078 | 100235.375 | 101403.1094 |
| 1067736.25 | 1398072 | 1321523.875 | 1434643.625 | 1725622.25 | 1611890.75 | 1598356.625 |
| 127130.7734 | 166886.4844 | 173960.2344 | 185357.3438 | 154826.875 | 162863.375 | 139169.6875 |
| 17752.29492 | 25661.50977 | 21484.10352 | 19206.17773 | 31695.57422 | 27182.73047 | 28751.16602 |
| 73462.35156 | 84901.52344 | 87338.01563 | 86470.61719 | 40454.90625 | 37791.61328 | 35305.51953 |
| 47985.46094 | 58516.89453 | 61152.77344 | 62873.82422 | 43742.10547 | 52314.11719 | 47555.39453 |
| 164291.875 | 205166.7188 | 168576.7969 | 170699.8125 | 203097.1875 | 217413.4844 | 159456.2188 |
| 18852.05273 | 23644.26172 | 19706.64844 | 25392.2207 | 23076.21289 | 20469.53125 | 23728.83594 |

| | | | | | | |
|---|---|---|---|---|---|---|
| 679624.3125 | 830933.0625 | 766241.9375 | 832284.6875 | 711972 | 708759.625 | 691691.3125 |
| 112455.5996 | 125084.044 | 129300.811 | 137285.7168 | 124239.5552 | 115559.6711 | 123611.6245 |
| 93914.41406 | 118095.2813 | 117065.0703 | 113583.8594 | 98978.16406 | 111302.6563 | 114480.2422 |
| 41205.55859 | 48782.35547 | 51536.28125 | 55237.91406 | 51184.04297 | 54735.41016 | 53920.03516 |
| 64792.02344 | 77915.5625 | 69745.32813 | 66385.36719 | 72310.09375 | 72524.82813 | 68388.52344 |
| 214364.8594 | 247123.6719 | 362976.5625 | 357247.75 | 466753.0313 | 514568.75 | 499074.1875 |
| 17999.91797 | 25877.38086 | 23521.625 | 21899.27344 | 21625.67969 | 24297.36719 | 22808.32227 |
| 5496.721191 | 8279.504883 | 7677.476074 | 7165.305664 | 9159.334961 | 7945.63623 | 9296.016602 |
| 253242.1563 | 296877.9063 | 312469.3438 | 387033.5625 | 605159.625 | 431426.9688 | 499842.9375 |
| 2337.421875 | 2937.0896 | 2297.841553 | 2877.838379 | 2575.156982 | 2934.588623 | 2868.193115 |
| 558444.125 | 693599.625 | 739749.375 | 710793.25 | 662514.25 | 714221.0625 | 710359.3125 |
| 162055.3906 | 204188.8594 | 206107.2969 | 190387.5625 | 178341.5938 | 196929.4375 | 182083.3438 |
| 50979.01172 | 66584.28906 | 62170.92578 | 57261.24219 | 37170.51563 | 41771.51172 | 38687.24219 |
| 2753.423096 | 5214.021484 | 4191.380371 | 4538.836914 | 10322.0791 | 8118.516113 | 8565.911133 |
| 130182.2188 | 159091.1875 | 147199.2344 | 162252.6563 | 158121.1719 | 173019.2969 | 167197.5781 |
| 13750.65332 | 14683.63867 | 18507.53711 | 15096.73828 | 21424.95703 | 30265.60547 | 22204.99414 |
| 77888.3125 | 100368.5625 | 97257.89844 | 108905.1563 | 197050.2344 | 148758.8281 | 171072.3906 |
| 4467.032227 | 7902.425781 | 6434.135742 | 7206.223145 | 6745.625488 | 6869.907227 | 7175.874512 |
| 14314.53516 | 17895.25586 | 17985.05469 | 17439.34766 | 28566.38477 | 25797.19141 | 26629.32031 |
| 179658.4219 | 265176.1563 | 202469.5781 | 169122.5156 | 127170.0078 | 155771.6563 | 153270.0313 |
| 33508.02344 | 34706.14844 | 16561.84961 | 29038.32617 | 41611.76953 | 62212.35547 | 26112.73047 |
| 15504.16504 | 14774.52734 | 17555.07422 | 19733.16211 | 15937.63184 | 13488.13867 | 15289.4209 |
| 7386.889648 | 10469.45605 | 9467.712891 | 9777.544922 | 9039.65332 | 11368.94238 | 9775.472656 |
| 41479.23438 | 52559.1875 | 51765.35156 | 46909.28516 | 45659.3125 | 44241.74219 | 45624.05859 |
| 94382.40625 | 118642.7031 | 115403.25 | 109871.8125 | 75140.55469 | 70554.4375 | 67942.14063 |
| 59329.32031 | 72266.84375 | 72042.71094 | 72569.51563 | 43635.01172 | 45618.33594 | 44593.57422 |
| 7378.540527 | 8153.321777 | 8214.111328 | 7459.197266 | 9990.185547 | 10128.95313 | 10936.875 |
| 1021592.625 | 1104006.5 | 1177142.5 | 1391778.125 | 1280394.875 | 1108790.375 | 1235755.625 |
| 80799.15625 | 103136.125 | 90179.54688 | 66823.55469 | 63668.27344 | 105932.6641 | 76693.48438 |
| 56536.76172 | 64129.71875 | 82218.29688 | 73238.67969 | 52466.80859 | 63387.95703 | 58252.14063 |
| 41042.23047 | 48324.71094 | 46296.55859 | 48906.42188 | 59998.67969 | 62795.62109 | 63313.66406 |
| 22740.1543 | 19006.68945 | 32860.07813 | 36604.66406 | 39243.67969 | 35831.39844 | 36941.12109 |
| 5857.806641 | 7563.552246 | 6889.188965 | 6789.395996 | 4426.821777 | 4827.851074 | 5421.609375 |

| | | | | | | |
|---|---|---|---|---|---|---|
| 74769.39063 | 98742.47656 | 87451.48438 | 89117.30469 | 115301.5703 | 115010.2813 | 112818.5 |
| 29381.71289 | 39696.65234 | 34896.22266 | 36676.40234 | 36672.19531 | 38912.95313 | 35971.28906 |
| 2505.129883 | 4348.92334 | 4166.900391 | 3427.554932 | 7696.972168 | 11654.96582 | 8129.039063 |
| 567002.25 | 683147.3125 | 653891.8125 | 682764.375 | 807714.5625 | 792837.875 | 758358 |
| 26267.60742 | 31165.83984 | 32411.61328 | 29413.17773 | 21722.41992 | 23648.48633 | 18943.50781 |
| 34412.39063 | 49156.95313 | 43408.75391 | 44120.55469 | 62060.28125 | 60669.75 | 60324.39844 |
| 13165.26172 | 17070.02148 | 15493.11816 | 16176.33789 | 12886.1748 | 14745.53906 | 13284.05273 |
| 143236.6406 | 178833.0156 | 169567.9063 | 167297.4844 | 177347.9375 | 183565.2344 | 179498.9219 |
| 20902.00195 | 142797.0781 | 49863.82422 | 74388.625 | 212596.1406 | 140019.9219 | 132256.25 |
| 28189.44141 | 44320.05078 | 34111.63281 | 37295.91406 | 41201.78906 | 56112.76172 | 40561.58203 |
| 62480.02344 | 74483.59375 | 73476.80469 | 74259.13281 | 85661.75781 | 75377.63281 | 77089.875 |
| 7648.006348 | 14745.22363 | 11917.11426 | 11855.74316 | 9548.523438 | 4769.596191 | 7742.853516 |
| 11505.58203 | 17167.41992 | 13995.37207 | 12660.14941 | 13066.04492 | 12363.94824 | 12181.36816 |
| 63043.71094 | 65465.64844 | 78805.30469 | 72251.57031 | 47131.82422 | 54716.53906 | 49244.13281 |
| 19664.73828 | 23563.87109 | 27471.9375 | 27389.5293 | 28446.00977 | 31299.83594 | 27212.12305 |
| 25751.45703 | 31282.39844 | 32441.05859 | 32120.96289 | 33982.37109 | 34974.5625 | 32229.68945 |
| 274284.5313 | 288249.2188 | 342789.9688 | 359965.0313 | 289667.25 | 289510.6563 | 327619.0625 |
| 47186.39453 | 52292.24219 | 58367.87109 | 62401.50391 | 75137.98438 | 63147.48828 | 68539.85156 |
| 75326.89844 | 96593.45313 | 91235.60938 | 110858.9531 | 108992.1953 | 93918.71094 | 102937.1875 |
| 229071.125 | 300576.7188 | 291448.5625 | 297986.3125 | 359251.125 | 364430.8438 | 365226.1563 |
| 12350.08984 | 15264.33301 | 14685.63574 | 15892.83105 | 16854.50781 | 16121.46094 | 15592.34473 |
| 54410.23828 | 71728.90625 | 70772.42188 | 64332.23438 | 36142.76563 | 44979.18359 | 41219.50781 |
| 139096.3125 | 176985.1875 | 129333.9609 | 153233.7188 | 109803.7422 | 152585.2031 | 121360.1406 |
| 35223.13281 | 43577.39453 | 46523.75 | 43638.57031 | 31209.35156 | 34587.41797 | 33995.11719 |
| 13238.75977 | 19396.31836 | 12897.5625 | 12467.27637 | 10509.38574 | 11595.57129 | 10170.12109 |
| 75741.82031 | 109399.3672 | 94508.14063 | 81101.47656 | 50400.35156 | 53426.59375 | 53691.35547 |
| 86262.86816 | 96951.22852 | 113784.5371 | 117113.167 | 129412.5488 | 122117.9238 | 125911.8682 |
| 12184.72461 | 15076.04688 | 13473.19141 | 17032.24805 | 13803.06738 | 13762.6875 | 14726.81934 |
| 25460.86719 | 33953.04297 | 26691.08984 | 25757.46289 | 26857.28516 | 29327.66797 | 31519.29297 |
| 3789.335938 | 3348.399902 | 3396.992432 | 3421.741455 | 2521.679443 | 2360.876221 | 2883.004883 |
| 17654.06836 | 21012.99805 | 20333.93945 | 19435.56641 | 16436.9043 | 19573.22852 | 16056.04883 |
| 46506.32422 | 56679.10938 | 54862.33203 | 54632.85938 | 74841.58594 | 77477.28906 | 74941.99219 |
| 1083281.672 | 1372162.031 | 1346293.328 | 1393487.781 | 1242575.594 | 1311770.953 | 1201038.766 |

| | | | | | | |
|---|---|---|---|---|---|---|
| 184854.6563 | 260017.7031 | 237960.4063 | 195834.4688 | 195111.0469 | 218077.7969 | 199479.8281 |
| 24142.30859 | 37036.35156 | 27891.16797 | 27131.27148 | 17448.25781 | 21972.58398 | 19909.07227 |
| 47195.89063 | 90913.1875 | 43778.67969 | 51336.63672 | 69915.39063 | 70880.53125 | 72861.57031 |
| 339841.125 | 450767.625 | 463404.5938 | 436394.25 | 350325.6563 | 335449.3125 | 356765.2813 |
| 310927.1875 | 405896.5938 | 394465.9375 | 412362.625 | 459093.875 | 409630.4688 | 421437.25 |
| 24294.26563 | 37538.82031 | 30888.05078 | 31237.5918 | 21571.11523 | 25668.19141 | 20887.05469 |
| 10010.02148 | 12178.34961 | 13828.30371 | 12474.20313 | 10937.29004 | 8273.199219 | 9353.888672 |
| 44082.16797 | 65618.80469 | 61673.11719 | 59197.47266 | 56530.20313 | 56230.19922 | 53395.38672 |
| 28357.51563 | 39790.09375 | 32352.0918 | 35842.60547 | 35384.78516 | 37985.13672 | 34654.97656 |
| 270638.6563 | 364743.0938 | 319794.7813 | 299874.375 | 326198.9063 | 382789.1875 | 334330.9688 |
| 413355.5313 | 513430.0938 | 502564.875 | 515414.5313 | 454273.8125 | 468608.0938 | 479891.4688 |
| 11591.83691 | 19359.05859 | 11923.41797 | 10900.42383 | 18489.5 | 14700.53223 | 14387.66016 |
| 41963.125 | 53620.71484 | 58298.41797 | 56953.89844 | 63467.83594 | 67540.85938 | 71294.35156 |
| 10720.09961 | 9463.049805 | 12750.94238 | 11504.39844 | 7844.52002 | 6892.106934 | 6718.932129 |
| 4366.17627 | 4387.397949 | 5936.342285 | 3932.920898 | 3225.093994 | 3530.649902 | 3867.115723 |
| 21815.13477 | 27618.31445 | 27136.19922 | 28974.68555 | 30975.76367 | 30060.41992 | 32704.19336 |
| 46179.09766 | 57449.03125 | 49140.24219 | 50842.05469 | 48248.13672 | 52618.89844 | 48096.04688 |
| 8461.230469 | 19159.86914 | 8221.953125 | 7612.050781 | 16828.12891 | 19438.49609 | 17712.58203 |
| 66418.05469 | 84820.71094 | 79024.1875 | 83341 | 80264.3125 | 79267.32813 | 77680.50781 |
| 33053.21875 | 47050.21875 | 33588.97266 | 36178.40234 | 37854.15625 | 35330.24219 | 37914.82422 |
| 50260.26563 | 66180.21094 | 68277.75 | 63918.96484 | 87584.77344 | 75468.94531 | 77094.85938 |
| 737078.8125 | 845735.625 | 856042.1875 | 963638 | 817436.875 | 916178.125 | 855472.5625 |
| 386251.875 | 504596.375 | 490743.6875 | 525494.9375 | 473447.1563 | 463623.3125 | 475323.625 |
| 59463.53516 | 79531.49219 | 79076.76563 | 76330.29688 | 89586.03125 | 76443.375 | 85179.16406 |
| 91357.4375 | 114423.2109 | 105368.0234 | 124227.3281 | 109527.5859 | 99786.78906 | 105595.5625 |
| 20806.7168 | 25513.71094 | 26247.28711 | 27049.26953 | 22246.16016 | 26287.77344 | 23991.20898 |
| 24768.0957 | 30560.53516 | 32796.02344 | 33556.81641 | 33203.16016 | 33365.80469 | 34069.79297 |
| 31064.4707 | 38734.37109 | 36089.62109 | 31405.98828 | 12376.11719 | 12834.02441 | 11345.95996 |
| 135536.3125 | 157540.3594 | 179142.875 | 171650.1094 | 194538.2656 | 178011.4219 | 204983.2031 |
| 2439.852783 | 2964.915771 | 2119.92041 | 1584.832764 | 5151.526855 | 6294.57373 | 5044.511719 |
| 41174.20703 | 48322.29297 | 43255.55078 | 45284.52734 | 48923.22266 | 43401.96875 | 44635.71484 |
| 109960.7969 | 157896.3906 | 152791.8281 | 164163.4844 | 80767.28125 | 79654.75781 | 83878.875 |
| 53415.12891 | 67614.42188 | 38524.00781 | 58055.38672 | 87551.67969 | 209933.3125 | 91587.07031 |

| | | | | | | |
|---|---|---|---|---|---|---|
| 649213.875 | 783208.8125 | 788575.4375 | 808078.25 | 720572.8125 | 772568.5 | 685107.3125 |
| 70973.92969 | 78298.34375 | 91176.77344 | 90088.5 | 102231.5547 | 114927.9922 | 118818.5781 |
| 14120.35352 | 20918.39844 | 17242.41992 | 15485.94043 | 17267.19141 | 14911.70996 | 14778.75586 |
| 1090163.625 | 1370661.25 | 1335381.375 | 1366219.75 | 688822.0625 | 611444 | 655832.125 |
| 101339.2734 | 115953.4141 | 117306.9844 | 117201.5156 | 112293.1875 | 111388.6953 | 123941.1875 |
| 4066.732178 | 6512.153809 | 3620.756836 | NA | 4447.414063 | 3599.309082 | NA |
| 43958.16016 | 62767.78125 | 12984.42773 | 43530.38281 | 74590.46875 | 155183 | 108934.6328 |
| 6644.999023 | 8664.388672 | 8532.181641 | 9708.303711 | 8060.75293 | 7781.532227 | 6560.005371 |
| 39056.59766 | 49062.09375 | 41455.0625 | 45762.75781 | 49378.71484 | 51637.30859 | 54410.40234 |
| 5704.523438 | 10005.80664 | 7402.547363 | 10625.0752 | 9592.658203 | 6819.123047 | 8037.544922 |
| 44297.92578 | 53152.21484 | 57968.89063 | 51186.91406 | 63078.65234 | 68563.14844 | 70288.27344 |
| 59724.64355 | 87682.08789 | 70061.43359 | 77171.64795 | 87158.86133 | 99237.0498 | 88469.85156 |
| 6087.223633 | 12414.75879 | 2083.991943 | 7149.393066 | 8142.292969 | 8921.347656 | 10828.00879 |
| 28336.26172 | 20538.52344 | 35967.75 | 44088.57813 | 24343.38086 | 21218.02344 | 23868.71484 |
| 65128.87891 | 83382.14063 | 81571.27344 | 70694.38281 | 72935.16406 | 80880.70313 | 79692.94531 |
| 1025361.625 | 1283633.25 | 1272177 | 1320877.625 | 1369102.75 | 1168504.875 | 1310612.625 |
| 7462.905273 | 9120.662109 | 9026.394531 | 10398.20117 | 11122.98535 | 12435.625 | 14536.34863 |
| 24178.61328 | 28096.24414 | 27407.88867 | 26934.71094 | 27596.06445 | 29028.86719 | 25459.73828 |
| 416677.5313 | 575863.125 | 508083.25 | 465972.25 | 466610.9375 | 515493.5625 | 474345.625 |
| 3150.022217 | 9102.354492 | 12183.65625 | 6153.679199 | 21225.97266 | 39363.03906 | 10300.74121 |
| 192986.2813 | 243627.2344 | 243208.0625 | 242204.125 | 208441.3438 | 209775.1875 | 216571.9063 |
| 424237.25 | 405616.7813 | 646516.6875 | 549123 | 390748.4375 | 418759.2813 | 477036.5938 |
| 121295.0391 | 125845.3828 | 153486.9219 | 112166.4141 | 69453.54688 | 68903.05469 | 64600.12109 |
| 14457.7207 | 18477.73828 | 18249.58008 | 15835.07324 | 14157.11523 | 12850.26367 | 13194.69336 |
| 183795.5 | 210840.5625 | 205125.6719 | 242187.1406 | 275244.125 | 251413.7969 | 272545.0625 |
| 217154.2969 | 283718.0625 | 274574.0938 | 256769.3594 | 272347.0625 | 282131 | 268937.0313 |
| 35807.88281 | 40624.17969 | 43068.66406 | 43060.08984 | 24139.57031 | 31159.51953 | 34028.82422 |
| 62997.97656 | 78238.45313 | 73037.42969 | 70833.60938 | 59404.88281 | 53028.16406 | 61105.53516 |
| 25455.06055 | 26736.51953 | 31441.11719 | 27511.04297 | 42231.77734 | 35070.52344 | 38404.5 |
| 292343.6563 | 451432 | 381869.3125 | 264724.4063 | 292809.6875 | 434347.875 | 403466.9063 |
| 130656.4844 | 201067.875 | 154057.5625 | 153636.2344 | 196578.6719 | 185274.1563 | 190510.5625 |
| 125816.3906 | 135968.7344 | 163292.4219 | 147426.4844 | 113338.9141 | 119630.0469 | 120354.125 |
| 23328.8457 | 33426.22266 | 24072.64063 | 27148.51953 | 38881.82422 | 35060.53906 | 38729.54297 |

| | | | | | | |
|---|---|---|---|---|---|---|
| 14043.7627 | 18599.18164 | 17586.29492 | 16286.54785 | 22939.71094 | 22160.30273 | 22756.54492 |
| 160972.9375 | 190834.1719 | 198632.5469 | 197252.6406 | 166997.7813 | 171861.5938 | 166999.6406 |
| 1243.72522 | 2786.781006 | 1986.262085 | 1690.528809 | 2717.635498 | 3688.059082 | 2926.524658 |
| 180107.4219 | 231980.7188 | 198649.5781 | 216983.125 | 236585.8594 | 277692.5938 | 243035.1875 |
| 55209.30859 | 61937.57813 | 66781.86719 | 62264.75 | 18370.31055 | 25166.79492 | 15836.63086 |
| 11004.09375 | 14094.21875 | 12288.61719 | 12264.74902 | 14423.3252 | 14787.49414 | 13977.18945 |
| 36204.66406 | 41281.21094 | 42892.87891 | 35558.57813 | 29228.64648 | 28873.30273 | 28777.98242 |
| 15099.03027 | 27483.875 | 14247.05176 | 11346.16406 | 43672.15625 | 25551.44141 | 24948.70898 |
| 7416.795898 | 9762.266602 | 8709.427734 | 8642.53418 | 17888.59961 | 16442.64258 | 17903.07227 |
| 66966.28125 | 89472.11719 | 73936.19531 | 77200.30469 | 50809.05859 | 50954.32031 | 48260.66406 |
| 12362819 | 15023258 | 14652141 | 15552506 | 12981175 | 11700123 | 12882695 |
| 45455.12109 | 69008.14844 | 65439.28516 | 72313.55469 | 105817.1953 | 101714.8203 | 96007.94531 |
| 56186.80859 | 87439.21094 | 57842.96484 | 56540.33984 | 71005.70313 | 66367.29688 | 74497.25781 |
| 58981.29688 | 72999.49219 | 69551.94531 | 68167.01563 | 75598.07031 | 76050.85938 | 79332.38281 |
| 104243.9063 | 165071.7344 | 123784.3281 | 144780.4844 | 246350.6875 | 198070.7031 | 202446.0781 |
| 335028.5938 | 406741.75 | 398970.9375 | 408660.5938 | 345085.4375 | 365092.7188 | 341955.0625 |
| 861507.5 | 1044128.625 | 936841 | 993768.875 | 1049586.375 | 1080742.25 | 973645.75 |
| 23637.17969 | 36872.80859 | 26482.33008 | 22149.57227 | 16620.71289 | 17033.70898 | 15827.9209 |
| 34558.35156 | 43407.27734 | 42358.42188 | 37986.30859 | 36551.34766 | 36499.13281 | 36796.03906 |
| 61172.05469 | 93901.88281 | 72753.50781 | 84297.23438 | 94640.3125 | 75130.65625 | 75223.51563 |
| 3059.813232 | 5075.711426 | 4205.416504 | 3320.562012 | 4431.22998 | 5834.28418 | 6353.829102 |
| 13277.52344 | 15030.58008 | 15631.70117 | 13209.97168 | 13194.94629 | 10211.46289 | 6284.372559 |
| 77661.54688 | 101285.4922 | 84704.96875 | 90639.82813 | 74996.8125 | 70884.10938 | 75341.74219 |
| 19346.12109 | 24213.83008 | 18272.83984 | 24409.22656 | 26687.3457 | 25092.25195 | 25885.91797 |
| 285323.0625 | 356480.9063 | 335265.9688 | 345755.5 | 370360.5625 | 373455.0938 | 391237.625 |
| 36443.09375 | 38435.33984 | 46929.12109 | 51189.625 | 55975.42969 | 56074.92969 | 55776.77734 |
| 25927.85352 | 26260.32617 | 28436.65234 | 30855.93945 | 28837.12109 | 33337.81641 | 31600.16602 |
| 144898.3125 | 144194.8438 | 186945.7656 | 192155.7344 | 213295.25 | 197756.8438 | 225248.5469 |
| 55973.46875 | 70622.70313 | 72982.61719 | 72978.94531 | 79203.90625 | 78145.03906 | 79845.61719 |
| 3193.196533 | 3808.662109 | 3097.571777 | 3293.672607 | 3207.676758 | 3742.685303 | 3385.268799 |
| 141101.9219 | 209296.6563 | 142615.3594 | 136203.875 | 118461.9688 | 130658.3438 | 128532.8828 |
| 870632.5625 | 1056426.5 | 1029723.313 | 1039906.063 | 1089802.75 | 1025552.938 | 1034712.125 |
| 62034.59766 | 72532.15625 | 70719.72656 | 71579.64844 | 61429.78906 | 61655.61719 | 57824.80859 |

| | | | | | | |
|---|---|---|---|---|---|---|
| 174398.6094 | 228925.5938 | 222424.0938 | 236284.0469 | 133418.7813 | 116004.4609 | 130775.5625 |
| 298353.1563 | 439268.3438 | 384236.875 | 378307.0625 | 563291.8125 | 543675.5 | 553282 |
| 16256.5752 | 21419.67773 | 17629.83594 | 16108.35059 | 16412.82813 | 15990.28516 | 15612.23047 |
| 85128.67969 | 121049 | 111142.2266 | 106363.7891 | 100781.0313 | 110260.5703 | 115171.0156 |
| 303957.3125 | 610513.3125 | 451364.375 | 331328.9063 | 367999.1563 | 507077.4063 | 394164.2188 |
| 3615.427246 | 3886.385986 | 3073.226074 | 3441.095703 | 3978.245361 | 3971.013428 | 3822.611328 |
| 10777.3252 | 12855.13672 | 12321.68652 | 17516.0918 | 21073.96875 | 17136.38281 | 18771.27734 |
| 92108.17188 | 110900.6641 | 119925.6172 | 122167.7344 | 134588.75 | 154328.9531 | 146307.5469 |
| 42660.78906 | 46886.42188 | 54146.48438 | 50620.08984 | 57358.55078 | 52766.99219 | 62017.57031 |
| 9064.076172 | 4730.843262 | 12567.02051 | 11581.18164 | 5514.906738 | 4309.234863 | 7079.097168 |
| 12488.57031 | 16205.00586 | 15152.32422 | 12304.06934 | 13323.79492 | 14287.61719 | 12969.42773 |
| 29138.51953 | 27353.8125 | 20377.36914 | 37300.89844 | 30143.06055 | 40954.86719 | 30833.375 |
| 38992.98438 | 44368.26172 | 52259.94531 | 51468.12891 | 50302.30469 | 46272.38672 | 49561.53516 |
| 24389.54688 | 33310.01953 | 28848.23047 | 28187.49805 | 30450.5 | 30161.34961 | 28052.6875 |
| 461079.2813 | 520433.2813 | 582490.3125 | 555040.25 | 477030.2813 | 540741.1875 | 527750.5 |
| 9497.081055 | 12579.2373 | 10197.11426 | 8411.00293 | 5744.597656 | 9833.519531 | 6900.277344 |
| 708957.8125 | 913691.875 | 859787.875 | 904621.875 | 943972.75 | 917461.625 | 904649.4375 |
| 97818.96094 | 105310.7266 | 104345.5156 | 121148.3672 | 115945.0859 | 134481.8438 | 120245.2891 |
| 15005.44629 | 39590.28516 | 4998.132324 | 13686.45801 | 39437.5625 | 37587.03125 | 35348.59375 |
| 23299.35156 | 29972.77148 | 33505.13672 | 29986.05078 | 34694.00781 | 38162.5625 | 34425.75 |
| 81145.66406 | 94240.34375 | 106911.9766 | 100626.5 | 106045.7422 | 107140.9297 | 106693.5703 |
| 6094.096191 | 6075.876465 | 7851.553223 | 7387.409668 | 6870.328613 | 7240.040039 | 6284.510742 |
| 68998.00781 | 79606.28906 | 68340.42188 | 69048.92188 | 77982.32813 | 86386.10156 | 65839.83594 |
| 65161.59375 | 79048.05469 | 81917.79688 | 80790.52344 | 88862.89063 | 89373.14063 | 92936.70313 |
| 177928.1094 | 197901.1875 | 210419.7188 | 198716.9688 | 292173.0625 | 297918.6875 | 301528.7813 |
| 2975.186523 | 3180.523682 | 3490.697998 | 4115.943848 | 4440.558105 | 5379.656738 | 6504.70166 |
| 135556.5469 | 150931.7813 | 167074.8438 | 157627.8125 | 153989.7031 | 155850.0625 | 164004.625 |
| 27791.9375 | 26252.92773 | 30060.44727 | 25979.10938 | 35283.75781 | 38056.15625 | 41345 |
| 16051.96582 | 26378.17578 | 19354.82031 | 18536.62695 | 19505.15625 | 16878.51953 | 18245.25195 |
| 9689.229492 | 13208.72363 | 10266.37695 | 11165.31738 | 10068.0957 | 8427.539063 | 9376.169922 |
| 111906.4375 | 143591.0313 | 131856.6406 | 130625.8828 | 91441.28906 | 82882.88281 | 88414.99219 |
| 47337.48438 | 57418.71094 | 56223.39063 | 50385.32422 | 24096.9375 | 21755.64453 | 22216.85156 |
| 7880.963379 | 11125.53809 | 9830.733398 | 10635.9873 | 9817.609375 | 9734.78418 | 10780.78516 |

| | | | | | | |
|---|---|---|---|---|---|---|
| 57475.66016 | 58021.10156 | 72731.9375 | 70779.52344 | 56170.60938 | 45271.23828 | 55065.41406 |
| 4172.336914 | 5748.402832 | 5675.766113 | 5785 | 6754.321777 | 6686.096191 | 6717.675781 |
| 15456.3877 | 16344.74609 | 17412.7207 | 14803.88965 | 10167.54297 | 13508.43652 | 11323.22949 |
| 48939.96484 | 60183.29297 | 70833.65625 | 53111.37109 | 89483.23438 | 123399.0078 | 97759.46094 |
| 737122.25 | 774279.8125 | 943164.75 | 934012.875 | 918248.25 | 943825.6875 | 922428.4375 |
| 66633.85156 | 82231.60156 | 81233.73438 | 83305.8125 | 101096.9531 | 104759.9453 | 103603.0938 |
| 290420.4375 | 330037.1875 | 363553.375 | 386946.375 | 312288.25 | NA | 284920.25 |
| 604647.4375 | 724265.9375 | 750960.6875 | 749592.9375 | 763049.4375 | 748190.8125 | 782190.25 |
| 125234.6719 | 159670.4844 | 149687.4844 | 150326.7188 | 179658.125 | 177358.2188 | 181449.7188 |
| 9427.643555 | 7030.88623 | 18822.97852 | 18184.39258 | 27008.26563 | 27035.17383 | 28997.88867 |
| 228396.125 | 272062.5625 | 298882.5625 | 222114.875 | 248540.8125 | 500705.7188 | 399464.3438 |
| 55103.11719 | 63679.79688 | 69315.71094 | 69824.67188 | 76324.00781 | 76051.04688 | 79366.33594 |
| 15417.19336 | 21876.01367 | 16639.74414 | 15132.72949 | 20066.25 | 19911.73242 | 17721.53125 |
| 21766.70508 | 26967.91797 | 24198.73633 | 22627.66602 | 25064.99609 | 23315.63281 | 23229.43945 |
| 68757.65625 | 71672.71094 | 67539.82031 | 85747.5 | 104279.9219 | 104066.5313 | 101476.1875 |
| 17511.25977 | NA | 14109.64648 | 28732.97461 | 23516.33984 | 5652.789063 | 23144.77539 |
| 24056122 | 97267368 | 30837412 | 50603740 | 92508864 | 57526936 | 68505184 |
| 275645.5938 | 311293 | 328140.9375 | 350748.5625 | 354349.75 | 325353.0938 | 360451.7813 |
| 25043.92773 | 23625.91016 | 28603.64063 | 31043.57813 | 31476.00195 | 30208.61133 | 30239.45117 |
| 16311.55273 | 21102.12891 | 14716.37012 | 17966.49414 | 24924.80859 | 23816.99609 | 25108.99805 |
| 228497.3594 | 282801.5625 | 272507.9063 | 259639.7813 | 203571.25 | 222058.8438 | 216377.5313 |
| 44317.27344 | 54917.92188 | 58053.50391 | 57885.44531 | 40670.85938 | 45460.49609 | 44232.13281 |
| 12796.56738 | 14651.73145 | 18335.6543 | 19831.69336 | 24298.43945 | 18451.73828 | 19485.95703 |
| 29634.65039 | 39763.38672 | 40404.76563 | 39895.76953 | 36265.56641 | 35069.34766 | 34993.80469 |
| 5097.528809 | 8675.880859 | 5699.708008 | 7065.65332 | 12060.65137 | 10907.79102 | 7152.391113 |
| 208970.75 | 282900.7813 | 238943.4844 | 255687.7188 | 263086.8438 | 273030.8438 | 295828.0156 |
| 820060.5625 | 1061121.375 | 1026588.063 | 1084358.125 | 748307.4375 | 733282 | 738281.5625 |
| 19073.80469 | 22131.02539 | 23335.56641 | 23565.74609 | 22332.00195 | 23216.10742 | 24230.30273 |
| 392849.4375 | 527406.625 | 515472.125 | 495084.6563 | 518931.0938 | 451844.7813 | 472301.3125 |
| 18223.28516 | 18396.78125 | 19469.66992 | 18559.98438 | 15528.83496 | 16073.52832 | 15447.16797 |
| 526925.0625 | 634798.75 | 666492.25 | 575187.75 | 267792.2813 | 400840.3125 | 317686.0938 |
| 14439.16797 | 21095.11719 | 18735.79297 | 19865.04688 | 22162.49414 | 23415.38086 | 18230.04297 |
| 19410.57422 | 24289.67188 | 22495.73828 | 23070.30078 | 26395.01563 | 25689.07422 | 25282.28125 |

| | | | | | | |
|---|---|---|---|---|---|---|
| 25382.51172 | 27778.29883 | 37677.98438 | 66569.42188 | 45244.77344 | 90912.11719 | 111556.0625 |
| 111604.3594 | 120262.2734 | 137688.7344 | 137293.4688 | 125213.3438 | 108932.6094 | 122855.1484 |
| 559435.25 | 674319.3125 | 710070.9375 | 545523.625 | 523182.9688 | 544303.75 | 536464 |
| 36175.22266 | 35384.23047 | 42689.14063 | 44458.39453 | 25175.11914 | 29898.7832 | 26627.72656 |
| 136157.2656 | 165001.0469 | 153713.7656 | 163753.7031 | 188165.375 | 188603.5625 | 178761.9531 |
| 501871.9375 | 582207.625 | 627352.5625 | 591446.9375 | 531996.625 | 582866.1875 | 564148.8125 |
| 143614.5625 | 165617.4531 | 168466.2813 | 167416.0781 | 111233.6016 | 112804.7891 | 115349.8125 |
| 7992.055176 | 9471.061523 | 4407.985352 | NA | 9388.145508 | 8491.164063 | 9280.74707 |
| 18811.66797 | 19796.29297 | 24139.11133 | 25703.53711 | 28352.13086 | 28263.48438 | 31598.31445 |
| 16198.5498 | 17919.51367 | 17342.01367 | 20666.9668 | 21080.31445 | 21419.30078 | 22207.74414 |
| 649774.375 | 793116.625 | 745315.5 | 776241.3125 | 687218.125 | 638608.625 | 698676.75 |
| 142340.9063 | 174411.9063 | 181284.6875 | 186664.4688 | 193717.7969 | 197689.2969 | 195572.0469 |
| 700959.0625 | 1098178.875 | 1101495.25 | 1231461.5 | 1450354.875 | 1401589.625 | 1428867 |
| 7875.01123 | 10162.91895 | 8002.729492 | 10129.68164 | 10218.29395 | 10485.92676 | 9684.603516 |
| 75830.25 | 101350.4141 | 85012.21094 | 89352.75781 | 83107.875 | 73793.40625 | 80209.01563 |
| 400454.9375 | 502525.5 | 502684.3438 | 487906.9688 | 444875.375 | 435161.7813 | 447401.9688 |
| 9528.37207 | 11468.76758 | 10196.50195 | 9875.986328 | 7946.640137 | 7151.775391 | 7462.390137 |
| 36734.69922 | 35916.42578 | 39688.67969 | 40611.23828 | 33403.14844 | 37877.19922 | 34601.05078 |
| 16578.21484 | 12163.3916 | 7889.885742 | 18453.5332 | 26089.26758 | 26516.50586 | 18169.44531 |
| 6904.944824 | 7495.677734 | 7617.242676 | 7908.56543 | 8140.916016 | 8101.473145 | 7204.383301 |
| 9048.367188 | 14439.12695 | 11419.49512 | 13006.54492 | 28173.76758 | 17500.54688 | 23288.73242 |
| 765667.125 | 880586.625 | 1049147.625 | 917172.6875 | 717596.0625 | 707135.9375 | 734400 |
| 163967.2031 | 167244.9219 | 208901.125 | 210634.0781 | 195749.5938 | 236543.5 | 214433.4063 |
| 8969.476563 | 10729.58691 | 11333.4707 | 11153.54492 | 6924.519043 | 7709.812012 | 6948.503418 |
| 122088.6328 | 138154.1875 | 150120.9531 | 135283.8906 | 132080.4375 | 137323.4531 | 135610.875 |
| 13957.53613 | 13738.33984 | 20813.60742 | 34524.52734 | 51541.44922 | 50843.35938 | 69009.96875 |
| 206559.7031 | 227991.7656 | 257252.1563 | 235456.4531 | 187584.25 | 200606.875 | 193570.0313 |
| 73436.80469 | 86020.96875 | 84316.14063 | 91118.14063 | 97864.79688 | 106962.1172 | 100553.125 |
| 30445.93359 | 41589.21875 | 31909.17773 | 25836.69922 | 17840.69141 | 22031.60547 | 19956.11914 |
| 17876.92969 | 25333.89844 | 22393.38281 | 26626.12109 | 33861.25391 | 32119.5957 | 33882.83984 |
| 44969.48828 | 58606.23828 | 41935.44922 | 42977.30469 | 56934.08203 | 51862.53125 | 52955.63281 |
| 53184.39453 | 59595.66797 | 66169.50781 | 57911.59375 | 41055.19141 | 49216.77734 | 50074.13281 |
| 180527.3281 | 207503.6406 | 209225.8281 | 204770.2813 | 216274.9688 | 208567.375 | 217433.8281 |

| | | | | | | |
|---|---|---|---|---|---|---|
| 253818.8438 | 302726.0313 | 284616.8125 | 302521.8125 | 354691.5 | 344708.5625 | 332175.4063 |
| 18268.34766 | 21775.90039 | 23282.24219 | 24895.46289 | 37512.66797 | 35180.53906 | 35660.40234 |
| 10522.79199 | 12800.44531 | 14006.31738 | 13913.73438 | 18698.88086 | 15817.61426 | 19857.29297 |
| 154820.6094 | 193952.4688 | 182070.3906 | 182854.2813 | 155554.4688 | 164365.0469 | 157786.0781 |
| 4861.946777 | 6300.62793 | 6130.601074 | 4722.279297 | 10569.83789 | 9616.842773 | 8293.850586 |
| 50709.33203 | 61691.93359 | 56720.28125 | 57465.875 | 70004.78906 | 60161.92969 | 70540.51563 |
| 70825.85156 | 58009.73047 | 92917.09375 | 77506.22656 | 70415.71094 | 70105.58594 | 65865.00781 |
| 203872.875 | 248626.1875 | 229303.2344 | 232268.7969 | 232936.6094 | 254107.875 | 234895.0781 |
| 365409.8438 | 339239.7188 | 319317.625 | 158634.6875 | 99528 | 886744 | 93055.15625 |
| 71857.44531 | 87703.1875 | 78279.42188 | 79274.22656 | 100382.3125 | 98826.00781 | 101827.7344 |
| 33516.85156 | 39909.66016 | 36414.82813 | 39224.62109 | 36081.54297 | 36902.04297 | 33768.83594 |
| 169378.9844 | 203619.375 | 194955.3594 | 233931.0781 | 294687.1875 | 248300.1719 | 272030.8125 |
| 141957.7031 | 143021.625 | 169128.1094 | 181243.1563 | 170735.4219 | 174459.7344 | 174447.8594 |
| 171204.9219 | 191725.5938 | 211488.3906 | 211028.8438 | 217547.3125 | 208411.0313 | 218876.2031 |
| 8492.765625 | 9508.450195 | 10034.24512 | 9173.041016 | 8615.100586 | 9508.988281 | 11085.77246 |
| 174169.1719 | 241466.3438 | 204677.3438 | 204046.7188 | 261090.2656 | 240873.1875 | 283971.875 |
| 18121.93359 | 20599.6543 | 22014.67188 | 20505.6582 | 24122.75 | 22498.32617 | 24308.32227 |
| 88140.28125 | 110669.5859 | 97953.91406 | 94138.14063 | 114196.1797 | 112850.2891 | 126972.4063 |
| 12294.93848 | 18187.72266 | 13667.5 | 15138.24219 | 20230.10352 | 15657.87109 | 15694.66309 |
| 35286.74609 | 45357.25781 | 40158.91016 | 36759.15625 | 42389.35547 | 43573.58594 | 44387.93359 |
| 130381.6484 | 142860.5938 | 143504.2813 | 175372.8906 | 361934.5313 | 290983.125 | 320190.875 |
| 30317.46875 | 40288.26172 | 40773.16797 | 35860.77734 | 29699.22461 | 24803.77148 | 27449.625 |
| 81368.5625 | 81683.46094 | 86582.82031 | 82751.07813 | 46082.98047 | 45316.52734 | 44717.45313 |
| 11513.4541 | 9303.100586 | 15846.84082 | 16251.62207 | 22712.23242 | 15008.93066 | 26668.60547 |
| 6082.181641 | 5585.55957 | 6822.267578 | 6188.963867 | 7094.993652 | 9267.473633 | 7873.741699 |
| 4910.922363 | 6377.404297 | 3217.670654 | 5103.487305 | 6982.774414 | 8597.401367 | 3421.541504 |
| 95122.95313 | 114094.4844 | 111316.8359 | 115906.5313 | 121124.2969 | 119321.6875 | 116121.9297 |
| 28573.33789 | 33804.21875 | 33101.56641 | 29064.06445 | 28287.00586 | 30454.06055 | 30182.23438 |
| 9777.953125 | 9247.655273 | 11964.67578 | 9186.310547 | 9833.740234 | 12411.08789 | 11837.73242 |
| 66975.52344 | 83457.73438 | 86017.44531 | 78850.16406 | 67070.35156 | 68129.50781 | 67394.6875 |
| 8832.165039 | 13233.50977 | 7645.585449 | 7943.571777 | 8049.943359 | 9939.682617 | 9371.103516 |
| 265124.3125 | 273525.5625 | 341563.4688 | 341737.4375 | 170266.4531 | 148517.1406 | 161931.3125 |
| 14848.76953 | 15103.44531 | 16991.05859 | 20818.20508 | 17817.57617 | 23203.96875 | 20254.76563 |

| | | | | | | |
|---|---|---|---|---|---|---|
| 149047.0781 | 208094.6719 | 158564.3125 | 155294.7969 | 274868.9375 | 249294.5 | 265138.9688 |
| 74271.14844 | 97634.35938 | 89741.01563 | 83954.32031 | 87313.64063 | 83063.25 | 85679.82031 |
| 11668.08594 | 16437.38672 | 13858.65137 | 11298.6377 | 18911.22656 | 17435.96094 | 17860.66797 |
| 121503.4688 | 147748.7188 | 147358.125 | 158050.5313 | 155231.7813 | 164580.5781 | 156529.9531 |
| 11052.98535 | 15606.99902 | 15118.71094 | 14051.28223 | 16671.47852 | 14967.03516 | 15813.91895 |
| 111590.1777 | 142493.5859 | 128299.543 | 133352.0723 | 185728.2656 | 178818.3086 | 185355.0039 |
| 51582.07813 | 70358.34375 | 60602.17188 | 58222.20313 | 64576.56641 | 66756.67188 | 64236.78516 |
| 35520.78125 | 41957.32422 | 42395.24609 | 39220.94922 | 31094.82422 | 33049.35938 | 32168.30273 |
| 4720.144531 | 9235.46582 | 4673.562012 | 4251.229004 | 15093.50293 | 19165.29102 | 17210.76953 |
| 60754.18359 | 97196.47656 | 69609.71094 | 75407.94531 | 87319.625 | 88293.625 | 90274.25 |
| 129106.5469 | 158518.7813 | 163643.4844 | 162219.625 | 169145.9375 | 141767.7344 | 153934.2188 |
| 109545.375 | 126403.4297 | 134617.3438 | 144583.3594 | 220997.1406 | 169184.4063 | 195271.1094 |
| 11795.86426 | 15426.47754 | 15991.00781 | 12201.17676 | 19365.12891 | 19595.40234 | 22512.41992 |
| 139388.3438 | 145049.9844 | 175909.7188 | 183446.4531 | 170514.6563 | 164110.0781 | 165480.7656 |
| 18203.63672 | 23001.20313 | 20179.27344 | 20222.18164 | 27746.79492 | 23916.33008 | 25153.13672 |
| 15941.60645 | 19356.52148 | 17877.30859 | 15061.94531 | 16373.85059 | 17919.12695 | 19112.45508 |
| 16929.20508 | 21980.52344 | 19473.61523 | 18839.16602 | 15931.56641 | 19935.625 | 15937.77734 |
| 79492.36719 | 94355.10156 | 96850.99219 | 94982.78125 | 61307.50781 | 63813.64453 | 63329.96875 |
| 251680.2813 | 271882.75 | 305596.625 | 266832.7188 | 499484.5313 | 486619.9063 | 543195.4375 |
| 70598.20313 | 96273.49219 | 89078.20313 | 82021.25781 | 95975.96094 | 96939.97656 | 92900.26563 |
| 7377.637695 | 10003.18359 | 6572.51123 | 7318.275879 | 11675.02637 | 14315.56055 | 12877.01758 |
| 26794.86914 | 29578.01758 | 29069.45117 | 29022.79102 | 15740.39746 | 16624.63477 | 14382.71875 |
| 62394.75781 | 99638.35156 | 65979.91406 | 68420.67969 | 92652.41406 | 76786.71094 | 85474.39844 |
| 101086.0078 | 123787.3438 | 118320.7969 | 116794.5 | 96800.1875 | 98914.96094 | 97199.47656 |
| 289212.4375 | 373443.4063 | 341476.3125 | 361493.4375 | 495807.875 | 438166.625 | 501804.6875 |
| 14804.63965 | 17936.63281 | 15664.04004 | 16953.26953 | 20434.13477 | 24969.05859 | 26357.69141 |
| 2284.375977 | 2855.718018 | 2865.161621 | 2516.337402 | 1846.698975 | 1322.334106 | 1805.252197 |
| 895399.0625 | 1028230.688 | 1025325.875 | 1095862.25 | 1446808.75 | 1440628.25 | 1531712.5 |
| 26512.19629 | 36163.86426 | 31095.88086 | 30037.69824 | 32735.76904 | 33214.12915 | 35788.55811 |
| 93020.20313 | 110968.4141 | 112646.1563 | 107157.3359 | 79535.02344 | 86217.85156 | 82721.73438 |
| 23135.17383 | 28110.34961 | 25746.74023 | 26856.94141 | 17740.67578 | 15534.77832 | 16431.4668 |
| 43396.72656 | 56864.19531 | 48216.44922 | 43336.63672 | 27290.76172 | 28575.67578 | 25042.99414 |
| 237073.6094 | 284708.4688 | 265475.4688 | 272558.7188 | 127067.7891 | 136348.4375 | 124865.9297 |

| | | | | | | |
|---|---|---|---|---|---|---|
| 856323.9375 | 1056740.875 | 887872.9375 | 886690.8125 | 885803.6875 | 811205.375 | 833580 |
| 21187.65625 | 24910.85352 | 19754.69336 | 22339.90234 | 25059.66016 | 34270.25 | 29496.91797 |
| 26853.07227 | 57916.10156 | 16014.87109 | 15126.72852 | 55659.3125 | 40599.85156 | 44210.67188 |
| 7271.225586 | 8852.293945 | 10913.79199 | 9711.132813 | 23540.98828 | 20242.375 | 22404.24805 |
| 8611.863281 | 10655.51953 | 11597.89844 | 11742.30664 | 17806.12109 | 11960.84375 | 13003.83789 |
| 512785.3438 | 569807.9375 | 701811.6875 | 604221.5625 | 515804.2813 | 510687.1875 | 520475.7188 |
| 150198.9375 | 203487.5938 | 166133.9063 | 172255.0938 | 154459.1563 | 169337.0781 | 173600.9219 |
| 148546.3906 | 159060.4375 | 168292.2188 | 177602.9375 | 174155.8906 | 192192.7656 | 198629.1719 |
| 14622.33398 | 17243.81055 | 15898.79492 | 14785.93262 | 13669.91992 | 13934.09863 | 13591.12695 |
| 10024.70801 | 13404.05176 | 12249.06055 | 13433.43359 | 13556.21777 | 10855.40918 | 11617.14941 |
| 58698.98828 | 46991.49609 | 25017.79688 | 56606.10938 | 46659.23828 | 63636.19922 | 49251.60156 |
| 7990.208984 | 7259.836426 | 10735.90723 | 11814.83008 | 16622.52734 | 12915.45313 | 16141.2334 |
| 152185.1563 | 164980.8438 | 182162.9531 | 195732.5781 | 208883.2969 | 205196.0625 | 205537.6563 |
| 79188.5625 | 92643.95313 | 99689.65625 | 92033.46094 | 86865.125 | 74054.25 | 94192.23438 |
| 4033.607422 | 7953.859375 | 4851.799805 | 4475.571777 | 6983.711426 | 5904.503418 | 7007.406738 |
| 227705.75 | 235244.6563 | 268895.7813 | 259154.9219 | 263278.6563 | 212907.2813 | 252140.2344 |
| 6745.855957 | 8690.205078 | 7798.228027 | 10337.80078 | 25304.5957 | 10404.24121 | 13398.84766 |
| 114605.6953 | 122562.8359 | 133613.1406 | 121630.1328 | 113107.1172 | 117976.8047 | 116907.7109 |
| 693084.75 | 812103.875 | 730217.25 | 816349 | 787616.625 | 768813.375 | 750030 |
| 17048.85156 | 28347.43945 | 14972.0293 | 14848.10254 | 34898.91406 | 41208.875 | 37128.64063 |
| 7566.647461 | 11478.31641 | 7214.414063 | 7932.828125 | 10074.80273 | 12063.58789 | 9610.791992 |
| 48439 | 55182.96875 | 56807.65234 | 57818.33594 | 47849.67188 | 54285.85938 | 50450.71875 |
| 23606.77344 | 32150.12695 | 27625.90234 | 26608.22656 | 38733.84375 | 36811.1875 | 32499.14453 |
| 55216.14844 | 61139.05469 | 57913.32031 | 62278.10547 | 63744.97266 | 66119.66406 | 65811.26563 |
| 762961.6875 | 1133915.125 | 887863 | 366557.75 | 256799.625 | 757041.8125 | 393530.1563 |
| 8058.571289 | 9444.481445 | 8953.422852 | 8546.067383 | 9529.583984 | 8196.84668 | 9311.550781 |
| 364798.125 | 426869.6563 | 459507.5625 | 445373.0938 | 387900 | 449459.5625 | 393301.375 |
| 28194.98828 | 27657.9082 | 32213.0625 | 30653.76953 | 27083.74219 | 32826.19141 | 31109.79492 |
| 16506.12305 | 16569.57813 | 22133.05664 | 16775.66992 | 11604.42969 | 10137.03125 | 9859.25293 |
| 20798.35352 | 24139.4707 | 24289.16992 | 24298.14258 | 27445.38477 | 23740.69922 | 25333.58594 |
| 31405.57813 | 43147.12109 | 29800.63281 | 32624.47461 | 54254.69531 | 43931.33594 | 49683.63672 |
| 34445.01172 | 58320.22656 | 43958.14844 | 39816.75391 | 55308.77734 | 58228.29297 | 57883.99609 |
| 41634.80859 | 51766.08203 | 50969.64063 | 47890.73047 | 39263.11328 | 47845.33594 | 42502.34766 |

| | | | | | | |
|---|---|---|---|---|---|---|
| 264794.375 | 305066.1875 | 326502.4688 | 323994.8125 | 289252.1563 | 284416.75 | 303506.5938 |
| 4396.751953 | 9189.890625 | 7334.441406 | 5595.222168 | 4704.333984 | 4266.251465 | 4179.099121 |
| 3975.30542 | 2716.946045 | 6511.720215 | 6641.557129 | 3203.864746 | 2443.322754 | 3976.372559 |
| 155951.4063 | 159198.125 | 194229.6875 | 188226.0313 | 138781.2969 | 154135.6719 | 153642.5156 |
| 49465.29297 | 54069.77734 | 62876.69922 | 59434.46484 | 69938.16406 | 67233.71875 | 73282.24219 |
| 9743.787109 | 10322.80469 | 8606.182617 | 7731.478516 | 5383.070313 | 6935.951172 | 8639.75293 |
| 12082.3457 | 14683.55664 | 13873.3623 | 13759.17969 | 15495.47461 | 14888.18457 | 12782.22461 |
| 708204.0625 | 793254.5625 | 785177.4375 | 806878.6875 | 398147.2188 | 404388.0625 | 404747.9688 |
| 6925.436523 | 10739.58398 | 9744.020508 | 9812.243164 | 12820.72852 | 12702.19531 | 12958.79004 |
| 9332.790039 | 10345.80371 | 11404.84961 | 8905.008789 | 6250.580078 | 4682.007813 | 4560.768066 |
| 75084.88281 | 82679.72656 | 91223.65625 | 88306.70313 | 94036.28906 | 81695.92969 | 86536.46094 |
| 72722.67188 | 75263.71875 | 81624.49219 | 81162.82031 | 40101.50391 | 41773.14844 | 37522.39844 |
| 181101.2656 | 231355.1406 | 216346.375 | 217843.0156 | 242006.9844 | 227331.4531 | 212613.8125 |
| 751161.375 | 882615.75 | 874006.875 | 857891 | 718834.9375 | 754380.3125 | 739499.5625 |
| 25373.78516 | 27965.80469 | 31049.13672 | 25097.32617 | 35199.69141 | 74330.21094 | 29618.86719 |
| 501910.2813 | 550456.6875 | 587266.75 | 596642.375 | 603383.75 | 649439.875 | 650939.5625 |
| 131866.0938 | 157032.375 | 141673.8594 | 146403.8906 | 119492.7969 | 110575.0313 | 115432.3594 |
| 25439.43359 | 32551.45313 | 29511.18359 | 28759.28125 | 22460.29883 | 24213.15039 | 23154.11523 |
| 79912.49219 | 101510.8203 | 92063.42188 | 88767.44531 | 122336.5 | 116055.6953 | 115499.3672 |
| 8891.807617 | 11165.76563 | 7719.646484 | 8535.229492 | 11312.56543 | 8338.673828 | 14865.62793 |
| 9036.931641 | 10638.61621 | 11856.94531 | 13437.45215 | 11950.875 | 11252.41699 | 11145.25391 |
| 11879.38965 | 13357.44336 | 11915.2207 | 11430.46289 | 11345.98242 | 11658.91797 | 10872.14551 |
| 21468.46094 | 5861.657227 | 40141.15234 | 45357.75 | 11973.22656 | 16305.97949 | 13241.84082 |
| 102666.6094 | 115407.25 | 122266.3281 | 116278.7656 | 103030.5469 | 101343.75 | 99893.19531 |
| 8626.962891 | 10399.50195 | 8965.058594 | 8997 | 10078.0625 | 9845.977539 | 9012.699219 |
| 290494.4063 | 340243.6563 | 367720.4063 | 361437.6875 | 432076.25 | 443092.125 | 444034.4375 |
| 32681.65234 | 44395.10547 | 38079.12109 | 42360.08984 | 37538.84375 | 37029.33594 | 34419.69922 |
| 34520.17969 | 37319.07813 | 38586.81641 | 30876.31445 | 27828.94727 | 26786.0918 | 27861.85352 |
| 28812.17578 | 35403.95703 | 32888.69141 | 31732.77734 | 23040.03711 | 28009.16992 | 25373.55078 |
| 4276623 | 6205897 | 4242557 | 3786623.75 | 1809036.25 | 2335491.75 | 1946003 |
| 169680.9531 | 231333.6094 | 169868.9844 | 178924.5313 | 269609.6563 | 245375.8594 | 252875.4531 |
| 131841.9688 | 153384.6563 | 147122.0781 | 147278.9063 | 130865.1875 | 121993.4453 | 126497.4922 |
| 196176.9688 | 221158.7344 | 227796.0781 | 228058.1719 | 175169.7188 | 167625.5781 | 170581.8906 |

| | | | | | | |
|---|---|---|---|---|---|---|
| 130913.5313 | 147893.75 | 153618.9063 | 150971.7031 | 126959.8438 | 120706.6328 | 135737.6406 |
| 728.9371948 | NA | 2407.536377 | 560.5797119 | 9099.671875 | NA | 4466.320801 |
| 16013.01465 | 14362.07422 | 22411.41211 | 21575.875 | 24792.45508 | 21505.4043 | 23918.93359 |
| 13304.37793 | 12623.09375 | 15293.74707 | 16080.91504 | 10614.7002 | 11011.75293 | 11720.27344 |
| 12196.34863 | 16485.74609 | 13939.10352 | 11307.11719 | 14404.57813 | 20063.41797 | 16902.2793 |
| 464791.8125 | 557141.75 | 569911.9375 | 535309.75 | 459371.8125 | 487970.3125 | 468147.4063 |
| 45507.48828 | 49722.05859 | 49093.76172 | 50355.60547 | 52874.72266 | 59204.01953 | 55200.0625 |
| 17828.87305 | 26996.5 | 28811.56836 | 21293.93164 | 19560.82227 | 18592.26953 | 20349.79883 |
| 34431.07422 | 37950.05469 | 37746.09766 | 37635.57422 | 34496.84375 | 34593.23047 | 35152.95313 |
| 17288.98633 | 25644.32422 | 21115.12305 | 20824.41992 | 23532.10352 | 20994.61328 | 20239.08398 |
| 171715.0781 | 199631.0156 | 203403.5781 | 214966.75 | 362375.2188 | 368844.7188 | 391170.25 |
| 307701.5625 | 333844.5313 | 369864.7188 | 321622.1563 | 267500.6875 | 260640.375 | 305505 |
| 162963.3281 | 185861.75 | 170020.1875 | 148798.9531 | 125254.0234 | 124517.6328 | 124273 |
| 1606.046265 | 1586.324463 | NA | NA | 1750.36853 | 1321.94751 | 1434.137085 |
| 48323.52734 | 57766.49609 | 54705.00781 | 54324.61328 | 60410.49609 | 63293.18359 | 63132.96484 |
| 22748.58398 | 25349.46289 | 26730.73047 | 25619.01563 | 23765.4043 | 23856.15234 | 25234.04492 |
| 11274.00684 | 16240.14355 | 14702.13281 | 14746.63379 | 30377.0293 | 22245.99414 | 24113.60156 |
| 21628.24609 | 25045.54883 | 22967.83594 | 25116.27734 | 24665.93945 | 26070.89844 | 24746.08008 |
| 38931.55078 | 55479.00781 | 45467.22266 | 49633.97266 | 57892.76563 | 56369.85938 | 53542.26953 |
| 444265.7188 | 534155.625 | 509290.75 | 552620 | 679019.625 | 650404.1875 | 590105.125 |
| 65529.72266 | 114538.3828 | 48556.23438 | 37096.84766 | 76782.14063 | 58430.55078 | 66477.01563 |
| 11328.33691 | 11438.83008 | 13454.03125 | 13848.61914 | 11177.50391 | 10736.79199 | 9992.336914 |
| 478346.6836 | 756197.1328 | 516491.4883 | 484438.6563 | 519732.6172 | 787402.5117 | 562069.3789 |
| 402411.4688 | 517858.9063 | 276888.125 | 374436.5313 | 296488.3438 | 327982.5313 | 176759.9219 |
| 333318.7422 | 399389.5781 | 379050.0156 | 383352.2969 | 454129.1875 | 412616.5469 | 416895.7813 |
| 16996.66992 | 21895.66602 | 16432.32422 | 14497.18652 | 21618.52539 | 24580.51563 | 23478.12109 |
| 165769.2344 | 211313.6406 | 192723.8906 | 218245.5313 | 184758.5781 | 163321.7031 | 178494.4375 |
| 22749.69531 | 29055.18359 | 29648.98438 | 25364.99023 | 38004.06641 | 33794.98047 | 39143.79688 |
| 15779.24512 | 20208.19141 | 16430.28516 | 13157.13184 | 11687.72363 | 9622.373047 | 9375.500977 |
| 4063.437134 | 4536.251709 | 6155.968018 | 4198.532471 | 17559.75439 | 13782.89648 | 14967.573 |
| 13134.85254 | 14703.29297 | 13127.84473 | 12660.81055 | 10228.82715 | 10698.15527 | 10355.75586 |
| 29703.48047 | 32501.27539 | 32094.37695 | 29307.85547 | 39111.74219 | 34665.32813 | 32182.23438 |
| 180577.2344 | 215754.8281 | 202917.4688 | 205758.5625 | 197950.3125 | 247671.0938 | 222770.3906 |

| | | | | | | |
|---|---|---|---|---|---|---|
| 115428.0469 | 126799.4844 | 141583.75 | 133496.6875 | 117869.6719 | 138844.7969 | 130092.7734 |
| 37151.39453 | 43599.95313 | 43954.69141 | 40783.65625 | 31734.43555 | 31049.12695 | 27489.26953 |
| 293312.375 | 337135.6563 | 372744.5938 | 354701.25 | 308976.125 | 328538.5938 | 316416.8125 |
| 20570.9375 | 24156.43164 | 25746.69531 | 26615.03906 | 30154.73633 | 26217.23047 | 26415.17773 |
| 46906.53516 | 57850.75781 | 53428.61328 | 52534.55859 | 60129.77734 | 60989.80859 | 61682.82813 |
| 74256.53906 | 95562.11719 | 83024.19531 | 96082.46875 | 131894.6719 | 113351.9063 | 113805.4219 |
| 1409199 | 1649519.125 | 1644276.25 | 1784073.875 | 1593545.25 | 1595081 | 1672689.125 |
| 30648.71875 | 35685.52734 | 40124.46484 | 40730.64453 | 50503.91016 | 41796.58203 | 45604.66797 |
| 7618.529785 | 12090.2002 | 6921.169434 | 7585.821777 | 9402.484375 | 8174.959961 | 9190.21875 |
| 48260.83594 | 55932.78125 | 55596.70313 | 56658.94922 | 35244.86719 | 35998.01563 | 38877.32422 |
| 390782.0625 | 398363.7813 | 464965.1563 | 513024.5625 | 367539.5625 | 355196.2188 | 407313.5 |
| 611594.0625 | 709422 | 633881.1875 | 681147.5 | 782340.4375 | 634439.375 | 746654.3125 |
| 7411.531738 | 6906.885254 | 6317.591797 | 9472.216797 | 6745.491211 | 6603.842285 | 8804.932617 |
| 20276.03516 | 21798.38477 | 29837.29883 | 22059.71094 | 18542.51367 | 18933.24414 | 22397.09766 |
| 131658.5313 | 140375.2656 | 164587.0156 | 145931.3125 | 117606.7031 | 134424.3438 | 128374.2578 |
| 53359.89453 | 66113.50781 | 65296.01953 | 64760.98828 | 79679.95313 | 83173.91406 | 74665.28906 |
| 38012.75391 | 54020.10938 | 50537.65234 | 56821.68359 | 52760.75 | 51823.85938 | 55112.30469 |
| 581828.125 | 708991.625 | 681534.125 | 619580.375 | 618883 | 607736.75 | 620734.8125 |
| 712237.6875 | 796974.3125 | 825500.375 | 842841.875 | 901890.6875 | 918920.5625 | 908944.9375 |
| 1336952.5 | 1643161.625 | 1546268.75 | 1498339 | 1585823.125 | 1612427.5 | 1617108.5 |
| 13424.32617 | 23590.68555 | 10117.95605 | 8891.131836 | 11096.67969 | 8994.451172 | 10460.9248 |
| 41911.35547 | 47643.16016 | 47929.11719 | 48416.86719 | 47137.00781 | 47174.40234 | 45687.94922 |
| 71729.50781 | 93253.92969 | 74943.00781 | 75904.44531 | 82003.67969 | 80193.75781 | 82210.72656 |
| 42783.74219 | 47815.71484 | 45915.88281 | 45457.17188 | 49582.14453 | 49044.32813 | 47944.02344 |
| 1056.884033 | 2316.026611 | 2066.032471 | 2187.621094 | 53938.44531 | 6199.236816 | 20158.41797 |
| 65472.32422 | 79498.50781 | 86417.33594 | 78950.57031 | 73230.23438 | 84652.09375 | 75505.73438 |
| 114908.6953 | 131130.8438 | 133996.9844 | 134289.7344 | 120685.7188 | 121277.8125 | 116714.7969 |
| 618865.9375 | 787273.1875 | 610663.4375 | 720046.5625 | 877270.25 | 719455.6875 | 796163.75 |
| 146975.2344 | 244012.0938 | 131645.875 | 139027.9219 | 197718.9844 | 238597.2969 | 233047.1875 |
| 139105.625 | 167783.7813 | 177316.0938 | 171758.9531 | 160676.8906 | 154464.75 | 162384.1406 |
| 7161.447754 | 9087.808594 | 8508.932617 | 8124.062012 | 4905.743164 | 5518.730469 | 5115.625488 |
| 6737.641602 | 6774.984863 | 7797.760742 | 7221.88916 | 13583.93652 | 10843.8457 | 11394.58301 |
| 37538.03125 | 41033.3125 | 39213.70313 | 37615.64063 | 34623.56641 | 39330.95703 | 33610.78906 |

| | | | | | | |
|---|---|---|---|---|---|---|
| 56890.37109 | 60763.04297 | 61913.17969 | 67618.69531 | 114433.3672 | 93051.17188 | 99644.85156 |
| 56259.52344 | 60498.37891 | 55743.36719 | 50080.33203 | 63850.57422 | 56559.64844 | 65078.96875 |
| 210339.2344 | 245209.5156 | 242075.4375 | 246580.75 | 222247.4531 | 230004.7656 | 225083.9531 |
| 54478.03125 | 60462.94531 | 78936.80469 | 72249.36719 | 46589.91406 | 43928.65625 | 41362.32813 |
| 10544.69336 | 10714.40234 | 12500.84375 | 10869.27246 | 9602.838867 | 10147.64941 | 9455.041992 |
| 41646.73828 | 45945.1875 | 46762.45703 | 43375.39453 | 53383.61719 | 50783.31641 | 53143.59766 |
| 31132.24805 | 32477.32031 | 34604.90234 | 32610.58203 | 27915.36133 | 30085.27734 | 28190.5293 |
| 2802.113281 | 3699.534912 | 3441.951904 | 4556.922363 | 2268.81665 | 2138.321533 | 2947.965332 |
| 643629.125 | 832083.4375 | 693260.8125 | 829089.25 | 1525994.75 | 1179613 | 1229911.5 |
| 92946.03125 | 110543.3438 | 110063.1719 | 107953 | 81288.59375 | 76639.19531 | 81279.32031 |
| 5460.441406 | 6494.066406 | 6326.656738 | 6649.841797 | 5238.25 | 6207.70166 | 5498.577637 |
| 34956.43359 | 60270.64063 | 32638.53125 | 25242.44141 | 47618.96094 | 38178.05078 | 44837.40625 |
| 188833.1719 | 240147.8438 | 227180.5781 | 253845.1406 | 373597.3125 | 319992.7188 | 352488.0313 |
| 140609.1875 | 159328.5938 | 158036.8594 | 158582.2031 | 157974.75 | 158000.2344 | 162103.2813 |
| 29766.63672 | 30412.85742 | 41895.13672 | 35752.33203 | 45898.90625 | 63472.80859 | 48964.92578 |
| 59514.68359 | 78178.64844 | 63383.55078 | 33355.58984 | 45865.12891 | 80981.61719 | 51060.42969 |
| 664459.6875 | 725301.625 | 848282.9375 | 799130.6875 | 752506.0625 | 739972.75 | 771621.0625 |
| 13185.70996 | 14503.66992 | 13231.71777 | 14705.55371 | 16351.53125 | 15146.11328 | 14138.19238 |
| 97547.96875 | 122886.9609 | 113444.6016 | 121373.8516 | 144418.3281 | 169274.6563 | 167613.3281 |
| 14056.71387 | 15409.26953 | 15918.24414 | 16685.4043 | 17034.76172 | 17729.79883 | 17862.875 |
| 184354.2344 | 198326.2344 | 222612.875 | 219988.9531 | 208296.7344 | 172026.7656 | 196086.9844 |
| 133512.5781 | 177143.3594 | 149236.0625 | 149979.4063 | 170773.7656 | 229129.0781 | 146277.1406 |
| 22763.62109 | 28815.0332 | 26983.61523 | 27238.9707 | 42532.49609 | 34547.48828 | 37855.71094 |
| 60875.05859 | 58512.60547 | 85836.80469 | 69531.75 | 67103.22656 | 67306.84375 | 70293.71094 |
| 150315.5625 | 168524 | 176874.0938 | 172506.4219 | 170621.6094 | 173258.5625 | 171786.5 |
| 6804.737305 | 6517.611328 | 6318.402344 | 6899.168457 | 5638.061035 | 7515.189941 | 7044.65625 |
| 34754.95703 | 42507.8125 | 39864.17578 | 40009.67969 | 43121.38281 | 42320.17188 | 39849.50781 |
| 40026.27734 | 47068.71875 | 44712.6875 | 51734.39063 | 49403.65625 | 55664.80078 | 44829.87891 |
| 852841.125 | 967246.1875 | 1046679.875 | 1072025.25 | 1001484.25 | 1032100.438 | 1028766.625 |
| 27758.625 | 31437.50586 | 30130.16016 | 31052.44727 | 28392.30859 | 25751.8457 | 33423.44922 |
| 113314.8047 | 119339.25 | 127345.6328 | 133353.8906 | 132263.6094 | 131781.8125 | 135550.7188 |
| 65794.96875 | 78308.125 | 84996.8125 | 76936.84375 | 94166.58594 | 97425.64844 | 94100.66406 |
| 63709.01758 | 82638.29297 | 51510.06445 | 59602.84766 | 73782.97266 | 67545.66797 | 70233.51172 |

| | | | | | | |
|---|---|---|---|---|---|---|
| 120823.5234 | 141083.1094 | 123218.7578 | 119310.4375 | 132109.6719 | 117099.9844 | 152465.5 |
| 60927.77734 | 66336.17188 | 66796.5625 | 68521.04688 | 66014.9375 | 60860.84766 | 64046.82422 |
| 644659.4375 | 835907.125 | 767812.625 | 782004.125 | 699126.75 | 712170.75 | 711761.625 |
| 21900.79102 | 24201.1543 | 24424.00195 | 23728.6582 | 15689.47852 | 14485.42578 | 13649.68262 |
| 68122.5 | 71612.00781 | 84345.17969 | 91585.14844 | 163351.2031 | 152996.1719 | 158439.2813 |
| 48557.65625 | 55810.22656 | 47568.31641 | 53878.52734 | 27818.52734 | 28696.53516 | 24565.375 |
| 212313.7656 | 216828.0625 | 215906.6094 | 242160.8594 | 196141.5313 | 144387.0156 | 183679.6875 |
| 41897.63672 | 48811.40234 | 41314.69141 | 44961.05469 | 26118.22266 | 27436.43945 | 26198.9375 |
| 666144.75 | 725618.5625 | 789749.3125 | 803102.5 | 599039.8125 | 614876.6875 | 686030 |
| 11036.56738 | 11739.43848 | 12276.27344 | 10335.99219 | 9077.392578 | 10194.96094 | 11108.76563 |
| 18531.99805 | 23763.64258 | 18098.17969 | 17761.18555 | 13153.68457 | 12045.25977 | 11926.625 |
| 131556.4063 | 160708.4375 | 158197.9688 | 158118.2656 | 163134.9531 | 196345.0313 | 184314.9531 |
| 392145.5938 | 505954.7813 | 465009.5938 | 467811.5 | 502366.7813 | 520093.25 | 470800.1563 |
| 54551.53906 | 65197.73047 | 66187.45313 | 66310.89063 | 72629.46875 | 74391.42188 | 73917.35156 |
| 15651.69141 | 21416.94727 | 19638.62891 | 12911.47266 | 18512.87695 | 26514.16992 | 23881.13477 |
| 61070.70703 | 71127.07031 | 61499.19141 | 71321.65625 | 141547.3125 | 114802.7109 | 127791.4375 |
| 15297.53125 | 14657.49121 | 14812.23828 | 15442.25488 | 21202.09961 | 22253.45313 | 22392.62891 |
| 526358.75 | 675801 | 652629.9375 | 685976.3125 | 563385.0625 | 540647.8125 | 505687.9375 |
| 325652.3125 | 339548.3438 | 393691.0938 | 347990.375 | 309625.9688 | 315911.6875 | 327072.25 |
| 41598.79297 | 49577 | 56894.67188 | 59580.70703 | 41350.10156 | 45004.25781 | 44526.5 |
| 13216.48633 | 13926.80566 | 20487.00195 | 14573.76855 | 19341.97656 | 18956.77148 | 16751.49219 |
| 2492920.25 | 2970559.75 | 2909330.5 | 2480023.5 | 2299481.25 | 2157676.5 | 2125254.5 |
| 111242.9063 | 111545.6641 | 121459.2109 | 127460.3203 | 126021.1641 | 124397.4453 | 132941 |
| 15992.20117 | 18130.9375 | 19163.85742 | 17966.01758 | 12725.19043 | 11777.2041 | 12066.46875 |
| 23132.37305 | 25001.23633 | 25450.23047 | 26407.99023 | 29400.12891 | 32643.10938 | 27178.96875 |
| 35227.49609 | 41058.98047 | 36084.42578 | 47049 | 69695.72656 | 70441.02344 | 71818.5 |
| 145438.5 | 170639.9688 | 135026.0781 | 123759.0078 | 73805.19531 | 92713.08594 | 78339.125 |
| 309883.9688 | 352275.8125 | 342089.0938 | 366402.9688 | 370097.0313 | 377585.7188 | 362000.4375 |
| 450279 | 527646.9375 | 495052.1875 | 502042.9375 | 519225.6563 | 520248.2188 | 519578.875 |
| 324000.0313 | 385367.4688 | 375983.4063 | 369558.125 | 274444.6563 | 278776.5625 | 287185.4375 |
| 367000.75 | 395919.6563 | 402859.2188 | 464630.4375 | 333904.8125 | 302396.2188 | 339749.3438 |
| 483451.625 | 546795.0625 | 555050.1875 | 578312.4375 | 514464.2813 | 491592.25 | 496525.5938 |
| 131142.4492 | 143274.5352 | 139923.085 | 146739.6797 | 157103.2441 | 150751.1807 | 158665.1172 |

| | | | | | | |
|---|---|---|---|---|---|---|
| 2869015.75 | 3401642 | 3004957.25 | 3140420.5 | 2991443.25 | 2860915 | 2827329.5 |
| 114396.1406 | 146296.3125 | 128565.3281 | 175961.625 | 300979.8125 | 316500.1563 | 338306.3438 |
| 4636.381348 | 6149.052734 | 6176.586914 | 3715.587646 | 4175.041016 | 4858.856445 | 4614.979004 |
| 236318.6875 | 281882.8125 | 301051.125 | 265334.25 | 162171.7031 | 66412.75 | 134529.0781 |
| 5437587 | 7433795.5 | 4351599 | 4390990 | 5655961.5 | 3886203.25 | 2962364 |
| 31480.61133 | 32745.87305 | 32276.83594 | 39599.77734 | 39035.05078 | 41366.11328 | 38328.78906 |
| 95302.10938 | 105537.6641 | 115069.375 | 113646.1719 | 85069.85156 | 90611.57813 | 91312.97656 |
| 263723.8438 | 286041.4375 | 311993.2188 | 324298.7188 | 452948.7188 | 417172.6875 | 429943.2813 |
| 104415.7422 | 113504.5 | 116801.7969 | 125339.1406 | 147094.2344 | 137148.4063 | 147811.6563 |
| 5459.279297 | 6464.389648 | 6501.254395 | 6794.599121 | 8516.928711 | 9031.209961 | 8820.930664 |
| 73107.32813 | 91658.01563 | 69933.10938 | 80799.89844 | 87968.65625 | 111058.125 | 86920.875 |
| 165813.1094 | 192602.2344 | 186194.875 | 197223.3281 | 190298.1406 | 188414.7031 | 190639.0781 |
| 117366.8047 | 171430.2969 | 147511.2969 | 138476.875 | 164733.1406 | 159265.2969 | 156262.6094 |
| 2316950.938 | 2734799.621 | 2515885.422 | 2621788.254 | 2315707.74 | 2062022.709 | 2137080.473 |
| 26234.91406 | 34091.89844 | 31301.1543 | 32191.29102 | 42302.03125 | 38095.63281 | 38872.26953 |
| 73658.64063 | 74865.375 | 86409.91406 | 95579.46875 | 115039.7578 | 106899.7109 | 99710.33594 |
| 135485.7031 | 150597.2813 | 140444.625 | 150896.0625 | 225663.0469 | 232976.1875 | 225381.1719 |
| 30611.19141 | 37292.52344 | 35248.98438 | 39377.77734 | 43548.86719 | 38368.03125 | 47497.34766 |
| 60785.37109 | 78754.51465 | 66015.64063 | 58645.48438 | 84162.77734 | 82318.03516 | 87036.15234 |
| 220744.6406 | 245777.9375 | 238807.2344 | 242662.0781 | 256526.7188 | 250133.6563 | 255874.9688 |
| 611029.125 | 659452.25 | 750878.5625 | 747680.3125 | 706511.875 | 731583.1875 | 742526.9375 |
| 336912.5625 | 427051.75 | 372131.9063 | 144580.4688 | 135198.4219 | 493596.1875 | 239751.5625 |
| 517388.4688 | 570177.625 | 625192.0625 | 640617.3125 | 680404.6875 | 658470.6875 | 658633.4375 |
| 13608.59375 | 15216.95898 | 14684.24414 | 14605.40332 | 12113.4668 | 13618.82715 | 10623.07031 |
| 23092.68164 | 19319.67578 | 18378.11914 | 32894.24219 | 98238.74219 | 67333.82031 | 110683.2578 |
| 21094.94141 | 24956.96484 | 23502.92773 | 23277.98242 | 22093.88281 | 25303.05469 | 24665.27344 |
| 55418.08984 | 86645.65625 | 46294.69922 | 49356.46094 | 98035.96875 | 102561.0469 | 100321.6953 |
| 396185.6875 | 430561.4063 | 400217.875 | 447223.7188 | 424949.5938 | 464659.75 | 424582.8125 |
| 39548.71875 | 39368.35547 | 47658.23438 | 47608.67578 | 51966.70703 | 55777.51172 | 54688.53516 |
| 91575.28125 | 93864.42188 | 117887.8047 | 140032.6406 | 124054.5156 | 114825.5625 | 124304.7031 |
| 185934.8281 | 178971.7969 | 243121.4531 | 198800.875 | 175133.5781 | 178737.375 | 181489.4375 |
| 253663.9736 | 279873.6797 | 318740.7012 | 329909.7598 | 400180.0996 | 395355.8379 | 460924.498 |
| 4123.38916 | 6678.722168 | 4727.59375 | 4688.954102 | 5496.357422 | 4990.058594 | 4407.210449 |

| | | | | | | |
|---|---|---|---|---|---|---|
| 36540.9375 | 40070.67969 | 46723.95313 | 37813.84375 | 27185.4375 | 34366.97266 | 31112.49609 |
| 42368.30078 | 66590.8125 | 47004.07813 | 54072.89063 | 111085.0781 | 106242.0938 | 102412.4297 |
| 331718.9063 | 388837.7188 | 372793.2188 | 396261.5313 | 405243.8438 | 419916.5625 | 400945.25 |
| 422386.5 | 507485.6875 | 447394.1563 | 471745.0313 | 324287.9688 | 300021.4063 | 298750.375 |
| 322121.8438 | 352248.0625 | 360769.9063 | 338209.5313 | 377233.125 | 345467.5 | 394138.25 |
| 108521.2188 | 120966.3359 | 117581.0781 | 128445.8828 | 129191.75 | 122284.625 | 126095.2734 |
| 22365.54102 | 24658.4375 | 35528.93359 | 35556.83984 | 83734.61719 | 90626.69531 | 85211.74219 |
| 745620.1875 | 905220.25 | 924271.75 | 865467.5 | 1111126.25 | 1181970.625 | 1141388.125 |
| 68144.96875 | 81124.32031 | 76270.82813 | 82750.54688 | 93956.71875 | 95984.35156 | 98771.28906 |
| 15301.24512 | 18121.99805 | 16635.06641 | 16344.05664 | 19254.26758 | 15976.57324 | 16283.07031 |
| 1307996.25 | 1447062.75 | 1493207.25 | 1517162 | 1659692.75 | 1657848.375 | 1614259.125 |
| 18956.61523 | 20342.01953 | 22011.41992 | 19354.1875 | 19895.44336 | 19208.66602 | 19424.49023 |
| 10632.9502 | 12348.41895 | 12962.32227 | 11524.48242 | 17142.38867 | 15019.09863 | 16139.61914 |
| 9854.34668 | 11585.55957 | 15105.71289 | 13938.14453 | 10562.20898 | 10630.05078 | 12359.54883 |
| 9220.833008 | 10855.08691 | 9698.271484 | 10352.73047 | 12680.16602 | 11788.53418 | 10416.9668 |
| 483904.5938 | 596305.25 | 529951.875 | 550269.875 | 504831.8438 | 559011.3125 | 515601.125 |
| 24780.34961 | 27040.4375 | 24521.44727 | 25092.65039 | 23397.15234 | 27353.92773 | 24879.55078 |
| 20017.37109 | 22919.42188 | 19926.14453 | 20034.05078 | 11532.35449 | 10638.95898 | 10355.65332 |
| 674938.125 | 751771 | 808454.75 | 778967.0625 | 762528.375 | 781926.3125 | 787318.3125 |
| 43956.53125 | 52870.42578 | 53036.35938 | 50277.24609 | 42917.45313 | 46127.41406 | 43453.16016 |
| 19063.03125 | 26283.39648 | 18672.27344 | 19083.21289 | 20998.14844 | 20690.02539 | 19672.20313 |
| 6497.183594 | 7128.803711 | 6560.946289 | 6450.525391 | 8249.714844 | 7316.626465 | 8000.196777 |
| 95030.15625 | 105769.375 | 115143.4844 | 113089.3359 | 105387.5313 | 103605.7969 | 108457.2969 |
| 2005.559082 | 1831.111328 | 2799.874512 | 2106.105957 | 2227.758301 | 3168.940186 | 2988.998779 |
| 48346.01953 | 49675.32813 | 55997.36328 | 56017.49219 | 38696.12891 | 38168.05078 | 38413.76953 |
| 48629.09375 | 77315.39844 | 42466.46484 | 51935.89844 | 52606.63672 | 67973.36719 | 56252.09375 |
| 82840.41406 | 101075.6875 | 84371.875 | 105122.9922 | 119337.8125 | 102836.2656 | 113224.8672 |
| 20262.0332 | 28428.98438 | 23311.53125 | 23056.59375 | 30914.85156 | 34559.67578 | 29443.81836 |
| 103046.7813 | 98744.79688 | 110807.7109 | 115875.3203 | 98969.27344 | 107186.0547 | 105126.6172 |
| 144915.8438 | 144050.7813 | 185818.2813 | 155935.4219 | 106018.5938 | 132752.8281 | 117219.8906 |
| 26817.72852 | 28115.73438 | 27381.83789 | 25378.9668 | 21652.34766 | 26168.17188 | 25435.10742 |
| 179060.125 | 224108.9844 | 192767.8281 | 185600.5313 | 181558.2344 | 188775.4531 | 179484.3125 |
| 23194.92969 | 25762.44727 | 25111.82617 | 24693.18359 | 23619.54883 | 24404.87695 | 21181.27148 |

| | | | | | | |
|---|---|---|---|---|---|---|
| 10653.98242 | 9191.739258 | 12255.5 | 11180.66406 | 6637.196289 | 7115.098633 | 4600.673828 |
| 21282.0332 | 24524.31641 | 24816.89453 | 21852.33398 | 17686.41016 | 19311.49414 | 20330.50977 |
| 173596.6875 | 187639.9844 | 188607.8438 | 199871.0625 | 180752.3281 | 161683.6094 | 173929 |
| 493684.8438 | 571061.25 | 549612.9375 | 463123.0313 | 432353.75 | 456526.0313 | 457670.1563 |
| 6198.441406 | 5130.735352 | 6621.908691 | 5693.342773 | 9986.019531 | 10734.37207 | 12161.22363 |
| 3421967.301 | 3689960.293 | 3785924.66 | 3569137.477 | 3317494.82 | 3453932.367 | 3433961.563 |
| 20071.13086 | 20588.56445 | 23598.7168 | 23080.12109 | 16688.88672 | 18366.85547 | 16790.88672 |
| 37665.70703 | 36380.27734 | 49119.61719 | 44159.88281 | 43079.03125 | 44551.97266 | 41139.09766 |
| 1974725.926 | 2218045.814 | 2091606.011 | 2229844.801 | 1717870.744 | 1774241.393 | 1899870.24 |
| 45041.24609 | 14229.02051 | 104249.5 | 17904.9375 | 14822.41699 | 32009.09961 | 54788.89063 |
| 12651.87695 | 14740.2041 | 14293.44531 | 10361.74707 | 10560.60547 | 11238.30957 | 11166.7334 |
| 40372.26172 | 40856 | 45990.85547 | 48056.71094 | 44523.77734 | 42993.60547 | 47266.30469 |
| 49258.16797 | 50831.28125 | 68248.5625 | 59211.46094 | 52267.78125 | 55120.88281 | 56281.71484 |
| 623785.25 | 681832.0625 | 683105.5625 | 758883.1875 | 860523.5625 | 675390.125 | 720065.0625 |
| 8262280.5 | 10058925 | 6411738 | 8324991.5 | 7469062 | 9507423 | 5830732.5 |
| 658091.1875 | 673969.3125 | 758767.4375 | 765009.1875 | 745551.9375 | 705794.5 | 738825.75 |
| 59145.35156 | 66885.74219 | 63215.66797 | 66890.49219 | 65023.39063 | 60714.19922 | 59014.34766 |
| 521020.3438 | 599580.5625 | 561880.6875 | 530790.25 | 446775.1875 | 466973.875 | 450499.1875 |
| 7389.283203 | 9853.298828 | 6300.604004 | 6125.098145 | 14910.44336 | 11342.0293 | 12686.58984 |
| 16448.3418 | 22961.99023 | 14603.48828 | 15389.46094 | 17658.81836 | 15150.63184 | 17551.47461 |
| 76006.46094 | 78902.03906 | 82177.96875 | 86350.82813 | 66472.89063 | 71296.95313 | 60962.6875 |
| 12007.49707 | 13717.39453 | 13511.48242 | 12917.58008 | 16790.62891 | 17946.96289 | 16430.12891 |
| 16181.36621 | 16896.07813 | 17282.72656 | 18205.12695 | 16645.39648 | 20443.50195 | 17301.6543 |
| 22612.28906 | 26599.21289 | 31490.60547 | 31250.94922 | 32081.0918 | 28709.8457 | 31728.06445 |
| 8234.744141 | 9102.792969 | 8194.071289 | 7704.633789 | 9606.979492 | 9492.041016 | 10299.82617 |
| 3496.82251 | 2443.603271 | 4451.571777 | 3638.224365 | 4660.370117 | 5114.76709 | 5557.223633 |
| 159299.1094 | 194940.4531 | 166461.7188 | 179436.8594 | 231481.9531 | 221820.5625 | 239374.8594 |
| 14790.92871 | 13743.98926 | 17032.57031 | 17419.2207 | 9685.298828 | 10795.59766 | 10989.36816 |
| 53492.01563 | 64730.09766 | 56239.35156 | 58967.14453 | 47749.64063 | 45697.16016 | 42872.72656 |
| 286933 | 324674.2188 | 347598.8125 | 336986.5938 | 308855.875 | 327099.4375 | 317527.1875 |
| 248052.2813 | 271023.5625 | 275031.8438 | 274261.0625 | 227761.7813 | 238147.1875 | 222141.2188 |
| 351286.1563 | 389205.9688 | 399615.5 | 415220.2188 | 446756.0313 | 446786.5938 | 482245.7188 |
| 105332.7188 | 125287.9844 | 123382.9297 | 113106.7813 | 80943.08594 | 85539.97656 | 83901.1875 |

| | | | | | | |
|---|---|---|---|---|---|---|
| 320774.0625 | 332297.1875 | 379293.1875 | 380441.4688 | 392889.625 | 379543.5 | 379213.875 |
| 66219.35938 | 69252.71875 | 76645.32031 | 71574.75781 | 59920.45313 | 67148.4375 | 67257.76563 |
| 30243.56055 | 38236.73047 | 34076.30078 | 38373.55469 | 92503.29688 | 65757.16406 | 72383.89063 |
| 29046.24805 | 31962.25391 | 31899.64844 | 29842.02734 | 29726.74414 | 27918.03906 | 29715.73828 |
| 6744.156738 | 5904.508789 | 7770.466797 | 6699.164063 | 6561.728027 | 6043.554199 | 4420.23584 |
| 76439.77344 | 110078.8281 | 84550.07031 | 82576.32813 | 55328.53516 | 56010.98047 | 52128.44141 |
| 3338.779541 | 4426.334961 | 5990.342285 | 2527.94873 | 4485.382324 | 4299.390137 | 3693.821777 |
| 608615.5625 | 616887 | 681897.1875 | 715426.25 | 680649.9375 | 643719.625 | 697649.1875 |
| 26563.99414 | 30409.17969 | 30257.56055 | 28150.93945 | 20883.88672 | 21186.05469 | 19242.7168 |
| 53978.19531 | 53385.76172 | 60784.99609 | 60781.67578 | 59455.95703 | 56902.02344 | 56729.1875 |
| 2095.868408 | 5887.697266 | 4819.849609 | 7404.706055 | 7433.822266 | 7447.006836 | 4925.104004 |
| 48173.51563 | 50105.73828 | 52753.04297 | 51176.32031 | 44960.48828 | 46891.875 | 47973.93359 |
| 159274.7813 | 169997 | 178678.6563 | 177596.3906 | 179325.2344 | 186987.9375 | 187678.8281 |
| 740686.375 | 731771.4375 | 816315 | 899490.5625 | 967372 | 973651.6875 | 1058853.125 |
| 69417.49219 | 70665.46875 | 74048.46094 | 73618.8125 | 80326.50781 | 85548.92969 | 84440.53125 |
| 116146.3281 | 121623.0703 | 138542 | 128373.6172 | 70850.71094 | 80391.42969 | 77289.32813 |
| 11800.61621 | 17207.37305 | 6371.54541 | 11838.71094 | 17948.50781 | 13810.05566 | 17495.40039 |
| 22571.54883 | 26816.8418 | 26546.80664 | 27832.96484 | 33542.10156 | 39547.05469 | 29768.87109 |
| 225866.6563 | 253863.75 | 276318.875 | 250866.7188 | 228706.3125 | 252445.5625 | 239867.6875 |
| 24081.45703 | 35281.85547 | 27859.66602 | 27191.38281 | 43696.76953 | 37052.41016 | 39256.41016 |
| 202190.4844 | 220730.8125 | 222011.1094 | 205123.2969 | 196334.4375 | 202884.0938 | 198560.3125 |
| 16655.20898 | 17148.49805 | 17728.375 | 19985.74414 | 20515.62109 | 18971.43555 | 21387.43555 |
| 30326.73633 | 26534.18555 | 30990.25 | 32100.70313 | 32147.32813 | 35743.96875 | 40722.92188 |
| 884862.6875 | 752790.0625 | 1047431.313 | 1112064.625 | 1262243.125 | 1195906.375 | 1233500.875 |
| 93629.64844 | 106665.0391 | 94071.64844 | 103876.0469 | 98901.32813 | 86435.97656 | 93561.3125 |
| 112920.6172 | 157435.6406 | 95539.25 | 99360.11719 | 128802.9688 | 153293.9063 | 111479.9219 |
| 39671408 | 53684820 | 31355682 | 38799624 | 38038396 | 39794112 | 28365704 |
| 172775.7344 | 228906.6094 | 182598.5625 | 201716.2344 | 224164.4688 | 257670.7813 | 237771.9375 |
| 320.3592224 | 427.0007935 | 552.968811 | 458.3832397 | 924.6676636 | NA | 793.6256714 |
| 67770.44531 | 77890.4375 | 78916.32813 | 79069.82031 | 79054.66406 | 68090.28906 | 77114.14063 |
| 149086.6406 | 148527.2031 | 167275.1406 | 176793.1406 | 228594.25 | 168922.0781 | 194509.0156 |
| 81384.71094 | 92272.28125 | 89978.25 | 96652.125 | 72260.63281 | 97328.40625 | 82526.99219 |
| 5306301.5 | 4446254.5 | 5990265 | 6791680 | 7429259.5 | 6357846 | 8884472 |

| | | | | | | |
|---|---|---|---|---|---|---|
| 41313.13281 | 40255.78516 | 45075.80469 | 48547.50781 | 61929.44141 | 62611.77734 | 69534.64844 |
| 33927.16016 | 40178.10938 | 36232.9375 | 39102.49219 | 49858.97266 | 46821.94922 | 50363.21484 |
| 243328.7031 | 290290.1875 | 277125.625 | 302859.7813 | 251947.3906 | 235667.0313 | 257942.5 |
| 878096.4375 | 964063.6875 | 941689.375 | 932159.625 | 925246.5 | 974767.25 | 896052.875 |
| 33019.15625 | 39169.87109 | 45499.14063 | 48730.16406 | 39948.18359 | 32888.87109 | 37364.32031 |
| 45440.30469 | 54470.1123 | 50212.34961 | 49738.79297 | 42592.02051 | 46046.6377 | 41810.1709 |
| 498976.7813 | 540576.875 | 492852.9688 | 535317.875 | 523113.8438 | 547792.125 | 514691.0625 |
| 105161.1094 | 122158.3359 | 124173.2422 | 122681.125 | 102761.4219 | 109505.3438 | 107354.2656 |
| 82512.52344 | 91227.33594 | 84054.94531 | 83009.90625 | 50765.66016 | 46023.44922 | 45329.03516 |
| 3125.726074 | NA | 2533.837891 | 5269.361816 | 8036.825195 | 6511.155762 | 8399.936523 |
| 37046.94922 | 47810.70703 | 40086.95313 | 40927.09766 | 41026.77344 | 37802.125 | 39140.94922 |
| 23154.47656 | 29210.01953 | 26887.85352 | 34036.69922 | 39665.05859 | 31548.29102 | 35261.75 |
| 115664.6875 | 125391.1641 | 132408.8281 | 135535.9375 | 151851.2969 | 145051.0781 | 150706.1406 |
| 56250.38281 | 64142.86719 | 65427.57031 | 57691.76953 | 64433.71484 | 67025.64063 | 64551.05859 |
| 36496.82422 | 63494.50391 | 48666.90234 | 56959.97266 | 72422.23438 | 96083.64063 | 101050.5078 |
| 99355.67188 | 98831.32031 | 105332.8984 | 85071.67969 | 75493.04688 | 96991.61719 | 74687.69531 |
| 23524.66992 | 24265.82031 | 25812.19922 | 23685.52344 | 20506.9668 | 21733.54883 | 20006.58789 |
| 64381.66406 | 61396.80078 | 64401.58203 | 62580.37891 | 60373.08203 | 55401.89844 | 52172.84375 |
| 671801.6875 | 734093.8125 | 792622 | 687249.375 | 595554.625 | 612243.3125 | 607777.8125 |
| 18225.45898 | 20927.57227 | 20063.53906 | 27479.20117 | 35978.55859 | 38910.75391 | 37104.04688 |
| 2034.05188 | 2575.428467 | 3135.61499 | 1259.314087 | 3335.739258 | 1490.272827 | 1576.01416 |
| 4734.256348 | 5258.282715 | 4885.703125 | 3519.096191 | 2757.06543 | 2125.910889 | 2654.131348 |
| 86461.24219 | 91228.07813 | 101568.3906 | 93709.42188 | 83474.79688 | 97438.21094 | 91931.90625 |
| 15047.48535 | 24777.66797 | 20650.02734 | 15342.3252 | 32357.54297 | 21496.70898 | 24638.46094 |
| 79199.27344 | 82295.39844 | 84446.26563 | 85383.8125 | 54268.98047 | 56872.27344 | 60199.53125 |
| 31863.0332 | 37232.25391 | 42388.10938 | 39796.69141 | 50475.73047 | 51669.35938 | 47128.53516 |
| 628571.125 | 655465.375 | 729576.5 | 720092.5 | 659467.8125 | 677530 | 663482 |
| 38058.5 | 31605.93164 | 41393.10156 | 46094.97656 | 47105.58984 | 40600.25 | 52173.46875 |
| 89132.99219 | 100244.1641 | 94881.50781 | 95696.58594 | 98910.125 | 106207.25 | 100031.1094 |
| 509505.5313 | 601469.1875 | 546024.1875 | 566848.875 | 492493.3438 | 480778.0938 | 487886.3125 |
| 64599.15625 | 75717.96094 | 70672.875 | 70791.58594 | 92075.28906 | 73796.70313 | 73572.39063 |
| 28219.23828 | 35174.76563 | 32186.22461 | 30451.14453 | 41903.92578 | 39341.20703 | 40977.30078 |
| 25683 | 29335.97656 | 32005.72461 | 34943.19141 | 33643.70313 | 29433.48828 | 29886.6875 |

| | | | | | | |
|---|---|---|---|---|---|---|
| 39364.75391 | 47985.95313 | 38534.00781 | 38298.68359 | 44679.81641 | 44212.52344 | 40662.01563 |
| 55941.63672 | 53339.64063 | 59178.90234 | 53539.56641 | 31296.05859 | 31729.67773 | 27350.80664 |
| 5883.50293 | 9759.021484 | 4945.434082 | NA | 799.9425659 | 8890.149414 | NA |
| 38118.01953 | 45184.82031 | 48668.56641 | 43780.29297 | 50380.04688 | 71472.64844 | 59212.25391 |
| 19331.25586 | 18865.24805 | 22855.59961 | 21504.0957 | 17899.25781 | 16222.40527 | 20909.96484 |
| 591844.375 | 651720.375 | 692155.125 | 682339.4375 | 670671.3125 | 667339 | 675743.75 |
| 478159.3438 | 650066.875 | 502388.5625 | 552572.0625 | 453452.7813 | 459161.4688 | 416891.9375 |
| 37204.29297 | 42295.38672 | 38327.38281 | 42691.39453 | 63393.96094 | 56798.88672 | 59460.92969 |
| 42321.59375 | 42376.48047 | 51045.125 | 43788.375 | 35679.17578 | 40471.82422 | 35232.41016 |
| 462964.5313 | 494853.625 | 476087.4375 | 462087.5313 | 522701.8125 | 501325.9375 | 535152.4375 |
| 10282.40039 | 12382.12695 | 10304.93848 | 8888.973633 | 12376.08398 | 10619.68457 | 12261.67285 |
| 487366.5 | 519223.3438 | 510627.75 | 543011 | 532451.4375 | 538513.9375 | 552945.5625 |
| 116587.6719 | 123837.0625 | 128912.0391 | 128007.1563 | 97505.33594 | 105576.9844 | 106228.8203 |
| 166749.7656 | 177464.3281 | 163763.4063 | 174952.5781 | 187101.7969 | 203584.5313 | 160058.4531 |
| 143541.1406 | 122517.1172 | 163641.3906 | 159483.0938 | 87422.32813 | 75853.91406 | 86958.28125 |
| 2755.291992 | 3359.979248 | 2191.346191 | 3849.370605 | 2146.951172 | 2689.060059 | 1885.734985 |
| 4745842 | 5365825.5 | 4980026 | 5602936.5 | 6698244 | 6043247 | 6433597.5 |
| 98767.46094 | 119390.7031 | 103391.375 | 120609.1641 | 138414.0156 | 129721.9375 | 132254.3281 |
| 982736.75 | 1123936.125 | 1043854.25 | 1073783.75 | 1010344.813 | 970108.1875 | 934445.4375 |
| 1655937.5 | 1557771 | 1760568.5 | 1740506.25 | 1512232.875 | 1325963.875 | 1927207 |
| 499037.6563 | 452510.2188 | 498864.6563 | 585278.5 | 385875.375 | 371450.2813 | 362577.25 |
| 69720.54688 | 80136.5 | 100072.5625 | 71885.67188 | 59888.63672 | 65815.08594 | 99533.35938 |
| 4859.418945 | 4951.200195 | 7460.967773 | 8343.456055 | 7131.855957 | 9671.635742 | 7142.584961 |
| 64952.53516 | 77945.05469 | 76711.72656 | 71420.36719 | 55224.13281 | 58467.07031 | 59960.96094 |
| 68464.35938 | 80241.67188 | 75203 | 75949.71094 | 90808.51563 | 84878.71094 | 78909.78906 |
| 10486.40918 | 2438.5271 | 13769.8291 | 12031.02637 | 18764.61914 | 14407.81738 | 18446.72266 |
| 126093.0938 | 140005.2031 | 137780.1719 | 126694.4609 | 96775.67188 | 110653.7422 | 108880.75 |
| 298375.7813 | 361761.125 | 343394.25 | 383189.4063 | 377485.5938 | 338099.9688 | 362043.375 |
| 12947.29395 | 14342.53906 | 13846.87402 | 12332.68262 | 11125.24707 | 10777.14258 | 10620.26758 |
| 91901.00781 | 121671.6563 | 88419.25 | 71351.57031 | 57651.65234 | 79227.88281 | 61847.25 |
| 9759.473633 | 16773.09375 | 12674.80859 | 16964.69531 | 46478.32422 | 39348.80469 | 41136.82031 |
| 327319.9375 | 352535.8125 | 359735.6563 | 358841 | 350385.9375 | 338646.4063 | 354766.5938 |
| 12595.6377 | 14067.33594 | 14183.49023 | 11851.51172 | 12633.33301 | 11628.62109 | 12326.52637 |

| | | | | | | |
|---|---|---|---|---|---|---|
| 257426.6094 | 280753.5313 | 282522.0938 | 265137.5938 | 270589.2188 | 310293.4688 | 277401.9063 |
| 38769.13672 | 45656.29688 | 47947.71094 | 37444.67188 | 37277.13672 | 45179.89453 | 40540.65625 |
| 8956.131836 | 9666.969727 | 9162.568359 | 9334.974609 | 7847.949219 | 8291.025391 | 8632.498047 |
| 39527.94531 | 47979.85938 | 46887.59375 | 45825.61719 | 30051.26367 | 30816.0332 | 30768.98438 |
| 7265.569336 | 11014.29492 | 14010.21191 | 12982.91699 | 30282.16406 | 23314.42188 | 24828.41797 |
| 3829.43457 | 3967.887207 | 3933.021729 | 3104.552734 | 2822.144043 | 2979.553711 | 3357.162842 |
| 104479.9844 | 110792.0313 | 125822.7734 | 120684.9297 | 126981.3672 | 141191.125 | 129604.7891 |
| 41342.81641 | 33785.28516 | 47962.69922 | 53583.03125 | 35559.78906 | 34601.60938 | 39232.39844 |
| 46263.96875 | 58285.67188 | 44308.85156 | 45267.99609 | 36755.83594 | 37339.96875 | 40312.80859 |
| 17566.72461 | 19041.24023 | 18475.81641 | 21279.36133 | 19279.98438 | 20890.5625 | 19172.42383 |
| 15820.14648 | 17780.52344 | 17935.3418 | 18822.13477 | 15308.22852 | 15182.72168 | 14870.28906 |
| 49308.88672 | 51707.01172 | 53099.50781 | 48719.61719 | 32826.39063 | 33659.71484 | 31625.28711 |
| 146453.6719 | 162506.375 | 155935.7188 | 151538.5781 | 170919.6875 | 151105.5313 | 157968 |
| 169197.1094 | 209112.1094 | 201402.625 | 221978.125 | 202083.1719 | 168940.5938 | 181626.6406 |
| 118538.2656 | 132853.8125 | 115496.9141 | 132488.9531 | 193225.0625 | 197789.8125 | 213263.2031 |
| 48258.17188 | 47490.50391 | 58904.69531 | 59726.8125 | 70793.80469 | 60086.62109 | 62482.68359 |
| 578572.4375 | 599429.75 | 668954.25 | 671002.5 | 596405.75 | 670878.8125 | 673792.75 |
| 26707.97656 | 28990.8418 | 27348.2168 | 29856.87305 | 29558.83984 | 29773.9043 | 27820.5957 |
| 26389.00977 | 37254.65234 | 29940.80078 | 30411.57031 | 42620.07031 | 22602.85938 | 40496.04688 |
| 36777.55078 | 44525.71094 | 32107.54102 | 37449.81641 | 43107.54688 | 35970.00781 | 39801.46484 |
| 324054.4688 | 330524.4375 | 372092.6563 | 391814.7188 | 419595.6563 | 428140.6875 | 486930.1563 |
| 28564.87305 | 33459.4375 | 27951.09961 | 37699.07031 | 22729.51563 | 35438.39844 | 32674.20703 |
| 60477.25 | 66590.38281 | 71663.58594 | 68749.88281 | 45280.34375 | 48675.17969 | 45962.05078 |
| 448018.0742 | 484106.0254 | 534750.0488 | 506710.1836 | 438680.3125 | 477579.8691 | 472777.9492 |
| 9007.245117 | 14436.68555 | 11590.78223 | 12844.30664 | 18871.31055 | 19214.20898 | 14416.47656 |
| 527071.8125 | 608029.9375 | 614255.0625 | 577437.9375 | 452178.8125 | 473860.75 | 468406 |
| 370835.9688 | 416117.4375 | 397201.7188 | 436753.25 | 446136.6563 | 436156 | 442167.25 |
| 10339.45215 | 12356.10059 | 12454.15527 | 10837.875 | 5835.312988 | 6941.649414 | 7727.115234 |
| 211558.4375 | 232453.7969 | 238160.0938 | 230592.4063 | 215220.1406 | 229079.4375 | 209217.7344 |
| 34640.55469 | 33846.66016 | 36957.28125 | 41406.57813 | 32960.44922 | 27221.30664 | 28490.60938 |
| 112001.1406 | 112650.9844 | 124078.4453 | 123129.8984 | 116231.4531 | 114973.9766 | 119112.4141 |
| 14596.1582 | 15435.59473 | 14016.74609 | 14291.93848 | 16255.78711 | 16813.30078 | 15179.30273 |
| 18687.84766 | 24860.45117 | 21093.30469 | 19982.28516 | 19642.44922 | 21146.22461 | 18958.66406 |

| | | | | | | |
|---|---|---|---|---|---|---|
| 92483.44531 | 80328.3125 | 107846.6406 | 113540.9141 | 99617.34375 | 100321.25 | 110594.5078 |
| 209328.9531 | 226144.9219 | 220257.7031 | 238566.1719 | 237076.7344 | 246053.8594 | 238268.8281 |
| 146721.3594 | 165907.0313 | 165157.6406 | 176871.6094 | 200175.6875 | 187021.3906 | 190435.3281 |
| 23633.5918 | 27361.17383 | 27500.54492 | 26688.67969 | 22274.13086 | 25148.31445 | 22119.75391 |
| 3968296.25 | 4640912.5 | 4861575.5 | 4129001.5 | 3434330.5 | 3238294 | 3353540 |
| 99073.10156 | 112889.6172 | 112798.8828 | 116197.2266 | 109764.375 | 118561.7734 | 120328.9688 |
| 430363.7813 | 418480.4063 | 485143.125 | 502179.2188 | 561720.1875 | 436008.7188 | 487089.1563 |
| 15133.22363 | 13871.08398 | 9759.09668 | 14843.35742 | 28735.45313 | 23539.14453 | 25746.375 |
| 552483.125 | 605934.3125 | 637711.9375 | 595509.5 | 553011.125 | 675479.8125 | 651127.375 |
| 7133.524414 | 6568.933105 | 8248.030273 | 6571.086426 | 6932.455078 | 6902.652344 | 6458.694824 |
| 15321.98828 | 13778.15137 | 17798.08789 | 17557.32227 | 16131.11621 | 15149.10645 | 16004.12793 |
| 193213.8906 | 207449.2969 | 212871.4375 | 223821.875 | 180890.5781 | 222962.8594 | 210445.5313 |
| 55827.67969 | 56980.09766 | 58849.97266 | 53450.96484 | 46963.08984 | 51242.92578 | 52540.57031 |
| 13766.17871 | 17823.4668 | 14852.5 | 14071.81836 | 16883.45703 | 13049.57617 | 13777.62207 |
| 69987.02344 | 79349.42188 | 81011.70313 | 83846.92969 | 77988.28125 | 83984.84375 | 80487.35938 |
| 41396.76563 | 46920.26172 | 47131.47656 | 50365.50781 | 72773.35156 | 69305.44531 | 66721.98438 |
| 20061.88086 | 14302.83008 | 18931.77344 | 22591.56836 | 37858.55469 | 51473.19922 | 44449.64844 |
| 26876.02148 | 30168.49805 | 27039.17773 | 29041.34961 | 27475.5 | 27834.03711 | 27494.41797 |
| 277899.6875 | 290655.9688 | 296001.7188 | 308368.4375 | 445255.375 | 375188.1563 | 405084.875 |
| 397215.1563 | 457239.3125 | 416944.875 | 404204.9063 | 299723.4688 | 320215.5938 | 273811.75 |
| 11255.2959 | 17798.55859 | 14343.21484 | 16496.6582 | 26104.57813 | 23480.28906 | 23052.34766 |
| 732746.8125 | 1050387.25 | 872243.0625 | 827269.375 | 1224433.625 | 1310084.25 | 1289442.625 |
| 77197.17188 | 97431.875 | 83673.85938 | 79325.88281 | 92420.27344 | 100568.4766 | 95524.375 |
| 18069.79297 | 20923.4082 | 22709.36523 | 19076.53516 | 15447.55371 | 17769.04297 | 16438.05469 |
| 26459.28516 | 36222.14844 | 27151.26758 | 23255.66211 | 20879.53906 | 21829.24609 | 24659.95313 |
| 16009.5957 | 19586.75195 | 18325.51563 | 19443.33203 | 16410.29688 | 17436.50391 | 18153.32813 |
| 271416.1875 | 302883.1875 | 262844.0313 | 306307.4063 | 405988.4063 | 380139.2188 | 402186.5938 |
| 21056.47656 | 21211.87305 | 23352.48438 | 25368.15625 | 26537.9375 | 21023.85938 | 22875.97852 |
| 186.0257874 | 4095.876221 | 379.318512 | 2299.863281 | NA | 12925.40625 | 6001.770508 |
| 72430.16406 | 87169.88281 | 78248.11719 | 82399.3125 | 183855.2031 | 157951.1563 | 155005.375 |
| 75782.39844 | 73972.01563 | 76270.1875 | 82195.375 | 85571.89063 | 93649.14844 | 90348.48438 |
| 56720.55078 | 60071.42578 | 61564.51563 | 55793.33984 | 56137.39453 | 61863.55859 | 60084.55469 |
| 83204.83594 | 87886.74219 | 92210.29688 | 91423.17969 | 92935.42969 | 99319.96094 | 100020.2891 |

| | | | | | | |
|---|---|---|---|---|---|---|
| 56743.58984 | 73652.28906 | 59926.38281 | 67176.91406 | 70550.73438 | 71655.58594 | 74245.625 |
| 18411.63672 | 23151.45313 | 16423.77148 | 15449.79688 | 24270.8418 | 32325.74805 | 24374.57617 |
| 140743.6719 | 156789.5781 | 144250.5469 | 161395.4844 | 158759.0781 | 148333.4688 | 152422.4844 |
| 2904.776123 | 4609.721191 | 3797.783203 | 2165.569092 | 2439.773438 | 2140.579102 | 2266.974854 |
| 106837.6484 | 109618.3047 | 112338.7969 | 111477.0078 | 79108.98438 | 83878.55469 | 83656.21094 |
| 33154.71094 | 43904.74219 | 35351.83203 | 36895.25391 | 72131.35938 | 69264.19531 | 67443.36719 |
| 133324.8281 | 133209.5469 | 149017 | 143871.6875 | 129676.1563 | 131090.9844 | 150386.4844 |
| 30535.95508 | 30822.99414 | 31889.55469 | 38857.75391 | 42242.72656 | 30589.9707 | 34007.58984 |
| 595317.3125 | 519658.5 | 701925.1875 | 654003.4375 | 593213.625 | 530153.625 | 562413.125 |
| 334767.125 | 314477.6563 | 359783.5313 | 404821.5 | 507259.5 | 510020.75 | 568833.5 |
| 60291.61719 | 67161.32031 | 65239.60156 | 67773.42969 | 65462.12109 | 63539.39844 | 63367.4375 |
| 9122.18457 | 11129.98926 | 10816.44336 | 10306.9248 | 11414.27051 | 9873.304688 | 11360.24023 |
| 73363.20313 | 84440.65625 | 82217.69531 | 81938.80469 | 65484.53125 | 69096.02344 | 62525.33594 |
| 13240.76758 | 14732.0166 | 14415.5791 | 14919.94336 | 18460.51367 | 16415.51367 | 15063.85156 |
| 10611.49707 | 13954.52051 | 14021.98145 | 12166.19238 | 12649.78809 | 14321.28516 | 11215.63867 |
| 3781.576416 | 3790.138184 | 3381.964844 | 3932.9375 | 11204.22852 | 6171.105957 | 9714.092773 |
| 2687080.5 | 3086427.25 | 2747220.75 | 2688715.25 | 2877549.75 | 2964489.75 | 2873106 |
| 18793.99609 | 20721.3418 | 19919.45508 | 21412.57813 | 50731.94531 | 34420.02344 | 42025.38281 |
| 27008.26172 | 30930.41016 | 30408.87891 | 29488.29492 | 34263.41406 | 32925.87109 | 31238.79297 |
| 14610.56445 | 16485.66016 | 15634.96582 | 15309.35059 | 15343.50293 | 14802.64355 | 13553.28125 |
| 103277.0391 | 111009.6563 | 107103.9844 | 111378.1016 | 131314.1094 | 132601.875 | 134725.8906 |
| 209459.1094 | 209018.8438 | 250312.6406 | 292264.7188 | 429474.9375 | 316126.4063 | 348430.875 |
| 35679.40625 | 17296.29102 | 44329.09766 | 43979.92578 | 32883.52344 | 28746.24609 | 39476.90625 |
| 309501.8438 | 324232.25 | 313055.125 | 342439.9375 | 320854.7813 | 312779.6875 | 309842.9375 |
| 123259.0547 | 141996.4844 | 149718.3906 | 130678.6484 | 119751.4375 | 105604.4297 | 94296.90625 |
| 123347.7344 | 130912.1641 | 134852.6719 | 144101.5156 | 116868.4922 | 127069.1406 | 118864.3438 |
| 25753.92188 | 37973.61328 | 36432.38281 | 35276.76563 | 32383.44727 | 36316.53125 | 32189.61719 |
| 667961.9297 | 758842.4805 | 723944.1211 | 746279.4688 | 738164.4688 | 691213.7461 | 711267.707 |
| 35283.57813 | 52494.69141 | 33625.51953 | 36991.25781 | 54551.25781 | 45428.08203 | 46228.83203 |
| 35203.76563 | 36101.24219 | 38609.44922 | 38052.38281 | 40278.24609 | 44562.875 | 41066.89844 |
| 31911.29492 | 42297.03125 | 35868.89844 | 35440.02344 | 66381.14063 | 57684.375 | 57198.87891 |
| 28175.8418 | 31444.8418 | 30510.43555 | 33493.10156 | 35651.83984 | 38619.70703 | 39154.61328 |
| 103239.6875 | 120770.1875 | 116894.0156 | 113992.4063 | 103981.2578 | 101907.125 | 99286.21875 |

| | | | | | | |
|---|---|---|---|---|---|---|
| 128477.3516 | 144106.0781 | 107203.2344 | 107762.1953 | 91470.35156 | 128897.1719 | 106905.8516 |
| 129091.6406 | 129303.4297 | 140295.625 | 146760.7344 | 124083.0469 | 122618.6719 | 123184.2578 |
| 47017.55469 | 46128.50391 | 52486.15234 | 57168.46484 | 50673.11719 | 52895.59375 | 54578.45313 |
| 1863478 | 2347955.25 | 2139435.25 | 2152179.5 | 1927984.625 | 1885438.75 | 1720560.875 |
| 287700.75 | 340061.1875 | 344404.1563 | 317208.375 | 318635.7813 | 304613.0313 | 300798.1875 |
| 113295.125 | 123397.9375 | 143295.6406 | 143358.5625 | 165002.6094 | 146720.1094 | 142181.9844 |
| 56610.5625 | 64040.41406 | 56535.46094 | 56984.58594 | 72708.44531 | 74925.70313 | 77173.82031 |
| 23942.97852 | 29150.54102 | 26375.88672 | 25902.50195 | 29095.29102 | 29880.33984 | 29235.7207 |
| 207521.0781 | 199236.6563 | 210318.5 | 226276.3125 | 232326.125 | 214396.7656 | 220011.2344 |
| 54916.58594 | 55018.46875 | 60015.29688 | 61430.17969 | 84042.36719 | 83206.84375 | 82525.92969 |
| 71580.05469 | 78846.46875 | 81933.21094 | 80948.85938 | 83988.5 | 78343.78125 | 84336.42969 |
| 4247.353027 | 2740.349121 | 5458.556152 | 3724.50293 | 3757.758057 | 6947.416504 | 4640.446289 |
| 21811.88086 | 28259.20898 | 25546.77148 | 24571.32813 | 17412.43555 | 20758.60938 | 18670.06836 |
| 132419.9063 | 124181.8438 | 147333.4375 | 156366.4688 | 183279.6719 | 176176.4531 | 177638.5313 |
| 211510.0156 | 209509.6719 | 226813.2813 | 245427.1563 | 260304.8594 | 258575.4375 | 266209.5 |
| 217058.6563 | 240347.3594 | 254425.0938 | 278667.2188 | 309426.375 | 317189.3438 | 316076.8125 |
| 386083.1563 | 439550.7188 | 462078.375 | 503520.4063 | 600076.125 | 569161.9375 | 577974.375 |
| 9907.415039 | 11370.60645 | 12450.73926 | 12615.68555 | 19999.25 | 17731.47266 | 22490.51367 |
| 15746.67188 | 17454.82617 | 12374.36133 | 14111.875 | 18469.82031 | 21476.9707 | 23203.58203 |
| 30045.73633 | 30091.05859 | 32443.84766 | 30217.29102 | 26772.83984 | 22585.2832 | 24742.29297 |
| 913974.0625 | 960204 | 935318.0625 | 902698.8125 | 837115.9375 | 852292.625 | 833227.625 |
| 10930.22363 | 11979.08398 | 12388.15625 | 11132.2959 | 9714.693359 | 9689.725586 | 8950.274414 |
| 34556.41797 | 32081.4043 | 36221.93359 | 36900.25 | 28074.76758 | 32298.29297 | 30852.51172 |
| 29768.67383 | 34861.64453 | 31032.38086 | 33617.05859 | 34058.26953 | 33445.97656 | 34430.27344 |
| 175927.9063 | 178858.75 | 204149.875 | 188178.3906 | 174046.8281 | 175042.6563 | 172683.6719 |
| 23835.21094 | 26404.92773 | 26010.79688 | 27325.23633 | 31509.54492 | 29455.36719 | 29498.64258 |
| 6114.162598 | 12701.1543 | 2099.237061 | 2930.281494 | 4056.556641 | 3828.743164 | 4047.30835 |
| 51444.99219 | 56000.01563 | 57885.40625 | 55732.35938 | 62348.59766 | 63859.68359 | 62284 |
| 242317.9463 | 305848.2695 | 265401.4248 | 279903.1846 | 336462.6943 | 387942.4746 | 355945.9629 |
| 73346.67969 | 82661.39063 | 80890.26563 | 81051.28125 | 93989.125 | 92211.32031 | 91849.40625 |
| 24023.49414 | 24442.07031 | 25248.17578 | 26911.40234 | 27538.09766 | 26496.88281 | 27309.19531 |
| 219466.1094 | 216821.3125 | 236025.1406 | 234262.3438 | 233011.0156 | 210878.375 | 218132.3438 |
| 144041.4219 | 219667.3438 | 143453.5469 | 165960.1875 | 233735.625 | 186566.2813 | 223286.7344 |

| | | | | | | |
|---|---|---|---|---|---|---|
| 4873840.5 | 14799151 | 8049105 | 11740045 | 17561346 | 15709831 | 14249826 |
| 722353.8125 | 696393.6875 | 287352.4063 | 598276 | 623598.5625 | 675969.75 | 654008.5625 |
| 33726.625 | 33563.80078 | 27517.57227 | 39679.74609 | 50687.93359 | 41573.89063 | 39653.0625 |
| 117703.9609 | 139612.0938 | 120478.8438 | 121217.2188 | 145016.0469 | 137683.8125 | 142796.75 |
| 451708.5 | 504224 | 547874.0625 | 528600.6875 | 690377.25 | 663606.25 | 729197.4375 |
| 125365.1484 | 125270.3359 | 142064.6563 | 136584.0313 | 150658.6875 | 165223.4844 | 184249.2656 |
| 89466.75 | 107732.2344 | 102232.8828 | 77922.23438 | 54286.34766 | 48358.16797 | 49496.05078 |
| 3470.649902 | 6596.521973 | 5186.907227 | 3567.416748 | 8301.652344 | 7429.989258 | 4352.594238 |
| 289522.125 | 289204.75 | 331024.7188 | 336364.9063 | 276620.5313 | 297158.75 | 319329.2813 |
| 69108.67188 | 46856.67578 | 94459.75781 | 51372.76953 | 42773.92578 | 86508.88281 | 74025.69531 |
| 121195.3125 | 131186.3906 | 161733.6719 | 177452.1094 | 213199.2031 | 230550.9688 | 247561.5469 |
| 80224.64844 | 100446.6094 | 83248.625 | 82297.29688 | 98743.5625 | 102008.9219 | 91819.46094 |
| 34550.43359 | 36587.20703 | 37151.54688 | 38396.99609 | 43942.55859 | 41811.75781 | 38326.67578 |
| 7221.576172 | 9832.716797 | 6728.171875 | 9122.668945 | 6902.211914 | 12376.33496 | 7810.673828 |
| 432417 | 500054.75 | 433148.1875 | 477092.5 | 449212.4375 | 445046.1875 | 437388 |
| 46769.71875 | 48974.54688 | 44689.68359 | 48099.69141 | 46315.71094 | 39416.07422 | 35837.63672 |
| 12937.32617 | 11612.27539 | 13406.58984 | 11983.53906 | 15005.11328 | 12360.35156 | 14553.96582 |
| 2602.956055 | 2314.218262 | 3267.476807 | 5139.690918 | 50773.46484 | 3970.094238 | 5956.161133 |
| 500195.75 | 438978.1875 | 536269.6875 | 557215.0625 | 582357.3125 | 614729 | 606010.9375 |
| 91754.14063 | 89581.19531 | 95205.92969 | 95292.99219 | 69276.55469 | 71757.44531 | 73944.9375 |
| 8541.491211 | 10090.78418 | 6574.779297 | 6123.165527 | 2283.563232 | 2494.141602 | 2298.073975 |
| 46019.09375 | 52757.15625 | 49772.35938 | 52019.29688 | 46902.14453 | 44320.99609 | 43420.09766 |
| 101636.4766 | 108156.1797 | 113187.1563 | 105457.8203 | 86724.875 | 91894.40625 | 94424.49219 |
| 5621.36084 | 8412.617188 | 6637.882813 | 7585.76123 | 9551.303711 | 8620.727539 | 9032.678711 |
| 69435.96875 | 84210.10938 | 76730.49219 | 69918.23438 | 75073.125 | 75625.95313 | 70386.0625 |
| 672491.5 | 798545.4375 | 716829.625 | 708517.125 | 827273.8125 | 775283.0625 | 760049.625 |
| 10032.59277 | 9714.277344 | 11204.76953 | 10538.06543 | 9960.089844 | 10972.85059 | 9114.645508 |
| 6043.143555 | 9238.990234 | 7588.493164 | 6094.433594 | 10923.83984 | 12385.54102 | 11453.96191 |
| 44952.47656 | 44058.42969 | 47656.64453 | 46377.74219 | 45774.32422 | 47706.32422 | 49382.66797 |
| 195815.7656 | 219227.6563 | 219346.7344 | 217545.3594 | 198777.5 | 235252.0781 | 232921.9844 |
| 196514.1875 | 220026.875 | 205388.1719 | 209670.8281 | 196373.4219 | 242143.1719 | 199018.0469 |
| 54906.46484 | 60767.25391 | 64970.36328 | 72133.53125 | 114389.3984 | 97562.39063 | 112072.0859 |
| 126605.6875 | 123568.0781 | 147825.0625 | 158318.2188 | 171709.6875 | 156887.3125 | 169841.2656 |

| | | | | | | |
|---|---|---|---|---|---|---|
| 43971.95703 | 53527.96484 | 47157.40234 | 52110.38672 | 48322.82031 | 39706.38281 | 42240.55469 |
| 921960.5 | 1098989 | 945052.4375 | 897983.6875 | 933832.125 | 936254.5 | 872263.8125 |
| 103805.9922 | 132454.875 | 102270 | 105919.6094 | 130836.9297 | 144014.3125 | 136809.7969 |
| 58333.87109 | 69750.46094 | 58527.98828 | 49127.35938 | 31969.53125 | 31698.39258 | 29613.56641 |
| 32684.01172 | 34486.52734 | 38656.05859 | 37847.22656 | 25500.37695 | 25033.5625 | 25822.02148 |
| 15931.3584 | 19978.07813 | 17603.82031 | 15102.71875 | 14917.24316 | 16044.17188 | 13765.0293 |
| 46343.92188 | 56306.71094 | 51658.64453 | 50357.39063 | 47271.74609 | 37173.31641 | 40715.64063 |
| 267664.5938 | 260281.3438 | 270311.125 | 281253.5313 | 282565 | 274883.625 | 272531.6563 |
| 14718.87207 | 12753.72168 | 14676.91699 | 15140.03809 | 15936.2998 | 16276.50781 | 16580.07227 |
| 10494.36328 | 9618.711914 | 10800.02246 | 11099.03711 | 10745.0752 | 10092.07422 | 9341.729492 |
| 5232.58252 | 6344.783203 | 5353.041992 | 5904.401855 | 4011.239746 | 4373.704102 | 3780.283203 |
| 40584.61328 | 43028.50781 | 41652.96094 | 44771.38281 | 42365.55078 | 46640.34375 | 43005.41406 |
| 26682.68945 | 28694.82031 | 28571.37891 | 27737.07422 | 27390.79492 | 28046.28906 | 26822.55859 |
| 59551.08203 | 62795.07031 | 60359.38281 | 58884.60156 | 65007.11328 | 66387.39063 | 67955.78906 |
| 415111.4375 | 633513.5 | 437902.0938 | 376723.625 | 433990.7188 | 353419.4688 | 402622 |
| 102648.4453 | 113653.0078 | 112935.4297 | 115802.8281 | 123340.8047 | 111585.1484 | 119412.6016 |
| 916093.5 | 997394.5 | 944878.1875 | 1009285.375 | 937306.125 | 844060.5 | 902060.8125 |
| 30418.06055 | 31864.36133 | 37962.25391 | 31355.20117 | 33990.41406 | 44092.63672 | 41060.24219 |
| 53801.76953 | 52076.37109 | 53390.08984 | 76488.3125 | 78377.67188 | 84967 | 79564.10156 |
| 34695.24609 | 40997.86719 | 41549.45313 | 37564.71484 | 27240.9043 | 27910.31445 | 27556.79492 |
| 163082.8281 | 176505.6094 | 176309.25 | 169469.1875 | 117516.9531 | 116286.3203 | 113565.8828 |
| 24663.49805 | 26558.17773 | 25036.89063 | 26200.26563 | 25555.56641 | 27998.42773 | 27483.76758 |
| 50341.55469 | 45488.62891 | 58787.85156 | 49306.42578 | 50152.63281 | 47999.53516 | 50141.73438 |
| 437218.3438 | 424798.9688 | 512337.1875 | 480705.375 | 452356.7813 | 490781.9375 | 500891.1563 |
| 9427.733398 | 35353.5 | 14915.12207 | 7226.582031 | 86750.96875 | 33550.61719 | 70588.8125 |
| 15563.07227 | 14009.03711 | 17779.06641 | 18906.50195 | 27804.63281 | 29406.04688 | 31201.85547 |
| 27076.17578 | 49342.35547 | 23544.81836 | 22326.41211 | 42138.05469 | 44916.63672 | 38222.36719 |
| 446919.25 | 480495.4375 | 470500.1875 | 465510.875 | 456523.0938 | 515148.125 | 474941.3438 |
| 44563.91406 | 37629.10156 | 51048.79297 | 47517.57813 | 50871.83594 | 49732.25781 | 51876.62891 |
| 20353.5957 | 18227.77734 | 19648.32422 | 19993.54102 | 28460.08789 | 22288.66211 | 25019.33203 |
| 438310.125 | 456602 | 465959.9375 | 458163.9375 | 363383.2813 | 342099.7188 | 364694.4063 |
| 20400.74805 | 25438.4707 | 18292.19922 | 17828.22266 | 23593.32617 | 21768.63281 | 21690.16992 |
| 15674017 | 17416974 | 16277351 | 18200956 | 31301948 | 29316260 | 30261594 |

| | | | | | | |
|---|---|---|---|---|---|---|
| 24154.17578 | 23900.93164 | 26613.29102 | 26398.97266 | 24458.85352 | 22566.89063 | 23764.87891 |
| 4878.277832 | 5317.791504 | 3480.640869 | 4486.443359 | 3791.297852 | 3761.223145 | 3116.511475 |
| 44919.71484 | 41082.24219 | 42686.8125 | 40560.19141 | 12511.16504 | 25844.94336 | 30949.8418 |
| 77382.16406 | 87655.01563 | 84443.35156 | 87738.25 | 100727 | 106764.7188 | 103303.1484 |
| 73564.21094 | 75386.54688 | 77407.375 | 73790.40625 | 79512.52344 | 77709.125 | 76295.36719 |
| 60442.12109 | 75908.64063 | 67397.66406 | 73902.40625 | 153080.3281 | 123330.0156 | 136439.0469 |
| 60417.44922 | 71287.28125 | 63904.6875 | 56419.44531 | 76944.02344 | 75918.60938 | 75270.24219 |
| 30428.85156 | 31117.5 | 33788.57422 | 32897.57813 | 27198.68945 | 28441.45703 | 27156.78906 |
| 865345.375 | 960586.75 | 841424.5 | 801943.6875 | 792256.5 | 819139.625 | 756164.5625 |
| 87796.19531 | 91201.03906 | 96702.64063 | 87854.0625 | 42605.38281 | 43692.92578 | 39820.02734 |
| 425805.375 | 443501.8125 | 461027.75 | 456913.6563 | 438857.2188 | 455605.8438 | 463156.9375 |
| 11802.30371 | 12468.66113 | 13116.71289 | 13837.10352 | 11306.05273 | 8811.049805 | 10205.28711 |
| 162673.8125 | 196983.2813 | 204297.7188 | 281756.0625 | 751922.1875 | 403639.4688 | 520720.375 |
| 31527.5957 | 32889.6875 | 33866.71484 | 30360.91602 | 31345.22266 | 38553.73828 | 34874.64063 |
| 69392.48438 | 63027.03906 | 79457.55469 | 72216.79688 | 69630.03125 | 65799.58594 | 70472.125 |
| 317734.5938 | 383499 | 372112.6563 | 385157.1563 | 349784.2188 | 369687.1563 | 370469.7188 |
| 37047.26953 | 37278.14844 | 47856.17969 | 48137.25 | 49757.21484 | 48603.74609 | 46357.35547 |
| 21108.82422 | 30398.74805 | 21610.36523 | 20899.48047 | 31727.91797 | 29851.41992 | 24878.98633 |
| 166337.1094 | 199670.1563 | 174040.3438 | 183289.6094 | 159267.7656 | 166077.1563 | 178926.3125 |
| 771348.25 | 808675.75 | 750779.125 | 762597.3125 | 751194.1875 | 809605.0625 | 778883.5625 |
| 115285.7109 | 131359.125 | 98240.39844 | 102834.9063 | 127564.2188 | 154364.875 | 120838.5078 |
| 53669.64453 | 52872.86719 | 58571.76563 | 58256.21875 | 52517.46875 | 53652.08203 | 55977.64063 |
| 62128.52344 | 68796.15625 | 33345.75781 | 55602.61328 | 72259.96094 | 83217.52344 | 84158.53125 |
| 96764.89844 | 104947.3359 | 103153.9297 | 100262.8828 | 81362.88281 | 84182.53906 | 82959.32813 |
| 49065.6875 | 55648.10156 | 51142.14844 | 52290.79688 | 81965.05469 | 70410.39063 | 73408.46875 |
| 63716.39844 | 70816.96094 | 66238.25781 | 70251.80469 | 88998.90625 | 87520.64844 | 83230.20313 |
| 8050.303223 | 7726.899902 | 9542.861328 | 9087.816406 | 9598.393555 | 10302.16309 | 9456.692383 |
| 75598.96875 | 79804.25781 | 78058.10938 | 74423.5 | 51468.28125 | 49515.35547 | 53291.89844 |
| 71010.79688 | 84432.41406 | 73047.36719 | 81313.08594 | 96029.78906 | 85986.91406 | 93288.9375 |
| 17016.50195 | 20703.82617 | 17569.42188 | 19165.57227 | 20497.52539 | 19006.41797 | 17714.48438 |
| 6433.518066 | 6518.024414 | 7175.51416 | 6922.927734 | 16167.31152 | 14963.72266 | 13028.43262 |
| 782856.3047 | 896166.3359 | 797856.2656 | 857645.7266 | 911296.9922 | 920677.8906 | 895057.1563 |
| 67056.59375 | 80181.5 | 69506.34375 | 66006.14063 | 64521.32031 | 67509.375 | 52556.89063 |

| | | | | | | |
|---|---|---|---|---|---|---|
| 8066.65332 | 9522.963867 | 8862.171875 | 7663.962402 | 7161.022949 | 7521.700684 | 6911.164063 |
| 185426.4531 | 231695.0938 | 190537.2031 | 200589.5156 | 171870.7031 | 188948.125 | 171511 |
| 25976.79492 | 30289.79492 | 27106.25391 | 28977.97461 | 29300.14844 | 28446.25195 | 26596.55078 |
| 14321.33691 | 17914.0957 | 17029.22461 | 20398.75195 | 14777.78516 | 16621.61914 | 13534.24707 |
| 51135.81641 | 68865.19531 | 54153.67188 | 62200.62109 | 90373.70313 | 99133.97656 | 83615.47656 |
| 13803.02148 | 17782.96875 | 19274.21094 | 14185.66309 | 26569.20313 | 26795.47852 | 28150.82813 |
| 99286.21875 | 102572.6406 | 91806.71875 | 83604.82031 | 99986.3125 | 106517.1953 | 92233.52344 |
| 30365.03125 | 29305.7207 | 30420.125 | 32729.76172 | 51881.96875 | 51353.15234 | 52997.25 |
| 1033102.563 | 982181.8125 | 1209797.375 | 1209514.125 | 782920.0625 | 782185.625 | 831137.8125 |
| 7692.642578 | 6794.802246 | 6002.693848 | 7207.169434 | 5172.421387 | 6343.661621 | 5989.967285 |
| 23574.45508 | 26005.91797 | 26666.97852 | 27085.81836 | 59869.21875 | 46792.12109 | 55417.875 |
| 41536.44922 | 47664.51953 | 52933.14453 | 48496.38281 | 29747.34961 | 35747.10156 | 35140.69141 |
| 40590.24609 | 41365.48047 | 41595.73828 | 45745.875 | 33078.33594 | 35158.625 | 35658.32813 |
| 59767.07031 | 59904.98047 | 60645.80078 | 69336.01563 | 80551.57031 | 85412.61719 | 83708.17969 |
| 1164484 | 1187069.75 | 1288555.25 | 1256476.125 | 903926.25 | 912576.8125 | 930329.8125 |
| 413476.9688 | 533536.625 | 487269.5625 | 452185.0625 | 303613.2188 | 401122.4063 | 350922.0938 |
| 50864.43359 | 53698.28516 | 44055.43359 | 46237.59766 | 39411.77734 | 44559.28516 | 38751.21875 |
| 51028.20703 | 55476.58984 | 53372.30859 | 54584.65625 | 50656.37891 | 49598.00781 | 52010.63281 |
| 601.6674805 | 427.2176514 | 583.741272 | 498.5734253 | 104.0494843 | NA | NA |
| 33992.29297 | 51180.62109 | 40331.11719 | 40789.76563 | 43363.50391 | 35115.25781 | 39754.78516 |
| 36915.17969 | 39253.82422 | 40235.23047 | 44026.3125 | 68940.82031 | 55930.05859 | 54640.41406 |
| 63758.12891 | 55608.6875 | 71435.60156 | 61606.46094 | 50976.64453 | 49552.16016 | 53385.06641 |
| 126687.8359 | 133766.5938 | 134660.3594 | 130120.7656 | 142349.9219 | 143227.3125 | 148666.9375 |
| 11071.11426 | 10431.08203 | 10991.10059 | 10610.16016 | 9761.029297 | 9719.195313 | 8804.485352 |
| 510948.6563 | 462128.375 | 623376.25 | 628024.0625 | 487676.5313 | 512234.5313 | 530205.4375 |
| 21279.38867 | 31003.16211 | 25467.26953 | 25563.42969 | 22386.21289 | 18042.24609 | 19412.18555 |
| 185386.4688 | 168511.8594 | 217562.4063 | 213716.7813 | 270424.1563 | 261756.875 | 288301.3438 |
| 41554.59375 | 43015.26563 | 42304.4375 | 40717.72656 | 37422.71484 | 40774.66406 | 40259.05859 |
| 250152.4375 | 285826.2188 | 252754.6719 | 266120.875 | 250177.625 | 231854.9219 | 265267.7813 |
| 440361.3438 | 467660.9375 | 472648.7188 | 488331.4063 | 469233.625 | 403236.3438 | 433055.875 |
| 14987.44141 | 17778.91602 | 16725.12695 | 14952.71777 | 15667.27832 | 13235.58398 | 15337.34668 |
| 6534.776367 | 5138.190918 | 5119.632813 | 6275.068848 | 6596.480957 | 5862.043945 | 7602.24707 |
| 9998.073242 | 10380.57617 | 8428.248047 | 8684.550781 | 12769.84082 | 10390.99121 | 11054.78223 |

| | | | | | | |
|---|---|---|---|---|---|---|
| 30836.7793 | 34360.51172 | 28676.83398 | 34094.47656 | 31706.87695 | 31339.6875 | 29839.24023 |
| 29156.76953 | 24415.53125 | 29151.03711 | 31450.36328 | 26628.02539 | 28948.99609 | 27388.78906 |
| 40632.0625 | 28627.47461 | 49533.46484 | 44246.09375 | 39113.53906 | 35642.26172 | 42099.05469 |
| 112592.8828 | 112287.7422 | 123944.4844 | 124099.7656 | 115064.2109 | 119304.0234 | 117039.75 |
| 89713.26563 | 116586.9375 | 78428.19531 | 62569.46094 | 74798.91406 | 117219.1172 | 89401.16406 |
| 64782.125 | 67446.20313 | 69242.73438 | 68067.09375 | 97224.13281 | 82266.72656 | 90161.53906 |
| 6608.338867 | 6189.974121 | 8759.357422 | 8824.673828 | 1127.016846 | 3093.530518 | 6951.454102 |
| 445218.5625 | 494794.5938 | 482493.9375 | 529141.9375 | 700422.125 | 666385.5 | 708121.625 |
| 133250.2188 | 123726.2422 | 137947.5156 | 144243.8438 | 116984.3516 | 145185.75 | 125655.1875 |
| 11121.95605 | 11325.36328 | 12614.61621 | 13548.92383 | 26308.91016 | 22131.2832 | 26340.94141 |
| 406387.5879 | 485461.1094 | 405145.6641 | 314018.5469 | 364649.332 | 549243.5234 | 409691.8242 |
| 972724.8125 | 966742.25 | 1069494.125 | 1010926.25 | 876294.8125 | 917584.75 | 910606.625 |
| 17248.02734 | 18959.5332 | 18939.3418 | 18570.57031 | 22219.88281 | 20335.41016 | 22327.74219 |
| 83785.33594 | 85451.82031 | 93973.82813 | 92978.16406 | 84350.23438 | 100245.3281 | 85704.40625 |
| 554452.8125 | 645894.75 | 550395.375 | 577663.5 | 553392.5 | 526925 | 541404.5625 |
| 384459.5 | 391456 | 452985.9063 | 438221.1875 | 308912.0625 | 299520.5313 | 307718.6563 |
| 8603.822266 | 7735.960449 | 8813.697266 | 7203.519531 | 7501.444336 | 7039.016113 | 6328.794922 |
| 1645714.625 | 1969080.5 | 1383378 | 999582.4375 | 777647.875 | 1049775.125 | 635013.3125 |
| 71304.21094 | 82542.17188 | 77387.52344 | 77497.25 | 83895.32031 | 82004.08594 | 79155.77344 |
| 15834.91992 | 17470.97461 | 17446.57617 | 17362.49023 | 16000.89844 | 15391.63477 | 15628.77539 |
| 8670.618164 | 10774.92676 | 7616.810059 | 6450.293945 | 4924.026367 | 5956.937988 | 4926.178223 |
| 35347.08984 | 38676.13672 | 41535.51172 | 45019.37109 | 66925.21094 | 67188.64063 | 67919.75781 |
| 149135.875 | 150324.4844 | 156928.4688 | 161257.125 | 97325.80469 | 97259.99219 | 100075.1719 |
| 335823.4688 | 337756.5 | 379794.625 | 387088 | 196245.4531 | 200264.2188 | 203354.0313 |
| 69177.35938 | 76945.16406 | 76718.89063 | 78720.20313 | 77739.35156 | 70197.32813 | 74560.58594 |
| 80104.32031 | 127721.0391 | 79212.07031 | 91930.60938 | 136416.6719 | 134045.2188 | 109422.2578 |
| 218533.2969 | 219647.75 | 233667.9531 | 223824.5156 | 204495.1875 | 209085.2188 | 224406.25 |
| 31907.77344 | 24300.05664 | 33728.83203 | 41490.38672 | 27924.09375 | 30642.10352 | 29857.50977 |
| 42144.96875 | 43888.42578 | 46481.19531 | 41097.07031 | 35422.27734 | 34529.59766 | 34291.33203 |
| 200291.125 | 187505.5313 | 220999.4063 | 211113.9375 | 129848.7109 | 163180.9375 | 152150.3281 |
| 154624.6875 | 169560.4375 | 161560.3125 | 158255.0313 | 171271.5625 | 153440.7188 | 173885.5313 |
| 350120.5313 | 334746.25 | 352146.25 | 379204.5625 | 331000.5313 | 345323.7813 | 329136.5938 |
| 11653.78809 | 16614.12891 | 14767.12012 | 10546.24316 | 6333.462402 | 8361.945313 | 6903.304688 |

| | | | | | | |
|---|---|---|---|---|---|---|
| 79418.96094 | 79837.85938 | 88822.44531 | 88352.99219 | 105978.0156 | 101654.5703 | 99420.44531 |
| 4683.894043 | 3777.908936 | 3716.498535 | 4546.280762 | 2181.749023 | 3958.805908 | 2496.314453 |
| 629195.125 | 670974.375 | 665069.1875 | 674500.9375 | 649436.5625 | 602442.4375 | 670224.875 |
| 25367.64844 | 30756.40234 | 24985.54883 | 22994.76563 | 23390.41016 | 25109.55859 | 23570.60156 |
| 46793.37891 | 53536.64063 | 48505.54688 | 45959.11328 | 23496.43555 | 24479.29102 | 22280.9043 |
| 303269.125 | 398128.4375 | 346907.5938 | 366499.4688 | 734179.1875 | 554932.5 | 590578.5625 |
| 155321.5469 | 178738.0313 | 163706.6719 | 182298.3125 | 229342.0469 | 211467.5313 | 218916.6406 |
| 76004.29688 | 80733.77344 | 84944.35938 | 82078.94531 | 64987.73438 | 67346.35156 | 63808.25391 |
| 10015.94336 | 13131.00391 | 12429.81934 | 13235.45898 | 16524.79688 | 16603.56445 | 18019.99414 |
| 9424.054688 | 8629.283203 | 8938.731445 | 9280.935547 | 6734.349609 | 7433.783691 | 7415.818848 |
| 2951.559814 | NA | 3459.843262 | 3451.634766 | 11471.2666 | 5085.006348 | 7620.787598 |
| 15307.00879 | 14980.25977 | 13656.66699 | 14612.82324 | 9088.757813 | 9429.089844 | 7746.987793 |
| 14455.18164 | 16992.78906 | 13525.84375 | 14909.94531 | 16489.18555 | 15028.46777 | 15980.96875 |
| 16695.30859 | 19803.79883 | 18590.57227 | 18116.47266 | 21897.6543 | 22834.61523 | 21019.07031 |
| 40815.82031 | 38805.87891 | 44391.01563 | 38087.44531 | 32230.72852 | 34993.22266 | 31381.07813 |
| 257850.875 | 272045.0313 | 283783.25 | 279834.8125 | 236396.0781 | 253927.9531 | 254322.5 |
| 21141.39063 | 25527.3457 | 27193.51172 | 27209.93555 | 32213.77539 | 26805.60938 | 27443.0625 |
| 256829.7813 | 288044.0938 | 267907.5 | 273814.0313 | 211339.3594 | 194614.2813 | 203966.9063 |
| 8634.430664 | 1759.410645 | 11386.50977 | 12023.67773 | 19778.66406 | 24907.07617 | 18124.51367 |
| 2567841.5 | 2453413.75 | 2628069.5 | 2972095.5 | 3914226 | 3603560 | 3854289.5 |
| 275178.7813 | 282930.125 | 305667.5938 | 307154.1875 | 307900.3125 | 294542.4063 | 306686.8125 |
| 20926.20313 | 27511.64844 | 19500.35742 | 17131.4043 | 19956.10742 | 23387.14453 | 22230.7832 |
| 154290.6973 | 166927.9121 | 173493.793 | 166144.8965 | 136905.4434 | 128295.6094 | 140592.9414 |
| 27174.98242 | 33044.32813 | 30721.35742 | 27669.57031 | 20344.93359 | 18243.3457 | 19803.39844 |
| 267478 | 297329.875 | 287028.5625 | 314029.4375 | 259772 | 247956.6875 | 256892.7344 |
| 11667.7959 | 13842.32031 | 11647.21875 | 10482.86133 | 11991.82422 | 13095.50293 | 9829.817383 |
| 243471.1719 | 282749.8438 | 256409.2969 | 287268.375 | 394637.5 | 351423.9375 | 356433.25 |
| 759848.8125 | 873337.25 | 746825.875 | 783897.875 | 935631 | 872714.1875 | 925019.3125 |
| 84188 | 82330.32813 | 88279.75781 | 92555.85156 | 92388.04688 | 105709.7188 | 101340.7813 |
| 44977.68359 | 46040.76953 | 46661.06641 | 53247.94922 | 60587.70703 | 55973.84375 | 58748.55469 |
| 230334.4688 | 231896.4063 | 232054.1563 | 226785.1719 | 242992.375 | 282852.3438 | 274245.25 |
| 64692.11328 | 54742.67578 | 64871.84375 | 70946.79688 | 74464.65625 | 62741.14063 | 73215.00781 |
| 4566.387695 | 5564.231934 | 4936.611328 | 4708.755859 | 3985.871582 | 4502.749512 | 4633.095703 |

| | | | | | | |
|---|---|---|---|---|---|---|
| 13632.75488 | 20613.65234 | 14708.57324 | 15652.89648 | 15883.34766 | 14504.87305 | 15258.10547 |
| 16981.76367 | 14298.61426 | 18005.44141 | 20300.4707 | 22864.0293 | 22204.97656 | 18606.30664 |
| 91899 | 93410.57031 | 95897.625 | 94253.08594 | 105481.6563 | 107701.6484 | 110928.0078 |
| 76351.33594 | 87568.65625 | 74046.73438 | 78439.5 | 85366.73438 | 78166.33594 | 80880.32031 |
| 49754.21094 | 50794.80078 | 53710.32422 | 53991.5 | 49532.49609 | 49319.46094 | 49507.53125 |
| 194334.0625 | 190326.5625 | 203263.625 | 197342.7656 | 191759.0625 | 196075.9688 | 202313.6094 |
| 7265.996094 | 7475.191895 | 7958.384277 | 8210.151367 | 6799.037598 | 6032.740234 | 6512.846191 |
| 165300.9219 | 166205.9531 | 180450.0156 | 162522.9375 | 135012.9688 | 145472.8594 | 152254.1875 |
| 311720.75 | 315159.2813 | 328066.6875 | 331250.25 | 263127.5 | 280244.0938 | 282737.9063 |
| 10111.65918 | 14999.54688 | 10938.06641 | 10764.86914 | 12270.51563 | 22512.5918 | 18578.67969 |
| 46465.48438 | 48019.63672 | 46611.44141 | 53490.28906 | 39486.94141 | 38876.58203 | 38345.21094 |
| 11199.61133 | 6919.130371 | 6852.007813 | 9927.25293 | 40008.86328 | 24475.95117 | 35226.95703 |
| 1502471.375 | 1571814.5 | 1699256.5 | 1670539.875 | 1575909.625 | 1589798.25 | 1594301 |
| 543822.625 | 614562.9375 | 587656.5 | 582783.3125 | 513295.5625 | 499161.25 | 451075.8438 |
| 5704.407227 | 5895.908203 | 8896.727539 | 7976.290527 | 1714.124146 | 766.0892944 | 9218.859375 |
| 3515.605957 | 4205.635742 | 3406.068359 | 4269.956055 | 4248.944336 | 4114.047363 | 4144.505371 |
| 265714.2188 | 289999.6563 | 292868.3438 | 289453.3125 | 281002.8125 | 300645.625 | 293028.1875 |
| 190741.9375 | 148880.8125 | 229371.7813 | 217589.8906 | 235482.625 | 200994.3281 | 245178.1719 |
| 10294.87988 | 9854.925781 | 10520.1582 | 11648.29297 | 6126.054688 | 4608.909668 | 5418.63916 |
| 179040.5 | 187319.6406 | 175537.3125 | 179998.9063 | 187010.7813 | 199412.8594 | 209336.3438 |
| 4511.791992 | 3775.634277 | 6181.33252 | 5950.882324 | 5375.805176 | 4360.944336 | 4736.981934 |
| 156338.2031 | 160132.9375 | 157590.4375 | 141693.9063 | 108251.9688 | 110952.8281 | 87264.39844 |
| 1421164.75 | 1755633 | 1267628.125 | 1208581 | 1645828.5 | 1598312.375 | 1541128.125 |
| 101986.3281 | 106960.3906 | 111552.6719 | 119657.7734 | 209159.5938 | 196429 | 200924.4531 |
| 237548.0156 | 216431.0156 | 255223.4844 | 263285.3438 | 157684.9063 | 167901.6875 | 171027.25 |
| 11194.25977 | 12228.21289 | 12199.36035 | 10521.20508 | 12332.70898 | 12906.06738 | 13007.58105 |
| 306909.1875 | 335199.9688 | 312726.1875 | 327592.7813 | 331427.1563 | 325025.2188 | 320203.875 |
| 16084.85352 | 17303.62305 | 18746.32617 | 16131.83105 | 21544.53711 | 20287.20117 | 22541.01758 |
| 129468 | 134016.4688 | 137037.5938 | 138656.2813 | 139663.6406 | 138950.3125 | 141941.2031 |
| 5898.853516 | 7875.646973 | 5692.899414 | 5540.106445 | 7366.657715 | 6947.330078 | 6320.577637 |
| 42218.50781 | 50628.08594 | 40080.10938 | 44077.38281 | 45389.85547 | 37365.13672 | 46329.19141 |
| 849439.5625 | 737207.5625 | 864669.4375 | 848226.1875 | 883585.75 | 1031210.438 | 994827.6875 |
| 24123.11328 | 12274.94824 | 37950.28516 | 42086.44922 | 17898.89063 | 19091.01758 | 16925.2793 |

| | | | | | | |
|---|---|---|---|---|---|---|
| 35377.07813 | 42582.51953 | 24939.17383 | 37727.89063 | 15856.00879 | 16142.43262 | 15093.99902 |
| 1780995.625 | 1954460.375 | 1723828.125 | 1862774.5 | 1933484.875 | 1763263.375 | 1809071.375 |
| 6444.444336 | 8124.836914 | 7475.013672 | 8157.129883 | 10290.04395 | 11795.89648 | 9912.347656 |
| 16609.96094 | 18369.08984 | 16350.29688 | 14784.22852 | 16270.49023 | 15690.10156 | 16114.46777 |
| 52404.83984 | 52414.19141 | 58429.98438 | 62006.75781 | 65549.73438 | 62632.10156 | 60432.64844 |
| 134741.75 | 157838.25 | 131163.5781 | 121720.3516 | 128426.1875 | 142745.4688 | 111735.7969 |
| 141148.8125 | 177736.2813 | 113023.6797 | 120171.9922 | 149832.6719 | 148380.2969 | 152355.0625 |
| 10131.25684 | 14240.97559 | 12661.4209 | 11192.84961 | 16415.83984 | 19586.4043 | 14445.39551 |
| 11708.40234 | 12998.36523 | 13408.57422 | 12304.47266 | 7223.751465 | 7799.246094 | 7773.618164 |
| 23423.60742 | 25971.48633 | 25620.14063 | 25703.01172 | 27187.875 | 25437.8125 | 27867.50195 |
| 110590.7188 | 94095.91406 | 111635.0234 | 117859.3203 | 124535.5 | 139802.5313 | 142110.8906 |
| 1475990.75 | 1381347.875 | 1629002.375 | 1631386.5 | 2191610 | 1975406.625 | 2089862.75 |
| 77690.33594 | 77175.72656 | 77205.82031 | 66713.69531 | 85026.29688 | 84894.03906 | 91772.52344 |
| 22917.41211 | 25702.06445 | 24223.07227 | 29732.53906 | 33980.78906 | 30213.60352 | 36567.55859 |
| 21787.18359 | 21371.36523 | 21672.44727 | 24086.33398 | 27482.48633 | 27555.06445 | 28263.50977 |
| 4871.52002 | 4104.10791 | 4738.280273 | 4598.19873 | 1227.064941 | 1273.806519 | 1750.841675 |
| 53252.98828 | 58580.75391 | 52967.53516 | 55594.58594 | 61523.27344 | 60481.92188 | 60870.60938 |
| 10630.0752 | 10817.59961 | 11157.32129 | 9719.989258 | 10841.61914 | 8429.700195 | 9833.882813 |
| 395733.9688 | 522543.6875 | 373456.125 | 370505.3438 | 422477.75 | 416146.1875 | 410130.6875 |
| 83203.73438 | 93903.51563 | 88183 | 72287.1875 | 44113.22266 | 50127.55859 | 46106.10938 |
| 32763.29297 | 33005.85547 | 28497.3125 | 33559.24609 | 38794.77734 | 38817.14063 | 38282.77344 |
| 45705.14844 | 57361.67188 | 43201.35938 | 41427.61328 | 57075.25 | 58774.02734 | 57183.26953 |
| 85001.91406 | 89002.70313 | 85512.88281 | 82681.92188 | 94455.63281 | 91892.46094 | 95483.88281 |
| 35021.12891 | 38732.41016 | 35802.01953 | 39826.73828 | 51781.61719 | 45644.56641 | 50131.66797 |
| 85840.71094 | 89836.41406 | 95600.14844 | 88503.39063 | 55425.21875 | 64985.26953 | 56083.69531 |
| 6804.30957 | 10624.27734 | 8210.206055 | 6311.436523 | 9003.573242 | 8079.456055 | 7545.929688 |
| 51453.31641 | 37339.94141 | 53490.50391 | 58938.83203 | 76482.28125 | 67319.8125 | 60211.04688 |
| 9411.179688 | 9883.03418 | 10411.39551 | 9321.055664 | 10588.19336 | 8983.704102 | 10019.49609 |
| 7055.799316 | 6178.392578 | 11661.49121 | 10541.75 | 11233.3623 | 11411.80273 | 15193.36816 |
| 11445.625 | 13461.76465 | 11621.19727 | 4284.70752 | 8993.890625 | 8972.449219 | 1973.280273 |
| 7843.570313 | 9593.193359 | 8203.915039 | 8608.800781 | 8307.821289 | 9892.269531 | 8657.94043 |
| 19096.04492 | 18169.27148 | 23328.08008 | 23760.04883 | 23438.67188 | 15768.61426 | 22750.26172 |
| 255874.8906 | 323414.6563 | 277840.0938 | 255870.5156 | 242245.25 | 257146 | 230574.1875 |

| | | | | | | |
|---|---|---|---|---|---|---|
| 23087.40234 | 25275.0293 | 25398.75977 | 25157.93555 | 21754.24609 | 21359.13086 | 21692.94336 |
| 150784.7656 | 148412.2188 | 154917.9219 | 170021.9063 | 195549.25 | 194566.5313 | 202230.0938 |
| 4908.875 | 7449.677246 | 3547.42334 | 2222.766113 | 13212.51465 | 5850.607422 | 13205.98828 |
| 7122.628906 | 8796.165039 | 7753.014648 | 8840.4375 | 8445.739258 | 7374.93457 | 7212.069336 |
| 72639.89063 | 73029.85156 | 82372.01563 | 99582.74219 | 144379.0625 | 129485.625 | 134649.0156 |
| 21141.79492 | 25382.14844 | 28006.34961 | 22353.19922 | 30330.33398 | 23552.34961 | 27176.60547 |
| 15052.18457 | 6862.068848 | 11435.58984 | 24812.20313 | 24690.09766 | 75630.80469 | 19406.73633 |
| 152807.7188 | 155262.4688 | 165031.6719 | 181580.4375 | 206169.5469 | 200328.6563 | 196921.375 |
| 101740.0234 | 120075.5469 | 100437.5781 | 92675.49219 | 157321.2969 | 167102.8594 | 167931.4531 |
| 47610.22656 | 63698.69922 | 40124.72656 | 54111.62109 | 96018.54688 | 75079.29688 | 68942.90625 |
| 275251.4063 | 302684.8438 | 310031.4375 | 286108.0938 | 260280.8906 | 275713.9063 | 247281.9063 |
| 21510.08789 | 18173.97852 | 23038.33984 | 20574.06836 | 30984.09961 | 25440.23633 | 28804.9082 |
| 3381923.164 | 3684793.125 | 3376750.859 | 3279547.313 | 3133519.625 | 3280555.484 | 3159942.867 |
| 678755.3125 | 708092.5625 | 661883.75 | 700464.8125 | 715778.625 | 610054.1875 | 647388.9375 |
| 10251.17871 | 11359.87891 | 9915.34082 | 11424.84766 | 9352.930664 | 8138.666016 | 8716.553711 |
| 24052.99414 | 28216.91602 | 27041.87695 | 27455.19531 | 40599.94531 | 30728.46094 | 34172.13672 |
| 17141.44727 | 17625.92578 | 16492.30078 | 17412.5332 | 23387.08594 | 22137.17188 | 22889.24219 |
| 95788.21094 | 97790.70313 | 99758.09375 | 104673.7188 | 124390.1719 | 103273.6953 | 111194.5703 |
| 5850.552246 | 6495.241699 | 5653.195801 | 4872.094727 | 7911.543457 | 12450.18164 | 10380.36328 |
| 8373.498047 | 11115.09082 | 16167.92383 | 11540.99609 | 18778.4668 | 11352.81445 | 14867.8916 |
| 59000.57813 | 55596.84766 | 65612.92188 | 57599.04297 | 53637.9375 | 58263.21094 | 56717.58594 |
| 280000.7188 | 302642.2813 | 288958.1563 | 330233.5 | 455576.3438 | 402893.8125 | 418279.8125 |
| 66258.9375 | 90643.83594 | 58686.29688 | 50134.02734 | 101130.1172 | 101644.5547 | 106858.125 |
| 2087.200439 | 2323.148682 | 1995.226074 | 1913.349121 | 2002.39209 | 2047.4646 | 2172.78125 |
| 6136.110352 | 6655.455078 | 7189.620605 | 6259.865723 | 3823.897217 | 4144.29834 | 4210.20166 |
| 236572.1875 | 291350.5938 | 269027.6875 | 293528.2813 | 332688.625 | 378081.0625 | 341120.375 |
| 38707.20313 | 36014.11328 | 39358.40625 | 40612.76172 | 67067.28125 | 62483.03906 | 64873.72266 |
| 79657.6875 | 94172.39844 | 78071.03125 | 80748.26563 | 107074.6563 | 102719.9688 | 102345.1406 |
| 142877.8125 | 152756.125 | 149367.8906 | 143070.125 | 126591.5703 | 112722.3047 | 120879.9063 |
| 22383.56445 | 25910.70898 | 19700.40234 | 16670.7168 | 8690.567383 | 10345.85742 | 10184.95996 |
| 96621.33594 | 89230.26563 | 98233.92969 | 98821.41406 | 109955.8594 | 106714.4766 | 107547.1172 |
| 19248.91211 | 21311.71094 | 18065.85742 | 18320.75977 | 23738.70898 | 22804.64453 | 20431.91797 |
| 1839.94397 | 2283.442871 | 1696.676025 | 1579.140137 | 1404.029663 | 1605.315063 | 1643.22168 |

| | | | | | | |
|---|---|---|---|---|---|---|
| 9895.914063 | 10042.93848 | 8687.421875 | 10387.55859 | 14989.21875 | 14412.45703 | 18029.03906 |
| 3598.38623 | 4525.622559 | 3335.252197 | 3281.984131 | 2333.678711 | 3926.821533 | 3799.456787 |
| 51482.41016 | 43943.14453 | 64147.29297 | 57692.82813 | 36573.56641 | 32951.79297 | 36893.82813 |
| 113298.5078 | 124594.4219 | 116739.4922 | 122911.4609 | 128334.4766 | 122192.8672 | 123492.7734 |
| 16758.72852 | 19269.7832 | 17477.33984 | 18761.92188 | 11733.50391 | 11301.26758 | 11102.66699 |
| 8378.056641 | 5668.642578 | 10604.64648 | 9777.573242 | 5454.235352 | 4949.415039 | 4007.749268 |
| 5624.083008 | 5847.915527 | 5551.491699 | 8104.564941 | 7156.316406 | 5106.218262 | 5108.395508 |
| 36148.45703 | 36401.67578 | 40547.99609 | 34812.38672 | 23871.09766 | 25223.47461 | 28383.625 |
| 23033.75195 | 24545.51172 | 23847.37695 | 26875.25977 | 27524.15234 | 28608.78711 | 28647.23633 |
| 395421.75 | 351638.3438 | 401100.9375 | 427576.6563 | 376581.0313 | 384362.2188 | 403289.1563 |
| 309021.75 | 310188.6563 | 332000.0313 | 321616.3438 | 301083.1563 | 317505.3125 | 333888.0625 |
| 71757.10156 | 83414.10938 | 64944.62891 | 71674.77344 | 92514.70313 | 92179.60938 | 87617.70313 |
| 495751.7813 | 505541.8438 | 526825.75 | 502910.3125 | 469177.9375 | 497717.9375 | 490847.7813 |
| 28183.92383 | 30128.52539 | 25962.67773 | 27477.20117 | 20921.66602 | 23692.79883 | 22819.41602 |
| 473126.9688 | 500838.2188 | 474784.8125 | 458010.3438 | 341112.875 | 362067.5625 | 314278.9375 |
| 31831.26953 | 39172.07813 | 28134.61133 | 22251.12891 | 22911.05664 | 61500.29297 | 44117.92188 |
| 39696.82031 | 39583.07813 | 45256.47656 | 45708.87891 | 52441.64063 | 50453.87891 | 51193.71094 |
| 26549.92969 | 28160.28906 | 24726.12891 | 24325.25781 | 26722.24805 | 23936.22266 | 22438.49219 |
| 45276.86328 | 50627.22266 | 46260.48047 | 50059.77344 | 42061.39453 | 38625.96094 | 38239.05469 |
| 80779.6875 | 85739.65625 | 85595.91406 | 89242.6875 | 74053.42188 | 77930.15625 | 77129.47656 |
| 41090.74219 | 46780.19922 | 40830.89453 | 40693.97656 | 62938.19141 | 55699.69141 | 54052.96094 |
| 59464.99219 | 42542.24219 | 75018.83594 | 85529.30469 | 130127.2031 | 134932.4375 | 145775.25 |
| 316043.875 | 380764.5313 | 350551 | 414823.4375 | 323186.5 | 290578.125 | 306533.9688 |
| 434178.9063 | 461996.375 | 439168 | 425111.625 | 369447.8438 | 359994.7188 | 368726.7188 |
| 32661.64844 | 39893.17188 | 36462.61719 | 38616.86719 | 61062.82031 | 58813.88672 | 63539 |
| 11493.76074 | 12536.35547 | 12649.42188 | 11170.19434 | 12344.16602 | 11457.89063 | 11809.86816 |
| 184154.6719 | 160174.75 | 171259.5313 | 200200.6094 | 202685.0469 | 178376.4375 | 199451.9219 |
| 8650.77832 | 9996.03125 | 8145.156738 | 9993.860352 | 11628.1748 | 12040.58594 | 11635.66309 |
| 2360413 | 2470710.5 | 2424650.75 | 2596711.5 | 2348290.5 | 1934929.625 | 2164863 |
| 48965.875 | 53990.08203 | 50810.90234 | 54503.18359 | 41223.63281 | 36986.5625 | 38610.46484 |
| 489143.2813 | 474796.0938 | 565192.125 | 564251.75 | 524293.0625 | 558482 | 558261.8125 |
| 41506.60547 | 40488.78125 | 37929.82813 | 36098.375 | 25763.76563 | 27752.58984 | 23468.76367 |
| 9794.617188 | 13131.89453 | 10388.16797 | 10999.73828 | 12922.9043 | 11249.97754 | 10691.34961 |

| | | | | | | |
|---|---|---|---|---|---|---|
| 27521.28125 | 24815.86719 | 25081.9707 | 25198.23242 | 26019.27734 | 25903.87305 | 27824.11523 |
| 27619.92578 | 29621.91992 | 24095.75781 | 20203.89453 | 28724.71289 | 32195.63672 | 33792.46094 |
| 173025.6094 | 176856.2969 | 190085.7344 | 188527.5625 | 177014.8906 | 183014.875 | 181462.6875 |
| 31715.88477 | 31035.79492 | 35956.83984 | 35335.48438 | 32879.91406 | 32256.46289 | 34306.63672 |
| 22234.55078 | 22774.12891 | 19673.17188 | 20253.33594 | 25421.80078 | 28984.70313 | 27513.83984 |
| 419756.5 | 461059.7109 | 441927.1172 | 484512.5527 | 572107.1611 | 506634.4609 | 560231.7422 |
| 20226.83203 | 20214.08398 | 20211.99414 | 20265.04492 | 13937.03418 | 14269.22656 | 12396.66504 |
| 87626.17969 | 85207.54688 | 90386.78906 | 81835.85156 | 98960.21094 | 99438.41406 | 92489.03125 |
| 33524.83984 | 41655.77734 | 38580.56641 | 38907.17188 | 43244.32813 | 40079.79297 | 41649.14844 |
| 52544.69922 | 69001.28125 | 44341.51563 | 43379.34375 | 63604.14063 | 58908.3125 | 59637.31641 |
| 1521300.375 | 1432985.5 | 1691915.25 | 1697511.75 | 1393259 | 1566968.25 | 1603216.625 |
| 174013.0781 | 173071.4063 | 161372.2031 | 177053.5938 | 209136.0469 | 216845.5625 | 210764.9375 |
| 182618.7969 | 187494.7344 | 179646.0625 | 175597.9219 | 158270.1563 | 151719.7969 | 155006.3438 |
| 27723.88867 | 26620.68945 | 28279.25977 | 29222.61523 | 38491.60156 | 34910.49609 | 35159.5625 |
| 72894.38281 | 79724.78906 | 72258.0625 | 72939.67188 | 65796.21094 | 68976.66406 | 63072.40234 |
| 166551.6406 | 158937.3281 | 167205.25 | 177154.1719 | 162685.7969 | 153878.0938 | 163661.3125 |
| 8065.055176 | 7536.21875 | 7268.737305 | 6695.612305 | 3129.678223 | 3687.596191 | 3086.101318 |
| 44706.71484 | 59858.81641 | 52488.44531 | 65650.83594 | 77199.375 | 61558.28516 | 61599.59375 |
| 185671.2344 | 192642.0781 | 176833.4219 | 196758.5938 | 155141.8438 | 135667.4063 | 160625.25 |
| 20704.11523 | 18663.875 | 19381.54297 | 21372.6875 | 22524.7793 | 19925.44727 | 22052.70508 |
| 8147.755371 | 8288.742188 | 7224.810059 | 7571.908691 | 5076.54248 | 5391.858398 | 5774.837891 |
| 40800.72266 | 39037.10547 | 43958.17578 | 43588.86328 | 35810.19141 | 43817.46484 | 39980.39063 |
| 1840345.375 | 1936111.375 | 1893369.5 | 1989978 | 1881230.25 | 1794509.125 | 1828509.25 |
| 4812.341309 | 1970.445923 | 8699.477539 | 1194.246338 | 1875.708008 | 7548.241699 | 3250.330566 |
| 539686.1875 | 532795.8125 | 592270.4375 | 631263.875 | 711131.9375 | 693497.5 | 734367.625 |
| 6934.422852 | 7586.336426 | 7899.685547 | 8281.154297 | 7961.877441 | 7778.598633 | 7905.615723 |
| 56097.75977 | 53020.92188 | 72277.70801 | 59885.30957 | 71734.85059 | 63171.45898 | 74893.57129 |
| 84005.65625 | 81236.82813 | 81169.85938 | 75021.98438 | 89948.75 | 92041.63281 | 101786.9219 |
| 6939.263672 | 6554.098633 | 6958.309082 | 6629.678223 | 5341.15332 | 6947.063965 | 5558.615234 |
| 14459.68945 | 21214.16406 | 13874.24609 | 14739.26367 | 20226.79688 | 22670.68555 | 21047.90625 |
| 331605.125 | 349362.4375 | 317097.25 | 384871.1875 | 466638.0625 | 408941.1563 | 431997.0938 |
| 33850.03516 | 28065.57227 | 25460.35156 | 31893.41406 | 23425.02539 | 27438.41016 | 22028.3125 |
| 100703.3359 | 97027.375 | 99896.14063 | 101954.0859 | 100698.7344 | 99706 | 99290.01563 |

| | | | | | | |
|---|---|---|---|---|---|---|
| 10725.03711 | 11450.98047 | 9667.365234 | 12252.47656 | 46580.64844 | 39562.05469 | 36987.21484 |
| 99270.04688 | 90220.53906 | 100630.6719 | 109597.5859 | 129974.875 | 118840.1797 | 120592.9922 |
| 134649.9219 | 110117.5234 | 139577.125 | 146925.25 | 141076.6406 | 145937.2188 | 167061.2031 |
| 16068.49902 | 17658.67578 | 16742.54492 | 16925.99023 | 15368.90918 | 12732.1416 | 13216.88184 |
| 440799.375 | 492022.2344 | 439481.5781 | 424922.3125 | 462214.1094 | 438955.625 | 477390.8125 |
| 454443.6563 | 411756.6875 | 484466.25 | 520267.6875 | 562739.5625 | 597127.125 | 634821.1875 |
| 164647.6406 | 190594.3438 | 173988.0313 | 167009.1406 | 149883.625 | 148882.4844 | 147795.6094 |
| 774714.125 | 787203.0625 | 799127 | 740387.75 | 810343.125 | 845868.5625 | 806365.125 |
| 527.0053711 | 543.4902344 | NA | NA | 494.9844666 | 492.0904236 | 437.8100586 |
| 154022.125 | 179756.625 | 178201.3281 | 164476.5938 | 266458.9063 | 299022.875 | 277468.4063 |
| 23790.23242 | 23762.89844 | 25208.5 | 18863.33008 | 12519.48242 | 13596.21387 | 12498.00977 |
| 519852.5625 | 417141.3125 | 585432.125 | 632892.8125 | 806603.25 | 741195.1875 | 771928.4375 |
| 85988.86719 | 97343.10156 | 91166.51563 | 92443.75 | 78229.65625 | 84222.25781 | 78900.54688 |
| 197121.7813 | 229582.7344 | 195831.8594 | 184157.125 | 152361.7344 | 156921.7969 | 163547.2656 |
| 1210754.125 | 1414898.625 | 896506.5625 | 1267877.625 | 1316783.5 | 1213408.625 | 1292185.625 |
| 14449.47461 | 18304.39063 | 16269.75 | 17641.91406 | 30188.89063 | 25870.61719 | 27068.63086 |
| 412900.4375 | 422287.0938 | 434884.8438 | 449382.8438 | 443238.3125 | 473241.9063 | 480764.25 |
| 210356.4219 | 180799.4688 | 218239.7031 | 215193.2188 | 432520.3438 | 369539.125 | 466219.5625 |
| 56802.54297 | 53137.43359 | 60856.58203 | 54575.78125 | 32099.10742 | 31511.38086 | 32914.78906 |
| 43755.99219 | 42356.97656 | 46855.32813 | 40835.61328 | 51213.33203 | 54675.4375 | 58644.53516 |
| 712787.625 | 794423.8125 | 682097.5625 | 725963.375 | 829786.3125 | 678594.4375 | 728893.875 |
| 30258.89844 | 38475.72656 | 36852.09375 | 16815.94922 | 6879.620605 | 14585.2998 | 14384.23633 |
| 92711.25781 | 95185.9375 | 99702.21875 | 98665.125 | 108174.1953 | 103616.625 | 109755.2344 |
| 753515.5 | 789090.375 | 769812.4375 | 803158.6875 | 772944.9375 | 814137 | 859900.5 |
| 125108.625 | 125947.3828 | 130903.0156 | 127802.8125 | 112704.3047 | 118585.9219 | 117699.1719 |
| 782127.9414 | 820057.707 | 850282.3418 | 778167.4844 | 789918.4336 | 794516.4258 | 781622.0469 |
| 102826.8203 | 105340 | 109209.5781 | 107050.5625 | 100568.4453 | 115882.1016 | 103052.1953 |
| 12069.68555 | 13516.25391 | 10323.64063 | 12159.49219 | 8100.111816 | 11005.35254 | 8829.860352 |
| 593015.625 | 613404.0625 | 672114 | 670100 | 621874.625 | 617347.4375 | 646079.375 |
| 1021643.125 | 1055115.75 | 1161680.25 | 1000371.313 | 737395.1875 | 730875.25 | 752826.0625 |
| 91588.50781 | 90181.03906 | 93772.21875 | 85369.78906 | 67703.66406 | 66568.98438 | 70240.75781 |
| 50254.26953 | 47807.66406 | 51793.55469 | 48621.83203 | 49323.79688 | 46394.46484 | 45168.12891 |
| 159966.7031 | 124384.125 | 172939.4063 | 137227.1875 | 118644.2031 | 135136.2344 | 122838.1953 |

| | | | | | | |
|---|---|---|---|---|---|---|
| 175199.625 | 172116.5625 | 187263.2344 | 189852.5625 | 136887.5313 | 144475.8906 | 144642.4219 |
| 13270.18066 | 19268.31836 | 12154.8252 | 10187.88477 | 9639.698242 | 8603.84375 | 9304.811523 |
| 2640.025879 | 4981.309082 | 801.9493408 | 810.9163818 | 4399.336426 | 3231.46875 | 4839.636719 |
| 88197.41406 | 95578.02344 | 86213.42969 | 84158.52344 | 79484.05469 | 83325.82031 | 69508.65625 |
| 43902.19141 | 43697.86328 | 42361.07031 | 43168.11719 | 37338.22266 | 66895.41406 | 48685.78516 |
| 133375.2344 | 126758.6328 | 158432.5938 | 146752.8438 | 132643.1875 | 133198.2188 | 125353.2734 |
| 58591.30469 | 78489.6875 | 49790.94141 | 45634.65234 | 51006.67578 | 41296.66797 | 49346.77344 |
| 69683.91406 | 56362.68359 | 75506 | 66776.34375 | 46541.45703 | 51649.22266 | 49990.59375 |
| 186112.1563 | 189277.125 | 194559.9063 | 213818.5156 | 199945.6875 | 187167.6563 | 196201.0781 |
| 2723327 | 2938936.5 | 2706052.5 | 2962056.25 | 3088932.5 | 3401503.5 | 3141967.25 |
| 16505.34961 | 17969.59375 | 15160.90234 | 16868.0957 | 13122.69043 | 12391.20996 | 11739.4873 |
| 14398.78516 | 15892.49707 | 13067.81055 | 13554.55078 | 29051.07422 | 22811.39844 | 29901.33984 |
| 7254.845215 | 6733.089355 | 5842.817383 | 5471.33252 | 7530.76416 | 8683.576172 | 9418.149414 |
| 35554.05078 | 37990.71875 | 39630.98047 | 33263.17969 | 43068.66016 | 41487.67188 | 39846.55078 |
| 616163.6875 | 656113.5625 | 658633.5625 | 711863.375 | 874587.625 | 873020.6875 | 909167.6875 |
| 1027903 | 1032744.875 | 945381.0625 | 825503.5625 | 1327750.875 | 1582505.125 | 1562381.75 |
| 14009.69141 | 16782.19531 | 13587.90137 | 13580.41406 | 18009.375 | 16359.74121 | 16764.66797 |
| 3978.078125 | 2364.696533 | 11897.84668 | 6918.692871 | 8875.21582 | 3626.915771 | 27288.29492 |
| 83111.90625 | 89013.98438 | 84187.15625 | 79835.79688 | 90404.28125 | 95919.8125 | 95175.95313 |
| 40653.1875 | 41741.38672 | 38646.51563 | 40931.25391 | 43380.33984 | 43015.25391 | 43957.01953 |
| 6907.004395 | 6001.946289 | 7297.883301 | 5027.644043 | 2584.769775 | 4795.216309 | 3140.477295 |
| 15032.95605 | 14792.64844 | 16165.1084 | 15460.29004 | 15321.39941 | 13115.16406 | 13240.02148 |
| 2051441.375 | 2075792.25 | 2179381 | 2300506.25 | 2572278.5 | 2504275 | 2682264 |
| 11363.38867 | 12600.91699 | 10926.24316 | 9700.225586 | 11952.27148 | 12096.35449 | 12686.86328 |
| 11443.84766 | 9820.795898 | 9827.003906 | 11477.33105 | 7122.251953 | 7994.631836 | 9550.119141 |
| 65434.98828 | 70766.89063 | 66126.625 | 66672.375 | 66389.94531 | 66738.78906 | 64199.32813 |
| 12960.9502 | 11276.40625 | 14921.99316 | 11676.27539 | 18516.82617 | 25944.82422 | 22138.41016 |
| 9809.486328 | 11681.65137 | 9047.087891 | 9278.714844 | 9785.956055 | 9761.194336 | 8166.194336 |
| 55517.38281 | 54109.55469 | 54736.46875 | 55304.75 | 59940.66016 | 56725.12891 | 61712.96875 |
| 12662.22949 | 13616.23633 | 11612.04395 | 11723.71777 | 15643.42969 | 15796.09082 | 17460.51758 |
| 21249.86719 | 21106.10938 | 23137.83398 | 20047.25977 | 23615.61523 | 23235.47266 | 21617.55273 |
| 9951.357422 | 10188.3623 | 8276.602539 | 12780.22461 | 12272.57227 | 11370.69336 | 10301.32129 |
| 19789.85156 | 24853.29292 | 23384.54492 | 19940.37891 | 44525.22321 | 52995.05747 | 46774.76022 |

| | | | | | | |
|---|---|---|---|---|---|---|
| 10325.57324 | 11487.80566 | 11510.94336 | 11686.88672 | 12715.02734 | 14795.83496 | 12903.28418 |
| 5588.530762 | 5444.893555 | 5194.408691 | 5004.085449 | 5199.412109 | 6838.71582 | 6737.463379 |
| 56912.93359 | 66198.57813 | 52918.4375 | 60857.06641 | 57415.18359 | 38563.21875 | 50229.59766 |
| 15153.71777 | 14645.91699 | 19819.53516 | 24052.54492 | 33709.19141 | 21253.78125 | 21526.41016 |
| 26477.81055 | 27957.09766 | 28468.33984 | 25636.89648 | 43845.50391 | 46053.24609 | 46535.88672 |
| 21411.87891 | 24130.00781 | 21575.53711 | 22238.49609 | 19200.43359 | 18074.14648 | 18449.80859 |
| 508437.5625 | 541373.5 | 494530.1563 | 477268.875 | 428844.7188 | 420133.9063 | 423803.375 |
| 89993.23438 | 102366.4766 | 92990.38281 | 94139.05469 | 122491.3359 | 103254.3672 | 110834.5313 |
| 71882.90625 | 77252.72656 | 75105.6875 | 73091.30469 | 49596.21875 | 53446.14844 | 49796.38281 |
| 143540.5938 | 140247.5625 | 152372.7188 | 155201.9688 | 176414.5938 | 169121.5938 | 178111.3594 |
| 214065.9063 | 250999.75 | 205983.6563 | 218369.0469 | 360767.2188 | 381131.875 | 369583.875 |
| 4778.788086 | 2525.977539 | 4475.981445 | 4239.860352 | 5995.228516 | 8420.34668 | 7409.425781 |
| 273507.5938 | 275398.375 | 299356.3125 | 294347.1563 | 211038.625 | 213599.9531 | 211189.2188 |
| 185002.8125 | 227676.5938 | 207274.8125 | 228485.9531 | 258251.9531 | 219299.0156 | 199675.875 |
| 595851 | 677339.8125 | 623979.6875 | 552639.875 | 602047.25 | 628702.125 | 619049.1875 |
| 6072.307129 | 7115.620117 | 5002.429199 | 5775.931641 | 5490.431152 | 5359.295898 | 5233.402344 |
| 355893.1563 | 373292.125 | 351897.9063 | 336643.5313 | 355679.4063 | 333073.7188 | 350830.1563 |
| 86812.86719 | 92710.5 | 88114.39844 | 81281.14844 | 70237.19531 | 61835.89453 | 64848.41797 |
| 30760.79688 | 30335.98438 | 30883.57031 | 38738.60156 | 69903.47656 | 52132.3125 | 57746.87109 |
| 1760.361694 | 2969.689941 | 1992.146118 | 3366.62793 | 2991.016113 | 2534.290283 | 2623.286133 |
| 1470659.875 | 1675935.375 | 1230853.375 | 1423798.5 | 1489818.875 | 1403072.625 | 1200035.25 |
| 7143761 | 6375264 | 7272399.5 | 7901351.5 | 5782542.5 | 5078779 | 5814789 |
| 62745.26563 | 67599.59375 | 66408.5625 | 65439.81641 | 40853.35156 | 32001.88281 | 32439.89844 |
| 40534.375 | 49457.65625 | 41140.75781 | 37143.08594 | 78784.38281 | 65208.66406 | 74836.9375 |
| 163369.9063 | 149646.6719 | 170755.875 | 160923.5781 | 155253.0156 | 140019.9688 | 157814.4844 |
| 802408.0625 | 879969.8125 | 810751.25 | 785903.5 | 801272.625 | 785474.4375 | 753766.5 |
| 227074.5625 | 207863.7969 | 269133.4063 | 229983.1563 | 194350.7656 | 191359.3906 | 253460.3281 |
| 69903.10938 | 62783.59766 | 75096.10156 | 69133.92188 | 63753.98828 | 82677.5625 | 64686.86719 |
| 56179.85156 | 55002.80859 | 52120.41797 | 57682.98438 | 53678.29297 | 51598.65234 | 51408.73828 |
| 47924.46094 | 52965.90234 | 50729.05859 | 46337.01953 | 73533.86719 | 68347.27344 | 71040.52344 |
| 137285.8906 | 120982.6484 | 141299.4844 | 147286.2656 | 193143.375 | 195513.5781 | 212839.7813 |
| 61370.65234 | 67549.84375 | 61417.23438 | 68079.30469 | 84359.16406 | 84683.41406 | 83802.75781 |
| 511268.1836 | 525135.2617 | 498588.5859 | 512843.4336 | 675847.6719 | 816475.4922 | 798601.5586 |

| | | | | | | |
|---|---|---|---|---|---|---|
| 127114.4609 | 149854.6094 | 161451.8906 | 153506.0313 | 159351 | 137030.5781 | 127590.7813 |
| 1346.880371 | 1953.883179 | 1349.555542 | 1325.781494 | 2908.723389 | 2359.739746 | 1764.863892 |
| 105771.8594 | 105039.3984 | 106229.4922 | 109344.2656 | 109859.25 | 98665.5 | 103355.4688 |
| 161197.1875 | 159178.5156 | 160728.7344 | 175105.7656 | 178516.8281 | 194557.3281 | 185735.7656 |
| 39570.52344 | 43609.55469 | 34392.71875 | 37020.21094 | 78525.32031 | 54942.07813 | 71338.53125 |
| 24936.55664 | 22962.94141 | 24379.32617 | 23067.20117 | 32076.04492 | 30586.94531 | 28332.09375 |
| 12233.86816 | 12763.79883 | 12276.3457 | 11954.18262 | 16138.18652 | 15856.75293 | 13745.81738 |
| 246541.6563 | 279124.5 | 219834.2813 | 214480.2188 | 149139.1406 | 156584.1875 | 131378.1094 |
| 197430.25 | 189531.9844 | 201164.9375 | 219997.4063 | 316634.75 | 270077.0625 | 300759.4063 |
| 496377.2188 | 517585.6563 | 471276.8438 | 461782.4063 | 359482.4688 | 394801.5313 | 332665.9063 |
| 167460.8906 | 168900.9844 | 160123.9375 | 164426.1563 | 169636.4844 | 172388.0313 | 173997.4844 |
| 127124.0781 | 146786.3281 | 142277.8594 | 139543.4375 | 142941.2188 | 130275.3359 | 117286.7188 |
| 231019.3906 | 229249.0781 | 231545.5469 | 223852.6406 | 244763.5625 | 288885.875 | 276997.9063 |
| 37864.29688 | 35476.20313 | 36911.48828 | 36621.19531 | 32755.26172 | 34386.02344 | 35965.60938 |
| 24168.13672 | 22855.10938 | 26099.7832 | 22177.27734 | 28252.79492 | 26046.5 | 30059.22461 |
| 20345.95508 | 16263.5625 | 23739.25977 | 22752.74219 | 25990.1875 | 22263.04883 | 22818.33203 |
| 45887.11719 | 80934.36719 | 43433.65625 | 58872.01172 | 42568.67188 | 50362.83594 | 57686.09766 |
| 111418.4609 | 123565.1719 | 117112.6172 | 104500.3125 | 120255.9375 | 117886 | 114630.0781 |
| 1714.760986 | 2317.804199 | 732.8544312 | 873.1708374 | 1582.530273 | 2082.27002 | 2200.370117 |
| 50243.92578 | 91944.875 | 43696.04297 | 32828.90234 | 72553.73438 | 81743.97656 | 59300.55859 |
| 15018.66406 | 13535.62988 | 14996.93652 | 17090.1582 | 12534.16406 | 12291.00488 | 12589.97168 |
| 29996.76855 | 33684.6543 | 32219.77197 | 30005.76123 | 36300.91699 | 35486.85156 | 34297.36133 |
| 16089.95703 | 18786.49609 | 17274.55273 | 14794.70605 | 18391.57617 | 18887.05664 | 16703.83398 |
| 19076.6543 | 18612.84375 | 21204.64258 | 19365.70313 | 13744.72266 | 11522.08105 | 12726.69043 |
| 17751.7793 | 18883.01953 | 18961.08984 | 16744.75195 | 17634.87695 | 18488.19141 | 18666.15039 |
| 292087.2188 | 315211.4688 | 302992.6563 | 305196.5938 | 321975.4063 | 318240.2813 | 340068.1875 |
| 90133.95313 | 91952.19531 | 94904.25 | 95444.86719 | 93601.76563 | 97946.17969 | 99795.9375 |
| 48596.49219 | 47769.26953 | 47981.77734 | 54175.50781 | 54172.12109 | 44840.3125 | 50759.11328 |
| 48695.04297 | 44966.43359 | 45589.33984 | 47167.44141 | 46102.90234 | 50215.77734 | 47056.85547 |
| 146141.0313 | 138686.4375 | 143287.8594 | 160123.0625 | 168330.0938 | 161242.9688 | 165473.1875 |
| 510191.5625 | 524795.9375 | 474548.125 | 556480.5625 | 849060.1875 | 685984.9375 | 777892 |
| 994021.9375 | 1024492.625 | 956274.875 | 1006433.375 | 977601.1875 | 893529.5 | 916477.375 |
| 17184.14453 | 12332.22754 | 21914.51758 | 18631.16211 | 18138.99805 | 14059.98633 | 18233.34766 |

| | | | | | | |
|---|---|---|---|---|---|---|
| 101943.3359 | 113566.8203 | 103906.4219 | 102452.1328 | 129413.6563 | 124809.0234 | 122755.125 |
| 164174.125 | 229160.0625 | 129674.2656 | 113073.9766 | 205151.25 | 195374.9063 | 218444.0625 |
| 57002.42188 | 47700.125 | 55889.67578 | 52107.49219 | 64207.15625 | 72451.125 | 61374.49219 |
| 242823.1719 | 294970.0313 | 217037.5 | 207687.7188 | 267181.9375 | 269101.8125 | 278170.4375 |
| 185396.6875 | 185524.4063 | 192488.9375 | 188855.0313 | 133805.2969 | 137879.4063 | 138016.1563 |
| 21643.4707 | 16966.24219 | 21932.44141 | 23065.05469 | 21378.875 | 19560.27148 | 23653.35156 |
| 6037.135742 | 5845.390137 | 5225.381348 | 7202.673828 | 3820.731201 | 4583.959473 | 5044.193848 |
| 63929.10547 | 65869.10156 | 60628.20313 | 72747.11719 | 107093.4219 | 75022.10938 | 89244.02344 |
| 13819.19336 | 19427.83203 | 12417.28613 | 11372.26172 | 12318.28906 | 11223.93359 | 12313.53711 |
| 54444.80859 | 62811.76953 | 71171.5 | 71426.23438 | 56755.62891 | 65092.82422 | 71331.34375 |
| 26448.36133 | 24521.22266 | 24378.38086 | 25553.22656 | 19242.51367 | 27340.3418 | 22619.14453 |
| 7496.304688 | 6126.112305 | 7497.748047 | 8837.140625 | 4584.481445 | 6912.508789 | 5925.586914 |
| 11015.48535 | 22193.71875 | 7967.588867 | 6423.908691 | 10478.92969 | 2399.845215 | 6038.003418 |
| 15607.18262 | 14746.45996 | 13760.17969 | 13555.48438 | 12653.32129 | 14801.08496 | 13333.47949 |
| 19423.24805 | 18747.97461 | 19807.55469 | 15174.21875 | 18040.8418 | 22571.13867 | 19816.1582 |
| 10678.38672 | 12101.12402 | 9746.766602 | 11841.40527 | 31553.64844 | 22288.70703 | 25094.75391 |
| 217277.25 | 218203.5313 | 198749.1094 | 220996.1875 | 395679.7813 | 324510.5625 | 351874.2813 |
| 230209.1563 | 232240.6875 | 231284.3281 | 233120.125 | 229686.1875 | 251790.8125 | 233441.1719 |
| 64957.52734 | 68946.89844 | 70883.50781 | 74976.97656 | 77632.97656 | 77852.07813 | 75660.33594 |
| 138759.0156 | 131839.1719 | 148561.9531 | 148232.8438 | 129961.4766 | 133061.8438 | 130440.7734 |
| 1330124.125 | 1447995.625 | 1188971.75 | 1169054.875 | 1237338.75 | 1320412.625 | 1346888.125 |
| 21425.26563 | 22665.01758 | 17751.61719 | 20493.49805 | 21841.94531 | 21068.76953 | 23014.44727 |
| 87912.25 | 76761.21875 | 114044.6328 | 125047.0859 | 86949.47656 | 72220.76563 | 73445.39844 |
| 27869.87109 | 28719.86523 | 28836.81641 | 30038.16797 | 38958.70313 | 37141.14844 | 39296.79688 |
| 41223.88672 | 41639.95703 | 43589.06641 | 45397.98438 | 44678.02344 | 46584.74219 | 46107.60156 |
| 28831.3457 | 34526.90625 | 28863.41504 | 27163.21973 | 27979.11621 | 33934.86426 | 28254.67578 |
| 47826.83984 | 66585.66016 | 45706.13867 | 43605.00977 | 44558.33203 | 127196.3672 | 58392.94922 |
| 51227.65625 | 54393.53125 | 53185.17188 | 59748.33203 | 51042.40234 | 48663.41797 | 46643.46094 |
| 152756.5781 | 174601.3438 | 132596.4531 | 133694.9844 | 62023.80078 | 63081.56641 | 57696.94141 |
| 11797.87793 | 18143.50195 | 11529.5 | 10680.77148 | 38244.80078 | 23163.72266 | 31565.87891 |
| 348723.9375 | 410839.0938 | 307387.6875 | 282351.6563 | 277461.4688 | 303893.9688 | 289789.5625 |
| 52391.01953 | 56321.4375 | 52020.80469 | 56770.46875 | 68545.28906 | 58614.33594 | 64073.15625 |
| 324160.3438 | 317432.2188 | 339239.8438 | 337873.5938 | 331233 | 331524.5313 | 344724.8125 |

| | | | | | | |
|---|---|---|---|---|---|---|
| 5299.711426 | 6977.218262 | 4981.148438 | 4961.475586 | 5985.11377 | 6952.994629 | 5942.036133 |
| 82883.32031 | 79992.08594 | 82966.79688 | 81614.03906 | 76250.70313 | 81446.11719 | 84748.25 |
| 28320.71484 | 26667.30664 | 32948.83203 | 35034.42188 | 35680.75391 | 37613.375 | 37724.1875 |
| 1215448.875 | 1171200.875 | 1162972.375 | 1023679.5 | 952714.6875 | 1031184.875 | 966372.5 |
| 21747.04688 | 26875.65234 | 23364.95703 | 23117.24219 | 25024.08984 | 23127.34961 | 24115.46484 |
| 11241.00879 | 12860.69629 | 10456.16992 | 9511.341797 | 15818.08203 | 14428.71387 | 15339.03125 |
| 3048.133789 | 3461.61792 | 2574.97583 | 2519.87207 | 2032.621582 | 3201.895264 | 1736.378174 |
| 105756.6797 | 94432.20313 | 96991.17969 | 114506.2891 | 110787.5313 | 109910.7891 | 112207.6797 |
| 16943.05078 | 21090.51758 | 15453.78906 | 11732.58691 | 19811.7168 | 25581.65039 | 18437.18164 |
| 111418.4219 | 114568.6953 | 120078.4844 | 114408.6563 | 84261.30469 | 91005 | 83124.95313 |
| 166895.5313 | 153845.1875 | 160336.8438 | 157769.6406 | 173502.2813 | 163238.8438 | 171533.1563 |
| 1054036.375 | 1124135.625 | 924148.3125 | 920837.125 | 921068.8125 | 1010902.5 | 934435.6875 |
| 89768.5625 | 96580.16406 | 93224.03906 | 85382.85156 | 113230.4922 | 100801.7969 | 108533.2422 |
| 25510.01758 | 27116.36523 | 24174.08008 | 27643.47461 | 33428.96875 | 33648.30859 | 31459.5293 |
| 48261.71094 | 46621.11328 | 45604.74219 | 54729.90234 | 64937.94922 | 63076.29297 | 62494.10547 |
| 116659.1172 | 87172.80469 | 140065.8906 | 135283.1094 | 133011.0313 | 114878.9531 | 130598.0313 |
| 13860.33398 | 8969.525391 | 13430.95215 | 12517.67676 | 16521.61719 | 19197.46289 | 20033.24219 |
| 223615.8438 | 243880.0313 | 225423.6719 | 202487.6875 | 160668.5313 | 148063.2969 | 155046.1094 |
| 96559.72656 | 88229.32813 | 90897.96094 | 93768.67188 | 62510.93359 | 69719.86719 | 63778.45313 |
| 19072.82422 | 20500.15625 | 19355.20508 | 20089.98633 | 17013.06641 | 18535.18945 | 15307.8418 |
| 1287119.672 | 1217107.484 | 1171746.109 | 1174158.188 | 1228181.453 | 1265276.016 | 1241158.656 |
| 7691.489258 | 9771.865234 | 6484.391602 | 8372.341797 | 7973.86084 | 6862.543457 | 9190.355469 |
| 10865.19434 | 12119.36426 | NA | 11016.28809 | 9300.806641 | NA | 8694.532227 |
| 302444.625 | 228282.3125 | 309638.2188 | 302067.5938 | 308725.1563 | 331087.0938 | 311246.9375 |
| 21246.6543 | 25757.91797 | 22644.83984 | 20481.45117 | 19005.25195 | 20632.89844 | 20940.67578 |
| 92975.17188 | 102343.0938 | 86878.36719 | 75955.55469 | 137191.0313 | 134320.9219 | 129742.7656 |
| 7872.658691 | 7061.820313 | 7813.516602 | 7797.158203 | 6305.37207 | 6663.574707 | 7128.20459 |
| 50726.53906 | 47634.125 | 44383.62109 | 43675.32031 | 61405.75781 | 59373.66406 | 59209.92188 |
| 31907.17383 | 33212.16797 | 32583.91797 | 31794.38086 | 39186.47656 | 39541.79688 | 38579.55469 |
| 586930 | 563636.625 | 575981.125 | 608329.8125 | 844925.5625 | 755277.1875 | 789839.5 |
| 14875.44629 | 15629.91309 | 15399.68652 | 16222.07031 | 16144.11914 | 15058.74512 | 15215.20801 |
| 99515.17188 | 90090.96875 | 105645.9531 | 109171.8281 | 131483.0156 | 110211.1484 | 128450.2578 |
| 51544.66406 | 45515.51172 | 51488.17578 | 57859.29688 | 48266.67578 | 51783.53906 | 54947.13281 |

| | | | | | | |
|---|---|---|---|---|---|---|
| 18247.10352 | 20488.90234 | 18641.26563 | 18219.66602 | 15987.88379 | 17646.42773 | 16857.11719 |
| 19866.98047 | 18821.35742 | 20654.89648 | 18227.88477 | 11859.5293 | 13511.5957 | 10188.8623 |
| 182392.7813 | 195206.9375 | 172297.0313 | 172936.25 | 201658.2031 | 199634.5781 | 198279.9219 |
| 218733.2031 | 205903.2031 | 235129.625 | 223635.6719 | 231689.2188 | 235214.0313 | 243869.2656 |
| 5703.31543 | 3779.575684 | 6080.861328 | 8457.269531 | 3649.196777 | 3779.760742 | 3743.715332 |
| 88386.80469 | 94362.41797 | 82848.99219 | 82149.61914 | 83738.70313 | 73316.53125 | 78004.52344 |
| 129283.0859 | 128075.0156 | 131993.2969 | 126940.5547 | 126828.2266 | 134354.0469 | 132560.5781 |
| 5177.545898 | 5245.04541 | 4911.701172 | 5425.049316 | 3634.332275 | 3203.19458 | 3543.499268 |
| 8771.030273 | 7521.599609 | 9953.594727 | 9394.731445 | 14062.98047 | 11270.81836 | 12324.75195 |
| 155417.5313 | 144051.75 | 158480.7813 | 159081.5469 | 211354.1094 | 215591.9219 | 230783.5625 |
| 119078.75 | 123711.9688 | 114531.6719 | 120581.2734 | 125750.7578 | 137182.1406 | 135198.4688 |
| 217404.6094 | 200226.1875 | 220837.875 | 221413.9375 | 135875.9219 | 148870.2188 | 139688.9375 |
| 323664.9375 | 378377.7813 | 340927.9063 | 354876.8438 | 329470.0625 | 467351.9375 | 371223.9688 |
| 1402858 | 1553210.25 | 1419101.875 | 1358922.5 | 1400390.25 | 1484062.5 | 1309714.125 |
| 158980.9316 | 160007.0234 | 176440.3613 | 175338.8965 | 155396.3887 | 141766.1221 | 151662.2861 |
| 107867.3438 | 100341.5234 | 115990.0938 | 115533.8828 | 119199.9453 | 103738.9219 | 105700.3125 |
| 345469.375 | 347191.2188 | 342918.5938 | 367597.0625 | 425538.7188 | 524682.0625 | 462158.7813 |
| 13783.2832 | 12249.8291 | 13923.71777 | 14832.27051 | 17079.22852 | 15238.90137 | 16466.90039 |
| 4960.156738 | 4510.936035 | 4801.438477 | 4632.245117 | 8227.729492 | 4113.797852 | 4476.706055 |
| 50359.44531 | 93875.28125 | 27457.38672 | 39806.33984 | 97194.82031 | 84072.26563 | 92548.25781 |
| 80049.11719 | 69550.48438 | 72024.77344 | 71464.11719 | 66409.30469 | 66334.75 | 69967.55469 |
| 60015.26172 | 66324.24219 | 58780.86719 | 60361.84375 | 62238.98828 | 67658.14844 | 63245.20313 |
| 333015.3125 | 287475.1875 | 334811.5 | 321733.9688 | 238253.6875 | 254265.9531 | 284856.9375 |
| 12249.06836 | 16827.1582 | 14273.22559 | 11528.43262 | 9258.960938 | 7216.268555 | 8579.686523 |
| 913447.25 | 898884.6875 | 895723.9375 | 949466.5625 | 1124961.25 | 1106055.875 | 1194584.875 |
| 32327.32813 | 27457.25 | 33515.19922 | 31074.41797 | 29917.86133 | 29415.10156 | 33528.44922 |
| 55103.92188 | 57760.02344 | 54810.52734 | 58806.77734 | 40332.04688 | 46524.80469 | 43720.28516 |
| 594238.375 | 596436.4375 | 590531.25 | 601935.1875 | 602518.5625 | 560653.875 | 575937.3125 |
| 736238.5 | 834436.875 | 683430.0625 | 629009.4375 | 571629.125 | 605888.125 | 621271.1875 |
| 170746.4844 | 181560.9063 | 164263.6406 | 174603.0469 | 145850.25 | 122110.6719 | 123644.1172 |
| 4999.834473 | 6132.117188 | 3497.619873 | 4663.277832 | 2843.793213 | 2197.758301 | 2765.324707 |
| 60574.67188 | 70948.38281 | 55236.20703 | 61642.81641 | 105751.9297 | 94010.78125 | 85585.1875 |
| 116293.2344 | 125610.6016 | 98644.19531 | 100470.5703 | 50418.88281 | 51546.58203 | 42827.71094 |

| | | | | | | |
|---|---|---|---|---|---|---|
| 40841.81641 | 37938.40234 | 40331.74609 | 42837.625 | 50995.83594 | 56700.37109 | 51811.34766 |
| 226515.5313 | 226979.6563 | 203732.6094 | 211630.4688 | 177123.1406 | 175954.0781 | 165579.125 |
| 13340.2793 | 14057.16699 | 14177.1084 | 11378.38867 | 15915.21191 | 14425.78809 | 14012.23828 |
| 11759.97168 | 13385.56445 | 10991.89063 | 11876.43457 | 13854.86035 | 12190.61133 | 10895.82813 |
| 168287.8281 | 172416.6875 | 167986.2656 | 163797.5469 | 190571.2813 | 176117.6875 | 174344.7813 |
| 102736.8359 | 95921.67188 | 106843.1328 | 105627.0469 | 138672.0313 | 123411.6953 | 123874.8672 |
| 48234.37109 | 49899.64844 | 46043 | 45992.75391 | 44356.83594 | 48549.125 | 46753.04297 |
| 7661.370605 | 8077.612793 | 8097.217285 | 8617.484375 | 10370.83887 | 8901.325195 | 7968.536133 |
| 848273.6875 | 823511.625 | 842930.0625 | 874295.8125 | 920508.25 | 789991.25 | 835117.875 |
| 54772.52344 | 45608.19922 | 51925.64844 | 62839.67969 | 66753.46094 | 96244.71875 | 67771.71875 |
| 550231.625 | 619393.625 | 473151.5 | 507664.0313 | 516898.125 | 483696.0938 | 485191.7813 |
| 305429.375 | 275060.6875 | 294848.2188 | 317959.1563 | 275085.625 | 288326.4688 | 301626.8125 |
| 110519.4141 | 119449.9297 | 107805.3906 | 113413.6172 | 134707.125 | 142007.9219 | 138648.125 |
| 40508.08984 | 37384.57031 | 38430.98047 | 41160.79688 | 26844.55469 | 27884.37109 | 25936.62891 |
| 6435.215332 | 7617.23291 | 6119.971191 | 5554.993164 | 4447.395996 | 6384.227051 | 5744.070313 |
| 4355.321777 | 7589.733887 | 5072.050293 | 4884.955078 | 9051.547852 | 6396.144043 | 8219.140625 |
| 4647.276855 | 4524.126465 | 5195.529297 | 4958.547852 | 3675.970215 | 5287.918457 | 3851.351807 |
| 6716.822266 | 8052.568359 | 6638.092773 | 8452.588867 | 8126.918457 | 9395.657227 | 9175.832031 |
| 109058.4453 | 98433.57031 | 117873.8906 | 113763.7422 | 120677.1797 | 111765.6328 | 117509.7266 |
| 1063117 | 996107.9375 | 1047292.938 | 1025034.313 | 935980.625 | 872049.9375 | 881198.6875 |
| 41191.91016 | 42620.39063 | 44840.05078 | 36765.05469 | 28969.42188 | 28963.02734 | 31565.70898 |
| 42725.0625 | 44224.0625 | 46860.07422 | 43012.00781 | 45789.65234 | 59484.67188 | 54641.62891 |
| 21320.35547 | 21910.67383 | 20221.11914 | 21148.75781 | 25567.91016 | 27461.59961 | 25497.97461 |
| 5731.457031 | 5863.062988 | 4924.921387 | 4897.119629 | 4097.455566 | 4161.938477 | 3299.596924 |
| 7552.82666 | 8723.338867 | 4675.246582 | 2667.73999 | 3395.342773 | 3522.801025 | 3397.198242 |
| 181562.6719 | 180505.75 | 177545.8594 | 171526.9063 | 144217.2813 | 156800.3281 | 139128.5938 |
| 10024.22656 | 8620.719727 | 7576.833008 | 9341.238281 | 4100.205566 | 4652.700195 | 5615.237793 |
| 3540.229492 | 3762.049805 | 3641.508301 | 3482.625732 | 2590.387939 | 1473.63623 | 1171.02771 |
| 73467.32813 | 84716.40625 | 72053.63281 | 71782.69531 | 71639.4375 | 67845.40625 | 67542.67969 |
| 147176.2188 | 151800.5469 | 156034.0781 | 148683.5625 | 147826.3281 | 177505.0781 | 151686.3594 |
| 23062.08203 | 30091.35352 | 27325.30859 | 26491.42773 | 30963.57422 | 30659.95117 | 29498.66406 |
| 16319.52441 | 24710.21484 | 26958.24414 | 12883.50098 | 11978.30859 | 23837.98047 | 15413.9248 |
| 13057.45508 | 12743.08398 | 10061.61914 | 12282.42773 | 16127.83008 | 15954.33105 | 14459.78223 |

| | | | | | | |
|---|---|---|---|---|---|---|
| 30516.62695 | 26440.71875 | 29355.27148 | 31456.54492 | 34015.625 | 31844.38477 | 35286.23438 |
| 12550.41113 | 12504.03906 | 15718.33203 | 17113.3457 | 19160.7168 | 17846.80273 | 17020.66406 |
| 45758.81641 | 46404.64063 | 42220.77734 | 48765.06641 | 43529.33984 | 42263.74609 | 41655.52734 |
| 128138.5078 | 152173.9219 | 114874.6328 | 76880.67969 | 114827.5469 | 213931.6875 | 157695.5 |
| 1759.913208 | 779.3104858 | 716.5263672 | 1350.344971 | 7288.137695 | 9898.214844 | 11952.96387 |
| 299091.9063 | 295184.625 | 257815.5 | 287856.9375 | 262753 | 266084.2188 | 264312.0313 |
| 72369.01563 | 86139.46875 | 64623.63672 | 62843.89063 | 55355.94922 | 48903.32813 | 42642.20313 |
| 170928.5938 | 207097.0313 | 183676.6875 | 178508.9219 | 208912.2344 | 211131.125 | 245386.8906 |
| 11373.12012 | 12256.55566 | 11035.49414 | 6046.226563 | 11561.7373 | 14903.83105 | 11355.43652 |
| 16634.56836 | 16119.5293 | 16190.85645 | 15196.27246 | 12883.33594 | 14870.57715 | 13567.13379 |
| 27834.70117 | 27015.06055 | 28659.95117 | 29636.27344 | 34465.42188 | 32246.19141 | 34476.84766 |
| 11771.73926 | 12065.05273 | 11229.1377 | 8882.091797 | 12073.77148 | 11528.08594 | 11774.32422 |
| 8253.554688 | 8714.962891 | 7414.394531 | 8600.9375 | 12469.28027 | 10832.56445 | 11904.21387 |
| 6788.389648 | 6625.428711 | 5859.068848 | 5415.403809 | 5456.671875 | 5381.091309 | 5207.606445 |
| 518140.3438 | 453501.8125 | 435755.2813 | 461409.1563 | 521428.8438 | 522342.4375 | 524016.375 |
| 14482.39355 | 14917.40039 | 15059.5293 | 14968.81348 | 19383.56641 | 23038.36914 | 20158.26172 |
| 93184.47656 | 91242.46094 | 93867.23438 | 91750.53125 | 90030.57813 | 98555.53906 | 93140.75781 |
| 24834.51563 | 22335.12695 | 22690.875 | 32333.57813 | 53086.92578 | 47173.44922 | 50130.25 |
| 27887.58594 | 26075.93945 | 26295.02344 | 30013.96484 | 35430.69531 | 32767.9707 | 33909.625 |
| 2554.155029 | 3037.47998 | 2267.774902 | 2630.333984 | 5326.996582 | 4163.182617 | 4183.665527 |
| 91343.71094 | 97184.65625 | 90465.8125 | 97327.75 | 87655.03906 | 88204.27344 | 90121.0625 |
| 18233.07031 | 16146.63184 | 18701.27734 | 19386.89453 | 14971.00586 | 13959.84375 | 14098.9707 |
| 98142.35938 | 117360.4922 | 95936.60938 | 86568.05469 | 99095.99219 | 99737.52344 | 93935.51563 |
| 1603303.625 | 1648663.125 | 1703437.25 | 1715302.75 | 1439371.5 | 1454768.375 | 1546286.5 |
| 14829.28809 | 14836.74902 | 16274.58105 | 14411.41797 | 12755.08691 | 13803.68848 | 12643.27832 |
| 13763.74023 | 16997.89258 | 12693.45996 | 10701.65332 | 8050.12793 | 6610.138184 | 7750.900879 |
| 71191.25 | 55110.71484 | 52784.62891 | 64781.36328 | 97599.98438 | 97963.46875 | 89794.19531 |
| 165285.1094 | 166990.9219 | 165320.5625 | 167543.1406 | 104229.3125 | 99056.11719 | 102909.8594 |
| 95848.74219 | 100241.6094 | 86719.69531 | 92012.27344 | 87354.91406 | 83649.51563 | 87583.42188 |
| 16506.52148 | 21234.41016 | 16026.4375 | 14276.36816 | 15893.84863 | 18866.32422 | 14396.84375 |
| 42326.98438 | 39974.28906 | 37732.14063 | 44684.03516 | 29684.85156 | 34836.77734 | 33156.24219 |
| 195500.9531 | 211146.7344 | 205339.25 | 213550.875 | 277791.0313 | 245516.5313 | 246935.1094 |
| 38994.49219 | 37175.14063 | 35225.73828 | 33928.66797 | 22072.28906 | 22195.17578 | 21618.125 |

| | | | | | | |
|---|---|---|---|---|---|---|
| 101602.3984 | 95819.60156 | 102281.0469 | 103695.1484 | 87452.14063 | 88364.53125 | 87898.85938 |
| 1175861.875 | 1145130.5 | 1158679.625 | 1111973.375 | 1188642.25 | 1220317.125 | 1273608 |
| 17285.51172 | 17344.82422 | 17109.47852 | 15589.73633 | 10263.31934 | 10250.43652 | 9562.375977 |
| 39954.76953 | 39623.89063 | 40334.03516 | 44096.90625 | 34851.66406 | 36589.03906 | 35476.13672 |
| 9393.865234 | 8849.952148 | 8923.182617 | 9848.740234 | 4248.820801 | 6379.896484 | 5905.801758 |
| 608984.375 | 645390.5625 | 592237.9375 | 624474.4375 | 579527.5 | 496834.9063 | 512954.7813 |
| 17762.5918 | 20194.57031 | 16162.06445 | 16862.73828 | 14854.41992 | 13883.28418 | 14557.74512 |
| 150911.5 | 169687.75 | 157886.375 | 156343.4063 | 172721.8906 | 140397.9688 | 147548.8438 |
| 474257.375 | 468063.5313 | 469427.7188 | 445971.5938 | 458177.6563 | 457934.0625 | 434223.0938 |
| 2066.234131 | 2230.29248 | 2367.955078 | 2407.955566 | 2321.960449 | 1830.574341 | 1920.724854 |
| 100041.8828 | 86309.23438 | 121705.5781 | 117312.4141 | 88086.25 | 70332.89063 | 76214.05469 |
| 459728.6875 | 455439.1875 | 446964.25 | 465587.4063 | 467089.4688 | 477765.5625 | 477986.625 |
| 4934.055664 | 5613.851074 | 5522.816895 | 4853.458496 | 5712.974609 | 4767.319824 | 3615.413086 |
| 367578.1875 | 385074.25 | 354291.9688 | 318742.6563 | 290775.0625 | 275273.6563 | 292954.6563 |
| 5397.776855 | 7663.132813 | 4307.116699 | 5110.080078 | 8587.798828 | 8491.755859 | 8838.944336 |
| 44416.16016 | 40265.97656 | 46757.58984 | 49429.23828 | 45290.23828 | 45381.38281 | 52833.74609 |
| 115811.1094 | 119848.2344 | 115099.7734 | 117359.3047 | 114389.8906 | 109029.5859 | 105989.2109 |
| 15092.98633 | 15055.37988 | 13501.66992 | 12640.0498 | 7211.536133 | 9168.523438 | 7181.130371 |
| 98794.94531 | 103595.2109 | 94495.24219 | 104323.5859 | 181334.5781 | 161021.3906 | 176274.1875 |
| 10457.52734 | 10176.74805 | 9740.681641 | 11084.78906 | 9203.768555 | 8590.880859 | 11024.94531 |
| 17885.90625 | 15625.04199 | 15920.97949 | 17696.36133 | 15140.96387 | 10413.88477 | 13217.90527 |
| 5786.776855 | 5106.864746 | 6155.919434 | 6536.131836 | 3991.903564 | 3739.319336 | 4091.370605 |
| 316703.9063 | 349292.125 | 300053.9063 | 277253.8438 | 328454.75 | 359305.6563 | 346935.8125 |
| 3158.031738 | 1953.856445 | 3026.695801 | 2489.225586 | 4530.567383 | 4203.586426 | 4707.347656 |
| 22173.28906 | 9280.955078 | 22398.43359 | 33516.06641 | 30592.5918 | 34536.98047 | 34420.82031 |
| 116487.9688 | 136468.4219 | 117308.9297 | 110701.25 | 172625.4844 | 201502.4063 | 171358.3594 |
| 85923.82031 | 96366.26563 | 85570.27344 | 79875.67969 | 84978.86719 | 93201.39844 | 89020.45313 |
| 16378.67773 | 17425.97656 | 16760.86719 | 18754.63086 | 11858.72363 | 8962.765625 | 11950.22656 |
| 10225.88574 | 9513.62793 | 7224.9375 | 5842.583496 | 3873.640625 | 5553.054688 | 3104.396484 |
| 44257.25391 | 43203.40625 | 48063.21094 | 47133.27344 | 46758.71484 | 50752.26953 | 48907.10938 |
| 1184.28833 | 1898.206787 | 666.18396 | 943.005188 | 1822.294556 | 2185.87793 | 2478.329346 |
| 801535.25 | 775155.4375 | 822133.1875 | 772591.375 | 654087.75 | 695309.1875 | 666263.125 |
| 72759.14844 | 73168.63281 | 72730.19531 | 72314.82031 | 70990.39063 | 65105.35938 | 67871.63281 |

| | | | | | | |
|---|---|---|---|---|---|---|
| 22573.02734 | 27751.04492 | 18850.69922 | 23598.23438 | 18913.43164 | 13529.23145 | 13128.9375 |
| 135598.8594 | 118295.0781 | 137085.1563 | 124149.0234 | 124465.9453 | 142283 | 128780.7109 |
| 92815.125 | 100988.7109 | 85088.40625 | 85111.57031 | 78536.86719 | 74445.4375 | 65094.32031 |
| 4881.849121 | 4779.194824 | 4718.506348 | 4158.970215 | 3431.372559 | 3014.310791 | 2609.174561 |
| 25341.94531 | 17227.77148 | 24117.75586 | 28340.1875 | 19994.61523 | 15721.62793 | 20141.6582 |
| 21803.33594 | 24206.4668 | 19713.67383 | 19515.05664 | 21719.42969 | 20428.29297 | 21947.2168 |
| 13749.29004 | 12531.4834 | 11346.65137 | 13042.49707 | 17636.75391 | 18966.42773 | 18682.9668 |
| 1175063.992 | 1102368.869 | 1064397.336 | 1007028.449 | 885886.4727 | 960604.4375 | 841678.7852 |
| 20176.42578 | 18276.19336 | 20344.71094 | 18359.04492 | 31372.29102 | 29473.91602 | 29533.73438 |
| 820295.125 | 1025990.938 | 759963 | 663957.3125 | 909913.8125 | 1044011.313 | 853051.5 |
| 22302.34766 | 27869.02539 | 17686.23828 | 10619.60156 | 20292.27344 | 16272.96484 | 17866.99414 |
| 130696.3594 | 129425.1484 | 129072.5469 | 124196.8359 | 125487.7656 | 128966.5469 | 122952.3906 |
| 14679.10938 | 13398.64258 | 14100.31152 | 13152.2002 | 10806.77344 | 11636.39063 | 11658.33691 |
| 73487.98438 | 44340.45313 | 70024.96875 | 73474.25781 | 86054.29688 | 84512.07813 | 89073.91406 |
| 60066.95313 | 39661.11719 | 68650.49219 | 78671.28906 | 94883.32031 | 88044.125 | 92113.125 |
| 642621.75 | 665641.6875 | 576157 | 617772.6875 | 599591.8125 | 536532.1875 | 576528.875 |
| 3497.370605 | 3994.464355 | 3351.791748 | 3482.360596 | 4563.732422 | 5755.150879 | 4982.354004 |
| 45086.98047 | 35100.89844 | 44356.10156 | 57702.07813 | 113374.2031 | 74668.125 | 86361.6875 |
| 290376.3125 | 339751.5 | 262909.625 | 283913.9375 | 394518.0938 | 337315.8438 | 350548.1875 |
| 57073.30859 | 53829.99219 | 50575.71094 | 50118.88672 | 47132.03906 | 47699.35547 | 48399.72656 |
| 11999.4707 | 11887.85645 | 9506.911133 | 6878.329102 | 8287.441406 | 4608.130859 | 9725.8125 |
| 124026.2188 | 121007.8516 | 115052.7031 | 112629.3984 | 87410.96875 | 88275.86719 | 87190.21094 |
| 2756067 | 2618934.25 | 2645317 | 2413230.25 | 2269327.5 | 2400404.5 | 2203433 |
| 13867.71973 | 14460.82813 | 4896.01416 | 9556.851563 | 54014.54297 | 44905.58984 | 43755.50781 |
| 190129.4375 | 157927.1719 | 192392.9844 | 196397.8438 | 168457.6094 | 166514.1406 | 180561.8438 |
| 39615.04688 | 39310.35547 | 40367.99219 | 40062.3125 | 34296.51172 | 31293.96289 | 37781.62109 |
| 103794.2891 | 102308.875 | 107635.0391 | 96867.14844 | 109691.3203 | 90080.03906 | 85048.14844 |
| 13297.45703 | 12680.88965 | 12394.87793 | 14247.99512 | 16433.23828 | 14903.15527 | 14316.39648 |
| 428822.4375 | 422094.1563 | 427753.5 | 426349.3438 | 414165.4063 | 395856.5 | 409675.7813 |
| 15832.97949 | 13627.42383 | 16233.42773 | 16383.58887 | 16067.34082 | 16496.95703 | 15308.75488 |
| 34320.12891 | 31350.50195 | 36337.26953 | 36655.12891 | 44210.34375 | 50246.38672 | 46343.15625 |
| 39545.31641 | 33729.07813 | 40040.55469 | 39733.67188 | 40379.26172 | 49705.99219 | 49499.69531 |
| 5717.046387 | 3605.55127 | 4581.269531 | 4078.09375 | 6439.594727 | 1829.20752 | 5213.893066 |

| | | | | | | |
|---|---|---|---|---|---|---|
| 98367.80469 | 92429.46875 | 85494.97656 | 90626.49219 | 75636.64063 | 80274.74219 | 73430.32031 |
| 2123120.828 | 1999185.063 | 2107008.273 | 2054043.305 | 1731032.82 | 1754476.656 | 1779583.133 |
| 5338.69043 | 6134.492188 | 5743.071289 | 5389.252441 | 10137.72168 | 8845.978516 | 10096.70703 |
| 100927.7969 | 100018.5547 | 103634.7891 | 98000.22656 | 87850.57031 | 87842.04688 | 83548.32813 |
| 56291.13672 | 51905.09766 | 57002.56641 | 58261.01563 | 60824.20313 | 73724.21875 | 70092.75781 |
| 18221.26367 | 12312.14258 | 24535.7168 | 27404.03125 | 20681.61133 | 21143.0332 | 24026.89063 |
| 386789.5938 | 418204.5938 | 388331.375 | 390323.1563 | 388409.1875 | 399666.1563 | 409260.875 |
| 867901.9375 | 851746.5625 | 774415 | 848801.4375 | 867003.625 | 828441.375 | 808475.875 |
| 9855.425781 | 11690.55176 | 8684.25293 | 8682.589844 | 11062.94141 | 8685.129883 | 9937.267578 |
| 29739.81445 | 29935.85938 | 29862.74609 | 29315.21289 | 23693.58398 | 21847.66992 | 23394.41211 |
| 264148.375 | 269856.7813 | 255713.6719 | 262607.7188 | 192971.5781 | 181179.375 | 186480.4844 |
| 20109.43945 | 19802.75977 | 18200.70117 | 14511.53223 | 15673.24805 | 22366.24609 | 18159.83984 |
| 20445.9707 | 23773.56055 | 20516.77539 | 23318.39453 | 25391.60352 | 22552.13672 | 22043.17969 |
| 82873.14844 | 67277.78125 | 78750 | 84882.08594 | 80295.21094 | 89155.71875 | 88572.3125 |
| 90475.61719 | 88799.89844 | 86217.94531 | 92106.96875 | 86199.32813 | 102517.8047 | 99494.07031 |
| 1738804 | 1596981.875 | 1654393.875 | 1807518 | 2154087.5 | 1979233.75 | 2222579.5 |
| 335025 | 310450.6563 | 301087.7188 | 283761 | 254598.9375 | 249556.7813 | 256328.5313 |
| 319497.125 | 344595.3438 | 300818.2813 | 300256.9688 | 257131.6094 | 244653.3594 | 253272.4375 |
| 25013.35156 | 27097.50586 | 21954.83789 | 23006.19531 | 20130.25586 | 23604.54297 | 19796.70703 |
| 314718.5176 | 310820.2017 | 314243.7847 | 291297.3291 | 280396.6328 | 276713.0967 | 286962.6973 |
| 99374.57813 | 75396.42969 | 100406.5313 | 89952.95313 | 122722.7656 | 102590.1484 | 119565.3594 |
| 5946.420898 | 5831.339355 | 5956.3125 | 6290.012207 | 3220.418213 | 4350.924805 | 3498.586426 |
| 3847.51709 | 708.4369507 | 5388.202148 | 5371.283691 | 8519.197266 | 10694.84961 | 11429.94629 |
| 20399.92969 | 20270.375 | 19058.51953 | 19111.9707 | 20729.71289 | 19627.65234 | 19880.69531 |
| 292833.9688 | 284145.3125 | 285135.9375 | 309160.875 | 265924.7188 | 248019.75 | 272472.7813 |
| 190153.8125 | 193648.8906 | 185054.9844 | 184450.6875 | 127845.2031 | 123892.3594 | 114294.3906 |
| 283326.625 | 283954.125 | 278907 | 268535.4375 | 205509.1094 | 211469.875 | 200340.0625 |
| 3821.901611 | 3196.850098 | 2546.545898 | 3997.179932 | 3360.865479 | 4234.369629 | 4014.018555 |
| 81735.6875 | 88549.625 | 76049.21094 | 77819.71875 | 55028.69141 | 62738.01563 | 60724.53516 |
| 13741.11719 | 15370.19727 | 13222.92578 | 12871.06543 | 22916.4043 | 20221.57227 | 20560.72656 |
| 161172.25 | 138890.5781 | 142708.1563 | 146329.625 | 152873.375 | 149414.6719 | 151589.625 |
| 44362.98438 | 45591.23047 | 45749.36719 | 47749.1875 | 65696.10938 | 58456.95313 | 64141.5 |
| 43421.19141 | 39325.48047 | 44954.34766 | 42658.10938 | 31532.04102 | 32881.83984 | 34434.42188 |

| | | | | | | |
|---|---|---|---|---|---|---|
| 118152.2891 | 131583.9375 | 115730.4453 | 119907.9922 | 131566.875 | 122517.8828 | 132209.0781 |
| 1622022.75 | 1892253.75 | 1458179.375 | 1524614.875 | 1262592.875 | 1324583 | 1271926.25 |
| 1083275.25 | 1474432.5 | 1369534.125 | 276685.875 | 526379.5625 | 844906.8125 | 342271.5625 |
| 31637.50586 | 29962.58008 | 57735.25781 | 53473.25781 | 32968.24609 | 33724.10938 | 38985.66016 |
| 94402.71094 | 94671.23438 | 93020.58594 | 89632.125 | 88398.92969 | 82992.07031 | 87311.89063 |
| 76936.92188 | 73722.51563 | 72201.29688 | 65852.14063 | 73504.64063 | 77622.32031 | 78722.92969 |
| 821951.5 | 705979.125 | 899343.4375 | 787896.8125 | 688773.3125 | 795008.5625 | 773940.875 |
| 11902.07031 | 10161.22168 | 11367.34375 | 10985.73438 | 9735.234375 | 9227.322266 | 10612.96484 |
| 226477.4688 | 183881.0781 | 209839.125 | 209650.2344 | 155912.0313 | 191833.2813 | 162532.0781 |
| 54852.89063 | 64066.02734 | 46441.98828 | 46970.64453 | 47240.07813 | 49673.71875 | 47112.94531 |
| 102818.9922 | 99990.25781 | 99201.49219 | 105132.0078 | 92948.35156 | 101126.9609 | 97121.10938 |
| 58167.03125 | 54509.21094 | 55695.35156 | 52266.54688 | 51064.17578 | 50778.53906 | 48218.09766 |
| 7753.755859 | 6410.538574 | NA | 8297.201172 | 7269.750488 | 5931.363281 | 7546.385742 |
| 41600.19531 | 44328.89453 | 33272.79688 | 33318.54688 | 31492.69531 | 29476.15625 | 44480.93359 |
| 8746.241211 | 7162.039063 | 8220.707031 | 6610.361816 | 6589.194336 | 7600.05127 | 6626.448242 |
| 445673.4688 | 419160.75 | 356252.5313 | 285419.3438 | 284113.625 | 449738.625 | 324549.4688 |
| 208131.3438 | 210058.4219 | 187293.1406 | 193345.3281 | 164001.6875 | 176170.3438 | 166660.3438 |
| 263276.125 | 268667.6875 | 272899.25 | 274030.0313 | 314328.4375 | 275062.5 | 290862.0625 |
| 899088.5625 | 868402.3125 | 859907.75 | 810652.125 | 806970.25 | 922858.4375 | 773087.5 |
| 252162.4063 | 296458.5 | 266738.2188 | 231550.2188 | 180083.25 | 188002.3438 | 159322.6875 |
| 214943.9688 | 191742.9375 | 211133.1406 | 226147.3438 | 147609.4375 | 133863.9063 | 139973.625 |
| 124266.5156 | 134390.9063 | 99136.625 | 84767.875 | 111482.8594 | 110673.0391 | 121168.875 |
| 524325.625 | 675646.3477 | 345453.125 | 319037.7441 | 325616.6655 | 795991.0186 | 386968.6445 |
| 147232.1094 | 169900.8438 | 114975.9609 | 113274.0156 | 152361.2969 | 148083.9844 | 160323.1563 |
| 204651.6719 | 216244.3125 | 198033.5469 | 200553.375 | 225922.9688 | 219102.2344 | 210701.4219 |
| 192621.4063 | 184195.5156 | 176047.1563 | 187093.5156 | 196989.9063 | 187537.3438 | 185599.4375 |
| 59395.42578 | 59213.54297 | 53315.00781 | 49306.35156 | 52883.28125 | 54716.74219 | 57554.34375 |
| 2194411.25 | 2035675.875 | 2081052.125 | 1909283.25 | 1605418.25 | 1746379.875 | 1631092.625 |
| 22366.82227 | 21144.25781 | 24612.86719 | 24821.98828 | 24605.33398 | 32675.35742 | 23180.65234 |
| 26645.5293 | 29088.50391 | 28112.32422 | 25155.51953 | 25884.94141 | 25115.86133 | 25599.60352 |
| 10223.33203 | 9821.486328 | 10677.18555 | 9901.180664 | 13462.54688 | 12376.14453 | 11539.67773 |
| 15408.27051 | 14138.89648 | 16014.46191 | 13744.87109 | 10071.27539 | 11405.52441 | 9701.255859 |
| 41306.98047 | 39934.24219 | 38234.51563 | 34217.89844 | 25511.87891 | 28601.80859 | 21865.00391 |

| | | | | | | |
|---|---|---|---|---|---|---|
| 436177.1875 | 380498.5625 | 419765.8125 | 413986.2188 | 403221.6875 | 371183.2813 | 393201.5938 |
| 9579.59668 | 8696.232422 | 9033.99707 | 7386.787109 | 7766.052246 | 8406.330078 | 7220.101563 |
| 525169.6875 | 569642.6875 | 516820.4688 | 486565.9688 | 632689.5 | 711375.5 | 717005.3125 |
| 1354291.5 | 1168917.125 | 1273319 | 1140901.625 | 981758.375 | 1070177.375 | 1007047.938 |
| 7304.015137 | 9232.845703 | 8681.52832 | 8860.701172 | 9906.049805 | 8335.679688 | 8920.1875 |
| 201474.3125 | 179529.9844 | 188886.0781 | 195238.75 | 202943.5625 | 213391.4844 | 207977.8594 |
| 122330.3828 | 126650.2188 | 117777.25 | 108863.2031 | 108252.0703 | 110638.6563 | 108953.6172 |
| 62682.44141 | 66483.39063 | 59592.94922 | 53799.23438 | 59110.4375 | 69684.67969 | 64934.02734 |
| 6853.125 | 6396.987305 | 3313.454346 | 5062.156738 | 5498.335449 | 5679.667969 | 5187.571777 |
| 20992.90039 | 22043.59375 | 17385.57813 | 18259.76367 | 20565.65625 | 19552.75195 | 20916.35742 |
| 59059.94531 | 58808.96094 | 51957.83594 | 50869.06641 | 47646.00391 | 36907.60547 | 43822.41797 |
| 1583560 | 1917005.625 | 1567167.625 | 1646183 | 2386019.75 | 1864765.875 | 1957722.125 |
| 8632.258789 | 14211.53906 | 10764.61719 | 2652.526123 | 2847.522461 | 11497.05859 | 1707.846191 |
| 16689.76172 | 19463.57031 | 17525.25586 | 20499.59961 | 27525.11133 | 18754.31641 | 23721.33203 |
| 52724.63281 | 58134.51563 | 49233.36328 | 53065.66797 | 54184.16797 | 47428.54688 | 48158.87891 |
| 37703.76172 | 34363.94141 | 38623.41797 | 38971.43359 | 41076.20313 | 34866.13281 | 35422.01953 |
| 38254.01172 | 38198.89063 | 34084.63672 | 42642.51953 | 64263.69141 | 63487.82031 | 57151.98047 |
| 101333.6875 | 96232.07813 | 99490.67188 | 99162.90625 | 114484.1094 | 113948.0469 | 111051.0625 |
| 11301.27832 | 9063.220703 | 9650.62793 | 7317.600586 | 5098.977539 | 6514.354004 | 2751.779297 |
| 1447649.5 | 1360781 | 1392498.25 | 1368179.5 | 1497236.75 | 1532603 | 1561710.5 |
| 45761.65234 | 41998.49609 | 41565.84375 | 42139.28125 | 22118.75195 | 27899.14453 | 24402.79102 |
| 13129.88672 | 9982.478516 | 16970.40039 | 15604.27344 | 21982.7168 | 19587.63086 | 21601.54297 |
| 238184.9063 | 204137.5781 | 218352.1406 | 213316.4063 | 192888.0469 | 216088.4688 | 196101.0938 |
| 22192.0332 | 24824.09961 | 20752.24219 | 15402.0332 | 21377.72461 | 26336.3418 | 22911.93555 |
| 433.1914063 | 444.906311 | 549.0179443 | 477.0780945 | 868.838501 | 384.0444031 | 573.2738037 |
| 11873.08301 | 11024.04883 | 10215.55762 | 11280.99414 | 9349.326172 | 7063.017578 | 7795.932617 |
| 14722.36133 | 12937.60645 | 16372.50488 | 15574.98047 | 16287.6709 | 18437.69531 | 16697.20508 |
| 15277.26563 | 14077.17676 | 17646.41797 | 17723.21289 | 17202.75781 | 16503.37891 | 21125.83203 |
| 11537.92969 | 11265.0957 | 14205.83008 | 12332.59082 | 13739.38965 | 14384.2373 | 13672.86523 |
| 154415.8125 | 155340.1563 | 143078.4844 | 154302.1406 | 112406.9844 | 119439.5859 | 119164 |
| 66582.36719 | 66714.25781 | 57995.125 | 65915.20313 | 75604.71094 | 78380.41406 | 79152.54688 |
| 116678.2734 | 121407.8438 | 112918.6328 | 118529.6172 | 151933.9063 | 139491.8438 | 139446.0938 |
| 11989.91406 | 11042.7998 | 11832.74609 | 11298.26074 | 11556.04883 | 10882.16797 | 12112.41016 |

| | | | | | | |
|---|---|---|---|---|---|---|
| 6936.066895 | 6717.006348 | 6163.308105 | 7117.211914 | 8209.551758 | 9102.49707 | 8930.824219 |
| 81539.47656 | 70172.30469 | 82926.70313 | 80590.85156 | 65540.09375 | 72966.94531 | 73307.88281 |
| 133878.25 | 127968.9609 | 126052.0234 | 122472.0547 | 162096.1094 | 145919.4375 | 162506.1094 |
| 39254.66016 | 41561.99609 | 35978.60938 | 36502.83984 | 50437.71875 | 52903.69141 | 55777.64453 |
| 43468.40625 | 36757.92578 | 37490.28125 | 38665.85156 | 34585.49609 | 34968.44141 | 35136.43359 |
| 30892.75 | 30930.15039 | 31111.08984 | 29945.37109 | 26889.28906 | 28953.07031 | 30945.77148 |
| 273995.8438 | 252976.5313 | 265475.0313 | 273984.5313 | 283493.3125 | 290854.5313 | 277416.3438 |
| 26654.28906 | 23381.54492 | 23663.8125 | 28026.44141 | 60264.19922 | 29223.91992 | 47138.71484 |
| 140367.9375 | 138601.9688 | 133841.7188 | 140455.8906 | 126586.7656 | 109399.75 | 127657.6484 |
| 211196.5781 | 158884.7656 | 191682.1719 | 199574.2188 | 110982.5781 | 138205.1719 | 122496.8125 |
| 1880732.625 | 2009376 | 1616496.625 | 1683208.875 | 1800512 | 1943990.125 | 1911524.625 |
| 122133.3281 | 98570.9375 | 124151.875 | 120311.6953 | 80177.89063 | 70459.71875 | 73411.17969 |
| 61393.5 | 55211.14063 | 54437.90625 | 62697.375 | 76061.54688 | 60006.54688 | 64388.81641 |
| 166982.3906 | 153636.8438 | 163775.1719 | 138233.9688 | 128034.0469 | 163689.4844 | 121582.9844 |
| 43022.94922 | 37066.67969 | 44149.05859 | 46291.48438 | 48766.94922 | 41689.65234 | 46353.59375 |
| 276280.3125 | 264596 | 271922.25 | 269068.25 | 289791.9688 | 324809.9375 | 292844.2813 |
| 65316.125 | 68561.42188 | 59352.97656 | 59245.5 | 81591.875 | 72263.82813 | 77222.21875 |
| 14498.4873 | 12727.87598 | 15059.95801 | 16460.89844 | 19990.71289 | 20789.10742 | 18910.85938 |
| 4710694.5 | 4336810 | 4591548.5 | 4649710.5 | 3795077.75 | 3962240.25 | 3830570 |
| 103804.8672 | 86357.00781 | 100988.3125 | 91892.38281 | 71289.07813 | 70352.33594 | 69765.99219 |
| 68431.14844 | 65425.67969 | 65307.39063 | 68440.97656 | 70992.85156 | 63143.14844 | 66511.63281 |
| 10896.5625 | 8116.012207 | 11604.14648 | 13968.88867 | 18362.61914 | 18634.89258 | 21065.0918 |
| 9122.970703 | 8947.120117 | 8479.391602 | 7088.219238 | 5890.758789 | 6498.970215 | 6003.729492 |
| 31564.85156 | 26174.57031 | 35028.06641 | 34920.87891 | 30760.73633 | 30184.33008 | 33686.32813 |
| 199814.2227 | 203755.1953 | 199858.2637 | 179713.4648 | 90858.21289 | 84983 | 90155.09766 |
| 441974.5313 | 461497.3438 | 447889.4688 | 519034.1563 | 888044.8125 | 707673.0625 | 779257.75 |
| 151174.3125 | 220372.8281 | 165695.4375 | 104188.7266 | 287386.0625 | 237677.8281 | 248708.4844 |
| 229500.7656 | 250991.2344 | 240510.4063 | 188824.0469 | 264218.6875 | 47765.87109 | 93238.63281 |
| 4483378.5 | 5900467.5 | 2201596.75 | 2306485.75 | 3044482.25 | 4812904.5 | 1139214.125 |
| 284938.25 | 291588.1875 | 267152.5313 | 262971.5938 | 389416.6563 | 366834.9688 | 370969.75 |
| 13452.45996 | 11998.01563 | 11418.15527 | 14880.74707 | 19427.54492 | 13451.51758 | 13938.35938 |
| 105542.7969 | 115901.0547 | 100931.8984 | 113850.4141 | 138952.625 | 122991.5234 | 131339.1563 |
| 80839.21094 | 89815.08594 | 73295.55469 | 83000.14063 | 81984.35938 | 79085.32031 | 76382.08594 |

| | | | | | | |
|---|---|---|---|---|---|---|
| 16543.70703 | 15743.77539 | 15456.96973 | 16356.20215 | 21063.38867 | 19067.99805 | 18432.31055 |
| 27772.58008 | 30288.1582 | 26801.57617 | 23905.91602 | 41015.11328 | 40492.01563 | 40349.67578 |
| 26154.02148 | 24598.66992 | 24861.94531 | 24798.66016 | 23196.41016 | 24194.06445 | 24079.42969 |
| 4698.520996 | 3954.200684 | 4668.714355 | 5074.868652 | 3483.00708 | 3795.522705 | 3024.712402 |
| 40144.16406 | 41708.49512 | 41509.12598 | 41369.90332 | 42491.63965 | 41817.49414 | 42238.1377 |
| 270439.0938 | 257881.4531 | 209174.4063 | 240041.0313 | 163875.4219 | 168849.1094 | 157710.9219 |
| 80311.35938 | 109889.1563 | 54570.71484 | 33642.36328 | 148721.625 | 107199.6953 | 140913.9688 |
| 13950.70996 | 13446.25488 | 13746.60059 | 14299.13281 | 22072.48633 | 26347.85352 | 24618.875 |
| 465949.375 | 502432.7188 | 441288.4375 | 445983.0938 | 472202.125 | 476406.0938 | 459551.1875 |
| 2363.335449 | 2530.984619 | NA | 1872.247314 | 1880.123901 | 1839.337402 | 3378.109131 |
| 133258.4219 | 131161.625 | 139392.6719 | 134052.0156 | 110613.1953 | 111050.5078 | 112806.7344 |
| 24411.58008 | 19979.66406 | 20839.32422 | 23210.46289 | 27890.71094 | 24048.06641 | 22288.51563 |
| 76654.88281 | 73991.42188 | 67861.48438 | 65828.82813 | 82871.50781 | 87988.49219 | 91141.32813 |
| 19960.36133 | 17458.75586 | 17161.66602 | 18088.17578 | 17800.59766 | 19510.43359 | 11254.98535 |
| 3049703 | 2570320.5 | 2800778 | 2406527 | 2156746.25 | 2321808.25 | 2170573.75 |
| 16497.16602 | 17426.48633 | 14721.5127 | 14428.6748 | 19505.86328 | 22764.73828 | 19794.25391 |
| 1172475 | 1136796.25 | 1023611.75 | 940158.0625 | 945359.3125 | 1002958.125 | 914277 |
| 48416.10156 | 48225.42969 | 45252.58203 | 48582.39453 | 50566.35547 | 51136.96484 | 49090.30859 |
| 2102881.5 | 1900993 | 2022309.125 | 1841479.25 | 1714941.25 | 1787361.625 | 1689880.75 |
| 274402.2188 | 256071.5781 | 254429.7031 | 239681.2031 | 155284.2188 | 145882.8125 | 145340.7813 |
| 31238.88281 | 36371.50781 | 20908.89063 | 18146.84961 | 27288.59375 | 26262.63086 | 31168.46094 |
| 248417.5625 | 284628.625 | 217824.1094 | 180614.8125 | 181524.1875 | 189076.875 | 237273.6719 |
| 459.8362122 | 686.4708862 | 409.89328 | 589.3718872 | 1385.615356 | 1472.968506 | 1298.821045 |
| 16796.71289 | 16122.48828 | 13907.08789 | 13251.77344 | 15881.75098 | 16319.75098 | 14795.18848 |
| 8065839.5 | 7043161 | 7231663.5 | 6203435 | 6361601.5 | 7726558 | 5219509.5 |
| 6530.723145 | 6849.074219 | 6488.314941 | 6530.921875 | 7214.354492 | 7492.097168 | 7819.64502 |
| 45847.66016 | 44104.04297 | 39160.66797 | 43186.70703 | 47609.57813 | 45611.8125 | 46330.78516 |
| 68770.25 | 71861.42969 | 63480.63672 | 57969.21875 | 47097.08594 | 50127.46484 | 46713.98438 |
| 39477.80469 | 36044.27344 | 40873.52734 | 34395.23828 | 25432.81836 | 29132.83008 | 31496.55273 |
| 15306.57227 | 10567.62988 | 13876.57324 | 10007.04102 | 10438.96484 | 7368.148926 | 10158.88379 |
| 495110.0313 | 482068.2188 | 477693.0938 | 494669.5625 | 504166.1563 | 540738.75 | 532459.8125 |
| 22744.03125 | 18148.57617 | 23463.14063 | 19243.18164 | 20065.19727 | 20861.31641 | 22823.26367 |
| 5540416.5 | 5227034.5 | 4884216.5 | 4770339 | 4214522.5 | 4707741.5 | 4370947.5 |

| | | | | | | |
|---|---|---|---|---|---|---|
| 585126.75 | 566409.375 | 540977.125 | 574494.125 | 490587 | 436781.9688 | 492404.0313 |
| 118145.0938 | 116011.4531 | 111080.2578 | 110922.4063 | 99748.17969 | 90443.17969 | 97288.72656 |
| 23679.47461 | 20196.38867 | 23635.08398 | 22699.78125 | 19784.43945 | 19733.55859 | 18513.05078 |
| 9473.90918 | 9774.634766 | 8302.709961 | 10250.78125 | 12034.28906 | 13864.94824 | 12907.05273 |
| 336215.1875 | 238043.0781 | 363965.8438 | 291919.8438 | 277692.625 | 285914.875 | 349078.4688 |
| 184007.8594 | 197912.9063 | 174465.4688 | 166350.0781 | 157553.5469 | 152908.5781 | 156604.125 |
| 6388.083984 | 3872.420898 | 5422.853027 | 6379.463379 | 4095.553711 | 4836.152344 | 3386.208496 |
| 25363.50391 | 19912.05078 | 20961.14063 | 24943.91211 | 23216.81445 | 26791.25586 | 29056.24414 |
| 214526.7188 | 251802.2109 | 212130.6094 | 199389.9453 | 265905.1484 | 279706.8359 | 340144.5469 |
| 2046651.5 | 1825364 | 1855913.5 | 1808953.375 | 1665266.375 | 1601669.75 | 1490016.5 |
| 1409356.75 | 1240035.125 | 1278287.125 | 1174357.5 | 1110572.625 | 1196331.75 | 1075243.25 |
| 984122.125 | 1196310.875 | 952187.875 | 903911.8125 | 915634.125 | 1026725.063 | 915187.4375 |
| 122829.9531 | 118284.2344 | 109256.6719 | 115137.5625 | 102241.9063 | 104257.8516 | 95806.8125 |
| 45162.03906 | 36628.74219 | 43334.31641 | 37291.41406 | 18874.37305 | 20311.54688 | 18673.58594 |
| 3169.342773 | 3141.651123 | 1523.381348 | 1803.981201 | 7980.51416 | 4865.673828 | 5387.766602 |
| 6743.021484 | 5397.513672 | 5849.742188 | 5495.074219 | 5650.279297 | 6075.93457 | 5327.286621 |
| 2297040.5 | 2164390.25 | 2114019.75 | 1941502.75 | 1748256.25 | 1825279.125 | 1804584.75 |
| 141418.0781 | 141818.9063 | 142071.9844 | 127996.9375 | 89032.25781 | 92826.09375 | 89100.54688 |
| 8443.203125 | 8116.187012 | 11122.14648 | 9472.670898 | 9046.703125 | 6368.953613 | 7182.818848 |
| 52847.59766 | 62770.97266 | 47642.11719 | 43620.60547 | 78111.52344 | 72973.47656 | 79907.89063 |
| 14879.71582 | 14351.38867 | 15816.78809 | 14785.56836 | 16594.02148 | 14825.6582 | 14244.02148 |
| 40052.76563 | 35652.49219 | 34325.54297 | 31975.80078 | 31436.77344 | 36947.97656 | 32781.71094 |
| 93453.24219 | 95496.58594 | 93123.95313 | 91126.53906 | 97400.5 | 95438.90625 | 101332.2656 |
| 19715.88867 | 21739.21484 | 18261.98438 | 17055.60938 | 20669.14844 | 21722.91797 | 23035.30273 |
| 8083.856445 | 9713.804688 | 7486.073242 | 7643.050781 | 8132.848633 | 6479.564453 | 7384.631348 |
| 43691.86719 | 45501.37891 | 42316.56641 | 43783.97656 | 43341.26953 | 44957.69141 | 47386.41406 |
| 13692.7832 | 13032.39746 | 11273.34668 | 9983.128906 | 8240.945313 | 10213.52539 | 10683.05273 |
| 1512356.625 | 1451616.875 | 1486368 | 1393384.875 | 1287718 | 1390417.5 | 1208365.375 |
| 55883.40625 | 52308.20313 | 58940.10156 | 50471.36719 | 39389.03516 | 41949.31641 | 45673.5625 |
| 169405.875 | 182306.2188 | 163702.8438 | 189352.125 | 238266.8125 | 257289.375 | 228970.6406 |
| 63960.12109 | 66748.26563 | 64066.43359 | 63431.95703 | 94254.90625 | 79271.14844 | 85898.03125 |
| 77949.33594 | 72603.60938 | 85108.1875 | 75513.42188 | 77180.35938 | 74623.28906 | 74407.25781 |
| 18111.09961 | 17700.75391 | 17012.38672 | 15498.03906 | 15199.57715 | 15022.37207 | 15033.50684 |

| | | | | | | |
|---|---|---|---|---|---|---|
| 19837.46289 | 14952.17188 | 19289.08789 | 17018.87109 | 33166.05078 | 24961.25977 | 28695.33203 |
| 137354.3408 | 139272.9385 | 143943.666 | 146705.3633 | 111235.3623 | 114548.5264 | 121883.1084 |
| 227305.4375 | 167786.3438 | 226947.6875 | 230242.5625 | 214146.7188 | 219893.6719 | 204770.7969 |
| 2080.023438 | 2152.422119 | 1906.047974 | 1661.097656 | 2523.82373 | 1893.012085 | 2562.157959 |
| 290338.6563 | 275259.2188 | 287843.5625 | 272392.7813 | 169884.5469 | 182365.6563 | 183874.4531 |
| 48753.37891 | 45184.72656 | 45084.68359 | 43530.47266 | 48309.91016 | 49762.34766 | 48873.40625 |
| 72991.07031 | 75423.10938 | 69805.74219 | 73209.53906 | 76366.3125 | 76328.27344 | 72467.21094 |
| 113451.3828 | 108176.375 | 113139.3906 | 120247.4063 | 153574.5469 | 137991.625 | 159063.1406 |
| 157874.0938 | 133914.4063 | 148144.4688 | 130979.8594 | 132746.4531 | 142462.9219 | 144459.1719 |
| 70196.53906 | 71282.78906 | 69637.26563 | 60890.47266 | 81745.75 | 90001.65625 | 82949.20313 |
| 101110.7344 | 99032.32031 | 86352.21875 | 88147.10156 | 75829.92969 | 70384 | 79336.5 |
| 229995.7813 | 186005.0313 | 222085.4688 | 235609.2813 | 237511.4375 | 226396.9531 | 235205.5469 |
| 9332.660156 | 9039.367188 | 9791.84668 | 8170.569336 | 8794.171875 | 8650.352539 | 8764.404297 |
| 184232.3125 | 183810.2344 | 176812.2031 | 180835.9063 | 176807.4063 | 173963.9531 | 178686.4844 |
| 7306.537598 | 10495.40723 | 9382.34375 | 7674.86084 | 9562.004883 | 8417.264648 | 7872.714355 |
| 9249.391602 | 7290.730957 | 8192.094727 | 9977.556641 | 14229.26074 | 14118.60742 | 17276.54102 |
| 60604.75391 | 78436.47656 | 48402.83984 | 53629.21094 | 56163.38281 | 70641.98438 | 80351.78125 |
| 29233.51367 | 30992.26953 | 28471.07031 | 26522.12891 | 23811.06836 | 28253.05664 | 24659.43945 |
| 50343.04297 | 48462.57422 | 50636.83203 | 53846.88281 | 38725.69531 | 103214.5156 | 43590.98047 |
| 70027.83594 | 57442.09375 | 73679.25 | 75368.30469 | 64905.34375 | 59541.48047 | 60754.97266 |
| 2871.729492 | 2562.150391 | 2464.143066 | 2087.77002 | 1043.719971 | 2209.404053 | 2525.038086 |
| 238486.1875 | 196228.2969 | 221982.3438 | 233242.0625 | 275572.8438 | 256843.4844 | 269967.3125 |
| 41356.32813 | 42328.73047 | 37666.19531 | 37146.92578 | 33654.94922 | 34868.17578 | 34117.53516 |
| 272755.75 | 369325.375 | 228557.4375 | 31711.01172 | 493784.9375 | 703428.0625 | 500673.5313 |
| 959497 | 846183.3125 | 836204.8125 | 765782.3125 | 678510 | 749423.75 | 684876.5625 |
| 4045.124756 | 5000.09668 | 4544.198242 | 4365.667969 | 4940.759277 | 4057.744141 | 5064.560059 |
| 66183.86719 | 59249.83984 | 57780 | 48463.77734 | 57652.13281 | 51464.1875 | 49271.98047 |
| 4441051.625 | 3825379.75 | 4233789.75 | 4379779 | 4684186.125 | 4023338.375 | 4920008.375 |
| 36658.94922 | 33955.59375 | 38783.8125 | 39017.41797 | 58320.99219 | 51434.72266 | 55798.52344 |
| 286232.9688 | 289187.2813 | 284001 | 289851.6875 | 246280.2813 | 231715.4531 | 249488.75 |
| 35665.67969 | 38781.28125 | 31196.95703 | 40298.23438 | 50456.13281 | 52576.64844 | 51079.76563 |
| 124677.9609 | 151187.0469 | 129512.9141 | 127124.4531 | 145027.2188 | 166903.8281 | 135437.0156 |
| 31545.95898 | 28436.99805 | 33860.11328 | 29746.40039 | 29674.50781 | 28653.3457 | 28792.58398 |

| | | | | | | |
|---|---|---|---|---|---|---|
| 55977.86719 | 53964.54297 | 56796.57813 | 56973.30469 | 48372.63672 | 54584.22266 | 50005.71875 |
| NA | 1758.062256 | 1849.000854 | NA | 1129.775879 | 2233.575195 | 1782.545288 |
| 80592.29688 | 72163.35156 | 80347.05469 | 82627.83594 | 38363.35938 | 40408.54297 | 42145.34766 |
| 5942.146973 | 4507.887207 | 4805.591797 | 4478.724121 | 1313.89563 | 2619.961182 | 2755.0625 |
| 562604.1875 | 527688.3125 | 550566.9375 | 531548.5 | 526680.25 | 523132.6563 | 545987.75 |
| 14088.6377 | 14881.24512 | 14521.72559 | 16812.62109 | 18348.75586 | 19652.51953 | 19321.5 |
| 70763.39844 | 64610.48828 | 79191.14063 | 70440.9375 | 63291.96484 | 62595.97266 | 65364.45313 |
| 25341.22461 | 30620.84375 | 21890.85742 | 19509.41211 | 10279.06152 | 14710.85352 | 11662.02441 |
| 2552362.5 | 2331578.5 | 2356483 | 2260916.5 | 2141594.75 | 2263944 | 2080906.25 |
| 49038.84766 | 48853.82422 | 45587.50391 | 46744.41406 | 56906.66016 | 55114.98047 | 55466.43359 |
| 67092.6875 | 66677.73438 | 67099.16406 | 78811.32031 | 196993.2813 | 171927.25 | 179304.3125 |
| 23997.80664 | 24170.24414 | 19134.77148 | 27424.18164 | 34646.53516 | 36756.39453 | 35466.60938 |
| 13010.74316 | 11172.57715 | 15684.95703 | 13490.8877 | 6285.935547 | 10049.09668 | 10007.17578 |
| 768679 | 754120.5 | 735533.5625 | 685364.6875 | 371125.2188 | 401909.9375 | 384137.6875 |
| 41567.01953 | 37510.24609 | 40758.77344 | 36305.73047 | 51586.72266 | 44010.00391 | 49989.32813 |
| 22451.28906 | 21929.44922 | 18975.36523 | 18628.81836 | 15409.20703 | 15173.2793 | 14949.84375 |
| 23818.57031 | 26450.47461 | 19569.22656 | 21632.24805 | 23273.58203 | 19096.5 | 20513.79883 |
| 528702 | 476209.4375 | 505925.6875 | 521992.0938 | 428345.4688 | 397743.375 | 436678.5625 |
| 45850.08594 | 37466.98047 | 45076.48047 | 44047.14453 | 47906.13672 | 48464.63672 | 50263.42578 |
| 93053.92188 | 100382.9219 | 81340.69531 | 76635.54688 | 107365.6953 | 111962.4375 | 109807.1016 |
| 18963.79297 | 15881.44922 | 15787.3877 | 16399.91992 | 10373.28027 | 8410.953125 | 11797.44629 |
| 4336087.5 | 3753529 | 3737343 | 4374741 | 4544866.5 | 3860422.25 | 4453841.5 |
| 635285.3125 | 512228.0625 | 498095.4375 | 648043.5 | 599265.75 | 669891.625 | 620221.625 |
| 146756.125 | 130092.4688 | 124011.2969 | 126567.3828 | 159265.7031 | 162355.7969 | 147452.3281 |
| 2768.595215 | 2260.004395 | 1893.227051 | 2442.364746 | 1145.954834 | 1607.367065 | 1667.010376 |
| 202713.3438 | 184573.0313 | 199766.4844 | 216911.75 | 247655.5 | 249361.1406 | 259998.5469 |
| 29112.71289 | 27301.86719 | 25748.88086 | 31809.94531 | 36364.54688 | 30196.43555 | 33154.37109 |
| 1135891 | 1028030.438 | 1125658.875 | 1149992.125 | 1513261.25 | 1445044.75 | 1440152.375 |
| 16617.22266 | 14168.23926 | 16608.36328 | 16467.93164 | 18357.24414 | 18667.6543 | 19131.03711 |
| 124100.6328 | 130261.5781 | 116938.5078 | 124536.5313 | 145992.1719 | 146529.9688 | 143504.375 |
| 56748.90234 | 54076.62891 | 47840.67578 | 50235.5625 | 61529.79297 | 61542.55078 | 65523.89063 |
| 1147572.375 | 999210.1875 | 1070582.75 | 989017.9375 | 937055.9375 | 941773.9375 | 884312.0625 |
| 41924.78906 | 51685.79688 | 36769.6875 | 33679.90625 | 57800.51172 | 61744.42578 | 59581.53516 |

| | | | | | | |
|---|---|---|---|---|---|---|
| 693390.4375 | 689183.25 | 633736.6875 | 574483.5625 | 621041.8125 | 646953.5625 | 592381.125 |
| 194067.8281 | 172524.9063 | 187449.2813 | 176767.8281 | 179464.5781 | 189878.6406 | 195818.0625 |
| 296641.125 | 293751.6563 | 252460.3281 | 271778.8438 | 320840.75 | 303340.0625 | 315035.875 |
| 4814.235352 | 3548.01123 | 7111.921387 | 3931.573486 | 3228.745117 | 3075.401611 | 3700.38208 |
| 17616.57617 | 19832.51563 | 16779.43359 | 14384.29297 | 17689.8125 | 26506.64648 | 17192.93945 |
| 187176.1719 | 188396.25 | 179064.7031 | 177801.6094 | 186404.6875 | 170786.3281 | 185481.3281 |
| 240116.2188 | 237435.75 | 206668.7031 | 146359.7969 | 147749.6875 | 254521.7656 | 172620.2188 |
| 30014.45703 | 22267.20898 | 26390.10742 | 28656.49316 | 21165.66455 | 23520.02832 | 24711.34424 |
| 13210.28516 | 11599.64258 | 12301.85254 | 13331.56445 | 13981.74414 | 14331.80176 | 15356.43457 |
| 365194.9688 | 434395.9688 | 347773.0938 | 325434.8438 | 349479.75 | 294260.2813 | 828078.3125 |
| 3467.867432 | 5486.072266 | 4771.584473 | 5015.006348 | 4655.865234 | 5400.623535 | 4941.699219 |
| 321670.9063 | 317341.4375 | 288955.5 | 292368.6875 | 352036.875 | 345308.125 | 359231.25 |
| 13545.05273 | 11992.83398 | 11737.12207 | 14344.32227 | 11358.28613 | 10696.49609 | 10734.82422 |
| 16864.13086 | 29479.28906 | 8809.609375 | 3829.83252 | 40994.40234 | 21566.13086 | 46552.87891 |
| 28502.55273 | 21412.18164 | 25680.08008 | 27907.07422 | 22360.55664 | 23476.46484 | 20281.49805 |
| 51014.01953 | 43172.97266 | 50199 | 51910.58984 | 59932.50781 | 49216.34375 | 46659.03125 |
| 174323.9844 | 165361.9844 | 152586.9375 | 165505.7969 | 223820.0156 | 189678.2656 | 211208.5781 |
| 40570.63281 | 46331.32422 | 35561.13281 | 39501.57813 | 40482.54688 | 45858.27734 | 45672.57813 |
| 11280.79883 | 8664.597656 | 11390.18359 | 10513.85059 | 13567.7666 | 12490.76465 | 13032.30078 |
| 62855.85547 | 64745.97266 | 59723.75391 | 63862.96484 | 78104.58594 | 72901.82031 | 73039.17969 |
| 455315.7188 | 445900.3125 | 495462.5938 | 471516.4688 | 279640.7813 | 283986.1875 | 292335.3438 |
| 47923.12891 | 45443.38281 | 43514.97656 | 38396.15234 | 35080.53125 | 38989.59375 | 35782.34766 |
| 64916.94922 | 69926.96094 | 65150.14063 | 61704.67188 | 70148.1875 | 64071.25391 | 62703.65625 |
| 181064.7813 | 174548.6094 | 162763.8125 | 175291.4688 | 192347.4531 | 199801.6875 | 189496.7656 |
| 117474.5391 | 110449.0313 | 122926.6797 | 112515.8203 | 111715.3438 | 116677.0938 | 119178.8516 |
| 55918.24219 | 58859.85938 | 54553.74609 | 62961.26172 | 76393.75781 | 67152.8125 | 62829.01172 |
| 97073.19531 | 100239.3281 | 88046.00781 | 92522.29688 | 166334.6563 | 146602.2656 | 140733.5156 |
| 80932.25 | 60064.91797 | 83947.14063 | 84736.07813 | 118623.0703 | 106342.9297 | 114267.9531 |
| 31348.67383 | 24956.19727 | 27378.28516 | 28551.08008 | 27813.75586 | 24273.58398 | 27163.16016 |
| 139934.7656 | 129947.4688 | 120444.2578 | 130313.0313 | 131321.625 | 131501.7656 | 128504.4922 |
| 15836.17188 | 15250.01953 | 12724.00098 | 14234.26855 | 15324.82813 | 15602.40918 | 14723.5127 |
| 101962.7734 | 79717.5625 | 95960.53125 | 89340.4375 | 65749.99219 | 72222.19531 | 71163.92969 |
| 39628.88672 | 45218.69531 | 41321.00781 | 40229.60156 | 57192.72266 | 57176.18359 | 54541.44922 |

| | | | | | | |
|---|---|---|---|---|---|---|
| 30902.1875 | 31350.64258 | 24104.71094 | 18960.29883 | 19332.99219 | 21838.68164 | 20539.24023 |
| 49189.16797 | 47078.34375 | 44333.67969 | 47515.09375 | 61872.67578 | 57908.52734 | 58452.39453 |
| 49218.875 | 50152.89453 | 49997.50391 | 50067.76953 | 71714 | 68524.07031 | 68176.59375 |
| 1623647.25 | 1461574.625 | 1461469 | 1314293.5 | 1211880 | 1258802.375 | 1154459.125 |
| 42874.82813 | 38794.26953 | 41627.78906 | 40313.66016 | 45404.55469 | 45321.29297 | 45248.82813 |
| 62015.88281 | 46160.47266 | 80291.22656 | 35921.76953 | 19255.21484 | 20735.88281 | 33234.26563 |
| 1918460.25 | 1815208.25 | 1708235.5 | 1603849.375 | 1560964.125 | 1554773.5 | 1355507.25 |
| 79221.66406 | 67193.35938 | 69970.53125 | 76384.9375 | 58677.11328 | 67033.84375 | 56978.53906 |
| 49293.09375 | 45493.21094 | 42821.60547 | 36785.51953 | 32660.71484 | 36641.53125 | 29761.80664 |
| 425764.7813 | 250576.8281 | 55173.30859 | 632922.4375 | 1556488.125 | 1811210.5 | 743609.1875 |
| 6435.71582 | 9113.878906 | 5313.152344 | 4888.712891 | 4539.378906 | 5024.55127 | 5805.879883 |
| 407855.9688 | 437719.5625 | 395978.2188 | 321905.5313 | 264234.5625 | 333931.9688 | 343436.3125 |
| 770236.1875 | 729598.6875 | 745268 | 725141.8125 | 765677.875 | 877626.6875 | 810454.5625 |
| 22697.68359 | 19852.57617 | 19858.75586 | 19931.15039 | 16507.26367 | 12231.35449 | 16226.38379 |
| 42203.62891 | 42953.5625 | 40997.39063 | 40398.64063 | 36812.07031 | 37217.73047 | 35939.69922 |
| 97948.96875 | 107623.2969 | 87161.95313 | 86119.01563 | 98815.71094 | 90421.28125 | 101928.2188 |
| 23194.91406 | 29602.26367 | 20241.9043 | 23129.82813 | 27247.17773 | 27614.20117 | 22981.76758 |
| 80614 | 94982.5 | 72843.95313 | 82489.67969 | 123497.4844 | 109948.5156 | 111609.1094 |
| 7226.013672 | 8563.412109 | 6755.661133 | 7260.204102 | 9520.180664 | 9395.038086 | 9880.532227 |
| 85474.78125 | 78595.51563 | 84897.04688 | 88922.6875 | 67109.375 | 86336.03125 | 78160.40625 |
| 31768.13672 | 26429.95117 | 31966.89648 | 28955.91016 | 32408.10547 | 36483.07031 | 38175.18359 |
| 41624.70313 | 46374.60547 | 35417.78906 | 40251.57813 | 75080.35938 | 57953.40234 | 65341.03125 |
| 7339.850098 | 8914.555664 | 7549.172852 | 8709.398438 | 8005.856445 | 17648.5918 | 9606.119141 |
| 2117318 | 2000310.625 | 1972527.125 | 2156122.75 | 2310761.25 | 1990992.375 | 2206750.75 |
| 1245429.625 | 1251677.875 | 1221875.5 | 1061959.125 | 1087046.875 | 1246141 | 1086090.5 |
| 11258.7959 | 11383.2334 | 12731.92188 | 11603.9707 | 9819.239258 | 7909.527832 | 7984.200684 |
| 41674.67188 | 44933.45313 | 44620.03906 | 35492.09375 | 39461.38281 | 44487.04688 | 43037.47656 |
| 440206.3125 | 436652.5938 | 457458.5313 | 407974.1875 | 367758.9063 | 389200.4375 | 361720.125 |
| 97772.61719 | 86974.90625 | 88336.53906 | 84920.89063 | 67094.91406 | 66496.74219 | 66613.15625 |
| 7546.925781 | 7124.594727 | 5764.561035 | 8174.60791 | 7502.36084 | 6726.04541 | 7069.783691 |
| 37027.1875 | 34396.57813 | 33257.38672 | 41517.14453 | 59449.5625 | 60109.27734 | 60679.27734 |
| 519797.9688 | 534759.875 | 449145.625 | 458598.0313 | 541879.375 | 530072.625 | 517579.8125 |
| 96957.96875 | 103975.8594 | 83384.07813 | 77325.17969 | 87630.14844 | 92630.02344 | 87978.07031 |

| | | | | | | |
|---|---|---|---|---|---|---|
| 135920.0156 | 111297.5156 | 121655.1172 | 118635.6484 | 88717.46875 | 89475.33594 | 94554.375 |
| 631644.4375 | 571927.25 | 594286.4375 | 582910.0625 | 548154.25 | 506606.25 | 546106.875 |
| 12909.63281 | 16535.30664 | 5730.689453 | 6849.769043 | 14614.54004 | 12508.60352 | 13506.07031 |
| 227647.3125 | 214318.1563 | 209208.5 | 211395.1719 | 237397.2031 | 237130.4688 | 239380.2813 |
| 1332062.5 | 1562656.25 | 959477.25 | 952708.25 | 608383.375 | 1058051.25 | 661717.5 |
| 225151.9688 | 199069.0625 | 216720.1563 | 216293.375 | 247660.7813 | 240869.2188 | 243900.4219 |
| 333661.5313 | 334520.9063 | 314473.9063 | 329492.9063 | 352931.8438 | 320034.8438 | 339012.9688 |
| 12711.58203 | 12196.81445 | 11825.59961 | 14284.18262 | 10415.40527 | 9193.207031 | 10082.63281 |
| 94113.35156 | 85580.5625 | 94236.17969 | 100871.6953 | 115925.4297 | 92494.03906 | 96615.94531 |
| 14175.63574 | 18261.45117 | 14820.76172 | 15593.05078 | 18681.13086 | 23862.0293 | 17332.87891 |
| 870448.5625 | 777779.9375 | 775743.375 | 689267.875 | 631771.75 | 702581.875 | 562143.25 |
| 27965.55469 | 29121.33203 | 20955.25586 | 24239.40625 | 33580.82422 | 33564.09766 | 34609.44141 |
| 58901.27734 | 57182.81641 | 53843.28125 | 49919.82031 | 40726.10547 | 40008.88672 | 40341.9375 |
| 22314.25781 | 17794.41406 | 23866.42578 | 21082.90625 | 16756.06445 | 16935.99609 | 15157.45801 |
| 121376.7578 | 130057.5859 | 110155.0547 | 116073.1328 | 130451.6797 | 119164.5313 | 124159.0547 |
| 1309886.375 | 1175250.625 | 1185055.75 | 1132239.75 | 1255165.25 | 1245291.125 | 1207048.625 |
| 357682.8438 | 345493.8438 | 320352.5625 | 308202.3125 | 294799.0313 | 293015.6875 | 279388 |
| 90135.96875 | 98774.03125 | 76742.98438 | 80745.78906 | 105188.375 | 101732.3984 | 103388.25 |
| 598853.25 | 537789 | 557124.5625 | 587506.5625 | 497622 | 461103.6563 | 495459 |
| 223522.3594 | 199342.8594 | 191739.2031 | 197311.2344 | 255054.625 | 253067.9219 | 240531.9219 |
| 3934.491943 | 4005.014404 | 3299.909912 | 3721.065674 | 4497.668945 | 4714.368164 | 6036.677734 |
| 33858.35156 | 28310.01172 | 38254.65625 | 29758.96094 | 37674.43359 | 29627.48242 | 32398.00977 |
| 6465.087402 | 8495.798828 | 4602.539063 | 3915.219727 | 5781.604004 | 14487.08691 | 12425.05957 |
| 2314649.25 | 2244268.75 | 2194450.5 | 2120180 | 1797318.5 | 1937662.375 | 1876995.875 |
| 24867.72461 | 23735.47461 | 25434.23828 | 22210.35938 | 20774.58789 | 16789.63281 | 19736.05859 |
| 119304.4063 | 92372.71875 | 124125.2031 | 116886.2578 | 85315.20313 | 86205.82813 | 76150.27344 |
| 28282.87109 | 27777.04102 | 27079.92578 | 26702.48242 | 26295.72461 | 29754.52539 | 27912.99609 |
| 33048.57031 | 32468.99023 | 29404.0293 | 28252.75 | 35003.73438 | 36973.58594 | 35525.97266 |
| 3216974.5 | 2804585.5 | 3067928.5 | 3016940.75 | 3196448 | 3311802.75 | 3497421.75 |
| 57173.8125 | 56295.42188 | 44537.36328 | 47588.95313 | 41803.85156 | 42892.73438 | 48362.87109 |
| 294960.9063 | 277591.4688 | 236253.5156 | 253384.8281 | 276004.1563 | 276201.125 | 246986.6719 |
| 352515.8438 | 340538.5 | 344769.125 | 328800.0313 | 345499.5313 | 368282.4375 | 361291.125 |
| 1343022.375 | 1257499.625 | 1365064.25 | 1308728.375 | 1237610.625 | 1206875.25 | 1303253.625 |

| | | | | | | |
|---|---|---|---|---|---|---|
| 114975.2031 | 102468.0078 | 112272.75 | 112305.3516 | 108650.1172 | 106311.0234 | 110587.8984 |
| 9585.844727 | 8969.892578 | 9847.026367 | 9063.37207 | 9076.626953 | 9007.749023 | 9160.56543 |
| 2066569.25 | 1751240.5 | 1947277.625 | 1825905.25 | 1916400.25 | 1926199.875 | 2063712.125 |
| 35083.19922 | 33832.07813 | 31397.67383 | 31178.22461 | 18712.76172 | 22305.18555 | 20094.43945 |
| 5886.651855 | 5139.529785 | 4888.936523 | 4428.484375 | 7603.620117 | 5257.220215 | 5575.018066 |
| 80707.96094 | 82202.66406 | 67846.02344 | 67043.3125 | 75174.14844 | 71995.98438 | 70145.24219 |
| 119751.6484 | 136742.6719 | 105297.6094 | 98578.13281 | 110164.2031 | 88014.55469 | 111352.7891 |
| 113540.5703 | 107675.4688 | 107981.9219 | 108893.1484 | 100372.8984 | 104664.7344 | 103499.6016 |
| 9553833 | 13238810 | 6014681 | 4951888 | 6896253 | 6574179 | 6757241 |
| 116010.2109 | 121531.2656 | 97334.86719 | 94155.98438 | 86631.09375 | 88946.42188 | 91222.28125 |
| 199987.1563 | 183230.5313 | 199988.2813 | 200445.7188 | 211238.8594 | 204540.2031 | 209110.0313 |
| 196897.0781 | 175281.1563 | 172382.6563 | 181081.1875 | 161548.1719 | 170876.3281 | 167231.0469 |
| 60651.45313 | 64042.35156 | 50808.32031 | 51132.98047 | 57198.47266 | 63996 | 61345.46484 |
| 73544.28125 | 71505.82031 | 66728.04688 | 64381.23828 | 79513.20313 | 75317.30469 | 75664.60938 |
| 25321.19531 | 28584.78125 | 19759.98438 | 18067.08594 | 32912.87109 | 29678.84766 | 28180.1543 |
| 206265.0313 | 191570.75 | 205351.6563 | 209627.4375 | 270916.0625 | 212294.4688 | 227924.5156 |
| 1593694.75 | 1370499.25 | 1456254 | 1281876.75 | 1126218.875 | 1263739.125 | 1094126 |
| 21558.73242 | 17935.4082 | 18706.48438 | 18744.92773 | 16289.9541 | 18301.50586 | 17899.13672 |
| 289409.0938 | 243913.8438 | 267179.5 | 302898.9063 | 289893.7188 | 280001.25 | 296612.125 |
| 13916.58105 | 12713.41406 | 11875.95605 | 10639.33691 | 12084.4043 | 7818 | 8895.265625 |
| 104048.1719 | 95004.48438 | 94103.57813 | 103845 | 59299.34375 | 61334.85938 | 61089.99609 |
| 12708.44727 | 12980.89258 | 13600.15137 | 14673.43555 | 18403.00586 | 15209.89648 | 18556.5918 |
| 826860.5 | 754054.8125 | 796076.4375 | 683703.6875 | 634918.125 | 711114.4375 | 594201 |
| 23164.11133 | 17727.7832 | 22279.0332 | 20823.85156 | 13105.73926 | 14365.87988 | 13769.12305 |
| 441034.0313 | 409162.9688 | 408173.7188 | 441494.2813 | 586586.3125 | 473410.3438 | 502578.5938 |
| 577664.6875 | 539308.8125 | 570834.0625 | 608704.5625 | 436755.2188 | 430115.4375 | 459961.7813 |
| 487876.3438 | 441040.7188 | 472699.6875 | 431660.1563 | 361211.6875 | 410544.75 | 390936.125 |
| 9774.837891 | 12671.125 | 8265.920898 | 10531.69727 | 9267.735352 | 8789.353516 | 8947.59668 |
| 5182.384766 | 6251.513184 | 4174.428223 | 4431.312988 | 3879.816162 | 6276.55127 | 4194.407715 |
| 3758.032471 | 3595.942871 | 2986.248779 | 3104.687988 | 2951.497803 | 3436.469971 | 2932.288574 |
| 187751.5625 | 134031.2188 | 186166.4688 | 170773.0781 | 220493.6406 | 214199.1563 | 246693.7031 |
| 25549.78125 | 23511.66797 | 25710.27734 | 26685.18359 | 17838.88672 | 15024.02344 | 18098.98242 |
| 49557.94922 | 44460.79297 | 48472.62891 | 50639.57813 | 40381.24219 | 41294.07031 | 39902.73047 |

| | | | | | | |
|---|---|---|---|---|---|---|
| 42057.23438 | 44246.86426 | 39274.50586 | 44949.01172 | 45562.75 | 44442.54102 | 45301.0293 |
| 114635.7344 | 116114.8594 | 108763.8438 | 103966.3359 | 79348.17969 | 85766.61719 | 83463.57813 |
| 70280.33594 | 63972.03906 | 67880.99219 | 67179.80469 | 58842.60938 | 56873.98047 | 55512.92188 |
| 216521.5313 | 186995.4688 | 191605.25 | 213746.375 | 287043.6875 | 269513.3125 | 287901.125 |
| 4724.721191 | 5168.812012 | 4831.573242 | 6783.854004 | 3702.303467 | 7221.682617 | 6742.445313 |
| 94936.58594 | 94672.24219 | 83629.83594 | 86223.76563 | 132120.5469 | 117449.4375 | 125201.0469 |
| 605829.75 | 613878 | 506054.2188 | 502563.5 | 635499.0625 | 680868.375 | 594043.3125 |
| 129559.3281 | 125126.2266 | 118877.2969 | 128544.2266 | 167547.3281 | 179182.75 | 171324.9688 |
| 90141.51563 | 76637 | 84718.10938 | 84188.33594 | 77085.21094 | 75482.16406 | 83334.84375 |
| 25984.77539 | 28722.50977 | 20713.79297 | 22181.74219 | 21303.6543 | 22087.52539 | 21121.27734 |
| 23354.84766 | 22894.29102 | 23843.375 | 21968.60547 | 29776.49609 | 31398.8457 | 27610.19141 |
| 5968481 | 6657369 | 4488436 | 5638877 | 5650151 | 6515823 | 4565294 |
| 72251.03125 | 69354.92188 | 65160.83984 | 67008.67188 | 76880.89063 | 73966.64844 | 75689.125 |
| 36926.14453 | 29977.85938 | 31468.41211 | 31019.72461 | 28761.41016 | 29707.57422 | 29122.2168 |
| 8944.041016 | 8797.039063 | 7283.154297 | 7203.092773 | 10488.69531 | 8365.891602 | 9712.577148 |
| 10793.1875 | 7997.703125 | 2656.545654 | 9873.393555 | 6741.907715 | 12185.97949 | 7073.357422 |
| 49379.28516 | 43952.66016 | 43870.25781 | 44902.56641 | 56090.42578 | 58463.3125 | 55918.10547 |
| 74878.28125 | 61700.88672 | 70059.85156 | 74155.10156 | 112469.125 | 109988.4766 | 111833.0547 |
| 42702.03906 | 34700.51172 | 34124.10156 | 35030.00391 | 40375.03125 | 37344.89063 | 39739.21484 |
| 7030.687988 | 6789.215332 | 5743.47168 | 5270.955566 | 4033.625977 | 5034.317871 | 5322.328613 |
| 101605.9375 | 116338.6719 | 99244.47656 | 98927.84375 | 84430.29688 | 79296.60938 | 79546.48438 |
| 338791.9063 | 305906.6563 | 311151.9688 | 351629.7813 | 442837.2813 | 383058.625 | 396315.3438 |
| 52320.375 | 46529.97656 | 51606.90234 | 47744.94922 | 26546.19336 | 25925.60156 | 25266.62891 |
| 56054.14063 | 53197.625 | 52832.47266 | 51001.63672 | 30441.00781 | 36640.875 | 29266.04688 |
| 11227060 | 9648921 | 10928727 | 9812334 | 9542936 | 10376309 | 9604575 |
| 712935.125 | 594824.75 | 638340.0625 | 610230.25 | 603101.625 | 570066.5625 | 558408.25 |
| 6605.496582 | 7630.375488 | 6155.42627 | 4051.687988 | 5781.17041 | 5369.439941 | 5364.253906 |
| 35278.63281 | 31613.04492 | 33373.60156 | 30850.50391 | 45283.17969 | 45153.53906 | 52947.36719 |
| 117745.5547 | 98284.55469 | 113701.1172 | 118041.1484 | 106675.7031 | 106199.5547 | 111542.1953 |
| 730513.9375 | 646964.1875 | 646237.9375 | 690236.875 | 1020904.75 | 838966.4375 | 940184.1875 |
| 50722.34766 | 50513.44922 | 59415.14844 | 58977.94922 | 95285.26563 | 66910.21875 | 74282.07813 |
| 48554.5625 | 44996.53516 | 42382.47266 | 40600.06641 | 47033.82031 | 49240.51563 | 49844.375 |
| 118365.6484 | 110648.2656 | 115459.625 | 118643.2813 | 86910.36719 | 82349.70313 | 87692.21094 |

| | | | | | | |
|---|---|---|---|---|---|---|
| 9029.588867 | 8886.599609 | 11153.38379 | 8847.254883 | 10545.95215 | 11537.58301 | 8336.585938 |
| 14077.85547 | 15090.50977 | 10540.59863 | 12057.84961 | 10464.49023 | 10427.60938 | 11436.01855 |
| 166402.4219 | 147025.6563 | 143241.8438 | 116386.3359 | 168617.7813 | 208434.8438 | 167796.0313 |
| 483444.2813 | 405179.1563 | 500934.5313 | 546398.75 | 638370.75 | 548091.1875 | 577065.375 |
| 24526.48438 | 23335.58594 | 20885.77539 | 20417.01172 | 19019.85742 | 20001.3125 | 19113.60156 |
| 5534824 | 4867290 | 4544636 | 4421978 | 4085075.25 | 4511528 | 3932346 |
| 8096.321289 | 9665.308594 | 6052.252441 | 6072.397461 | 14468.06348 | 12563.15625 | 12348.66504 |
| 21975.46094 | 18821.27734 | 21457.23438 | 21058.16602 | 19677.59961 | 19588.82422 | 19503.41602 |
| 43010.00391 | 41932.11719 | 41132.64453 | 40766.84766 | 85086.4375 | 85329.11719 | 85280.76563 |
| 6940.743164 | 7221.30127 | 7146.715332 | 6567.163574 | 8144.961914 | 7455.393066 | 6670.395508 |
| 174857.7813 | 164725.4375 | 160227.0938 | 148214.5156 | 145528.6719 | 132200.0625 | 151279.875 |
| 102073.3984 | 96645.8125 | 93257.96875 | 94269.24219 | 84989.07031 | 85385.82813 | 84592.125 |
| 97100.75 | 75277.72656 | 83992.71875 | 88029.90625 | 120283.5313 | 115991.2266 | 128075.5156 |
| 26879.98438 | 23498.65625 | 27658.32617 | 28633.48242 | 32077.19531 | 32764.52148 | 35146.59766 |
| 2564431 | 2180771 | 2330336 | 1994968.375 | 1913003.25 | 1948364.625 | 1672718.125 |
| 15203.00195 | 13280.4375 | 10412.4375 | 13554.53809 | 16270.21973 | 21597.87891 | 19074.94727 |
| 34578.80469 | 29005.35352 | 30653.16211 | 40152.27344 | 51127.21094 | 31808.58398 | 47417.00781 |
| 56032.3125 | 47431.10156 | 51570.50781 | 51451.82031 | 50194.03125 | 45071.00391 | 49963.68359 |
| 2037.790283 | 4216.989258 | 2040.553223 | 1786.570557 | 4918.709961 | 1830.691528 | 1587.8927 |
| 612100.875 | 603586.0625 | 566234 | 639191.75 | 685293.5 | 663149.0625 | 644704.0625 |
| 13580.87695 | 11613.84766 | 14334.03125 | 12682.2959 | 10574.32324 | 11727.24414 | 12441.14746 |
| 2388.834229 | 1926.001953 | 2623.237305 | 2425.646973 | 2462.462158 | 3024.904053 | 2442.742188 |
| 6958.383301 | 7901.807129 | 6241.044434 | 4508.852051 | 4840.207031 | 5738.141113 | 5438.685059 |
| 29188.25 | 26299.35938 | 24927.5293 | 24543.71484 | 16819.17578 | 19301.4707 | 18050.31641 |
| 4456.943848 | 4089.018799 | 3544.671387 | 4713.67041 | 9198.470703 | 5821.732422 | 7187.304688 |
| 30697.14063 | 34211.75 | 34622.20703 | 29730.97461 | 31587.66211 | 29987.5293 | 25995.21875 |
| 107209.1406 | 100247.125 | 90232.22656 | 78844.92969 | 49328.57813 | 47463.08203 | 41712.55469 |
| 185976.7031 | 185863.7031 | 174291.2813 | 171799.2344 | 105940.3047 | 108104.1406 | 98652.02344 |
| 15482.52148 | 14179.38379 | 18450.17383 | 17089.9375 | 12155.56543 | 13040.53027 | 9776.454102 |
| 128140.1406 | 116379.7266 | 111969.7422 | 106597.4688 | 64055.07031 | 62705.27734 | 62663.4375 |
| 285852.9688 | 241142.0313 | 261058.875 | 274310.7813 | 227266.875 | 255545.1719 | 247081.7344 |
| 13945.22754 | 13471.37305 | 10236.84863 | 11196.75488 | 11725.46777 | 13000.82813 | 11084.54492 |
| 2302782.25 | 2116149.75 | 2177027.25 | 1958915.875 | 1727283.25 | 1916913.75 | 1654113 |

| | | | | | | |
|---|---|---|---|---|---|---|
| 16995.9082 | 16170.48047 | 18816.28906 | 17270.38086 | 12988.14648 | 8456.064453 | 10587.00391 |
| 42610.29688 | 41927.25781 | 39232.09375 | 39480.32031 | 32650.74414 | 34398.39063 | 33384.03516 |
| 148523.4063 | 137437.0781 | 137054.7188 | 140815.8438 | 152684.8438 | 144400.6563 | 154148.4844 |
| 23192.13281 | 20778.26172 | 20912.90234 | 17668.47461 | 18808.15625 | 20192.13086 | 18922.21875 |
| 34213.66406 | 53322.98438 | 21231.4043 | 20252.67773 | 65701.78125 | 44651.47656 | 64813.70703 |
| 163511.0127 | 149093.3809 | 153407.0176 | 153100.5547 | 145082.5615 | 163529.2744 | 159500.3428 |
| 173381.6094 | 146890.4219 | 170971.25 | 162320.75 | 208946.9219 | 190864.8281 | 189456.0469 |
| 2000257.625 | 1841282 | 1821824.125 | 1642125.875 | 1546247.625 | 1636040.5 | 1477896.75 |
| 122899.2109 | 114735.0391 | 112950.3984 | 111230.3203 | 106961.9609 | 108154.2656 | 105454.1484 |
| 13829.66797 | 14069.40723 | 12363.44238 | 11467.08008 | 9627.978516 | 11468.91797 | 10533.80762 |
| 14959.9873 | 14516.79199 | 13817.66797 | 14091.4707 | 13761.28125 | 13000.0498 | 13935.90234 |
| 69576.41406 | 69993.72656 | 62443.14844 | 66647.75 | 70102.5625 | 72596.84375 | 71863.32813 |
| 36904.48828 | 30973.75195 | 35016.18359 | 33594.29297 | 27575.08594 | 30696.23633 | 27268.17969 |
| 3020408.75 | 2666849.25 | 2537878.25 | 2448037.75 | 2170022.25 | 2408614.25 | 2121824 |
| 287338.1563 | 294467.375 | 251773.4063 | 223439.2813 | 224301.9688 | 242663.2188 | 229771.7188 |
| 2618.571289 | 3191.27417 | 2297.680176 | 1437.508789 | 3640.923096 | 3844.18457 | 4124.025391 |
| 105233.5391 | 93065.24219 | 97956.46875 | 90909.82813 | 125340.9688 | 121474.2188 | 122182.6172 |
| 23855.5918 | 20178.85156 | 21298.57227 | 21751.66211 | 26136.66992 | 23146.64063 | 26151.13672 |
| 38725.17578 | 38151.92188 | 37327.83203 | 31742.23438 | 33643.76563 | 37248.02734 | 37235.18359 |
| 1516940.875 | 1269961.625 | 1322704.875 | 1234491.25 | 1088042.875 | 1211932.125 | 1055302.375 |
| 7270.233887 | 7433.554199 | 6453.121582 | 6734.666016 | 3748.733398 | 5178.781738 | 5219.204102 |
| 61869.74609 | 62004.32813 | 63626.82422 | 68270.44531 | 75971.73438 | 78942.61719 | 76650.1875 |
| 26065.76563 | 26199.00977 | 28491.66992 | 26043.1875 | 25739.37305 | 25306.67188 | 26290.18359 |
| 3768.551758 | 2086.982666 | 4400.48291 | 4463.312988 | 2230.405273 | 3955.625488 | 2171.156738 |
| 16440.51172 | 17480.7207 | 15682.21191 | 16902.58398 | 17670.77344 | 17116.06055 | 16446.83008 |
| 4636.102539 | 5273.101074 | 5425.13623 | NA | 4451.82373 | 2106.896484 | 4819.965332 |
| 296591.5625 | 286427.5938 | 287583 | 290421.5625 | 293343 | 246031 | 263416.9375 |
| 1039922.75 | 789431.25 | 934094.5625 | 942410.75 | 1022630.313 | 852690.1875 | 968781.375 |
| 2042570.75 | 1835369 | 1906123 | 1723115.5 | 1655877.375 | 1657293.625 | 1460997.125 |
| 2142866.5 | 1995724.625 | 2018955.125 | 1875839.375 | 2026753.625 | 2015680.5 | 2074412.375 |
| 35931.55859 | 36140.20703 | 29308.01563 | 32159.86523 | 24553.17383 | 27770.49219 | 24188.79883 |
| 47868.05078 | 60011.71094 | 64394.64844 | 57949.58203 | 52450.74219 | 76613.41406 | 65714.59375 |
| 3951702.25 | 3784427.25 | 3311593 | 2860736 | 3307462.75 | 3545350.75 | 3244948.5 |

| | | | | | | | |
|---|---|---|---|---|---|---|---|
| | 2688469.75 | 2497965.5 | 2367925.75 | 2160570.5 | 1993542.375 | 1950909.625 | 1808103.25 |
| | 222026.8125 | 237962.1875 | 177227.4375 | 160324.0625 | 187339.875 | 162741 | 273253.6563 |
| | 35397.375 | 34691.75 | 33665.95313 | 35302.21094 | 28367.24023 | 31019.64844 | 30822.29883 |
| | 5465.27002 | 4574.85498 | 5232.30127 | 4430.120605 | 2865.007324 | 3151.560547 | 3641.94165 |
| | 3042830.75 | 3032568.25 | 3148708.5 | 3087407.75 | 3962642.5 | 3559174.25 | 3864214 |
| | 1931695 | 1652510 | 1778360.5 | 1574487.875 | 1489278.625 | 1490259.25 | 1407090.375 |
| | 132290.542 | 142774.0605 | 115035.1543 | 119038.1611 | 131571.3711 | 137144.2227 | 142509.0986 |
| | 63235.58203 | 63825.79297 | 61356.27734 | 62750.51563 | 86415.94531 | 102113.7266 | 101492.5859 |
| | 365184.1489 | 347415.1294 | 333308.6069 | 397703.9385 | 476501.6318 | 421751.4297 | 421999.3335 |
| | 129642.1719 | 149476.9844 | 116069.7344 | 127545 | 116807.3281 | 110120.3125 | 110911.4063 |
| | 6158.922363 | 6600.702148 | 6911.07666 | 6233.946289 | 7728.737305 | 8010.346191 | 6850.908203 |
| | 100372.0938 | 88124.95313 | 88227.09375 | 84288.80469 | 65518.3125 | 66213.58594 | 64623.18359 |
| | 54127.70313 | 46773.76172 | 53099.05469 | 57066.82422 | 82525.90625 | 73320.39063 | 73805.64063 |
| | 84951.22656 | 77173.48438 | 78033.25781 | 78307.28906 | 84838.67188 | 83118.23438 | 91459.125 |
| | 35965.80469 | 33762.27734 | 29088.76953 | 30361.42773 | 24365.91992 | 26016.30859 | 25737.07227 |
| | 77049.55469 | 82841.08594 | 72528.97656 | 67623.41406 | 62760.60547 | 65943.28906 | 61652.96484 |
| | 174530.875 | 171897.9219 | 173078.8125 | 151742.2344 | 151508.0469 | 138507.7188 | 166588.7188 |
| | 46003.65234 | 38250.09375 | 41130.08594 | 49569.14453 | 59038.93359 | 55777.81641 | 56032.03906 |
| | 206399.7813 | 246953.1875 | 165701.6719 | 170822.7813 | 140694.2344 | 208713.125 | 120210.4453 |
| NA | | 948.5690308 | 1430.005981 | 1495.492432 | 2803.395996 | 3406.880127 | 3957.830322 |
| | 18146.5918 | 14467.53906 | 16022.43848 | 15124.65234 | 13240.78613 | 13631.90918 | 11466.81543 |
| | 9301.805664 | 11446.14258 | 7709.589355 | 7611.64502 | 12061.63867 | 13472.58984 | 12718.38965 |
| | 96407.8125 | 76910.11719 | 98104.39844 | 87764.5 | 72395.82031 | 69243.17969 | 76535.22656 |
| | 12911.22559 | 15649.3125 | 11624.60449 | 9854.999023 | 10948.87695 | 9854.22168 | 10249.41016 |
| | 38258.57031 | 40205.94141 | 38622.33984 | 40154.55078 | 63738.98438 | 62854.89844 | 63290.03125 |
| | 98079.65625 | 87173.40625 | 86093.75781 | 78680.61719 | 84271.3125 | 78240.01563 | 81841.67969 |
| | 107347.6875 | 92333.49219 | 108557.1563 | 78199.08594 | 73139.64063 | 91789.29688 | 89274.64844 |
| | 174477.4219 | 177556.125 | 155893.7031 | 154605.4531 | 157753.3281 | 171044.5625 | 164469.5781 |
| | 39027.38672 | 33777.03125 | 37090.92578 | 35949.49609 | 32716.20313 | 30222.11523 | 28514.93555 |
| | 5315.382813 | 5214.532227 | 4628.15625 | 4958.029785 | 6847.347168 | 6511.618652 | 7936.314941 |
| | 159512.6406 | 156397.125 | 156520.4063 | 154853.2344 | 112361.3516 | 106230.2188 | 113997.1094 |
| | 80515.0625 | 62046.34375 | 77079.42969 | 71043.98438 | 62375.64844 | 62915.96094 | 68429.375 |
| | 9584.817383 | 8498.430664 | 9917.727539 | 8819.805664 | 14272.84668 | 11136.05664 | 16770.2207 |

| | | | | | | |
|---|---|---|---|---|---|---|
| 26840.09961 | 23755.00586 | 24992.63086 | 23212.33789 | 21609.55469 | 20530.9668 | 21028.89648 |
| 185218.7969 | 174527.625 | 171431.3125 | 177410.5313 | 222168.6563 | 196083.125 | 216046.5781 |
| 1151893.5 | 989347.25 | 1041381.063 | 1006128.188 | 1211811.875 | 1124820 | 1120502.875 |
| 176223.4844 | 163760.1406 | 160394.7031 | 146706.1094 | 112291.2266 | 104174.6563 | 107420.1484 |
| 656748.5625 | 671903.75 | 619302.75 | 649961.25 | 787712.4375 | 726563.5 | 753983.625 |
| 50411.70313 | 52749.67578 | 43101.68359 | 41614.24219 | 47711.62891 | 50011.46094 | 43142.35547 |
| 43811.94141 | 45324.55469 | 38871.08984 | 32951.94531 | 21247.32813 | 20993.26563 | 22105.81836 |
| 358750.8438 | 346029.0313 | 312437.3438 | 341642.3438 | 391701 | 410987.0625 | 409345.4063 |
| 43965.57813 | 35183.54688 | 47653.32031 | 47043.76172 | 42499.0625 | 59421.39063 | 53470.47656 |
| 5255511 | 4367630 | 4738369 | 4319654 | 3794805.75 | 3907040 | 3618790.25 |
| 222795.5156 | 210125.2031 | 212162.4844 | 214432.1719 | 222688.8906 | 252142.1563 | 230871.8281 |
| 78172.625 | 64782.41406 | 73777.60156 | 69257.10938 | 47855.05469 | 49819.94922 | 45882.52344 |
| 68312.50781 | 68310.21094 | 46588.39453 | 51621.88281 | 112211.625 | 143782.6875 | 86018.29688 |
| 6401.29248 | 6821.702637 | 6665.666992 | 5558.876465 | 9362.451172 | 9294.625977 | 8159.803711 |
| 86523.55469 | 78947.71094 | 77661.97656 | 73617.96094 | 35072.09766 | 40464.09375 | 38648.92969 |
| 286616.2188 | 236091.3438 | 281541.0625 | 265708.5625 | 245806.7969 | 256805.7813 | 270651.6875 |
| 103114.5547 | 109800.8672 | 96701.45313 | 97577.08594 | 105177.8203 | 94240.42188 | 97273.34375 |
| 3320.559326 | 3944.770996 | 3346.972412 | 3510.578125 | 3124.271973 | 3196.413574 | 3311.809814 |
| 861093.4375 | 753830.6875 | 737439.625 | 771323.3125 | 733750.4375 | 706380.9375 | 702105 |
| 114418.2734 | 118797.9688 | 103973.3594 | 97806.67188 | 85874.32813 | 67288.78906 | 62951.08984 |
| 171849.0469 | 155674.7031 | 165465.5469 | 151461.2344 | 157180.7344 | 182681.5469 | 180353.25 |
| 4632.357422 | 10882.67578 | 927.5062866 | 2728.771729 | 9709.459961 | 5647.861816 | 9661.400391 |
| 39378.59766 | 24126.06445 | 41062.71484 | 39492.03906 | 33004.99219 | 29385.94141 | 31147.70313 |
| 137664.3125 | 135272 | 118897.4297 | 114136.0781 | 108422.6797 | 92830.02344 | 108773.5469 |
| 26309.82617 | 24116.97656 | 24103.85156 | 23906.77148 | 20792.42383 | 21360.23047 | 20937.41992 |
| 23980.58398 | 19906.46875 | 22338.92578 | 17809.50977 | 37119.60938 | 37195.29297 | 40717.19141 |
| 176360.9688 | 156070.9063 | 148107.1875 | 154814.7031 | 236137.9531 | 243881.1719 | 247415.5625 |
| 146671.5156 | 144465.4844 | 121718.7344 | 134614.8438 | 133477.0938 | 135939.8125 | 137219.7969 |
| 3332.495605 | 2496.446533 | 3155.536133 | 2333.470215 | 1346.0802 | 1908.232788 | 1576.764404 |
| 21060.0293 | 46870.76953 | 30062.9043 | 40976.09766 | 23065.67969 | 11458.93945 | 44714.34375 |
| 31427.26563 | 24336.35742 | 30251.98047 | 30605.62695 | 32482.46094 | 31052.97656 | 29966.10352 |
| 6896.778809 | 7573.554199 | 4360.596191 | 4757.253418 | 8485.001953 | 8210.336914 | 7363.625 |
| 712191.5625 | 571589.6875 | 666767 | 645567.9375 | 564171.5625 | 602448.5625 | 631351.125 |

| | | | | | | |
|---|---|---|---|---|---|---|
| 83061.67969 | 52920.92578 | 72892.46094 | 80825.11719 | 39533.32031 | 37471.04688 | 42950.5625 |
| 1688458.75 | 1424111.5 | 1471701.625 | 1365000.5 | 1188996.25 | 1350640.25 | 1162851.625 |
| 17593.44922 | 10346.59766 | 17257.08984 | 14091.90918 | 10693.35938 | 8986.085938 | 9947.381836 |
| 128613 | 135818.4219 | 97516.73438 | 97221.40625 | 128532.4297 | 113583.4063 | 109799.0781 |
| 1218299.125 | 1291809.75 | 1088470.375 | 1106787.25 | 1353293 | 1342160.375 | 1379517 |
| 36117.94141 | 33684.01953 | 35204.96094 | 34073.01563 | 32672.37695 | 36478.69922 | 35373.41016 |
| 1849508.25 | 1464550.625 | 1624919.75 | 1479905.125 | 1377737.75 | 1545269.625 | 1368577.125 |
| 279659.1875 | 257048.3594 | 246890.6094 | 230066.4063 | 211256.4063 | 210657.5781 | 209947.375 |
| 8210.110352 | 7568.579102 | 7013.98291 | 6542.216797 | 6659.067383 | 7035.714355 | 6903.074707 |
| 166674.9375 | 138894.0781 | 160563.4375 | 152819.75 | 144214.2813 | 159531.6406 | 161294.6719 |
| 93424.59375 | 101049.4297 | 69451.75 | 88868.55469 | 89715.71094 | 94956.21094 | 87846.96094 |
| 34365.48047 | 29171.00195 | 32398.16211 | 33434.0625 | 26056.26367 | 28996.75781 | 29374.91602 |
| 11018.56934 | 6731.138184 | 10509.28906 | 4695.870117 | 5424.114746 | 29744.36914 | 14604.91211 |
| 173072.3281 | 154469.7656 | 154533.6406 | 153511.5781 | 128473.1016 | 119848.4688 | 123753.9297 |
| 16308.1875 | 16944.95898 | 17679.15625 | 15107.24121 | 15476.65625 | 15560.28125 | 16701.13867 |
| 19977.14258 | 18604.12695 | 17918.25586 | 17335.98633 | 19181.51172 | 17491.60742 | 18767.32422 |
| 367241.6875 | 311174.0625 | 332512.5 | 338771.25 | 307750.4063 | 300444.0313 | 294880.8125 |
| 6720.172363 | 5189.657227 | 5761.719727 | 6269.221191 | 3699.449463 | 4050.361084 | 4507.717285 |
| 188403.0625 | 177741.5 | 170923.125 | 172569.5938 | 189507.7656 | 172961.625 | 195770.6875 |
| 75001.25781 | 69904.76563 | 66784.99219 | 72857.4375 | 141043.625 | 148484.7813 | 152081.7188 |
| 424495.375 | 334085.8125 | 365791.1875 | 322877.625 | 281720.9688 | 331047.3438 | 291220.5938 |
| 329747.3398 | 318678.291 | 298878.9082 | 303171.7676 | 225770.8555 | 238914.8828 | 243658.5234 |
| 28372.66016 | 26622.83008 | 27361.98242 | 26434.80664 | 30693.42383 | 27097.16797 | 30489.49414 |
| 115467.2344 | 90847.14063 | 111078.125 | 86947.97656 | 60169.44922 | 75009.72656 | 62147.53906 |
| 8187.919434 | 8482.072266 | 5994.005371 | 7847.155762 | 10922.3418 | 9857.561523 | 10090.38379 |
| 3031288 | 2541221.75 | 2586318 | 2523920.25 | 2519835 | 2591841.75 | 2409607.5 |
| 19487.60938 | 16691.4375 | 15737.88574 | 16154.47461 | 17757.33984 | 17098.68945 | 17324.86719 |
| 15013.92188 | 14031.24707 | 11805.4082 | 12872.92773 | 16433.70508 | 17591.90234 | 14932.55078 |
| 178414.0469 | 142829.0156 | 163980.5781 | 169000.7969 | 194264.75 | 194602.4844 | 207704.2031 |
| 4715764 | 4154979.25 | 4103016.75 | 3674552.75 | 3216031 | 3486793.5 | 3116072.25 |
| 62091.83594 | 67352.41406 | 50024.57422 | 47869.13672 | 57069.10938 | 55480.03516 | 59817.5 |
| 1105477.875 | 2460191 | 1252306.375 | 403971.5313 | 375620.625 | 1083146.625 | 385376.0938 |
| 62176.22656 | 57436.58594 | 63372.51172 | 57215.53125 | 53320.88281 | 66903.86719 | 51422.40625 |

| | | | | | | |
|---|---|---|---|---|---|---|
| 371010.4063 | 309893.875 | 350796.6563 | 361913.3125 | 142810.1406 | 140894.6094 | 156725.6094 |
| 1244673.875 | 1044008.063 | 1037004.063 | 965132.5 | 876734.5 | 949378.75 | 869217 |
| 30740.69141 | 27630.01563 | 26359.07813 | 28842.74219 | 31319.94727 | 33901.16016 | 32836.33203 |
| 353201.9688 | 310578.5625 | 286145.4688 | 286542.0938 | 389098.2813 | 359419.6875 | 372830.875 |
| 12690.46191 | 12998.12988 | 11528.39551 | 9980.404297 | 7577.64209 | 7457.065918 | 6801.10791 |
| 924654.875 | 763050.1875 | 802437.0625 | 823362.875 | 705508.9375 | 721574.125 | 677342.875 |
| 2358903.25 | 2047123.375 | 2027925.875 | 1916700.375 | 1714985.5 | 1815679.375 | 1493587.5 |
| 7310.902832 | 7606.246094 | 5295.90625 | 8477.191406 | 10257.04688 | 7938.76709 | 8454.344727 |
| 261946.25 | 269235.0938 | 231141.2969 | 252627.4219 | 301348.125 | 304629.25 | 281813.5313 |
| 203859.625 | 187839.625 | 160918.5313 | 182162.5313 | 281743.3438 | 244420.1406 | 239081.9375 |
| 242846.5156 | 218576.9219 | 227035.9688 | 222339.2969 | 245948 | 244407.9375 | 246241.9844 |
| 1171263.25 | 1150326 | 1154502.875 | 1320645.625 | 984882.8125 | 852433.75 | 1012422.938 |
| 666822.5 | 584363 | 641066 | 586570.875 | 470419.75 | 510854.5625 | 505973.875 |
| 32600.71875 | 33044.53906 | 24744.29688 | 25924.78125 | 24545.15234 | 22007.78516 | 26471.23047 |
| 24049.54883 | 30284.35156 | 21632.8457 | 20588.24609 | 30899.35156 | 29659.78516 | 27284.67969 |
| 34370.28516 | 41485.82031 | 26140.625 | 17215.48633 | 20182.83398 | 44581.44922 | 22626.15234 |
| 6139.675293 | 9773.326172 | 6019.355957 | 5201.747559 | 6891.91748 | 22962.32422 | 7060.046875 |
| 8664.950195 | 8370.758789 | 8701.141602 | 8218.52832 | 7494.04541 | 7301.736328 | 7604.214355 |
| 176349.25 | 173110.9688 | 174860.5156 | 173020.5938 | 163752.2344 | 172624.7188 | 167818.1406 |
| 20309.19141 | 31954.2168 | 17875.21289 | 19572.83203 | 96402.92969 | 46081.42969 | 54496.8125 |
| 47229.48047 | 56905.89063 | 30202.56055 | 29015.74023 | 31326.73242 | 32466.01367 | 33307.42188 |
| 3765533.75 | 3271920.75 | 3340357.5 | 2983552.25 | 2943837.75 | 3043212.75 | 2611564.25 |
| 62781.14844 | 61170.98438 | 61357.65234 | 52468.41406 | 54068.86719 | 49734.58594 | 47024.71484 |
| 137711.7969 | 126094.9766 | 128503.7031 | 136843.6094 | 163310.1094 | 149654.7969 | 162768.0313 |
| 1044883.75 | 755191.8125 | 876668.5625 | 950414.8125 | 897689.625 | 983397.875 | 1032509.313 |
| 112946.6719 | 93599.90625 | 95093.72656 | 90990.72656 | 59908.91406 | 74750.88281 | 63054.12109 |
| 29959.1543 | 27244.89453 | 28484.58594 | 27327.43164 | 25464.80273 | 28781.79492 | 26276.30469 |
| 56643.78125 | 54552.76563 | 62998.80078 | 45476.74219 | 33301.60156 | 39404.69141 | 36702.27344 |
| 255446.5313 | 249641.7656 | 252959.6563 | 234182.0469 | 224399.4688 | 240835.0625 | 244032.4688 |
| 863394.375 | 755243.5625 | 748689.3125 | 692606.375 | 648596.6875 | 656011.1875 | 614060.8125 |
| 111677.8125 | 83663.32813 | 105140.2266 | 89807.22656 | 67856.77344 | 74638.16406 | 66754.50781 |
| 578311 | 489849.3438 | 550952.5625 | 556018.75 | 431728.9375 | 441682.0938 | 452456.9688 |
| 24994.12109 | 21692.70117 | 23270.94141 | 22379.62891 | 20351.14648 | 19242.99219 | 20726.49023 |

| | | | | | | |
|---|---|---|---|---|---|---|
| 122520.0391 | 102930.7109 | 112971.3828 | 110798.4844 | 73461.45313 | 83851.0625 | 72592.75781 |
| 69812.89063 | 58508.59766 | 64929.41797 | 69780.95313 | 40489.81641 | 42651.4375 | 42537 |
| 33473.47266 | 28977.54297 | 32558.0332 | 31025.22266 | 34528.1875 | 33444.05469 | 31013.58008 |
| 214024.207 | 240561.3438 | 159831.3301 | 166191.6992 | 209472.3594 | 203094.2852 | 233415.6641 |
| 58092.88672 | 45675.81641 | 40734.96484 | 51346.78125 | 48933.04297 | 53996.71875 | 60760.82813 |
| 18721.04883 | 16013.18555 | 13753.72754 | 17135.42969 | 14231.52246 | 14001.14551 | 14457.6709 |
| 72279.84375 | 68110.41406 | 50537.70313 | 39771.64453 | 46340.64844 | 79930.6875 | 58063.89063 |
| 2937.373291 | 2579.523438 | 2399.19043 | 1815.553101 | 3735.795654 | 4206.309082 | 3908.812744 |
| 17505.13086 | 18165.20313 | 15702.87793 | 13735.60547 | 35390.26172 | 19028.1582 | 20915.94727 |
| 193710.8281 | 217909.625 | 247764.6406 | 157689.6406 | 156956.7344 | 236060.4063 | 177239.3125 |
| 25356.94141 | 19055.44922 | 27172.96484 | 21008.05859 | 42403.45703 | 43861.43359 | 33766.86328 |
| 47476.35156 | 42646.72266 | 52132.46094 | 60891.26563 | 65496.83203 | 78053.01563 | 80334.75 |
| 180213.4219 | 160416.3281 | 163596.1563 | 154426.3125 | 70396.4375 | 90811.45313 | 73281.9375 |
| 45578.50391 | 47769.59766 | 40594.33984 | 39319.06641 | 33321.36719 | 32278.28711 | 30861.64648 |
| 1440960.75 | 1238401.25 | 1336835 | 1191574.125 | 1137683 | 1139491.125 | 1005858.313 |
| 33912.25 | 23642.80664 | 39622.67188 | 26084.46875 | 10873.84766 | 12611.21289 | 8781.640625 |
| 1750402.5 | 1580162.125 | 1577982.5 | 1544911.375 | 1339060 | 1389459.5 | 1379805.5 |
| 230386.3281 | 158890.3281 | 236639.6094 | 180825.4688 | 248013.2656 | 236793.375 | 278891.1563 |
| 49720.36328 | 46860.97266 | 43298.99609 | 38948.72656 | 58932.03906 | 61628.76172 | 64039.14844 |
| 3155.125 | 2412.798096 | 3152.943115 | 3411.61792 | 5200.272461 | 6459.845703 | 6329.114258 |
| 20825.18555 | 15039.49219 | 20294.90234 | 15851.16797 | 16432.98242 | 20391.67773 | 14944.8252 |
| 204317.0625 | 182803.6406 | 182020.2031 | 187009.9688 | 177839.7188 | 186160.7344 | 187186.9063 |
| 175887.3594 | 163102.0313 | 164009.2813 | 179668.6719 | 125266.1797 | 114001.2891 | 120646.7422 |
| 52739.27344 | 58114.60547 | 40255.03125 | 36487.89844 | 30621.38672 | 30986.75977 | 31796.87695 |
| 69723.53906 | 61837.99609 | 56122.82422 | 60163.75391 | 105818.1016 | 101339.6172 | 97365.96875 |
| 102692.9531 | 98518.35156 | 91016.63281 | 102931.5156 | 120917.5625 | 118264.0391 | 121403.1953 |
| 39010.36328 | 41012.24609 | 38581.61328 | 31882.86523 | 46709.76172 | 51327.74219 | 51392.25781 |
| 31533.01758 | 29589.56836 | 27863.02148 | 24317.07227 | 28838.58789 | 24669.62109 | 26696.99609 |
| 3590.057861 | 5202.904785 | 2626.425293 | 5465.075195 | 6862.803223 | 4275.682617 | 4490.953613 |
| 107697.9297 | 104211.5547 | 91237.67188 | 95636.92969 | 104122.1641 | 106791.2422 | 103442.5 |
| 3376.719727 | 4172.927246 | 2273.347412 | 3710.574707 | 5739.53418 | 7606.869141 | 5904.362305 |
| 90536.59375 | 88658.41406 | 83259.67188 | 89758.86719 | 134849.0469 | 110242.1953 | 115646.1875 |
| 34751.62891 | 35894.85156 | 32615.76953 | 32150.14063 | 26679.95508 | 30242.34766 | 28232.63477 |

| | | | | | | |
|---|---|---|---|---|---|---|
| 934334.4375 | 806761.0625 | 819918.5625 | 747517.625 | 641826 | 683971.6875 | 619775.875 |
| 25939.58789 | 19807.65234 | 26461.95703 | 23994.16602 | 12511.61328 | 13841.32715 | 14854.96582 |
| 104479.7109 | 84136.61719 | 102170.7969 | 86322.6875 | 68958.15625 | 70932.21094 | 65878.75 |
| 37258.64844 | 36539.38672 | 31304.86914 | 34696.37891 | 68203.82031 | 56908.28906 | 56106.49609 |
| 57921.04297 | 116477.5234 | 28167.11133 | 15564.18945 | 97667.26563 | 64170.46094 | 89222.71875 |
| 48997.19141 | 48823.96094 | 52549.12891 | 46993.42188 | 59887.36328 | 52849.38281 | 63196.70313 |
| 7705.039551 | 6767.418945 | 5874.258301 | 5634.400391 | 5379.588379 | 8773.746094 | 5539.867188 |
| 23969.3125 | 22367.71094 | 23653.26758 | 23152 | 30747.51953 | 28062.66016 | 28252.86914 |
| 96971.125 | 91004.65625 | 88053.03906 | 82949.03125 | 88915.96094 | 81792.73438 | 87898.82031 |
| 121781 | 116232.4766 | 107383.9375 | 112481.4453 | 98451.14844 | 107945.1094 | 102377.6172 |
| 68738.53906 | 64244.70313 | 59400.71094 | 60968.58984 | 50888.04688 | 52106.35938 | 49229.34375 |
| 20713.21289 | 15679.3584 | 18062.45117 | 19216.38867 | 21967.10742 | 22423.83203 | 21761.50977 |
| 410750.9375 | 353780.25 | 377570.125 | 327786.25 | 279440.5625 | 324832.7188 | 290099.5 |
| 77299.76563 | 60182.05469 | 72138.55469 | 64761.93359 | 70316.17969 | 67140.90625 | 75191.49219 |
| 140126.7188 | 119242.1406 | 135552.5781 | 137726.5 | 209201.9844 | 191949.6406 | 203075.0781 |
| 5925.912598 | 5423.276855 | 5675.416016 | 4913.560059 | 4283.272461 | 5977.386719 | 4779.336914 |
| 54963.55859 | 55312.25 | 40380.53125 | 42783.50781 | 53553.64063 | 38800.16406 | 47428.84375 |
| 102611.3672 | 95028.19531 | 102787.4375 | 105785.9141 | 107100.375 | 88736.70313 | 104326.0234 |
| 11474.90332 | 10346.47363 | 12207.72754 | 11140.1709 | 11693.03809 | 11075.90625 | 12847.7334 |
| 19084.09375 | 16824.20117 | 16647.48047 | 16500.7207 | 26563.51953 | 26894.45898 | 25495.99023 |
| 4612.155762 | 4421.118652 | 4350.746094 | 3950.171631 | 5230.754883 | 5111.880859 | 4962.519531 |
| 237429.6563 | 210564.5625 | 212553.2813 | 194805.3438 | 138185.1094 | 172193.3594 | 155175.9531 |
| 5392.607422 | 10144.88672 | 1620.176147 | 1290.474731 | 5598.467773 | 6717.985352 | 7682.180176 |
| 16784656.5 | 15590911.5 | 14838521.94 | 14376283.44 | 11313175.38 | 11540228.97 | 11940522.53 |
| 123072.9922 | 104294.8359 | 133673.75 | 121867.5078 | 135497.4688 | 115177.2109 | 146271.4063 |
| 71756.32813 | 61959.64453 | 67228.01563 | 67564.53125 | 64287.79688 | 70614.25 | 66293.33594 |
| 120948.0469 | 112227.7852 | 114534.7773 | 102567.0742 | 100736.7266 | 98483.79297 | 97620.39063 |
| 235185.125 | 216910.4063 | 232448.8906 | 221289.3906 | 178402.4375 | 223888.2188 | 202912.2969 |
| 96394.22656 | 88177.07813 | 83388.625 | 85391.75 | 89513.10156 | 93368.02344 | 93266.96875 |
| 20181.01758 | 9189.807617 | 35880.70703 | 35685.38672 | 39247.28906 | 44704.92969 | 53094.45703 |
| 962017.5 | 737676.1875 | 910380.75 | 770869 | 704755.25 | 868182.875 | 738396.6875 |
| 2392168 | 2059635.125 | 2013361.5 | 1925049.5 | 1915918 | 1897187.375 | 1756972.125 |
| 7071.966309 | 5046.334473 | 6707.223145 | 6721.870605 | 6299.167969 | 6001.164063 | 6150.79541 |

| | | | | | | |
|---|---|---|---|---|---|---|
| 5961.530273 | 4326.736328 | 6882.921387 | 3838.546875 | 5784.722168 | 7624.452148 | 8886.90332 |
| 44966.16016 | 40873.78906 | 40483.27734 | 40808.52344 | 37476.81641 | 37254.07813 | 38999.24219 |
| 61720.125 | 45814.34766 | 59730.95313 | 54688.78906 | 51476.03906 | 54853.99219 | 57930.77344 |
| 40650.10938 | 49608.36719 | 27224.73047 | 28293.91211 | 38722.09766 | 33263.19922 | 26574.11719 |
| 5894.648438 | 5799.666504 | 4498.644043 | 5014.499512 | 6346.016113 | 4681.219238 | 5556.70459 |
| 381069.6875 | 324664.7813 | 358942.9375 | 376506.6563 | 307265.125 | 338595.75 | 309055.375 |
| 132968.5625 | 148164.7031 | 106826.8125 | 97546.5625 | 100132.8906 | 104004.8203 | 90949.4375 |
| 28802.03516 | 27068.20898 | 26076.47852 | 25979.83008 | 16026.37793 | 16499.63086 | 16180.68359 |
| 35714.77734 | 32082.39453 | 31196.10547 | 27838.47656 | 23260.18945 | 26911.80078 | 24879.15234 |
| 118997.3438 | 94130.72656 | 104989.2422 | 106792.6563 | 115518.625 | 115880.3125 | 116533.6484 |
| 113071.7734 | 101646.7031 | 104782.4688 | 116282.3359 | 153282 | 131578.5469 | 146422.5781 |
| 18107.82813 | 17726.04688 | 12445.9668 | 20993.27734 | 18973.69141 | 20768.38672 | 23682.19531 |
| 59911.38281 | 47697.71484 | 56246.13281 | 63672.70703 | 54565.76563 | 56350.55469 | 54900.87109 |
| 180412.1719 | 124452.9609 | 189050.4531 | 186021.1094 | 198912.6563 | 198076.2188 | 195217.4219 |
| 115562.2422 | 88719.60156 | 109563.6641 | 111518.8438 | 99091.00781 | 98520.14063 | 108166.8984 |
| 75915.76563 | 61718.54688 | 75073.82031 | 64098.04688 | 47017.36719 | 47141.12109 | 48464.35156 |
| 331813.2813 | 220820.3281 | 340919.3125 | 332935.25 | 362769.1875 | 348959.6563 | 387447.8125 |
| 32905.32422 | 27332.57813 | 27316.86523 | 23280.24219 | 20494.5918 | 36978.54688 | 25916.62695 |
| 104669.6406 | 107259.6953 | 75625.66406 | 73234.54688 | 124111.1406 | 114236.9375 | 118973.9766 |
| 131164.1875 | 115778.6719 | 105244.1484 | 123088.9688 | 123756.3281 | 122236.7813 | 131443.5625 |
| 982563.1875 | 893072.0625 | 816974.875 | 783825.4375 | 763102.0625 | 775746.625 | 686455.4375 |
| 69867.01563 | 58337.73438 | 65115.50391 | 58475.53516 | 59860.44922 | 57561.92188 | 63195.30859 |
| 3781.018311 | 3261.735596 | 4419.300293 | 3845.310791 | 4143.995117 | 3475.320557 | 3146.483398 |
| 95931.11719 | 85427.26563 | 93387.14844 | 89065.92188 | 92787.00781 | 82270.25781 | 93608.77344 |
| 296138.125 | 243635.3281 | 257816.0156 | 243522.7031 | 227735.9844 | 234703.7969 | 242722.875 |
| 503073.375 | 489375.7813 | 416475.8125 | 429861.9375 | 522311.6563 | 516344.125 | 544643.75 |
| 2338.791504 | 2814.368408 | 2422.279053 | 538.5976563 | 3269.750732 | 3472.450195 | 3585.383301 |
| 39550.96875 | 32613.71289 | 35601.08203 | 35054.71094 | 53156.36719 | 49421.29688 | 53098.17578 |
| 1056617.125 | 879987.1875 | 922030.5625 | 937720.875 | 916163.9375 | 948686.625 | 962724 |
| 41143.98438 | 35073.61328 | 36597.16797 | 41622.88672 | 28037.24805 | 22435.2793 | 25287.30469 |
| 228105.3125 | 215583.3906 | 208771.1563 | 205292.1094 | 209993.7813 | 224328.125 | 209680.2031 |
| 11446.92578 | 9907.851563 | 10296.28711 | 10951.57617 | 12393.48145 | 13115.4873 | 11690.99707 |
| 6138.191895 | 5790.341309 | 5822.783203 | 6001.419922 | 6701.914063 | 5391.487793 | 5512.386719 |

| | | | | | | |
|---|---|---|---|---|---|---|
| 69649.75 | 67986.22656 | 63466.66797 | 60042.46484 | 43922.90234 | 50769.48828 | 41412.25781 |
| 50402.91797 | 46154.22656 | 42619.09375 | 45764.18359 | 62024.34766 | 65476.49609 | 62378.94141 |
| 142497.2344 | 83853.125 | 97305.375 | 57334.08203 | 36469.24219 | 43673.5625 | 38760.95313 |
| 2127.225586 | 2246.583252 | 798.7106323 | 3285.710205 | 3337.53418 | 5703.651855 | 7074.963379 |
| 316935.125 | 298303.5938 | 265875.4688 | 283503.6875 | 304216.875 | 311286.25 | 302682.75 |
| 12881202 | 11032108 | 11683873 | 12683602 | 10378490 | 10057241 | 10239384 |
| 23821.13672 | 16859.77734 | 23868.13477 | 28865.1543 | 30361.11523 | 28733.06836 | 31168.9375 |
| 110019.5938 | 95651.21094 | 107674.0391 | 114703.0625 | 140977.8438 | 120205.1406 | 141278.2344 |
| 5144.724609 | 4792.822266 | 4838.808594 | 6534.779297 | 7080.29248 | 6737.432617 | 7533.910645 |
| 30054.17188 | 22703.47852 | 27620.5957 | 22490.96484 | 20300.74609 | 31229.6582 | 22055.06055 |
| 122604.2969 | 125607.5703 | 113870.4922 | 111234.9297 | 130833.3125 | 132797.6094 | 115120.4063 |
| 325464.25 | 283954.9375 | 293659.2813 | 306092.5938 | 215539.1563 | 199120.1406 | 206954.0156 |
| 57537.77734 | 49359.30078 | 64493.28516 | 62209.73047 | 53090.42969 | 100038.6484 | 103542.4688 |
| 324071.0313 | 259123.2969 | 304193.875 | 278733.7813 | 239725.7031 | 244349.6563 | 248116.125 |
| 20655.45898 | 16216.68164 | 21044.51563 | 23005.06641 | 21985.91406 | 22265.05273 | 21753.56641 |
| 9158.811523 | 10448.08594 | 6012.09082 | 5991.253906 | 4544.925781 | 4158.010742 | 4213.515137 |
| 15264.38477 | 13318.14453 | 18235.07617 | 14297.68652 | 17543.92188 | 12337.46875 | 16639.88477 |
| 132822.2969 | 156279.1406 | 111922.0781 | 100199.3594 | 144545.5313 | 149571.625 | 156475.4688 |
| 256695.4219 | 283324 | 242534.9063 | 226154.25 | 221195.0781 | 232288.4219 | 209872.1094 |
| 77869.55469 | 88982.32031 | 57586.47656 | 59097.05469 | 99949.125 | 95355.96875 | 100039.0391 |
| 158070.1406 | 127117.9375 | 136154.5 | 132876.5938 | 81149.125 | 84070.92188 | 83544.15625 |
| 43870.28516 | 35014.58203 | 42602.51563 | 42611.54688 | 55677.41797 | 84106.07031 | 56506.625 |
| 109845.7031 | 86511.70313 | 99702.92188 | 94786.35938 | 74747.69531 | 76051.92188 | 77159.57031 |
| 134245.4688 | 117981.1094 | 118426.5078 | 121352.9922 | 100399.1875 | 103788.5547 | 99399.94531 |
| 70302.15625 | 46421.37109 | 66576.45313 | 69861.38281 | 95305.35156 | 74595.40625 | 88673.53906 |
| 55049.87891 | 59505.73828 | 42250.21484 | 37789.78906 | 46434.59766 | 50055.48828 | 45712.98828 |
| 39827.15625 | 37557.34375 | 34310.14063 | 33545.86328 | 25796.64844 | 28499.46875 | 31374.44141 |
| 96294.5625 | 105532.25 | 83578.28125 | 82022.78906 | 91080.125 | 90981.26563 | 81731.27344 |
| 96429.88281 | 77574.21094 | 91507.28906 | 86871.53906 | 82953.96094 | 76868.89844 | 78020.125 |
| 167103.5625 | 154027.8906 | 140478.1875 | 139827.75 | 149257.7969 | 134166.0156 | 140437.5938 |
| 43336.64063 | 40457.36719 | 37408.91797 | 34785.14063 | 40945.59375 | 46844.65234 | 47894.77734 |
| 88275.96094 | 88457.33594 | 74650.60156 | 76708.875 | 96814.5625 | 98203.20313 | 98748.55469 |
| 2916474.75 | 2415865.75 | 2587681.5 | 2283348.75 | 2033029.375 | 2143216.25 | 1944604.75 |

| | | | | | | |
|---|---|---|---|---|---|---|
| 4010016 | 3388488.5 | 3389558 | 3066249.25 | 2874298.5 | 2853073.5 | 2650283.75 |
| 11362.44629 | 9232.52832 | 10197.69434 | 9762.673828 | 5323.498535 | 7015.536133 | 7102.639648 |
| 41760.51953 | 48912.54688 | 28399.70117 | 30757.61133 | 52911.17188 | 49825.37109 | 56095.89844 |
| 581135.25 | 501514.5625 | 525225.4375 | 564072 | 670893.625 | 589776.5 | 675191.9375 |
| 39401.55859 | 34014.11328 | 40774.46484 | 36807.86719 | 38384.53125 | 53317.06641 | 43789.77734 |
| 434575.0313 | 350536.8438 | 398301.3438 | 384489.8438 | 383477.7813 | 397286.9063 | 427184.625 |
| 156647.7969 | 150091.0781 | 142058.2813 | 143814.4531 | 148309.5469 | 140976.4688 | 140236.7031 |
| 768809.3125 | 721690.375 | 633342.8125 | 555681.5 | 489450.3125 | 617523.125 | 483294.7188 |
| 9886.451172 | 8920.054688 | 11178.52832 | 8826.871094 | 15716.47559 | 7637.709961 | 15958.68848 |
| 148281.625 | 138544.7969 | 130368.8359 | 139420.4531 | 162139.2188 | 161843.3438 | 154190.4375 |
| 286253.5313 | 265686.8438 | 255370.1563 | 250590.5 | 264520.1563 | 269524.25 | 276290.0625 |
| 97283.64844 | 107728.0469 | 98291.42188 | 109764.2656 | 145876.9375 | 171957.2188 | 166706.1719 |
| 658244.9375 | 736708.3125 | 512177.1875 | 433205.9375 | 426566.0938 | 506011.6563 | 381782.0625 |
| 152916.6406 | 146507.5781 | 130304.6016 | 134023.8594 | 264767.375 | 230163.6094 | 257541.2813 |
| 636493.1875 | 742147.375 | 599297.1875 | 431776.6563 | 395674.0313 | 539510.3125 | 1161794.875 |
| 135891.6406 | 139074.4531 | 120553.0156 | 122467.7891 | 137581.5938 | 117045.0078 | 136479.4063 |
| 38081.98828 | 25909.86914 | 43092.0625 | 38398.89063 | 30381.08398 | 35769.71094 | 39111.03516 |
| 115107.3535 | 131190.0117 | 117306.0723 | 99963.50781 | 141559.8047 | 127553.2773 | 121519.6426 |
| 25553.04883 | 24486.9082 | 23724.24805 | 21960.04297 | 17565.3418 | 19750.70898 | 18920.48047 |
| 255719.8438 | 214303 | 234070.6406 | 233082.6719 | 241272.8906 | 247981.7969 | 225014.3281 |
| 61154.70313 | 61002.57422 | 58777.47656 | 64083.83203 | 46219.07031 | 42769.21094 | 45890.28516 |
| 350071 | 278463.25 | 341177.5938 | 318040.1875 | 364342.5625 | 343307.8438 | 346504.1563 |
| 1973.173218 | 3689.674805 | 772.1557617 | 686.7589722 | 5596.158203 | 5602.117188 | 5466.712402 |
| 19702.68945 | 4240.094238 | 17973.10742 | 18116.82813 | 20866.4082 | 27061.00781 | 24101.99219 |
| 22674.45508 | 19233.92773 | 22952.62109 | 21162.55078 | 22072.93555 | 20715.42773 | 20745.55664 |
| 20570.32813 | 18201.6582 | 16328.6748 | 17407.82227 | 28442.95703 | 28941.50586 | 29074.59766 |
| 50895.07813 | 53393.71875 | 42048.54297 | 37855.30859 | 43285.78906 | 41228.13672 | 44940.31641 |
| 51231.53516 | 44942.33594 | 44642.22266 | 47519.36328 | 44018.91016 | 43749.56641 | 46769.83203 |
| 259454 | 270486.9063 | 200316.5938 | 183094.8906 | 361604.7188 | 374563.0625 | 401754.9375 |
| 122181.5078 | 113085.3438 | 106177.5938 | 95691.74219 | 63082.13672 | 68462.98438 | 63075.89844 |
| 28454.61719 | 28461.92188 | 22275.37695 | 25361.6543 | 30485.00586 | 21621.76563 | 31070.58008 |
| 1104565.125 | 836461.4375 | 932801.5625 | 940019.3125 | 850876.9375 | 950798.375 | 852893.25 |
| 22576.87695 | 22338.08789 | 19327.26172 | 19535.50781 | 20560.67773 | 24302.96484 | 21510.8418 |

| | | | | | | |
|---|---|---|---|---|---|---|
| 1915763.875 | 1631955.75 | 1896426.5 | 2125596.75 | 2406733.75 | 2243581.25 | 2369652.25 |
| 3922927.75 | 3052582.5 | 3253209.75 | 3175111.5 | 2912360 | 3149865.25 | 2959860.5 |
| 5150.099121 | 4625.967773 | 5618.033691 | 4960.275391 | 4899.538574 | 4670.568848 | 5066.662598 |
| 17009.40625 | 18730.85156 | 13490.24609 | 12739.11914 | 13677.49414 | 13974.52148 | 14528.34863 |
| 57768.07813 | 54565.57422 | 49240.76953 | 49369.70703 | 47320.80078 | 41352.81641 | 44187.90625 |
| 65355.53516 | 60353.01953 | 56731.75 | 63095.05469 | 70093.875 | 65329.61719 | 69259.89844 |
| 28113.4043 | 31852.14648 | 26292.13477 | 21563.60156 | 28882.99609 | 36868.65625 | 30053.4668 |
| 2567109.5 | 2004686 | 2288840 | 2038555.625 | 1954163.25 | 1981726.75 | 1854806.125 |
| 16251.0332 | 11305.13574 | 12148.68652 | 14108.67188 | 12427.05273 | 12153.92285 | 9748.199219 |
| 90697.13281 | 83235.85938 | 84902.75781 | 87195.25 | 83892.29688 | 89433.42969 | 88697.46875 |
| 93661.27344 | 78915.60938 | 79627.07031 | 78275.01563 | 58839.20703 | 70350.10938 | 58044.55859 |
| 380866.7188 | 303454.25 | 345886.3125 | 341580 | 310764.1875 | 277831.5938 | 308756.875 |
| 112170.2422 | 102478.8672 | 97752.78125 | 95706.15625 | 90245.3125 | 85780.00781 | 84300.28125 |
| 938671.8672 | 698624.5615 | 856305.8701 | 1024628.288 | 1400268.107 | 1431004.797 | 1408556.758 |
| 75049.78125 | 69016.78906 | 62541.64453 | 70869.49219 | 71787.48438 | 63756.83594 | 68101.75 |
| 37343.64453 | 32903.30859 | 31921.86719 | 32194.61328 | 33998.91406 | 34988.68359 | 33360.21094 |
| 151140.9531 | 135650.4219 | 134920.3281 | 136970.5781 | 61757.29688 | 53252.9375 | 55975.82031 |
| 102161.0625 | 79054.84375 | 93720.82031 | 81537.82031 | 66906.16406 | 67823.16406 | 67409.0625 |
| 21380.91992 | 18638.83398 | 20359.96484 | 19596.94922 | 18014.27148 | 24501.27734 | 20647.66602 |
| 2597417.75 | 2145247.75 | 2196975.25 | 1982595.375 | 1913981.625 | 1953822.125 | 1785681.375 |
| 85352.53906 | 64474.72656 | 61123.72266 | 71406.27344 | 110079.9141 | 86499.28906 | 85938.05469 |
| 83590.17969 | 72929.35938 | 76552.22656 | 85315.40625 | 62905.14453 | 60776.69141 | 60405.02734 |
| 40739.23828 | 35590.49219 | 33467.79688 | 36485.73438 | 29929.61719 | 33462.72656 | 29624.51367 |
| 12154.08594 | 10688.23535 | 10161.39648 | 10617.9502 | 17209.51563 | 19455.87305 | 17732.38086 |
| 40485.59375 | 37077.42188 | 31902.02539 | 36841.16016 | 35946.89063 | 28328.66602 | 26857.38086 |
| 75041.78125 | 60417.46094 | 61677.55078 | 59886.29297 | 40840.52344 | 45350.70313 | 43566.03125 |
| 43863.74219 | 37629.26563 | 41025.67578 | 41941.58984 | 45098.36719 | 43219.18359 | 44088.33594 |
| 8226074 | 7301918.5 | 7110983 | 6768770.5 | 5458748.5 | 5551542 | 5575190 |
| 33763.01172 | 30247.98047 | 27681.26563 | 33079.95313 | 39345.50391 | 50720.17578 | 42891.87891 |
| 66360.05469 | 61717.22266 | 56230.83984 | 55373.44531 | 53659.375 | 55423.17188 | 55517.61328 |
| 55595.07813 | 42326.57031 | 46601.75 | 51855.01563 | 55618.5 | 59118.72656 | 60424.70703 |
| 9592.473633 | 11271.25879 | 7067.831543 | 6196.884277 | 8548.118164 | 7607.154297 | 9967.339844 |
| 360265.8438 | 343428.0625 | 318342.2813 | 359474.7188 | 378723.9688 | 381574.625 | 345177.8125 |

| | | | | | | |
|---|---|---|---|---|---|---|
| 265681.1563 | 218951.7813 | 247035.6563 | 232100.4219 | 232963.4375 | 272132.3438 | 251135.1875 |
| 42211.75 | 30815.20898 | 29166.98438 | 34187.89453 | 59971.48828 | 55537.14844 | 55636.30859 |
| 52950.6875 | 49081.48047 | 45572.85547 | 44545.60938 | 50050.55469 | 50410.57422 | 51224.39844 |
| 22972.22266 | 29225.9043 | 20185.2832 | 16468.38672 | 13253.66016 | 15764.24512 | 15362.51758 |
| 124452.9688 | 105031.4766 | 104110.5547 | 121311.4844 | 127842.8125 | 123195.7891 | 125526.2969 |
| 3342548 | 2846435.75 | 2827878.5 | 2668634.5 | 2567120.25 | 2712277.5 | 2421055.75 |
| 126385.9219 | 117907.0938 | 111751.5547 | 120025.6328 | 99318.07031 | 110386.7656 | 100246.9219 |
| 75291.27344 | 70123.07031 | 67019.34375 | 61964.16406 | 84656.28125 | 81687.60938 | 80175.39063 |
| 2983.4375 | NA | 2264.876221 | 3863.659668 | NA | 6221.76416 | 5443.339844 |
| 18800.93164 | 18117.61328 | 15804.55371 | 18114.62109 | 13929.59277 | 12050.64746 | 13466.36816 |
| 1362873.375 | 1028554.063 | 1319388.75 | 1279513 | 1274461.25 | 1378837 | 1330799.25 |
| 15840.53613 | 13993.36426 | 18402.75781 | 14573.91211 | 22593.84961 | 15339.24023 | 19164.25781 |
| 10606.59668 | 9886.674805 | NA | NA | 10506.75293 | 10405.06934 | 14450.82617 |
| 97432.9375 | 78188.96875 | 88644.76563 | 85489.24219 | 79131.28906 | 151593.875 | 76552.76563 |
| 81010.75 | 60266.48438 | 92517.19531 | 80152.03125 | 76019.57813 | 64880.11719 | 73724.53906 |
| 46998.21484 | 43472.17188 | 47310.55859 | 40473.51953 | 24114.26563 | 28016.65234 | 26442.65234 |
| 3019370 | 2406052.5 | 2581985.5 | 2326525.25 | 2207789.75 | 2175686.75 | 1960945 |
| 86028.59375 | 83822.52344 | 76534.01563 | 75271.40625 | 77821.15625 | 67757.30469 | 67998.72656 |
| 4287.621582 | NA | NA | 3136.205811 | NA | 3970.565186 | 4382.766602 |
| 1549706.578 | 1257503.047 | 1431039.594 | 1437439.965 | 1319872.191 | 1423213.961 | 1520623.094 |
| 65044.39453 | 64882.91406 | 60623.78516 | 46995.84766 | 39078.05078 | 52621.1875 | 43974.40625 |
| 44548.46875 | 41582.68359 | 41728.26563 | 45713.47266 | 77788.11719 | 62913.28125 | 67755.28906 |
| 60610.70313 | 32827.47266 | 58843.46484 | 57070.96484 | 36009.30469 | 36404.11328 | 39184.74609 |
| 74808.32813 | 64726.82031 | 63099.125 | 54174.875 | 49669.5 | 52473.98828 | 44894.65234 |
| 172671.7656 | 148959.8438 | 158582.2656 | 160006.4219 | 180197.5938 | 171129.1563 | 182074.9375 |
| 686502.625 | 654886.25 | 564648.125 | 632936.4375 | 984483.6875 | 879523.3125 | 902087.8125 |
| 69052.65625 | 62035.88281 | 64006.85156 | 62066.85156 | 62371.3125 | 66411.83594 | 67775.57813 |
| 140910.6406 | 128317.4375 | 128846.2734 | 110965.5781 | 126825.2578 | 129605.8281 | 127460.5 |
| 36109.28516 | 37392.19531 | 33527.21484 | 35113.52344 | 46577.64844 | 27398.875 | 40741.69531 |
| 39326.87891 | 34925.94141 | 29904.71484 | 30216.08594 | 22896.36914 | 23299.57031 | 16135.61621 |
| 124297 | 92495.74219 | 112207.3984 | 110457.9766 | 107956.2344 | 111727.5391 | 110775.2891 |
| 30607.45117 | 28149.72461 | 26779.73047 | 25424.95703 | 29808.13672 | 25681.56445 | 28951.11133 |
| 5431.84375 | 6003.461426 | 3201.105225 | 2533.636963 | 5377.61084 | 5511.206543 | 4743.302734 |

| | | | | | | |
|---|---|---|---|---|---|---|
| 76891.53906 | 66654.35156 | 65989.6875 | 69314.39063 | 78100.36719 | 72163.10938 | 69945.78906 |
| 677026.125 | 610508.4375 | 597775.5 | 611709.5625 | 521281.8125 | 502567.5625 | 510462.4375 |
| 12326.70508 | 18160 | 6013.391602 | 5723.476074 | 11581.625 | 9909.128906 | 11336.48828 |
| 2308.436279 | 1370.158936 | 2745.523926 | 2689.625732 | 2364.06958 | 2771.151367 | 2790.476807 |
| 388948.8438 | 339925.0938 | 335305.375 | 363048 | 323753.375 | 344803.2188 | 296850.4063 |
| 202000.7031 | 201410.7188 | 174055.9844 | 179552.8438 | 183521.2188 | 194579.375 | 186145.375 |
| 239419.5781 | 184869.5313 | 206935.4219 | 201227.8438 | 124180.4453 | 132952.7031 | 127952.7656 |
| 75235.07031 | 62873.13672 | 65439.21094 | 70344.78906 | 72801.82031 | 66255.07031 | 69895.60156 |
| 899903.625 | 779279.6875 | 831351.875 | 754235.875 | 473237 | 464467.8438 | 493193.625 |
| 32002.89063 | 29141.00195 | 28936.01563 | 28742.07617 | 19764.56836 | 20462.92383 | 18330.71289 |
| 242960.1094 | 208698.7969 | 205063.25 | 215229.8125 | 223915.5313 | 192762.0469 | 206962.2813 |
| 1609188.75 | 1922599.25 | 1517506.625 | 1687408.375 | 1700105.875 | 1625897 | 11923357 |
| 228363.2813 | 199499.8906 | 210367.8281 | 206594 | 192281.9688 | 203209.75 | 209088.8906 |
| 89329.58203 | 88904.59375 | 69974.01953 | 75732.02734 | 89787.87109 | 86189.25781 | 88364.16016 |
| 1933195.75 | 1775553.5 | 1310149.75 | 1651262.625 | 1571199 | 1455944.75 | 1324868.5 |
| 2219662.5 | 2020091.125 | 1951993.375 | 2062768.75 | 2003841 | 1979250.75 | 1975205.125 |
| 6396.57373 | 5289.319824 | 6289.465332 | 8170.566895 | 7563.158691 | 5275.347656 | 7100.811523 |
| 62877.01953 | 58893.24609 | 58858.375 | 56043.26172 | 71110.32031 | 66092.71875 | 74594.4375 |
| 5030.288086 | 5540.852051 | 4256.765625 | 4384.995117 | 4945.826172 | 4493.730469 | 3957.37207 |
| 259863.4219 | 204193.625 | 217564.2188 | 198633.9844 | 203841.9688 | 175323.7344 | 192910.9219 |
| 41884.125 | 41240.3125 | 32044.90625 | 28184.16016 | 45394.46484 | 43935.59766 | 45780.76563 |
| 64139.32813 | 55007.80078 | 55728.36719 | 58304.73047 | 53444.26563 | 49896.92969 | 55520.10938 |
| 70490.42188 | 61061.00781 | 62630.39453 | 64639.27344 | 63718.41016 | 54602.66406 | 61010.03516 |
| 451394.9688 | 364678.9375 | 380475.875 | 376332.0938 | 264103.5938 | 258733.6563 | 276066.125 |
| 122927.4219 | 96046.05469 | 111555.8125 | 104661.2734 | 81370.5625 | 84904.60156 | 84802.26563 |
| 99423.36719 | 77960.85156 | 96341.78906 | 83945.42188 | 70400.28125 | 77279.44531 | 75067.64844 |
| 855939.4102 | 908323.2969 | 698402.9004 | 571141.9219 | 561075.5801 | 727848.625 | 502056.0117 |
| 19994.22266 | 15297.22852 | 18763.125 | 17942.06445 | 15612.93652 | 16640.24805 | 14744.63672 |
| 89361.32031 | 86012.63281 | 63913.43359 | 61434.91016 | 67624.09375 | 59081.52734 | 51185.38672 |
| 12520.72559 | 14518.97559 | 9070.43457 | 9129.850586 | 13963.86328 | 14336.77734 | 14537.96777 |
| 5312.862305 | 5555.740723 | 4378.751465 | 3833.463623 | 4045.289063 | 3773.608887 | 4042.056885 |
| 56381.69922 | 44094.87891 | 52132.53125 | 46449.82422 | 38382.63281 | 35836.32422 | 35853.00391 |
| 786318.9375 | 745164.5625 | 752060.1875 | 619148.25 | 740303.1875 | 713191.4375 | 645959.6875 |

| | | | | | | |
|---|---|---|---|---|---|---|
| 118490.4531 | 109863.3828 | 104374.1094 | 118917.4141 | 182850.0938 | 176593.4219 | 171969.0469 |
| 30402.45508 | 32097.45313 | 25891.77344 | 28821.8125 | 26276.96094 | 34060.85156 | 28427.32227 |
| 87048 | 87923.32813 | 83715.59375 | 72271.14063 | 68299.33594 | 77205.11719 | 62751.81641 |
| 27467.71875 | 34304.23438 | 27225.1875 | 26336.36523 | 13704.52051 | 12228.70117 | 13534.1748 |
| 38488.8125 | 30761.38867 | 35044.52734 | 35959.21484 | 39123.23828 | 37386.89063 | 39292.83984 |
| 480445.875 | 418908.9688 | 392921.9375 | 387453.9688 | 370862.7188 | 402019.125 | 403857.375 |
| 67861.64063 | 51137.50781 | 59422.58594 | 49069.42969 | 37746.41016 | 43578.35938 | 39888.60547 |
| 948735.25 | 780470.375 | 803914.1875 | 747961.125 | 693887.9375 | 782996.6875 | 632639.4375 |
| 99218.75781 | 90497.17188 | 81614.71094 | 83815.42188 | 73518.55469 | 81422.75781 | 73025.94531 |
| 341426.2188 | 354639.8438 | 234621.1875 | 194105.7813 | 240984.6406 | 203980.9688 | 501137.8438 |
| 1806050.625 | 1374372.375 | 1627207.625 | 1420737.875 | 1430475.25 | 1478987.875 | 1294817.875 |
| 95354.29688 | 79216.35156 | 88228.88281 | 90113.49219 | 92592.10156 | 84683.54688 | 92653.96875 |
| 12775.15332 | 10866.8584 | 10423.88086 | 8968.125 | 8238.224609 | 7648.595215 | 6942.603516 |
| 104687.5234 | 94242.50781 | 97273.5 | 90251.53906 | 106778.0469 | 131425.5781 | 133332.25 |
| 220727.0781 | 192642.2188 | 186895.1406 | 180092.9844 | 186582.7344 | 193458.5 | 180439.3906 |
| 55145.77344 | 41079.47266 | 47460.91797 | 46373.16016 | 47225.87891 | 41189.40234 | 48463.11328 |
| 39741.45313 | 29355.69141 | 36456.73047 | 25911.55469 | 34098.12891 | 29443.07422 | 33047.53516 |
| 132909.8594 | 103439.3828 | 121297.3125 | 115095.1563 | 128984.9453 | 137013.1406 | 128103.3516 |
| 332831.5 | 210833.9688 | 348317.9063 | 391335.7188 | 362177.2813 | 288720.4375 | 335322.5938 |
| 214799.125 | 174780.2344 | 183127.8594 | 195115.0938 | 140938.1563 | 138577.4375 | 151126.625 |
| 14016.7959 | 15779.70313 | 9087.041992 | 12635.78223 | 12912.24414 | 11214.55078 | 11109.10254 |
| 548183.5 | 520542.2188 | 485607.0625 | 504833.25 | 515154.375 | 545352.8125 | 534543.9375 |
| 85284.125 | 78035.27344 | 76322.03125 | 74139.875 | 70443.54688 | 71080.35156 | 68342.71094 |
| 1232245.875 | 924001.4375 | 1097840.625 | 1103921.75 | 783689.25 | 842521.625 | 899712.25 |
| 306411.7813 | 274792.75 | 251886.2969 | 282122.3125 | 253056.4219 | 255664.7031 | 263105.4688 |
| 11887.1875 | 16413.46484 | 11896.20215 | 10813.25977 | 40121.83203 | 23997.4043 | 33422.01563 |
| 51503.5625 | 44761.47656 | 49932.23828 | 41736.66016 | 45665.46484 | 52664.39844 | 46136.81641 |
| 189510.4688 | 174052.7813 | 178271.2813 | 173764.4531 | 181076.0469 | 185366 | 181288.0625 |
| 658587.0439 | 768701.25 | 456075.5215 | 456995.4023 | 660305.2402 | 595716.5313 | 666367.1758 |
| 392123.375 | 349258.6875 | 335236.3438 | 357981.3125 | 315769.875 | 299903.7188 | 322088.7813 |
| 11658.03711 | 10959.63965 | 8357.511719 | 10469.7373 | 12978.75879 | 16859.47656 | 14083.95801 |
| 29425.88281 | 31857.5293 | 18115.71094 | 21921.30664 | 32520.67578 | 28175.12695 | 30586.7168 |
| 31240.125 | 23352.05664 | 23789.72852 | 29542.26367 | 25875.38477 | 26385.6582 | 26379.2832 |

| | | | | | | |
|---|---|---|---|---|---|---|
| 291742.1875 | 245243.9219 | 307695.7813 | 308474 | 341301.2188 | 320804.5938 | 339525.6875 |
| 142095.0313 | 103712.7734 | 125247.0313 | 93457.34375 | 138629.5313 | 173705.25 | 181443.6563 |
| 18259.94141 | 16948.22656 | 15965.86328 | 15809.14453 | 14012.10449 | 12800.1748 | 12388.90527 |
| 171812.4531 | 158854.8125 | 136173.4063 | 154942.1094 | 206259.6406 | 207538.9063 | 208861.9844 |
| 25221.80469 | 23647.70313 | 16796.90039 | 22071.2207 | 25136.04492 | 23473.62891 | 22585.58203 |
| 8802.128906 | 9676.612305 | 9135.111328 | 7953.524902 | 8103.512695 | 5580.201172 | 10077.85449 |
| 107178.5156 | 96655.07031 | 96045.5 | 90843.10156 | 68241.3125 | 67400.61719 | 68067.05469 |
| 2059729.25 | 2164045.75 | 1801492.25 | 1822418.125 | 786180.4375 | 701839.9375 | 778653.3125 |
| 39396.58203 | 39674.51953 | 27973.75 | 27496.51367 | 20240.13672 | 22272.62305 | 20753.54102 |
| 4663.180176 | 4219.727539 | 3756.759277 | 4794.34082 | 4325.018066 | 3898.448242 | 3991.878906 |
| 6211.814941 | NA | NA | 4174.398438 | NA | NA | 3006.145264 |
| 34504.79688 | 33945.21094 | 29867.48047 | 31215.22461 | 38510.23438 | 38161.66016 | 38662.84766 |
| 146685.2969 | 126670.3516 | 133322.6719 | 124916.9922 | 107705.3906 | 112175.0156 | 108671.6484 |
| 32933.96094 | 27358.05078 | 30752.1543 | 28479.70508 | 26715.73242 | 29879.80273 | 29239.38477 |
| 307556.625 | 263966.8438 | 273569.4063 | 273008.3125 | 276113.5313 | 249142.3281 | 279634.375 |
| 217570.0469 | 218308.5938 | 172433.3594 | 175854.9063 | 203367.375 | 204497.2188 | 230590.5156 |
| 6344.231445 | 7626.200684 | 5214.156738 | 5812.155762 | 4438.775391 | 4621.98584 | 5070.491211 |
| 109516.7188 | 98117.02344 | 95018.39063 | 88720.85938 | 82319.47656 | 84393.19531 | 87787.8125 |
| 23045.14258 | 14791.16406 | 26013.16797 | 22952.21484 | 19795.2207 | 17152.66211 | 22870.41406 |
| 51537.86328 | 38761.42578 | 45816.19531 | 44292.85156 | 46672.18359 | 49148.44141 | 50630.35938 |
| 9326.912109 | 9247.745117 | 8409.574219 | 7668.330566 | 7162.94873 | 7474.004883 | 6996.299316 |
| 159188.9844 | 163772.875 | 161729.2813 | 107383.7031 | 71446.48438 | 110415.3359 | 69428.92969 |
| 308644.5625 | 291650.4688 | 257381.4063 | 267161.5 | 229607.3125 | 214021.25 | 226689.3125 |
| 12699.48145 | 10104.66602 | 12819.79102 | 10010.08691 | 12098.88477 | 15265.3877 | 13164.22949 |
| 56653.36328 | 58030.80078 | 46943.375 | 49448.90625 | 53788.84375 | 50957.19141 | 56806.84766 |
| 571571.9375 | 483592.5 | 504978.6875 | 543658.5625 | 651925.875 | 622460.6875 | 612721.8125 |
| 8825.207031 | 10180.39844 | 7613.139648 | 9496.519531 | 10180.13086 | 11463.24219 | 8927.37207 |
| 13914.62402 | 10719.13281 | 12863.69727 | 10931.33398 | 19167.64258 | 15230.62891 | 16820.91797 |
| 34730.03906 | 28782.52539 | 29867.2207 | 31559.24609 | 21057.14648 | 25635.4707 | 22260.04102 |
| 36030.17969 | 34648.5625 | 32361.37891 | 37713.37891 | 54907.86719 | 47458.79688 | 53772.1875 |
| 258700.7656 | 176504.3906 | 251547.6719 | 233163.3594 | 190898.7344 | 210406.9063 | 201338.1563 |
| 206594.0781 | 159763.7969 | 181366.3438 | 178436.4375 | 178090.25 | 186707.9844 | 178724.5313 |
| 43665.92188 | 42517.30078 | 39854.28125 | 39206.55078 | 42739.27734 | 44443.76953 | 44766.51953 |

| | | | | | | |
|---|---|---|---|---|---|---|
| 892169.75 | 776992.375 | 740823.5625 | 507813.4688 | 499476.875 | 914738.1875 | 668713.875 |
| 279374.9063 | 192357.2813 | 262042.1406 | 239577.9844 | 238793.375 | 259821.8594 | 233160.6406 |
| 104874.9766 | 93886.41406 | 92249.51563 | 91575.53125 | 90980.21094 | 99362.125 | 89127.85938 |
| 1110663.25 | 866158.5625 | 929613.875 | 862262 | 883311.1875 | 907102.4375 | 731767.4375 |
| NA | NA | 1894.911499 | 1081.244995 | 2181.37207 | NA | NA |
| 410701.9297 | 358002.9297 | 377123.9063 | 275800.5469 | 331071.6602 | 419869.9922 | 346143.0234 |
| 12783.42871 | 12051.99316 | 12728.38574 | 12905.78906 | 10687.20703 | 14658.50098 | 15633.95898 |
| 64315.20313 | 57950.79688 | 57784.33203 | 58135.37109 | 53651.10156 | 58647.45703 | 54203.85547 |
| 708115.6875 | 603012.8125 | 605089.375 | 554823.8125 | 297232.2188 | 278058.2813 | 277926.75 |
| 49596.59375 | 41643.23438 | 44740.33203 | 43900.60938 | 53174.75781 | 51519.875 | 53684.20703 |
| 11226.59961 | 12117.53125 | 10620.7168 | 9140.027344 | 13294.19238 | 14285.15137 | 14763.41895 |
| 95229.57031 | 73769.98438 | 86718.78125 | 87152.17969 | 70009.15625 | 65699.32031 | 71073.89844 |
| 397156.6875 | 301916.125 | 371116.75 | 387259.0938 | 278675.1563 | 303374.5313 | 314898.7188 |
| 201380.6563 | 173661.5156 | 181225.0625 | 177718.5156 | 144921.5313 | 135661.625 | 138563.2656 |
| 61055.74609 | 51594.07422 | 56255.95703 | 55708.25391 | 50541.83594 | 52952.06641 | 52880.55469 |
| 43898.46875 | 40902.37891 | 35575.68359 | 39510.72656 | 60497.77734 | 52205.17188 | 56261.25391 |
| 9214.005859 | 12112.45898 | 7901.862793 | 6351.926758 | 4126.418457 | 4713.046875 | 3371.541992 |
| 25074.49414 | 20232.66797 | 24263.82031 | 22502.44141 | 24437.22461 | 25467.21289 | 25833.71289 |
| 725528.7773 | 601420.2148 | 611934.1563 | 549161.4102 | 657037.875 | 705222.9258 | 730840.1406 |
| 699338.8125 | 564392.625 | 609517.3125 | 566020 | 438979.8438 | 467729.6875 | 463465.2813 |
| 24766.45313 | 23454.17578 | 19396.88281 | 19692.2793 | 16500.94141 | 18330.8418 | 16948.73633 |
| 241024.375 | 180731.3125 | 215017.5 | 198105.7188 | 179877.8125 | 180129.5156 | 178039.1563 |
| 149245.4219 | 113912.0625 | 124564.0469 | 124579.5859 | 71197.45313 | 67790.47656 | 68002.79688 |
| 48243.48828 | 46909.76172 | 35344.82422 | 39984.46094 | 42658.24219 | 37311.57813 | 38089.73828 |
| 28828.33398 | 35328.08984 | 30245.54297 | 32310.44336 | 49217.26563 | 49928.46484 | 50900 |
| 341412.0625 | 346413.4063 | 276955.9375 | 327710.25 | 497794.9375 | 509445.1875 | 512762.5625 |
| 2730.14624 | 2391.503174 | 2989.655273 | 3392.380371 | 3360.101807 | 2715.375244 | 3630.381836 |
| 464271.2813 | 382313.0625 | 378255.0625 | 350662.4375 | 269862.0313 | 293597.5625 | 281846.7188 |
| 7287.446777 | 2948.753906 | 7016.572754 | 6652.917969 | 8179.927246 | 8714.006836 | 7927.610352 |
| 111415.9297 | 93972.53125 | 101345.2188 | 105550.8203 | 88597.42188 | 85229.50781 | 85959.42188 |
| 47319.19141 | 50662.48828 | 30875.99414 | 25291.1582 | 20698.38672 | 22866.94922 | 20591.25781 |
| 557681.0313 | 513515.3203 | 445577.0469 | 343726.1563 | 369332.0078 | 562204.5234 | 419568.5469 |
| 22487.00977 | 16179.03027 | 19164.71289 | 18082.50391 | 13735.11914 | 14336.10645 | 15160.54785 |

| | | | | | | |
|---|---|---|---|---|---|---|
| 51599.91406 | 47811.00391 | 43563.73438 | 45489.02734 | 65483.57813 | 55610.86328 | 55392.99219 |
| 1730.990112 | 1275.727661 | NA | 2292.864014 | 2264.056885 | 1392.846191 | 899.4387817 |
| 3972929 | 3161591.75 | 3652439.5 | 3912714.75 | 3408670.75 | 2988955 | 3483353.25 |
| 1431889.75 | 1166025.375 | 1207367.875 | 1125257.125 | 1029183.563 | 1080709.875 | 996519.0625 |
| 198459.0781 | 205145.6719 | 169757.3906 | 179142.6563 | 208504.0156 | 208306.9531 | 188032.2344 |
| 23984.01758 | 20623.38672 | 19459.44141 | 15901.84473 | 10922.79688 | 12607.79395 | 11808.9082 |
| 57855.27734 | 52234.03125 | 45040.80859 | 44597.69922 | 67998.30469 | 63915.65625 | 71468.47656 |
| 220370.8906 | 187583.9375 | 212958.9063 | 191531.3906 | 218033.4688 | 270962.375 | 223246.4531 |
| 3703.804199 | 3042.224121 | 3577.511963 | 3002.703369 | 5015.809082 | 3221.671387 | 4556.416016 |
| 144982.7813 | 158752.7188 | 99825 | 102873.3672 | 137919.7813 | 143542.5781 | 142000.4531 |
| 85617.42188 | 76761.22656 | 74728.02344 | 71072.92188 | 47608.33594 | 48279.53516 | 45922.22266 |
| 865705.8125 | 770848.75 | 740243.5625 | 642507.75 | 605699 | 579019.5 | 630739.4375 |
| 39628.63281 | 34253.62109 | 39242.32813 | 36487.85547 | 46476.51953 | 48080.27344 | 48928.33203 |
| 76142.33594 | 57084.81641 | 69000.32031 | 68716.82031 | 53685.55078 | 48289.34766 | 54184.58594 |
| 14707287 | 11674362 | 13807372 | 12708783 | 11756988 | 11814809 | 11630790 |
| 3010.371338 | 3775.243896 | 2836.230225 | 3470.916748 | 5212.452637 | 4596.262207 | 4790.233887 |
| 96589.50781 | 79091.35938 | 85291.42969 | 78848.32813 | 76215.91406 | 78211.89844 | 82581.99219 |
| 29042.01563 | 24436.62695 | 24291.02148 | 28450.95313 | 47521.55859 | 37257.26563 | 38257.38672 |
| 13765.53516 | 14116.72754 | 12921.48145 | 13228.91602 | 17247.25977 | 15853.50977 | 15899.69629 |
| 3613803.25 | 2661960.25 | 3072836.75 | 2710376.75 | 2422104.75 | 2679373 | 2355085 |
| 25946.42383 | 19656.92578 | 22940.25977 | 22246.7207 | 26232.30078 | 27305.28906 | 26999.45898 |
| 599717.0625 | 491541.6875 | 520401.4063 | 542028.4375 | 439452.1875 | 496079.1875 | 497320.8125 |
| 59642.22266 | 45590.84766 | 51305.55469 | 54616.27344 | 49457.75391 | 45916.03516 | 47794.76953 |
| 70602 | 67724.27344 | 52875.81641 | 54248.19922 | 83502.5 | 89047.38281 | 86577.23438 |
| 203214.9063 | 115482.6641 | 211143.7813 | 169202.6719 | 131598.4063 | 190534.9219 | 186912.0938 |
| 71800.72656 | 66884.16406 | 60102.99219 | 56950.46875 | 65213.13281 | 62720.80078 | 62177.88672 |
| 624746.125 | 515199.2813 | 565729.125 | 608815.25 | 761362.125 | 698611.8125 | 708217.875 |
| 17038.75 | 22168.39063 | 9363.512695 | 6112.006348 | 13517.69629 | 15344.24121 | 17157.04102 |
| 310351.9375 | 277162.1875 | 262940.5938 | 284235.1563 | 386470.5938 | 338958.4375 | 352387.875 |
| 9031.262695 | 8082.075195 | 7865.191895 | 7951.717285 | 7226.48291 | 8597.364258 | 8231.84668 |
| 66678.13281 | 55241.26953 | 61705.78516 | 58060.06641 | 62562.90234 | 65942.45313 | 66977.54688 |
| 15575.80957 | 16711.93945 | 11666.11328 | 10985.65234 | 16881.68555 | 14108.5 | 15225.40918 |
| 41831.67969 | 31961.62891 | 34960.72266 | 35653.75 | 25748.2832 | 27983.22461 | 28023.04688 |

| | | | | | | |
|---|---|---|---|---|---|---|
| 1149015.375 | 921005.8125 | 969466.0625 | 981088.4375 | 938416.75 | 1009964.75 | 1021695.438 |
| 200494.5156 | 201360 | 158937.8594 | 171378.8281 | 178760.875 | 148618.8438 | 166080.125 |
| 39745.46484 | 34606.83594 | 34112.67188 | 35057.76953 | 30427.95313 | 31316.78516 | 31624.07031 |
| 67348.71094 | 50289.77344 | 70104.0625 | 76612.80469 | 119690.8203 | 77436.52344 | 107472.4453 |
| 849341.6875 | 717138.875 | 747150.5625 | 694301.875 | 487680.4063 | 500684.9688 | 465088.0625 |
| 10432.23438 | 10161.40918 | 10535.83789 | 8574.083984 | 14369.0127 | 12198.41504 | 9856.213867 |
| 326741.8125 | 280205.1875 | 308693.4688 | 313313.5938 | 353986.8125 | 344598.6563 | 359690.1875 |
| 303264.0938 | 248475.6875 | 266528.4063 | 282582.25 | 324503.5938 | 278559 | 298219.1563 |
| 122380.8047 | 108411.2031 | 98753.70313 | 101468.8125 | 99049.47656 | 108770.7266 | 97521.96875 |
| 150277.4063 | 128978.8203 | 125824.3672 | 127728.1328 | 126432.8594 | 125223.2109 | 124736.875 |
| 111512.0859 | 99675.21094 | 98696.23438 | 92776.75781 | 97070.0625 | 104881.4531 | 98629.20313 |
| 91547.67969 | 77821.02344 | 76872.44531 | 76661.45313 | 74839.8125 | 75467.22656 | 74111.19531 |
| 9623.935547 | 8965.114258 | 6508.669922 | 5529.518066 | 3558.217285 | 4018.794922 | 4035.57666 |
| 51514.41406 | 43855.34375 | 46826.48047 | 41288.56641 | 43675.51953 | 46978.07813 | 43172.48828 |
| 245329.1719 | 214065.4531 | 220789.9844 | 206809.1094 | 93081.34375 | 81537.03906 | 88605.25781 |
| 9912.229492 | 6482.185547 | 6898.672363 | 7612.988281 | 5944.69043 | 5047.806641 | 5566.96582 |
| 111780.9766 | 88911.28906 | 82821.88281 | 89967.75 | 87799.50781 | 93005.625 | 89262.74219 |
| 70469.23438 | 59718.5625 | 55703.75 | 55510.66016 | 79217.35938 | 98016.82031 | 85477.84375 |
| 43510.47656 | 48651.56641 | 33796.07031 | 40271.10938 | 48022.62109 | 52163.25 | 46337.95703 |
| 93961.03906 | 77512.44531 | 86235.41406 | 96489.90625 | 127970.7109 | 122649.4922 | 121852.3281 |
| 15901.05859 | 14244.06641 | 11902.67676 | 14188.83203 | 21186.55078 | 20852.25586 | 19926.04492 |
| 6543.09668 | 4996.731934 | 5457.562012 | 6875.34375 | 7502.030762 | 8118.032227 | 6193.313965 |
| 20300.00391 | 20459.67969 | 16191.6416 | 16969.35156 | 18084.31445 | 17842.26953 | 18811.29688 |
| 793.1764526 | 770.3460693 | 550.1833496 | 629.2761841 | 860.9059448 | 1188.286499 | 1006.64386 |
| 1728984.5 | 1393899.375 | 1469213.25 | 1371955.625 | 1281812.625 | 1302584.5 | 1253823 |
| 263221.4231 | 216947.4844 | 219813.9329 | 210980.1548 | 135725.8557 | 144809.7753 | 155352.2996 |
| 99714.00781 | 77779.60938 | 83586.89844 | 88817.04688 | 92787.875 | 100551.9453 | 100045.3359 |
| 6991.074707 | 7193.785645 | 8291.297852 | 3041.005615 | 11874.19922 | 9522.75 | 11696.0791 |
| 177225 | 132512.6719 | 158849.8125 | 150272.9844 | 116683.6719 | 131906.3594 | 122339.0938 |
| 58403.00781 | 47484.98047 | 49876.88281 | 51224.69141 | 46993.96094 | 46253.69531 | 45225.35547 |
| 36561.94531 | 28644.52344 | 33158.39844 | 33474.46875 | 29849.26172 | 25490.87891 | 28567.91406 |
| 978358.1328 | 951915.1953 | 741674.2031 | 699796.4844 | 737356.3906 | 757609.3906 | 726046.1172 |
| 338113.1563 | 298442.5313 | 293460.625 | 274308.2813 | 432646.0625 | 419214.9375 | 440801.125 |

| | | | | | | |
|---|---|---|---|---|---|---|
| 1477251.625 | 1105995.25 | 1286500.75 | 1288824.25 | 1456590.125 | 1528412 | 1576386.125 |
| 144027.6875 | 120138.9922 | 110681.7969 | 114735.9141 | 113406.5156 | 113137.7734 | 112438.1797 |
| 824905.625 | 615821.625 | 723975.125 | 620731.875 | 567580.5 | 706086.875 | 571221.375 |
| 83409.30469 | 75008.38281 | 68493.40625 | 65831.08594 | 71559.03906 | 63144.80078 | 68121.78906 |
| 137887.2813 | 112976.7891 | 113720.2656 | 113335.3828 | 94304.39063 | 87626.39063 | 92201.4375 |
| 27101.45508 | 24918.77148 | 19256.91602 | 17245.63672 | 29058.34375 | 30706.28125 | 32777.35938 |
| 957378.3125 | 1299505 | 724603.0625 | 784974.5 | 1022401.688 | 1387720 | 1034884.938 |
| 44072.88672 | 38042.61328 | 35694.5625 | 35294.31641 | 37235.78906 | 36034.55078 | 40394.88281 |
| 701448.375 | 594890.375 | 598613.1875 | 609111.75 | 497846.9375 | 479066.3125 | 455606.4375 |
| 34271.53516 | 26971.91016 | 29425.23438 | 30947.89063 | 22558.93945 | 26076.33008 | 23311.29297 |
| 114718.6719 | 93737.24219 | 95965.57813 | 91845.96875 | 116441.4219 | 159724.4375 | 114922.3906 |
| 1430290.5 | 1171552.375 | 1204405.75 | 1046176.438 | 1060757.625 | 1067783.75 | 1028626.938 |
| 142099.375 | 125381.9297 | 117612.0859 | 120637.3516 | 150650.0156 | 127959.8438 | 134792.2344 |
| 256495.5 | 266246.875 | 204420.2344 | 233376.7813 | 222705.7344 | 218023.5625 | 220062.4688 |
| 353226.0625 | 296846.875 | 309652.0313 | 288698.1563 | 304740.375 | 288573.7188 | 299719.8125 |
| 11705.84277 | 11904.0957 | 9352.828125 | 10539.62598 | 12794.25977 | 10486.59766 | 10047.86523 |
| 18597.52344 | 18818.99414 | 16594.44141 | 15421.75879 | 20563.09961 | 22112.80859 | 19493.80273 |
| 32454.91211 | 22889.37891 | 29219.65625 | 29202.16016 | 31906.21094 | 30388.18945 | 32122.79492 |
| 132126.8438 | 107654.4766 | 109781.9922 | 94441.125 | 107716.2891 | 133388.2813 | 111350.3203 |
| 95150.39844 | 155093.2969 | 103745.3047 | 110749.2969 | 110014.9609 | 269367.8438 | 125431.1563 |
| 52946.78125 | 54341.25781 | 38183.24219 | 38677 | 35122.19922 | 33742.48047 | 35474.25781 |
| 198966.6719 | 185171.4531 | 171722.9063 | 170148.0781 | 195917.7344 | 196804.625 | 193838.6875 |
| 20886.58984 | 20213.69531 | 19048.10938 | 18357.10938 | 18823.72266 | 17929.72852 | 18051.51367 |
| 33986.82422 | 26781.9082 | 26075.63086 | 26428.84766 | 25862.13281 | 20331.65625 | 22351.0625 |
| 140791.2344 | 130652.4453 | 126903.5469 | 123699.9531 | 134329.7969 | 134406.0313 | 137102.8438 |
| 15990.11914 | 13571.01953 | 14156.76465 | 13397.52051 | 11541.14453 | 11893.74512 | 10722.90039 |
| 24578.23828 | 17841.30078 | 20433.25586 | 21454.08203 | 23025.72852 | 16980.93945 | 19956.09766 |
| 2410.563721 | 4610.431152 | 2389.463135 | 2669.721436 | 5288.178223 | 16959.48438 | 7268.31543 |
| 526872.375 | 502116.3438 | 376463.8125 | 267203.3125 | 382214.0313 | 588494.4375 | 477455.7188 |
| 584217.8125 | 448965.8125 | 490802.2188 | 503289.9063 | 381905.875 | 399145.5313 | 410495.0313 |
| 60525.09375 | 54662.54688 | 49806.51563 | 52770.98438 | 41065.30078 | 45541.05469 | 43433.28125 |
| 26489.16992 | 28343.02148 | 25045.84375 | 22884.23633 | 26973.85547 | 36158.71484 | 22165.14063 |
| 1653656.25 | 1125627.75 | 1358244.25 | 1405582.75 | 1275287 | 1374690.125 | 1346509.625 |

| | | | | | | |
|---|---|---|---|---|---|---|
| 316898.125 | 298293.125 | 253828.0469 | 287511.125 | 434613.0938 | 360488.4375 | 394415.9375 |
| 15992.99707 | 15301.33691 | 12269.27539 | 11574.46582 | 2581.141602 | 6923.419434 | NA |
| 153148.3125 | 130068.4453 | 135273.125 | 131296.2969 | 128866.1641 | 123700.3906 | 125830.6484 |
| 7181.263184 | 7438.918945 | 5848.756348 | 3490.600098 | 4948.518066 | 4105.800781 | 4201.328125 |
| 18286.05078 | 17226.15039 | 13036.11426 | 14125.77637 | 17852.58984 | 17427.94727 | 17555.5918 |
| 23997.91602 | 22448.10156 | 18700.83203 | 20271.75586 | 34057.59766 | 31803.92969 | 32856.51172 |
| 38513.94922 | 31592.21289 | 32822.46484 | 34880.79688 | 41406.07813 | 40536.21875 | 39485.85156 |
| 203448.9531 | 99891.59375 | 188585.6406 | 200736.9531 | 240264.4531 | 213814.4063 | 254685.7188 |
| 44306.625 | 35069.20313 | 40677.85547 | 41029.50391 | 52713.32031 | 51025.22266 | 53542.10156 |
| 12157.00488 | 9642.287109 | 9877.699219 | 9658.392578 | 6516.651367 | 5529.287109 | 5506.583008 |
| 3020911 | 2506083.5 | 2439636 | 2242690 | 2123833.25 | 2153509.75 | 1948217.375 |
| 1072219.5 | 913948.5625 | 944813.3125 | 994068.5625 | 1227743.875 | 1096732.5 | 1188289.25 |
| 123998.1094 | 116472.5469 | 107054.9766 | 91165.42969 | 99355.04688 | 101437.5234 | 96626.92188 |
| 712497.75 | 513145.875 | 490320.9063 | 448750.6875 | 399508.8438 | 425039.5938 | 363840.4688 |
| 75677.71875 | 67612.17969 | 57306.53906 | 64924.09766 | 86748.38281 | 97545.09375 | 85321.97656 |
| 47889.94922 | 45277.09375 | 39760.65625 | 39758.65234 | 40234.03125 | 41440.875 | 40084.97266 |
| 3668.98877 | 3629.098389 | 2537.690918 | 2231.700439 | 1607.692871 | 2000.733765 | 2495.924805 |
| 25011.39844 | 20424.37891 | 19143.9707 | 20802.49414 | 13746.92578 | 12937.56934 | 13011.3584 |
| 23802.33789 | 17290.58203 | 19915.58789 | 17771.8457 | 10323.16797 | 13026.87012 | 12713.5293 |
| 100583.2422 | 77537.16406 | 80306.8125 | 88985.34375 | 106141.8672 | 103205.7656 | 108881.1719 |
| 88628.60156 | 68531.67969 | 71502.22656 | 68077.76563 | 71975.17969 | 78305.55469 | 81908.875 |
| 132051.625 | 115160.1719 | 99819.21875 | 112482.4375 | 180492.3438 | 159077.7969 | 188226.5781 |
| 25566.42578 | 21900.52148 | 20810.07813 | 23034.26367 | 23293.98242 | 21189.00586 | 21598.23633 |
| 616563.1563 | 536120.8438 | 506175.5078 | 497287.2422 | 526598.3203 | 524859.9063 | 523163.5 |
| 8707.361328 | 11328.9043 | 5078.619629 | 6427.165527 | 4750.212891 | 7439.311035 | 7874.884766 |
| 539480.6875 | 473337.6563 | 417273.3125 | 434465.4063 | 607349.8125 | 552898.0625 | 610591.75 |
| 729263.3125 | 631984.125 | 654528.5625 | 559241.75 | 646873.5625 | 672478.5625 | 601615.3125 |
| 16349.43945 | 11431.64355 | 13424.74316 | 15878.01074 | 11187.1582 | 13599.33594 | 11793.55664 |
| 184249.6563 | 153217.2031 | 165083.3438 | 163892.875 | 143123.2969 | 141187.9063 | 145320.6875 |
| 74021.24219 | 43060.82031 | 64049.50781 | 61456.35156 | 53185.96484 | 51030.98438 | 60105.17578 |
| 11583.67578 | 10433.73633 | 10042.22266 | 8376.120117 | 7385.619141 | 8788.677734 | 9185.348633 |
| 256379.1719 | 211313.2969 | 204462.5469 | 207668.3438 | 213110.7188 | 203907.5781 | 211425.3281 |
| 58012.48047 | 51284.26172 | 48948.52734 | 50825.52344 | 47604.91406 | 54096.22266 | 51572.59766 |

| | | | | | | |
|---|---|---|---|---|---|---|
| 99373.5 | 98552.92188 | 89782.125 | 83314.71875 | 87934.875 | 80607.03906 | 80217.64844 |
| 1370870.867 | 1118065.57 | 1089241.281 | 1181400.5 | 1572652.148 | 1491563.961 | 1696836.82 |
| 32925.21875 | 30518.47656 | 25949.08008 | 27750.58984 | 29460.125 | 28345.07617 | 27120.00195 |
| 138648.8125 | 124265.5859 | 106983.6641 | 96608.76563 | 52889.19531 | 52191.72656 | 47821.82031 |
| 65151.26953 | 62839.17969 | 57814.27734 | 60074.25 | 68464.77344 | 70659.03906 | 67421.92969 |
| 91397.72656 | 71740.19531 | 82804.46094 | 83154.9375 | 73581.25 | 72791.83594 | 76144.74219 |
| 35100.3125 | 35203.25 | 26052.17383 | 23772.28906 | 21401.91797 | 23167.53711 | 22805.58008 |
| 248226.75 | 167696.6094 | 226131.7031 | 203216.4219 | 250807.4531 | 214526.6875 | 250760.9375 |
| 5350.457031 | 3905.700195 | 4495.737305 | 4359.252441 | 7004.370605 | 7630.443359 | 7320.751953 |
| 16713.68164 | 15794.41699 | 12003.97461 | 14253.68652 | 18056.46484 | 16474.1543 | 15522.54688 |
| 53855.94141 | 36913.74609 | 46153.26563 | 49823.21094 | 44001.77344 | 40370.55859 | 44354.90625 |
| 66455.02344 | 54427.375 | 54054.99219 | 58396.66406 | 62861.58984 | 62289.91406 | 62102.77344 |
| 22679.12305 | 16453.15039 | 23410.57227 | 21497.46094 | 18463.37891 | 13880.12402 | 17164.35156 |
| 84132.3125 | 64511.55469 | 74994.35938 | 67984.19531 | 82886.94531 | 89835.70313 | 81306.74219 |
| 34010.41992 | 29415.45996 | 24386.46777 | 26986.18359 | 39264.36719 | 39725.34766 | 44940.59082 |
| 70209.69531 | 67765.11719 | 63716.64063 | 66126.82031 | 73669.1875 | 84193.70313 | 71039.35156 |
| 9316.697266 | 9040.546875 | 9123.043945 | 7850.216797 | 8721.566406 | 16929.05664 | 14103.71582 |
| 35705.57031 | 30998.19922 | 27057.60547 | 30116.60547 | 26701.57227 | 33431.94531 | 28668.84961 |
| 65092.91406 | 57374.0625 | 59405.36719 | 33795.36328 | 30015.62305 | 55334.14063 | 37025.67969 |
| 11289.00098 | 9078.837891 | 8851.488281 | 9248.349609 | 10036.50391 | 12582.18262 | 8222.03418 |
| 160166.4688 | 141674.7344 | 113972.8047 | 113537.6563 | 128361.6016 | 121849.7734 | 132060.2813 |
| 353763.9063 | 301503.875 | 296021.8438 | 264911.1875 | 268392.6875 | 251697.7188 | 248780.2656 |
| 132911.4688 | 127512.7969 | 108798.9063 | 126358.8984 | 151175.9844 | 150287.6094 | 132146.5156 |
| 29142.99414 | 31733.22266 | 22288.28516 | 20682.875 | 25108.10547 | 23344.41992 | 25956.78711 |
| 240485.4375 | 157006.4844 | 244151.4375 | 209392.3594 | 252760.1406 | 237617.2969 | 225247.7656 |
| 2812259.5 | 2215665 | 2329544.75 | 2281021 | 2010094.5 | 2167137.5 | 1904366.75 |
| 450772.7109 | 378574.498 | 384225.1328 | 374834.127 | 352639.5615 | 380389.3994 | 388989.5234 |
| 29459.78906 | 28413.63867 | 23201.30859 | 27245.68359 | 30592.51953 | 26054.80078 | 30115.56445 |
| 2291392.5 | 2102349.25 | 1734954.625 | 1812053.625 | 1841662.125 | 1646372.625 | 1671801 |
| 258917.4219 | 284920.5 | 231360.7813 | 245237.4219 | 322353.75 | 336505.9375 | 313829.4063 |
| 88920.9375 | 80661.24219 | 73180.54688 | 71962.60156 | 71098.26563 | 68855.07813 | 69717.01563 |
| 49632.82813 | 40430.61719 | 41190.05469 | 36274.6875 | 29053.32617 | 33921.64844 | 26749.99805 |
| 126833.6875 | 86544.0625 | 112261.7109 | 102947.7344 | 69380.39844 | 66520.60938 | 71058.70313 |

| | | | | | | |
|---|---|---|---|---|---|---|
| 42136.91797 | 37249.41797 | 37355.57813 | 36145.39063 | 39262.14844 | 40227.63281 | 36575.19141 |
| 25390.08398 | 18863.69336 | 28750.75781 | 19820.2168 | 24773.7793 | 21796.57422 | 3644.015869 |
| 17551.61523 | 14396.19922 | 15174.53027 | 15622.97363 | 17418.51367 | 14699.17773 | 15468.83789 |
| 421464.4063 | 344973.625 | 380616.7813 | 355199.5625 | 287451.7813 | 347271.0938 | 309598.4688 |
| 149577.2656 | 137723.1875 | 130711.6953 | 128890.1875 | 123780.3047 | 121988.8438 | 129089.9844 |
| 16596.82031 | 12099.88184 | 13964.79492 | 11605.72852 | 8716.310547 | 10460.25586 | 10723.35449 |
| 8912.428711 | 6970.083984 | 6918.891602 | 7209.093262 | 9276.948242 | 8703.930664 | 8962.024414 |
| 7056.345215 | 5361.257324 | 6178.87207 | 6055.043457 | 13959.15332 | 10394.7207 | 11372.2959 |
| 205279.6094 | 157943.2969 | 169673.125 | 164466.8594 | 171864.9844 | 176005.1719 | 175316.4375 |
| 258287.3906 | 198123.5938 | 246479.7969 | 230613.125 | 219177.7031 | 202328.7656 | 215835.8438 |
| 115549.7266 | 76123.83594 | 115165.3672 | 112789.5781 | 141155.0313 | 132056.0156 | 137404.2344 |
| 2912.474854 | 1179.295166 | 3352.159912 | 2639.648682 | 2592.520264 | 2229.016113 | 2651.437012 |
| 1887528.625 | 1438254.125 | 1619099.5 | 1470420.375 | 1439131.25 | 1465598.5 | 1387527.75 |
| 641570.625 | 536892.875 | 546180.4375 | 511300.0625 | 458595.625 | 464668.0625 | 474248.0938 |
| 706801.8125 | 652565.0625 | 568816.25 | 584018.4375 | 422630.6875 | 437811.3125 | 432049.6875 |
| 199146.8594 | 142368.1875 | 173565.7656 | 188693.8125 | 215776.9844 | 233102.8281 | 238510.4063 |
| 1011784.625 | 881879.25 | 809717.375 | 817182.375 | 775741.8125 | 792547.375 | 753922.875 |
| 209075.8281 | 181031.375 | 162322.8594 | 178428.1875 | 258614.125 | 222983.4375 | 235141.2813 |
| 22284.50781 | 17931.57422 | 17642.35547 | 16688.08008 | 17978.35352 | 21302.03711 | 19022.28711 |
| 1321518.875 | 1330311.375 | 1076859.375 | 1142812.5 | 1781663.5 | 1828303.625 | 1679098.125 |
| 44246.58984 | 56764.96484 | 29423.58008 | 27374.81445 | 58737.875 | 51630.49219 | 58604.31641 |
| 76657.86719 | 58947.77344 | 65962.53125 | 64793.8125 | 52278.9375 | 50183.75 | 47558.75 |
| 37558.24219 | 31066.54102 | 29236.65039 | 26053.17578 | 32803.57422 | 29988.47656 | 35352.65234 |
| 85832.63281 | 74155.57813 | 73730.07813 | 74268.50781 | 76010.53906 | 72687.53906 | 71521.625 |
| 279791.875 | 230232.7031 | 226818.7656 | 254309.6563 | 275808.2813 | 338071.3125 | 346147.25 |
| 198723.8906 | 170214.625 | 161054.5625 | 159017.4531 | 107457.6094 | 109520.9375 | 106629.5703 |
| 46824.45703 | 46285.41016 | 42840.23438 | 35369.03906 | 42617.52344 | 45831.60547 | 43661.54688 |
| 393997.8125 | 326604.6563 | 325720.6875 | 311983.9688 | 306552.9063 | 326135.5625 | 307821 |
| 20359.38867 | 16200.94434 | 16717.16602 | 15680.50879 | 16890.49219 | 15081.50098 | 17291.74805 |
| 21752.25195 | 20845.27148 | 15795.14746 | 15720.09863 | 23462.55078 | 22700.16797 | 24186.95703 |
| 127164.5859 | 106679.3203 | 102543.9844 | 103610.3047 | 94864.22656 | 101288.375 | 93175.11719 |
| 41333.01953 | 33615.23828 | 35455.40234 | 32007.61328 | 31233.9375 | 34924.50391 | 33433.54297 |
| 5680.349121 | 4474.865234 | 5364.995117 | 5068.547363 | 5668.237793 | 4119.539063 | 4764.440918 |

| | | | | | | |
|---|---|---|---|---|---|---|
| 38261.01563 | 30028.45117 | 30257.02148 | 31591.56641 | 26921.29883 | 28636.76953 | 28453.39844 |
| 359778.625 | 216432.3125 | 415144.5313 | 395131.75 | 247094.5313 | 409267.6875 | 234124.7344 |
| 10405.51367 | 9142.362305 | 9208.87793 | 6991.803223 | 7817.104004 | 10317.03418 | 5732.574219 |
| 3473.150146 | 3956.307373 | 2415.564941 | 2157.588623 | 3842.099365 | 3591.871094 | 3960.095947 |
| 58774.98438 | 49842.94141 | 44274.97656 | 47510.96094 | 113771.1484 | 113947.0234 | 102684.3828 |
| 273520.0313 | 257057.3906 | 246217.4531 | 259930.5156 | 275913.0938 | 277749.2188 | 255297.4844 |
| 9367.911133 | 8597.392578 | 8698.254883 | 6243.607422 | 8909.640625 | 7900.066406 | 9960.714844 |
| 63784.60547 | 54407.64844 | 48258.57422 | 50876.05078 | 39747.5625 | 54868.50391 | 44841.17969 |
| 192377.8906 | 156684.1406 | 156145.9844 | 156219.875 | 134942.5938 | 133064.0156 | 127663.3828 |
| 10672.87988 | 10898.7334 | 9782.250977 | 9309.893555 | 10821.69336 | 9983.511719 | 10937.62012 |
| 16360.00781 | 13905.57715 | 12377.50391 | 12949.51758 | 12917.61426 | 12009.85254 | 12064.80469 |
| 1183258.125 | 871866.3125 | 1060123 | 1095209.5 | 948800.4375 | 1065148.75 | 1069156.875 |
| 422311.9375 | 293991.0625 | 346155.9063 | 378925.2188 | 359696.5313 | 350139.3125 | 373544.0938 |
| 66636.92188 | 63959.18359 | 62538.03516 | 65732.67188 | 79224.88281 | 68239.375 | 83790.75781 |
| 1901677.625 | 1625462 | 1681645.375 | 1587970.75 | 1761569.25 | 1801404.125 | 1729250.75 |
| 6018.077637 | 5255.491211 | 5961.850586 | 6966.745605 | 7752.580078 | 7452.829102 | 7295.10498 |
| 50634.44141 | 71596.21875 | 39877.16797 | 32647.4043 | 31817.33594 | 47751.22266 | 40212.78125 |
| 1054197.75 | 782163.0625 | 917832 | 864083 | 864143.9375 | 916134.5 | 918600.875 |
| 2292.6875 | 3305.271973 | 1285.526733 | 1563.731201 | 2315.390625 | 2960.510742 | 2390.386719 |
| 11010.32129 | 10385.92285 | 8916.094727 | 10771.12109 | 10974.2373 | 15033.38574 | 13216.64941 |
| 147533.9063 | 115357.9844 | 109028.0781 | 97046.13281 | 90595.82813 | 87749.625 | 78549.90625 |
| 101340.2734 | 94988.48438 | 90622.51563 | 94223.46875 | 95957.5 | 92610.32031 | 97152.125 |
| 21321.5918 | 15590.28418 | 13721.86816 | 9465.585938 | 9512.898438 | 18214.77344 | 10828.72852 |
| 35684.13281 | 32420.64648 | 27186.61719 | 31904.89453 | 39596.00391 | 29281.41992 | 32653.40625 |
| 73451.69531 | 57842.11719 | 65241.23047 | 66842.85156 | 79046.32031 | 55104.57422 | 64136.71484 |
| 44785.07031 | 35184.84375 | 41813.58984 | 36666.85547 | 40013.62891 | 37437.21875 | 40664.77344 |
| 57913.50391 | 39033.16797 | 45449.14063 | 48539.62891 | 35906.73438 | 30847.77734 | 33962.79688 |
| 39322.31641 | 35451.42578 | 31646.86133 | 29237.66602 | 43384.94922 | 38815.60547 | 42201.51953 |
| 2762018.75 | 2408884 | 2196989 | 2471939.25 | 2985034 | 2703954 | 2968569.75 |
| 78040.99219 | 64315.44922 | 65338.46484 | 68542.28906 | 56659.54688 | 59956.20703 | 61608.30078 |
| 1493368.375 | 1135599.25 | 1212608.75 | 1342228.875 | 1396761.25 | 1579183.25 | 1668994.125 |
| 6362.924316 | 5919.026855 | 6207.567383 | 6310.489746 | 9749.286133 | 6840.973633 | 10068.35645 |
| 15244.05371 | 12683.41211 | 12764.27051 | 13576.08301 | 14064.64746 | 16008.62988 | 14141.92383 |

| | | | | | | |
|---|---|---|---|---|---|---|
| 29417.95508 | 29798.68164 | 22795.76172 | 17014.8125 | 30718.35742 | 24824.54102 | 31448.31641 |
| 4788347.5 | 4342924 | 3793347 | 3606630 | 5659958 | 5325451 | 5518587 |
| 336939.375 | 288144.7813 | 267622.7188 | 258219.3438 | 224186.6719 | 272773.9688 | 273542.4688 |
| 265541.5938 | 222082.8125 | 215183.2813 | 220592.9219 | 169601.2344 | 164577.0781 | 167169.3125 |
| 76326.08594 | 63581.71094 | 65046.37109 | 56037.65625 | 70926.64844 | 72855.36719 | 70524.11719 |
| 15034.97754 | 12984.70117 | 13221.24805 | 11551.15039 | 10902.6377 | 10783.78809 | 11310.07617 |
| 58606.00391 | 53948.20313 | 44227.93359 | 44339.17188 | 48440.03516 | 49368.51953 | 47862.85938 |
| 183902.8438 | 207939.8281 | 161524.5625 | 174300.8906 | 250959.7656 | 206223.125 | 212070.2188 |
| 184645.2969 | 177313.0625 | 143524.5469 | 173303.3438 | 156212.5625 | 129909.1875 | 131938.8125 |
| 41312.45313 | 30192.74805 | 37603.54688 | 38423.27734 | 38258.26953 | 45095.78125 | 47687.46484 |
| 1893.414795 | 1082.387695 | 2344.975098 | 2608.720215 | 5881.677246 | 6263.271973 | 4762.007324 |
| 54626.38672 | 47379.32422 | 47759.37891 | 47566.17969 | 46511.15234 | 51796.98438 | 49492.53516 |
| 243512.0625 | 229556.9531 | 211142.1563 | 222882.4219 | 260064.7344 | 260171.5781 | 249978.2344 |
| 766499.5 | 656771.75 | 614734.3125 | 640974.1875 | 574424.5 | 603776.6875 | 545447.6875 |
| 28275.09375 | 22355.20117 | 22060.47656 | 24208.67773 | 26374.20898 | 26557.20313 | 27254.9043 |
| 47930.60938 | 45344.82031 | 38323.35938 | 34935.46094 | 42014.06641 | 40212.54688 | 46304.57422 |
| 41720.14453 | 32466.96094 | 33304.80078 | 30659.44141 | 27166.23242 | 29624.44336 | 28228.75391 |
| 469404.3125 | 379435.625 | 425138.2813 | 434544.1875 | 334024.6875 | 357026.625 | 381923.6563 |
| 11828.93066 | 10783.87207 | 9248.338867 | 9443.067383 | 12389.08301 | 27125.72461 | 10940.20117 |
| 1425056.125 | 1037612.563 | 1084209.75 | 1041346.313 | 914098.75 | 965596.375 | 925533.9375 |
| 15098.59277 | 11587.73926 | 10633.53809 | 9845.899414 | 17899.18359 | 17029.58398 | 17309.30078 |
| 1415016.25 | 981444.625 | 1176239.5 | 1041804.438 | 949272.8125 | 1046209.625 | 957663 |
| 84773.42969 | 72015.32813 | 71295.53125 | 64968.70313 | 67412.5 | 74620.80469 | 70428.71875 |
| 474848.0313 | 390493.6563 | 419250.125 | 409039.5 | 414813.9063 | 389084.375 | 389511.1875 |
| 109791.2578 | 102242.1328 | 86555.50781 | 78269.11719 | 111637.1641 | 108758.2891 | 105949.0859 |
| 60675.89453 | 36514.78516 | 49652.375 | 50402.95703 | 49103.48047 | 36964.12109 | 52122.29297 |
| 16036.85742 | 14426.95898 | 11720.37402 | 11848.83594 | 19182.67773 | 23606.42578 | 20776.11133 |
| 40827.58984 | 27609.84961 | 35951.85156 | 36690.19141 | 34681.49219 | 37444.94141 | 39995.18359 |
| 164146.4531 | 147713.4844 | 117080.4063 | 112556.7344 | 161589.4375 | 168783.9688 | 156011.4531 |
| 31895.83008 | 30796.60547 | 26700.76172 | 27516.30859 | 18988.85547 | 16382.41992 | 14380.4502 |
| 159293.4531 | 120323.6953 | 130273.6953 | 145775.4531 | 99922.07031 | 88152.75 | 97651.48438 |
| 14522.08789 | 13977.62598 | 12167.97168 | 8883.186523 | 16160.52637 | 14110.15625 | 15990.57422 |
| 40817.75391 | 40678.76172 | 32258.0957 | 36179.75 | 38993.41797 | 42750.46875 | 37436.03125 |

| | | | | | | |
|---|---|---|---|---|---|---|
| 29309.50781 | 47112.21875 | 17334.02539 | 10595.97949 | 31870.78906 | 26913.29492 | 31449.30664 |
| 7311.869141 | 5773.75 | 6886.769043 | 7475.04541 | 9371.597656 | 11243.74707 | 11061.73633 |
| 30891.25781 | 19823.48047 | 26767.86719 | 26568.73828 | 34457.09766 | 31913.0957 | 34382.14063 |
| 91351.60156 | 74415.79688 | 73906.38281 | 65665.33594 | 47575.03906 | 49004.625 | 48967.25781 |
| 24255.62891 | 23954.82422 | 23614.42383 | 23130.30273 | 30105.41797 | 22719.39648 | 22863.46094 |
| 299470.25 | 241314.8125 | 234462.9219 | 220834.3594 | 218183 | 223206.6406 | 230316.6094 |
| 180455.0781 | 136289.5 | 138537.8281 | 150211.3438 | 176375 | 172524.3438 | 187068.0313 |
| 410266.6563 | 343298.8125 | 351153.9375 | 371701.1563 | 433700.3125 | 414490.0938 | 459671.75 |
| 11106.24121 | 11010.51172 | 6248.523926 | 6236.60498 | 6481.629883 | 6110.529785 | 7910.21582 |
| 122069.6016 | 104213.6406 | 100707.6875 | 92365.77344 | 120087.9063 | 118566.6953 | 118289.3594 |
| 5873.476563 | 6885.578613 | 3549.990967 | 3116.299561 | 2421.022949 | 2331.370605 | 2320.327148 |
| 19973.90234 | 17751.15039 | 19261.66797 | 17495.50586 | 31764.25586 | 31425.05273 | 30688.32617 |
| 17000.89453 | 15139.29492 | 16553.33203 | 16120.58984 | 14814.88379 | 16130.56348 | 16130.14941 |
| 264295.125 | 233328.0313 | 211633.5313 | 220914.625 | 274537.125 | 267384.25 | 292414.625 |
| 165995.9688 | 134692.2656 | 136454.0781 | 146464.125 | 138340.2031 | 150408.4219 | 155360.6563 |
| 146243.9375 | 129732.5 | 133325.3594 | 99921.14063 | 77698.04688 | 75868.02344 | 77610.86719 |
| 32570.20117 | 30830.99023 | 26175.05859 | 32130.48828 | 29752.87109 | 30219.62305 | 27119.15234 |
| 249672.25 | 235494.875 | 205889.1094 | 203802.3438 | 251183.8125 | 271919.0938 | 274143.8438 |
| 147050.7031 | 122039.0469 | 125182.6875 | 122062.3672 | 105767.1875 | 106615.3516 | 110885.9063 |
| 58114.05078 | 48751.89844 | 48229.10938 | 51447.56641 | 68335.11719 | 62884.30078 | 72090.96875 |
| 35031.47656 | 29389.91016 | 33919.42578 | 29209.07422 | 19644.34375 | 18352.70508 | 18649.42383 |
| 9147.887695 | 7044.524902 | 7203.251953 | 7542.091797 | 5779.840332 | 6842.092285 | 5760.196777 |
| 75858.86719 | 69472.16406 | 60129.50781 | 60547.21875 | 73568.78125 | 77923.96094 | 72618.95313 |
| 16666.61523 | 14920.89551 | 15031.9375 | 12901.24902 | 17198.5 | 16193.33691 | 18953.10742 |
| 43816.45313 | 38955.82031 | 37073.57422 | 33917.34766 | 48723.15234 | 43125.38281 | 46036.58984 |
| 78520.07031 | 73698.94531 | 71042.99219 | 63756.05859 | 52415.44922 | 58438.72656 | 54074.49609 |
| 39401.57813 | 28911.69922 | 34152.33594 | 35824.60156 | 29145.42969 | 19068.42383 | 31668.92383 |
| 973599.6875 | 735210.25 | 799119.375 | 689471.4375 | 681851.5625 | 845284.6875 | 653062.375 |
| 18429.55273 | 15741.48145 | 14583.87598 | 13266.44043 | 16982.2168 | 21844.93359 | 17276.47656 |
| 1604386.375 | 902924.6875 | 1571712 | 1184129 | 981192.0625 | 1296311.875 | 1299109.25 |
| 46707.86328 | 50735.23438 | 31407.08398 | 27524.35352 | 42996.67969 | 43035.29688 | 44221.38672 |
| 99201.9375 | 81348.42969 | 79867.88281 | 81496.30469 | 82585.08594 | 77145.09375 | 82950.63281 |
| 66853.50781 | 56504.48438 | 52904.00391 | 49962.44141 | 53889.42188 | 58541.51172 | 53091.85938 |

| | | | | | | |
|---|---|---|---|---|---|---|
| 4288.172363 | 3891.915283 NA | | 3652.471924 | 4500.766113 | 13293.68164 | 7277.951172 |
| 30648.19531 | 24508.19141 | 25527.57813 | 23998.10938 | 30609.73633 | 31067.54688 | 31565.41992 |
| 22286.45313 | 18453.19531 | 20377.72461 | 14106.75684 | 16566.13867 | 16125.41895 | 25824.09766 |
| 696495.0625 | 916468.9375 | 466382.25 | 356768.0313 | 890781.6875 | 764676.625 | 905540.25 |
| 16497.32227 | 15050.9707 | 14184.14063 | 11143.89746 | 9462.469727 | 12754.62402 | 10339.27344 |
| 154880.0781 | 131738.2656 | 136801.7656 | 129623.4844 | 93600.99219 | 93111.74219 | 98998.26563 |
| 23706.35742 | 16903.49219 | 16890.25586 | 17888.00391 | 25738.375 | 26863.27539 | 23013.07031 |
| 2190487.75 | 1749009.375 | 1816061 | 1773395 | 1296071.625 | 1465575 | 1368597.125 |
| 2328.649902 | 1745.310059 | 1629.102295 | 1425.02832 | 2430.525391 | 1817.183472 | 2397.538086 |
| 43336.45313 | 34853.5 | 38743.16016 | 34484.23438 | 32691.4043 | 33611.49609 | 35710.03906 |
| 12756.28809 | 8745.147461 | 8894.762695 | 8710.541992 | 4670.672852 | 5909.191406 | 5101.397461 |
| 509651.75 | 380027.1875 | 406976.125 | 415070.6563 | 411684.5 | 385289.3438 | 420399.375 |
| 2107609.25 | 1613341.25 | 1797121.625 | 1745100.125 | 1634624.5 | 1804756.625 | 1737573.875 |
| 23036.86133 | 19259.20898 | 17594.41016 | 16666.51563 | 17985.08594 | 18329.58008 | 18522.04492 |
| 53962.84766 | 51161.9375 | 34711.76563 | 46463.03906 | 74697.47656 | 70305.15625 | 70703.24219 |
| 96014.60938 | 81646.92188 | 74012.21875 | 67226.71875 | 76599.94531 | 77540.73438 | 79964.89063 |
| 617890.25 | 511331.125 | 505433.0313 | 543662.8125 | 610498.1875 | 504073.9063 | 546675.9375 |
| 85252.74219 | 70346.55469 | 72908.19531 | 76558.10156 | 66690.69531 | 72629.16406 | 67859.25781 |
| 676.7974854 | 384.793457 | 480.8236694 | 1179.286987 | 5985.67627 | 2308.455811 | 3123.042236 |
| 31966.5918 | 31777.01367 | 21186.61719 | 20134.81836 | 22324.54883 | 23366.03711 | 25160.21484 |
| 135135.7969 | 116835.8359 | 120239.3828 | 113110.9844 | 120364.9844 | 120005.2656 | 124110.75 |
| 463328.125 | 273265.9063 | 379731.9063 | 391287.1563 | 443308.9063 | 416483.7813 | 480492.9688 |
| 10931.11133 | 10397.66699 | 7636.908691 | 10138.33105 | 9852.18457 | 9215.192383 | 9356.53125 |
| 11872.23926 | 8335.459961 | 12408.45703 | 12054.45313 | 11660.97656 | 9945.032227 | 11137.36523 |
| 11295.69531 | 9309.00293 | 9972.354492 | 9908.429688 | 8858.521484 | 7829.05127 | 8630.529297 |
| 299145.1563 | 280221.5 | 242522.2344 | 214529.2188 | 261661.5781 | 258123.8906 | 250959.4375 |
| 24469.94531 | 17316.2793 | 17535.29297 | 18231.25 | 15229.81836 | 15524.59082 | 12552.26367 |
| 49913.06445 | 37540.09082 | 37368.44287 | 41611.50098 | 46099.26074 | 53199.625 | 45874.94824 |
| 31646.09961 | 24426.1543 | 28988.55859 | 25819.32422 | 23688.36133 | 22234.58398 | 21744.12891 |
| 65190.05859 | 52002.98438 | 56323.42578 | 52837.83594 | 47377.32031 | 46167.16406 | 44951.43359 |
| 129144.8203 | 123763.5078 | 111643.8984 | 95448.8125 | 95289.23438 | 100372.6875 | 119236.6641 |
| 40741.12891 | 37456.40234 | 32356.73633 | 34245.12109 | 35877.26953 | 34587.08203 | 33533.05859 |
| 27276.81445 | 24900.6582 | 23012.60938 | 21488.35938 | 20084.43359 | 16913.93164 | 18092.4043 |

| | | | | | | |
|---|---|---|---|---|---|---|
| 49426.04297 | 38512.19141 | 51481.5 | 53687.53516 | 38177.14063 | 39798.14844 | 39094.03516 |
| 173528.5469 | 155875.8906 | 142964.2813 | 141931.8438 | 167992.3438 | 168489.8594 | 151980.0469 |
| 286226.9688 | 239511.5469 | 248175.25 | 254763.7031 | 214743.3594 | 209621.5313 | 212410.3438 |
| 385582.7813 | 323661.8125 | 321978.875 | 348821.8125 | 467445.75 | 448032.9688 | 467929.3125 |
| 57506.54297 | 86573.73438 | 22974.77539 | 12057.17871 | 49510.59375 | 50333.60938 | 53346.4375 |
| 5169.459961 | 4731.300781 | 5212.98291 | 4310.938965 | 4621.961914 | 4344.18457 | 4681.452148 |
| 4501.213379 | 3044.142334 | 3652.643311 | 4000.553711 | 3644.37793 | 4129.72998 | 3676.180908 |
| 190269.9219 | 187193.125 | 131429.5625 | 156063.1719 | 200993.2656 | 201063.6875 | 221258.4688 |
| 70908.34375 | 76840.29688 | 47797.65625 | 47865.25391 | 83638.98438 | 84851.91406 | 84919.33594 |
| 167680.5781 | 161176 | 133985.7031 | 153869.9531 | 162218.8906 | 163865.0625 | 148622.2188 |
| 8351.961914 | 12885.13965 | 5050.537598 | 6769.402344 | 11802.36523 | 5465.383301 | 15907.875 |
| 530753.375 | 375882.4063 | 488947.625 | 519569.625 | 577850.9375 | 624965.6875 | 590255.5 |
| 348452.6563 | 295467.75 | 274966 | 242462.7188 | 226512.2188 | 215138.9688 | 215520.7969 |
| 825992.25 | 658391.3125 | 636062.4375 | 691314.9375 | 882336.0625 | 821527.875 | 844560.3125 |
| 104747.0625 | 85598.26563 | 85623.82813 | 96111.17188 | 139995.9219 | 129438.6406 | 128421.8047 |
| 34636.00391 | 32346.625 | 22501.68945 | 22322.53906 | 21094.42773 | 11470.33496 | 16122.65527 |
| 186981.3281 | 134608.6406 | 150739.5313 | 152408.75 | 173493.6563 | 186769.7344 | 196172.3906 |
| 605827.375 | 443389.1875 | 526544.6875 | 528626.125 | 551628.1875 | 596659.0625 | 619898.125 |
| 295291.9063 | 245051.375 | 239355.2813 | 234037.8125 | 200528.1563 | 216375.6563 | 210507.1563 |
| 225721.2656 | 190062.6875 | 187018.0156 | 182583.6406 | 201359.5 | 215125.9219 | 195473.5156 |
| 235988.8281 | 132577.8438 | 207692.2031 | 218095.0938 | 259327.1406 | 240809.3281 | 291164.1875 |
| 31686.03711 | 27927.79688 | 24710.69141 | 25879.60938 | 27071.08789 | 27110.95508 | 26595.74414 |
| 106839.0156 | 75760.24219 | 93307.39063 | 82350.8125 | 68971.125 | 83472.77344 | 74174.32031 |
| 501006.1055 | 404526.4297 | 406815.0879 | 391529.6758 | 367086.7148 | 354875.5781 | 376375.8008 |
| 14388.02148 | 13361.98242 | 13962.78711 | 8155.547363 | 17600.50977 | 13191.29395 | 15267.72168 |
| 58995.44531 | 50010.66016 | 49886.52734 | 46625.66016 | 54934.83984 | 54684.64063 | 52756.69141 |
| 13100.48926 | 14253.8584 | 9346.370117 | 10061.79785 | 13508.45703 | 10015.46191 | 12672.31055 |
| 109669.4961 | 99583.30859 | 79580.96875 | 93025.92188 | 140506.4141 | 138073.3594 | 147287.4766 |
| 69271.27344 | 72787.85938 | 76066.61719 | 70190.92188 | 51577.28516 | 70120.55469 | 64775.11719 |
| 8045.40918 | 10697.50098 | 3145.283203 | 3363.152588 | 5827.940918 | 5092.316895 | 6391.293945 |
| 15658.25391 | 14506.21094 | 11423.96484 | 12035.44824 | 14822.08105 | 14177.13867 | 14554.01172 |
| 585623.8125 | 536649.75 | 383727.125 | 348052.2813 | 210385.6719 | 254424.1719 | 208214.5938 |
| 1149189.5 | 940199.875 | 938117.125 | 996584.8125 | 1315591.25 | 1268639.375 | 1296412.75 |

| | | | | | | |
|---|---|---|---|---|---|---|
| 584299.625 | 476551.5625 | 497438.9375 | 548998.5 | 564894.0625 | 657374.125 | 685117.125 |
| 180405.0313 | 138036.3906 | 158327.3438 | 104984.8516 | 117274.8281 | 111799.2188 | 120848.1797 |
| 29454.02734 | 25834.49414 | 25206.73633 | 24397.69531 | 19012.35352 | 19830.79102 | 19473.29297 |
| 7279.462891 | 10021.13086 | 11147.89941 | 8309.238281 | 8067.430664 | 9427.959961 | 9082.586914 |
| 59275.80469 | 49181.34766 | 47528.57813 | 47918.59766 | 49317.625 | 49254.07422 | 49063.98438 |
| 9576.34668 | 7103.581543 | 6457.212891 | 6762.084961 | 5069.087891 | 6199.092285 | 7146.245605 |
| 153788.0313 | 132178.3594 | 115492.7891 | 111483.1641 | 148302.1719 | 145876.0313 | 143837.9063 |
| 848016.25 | 688114.875 | 722105.4375 | 787157.8125 | 947781.4375 | 900448.0625 | 876041.125 |
| 360328.9063 | 268351.75 | 313038.8438 | 300316.125 | 258657.3125 | 250906.625 | 253986.5 |
| 427999.0313 | 273872.7188 | 355134.9063 | 341098.2813 | 273153.2813 | 249798.7969 | 258771.9688 |
| 1725150.125 | 1197546.125 | 1433993.5 | 1459934.75 | 1395666.125 | 1495786.125 | 1514636.875 |
| 129591.0078 | 111365.0781 | 93864.25781 | 90593.625 | 82064.22656 | 88567.35156 | 81732.15625 |
| 112765.9375 | 97159.88281 | 89430.75781 | 93606.82813 | 87561.71875 | 88692.45313 | 80947.47656 |
| 46756.05859 | 42091.87109 | 39967.875 | 36658.40234 | 39685.11328 | 35293.35938 | 34514.57813 |
| 4309.884766 | 3761.558594 | 3452.54248 | 4071.443115 | 3356.44873 | 5458.070801 | 2192.2854 |
| 135455.5313 | 90967.75 | 103014.1094 | 121996.1016 | 94710.80469 | 99108.98438 | 105975.1094 |
| 11919.13672 | 8617.852539 | 10193.02246 | 7823.181152 | 8819.056641 | 6416.810059 | 9937.52832 |
| 70408.10938 | 65324.56641 | 59661.69531 | 64064.95313 | 95563.99219 | 80266.44531 | 88140.10938 |
| 34610.41797 | 30634.8418 | 25230.66602 | 25473.02734 | 29132.30469 | 30820.07617 | 29294.25391 |
| 10753.35352 | 9480.551758 | 7635.333984 | 7898.522461 | 6266.005859 | 5689.609375 | 7620.421387 |
| 79129.25 | 63091.69531 | 59598.77734 | 59754.66016 | 50523.05078 | 49423.40234 | 49361.49219 |
| 484806.2188 | 366084.0313 | 403959.4063 | 454357.25 | 272685.9375 | 264994.6875 | 268338.5938 |
| 55362.21875 | 37557.28125 | 45235.89453 | 36872.92188 | 28640.98633 | 43114.97656 | 26942.95117 |
| 191140.0625 | 175307.3125 | 138065.7656 | 148692.7656 | 238856.0625 | 232135.2969 | 252901.9531 |
| 27400.19336 | 23156.7168 | 25930.81641 | 27799.12891 | 32140.93945 | 27039.77344 | 30304.76367 |
| 172793.4844 | 158881.5625 | 152889.4531 | 150530.9375 | 160722.375 | 185687.7969 | 149274.375 |
| 30311.52539 | 27374.08594 | 26934 | 27027.20703 | 38664.28906 | 37398.20313 | 36765.23828 |
| 31513.92188 | 27368.60352 | 26536.83398 | 28009.99805 | 31152.88281 | 37933.73047 | 38507.86719 |
| 170569.8125 | 131418.7246 | 135881.2715 | 126951.4902 | 150147.3691 | 172433.6504 | 156457.5918 |
| 172040.8906 | 160930.7656 | 148364.7813 | 149376.7188 | 183149.5781 | 174270.8125 | 173078.2656 |
| 128460.7656 | 95061.50781 | 104094.7734 | 102645.4375 | 57295.66016 | 67314.64844 | 59269.89844 |
| 21783.54883 | 25143.65234 | 14022.00586 | 12508.44434 | 25948.58398 | 23859.91211 | 23639.04297 |
| 16336.04883 | 14369.5625 | 14934.65039 | 14863.39453 | 15439.98535 | 17935.91016 | 14159.30176 |

| | | | | | | |
|---|---|---|---|---|---|---|
| 253860.25 | 226468.875 | 213990.1875 | 227835.125 | 241658.0313 | 243094.4688 | 231768.875 |
| 9254.772461 | 8305.929688 | 7588.000488 | 5625.508301 | 6819.678711 | 8237.519531 | 8447.908203 |
| 102092.3984 | 80649.19531 | 87463.04688 | 80895.83594 | 63085.09375 | 71750.38281 | 71473.94531 |
| 81082.25 | 64584.48438 | 59209.97656 | 71517.40625 | 61213.85547 | 56727.33594 | 58299.98828 |
| 1117.76001 | 1169.285034 | 937.9546509 | 825.7650757 | 875.0909424 | 911.3412476 | 839.6447754 |
| 157774.3438 | 124492.5859 | 135896.7344 | 133214.7969 | 149325.75 | 138732.75 | 156932.7188 |
| 31893.92383 | 24810.97656 | 26561.92773 | 21685.85742 | 19829.12305 | 20539.71289 | 21260.68164 |
| 44238.41016 | 33903.86719 | 35731.92969 | 36848.42578 | 39311.05469 | 40292.95313 | 41410.49609 |
| 610302.5 | 499898.0313 | 477773.5 | 532712.75 | 689676.75 | 657640.125 | 674062.375 |
| 83026.65625 | 61013.54297 | 68014.59375 | 67852.3125 | 69243.29688 | 70617.17188 | 68648.85938 |
| 76192.01563 | 61957.41797 | 61025.125 | 64309.62109 | 61351.48828 | 67209.39844 | 65279.41797 |
| 12745.84766 | 3950.143799 | 7533.26123 | 8960.142578 | 4407.872559 | 5258.239746 | 6222.150391 |
| 183277.7656 | 167825.4375 | 171250.4688 | 142218.0625 | 132174.9219 | 148459.5156 | 122582.3125 |
| 65591.78125 | 48554.84375 | 54739.98828 | 52773.67969 | 51576.52344 | 57078.38672 | 58599.92969 |
| 5337.351563 | 3468.668945 | 4902.499512 | 4676.715332 | 7684.428711 | 6211.890625 | 6995.861816 |
| 213385.25 | 161885.5313 | 173368.25 | 183295.9844 | 177481.9688 | 196974.2969 | 196944 |
| 7033.474121 | 6843.671875 | 3926.347168 | 3737.141113 | 6361.026367 | 6557.855469 | 5714.166992 |
| 119233.4219 | 98494.8125 | 93354.71094 | 90450.375 | 93329.03125 | 103582.6016 | 95808.16406 |
| 144254.6406 | 117073.6641 | 126110.3906 | 129274.2422 | 138329.5938 | 170672.4063 | 182415.5469 |
| 754778.4375 | 659131.5 | 451337.1563 | 534491.125 | 463391.8125 | 426825.5 | 451685.1563 |
| 1280417.875 | 902890.3125 | 1075049.5 | 1038388 | 994968.625 | 1009064 | 976537.375 |
| 75983.00781 | 66049.29688 | 63433.92969 | 63299.27344 | 64454.58594 | 59657.16016 | 57783.08594 |
| 378857.7813 | 248594.625 | 329789.625 | 337785.0938 | 345759.9375 | 339847.25 | 366966.0625 |
| 13634.88672 | 10348.33887 | 10191.25977 | 11952.93945 | 9574.299805 | 7578.411133 | 10593.43457 |
| 812172.5625 | 704395.25 | 640283.8125 | 694462.125 | 722547.4375 | 672786.75 | 703019.1875 |
| 16136.86133 | 13207.22168 | 11926.52832 | 14198.72168 | 12716.65332 | 11815.26074 | 12758.37793 |
| 732292.7422 | 591977.2344 | 630204.6445 | 658354.7188 | 686758.9961 | 599684.2656 | 626814.0352 |
| 106640.125 | 81396.84375 | 87757.79688 | 89723.26563 | 104911.2031 | 88308.55469 | 102494.8984 |
| 306040.9063 | 260484.3125 | 240920.5938 | 242920.4063 | 205466.1563 | 216347.2813 | 204754.2656 |
| 5682497 | 4321060 | 5059314 | 5458751 | 6555202 | 6704203 | 6837990.5 |
| 41393.71484 | 29228.60156 | 36177.02734 | 36964.77734 | 25429.66406 | 24846.41211 | 25489.74609 |
| 93871.25 | 74150.84375 | 72245.1875 | 75361.90625 | 42437.76953 | 47147.82813 | 39591.97266 |
| 56533.75391 | 46235.28125 | 39791.46484 | 42461.29688 | 36995.82031 | 39424.8125 | 37372.11328 |

| | | | | | | |
|---|---|---|---|---|---|---|
| 18883.79297 | 13876.36914 | 13513.87598 | NA | 12142.43652 | NA | NA |
| 18714.73828 | 10430.42383 | 21532.85938 | 14210.47852 | 11417.66992 | 17800.16797 | 12144.57227 |
| 84601.42188 | 65782.44531 | 62636.83594 | 66304.09375 | 57536.17188 | 54330.68359 | 54439.39844 |
| 119211.2813 | 120671.0547 | 98406.96875 | 102632.4531 | 98954.89063 | 108640.9297 | 91565.41406 |
| 93578.1875 | 76436.46875 | 99440.09375 | 69033.8125 | 61830.07422 | 68240.29688 | 65920.375 |
| 66599.9375 | 48585.57422 | 55368.64453 | 42465.03125 | 39716.17969 | 58168.55859 | 42052.60547 |
| 426999.0938 | 353146.9688 | 377160.5313 | 345513.4688 | 281054.4375 | 292092.2813 | 273799.0313 |
| 34540.80859 | 33013.84766 | 27879.04492 | 25028.98633 | 19992.20508 | 21985.85742 | 20818.83008 |
| 34469.12891 | 27846.16992 | 28420.94531 | 26129.23242 | 12916.59277 | 17166.91016 | 16084.54297 |
| 45690.02344 | 37198.9375 | 35671.07031 | 32082.76563 | 40009.83203 | 46692.56641 | 35159.39844 |
| 24017.61328 | 24834.11133 | 16837.86328 | 14000.29883 | 30380.33594 | 28544.41406 | 30992.34766 |
| 228.1022186 | 322.2366028 | 114.8527527 | 158.506073 | NA | NA | 91.45020294 |
| 11476.62207 | 8988.151367 | 7666.794922 | 9053.076172 | 12550.17871 | 11833.00977 | 10645.20898 |
| 116782.7031 | 88283.92188 | 98549.83594 | 89903.82813 | 83127.79688 | 86675.04688 | 88520.33594 |
| 544345.75 | 420777.5625 | 414079.1875 | 433635.625 | 462386.375 | 465767.8438 | 472440.5625 |
| 128160.1797 | 82792.50781 | 109171.9922 | 82076.04688 | 90817.47656 | 113289.7109 | 101125.375 |
| 66563.15625 | 60479.47656 | 52069.70313 | 48144.11719 | 43434.8125 | 46662.45313 | 46636.13672 |
| 38366.42969 | 29755.47266 | 30697.57617 | 24722.37109 | 14966.5791 | 15177.69238 | 16040.97559 |
| 18559.66211 | 15216.18945 | 13027.93652 | 14046.64844 | 13146.20313 | 12507.73828 | 11492.51563 |
| 9505.931641 | 9448.541992 | 7128.988281 | 8447.931641 | 15402.26563 | 15381.34473 | 15278.48145 |
| 444230.8125 | 307887.875 | 393758.7813 | 384101.7188 | 428064.4375 | 452825.625 | 453295.0313 |
| 138672.5625 | 107935.4063 | 109099.9453 | 118610.0156 | 105007.6797 | 115195.9688 | 111434.7266 |
| 52300.4209 | 46050.10352 | 47416.88574 | 46625.86816 | 17127.59375 | 18002.17896 | 21154.14551 |
| 241177.5469 | 204656.625 | 182832.1094 | 177481.1875 | 174408.9844 | 156625.125 | 167031.6094 |
| 63242.35156 | 52547.41211 | 52658.23584 | 46450.2085 | 58552.30957 | 54819.54932 | 56616.01563 |
| 8454.118164 | 5750.080566 | 7405.783691 | 7289.911133 | 6005.477051 | 5363.129395 | 7543.865723 |
| 117067.5703 | 84930.8125 | 116510.4922 | 151538.9219 | 173770.4375 | 156176.875 | 200692.0156 |
| 23960.66406 | 22556.48242 | 17336.51172 | 18048.23242 | 20750.79688 | 17374.79492 | 15682.12109 |
| 114126.9453 | 81446.45313 | 90025.0625 | 88913.60938 | 142982.7344 | 138181.4063 | 142430.8281 |
| 59220.29688 | 52966.06641 | 52016.25391 | 37651.58203 | 58457.5625 | 50465.66797 | 53680.87109 |
| 13235.06348 | 11758.21387 | 9807.256836 | 12294.75391 | 16195.51465 | 19282.25781 | 13940.18652 |
| 254204.5313 | 220897.3594 | 204057.7969 | 196039.4219 | 212468.7031 | 222745.4844 | 215772.7969 |
| 29398.52344 | 21831.8418 | 26784.22266 | 24641.0332 | 25125.69141 | 24772.54102 | 27192.40625 |

| | | | | | | |
|---|---|---|---|---|---|---|
| 242471.9531 | 196257.4531 | 203184.4375 | 181285.7656 | 181779.7656 | 188460.4531 | 186335.8906 |
| 195331.3281 | 167832.7656 | 163368.7813 | 174289.1875 | 175372.2031 | 146314.0156 | 176901.7813 |
| 25239.25977 | 21343.31641 | 23600.50586 | 22279.43359 | 18750.08984 | 17025.93555 | 19717.49609 |
| 515810.0625 | 397818.7188 | 438197.25 | 435855.9063 | 434884.0938 | 437019.9375 | 442222.75 |
| 139930.0625 | 112773.8359 | 120786.375 | 136679.4688 | 186299.8438 | 223435.2813 | 189010.9531 |
| 532841.9375 | 360245.875 | 438272.5938 | 370553.4063 | 346412.4688 | 417170.7813 | 339712.9375 |
| 12867.0625 | 11787.48828 | 11651.83691 | 12125.97168 | 13986.36914 | 13090.97949 | 14572.3291 |
| 28066.32813 | 23509.31641 | 15195.70313 | 23583.39453 | 27079.20898 | 18703.87305 | 25011.65625 |
| 208630.9531 | 190323.0313 | 195865.0938 | 124321.9531 | 110898.9219 | 133116.1875 | 107850.1016 |
| 1380.439941 | 1265.348145 | 1058.704346 | 975.5413208 | 1370.642334 | 1521.025146 | 1479.286011 |
| 70742.875 | 56896.44141 | 54935.15234 | 50796.63672 | 45245.26563 | 46105.375 | 46822.72266 |
| 492562.6641 | 191109.9063 | 544450.9688 | 600104.9688 | 307359.1992 | 322979.9883 | 347660.2539 |
| 16904.86719 | 15522.20996 | 14179.16992 | 12013.2002 | 16654.03711 | 16671.21094 | 19436.02734 |
| 49852.92969 | 45736.55078 | 40934.29297 | 43946.64063 | 58201.26563 | 48916.43359 | 51177.23438 |
| 78908.64063 | 65414.71094 | 63125.42578 | 56264.29297 | 58953.76563 | 56954.33594 | 52646.4375 |
| 262385.4375 | 212354.375 | 216900.2813 | 215960.2813 | 297571.9063 | 284595.4688 | 300774.7813 |
| 715252.4375 | 605678.8125 | 601597 | 516296.0313 | 454289.8125 | 505291.875 | 518490.9063 |
| 3220849.75 | 2540045.25 | 2456942 | 2585892.5 | 2826979 | 2695828.5 | 2990957.75 |
| 184328.2031 | 172045.7813 | 146875.6406 | 157378.6406 | 202699.3594 | 194501.3906 | 180961.9219 |
| 465693.125 | 339416.6875 | 383156.7813 | 372833.4063 | 300960.6875 | 315623.4375 | 307815.125 |
| 84850.20313 | 69487.55469 | 66276.24219 | 67059.91406 | 71465.20313 | 73136.79688 | 68741.35938 |
| 67050.11719 | 58268.48828 | 54518.97266 | 52518.63672 | 54030.25 | 56711.53125 | 53236.35156 |
| 52621.11328 | 43185.22266 | 35639.82031 | 38267.95703 | 47758.26563 | 42007.33984 | 49785.35156 |
| 101881.6406 | 83363.00781 | 78221.09375 | 74973.04688 | 63844.55078 | 60254.49609 | 61487.52734 |
| 32925.88672 | 17934.60547 | 29499.66602 | 28645.28516 | 28536.22656 | 23838.08984 | 27588.90234 |
| 13606.51953 | 11907.30664 | 10076.88574 | 9532.264648 | 7372.32666 | 9439.3125 | 7765.767578 |
| 120287.0859 | 93511.42188 | 90958.21875 | 76323.47656 | 40673.875 | 41369.875 | 40362.51563 |
| 303968.9688 | 265480.0625 | 247275.5313 | 254886.5313 | 293015.375 | 257181.625 | 284444.7188 |
| 419248.0625 | 367871.1875 | 333126.4688 | 278228.4375 | 298082.5625 | 340381.1563 | 276375.4063 |
| 36110.39844 | 25956.31055 | 27423.74805 | 24147.1582 | 19372.34766 | 21929.15234 | 21604.13086 |
| 90271.89063 | 76539 | 74976.77344 | 68972.86719 | 73570.44531 | 81699.11719 | 80686.60938 |
| 78517.1875 | 63407.02344 | 75412.67969 | 61313.51563 | 50390.05859 | 44673.06641 | 45400.40234 |
| 143034 | 113771.7344 | 118341.5156 | 107887.4063 | 101853.1016 | 110265.2188 | 104376.2969 |

| | | | | | | |
|---|---|---|---|---|---|---|
| 459161.125 | 275288.0938 | 429103.1875 | 430006.9063 | 440325.625 | 398826.0313 | 427428.25 |
| 220478.4688 | 177262.6719 | 177088.9375 | 165906.1875 | 155386.2969 | 162917.8906 | 155169 |
| 40346.51953 | 33508.70703 | 34662.27344 | 38650.22656 | 44497.21484 | 54587.37891 | 48818.45703 |
| 19505.37109 | 12828.57422 | 15989.75488 | 17853.38477 | 11401.46777 | 12599.60352 | 13549.66309 |
| 425020.4063 | 366204.8438 | 312074.1563 | 246774.1563 | 298631.7188 | 360244.6875 | 298039.25 |
| 71894.33594 | 60400.50781 | 52648.20703 | 51811.44922 | 72150.75 | 68384.10938 | 69526.64844 |
| 1384923.75 | 1352639.625 | 1183926.875 | 1341085.125 | 1317530.875 | 1070567.875 | 1079509.75 |
| 43615.85938 | 17056.80859 | 43718.92578 | 56659.22656 | 117698.6875 | 84193.64844 | 107457.0156 |
| 136333.2188 | 96316.08594 | 116285.0547 | 116277.5938 | 79619.27344 | 82331.80469 | 89992.46875 |
| 14221.45605 | 13802.47363 | 10281.42871 | 10282.31152 | 20959.76758 | 17293.82422 | 19373.44727 |
| 6787.377441 | 6541.135254 | 6375.192871 | 6641.460449 | 6473.911621 | 7727.844238 | 6262.133789 |
| 54260.75391 | 49705.90625 | 39263.01563 | 38032.78906 | 32327.69141 | 29572.01172 | 28804.61328 |
| 8906.209961 | 4394.725586 | 7954.100098 | 8755.791992 | 7041.945801 | 7010.125 | 8463.900391 |
| 78648.04688 | 62631.03125 | 62165.57031 | 56648.15625 | 71068.28125 | 69867.64844 | 74729.42188 |
| 54776.64844 | 47885.01172 | 43863.32813 | 42683.87109 | 43782.69141 | 43181.51172 | 43239.46875 |
| 7383.788086 | 6758.40332 | 6574.157715 | 4987.709473 | 9245.008789 | 6922.66748 | 7809.76416 |
| 137168.3594 | 93531.375 | 129495.1953 | 113952.3047 | 166066.2656 | 180601.875 | 181606.8125 |
| 16003.82324 | 12023.50586 | 13723.74805 | 11277.58984 | 13208.6875 | 13778.7207 | 13284.35742 |
| 14657.28711 | 11823.94238 | 6079.423828 | 11907.08203 | 10167.35254 | NA | NA |
| 5671.575195 | 5388.14209 | 3829.354248 | 4308.890625 | 7110.980469 | 6105.020996 | 5995.078613 |
| 40790.30078 | 31096.73047 | 35301.61328 | 34671.96484 | 27759.01563 | 26299.9043 | 26939.05664 |
| 28555.48633 | 22936.90234 | 20035.19922 | 20037.45117 | 13862.39941 | 19939.54688 | 15899.91895 |
| 17758.24023 | 17121.41016 | 13279.72363 | 14543.9873 | 17129.10938 | 16115.69824 | 16244.47266 |
| 441131.5313 | 296435.2813 | 356016.6563 | 294266.4063 | 299311.8438 | 345447.375 | 274799.1563 |
| 26466.72656 | 18309.20898 | 21481.63281 | 18707.17383 | 15040.33496 | 16659.86133 | 16422.13281 |
| 14473.51758 | 12342.01465 | 12295.04883 | 14032.29492 | 15378.74512 | 16396.40234 | 15581.99609 |
| 632663.8828 | 454069.8594 | 492566.7031 | 522343.1484 | 529851.0625 | 493160.5625 | 538155.8828 |
| 380641.5313 | 304853.8438 | 323063.1563 | 289551 | 237801.1563 | 242315.8438 | 241285.8125 |
| 2147.046387 | 1541.095581 | 1753.634277 | 2441.886475 | 4900.341309 | 3326.004883 | 3868.26001 |
| 7531.39502 | 6078.771973 | 4638.322266 | 4486.889648 | 3473.971924 | 3930.851807 | 4051.534912 |
| 47905.64063 | 31143.86523 | 41656.09375 | 38208.31641 | 28042.50195 | 24171.27734 | 21833.98047 |
| 2753453.75 | 2308669 | 2102807.25 | 2172491.25 | 2541708.5 | 2677498.75 | 2585536.5 |
| 126106.4688 | 103591.5234 | 105932.7109 | 93837.64844 | 80857.33594 | 92166.44531 | 84627.88281 |

| | | | | | | |
|---|---|---|---|---|---|---|
| 555850.625 | 447613.4375 | 408920.75 | 434934.6563 | 390779.875 | 425228.4063 | 442481.6563 |
| 23944.41602 | 15443.91895 | 18350.22461 | 17526.79883 | 9546.194336 | 13113.24316 | 12176.66504 |
| 82805.28906 | 66913.55469 | 62953.88672 | 62448.22656 | 63890.28125 | 68078.54688 | 63529.92969 |
| 62236.64453 | 49706.82813 | 52528.78125 | 60035.54688 | 88421.80469 | 81899.96094 | 81990.71094 |
| 90130.38281 | 70314.0625 | 76967.51563 | 81891.8125 | 91898.63281 | 89512.22656 | 102180.2344 |
| 210744.7656 | 171193.4531 | 155738.7813 | 147266.6875 | 65544.28125 | 63445.43359 | 60085.74609 |
| 256916.5938 | 203959.2656 | 215826.7969 | 198370.4531 | 150996.8281 | 172894.9531 | 152885.9688 |
| 3968146 | 3454692.5 | 3131293 | 3129150.5 | 3231968 | 3317754 | 3181839 |
| 13513.93066 | 12370.42188 | 9865.302734 | 8833.813477 | 10490.16504 | 11464.75488 | 11876.90527 |
| 40213.27734 | 28302.35352 | 32837.43359 | 32207.00586 | 29941.43359 | 27912.9707 | 29567.77539 |
| 283785.625 | 223370.4688 | 230179.6563 | 219948.7031 | 239561.7969 | 235058.9844 | 246912.5625 |
| 3391.081299 | 3303.901855 | 2718.280273 | 2075.758057 | 2048.788086 | 1547.764038 | 1966.411011 |
| 361063.1414 | 297191.3708 | 254206.0515 | 324661.4141 | 354493.9739 | 358584.9924 | 376327.1509 |
| 5174.254883 | 4386.894531 | 4006.681641 | 4778.418945 | 5269.270508 | 4817.158691 | NA |
| 17136.36133 | 13331.81738 | 10965.16797 | 11928.70508 | 8382.214844 | 10866.62012 | 9965.681641 |
| 208636.5469 | 152851.4063 | 176405.5 | 164405.8438 | 128116.5625 | 149437.7188 | 143709.8594 |
| 12295.50977 | 8651.134766 | 11544.50977 | 9204.005859 | 6828.057129 | 8362.314453 | 6774.598633 |
| 58474.54688 | 47112.36719 | 47060.58984 | 51554.84375 | 82742.625 | 62518.05078 | 63276.33594 |
| 1391095.25 | 815700.625 | 1193170.25 | 1142309 | 1079877 | 1104048.5 | 1189710.875 |
| 476566.5625 | 366014.7188 | 363944.2813 | 373493.5 | 424151.3438 | 376138.2813 | 395222.25 |
| 123438.5703 | 91276.92188 | 94259.35156 | 113264.5547 | 168533.4531 | 162560.8281 | 162450.6875 |
| 96474.08594 | 102446.875 | 89825.09375 | 92920.53906 | 119101.5703 | 100380.7422 | 113892.1719 |
| 129240.9648 | 123502.7461 | 91177.33008 | 84089.9873 | 79431.23828 | 78011.89258 | 77463.13867 |
| 76488.86719 | 64891.40625 | 69920.14844 | 58837.80859 | 48969.01563 | 46909.10938 | 47532.53516 |
| 106308.946 | 112622.501 | 86243.35645 | 98129.73584 | 250944.457 | 125387.7705 | 160677.3535 |
| 1797892.625 | 1206170 | 1465918 | 1422240.75 | 1404759.375 | 1485894 | 1426247.5 |
| 87198.33594 | 72674.85156 | 51254.53125 | 53265.21094 | 70254.375 | 109471.5391 | 82368.77344 |
| 1360556.125 | 864845.7813 | 1101579.813 | 1098188.438 | 2044938.688 | 2126762.438 | 2120423.688 |
| 95906.39844 | 80958 | 76583 | 82239.66406 | 74619.07813 | 68652.53906 | 73061.72656 |
| 43214.84766 | 43120.66016 | 30707.96289 | 30442.42188 | 61488.03516 | 49259.91406 | 49611.81641 |
| 44278.51563 | 38644.94531 | 31978.55469 | 30496.13477 | 53541.37891 | 43536.97656 | 48989.64063 |
| 62635.48828 | 71914.66406 | 37869.97266 | 34614.85547 | 57137.58594 | 48914.33594 | 52215.03906 |
| 627434.875 | 498410.1875 | 539462.9375 | 511527.9688 | 505441.4063 | 497887.0313 | 532938.4375 |

| | | | | | | |
|---|---|---|---|---|---|---|
| 168683.2813 | 121527.375 | 140685.0313 | 144008.7344 | 194006 | 199676.1406 | 203977.9219 |
| 13388.77344 | 13505.97559 | 7825.669434 | 11066.36133 | 11428.02051 | 10632.37109 | 11803.5293 |
| 101938.0469 | 73767.26563 | 81228.98438 | 99406.07031 | 83092.17188 | 79483.07813 | 79223.9375 |
| 29243.55859 | 29202.10352 | 17633.95703 | 17644.58594 | 18133.88672 | 17200.60156 | 18086.48633 |
| 30266.72852 | 27820.20117 | 26153.10156 | 21828.85352 | 38256.32031 | 32008.17773 | 32948.55469 |
| 36201.42578 | 27234.09375 | 28057.28906 | 28304.06445 | 27969 | 32661.73047 | 30186.02148 |
| 616174.9375 | 520052.8125 | 505300.2813 | 466968.3125 | 399213.8438 | 453621.0938 | 406199.3438 |
| 245170.7344 | 206171.3281 | 200198.8438 | 203954.7188 | 242014.7813 | 244927.125 | 240614.6563 |
| 211384.0938 | 161607.1094 | 160475.5781 | 112324.1016 | 141265.0781 | 198214.7344 | 124325.7734 |
| 136889.6875 | 114907.7656 | 98818.36719 | 94500.98438 | 140634.7969 | 181163.6563 | 161883.5 |
| 187395.3594 | 140976.7031 | 162846.2188 | 157211.0313 | 153746.1094 | 150814.9688 | 144843.0313 |
| 41090.35156 | 37493.50391 | 28923.78711 | 29066.14063 | 55216.62109 | 51728.125 | 57210.58594 |
| 6959.277832 | 5137.129395 | 7077.128906 | 5965.375488 | 9270.421875 | 8887.385742 | 9774.010742 |
| 52505.11719 | 38019.39453 | 39989.79688 | 37925.46094 | 34122.00391 | 35581.44922 | 36535.30469 |
| 413630.2813 | 309140.5313 | 285565.375 | 309384.5313 | 295334.4063 | 249384.125 | 237302.25 |
| 24998.93164 | 18889.82422 | 20369.63477 | 16185.66113 | 12050.10938 | 13508.36914 | 12564.6875 |
| 10136.50684 | 6202.73291 | 8385.854492 | 9205.325195 | 3364.197754 | 6137.020508 | 4557.417969 |
| 234693.4063 | 171492.6406 | 193102.6719 | 190497.2656 | 210653.4844 | 214103.75 | 202921.0313 |
| 77889.07031 | 54947.51563 | 62708.60547 | 62615.44141 | 75985.44531 | 66295.10938 | 72427.10938 |
| 61328.21875 | 45262.16016 | 51347.54297 | 48019.67969 | 37697.26953 | 37479.79688 | 42322.58203 |
| 132362.6094 | 90621.10156 | 100978.5 | 106297.8516 | 131325.2031 | 115451.5234 | 130729.5625 |
| 13324.04395 | 9700.707031 | 7577.500977 | 8200.833984 | 9149.999023 | 4696.51416 | 5564.453613 |
| 12503.63184 | 8927.984375 | 9902.519531 | 9984.148438 | 11507.16309 | 11942.64746 | 13129.20313 |
| 130136.8672 | 94923.78125 | 97720.00781 | 88181.69531 | 87960.44531 | 96121.47656 | 92621.94531 |
| 5468.45459 | 3570.143311 | 4952.637207 | 6816.166992 | 6889.174316 | 8064.520996 | 8146.338867 |
| 10933.28027 | 8145.083496 | 8055.649902 | 8855.705078 | 5530.740234 | 8272.959961 | 8423.441406 |
| 940413.875 | 991513.5 | 556859.1875 | 593849.0625 | 579732.0625 | 593858.1875 | 523794.8125 |
| 71720.39063 | 54191.94141 | 61562.82031 | 56527.81641 | 52785.34766 | 50447.94531 | 50898.40234 |
| 75998.51563 | 60872.05469 | 54377.96875 | 52962.51563 | 36591.37891 | 32297.10352 | 33116.07031 |
| 57206.46094 | 43994.67188 | 45541.01172 | 47006.91797 | 55883.4375 | 59515.38281 | 58672.53125 |
| 34616.36328 | 30316.82227 | 25375.33789 | 27026.78516 | 28076.80078 | 27606.33594 | 27692.32031 |
| 19153.48047 | 17687.51758 | 13461.77734 | 13795.97559 | 20152.38867 | 14867.93555 | 18881.35352 |
| 43389.63281 | 33014.34766 | 32849.03516 | 29090.86914 | 29961.14844 | 33851.75391 | 39009.56641 |

| | | | | | | |
|---|---|---|---|---|---|---|
| 54268.40234 | 45448.77344 | 46373.88281 | 47836.24219 | 38536.19922 | 38240.00781 | 37052.22266 |
| 55345.79297 | 44353.91016 | 43942.14453 | 47408.56641 | 60289.19922 | 57145.28906 | 57734.80469 |
| 35306.81641 | 25422.23047 | 28587.50586 | 38944.78516 | 40995.23047 | 50404.875 | 54424.07031 |
| 23342.06641 | 23073.95898 | 19847.33203 | 20901.92969 | 32759.02734 | 30333.63281 | 29552.95313 |
| 65833.84375 | 61695.88281 | 50870.96484 | 51748.1875 | 59199.30469 | 56844.64063 | 53794.01953 |
| 116074.8438 | 147293.7344 | 130711.5547 | 5494.050293 | 154620.1875 | 175171.9844 | 176182.25 |
| 63160.15625 | 49179.80078 | 52273.41016 | 54697.33984 | 51519.11719 | 44980.875 | 47681.46875 |
| 165756.4688 | 115486.625 | 128208.6953 | 122857.9297 | 125587.25 | 122670.875 | 124143.7031 |
| 14922.04639 | 13494.70361 | 11216.23633 | 10491.76172 | 12947.79785 | 11536.38672 | 11608.89941 |
| 753085.2969 | 479337.8516 | 595534.9688 | 614571.0469 | 686848.625 | 663650.6875 | 675880.8672 |
| 26786.66992 | 29053.81445 | 18165.625 | 14683.52637 | 18220.04297 | 17496.3125 | 20763.95898 |
| 93649.25781 | 76509.53906 | 75050.92188 | 70930.76563 | 63249.01172 | 68739.20313 | 61138.73828 |
| 278684.3125 | 233108.6406 | 215410.1406 | 222484.8125 | 226055.3594 | 232767.2969 | 225467.6406 |
| 589116.8125 | 469838.0625 | 486979.9688 | 452314.3125 | 324446.0625 | 344036.1875 | 324074.2188 |
| 70432.91406 | 59838.94141 | 47452.10547 | 54130.77734 | 49961.39063 | 43887.25391 | 48073.08203 |
| 687485.0625 | 535131.5 | 544581.4375 | 541869.125 | 457107.0625 | 485729.8438 | 489531.5 |
| 157798.5469 | 106453.4531 | 125887.2578 | 119750.0547 | 136135.2188 | 148569.6875 | 156615.2969 |
| 63806.41797 | 41416.58594 | 48224.48828 | 50324.04688 | 59958.85156 | 62059.73438 | 62884.9375 |
| 4332.166016 | 3817.604248 | 3256.036133 | 2673.43335 | 2797.527344 | 2896.441406 | 2436.883545 |
| 144733.2969 | 126050.9141 | 121265.0625 | 123279.7969 | 165306.6406 | 168936.2344 | 171941.9219 |
| 32075.08398 | 22815.66406 | 25836.50391 | 27214.49609 | 31388.48242 | 27649.9375 | 29860.42383 |
| 24847.97852 | 21634.41602 | 23334.71484 | 19257.4668 | 19659.96875 | 20607.35742 | 20928.73438 |
| 77075.6875 | 63179.35938 | 62049.13281 | 57585.44531 | 84415.52344 | 66187 | 63758.75391 |
| 44976.14063 | 37208.71484 | 40412.52344 | 40521.07031 | 64282.49609 | 60459.9375 | 62823.40234 |
| 88852.03125 | 54167.48438 | 68324.74219 | 71937.27344 | 101760.6641 | 71478.78906 | 99408.35156 |
| 29829.19141 | 26390.37891 | 24269.06445 | 20484.99219 | 24737.25195 | 22103.77539 | 24399.89453 |
| 475467.625 | 354385.625 | 383607.7813 | 390084.5313 | 367893.0313 | 353506.7813 | 384576.5938 |
| 8592.176758 | 6817.52002 | 6891.756348 | 7622.470215 | 6661.423828 | 6469.04541 | 6000.17041 |
| 23821.60352 | 22850.93359 | 18120.48633 | 14796.95801 | 22133.76563 | 16778.52734 | 18360.27148 |
| 77239.96875 | 38165.80078 | 66598.52344 | 48355.37891 | 30329.32617 | 58990.58594 | 24997.10742 |
| 75685.16406 | 61911.64844 | 57779.91016 | 52060.16406 | 67195.92969 | 68940.61719 | 71223.52344 |
| 27754.12891 | 23713.25977 | 22245.80469 | 21394.82422 | 28914.24805 | 28346.58984 | 26427.91992 |
| 1358815.5 | 1052331.875 | 1022930.813 | 920722.25 | 996678.8125 | 1059785.125 | 1075012.125 |

| | | | | | | |
|---|---|---|---|---|---|---|
| 107901.9219 | 79793.46094 | 81032.42969 | 86310.86719 | 100683.8828 | 104125.7891 | 106492.6328 |
| 931075.3125 | 612922.1875 | 745915.375 | 808834.8125 | 841035.875 | 824268.5625 | 763165.25 |
| 97119.3125 | 79857.85938 | 75897.25 | 70956.21875 | 65118.42969 | 68326.05469 | 62918.21484 |
| 14066.66699 | 10129.20801 | 11419.42285 | 11920.80469 | 12734.53027 | 12280.48633 | 13010.22363 |
| 35693.67578 | 28452.59961 | 27575.76953 | 25426.66602 | 21337.14453 | 20449.39844 | 19312.51953 |
| 322991.8125 | 290828.9375 | 292692.5938 | 288096.5 | 348769.9688 | 318000.6875 | 337929.4063 |
| 19377.74023 | 15782.08105 | 11675.70605 | 14257.9248 | 10368.35254 | 12594.14355 | 11495.27539 |
| 25567.15039 | 23577.44141 | 18874.02539 | 17187.28711 | 29968.6875 | 24885.77734 | 25711.2207 |
| 1037335.563 | 721100.125 | 844819.125 | 808632.1875 | 780567.125 | 745356.3125 | 754385.5625 |
| 262154.0938 | 206989.75 | 203626.125 | 197174.9688 | 207987.9844 | 176991.4375 | 221963.0625 |
| 535458.75 | 398101.9063 | 455065.6563 | 455539.2813 | 463222.4063 | 450983.4063 | 486966.25 |
| 114570.6641 | 71610.75 | 116869.2344 | 113297.4922 | 152749.0313 | 133157.5156 | 123231.7656 |
| 4065.445557 | 7083.804688 | 1739.9021 | 3754.311279 | 7308.192871 | 7526.662109 | 7647.54834 |
| 150918.4688 | 120132.6484 | 111438.8828 | 115704.0156 | 130731.6563 | 129267.75 | 122798.9531 |
| 305745.5938 | 249758.5469 | 234725.7969 | 242496.2813 | 140772.375 | 128701.6797 | 129277.4844 |
| 3234114 | 2176423.25 | 2557428.5 | 2483381.5 | 2383585.75 | 2434960.75 | 2336461.5 |
| 9002.375 | 6963.735352 | 6848.116211 | 6374.574219 | 6347.513672 | 5631.941895 | 5986.671387 |
| 22636.88086 | 19691.1543 | 16169.72754 | 15060.0293 | 18548.29297 | 10893.34961 | 14080.16211 |
| 17914.89453 | 12359.48535 | 12760.77344 | 13737.23047 | 27971.95508 | 25707.89063 | 29310.81055 |
| 29832.16016 | 24073.95117 | 20185.89258 | 24736.30078 | 34180.61328 | 32826.07422 | 32876.33984 |
| 75810.05469 | 65129.29297 | 61964.63672 | 69752.52344 | 126267.4297 | 97005.82813 | 101653.1875 |
| 319485.1563 | 236816.1563 | 266730.6875 | 277272.2813 | 285227.7813 | 330477 | 307884.375 |
| 88814.63281 | 99035.0625 | 73048.10938 | 59553.125 | 100974.2969 | 83710.17969 | 81222.48438 |
| 570580.125 | 488640.2188 | 511007.625 | 536013.4375 | 511330.875 | 484924.8125 | 438691.7813 |
| 528277.75 | 354895 | 426541.25 | 418266.3438 | 356223.4063 | 360351.8125 | 353660.0625 |
| 213452.8438 | 152841.3906 | 174944.9219 | 167069.1719 | 167130.3906 | 178117.2969 | 169152.5781 |
| 35991.92969 | 29898.62695 | 24065.57617 | 21269.20898 | 11735.65234 | 15658.78613 | 10316.75098 |
| 5531.381348 | 4866.46582 | 3833.018311 | 3564.738037 | 1945.640381 | 1959.572021 | 2172.57251 |
| 2095890.25 | 1430773.625 | 1780736.375 | 1783808.5 | 1247793 | 1244752.75 | 1321864.125 |
| 164513.7656 | 131145.3594 | 132437.1406 | 144200.0781 | 174706.875 | 172821.5156 | 182306.5469 |
| 22898.85938 | 16483.71094 | 18571.06641 | 19295.9043 | 21014.87891 | 21365.00586 | 19007.23242 |
| 885904.9375 | 660635.5 | 715113 | 671492.4375 | 692729.625 | 686943.5 | 708906.4375 |
| 735452.1875 | 531877.8125 | 592238.5 | 635065.375 | 925893.125 | 778933.1875 | 918397.5 |

| | | | | | | |
|---|---|---|---|---|---|---|
| 1717003.5 | 1405552.25 | 1195654.875 | 1247759.125 | 1986467.125 | 2029504.25 | 2079110 |
| 33424.50391 | 31332.37891 | 21931.95703 | 27219.82031 | 43228.23828 | 34204.73047 | 42170.87109 |
| 88091.91406 | 69930.14063 | 64753.63281 | 65279.24609 | 67137.78906 | 66696.82813 | 65295.51172 |
| 8961.378906 | 6792.656738 | 7700.02002 | 7678.435059 | 7865.189941 | 4180.978516 | 5472.937988 |
| 5314.669922 | 7368.342285 | 2283.566406 | 2293.050537 | 7735.600098 | 6914.530762 | 6613.750977 |
| 459563.9063 | 375998.7188 | 397499.0313 | 377445.4688 | 438978.375 | 401528.0938 | 422810.2188 |
| 778933.0625 | 609620.25 | 619684.5 | 627068.6875 | 662703.75 | 736844.1875 | 745338.0625 |
| 15860.23047 | 12263.85059 | 13109.97363 | 11961.96875 | 12305.01172 | 11600.2959 | 11833.14453 |
| 34159.76953 | 32268.62891 | 29649.60352 | 29184.00781 | 30823.05078 | 31245.67773 | 30287.84766 |
| 60449.27734 | 44362.78125 | 45477.48828 | 53201.5625 | 75317.74219 | 63399.83594 | 68817.71094 |
| 88157.0625 | 73289.67969 | 73082.78906 | 67443.35156 | 73589.01563 | 75643.17969 | 73153.72656 |
| 14446.50391 | 11648.4541 | 10392.13086 | 11721.86719 | 12821.9668 | 13969.11719 | 12692.79102 |
| 87765.35156 | 73501.46094 | 71424.79688 | 60410.20703 | 90000.35938 | 94979.77344 | 104499.2031 |
| 498705.0313 | 379855.1563 | 387656.0938 | 379338.75 | 409947.1875 | 442896.1875 | 438955.7813 |
| 108505.3125 | 67416.52344 | 85352.875 | 112801.2734 | 150023.2813 | 125148.5781 | 147635.3438 |
| 20989.41992 | 13276.08398 | 19943.83398 | 19262.40234 | 30719.46484 | 23986.67383 | 25232.10156 |
| 40842.51563 | 34068.21875 | 32298.3418 | 32996.65625 | 34779.10156 | 32659.3457 | 33682.56641 |
| 65378.85156 | 45655.51953 | 52236.77734 | 45642.99609 | 43462.92188 | 43053.73438 | 42707.125 |
| 49697.49219 | 34427.73047 | 40700.16016 | 33312.81641 | 41342.79297 | 42188.69141 | 41948.20703 |
| 18801.9707 | 13243.51953 | 14219.28906 | 12235.42871 | 12731.00098 | 13345.58398 | 12956.52344 |
| 4265.891602 | 3455.352783 | 3392.814941 | 3912.060303 | 6231.88916 | 5998.835449 | 7102.431152 |
| 722857.6875 | 606343.625 | 601148.1875 | 534899.4375 | 421520.125 | 433038.9375 | 420451.9063 |
| 10855.36133 | 9594.635742 | 8966.407227 | 8119.214844 | 8520.532227 | 7726.135742 | 9204.950195 |
| 6704.964355 | 5937.336914 | 3958.093262 | 6411.004395 | 12849.38867 | 11985.14551 | 12073.4248 |
| 63525.49219 | 53720.58594 | 47044.21094 | 43751.48828 | 43421.72656 | 45451.91406 | 42875.35938 |
| 177275.6563 | 137981.0625 | 153175.75 | 137631.9063 | 98173.5625 | 93861.67969 | 94736.72656 |
| 54259.22656 | 41951.43359 | 41133.48438 | 43185.85938 | 44772.19922 | 44447.61719 | 39274.24219 |
| 447643.0938 | 330970.5625 | 354334.9375 | 308653.0625 | 350408.6875 | 339846.5 | 358608.7813 |
| 3556828 | 2724853.5 | 2723345.25 | 2744021.75 | 2939645 | 3024580 | 3078275.75 |
| 113783.0234 | 94830.89844 | 87220.78125 | 79598.29688 | 45536.60938 | 44674.96094 | 44797.53125 |
| 27108.15625 | 22414.07813 | 22109.98047 | 23843.16602 | 23333.16016 | 22758.68359 | 22678.74023 |
| 17104.73633 | 14597.00293 | 11649.72852 | 11770.75098 | 15478.81152 | 13890.62695 | 13371.96289 |
| 6844.992676 | 4768.574219 | 5284.991211 | 5458.274414 | 6284.297363 | 5521.154785 | 4933.189941 |

| | | | | | | |
|---|---|---|---|---|---|---|
| 334093.8125 | 272866.8438 | 245099.6563 | 239610.4531 | 279990.8125 | 273570.5 | 303173.375 |
| 20255.87305 | 18039.72461 | 13462.37793 | 11245.12891 | 15383.77246 | 12289.75391 | 14046.47559 |
| 15807.29004 | 16729.67578 | 12918.66602 | 14450.11621 | 27772.58203 | 25624.39453 | 25671.68164 |
| 15333.76855 | 14108.71191 | 11640.94043 | 11842.82227 | 23461.54492 | 19701.49414 | 20066.05078 |
| 4982520 | 7156267.5 | 4195781.5 | 1508882 | 1329167.625 | 6100814 | 1050197.25 |
| 117456.6016 | 77037.6875 | 96945.51563 | 105644.0078 | 84155.52344 | 86431.14063 | 89925.09375 |
| 56173.37891 | 48309.55859 | 41950.46875 | 37462.08594 | 33971.6875 | 35485.29688 | 36712.9375 |
| 4952.60791 | 4458.626953 | 4847.674805 | NA | 5076.477539 | 6723.751465 | 4315.860352 |
| 1209505.25 | 1076534 | 883769.6875 | 408163.625 | 567880.75 | 1368466.125 | 788481.25 |
| 266065.1875 | 204106.4688 | 219977.6094 | 201993.8594 | 250353.0156 | 227309.4063 | 251195.5 |
| 500642.0313 | 403008.9063 | 369523 | 427503 | 452838.2188 | 598578.75 | 450819.5625 |
| 79365.11719 | 63087.98438 | 57490.96094 | 53711.69531 | 55368.64844 | 51649.66016 | 60383.46094 |
| 16133.29004 | 11009.21582 | 14729.26855 | 13614.37012 | 9788.669922 | 9283.004883 | 9923.578125 |
| 1665739 | 1142043.875 | 1338768.375 | 1373084.125 | 1211200.875 | 1351143.375 | 1287039 |
| 9310.411133 | 7166.527832 | 6767.978027 | 6539.272949 | 5696.911621 | 5980.178223 | 5511.399902 |
| 26832.49219 | 23682.16992 | 21606.02344 | 17209.73828 | 22888.51758 | 23975.10352 | 20988.50977 |
| 476739.0313 | 299137.1563 | 339428.3125 | 369096.0625 | 421002.5625 | 392097.3125 | 439977.125 |
| 71924.10938 | 49234.80469 | 54323.89063 | 55518.73438 | 62761.51172 | 69165.60156 | 65424.65625 |
| 295805 | 245538.9844 | 217434 | 211235.8281 | 180404.1563 | 175496.1406 | 173717.875 |
| 10727.36328 | 8860.694336 | 8184.925781 | 8599.381836 | 16805.30469 | 12945.8418 | 13861.78809 |
| 324322.5 | 224429.0781 | 254538.1875 | 251545.0781 | 236966.0469 | 247811.3906 | 265616.8438 |
| 81405.80469 | 58505.09766 | 68086.25 | 62708.39063 | 59167.40625 | 98821.57813 | 80429.76563 |
| 56895.65234 | 42455.98828 | 43113.50391 | 42534.59766 | 24832.22852 | 23629.50195 | 25389.33398 |
| 80453.53125 | 59327.85156 | 60335.08594 | 53352.82422 | 53847.88281 | 61633.80078 | 56462 |
| 167719.6719 | 132228.7813 | 124209.5078 | 137609.3906 | 177578.4531 | 166102.1094 | 165487.6563 |
| 73850.46875 | 49474.15234 | 59848.16797 | 66752.07031 | 61281.94141 | 63839.16797 | 64319.44141 |
| 20849.18359 | 15765.15234 | 15676.51465 | 18674.42383 | 31865.88672 | 27427.33594 | 28353.10547 |
| 17354.53906 | 14666.7998 | 9406.632813 | 10881.1123 | 9463.757813 | 10938.43945 | 8627.713867 |
| 217543.1563 | 157632.9688 | 174117 | 168680.4844 | 164413.7969 | 145829.2188 | 162590.6094 |
| 128995.8516 | 86221.07813 | 104724.6563 | 102634.9375 | 108819.8828 | 107696.1328 | 113137.4219 |
| 43109.33984 | 31096.00391 | 31840.59766 | 53289.42969 | 103923.9844 | 61401.44141 | 97993.32031 |
| 77330.0625 | 67604.38281 | 55485.40625 | 61043.38672 | 88924.39844 | 75139.9375 | 74035.9375 |
| 832917.875 | 712852.875 | 734797 | 800231.4375 | 1110422.375 | 937092.25 | 990475.5 |

| | | | | | | |
|---|---|---|---|---|---|---|
| 4872638.5 | 8569838 | 5014630.5 | 1569354.875 | 1314924.75 | 4036428.5 | 1263348.875 |
| 752.3095703 | 753.0222778 | 577.6083984 | 708.5894165 | 887.8482056 | 730.2406006 | 600.345459 |
| 70445.74219 | 58334.77734 | 55793.20703 | 64410.9375 | 67759.8125 | 58106.06641 | 66908.07813 |
| 24383.00977 | 18269.01172 | 18028.56055 | 16129.32227 | 15449.4834 | 14725.48145 | 13280.35742 |
| 5132.605469 | 4554.713867 | 4015.88208 | 3719.747314 | 4604.406738 | 5446.979492 | 5912.271973 |
| 104281.9531 | 78816.79688 | 79919.98438 | 82010.89844 | 87129.77344 | 81277.74219 | 90119.57031 |
| 86978.76563 | 66270.67969 | 65193.73438 | 62082.07031 | 49134.74609 | 46703.80469 | 47258.08203 |
| 13339.58203 | 12392.70215 | 11380.74609 | 11738.07129 | 20440.75195 | 18102.3418 | 17788.13477 |
| 479673.25 | 393671.9688 | 369049.3438 | 406078 | 438840.875 | 397135.375 | 409814.875 |
| 11667.83789 | 6824.38916 | 8319.302734 | 7640.812988 | 8748.049805 | 9286.097656 | 7698.852051 |
| 46822.21875 | 38116.98047 | 39131.60547 | 42942.48438 | 35205.30859 | 33964.27734 | 39096.39844 |
| 95925.9375 | 69289.32813 | 79829.34375 | 81382.71094 | 92703.03906 | 99875.92188 | 100985.8281 |
| 206242.4063 | 148247.6875 | 159878.4688 | 144269.3281 | 157862.0313 | 176421.6406 | 172500.9375 |
| 103758.1953 | 82390.08594 | 70728.92188 | 61018.63281 | 43823.73438 | 50888.92969 | 45443.41016 |
| 12204.0293 | 8111.300293 | 9035.560547 | 10427.53223 | 9743.837891 | 10632.55762 | 10680.01953 |
| 2840068 | 1587616.25 | 2173697 | 1966452.375 | 2107112.5 | 2126063.75 | 2029894.875 |
| 155976.9844 | 118713.7188 | 117494.8438 | 111685.7813 | 98219.6875 | 93504.65625 | 99142.29688 |
| 31503.45898 | 22853.01953 | 23540.53711 | 23832.09961 | 12955.29883 | 14808.39844 | 14339.12988 |
| 2174244.75 | 1398297.25 | 1762313.5 | 1594678.25 | 1560105.875 | 1913902 | 1791789.875 |
| 21315.0332 | 17285.19141 | 15357.58691 | 13083.28613 | 13038.4707 | 12439.79102 | 12125.31738 |
| 107567.7969 | 75221.52344 | 91556.53906 | 67833.90625 | 95223.91406 | 113187.4375 | 108173.2734 |
| 83214.85156 | 60346.50391 | 65392.04102 | 63175.5957 | 54574.11523 | 53031.41797 | 54312.99023 |
| 27023.13281 | 21985.87891 | 20504.24414 | 22393.26563 | 43037.60547 | 45077.52734 | 40886.36719 |
| 155438.3906 | 116927.0703 | 107113.0313 | 91888.70313 | 109473.9141 | 107160.9297 | 87181.91406 |
| 2520547.75 | 1648121.625 | 1917600.125 | 1786927 | 1848145.75 | 1880106.125 | 1810973.875 |
| 112287.2422 | 73358.90625 | 89545.78906 | 90062.00781 | 72070.67969 | 82662.64844 | 75372.875 |
| 71455.21094 | 51952.24609 | 66107.14063 | 61263.33984 | 73157.96875 | 68044.51563 | 72853.82031 |
| 10586.24414 | 8939.682617 | 8781.929688 | 7584.937988 | 14768.93652 | 12946.98633 | 14087.02637 |
| 26439.66797 | 22007.68945 | 19594.05078 | 22999.64258 | 29713.58594 | 25626.16406 | 26381.18359 |
| 19709.39453 | 16769.62305 | 13757.50391 | 11171.50781 | 14399.17773 | 14558.89844 | 14257.15918 |
| 186677.7461 | 184583.4023 | 120657.0781 | 126780.0547 | 158411.7344 | 162739.6338 | 159453.5352 |
| 17755.80469 | 16819.67969 | 16192.89551 | 17291.83789 | 20053.15039 | 19000.13672 | 18532.28906 |
| 599377.9375 | 476882 | 461185.375 | 448373.4688 | 486170.0313 | 501609.4063 | 491145.625 |

| | | | | | | |
|---|---|---|---|---|---|---|
| 271737.1563 | 246881.9063 | 214300.9063 | 235502.0625 | 257265.8125 | 268546.3438 | 239874.5313 |
| 4105325.75 | 3160155 | 3215428.5 | 3013755.5 | 2680673 | 2837673 | 2745134 |
| 7428.973145 | 5317.210938 | 5622.07959 | 5447.665039 | 6550.01123 | 5454.102539 | 5723.176758 |
| 36649.07031 | 42255.50781 | 23826.0332 | 21181.12695 | 48199.22266 | 42976.91406 | 46621.03125 |
| 16063.40723 | 11016.91797 | 12275.60254 | 13697.61816 | 12562.52148 | 14873.89648 | 12215.20117 |
| 27160.66211 | 20579.11523 | 18815.37109 | 19499.96875 | 23833.62109 | 20331.90039 | 19922.09961 |
| 10429.16309 | 9456.392578 | 8694.288086 | 8721.609375 | 11231.50293 | 8505.007813 | 13502.96387 |
| 33271.29688 | 28950.55273 | 30071.67188 | 26226.56055 | 39452.25 | 45286.69531 | 42883.09375 |
| 1259236.875 | 1102907.375 | 965141.8125 | 1088160.875 | 1210118.5 | 990682.9375 | 1063430.125 |
| 175120.1563 | 165266.5313 | 142654.875 | 127306.3125 | 213409.75 | 217765.75 | 191574.75 |
| 29046.18945 | 10891.76465 | 27482.90625 | 29061.63281 | 32711.98633 | 29170.88867 | 33003.71875 |
| 384097.9063 | 312279.0313 | 313761.625 | 287615.1875 | 321650.875 | 336030.4063 | 320087.125 |
| 38200.72656 | 29413.51367 | 30106.13281 | 28947.19336 | 25865.67773 | 26221.90625 | 27822.44922 |
| 103349.4766 | 77291.75781 | 73747.84375 | 80652.35938 | 81131.65625 | 86060.1875 | 80727.48438 |
| 32090.56641 | 23656.98242 | 23712.94531 | 25114.39063 | 26454.89258 | 29868.89844 | 28732.41602 |
| 134343.1094 | 88609.40625 | 86237.44531 | 93093.57031 | 109328.3516 | 101823.5156 | 108801.3281 |
| 218631.1094 | 133255.5625 | 160313.75 | 174179.7813 | 156115.4844 | 138798.875 | 161962.2344 |
| 106197.4063 | 80666.61719 | 83612.32813 | 75683.8125 | 96878.32031 | 93849.77344 | 94854.74219 |
| 4723.778809 | 4509.096191 | 2815.960938 | 3486.211426 | 4933.89209 | 3699.02002 | 4031.088135 |
| 250773.6406 | 167975.9219 | 195398.8125 | 182621.4844 | 151600.7188 | 181080.1094 | 143020.4375 |
| 319432.5313 | 259183.4688 | 256362.1563 | 264444.5 | 372294.7813 | 383148.2813 | 403520.1563 |
| 7348.946289 | 6698.262695 | 4873.047363 | 6332.084961 | 5039.445801 | 5024.584961 | 4294.098633 |
| 124343.9219 | 102480.5078 | 98061.5625 | 75567.28125 | 72774.97656 | 84848.90625 | 74329.42969 |
| 42205.28516 | 34008.20313 | 31023.36719 | 32386.00586 | 33678.86328 | 31877.88281 | 32110.50977 |
| 359150.3438 | 277594.7813 | 296396.4063 | 278842.6563 | 252608.2031 | 208732.9844 | 221457 |
| 275250 | 199831.9844 | 207695.0625 | 212730.8281 | 228325.3594 | 228659.5156 | 238972.0938 |
| 5338.731934 | 3405.235596 | 5033.735352 | 3932.467773 | 7237.609863 | 7356.328125 | 6403.152832 |
| 24579.92773 | 16942.25 | 18883.32422 | 18114.10547 | 23451.16211 | 24888.82422 | 23529.50195 |
| 79557.17969 | 65851.96875 | 62417.13672 | 60602.53516 | 61457.14453 | 64068.52734 | 60772.4375 |
| 819160.4375 | 836046.9375 | 587291.9375 | 680562.25 | 942028.25 | 1007924.625 | 789933.1875 |
| 168688.0469 | 128207.5703 | 138717.3438 | 129611.4063 | 164786.0156 | 171701.1406 | 171623.1875 |
| 442922.375 | 351364.2813 | 333362.75 | 311854.1875 | 334687.5313 | 356156.6875 | 324032.0313 |
| 18136.1875 | 17097.85156 | 7679.55127 | 14184.22168 | 15400.07324 | 15963.36523 | 18237.97461 |

| | | | | | | |
|---|---|---|---|---|---|---|
| 113752.4766 | 98745.0625 | 84871.58594 | 83672.15625 | 64239.20703 | 71949.32031 | 58341.89844 |
| 359657.0625 | 280227.375 | 262215.5313 | 261217.5469 | 302124.3438 | 316708.4063 | 319766.3438 |
| 8895.682617 | 9486.507813 | 5007.223633 | 4128.270508 | 4904.458496 | 2584.342285 | 5460.317383 |
| 109990.9844 | 79899.9375 | 79762.82031 | 81371.90625 | 80176.89844 | 71475.76563 | 80212.78906 |
| 2858.678711 | 2119.038818 | 3645.886475 | 1965.981567 | 2389.132813 | 2311.342773 | 2008.425171 |
| 103512.7813 | 79336.50781 | 82006.5 | 61051.42578 | 71092.90625 | 80054.39063 | 66716.17969 |
| 390815.4375 | 259578.7969 | 333692.75 | 335234.7188 | 219150.5469 | 282358.3438 | 266536.9375 |
| 38360.89453 | 32126.28711 | 29149.40234 | 33421.44922 | 42849.33594 | 51207.27734 | 41383.59375 |
| 746028.75 | 565785.375 | 545190.4375 | 553948.6875 | 579212.5 | 583526 | 591731.1875 |
| 288521.0938 | 223583.0313 | 206245.3125 | 209080.6719 | 211625.5156 | 227932.9219 | 214790.8281 |
| 61702.58203 | 57777.29297 | 41869.03516 | 52118.36328 | 67288.92969 | 62842.02734 | 55675.14844 |
| 26812.20313 | 20356.51367 | 18642.0332 | 21018.90039 | 19924.67969 | 19520.45313 | 18762.7168 |
| 667181.875 | 450140.7188 | 539892.125 | 408124.2188 | 474136.9063 | 525240.25 | 499562.625 |
| 36842.66406 | 31935.45313 | 28290.20898 | 28629.68164 | 28142.1582 | 27709.4668 | 25991.85352 |
| 22205.34961 | 10000.84863 | 18400.26758 | 20333.95703 | 23264.94336 | 23850.62891 | 19503.91211 |
| 7372.433105 | 8438.352539 | 3602.380371 | 3796.384766 | 7543.242676 | 7472.986816 | 7441.688477 |
| 269757.751 | 258516.1533 | 181867.9014 | 183014.4395 | 248179.458 | 257126.0752 | 254752.1992 |
| 11303.03223 | 5430.726074 | 8638.788086 | NA | 5557.908691 | 5022.360352 | 5408.947266 |
| 34232.80859 | 29930.2207 | 21996.99805 | 23492.2832 | 28995.9707 | 32766.54492 | 27322.03906 |
| 4397.28125 | 3696.939697 | 2945.704102 | 2933.112549 | 6258.824219 | 6346.203613 | 6701.463867 |
| 267609.0313 | 210277.8281 | 249511.6875 | 223714.4219 | 201234.1875 | 230783.5156 | 202997.1094 |
| 28153.2168 | 20606.06836 | 19913.99219 | 21512.43945 | 17192.15625 | 18498.94336 | 22588.95313 |
| 32698.07617 | 21719.2168 | 24370.04297 | 25710.56445 | 42325.35547 | 39014.0625 | 38659.05469 |
| 126462.6016 | 96234.35938 | 103800.8906 | 88630.35156 | 60176.55078 | 57670.77734 | 63727.38281 |
| 167483.7031 | 120263.5156 | 118110.8047 | 121741.6172 | 90391.57031 | 84644.4375 | 87299.46094 |
| 146886.5469 | 105862.2891 | 124728.8516 | 114229.2656 | 124457.0313 | 115414.8906 | 124073.0156 |
| 63885.09766 | 47270.47266 | 45967.89063 | 43415.5625 | 49671.06641 | 55328.07031 | 49374.23047 |
| 191714.8438 | 150307.3438 | 155093.875 | 136095.3281 | 144868.9844 | 143088.1875 | 148290.8438 |
| 403520.5625 | 286528.0938 | 308745.8438 | 286662.8125 | 312646.3438 | 331597.375 | 324160.1563 |
| 5870.15332 | 4419.913086 | 4322.432129 | 4997.770508 | 4546.4375 | 3725.532227 | 4026.11499 |
| 826871.8125 | 522816.9375 | 721746.9375 | 740542.1875 | 935377.6875 | 790062.5 | 897169.6875 |
| 27377.6875 | 20164.83008 | 20714.58008 | 25073.07813 | 25561.80078 | 23724.33594 | 24668.02734 |
| 50522.25977 | 38312.73779 | 41724.43506 | 40644.46875 | 49028.59326 | 49152.79395 | 47589.34326 |

| | | | | | | |
|---|---|---|---|---|---|---|
| 393770.4063 | 279616.5625 | 293291.8438 | 311792 | 209783.3594 | 212412.5781 | 215972.3281 |
| 8596.546875 | 5369.459473 | 5568.019531 | 7648.32959 | 8304 | 7767.271484 | 10588.91504 |
| 350044.2813 | 259736.125 | 277889 | 258630.0781 | 240061.875 | 248978.9688 | 239839.4219 |
| 196226.3906 | 126293.0078 | 151431.3594 | 120012.1484 | 105037.3516 | 141710.0625 | 102879.5156 |
| 12218.4082 | 12596.73242 | 8391.953125 | 8484.064453 | 10884.32031 | 9895.109375 | 10995.81543 |
| 146759.0469 | 118764.25 | 116673.4219 | 118634.6328 | 139235.7031 | 136439.875 | 138381.9531 |
| 175747.1328 | 137488.1455 | 132243.6953 | 133940.5918 | 154742.6465 | 162169.3809 | 164365.3594 |
| 188522.2656 | 295341.4063 | 241230.0781 | 261977.3438 | 722062.125 | 190648.1875 | 250187.9531 |
| 57750.41016 | 55714.44727 | 37768.05469 | 31426.61865 | 66266.80859 | 69444.14063 | 68966.85938 |
| 40324.89844 | 33309.07422 | 33464.34766 | 29128.24609 | 31973.93945 | 33387.84766 | 32483.41992 |
| 1563926.857 | 996311.4189 | 1284327.049 | 1290074.922 | 1176113.553 | 1141554.523 | 1257074.199 |
| 834830.625 | 598109.625 | 695268.9375 | 664250.3125 | 1093029 | 1036266.063 | 1067476.375 |
| 241560.2813 | 256859.0781 | 135689.6094 | 170436.7813 | 308366.7813 | 371732.1875 | 287860.9688 |
| 58765.21094 | 47690.36719 | 42287.29688 | 43193.08594 | 36248.36328 | 38340.6875 | 35530.88672 |
| 51580.55469 | 37519.60156 | 42406.21094 | 43921.55469 | 45820.81641 | 39409.75391 | 42273.63672 |
| 26752.26953 | 20565.64258 | 18454.31641 | 18998.83203 | 15062.3252 | 16637.14844 | 16357.73438 |
| 88804.24219 | 80072.23438 | 73157.17188 | 66652.80469 | 65361.32031 | 84162.74219 | 61010.59375 |
| 239111.041 | 195614.3232 | 164107.6406 | 177201.4424 | 369619.5625 | 344574.6035 | 381248.6348 |
| 16623.12305 | 14954.90137 | 13505.32617 | 8285.662109 | 7530.542969 | 7141.84375 | 7487.681641 |
| 769634.5 | 610842.8125 | 599598.5625 | 578469.1875 | 565727.5 | 621908.125 | 608665 |
| 8993.780273 | 7825.481445 | 6961.707031 | 7920.163086 | 14880.27051 | 13223.61523 | 10989.57129 |
| 35781.52734 | 30502.77148 | 25402.93359 | 25444.71289 | 27526.86719 | 30705.6582 | 27306.125 |
| 35589.38281 | 28828.45703 | 25604.98242 | 21361.08984 | 14121.19141 | 17031.71094 | 15469.9668 |
| 117844.6016 | 93842.45313 | 78220.92969 | 83185.71875 | 88049.84375 | 89491.50781 | 90993.70313 |
| 112680.9219 | 98473.3125 | 85287.28125 | 86247.07813 | 90825.32813 | 85110.00781 | 87117.07813 |
| 2514836.75 | 2018915 | 1699786 | 1537611.5 | 1443296.5 | 1586516.375 | 1800708.875 |
| 129340.5156 | 107580.8984 | 84602.65625 | 87454.375 | 65905.89063 | 69837.625 | 66677.22656 |
| 64346.31641 | 52752.25781 | 47610.35938 | 48390.33984 | 60838.17188 | 53734 | 58629.83984 |
| 62985.14063 | 52856.71484 | 42647.84375 | 40280.21094 | 49952.0625 | 44977.21094 | 42742.57422 |
| 160862.6563 | 116164.6797 | 123210.1641 | 122717.4766 | 119051.8438 | 120511.9609 | 120964.2734 |
| 590057.4375 | 429059.75 | 465471.2188 | 423680.7813 | 330189.7813 | 367103.4688 | 335081.1875 |
| 122756.7188 | 91790.07031 | 90294.3125 | 87447.24219 | 92342.82031 | 94339.08594 | 96835.76563 |
| 113455.1641 | 77329.5625 | 103454.625 | 103027.3828 | 106950.4922 | 99961.30469 | 108509.3516 |

| | | | | | | |
|---|---|---|---|---|---|---|
| 54963.74609 | 43037.17188 | 41031.11719 | 45256.77734 | 45224.12891 | 39699.76172 | 42346.28125 |
| 138838.4375 | 17654.68164 | 153322.9531 | 147763.1094 | 162848.9531 | 182156.3281 | 151629.9063 |
| 57583.23047 | 37372.27734 | 42393.35938 | 43015.65234 | 52449.64453 | 57404.89063 | 62054.75391 |
| 61679.03125 | 37625.6875 | 55468.15625 | 42851.09375 | 33926.40625 | 49376.53516 | 46740.0625 |
| 49815.03906 | 45847.98047 | 37402.37891 | 38191.14453 | 44580.08984 | 43197.68359 | 40329.72656 |
| 150150.25 | 120187.4063 | 121334.5938 | 102459.5 | 167266.9375 | 181322.125 | 180017.1563 |
| 44362.19922 | 30824.33789 | 31453.98438 | 31526.41016 | 30262.20898 | 34311.6875 | 28282.42383 |
| 28205.98242 | 26360.24219 | 21130.35938 | 19522.76563 | 35833.9375 | 43479.32422 | 37323.90625 |
| 6617.345215 | 5243.850586 | 4654.041016 | 4529.423828 | 7073.721191 | 6480.625977 | 7350.17041 |
| 114575.9688 | 80562.57031 | 77813.21094 | 87547.09375 | 104824.2656 | 89133.69531 | 96970.58594 |
| 14670.87988 | 9552.039063 | 12682.68164 | 13107.4043 | 11614.48926 | 10046.75195 | 9534.780273 |
| 11076.43066 | 7234.168945 | 13182.09863 | 13466.41699 | 13811.2002 | 19786.00195 | 19026.81836 |
| 487.7626648 | 337.9053345 | 404.0436401 | 347.7467957 | NA | NA | NA |
| 6564.494629 | 4284.601074 | 4348.230957 | 5448.709961 | 2892.789063 | 2694.123047 | 3246.356689 |
| 103580.7422 | 81869.53906 | 80052.61719 | 77005.20313 | 86930.57031 | 89333.85156 | 92643.91406 |
| 35433.16406 | 22662.5625 | 26775.58789 | 26440.14844 | 33673.66016 | 28128.94922 | 36893.03125 |
| 67670.21094 | 46348.68359 | 54975.57813 | 49943.39063 | 54234.53516 | 44244.09375 | 49672.20313 |
| 136282 | 99315.70313 | 104099.1563 | 93858.75781 | 110833.8281 | 117232.0313 | 128168.6406 |
| 21240.41992 | 13628.03809 | 14614.7168 | 16343.52051 | 11172.21484 | 11089.29004 | 11859.01563 |
| 517887.6875 | 403693.8125 | 366630.3125 | 363134.2188 | 394198.1875 | 424696.5 | 331743.125 |
| 46091.79297 | 35884.5625 | 38883.38281 | 41419.33984 | 48554.57422 | 48272.51563 | 45700.69531 |
| 225891.9063 | 164407.2344 | 159065.6719 | 161579.4531 | 191225.2813 | 180849.2656 | 186664.0156 |
| 428232.5625 | 355567.1563 | 311502.8438 | 289779.75 | 307669.9688 | 343512 | 324281.6563 |
| 220156.7813 | 148545.8125 | 161511.3438 | 161050.8438 | 146864.9688 | 152705.6406 | 156315.1406 |
| 1230.445557 | 22059.49609 | 5734.933594 | 33108.38672 | 35667.42969 | 51376.99609 | 111645.7578 |
| 191878.6641 | 152145.75 | 140175.084 | 132949.0547 | 135572.4609 | 128539.0293 | 129946.5703 |
| 94371.98438 | 65949.17188 | 71999.64063 | 76234.8125 | 71837.17188 | 71620.27344 | 77996.61719 |
| 131381.6875 | 75637.41406 | 112154.2188 | 69095.89063 | 68790.46875 | 106788.8359 | 68888.71094 |
| 66008.83594 | 54722.9375 | 48090.14844 | 52118.13672 | 70408.85938 | 69137.20313 | 62121.23047 |
| 170504.8438 | 117428.8672 | 127025.125 | 117283.6328 | 90081.23438 | 92606.82031 | 95455.64063 |
| 48131148 | 36763844 | 35228656 | 33729236 | 32268168 | 32958884 | 34426236 |
| 14301.57422 | 12291.7793 | 8705.400391 | 9194.851563 | 7884.307129 | 9710.139648 | 7974.942871 |
| 52646.49219 | 35201.03516 | 41898.61719 | 36998.1875 | 35751.32031 | 37819.65625 | 37698.36719 |

| | | | | | | |
|---|---|---|---|---|---|---|
| 187205.9844 | 150704.7656 | 144397.1719 | 141868.9375 | 112627.1953 | 118822.9844 | 117504.3281 |
| 154585.8281 | 125921.6484 | 106316.4141 | 98998.24219 | 116234.7578 | 115529.4844 | 120738.9844 |
| 298137.3125 | 223786.9375 | 218219.8594 | 219165.5625 | 196709.2969 | 200816.7031 | 199220.8438 |
| 23619.60742 | 17567.2832 | 16024.09473 | 15004.68555 | 16165.26465 | 17284.54883 | 15422.30762 |
| 37520.80078 | 21647.4375 | 33581.30859 | 35871.55078 | 46482 | 49310.74219 | 60237.80469 |
| 266390.8438 | 155910.9531 | 197825.5469 | 127287.0469 | 158352.4063 | 311993 | 129582.5156 |
| 132475.8125 | 82768.75781 | 103636.3125 | 100237.5156 | 106484.0859 | 113148.8438 | 82493.76563 |
| 151234.6094 | 94168.32031 | 117610.9922 | 96281.14063 | 70601.48438 | 97503.82031 | 79242.95313 |
| 4752.162598 | 3545.945801 | 8375.21582 | 4912.216309 | 9533.643555 | 7701.170898 | 17332.74805 |
| 135873.0625 | 116956.9063 | 96990.95313 | 97627.67188 | 144881.8594 | 139776.2188 | 155323.8594 |
| 29296.17578 | 20979.99609 | 22904.22461 | 22019.91016 | 24682.08203 | 23903.28711 | 23019.17578 |
| 202624.6875 | 145704.9219 | 127448.6875 | 141567.6719 | 140839.7813 | 127675.8281 | 142516.2656 |
| 6089.275879 | 7076.594727 | 2586.116699 | 1606.583862 | 6416.256348 | 6781.46582 | 5092.290527 |
| 31610.00977 | 26696.13672 | 24162.42188 | 21080.71289 | 24538.46484 | 26775.31641 | 25261.71094 |
| 10050.05957 | 1756.389282 | 10851.12305 | 8513.478516 | 15516.79102 | 11581.12207 | 13679.30664 |
| 54376.75781 | 38688.47656 | 38350.52344 | 38200.42969 | 41664.74609 | 39139.49609 | 42975.55469 |
| 9634.489258 | 7989.046387 | 7686.972168 | 7287.486816 | 10904.10547 | 11054.29785 | 11143.83203 |
| 52987.42578 | 43268.35547 | 41417.14844 | 45457.79688 | 53546.36719 | 64369.3125 | 66956.41406 |
| 32153.8125 | 24288.36914 | 27578.60352 | 23073.44922 | 53023.93359 | 24017.27734 | 35514.15625 |
| 214684.1563 | 168718.9375 | 178512.5 | 187066.5469 | 224145.2969 | 197541.1406 | 210599.2344 |
| 27266.92578 | 22527.94336 | 19293.40039 | 17186.23242 | 24786.11133 | 18459.80469 | 25031.31055 |
| 15282.46387 | 15695.73926 | 9701.5625 | 10470.38379 | 11691.56934 | 14651.35059 | 11060.30469 |
| 134433.7656 | 101567.1953 | 100143.8828 | 107252.5781 | 129503.3359 | 133730.0313 | 118964.8906 |
| 228350.7188 | 163198.0156 | 180804.9219 | 166838.7656 | 205721.3438 | 208390.1875 | 220892.0938 |
| 21889.10352 | 18379.90625 | 15427.0752 | 15854.43945 | 15730.00879 | 16059.29395 | 12509.44336 |
| 241939.125 | 169072.0156 | 179086.4844 | 161154.7031 | 143802.3125 | 156931 | 144326.75 |
| 15900.73828 | 10174.30762 | 12005.02148 | 11030.63184 | 8039.100586 | 8532.773438 | 8152.504883 |
| 45151.74609 | 31203.92773 | 35136.03906 | 36187.37109 | 52484.05469 | 46240.66406 | 55138.875 |
| 66083.44531 | 47131.62109 | 52806.20703 | 56899.43359 | 61250.25781 | 59202.62891 | 63980.42969 |
| 1684802.375 | 1480383 | 1274799.25 | 1095176.875 | 586007.8125 | 651276.4375 | 714873.125 |
| 53224.40234 | 39842.80859 | 40284.37891 | 41887.32031 | 55329.58203 | 66921.14844 | 53017.91016 |
| 30631.99805 | 22995.75977 | 22285.99414 | 21881.72266 | 24148.66797 | 23597.10547 | 23540.26563 |
| 19439.92578 | 15585.59473 | 16296.96191 | 12558.27441 | 13640.4209 | 15208.48633 | 13883.91016 |

| | | | | | | |
|---|---|---|---|---|---|---|
| 189754.9063 | 137909.4063 | 146947 | 137897.0781 | 129496.2656 | 137004.3281 | 135603.9531 |
| 5217.646484 | 4424.21875 | 4908.090332 | 4442.612305 | 5583.403809 | 4879.524902 | 5942.849609 |
| 280979.8984 | 259254.9453 | 169094.0723 | 179946.2148 | 278556.3281 | 234465.5664 | 256996.2266 |
| 2146.868896 | 1768.589233 | 1755.740601 | 1813.144043 | 12222.68945 | 12504.08887 | 11414.46289 |
| 17075.42188 | 10454.84863 | 13835.9834 | 16357.50781 | 21254.27148 | 17323.80859 | 18247.50977 |
| 53971.14844 | 32653.22266 | 56669.10938 | 67382.78906 | 92030.94531 | 103574.1406 | 137268.7344 |
| 180013.5 | 132850.4531 | 128345.6016 | 133907.1406 | 154789.5 | 142930.6719 | 140507.8438 |
| 190338.6406 | 128277.6641 | 139392.2031 | 134473.1094 | 117010.1172 | 118386.0391 | 113575.7031 |
| 77818.84375 | 64792.92578 | 56636.63281 | 53085.99219 | 63922.24609 | 55524.41406 | 61627.66797 |
| 149996.2031 | 128620.7344 | 107746.7109 | 104968.4844 | 107113.5078 | 96405.94531 | 101845.1719 |
| 46071.69531 | 40582.08203 | 32034.62109 | 31887.41992 | 46217.59375 | 39881.99219 | 37298.05078 |
| 121891.4609 | 93074.78125 | 101084.3906 | 89620.46875 | 100512.8828 | 99789.90625 | 104541.4219 |
| 161335.4219 | 121346.6797 | 139842.4063 | 129588.6875 | 101113.625 | 114805.4219 | 113963.7813 |
| 33829.64063 | 22267.07227 | 23119.0293 | 24910.43359 | 36618.85547 | 36625.18359 | 37736.34766 |
| 68604.97656 | 48887.96484 | 48799.15234 | 52673.01563 | 68115.78906 | 64008.60156 | 68975.28125 |
| 237704.5625 | 165405.0625 | 152714.0156 | 165425.0469 | 186013.4688 | 179558.8438 | 181213.3125 |
| 68642.97656 | 45938.96094 | 53843.69141 | 48316.20703 | 41607.96484 | 48202.57422 | 44110.42578 |
| 2166.97876 | NA | 2343.276123 | 918.532959 | 1455.000366 | NA | 1428.667603 |
| 12489.69043 | 10806.58984 | 10350.76563 | 3390.911133 | 15642.33398 | 13641.89648 | 11059.08496 |
| 105840.875 | 77177.07031 | 86476 | 82117.23438 | 74441.34375 | 72638.45313 | 80373.47656 |
| 185045.0781 | 132255.9219 | 137377.7188 | 131519.8594 | 133302.0938 | 147070.7344 | 145563.0938 |
| 69097.78125 | 50862.37891 | 45154.84766 | 38655.78516 | 26595.16406 | 25735.15234 | 25486.29492 |
| 517664.4063 | 383533.3438 | 387330 | 363865.0313 | 394554.7188 | 422513.5313 | 382041.6563 |
| 36252.02344 | 29030.79297 | 28774.53125 | 27027.80664 | 36300.21484 | 39219.79688 | 37526.49219 |
| 25749.7207 | 14006.4873 | 15756.09277 | 20036.13672 | 23009.00781 | 22034.32813 | 25782.11523 |
| 14538.68945 | 10815.37012 | 9379.597656 | 7981.552734 | 8721.510742 | 8868.572266 | 8293.990234 |
| 19114.54688 | 14276.11523 | 13978.10449 | 13255.35156 | 12284.39648 | 14817.91797 | 13379.44043 |
| 5125.466309 | 4186.272949 | 3743.565186 | 4374.359863 | 4687.621094 | 3842.334473 | 4080.201172 |
| 27710.98047 | 21907.79883 | 22888.18359 | 19669.15234 | 24125.67969 | 20987.94922 | 20838.45508 |
| 106812.8203 | 79946.6875 | 72608.10156 | 78367.46875 | 83192.17969 | 79902.25 | 73645.69531 |
| 46795.73584 | 33288.88501 | 34144.26392 | 29453.56396 | 35168.18164 | 36620.44629 | 37463.53247 |
| 21948558 | 16624383 | 15675626 | 15336113 | 17361630 | 15990153 | 16243003 |
| 9660.573242 | 8204.52832 | 7316.104004 | 7925.972168 | 8503.313477 | 9000.735352 | 7010.669922 |

| | | | | | | |
|---|---|---|---|---|---|---|
| 75706.49219 | 45106.23828 | 58162.44531 | 60710.27734 | 54142.46094 | 57424.13672 | 58276.12109 |
| 24809.32617 | 20907.73633 | 18250.95508 | 17371.58789 | 23273.16992 | 21591.45117 | 23600.31445 |
| 5954463 | 7090127.5 | 5012707.5 | 1394348.625 | 1430798.875 | 6732842 | 1235098.25 |
| 477354.4688 | 348508.875 | 364917.4063 | 365590.0313 | 386841.9375 | 362881.4688 | 384208.875 |
| 24265.74219 | 14237.79492 | 16459.21484 | 15854.37402 | 20234.64844 | 8037.405273 | 7866.625488 |
| 383557.5 | 283618.6875 | 277356.875 | 274982.9063 | 269539.4375 | 265132.5625 | 268120.375 |
| 114378.9688 | 86989.42969 | 87886.39844 | 87631.42969 | 87912.89063 | 93925.32813 | 95741.79688 |
| 271982.7813 | 148560.6406 | 223302.2656 | 207877.4375 | 263737.3125 | 247251.4531 | 265409.2188 |
| 63785.87891 | 51869.02734 | 47454.13672 | 50124.10547 | 56910.64453 | 46071.14453 | 52398.50781 |
| 12454.63965 | 8839.522461 | 8925.211914 | 9279.542969 | 11103.49902 | 10199.32031 | 10153.72754 |
| 55937.28125 | 47165.19531 | 37787.57422 | 40252.24219 | 70506.4375 | 72596.91406 | 72963.17969 |
| 12135.27637 | 7720.258789 | 7386.289063 | 8267.033203 | 8377.226563 | 9710.858398 | 9290.291016 |
| 17957.29883 | 13571.55859 | 14039.63086 | 16617.58008 | 18819.68945 | 19045.82422 | 15890.10645 |
| 22281.01953 | 17400.76563 | 15243.82031 | 16222.61719 | 17217.4043 | 18415.48828 | 15849.87402 |
| 187581.4063 | 125042.8672 | 151715.7656 | 135503.4375 | 98487.98438 | 109046.2969 | 107412.9375 |
| 103312.1016 | 68728.14844 | 85634.03125 | 77571.01563 | 67399.51563 | 60017.25391 | 66647.28906 |
| 82789.48438 | 63591.74609 | 65901.4375 | 64397.67188 | 79320.85156 | 65132.19531 | 72680.84375 |
| 8477.195313 | 5149.299316 | 7651.016113 | 6900.020996 | 10130.41504 | 5119.559082 | 9362.003906 |
| 30522.92188 | 22138.67188 | 21858.83984 | 22245.16602 | 38602.60156 | 45101.54297 | 45521.84375 |
| 20186.19336 | 13320.55273 | 15378.43164 | 15322.62305 | 17121.71289 | 17660.89258 | 16805.65625 |
| 11517.94922 | 10542.05371 | 13320.88281 | 9650.09082 | 10520.18262 | 12473.09375 | 8881.623047 |
| 4642.955566 | 4419.922363 | 3564.137451 | 2478.623535 | 3220.155518 | 3920.492188 | 2929.17334 |
| 695091.9375 | 412336.3438 | 510204.0938 | 385536.625 | 498132.3438 | 763197.75 | 442938.875 |
| 6634.73584 | 4284.617676 | 10251.78809 | 11021.96289 | 9838.573242 | 1726.425171 | 12389.63281 |
| 108688.9297 | 80793.25781 | 85744.07813 | 80236.13281 | 87323 | 86363.58594 | 85935.59375 |
| 8624.663086 | 4919.136719 | 6174.32373 | 6866.630859 | 5284.398926 | 7187.945313 | 7421.245605 |
| 297687.5938 | 177090.1563 | 193657.7188 | 175700.8281 | 98911.44531 | 98982.73438 | 92681.53906 |
| 329513.4688 | 186306.5469 | 267343 | 281863.3125 | 472529.875 | 419359.7813 | 461458 |
| 24921.25391 | 15210.34277 | 16140.34668 | 16432.83594 | 26585.85547 | 23230.74219 | 24266.42969 |
| 9163.899414 | 7881.83252 | 6247.343262 | 6228.650879 | 5593.214355 | 5574.472656 | 6382.550293 |
| 3598.962158 | 2617.486816 | 2339.164063 | 3006.10083 | 4043.453125 | 5404.857422 | 3150.369141 |
| 259532.6719 | 198903.4531 | 188709.3281 | 205029.9688 | 232842.3438 | 217546.625 | 219403 |
| 173280.1406 | 143725.9844 | 140708.2656 | 176798.4375 | 258393.2188 | 259966.7344 | 303983.25 |

| | | | | | | |
|---|---|---|---|---|---|---|
| 11811.41699 | 10550.41211 | 10350.05664 | 11320.63477 | 29537.74219 | 28050.79102 | 26476.97656 |
| 13232.63965 | 9337.882813 | 8553.929688 | 9844.811523 | 12367.23926 | 13622.77344 | 14683.72168 |
| 30100.37891 | 20587.98242 | 18422.66211 | 19148.39648 | 18965.05273 | 19269.54102 | 17851.33398 |
| 9895.495117 | 7411.713379 | 6897.713867 | 7743.038574 | 9612.208008 | 9737.498047 | 9646.514648 |
| 21458.29688 | 17885.26563 | 14272.76855 | 15354.82715 | 20279.86133 | 18448.66016 | 18836.375 |
| 16645.71289 | 12435.00195 | 12832.55078 | 11271.10645 | 17006.22852 | 16448.31641 | 16733.0293 |
| 2237158.25 | 1518736.125 | 1709905.375 | 1734290.875 | 1836522.125 | 2055065.25 | 1973778.125 |
| 77267.65625 | 43549.19922 | 64816.03906 | 40397.48047 | 37640.51563 | 70736.10156 | 38685.85156 |
| 41805.03516 | 29185.09766 | 27985.73828 | 28252.91211 | 26761.68945 | 29699.52148 | 27484.08008 |
| 32264.22266 | 22632.1543 | 22737.77344 | 19409.54492 | 21380.37305 | 23456.56055 | 24420.20313 |
| 5808.20166 | 3606.366455 | 4015.657715 | 3279.508301 | 11146.25879 | 6972.189941 | 9180.068359 |
| 14447.05273 | 11832.5752 | 9838.131836 | 10869.95313 | 9268.104492 | 8326.166992 | 8256.924805 |
| 45304.97266 | 40419.57422 | 37566.42188 | 39320.75 | 72539.96094 | 75171.61719 | 65638.10938 |
| 109082.0938 | 96912 | 70110.55469 | 71947.91406 | 93506.6875 | 103363.9766 | 115338.3984 |
| 230502 | 179541.7969 | 177510.1563 | 180001.375 | 177617.0781 | 231174.75 | 197396.7344 |
| 131210.9688 | 100014.7344 | 95392.30469 | 108147.1172 | 146675.9688 | 151288.875 | 142500.7344 |
| 74511.875 | 166070.8438 | 69311.9375 | 16501.90039 | 20503.33203 | 70111.02344 | 17784.43164 |
| 24254.31055 | 16520.86523 | 19747.33984 | 17022.71875 | 16461.30664 | 17951.48242 | 16735.87305 |
| 245533.2446 | 195319.2698 | 175252.3857 | 168927.0508 | 171913.1313 | 179439.2568 | 185090.04 |
| 7247770 | 5205628 | 5400207.5 | 5604825 | 6853054.5 | 6581272 | 6472593.5 |
| 172916.25 | 94517.73438 | 122085.6406 | 144444.3281 | 201566.6719 | 176858.6563 | 199478.8125 |
| 13246.8877 | 8686.510742 | 9130.847656 | 9820.179688 | 7229.105469 | 7884.739746 | 3963.619385 |
| 219714.9844 | 147407.9375 | 157266.0938 | 156165.3125 | 122951.8516 | 126313.2109 | 125277.3516 |
| 83627.39063 | 61624.10938 | 59466.9375 | 59507.5 | 70326.00781 | 60897.32422 | 67194.25781 |
| 298142.0625 | 190589.2969 | 239787.1563 | 242819.9219 | 228962.1563 | 235180.3125 | 231440.5156 |
| 1163847.75 | 786446.0625 | 856493.3125 | 871435.9375 | 857919.9375 | 915446.3125 | 893837.125 |
| 156580.4844 | 119128.5625 | 117072.7734 | 113476.5703 | 122381.7813 | 120621.6719 | 115232.2734 |
| 143714.5 | 96468.49219 | 104420.2578 | 102840.3359 | 117674.2656 | 116080.125 | 128711.9297 |
| 32507.47266 | 24375.31445 | 20962.50977 | 23538.78711 | 28873.58594 | 26652.90625 | 26278 |
| 117969.4297 | 88191.67188 | 99558.42969 | 89322.49219 | 92619.96094 | 104502.6719 | 99946.21094 |
| 44733.38672 | 33846.63672 | 29617.88477 | 32281.16211 | 40799.17969 | 42449.55469 | 40955.02734 |
| 11636.52148 | 12236.7002 | 9637.31543 | 7701.743164 | 10847.18262 | NA | 10616.10059 |
| 9172.291992 | 7473.285645 | 5338.102539 | 4885.841797 | 13591.17871 | 9079.427734 | 10848.67383 |

| | | | | | | |
|---|---|---|---|---|---|---|
| 14944.41602 | 12254.59766 | 10933.14355 | 11752.64258 | 12826.87988 | 13086.79785 | 15072.4082 |
| 21433.57422 | 15842.11816 | 15414.66699 | 14442.07129 | 10246.17773 | 9403.548828 | 9527.146484 |
| 117756.6797 | 81825.24219 | 98040.21875 | 101059.6016 | 107348.0391 | 125497.4063 | 111676.7266 |
| 8176.170898 | 5226.037109 | 5265.584961 | 5922.600098 | 10423.6875 | 6775.749023 | 6956.517578 |
| 69875.05762 | 49433.37354 | 54771.43457 | 48590.97314 | 49668.4209 | 54249.33887 | 46270.96094 |
| 37242.15234 | 27405.58984 | 27077.57227 | 34465.80469 | 38691.14063 | 29494.13672 | 39143.35547 |
| 14346.79004 | 10369.51172 | 10856.31055 | 6296.353027 | 6331.095215 | 12473.20996 | 8171.324219 |
| 173540.0781 | 125750.8047 | 129726.2422 | 127100.9063 | 157261.6563 | 145826.5781 | 160036.0156 |
| 81533.84375 | 58031.07031 | 63424.92969 | 64237.07031 | 67291.47656 | 66091.14844 | 63274.19141 |
| 378277.0313 | 277438.4063 | 315749.4063 | 315150.7813 | 434673.1563 | 395995.5313 | 424097.6563 |
| 52742.67578 | 32363.45313 | 41960.92188 | 40435.93359 | 45843.17969 | 45635.98828 | 48941.49609 |
| 8933.520508 | 7342.285645 | 7150.052734 | 6011.769043 | 4437.387695 | 6823.541504 | 4436.849609 |
| 1665919.75 | 1092033.875 | 1195927.375 | 1132830.875 | 1092475 | 1196183.25 | 1169007.875 |
| 81621 | 62030.65625 | 57990.85547 | 63312.32031 | 89569.71094 | 80999.9375 | 86490.14063 |
| 2082381.25 | 1406848 | 1410294.75 | 1502921.375 | 1833780.375 | 1881144.25 | 1937996.125 |
| 300522.1563 | 216734.625 | 218551.8281 | 222395.7188 | 198235.7031 | 188845.0938 | 201163.7969 |
| 82130.02344 | 51844.75781 | 56667.66797 | 57513.53125 | 64112.80859 | 71985.82813 | 62437.34766 |
| 11272.31445 | 8922.102539 | 6655.599121 | 9314.308594 | 9416.415039 | 9687.969727 | 10109.58301 |
| 87709.50781 | 74672.08594 | 66723.09375 | 63937.58984 | 48244.51172 | 46856.16797 | 48501.87109 |
| 186704.2188 | 141267.6719 | 143085.125 | 142681.8438 | 148441.8594 | 139363.2344 | 148744.0469 |
| 24906.5332 | 18192.125 | 16972.32227 | 17359.11914 | 17779.6582 | 16922.31641 | 18354.26563 |
| 83633.66406 | 60470.48828 | 56502.85938 | 58281.96484 | 59261.30469 | 61069.71484 | 59333.19922 |
| 121583.1953 | 92343.82031 | 93417.6875 | 85095.96875 | 95266.73438 | 84995.28125 | 94813.55469 |
| 99868.35156 | 65108.51172 | 75314.07813 | 71489.70313 | 58073.60938 | 62278.91797 | 64463.44531 |
| 73268.71875 | 47212.19141 | 58432.10547 | 59926.61328 | 48309.15625 | 53960.94531 | 48182.40625 |
| 24862.29492 | 20022.55078 | 17850.75195 | 19739.6875 | 42027.83203 | 36931.07813 | 35909.53516 |
| 8475.604492 | 9495.944336 | 3727.586914 | 3730.311768 | 11056.15625 | 19005.64453 | 14177.62891 |
| 60435.77734 | 51432.70703 | 36501.07813 | 31113.01953 | 22417.75391 | 22092.19531 | 21113.33008 |
| 9595.25293 | 7124.753906 | 6798.39502 | 6950.03125 | 6626.501465 | 6896.981445 | 7416.655762 |
| 7212660 | 5122741.5 | 5103566 | 5726147 | 5773036 | 5371308.5 | 5862461.5 |
| 15451.7959 | 18127.5918 | 12647.42285 | 9457.585938 | 20674.58203 | 23217.27539 | 18282.92969 |
| 144447.7031 | 114847.0938 | 96973.67969 | 115812.7188 | 126892.8984 | 115505.7031 | 119958.9922 |
| 8806.273438 | 6766.286621 | 6696.202148 | 4245.755859 | 3341.220947 | 1439.079346 | 2394.540771 |

| | | | | | | |
|---|---|---|---|---|---|---|
| 27201.5332 | 20577.32617 | 18859.76758 | 20771.46289 | 20688.95508 | 19277.07617 | 22324.88086 |
| 20780.63477 | 13739.5625 | 16733.20313 | 11325.46777 | 8627.743164 | 11956.94727 | 12220.05664 |
| 50493.41016 | 31354.25586 | 45792.72266 | 30943.51172 | 42139.91797 | 97387.28125 | 42730.89844 |
| 87563.85938 | 56917.96875 | 67638.57813 | 65243.04688 | 66110.78906 | 65553.04688 | 68410.84375 |
| 2892.712891 | 2418.079346 | 2015.291382 | 1491.96814 | 2140.735596 | 1491.537842 | 2291.744629 |
| 273195.625 | 231504.7344 | 205874.0156 | 240670.2969 | 458431.875 | 384134.5313 | 372332.625 |
| 53844.93359 | 38687.125 | 38138.95703 | 39733.95703 | 61982.15625 | 55855.03516 | 56392.03125 |
| 18581.8418 | 21667.49609 | 9686.50293 | 9044.379883 | 30998.86914 | 33506.77344 | 31371.0918 |
| 577598.625 | 382909.9063 | 410589.3125 | 346703.7813 | 339608.75 | 408871.25 | 347708.75 |
| 193662.3281 | 133765.5469 | 146358.7188 | 176518.4375 | 344721.9375 | 223773.9219 | 264437.0625 |
| 131462.1719 | 98778.70313 | 96224.30469 | 96950.10156 | 110800.9219 | 110156.5859 | 109657.7578 |
| 61413.84375 | 53278.38281 | 43941.08203 | 45774.25391 | 47602.05078 | 49757.87109 | 47007.87109 |
| 2233123.25 | 2128190.25 | 1512697.5 | 1557280.25 | 1446682.875 | 1341159.75 | 1127706.375 |
| 876295.1875 | 623626.9375 | 614617.25 | 600052.5625 | 565909.3125 | 505253.6563 | 529000.6875 |
| 54837.83594 | 36919.29297 | 44836.84766 | 38418.11719 | 41577.11719 | 42945.34766 | 46935.92969 |
| 103978.3516 | 75210.125 | 81773.82813 | 85235.20313 | 117410.625 | 127507.3125 | 124122.6172 |
| 18855.67773 | 11756.27539 | 17797.62305 | 12677.75293 | 12518.59766 | 12914.62891 | 7798.552734 |
| 16136.1582 | 12334.76172 | 11755.52539 | 12731.11035 | 21340.85156 | 15917.6377 | 17880.03125 |
| 10291.5791 | 11676.09473 | 6773.223633 | 5834.65918 | 5852.010742 | 4841.26123 | 5970.299805 |
| 8150.510742 | 7341.646973 | 5247.762207 | 5371.136719 | 3406.924072 | 5067.80957 | 4132.868164 |
| 12133.11035 | 12081.16504 | 6973.509277 | 9109.288086 | 12597.91504 | 11700.35059 | 11364.83496 |
| 27817.65234 | 20849.16797 | 22925.72461 | 19503.67773 | 18537.625 | 18408.18359 | 19493.18945 |
| 22403.75195 | 18793.26563 | 12867.93359 | 12584.12988 | 22149.10156 | 15499.41602 | 18794.98438 |
| 104563.9531 | 75990.60938 | 65061.85938 | 74321.28125 | 67388.39063 | 75766.32813 | 62103.34766 |
| 168448.5156 | 129726.6875 | 115242.1484 | 121124.2578 | 169283.125 | 147530.0469 | 160343.6563 |
| 17021.5 | 13310.15039 | 15594.46582 | 16332.67285 | 17905.2168 | 16194.43164 | 12731.53613 |
| 55124.05859 | 41246.41797 | 38699.64453 | 38783.08984 | 39710.32813 | 37258 | 37655.80469 |
| 79864.42969 | 65256.84766 | 54145.07422 | 53494.95313 | 55249.36328 | 51149.74609 | 49069.41016 |
| 508327.1875 | 358600.125 | 362899.375 | 272178.1875 | 266672.4063 | 441569.2813 | 317633.4688 |
| 96915.22803 | 113524.4009 | 61274.0354 | 82343.61035 | 601822.0859 | 201204.3311 | 316797.5156 |
| 866696.4375 | 695958.4375 | 562625.8125 | 623468.5 | 928243.625 | 899928.9375 | 886655.3125 |
| 211625.1094 | 141002.9531 | 148782.7344 | 119449.9766 | 133907.2656 | 162688.2969 | 135263.1563 |
| 654117.0625 | 534941.25 | 435906.3438 | 339423.2813 | 483987.8125 | 532984.6875 | 463686.8125 |

| | | | | | | |
|---|---|---|---|---|---|---|
| 64631.42969 | 45263.16406 | 42327.125 | 48845.11719 | 41077.56641 | 39624.23438 | 39246.13672 |
| 29423.15234 | 15337.6748 | 19499.85547 | 23113.09766 | 18554.39453 | 17545.37305 | 16881.51367 |
| 38898.36328 | 31171.68164 | 29185.20703 | 28817.1875 | 30700.23633 | 29601.2207 | 31634.47656 |
| 211068.4063 | 156678.4219 | 148124.2344 | 138422.7344 | 153564.4688 | 180498.5938 | 159619 |
| 435970.1563 | 322009.0625 | 309227.4063 | 291975.7813 | 290133 | 299604.8125 | 297508.3125 |
| 383131.4063 | 270625.9063 | 277135.5313 | 246781.5469 | 219087.4844 | 231861.5938 | 207271.5938 |
| 83460.71875 | 59295.83594 | 58993.8125 | 55729.22656 | 48496.49219 | 48914.60547 | 53264.63281 |
| 105519.0859 | 79856.82031 | 86011.69531 | 101892.5469 | 118924.5 | 104790.7656 | 109199.4063 |
| 26853.63477 | 20422.85742 | 15313.80859 | 14394.24316 | 11500.94531 | 12266.4502 | 10979.98438 |
| 13928.76172 | 7032.287598 | 8589.625 | 10302.37793 | 8345.254883 | 7594.783691 | 10547.69531 |
| 7100.087891 | 6830.554199 | 6118.750488 | 3656.986084 | 4793.666992 | 5619.992188 | 4219.143555 |
| 128340.9297 | 82422.92188 | 92919.14844 | 95864.65625 | 119606.8281 | 127250.6328 | 124126.7109 |
| 168280.5313 | 96890.57813 | 124239.3359 | 124440.0625 | 134803.0781 | 133415.0313 | 141326.3438 |
| 949080.75 | 627700 | 698516.0625 | 765089 | 578062.125 | 609361.8125 | 636531.0625 |
| 71838.92969 | 54426.74219 | 53012.59375 | 55611.79688 | 59161.76172 | 56467.5 | 55713.80469 |
| 137142.3594 | 100377.0625 | 107527.1406 | 97555.14063 | 109214.6016 | 109712.3047 | 109751.2031 |
| 12703.4375 | 8244.360352 | 7502.636719 | 6824.443848 | 14039.21973 | 12622.03516 | 17606.5918 |
| 92465.03125 | 64399.03516 | 70866.82813 | 69707.38281 | 43031.19531 | 44796.87109 | 40128.44141 |
| 18846.04102 | 13674.99414 | 14082.7959 | 13459.90039 | 18203.74609 | 14373.17578 | 15575.63867 |
| 215048.9531 | 147206.5938 | 159667.0781 | 158648.7813 | 151560.9688 | 149765.5938 | 153042.9219 |
| 42622.91016 | 34679.86719 | 31765.3457 | 31186.43164 | 65836.46094 | 59587.05078 | 60944.73828 |
| 9237.584961 | 5945.930176 | NA | 6699.339844 | 5705.041992 | NA | 7810.803711 |
| 17785.99414 | 14695.60059 | 12515.58398 | 12571.75977 | 19152.14648 | 23294.75977 | 23369.36328 |
| 15789.44141 | 15099.31934 | 10736.4043 | 8678.581055 | 14635.72949 | 13382.4668 | 15150.90332 |
| 43148.36719 | 37133.52344 | 27967.94922 | 26197.67578 | 54968.80078 | 50803.30469 | 46263.4375 |
| 37168.64453 | 29220.96289 | 26943.05664 | 28697.07422 | 35873.09766 | 31864.66602 | 33959.17969 |
| 112732.4141 | 74298.60938 | 74837.32813 | 73803.5 | 57223.23438 | 61034.05469 | 62616.54297 |
| 139988.5 | 101779.1953 | 99246.26563 | 95989.64063 | 97018.41406 | 98911.64844 | 95920.92188 |
| 575.9699707 | 336.5834351 | 235.6847839 | NA | 331.4283447 | 418.3752441 | 696.2949829 |
| 158219.6094 | 130246.8047 | 105524.5625 | 107882.7422 | 88876.74219 | 87189.27344 | 81625.05469 |
| 46666.13672 | 35451.12109 | 30303.52148 | 32562.81055 | 33159.07422 | 30675.92578 | 30387.61523 |
| 60943.12109 | 43724.45703 | 40155.00781 | 39148.86328 | 38299.69531 | 39047.16016 | 37481.72656 |
| 21897.60547 | 11405.36914 | 18183.30273 | 19283.96484 | 20055.63672 | 22132.05078 | 25682.01953 |

| | | | | | | |
|---|---|---|---|---|---|---|
| 149999.5156 | 116219.8906 | 112526.8281 | 100782.1875 | 120329.8438 | 123581.9141 | 119173.1953 |
| 199179 | 147286.8125 | 137912.375 | 144537.75 | 120997.1406 | 125068.4453 | 122233.5625 |
| 395139.1875 | 286580.0313 | 269619.4063 | 271317.1563 | 286908.9063 | 259406.8594 | 270252.4063 |
| 1125536.875 | 745082 | 945019.1875 | 827266.8125 | 1256178 | 1367019 | 1303181.25 |
| 179294.1094 | 126050.1875 | 134759.8594 | 137263.4375 | 163449.2188 | 179310.8281 | 154624.125 |
| 19114.58398 | 11295.43359 | 12395.39063 | 10948.05957 | 21033.72461 | 17852 | 21315.54688 |
| 44789.32422 | 31364.53906 | 37667.76563 | 14770.14844 | 18575.19531 | 35493.58203 | 28855.60156 |
| 18758.07422 | 16438.13281 | 12354.70313 | 13423.13477 | 16231.10254 | 14885.89746 | 11331.58887 |
| 73217.41406 | 49999.96484 | 53408.83594 | 53880.04297 | 68605.50781 | 67683.23438 | 64183.12891 |
| 14911.00098 | 12038.14453 | 10565.63867 | 10683.2334 | 9395.116211 | 10581.74121 | 11665.08496 |
| 102663.9609 | 68604.60156 | 67859.38281 | 81645.00781 | 111930.4688 | 105156.5078 | 101973.3281 |
| 47307.4375 | 43927.03516 | 39094.32813 | 35865.90234 | 48309.69531 | 63299.10547 | 45756.8125 |
| 3697215.25 | 2413471.75 | 2511299.5 | 2484980.25 | 2328819.5 | 2755230.5 | 2614430 |
| 213347.9375 | 174965.125 | 133383.9688 | 132379.75 | 138724.5 | 126700.7344 | 141528.4531 |
| 2622.338379 | 2188.673096 | 3845.119385 | 1199.13855 | 4705.92627 | 11689.9248 | 10911.81641 |
| 51136.15625 | 33074.97656 | 35423.52734 | 36120.22266 | 34390.76172 | 33856.6875 | 34361.09375 |
| 22816.81055 | 10539.0918 | 18761.75977 | 20849.31641 | 43543.15234 | 33566.69531 | 41856.36328 |
| 485240 | 299718.4688 | 365076.25 | 365729.5 | 355847.125 | 388299.375 | 381816.125 |
| 3050.251709 | 2243.320801 | 2011.364868 | 1707.567871 | 1924.819702 | 1686.468384 | 1640.433716 |
| 154722.9531 | 113967.2344 | 106484.9375 | 108819.9063 | 127406.4375 | 125968.3281 | 120843.0156 |
| 160667.7969 | 122127.5938 | 111102.5078 | 123202.5547 | 157008.5156 | 145535 | 143829.3906 |
| 3734302 | 2553271 | 2320254.75 | 2406928.5 | 2624792.25 | 2576263.25 | 2564071 |
| 281268.4063 | 202468.3125 | 206356.4844 | 221867.0469 | 279690.0938 | 283419.0938 | 275667.8438 |
| 58918.33203 | 49747.40625 | 42319.73047 | 36665.27734 | 40706.98828 | 35894.27344 | 33606.58594 |
| 129810.6875 | 84686.625 | 88503.42188 | 92404.52344 | 96657.78125 | 95868.04688 | 99328.07813 |
| 2170162.25 | 1258159.5 | 1541649.875 | 1660899.375 | 1106963.875 | 1091468 | 1343974.75 |
| 128380.4609 | 81544.79688 | 96816.90625 | 87917.9375 | 67083.36719 | 62674.01953 | 71109.57813 |
| 24346.15625 | 16054.85254 | 19182.22266 | 16458.55664 | 16051.88379 | 17913.72461 | 17222.72656 |
| 24053.79102 | 19011.14648 | 16008.28223 | 17084.17969 | 23023.58594 | 18651.98047 | 19802.93555 |
| 56286.01953 | 51266.53906 | 32844.37109 | 33999.67188 | 61006.00391 | 55098.17969 | 64409.92578 |
| 76863.28125 | 57819.87891 | 52035.03906 | 51831.46094 | 45318.08984 | 52184.83594 | 48712.30469 |
| 91416.77344 | 58049.22656 | 66394.20313 | 70255.53906 | 58765.24609 | 66810.60156 | 64940.64844 |
| 27870.34375 | 18429.18945 | 17712.43164 | 20921.47461 | 26440.07422 | 24495.33789 | 25073.56055 |

| | | | | | | |
|---|---|---|---|---|---|---|
| 178313.125 | 130656.0625 | 129919.5078 | 123354.1484 | 136499.7188 | 141963.4531 | 139181.375 |
| 5136.373535 | 3146.460205 | 3892.299072 | 3445.602051 | 6965.196777 | 6653.458984 | 6140.410156 |
| 147780.3281 | 100905.5469 | 113033.6719 | 100826.9453 | 97906.59375 | 99365.86719 | 107686.5625 |
| 275247.3438 | 184970.4531 | 191232.5625 | 189394.4063 | 171529.7031 | 169782.0625 | 164202.25 |
| 24818.33984 | 22472.03906 | 17854.27734 | 15312.42871 | 14762.75391 | 33187.15234 | 18525.01563 |
| 329453.75 | 236345.8906 | 279799.6563 | 243462.7813 | 306777 | 347524.5938 | 359440.4375 |
| 129104.5469 | 101713.6094 | 106627.875 | 91914.34375 | 105929.6797 | 99950.04688 | 101308.2344 |
| 120286.5547 | 61350.83984 | 105734.6172 | 101176.3516 | 68672.64063 | 70376.38281 | 71737.79688 |
| 15805.47559 | 12790.29688 | 11561.91406 | 11040.0625 | 17511.46875 | 14973.93457 | 14410.24023 |
| 12424.99609 | 14592.9541 | 9696.03418 | 4312.606445 | 11866.14941 | 11483.5957 | 9742.191406 |
| 89338.64844 | 66186.75781 | 69590.29688 | 63203.39453 | 74738.95313 | 70162.375 | 71162.71875 |
| 121934.4063 | 96554.13281 | 89251.89063 | 88884.09375 | 100278.0234 | 101047.8438 | 93801.20313 |
| 59659.57813 | 42559.73047 | 33457.80469 | 42016.10938 | 27061.13281 | 30305.17969 | 32242.83398 |
| 48131.46875 | 33864.80469 | 33689.66016 | 32348.85938 | 37918.07031 | 38993.79688 | 37008.23828 |
| 10327.73145 | 7587.261719 | 7045.244629 | 6376.538574 | 7051.745117 | 6077.3125 | 6760.94043 |
| 3486.19165 | 2639.511475 | 6551.473633 | 1412.286987 | 34116.40625 | 3480.760986 | 4902.328613 |
| 20322132 | 13374005 | 13647942 | 14173193 | 15766739 | 15398876 | 15622709 |
| 26022.66602 | 19507.07422 | 16888.79492 | 16715.99609 | 16910.05469 | 17197.09766 | 16237.37305 |
| 6448.645508 | 5164.902832 | 4433.885742 | 4372.45459 | 5204.966309 | 4301.020996 | 5462.677246 |
| 16838.71289 | 7841.617188 | 9827.754883 | 9934.990234 | 7490.466309 | 10719.6543 | 13070.41699 |
| 14970.22363 | 12868.25098 | 9558.681641 | 8010.258301 | 11405.52441 | 11530.32227 | 9819.133789 |
| 104483.6953 | 68774.09375 | 75148.78125 | 84051.17969 | 99825.53125 | 103701.5313 | 109929.1797 |
| 803776.3877 | 700258.1104 | 519743.8311 | 498115.793 | 459859.3291 | 662291.5313 | 424336.3535 |
| 65112.86328 | 38839.84766 | 51973.08984 | 48081.35547 | 53712.96875 | 56173.05859 | 57727.82031 |
| 269392.4375 | 195244.1406 | 196968.5781 | 182982.625 | 206832.2656 | 200462.5156 | 200942.2188 |
| 1249213.5 | 900887.75 | 875148.9375 | 866256.9375 | 600692 | 599329.0625 | 615704.8125 |
| 17632.04883 | 11442.60547 | 10495.05566 | 9945.707031 | 7242.125977 | 9399.451172 | 6367.653809 |
| 21222.26172 | 14456.38184 | 14796.0459 | 14917.3584 | 14655.45215 | 13634.1875 | 14262.76953 |
| 73399.875 | 46352.41797 | 63545.91406 | 62672.44531 | 82536.65625 | 81357.04688 | 77525.91406 |
| 967994.1875 | 633495.9375 | 663566.25 | 698600.75 | 685427.625 | 744691.4375 | 757910.0625 |
| 50980.59766 | 35574.85547 | 33021.48047 | 27676.70313 | 15661.9873 | 17867.69141 | 15838.76758 |
| 18627.45117 | 14036.47949 | 12580.3877 | 13692.63574 | 14414.6123 | 13815.37402 | 13492.34863 |
| 115381.6328 | 70723.19531 | 82755.58594 | 68184.29688 | 48864.44922 | 45375.16406 | 48544.71484 |

| | | | | | | |
|---|---|---|---|---|---|---|
| 493327.2188 | 363787.4375 | 347213.625 | 342964.0938 | 422489.1563 | 383842.9375 | 396724.6563 |
| 72757.91406 | 50063.43359 | 50191.55859 | 69868.69531 | 173905.5469 | 80534.52344 | 104097.9453 |
| 675249.875 | 442398.2188 | 486082.25 | 484812.2813 | 412229.7188 | 381297.9688 | 401055.375 |
| 30928.72852 | 16416.76367 | 22907.73633 | 22589.43555 | 23157.60352 | 28928.44531 | 27497.94922 |
| 2140.585205 | 1142.055298 | 1575.11792 | 717.3312988 | 424.9326172 | 2623.486328 | 325.8370056 |
| 121109.0625 | 86915.42188 | 85755.28906 | 81624.60938 | 101431.9609 | 96059.9375 | 100553.0703 |
| 248607.9063 | 186551.9219 | 190843.1094 | 168047.7813 | 166835.5781 | 173512.5625 | 172733.1875 |
| 2057796.625 | 1272445 | 1429191.875 | 1334546.625 | 1389374.625 | 1492933.25 | 1456590.25 |
| 22321.02539 | 14148.83496 | 15725.28711 | 14798.09277 | 9707.478516 | 8681.642578 | 10162.05762 |
| 38066.17188 | 24380.44922 | 15673.07129 | 14638.0752 | 34649.30859 | 16733.30469 | 39465.84766 |
| 73242.64063 | 52120.09766 | 46452.66406 | 52029.03906 | 64485.01172 | 60587.05859 | 64019.0625 |
| 60756.85547 | 52813.29688 | 42982.46094 | 40887.36719 | 45539.28125 | 47335.91016 | 46707.73047 |
| 23864.1543 | 17805.85547 | 16636.45313 | 15372.47559 | 16541.88672 | 16208.25879 | 15880.66016 |
| 2396849.063 | 1855719.75 | 1436980.922 | 1712546.922 | 1762632.922 | 1747604.953 | 1712185.141 |
| 40028.74219 | 25425.60352 | 35777.94922 | 24988.35742 | 87875.80469 | 53206.35156 | 82783.39844 |
| 111804.9844 | 79890.85156 | 85248.89844 | 82837.86719 | 87107.89844 | 89324.66406 | 84905.57813 |
| 48743.48047 | 37163.73828 | 28105.01953 | 28849.9375 | 42206.90625 | 32688.77734 | 25955.16406 |
| 58801.91797 | 45687.64453 | 38833.34375 | 41157.07813 | 41244.85938 | 44792.53906 | 40862.17969 |
| 1999522.125 | 1198195.25 | 1461612.125 | 1501302 | 947862.1875 | 863293.125 | 883012.8125 |
| 16150.01465 | 12834.87793 | 9670.495117 | 10425.15137 | 9239.429688 | 8633.490234 | 8336.076172 |
| 69678.17188 | 44222.34375 | 47219.16016 | 54513.87891 | 56553.64844 | 60871.59375 | 58760.88672 |
| 6506726.5 | 11964986 | 7467998 | 11242251 | 20452300 | 12530461 | 20359414 |
| 164412.6875 | 110055.3906 | 129948.1094 | 122354.1016 | 106812.4531 | 114188.2969 | 110021.1641 |
| 75805.17188 | 48405.39453 | 55888.83203 | 55307.41797 | 66134.10938 | 58067.83203 | 68421.94531 |
| 46037.92578 | 41163.67969 | 27640.50977 | 27698.52344 | 34540.86328 | 36372.30078 | 33295.51563 |
| 27921.64844 | 25198.79883 | 13374.12012 | 13279.24805 | 25028.62109 | 28250.5918 | 28898.90039 |
| 402184.0938 | 270152.4063 | 289112.9063 | 342544.9375 | 388454.2188 | 425222.5 | 463183.9375 |
| 58998.58984 | 41875.19531 | 43975.92188 | 40788.07813 | 46221.09766 | 39026.79688 | 40766.60547 |
| 4460.669922 | 4400.402344 | 3047.598389 | 2896.493652 | 3603.239746 | 4026.825439 | 4017.056396 |
| 61855.28906 | 45297.01563 | 45475.52344 | 42740.45703 | 50882.51563 | 50578.59375 | 48673.07031 |
| 22896.77734 | 12660.45703 | 18176.87109 | 16825.43555 | 24389.09766 | 20781.74609 | 24895.55273 |
| 63108.48828 | 45964.84375 | 42541.96484 | 46257.26953 | 44795.50391 | 43430.35938 | 42458.94141 |
| 268219.375 | 175437.7188 | 183734.3438 | 183784.7969 | 220543.2031 | 209190.2969 | 214548.0625 |

| | | | | | | |
|---|---|---|---|---|---|---|
| 3711.473877 | 2407.195801 | 2218.553711 | 2849.551025 | 4412.07666 | 4214.040527 | 3894.223389 |
| 40796.65625 | 33453.54297 | 25747.21484 | 26936.84766 | 37082.71094 | 32743.24219 | 32455.69922 |
| 24901.01172 | 16410.93945 | 19730.17188 | 16851.97852 | 14514.99219 | 15013.25293 | 16259.14453 |
| 39708.625 | 32054.39453 | 28479.76563 | 27552.37305 | 33194.19531 | 34481.42188 | 40178.52344 |
| 226919.6406 | 151960 | 157861.0938 | 155121.5 | 179293.7031 | 168721.9531 | 170194.3125 |
| 5028.535156 | 4807.855957 | 3227.810303 | 2895.05127 | 7352.927734 | 6052.981934 | 5998.243652 |
| 7903.240234 | 7309.803223 | 3750.727783 | 4446.859375 | 3968.689209 | 5137.646973 | 4510.411133 |
| 476522.2813 | 348875.4375 | 346467.375 | 343427.625 | 416751.4375 | 421376.3438 | 425716.375 |
| 1029830.125 | 615353.1875 | 732132.875 | 645136.6875 | 659536.875 | 699813.9375 | 617773 |
| 32089.83594 | 24547.02148 | 22355.03711 | 25931.28125 | 27241.98828 | 23180.0918 | 24194.99219 |
| 6912.030762 | 4801.845703 | 5182.690918 | 5276.905273 | 3556.960938 | 4032.369385 | 3456.885986 |
| 131666.3906 | 92786.07813 | 92650.48438 | 86816.26563 | 96920.21875 | 94009.09375 | 97567.83594 |
| 1248.984619 | 1054.209473 | 642.6262817 | 617.512207 | 981.7978516 | 879.3726196 | 918.7026367 |
| 3039813.5 | 2164467.5 | 2088377.5 | 1988084.125 | 1835077.25 | 1875442.75 | 1822732.875 |
| 24302.32422 | 20068.83789 | 14074.14258 | 18976.10547 | 19704.61719 | 24455.34961 | 21191.52148 |
| 10179.05176 | 9862.423828 | 5890.557617 | 6347.61377 | 8693.338867 | 11306.00977 | 6417.05127 |
| 358147.8438 | 244946.3281 | 259877.2188 | 242534 | 311222.5313 | 298560.375 | 321807.0625 |
| 135431.8125 | 80518.39844 | 94717.60156 | 98956.26563 | 108512.375 | 119667.1094 | 118646.4531 |
| 136634.625 | 114903.5078 | 102588.0469 | 114669.8828 | 163274.9375 | 166022.625 | 156029.7969 |
| 31593.09375 | 22122.13086 | 23372.5332 | 20310.63281 | 11272.57324 | 17768.55469 | 12654.61914 |
| 10915.51563 | 8030.375 | 6826.218262 | 7376.853516 | 8728.801758 | 8198.44043 | 8638.5 |
| 10401.46973 | 8109.282227 | 7719.241211 | 7707.54541 | 6966.902344 | 6003.727051 | 4534.224121 |
| 1061233 | 686212.125 | 740704.875 | 768788.75 | 715354.6875 | 687494.375 | 727121.0625 |
| 34515.19531 | 26015.8457 | 23743.40625 | 26114.43555 | 30556.33594 | 30808.97852 | 30123.80273 |
| 1528001 | 933995.3125 | 1094478.375 | 1005904.25 | 968696.3125 | 1021016.188 | 1042372.563 |
| 130309.2813 | 91197.90625 | 87635.13281 | 82329.08594 | 99338.27344 | 95063.65625 | 91140.4375 |
| 67094.17969 | 44376.27344 | 44816.76172 | 46265.08203 | 33795.48828 | 28568.06445 | 29948.66016 |
| 188798.4219 | 141695.5313 | 136602.6563 | 132609.3281 | 150919.9219 | 150716.7188 | 146195.8438 |
| 143462.5781 | 85777.32031 | 111736.7578 | 109897.4766 | 144328.75 | 116718.6484 | 133690.9375 |
| 6151.547852 | 4109.032715 | 3905.771729 | 3128.042236 | 2473.040283 | 3179.071045 | 2416.547363 |
| 155462.7031 | 105557.5313 | 99246.35156 | 105298.0781 | 111383.1172 | 123205.6563 | 116248.2969 |
| 558350.9375 | 485379.8438 | 374642.2188 | 386840.1875 | 758041.8125 | 639841.4375 | 745389.25 |
| 21722.70313 | 14440.48242 | 12000.92188 | 14379.71191 | 14131.22168 | 18594.87891 | 15019.80566 |

| | | | | | | |
|---|---|---|---|---|---|---|
| 131644.4375 | 85356.94531 | 94646.03125 | 83241.42188 | 84115.42969 | 91316.5625 | 77282.6875 |
| 32115.83008 | 27001.63281 | 23099.1582 | 24960.37305 | 34432.52344 | 32133.50586 | 34487.58594 |
| 30849.46289 | 16534.37109 | 20119.30469 | 21187.81055 | 13775.70313 | 13345.13672 | 12923.52148 |
| 19927.375 | 13418.21582 | 11866.80762 | 12976.04785 | 23055.36328 | 25746.32813 | 23018.12891 |
| 178602.875 | 99500.875 | 115919.6484 | 131988.9844 | 178892.9375 | 159140.25 | 208498.4531 |
| 58288.37109 | 43716 | 40686.06641 | 37775.20313 | 56713.99219 | 59863.58594 | 64371.95313 |
| 155216.1875 | 111407.0078 | 113638.5938 | 125795.8672 | 131264.0313 | 137942.6094 | 125557.3828 |
| 36555.91406 | 25629.95508 | 26690.00391 | 24918.94727 | 21765.56055 | 22232.63086 | 22399.62305 |
| 82956.13281 | 60945.82031 | 59694.3125 | 57679.48047 | 87557.99219 | 73935.51563 | 82139.63281 |
| 29278.89648 | 23083.29102 | 11403.27148 | 5862.179199 | 19637.21094 | 40017.62109 | 39572.81641 |
| 39755.09375 | 39628.63672 | 22518.61523 | 22813.07031 | 39499.03125 | 31449.12695 | 37275.28906 |
| 61469.26172 | 40924.32813 | 42531.41797 | 40422.82031 | 43072.90625 | 44868.375 | 41533.57813 |
| 8812.347656 | 3830.273682 | 8455.885742 | 4524.064941 | 5176.530762 | 9014.524414 | 5315.080566 |
| 284515.25 | 193847.2656 | 187179.1875 | 190558.875 | 181984.5469 | 170931.8438 | 185703.4063 |
| 47827.64453 | 38315.84766 | 38880.33594 | 34333.92969 | 38240.54688 | 33507.53906 | 37536.73828 |
| 5118.722168 | 3659.605225 | 3366.419922 | 2449.944336 | 5802.808594 | 4748.391113 | 4560.410156 |
| 6039.818848 | 2031.827881 | 5127.391113 | 3014.057617 | 2343.001465 | 2843.376465 | 2765.78418 |
| 760904.3125 | 359153.1875 | 532966.375 | 628103.4375 | 1074892 | 1016253.875 | 1087852.125 |
| 16921.96484 | 12737.99707 | 10933.73828 | 11281.59277 | 12423.35547 | 14908.61719 | 12548.6377 |
| 39627.59375 | 33866.19531 | 27542.38086 | 30589.82813 | 45382.17578 | 60252.91016 | 42305.57813 |
| 12343.18066 | 7511.518066 | 8191.978516 | 7212.239746 | 10816.35352 | 13416.98047 | 11676.31348 |
| 17010.39453 | 11198.33496 | 10222.15332 | 11786.83105 | 13565.44824 | 14918.2041 | 14396.6084 |
| 32238.8418 | 21336.29102 | 20735.12695 | 22303.82422 | 22149.51172 | 20843.47266 | 20050.77734 |
| 3760.612549 | 2773.223633 | 3156.433838 | 3117.653564 | 3611.004639 | 5710.961914 | 4399.334473 |
| 6688.760254 | 4638.71875 | 5464.512207 | 4746.336914 | 5379.392578 | 3796.238281 | 4037.712402 |
| 774626.6875 | 563740.6875 | 509177.4688 | 513654.8438 | 563579.9375 | 614456.5 | 575869.5625 |
| 120519.0391 | 72994.25781 | 92563.55469 | 92249.78906 | 115804.8047 | 107554.1328 | 113778.2109 |
| 1624792.625 | 1072642.5 | 1093637 | 996361.5625 | 905762.625 | 952176 | 906473.875 |
| 9019.697266 | 6768.263672 | 5009.251953 | 7000.129395 | 17882.73438 | 8599.424805 | 9366.09375 |
| 19397.52734 | 15145.54297 | 12410.43848 | 16861.28125 | 24381.30859 | 24930.82422 | 21574.57422 |
| 29336.63672 | 28483.08008 | 17191.74805 | 16134.32227 | 25394.67188 | 22171.99219 | 27084.37695 |
| 154900.7031 | 91850.14844 | 113476.7188 | 121727.4375 | 84550.99219 | 80921.85938 | 85865.49219 |
| 102857.9453 | 84066.71094 | 67194.32813 | 63412.96094 | 44999.24219 | 58336 | 47502.63672 |

| | | | | | | |
|---|---|---|---|---|---|---|
| 619666.3125 | 498930.2813 | 435509.4063 | 459700.625 | 524038.3125 | 476220.0938 | 507555.9688 |
| 57956.50781 | 30763.94141 | 30197 | 46615.50781 | 51168.40234 | 45521.09766 | 46254.21484 |
| 27722.29297 | 22177.25195 | 18978.07227 | 17276.68555 | 23714.58398 | 24632.61719 | 20460.51758 |
| 86911.05469 | 62019.96875 | 61018.03516 | 61998.23438 | 68223.71094 | 64742.15625 | 64346.60547 |
| 29813.46289 | 25948.94727 | 14020.92383 | 15105.94824 | 24077.77734 | 20436.81836 | 23183.92578 |
| 113633.9844 | 72288.6875 | 81433.91406 | 86915.77344 | 120899.6953 | 89815.00781 | 119252.3203 |
| 19619.21875 | 12683.63965 | 12037.31738 | 9956.411133 | 9337.737305 | 9725.165039 | 9603.166992 |
| 56861.25 | 36016.9375 | 38997.67188 | 32591.57227 | 49816.29297 | 51640.62109 | 51662.72656 |
| 72556.21094 | 54960.52734 | 43541.73047 | 44373.17188 | 47437.35156 | 44830.16406 | 47530.53906 |
| 201884.75 | 118130.6016 | 149143.0781 | 169583.3281 | 224622.125 | 184165.75 | 212111.125 |
| 165108.1719 | 121433.3906 | 120103.7813 | 115904.8984 | 120834.0703 | 121266.8047 | 120235.5234 |
| 260739.7969 | 177879.4688 | 175062.9688 | 178051.8594 | 157234.3594 | 158939 | 158198.3906 |
| 36781.44531 | 27545.41211 | 28438.22852 | 21287.23242 | 28659.19141 | 33731.1875 | 33588.29297 |
| 10772.91895 | 9160.822266 | 6339.740234 | 5596.787598 | 9203.212891 | 10098.03613 | 9422.05957 |
| 10693.48145 | 6263.722168 | 7643.54248 | 8070.618164 | 12275.83984 | 9483.583008 | 20599.00586 |
| 76972.36719 | 46131.10156 | 62969.48828 | 58583.35938 | 49638.16016 | 50179.00391 | 53533.12891 |
| 183070.25 | 118882.6719 | 120379.2422 | 122128.0625 | 124575.5313 | 131244.4688 | 115539.9531 |
| 121312.8984 | 84576.63281 | 91578.125 | 83604.76563 | 93713.82813 | 96757.08594 | 90327.29688 |
| 193462.3281 | 112167.9531 | 132583.8281 | 138099.9219 | 116779.7969 | 129383.8281 | 138798.5781 |
| 72409.73438 | 47849.35156 | 50134.59766 | 47567.89063 | 43734.42188 | 45404.91016 | 42916.38672 |
| 15981.9043 | 12102.48535 | 10340.59473 | 9051.260742 | 10079.61426 | 9211.283203 | 10545.20215 |
| 10204.05371 | 7796.312988 | 6353.035645 | 6592.874512 | 8119.824707 | 7973.503418 | 7232.331543 |
| 371522.1563 | 276181.9063 | 258097.0938 | 269578.4063 | 400745.5313 | 349090.1563 | 369243.75 |
| 23217.55469 | 29809.53906 | 8511.448242 | 7310.816895 | 31074.67578 | 23020.78711 | 29119.82813 |
| 83005.60938 | 53135.45313 | 53829.28906 | 39818.45703 | 39153.58984 | 55669.66797 | 52072.40234 |
| 55086.62109 | 42518.33594 | 42558.78906 | 40230.39453 | 46998.89844 | 49955.28906 | 50520.00781 |
| 116173.4375 | 71801.375 | 84445.92188 | 83837.84375 | 96945.26563 | 86371.98438 | 95791.3125 |
| 211452.9688 | 144051.4219 | 156007.0625 | 148266.5156 | 156370.8594 | 163324.5313 | 158763.6094 |
| 278875.5 | 199726.9219 | 183101.5156 | 187358.4844 | 179837.1563 | 187632.7031 | 185061.5625 |
| 8607.408203 | 5199.557617 | 5693.805176 | 5664.229004 | 7042.125977 | 5983.160645 | 6551.634766 |
| 9143.023438 | 9803.758789 | 7489.521484 | 4335.490234 | 8371.200195 | 11518.95215 | 8840.920898 |
| 189609.6094 | 112728.4297 | 142014.8125 | 131790.0313 | 148245.6094 | 158764.1875 | 122051 |
| 359329.1563 | 185869.4531 | 266725.4688 | 252496.0313 | 197109.5469 | 203209.4844 | 202668.5313 |

| | | | | | | |
|---|---|---|---|---|---|---|
| 5581644 | 3756895.5 | 3872354.5 | 3833637.25 | 4014365.75 | 3882628 | 4115797.75 |
| 243940.1406 | 155349.7344 | 168008.2656 | 168104.7188 | 184862.5156 | 162093.4844 | 174973.5625 |
| 258430.2656 | 163177.7188 | 165936.7188 | 156693.5313 | 166393.7813 | 178099.8594 | 156978.7656 |
| 2154215 | 1497797.875 | 1386968.75 | 582634.8125 | 398707.0938 | 1450722.75 | 714314.5 |
| 135576.1094 | 95224.50781 | 85444.84375 | 82389.26563 | 109055.9297 | 99118.36719 | 109775.1094 |
| 14628.27441 | 10695.80859 | 10285.84277 | 10850.31836 | 15125.16992 | 14895.99512 | 13339.23633 |
| 51513.32031 | 49715.30469 | 31777.67773 | 28006.32031 | 34417.51172 | 33373.86328 | 36759.46484 |
| 79543.25781 | 63854.49219 | 48933.66016 | 58536.87109 | 54572.14063 | 52484.62891 | 51819.19922 |
| 11962.75781 | 6083.762207 | 6894.131836 | 9485.613281 | 11330.75488 | 13014.09277 | 11438.40137 |
| 63537.49609 | 42660.46484 | 40006.9375 | 39647.59766 | 66229.39844 | 66783.80469 | 68909.46875 |
| 16162.9873 | 10364.68359 | 11110.98535 | 9922.134766 | 10612.35742 | 10788.57227 | 11320.23633 |
| 68937.23438 | 73121.55469 | 68679.02344 | 38384.84375 | 17848.3418 | 33617.44531 | 46818.375 |
| 325623.1875 | 222312.1563 | 238204.2344 | 218187.4375 | 274663.9063 | 258230.1563 | 287525.5313 |
| 145246.125 | 98126.98438 | 95929.46875 | 93689.48438 | 113586.1563 | 101499.4453 | 101395.0469 |
| 111739.5547 | 76367.32031 | 70659.03125 | 69329.99219 | 84752.63281 | 90673.75 | 82459.4375 |
| 95642.8125 | 64099.41406 | 64752.48438 | 63017.96484 | 65653.79688 | 65037.16406 | 72429.45313 |
| 279764.7813 | 197885.875 | 200058.7656 | 184797.9063 | 218832.9688 | 215143.2813 | 219651.7188 |
| 2193.051025 | 2054.438721 | 1299.93335 | 1203.871582 | 1464.848755 | 615.4416504 | 1445.017456 |
| 180944.2656 | 118671.7891 | 121953.8828 | 121355.3906 | 135010.875 | 122267.8672 | 124528.1719 |
| 210744.875 | 149657.8906 | 146427.2031 | 153490.5469 | 203478.1719 | 196656.7188 | 200005.2813 |
| 26048.32227 | 17731.19727 | 16418.6543 | 16982.85547 | 19731.20898 | 16382.69434 | 15307.29004 |
| 153750.0938 | 101804.2109 | 105806.5078 | 105354.9297 | 112794.1719 | 111352.5 | 120893.7344 |
| 42485.19922 | 34701.96484 | 19061.27148 | 21060.35156 | 31657.7207 | 31893.08398 | 28960.46875 |
| 11933.11328 | 7980.535156 | 8165.060059 | 8937.351563 | 8455.897461 | 8171.811035 | 8292.6875 |
| 29336.07617 | 20434.84961 | 21585.43555 | 18142.68555 | 12918.11523 | 14346.24414 | 12947.21973 |
| 165412.7969 | 125312.875 | 118646.7109 | 111720.7891 | 138161.625 | 130154.8438 | 122440.4609 |
| 28247.55469 | 18758.12695 | 21318.67773 | 22548.23242 | 27761.37695 | 25156.79688 | 29399.85547 |
| 224126.3438 | 142546.8281 | 145715.6875 | 129772.8438 | 165170.5938 | 164010.2344 | 168401.8438 |
| 58424.76172 | 35142.92188 | 38573.20703 | 36348.37109 | 27110.56055 | 33004.27734 | 34544.76563 |
| 114303.0938 | 73095.6875 | 74070.07031 | 62364.12891 | 69302.46094 | 68492.15625 | 72796.61719 |
| 34021.41016 | 23417.05859 | 23259.12109 | 26702.75 | 35113.76172 | 39112.20313 | 49567.80859 |
| 57831.60938 | 42081.44531 | 35831.96094 | 37018.32422 | 59625.89844 | 60315.38672 | 59700.74609 |
| 44360.58594 | 32642.16211 | 34162.64844 | 29552.75977 | 38619.68359 | 40491.33594 | 38616.25391 |

| | | | | | | |
|---|---|---|---|---|---|---|
| 2504.001221 | 1895.066528 | 1473.099609 | 1563.445435 | 3012.948242 | 3073.158203 | 2878.102051 |
| 355553.0938 | 232412.0938 | 237773.9219 | 243443.1406 | 264783.375 | 258168.0938 | 268052.1563 |
| 6310.437988 | 4031.089355 | 4199.931152 | 4695.58252 | 5075.765625 | 6362.418457 | 5332.506836 |
| 247143.5156 | 147874.8281 | 166510.875 | 151738.8125 | 140812.8906 | 158759.6875 | 148664.9531 |
| 9415.898438 | 5341.251465 | 6049.813477 | 5940.078125 | 3562.884766 | 6010.376953 | 5355.440918 |
| 30636.14453 | 19873.01172 | 20828.33203 | 20936.27148 | 17537.73828 | 15316.51953 | 19975.10156 |
| 34269.57813 | 21510.43164 | 24123.41602 | 20646.6875 | 20189.83008 | 20696.7168 | 19583.89648 |
| 66772.19531 | 50123.57422 | 45762.75391 | 43004.97266 | 46740.19531 | 45067.68359 | 48260.61328 |
| 83967.92969 | 51190.52344 | 63788.84375 | 65130.29297 | 122076.9844 | 132506.5625 | 134030.9844 |
| 11165.85645 | 7783.041992 | 8411.697266 | 5501.59082 | 6079.65625 | 6324.995117 | 6162.23584 |
| 26393.7832 | 31006.69531 | 18286.68359 | 25973.2168 | 40775.94141 | 37216.24609 | 34969.78125 |
| 109963.1797 | 87587.76563 | 68437.55469 | 77735.24219 | 98380.45313 | 92182.75 | 87890.85938 |
| 17033.26563 | 8185.496582 | 10933.37598 | 10645.21289 | 9245.59082 | 9789.830078 | 10559.30762 |
| 20955.25977 | 17124.80078 | 12700.09863 | 11395.19141 | 9768.105469 | 9417.180664 | 10257.62109 |
| 133870.0625 | 93337.54688 | 93724.96875 | 92514.15625 | 96547.73438 | 101914.3125 | 90398.875 |
| 182740.0313 | 126439.7188 | 122131.7656 | 138734.6563 | 235723.5313 | 201359.7656 | 223003.9375 |
| 17693.04883 | 10302.0293 | 11903.91504 | 9756.819336 | 12422.66211 | 12861.8584 | 13223.03711 |
| 532524.875 | 380885.4688 | 361977 | 381331.5313 | 567286.8125 | 470390.4063 | 500543.4375 |
| 22532.29883 | 17976.61719 | 16480.45898 | 14210.59375 | 17285.75195 | 16664.83203 | 16164.38086 |
| 90044.32031 | 48207.98828 | 63732.83984 | 43243.78125 | 43732.64453 | 70142.17188 | 40054.92578 |
| 54380.69141 | 19525.40234 | 43046.79688 | 44685.32422 | 38575.71484 | 35339.67188 | 42557.05859 |
| 5337.458496 | 2464.156982 | 4119.941895 | 4919.213379 | 5040.420898 | 5508.800781 | 5140.163086 |
| 1093536.625 | 753249.25 | 750532.9375 | 689279.125 | 642125.625 | 669595.0625 | 649672.875 |
| 46443.70313 | 31117.88086 | 31120.80859 | 34219.51172 | 52373.63672 | 42091.86719 | 45544.67969 |
| 1006783.875 | 728677.5625 | 736302.6875 | 722103.0625 | 667291.4375 | 600113.375 | 679015.25 |
| 1533.568481 | 2926.377197 | 1046.662476 | 1387.955933 | 939.2208862 | NA | NA |
| 22585.11523 | 25670.85742 | 9526.480469 | 8607.532227 | 19137.03906 | 15320.72559 | 17082.66602 |
| 58007.29688 | 41168.69141 | 43442.67969 | 45284.06641 | 42172.13281 | 45201.08203 | 44567.21094 |
| 149751.3594 | 99169.90625 | 110361.3828 | 103535.8281 | 97157.0625 | 92626.625 | 93053.35156 |
| 466257.625 | 309352.9375 | 358216.5938 | 363700.0313 | 428181.1563 | 427846.0625 | 410173.0625 |
| 119427.0625 | 67573.03125 | 86810.39063 | 96603.05469 | 103586.3359 | 117767.6719 | 121963.0547 |
| 2909490.594 | 2011343.219 | 2412720.063 | 1570371.375 | 1581217.109 | 2798200.594 | 1746976.719 |
| 94560.92188 | 63148.87891 | 62427.64844 | 64282.20703 | 75062.29688 | 80993.125 | 76099.78906 |

| | | | | | | |
|---|---|---|---|---|---|---|
| 27936.56641 | 13263.55078 | 18301.66016 | 14405.97754 | 14245.55859 | 19809.71484 | 9517.105469 |
| 1011727.063 | 698187.125 | 639349.9375 | 667375.9375 | 462168.125 | 419861.4063 | 402465.7188 |
| 600150.75 | 363845.625 | 369761.7813 | 489675.375 | 337545.6875 | 324858.125 | 349313.1875 |
| 91458.71875 | 72933.89844 | 54597.02344 | 53387.6875 | 48135.79297 | 49273.86328 | 47264.19922 |
| 102379.2891 | 84367.53125 | 58762.04297 | 60827.78906 | 103768.7578 | 97081.55469 | 105982.0469 |
| 61383.91016 | 33493.63281 | 41635.20313 | 46237.81641 | 42552.43359 | 46941.59375 | 48099.125 |
| 668582.1875 | 470672.125 | 386448.9375 | 457148.1563 | 325981.625 | 414358.5 | 399322.9063 |
| 116594.0781 | 81719.67969 | 66441.375 | 71428.21875 | 63458.05469 | 65394.10156 | 62523.09766 |
| 99855.96875 | 60443.41797 | 63599.83984 | 64337.24609 | 57874.875 | 58893.26953 | 55657.20703 |
| 499578.5313 | 365491.6875 | 327425.5 | 305927.8438 | 354712.4375 | 364701.6875 | 384473.8125 |
| 849610 | 615736.625 | 694257.1875 | 656171.1875 | 667103.4375 | 519555.0313 | 580785.625 |
| 38386.87109 | 25396.49414 | 24502.51953 | 25721.17773 | 23560.81641 | 24371.51563 | 22508.48633 |
| 64340.75781 | 37300.64453 | 44718.57422 | 44400.01953 | 47324.20313 | 48404.08203 | 45449.0625 |
| 36071.5625 | 19845.96484 | 26612.44141 | 22668.29297 | 22027.93945 | 21388.78125 | 22631.4668 |
| 257693.1719 | 168589.7656 | 166322.0313 | 164906.6563 | 183369.7188 | 188122.3906 | 175041.375 |
| 10916.13281 | 12341.4834 | 6166.817871 | 10612.76465 | 19940.41211 | 13032.10156 | 13769.14551 |
| 158191.625 | 109663.1484 | 107545.6172 | 115827.5547 | 135408.8125 | 126577.8828 | 120508.5 |
| 68307.78906 | 44648.72656 | 52379.93359 | 49012.84375 | 43818.18359 | 49216.1875 | 47591.26953 |
| 16166.39063 | 11585.83887 | 11011.90723 | 9708.266602 | 11385.24219 | 11232.66016 | 10941.07813 |
| 42504.35547 | 47532.69531 | 21209.5625 | 20559.76953 | 6527.141113 | 17760.01367 | 10652.23242 |
| 8064.103027 | 5986.818359 | 5261.007813 | 6357.433594 | 9000.910156 | 9420.708008 | 8398.953125 |
| 116787.6016 | 64819.86328 | 82192.46094 | 56027.0625 | 55915.40234 | 95442.89063 | 51623.25 |
| 10869.22754 | 8341.24707 | 9103.545898 | 7984.138672 | 4585.802246 | 4914.995605 | 4140.629395 |
| 143278.5 | 109320.9219 | 103289.7656 | 118871.5547 | 120608.1875 | 127804.1953 | 120596.9219 |
| 284827.7188 | 172047.4219 | 186065.9063 | 193857.375 | 162954.75 | 149598.0156 | 157345.3281 |
| 1594027.489 | 1012095.399 | 1014402.57 | 1045406.854 | 1424770.159 | 1222668.047 | 1383488.955 |
| 2880.151123 | 2135.706055 | 2028.641479 | 1860.112671 | 3073.28418 | 1870.943604 | 2634.442383 |
| 18136.33008 | 11944.72168 | 12059.33203 | 12790.0459 | 15265.44336 | 11035.80762 | 14645.36816 |
| 516830.5938 | 383786.0938 | 408579.4375 | 456148.9688 | 357402.7813 | 358617.4375 | 366190.9688 |
| 361754.9063 | 225578.1094 | 231865.7344 | 222703.2188 | 173849.8281 | 217262.2813 | 167827.5313 |
| 94143.91406 | 56410.57813 | 63021.65234 | 60680.14844 | 74105.60156 | 78643.72656 | 66947.16406 |
| 296554.5 | 233870.2031 | 190752.0938 | 196729.8438 | 248496.0938 | 193058.5469 | 211734.125 |
| 3020.748779 | 1823.658569 | 1760.811401 | 1817.75 | 1890.305542 | 1988.949585 | 2241.375244 |

| | | | | | | |
|---|---|---|---|---|---|---|
| 154120.0469 | 101232.5 | 95652.26563 | 102437.2734 | 108875.9531 | 118694.2188 | 111646.4141 |
| 60611.88672 | 33265.01953 | 41828.29297 | 48832.44531 | 59205.14063 | 54483.0625 | 53827.67188 |
| 148869.2969 | 71700.88281 | 97077.49219 | 98808.0625 | 145583.5156 | 111375.2266 | 162932.7188 |
| 10927.22363 | 7902.188477 | 7069.326172 | 5386.705566 | 9299.543945 | 6796.489746 | 8660.774414 |
| 8556.077148 | 8289.958008 | 5619.179199 | 4739.422363 | 2758.919189 | 3781.64917 | 2621.918945 |
| 273339.875 | 166688.9844 | 189698.0469 | 128857.375 | 126093.8828 | 202861.6094 | 122872 |
| 52814.24609 | 28545.94141 | 32844.21094 | 31290.36523 | 40095.44141 | 38421.26953 | 43754.45313 |
| 77237.69531 | 44100.80859 | 59100.87891 | 54755.14453 | 65999.16406 | 49610.625 | 71346.25781 |
| 105522.3906 | 70453.26563 | 70591.11719 | 68854.97656 | 72721.14844 | 69820.42969 | 72443.90625 |
| 44773.32813 | 31856.70703 | 33659.69531 | 32153.65039 | 40152.38672 | 41120.88672 | 36831.03906 |
| 4125287.75 | 2444113.75 | 2749026.25 | 2970120 | 3060557.25 | 3001523.75 | 3236879.75 |
| 32169.11328 | 16931.35156 | 20123.76563 | 18925.60156 | 16433.97656 | 14564.6123 | 17813.9082 |
| 193076.5469 | 131598.5938 | 135212.4063 | 128827.1172 | 153028.2969 | 150561.2656 | 147118.9531 |
| 39937.85938 | 24168.09961 | 24416.09766 | 25466.22266 | 50706.08203 | 58053.55078 | 61557.5625 |
| 29816.08203 | 19424.41211 | 19662.83398 | 21411.48828 | 23505.41016 | 22339.46289 | 26125.01172 |
| 41140.9375 | 22152.75195 | 32183.63281 | 35797.64063 | 43667.79688 | 30197.17773 | 38135.92188 |
| 45794.95703 | 44123.37891 | 27795.13281 | 25726.86328 | 54462.51172 | 41886.36719 | 39364.32031 |
| 8433.355469 | 7488.016602 | 2793.398193 | 4545.489746 | 6392.91748 | 6443.697266 | 6949.71582 |
| 186231.4063 | 133492.8906 | 131456.1094 | 131354.6406 | 146919.5469 | 145197.3906 | 144529.5469 |
| 31219.1875 | 25082.83398 | 19030.25195 | 16247.17383 | 24530.57422 | 23365.71875 | 22005.79883 |
| 179313.5938 | 120796.5547 | 128161.3203 | 120699.1719 | 142774.0781 | 131525.6094 | 142539.125 |
| 24589.26172 | 20750.95703 | 13539.48145 | 13753.79004 | 25708.64648 | 25651.55273 | 25013.12891 |
| 68390.91406 | 48472.97656 | 46523.40234 | 43572.42188 | 65435.05078 | 58284.78125 | 61283.05078 |
| 186491.5156 | 135248.4375 | 123079.0625 | 135087.6094 | 141123.7031 | 136492.1406 | 133088.7188 |
| 17395.25391 | 11463.31348 | 10994.96289 | 10643.09375 | 14597.1875 | 14073.67578 | 15110.55957 |
| 20123.40234 | 13737.0752 | 14616.44531 | 12472.80371 | 12863.26172 | 14238.00977 | 14112.0957 |
| 80624.50781 | 45589.99609 | 63019.58594 | 36785.78516 | 40888.41016 | 58954.87891 | 42745.40625 |
| 7165.368164 | 5760.245117 | NA | 4251.561523 | 11803.23145 | 7160.193848 | 9374.076172 |
| 40715.54297 | 31242.2793 | 24740.70313 | 22468.53125 | 25081.79492 | 24888.77539 | 24159.6543 |
| 34997.35938 | 18666.67188 | 21528.85547 | 20961.24414 | 16089.21094 | 16774.84375 | 20808.45703 |
| 4288.166016 | 1903.109619 | 3506.680176 | 3090.897217 | 5143.677734 | 5149.761719 | 6899.239746 |
| 15928.78613 | 8848.913086 | 11681.10449 | 10828.7793 | 7144.526367 | 7379.03125 | 7868.767578 |
| 62556.01172 | 39437.75781 | 44404.19922 | 40621.36328 | 37628.28906 | 41092.20703 | 39794.86719 |

| | | | | | | |
|---|---|---|---|---|---|---|
| 87611.82031 | 79509.64844 | 46111.17578 | 46430.98438 | 80328.34375 | 60364.57422 | 79421.48438 |
| 13843.54688 | 6888.195313 | 12418.34863 | 6456.93457 | 15327.46875 | 13896.2832 | 15136.31738 |
| 230300.6875 | 150609.625 | 156552.2031 | 156792.6719 | 173861.8906 | 171470.375 | 167418.7969 |
| 71026.92188 | 44147.10938 | 48186.22266 | 35689.05469 | 33365.31641 | 50529.58594 | 31101.81641 |
| 92893.74219 | 76221.82031 | 56941.95703 | 52286.87891 | 76233.65625 | 75493.53906 | 80788.23438 |
| 485208.1875 | 314771.5625 | 310982.125 | 312637.4688 | 229663.2031 | 234801.1719 | 240701.9219 |
| 1318549 | 771525.6875 | 933658.25 | 915752.4375 | 824972.1875 | 848838.1875 | 865874.625 |
| 10652.23242 | 7298.199707 | 6186.797852 | 5681.372559 | 6649.807129 | 7150.611328 | 6423.504883 |
| 66305.04688 | 45044.79688 | 38123.57031 | 38091.98438 | 45176.43359 | 37528.05078 | 38730.00781 |
| 13508.19531 | 8958.081055 | 7890.777832 | 8070.833984 | 7158.955078 | 6923.710449 | 6014.61377 |
| 86640.74219 | 60854.53125 | 56319.68359 | 56640.9375 | 71998.39063 | 66267.35156 | 64860.23438 |
| 24020.83594 | 17765.82031 | 15797.95508 | 19484.81055 | 19633.16797 | 20975.81836 | 19222.67578 |
| 432664.0391 | 324625.0508 | 306678.1523 | 302876.0977 | 521186.1797 | 440210.3516 | 463823.9219 |
| 165738.3281 | 121210.9844 | 89939.96875 | 95560.39844 | 112250.5938 | 97436.39844 | 108696.4609 |
| 2478.221924 | 1412.819824 | 1710.28894 | NA | 15864.96875 | 12881.02734 | 13468.4502 |
| 2460162.5 | 1593099.125 | 1649329.375 | 1658668.25 | 1614904.625 | 1563995.125 | 1743882.875 |
| 43759.12109 | 35924.12891 | 26202.62109 | 23792.78906 | 33769.76172 | 31764.25977 | 34326.02734 |
| 47501.05469 | 33926.74219 | 30332.36133 | 27739.33398 | 23858.36328 | 28289.13477 | 25325.51172 |
| 161100.0938 | 96838.82813 | 109805.375 | 104346.1484 | 92243.99219 | 100451.0547 | 92690.76563 |
| 386150.8125 | 243947.5625 | 247791.3125 | 268370.6563 | 375967.4375 | 266113.8125 | 317756.3438 |
| 13414729 | 10857638 | 8750706 | 8575897 | 28592234 | 27057990 | 33943812 |
| 977440.75 | 514772.9688 | 809589.0625 | 174196.5938 | 138622.3125 | 1061280.375 | 116818.6953 |
| 86619.50781 | 61016.19531 | 58279.5 | 58970.96484 | 45599.77734 | 56030.50781 | 49685.24219 |
| 26244.43359 | 26301.875 | 17219.13867 | 13706.77441 | 23190.59766 | 17743.17578 | 25021.98828 |
| 46950.51172 | 22261.83398 | 35573.31641 | 24953.45117 | 27744.83203 | 43769.6875 | 40135.84375 |
| 31424.63867 | 23485.22266 | 21467.81055 | 21789.87695 | 33557.89453 | 29271.06055 | 29045.57031 |
| 426546.25 | 261984.0938 | 273085.6875 | 277641.5625 | 311659.4063 | 295163.2188 | 296133.2188 |
| 29398.51758 | 18106.27344 | 21225.35742 | 18096.875 | 20970.0918 | 20450.25781 | 20337.56836 |
| 16529.62695 | 11915.1709 | 8403.461914 | 8783.067383 | 7174.843262 | 9325.34375 | 8072.248535 |
| 16300.76172 | 15401.9834 | 10370.62207 | 7895.01709 | 12799.99414 | 12588.00391 | 12753.3125 |
| 102578.3438 | 68155.375 | 69919.05469 | 62569.96875 | 44576.125 | 53749.92969 | 46064.15625 |
| 993138.5 | 698350.8125 | 707687.6875 | 726836.0625 | 1147369 | 897594.125 | 962867.8125 |
| 205827.1094 | 138630.0781 | 151128.8906 | 130405.9453 | 158122.1094 | 163662.9844 | 148841.3438 |

| | | | | | | |
|---|---|---|---|---|---|---|
| 3825.615723 | 2473.845459 | 2137.040039 | 2728.214355 | 3383.750488 | 2733.054443 | 3159.443359 |
| 342430.1641 | 232457 | 237808.1738 | 218773.5605 | 219356.8809 | 248174.9336 | 230526.0508 |
| 192667.9688 | 146228.0313 | 111745.1641 | 112251.3828 | 162662.2344 | 179612.4688 | 155985.0625 |
| 342006.5938 | 189492.75 | 225897.5625 | 156650.1875 | 199563.9844 | 349814.8438 | 162829.5 |
| 90408.5 | 58210.31641 | 67690.14844 | 58176.11719 | 49090.46484 | 52769.46094 | 52402.47656 |
| 58586.51074 | 47476.03467 | 35030.63818 | 34161.34668 | 71755.49414 | 76478.08887 | 74556.08789 |
| 508866.4375 | 301980.8125 | 337847.2188 | 334516.7813 | 309312.1875 | 375000.75 | 373067.9063 |
| 79102.42969 | 45682.20703 | 52708.32422 | 57391.14063 | 55822.58594 | 60289.85938 | 61186.44531 |
| 230745.5 | 152111.1563 | 155485.6563 | 149809.9688 | 171062.125 | 174596.75 | 170580.3125 |
| 58209.39453 | 37269.76172 | 38689.86328 | 37968.51563 | 46660.67188 | 44141.30469 | 47511.99219 |
| 165797.9048 | 121787.4658 | 106224.2939 | 113141.355 | 109076.0293 | 103510.3799 | 103476.7368 |
| 1107315.875 | 756954.1875 | 725784.8125 | 633588.6875 | 429316.4375 | 544974 | 468418.4063 |
| 15385.44727 | 13088.50879 | 7443.078613 | 11377.59277 | 23723.61719 | 17882.875 | 17270.38867 |
| 108080.8281 | 76040.25781 | 68638.30469 | 73103.625 | 88949.91406 | 93637.45313 | 90412.53125 |
| 84618.94531 | 53715.46094 | 46275.05078 | 49390.41797 | 46159.39453 | 47337.56055 | 47201.08203 |
| 93137.13867 | 66129.50977 | 62775.02344 | 53733.25781 | 59547.64844 | 67028.04492 | 38984.65625 |
| 148632.1563 | 105337.1797 | 98981.59375 | 93455.94531 | 111492.2656 | 108706.6484 | 106854.8906 |
| 14435.14648 | 10401.72754 | 8737.198242 | 10289.56152 | 12216.41211 | 11320.64746 | 11127.69238 |
| 153748.625 | 94378.61719 | 76349.11719 | 72120.90625 | 128334.6484 | 93604.82031 | 144110.125 |
| 85066.94531 | 58594.24609 | 58077.92188 | 56404.10938 | 114819.2422 | 96381.11719 | 94444.17188 |
| 4068911.25 | 2583681.25 | 2675200 | 2752085 | 5359260.5 | 4895855 | 5217474.5 |
| 137247.25 | 92131.67188 | 93541.25 | 99616.14063 | 129211.2891 | 127310.7344 | 127021.7813 |
| 98277.45313 | 66539.86719 | 64675.4375 | 65143.79688 | 50383.29688 | 49264.76953 | 48729.61328 |
| 14141.05664 | 9959.683594 | 8892.241211 | 9046.621094 | 9443.779297 | 8203.650391 | 8194.305664 |
| 72342.07031 | 46151.78516 | 46530.69141 | 44885.04297 | 38591.89453 | 41628.61719 | 40570.77734 |
| 7216.447754 | 4602.741699 | 4706.063965 | 4470.354492 | 5088.77832 | 6255.119629 | 5255.494629 |
| 147202.625 | 95500.8125 | 99931.34375 | 97245.8125 | 82624.60938 | 78182.27344 | 81620.54688 |
| 3590285 | 2518221.5 | 2501920.75 | 2618866.5 | 2082339.25 | 1998214.75 | 2057977.5 |
| 63733.25391 | 38279.82422 | 42375.67578 | 42761.56641 | 49363.44922 | 44496.53125 | 52481.6875 |
| 448807.3438 | 249006.875 | 303565.2188 | 252105.8281 | 166974.7188 | 225671.875 | 163619.0781 |
| 129047.4844 | 90731.75781 | 91368.72656 | 98583.00781 | 136259.6406 | 149219.5469 | 153749.8281 |
| 101429.5391 | 71342.0625 | 70945.61719 | 73727.82813 | 322639.3438 | 165049.2813 | 191677.5313 |
| 18871.45313 | 14650.29297 | 12771.44531 | 11716.90625 | 12329.14551 | 10651.24902 | 11571.45508 |

| | | | | | | |
|---|---|---|---|---|---|---|
| 213602.6875 | 142049.2969 | 143349.125 | 134685.6563 | 149054.5156 | 150433.7813 | 153580.5 |
| 87108.5625 | 58478.79297 | 61777.89844 | 53563.49219 | 48802.47656 | 53094.19922 | 52627.72656 |
| 54952.35938 | 36072.73438 | 30327.99805 | 29902.99609 | 22446.74609 | 23002.01563 | 22069.52734 |
| 12047.48438 | 10124.54199 | 7685.665527 | 6593.98877 | 6639.538086 | 5495.247559 | 5984.838379 |
| 30358.69727 | 21814.07617 | 19265.84961 | 17406.82227 | 15596.42578 | 12927.63477 | 12349.09766 |
| 4635.932617 | 2707.036621 | 2865.221924 | 2902.067871 | 2960.462158 | 4116.083496 | 4272.115234 |
| 98681.21094 | 64555.51563 | 66538.67969 | 62483.11719 | 56327.40234 | 56451.23047 | 55357.55859 |
| 13399.50879 | 9754.743164 | 7844.370117 | 7065.333984 | 8319.138672 | 6784.055664 | 6496.100098 |
| 53929.13281 | 34473.52344 | 37628.35156 | 33948.41797 | 32006.15234 | 38098.03516 | 37142.92578 |
| 16130.33105 | 12127.39648 | 8033.772949 | 9696.817383 | 13333.09961 | 10571.4668 | 11452.29492 |
| 135776.6563 | 94889.98438 | 93490.78125 | 69243.40625 | 64941.73047 | 90372.59375 | 66425.14844 |
| 31814.58008 | 22625.88672 | 20882.26367 | 20366.83398 | 27276.47852 | 26870.92383 | 25825.73047 |
| 32023.07227 | 18981.17188 | 21335.80078 | 19222.75 | 16088.55078 | 14495.94043 | 14832.04883 |
| 76529.39844 | 46393.10547 | 42626.44141 | 46376.64453 | 113281.3516 | 77921.92188 | 110675.2734 |
| 89800.39844 | 55511.42578 | 61883.89453 | 59536.96094 | 60945.84766 | 62232.67578 | 62330.77734 |
| 73691.80469 | 48718.32422 | 50179.02344 | 52597.67969 | 56768.80078 | 58944.24219 | 55054.79688 |
| 55783.89063 | 36132.14844 | 34947.51563 | 33984.04297 | 41757.89453 | 39438.8125 | 40013.71484 |
| 15072.33984 | 8670.574219 | 8221.631836 | 9268.235352 | 6615.497559 | 7911.385742 | 6869.57959 |
| 3206.508057 | 1514.458862 | 2434.630859 | 672.9390259 | 2492.766113 | 855.9961548 | 2918.300293 |
| 10761.10938 | 4576.610352 | 7852.659668 | 6098.532715 | 9543.397461 | 9083.981445 | 10175.75488 |
| 585821 | 333838.3125 | 365824.4063 | 233955.9844 | 290454.1563 | 552999.6875 | 236930.8438 |
| 334632.5625 | 228637.0625 | 221338.9219 | 246025.0469 | 259130.4219 | 229636.2656 | 254332.2344 |
| 89319.86719 | 61262.64063 | 61143.82031 | 61049.45313 | 60797.89844 | 55505.76563 | 54430.89453 |
| 225269.3281 | 165736.2969 | 144193.7031 | 144073.2656 | 171410.2344 | 169268.1875 | 166218.5469 |
| 55278.53125 | 37411.46484 | 34191.17969 | 26670.57227 | 29199.08984 | 35046.42969 | 25484.17773 |
| 14548.19727 | 9554.669922 | 8214 | 7838.279297 | 12853.1084 | 14204.37109 | 14495.07813 |
| 6925.043457 | 6128.691406 | 3632.068604 | 4406.01709 | 7149.225098 | 6045.391602 | 7071.404785 |
| 308076.7188 | 189150.6563 | 203047.3281 | 220361.2813 | 287050.8125 | 270248.6563 | 265690.5 |
| 39695.99609 | 36134.10156 | 19350.79883 | 19906.89648 | 38645.27344 | 39434.02734 | 34196.21094 |
| 175802.9219 | 116001.1953 | 117783.3984 | 115309.7109 | 125794.9453 | 122094.5547 | 125520.9297 |
| 245919.0439 | 153637.3193 | 164876.1265 | 157832.9121 | 152078.4805 | 139837.3218 | 157887.5615 |
| 15893.30371 | 11068.20508 | 11004.18359 | 9818.546875 | 21756.13672 | 30816.20313 | 30322.55078 |
| 231995.6094 | 201816.375 | 124242.0938 | 119652.0859 | 160751.5625 | 136433.4688 | 155253.9531 |

| | | | | | | |
|---|---|---|---|---|---|---|
| 31236.15234 | 18799.87109 | 19864.14258 | 18300.89648 | 20130.84961 | 20944.76758 | 19936.51758 |
| 120103.4844 | 85423.8125 | 81070.49219 | 75229.02344 | 98649.79688 | 100077.6016 | 104762.3203 |
| 7292.642578 | 8545.838867 | 4032.580322 | 6634.534668 | 8776.205078 | 27245.59766 | 12606.46387 |
| 74204.07031 | 55332.76953 | 51724.17188 | 44053.23438 | 38872.65625 | 39869 | 38247.52734 |
| 27404.12695 | 15475.29395 | 18648.64063 | 16181.45898 | 24169.13086 | 18071.19531 | 22389.74414 |
| 411457.5 | 213927.3281 | 315089.5625 | 216441.1094 | 172621.7188 | 248972.9531 | 175296.6719 |
| 172993.1406 | 123129.1797 | 92118.92188 | 67663.8125 | 61037.60156 | 79479.79688 | 45491.08203 |
| 1573124.5 | 1076053 | 1046606.25 | 987977.6875 | 884202.875 | 1018189.125 | 918573.5 |
| 74150.28125 | 51863.13281 | 50524.14844 | 54922.68359 | 114512.5781 | 94761.23438 | 98661.95313 |
| 259454.8906 | 155330.6875 | 160898.1563 | 167318.8125 | 182327.3906 | 188923.6875 | 195910.5625 |
| 1260.161865 | 789.1400757 | 778.2171021 | 766.7327271 | 1198.004272 | 1084.689697 | 909.5095215 |
| 1287.66687 | 1057.659058 | 703.3370972 | 698.9483032 | 2432.404785 | 2822.923584 | 1584.436768 |
| 12851.03418 | 11687.125 | 8979.740234 | 6536.817383 | 5109.209473 | 7310.043945 | 5272.666992 |
| 108115.7813 | 59017.18359 | 70981.85938 | 70293.86719 | 51717.22266 | 52676.80469 | 52158.44141 |
| 41208.38672 | 32684.64258 | 23127.01953 | 23869.72656 | 23918.79102 | 20395.14453 | 22108.43555 |
| 130460.2813 | 63263.11719 | 89810.64063 | 89693.94531 | 92616.85938 | 94951.67969 | 95586.35156 |
| 1311690.625 | 801636.375 | 849574.1875 | 842128.4375 | 838390.875 | 773310.625 | 819100.0625 |
| 11269.1582 | 10031.11328 | 6698.876465 | 4524.877441 | 12605.47168 | 8987.900391 | 10463.69531 |
| 58879.46484 | 75101.4375 | 14078.89063 | 17415.21289 | 65611.65625 | 57725.31641 | 68856.35156 |
| 10921.01465 | 12234.84375 | 5506.298828 | 3910.412354 | 12694.34863 | 9666.321289 | 13077.69336 |
| 27010.94531 | 11812.74707 | 19013.89453 | 19321.0625 | 11998.79395 | 11887.5127 | 10870.73242 |
| 26159.29688 | 13531.24902 | 15370.54199 | 17441.58203 | 19488.19336 | 23893.43945 | 21497.87109 |
| 10701.64453 | 6335.063965 | 9087.833984 | 3085.595947 | 5411.791504 | 5570.702637 | 7902.892578 |
| 3023776 | 2508040.25 | 1792127.5 | 1067288.375 | 1049861.625 | 2159921.75 | 1718691.375 |
| 33671.44141 | 19979.49805 | 19725.98438 | 16478.95313 | 27404.77344 | 25483.80273 | 24424.41797 |
| 55259.54297 | 50663.08984 | 31157.06641 | 28045.48242 | 49329.80469 | 48405.76563 | 49814.75 |
| 21723.95313 | 12608.51367 | 17056.60547 | 14377.14453 | 12833.72949 | 18888.92969 | 15421.39551 |
| 33575.32422 | 23323.51367 | 19250.95898 | 23489.78125 | 35252.74219 | 32639.20703 | 31691.19336 |
| 56625.26563 | 41634.41797 | 28622.90039 | 30864.75586 | 40411.89063 | 36243.97656 | 40546.50391 |
| 78719.1875 | 46863.87891 | 44223.25 | 53742.92188 | 34470.88281 | 45332.63281 | 40554.89063 |
| 13823.09863 | 8601.620117 | 9181.237305 | 9096.594727 | 12640.18945 | 13876.37207 | 13400.89453 |
| 10438.7207 | 8661.336914 | 7284.205566 | 7780.20166 | 18825.58203 | 16342.94727 | 15958.4248 |
| 504111.7188 | 269929.2813 | 321965.5938 | 276651.7188 | 298853.4063 | 338415.2188 | 300772.75 |

| | | | | | | |
|---|---|---|---|---|---|---|
| 2515.185303 | 2153.060791 | 1697.820801 | 1598.802124 | 2399.029785 | 1659.354248 | 1876.649048 |
| 126309.1328 | 75391.07813 | 88642.84375 | 81910.97656 | 95114.01563 | 91526.32813 | 95392.64063 |
| 11790.90918 | 10042.65625 | 6429.120117 | 6381.608887 | 4035.797607 | 5546.568848 | 4637.623535 |
| 146418.1875 | 124702.4453 | 84734.58594 | 64817.71094 | 49183.04688 | 70359.39063 | 49213.10938 |
| 58551.77344 | 41092.03125 | 40222.92969 | 32303.60352 | 44898.88672 | 52707 | 45002.01172 |
| 405149.0313 | 311792.2813 | 383015.25 | 378697.2813 | 448605.7188 | 415311.6563 | 708917.625 |
| 176974.25 | 124138.8281 | 112742.4922 | 99084.05469 | 83653.94531 | 86815.5625 | 82821.10938 |
| 345028.875 | 224764.2813 | 201906.9531 | 219630.5625 | 143613.5156 | 147918.9375 | 132138.0469 |
| 130508.7188 | 81578.29688 | 81085.90625 | 84730.19531 | 64091.49219 | 72416.71094 | 68311.97656 |
| 19543.92578 | 15351.77734 | 12421.28027 | 11047.2334 | 12362.9248 | 10949.625 | 11428.69434 |
| 180211.7813 | 65157.83984 | 111300.6641 | 94321.36719 | 132621.625 | 122132.4453 | 191763.1875 |
| 50327.14844 | 34433.92188 | 30871.11719 | 23113.48828 | 25982.19141 | 35652.04688 | 25087.06445 |
| 26399.79883 | 15719.1875 | 15078.62012 | 16724.64648 | 28854.65039 | 23997.16797 | 26251.82031 |
| 80253.34375 | 51993.58984 | 49787.83984 | 51098.13281 | 97941.61719 | 108450.1328 | 102562.5234 |
| 61602.97266 | 39277.84766 | 37969.20703 | 35392.82422 | 15437.33496 | 17448.26953 | 16225.99219 |
| 4063.737549 | 963.2142944 | 2453.548828 | 3831.466797 | 3468.322021 | 2150.173096 | 3010.095459 |
| 20548.3418 | 14829.43555 | 12820.9541 | 12965.8457 | 6518.863281 | 7307.58252 | 6111.89502 |
| 2859698 | 1580704 | 1919440.375 | 1841655.25 | 2338681 | 2132367.25 | 2283459 |
| 652169.4375 | 408730.625 | 370941.7188 | 370829.9375 | 337249.375 | 345700.25 | 310664.5938 |
| 485854.125 | 373018.3438 | 312435.6875 | 319700.1563 | 447922.75 | 374867.9063 | 401918.2188 |
| 62939.3125 | 35392.81641 | 42577.09375 | 40883.88281 | 45068.66016 | 45055 | 42619.22266 |
| 42587.89453 | 25527.28711 | 26532.375 | 23100.16992 | 17077.94922 | 14411.97754 | 11778.64355 |
| 221547.2344 | 141487.7969 | 140340.3281 | 128564.1484 | 109880.4297 | 119442.4141 | 103126.9922 |
| 229547.0313 | 151591.1875 | 155709.5313 | 146123.9688 | 166183.9063 | 166104.8281 | 172928.4063 |
| 7586.282227 | 4996.610352 | 4040.885742 | 1565.838379 | 2158.931152 | 2035.810181 | 1872.166138 |
| 66332.92188 | 32221.23438 | 45034.16406 | 35540.58984 | 30665.10938 | 47856.23047 | 32308.25195 |
| 11117.73145 | 9245.029297 | 5646.367676 | 6687.035645 | 10295.13184 | 10419.33496 | 8301.392578 |
| 244704.4688 | 121472.5938 | 190273.5938 | 137058.6094 | 51117.0625 | 76644.21094 | 64199.51953 |
| 12513.0918 | 9148.155273 | 6440.573242 | 6890.393066 | 10063.26953 | 8462.414063 | 9982.301758 |
| 17850.05469 | 13831.75586 | 9083.264648 | 8886.095703 | 16654.41797 | 5599.885254 | 11721.64941 |
| 18352.5293 | 10786.05859 | 10539.56934 | 11637.59766 | 13617.56445 | 16925.20703 | 14463.07813 |
| 64797.4375 | 41602.10547 | 40216.26953 | 35378.27344 | 42139.62891 | 42963.96875 | 43017.67969 |
| 65788.82031 | 60454.33984 | 52468.55469 | 43403.41016 | 104160.2109 | 57780.47656 | 67191.57031 |

| | | | | | | |
|---|---|---|---|---|---|---|
| 1575.747803 | 1023.785095 | 1008.79718 | 905.0951538 | 1129.158447 | 1408.247192 | 1040.278442 |
| 16979.42969 | 12063.94043 | 10603.83594 | 11679.92188 | 12891.23828 | 11038.87207 | 11683.06348 |
| 395096.875 | 285031.625 | 201231.75 | 238825.4688 | 158231.2188 | 141379.9531 | 132956.5781 |
| 77437.52344 | 39899.51172 | 55488.92188 | 51547.24609 | 54242.32813 | 59902.99219 | 57924.47656 |
| 17837.61719 | 10620.4834 | 9021.182617 | 11219.99512 | 13124.31543 | 11387.13672 | 10558.07324 |
| 98134.08594 | 65769.21094 | 59373.75 | 60033.11719 | 66646.57031 | 77016.15625 | 76576.45313 |
| 268676.2813 | 164210.4375 | 189393.0781 | 180016.8438 | 203422.6719 | 209257.2813 | 208550.7969 |
| 64280.56641 | 50306.38281 | 44897.85156 | 40505.42969 | 45210.95703 | 63515.92578 | 126371.5469 |
| 24649.96289 | 13356.58789 | 14682.76172 | 13251.91797 | 12656.24219 | 14749.56836 | 12356.91602 |
| 59907.96484 | 37085.70313 | 39047.30859 | 37949.10156 | 62008.73047 | 57038.98438 | 63692.49609 |
| 392846.0313 | 236424.8594 | 263028.0313 | 246318.2656 | 284571.8125 | 265659 | 284449.8438 |
| 7516.998535 | 6226.140137 | 2960.90332 | 5280.765137 | 6606.599609 | 6063.450195 | 5958.509277 |
| 55668.33984 | 37320.90234 | 33394.79688 | 26351.00195 | 28644.63477 | 34766.50391 | 26864.52734 |
| 430346.8125 | 324127.0625 | 311396.75 | 265184.1563 | 323343.375 | 311099.0938 | 316185.375 |
| 27879.68359 | 15764.66699 | 17588.66211 | 16000.7168 | 18546.68359 | 20866.72852 | 16619.78906 |
| 133888.1563 | 77810.4375 | 80210.25 | 85773.625 | 97152.04688 | 83203.77344 | 91333.46094 |
| 27487.71289 | 18563.01367 | 15707.39551 | 17649.41406 | 12381.14453 | 13177.59082 | 12670.05371 |
| 54705.9375 | 30645.95898 | 36673.75391 | 33844.96484 | 36644.14844 | 38368.24219 | 37784.87891 |
| 2784.594482 | 2415.584961 | 1356.179321 | 1782.299194 | 2254.458984 | 1180.814209 | 1309.588745 |
| 12320.16309 | 8974.995117 | 6224.237305 | 4981.429688 | 3400.836914 | 4064.572021 | 3305.245117 |
| 418781.4688 | 241986.2344 | 291731.625 | 302110.5625 | 402756.5313 | 391482.4375 | 400985.4375 |
| 2831.953613 | 1651.80127 | 1734.880493 | 1676.860229 | 4226.064453 | 3080.878906 | 3605.124512 |
| 53158.73047 | 28195.48242 | 39879.44141 | 25365.06055 | 33577.70703 | 28764.91797 | 18371.6875 |
| 11515.6748 | 6184.605469 | 8160.459473 | 7170.969727 | 8798.375977 | 6033.791016 | 7737.078613 |
| 520011.0508 | 400186.3789 | 349199.8203 | 352361.7148 | 564111.2969 | 504748.7383 | 507526.6016 |
| 9019.0625 | 5493.95752 | 4585.756348 | 6986.897461 | 7285.597656 | 6071.415527 | 6984.456055 |
| 3298.894043 | 1948.533691 | 2024.970093 | 2233.6604 | 2592.211914 | 1784.181152 | 2538.133057 |
| 39915.26953 | 18943.05469 | 36130.68359 | 20207.71484 | 35238.65625 | 44081.61328 | 34731.73438 |
| 196032.8906 | 147043.1875 | 118147.5234 | 96528.73438 | 49213.58984 | 67822.74219 | 57233.67188 |
| 41122.9375 | 28738.92578 | 23968.05664 | 23818.72852 | 29071.61914 | 30068.29102 | 28485.61523 |
| 440749.6875 | 278052.5625 | 272310.4375 | 236947.0469 | 390749.2188 | 378163.625 | 381316.7188 |
| 15879.76855 | 7823.509766 | 7647.348145 | 9763.132813 | 8776.669922 | 10178.4082 | 5078.751465 |
| 19889.56055 | 11906.21875 | 12807.19336 | 13781.17676 | 20778.08789 | 17434.2793 | 20023.28906 |

| | | | | | | |
|---|---|---|---|---|---|---|
| 359864.4063 | 208381.2656 | 236512.8906 | 220816.75 | 242290.6563 | 258504.9375 | 255334.0938 |
| 320079.4688 | 111433.8438 | 234437.6094 | 246307.3281 | 250105.9063 | 244681.2813 | 250093.7656 |
| 51442.22656 | 36946.375 | 32776.70703 | 30681.80273 | 28114.04102 | 27826.9375 | 24976.07617 |
| 116316.9141 | 66147.13281 | 66707.125 | 66302 | 70692.96094 | 72044.5625 | 63239.26953 |
| 41668.97656 | 28933.16406 | 25081.22461 | 25343.91211 | 24204.25977 | 21995.58008 | 27365.33203 |
| 29741.75781 | 17314.10352 | 19630.00391 | 19243.4043 | 47931.39844 | 41393.10156 | 38793.26172 |
| 206052.6719 | 134142.5 | 121843.0078 | 116690.4219 | 134555.8125 | 142729.2656 | 135806.3906 |
| 22084.875 | 14250.9043 | 15483.17676 | 10759.19238 | 11345.49805 | 19247.38672 | 11191.63281 |
| 563065.3125 | 556719.875 | 409654.6875 | 121004.4766 | 114235.6406 | 630668.875 | 99112.22656 |
| 58265.27734 | 37558.89453 | 34497.03906 | 33582.80859 | 50016.08203 | 50669.77734 | 50089.71094 |
| 12102.7168 | 8277.53418 | 6881.90625 | 7115.99707 | 5714.513672 | 5643.705566 | 4652.362305 |
| 256205.125 | 153297.9375 | 165193.5781 | 188343.4063 | 245584.5938 | 259193.1094 | 253843.5781 |
| 33751.26563 | 23637.19336 | 26444.15625 | 18472.28125 | 19600.63477 | 50880.32422 | 27485.84375 |
| 68189.90625 | 45369.24219 | 44504.27734 | 39776.29688 | 39700.47656 | 38616.01563 | 36619.64063 |
| 35247.00781 | 12396.66504 | 20313.20703 | 27022.38281 | 85033.21094 | 63654.33203 | 80598.89844 |
| 38303.13281 | 21753.90234 | 26973.57617 | 39044.61328 | 35279.98438 | 37087.95703 | 44247.125 |
| 55556.39453 | 36619.69141 | 37164.62109 | 33399.64063 | 36404.92578 | 39860.57031 | 35080.14063 |
| 91489.08594 | 43934.125 | 61249.06641 | 29896.20117 | 26785.68945 | 63203.91797 | 28381.42969 |
| 8522.891602 | 7283.695313 | 6578.547852 | 6117.305176 | 7843.819336 | 8317.587891 | 8564.55957 |
| 62421.79688 | 35550.97266 | 38408.92578 | 39775.59375 | 36654.69531 | 34893.00781 | 41561.92188 |
| 2127781.5 | 1171937.875 | 1297299.625 | 1326619.5 | 1466811.625 | 1460683.375 | 1530648.375 |
| 442116.9063 | 286939.3438 | 273323.8438 | 256874.6563 | 235671.875 | 249775.8906 | 251769.5781 |
| 196887.5938 | 134048.25 | 130899.7734 | 125356.2031 | 152741.5469 | 137884.3438 | 142133.0156 |
| 15181.84863 | 12291.5791 | 8277.641602 | 7843.800293 | 9791.563477 | 8816.4375 | 8482.987305 |
| 17172.14063 | 10257.37598 | 11982.11523 | 11765.40527 | 18719.79883 | 17914.68945 | 23579.25 |
| 18361.10547 | 17624.02344 | 13157.78906 | 15940.37598 | 24401.00977 | 18815.02539 | 22497.07617 |
| 2205.55957 | 1132.44397 | 1839.907837 | 1630.277466 | 2635.376953 | 2447.624756 | 2201.788818 |
| 11088.18848 | 5800.946777 | 5936.13623 | 6852.184082 | 4158.585449 | 4430.535156 | 4457.765625 |
| 105759.6641 | 41951.52734 | 70012.9375 | 72104.21094 | 72382.29688 | 63422.15234 | 55317.33984 |
| 45073.83984 | 28581.2168 | 30134.61133 | 20536.94336 | 14521.29688 | 16754.81055 | 14389.5459 |
| 35865.22266 | 24536.58398 | 20824.36523 | 20387.15039 | 12245.09668 | 17861.0293 | 13607.38379 |
| 82677.05469 | 38919.19922 | 48617.61328 | 46995.66797 | 49926.33203 | 52393.95703 | 56013.67578 |
| 3987.533936 | 3913.291016 | 3900.908691 | 3480.919189 | 31129.30469 | 11015.00293 | 15845.07031 |

| | | | | | | |
|---|---|---|---|---|---|---|
| 91941.875 | 52518.03516 | 58559.60938 | 58409.09766 | 68949.64844 | 65394.01563 | 66453.82813 |
| 43960.52734 | 26323.28711 | 26677.73438 | 27054.85547 | 35816.06641 | 34369.95313 | 31647.37109 |
| 3095.744141 | 3250.434326 | 2015.04126 | 2202.156738 | 1273.260498 | 2109.514648 | 1987.615234 |
| 779994.1875 | 483951.0313 | 424520.4375 | 469745.375 | 466072.5625 | 466966.4375 | 468688.125 |
| 140654.5625 | 90312.49219 | 87558.86719 | 66605.85156 | 75547.66406 | 92799.42188 | 66758.91406 |
| 212643.75 | 146996.1875 | 146748.25 | 158094.6719 | 184494.8594 | 172813.9063 | 170582.6719 |
| 148666.2656 | 104012.4609 | 138180.1563 | 173104.2031 | 568901 | 108350.3203 | 198039.9531 |
| 15001.52344 | 9462.018555 | 9709.704102 | 8881.514648 | 11703.43066 | 11636.6377 | 9947.77832 |
| 62726.86328 | 42049.64063 | 41440.55078 | 34320.24609 | 41908.62891 | 51895.30859 | 43103.33594 |
| 15704.15039 | 7957.476074 | 11556.44043 | 9515.714844 | 12474.49316 | 8592.606445 | 8366.506836 |
| 288468.5625 | 154999.9375 | 162336.4531 | 146455.9219 | 159071.9688 | 183070.9844 | 160018.4219 |
| 41155.27734 | 22861.57031 | 25984.12109 | 25183.32617 | 34373.69141 | 30537.12891 | 33906.26563 |
| 196530.4844 | 100108.8281 | 141608.1094 | 133058.2344 | 138983.375 | 141614.8906 | 156542 |
| 9858.251953 | 4071.734863 | 6489.064453 | 7171.875 | 9322.56543 | 8602.749023 | 7653.029785 |
| 106782.3594 | 66799.25781 | 65932.82813 | 59150.19922 | 50043.625 | 47334.36719 | 45483.51953 |
| 805894.75 | 635503.5625 | 414867.3438 | 394642.4375 | 689353.8125 | 627353.8125 | 685303.875 |
| 208989.4219 | 114945.9688 | 129213.3984 | 135103.125 | 131699.2188 | 123790.1094 | 138130.7188 |
| 22537.37695 | 12410.09277 | 13623.01758 | 12668.39844 | 12315.50195 | 13138.57324 | 12838.5957 |
| 13887.54199 | 8566.512695 | 8593.931641 | 8707.067383 | 17391.01367 | 13686.68164 | 15929.10449 |
| 6972.822266 | 3799.039063 | 5581.026367 | 3842.62915 | 4442.532715 | 6519.506348 | 5952.374023 |
| 4797.967773 | 3139.104736 | 2658.681885 | 3577.695557 | 4537.891113 | 4772.959473 | 4857.592285 |
| 13807.51855 | 9455.571289 | 8706.366211 | 9867.491211 | 21727.42969 | 19410.55859 | 23649.72852 |
| 15985.08789 | 9612.333008 | 10400.39453 | 11820.39551 | 15683.12305 | 14076.19531 | 17030.70313 |
| 49250.92578 | 36532.74219 | 32507.80859 | 28509.05664 | 54558.19141 | 82564.28125 | 81445.96875 |
| 297169.4688 | 190319.1719 | 179796.5469 | 170647.5469 | 199181.6406 | 202440.5156 | 192749.6563 |
| 38094.65625 | 28263.125 | 24157.72461 | 25835.54688 | 18103.92188 | 20919.66016 | 17096.0918 |
| 93240.14063 | 55657.65625 | 57645.51172 | 64564.71875 | 105643.2734 | 123927.4453 | 116896.9688 |
| 7162.04541 | 3541.428223 | 3419.916992 | 3769.828613 | 3708.279053 | 3130.739014 | 3058.039795 |
| 99983.57422 | 69951.49609 | 64179.49219 | 71752.37109 | 133417.5898 | 100897.9395 | 105422.1953 |
| 330210.7813 | 190743.1563 | 204271.8594 | 192846.0625 | 195812.0781 | 205961.0313 | 207957.8125 |
| 24843.75391 | 16380.90527 | 18543.26172 | 15460.88281 | 17742.35742 | 18301.63477 | 14886.14258 |
| 698069.625 | 463947.6563 | 369036.1875 | 465567.7188 | 494598.5938 | 419325.0938 | 464637.5313 |
| 26864.11133 | 16426.66992 | 15608.53906 | 16315.28027 | 17073.9043 | 12649.4873 | 15403.68945 |

| | | | | | | |
|---|---|---|---|---|---|---|
| 59559.74219 | 36010.94141 | 36525.58594 | 33238.41406 | 38848.24219 | 47046.02344 | 42120.76172 |
| 2088.730225 | 2224.395752 | 735.5831299 | 2091.512695 | 2016.507568 | 2320.888428 | 1695.940063 |
| 14196.13867 | 10355.77734 | 8898.825195 | 11003.09668 | 13138.20996 | 12951.25 | 13011.82324 |
| 11582.97363 | 10733.87793 | 8126.153809 | 10235.41797 | 12232.80566 | 22752.58594 | 13096.64453 |
| 12474.84473 | 6480.155762 | 6456.57373 | 6247.662598 | 10749.27246 | 16377.0498 | 15825.47949 |
| 14818.64453 | 13072.57422 | 7324.422852 | 9906.75 | 8905.385742 | 4510.889648 | 7043.68457 |
| 90617.99219 | 53224.46094 | 52684.8125 | 52206.78516 | 47977.91797 | 55667.82813 | 50986.30078 |
| 24609.88477 | 30938.27344 | 14990.2627 | 24989.14844 | 16552.22656 | 36103.10156 | 24463.86719 |
| 178060.2031 | 109480.4531 | 113369.2578 | 107829.75 | 123864.7656 | 122013.9375 | 125012.0313 |
| 16114.68066 | 12041.27344 | 8629.55542 | 8502.738281 | 17909.49829 | 17827.73291 | 19038.56044 |
| 137019.3125 | 81727.17188 | 82590.73438 | 83672.32813 | 80673.63281 | 86351.51563 | 90260.60938 |
| 33655.32422 | 20139.26953 | 23781.37109 | 22736.36523 | 28540.67188 | 25138.2207 | 26329.42188 |
| 112091.4297 | 58711.09766 | 75466.92969 | 71328.35938 | 54639.58594 | 57003.65625 | 56189.61719 |
| 10634.55664 | 6136.984375 | 6677.768555 | 7974.008789 | 7194.445801 | 6097.440918 | 5271.462891 |
| 89419.83594 | 61413.26953 | 49493.70703 | 50147.01172 | 101680.2969 | 98608.89063 | 94574.57813 |
| 29496.42969 | 21127.33984 | 18359.17383 | 17294.22852 | 20550.12305 | 19898.72461 | 19998.84375 |
| 14482.96582 | 8088.962891 | 7505.178223 | 6686.405762 | 4943.824219 | 6110.166992 | 6243.816406 |
| 16719.54492 | 11480.69824 | 9176.901367 | 8860.613281 | 5200.375488 | 6088.901367 | 4618.649902 |
| 26501.51563 | 14596.8125 | 16491.89453 | 15360.9043 | 17109.88867 | 17693.6543 | 18709.70117 |
| 88129.39844 | 52951.75781 | 52579.99609 | 61695.83203 | 88970.35156 | 87504.02344 | 94879.75 |
| 492217.875 | 252609.8125 | 307945.4688 | 336774.8438 | 447558.2813 | 444157.4063 | 482573.5 |
| 1116623.875 | 700298.75 | 697205.875 | 723321.8125 | 842646.625 | 815902.375 | 875586.5 |
| 73253.98438 | 41818.01563 | 45316.74609 | 48849.14063 | 40346.92578 | 37835.83594 | 39245.19531 |
| 377897.5 | 234724.8594 | 241033.625 | 219397.1875 | 234161.8906 | 236634.3281 | 238276.2344 |
| 19469.54102 | 10160.10254 | 13808.65625 | 12593.7002 | 14939.75391 | 14913.35645 | 13570.33887 |
| 63280.01172 | 50983.91406 | 36530.22266 | 30891.7168 | 59907.10938 | 55131.69531 | 65221.00391 |
| 14387.32422 | 9559.913086 | 7978.966309 | 9045.083984 | 12290.84961 | 11221.52051 | 9535.200195 |
| 23095.83594 | 12268.92285 | 15000.93945 | 11918.75391 | 12704.29492 | 13445.40918 | 10681.07617 |
| 97270.61914 | 48453.10547 | 40157.63721 | 75121.58301 | 228684.9492 | 129532.4844 | 143020.0215 |
| 4373.711426 | 676.100769 | 4463.382813 | 3427.896973 | 13255.21875 | 9946.021484 | 8725.03418 |
| 105592.5859 | 60080.16406 | 67484.875 | 70478.52344 | 68365.35938 | 68329.8125 | 70349.71094 |
| 85956.99219 | 48032.70313 | 52991.95703 | 60321.38281 | 86499.55469 | 64226 | 73176.03906 |
| 164480.8906 | 94956.17188 | 99360.39063 | 96342.65625 | 119115.9844 | 129831.6172 | 124150.2969 |

| | | | | | | |
|---|---|---|---|---|---|---|
| 42065.96484 | 22740.20117 | 28190.11523 | 28993.85938 | 29909.91211 | 43510.77344 | 34751.89453 |
| 15027.5459 | 9721.867188 | 9851.170898 | 9576.450195 | 8984.049805 | 9773.400391 | 10228.3418 |
| 3753.810547 | 1698.012817 | 3139.989014 | 2071.76709 | 2948.543945 | 2376.895508 | 2744.272217 |
| 15992.80078 | 12730.4375 | 13023.86523 | 11316.61035 | 17998.77539 | 20259.30273 | 20188.84961 |
| 12070.45313 | 5944.822266 | 7025.192383 | 8250.001953 | 8349.461914 | 11174.30664 | 12071.14453 |
| 85843.54688 | 51723.66797 | 49724.66406 | 44011.67578 | 21387.11523 | 31070.17188 | 21952.64258 |
| 350209.1563 | 177253.5938 | 204910.6875 | 198020.7656 | 259430.3125 | 233193.4375 | 229101.4531 |
| 24963.17969 | 10723.39844 | 14665.22949 | 13880.35352 | 26663.57422 | 14079.07031 | 17799.98438 |
| 112436.0156 | 68423.39063 | 69608.28125 | 59086.09375 | 52812.19922 | 59182.61328 | 53265.14844 |
| 20258.97656 | 13860.39063 | 13148.74414 | 14107.28809 | 13800.74512 | 13164.90234 | 14001.93262 |
| 178214.7813 | 99877.53125 | 99487.40625 | 102099.2109 | 148259.9375 | 139186.5313 | 147017 |
| 30221.64063 | 23670.60938 | 22137.19727 | 21327.26563 | 36875.63672 | 54239.91016 | 36450.89844 |
| 231655.4531 | 148398.1563 | 148486.0781 | 141899.5 | 114872.1875 | 100918.7344 | 105790.4922 |
| 114626.7969 | 46268.07813 | 98006.67969 | 107044.1172 | 144760.2188 | 140835.4219 | 145863.9688 |
| 2270.055664 | 939.5684814 | NA | 1388.634277 | 6922.317871 | 2771.658691 | 3472.645508 |
| 4941.408691 | 5331.949219 | 2724.505615 | 2862.092773 | 5346.867188 | 5723.071777 | 5287.86377 |
| 1508677.625 | 903634.625 | 933704.25 | 903873.875 | 1334873.375 | 1391928.875 | 1509663.5 |
| 142723.4688 | 78087.55469 | 82739.27344 | 84748.44531 | 196864.8281 | 185450.375 | 240357.1563 |
| 866259.375 | 535224.5 | 589790.625 | 566678.75 | 603980.4375 | 609478.25 | 610787.125 |
| 28360.36328 | 15023.55469 | 15285.84375 | 15638.62891 | 31260.54688 | 34420.02734 | 32419.77344 |
| 111643.9531 | 65422.28125 | 60747.75391 | 54173.30859 | 34066.39453 | 34285.13281 | 106183.5859 |
| 542575.1719 | 312116.0859 | 309127.7031 | 307545.8125 | 269584.7109 | 292961.25 | 266892.3281 |
| 134297.2813 | 78167.20313 | 81296.53125 | 86289.53125 | 128995.1406 | 126958.2344 | 128077.2969 |
| 311937.5313 | 171757.375 | 198476.1719 | 142909.1406 | 139218.7344 | 228136.7188 | 152497.4219 |
| 875479.5625 | 461914.75 | 525162.5625 | 600664.6875 | 1229738.625 | 763672.4375 | 1091580 |
| 15378.16211 | 10122.96094 | 8878.047852 | 10226.87402 | 10424.28809 | 8509.270508 | 9165.358398 |
| 39989.15625 | 19145.43359 | 24211.11719 | 28085.09375 | 28439.04492 | 34483.46094 | 32644.4707 |
| 21777.11133 | 20377.64063 | 10183.98438 | 10616.41016 | 26341.8457 | 25719.625 | 28560.74414 |
| 12983.18945 | 7459.331055 | 7743.444824 | 10649.36426 | 26664.25391 | 25787.51563 | 24609.68359 |
| 159292.9063 | 28606.89453 | 103895.1797 | 122065.1953 | 185235.8281 | 79899.77344 | 183665.9531 |
| 12420.08496 | 10208.46094 | 6820.247559 | 6812.577148 | 4218.640137 | 4951.307617 | 3795.003174 |
| 123654.75 | 71305.25781 | 69997.76563 | 66483.46875 | 60838.01172 | 69178.61719 | 63012.05469 |
| 25584.49023 | 15343.98145 | 16386.25977 | 14352.49414 | 17170.73828 | 18223.7832 | 17879.36133 |

| | | | | | | |
|---|---|---|---|---|---|---|
| 176846.0469 | 101831.4063 | 105442.2891 | 114708.5156 | 108464.0703 | 107569.2656 | 106948.3203 |
| 12651.85352 | 6602.270996 | 8258.032227 | 7595.629395 | 6424.985352 | 6278.168945 | 6184.45459 |
| 172349.125 | 99567.58594 | 100861.9141 | 113728.4922 | 53860.45313 | 53001.78906 | 58894.03906 |
| 813938.875 | 518447.7188 | 453773.625 | 492942.7188 | 505304.0313 | 487869.1875 | 495274.0938 |
| 24220.33203 | 12892.32715 | 14896.30371 | 18298.98633 | 12016.19531 | 11227.39453 | 12526.31348 |
| 360152.1563 | 236288.2813 | 188412.2969 | 239299.0781 | 249485.7031 | 216865.2031 | 222568.7813 |
| 32032.23242 | 15585.85938 | 18972.46289 | 17369.29883 | 19969.90234 | 22355.26172 | 22366.57227 |
| 21824.04688 | 16702.25977 | 14855.05566 | 15034.20898 | 17129.98438 | 16611.86719 | 15051.03613 |
| 25054.54102 | 14456.3252 | 16326.6123 | 14745.71191 | 13598.89258 | 12569.68164 | 13165.2002 |
| 15433.50293 | 12871.2168 | 6830.505371 | 6668.31543 | 4492.743652 | 3223.751953 | 2342.437012 |
| 31512.91602 | 18797.62891 | 19824.63672 | 20876.98242 | 25382.90039 | 28590.24609 | 24241.19531 |
| 190101.9063 | 106411.2188 | 116334.5234 | 100046.8359 | 94664.6875 | 121506.5938 | 97329.46094 |
| 7865.802246 | 5982.029297 | 5198.07959 | 4097.307617 | 2589.306885 | 1557.040649 | 4231.203125 |
| 27669.12695 | 10288.27734 | 14333.79688 | 17397.78906 | 17346.74219 | 18042.74414 | 28586.17383 |
| 444125.375 | 254045.2188 | 257582.4063 | 306246.6875 | 278890.75 | 238814.9844 | 266822.0313 |
| 114508.1172 | 61250.25391 | 72241.34375 | 65696.94531 | 73731.60156 | 83638.55469 | 78716.51563 |
| 32940.9375 | 26237.20508 | 15912.93848 | 13697.66016 | 35123.79688 | 30289.96094 | 34240.49219 |
| 283879.4375 | 177667.5625 | 154986 | 162247.5313 | 163388.2969 | 152829.7813 | 160473.2969 |
| 612528.6875 | 341062.9063 | 354663.7188 | 361414.6875 | 443862.7188 | 433409.7813 | 474413.9375 |
| 82767.83594 | 56597.35156 | 41790.41406 | 40853.42188 | 43671.84375 | 47922.34766 | 37751.41797 |
| 115326.5938 | 76232.29688 | 84828.71094 | 79128.92188 | 62769.47656 | 113712.4922 | 97024.27344 |
| 10956.53418 | 8115.101074 | 5140.583008 | 3821.322266 | 5634.97998 | 4881.863281 | 3721.063477 |
| 68130.05469 | 59170.21094 | 36471.13672 | 43849.58984 | 43815.4375 | 46062.92969 | 66724.63281 |
| 8259.384766 | 4961.362793 | 5994.919434 | 5252.483398 | 5414.702148 | 5948.23584 | 4691.116211 |
| 63504.82031 | 50242.93994 | 30053.20142 | 37694.63672 | 52190.44727 | 53544.56152 | 58568.21289 |
| 122034.1016 | 70248.125 | 67251.8125 | 68561.4375 | 64640.54688 | 67699.79688 | 62871.30859 |
| 27571.61133 | 14333.04297 | 17131.8418 | 18576.02148 | 15657.37402 | 13920.92383 | 14109.21875 |
| 22350.5918 | 10798.86719 | 17929.05664 | 12576.40527 | 15983.46777 | 24900.63281 | 16977.48047 |
| 9982.771484 | 6403.132813 | 5863.92334 | 5483.631348 | 7049.814941 | 6116.181152 | 5842.517578 |
| 30422.03516 | 18275.82227 | 17953.53906 | 20437.05859 | 32024.55664 | 28316.62109 | 27379.96094 |
| 193018.5313 | 101044.2422 | 121875.875 | 90600.11719 | 93557.5 | 140613.3906 | 92408.17969 |
| 21787.75586 | 19288.20117 | 16111.44043 | 20152.90039 | 47133.30469 | 16086.43945 | 47123.39063 |
| 69304.79688 | 38898.88672 | 36934.65234 | 33654.25781 | 26818.05664 | 25702.50977 | 20620.74023 |

| | | | | | | |
|---|---|---|---|---|---|---|
| 854902.625 | 478937.8438 | 546136.6875 | 526502.625 | 457046.5938 | 503290.0625 | 490738.5313 |
| 12894.58203 | 7971.539551 | 8372.030273 | 9180.162109 | 9218.714844 | 9376.139648 | 9053.249023 |
| 6032.383301 | 2918.777344 | 3456.73584 | 3685.952881 | 6915.592285 | 5599.375 | 4990.650391 |
| 52608.63281 | 28974.93359 | 39224.90625 | 39320.91797 | 37563.47266 | 40382.25391 | 43483.26172 |
| 53254.08203 | 30033.71094 | 31740.03711 | 28911.16016 | 30625.45313 | 32483.76172 | 31804.88867 |
| 52019.47266 | 33336.26953 | 26745.08984 | 19650.51758 | 31974.14453 | 51836.83984 | 40496.57422 |
| 3217.302246 | 1007.58136 | 3244.07666 | 3069.883789 | 4267.693848 | 4756.714355 | 5564.855957 |
| 1958314.625 | 1056091.75 | 1232128 | 1419133.125 | 2229821 | 1468591.375 | 1914097.875 |
| 3468.262939 | 3695.006592 | 1221.301392 | 1066.833496 | 5877.454102 | 3414.888184 | 5536.450195 |
| 16123.92188 | 16287.47852 | 15048.32617 | 13857.69727 | 17163.44141 | 10611.31445 | 11573.85645 |
| 119041 | 133067.1094 | 95576.80469 | 25527.14258 | 25405.83789 | 169378.2188 | 22855.37305 |
| 401636.5 | 296349.5938 | 194634.6094 | 196202.0625 | 293903.4375 | 285693.0625 | 314815.9375 |
| 37928.78516 | 22465.32422 | 20443.6543 | 19184.30859 | 30364.85742 | 32735.23828 | 32856.11328 |
| 33003.55859 | 13707.57617 | 19526.17578 | 27005.81055 | 31873.59961 | 31862.875 | 33874.12891 |
| 7397.845703 | 5847.9375 | 4137.368164 | 3905.975098 | 6822.890137 | 5016.02832 | 4692.345215 |
| 91167.51563 | 52432.54297 | 54716.59766 | 48813.10547 | 53399.14844 | 54058.875 | 55791.25 |
| 196631.6094 | 105758.6719 | 119877.2813 | 126094.6484 | 149691.9219 | 151192.9063 | 151469.6406 |
| 109687.75 | 65883.25 | 72809.83594 | 72085.83594 | 134679.2188 | 147584.6563 | 141564.0625 |
| 28431.25 | 17027.46875 | 15535.59082 | 11952.74316 | 7515.055176 | 8821.494141 | 9723.825195 |
| 184067.3125 | 71671.16406 | 103739.0859 | 108887.7266 | 111428.4453 | 113431.5781 | 120492.6641 |
| 10432.75293 | 6617.850586 | 5269.803711 | 6047.9375 | 6345.056641 | 5887.856934 | 6188.987793 |
| 103799.9453 | 65778.82031 | 63161.35547 | 59130.76953 | 56620.73047 | 72417.97656 | 63988.83984 |
| 2400.473633 | NA | 1267.37915 | 1270.289307 | NA | 1107.961426 | 840.3285522 |
| 14553.60449 | 10268.89258 | 9165.777344 | 10350.87598 | 9733.432617 | 10395.92969 | 14081.1416 |
| 85860.64063 | 50607.46484 | 49308.05469 | 52788.14063 | 60245.36719 | 56934.16406 | 54764.59375 |
| 80881.19531 | 53331.05078 | 48029.89453 | 44052.75781 | 53432.82422 | 53431.60547 | 58572.17578 |
| 85071.59375 | 52253.47656 | 44672.74609 | 41886.51563 | 42615.40625 | 59845.84766 | 42489.5625 |
| 168958.7188 | 119615.3906 | 94511.32031 | 86935.55469 | 146973.8438 | 153749.375 | 165873.6719 |
| 2856249.75 | 1300094.5 | 1666535.125 | 1761932.75 | 2001643.25 | 1649265.25 | 2077292.5 |
| 1530772.25 | 930668 | 955509.5625 | 847815.75 | 1027214.875 | 1016873.25 | 1052155 |
| 6528.312012 | 3410.454346 | 3622.14209 | 2782.762939 | 1834.572266 | 2584.907471 | 1221.940186 |
| 75591.97656 | 46028.80859 | 46448.46875 | 43030.69141 | 45261.55859 | 47824.86719 | 46491.28906 |
| 6079859.5 | 3205347.75 | 3680548.25 | 3798816.5 | 3919991.75 | 4034642.25 | 4004358.5 |

| | | | | | | |
|---|---|---|---|---|---|---|
| 19647.59375 | 21537.10156 | 6768.160645 | 14975.08301 | 22869.59766 | 40890.26563 | 29381.92383 |
| 54307.89063 | 30706.63867 | 30333.96289 | 29267.93945 | 32838.21484 | 29962.19922 | 31092.50195 |
| 782436.4375 | 452174 | 469886.0625 | 505386.8438 | 447408.0625 | 401981.875 | 422691.9688 |
| 38722.79688 | 28479.4375 | 23428.56055 | 21241.83203 | 16833.79688 | 14217.97363 | 12843.61523 |
| 2311.355225 | 2065.95166 | 1719.744629 | 1486.063721 | 1720.92627 | 1619.429199 | 1355.885742 |
| 3889.095947 | 2779.837891 | 1646.690308 | 2465.104248 | 3101.047119 | 3705.878174 | 3668.18335 |
| 181772.5 | 110027.625 | 117150.2188 | 135487.3594 | 149331.0156 | 147162.5625 | 180665.6563 |
| 24659.11914 | 20009.88672 | 13808.51465 | 11457.01953 | 25852.7832 | 20250.32617 | 22311.24023 |
| 192283.0313 | 113701.5859 | 104573.9219 | 104112.4609 | 93791.16406 | 110372.6641 | 92185.49219 |
| 52589.70313 | 32008.4375 | 31447.6875 | 29708.11328 | 53496.51953 | 49340.48047 | 51611.65625 |
| 123458.4219 | 64601.19531 | 87850.71875 | 70105.46094 | 91427.48438 | 99566.83594 | 110365.6094 |
| 40539.73438 | 28338.18359 | 26497.41016 | 32063.83008 | 56866.44531 | 22069.39648 | 27499.03516 |
| 82462 | 40363.73438 | 47983.76953 | 43103.35547 | 32463 | 36491.75781 | 37922.22656 |
| 53922.52344 | 24488.87891 | 31287.10938 | 34545.95313 | 35143.45703 | 33128.54688 | 33649.70313 |
| 9823.59375 | 6565.083008 | 5658.464844 | 6370.874023 | 6117.481934 | 10620.04492 | 9218.499023 |
| 32805.79688 | 22639.81641 | 15881.31055 | 16521.05664 | 21692.50781 | 19723.98047 | 21762.97852 |
| 122928.6641 | 73452.28125 | 79425.47656 | 66423.92188 | 58319.64453 | 69278.05469 | 67895.39063 |
| 91419.78125 | 53412.51953 | 56748.91406 | 60066.53516 | 71778.09375 | 80551.08594 | 74966.26563 |
| 17993.30273 | 10405.13281 | 10349.46973 | 9626.759766 | 11870.64551 | 12185.54883 | 12015.64355 |
| 136431.5313 | 78150.17969 | 79323.96875 | 78832.08594 | 68611.04688 | 66394.21094 | 74912.28906 |
| 26232.39648 | 17744.96289 | 11595.78711 | 18495.40625 | 28662.79102 | 23121.41992 | 18030.13867 |
| 10940.86328 | 11020.24316 | 5279.103516 | 4416.129395 | 13497.65918 | 13020.45801 | 12929.50488 |
| 549476.625 | 285212.1875 | 340818.5625 | 337025.875 | 264193.8125 | 254174.3438 | 273673.0625 |
| 2003840.625 | 1304506.25 | 1096687.25 | 1141209.25 | 1285553.875 | 1220262.25 | 1265518.5 |
| 6088.903809 | 3403.94873 | 3358.554199 | 3993.304443 | 5419.936523 | 5325.308105 | 5253.912598 |
| 43899.43555 | 31328.57617 | 25905.52344 | 24206.12549 | 18666.8147 | 18855.91577 | 17305.42969 |
| 27244.99219 | 16014.82129 | 14807.0166 | 13165.12598 | 16418.11914 | 18366.54688 | 19042.07227 |
| 14781.38281 | 10704.82617 | 8378.799805 | 7395.07373 | 6548.315918 | 7696.667969 | 6243.414551 |
| 95701.58594 | 54262.07813 | 55018.45313 | 52833.42969 | 49514.54688 | 47761.23828 | 51271.08594 |
| 5039.841309 | NA | 3052.160645 | 2983.581055 | 16177.92383 | 14560.73535 | 11916.83301 |
| 382520.2188 | 225763.7656 | 243685.8125 | 255868.8281 | 250221.4063 | 256544.0469 | 264471.5938 |
| 226677.8594 | 129992.8359 | 131975.9063 | 132711.4531 | 177845.2344 | 176154.5469 | 174985.5938 |
| 910304.6411 | 556738.1147 | 529674.0896 | 552424.3259 | 499554.9521 | 663481.6782 | 623332.5615 |

| | | | | | | |
|---|---|---|---|---|---|---|
| 162306.1563 | 93378.57813 | 80989.03125 | 71061.22656 | 56832.90625 | 57866.90234 | 55725.99219 |
| 15022.33887 | 7996.862305 | 8818.660156 | 7093.713379 | 10230.75977 | 8625.079102 | 7665.735352 |
| 42540.39063 | 12337.79199 | 20111.85742 | 37741.84766 | 71636.45313 | 78031.25 | 99235.63281 |
| 20336.88672 | 10930.80957 | 12131.13672 | 10868.94043 | 9596.550781 | 12197.40625 | 12083.34277 |
| 188256.625 | 111232.7109 | 105020.9063 | 114311.8125 | 118071.3594 | 112358.7344 | 118136.1641 |
| 15197.30762 | 9891.085938 | 8303.423828 | 7408.67334 | 9046.509766 | 9947.121094 | 9459.985352 |
| 44168.35156 | 29368.35938 | 20711.26953 | 22754.51758 | 36245.85938 | 34890.60938 | 35514.73828 |
| 37326.14453 | 23532.41211 | 22619.875 | 20480.97852 | 17755.97656 | 19340.35352 | 17068.75586 |
| 285320.6875 | 161867.8906 | 164917.7813 | 186488.2969 | 433100.7188 | 352581.5313 | 402014.8125 |
| 15127.63379 | 6618.661621 | 10436.30762 | 10202.42285 | 11976.93652 | 10326.89941 | 10323.60742 |
| 95458.46094 | 57777.62891 | 55521.18359 | 46938.21875 | 60394.78906 | 62572.48828 | 60314.59375 |
| 6525.498535 | 4181.080078 | 3159.02002 | 5670.170898 | 28692.67773 | 10896.73438 | 15342.78906 |
| 121645.5078 | 55329.27344 | 73513.67188 | 74978.71094 | 74944.75781 | 84326.74219 | 90042.875 |
| 8383.825195 | 4226.378906 | 5287.28418 | 6015.589844 | 5995.700684 | 5135.995117 | 4745.835938 |
| 241217.4688 | 143961.5469 | 126295.4531 | 130021.3906 | 154167.4375 | 149890.2969 | 142810.6406 |
| 406687.75 | 225421.4375 | 246262.3594 | 228381.5938 | 232177.0625 | 212884.625 | 213883.3125 |
| 117830.5625 | 70417.92969 | 69280.03125 | 61775.6875 | 66128.35156 | 66967.67188 | 62450.97656 |
| 79583.57031 | 46222.30078 | 39459.35156 | 42355.12891 | 37817.35156 | 35598.05859 | 35645.84766 |
| 19864.36328 | 11513.85547 | 13696.31641 | 8826.365234 | 17739.06055 | 14965.36621 | 20389.27734 |
| 2969524.25 | 1732769.625 | 1657821.125 | 1755140.125 | 1999423.875 | 1840204.625 | 2035308.25 |
| 3839.161133 | 3731.100586 | 2061.472656 | 1603.445557 | 1738.796021 | 1887.465942 | 1219.178711 |
| 27964.52148 | 9398.754883 | 20579.79102 | 9561.428711 | 8733.959961 | 24287.02148 | 9257.472656 |
| 48639.44922 | 15980.9873 | 30233.52734 | 45996.09766 | 72714.72656 | 19691.41406 | 70555.42188 |
| 5125.160645 | 1889.325317 | 3584.930176 | 3209.904053 | 1904.250854 | 3196.383301 | 2752.812012 |
| 74238.39844 | 60267.51563 | 38424.25 | 36300.57813 | 79260.95313 | 67493.89844 | 51450.87891 |
| 38563.17578 | 20681.25 | 18798.36133 | 17828.80469 | 12652.11133 | 12137.58984 | 10901.75293 |
| 1218578.875 | 639638.875 | 641116.875 | 913104.1875 | 2105575.25 | 1392416.625 | 1803188.75 |
| 128060.7344 | 73895 | 77145.625 | 74220.02344 | 92592.67969 | 93276.80469 | 92690.66406 |
| 40780.79688 | 26827.55859 | 26922.9082 | 23229.25977 | 33447.84375 | 29001.82813 | 34652.89453 |
| 78898.53906 | 42693.51953 | 42176.51172 | 39392.53906 | 41407.75781 | 48911.91406 | 39673.875 |
| 48015.03906 | 23251.27539 | 30734.49609 | 16561.55859 | 18564.21875 | 30966.72656 | 18168.05664 |
| 62063.26563 | 34586.63672 | 36622.86719 | 37171.64844 | 43308.9375 | 47316.57422 | 46276.09375 |
| 56286.64063 | 31651.33789 | 36985.51172 | 41181.4375 | 49257.66406 | 38449.03906 | 46688.47656 |

| | | | | | | |
|---|---|---|---|---|---|---|
| 390745.6875 | 237774.0938 | 214683.75 | 207298.4531 | 219734.7813 | 201875.3281 | 204065.5938 |
| 8588.535156 | 5420.768066 | 5363.871582 | 4382.111328 | 7300.717773 | 6180.625977 | 6431.266113 |
| 178149.2656 | 62304.79297 | 106241.2578 | 86711.71875 | 86134.60156 | 153087.9531 | 147946.9531 |
| 730859.5625 | 401645.375 | 434687.2188 | 444210.9375 | 497094.1875 | 498349.3125 | 514218.1875 |
| 26749.43555 | 16536.20898 | 13625.74121 | 14058.83398 | 15227.7959 | 11153.52441 | 11424.66504 |
| 224723.0938 | 104369.1719 | 135997.4375 | 138700.7188 | 163502.1406 | 150005.8281 | 175273.0156 |
| 15976.1123 | 7978.717773 | 10192.05273 | 11193.90332 | 16576.95508 | 14979.86035 | 18959.38867 |
| 39465.28125 | 20297.8457 | 21841.85742 | 29876.05664 | 31358.70313 | 30776.36719 | 32473.23242 |
| 788842 | 438809.7188 | 464611.6875 | 472450.25 | 584919.5 | 568619.3125 | 608553.0625 |
| 26190.26367 | 14426.58496 | 13994.51172 | 16324.50977 | 24506.45508 | 24279.06445 | 25440.3125 |
| 6773.616699 | 4893.205078 | NA | NA | 2304.438232 | 4665.288574 | 5650.48291 |
| 16117.4502 | NA | 12268.89258 | NA | 30701.75391 | 8009.614746 | 66153.04688 |
| 5924.937988 | 3770.711182 | 3455.277588 | 4502.450684 | 3113.817383 | 2289.821289 | 2628.182129 |
| 14351.33691 | 6254.643066 | 8460.283203 | 8025.991211 | 6607.251953 | 9222.592773 | 8402.012695 |
| 7982.66748 | 7017.112793 | 3064.994629 | 2224.319824 | 17073.56641 | 9377.166992 | 6505.330078 |
| 38833.54297 | 20468.24609 | 21650.84961 | 22606.76953 | 25087.69531 | 27702.50977 | 26329.11523 |
| 182486.8594 | 101232.5859 | 106768.6328 | 109359.4141 | 123599.1797 | 145463.4688 | 127734.3672 |
| 39751.01172 | 22064.30664 | 20388.18359 | 19245.88867 | 19606.53906 | 19189.62695 | 19003.15234 |
| 38561.44531 | 20114.06445 | 23334.38281 | 22704.20703 | 30590.875 | 27218.89453 | 30494.28711 |
| 13682.70898 | 5829.308594 | 7287.339355 | 13125.57129 | 10824.28906 | 20549.82031 | 17118.19922 |
| 156906.2656 | 68491.48438 | 90024.67188 | 91075.42188 | 77851.20313 | 81325.17969 | 81788.70313 |
| 1175579 | 661533.125 | 683194.4375 | 660817.9375 | 780045.875 | 855400.25 | 835875.375 |
| 55592.79297 | 28802.99414 | 30333.08008 | 28435.58789 | 33678.125 | 31981.25 | 31335.53711 |
| 12394.00781 | 11260.02539 | 5574.449219 | 9126.084961 | 7892.182129 | 16449.00977 | 12188.34473 |
| 458105.5938 | 206137.1094 | 335514.5938 | 378395.6875 | 265609.7813 | 270613.8438 | 272980.0625 |
| 60935.24219 | 32489.73633 | 31945.20117 | 32859.89844 | 38864.49219 | 39641.22656 | 37825.10156 |
| 205195.2969 | 115368.9766 | 110361.0469 | 114128.2656 | 119105.1172 | 124791.9766 | 119044.3125 |
| 33408.19531 | 18741.30469 | 19957.55273 | 20031.57031 | 24319.80273 | 24456.33984 | 22432.9668 |
| 44960.66797 | 27437.0918 | 24883.59961 | 25136.92969 | 48463.54297 | 50581.82813 | 49391.50781 |
| 34501.22266 | 18618.91797 | 19335.96484 | 19636.70313 | 25032.88672 | 20827.12109 | 24164.46289 |
| 71065.96875 | 37784.67188 | 41296.39063 | 29361.07031 | 27613.18945 | 43872.99609 | 24725.41797 |
| 173250.8125 | 92881.40625 | 98103.00781 | 95332.08594 | 91396.96875 | 93032.32813 | 96543.36719 |
| 1169981.875 | 829017.5 | 575351.25 | 482043.1875 | 1103249.25 | 1135204.125 | 1218461.75 |

| | | | | | | |
|---|---|---|---|---|---|---|
| 26632.85938 | 12938.47266 | 13387.75391 | 14920.95313 | 15166.47656 | 15046.05859 | 14284.42285 |
| 746773.375 | 522787.4688 | 487010.875 | 434998.5625 | 352655.9063 | 362249.9063 | 366748.125 |
| 133439.2188 | 75425.60938 | 75005.58594 | 84767.92188 | 179383.25 | 147987.7969 | 164405.3125 |
| 103808.6797 | 58968.77734 | 55952.59766 | 55532.40625 | 43634.77344 | 42082.63281 | 39834.91016 |
| 50974.97266 | 29808.23828 | 25696.16406 | 26151.96484 | 32200.26563 | 30460.29297 | 34938.51953 |
| 250742.5313 | 117748.0547 | 143614.875 | 133307.4063 | 106631.2813 | 104790.7109 | 111508.9063 |
| 11913.29297 | 7659.401367 | 6030.418945 | 6302.624023 | 6535.662109 | 6816.449707 | 7686.61084 |
| 43978.71875 | 21157.4043 | 24682.96289 | 24125.83398 | 21652.8125 | 22916.29688 | 25285.51953 |
| 3473.993408 | 1713.900391 | 2618.209473 | 2367.384766 | 2610.437988 | 2073.377441 | 2521.281006 |
| 42409.60547 | 29222.09961 | 22359.35938 | 24678.6543 | 22600.76172 | 21799.8125 | 22326.52539 |
| 24978.58984 | 12147.19336 | 12449.56445 | 11858.87109 | 8932.173828 | 12223.04297 | 11517.35449 |
| 79016.40625 | 38941.58594 | 46725.36719 | 45223.76953 | 41473.25781 | 39657.64453 | 41038.89063 |
| 100075.8047 | 54690.23438 | 59334.54688 | 54735.29688 | 54509.05469 | 55641.86328 | 62094.31641 |
| 6354.13916 | 4407.101074 | 2650.757568 | 2898.407227 | 6776.979492 | 8375.87793 | 10112.8125 |
| 77117.38281 | 43422.66406 | 56457.45313 | 58732.41016 | 41041.13672 | 42249.48438 | 42135.74219 |
| 1980.684082 | 1365.449707 | 902.2043457 | 1312.352783 | NA | 1281.452026 | 1080.980225 |
| 150653.3438 | 60642.99219 | 105265.5703 | 39993.52344 | 44972.32031 | 109660.0859 | 46082.68359 |
| 108053.3711 | 55105.33398 | 58830.86523 | 58231.37891 | 51886.51367 | 57663.67773 | 51306.12695 |
| 70071.875 | 33089.32813 | 41521.97656 | 36732.70703 | 37528.28906 | 35209.6875 | 34252.22266 |
| 5837.091797 | 3440.738525 | 3476.593506 | 3549.187012 | 5207.291992 | 7144.495605 | 5892.62207 |
| 39461.54297 | 19661.33984 | 24820.06641 | 20616.18555 | 29536.82813 | 28763.96875 | 32025.85352 |
| 172424.3594 | 91967.51563 | 90678.52344 | 96158.99219 | 112512.4219 | 102008.5469 | 106139.8984 |
| 23159.24805 | 11873.76855 | 14101.02051 | 13368.71582 | 9067.635742 | 10962.99805 | 9584.430664 |
| 10063.35645 | 3157.86499 | 6776.510254 | 8632.928711 | 18163.7832 | 14496.68457 | 17468.61133 |
| 119953.3203 | 74936.71875 | 68422.03906 | 70465.64063 | 97601.17188 | 106986.5313 | 101603 |
| 18833.92773 | 10386.14746 | 9844.818359 | 10126.09375 | 13560.73047 | 12237.30078 | 12182.26758 |
| 1874.482544 | NA | 2191.225586 | 1020.595154 | 5175.054688 | NA | 6420.026855 |
| 1713739.5 | 1227224 | 1035360.688 | 863402.625 | 1102181.5 | 1335343.125 | 1607144.125 |
| 47138.31641 | 34982.98438 | 24040.66016 | 23662.67578 | 47278.47266 | 37041.33984 | 34326.72266 |
| 8889.78125 | 6014.361816 | 3643.025635 | 4933.335449 | 7812.081543 | 7893.709473 | 7806.521484 |
| 30220.29102 | 16915.35938 | 18904.44531 | 17280.79297 | 23607.6875 | 20635.36523 | 23601.71094 |
| 86525.08594 | 51184.61328 | 51361.17188 | 51668.51172 | 52048.24609 | 58485.82031 | 54693.39063 |
| 11836.14746 | 7271.566406 | 4596.862305 | 6514.240234 | 20999.55664 | 15840.84082 | 21579.52734 |

| | | | | | | |
|---|---|---|---|---|---|---|
| 122285.9609 | 59444.0625 | 77195.09375 | 71957.53906 | 87957.67188 | 85492.64844 | 88877.70313 |
| 27429.73828 | 15184.59375 | 13889.05273 | 12144.65137 | 10942.6748 | 11254.33887 | 10250.50195 |
| 12194.2832 | 4666.835449 | 6205.19873 | 7042.62793 | 1131.103516 | 4633.771973 | 3316.488037 |
| 38839.61328 | 17931.01758 | 21245.28516 | 19465.77148 | 22550.23828 | 27762.00586 | 24085.4375 |
| 137515.1094 | 87119.41406 | 71408.73438 | 91754.35938 | 95352.10156 | 81243.28906 | 89594.60938 |
| 36902.375 | 24757 | 18855.59766 | 18779.97461 | 37105.64844 | 32249.36523 | 28414.75391 |
| 9210.219727 | 3557.169434 | 5816.830566 | 5296.693848 | 2624.077393 | 1943.028809 | 2208.985596 |
| 57675.41797 | 35953.42578 | 26764.17383 | 24017.31445 | 25525.08398 | 31262.48047 | 22061.43359 |
| 27952.54102 | 22630.04688 | 14060.2207 | 9123.827148 | 19750.30664 | 21398.55469 | 22031.1582 |
| 18511.58594 | 9715.878906 | 10132.59473 | 9235.405273 | 13383.28906 | 14039.2334 | 15138.00391 |
| 52619.78125 | 38240.58984 | 34206.36719 | 42582.73828 | 43737.74609 | 46186.4375 | 43561.46094 |
| 114112.7031 | 78303.65625 | 55209.42578 | 72480.0625 | 120302.7344 | 143259.375 | 137764.8438 |
| 35550.09375 | 21743.55664 | 17941.07031 | 19257.08984 | 40245.97266 | 29893.00977 | 32016.73828 |
| 709794.625 | 404137.0938 | 337936.0938 | 400198.5625 | 473504.2188 | 431307.2188 | 437084.3125 |
| 32465.53516 | 20620.98438 | 18226.35352 | 16453.27344 | 21087.57031 | 19828.3418 | 20079.54102 |
| 24148.9082 | 16672.24414 | 11898.83887 | 10895.66602 | 9346.678711 | 9846.858398 | 6040.749512 |
| 11865.07715 | 5234.665039 | 5971.224121 | 8480.957031 | 13316.86719 | 9343.301758 | 8814.126953 |
| 20085.65625 | 13629.39063 | 8656.490234 | 14540.1582 | 19370.95508 | 23121.87305 | 18936.43945 |
| 33690.19141 | 21721.39453 | 16733.11133 | 15309.21973 | 25793.66211 | 27174.76758 | 33361.45703 |
| 265365.8438 | 139136.0313 | 145884.9063 | 128064.3984 | 160916.9219 | 177073.8125 | 176725.5 |
| 96817.20313 | 52902.46094 | 52552.875 | 55854.08984 | 110653.1719 | 96241.85156 | 113038.1875 |
| 573547.8125 | 338683.9063 | 290104.5938 | 334775.4063 | 636611.5625 | 575513.4375 | 606350.25 |
| 34260.98438 | 21671.3125 | 18106.55469 | 15736.68457 | 9392.219727 | 11748.69727 | 8517.792969 |
| 20526.85742 | 14828.28613 | 10409.48145 | 11262.7373 | 15816.95117 | 15957.50293 | 13265.83496 |
| 14068.94336 | 8734.77832 | 6777.189453 | 6094.041992 | 9458.849609 | 13287.22363 | 8186.25293 |
| 72144.21875 | 42303.48047 | 43298.23047 | 36182.23047 | 44118.98438 | 47582.24219 | 43110.98828 |
| 49105.32031 | 27054.13672 | 26273.19727 | 19988.61914 | 19745.25391 | 25050.01172 | 21604.98438 |
| 66768.35938 | 49521.04297 | 25784.49609 | 29660.54102 | 45064.08594 | 40772.71875 | 45917.79688 |
| 163843.1094 | 88366.49219 | 83203.8125 | 71562.9375 | 47601.94141 | 45654.44922 | 40417.40234 |
| 75013.29688 | 51614.23828 | 38142.78516 | 39101.93359 | 88539.79688 | 70680.66406 | 85223.52344 |
| 190436.5938 | 107946.8281 | 106215.125 | 98187.95313 | 111505.2109 | 111393.875 | 108646.1094 |
| 305889.5 | 197605.25 | 151727.7188 | 144261.7188 | 310778.4688 | 303658.6875 | 260219.5938 |
| 24390.09961 | 10776.56445 | 10847.37109 | 10741.16504 | 11282.2041 | 10748.80371 | 10728.02148 |

| | | | | | | |
|---|---|---|---|---|---|---|
| 26971.19141 | 13541.50781 | 14114.32715 | 11993.72852 | 10376.33398 | 11023.18262 | 10010.74609 |
| 81651.79688 | 54551.05078 | 36020.5625 | 31128.85352 | 33960.42188 | 20426.58203 | 13098.79883 |
| 54152.01953 | 34342.80859 | 34216.64844 | 30764.03516 | 48192.52734 | 34553.78125 | 42451.65625 |
| 141069.3281 | 81968.63281 | 80645.95313 | 69770.29688 | 77855.52344 | 74314.67188 | 84128.125 |
| 27747.49609 | 18024.80273 | 13864.88574 | 14771.17188 | 22651.5918 | 20300.59961 | 20858.25781 |
| 97731.60156 | 60237.87109 | 50440.19531 | 41344.3125 | 47957.17188 | 58875.26563 | 50083.8125 |
| 526093.1533 | 257312.2656 | 296110.1624 | 283361.4364 | 294686.5906 | 292165.0725 | 307430.4333 |
| 483731.7813 | 260279.5781 | 283087.9063 | 268127.9688 | 260141.9375 | 256966.0156 | 246122.6875 |
| 25348.35156 | 12824.95508 | 18604.57227 | 14004.57617 | 21385.53516 | 20765.35352 | 20073.19141 |
| 90103.35938 | 47519.00391 | 50848.9375 | 47643.29688 | 55245.11719 | 57252.35156 | 55644.37109 |
| 72674.85938 | 36814.95703 | 40572.51563 | 38217.375 | 35283.56641 | 34505.30078 | 33051.37891 |
| 19411.89258 | 10649.00098 | 13820.04883 | 7000.061035 | 6568.563965 | 17582.03516 | 7854.25293 |
| 4720.359375 | 2642.176514 | 3083.490479 | 3183.873535 | 9303.249023 | 7345.8125 | 5032.825195 |
| 6542.124512 | 5002.47168 | 2016.493652 | 3205.076172 | 4089.592773 | 4528.317383 | 6668.935059 |
| 403117.2188 | 275533.7188 | 222544.5781 | 201550.5781 | 130352.4453 | 127001.5703 | 109876.0469 |
| 81264.69531 | 41855.90234 | 41185.20313 | 40870.45313 | 39833.23438 | 43407.53906 | 36019.08984 |
| 12654.77148 | 6459.386719 | 6480.496094 | 7799.85791 | 7814.881836 | 8071.947266 | 6905.05127 |
| 50924.18359 | 33659.91797 | 30752.49023 | 31889.93555 | 30772.20898 | 25047.9375 | 28004.00195 |
| 17252.87109 | 9662.361328 | 8674.777344 | 8848.949219 | 8681.832031 | 7995.394531 | 9052.875 |
| 42341.44141 | 24982.75781 | 22852.86133 | 22329.34766 | 27673.90234 | 27043.25781 | 26250.01758 |
| 34289.64453 | 18003.2207 | 15046.41113 | 14627.71094 | 10186.84668 | 9376.912109 | 8616.867188 |
| 38940.82422 | 22063.9707 | 18686.01953 | 16749.70898 | 22099.09961 | 25276.0332 | 14576.20117 |
| 598808.5 | 270029.875 | 354297.2813 | 424027.875 | 270787.8125 | 231740.1719 | 268338 |
| 47193.67578 | 26523.89648 | 25414.32031 | 22025.43555 | 16640.08008 | 14828.68457 | 11295.44141 |
| 19648.47852 | 11785.88477 | 13998.13184 | 11266.08301 | 32224.60156 | 27011.34375 | 25338.70898 |
| 42230.50781 | 20417.37891 | 20428.41016 | 20885.21289 | 28640.67578 | 28093.375 | 29843.33984 |
| 47798.53125 | 42183.92188 | 14728.93652 | 14588.73438 | 41282.16797 | 36661.40234 | 40213.60938 |
| 4976.939453 | 3018.080078 | 2082.383789 | 3434.592529 | 5373.529297 | 5054.678223 | 4627.383301 |
| 14783.41602 | 14509.82715 | 6912.223633 | 5532.51123 | 16553.68359 | 18821.01172 | 15802.40527 |
| 157011.4063 | 128918.3047 | 74063.08594 | 81238.07813 | 116688.6094 | 139257.3281 | 140989.9531 |
| 3436.784912 | 2855.581299 | 885.5490723 | 1706.748291 | 5236.982422 | 6195.358887 | 5793.679688 |
| 31834.21289 | 11229.37598 | 20834.1582 | 23225.95703 | 25897.21875 | 29025.21094 | 32221.79883 |
| 533357.5 | 213866.2188 | 305672.9063 | 303099.625 | 265722.6875 | 306482.4063 | 333268.4375 |

| | | | | | | |
|---|---|---|---|---|---|---|
| 27353.79688 | 20509.83594 | 13753.69336 | 10580.81836 | 8980.00293 | 10884.89453 | 10301.52051 |
| 11557.51953 | 5808.126953 | 7128.234375 | 4767.932129 | 5183.801758 | 7038.400879 | 9444.881836 |
| 14848.24316 | 8959.765625 | 7297.577148 | 6137.944336 | 5216.61084 | 5198.414063 | 5461.078125 |
| 91558.00781 | 47002.48828 | 51552.81641 | 43017.59375 | 27397.08984 | 45285.34375 | 23621.71484 |
| 9574.728516 | 4403.225098 | 4003.363037 | 5436.686035 | 8656.777344 | 5142.283691 | 5226.251953 |
| 95570.51563 | 48963.52734 | 54880.01953 | 36545.53125 | 37668.45313 | 57101.88281 | 46981.24609 |
| 2520.435059 | 1529.234619 | 881.7373047 | 1212.779419 | 6057.894043 | 3634.566895 | 3054.619385 |
| 24134.58398 | 13324.11328 | 15982.94629 | 12834.125 | 8643.549805 | 9224.998047 | 9455.985352 |
| 297128.4063 | 153313.125 | 136475.1406 | 157682.1563 | 293936.7188 | 272986.8125 | 289806.5625 |
| 42908.42969 | 23488.79492 | 23342.42969 | 22992.69922 | 21840.05078 | 24502.27344 | 23524.08984 |
| 389.4604492 | NA | NA | 394.5737305 | NA | NA | NA |
| 84000.09375 | 54819.55469 | 43897.49609 | 43319.37891 | 48203.05078 | 45983.45313 | 44800.36328 |
| 10661.22949 | 8787.376953 | 5164.614258 | 3411.012451 | 9819.944336 | 7688.39502 | 10426.90723 |
| 9400.217773 | 5388.266602 | 4147.110352 | 4365.962402 | 3373.121582 | 4469.817383 | 3839.427002 |
| 3568.558594 | 3844.492676 | 591.5324707 | 554.4711914 | 3575.477051 | 2136.946777 | 3117.291748 |
| 112055.3203 | 86584.49219 | 43689.21875 | 45127.22656 | 112269.9063 | 100908.1172 | 98689.63281 |
| 5513.932617 | 3884.279297 | 2581.064453 | 3319.051758 | 3306.275391 | 3566.935059 | 3350.568604 |
| 107812.9609 | 70722.92188 | 57015.84375 | 59483.91406 | 97607.05469 | 100812.2109 | 85334.28906 |
| 8178.908691 | 4358.425781 | 4052.151367 | 3337.972412 | 4230.020996 | 4498.425781 | 3886.99707 |
| 8449.742188 | 4583.903809 | 4136.521484 | 5109.150391 | 8359.241211 | 8618.847656 | 7696.461426 |
| 5859462 | 3197588 | 3248297 | 3013670.75 | 2519062.25 | 2758875.75 | 2669160.5 |
| 29455.41211 | 14123.3457 | 17209.5957 | 15375.13184 | 12789.80566 | 16355.73242 | 13098.69043 |
| 26358.41797 | 17680.55664 | 12363.80078 | 10746.15723 | 24937.90039 | 20223.80078 | 25176.51758 |
| 51726.87109 | 28572.33203 | 31709.19141 | 30348.11328 | 24660.35352 | 19471.25586 | 19626.96094 |
| 26363.4668 | 13578.17969 | 11603.51367 | 14470.08496 | 22094.75 | 21780.09961 | 18749.61914 |
| 142788.7188 | 83285.8125 | 78166.63281 | 79510.82031 | 126053.3359 | 112447.8281 | 114637.5703 |
| 123000.6484 | 58062.25781 | 63162.29688 | 57401.55469 | 48666.57422 | 58067.76172 | 46770.94922 |
| 30145.85742 | 14041.91992 | 18022.75586 | 15640.14453 | 9915.46875 | 11681.49023 | 10549.77246 |
| 246973.75 | 130533.0391 | 142342.0313 | 129917.7969 | 133093.9531 | 140324.0313 | 135773.4375 |
| 482533.0625 | 287370.9688 | 267249.5 | 248528.7031 | 255717.9219 | 227224.3438 | 235048.7656 |
| 67574.03906 | 39854.59766 | 36581.5625 | 36540.99219 | 28817.64258 | 30521.57617 | 31294.10547 |
| 25360.74414 | 15317.00488 | 16263.45801 | 16605.63867 | 30372.2168 | 16867.58984 | 21864.62695 |
| 254757.7031 | 137829.3594 | 131673.2969 | 138600.8594 | 179775.7813 | 147882.5313 | 161897.0781 |

| | | | | | | |
|---|---|---|---|---|---|---|
| 386505.5 | 181410.9297 | 203884.9453 | 196733.9219 | 258726.4531 | 291307.2813 | 304074.2813 |
| 8364.369141 | 3783.284912 | 4128.033691 | 3905.867188 | 4226.432129 | 3686.98291 | 3059.959229 |
| 349944.25 | 294999.3438 | 144159.7188 | 136479.7031 | 313475.4063 | 277727.3438 | 313979.4688 |
| 80734.17969 | 36787.81641 | 52806.19141 | 32347.88477 | 30696.25 | 40686.89844 | 28458.00781 |
| 30220.68164 | 12556.21777 | 15531.11035 | 15998.78809 | 12225.61426 | 10305.37988 | 10290.64355 |
| 17293.64648 | 8147.719727 | 8269.575195 | 10828.05566 | 6613.008789 | 8273.636719 | 6826.522949 |
| 34193.4375 | 21356.68555 | 17502.85156 | 17634.75586 | 12156.65039 | 15470.56641 | 14462.67383 |
| 6020.228516 | 2429.598877 | 3982.383789 | 1435.733521 | NA | 3089.714844 | 2238.884033 |
| 28061.71875 | 13281.23145 | 12380.01074 | 17520.9375 | 21456.1543 | 20400.42383 | 22125.43359 |
| 78063.22656 | 40233.51172 | 44586.89844 | 37972.76172 | 66703.76563 | 60088.51563 | 63020.48828 |
| 5408.200195 | NA | 2470.468018 | NA | 6862.607422 | 3874.121094 | 14038.24023 |
| 254274.9219 | 104801.4141 | 139517.125 | 143037.4531 | 187467.5781 | 158709.8438 | 180147.4531 |
| 327583.375 | 160917.7188 | 179519.5625 | 176459.9375 | 194963.0781 | 174852.75 | 188980 |
| 899974.3125 | 466786.375 | 546575.6875 | 543926.9375 | 593809.6875 | 592726.5625 | 581108.0625 |
| 3189.38208 | 1671.30957 | 1946.699707 | 1728.738525 | 3433.318604 | 2935.618896 | 2884.538818 |
| 146390.0469 | 90599.74219 | 93932.47656 | 91301.35938 | 72098.125 | 81992.28125 | 75942.38281 |
| 43438.51172 | 18190.47461 | 22158.31445 | 19209.78516 | 23082.01758 | 20610.30859 | 20651.00391 |
| 133632.5938 | 78720.88281 | 61931.32422 | 88398.83594 | 114938.6563 | 91676.92969 | 97785.52344 |
| 6306.841797 | 4415.156738 | 3093.125977 | 2595.786865 | 2797.526855 | 3330.642822 | 2688.596436 |
| 53021.40625 | 22506.75 | 29536.16992 | 29027.98242 | 31382.61328 | 32831.55078 | 33564.36719 |
| 8611.999023 | 9609.269531 | 5680.241211 | 7166.166504 | 9639.054688 | 7017.061523 | 6737.513672 |
| 4209.793945 | 2248.476563 | 2424.398926 | 2301.947021 | 2349.653076 | 2902.2771 | 2760.833252 |
| 32179.70117 | 19600.02344 | 14151.95801 | 11546.16016 | 9967.243164 | 10707.82129 | 10793.52344 |
| 39343.83984 | 19046.74414 | NA | 23560.17578 | 47714.04297 | 26836.04492 | 20708.81641 |
| 129347.2422 | 72211.71875 | 62458.42188 | 55795.23828 | 71007.28125 | 89326.51563 | 57936.97266 |
| 52085.53516 | 22901.9707 | 28747.49609 | 26965.25977 | 30676.11914 | 34828.17578 | 33597.22266 |
| 421701.0625 | 220017.9531 | 190902.3281 | 212876.4531 | 174647.125 | 194110.7031 | 171898.5 |
| 18624.01563 | 10583.92285 | 10982.7666 | 7475.682129 | 14604.50781 | 13535.73242 | 12728.84668 |
| 26588.50391 | 11168.98242 | 15277.87012 | 12497.72363 | 9736.610352 | 10253.76758 | 12325.39746 |
| 124663.2266 | 55443.13672 | 59108.34766 | 51063.01953 | 47821.50391 | 50171.19141 | 31961.75 |
| 19066.88086 | 11268.84961 | 9429.905273 | 10816.99121 | 36235.57422 | 30357.66797 | 30791.80078 |
| 5860.219727 | 2656.436523 | 3266.973389 | 2636.840576 | 2042.762573 | 3125.990234 | 1680.650757 |
| 77278.47656 | 39904.72266 | 39624.80859 | 33753.69141 | 22128.23242 | 25177.23242 | 21005.38672 |

| | | | | | | |
|---|---|---|---|---|---|---|
| 9714.902344 | 5790.68457 | 4969.769531 | 6067.461914 | 8899.254883 | 11615.25684 | 11102.48926 |
| 168390.2813 | 127336.6016 | 66276.73438 | 60391.41016 | 130154.5547 | 93450.10156 | 117372.4609 |
| 36588.00781 | 20338.07227 | 19228.37695 | 19723.29883 | 18371.31055 | 16675.89063 | 17700.80664 |
| 84444.35938 | 44702.88672 | 35319.45703 | 40394.98828 | 38682.47656 | 37460.13281 | 35544.99219 |
| 813.0259399 | 674.888916 | 546.7220459 | 667.3352051 | 909.0834961 | 1947.632446 | 1052.914795 |
| 99221.39844 | 45729.32422 | 52108.41797 | 43164.49219 | 57459.39453 | 66031.625 | 67976.41406 |
| 212100.1563 | 103918.4219 | 119462.8984 | 125150.3516 | 135943.3438 | 141208.2969 | 146136.7188 |
| 53165.28125 | 17033.60938 | 28672.45117 | 14035.19727 | 18649.8125 | 44686.54297 | 17884.77344 |
| 29952.55078 | 12007.2334 | 13520.3916 | 14210.9668 | 58351.49219 | 65268.01172 | 64824.42969 |
| 10605.05566 | NA | 5550.473145 | NA | 9198.670898 | 6677.529297 | 5862.438477 |
| 85067.96875 | 44469.41016 | 44156.52344 | 42126.41016 | 74807.35938 | 70901.55469 | 80981.1875 |
| 7639.577637 | 3689.471191 | 4658.556152 | 5853.04541 | 3027.81958 | 7099.054688 | 4783.854004 |
| 15909.00879 | 11538.66992 | 9798.257813 | 11977.06738 | 18747.125 | 9387.430664 | 10343.35156 |
| 68487.23438 | 37392.91797 | 37657.76172 | 32569.12109 | 34812.83203 | 42334.17188 | 35285.01172 |
| 21879.32813 | 13255.66797 | 10766.75098 | 9606.658203 | 21102.78125 | 17305.28906 | 21374.06055 |
| 79971.75 | 40655.70313 | 39952.33203 | 33924.14844 | 41642.35938 | 48287.44141 | 42459.17578 |
| 9323.62793 | 3576.716309 | 5852.628418 | 4040.667236 | 4142.04834 | 4657.137207 | 3739.19873 |
| 11118.70508 | 5022.71875 | 5441.722168 | 5705.358398 | 5131.657715 | 4394.071289 | 4215.818359 |
| 69040.71875 | 37914.82031 | 38284.88672 | 37437.9375 | 31872.48438 | 34532.375 | 28050.78906 |
| 50882.62109 | 22716.72852 | 29358.99805 | 20731.58203 | 22630.84375 | 35758.33984 | 25146.33398 |
| 70518.79688 | 41886.55469 | 33479.33203 | 35558.34375 | 45914.54688 | 40767.78516 | 42203.34375 |
| 2181975.5 | 883684.875 | 1102652.625 | 1129350.375 | 1116929.375 | 1184772.375 | 1266779.625 |
| 10803.04395 | 8185.04834 | 6039.17334 | 4664.546387 | 5717.164063 | 7416.53418 | 5323.776855 |
| 13806.97461 | 11551.3916 | 6202.554199 | 5225.344238 | 7926.928711 | 7411.578613 | 8991.255859 |
| 67058.10938 | 31655.75781 | 40928.36719 | 28412.01367 | 30873.33984 | 29251.66602 | 31043.44531 |
| 284576.7813 | 213850.125 | 108035.9688 | 108006.6875 | 334059.7188 | 315186.4688 | 351740.2813 |
| 12200.75098 | 4990.233398 | 5595.806152 | 6784.63916 | 7741.90918 | 17635.57617 | 9405.986328 |
| 11148.1084 | 5828.697266 | 6597.217285 | 4961.263184 | 9097.515625 | 9104.022461 | 6852.187012 |
| 9857.706055 | 5911.778809 | 4303.25293 | 4470.476563 | 6311.131348 | 6123.522949 | 6228.371582 |
| 34083.89063 | 15957.62305 | 15540.34082 | 18072.69336 | 27144.53711 | 29337.46094 | 30430.42188 |
| 192780.25 | 117109.6875 | 81063.32813 | 77052.96094 | 173615.7969 | 167521.2969 | 161383.3438 |
| 32629.77734 | 22415.14648 | 19033.81836 | 23800.01172 | 81834.11719 | 51931.4375 | 55289.93359 |
| 38786.74609 | 17561.75781 | 19169.19727 | 17647.48438 | 20101.40039 | 20835.79492 | 18816.88086 |

| | | | | | | |
|---|---|---|---|---|---|---|
| 28263.47852 | 16438.79297 | 6109.736816 | 16358.52734 | 15422.64746 | 15924.47168 | 17193.56445 |
| 233919.8594 | 113921.9453 | 113005.6016 | 70301.07813 | 84743.71094 | 144639.2031 | 76900.78906 |
| 34374.11719 | 17367.88672 | 18015.30273 | 13733.6416 | 20603.64648 | 18642.83398 | 22263.77344 |
| 14301.45996 | 6917.146973 | 5474.129395 | 5926.395996 | 18367.55078 | 13358.3125 | 15065.35547 |
| 18913.55078 | 9718.860352 | 9196.327148 | 8583.878906 | 9759.301758 | 9292.191406 | 10744.28027 |
| 27927.45313 | 14169.95605 | 15777.76172 | 15771.09961 | 19595.04688 | 20672.00586 | 19329.67773 |
| 167625.4844 | 90180.125 | 67769.27344 | 109406.5703 | 186762.2344 | 158821.9688 | 187047.3594 |
| 6129.338867 | 3946.988281 | 1854.364136 | 2233.608643 | 5055.509277 | 4947.374023 | 5553.208008 |
| 34975.00391 | 20878.83594 | 16234.89258 | 16871.88867 | 16384.10938 | 15195.91016 | 15398.44727 |
| 20526.22656 | 11602.05859 | 11064.24219 | 11295.43457 | 12764.4502 | 14018.27344 | 11870.21094 |
| 69242.07031 | 32996.01953 | 34612.03125 | 38372.66406 | 39990.52734 | 32385.57813 | 31727.06445 |
| 779.7069702 | 317.8798523 | 320.1531982 | 537.3098145 | NA | NA | 245.7769928 |
| 5788.338379 | 1656.930664 | 990.7208862 | 3186.348389 | 7476.996582 | 8497.296875 | 7401.398438 |
| 26134.77539 | 12513.07617 | 11563.8418 | 12724.37598 | 11679.58984 | 14346.91895 | 13692.47852 |
| 39922.60547 | 18435.45703 | 17348.28906 | 15837.49219 | 10899.34277 | 9105.745117 | 10890.87402 |
| 6547.824707 | NA | 2921.549072 | 3401.128662 | NA | 4881.303711 | 4519.438965 |
| 288858.9375 | 161875.375 | 157270.9844 | 147342.7656 | 203636.1875 | 176856.6406 | 177217.9844 |
| 176905.2969 | 82234.35938 | 111359.7031 | 108987.5313 | 86388.73438 | 110740.6953 | 108638 |
| 125735.2891 | 62788.33203 | 60591.85938 | 50798.53516 | 30683.19141 | 50648.61328 | 28285.24023 |
| 155336.4844 | 68168.46094 | 82605.52344 | 75896.00781 | 78334.53906 | 74333 | 72055.82813 |
| 418283.375 | 202485.875 | 195662.7344 | 198213.5625 | 133204.9219 | 141940.9375 | 120617.8672 |
| 8686.326172 | 3841.107422 | 4220.424316 | 4156.331055 | 4411.650879 | 5643.26123 | 4654.106445 |
| 17641.25 | 9889.573242 | 8401.990234 | 7571.337891 | 9630.431641 | 11812.98145 | 9539.852539 |
| 64309.09375 | 42403.24609 | 30073.34766 | 28408.15234 | 44933.89453 | 65646.25 | 55578.35156 |
| 72123.24219 | 59741 | 54753.05078 | 48168.23438 | 83792.41406 | 42215.78516 | 54319.60547 |
| 4890.606445 | 3488.939209 | 4411.340332 | 4154.919922 | 2108.585938 | NA | 1904.486084 |
| 153564.6406 | 75175.42383 | 75485.16406 | 66202.44922 | 57054.67773 | 71610.98828 | 64291.59375 |
| 28259.94727 | 12434.84277 | 14700.80664 | 13703.17969 | 22239.06836 | 20018.86328 | 24591.08594 |
| 176398.0469 | 62007.52734 | 89317.92188 | 97221.5625 | 117057.7734 | 96945.10938 | 118293.0234 |
| 53635.32031 | 25915.08008 | 26503.44727 | 24707.92773 | 25674.74023 | 30944.20703 | 35308.6875 |
| 24277.10547 | 13220.41406 | 11443.0127 | 10849.63965 | 13787.32617 | 11777.29688 | 11761.25977 |
| 49220.19922 | 25880.57813 | 22763.45117 | 21146.88867 | 32984.21484 | 30722.50586 | 31789.20313 |
| 13562.83105 | 7086.008301 | 5630.450684 | 4277.370605 | 4933.350586 | 3468.140869 | 5125.334473 |

| | | | | | | |
|---|---|---|---|---|---|---|
| 96638.67969 | 52426.97656 | 44537.73438 | 45057.54688 | 50202.43359 | 40065.91797 | 52767.96094 |
| 98888.78906 | 49815.04688 | 47333.80859 | 44731.13672 | 54525.96484 | 56888.98828 | 57905.61328 |
| 1605686.125 | 825632.25 | 758381 | 945893.875 | 1760170.25 | 1312440.75 | 1435308 |
| 19520.37109 | 17117.81641 | 5736.154785 | 3768.664307 | 21565.81055 | 19980.84375 | 21636.25586 |
| 18467.95898 | 5085.483887 | 7224.365723 | 14098.45801 | 25470.32227 | 13635.92773 | 15004.12793 |
| 112273.7344 | 45630.23438 | 55731.18359 | 50715.40625 | 78818.125 | 63759.39453 | 70018.39844 |
| 6298.083008 | 3294.899414 | 3113.170166 | 3312.886963 | 5583.495605 | 7593.464844 | 7097.642578 |
| 9948.328125 | 6140.753906 | 2760.317139 | 6028.101563 | 11907.86133 | 7939.412598 | 10720.50195 |
| 10778.30957 | 7276.485352 | 5192.72998 | 4181.927246 | 8911.90625 | 8750.052734 | 8780.660156 |
| 8011.680664 | 4474.406738 | 4338.045898 | 4345.333008 | 4955.384766 | 3970.011963 | 1486.121948 |
| 34276.30078 | 17224.94531 | 16502.41406 | 17368.42773 | 31472.78125 | 26294.1543 | 27346.22266 |
| 272865.25 | 141641.5938 | 143225.2188 | 138926.9688 | 178533.1094 | 161555.4844 | 142797.5 |
| 3753.097412 | 2402.579102 | 2322.840088 | 1419.285034 | 1378.60437 | 886.3158569 | 697.1279297 |
| 29236.69336 | 13675.6582 | 15151.60352 | 15493.54395 | 16649.33789 | 18329.78711 | 14438.91113 |
| 364921.4063 | 171779.2344 | 187164.3906 | 165364.5469 | 143249.3906 | 178301.4063 | 169053.6719 |
| 25288.05859 | 12771.99023 | 11061.39746 | 11638.21777 | 16996.375 | 15286.81738 | 16271.69531 |
| 100271.1953 | 44086.51172 | 47751.04688 | 48440.82422 | 39167.08594 | 41419.82422 | 42655.89453 |
| 31798.86133 | 11491.11914 | 16894.04492 | 21812.56641 | 17182.66797 | 23694.4668 | 20629.12109 |
| 134929.9063 | 65054.56641 | 64401.85938 | 45958.08984 | 56754.16016 | 86913.78906 | 72466.61719 |
| 305769.5313 | 147111.5313 | 159728.7969 | 150887.4688 | 161457.6875 | 156478.1719 | 162813.8281 |
| 272964.6758 | 158613.1943 | 108980.3086 | 113769.4746 | 166765.752 | 160151.0225 | 170749.9609 |
| 148224.8594 | 70750.54688 | 75475.00781 | 73757.14844 | 56911.20313 | 55535.22656 | 56682.01953 |
| 24365.77734 | 11750.79004 | 12982.3291 | 10435.68262 | 16282.38281 | 18866.54297 | 18290.79492 |
| 33218.84375 | 17310.89258 | 16509.46484 | 13908.80176 | 19172.37695 | 20773.65625 | 23478.69922 |
| 100116.6484 | 62935.90625 | 44827.98047 | 39913.35547 | 61154.19141 | 58633.93359 | 67235.04688 |
| 250054.6563 | 104245.4063 | 124890.0469 | 106209.7031 | 118603.0313 | 145880.2188 | 123248.4297 |
| 1693348.5 | 851581.8125 | 759859.9375 | 681563.625 | 596851.9375 | 565653.0625 | 515932.1563 |
| 22316.42188 | 9402.286133 | 9213.912109 | 10050.61328 | 22481.51367 | 15927.96484 | 19669.9375 |
| 327321.6875 | 141313.5 | 152860.8125 | 100869.2734 | 106891.1875 | 172726.4531 | 83303.53906 |
| 152023.3438 | 64903.55078 | 74616.01563 | 73608.98438 | 94694.98438 | 107317.2266 | 118546.0703 |
| 194327.7344 | 76554.86719 | 48864.19922 | 73597.6875 | 74847.6875 | 95243.46094 | 91846.17969 |
| 49737.42969 | 25263.23047 | 20449.63281 | 22662.87695 | 39356.87109 | 32173.16211 | 27824.75586 |
| 483159.8438 | 221933.5625 | 240051.75 | 239450.8281 | 191566.0469 | 240185.9375 | 245965.4375 |

| | | | | | | |
|---|---|---|---|---|---|---|
| 18661.63477 | 10173.51953 | 6774.34082 | 7754.43457 | 10351.6875 | 9213.210938 | 10666.76367 |
| 346985.7188 | 168665.2656 | 144973.5625 | 114837.2188 | 99203.61719 | 163920.2969 | 105319.5391 |
| 6304.307617 | 3441.211914 | 3524.611328 | 3610.742676 | 4663.643066 | 6347.085938 | 5757.807129 |
| 6385.563477 | 2859.483398 | 1451.38269 | 2154.788818 | 3428.096436 | 4089.078613 | 2750.748291 |
| 10845.64941 | 5310.04541 | 5380.183594 | 5866.373535 | 9815.947266 | 7113.427246 | 7182.635254 |
| 418522.6426 | 194922.1636 | 205540.6543 | 183017.9707 | 196547.3125 | 197675.2002 | 188452.9834 |
| 6191.931641 | 3697.871826 | 3959.908447 | 2535.568848 | 3304.371338 | 3080.51416 | 2400.842285 |
| 60490.99609 | 27449.07813 | 29723.02148 | 30749.6543 | 26196.74219 | 26550.16016 | 27557.22852 |
| 24867.73828 | 24190.91602 | 215.4902039 | NA | 15003.625 | 10327.1709 | 13810.18945 |
| 56010.67578 | 31972.49219 | 25174.36523 | 24400.8125 | 65709.23438 | 66991.46094 | 57813.32031 |
| 2137847.5 | 1103011.125 | 1013667.688 | 1078112.375 | 1997135.25 | 1865852.625 | 1898017.875 |
| 65033.28125 | 39294.69141 | 23649.87695 | 31529.01758 | 49949.72266 | 48255.42188 | 42965.67578 |
| 87446.96875 | 49289.25781 | 33751.98828 | 31471.01758 | 35607.96875 | 34719.54297 | 35197.82422 |
| 296496.1064 | 258562.438 | 66052.58154 | 64739.30859 | 356535.1592 | 285307.4648 | 348669.4058 |
| 299397.75 | 140932.2188 | 142913.7813 | 127955.2031 | 135025.5 | 147512.1406 | 132237.2969 |
| 78454.39844 | 41500.07813 | 33430.56641 | 32518.55859 | 75421.76563 | 76411.17969 | 80822.84375 |
| 2.728205442 | NA | NA | 1.614615321 | NA | NA | NA |
| 4232.027344 | 2770.226074 | 1303.903076 | 1178.316284 | 2055.426025 | 1944.785034 | 2489.510498 |
| 33013.12891 | 13363.58008 | 15151.64941 | 16427.52148 | 11715.03809 | 12017.57813 | 13283.69141 |
| 105619.1494 | 62847.72656 | 45807.66846 | 40173.6355 | 51444.62402 | 52633.00732 | 51809.67773 |
| 92659.21875 | 44025.91406 | 40989.51563 | 26358.0625 | 26114.33008 | 51889.53906 | 26226.49805 |
| 39506.30078 | 12210.43555 | 17644.64063 | 14738.04688 | 19345.44727 | 17678.55078 | 20213.94531 |
| 54917.98047 | 24165.92969 | 28855.60938 | 14156.06055 | 16388.42188 | 46783.78516 | 19529.26953 |
| 304928.3936 | 214456.9932 | 156702.3193 | 108194.082 | 321529.9414 | 308772.8672 | 245448.7695 |
| 417455.4688 | 172816.375 | 182975.375 | 241474.2656 | 227497.2656 | 205264.9063 | 232525.125 |
| 13366.04199 | 6674.862305 | 3981.325439 | 2664.095459 | 5540.729492 | 3672.71875 | 3540.562988 |
| 150484.7188 | 73165.82813 | 63567.89063 | 55117.76953 | 66170.03125 | 85431.03906 | 65451.41797 |
| 21482.98047 | 13005.59961 | 8519.212891 | 8303.94043 | 8646.108398 | 7062.048828 | 6521.037598 |
| 8990.601563 | 9315.495117 | 7310.921875 | 6541.120117 | 16803.79102 | 6858.111816 | 6417.904785 |
| 31455.0625 | 18444.71875 | 15355.71387 | 14303.86914 | 13290.78809 | 15932.80566 | 15261.39844 |
| 35382.76172 | 17574.81445 | 15986.62988 | 14065.3418 | 12870.57031 | 14446.19824 | 11614.37793 |
| 38307.40625 | 20893.40234 | 13352.24414 | 15226.41797 | 26453.48438 | 24319.9707 | 24222.02344 |
| 1801.672729 | 4881.494141 | 3432.051025 | 193.6780701 | 14058.18652 | 1687.925659 | 4830.236328 |

| | | | | | | |
|---|---|---|---|---|---|---|
| 21288.44141 | 9681.902344 | 8424.541016 | 9323.816406 | 11982.9043 | 9636.757813 | 11409.34277 |
| 88539.44531 | 42142.6875 | 38328.79688 | 38160.26172 | 40087.33594 | 48594.83203 | 47608.48047 |
| 4657.394043 | 2446.909668 | 5003.79248 | 2692.166992 | 19335.45508 | 9204.616211 | 10800.00293 |
| 88489.05469 | 41359.3125 | 44599.17578 | 32886.91797 | 34623.22266 | 58285.17188 | 32273.8125 |
| 14709.80469 | 6658.191406 | 6263.547363 | 5823.65332 | 11929.09473 | 10437.16699 | 8565.962891 |
| 20953.83594 | 10281.38867 | 10614.19922 | 11000.59082 | 16504.42383 | 18190.91406 | 16725.82813 |
| 17452.75195 | 8500.370117 | 7218.799805 | 7920.004883 | 13754.12402 | 11525.44922 | 10400.61328 |
| 92781.03906 | 35378.03125 | 42235.14844 | 41464.71875 | 35029.41016 | 35423.50391 | 35325.3125 |
| 128527.5313 | 48684.3125 | 63626.19141 | 34400.28125 | 36630.52344 | 82721.17969 | 36962.49219 |
| 34611.24609 | 18636.29492 | 16303.07324 | 15060.60938 | 14587.90234 | 16202.69629 | 13556.21289 |
| 5712.93457 | 1828.333862 | 3280.250977 | 1783.237183 | 4088.58252 | 2669.069336 | 2195.352539 |
| 133482.1563 | 64457.27344 | 61966.64453 | 55417.97266 | 65255.96875 | 68614.29688 | 68939.4375 |
| 40711.05859 | 21719.4082 | 16764.27734 | 17168.98438 | 28239.49805 | 31278.24609 | 23176.90039 |
| 36955.35938 | 15423.98535 | 19104.76758 | 17555.19336 | 20800.05469 | 14081.46973 | 13566.18359 |
| 460571.375 | 221556.1094 | 184081.8906 | 184683.6875 | 231524.3438 | 244157.3125 | 254130.25 |
| 24246.2832 | 13803.44336 | 9992.40332 | 8632.78418 | 16178.34668 | 15327.27539 | 16212.77344 |
| 20819.84375 | 7369.613281 | 10026.99707 | 8998.930664 | 5910.244629 | 4961.631348 | 5718.886719 |
| 52107.75 | 32568.36914 | 16123.78027 | 29524.68164 | 47512.14844 | 25709.71875 | 28925.99414 |
| 31133.28125 | 11811.49512 | 13227.19922 | 16512.02539 | 32682.44727 | 24361.36523 | 29999.64648 |
| 609597.8125 | 257374.1406 | 286889.5 | 252447.8281 | 232018.0625 | 277967.3438 | 248594.5625 |
| 89299.05469 | 30511.95313 | 39016.73828 | 40209.70703 | 23475.47461 | 24408.36523 | 25843.08984 |
| 59546.80859 | 26927.05664 | 27433.86719 | 27541.63867 | 97559.50781 | 98347.40625 | 98121.24219 |
| 55845.20703 | 15485.98242 | 25183.68359 | 27027.29297 | 47973.41016 | 41627.56641 | 49245.88672 |
| 5958.28418 | 2640.157959 | 4109.451172 | NA | 13999.30762 | 11053.1875 | 9566.580078 |
| 404455.3125 | 189311.9219 | 176825.9688 | 173882.9844 | 160857.7188 | 158677.1406 | 153888.5938 |
| 9088.938477 | 3883.975098 | 2557.627441 | 3276.13208 | 2085.926758 | NA | 2062.47998 |
| 363301.3125 | 186392.4375 | 165978.4219 | 196072.375 | 94677.76563 | 88535.02344 | 77518.54688 |
| 10789.93262 | 6105.953125 | 8485.693359 | 3946.522705 | 8173.346191 | 5743.204102 | 8621.238281 |
| 237438.4063 | 132468.5781 | 92388.15625 | 147308.6406 | 123792.2969 | 183913 | 150520.375 |
| 5534.157715 | 2585.637207 | 3936.355957 | 2814.023926 | 4562.838867 | 4015.23999 | 4565.499023 |
| 2250359 | 746399.125 | 1086274.25 | 643098.6875 | 585120.75 | 1279346.375 | 617197.5 |
| 94338.3125 | 46976.59375 | 44339.56641 | 42059.77344 | 41185.39453 | 45579.42578 | 43798.71094 |
| 89140.32813 | 38191.71875 | 43159.55859 | 38149.70703 | 48139.88281 | 52123.47266 | 51658.9375 |

| | | | | | | |
|---|---|---|---|---|---|---|
| 11294.98438 | 4341.445313 | 11176.65723 | 4045.691162 | 20116.55859 | 6285.748047 | 58216.71094 |
| 256867.6563 | 156697.75 | 135134.0469 | 137485.4219 | 360164.6875 | 169462.7813 | 198798.5469 |
| 8866.43457 | 4339.76123 | 3869.452637 | 3575.997559 | 3550.906738 | 5221.617188 | 3239.875488 |
| 200933.125 | 73119.29688 | 97391.77344 | 65186.21875 | 64377.09766 | 117675.4922 | 66137.73438 |
| 3905.619629 | 1340.609131 | 2231.204102 | 2301.178711 | 1975.032349 | 2721.305664 | 2515.875244 |
| 40869.78906 | 18208.96875 | 19325.12695 | 20036.17773 | 45114.19141 | 38273.29297 | 49238.46875 |
| 28033.75977 | 16592.33203 | 13918.42969 | 15485.0957 | 18677.21484 | 14426.45508 | 12259.74219 |
| 83652.92188 | 37212.125 | 40176.53906 | 48401.15625 | 72209.35156 | 60997.72266 | 64148.98828 |
| NA | 1112.914185 | 709.6860352 | NA | 38500.39453 | 42146.43359 | 47435.30078 |
| 56407.19922 | 30305.15039 | 20859.22656 | 16411.04883 | 18124.96094 | 26101.37109 | 15946.39844 |
| 58298.67188 | 26247.5 | 27968.02539 | 20070.04492 | 22974.00195 | 32159.08789 | 31521.42773 |
| 21076.375 | 7172.081055 | 8883.44043 | 10759.17383 | 26893.99023 | 16683.27344 | 22487.51758 |
| 4116.856445 | 3167.103271 | 2418.939697 | 3610.648926 | 10263.00977 | 4841.19873 | 6444.020508 |
| 111563.1719 | 36155.37109 | 54572.51563 | 32295.97656 | 34206.19531 | 68550.96094 | 32099.08008 |
| 573112.625 | 284255.3438 | 256650.9063 | 166007.1406 | 95434.875 | 152524.0156 | 104900.6875 |
| 7448.371094 | 3102.385986 | 4431.714844 | 4509.02002 | 6292.93457 | 3218.701172 | 4369.631836 |
| 6911.972168 | 3900.956299 | 2947.274658 | 1795.818237 | 3424.791504 | 3262.837891 | 3815.092285 |
| NA | NA | 3368.864258 | NA | NA | 3167.103516 | 18568.18555 |
| 36572.40625 | 13535.22461 | 18359.64258 | 18885.71875 | 19383.08008 | 27577.25391 | 23231.9668 |
| 133845.875 | 60861.64453 | 64255.24609 | 65311.79297 | 83655.24219 | 108767.1953 | 87671.11719 |
| 35181.55859 | 10270.88672 | 15095.96582 | 20283.97266 | 34291.69922 | 33649.4375 | 37539.91406 |
| 5146.436035 | 2727.894043 | 3120.94043 | 1968.177979 | 2260.278564 | 2903.312012 | 2408.745117 |
| 3678.827881 | 1544.734131 | NA | NA | NA | 393.0088806 | NA |
| 210778.1563 | 80125.42188 | 101629.5391 | 93660.03125 | 80492.78906 | 106892.0156 | 90367.86719 |
| 2061.737793 | 1236.730957 | 871.1715698 | 1073.012573 | 1988.397949 | 1453.54248 | 1107.901733 |
| 20933.99414 | 5843.562988 | 17118.37305 | 24029.02344 | 36479.61328 | 30329.02734 | 76975.625 |
| 396798.125 | 176700.8594 | 151372.5781 | 149144.2656 | 154802.4844 | 135354.4531 | 144056.75 |
| 55913.44922 | 24559.85156 | 23273.26563 | 17192.45117 | 20594.13086 | 30400.61914 | 21312.98047 |
| 56626.83984 | 18389.94922 | 22343.9668 | 21796.45117 | 13197.24121 | 17214.61133 | 13699.66504 |
| 125685.7734 | 52397.97266 | 63829.27734 | 51744.49609 | 63041.76953 | 56936.89063 | 54146.85938 |
| 37525.17188 | 15732.27539 | 15309.98242 | 15476.85254 | 19078.57813 | 18662.99219 | 20342.64648 |
| 204453.625 | 81594.85938 | 85657.25781 | 77382.82813 | 74590.79688 | 82558.60156 | 78957.25781 |
| 43521.92188 | 15009.05566 | 21799.94141 | 13789.02832 | 15770.31348 | 25476.69336 | 13830.61133 |

| | | | | | | |
|---|---|---|---|---|---|---|
| 83228.16406 | 40106.02344 | 33412.01172 | 27866.9375 | 30678.64258 | 32231.6543 | 33144.90234 |
| 43875.35156 | 20680.4668 | 18143.19141 | 15946.74512 | 19458.11719 | 20536.46875 | 16143.46289 |
| 19463.62695 | 8512.193359 | 7680.316895 | 7308.97998 | 8177.4375 | 11240.95898 | 10463.72363 |
| 30532.18945 | 13163.20508 | 10377.31738 | 14753.2998 | 27785.63086 | 32416.79492 | 28612.2168 |
| 541986.875 | 206815.6563 | 234706.0781 | 220510.9219 | 207874.375 | 205801.1875 | 209413.7031 |
| 6989.490234 | 2471.215088 | 2964.251465 | 2992.651855 | 3574.049316 | 3744.884766 | 3651.840088 |
| 151210.875 | 89362.96875 | 156084.0469 | 76849.21875 | 185818.0313 | 60096.07813 | 733395.4375 |
| 2329.995605 | 1417.838989 | 801.8303223 | 1169.027344 | 3724.438721 | 2507.015381 | 2704.426758 |
| 112427 | 47271.87891 | 46265.33594 | 44092.10156 | 68268.80469 | 79416.41406 | 73651.89844 |
| 50038.72266 | 25934.91016 | 19304.39258 | 23079.70898 | 60411.35156 | 63531.85156 | 54831.31641 |
| 1975.412109 | 700.6365356 | 456.9185791 | 1264.379761 | 1468.653809 | 1167.390991 | 1551.332397 |
| 84404.28125 | 28114.82031 | 39431.28906 | 35177.53906 | 26035.50391 | 28718.2207 | 28549.84961 |
| 39612.62891 | 15138.50781 | 14840.86133 | 14340.6123 | 20737.98438 | 23207.4668 | 22300.83789 |
| 4962.039063 | 3491.681152 | 2326.09668 | 4167.348145 | 10618.05566 | 3102.193359 | 4830.071289 |
| 11116.76074 | 6837.102539 | 4144.166504 | 3407.206543 | NA | 2900.332275 | 3318.077393 |
| 102592.1094 | 41110.46484 | 41937.01563 | 38357.0625 | 44230.24609 | 40883.00391 | 42815.18359 |
| 78522.35156 | 41090.78906 | 38120.35547 | 28535.41602 | 22202.14844 | 29254.67188 | 21319.86719 |
| 435806.875 | 161159.2813 | 164995.875 | 178546.5313 | 630544.375 | 434608.0313 | 505693.0938 |
| 86590.46875 | 37009.21094 | 33831.09375 | 31076.23633 | 30076.13281 | 31369.41797 | 28694.15039 |
| 7226216 | 3875915.5 | 2685549.75 | 2257462.75 | 3356568.75 | 3650901.25 | 3638649.75 |
| 28103.90381 | 13798.96948 | 10722.73535 | 10250.06946 | 16551.9519 | 15339.1084 | 15367.80322 |
| 386366.0313 | 130019.7891 | 142811.6094 | 146514.9375 | 138156.6094 | 112816.0391 | 132456.5 |
| 29084.0957 | 11477.96094 | 9810.024414 | NA | 13534.6748 | 8660.873047 | 10580.88184 |
| 566784.3125 | 238834.4063 | 239119.9375 | 209748.9219 | 358985.9688 | 349773 | 347220.9063 |
| 6226.050293 | 4441.484375 | 1985.123779 | 1980.980347 | NA | NA | NA |
| 371300.7813 | 147988.8281 | 142036.2344 | 152698.75 | 222389.7344 | 222722.3125 | 265624.875 |
| 76827.42188 | 28105.85938 | 32626.92969 | 25591.85742 | 24964.64063 | 47690.90625 | 27180.81055 |
| 6811.970703 | 3518.680908 | 1193.787964 | 1406.015747 | 6459.128906 | 6388.270508 | 6402.006348 |
| 9991.041016 | 4607.51123 | 5472.914551 | 4803.925781 | 17162.91016 | 6448.082031 | 6414.655762 |
| 37890.41016 | 15816.39258 | 16640.14258 | 14204.52051 | 12544.44824 | 13895.37402 | 13546.13867 |
| 611490.9375 | 263442.0625 | 251208.3281 | 258170.1875 | 176454.5156 | 162707.6563 | 165523.7656 |
| 1669.09314 | 2358.21167 | 568.7139893 | 1100.644531 | 2036.202637 | 1495.736084 | 2087.243896 |
| 30696.08789 | 14459.58984 | 14053.80078 | 11135.57715 | 13080.38184 | 12769.72656 | 12106.07617 |

| | | | | | | |
|---|---|---|---|---|---|---|
| 7089.712402 | 2931.231445 | 2503.191162 | 3060.267822 | 5206.473145 | 5042.455566 | 5598.492676 |
| 7206.126465 | 2136.203613 | 3679.187256 | 2171.150879 | 3046.51416 | 2912.299561 | 3422.94043 |
| 39780.04297 | 16439.97461 | 16161.80176 | 14743.9502 | 11153.99609 | 15275.87305 | 13306.9707 |
| 54835.09375 | 19763.02539 | 21493.44922 | 22986.90039 | 18645.00586 | 21497.75586 | 20789.01172 |
| 105195.9375 | 38356.34766 | 39989.55469 | 42995.46484 | 87370.24219 | 76897.45313 | 88788.39844 |
| 155272.0156 | 75003.50781 | 47476.32813 | 52153.82031 | 146130.0469 | 129344.9609 | 153051.5469 |
| 59301.47656 | 21314.4668 | 19367.80078 | 24696.29492 | 29101.88281 | 23569.91602 | 28846.95898 |
| 334669.6563 | 134146.6406 | 133872.7188 | 146842.6406 | 206335.5625 | 163814.5938 | 181171 |
| 317361.8438 | 108491.1797 | 122209.3125 | 120377.2578 | 138333.1094 | 138111.3438 | 146361.2656 |
| 10871.67383 | 4449.813965 | 3731.518555 | 4422.784668 | 4358.65625 | 4208.337891 | 4582.701172 |
| 233378.7305 | 93498.70166 | 78167.72607 | 78595.20166 | 24962.51903 | 26505.34961 | 29015.62958 |
| 69456.41406 | 21830.60156 | 29849.35352 | 14702.7334 | 21426.76172 | 43932.97266 | 22373.91797 |
| 39887.97266 | 13904.57715 | 20540.20117 | 15523.17969 | 19663.22461 | 28055.66016 | 20998.41992 |
| 14370.99902 | 5185.753906 | 4562.63623 | 4665.853516 | 5869.512207 | 7017.266113 | 6706.501953 |
| 266508.3438 | 116146.3672 | 107802.9063 | 124325.8359 | 226199.2344 | 210769.7813 | 216788.3125 |
| 30881.20703 | 14080.93359 | 11596.47754 | 9328.310547 | 16746.98047 | 13501.65039 | 9434.564453 |
| 8850.978516 | 4539.789063 | 3506.113525 | 4079.055664 | 5151.76123 | 5476.771484 | 4756.485352 |
| 49589.4375 | 17899.57813 | 13608.25977 | 20356.5957 | 41105.16797 | 35113.28125 | 41978.88281 |
| 84734.96875 | 40030.05469 | 13378.28223 | 18546.58984 | 18474.48633 | 16914.01367 | 16367.37207 |
| 21386.01953 | 6535.232422 | 8728.055664 | 7042.499023 | 7360.539063 | 7034.985352 | 6574.275879 |
| 4266.890625 | 2003.751831 | 1336.134155 | 2064.221436 | 2218.105957 | 2856.482666 | 2425.039795 |
| 37167.53125 | 7556.203613 | 4347.858887 | 12227.51758 | 14795.42383 | 22375.91016 | 17276.02734 |
| 74386.19531 | 27098.63965 | 28670.88843 | 32823.89893 | 58065.98926 | 57170.60352 | 53548.57227 |
| 120435.0156 | 44127.16016 | 48227.99219 | 46503.41016 | 58751.19141 | 65830.39844 | 59325.48828 |
| 4417.45166 | 691.5117188 | 2027.489502 | 2247.603271 | 2554.737549 | 2347.866699 | 2264.5979 |
| 71352.60156 | 34778.04688 | 18093.25 | 14375.35449 | 47177.51172 | 47249.53125 | 51373.8125 |
| 219281.7031 | 92168.64063 | 80627.21094 | 88513.07031 | 131298.2813 | 126471.9453 | 118828.6406 |
| 60141.86719 | 38802.42578 | 35481.35938 | 22902.53516 | 44422.10547 | 22180.83008 | 208105.4219 |
| 3032.411621 | 1376.910156 | 1714.496948 | 776.2557983 | 1820.709839 | 705.5383911 | 1111.666138 |
| 133835.3906 | 66770.24219 | 46293.50391 | 49556.79688 | 92196.49219 | 89128.8125 | 84873.03906 |
| 13854.41797 | 5501.675781 | 5675.392578 | 5479.940918 | 8074.412109 | 8848.274414 | 7542.76709 |
| 18846.49219 | 5530.015137 | NA | 6122.43457 | 4479.190918 | 8649.77832 | 6019.77002 |
| 32823.39453 | 11586.38281 | 12281.13379 | 13088.29004 | 22613.01172 | 21324.21875 | 24659.26563 |

| | | | | | | |
|---|---|---|---|---|---|---|
| 10951.14648 | 4139.783203 | 4449.016602 | 4855.402832 | 3160.07373 | 4725.163086 | 4367.525391 |
| 15681.20605 | NA | 4443.211426 | NA | NA | NA | 3988.784912 |
| 30400.44336 | 11925.70898 | 10065.59766 | 9135.081055 | 12617.85449 | 12756.49805 | 14529.70801 |
| 57220.26172 | 21656.65234 | 23514.31055 | 25403.74414 | 35379.25 | 38860.49609 | 44098.94141 |
| 89421.42969 | 35388.42578 | 28179.91211 | 32383.70313 | 67700.35156 | 56570.77734 | 52942.86328 |
| 31061.70508 | 17959.7207 | 17841.01953 | 22219.65039 | 65924.98438 | 22778.37695 | 27395.14258 |
| 17229.53906 | 6718.133301 | NA | NA | 25158.44531 | NA | 10875.625 |
| 70802.67969 | 29590.70313 | 27465.23242 | 30457.82227 | 19752.52148 | 22511.51367 | 19677.10742 |
| 12395.96387 | 5605.19043 | 4345.320801 | 4307.998535 | 3338.009521 | 4785.314453 | 4450.092773 |
| 458.1629333 | 203.5567627 | 116.585434 | NA | NA | NA | NA |
| 23919.10352 | 4517.598633 | 8478.516602 | 10857.50293 | 8995.748047 | 7397.437988 | 5498.460449 |
| 13200.54102 | 5740.359375 | 5113.19873 | 2674.200195 | 3657.89917 | NA | 3073.406738 |
| 8111.253906 | 4227.786133 | 3193.530029 | 2736.572754 | 2889.750488 | 3082.298828 | 2623.709717 |
| 32704.46875 | 12720.79004 | 9464.170898 | 10648.08398 | 25173.07227 | 26177.52148 | 27828.37305 |
| 14545.72461 | 5813.692383 | 5273.719727 | 4602.379395 | 5644.123047 | 4599.424805 | 4655.31543 |
| 258862.1406 | 126796.0156 | 100451.6875 | 106363.0313 | 38051.43359 | 35347.32031 | 28417.11523 |
| 10300.96777 | 3446.327393 | 2911.75 | 3687.524902 | 3023.040039 | 2849.733154 | 1745.902954 |
| 15722.80664 | 6692.313965 | 6738.802734 | 4246.897949 | 13222.48438 | 8632.96875 | 10536.91309 |
| 1513719.977 | 492997.9648 | 568772.1465 | 507412.543 | 270718.0986 | 382277.8457 | 398998.5645 |
| 4578.140137 | 514.9755859 | 3436.967773 | 184.5651398 | 26508.39258 | 3533.460693 | 6833.854492 |
| 25047.52734 | 11203.13281 | 6624.37207 | 6018.370117 | 7871.803711 | 8660.400391 | 7858.118652 |
| 29813.16016 | 11891.7207 | 16219.78125 | 15364.14648 | 60733.14453 | 11875.43457 | 20582.16797 |
| 3065.7146 | 1442.374268 | 998.84375 | 1027.484009 | 1194.610229 | 560.7443848 | NA |
| 37183.75 | 12280.14746 | 13146.12793 | 12867.86621 | 16184.67773 | 16163.5498 | 15368.80859 |
| 24678.85742 | 11970.00488 | 15517.05566 | 10301.23828 | 30045.74023 | 13813.46387 | 85993.21875 |
| 18278.23633 | 1614.96106 | 9695.036133 | 5973.578613 | 6535.259277 | 13834.07129 | 13063.36914 |
| 73723.42188 | 35790.66016 | 24611.57031 | 22876.31641 | 37925.5625 | 35721.79688 | 28701.28906 |
| 116916.4219 | 34446.63672 | 40362.34375 | 41453.52734 | 38036.03906 | 34318.66406 | 38383.52344 |
| 3726266 | 1397093.375 | 1130507.375 | 1259555.125 | 3084298.75 | 2564123 | 3258055 |
| 181085.4688 | 62972.91797 | 69018.28906 | 46450.24609 | 55286.72266 | 91281.71875 | 52165.9375 |
| 37519.90234 | 18619.80469 | 10134.4082 | 11578.24512 | 24096.11328 | 5175.565918 | 15808.61328 |
| 72006.17969 | 24981.71094 | 29101.22266 | 18569.78516 | 20444.79688 | 23259.08984 | 21496.93164 |
| 45814.31641 | 19535.03906 | 15773.89844 | 15889.12891 | 66447.92969 | 65109.60938 | 64851.66406 |

| | | | | | | |
|---|---|---|---|---|---|---|
| 320975.0313 | 82042.28125 | 131655.1406 | 89570.89063 | 101558.5078 | 170617.7188 | 102324.3047 |
| 134626.6875 | 56505.50391 | 41867.26953 | 28433.08398 | 51486.22656 | 70774.42969 | 34648.28906 |
| 9346.647461 | 2659.275879 | 2501.974854 | 4768.40918 | 4128.054688 | 2265.619629 | 3097.743164 |
| 1233.98877 | 419.8770752 | NA | NA | 839.4413452 | 375.6622009 | 600.1731567 |
| 28566.90039 | 23830.07617 | 13803.56787 | 9256.46875 | 36464.46582 | 12376.58569 | 16152.68457 |
| 12750.875 | 5746.513184 | 3465.471924 | 3693.916504 | 4249.69043 | 5316.785156 | 4316.327148 |
| 29059.06055 | 5896.691406 | 11755.03711 | 10517.05762 | NA | 1336.664429 | 23476.36328 |
| 34983.16797 | 25321.76172 | 7543.659668 | 7539.304199 | 14445.96484 | 14898.31152 | 9147.369141 |
| 7222.449707 | 3604.731934 | NA | 2885.831055 | 5668.398438 | 4646.44043 | NA |
| 15655.34961 | 5219.727051 | 3706.989014 | 4666.657715 | 22319.63672 | 13016.01172 | 16300.99316 |
| 81263.39063 | 36768.45703 | 38621.90625 | 32362.1543 | 108502.8906 | 24219.75 | 249282.3281 |
| 941742.25 | 270037.9375 | 374289.3125 | 374292.3125 | 598077.0625 | 594608.8125 | 724109.6875 |
| 365793.4063 | 75955.21094 | 226826.7813 | 49602.41797 | 300866.8125 | 222292.7813 | 850219.3125 |
| 49011.19922 | 21242.12305 | 13859.1084 | 14753.49219 | 24276.43555 | 21761.81641 | 23013.64063 |
| 244395.8906 | 87878.17969 | 80185.64844 | 90334.46875 | 75187.10156 | 69733.4375 | 72926.14063 |
| 197911.9375 | 58046.10938 | 65375.84375 | 61738.16797 | 60693.07813 | 67110.76563 | 65263.60938 |
| 173905.9688 | 43077.18359 | 41607.125 | 74365.1875 | 114327.8984 | 141913.1719 | 124412.3203 |
| 253755.2344 | 59826.33594 | 190696.25 | 38673.14844 | 693990.4375 | 204166.6406 | 958818.0625 |
| 8304.671875 | 3636.551514 | 1402.201904 | 4439.913574 | 6352.107422 | 4976.624023 | 6120.171387 |
| 9833.055664 | 4613.081055 | 1496.39624 | NA | 6174.026367 | 6355.925781 | 6951.632813 |
| 15137.91113 | 4438.964844 | 4642.130371 | 7425.400391 | 19496.94531 | 11617.22754 | 16807.18164 |
| 231595.5313 | 82936.45313 | 76001.05469 | 69784.21094 | 97359.35156 | 94694.22656 | 100597.8047 |
| 35100.65625 | 13111.03027 | 9850.435547 | 13818.48438 | 8671.954102 | 9557.422852 | 6707.081055 |
| 169722.5625 | 52870.73438 | 47161.25781 | 40963.09766 | 41402.46875 | 58239.22266 | 39673.16406 |
| 283573.5938 | 104422.6953 | 82968.97656 | 75636.36719 | 98818.83594 | 120892.6484 | 81724.49219 |
| 254005.1094 | 83023.32031 | 72260.5625 | 74162.92969 | 75193.48438 | 71285.09375 | 78165.67969 |
| 11823.16211 | 4260.114746 | 5339.046875 | 3381.790039 | 6602.251465 | 4043.480713 | 6882.594238 |
| 142382.75 | 50019.26563 | 43047.97656 | 40815.60547 | 26014.3125 | 30909.16211 | 28669.14648 |
| 9985.235352 | 3364.037109 | 3374.391846 | 3088.419189 | 4075.871094 | 4409.460938 | 4708.352051 |
| 133554.125 | 22628.05664 | 61815.65625 | 8608.267578 | 77395.85938 | 80613.44531 | 220734.4688 |
| 12869.74805 | 5043.039551 | 4361.936523 | 6100.240723 | 12587.05566 | 9853.155273 | 8365.883789 |
| 9183.061523 | 2723.395264 | 2801.684326 | 3817.98999 | 6108.575684 | 10940.45508 | 7126.274902 |
| 32275.91211 | 8786.009766 | 10565.36133 | 12398.91699 | 26524.52344 | 15554.35352 | 18366.4043 |

| | | | | | | |
|---|---|---|---|---|---|---|
| 29298.24805 | 17823.6875 | 7990.833984 | 9586.664063 | 29288.82227 | 32240.17188 | 25785.00391 |
| 504320.5938 | 144961.0625 | 123375.4531 | 186486 | 409615.5313 | 249823.5 | 286154.0313 |
| 78806.6875 | 21382.78516 | 24079.73633 | 23681.33203 | 16726.82031 | 23169.4082 | 18542.66992 |
| 86765.05469 | 33501.44141 | 23809.01563 | 25299.27734 | 64214.83203 | 59106.5 | 56889.85547 |
| 19357.87891 | 11499.00098 | 5505.762695 | 7990.748535 | 14936.88574 | 7834.850586 | 9852.686523 |
| 388545.75 | 230127.2656 | 120760.3125 | 155305.6875 | 73253.82813 | 52605.79688 | 50016.94141 |
| 108333.25 | 23062.91602 | 31383.18555 | 38692.4375 | 98331.49219 | 74119.82031 | 86464.1875 |
| 24622.20898 | 8949.69043 | 7379.490234 | 5572.174805 | 13453.66309 | 11925.83008 | 12424.69141 |
| 115590.4453 | 27042.29688 | 38051.34375 | 35375.33594 | 32327.39258 | 33830.70703 | 38183.89453 |
| 25013.16602 | 8967.140625 | 5979.795898 | 7019.962891 | 7702.060547 | 6733.00293 | 6512.853027 |
| 22042.86914 | 9490.355469 | NA | 10919.17676 | 20719.33008 | 31652.34375 | 15250.29297 |
| 64460.53516 | 26242.86719 | 16098.31445 | 21884.92969 | 39995.19141 | 35568.80859 | 35094.98047 |
| 168767.4688 | 63921.24609 | 54914.94531 | 49610.08984 | 59836.04297 | 53177.29297 | 50876.0625 |
| 55778.36328 | 20862.08203 | 15168.83984 | 20072.76563 | 38551.76953 | 46883.44141 | 46897.43359 |
| 793.8706055 | 222.6809845 | NA | NA | NA | NA | NA |
| 118887.1719 | 38723.26953 | 30251.18164 | 59351.57813 | 98751.42188 | 25020.65039 | 30996.52148 |
| 40756.69531 | 14953.83496 | 10863.27148 | 9758.666016 | 13622.00586 | 17327.99609 | 13494.6709 |
| 39369.55859 | 11435.33398 | 12750.72266 | 11198.91797 | 10953.64746 | 10349.1543 | 11410.48438 |
| 150179.9063 | 48910.96094 | 38369.95313 | 43174.01563 | 41859.90234 | 27589.05859 | 30201.35156 |
| 12659.13086 | 3336.275879 | NA | 3350.035645 | NA | 2783.248047 | 2164.859131 |
| 60226.83203 | 18756.74609 | 24145.38672 | 22805.43359 | 3953.898193 | 3208.811279 | 4602.339355 |
| 13460.8916 | 4749.716797 | 2461.930176 | 2887.973389 | 4882.053711 | 4605.363281 | 5055.283203 |
| 32437.32617 | 9759.186523 | 5010.899902 | 9896.773438 | 9751.774414 | 10939.32227 | 7340.407227 |
| 2447163.75 | 794574.8516 | 849427.1406 | 186228.5195 | 151409.5781 | 50468.7334 | 73839.89941 |
| 2137.461914 | 908.8108521 | 602.0793457 | 352.1578064 | 311.6470947 | 558.8145142 | NA |
| 71987.21875 | 14295.55469 | 16890.20313 | 19428.43945 | 21113.72852 | 19933.18945 | 20523.61133 |
| 42799.51172 | 12620.31836 | 13669.77832 | 7825.708984 | 10435.91211 | 18321.32422 | 22313.75391 |
| 146978.4219 | 41750.72656 | 53891.69922 | 51827.67578 | 123504.5156 | 54389.625 | 63572.66406 |
| 122248.3359 | 27743.39063 | 34196.35156 | 35582.59375 | 25344.32422 | 22438.5293 | 27415.38281 |
| 192394.9375 | 40220.57813 | 54847.14844 | 48670.90625 | 58385.92578 | 49461.88672 | 49067.78516 |
| 184026.2031 | 51556.32813 | 39349.16406 | 30980.99805 | 29222.45508 | 23060.07227 | 22528.00195 |
| 14606.39648 | 5838.934082 | 2807.887695 | 4071.083496 | 10285.7334 | 6188.86377 | 6668.282227 |
| 35425.00391 | 8297.648438 | 10308.52539 | 9331.235352 | 20940.24219 | 18742.29883 | 21345.47266 |

| | | | | | | |
|---|---|---|---|---|---|---|
| 27662.20703 | 4456.666992 | 10384.78613 | 7119.540527 | 11643.68652 | 12549.62012 | 12448.70703 |
| 1961326.125 | 963360.25 | 848438.25 | 742354.0625 | 1575603.875 | 857266.5 | 3595037 |
| 164675.125 | 42529.85547 | 49818.51563 | 45437.03516 | 58533.89453 | 52616.78906 | 52869.15234 |
| 18181.86719 | 3826.171631 | 5156.592773 | 3918.773193 | 5838.035645 | 4056.819336 | 5775.52832 |
| 14068.44043 | 3165.039063 | 4190.787109 | 3922.905273 | 3868.303223 | 4999.298828 | 3655.492432 |
| 13127.5293 | 3546.839844 | 3396.873047 | 3726.823975 | 5013.173828 | 2879.489014 | 3898.500732 |
| 426534.7344 | 173935.9434 | 68847.19629 | 62466.87598 | 95638.01172 | 92559.1875 | 97619.69336 |
| 12675.59277 | 3887.964111 | 3038.575684 | 3637.202148 | 14253.48145 | 10892.05859 | 12909.2168 |
| 17073.96484 | 4896.16748 | 3771.43457 | 3696.535889 | 5626.914063 | 6437.519043 | 6862.832031 |
| 21481.64648 | 3810.037109 | 7691.391602 | 5307.332031 | 8880.509766 | 8511.852539 | 3430.371582 |
| 8303672.891 | 2032747.264 | 2140890.421 | 1960763.611 | 2472101.91 | 3499078.242 | 3374122.391 |
| 74084.96094 | 22153.17188 | 19071.12891 | 15050.62012 | 25404.93945 | 24824.96875 | 24394.60938 |
| 7043.188965 | NA | 3763.225098 | 1319.800293 | 4857.989746 | 9508.155273 | 7166.570801 |
| 122347.9766 | 34009.39453 | 30311.70898 | 20928.18555 | 22289.98633 | 30313.52734 | 25285.85547 |
| 36551.30859 | 9129.295898 | 7490.219727 | 6841.61084 | 10194.13672 | 8875.792969 | 10456.98633 |
| 8600.207031 | NA | 1872.685669 | 1800.500854 | NA | 660.9545898 | 1870.352661 |
| 113349.5078 | 31103.31445 | 14128.25293 | 28192.39844 | 18777.34375 | 27530.98242 | 22410.01563 |
| 15680.49414 | 8446.844727 | 5318.685059 | 4093.228027 | 6185.891602 | 11973.20313 | 9485.601563 |
| 1950044.5 | 494819.9375 | 505962 | 409849.0938 | 527521.1875 | 905935.25 | 763518.625 |
| 26382.10938 | 6101.625977 | 5378.242676 | 6087.145508 | 36590.17578 | 21354.86523 | 29242.87891 |
| 3672.655762 | 1235.782349 | 834.3313599 | 542.1774902 | 1444.71875 | 4248.187012 | 1524.895386 |
| 182669.0518 | 48550.64917 | 51992.38086 | 50851.05078 | 68536.5752 | 64607.10986 | 62987.62695 |
| 83029.57813 | 19892.69922 | 17029.01758 | 24853.61133 | 78325.0625 | 51954.34375 | 62118.13281 |
| 3777.901367 | 1143.226318 | NA | NA | 4399.904297 | 963.5912476 | 1709.60083 |
| 54514.36328 | 12197.00195 | 13557.4668 | 11244.77734 | 37556.57422 | 27604.68359 | 32538.64453 |
| 242604.6875 | 49017.45313 | 34415.17188 | 62449.33594 | 195066.125 | 122164.1797 | 168978.7344 |
| 21881.44727 | 5581.761719 | 3892.270996 | 4815.335938 | 5025.552246 | 6273.069336 | 5993.091797 |
| 497389.5 | 69118.99219 | 96469.89844 | 115591.4453 | 60053.41406 | 82897.50781 | 69296.27344 |
| 2392305 | 485730.5938 | 393746.4688 | 544778.75 | 684691.6875 | 914143.0625 | 989628.875 |
| 32715.86914 | 6253.205566 | NA | NA | 5420.213379 | NA | 4032.689697 |
| 3581.5979 | 3169.794678 | 1845.515015 | 2564.587891 | 36523.40625 | 5887.282715 | 12229.40527 |
| 5665.520996 | 2544.437744 | 2143.656494 | 3321.863525 | 23528.44922 | 5321.946777 | 7357.362305 |
| 43972.66016 | 23355.95703 | 17233.50195 | 19522.71875 | 200220.9219 | 48610.50781 | 69754.67969 |

| | | | | | | |
|---|---|---|---|---|---|---|
| 6029.40332 | NA | 1630.989746 | NA | NA | NA | 1364.786011 |
| 199901.875 | 47151.85938 | 34968.67578 | 44616.07031 | 47838.12891 | 33253.73438 | 35552.88672 |
| 15188.88672 | 8752.110352 | 3550.537842 | 5699.006836 | 33984.17578 | 6839.425781 | 11630.57715 |
| 176066.3281 | 43091.21875 | 42918.83984 | 25592.01172 | 18509.15625 | 31727.75 | 24480.23242 |
| 1046.515869 | 167.2893372 | NA | 290.7576904 | 179.2479095 | 166.9843292 | NA |
| 13047.73828 | 3223.691406 | 1386.012695 | 3107.760742 | 1337.769775 | 2953.990967 | 1961.876099 |
| 9116.694336 | 2283.544434 | 519.9208374 | 460.5877991 | NA | 976.8382568 | NA |
| 14004.78906 | 2956.536865 | 2118.437256 | 4315.706055 | 3807.145508 | 4014.032959 | 3478.832031 |
| 121592.875 | 24368.71289 | 32496.79297 | 26304.80078 | 61303.62109 | 20023.95117 | 22690.34961 |
| 34225.94531 | 6341.063477 | 6592.627441 | 5258.757813 | 5648.069824 | 6552.53125 | 5251.982422 |
| 112155.1641 | 34123.125 | 18773.22266 | 22802.47656 | 27382.61523 | 24483.84375 | 29591.18555 |
| 20187.85547 | 2504.21875 | NA | 3329.651367 | 21650.55469 | 14612.82129 | 29901.44727 |
| 1790.361694 | NA | 403.0919495 | 1061.94397 | 36302.68359 | 3620.372559 | 22695.31055 |
| 31418.87305 | 5517.646484 | NA | NA | NA | NA | NA |
| 9921.275391 | 1819.834106 | 1881.810791 | 1178.579468 | 483.7142334 | NA | 379.1604614 |
| 87426.47656 | 14555.49512 | 13133.33105 | 15430.07813 | 59499.94141 | 53578.96484 | 69403.05469 |
| 102118.6563 | 18469.25781 | 19759.41016 | 17035.6875 | 17411.35547 | 21817.65234 | 20121.90625 |
| 54543.10156 | 7079.758301 | 9421.482422 | 12029.7334 | 20279.11719 | 20814.93945 | 22312.73828 |
| 32197.10938 | 10402.72363 | 5404.543945 | 9576.692383 | 28958.83398 | 12327.72168 | 17252.23438 |
| 5093.134766 | 1096.679565 | NA | 1108.305786 | 2795.033936 | 2929.374023 | 2593.82251 |
| 3767.675049 | NA | 1044.953003 | 749.5318604 | 4419.780273 | 4070.379639 | NA |
| 16899.08008 | 3073.712158 | 3530.133301 | 2232.711182 | NA | NA | NA |
| 1290227.875 | 425004.4063 | 119832.9375 | 109953.3516 | 94849.36719 | 90780.09375 | 67808.41406 |
| 5734.08252 | 2056.968506 | 444.9511108 | 1234.842773 | NA | 714.6929321 | 1234.431763 |
| 7471.461914 | 1164.235596 | 1130.268433 | 1488.025269 | 818.8373413 | NA | 929.5625 |
| 134048.9531 | 28009.72656 | 17098 | 24837.91602 | 50033.98438 | 38443.52734 | 41666.69141 |
| 25207.60352 | 3036.212891 | 3472.749023 | 4303.373047 | 12112.5918 | 7743.623535 | 15293.48828 |
| 49020320 | 12908465 | 7881535 | 10543970 | 25647116 | 42587300 | 33052318 |
| 109470.6328 | 13512.77148 | 23206.92969 | 24976.41211 | 26058.17969 | 36767.66406 | 30391.65625 |
| 29173.42969 | 1079.954834 | 4949.150391 | 7327.144531 | NA | NA | NA |
| 33771.69922 | 4901.055176 | 4631.125977 | 3812.524414 | 2962.953857 | 4178.45166 | 2492.36084 |
| 5559.202637 | 1271.809326 | 857.7078247 | 778.59021 | 2950.889893 | 3289.643066 | 2364.744141 |
| 84231.21094 | 14076.56543 | 12783.00781 | 9367.258789 | 23785.90625 | 19023.0332 | 26231.2168 |

| | | | | | | |
|---|---|---|---|---|---|---|
| 87359.39844 | 18560.33789 | 19030.41016 | 8659.288086 | 23717.79102 | 17523.52539 | 17529.92578 |
| 65218.11328 | 35439.30078 | 34682.45703 | 21491.5293 | 70428.5 | 24808.44727 | 29987.88477 |
| 108060.1484 | 16350.67871 | 15306.00098 | 13415.13184 | 12337.50391 | 14086.42188 | 11593.21094 |
| 2974852.75 | 460559.1875 | 353204.5313 | 385900.5 | 419016 | 609924.125 | 396720.5313 |
| 52924.61719 | 10959.97363 | 8262.776367 | 8014.435547 | 12249.66895 | 9812.229492 | 9590.257813 |
| 165657.4063 | 26721.2168 | 25472.45508 | 16190.68066 | 15867.58203 | 16674.57617 | NA |
| 24749.00781 | 2068.346191 | 1360.462402 | 2373.28418 | 14913.33008 | 2376.922607 | 5311.943848 |
| 15728.17871 | 3516.374512 | 1074.097778 | 3287.586426 | 3759.931396 | 3054.837402 | 3675.218262 |
| 33357.42188 | 3310.916504 | 5143.875977 | 2765.162842 | 3435.650635 | 3558.98291 | 4313.96582 |
| 435124.7891 | 58618.20313 | 47199.64746 | 50932.99805 | 61283.65234 | 62301.2793 | 58762.5 |
| 3326343 | 282291.1875 | 412904.5313 | 370753.0313 | 532660.25 | 1387784.125 | 893214.1875 |
| 166027.2188 | 21379.58594 | 22842.61719 | 26546.12305 | 37902.07813 | 30248.35547 | 26547.13281 |
| 89387.17969 | 21088.30273 | 9852.580078 | 9483.393555 | 64513.96875 | 30172.83203 | 37522.69141 |
| 148497.2344 | 18359.01563 | 17709.25391 | 16525.5625 | 53438.09766 | 47595.75781 | 50141.71484 |
| 79005.11719 | 15519.20313 | 10925.24414 | 11462.03027 | 31925.44727 | 31340.98828 | 20516.92383 |
| 740878.8125 | 81464.17773 | 88286.68555 | 92397.23291 | 120120.2969 | 102435.1738 | 109699.3516 |
| 22381.3457 | 3170.942627 | 3398.078613 | 3758.074463 | 29330.23438 | 6525.88916 | 10074.42773 |
| 68850.1875 | 6591.71582 | 7998.559082 | 8431.935547 | 79801.78906 | 51163.71875 | 60118.16016 |
| 17298.56445 | 825.6362305 | 1300.36145 | 3229.327393 | 10630.00293 | 8798.039063 | 10318.10156 |
| 4260.345703 | NA | NA | 460.8201904 | 3719.317383 | 1057.588501 | NA |
| 34558.76953 | 2968.157227 | 2970.941162 | 3219.257324 | 11650.26855 | 9080.920898 | 10595.08496 |
| 241200.7656 | 41170.58203 | 31311.53711 | 37027.94531 | 187624.8906 | 39062.33984 | 75633.03125 |
| 8959.797852 | 1162.423828 | 833.2860718 | 751.3850098 | 1157.314819 | 954.1676025 | 606.4274292 |
| 54550.04688 | 6673.244629 | 7607.104492 | 8590.817383 | 2294.602295 | 4995.119629 | 5303.40332 |
| 13074.66797 | NA | NA | 1173.389771 | 7377.007813 | 5738.087891 | 4159.017578 |
| 61858.22266 | 5360.647949 | 7148.758301 | 5025.93457 | 20570.72461 | 19187.19141 | 21803.36719 |
| 1789.514038 | 817.8673096 | NA | NA | 10248.78906 | NA | NA |
| 18611.99805 | 1710.777222 | NA | NA | 7611.310059 | 5405.330078 | 2925.045898 |
| 2166.494141 | NA | 351.4532471 | 773.3787842 | 7712.673828 | NA | 2806.717041 |
| 271270.2813 | 16686.37891 | 26291.93555 | 17166.25195 | 12000.94824 | 17442.17188 | 15828.20996 |
| 275498.1875 | 27489.10742 | 16562.125 | 15641.64648 | 44164.51172 | 67603.50781 | 59125.08203 |
| 67147.6875 | 7030.563477 | 6050.828125 | NA | 8906.089844 | 6217.557617 | 6882.315918 |
| 53948.00391 | 4450.89209 | 4051.542236 | 5193.222168 | 25962.81445 | 33010.60547 | 27659.59961 |

| | | | | | | |
|---|---|---|---|---|---|---|
| 92531.49219 | 6898.530273 | 4664.958984 | NA | 5954.634277 | 3516.69873 | 5279.274902 |
| 18207.86523 | NA | NA | 2608.535889 | 65414.36719 | 8076.940918 | 14309.39453 |
| 301828.8438 | 18681.1875 | 18570.23242 | 20309.91992 | 28161.32422 | 20439.24219 | 23673.29492 |
| 287757.1563 | 6402.249512 | 13701.49902 | 17800.30469 | 11503.84277 | 10422.15625 | 11746.84863 |
| 335423.3125 | 11351.37012 | 13556.68359 | 20905.4043 | 93270.65625 | 55319.19922 | 71057.61719 |
| 48857.36328 | 1775.850098 | 3077.706299 | 2846.626465 | 10147.87695 | 11488.58008 | 11987.875 |
| 732.9219971 | NA | 127.1963654 | NA | 4243.45459 | NA | 1255.153564 |
| 99948.96094 | 4780.639648 | 5304.135254 | 4134.541504 | 4613.039551 | 4014.527344 | 5153.63916 |
| 2726.498535 | 86.19256592 | NA | 129.1618805 | 159.429657 | 47.48683167 | 112.4372025 |
| 172177.0469 | 6875.467285 | 4825.227539 | 9013.181641 | 5330.358398 | 11356.72656 | 8475.774414 |
| 29328.54883 | 1218.462036 | NA | NA | NA | NA | NA |
| 31106.42969 | NA | NA | 742.09021 | 1132.143799 | NA | NA |
| 871451.75 | 15327.57422 | 10155.84473 | 16962.88672 | 29821.75391 | 18451.39648 | 32325.04688 |

| KO3_r1 | KO3_r2 | KO3_r3 | FC_LTED_vs_parental | padj_LTED_vs_ | FC_KO1_vs_LT | padj_KO1_vs_ | FC_KO2_vs_LT | padj_KO2_vs_ |
|---|---|---|---|---|---|---|---|---|
| 14956.88379 | 19823.95117 | 15750.14844 | NA | NA | 3.624389974 | NA | 3.564828504 | NA |
| 7269.272461 | NA | 4234.811523 | NA | NA | 1.381407852 | NA | 1.369654477 | NA |
| NA | NA | NA | NA | NA | NA | NA | NA | NA |
| 717.2216797 | 984.6264038 | 926.3806763 | NA | NA | 1.531014727 | NA | 1.50537061 | NA |
| 8062.212891 | 4479.221191 | 819.746582 | NA | NA | NA | NA | NA | NA |
| 283.962616 | 210.1411591 | 588.7685547 | NA | NA | NA | NA | NA | NA |
| NA | 0.799118042 | NA | NA | NA | -2.249970172 | NA | -3.803628113 | NA |
| NA | NA | NA | NA | NA | -0.418489179 | NA | NA | NA |
| 67208.625 | 38893.88281 | 19648.375 | NA | NA | NA | NA | NA | NA |
| NA | NA | NA | NA | NA | NA | NA | NA | NA |
| 2499.503418 | 3159.428467 | 2716.894043 | NA | NA | NA | NA | NA | NA |
| NA | NA | NA | NA | NA | NA | NA | NA | NA |
| 2121.79126 | 2124.632813 | 1411.89209 | NA | NA | -0.694237474 | 0.46953657 | 0.84770192 | 0.118373568 |
| NA | NA | NA | NA | NA | NA | NA | NA | NA |
| 9008.729492 | 11157.09082 | NA | NA | NA | NA | NA | NA | NA |
| 1045.766357 | 922.8390503 | 1376.150024 | NA | NA | NA | NA | NA | NA |
| NA | NA | NA | NA | NA | NA | NA | NA | NA |
| 689.7990112 | 251.2953949 | NA | NA | NA | -0.153256997 | NA | 0.636363406 | NA |
| 581.3278809 | NA | NA | NA | NA | NA | NA | NA | NA |
| NA | NA | NA | NA | NA | NA | NA | NA | NA |
| 2042.495728 | 2350.213379 | NA | NA | NA | 1.341499318 | 0.097594832 | 0.682778057 | NA |
| 4906.279297 | 6037.960938 | 3535.258545 | NA | NA | 1.103778706 | NA | 1.581001009 | NA |
| NA | NA | NA | NA | NA | NA | NA | NA | NA |
| NA | NA | NA | NA | NA | 2.07437308 | NA | NA | NA |
| 7754.713379 | 6995.654785 | 10425.25684 | NA | NA | -0.42204452 | 0.374500771 | 2.697722325 | 0.065467296 |
| 1608.075928 | 1331.665283 | NA | NA | NA | NA | NA | NA | NA |
| 13910.59863 | 9570.53418 | 12636.02051 | NA | NA | NA | NA | NA | NA |
| 4194.758301 | 3602.245605 | 4016.994385 | NA | NA | 1.980087703 | NA | 2.849575595 | NA |
| 1372.526978 | NA | 399.3907776 | NA | NA | NA | NA | NA | NA |
| NA | NA | NA | NA | NA | NA | NA | NA | NA |
| 12789.26367 | 18997.96875 | 19122.85547 | NA | NA | 0.274190327 | 0.64559659 | 1.724045679 | 0.061996002 |
| NA | NA | NA | NA | NA | NA | NA | NA | NA |

| | | | | | | | | |
|---|---|---|---|---|---|---|---|---|
| 2635.561279 | 3294 | 1922.567871 | NA | NA | NA | NA | NA | NA |
| NA | NA | NA | NA | NA | NA | NA | NA | NA |
| NA | 814.9692383 | NA | NA | NA | NA | NA | NA | NA |
| 13219.84668 | 15014.91797 | 16738.00391 | NA | NA | 0.839761221 | 0.205381063 | 0.965947318 | 0.138038355 |
| 5476.356934 | 6590.447754 | 4614.768555 | NA | NA | NA | NA | NA | NA |
| NA | NA | NA | NA | NA | NA | NA | NA | NA |
| NA | NA | NA | NA | NA | NA | NA | NA | NA |
| 1740.327393 | 1312.567871 | 3038.701904 | NA | NA | NA | NA | NA | NA |
| 2179.153809 | NA | NA | NA | NA | NA | NA | NA | NA |
| 8646.071289 | 7844.679688 | 10156.2627 | NA | NA | 2.259825982 | NA | 4.410325844 | NA |
| NA | NA | 440.7835693 | NA | NA | NA | NA | NA | NA |
| 1704.958496 | 2700.479736 | 2833.741943 | NA | NA | 2.228071084 | NA | 1.480437125 | NA |
| 4193.542969 | NA | NA | NA | NA | -0.710880994 | 0.54034971 | -1.712595438 | NA |
| NA | NA | NA | NA | NA | NA | NA | NA | NA |
| NA | NA | NA | NA | NA | NA | NA | NA | NA |
| NA | NA | NA | NA | NA | NA | NA | NA | NA |
| 9336.292969 | 9501.042969 | 8232.431641 | NA | NA | 0.227700846 | 0.603053058 | 1.282390597 | 0.041088687 |
| 599.7752075 | 1202.348633 | NA | NA | NA | -1.432858717 | 0.341543595 | -2.40291708 | NA |
| 3258.72876 | 3519.009277 | 1739.812622 | 5.565766885 | NA | -0.37250222 | 0.245897577 | -0.93402179 | 0.084858149 |
| 16724.45703 | 15202.73926 | 14972.13965 | 5.506679554 | NA | -0.348934117 | 0.478993548 | 0.383445424 | 0.321105038 |
| 2119056.25 | 1835543.375 | 1778667.875 | 5.014520385 | 0.130973753 | -1.53869059 | 0.22025368 | 0.252411747 | 0.646788596 |
| 10342.63477 | 11983.69043 | 14096.67969 | 4.903498054 | 0.077066654 | -0.926000648 | 0.215727673 | 0.530377846 | 0.209736296 |
| 815268.2246 | 878093.1582 | 680586.6079 | 4.728329969 | 0.024701492 | -0.905691809 | 0.07388302 | -0.950835514 | 0.044841696 |
| 646724.9375 | 652366.625 | 644165.5 | 4.666054523 | 0.136409711 | -0.955907292 | 0.331953431 | -0.455657555 | 0.534750592 |
| 13979.94434 | 8180.976074 | 9993.020508 | 4.480826531 | NA | -2.811988444 | NA | -2.41728292 | NA |
| 3811.105957 | 2654.672852 | 1033.154053 | 4.475687577 | NA | -1.709880727 | 0.052476573 | -1.497389667 | 0.056570421 |
| 206688.6875 | 241332.3438 | 216232.8281 | 4.344004223 | 0.047406642 | -1.564353903 | 0.094955363 | -1.268626374 | 0.091928987 |
| 25966.63867 | 24896.33203 | 26342.47461 | 4.297420393 | 0.029811229 | -1.399694465 | 0.05818214 | -1.558622687 | 0.05270761 |
| 24880.06641 | 19629.13477 | 23889.55859 | 4.173591601 | 0.068365552 | -1.418780553 | 0.138660529 | 0.219418677 | 0.529273607 |
| 983.9118652 | 1089.377808 | NA | 4.066666749 | NA | 0.818796286 | 0.649333082 | -2.453550805 | NA |
| 261546.9219 | 296164.625 | 278047.8438 | 3.947452809 | 0.013439299 | -1.480727861 | 0.024557735 | -1.364787694 | 0.011939492 |
| 919418.4375 | 1180221.5 | 1237017.25 | 3.922218148 | 0.04179011 | 0.957915182 | 0.166172033 | 2.067400619 | 0.047850077 |
| 18425.92578 | 25298.89648 | 17066.35547 | 3.899840976 | 0.06543655 | -1.301651761 | 0.136651367 | -1.554678675 | 0.103046053 |

| | | | | | | | | |
|---|---|---|---|---|---|---|---|---|
| 123882.8594 | 127769.2422 | 83301.1875 | 3.867410951 | 0.015044187 | 0.149950359 | 0.356748392 | 0.217481924 | 0.503548482 |
| 1018056.719 | 771709.6323 | 711702.2275 | 3.789834623 | 0.032652748 | -1.31913234 | 0.069086457 | -1.796778857 | 0.036229875 |
| 263452.125 | 260167.5313 | 248894.8906 | 3.780186663 | 0.032339427 | -1.34045592 | 0.054639347 | -1.774933608 | 0.052349495 |
| 82102.64063 | 73835.35156 | 80612.3125 | 3.733193305 | 0.02930047 | -0.653610825 | 0.096995825 | -0.373887693 | 0.157967349 |
| 28628.73047 | 32580.48633 | 26972.66211 | 3.704434594 | 0.025187448 | -0.674394176 | 0.063686231 | 0.058161733 | 0.733558803 |
| 150190.6406 | 147496.8125 | 143331.4219 | 3.692609228 | 0.026154212 | -0.943916225 | 0.071811131 | -0.819778773 | 0.069658709 |
| 35663.12109 | 41558.20313 | 35224.58203 | 3.686136571 | 0.038502747 | -0.186644457 | 0.508456536 | 0.163849539 | 0.454451558 |
| 888574.5 | 987283.8125 | 1029909.875 | 3.677494016 | 0.004709568 | 0.979370119 | 0.028434447 | 2.119911314 | 0.029219266 |
| 8980.504883 | 11199.9873 | 8499.489258 | 3.654761095 | 0.067708966 | -1.508774575 | 0.120245081 | -2.382088896 | 0.091109394 |
| 18505.97656 | 20760.29102 | 19865.79883 | 3.641816159 | NA | -0.74977231 | 0.371581571 | 0.785292884 | 0.159646627 |
| 5531.303711 | 7883.757813 | 4907.563965 | 3.629869754 | 0.017866665 | -2.032449635 | 0.02118733 | -2.463820215 | 0.020008995 |
| 13500.03711 | 15172.88379 | 11580.93848 | 3.629262379 | 0.028428652 | -1.659847933 | 0.050817911 | -0.700489342 | 0.057515439 |
| 2007.959717 | 2350.345459 | 3512.807373 | 3.593067582 | NA | NA | NA | -2.221467627 | 0.105746635 |
| 2557.478271 | 5032.979004 | 1892.476563 | 3.558661036 | 0.295346854 | -3.176717568 | 0.337374554 | -3.637419136 | 0.315909231 |
| 161854.8281 | 177206.5781 | 173672.1875 | 3.528364468 | 0.030959545 | -0.509561427 | 0.117302538 | -0.160858601 | 0.395608126 |
| 12950.89941 | 16084.99316 | 12468.46777 | 3.439666322 | 0.03740048 | -1.365291925 | 0.078527985 | -1.469406837 | 0.053142343 |
| 74491.25 | 56774.90234 | 42561.53516 | 3.402717698 | 0.029514505 | 0.392281085 | 0.108875676 | -0.697488839 | 0.101483387 |
| 12195.01953 | 9105.020508 | 10849.04395 | 3.380211523 | 0.219971212 | 2.958120285 | 0.137910094 | 2.793158063 | 0.024451141 |
| 197283.2656 | 213266.625 | 190910.0469 | 3.340993772 | 0.023710065 | -0.293074462 | 0.25313315 | -0.856695058 | 0.084180144 |
| 24446.38086 | 24817.23828 | 23687.96094 | 3.254395643 | NA | -0.588191442 | 0.252436809 | 0.260925302 | 0.415586176 |
| 41705.66406 | 54154.36719 | 52647.19141 | 3.231119014 | 0.011814081 | -2.117884867 | 0.01179292 | -1.960347777 | 0.007057904 |
| 20036.54688 | 18939.24805 | 22792.23828 | 3.204449595 | NA | -0.214638491 | 0.393273993 | 0.615816108 | 0.058215835 |
| 632777.7412 | 764428.3164 | 735371.6299 | 3.174376848 | 0.111362172 | 1.075398751 | 0.277660269 | 3.031298717 | 0.020008995 |
| 108479.9219 | 111626.7344 | 106921.7734 | 3.165066512 | 0.026153568 | -1.423345864 | 0.047452686 | -2.199813802 | 0.036848746 |
| 7317.398438 | 10948.40625 | 14108.14746 | 3.162208504 | NA | -0.050770197 | NA | 1.43827164 | NA |
| 23021.04297 | 19714.88281 | 28591.88477 | 3.142888371 | NA | 0.106687829 | 0.70926014 | 1.057278469 | 0.074615902 |
| 833840.9375 | 893946.625 | 1091408.375 | 3.075042636 | 0.046448757 | -1.405592078 | 0.085927719 | 0.424058889 | 0.169381511 |
| 1964316.625 | 2320130.25 | 2237203.5 | 3.07214228 | 0.046945973 | -1.414969926 | 0.078283911 | 0.494438912 | 0.108322697 |
| 6125.019531 | 9901.9375 | 10627.66016 | 3.063107182 | NA | -1.400676541 | 0.063663792 | -1.013967905 | 0.074812764 |
| 130539.2734 | 175906.9688 | 118139.9609 | 3.036628914 | 0.009008041 | -1.325775241 | 0.042608596 | -1.357502334 | 0.027533174 |
| 1320039 | 3057658.5 | 1492136.25 | 3.012902462 | 0.009974872 | 0.260640829 | 0.137613908 | -0.328121214 | 0.564224231 |
| 11231.69434 | 14948.03613 | 11151.03223 | 3.001789511 | 0.053170792 | -0.864742252 | 0.130039916 | -3.171895132 | 0.065303359 |
| 84754.92969 | 84249.53906 | 80645.30469 | 2.986408033 | 0.021326467 | -0.262446018 | 0.152134335 | 0.051858748 | 0.645234682 |

| | | | | | | | | |
|---|---|---|---|---|---|---|---|---|
| 196199.7813 | 203372.8594 | 248443.4688 | 2.936808706 | 0.085294501 | -2.109137923 | 0.121847149 | 0.949328239 | 0.062080903 |
| 6516.683594 | 9372.423828 | 8608.152344 | 2.923409344 | 0.08963545 | -1.481886098 | 0.170410796 | 0.720524616 | 0.141604552 |
| 31610.95508 | 24796.75391 | 25976.02344 | 2.921103024 | 0.005790576 | 0.56153997 | 0.306401385 | 0.582137116 | 0.116261556 |
| 3221.07959 | 3537.566162 | 2495.318604 | 2.896537008 | NA | -0.369312256 | NA | -0.787787042 | NA |
| 83019 | 86486.42969 | 83752.14063 | 2.878315925 | 0.018311389 | -1.042337124 | 0.021906439 | -1.653603313 | 0.030832901 |
| 22933.04688 | 28804.01563 | 30969.38086 | 2.877956976 | 0.039205423 | -1.983557345 | 0.067039668 | -0.496919365 | 0.137726349 |
| 7116.710938 | 6097.662109 | 6781.689453 | 2.870431585 | 0.030271458 | -2.048978309 | 0.055241445 | -2.034189312 | 0.051065331 |
| 103176.0391 | 110239.7344 | 106126.8359 | 2.838249826 | 0.034713383 | -0.669528203 | 0.092696398 | 0.205253351 | 0.302562509 |
| 29709.18945 | 24906.6582 | 21775.31641 | 2.831651792 | 0.029750807 | 0.049147794 | 0.773923011 | 0.278392834 | 0.290771554 |
| 26224.45313 | 21800.70898 | 18512.48828 | 2.826514633 | 0.034890445 | 0.56726108 | 0.059340088 | 0.418108838 | 0.203156123 |
| 53780.83594 | 51867.00391 | 41443.28125 | 2.795676316 | 0.045233346 | 0.911999229 | 0.089716327 | -1.676586453 | 0.062345205 |
| 6283.106445 | 5871.151855 | 5811.892578 | 2.769171271 | 0.019995793 | -0.002474588 | 0.984856255 | 0.369505069 | 0.05747179 |
| 3278.577881 | 3250.151123 | 3912.050049 | 2.752690459 | NA | -1.446828667 | 0.089889925 | -1.056381388 | 0.126782073 |
| 477169.9063 | 351721.25 | 550235.625 | 2.746057549 | 0.015698773 | -0.709417697 | 0.05253541 | 0.336212764 | 0.346756059 |
| 155603.3281 | 154548.9844 | 136720.5938 | 2.734889865 | 0.004709568 | -0.914276783 | 0.012690505 | -0.53081385 | 0.02183874 |
| 29717.11719 | 28061.44141 | 25024.44336 | 2.732456079 | 0.238382947 | 1.19053549 | 0.146807068 | 1.734287797 | 0.061962869 |
| 8012.44873 | 9848.798828 | 5020.87793 | 2.729481565 | 0.109755066 | -1.334301515 | 0.197213027 | -1.73263377 | 0.154013101 |
| 1175.999512 | 1709.026489 | 946.2931519 | 2.72496732 | 0.019286562 | -2.014370677 | 0.034480785 | -2.831866412 | 0.026764821 |
| 2294.686768 | 1518.104126 | 1823.980957 | 2.703205092 | NA | 0.517826423 | 0.155173707 | -1.504456121 | 0.091928987 |
| 1415842.625 | 1763120.07 | 2019563.109 | 2.695394849 | 0.027405706 | -1.540684345 | 0.048024604 | -0.577117657 | 0.072839808 |
| 41947.05469 | 40337.79297 | 43708.96875 | 2.685976422 | 0.019182942 | -1.210852615 | 0.021906439 | -1.548765702 | 0.027949491 |
| 77501.63281 | 87109.5625 | 85819.30469 | 2.677755784 | 0.074273374 | -0.29286909 | 0.478541077 | -0.352010705 | 0.397891679 |
| 15934.10938 | 14787.35449 | 29674.93359 | 2.675151805 | 0.077217864 | -0.144360502 | 0.726897808 | 1.118026051 | 0.235836583 |
| 747.901123 | 538.6050415 | NA | 2.660303767 | NA | -0.225255035 | 0.498804814 | -1.156557015 | NA |
| 223245.0781 | 252088.7188 | 279139.9063 | 2.659416273 | 0.02054676 | 0.534213575 | 0.280688037 | 0.845902163 | 0.045887602 |
| 6095.309082 | 6902.674316 | 6866.533691 | 2.658757306 | NA | 0.764029106 | 0.207303274 | 1.083596804 | 0.017279709 |
| 3452.644287 | 3697.217285 | 1991.550781 | 2.655225701 | 0.080233873 | 0.115295562 | 0.788500178 | -0.775777489 | 0.224000425 |
| 22891.13867 | 28851.35742 | 27149.82031 | 2.64799627 | 0.013585622 | -0.642574533 | 0.191112268 | 0.528112535 | 0.094991118 |
| 12274.7168 | 14142.6582 | 9255.323242 | 2.642671583 | 0.029239337 | 2.608533845 | 0.11146465 | 0.842925968 | 0.133481008 |
| 131065.8438 | 142835.8125 | 149700.7656 | 2.640730699 | 0.025086961 | -1.680378227 | 0.038037652 | -1.84037816 | 0.037292993 |
| 6085.197754 | 33662.64844 | 6725.646973 | 2.631380292 | 0.307746917 | -2.727603392 | 0.333170371 | -3.474969515 | 0.301801908 |
| 85302.67969 | 83934.11719 | 93685.32813 | 2.630543809 | 0.022440374 | 0.378919513 | 0.112102762 | 0.000391356 | 0.997605405 |
| 4790.375 | 5884.520508 | 3691.384277 | 2.618297486 | NA | -2.316823389 | NA | 2.464777218 | 0.058909257 |

| | | | | | | | | |
|---|---|---|---|---|---|---|---|---|
| 8596.27832 | 6598.895508 | 9987.162109 | 2.610169884 | NA | 0.825971438 | 0.112543356 | 1.476173978 | 0.061706361 |
| 54823.9375 | 63847.23438 | 30051.44922 | 2.609255002 | 0.109761076 | -0.662346709 | 0.328222763 | 0.135947951 | 0.820784824 |
| 39473.79297 | 43393.36719 | 47994.59375 | 2.599008736 | 0.01455479 | -0.690547089 | 0.058877608 | 0.267370584 | 0.166665266 |
| 302061.3438 | 351158.9375 | 453404.375 | 2.593656938 | 0.089820226 | -0.951714288 | 0.20864513 | 0.465880695 | 0.30221168 |
| 39583.80078 | 41947.4375 | 79855.01563 | 2.584474981 | 0.118360556 | -2.972477962 | 0.13681424 | 0.12572526 | 0.852856834 |
| 27425.86914 | 35876.48438 | 33201.44141 | 2.583728991 | 0.008009891 | -0.35583886 | 0.404718985 | 0.065307173 | 0.716011038 |
| 18709.66016 | 18451.71094 | 16759.63281 | 2.576160664 | 0.024583014 | 0.651504228 | 0.041310052 | 0.616228003 | 0.044156984 |
| 38916.98535 | 41974.10059 | 47150.72363 | 2.57022753 | 0.019930448 | -0.29847729 | 0.187014813 | -0.000834051 | 0.997178143 |
| 32535.91016 | 29949.57227 | 30457.97852 | 2.568023468 | 0.016050563 | 0.181902784 | 0.420257264 | 0.866198879 | 0.011850971 |
| 2091.352783 | 5953.562988 | 3292.49707 | 2.563292435 | NA | 1.438678389 | 0.097722369 | -1.008683073 | 0.128960705 |
| 8221.822266 | 9257.621094 | 7149.461426 | 2.550757977 | 0.02930047 | -1.488917384 | 0.05467585 | -1.526498145 | 0.049557263 |
| 5377.521484 | 2175.088135 | NA | 2.549372881 | NA | -0.620747265 | NA | 0.046896542 | NA |
| 6209.075195 | 7550.969238 | 5403.974609 | 2.544207428 | 0.016105265 | -0.693137538 | 0.093751816 | -1.157956689 | 0.03119922 |
| 5240.17627 | 6196.45459 | 11191.14941 | 2.541223711 | 0.008009891 | -1.655102732 | 0.016244911 | -2.053740329 | 0.01858328 |
| 13018.48242 | 12918.78418 | 11460.32422 | 2.5332639 | NA | 0.31814643 | 0.262820793 | 0.206587953 | 0.403921495 |
| 11210.56738 | 8989.370117 | 8465.136719 | 2.518975632 | NA | 1.302039979 | 0.023497318 | 0.663324828 | 0.09816938 |
| 53973.40625 | 57642.24609 | 53329.28516 | 2.510612544 | 0.033688384 | 0.163962793 | 0.601428216 | 0.619372581 | 0.101451879 |
| 16903.74414 | 18197.46289 | 17534.21094 | 2.485752684 | 0.068494602 | -1.043327302 | 0.135908153 | -0.918681283 | 0.153886288 |
| 127390.1875 | 148359.8125 | 147667.375 | 2.448422461 | 0.032646789 | 0.005915543 | 0.976440441 | -0.00287874 | 0.989480447 |
| 6412.82959 | 1594.251343 | 6946.900391 | 2.431282526 | 0.240260695 | 0.369414886 | 0.648824794 | 0.026290314 | 0.982802241 |
| 373282.2813 | 436056.5 | 408532.3438 | 2.427795503 | 0.037600114 | -0.797721072 | 0.107063979 | -0.994178454 | 0.070421533 |
| 1872.223022 | 1139.498657 | 1137.320068 | 2.416701871 | 0.029476989 | -1.854446576 | 0.038454242 | -0.368559081 | 0.356361596 |
| 5499.573242 | 5071.267578 | 3369.152344 | 2.388584796 | 0.071859228 | -1.135106163 | 0.152452638 | -0.451353783 | 0.340777735 |
| 15309.19824 | 20922.03906 | 12447.85254 | 2.386738714 | NA | 1.650598507 | 0.193587721 | 0.894820549 | 0.15166044 |
| 695536.625 | 708846.0625 | 730400.4375 | 2.361178766 | 0.037022948 | -0.698152562 | 0.135549097 | 0.094202408 | 0.663092542 |
| 36887.67969 | 36172.64063 | 37560.08984 | 2.335217133 | 0.059344159 | -1.455615972 | 0.104928475 | 0.016124064 | 0.957668955 |
| 8071.121582 | 8691.388672 | 7659.907715 | 2.307270422 | 0.07101293 | -1.158117859 | 0.137613908 | -0.602732335 | 0.221788957 |
| 3605720 | 3747985.5 | 3946809.75 | 2.29784649 | 0.005752145 | -0.208204419 | 0.095980292 | 0.022323945 | 0.77681132 |
| 60348.02344 | 45199.50781 | 68918.33594 | 2.277811628 | 0.16335134 | -0.15899495 | 0.805534524 | 1.008621132 | 0.114669448 |
| 9564.300781 | 9886.279297 | 8852.375977 | 2.268949002 | NA | 0.870050306 | NA | 1.533735202 | NA |
| 836.3778076 | 1549.05481 | NA | 2.228274178 | 0.168887111 | -0.720663389 | 0.429618137 | -0.817416695 | NA |
| 46440.49609 | 44185.71484 | 44063.51172 | 2.223961497 | 0.014306629 | -0.98566506 | 0.015625705 | -1.325955012 | 0.021984636 |
| 1614567.25 | 1633095.25 | 1333643 | 2.222586656 | 0.033688384 | 1.236409555 | 0.034038555 | 0.707228016 | 0.049493862 |

| | | | | | | | | |
|---|---|---|---|---|---|---|---|---|
| 68326.49219 | 61497.02344 | 57854.65625 | 2.211684744 | 0.037492489 | -0.865561153 | 0.096609145 | 0.050571268 | 0.793906739 |
| 6662.57373 | 6475.686523 | 7433.756348 | 2.201487932 | 0.023479107 | 0.535560363 | 0.047347558 | -0.436848251 | 0.06365738 |
| 475264.7188 | 466541.5625 | 439302.875 | 2.201369796 | 0.016885305 | 0.095039856 | 0.431724911 | 0.26222234 | 0.102433019 |
| 47212.17578 | 48580.48438 | 53468.19922 | 2.184338689 | 0.054728089 | 1.398124469 | 0.249900189 | 2.835608899 | 0.014469072 |
| 2204984.75 | 2274343.25 | 1845132.875 | 2.180315952 | 0.029239337 | 1.189556256 | 0.015625705 | 0.668858201 | 0.052657999 |
| 4249.371094 | 3954.668945 | 1782.866821 | 2.173815718 | 0.135173059 | -0.728821091 | 0.356945511 | -2.983910104 | 0.141055321 |
| 91312.79688 | 133629.0313 | 165836.75 | 2.168374146 | 0.375561534 | -1.159175418 | 0.530091357 | -1.291302626 | 0.497023216 |
| 4948.146973 | 5873.489258 | 4073.72876 | 2.167909991 | NA | 0.076315822 | 0.721558875 | -0.332124274 | 0.24971241 |
| 10915.84375 | 15621.0498 | 13906.09375 | 2.166818805 | 0.075944008 | -0.154863073 | 0.703049798 | 0.433829085 | 0.261028666 |
| 48359.8125 | 46568.55859 | 46532.48047 | 2.158827519 | 0.029603305 | 0.265152628 | 0.276749208 | 0.356740845 | 0.174451245 |
| 229297.2813 | 226929.9844 | 187856.0781 | 2.158815538 | 0.009974872 | -0.204788069 | 0.150271527 | -0.900798623 | 0.01865088 |
| 13010.7373 | 11985.80273 | 10647.87305 | 2.156823527 | 0.047406642 | -0.004013722 | 0.989771736 | -0.941876993 | 0.098799268 |
| 202447.5 | 210252.2656 | 227772.4375 | 2.154982519 | 0.006735623 | -0.157667121 | 0.218573457 | -0.113115901 | 0.303769038 |
| NA | 287.3918457 | NA | 2.153769362 | NA | -0.540665122 | 0.358946075 | -2.114863026 | NA |
| 7605.976563 | 8947.181641 | 7614.87793 | 2.153140953 | 0.133581817 | -0.125524746 | 0.852417959 | 0.949503366 | 0.107554917 |
| 13851.5752 | 10830.53125 | 13016.13574 | 2.146312134 | 0.042134056 | -0.542687094 | 0.175245919 | 0.01025493 | 0.970918035 |
| 16276.82715 | 18788.98242 | 17413.56445 | 2.144058693 | 0.01031238 | -0.515130646 | 0.05467585 | -0.95768261 | 0.009714527 |
| 88443.85156 | 81449.29688 | 55981.38281 | 2.13892375 | 0.008854154 | 0.464323603 | 0.055340142 | -0.088730263 | 0.759571084 |
| 22414.69727 | 25709.66992 | 21558.81055 | 2.126245938 | 0.044061266 | -1.204782998 | 0.065424224 | -0.944275787 | 0.07580998 |
| 11355.10938 | 9559.884766 | 9758.533203 | 2.124096533 | 0.029199743 | 1.912209659 | 0.050817911 | 0.551971156 | 0.063818875 |
| 18359.49219 | 19082.57422 | 17585.91016 | 2.11533594 | 0.004709568 | 0.259209747 | 0.063918451 | -0.369924423 | 0.020008995 |
| 339700.2188 | 349243.7813 | 338471 | 2.113665803 | 0.026154212 | -0.739144978 | 0.068524409 | -0.634421487 | 0.068838627 |
| 73587.26563 | 74598.36719 | 69324.3125 | 2.094506846 | 0.028932331 | -0.14712693 | 0.444085145 | 0.33511629 | 0.08152982 |
| 9109.792969 | 11151.94141 | 12600.40234 | 2.089887102 | 0.022960243 | -0.42623793 | 0.134788265 | -0.011558686 | 0.960764822 |
| 3945.904785 | 5969.913574 | 4091.224609 | 2.083581652 | 0.101863182 | -1.147000121 | 0.203239255 | -0.854932167 | 0.233347919 |
| 4606.390137 | 3276.397949 | 4754.529785 | 2.076708114 | 0.165760942 | -0.447220736 | 0.548184123 | -0.677566674 | 0.412823101 |
| 32169.49414 | 37920.38672 | 40001.26563 | 2.065410907 | 0.01516079 | 0.089088592 | 0.355229313 | 0.569822805 | 0.082189314 |
| 73164.78125 | 69927.99219 | 74194.80469 | 2.061622469 | 0.054134182 | -1.234655865 | 0.073326865 | -0.164358146 | 0.5195709 |
| 2772.375244 | 2226.044678 | 1490.579346 | 2.052597178 | 0.205289963 | -0.618100486 | 0.528875081 | -1.923301631 | 0.230308748 |
| 90700.39844 | 88612.35938 | 82782.53125 | 2.050743822 | 0.058363046 | -0.310761791 | 0.343549377 | -0.437735666 | 0.239208062 |
| 36622.38672 | 29728.56641 | 32974.33984 | 2.046900245 | NA | 2.4155011 | 0.342966569 | 1.926083642 | 0.063818875 |
| 9486.325195 | 7889.562012 | 6260.166992 | 2.045928574 | 0.160984058 | 1.18454159 | 0.087018661 | 0.713503458 | 0.210955784 |
| 4163.681641 | 4521.039063 | 3833.483887 | 2.045891597 | 0.011384748 | 0.071239834 | 0.675218019 | 0.591819736 | 0.052439423 |

| | | | | | | | | |
|---|---|---|---|---|---|---|---|---|
| 1081447.125 | 1302202 | 1502129.75 | 2.040654448 | 0.030348789 | -0.047151135 | 0.818858061 | 0.477621125 | 0.151966642 |
| 519898.4688 | 648449 | 580697.1875 | 2.03349455 | 0.017533506 | -0.927436674 | 0.042887894 | -0.907338207 | 0.021332797 |
| 82362.77344 | 95336.125 | 95759.33594 | 2.032079617 | 0.005752145 | 0.17397599 | 0.100528183 | 0.12802768 | 0.322383021 |
| 19451.69922 | 36718.68359 | 24723.87305 | 2.030803822 | NA | 0.198111024 | 0.723454571 | 0.632522634 | 0.348056184 |
| 3097.9646 | 3714.605225 | 6134.746582 | 2.014602287 | 0.2355857 | 0.865678358 | 0.342886559 | -0.121481781 | 0.890142907 |
| 11730.16602 | 12808.6582 | 10116.50195 | 2.008036972 | 0.010297098 | -0.094846047 | 0.432898177 | -0.504182111 | 0.078680305 |
| 38423.01563 | 29273.57422 | 35805.51563 | 2.002506223 | 0.018050472 | -0.404703569 | 0.094777667 | -0.629462749 | 0.04784105 |
| 12597.29102 | 7439.78125 | 5837.782715 | 1.994166408 | 0.068289248 | -0.618517162 | 0.272720515 | -0.303999162 | 0.573204391 |
| 350381.9688 | 261707.6875 | 290285.5 | 1.987652933 | 0.047406642 | -1.2295655 | 0.087594938 | -1.766605367 | 0.064726471 |
| 1276205.625 | 1448092.375 | 1602197.375 | 1.986987835 | 0.072163652 | -0.047782398 | 0.885440998 | 0.531197914 | 0.126366029 |
| 30412.19922 | 29400.40039 | 23884.46094 | 1.986165411 | 0.007997389 | -0.251303775 | 0.140039224 | -0.47811935 | 0.060539652 |
| 2709.51709 | 1945.779907 | 1907.186523 | 1.978810243 | 0.040923714 | 0.448688286 | 0.197242348 | -0.247699708 | 0.438271266 |
| 40777.22266 | 39559.77734 | 36119.17188 | 1.978444664 | 0.016115022 | 0.756215194 | 0.135901972 | -0.342838694 | 0.071151453 |
| 61690.33984 | 51648.21875 | 51407.75391 | 1.977065527 | 0.108283439 | 0.025544339 | 0.952846884 | -0.44641914 | 0.397891679 |
| 7983.939941 | 8445.334961 | 8285.261719 | 1.970404189 | 0.02753514 | -0.395839138 | 0.129506032 | 0.200475345 | 0.233299785 |
| 54219.58203 | 46949.57813 | 39503.75 | 1.967086815 | 0.052665954 | -0.261960774 | 0.395857551 | 0.19491886 | 0.507600785 |
| 12286.59375 | 12342.19922 | 4347.39209 | 1.960441134 | 0.101401941 | 1.762038379 | 0.089248231 | -1.248834811 | 0.136000191 |
| 13232.15625 | 11958.21973 | 8264.97168 | 1.957120526 | 0.068084878 | -0.264715004 | 0.428256136 | -0.46909836 | 0.254146446 |
| 3292.571533 | 4693.902832 | 6940.007813 | 1.932398505 | NA | -0.236620823 | 0.529223913 | 0.638014367 | 0.343211558 |
| 2812.043701 | 4148.836426 | 2423.635986 | 1.926202356 | 0.125428544 | -1.676619563 | NA | -1.063663441 | 0.218872239 |
| 29590.13281 | 11763.49121 | 16902.30078 | 1.918783787 | 0.04179011 | -0.058889988 | 0.843871989 | -0.591393012 | 0.316138998 |
| 76147.94531 | 74497.11719 | 57017.47266 | 1.914220799 | 0.05049279 | 0.583851328 | 0.104273255 | 0.510738155 | 0.137834136 |
| 164813.1406 | 160272.9844 | 176052.625 | 1.902004353 | 0.08654621 | -0.192417515 | 0.624832492 | 0.447656747 | 0.226987124 |
| 5133.119629 | 5159.430176 | 4321.158691 | 1.900783441 | 0.019610279 | -0.625824842 | 0.024404888 | -0.326865581 | 0.08376499 |
| 6456.793945 | 4627.48291 | 5370.526855 | 1.899108929 | 0.006395507 | 0.222635806 | 0.289867338 | -0.3096326 | 0.208408498 |
| 3010.359375 | 2698.38623 | 2500.048828 | 1.887692016 | 0.030009295 | -1.371682156 | 0.042884533 | -0.781123435 | 0.070464402 |
| 39130.28516 | 33520.86328 | 28117.5 | 1.887300896 | 0.036590352 | -0.324516311 | 0.18152121 | -0.206886523 | 0.397327146 |
| 191703.3906 | 197416.625 | 248443.5156 | 1.879596685 | 0.132098822 | 1.444584924 | 0.182060286 | 3.85834447 | 0.042310808 |
| 23550.78125 | 43140.17188 | 27618.00781 | 1.869554628 | 0.005752145 | -0.712840438 | NA | 0.006858399 | 0.988723353 |
| 15835.44043 | 23011.79297 | 15954.08887 | 1.867980252 | 0.263341433 | -0.256884065 | 0.779772909 | -0.048166172 | 0.957356765 |
| 349918.9688 | 398541.5938 | 389996.9688 | 1.865066575 | 0.012256654 | -0.379885846 | 0.068255066 | -0.937129568 | 0.02146481 |
| 13438.01172 | 11641.76367 | 10092.47266 | 1.863909559 | 0.103160411 | -0.91222079 | 0.200310376 | -1.054963511 | 0.173821986 |
| 7606.444824 | 10254.57422 | 7813.299805 | 1.856477346 | 0.056696683 | 0.027109718 | 0.814025375 | 0.196441545 | 0.441920881 |

| | | | | | | | | |
|---|---|---|---|---|---|---|---|---|
| 80326.375 | 77938.11719 | 76173.13281 | 1.851546347 | 0.009974872 | -1.055711802 | 0.0155094 | 0.170549188 | 0.086851601 |
| 174708.9531 | 180965.4688 | 208665.9531 | 1.848126047 | 0.094031642 | -0.087764379 | 0.813774916 | 0.446170995 | 0.224000425 |
| 413172.4688 | 450997.9375 | 469276.5938 | 1.84487272 | 0.011206547 | -0.272072438 | 0.054772146 | 0.524201351 | 0.037571191 |
| 86098.5625 | 63034.35938 | 81493.57813 | 1.837669112 | 0.044205353 | -0.571436447 | 0.135790392 | 0.182283506 | 0.478935691 |
| 971641.25 | 927295.125 | 945981.6875 | 1.837278309 | 0.026565565 | 0.035271812 | 0.831728137 | 0.089580564 | 0.543506636 |
| 63121.95313 | 62309.98438 | 63979.78516 | 1.835359332 | 0.024554363 | -0.414104785 | 0.166408179 | 0.189077555 | 0.311264669 |
| 4213.664551 | NA | 690.1661987 | 1.835054463 | NA | NA | NA | 0.514696772 | NA |
| 855.550415 | 3601.210693 | NA | 1.833211229 | NA | -0.311228091 | 0.8496215 | -0.941831816 | NA |
| 9366.086914 | 9442.299805 | 8221.729492 | 1.826342103 | 0.008503176 | -0.501100284 | 0.048271656 | -0.236916386 | 0.117459662 |
| 2046.733765 | NA | 1232.362061 | 1.824623897 | NA | NA | NA | -2.368813713 | NA |
| 78668.07031 | 74007.30469 | 86242.46875 | 1.803013539 | 0.089436106 | 0.260215249 | 0.448317772 | 0.281318563 | 0.396993313 |
| 19924.59375 | 22655.21289 | 19759.09766 | 1.797582901 | 0.052312419 | 0.364842105 | 0.2482979 | 0.144917059 | 0.538145904 |
| 366242.2188 | 382479 | 340855.5625 | 1.796265849 | 0.017355582 | -0.861243495 | 0.01915096 | -1.328786487 | 0.015632484 |
| 2675093.25 | 2871260.25 | 3334697.5 | 1.795575974 | 0.020823572 | -0.301153143 | 0.226695454 | 0.051496786 | 0.804302828 |
| 8423.37793 | 18101.74414 | 18180.95313 | 1.792159984 | 0.542216889 | -0.14479268 | 0.936763383 | -0.640484186 | 0.763776161 |
| 7929.402832 | 10596.0127 | 9333.546875 | 1.787779252 | 0.081090664 | 0.084001346 | 0.852236385 | 0.040879346 | 0.922299095 |
| 156098.0469 | 141045.2031 | 138111.5938 | 1.786183525 | 0.029183044 | -0.62579809 | 0.064535327 | -0.896319263 | 0.041386295 |
| 16643.00781 | 11928.33691 | 13835.0498 | 1.779160032 | 0.023619216 | 0.634773244 | 0.051924975 | 0.958960191 | 0.076969062 |
| 4070.924072 | 5786.203125 | 5209.880371 | 1.77666171 | 0.024701492 | -5.110569779 | NA | -2.899318699 | 0.039446558 |
| 3283.083984 | 2056.027344 | 3318.908447 | 1.77033582 | 0.042872978 | 0.300468951 | 0.311163334 | 0.583517397 | 0.225747871 |
| 29355.12891 | 28677.94531 | 29330.58203 | 1.770255042 | 0.006970105 | -0.209513037 | 0.055241445 | -0.286024219 | 0.04905244 |
| 51230.39063 | 65823.36719 | 79965.04688 | 1.766416012 | 0.044205353 | -1.676984335 | 0.090523063 | 0.707646327 | 0.147429332 |
| 53905.50391 | 14020.51758 | 24445.39746 | 1.763239611 | 0.035482115 | 0.028041054 | 0.967598067 | -0.524724956 | 0.487908677 |
| 4417.962891 | 3303.668213 | 2609.528564 | 1.762388561 | 0.009008041 | -1.223367048 | 0.020431201 | -0.73936496 | 0.103879505 |
| 51155.04297 | 42151.07813 | 43031.47266 | 1.761891095 | 0.018795991 | -0.544302051 | 0.068255066 | -0.34809218 | 0.109149667 |
| 8918.841797 | 7306.935059 | 6896.553711 | 1.756391612 | 0.006395507 | -1.146374185 | 0.013885891 | -1.072558556 | 0.017015337 |
| 50259.44141 | 51642.18359 | 51583.00391 | 1.752713343 | 0.009015232 | -0.675909382 | 0.039582042 | -0.489260278 | 0.065490138 |
| 17089.70898 | 21067.19727 | 19476.24609 | 1.748562119 | 0.036547632 | 0.172900459 | 0.497082817 | 0.474361617 | 0.110105852 |
| 20632.87891 | 20363.92969 | 18881.65234 | 1.741931602 | 0.033894086 | 0.068933397 | 0.732942123 | 0.385683035 | 0.089381209 |
| 22921.78516 | 23611.83984 | 22287.83594 | 1.734162497 | 0.011814081 | -1.123164943 | 0.054857325 | -1.549096352 | 0.034655694 |
| 27760.01367 | 20165.5 | 18597.32227 | 1.733728697 | 0.010297098 | -2.424501327 | 0.030265482 | -2.360115549 | 0.01123893 |
| 18516.94727 | 18737.28906 | 14917.20996 | 1.731589049 | 0.004709568 | -0.656813634 | 0.013910966 | -0.985032669 | 0.020008995 |
| 34632.00391 | 37252.67578 | 31724.04688 | 1.729434574 | 0.048391875 | -1.170849033 | 0.086540831 | -0.839444322 | 0.095685399 |

| | | | | | | | | |
|---|---|---|---|---|---|---|---|---|
| 62480.69141 | 53851.67188 | 60383.19141 | 1.728637977 | 0.006564049 | -0.485471678 | 0.022836499 | -0.208975636 | 0.128367922 |
| 8336.794922 | 7138.744141 | 2595.60376 | 1.724051623 | 0.079009944 | -1.475268507 | 0.101856807 | -1.991905133 | 0.075009766 |
| 583605.875 | 620007.1875 | 601823.5 | 1.720771817 | 0.026892803 | 0.115327394 | 0.633575766 | 1.15268414 | 0.020008995 |
| 29211.2793 | 33151.58594 | 31265.7793 | 1.713128459 | 0.041314706 | 0.546457331 | 0.075404888 | 0.971494678 | 0.020008995 |
| 57972.83594 | 58193.54297 | 54641.22266 | 1.713072101 | 0.010297098 | 0.114367592 | 0.230859884 | 0.38332079 | 0.02146481 |
| 48496.09766 | 45131.58203 | 42329.94141 | 1.707503004 | 0.036830197 | 0.060178666 | 0.724165773 | -0.1109362 | 0.536607309 |
| 100876.6953 | 89291.58594 | 95035.65625 | 1.705104328 | 0.026183333 | -0.572875595 | 0.070271638 | -0.498792819 | 0.084308715 |
| 70406.55469 | 78603.45313 | 62563.17969 | 1.699292478 | 0.013723547 | -1.284018082 | 0.022702163 | -1.341939222 | 0.011394883 |
| 386407.125 | 395192.4375 | 401579.0625 | 1.697612869 | 0.037380825 | 0.291481066 | 0.162317825 | 0.442789448 | 0.093609212 |
| 25542.5918 | 26069.5625 | 24021.59961 | 1.696036354 | 0.018311389 | -0.057642604 | 0.741853858 | 0.202975818 | 0.244006157 |
| 746826.25 | 760454.1875 | 809882.875 | 1.676118612 | 0.008009891 | 0.379477274 | 0.018539883 | 0.473996119 | 0.024074394 |
| 9218.174805 | 10443.71191 | 10233.01465 | 1.672505566 | 0.033123824 | -0.99412821 | 0.069174232 | -0.689153426 | 0.064893735 |
| 79551.84375 | 77527.0625 | 94927.02344 | 1.664890813 | 0.01904327 | -0.084868755 | 0.610945492 | 0.671747908 | 0.074184841 |
| 93803.67969 | 104352.1406 | 98366.55469 | 1.664816344 | 0.037600114 | 0.48451076 | 0.196976249 | 0.941161692 | 0.019022762 |
| 47322.26172 | 52208.08203 | 43772.41406 | 1.663531531 | 0.028196123 | -0.318883934 | 0.126441864 | -1.167833465 | 0.027949491 |
| 46643.21484 | 48544.60547 | 43651.36328 | 1.663124 | 0.011979502 | -0.465598185 | 0.06794739 | -0.259373415 | 0.110105852 |
| 1103278.75 | 1055090.5 | 994527.875 | 1.655669554 | 0.048937447 | 0.174555108 | 0.464701252 | 0.56447878 | 0.071851903 |
| 55133.10156 | 57417.55859 | 58539.60156 | 1.654391458 | 0.007467253 | -0.781919492 | 0.046804468 | -0.914376441 | 0.017279709 |
| 36273.89844 | 40115.83984 | 39859.12109 | 1.65423794 | 0.006735623 | 0.264687635 | 0.055241445 | 0.073378287 | 0.436948967 |
| 3855.122559 | 4154.690918 | 5331.193359 | 1.643134891 | NA | -0.738167331 | 0.102883408 | -0.442656064 | 0.173821986 |
| 4518.973145 | 5373.004395 | 5555.788574 | 1.642662443 | 0.061766802 | -0.470602923 | 0.223159833 | -0.505366883 | 0.181045061 |
| 223928.4688 | 210257.3906 | 200542.7656 | 1.635221701 | 0.013585622 | -0.553590954 | 0.041367426 | -0.721637828 | 0.015734747 |
| 50134.67188 | 57130.21094 | 45461.32813 | 1.634511697 | 0.012774217 | 0.707626102 | 0.059057385 | 0.183435659 | 0.331318816 |
| 55509.73828 | 67291.45313 | 69048.39844 | 1.632688959 | 0.135952427 | -0.45102352 | 0.422801 | 0.418846166 | 0.329050274 |
| 152937.6484 | 165553.2266 | 142085.4531 | 1.626363628 | 0.006351124 | -0.663687225 | 0.031437224 | -0.488462274 | 0.027982615 |
| 13039.6875 | 5455.166016 | 15070.64551 | 1.621997898 | 0.027973416 | 0.198664947 | 0.354607038 | -0.493634278 | 0.367550122 |
| 395811.9375 | 411166.8438 | 433509.8438 | 1.615088739 | 0.035879745 | -0.166405494 | 0.416045961 | 0.568723183 | 0.035693643 |
| 1332.43103 | 3880.656982 | 16950.32813 | 1.614748255 | 0.329662688 | -0.415074961 | 0.795956929 | -0.700261895 | 0.635657892 |
| 42460.05859 | 39513.05078 | 31782.69727 | 1.613484223 | 0.015034245 | -0.064141486 | 0.532402501 | 0.325261329 | 0.215760001 |
| 23503.00977 | 21368.63672 | 19396.64648 | 1.611814927 | 0.09428407 | 1.652209974 | 0.025824895 | 2.078396387 | 0.017083334 |
| 8195.874023 | 10780.69922 | 8163.573242 | 1.606473218 | NA | -0.246186787 | NA | -0.228077454 | 0.405108106 |
| 8605.299805 | 6988.572266 | 7440.339844 | 1.602065614 | 0.008705223 | -0.046024223 | 0.490854054 | 0.24988137 | 0.203834997 |
| 53607.22656 | 57696.63672 | 55201.20703 | 1.600676845 | 0.006930304 | 0.095729282 | 0.264081584 | -0.106660892 | 0.239208062 |

| | | | | | | | | |
|---|---|---|---|---|---|---|---|---|
| 12820.31934 | 13691.43652 | 13232.78906 | 1.600256712 | 0.013723547 | -1.032975358 | 0.0155094 | -0.048320756 | 0.436305727 |
| 17636.11719 | 13616.71875 | 17151.72656 | 1.600040843 | NA | -0.599657697 | NA | -1.123353971 | NA |
| 28869.23242 | 20825.13477 | 22101.96484 | 1.59630271 | 0.006735623 | -0.801757773 | 0.022035738 | -1.545740072 | 0.011046095 |
| 4568.981934 | 7752.279785 | 6606.121582 | 1.595306316 | 0.2756587 | 0.327977123 | 0.648119697 | -0.574730058 | 0.558686385 |
| 4439.302734 | 5501.991211 | 5496.595703 | 1.592337328 | 0.137237024 | 2.16833663 | 0.034038555 | 1.30179628 | 0.043404927 |
| 28027.44141 | 31297.5918 | 27362.68945 | 1.591794769 | 0.232095578 | -0.1883021 | 0.818237711 | 1.802402812 | 0.040645703 |
| 13805.04102 | 13870.60938 | 18831.34766 | 1.590884038 | 0.076933208 | 1.07179874 | 0.14734576 | 0.883698598 | 0.086952277 |
| 72227.51563 | 72954.14844 | 61544.82031 | 1.589795714 | 0.00799298 | -1.195983708 | 0.018539883 | 0.223160417 | 0.181929698 |
| 9334959 | 10618336 | 11025278 | 1.584399479 | 0.01031238 | 0.186553047 | 0.181649566 | 0.645041663 | 0.041258108 |
| 762081.875 | 743354.875 | 795514.6875 | 1.582607745 | 0.081796003 | -1.186453083 | 0.129928099 | -0.1635853 | 0.611068784 |
| 25813.1543 | 25830.61328 | 31466.56641 | 1.581368791 | 0.061715176 | 1.029415101 | 0.107505082 | 1.11926778 | 0.032840382 |
| 26244.05859 | 13765.23047 | 10213.47559 | 1.572112978 | 0.329857534 | -1.671586676 | 0.350656099 | -1.541693002 | 0.352451091 |
| 12328.09961 | 12246.84961 | 11935.7666 | 1.570397482 | 0.061681941 | 0.321480974 | 0.370232693 | 0.197854875 | 0.510430611 |
| 118364.2813 | 101233.9297 | 109252.6328 | 1.569978641 | 0.036762746 | 0.472973747 | 0.134788265 | 0.69174724 | 0.041258108 |
| 56193.14453 | 29225.00977 | 42495.64453 | 1.568908426 | 0.145132831 | 1.765337577 | 0.419290117 | 0.492178396 | 0.410161823 |
| 556978.3125 | 574579.6875 | 511262.3125 | 1.566393026 | 0.027921552 | -0.412383091 | 0.116553799 | -0.523329785 | 0.065399262 |
| 54490.10156 | 58064.66016 | 56026.32031 | 1.56019129 | 0.025025466 | -1.497295022 | 0.041910161 | -1.450076853 | 0.040934649 |
| 3505373 | 3834570.75 | 3355666 | 1.559454923 | 0.036480793 | -0.039027827 | 0.847283599 | 0.218769215 | 0.297710892 |
| 106522.4531 | 105848.1484 | 106600.4453 | 1.5548222 | 0.003149457 | 0.100848283 | 0.24069685 | 0.613463614 | 0.009587646 |
| 68202.61719 | 60288.08203 | 83164.21094 | 1.552712224 | 0.011063961 | 0.677401408 | 0.05467585 | 0.665019438 | 0.119166108 |
| 4045.042725 | 4693.470703 | 4922.640625 | 1.550492871 | 0.016189975 | -0.696972376 | 0.048593497 | -0.427549871 | 0.062822 |
| 10177.18457 | 12189.96387 | 11489.29102 | 1.549989494 | 0.338882582 | 0.648645793 | 0.423723931 | 0.253480122 | 0.763765593 |
| 96234.78906 | 95602.53125 | 92031.03125 | 1.549292402 | 0.021798208 | -0.454038214 | 0.099830362 | -0.699032438 | 0.02183874 |
| 109087.0313 | 110939.3984 | 122798.1797 | 1.547095888 | 0.018050472 | -0.01241963 | 0.903989395 | 0.320547961 | 0.062345205 |
| 20768.33398 | 22769.49609 | 15820.5332 | 1.545490981 | 0.089040397 | 0.804984282 | 0.349008413 | 0.709183097 | 0.115275002 |
| 11816.89648 | 10804.29395 | 9247.512695 | 1.543392991 | 0.140640517 | 1.902359792 | 0.03520579 | -0.696874797 | 0.298016429 |
| 16269.47754 | 14808.9502 | 9451.922852 | 1.539255991 | 0.009011817 | 0.536382447 | 0.235640931 | -1.945984898 | 0.010623938 |
| 309878.0625 | 300994.0938 | 213303.7969 | 1.531749557 | 0.170961164 | 1.647670745 | 0.30612213 | -0.414244024 | 0.531405981 |
| 11828.69824 | 12777.52344 | 10042.61621 | 1.52839516 | NA | 0.567368646 | NA | 1.856536482 | NA |
| 9997.882813 | 9578.055664 | 10060.06641 | 1.522763635 | 0.005752145 | -0.007636921 | 0.933858968 | -0.182678504 | 0.040645703 |
| 309749.3438 | 248026.1406 | 155456.5156 | 1.52162043 | 0.029355394 | 0.098695608 | 0.539179679 | -0.25573556 | 0.527330576 |
| 19634.22461 | 21266.21875 | 18742.11914 | 1.521618318 | 0.074010001 | 0.158083785 | 0.695596157 | 0.808296776 | 0.063440273 |
| 20181.28516 | 22966.19727 | 15790.83203 | 1.51914428 | 0.072333886 | 0.015062702 | 0.955379211 | 0.061555543 | 0.850636261 |

| | | | | | | | | |
|---|---|---|---|---|---|---|---|---|
| 28449.90234 | 19641.65234 | 28916.05078 | 1.517405428 | 0.169115491 | 0.243498157 | 0.611644229 | 0.18342924 | 0.719887357 |
| 260368.8906 | 278359.9375 | 306131.375 | 1.514438929 | 0.006735623 | 0.158525527 | 0.385120531 | 0.210233752 | 0.179985011 |
| 27510.37109 | 31755.43945 | 25359.79102 | 1.513507582 | 0.016596319 | 0.057574833 | 0.54213578 | -0.274657067 | 0.167717489 |
| 1343919.375 | 1046820.188 | 1240942.625 | 1.512970056 | 0.049763403 | 0.169297154 | 0.45507774 | 0.006857133 | 0.98127134 |
| 89702.51563 | 97742.04688 | 108364.2734 | 1.511580191 | 0.009974872 | -0.205854498 | 0.144798646 | 0.100727887 | 0.484109463 |
| 16582.22266 | 18443.22461 | 14676.41699 | 1.506860376 | 0.039721652 | -1.812572626 | 0.053336389 | -2.041133846 | 0.045975624 |
| 331747.4375 | 342822.3125 | 362659 | 1.505033952 | 0.005752145 | -0.40286053 | 0.049184287 | -0.389763318 | 0.021836949 |
| 158925.875 | 151945.5625 | 162879.0313 | 1.504500843 | 0.004709568 | -0.299018615 | 0.033056067 | -0.23266755 | 0.031785503 |
| 62362.29688 | 67952.75 | 57600.15625 | 1.494885847 | 0.02624826 | 0.462543626 | 0.102526216 | -0.175932005 | 0.249497567 |
| 13375.13965 | 15211.28809 | 12702.74316 | 1.491617234 | 0.012414001 | -0.006543743 | 0.954690016 | 0.127786184 | 0.396993313 |
| 4832.35791 | 4289.212402 | 6267.69043 | 1.486169583 | 0.127138267 | 0.59385969 | 0.192808829 | 0.890838041 | 0.102257588 |
| 24855.64453 | 22418.96875 | 27977.74609 | 1.484902851 | 0.153739559 | 0.047782056 | 0.922182791 | 1.369656017 | 0.036042199 |
| 7405.083984 | 8111.032227 | 15869.38965 | 1.482152357 | NA | 2.5937129 | 0.012254503 | 1.518473225 | 0.218219327 |
| 149451.5625 | 171395.9531 | 160591.1563 | 1.478436281 | 0.014003671 | 0.351524211 | 0.088308366 | 0.51294307 | 0.035414333 |
| 82241.92969 | 116714.9688 | 65545.6875 | 1.477651836 | 0.015030322 | 0.049344537 | 0.848507078 | -0.00779349 | 0.986294081 |
| 21158.12305 | 20655.41992 | 20979.45313 | 1.463283702 | 0.012414001 | 0.037259871 | 0.826734157 | 0.034807088 | 0.258294119 |
| 212407.25 | 204540.4063 | 237426.4688 | 1.461247635 | 0.01508889 | -0.216012773 | 0.11146465 | 0.364505233 | 0.070481383 |
| 48661.27344 | 36134.46094 | 39470.77734 | 1.460333608 | 0.009974872 | -0.454554644 | 0.052457323 | -0.611022835 | 0.056222926 |
| 116960.6406 | 107838.3984 | 110421.1094 | 1.459468253 | 0.01391113 | 0.3413959 | 0.216140905 | 0.190532368 | 0.181045061 |
| 9812.061523 | 9612.726563 | 6423.804199 | 1.454500818 | 0.436947846 | 1.543285256 | 0.131789366 | 0.139008299 | 0.903232066 |
| 56433.17969 | 59224.37891 | 52494.19531 | 1.453732189 | 0.013723547 | -0.271038064 | 0.107325642 | -0.28969758 | 0.046487363 |
| 82456.98438 | 81301.92188 | 88530.4375 | 1.453234246 | 0.029603305 | -0.169097006 | 0.358946075 | -0.434827781 | 0.086952277 |
| 19354.88281 | 22452.07422 | 25093.55469 | 1.452505312 | 0.080233873 | -0.937494482 | 0.125467937 | 0.014631138 | 0.963218346 |
| 18850.96875 | 18322.68359 | 20849.56055 | 1.451976885 | 0.037492489 | 0.168355711 | 0.501923815 | -0.160928553 | 0.518850652 |
| 180715.6406 | 92951.15625 | 127047.6172 | 1.447863468 | 0.109140206 | 0.629403267 | 0.178660809 | 0.925405231 | 0.205642234 |
| 9516.075195 | 9763.667969 | 7705.566895 | 1.447671161 | 0.042404592 | 0.060294609 | 0.704376222 | -0.43074684 | 0.118706964 |
| 16105.7832 | 15700.63184 | 14803.39453 | 1.447637664 | 0.03211333 | 0.546762024 | 0.050817911 | 0.358690308 | 0.083362426 |
| 391241.3438 | 395218.6563 | 408279.0625 | 1.446618364 | 0.094885214 | -0.52063585 | 0.278265841 | -0.467744049 | 0.297215475 |
| 50069.44141 | 43211.50781 | 40679.83203 | 1.445331978 | 0.011269714 | 0.539144559 | 0.356546603 | 1.72162961 | 0.043050202 |
| 15452.65918 | 11123.75098 | 16102.35938 | 1.44238462 | 0.005713167 | -0.260312777 | 0.17813923 | -0.163168428 | 0.500573086 |
| 4203.561523 | 3280.852295 | 2813.083496 | 1.441018611 | NA | 0.101651416 | 0.773721709 | 0.306781177 | 0.402996452 |
| 7415.19873 | 6689.601074 | 7863.60498 | 1.440752491 | 0.021027835 | 0.011726577 | 0.896060924 | -0.20085453 | 0.155520889 |
| 695088.375 | 751321.5625 | 733489.125 | 1.439817088 | 0.219971212 | 1.086461177 | 0.420830585 | 2.530688636 | 0.017279709 |

| | | | | | | | | |
|---|---|---|---|---|---|---|---|---|
| 69106.60938 | 59915.25 | 51388.375 | 1.437924899 | 0.027158619 | -0.311073586 | 0.195590467 | -0.000271645 | 0.998983823 |
| 79510.38281 | 74028.52344 | 74827.36719 | 1.437427943 | 0.004709568 | -0.467323667 | 0.028434447 | -0.387486302 | 0.017763718 |
| 7501.763672 | 9972.819336 | 3981.326904 | 1.435517106 | 0.06480491 | -0.318440799 | 0.369927787 | -1.698691912 | 0.052967087 |
| 52908.57422 | 24569.32227 | 38082.40234 | 1.427888828 | 0.046236587 | -0.231489371 | 0.687518345 | 0.15173695 | 0.765741675 |
| 15696.81348 | 11774.85352 | 13745.63281 | 1.427728236 | 0.040326505 | 0.216264155 | 0.467062508 | -0.05514219 | 0.840720144 |
| 50789.29688 | 68928.41406 | 40541.83203 | 1.423597778 | 0.044665667 | 0.27205209 | 0.320787116 | -0.410906525 | 0.29061103 |
| 196551.75 | 212821.0313 | 218508.7031 | 1.421327767 | 0.013186491 | 0.252939453 | 0.111308998 | 0.264800759 | 0.080462751 |
| 6366.512695 | 3334.419434 | 4898.039063 | 1.420325085 | 0.066639151 | -1.440284176 | 0.086089321 | -1.379219878 | 0.054887084 |
| 204032.4219 | 156061.6406 | 160163.5 | 1.419474919 | 0.06543655 | -0.292633291 | 0.555335026 | -0.236459768 | 0.510714902 |
| 18658.98633 | 19653.60547 | 17876.22266 | 1.41752698 | 0.025818052 | 1.178802989 | 0.036071705 | 1.842479192 | 0.009026966 |
| 17467.74805 | 18976.83789 | 17920.75195 | 1.416435418 | 0.011062932 | -0.312733988 | 0.087295253 | -0.826926216 | 0.034677634 |
| 203665.25 | 147633.2969 | 143760.3281 | 1.415337366 | 0.035776373 | -0.779412858 | 0.112996996 | 3.059302619 | 0.06365738 |
| 55521.03125 | 57501.83984 | 51419.51563 | 1.41532247 | 0.037709795 | -0.053763098 | 0.729706165 | 0.005129055 | 0.975148062 |
| 27456.26563 | 40940.17188 | 29240.29297 | 1.413320607 | 0.041542612 | -0.513617773 | 0.167753597 | 0.132862673 | 0.689648336 |
| 2757.640869 | 3265.415283 | 3535.725098 | 1.410535406 | 0.238909917 | -0.467619956 | 0.601291377 | -0.475381398 | 0.544630143 |
| 154207.7656 | 169816.5781 | 185648.5313 | 1.409454906 | 0.037729096 | 1.377243702 | 0.158510575 | 2.057843663 | 0.026553937 |
| 3309.731689 | NA | NA | 1.40916133 | 0.401602606 | 0.359925876 | 0.723122465 | -0.766627886 | NA |
| 48549.34766 | 52791.27734 | 47214.98047 | 1.408600318 | 0.018497551 | -0.762609166 | 0.067039668 | -0.58905795 | 0.065367829 |
| 51668.5625 | 54681.89453 | 56378.89307 | 1.404227705 | 0.041945948 | 0.392866046 | 0.147323941 | 1.041919412 | 0.020008995 |
| 31744.99805 | 30856.41602 | 31010.89844 | 1.396568161 | 0.029183044 | -0.288623022 | 0.207242983 | 0.059909646 | 0.697131411 |
| 16322.95703 | 10908.74219 | 17177.63867 | 1.396069274 | 0.038727982 | 0.102112359 | 0.642450324 | 0.456638175 | 0.265688274 |
| 3149092.75 | 2798566.25 | 3493532.75 | 1.38956938 | 0.009974872 | 0.366733273 | 0.039699307 | 1.118099308 | 0.049973194 |
| 297543.5938 | 288526.7188 | 303598.875 | 1.383796308 | 0.026880281 | 0.446348389 | 0.174475307 | 1.160958325 | 0.006394096 |
| 6940.299805 | 6302.861816 | 7066.305664 | 1.371077484 | 0.028582048 | -0.261549532 | 0.23287903 | -1.29826402 | 0.053053903 |
| 8029.554688 | 7439.63916 | 7959.904785 | 1.366670177 | 0.282458905 | -0.290212216 | 0.726796503 | 0.360205789 | 0.580443707 |
| 31152.52148 | 23672.81055 | 27950.9043 | 1.36647547 | 0.009015232 | 0.226124162 | 0.328806215 | -0.027390573 | 0.882286856 |
| 199680.3281 | 197217.375 | 227529.5938 | 1.366334257 | 0.029239337 | 0.15730142 | 0.348633551 | 0.699864401 | 0.031700361 |
| 7814.35791 | 7248.726074 | 7720.365723 | 1.366261406 | 0.036480793 | -1.033875826 | 0.047212306 | -0.310147108 | 0.191124445 |
| 54366.80859 | 69716.77344 | 49105.3125 | 1.365627192 | 0.055439293 | 0.923863319 | 0.055375611 | -0.315873024 | 0.372845622 |
| 33250.08594 | 28464.9668 | 22799.9707 | 1.361787091 | 0.169788346 | -0.466087559 | 0.437151431 | -0.444941803 | 0.444894646 |
| 131675.5781 | 98402.14063 | 112227.1953 | 1.361519532 | 0.032646789 | -0.225829647 | 0.354607038 | 0.403361815 | 0.170776308 |
| 9136.714844 | 6976.858398 | 10220.48535 | 1.360772539 | NA | 0.067376288 | 0.80599196 | -0.563470507 | 0.123837484 |
| 8545.073242 | 8816.609375 | 7174.274414 | 1.35391828 | 0.036547632 | 0.086780035 | 0.677878706 | 0.15531425 | 0.469755826 |

| | | | | | | | | |
|---|---|---|---|---|---|---|---|---|
| 17683.53906 | 16875.7832 | 13563.04004 | 1.353817688 | 0.027973599 | -1.695726569 | 0.007509388 | -1.646788154 | 0.010654184 |
| 6360.936523 | 5618.741699 | 6919.117188 | 1.353115375 | 0.293892903 | -0.139715625 | 0.861195218 | 0.010392801 | 0.989480447 |
| 26960.31445 | 28017.24414 | 26770.19336 | 1.352799007 | 0.013964852 | -0.532590369 | 0.028138975 | -0.456233429 | 0.021332797 |
| 63695.91016 | 72420.82813 | 87537.46094 | 1.348827908 | 0.066869554 | 0.31933087 | 0.257671339 | 1.186845246 | 0.056004888 |
| 247442.4688 | 253711.2031 | 289664.375 | 1.34413919 | 0.026670912 | 0.464726463 | 0.042884533 | 1.138258155 | 0.031136253 |
| 48437.96484 | 54978.69531 | 58960.6875 | 1.343592544 | 0.006564049 | -0.088019644 | 0.439995312 | -0.240663101 | 0.142837957 |
| 13237.6709 | 16701.625 | 19852.51953 | 1.337400132 | 0.009974872 | -0.643210389 | 0.080440542 | -1.119272815 | 0.023855453 |
| 35608.72656 | 40082.76172 | 40759.14844 | 1.336734115 | 0.036534375 | -1.542171579 | 0.039887225 | -0.697907068 | 0.063474265 |
| 33271.20703 | 28637.47461 | 29379.70703 | 1.336666822 | 0.038395823 | -0.522203331 | 0.105321264 | -0.497358384 | 0.082931684 |
| 408535.1875 | 452075.9063 | 540900.75 | 1.33549175 | 0.099406137 | -0.783049044 | 0.204902073 | -0.350016353 | 0.426736751 |
| 15826.08008 | 13230 | 10119.55469 | 1.32805326 | 0.023527743 | 0.144993079 | 0.714831692 | -0.380514048 | 0.220732169 |
| 9470.395508 | 11625.81836 | 11585.20605 | 1.327453982 | 0.085292813 | 0.773416145 | 0.054768492 | 0.326425644 | 0.145071579 |
| 2282540 | 1177707.625 | 2652513 | 1.325486063 | 0.425266977 | -1.447795505 | 0.433110489 | -0.388427295 | 0.753922782 |
| 20351.89063 | 20846.91797 | 27595.7832 | 1.316005081 | 0.036830197 | -1.081919958 | 0.032229098 | 0.574022503 | 0.16200644 |
| 9853.101563 | 11311.18262 | 11181.63574 | 1.315200674 | 0.034381342 | -0.69476094 | 0.053389337 | -0.809824128 | 0.04324216 |
| 137152.6406 | 137579.4531 | 143388.2031 | 1.313595512 | 0.046112206 | 0.435134905 | 0.111891262 | 0.692660157 | 0.05953462 |
| 59747.25 | 66749.42188 | 51672.27734 | 1.312957829 | 0.026435669 | -0.562529354 | 0.054857325 | -0.171690452 | 0.340584009 |
| 17146.04492 | 18934.79883 | 16342.99707 | 1.309861371 | 0.081999941 | 0.639651004 | 0.169437739 | 1.105360851 | 0.031308498 |
| 239995.5156 | 231587.9375 | 224198.0781 | 1.308330913 | 0.099542225 | -0.128639452 | 0.703047486 | -0.106353575 | 0.747473963 |
| 85114.44531 | 93610.54688 | 87175.40625 | 1.307945568 | 0.024701492 | -0.45979981 | 0.071542761 | -0.542614278 | 0.062063322 |
| 53967.44922 | 56234.98828 | 55812.72656 | 1.30393197 | 0.011946285 | -0.555131347 | 0.020431201 | -0.393762285 | 0.027982615 |
| 102701.3828 | 69867.17188 | 85474.96094 | 1.29889585 | 0.05267493 | 0.536153966 | 0.162537117 | 0.875197258 | 0.101487706 |
| 1028.976807 | 1729.989502 | 716.0490723 | 1.297777945 | NA | -1.020440809 | 0.175968875 | -1.117664602 | 0.148086138 |
| 16791.79883 | 22126.31445 | 17903.75977 | 1.296477067 | NA | 0.669454192 | 0.444175174 | 0.905804296 | 0.214490455 |
| 28035.93945 | 26201.27539 | 25914.14258 | 1.295849625 | 0.06007694 | 0.055354918 | 0.852417959 | 0.792342777 | 0.053012031 |
| 5935.796387 | 5816.563477 | 6077.152832 | 1.293580915 | 0.06329053 | -0.493315022 | 0.224985976 | -0.630574491 | 0.170476688 |
| 22660.35938 | 21039.58203 | 20608.05469 | 1.292919387 | 0.1622947 | -0.212776397 | 0.097793713 | -0.179564618 | 0.114252413 |
| 156633.2969 | 146668.25 | 116874.3672 | 1.29068416 | 0.007485535 | 0.526723046 | 0.05283098 | -0.05928503 | 0.753859857 |
| 14885.2041 | 10676.78906 | 12755.50879 | 1.289315368 | 0.05239238 | -0.093102278 | 0.394278648 | -0.228428185 | 0.31203033 |
| 207752.8125 | 235481.9063 | 201110.3281 | 1.288181259 | 0.030839901 | 1.36848212 | 0.013579478 | 1.49090671 | 0.020008995 |
| 6856.706055 | 7480.774902 | 3759.794434 | 1.283501909 | 0.039762681 | -1.253887888 | 0.054857325 | -1.683118921 | 0.0236713 |
| 44429.48047 | 46685.55859 | 40223.83594 | 1.278834408 | 0.027671686 | -0.127289763 | 0.620414583 | 0.130195213 | 0.441242238 |
| 991808.6875 | 1082831.375 | 1140739 | 1.2786942 | 0.025086961 | 0.754111893 | 0.058077004 | 0.542567098 | 0.034567592 |

| | | | | | | | | |
|---|---|---|---|---|---|---|---|---|
| 4744.948242 | 3943.174072 | 1446.005493 | 1.278311783 | 0.085560167 | -1.064884417 | 0.096457635 | -1.376314858 | 0.069955133 |
| 60737.21484 | 53161.53516 | 65623.5 | 1.277293621 | 0.066082895 | -0.526523867 | 0.177852058 | -0.420422886 | 0.185585346 |
| 49609.94141 | 37498.88672 | 48694.83594 | 1.272329453 | 0.026764186 | -0.167595495 | 0.61125836 | 0.477788636 | 0.134550901 |
| 109627.5547 | 111523.4219 | 114796.6406 | 1.267833524 | 0.012414001 | 0.079046459 | 0.42300023 | 0.348281678 | 0.054887084 |
| 33281.125 | 33265.28516 | 31058.04102 | 1.267617816 | 0.006564049 | -0.437615052 | 0.03633877 | -0.698327978 | 0.020157267 |
| 1410677.625 | 1576573.75 | 1645282.75 | 1.267294271 | 0.058208305 | 0.165434907 | 0.462352123 | 0.242317273 | 0.297701087 |
| 14218.70898 | 13679.55957 | 15579.69727 | 1.266396602 | 0.004709568 | -0.275185839 | 0.036006083 | -0.653608519 | 0.016463359 |
| 162474.7656 | 165607.2656 | 180721.0938 | 1.263096006 | 0.029061881 | -0.595859587 | 0.052305796 | -0.998577045 | 0.033709967 |
| 491289.4688 | 502921.875 | 487735.4688 | 1.262477963 | 0.033748356 | 0.430973699 | 0.083003488 | 1.118052338 | 0.024273435 |
| 12386.00195 | 12568.10547 | 8948.769531 | 1.26062324 | 0.186721358 | 0.508359796 | 0.308832804 | -0.035202582 | 0.950211597 |
| 17501.60938 | 16171.23438 | 31085.66602 | 1.258905751 | 0.153779868 | -0.620621559 | 0.332451144 | -0.183932688 | 0.762717902 |
| 92942.66406 | 101991.4766 | 95871.32813 | 1.25661611 | 0.022480145 | -0.565388208 | 0.076036407 | -0.780766091 | 0.063099035 |
| 20732.30469 | 21102.99609 | 24583.04297 | 1.254964968 | 0.054407811 | 1.598434953 | 0.306289709 | 2.72577361 | 0.026764821 |
| 11068.22559 | 9714.166016 | 9494.509766 | 1.253597511 | 0.04646045 | 0.638399362 | 0.07908104 | 0.461257554 | 0.110277386 |
| 19212.60547 | 18477.76953 | 14985.36621 | 1.252466468 | 0.005752145 | -0.140011464 | 0.076552776 | -0.539957654 | 0.065367829 |
| 41055.85156 | 47704.22656 | 43113.34766 | 1.249996592 | 0.039721652 | -0.003834306 | 0.977473209 | 0.063847357 | 0.676744371 |
| 28575.13281 | 32837.78125 | 30767.23633 | 1.248299722 | 0.182074985 | -0.145252334 | 0.808922116 | 1.020603577 | 0.085236358 |
| 17592.08203 | 15910.7832 | 16053.07422 | 1.247867263 | 0.072163652 | 0.453904604 | 0.212030117 | 0.639451961 | 0.092209064 |
| 114052.4844 | 113371.7734 | 118310.7188 | 1.239615417 | 0.017729013 | 0.926687832 | 0.123324423 | 2.104628345 | 0.00797582 |
| 20149.4082 | 22837.15234 | 19229.46289 | 1.239299102 | 0.028047191 | -0.777065094 | 0.026826422 | -0.616768323 | 0.027559784 |
| 36131.64063 | 36764.78516 | 40708.11328 | 1.237761435 | 0.010297098 | 0.331160933 | 0.260639434 | 1.518357256 | 0.021984636 |
| 22987.33594 | 31622.17383 | 38834.67578 | 1.236175734 | 0.178859946 | -1.085415676 | 0.245391836 | 0.785321541 | 0.185585346 |
| 2182.114014 | 3767.961182 | 3427.406494 | 1.235255532 | 0.132688125 | 0.725055167 | 0.110657758 | 1.50703575 | 0.101659771 |
| 12219.32617 | 6569.59082 | 7468.862305 | 1.234549482 | 0.033433262 | -0.474161676 | 0.220151814 | -0.897290993 | 0.086503564 |
| 5877.722656 | 8204.59082 | 5639.906738 | 1.234283134 | 0.017627122 | -0.806827606 | 0.051888756 | -1.1219488 | 0.027087229 |
| 1542.912231 | 1833.136353 | 383.8429871 | 1.231489027 | 0.406258935 | -0.068273185 | 0.947443559 | -1.513088395 | 0.377816458 |
| 2073.64917 | 4308.822266 | NA | 1.229574426 | 0.09808182 | 0.176730926 | 0.602152681 | -0.541998683 | NA |
| 203743.1406 | 247699.1719 | 188016.1875 | 1.22837747 | 0.069662211 | 0.146999268 | 0.521939419 | 0.10195133 | 0.702969142 |
| 815353.6875 | 1103478.75 | 1280905.875 | 1.228111269 | 0.048232496 | -0.444159652 | 0.208267253 | 0.338170034 | 0.356805712 |
| 240284.4688 | 241758.9844 | 226349.7344 | 1.226616576 | 0.009008041 | -0.356105942 | 0.230974993 | -0.684205435 | 0.020157267 |
| 18870.80469 | 28970 | 10344.68945 | 1.21965935 | 0.188632173 | 0.262716931 | 0.614330411 | -0.553745943 | 0.457083172 |
| 45317.17969 | 50170.4375 | 50847.60547 | 1.218794966 | 0.016511099 | 0.601582443 | 0.029364116 | 0.771697735 | 0.020008995 |
| 140490.4844 | 132214.75 | 126676.7188 | 1.215838007 | 0.018238823 | 0.576068519 | 0.060391495 | 0.942659625 | 0.016338114 |

| | | | | | | | | |
|---|---|---|---|---|---|---|---|---|
| 5640.438965 | 8207.512695 | 7632.932617 | 1.214010717 | 0.13260851 | -0.546822918 | 0.327559963 | -1.023946824 | 0.168508025 |
| 29047.4707 | 25921.68555 | 21623.35156 | 1.21002406 | 0.010025291 | -0.233008278 | 0.100636165 | -0.36471532 | 0.129065826 |
| 645063.375 | 682361.375 | 632936.3125 | 1.208127233 | 0.024149941 | 0.031992434 | 0.842416661 | 0.653395158 | 0.021366148 |
| 37835.82813 | 31039.11719 | 34069.05469 | 1.207793498 | 0.009316552 | -0.557964227 | 0.058077004 | 1.074736281 | 0.049145076 |
| 72965.125 | 102925.75 | 87149.46094 | 1.202933748 | 0.146342609 | -0.040428421 | 0.944504375 | 0.261760284 | 0.52683785 |
| 6680.02832 | 4095.674561 | 4779.875 | 1.199676631 | 0.180746458 | -0.167834731 | 0.750780433 | -0.244698879 | 0.668945234 |
| 15426.85156 | 19003.15625 | 17037.35547 | 1.197395197 | 0.048531905 | 0.656382815 | 0.126364459 | -0.031764414 | 0.856578816 |
| 6397.613281 | 7639.38623 | 7881.663086 | 1.197327677 | 0.010484689 | -0.624585315 | 0.042044993 | -1.187220873 | 0.014469072 |
| 32977.625 | 33704.65625 | 34020.88281 | 1.196240864 | 0.05114005 | -1.097508971 | 0.069257825 | -0.720656052 | 0.102257588 |
| 6911.306641 | 6636.811523 | 5047.71875 | 1.195140336 | 0.005752145 | -0.903076138 | 0.037014778 | -1.785028755 | 0.009587646 |
| 3268.83252 | 2774.675049 | 1495.742676 | 1.193333117 | 0.235000038 | 0.405076177 | 0.597251041 | -0.367740454 | 0.606124172 |
| 12004.75391 | 15256.42969 | 12208.21387 | 1.192713974 | 0.119023657 | -0.385490216 | 0.379832076 | 0.277540075 | 0.423557711 |
| 19919.19141 | 20374.68945 | 16800.51953 | 1.190642753 | NA | 0.981497239 | 0.049358751 | 1.926631701 | 0.025120016 |
| 20153.24219 | 36253.14844 | 30665.63867 | 1.189462194 | 0.031442732 | 0.691666186 | 0.210646218 | -0.047843732 | 0.894865646 |
| 17006.76953 | 15931.86426 | 16092.31055 | 1.188827228 | 0.010297098 | -0.447344016 | 0.05467585 | -0.559172121 | 0.040786554 |
| 7419.416016 | 10264.2334 | 8365.442383 | 1.183634575 | 0.024111712 | -0.005580976 | 0.955379211 | -0.434127605 | 0.112907293 |
| 20940.07031 | 24476.42578 | 25470.83984 | 1.183486765 | 0.083609247 | 0.389095729 | 0.220221768 | 0.451583818 | 0.145687527 |
| 14991.59863 | 13692.55664 | 12722.61328 | 1.18344464 | 0.029906057 | -0.331261469 | 0.100538317 | -0.133051146 | 0.329050274 |
| 29848.40039 | 8173.754395 | 9877.962891 | 1.181149457 | 0.026698267 | -1.730430675 | 0.028434447 | -1.298079406 | 0.14314135 |
| 46336.77344 | 35303.70313 | 31451.59375 | 1.181126568 | 0.032172412 | -0.164554378 | 0.446654651 | -1.065258337 | 0.044248771 |
| 12952.76465 | 17039.96289 | 12969.92871 | 1.180738529 | 0.124822319 | -0.246433464 | 0.586104878 | 0.355868038 | 0.371124353 |
| 6012.884277 | 3938.21167 | 4387.619141 | 1.180473352 | 0.020740891 | -0.554166536 | 0.066354581 | -0.678294606 | 0.081022357 |
| 13004.53613 | 20840.30273 | 14308.77246 | 1.179134885 | NA | 0.402029368 | 0.135823019 | 0.545652504 | 0.263528267 |
| 77001.79688 | 75487.38281 | 64007.22266 | 1.176080483 | 0.134042679 | 1.510432734 | 0.033067345 | 0.109009372 | 0.771355563 |
| 53606.98438 | 48341.19922 | 56937.34766 | 1.175929945 | 0.006351124 | 0.763716128 | 0.013579478 | 1.049729786 | 0.039446558 |
| 63473.46875 | 57551.53906 | 80953.55469 | 1.175641956 | 0.012774217 | 0.013127058 | 0.941915718 | -0.248414435 | 0.312919238 |
| 7951.791992 | 8405.630859 | 8584.006836 | 1.174298428 | 0.211214576 | 0.542353598 | 0.312626085 | 0.28331679 | 0.563915247 |
| 7072.018555 | 3328.143799 | NA | 1.171936364 | NA | 0.354477489 | 0.697793261 | 1.193027787 | NA |
| 68227.35156 | 65163.37109 | 70960.24219 | 1.170939877 | 0.037138683 | 0.58436705 | 0.14131635 | 1.468853386 | 0.009026966 |
| 98094.16406 | 95533.17188 | 94508.39844 | 1.170545087 | 0.029016212 | 0.55382107 | 0.06919378 | 0.851365296 | 0.034518294 |
| 84494.39063 | 85606.5625 | 95783.28906 | 1.169165683 | 0.022444544 | -0.756224818 | 0.033121608 | 0.371733188 | 0.057068534 |
| 40704.5625 | 44010.74219 | 48228.24609 | 1.168633467 | 0.063885064 | -0.152089479 | 0.494663499 | -0.290101486 | 0.259190799 |
| 13414.6709 | 13533.98633 | 14819.60742 | 1.167829952 | 0.020740891 | -0.581377742 | 0.047896832 | -0.331596268 | 0.068248139 |

| | | | | | | | | |
|---|---|---|---|---|---|---|---|---|
| 51736.27344 | 44698.05469 | 45063.65234 | 1.166257553 | 0.037309446 | -0.080373852 | 0.77771169 | 0.077191064 | 0.692619488 |
| 148576.5781 | 162958.5938 | 160467.2344 | 1.166224478 | 0.019286562 | -0.339833793 | 0.114739915 | 0.443276764 | 0.044885273 |
| 8291.416992 | 11367.9502 | 9627.420898 | 1.166030065 | 0.036480793 | -0.024056379 | 0.943661372 | 0.536134252 | 0.126366029 |
| 4999.904785 | 5683.711914 | 4902.153809 | 1.165503301 | 0.052312419 | -0.150031665 | 0.659670874 | -1.208030043 | 0.086843504 |
| 64019.46484 | 66426.20313 | 66106.39063 | 1.164847257 | 0.01488475 | -1.041093625 | 0.024042158 | 0.215758837 | 0.082930409 |
| 18860.91406 | 17608.62305 | 18829.38672 | 1.163599052 | 0.026425936 | 0.296453639 | 0.272913308 | 0.952682958 | 0.009587646 |
| 66967.625 | 129434.4297 | 116455.9141 | 1.161510391 | 0.415082806 | -0.061313999 | 0.953723766 | 0.49726878 | 0.565082073 |
| 68251.16406 | 63914.11719 | 68497.10156 | 1.160272647 | 0.048596967 | -0.014431174 | 0.938183324 | 0.093185703 | 0.583521768 |
| 9385.25 | 10801.60742 | 18094.5293 | 1.158160762 | 0.048820442 | -0.774461662 | 0.108962882 | 0.586795513 | 0.360914175 |
| 629441.625 | 666295.375 | 725805.125 | 1.157640797 | 0.006395507 | -1.572245677 | 0.022021696 | -0.776353117 | 0.012551348 |
| 74761.94531 | 77240.46094 | 52133.80078 | 1.15535076 | 0.006789933 | -1.727700546 | 0.013427556 | -1.752828308 | 0.010236624 |
| 133691.4063 | 126862.3125 | 132841.7813 | 1.154319801 | 0.013723547 | -1.057230623 | 0.021408573 | -0.84903068 | 0.036013501 |
| 24774.98438 | 27412.33789 | 25762.00195 | 1.1541967 | 0.26557175 | -0.104379503 | 0.881715975 | -0.059281755 | 0.929695889 |
| 29514.06836 | 30725.22852 | 30729.25195 | 1.152433155 | 0.031396254 | 0.46757548 | 0.081663959 | 0.023225887 | 0.894936858 |
| 95688.15625 | 105179.2969 | 119350.4375 | 1.152307246 | 0.063774293 | -0.11813264 | 0.629009067 | 0.995969472 | 0.034457303 |
| 3534.907227 | 7958.088867 | 7796.617188 | 1.149485698 | 0.238398694 | -1.214964883 | 0.093818896 | -1.019585895 | 0.10057243 |
| 88145.79688 | 134721.2969 | 129874.7891 | 1.149191844 | 0.041722383 | 0.026921142 | 0.858056341 | 0.289520105 | 0.394131236 |
| 172158.5938 | 173300.7969 | 173320.6406 | 1.148733794 | 0.010297098 | -0.056955559 | 0.258883868 | -0.21358082 | 0.042737273 |
| 31122.30469 | 31702.31055 | 28308.62695 | 1.14721627 | 0.037600114 | -0.374747285 | 0.015829308 | -1.034217307 | 0.007057904 |
| 318208.125 | 357985.5 | 334882.75 | 1.144707602 | 0.18659711 | -0.367681973 | 0.517875146 | -2.012120252 | 0.145687527 |
| 47879.68359 | 50566.12891 | 49231.39063 | 1.143398617 | 0.193611705 | -0.540209471 | 0.398870378 | -0.078309885 | 0.871042009 |
| 1151392 | 1220465.375 | 1297411.75 | 1.1417385 | 0.014003671 | -0.372340779 | 0.029435257 | -0.535894742 | 0.018257145 |
| 52876.57813 | 67702.72656 | 46130.20313 | 1.141709123 | 0.122520185 | -0.434861554 | 0.382214644 | 0.645937419 | 0.163777675 |
| 547407.0625 | 508895 | 561232.4375 | 1.139980473 | 0.005752145 | 0.293131906 | 0.033067345 | 0.017859655 | 0.78762586 |
| 199609.6719 | 234202.3906 | 254941.0313 | 1.137697465 | 0.006395507 | -0.234659941 | 0.124127483 | 0.135597151 | 0.440211328 |
| 666995.3125 | 670659.1875 | 678779 | 1.136765139 | 0.030049926 | -0.546448847 | 0.074598722 | -0.058314773 | 0.621327615 |
| 232203.2031 | 171056.6875 | 391250.5625 | 1.134348049 | 0.542216889 | -2.309053799 | 0.428256136 | -0.61485668 | 0.706396046 |
| 4042.189453 | 4221.57666 | 3705.204102 | 1.130325105 | 0.078894657 | -0.098134135 | 0.703427718 | -0.063203407 | 0.801045589 |
| 3980.297119 | 11391.375 | 4813.884277 | 1.128769641 | 0.37697881 | -1.118666856 | 0.411843428 | -0.342385756 | 0.753625984 |
| 48713.11328 | 59770.33984 | 73558.31641 | 1.128086845 | 0.171423534 | 0.581002403 | 0.204821604 | 0.408660576 | 0.368735396 |
| 27739.92773 | 28607.33398 | 25178.94141 | 1.126959165 | 0.013618049 | -0.212401373 | 0.126224068 | -0.273402265 | 0.07988042 |
| 40212.98047 | 40413.81641 | 33191.92578 | 1.124867294 | 0.025437157 | 0.648230284 | 0.043921768 | 0.474667951 | 0.080533614 |
| 15920.2041 | 13972.58691 | 14635.28906 | 1.124369425 | 0.036275447 | -0.898142517 | 0.088587863 | -1.0176655 | 0.064726471 |

| | | | | | | | | |
|---|---|---|---|---|---|---|---|---|
| 12495.25 | 12349.26758 | 12774.00684 | 1.12160131 | 0.022906187 | -0.204425338 | 0.261269703 | -0.115428987 | 0.441678579 |
| 20046.41211 | 16296.70508 | 22014.76758 | 1.117462831 | 0.109739471 | -0.378431955 | 0.353604166 | -0.024855974 | 0.946678732 |
| 4012213.75 | 4032679.25 | 4446028 | 1.117088559 | 0.080764036 | 0.141631565 | 0.580580715 | 0.49413968 | 0.118023574 |
| 8824.113281 | 8764.856445 | 6752.836426 | 1.117013938 | 0.010782008 | 0.076077263 | 0.534566787 | 0.344947151 | 0.205451497 |
| 350783.2188 | 320133.3438 | 321257.3438 | 1.116203897 | 0.059622902 | -0.256309167 | 0.3150978 | 0.236020441 | 0.26481418 |
| 603716.375 | 589077.75 | 609794.375 | 1.115844314 | 0.014444503 | -0.112546735 | 0.253723346 | -0.130947748 | 0.123837484 |
| 10238.40723 | 10595.05078 | 14650.91699 | 1.114711545 | NA | -0.229106645 | 0.607218021 | 1.588518587 | 0.063937169 |
| 72498.03125 | 81580.60156 | 84505.96875 | 1.112900678 | 0.045092515 | 0.488886681 | 0.152415276 | 0.894383302 | 0.023920487 |
| 44113.16406 | 49305.93359 | 43593.39844 | 1.106003905 | 0.130732496 | 0.100324897 | 0.761592516 | -0.765322703 | 0.199326473 |
| 42041.60156 | 38597.61328 | 37609.80078 | 1.104072435 | 0.004709568 | 0.292255241 | 0.044006658 | 0.037345697 | 0.639103574 |
| 71191.64844 | 70493.66406 | 68932.60156 | 1.103515265 | 0.058893147 | -0.01138871 | 0.954509825 | 0.497308321 | 0.089275366 |
| 16022.39551 | 15549.21094 | 17658.58594 | 1.102511071 | 0.015034245 | -0.31781732 | 0.080311688 | -0.95164357 | 0.021984636 |
| 3575.043457 | 6504.27832 | 4577.529297 | 1.102303066 | 0.036971527 | 0.542562998 | 0.13460208 | -0.088316448 | 0.827301412 |
| 321798.9063 | 342654.1875 | 361793.0313 | 1.100697916 | 0.054961851 | 0.165902405 | 0.393083848 | 0.389756286 | 0.107612185 |
| 4012.509521 | 3123.991943 | 3597.11084 | 1.097880887 | 0.088657003 | -0.491148329 | 0.242583672 | -0.488219926 | 0.233037555 |
| 17786.30859 | 17551.40234 | 16002.79297 | 1.09709708 | 0.035879745 | 0.033439644 | 0.531229987 | -0.436074813 | 0.021836949 |
| 28508.36719 | 29343.05664 | 27187.25781 | 1.09540706 | 0.013723547 | -0.480072781 | 0.065813387 | -0.756143537 | 0.02909479 |
| 237155.9297 | 242973.5195 | 252930.5352 | 1.092659993 | 0.014615724 | 0.684187205 | 0.090322517 | 0.870048302 | 0.009014131 |
| 609275.0625 | 621748.9375 | 646906.3125 | 1.09053186 | 0.105696928 | 0.012432479 | 0.968951556 | -0.030797261 | 0.923778828 |
| 41753.26953 | 37112.42969 | 43260.08984 | 1.090415318 | 0.009974872 | -0.764399216 | 0.047452686 | -0.52800936 | 0.027783576 |
| 21172.69336 | 21872.80469 | 19035.96094 | 1.087584651 | 0.016471012 | -0.01837012 | 0.795956929 | -0.409244794 | 0.039195867 |
| 2293.60376 | 3352.81543 | NA | 1.087573987 | 0.047817825 | -1.032818228 | 0.078077873 | -0.921684829 | NA |
| 64551.60205 | 78945.65234 | 99304.52051 | 1.08740473 | 0.482548201 | 0.830691262 | 0.432802496 | 0.808728883 | 0.371497539 |
| 6653.658691 | 6011.80957 | 5862.706055 | 1.084928952 | 0.04186746 | -0.219146236 | 0.23287903 | 0.072440043 | 0.45718387 |
| 12781.06641 | 11182.79102 | 10445.19824 | 1.083414793 | 0.170961164 | -0.860508865 | 0.098459633 | -1.603574214 | 0.064726471 |
| 52647.375 | 48560.35938 | 41248.46094 | 1.081535996 | 0.024701492 | -0.00218985 | 0.988847797 | -0.234824832 | 0.243417346 |
| 17705.58398 | 17102.69531 | 14577.20313 | 1.07917626 | 0.023448038 | -0.095669272 | 0.480429158 | 0.055593912 | 0.731461707 |
| 1009.104553 | 1273.513794 | 1236.276978 | 1.07841232 | 0.046351003 | 0.345812564 | 0.218573457 | 0.17316452 | 0.443995111 |
| 7299.49707 | 7942.35498 | 6722.050781 | 1.077153263 | 0.048752402 | 0.511128966 | 0.059562109 | -0.315787955 | 0.17086882 |
| 4646.802246 | 4618.116211 | 6570.645508 | 1.076167838 | 0.123070256 | -0.522957662 | NA | -0.574114085 | 0.232012235 |
| 132429.6719 | 145002.1406 | 161315.6563 | 1.073658761 | 0.037380825 | 0.773722736 | 0.021906439 | 0.79711045 | 0.04174272 |
| 10618.4668 | 10121.68652 | 6548.87207 | 1.072187636 | 0.013723547 | -0.930585614 | 0.03588199 | -1.096878343 | 0.033620828 |
| 445187.875 | 455013.3125 | 467258.8125 | 1.069066839 | 0.037600114 | 0.194466764 | 0.216856868 | 0.541208488 | 0.046593982 |

| | | | | | | | | |
|---|---|---|---|---|---|---|---|---|
| 17206.82617 | 28926.10938 | 15309.1709 | 1.068963133 | 0.029183044 | -0.425861827 | 0.14231796 | 0.005203138 | 0.99140824 |
| 30645.29883 | 30128.10156 | 30699.4668 | 1.068324465 | 0.0224814 | -0.040074272 | 0.876926289 | 0.621462511 | 0.049256739 |
| 270545.4063 | 270855.7188 | 256029.2656 | 1.065917083 | 0.027379407 | -0.215288087 | 0.223895569 | -0.082194652 | 0.538909094 |
| 28985.67578 | 28998.14648 | 27544.52734 | 1.062354884 | 0.047406642 | -0.498796561 | 0.13812939 | -0.523806996 | 0.134445271 |
| 28901.89648 | 29219.12305 | 32077.10156 | 1.060922923 | 0.005752145 | 0.290028006 | 0.098036049 | 0.661883615 | 0.033400717 |
| 7667008.5 | 10552360 | 11159200 | 1.060269437 | 0.113315536 | -0.49150753 | 0.265441238 | -0.175482606 | 0.617864353 |
| 15978.5791 | 17508.5625 | 7183.218262 | 1.059492681 | 0.634817545 | 0.181974751 | 0.911966349 | 0.298548419 | 0.845265499 |
| 36508.08984 | 32895.11328 | 31872.54297 | 1.059284546 | 0.007626474 | -0.121950938 | 0.306017296 | -0.07072149 | 0.472928318 |
| 166597.1563 | 143338.375 | 140306.9531 | 1.057727104 | 0.005944135 | -0.28549073 | 0.122851676 | 0.264105067 | 0.167717489 |
| 104270.9219 | 103170.9609 | 92325.32031 | 1.054693471 | 0.017507272 | -0.093466267 | 0.436667107 | -0.113606131 | 0.339165145 |
| 854831.3125 | 917941.0625 | 919574.75 | 1.053355658 | 0.012731179 | 0.24371774 | 0.168900641 | 0.222542143 | 0.075895689 |
| 3654.699707 | 3909.450195 | 2760.237305 | 1.051421563 | 0.02402416 | -1.223987737 | 0.015993065 | -0.474853638 | 0.107554917 |
| 8325.78125 | 7978.071289 | 7336.702637 | 1.050545824 | 0.048656739 | -0.13318817 | 0.306017296 | -0.161649545 | 0.138029431 |
| 12337.88574 | 16467.46484 | 4322.126465 | 1.050247134 | 0.289025013 | -0.403125282 | 0.591184368 | -0.263876598 | 0.763128234 |
| 919.3852539 | 1668.737305 | 884.8944702 | 1.04887208 | 0.061954848 | -0.04754838 | 0.897511445 | -0.71261316 | 0.162108808 |
| 3707461.75 | 4089129.5 | 5008816.5 | 1.044036132 | 0.061375594 | 0.753738471 | 0.054857325 | 0.159048965 | 0.574930993 |
| NA | 3372.234863 | 2324.192383 | 1.043320934 | 0.03927615 | -0.039639916 | 0.837024526 | -1.602937553 | NA |
| 66782.52344 | 70028.24219 | 73068.78125 | 1.041914149 | 0.036971527 | -1.015372796 | 0.055241445 | -1.53531803 | 0.038631271 |
| 74307.65625 | 81391.59375 | 77888.71875 | 1.040556618 | 0.011206547 | 0.289199263 | 0.028354175 | -0.271551724 | 0.040133731 |
| 2666.323486 | 2841.099609 | 1762.457397 | 1.039242473 | NA | 0.220077445 | 0.345103309 | -0.272063419 | 0.409848335 |
| 990773.4375 | 996836.1875 | 1097432.5 | 1.037930715 | 0.025437157 | -0.080380052 | 0.414187935 | 0.374999162 | 0.04296087 |
| 73047.17969 | 69334.60156 | 73193.89063 | 1.037431941 | 0.065789073 | -0.743115866 | 0.100538317 | -0.36322999 | 0.201736266 |
| 35183.45313 | 36175.14844 | 38148.64453 | 1.035933789 | 0.023465918 | -0.218762403 | 0.288975313 | 0.452342824 | 0.045009854 |
| 114002.3906 | 103863.9141 | 110346.0859 | 1.03539501 | 0.013618049 | -0.011752567 | 0.834946164 | -0.061753022 | 0.408181376 |
| 18593.54492 | 22418.28125 | 19302.36133 | 1.035190515 | 0.024610025 | -0.733175841 | 0.067911877 | -0.288011088 | 0.157477573 |
| 28548.15234 | 26183.55859 | 32522.25586 | 1.035013036 | 0.155839329 | -0.593183943 | 0.284889693 | 0.132514028 | 0.713705355 |
| 88899.17969 | 59636.22266 | 69051.96875 | 1.03490037 | 0.012028293 | -0.405386852 | 0.04667516 | -0.247951269 | 0.361492414 |
| 85104.82031 | 91197.64063 | 88842.40625 | 1.033102949 | 0.006351124 | -0.208200817 | 0.051593867 | -0.304910895 | 0.020351059 |
| 13497.57422 | 14481.93457 | 16008.74219 | 1.03292511 | 0.011411241 | -0.031006349 | 0.786431195 | 0.29823996 | 0.123837484 |
| 116027.3984 | 99701.00781 | 78346.58594 | 1.032792841 | 0.016551837 | -0.008634704 | 0.929570723 | -0.15620481 | 0.527185627 |
| 4136.76709 | 5937.595215 | 4372.237793 | 1.03117756 | 0.007922417 | -0.417886893 | 0.064324132 | -0.601446287 | 0.09942999 |
| 130163.75 | 124924.9141 | 112924.7422 | 1.03105614 | 0.012414001 | -0.069354988 | 0.532402501 | 0.386821088 | 0.056482153 |
| 49870.51172 | 44040 | 39639.83203 | 1.030246381 | 0.104757749 | -0.164521567 | 0.604199716 | -0.064048989 | 0.836731763 |

| | | | | | | | | |
|---|---|---|---|---|---|---|---|---|
| 217388.6875 | 156854.5313 | 174318.7188 | 1.027764547 | 0.018311389 | 0.419390899 | 0.100538317 | 0.014811137 | 0.952860484 |
| 7838.630371 | 3179.076172 | 8845.275391 | 1.026263243 | 0.161921798 | -0.491231666 | 0.137613908 | -0.274478866 | 0.616310209 |
| 70178.66406 | 56805.35938 | 66494.55469 | 1.024167076 | 0.024259746 | 0.538214274 | 0.011710842 | 0.314088633 | 0.160764344 |
| 547.788147 | 1499.522705 | 1779.828979 | 1.023502368 | 0.271839224 | -0.59560807 | 0.466149389 | -0.667524152 | 0.434631993 |
| 17948.55078 | 15727.22266 | 16041.44629 | 1.01932183 | 0.015044187 | -0.370956133 | 0.012254503 | -1.071946676 | 0.014388274 |
| 45707.75 | 45647.90625 | 67486.84375 | 1.019279239 | 0.164028708 | 0.383513848 | 0.340389871 | 1.595096243 | 0.080462751 |
| 359664.0938 | 319434.0938 | 353140.9063 | 1.017050429 | 0.019930448 | 0.150325645 | 0.284636711 | 0.586966082 | 0.027982615 |
| 27166.0625 | 27741.91797 | 32637.25391 | 1.016843378 | 0.04765314 | -0.090625931 | 0.679510784 | 1.249063017 | 0.030633392 |
| 122164.5859 | 130338.3594 | 132428.8438 | 1.016463001 | 0.005752145 | -0.010648582 | 0.722366426 | -0.144777002 | 0.08767208 |
| 81059.51563 | 74107.14063 | 66074.625 | 1.015941184 | 0.008731287 | -0.260288821 | 0.068483508 | -0.454593204 | 0.056570421 |
| 104617.1406 | 107408.3984 | 114740.4688 | 1.014810617 | 0.015044187 | 0.015934106 | 0.858056341 | 0.193999361 | 0.08376499 |
| 9757.486328 | 8868.397461 | 7020.625977 | 1.014432466 | NA | 0.842906828 | NA | 1.27457696 | NA |
| 65953.10938 | 71466.0625 | 78299.875 | 1.012884665 | 0.063399586 | 0.406447918 | 0.140334789 | 0.913072968 | 0.026321276 |
| 10277.11133 | 11458.12695 | 10541.44336 | 1.012541293 | 0.333686137 | 0.395695581 | 0.536301388 | 0.565171127 | 0.375359006 |
| 77864.17188 | 50053.72266 | 66759.46875 | 1.012011151 | 0.044763187 | 0.204690233 | 0.256913558 | -0.486314039 | 0.138525852 |
| 39883.05164 | 41721.24829 | 35184.27954 | 1.010950652 | 0.029016212 | -0.027915495 | 0.855467828 | -0.551477166 | 0.076144169 |
| 49008.48047 | 44790.91797 | 37764.25781 | 1.010619426 | 0.128084307 | 0.450535877 | 0.275021879 | 1.699019776 | 0.029628992 |
| 857724.6875 | 815102.875 | 753615.625 | 1.008657508 | 0.01031238 | 0.027668837 | 0.781053515 | -0.025805397 | 0.788453891 |
| 9525.464844 | 12237.0918 | NA | 1.008284946 | NA | NA | NA | 1.594544259 | NA |
| 11997.2832 | 11762.11914 | 13218.14844 | 1.007585661 | 0.3674342 | 1.217106202 | 0.133233444 | 1.108322905 | 0.145132006 |
| 16246.2002 | 16338.04102 | 13108.59766 | 1.004264844 | 0.075499022 | -1.075890104 | 0.105604495 | 1.008319238 | 0.038631271 |
| NA | 7731.489258 | 9312.990234 | 1.002896139 | NA | -0.813000466 | NA | -1.2227342 | NA |
| 22891.24805 | 23703.32617 | 21050.35156 | 1.000794174 | 0.111176969 | 0.389980919 | 0.242732622 | 0.332196141 | 0.273802519 |
| 18291.76953 | 23934.29492 | 19415.73047 | 1.000721294 | 0.02600233 | -0.628189364 | 0.068436129 | -0.899777845 | 0.025666912 |
| 24076.72266 | 24177.72656 | 22788.00391 | 0.999756117 | 0.037210637 | -0.000358868 | 0.998405218 | 0.310100036 | 0.16248539 |
| 34546.58984 | 26527.60547 | 30332.1875 | 0.99893678 | 0.046945973 | 0.605038022 | 0.10108711 | -0.049732794 | 0.840549457 |
| 17098.07617 | 17810.15234 | 14981.88965 | 0.998416397 | 0.128931164 | 1.015161595 | 0.082833891 | 1.238245146 | 0.029219266 |
| 56728.70313 | 57520.00391 | 59930.28516 | 0.997017426 | 0.021353118 | 0.74920868 | 0.152621595 | 0.763969686 | 0.017279709 |
| 125032.7891 | 125771.3438 | 132043.25 | 0.99668729 | 0.006351124 | 0.619248598 | 0.01179292 | -0.17922079 | 0.053012031 |
| 27374.21484 | 31361.61133 | 29616.50391 | 0.996069383 | 0.011946285 | -0.140623834 | 0.497760151 | 0.175983374 | 0.170102465 |
| 186295.7656 | 181800.4219 | 186647.4375 | 0.995119074 | 0.072163652 | -0.062048629 | 0.847335172 | 0.487085851 | 0.117181297 |
| 56535.59375 | 48269.78516 | 58382.04688 | 0.992571689 | 0.018050472 | 0.003279482 | 0.978921212 | -0.242876431 | 0.15808356 |
| 14388.47168 | 17013.53125 | 17814.44922 | 0.992114619 | 0.051750296 | -0.295718354 | 0.271060093 | 0.540577347 | 0.076362612 |

| | | | | | | | | |
|---|---|---|---|---|---|---|---|---|
| 22164.24609 | 21737.83203 | 20605.3418 | 0.991265009 | 0.103527693 | -0.63893628 | 0.20864513 | 0.29078314 | 0.336101329 |
| 13210.25977 | 12334.76074 | 15506.92871 | 0.989301528 | 0.020318767 | -0.478860169 | 0.038159713 | 0.35713349 | 0.161931175 |
| 41825.96094 | 37193.36328 | 26249.12305 | 0.988866961 | 0.029016212 | -0.788658612 | 0.044099436 | -1.05730543 | 0.036624925 |
| 84879.10938 | 83868.0625 | 82156.51563 | 0.988667645 | 0.004709568 | -0.138644877 | 0.096169757 | -0.309864803 | 0.010654184 |
| 1315391.375 | 1307239.75 | 1225376.5 | 0.988500935 | 0.015716954 | 0.360174116 | 0.092495024 | 0.300892761 | 0.063869245 |
| 3118.387451 | 3519.312988 | 2463.333984 | 0.987488977 | 0.09684552 | 0.164943657 | 0.566154586 | -0.328032752 | 0.371644128 |
| 69673.52344 | 68047.85938 | 76277.47656 | 0.985843649 | 0.025300184 | -0.371583379 | 0.10117671 | -0.641155916 | 0.040915765 |
| 45711.82422 | 44368.11328 | 33788.63281 | 0.985365248 | 0.050774241 | 0.11732532 | 0.500982577 | -0.113092775 | 0.639103574 |
| 1203171 | 1405844.125 | 1273921.375 | 0.981489813 | 0.039633894 | -0.634564525 | 0.073324814 | -0.430363252 | 0.072839808 |
| 83899.64063 | 64964.17188 | 83094.30469 | 0.980672578 | 0.012149163 | -0.238983145 | 0.128915752 | -0.316603807 | 0.147031451 |
| 27463.8457 | 26047.94141 | 25839.85352 | 0.980080599 | 0.068494602 | -0.547806663 | 0.143106258 | -0.156883171 | 0.50437262 |
| 26588.42383 | 18416.68359 | 18983.42188 | 0.979634057 | 0.013723547 | -0.020054739 | 0.919703825 | -0.208925554 | 0.445667792 |
| 537000.25 | 558035.625 | 596112 | 0.978201792 | 0.001230443 | -0.07840136 | 0.088294231 | -0.000668771 | 0.992472642 |
| 28813.48242 | 26762.69727 | 24791.24219 | 0.974605742 | 0.024701492 | -0.075640072 | 0.330821995 | -0.25728831 | 0.107554917 |
| 140545.9063 | 86934.64063 | 141584.2813 | 0.973965873 | 0.061354438 | 0.047582042 | 0.832010656 | -0.075651248 | 0.833116553 |
| 11561.85547 | 10324.54297 | 9489.660156 | 0.973668889 | 0.012291622 | -0.155523735 | 0.355609923 | -0.119136389 | 0.380984766 |
| 2201649.5 | 2211416 | 2276437.75 | 0.973076641 | 0.046236587 | 0.431164402 | 0.231823309 | 0.58553317 | 0.050623317 |
| 164368.2813 | 140827.8125 | 139253.2813 | 0.972329864 | 0.011384748 | -0.084499484 | 0.31383898 | -0.27251481 | 0.106144265 |
| 4063.333008 | 2944.724609 | 2090.349121 | 0.972245142 | 0.166935656 | 2.462797959 | 0.01179292 | -0.076032683 | 0.885245898 |
| 26688.08594 | 18572.05664 | 23381.97266 | 0.970936443 | 0.010484689 | -0.120805496 | 0.223235675 | -0.150506572 | 0.511169671 |
| 27258.94336 | 27848.22461 | 25174.67188 | 0.970354109 | 0.022724126 | 0.510956274 | 0.090033554 | 0.397176295 | 0.04761948 |
| 450126.375 | 431305.7188 | 495922.5625 | 0.968753647 | 0.022729015 | -1.089486327 | 0.028434447 | -0.098956561 | 0.353518154 |
| 20067.78906 | 15192.00195 | 15411.3623 | 0.968580476 | 0.064151421 | -0.399323553 | 0.200881472 | -0.512140171 | 0.132636805 |
| 10415.28027 | 11340.97754 | 10283.62793 | 0.96748836 | 0.021667504 | -0.025696668 | 0.864987005 | -0.15645846 | 0.282951822 |
| 8945.990234 | 12576.9043 | 11245.28418 | 0.966793198 | NA | -0.159625359 | NA | -1.818760281 | NA |
| 78507.03906 | 82038.25781 | 75508.23438 | 0.966608318 | 0.016471012 | -1.083203671 | 0.018539883 | -0.70445964 | 0.009587646 |
| 64806.02637 | 61878.86719 | 38645.29541 | 0.965279668 | 0.0424199 | -0.41868731 | 0.115660935 | -0.565436978 | 0.138136273 |
| 73652.17969 | 74183.00781 | 73342.53906 | 0.965081801 | 0.031883977 | -0.299986476 | 0.170360288 | 0.635799167 | 0.055746166 |
| 86337.13281 | 88281.9375 | 111286.2969 | 0.963170964 | 0.038217062 | -0.009128721 | 0.945348035 | -0.017850826 | 0.935380368 |
| 37740.15234 | 40525.80859 | 37683.29688 | 0.962358599 | 0.016596319 | 0.176407969 | 0.117794934 | 0.111984063 | 0.233347919 |
| 3537.334473 | 4012.608887 | 3244.685791 | 0.962026009 | 0.024701492 | 0.146774819 | 0.240374068 | -0.397106464 | 0.070968049 |
| 111627.9453 | 120408.4766 | 113378.3672 | 0.96146645 | 0.014324326 | -1.060194675 | 0.01179292 | -1.238676112 | 0.009587646 |
| 392035.2188 | 434647.5625 | 380841.5313 | 0.961216812 | 0.015034317 | -0.319160203 | 0.034038555 | -0.561803655 | 0.02354608 |

| | | | | | | | | |
|---|---|---|---|---|---|---|---|---|
| 13951.12305 | 14167.37305 | 9649.330078 | 0.960979359 | 0.045484912 | 0.520908462 | 0.073324814 | 0.48269913 | 0.216424718 |
| 3622.804932 | 4468.04834 | 3209.89209 | 0.960755239 | 0.013583579 | -0.052071367 | 0.590470503 | -0.355856797 | 0.167225632 |
| 133725.8438 | 136047.1094 | 149916.6875 | 0.96071295 | 0.043150956 | 0.038817277 | 0.816076757 | 0.808898083 | 0.020849231 |
| 103652.6797 | 93480.16406 | 92559.36719 | 0.95944817 | 0.045160985 | 0.873303649 | 0.008408055 | 0.579003727 | 0.049693654 |
| 195725.9688 | 205231.7656 | 205491.9219 | 0.958980858 | 0.012364839 | -0.227158457 | 0.082763602 | 0.391549162 | 0.031308498 |
| 7572.282227 | 9119.551758 | 9444.842773 | 0.958795799 | 0.219449972 | -0.112064726 | 0.821958214 | -0.032157644 | 0.950160109 |
| 19965.9668 | 13630.60156 | 17588.2793 | 0.958607789 | 0.292353414 | 0.573637521 | 0.118726695 | 0.062183468 | 0.857977393 |
| 36135.01563 | 43418.32813 | 33616.02734 | 0.957760594 | 0.04950513 | 0.388461971 | 0.081503088 | 0.085313372 | 0.681870325 |
| 31100.4375 | 24018.37891 | 23426.80859 | 0.95286194 | 0.034061991 | -0.816101084 | 0.042884533 | -0.859365501 | 0.052349495 |
| 86741.19531 | 76768.21094 | 94886.19531 | 0.951769565 | 0.015044187 | 0.246110519 | 0.094508793 | 0.31166171 | 0.138522639 |
| 30411.04883 | 32362.09961 | 33085.97656 | 0.951214411 | 0.290808764 | -0.308389954 | 0.64026592 | -0.154538512 | 0.800455624 |
| 9588.151367 | 10483.61621 | 5847.990234 | 0.951169254 | NA | 1.15738769 | NA | 1.881189759 | NA |
| 32849.21875 | 34949.37109 | 36018.38281 | 0.949932531 | 0.006735623 | -0.416077829 | 0.029180754 | -0.113825471 | 0.15164826 |
| 25443.47852 | 44407.46875 | 35438.59375 | 0.94403821 | 0.289811436 | -0.133171941 | 0.836602453 | 0.117429306 | 0.852652233 |
| 1735196.75 | 2011771.938 | 1773743.063 | 0.942867429 | 0.010297098 | -0.108834434 | 0.292995226 | -0.042659233 | 0.713487699 |
| 6751.438477 | 6893.837402 | 6104.108398 | 0.939452827 | 0.035947727 | 0.136621353 | 0.21109765 | -0.301184861 | 0.06710618 |
| 36202.41797 | 36294.39453 | 31389.03125 | 0.936538444 | 0.042404592 | 0.099345094 | 0.575891963 | 0.385405572 | 0.078910724 |
| 3733.998047 | 3915.67627 | 3134.985107 | 0.934581917 | 0.063399586 | 0.102417698 | 0.73629763 | -0.559417065 | 0.135721265 |
| 10059.10742 | 11499.72363 | 7834.945801 | 0.9336203 | 0.161511845 | 0.010258264 | 0.977833898 | 0.137744684 | 0.728591204 |
| 925851.625 | 931287.125 | 942908.4375 | 0.931839654 | 0.007230739 | 0.04497789 | 0.782907118 | 0.044311743 | 0.393050062 |
| 201858.6563 | 202194.3594 | 207134.6094 | 0.929282353 | 0.018312909 | -0.864619875 | 0.031437224 | -0.490522373 | 0.048302686 |
| 10030.44141 | 10287.86426 | 8574.197266 | 0.929087387 | 0.019930448 | 0.277472734 | 0.243646891 | -1.269405792 | 0.0236713 |
| 10142.49805 | 9905.078125 | 10322.48633 | 0.928923326 | 0.028143177 | 0.462258854 | 0.135019733 | 0.727108845 | 0.026553937 |
| 10274.61621 | 10779.66797 | 7829.139648 | 0.928575506 | 0.048531905 | 0.041769757 | 0.788148361 | -1.117464116 | 0.033327007 |
| 4072.962646 | 3621.073975 | 1876.744141 | 0.927981016 | 0.028196123 | -0.605568995 | 0.052953537 | -0.916639552 | 0.120481022 |
| 1360602.625 | 1437915.625 | 1415791.25 | 0.927151909 | 0.039517869 | 0.443509311 | 0.124688551 | 0.750822983 | 0.025930246 |
| 113297.8516 | 95734.89063 | 95835.58594 | 0.925653713 | 0.062609 | 0.046649711 | 0.771263379 | -0.477332413 | 0.103328406 |
| 82232.21094 | 71756.88281 | 72835.47656 | 0.9256273 | 0.05239238 | -0.05555995 | 0.801215147 | 0.055403586 | 0.785749164 |
| 4978.655273 | 6127.670898 | 5294.509277 | 0.924477615 | 0.057923073 | 0.145838898 | 0.677152193 | -0.387208801 | 0.101363503 |
| 291732.4063 | 266935.2188 | 264694.5 | 0.923580446 | 0.009974872 | -0.355224817 | 0.098459633 | -0.121430635 | 0.213539875 |
| 1614177.75 | 1798992.625 | 1483926.125 | 0.92345387 | 0.028368096 | 0.157276311 | 0.535392728 | -0.625375797 | 0.056482153 |
| 26627.43359 | 23703.12109 | 27243.67578 | 0.922874622 | 0.059894429 | 0.282394991 | 0.015395012 | 0.864439259 | 0.04403672 |
| 16073.8916 | 15573.4873 | 13595.12012 | 0.922473929 | 0.042146623 | -0.253464064 | 0.197534048 | 0.336421714 | 0.096134514 |

| | | | | | | | | |
|---|---|---|---|---|---|---|---|---|
| 8194.56543 | 10264.88184 | 7182.530762 | 0.920135151 | 0.009974872 | 0.062225382 | 0.556099514 | -0.185965272 | 0.434321145 |
| 231612.0156 | 232729.4531 | 226897.8125 | 0.919642397 | 0.076390655 | -1.128889101 | 0.082612243 | -0.453075498 | 0.176516532 |
| 17234.63086 | 16294.77637 | 9885.374023 | 0.91893812 | NA | 0.580735128 | 0.205468655 | 0.07526922 | 0.840910557 |
| 22736.68555 | 20510.11523 | 19726.59961 | 0.916960997 | 0.018312909 | 0.823836379 | 0.013885891 | 0.296257323 | 0.085185093 |
| 12519.48828 | 12532.06543 | 8846.113281 | 0.916762776 | 0.038502747 | 0.519636539 | 0.114863889 | 0.224488008 | 0.431199487 |
| 58304.68359 | 55006.32813 | 58872.96484 | 0.916610274 | 0.038176133 | -0.221215121 | 0.175682075 | 0.079257374 | 0.481104638 |
| 30471.17188 | 29460.53125 | 30207.40625 | 0.916477502 | 0.051555299 | 0.458037821 | 0.111349456 | 0.214625847 | 0.304878528 |
| 1698326.875 | 1689008.5 | 1667560.125 | 0.916291649 | 0.041945948 | 0.013157964 | 0.966180385 | 0.827927858 | 0.035986636 |
| 874283.0078 | 1021112.961 | 1120770.492 | 0.914913661 | 0.025478708 | -0.368733682 | 0.078437749 | -0.169416067 | 0.329172546 |
| 10847.23242 | 12574.36523 | 12907.78613 | 0.914331423 | 0.07166358 | -0.453277098 | 0.178660809 | -1.059969196 | 0.07832051 |
| 159645.7188 | 161501.9531 | 183827.9531 | 0.913692351 | 0.047406642 | -0.076279263 | 0.735510343 | 0.600879132 | 0.050613123 |
| 333056.5313 | 350358.0625 | 363887 | 0.913678052 | 0.039594932 | -0.359086128 | 0.159217919 | -0.145921094 | 0.400680917 |
| 46120.24219 | 43655.10156 | 43573.55859 | 0.913555916 | 0.013585622 | 0.258031503 | 0.057189248 | -0.160300964 | 0.135536925 |
| 29389.06445 | 20747.50586 | 21340.96484 | 0.913048755 | 0.132688125 | -0.486473782 | 0.302138556 | -0.357556772 | 0.397327146 |
| 13951.29297 | 14469.98828 | 13133.98047 | 0.912028219 | 0.047817825 | 0.085337037 | 0.713585003 | -0.319157991 | 0.227680787 |
| 30092.28906 | 24283.38477 | 34591.62891 | 0.911786524 | 0.139794213 | -0.377463721 | 0.395202582 | -0.297759002 | 0.474317331 |
| 25386.3125 | 20019.66797 | 20804.79102 | 0.911707118 | 0.032693403 | 0.365725646 | 0.108508712 | -0.476229628 | 0.091108554 |
| 2174.324463 | 2036.88269 | 1951.815674 | 0.911679669 | 0.136265249 | -0.95054204 | 0.144097346 | -1.365781646 | 0.115681603 |
| 118547.4375 | 101218.1172 | 90554.95313 | 0.910808456 | 0.075590877 | 0.261256161 | 0.416577532 | -0.00087303 | 0.997782424 |
| 6744.458496 | 7221.995117 | 7566.344238 | 0.910529877 | 0.029514505 | -0.822156765 | 0.052965881 | -0.269774893 | 0.174083052 |
| 29269.75 | 28434.75195 | 26484.96289 | 0.910113886 | 0.013596234 | 0.507318788 | 0.034038555 | 0.33411362 | 0.044248771 |
| 4177.705566 | 2982.415527 | 1986.536255 | 0.909908957 | 0.034127829 | -0.287871501 | 0.208496317 | -0.612858988 | 0.196956833 |
| 55935.36719 | 48748.17188 | 50245.05469 | 0.909674355 | 0.03211333 | 0.034169445 | 0.534693749 | -0.278396843 | 0.094664375 |
| 7352.210938 | 6118.799805 | 5488.257813 | 0.909659633 | 0.30624524 | -0.420650527 | 0.621458945 | -1.144417302 | 0.249456378 |
| 57753.17578 | 47197.96875 | 51553 | 0.90921634 | 0.166054997 | 0.79229454 | 0.100849207 | 0.411331424 | 0.268750571 |
| 14892.95508 | 14000.97949 | 7775.054199 | 0.906769971 | 0.029571758 | 0.003311859 | 0.971700493 | -0.300803185 | 0.442890403 |
| 15919.07129 | 18017.9043 | 16890.18359 | 0.906768447 | 0.199351416 | -0.305510732 | 0.529223913 | -0.933146993 | 0.214658701 |
| 899.1339111 | 964.3866577 | 1361.223633 | 0.906410628 | 0.093835222 | -0.171372949 | 0.606716614 | -0.434033286 | 0.282508948 |
| 121126.7109 | 152069.1094 | 129577.6406 | 0.906270734 | 0.016471012 | -0.139986911 | 0.561472953 | 0.382251234 | 0.153102212 |
| 137058.3594 | 140727.4219 | 152261.0938 | 0.906093063 | 0.035524295 | -0.284183458 | 0.112996996 | -0.35666339 | 0.06471715 |
| 350401.8438 | 366820.8438 | 391244.4375 | 0.904288068 | 0.053022874 | 0.072788742 | 0.744462234 | 0.90357377 | 0.020008995 |
| 5609.906738 | 4760.69043 | 3925.338623 | 0.903613368 | 0.290126842 | -2.270088233 | 0.183605997 | -1.45576702 | 0.21989921 |
| 52889.08594 | 45801.46094 | 52796.57422 | 0.899448395 | 0.012665235 | -0.192980067 | 0.065871863 | 0.375374278 | 0.082931684 |

| | | | | | | | | |
|---|---|---|---|---|---|---|---|---|
| 17233.32031 | 20724.24414 | 24430.79102 | 0.899261824 | 0.362732802 | 0.442468835 | 0.497000891 | 0.366158294 | 0.560867903 |
| 15274.53809 | 13014.45508 | 13120.37891 | 0.898384364 | 0.036830197 | 1.148703439 | 0.050235996 | 0.50116484 | 0.063283918 |
| 27480.9082 | 28497.99414 | 28582.26367 | 0.897078204 | 0.029603305 | 0.137033906 | 0.230523816 | 0.272819328 | 0.077004386 |
| 31046.89453 | 35669.30859 | 32888.3125 | 0.896884136 | 0.106077729 | 0.384133644 | 0.191184742 | 0.365389154 | 0.191055359 |
| 9037.727539 | 8987.242188 | 9322.921875 | 0.89662618 | 0.019930448 | -0.739241945 | 0.041058966 | 0.161658361 | 0.080462751 |
| 8019.954102 | 7454.562012 | 10638.93262 | 0.896329684 | 0.061674089 | -0.290388079 | 0.313982423 | -0.338167668 | 0.291228946 |
| 102646.7422 | 101876.7188 | 99928.46875 | 0.894774931 | 0.035947727 | -1.024184893 | 0.058122192 | -0.654594689 | 0.069955133 |
| 10553.64648 | 9141.271484 | 6410.186523 | 0.894483037 | 0.060886955 | 1.233429085 | 0.021906439 | -1.972855503 | 0.016313136 |
| 12628.00293 | 9624.538086 | 10862.53516 | 0.894179819 | 0.053770474 | 0.084011196 | 0.335018067 | -0.306971321 | 0.155392782 |
| 32751.91797 | 26730.22266 | 32958.66016 | 0.894164193 | 0.018312909 | -0.299607677 | 0.014951154 | -0.560795533 | 0.060937162 |
| 39217.96094 | 39662.95313 | 39192.35938 | 0.890526682 | 0.025587445 | 0.523720871 | 0.040327952 | 0.806792914 | 0.03405215 |
| 8593.668945 | 6828.260742 | 7360.814453 | 0.888615029 | 0.094031642 | 0.052563984 | 0.851894319 | 0.096282118 | 0.731115724 |
| 33092.16016 | 30924.37891 | 36132.63672 | 0.887865152 | 0.035393978 | 0.050515845 | 0.800958637 | 0.489542057 | 0.054887084 |
| 2932.739258 | 2545.713623 | 2418.704102 | 0.887791959 | 0.094792234 | -0.237030322 | 0.419309963 | -0.344982771 | 0.283693593 |
| 8495.881836 | 6645.413086 | NA | 0.886381724 | 0.031284382 | -0.797406595 | 0.047347558 | -1.007010586 | NA |
| 162806.8906 | 100797.7188 | 143933.75 | 0.886157969 | 0.050822105 | 0.124339267 | 0.569857055 | -0.207253929 | 0.5195709 |
| 21419.51953 | 13384.51465 | 12865.80664 | 0.88591665 | 0.042396393 | -0.176844043 | 0.122107272 | -0.215112936 | 0.557456388 |
| 109998.4609 | 102828 | 108706.9219 | 0.885018304 | 0.009974872 | -0.443864609 | 0.042199929 | -1.079898748 | 0.020008995 |
| 301833.4375 | 298386.4063 | 291164.3438 | 0.884039048 | 0.008731287 | -0.44900903 | 0.015625705 | -0.355367497 | 0.02163333 |
| 497424.3125 | 498602.0938 | 444288.9688 | 0.882694387 | 0.015175173 | 0.278984077 | 0.152866766 | -0.203304732 | 0.126064355 |
| 7519.217285 | 5178.223633 | 6907.868652 | 0.88101744 | 0.088231376 | -0.246127848 | 0.431639915 | -0.595424486 | 0.16248539 |
| 379122.7188 | 367131 | 399582.8438 | 0.880613233 | 0.021667504 | -0.257499476 | 0.152585278 | -0.460626194 | 0.068784766 |
| 222792.3906 | 213334.1406 | 208665.6094 | 0.879936557 | 0.137626056 | -0.62946509 | 0.215277105 | -0.482481346 | 0.263394123 |
| NA | NA | NA | 0.878258506 | NA | 2.648333902 | NA | NA | NA |
| 240420.3438 | 205960.375 | 205926.0625 | 0.877879092 | 0.011817481 | -0.69246029 | 0.026826422 | -0.102051658 | 0.433601327 |
| 7552.272461 | 8366.416992 | 4446.962891 | 0.876246317 | 0.057336343 | 0.449206679 | 0.096243216 | -0.112048931 | 0.778561634 |
| 24090.08594 | 16230.2334 | 19115.12109 | 0.875182717 | 0.075030284 | 0.845332841 | 0.036154671 | 0.394752081 | 0.263528267 |
| 3367638.5 | 3554675.75 | 3264665.75 | 0.874450451 | 0.009905914 | 0.11951061 | 0.157905281 | 0.162248722 | 0.101910514 |
| 29705.63281 | 29702.79688 | 29994.38477 | 0.869953055 | 0.049763403 | 0.126341668 | 0.582568898 | 0.858981479 | 0.048083719 |
| 87025.79688 | 85683.57813 | 70145.00781 | 0.869906035 | 0.125805569 | 0.733377438 | 0.071050221 | 1.368384154 | 0.027783576 |
| 73649.14844 | 74195.05469 | 71385.80469 | 0.867045738 | 0.046600136 | 0.161010073 | 0.343475536 | 0.627078872 | 0.053012031 |
| 26961.69336 | 29403.77734 | 23985.23438 | 0.866695662 | 0.050055523 | 0.476016158 | 0.017037525 | -0.092401583 | 0.493938048 |
| 267832.9688 | 263127.4063 | 239800.4844 | 0.866064633 | 0.061344134 | 0.590774735 | 0.065362897 | 0.68157087 | 0.032502 |

| | | | | | | | | |
|---|---|---|---|---|---|---|---|---|
| 28777.95703 | 28957.79688 | 23933.30078 | 0.865691652 | 0.071921738 | -0.031637713 | 0.909428871 | 0.139972192 | 0.405672325 |
| 8414.376953 | 9327.235352 | 11160.88281 | 0.865633304 | 0.046564243 | 0.382709036 | 0.046794787 | 0.497940067 | 0.132715015 |
| 15620.20898 | 10405.49316 | 13219.49023 | 0.86525217 | 0.063613605 | -1.291487223 | 0.057171461 | -1.325965891 | 0.047120928 |
| 23354.68359 | 26174.3125 | 24384.74023 | 0.864617151 | 0.047290455 | -0.365071515 | 0.080440179 | -0.705853227 | 0.020337832 |
| 10607.26855 | 9465.638672 | 10076.3418 | 0.863055644 | 0.264058004 | 1.096140946 | 0.095527512 | 1.234893056 | 0.062318926 |
| 15295.4502 | 15724.50586 | 12865.92676 | 0.861001225 | 0.059977515 | 0.133870384 | 0.54678916 | -0.177500491 | 0.449559439 |
| 92709.79688 | 88956.36719 | 96764.69531 | 0.860696174 | 0.03228159 | -0.014921748 | 0.888375183 | 1.080654619 | 0.009587646 |
| 17093.91797 | 20648.16797 | 21142.0957 | 0.860577032 | 0.093428026 | 0.206004836 | 0.47004298 | -0.083604578 | 0.768641076 |
| 33058.96484 | 34103.15234 | 40098.34766 | 0.859633704 | 0.011814081 | -0.376750386 | 0.063325993 | 0.259846248 | 0.185585346 |
| 7376.933594 | 1759.08374 | 6060.464355 | 0.858037637 | 0.080149199 | -0.139898733 | 0.536495 | -0.443233696 | 0.51004668 |
| 76062.14844 | 71192.25781 | 66640.75 | 0.857396066 | 0.014676535 | 0.292311675 | 0.059937144 | 0.413955634 | 0.045009854 |
| 7450.263184 | 8953.391602 | 6046.279785 | 0.856562264 | 0.035947727 | 0.581366406 | 0.05467585 | 0.18813941 | 0.514699959 |
| 19900.0332 | 13259.41406 | 14319.38379 | 0.854714127 | 0.00953954 | -0.131691252 | 0.677701081 | -0.579097628 | 0.123837484 |
| 26493.65234 | 17973.37695 | 24178.54883 | 0.852403389 | 0.010297098 | -0.117978968 | 0.360253167 | -0.450244215 | 0.143757509 |
| 8725.113281 | 6850.588379 | 6170.481445 | 0.852350288 | 0.047406642 | 0.912670015 | 0.1131055 | 0.704354238 | 0.122834452 |
| 51999.85156 | 54667.21094 | 25291.85547 | 0.852280941 | 0.097612498 | -1.390504967 | 0.10111096 | -1.456555973 | 0.057576847 |
| 65345.83594 | 65117.36719 | 55305.06641 | 0.852088259 | 0.07291087 | -0.107287041 | 0.437441812 | 0.357928123 | 0.092731087 |
| 4940.841309 | 6571.239746 | 5243.443359 | 0.851823818 | 0.133433633 | 0.235549135 | 0.591244008 | 0.664201585 | 0.103879505 |
| 434239.9063 | 446702.1875 | 456617.7813 | 0.849975358 | 0.044763187 | -0.453845479 | 0.137613908 | -0.15887945 | 0.392157851 |
| 45343.58203 | 50083.55078 | 40328.27344 | 0.849847866 | 0.034518918 | 0.409086945 | 0.09678603 | 0.412534429 | 0.096952234 |
| 8576.243164 | 10879.79297 | 10382.2002 | 0.849635736 | 0.159577005 | 0.629185094 | 0.138525684 | 0.869027934 | 0.065367829 |
| NA | 409.8201904 | NA | 0.849584383 | NA | 0.052643939 | NA | -2.701266306 | NA |
| 121732.5781 | 184062.4844 | 113142.4531 | 0.849325208 | 0.168226751 | 0.518951418 | 0.198890201 | 0.174140812 | 0.711855361 |
| 191874.6406 | 198350.2656 | 204796.3125 | 0.849035617 | 0.010297098 | 0.060353355 | 0.337199249 | -0.005232056 | 0.938347052 |
| 18781.48828 | 16981.68164 | 18198.61719 | 0.848269657 | 0.02883182 | 0.352412689 | 0.320787116 | 0.515732649 | 0.039032326 |
| 62533.46094 | 66957.17969 | 70268.85156 | 0.846978458 | 0.030955339 | -0.321259155 | 0.137939807 | 0.2053659 | 0.177455256 |
| 11028.3623 | 10136.4248 | 8586.68457 | 0.846756205 | 0.154102524 | 0.056063165 | 0.875125893 | -0.290551863 | 0.448196169 |
| 818231.5 | 977777.6875 | 1051229.875 | 0.846381006 | 0.079098326 | -0.27736034 | 0.298413403 | 0.191357184 | 0.420966129 |
| 192335.9063 | 201279.3438 | 208393.2031 | 0.845953856 | 0.012925173 | -0.004736581 | 0.947346059 | 0.050954115 | 0.490058892 |
| 1252.967041 | 1333.309448 | NA | 0.845937944 | 0.346333781 | 0.565693817 | 0.464869498 | -1.470217847 | NA |
| 1722737.75 | 2001864.5 | 2087121.25 | 0.845670001 | 0.020807424 | -0.217798021 | 0.149035707 | 0.122012926 | 0.421123997 |
| 2332567.5 | 2284935.75 | 1767800.375 | 0.844551421 | 0.193810113 | -0.258740317 | 0.43114209 | -0.248607874 | 0.268450393 |
| 13230.53125 | 19583.50195 | 18552.33008 | 0.844470085 | 0.235575859 | -0.813526041 | 0.280400081 | -0.048702566 | 0.926262361 |

| | | | | | | | | |
|---|---|---|---|---|---|---|---|---|
| 24892.80664 | 21886.14063 | 32111.74023 | 0.844241009 | 0.064774085 | -0.220834338 | 0.422801 | -0.006331447 | 0.985804247 |
| 5379.730957 | 6178.864258 | 6030.327148 | 0.843048684 | 0.059460749 | 0.05239509 | 0.760166132 | -0.704218378 | 0.08376499 |
| 44046.57422 | 44487.44531 | 46565.38281 | 0.842825365 | 0.01391113 | 0.187114307 | 0.215727673 | 0.274913626 | 0.058410816 |
| 47209.08984 | 57063.48828 | 33671.94531 | 0.84242075 | 0.103174294 | -0.232040286 | 0.41315045 | -0.725927917 | 0.112503484 |
| 4073.857422 | 4212.445801 | 2901.317871 | 0.842261149 | 0.039022378 | -0.468495258 | 0.130087098 | -0.466609955 | 0.136000191 |
| 63623.6875 | 63093.19141 | 64523.19141 | 0.842158886 | 0.064507015 | -0.215361234 | 0.431374349 | -0.059709272 | 0.714462192 |
| 62526.20313 | 65065.24609 | 62014.05859 | 0.841803014 | 0.010297098 | 0.042741843 | 0.660718628 | 0.029620854 | 0.62206986 |
| 95996.33594 | 152899.6719 | 64697.42969 | 0.841614799 | 0.2756587 | 0.258128498 | 0.629908746 | -0.447270746 | 0.506703416 |
| 7380.677246 | 8987.628906 | 8222.5 | 0.841058817 | 0.037492489 | -0.559096445 | 0.124774134 | -0.660449162 | 0.052768746 |
| 728104.125 | 673907.8125 | 806570.9375 | 0.840942105 | 0.13224933 | 0.500745855 | 0.502287468 | 1.246704834 | 0.023619478 |
| 144990.1406 | 143614.0625 | 129000.6094 | 0.838342407 | 0.020740891 | 0.070655693 | 0.529223913 | 0.038216913 | 0.758482113 |
| 18613.73047 | 18236.72852 | 16857.20703 | 0.837424151 | 0.014038682 | 0.119157664 | 0.093561793 | -0.328093695 | 0.02953449 |
| 10264.16992 | 9909.241211 | 10188.25488 | 0.835782929 | 0.068191065 | -0.118306432 | 0.52231553 | 0.567240797 | 0.057068534 |
| 8743.854492 | 8306.170898 | 5078.556641 | 0.835557327 | 0.148006693 | 0.614251842 | 0.181757098 | -0.206760714 | 0.620264646 |
| 5334.771484 | 4800.266113 | 5181.891602 | 0.835457718 | 0.043773448 | -0.672091189 | 0.05818214 | -0.824567673 | 0.033327007 |
| 9409.25 | 9531.003906 | 5240.992676 | 0.83482933 | 0.038502747 | -0.113541633 | 0.505325243 | -1.45354697 | 0.04007821 |
| 71080.10938 | 73452.1875 | 79017.35938 | 0.834660206 | 0.033164349 | -0.244876641 | 0.158752643 | -0.243622388 | 0.136000191 |
| 11671.68164 | 8449.209961 | 10322.03906 | 0.834452562 | 0.019078369 | -0.113060408 | 0.516940522 | -0.248180082 | 0.269198637 |
| 2911.864746 | 3416.363037 | 2766.639404 | 0.834302259 | 0.083966614 | -1.47137732 | 0.076036407 | -1.629557006 | 0.074096262 |
| 35845.52344 | 42803.35156 | 33836.14063 | 0.834128494 | 0.023527734 | -0.099098673 | 0.401614835 | 0.509017303 | 0.100909111 |
| 37911.96484 | 51715.94141 | 47675.625 | 0.832757429 | 0.073001312 | -0.00357739 | 0.991037626 | -0.779310719 | 0.082930409 |
| 9759.800781 | 10320.41309 | 8062.331055 | 0.832049415 | NA | -0.307439569 | 0.541557838 | 1.535264661 | 0.027949491 |
| 80148.44531 | 76708.95313 | 16675.1582 | 0.831882673 | 0.170437625 | -0.306062116 | 0.163662129 | -0.504531426 | 0.471646181 |
| 90043.67188 | 89635.21094 | 81207.875 | 0.831830342 | 0.016596319 | 1.088576073 | 0.13262544 | 1.289774125 | 0.017112805 |
| 8482.598633 | 11043.28809 | 5312.689941 | 0.831343364 | 0.071197426 | 0.415772001 | 0.306000099 | 0.176771848 | 0.720196825 |
| 32428.85352 | 28807.38281 | 28917.32617 | 0.83072996 | 0.029061881 | 0.078828034 | 0.553204641 | 0.64874517 | 0.027629097 |
| 64071.6875 | 52925.96484 | 51950.39063 | 0.82990605 | 0.243383004 | 0.073786991 | 0.887113987 | 0.314178251 | 0.484259271 |
| 15216.32227 | 15032.19434 | 13277.05859 | 0.829513415 | 0.028446198 | -0.303979227 | 0.033670234 | -1.632891886 | 0.005561476 |
| 12706.79785 | 7476.990723 | 8755.5 | 0.82905335 | 0.10748156 | 0.864646299 | 0.115260875 | 1.306302355 | 0.1278105 |
| 39643.78516 | 38649.61719 | 38467.37891 | 0.828009567 | 0.031115452 | 0.802565267 | 0.039856414 | 0.551719471 | 0.025666912 |
| 13869.84668 | 12381.17188 | 15942.4541 | 0.827449476 | 0.193611705 | -0.243364115 | 0.565659073 | -0.472226252 | 0.339927953 |
| 3302005 | 3740630.25 | 3943542 | 0.826687042 | 0.037600114 | 0.940395046 | 0.087781599 | 1.674204552 | 0.029120377 |
| 9038.025391 | 15723.61035 | 12361.58105 | 0.826461555 | 0.26557175 | 1.43608323 | 0.044099436 | 1.943836333 | 0.086503564 |

| | | | | | | | | |
|---|---|---|---|---|---|---|---|---|
| 2318.162354 | 1726.895874 | 1559.104858 | 0.826107412 | 0.009974872 | -0.61310158 | 0.041554876 | -0.380692621 | 0.216674879 |
| 560825.125 | 593839.5 | 653388.875 | 0.82519942 | 0.024701492 | -0.212397722 | 0.233503043 | 0.112176605 | 0.380575378 |
| 227667.5469 | 214951.7188 | 217152.7031 | 0.824888983 | 0.036480793 | 0.436503807 | 0.01179292 | 0.39582715 | 0.021332797 |
| 8076.253418 | 7115.469238 | 6426.277832 | 0.823745341 | 0.333115267 | 1.576651945 | 0.279957252 | 0.805642453 | 0.166647679 |
| 11562.26758 | 12150.61719 | 12347.2627 | 0.823221865 | 0.04179011 | -0.32116209 | 0.182060286 | -0.183162174 | 0.334263521 |
| 562237.8125 | 291599.1563 | 527155.75 | 0.822849673 | 0.061766802 | -0.057583524 | 0.693159679 | -0.430056355 | 0.298904019 |
| 9329.419922 | 7159.908203 | 7815.875 | 0.822790562 | 0.025437157 | 0.420021538 | 0.111298502 | -0.12955283 | 0.509701467 |
| 16793.20313 | 17245.98633 | 13310.10938 | 0.821992631 | 0.027671686 | -0.990895561 | 0.02383791 | -0.918412101 | 0.021984636 |
| 1433.125244 | 1691.221436 | 1111.557617 | 0.821067075 | NA | -0.207491897 | 0.344433056 | -0.324121717 | 0.282631166 |
| 57831.42188 | 54190.87109 | 53451.13281 | 0.820670663 | 0.061350067 | 0.380399811 | 0.048024604 | 0.519977077 | 0.019022762 |
| 519543.375 | 543154.6875 | 577538.8125 | 0.820605467 | 0.016090856 | 0.049390935 | 0.463393444 | 0.40965713 | 0.061031191 |
| 72415.85938 | 76417.82813 | 60424.93359 | 0.820424172 | 0.083033915 | -1.393633215 | 0.084935024 | -0.807802887 | 0.088049383 |
| 8292171.5 | 10305612 | 8739521 | 0.820212737 | 0.205209259 | -0.729563105 | 0.236672563 | -0.490330475 | 0.335259007 |
| 236882.7813 | 199738.375 | 233528.7031 | 0.819812986 | 0.023465918 | -0.11765221 | 0.318820359 | -0.349483718 | 0.074385174 |
| 17698.63281 | 18412.5332 | 14030.7041 | 0.819742398 | 0.065883116 | 0.175907824 | 0.42765344 | 0.204234532 | 0.405343283 |
| 18227.9668 | 14784.24707 | 12645.5498 | 0.817622209 | 0.022260115 | -0.156752447 | 0.022454421 | -0.380678206 | 0.190976702 |
| 53241.50781 | 49998.77344 | 43016.24609 | 0.81609652 | 0.080233873 | -0.930942697 | 0.086540831 | -0.816958912 | 0.079927965 |
| 2834.612549 | 1767.470215 | 1859.236084 | 0.815834109 | 0.073576137 | -1.653965402 | NA | -1.843679097 | 0.029219266 |
| 48296.83984 | 45781.47656 | 45412.15625 | 0.814129855 | 0.019930448 | -0.058626672 | 0.502084571 | 0.013758482 | 0.763946881 |
| 26456.12109 | 23038.10742 | 24522.95703 | 0.812899516 | 0.01111345 | 0.141657618 | 0.144376338 | 0.160054411 | 0.199393614 |
| 8570.601563 | 6699.781738 | 6264.218262 | 0.812875395 | 0.549397113 | -0.052080666 | 0.966757644 | 0.56968582 | 0.526752622 |
| 42943.85156 | 38037.66797 | 27783.9668 | 0.811261023 | 0.195658833 | 0.133111564 | 0.715488289 | -0.283783421 | 0.531405981 |
| 70976.28125 | 64370.23828 | 77333.125 | 0.810155778 | 0.010510896 | -0.418377496 | 0.042887894 | -0.35849324 | 0.061382227 |
| 35211.28516 | 28907.24805 | 32483.69336 | 0.809482003 | 0.043802925 | -0.680548193 | 0.105345406 | -1.204685358 | 0.050138206 |
| 151053.75 | 133668.1563 | 163965.7813 | 0.80911981 | 0.021985747 | -0.011729224 | 0.948117667 | 0.630167848 | 0.056570421 |
| 55677.53906 | 61546.66016 | 43508.12109 | 0.808844047 | 0.016885305 | -0.79226094 | 0.013885891 | -0.841991483 | 0.044248771 |
| 280318.1875 | 96026.92969 | 192637.125 | 0.808808674 | 0.084263766 | 1.337891235 | 0.055083913 | 0.127202065 | 0.848147401 |
| 104449.6953 | 98637.5 | 86742.57813 | 0.804855632 | 0.028820473 | -0.561069409 | 0.047212306 | -0.735804779 | 0.020008995 |
| 291222.125 | 279886.8438 | 308419.9688 | 0.804465468 | 0.029801769 | 0.286018187 | 0.079287615 | 0.983529457 | 0.011723127 |
| 12159189 | 13490060 | 13500794 | 0.802769639 | 0.036971527 | -0.121663177 | 0.35047273 | -0.081994758 | 0.512810702 |
| 3489.508789 | 3664.177002 | 4292.548828 | 0.801933959 | 0.020940469 | -0.902782431 | 0.028573694 | -1.102817188 | 0.021134419 |
| 11324.28711 | 11087.57129 | 10171.50391 | 0.801551373 | 0.009194956 | -0.056441533 | 0.132973237 | -0.120355776 | 0.187440064 |
| 12278.54004 | 13239.87207 | 12771.35156 | 0.80103217 | 0.039409697 | -0.590002851 | 0.03062316 | -0.301897412 | 0.056570421 |

| | | | | | | | | |
|---|---|---|---|---|---|---|---|---|
| 48299.16406 | 50063.94531 | 50069.39063 | 0.800500418 | 0.005752145 | -0.215985516 | 0.095375186 | -0.034360076 | 0.435531711 |
| 80431.33594 | 76635.21875 | 79842.65625 | 0.799559985 | 0.036160457 | -0.162496261 | 0.015395012 | 0.238189343 | 0.044885555 |
| 14607.37988 | 15583.50684 | 10862.63965 | 0.799015517 | 0.047406642 | -0.331257957 | 0.130039916 | -0.481646552 | 0.113248184 |
| 68098.91406 | 64998.98438 | 67767.03125 | 0.798002873 | 0.01031238 | -0.208544198 | 0.019456985 | -0.290656931 | 0.014469072 |
| 1520163 | 1858399.875 | 1744073.5 | 0.797813907 | 0.023774301 | -0.054036476 | 0.630875263 | 0.255324229 | 0.174066419 |
| 5677.504395 | 6171.32959 | 4620.869629 | 0.796846572 | 0.016132048 | 0.889048001 | 0.555335026 | -0.002217687 | 0.991173771 |
| 238107.4375 | 226184.1094 | 240887.2188 | 0.795364156 | 0.01508889 | -0.099830676 | 0.111472792 | -0.117710025 | 0.090508565 |
| 72682.03906 | 72611.03906 | 54198.80469 | 0.793796338 | 0.029508458 | -0.800481224 | 0.042044993 | -0.841501743 | 0.033709967 |
| 6281.6875 | 5052.941406 | 5316.315918 | 0.793341398 | 0.034514668 | -0.666055688 | 0.036730784 | -0.580171863 | 0.039179608 |
| 136598.0313 | 130826.4609 | 139894.75 | 0.793026171 | 0.137409996 | 0.910053697 | 0.074598722 | 1.299738145 | 0.031136253 |
| 388923.8125 | 263232.6875 | 355384.9375 | 0.792777307 | 0.187040833 | -0.314325971 | 0.515992993 | 0.08140267 | 0.845265499 |
| 16283.90723 | 19161.02734 | 16842.48242 | 0.791356331 | 0.052671628 | 0.054423255 | 0.495702417 | 0.374445379 | 0.099306515 |
| 131674.2813 | 63877.33984 | 95674.67969 | 0.79129021 | 0.047058169 | -0.46855207 | 0.044605958 | -0.129249114 | 0.762218779 |
| 49417.71484 | 31965.21094 | 42642.42578 | 0.789435215 | 0.054407811 | 0.055664355 | 0.931623042 | -0.242937191 | 0.381068951 |
| 358508.7188 | 250694.2656 | 293093.8125 | 0.789008973 | 0.093577918 | -0.303799576 | 0.170410796 | 0.297266755 | 0.319455937 |
| 52720.22656 | 57925.67188 | 63911.1875 | 0.788295525 | 0.059151449 | 0.568195167 | 0.048530044 | 0.854828792 | 0.033327007 |
| 59759.54688 | 60210.01172 | 61319.12891 | 0.787997185 | 0.018999567 | -0.253755986 | 0.092746718 | -0.034174893 | 0.664545179 |
| 86185.38281 | 76350.14844 | 84893.13281 | 0.787442189 | 0.07827194 | -0.226710791 | 0.369408443 | -0.707619573 | 0.106167384 |
| 40336.09375 | 34972.66406 | 36241.53516 | 0.787198248 | 0.138753516 | -0.059959522 | 0.850899272 | -0.383875091 | 0.315117797 |
| 65906.25 | 59986.83203 | 51859.29688 | 0.786322001 | 0.073155658 | 0.163773606 | 0.45473591 | -0.149681398 | 0.546493907 |
| 4343.123047 | 6556.122559 | 6575.910156 | 0.784959028 | 0.113673327 | 0.261577035 | 0.252424496 | -0.374619415 | 0.22851844 |
| 55954.36719 | 60907.92578 | 63173.51172 | 0.78486039 | 0.006888749 | 0.019557252 | 0.702653772 | 0.185619499 | 0.137293079 |
| 54052.94531 | 53985.15625 | 61550.51563 | 0.784610272 | 0.017824776 | -0.572380652 | 0.034038555 | 0.164328957 | 0.215749573 |
| 12922.7793 | 14553.42773 | 13847.68066 | 0.784392984 | 0.358311233 | -0.167738407 | 0.591597867 | -0.104412756 | 0.243466924 |
| 60951.96484 | 55219.82422 | 60545.44922 | 0.784294559 | 0.040005201 | 0.535963215 | 0.053185315 | 1.092601708 | 0.012256037 |
| 11880.0498 | 13825.96289 | 11119.20996 | 0.784009159 | 0.030838583 | 0.293107705 | 0.111882773 | 0.373132628 | 0.121714657 |
| 52277.61719 | 53025.55078 | 56939.95313 | 0.783107356 | 0.093835222 | 0.144197734 | 0.491759763 | 0.156632172 | 0.449699841 |
| 14675.05176 | 13102.24609 | 7398.54248 | 0.782935307 | 0.040958886 | 0.272122791 | 0.208500335 | 0.046952972 | 0.916832689 |
| 19210.34766 | 20247.85938 | 23441.68164 | 0.782055424 | 0.04977426 | -0.240125227 | 0.185493647 | -0.128900885 | 0.441678579 |
| 27662.53906 | 36741.73047 | 34191.45703 | 0.781361173 | 0.117038402 | -0.598432145 | 0.061247369 | -0.450613011 | 0.085236358 |
| 49382.35547 | 47000.80859 | 44114.33984 | 0.78043395 | 0.067459302 | 0.46395775 | 0.104410984 | 0.73639169 | 0.040786554 |
| 1742447.5 | 1779760 | 1479525.875 | 0.778450142 | 0.072487978 | 0.474716864 | 0.183009351 | -0.415928942 | 0.135824856 |
| 162530.4063 | 39031.36719 | 133571.1719 | 0.778297092 | 0.295286026 | -2.35712938 | 0.141443989 | -0.984657345 | 0.263394123 |

| | | | | | | | | |
|---|---|---|---|---|---|---|---|---|
| 27595.62695 | 25881.51563 | 27681.2793 | 0.778286831 | 0.037600114 | 0.19955329 | 0.201166198 | 0.515516527 | 0.04016603 |
| 342071.1563 | 334504.9063 | 379707.125 | 0.778258176 | 0.06853067 | 0.079481342 | 0.67818072 | 0.519700931 | 0.068366118 |
| 62124.78516 | 66743.84375 | 64607.09375 | 0.77804456 | 0.017716941 | 0.437768314 | 0.059039944 | 0.540051765 | 0.021134419 |
| 40294.54297 | 31834.42773 | 36419.86719 | 0.777735784 | 0.009974872 | -0.23748269 | 0.055605274 | -0.567446455 | 0.045454735 |
| 26556.49609 | 22580.62695 | 24687.11914 | 0.777596947 | 0.04541773 | -0.431287389 | 0.111429162 | -0.380413934 | 0.087278746 |
| 47530.30859 | 41288.09766 | 41185.38672 | 0.777517434 | 0.039704162 | 0.321637211 | 0.097877924 | 0.496205196 | 0.051408747 |
| 344851.125 | 343972.3125 | 339304.5 | 0.777088686 | 0.074826564 | -0.416019225 | 0.155418267 | -0.381187708 | 0.178198454 |
| 108925.2109 | 125416.4688 | 133388.8906 | 0.775871457 | 0.016105265 | 0.035501857 | 0.567693144 | 0.22252734 | 0.215578616 |
| 26539.62891 | 30900.375 | 27550.25 | 0.775711097 | 0.04646045 | -0.692732632 | 0.01179292 | -0.354084962 | 0.063283918 |
| 25815.45313 | 25615.66797 | 29941.18164 | 0.77379099 | 0.006395507 | -0.239338859 | 0.160920103 | -0.02504956 | 0.828430667 |
| 3362.429932 | 4099.690918 | 3560.097656 | 0.772052419 | NA | 0.245339146 | NA | -0.096726359 | NA |
| 65822.01563 | 61051.89844 | 65992.5 | 0.771353163 | 0.020732204 | -0.284752895 | 0.116079932 | -0.149171376 | 0.237901119 |
| 48487.42188 | 54629.65234 | 44179.33203 | 0.770705253 | 0.254204903 | -0.398971562 | 0.469724355 | -0.162800985 | 0.732034702 |
| 1345341.875 | 1213331.125 | 1111213.875 | 0.770080145 | 0.029603305 | -0.429588547 | 0.091850418 | -0.746936731 | 0.042737273 |
| 10395.31055 | 7424.90625 | 9334.795898 | 0.769952062 | 0.032646789 | -0.581972428 | 0.024656861 | -0.900683174 | 0.03161115 |
| 83740.08594 | 65263.08984 | 49291.65234 | 0.76922069 | 0.02242334 | -0.497848053 | 0.041240492 | -0.815062911 | 0.087262349 |
| 61911.64844 | 69880.78125 | 71577.14844 | 0.768602924 | 0.102272218 | 0.141220543 | 0.541058779 | -0.047152357 | 0.842303553 |
| 79165.17188 | 88054.02344 | 89750.76563 | 0.768227208 | 0.019149891 | 0.564272778 | 0.017786674 | 0.381822721 | 0.057544683 |
| 119889.8984 | 129410.0469 | 119278.3594 | 0.766971466 | 0.011063961 | -0.828141625 | 0.017037525 | -0.218348367 | 0.069955133 |
| 40268.90234 | 42651.37891 | 36770.51172 | 0.766339457 | 0.021027835 | 0.058806377 | 0.628940433 | -0.016878867 | 0.893240084 |
| 26816.6875 | 22435.95508 | 24290.99609 | 0.765122787 | 0.063370218 | 0.389911487 | 0.155137511 | -0.408266588 | 0.156978352 |
| 230785.5469 | 262895.5625 | 283977.2188 | 0.764976196 | 0.075602358 | -0.234290049 | 0.302384728 | 0.467095036 | 0.084308715 |
| 28832.81055 | 30303.71875 | 25311.39063 | 0.764073923 | 0.018050472 | -0.193508682 | 0.295602028 | -0.721951946 | 0.021134419 |
| 16355.31152 | 16696.89258 | 15490.18262 | 0.76350843 | 0.028196123 | 0.377957444 | 0.089716327 | 0.034220403 | 0.776446789 |
| 189758.2188 | 179415 | 175141.8438 | 0.76290359 | 0.006614001 | -0.083187071 | 0.655353432 | 0.646274574 | 0.02183874 |
| 2242650 | 2359024.5 | 2362124.5 | 0.762094537 | 0.036547632 | 0.115561035 | 0.118728218 | 0.17879939 | 0.057402586 |
| 23071.29883 | 23100.25195 | 17819.32813 | 0.761871435 | 0.080105106 | 0.509152448 | 0.112520457 | -0.40787099 | 0.207497392 |
| 11746.39941 | 9793.03418 | 7519.845215 | 0.761561496 | 0.100751002 | -0.797898666 | 0.124487169 | -0.469049425 | 0.190240855 |
| 139.5052032 | 174.6486816 | NA | 0.761407803 | NA | -0.969578147 | NA | -1.256749749 | NA |
| 63752.24219 | 53746.77344 | 61338.44141 | 0.761313346 | 0.028446198 | -0.19234915 | 0.453596772 | -0.020513248 | 0.868514218 |
| 16537.80664 | 17418.88672 | 14185.28711 | 0.761148652 | 0.037600114 | -0.455290573 | 0.105107468 | -0.196524337 | 0.31203033 |
| 37835.79297 | 38040.16016 | 32486.83789 | 0.761114576 | 0.142870238 | -1.50430097 | 0.111349456 | -1.143494231 | 0.118373568 |
| 32868.73438 | 27402.82617 | 24619.24219 | 0.760806184 | 0.153739559 | 0.277507379 | 0.585714677 | -0.010155492 | 0.979696342 |

| | | | | | | | | |
|---|---|---|---|---|---|---|---|---|
| 642023.375 | 559687.3125 | 567450.9375 | 0.759906355 | 0.116764456 | -0.278548445 | 0.089463488 | 0.343448701 | 0.079165593 |
| 39496.86719 | 39210.46484 | 35601.98047 | 0.758367436 | 0.028047191 | 0.197975991 | 0.196035101 | -0.266124638 | 0.138724083 |
| 79589.29688 | 88513.28906 | 93129.35156 | 0.758303645 | 0.018311389 | -0.387145084 | 0.087791427 | -0.018927691 | 0.882379732 |
| 21748.7207 | 15415.45703 | 15681.36426 | 0.757797568 | 0.062556877 | 0.82380352 | 0.129412395 | -0.085141367 | 0.768663119 |
| 943889.8125 | 979036.625 | 798616.9375 | 0.757567194 | 0.015034317 | -0.224538567 | 0.259843823 | -0.492282581 | 0.043327982 |
| 79464.14844 | 90435.8125 | 97016.6875 | 0.755480301 | 0.541961432 | 0.509788199 | 0.553204641 | 1.059098581 | 0.217909765 |
| 160954.5156 | 142347.9688 | 177399.6563 | 0.755227084 | 0.029183044 | 0.382301827 | 0.059937144 | 0.090747496 | 0.588834899 |
| 231625.1563 | 263216.9375 | 211654.8438 | 0.754072969 | 0.066355015 | -0.035184757 | 0.876081794 | -0.432825359 | 0.140150822 |
| 456188.6563 | 407033.75 | 480547.9688 | 0.753592207 | 0.066744288 | 0.999107729 | 0.053882844 | 1.160163319 | 0.020256835 |
| 66468.58594 | 86099.78125 | 107532.1484 | 0.752943389 | 0.070275485 | -0.412741942 | 0.119711295 | -0.099062481 | 0.744940584 |
| 5211.634277 | 8016.600586 | 6338.224121 | 0.752196681 | 0.065637909 | 0.205409112 | 0.185362734 | -0.045083339 | 0.875605333 |
| 99406.99219 | 45215.17578 | 82714.80469 | 0.752155644 | 0.175388722 | -0.453685119 | 0.155539449 | -0.92075 | 0.107557275 |
| 1203861 | 1154640 | 1180466.5 | 0.752031334 | 0.032189911 | 0.231093405 | 0.231386286 | 0.336225752 | 0.042175609 |
| 635157.0625 | 724121.25 | 557669.4375 | 0.751673514 | 0.37942274 | 0.164697782 | 0.789721093 | -0.007989897 | 0.990553075 |
| 441744.4688 | 354764.2813 | 373445.8438 | 0.751607074 | 0.035361197 | -0.287675901 | 0.106271335 | -0.363918064 | 0.091480899 |
| 18649.38086 | 19061.25 | 16242.83203 | 0.751013741 | 0.072479892 | 0.563997869 | 0.057524181 | 0.345663582 | 0.127304768 |
| 268539.9297 | 299474.4688 | 319989.3691 | 0.751012549 | 0.014306629 | 0.387374292 | 0.03633877 | 1.385552142 | 0.040645703 |
| 49167.69141 | 50975.125 | 43582.11328 | 0.750966932 | 0.04186746 | -0.179169745 | 0.082612243 | -0.294157176 | 0.097178029 |
| 30650.69922 | 38827.86328 | 37361.92969 | 0.749368915 | 0.029390153 | 0.258010083 | 0.148589158 | -0.116968159 | 0.522132406 |
| 4108.199219 | 4336.62207 | 2435.924316 | 0.748668422 | 0.093994529 | -0.143921071 | 0.579418371 | -0.6527856 | 0.147524027 |
| 7649.025391 | 9601.864258 | 7442.879395 | 0.748542692 | 0.063167428 | 0.607506984 | 0.055241445 | 0.994302286 | 0.060012809 |
| 13230.16406 | 12408.96582 | 10198.98047 | 0.747154692 | 0.083669663 | -0.328419806 | 0.284636711 | -0.488517204 | 0.166853085 |
| 328307.7813 | 342117.75 | 349030.125 | 0.745031 | 0.025086961 | -0.354965252 | 0.113912361 | -0.120747274 | 0.331779341 |
| 111281.9966 | 127931.0127 | 153406.9834 | 0.744642368 | 0.123186057 | 0.284613104 | 0.429618137 | -0.429265774 | 0.225935566 |
| 8801.858398 | 8521.630859 | 6767.983887 | 0.744491084 | 0.04950513 | -0.357839449 | 0.148476207 | -0.623806439 | 0.062985995 |
| 659465.8125 | 721102.8125 | 729763.125 | 0.744478583 | 0.02237899 | 0.101512794 | 0.105216387 | 0.373722349 | 0.053570169 |
| 9336.74707 | 9662.716797 | 9737.387695 | 0.7431278 | 0.037600114 | 0.265203829 | 0.147710173 | 0.333161691 | 0.111646722 |
| 1081899.5 | 1075554.375 | 1058154 | 0.742728297 | 0.012750417 | 0.169905295 | 0.435676906 | 0.067218576 | 0.122834452 |
| 25026.4043 | 27491.56055 | 26842.61133 | 0.74266068 | 0.055757407 | 0.002155891 | 0.991322905 | 0.666191217 | 0.025404397 |
| 18708.27344 | 22781.54492 | 24533.52148 | 0.742148701 | 0.113673327 | 0.32245787 | 0.408992835 | 0.070886609 | 0.701341354 |
| 55465.51953 | 56633.46484 | 56475.54688 | 0.740944745 | 0.021745834 | -0.143905943 | 0.273114108 | 0.015327269 | 0.877391863 |
| 249134.9531 | 219297.2188 | 225336.4219 | 0.740731323 | 0.010297098 | -0.706585409 | 0.016400229 | -0.442457966 | 0.027800907 |
| 6514.108887 | 7624.757324 | 5263.305664 | 0.739020699 | 0.405460335 | 0.21241321 | 0.734206841 | 0.275153151 | 0.664000423 |

| | | | | | | | | |
|---|---|---|---|---|---|---|---|---|
| 15013.60254 | 14801.25781 | 16166.53613 | 0.73842191 | 0.043122548 | 0.176430977 | 0.449457465 | 0.810574275 | 0.02163333 |
| 240653.0938 | 317238.875 | 284486.0625 | 0.738381134 | 0.052969391 | -0.272499086 | 0.610732715 | 1.603899915 | 0.049905588 |
| 8286.822266 | 7849.261719 | 6216.095215 | 0.737560654 | 0.113551189 | 0.148228392 | 0.623543083 | 0.301885623 | 0.31013683 |
| 172057.3125 | 170626.5469 | 162125.5156 | 0.7354495 | 0.016471012 | 0.104854824 | 0.214737255 | 0.366923386 | 0.024573227 |
| 445.2478638 | 721.8308716 | 395.803894 | 0.734386702 | 0.204825812 | -0.34427245 | NA | -0.262295284 | 0.565766815 |
| 29614.36523 | 30929.66016 | 26604.60938 | 0.733277166 | 0.072220768 | 0.023045251 | 0.681696486 | -0.663275717 | 0.017279709 |
| 11870.01563 | 10460.47754 | 5867.735352 | 0.733040958 | 0.033164349 | -0.472712054 | 0.072901076 | -0.790954154 | 0.114763272 |
| 43790.89063 | 42720.69141 | 43831.15234 | 0.732430783 | 0.023401535 | -0.093627117 | 0.268936384 | -0.265982732 | 0.078781357 |
| 159376.4063 | 164965.9063 | 164486.6094 | 0.731508455 | 0.030661161 | 0.174231225 | 0.242733442 | 0.297189472 | 0.107612185 |
| 466344.8203 | 429325.6055 | 431463.3594 | 0.730269247 | 0.02423784 | -0.410087913 | 0.105099839 | -0.271206038 | 0.118841505 |
| 182619.7188 | 182180.9375 | 180456.6719 | 0.729155515 | 0.005752145 | 0.001874329 | 0.975901807 | 0.265646774 | 0.023137414 |
| 72188.15625 | 43499.96484 | 47134.94531 | 0.728461945 | 0.11294756 | 1.013175396 | 0.038454242 | 0.451883683 | 0.35314512 |
| 472649.4375 | 474963.75 | 455806.5313 | 0.727327819 | 0.025818052 | -0.413280336 | 0.07898558 | 0.124585939 | 0.306287848 |
| 31251.03711 | 34114.09766 | 35023.40234 | 0.725779832 | 0.115609849 | -0.68725652 | 0.153115547 | 0.132140619 | 0.604993935 |
| NA | 13938.24023 | 4300.942383 | 0.725672225 | 0.284469204 | 0.226837464 | 0.62182213 | -0.492887546 | NA |
| 24003.33008 | 20394.23633 | 20577.06836 | 0.724238497 | 0.023132924 | -0.041187156 | 0.702514594 | -0.216679816 | 0.179657466 |
| 7714.158691 | 8484.569336 | 7389.011719 | 0.723821239 | 0.068352646 | 0.674079141 | 0.095980292 | 0.568841316 | 0.033916706 |
| 12645.74805 | 14827.99707 | 15582.2041 | 0.723635813 | 0.026440649 | 0.002619362 | 0.985576344 | -0.351784992 | 0.09000305 |
| 238307.0625 | 252110.2813 | 271325.0313 | 0.723066393 | 0.060886955 | -0.869260865 | 0.069037719 | -0.544009829 | 0.072610739 |
| 8919.59082 | 8439.063477 | 9537.070313 | 0.722976297 | 0.089303227 | 0.046126454 | 0.717559738 | 0.396729928 | 0.056004888 |
| 352908.3438 | 376573.625 | 364701.9375 | 0.722837874 | 0.016859654 | -0.259948524 | 0.089889925 | -0.17242436 | 0.142748783 |
| 401128.125 | 441960.75 | 409344.9375 | 0.72280312 | 0.017982551 | -0.232110839 | 0.125646125 | 0.197555505 | 0.088892469 |
| 16993738 | 18956572 | 17994124 | 0.722384883 | 0.01391113 | 0.108942317 | 0.220151814 | -0.031569322 | 0.72315966 |
| 53191.08203 | 55108.76172 | 61450.84375 | 0.721833088 | 0.047406642 | -0.134335889 | 0.473820477 | 0.159163692 | 0.370576128 |
| 883036.3125 | 943359.25 | 976940.6875 | 0.721830441 | 0.027973416 | -0.038309393 | 0.687866158 | 0.552792723 | 0.044885273 |
| 15462.95703 | 23768.05078 | 35379.44141 | 0.71929026 | 0.143627731 | 0.728411836 | 0.058077004 | 0.594378161 | 0.391006453 |
| 12934.15234 | 13157.94043 | 12534.15234 | 0.71917468 | NA | 0.9396754 | 0.114739915 | 1.68110599 | 0.022221637 |
| 11988.08301 | 13626.16602 | 12187.50977 | 0.719151851 | 0.027230193 | -0.136273337 | 0.266303698 | -0.061950235 | 0.578085212 |
| 14750.62207 | 16144.44043 | 7589.886719 | 0.71818387 | 0.064239122 | 0.421449836 | 0.100046608 | 0.418797417 | 0.456912711 |
| 31265.20313 | 28734.62695 | 26981.72656 | 0.717937909 | 0.034461901 | -1.205293895 | 0.036154671 | -0.38555932 | 0.046772816 |
| 20534.36328 | 18408.70898 | 20552.97656 | 0.716688043 | 0.049438736 | 0.546493544 | 0.053336389 | -0.104092335 | 0.45869086 |
| 164801.8906 | 173368.6719 | 197878.7656 | 0.716524331 | 0.046689196 | -0.092994134 | 0.534139527 | 0.010158131 | 0.955025524 |
| 8748.216797 | 5307.743164 | 2323.11377 | 0.71623741 | NA | 0.255680234 | NA | -0.746704255 | NA |

| | | | | | | | | |
|---|---|---|---|---|---|---|---|---|
| 62665.0625 | 65899.3125 | 68660.78125 | 0.716193631 | 0.005752145 | 0.701668929 | 0.048221719 | 0.255288562 | 0.07832051 |
| 10237.69531 | 12483.18555 | 13679.19727 | 0.715701198 | 0.208851744 | 0.523300741 | 0.161939384 | 1.326689045 | 0.039195867 |
| 12839.68945 | 10835.82813 | 9121.724609 | 0.715575957 | 0.295928701 | 3.668088845 | 0.205304317 | 0.314196758 | 0.510714902 |
| 61807.67188 | 81634.04688 | 67627.30469 | 0.715429959 | 0.127230164 | -0.487445503 | 0.231498847 | 0.307392402 | 0.317865083 |
| 24989.94922 | 23875.85352 | 20115.09961 | 0.714672206 | 0.012750417 | 0.13415394 | 0.311163334 | -0.325205628 | 0.101436425 |
| 76237.01563 | 81574.02344 | 82444.21094 | 0.713813473 | 0.094792234 | 0.018401962 | 0.941915718 | -0.188838749 | 0.439110295 |
| 51944.03906 | 55892.51953 | 50316.95313 | 0.71313479 | 0.026775338 | -0.057568305 | 0.63348405 | 0.005919698 | 0.960748672 |
| 29704.95117 | 30695.18359 | 32118.32422 | 0.711622644 | 0.019930448 | 0.244439343 | 0.094508793 | -0.074847272 | 0.452220015 |
| 10688.4668 | 10237.56348 | 10874.71191 | 0.709448402 | 0.223885477 | -0.293629392 | 0.525949062 | -0.159393517 | 0.700372945 |
| 25232.39453 | 30002.70703 | 26117.68359 | 0.707783942 | 0.029508458 | -0.217137357 | 0.064039072 | -0.633668848 | 0.024070367 |
| 203436.3018 | 193683.5313 | 175509.7256 | 0.707480659 | 0.034890445 | -0.008776138 | 0.949050855 | 0.215019994 | 0.163723435 |
| 24139.51367 | 33260.98828 | 30574.48828 | 0.707480179 | 0.070275485 | -0.029992673 | 0.876454953 | -0.153812262 | 0.520924639 |
| 12194.83594 | 15394.95898 | 8499.254883 | 0.706944259 | NA | 2.050313487 | 0.039567204 | 1.666812336 | 0.100522212 |
| 4865.036621 | 6201.434082 | 8486.075195 | 0.705959206 | 0.179094466 | -0.125053696 | 0.754273728 | 1.096988707 | 0.13790491 |
| 26894.02344 | 29049.78516 | 28768.02539 | 0.70573126 | 0.033123824 | 0.009296633 | 0.922008276 | -0.231117258 | 0.105819633 |
| 170131.625 | 153220.4844 | 167023.4688 | 0.70473662 | 0.04113039 | 0.203981562 | 0.163834679 | 0.838183914 | 0.016683666 |
| 32162.17383 | 31572.16797 | 24265.41797 | 0.70365481 | 0.013660857 | 0.139700594 | 0.240911037 | -0.356891023 | 0.138562169 |
| 4943.088379 | 5743.603516 | 3319.480225 | 0.702703068 | 0.083476795 | -0.40287889 | 0.173549208 | -0.276631576 | 0.426621147 |
| 85235.88281 | 84812.35156 | 82031.29688 | 0.702329959 | 0.013596234 | 0.316902705 | 0.159703167 | 0.313177217 | 0.028161099 |
| 126997.9922 | 138784.125 | 123916.2656 | 0.702114873 | 0.021798208 | 0.176576706 | 0.127892721 | 0.32052027 | 0.066077335 |
| 33986.25391 | 37369.10938 | 34959.45703 | 0.701584728 | 0.026045423 | -0.273115211 | 0.099431528 | 0.45755108 | 0.027982615 |
| 12047.12402 | 12632.8457 | 10427.97168 | 0.701351114 | 0.177388967 | 0.07978827 | 0.808336452 | 0.220340696 | 0.502653814 |
| 718709.25 | 745334.375 | 742192.9375 | 0.699964543 | 0.068463493 | 0.0286952 | 0.89833266 | 0.404322704 | 0.089218964 |
| 61378.70313 | 54938.82031 | 56219.65234 | 0.699398833 | 0.027470745 | 0.510460676 | 0.052963377 | 0.431309965 | 0.042414121 |
| 233261.5469 | 200733.7656 | 187338.7031 | 0.698623231 | 0.033526967 | -0.105188217 | 0.409970395 | -0.237897449 | 0.200720922 |
| 58726.46484 | 59698.6875 | 57322.92969 | 0.697836857 | 0.0224814 | -0.368773329 | 0.017861237 | -0.260705909 | 0.017279709 |
| 7881.15332 | 10208.48438 | 11043.99414 | 0.697819438 | 0.212496914 | 0.731019774 | 0.335988204 | 0.867212962 | 0.077431074 |
| 27360.43945 | 28260.65039 | 23638.4043 | 0.697588055 | 0.026154212 | 0.188642733 | 0.079287989 | -0.733959055 | 0.039446558 |
| NA | 10720.68262 | NA | 0.697379286 | 0.173460215 | -0.160864676 | NA | 0.447241546 | NA |
| 164031.8594 | 165212.3281 | 165230.6406 | 0.696574859 | 0.005752145 | -0.226231043 | 0.069257825 | -0.453993199 | 0.020008995 |
| 84184.04688 | 75663.27344 | 77729.17969 | 0.696378354 | 0.046810226 | -0.658237911 | 0.052447104 | -0.342731509 | 0.112229745 |
| 126135.1475 | 100564.4619 | 94926.33887 | 0.695647983 | 0.060362902 | 0.883114744 | 0.137613908 | 1.201576982 | 0.077431074 |
| 7243.953125 | 6343.967773 | 7940.508301 | 0.695314916 | 0.054926056 | 0.068020024 | 0.76241249 | -0.202185113 | 0.296204413 |

| | | | | | | | | |
|---|---|---|---|---|---|---|---|---|
| 22149.9707 | 22532.16602 | 21629.77539 | 0.695016751 | 0.06611991 | 0.491758438 | 0.141443989 | 0.872782066 | 0.036066713 |
| 175093.6875 | 178540.0625 | 183629.1094 | 0.694637859 | 0.090037646 | 0.521384259 | 0.067104052 | 0.55026234 | 0.074131758 |
| 145485.2344 | 152482.5156 | 147265.7813 | 0.694269756 | 0.024442894 | 0.009389291 | 0.891327899 | -0.054407441 | 0.442628123 |
| 116922.2266 | 128939.9297 | 120614.1953 | 0.693564249 | 0.032155886 | 0.343587566 | 0.081173768 | 0.381556462 | 0.05899079 |
| 397485.1875 | 407189.375 | 362715.3125 | 0.691930699 | 0.178834579 | 0.325185681 | 0.321888877 | -0.084859117 | 0.787711053 |
| 54551.76563 | 58828.90625 | 53903.46484 | 0.690738692 | 0.024706248 | 0.409212046 | 0.045837958 | 0.518447685 | 0.02163333 |
| 16750.73438 | 15715.39258 | 18362.03516 | 0.689572157 | 0.013723547 | -0.51328336 | 0.047452686 | -0.31643694 | 0.060542864 |
| 7013.282227 | 3361.926514 | NA | 0.688136421 | 0.110984157 | 0.269430709 | 0.312626085 | -0.351016945 | NA |
| 1486010.75 | 1641506.875 | 1676192.125 | 0.687377698 | 0.023699066 | -0.239171165 | 0.048593497 | 0.182622596 | 0.141195154 |
| 29573.08594 | 33632.07031 | 35209.82813 | 0.68709348 | 0.046568492 | -0.036272448 | 0.638611855 | -0.218196973 | 0.138562169 |
| 187769.5625 | 205020.5156 | 197777.0313 | 0.686557042 | 0.053893741 | 0.182529698 | 0.323498608 | 0.217293991 | 0.237224538 |
| 32336.38672 | 25168.70117 | 30896.13086 | 0.684681227 | 0.052403056 | 0.268264684 | 0.179933858 | 0.582602977 | 0.081899796 |
| 7235.229492 | 6967.263184 | 6123.710938 | 0.684262928 | 0.330645004 | 0.821986065 | 0.168363676 | 0.059810667 | 0.909535698 |
| 56098.74219 | 67459.15625 | 69751.03906 | 0.68382123 | 0.05239238 | -0.120815823 | 0.41103358 | 0.26239809 | 0.205588881 |
| 95852.33594 | 95519.44531 | 68793.07813 | 0.683057076 | 0.030588379 | -1.036790322 | 0.031419447 | -0.312948743 | 0.224512305 |
| 4528.710449 | 8162.506836 | 7381.774414 | 0.682794158 | 0.510558024 | 0.804543058 | 0.440401575 | 0.505771519 | 0.512774056 |
| 58743.71094 | 56858.57031 | 55869.59375 | 0.682368352 | 0.025307586 | -0.457789268 | 0.087018661 | -0.558620062 | 0.059847517 |
| 5857.820313 | 6747.440918 | 6856.62207 | 0.681178733 | 0.040749235 | -0.286174247 | 0.173986552 | -0.365242783 | 0.114946634 |
| 200592.4063 | 197103.1406 | 206321.1563 | 0.680872247 | 0.024701492 | 0.148220585 | 0.146984394 | 0.011130288 | 0.861196469 |
| 7427.165527 | 6508.939941 | 7239.419922 | 0.680623985 | 0.095338477 | 1.216010565 | 0.051593867 | -0.313501737 | 0.276382075 |
| 1834.83313 | 2198.640381 | 1182.786987 | 0.680066583 | 0.180041386 | 0.448256253 | 0.601186127 | -1.073584914 | 0.115349249 |
| 34633.88672 | 53605.86719 | 23210.12695 | 0.679695565 | 0.305890648 | 0.158298628 | 0.75356214 | -0.58792374 | 0.385095259 |
| 45108.10938 | 25324.44336 | 34900.60547 | 0.678978263 | 0.065355995 | -0.174599607 | 0.390316657 | 0.051249682 | 0.895525205 |
| 5819.04834 | 4390.537109 | 3242.843994 | 0.678707926 | 0.240283975 | 0.021929856 | 0.964274443 | 0.671051881 | 0.24326618 |
| 24202.1582 | 23180.01172 | 25963.79297 | 0.678672957 | 0.076603897 | 0.221498869 | 0.314773504 | 0.416936429 | 0.111750711 |
| 55890.79297 | 89247.53125 | 22094.31836 | 0.678601504 | 0.337270865 | 0.182080631 | 0.724035141 | -0.616025703 | 0.449231325 |
| 1175.131592 | 1061.772583 | 1100.37439 | 0.678120749 | 0.033901565 | -0.560483412 | NA | -0.433034492 | 0.086651286 |
| 14606.78809 | 13092.39746 | 11538.97754 | 0.677951798 | 0.147715547 | -0.419436672 | 0.25313315 | -0.46512087 | 0.213264642 |
| 5883.716309 | 5089.527832 | 4742.033691 | 0.677040714 | 0.013180328 | 0.083313125 | 0.421355009 | -0.59185125 | 0.047120928 |
| 29693.89258 | 30718.41406 | 29821.83203 | 0.67670335 | 0.377796751 | 0.875734685 | 0.159217919 | 0.400777678 | 0.455335597 |
| 213128.4219 | 216521.2188 | 191912.0625 | 0.676261428 | 0.01031238 | 0.062522733 | 0.346783035 | -0.165850281 | 0.135536925 |
| 10278.58984 | 11444.70703 | 10466.12109 | 0.676129313 | 0.110302818 | -0.036953573 | 0.873585467 | -0.247206536 | 0.332650897 |
| 78031.46094 | 86645.04688 | 76821.52344 | 0.675956455 | 0.066744288 | -0.649658033 | 0.114739915 | -0.15024197 | 0.450472031 |

| | | | | | | | | |
|---|---|---|---|---|---|---|---|---|
| 238674.9844 | 262142.3594 | 222571.6406 | 0.675875829 | 0.016471012 | 0.274329961 | 0.059562109 | 0.412063212 | 0.066550319 |
| 117333.6094 | 107785.0781 | 114452.9063 | 0.67586077 | 0.091607129 | 0.079179626 | 0.700417395 | -0.199190115 | 0.389456703 |
| 234873.1563 | 268643.5 | 245112.5 | 0.675813182 | 0.010373168 | -0.172371994 | 0.108625401 | 0.139615637 | 0.227573791 |
| 12953.76172 | 18433.56641 | 8767.089844 | 0.6757102 | 0.259597065 | -0.087726322 | 0.848312804 | -0.755182756 | 0.234271542 |
| 27733.33984 | 73710.08594 | 20412.22266 | 0.675175372 | 0.494764936 | 1.094316769 | 0.303517158 | 0.403492251 | 0.722351194 |
| 102375.5234 | 110403.8594 | 119869.4766 | 0.674352507 | 0.029627098 | 0.363386042 | 0.070731667 | 0.644624212 | 0.034410747 |
| 30258.36914 | 38649.375 | 31496.18359 | 0.673691847 | 0.224955599 | -0.396091742 | 0.406536014 | -0.220375945 | 0.588834899 |
| 31841.29883 | 31423.10156 | 27202.01367 | 0.673639529 | 0.020015311 | 0.052517584 | 0.560671011 | -0.219095932 | 0.145662926 |
| 2758.900391 | 4742.577637 | 4678.458008 | 0.672864309 | 0.177171841 | 0.507033727 | 0.168359532 | 0.185615201 | 0.670590816 |
| 13162.01953 | 14192.26074 | 14615.48145 | 0.672516118 | 0.011814081 | 0.065273135 | 0.141443989 | 0.012719195 | 0.856578816 |
| 67115.79688 | 63552.04297 | 68641.66406 | 0.672292006 | 0.012750417 | -0.55239202 | 0.033121608 | -0.011810984 | 0.872236202 |
| 5126.666016 | 6244.547852 | 6546.996094 | 0.672198974 | 0.019694688 | 0.514227173 | 0.14785027 | 0.416518336 | 0.137062098 |
| 8955.727539 | 8681.194336 | 6939.319824 | 0.671487576 | 0.120016833 | -0.298761288 | 0.121769831 | -0.50392689 | 0.066427455 |
| 33468.58203 | 34955.32422 | 29661.66016 | 0.671428928 | 0.061350067 | 0.592697426 | 0.053185315 | 0.317401377 | 0.147103987 |
| 15052.88281 | 16464.08789 | 16120.03711 | 0.671235881 | 0.016471012 | 0.76634534 | 0.013906082 | 0.119474243 | 0.223290576 |
| 120750.7031 | 122926.8828 | 125898.3984 | 0.671205787 | 0.025086961 | -0.434179242 | 0.083003488 | -0.10521351 | 0.338338012 |
| 6424.320313 | 5629.843262 | 4147.643555 | 0.670763668 | 0.013660857 | -0.473874818 | 0.049277594 | -0.302280942 | 0.297300949 |
| 24916.34961 | 23433.56641 | 19839.00391 | 0.670582671 | 0.060362902 | 0.697565174 | 0.038647166 | 0.276100897 | 0.205588881 |
| 216074.1875 | 218027.125 | 200100.8594 | 0.668299992 | 0.042979653 | 1.165621676 | 0.103477445 | 1.237603237 | 0.009587646 |
| 240115.0938 | 227236.6875 | 226517.6094 | 0.668053727 | 0.060213519 | -0.199774873 | 0.313455345 | -0.191152173 | 0.262901299 |
| 120689.4609 | 127187.6016 | 122338.9063 | 0.667058389 | 0.023724131 | -0.154607172 | 0.215727673 | 0.084769774 | 0.383564774 |
| 39397.125 | 38251.32422 | 36828.47266 | 0.666928802 | 0.02194812 | 0.660700595 | 0.096721331 | -0.029629741 | 0.760630262 |
| 8283.404297 | 9892.209961 | 5879.515625 | 0.666734841 | 0.163314204 | -0.543897047 | 0.234571805 | -0.728348478 | 0.157477573 |
| 19305.56445 | 14484.65137 | 16683.94922 | 0.666087263 | 0.110453942 | 0.947980486 | 0.052292511 | 1.143987671 | 0.04728688 |
| 10565.97461 | 23503.86328 | 8686.939453 | 0.664863859 | 0.500705085 | 0.261905487 | 0.732502775 | -0.360889467 | 0.720547247 |
| 16328.05566 | 15168.43945 | 16497.58984 | 0.66305121 | 0.214500791 | 0.371095437 | 0.078083663 | 0.902197474 | 0.012256037 |
| 2825.288574 | 1824.968018 | 1832.128662 | 0.662907674 | 0.036480793 | 0.411134328 | 0.073326865 | 0.166784724 | 0.649231493 |
| 292085.5938 | 292336.1563 | 292756.3125 | 0.662754 | 0.010297098 | -0.419079224 | 0.042487934 | -0.565857539 | 0.029710029 |
| 10388.06348 | 8654.469727 | 6470.494141 | 0.662611145 | 0.103144215 | 0.177093453 | 0.431424722 | -0.129703495 | 0.695461646 |
| 141727.0938 | 137161.2031 | 153523.9688 | 0.66247411 | 0.056093567 | -0.203791493 | 0.26950249 | -0.061009007 | 0.67778571 |
| 146404.2813 | 181785.9375 | 101381.1172 | 0.662278271 | 0.031790847 | 0.177801055 | 0.219820279 | -0.325750032 | 0.365941752 |
| 65052.84766 | 56356.25781 | 51222.88672 | 0.662225449 | 0.100951177 | -0.376980433 | 0.227071428 | -0.653604676 | 0.095635733 |
| 35822.96094 | 41016.20703 | 38782.15234 | 0.662042793 | 0.143738987 | -0.365208135 | 0.289589686 | 0.266603909 | 0.306469701 |

| | | | | | | | | |
|---|---|---|---|---|---|---|---|---|
| 84708.35938 | 75477.8125 | 75342.21094 | 0.662016315 | 0.02237899 | 0.327713781 | 0.06919378 | -0.142211214 | 0.218029927 |
| 22617.11914 | 27421.41992 | 25651.69336 | 0.661071439 | 0.039111956 | 0.621508779 | 0.044791528 | 0.361746936 | 0.096997136 |
| 176866.0313 | 138153.1719 | 140568.2813 | 0.660590161 | 0.2891986 | 0.195199977 | 0.709101354 | -0.234145724 | 0.570514346 |
| 7238.279297 | 6292.110352 | 7497.319824 | 0.660298136 | 0.070477311 | 1.057161531 | 0.026826422 | 0.461035212 | 0.066274975 |
| NA | 808.980896 | 1528.368286 | 0.659636745 | 0.331299053 | -2.320875902 | 0.147323941 | -2.624943819 | NA |
| 14522.07227 | 11659.45215 | 11473.98438 | 0.659535039 | 0.063208632 | 0.502772769 | 0.497345646 | -0.16470333 | 0.473264096 |
| 23498.75195 | 42893.75 | 13928.73438 | 0.658610234 | 0.292211838 | 0.258976043 | 0.553733446 | -0.612354149 | 0.396866599 |
| 17271.17773 | 15860.98926 | 13988.3623 | 0.658523206 | 0.022477079 | -0.607272404 | 0.050235996 | -0.574173436 | 0.032218853 |
| 2350.870117 | 12899.14551 | 19626.80859 | 0.658294909 | NA | 1.764503497 | NA | 0.631925394 | NA |
| 32367.99023 | 36146.49609 | 31516.05078 | 0.658189098 | 0.024442894 | -0.023068646 | 0.786659698 | -0.258777471 | 0.088892469 |
| 828732.0625 | 803467.5 | 766857.375 | 0.658117811 | 0.016471012 | 0.195405758 | 0.091210216 | 0.183249823 | 0.084180144 |
| 103204.1953 | 93671.07813 | 100555.0547 | 0.657968911 | 0.016050563 | 0.423957364 | 0.093738971 | 0.277032412 | 0.054109248 |
| 2762.351074 | 1256.76709 | 767.9111938 | 0.657616432 | 0.248611406 | -0.583161099 | 0.319951282 | -1.099301565 | 0.182679461 |
| 5689.433105 | 6658.479004 | 6164.566406 | 0.656317268 | 0.096971085 | 0.856570353 | 0.050817911 | 1.128925741 | 0.019880539 |
| 1449000.25 | 1500237.875 | 1596722.25 | 0.654869608 | 0.029750807 | -0.161789914 | 0.22769056 | 0.400722577 | 0.029820656 |
| 12820.06836 | 14600.71777 | 11405.48535 | 0.654360551 | 0.044599418 | -0.425221463 | 0.125379248 | -0.418338406 | 0.075839948 |
| 13415.38574 | 12142.84863 | 12372.31152 | 0.653870325 | 0.434484496 | -0.794680881 | 0.399613451 | -1.072344064 | 0.317050668 |
| 257677.625 | 270926.7188 | 282118.0938 | 0.653643244 | 0.019286562 | -0.198342215 | 0.150992552 | 0.003293919 | 0.970723265 |
| 119159.5313 | 121195.7109 | 110219.1563 | 0.652912278 | 0.08654621 | -0.080668602 | 0.688199662 | 0.166854807 | 0.401391855 |
| 103644.1484 | 97385.42969 | 89407.52344 | 0.651965826 | 0.026784983 | 0.215915223 | 0.132973237 | 0.422475532 | 0.05183905 |
| 851715.25 | 826473.5 | 797763.3125 | 0.650756306 | 0.089295939 | 0.245944724 | 0.231359769 | -0.262120014 | 0.263528267 |
| 660008.5 | 671206.8125 | 672286.75 | 0.650148959 | 0.021100999 | -0.305259404 | 0.068255066 | -0.363746386 | 0.010654184 |
| 3136.733887 | 2510.318848 | 2414.719482 | 0.650034277 | 0.084617214 | 0.078145415 | 0.625385428 | 0.267582289 | 0.27322162 |
| 13417.33496 | 19013.41016 | 18354.44141 | 0.64993624 | 0.131566823 | -0.437614774 | 0.223404971 | -0.525072865 | 0.169075369 |
| 20707.20703 | 20577.62109 | 17130.22852 | 0.649758536 | 0.125553661 | 0.6670156 | 0.065871863 | 0.433195819 | 0.136000191 |
| 142155.5313 | 128784.9141 | 130296 | 0.649504589 | 0.176820059 | -0.124489344 | 0.690837397 | -0.070961297 | 0.813655766 |
| 175264.7031 | 169795.1406 | 173370.2188 | 0.64925463 | 0.023724131 | 0.653364848 | 0.091427444 | 1.727597401 | 0.002683747 |
| 21454.77734 | 18006.70508 | 17569.21484 | 0.648487396 | 0.197302391 | 0.507548448 | 0.190813605 | 0.018775152 | 0.959679156 |
| 17244.42188 | 20541.77148 | 15639.43848 | 0.648103749 | 0.148078396 | -1.189626364 | 0.103460535 | -1.015248403 | 0.110685143 |
| 23295.61914 | 38615.19141 | 17138.96289 | 0.646954869 | 0.329282319 | 0.050034738 | 0.922008276 | -0.362464098 | 0.584137475 |
| 124689.5703 | 133039.0781 | 122713.5313 | 0.64651298 | 0.009594272 | -1.150285756 | 0.021680265 | -1.295501848 | 0.01107691 |
| 11417.42383 | 12802.73828 | 9489.668945 | 0.645589324 | 0.051607729 | 0.292074975 | 0.225445341 | 0.511626032 | 0.12131068 |
| 55071.125 | 49801.56641 | 52438.41797 | 0.644730729 | 0.023401535 | -0.348085473 | 0.04667516 | -0.009779392 | 0.908710949 |

| | | | | | | | | |
|---|---|---|---|---|---|---|---|---|
| 32758.69922 | 33121.54688 | 39952.10156 | 0.64435439 | 0.026675593 | -0.605098617 | 0.047212306 | -0.36520745 | 0.088388282 |
| 30302.03711 | 32570.875 | 32543.81055 | 0.643993008 | 0.228635392 | -0.199306993 | 0.628159256 | -0.23634013 | 0.555307601 |
| 830147.75 | 721899.875 | 692481.5 | 0.643953584 | 0.025307586 | 0.055020489 | 0.683528942 | 0.424612458 | 0.081899796 |
| 38321.61719 | 32993.27344 | 35491.68359 | 0.643754277 | 0.013583579 | 0.010836599 | 0.888691051 | 0.134449212 | 0.266209604 |
| 329475.25 | 363908.7188 | 340184.375 | 0.643300411 | 0.031442732 | -0.313117641 | 0.117302538 | -0.37078034 | 0.080462751 |
| 32631.50586 | 36605.8125 | 35292.38672 | 0.642662221 | 0.014665071 | 0.316940151 | 0.078237525 | 0.307743822 | 0.05511506 |
| 82257.28906 | 89810.46875 | 79377.3125 | 0.64266088 | 0.015175173 | -0.363591041 | 0.05818214 | -0.012733724 | 0.902702952 |
| 289424.9688 | 268875.6563 | 293511.1875 | 0.642637886 | 0.025086961 | -0.025434819 | 0.807829651 | -0.343342177 | 0.078572249 |
| 19393.54883 | 16401.75781 | 11871.82227 | 0.642116577 | 0.033344102 | 0.020934896 | 0.84622109 | -0.276744973 | 0.370714373 |
| 6673.79541 | 3159.650146 | 2456.893555 | 0.641379509 | 0.145553706 | -1.61816794 | 0.01179292 | -2.1249279 | 0.036229875 |
| 193592.875 | 216475.3594 | 203525.9844 | 0.639838887 | 0.145580182 | -0.055074502 | 0.842124822 | 1.234353294 | 0.020920078 |
| 59268.57031 | 59255.02344 | 67673.03906 | 0.639467149 | 0.268867993 | 2.729969994 | 0.052702293 | 2.986110659 | 0.015632484 |
| 40343.78125 | 50348.94141 | 71857.53125 | 0.639131932 | 0.088231376 | -0.064371236 | 0.760749229 | 0.569014036 | 0.2939741 |
| 36436.41797 | 32636.55859 | 36898.28516 | 0.638858707 | 0.044205353 | 0.44855229 | 0.103460535 | 0.715385282 | 0.024635256 |
| 46512.60156 | 85960.10156 | 37273.35547 | 0.637816276 | 0.367769754 | 0.177018134 | 0.736533491 | -0.304562658 | 0.664329886 |
| 85886.64844 | 86723.08594 | 90823.60156 | 0.63777357 | 0.013777247 | -0.418729337 | 0.050817911 | -0.299091445 | 0.031136253 |
| 5334.826172 | 4370.11377 | 5428.939453 | 0.637613401 | 0.088240896 | -0.947039716 | 0.072465942 | -1.778260326 | 0.051065331 |
| 46674.80859 | 50001.60156 | 47720.71875 | 0.636688857 | 0.016885305 | -0.211522166 | 0.074808934 | -0.095450987 | 0.189363824 |
| 68426.25 | 65695.10938 | 55584.95313 | 0.635384074 | 0.016471012 | -0.40353301 | 0.01179292 | -0.53495729 | 0.060216651 |
| 35215.53125 | 37928.58594 | 24236.04688 | 0.635165909 | 0.11374283 | -0.363665404 | 0.24146716 | -0.200743081 | 0.537307257 |
| 74364.64844 | 73989.80469 | 62598.85547 | 0.634755446 | 0.013583579 | -0.738966905 | 0.047652737 | -0.076857914 | 0.546143784 |
| 86393.86719 | 94984.54688 | 89270.35938 | 0.634488831 | 0.010297098 | -1.083635471 | 0.017096626 | -0.465657881 | 0.019880539 |
| 14596.11328 | 18861.05469 | 18127.83398 | 0.634436321 | 0.172476895 | 0.279347951 | 0.378617903 | 0.280488753 | 0.379296337 |
| 140747.7031 | 139286.1875 | 140203.4531 | 0.634221216 | 0.145553706 | 0.112509444 | 0.689375816 | 0.87734256 | 0.062863768 |
| 994576.3125 | 1142530 | 1010349.563 | 0.634181047 | 0.044763187 | 0.943260485 | 0.018539883 | 1.158685153 | 0.021134419 |
| 20653.05273 | 17931.58398 | 13030.0332 | 0.634114557 | 0.036830197 | -0.857802407 | 0.035570159 | -0.994899553 | 0.062345205 |
| 181575.9375 | 190122.7695 | 192913.2578 | 0.634083956 | 0.039413354 | -0.162970523 | 0.313982423 | -0.115294418 | 0.423557711 |
| 879138.5625 | 944638.375 | 936472.4375 | 0.634079736 | 0.04251435 | 0.656872281 | 0.048820023 | 0.605835875 | 0.027087229 |
| 34768.66797 | 28215.20508 | 35363.29297 | 0.633763022 | 0.022729015 | -0.440883297 | 0.042753324 | -0.811963625 | 0.021723356 |
| 43697.98828 | 42819.71875 | 43083.89453 | 0.633539085 | 0.016596319 | -0.08161625 | 0.068834111 | -0.003813847 | 0.83964397 |
| 22468.54102 | 21049.31641 | 17443.37305 | 0.633493819 | 0.154381173 | 1.059180885 | 0.033232099 | 0.140410727 | 0.616686362 |
| 62806.58594 | 60936.3125 | 61777.91797 | 0.633024784 | 0.008009891 | -0.659173359 | 0.023578081 | -0.35748368 | 0.006394096 |
| 8377.382813 | 5888.323242 | 7927.808594 | 0.632858908 | 0.111187644 | -0.464430037 | 0.22425544 | -0.672098012 | 0.112503484 |

| | | | | | | | | |
|---|---|---|---|---|---|---|---|---|
| 25398.06445 | 26670.13281 | 25045.12891 | 0.632852157 | 0.121235391 | 0.267046452 | 0.295727216 | 0.13585569 | 0.54725722 |
| 210883.7813 | 243393.75 | 238976.1094 | 0.632397345 | 0.05464424 | 0.199835524 | 0.333199228 | -0.06694363 | 0.669239917 |
| 144756.4688 | 137047.875 | 144760.5469 | 0.631207228 | 0.010603888 | -0.385477511 | 0.021622074 | -0.471401683 | 0.015112322 |
| 89286.05469 | 82862.10156 | 91744.67969 | 0.630415781 | 0.020421119 | -0.319713924 | 0.243784441 | -0.038532636 | 0.6581689 |
| 22289.11328 | 24509.17578 | 19778.17969 | 0.628691923 | 0.018311389 | 0.073291404 | 0.393066742 | -0.10831698 | 0.461562059 |
| 234345.75 | 176953.875 | 216535.2344 | 0.628501167 | 0.014544337 | 0.00792501 | 0.873176331 | 0.003190518 | 0.988003013 |
| 161124.0313 | 158097.7344 | 155412.9375 | 0.628287028 | 0.016048565 | -0.231861353 | 0.050958871 | -0.118762016 | 0.110402248 |
| 6347.412598 | 5515.013672 | NA | 0.628274045 | 0.1694675 | -0.794823485 | 0.125579796 | -0.668907423 | NA |
| 77847.55469 | 69720.05469 | 78129.75 | 0.628114013 | 0.054690884 | -0.27468017 | 0.502558557 | 0.709466267 | 0.027783576 |
| 29017.00586 | 26498.51367 | 29411.99805 | 0.627658662 | 0.01446452 | -0.1707663 | 0.119643706 | -0.450404756 | 0.027868741 |
| 173808.2813 | 187445.2188 | 188696.7344 | 0.627455368 | 0.03777619 | -0.268115972 | 0.138466365 | -0.209175042 | 0.174083052 |
| 21298.98047 | 19876.43164 | 17047.375 | 0.626431869 | 0.029514505 | -0.442122144 | 0.011710842 | -0.009518271 | 0.950160109 |
| 127788.9453 | 134044.0156 | 120280.6641 | 0.626392946 | 0.011979502 | 0.365462136 | 0.065566999 | 0.434677829 | 0.035986636 |
| 31219.14453 | 28590.80078 | 27453.57617 | 0.626108063 | 0.005752145 | -0.193237451 | 0.068436129 | -0.221633635 | 0.097178029 |
| 26972.95117 | 22293.16602 | 27909.83398 | 0.626080725 | 0.057316826 | -0.141917697 | 0.200677134 | -0.299071067 | 0.145071579 |
| 9462.427734 | 13144.70215 | 9033.489258 | 0.62523717 | 0.199046058 | 0.030039534 | 0.852417959 | -0.261059375 | 0.362335792 |
| 69786.20313 | 63266.00391 | 67666.85156 | 0.625130429 | 0.015216041 | -0.088846138 | 0.661673786 | 0.313344356 | 0.053053903 |
| 27621.39063 | 26503.59766 | 27342.87695 | 0.624717146 | 0.045580081 | -0.236798935 | 0.127902424 | -0.386109459 | 0.081631245 |
| 103891.7891 | 178466 | 76569.95313 | 0.624628067 | 0.39571103 | 0.152996258 | 0.773923011 | -0.393995917 | 0.573204391 |
| 10999.27051 | 10979.43066 | 11226.21094 | 0.624573445 | 0.014676535 | 0.351077886 | 0.036088129 | -0.249322502 | 0.079165593 |
| 11635.51465 | 10215.0918 | 7518.181152 | 0.624421021 | 0.071134978 | -0.195855452 | 0.290124454 | -0.806566008 | 0.052349495 |
| 7464.009766 | 7015.503906 | 6066.282227 | 0.624045348 | 0.052584221 | 0.293981971 | 0.267197239 | -1.241436007 | 0.041386295 |
| 40192.42102 | 38561.85156 | 32666.45703 | 0.623178147 | 0.041945948 | 0.250970205 | 0.162041667 | -0.045395683 | 0.794795508 |
| 35890.42188 | 57671.98438 | 12444.1582 | 0.623156456 | 0.329719661 | 0.17411061 | 0.686901993 | -0.254929226 | 0.757455431 |
| 760805.1875 | 793673.8125 | 852241.8125 | 0.623067317 | 0.06257795 | -0.084716167 | 0.530684856 | 0.32050289 | 0.079165593 |
| 16716.05469 | 13839.11328 | 14407.50586 | 0.62270175 | 0.156126659 | 0.55034004 | 0.139057647 | 0.018922358 | 0.951388228 |
| 37783.30859 | 35690.04297 | 41890.51563 | 0.622688503 | 0.037309446 | 0.398264986 | 0.34166974 | 1.165634132 | 0.036229875 |
| 47483.44531 | 52200.51563 | 49630.8125 | 0.622622837 | 0.116546 | -0.295952714 | 0.330821995 | -0.458989503 | 0.202986451 |
| 5823.097656 | 6639.928711 | 6318.541504 | 0.620713196 | 0.068728608 | -0.097192715 | 0.612877707 | -0.606182768 | 0.095708676 |
| 164335.2969 | 175817.5938 | 171039.3125 | 0.619523098 | 0.057710534 | -0.089118307 | 0.605041973 | -0.105313416 | 0.520080533 |
| 34215.04297 | 36010.53516 | 34541.45313 | 0.618003637 | 0.028167774 | -0.567238545 | 0.034488442 | 0.160969205 | 0.102620951 |
| 10316326 | 11596725 | 11595104 | 0.617736716 | 0.153739559 | 0.15377661 | 0.618374257 | 1.28746055 | 0.019022762 |
| 139133.2813 | 135763.0156 | 131844.25 | 0.61731866 | 0.005752145 | -0.290683928 | 0.021906439 | -0.395256554 | 0.009824429 |

| | | | | | | | | |
|---|---|---|---|---|---|---|---|---|
| 30277.8457 | 27389.44922 | 28771.93555 | 0.61706209 | 0.05267493 | 0.540903004 | 0.082307494 | 0.54885511 | 0.045311345 |
| 10691.4834 | 10215.91211 | 4161.197266 | 0.616827225 | 0.034514668 | 0.099012001 | 0.465007104 | -0.025108071 | 0.966538084 |
| 111978.7344 | 126856.0391 | 105589.3047 | 0.616629851 | 0.032310891 | -0.743844412 | 0.024165741 | -0.726107843 | 0.020008995 |
| 25489.66406 | 25293.85938 | 21853.74609 | 0.616457058 | 0.032595469 | -0.087889637 | 0.471790095 | -0.017131305 | 0.906660092 |
| 545381.0625 | 554711.875 | 566321.0625 | 0.615091335 | 0.023132924 | -0.355654894 | 0.018539883 | -0.217087377 | 0.017279709 |
| 1274.115112 | 1199.78772 | 698.3597412 | 0.614933586 | 0.188824978 | -0.371138685 | 0.375150282 | -0.438043599 | 0.347817837 |
| 111175.3594 | 128344.4844 | 120338.7813 | 0.614812994 | 0.043150549 | -0.208158265 | 0.188231791 | 0.105537718 | 0.420194723 |
| 18096304 | 34193856 | 12226955 | 0.614675381 | 0.40453073 | 0.221580925 | 0.677447915 | -0.425394759 | 0.573238987 |
| 10658.23828 | 11237.50879 | 8065.429199 | 0.614639976 | 0.039715426 | -1.622935712 | 0.029419521 | -1.272600043 | 0.015734747 |
| 448272.5938 | 457434.5938 | 469473.25 | 0.613765509 | 0.14222539 | 0.079809203 | 0.751217547 | 0.491547598 | 0.129008887 |
| 1604047.25 | 1685634.125 | 1777039.375 | 0.613733292 | 0.028829273 | -0.199650791 | 0.166408179 | 0.145598293 | 0.219268191 |
| 14096.93945 | 15146.22266 | 12713.21777 | 0.613440371 | 0.055228595 | -0.404585721 | 0.129702828 | -0.623385593 | 0.06365738 |
| 21009.59961 | 26008.17578 | 22658.4707 | 0.612943894 | 0.279965855 | 0.731562348 | 0.160182447 | 0.275472397 | 0.493412337 |
| 85887.16406 | 147479.1406 | 96424.85156 | 0.612682569 | 0.686086009 | 0.711235183 | 0.507997025 | 0.522801323 | 0.640857701 |
| 41071.23047 | 42536.35547 | 42930.63281 | 0.612595777 | 0.079968566 | -0.138820653 | 0.175716398 | 0.114127713 | 0.107956786 |
| 19203.43359 | 16658.08398 | 14641.99414 | 0.611334033 | 0.027671686 | -0.809046734 | 0.040932722 | -0.660217167 | 0.036652305 |
| 564763.1875 | 529874.375 | 604521.6875 | 0.611111862 | 0.030009295 | 0.546032024 | 0.042480281 | 0.858423578 | 0.02183874 |
| 350283.2188 | 389109.8438 | 423619.7188 | 0.610405785 | 0.042621515 | -0.278116129 | 0.150271527 | 0.201399158 | 0.23571729 |
| 7220347.5 | 6010828 | 7388290.5 | 0.60994569 | 0.0140944 | -0.148865432 | 0.13812939 | -0.048449027 | 0.735424865 |
| NA | 2061.240967 | NA | 0.609935311 | NA | 0.48830112 | NA | -0.4634437 | NA |
| 38117.62891 | 37882.12109 | 34998.40234 | 0.609862252 | 0.018432262 | 0.454859838 | 0.042884533 | 0.061530174 | 0.410416651 |
| 34843.64844 | 36564.38281 | 29983.25195 | 0.60947085 | 0.055915605 | 0.031189688 | 0.833585594 | 0.006630887 | 0.973455039 |
| 1835.16687 | 2238.01709 | 2695.11377 | 0.609025674 | 0.727807099 | -1.076955732 | 0.504348764 | -2.003694686 | 0.363704844 |
| 12916 | 14098.00879 | 15302.53906 | 0.608975079 | 0.192973731 | 0.934570086 | 0.052817461 | 0.915020403 | 0.046624125 |
| 35972.89844 | 42420 | 27592.60547 | 0.608364411 | 0.177773716 | 0.752516244 | 0.065369282 | 0.029512563 | 0.926080265 |
| 118846.2969 | 105538.6094 | 127879.9375 | 0.607932166 | 0.506809817 | 0.284719756 | 0.808922116 | 3.042849072 | 0.014388274 |
| 9854.546875 | 23161.91406 | 9371.298828 | 0.607911069 | 0.298673579 | 0.031198828 | 0.963570702 | 0.225388901 | 0.784039301 |
| 59056.11719 | 54166.54688 | 47607.92578 | 0.607632744 | 0.083280787 | 0.051458644 | 0.795956929 | -0.284087876 | 0.262261709 |
| 40521.76953 | 39164.3125 | 42242.19531 | 0.607563611 | 0.081471147 | -0.535448735 | 0.110779853 | -0.040729191 | 0.812775264 |
| 3746.147705 | 3656.334229 | 3841.939697 | 0.607269611 | 0.048391875 | -0.924030419 | 0.039808229 | -0.342776579 | 0.101436425 |
| 17658.71094 | 21041.43555 | 16060.22754 | 0.607180699 | 0.269924077 | 0.630890794 | 0.166391835 | 0.841204234 | 0.082885342 |
| 58064.22656 | 58814.13672 | 57204.35938 | 0.606873904 | 0.01111345 | -0.497871004 | 0.021906439 | -0.157727052 | 0.086923184 |
| 65503.07813 | 68523.75781 | 66820.57813 | 0.606822458 | 0.027671686 | 6.30E-05 | 0.999222442 | 0.129420047 | 0.214088222 |

| | | | | | | | | |
|---|---|---|---|---|---|---|---|---|
| 19116.99805 | 17929.72656 | 18002.36523 | 0.606782832 | 0.028196123 | 0.043287082 | 0.631481364 | 0.054799668 | 0.359960679 |
| 5770.238281 | 8749.251953 | 8815.164063 | 0.606389209 | 0.067008234 | -0.654267636 | 0.093751816 | -0.596042937 | 0.106177326 |
| 1899.479614 | 2129.058594 | 1413.399414 | 0.606106197 | 0.150795791 | -0.25787723 | 0.410615137 | -0.825195535 | 0.106413249 |
| 69833.55469 | 66194.26563 | 70901.54688 | 0.605710505 | 0.036547632 | 0.080695608 | 0.50345588 | 0.620340214 | 0.02909479 |
| 98737.72656 | 100915.5313 | 82719.66406 | 0.605437803 | 0.064507716 | 0.65086778 | 0.055527809 | 0.534928004 | 0.065832543 |
| 29768.34961 | 30812.26172 | 28500.59961 | 0.604306232 | 0.042621515 | -0.127471819 | 0.364154084 | 0.04438298 | 0.707975512 |
| 60848.08984 | 66582.88281 | 66023.78125 | 0.603626465 | 0.006735623 | -0.383847679 | 0.017096626 | -0.066408103 | 0.341659762 |
| 116595.8584 | 126723.3174 | 115738.5459 | 0.603616036 | 0.012731179 | -0.112800749 | 0.179124475 | -0.022955886 | 0.777851309 |
| 23598.82422 | 14252.71387 | 14767.94043 | 0.603382655 | 0.048820442 | 0.078503212 | 0.809869006 | -0.148499383 | 0.692411172 |
| 83606.16406 | 87726.20313 | 92460.125 | 0.602689436 | 0.023401535 | -0.01300625 | 0.870533166 | 0.764914021 | 0.017279709 |
| 5522.57666 | 7192.562012 | 8321.00293 | 0.601716967 | 0.029166847 | -0.157202508 | 0.464222823 | -0.136284996 | 0.593820705 |
| 140143.7656 | 140023.2188 | 144846.875 | 0.601504995 | 0.024508165 | -0.154153895 | 0.192361772 | 0.053450423 | 0.529505889 |
| 4885.367676 | 3926.418945 | 5471.108887 | 0.601440908 | 0.096286953 | -0.821840824 | 0.09466445 | -1.848350465 | 0.051065331 |
| 439888.875 | 356038.1875 | 440014.5625 | 0.600725876 | 0.060329638 | -0.057224538 | 0.701829892 | 0.00270772 | 0.989480447 |
| 4448.650879 | 1223.769165 | 633.109314 | 0.599209843 | 0.111187644 | 0.081064097 | 0.643779366 | -0.885802631 | 0.379636564 |
| 200202.0313 | 202675.6875 | 215425.5313 | 0.598789801 | 0.017533506 | -0.43426828 | 0.024165741 | -0.126909803 | 0.100363691 |
| 49641.30859 | 47110.13672 | 39466.39063 | 0.598076146 | 0.0140944 | 0.289183024 | 0.260944141 | -0.906248649 | 0.017763718 |
| 8296.318359 | 10871.50586 | 21561.86523 | 0.597684071 | 0.668087122 | -0.381367567 | 0.781102846 | 0.209579983 | 0.866968244 |
| 2997.884521 | 3684.19751 | 4752.711426 | 0.596698735 | NA | -0.750023943 | 0.100027506 | -0.000586176 | 0.998511466 |
| 20710.30469 | 20058.71484 | 20906.97461 | 0.596499829 | 0.0958851 | 0.251436982 | 0.448541012 | 0.328273856 | 0.054887084 |
| 140321.0332 | 149981.7012 | 151996.6406 | 0.596448781 | 0.186426612 | 0.460469158 | 0.194056605 | 0.413652116 | 0.215760001 |
| 13643.5459 | 12599.73535 | 12605.54102 | 0.595926865 | 0.356928901 | 0.574059218 | 0.256651639 | 1.23673768 | 0.069207996 |
| 42998.66797 | 46882.98047 | 47836.61328 | 0.594929848 | 0.032646789 | 0.021337049 | 0.857985488 | 0.066176659 | 0.570887829 |
| 30565.9668 | 29802.61328 | 27416.37891 | 0.594289296 | 0.029933445 | -0.062623555 | 0.63851583 | -0.310758563 | 0.091523165 |
| 10476.2998 | 12757.13086 | 14948.44531 | 0.594241881 | 0.313971512 | 0.668532312 | 0.254161368 | 1.150497833 | 0.062635252 |
| 6613.336426 | 3919.422363 | 10825.40723 | 0.594050413 | 0.458148137 | 1.107872817 | 0.131425065 | 0.254114199 | 0.746847228 |
| 35207.02344 | 35345.1875 | 33729.36719 | 0.594043039 | 0.030417399 | 0.1954433 | 0.030506702 | 0.280279094 | 0.020008995 |
| 54571.10547 | 51181.00391 | 51996.41406 | 0.593606237 | 0.053826615 | 0.235720251 | 0.024165741 | 0.154187799 | 0.063283918 |
| 184090.5938 | 173713.875 | 171159.5156 | 0.592517949 | 0.034514668 | 0.051114871 | 0.754355045 | 0.561744283 | 0.017602156 |
| 127941.5283 | 121378.9282 | 116718.2378 | 0.591824337 | 0.056093567 | -0.372689739 | 0.120256734 | -0.14822061 | 0.274843527 |
| 39549.90625 | 36743.44922 | 34661.80078 | 0.591594364 | 0.016471012 | 0.232812149 | 0.021969813 | -0.053233352 | 0.535773657 |
| 52321.375 | 52802.38672 | 54629.01563 | 0.591432786 | 0.049249627 | -0.367140418 | 0.119643706 | -0.245014509 | 0.207846735 |
| 135657.1094 | 133127.2969 | 138684.1406 | 0.589998402 | 0.02406427 | -0.074215684 | 0.484330897 | 0.24019169 | 0.091915425 |

| | | | | | | | | |
|---|---|---|---|---|---|---|---|---|
| 211360.5469 | 182123.9531 | 178812.3281 | 0.589899153 | 0.020626238 | 0.348027711 | 0.046114261 | 0.360977348 | 0.098490466 |
| 21951.84766 | 22722.10742 | 25186.59961 | 0.589621531 | 0.055042495 | 0.218028038 | 0.191226807 | 0.510214104 | 0.04324216 |
| 90249.50781 | 98570.83594 | 103656 | 0.588925499 | 0.037346365 | -0.885479057 | 0.058077004 | -0.053011177 | 0.70128144 |
| 488345.5938 | 539303 | 466695.5 | 0.588893712 | 0.078489996 | -0.151522503 | 0.483003566 | 0.158726845 | 0.376493076 |
| 25554.40234 | 22826.68945 | 24490.39648 | 0.588753522 | 0.081796003 | -0.335541704 | 0.17536027 | -0.325544406 | 0.149748939 |
| 7674.302734 | 8075.963867 | 6309.87793 | 0.587478578 | 0.016471012 | -0.140104959 | 0.382940485 | -0.475007998 | 0.062838928 |
| 337.9955444 | NA | NA | 0.587393686 | NA | -0.245427482 | NA | -2.421053083 | NA |
| 4204.668945 | 4188.262207 | 3345.390869 | 0.587303321 | 0.070959976 | -0.195150639 | 0.337838294 | 0.005673188 | 0.981575243 |
| 1036372.867 | 995008.8711 | 953313.8086 | 0.586639847 | 0.025307586 | -0.056406092 | 0.420028809 | 0.005913025 | 0.941811211 |
| 8557.762695 | 8593.09668 | 8474.168945 | 0.585870867 | 0.100401711 | -0.358909139 | 0.21109765 | -0.087062277 | 0.644909155 |
| 48592.39453 | 53664.38281 | 23022.48438 | 0.585641654 | 0.260927802 | 0.94961499 | 0.47004298 | 0.702536586 | 0.335126348 |
| 29753.6582 | 30681.16406 | 30542.33789 | 0.58500529 | 0.097964968 | -0.034970784 | 0.860618813 | 0.000534494 | 0.997864759 |
| 4110.206055 | 3979.306396 | 2960.368896 | 0.584846729 | 0.092356264 | 0.134996634 | 0.440131477 | 0.161274363 | 0.50760018 |
| 309825.625 | 354842.875 | 262307.6563 | 0.584605878 | 0.093016235 | -0.141609911 | 0.552005408 | -0.134682646 | 0.591044383 |
| 130401.6875 | 125018.4766 | 123563.2891 | 0.583558248 | 0.004709568 | -0.067415099 | 0.490292673 | 0.333049406 | 0.026263251 |
| 69864.1875 | 55005.42969 | 58218.91406 | 0.583340355 | 0.139920717 | -0.013616855 | 0.953751847 | 0.006027916 | 0.983784117 |
| NA | 6943.477051 | 5203.473145 | 0.583228831 | 0.204593648 | -0.561569843 | 0.247130077 | -0.814926647 | NA |
| 14491.97363 | 17042.60742 | 15471.17871 | 0.583162467 | 0.313021144 | 0.062572289 | 0.885440998 | 1.451019788 | 0.029215308 |
| NA | 3285.168945 | 1839.88916 | 0.583068322 | 0.49108837 | -0.26063579 | 0.729919166 | -1.926865956 | NA |
| 93437.24219 | 91508.88281 | 99020.71875 | 0.582470301 | 0.029391485 | -0.149293202 | 0.047212306 | -0.359005277 | 0.020045376 |
| 1530.318726 | 4238.678223 | 3367.959717 | 0.582190887 | 0.296534519 | -1.803584247 | 0.12050319 | -1.223647529 | 0.126688319 |
| 19992.98047 | 16967.6582 | 18603.03906 | 0.58215642 | 0.032652748 | -0.112540294 | 0.321230521 | 0.128504715 | 0.343566025 |
| 293343.0313 | 283113.1563 | 314396.4063 | 0.582049315 | 0.055228595 | -0.145244159 | 0.401994763 | 0.129397634 | 0.402758761 |
| 59079.32813 | 72313.05469 | 65142.59375 | 0.581815119 | 0.039126605 | -0.573686027 | 0.065457698 | -1.131550297 | 0.02527614 |
| 584172.1875 | 595790.25 | 654346.8125 | 0.581458332 | 0.022137298 | 0.187359471 | 0.121571366 | 0.421989599 | 0.050623317 |
| 170893.375 | 178297.5 | 199993.8906 | 0.580860684 | 0.013964852 | 0.169604831 | 0.134634811 | 0.099831224 | 0.413077372 |
| 66851.15625 | 63502.70703 | 62227.8125 | 0.580138532 | 0.010881569 | 0.04063094 | 0.553393539 | -0.19071295 | 0.044524888 |
| 6869.80127 | 9105.948242 | 4130.515137 | 0.579763102 | 0.465993763 | 0.401595672 | 0.526766343 | -0.774555883 | 0.399380775 |
| 818008.5 | 853627.1875 | 838216.0625 | 0.579564536 | 0.046600136 | 0.304194643 | 0.100849207 | 0.391908809 | 0.051942155 |
| 10096.63281 | 5815.330078 | 9937.493164 | 0.579433168 | 0.231091988 | 0.026850057 | 0.945953011 | -0.067093712 | 0.88560307 |
| 33188.10938 | 24901.72852 | 15442.5 | 0.578381056 | 0.217831937 | 0.919515401 | 0.058077004 | 0.008238109 | 0.989076637 |
| 50220.02734 | 49079.19922 | 44876.54688 | 0.577590978 | 0.046025893 | 0.100589332 | 0.349103174 | -0.235434738 | 0.120850847 |
| 122519.9844 | 144002.9375 | 123939.875 | 0.577515816 | 0.025025466 | 0.031924958 | 0.735373025 | 0.130640222 | 0.363209271 |

| | | | | | | | | |
|---|---|---|---|---|---|---|---|---|
| 24602.92383 | 22977.36133 | 21097.04297 | 0.576070255 | 0.054033703 | 0.395594385 | 0.115283074 | 0.140452276 | 0.382150932 |
| 7596.118652 | 8313.435547 | 7150.355957 | 0.575933824 | 0.131546744 | 0.459227519 | 0.221929915 | -0.125569889 | 0.315117797 |
| 25794.91992 | 25028.44141 | 26428.41992 | 0.575772609 | 0.025939018 | -0.248277618 | 0.05253541 | -0.181283733 | 0.069519222 |
| 94431.53906 | 96873.92188 | 105874.3828 | 0.575756832 | 0.080105106 | -0.007512755 | 0.921811704 | -0.196819566 | 0.112528358 |
| 336118.5 | 320251.2813 | 270310.6563 | 0.575217884 | 0.096750655 | 0.639505381 | 0.109509233 | -0.400716724 | 0.176743286 |
| 12496.76367 | 8260.591797 | 11789.93066 | 0.574911077 | 0.121770031 | -0.492874132 | 0.085659818 | -1.101875018 | 0.025983861 |
| 1618.114258 | 1611.859985 | 1172.905762 | 0.574080405 | 0.360794803 | -0.365977032 | 0.382455158 | -0.083570625 | 0.824283778 |
| 182357.1094 | 316400.0938 | 128513.125 | 0.573796843 | 0.403414633 | 0.204354974 | 0.689502716 | -0.388903163 | 0.570887829 |
| 58929.00391 | 66196.01563 | 61989.89453 | 0.572309684 | 0.132673516 | 0.32162994 | 0.085927719 | 0.602739932 | 0.023614322 |
| 25712.08984 | 24800.97461 | 24815.37891 | 0.57226676 | 0.025086961 | -0.126320383 | 0.228110774 | -0.339066231 | 0.02527614 |
| 254547.3125 | 257758.25 | 235038.4375 | 0.572048197 | 0.041635583 | 0.551771066 | 0.094472675 | 0.694719672 | 0.020045376 |
| 597859.1875 | 603626.0625 | 632430.625 | 0.571781166 | 0.017243194 | 0.076343577 | 0.291070942 | -0.043061261 | 0.529445312 |
| 22126.38477 | 30608.26563 | 32473.64063 | 0.571204308 | 0.118081327 | 0.022116641 | 0.778964034 | 0.049474115 | 0.845545984 |
| 22398.38672 | 22009.57617 | 22853.01953 | 0.571098136 | 0.142471188 | 0.204489169 | 0.41941562 | 0.274809974 | 0.299059995 |
| 7241.716797 | 9706.078125 | 8265.376953 | 0.571025925 | 0.037314581 | -0.417595464 | 0.088587863 | -1.135836723 | 0.02183874 |
| 13848.01855 | 14079.21875 | 16215.55371 | 0.570955856 | 0.113673327 | 0.204726245 | 0.108625401 | 0.060024217 | 0.642883208 |
| 12064.6123 | 11198.86719 | 13998.71387 | 0.570132538 | NA | -0.26302383 | 0.365468558 | -0.414471071 | 0.102308303 |
| 110933.1563 | 114269.3047 | 113276.3047 | 0.569525332 | 0.010484689 | -0.253996763 | 0.052447104 | -0.062245927 | 0.28518673 |
| 255620.2969 | 258515.7344 | 276296.4688 | 0.569007899 | 0.06703077 | 0.139293629 | 0.407540737 | 0.133518758 | 0.31598483 |
| 698.0473633 | 1108.995117 | NA | 0.569007682 | 0.080487571 | -1.043472634 | 0.013427556 | -2.007909582 | NA |
| 65050.47266 | 62895.10547 | 66456.21094 | 0.568061339 | 0.064437538 | -0.522265422 | 0.092287364 | -0.951629653 | 0.05752489 |
| 129161.9844 | 142638.2969 | 140974.7969 | 0.567896409 | 0.031396254 | -0.17358075 | 0.167303467 | 0.320804022 | 0.048431709 |
| 31226.32031 | 35288.86328 | 36463.70703 | 0.56789366 | 0.153465292 | 0.096202782 | 0.571596501 | 0.615196391 | 0.042175609 |
| 13341.03906 | 21236.43555 | 9154.776367 | 0.567756349 | 0.400522684 | 0.297709112 | 0.535797708 | -0.146740976 | 0.820794202 |
| 12386.96875 | 10413.48145 | 10124.01074 | 0.566709692 | 0.063843436 | -0.581689291 | 0.098459633 | -0.40900461 | 0.102613568 |
| 34519.76172 | 21604.37305 | 44296.89844 | 0.566322953 | 0.531449704 | 0.271173454 | 0.55734223 | 1.178617776 | 0.169075369 |
| 5557.489258 | 13778.12598 | 3497.94458 | 0.566165722 | 0.46699162 | 0.263177482 | 0.693893294 | -0.207564972 | 0.833839827 |
| 337148.3125 | 369626.4375 | 356988.5938 | 0.564658106 | 0.031140524 | -0.223448322 | 0.109653398 | -0.228979678 | 0.09105947 |
| 84718.83594 | 79099.90625 | 87731.32031 | 0.564330624 | 0.029432233 | 0.061929534 | 0.374414438 | -0.137746159 | 0.15172347 |
| 12111.84961 | 14797.8252 | 16718.36914 | 0.564044499 | 0.090594806 | -0.237579202 | 0.299244779 | 0.211782126 | 0.38339742 |
| 346891.1875 | 489734.8438 | 132068.8125 | 0.563458716 | 0.368213807 | 0.472919776 | 0.356546603 | -0.187037116 | 0.808676121 |
| 12454.89941 | 16184.21289 | 14815.50586 | 0.563442643 | 0.318687987 | 1.043367629 | 0.104928475 | 1.110704014 | 0.059345754 |
| 61778.41797 | 58961.39844 | 61409.05078 | 0.562500264 | 0.037877771 | -0.163372295 | 0.259839565 | -0.192871832 | 0.206000009 |

| | | | | | | | | |
|---|---|---|---|---|---|---|---|---|
| 3798.896973 | 4123.262695 | 2905.286865 | 0.560881623 | 0.154273484 | -0.704011792 | 0.035493707 | -1.070940904 | 0.021836949 |
| 135191.0156 | 137313.9688 | 152237.25 | 0.560572422 | 0.036566441 | -0.267316816 | 0.142889936 | -0.433868466 | 0.064943902 |
| 38883.79688 | 54977.00781 | 51074.99219 | 0.560001759 | 0.114045475 | 0.232753354 | 0.312626085 | -0.060260694 | 0.832271436 |
| 31460.24805 | 31787.62305 | 37296.72656 | 0.559950864 | 0.271021856 | -0.497362966 | 0.344433056 | -0.314095684 | 0.477106314 |
| 17845.74805 | 18771.00391 | 18665.75195 | 0.559468761 | 0.195658833 | 0.316819716 | 0.303308952 | 0.094011976 | 0.730506011 |
| 86098.64844 | 103508.1094 | 98387.84375 | 0.559385812 | 0.047406642 | 0.489362214 | 0.031794134 | 0.300893584 | 0.120303611 |
| 131724.5 | 108662.1719 | 153438.9844 | 0.559174819 | 0.312637589 | -0.021801384 | 0.960979488 | 0.82965232 | 0.091108554 |
| 1116362.125 | 1110497.25 | 1028380.375 | 0.558667282 | 0.018432262 | -0.275179506 | 0.122107272 | 0.32586439 | 0.034567592 |
| 112171.6094 | 124866.8047 | 100279.4844 | 0.557611331 | 0.011979502 | -0.711551597 | 0.037351065 | -0.74892587 | 0.023855453 |
| 22123.99414 | 23126.3418 | 23195.57813 | 0.557476607 | 0.084021006 | 0.713036328 | 0.038407677 | 0.984862939 | 0.025917161 |
| 137796.9688 | 135562.5625 | 140536.2344 | 0.557257871 | 0.033384245 | -0.087186758 | 0.149058018 | 0.136999963 | 0.052505925 |
| 116393.0625 | 113279.5781 | 104282.1172 | 0.557172706 | 0.023105653 | -0.009242369 | 0.931745588 | -0.295626168 | 0.063283918 |
| 52898.94531 | 53603.82031 | 47930.12891 | 0.557062749 | 0.030421994 | 0.098047276 | 0.371100531 | 0.083587324 | 0.472735727 |
| 336611.0938 | 378485.875 | 361203.9063 | 0.556950685 | 0.010484689 | -0.039301736 | 0.239014656 | -0.206703706 | 0.089895191 |
| 280984.2188 | 300765.7813 | 290117.5313 | 0.556794096 | 0.050254511 | -0.10335513 | 0.290934437 | 0.255807745 | 0.057438754 |
| 186981.1719 | 191123.5938 | 184028.0156 | 0.556455948 | 0.040854235 | 0.228353355 | 0.243784441 | 0.041516056 | 0.212201362 |
| 70098.13281 | 69998.27344 | 51714.4375 | 0.556356023 | 0.214135066 | 0.823866658 | 0.171339044 | -0.729037742 | 0.090359716 |
| 25845.10547 | 22187.74414 | 21300.98438 | 0.55555208 | 0.116879496 | 0.062628458 | 0.756806073 | -0.42661555 | 0.171025182 |
| 250126.375 | 260239.8281 | 265905.7813 | 0.555354424 | 0.00762892 | 0.03721491 | 0.744303485 | 0.467571665 | 0.027868741 |
| 41984.41797 | 36171.85938 | 34567.05469 | 0.554323969 | 0.035293563 | -0.149649269 | 0.133875076 | -0.040261855 | 0.771014523 |
| 30519.79883 | 31440.39844 | 28684.61523 | 0.554238543 | 0.096314216 | 0.603051536 | 0.032745004 | 0.665548814 | 0.017015337 |
| 16324.66406 | 17618.3418 | 19003.25977 | 0.554200843 | 0.09675935 | 0.55199612 | 0.110860227 | 0.553169151 | 0.05167048 |
| 61667.90234 | 56757.77734 | 61475.95313 | 0.553974254 | 0.148322176 | -0.637377921 | 0.156070866 | -0.143537221 | 0.557456388 |
| 35485.32031 | 35586.32031 | 33563.02734 | 0.553969754 | 0.010946719 | -0.689242958 | 0.028434447 | -0.357384224 | 0.033053706 |
| 58064.15234 | 57639.92578 | 50019.60547 | 0.553936432 | 0.04179011 | 0.236310005 | 0.021906439 | 0.050914163 | 0.639552995 |
| 5430.555176 | 6170.276367 | 1683.326294 | 0.553900147 | 0.048001737 | -0.524218664 | 0.185953071 | -1.216611712 | 0.106472402 |
| 89817.99219 | 70409.82813 | 70340.60938 | 0.553885608 | 0.013660857 | 0.072345411 | 0.428256136 | 0.043046646 | 0.820784824 |
| 13710.97559 | 15873.97754 | 12395.62012 | 0.553607521 | 0.077857993 | -0.485032604 | 0.047347558 | -0.586611686 | 0.040959479 |
| 4948.459961 | 7041.890137 | 5654.967285 | 0.553223145 | 0.069367958 | -0.105675938 | 0.565383328 | -0.469052071 | 0.115074465 |
| 5076.168457 | 2909.725586 | 3722.804443 | 0.552562699 | 0.532420875 | 0.423090958 | 0.533779394 | 0.242463174 | 0.745482456 |
| 574557.5625 | 560508.4375 | 575281 | 0.551729917 | 0.035879745 | 0.601204977 | 0.149088844 | 0.889123738 | 0.006394096 |
| 4443.664551 | 5111.840332 | 2312.481934 | 0.551626738 | 0.368557413 | -0.18098705 | 0.734206841 | -1.066528922 | 0.21341354 |
| 60647.85938 | 59042.16406 | 53551.22266 | 0.551454375 | 0.006735623 | 0.040193749 | 0.747429573 | 0.250942414 | 0.101910514 |

| | | | | | | | | |
|---|---|---|---|---|---|---|---|---|
| 26089.67969 | 22117.86914 | 20546.81641 | 0.551302158 | 0.096314216 | 0.793944544 | 0.034840225 | 0.205698845 | 0.370368769 |
| 5420.860352 | 4696.373535 | 4788.910156 | 0.550980407 | 0.120063669 | 0.694570388 | 0.097232773 | 0.322258227 | 0.10621357 |
| 43543.30469 | 52276.83203 | 67061.57813 | 0.550747977 | 0.094031642 | 0.745894878 | 0.04128058 | 1.006197708 | 0.113248184 |
| 280187.125 | 259927.4063 | 288763.3125 | 0.550147581 | 0.02194812 | 0.108343639 | 0.246470557 | 0.159964931 | 0.136438488 |
| 67482.33594 | 71632.27344 | 69921.92969 | 0.549676153 | 0.045302846 | -0.110524537 | 0.124487169 | -0.428373641 | 0.017279709 |
| 96228.67188 | 93382.0625 | 94594.53906 | 0.549648892 | 0.05158177 | -0.36687798 | 0.077511889 | -0.245634785 | 0.142342008 |
| 106324.8984 | 111010.2109 | 99665.14063 | 0.549486801 | 0.036733943 | -0.1640273 | 0.226305841 | 0.872563825 | 0.017015337 |
| 12688.08594 | 9487.992188 | 8300.632813 | 0.548936784 | 0.140553499 | -1.062936473 | 0.05919192 | -1.174957834 | 0.037542695 |
| 96359.67969 | 101262.9063 | 100962.125 | 0.548756489 | 0.064726008 | 0.03847027 | 0.782663525 | 0.532006399 | 0.051133451 |
| 18263.14453 | 18535.17773 | 17694.40625 | 0.548689947 | 0.086055451 | 0.391890459 | 0.240361908 | 0.809979134 | 0.039408515 |
| 19105.88281 | 16918.94336 | 17243.04883 | 0.548037809 | 0.062398014 | 1.063161365 | 0.027435011 | 0.971611829 | 0.01858328 |
| 558381.3125 | 547058 | 513441.2813 | 0.547896407 | 0.018310596 | -0.098759361 | 0.485554805 | 0.09717889 | 0.194351963 |
| 39961.05859 | 48489.80078 | 42265.92188 | 0.547255612 | NA | -0.001559395 | NA | 0.151127213 | NA |
| 7999972.031 | 9143969.375 | 8656348.844 | 0.547209293 | 0.080266611 | 0.187995063 | 0.470566249 | 1.278581415 | 0.014469072 |
| 245025.7188 | 226487.3438 | 236321.5625 | 0.547032577 | 0.313971512 | 1.129632161 | 0.094777667 | 0.421536397 | 0.309038899 |
| 40278.00391 | 41094.76172 | 41636.36328 | 0.546783395 | 0.084240527 | 0.344609052 | 0.154104796 | 1.034442476 | 0.03161115 |
| 24692.83008 | 27531.49023 | 23105.32813 | 0.546590771 | 0.024311948 | -0.620414318 | 0.031738972 | -0.561399647 | 0.033243122 |
| 57338.62109 | 57313.26563 | 53254.86328 | 0.546025287 | 0.01751483 | -0.219892306 | 0.150271527 | -0.260549759 | 0.060096206 |
| 170445.6406 | 171651.1094 | 187322.0313 | 0.546004962 | 0.029593095 | -0.772050469 | 0.011710842 | -0.954588338 | 0.007057904 |
| 100745.8281 | 117682.7656 | 109737.0391 | 0.545810444 | 0.160003781 | 0.066288409 | 0.796141333 | 0.15258422 | 0.5195709 |
| 8484.508789 | 3977.574707 | 7130.958984 | 0.545728748 | 0.042621515 | -1.512801778 | 0.021622074 | -1.240246229 | 0.049415261 |
| 355161.7188 | 268690.4375 | 330883.0313 | 0.545656842 | 0.029812678 | -0.307635797 | 0.094163342 | -0.046676255 | 0.803948135 |
| 15964.59375 | 11425.96875 | 12575.14453 | 0.545325416 | NA | 1.50024073 | NA | 1.560026785 | NA |
| 119406.4219 | 113878.9375 | 116282.8672 | 0.545166276 | 0.098021382 | 0.470924772 | 0.096073212 | 0.394008924 | 0.106029703 |
| 283690.625 | 291115.6563 | 286809.6563 | 0.545080151 | 0.009974872 | 0.648330746 | 0.011710842 | 0.359974705 | 0.020746708 |
| 601162.1875 | 925550.8125 | 378068.8438 | 0.544718756 | 0.405865623 | 0.110045435 | 0.833336814 | -0.530921534 | 0.431987073 |
| 19954.50586 | 22436.20703 | 21876.00977 | 0.544576919 | 0.032759649 | -0.046510127 | 0.698316027 | -0.76676576 | 0.021984636 |
| 92653.46094 | 81538.22656 | 60046.09375 | 0.544529786 | 0.314910702 | 2.404833649 | 0.012254503 | 1.959971522 | 0.063283918 |
| 8858.44043 | 10843.46875 | 8076.397461 | 0.544061153 | 0.05343731 | 0.310576168 | 0.160918214 | 0.476692573 | 0.163723435 |
| 1599579.125 | 1462931 | 1486543.25 | 0.543870694 | 0.227207392 | -0.101171452 | 0.76362454 | 0.067788591 | 0.827301412 |
| 97619.10156 | 105154.5547 | 97742.21875 | 0.542769287 | 0.032929334 | -0.183660367 | 0.167303467 | -0.051204781 | 0.582846665 |
| 7577.54248 | 6836.854004 | 5227.380859 | 0.54276371 | 0.080015674 | -0.379491426 | 0.023005377 | -0.805593704 | 0.065367829 |
| 25539.81055 | 27248.50195 | 28739.45703 | 0.542524948 | 0.213902628 | 0.292614429 | 0.454547212 | 0.188695125 | 0.51695586 |

| | | | | | | | | |
|---|---|---|---|---|---|---|---|---|
| 616.9764404 | 772.5064697 | 384.4484863 | 0.542323283 | 0.39188184 | -0.227610767 | 0.640999613 | 0.110490617 | 0.846348122 |
| 18205.77148 | 16820.85547 | 14203.69141 | 0.541591364 | 0.051672933 | -0.399507881 | 0.057189248 | -0.856547197 | 0.021134419 |
| 72821.35938 | 76257.5 | 67915.99219 | 0.541326169 | 0.029016212 | 0.044850617 | 0.639796921 | 0.320956517 | 0.058141533 |
| 657013.625 | 646654.6875 | 663711 | 0.541293647 | 0.021498678 | 0.114254384 | 0.091981596 | 0.258819912 | 0.009587646 |
| 202086.9063 | 231814.9531 | 263564.7813 | 0.541100195 | 0.036218901 | 0.402029533 | 0.06824263 | 0.565543201 | 0.098737916 |
| 227664.1563 | 471767.9688 | 187841.9688 | 0.540782858 | 0.429715592 | 0.213182436 | 0.662482496 | -0.238434888 | 0.746069294 |
| 18763.45898 | 21364.66992 | 22453.52539 | 0.539603854 | 0.533078483 | 0.970953515 | 0.194685429 | 0.736640994 | 0.276064537 |
| 71878.16406 | 69822.88281 | 68740.58594 | 0.538837928 | 0.092740746 | 0.063599286 | 0.71916924 | -0.132531947 | 0.4918172 |
| 144716.7969 | 206863.5 | 164859.8438 | 0.538451357 | 0.014665071 | -0.422806731 | 0.034038555 | -0.554779417 | 0.097278088 |
| 66716.0625 | 91509.53125 | 44808.03125 | 0.538306646 | 0.421238544 | 0.119119704 | 0.808694324 | -0.241057177 | 0.695364867 |
| 10345.99121 | 9407.514648 | 8890.466797 | 0.537382381 | 0.136416657 | 0.965774322 | 0.138196844 | 0.737542276 | 0.029219266 |
| 68199.24219 | 60601.39844 | 50903.42188 | 0.536377384 | 0.049895713 | -0.638262335 | 0.082329519 | -0.448155938 | 0.094103554 |
| 2705311.75 | 2945328.25 | 3078228.25 | 0.536375565 | 0.030847101 | 0.214364183 | 0.071935047 | 0.346752258 | 0.056570421 |
| 40191.10156 | 40284.33594 | 46172.34766 | 0.535932179 | 0.029239337 | -0.367106452 | 0.037014778 | 0.218173819 | 0.156297214 |
| 2620.674316 | 3000.33252 | 1753.969604 | 0.535186502 | 0.19253889 | -0.315971739 | 0.296640926 | -0.125504591 | 0.731461707 |
| 26059.13281 | 22462.07422 | 23230.40625 | 0.535000322 | 0.008878158 | 0.06954249 | 0.414751124 | -0.185575597 | 0.174355634 |
| 101740.4844 | 114650.4141 | 102597.5625 | 0.534216091 | 0.399616017 | -0.243644494 | 0.691779308 | -0.61877943 | 0.376019906 |
| 7580.874023 | 6625.345703 | 5751.67041 | 0.533891779 | 0.138821389 | -0.40569045 | 0.129311198 | -0.550779979 | 0.053012031 |
| 33020.68359 | 33282.36328 | 32335.13477 | 0.533791399 | 0.043580626 | 0.09611113 | 0.47094807 | 0.328449556 | 0.095599918 |
| 55903.89453 | 56382.05078 | 50203.48438 | 0.532762317 | 0.010603888 | -0.149391833 | 0.097722369 | -0.02665236 | 0.758792713 |
| 36733.12109 | 33831.9043 | 30479.2417 | 0.532258803 | 0.026183333 | 0.119789353 | 0.214966186 | 0.017138449 | 0.894865646 |
| 28167.01758 | 22051.50781 | 22271.54102 | 0.531782947 | 0.181523185 | 0.793160314 | 0.07830547 | 0.885046219 | 0.075839948 |
| 37710.73438 | 33651.42578 | 38501.34375 | 0.5313069 | 0.056620854 | -0.530741695 | 0.107105567 | -0.735460901 | 0.058410816 |
| 25854.65234 | 28630.82031 | 33200.74609 | 0.531030286 | 0.048520642 | 0.815050694 | 0.078996408 | 0.77749995 | 0.063937169 |
| 8203.777344 | 9060.745117 | 14357.53027 | 0.530619606 | 0.071921738 | 0.225499993 | 0.408157212 | -0.137853544 | 0.72266742 |
| 1329568.125 | 1702478.125 | 1797442.75 | 0.530588913 | 0.022022602 | 0.217743828 | 0.038454242 | 1.598357259 | 0.06365738 |
| 28223.48828 | 35158.51953 | 29214.88281 | 0.530377335 | 0.084757955 | -0.193370593 | 0.355035916 | -0.380270731 | 0.146875127 |
| 46405.69531 | 44551.66016 | 43815.34375 | 0.530277253 | 0.021320954 | -0.246511982 | 0.083732049 | -0.426856547 | 0.045887602 |
| 404576.7188 | 440648.9688 | 399887.4375 | 0.530169273 | 0.073001312 | 0.289653662 | 0.131196907 | 0.220304526 | 0.190240855 |
| 109806.6797 | 125671.0234 | 120620.2109 | 0.529802655 | 0.065549785 | 0.075962748 | 0.61581529 | -0.678312461 | 0.061031191 |
| 1667980.375 | 2876366.75 | 1103304 | 0.529079841 | 0.398875467 | 0.250360935 | 0.560671011 | -0.304363294 | 0.654381087 |
| 69395.01563 | 72353.5625 | 60532.80859 | 0.528694676 | 0.018805749 | -0.581855022 | 0.047212306 | -0.617468362 | 0.041258108 |
| 9753.231445 | 17552.33398 | 16581.83594 | 0.528411718 | 0.122673099 | 0.578498825 | 0.126364459 | 0.259377257 | 0.547254221 |

| | | | | | | | | |
|---|---|---|---|---|---|---|---|---|
| 16151.21484 | 16679.42578 | 15838.94336 | 0.52819431 | 0.216793672 | 0.150402137 | 0.431724911 | 0.979425893 | 0.009026966 |
| 77414.89063 | 75875.64844 | 84665.53125 | 0.525914726 | 0.046915807 | 0.087021121 | 0.497760151 | 0.513595218 | 0.053337315 |
| 79282.58594 | 70948.32813 | 74479.96094 | 0.525529627 | 0.058893147 | -0.076915725 | 0.590283746 | -0.117435755 | 0.425145226 |
| 13346.85742 | 14999.93457 | 14207.34082 | 0.525069909 | 0.078653911 | 0.794433612 | 0.018539883 | 0.775050098 | 0.019022762 |
| 62953.87891 | 55874.26953 | 64720.80078 | 0.523764006 | 0.093428026 | -0.329921282 | 0.254426984 | -0.514408429 | 0.10119601 |
| 984428.75 | 1159729.625 | 1197925 | 0.523698176 | 0.032646789 | -0.073604548 | 0.456300228 | 0.172179332 | 0.294592752 |
| 25538.42773 | 21694.93945 | 24991.33008 | 0.523570309 | 0.05267493 | -0.406261338 | 0.115742717 | -0.118416858 | 0.432857345 |
| 25887.8457 | 22617.93945 | 21718.79883 | 0.523488844 | 0.023943571 | -0.211407999 | 0.150941204 | -0.085944478 | 0.5195709 |
| 44509.15234 | 40784.21875 | 41619.87109 | 0.523115246 | 0.022724126 | 0.387469564 | 0.128657577 | 0.521907596 | 0.020337832 |
| 136040.5625 | 134540.2344 | 153001.125 | 0.523036771 | 0.006735623 | -0.286233342 | 0.053543594 | -0.291017241 | 0.071851903 |
| 119425.4844 | 130708.9844 | 130214.0156 | 0.522956633 | 0.020651034 | 0.092421305 | 0.256229464 | 0.136857655 | 0.167717489 |
| 110667.4141 | 112329.9609 | 95052.48438 | 0.522841483 | 0.036547632 | 0.645203531 | 0.023497318 | -0.222147159 | 0.138917487 |
| 72702.17969 | 77934.05469 | 77087.25781 | 0.522526606 | 0.113673327 | -0.130190298 | 0.565383328 | -0.514621706 | 0.128484296 |
| 49617.34766 | 49783.58984 | 47575.40234 | 0.521938475 | 0.034061991 | -0.178795268 | 0.199670392 | -0.358800904 | 0.088892469 |
| 16256.16406 | 15600.53711 | 20042.41211 | 0.521898223 | 0.166415915 | -0.680813105 | 0.142579096 | -1.110974144 | 0.077950438 |
| 107879.1797 | 90628.70313 | 105429.5625 | 0.521650564 | 0.04045616 | 0.117451657 | 0.232501965 | 0.049649883 | 0.706124008 |
| 6226.579102 | 23134.16406 | 21574.87109 | 0.521353458 | 0.222058972 | -0.358782628 | 0.467297307 | -0.433246938 | 0.499893318 |
| 357357.7188 | 355070.1875 | 327443.875 | 0.520618678 | 0.027671686 | -0.301744335 | 0.041058966 | -0.324533171 | 0.029820656 |
| 4038.074463 | 4031.977295 | 2873.367188 | 0.519672057 | 0.181247944 | 0.083065806 | 0.619271606 | -0.458743701 | 0.133481008 |
| 35681.21094 | 44337.60156 | 18979.44336 | 0.519183198 | 0.423578336 | 0.034479688 | 0.947783674 | -0.064721048 | 0.92777397 |
| 112465.3828 | 113714.7031 | 115420.8047 | 0.519122174 | 0.025025466 | -0.430773342 | 0.011710842 | -0.313272788 | 0.009587646 |
| 122391.3359 | 114826.6094 | 121592.9531 | 0.518794576 | 0.039856301 | 0.151787528 | 0.081173768 | 0.168487983 | 0.06065526 |
| 420382.4688 | 418179.1875 | 414989.2188 | 0.518427196 | 0.010603888 | -0.46006405 | 0.038037652 | -0.403681921 | 0.039424444 |
| 13925.78418 | 11802.85449 | 11791.26953 | 0.518248667 | 0.115609849 | -0.336099592 | 0.108795517 | -0.750044661 | 0.034567592 |
| 68432.39063 | 71528.16406 | 65658.36719 | 0.517781884 | 0.006395507 | 0.049982384 | 0.189281807 | -0.15547103 | 0.079143743 |
| 40254.32422 | 32847.875 | 24384.37305 | 0.517559221 | 0.037043051 | -0.655416776 | 0.033067345 | -0.370969032 | 0.262168702 |
| 99839.67969 | 101954.4922 | 97340.72656 | 0.517226532 | 0.033529204 | -0.455921591 | 0.056328031 | -0.063113661 | 0.536804361 |
| 31494.52539 | 70872.91406 | 22046.67578 | 0.516871624 | 0.753421136 | -0.157275927 | 0.924427311 | -0.550340602 | 0.763019905 |
| 5881.743164 | 6758.825195 | 2108.54126 | 0.516839252 | 0.193611705 | 0.29964025 | 0.307600544 | -0.389789006 | 0.523326184 |
| 114597.1797 | 113771.3047 | 114837.1172 | 0.516688991 | 0.025680323 | -0.257404257 | 0.051743333 | -0.154214784 | 0.120615779 |
| 105063.8047 | 109771.2344 | 112653.9297 | 0.516658039 | 0.041945948 | -0.294970918 | 0.083773082 | -0.148582727 | 0.186660744 |
| 60674.32813 | 55345.21484 | 57860.3125 | 0.515913551 | 0.10884414 | -0.276357388 | 0.280941693 | -0.727654753 | 0.10188566 |
| 18487.12305 | 19030.51367 | 17932.51563 | 0.515072738 | 0.055041746 | -0.012797947 | 0.902074369 | 0.265077452 | 0.08152559 |

| | | | | | | | | |
|---|---|---|---|---|---|---|---|---|
| 23881.24219 | 28298.22461 | 26042.33008 | 0.514709182 | 0.123181907 | 0.559334547 | 0.085104375 | 0.394454753 | 0.105824879 |
| 8395.992188 | 8547.501953 | 8994.521484 | 0.514246035 | 0.022960243 | -0.006076324 | 0.93427043 | -0.145993573 | 0.130422289 |
| 623.0413208 | 1614.376587 | 1644.545044 | 0.513630179 | NA | -0.688541883 | NA | -0.912236672 | NA |
| 109670.3984 | 92232.28906 | 93282.84375 | 0.513607369 | 0.56288987 | 1.257787299 | 0.09678603 | 0.360838358 | 0.565669968 |
| 35655.53125 | 24838.73828 | 26312.71094 | 0.513432227 | 0.097252031 | 0.43983085 | 0.036154671 | -0.135195819 | 0.588834899 |
| 43760998.69 | 27112320.15 | 47482486.64 | 0.513375012 | 0.491789048 | 0.315598698 | 0.587139619 | 0.356864142 | 0.54725722 |
| 81264.29688 | 87214.0625 | 89134.77344 | 0.513306799 | 0.356678031 | -0.752327075 | 0.289589686 | -0.215923188 | 0.649485334 |
| 172067.2344 | 151170.2031 | 148007.3438 | 0.512746834 | 0.061786866 | -0.219295192 | 0.23288998 | -0.035158659 | 0.806396591 |
| 549050.5 | 555593.8125 | 637781 | 0.512691857 | 0.068444011 | 0.059550022 | 0.834109875 | 0.478853456 | 0.055019141 |
| 95990.29688 | 78179.98438 | 66100.05469 | 0.512481947 | 0.036160457 | 0.424521639 | 0.017096626 | 0.382678093 | 0.260272253 |
| 431137.875 | 381757.875 | 364789.25 | 0.512403509 | 0.097443336 | -0.038054147 | 0.698637979 | -0.345302743 | 0.064726471 |
| 320376.1875 | 309107.2188 | 280777.625 | 0.51171169 | 0.10058931 | 0.422689642 | 0.119643706 | -0.288893067 | 0.239535779 |
| 6574.709473 | 5848.709473 | 4263.078613 | 0.511434307 | 0.382591812 | -0.523588916 | 0.406651721 | -0.99876733 | 0.225772142 |
| 13037563 | 15199258 | 13490776 | 0.511426158 | 0.029627098 | 0.280513652 | 0.087467078 | 0.40996949 | 0.091954906 |
| 49236.82422 | 46961.38672 | 31581.26953 | 0.5106843 | 0.01904327 | -0.852569542 | 0.0155094 | -0.685503057 | 0.09236595 |
| 105752.5938 | 111425.3125 | 77648.53125 | 0.510583744 | 0.046681455 | 0.237838149 | 0.14150374 | 0.343687192 | 0.275100616 |
| 46315.91797 | 46252.875 | 52969.73047 | 0.51045368 | 0.028829273 | -0.470673421 | 0.018539883 | -0.079778049 | 0.451006888 |
| 15082.68652 | 20074.10742 | 13459.31934 | 0.510372564 | 0.215660117 | -0.576154548 | 0.234710203 | -0.500576516 | 0.2429355 |
| 68549.29688 | 55149.74219 | 66410.28906 | 0.510170334 | 0.226802749 | -1.199252597 | 0.193488746 | -0.185863513 | 0.265248575 |
| 76780.40625 | 81508.91406 | 89242.67188 | 0.509633267 | 0.068444011 | -0.279382658 | 0.039856414 | 0.029281239 | 0.771252617 |
| 14870.19434 | 15535.23438 | 10190.09082 | 0.509427749 | 0.087043567 | -0.019289514 | 0.786174208 | 0.02951876 | 0.918471363 |
| 27640.67383 | 33396.98438 | 31629.55859 | 0.509087524 | 0.019078369 | 0.353214004 | 0.047802304 | 0.320696211 | 0.120554018 |
| 6298.312012 | 5721.209473 | NA | 0.509082725 | 0.321360807 | 2.205445886 | 0.401614835 | -0.061016467 | NA |
| 42543.71094 | 40712.55078 | 36290.95703 | 0.50903312 | 0.15400688 | -0.530506772 | 0.178660809 | -0.539136771 | 0.167298956 |
| 5614.098145 | 6107.666016 | 7154.131348 | 0.508604704 | 0.047299472 | -0.36462256 | 0.387322672 | -0.239968992 | 0.227629112 |
| 140069.6875 | 145359.8594 | 140444.1875 | 0.507275021 | 0.015638289 | 0.130058447 | 0.057893293 | 0.146483161 | 0.032577228 |
| NA | NA | NA | 0.507243663 | NA | -0.243124701 | NA | NA | NA |
| 46804.33984 | 41079.15625 | 40657.98047 | 0.507085946 | 0.010603888 | 0.165124535 | 0.168058041 | -0.107664762 | 0.340314063 |
| 55639.48047 | 55062.05469 | 48529.98047 | 0.506643502 | 0.040928921 | 0.016815544 | 0.57418887 | 0.095136564 | 0.382649966 |
| 48070.55859 | 44142.33984 | 37362.21875 | 0.506271344 | 0.044605998 | -0.508109197 | 0.096829709 | -0.410110905 | 0.088296374 |
| 29483.69922 | 32174.11523 | 30975.86719 | 0.50600025 | 0.029391485 | 0.285779232 | 0.05467585 | 0.569597114 | 0.019816721 |
| 6822.824707 | 8273.169922 | 5570.923828 | 0.505841363 | 0.174729987 | -0.118191497 | 0.681896541 | -0.262679794 | 0.436076836 |
| 17571.9707 | 15562.2832 | 13293.12598 | 0.505831116 | 0.021027835 | -0.041727489 | 0.782928238 | 0.002978491 | 0.98861914 |

| | | | | | | | | |
|---|---|---|---|---|---|---|---|---|
| 21642.10352 | 17919.14648 | 20027.08008 | 0.505697586 | 0.016315242 | -0.139721045 | 0.246891752 | -0.333679606 | 0.0741034 |
| 15294.21289 | 12946.69629 | 9244.467773 | 0.505679001 | 0.062398014 | -0.590550631 | 0.089737108 | -0.849828739 | 0.055868946 |
| 20689.82422 | 19433.41797 | 23145.68164 | 0.505612589 | 0.025187448 | -0.056681765 | 0.539179679 | 0.130731164 | 0.35136918 |
| 13883.2832 | 11839.30859 | 7527.523926 | 0.505396356 | 0.147715547 | -1.017800075 | 0.105099839 | -0.992418768 | 0.068375475 |
| 430494.9375 | 395251.3125 | 416420.7813 | 0.505322788 | 0.047284259 | -0.400478881 | 0.087630144 | -0.064350515 | 0.528908482 |
| 112419.2422 | 112181.8438 | 105500.4922 | 0.505284412 | 0.013723547 | -0.577456091 | 0.028434447 | -0.533475116 | 0.018257145 |
| 3892.377197 | 3809.536377 | 3053.467529 | 0.50459732 | 0.033688384 | -0.248425391 | 0.126158269 | -0.186503014 | 0.314984889 |
| 9056.248047 | 8528.624023 | 8043.874512 | 0.504340378 | 0.037138683 | -0.124813097 | 0.219605065 | -0.346675032 | 0.044854747 |
| 23192.91406 | 24170.46289 | 38978.40234 | 0.503480675 | 0.257974029 | 0.630155003 | 0.05818214 | 0.676923682 | 0.258591127 |
| 155601.2969 | 164447.0625 | 190432.1875 | 0.503453183 | 0.041269027 | -0.196375875 | 0.21691719 | 0.410580999 | 0.095599918 |
| 216875.9688 | 244804.0938 | 105306.5156 | 0.503172624 | 0.423302296 | 0.182495174 | 0.708708221 | -0.652675808 | 0.342246912 |
| 4706.611816 | 4511.061523 | 2978.497803 | 0.502833386 | 0.107769589 | -0.255001376 | 0.292995226 | -0.418926147 | 0.202896674 |
| 27078.78711 | 22729.24805 | 19577.67383 | 0.502677993 | 0.053893741 | -0.583625982 | 0.048370376 | -1.066000393 | 0.034567592 |
| 12753.36621 | 11069.99219 | 9960.317383 | 0.502426734 | 0.123102055 | 0.150311791 | 0.609951224 | 0.609116929 | 0.074322497 |
| 20342.18945 | 22566.30469 | 16349.18359 | 0.501407176 | 0.015391678 | -0.175518588 | 0.16564489 | -0.6007014 | 0.071151453 |
| 10981.66504 | 19195.4707 | 15600.43945 | 0.50127982 | 0.144037739 | -0.080805697 | 0.66660509 | -0.413525973 | 0.252366687 |
| 7613.912598 | 5025.59668 | 6215.053223 | 0.49993849 | 0.130960943 | 0.581607852 | 0.181761214 | 0.327260888 | 0.339927953 |
| 139070.9688 | 148801.0469 | 146868.7031 | 0.499495094 | 0.065880743 | -0.088615075 | 0.140163976 | -0.319260674 | 0.020008995 |
| 108356.1016 | 110348.9141 | 98734.79688 | 0.499110183 | 0.025025466 | -0.201541594 | 0.096073212 | -0.219857971 | 0.079168003 |
| 531807.9375 | 729229.375 | 426650.3125 | 0.499091856 | 0.713928646 | -0.463936077 | 0.744419479 | -0.34938637 | 0.780623364 |
| 72120.61719 | 69803.67969 | 55688.97656 | 0.498948945 | 0.35114767 | 0.800801856 | 0.123184236 | 0.767056874 | 0.111523548 |
| 372995.4375 | 391131.2813 | 381997.8125 | 0.498898913 | 0.111787093 | -0.099963595 | 0.658567978 | -0.096834232 | 0.619154814 |
| 112952.7109 | 112588.0547 | 126006.6406 | 0.497338704 | 0.016885305 | -0.096435147 | 0.089931608 | -0.330418407 | 0.061106674 |
| 73823.79688 | 71485.17188 | 71925.26563 | 0.497256532 | 0.05114005 | 0.41877984 | 0.028434447 | 0.55113529 | 0.018257145 |
| 39887.16797 | 41495.76172 | 41261.71484 | 0.496546768 | 0.064978842 | -0.365500202 | 0.142833812 | -0.614646476 | 0.085665996 |
| 7687.67041 | 7295.629395 | 8709.256836 | 0.496219522 | 0.337270865 | 0.127523279 | 0.841310571 | 0.609133084 | 0.113248184 |
| 44003.12891 | 44944.76563 | 43481.35547 | 0.495982867 | 0.16947575 | 0.329728191 | 0.210455828 | 0.513492413 | 0.116344276 |
| 48251.74219 | 44498.28906 | 42462.48828 | 0.495250994 | 0.070959976 | -0.493254546 | 0.09861697 | -0.014017156 | 0.905572751 |
| 4410.421387 | 10896.82031 | 9569.71875 | 0.494768485 | 0.485058336 | -0.156818734 | 0.893894927 | -0.198022269 | 0.791258063 |
| 19782.23633 | 15171.8252 | 16818.04688 | 0.494673981 | 0.036480793 | -0.011015118 | 0.948953473 | -0.1677974 | 0.371644128 |
| 207113.7969 | 198348.5781 | 181318.7188 | 0.494490368 | 0.058238988 | -0.266246216 | 0.184581496 | -0.235950161 | 0.19608824 |
| 399093.125 | 381185.9375 | 391294.5625 | 0.494400981 | 0.008762412 | -0.240643807 | 0.189538652 | -0.121808946 | 0.049415261 |
| 6218.497559 | 16312.21191 | 5124.211426 | 0.494398176 | 0.485046454 | 1.011829494 | 0.085565133 | -0.002410972 | 0.998066098 |

| | | | | | | | | |
|---|---|---|---|---|---|---|---|---|
| 24790.41797 | 26451.93945 | 24322.41992 | 0.493379505 | 0.03438191 | -0.308133375 | 0.096106105 | -0.301528312 | 0.084613954 |
| 11049.11621 | 16246.62402 | 11604.97656 | 0.492546506 | 0.058363046 | -0.148018327 | 0.555335026 | -0.227475464 | 0.423206002 |
| 539056.375 | 556360.75 | 542134.8125 | 0.492462003 | 0.065975508 | 0.200387926 | 0.134028069 | 0.144630856 | 0.217783381 |
| 46984.44141 | 41600.35547 | 41978.40234 | 0.492101495 | 0.153804457 | 0.111447021 | 0.617383969 | 0.184770238 | 0.395364978 |
| 143561.2344 | 139901.6563 | 79379.57031 | 0.491812491 | 0.219971212 | -0.569676839 | 0.224179033 | -0.648027591 | 0.185378125 |
| 62774.84375 | 63309.46875 | 63042.54297 | 0.491636714 | 0.085777338 | 0.1508948 | 0.378345732 | -0.012427583 | 0.940095284 |
| 364129.5625 | 373627.4375 | 394976.0938 | 0.491464538 | 0.056312464 | 0.291929367 | 0.075842188 | 0.379498507 | 0.027391709 |
| 117239.2109 | 113247.7734 | 96071.5 | 0.491012684 | 0.021539691 | 0.000871196 | 0.989874203 | 0.330033542 | 0.135536925 |
| 26704.84766 | 28253.35742 | 23969.32227 | 0.490382222 | 0.066846369 | -0.196496701 | 0.098408786 | -0.066440957 | 0.557774325 |
| 207402.6406 | 209809.5781 | 204737.6563 | 0.490348986 | 0.108716757 | -0.350021254 | 0.196499103 | -0.379045525 | 0.174887902 |
| 30276.33984 | 28579.01172 | 29265.65039 | 0.489920446 | 0.014544337 | -0.136394281 | 0.580924969 | -0.246510899 | 0.029820656 |
| 7750.953125 | 6994.77002 | 5533.541992 | 0.48975566 | 0.081289628 | -0.788413725 | 0.042887894 | -1.213192337 | 0.017015337 |
| 227428.8125 | 215406.7813 | 226843.7813 | 0.489263286 | 0.027230193 | 0.434406508 | 0.26604978 | 0.861899789 | 0.009587646 |
| 33466.03125 | 28321.88086 | 26032.38672 | 0.48845604 | 0.019610279 | -0.15743764 | 0.170357294 | -0.32510053 | 0.151772717 |
| 167853.6523 | 174639.2852 | 148877.1914 | 0.488391763 | 0.025419349 | -0.423676059 | 0.050410119 | -0.149193 | 0.249456378 |
| 56098.92188 | 56655.46094 | 55386.84766 | 0.487786204 | 0.124546367 | -0.084684633 | 0.709967054 | 0.995884837 | 0.042414121 |
| 12354.73145 | 11140.875 | 6653.484863 | 0.487765555 | 0.066744288 | -0.354245116 | 0.135938622 | -0.346667833 | 0.363820887 |
| 37229.71094 | 41512.88281 | 39832.87109 | 0.486675516 | 0.025437157 | -0.207900267 | 0.127763601 | -0.497808266 | 0.026764821 |
| 42912.07031 | 75755.61719 | 49013.35938 | 0.48653419 | 0.161230279 | -0.565737931 | 0.175868431 | -0.779339718 | 0.113039242 |
| 22583.10156 | 21530.61914 | 19945.74609 | 0.486335352 | 0.103546806 | 0.196389715 | 0.109653398 | 0.046149519 | 0.650869865 |
| 11238.10547 | 8469.470703 | 4590.987305 | 0.485687577 | 0.037600114 | -1.230064123 | 0.039752718 | -1.768146885 | 0.02527614 |
| 21361.1875 | 22032.25195 | 20367.16797 | 0.485416908 | 0.176268956 | -0.271384544 | 0.34708379 | -0.254807946 | 0.358664612 |
| 7541.827637 | 10285.55176 | 7290.873047 | 0.485280251 | 0.141838437 | 0.160351357 | 0.529211919 | 0.070636127 | 0.803948135 |
| 99231.79688 | 95425.27344 | 96398.9375 | 0.485039778 | 0.039413354 | 0.008135729 | 0.922008276 | 0.064873469 | 0.405343283 |
| 112743.5391 | 105868.2422 | 117017.7422 | 0.484966682 | 0.05315306 | 0.065577764 | 0.412125285 | -0.035992494 | 0.6407955 |
| 22053.60938 | 23139.67188 | 23427.56445 | 0.484699913 | 0.239968322 | 0.835026552 | 0.061247369 | 1.145703358 | 0.043437905 |
| 459730.9141 | 430782.9531 | 415904.5391 | 0.484686792 | 0.060329638 | 0.72639189 | 0.032675377 | 0.44820085 | 0.046487363 |
| 16958.09375 | 12109.05273 | 18014.89453 | 0.484027511 | 0.42956317 | 0.696106618 | 0.401614835 | 0.882270286 | 0.118341115 |
| 50041.3125 | 50498.56641 | 43909.86328 | 0.483549537 | 0.013723547 | 0.206192418 | 0.052447104 | -0.116807112 | 0.294592752 |
| 211191 | 217617.6406 | 213990.6563 | 0.482827999 | 0.028462306 | 0.110136552 | 0.534862174 | -0.154144794 | 0.127051426 |
| 27480.24219 | 25521.47266 | 25068.42773 | 0.481350835 | 0.103193326 | 0.526553355 | 0.073324814 | 0.107154244 | 0.541582817 |
| 951577.625 | 990190 | 933007.9375 | 0.481046439 | 0.020740891 | 0.179469979 | 0.065813387 | -0.051851172 | 0.401179264 |
| 39295.75391 | 39268.61328 | 35232.03906 | 0.480739354 | 0.057497846 | 0.49910477 | 0.04780612 | 0.304399538 | 0.093070634 |

| | | | | | | | | |
|---|---|---|---|---|---|---|---|---|
| 65032.90234 | 62351.98047 | 62107.37891 | 0.480489108 | 0.054546889 | -0.497658774 | 0.071840717 | -0.28757843 | 0.099486344 |
| 125960.6172 | 115847.4453 | 114013.8281 | 0.480205971 | 0.016885305 | -0.012847428 | 0.926516232 | 0.678984986 | 0.034722024 |
| 39029.98438 | 42238.65625 | 45258.54297 | 0.480013074 | 0.078148867 | 0.594516684 | 0.052447104 | 0.633553423 | 0.031173695 |
| 38217.54297 | 37021.85547 | 34051.88281 | 0.479143478 | 0.040558223 | 0.549642951 | 0.027445011 | 0.293931508 | 0.066134523 |
| 256360.0781 | 274234.3125 | 275586.625 | 0.478844946 | 0.04186746 | 0.079598783 | 0.433946383 | 0.152265382 | 0.091928987 |
| 866582.4375 | 927027.5 | 954819.125 | 0.47862876 | 0.016859654 | -0.221560008 | 0.053185315 | -0.325364058 | 0.027982615 |
| 258396.0156 | 272312.2188 | 281215.0625 | 0.478519252 | 0.057497846 | 0.338274871 | 0.095375186 | 0.445734009 | 0.035626762 |
| 4298898.5 | 3940121.5 | 4128587.5 | 0.478388794 | 0.022960243 | 0.31520137 | 0.055701361 | 0.290980346 | 0.038794276 |
| 7311879 | 8975396 | 8874464 | 0.478145763 | 0.030588379 | 0.093332576 | 0.06794739 | 0.234241364 | 0.239974773 |
| NA | 3204.865234 | NA | 0.477983422 | NA | -1.772673961 | NA | -0.766246179 | NA |
| 23798.86133 | 21975.78711 | 18888.97461 | 0.477722962 | 0.038597679 | -0.065066249 | 0.573817751 | -0.303757775 | 0.126366029 |
| 196022.3594 | 178431.9531 | 153532.2188 | 0.476432722 | 0.412203043 | 0.083927779 | 0.858056341 | 0.172803004 | 0.708835598 |
| 703041.3125 | 685256.3125 | 703479.6875 | 0.47639646 | 0.042621515 | 0.334897114 | 0.048107216 | 0.297241368 | 0.052503815 |
| 30251.52344 | 31118.87109 | 30731.88672 | 0.476316891 | 0.034518918 | -0.722090554 | 0.055241445 | -0.860239997 | 0.042262939 |
| 409485.1563 | 346037.7813 | 411470.0625 | 0.47626319 | 0.040558223 | 0.04768767 | 0.489969083 | 0.044044024 | 0.733336166 |
| 25131.63818 | 30886.97461 | 23797.60059 | 0.476211221 | 0.211438669 | 0.724674811 | 0.274382338 | 0.254444332 | 0.399077635 |
| 77266.28906 | 81840.73438 | 91759.14063 | 0.475271146 | 0.201802341 | 0.321664952 | 0.269442213 | 0.248100295 | 0.363820887 |
| 19584.04492 | 23948.34961 | 26570.02344 | 0.474807381 | 0.326887873 | 0.20793661 | 0.568926403 | 0.68207481 | 0.118023574 |
| 287210.8125 | 265325.0313 | 240079.2344 | 0.474369469 | 0.037913948 | 0.168772465 | 0.157674375 | -0.079084206 | 0.548141599 |
| 181597.2031 | 192484.3281 | 187153.6094 | 0.474307341 | 0.045092515 | 0.712056106 | 0.068255066 | 1.365840181 | 0.006427997 |
| 279531.25 | 286531.0625 | 268249.25 | 0.474025832 | 0.129985711 | 0.371062301 | 0.142833812 | 0.541710845 | 0.081899796 |
| 319285.1875 | 270272.875 | 306361.1563 | 0.473407035 | 0.045675045 | -0.086657961 | 0.507917236 | 0.169698738 | 0.261028666 |
| 58455.45703 | 46751.41797 | 84145.32813 | 0.473359442 | NA | 1.139045373 | NA | 2.228083091 | NA |
| 44072.96094 | 43032.03125 | 43527.68359 | 0.472192151 | 0.040928921 | -0.275186665 | 0.112184087 | -0.367077854 | 0.077710677 |
| 371642.7188 | 386828.625 | 407938.5938 | 0.47172505 | 0.023770785 | 0.20562725 | 0.253688447 | 0.796779542 | 0.016395134 |
| 10244.38574 | 13870.79492 | 11935.05273 | 0.471715331 | 0.358556216 | -0.031540385 | 0.940558645 | -0.051997728 | 0.909137956 |
| 49114.88672 | 47884.40625 | 51840.22266 | 0.471424827 | 0.152253903 | 0.190347143 | 0.1315555 | 0.48739035 | 0.019880539 |
| 112367.6641 | 112739.3203 | 111664.625 | 0.471167999 | 0.008878158 | 0.04998136 | 0.59453601 | -0.155957707 | 0.057068534 |
| 405754.0625 | 405051.4688 | 408949.2188 | 0.470652124 | 0.007152279 | -0.519432374 | 0.036730784 | 0.027841583 | 0.368053981 |
| 5308.120117 | 3359.191895 | 3890.155518 | 0.470596228 | 0.483182747 | 0.358938278 | 0.588909408 | -0.855511543 | 0.231918617 |
| 118800.1797 | 113198.1797 | 124882.5234 | 0.470259947 | 0.047406642 | 0.124103027 | 0.302578553 | 0.077216542 | 0.496281651 |
| 112893.2734 | 102484.0703 | 102984.3125 | 0.470249601 | 0.20270675 | 0.176600549 | 0.573423422 | 0.300181782 | 0.274800325 |
| 2670.158691 | 1607.815186 | 2146.773682 | 0.469896118 | 0.332613034 | -0.312855379 | 0.517875146 | -0.462407291 | 0.210758021 |

| | | | | | | | | |
|---|---|---|---|---|---|---|---|---|
| 15353.375 | 15123.12793 | 13122.25586 | 0.469860795 | 0.138753516 | 0.207592208 | 0.245788263 | 0.009018798 | 0.948081141 |
| 31629.49219 | 36903.33203 | 35541.76172 | 0.469646054 | 0.056174806 | -0.107744255 | 0.437955083 | 0.281329195 | 0.118373568 |
| 417592.0313 | 453151.7188 | 443155.1563 | 0.469527454 | 0.029603305 | -0.647408598 | 0.028434447 | -0.558789231 | 0.03581165 |
| 76133.80469 | 69426.91406 | 76581.76563 | 0.468986673 | 0.019078369 | 0.195921375 | 0.08416302 | -0.011906069 | 0.88560307 |
| 3442.775879 | 3832.937256 | 3360.768555 | 0.468788202 | 0.049995393 | -0.480218888 | 0.082763602 | -0.452690306 | 0.059770277 |
| 43438.66797 | 38191.65625 | 38187.72266 | 0.468457202 | 0.024243002 | 0.094849233 | 0.28831421 | -0.090537703 | 0.426794374 |
| 6158.952637 | 17959.16992 | 4688.475586 | 0.468157288 | 0.431148265 | 0.508862366 | 0.393711547 | -0.521505987 | 0.55314944 |
| 6270089 | 6958614.5 | 6699089.5 | 0.467019521 | 0.103338591 | -0.021186878 | 0.865351603 | 0.040740854 | 0.734524444 |
| 96360.20313 | 109416.1563 | 109323.5781 | 0.466652478 | 0.090202714 | -0.511341166 | 0.1268046 | -0.365378182 | 0.148445737 |
| 34978.64844 | 37500.82422 | 45862.98047 | 0.465666218 | 0.257894491 | 0.036845764 | 0.773923011 | 0.115412681 | 0.578503308 |
| 15101.92871 | 16397.86133 | 12813.23535 | 0.465631919 | 0.184517222 | 0.289139634 | 0.26604978 | 0.309197237 | 0.25417709 |
| 166985.7383 | 157314.4902 | 167107.5293 | 0.465532566 | 0.031284382 | 0.071593184 | 0.444741164 | 0.029438904 | 0.749715423 |
| 9779.582031 | 10101.59473 | 7842.015625 | 0.465429639 | 0.230116038 | -0.396900127 | 0.305910057 | -0.298052474 | 0.380080047 |
| 471071.25 | 515158.375 | 536588.5 | 0.465283321 | 0.073566057 | -0.601642288 | 0.068812228 | -0.453230606 | 0.062863768 |
| 5021.018066 | 7093.005859 | 6570.79834 | 0.464686819 | 0.289991018 | 0.396214335 | 0.122669631 | 0.522856803 | 0.149640357 |
| 9489828 | 9109225 | 10892958 | 0.464646559 | 0.078182482 | -0.481654057 | 0.084356576 | -0.174440082 | 0.312849579 |
| 56448.19531 | 54996.07422 | 52997.64453 | 0.464386645 | 0.184517222 | -0.30178137 | 0.329687018 | 0.16907567 | 0.473225696 |
| 143107.1406 | 118499.0703 | 132600.75 | 0.46352972 | 0.041912796 | 0.251004995 | 0.094211938 | 0.541075285 | 0.059641593 |
| 31954.37891 | 29391.26367 | 22830.44141 | 0.462674086 | 0.329662688 | 0.380868325 | 0.329328421 | 0.269670963 | 0.491112474 |
| 35033.29297 | 39782.69531 | 44262.94141 | 0.462357535 | 0.033348676 | 0.111002055 | 0.332606573 | 0.297947957 | 0.186529757 |
| 62799.00781 | 53468.17578 | 56827.80859 | 0.461953956 | 0.028302957 | -0.014184465 | 0.875616587 | 0.019773332 | 0.871225252 |
| 6306.99707 | 7172.974609 | 8040.206055 | 0.461834519 | 0.075287012 | 0.524371572 | 0.042044993 | -0.015403234 | 0.934898017 |
| 115872.3047 | 118237.4375 | 94409.69531 | 0.461719359 | 0.021027835 | 0.186496201 | 0.070283935 | 0.112304101 | 0.505676351 |
| 100397.0547 | 160153.9688 | 166278.75 | 0.461365845 | 0.153537772 | -0.104306089 | 0.610969415 | 0.610656318 | 0.229212907 |
| 174094.8125 | 178963.4844 | 164228.9063 | 0.461231894 | 0.053873527 | -0.458458066 | 0.053185315 | -0.664322436 | 0.021984636 |
| 44381.31641 | 44480.5625 | 45251.95703 | 0.460865335 | 0.021027835 | 0.078822184 | 0.291253414 | -0.116652439 | 0.194140122 |
| 9415.857422 | 11132.31836 | 9789.432617 | 0.460791812 | 0.054407811 | -0.454714494 | 0.096721331 | -0.783674854 | 0.034761591 |
| 29357.37891 | 27956.84375 | 29695.96484 | 0.460781577 | 0.054479995 | -0.574846563 | 0.055241445 | -0.292306359 | 0.116465028 |
| 19905.26172 | 19372.38672 | 19605.38672 | 0.460776845 | 0.085638558 | 0.487484226 | 0.053827713 | 0.066805473 | 0.394165962 |
| 76720.07031 | 77288.67969 | 73866.45313 | 0.460520384 | 0.011814081 | 0.453718255 | 0.090139838 | 0.206772929 | 0.028738995 |
| 61528.82813 | 71223.60938 | 64096.23828 | 0.45921453 | 0.088240896 | 0.078932753 | 0.629336543 | -0.483065983 | 0.095599918 |
| 699421.3125 | 662545.4375 | 680915.3125 | 0.458932783 | 0.037600114 | -0.07053497 | 0.419453192 | 0.082247892 | 0.334926612 |
| 164890.6563 | 167663.625 | 157035.6563 | 0.458587626 | 0.046945973 | 0.178583419 | 0.185649792 | 0.64964706 | 0.012394221 |

| | | | | | | | | |
|---|---|---|---|---|---|---|---|---|
| 105948.2422 | 105579.2969 | 89281.0625 | 0.458220484 | 0.047058169 | -0.132598076 | 0.323674771 | -0.416205859 | 0.065710106 |
| 124029.4609 | 111527.1563 | 87190.46875 | 0.457875393 | 0.320614099 | 0.30507348 | 0.567725619 | -0.122125089 | 0.7618552 |
| 150753.2188 | 156718.8125 | 153010.25 | 0.457423862 | 0.027671686 | -0.166940758 | 0.225945972 | 0.494039562 | 0.009587646 |
| 761070.25 | 921129.125 | 878490.375 | 0.456907377 | 0.224955599 | 0.136178173 | 0.63348405 | 0.622575982 | 0.081899796 |
| 564584.25 | 625489.125 | 655542.875 | 0.456863687 | 0.018311389 | 0.180053842 | 0.07908104 | 0.306746388 | 0.088642968 |
| 27479.78516 | 27191.0293 | 27111.04883 | 0.456605922 | 0.027158619 | -0.11922332 | 0.260384376 | -0.347088448 | 0.078910724 |
| 1206264.875 | 1269251.375 | 1351443.125 | 0.456602076 | 0.179549142 | 0.081172096 | 0.765031984 | 0.195744134 | 0.414098843 |
| 74583.39844 | 79674.03125 | 42125.05859 | 0.456399705 | 0.076933208 | -0.964360944 | 0.057530184 | -0.923761542 | 0.067488244 |
| 129025.8203 | 120743.6875 | 120927.9922 | 0.456292878 | 0.017524813 | -0.084329623 | 0.26604978 | 0.1323315 | 0.124509143 |
| 150822.6563 | 147211.25 | 156880.1719 | 0.455654905 | 0.061707718 | 0.068665513 | 0.436953642 | 0.714872337 | 0.010654184 |
| 684703.8125 | 711408.75 | 678549.5625 | 0.455483106 | 0.250607381 | 0.215155034 | 0.456419987 | 0.324219136 | 0.289430057 |
| 206132.5625 | 207277.4063 | 205529.8281 | 0.45503124 | 0.038651604 | -0.173987447 | 0.222006972 | 0.311711008 | 0.0890834 |
| 14455.92969 | 12905.87695 | 10699.45605 | 0.454981634 | 0.060858186 | -0.07700613 | 0.693593538 | -0.081957346 | 0.69525709 |
| 765300.5938 | 710787.3633 | 786857.8633 | 0.454449888 | 0.062785336 | -0.105942353 | 0.541557838 | 0.173238875 | 0.237224538 |
| 54976.40625 | 55238.17969 | 52896.82031 | 0.45420285 | 0.015840964 | 0.412509262 | 0.019456985 | 0.433967968 | 0.012701134 |
| 202580.8594 | 204415.6406 | 201502.7344 | 0.454053562 | 0.02237899 | -0.04466388 | 0.46485074 | 0.141647785 | 0.116993358 |
| 14409.14355 | 13103.7666 | 7551.053223 | 0.453367955 | 0.063399586 | -0.439812253 | 0.079837944 | -0.552088901 | 0.194516083 |
| 1969.851563 | 2961.826172 | 2517.702393 | 0.453206605 | NA | -0.078144532 | NA | -0.646239818 | NA |
| 129995.0703 | 123382.5234 | 112724 | 0.452542735 | 0.062398014 | 0.409683485 | 0.063199811 | 0.326442599 | 0.087771237 |
| 281075.5 | 311191.5938 | 290637.5625 | 0.451726904 | 0.029603305 | -0.390662621 | 0.048593497 | -0.563032803 | 0.016395134 |
| 23625.71094 | 26087.37891 | 23705.75781 | 0.451490573 | 0.177388967 | -0.018447321 | 0.929570723 | -0.356147018 | 0.210335561 |
| 191458.8281 | 204720.6094 | 218566.1563 | 0.451419795 | 0.028196123 | 0.70329929 | 0.044099436 | 1.010140765 | 0.030482431 |
| 207863.875 | 196858.2031 | 211398.9375 | 0.450971571 | 0.042240011 | 0.431258432 | 0.0155094 | 0.157890314 | 0.070298132 |
| 3143.061523 | 2721.150635 | 1262.312744 | 0.450620059 | 0.171291623 | 0.517335897 | 0.098225909 | -0.724242874 | 0.17916788 |
| 153183.7813 | 110983.3359 | 164049 | 0.450251546 | 0.013596234 | 0.00864576 | 0.80108348 | 0.089430515 | 0.730529882 |
| 885708.0625 | 1010653 | 1052798.375 | 0.448683478 | 0.011814081 | -0.138520956 | 0.092018139 | 0.234588144 | 0.168508025 |
| 27109.57031 | 24899.12891 | 32268.35352 | 0.447929779 | 0.032969545 | 0.144950308 | 0.19372563 | 0.529546488 | 0.112639617 |
| 46731.5 | 44865.80469 | 45914.19141 | 0.447865512 | 0.016105265 | -0.632820608 | 0.031437224 | -0.691165584 | 0.021553947 |
| 154994.8438 | 161744.9531 | 161834.5938 | 0.447681418 | 0.187040833 | 0.244975634 | 0.321888877 | -0.147818557 | 0.544452231 |
| 10061.45801 | 6920.019531 | 9839.517578 | 0.447616659 | 0.149894467 | 0.358615068 | 0.192808829 | -0.86904748 | 0.05519156 |
| 20789.62305 | 23462.50977 | 22986.68359 | 0.447039407 | 0.214500791 | 0.412308071 | 0.181752348 | 0.620936866 | 0.083808454 |
| 46667.82422 | 49182.25781 | 25902.22852 | 0.446990345 | 0.045880156 | -0.543546265 | 0.076429813 | -0.778432587 | 0.108322697 |
| 31304.69336 | 18965.92578 | 23443.43945 | 0.446874074 | 0.505332989 | 0.951471976 | 0.06863053 | 0.917159108 | 0.145702448 |

| | | | | | | | | |
|---|---|---|---|---|---|---|---|---|
| 288165.5 | 281586.2813 | 249616.0469 | 0.446715976 | 0.054037379 | 0.129442189 | 0.374048576 | 0.089456991 | 0.504964568 |
| 210545.4844 | 213666.5938 | 220086.9375 | 0.446473642 | 0.018937861 | -1.208294393 | 0.007509388 | -1.607750848 | 0.005647029 |
| 23274.72656 | 16024.64746 | 15455.7168 | 0.446253754 | NA | 1.949843084 | NA | 2.465056433 | NA |
| NA | 50.75479889 | NA | 0.445816754 | NA | -0.559492563 | NA | -2.203399854 | NA |
| 12661.89844 | 16791.25977 | 13522.58691 | 0.445727017 | 0.046112206 | -0.203686968 | 0.210052496 | -0.290083763 | 0.199188148 |
| 86304.875 | 81263.89844 | 91279.17188 | 0.445346506 | 0.015034317 | 0.080731985 | 0.530165229 | 0.095399079 | 0.299167804 |
| 50039.91797 | 38001.95313 | 47494.78516 | 0.445169923 | 0.040287826 | 0.121368287 | 0.292396668 | -0.077945258 | 0.674949057 |
| 78407.125 | 83013.24219 | 77496.33594 | 0.445005421 | 0.036480793 | -0.115034605 | 0.319523239 | -0.052560439 | 0.594337835 |
| 56386.83594 | 61833.41406 | 61757.1875 | 0.444688926 | 0.39188184 | 0.527028631 | 0.245062116 | 1.055569893 | 0.073130408 |
| 76714.01563 | 77706.53906 | 90386.97656 | 0.444624134 | 0.081385334 | -0.953230833 | 0.021906439 | -0.217074679 | 0.15166044 |
| 22102.84766 | 3789.855469 | 3645.382568 | 0.444551604 | 0.487751762 | 0.251907405 | 0.786782444 | -0.343059946 | 0.787711053 |
| 182536.9531 | 191814.3906 | 182036.5 | 0.444165359 | 0.043033157 | 0.147024706 | 0.054285958 | 0.049252787 | 0.297195194 |
| 39829.36328 | 44470.14063 | 39049.49219 | 0.443841359 | 0.072625307 | 0.334833678 | 0.142579096 | -0.536105728 | 0.067380632 |
| 313373.9688 | 281643.3438 | 316244.2813 | 0.443517241 | 0.011471164 | -0.604805107 | 0.037014778 | -0.863297281 | 0.009587646 |
| 2253235.852 | 2991194.776 | 2583040.255 | 0.443451494 | 0.063436145 | 0.187402796 | 0.235421907 | 0.643978623 | 0.092909047 |
| 31257.71094 | 37716.17578 | 42703.55859 | 0.443437208 | 0.542216889 | -0.065744153 | 0.922226898 | 0.057383256 | 0.931717307 |
| 16253714.2 | 16507398.94 | 17097566.22 | 0.443144064 | 0.027230193 | 0.029598991 | 0.675625552 | -0.034910895 | 0.610401648 |
| 290661.3125 | 299279.1563 | 320917.9063 | 0.44259382 | 0.011979502 | -0.074181216 | 0.54074752 | 0.24348809 | 0.067849371 |
| 15616.62988 | 17215.09375 | 13557.79688 | 0.442230479 | 0.161540693 | -1.92214896 | 0.026024574 | -1.000814828 | 0.023619478 |
| 157211.75 | 183391.6875 | 171358.7031 | 0.442125211 | 0.025437157 | -0.048701844 | 0.378617903 | 0.113455342 | 0.334263521 |
| 98672.48438 | 89598.78125 | 85195.875 | 0.441945418 | 0.015216041 | -0.602558757 | 0.011710842 | -0.573520782 | 0.032510751 |
| 481145.125 | 513240.8438 | 465395.75 | 0.441906494 | 0.13516676 | 0.092192459 | 0.595824479 | 0.182123504 | 0.326913155 |
| 4689.526367 | 4501.874023 | 3803.665283 | 0.441716107 | 0.346157494 | 0.31758327 | 0.426191228 | -0.358129858 | 0.432150607 |
| 11085.88574 | 8965.688477 | 10766.24219 | 0.441667341 | 0.122108564 | -0.640949596 | 0.124127483 | -0.471576893 | 0.121413405 |
| 80446.5 | 85765.875 | 84166.14063 | 0.440730944 | 0.020940469 | -0.112377363 | 0.198281858 | 0.154703281 | 0.089665148 |
| 7132.000977 | 16584.95313 | 28146.5 | 0.440627192 | 0.611465565 | 1.112514166 | 0.142476036 | 0.736782091 | 0.504851871 |
| 149001.1094 | 136336.6719 | 158176.1406 | 0.440276261 | 0.083280787 | 0.030505536 | 0.532402501 | -0.031867792 | 0.737230437 |
| 13503348 | 14101079 | 15278966 | 0.440124025 | 0.11338479 | -0.289656401 | 0.148589158 | -0.339012487 | 0.0890834 |
| 83283.29688 | 89693.6875 | 76676.11719 | 0.439524685 | 0.081995281 | 0.605296532 | 0.048138491 | 0.452904717 | 0.076783998 |
| 29135.63477 | 28296.15625 | 27558.3418 | 0.439017417 | 0.025307586 | -0.701250102 | 0.019428587 | -0.181181876 | 0.092205529 |
| 405083 | 374359.1875 | 397419.9375 | 0.438666218 | 0.004709568 | -0.368480999 | 0.078617016 | -0.090080096 | 0.187991349 |
| 238913.8125 | 259761.5469 | 251668.4375 | 0.438657623 | 0.139325834 | 0.060056516 | 0.741007407 | 0.406977465 | 0.110105852 |
| 48436.79883 | 48194.56445 | 45401.38477 | 0.43817363 | 0.119451239 | 0.189782506 | 0.284636711 | -0.278662326 | 0.210821888 |

| | | | | | | | | |
|---|---|---|---|---|---|---|---|---|
| 50295.32031 | 50032.76563 | 65907.4375 | 0.436500117 | 0.050592788 | -0.037295012 | 0.801584301 | -0.259889846 | 0.261699898 |
| 17929.34961 | 18562.21289 | 17821.71289 | 0.436367345 | 0.087597224 | 0.322405569 | 0.075228015 | -0.254444109 | 0.022638191 |
| 366868.9375 | 359824.5938 | 316319.1563 | 0.436175159 | 0.021539691 | -0.073882017 | 0.634892576 | -0.091451419 | 0.39677885 |
| 5683.449219 | 5081.98584 | 4300.020508 | 0.436123359 | 0.078758366 | -0.177125115 | 0.279104324 | -0.605612228 | 0.050514656 |
| 546474 | 534554.0625 | 489135.9688 | 0.436037139 | 0.019930448 | 0.560961582 | 0.015625705 | 0.820198387 | 0.023619478 |
| 11735.90918 | 9286.551758 | 10876.01074 | 0.434670131 | 0.118201067 | 0.097803226 | 0.571319729 | -0.395429529 | 0.076647788 |
| 11094.23828 | 10310.47266 | 8135.086914 | 0.433957348 | 0.204291739 | 0.325469721 | 0.23374866 | 0.212469765 | 0.454490742 |
| 103073.5313 | 93754.42969 | 93267.40625 | 0.433743059 | 0.050577071 | -0.988727614 | 0.024018676 | 0.077894013 | 0.477266228 |
| 33500.18359 | 26975.91406 | 20332.88086 | 0.433723175 | 0.012414001 | -0.034179723 | 0.6777223 | -0.609208541 | 0.136114629 |
| 902455.875 | 926144.125 | 926418.875 | 0.433059227 | 0.038901139 | 0.079156685 | 0.524482286 | 0.326225463 | 0.071891702 |
| 210289.4531 | 214437.0781 | 219249.125 | 0.432938459 | 0.025187448 | -0.216370095 | 0.038286427 | -0.197166748 | 0.02592872 |
| 11114.67969 | 9848.151367 | 8397.900391 | 0.432656558 | 0.068921342 | 0.299607395 | 0.065813387 | -0.471045618 | 0.071876894 |
| 179184.7969 | 185588.3906 | 187968.9375 | 0.43231748 | 0.024810615 | 0.362843074 | 0.033121608 | 0.540748903 | 0.016458149 |
| 29430.48047 | 26417.91797 | 23080.41211 | 0.432304641 | 0.10217612 | -0.293179104 | 0.215436149 | -0.392838545 | 0.125742768 |
| 8791.924805 | 11412.18164 | 4159.635254 | 0.432029263 | 0.515727508 | 2.455970353 | 0.013427556 | 0.773140901 | 0.36320612 |
| 2301.311523 | 2804.574707 | 3730.909912 | 0.431468269 | 0.190478218 | 0.500524149 | 0.089993802 | 1.136879118 | 0.121525007 |
| 29052.46484 | 29705.58008 | 27475.26172 | 0.431277103 | 0.054063082 | 0.125865685 | 0.575220987 | -0.108006934 | 0.26303995 |
| 6114215 | 6055723 | 6744066 | 0.431196574 | 0.025955593 | 0.076113845 | 0.225445341 | 0.179078058 | 0.14578286 |
| 3548.269775 | 3972.555176 | 1278.037964 | 0.431144815 | 0.338818972 | -1.258333824 | 0.120977365 | -1.261361474 | 0.108298926 |
| 23877.91797 | 25295.15234 | 23397.38477 | 0.430942024 | 0.111903085 | -0.244368225 | 0.118156126 | -0.204390411 | 0.09236595 |
| 554769.5 | 589402.375 | 547256.75 | 0.430268596 | 0.027158619 | 0.045489942 | 0.521166402 | -0.022552292 | 0.773298647 |
| 554072.8125 | 581112.6875 | 613611.5 | 0.430141672 | 0.041446273 | 0.129811925 | 0.297944313 | 0.404908685 | 0.030831306 |
| 2785.271484 | 1205.144775 | 668.3234863 | 0.429800091 | 0.582738039 | 0.356499685 | 0.629497737 | -0.127417605 | 0.906660092 |
| 43069.48828 | 37052.37109 | 31740.72461 | 0.429721051 | 0.047549099 | 0.063128486 | 0.630482257 | -0.454195403 | 0.088892469 |
| 9896704 | 7955321 | 8842910 | 0.429362334 | 0.48584679 | 0.501045231 | 0.313122839 | 0.476416993 | 0.308503071 |
| 29945.61133 | 31750.60156 | 28432.13477 | 0.42919005 | 0.2756587 | -0.24497816 | 0.491319916 | -0.066682158 | 0.831540545 |
| 1335022.875 | 1148371.875 | 1072518.75 | 0.429139381 | 0.277102683 | 0.56107181 | 0.134788265 | 1.256163088 | 0.028449017 |
| 931164.1875 | 1310967.75 | 668093.25 | 0.428989056 | 0.212061418 | 0.064550717 | 0.737109807 | -0.444913827 | 0.304108984 |
| 322790.375 | 586736.3125 | 196690.0938 | 0.42830758 | 0.543230636 | 0.223630032 | 0.660062477 | -0.476098823 | 0.528571712 |
| 26113.59766 | 26461.03516 | 24730.33203 | 0.428243117 | 0.071921738 | 0.321990785 | 0.271315379 | 0.285431981 | 0.032836911 |
| 405942.3438 | 439012.7188 | 424036.9375 | 0.428197597 | 0.037152165 | 0.390160227 | 0.042887894 | 0.22628718 | 0.065059183 |
| 126475.6484 | 136316.7031 | 132767.1094 | 0.42745226 | 0.012774217 | -0.028843258 | 0.471790095 | -0.212020989 | 0.037907761 |
| 18095.78516 | 21243.19727 | 21321.32227 | 0.427238503 | 0.031284382 | -0.067835182 | 0.475179451 | 0.625794952 | 0.052349495 |

| | | | | | | | | |
|---|---|---|---|---|---|---|---|---|
| 7193.901367 | 7169.606934 | 8469.038086 | 0.426931937 | 0.282458905 | 0.295181078 | 0.422678148 | 0.291744812 | 0.349607929 |
| 112394.6328 | 118645.7813 | 117089.5469 | 0.426837553 | 0.019286562 | 0.09096881 | 0.211541758 | -0.005853023 | 0.90315795 |
| 96523.58594 | 117430.1016 | 88499.09375 | 0.426800452 | 0.381106207 | 0.411501461 | 0.275888795 | 0.550414589 | 0.169075369 |
| 77317.40625 | 71105.15625 | 67817.0625 | 0.426701677 | 0.006351124 | -0.226284856 | 0.110625004 | 0.115346694 | 0.280633075 |
| 10173.06738 | 14749.82715 | 13923.1416 | 0.426488477 | 0.292515286 | -0.752256623 | 0.189272555 | -0.671806344 | 0.190517393 |
| 159611.5313 | 164196.0469 | 158058.9688 | 0.426189636 | 0.059865575 | -0.185357999 | 0.242699931 | -0.12169475 | 0.371124353 |
| 23333.04883 | 28462.27539 | 28015.89258 | 0.425682296 | 0.056696683 | 0.198635219 | 0.178579756 | 0.104847908 | 0.484439802 |
| 39315.60156 | 37393.26563 | 30158.1875 | 0.424379823 | 0.057184572 | 0.45372839 | 0.192808829 | 0.131691604 | 0.505012783 |
| 41463.49219 | 39152.10156 | 44551.63281 | 0.424349377 | 0.076604576 | -0.676344067 | 0.054857325 | -0.423102259 | 0.068366118 |
| 15914.16602 | 8815.548828 | 15684.2959 | 0.423692124 | 0.368167651 | -0.090895387 | 0.898502604 | 0.165020073 | 0.713483095 |
| 541744.5 | 528704.6875 | 489978.5938 | 0.423518278 | 0.018050472 | -0.678404309 | 0.019428587 | -0.451318812 | 0.02183874 |
| 27602.6543 | 30113.37305 | 29089.42969 | 0.423452677 | 0.226456157 | 0.674601515 | 0.10108711 | 0.516400233 | 0.026553937 |
| 30432.37598 | 28808.91699 | 29642.03418 | 0.422702556 | 0.095278437 | 0.32243594 | 0.027730297 | 0.571753048 | 0.009587646 |
| 38991.07813 | 28122.6582 | 36741.94922 | 0.422440126 | 0.060329638 | -0.375238549 | 0.030387624 | -0.467996547 | 0.112503484 |
| 10620.18848 | 8726.124023 | 7625.081055 | 0.421877665 | 0.471239653 | 0.196000578 | 0.663045481 | 0.358242293 | 0.433601327 |
| 195214.1406 | 231029.625 | 253467.6719 | 0.421695555 | 0.035276296 | 0.218363927 | 0.032817111 | 0.479814668 | 0.132636805 |
| 307294.4375 | 304406.3125 | 324232.5 | 0.421193733 | 0.084757955 | 0.135406562 | 0.351133652 | 0.426082417 | 0.060937162 |
| 114968.1406 | 114522.4531 | 120853.8906 | 0.4204674 | 0.073255783 | 0.730388029 | 0.041554876 | 1.428850078 | 0.007609795 |
| 8866.93457 | 13050.19238 | 7163.076172 | 0.418853912 | 0.41215279 | 0.349090152 | 0.401614835 | -0.429261459 | 0.442993963 |
| 1100933.25 | 1103549.5 | 1152225.625 | 0.418851649 | 0.039781174 | -1.36411719 | 0.007509388 | -0.44267664 | 0.011811065 |
| 52573.83984 | 52286.49219 | 49568.92578 | 0.418573497 | 0.02227739 | -0.368793976 | 0.042884533 | -0.490415803 | 0.027949491 |
| 60451.12109 | 64336.02344 | 65746.89063 | 0.418118038 | 0.045477314 | 0.645077021 | 0.019592862 | 0.476011263 | 0.02163333 |
| 7845.027832 | 22688.45313 | 28854.45117 | 0.417517662 | 0.405753158 | 0.708185986 | 0.147248245 | -0.202863098 | 0.778343026 |
| 3153.32373 | 4910.677246 | 2635.630127 | 0.41730635 | 0.498556044 | 0.558392461 | 0.355229313 | -0.324831486 | 0.531726747 |
| 28557.01367 | 30890.98828 | 24465.76367 | 0.415902598 | 0.197199319 | -0.498787037 | 0.171581754 | -0.753543516 | 0.104391241 |
| 122707.6719 | 127551.4375 | 145786.125 | 0.415621801 | 0.091129794 | -0.129123556 | 0.087322601 | -0.013626956 | 0.910023272 |
| 60448.125 | 59009.89063 | 58198.24219 | 0.414254574 | 0.062373059 | 0.49054941 | 0.106670824 | 0.834525386 | 0.021134419 |
| 274994.75 | 308196.9063 | 295894.2188 | 0.413518234 | 0.084019131 | 0.056468495 | 0.708396864 | 0.003738824 | 0.982235546 |
| 94153.07813 | 109273.5547 | 99382.55469 | 0.413468921 | 0.04765314 | -0.370507972 | 0.015540203 | -0.048288504 | 0.619317411 |
| 56766.82031 | 70002.36719 | 72598.45313 | 0.413287133 | 0.524294855 | 0.549146602 | 0.132025913 | 0.185040444 | 0.475738237 |
| 46984.71875 | 48374.00391 | 37818.03125 | 0.412622128 | 0.081471147 | -0.059269763 | 0.320787116 | -0.222187216 | 0.240299186 |
| 481604.5 | 550247.5 | 599074.4375 | 0.412511451 | 0.024706248 | -0.124774715 | 0.270028971 | 0.314482704 | 0.153543767 |
| 39675.98047 | 38496.91016 | 36587.53516 | 0.412385391 | 0.147715547 | 0.070183779 | 0.554396013 | 0.177292057 | 0.168508025 |

| | | | | | | | | |
|---|---|---|---|---|---|---|---|---|
| 35776.67578 | 32082.55469 | 24979.59375 | 0.412114688 | 0.063544728 | 0.576045805 | 0.046373997 | 0.174827549 | 0.499893318 |
| 10459.57715 | 18223.66406 | 10853.57715 | 0.411875177 | 0.216609416 | -0.957903686 | 0.095165103 | -0.639935285 | 0.180606854 |
| 41491.98828 | 37506.07422 | 33658.88281 | 0.411316764 | 0.016471012 | -0.295519669 | 0.079287615 | -0.285876995 | 0.122550756 |
| 1734815.25 | 1676897.375 | 1562214.75 | 0.411015387 | 0.069867455 | -0.017797577 | 0.897511445 | 0.047569973 | 0.687539514 |
| 5680.199707 | 7817.586914 | 8650.280273 | 0.410534953 | 0.166087729 | 0.284887711 | 0.230659563 | 0.394512757 | 0.27677195 |
| 1109767.5 | 1133339.875 | 1110648.5 | 0.409759541 | 0.041945948 | -0.401565282 | 0.058754959 | -0.207413097 | 0.108368576 |
| 14491.49316 | 26223.28711 | 10217.09277 | 0.408768335 | 0.515809019 | -0.047405676 | 0.922008276 | -0.359528429 | 0.596690124 |
| 2988.69458 | 4028.063965 | 680.920105 | 0.408572346 | 0.427440307 | 0.974885622 | 0.048024604 | -1.58782526 | 0.076969062 |
| 597276.1875 | 627583.8125 | 601582.1875 | 0.408470814 | 0.344637096 | 0.285557626 | 0.41896461 | 0.210793695 | 0.529641658 |
| 17688.2793 | 16179.50684 | 11685.98633 | 0.408378474 | 0.206147441 | 0.065020368 | 0.661192958 | 0.101373357 | 0.725853661 |
| 21971.18164 | 23439.0957 | 25300.37695 | 0.408357809 | 0.05132168 | -0.095430803 | 0.380876678 | -0.348172021 | 0.064571582 |
| 73500.53125 | 76419.05469 | 68811.76563 | 0.408161318 | 0.023105653 | -0.071608061 | 0.078996408 | 0.09291958 | 0.268450393 |
| 44620.92188 | 43872.86328 | 46069.55078 | 0.407843166 | 0.14905608 | 0.098442321 | 0.604923637 | 0.118910709 | 0.528938224 |
| 140618.7813 | 132805.6563 | 95470.09375 | 0.407705464 | 0.080784362 | -0.458793141 | 0.015540203 | -0.545827816 | 0.1187593 |
| 83129.38281 | 75323.39063 | 67353.51563 | 0.407355617 | 0.03279857 | -0.918903219 | 0.007509388 | -0.80749052 | 0.037571191 |
| 12419.44238 | 12778.43164 | 11357.5498 | 0.407292919 | 0.469297461 | 1.60308606 | 0.079283683 | 1.712666474 | 0.010178497 |
| 2614760.5 | 2873844.75 | 2856102 | 0.406806555 | 0.021024291 | -0.839580582 | 0.033738109 | -0.337818201 | 0.034677634 |
| 125480.8438 | 134671.7656 | 129857.5391 | 0.406581357 | 0.1377304 | 0.353163215 | 0.150271527 | 0.306140368 | 0.171025182 |
| 29822.21289 | 26336.01563 | 27850.79688 | 0.406535801 | 0.084819075 | 0.172550048 | 0.348633551 | 0.376750179 | 0.070873774 |
| 52778.40234 | 38793.58203 | 48091.61328 | 0.40631864 | 0.021353118 | -0.192333212 | 0.086634371 | -0.615424628 | 0.059641593 |
| 476939.3125 | 481597 | 455922.7813 | 0.406198884 | 0.01488475 | 0.460393891 | 0.013885891 | 0.464252592 | 0.012602644 |
| 10662.64844 | 12018.39258 | 9490.274414 | 0.406054561 | 0.222830607 | 0.126349744 | 0.610552707 | -0.026041729 | 0.9274676 |
| 28428.12305 | 26828.45508 | 17809.04688 | 0.405896462 | 0.046600136 | -0.036715335 | 0.751608736 | -0.706932268 | 0.085236358 |
| 23384.00977 | 15421.79199 | 17643.46875 | 0.405222965 | 0.074930198 | -0.192031085 | 0.069370812 | -0.813170881 | 0.076783998 |
| 207714.2969 | 208494.8594 | 177470.2188 | 0.405216348 | 0.042192618 | -0.731984476 | 0.096721331 | 0.14797191 | 0.307275286 |
| 101539.1953 | 97500.71875 | 92948.03125 | 0.405057397 | 0.02624826 | 0.317186646 | 0.055340142 | 0.275565214 | 0.046624125 |
| 592953.2188 | 603969.875 | 591070.4063 | 0.404929328 | 0.062989607 | -0.10632189 | 0.411920724 | -0.118037546 | 0.366246911 |
| 4676.269043 | 7592.845215 | 2907.759521 | 0.404620547 | 0.53362741 | 0.235433981 | 0.623490604 | -0.563586788 | 0.433773113 |
| 364861.6563 | 110449.0781 | 383008.75 | 0.404054474 | 0.563894297 | -0.417896019 | 0.595637404 | -0.340889232 | 0.627065516 |
| 3252.702393 | 4623.45459 | 2672.213135 | 0.403943511 | 0.307652251 | -0.959559521 | 0.166004104 | -0.223831456 | 0.609648287 |
| 6596.349609 | 5751.208008 | 4970.515625 | 0.403712038 | 0.121041004 | -0.248352527 | 0.362149312 | 0.000774813 | 0.997782424 |
| 631215.3125 | 572058.25 | 649460.1875 | 0.403522682 | 0.113673327 | 0.100638807 | 0.591189473 | -0.08891112 | 0.601247781 |
| 1522.269775 | 1240.170898 | 1030.889038 | 0.403407804 | 0.047290455 | 0.505341766 | 0.227071428 | -0.554831367 | 0.093070634 |

| | | | | | | | | |
|---|---|---|---|---|---|---|---|---|
| 741322.9375 | 723321 | 729968.5 | 0.402726577 | 0.00799298 | 0.112181593 | 0.215335332 | 0.033901021 | 0.301755353 |
| 29370.52344 | 40329.75 | 25863.34766 | 0.402467365 | 0.220149573 | 0.193861836 | 0.408157212 | -0.01652519 | 0.965408768 |
| 50248.10156 | 46657.41406 | 48542.61328 | 0.402419214 | 0.034713383 | 0.008629556 | 0.926516232 | 0.045503551 | 0.471734685 |
| 11465.96289 | 14104.58789 | 15653.10938 | 0.401968991 | 0.193815098 | 0.402853101 | 0.151717878 | 0.995272629 | 0.060920196 |
| 163977.7813 | 169252.0156 | 151722.4063 | 0.401951944 | 0.123070256 | 0.126613808 | 0.532853413 | 0.247981354 | 0.142342008 |
| 22780.04688 | 24846.60547 | 18251.15234 | 0.401945754 | 0.085283871 | -1.047082223 | 0.031437224 | -0.865629244 | 0.027533174 |
| 492503.5 | 535142.125 | 503990.1875 | 0.40176362 | 0.0140944 | -0.111397341 | 0.086089321 | -0.001074843 | 0.987293677 |
| 93978.69141 | 88410.63135 | 88299.93164 | 0.40164415 | 0.028452905 | 0.065592962 | 0.490020706 | -0.134037972 | 0.07481103 |
| 56207.92188 | 54537.08594 | 57044.30078 | 0.401422773 | 0.087043567 | 0.332805465 | 0.099830362 | 0.314960478 | 0.0890834 |
| NA | 2346.926025 | NA | 0.401040597 | 0.081289628 | 0.688562565 | 0.242349552 | -0.352764774 | NA |
| 397988.4688 | 389479.9688 | 368431.9063 | 0.401031504 | 0.042134056 | -0.459085 | 0.043644056 | -0.402404828 | 0.031785503 |
| 43807.26953 | 42686.04688 | 41856.65234 | 0.400977108 | 0.045667508 | -0.123179776 | 0.14824422 | -0.125716101 | 0.129731124 |
| 135.1269226 | 89.0185318 | NA | 0.40039442 | 0.103953305 | -2.002244828 | NA | -3.251246704 | NA |
| 40328.94922 | 41926.93359 | 43594.58984 | 0.399971575 | 0.067188303 | 0.193894745 | 0.175868431 | 0.315682728 | 0.063937169 |
| 1100111.75 | 1104834.5 | 1111672.25 | 0.399784348 | 0.02187351 | -1.133930589 | 0.005312664 | -0.538660758 | 0.014388274 |
| 705243.8125 | 755846.875 | 702797.9375 | 0.39947272 | 0.055228595 | -0.36411908 | 0.069743814 | -0.569597157 | 0.035868549 |
| 115417.8281 | 121233.6484 | 112794.5 | 0.399421804 | 0.049895713 | 0.350217758 | 0.092761595 | 0.154320743 | 0.205514355 |
| 238040.2344 | 239568.1094 | 264917.6875 | 0.39941244 | 0.075602358 | 0.220144963 | 0.125004947 | 0.515638123 | 0.03009785 |
| 10166786 | 10719842 | 9909200 | 0.398578601 | 0.005752145 | -0.147399377 | 0.08053869 | -0.089289438 | 0.185596413 |
| 367064.6875 | 341567.5 | 294676.0313 | 0.398351206 | 0.043802258 | -0.039280491 | 0.848507078 | 0.015873862 | 0.920066992 |
| 2668.426758 | 2121.13501 | 1426.746216 | 0.397564958 | 0.195658833 | 0.426311831 | 0.133875076 | -0.033591929 | 0.939096394 |
| 1743707.125 | 1676940.25 | 1437416.625 | 0.397409408 | 0.031387038 | 0.013278068 | 0.807635266 | 0.099166851 | 0.477536926 |
| 287355.4688 | 275342.8438 | 271012.125 | 0.3969791 | 0.089040397 | 0.086393931 | 0.555335026 | 0.045598203 | 0.74108401 |
| 4656.214844 | 4973.093262 | 6846.311035 | 0.39671138 | 0.234602879 | 0.023413665 | 0.927376589 | -0.261456981 | 0.437260941 |
| 343066.0313 | 620703.25 | 221903.0938 | 0.396639297 | 0.594734687 | 0.378531025 | 0.488412384 | -0.256877492 | 0.74476335 |
| 1289.655518 | 2074.65332 | 2180.108643 | 0.396195418 | 0.138806746 | 0.550527087 | 0.328806215 | -0.058610549 | 0.866540579 |
| 22689.48828 | 19878.51953 | 20192.10352 | 0.396174646 | 0.024042097 | 0.063321412 | 0.393210193 | -0.136763887 | 0.23354293 |
| 137925.375 | 135420.2344 | 120392.6797 | 0.395882078 | 0.193810113 | 1.281372968 | 0.021906439 | 1.092253197 | 0.020337832 |
| 32974.08984 | 32630.94141 | 29320.88477 | 0.395534948 | 0.041313942 | -0.724538966 | 0.042887894 | -0.538888988 | 0.031576918 |
| 124640.9453 | 124476.0547 | 130202.5625 | 0.395426778 | 0.036590352 | 0.152425963 | 0.169499862 | 0.140732046 | 0.179439731 |
| 40610.88672 | 44200.38672 | 41467.36719 | 0.395268132 | 0.087885618 | -0.092911987 | 0.467378475 | -0.417301182 | 0.077431074 |
| 14232.79688 | 14810.16699 | 15331.21289 | 0.395109953 | 0.110302818 | 0.466105085 | 0.068255066 | 0.645717967 | 0.030633392 |
| 13862.05273 | 14118.16113 | 12909.19824 | 0.395030052 | 0.081999941 | 0.008455251 | 0.88033198 | 0.15293781 | 0.128367922 |

| | | | | | | | | |
|---|---|---|---|---|---|---|---|---|
| 74299.08594 | 71659.03906 | 62624.61328 | 0.394526448 | 0.216609416 | 0.133650449 | 0.569857055 | -0.029947941 | 0.906660092 |
| 43397.10156 | 44776.87891 | 44041.0625 | 0.394346185 | 0.060975436 | -0.119604506 | 0.1705284 | -0.084707138 | 0.166647679 |
| 10151.33691 | 12278.63574 | 10639.74316 | 0.394160905 | 0.437128624 | 0.438290952 | 0.321888877 | 0.312865123 | 0.454130357 |
| 28216.4375 | 28710.42773 | 31977.01758 | 0.394147138 | 0.153739559 | 0.040094192 | 0.861911572 | 0.21192303 | 0.190521847 |
| 151126.1563 | 152450.625 | 155788.0938 | 0.393607629 | 0.033282394 | -0.035174335 | 0.748822862 | 0.084500165 | 0.295964678 |
| 10748.20508 | 11197.2168 | 5701.320801 | 0.393395132 | 0.136902686 | -0.787872981 | 0.100528183 | -0.465193245 | 0.277618275 |
| 338492.8125 | 326053.1875 | 297377.2188 | 0.393205655 | 0.056599263 | -0.618492143 | 0.038184043 | -0.679859991 | 0.031082912 |
| 22741.84961 | 22801.26367 | 29111.67773 | 0.392964547 | 0.183506434 | 0.270691492 | 0.111472792 | 0.444186513 | 0.150131299 |
| 28896.51563 | 29403.5918 | 27728.39453 | 0.392426864 | 0.019930448 | -0.649607369 | 0.047136323 | 0.199372652 | 0.040645703 |
| 119828.5313 | 109242.4531 | 120357.4297 | 0.392127335 | 0.040854235 | 0.465660543 | 0.08608279 | 0.285100344 | 0.069238464 |
| 22032.32031 | 24210.51953 | 20781.7168 | 0.392092582 | 0.08816872 | -0.215452543 | 0.440401575 | 0.738409772 | 0.027949491 |
| 332202.0625 | 361711.5313 | 347624.0313 | 0.391092283 | 0.034890445 | -0.194607685 | 0.112543356 | -0.553482057 | 0.02354608 |
| 209804.1406 | 273239.2188 | 250636.2813 | 0.390769619 | 0.064026439 | -0.152085313 | 0.415383613 | -0.347668893 | 0.121306313 |
| 9402.198242 | 7310.367188 | 8256.853516 | 0.39063312 | 0.033123824 | 1.002958757 | 0.007509388 | -0.137338832 | 0.411945771 |
| 69815.55469 | 64281.80078 | 61399.58984 | 0.390175448 | 0.037393884 | 0.112812143 | 0.23786621 | 0.205017638 | 0.124211802 |
| 153521.4688 | 156562.7344 | 151145.6406 | 0.389145094 | 0.025740944 | -1.076169421 | 0.011710842 | -1.563505117 | 0.005561476 |
| 28159.2207 | 27031.83789 | 29801.25977 | 0.389137951 | 0.050166486 | 0.214396642 | 0.03376576 | -0.060395002 | 0.394060816 |
| 15772.53613 | 17804.34766 | 11135.39453 | 0.388802947 | 0.541808174 | -0.696869301 | 0.27704244 | -0.902741077 | 0.116993358 |
| 1176088.75 | 1285209.5 | 1121726.625 | 0.388758069 | 0.091259043 | -0.016833565 | 0.885440998 | 0.120881615 | 0.371644128 |
| 15440.64258 | 31066.77539 | 25929.10938 | 0.388132459 | 0.422930592 | 0.761480503 | 0.129928099 | 0.189471538 | 0.728068597 |
| 129654.6563 | 132674.2031 | 127181.2656 | 0.387650165 | 0.103960261 | -0.102079584 | 0.506862757 | -0.144612417 | 0.364029016 |
| 552.0963135 | 1287.227905 | NA | 0.387600954 | 0.686880627 | -0.296925404 | 0.763032066 | -0.778879044 | NA |
| 10806.05273 | 8949.738281 | NA | 0.387362297 | 0.160645659 | -0.328454688 | 0.230789198 | -0.254662318 | NA |
| 1205077.25 | 1331968.875 | 1386544.5 | 0.386601693 | 0.414640258 | 0.232739558 | 0.593477001 | 1.89746847 | 0.014665336 |
| 1823846 | 1867211 | 1859254.375 | 0.386015623 | 0.03777619 | 0.069064894 | 0.570522708 | 0.323229585 | 0.025878597 |
| 85049.0625 | 84646.22656 | 80329.3125 | 0.385720732 | 0.065091163 | 0.075829992 | 0.571319729 | 0.426628191 | 0.045697257 |
| 11684.42578 | 9686.428711 | 4185.666016 | 0.385310517 | 0.109140206 | 0.910577879 | 0.087142005 | 0.55890933 | 0.456448725 |
| 2518.855713 | 2703.913574 | 3208.993896 | 0.385075335 | 0.160518598 | -0.920922726 | 0.056841901 | 0.138849854 | 0.525435103 |
| 63463.27734 | 61858.11328 | 57858.19922 | 0.384778698 | 0.019376457 | 0.269602654 | 0.108786013 | -0.148380803 | 0.098401903 |
| 76070.5625 | 71437.78125 | 80677.48438 | 0.384777189 | 0.012414001 | -0.260733188 | 0.040932722 | -0.231508149 | 0.076144169 |
| 28714.78516 | 31776.91797 | 23716.87891 | 0.384607876 | 0.014038682 | -0.879563836 | 0.02118733 | -0.664567907 | 0.058506939 |
| 353431.7813 | 355766.4688 | 322216.0938 | 0.384416933 | 0.021027835 | -0.053217551 | 0.68225883 | -0.167404719 | 0.111473781 |
| 52751.28125 | 44764.19141 | 37735.58984 | 0.384401064 | 0.103160411 | -0.343838487 | 0.121179966 | -0.95475911 | 0.030225962 |

| | | | | | | | | |
|---|---|---|---|---|---|---|---|---|
| 33890.31641 | 32839.46094 | 33306.12891 | 0.383485233 | 0.010893509 | 0.293709502 | 0.048024604 | 0.071748044 | 0.134259503 |
| 37214.86719 | 38493.84375 | 31017.68164 | 0.383143354 | 0.031790847 | -0.043265559 | 0.684055965 | -0.207834755 | 0.215760001 |
| 45952.21484 | 35567.82422 | 35765.77344 | 0.383083891 | 0.578855534 | -0.025511835 | 0.954688064 | 1.952551117 | 0.043034683 |
| 29937.05078 | 29548.64844 | 24166.73633 | 0.382848123 | 0.016885305 | 0.203111052 | 0.129824422 | -0.398763468 | 0.081899796 |
| 285928.9688 | 268538.6563 | 299698.4688 | 0.382460427 | 0.156655374 | -0.08448898 | 0.684055965 | -0.163559298 | 0.436569813 |
| 7892.768555 | 6356.164063 | 9081.338867 | 0.382256965 | 0.234602879 | -0.964004442 | 0.110625004 | -1.612266019 | 0.041088687 |
| 150629.7344 | 146543.3906 | 146266.0313 | 0.381480576 | 0.030839901 | -0.317865442 | 0.036918072 | 0.14318106 | 0.073672213 |
| 26370.10352 | 23876.48633 | 25427.56836 | 0.380801626 | 0.037913948 | -0.186856644 | 0.151284841 | 0.072575032 | 0.366486609 |
| 240588.8906 | 236391.9219 | 232155.5469 | 0.379906184 | 0.091934878 | 0.216420078 | 0.052817461 | 0.0666022 | 0.29000436 |
| 8172.397461 | 9090.574219 | 6474.928223 | 0.37986727 | 0.048531905 | -0.244425984 | 0.345066656 | -0.934134003 | 0.026263251 |
| 243191.9375 | 244135.1875 | 226785.7813 | 0.379251801 | 0.151189017 | 0.076256309 | 0.63348405 | 0.228568837 | 0.214658701 |
| 23419.83789 | 20739.76172 | 17444.36133 | 0.379024996 | 0.238040125 | -0.002609801 | 0.99201113 | 0.075745803 | 0.787851276 |
| 4949.061523 | 5606.442871 | 358.6928711 | 0.378945597 | 0.358049798 | 0.473564218 | 0.174452979 | -0.042548974 | 0.969402997 |
| 60437.29297 | 63430.73438 | 51081.31641 | 0.378432877 | 0.13131902 | 0.306926402 | 0.045744926 | -0.084659769 | 0.547829996 |
| 733105 | 762019.8125 | 695584 | 0.378283494 | 0.064047558 | -0.654664076 | 0.047212306 | -0.445445013 | 0.047526425 |
| 30629.89063 | 34161.44531 | 33799.65625 | 0.378138255 | 0.056932452 | 0.078023667 | 0.414751124 | -0.070171001 | 0.515665974 |
| 7314.847656 | 7733.936035 | 7686.51416 | 0.378080158 | 0.412066495 | -0.124420826 | 0.765555194 | -0.48992561 | 0.33090215 |
| 253185.0156 | 344090 | 326755.6563 | 0.377818505 | 0.494503803 | 0.081254692 | 0.865919704 | -0.017105592 | 0.975905464 |
| 158054.7188 | 176190.0313 | 171720.6094 | 0.377628242 | 0.329719661 | -0.019277342 | 0.955667011 | -0.314827675 | 0.406218733 |
| 118141.6406 | 104320.4922 | 116581.8984 | 0.377403483 | 0.084596054 | -0.197664254 | 0.217192228 | -0.41908188 | 0.070587266 |
| 15356.07227 | 16741.49219 | 15220.36719 | 0.376851432 | 0.048340163 | 0.336193808 | 0.027675546 | -0.159970909 | 0.10853769 |
| 27909.38086 | 23942.9375 | 14959.93945 | 0.37618186 | 0.17760319 | -0.281441419 | 0.289050544 | 0.163376607 | 0.688894986 |
| 471656.5 | 469531.375 | 530055 | 0.375871594 | 0.064338292 | -0.487802236 | 0.054504325 | 0.835394627 | 0.0236713 |
| 345817.2813 | 343500.3125 | 399490.4063 | 0.374712394 | 0.046164606 | -0.06590286 | 0.502245514 | 0.378486523 | 0.078691633 |
| 130819.4844 | 138567.9063 | 88085.49219 | 0.37442909 | 0.239971035 | 0.68496274 | 0.17671494 | 0.466816578 | 0.262180774 |
| 36180.01172 | 38048.47656 | 36279.11328 | 0.374364844 | 0.021027835 | -0.19799689 | 0.08026683 | -0.605578801 | 0.021984636 |
| 31324.27539 | 32940.45313 | 34576.62109 | 0.374223322 | 0.05267493 | -0.173174973 | 0.206042711 | -0.152021321 | 0.231918617 |
| 42726.65234 | 46371.25781 | 54163.40625 | 0.374198845 | 0.271312801 | 0.429373935 | 0.32242293 | 0.03986343 | 0.832420876 |
| 91046.80469 | 96198.91406 | 68669.75781 | 0.374129102 | 0.080007818 | -0.460632942 | 0.061296684 | -0.443581872 | 0.11327823 |
| 39274.76563 | 42558.20313 | 36185.21484 | 0.373908847 | 0.037562551 | -0.258677252 | 0.07650431 | -0.427318004 | 0.037772514 |
| 55604.05469 | 61923.10547 | 46657.35938 | 0.373689268 | 0.160003781 | 0.385326453 | 0.084352289 | -0.11643005 | 0.54725722 |
| 175499 | 173461.4063 | 165969.7813 | 0.373637002 | 0.329558949 | 0.054344726 | 0.863198658 | 0.152400707 | 0.609506134 |
| 15663.67871 | 15225.82813 | 13991.81055 | 0.373416181 | 0.154178973 | 0.023930505 | 0.899247296 | -0.013257993 | 0.942386927 |

| | | | | | | | | |
|---|---|---|---|---|---|---|---|---|
| 12281.72949 | 6392.437988 | 11948.30664 | 0.373312824 | 0.319021215 | -0.286192687 | 0.532402501 | -0.460855475 | 0.299628534 |
| 11329.67773 | 7477.936523 | 11759.00781 | 0.373070188 | 0.054063082 | -0.311190511 | 0.227457591 | -0.469961316 | 0.173821986 |
| 24931.14453 | 23009.18164 | 27135.38867 | 0.372911654 | 0.018432262 | 0.07524947 | 0.326455795 | 0.10652185 | 0.381812902 |
| 35288.9375 | 37024.96484 | 36337.62109 | 0.372785069 | 0.1214887 | -0.146387609 | 0.127011012 | 0.051242501 | 0.405343283 |
| 83257.86719 | 83238.22656 | 87338.04688 | 0.372769607 | 0.014862855 | -0.044974077 | 0.356546603 | -0.101345999 | 0.116465028 |
| 53768.48828 | 59770.61328 | 55640.12891 | 0.372739663 | 0.10217612 | -0.083888826 | 0.407072973 | -0.286176059 | 0.070298355 |
| 9572.074219 | 8942.686523 | 7668.315918 | 0.372722615 | 0.283428713 | 0.424895102 | 0.179324339 | -0.08805809 | 0.763776161 |
| 1481639.75 | 1706142.875 | 1633108.75 | 0.372591733 | 0.155196387 | -0.153134658 | 0.447758985 | -0.049516749 | 0.790960569 |
| 2623534 | 2839255.5 | 2799152 | 0.372314793 | 0.163314204 | 0.278830347 | 0.192808829 | 0.292330802 | 0.162108808 |
| 811995.2852 | 730083.3789 | 648655.5781 | 0.372109496 | 0.239008795 | -0.58135567 | 0.185493647 | -0.467776301 | 0.196239339 |
| 24397.44922 | 23244.41406 | 33192.6875 | 0.372008394 | 0.227705977 | -0.926543099 | 0.093751816 | -0.251395595 | 0.434321145 |
| 51670.47656 | 56730.75781 | 49310.1875 | 0.371929721 | 0.060311673 | -0.18622479 | 0.208057773 | -0.511483085 | 0.04557602 |
| 800023.1875 | 815731.0625 | 784267.125 | 0.371720916 | 0.050577071 | 0.137818802 | 0.213352365 | 0.385153787 | 0.048421191 |
| 92319.96094 | 96923.57031 | 94102.59375 | 0.371361493 | 0.0623818 | -0.098718932 | 0.333718404 | 0.161354546 | 0.147916631 |
| 199690.625 | 188233.7813 | 212994.3125 | 0.371262147 | 0.038502747 | -0.19774611 | 0.029180754 | -0.296339796 | 0.056482153 |
| 50764.08594 | 50254.75781 | 50687.96094 | 0.371260315 | 0.058516905 | -0.110497592 | 0.101716174 | 0.24272426 | 0.023619478 |
| 148739.1875 | 140797.7344 | 131282.0156 | 0.371225886 | 0.029716072 | 0.10418892 | 0.147577182 | 0.131658861 | 0.212405887 |
| 37200.95703 | 36799.94531 | 32243.5625 | 0.371165513 | 0.039443869 | -0.567415933 | 0.036485471 | -0.007892075 | 0.950211597 |
| 19856.1543 | 21701.86328 | 25100.11719 | 0.370962845 | 0.081090664 | -0.538233193 | 0.089100892 | -0.34340338 | 0.112223514 |
| 96437.52344 | 97518.19531 | 84860.625 | 0.370875223 | 0.065637909 | 0.204109374 | 0.166004104 | -0.138506618 | 0.315738114 |
| 27090.92188 | 41219.75 | 15038.79102 | 0.370800253 | 0.486883738 | -0.033758706 | 0.934339033 | -0.550498203 | 0.394984688 |
| 101263.3281 | 85017.50781 | 90613.82031 | 0.370507544 | 0.024973429 | -0.401537654 | 0.037138856 | -0.559533576 | 0.028173069 |
| 77416.15625 | 83902.10938 | 76130.1875 | 0.370374116 | 0.01802855 | -0.686593598 | 0.014340141 | -0.508262454 | 0.016683666 |
| 18463.20508 | 14190.92578 | 17461.85547 | 0.369629269 | 0.074738377 | 0.403441849 | 0.065422444 | -0.276950381 | 0.182917302 |
| 46158.05859 | 43488.13281 | 35530.49219 | 0.368953892 | 0.109154846 | 0.688992717 | 0.040327952 | 0.452728512 | 0.119204395 |
| 341403.3125 | 359061.3125 | 355928.2813 | 0.368870634 | 0.038502747 | -0.630023931 | 0.023005377 | -0.202688206 | 0.038798046 |
| 293514.0938 | 291550.1875 | 245068.7969 | 0.36777365 | 0.076438569 | -0.398018702 | 0.028434447 | -0.77031022 | 0.039195867 |
| 64746.09766 | 60944.02344 | 81884.1875 | 0.367596461 | 0.084004676 | -0.053902938 | 0.398049561 | 0.61574148 | 0.136000191 |
| 115517.4844 | 123310.3047 | 127258.2109 | 0.366571517 | 0.057292418 | -0.144355851 | 0.26192413 | -0.331986732 | 0.052768746 |
| 789.7149048 | 348.7382813 | NA | 0.366128469 | NA | -0.735726369 | NA | -1.587211343 | NA |
| 303324.0625 | 335299.5313 | 344824.4688 | 0.365968587 | 0.063237602 | -0.389716306 | 0.062122829 | -0.443816952 | 0.045495106 |
| 49608.75781 | 54157.625 | 59892.68359 | 0.365863962 | 0.143584671 | 0.53162579 | 0.07468596 | 1.168675469 | 0.025966989 |
| 96829.57813 | 98674.1875 | 101441.3359 | 0.365289089 | 0.071903471 | -0.113818697 | 0.385249231 | -0.102441183 | 0.382150932 |

| | | | | | | | | |
|---|---|---|---|---|---|---|---|---|
| 5569.926758 | 5803.826172 | 6364.26123 | 0.36510085 | 0.418203491 | 0.466351893 | 0.247188011 | 0.541014019 | 0.189440449 |
| 64022.24609 | 60332.10938 | 66933.94531 | 0.36499082 | 0.037397136 | -0.572181208 | 0.052437153 | -0.142976515 | 0.148438072 |
| 298796.1563 | 293545.7188 | 292660.8438 | 0.364504189 | 0.181816436 | -0.157606547 | 0.470469741 | -0.033823465 | 0.860778751 |
| 23964.46484 | 11227.15723 | 1597.129517 | 0.364402234 | 0.056818188 | 1.329801962 | 0.137284361 | -0.102017313 | 0.931604241 |
| 68637.54688 | 54921.16797 | 58552.65625 | 0.364194263 | 0.035776373 | 0.302744533 | 0.023592908 | 0.147336054 | 0.395484708 |
| 13945.62988 | 12814.80762 | 14313.52734 | 0.363874344 | 0.010297098 | -0.260668961 | 0.024404888 | -0.074066452 | 0.358094258 |
| 81756.58594 | 77006.10938 | 76265.39844 | 0.362953741 | 0.154530346 | 0.121996995 | 0.460748219 | 0.171700248 | 0.317427062 |
| 140361.6875 | 135610.75 | 121152.2109 | 0.362457774 | 0.037492489 | -0.469949682 | 0.039699307 | -0.595415526 | 0.021195162 |
| 242959.4531 | 250900.2344 | 300194.4063 | 0.362277177 | 0.09706261 | 0.148049205 | 0.280546656 | -0.143763136 | 0.374030298 |
| 62137.75391 | 56383.76953 | 53258.82422 | 0.362222509 | 0.005752145 | -0.071013109 | 0.669186511 | 0.49365992 | 0.074385174 |
| 155293 | 145404.6094 | 138664.2031 | 0.362049348 | 0.059440088 | 0.831447762 | 0.015395012 | 0.811581798 | 0.017622287 |
| 26407.25195 | 28173.08203 | 22166.1543 | 0.361770582 | 0.065975508 | -0.356142836 | 0.097793713 | -0.61193721 | 0.036566175 |
| 674187.25 | 958339.625 | 915751.6875 | 0.361206425 | 0.177108689 | -0.20245173 | 0.388239791 | 0.346456515 | 0.277706206 |
| 6733.835938 | 7919.437012 | 9420.27832 | 0.361196737 | 0.749455721 | 0.124318134 | 0.903989395 | 0.035830303 | 0.975159144 |
| 58485.04688 | 63463.53125 | 61136.81641 | 0.360609471 | 0.02753514 | 0.062874482 | 0.382650392 | 0.285723821 | 0.036224874 |
| 239046.2813 | 79088.75 | 261679.4844 | 0.360549763 | 0.601468792 | -0.455125335 | 0.553393539 | -0.379498739 | 0.573238987 |
| 59648.58594 | 50404.27344 | 24875.78125 | 0.360063032 | 0.542216889 | 0.807637021 | 0.143959847 | 0.194470149 | 0.767152117 |
| 26757 | 27906.78516 | 34270.29297 | 0.360046095 | 0.318215645 | 0.09715089 | 0.7417052 | -0.40044109 | 0.303269839 |
| 8883.168945 | 9066.916016 | 9865.375977 | 0.359992262 | 0.243529133 | -0.11465242 | 0.667331576 | -0.481731672 | 0.198223724 |
| 22568.03906 | 18920.78906 | 13858.7832 | 0.359925559 | 0.095278437 | -0.391295282 | 0.122113311 | -0.870699858 | 0.050460516 |
| 25808.81055 | 28669.28125 | 28844.17188 | 0.35965888 | 0.020732204 | 0.019650873 | 0.848041402 | 0.235965554 | 0.0890834 |
| 75354.14844 | 73830.54688 | 60531.20313 | 0.35911578 | 0.161511845 | 0.406462782 | 0.202608385 | 1.043864864 | 0.043108871 |
| 55986.92188 | 65096.13281 | 63940.49609 | 0.359015286 | 0.214335473 | 0.167480674 | 0.439143602 | 0.456521572 | 0.09843293 |
| 276101.8125 | 259910.2031 | 262036.3125 | 0.358947653 | 0.106930422 | 0.46539121 | 0.079547946 | 0.827020417 | 0.012701134 |
| 82855.67188 | 82758.84375 | 65567.73438 | 0.358457092 | 0.015030322 | -0.267073457 | 0.02118733 | -0.222189395 | 0.246737577 |
| 75487.78906 | 69703.02344 | 73281 | 0.358353708 | 0.145754807 | 0.557044565 | 0.063199811 | 0.389784962 | 0.02411991 |
| 24894.48828 | 22616.20117 | 20068.26563 | 0.358206167 | 0.19452447 | -0.116516938 | 0.6115573 | -0.396073741 | 0.16847165 |
| 280369.0625 | 291771.4063 | 293828.9375 | 0.357648398 | 0.033526967 | 0.056784772 | 0.526766343 | 0.173342833 | 0.095206618 |
| 42386.86328 | 49422.10938 | 19528.47656 | 0.357560578 | 0.610763682 | 0.181275631 | 0.717802034 | -0.40695888 | 0.560298855 |
| 100687.1406 | 111296.2578 | 119612.0625 | 0.357067388 | 0.440821363 | 0.429445123 | 0.303308952 | 0.587063999 | 0.172011685 |
| 28381.32227 | 30106.51367 | 33722.9541 | 0.357050773 | 0.089436106 | 0.399655938 | 0.130757652 | -0.397488326 | 0.082387931 |
| 102284.1875 | 97885.91406 | 60912.09766 | 0.356727306 | 0.096833175 | -0.868346753 | 0.064138817 | -0.474444796 | 0.192404323 |
| 143604.4219 | 152949.9219 | 148056.4063 | 0.35666169 | 0.061588651 | 0.386331002 | 0.05467585 | 0.459693727 | 0.036016754 |

| | | | | | | | | |
|---|---|---|---|---|---|---|---|---|
| 46438.6875 | 48946.58203 | 42697.85938 | 0.356661198 | 0.102165776 | -0.628544764 | 0.088587863 | -0.515707312 | 0.07755543 |
| 128551.5 | 127076.2422 | 120941.5781 | 0.356595494 | 0.132174911 | 0.409322295 | 0.112102762 | 0.340324559 | 0.120695883 |
| 10923.49707 | 5894.294922 | 8847.108398 | 0.356290152 | 0.185482817 | 0.56488227 | 0.231558384 | 0.491628367 | 0.340584009 |
| 13406.6709 | 18109.73242 | 11454.74219 | 0.356106309 | 0.228860613 | -0.269400745 | 0.467062508 | -0.30126149 | 0.343143637 |
| 207666.8281 | 199883.9844 | 200783.625 | 0.356099341 | 0.026764186 | -0.257323954 | 0.104959595 | -0.035749492 | 0.582848072 |
| 35565.58203 | 37061.01563 | 31062.07031 | 0.355219624 | 0.049996115 | -0.441646136 | 0.061625352 | -0.563424882 | 0.033928174 |
| 142481.625 | 137814.0156 | 147502.0625 | 0.354894217 | 0.025960199 | -0.100068316 | 0.231229095 | -0.250501958 | 0.055868946 |
| 107865.7422 | 63291.46875 | 107478.9219 | 0.354775125 | 0.404501862 | 1.070133986 | 0.30772129 | 2.377029291 | 0.091109394 |
| 5882.183105 | 7656.061035 | 6177.826172 | 0.354668346 | 0.051358293 | -0.479498022 | 0.048593497 | -0.417714495 | 0.095599918 |
| 19040.58398 | 18879.99023 | 17503.22852 | 0.354430747 | 0.046236587 | -0.20260346 | 0.175868431 | -0.236726543 | 0.095599918 |
| 74933.25 | 71665.40625 | 96566.71094 | 0.353843822 | 0.08771097 | -0.064737783 | 0.689502716 | 0.460814628 | 0.178411577 |
| 153845.5625 | 156800.3906 | 159418.6719 | 0.353580718 | 0.010297098 | -0.463954428 | 0.014298932 | -0.144163659 | 0.03286768 |
| 4918.030273 | 5687.469727 | 4367.414551 | 0.353514091 | 0.201160451 | 0.004460751 | 0.984569317 | -0.075096694 | 0.656039733 |
| 564177.5625 | 639803.4375 | 679454.125 | 0.353476355 | 0.482200649 | 0.748281119 | 0.149058018 | 1.057132985 | 0.060016882 |
| 178444.8906 | 186532.6406 | 160557.0781 | 0.353072468 | 0.168887111 | -0.469339953 | 0.125467937 | -1.290987385 | 0.056756997 |
| 74880.07031 | 74933.21875 | 73921.00781 | 0.352151667 | 0.296551702 | -0.183342682 | 0.534935991 | -0.521887115 | 0.21485344 |
| 193956.4063 | 200258.5156 | 190070.0313 | 0.351664663 | 0.033353006 | 0.10088135 | 0.058154495 | 0.165882915 | 0.04016603 |
| 17822.99805 | 18920.5 | 25844.45117 | 0.351355343 | 0.143267795 | 0.064854601 | 0.808781142 | 0.35650605 | 0.308214918 |
| 10871.31738 | 10170.60938 | 11482.92969 | 0.350838381 | 0.154102524 | -0.156011233 | 0.498876186 | 0.10741808 | 0.528968658 |
| 79944.53125 | 91782.19531 | 83190.92188 | 0.35076699 | 0.216909187 | 0.385024045 | 0.193310462 | 0.762462239 | 0.037156974 |
| 49371.5 | 45969.0625 | 58415.62891 | 0.35072141 | 0.050928677 | -0.006744787 | 0.936763383 | 0.617548571 | 0.097707274 |
| 32635.22461 | 32846.96875 | 26560.75586 | 0.350506714 | 0.255136449 | -0.463733285 | 0.090033554 | -0.691839892 | 0.036013501 |
| 74834.65625 | 74726.92969 | 76805.375 | 0.349544227 | 0.068715601 | 0.209397918 | 0.098459633 | 0.107189246 | 0.104472821 |
| 7514.714844 | 7462.358398 | 6725.82666 | 0.349438349 | 0.13402188 | -0.42447154 | 0.100528183 | -0.490158248 | 0.08197271 |
| 14951.98828 | 13282.30566 | 14552.56445 | 0.349412935 | 0.435526499 | 0.792846666 | 0.107845807 | 0.730218404 | 0.121306313 |
| 62050.42188 | 57579.52734 | 72593.00781 | 0.348945432 | 0.187040833 | 0.831226194 | 0.060145223 | 1.181870558 | 0.036848746 |
| 75343.80469 | 96825.71094 | 92484.13281 | 0.348453971 | 0.414254319 | 0.124945495 | 0.7377768 | 0.451978423 | 0.240161898 |
| 133311.1563 | 89565.49219 | 106208.7422 | 0.348294865 | 0.158200263 | 0.10796575 | 0.472698462 | -0.273232099 | 0.326461577 |
| 95469 | 92950.13281 | 92209.23438 | 0.348281218 | 0.067008234 | -0.32606288 | 0.061296684 | -0.40852741 | 0.020159082 |
| 38332.11328 | 42346.77344 | 38115.32422 | 0.348232812 | 0.037238828 | 0.00132566 | 0.985576344 | -0.001437552 | 0.989140123 |
| 4164.018555 | 12981.52832 | 14839.20703 | 0.348080735 | 0.508061586 | 1.300089096 | 0.284636711 | 0.600668275 | 0.504418582 |
| 161857.375 | 171494.125 | 180241.5469 | 0.347858071 | 0.017824776 | -0.176937751 | 0.048325476 | 0.329801309 | 0.056570421 |
| 549080.25 | 623489.375 | 598140.375 | 0.347559632 | 0.155053391 | 0.515996149 | 0.058070756 | 0.488437467 | 0.054109248 |

| | | | | | | | | |
|---|---|---|---|---|---|---|---|---|
| 988265.875 | 953433.6875 | 991874.0625 | 0.347321262 | 0.268922439 | 0.149353488 | 0.555335026 | 0.218650778 | 0.381867669 |
| 5493.276367 | 4631.906738 | 4658.774414 | 0.347256192 | 0.131546744 | 0.089953531 | 0.807635266 | -0.532753585 | 0.06688778 |
| 24277.67578 | 25840.27148 | 21295.08008 | 0.347196047 | 0.071472027 | -0.327462612 | 0.076429114 | -0.695528011 | 0.022925497 |
| 1840684.828 | 1647092.07 | 1818112.266 | 0.346595006 | 0.048840281 | -0.062289689 | 0.520920322 | 0.022853594 | 0.833250417 |
| 43687.8125 | 42438.92969 | 40722.63281 | 0.346572072 | 0.047406642 | 0.118795198 | 0.292995226 | 0.131124042 | 0.199326473 |
| 2859.173584 | 996.5511475 | 2435.207764 | 0.346487222 | 0.794392599 | 0.221328396 | 0.867720056 | 0.045860916 | 0.972131458 |
| 26593.24609 | 26670.31055 | 24280.19727 | 0.346178278 | 0.011814081 | -0.075216951 | 0.475752829 | -0.2334241 | 0.059258927 |
| 140897.7188 | 251852.1719 | 92367.71875 | 0.34602411 | 0.633323088 | 0.254485811 | 0.686661605 | -0.534832661 | 0.476446871 |
| 11985.26953 | 10524.94824 | 7501.614258 | 0.346003677 | 0.425653141 | -0.205511074 | 0.272117866 | -0.217752042 | 0.466024706 |
| 212138.8281 | 203253.4688 | 189211.6719 | 0.345753573 | 0.276360085 | 0.646482148 | 0.091622784 | 0.32895373 | 0.236918778 |
| 281382.3125 | 278006.1875 | 263426.3438 | 0.345729769 | 0.058810299 | 0.076091099 | 0.330702401 | -0.009082059 | 0.908220353 |
| 193731.7188 | 196233.0469 | 199825.0313 | 0.34533141 | 0.058363046 | 0.283767621 | 0.047212306 | 0.309722417 | 0.04174272 |
| 2378.287598 | 2081.565674 | 1249.652466 | 0.344983363 | 0.252381971 | -0.274064388 | 0.34315805 | -0.901659433 | 0.085314833 |
| 64288.76563 | 61444.07031 | 64114.91797 | 0.344669157 | 0.073614057 | 0.065205417 | 0.26604978 | -0.010258221 | 0.799884139 |
| 11497.08984 | 13246.31055 | 10634.82324 | 0.344648202 | 0.031883977 | -0.069662736 | 0.491534808 | -0.54938816 | 0.04014804 |
| 78239.65625 | 82836.03906 | 61404.13281 | 0.344627193 | 0.071472027 | -0.176018231 | 0.198421115 | -0.346067729 | 0.149087782 |
| 419233.5781 | 470379.1875 | 463590.9531 | 0.344007315 | 0.18705416 | 0.310847808 | 0.445860571 | 0.184483254 | 0.225631685 |
| 163448.5 | 171835.3594 | 152487.1875 | 0.34323005 | 0.031155519 | -0.413943889 | 0.028138975 | -0.141283592 | 0.159743169 |
| 23571.20703 | 22857.57813 | 20973.98828 | 0.343003132 | 0.387552052 | -0.601584242 | 0.262686667 | -0.5653873 | 0.261882466 |
| 84967.66406 | 104072.3359 | 90200.3125 | 0.342813937 | 0.068718798 | -0.35576706 | 0.220404142 | -0.186577527 | 0.266042939 |
| 48173.56641 | 49572.27734 | 60044.55859 | 0.342725213 | 0.087469248 | 0.811218147 | 0.035356834 | 1.008941973 | 0.053570169 |
| 792512.5625 | 772812.1875 | 832684.1875 | 0.342331545 | 0.010893509 | -0.066666167 | 0.310062196 | -0.187136841 | 0.047467226 |
| 273535 | 284301.8125 | 257189.5 | 0.342061543 | 0.057009196 | -0.087058916 | 0.332936378 | -0.330410597 | 0.048590503 |
| 11809.60742 | 11788.54883 | 12263.61816 | 0.34190474 | 0.058238988 | -0.46886528 | 0.066565 | -0.428788556 | 0.072139147 |
| 66111.78125 | 45037.12109 | 62496.30859 | 0.341824718 | 0.290356328 | -0.23688651 | 0.082329519 | -0.051652675 | 0.835784721 |
| 30972.40625 | 33876.74609 | 29799.03125 | 0.341800218 | 0.056312464 | -0.070837392 | 0.436953642 | -0.09397029 | 0.395734266 |
| 72682.82031 | 97374.0625 | 87173.60156 | 0.340573554 | 0.420601191 | -0.605520522 | 0.06106734 | 0.694094195 | 0.088845579 |
| 4332.522461 | 2750.813721 | 4576.067383 | 0.339896591 | 0.717665729 | 0.879498456 | 0.266525019 | -0.191506637 | 0.841765657 |
| 50914.93359 | 55722.33594 | 63497.16797 | 0.339045166 | 0.1694675 | 0.72281903 | 0.052036261 | 0.285806326 | 0.212405887 |
| 444131.875 | 424378.4688 | 416271.4063 | 0.338872329 | 0.089644612 | 0.686175124 | 0.050410119 | 1.136616886 | 0.009587646 |
| 3353.26001 | 5083.814941 | 4227.446289 | 0.338600069 | 0.227207392 | 0.20869396 | 0.159703167 | -0.169941469 | 0.51134221 |
| 3348.999023 | 3547.334717 | 1841.407471 | 0.338501846 | 0.371974804 | -0.37710438 | 0.366494549 | -0.622849374 | 0.23126742 |
| 196510.8125 | 207641.7969 | 197874.8281 | 0.337721441 | 0.026183333 | 0.423774169 | 0.021906439 | -0.040527557 | 0.538909094 |

| | | | | | | | | |
|---|---|---|---|---|---|---|---|---|
| 24372.45313 | 23392.35742 | 25426.85742 | 0.337600225 | 0.052921297 | 0.171187733 | 0.118137091 | 0.087672213 | 0.235870699 |
| 82462.29688 | 83014.90625 | 81801.58594 | 0.33746528 | 0.061050189 | -0.245955816 | 0.127902424 | -0.047288593 | 0.634317573 |
| 58181.09766 | 33372.88672 | 45153.46484 | 0.337445189 | 0.374803124 | -0.8013339 | 0.244443156 | 1.016614683 | 0.140858951 |
| 307084.5625 | 314756.875 | 291197.5625 | 0.337374192 | 0.111176969 | 0.016526492 | 0.93172794 | -1.301386828 | 0.04728688 |
| 12325.71191 | 15970.62012 | 11252.2373 | 0.337290476 | 0.177154 | -0.108275149 | 0.245062116 | -0.512736423 | 0.110015219 |
| 1776.759888 | 1424.96228 | 1825.449219 | 0.33634299 | 0.340812741 | -0.116018925 | 0.623490604 | -0.415815457 | 0.169029222 |
| 60330.6875 | 71005.57031 | 70407.27344 | 0.336263891 | 0.096429314 | -0.199061988 | 0.108400844 | -0.619551452 | 0.035626762 |
| 137490.6719 | 141817.0313 | 130898.2031 | 0.335844888 | 0.208728742 | -0.377181685 | 0.211473837 | -0.730238815 | 0.110105852 |
| 50088.37109 | 50388.85938 | 48320.60938 | 0.335597314 | 0.046747318 | -0.063761512 | 0.472503115 | -0.222036716 | 0.112926052 |
| 183051.9063 | 192297.625 | 175301.1094 | 0.335227019 | 0.063867921 | -0.223137696 | 0.134845721 | -0.199435526 | 0.122834452 |
| 15979.80469 | 16820.37891 | 15555.09766 | 0.334417298 | 0.248482121 | 0.298973482 | 0.031437224 | 0.35440192 | 0.026263251 |
| 49867.93359 | 45367.01172 | 47285.81641 | 0.334363936 | 0.234680715 | 0.142127652 | 0.514186354 | 0.092714286 | 0.661123692 |
| 60004.27295 | 54449.271 | 56582.3833 | 0.33428108 | 0.045672283 | -0.705239142 | 0.033121608 | -0.77924703 | 0.027868741 |
| 2188788 | 2398168.75 | 2645264.25 | 0.33381848 | 0.036692024 | 0.121595115 | 0.16329317 | 0.217709432 | 0.199146446 |
| 40721.76172 | 26419.79688 | 33040.95313 | 0.333629328 | 0.107630582 | -0.345822312 | 0.120427714 | -0.735251572 | 0.060937162 |
| 115905.6328 | 123930.4297 | 141255.375 | 0.333319015 | 0.038541031 | -0.165090379 | 0.058877608 | -0.082653476 | 0.529073522 |
| 198160.7344 | 178598.8281 | 158990.8438 | 0.332551714 | 0.080233873 | -0.655991192 | 0.048593497 | -0.062104457 | 0.696406851 |
| 17383.68164 | 12768.37402 | 12620.93164 | 0.332454245 | 0.105326647 | 0.35139508 | 0.086219414 | 0.171882457 | 0.531726747 |
| 4302.278809 | 4850.217285 | 2801.355713 | 0.332282936 | 0.06329053 | -0.188400439 | 0.098811849 | -0.630123768 | 0.133712681 |
| 24810.77539 | 20802.12891 | 21039.44336 | 0.33199515 | 0.134688431 | -0.475743076 | 0.109341875 | -0.420443615 | 0.095160157 |
| 123477.6328 | 130127.3906 | 134452.0156 | 0.331748558 | 0.189940826 | -0.299615529 | 0.216934321 | -0.043490393 | 0.801698552 |
| 442834.5938 | 398318.1563 | 428822.2188 | 0.331365831 | 0.024701492 | -0.134798575 | 0.108469073 | -0.241606995 | 0.053012031 |
| 328228.2383 | 386648.8516 | 369479.1719 | 0.331247866 | 0.249581627 | 0.355810421 | 0.191184742 | 0.218700851 | 0.358051516 |
| 1322.047119 | 1519.376099 | 1414.453003 | 0.331191429 | 0.464094229 | -0.059467653 | 0.893931229 | -0.794354897 | 0.235469026 |
| 71137.78906 | 69623.85938 | 63016.98047 | 0.33078269 | 0.144037739 | -0.142148037 | 0.327625892 | -0.463922908 | 0.023357745 |
| 49475.8125 | 54563.30078 | 49979.53516 | 0.33051026 | 0.028302957 | -0.144194948 | 0.116818656 | 0.266013189 | 0.069207996 |
| 8957.662109 | 11744.71777 | 7848.406738 | 0.329752173 | 0.478874032 | 0.673423142 | 0.178610045 | 0.746814512 | 0.14542672 |
| 58036.24609 | 64735.23047 | 60770.24609 | 0.329509525 | 0.265636509 | 0.013844566 | 0.95424101 | -0.180912027 | 0.484259271 |
| 57449.75391 | 62754.76172 | 55258.65625 | 0.328658683 | 0.138177771 | 0.269599028 | 0.166004104 | 0.277170286 | 0.139519246 |
| 51433.55859 | 57103.09375 | 54057.3125 | 0.328641218 | 0.019930448 | 0.123558738 | 0.078996408 | -0.021776482 | 0.764349564 |
| 136590.4688 | 142626.4688 | 132409.4688 | 0.328360713 | 0.070275485 | 0.284081465 | 0.064588354 | 0.120534904 | 0.208720864 |
| 48728.87109 | 49557.79297 | 52483.30469 | 0.328084272 | 0.094741101 | -0.114918415 | 0.414587246 | -0.340245081 | 0.118023574 |
| 150430.125 | 154864.1406 | 156942.0781 | 0.32776342 | 0.021024291 | -0.072491576 | 0.221991685 | -0.096736873 | 0.132673685 |

| | | | | | | | | |
|---|---|---|---|---|---|---|---|---|
| 40321.14453 | 39071.97266 | 43416.51953 | 0.327718348 | 0.197651318 | 1.14403147 | 0.055241445 | 2.121496517 | 0.009714527 |
| 11254.13867 | 10762.24023 | 7790.719238 | 0.327333837 | 0.271021856 | -0.331574868 | 0.305109592 | -0.32794744 | 0.322805024 |
| 25197.1582 | 21961.7832 | 21724.61328 | 0.327306182 | 0.029075736 | -0.203256502 | 0.080311688 | -0.6949374 | 0.018257145 |
| 199124.7813 | 204352.2813 | 190494.1719 | 0.327278055 | 0.10745682 | -0.080121506 | 0.328944751 | -0.024284313 | 0.716340393 |
| 13539.44336 | 15591.09766 | 10734.92969 | 0.327232623 | 0.316823258 | 0.760796464 | 0.047212306 | 0.243814241 | 0.423557711 |
| 309485.5625 | 308014.9688 | 332438.6875 | 0.326708753 | 0.027158619 | -0.117408357 | 0.166004104 | -0.196402942 | 0.071696088 |
| 993920.875 | 1065471.75 | 1067090.875 | 0.325763023 | 0.046600136 | 0.129418465 | 0.111429162 | 0.253338393 | 0.041088687 |
| 81744.70313 | 88544.91406 | 92147.75 | 0.325653941 | 0.077055795 | -0.569995667 | 0.060234163 | -0.248892768 | 0.119004686 |
| 352735.4688 | 343496.9375 | 385036.0938 | 0.325077206 | 0.142471188 | 0.078065627 | 0.656644615 | 0.285155491 | 0.136201857 |
| 30622.89063 | 31197.29102 | 28978.70898 | 0.324846943 | 0.110302818 | 0.109718773 | 0.137806142 | -0.372235922 | 0.021134419 |
| 65725.97656 | 40598.80859 | 44673.97656 | 0.324797741 | 0.024701492 | -0.09368125 | 0.628940433 | -0.298522977 | 0.383761435 |
| 1813717.51 | 1806528.906 | 1731676.032 | 0.324714594 | 0.03666886 | 0.171512343 | 0.296013798 | -0.008650267 | 0.811284277 |
| 229467.3594 | 249003.6406 | 238980.0781 | 0.323903602 | 0.038502747 | -0.419626178 | 0.021906439 | -0.116378756 | 0.118023574 |
| 21224.39648 | 40194.07031 | 13713.59473 | 0.322707423 | 0.651521079 | 0.350365369 | 0.510981852 | -0.359770357 | 0.639103574 |
| 71694.35156 | 73856.92188 | 71509.00781 | 0.322706598 | 0.030588379 | -0.481190115 | 0.01179292 | -0.317230339 | 0.012551348 |
| 396205.8125 | 378716.2813 | 388927.375 | 0.322515132 | 0.044225724 | -0.046016503 | 0.492413341 | -0.051461839 | 0.377185359 |
| 63533.25 | 64687.57031 | 67019.55469 | 0.322085398 | 0.052584221 | -0.124744226 | 0.23288998 | -0.131447863 | 0.204241692 |
| 3653587 | 4131118.5 | 3888728.75 | 0.321920388 | 0.060948791 | 0.403861453 | 0.007509388 | 0.131165555 | 0.221769091 |
| 22073.26953 | 18690.49023 | 17592.01172 | 0.321274176 | 0.035361197 | 0.252587829 | 0.107911419 | 0.00054187 | 0.997782424 |
| 9216.46582 | 10056.59863 | 7919.338867 | 0.321100447 | 0.034061991 | -0.38867477 | 0.042884533 | -0.326889437 | 0.121306313 |
| 140990.3438 | 165938.2188 | 182363.4375 | 0.320613716 | 0.129251227 | 0.205434006 | 0.300646342 | 0.262036073 | 0.241606904 |
| 3768.961426 | 4328.067871 | 3540.578369 | 0.320440005 | 0.200756526 | 0.38079224 | 0.122107272 | -0.021153548 | 0.922587594 |
| 783286.5273 | 817066.2148 | 821197.3359 | 0.320034782 | 0.087656352 | -0.552547448 | 0.076930836 | -0.467931552 | 0.075097442 |
| 119153.875 | 117487.3438 | 113844.3047 | 0.319878393 | 0.017824776 | -0.351821123 | 0.029468331 | -0.054220739 | 0.233050473 |
| 28042.45898 | 28579.77734 | 25520.76172 | 0.319503807 | 0.052584221 | -0.02612919 | 0.762525545 | -0.23670095 | 0.06365738 |
| 21963.83203 | 22786.11523 | 26682.36914 | 0.318901993 | 0.307058839 | 0.058874407 | 0.786782444 | 0.345881285 | 0.183356293 |
| 25630.08203 | 26574.5918 | 25472.65625 | 0.318648711 | 0.037600114 | -0.422359539 | 0.027031346 | -0.510594945 | 0.018947576 |
| 18570.35742 | 22365.45508 | 20825.51953 | 0.318420018 | 0.63242956 | 0.106292336 | 0.861195218 | -0.135428127 | 0.834815346 |
| 83694.26563 | 84082.03906 | 45647.81641 | 0.318395017 | 0.444866261 | 0.762111632 | 0.100464534 | -0.135520504 | 0.784381383 |
| 60148.42578 | 60657.94141 | 62074.91406 | 0.318134333 | 0.090360627 | -0.062061344 | 0.277926385 | -0.313681293 | 0.034722024 |
| 110427.3438 | 104482.4453 | 95200.21875 | 0.31813052 | 0.076170539 | 0.56103403 | 0.03588199 | 0.855809983 | 0.02592872 |
| 104587.4375 | 103520.4531 | 117727.1484 | 0.318021437 | 0.030693602 | 0.171566904 | 0.184317721 | 0.787266125 | 0.048710777 |
| 3563.170654 | 3933.726563 | 2746.958008 | 0.3179656 | 0.219433007 | 0.337423477 | 0.289417877 | 0.297692598 | 0.32247649 |

| | | | | | | | | |
|---|---|---|---|---|---|---|---|---|
| 96606.82031 | 79816.90625 | 107406.2891 | 0.317856555 | 0.581827098 | 0.739028427 | 0.183896983 | 0.1007506 | 0.849076179 |
| 1193505.25 | 1186705.25 | 1303962.5 | 0.317341594 | 0.301134416 | 0.052214293 | 0.84401622 | 0.253292359 | 0.323998345 |
| 69526.3125 | 55542.82422 | 40290.875 | 0.317188643 | 0.044061266 | -0.145003388 | 0.154482527 | -0.06847007 | 0.837615794 |
| 90678.50781 | 84177.35156 | 75980.38281 | 0.316878891 | 0.05963854 | -0.339230983 | 0.078514758 | -0.564476984 | 0.03271316 |
| 11475.31152 | 17114.86328 | 10325.71484 | 0.316593271 | 0.59905868 | -0.236737789 | 0.417514232 | 0.419312928 | 0.366486609 |
| 35654.84375 | 40600.19922 | 34507.34375 | 0.316545119 | 0.180233605 | -0.197687479 | 0.330185027 | -0.045521483 | 0.803981249 |
| 1263702.125 | 1418371 | 1406332.125 | 0.316220879 | 0.058760777 | -0.039771227 | 0.445857703 | 0.026696 | 0.748712625 |
| 72056.54688 | 66459.48438 | 71637.67188 | 0.315624938 | 0.064507716 | 0.019557337 | 0.676596796 | 0.244353957 | 0.063419062 |
| 46175.61328 | 33548.16016 | 30352.13867 | 0.314665975 | 0.232084106 | 0.325897254 | 0.392329965 | -0.74058757 | 0.084141795 |
| 104378.6797 | 106514.8203 | 96823.42188 | 0.314300723 | 0.312447294 | -0.500213065 | 0.223159833 | -0.737505229 | 0.141180478 |
| 6753.605957 | 6323.369629 | 6388.231934 | 0.313700582 | 0.062978922 | 0.084217507 | 0.32242293 | -0.08592947 | 0.279579647 |
| 251066.7656 | 238142.875 | 261831.5 | 0.313233122 | 0.155839329 | 1.143231499 | 0.284889693 | 2.456076205 | 0.010654184 |
| 110734.3984 | 108452.1641 | 95890.08594 | 0.313210904 | 0.015698773 | -0.247648998 | 0.019141974 | -0.593541628 | 0.029820656 |
| 62050.34375 | 64060.15625 | 67399.78906 | 0.312691202 | 0.199404645 | -0.244292359 | 0.259078883 | -0.071500034 | 0.656827719 |
| 246915.8125 | 244644.3438 | 256457.0625 | 0.312681249 | 0.019930448 | -0.017675292 | 0.68780961 | -0.139052654 | 0.042737273 |
| 60637.48047 | 57911.94141 | 61829.07813 | 0.312143461 | 0.124156216 | -0.366702713 | 0.134252261 | -0.610278884 | 0.071851903 |
| 335273.9375 | 355988.125 | 380574.3438 | 0.311884856 | 0.033350701 | 0.310651681 | 0.050817911 | 0.203811374 | 0.116064096 |
| 120850.7578 | 116317.1563 | 127041.4141 | 0.311464259 | 0.130768956 | -0.160816311 | 0.13871131 | -0.715440524 | 0.023619478 |
| 18138.88672 | 17125.8457 | 17147.21289 | 0.311401302 | 0.231558429 | 0.683208815 | 0.051075754 | 0.510726897 | 0.090965556 |
| 733032.5 | 774439.8125 | 740148.0625 | 0.310366672 | 0.03951042 | -0.493184134 | 0.037014778 | -0.394449082 | 0.044743801 |
| 6284.992676 | 5993.526367 | 4882.495117 | 0.310121145 | 0.293669001 | 0.49722506 | 0.307600544 | 0.130999316 | 0.592495191 |
| 96790.42188 | 119418.6094 | 103212.1016 | 0.309899425 | 0.042404592 | 0.069762801 | 0.374952259 | 0.114005121 | 0.470065733 |
| 257928.8906 | 276856.7188 | 278317.1875 | 0.309784948 | 0.086929104 | -0.092336806 | 0.437819947 | 0.038260125 | 0.726545085 |
| 39443.58984 | 41390.15625 | 43415.98438 | 0.30939335 | 0.067532444 | 0.098264974 | 0.259391884 | 0.033701596 | 0.615352664 |
| 36112.80859 | 39624.50781 | 35612.44531 | 0.309384037 | 0.06480491 | -0.877960673 | 0.027730297 | -0.844808681 | 0.015239677 |
| 89617.82813 | 89427.70313 | 84831.92969 | 0.309267852 | 0.024111712 | 0.396993383 | 0.035031556 | 0.202494541 | 0.040959667 |
| 2091712.125 | 1750859.125 | 2067830.875 | 0.309205918 | 0.025773059 | 0.248719895 | 0.053185315 | 0.508671897 | 0.079566863 |
| 185151.0938 | 178552.3906 | 188433.2344 | 0.308380608 | 0.07907385 | -0.20387045 | 0.147248245 | 0.069403161 | 0.360575146 |
| 22029.78125 | 22754.56055 | 23624.63672 | 0.308320323 | 0.224368093 | 0.401280736 | 0.119067553 | 0.044050279 | 0.815621633 |
| 35856.28516 | 42861.75391 | 16161.27539 | 0.307385942 | 0.033123329 | -1.187983079 | 0.012254503 | -1.4471712 | 0.069955133 |
| 38210.83203 | 44870.80859 | 31490.27148 | 0.307062899 | 0.078882271 | -0.346067779 | 0.07908104 | -0.671986833 | 0.069207996 |
| 228923.4531 | 215586.2031 | 152580.0313 | 0.306799651 | 0.47594001 | 0.091188657 | 0.698316027 | 0.133165767 | 0.647811777 |
| 23645.83203 | 23713.69922 | 23652.7793 | 0.306604098 | 0.173810033 | -0.031154079 | 0.866033702 | 0.046853589 | 0.76812776 |

| | | | | | | | | |
|---|---|---|---|---|---|---|---|---|
| 792016.3125 | 800879.3125 | 809498.25 | 0.306246006 | 0.037309446 | -0.201736782 | 0.101742955 | -0.016162792 | 0.789122922 |
| 126301.6724 | 113416.1421 | 116823.2039 | 0.306164463 | 0.034240061 | -0.108039239 | 0.211934309 | -0.135572674 | 0.172385547 |
| 115117.4141 | 126834.6094 | 118240.375 | 0.305789812 | 0.021353118 | -0.102790599 | 0.346664849 | 0.046598466 | 0.505506227 |
| 57665.28516 | 56603.44922 | 57610.95313 | 0.305658885 | 0.136228366 | 0.039184719 | 0.651152065 | 0.143959908 | 0.179875711 |
| 61670.52734 | 66847.58594 | 56187.48828 | 0.305444801 | 0.180541072 | -0.005556532 | 0.959223724 | -0.212694993 | 0.187569108 |
| 436275.9063 | 266708.4063 | 327711.75 | 0.305242423 | 0.382591812 | 0.613876243 | 0.102963557 | 0.091512096 | 0.813918003 |
| 31217.74609 | 29999.67188 | 30759.93555 | 0.305006331 | 0.08771097 | -0.052898561 | 0.672900092 | 0.367411428 | 0.069207996 |
| 8839.729492 | 8609.911133 | 9285.359375 | 0.304899672 | 0.137237024 | 0.191307832 | 0.219535071 | 0.2094456 | 0.108339667 |
| 458901.7813 | 444727.9375 | 420633 | 0.304620722 | 0.215566621 | 0.624812454 | 0.129412395 | 0.41042022 | 0.110006949 |
| 2587.102295 | 2606.455322 | 2260.899902 | 0.304381338 | 0.285271062 | 0.046400827 | 0.797265661 | -0.122091137 | 0.520924639 |
| 759486.0625 | 713396 | 716066.3125 | 0.304341436 | 0.018310596 | -0.038904611 | 0.54034971 | 0.029837307 | 0.597007015 |
| 189753.8438 | 200269.5313 | 203638.3906 | 0.303820368 | 0.035701769 | -0.108010652 | 0.233503043 | -0.01696436 | 0.801045589 |
| 45863.91406 | 50117.61719 | 50536.50391 | 0.303746995 | 0.097875287 | -0.661183175 | 0.045064018 | -0.344352004 | 0.072839808 |
| 8223.479492 | 6295.677246 | 8209.097656 | 0.30358871 | 0.450224274 | 0.953637842 | 0.062261238 | 0.704817706 | 0.084504925 |
| 173563.1406 | 169115.75 | 165704.6406 | 0.303490669 | 0.045092515 | 0.088943028 | 0.307103937 | 0.1177356 | 0.162206264 |
| 10856.36816 | 9857.850586 | 7820.635742 | 0.303463402 | 0.29607963 | 0.613825474 | 0.168359532 | -0.758936937 | 0.060012809 |
| 217426.0781 | 203621.0156 | 219186.5156 | 0.303358146 | 0.050082631 | 0.753797548 | 0.100538317 | 1.062562526 | 0.009587646 |
| 9513.680664 | 10036.35449 | 8694.003906 | 0.303331264 | 0.27979599 | -0.05121734 | 0.708708221 | 0.390741669 | 0.066292531 |
| 27887.08398 | 27122.28125 | 24979.88086 | 0.303010968 | 0.11689084 | 0.603127849 | 0.04780226 | 0.585138522 | 0.044163517 |
| 164071.5938 | 170175.8594 | 166999.5781 | 0.302750817 | 0.474195126 | -0.545739953 | 0.23503716 | -0.345246616 | 0.359762084 |
| 15353.52344 | 30359.14063 | 11409.42773 | 0.302409791 | 0.639681593 | 0.694225186 | 0.374500771 | -0.491464795 | 0.49278786 |
| 13701.69238 | 13754.12598 | 15457.60059 | 0.301938278 | 0.211214576 | -0.219486845 | 0.344957982 | -0.278823051 | 0.253361125 |
| 8854.990234 | 9185.452148 | 8207.263672 | 0.301903771 | 0.058051058 | 0.022609784 | 0.88997349 | -0.178986146 | 0.117034613 |
| 47110.60938 | 45885.09375 | 42417.00781 | 0.301879989 | 0.07316401 | -0.158220629 | 0.183896983 | -0.159417673 | 0.16248539 |
| 86701.08594 | 88833.35156 | 88890.39063 | 0.301685047 | 0.023818772 | -0.686901215 | 0.0155094 | -0.379205998 | 0.039508617 |
| 62837.94922 | 64789.63672 | 72105.14844 | 0.301587025 | 0.041912796 | -0.696306873 | 0.014164027 | -0.118820037 | 0.281503704 |
| 12939.55273 | 15643.15234 | 15010.6582 | 0.301573214 | 0.360794803 | 0.382297722 | 0.039582042 | 0.871533578 | 0.050152521 |
| 1228481.75 | 1312185.5 | 1316759.5 | 0.301563569 | 0.182751618 | -0.018972808 | 0.917056233 | 0.07070849 | 0.651680124 |
| 63141.11328 | 92307.53906 | 53299.72656 | 0.301345262 | 0.427440307 | -0.078900007 | 0.847462119 | -0.317518967 | 0.450252535 |
| 39323.67578 | 39414.35547 | 37831.4375 | 0.301256655 | 0.173810033 | -0.334817213 | 0.174050429 | -0.913600669 | 0.072468598 |
| 48049.69922 | 47662.89063 | 41501.85156 | 0.301084197 | 0.125127995 | 0.374810691 | 0.019428587 | -0.06489678 | 0.537160122 |
| 27802.04688 | 23524.69531 | 21574.12891 | 0.301001676 | 0.498556044 | 0.340423809 | 0.400065651 | -0.279275615 | 0.544452231 |
| 6456.130859 | 6997.980469 | 6394.192383 | 0.30042151 | 0.064437538 | -0.533442372 | 0.043909427 | -0.097913209 | 0.317922761 |

| | | | | | | | | |
|---|---|---|---|---|---|---|---|---|
| 103401.7969 | 95204.17188 | 104831.3516 | 0.29989991 | 0.065355995 | 0.317693099 | 0.079859428 | 0.140334891 | 0.204241692 |
| 55913.57031 | 51521.84375 | 55847.42969 | 0.299879581 | 0.119441524 | 0.003718465 | 0.967607137 | 0.55331791 | 0.020008995 |
| 6677.487305 | 10563.65234 | 5284.93457 | 0.299486064 | 0.543069772 | 1.20222228 | 0.126413717 | 0.915384888 | 0.247912817 |
| 820915.875 | 938480.875 | 937426.75 | 0.298910218 | 0.104525269 | 0.223905699 | 0.043830603 | 0.417046159 | 0.075839948 |
| 25746.48828 | 24223.55469 | 22860.88281 | 0.298856764 | 0.130732496 | -0.531943258 | 0.049696115 | -0.352534183 | 0.036383418 |
| 44371.35156 | 42490.76172 | 45829.19531 | 0.298730571 | 0.081990933 | 0.42140444 | 0.05467585 | -0.04279437 | 0.655615126 |
| 15811.97266 | 18084.37891 | 13240.1123 | 0.2986577 | 0.052520339 | -0.252433904 | 0.090012942 | -0.048247253 | 0.807833796 |
| 183337.3438 | 199896.9531 | 198512.1406 | 0.298581476 | 0.024227068 | 0.067532374 | 0.234571805 | 0.173862884 | 0.090736213 |
| 212011.2031 | 142480.9531 | 180490.8594 | 0.298459796 | 0.761377655 | 0.860497544 | 0.231902804 | 1.002385759 | 0.139434127 |
| 37251.77344 | 29246.25195 | 34494.15234 | 0.298184539 | 0.214698397 | 0.252639793 | 0.415638095 | -0.196489392 | 0.382092385 |
| 86861.00781 | 90444.10938 | 90921.00781 | 0.29711992 | 0.048820442 | 0.099759313 | 0.355229313 | 0.271464138 | 0.035868549 |
| 23487.10742 | 31614.12695 | 29843.59961 | 0.297000664 | 0.514540705 | -0.804039399 | 0.110375785 | 1.14099038 | 0.060920936 |
| 16345.41016 | 18735.54883 | 16814.79688 | 0.296934463 | 0.258416959 | -0.220517797 | 0.379832076 | 0.243930235 | 0.27345257 |
| 38432.69531 | 31855.0957 | 32224.89648 | 0.296564037 | 0.110302818 | -0.519085095 | 0.066565 | -1.078714631 | 0.02852503 |
| 23143.28125 | 25405.54492 | 26255.72266 | 0.29650957 | 0.100563497 | 0.148998401 | 0.298454935 | -0.068193837 | 0.592078283 |
| 34603.42188 | 35819.46094 | 34167.22656 | 0.295927542 | 0.010297098 | 0.078252232 | 0.252331929 | 0.125980083 | 0.052598684 |
| 288652.7813 | 286931.625 | 280978.5625 | 0.295927297 | 0.155196387 | -0.128111631 | 0.459072637 | -0.210331892 | 0.273434049 |
| 68272.8125 | 68167.38281 | 65883.23438 | 0.295424128 | 0.115723457 | 0.257127055 | 0.158732431 | 0.225377609 | 0.146976714 |
| 137619.7656 | 127985.5156 | 134917.5625 | 0.294917465 | 0.166443617 | 0.034175931 | 0.824276343 | 0.423245176 | 0.060920196 |
| 395392.8438 | 428646.5938 | 437221.25 | 0.294722745 | 0.031284382 | 0.290986199 | 0.01179292 | 0.50297046 | 0.04559071 |
| 16924.85742 | 19648.69141 | 20203.72852 | 0.294415397 | 0.087656352 | 0.083320232 | 0.255783923 | 0.308618511 | 0.1278105 |
| 45344.05078 | 51464.71875 | 43607.71094 | 0.294109987 | 0.056312464 | -0.757556945 | 0.025293799 | -0.558758036 | 0.027949491 |
| 117888.1406 | 199823.7813 | 187409.1406 | 0.293998834 | 0.302727482 | -0.260067525 | 0.369856356 | 0.136398361 | 0.718252335 |
| 30640.32813 | 32508.02539 | 42658.39453 | 0.293724261 | 0.030755796 | -0.422433515 | 0.026392799 | -0.337996305 | 0.213824986 |
| 9482.00293 | 13847.91113 | 8373.832031 | 0.2933972 | 0.436175859 | -0.471826748 | 0.318729418 | -0.497594017 | 0.305016128 |
| 63101.5 | 54550.18359 | 50590.35547 | 0.29332402 | 0.239971035 | -0.855487945 | 0.103637757 | -0.760469158 | 0.081190238 |
| 131583.3721 | 127225.373 | 115908.1406 | 0.293206039 | 0.132098822 | 0.203225073 | 0.205468655 | 0.192769793 | 0.212530411 |
| 13347.14063 | 13140.77734 | 9729.104492 | 0.293026128 | 0.182944903 | -0.108043594 | 0.518876214 | -0.331780027 | 0.207777277 |
| 34924.08203 | 40327.875 | 40467.35156 | 0.292743773 | 0.247825184 | 0.021588734 | 0.922415374 | 0.421499746 | 0.107540112 |
| 1957.71875 | 2993.981934 | 1784.617798 | 0.292417998 | 0.48317398 | -0.388751055 | 0.103460535 | -0.593880774 | 0.176516532 |
| 20958.99023 | 18925.27148 | 16652.48242 | 0.292380037 | 0.115723457 | -0.223309499 | 0.205568001 | -0.104467304 | 0.487243302 |
| 62959.92969 | 56752.75781 | 57877.19531 | 0.292319712 | 0.094031642 | 0.451652 | 0.01179292 | 0.095852111 | 0.276951269 |
| 1401554.938 | 1499814.594 | 1448992.359 | 0.2923121 | 0.014306629 | -0.130859376 | 0.129861867 | 0.081315103 | 0.171025182 |

| | | | | | | | | |
|---|---|---|---|---|---|---|---|---|
| 273256.125 | 334888.2188 | 366894.875 | 0.292309926 | 0.221505574 | -0.179438913 | 0.40775843 | 0.490914254 | 0.125742768 |
| 8016.913574 | 9735.227539 | 11393.30371 | 0.292019904 | 0.491789048 | -0.633795803 | 0.143043237 | -1.659285076 | 0.062345205 |
| 97514.3125 | 114534.3125 | 124596.5313 | 0.291965664 | 0.603255079 | 0.199776215 | 0.686661605 | 0.855705014 | 0.124886255 |
| 328435.0313 | 328286.8125 | 352977.6563 | 0.291926884 | 0.044665667 | -0.373461561 | 0.020431201 | -0.419638653 | 0.017015337 |
| 475458.75 | 479604.5313 | 462984.2188 | 0.291713628 | 0.033164349 | 0.089299151 | 0.337901514 | 0.225653234 | 0.017983315 |
| 21462.20898 | 23973.84961 | 22163.56055 | 0.291661349 | 0.195658833 | -0.548866025 | 0.081663959 | -0.56006406 | 0.084114644 |
| 9186.521484 | 9219.600586 | 5819.626953 | 0.291604356 | 0.065637909 | -0.429923797 | 0.098459633 | -0.667599636 | 0.095764715 |
| 54468.75391 | 51960.12891 | 51375.83984 | 0.291557687 | 0.120011001 | -0.166557029 | 0.126731731 | -0.240953206 | 0.070298132 |
| 34043.49219 | 37006.57813 | 36180.48438 | 0.291417067 | 0.136407599 | 0.000535742 | 0.996723914 | -0.010112081 | 0.945888168 |
| 330487.6875 | 383090.6875 | 375410.9375 | 0.291195296 | 0.130960943 | 0.083845907 | 0.60181352 | 0.145655279 | 0.350351437 |
| 451039.375 | 464872.875 | 479796.6563 | 0.290742766 | 0.006395507 | -0.126578159 | 0.066788385 | -0.133862033 | 0.065710106 |
| 16600.38867 | 17226.31836 | 16692.08789 | 0.29059896 | 0.527167215 | 0.173627005 | 0.683405915 | 0.260161931 | 0.515590007 |
| 73783.71875 | 62658.80469 | 70366.08594 | 0.290568374 | 0.058516905 | 0.260579098 | 0.081402616 | 0.292357295 | 0.111977423 |
| 5587.599609 | 4973.575684 | 3908.46582 | 0.290270518 | 0.258416959 | -0.652184183 | 0.111349456 | -1.220506539 | 0.049004518 |
| 3910.469238 | 4113.328125 | 1931.030762 | 0.289995469 | 0.380762215 | -0.424463962 | 0.289928504 | -0.518167834 | 0.301475535 |
| 44546.37109 | 43107.52734 | 41740.70703 | 0.289866143 | 0.02930047 | 0.162939769 | 0.097735536 | 0.627974739 | 0.011671599 |
| 54725.93359 | 58146.02734 | 57464.60547 | 0.289545875 | 0.097116923 | -0.079767832 | 0.520920322 | 0.113665515 | 0.334263521 |
| 17341.33398 | 18000.46484 | 23062.94922 | 0.288567946 | 0.701331836 | 0.625301436 | 0.349097027 | 0.738983311 | 0.260165601 |
| 92561.54688 | 89538.61719 | 89484.25781 | 0.288440111 | 0.027581684 | -0.059418545 | 0.302578553 | 0.135804287 | 0.076920623 |
| 33030.73828 | 33911.35547 | 34778.15625 | 0.288217033 | 0.311243541 | -0.072408851 | 0.767412356 | -0.19965066 | 0.45643538 |
| 56924.84375 | 57863.01563 | 54845.06641 | 0.287843248 | 0.04127274 | 0.275683038 | 0.136994383 | -0.225828004 | 0.035612683 |
| 800293.8125 | 925218.625 | 822175.125 | 0.287826736 | 0.079968566 | -0.041916857 | 0.70926014 | -0.065172253 | 0.604668962 |
| 456162.8438 | 494359.9063 | 495896.1563 | 0.287607623 | 0.156419108 | -0.106720946 | 0.138536351 | -0.072378199 | 0.317050668 |
| 84219.63281 | 83292.96094 | 89206.17188 | 0.286742938 | 0.073571365 | 0.096602198 | 0.415338319 | 0.127905271 | 0.101114032 |
| 141969.6094 | 135403.9063 | 142587.5156 | 0.286637979 | 0.105830502 | -0.127547127 | 0.329940419 | 0.287769053 | 0.085430016 |
| 21989.3457 | 25284.70508 | 23288.72461 | 0.285784922 | 0.016596319 | -0.119902373 | 0.325827927 | -0.159476293 | 0.166647679 |
| 30137.94727 | 28586.12109 | 27615.01758 | 0.285738834 | 0.084947579 | 0.05441833 | 0.4202179 | -0.166681966 | 0.104365856 |
| 19990.35742 | 17992.03516 | 14019.5752 | 0.285726021 | 0.141949735 | -1.539006853 | 0.048820023 | -1.030552863 | 0.030007011 |
| 188970.5781 | 206579.4375 | 211764.7969 | 0.285652344 | 0.065789073 | 0.184128244 | 0.181741873 | 0.256669691 | 0.07832051 |
| 3754.25708 | 5109.105469 | 5077.486328 | 0.285431517 | 0.687239876 | 1.305957909 | 0.042884533 | 1.063631353 | 0.062838928 |
| 49388.45313 | 45512.84766 | 49664.78906 | 0.284622596 | 0.072654291 | 0.001038306 | 0.99201113 | 0.079003339 | 0.37741689 |
| 168738.0781 | 173031.25 | 181108.2969 | 0.284579693 | 0.163060402 | -0.958817767 | 0.01915096 | 0.138996308 | 0.0890834 |
| 109096.0156 | 88046.44531 | 73829.625 | 0.284574366 | 0.573143374 | 1.244637595 | 0.315613662 | 0.722744445 | 0.129291758 |

| | | | | | | | | |
|---|---|---|---|---|---|---|---|---|
| 815969.6875 | 916029.8125 | 932200.4375 | 0.284556016 | 0.017029724 | -0.12770959 | 0.208496317 | 0.162824133 | 0.205658715 |
| 65178.40625 | 65311.41016 | 66845.03906 | 0.28446857 | 0.093994529 | 0.372279128 | 0.078053315 | -0.395442373 | 0.095599918 |
| 16666.22852 | 13139.05762 | 12445.38477 | 0.284443505 | 0.254204903 | -0.192130577 | 0.423580857 | -0.344516759 | 0.237683126 |
| 1221109.375 | 1169674 | 1230053.875 | 0.283681176 | 0.030223639 | -1.057851728 | 0.012254503 | -0.169507185 | 0.03581165 |
| 93998.25781 | 109281.8594 | 128853.8984 | 0.283049973 | 0.192885661 | -0.011733839 | 0.880969998 | -0.077492185 | 0.69422618 |
| 3655.755127 | 5372.608887 | 3803.970947 | 0.282972623 | NA | -0.332575342 | NA | -0.244227438 | NA |
| 33371.66016 | 35192.25781 | 20367.98242 | 0.282268308 | 0.748348017 | 1.505658985 | 0.155243406 | -0.42361059 | 0.658231333 |
| 9339.113281 | 7308.31543 | 7494.53418 | 0.282088692 | 0.092864964 | -0.264221303 | 0.153891574 | -0.156324571 | 0.410415738 |
| 43003.67578 | 46156.10156 | 46367.81641 | 0.281884297 | 0.133144067 | 0.18965887 | 0.175752242 | -0.007986325 | 0.948264111 |
| 10601.80566 | 9456.40625 | 7750.683594 | 0.281270361 | 0.417614366 | -0.197353529 | 0.520977849 | -0.011601787 | 0.9709651 |
| 47793.39844 | 50637.97266 | 52401.23828 | 0.281234598 | 0.074984572 | 0.315121509 | 0.05975961 | -0.105785917 | 0.304244609 |
| 95023.48535 | 104666.0391 | 105670.8364 | 0.28082066 | 0.196593962 | 0.226587339 | 0.212463985 | 0.378372919 | 0.077576039 |
| 5952.486816 | 7664.399902 | 2450.323486 | 0.280636573 | 0.767721713 | 0.365589925 | 0.660062477 | -0.430123876 | 0.703463561 |
| 26245.75781 | 23263.14453 | 18436.93555 | 0.280587734 | 0.566690873 | -0.534922918 | 0.392329965 | -0.566099385 | 0.361751877 |
| 88100.04688 | 95891.17188 | 96265.82813 | 0.279788757 | 0.133330992 | -0.013155196 | 0.920451359 | 0.250118456 | 0.107790858 |
| 1360841.125 | 1284486 | 1194265.5 | 0.279716514 | 0.015440135 | -0.010633211 | 0.922008276 | -0.013871332 | 0.870814221 |
| 4179.433594 | 4898.986328 | 3609.944092 | 0.27944495 | 0.318687987 | 0.41634901 | 0.154889752 | -1.169745026 | 0.019880539 |
| 33649.85547 | 30176.40234 | 27980.0918 | 0.279389907 | 0.062609 | -0.00621146 | 0.943583387 | 0.155269668 | 0.299587425 |
| 524466.875 | 647881 | 605332.5625 | 0.27936341 | 0.166443617 | -0.089736174 | 0.559205077 | 0.197803531 | 0.296423725 |
| 21039.64844 | 5803.372559 | 19535.08398 | 0.279215595 | 0.709777983 | 1.369312922 | 0.344912699 | 0.757179693 | 0.433601327 |
| 211981.5313 | 213844.3594 | 218626.2188 | 0.278619498 | 0.035879745 | -0.199720978 | 0.045660829 | -0.177923676 | 0.031136253 |
| 248134.1719 | 277883.4063 | 229101.3594 | 0.278251037 | 0.404110576 | -0.31570325 | 0.391636489 | -1.084428595 | 0.120108192 |
| 74476.64063 | 83351.8125 | 69133.39844 | 0.278212196 | 0.280139443 | -0.947835458 | 0.104541915 | -0.786557761 | 0.094638121 |
| 15930.48438 | 15845.4541 | 14994.79199 | 0.277995579 | 0.166243149 | -0.386761051 | 0.089652719 | -0.168424959 | 0.233037555 |
| 318534.2813 | 306399.125 | 295493.75 | 0.277983047 | 0.141002246 | 0.280138144 | 0.104928475 | 0.483879723 | 0.039195867 |
| 288466.5313 | 292493.9063 | 308679.1875 | 0.277626404 | 0.046688926 | 0.014710946 | 0.834026179 | 0.126312099 | 0.147966231 |
| 29277.27148 | 24849.51953 | 21143.53125 | 0.277532875 | 0.024037284 | -0.504836159 | 0.118860675 | -0.751866084 | 0.053012031 |
| 70971.02344 | 81648.625 | 86417.60938 | 0.277065186 | 0.124546636 | -0.356014604 | 0.052965881 | 0.105964453 | 0.459242178 |
| 26200.77539 | 23888.57813 | 24434.62695 | 0.276730334 | 0.110984157 | 0.433297135 | 0.082329519 | -0.201399889 | 0.196327908 |
| 306563.8438 | 417690.625 | 326856.6875 | 0.276434202 | 0.624643671 | 0.042207978 | 0.917495271 | -0.062996603 | 0.867909164 |
| 159081.2969 | 129038 | 145289.3594 | 0.27639597 | 0.2756587 | 0.16994141 | 0.429253521 | -0.23126202 | 0.363640088 |
| 106105.2188 | 109053.9453 | 128494.3047 | 0.276323288 | 0.122520185 | -0.33827832 | 0.129745502 | -0.378311816 | 0.09056925 |
| 36738.16797 | 40760.34375 | 40025.14844 | 0.276222188 | 0.298788798 | 0.412590469 | 0.142579096 | 0.473413865 | 0.104817936 |

| | | | | | | | | |
|---|---|---|---|---|---|---|---|---|
| 19781.70703 | 19763.5918 | 19301.375 | 0.275061365 | 0.083539806 | 0.370939813 | 0.056841901 | 0.165412421 | 0.157967349 |
| 166839.4844 | 158370.3438 | 146532.1406 | 0.274981948 | 0.011384748 | -0.213935238 | 0.026171282 | -0.314672639 | 0.056784127 |
| 4804.95166 | 2999.57959 | 2695.384033 | 0.274444727 | 0.53531189 | 0.529888553 | 0.193402284 | 0.699974135 | 0.266209604 |
| 292580.5938 | 288872.3125 | 296514.3125 | 0.274347133 | 0.142694155 | 0.225758029 | 0.184317721 | 0.439034188 | 0.055019141 |
| 46730.59766 | 40325.40234 | 42428.32031 | 0.274345544 | 0.205630433 | -1.685556446 | 0.020651809 | -0.560675847 | 0.02163333 |
| 13468.72168 | 12165.07031 | 12736.53613 | 0.274199113 | 0.113705211 | 0.160252537 | 0.218789993 | -0.010387163 | 0.932859665 |
| 33182.58984 | 31438.40234 | 34982.53906 | 0.274177598 | 0.136100201 | -0.462721959 | 0.11146465 | -0.265548036 | 0.154013101 |
| 13450.95117 | 21959.16406 | 1365.398071 | 0.274116908 | 0.694286699 | 0.827213569 | 0.273114108 | -0.52934382 | 0.620264646 |
| 18107.25195 | 19040.83008 | 12919.44629 | 0.27408699 | 0.084757955 | 0.945947751 | 0.015540203 | 0.884825114 | 0.121306313 |
| 66944.89063 | 65315.11328 | 60188.30859 | 0.27354269 | 0.159684245 | -0.681494586 | 0.082612243 | -0.322217291 | 0.132620714 |
| 11979245 | 12408517 | 14130602 | 0.27304414 | 0.059908598 | -0.267862743 | 0.056841901 | -0.231666742 | 0.134102681 |
| 79680.72656 | 63011.35547 | 69993.29688 | 0.272697739 | 0.252715703 | 0.553922456 | 0.024939682 | 0.040756855 | 0.798323297 |
| 58932 | 62782.22656 | 61127.27734 | 0.272313447 | 0.461795637 | 0.070093945 | 0.834109875 | -0.142493542 | 0.684678501 |
| 73757.53125 | 74744.8125 | 78714.0625 | 0.272033495 | 0.117342707 | 0.132459475 | 0.08807311 | 0.108750349 | 0.129572086 |
| 312187.75 | 321873.2813 | 341495.2188 | 0.271970637 | 0.249779609 | 0.576983809 | 0.089359332 | 1.169738708 | 0.015632484 |
| 335921.5313 | 353119.8438 | 349034.0938 | 0.271519178 | 0.021745834 | -0.206894567 | 0.054857325 | -0.226300622 | 0.0236713 |
| 991146.875 | 1021900.188 | 1029383.375 | 0.270809566 | 0.064028623 | 0.061353796 | 0.506109273 | 0.032461426 | 0.646553159 |
| 16215.99902 | 15774.52148 | 12753.79492 | 0.27074314 | 0.471239653 | -0.789090101 | 0.210737089 | -0.934299503 | 0.161199638 |
| 38192.24219 | 45258.78516 | 39856.64453 | 0.270516998 | 0.090992566 | -0.171962773 | 0.207662416 | -0.005189369 | 0.972680155 |
| 115755.9453 | 115584.5313 | 105649.2813 | 0.270397259 | 0.260794226 | -0.034665767 | 0.878990407 | 0.425289707 | 0.077915852 |
| 5754.146484 | 4316.95166 | 3384.361328 | 0.269866799 | 0.388207029 | 0.399246155 | 0.267197239 | 0.094574438 | 0.817449237 |
| 15492.82324 | 12216.41797 | 9914.731445 | 0.269324826 | 0.152033568 | -0.563293715 | 0.234571805 | -0.221656605 | 0.441917671 |
| 78481.20313 | 82757.5625 | 79103.07813 | 0.269213008 | 0.136902686 | -0.322459934 | 0.133305993 | -0.202871285 | 0.213076586 |
| 24665.55078 | 23830.24805 | 23015.90234 | 0.269079011 | 0.274371306 | 0.215356452 | 0.330637946 | 0.096261608 | 0.623632122 |
| 472976.5625 | 462536.1563 | 444256.1563 | 0.269021432 | 0.028316583 | 0.129645383 | 0.085412183 | 0.411311982 | 0.017279709 |
| 58078.78906 | 59607.78906 | 56464.5 | 0.268143783 | 0.246087007 | 0.297503528 | 0.208916626 | 0.350866922 | 0.14724171 |
| 30867.78125 | 29771.07227 | 25130.51953 | 0.267942833 | 0.182780868 | 0.132387889 | 0.333825431 | 0.003645513 | 0.984470463 |
| 186900.0156 | 180164.8281 | 175755.4375 | 0.267749763 | 0.265384605 | 0.282080395 | 0.217458834 | 0.052846446 | 0.782159539 |
| 70269.80469 | 71983.8125 | 71353.88281 | 0.26764692 | 0.081995281 | 0.1311425 | 0.038246759 | -0.019966136 | 0.466873769 |
| 3149.414063 | 4351.261719 | 3851.338135 | 0.267354461 | 0.50068146 | 0.019070523 | 0.909158577 | 0.154392336 | 0.518914722 |
| 122766.3125 | 155289.9531 | 139026.2188 | 0.267319496 | 0.463957081 | -0.370161726 | 0.374447663 | -0.226890993 | 0.531406693 |
| 1039203.75 | 1039854.25 | 1080002.125 | 0.266966291 | 0.053424232 | 0.011039321 | 0.805915462 | 0.01515183 | 0.623768537 |
| 65122.74609 | 66816.79688 | 65721.65625 | 0.266782558 | 0.066021112 | -0.247931886 | 0.045588559 | -0.120176057 | 0.02183874 |

| | | | | | | | | |
|---|---|---|---|---|---|---|---|---|
| 282885.3125 | 274307.1563 | 279450.4063 | 0.266395557 | 0.132174911 | -0.854886327 | 0.014119111 | 0.28297161 | 0.020351204 |
| 563995.25 | 571741.625 | 552201.6875 | 0.266369391 | 0.162383379 | 0.466188331 | 0.058498389 | 0.490050824 | 0.048505224 |
| 17527.70508 | 18672.25781 | 15820.97656 | 0.265806783 | 0.261463595 | -0.20007109 | 0.391455472 | -0.084474143 | 0.687534966 |
| 80332.08594 | 89668.71875 | 95923.67969 | 0.26577598 | 0.085651859 | -0.053577709 | 0.63549743 | -0.348373677 | 0.066292531 |
| 208081.625 | 215805.9063 | 203520.6094 | 0.265732136 | 0.540918348 | -0.134443424 | 0.758616558 | -1.150933053 | 0.16847165 |
| 3466.628418 | 3695.234131 | 2890.46167 | 0.265540149 | 0.351526078 | 0.178661823 | 0.302578553 | -0.049152598 | 0.808452938 |
| 17363.33398 | 20522.3125 | 17118.98828 | 0.264531604 | NA | 0.416500192 | 0.17813923 | 0.365556515 | 0.207863504 |
| 162035.1406 | 167757.9063 | 155189.7813 | 0.264290908 | 0.045235089 | 0.302118533 | 0.085441567 | 0.458289941 | 0.020008995 |
| 54401.75781 | 51002.5 | 50183.03125 | 0.264285739 | 0.106647829 | 0.182834465 | 0.221183294 | 0.036950202 | 0.712863184 |
| 8061.997559 | 10049.02832 | 10341.80176 | 0.263683708 | 0.670827091 | -0.772723352 | 0.361167013 | -0.021451021 | 0.973455039 |
| 10867.30371 | 11595.81055 | 11539.02148 | 0.263521476 | 0.240223025 | -0.105559649 | 0.587433369 | -0.360733004 | 0.191130479 |
| 21168.12109 | 32453.35156 | 12261.4209 | 0.26295721 | NA | 0.261518031 | 0.52544407 | -0.368103311 | 0.553731254 |
| 47773.45313 | 45732.13672 | 43300.22656 | 0.262878336 | 0.120564107 | -0.019222087 | 0.876081794 | -0.114406378 | 0.3849966 |
| 25999.24023 | 23590.40039 | 22519.53711 | 0.262857209 | 0.148857765 | -0.027099515 | 0.834109875 | -0.325273814 | 0.106751573 |
| 390518.75 | 422782.7188 | 420757.5938 | 0.262734596 | 0.060388606 | -0.101318384 | 0.348633551 | -0.426001101 | 0.036848746 |
| 5960.856445 | 6438.535156 | 3732.284668 | 0.262277412 | 0.404110576 | -0.472422181 | 0.278495147 | -0.95106482 | 0.09056925 |
| 945792.9375 | 963851.5625 | 979702.375 | 0.261296796 | 0.057894371 | 0.046634164 | 0.348677945 | 0.109532707 | 0.089381209 |
| 130249.9688 | 127188.1328 | 130282.4297 | 0.261248468 | 0.1152822 | 0.164164655 | 0.276825942 | 0.229037796 | 0.143658547 |
| 25918.75 | 27191.00977 | 25726.04688 | 0.261245307 | 0.832358028 | 0.947351924 | 0.342370709 | 0.435977023 | 0.669450319 |
| 37151.74609 | 30986.46484 | 33774.23828 | 0.261207025 | 0.134144764 | 0.198930292 | 0.132066965 | 0.124848242 | 0.377609807 |
| 88242.70313 | 95430.45313 | 86781.25 | 0.26038385 | 0.07362475 | 0.084040328 | 0.359166221 | -0.158106155 | 0.170158914 |
| 6131.96875 | 5900.484863 | 3827.7854 | 0.260234206 | 0.247683013 | -0.063651206 | 0.727571165 | -0.42644367 | 0.213264642 |
| 87630 | 90365.67188 | 61450.82422 | 0.260221521 | 0.54532205 | 0.085273627 | 0.666605955 | 0.142037317 | 0.609610248 |
| 85172.07031 | 75714.35938 | 78970.82031 | 0.260110126 | 0.043122548 | 0.165658194 | 0.044791528 | -0.011377886 | 0.885245898 |
| 230306.9063 | 235055.7969 | 238956.8906 | 0.259515103 | 0.056527689 | 0.554643304 | 0.012254503 | 0.214443685 | 0.034677634 |
| 3432.800781 | 3402.88208 | 2435.026611 | 0.259480093 | 0.283623624 | 0.597759786 | 0.161989613 | -0.218564172 | 0.414213561 |
| 145980.5156 | 148602.7031 | 171488.5156 | 0.258632058 | 0.046164606 | -0.005439815 | 0.942648968 | -0.029301154 | 0.815420178 |
| 43535.47266 | 41005.25781 | 38874.30078 | 0.258544823 | 0.296551702 | 0.478843045 | 0.055527809 | 0.584685534 | 0.024635256 |
| 21569.96484 | 17692.66211 | 16532.06641 | 0.258513512 | 0.401743865 | -0.234471905 | 0.444625358 | -0.204009097 | 0.508210416 |
| 11446.53613 | 10686.90332 | 11603.80273 | 0.25773209 | 0.240333703 | -0.313649918 | 0.204377767 | -0.038114291 | 0.826852205 |
| 113568.25 | 109087.4688 | 116325.6172 | 0.257309989 | 0.054728089 | -0.62810981 | 0.027031346 | -0.260535234 | 0.06479708 |
| 30391.70508 | 26292.90625 | 19672.88867 | 0.25727709 | 0.088231376 | -1.268858047 | 0.032222933 | -1.103095283 | 0.027331143 |
| 13768.81152 | 13164.88867 | 14071.19336 | 0.257105322 | 0.102165776 | -0.0586744 | 0.564994633 | 0.376225001 | 0.031136253 |

| | | | | | | | | |
|---|---|---|---|---|---|---|---|---|
| 58073.74609 | 63112.16797 | 66025.23438 | 0.256908332 | 0.210744849 | -0.364783362 | 0.150271527 | -0.10634661 | 0.530924104 |
| 5929.542969 | 6766.071289 | 3760.223877 | 0.256656525 | 0.164604942 | 0.228181788 | 0.01179292 | -0.064577921 | 0.854507456 |
| 10730.53906 | 9483.53418 | 8855.924805 | 0.256418032 | 0.313186894 | -0.472486404 | 0.089248231 | -0.740277746 | 0.029738889 |
| 86220.47656 | 78381.73438 | 97581.67969 | 0.256149567 | 0.267565535 | 0.754539981 | 0.107208456 | 0.509867504 | 0.082022058 |
| 791764.3125 | 635534.0625 | 750117.5 | 0.256040186 | 0.192968202 | 0.070633933 | 0.61125836 | -0.284168609 | 0.178411577 |
| 108936.9063 | 102675.6953 | 95542.46094 | 0.255821132 | 0.072163652 | 0.326579176 | 0.017037525 | 0.315793432 | 0.090965556 |
| 321006.75 | 240664.3281 | 265057.9688 | 0.255691094 | 0.112665115 | -0.270479286 | NA | -0.386261909 | 0.118033234 |
| 812326.875 | 850914.375 | 784426.9375 | 0.255679323 | 0.030661161 | 0.043818853 | 0.268818061 | 0.13771966 | 0.109799791 |
| 157138.2813 | 171459.2031 | 175063.4063 | 0.255503039 | 0.056964374 | 0.22821081 | 0.052965881 | 0.1318081 | 0.182273504 |
| 25145.72266 | 25583.93359 | 32981.42578 | 0.255489247 | 0.697160003 | 0.915072138 | 0.162317825 | 0.926661246 | 0.122834452 |
| 278299.5625 | 267919.4063 | 250778.6563 | 0.255127134 | 0.270976146 | 0.5345138 | 0.361768019 | 0.007145406 | 0.973455039 |
| 73624.84375 | 74081.76563 | 74285.47656 | 0.25511551 | 0.069867455 | 0.192314541 | 0.084935024 | 0.130307044 | 0.161137259 |
| 23740.55664 | 24866.48828 | 24644.12109 | 0.25485447 | 0.380762215 | 0.105021666 | 0.681538923 | 0.449314232 | 0.160773455 |
| 27639.31445 | 26535.4707 | 23767.99609 | 0.254139864 | 0.131627208 | -0.043347337 | 0.725035302 | 0.078905631 | 0.566086724 |
| 100501.7422 | 130613.6875 | 143100.8438 | 0.253043614 | 0.268099056 | 0.461773875 | 0.103398614 | 0.734203598 | 0.107554917 |
| 19158.81445 | 12668.93555 | 29841.30664 | 0.252891961 | NA | -0.296814802 | NA | -0.05946247 | NA |
| 105691912 | 85314696 | 141416784 | 0.252170655 | 0.767721713 | 0.290295443 | 0.686184966 | 0.895413488 | 0.209039618 |
| 396995.2813 | 427369 | 430931.2188 | 0.252152626 | 0.070360777 | 0.071031652 | 0.469567595 | 0.342260661 | 0.036013501 |
| 24826.17773 | 25800.86328 | 23648.55664 | 0.25152484 | 0.248943392 | 0.142591556 | 0.435555266 | -0.164962724 | 0.410938317 |
| 19072.46094 | 25188.88477 | 25241.18359 | 0.251295293 | 0.480710218 | 0.457409908 | 0.147323941 | 0.369861786 | 0.224000425 |
| 238457.1875 | 248210.4375 | 233826.1406 | 0.251113436 | 0.044665667 | -0.344119788 | 0.036578039 | -0.177724273 | 0.075097442 |
| 58167.12891 | 58996.67188 | 58177.58594 | 0.251051534 | 0.056093567 | -0.390248406 | 0.032996993 | 0.037378295 | 0.387223764 |
| 27507.02734 | 23226.82227 | 24376.76953 | 0.250856052 | 0.314306229 | 0.236693315 | 0.378072237 | 0.507957742 | 0.073175112 |
| 39685.8125 | 42927.28906 | 43927.76172 | 0.250499988 | 0.151648018 | -0.175271387 | 0.034038555 | 0.075800644 | 0.340777735 |
| 11367.42676 | 4987.752441 | 9034.053711 | 0.250458836 | 0.476745676 | 0.490373202 | 0.299034856 | 0.243828317 | 0.663092542 |
| 250835.3906 | 234138.6875 | 234391.6328 | 0.250272067 | 0.158200263 | 0.097587351 | 0.445857703 | -0.112176629 | 0.377470755 |
| 1009426.75 | 976869.1875 | 1042336.813 | 0.249716711 | 0.079619196 | -0.514947658 | 0.028225257 | -0.066756851 | 0.225833549 |
| 17783.57031 | 19253.24219 | 15861.54492 | 0.249316426 | 0.026060442 | 0.015508423 | 0.808922116 | -0.384049419 | 0.066846273 |
| 525781 | 492068.4375 | 521904.375 | 0.248841914 | 0.130960943 | -0.091872708 | 0.369856356 | 0.001678524 | 0.979696342 |
| 15509.76758 | 13489.02832 | 14705.0332 | 0.248709251 | 0.195658833 | -0.262190868 | 0.036355545 | -0.368611531 | 0.042175609 |
| 404796.8125 | 551773.4375 | 568572.4375 | 0.248707706 | 0.194423598 | -0.927902179 | 0.043906958 | -0.29908373 | 0.233826063 |
| 24748.83398 | 23485.75 | 22569.88086 | 0.248055888 | 0.196367569 | 0.096102233 | 0.594628825 | 0.246207111 | 0.063450205 |
| 34845.55078 | 36288.51563 | 39377.03906 | 0.247208055 | 0.038502747 | 0.14732854 | 0.086431911 | 0.661741385 | 0.03418844 |

| | | | | | | | | |
|---|---|---|---|---|---|---|---|---|
| 81454.72656 | 12604.08691 | 53989.20313 | 0.246769164 | 0.719961518 | 0.907850464 | 0.297032792 | 0.165246317 | 0.875423733 |
| 129712.5469 | 140386.2813 | 139488.6719 | 0.245844416 | 0.109833334 | -0.146816776 | 0.286559326 | 0.051426386 | 0.618929829 |
| 422202.5313 | 474968.25 | 485928.5 | 0.245429038 | 0.258416959 | -0.266906651 | 0.27175302 | -0.480631747 | 0.121683988 |
| 24776.08008 | 24599.14063 | 15695.77637 | 0.245016149 | 0.228580711 | -0.584719061 | 0.086909686 | -0.913069243 | 0.048710777 |
| 139593.7031 | 153216.75 | 140215.0156 | 0.244901056 | 0.03279857 | 0.203432336 | 0.051411338 | -0.155982919 | 0.113977669 |
| 510492.4375 | 547833.875 | 550674.0625 | 0.244889123 | 0.092740746 | -0.10119167 | 0.218095208 | -0.162639226 | 0.083625755 |
| 113882.6406 | 113866.6406 | 106973.0859 | 0.244705966 | 0.063884892 | -0.563312743 | 0.007509388 | -0.583284202 | 0.017015337 |
| 9080.695313 | 11817.55371 | 5443.452148 | 0.244361338 | NA | 0.383615491 | NA | 0.339477522 | NA |
| 30821.13672 | 30664.16016 | 37637.63281 | 0.244016684 | 0.25332178 | 0.341112214 | 0.13116248 | 0.509324556 | 0.087970617 |
| 19502.26172 | 17727.32031 | 17514.16992 | 0.243884456 | 0.166087729 | 0.210346317 | 0.188452025 | -0.030889171 | 0.821324688 |
| 764905 | 764126.8125 | 683138.875 | 0.243862385 | 0.039781174 | -0.19324053 | 0.072221754 | -0.065345961 | 0.454490742 |
| 251017.25 | 253828.125 | 243910.5 | 0.243820568 | 0.051610988 | 0.114055628 | 0.116818656 | 0.46524181 | 0.010236624 |
| 764139.125 | 823706.5625 | 756813.8125 | 0.243790641 | 0.405810916 | 0.319198156 | 0.071935047 | -0.549307729 | 0.038904509 |
| 10307.79492 | 11481.75977 | 9717.649414 | 0.243415975 | 0.271759161 | 0.102976889 | 0.550128659 | 0.15511777 | 0.40758312 |
| 80247.57031 | 86289.47656 | 92559.97656 | 0.243302461 | 0.164136479 | -0.217621446 | 0.199074765 | -0.089687319 | 0.523666735 |
| 412061.5 | 454405.0625 | 437680.6563 | 0.243301914 | 0.110984157 | -0.169681336 | 0.024413553 | -0.1952203 | 0.076969062 |
| 10068.79199 | 8102.614258 | 8088.076172 | 0.242937868 | 0.48584679 | -0.483421516 | 0.065266848 | -0.264401623 | 0.190240855 |
| 31383.19727 | 36444.81641 | 29473.31641 | 0.242647505 | 0.296437226 | -0.134363818 | 0.265347062 | -0.256281541 | 0.151776851 |
| 11469.37012 | 20805.19922 | 18626.29688 | 0.242560601 | 0.725013942 | 0.878130671 | 0.142784851 | 0.402576576 | 0.488353105 |
| 4866.26709 | 5952.493652 | 3634.628174 | 0.242434489 | 0.172140645 | 0.026408394 | 0.77405498 | -0.671573146 | 0.113641677 |
| 31613.0918 | 26846.80664 | 25812.23242 | 0.242278154 | 0.281112501 | 0.827345801 | 0.162826583 | 1.11657789 | 0.03695751 |
| 699885.875 | 769893.375 | 740779.0625 | 0.242073596 | 0.155789183 | -0.398943887 | 0.119711295 | -0.364984542 | 0.102257588 |
| 226212.3438 | 278198.4063 | 256509.4688 | 0.241872422 | 0.233064592 | 0.140335721 | 0.461673965 | 0.374925192 | 0.117034613 |
| 7038.521484 | 6646.969238 | 4366.983398 | 0.241871518 | 0.066744288 | -0.622020173 | 0.018539883 | -0.879707945 | 0.073672213 |
| 113318.5781 | 110511.1328 | 110368.5391 | 0.241508132 | 0.066744288 | -0.064588546 | 0.426150538 | -0.341858873 | 0.067792711 |
| 41845.20703 | 36226.28906 | 24316.71094 | 0.240850694 | 0.705950888 | 1.311056789 | 0.072360314 | 0.567783196 | 0.348056184 |
| 181565.0469 | 180719.9219 | 177220.5625 | 0.240809361 | 0.08662098 | -0.308972624 | 0.083003488 | -0.417761845 | 0.067125324 |
| 81401.3125 | 79687.21875 | 72900.03906 | 0.240506653 | 0.086055451 | 0.224041725 | 0.068282276 | -0.16012598 | 0.122834452 |
| 20875.53906 | 26413.48828 | 31294.76367 | 0.240066387 | 0.47570742 | -0.731472653 | 0.191184742 | -0.338071753 | 0.396137806 |
| 34032.41406 | 31492.73828 | 33616.30078 | 0.239945228 | 0.240263529 | 0.425561448 | 0.05746166 | 0.415089644 | 0.045861991 |
| 42915.23828 | 51063.19922 | 53835.30469 | 0.239579946 | 0.407882687 | 0.172544081 | 0.497760151 | 0.042538738 | 0.877555686 |
| 47569.96484 | 50904.65625 | 47988.05859 | 0.239122206 | 0.09670005 | -0.388180183 | 0.081402616 | -0.326636077 | 0.079780753 |
| 177175.7344 | 184006.5 | 178609.9531 | 0.239030151 | 0.017850787 | 0.047439963 | 0.217021068 | -0.203349714 | 0.020008995 |

| | | | | | | | | |
|---|---|---|---|---|---|---|---|---|
| 354202.4375 | 360542.4063 | 358731.3438 | 0.23899922 | 0.248502527 | 0.213191539 | 0.04780226 | 0.270632357 | 0.038294126 |
| 27843.16211 | 31257.875 | 27814.10156 | 0.238908679 | 0.249779609 | 0.63127689 | 0.018539883 | 0.313209067 | 0.069238464 |
| 13648.48828 | 5515.290039 | 8784.291016 | 0.238212597 | 0.081030942 | 0.417156168 | 0.131985724 | -0.543006548 | 0.317399806 |
| 148089.3125 | 142716.0938 | 134214.8125 | 0.237778035 | 0.034345213 | -0.226409396 | 0.048107216 | -0.394999693 | 0.021836949 |
| 9399.897461 | 8977.774414 | 8170.738281 | 0.237517926 | 0.310484451 | 0.731472385 | 0.062630775 | 0.630121796 | 0.047837702 |
| 114964.1016 | 80758.5 | 96938.57031 | 0.23701971 | 0.053733207 | 0.19051685 | 0.22093565 | 0.734655583 | 0.126064355 |
| 61035.76953 | 70716.69531 | 68543.13281 | 0.236940707 | 0.469807364 | -0.146424162 | 0.643665701 | -0.18964069 | 0.55724175 |
| 252467.4688 | 223296.7031 | 252287.5 | 0.236544905 | 0.102216965 | 0.023656324 | 0.76362454 | 0.035819104 | 0.717271407 |
| 102802.8047 | 480650.5 | 148830.0469 | 0.23637928 | 0.773313113 | 0.401385187 | 0.834109875 | -0.158273139 | 0.885245898 |
| 90802.38281 | 85499.96094 | 86668.17188 | 0.23597458 | 0.22713782 | 0.295642938 | 0.078077873 | 0.100607704 | 0.279983734 |
| 30918.19922 | 29129.45508 | 26591.5957 | 0.235454953 | 0.05642147 | -0.114237339 | 0.210255356 | -0.415413481 | 0.034722024 |
| 238624.1875 | 227661.2188 | 221065.9375 | 0.235289941 | 0.168081422 | 0.365753489 | 0.095325195 | 0.119968992 | 0.369415108 |
| 183882.1094 | 189884.6563 | 161602.2813 | 0.235147899 | 0.231728533 | 0.074783933 | 0.624370758 | 0.117796802 | 0.493707745 |
| 175784.9531 | 182160.7813 | 181735.9063 | 0.234938051 | 0.143726661 | 0.07011993 | 0.378422548 | -0.186700499 | 0.120295504 |
| 6360.711914 | 6566.052246 | 6135.167969 | 0.234502536 | 0.093577918 | 0.024613953 | 0.88755067 | -0.591147197 | 0.022925497 |
| 253129.5469 | 336962.0625 | 289653.1875 | 0.234201096 | 0.183818223 | 0.273548334 | 0.146155799 | 0.436222808 | 0.142665954 |
| 21069.82227 | 21153.19336 | 20562.60547 | 0.234089065 | 0.036275447 | 0.168286864 | 0.094230752 | -0.007662651 | 0.886748012 |
| 98963.65625 | 96407.17969 | 105247.1953 | 0.234019447 | 0.145925272 | 0.225643852 | 0.150271527 | -0.010250884 | 0.934841883 |
| 25608.87891 | 21613.98633 | 17847.44727 | 0.232731906 | 0.42340664 | 0.134425406 | 0.600442096 | 0.46953934 | 0.181045061 |
| 47596.40234 | 48213.57031 | 49097.76172 | 0.232325448 | 0.198269969 | 0.092266977 | 0.513385141 | 0.24500131 | 0.172393686 |
| 309169.6563 | 239740.3594 | 240947.5781 | 0.231775758 | 0.378842436 | 1.075526983 | 0.045329621 | 0.774518833 | 0.080171572 |
| 18505.65039 | 14291.4541 | 22092.57617 | 0.231685083 | 0.161527162 | -0.51268697 | 0.045976225 | -1.090942116 | 0.027868741 |
| 48146.46875 | 48428.82422 | 41548.44531 | 0.231449928 | 0.367030807 | -0.882940293 | 0.021906439 | -0.861825853 | 0.014448708 |
| 34522.25 | 38224.82813 | 39295.05859 | 0.230890039 | 0.547458136 | 0.637146214 | 0.254631118 | 1.436284247 | 0.02425784 |
| 11658.90918 | 12051.44824 | 10146.87988 | 0.230603657 | 0.269289238 | 0.382110353 | 0.1687782 | 0.864410559 | 0.030832901 |
| 3774.355469 | 6838.434082 | 3147.384521 | 0.23031882 | 0.64719338 | 0.370454764 | 0.585714677 | -0.095176323 | 0.883557065 |
| 126967.6484 | 129775.2813 | 130377.2969 | 0.230308018 | 0.073703861 | 0.063061015 | 0.147741919 | 0.181665817 | 0.020337832 |
| 21551.24023 | 24996.69141 | 22287.17773 | 0.230144811 | 0.127230164 | -0.110019731 | 0.369927787 | -0.479436633 | 0.049415261 |
| 6800.830566 | 8675.912109 | 8955.295898 | 0.230052815 | 0.361664975 | 0.165026869 | 0.486706805 | -0.315232612 | 0.260534667 |
| 87564.52344 | 86841.48438 | 87075.92969 | 0.230042967 | 0.106925332 | -0.293636166 | 0.073861046 | 0.074479784 | 0.270400252 |
| 7183.719727 | 9568.445313 | 8367.478516 | 0.229983267 | 0.611412475 | -0.07509714 | 0.866635988 | -0.198387843 | 0.669288977 |
| 157300.2031 | 161089.3594 | 170494.0469 | 0.229832689 | 0.252797361 | -0.993075759 | 0.068436129 | -0.968766262 | 0.066292531 |
| 25786.81445 | 26062.6543 | 26208.77734 | 0.22926239 | 0.384199211 | 0.211715153 | 0.413443071 | 0.560936765 | 0.098825691 |

| | | | | | | | | |
|---|---|---|---|---|---|---|---|---|
| 220040.0938 | 205015.2344 | 214853.75 | 0.229012079 | 0.360108859 | 0.596656085 | 0.083773082 | 0.293944871 | 0.235404176 |
| 112964.5859 | 116704.6484 | 141113.6094 | 0.228973169 | 0.125515217 | -0.083583579 | 0.447004412 | 0.450527541 | 0.113848811 |
| 15518.09863 | 12981.44043 | 14770.01465 | 0.228479443 | 0.418329472 | 0.382103061 | 0.191849347 | 0.056953369 | 0.819104556 |
| 165494.2969 | 169851.3594 | 161812.9063 | 0.228209921 | 0.111362172 | 0.071986542 | 0.280688037 | 0.13369389 | 0.089383058 |
| 15815.95801 | 15633.44336 | 16021.9082 | 0.228134973 | 0.190614833 | 0.083724427 | 0.376466201 | 0.084298228 | 0.287067716 |
| 209208.832 | 232128.9805 | 217439.7461 | 0.227889708 | 0.080663793 | 0.44429971 | 0.029156106 | 0.704917693 | 0.020157267 |
| 57165.17578 | 52371.32813 | 51560.59375 | 0.227741791 | 0.217435865 | 0.047904945 | 0.713585003 | -0.231849806 | 0.200717552 |
| 31221.72656 | 33217.23828 | 25638.19531 | 0.227641409 | 0.060903498 | -0.35957487 | 0.03821916 | -0.456136373 | 0.07832051 |
| 13059.81348 | 14248.18848 | 13221.40039 | 0.227479463 | 0.705097252 | 1.50293494 | 0.048370376 | 1.158184295 | 0.098401903 |
| 68670.75 | 71014.67188 | 70232.30469 | 0.227423463 | 0.398875467 | 0.134532908 | 0.559205077 | -0.206459046 | 0.438757478 |
| 167443.7188 | 163770.7031 | 162944.2188 | 0.226766502 | 0.065789073 | -0.05938622 | 0.606716614 | 0.028829338 | 0.293588797 |
| 284243 | 250566.6875 | 261982.1563 | 0.226220645 | 0.0958851 | 0.529480157 | 0.121581067 | 0.974130527 | 0.02146481 |
| 12237.87988 | 17108.30859 | 14249.08105 | 0.225870856 | 0.317987095 | 0.495006227 | 0.08026683 | -0.000773921 | 0.997969366 |
| 162775.4375 | 181715.1406 | 170941.8438 | 0.225678693 | 0.247714225 | -0.012353412 | 0.937984911 | 0.031197405 | 0.847249863 |
| 25249.53906 | 25051.00586 | 23531.57422 | 0.225584037 | 0.176366736 | 0.276868429 | 0.11332333 | 0.21970524 | 0.109873058 |
| 19142.67969 | 15474.57617 | 17951.21289 | 0.225388434 | 0.254204903 | 0.0302921 | 0.866635988 | 0.007503306 | 0.972680155 |
| 10410.90723 | 11074.26953 | 12973.04297 | 0.225199335 | 0.143584671 | -0.218907329 | 0.280400081 | -0.807149639 | 0.027868741 |
| 55450.29297 | 54131.21484 | 52966.23438 | 0.225056736 | 0.028143177 | -0.602777214 | 0.007509388 | -0.816104074 | 0.004289827 |
| 456643.75 | 488719.0938 | 467667.6563 | 0.224598248 | 0.118081327 | 0.857023043 | 0.02118733 | 0.742944322 | 0.014388274 |
| 84275.42969 | 79366.89844 | 86277.45313 | 0.224252823 | 0.135465479 | 0.096234173 | 0.39272617 | -0.097388437 | 0.406371799 |
| 10835.40527 | 14059.82324 | 8863.926758 | 0.224233343 | 0.477728118 | 0.701921573 | 0.083527411 | 0.498632184 | 0.254951037 |
| 19054.16016 | 20566.91992 | 16988.10547 | 0.223730407 | 0.089665988 | -0.907190564 | 0.024921531 | -0.631051355 | 0.046124274 |
| 88393.51563 | 86643.34375 | 78911.8125 | 0.223497714 | 0.500701937 | 0.12325929 | 0.691779308 | 0.117788307 | 0.69501344 |
| 100648.1641 | 97904.90625 | 102597.4922 | 0.223475739 | 0.099542225 | -0.29311231 | 0.046373997 | -0.253107584 | 0.029219266 |
| 424824.7188 | 374744.4688 | 399340.2188 | 0.223440999 | 0.061425128 | 0.415601961 | 0.06919378 | 0.155490719 | 0.168917579 |
| 27901.79883 | 32287.61719 | 31070.7793 | 0.222978354 | 0.640030711 | 0.505374075 | 0.122318166 | 0.852162138 | 0.028738995 |
| 2589.583496 | 3492.845215 | 2072.554688 | 0.222511319 | 0.131501491 | -0.727667753 | 0.053327615 | -0.014475063 | 0.969282881 |
| 1150544.125 | 1212157.5 | 1100159 | 0.222277003 | 0.036160457 | 0.488683117 | 0.018539883 | 0.136878701 | 0.138109111 |
| 27713.16406 | 34222.45361 | 29295.90527 | 0.221824404 | 0.191834923 | 0.064391308 | 0.636784361 | -0.092869496 | 0.596749137 |
| 79257.125 | 85348.54688 | 83649.58594 | 0.221505795 | 0.109154846 | -0.41273827 | 0.018539883 | -0.414012433 | 0.0146504 |
| 19487.39258 | 19524.70117 | 18558.33594 | 0.221478476 | 0.329618532 | -0.699372754 | 0.017297259 | -0.487491512 | 0.027868741 |
| 30784.91016 | 25803.54297 | 25137.30859 | 0.221446853 | 0.298390687 | -0.875279292 | 0.088108869 | -0.860796261 | 0.064734893 |
| 171989.3594 | 122408.0781 | 130404.3438 | 0.221211421 | 0.075807406 | -1.083335835 | 0.012254503 | -0.953651428 | 0.042860672 |

| | | | | | | | | |
|---|---|---|---|---|---|---|---|---|
| 1112790.375 | 1228512.875 | 1100666.5 | 0.220905049 | 0.20270675 | -0.16199373 | 0.328806215 | 0.281767439 | 0.116465028 |
| 20107.2168 | 24918.92188 | 18780.02734 | 0.2207446 | 0.629100374 | 0.406717048 | 0.189627399 | -0.07058258 | 0.754978058 |
| 39434.8125 | 43994.44531 | 37249.21875 | 0.220629804 | 0.837136363 | 0.657448086 | 0.469445813 | 0.438356112 | 0.629670066 |
| 20342.14258 | 18099.23047 | 16013.40625 | 0.220319776 | 0.504596086 | 1.166960797 | 0.023661483 | 0.885458266 | 0.051065331 |
| 14527.78613 | 14026.21387 | 14975.22852 | 0.220292711 | 0.176820059 | 0.331272964 | 0.362329238 | 0.35663113 | 0.026553937 |
| 463736.4375 | 448899.7813 | 460414.5 | 0.220256947 | 0.219471631 | -0.278095095 | 0.208744897 | -0.450152774 | 0.11805192 |
| 147748.125 | 155035.625 | 149382.3594 | 0.22021965 | 0.223885477 | -0.123566072 | 0.456639742 | -0.261111468 | 0.208408498 |
| 224730.8281 | 245390.1719 | 245973.4531 | 0.220180845 | 0.076438569 | 0.16203773 | 0.195611454 | 0.503993371 | 0.0236713 |
| 13051.73438 | 12913.33984 | 12141.63086 | 0.219932419 | 0.131122522 | -0.218410597 | 0.177893197 | -0.330840066 | 0.088350903 |
| 18594.17773 | 20491.00391 | 20955.42773 | 0.219721214 | 0.216496858 | -0.117522444 | 0.459712252 | 0.619266521 | 0.029653052 |
| 27743.21484 | 51547.89453 | 17477.39453 | 0.219069228 | 0.757133796 | 0.310918412 | 0.522105467 | -0.410453961 | 0.582899134 |
| 13161.48828 | 12135.53613 | 10268.21191 | 0.21905533 | 0.565062132 | 0.615713681 | 0.120506435 | 0.254643609 | 0.423206002 |
| 178779.1094 | 200898.7813 | 197640.375 | 0.219023483 | 0.160761277 | 0.190753062 | 0.198186577 | 0.088743169 | 0.478935691 |
| 81139 | 66002.20313 | 74998.67188 | 0.218776985 | 0.047817825 | -0.156625857 | 0.353137578 | -0.356286025 | 0.076783998 |
| 4799.848633 | 5179.045898 | 4797.041016 | 0.218255057 | 0.638465231 | 0.203244987 | 0.623543083 | -0.225956507 | 0.624114216 |
| 246048.5938 | 233643.1563 | 270809.2813 | 0.217881694 | 0.107059407 | -0.067656732 | 0.660718628 | -0.024387366 | 0.840929596 |
| 20524.22461 | 18895.88867 | 23292.38086 | 0.217735662 | 0.375561534 | 0.872304271 | 0.361561972 | 1.225108368 | 0.029653052 |
| 87582.07813 | 108282.4922 | 108052.4922 | 0.217712281 | 0.072387792 | -0.118593413 | 0.209677596 | -0.313968358 | 0.117048888 |
| 760049.5625 | 839276.4375 | 801704.625 | 0.217346345 | 0.087648224 | -0.03229339 | 0.69897549 | 0.025680143 | 0.785580722 |
| 6846.950684 | 8620.523438 | 10666.61914 | 0.21727254 | 0.680305764 | 0.961051195 | 0.113261078 | -1.154281963 | 0.226153996 |
| 7064.166504 | 6739.033203 | 5783.797852 | 0.217249371 | 0.534135477 | 0.253907657 | 0.458289776 | -0.442915525 | 0.317582832 |
| 44343.40234 | 47299.03125 | 46458.67578 | 0.217151227 | 0.029811229 | -0.154287597 | 0.165945937 | -0.298187651 | 0.019880539 |
| 42407.79688 | 46484.79688 | 43771.22266 | 0.216780192 | 0.427440307 | 0.322781004 | 0.122695009 | 0.618934634 | 0.027982615 |
| 61554.97266 | 63425.21094 | 64734.83203 | 0.216479012 | 0.07321371 | 0.109844645 | 0.124127483 | 0.065212417 | 0.262465444 |
| 488119.5938 | 1351472.875 | 539284.875 | 0.216424908 | 0.765814241 | -0.76300244 | 0.423723931 | -0.005724852 | 0.995859506 |
| 9057.618164 | 8752.157227 | 9684.209961 | 0.216306889 | 0.059865575 | 0.005024922 | 0.964381518 | 0.029153519 | 0.732715412 |
| 375870.7188 | 385640.8438 | 378305.8438 | 0.216306194 | 0.078894657 | -0.113890283 | 0.343070048 | -0.22452089 | 0.060470082 |
| 17603.31641 | 19364.24219 | 18683.1875 | 0.215600595 | 0.41215279 | 0.007867151 | 0.957006417 | -0.701911102 | 0.041386295 |
| 10773.24121 | 10498.35645 | 9529.574219 | 0.215509359 | 0.426576337 | -0.811966553 | 0.112885657 | -0.848938461 | 0.104803909 |
| 26469.60547 | 26840.66797 | 23473.66016 | 0.215408041 | 0.005752145 | 0.073343885 | 0.479572156 | 0.078317705 | 0.44269115 |
| 39781.35938 | 36419.67969 | 37277.46484 | 0.215313802 | 0.490789553 | 0.486095773 | 0.135029738 | 0.104188661 | 0.6771622 |
| 51398.74609 | 49134.33594 | 48219.35156 | 0.215286507 | 0.477538026 | 0.270687318 | 0.339659199 | 0.066056172 | 0.797491208 |
| 32525.13672 | 46327.4375 | 36132.53125 | 0.214082838 | 0.155490733 | -0.216789256 | 0.184954137 | -0.389528165 | 0.17603045 |

| | | | | | | | | |
|---|---|---|---|---|---|---|---|---|
| 296315.9688 | 299960.5313 | 305875.375 | 0.21360631 | 0.07362475 | -0.123486183 | 0.119711295 | -0.082981373 | 0.190591136 |
| 5376.252441 | 4095.223877 | 3002.79834 | 0.213447635 | 0.609001008 | -0.750294116 | 0.196521734 | -0.82636638 | 0.145687527 |
| 4696.594238 | 3600.167236 | 3569.281006 | 0.212780473 | 0.710608258 | -0.721679839 | 0.361561972 | -0.419483583 | 0.519508074 |
| 169534.4531 | 185407.0156 | 193409.0469 | 0.212588952 | 0.211578922 | -0.278518795 | 0.174050429 | 0.017727226 | 0.908031064 |
| 57167.19922 | 58910.46875 | 57075.96484 | 0.212430024 | 0.131501491 | 0.254811107 | 0.093818896 | -0.026642025 | 0.786883312 |
| 5691.061523 | 5535.584473 | 4737.369629 | 0.212309735 | 0.554251928 | -0.347147632 | 0.31156031 | -0.739878392 | 0.086952277 |
| 14013.42871 | 13313.47852 | 9522.871094 | 0.212261792 | 0.04251435 | 0.028684904 | 0.831728137 | -0.199550683 | 0.431199487 |
| 629485.5 | 657442.25 | 552637.0625 | 0.212147179 | 0.129251227 | -0.982412992 | 0.01179292 | -0.374812683 | 0.07832051 |
| 11007.92871 | 13615.98535 | 8375.347656 | 0.211631513 | 0.361920409 | 0.345052982 | 0.058077004 | 0.123313676 | 0.700974317 |
| 2886.206543 | 4492.185059 | NA | 0.21148265 | 0.268379972 | -0.98450389 | 0.048146568 | -1.469812798 | NA |
| 87622.89063 | 81719.85938 | 74620.5625 | 0.210918465 | 0.088231376 | 0.00032235 | 0.997242283 | -0.10405897 | 0.369265324 |
| 40832.625 | 42716.29297 | 35934.03125 | 0.21078925 | 0.334961277 | -0.995503678 | 0.015625705 | -0.994466125 | 0.01054584 |
| 265756.1563 | 267171.25 | 263419.4063 | 0.210753563 | 0.088240896 | 0.035135527 | 0.70134953 | 0.258861668 | 0.047558276 |
| 762149.1875 | 839385.75 | 799548.0625 | 0.210705491 | 0.019078369 | -0.240725068 | 0.021776816 | -0.122857313 | 0.150169758 |
| 26690.83594 | 28786.73828 | 25528.16602 | 0.209983559 | 0.308539067 | 0.726240017 | 0.440449259 | -0.054292086 | 0.716525566 |
| 547805.8125 | 594193.5625 | 552334.4375 | 0.209432829 | 0.05144423 | 0.134445806 | 0.137540222 | -0.033690086 | 0.64388549 |
| 182880.875 | 176064.3125 | 184701.5625 | 0.209261717 | 0.071158565 | -0.365475825 | 0.049184287 | 0.288507205 | 0.041664392 |
| 39260.57813 | 39526.16797 | 36462.66797 | 0.209213043 | 0.160946254 | -0.379243851 | 0.06863053 | 0.343647007 | 0.040520608 |
| 90918.16406 | 101125.0625 | 99768.74219 | 0.209198733 | 0.116674977 | 0.325865208 | 0.058077004 | 0.0475969 | 0.655621396 |
| 10821.25195 | 9772.442383 | 8764.480469 | 0.208503028 | 0.458489299 | 0.3320392 | 0.469555906 | 0.09849995 | 0.710273063 |
| 9212.588867 | 11236.46387 | 6483.479004 | 0.208496112 | 0.337496614 | -0.065060912 | 0.682560717 | -0.415960084 | 0.246736339 |
| 11697.63672 | 14475.67285 | 15042.29395 | 0.208132283 | 0.448932809 | -0.115594904 | 0.359216188 | 0.16728764 | 0.399866811 |
| 17131.10938 | 10328.91504 | 10366.6543 | 0.207653091 | 0.841038496 | -1.137728668 | 0.431374349 | -1.272167378 | 0.395364978 |
| 82019.25 | 81737.45313 | 85452.30469 | 0.207451439 | 0.052854398 | -0.218214923 | 0.052965881 | -0.506198912 | 0.017279709 |
| 12375.9834 | 11553.27148 | 11316.73828 | 0.206904169 | 0.168060614 | 0.028965444 | 0.820444407 | 0.313522342 | 0.070587266 |
| 330229.6563 | 352816.6563 | 357876.3438 | 0.206310118 | 0.060886955 | 0.302862437 | 0.0395519 | -0.038941067 | 0.570103979 |
| 32021.29492 | 31187.61328 | 24834.76172 | 0.20629533 | 0.187660694 | -0.195846885 | 0.18008437 | -0.503723392 | 0.061106674 |
| 21449.63672 | 23827.62891 | 24707.80859 | 0.206236467 | 0.414321081 | -0.372609416 | 0.163673552 | -0.609552124 | 0.075895689 |
| 24949.01172 | 25292.38672 | 25290.50977 | 0.205408338 | 0.102981042 | -0.388292545 | 0.06723286 | -0.40520863 | 0.063283918 |
| 3767479 | 5578141.5 | 4349435 | 0.205381027 | 0.670294273 | -1.224810454 | 0.146345919 | -0.05579526 | 0.893568216 |
| 360967.3438 | 410159.4063 | 377597.7813 | 0.205110652 | 0.404110576 | 0.404476039 | 0.155243406 | 0.985591893 | 0.024451141 |
| 134448.4844 | 132850.3125 | 117185.5078 | 0.204978906 | 0.212024069 | -0.239255434 | 0.034480785 | -0.219883559 | 0.115929208 |
| 172240.3906 | 177998.3281 | 168548.4844 | 0.204948117 | 0.096004496 | -0.399164314 | 0.01179292 | -0.38404061 | 0.010418649 |

| | | | | | | | | |
|---|---|---|---|---|---|---|---|---|
| 121881.8438 | 123718.9688 | 114534.3594 | 0.2047849 | 0.020651493 | -0.239002095 | 0.086909686 | -0.32932942 | 0.024074394 |
| NA | NA | 2495.671387 | 0.204596722 | NA | 2.192375194 | NA | 0.749880463 | NA |
| 21775.83984 | 23618.80273 | 24662.2793 | 0.203881635 | 0.533024504 | 0.26709924 | 0.378617903 | 0.263810736 | 0.37239437 |
| 10261.25781 | 7858.346191 | 7155.637695 | 0.202939939 | 0.285438689 | -0.399884792 | 0.152415276 | -0.799705052 | 0.053854543 |
| 11940.2627 | 13625.39844 | 15388.71973 | 0.202813467 | 0.437339197 | 0.299780892 | 0.340114217 | -0.027135131 | 0.923992665 |
| 456385.6563 | 456995 | 477774.8438 | 0.202396175 | 0.089123154 | -0.231934749 | 0.03968601 | -0.256952139 | 0.028663646 |
| 51506.53906 | 52100.70313 | 49903.62891 | 0.202185359 | 0.060886955 | 0.165283448 | 0.167865792 | 0.041369616 | 0.241289486 |
| 45193.07031 | 38290.97266 | 45649.34766 | 0.20197291 | 0.474195126 | -0.398260374 | 0.204833051 | 0.744021905 | 0.040202519 |
| 34319.04688 | 34913.03125 | 32926.52344 | 0.201777164 | 0.136222118 | -0.120600905 | 0.025293799 | -0.14974111 | 0.073390601 |
| 23908.64258 | 29898.11133 | 23363.85156 | 0.2014954 | 0.322257553 | -0.061446698 | 0.726432734 | 0.19137252 | 0.403026694 |
| 253072.125 | 247881.9375 | 247422.6406 | 0.200946018 | 0.043802925 | 0.860892421 | 0.017786674 | 0.276154673 | 0.037156974 |
| 233443.2188 | 231434.5938 | 240316.2031 | 0.20092282 | 0.14383958 | -0.298583391 | 0.104541915 | -0.539998154 | 0.061900682 |
| 128428.7969 | 138814.375 | 118440.9297 | 0.200800673 | 0.274703862 | -0.432160941 | 0.137613908 | -0.387951784 | 0.11222326 |
| 1190.620117 | 2143.715088 | 1360.591064 | 0.200742655 | NA | -0.078658003 | NA | -0.019547886 | NA |
| 52580.60156 | 47680.60156 | 55712.59375 | 0.200614243 | 0.147312752 | 0.163691736 | 0.05744085 | -0.096782024 | 0.371139574 |
| 30055.25 | 31442.83594 | 30377.94336 | 0.200588623 | 0.036480793 | -0.092860012 | 0.150726837 | 0.241788596 | 0.02183874 |
| 25323.3457 | 24654.98242 | 25636.68555 | 0.200238714 | 0.258756218 | 0.748071285 | 0.125154882 | 0.726828686 | 0.013271537 |
| 33179.11328 | 29133.67578 | 30031.43359 | 0.200000801 | 0.084947579 | 0.045693499 | 0.528259024 | 0.336565008 | 0.065367829 |
| 63236.53125 | 59828.92188 | 70872.55469 | 0.199701988 | 0.253322022 | 0.156252686 | 0.290374442 | 0.365063459 | 0.087771237 |
| 652369.875 | 659506.0625 | 530042.4375 | 0.199416274 | 0.338882582 | 0.26623166 | 0.109083959 | 0.20668848 | 0.28577659 |
| 123538.5313 | 150853.2031 | 144264.8906 | 0.199136995 | 0.807911996 | 0.010756994 | 0.990463209 | 1.064386989 | 0.161137259 |
| 12574.08008 | 12180.10254 | 14484.69336 | 0.198980814 | 0.402084008 | -0.280022721 | 0.164383952 | 0.018404629 | 0.914110816 |
| 796466.6406 | 836397.6914 | 897544.3945 | 0.198940452 | 0.594734687 | 0.089205725 | 0.831472283 | 0.52615174 | 0.161930117 |
| 351489.1875 | 275635.5625 | 252963.6406 | 0.198878089 | 0.737915376 | -0.545211671 | 0.346376894 | -0.40978112 | 0.413077372 |
| 403398.5625 | 331961.0781 | 324908.9688 | 0.198684722 | 0.148988393 | 0.14389069 | 0.162317825 | -0.13192186 | 0.418228234 |
| 11359.58301 | 15458.90625 | 11355.46973 | 0.198584003 | 0.522306269 | 0.399460176 | 0.214766804 | -0.468636875 | 0.244006157 |
| 233376.7969 | 220039.5781 | 172116.9219 | 0.198345593 | 0.134823843 | -0.240932721 | 0.104928475 | 0.007515701 | 0.973660041 |
| 37936.87891 | 36309.87109 | 27219.69141 | 0.198056844 | 0.180419232 | 0.400168079 | 0.06794739 | 0.271354123 | 0.325000572 |
| 8671.619141 | 9093.202148 | 5870.923828 | 0.19745039 | 0.504874232 | -0.69845683 | 0.165292229 | -1.075048167 | 0.091109394 |
| 10168.66919 | 9958.604004 | 8531.105225 | 0.19726076 | 0.497120042 | 1.636914099 | 0.038184043 | 0.944540347 | 0.035612683 |
| 9718.49707 | 8212.536133 | 8410.569336 | 0.197186494 | 0.247616465 | -0.372268257 | 0.103004848 | -0.620291942 | 0.034722024 |
| 46501.90625 | 46693.625 | 42393.86719 | 0.197134641 | 0.16722446 | 0.174259325 | 0.289928504 | 0.529993424 | 0.024912072 |
| 198918.6406 | 207975.7031 | 204587.0625 | 0.197009494 | 0.156500695 | 0.09815219 | 0.532853413 | -0.030233207 | 0.525214393 |

| | | | | | | | | |
|---|---|---|---|---|---|---|---|---|
| 118535.2188 | 114331.6797 | 81827.14844 | 0.196643546 | 0.109633261 | -0.055149695 | 0.640666998 | -0.352814614 | 0.206336913 |
| 36977.59375 | 25549.92383 | 29872.68555 | 0.196127338 | 0.202541133 | -0.507588014 | 0.031120558 | -0.473983859 | 0.131166582 |
| 287207.4375 | 298683.375 | 299685.4375 | 0.196010465 | 0.136265249 | -0.158328776 | 0.127902424 | -0.265597964 | 0.06578999 |
| 25932.82422 | 23425.91406 | 18985.88867 | 0.19597752 | 0.271759161 | 0.113604463 | 0.365129709 | -0.162974374 | 0.421974631 |
| 54980.40625 | 48826.23438 | 50775.17969 | 0.19472801 | 0.097116923 | 0.158227083 | 0.133190834 | -0.08368738 | 0.387952458 |
| 117007.2656 | 112722.1328 | 98350.3125 | 0.19469841 | 0.178228644 | 0.386499855 | 0.088587863 | 0.256353434 | 0.14559756 |
| 1486355.625 | 1712597.25 | 1791569.25 | 0.194401841 | 0.063885064 | -0.062876516 | 0.37992376 | -0.025032485 | 0.846177482 |
| 44909.10547 | 45087.71875 | 49530.69922 | 0.194155739 | 0.172511971 | 0.242843343 | 0.181741873 | 0.259716641 | 0.079165593 |
| 9872.813477 | 7219.50293 | 7180.97998 | 0.193644549 | 0.642613461 | 0.009217283 | 0.98257662 | -0.131903865 | 0.773308828 |
| 59843.28125 | 58010.39453 | 59640.71094 | 0.193530251 | 0.042192618 | -0.610999267 | 0.029180754 | 0.077694906 | 0.064597187 |
| 370122.3125 | 369387.4063 | 343469.9063 | 0.193526173 | 0.303961749 | -0.284465367 | 0.219022122 | -0.345844956 | 0.172176377 |
| 827221.125 | 926120.3125 | 947852.625 | 0.193380635 | 0.096323959 | 0.095792691 | 0.536495 | 0.41606679 | 0.052545194 |
| 7642.000977 | 9537.899414 | 10331.62891 | 0.193346063 | 0.529580181 | -0.034897624 | 0.921217151 | 0.277554177 | 0.382150932 |
| 18745.56641 | 22400.50195 | 21332.83789 | 0.193129404 | 0.455517192 | -0.299672386 | 0.335186404 | -0.238205265 | 0.401391855 |
| 114298.4375 | 115794.2813 | 117370.4531 | 0.192706429 | 0.219131856 | -0.245249648 | 0.150271527 | -0.37592694 | 0.104282201 |
| 72966.57031 | 84016.22656 | 78726.54688 | 0.192566793 | 0.186686406 | 0.275935652 | 0.095980292 | 0.264900728 | 0.12089792 |
| 56002.5625 | 58453.52344 | 44013.46484 | 0.192434001 | 0.462864489 | -0.015120401 | 0.855436492 | -0.026251177 | 0.893928054 |
| 544664.875 | 601508.9375 | 615727.0625 | 0.192207578 | 0.160107556 | -0.121810882 | 0.287985731 | -0.190138911 | 0.152064039 |
| 954121 | 992528.8125 | 1013514 | 0.191642269 | 0.037492489 | 0.14699919 | 0.066380875 | 0.263904106 | 0.02616038 |
| 1671410.25 | 1707350.5 | 1746961.25 | 0.191512344 | 0.071134978 | 0.038741802 | 0.539128604 | 0.128853639 | 0.126366029 |
| 8365.199219 | 7909.624023 | 8645.412109 | 0.191183144 | 0.79642005 | -0.479576304 | 0.593832889 | -0.773528064 | 0.447937242 |
| 50411.89844 | 48068.03516 | 47148.77344 | 0.191153457 | 0.04382682 | -0.040539824 | 0.192808829 | 0.016334725 | 0.717031719 |
| 109513.6875 | 102208.8906 | 122913.9531 | 0.191132793 | 0.312750899 | 0.001812009 | 0.991529214 | 0.455114481 | 0.076783998 |
| 61304.33594 | 53453.65625 | 57466.29297 | 0.190818861 | 0.122096077 | 0.074551914 | 0.137176808 | 0.307245778 | 0.087971481 |
| 4714.570313 | 2237.836182 | 4210.131348 | 0.190685566 | 0.825752436 | 3.611434875 | 0.343658822 | 0.764770022 | 0.280633075 |
| 60594.91406 | 67043.35938 | 55374.77344 | 0.190653295 | 0.210383422 | -0.069264136 | 0.538096429 | -0.420048406 | 0.049492888 |
| 149156.5469 | 142944.1406 | 153028.0625 | 0.19018663 | 0.110631525 | -0.155207192 | 0.032675377 | 0.156324915 | 0.079264515 |
| 997897.3125 | 978775.4375 | 999561.8125 | 0.190142492 | 0.316823258 | 0.176062753 | 0.374500771 | 0.490797095 | 0.085652968 |
| 234718.9063 | 349071.6563 | 338108.7813 | 0.19011148 | 0.717472921 | 0.379097583 | 0.411843428 | 0.840917007 | 0.128960705 |
| 142822.2813 | 149352.7813 | 158418.8125 | 0.190061196 | 0.107769315 | -0.11419166 | 0.088709063 | -0.197942582 | 0.070968049 |
| 6409.334473 | 5426.540039 | 3386.697998 | 0.189719891 | 0.198381967 | -0.726939991 | 0.024165741 | -0.756723512 | 0.116453865 |
| 12328.59766 | 10937.13184 | 10926.92969 | 0.18964735 | 0.171512612 | 0.71688757 | 0.082791584 | 0.649713599 | 0.029820656 |
| 28589.56641 | 29099.54883 | 26154.9707 | 0.189507962 | 0.132446456 | -0.131893771 | 0.298829753 | -0.491325756 | 0.020351059 |

| | | | | | | | | |
|---|---|---|---|---|---|---|---|---|
| 96152.625 | 89556.25781 | 98908.99219 | 0.189497632 | 0.103527693 | 0.690609544 | 0.075404888 | 0.580789238 | 0.020746708 |
| 44238.89844 | 53185.77734 | 42120.16016 | 0.189240234 | 0.332739002 | 0.157355433 | 0.303929178 | -0.253250953 | 0.212947102 |
| 290805.5625 | 292591.6563 | 321638.5938 | 0.189196561 | 0.031284382 | -0.115644033 | 0.042239422 | 0.302458806 | 0.083243599 |
| 59312.26172 | 39941.85547 | 49795.98828 | 0.189114898 | 0.358576039 | -0.682438851 | 0.100636165 | -0.505877075 | 0.125264621 |
| 8192.253906 | 8298.435547 | 8690.355469 | 0.188893328 | 0.194985779 | -0.222875026 | 0.185953071 | -0.436778454 | 0.084522976 |
| 48686.89844 | 49023.29688 | 50732.46094 | 0.188737198 | 0.183352415 | 0.209127991 | 0.048593497 | 0.125418457 | 0.095239671 |
| 16017.09766 | 18199.57617 | 17320.59375 | 0.188571304 | 0.262236638 | -0.209949339 | 0.05610328 | -0.951873332 | 0.009026966 |
| NA | NA | NA | 0.188543765 | 0.616828703 | -0.669494806 | 0.09842296 | NA | NA |
| 1949985.25 | 1967687.625 | 2064647.125 | 0.188442223 | 0.309753125 | 0.741174022 | 0.092018139 | 1.345325109 | 0.008387835 |
| 101191.9688 | 100577.5938 | 91192.60156 | 0.188044949 | 0.039043192 | -0.457894458 | 0.018539883 | -0.165440372 | 0.131961597 |
| 4623.029297 | 5330.467285 | 3620.553711 | 0.187883059 | 0.101446672 | -0.200475184 | 0.184317721 | -0.520443481 | 0.12348787 |
| 36332.73047 | 45929.94922 | 42542.21484 | 0.187576146 | 0.762776035 | 0.144895722 | 0.805890965 | 0.079035186 | 0.895597858 |
| 445989.2188 | 439800.2813 | 436170.5625 | 0.187450881 | 0.214939184 | 0.5365721 | 0.05468055 | 0.874260177 | 0.015712916 |
| 144552.0938 | 116788.8594 | 138104.9531 | 0.187131971 | 0.25252148 | 0.006443902 | 0.741007407 | -0.252802774 | 0.17603045 |
| 35772 | 30430.62305 | 22684.07031 | 0.186946547 | 0.639875121 | 0.551158206 | 0.158732431 | -0.281797578 | 0.385095259 |
| 49481.03906 | 82578.49219 | 69685.52344 | 0.186905579 | 0.717472921 | 0.024447684 | 0.967442364 | 0.205856291 | 0.701495778 |
| 711450.3125 | 740004.8125 | 769703.5 | 0.186764975 | 0.167516948 | -0.067601372 | 0.534862174 | -0.095226546 | 0.405108106 |
| 14376.59277 | 19838.28125 | 16106.12207 | 0.186193513 | 0.161072919 | 0.104710531 | 0.374951761 | 0.245703823 | 0.349498429 |
| 113611.1328 | 88839.125 | 110796.9688 | 0.186161667 | 0.14802893 | 0.42818339 | 0.090297888 | -0.191470218 | 0.294592752 |
| 14345.58398 | 14336.59082 | 14464.35156 | 0.185580568 | 0.052671628 | 0.132392478 | 0.102526216 | -0.154178202 | 0.113977669 |
| 194887.8438 | 189530.0938 | 177344.8125 | 0.185388159 | 0.116084643 | -0.153065861 | 0.299255323 | -0.190201475 | 0.124439663 |
| 129908.6641 | 119323.8125 | 128571.375 | 0.185189191 | 0.271858743 | 0.197327719 | 0.562301807 | -0.334411316 | 0.127304768 |
| 38900.87109 | 41159.83203 | 37594.86328 | 0.184917297 | 0.137796298 | 0.468989938 | 0.110712084 | 0.502732593 | 0.02527614 |
| 54577.54297 | 56335.87891 | 54629.15625 | 0.18444747 | 0.488966028 | -0.063271656 | 0.805890965 | -0.369607077 | 0.280586597 |
| 150923.3906 | 159090.7344 | 163249.375 | 0.184213602 | 0.035776373 | -0.006247332 | 0.847098346 | -0.130042478 | 0.091928987 |
| 6573.183594 | 9184.195313 | 7675.079102 | 0.183705083 | 0.449952851 | 0.033435985 | 0.863868059 | 0.247738464 | 0.351032927 |
| 51164.09375 | 43777.75 | 44814.90234 | 0.183637988 | 0.044714174 | 0.033896038 | 0.607218021 | 0.191530457 | 0.206000009 |
| 46414.98438 | 50133.73828 | 47461.32813 | 0.183356829 | 0.158621062 | 0.062774824 | 0.696860822 | 0.00495996 | 0.964490899 |
| 921471.25 | 985962.875 | 996669 | 0.183294122 | 0.143584671 | -0.011075502 | 0.876081794 | -0.087622616 | 0.309561656 |
| 45090.89453 | 43228.375 | 42944.48438 | 0.183277992 | 0.02998338 | -0.080928292 | 0.635648296 | 0.503071171 | 0.013917035 |
| 112725.9063 | 116382.7969 | 110419.6719 | 0.183116872 | 0.137796298 | 0.072396032 | 0.364459731 | -0.162614469 | 0.136000191 |
| 78308.42188 | 85833.86719 | 82772.13281 | 0.182952776 | 0.110302818 | 0.249978098 | 0.068255066 | 0.039524028 | 0.63489244 |
| 65874.30273 | 60991.58203 | 55891.19922 | 0.182701585 | 0.593565321 | 0.126876274 | 0.685914317 | -0.084277955 | 0.801698552 |

| | | | | | | | | |
|---|---|---|---|---|---|---|---|---|
| 95931.82813 | 95757.60938 | 102957.5 | 0.182599107 | 0.300449017 | 0.066380236 | 0.734206841 | -0.380661726 | 0.095957305 |
| 71411.55469 | 73474.46875 | 69020.5625 | 0.18255555 | 0.02237899 | -0.078892499 | 0.229616344 | 0.085100444 | 0.141028512 |
| 778240.4375 | 857211.4375 | 820805.0625 | 0.182089363 | 0.215236407 | -0.168282525 | 0.124656054 | 0.042034408 | 0.587409713 |
| 11619.77637 | 11607.86133 | 8609.878906 | 0.181906531 | 0.255554892 | -0.723328584 | 0.028434447 | -1.18434127 | 0.033327007 |
| 145931.2969 | 160472.8906 | 149258.4219 | 0.181877077 | 0.360794803 | 0.939604106 | 0.026826422 | 0.880290708 | 0.020008995 |
| 37073.12109 | 38400.46484 | 30239.33398 | 0.181795631 | 0.239106331 | -0.955699104 | 0.036240475 | -0.572973173 | 0.046092295 |
| 218387.6719 | 195299.75 | 196514.0938 | 0.181714788 | 0.171522145 | -0.364524185 | 0.149301011 | -0.145378438 | 0.226466969 |
| 39167.3125 | 34884.64063 | 35791.79688 | 0.181433139 | 0.2756587 | -0.76026889 | 0.059340088 | -0.298437646 | 0.101959807 |
| 522749.5938 | 525112.875 | 514502.5625 | 0.181351888 | 0.10610949 | -0.287214498 | 0.077511889 | -0.569441728 | 0.045096398 |
| 6841.45166 | 8657.431641 | 6552.000488 | 0.181312567 | 0.304406879 | -0.177206654 | 0.298829753 | -0.63954519 | 0.049415261 |
| 16010.93359 | 17170.70508 | 10863.36133 | 0.181237309 | 0.46699162 | -0.683456007 | 0.138108349 | -0.436890904 | 0.225155825 |
| 178218.3281 | 201158.2188 | 193515.0156 | 0.181236917 | 0.174127934 | 0.188998836 | 0.256838624 | 0.264198205 | 0.103046053 |
| 627320.6875 | 581993.625 | 533424.6875 | 0.180825086 | 0.147088229 | 0.05362369 | 0.523473925 | 0.276514647 | 0.112639617 |
| 54425.74219 | 59925.24219 | 53506.11328 | 0.179717609 | 0.161932491 | 0.160358934 | 0.019268783 | -0.236050682 | 0.094923927 |
| 15339.61816 | 19519.45313 | 21743.35156 | 0.179695959 | 0.591325936 | 0.352596492 | 0.34708379 | 0.068785012 | 0.848721507 |
| 124780.4141 | 132096.9844 | 129612.5391 | 0.178042358 | 0.196593962 | 0.913436905 | 0.052756858 | 0.922230074 | 0.011850971 |
| 15586.05762 | 18840.65039 | 18393.34375 | 0.177831153 | 0.377103325 | 0.552043135 | 0.015625705 | 0.233985248 | 0.205896969 |
| 573250.9375 | 514019.7813 | 462689.75 | 0.177330367 | 0.219608402 | -0.323544889 | 0.043906958 | -0.37812396 | 0.088892469 |
| 369901.5 | 399863.8438 | 461380.1563 | 0.177292563 | 0.196492036 | -0.18271574 | 0.218053066 | 0.187328077 | 0.309561656 |
| 36210.57422 | 44029.36328 | 36806.59766 | 0.177031276 | 0.468515683 | -0.343384265 | 0.122322519 | -0.504556163 | 0.06065526 |
| 16626.54297 | 14450.75 | 11125.57715 | 0.175888863 | 0.556962894 | 0.168332948 | 0.555335026 | -0.215074795 | 0.545479767 |
| 2229584.25 | 2268906.25 | 2250745.5 | 0.175310262 | 0.243506704 | -0.344871674 | 0.128244359 | -0.308763857 | 0.148715498 |
| 116148.7266 | 112864.0703 | 115544.5781 | 0.175185033 | 0.142526436 | 0.088838376 | 0.362329238 | -0.065115729 | 0.481725146 |
| 21371.49219 | 19414.51953 | 19818.0332 | 0.175096438 | 0.045230483 | -0.595640992 | 0.015625705 | 0.133157313 | 0.153754411 |
| 27020.66406 | 25112.96875 | 24651.40234 | 0.174520528 | 0.052665954 | 0.215180051 | 0.215799184 | -0.001397617 | 0.986240204 |
| 84916.83594 | 91175.07031 | 89237.99219 | 0.174358416 | 0.41183809 | 0.771182739 | 0.055241445 | 1.095210291 | 0.018192328 |
| 78234.22656 | 97402.625 | 74531.89844 | 0.174123432 | 0.506010705 | -0.810464533 | 0.106102514 | -0.779504782 | 0.092242533 |
| 384035.5938 | 410368.5625 | 443605.0625 | 0.173468502 | 0.05059713 | 0.065038859 | 0.24184923 | 0.222913068 | 0.132010281 |
| 485162.3438 | 532226.3125 | 508939.7188 | 0.17306672 | 0.066021112 | 0.032104536 | 0.487594397 | 0.001500172 | 0.985129662 |
| 338730.6875 | 344232.7188 | 363580.4375 | 0.17265394 | 0.118163053 | -0.428323352 | 0.012254503 | -0.111850093 | 0.10883673 |
| 325977 | 325868.4688 | 280465.75 | 0.17261973 | 0.234724186 | -0.372294598 | 0.09842296 | -0.438438579 | 0.066134523 |
| 613323.25 | 602717.4375 | 628130.125 | 0.171953184 | 0.220198025 | -0.161152795 | 0.054678041 | 0.134375648 | 0.054109248 |
| 153390.2363 | 157176.1738 | 154035.0889 | 0.17184952 | 0.240263529 | 0.117811237 | 0.078888051 | 0.11186753 | 0.05536545 |

| | | | | | | | | |
|---|---|---|---|---|---|---|---|---|
| 3222150.25 | 2991704 | 3468677.5 | 0.171797328 | 0.194241398 | -0.137407301 | 0.218612366 | 0.020333883 | 0.861875296 |
| 239896.25 | 233448.9063 | 232895.8906 | 0.171077463 | 0.579372588 | 1.084126137 | 0.022021696 | 0.647598703 | 0.075467557 |
| 3949.199463 | 4012.912598 | 4331.124512 | 0.171061789 | 0.631616779 | -0.233002327 | 0.54685652 | -0.383919725 | 0.374030298 |
| 259391.1875 | 229995.4531 | 260783.8438 | 0.171034592 | 0.141771455 | -1.224099765 | 0.080723218 | -0.177301957 | 0.166774963 |
| 4993715 | 4932803 | 6319634 | 0.17075316 | 0.702894326 | -0.371438506 | 0.517875146 | 0.006208853 | 0.989480447 |
| 49303.91406 | 48382.72656 | 43955.5 | 0.170258605 | 0.327649439 | 0.182490982 | 0.289589686 | 0.437057591 | 0.063419062 |
| 96492.9375 | 100420.2188 | 97423.51563 | 0.170153903 | 0.102272218 | -0.324131903 | 0.045329621 | -0.183474118 | 0.091109394 |
| 395400.9688 | 399367.2188 | 419711.5938 | 0.170059704 | 0.137336288 | 0.495223201 | 0.024921531 | 0.396978264 | 0.030321148 |
| 146283.625 | 153620.1094 | 153242.1094 | 0.169665453 | 0.095307674 | 0.280772974 | 0.049184287 | 0.349535797 | 0.027949491 |
| 7743.424316 | 6528.556152 | 7719.79248 | 0.169645528 | 0.12131353 | 0.416245954 | 0.01878777 | 0.154363217 | 0.303928412 |
| 72026.17188 | 73120.35156 | 68335.28125 | 0.169593644 | 0.378186742 | 0.238414809 | 0.344752122 | -0.183223159 | 0.371142525 |
| 224291.3281 | 214356.9531 | 228540.8125 | 0.169330186 | 0.052584221 | -0.016799328 | 0.660053795 | 0.211975706 | 0.038131463 |
| 193618.6563 | 186439.8594 | 223572.3438 | 0.169010451 | 0.397275895 | 0.070304182 | 0.627813772 | 0.400151945 | 0.091108554 |
| 2459800.398 | 2547836.609 | 2760731.551 | 0.168826193 | 0.104262569 | -0.27309365 | 0.05818214 | -0.01920531 | 0.826852205 |
| 64322.97656 | 59993.89063 | 62829.89063 | 0.168294455 | 0.17437849 | 0.289508796 | 0.066354581 | 0.939448447 | 0.009714527 |
| 145770.6875 | 132756.0469 | 147618.2813 | 0.168060637 | 0.345950084 | 0.324538015 | 0.127872116 | 0.73039166 | 0.026263251 |
| 254913.6719 | 263799.5625 | 264109.3125 | 0.1679599 | 0.064047558 | 0.630195481 | 0.01179292 | 0.824841425 | 0.006394096 |
| 40882.52344 | 33728.18359 | 42990.99219 | 0.167837183 | 0.273264767 | 0.209537787 | 0.253751108 | 0.071450253 | 0.673365544 |
| 91196.3457 | 94646.04102 | 95289.28711 | 0.167804825 | 0.42324628 | 0.317651687 | 0.194685429 | 0.466815399 | 0.107440971 |
| 205364.9844 | 181249.25 | 191992.5469 | 0.167802115 | 0.018311389 | 0.068358273 | 0.068766483 | -0.329862704 | 0.058506939 |
| 676922.5 | 694969.25 | 703099.875 | 0.167744855 | 0.197302391 | 0.015037405 | 0.876081794 | -0.056596789 | 0.557774325 |
| 187607.2656 | 458144.25 | 198895.375 | 0.167641632 | 0.824285485 | -0.119823787 | 0.902675393 | -0.160077991 | 0.852108327 |
| 681810.8125 | 698500.4375 | 707683.3125 | 0.167482649 | 0.171503557 | 0.121646024 | 0.218950662 | 0.185562111 | 0.113248184 |
| 13040.80371 | 13406.11426 | 9141.892578 | 0.167434889 | 0.129225288 | -0.291851251 | 0.174452979 | -0.322595786 | 0.256923293 |
| 31401.33008 | 33970.86328 | 34050.25391 | 0.167430109 | 0.714543616 | 1.968427501 | 0.087322601 | 0.494066161 | 0.274194007 |
| 21768.49219 | 21816.22461 | 21382.11719 | 0.167241284 | 0.096730498 | 0.006507905 | 0.947783674 | -0.14303161 | 0.107790858 |
| 86006.07031 | 89691.17969 | 75984.26563 | 0.167001663 | 0.727807099 | 0.723084449 | 0.187316107 | 0.465309897 | 0.316722571 |
| 422751.5313 | 426530.5313 | 424544.8438 | 0.167001163 | 0.150947498 | 0.040285014 | 0.660613314 | -0.004721972 | 0.950211597 |
| 45818.65625 | 44261.23828 | 36976.11328 | 0.166888903 | 0.298674624 | 0.270786224 | 0.147248245 | -0.083591757 | 0.644791506 |
| 144201.5469 | 134241.9688 | 108611.0234 | 0.166870232 | 0.553168313 | 0.046010326 | 0.856816392 | 0.137843445 | 0.625396737 |
| 158323.9219 | 173775.9063 | 150369.5625 | 0.166834979 | 0.446293458 | -0.213836878 | 0.393001672 | -0.363909865 | 0.214765519 |
| 338555.3037 | 352063.3438 | 353164.4453 | 0.166710114 | 0.238945313 | 0.436353919 | 0.073853963 | 0.168810614 | 0.20663369 |
| 6479.492188 | 6791.182129 | 5586.358398 | 0.166612487 | 0.582738039 | -0.1119417 | 0.709576057 | 0.228465896 | 0.413778532 |

| | | | | | | | | |
|---|---|---|---|---|---|---|---|---|
| 12518.18848 | 13711.71875 | 11236.06934 | 0.166382171 | 0.318215645 | -0.427307237 | 0.110620378 | -1.73374925 | 0.038147816 |
| 104789.2578 | 107774.1328 | 99687.42969 | 0.166016692 | 0.535047385 | 0.931291951 | 0.046114261 | 0.897099872 | 0.041386295 |
| 461866.2188 | 489783.9375 | 500464.75 | 0.165705498 | 0.0424199 | 0.082582025 | 0.153662285 | 0.326654339 | 0.032474855 |
| 436256.3438 | 492902.2188 | 450872.125 | 0.165674171 | 0.148006693 | -0.628110092 | 0.039922031 | -0.047905686 | 0.652263825 |
| 332103.4063 | 307525.75 | 282656.7188 | 0.164829756 | 0.12557053 | 0.087346143 | 0.384749949 | -0.188789014 | 0.154329596 |
| 115369.1719 | 108631.5781 | 111002.5234 | 0.164712473 | 0.087043567 | 0.040996725 | 0.53018049 | -0.13157852 | 0.13062922 |
| 61990.08984 | 62881.83984 | 58109.16797 | 0.164626535 | 0.613531514 | 1.438883466 | 0.021204995 | 0.934437459 | 0.044458414 |
| 1002112.688 | 977685 | 977943.3125 | 0.164614868 | 0.230892382 | 0.349830191 | 0.022353303 | 0.134232133 | 0.073672213 |
| 82679.09375 | 93092.1875 | 89440.48438 | 0.164500223 | 0.17113456 | 0.265722921 | 0.039038389 | 0.143234831 | 0.177030473 |
| 17403.95703 | 18383.55469 | 17998.09375 | 0.1640394 | 0.122096077 | 0.011607137 | 0.937984911 | 0.073865177 | 0.354260455 |
| 1756447.625 | 1642840.875 | 1753592.125 | 0.163611882 | 0.157447235 | 0.145901579 | 0.042887894 | 0.209166381 | 0.052693084 |
| 16577.32422 | 20514.51367 | 16373.28027 | 0.163593114 | 0.370772663 | -0.076307055 | 0.423723931 | -0.206850863 | 0.269608292 |
| 13139.01367 | 14561.48438 | 12507.70508 | 0.163399598 | 0.143252431 | 0.390970236 | 0.055241445 | 0.126403863 | 0.317055242 |
| 11139.06348 | 11497.80078 | 16226.07617 | 0.163337509 | 0.495364972 | -0.276134127 | 0.243612448 | -0.064129601 | 0.834160705 |
| 11011.89453 | 10008.58887 | 9763.587891 | 0.163317981 | 0.159163342 | 0.174743323 | 0.284802359 | -0.005707049 | 0.958693133 |
| 648787.75 | 565524.3125 | 553922.375 | 0.163257607 | 0.137237024 | -0.086058674 | 0.399844084 | 0.076833846 | 0.550150503 |
| 27732.68359 | 28221.8125 | 23765.8125 | 0.162675898 | 0.200137373 | -0.019400496 | 0.867434697 | 0.056576145 | 0.661949635 |
| 10345.36133 | 9554.602539 | 8051.805664 | 0.162662108 | 0.327649439 | -0.95095614 | 0.043360794 | -1.169652816 | 0.020337832 |
| 726655.0625 | 769745.5625 | 889385.6875 | 0.162504562 | 0.074211155 | -0.004583436 | 0.928153584 | 0.028454048 | 0.839985854 |
| 44373.35156 | 47253.96484 | 47377.19922 | 0.162430505 | 0.196492036 | -0.237276075 | 0.041910161 | -0.168115205 | 0.056482153 |
| 22159.19922 | 25895.27539 | 23207.52344 | 0.162342171 | 0.547172327 | -0.061640794 | 0.817761485 | 0.154184795 | 0.547848654 |
| 4951.022949 | 5618.477539 | 6149.519043 | 0.161997121 | 0.294005007 | 0.226656433 | 0.105235029 | -0.268593166 | 0.132786986 |
| 92813.64844 | 99800.21875 | 90538.20313 | 0.161888943 | 0.257609696 | -0.073325367 | 0.298070114 | -0.238280507 | 0.057068534 |
| 1874.760986 | 2435.970215 | 2349.883789 | 0.16184839 | 0.588172074 | 0.315804791 | 0.368727629 | -0.016470566 | 0.961484633 |
| 43787.52344 | 42134.08203 | 46951.46094 | 0.161396802 | 0.165729568 | -0.488115518 | 0.076429114 | -0.283183396 | 0.078572249 |
| 51517.76172 | 73207.07031 | 55793.82422 | 0.160637744 | 0.695559783 | 0.042340896 | 0.922415374 | 0.072108672 | 0.870814221 |
| 257206.8125 | 227171.3594 | 224828.1875 | 0.160436021 | 0.333115267 | 0.206989639 | 0.24635716 | 1.28731696 | 0.019880539 |
| 34278.06641 | 41152.61719 | 37316.59766 | 0.160410758 | 0.371068627 | 0.34370444 | 0.122669631 | 0.592038224 | 0.047847186 |
| 128968.8516 | 140938.6406 | 142051.2813 | 0.160274514 | 0.228724059 | -0.064115723 | 0.577752036 | 0.340162343 | 0.056570421 |
| 131731.25 | 136403.0625 | 134940.2656 | 0.160259728 | 0.396209127 | -0.448533773 | 0.132089414 | -0.269329012 | 0.25220534 |
| 19975.16797 | 19747.56445 | 15287.43164 | 0.16023523 | 0.294591633 | -0.142781681 | 0.32677734 | -0.556023056 | 0.060841062 |
| 206090.7813 | 204720.6563 | 216342.3906 | 0.160053383 | 0.296394035 | -0.131952851 | 0.395285409 | 0.057912361 | 0.658833156 |
| 20049.52734 | 18204.33398 | 19860.5 | 0.159932282 | 0.216596761 | -0.126874264 | 0.260639434 | -0.378870345 | 0.029219266 |

| | | | | | | | | |
|---|---|---|---|---|---|---|---|---|
| 3911.94043 | 5107.575195 | 4012.13623 | 0.159782697 | 0.46699162 | -0.830092832 | 0.073442029 | -1.324086315 | 0.048299571 |
| 17980.56641 | 17930.73438 | 15026.80371 | 0.159531991 | 0.232198861 | -0.31249607 | 0.082329519 | -0.483001164 | 0.044191148 |
| 166375.3281 | 156144.9219 | 146477.2813 | 0.159468261 | 0.057845696 | -0.157975522 | 0.128476695 | -0.296786235 | 0.048590503 |
| 366999.8125 | 447974.0938 | 447315.3125 | 0.159404073 | 0.329576364 | -0.234119898 | 0.236672563 | -0.32734513 | 0.142656456 |
| 8668.637695 | 10168.93945 | 11687.49121 | 0.159204596 | 0.729581224 | 0.914385965 | 0.041250713 | 0.807099321 | 0.072978884 |
| 3363732.344 | 3669114.359 | 3838524.602 | 0.158961409 | 0.049995393 | -0.114065209 | 0.092926111 | -0.022862416 | 0.796833165 |
| 16322.61328 | 13350.91504 | 7226.608887 | 0.158948363 | 0.199046058 | -0.37565736 | 0.068812228 | -0.866281366 | 0.1193758 |
| 47090.98438 | 47834.64453 | 54986.80078 | 0.158533374 | 0.439269874 | -0.00993334 | 0.958320252 | 0.209389006 | 0.322805024 |
| 1725323.382 | 1810269.961 | 1725666.614 | 0.158440156 | 0.054063082 | -0.278363808 | 0.047347558 | -0.313771255 | 0.027391709 |
| 14312.08301 | 29729.31836 | 8318.290039 | 0.158107502 | 0.914068274 | -0.424478519 | 0.808336452 | -1.38114017 | 0.546828747 |
| 11093.37402 | 13429.20801 | 8411.194336 | 0.158080195 | 0.557959822 | -0.257063595 | 0.382455158 | -0.258459106 | 0.454419927 |
| 46419.72266 | 42990.07813 | 48408.41797 | 0.158024983 | 0.230656554 | -0.001282587 | 0.991588289 | 0.030838127 | 0.799843956 |
| 64589.33594 | 72635.82813 | 85473.21094 | 0.157928768 | 0.499210684 | -0.123443098 | 0.550165596 | 0.320854687 | 0.23049762 |
| 900424.4375 | 969604.375 | 999099.5625 | 0.157919423 | 0.150795791 | 0.087091428 | 0.630875263 | 0.43395047 | 0.030007011 |
| 8340001.5 | 10692444 | 9821178 | 0.157865265 | 0.600673217 | -0.12059686 | 0.757893323 | 0.2186652 | 0.461197364 |
| 680930.25 | 674846.375 | 688514.125 | 0.157797773 | 0.19588088 | -0.004980322 | 0.956378635 | -0.10442422 | 0.329987935 |
| 72155.03906 | 64027.60156 | 73776.38281 | 0.157209208 | 0.047979477 | -0.092546833 | 0.25003719 | 0.091971466 | 0.397327146 |
| 478367.75 | 551144.3125 | 503194.0625 | 0.157126834 | 0.207480826 | -0.310837803 | 0.087781599 | -0.142862889 | 0.237901119 |
| 17249.1543 | 17437.0918 | 17005.19922 | 0.156882201 | 0.683124393 | 0.805533608 | 0.093584 | 1.214241 | 0.059847517 |
| 21158.24414 | 21419.6875 | 19909.30078 | 0.156629692 | 0.64439335 | -0.072460484 | 0.839865475 | 0.238796184 | 0.465027161 |
| 60691.15234 | 48678.75 | 29310.62695 | 0.156301244 | 0.288211918 | -0.316197255 | 0.06824263 | -0.835260059 | 0.120303611 |
| 21260.55664 | 24407.07422 | 21778.79688 | 0.156194386 | 0.047692642 | 0.349961295 | 0.046508254 | 0.74846936 | 0.045697257 |
| 16007.35254 | 14865.16406 | 14463.31934 | 0.156114587 | 0.289180273 | 0.054231743 | 0.718681361 | -0.208472458 | 0.067380632 |
| 28871.12305 | 26306.80078 | 29105.41797 | 0.156030125 | 0.460141058 | 0.050431053 | 0.703047486 | -0.084071132 | 0.528571712 |
| 7879.032715 | 8344.961914 | 6020.011719 | 0.156019186 | 0.314910702 | 0.233745018 | 0.127335659 | -0.168597887 | 0.449433318 |
| 3204.984619 | 5682.583984 | 3046.680176 | 0.155517649 | 0.68199208 | 0.541608748 | 0.191226807 | 0.180136648 | 0.753859857 |
| 223013.5938 | 233062.2188 | 206396.5938 | 0.155486 | 0.217728505 | 0.35698428 | 0.058070756 | 0.292661092 | 0.075911738 |
| 6978.4375 | 7886.605957 | 6310.728516 | 0.155273074 | 0.478603334 | -0.614917527 | 0.099166239 | -1.186492274 | 0.047120928 |
| 69102.21875 | 62301.60547 | 69264.82813 | 0.155229344 | 0.336825283 | -0.400496379 | 0.065514518 | 0.157326645 | 0.201174116 |
| 285930.5313 | 304146.7813 | 298537.1875 | 0.155162537 | 0.227260721 | -0.08201898 | 0.193262335 | -0.183667758 | 0.050501801 |
| 304079.25 | 298925.3438 | 310064.4688 | 0.155116171 | 0.089040397 | -0.253666791 | 0.052457323 | 0.154543394 | 0.036848746 |
| 458415.75 | 495705.25 | 508685.2188 | 0.155078972 | 0.048531905 | 0.192373194 | 0.082763602 | 0.280853322 | 0.061106674 |
| 77829.33594 | 78743.42969 | 78063.625 | 0.154335848 | 0.127443855 | -0.530959649 | 0.046068794 | -0.624676681 | 0.044156984 |

| | | | | | | | | |
|---|---|---|---|---|---|---|---|---|
| 406845.9688 | 378511.0625 | 384582.6563 | 0.154082869 | 0.208916043 | 0.076683647 | 0.45682574 | 0.099419232 | 0.352925267 |
| 55000.76953 | 60695.53125 | 53540.39844 | 0.153864766 | 0.114738011 | -0.162351137 | 0.168900641 | -0.361792496 | 0.040959479 |
| 106878.5078 | 103020.8906 | 104307.4609 | 0.153502245 | 0.327649439 | 1.059189568 | 0.106169282 | 1.505234283 | 0.005561476 |
| 55176.85156 | 21607.35352 | 26393.74609 | 0.153116441 | 0.167171964 | -0.101128084 | 0.17813923 | 0.138953247 | 0.844045457 |
| 6047.794434 | 6749.9375 | 4229.01416 | 0.153084982 | 0.530716567 | -0.259040441 | 0.378072237 | -0.258936385 | 0.410161823 |
| 90581.61719 | 84696.60938 | 82914.6875 | 0.153024332 | 0.543069772 | -0.761946592 | 0.126731731 | -0.102505052 | 0.646918782 |
| 3549.496094 | 4356.963867 | 3618.346924 | 0.152825856 | 0.776365661 | -0.052897862 | 0.923262896 | -0.167610704 | 0.761804427 |
| 618848.875 | 585641.75 | 564030.0625 | 0.1528036 | 0.207403547 | 0.005581956 | 0.956152889 | -0.187671324 | 0.167349766 |
| 17727.75781 | 18248.35742 | 10956.36426 | 0.152727027 | 0.080486002 | -0.534661928 | 0.021622074 | -0.920260201 | 0.069238464 |
| 58772.77734 | 54975.99219 | 58051.71875 | 0.152654767 | 0.203641334 | -0.01546396 | 0.876081794 | -0.026228596 | 0.793189509 |
| 14780.4502 | 13582.70996 | 19904.62891 | 0.152564866 | 0.815681519 | 0.128966662 | 0.725923812 | 1.414094735 | 0.07648851 |
| 40935.44922 | 42349.33594 | 37419.95313 | 0.152391217 | 0.074826564 | -0.139623439 | 0.072240849 | -0.35177684 | 0.041176746 |
| 161460.3125 | 180112.3594 | 177168.8906 | 0.152264159 | 0.096483653 | 0.074056375 | 0.162505567 | -0.020792802 | 0.789122922 |
| 857651 | 849587.125 | 836784.3125 | 0.152128019 | 0.318687987 | 0.293549012 | 0.116475447 | 0.055757148 | 0.668657953 |
| 71790.96875 | 67056.48438 | 62452.16016 | 0.15198674 | 0.232084106 | 0.19722183 | 0.051888756 | -0.117184099 | 0.256723819 |
| 91128.70313 | 94625.02344 | 82040.92188 | 0.151903912 | 0.195925 | -0.76566542 | 0.030694475 | -0.536931127 | 0.031136253 |
| 10358.31152 | 10367.16406 | 6254.973145 | 0.151641823 | 0.80060403 | 0.475772183 | 0.398951669 | -0.392553309 | 0.575306589 |
| 24425.81641 | 24466.6875 | 26944.75 | 0.151565571 | 0.200813951 | 0.341162932 | 0.228114774 | -0.098512829 | 0.249396712 |
| 240527.5313 | 197507.6094 | 251391.5938 | 0.151339026 | 0.134574193 | -0.115375292 | 0.238316696 | -0.180016451 | 0.301755353 |
| 50229.32813 | 43745.16797 | 53832.12891 | 0.151145929 | 0.494764936 | 0.409778109 | 0.117302538 | 0.710387437 | 0.046487363 |
| 166913.2969 | 194144.1875 | 147728.6094 | 0.150059108 | 0.28738368 | -0.116081869 | 0.168900641 | -0.348634468 | 0.131531367 |
| 20356.78906 | 23193.53906 | 20369.28906 | 0.149868767 | 0.240260695 | 0.150014386 | 0.25963333 | 0.220435321 | 0.156337302 |
| 26347.81641 | 27383.17578 | 23453.62305 | 0.149838224 | 0.380648504 | 0.277237628 | 0.214649734 | -0.215590134 | 0.215760001 |
| 1245879 | 1329868.25 | 1013259.75 | 0.14958976 | 0.565985157 | 0.342113884 | 0.246468737 | 0.301432695 | 0.302401454 |
| 79307.76563 | 80311.53906 | 81478.375 | 0.149251305 | 0.183137723 | -0.127235474 | 0.29276405 | -0.337358489 | 0.084288402 |
| 117372 | 116875.3125 | 52969.37891 | 0.149089791 | 0.759466989 | 0.159697342 | 0.687745186 | -0.294808234 | 0.597655284 |
| 43262144 | 57445528 | 60241960 | 0.148841522 | 0.688031657 | -0.221719368 | 0.596219861 | 0.378130434 | 0.326227143 |
| 199590.2031 | 195369.0625 | 186733.0781 | 0.148551567 | 0.370458663 | 0.230801537 | 0.205568001 | -0.076151744 | 0.617308452 |
| NA | 744.7860718 | 818.4251709 | 0.148517118 | NA | 0.841521216 | NA | 0.70505762 | NA |
| 87541.1875 | 87356.47656 | 87725.82813 | 0.147616308 | 0.135239277 | -0.07286573 | 0.490020706 | 0.154963756 | 0.019260832 |
| 147980.0469 | 155678.7344 | 153142.5156 | 0.147427044 | 0.268922439 | 0.265255534 | 0.301926295 | -0.108836656 | 0.395364978 |
| 82600.1875 | 71763.52344 | 76521.47656 | 0.14724577 | 0.06931047 | -0.145673817 | 0.464654667 | -0.272586055 | 0.066134523 |
| 5948726 | 6793318.5 | 6492096.5 | 0.147113515 | 0.594192447 | 0.396112843 | 0.254161608 | 0.158897465 | 0.544452231 |

| | | | | | | | | |
|---|---|---|---|---|---|---|---|---|
| 34355.73438 | 41465.91016 | 36323.74609 | 0.147038646 | 0.294587393 | 0.535690002 | 0.04183385 | -0.255560408 | 0.169021874 |
| 43764.64063 | 45369.21484 | 46866.64063 | 0.146970553 | 0.352815992 | 0.348187303 | 0.036088129 | 0.235549958 | 0.053854543 |
| 275057.125 | 282744.9063 | 275089.7188 | 0.146645534 | 0.249807798 | -0.223153834 | 0.068255066 | -0.063343335 | 0.339006931 |
| 948180.4375 | 951811.875 | 1152976.5 | 0.146616294 | 0.167251239 | -0.021431466 | 0.706433309 | 0.105382648 | 0.510430611 |
| 38279.49609 | 36351.19141 | 36208.91016 | 0.146576169 | 0.49237983 | -0.275607528 | 0.193488746 | -0.267276374 | 0.196593026 |
| 43707.85254 | 37212.16699 | 40196.71631 | 0.146466439 | 0.132446456 | -0.243387347 | 0.073442029 | -0.350473112 | 0.052503815 |
| 515810.4375 | 551627.1875 | 498166.0625 | 0.146184956 | 0.150795791 | 0.015412805 | 0.834026179 | -0.002894302 | 0.975957362 |
| 103256.3438 | 103092.25 | 102048.7031 | 0.146153827 | 0.152553608 | -0.20730814 | 0.055241445 | -0.258880361 | 0.00810066 |
| 49745.21094 | 55307.23828 | 22941.26758 | 0.146113057 | 0.322065711 | -0.861913249 | 0.019141974 | -1.012931017 | 0.106375556 |
| 3964.844727 | 2643.042969 | 1699.761475 | 0.145808742 | NA | 0.971263006 | NA | -0.494588031 | NA |
| 40355.42188 | 42842.84375 | 43820.94531 | 0.145707758 | 0.314260232 | -0.126991717 | 0.39679721 | -0.020363167 | 0.879736047 |
| 48516.24219 | 41265.1875 | 36429.19531 | 0.14521673 | 0.47152965 | 0.240363598 | 0.271818262 | 0.485680887 | 0.124439663 |
| 142807.2344 | 146404.1875 | 139245.5938 | 0.145211024 | 0.076923702 | 0.186475507 | 0.057917515 | 0.123388498 | 0.101150402 |
| 47760.15625 | 43574.42578 | 49617.01172 | 0.145151748 | 0.194454556 | 0.065870541 | 0.468709868 | -0.409859976 | 0.045593044 |
| 73600.96094 | 74957.78125 | 59941.67969 | 0.144945918 | 0.762776035 | 0.672528027 | 0.1214746 | 0.301991258 | 0.200766379 |
| 118349.6953 | 137779.3125 | 126364.0625 | 0.144754725 | 0.673150712 | -0.226729204 | 0.332451144 | 0.403186987 | 0.069042103 |
| 20229.16211 | 25544.97656 | 21491.77734 | 0.144698626 | 0.460627897 | -0.244901245 | 0.057753648 | -0.133032283 | 0.420233357 |
| 72574.78125 | 77855.17188 | 63490.07813 | 0.144412472 | 0.433494258 | -0.165623216 | 0.185475723 | 0.18343526 | 0.276254237 |
| 544512.375 | 567158.9375 | 583315.3125 | 0.144246401 | 0.215495312 | -0.286205413 | 0.122851676 | -0.385358648 | 0.069207996 |
| 40252.11328 | 38577.59766 | 47303.30859 | 0.144135723 | 0.653484121 | 0.709862688 | 0.072221754 | 0.881395497 | 0.039446558 |
| 1413.523926 | 1285.739258 | 950.303894 | 0.143917757 | 0.783418048 | -0.12270407 | 0.875125893 | -0.933507324 | 0.280633075 |
| 2195.797607 | 3621.500244 | NA | 0.143783566 | 0.616828703 | -0.858200048 | 0.125646125 | -0.646899375 | NA |
| 79327.60938 | 84077.84375 | 78542.59375 | 0.143720191 | 0.148202641 | -0.07048365 | 0.540771148 | -0.243867508 | 0.075839948 |
| 20805.60938 | 23630.48438 | 17711.64063 | 0.143596906 | 0.663538898 | 0.369198945 | 0.355229313 | 0.032342019 | 0.925417836 |
| 60388.55469 | 60876.80078 | 65940.07813 | 0.143458557 | 0.069867455 | -0.557273301 | 0.021622074 | -0.429519596 | 0.023619478 |
| 44268.51172 | 50916.22656 | 41973.5625 | 0.143150504 | 0.372680569 | 0.321950388 | 0.052756858 | 0.199833045 | 0.24120794 |
| 670812.3125 | 649280.5625 | 640541.8125 | 0.142884965 | 0.153864436 | -0.073566256 | 0.392329965 | -0.102591574 | 0.262772885 |
| 53800.64453 | 52307.80469 | 51290.93359 | 0.142859353 | 0.566211171 | 0.23208172 | 0.39602624 | 0.402329034 | 0.174355634 |
| 89763.625 | 80868.44531 | 88638.29688 | 0.142806863 | 0.174750491 | 0.069373736 | 0.284902041 | -0.165680363 | 0.107790858 |
| 587676.5625 | 647904.3125 | 688114.1875 | 0.142780608 | 0.134417501 | -0.230543213 | 0.099454811 | 0.166224969 | 0.215760001 |
| 100988.8125 | 92387.11719 | 93315.86719 | 0.142734108 | 0.12643787 | 0.140783251 | 0.469555906 | 0.400593285 | 0.032277182 |
| 42256.34766 | 39753.09375 | 42991.77344 | 0.142471309 | 0.252399814 | 0.321423747 | 0.065260161 | 0.353856751 | 0.044724508 |
| 58984.39063 | 44748.39844 | 65853.32031 | 0.142271654 | 0.413482116 | -0.050639186 | 0.713335915 | 0.816636648 | 0.119708622 |

| | | | | | | | | |
|---|---|---|---|---|---|---|---|---|
| 42439.66406 | 45950.60547 | 45443.86719 | 0.142014846 | 0.439671939 | 0.053724068 | 0.758181615 | 0.100612697 | 0.553731254 |
| 35975.20313 | 35562.41016 | 27109.02539 | 0.141909713 | 0.31287003 | -0.8776679 | 0.021204995 | -0.75134638 | 0.035174528 |
| 1561.970825 | 21176.15039 | 3834.239502 | 0.141809219 | NA | -0.601671104 | NA | 0.268710534 | NA |
| 54098.07813 | 59049.59375 | 61857.79297 | 0.141790977 | 0.268790045 | 0.395673746 | 0.241244667 | 0.346566425 | 0.060012809 |
| 34027.87109 | 37809.69922 | 39372.87109 | 0.141676312 | 0.316898283 | -0.200232833 | 0.323640378 | 0.814726503 | 0.023855453 |
| 641570.9375 | 670239.125 | 745426.0625 | 0.141632879 | 0.115980925 | -0.008899722 | 0.82951387 | 0.021920172 | 0.834160705 |
| 456892.5313 | 404229.1875 | 398085.6563 | 0.141502499 | 0.442408553 | -0.358904517 | 0.187014813 | -0.437279113 | 0.130599473 |
| 49931.34766 | 46453.73828 | 44908.12891 | 0.141461279 | 0.180251704 | 0.54288075 | 0.03062316 | 0.196353667 | 0.107554917 |
| 30523.91406 | 30077.26172 | 24998.12109 | 0.141288997 | 0.336825283 | -0.300851057 | 0.149895744 | -0.680714574 | 0.047120928 |
| 559791.1875 | 644812.625 | 557555.6875 | 0.141283234 | 0.314306229 | 0.121720267 | 0.110122293 | 0.298277118 | 0.119027258 |
| 11316.87598 | 11997.78027 | 11631.19336 | 0.141025612 | 0.530730634 | 0.159097575 | 0.485213684 | 0.146290913 | 0.493991598 |
| 460847.5313 | 490754.5625 | 478922.8438 | 0.140626757 | 0.055345971 | 0.046080328 | 0.342664252 | -0.136847535 | 0.074131758 |
| 93070.45313 | 102006.7656 | 101456.7422 | 0.140447021 | 0.032875666 | -0.299808951 | 0.042487934 | -0.360670407 | 0.029492107 |
| 211610.125 | 194434.5313 | 146325.8281 | 0.140366195 | 0.746281035 | 0.093508802 | 0.576394593 | 0.097761115 | 0.690688311 |
| 113890.5078 | 99368.21094 | 117254.375 | 0.14017439 | 0.502961317 | -0.832603074 | 0.098835098 | -0.43117649 | 0.16241362 |
| 3264.509277 | 2417.931396 | 1468.039429 | 0.140147102 | 0.74277824 | -0.483931507 | 0.321888877 | -0.394727469 | 0.463215413 |
| 6032878.5 | 6318787.5 | 6467872 | 0.139894209 | 0.169115491 | 0.265786443 | 0.069257825 | 0.238784419 | 0.062565713 |
| 164221.4688 | 136129 | 131708.1406 | 0.139422615 | 0.274371306 | 0.22155371 | 0.139976934 | 0.331373784 | 0.160773455 |
| 1068639.75 | 1146071.75 | 1026978.938 | 0.139381316 | 0.413987351 | -0.153248946 | 0.094777667 | 5.18E-05 | 0.999456208 |
| 1737293.125 | 1707527.75 | 969148.8125 | 0.139229694 | 0.642217762 | -0.086209798 | 0.729706165 | -0.196731528 | 0.584551698 |
| 417623.4688 | 450772.2813 | 314979.2188 | 0.139154324 | 0.449151467 | -0.456418113 | 0.152460574 | -0.376884487 | 0.192235011 |
| 54936.05469 | 55905.25391 | 34249.64063 | 0.139127937 | 0.598534405 | -0.162521618 | 0.680064291 | -0.797008457 | 0.090965556 |
| 6673.439453 | 7425.114258 | 6585.299316 | 0.13910002 | 0.719910414 | 0.206286996 | 0.578054117 | -0.004997349 | 0.989140123 |
| 71999.57813 | 64498.31641 | 67328.47656 | 0.138960853 | 0.120050919 | -0.380614784 | 0.036730784 | -0.149474454 | 0.141398238 |
| 118826.8359 | 121263.6172 | 120213.0313 | 0.138872227 | 0.06853067 | 0.137861665 | 0.246470557 | 0.638858761 | 0.011811065 |
| 17013.15234 | 14038.03516 | 18313.14258 | 0.13876178 | 0.908725451 | 0.870198096 | 0.253993044 | 0.805760403 | 0.275313311 |
| 102514.0234 | 120579.3438 | 105684.8281 | 0.138677565 | 0.193815098 | -0.354727868 | 0.052817461 | -0.298953161 | 0.079203886 |
| 445515.125 | 403240.4375 | 449231.5313 | 0.138545566 | 0.263336537 | -0.014275169 | 0.88053234 | 0.254140372 | 0.074184841 |
| 9719.010742 | 10189.75488 | 10056.0332 | 0.138432615 | 0.250607381 | -0.317263653 | 0.115650681 | -0.4354401 | 0.076362612 |
| 85647.91406 | 112280.3281 | 101839.3047 | 0.138388892 | 0.705229337 | -0.502053745 | 0.31156031 | 0.091004121 | 0.801773096 |
| 45191.01563 | 43670.67578 | 41761.08594 | 0.138366178 | 0.774038969 | 1.451830558 | 0.023005377 | 1.492818138 | 0.01054584 |
| 319049.25 | 299899.5 | 277518.0625 | 0.138355997 | 0.085338824 | -0.037266135 | 0.306709414 | -0.256787893 | 0.096631602 |
| 9099.672852 | 9945.96875 | 11021.39551 | 0.137812446 | 0.332906865 | -0.132296844 | 0.379832076 | -0.415503638 | 0.082912076 |

| | | | | | | | | |
|---|---|---|---|---|---|---|---|---|
| 266915.9063 | 281634.875 | 297387.4688 | 0.137690711 | 0.066736762 | 0.051105528 | 0.621972901 | 0.030201782 | 0.695793958 |
| 30024.99023 | 29105.99805 | 32521.73633 | 0.137656645 | 0.483402665 | -0.091471627 | 0.637007251 | -0.515853144 | 0.10883673 |
| 5670.723145 | 5740.270508 | 5096.727539 | 0.137436086 | 0.152416319 | -0.185198871 | 0.079287989 | -0.770737533 | 0.010654184 |
| 30847.93359 | 26414.41797 | 29448.96289 | 0.137207929 | 0.438477855 | -0.618560454 | 0.017037525 | -0.698259112 | 0.021984636 |
| 37093.86719 | 34967.34375 | 36528.61328 | 0.136750694 | 0.721385477 | 1.045032454 | 0.065909751 | 1.51453577 | 0.008614663 |
| 2181.763672 | 2643.061035 | 2826.696777 | 0.136733604 | 0.721394761 | -0.26497963 | 0.258368613 | -0.524401109 | 0.09810161 |
| 129639.5547 | 128774.1797 | 134346.4688 | 0.136545555 | 0.192885661 | 0.154826039 | 0.192295741 | 0.136513913 | 0.186667547 |
| 32392.74609 | 31450.48828 | 26880.93945 | 0.136348884 | 0.639887147 | -0.306961586 | 0.399403845 | -0.5769336 | 0.208663062 |
| 39322.83984 | 39802.76953 | 41532.33203 | 0.136300198 | 0.537423719 | -0.370060732 | 0.228822955 | -0.293333528 | 0.281316231 |
| 19492.53125 | 21792.01563 | 20825.3457 | 0.136267588 | 0.246849543 | 0.013348885 | 0.90544874 | 0.079094808 | 0.479451609 |
| 18609.42969 | 14070.44824 | 15883.88477 | 0.136153016 | 0.604697925 | -0.26580174 | 0.045329621 | -0.16738159 | 0.377470755 |
| 29776.31836 | 27645.625 | 26017.32617 | 0.136136169 | 0.175245521 | -0.64599171 | 0.026250916 | -0.879685852 | 0.010654184 |
| 148433.5469 | 152956.7188 | 155804.7969 | 0.13606747 | 0.07312577 | 0.030412597 | 0.73064637 | -0.039791603 | 0.448159735 |
| 151455.0156 | 167568.1719 | 161177.4219 | 0.135751832 | 0.328216285 | -0.194682039 | 0.18823259 | -0.397411958 | 0.029820656 |
| 140600.3594 | 131767.8438 | 96325.03906 | 0.135663983 | 0.255950078 | 0.666028804 | 0.022223954 | -0.046762827 | 0.852108327 |
| 61745.75391 | 47288.19531 | 53294.03125 | 0.135636177 | 0.426378548 | 0.21906933 | 0.263424805 | -0.033331549 | 0.879817022 |
| 554743.6875 | 691965.8125 | 683051.375 | 0.135616685 | 0.199046058 | 0.001257235 | 0.991529214 | -0.007178259 | 0.96818931 |
| 27290.28516 | 34422.95703 | 34636.83594 | 0.135446511 | 0.107464969 | 0.015936174 | 0.820507974 | 0.160666059 | 0.403879392 |
| 37574.14453 | 24611.78906 | 31541.79688 | 0.135343011 | 0.625707737 | 0.115177496 | 0.791291884 | -0.058509013 | 0.840720144 |
| 48731.07031 | 47293.44922 | 46414.38281 | 0.135304356 | 0.536898334 | 0.059409432 | 0.794486514 | 0.320258533 | 0.204994112 |
| 327667.3125 | 332449.625 | 281183.5 | 0.135286645 | 0.294498236 | 0.286297222 | 0.124254466 | -0.217454901 | 0.191706351 |
| 37199.93359 | 47175.61719 | 37352.44141 | 0.135286044 | 0.498332261 | -0.125663523 | 0.689773477 | 0.296564144 | 0.257497935 |
| 39254.94141 | 34778.54688 | 40816.17969 | 0.135284559 | 0.156854993 | -0.565080429 | 0.018539883 | -0.849910925 | 0.011923035 |
| 423200.6387 | 446655.959 | 437892.6797 | 0.135281764 | 0.132311059 | -0.135258636 | 0.168390259 | -0.222258867 | 0.07588923 |
| 16825.8418 | 20373.38281 | 20488.18945 | 0.135194148 | 0.608614091 | 0.433649309 | 0.170700977 | 0.569533628 | 0.063419062 |
| 450626.5938 | 491197.5 | 441870.5 | 0.134949728 | 0.06480491 | -0.36808321 | 0.027435011 | -0.379249275 | 0.027629097 |
| 475778.7813 | 496872.5625 | 509569.5 | 0.134670436 | 0.122437469 | 0.083392511 | 0.26604978 | 0.245748758 | 0.04296087 |
| 5287.922852 | 6796.166016 | 6699.185547 | 0.134643465 | 0.252872698 | -0.797915587 | 0.034480785 | -0.924377849 | 0.021984636 |
| 197417.9531 | 180747.875 | 188675.3281 | 0.13447785 | 0.064774085 | -0.101613496 | 0.197313963 | -0.306894441 | 0.033928174 |
| 29537.55664 | 29614.58203 | 27175.4043 | 0.134458326 | 0.355407665 | -0.339651469 | 0.131554483 | -0.3783151 | 0.112423117 |
| 145672.25 | 153531.6406 | 155402.25 | 0.134427174 | 0.146737393 | -0.038768581 | 0.592296138 | 0.337184116 | 0.029219266 |
| 15939.52051 | 16241.29004 | 16223.7666 | 0.133871289 | 0.404498232 | 0.141386461 | 0.174452979 | 0.146049116 | 0.136989085 |
| 29423.75586 | 30182.55273 | 10121.41016 | 0.133697352 | 0.434484496 | -0.142192833 | 0.419274561 | 0.080665016 | 0.903657204 |

| | | | | | | | | |
|---|---|---|---|---|---|---|---|---|
| 75483.74219 | 70352.96094 | 75117.27344 | 0.13369522 | 0.563894297 | 0.041556492 | 0.858230595 | -0.449444674 | 0.206265186 |
| 210048.2031 | 210812.625 | 209533.1406 | 0.132816969 | 0.097443336 | 0.074759998 | 0.246470557 | -0.119784537 | 0.154716822 |
| 248087.2813 | 251062.25 | 241154.125 | 0.132704981 | 0.187635714 | 0.185504155 | 0.065557261 | 0.543469626 | 0.009824429 |
| 20106.78516 | 23004.73438 | 19683.78906 | 0.132672093 | 0.155497883 | -0.229803234 | 0.126707606 | -0.377035146 | 0.073130408 |
| 2278927.75 | 2610959.75 | 2118597 | 0.132671931 | 0.48022212 | -0.443173403 | 0.090276742 | -0.959768795 | 0.024715562 |
| 85034.53125 | 82945.67188 | 79675.27344 | 0.13239501 | 0.208744132 | 0.028286463 | 0.66660509 | -0.465179668 | 0.01054584 |
| 346854.5 | 307943.0938 | 342537.5625 | 0.132326772 | 0.335712418 | 0.078891984 | 0.683876356 | -0.495243871 | 0.067667169 |
| 59621.625 | 59390.65625 | 54560.00391 | 0.13192063 | 0.732781314 | 1.019995476 | 0.03968601 | 2.173598193 | 0.008041683 |
| 497040.5313 | 551645.375 | 573431.625 | 0.131606202 | 0.080486002 | 0.031396068 | 0.822759557 | -0.181165306 | 0.139201262 |
| 8845.479492 | 7841.199219 | 9027.047852 | 0.131578402 | 0.524853127 | -0.075765748 | 0.68122239 | 0.265733767 | 0.196239339 |
| 15953.43066 | 18643.54297 | 13910.87109 | 0.130553678 | 0.562059843 | -0.055346036 | 0.762412522 | -0.018490739 | 0.942386927 |
| 176637.8906 | 207478.4219 | 170614.3281 | 0.130547636 | 0.10058931 | -0.068439166 | 0.623543083 | -0.215592713 | 0.192235011 |
| 47938.07813 | 52457.33594 | 48987.55859 | 0.130401022 | 0.245093451 | -0.167295026 | 0.137745823 | -0.180404642 | 0.094664375 |
| 15661.22266 | 15115.04004 | 8850.298828 | 0.130068125 | 0.743680462 | -0.096913008 | 0.667331576 | -0.238430227 | 0.515342263 |
| 96982.60938 | 99408.09375 | 91079.35156 | 0.129644743 | 0.274068475 | -0.010361146 | 0.855467828 | 0.235300895 | 0.058410816 |
| 52781.69531 | 50986.48438 | 61352.55078 | 0.129155492 | 0.370141645 | 0.531884073 | 0.024165741 | 0.193278221 | 0.249879244 |
| 14522.04688 | 18089.76758 | 12435.69922 | 0.128941889 | 0.691165074 | 1.260864037 | 0.053365234 | -0.30949415 | 0.402592316 |
| 26880.75586 | 24488.37695 | 26305.78125 | 0.128247917 | 0.284352047 | -0.058808476 | 0.456340808 | -0.151059414 | 0.149855034 |
| 376321.0313 | 365474.1875 | 367077.5 | 0.128233918 | 0.083280787 | 0.453402225 | 0.092766836 | 0.309092077 | 0.01858328 |
| 308323.4688 | 311252.9063 | 277990.125 | 0.127860092 | 0.217906015 | -0.516382452 | 0.040327952 | -0.510236269 | 0.034200633 |
| 23077.55859 | 21603.54102 | 23833.30078 | 0.127458759 | 0.63242956 | 0.578612176 | 0.042884533 | 0.494310537 | 0.045495106 |
| 1117179.5 | 1327112.375 | 1113621.125 | 0.1273346 | 0.676220544 | 0.475688629 | 0.087437274 | 0.371652241 | 0.120554018 |
| 86522.50781 | 91428.24219 | 87564.65625 | 0.126863499 | 0.407278175 | 0.147732345 | 0.341247073 | 0.027890968 | 0.842137752 |
| 14961.33887 | 20474.05273 | 14681.19922 | 0.126771971 | 0.362249448 | -0.336750746 | 0.102526216 | -0.323391334 | 0.238240967 |
| 11801.18555 | 10904.6123 | 10699.07422 | 0.126702996 | 0.670886568 | -0.362772085 | 0.350428319 | -1.374792831 | 0.107874808 |
| 16477.39648 | 17288.14453 | 20250.41406 | 0.12656743 | 0.295415945 | -0.141419143 | 0.134227499 | -0.086548734 | 0.544452231 |
| 381869.4063 | 390898.7188 | 376632.3125 | 0.12638996 | 0.301892106 | 0.446459045 | 0.047347558 | 0.398424179 | 0.062345205 |
| 23297.33594 | 20730.85547 | 19375.10938 | 0.126036713 | 0.328932057 | 0.010385983 | 0.95424101 | -0.141405414 | 0.370576128 |
| NA | 4806.878906 | NA | 0.125846352 | 0.925171957 | 2.067116525 | NA | 1.089829661 | NA |
| 173084.4063 | 167093.0469 | 158264.6875 | 0.125444619 | 0.20270675 | 1.003422272 | 0.052817461 | 1.008149068 | 0.012701134 |
| 70040.34375 | 68102.08594 | 67428.46094 | 0.125173585 | 0.207247298 | 0.213814072 | 0.084811505 | -0.177207338 | 0.126766337 |
| 55893.84375 | 61738.73438 | 62895.58203 | 0.125138598 | 0.163060402 | 0.005326 | 0.95057864 | 0.024979779 | 0.794935317 |
| 83308.86719 | 88942.625 | 80082.38281 | 0.125106688 | 0.245606762 | 0.106270155 | 0.160437134 | -0.105725736 | 0.20605118 |

| | | | | | | | | |
|---|---|---|---|---|---|---|---|---|
| 66728.30469 | 74148.89063 | 70230.46875 | 0.125034626 | 0.420245262 | 0.108606636 | 0.435753831 | 0.072538825 | 0.596258185 |
| 18411.19336 | 24539.96289 | 18488.18945 | 0.124664732 | 0.686369329 | 0.55732045 | 0.16494964 | 0.159075049 | 0.632470042 |
| 130185.2813 | 125395.5391 | 113188.0469 | 0.124528015 | 0.236422852 | -0.009140441 | 0.914247671 | -0.326535632 | 0.055401366 |
| 2858.407715 | 2801.62207 | 2129.661377 | 0.124469797 | 0.797520944 | -0.626781916 | 0.343585509 | -0.440756779 | 0.436300539 |
| 78017.65625 | 85182.28906 | 79544.55469 | 0.124155494 | 0.227575325 | -0.434972966 | 0.018539883 | -0.457963087 | 0.026263251 |
| 60215.8125 | 66001.0625 | 63918.28125 | 0.123971331 | 0.458032129 | 0.846378755 | 0.029594208 | 0.711013452 | 0.027949491 |
| 113551.9141 | 159640.2031 | 145871.6719 | 0.123968404 | 0.186588896 | -0.051508525 | 0.674335952 | -0.024016209 | 0.916489217 |
| 50715.22656 | 39119.63281 | 36778.25391 | 0.123955253 | 0.474480685 | 0.072977111 | 0.77771169 | 0.317948185 | 0.290063497 |
| 640580.0625 | 627460.625 | 536732.0625 | 0.123699565 | 0.542678823 | -0.153925688 | 0.487787786 | -0.055525091 | 0.798118541 |
| 492922.875 | 506123.3438 | 503874.625 | 0.123500801 | 0.46699162 | 0.55569082 | 0.052817461 | 0.477963608 | 0.086952277 |
| 61988.04688 | 64888.30469 | 65672.46875 | 0.123264205 | 0.222800905 | -0.057381126 | 0.149058018 | -0.056032849 | 0.23873122 |
| 11661.58398 | 10811.64648 | 8424.427734 | 0.123195762 | 0.377522736 | 0.017537114 | 0.875773261 | -0.061951849 | 0.763776161 |
| 71141.67188 | 80442.5 | 71436.42188 | 0.123176564 | 0.156563923 | -0.3348389 | 0.046285147 | -0.156632858 | 0.185030852 |
| 16658.54297 | 13754.67871 | 13587.27148 | 0.122982583 | 0.386749047 | 0.180475991 | 0.298644378 | -0.002196638 | 0.989329895 |
| 15303.41699 | 14454.57422 | 13730.25098 | 0.122587898 | 0.766981151 | -0.072066751 | 0.679388407 | 0.115487944 | 0.329050274 |
| 4345.554688 | 3474.365479 | 4740.529785 | 0.122488811 | 0.605864892 | 1.286515233 | 0.156220504 | 0.177673489 | 0.438271266 |
| 2816229.75 | 3284102 | 3513157.5 | 0.122464799 | 0.318687987 | 0.032271205 | 0.739638889 | 0.173806612 | 0.326227143 |
| 43416.26563 | 42745.01563 | 44760.20313 | 0.122385772 | 0.130461034 | 1.035260187 | 0.119032068 | 1.077120281 | 0.006394096 |
| 24603.12695 | 26950.32617 | 22366.66797 | 0.1222739 | 0.296437226 | 0.11593942 | 0.176130095 | -0.297163385 | 0.104365856 |
| 13562.24023 | 14786.13477 | 13086.90234 | 0.122241749 | 0.110478433 | -0.118184768 | 0.238358909 | -0.194939577 | 0.098799268 |
| 122895.3516 | 111915.3672 | 112538.2031 | 0.122152168 | 0.040749235 | 0.274850956 | 0.015625705 | 0.076143311 | 0.339927953 |
| 369481.8438 | 296740.9063 | 262116.2656 | 0.121959554 | 0.59228898 | 0.541625542 | 0.13928207 | 0.304693974 | 0.324580717 |
| 38087.58984 | 34383.75 | 31053.60156 | 0.121855667 | 0.827936728 | -0.062804181 | 0.919703825 | -0.028704052 | 0.963218346 |
| 321322.0625 | 322002.0625 | 328395.6875 | 0.121747309 | 0.137440936 | -0.05439229 | 0.414751124 | -0.011839875 | 0.841259847 |
| 125392.1094 | 118816.125 | 128846.4219 | 0.121551684 | 0.586014066 | -0.402082103 | 0.087875794 | -0.179200699 | 0.154835571 |
| 109576.9766 | 119057.4297 | 116104.8047 | 0.121271705 | 0.163416464 | -0.175971239 | 0.110122293 | -0.249648154 | 0.05752489 |
| 46433.30078 | 40716.28516 | 39398.72656 | 0.121172435 | 0.635577559 | -0.120559399 | 0.259300939 | 0.206351467 | 0.19608824 |
| 655304.8789 | 685617.4453 | 619443.3398 | 0.12051256 | 0.147312752 | -0.058393196 | 0.276203697 | -0.185316596 | 0.078691633 |
| 67617.52344 | 63959.62891 | 64773.90625 | 0.1204574 | 0.706689159 | 0.248058783 | 0.442202166 | 0.67347021 | 0.11550321 |
| 42367.84375 | 40896.40234 | 37773.72266 | 0.120434215 | 0.184741724 | 0.159075461 | 0.142053668 | 0.102164966 | 0.269349179 |
| 35024.39063 | 33967.25 | 33726.58984 | 0.120112891 | 0.429442948 | 0.67405713 | 0.043906958 | -0.145346182 | 0.350351437 |
| 39793.9375 | 39384.39453 | 40003.14844 | 0.120071346 | 0.264863333 | 0.248960816 | 0.067679724 | 0.320367474 | 0.049415261 |
| 121759.5938 | 119933.0391 | 121985.1719 | 0.120024064 | 0.098618413 | -0.204532435 | 0.042222958 | 0.048493548 | 0.268750571 |

| | | | | | | | | |
|---|---|---|---|---|---|---|---|---|
| 98183.42969 | 144638.0938 | 80005 | 0.119735567 | 0.682301096 | -0.133774942 | 0.650953861 | -0.153512065 | 0.713487699 |
| 122279.0859 | 120897.375 | 124579.3281 | 0.119667123 | 0.232568235 | -0.170750139 | 0.188452025 | -0.179082692 | 0.159389389 |
| 39015.26563 | 40587.00391 | 37765.375 | 0.119236123 | 0.435137799 | 0.021728758 | 0.878651713 | -0.408504181 | 0.118023574 |
| 2193307.25 | 2297509.75 | 2350091.75 | 0.119201633 | 0.329483256 | -0.262771273 | 0.073326865 | 0.043098175 | 0.564224231 |
| 334747.7813 | 350612.4688 | 341780.5938 | 0.119068716 | 0.422797031 | -0.116374509 | 0.160926773 | 0.036221373 | 0.544630143 |
| 224932.1719 | 206609.1719 | 181270.2031 | 0.119030841 | 0.468515683 | 0.146582068 | 0.321888877 | 0.579636113 | 0.065367829 |
| 80541.53906 | 82734.10938 | 74813.5625 | 0.118738285 | 0.295346854 | 0.340382817 | 0.05744085 | 0.423191887 | 0.032840382 |
| 27870.74219 | 26564.53711 | 26949.33398 | 0.118557222 | 0.237804355 | 0.115422883 | 0.258368613 | -0.000785378 | 0.992953033 |
| 282176.4375 | 293292.1875 | 290805.4688 | 0.118473212 | 0.401593012 | 0.068467149 | 0.465719209 | 0.446179153 | 0.033948107 |
| 85387.63281 | 89262.1875 | 74431.78125 | 0.117991879 | 0.200486826 | 0.501256451 | 0.042884533 | 0.497245015 | 0.068366118 |
| 93489.29688 | 86462.04688 | 95448.42969 | 0.117616927 | 0.064954349 | 0.029186915 | 0.609108121 | 0.188139561 | 0.107956786 |
| 2570.472656 | 6556.142578 | 3367.818848 | 0.117562057 | 0.807112963 | 0.364030339 | 0.52823365 | 0.067488881 | 0.931717307 |
| 16901.73047 | 45794.5 | 30521.58398 | 0.117359511 | 0.405410011 | -0.463501222 | 0.053580444 | 0.250169655 | 0.7008148 |
| 153440.25 | 117621.7578 | 130858.9688 | 0.117294418 | 0.527524779 | 0.327964221 | 0.138525684 | -0.090300272 | 0.667945505 |
| 225592.7188 | 211247.5 | 204359.25 | 0.117270565 | 0.312095965 | 0.203614649 | 0.163084143 | -0.088469796 | 0.449433318 |
| 272130.0313 | 232076.7344 | 235825.8906 | 0.117228872 | 0.346157494 | 0.285498532 | 0.102497957 | -0.063699831 | 0.643399583 |
| 588081.3125 | 586697.3125 | 624627.875 | 0.117000714 | 0.531144616 | 0.314331398 | 0.065266848 | 0.356797506 | 0.041088687 |
| 24447.70117 | 23461.68945 | 22919.32617 | 0.116782215 | 0.40453073 | 0.724866908 | 0.085130535 | 0.958928987 | 0.008387835 |
| 18908.37695 | 25400.47656 | 24804.91211 | 0.116709767 | 0.653205948 | 0.523221502 | 0.10623247 | 0.653403331 | 0.094923927 |
| 21048.13477 | 20571.78711 | 20811.69727 | 0.116528168 | 0.221523033 | -0.323899817 | 0.076036407 | -0.571104577 | 0.034677634 |
| 714699.75 | 794811.5 | 794860.5 | 0.116497817 | 0.167007229 | -0.149577688 | 0.084443256 | -0.280136368 | 0.050613123 |
| 8280.967773 | 7706.046875 | 8600.526367 | 0.116446033 | 0.189747121 | -0.324212619 | 0.053543594 | -0.529872714 | 0.02527614 |
| 32072.73242 | 33960.76563 | 31335.78711 | 0.116200857 | 0.442980522 | -0.205673706 | 0.168363676 | -0.111645252 | 0.329172546 |
| 38501.73438 | 40954.64844 | 39355.11719 | 0.115676999 | 0.222892493 | 0.034713561 | 0.650768422 | 0.255745351 | 0.056570421 |
| 190327.5156 | 182781.8125 | 181836.6875 | 0.11562175 | 0.256643853 | -0.130540461 | 0.265347062 | -0.041615749 | 0.643774919 |
| 35047.69531 | 36206.45313 | 34798.21094 | 0.11546596 | 0.056649614 | 0.182015696 | 0.075045269 | 0.411383821 | 0.009740856 |
| 5028.649414 | 7359.536133 | 5345.07666 | 0.115395445 | 0.929305291 | -0.571337873 | 0.713789131 | 0.000210673 | 0.999822993 |
| 46526.82031 | 49035.77344 | 44395.78516 | 0.115333375 | 0.347753922 | 0.152218029 | 0.032914334 | -0.277289558 | 0.043517371 |
| 249155.6572 | 274730.123 | 244283.9551 | 0.115153234 | 0.400305876 | 0.3440101 | 0.077029113 | -0.147993158 | 0.240458637 |
| 85902.53906 | 85869.0625 | 83910.52344 | 0.114613322 | 0.344015413 | 0.184901837 | 0.0155094 | 0.063909574 | 0.058589883 |
| 22363.63281 | 23398.43945 | 18857.46094 | 0.114416264 | 0.224060759 | 0.086663614 | 0.2584075 | -0.24540511 | 0.160764344 |
| 207776.8281 | 232228.375 | 241653.9531 | 0.114266077 | 0.393813191 | -0.053659972 | 0.520977849 | -0.011488012 | 0.918818285 |
| 197277.3906 | 199610.9688 | 212173.7656 | 0.113947342 | 0.705880057 | 0.28265004 | 0.357910134 | 0.203100569 | 0.469755826 |

| | | | | | | | | |
|---|---|---|---|---|---|---|---|---|
| 24829292 | 18422052 | 25063712 | 0.11380507 | 0.861446729 | 0.458281137 | 0.25313315 | 0.981919421 | 0.067488244 |
| 408358.6875 | 671504.6875 | 270965.5313 | 0.113760693 | 0.863810045 | 0.30434828 | 0.536150475 | -0.227924791 | 0.751199361 |
| 96740.1875 | 102619.9375 | 119524.1953 | 0.113609247 | 0.650027478 | 0.388668332 | 0.198917031 | 1.66209411 | 0.027800907 |
| 142772 | 148185.3281 | 145375.1719 | 0.113425072 | 0.351588472 | 0.158190245 | 0.236628268 | 0.194470545 | 0.174066419 |
| 681593.4375 | 709706.5625 | 666527.5625 | 0.113310827 | 0.228860613 | 0.398225707 | 0.037014778 | 0.380559651 | 0.019816721 |
| 124967.5703 | 126775.7266 | 116021.0078 | 0.113274109 | 0.25326502 | 0.308241199 | 0.14612569 | -0.135284636 | 0.21485344 |
| 118186.4531 | 106252.6172 | 113325.8516 | 0.113096344 | 0.596723648 | -0.920099524 | 0.105359552 | 0.230515102 | 0.309106111 |
| 11650.58887 | 6182.517578 | 4651.637207 | 0.112934766 | 0.807212628 | 0.38774541 | 0.471790095 | 0.550628275 | 0.495843688 |
| 236140.1719 | 248757.3594 | 247898.9375 | 0.112872586 | 0.332613034 | -0.099071754 | 0.457960453 | -0.384494757 | 0.088049383 |
| 37109.16797 | 59040.55469 | 53529.91406 | 0.112691307 | 0.831801391 | 0.077394825 | 0.90544874 | -0.364397774 | 0.557769502 |
| 175247.125 | 409732.9063 | 205993.1406 | 0.112545326 | 0.641237347 | 0.555533672 | 0.068282276 | 0.749826052 | 0.392418029 |
| 135776.9844 | 136662.7188 | 136294.2813 | 0.11243871 | 0.470739135 | 0.137405699 | 0.395790227 | 0.619776465 | 0.060920196 |
| 40945.95313 | 48201.34766 | 47783.48828 | 0.112265331 | 0.468715163 | 0.146035831 | 0.228751244 | 0.288200578 | 0.139434127 |
| 4906.524414 | 10768.50098 | 14274.42285 | 0.112030752 | 0.773960816 | 0.076873836 | 0.87125384 | 0.22168437 | 0.742102915 |
| 403402.9063 | 441073.3125 | 430617.375 | 0.1115607 | 0.294591633 | -0.082786129 | 0.431724911 | -0.145394272 | 0.228837085 |
| 67039.65625 | 70775.83594 | 78198.4375 | 0.111481279 | 0.262755123 | -0.221710025 | 0.256398999 | 0.607633926 | 0.047847186 |
| 15948.16406 | 13429.80371 | 16136.58887 | 0.111296451 | 0.639361146 | 0.180000107 | 0.296013798 | 0.298709012 | 0.135230847 |
| 5427.725586 | 1830.127808 | 3399.092285 | 0.111014817 | 0.838790359 | 2.501198447 | 0.508238097 | -0.008697404 | 0.991353382 |
| 502347.4688 | 478107.25 | 449150.625 | 0.110942804 | 0.508098633 | 0.234625028 | 0.227543266 | -0.100235251 | 0.557774325 |
| 144358.9688 | 131330.1719 | 148075.5469 | 0.11088327 | 0.574372824 | -0.381644251 | 0.024404888 | 0.597423802 | 0.038522104 |
| 2703.691895 | NA | 1365.729614 | 0.110735567 | 0.779387269 | -1.687359609 | 0.132821529 | -1.900462336 | NA |
| 51846.80859 | 50734.04688 | 48081.36719 | 0.110588283 | 0.085651859 | -0.198920789 | 0.054678041 | -0.036744851 | 0.52683785 |
| 80165.6875 | 85943.89063 | 77082.29688 | 0.110514583 | 0.163314204 | -0.25928083 | 0.044791528 | -0.426318058 | 0.02183874 |
| 5675.003906 | 4746.088379 | 5189.027832 | 0.110439983 | 0.639033033 | 0.265220756 | 0.168370493 | -0.536154053 | 0.082022058 |
| 70635.57031 | 66324.13281 | 76134.8125 | 0.110198935 | 0.395528635 | -0.062408888 | 0.628159256 | -0.115517488 | 0.420915559 |
| 720981.375 | 553011.875 | 635544.5625 | 0.109870274 | 0.273669344 | 0.087292515 | 0.387915664 | -0.219863361 | 0.249456378 |
| 8534.477539 | 7038.867676 | 7366.954102 | 0.109798094 | 0.346510839 | -0.066137153 | 0.636049699 | -0.455502025 | 0.051354931 |
| 7549.793457 | 7591.654785 | 11101.18262 | 0.109556812 | 0.713928646 | 0.600839112 | 0.102454913 | 0.195184656 | 0.597778823 |
| 39667.10938 | 43657.15625 | 46927.69531 | 0.109354946 | 0.261806756 | 0.048997245 | 0.463776372 | -0.08433318 | 0.458182767 |
| 244776.3438 | 434043.1875 | 361861.2813 | 0.10912886 | 0.097194367 | 0.023622841 | 0.8446808 | 0.665496593 | 0.241197073 |
| 213030.1094 | 198022.9219 | 205046.4375 | 0.108968322 | 0.104313747 | 0.00555205 | 0.971700493 | -0.043788392 | 0.471357676 |
| 84842.96094 | 79538.8125 | 64374.10547 | 0.108939101 | 0.502730446 | 0.71153885 | 0.042884533 | 0.209247552 | 0.343143637 |
| 153358.3438 | 142317.6563 | 133935 | 0.108842157 | 0.511958098 | 0.214046915 | 0.250511865 | -0.000336982 | 0.998467793 |

| | | | | | | | | |
|---|---|---|---|---|---|---|---|---|
| 44490.66016 | 45648.09375 | 43240.62109 | 0.108788676 | 0.351147248 | -0.230102252 | 0.177376869 | -0.196068858 | 0.138522639 |
| 801033.8125 | 895279.1875 | 683034.0625 | 0.108572713 | 0.562587031 | -0.101396626 | 0.509176589 | -0.306243842 | 0.179221371 |
| 142972.2656 | 151374.8594 | 140893.9063 | 0.108548999 | 0.565200366 | 0.273190067 | 0.21914142 | 0.353547722 | 0.142342008 |
| 29837.85938 | 33677.73047 | 15086.65234 | 0.108377221 | 0.644961715 | -0.927390505 | 0.114739915 | -1.174410875 | 0.063089609 |
| 21629.56836 | 21112.66992 | 25087.49219 | 0.108159076 | 0.246012528 | -0.539614567 | 0.05744085 | -0.710437596 | 0.02163333 |
| 16135.9043 | 17448.19922 | 17067.96094 | 0.107864772 | 0.550359435 | -0.236253715 | 0.308059211 | -0.056760715 | 0.758466735 |
| 71358.52344 | 65924.04688 | 70230.19531 | 0.107678142 | 0.261850791 | -0.339086873 | 0.119643706 | 0.390331549 | 0.02796904 |
| 280928.8438 | 290907.1875 | 262389.6875 | 0.107676121 | 0.370662358 | 0.031870974 | 0.552589151 | 0.039231704 | 0.610401648 |
| 12214.27832 | 12020.52441 | 10445.64355 | 0.107359147 | 0.518212038 | 0.196810612 | 0.193691583 | -0.295737561 | 0.11805192 |
| 6755.067383 | 11332.92969 | 9339.121094 | 0.107258151 | 0.387027791 | -0.062626317 | 0.610888939 | -0.200562488 | 0.523801675 |
| 6309.753906 | 5535.861816 | 5857.261719 | 0.107165742 | 0.659279692 | -0.532994725 | 0.056230349 | 0.008225905 | 0.952354382 |
| 41609.86328 | 47374.97266 | 44439.77344 | 0.107096797 | 0.200303108 | 0.028234777 | 0.71382144 | 0.043598043 | 0.629083435 |
| 28056.89453 | 27425.54297 | 25073.5957 | 0.107041292 | 0.232132645 | -0.047333597 | 0.225638357 | -0.077525757 | 0.348056184 |
| 60290.35547 | 60821.80078 | 61525.47266 | 0.106660377 | 0.371155319 | 0.131057726 | 0.077209499 | 0.004736035 | 0.914274102 |
| 506411.5313 | 344460.4063 | 468468.4063 | 0.106410589 | 0.793038329 | -0.283199709 | 0.50345588 | -0.134383527 | 0.746141254 |
| 109902.5156 | 109450.3281 | 104981.5313 | 0.106364813 | 0.176256277 | 0.049482551 | 0.488418166 | -0.078163961 | 0.106159419 |
| 863168.0625 | 868107.875 | 925992.1875 | 0.106294486 | 0.314242842 | -0.137399846 | 0.147248245 | -0.151532916 | 0.079935933 |
| 32762.75 | 36070.33984 | 42186.61719 | 0.10620713 | 0.468442429 | 0.235747695 | 0.301284483 | 0.133865747 | 0.512102022 |
| 68236.08594 | 62091.94922 | 62985.38281 | 0.106104448 | 0.717472921 | 0.416834699 | 0.219332589 | 0.087361879 | 0.758792713 |
| 21339.04492 | 24068.16602 | 22825.00391 | 0.106042678 | 0.391494686 | -0.53828169 | 0.053882844 | -0.815855763 | 0.018546147 |
| 103625.8438 | 107024.3203 | 97809.52344 | 0.105764698 | 0.136100201 | -0.588364978 | 0.014983141 | -0.759752681 | 0.008361136 |
| 22793.08398 | 24804.88672 | 20201.20898 | 0.105648189 | 0.084250392 | 0.058910699 | 0.41771262 | -0.19841581 | 0.2081433 |
| 48819.03516 | 42948.28906 | 43048.5625 | 0.105632634 | 0.630765638 | -0.050558376 | 0.775185663 | -0.188027139 | 0.363209271 |
| 431214.25 | 425156.9375 | 487000.6875 | 0.105294 | 0.414354381 | 0.026404295 | 0.842902428 | -0.077837565 | 0.586390561 |
| 46380.55859 | 21758.63281 | 16176.19629 | 0.104534046 | 0.924854312 | 1.731231013 | 0.181437192 | 0.552354329 | 0.61132384 |
| 49467.22266 | 51362.16406 | 53951 | 0.104406842 | 0.597550855 | 0.802418695 | 0.039011452 | 1.610318529 | 0.009026966 |
| 48281.03516 | 40960.27344 | 38095.44141 | 0.104329729 | 0.866347862 | 0.395882895 | 0.493167957 | 0.419409498 | 0.459131485 |
| 489132.625 | 571354.1875 | 645166.125 | 0.103980797 | 0.417173199 | 0.030341363 | 0.711810406 | 0.267986898 | 0.263528267 |
| 39137.85938 | 41335.57031 | 30185.24805 | 0.103714697 | 0.605864892 | 0.162948115 | 0.428256136 | -0.299562157 | 0.286045361 |
| 27078.48242 | 29020.70117 | 29292.36914 | 0.103704779 | 0.481301871 | 0.388783495 | 0.156070866 | 0.561286626 | 0.017329006 |
| 285501.0625 | 307394.4688 | 294111.4688 | 0.103566571 | 0.122286474 | -0.367576944 | 0.029468331 | -0.638409525 | 0.012551348 |
| 25211.77734 | 25780.01758 | 24258.74023 | 0.103322138 | 0.69868807 | 0.12331586 | 0.639324884 | 0.289734564 | 0.307275286 |
| 22803608 | 26190644 | 24967644 | 0.103205354 | 0.238040125 | 0.808356336 | 0.01179292 | 0.511178853 | 0.040786554 |

| | | | | | | | | |
|---|---|---|---|---|---|---|---|---|
| 23032.35742 | 21850.40039 | 23938.92383 | 0.103076634 | 0.542532097 | -0.119672857 | 0.236744048 | -0.160367985 | 0.146688991 |
| 6090.983398 | 5298.055664 | 2975.298584 | 0.102962923 | 0.736907454 | -0.316355383 | 0.356068496 | 0.112706738 | 0.818639379 |
| 15417.02832 | 10204.44922 | 5092.884277 | 0.102878624 | 0.641084841 | -0.843114579 | 0.168058041 | -2.017180391 | 0.042200911 |
| 104409.4219 | 94591.78125 | 89890.85938 | 0.102867307 | 0.265542969 | 0.258357922 | 0.033504796 | 0.15292581 | 0.240117089 |
| 97180.07031 | 98161.23438 | 91597.85938 | 0.102655787 | 0.129268905 | 0.043479599 | 0.289435934 | 0.340696814 | 0.029820656 |
| 164024.7344 | 149055.8906 | 146747.9375 | 0.102349571 | 0.551436289 | 0.926533614 | 0.060519327 | 1.08201412 | 0.020560319 |
| 70263.46094 | 78648.65625 | 87711.92969 | 0.101981247 | 0.542216889 | 0.251690857 | 0.202104071 | 0.304413191 | 0.169021874 |
| 28083.10547 | 30276.14063 | 29737.0625 | 0.101713264 | 0.173993181 | -0.24031097 | 0.058077004 | -0.150806773 | 0.096058082 |
| 790697.0625 | 879268.9375 | 874122.5 | 0.101702919 | 0.589921226 | -0.137303076 | 0.353741919 | -0.033555624 | 0.804711404 |
| 50445.19531 | 44990.29688 | 37532.11328 | 0.101354265 | 0.201158113 | -1.128623352 | 0.020431201 | -1.052326583 | 0.019577244 |
| 426900.9063 | 403241.7813 | 401131.875 | 0.101141515 | 0.240800716 | -0.00405709 | 0.924427311 | -0.144984309 | 0.063419062 |
| 12597.1084 | 11362.36426 | 11609.19727 | 0.100577783 | 0.218653229 | -0.378634993 | 0.102892509 | -0.14841142 | 0.156588849 |
| 697755.5625 | 724705.5 | 593568.1875 | 0.100233941 | 0.757837592 | 1.295229132 | 0.165197811 | 1.561480802 | 0.021863066 |
| 37947.55078 | 41483.34375 | 40316.75781 | 0.100185875 | 0.396119538 | 0.109474775 | 0.473742203 | 0.302196881 | 0.042175609 |
| 56512.15234 | 53407.51172 | 52698.72266 | 0.100061303 | 0.513130981 | -0.060375499 | 0.697163698 | -0.400841148 | 0.132636805 |
| 537924.125 | 483589.1875 | 549915 | 0.099654066 | 0.40512287 | -0.065756279 | 0.215837595 | 0.462070013 | 0.063474265 |
| 40966.11328 | 40720.75391 | 42705.44141 | 0.099643344 | 0.609089925 | 0.118878389 | 0.517397569 | -0.099471856 | 0.594012413 |
| 34243.38672 | 35863.94531 | 39774.40234 | 0.09959297 | 0.718505305 | 0.245915998 | 0.417167466 | 0.591790807 | 0.097707274 |
| 240708.3594 | 255596.6094 | 245864.4219 | 0.099497135 | 0.455514623 | -0.14347768 | 0.248672029 | 0.41407089 | 0.034593218 |
| 771260.6875 | 907301.6875 | 921035.125 | 0.099405761 | 0.441018848 | 0.010912562 | 0.869754532 | 0.162887869 | 0.286326519 |
| 134231.9688 | 138990.0781 | 156488.5625 | 0.099171991 | 0.689568168 | 0.276877831 | 0.297944313 | 0.37029559 | 0.143217293 |
| 54021.67578 | 54797.125 | 58358.04297 | 0.099009323 | 0.254836953 | -0.06568977 | 0.435133424 | -0.021618776 | 0.797672854 |
| 104370.6172 | 84328.82813 | 120007.6953 | 0.098946004 | 0.839961736 | 0.603254797 | 0.204833051 | 0.9686489 | 0.081095782 |
| 75827.91406 | 76084.39844 | 72703.38281 | 0.098929494 | 0.074071792 | -0.311361614 | 0.021217492 | -0.4571763 | 0.009740856 |
| 71574.73438 | 77880.80469 | 79289.97656 | 0.098845337 | 0.40427953 | 0.505180744 | 0.067039668 | 0.523981499 | 0.028173069 |
| 94638.70313 | 92636.875 | 106997.0156 | 0.098725672 | 0.127230164 | 0.325353409 | 0.028434447 | 0.505384192 | 0.062822 |
| 6047.15625 | 5498.978027 | 3246.658936 | 0.098675497 | 0.654529515 | 0.155498997 | 0.318694029 | -0.833323296 | 0.080462751 |
| 38286.50781 | 37018.44141 | 33820.39844 | 0.098449023 | 0.210699315 | -0.590392081 | 0.017029168 | -1.08991513 | 0.008387835 |
| 98190.76563 | 99452.29688 | 111759.1172 | 0.098400377 | 0.363908989 | 0.205274426 | 0.137613908 | 0.373723617 | 0.060171057 |
| 18507.95313 | 15559.20313 | 13449.46582 | 0.098204375 | 0.41842039 | -0.005546264 | 0.966180385 | -0.27359384 | 0.236411423 |
| 8964.260742 | 6469.074219 | 4117.923828 | 0.098031484 | 0.228317381 | 1.098925678 | 0.05818214 | -0.076535577 | 0.869766479 |
| 865537.0781 | 773351.6094 | 895656.8984 | 0.097548683 | 0.271101668 | 0.095890834 | 0.284677807 | -0.009713704 | 0.935329166 |
| 61083.34766 | 66734.50781 | 62980.67578 | 0.097352719 | 0.518013064 | -0.22468041 | 0.28645625 | -0.176935875 | 0.283755733 |

| | | | | | | | | |
|---|---|---|---|---|---|---|---|---|
| 7826.186035 | 7019.283203 | 7309.497559 | 0.097134642 | 0.517891679 | -0.27061042 | 0.198421115 | -0.233603261 | 0.217152139 |
| 151800.5938 | 151755.4375 | 145422.7344 | 0.097003663 | 0.512648148 | -0.226499064 | 0.223042946 | -0.472172257 | 0.102372782 |
| 28558.9707 | 29252.25781 | 27686.64453 | 0.096782699 | 0.26956076 | -0.034329992 | 0.699065886 | -0.014708964 | 0.846743679 |
| 15743.23535 | 17144.86133 | 15712.01758 | 0.096633966 | 0.695769702 | -0.300580364 | 0.14785027 | -0.187416961 | 0.23707058 |
| 77114.52344 | 82552.95313 | 81588.66406 | 0.096340184 | 0.634340373 | 0.560315733 | 0.054829172 | 0.38132976 | 0.094103554 |
| 25662.60938 | 18373.37891 | 18757.06836 | 0.096313758 | 0.732446514 | 0.66972407 | 0.068436129 | 0.293254528 | 0.363820887 |
| 89490.4375 | 87180.61719 | 75177.09375 | 0.096038218 | 0.723069509 | 0.103873304 | 0.494902041 | -0.142448656 | 0.397327146 |
| 48360.34375 | 47426.70703 | 54222.42188 | 0.095605781 | 0.332906865 | 0.756860682 | 0.017786674 | 0.698220879 | 0.034722024 |
| 760347.5625 | 744937.5625 | 798933.5625 | 0.095500448 | 0.538853002 | -0.505393734 | 0.117855098 | -0.561890607 | 0.095708676 |
| 4196.44873 | 6245.467285 | 3822.911377 | 0.095310635 | 0.888334398 | -0.192482887 | 0.280688037 | -0.487874217 | 0.198354379 |
| 40414.125 | 32309.11523 | 38599.08594 | 0.095269896 | 0.17785882 | 1.022985022 | 0.078098956 | 0.481028892 | 0.116344276 |
| 30239.54102 | 32182.60352 | 30607.97266 | 0.095082775 | 0.520362748 | -0.56708846 | 0.037811274 | -0.680452905 | 0.031136253 |
| 39022.60938 | 37514.28516 | 39327.85156 | 0.09491289 | 0.271519724 | -0.308961331 | 0.065678888 | -0.151649903 | 0.158426237 |
| 109757.6953 | 108445.2891 | 96288.16406 | 0.094844146 | 0.405753158 | 0.394896562 | 0.059039944 | 0.727879816 | 0.025945806 |
| 885309.0625 | 885032.5625 | 874289.375 | 0.094710431 | 0.17954942 | -0.442218942 | 0.046500865 | -0.496921638 | 0.04014804 |
| 336739.9063 | 391318 | 392894.25 | 0.094688803 | 0.489182982 | -0.480606722 | 0.081958166 | -0.394024157 | 0.068413268 |
| 43684.83984 | 46100.81641 | 40256.24219 | 0.094525409 | 0.643009308 | -0.230584605 | 0.225638357 | -0.147005288 | 0.366486609 |
| 48085.69531 | 47098.89453 | 44967.83594 | 0.094078911 | 0.050984999 | -0.102119671 | 0.073326865 | -0.2217075 | 0.031136253 |
| 435.6015625 | 534.5628052 | NA | 0.093984559 | 0.828897042 | -2.273797454 | NA | -0.052838075 | NA |
| 47619.60938 | 33136.80859 | 39095.57031 | 0.093929752 | 0.688811545 | -0.162190026 | 0.462157906 | -0.142575618 | 0.591863123 |
| 73292.67969 | 66443.50781 | 69764.67188 | 0.09302585 | 0.314273917 | 0.53938406 | 0.121240742 | 0.762265616 | 0.016395134 |
| 43497.23047 | 43773.08594 | 38657.38281 | 0.092747949 | 0.638521857 | -0.293594579 | 0.225427842 | -0.583122181 | 0.092354827 |
| 109217.1953 | 112731.125 | 99669.10156 | 0.092641709 | 0.048963509 | 0.123754015 | 0.058011247 | -0.309406912 | 0.059847517 |
| 8967.457031 | 9402.006836 | 7340.650391 | 0.092605812 | 0.427440307 | -0.17950694 | 0.110620378 | -0.317193296 | 0.145132006 |
| 624537.0625 | 582647.75 | 525842.75 | 0.091782371 | 0.663551774 | -0.163328844 | 0.498850696 | 0.016324254 | 0.946678732 |
| 12696.95801 | 12314.86719 | 12788.59082 | 0.091759147 | 0.719961518 | -0.455093823 | 0.106309393 | -1.117817425 | 0.060457482 |
| 292812.8438 | 277736.8438 | 263814.4688 | 0.091423084 | 0.61953157 | 0.452012356 | 0.092696398 | 0.476217195 | 0.071994935 |
| 34511.73438 | 39836.40234 | 35572.52734 | 0.091332663 | 0.252282983 | -0.089495635 | 0.227457591 | -0.197389613 | 0.137812617 |
| 214454.5156 | 218686.2188 | 215397.5156 | 0.090658264 | 0.313243762 | -0.106766042 | 0.351590709 | -0.311262533 | 0.088176741 |
| 476282.5 | 441615.2813 | 436283.75 | 0.090565071 | 0.268941926 | -0.130012476 | 0.265924368 | -0.077222345 | 0.277706206 |
| 10640.33203 | 9852.876953 | 9840.273438 | 0.090147211 | 0.443522229 | -0.16080955 | 0.313455345 | -0.705256616 | 0.051121426 |
| 8090.687988 | 7601.359863 | 4977.633301 | 0.089383277 | 0.772850314 | 0.279037965 | 0.249697654 | 0.322176975 | 0.398529846 |
| 9032.964844 | 12100.26074 | 10336.79102 | 0.089097044 | 0.777698004 | 0.315570829 | 0.166818836 | 0.194893921 | 0.404479667 |

| | | | | | | | | |
|---|---|---|---|---|---|---|---|---|
| 30246.125 | 31920.52539 | 28179.65625 | 0.088669505 | 0.543069772 | -0.064485895 | 0.633452303 | -0.104478403 | 0.47454841 |
| 27449.66797 | 25446.81641 | 26686.96094 | 0.088569971 | 0.627975352 | -0.035233254 | 0.837247847 | -0.095281834 | 0.57744406 |
| 74089.76563 | 60236.99609 | 49322.30078 | 0.088301595 | 0.791331725 | -0.066968773 | 0.852971034 | 0.585265077 | 0.185480538 |
| 103503.375 | 96748.53906 | 99499.26563 | 0.087588276 | 0.353461258 | -0.036179776 | 0.634892576 | -0.265561316 | 0.069879424 |
| 117094.4844 | 141417.875 | 171529.1875 | 0.087572543 | 0.835052252 | 0.127674429 | 0.787778179 | 0.739429739 | 0.136000191 |
| 63765.60156 | 64787.98828 | 64933.64844 | 0.08756318 | 0.297441327 | 0.397194918 | 0.10820268 | -0.081667534 | 0.039602697 |
| 11996.21582 | 8076.512695 | 12841.79785 | 0.087396748 | 0.730913878 | -1.089497317 | 0.220094258 | 0.469339453 | 0.263631739 |
| 670310.75 | 668301.4375 | 593910.875 | 0.087321001 | 0.35982448 | 0.46192806 | 0.019645803 | 0.359351546 | 0.060161253 |
| 110271.7734 | 107699.0078 | 105070.5547 | 0.086970274 | 0.594194983 | -0.065780019 | 0.686644956 | -0.329468118 | 0.101027689 |
| 25622.82227 | 24130.28516 | 27277.09961 | 0.086674512 | 0.647456546 | 0.996210741 | 0.044747456 | 1.03896068 | 0.016683666 |
| 416107.6289 | 601080.0586 | 481093.8438 | 0.086464574 | 0.791693363 | 0.135866009 | 0.709967054 | 0.314724247 | 0.363680339 |
| 839084.125 | 886559.125 | 879480.625 | 0.085710766 | 0.260141689 | -0.172111968 | 0.122540819 | -0.226114523 | 0.074184841 |
| 23004.50195 | 22259.80664 | 23899.5918 | 0.085617958 | 0.086423195 | 0.200370849 | 0.121847149 | 0.29254877 | 0.046772816 |
| 69484.15625 | 48286.8125 | 56463.59375 | 0.084664874 | 0.641311793 | -0.01118555 | 0.935075242 | -0.644716059 | 0.070873774 |
| 602164.5 | 652100.25 | 596590.8125 | 0.084283703 | 0.464841982 | -0.129441709 | 0.328806215 | 0.061224019 | 0.592078283 |
| 332061.4375 | 328063.8438 | 308980.875 | 0.084176442 | 0.422516256 | -0.485484578 | 0.081451772 | -0.404415667 | 0.068838627 |
| 6798.600586 | 4929.111328 | 4909.843262 | 0.084149841 | 0.772117891 | -0.186741377 | 0.297032792 | -0.513677052 | 0.107933864 |
| 755220.4375 | 948650.5 | 564387.625 | 0.083871804 | 0.885903743 | -0.821605716 | 0.249697654 | -0.940107023 | 0.209718967 |
| 83162.3125 | 83879.99219 | 89070.04688 | 0.083851758 | 0.331236723 | 0.045622967 | 0.431724911 | 0.109293158 | 0.144559354 |
| 13041.6377 | 14804.35059 | 14630.36719 | 0.083811265 | 0.165631416 | -0.152945584 | 0.053185315 | -0.299600287 | 0.08376499 |
| 5274.566895 | 6117.964844 | 7123.677734 | 0.083629427 | 0.831656064 | -0.652206516 | 0.243784441 | -0.423994325 | 0.348589806 |
| 74981.52344 | 73295.27344 | 78891.21094 | 0.083486941 | 0.648602476 | 0.690003353 | 0.041910161 | 0.859167714 | 0.010623938 |
| 115314.5156 | 97457.21875 | 116032.4141 | 0.083109731 | 0.427440307 | -0.669024111 | 0.027859864 | -0.510851581 | 0.040786554 |
| 182874.2813 | 189347.0156 | 190145.6563 | 0.082901981 | 0.420983699 | -0.880868488 | 0.051593867 | -0.973991408 | 0.040202519 |
| 83274.67188 | 80891.53125 | 75507.14844 | 0.081987089 | 0.287229066 | -0.062724735 | 0.389120839 | 0.044557169 | 0.512102022 |
| 193755.0781 | 216834.6094 | 235579.2344 | 0.081751392 | 0.817346175 | 0.346071745 | 0.318729418 | 1.112423596 | 0.032840382 |
| 200389.0625 | 192565.8125 | 214890.6406 | 0.081599267 | 0.412066495 | -0.085928361 | 0.269262694 | -0.155749902 | 0.118486034 |
| 27749.22656 | 29284.53516 | 25618.95117 | 0.081588664 | 0.816015197 | -0.170542763 | 0.637204757 | -0.267913808 | 0.486026097 |
| 27379.75195 | 27526.56055 | 20849.29492 | 0.081577612 | 0.404888951 | -0.334743948 | 0.088308366 | -0.795272737 | 0.030633392 |
| 153519.0781 | 162018.3594 | 159608.5625 | 0.081424659 | 0.609981483 | -0.476992388 | 0.067579251 | -0.383010208 | 0.092205529 |
| 210075.7813 | 188311.3281 | 185545.6094 | 0.081283771 | 0.287500692 | 0.026933848 | 0.77405498 | 0.254859445 | 0.102613568 |
| 364991.7188 | 379224.7813 | 373667.7188 | 0.081238591 | 0.404110576 | -0.084481805 | 0.374951761 | 0.06843198 | 0.423267439 |
| 9243.342773 | 12668.30371 | 10849.74316 | 0.081236633 | 0.826782722 | -0.956951231 | 0.1315555 | -0.355899987 | 0.338959672 |

| | | | | | | | | |
|---|---|---|---|---|---|---|---|---|
| 103642.0781 | 108096.8281 | 117193.2813 | 0.08122509 | 0.437016098 | 0.256644845 | 0.063686231 | 0.355947188 | 0.050810933 |
| 2771.280518 | 36397.06641 | 2799.591553 | 0.081166573 | 0.893917074 | -0.47933748 | 0.275021879 | 1.80136974 | 0.567979851 |
| 609549 | 603534.5625 | 662094.5 | 0.080997743 | 0.051623428 | -0.064899963 | 0.399976168 | -0.100558671 | 0.226466969 |
| 16701.05078 | 20796.76563 | NA | 0.080637998 | 0.751304267 | -0.127626294 | 0.555300995 | -0.485267469 | NA |
| 27118.54883 | 21175.11914 | 15247.32617 | 0.079986604 | 0.45101217 | -1.074903557 | 0.042884533 | -1.219850288 | 0.032840382 |
| 748633.0625 | 829510.5625 | 896131.25 | 0.079770781 | 0.672418102 | 0.757940917 | 0.106881887 | 1.154451757 | 0.034567592 |
| 181876.7969 | 178066.4531 | 180935.0313 | 0.079392123 | 0.47429724 | 0.330256382 | 0.042608596 | 0.043692909 | 0.548836711 |
| 60477.34766 | 69252.53125 | 59985.19922 | 0.079307302 | 0.173500617 | -0.337025281 | 0.019141974 | -0.385091929 | 0.057347686 |
| 17555.52539 | 14786.08301 | 8968.03125 | 0.079287125 | 0.721643044 | 0.398769447 | 0.043916515 | 0.090560067 | 0.829646836 |
| 5996.254395 | 7888.698242 | NA | 0.079238141 | 0.530526722 | -0.314906624 | 0.043222528 | -0.366380923 | NA |
| 11834.02344 | 7624.024902 | 7689.016602 | 0.078903044 | NA | 1.221610181 | NA | 1.388767555 | NA |
| 12628.4541 | 13285.11621 | 10118.7041 | 0.078750348 | 0.627678094 | -0.719559247 | 0.024939682 | -0.263402074 | 0.206000009 |
| 13248.0127 | 12324.30664 | 8370.401367 | 0.078668771 | 0.6277698 | 0.064285511 | 0.674526148 | -0.420497941 | 0.199326473 |
| 16563.81445 | 15663.8125 | 13322.53711 | 0.07861968 | 0.420601191 | 0.218492563 | 0.065601851 | -0.31107148 | 0.109020087 |
| 37759.73828 | 40163.50391 | 35777.85547 | 0.078448054 | 0.661080189 | -0.298660145 | 0.118404 | -0.093147101 | 0.464555918 |
| 251937.4844 | 280554.125 | 254584.7188 | 0.078321506 | 0.457896539 | -0.166365652 | 0.077996184 | -0.086417872 | 0.317427062 |
| 28641.91992 | 23467.67773 | 22203.35938 | 0.07778702 | 0.780681403 | 0.11332227 | 0.450125745 | -0.105137624 | 0.548141599 |
| 246161.1719 | 244351.2344 | 244256.7031 | 0.077725871 | 0.240263529 | -0.444082573 | 0.021906439 | -0.175412916 | 0.091109394 |
| 19280.54492 | 19221.80664 | 16360.7627 | 0.077619846 | 0.929811497 | 1.319317919 | 0.117377154 | 1.124154335 | 0.167717489 |
| 4839231.5 | 4779418.5 | 4648205 | 0.076886218 | 0.543599714 | 0.497793664 | 0.048024604 | 0.824965423 | 0.026725136 |
| 274772.125 | 267965.1875 | 272837.9063 | 0.0766137 | 0.26038194 | 0.021386653 | 0.732837711 | -0.135281269 | 0.142772717 |
| 21433.36523 | 20493.27539 | 16785.19531 | 0.076427912 | 0.809956349 | 0.031823619 | 0.925830781 | -0.127649685 | 0.720547247 |
| 121431.0215 | 124016.2188 | 129431.4941 | 0.076100837 | 0.176121329 | -0.320004449 | 0.035356834 | -0.434327973 | 0.010654184 |
| 23538.52344 | 20482.42578 | 19641.36328 | 0.075849115 | 0.543350794 | -0.646987787 | 0.058077004 | -0.522310988 | 0.051693365 |
| 241325.0156 | 233262.1563 | 243310.8906 | 0.075611652 | 0.471770554 | -0.232592837 | 0.074419062 | -0.323559453 | 0.047629307 |
| 11084.05371 | 9164.97168 | 11219.14258 | 0.07543604 | 0.673603575 | -0.042954849 | 0.861048354 | -0.192997242 | 0.394165962 |
| 445383.4375 | 439600.4375 | 452718.7188 | 0.075028555 | 0.47594001 | 0.41581151 | 0.050921156 | 0.694797209 | 0.020134795 |
| 792798.6875 | 809297.5625 | 841756.5 | 0.07495397 | 0.586536454 | 0.185204356 | 0.172806689 | 0.023683858 | 0.825112663 |
| 85863.82031 | 143765.8125 | 104055.7266 | 0.074750143 | 0.398513138 | 0.186287184 | 0.164081629 | 0.342515621 | 0.411293423 |
| 57087.71875 | 57880.22656 | 57655.78516 | 0.074705935 | 0.493397529 | 0.264437073 | 0.100538317 | 0.242158717 | 0.132134669 |
| 190213.8906 | 191646.7031 | 183009.9063 | 0.07432486 | 0.302067193 | 0.212028376 | 0.188231791 | -0.290213703 | 0.016242079 |
| 99208.36719 | 88177.6875 | 99943.65625 | 0.074168261 | 0.722699186 | 0.143022069 | 0.456340808 | 0.592451896 | 0.041771828 |
| 3123.216309 | 4975.618652 | 3128.991699 | 0.074070888 | 0.591093347 | -0.213025639 | 0.200251432 | -0.437903432 | 0.248884848 |

| | | | | | | | | |
|---|---|---|---|---|---|---|---|---|
| 15029.10059 | 11633.04883 | 12545.07422 | 0.074020241 | 0.777698004 | -0.159294575 | 0.549561464 | -0.378673841 | 0.250975288 |
| 13820.53906 | 11032.27441 | 12273.0166 | 0.073953678 | 0.74192438 | 0.275547193 | 0.286605361 | -0.502763659 | 0.159338304 |
| 98468.41406 | 82420.41406 | 98657.00781 | 0.073871334 | 0.069880665 | 0.192828822 | 0.046373997 | -0.020575699 | 0.872236202 |
| 83531.82813 | 80537.42188 | 80483.30469 | 0.073774968 | 0.511747456 | 0.025959 | 0.829546664 | 0.026780207 | 0.808411584 |
| 51712.22266 | 54103.82031 | 51107.72266 | 0.072644292 | 0.187013619 | -0.095354923 | 0.171590168 | -0.01438826 | 0.78904813 |
| 182100.9844 | 190611.2969 | 196427.5938 | 0.072631427 | 0.21324318 | -0.001916081 | 0.969557622 | -0.054211192 | 0.374523462 |
| 7974.995117 | 6806.62207 | 4545.514648 | 0.072519994 | 0.58307644 | -0.289524812 | 0.056841901 | -0.290829945 | 0.397327146 |
| 137925.5156 | 147893.6094 | 156789.0469 | 0.07243606 | 0.355531488 | -0.234672121 | 0.093818896 | -0.202142566 | 0.111558718 |
| 277237.0938 | 295992.625 | 302188.875 | 0.072344417 | 0.401802803 | -0.238293956 | 0.040066052 | -0.154653951 | 0.082930409 |
| 10149.32031 | 9104.839844 | 9205.982422 | 0.072245804 | 0.783327271 | 0.539929339 | 0.312461016 | -0.366934756 | 0.281897639 |
| 44431.3125 | 44553.92578 | 39085.59375 | 0.072216754 | 0.460764722 | -0.343867236 | 0.106670824 | -0.20983783 | 0.151192408 |
| 29108.66797 | 20742.09766 | 28321.33594 | 0.071747813 | 0.896303807 | 2.072974715 | 0.093773752 | 1.721864706 | 0.055528595 |
| 1446136 | 1587426 | 1537049.25 | 0.071554042 | 0.327619892 | -0.054017147 | 0.376242308 | -0.112594294 | 0.177813604 |
| 653060.1875 | 900759.9375 | 848253.0625 | 0.070964446 | 0.386662122 | -0.286471359 | 0.068812228 | 0.428353812 | 0.198048338 |
| 2679.455078 | 1063.301147 | 498.8885803 | 0.070933908 | 0.941157339 | -0.960672062 | 0.420604095 | -2.424370539 | 0.040771421 |
| 4051.118164 | 4398.987305 | 3074.103027 | 0.070850272 | 0.746281035 | 0.074056685 | 0.634825579 | -0.044068763 | 0.865392889 |
| 330870.1563 | 305955.125 | 308395.9375 | 0.070756192 | 0.51034672 | 0.003890102 | 0.931230576 | 0.115792378 | 0.171126239 |
| 216504.3438 | 216667.625 | 239422.3438 | 0.070477442 | 0.794535026 | 0.194111017 | 0.506573912 | 0.174805586 | 0.52607612 |
| 10235.45508 | 9871.878906 | 10155.32227 | 0.070404209 | 0.531204155 | -0.987269418 | 0.028434447 | -0.081586829 | 0.489401961 |
| 182610.8906 | 181241.8906 | 179505.7969 | 0.070299475 | 0.231408883 | 0.134162044 | 0.190057963 | 0.001335408 | 0.977618946 |
| 4047.60376 | 5573.514648 | 4648.366211 | 0.070135522 | 0.833974595 | -0.136301876 | 0.697931686 | -0.156805555 | 0.670209128 |
| 106731.5469 | 105671.0156 | 102265.4766 | 0.069718236 | 0.816015197 | -0.584063057 | 0.049805123 | -0.545974474 | 0.05183905 |
| 1488228 | 1843531 | 2027702.375 | 0.069542584 | 0.809105534 | 0.17731422 | 0.517240964 | 0.340802211 | 0.288252409 |
| 141231.1719 | 150752.5781 | 140277.0469 | 0.069331167 | 0.471770554 | 0.842786434 | 0.01179292 | 0.354149753 | 0.030633392 |
| 142649.625 | 151202.1094 | 138524.1719 | 0.068897314 | 0.598139599 | -0.565501688 | 0.090012942 | -0.765340035 | 0.061479799 |
| 13297.54688 | 14494.79102 | 13084.19043 | 0.068654441 | 0.565062132 | 0.130080327 | 0.2967259 | 0.226030647 | 0.123837484 |
| 342283.1875 | 365719.7813 | 361793.0938 | 0.068576693 | 0.285547261 | 0.001680992 | 0.971700493 | 0.133094014 | 0.091854175 |
| 22586.91211 | 20805.58203 | 19377.02148 | 0.068180385 | 0.495678002 | 0.302904068 | 0.071815633 | 0.266517911 | 0.110015219 |
| 155502.4844 | 151649.9688 | 142511.0469 | 0.068176099 | 0.082006863 | 0.037690708 | 0.199074765 | 0.134241466 | 0.134259503 |
| 4433.049316 | 7597.870605 | 6954.500977 | 0.067869816 | 0.820454661 | 0.110836984 | 0.677602288 | -0.009334088 | 0.984083418 |
| 32828.99609 | 32668.3125 | 31241.41016 | 0.067538963 | 0.649999812 | -0.062353897 | 0.751608736 | -0.478500803 | 0.119146128 |
| 908749.875 | 938550.8125 | 998211.8125 | 0.06753426 | 0.722080384 | 0.247990159 | 0.148419129 | 0.215846007 | 0.132907135 |
| 28068.35352 | 13644.22461 | 14049.48145 | 0.067365145 | 0.930893729 | -0.775821515 | 0.423723931 | -0.727229393 | 0.439792148 |

| | | | | | | | | |
|---|---|---|---|---|---|---|---|---|
| 15353.02539 | 17031.58008 | NA | 0.067312141 | 0.85906459 | -1.160247138 | 0.152270472 | -1.115471957 | NA |
| 1937690.375 | 2319307.75 | 2388122.5 | 0.06723466 | 0.427440307 | -0.009205423 | 0.925830781 | 0.262132618 | 0.188666258 |
| 9235.05957 | 9025.702148 | 10326.77441 | 0.067228115 | 0.682393165 | 0.429643243 | 0.09678603 | 0.267034804 | 0.098920233 |
| 17938.64844 | 20506.92578 | 20742.13477 | 0.067091826 | 0.637916865 | -0.042245243 | 0.768316659 | 0.257763726 | 0.15813376 |
| 88560.78125 | 84340.03125 | 75787.85938 | 0.066671421 | 0.553467034 | 0.125912074 | 0.31156031 | 0.524812396 | 0.042737273 |
| 129143.0547 | 137217.2031 | 128437.4297 | 0.06657723 | 0.729581224 | -0.101167115 | 0.645770679 | -0.057049318 | 0.75291637 |
| 144177.6563 | 142317.9531 | 122605.6406 | 0.066267339 | 0.837586473 | 0.132845439 | 0.680399418 | -0.00644157 | 0.987223198 |
| 10830.92871 | 12678.57324 | 7369.598633 | 0.066054173 | 0.75710877 | 0.405175795 | 0.188306417 | -0.302980243 | 0.377029093 |
| 5598.686523 | 5972.070313 | 7308.227539 | 0.065889438 | 0.661080189 | -0.76393927 | 0.019141974 | -1.03597778 | 0.020018964 |
| 29832.25391 | 20852.83594 | 28699.51172 | 0.065600724 | 0.41874454 | 0.058498373 | 0.399377103 | 0.038490842 | 0.872413573 |
| 100155.9453 | 107514.2109 | 87406.30469 | 0.065560419 | 0.667416519 | 0.328905887 | 0.114739915 | -0.133079336 | 0.473466754 |
| 1877782.875 | 1924364.25 | 1644838.875 | 0.065338751 | 0.590640274 | 0.430778586 | 0.052817461 | 0.230793455 | 0.170228714 |
| 62363.43359 | 65211.76563 | 57692.42969 | 0.06519933 | 0.639033033 | 0.243206579 | 0.112543356 | -0.255057153 | 0.124509143 |
| 43259.85547 | 46788.5 | 38767.51953 | 0.064936158 | 0.755050946 | 0.339065673 | 0.125482692 | 0.693425108 | 0.04014804 |
| 24270.16992 | 24696.45117 | 24031.72461 | 0.064802844 | 0.508982648 | 0.311374073 | 0.077476727 | 0.120902979 | 0.240008368 |
| 3349.974121 | 3095.125732 | 2235.092529 | 0.064717704 | 0.694156601 | -1.660479972 | 0.015829308 | -0.630797215 | 0.076783998 |
| 51201.47266 | 47075.0625 | 41649.29688 | 0.064434647 | 0.359445722 | 0.129782356 | 0.170357294 | -0.25641953 | 0.133621587 |
| 9218.419922 | 8871.28418 | 9303.430664 | 0.064224561 | 0.613778303 | -0.122974141 | 0.484068607 | -0.210436806 | 0.145132006 |
| 470520.1875 | 560239.9375 | 479340.3438 | 0.063010918 | 0.804888159 | -0.020362929 | 0.939755302 | 0.253791586 | 0.359958496 |
| 52249.45703 | 55441.95703 | 32884.58984 | 0.062868772 | 0.706689159 | -0.857952433 | 0.084023448 | -0.855599184 | 0.063869245 |
| 72629.0625 | 76529.72656 | 94900.64063 | 0.062665262 | 0.627975352 | 0.285867541 | 0.126364459 | 1.36028555 | 0.06065526 |
| 49755.39063 | 54886.49609 | 52993.01563 | 0.062066906 | 0.784725637 | 0.285247553 | 0.28939927 | 0.150791124 | 0.520269912 |
| 108730.3359 | 114872.5938 | 122307.6563 | 0.061819929 | 0.3674342 | 0.131958645 | 0.093751816 | 0.427522502 | 0.044417141 |
| 58794.73828 | 56283.05469 | 57318.29297 | 0.061779028 | 0.60592196 | 0.367683429 | 0.058077004 | 0.592129761 | 0.016500036 |
| 49172.625 | 43365.01172 | 32542.67773 | 0.061630509 | 0.600673217 | -0.634239053 | 0.028019426 | -1.131004942 | 0.031136253 |
| 6458.361816 | 13053.05176 | 6559.133301 | 0.061596935 | 0.862871796 | -0.029968684 | 0.932304307 | 0.052096513 | 0.936056793 |
| 55353.44531 | 52639.99609 | 53858.98828 | 0.061517833 | 0.827859454 | 0.445920371 | 0.189981295 | 0.111937314 | 0.687539514 |
| 8193.389648 | 10794.36328 | 9718.295898 | 0.061311192 | 0.45827242 | -0.001174091 | 0.99201113 | -0.044996962 | 0.801698552 |
| 11797.81738 | 16772.24023 | 16111.49121 | 0.061168623 | 0.900733744 | 0.414898524 | 0.332662365 | 0.654721555 | 0.153907433 |
| 9336.03418 | 7912.589844 | 4397.535156 | 0.060873634 | 0.926993948 | -0.558590856 | 0.552697805 | -0.44011772 | 0.574930993 |
| 7196.771484 | 7408.839844 | 5311.263672 | 0.060852414 | 0.648910173 | 0.02449274 | 0.87125384 | -0.406869512 | 0.135536925 |
| 21366.08398 | 16504.70703 | 24876.01758 | 0.060814033 | 0.775458257 | -0.074861471 | 0.805534524 | -0.056599434 | 0.847405864 |
| 209234.6719 | 205862.8594 | 197053.1875 | 0.060567985 | 0.745023488 | -0.231677907 | 0.26626937 | -0.485619159 | 0.118785258 |

| | | | | | | | | |
|---|---|---|---|---|---|---|---|---|
| 18169.6875 | 15664.98926 | 14692.92285 | 0.060530398 | 0.29607963 | -0.226667013 | 0.01179292 | -0.643996034 | 0.055741156 |
| 192896.0625 | 212421.6875 | 195383.25 | 0.060476548 | 0.540500239 | 0.323526261 | 0.071187644 | 0.343733505 | 0.047120928 |
| 5700.841309 | 9874.236328 | 6259.070313 | 0.060473682 | NA | 1.287446174 | 0.206278656 | 0.723878603 | 0.343531881 |
| 7156.549805 | 5304.148926 | 5353.212402 | 0.060374731 | 0.680230584 | -0.140552399 | 0.327625892 | -0.511234341 | 0.091928987 |
| 227465.7031 | 246679.8438 | 238981.7656 | 0.060299003 | 0.81092893 | 0.679974303 | 0.054857325 | 1.48374949 | 0.010623938 |
| 34680.92578 | 24189.49805 | 23082.48242 | 0.059987852 | 0.837651404 | 0.097889772 | 0.630134812 | 0.113707361 | 0.727949232 |
| 11387.73828 | 39670.70703 | 28567.31445 | 0.059764772 | 0.963716179 | 1.473666388 | 0.410282257 | 0.885217295 | 0.403223622 |
| 182016.875 | 182924.4375 | 153648.6875 | 0.059593567 | 0.642613461 | 0.265834636 | 0.108508712 | 0.047267517 | 0.751918242 |
| 106607.5391 | 111261.2344 | 112978.8516 | 0.059118268 | 0.73774346 | 0.652668916 | 0.055241445 | 0.079135144 | 0.643512365 |
| 178495.5938 | 183609.3594 | 176283.5938 | 0.058687864 | 0.835403357 | 0.603947992 | 0.150108069 | 1.769316369 | 0.02183874 |
| 259857 | 260747.7813 | 285456.5938 | 0.058230059 | 0.645442215 | -0.1985172 | 0.089964733 | -0.157149557 | 0.118023574 |
| 29173.39844 | 20970.28906 | 25282.19922 | 0.057764877 | 0.757156456 | 0.464059531 | 0.08608279 | 0.287770745 | 0.294442518 |
| 3345232.125 | 3958108.633 | 3630378.203 | 0.057737746 | 0.650027478 | -0.111192021 | 0.273058477 | 0.080395757 | 0.524280218 |
| 711948.375 | 764481.1875 | 747282.4375 | 0.057542205 | 0.279048922 | -0.069385084 | 0.532402501 | 0.103031773 | 0.148543157 |
| 9671.943359 | 7893.186035 | 7969.593262 | 0.057135647 | 0.649999812 | -0.319278068 | 0.094800581 | -0.356833219 | 0.096997136 |
| 18667.49609 | 18868.61523 | 10075.18066 | 0.056897807 | 0.692885865 | 0.351047186 | 0.220247361 | -0.796827554 | 0.124271916 |
| 18706.60352 | 18761.64844 | 15730.25098 | 0.055929458 | 0.581907902 | 0.408847176 | 0.017037525 | 0.045951783 | 0.724760034 |
| 136208.5625 | 142353.5 | 149595.2813 | 0.055880915 | 0.309686498 | 0.165071549 | 0.291253414 | 0.502529925 | 0.026725136 |
| 4986.402344 | 6076.319824 | 5352.442383 | 0.055838331 | 0.762776035 | 0.852939024 | 0.143770745 | -0.052246938 | 0.805147292 |
| 12322.24121 | 7387.364258 | 6552.280762 | 0.055597905 | 0.8926662 | 0.212949319 | 0.598335471 | -0.563978624 | 0.257876793 |
| 45166.53906 | 45857.8125 | 46881.03125 | 0.05523938 | 0.652697474 | -0.084653005 | 0.509387885 | -0.374739071 | 0.109020087 |
| 648732.25 | 639601.3125 | 718596.75 | 0.055238522 | 0.693763698 | 0.469897237 | 0.043784011 | 1.122411735 | 0.020898684 |
| 63318.50781 | 74337.92969 | 71556.96875 | 0.054934977 | 0.8926662 | 0.634428747 | 0.190057963 | 0.068845756 | 0.86637512 |
| 1676.472168 | 1888.721191 | 1295.2854 | 0.054910247 | 0.683925485 | -0.002105004 | 0.989297728 | -0.358532883 | 0.193784104 |
| 5122.608398 | 5313.829102 | 5171.731934 | 0.054768851 | 0.749074148 | -0.723225857 | 0.041240492 | -0.365248787 | 0.081658507 |
| 332878.25 | 368678.625 | 322352.2188 | 0.054732239 | 0.74476981 | 0.300833812 | 0.08804938 | 0.261937515 | 0.096592219 |
| 58548.43359 | 52785.00391 | 65462.14453 | 0.054601116 | 0.653598418 | 0.745264901 | 0.013885891 | 0.608140487 | 0.070968049 |
| 217473.8438 | 214825.8594 | 203416.7656 | 0.054347438 | 0.674253346 | 0.303102136 | 0.126735053 | 1.329293435 | 0.008387835 |
| 182711.5 | 188599.1719 | 196252.1094 | 0.054309343 | 0.268099056 | -0.305661332 | 0.048024604 | 0.350345381 | 0.023204793 |
| 13636.43066 | 13603.53223 | 8868.3125 | 0.054307549 | 0.854804669 | -1.091786744 | 0.131086275 | -0.786481776 | 0.137554403 |
| 131527.6094 | 121650.2188 | 115829.0938 | 0.054274983 | 0.518013064 | 0.179506603 | 0.122107272 | 0.366192726 | 0.051133451 |
| 22125.27539 | 18642.41602 | 17338.7168 | 0.054236561 | 0.669094924 | 0.215099008 | 0.194228932 | 0.010167763 | 0.960748672 |
| 1871.113647 | 3136.97876 | 2395.487061 | 0.05412137 | 0.857725988 | -0.256865864 | 0.419290117 | 0.413330345 | 0.347944229 |

| | | | | | | | | |
|---|---|---|---|---|---|---|---|---|
| 12751.40332 | 13637.20703 | 8383.294922 | 0.053898041 | 0.662042929 | 0.703914362 | 0.078077873 | 0.25601487 | 0.477536926 |
| 3617.674805 | 2821.501953 | 3626.087891 | 0.05379097 | 0.817346175 | -0.147495285 | 0.700745399 | -0.146734322 | 0.604535078 |
| 24960.72852 | 24391.53906 | 22184.95117 | 0.053722854 | 0.816650151 | -0.639540093 | 0.163729974 | -1.212532452 | 0.090232821 |
| 109693.8359 | 115546.8047 | 124067.1094 | 0.053425591 | 0.682393165 | 0.038205216 | 0.466251631 | -0.060412002 | 0.482696616 |
| 14266.08691 | 14639.1543 | 11333.8252 | 0.053294166 | 0.553467034 | -0.701368062 | 0.034038555 | -0.464126037 | 0.075477785 |
| 6857.331543 | 7110.959961 | 2984.125977 | 0.053260245 | 0.887963154 | -0.854120669 | 0.220094258 | -0.619840106 | 0.3169493 |
| 8003.599121 | 8412.455078 | 7693.493164 | 0.053227958 | 0.849879367 | -0.167092963 | 0.639466845 | 0.305836593 | 0.29550852 |
| 30986.25977 | 30589.32422 | 27918.37109 | 0.053082947 | 0.598139599 | -0.528568208 | 0.051139483 | -0.320568357 | 0.086777585 |
| 28470.39258 | 28436.59766 | 29911.58594 | 0.052924559 | 0.685684032 | 0.171687475 | 0.141906061 | 0.20596428 | 0.091810798 |
| 340820.5625 | 371071.2188 | 277683.9688 | 0.052547299 | 0.686476859 | -0.019794 | 0.881315433 | -0.254291028 | 0.246207108 |
| 270562.0625 | 305968.9063 | 294893.0313 | 0.052125832 | 0.293018436 | -0.017057787 | 0.821958214 | -0.145366485 | 0.15696864 |
| 82984.47656 | 80845.14063 | 78598.42188 | 0.052049787 | 0.746135232 | 0.307537375 | 0.163673552 | 0.139833303 | 0.408763244 |
| 433643.9063 | 456336.8125 | 480069.0938 | 0.05182831 | 0.280015719 | -0.074762782 | 0.183057628 | -0.164271479 | 0.093865648 |
| 18226.29297 | 17898.5332 | 20535.3457 | 0.051762347 | 0.626429783 | -0.30948396 | 0.094128959 | -0.56062261 | 0.037772514 |
| 364374.375 | 353425 | 316788.8125 | 0.051624309 | 0.476745676 | -0.494704937 | 0.029364116 | -0.470619504 | 0.027087229 |
| 31815.58398 | 42106.0625 | 43376.23828 | 0.051369349 | 0.888900123 | 0.521205669 | 0.491319916 | 0.389285708 | 0.320774355 |
| 50538.27344 | 49737.10156 | 47641.1875 | 0.05130213 | 0.630765638 | 0.239180886 | 0.126731731 | 0.180198455 | 0.161358011 |
| 17832.33789 | 19832.05273 | 19781.53711 | 0.051184583 | 0.819127104 | -0.07900756 | 0.586831771 | -0.426614399 | 0.063474265 |
| 35881.79297 | 39878.53125 | 37427.96875 | 0.051167105 | 0.62114954 | -0.305231453 | 0.051111089 | -0.376575836 | 0.031136253 |
| 85538.61719 | 89354.25781 | 89586.55469 | 0.051136981 | 0.415576072 | -0.185657048 | 0.038133348 | 0.021438806 | 0.602118291 |
| 69269.30469 | 70929.07031 | 74319.25781 | 0.051130841 | 0.619539817 | 0.428613408 | 0.068282276 | 0.741518055 | 0.016560894 |
| 125664.2422 | 146974.7656 | 152209.0469 | 0.051051519 | 0.924866197 | 1.016436786 | 0.087142005 | 1.064825035 | 0.056570421 |
| 400716 | 367109.9063 | 328998.7188 | 0.050976604 | 0.828897042 | -0.316608052 | 0.109650618 | -0.063449114 | 0.677060914 |
| 399756.4063 | 418248.6875 | 403392.0938 | 0.050883461 | 0.379981327 | -0.272280573 | 0.067403223 | -0.118848605 | 0.136000191 |
| 80624.20313 | 62611.90234 | 56055.39844 | 0.050610689 | 0.750995763 | 0.673826403 | 0.015625705 | 0.793589419 | 0.131886359 |
| 8778.818359 | 11177.2207 | 10851.7959 | 0.050548977 | 0.561698942 | -0.02983194 | 0.754243703 | -0.238895055 | 0.209261477 |
| 238503.375 | 223226.4531 | 234148.625 | 0.050459018 | 0.746281035 | 0.126893595 | 0.443820431 | 0.388399547 | 0.089733184 |
| 7627.755371 | 9478.855469 | 6445.206543 | 0.050346928 | 0.73314776 | 0.32748053 | 0.13610541 | -0.256530052 | 0.345946596 |
| 2280052.5 | 2585739 | 2483755.75 | 0.049967954 | 0.326655688 | -0.216494566 | 0.177376869 | -0.027709529 | 0.752300704 |
| 44992.94141 | 44921.70703 | 47624.62891 | 0.049813558 | 0.436807126 | -0.447488571 | 0.024165741 | -0.211940343 | 0.047514693 |
| 499443.4063 | 489528.0625 | 506223.4688 | 0.049758518 | 0.708474379 | 0.032717695 | 0.800473631 | -0.101556373 | 0.45861981 |
| 20252.17969 | 21071.27148 | 8326.828125 | 0.049454824 | 0.828467897 | -0.572910073 | 0.031635436 | -1.205687846 | 0.080646782 |
| 13398.96582 | 9061.709961 | 10565.2666 | 0.049409464 | 0.788314219 | 0.01432354 | 0.941883073 | -0.06382441 | 0.820959074 |

| | | | | | | | | |
|---|---|---|---|---|---|---|---|---|
| 24576.09766 | 28545.98633 | 24202.98828 | 0.049369465 | 0.746281035 | 0.086697637 | 0.230183652 | 0.042198861 | 0.728022021 |
| 27394.27148 | 29236.69531 | 32232.70117 | 0.049240643 | 0.880090511 | 0.357564131 | 0.210538099 | 0.265598249 | 0.309518203 |
| 162222.25 | 154510.7656 | 157916.4375 | 0.048878711 | 0.435526499 | -0.03676671 | 0.506573912 | -0.22684552 | 0.050533787 |
| 28096.53125 | 29347.85547 | 27364.66797 | 0.048782732 | 0.636489801 | -0.04126072 | 0.69897549 | -0.270912435 | 0.118033234 |
| 24240.34375 | 28585.91016 | 31902.62695 | 0.04867555 | 0.779670979 | 0.385741676 | 0.058077004 | 0.43437372 | 0.136837137 |
| 521759.8301 | 480556.0176 | 453531.6602 | 0.048112875 | 0.631078063 | 0.240305281 | 0.102526216 | 0.069372121 | 0.504948403 |
| 17796.94141 | 13426.96094 | 14275.12598 | 0.048066787 | 0.66030774 | -0.579901827 | 0.051139483 | -0.415649782 | 0.136356438 |
| 140333.8906 | 149588.875 | 145630.9375 | 0.047808556 | 0.562292487 | 0.176280836 | 0.098228912 | 0.758669371 | 0.009587646 |
| 45346.08984 | 39611.46875 | 42019.03125 | 0.047139159 | 0.762776035 | 0.068919115 | 0.338377194 | 0.091862103 | 0.372162147 |
| 55795.83203 | 54658.1875 | 54680.35156 | 0.047097331 | 0.890671487 | 0.216916161 | 0.534693749 | 0.075431426 | 0.827620964 |
| 1393175 | 1610908.25 | 1540482.875 | 0.046973526 | 0.715015377 | -0.079632074 | 0.585401205 | -0.085612496 | 0.547919535 |
| 230580.3438 | 251568.0156 | 250524.625 | 0.046916573 | 0.764304977 | 0.315992879 | 0.042884533 | 0.518443069 | 0.022340012 |
| 152945.0625 | 162855.8438 | 154047.2188 | 0.046781697 | 0.423302296 | -0.223038607 | 0.04939536 | -0.208063342 | 0.041803864 |
| 34959.51563 | 34044.82422 | 30262.94922 | 0.046726206 | 0.539642904 | 0.367949952 | 0.048593497 | 0.238825563 | 0.121306313 |
| 60146.66016 | 63038.82813 | 58416.35156 | 0.046445091 | 0.515809019 | -0.18505551 | 0.117476758 | -0.308649319 | 0.053823402 |
| 187678.2969 | 196243.125 | 190266.7031 | 0.046349004 | 0.51585406 | -0.067698161 | 0.414034856 | 0.190114212 | 0.086224164 |
| 4720.783691 | 5129.735352 | 1790.221924 | 0.046301469 | 0.835128657 | -1.118382516 | 0.017018475 | -0.885191905 | 0.161137259 |
| 72536.16406 | 76151.66406 | 83667.01563 | 0.046154482 | 0.863810045 | 0.170712921 | 0.436924695 | 0.384467898 | 0.076362612 |
| 157413.7969 | 139134.4531 | 137959.2813 | 0.045853695 | 0.525725981 | -0.326881941 | 0.078206113 | -0.382017435 | 0.043437905 |
| 16184.82129 | 17447.42773 | 14942.28125 | 0.045850651 | 0.743096884 | 0.118462126 | 0.328806215 | -0.290702561 | 0.086843504 |
| 5516.353516 | NA | 2481.508545 | 0.045709306 | 0.704271223 | -0.507145216 | 0.047347558 | -0.944335733 | NA |
| 37988.30469 | 33935.65625 | 38333.64844 | 0.045588394 | 0.690885405 | -0.081782256 | 0.57025817 | -0.199218483 | 0.136049841 |
| 1704918.875 | 1726837.5 | 1743016.75 | 0.04556902 | 0.72050444 | -0.080339372 | 0.129311198 | -0.169389397 | 0.047120928 |
| 3023.249268 | 5216.123047 | 336.0046692 | 0.045441882 | 0.976103645 | 0.095292673 | 0.950005687 | -0.468339151 | 0.785580722 |
| 630899.0625 | 621078.75 | 576944.875 | 0.044710279 | 0.69431535 | 0.284370458 | 0.102618064 | 0.058430047 | 0.611231378 |
| 9995.639648 | 10802.15625 | 11411.88281 | 0.044517415 | 0.694156601 | -0.007368754 | 0.901034011 | 0.43852376 | 0.048431709 |
| 52221.77246 | 45979.85547 | 53124.97266 | 0.044187424 | 0.825031307 | 0.180054876 | 0.417514232 | -0.291293372 | 0.273802519 |
| 72544.51563 | 77111.21094 | 85703.98438 | 0.04401044 | 0.730858472 | 0.257265048 | 0.101338323 | -0.012626785 | 0.917138704 |
| 5088.34082 | 6675.554199 | 4430.334961 | 0.043896089 | 0.832358028 | -0.174545053 | 0.38601673 | -0.314733227 | 0.286081151 |
| 19090.85742 | 23158.10156 | 18537.01172 | 0.043623025 | 0.889653494 | 0.359893101 | 0.284636711 | 0.286791166 | 0.374943822 |
| 527574.0625 | 532978.5625 | 481050.5625 | 0.043559545 | 0.77009062 | 0.314678361 | 0.099835584 | 0.552214674 | 0.030633392 |
| 25850.37305 | 20286.89258 | 22970.89648 | 0.043506516 | 0.912540692 | -0.228807217 | 0.280916524 | -0.305706603 | 0.176540122 |
| 97746.75781 | 103767.5313 | 102165.8516 | 0.043389604 | 0.573636658 | 0.003939028 | 0.906346526 | 0.022997783 | 0.622107185 |

| | | | | | | | | |
|---|---|---|---|---|---|---|---|---|
| 61738.01953 | 60600.87891 | 57164.51953 | 0.043320397 | 0.766981151 | 1.883522208 | 0.047896832 | 2.427352175 | 0.008614663 |
| 132250.9844 | 126310.4063 | 145383.8438 | 0.043029903 | 0.722598037 | 0.298096148 | 0.104431683 | 0.427040565 | 0.055298525 |
| 157846.8281 | 152969.8594 | 166413.7656 | 0.042865767 | 0.825273741 | 0.195173748 | 0.363796172 | 0.266929048 | 0.22024877 |
| 13218.42969 | 13541.48535 | 12264.37305 | 0.042304301 | 0.728494815 | -0.312955799 | 0.117831745 | -0.395351555 | 0.021984636 |
| 519657.6563 | 511578.7969 | 541685.5625 | 0.042281519 | 0.664936285 | 0.023352156 | 0.832010656 | 0.213636668 | 0.123837484 |
| 516610.1875 | 528500.125 | 533696.8125 | 0.041633591 | 0.794535026 | 0.341411905 | 0.109845349 | 0.156513882 | 0.339148153 |
| 159513.4688 | 169952.6563 | 156130.7031 | 0.041572992 | 0.741457034 | -0.251458497 | 0.136651367 | -0.130559081 | 0.2446185 |
| 845639.1875 | 911219.5625 | 973629.8125 | 0.041263595 | 0.531679333 | 0.081872476 | 0.218053066 | 0.230862864 | 0.11743518 |
| 188.2764435 | 298.4723206 | NA | 0.040814289 | NA | -0.194443084 | NA | -1.159076797 | NA |
| 274165.4688 | 313015.7813 | 312806.9688 | 0.040771293 | 0.700025048 | 0.69019708 | 0.032593212 | 0.784655815 | 0.036652305 |
| 19876.91016 | 21517.46484 | 14488.80078 | 0.040606498 | 0.835052252 | -0.812911034 | 0.103128859 | -0.279610052 | 0.346059704 |
| 770873.5 | 689746.5625 | 617439.375 | 0.040345777 | 0.874213658 | 0.504252964 | 0.149058018 | 0.345534932 | 0.240458637 |
| 86743.79688 | 82621.19531 | 90388.07813 | 0.040333726 | 0.678408917 | -0.219189154 | 0.052457323 | -0.113189889 | 0.151995 |
| 150379.2188 | 118397.1875 | 123875.8125 | 0.0401139 | 0.780681403 | -0.366471965 | 0.160918214 | -0.634543921 | 0.061962869 |
| 1468362.625 | 995224.4375 | 1153819.375 | 0.039999953 | 0.885533339 | 0.09479982 | 0.735373025 | 0.015285162 | 0.969246541 |
| 20737.00586 | 21739.45117 | 20758.89453 | 0.039939439 | 0.81532173 | 0.670843453 | 0.048787226 | 0.276237864 | 0.053375696 |
| 410444.0625 | 425611.3438 | 411557.5313 | 0.03989631 | 0.377880293 | 0.096816849 | 0.203239255 | -0.066597197 | 0.203158741 |
| 256374.1406 | 263304.0938 | 251922.3594 | 0.039894926 | 0.761691235 | 1.046015687 | 0.052874011 | 0.329069661 | 0.104116883 |
| 31946.25977 | 29977.74414 | 30532.66406 | 0.03977259 | 0.674529959 | -0.804367368 | 0.05283098 | -0.866496792 | 0.039966686 |
| 39551.42188 | 37582.16016 | 42162.76563 | 0.039444858 | 0.791670943 | 0.33933634 | 0.071544796 | -0.124493428 | 0.29550852 |
| 790736.5625 | 883202.4375 | 857315.125 | 0.039227975 | 0.68087566 | 0.022610328 | 0.886633393 | 0.200720324 | 0.137551235 |
| 8210.621094 | 11177.28809 | 5082.970703 | 0.038587654 | 0.939941347 | -1.361947435 | 0.199074765 | -1.912820642 | 0.152372879 |
| 103047.4063 | 101244.6719 | 105817.5391 | 0.038313662 | 0.419717719 | 0.131403005 | 0.07692885 | 0.079155885 | 0.110105852 |
| 949539.5625 | 1026245.313 | 1045906.188 | 0.038220033 | 0.481848069 | 0.050957219 | 0.500849741 | 0.355309639 | 0.05263698 |
| 173212.2031 | 169282.9688 | 182437.9531 | 0.038132354 | 0.217999855 | -0.14037514 | 0.052817461 | 0.448575288 | 0.029710029 |
| 672307.7012 | 647507.125 | 652747.9844 | 0.037845798 | 0.540918348 | -0.049417862 | 0.437151431 | -0.311831395 | 0.049651271 |
| 97000.22656 | 112490.9609 | 93146.74219 | 0.037813676 | 0.27685626 | -0.009439724 | 0.926516232 | -0.087675225 | 0.507481359 |
| 3660.917969 | 3939.213867 | 3412.979736 | 0.037550555 | 0.866400821 | -0.36588174 | 0.200649535 | -1.708750193 | 0.051133451 |
| 514435.4375 | 466228.4375 | 479706.0625 | 0.037414948 | 0.665543406 | -0.052829424 | 0.473700287 | -0.42129076 | 0.029653052 |
| 776402.4375 | 767938.375 | 797038.4375 | 0.037361592 | 0.710150856 | -0.534518892 | 0.077625195 | -0.45843233 | 0.074184841 |
| 69333.78125 | 76574.52344 | 72053.32813 | 0.037244977 | 0.755926324 | -0.397142274 | 0.045536435 | -0.305263488 | 0.040766293 |
| 47678.38672 | 42559.20313 | 43161.4375 | 0.037098696 | 0.641235022 | -0.073237568 | 0.349103174 | -0.152021677 | 0.152597158 |
| 117365.7422 | 176131.25 | 164123.4844 | 0.037070361 | 0.896724734 | -0.206420194 | 0.433732426 | 0.074627036 | 0.816577853 |

| | | | | | | | | |
|---|---|---|---|---|---|---|---|---|
| 119365.2266 | 125991.8594 | 119450.0234 | 0.036987871 | 0.636286379 | -0.366543407 | 0.052817461 | -0.590282783 | 0.030727383 |
| 7850.564453 | 8107.779297 | 5705.274902 | 0.036923365 | 0.932924274 | -0.594999846 | 0.34872471 | -0.941691693 | 0.221789362 |
| 3263.468018 | 5494.732422 | 5578.171387 | 0.036824422 | 0.979227978 | 0.919248552 | 0.414751124 | 1.120415924 | 0.303877641 |
| 80301.98438 | 86278.02344 | 78345.15625 | 0.036776836 | 0.72315698 | -0.195050672 | 0.209379283 | -0.118813903 | 0.286659111 |
| 33641.03125 | 29530.24023 | 24487.42969 | 0.036382525 | 0.833974595 | 0.242863554 | 0.567455459 | -0.55993889 | 0.090708074 |
| 115932.8125 | 115045.5625 | 111127.625 | 0.036292469 | 0.80960355 | -0.14295777 | 0.398951669 | -0.336401112 | 0.158868941 |
| 48696.02344 | 54662.26172 | 51762.68359 | 0.036265313 | 0.926756367 | -0.296052907 | 0.526766343 | -0.164990984 | 0.695058584 |
| 24774.83594 | 31251.37109 | 22152.41602 | 0.035832242 | 0.863008177 | -0.422829542 | 0.177893197 | -1.345346599 | 0.045311345 |
| 262295.5938 | 235378.8438 | 212529.1875 | 0.035459932 | 0.745111336 | -0.035040496 | 0.698316027 | 0.248918484 | 0.189972522 |
| 3388315 | 3925577 | 3995750 | 0.034961108 | 0.595239978 | 0.162377737 | 0.137613908 | 0.393963304 | 0.084223187 |
| 14510.30273 | 12740.10059 | 12726.45898 | 0.03456221 | 0.783327271 | -0.424515837 | 0.081070686 | -0.32272222 | 0.093202677 |
| 27388.08594 | 31114.77734 | 31467.41992 | 0.033778387 | 0.819276169 | 0.943495335 | 0.076036407 | 1.081481431 | 0.020159082 |
| 8363.322266 | 9110.921875 | 9564.755859 | 0.033687809 | 0.924956664 | 0.506195457 | 0.088618776 | 0.583263636 | 0.034111087 |
| 46301.20313 | 46717.46094 | 44406.26563 | 0.03279859 | 0.777598159 | 0.165957276 | 0.231386286 | 0.309581099 | 0.089218964 |
| 1306264.375 | 1299126.625 | 1223646.625 | 0.032700594 | 0.786345636 | 0.390607756 | 0.025489497 | 0.917912997 | 0.009587646 |
| 916723.5 | 1188827.25 | 1105360 | 0.032551571 | 0.87622214 | 0.673830206 | 0.050075499 | 0.195686855 | 0.363209271 |
| 19736.3125 | 13209.44531 | 17226.25391 | 0.032389995 | 0.85372591 | 0.218396866 | 0.24706799 | 0.191002855 | 0.51721563 |
| 5368.54834 | 3712.825195 | 2868.753906 | 0.031904186 | 0.979395647 | 0.90963458 | 0.596459242 | -0.82576078 | 0.49125304 |
| 91593.47656 | 93645.76563 | 92545.76563 | 0.031855266 | 0.70461778 | 0.153787162 | 0.127225665 | 0.134620738 | 0.162707853 |
| 48828.46875 | 48550.82422 | 49750.06641 | 0.031758507 | 0.658737241 | 0.103612137 | 0.162317825 | 0.278277793 | 0.037962113 |
| 3634.799072 | 5446.303223 | 2842.738037 | 0.031593154 | 0.951462775 | -0.80080968 | 0.128915752 | -0.620158968 | 0.191130479 |
| 8127.887695 | 7975.758301 | 7370.767578 | 0.031579017 | 0.672802027 | -0.155448754 | 0.253983381 | -0.983596702 | 0.013391184 |
| 2317886.5 | 3013364 | 2656989 | 0.031510371 | 0.661967827 | 0.243091468 | 0.055581979 | 0.285132263 | 0.223102126 |
| 9892.37793 | 10890.27246 | 9525.089844 | 0.030986743 | 0.861468351 | 0.1448017 | 0.418226191 | -0.132686502 | 0.487247142 |
| 7455.242188 | 8449.432617 | 5616.986816 | 0.030788524 | 0.849614988 | -0.335497466 | 0.169811009 | -0.532290434 | 0.102620951 |
| 72350.55469 | 65338.28906 | 62418.11328 | 0.030777707 | 0.604697925 | -0.044899705 | 0.419290117 | -0.024724519 | 0.811284277 |
| 18777.98242 | 29431.49023 | 20330.73047 | 0.030484544 | NA | 0.814289987 | 0.095989926 | 0.855716966 | 0.156467569 |
| 16310.66406 | 16078.4834 | 16923.34961 | 0.030244285 | 0.868395759 | -0.114740082 | 0.601692147 | 0.716632387 | 0.053053903 |
| 58873.82031 | 60752.46875 | 60577.8125 | 0.030180654 | 0.46699162 | 0.119922275 | 0.16635868 | 0.134610301 | 0.031136253 |
| 15865.18457 | 14784.45605 | 12459.34961 | 0.029750444 | 0.801216945 | 0.404183214 | 0.058077004 | 0.222336395 | 0.274434424 |
| 20348.75391 | 19451.02344 | 18968.90039 | 0.029663793 | 0.853712828 | 0.09082202 | 0.402638339 | -0.12957391 | 0.268328876 |
| 7573.137207 | 11075.50586 | 10626.87695 | 0.029605843 | 0.940842952 | 0.119547441 | 0.673113251 | -0.09393929 | 0.787977398 |
| 38777.39075 | 43020.79437 | 45926.04297 | 0.029295357 | 0.873134516 | 1.081638948 | 0.033804616 | 0.905649447 | 0.026417259 |

| | | | | | | | | |
|---|---|---|---|---|---|---|---|---|
| 16589.57617 | 16961.29688 | 17277.97852 | 0.028951853 | 0.844972921 | 0.220522052 | 0.199414398 | 0.551309307 | 0.017015337 |
| 7357.955078 | 7645.078125 | 3585.770996 | 0.028950466 | 0.85906459 | 0.263305371 | 0.297572186 | 0.248880993 | 0.627065516 |
| 80069.32813 | 75452.125 | 76258.50781 | 0.028538832 | 0.835128657 | -0.299766907 | 0.291065687 | 0.36496668 | 0.093324351 |
| 39270.83594 | 28563.05469 | 22223.23633 | 0.028090271 | 0.941581581 | 0.386379117 | 0.420604095 | 0.621960062 | 0.255662977 |
| 56895.96875 | 47253.38281 | 50654.85156 | 0.02794382 | 0.686594537 | 0.733417856 | 0.012254503 | 0.91565255 | 0.046487363 |
| 16875.09961 | 18952.28711 | 14620.30371 | 0.027413315 | 0.728917468 | -0.286038142 | 0.082108346 | -0.429558804 | 0.076613575 |
| 444014.4375 | 507650.375 | 445110.0313 | 0.027379416 | 0.86454987 | -0.249591154 | 0.129928099 | -0.115477011 | 0.354400658 |
| 156400.7656 | 163009.8906 | 187652.625 | 0.027238539 | 0.726620132 | 0.217407471 | 0.183085834 | 0.80862282 | 0.052598684 |
| 46419.03906 | 51685.61328 | 44900.32031 | 0.026525628 | 0.674827859 | -0.560795347 | 0.013679803 | -0.656740396 | 0.020008995 |
| 151212.1563 | 141138.4531 | 137337.4844 | 0.026234431 | 0.695559783 | 0.225669889 | 0.06863053 | -0.059636408 | 0.49402204 |
| 313389.375 | 318989.875 | 323518 | 0.026191507 | 0.838790359 | 0.718773324 | 0.037014778 | 0.501214938 | 0.063438208 |
| 5515.644043 | 4617.049805 | 227.9144287 | 0.025852211 | 0.957800688 | 0.957106183 | 0.081451772 | -0.117766828 | 0.91311394 |
| 314874.2813 | 319004.8438 | 342688.8125 | 0.025214108 | 0.770955341 | -0.450889157 | 0.052817461 | 0.168195167 | 0.095653818 |
| 526986.4375 | 541325.6875 | 457713.7188 | 0.024611839 | 0.881107987 | 0.029678904 | 0.87026131 | 1.201747235 | 0.036848746 |
| 588589 | 585107.125 | 702341.375 | 0.024179433 | 0.846345659 | -0.003241457 | 0.980414266 | 0.017079035 | 0.926180903 |
| 4665.148926 | 4531.253906 | 2579.894287 | 0.023982092 | 0.920219187 | -0.153926238 | 0.54508114 | -0.60358841 | 0.171907805 |
| 285793.375 | 286040.9063 | 274147.375 | 0.023686714 | 0.755050946 | -0.030552311 | 0.687498922 | -0.327859362 | 0.056570421 |
| 57783.26563 | 50255.06641 | 53477.54688 | 0.023604167 | 0.772187194 | -0.412529946 | 0.052457323 | -0.698474632 | 0.02527614 |
| 79731.78125 | 76749.53125 | 76841.10156 | 0.023586595 | 0.908823352 | 0.846857685 | 0.081994347 | 1.22292871 | 0.022312939 |
| 1615.944702 | 2826.784668 | 6028.310547 | 0.023559605 | 0.964557468 | -0.031499555 | 0.928153584 | 0.330282378 | 0.729376986 |
| 1503325.25 | 1703507.875 | 1521998.625 | 0.023496412 | 0.916063354 | -0.081429828 | 0.723806745 | 0.126921137 | 0.531726747 |
| 5562791.5 | 6747017.5 | 6094592.5 | 0.023487127 | 0.857050688 | -0.369839089 | 0.124478178 | -0.227571091 | 0.233347919 |
| 34564.78125 | 24343.16602 | 40478.83594 | 0.023394411 | 0.742989299 | -0.921573708 | 0.048221719 | -1.004886544 | 0.066077335 |
| 80650.05469 | 84016.09375 | 100294.5781 | 0.023212161 | 0.90084342 | 0.776583135 | 0.045329621 | 1.052551238 | 0.038826476 |
| 145952.6875 | 150518.7188 | 148207.8594 | 0.023106683 | 0.794535026 | -0.087225141 | 0.437068655 | -0.114249594 | 0.280069359 |
| 830073.5 | 781732.4375 | 877625.0625 | 0.023023756 | 0.810385109 | -0.081550059 | 0.365797018 | 0.007440845 | 0.943605034 |
| 201562.1719 | 230304.2813 | 213556.8438 | 0.022893141 | 0.896724734 | -0.145469347 | 0.562301807 | -0.131424483 | 0.489970707 |
| 39909.32031 | 65540.03906 | 63995.84766 | 0.022703 | 0.840812438 | 0.028326748 | 0.89833266 | -0.288906868 | 0.378495398 |
| 61404.54688 | 63350.22656 | 57583.76563 | 0.022456239 | 0.791331725 | -0.072897247 | 0.347728415 | 0.145848908 | 0.140806999 |
| 57855.05469 | 60780.34766 | 63291.00781 | 0.022454102 | 0.858884107 | 0.505052692 | 0.026250916 | 0.278084938 | 0.060161253 |
| 150692.5 | 168869.7656 | 158971.5938 | 0.022437529 | 0.857050688 | 0.554452501 | 0.039936585 | 0.224516604 | 0.163299322 |
| 94115.80469 | 94133.25 | 101114.7734 | 0.021869147 | 0.835052252 | 0.3597199 | 0.062087755 | 0.554349323 | 0.017279709 |
| 617584.4492 | 610345.3672 | 628452.7461 | 0.021832412 | 0.896724734 | 0.576219115 | 0.093751816 | 0.272783112 | 0.017279709 |

| | | | | | | | | |
|---|---|---|---|---|---|---|---|---|
| 202067.4531 | 195704.75 | 209864.0156 | 0.021671709 | 0.90288932 | -0.132678752 | 0.405351976 | 0.386558902 | 0.020008995 |
| 3892.604492 | 4120.648438 | 2804.693359 | 0.021574485 | 0.951103717 | 0.603438105 | 0.225638357 | 1.22458554 | 0.070968049 |
| 109940.7422 | 106577.9297 | 108108.7813 | 0.021401191 | 0.527886081 | -0.039841576 | 0.58415851 | 0.017951384 | 0.557774325 |
| 179660 | 201696.5313 | 194832.0313 | 0.021306978 | 0.78527275 | 0.174891177 | 0.109341875 | 0.219074564 | 0.091928987 |
| 114382.0078 | 81719.30469 | 113128.7656 | 0.021226229 | 0.94882882 | 0.832341582 | 0.108875676 | 1.42676475 | 0.071043799 |
| 31419.93555 | 28038.88867 | 29551.64258 | 0.020584318 | 0.827859454 | 0.370019148 | 0.066504374 | 0.338205536 | 0.055355517 |
| 16284.2373 | 13400.32324 | 15457.4375 | 0.020003926 | 0.852675579 | 0.306176178 | 0.129928099 | 0.287166216 | 0.155548876 |
| 209778.125 | 239325.6406 | 126028.7188 | 0.019478619 | 0.920219187 | -0.70682196 | 0.106378122 | -0.310895757 | 0.433601327 |
| 304114.0625 | 303785.8438 | 281794.25 | 0.019477868 | 0.861910935 | 0.539249887 | 0.051888756 | 0.54285903 | 0.022233336 |
| 373362.9688 | 373743.5938 | 326193.3438 | 0.019454191 | 0.844315361 | -0.416408962 | 0.053116693 | -0.434641158 | 0.039408515 |
| 147483.7344 | 156798.875 | 155420.9063 | 0.019357396 | 0.688041815 | 0.064525515 | 0.168942668 | -0.10220336 | 0.110015219 |
| 157040.125 | 145297.6563 | 157601.3906 | 0.01929874 | 0.880982314 | -0.134322874 | 0.342068721 | 0.101785558 | 0.190246465 |
| 198756.8281 | 213405.0625 | 205120.6406 | 0.018952118 | 0.759466989 | 0.24371354 | 0.174475307 | -0.14942993 | 0.065461015 |
| 25379.01758 | 26360.06445 | 23376.64453 | 0.018860433 | 0.820538814 | -0.080304947 | 0.26626937 | -0.537258859 | 0.022750858 |
| 22561.39258 | 21844.46875 | 20625.94727 | 0.018629608 | 0.865523853 | 0.246031602 | 0.141443989 | -0.129356627 | 0.339568705 |
| 19519.2793 | 17669.28125 | 13131.76855 | 0.018434484 | 0.94279195 | 0.179529149 | 0.488682915 | -0.318602029 | 0.353909834 |
| 44118.28125 | 52877.82422 | 28108.31445 | 0.017972448 | 0.976602645 | -0.28284433 | 0.5443838 | -0.550601984 | 0.330633227 |
| 80652.85156 | 81821.30469 | 73070.90625 | 0.017918746 | 0.883803356 | 0.031395179 | 0.775164931 | -0.551337851 | 0.05183905 |
| 1219.381836 | 1578.536987 | 1337.746826 | 0.017825429 | 0.984192099 | 0.579910683 | 0.450893322 | 0.075857212 | 0.931717307 |
| 94909.80469 | 126031.1641 | 134458.4063 | 0.017645295 | 0.980860528 | 0.342409789 | 0.610324144 | 1.076950974 | 0.124211802 |
| 13926.72754 | 16741.14453 | 17001.625 | 0.017633392 | 0.919971045 | -0.28613041 | 0.21914142 | 0.063313797 | 0.73671998 |
| 25291.98682 | 24228.771 | 24029.73218 | 0.017600902 | 0.860320839 | 0.145466475 | 0.15267096 | -0.382949113 | 0.054887084 |
| 16936.125 | 18651.64063 | 18610.83594 | 0.017584783 | 0.907439401 | 0.086079864 | 0.5989997 | 0.091844588 | 0.557015205 |
| 16074.26172 | 17079.53906 | 16491.60742 | 0.017464013 | 0.834212312 | -0.639435049 | 0.033121608 | -0.253527161 | 0.104116883 |
| 14878.39063 | 14763.92285 | 11193.68457 | 0.017448768 | 0.835128657 | 0.005285428 | 0.954688064 | -0.418765116 | 0.106643735 |
| 321861.75 | 321894.6875 | 334397.9375 | 0.017426448 | 0.809956349 | 0.086242736 | 0.182014886 | 0.083105318 | 0.090074565 |
| 78471.20313 | 82518.75781 | 76029.89844 | 0.016994267 | 0.809956349 | 0.045487183 | 0.365051532 | -0.252227892 | 0.036342938 |
| 44600.53516 | 41437.87891 | 41960.62109 | 0.016767831 | 0.863810045 | -0.001492362 | 0.99201113 | -0.228122998 | 0.125742768 |
| 80058.86719 | 87485.21094 | 84649.13281 | 0.016345942 | 0.875079488 | 0.058027108 | 0.393273993 | 0.872757676 | 0.023204793 |
| 152568.5313 | 159333.5469 | 161698 | 0.016174304 | 0.875623063 | 0.163199199 | 0.200777525 | 0.099305202 | 0.353863449 |
| 860654.5 | 847667.75 | 861233.875 | 0.016111751 | 0.868882653 | 0.572046602 | 0.077376896 | 0.723839718 | 0.032502 |
| 974729 | 1030903.188 | 969302.75 | 0.015553465 | 0.872320507 | -0.099766523 | 0.213349541 | -0.005936139 | 0.923534132 |
| 16202.29395 | 15966.17285 | 13508.51074 | 0.015480707 | 0.965091546 | -0.068316142 | 0.861576827 | -0.21119789 | 0.586939472 |

| | | | | | | | | |
|---|---|---|---|---|---|---|---|---|
| 117154.0469 | 114958.4766 | 115081.5234 | 0.015377053 | 0.835112801 | 0.23674416 | 0.071544796 | 0.118006786 | 0.206000009 |
| 160746.1406 | 197022.5625 | 193632.6563 | 0.015341409 | 0.976602645 | 0.391363437 | 0.428210643 | 0.224596264 | 0.642626047 |
| 62971.71484 | 72953.47656 | 63201.08203 | 0.01504933 | 0.920899159 | 0.346995348 | 0.099547208 | 0.354939727 | 0.082931684 |
| 210771.875 | 228839.4375 | 241255.7969 | 0.015048502 | 0.957465821 | 0.178447414 | 0.491534808 | -0.080012894 | 0.765299476 |
| 140323.3438 | 137737.3125 | 130411.4922 | 0.015032663 | 0.809940214 | -0.468442866 | 0.01179292 | -0.472776073 | 0.012640854 |
| 19052.34766 | 17512.35156 | 17516.87891 | 0.014786714 | 0.946061916 | 0.059942431 | 0.787503164 | -0.196286934 | 0.40366518 |
| 3039.793701 | 6110.082031 | 1434.44751 | 0.014784628 | 0.948568397 | -0.442262151 | 0.18823259 | -0.787819641 | 0.298435811 |
| 119636.9609 | 120360.0938 | 128097.2031 | 0.014673381 | 0.896724734 | 0.445666373 | 0.208916626 | 0.88553591 | 0.013592103 |
| 14415.34668 | 20558.26367 | 17854.94531 | 0.014544289 | 0.973249983 | -0.269406721 | 0.548417707 | 0.289706358 | 0.475229703 |
| 51114.73828 | 38188.36719 | 36491.5 | 0.013679332 | 0.955213556 | -0.088558712 | 0.584095922 | -0.707432878 | 0.056570421 |
| 21721.42383 | 21957.86328 | 16580.32422 | 0.013631412 | 0.908823352 | -0.105512955 | 0.636784361 | -0.305135321 | 0.196593026 |
| 6518.59668 | 8729.743164 | 7420.94043 | 0.013397055 | 0.95401752 | -0.36646418 | 0.296007565 | 0.013316365 | 0.964697558 |
| 20028.7832 | 19403.60742 | 23612.64063 | 0.013045841 | 0.987952683 | -0.951594382 | 0.489603489 | 0.785121884 | 0.323066848 |
| 10800.03418 | 12660.93848 | 13187.9834 | 0.012935406 | 0.926116021 | -0.044380978 | 0.69180654 | -0.1987496 | 0.20370394 |
| 14454.67578 | 12119.7998 | 15846.15332 | 0.012919043 | 0.948108125 | 0.169499431 | 0.430750734 | -0.340955063 | 0.19182538 |
| 37909.65625 | 23369.64453 | 24496.7207 | 0.0129068 | 0.940208105 | 1.228413413 | 0.090276742 | 1.348284125 | 0.135185535 |
| 412544.5313 | 447341.6875 | 456452.125 | 0.012746924 | 0.884624934 | 0.748879265 | 0.064644006 | 1.045017748 | 0.017279709 |
| 213872.8438 | 232764.1719 | 204694.9063 | 0.012687384 | 0.874707705 | 0.037354165 | 0.577348282 | -0.097031023 | 0.315155469 |
| 83431 | 76631.47656 | 73728.89844 | 0.012312045 | 0.923831846 | 0.105757015 | 0.167463497 | 0.122178128 | 0.244446425 |
| 131265.0938 | 127159.9922 | 127364.8438 | 0.012224472 | 0.883803356 | -0.1235142 | 0.276759353 | -0.151930701 | 0.205642234 |
| 1178393.25 | 1396262.125 | 1397482 | 0.012151633 | 0.959677376 | 0.036905361 | 0.821248921 | 0.061631523 | 0.737230437 |
| 18691.94727 | 18466.62305 | 17499.85352 | 0.011419406 | 0.95007546 | 0.114146963 | 0.484644469 | -0.156238392 | 0.387879439 |
| 101000.7578 | 96161.10938 | 72124.71875 | 0.01117441 | 0.975513759 | -0.441304886 | 0.306709414 | -0.230110623 | 0.531726747 |
| 40487.14453 | 38847.67969 | 40213.16016 | 0.011128143 | 0.826465565 | 0.397683373 | 0.021622074 | 0.448671856 | 0.009587646 |
| 23953.24023 | 24007.23047 | 18916.27148 | 0.011118223 | 0.926993948 | 0.072617615 | 0.275840689 | -0.965876737 | 0.020008995 |
| 31081.14941 | 32451.52832 | 23937.13135 | 0.010279844 | 0.95078707 | -0.006145036 | 0.976298184 | -0.049985913 | 0.835784721 |
| 50375.73633 | 61578.04297 | 52082.89063 | 0.009914893 | 0.978177397 | 0.561968307 | 0.551698166 | 0.073426967 | 0.820784824 |
| 55189.93359 | 52330.38281 | 51685.41016 | 0.009182398 | 0.957465821 | -0.193254901 | 0.133190834 | -0.071778332 | 0.431147819 |
| 79555.01563 | 78704.5 | 41572.43359 | 0.009000064 | 0.968709187 | -1.270143566 | 0.088077008 | -1.141640596 | 0.056570421 |
| 32172.26758 | 24649.22852 | 32622.20117 | 0.008914622 | 0.982392172 | 1.204129984 | 0.109509233 | 1.14827624 | 0.049082028 |
| 299574.9063 | 351880.6563 | 321645.9063 | 0.008913669 | 0.978177397 | -0.199849491 | 0.506362982 | -0.040172115 | 0.885245898 |
| 60164.98438 | 61758.05469 | 67234.70313 | 0.008906514 | 0.896724734 | 0.211878664 | 0.163729974 | 0.196138639 | 0.108589973 |
| 295768.5313 | 282975.375 | 265909.8125 | 0.008856978 | 0.896724734 | 0.018645049 | 0.721955757 | -0.235677598 | 0.054887084 |

| | | | | | | | | |
|---|---|---|---|---|---|---|---|---|
| 6087.479492 | 5813.705078 | 4905.366211 | 0.008837519 | 0.973802851 | 0.158153688 | 0.559278148 | -0.009692451 | 0.975601748 |
| 69796.10938 | 78948.375 | 64764.14844 | 0.008658644 | 0.811382893 | -0.012606764 | 0.858230595 | -0.19597028 | 0.216343174 |
| 31223.33398 | 31423.89063 | 22957.49414 | 0.008642305 | 0.965031901 | 0.230114765 | 0.266009189 | -0.144920726 | 0.56808558 |
| 858873 | 894082.125 | 800879.625 | 0.00837112 | 0.95078707 | -0.186690966 | 0.180354112 | -0.39487379 | 0.060352444 |
| 32142.47266 | 28882.31641 | 29694.0625 | 0.008339701 | 0.974559706 | -0.021616292 | 0.855764638 | 0.30645099 | 0.074131758 |
| 12580.19043 | 12678.96094 | 10587.06836 | 0.008316913 | 0.971328374 | 0.473649354 | 0.108625401 | 0.126884983 | 0.558686385 |
| 1549.12915 | 1459.615845 | 1158.438599 | 0.008176115 | 0.976463232 | -0.295671125 | 0.509478737 | -1.037942364 | 0.084288402 |
| 104017.5156 | 107740.5469 | 94405.5625 | 0.008121272 | 0.957465821 | 0.121914833 | 0.411843428 | 0.001102848 | 0.994798118 |
| 14623.16992 | 20666.91797 | 15051.20605 | 0.008056867 | 0.984192099 | 0.402914206 | 0.333066325 | 0.060409452 | 0.88793846 |
| 89281.92188 | 95935.3125 | 96217.11719 | 0.007884765 | 0.919806511 | -0.433900572 | 0.021979749 | -0.310659399 | 0.023920487 |
| 155272.4844 | 143082.5313 | 111848.25 | 0.006624378 | 0.944301598 | 0.106968098 | 0.114271926 | -0.202300157 | 0.361200142 |
| 912001.9375 | 980691.1875 | 1006898.563 | 0.006339106 | 0.980529569 | -0.050792447 | 0.751608736 | -0.034186161 | 0.83049333 |
| 110408.9922 | 105690.3203 | 107182.5938 | 0.006139019 | 0.966967584 | 0.229179678 | 0.102916055 | 0.232380159 | 0.081660057 |
| 38107.44531 | 39846.60547 | 38562.50781 | 0.006079124 | 0.975647436 | 0.320017321 | 0.058077004 | 0.561817695 | 0.026321276 |
| 64431.9375 | 45109.59766 | 39151.68359 | 0.005827336 | 0.96605672 | 0.374472329 | 0.100464534 | 0.016956968 | 0.96468232 |
| 131647.9531 | 124295.0781 | 123056.8828 | 0.005161042 | 0.986291242 | 0.062179848 | 0.844829988 | 0.064129749 | 0.833116553 |
| 12517.64844 | 11293.28613 | 12667.72949 | 0.005079966 | 0.987633122 | 0.675054596 | 0.076429114 | 0.063075592 | 0.807600061 |
| 160699.7813 | 155278.8594 | 130647.7266 | 0.004789325 | 0.968716865 | -0.534579116 | 0.083003488 | -0.588944882 | 0.04761948 |
| 68015.86719 | 75255.22656 | 75934.98438 | 0.004770621 | 0.959116844 | -0.477429274 | 0.024165741 | -0.31606324 | 0.041386295 |
| 15548.74609 | 15923.4375 | 13400.75391 | 0.004743693 | 0.93322496 | -0.237227027 | 0.157633835 | -0.417802161 | 0.056962705 |
| 1483200.125 | 1500555.031 | 1325609.734 | 0.004706186 | 0.963648889 | 0.067862705 | 0.11330346 | 0.274378008 | 0.101895754 |
| 7446.780273 | 8085.276367 | 6473.010254 | 0.004574846 | 0.98594118 | -0.03569244 | 0.908970621 | -0.162498783 | 0.57265858 |
| 7964.194336 | 6170.699707 | 6306.529297 | 0.004213572 | NA | -0.362494509 | NA | -0.763584624 | NA |
| 330213.5313 | 342254.2188 | 300388.5625 | 0.004083269 | 0.987549311 | 0.179166195 | 0.409739927 | 0.211857807 | 0.333735773 |
| 18804.67578 | NA | NA | 0.003995643 | 0.979955316 | -0.185359674 | 0.322217412 | -0.288119364 | NA |
| 167979.0781 | 157213.1094 | 145775.3281 | 0.003767913 | 0.984457944 | 0.597562624 | 0.082612243 | 0.828671821 | 0.027949491 |
| 7333.206543 | 4910.861816 | 6271.576172 | 0.003613553 | 0.970475218 | -0.173952145 | 0.153863035 | -0.292198398 | 0.279338111 |
| 64595.58203 | 64126.73438 | 63706.125 | 0.003464223 | 0.983042221 | 0.40756449 | 0.026250916 | 0.503975044 | 0.029219266 |
| 38473.12891 | 42893.47656 | 36588.78125 | 0.003419478 | 0.98305119 | 0.265487161 | 0.017291214 | 0.273429175 | 0.135230932 |
| 631406 | 642800.625 | 588612 | 0.002784837 | 0.978177397 | 0.451374216 | 0.046114261 | 0.091825306 | 0.244370877 |
| 16951.92773 | 12470.0459 | 12227.6123 | 0.002580335 | 0.971664759 | -0.025678707 | 0.645544015 | -0.182063164 | 0.457652244 |
| 116836.75 | 128690.0625 | 135850.7969 | 0.002280345 | 0.98725301 | 0.279710735 | 0.160011478 | 0.322842594 | 0.093865648 |
| 46163.22266 | 50845.08594 | 46271.11719 | 0.002188249 | 0.988200136 | 0.001251178 | 0.993263115 | -0.112160889 | 0.510020897 |

| | | | | | | | | |
|---|---|---|---|---|---|---|---|---|
| 18433.37695 | 20195.06445 | 18009.73828 | 0.002155645 | 0.98594118 | -0.183750809 | 0.14612569 | -0.018014411 | 0.864527019 |
| 16250.31348 | 15344.49707 | 8098.785645 | 0.001973056 | 0.988474238 | -0.698419987 | 0.042884533 | -0.539768521 | 0.231001576 |
| 217623.0625 | 249218.8281 | 245631.5 | 0.001954382 | 0.98305119 | 0.149799661 | 0.220633017 | 0.398701092 | 0.054887084 |
| 230314.1563 | 237066.6406 | 251295.5625 | 0.001876608 | 0.98305119 | 0.096752913 | 0.322217412 | 0.112707247 | 0.268450393 |
| 3756.224121 | 3605.591553 | 3278.955811 | 0.001829089 | 0.996623135 | -0.713264518 | 0.338306497 | -0.783636129 | 0.302062013 |
| 87973.00391 | 83795.23438 | 99548.27344 | 0.001820073 | 0.988300607 | -0.14193415 | 0.299034856 | 0.065015128 | 0.647661418 |
| 124796.9531 | 120620.9922 | 128428.2109 | 0.001738264 | 0.948396286 | 0.02488711 | 0.575414488 | -0.049921922 | 0.29648883 |
| 1830.57959 | 1545.995605 | 1653.138306 | 0.000837292 | 0.991744627 | -0.585912467 | 0.023592908 | -1.631313393 | 0.011038949 |
| 11914.79199 | 10571.55566 | 10164.10449 | 0.000790703 | 0.995888772 | 0.486985145 | 0.100528183 | 0.281110634 | 0.19500804 |
| 203741.6406 | 184117.0781 | 167270.7344 | 0.000585043 | 0.992559074 | 0.510807361 | 0.024656861 | 0.266137016 | 0.158426237 |
| 117856.7578 | 130601.9297 | 123567.5859 | -2.60E-05 | 0.999548057 | 0.149964477 | 0.122695009 | 0.052124448 | 0.504948403 |
| 156770.6875 | 174152.5469 | 170005.0156 | -0.000168544 | 0.997914794 | -0.598103159 | 0.033067345 | -0.359043079 | 0.042310808 |
| 358608.6875 | 348514.75 | 281975.1563 | -0.000633619 | 0.995013394 | 0.120857888 | 0.630875263 | -0.119052923 | 0.477266228 |
| 1903061 | 1822204.25 | 2149736.25 | -0.001282413 | 0.98808961 | -0.046394062 | 0.677878706 | 0.439810894 | 0.065059183 |
| 142744.4512 | 130572.1826 | 123071.0195 | -0.001299895 | 0.988344101 | -0.189389091 | 0.111298502 | -0.368629449 | 0.045177596 |
| 136672.6875 | 117079.125 | 152169.1406 | -0.001372324 | 0.994161663 | -0.014094172 | 0.919703825 | 0.290600154 | 0.210614344 |
| 428192.8438 | 462280.0625 | 425588.9063 | -0.001449059 | 0.98305119 | 0.417187985 | 0.125406681 | 0.315287386 | 0.039446558 |
| 17181.39063 | 14293.28516 | 15987.58984 | -0.001579916 | 0.988446298 | 0.250609928 | 0.137136478 | 0.210952374 | 0.223102126 |
| 14185.45996 | 12678.88965 | 17828.49219 | -0.00166229 | 0.990641317 | 0.27031759 | 0.642450324 | 1.680335199 | 0.071956774 |
| 62733.51563 | 79085.13281 | 77704.71094 | -0.00193076 | 0.997813908 | 0.764897515 | 0.32159658 | 0.446068687 | 0.546828747 |
| 63885.54297 | 64288.10938 | 68791.67188 | -0.002059957 | 0.98808961 | -0.071691386 | 0.177221267 | -0.113178436 | 0.129065826 |
| 64532.92188 | 58676.02344 | 63964.33984 | -0.002257586 | 0.98594118 | 0.058502313 | 0.537359718 | 0.013212389 | 0.895629548 |
| 229272.125 | 235723.1094 | 233574.5156 | -0.002273717 | 0.985420105 | -0.280204787 | 0.125736204 | -0.434414256 | 0.081899796 |
| 10955.61035 | 8475.839844 | 7090.732422 | -0.002339858 | 0.991744627 | -0.766735297 | 0.125467937 | -0.684629494 | 0.121525007 |
| 1355689.5 | 1247183.375 | 1135234.125 | -0.002398313 | 0.976647012 | 0.320037531 | 0.03968601 | 0.445987854 | 0.083898963 |
| 23516.57617 | 18305.72656 | 22615.64844 | -0.002525749 | 0.98725301 | 0.012710599 | 0.933325598 | -0.514457997 | 0.066846273 |
| 45884.25 | 49467.53516 | 43410.52734 | -0.002582809 | 0.973379537 | -0.392273953 | 0.044791528 | -0.304560476 | 0.04905244 |
| 689664.375 | 641174.9375 | 615907.9375 | -0.002620156 | 0.923927259 | -0.040726082 | 0.426604342 | 0.121990527 | 0.2258352 |
| 655154.3125 | 704029.8125 | 641729.75 | -0.002748244 | 0.991744627 | -0.255213602 | 0.31156031 | -0.101580047 | 0.613827778 |
| 174393.7813 | 170923.5625 | 181211.0313 | -0.003021492 | 0.978177397 | -0.410298128 | 0.065813387 | 0.016813824 | 0.811284277 |
| 5915.48291 | 7440.411621 | 5445.592285 | -0.003089292 | 0.991744627 | -0.872492962 | 0.193244598 | 0.395536491 | 0.30530221 |
| 76261.71875 | 77592.625 | 76921.51563 | -0.003137619 | 0.984457944 | 0.603314353 | 0.065871863 | 0.297084732 | 0.168518633 |
| 61535.14453 | 58038.81641 | 35337.96484 | -0.003223512 | 0.986190374 | -1.165226482 | 0.067039668 | -1.067771878 | 0.047847186 |

| | | | | | | | | |
|---|---|---|---|---|---|---|---|---|
| 34379.15234 | 36570.57422 | 31661.79688 | -0.003371619 | 0.966009107 | 0.397333286 | 0.046756616 | -0.239098712 | 0.09843293 |
| 193126.0625 | 187028.6563 | 154746.0313 | -0.003991282 | 0.966093377 | -0.30856434 | 0.062911841 | -0.264072054 | 0.151734273 |
| 11683.23145 | 8810.96582 | 11547.58496 | -0.004429787 | 0.976647012 | 0.163077711 | 0.348633551 | -0.306007334 | 0.217783381 |
| 10458.85059 | 12919.49902 | 8758.807617 | -0.004458372 | 0.973249983 | 0.02709884 | 0.887811168 | -0.173893472 | 0.504418582 |
| 195557.3906 | 211730.3438 | 199933.8906 | -0.004778582 | 0.921340182 | 0.101721044 | 0.228162961 | 0.268225622 | 0.045311345 |
| 114518.1953 | 98200.50781 | 11929.93359 | -0.004846754 | 0.948568397 | 0.323681435 | 0.074598722 | -0.457094339 | 0.573238987 |
| 37595.41797 | 39659.92969 | 42265.64063 | -0.005093353 | 0.950135609 | -0.023325891 | 0.766656897 | -0.247970497 | 0.063283918 |
| 7869.240234 | 8792.835938 | 7701.756348 | -0.005180419 | 0.967067233 | 0.135870828 | 0.483792142 | -0.025151866 | 0.798323297 |
| 925550.875 | 856645.625 | 872280.1875 | -0.005212471 | 0.89622256 | 0.002768259 | 0.97912062 | 0.063180142 | 0.327695945 |
| 69089.71094 | 84887.95313 | 78003.89844 | -0.005341431 | 0.981575164 | 0.525019034 | 0.223159833 | 0.532574129 | 0.079082931 |
| 548425.25 | 596959.25 | 543857.0625 | -0.005555982 | 0.976602645 | -0.107034148 | 0.591184368 | 0.078115163 | 0.67247313 |
| 313522.4375 | 349160.0938 | 309170.5938 | -0.00572998 | 0.951925361 | -0.037580286 | 0.719242791 | 0.13039299 | 0.293399859 |
| 149249.375 | 185573.5313 | 185347.0313 | -0.005846412 | 0.963965997 | 0.286002898 | 0.047069121 | 0.610612616 | 0.088642968 |
| 28203.32617 | 26757.74414 | 24252.19531 | -0.006128379 | 0.941581581 | -0.53619219 | 0.034038555 | -0.56240289 | 0.021984636 |
| 5227.834473 | 3909.545166 | 2115.699463 | -0.0063467 | 0.976647012 | -0.218948248 | 0.461561539 | -0.777697657 | 0.156588849 |
| 6214.916992 | 7928.833008 | 6225.038574 | -0.006872391 | 0.988200136 | 0.431663677 | 0.269046934 | 0.215157222 | 0.531405981 |
| 4601.853027 | 4954.291992 | 3647.825928 | -0.007154394 | 0.97318379 | -0.195814864 | 0.470177395 | -0.152703546 | 0.454130357 |
| 7068.80957 | 9937.526367 | 7434.277832 | -0.007253231 | 0.984281881 | 0.206162227 | 0.274179898 | 0.07868904 | 0.777319405 |
| 88007.88281 | 89641.72656 | 87027.51563 | -0.007985615 | 0.947063588 | 0.084381999 | 0.506109273 | -0.318543763 | 0.137551235 |
| 790153.3125 | 841844.875 | 834629.4375 | -0.008139579 | 0.921630415 | -0.190310364 | 0.054285958 | -0.314963175 | 0.01910141 |
| 33236.54688 | 33631.46875 | 37655.49219 | -0.008251579 | 0.957465821 | -0.473031405 | 0.098005404 | -0.249133856 | 0.177115155 |
| 58743.69531 | 59918.25781 | 56588.86328 | -0.008380522 | 0.935529534 | 0.25404611 | 0.26833397 | 0.386153407 | 0.019816721 |
| 26191.6543 | 22476.20313 | 24259.17383 | -0.008484946 | 0.962345406 | 0.311435524 | 0.04130763 | 0.204691509 | 0.15573211 |
| 7982.706055 | 10499.47559 | 7408.467285 | -0.008789515 | 0.957800688 | -0.440379631 | 0.101658109 | 0.72303603 | 0.129065826 |
| 2822.537598 | 3956.266113 | 1378.029785 | -0.008820952 | 0.990287588 | -0.639248346 | 0.512809933 | -0.977958897 | 0.385275383 |
| 146608.8125 | 138894.3125 | 123282.8828 | -0.009689451 | 0.850358984 | -0.266861772 | 0.07036809 | -0.373498779 | 0.06527068 |
| 3713.882324 | 4676.461426 | 1713.974243 | -0.009869133 | 0.963965997 | -0.829816561 | 0.04667516 | -1.337718283 | 0.055298525 |
| 2837.856689 | 3216.624756 | 2358.30957 | -0.010091936 | 0.954854544 | -1.056222565 | 0.1214746 | -0.371841893 | 0.142776128 |
| 71318.38281 | 64338.17969 | 63410.25 | -0.010153679 | 0.939127804 | -0.142704499 | 0.348220981 | -0.199274358 | 0.228116839 |
| 173835.0625 | 167366.0156 | 147098.6094 | -0.010334913 | 0.925934822 | 0.06337066 | 0.642705727 | 0.097094517 | 0.428577571 |
| 34748.11719 | 40043.86328 | 41177.86719 | -0.01049219 | 0.960189057 | 0.1189925 | 0.253952505 | 0.466867934 | 0.063474265 |
| 13054.06055 | 17325.30664 | 6082.054688 | -0.01093472 | 0.990070637 | -0.333465986 | 0.585220616 | -0.824090077 | 0.266313453 |
| 12933.97949 | 14985.21191 | 3219.048096 | -0.011408486 | 0.95078707 | 0.407589471 | 0.088901574 | -0.172254601 | 0.815420178 |

| | | | | | | | | |
|---|---|---|---|---|---|---|---|---|
| 37931.30078 | 37614.42188 | 39310.32813 | -0.011595518 | 0.922134001 | 0.213173794 | 0.152585278 | 0.396557907 | 0.062037824 |
| 12462.90039 | 12921.52539 | 9566.134766 | -0.011860583 | 0.970845797 | 0.253064085 | 0.26626937 | -0.375332501 | 0.210234556 |
| 42807.45313 | 43017.74609 | 39796.14063 | -0.012033061 | 0.891067226 | -0.108366411 | 0.338611479 | -0.129200525 | 0.266221642 |
| 141207.3906 | 221012.3594 | 139840.8438 | -0.012035469 | 0.978177397 | 0.500193814 | 0.382836467 | 0.545749742 | 0.310381232 |
| 1283.501709 | 1625.782959 | 1309.18335 | -0.012081753 | 0.989743807 | 3.355867299 | 0.082846844 | 0.567691335 | 0.231918617 |
| 273288.8438 | 282711.2188 | 281921.25 | -0.01211543 | 0.909145206 | -0.084268208 | 0.419290117 | -0.005045808 | 0.959233168 |
| 55415.38672 | 57547.07031 | 50853.03125 | -0.012705173 | 0.956352726 | -0.540109964 | 0.169417269 | -0.382887133 | 0.245828736 |
| 176936.8438 | 206429.4688 | 190600.4844 | -0.012964052 | 0.946061916 | 0.225142344 | 0.198627661 | 0.011822174 | 0.933069277 |
| 7206.041504 | 12717.52246 | 11032.7373 | -0.013513833 | 0.978177397 | 0.366403581 | 0.407540737 | 0.077448991 | 0.882379732 |
| 9340.167969 | 10940.70605 | 10146.75781 | -0.013582367 | 0.874877324 | -0.20125292 | 0.130503277 | -0.642747672 | 0.020008995 |
| 30758.42188 | 29859.24023 | 27631.82031 | -0.013600304 | 0.845267263 | 0.246236279 | 0.052305796 | 0.04885122 | 0.564166547 |
| 9337.569336 | 8863.87207 | 9189.769531 | -0.013644429 | 0.948982367 | 0.136780697 | 0.498876186 | -0.232284594 | 0.343177751 |
| 8702.000977 | 9155.699219 | 7895.765625 | -0.013895556 | 0.897542844 | 0.509544278 | 0.04667516 | 0.058487154 | 0.665761001 |
| 4013.315186 | 4348.821777 | 3005.23584 | -0.013942636 | 0.938919112 | -0.157794613 | 0.337374554 | -0.655052777 | 0.062822 |
| 487908.6563 | 518217.5 | 481783.8438 | -0.014208443 | 0.937209936 | 0.215058943 | 0.057524181 | 0.139616036 | 0.093865648 |
| 21687.8125 | 28853.84961 | 24550.95898 | -0.014382312 | 0.885903743 | 0.477521722 | 0.104714354 | 0.740486666 | 0.103879505 |
| 70561.64063 | 69510.09375 | 67046.42188 | -0.014423367 | 0.604716028 | 0.025139382 | 0.683335905 | -0.418703785 | 0.009587646 |
| 47003.48438 | 41049.18359 | 45461.15234 | -0.015045456 | 0.960071315 | 0.959062767 | 0.050472986 | 0.787337217 | 0.059770277 |
| 34498.64453 | 34174.94922 | 37775.4375 | -0.015211087 | 0.875733524 | 0.309647689 | 0.076955097 | 0.369710494 | 0.049415261 |
| 7225.812988 | 6776.355469 | 6508.564453 | -0.01547043 | 0.958413739 | 0.785009759 | 0.081503088 | 1.369969856 | 0.010623938 |
| 91117.36719 | 93166.95313 | 85880.27344 | -0.015540966 | 0.896329828 | -0.099531854 | 0.192597059 | -0.077013041 | 0.294614421 |
| 18921.40234 | 21290.49414 | 21954.67383 | -0.015542067 | 0.891343761 | -0.333882143 | 0.132066965 | 0.196920029 | 0.213371407 |
| 107498.0078 | 108926.6016 | 94348.21875 | -0.015673944 | 0.940597933 | -0.034550938 | 0.869626054 | 0.051546476 | 0.807833796 |
| 1545797.25 | 1724353.375 | 1735051.5 | -0.01572511 | 0.853712828 | -0.190548391 | 0.056581751 | -0.017818273 | 0.830763714 |
| 16892.05273 | 15710.76758 | 15500.03711 | -0.015778643 | 0.85372591 | -0.215658415 | 0.119439839 | 0.079534945 | 0.406520313 |
| 8375.679688 | 7530.399414 | 4188.713379 | -0.015903231 | 0.960071315 | -0.849887734 | 0.167655329 | -1.00728383 | 0.11327823 |
| 152916.2813 | 182467.3906 | 166789.75 | -0.016311356 | 0.954854544 | 0.724697746 | 0.028686404 | 1.540112189 | 0.025666912 |
| 155159.5469 | 154917.3281 | 162166.0469 | -0.01648076 | 0.692932191 | -0.707056484 | 0.01179292 | -0.081979397 | 0.120554018 |
| 84798.35938 | 85894.76563 | 84465.89063 | -0.016633112 | 0.870845851 | -0.10947399 | 0.338882942 | -0.128731872 | 0.281582889 |
| 14856.47754 | 15364.97852 | 14707.6582 | -0.016666164 | 0.953849986 | -0.068217242 | 0.8286291 | -0.19796398 | 0.511872946 |
| 29080.34766 | 30239.05859 | 26649.51758 | -0.016759389 | 0.924337975 | -0.325387503 | 0.108075904 | -0.509604014 | 0.05752489 |
| 262222.9063 | 247402.0781 | 255207.5313 | -0.016942856 | 0.889858508 | 0.289876818 | 0.115989509 | 0.279707625 | 0.026321276 |
| 33769.8125 | 31500.10547 | 38718.76953 | -0.017101956 | 0.90656976 | -0.690507683 | 0.039887225 | -0.032115957 | 0.820784824 |

| | | | | | | | | |
|---|---|---|---|---|---|---|---|---|
| 86229.88281 | 85639.72656 | 91236.55469 | -0.017136649 | 0.733572873 | -0.194590174 | 0.103787337 | -0.197927665 | 0.061706361 |
| 1081518.25 | 1129088.5 | 1120358.875 | -0.017331193 | 0.817346175 | 0.108495384 | 0.127892721 | -0.036276071 | 0.342470888 |
| 17575.63086 | 17459.00781 | 16028.35156 | -0.017381614 | 0.813966042 | -0.734579204 | 0.033801995 | 0.02907979 | 0.748277205 |
| 45792.93359 | 43527.30078 | 42526.16406 | -0.017747838 | 0.854954458 | -0.214488829 | 0.11629032 | 0.087880232 | 0.314191738 |
| 6069.843262 | 6550.82959 | 3257.581543 | -0.017848558 | 0.809876851 | -0.740330174 | 0.069810176 | -0.798758939 | 0.120615779 |
| 853560.0625 | 837691.3125 | 854764.0625 | -0.018074721 | 0.794535026 | -0.228525749 | 0.126707606 | 0.451308339 | 0.02527614 |
| 17627.20703 | 16748.08398 | 16553.13281 | -0.018442492 | 0.922134001 | -0.297736123 | 0.211473837 | -0.063480446 | 0.69422618 |
| 181310.9844 | 179130.75 | 176863.8594 | -0.018711363 | 0.855390604 | -0.071031618 | 0.629234482 | 0.150981652 | 0.106909019 |
| 381293.3125 | 370203.4375 | 358769.5 | -0.018747197 | 0.759757801 | -0.034966733 | 0.491839365 | -0.31737821 | 0.019880539 |
| 1832.617798 | 1958.064575 | 909.0074463 | -0.018767502 | 0.935335256 | -0.206161767 | 0.278713542 | -0.576067274 | 0.237224538 |
| 114466.5938 | 107410.6797 | 103336.5 | -0.01922981 | 0.94210358 | -0.47148443 | 0.192808829 | -0.000503207 | 0.998369251 |
| 407996.625 | 390913.5313 | 375487.75 | -0.019488215 | 0.697888216 | 0.056716544 | 0.137700061 | -0.220137273 | 0.044994481 |
| 11170.02832 | 12392.05078 | 10448.68652 | -0.019547157 | 0.875891098 | -0.181925455 | 0.530684856 | 1.088810134 | 0.031136253 |
| 247134.3906 | 276865.9063 | 289965.5625 | -0.01964732 | 0.865799459 | -0.300752378 | 0.150108069 | -0.378447893 | 0.087771237 |
| 7410.646973 | 9526.077148 | 7068.275391 | -0.01977815 | 0.960102505 | 0.601646358 | 0.196535919 | 0.490999093 | 0.244446425 |
| 45543.91016 | 34909.18359 | 44154.77734 | -0.019838759 | 0.89487344 | 0.072702647 | 0.657948979 | -0.131006847 | 0.520924639 |
| 121350.1641 | 128332.3516 | 114628.1094 | -0.01991044 | 0.773960816 | -0.096955816 | 0.15765519 | 0.048334652 | 0.529505889 |
| 9113.797852 | 8908.163086 | 5313.01416 | -0.020820725 | 0.874117088 | -0.806130368 | 0.039779424 | -0.820048844 | 0.073672213 |
| 145172.1875 | 148274.2344 | 139063.4844 | -0.02112391 | 0.813574836 | 0.77818098 | 0.029439089 | 0.516208061 | 0.016913656 |
| 10102.33691 | 9481.607422 | 7728.560059 | -0.02169623 | 0.857904207 | -0.105321421 | 0.556770167 | -0.182809862 | 0.332189784 |
| 13235.74805 | 13087.64258 | 13977.54883 | -0.021801828 | 0.835128657 | -0.344857366 | 0.210578593 | -0.289087105 | 0.086503564 |
| 4227.921387 | 3761.100342 | 3218.98999 | -0.021922786 | 0.889892479 | -0.590242829 | 0.109199532 | -0.667259012 | 0.064582407 |
| 271088.3438 | 308062.3125 | 333806.75 | -0.022019921 | 0.877108871 | 0.15918887 | 0.344936914 | -0.021398934 | 0.91311394 |
| 7604.433594 | 9405.55957 | 13413.45313 | -0.022406798 | 0.954854544 | 0.847557062 | 0.060391495 | 2.026046347 | 0.107956786 |
| 26188.02539 | 26432.50195 | 27934.16211 | -0.022546209 | 0.975647436 | 0.610655611 | 0.360346489 | 0.305197183 | 0.6366273 |
| 113569.2891 | 101477.7578 | 106175.1484 | -0.022854832 | 0.884594659 | 0.581708714 | 0.056194202 | -0.182262535 | 0.299059995 |
| 91363.26563 | 120851.2891 | 130636.125 | -0.023012242 | 0.856898147 | 0.029391459 | 0.82401933 | 0.389047953 | 0.230299136 |
| 11244.90332 | 8562.730469 | 10476.37793 | -0.023403609 | 0.851662636 | -0.69194717 | 0.052305796 | -0.805842048 | 0.026321276 |
| 4218.67334 | 5984.263672 | 7707.377441 | -0.023737105 | 0.960072248 | -0.849606774 | 0.156712278 | -0.334328225 | 0.4589703 |
| 40007.86719 | 39899.6875 | 34149.25 | -0.023814772 | 0.8285165 | 0.081251044 | 0.352029077 | -0.279090282 | 0.09057246 |
| 1382.386353 | 2249.918213 | 2426.812988 | -0.024449127 | 0.97318379 | 0.887040343 | 0.196976249 | 0.788707396 | 0.26011348 |
| 672816.6875 | 708729.625 | 739998.8125 | -0.025031771 | 0.589921226 | -0.233561658 | 0.045660829 | -0.159698645 | 0.090232821 |
| 70856.89844 | 69516.69531 | 64737.83984 | -0.025364677 | 0.633668112 | -0.097402876 | 0.204546842 | -0.089333425 | 0.235430997 |

| | | | | | | | | |
|---|---|---|---|---|---|---|---|---|
| 25367.59961 | 20395.15625 | 21347.79102 | -0.025746551 | 0.921903697 | -0.623335547 | 0.15358024 | -0.064931076 | 0.812775264 |
| 122750.7109 | 153158.2188 | 180659.7656 | -0.025874916 | 0.831656064 | 0.059574763 | 0.630875263 | 0.266620837 | 0.370623248 |
| 88970.6875 | 90487.69531 | 82680.25781 | -0.026011208 | 0.834514511 | -0.314461858 | 0.144097346 | -0.048966968 | 0.723564444 |
| 4686.281738 | 4361.601074 | 4002.627441 | -0.026044715 | 0.82764206 | -0.592841983 | 0.053185315 | -0.065499676 | 0.60672012 |
| 22001.23828 | 20722.90625 | 22642.38281 | -0.026430454 | 0.940664982 | -0.319101571 | 0.414031192 | -0.092310888 | 0.772564043 |
| 17014.41797 | 19200.30078 | 16531.82227 | -0.026443966 | 0.860585436 | 0.014926907 | 0.929570723 | -0.266207059 | 0.224000425 |
| 16445.92188 | 15558.8252 | 14289.64941 | -0.026556935 | 0.866568928 | 0.582490826 | 0.021969813 | 0.326410336 | 0.068496057 |
| 827722.3359 | 905293.8906 | 872891.1758 | -0.026783376 | 0.819940924 | -0.239584575 | 0.088308366 | -0.284422968 | 0.041771828 |
| 24972.30273 | 22766.3125 | 29118.85156 | -0.027159198 | 0.853986673 | 0.665548286 | 0.015625705 | 0.43173109 | 0.128365896 |
| 761566.1875 | 927030.5625 | 709986.625 | -0.027277535 | 0.95462945 | 0.196287563 | 0.519529115 | -0.030546899 | 0.929695889 |
| 18299.63086 | 20156.94141 | NA | -0.027530424 | 0.965156909 | -0.045463552 | 0.941673059 | 0.03826792 | NA |
| 121086.0078 | 128046.7813 | 120567.4922 | -0.027762408 | 0.621067403 | -0.020073238 | 0.613687235 | -0.049837144 | 0.331161608 |
| 7609.012695 | 7245.554199 | 5670.348145 | -0.028406501 | 0.762776035 | -0.253461078 | 0.046114261 | -0.985920068 | 0.027510643 |
| 87010.89844 | 89201.17188 | 90501.36719 | -0.028736912 | 0.935335256 | 0.467012414 | 0.221069387 | 0.505788659 | 0.187797081 |
| 99275.55469 | 105993.3125 | 99357.02344 | -0.028998353 | 0.945598346 | 0.556738115 | 0.229062299 | 0.704132237 | 0.150025452 |
| 503206.4688 | 564271.75 | 586497 | -0.029160507 | 0.762776035 | -0.118737351 | 0.32242293 | -0.169032504 | 0.221789362 |
| 4611.856934 | 4273.150391 | 4799.682617 | -0.029191961 | 0.891343761 | 0.498799361 | 0.100528183 | 0.33771379 | 0.071151453 |
| 100250.9375 | 93624.32813 | 82938.36719 | -0.02922735 | 0.925435999 | 1.000451481 | 0.109509233 | 1.013064883 | 0.039446558 |
| 554282.375 | 569179.375 | 672234.9375 | -0.029310154 | 0.86018616 | 0.287895231 | 0.185863614 | 1.018228996 | 0.034045682 |
| 44779.42969 | 45323.99219 | 42012.58984 | -0.029314884 | 0.849597117 | -0.109491442 | 0.162505567 | -0.226031112 | 0.047526836 |
| 6759.091797 | 9191.730469 | 13025.90723 | -0.029597188 | 0.948108125 | -0.32174266 | 0.535797708 | 0.035464865 | 0.948114044 |
| 77100.25781 | 79797.02344 | 75839.66406 | -0.03075022 | 0.783327271 | -0.407556618 | 0.049358751 | -0.58324455 | 0.020045376 |
| 2085722.5 | 1876236.375 | 1850427.625 | -0.03176204 | 0.707693403 | -0.159658537 | 0.127187484 | -0.401502491 | 0.032840382 |
| 39557.36328 | 40814.33203 | 41432.80859 | -0.031947862 | 0.965156909 | 2.302914205 | 0.023068087 | 2.074742644 | 0.040202519 |
| 144351.0313 | 137953.6094 | 125297.6328 | -0.032102701 | 0.817518336 | -0.084730434 | 0.600800179 | -0.423634777 | 0.111646722 |
| 32898.8125 | 30176.21094 | 27689.68359 | -0.032197629 | 0.639820022 | -0.212066333 | 0.193488746 | -0.399709795 | 0.069586726 |
| 109628.0547 | 91569.64844 | 97732.14063 | -0.03249054 | 0.626970721 | -0.10730254 | 0.554396013 | -0.037543566 | 0.767517528 |
| 13338.34766 | 13452.57422 | 12322.22168 | -0.032696617 | 0.72050444 | 0.21530166 | 0.157674375 | -0.007747881 | 0.947307151 |
| 453287.5 | 446828.5313 | 435364.4375 | -0.033436576 | 0.534090887 | -0.06532744 | 0.15090616 | 0.065507829 | 0.12131068 |
| 18590.4043 | 16995.59766 | 14360.79492 | -0.033485854 | 0.801548444 | 0.049933872 | 0.705308598 | 0.111112227 | 0.56808558 |
| 37876.90625 | 39663.09766 | 35449.66406 | -0.033590234 | 0.749455721 | 0.432313732 | 0.05253541 | 0.114858414 | 0.360590046 |
| 42769.95703 | 48141.73047 | 34905.32031 | -0.033808497 | 0.848955033 | 0.298409088 | 0.182826975 | 0.148592655 | 0.522523924 |
| 3955.209961 | 1916.391113 | 2287.076416 | -0.033890777 | 0.95078707 | 0.13657175 | 0.8517042 | -0.588129965 | 0.243867938 |

| | | | | | | | | |
|---|---|---|---|---|---|---|---|---|
| 62180.23047 | 54188.95313 | 57928.89063 | -0.033975701 | 0.761377655 | -0.227697042 | 0.05818214 | -0.623639128 | 0.017015337 |
| 1814902.68 | 1980289.828 | 1832359.977 | -0.034389938 | 0.705450603 | -0.226526864 | 0.04183385 | -0.130478165 | 0.126779284 |
| 12529.23828 | 9167.405273 | 11169.5459 | -0.034784554 | 0.924337975 | 0.752045404 | 0.039922031 | 0.928602131 | 0.085058449 |
| 85648.60938 | 86473.35938 | 90158.92188 | -0.034977293 | 0.427440307 | -0.218599058 | 0.037014778 | -0.201779959 | 0.03286768 |
| 44991.70313 | 55217.43359 | 48940.40625 | -0.035000602 | 0.714199995 | 0.291791913 | 0.148709893 | -0.164545016 | 0.279512653 |
| 23375.33594 | 27798.12305 | 31302.94922 | -0.035400545 | 0.951248394 | 0.035478167 | 0.941883073 | 0.360242599 | 0.436830073 |
| 500458.4688 | 439337.125 | 488058.8125 | -0.035408344 | 0.769706422 | 0.000574975 | 0.99201113 | 0.254595528 | 0.095208361 |
| 818683.125 | 898714.9375 | 901435.25 | -0.035411372 | 0.607751104 | 0.016782016 | 0.831472283 | 0.081517278 | 0.376260129 |
| 12340.21777 | 12993.14746 | 12553.47656 | -0.035454113 | 0.870845851 | 0.030845234 | 0.908970621 | 0.382791536 | 0.177455256 |
| 33149.19141 | 32739.17773 | 32259.91992 | -0.035824786 | 0.386648818 | -0.37039856 | 0.039582042 | 0.139313959 | 0.020008995 |
| 175181.9531 | 182019.3594 | 191118.6719 | -0.035962384 | 0.42324628 | -0.491469149 | 0.013885891 | -0.52350374 | 0.011546635 |
| 11509.36328 | 14768.9043 | 9303.831055 | -0.036165428 | 0.906235705 | 0.097823659 | 0.732837711 | -0.561577771 | 0.138038355 |
| 20962.49609 | 21282.54688 | 20083.10352 | -0.036198549 | 0.854775418 | 0.04987576 | 0.703855443 | -0.117325747 | 0.327637601 |
| 64431.29297 | 84054.03906 | 70244.51563 | -0.036402513 | 0.80053545 | 0.160171203 | 0.33801894 | -0.078179721 | 0.713487699 |
| 73099.53906 | 72292.47656 | 66274.60156 | -0.036602667 | 0.486426842 | 0.109612426 | 0.415960806 | -0.335720236 | 0.02938023 |
| 1999165.875 | 2387958.25 | 2160801.25 | -0.036862667 | 0.642928108 | 0.329274657 | 0.059562109 | 0.372215941 | 0.085082134 |
| 224434.9531 | 269105.9375 | 239237.9219 | -0.036960516 | 0.835128657 | -0.23545181 | 0.087367756 | -0.288992389 | 0.092300731 |
| 287435.625 | 321084.625 | 308602.2813 | -0.037012612 | 0.707693403 | -0.324751401 | 0.118137091 | -0.044223239 | 0.707910779 |
| 17743.61328 | 20737.70703 | 15811.94434 | -0.037769921 | 0.808150017 | -0.181697187 | 0.348524737 | -0.408396167 | 0.118298395 |
| 260521.6777 | 282470.0923 | 243992.2498 | -0.038023411 | 0.466946593 | -0.118549759 | 0.14612569 | -0.219582215 | 0.102257588 |
| 85773.64844 | 90497.65625 | 94259.00781 | -0.038637428 | 0.845976666 | 0.375985448 | 0.132680703 | 0.025688457 | 0.890901767 |
| 4805.731934 | 4679.555176 | 3857.804199 | -0.038839402 | 0.74476981 | -0.707562243 | 0.06015933 | -0.438115323 | 0.069298557 |
| 4624.567871 | 5234.316895 | 4738.936035 | -0.039031899 | 0.965031901 | 1.418000208 | 0.105604495 | 0.348148919 | 0.665559747 |
| 19939.27734 | 18890.86719 | 18229.14453 | -0.039085631 | 0.415082806 | 0.043697783 | 0.432194541 | -0.034515766 | 0.608146408 |
| 259994.3281 | 277084.0625 | 248387.1719 | -0.039441651 | 0.535711942 | -0.159652272 | 0.129000147 | -0.161399215 | 0.1278105 |
| 140733.6563 | 128213.3125 | 81253.96094 | -0.039694591 | 0.764304977 | -0.621561341 | 0.015993065 | -0.685348023 | 0.126366029 |
| 211556.1875 | 217311.2969 | 230468.1563 | -0.03977626 | 0.654184037 | -0.429520478 | 0.017291214 | -0.334523714 | 0.023137414 |
| 1837.092773 | 3581.154297 | 2535.399902 | -0.04004739 | 0.930620943 | 0.253196245 | 0.410684021 | -0.292390548 | 0.526000405 |
| 60011.75391 | 54258.99609 | 54738.37109 | -0.040707333 | 0.719961518 | -0.441646838 | 0.070010854 | -0.520399019 | 0.055868946 |
| 12820.85156 | 12070.56836 | 13689.19824 | -0.040902283 | 0.766981151 | 0.619398144 | 0.036826559 | -0.103989597 | 0.473264096 |
| 136384.9531 | 145153.6406 | 141448.5469 | -0.040945981 | 0.839932706 | 0.084934226 | 0.112181717 | -0.016755487 | 0.728308808 |
| 53707.125 | 54311.33203 | 55552.29688 | -0.041156804 | 0.683040212 | 0.436974868 | 0.060391495 | 0.233898349 | 0.020008995 |
| 42385.96484 | 42248.75781 | 37064.125 | -0.041193354 | 0.70461778 | -0.360835207 | 0.068812228 | -0.060807799 | 0.604461912 |

| | | | | | | | | |
|---|---|---|---|---|---|---|---|---|
| 145609.9375 | 146170.7031 | 140828.0313 | -0.04131855 | 0.696706503 | 0.073044691 | 0.455342372 | 0.23640862 | 0.091109394 |
| 1351149.125 | 1734107.125 | 1655722.75 | -0.041948849 | 0.827859454 | -0.337151201 | 0.230523816 | -0.040231421 | 0.861875296 |
| 424719.625 | 655050 | 447033.4063 | -0.04202188 | 0.957975824 | -0.864852743 | 0.467532121 | -1.031333759 | 0.408044366 |
| 27786.31836 | 28246.94336 | 29606.02539 | -0.042158357 | 0.935975094 | -0.41776945 | 0.423207125 | -0.721100041 | 0.263528267 |
| 117266.4453 | 105408.2188 | 111855.8906 | -0.042260358 | 0.39188184 | -0.100276016 | 0.128915752 | 0.270565797 | 0.06664224 |
| 84002.95313 | 86045.39844 | 67705.75 | -0.042834648 | 0.624643671 | 0.118153197 | 0.218601949 | 0.166931447 | 0.382150932 |
| 684919.5 | 783312.8125 | 773842.4375 | -0.043179784 | 0.767820616 | -0.084084315 | 0.634960279 | -0.09411827 | 0.586390561 |
| 7089.258789 | 7371.943848 | 6808.255859 | -0.043484705 | 0.643493907 | -0.136670749 | 0.25313315 | -0.612291052 | 0.029820656 |
| 210335.9688 | 174766.8906 | 139981.4844 | -0.043528047 | 0.790845624 | -0.241762292 | 0.191836121 | -0.200494833 | 0.441242238 |
| 36944.98828 | 50794.88281 | 44946.44922 | -0.043818817 | 0.856398887 | -0.128819634 | 0.625385428 | -0.247136704 | 0.423557711 |
| 87487.21875 | 87474.50781 | 92018.07031 | -0.043912897 | 0.713238474 | -0.063614338 | 0.341543595 | -0.188876393 | 0.043108871 |
| 51021.44141 | 52050.44141 | 46556.07813 | -0.04391925 | 0.430563789 | -0.114635926 | 0.111349456 | -0.118803404 | 0.19853994 |
| 11117.9043 | 6539.447754 | NA | -0.044017172 | NA | -0.088600581 | NA | 0.263693455 | NA |
| 37005.71094 | 35431.34766 | 27167.01367 | -0.04411581 | 0.870421302 | -0.072966465 | 0.834946164 | -0.155245988 | 0.575112472 |
| 6403.208008 | 7005.093262 | 7882.535156 | -0.044392662 | 0.838790359 | -0.079379842 | 0.638775352 | -0.046818862 | 0.797672854 |
| 289251.25 | 423582.3438 | 288742.1875 | -0.044911097 | 0.919806511 | -0.003309741 | 0.992236761 | -0.082925468 | 0.816088807 |
| 190365.8281 | 181034.9688 | 176822.25 | -0.045317821 | 0.584096954 | -0.220909335 | 0.106309393 | -0.107655039 | 0.263142696 |
| 287485.3125 | 291216.4375 | 278526.1563 | -0.0455286 | 0.515727508 | 0.110061826 | 0.313982423 | 0.071822403 | 0.118033234 |
| 864279.0625 | 890753.875 | 729884.125 | -0.045619681 | 0.686476859 | -0.020628927 | 0.871137795 | -0.031041209 | 0.81963682 |
| 147286.0156 | 144485.0781 | 125505.2266 | -0.046720436 | 0.77354208 | -0.591575328 | 0.093078364 | -0.9294925 | 0.057347867 |
| 151715.4063 | 149042.2031 | 143971.6094 | -0.046848425 | 0.6412557 | -0.577762631 | 0.059562109 | -0.500186544 | 0.071353321 |
| 103735.6875 | 111827.4531 | 121194.6641 | -0.047069538 | 0.878992455 | 0.109207627 | 0.717241613 | 0.081345046 | 0.790960569 |
| 485538.6472 | 716944.8835 | 521603.6514 | -0.047150006 | 0.942308793 | 0.170806968 | 0.838031445 | 0.363452003 | 0.516064937 |
| 145752.8438 | 159157.8594 | 148181.7656 | -0.047324458 | 0.871817708 | 0.210726866 | 0.498192509 | 0.186490481 | 0.535773657 |
| 216254.4844 | 217800.0938 | 228865.2344 | -0.047387817 | 0.544660123 | 0.092904017 | 0.242122295 | 0.108643913 | 0.172011685 |
| 188361.8594 | 180859.1094 | 194324.8125 | -0.047470451 | 0.454648558 | 0.058855291 | 0.307456205 | 0.042105572 | 0.461115334 |
| 48084.25 | 52771.25 | 50032.88281 | -0.047943733 | 0.704147678 | 0.029292337 | 0.816502262 | -0.101041048 | 0.448929881 |
| 1401828.625 | 1952198.75 | 1865960.125 | -0.048393824 | 0.790141348 | -0.274220423 | 0.05087265 | -0.207157014 | 0.362743641 |
| 23222.53516 | 3892.656738 | 9412.087891 | -0.048497935 | 0.707088135 | 0.189054527 | 0.506109273 | -0.950266986 | 0.28527444 |
| 24176.91016 | 26131.86133 | 27272.98242 | -0.048567725 | 0.696577081 | -0.104527807 | 0.362858173 | -0.086162459 | 0.458699163 |
| 6043.092285 | 6829.008789 | 3469.948975 | -0.048606797 | 0.766771986 | 0.298139298 | 0.106271335 | -0.89547535 | 0.097083282 |
| 12916.75391 | 11303.77441 | 8873.133789 | -0.048789387 | 0.630910713 | -0.493631793 | 0.057239807 | -0.407607806 | 0.14724171 |
| 28989.59375 | 30465.88672 | 14467.73633 | -0.049047504 | 0.700814002 | -0.564803984 | 0.056128038 | -0.604371311 | 0.207777277 |

| | | | | | | | | |
|---|---|---|---|---|---|---|---|---|
| 431012.2813 | 507420.1563 | 493412.0938 | -0.049086007 | 0.694887813 | -0.056511967 | 0.494902041 | 0.237808662 | 0.146578308 |
| 8365.759766 | 8322.703125 | 7686.106934 | -0.049144718 | 0.881977802 | -0.102620051 | 0.52210667 | -0.043288354 | 0.759627144 |
| 636323.5625 | 918137.3125 | 988564.75 | -0.049275812 | 0.621182888 | 0.38984834 | 0.06015933 | 0.693000617 | 0.170158914 |
| 911288 | 1039113.25 | 968717.6875 | -0.049363459 | 0.685054567 | -0.228171094 | 0.096721331 | -0.295690514 | 0.062345205 |
| 9070.664063 | 9915.664063 | 9185.144531 | -0.049442079 | 0.825747772 | 0.020694677 | 0.861195218 | 0.073344356 | 0.330301431 |
| 262900.6563 | 269539.4688 | 271528.8438 | -0.049472999 | 0.492900774 | 0.147457377 | 0.093880063 | 0.512327905 | 0.015712916 |
| 94365.1875 | 100405.1953 | 101206.0078 | -0.050330763 | 0.630412128 | -0.107844653 | 0.357606274 | -0.255373558 | 0.117544052 |
| 53799.58984 | 57176.50781 | 54793.95703 | -0.050413956 | 0.690885405 | 0.107041716 | 0.508503755 | -0.117815876 | 0.438195855 |
| 2474.491455 | 5055.152344 | 1331.357544 | -0.050618174 | 0.921575258 | 0.147740702 | 0.703047486 | -0.737381962 | 0.348871035 |
| 23762.02344 | 23884.60352 | 25042.78711 | -0.051127041 | 0.77354208 | 0.081336598 | 0.626219461 | 0.333450633 | 0.136474059 |
| 54338.49219 | 58100.96875 | 59449.71094 | -0.051215814 | 0.777513485 | -0.332371513 | 0.134444814 | 0.088731331 | 0.429144799 |
| 3164054.5 | 4379210.5 | 4483530 | -0.051347735 | 0.748625706 | 0.275187541 | 0.25469106 | 1.229121318 | 0.081631245 |
| 5422.692383 | 15929.69824 | 3532.610107 | -0.05143209 | 0.968762606 | -0.783375301 | 0.562301807 | -0.150890331 | 0.904536989 |
| 33381.31641 | 35393.60547 | 36803.29688 | -0.051501358 | 0.793038329 | 0.284099055 | 0.355609923 | 0.8769688 | 0.012916212 |
| 64665.96875 | 67738.36719 | 67621.36719 | -0.051677446 | 0.634019081 | -0.09921182 | 0.47004298 | 0.318209554 | 0.077431074 |
| 51476.30078 | 48070.69141 | 47291.56641 | -0.051705404 | 0.589921226 | -0.007680297 | 0.955379211 | 0.391263019 | 0.03405215 |
| 73139.89844 | 68418.20313 | 80877.25781 | -0.051769649 | 0.762776035 | 0.686066665 | 0.032675377 | 0.952680336 | 0.025700816 |
| 113339.0469 | 119364.4063 | 110404.6797 | -0.051812495 | 0.613811441 | 0.203184612 | 0.019456985 | 0.218507674 | 0.062384795 |
| 4204.625488 | 3787.364014 | 1632.728271 | -0.052052282 | 0.863936575 | -0.857686507 | 0.129149835 | -1.435439489 | 0.042527001 |
| 1341613.625 | 1457143.625 | 1389244.375 | -0.052089408 | 0.374464563 | 0.155826279 | 0.044791528 | 0.023106946 | 0.672587482 |
| 21162.66406 | 20854.89844 | 17093.35938 | -0.052089726 | 0.76356807 | -0.756250586 | 0.05467585 | -1.088529584 | 0.029324108 |
| 21520.59766 | 15612.76074 | 16222.79492 | -0.052189616 | 0.883803356 | 0.569881132 | 0.160918214 | 0.326253042 | 0.385275383 |
| 186330.2188 | 180741.6563 | 171097.8594 | -0.052362664 | 0.701490098 | -0.071466715 | 0.423723931 | -0.240525656 | 0.039552949 |
| 26918.71875 | 32133.12695 | 23175.91602 | -0.052432634 | 0.87995149 | 0.211901725 | 0.493747329 | 0.431327854 | 0.226707644 |
| 325.0366516 | 192.2979889 | 232.6407166 | -0.052674657 | 0.909145206 | 0.312011021 | 0.600800179 | -0.971884454 | 0.049415261 |
| 9399.131836 | 8383.060547 | 8397.735352 | -0.053365861 | 0.64826161 | -0.425853538 | 0.109341875 | -0.312892786 | 0.051088524 |
| 13693.06543 | 13850.66602 | 12249.96973 | -0.053484668 | 0.807389856 | 0.196165406 | 0.276405455 | -0.173696257 | 0.350351437 |
| 18008.69336 | 20692.01953 | 19477.36914 | -0.053622733 | 0.81411754 | 0.149139894 | 0.512809933 | 0.234598355 | 0.215498909 |
| 16125.02148 | 10060.24316 | 8016.574219 | -0.053696162 | 0.762776035 | 0.144861423 | 0.37870309 | -0.144446551 | 0.755289431 |
| 143653.1563 | 155745.2969 | 143834.5 | -0.053990303 | 0.513455136 | -0.367107056 | 0.040066052 | -0.030556386 | 0.690662917 |
| 63755.14453 | 61893.31641 | 55134.26953 | -0.054010185 | 0.784001384 | 0.290447939 | 0.089889925 | -0.076477335 | 0.544387447 |
| 146034.2813 | 144603.6719 | 149056.5469 | -0.054061436 | 0.418600089 | 0.2881789 | 0.057189248 | 0.317421561 | 0.024517825 |
| 9343.413086 | 9138.695313 | 7587.748535 | -0.054159286 | 0.276360085 | 0.015820931 | 0.842124822 | -0.390508294 | 0.080600351 |

| | | | | | | | | |
|---|---|---|---|---|---|---|---|---|
| 8615.739258 | 9024.167969 | 8095.315918 | -0.054195935 | 0.680230584 | 0.39210412 | 0.047212306 | 0.363922738 | 0.044411216 |
| 64341.75391 | 68024.89844 | 53783.63281 | -0.0544971 | 0.626857395 | -0.141790904 | 0.32242293 | -0.328127306 | 0.121673185 |
| 185421.5781 | 188423.3906 | 183494.3281 | -0.05460301 | 0.288813407 | 0.321637935 | 0.084811505 | 0.565932687 | 0.006394096 |
| 41380.93359 | 42933.14453 | 40307.17969 | -0.054809135 | 0.721803576 | 0.480523081 | 0.038037652 | 0.127966654 | 0.277205993 |
| 23299.22656 | 25371.01563 | 23958.95117 | -0.054981974 | 0.719961518 | -0.109096367 | 0.099830362 | -0.636603661 | 0.009587646 |
| 17840.86719 | 20104.00195 | 21122.89063 | -0.055125347 | 0.775671966 | -0.083926139 | 0.395934625 | -0.639052995 | 0.031136253 |
| 249746.4531 | 254473.0156 | 223358.625 | -0.055160212 | 0.513194681 | 0.10415948 | 0.147710173 | -0.123192487 | 0.240299186 |
| 36251.13281 | 25986.37109 | 40121.94922 | -0.055241544 | 0.821952288 | 0.863897904 | 0.255621964 | 0.447301387 | 0.24971241 |
| 130168.2891 | 130893.6328 | 118024.6953 | -0.055417833 | 0.486072526 | -0.183263521 | 0.19372563 | -0.123263418 | 0.180958636 |
| 139102.5938 | 157418.9219 | 109482.3281 | -0.055751361 | 0.747160698 | -0.565723067 | 0.08472494 | -0.43830845 | 0.136000191 |
| 1494354.25 | 1773384.75 | 1692475.375 | -0.056110155 | 0.718505305 | 0.091326655 | 0.559568799 | -0.098059815 | 0.578032057 |
| 80937.95313 | 80914.33594 | 82929.51563 | -0.056193689 | 0.701595298 | -0.614540784 | 0.094097366 | -0.48685739 | 0.122834452 |
| 82571.28906 | 85304.60938 | 84864.8125 | -0.05674106 | 0.6244989 | 0.217980804 | 0.297032792 | 0.552346722 | 0.037883801 |
| 127899.7969 | 179195.6875 | 139923.3281 | -0.056742845 | 0.825031307 | -0.140701246 | 0.531076327 | -0.027577805 | 0.912637994 |
| 48545.61719 | 36253.32422 | 41407.25 | -0.057012312 | 0.687712573 | 0.101596775 | 0.540404601 | -0.014796283 | 0.950211597 |
| 282976.9063 | 277084.625 | 286049.25 | -0.057195686 | 0.444228235 | 0.171772585 | 0.183896983 | 0.070807061 | 0.065367829 |
| 96988.30469 | 89733.875 | 96432.20313 | -0.057242677 | 0.70596069 | 0.304108075 | 0.088915315 | 0.59731751 | 0.022968762 |
| 17683.24414 | 15451.99805 | 11278.52539 | -0.057428387 | 0.714543616 | 0.431869554 | 0.090723649 | 0.005370832 | 0.988179652 |
| 4921449 | 5164446.5 | 5144342 | -0.057571408 | 0.419590652 | -0.228660676 | 0.053580667 | 0.165660118 | 0.055516192 |
| 73749.59375 | 78860.72656 | 75838.94531 | -0.058116298 | 0.611465565 | -0.40146783 | 0.102709172 | -0.289619945 | 0.112298345 |
| 72606.55469 | 68353.48438 | 68157.1875 | -0.05861581 | 0.166325586 | 0.010634465 | 0.89718243 | 0.070282241 | 0.238701326 |
| 20250.62891 | 20388.18164 | 16238.80078 | -0.058693868 | 0.87137302 | 0.78532963 | 0.089431851 | 0.755581244 | 0.080292195 |
| 5697.678223 | 4507.270508 | 4515.693359 | -0.059045026 | 0.776365661 | -0.414456155 | 0.137282102 | -0.735808328 | 0.053053903 |
| 37085.99609 | 41844.44922 | 45256.14063 | -0.059106451 | 0.845668534 | -0.022570529 | 0.919069808 | 0.369548029 | 0.146345197 |
| 153366.8223 | 155477.3047 | 148310.2402 | -0.059594817 | 0.512681345 | -1.132898429 | 0.037760167 | -0.351623262 | 0.075542979 |
| 849333.1875 | 847955.1875 | 838900.375 | -0.059659456 | 0.698929048 | 0.733491489 | 0.070731667 | 0.828240901 | 0.029120377 |
| 290582.2813 | 293741.6563 | 357030.5938 | -0.059883852 | 0.889892479 | 0.658371016 | 0.184451984 | 0.941199964 | 0.080852625 |
| 304577.6875 | 155324.4688 | 286674.4375 | -0.060579568 | 0.72812367 | -0.747508819 | 0.418126829 | 0.134064716 | 0.76812776 |
| 3263023.75 | 4501628.5 | 2406438 | -0.060625236 | 0.957465821 | -0.210317177 | 0.842416661 | -0.033294771 | 0.97158091 |
| 331564.9375 | 338285.3438 | 328779.625 | -0.060697329 | 0.408583166 | 0.456065585 | 0.024018676 | 0.281316728 | 0.061583511 |
| 19301.85742 | 17455.66211 | 19329.8125 | -0.060777325 | 0.794535026 | 0.289817198 | 0.404875207 | 0.550446666 | 0.060841062 |
| 122415.4297 | 110451.1953 | 131514.6563 | -0.0609175 | 0.604697925 | 0.250118461 | 0.104557189 | 0.139998366 | 0.330512624 |
| 80513.22656 | 80296.69531 | 82086.375 | -0.061078016 | 0.640761405 | -0.051673375 | 0.704789036 | -0.018967363 | 0.885245898 |

| | | | | | | | | |
|---|---|---|---|---|---|---|---|---|
| 22709.26367 | 23061.76953 | 21749.55664 | -0.061096504 | 0.176366736 | 0.300350527 | 0.103460535 | 0.505671397 | 0.013843758 |
| 37530.62891 | 41056.62891 | 38166.83984 | -0.061115643 | 0.66651855 | 0.589270476 | 0.07457353 | 0.527556844 | 0.048228885 |
| 22457.43359 | 20040.05078 | 20513.53125 | -0.061615947 | 0.403377498 | -0.055235381 | 0.185649792 | -0.236967174 | 0.095647158 |
| 3485.575439 | 2918.628418 | 1513.597168 | -0.061766323 | 0.811151714 | -0.410844092 | 0.13460208 | -0.790770765 | 0.133621587 |
| 41783.87305 | 44990.03418 | 39276.17969 | -0.061789739 | 0.465702136 | 0.02251679 | 0.144451236 | 0.016837502 | 0.847227591 |
| 149904.2656 | 143300.1875 | 144594.5938 | -0.061893227 | 0.697103632 | -0.527844641 | 0.101045598 | -0.691639102 | 0.074185756 |
| 102286.5313 | 104019.9844 | 121759.0547 | -0.062481279 | 0.940664982 | 1.002295173 | 0.173056576 | 0.727739042 | 0.295039695 |
| 13765.33691 | 13628.95801 | 9309.106445 | -0.06262676 | 0.659279692 | 0.815838464 | 0.059562109 | -0.176919026 | 0.500106023 |
| 525113.75 | 524215.625 | 552427.9375 | -0.062798147 | 0.507584438 | 0.019032771 | 0.843999416 | 0.204877688 | 0.11945215 |
| NA | 1682.889648 | NA | -0.062830957 | NA | 0.103799922 | NA | -0.38762226 | NA |
| 119093.7656 | 107652.5234 | 106034.6328 | -0.062891819 | 0.445968126 | -0.274640126 | 0.050410119 | -0.281946101 | 0.056398361 |
| 26619.75977 | 32648.19531 | 33018.67188 | -0.062965719 | 0.63829945 | 0.213214111 | 0.280688037 | 0.527385961 | 0.085462263 |
| 70443.14844 | 79664.76563 | 78128.53125 | -0.063022745 | 0.494165102 | 0.335199788 | 0.048024604 | 0.136154846 | 0.239658156 |
| 17500.03711 | 18562.59375 | 18869.99414 | -0.063049537 | 0.80053545 | -0.118091148 | 0.723326996 | 0.059624925 | 0.328763572 |
| 1998131.375 | 2320994.25 | 2179021.25 | -0.063076885 | 0.739240209 | -0.226164564 | 0.148395417 | -0.259301417 | 0.11365489 |
| 18308.17383 | 23109.73828 | 11838.79199 | -0.063115762 | 0.639033033 | 0.41416895 | 0.085088535 | 0.193355531 | 0.664329886 |
| 839715.3125 | 825918.5 | 827843.0625 | -0.063128634 | 0.683282587 | -0.11520834 | 0.420908194 | -0.314372806 | 0.145574695 |
| 46339.09375 | 44323.8125 | 36436.89844 | -0.063148151 | 0.370851558 | 0.086070959 | 0.187408941 | -0.160542703 | 0.344155352 |
| 1542814.75 | 1511690 | 1477096.75 | -0.06339148 | 0.42956317 | -0.15092449 | 0.128081636 | -0.347244897 | 0.049004518 |
| 138284.2031 | 128000.2813 | 104264.8125 | -0.063534384 | 0.56388701 | -0.74855616 | 0.015540203 | -1.017576091 | 0.022744861 |
| 15462.53223 | 20463.28516 | 18578.54883 | -0.063588243 | 0.900759821 | 0.167611444 | 0.727571165 | -0.468713985 | 0.453876956 |
| 205133.4375 | 224334.2656 | 192898.6875 | -0.063813199 | 0.874874841 | -0.168254194 | 0.628159256 | -0.134264097 | 0.672052206 |
| 1122.116821 | 823.171875 | NA | -0.063883112 | 0.85702865 | 1.302304603 | 0.031437224 | 0.791568117 | NA |
| 13458.99805 | 11313.23047 | 9482.083984 | -0.063921488 | 0.691392083 | 0.118813678 | 0.416925425 | -0.337459856 | 0.198589173 |
| 5489070 | 6069830.5 | 5387102 | -0.064246611 | 0.706939023 | -0.084919103 | 0.738562038 | -0.273148115 | 0.109009095 |
| 7086.960938 | 6137.98291 | 4035.43457 | -0.064351943 | 0.538789485 | 0.181128108 | 0.077543002 | -0.203005143 | 0.542217075 |
| 46393.65234 | 45141.30078 | 45142.46094 | -0.064388703 | 0.495678002 | 0.142221403 | 0.182716493 | 0.112191643 | 0.241436458 |
| 46020.30469 | 40225.17969 | 38981.63281 | -0.064391753 | 0.652404567 | -0.42547298 | 0.125411718 | -0.626378859 | 0.061031191 |
| 21319.59766 | 21951.02344 | 18845.56445 | -0.064447303 | 0.56388701 | -0.371170953 | 0.103460535 | -0.841581454 | 0.037292993 |
| 9005.783203 | 6326.648438 | 4958.53418 | -0.064493108 | 0.87058201 | -0.300882291 | 0.358705938 | -0.763718547 | 0.118023574 |
| 549701.875 | 588361.875 | 568640.4375 | -0.064659115 | 0.271519724 | 0.117061548 | 0.126731731 | 0.230758296 | 0.044991933 |
| 10954.03516 | 18711.98242 | 11960.04102 | -0.064803144 | 0.711280719 | 0.067046908 | 0.717559738 | -0.547886434 | 0.190246465 |
| 3842834 | 4095209.5 | 4072323.75 | -0.064829592 | 0.697309158 | -0.16283899 | 0.139855885 | -0.309248411 | 0.043437905 |

| | | | | | | | | |
|---|---|---|---|---|---|---|---|---|
| 475781.9375 | 487269 | 492599.7188 | -0.06507447 | 0.180541072 | -0.244415028 | 0.087142005 | -0.208411148 | 0.043437905 |
| 105257.2266 | 99066.8125 | 94616.16406 | -0.065240863 | 0.504877484 | -0.23362148 | 0.059340088 | -0.17722657 | 0.091037344 |
| 24240.1875 | 22991.06055 | 17477.97852 | -0.065298498 | 0.602426846 | -0.19720722 | 0.191377912 | -0.040060791 | 0.858927248 |
| 12925.94824 | 14604.37207 | 10946.18945 | -0.065618312 | 0.745111336 | 0.454055351 | 0.06106734 | 0.441742772 | 0.138331403 |
| 236609.7969 | 296170.1563 | 282908.4375 | -0.065689054 | 0.80053545 | 0.02995868 | 0.923246587 | -0.132141753 | 0.661949635 |
| 124927.6563 | 130887.7344 | 117735.0156 | -0.06631143 | 0.607487192 | -0.205931063 | 0.228751244 | -0.528255333 | 0.064834271 |
| 5023.807617 | 4862.69873 | 3008.369141 | -0.066561362 | 0.883955097 | -0.34768324 | 0.393001672 | -0.281643422 | 0.506889831 |
| 29709.32031 | 35576.75781 | 24370.33398 | -0.066570507 | 0.716159442 | 0.264564146 | 0.230974993 | 0.445944229 | 0.2081433 |
| 139251.875 | 137906.9922 | 109648.75 | -0.066892845 | 0.688811545 | 0.417199157 | 0.147949281 | -0.778094836 | 0.055298525 |
| 1469855.625 | 1443565.125 | 1544935.875 | -0.067264607 | 0.579001766 | -0.206829161 | 0.104928475 | -0.300354796 | 0.027949491 |
| 1006160.875 | 1026417.125 | 1023638.75 | -0.06732749 | 0.599013051 | -0.126728517 | 0.191239401 | -0.272921285 | 0.063474265 |
| 915722.75 | 1105843 | 1070824 | -0.067587125 | 0.722927832 | -0.095171986 | 0.658237453 | 0.018773137 | 0.935152011 |
| 90870.14844 | 84830.75 | 81677.49219 | -0.067677837 | 0.274703862 | -0.180843212 | 0.08416302 | -0.412964616 | 0.02354608 |
| 19597.32813 | 21539.03906 | 11214.71875 | -0.067690266 | 0.64439335 | -1.019017048 | 0.053419675 | -1.163351586 | 0.042674406 |
| 5314.429199 | 5126.244629 | 1112.389526 | -0.068160517 | 0.947463938 | 1.495009821 | 0.108760394 | 0.836657773 | 0.453790172 |
| 5076.560059 | 7394.006348 | 3750.189697 | -0.068347882 | 0.787299254 | 0.026567579 | 0.782663525 | -0.045659256 | 0.918174481 |
| 1581808.125 | 1927098.125 | 2049441.5 | -0.068389544 | 0.675850705 | -0.209792364 | 0.116818656 | -0.162238246 | 0.364262686 |
| 102207.4375 | 109959.4297 | 111213.7031 | -0.068443736 | 0.404110576 | -0.604177466 | 0.048615365 | -0.349018493 | 0.048991451 |
| 7306.885742 | 3361.497559 | 5774.699707 | -0.068639422 | 0.825031307 | -0.345378344 | 0.270406284 | -0.804122978 | 0.119412835 |
| 81561.39063 | 83666.24219 | 86780.74219 | -0.068641488 | 0.770630681 | 0.584602471 | 0.108338517 | 0.710225701 | 0.077902102 |
| 17267.21875 | 15483.87109 | 15980.14746 | -0.068804118 | 0.478561925 | 0.022606486 | 0.847335172 | 0.116405541 | 0.2384734 |
| 34006.98438 | 35229.10156 | 30448.3418 | -0.06896046 | 0.639681593 | -0.011184975 | 0.929570723 | -0.032476001 | 0.7678184 |
| 95240.75 | 99706.50781 | 93204.40625 | -0.069156909 | 0.343316347 | 0.072535262 | 0.191362386 | 0.042705288 | 0.408925574 |
| 19875.38477 | 18029.95508 | 17461.17578 | -0.069456807 | 0.649999812 | 0.197495207 | 0.28939927 | -0.043385322 | 0.807597809 |
| 8228.350586 | 7285.556152 | 7635.672852 | -0.069457501 | 0.709777983 | -0.175525568 | 0.461684928 | -0.101849268 | 0.620264646 |
| 37815.89844 | 36748.17578 | 36501.73828 | -0.070091758 | 0.521511046 | 0.044084669 | 0.532402501 | -0.244765753 | 0.046581887 |
| 3465.795166 | 5836.305664 | 5397.619141 | -0.070092274 | 0.730344982 | -0.234862229 | 0.332429777 | -1.221951817 | 0.036848746 |
| 1203283.375 | 1149589.75 | 1027513.625 | -0.070262359 | 0.476002496 | -0.156351493 | 0.182543644 | -0.35763502 | 0.05536545 |
| 29188.23633 | 33245.45313 | 29552.60352 | -0.070421294 | 0.494764936 | -0.348531349 | 0.091790291 | -0.814004378 | 0.033709967 |
| 220695.1406 | 216726.75 | 167371.0156 | -0.070429637 | 0.603052824 | 0.436526561 | 0.04542656 | 0.175928667 | 0.423098337 |
| 91155.0625 | 90198.41406 | 93463.45313 | -0.070455082 | 0.432950373 | 0.417422667 | 0.101962771 | 0.500581118 | 0.008387835 |
| 91042.89063 | 80345.79688 | 83533.88281 | -0.070598159 | 0.624312645 | -0.044055309 | 0.693736734 | 0.128335316 | 0.326227143 |
| 16124.21094 | 14787.44238 | 13765.10352 | -0.07060565 | 0.420983699 | -0.149916885 | 0.229462977 | -0.168484159 | 0.213904657 |

| | | | | | | | | |
|---|---|---|---|---|---|---|---|---|
| 24243.23242 | 24680.62891 | 24413.76172 | -0.071100602 | 0.70461778 | 0.760234172 | 0.082888448 | 0.516716346 | 0.081899796 |
| 149087.4434 | 139606.7129 | 119731.6982 | -0.07118149 | 0.499820309 | -0.306368707 | 0.037014778 | -0.074000634 | 0.595667125 |
| 251400.4375 | 260563.5313 | 241908.4688 | -0.071349258 | 0.721803576 | 0.031587419 | 0.885440998 | 0.270518266 | 0.260165601 |
| 1410.5271 | 1003.241333 NA | | -0.071400673 | 0.664145118 | 0.287112924 | 0.308392763 | -0.659656344 | NA |
| 210812.2188 | 217903.6719 | 212893.0625 | -0.071566745 | 0.140783489 | -0.64006368 | 0.012254503 | -0.380937892 | 0.02354608 |
| 43104.62109 | 45089.76563 | 45533.97266 | -0.071574252 | 0.234431096 | 0.135205937 | 0.039557441 | -0.000771406 | 0.98785686 |
| 81317.39844 | 77852.53125 | 68454.32813 | -0.071584994 | 0.424834696 | 0.043735634 | 0.465719209 | 0.05942788 | 0.618473137 |
| 132399.7031 | 128439.8047 | 125848.0469 | -0.071621363 | 0.491610667 | 0.399788732 | 0.066380875 | 0.179015904 | 0.088892469 |
| 131820.875 | 163367.2656 | 116758.7969 | -0.071646895 | 0.764304977 | 0.022973258 | 0.815041523 | -0.003818565 | 0.988689536 |
| 67388.99219 | 60215.80859 | 55975.31641 | -0.071912854 | 0.489346028 | 0.33577832 | 0.06824263 | -0.136591619 | 0.370397499 |
| 93309.35938 | 62102.63672 | 95004.48438 | -0.072207138 | 0.655485241 | -0.278257691 | 0.106378122 | -0.127378213 | 0.649614705 |
| 240838 | 212703.7031 | 212712.5781 | -0.072386287 | 0.648910173 | 0.119139618 | 0.45811699 | 0.04968492 | 0.765741675 |
| 4109.120117 | 5965.076172 | 4602.385742 | -0.072778944 | 0.535047385 | -0.042996302 | 0.715505426 | -0.879538662 | 0.036848746 |
| 180417.75 | 185677 | 179011.7969 | -0.072799456 | 0.295346854 | -0.032334531 | 0.303308952 | 0.009687897 | 0.765721035 |
| 7590.341797 | 6026.433105 | 5882.852051 | -0.073009514 | 0.817376907 | -0.091914086 | 0.686521476 | -0.49874256 | 0.134304215 |
| 29166.07422 | 32627.38086 | 32319.64258 | -0.073035552 | 0.862495139 | 0.841551793 | 0.052756858 | 1.886141432 | 0.010236624 |
| 58823.49219 | 85200.63281 | 72068.91406 | -0.073136158 | 0.869533034 | 0.198978343 | 0.598729228 | 0.259905338 | 0.490425837 |
| 19166.00391 | 17919.50977 | 19663.91797 | -0.073249361 | 0.507457676 | -0.164423123 | 0.28983684 | -0.599486926 | 0.047847186 |
| 44173.49219 | 20201.90039 | 22546.58203 | -0.073336751 | 0.832358028 | 0.278636649 | 0.734444561 | -0.815232249 | 0.18058763 |
| 58303.73828 | 46617.00391 | 51017.34766 | -0.073420408 | 0.673063365 | -0.156971363 | 0.460233965 | -0.405096092 | 0.156588849 |
| 1094.623169 | 1789.680664 | 562.382019 | -0.073861101 | 0.707693403 | -0.300063167 | 0.544751146 | -1.045464037 | 0.131502216 |
| 266214.9375 | 251614.2813 | 243977.8594 | -0.073908962 | 0.507427207 | 0.300631752 | 0.094078555 | 0.225765253 | 0.128484296 |
| 31837.64063 | 31343.70508 | 30677.28516 | -0.07393 | 0.468038758 | -0.190654249 | 0.189627399 | -0.319695283 | 0.09942999 |
| 1253046 | 334094.125 | 162122.5469 | -0.074183264 | 0.944040927 | 1.431246753 | 0.124972519 | 1.474253648 | 0.497255892 |
| 633513.125 | 816424.0625 | 744646.9375 | -0.074370015 | 0.666442307 | -0.212540751 | 0.100046608 | -0.157756425 | 0.361281503 |
| 4274.59375 | 4529.120117 | 4249.480957 | -0.074431682 | 0.706689159 | 0.015792332 | 0.926516232 | -0.091715615 | 0.37957979 |
| 44405.84766 | 47130.24219 | 45429.17578 | -0.074567448 | 0.735207913 | -0.0633098 | 0.696860822 | -0.272965503 | 0.184624365 |
| 4002190.125 | 4266082.5 | 4180422.375 | -0.074638758 | 0.475746506 | 0.131659865 | 0.412292948 | 0.001129977 | 0.990842721 |
| 55131.71875 | 55287.77734 | 41026.86719 | -0.075199482 | 0.489291353 | 0.566941015 | 0.034480785 | 0.438444005 | 0.178181856 |
| 403837.5313 | 386759.9688 | 380366.2188 | -0.07521578 | 0.296107668 | -0.246510896 | 0.053882844 | 0.440197096 | 0.02896549 |
| 54967.68359 | 30829.15039 | 43655.83984 | -0.075413263 | 0.673593051 | 0.482859295 | 0.099165657 | 0.231299738 | 0.566209266 |
| 134763.1875 | 151793.1094 | 164854.4844 | -0.075857132 | 0.685054567 | 0.133514123 | 0.458093846 | 0.146492732 | 0.389796538 |
| 29707.46484 | 28505.38086 | 31349.98828 | -0.076011546 | 0.510801673 | -0.079304829 | 0.536495 | -0.039415848 | 0.758090194 |

| | | | | | | | | |
|---|---|---|---|---|---|---|---|---|
| 37395.84375 | 37391.89844 | 29298.28906 | -0.076273597 | 0.260122085 | -0.133000073 | 0.197991611 | -0.688402351 | 0.04784105 |
| 2579.243896 | NA | 3343.286133 | -0.07636236 | NA | -0.072364764 | NA | 0.715388943 | NA |
| 61014.13672 | 59416.65234 | 58829.85547 | -0.076589033 | 0.406891053 | -0.95948912 | 0.038454242 | -0.391450405 | 0.077163447 |
| NA | 611.8705444 | 1522.978271 | -0.077050679 | 0.825989203 | -1.04400788 | 0.099817974 | -2.106684615 | NA |
| 487519.5938 | 514357.9688 | 516691.6875 | -0.077296923 | 0.245930178 | -0.012604376 | 0.744632174 | -0.084172121 | 0.138522639 |
| 14862.07324 | 13368.81543 | 13969.10059 | -0.077417688 | 0.728917468 | 0.310728805 | 0.077848578 | -0.131137025 | 0.302562509 |
| 83701.4375 | 85870.41406 | 69995.0625 | -0.077938457 | 0.557959822 | -0.163767367 | 0.326871267 | 0.161183527 | 0.377470755 |
| 9101.649414 | 10699.13379 | 8812.768555 | -0.078069577 | 0.796679771 | -0.974530359 | 0.141443989 | -1.331721312 | 0.113631071 |
| 2042392.25 | 2305326 | 2272007 | -0.078322222 | 0.593204455 | -0.099372811 | 0.177834038 | -0.070029474 | 0.425809369 |
| 41965.37891 | 45932.67578 | 49785 | -0.079137079 | 0.161527162 | 0.24646397 | 0.038037652 | -0.036243393 | 0.747473963 |
| 185635.1875 | 201245.9688 | 200299.6875 | -0.079391725 | 0.646266596 | 1.36670606 | 0.021959652 | 1.465743255 | 0.009014131 |
| 35202.21484 | 34137.94141 | 43134.52344 | -0.07956838 | 0.705696339 | 0.595472909 | 0.100849207 | 0.669222466 | 0.071851903 |
| 7120.720703 | 12307.73242 | 6844.499512 | -0.079979021 | 0.770630681 | -0.61513597 | 0.145357352 | -0.618933903 | 0.193809689 |
| 391261.0625 | 428439.0625 | 412140.2188 | -0.080008828 | 0.25326502 | -0.910423466 | 0.023497318 | -0.820212554 | 0.019274211 |
| 31047.4668 | 26513.58203 | 24247.53711 | -0.080200456 | 0.648757503 | 0.345583032 | 0.094078555 | -0.485964968 | 0.060012809 |
| 17082.43164 | 15456.58887 | 16314.33398 | -0.080535286 | 0.528395401 | -0.386813642 | 0.121984799 | -0.285247304 | 0.131166582 |
| 30576.33984 | 32164.29492 | 34133.00391 | -0.080600773 | 0.665671044 | -0.105441247 | 0.64741062 | 0.517972629 | 0.077833101 |
| 445874.375 | 429260 | 413125.5313 | -0.081072556 | 0.230254167 | -0.252337667 | 0.059562109 | -0.223502909 | 0.060937162 |
| 48894.92578 | 48560.35547 | 44276.15625 | -0.081586713 | 0.521991297 | 0.212051276 | 0.193691583 | 0.162989455 | 0.259140096 |
| 105937.6016 | 113070.4297 | 109093.7266 | -0.081717222 | 0.664145118 | 0.34930242 | 0.169553637 | 0.344765266 | 0.152372879 |
| 8455.90918 | 8677.688477 | 10279.09082 | -0.082602589 | 0.579372588 | -0.652431844 | 0.088323423 | -0.810255666 | 0.031136253 |
| 4504378.5 | 4470189.5 | 4010897.75 | -0.082627395 | 0.51043185 | 0.116006432 | 0.451971433 | 0.130111125 | 0.343159458 |
| 565174.25 | 567977.625 | 454001.8125 | -0.082793851 | 0.662717815 | 0.188148815 | 0.364020618 | -0.063321479 | 0.785580722 |
| 146614.1719 | 180211.4375 | 214216.0156 | -0.082807864 | 0.444840895 | 0.301269778 | 0.066380875 | 0.507194362 | 0.198718671 |
| NA | 1842.119263 | NA | -0.082861872 | 0.724564525 | -0.577348253 | 0.106067796 | -0.255174106 | NA |
| 216602.8594 | 209794.0625 | 198648.9688 | -0.082989618 | 0.466759028 | 0.332354219 | 0.079209365 | 0.05599409 | 0.607583742 |
| 37702.41797 | 38372.17188 | 36829.83984 | -0.083245555 | 0.544919116 | 0.232719192 | 0.227656066 | 0.411933718 | 0.086952277 |
| 1182512.875 | 1118426.125 | 1010594.875 | -0.083537995 | 0.298117548 | 0.41292338 | 0.032914334 | 0.003425037 | 0.979696342 |
| 18588.04688 | 19735.68359 | 15330.15234 | -0.083561068 | 0.464841982 | 0.24929109 | 0.136651367 | 0.183535149 | 0.362075745 |
| 145968.0469 | 147377.4688 | 146577.8125 | -0.083689672 | 0.387895744 | 0.230135066 | 0.091622784 | 0.242971301 | 0.081660057 |
| 47424.77344 | 54184.55078 | 52072.67969 | -0.083700156 | 0.360867419 | 0.30977935 | 0.051139483 | 0.014426716 | 0.897530165 |
| 834995.875 | 859923.0625 | 757411.75 | -0.083712862 | 0.518212038 | -0.146661236 | 0.114739915 | -0.318817374 | 0.045009854 |
| 44710.26172 | 40967.52734 | 47583.55078 | -0.084200872 | 0.766848849 | 0.552497261 | 0.159217919 | 0.125777 | 0.672587482 |

| | | | | | | | | |
|---|---|---|---|---|---|---|---|---|
| 415942.2188 | 415196.5938 | 367344.6563 | -0.084221466 | 0.468441827 | -0.028431892 | 0.822508142 | -0.662816504 | 0.04728688 |
| 169698.1406 | 175242.7813 | 173346.3594 | -0.084248437 | 0.210412515 | 0.074433806 | 0.338904732 | -0.050476835 | 0.408792062 |
| 480016.5625 | 512278.0625 | 503275.875 | -0.084258723 | 0.398496937 | 0.199373375 | 0.144097346 | 0.870539348 | 0.01054584 |
| 6557.877441 | 5484.994629 | 6172.127441 | -0.084283602 | 0.861912685 | -0.544475582 | 0.428011055 | 0.319998154 | 0.508589617 |
| 15816.10352 | 15371.41602 | 17385.47266 | -0.084302122 | 0.6842726 | 0.267598591 | 0.503628106 | -0.07023658 | 0.747716162 |
| 174122.6406 | 168629.7656 | 167589.9375 | -0.084662651 | 0.228958194 | -0.006869718 | 0.924830326 | -0.095485755 | 0.121167761 |
| 156735.7188 | 219161.9375 | 143004.4063 | -0.084756768 | 0.853712828 | -0.038559582 | 0.933325598 | -0.186387443 | 0.635263628 |
| 20353.43555 | 24239.48389 | 26900.75098 | -0.084925943 | 0.74277824 | -0.155852059 | 0.422801 | -0.112910607 | 0.60486123 |
| 10715.42383 | 10356.02246 | 11652.40527 | -0.08499003 | 0.383125521 | 0.230057714 | 0.101338323 | -0.18624161 | 0.160292276 |
| 567617.125 | 400437.3125 | 445881.2813 | -0.085253549 | 0.859147033 | 0.410157574 | 0.64923649 | 0.352274486 | 0.275313311 |
| 4450.421387 | 4086.245117 NA | | -0.085346327 | 0.881702113 | -0.026163435 | 0.835648534 | -0.254244421 NA | |
| 303797.4063 | 349296.4375 | 353606.9688 | -0.085400196 | 0.249016055 | 0.233540563 | 0.07036809 | 0.16377866 | 0.233367123 |
| 12955.50195 | 12831.44336 | 12461.91992 | -0.085697197 | 0.559103651 | -0.215578105 | 0.273114108 | 0.006600255 | 0.966538084 |
| 16619.04883 | 19597.98047 | 22314.00586 | -0.085850905 | 0.946061916 | 1.373294574 | 0.21119114 | 0.474739825 | 0.654568601 |
| 20695.80078 | 21846.66797 | 15491.11621 | -0.08596052 | 0.650237824 | -0.1818234 | 0.380695769 | -0.369989769 | 0.194422602 |
| 37338.57813 | 40264.33594 | 31854.89648 | -0.086395563 | 0.652708799 | 0.100906671 | 0.62199953 | -0.40848665 | 0.088388282 |
| 212201.8281 | 190154.2656 | 207070.7031 | -0.086407964 | 0.37365573 | 0.3697986 | 0.095052628 | 0.334072167 | 0.048402783 |
| 40832.83984 | 52898.60156 | 51238.37891 | -0.086633554 | 0.692051141 | 0.120986868 | 0.514075591 | 0.256055058 | 0.290785919 |
| 10137.71289 | 13409.74023 | 11308.75098 | -0.08664166 | 0.617270607 | 0.354778309 | 0.138660529 | 0.189363483 | 0.4289352 |
| 85568.78906 | 84575.54688 | 84540.89844 | -0.086709478 | 0.330645004 | 0.250508854 | 0.050075499 | 0.435431871 | 0.031081866 |
| 296180.375 | 300154.125 | 302812.7188 | -0.086881844 | 0.553671904 | -0.723019118 | 0.039887225 | -0.652009046 | 0.04014804 |
| 38933.41406 | 38671.58203 | 30677.5957 | -0.086996928 | 0.736060839 | -0.213282727 | 0.185515134 | -0.234048745 | 0.236563713 |
| 93026.28125 | 90158.61719 | 78761.78125 | -0.087159488 | 0.730072928 | 0.001035739 | 0.99201113 | 0.412676575 | 0.070968049 |
| 199995.7813 | 189765.4844 | 196738.1094 | -0.087258135 | 0.171832714 | 0.182296709 | 0.063663792 | 0.194285142 | 0.049905588 |
| 99131.96875 | 91209.57031 | 96026.38281 | -0.087413088 | 0.291824215 | 0.006989223 | 0.933215071 | -0.272449796 | 0.06618804 |
| 83676.89063 | 86474.14063 | 76820.61719 | -0.087681636 | 0.613778303 | 0.226627287 | 0.216249752 | 0.485700438 | 0.031136253 |
| 235367.7344 | 222666.9375 | 241639.0469 | -0.087830022 | 0.429788696 | 0.692062502 | 0.052817461 | 1.317100151 | 0.009587646 |
| 117979.7578 | 124011.9375 | 133640.7969 | -0.087910906 | 0.719049006 | 0.568520581 | 0.098459633 | 0.71556187 | 0.054109248 |
| 25953.01367 | 24253.48633 | 26835.1875 | -0.088281372 | 0.517345912 | -0.029462177 | 0.805890965 | -0.070242911 | 0.496294 |
| 123655.4453 | 124270.1328 | 126989.3516 | -0.088320441 | 0.289614586 | 0.039705309 | 0.508171609 | -0.022109323 | 0.703463561 |
| 16006.05859 | 15126.11816 | 13614.94824 | -0.088320693 | 0.46970317 | 0.113112213 | 0.374500771 | 0.084268642 | 0.553097148 |
| 77490.70313 | 74236.60938 | 75264.92969 | -0.088493534 | 0.512681345 | -0.34165101 | 0.118324098 | -0.223450251 | 0.197962658 |
| 60441.98828 | 65882.08594 | 56208.46484 | -0.088524271 | 0.605477695 | 0.414052318 | 0.037014778 | 0.525948208 | 0.048228885 |

| | | | | | | | | |
|---|---|---|---|---|---|---|---|---|
| 21873.17969 | 25450.71094 | 21421.71289 | -0.089207154 | 0.801685883 | -0.270080441 | 0.479482194 | -0.114338653 | 0.739478415 |
| 68271.30469 | 71454.91406 | 75566.4375 | -0.089426936 | 0.150110417 | 0.359441089 | 0.026826422 | 0.631970883 | 0.025759614 |
| 68846.625 | 67888.35156 | 62031.80469 | -0.089496054 | 0.503296939 | 0.472397481 | 0.038037652 | 0.404017359 | 0.063886777 |
| 1124904.25 | 1329600.625 | 1140491.125 | -0.089512806 | 0.530526722 | -0.225120606 | 0.102596936 | -0.237167749 | 0.137812617 |
| 39989.00781 | 38316.52734 | 35863.78906 | -0.08953829 | 0.163060402 | 0.171485436 | 0.090012942 | -0.08067748 | 0.321989721 |
| 26779.4082 | 28763.70703 | 12455.78906 | -0.08970374 | 0.85906459 | -1.14890057 | 0.256063289 | -1.255732516 | 0.224000425 |
| 1361508.25 | 1532591.25 | 1421583 | -0.089709693 | 0.530096414 | -0.197520819 | 0.164575062 | -0.248608647 | 0.084223187 |
| 94346.59375 | 95523.50781 | 89453.45313 | -0.0897651 | 0.329628012 | -0.225172294 | 0.156170999 | 0.387371257 | 0.033096306 |
| 32081.94727 | 31141.55273 | 25493.76953 | -0.090040408 | 0.757252703 | -0.336652115 | 0.143502718 | -0.495798794 | 0.07561358 |
| 474285.125 | 1440175.75 | 1466894.125 | -0.090535544 | 0.95007546 | 2.130903477 | 0.14612569 | 1.848907576 | 0.20322079 |
| 5821.491699 | 5969.856934 | 5757.579102 | -0.09054507 | 0.835112801 | -0.329677922 | 0.54685652 | -0.138394376 | 0.774288368 |
| 294421 | 353047.9375 | 310743.125 | -0.090586432 | 0.826782722 | -0.295455662 | 0.284889693 | -0.270229364 | 0.285412368 |
| 702855.5625 | 749521.75 | 727613.0625 | -0.090616009 | 0.340812741 | 0.157484535 | 0.219649135 | -0.013187332 | 0.759571084 |
| 16989.16016 | 16727.99414 | 16003.91895 | -0.090860448 | 0.326540039 | -0.407537775 | 0.160175093 | -0.262482869 | 0.043783164 |
| 51251.53516 | 50943.38281 | 51968.53125 | -0.090951376 | 0.460125956 | -0.177298346 | 0.063274608 | 0.310058961 | 0.027087229 |
| 105694.9766 | 113668.5938 | 96933.39063 | -0.091056694 | 0.563894297 | 0.051759457 | 0.767012555 | 0.171201162 | 0.355811485 |
| 34603.68359 | 37621.88672 | 34977.78516 | -0.091128752 | 0.709892588 | 0.093187568 | 0.724165773 | 0.554895699 | 0.107323815 |
| 103784.3906 | 99511.07813 | 92699.64063 | -0.091208987 | 0.631078063 | 0.46307552 | 0.080440179 | 0.241822064 | 0.217152139 |
| 8149.491699 | 9876.333008 | 8353.230469 | -0.091688595 | 0.717102937 | 0.350856634 | 0.133621013 | 0.224393579 | 0.274556211 |
| 61751.98828 | 66314.92188 | 54315.61328 | -0.092292677 | 0.519382561 | -0.124127615 | 0.470792459 | -0.468831583 | 0.044133846 |
| 30580.01367 | 31597.17383 | 24432.27539 | -0.092913857 | 0.441018848 | 0.293580095 | 0.127885085 | -0.01232859 | 0.953554342 |
| 78791.70313 | 83691.95313 | 85033.92188 | -0.093030017 | 0.598139599 | 0.70082767 | 0.076177015 | 1.020129939 | 0.020008995 |
| 6236.12793 | 11125.75977 | 7136.765137 | -0.093117696 | 0.820454661 | 0.486171253 | 0.496669713 | -0.039182018 | 0.927109931 |
| 2256988.5 | 2218069.5 | 2147424 | -0.093175305 | 0.220006554 | 0.08668365 | 0.446737466 | 0.11172961 | 0.16248539 |
| 1213863.375 | 1252398 | 1278557.125 | -0.093299585 | 0.643395293 | -0.048227478 | 0.732942123 | 0.082976437 | 0.46957326 |
| 12864.09473 | 7832.901367 | 9943.526367 | -0.093390275 | 0.530730634 | -0.474200644 | 0.070271638 | -0.221255882 | 0.477106314 |
| 40396.91016 | 36863.85156 | 36270.35156 | -0.093871102 | 0.586822171 | 0.022214254 | 0.901964742 | -0.139366409 | 0.454451558 |
| 293137.2188 | 306344.0313 | 322231.9688 | -0.093900771 | 0.470928594 | -0.219027252 | 0.100389463 | -0.49843413 | 0.031136253 |
| 81728.75 | 79881.26563 | 78836.64063 | -0.093999292 | 0.34282486 | -0.378323586 | 0.029439089 | -0.114083376 | 0.044248771 |
| 7244.65625 | 7606.38916 | 6589.125488 | -0.094352899 | 0.677149842 | 0.015967586 | 0.944405642 | 0.025553165 | 0.914110816 |
| 60632.46875 | 66799.42969 | 55822.90234 | -0.094439988 | 0.663675216 | 0.723313031 | 0.058077004 | 0.747259876 | 0.034122066 |
| 522825.25 | 561179.875 | 593190.1875 | -0.094466889 | 0.461919485 | 0.14002704 | 0.324535676 | 0.217475853 | 0.168528497 |
| 69060.76563 | 72010.14063 | 68986.69531 | -0.09482587 | 0.624043697 | 0.019237857 | 0.926516232 | -0.333492082 | 0.247014704 |

| | | | | | | | | |
|---|---|---|---|---|---|---|---|---|
| 99560.35156 | 100988.5313 | 95764.28906 | -0.09565246 | 0.47152965 | -0.366322706 | 0.040327952 | -0.246764226 | 0.057202212 |
| 720132.8125 | 703966.6875 | 743185.875 | -0.095874346 | 0.111075047 | -0.127778571 | 0.127892721 | 0.309256768 | 0.021863066 |
| 8976.670898 | 14159.82617 | 14183.37891 | -0.096006713 | 0.896724734 | 0.480716918 | 0.499021027 | 0.35814366 | 0.624394829 |
| 207563.8594 | 207482.5313 | 209127.6719 | -0.096098547 | 0.02237899 | 0.169159097 | 0.021149096 | -0.02463057 | 0.215760001 |
| 915246.6875 | 1256331.375 | 997619.25 | -0.09613763 | 0.801548444 | -0.577761947 | 0.322217412 | -0.132829793 | 0.758897267 |
| 258390.5625 | 252743.4219 | 267226.25 | -0.096216924 | 0.279048922 | 0.212578629 | 0.081451772 | 0.30032492 | 0.030633392 |
| 336482.1875 | 362500.75 | 371739.75 | -0.096298106 | 0.150947498 | 0.048554639 | 0.506573912 | 0.129959228 | 0.159719054 |
| 9917.435547 | 11375.30957 | 10899.97559 | -0.096313223 | 0.508982648 | -0.367555262 | 0.123343898 | -0.250858348 | 0.183628222 |
| 117305.3438 | 116281.1016 | 102414.7813 | -0.096770922 | 0.693062064 | 0.120007111 | 0.520791665 | 0.259496864 | 0.115926052 |
| 17339.69531 | 23887.72266 | 16649.90625 | -0.096876991 | 0.670331039 | 0.298789993 | 0.31156031 | 0.249809431 | 0.4289352 |
| 594022.9375 | 681967.5625 | 769829.4375 | -0.096977426 | 0.563664711 | -0.241958312 | 0.183642732 | -0.132616134 | 0.449433318 |
| 25311.03906 | 29033.16797 | 18658.46875 | -0.097013222 | 0.688352406 | 0.453346016 | 0.141473046 | -0.025723394 | 0.940662021 |
| 48484.98438 | 47900.80469 | 49210.33203 | -0.097143128 | 0.323994033 | -0.41065199 | 0.087467078 | -0.144604066 | 0.224000425 |
| 19841.12305 | 17196.93359 | 13260.16797 | -0.097287211 | 0.579372588 | -0.361127324 | 0.207485122 | -0.31896419 | 0.288940511 |
| 114351.7734 | 110536.8047 | 119753.4688 | -0.097344728 | 0.619539817 | 0.069136225 | 0.560671011 | -0.047936154 | 0.683612427 |
| 1258324.25 | 1303222.625 | 1252948.875 | -0.09773535 | 0.659279692 | 0.086169469 | 0.100528183 | 0.127213186 | 0.047850077 |
| 299115.2188 | 294200.5938 | 310358.25 | -0.097751284 | 0.349823026 | -0.167624653 | 0.152229165 | -0.108191418 | 0.250851318 |
| 118227.7188 | 131789.2656 | 130649.9453 | -0.098205516 | 0.555128814 | 0.276081556 | 0.216410714 | 0.570905148 | 0.050626058 |
| 483215 | 496960.3125 | 478434.9063 | -0.098213757 | 0.167067405 | -0.210327535 | 0.073324814 | -0.205943606 | 0.077094395 |
| 227580.875 | 202463.3438 | 204106.25 | -0.098269709 | 0.373703142 | 0.347519205 | 0.030793508 | 0.108044387 | 0.299059995 |
| 6399.745117 | 6669.279297 | 9873.866211 | -0.098303563 | NA | 0.467779461 | 0.176036688 | 1.057138974 | 0.140158604 |
| 35244.21875 | 37333.17188 | 41092.78516 | -0.098396401 | 0.622840655 | 0.049702685 | 0.84440066 | 0.238892126 | 0.302401454 |
| 7061.196777 | 15256.47266 | 13501.11816 | -0.098507002 | 0.88860779 | 0.942330039 | 0.289971639 | 1.074031658 | 0.200746623 |
| 1714527.25 | 1759966.75 | 2030322 | -0.098538041 | 0.134212026 | -0.224944682 | 0.042884533 | -0.252759372 | 0.1278105 |
| 17888.14648 | 9412.730469 | 19793.40039 | -0.098813789 | 0.313971512 | -0.316979191 | 0.112543356 | -0.5999696 | 0.207463477 |
| 78042.85938 | 71136.60156 | 56895.72656 | -0.099166761 | 0.619539817 | -0.428758918 | 0.17813923 | -0.694014837 | 0.082528683 |
| 28082.93164 | 28508.92383 | 25495.36719 | -0.099174527 | 0.074128336 | 0.041905911 | 0.618328902 | 0.009305643 | 0.908522291 |
| 32619.53516 | 36653.41406 | 34443.44141 | -0.099240858 | 0.463954586 | 0.254369274 | 0.095792771 | 0.202632328 | 0.129291758 |
| 3266857.75 | 3765486.5 | 3698738.75 | -0.099844714 | 0.174418417 | 0.1706513 | 0.107845807 | 0.271628846 | 0.103046053 |
| 31769.70508 | 44220.23047 | 30499.47461 | -0.09993988 | 0.555933112 | -0.157631357 | 0.414380687 | -0.47899244 | 0.145702448 |
| 323864.6563 | 303962.2188 | 297844.0313 | -0.100093632 | 0.534135477 | 0.058883276 | 0.632281284 | 0.270840598 | 0.082912076 |
| 351363.5313 | 339142.0313 | 391613.9063 | -0.100208398 | 0.237119361 | 0.084223936 | 0.16720015 | 0.09364897 | 0.396866599 |
| 1270984.625 | 1474583.5 | 1533983.625 | -0.100222216 | 0.167028974 | -0.06898294 | 0.338904732 | 0.122456077 | 0.39090437 |

| | | | | | | | | |
|---|---|---|---|---|---|---|---|---|
| 99124.96094 | 91906.46875 | 89205.74219 | -0.100289766 | 0.230507258 | -0.006619172 | 0.92625747 | -0.222845718 | 0.078145737 |
| 8693.613281 | 8546.134766 | 6687.316406 | -0.100373581 | 0.391472313 | -0.033257232 | 0.630524877 | -0.220602213 | 0.266340398 |
| 1911858.875 | 1830069.125 | 1969413.625 | -0.100466901 | 0.277592349 | 0.096433742 | 0.276825942 | 0.048005795 | 0.534750592 |
| 17106.72461 | 14712.07715 | 13468.91504 | -0.10079968 | 0.293892903 | -0.657687691 | 0.024233498 | -1.090032709 | 0.013482424 |
| 3229.852295 | 4397.413086 | 4946.241699 | -0.101039025 | 0.556189585 | 0.350751306 | 0.324535676 | -0.201376174 | 0.454276381 |
| 64088.35547 | 69415.42969 | 73608.10938 | -0.101089721 | 0.573143374 | 0.001483686 | 0.99201113 | -0.067896152 | 0.684644513 |
| 134127.4531 | 141855.875 | 153534.7188 | -0.101138881 | 0.664737228 | -0.138069976 | 0.612901 | 0.33456232 | 0.200720922 |
| 96135.35938 | 94148.11719 | 100954.1563 | -0.101263378 | 0.064613771 | -0.072998548 | 0.121112119 | -0.156246364 | 0.084288402 |
| 4896735.5 | 5975408.5 | 2259407 | -0.101728942 | 0.87622214 | -0.258997322 | 0.741007407 | -0.882290357 | 0.401842443 |
| 80725.26563 | 88381.625 | 80270.91406 | -0.101747703 | 0.559820589 | -0.230507029 | 0.332666337 | -0.32793153 | 0.215498909 |
| 216247.9063 | 163073.8438 | 229785.5625 | -0.101973062 | 0.217097917 | 0.098460761 | 0.228320099 | 0.061557025 | 0.789122922 |
| 156304.2969 | 157745.4219 | 166629.6406 | -0.102120449 | 0.340812741 | -0.081638245 | 0.144097346 | -0.137497261 | 0.069042103 |
| 41618.67188 | 47410.95703 | 40485.84375 | -0.102586435 | 0.537230454 | 0.137170984 | 0.458269869 | -0.357916697 | 0.171907805 |
| 81518.5625 | 82297.78906 | 83042.39063 | -0.102998356 | 0.205289963 | 0.185994454 | 0.094078555 | 0.284944002 | 0.063283918 |
| 35409.75 | 39710.78125 | 47094.52734 | -0.103045696 | 0.73944516 | 0.450804606 | 0.218053066 | 0.87990943 | 0.064209546 |
| 268205.5 | 260587.5 | 259015.7656 | -0.103677427 | 0.45238856 | 0.229497353 | 0.291065687 | 0.377219264 | 0.026263251 |
| 1106668 | 1140143.875 | 1034621.813 | -0.103816719 | 0.423142746 | -0.237878008 | 0.122344477 | -0.324331137 | 0.060634835 |
| 13918.6543 | 12137.87305 | 12602.2168 | -0.103902819 | 0.337551113 | -0.07748372 | 0.374107188 | -0.518747946 | 0.028914056 |
| 249097.9688 | 238998.7344 | 244740.8906 | -0.103987481 | 0.449692985 | 0.090196502 | 0.533229239 | -0.151521545 | 0.363640088 |
| 9185.913086 | 8362.088867 | 4311.203613 | -0.104540599 | 0.466554528 | -0.290799441 | 0.350656099 | -0.688510556 | 0.161898002 |
| 59838.72266 | 54860.61719 | 43096.34766 | -0.104654029 | 0.203203033 | -0.688918984 | 0.045779673 | -0.892611754 | 0.029324108 |
| 12286.94238 | 4704.292969 | 13354.01465 | -0.104758005 | 0.595906478 | 0.338655666 | 0.146253591 | -0.443080072 | 0.423206002 |
| 606495.5625 | 609244.625 | 590828 | -0.104880941 | 0.42340664 | -0.203292254 | 0.208500335 | -0.306270877 | 0.111785532 |
| 23598.70898 | 24928.27734 | 18525.0625 | -0.104922188 | 0.490861401 | -0.56072855 | 0.102526216 | 0.140482569 | 0.54164829 |
| 560431.9375 | 574213.5 | 569264.125 | -0.105003108 | 0.150795791 | 0.311841092 | 0.17813923 | 0.436764188 | 0.026764821 |
| 407762.1875 | 333801.5625 | 401239.6875 | -0.105145227 | 0.431503081 | -0.373455311 | 0.061137147 | -0.588864218 | 0.031785503 |
| 343742.2188 | 348133.2188 | 373853.9375 | -0.105255649 | 0.180562259 | -0.210566095 | 0.104273255 | -0.336194631 | 0.035892596 |
| 8743.654297 | 11152.93262 | 7200.284668 | -0.105693745 | 0.709049574 | -0.220709811 | 0.482392757 | -0.215793262 | 0.54725722 |
| 3373.592041 | 3730.263916 | 3386.480957 | -0.105697557 | 0.71414429 | -0.050038867 | 0.909428871 | -0.502106519 | 0.248833486 |
| 3609.643311 | 2905.178223 | 3155.682129 | -0.105736508 | 0.605864892 | -0.05566238 | 0.732111347 | -0.002440985 | 0.989480447 |
| 216396.5 | 192658.1563 | 203554.4531 | -0.105778657 | 0.606039053 | 0.472836241 | 0.104110714 | 0.319329696 | 0.190246465 |
| 18597.54492 | 19927.90625 | 16718.30078 | -0.106024402 | 0.797091413 | -0.574816528 | 0.039582042 | -0.458424079 | 0.040959479 |
| 42973.94141 | 40623.74219 | 40495.50781 | -0.106065282 | 0.395528635 | -0.239901764 | 0.129294865 | -0.210360503 | 0.12131068 |

| | | | | | | | | |
|---|---|---|---|---|---|---|---|---|
| 55909.22461 | 54139.375 | 74200.83984 | -0.106122411 | 0.588172074 | 0.074793439 | 0.449267831 | 0.520224438 | 0.174355634 |
| 91790.0625 | 99343.41406 | 88277.20313 | -0.106153464 | 0.200542122 | -0.403707111 | 0.048593497 | -0.235020741 | 0.08213851 |
| 52449.83203 | 47734.43359 | 44708.62109 | -0.106230136 | 0.086055451 | -0.217075088 | 0.042884533 | -0.458019693 | 0.037772514 |
| 213783.3594 | 213549.0938 | 209900.2813 | -0.106238466 | 0.286127402 | 0.511583094 | 0.029012089 | 0.105377589 | 0.316079507 |
| 4569.989746 | 6461.837402 | 5128.614746 | -0.106274735 | 0.810385109 | 0.073903498 | 0.875616587 | -0.054640501 | 0.860919867 |
| 83787.32031 | 80461.51563 | 81623.52344 | -0.106417938 | 0.314376248 | 0.502600763 | 0.036154671 | -0.10549912 | 0.301475535 |
| 636515.5 | 628645.4375 | 551055 | -0.106468561 | 0.559171823 | 0.235668188 | 0.204833051 | 0.162721069 | 0.343877023 |
| 136975.4375 | 158405.4063 | 152199 | -0.107286055 | 0.671186503 | 0.475680052 | 0.018539883 | 0.264719718 | 0.10732392 |
| 71963.57031 | 74775.51563 | 77616.05469 | -0.10784425 | 0.195718606 | -0.057789323 | 0.532402501 | -0.13019407 | 0.179985011 |
| 22338.61914 | 23248.03906 | 20164.07813 | -0.107957722 | 0.612811759 | -0.150745371 | 0.551512542 | -0.123316085 | 0.616130362 |
| 21777.53516 | 28219.2207 | 22841.48633 | -0.10800033 | 0.620025412 | 0.369882828 | 0.064946832 | 0.084254312 | 0.659530356 |
| 5576157 | 7001865.5 | 7166897.5 | -0.108075595 | 0.736240107 | -0.004598418 | 0.989627785 | 0.234336318 | 0.399983931 |
| 61437.35938 | 63380.76953 | 64137.15234 | -0.108140686 | 0.10058931 | 0.168789796 | 0.049696115 | -0.092909925 | 0.122834452 |
| 31029.80664 | 32877.19922 | 29541.88867 | -0.108241017 | 0.533078483 | -0.078136966 | 0.122107272 | 0.015254678 | 0.834160705 |
| 9003.078125 | 8804.833984 | 6179.428711 | -0.108471367 | 0.685684032 | 0.295063133 | 0.19372563 | 0.042978543 | 0.879527307 |
| 2685.493652 | 15124.68262 | NA | -0.108483562 | NA | 0.341012745 | 0.649015214 | 0.380096357 | NA |
| 46752.04688 | 47428.58594 | 46154.41016 | -0.108663288 | 0.342326528 | 0.361088067 | 0.027031346 | 0.080429902 | 0.042723587 |
| 87268.64063 | 87465.6875 | 83877.14844 | -0.108940968 | 0.368167651 | 0.699048219 | 0.04667516 | 0.328731488 | 0.089275366 |
| 43834.54297 | 46013.97266 | 51777.20313 | -0.10897866 | 0.530730634 | 0.177593583 | 0.100528183 | 0.447517921 | 0.090703901 |
| 3873.24585 | 3580.162598 | 2531.249023 | -0.109224457 | 0.590927781 | -0.307078535 | 0.23446912 | -0.834387602 | 0.056579133 |
| 88772.10156 | 91775.45313 | 86390.88281 | -0.109283359 | 0.466759028 | -0.370531914 | 0.121848802 | -0.236603381 | 0.187314946 |
| 436989.9688 | 432369.2188 | 433592.5625 | -0.109291494 | 0.298390687 | 0.335389086 | 0.088308366 | 0.42767908 | 0.062345205 |
| 25917.49219 | 25416.69727 | 23397.99219 | -0.109333105 | 0.188598126 | -0.908100422 | 0.036485471 | -0.96499864 | 0.01987292 |
| 29284.92383 | 23235.13477 | 16706.05859 | -0.10945418 | 0.223624482 | -0.70473057 | 0.052953537 | -1.181667805 | 0.056534017 |
| 8645105 | 10594452 | 9619435 | -0.109783579 | 0.368847183 | -0.041716421 | 0.681696486 | -0.074574911 | 0.603485071 |
| 535465.875 | 548652.5 | 506644.8125 | -0.10981339 | 0.566211171 | -0.090279217 | 0.216249752 | -0.21264621 | 0.047790094 |
| 5846.397949 | 5872.548828 | 5600.366211 | -0.110069822 | 0.75519499 | -0.111147489 | 0.785487872 | -0.042530897 | 0.916489217 |
| 41426.0625 | 44345.45313 | 47861.63281 | -0.11007736 | 0.659937863 | 0.581227993 | 0.078996408 | 0.479621004 | 0.054745581 |
| 104341.9141 | 98846.10156 | 92549.42969 | -0.110225714 | 0.371068627 | -0.024731837 | 0.844905766 | -0.158266348 | 0.295151813 |
| 701605.375 | 670560.1875 | 591156.5625 | -0.110434984 | 0.127348903 | 0.49745133 | 0.096169757 | -0.014707131 | 0.90229668 |
| 88647.0625 | 72570.54688 | 109150.8125 | -0.110890893 | 0.580556008 | 0.485478055 | 0.205304317 | 0.678701411 | 0.153754411 |
| 33453.3125 | 38394.24609 | 36317.64063 | -0.110917737 | 0.270935076 | 0.191232996 | 0.085196118 | -0.24267153 | 0.08197271 |
| 98518.21875 | 106603.3828 | 97943.75 | -0.111343409 | 0.273267781 | -0.424054995 | 0.021906439 | -0.185926533 | 0.069876425 |

| | | | | | | | | |
|---|---|---|---|---|---|---|---|---|
| 6914.129883 | 6256.630371 | 5752.415039 | -0.111454049 | 0.687216284 | 0.074593612 | 0.771622264 | -0.610278092 | 0.091879002 |
| 11603.53418 | 15092.2666 | 12775.16309 | -0.111568838 | 0.614858464 | -0.221352379 | 0.431639915 | 0.066649841 | 0.806089164 |
| 178491.0156 | 207555.6406 | 197884.4375 | -0.111589148 | 0.714543616 | 0.422054322 | 0.136936932 | 0.521996912 | 0.051065331 |
| 542564.0625 | 563010.25 | 507471.1875 | -0.111681232 | 0.548913686 | 0.279913402 | 0.218601949 | 0.151236617 | 0.439514435 |
| 14927.46191 | 18168.52734 | 13032.80664 | -0.111740377 | 0.288647574 | -0.152989561 | 0.231614459 | -0.486723252 | 0.09236595 |
| 3718277.5 | 4216963.5 | 3870654.75 | -0.111802605 | 0.50652416 | -0.142942923 | 0.225621636 | -0.228700843 | 0.083348055 |
| 11417.95898 | 12637.87402 | 12727.37598 | -0.112002529 | 0.736208546 | 0.853795441 | 0.084215003 | 0.755383898 | 0.105346661 |
| 18619.50586 | 17785.83789 | 16910.94531 | -0.112234322 | 0.262311587 | -0.061674059 | 0.524069349 | -0.202173657 | 0.134550901 |
| 66222.38281 | 60989.89063 | 61012.625 | -0.112274733 | 0.129985841 | 1.046051774 | 0.011710842 | 0.604077855 | 0.036383418 |
| 5156.839355 | 5639.888184 | 4299.986328 | -0.112308153 | 0.358868205 | 0.089220841 | 0.534583821 | -0.471694785 | 0.072139147 |
| 148495.375 | 126319.2266 | 111012.8672 | -0.112382875 | 0.407059023 | -0.141343002 | 0.226271467 | -0.294393744 | 0.184452943 |
| 95863.91406 | 94058.95313 | 91025.09375 | -0.11259135 | 0.044365635 | -0.15645894 | 0.052691197 | -0.016466676 | 0.664794413 |
| 122923.5078 | 129783.0469 | 126734.1719 | -0.112800169 | 0.44802068 | 0.559061783 | 0.028434447 | 0.617610245 | 0.024151365 |
| 20025.12891 | 22161.55664 | 17600.98242 | -0.112930689 | 0.445541262 | 0.325543147 | 0.099684213 | -0.416368412 | 0.09149674 |
| 1777137.125 | 1816523.5 | 1718755.875 | -0.112948467 | 0.566211171 | -0.233442473 | 0.155441252 | -0.292419276 | 0.111646722 |
| 8520.601563 | 11631.08496 | 8397.694336 | -0.113219574 | 0.581635738 | 0.612379435 | 0.098656718 | -0.383680772 | 0.208408498 |
| 66841.92188 | 48530.64453 | 52774.42188 | -0.113240427 | 0.725362866 | 0.385153925 | 0.346969068 | 0.752455261 | 0.091941766 |
| 55258.49609 | 46746.09766 | 50793.07031 | -0.113349344 | 0.187640538 | -0.050990222 | 0.576955884 | 0.022305494 | 0.848721507 |
| 5478.445801 | 2991.389404 | 3618.561279 | -0.113424914 | 0.863810045 | 0.051645888 | 0.958449529 | 0.587617625 | 0.387761522 |
| 688728.125 | 714822.0625 | 676993.875 | -0.113771924 | 0.289180273 | 0.139846017 | 0.17294672 | 0.201759014 | 0.089218964 |
| 10639.9043 | 11259.9541 | 13424.4873 | -0.114447304 | 0.59265437 | -0.153017713 | 0.377381492 | -0.129065367 | 0.505254865 |
| 1587.647095 | 1238.377686 | 1727.743896 | -0.114522148 | 0.546269698 | 0.185170902 | 0.441278659 | -0.615108678 | 0.101686086 |
| 3777.54126 | 2599.840332 | 1345.053955 | -0.11459275 | 0.722927832 | -0.219700464 | 0.578271639 | -1.27217958 | 0.110105852 |
| 18023.41211 | 20120.17773 | 17527.86523 | -0.114713226 | 0.260684155 | -0.484118504 | 0.029180754 | -0.444700448 | 0.031136253 |
| 5176.261719 | 5035.652344 | 5383.780762 | -0.114884403 | 0.524853127 | 0.846844818 | 0.152229165 | 0.336945149 | 0.151280553 |
| 33643.04688 | 34409.88281 | 36705.85156 | -0.115173131 | 0.52911597 | -0.170630973 | 0.299315782 | 0.087924833 | 0.443421186 |
| 53425.07422 | 56500.03125 | 35231.47656 | -0.115573854 | 0.502981636 | -0.959414424 | 0.060391495 | -0.891734312 | 0.051065331 |
| 113003.9219 | 109342.8906 | 86464.16406 | -0.115680289 | 0.290808764 | -0.766539162 | 0.017029168 | -0.784578056 | 0.033213222 |
| 13466.01855 | 12069.29102 | 9395.660156 | -0.116015027 | 0.613763545 | -0.507588661 | 0.113609952 | -0.509304952 | 0.114083012 |
| 86928.07813 | 83169.63281 | 78620.25 | -0.116267172 | 0.221775028 | -0.822315066 | 0.03520579 | -0.429421867 | 0.02354608 |
| 213335.8125 | 203026.0313 | 208487.875 | -0.116508726 | 0.295023996 | -0.08932111 | 0.414279333 | -0.313498387 | 0.08082521 |
| 9426.185547 | 9317.18457 | 7173.920898 | -0.116639288 | 0.541659174 | 0.03696365 | 0.854867034 | -0.429517813 | 0.148333026 |
| 1590218.875 | 1880673.375 | 1716568 | -0.117009287 | 0.279124697 | -0.238806953 | 0.109641795 | -0.269310833 | 0.089724983 |

| | | | | | | | | |
|---|---|---|---|---|---|---|---|---|
| 9073.728516 | 8282.498047 | 15682.76855 | -0.117162582 | 0.391494686 | -0.70614994 | 0.077511889 | -0.661458648 | 0.18581375 |
| 34740.59766 | 37657.12891 | 36626.41016 | -0.117510145 | 0.084806455 | -0.264468585 | 0.04542656 | -0.146056843 | 0.091810798 |
| 141949.4063 | 138904.6875 | 153969.9531 | -0.117526617 | 0.316823258 | 0.119695313 | 0.120946561 | 0.066251313 | 0.400632465 |
| 16809.94922 | 16295.28906 | 14735.38574 | -0.117719347 | 0.46623402 | -0.035358292 | 0.780724906 | -0.311246102 | 0.114499805 |
| 47687.91797 | 44901.18359 | 33856.95313 | -0.118199513 | 0.8789414 | 0.88566422 | 0.221048551 | 0.415455524 | 0.545831185 |
| 131105.4448 | 128851.3481 | 143288.4741 | -0.118316485 | 0.02237899 | 0.039083534 | 0.638775352 | -0.176113165 | 0.118033234 |
| 145061.1094 | 127643.5313 | 132436.75 | -0.118357814 | 0.274017483 | 0.295340834 | 0.078527985 | -0.245157108 | 0.119204395 |
| 1403390 | 1682532.875 | 1516333.375 | -0.118466641 | 0.42065175 | -0.187028653 | 0.139250597 | -0.205074528 | 0.170506901 |
| 119335.0391 | 113951.5547 | 125980.5781 | -0.118547975 | 0.123080398 | -0.080285609 | 0.05744085 | 0.084130399 | 0.286081151 |
| 9081.676758 | 10883.84375 | 8873.083984 | -0.118950938 | 0.45533667 | -0.260869501 | 0.192808829 | -0.394193818 | 0.103569502 |
| 14907.98438 | 13682.61035 | 14457.14551 | -0.119488539 | 0.105673253 | -0.060015584 | 0.300186184 | 0.020991199 | 0.728465526 |
| 72160.03125 | 70407.82813 | 77141.71094 | -0.119502194 | 0.172104341 | 0.10801772 | 0.235937927 | 0.142215905 | 0.161733344 |
| 29377.32227 | 27377.22266 | 32323.67969 | -0.119583133 | 0.224685731 | -0.219326469 | 0.121179966 | -0.160844471 | 0.229590822 |
| 2049136.5 | 2113930.25 | 1982662.75 | -0.119608063 | 0.474195126 | -0.191720895 | 0.128223548 | -0.316396876 | 0.036013501 |
| 231671.3906 | 265168.8438 | 250617.8281 | -0.119982939 | 0.533024504 | -0.143644892 | 0.480060312 | -0.04226624 | 0.82622823 |
| 3408.218018 | 1725.931519 | 2193.273926 | -0.120140497 | 0.817518336 | 0.745069478 | 0.174475307 | 0.081187138 | 0.898154456 |
| 120863.2734 | 133794.125 | 151195.5781 | -0.120186663 | 0.311128404 | 0.388267355 | 0.022702163 | 0.525612316 | 0.097178029 |
| 21398.03125 | 21967.58203 | 21342.09375 | -0.120308302 | 0.14678023 | 0.25463512 | 0.11104965 | 0.033349192 | 0.519367772 |
| 32235.43164 | 31186.11719 | 27025.42383 | -0.120328507 | 0.60697985 | 0.012125712 | 0.93172794 | -0.245456673 | 0.183230134 |
| 1069822.5 | 1208608.25 | 1105592.375 | -0.120554721 | 0.429610905 | -0.189841332 | 0.144798646 | -0.177533941 | 0.118023574 |
| 4504.336914 | 4521.795898 | NA | -0.120594685 | 0.344779493 | -0.543670693 | 0.093960606 | -0.606995797 | NA |
| 76472.73438 | 84914.17969 | 80048.94531 | -0.120673923 | 0.424042107 | 0.256089651 | 0.057932046 | 0.316315278 | 0.045311345 |
| 16932.91602 | 21143.15625 | 15800.86035 | -0.120681612 | 0.383651804 | -0.062029589 | 0.409040093 | -0.583506283 | 0.06456861 |
| 569.4139404 | 3479.152832 | NA | -0.120987628 | NA | -0.389944035 | 0.54315586 | -0.850587767 | NA |
| 23311.88867 | 26600.1875 | 23815.99805 | -0.121008795 | 0.478101102 | 0.033274805 | 0.675218019 | 0.558396807 | 0.04066359 |
| 5151.40332 | 5854.046387 | 5628.514648 | -0.121021367 | NA | -0.496001703 | NA | 0.051796434 | NA |
| 330805.25 | 332046.5625 | 333753.5313 | -0.121289647 | 0.160984058 | -0.10672841 | 0.39679721 | 0.205269569 | 0.006394096 |
| 898690.25 | 708127.75 | 632110.875 | -0.121913532 | 0.414354381 | 0.093330692 | 0.559568799 | -0.251833753 | 0.322943908 |
| 1515933.125 | 1445439.75 | 1346045.25 | -0.122320212 | 0.216909187 | -0.194868504 | 0.135981012 | -0.343294503 | 0.040133731 |
| 1896915.125 | 1940802.625 | 1835974.75 | -0.122418005 | 0.110183943 | 0.05439329 | 0.343658822 | -0.054106821 | 0.368528672 |
| 23944.98242 | 23605.48047 | 22630.69336 | -0.12270044 | 0.371017022 | -0.351305896 | 0.12948378 | -0.475916979 | 0.102372782 |
| 46253.73047 | 46525.625 | 40496.875 | -0.122711067 | 0.7263536 | 0.095079085 | 0.706518199 | -0.452337704 | 0.032577228 |
| 3043051.25 | 4393343 | 3400218.5 | -0.122998591 | 0.691229987 | 0.020287862 | 0.914684262 | 0.12216616 | 0.672824986 |

| | | | | | | | | |
|---|---|---|---|---|---|---|---|---|
| 1823064.125 | 2128129.25 | 1968132.75 | -0.123292943 | 0.419717719 | -0.288595453 | 0.09861697 | -0.247365443 | 0.118033234 |
| 177076.6406 | 172174.6719 | 146784.125 | -0.123696488 | 0.666442307 | 0.115156396 | 0.789764311 | -0.214407048 | 0.493532407 |
| 30176.90625 | 30752.47461 | 27827.71875 | -0.12389043 | 0.196895924 | -0.200511823 | 0.076177015 | -0.223923674 | 0.062318926 |
| 3771.570068 | 4231.758789 | 2875.757813 | -0.123965721 | 0.348780804 | -0.559800752 | 0.058077004 | -0.388115787 | 0.169217885 |
| 4870403.5 | 5114814.5 | 5135290.5 | -0.124323193 | 0.333728081 | 0.296828404 | 0.085053588 | 0.70607016 | 0.016834605 |
| 1420564.375 | 1365309 | 1397530.875 | -0.124340463 | 0.487668097 | -0.190360919 | 0.137613908 | -0.258795952 | 0.097178029 |
| 117146.7793 | 128326.0537 | 124976.7627 | -0.124546113 | 0.417218549 | 0.125946548 | 0.435798634 | -0.024703088 | 0.878629235 |
| 63508.34375 | 56105.01172 | 53509.08203 | -0.125491276 | 0.228243249 | 0.625948385 | 0.081805946 | -0.118422523 | 0.346457757 |
| 272637.6989 | 236259.3457 | 239768.6558 | -0.12550545 | 0.337821825 | 0.291884334 | 0.112996996 | -0.52653586 | 0.059258927 |
| 125948.625 | 122473.7969 | 97013.07813 | -0.125848819 | 0.521368013 | -0.218529889 | 0.313455345 | -0.186449616 | 0.410930546 |
| 5521.244629 | 4596.791016 | 4258.753906 | -0.126021796 | 0.456360515 | 0.194143394 | 0.191362386 | -0.4577988 | 0.077912401 |
| 73774.60938 | 77299.125 | 76875.64844 | -0.12628659 | 0.247645442 | -0.408598284 | 0.027435011 | -0.19334972 | 0.030493787 |
| 78232.29688 | 84008.09375 | 92598.16406 | -0.126288011 | 0.338882582 | 0.549239153 | 0.042887894 | 0.699373759 | 0.034150933 |
| 80363.4375 | 84192.53906 | 89191.13281 | -0.126631002 | 0.00744439 | 0.151758397 | 0.164062441 | 0.119882147 | 0.206265186 |
| 21967.47656 | 21970.31445 | 21516.77148 | -0.126692475 | 0.263383916 | -0.292260734 | 0.117377154 | -0.510029285 | 0.071501012 |
| 78536.53906 | 71619.63281 | 70157.44531 | -0.126752649 | 0.364050541 | -0.228294949 | 0.231386286 | -0.017442838 | 0.908710949 |
| 151702.0156 | 141660.625 | 139870.8594 | -0.126762085 | 0.299735363 | -0.121484813 | 0.401614835 | -0.197285037 | 0.148459899 |
| 51830.91797 | 49482.31641 | 49451.62891 | -0.126801735 | 0.505886967 | 0.40591581 | 0.121179966 | 0.225496026 | 0.258294031 |
| 86548.28125 | 115030.2734 | 91988.79688 | -0.126868012 | 0.679427532 | -0.313190265 | 0.488418166 | -0.990985841 | 0.124736435 |
| NA | NA | NA | -0.126958779 | NA | 1.392130063 | 0.05975961 | NA | NA |
| 12887.92383 | 14040.03906 | 10251.43066 | -0.12705154 | 0.594714992 | -0.250664684 | 0.124326928 | -0.294993373 | 0.204994112 |
| 10541.44336 | 14164.89746 | 11746.20117 | -0.1271 | 0.716159442 | 0.515082725 | 0.174475307 | 0.445543653 | 0.215760001 |
| 60770.62109 | 44322.16797 | 47985.5 | -0.127732151 | 0.40695102 | -0.26836939 | 0.228114774 | -0.779580392 | 0.051785074 |
| 9766.694336 | 9393.799805 | 9438.888672 | -0.127827634 | 0.669212831 | -0.257833365 | 0.478541077 | -0.376559183 | 0.347934694 |
| 57652.06641 | 51727.48438 | 57896.34375 | -0.12832056 | 0.400644934 | 0.67436427 | 0.011710842 | 0.491476136 | 0.053012031 |
| 84518.20313 | 99803.67188 | 96816.8125 | -0.128346551 | 0.316823258 | -0.044157888 | 0.588901453 | 0.158157245 | 0.262261709 |
| 68998.03125 | 69365.77344 | 66574.5 | -0.128455922 | 0.611451298 | -0.134744677 | 0.591184368 | -0.445539398 | 0.191521117 |
| 181883.7969 | 189137.5 | 194294.2813 | -0.128695425 | 0.268099056 | 0.015325575 | 0.88997349 | 0.212011884 | 0.125876501 |
| 26926.48438 | 23771.02148 | 20384.48242 | -0.128910397 | 0.136902686 | -0.224040984 | 0.102825998 | -0.58759145 | 0.053012031 |
| 5613.6875 | 4878.256836 | 5173.890137 | -0.129112865 | 0.439947751 | 0.524866561 | 0.090297888 | 0.08195443 | 0.462677693 |
| 141029.7813 | 141883.75 | 136976.1406 | -0.129929028 | 0.189350067 | -0.492062636 | 0.029364116 | -0.155791398 | 0.034677634 |
| 54344.83594 | 47067.07813 | 41846.32422 | -0.129987327 | 0.358433598 | -0.11757485 | 0.461066015 | -0.552936983 | 0.066458638 |
| 9335.560547 | 9690.878906 | 6604.356445 | -0.13003294 | 0.263777163 | 0.631016209 | 0.177361642 | -0.087634646 | 0.732715412 |

| | | | | | | | | |
|---|---|---|---|---|---|---|---|---|
| 16974.41406 | 19922.3418 | 15262.10352 | -0.130151368 | 0.123324053 | -0.187968398 | 0.068282276 | -0.464282357 | 0.08331162 |
| 193927.2656 | 186085.5 | 181449.875 | -0.130230612 | 0.064338292 | 0.277331845 | 0.109975763 | 0.101360264 | 0.124736435 |
| 1042845.688 | 1050746.125 | 1128308.375 | -0.130328161 | 0.343477561 | 0.186998098 | 0.090297888 | 0.085333062 | 0.233299785 |
| 128684.9063 | 123511.7813 | 122997.8125 | -0.130553878 | 0.271140415 | -0.539814842 | 0.042753324 | -0.32766241 | 0.064495435 |
| 869594 | 922488.5 | 928309.25 | -0.131030259 | 0.297441327 | 0.224660944 | 0.057138779 | 0.486889625 | 0.016683666 |
| 51663.22656 | 53541.28125 | 51765.79297 | -0.131319645 | 0.427465872 | 0.035247104 | 0.849762732 | 0.191420969 | 0.305476348 |
| 21706.42969 | 23552.0332 | 15396.14453 | -0.131412669 | 0.497370191 | -0.86439566 | 0.110734991 | -0.949638129 | 0.062691233 |
| 387607.7813 | 407154.5938 | 326488.4688 | -0.131421492 | 0.144653292 | 0.277272698 | 0.053327615 | 0.164952231 | 0.342470888 |
| 30353.48047 | 31424.26758 | 13959.60742 | -0.131425555 | 0.636523029 | 0.258716019 | 0.378235281 | -0.778109323 | 0.140758419 |
| 3508913.75 | 3814175.5 | 3624527.25 | -0.131770316 | 0.352400798 | -0.246037248 | 0.072104237 | -0.294375458 | 0.048431709 |
| 228428.3281 | 219548.3906 | 214086.3594 | -0.131872692 | 0.408015654 | 0.148402125 | 0.220094258 | 0.056309914 | 0.273802519 |
| 55482.73047 | 61040.27344 | 45032.14844 | -0.132141955 | 0.178701914 | -0.533685564 | 0.048677238 | -0.363287781 | 0.133712681 |
| 85643.65625 | 30678.42773 | 43354.61719 | -0.132260164 | 0.613547565 | 1.038349822 | 0.140163976 | -0.060545873 | 0.934941973 |
| 8815.726563 | 7945.987305 | 9094.80957 | -0.132380133 | 0.409601323 | 0.493634765 | 0.056005515 | 0.441021631 | 0.057068534 |
| 35843.35547 | 36125.14063 | 34451.76563 | -0.132895904 | 0.173510342 | -1.011686427 | 0.01179292 | -1.113288243 | 0.017279709 |
| 276359.1875 | 272831.25 | 292701.875 | -0.132954945 | 0.274068475 | -0.018678869 | 0.881715975 | 0.103995295 | 0.39123981 |
| 111465.8516 | 116033.5078 | 135308.8281 | -0.132968117 | 0.253948329 | -0.0354841 | 0.748766608 | 0.2547589 | 0.181929698 |
| 2218.043213 | 3868.375732 | 2499.382568 | -0.13309035 | 0.700998197 | -0.165359905 | 0.261220712 | -0.33131667 | 0.392418029 |
| 743714.25 | 781765.375 | 784832.5625 | -0.133162143 | 0.106595519 | -0.078859815 | 0.122107272 | 0.030110322 | 0.503548482 |
| 85910.21875 | 81590.21875 | 83121.99219 | -0.133224065 | 0.337393214 | -0.568881506 | 0.083510216 | -0.355159887 | 0.131502216 |
| 137720.25 | 150797.8125 | 158877.0938 | -0.133745957 | 0.132174911 | 0.138485312 | 0.296689492 | -0.079074524 | 0.440211328 |
| 4003.628418 | 8600.186523 | 5885.234375 | -0.133831586 | 0.922117434 | 0.783084647 | 0.498930711 | 0.346747586 | 0.789122922 |
| 26660.71875 | 23410.89063 | 20650.26367 | -0.133836918 | 0.672468428 | -0.162362801 | 0.662482496 | -0.565768718 | 0.260953322 |
| 97894.57031 | 84731.83594 | 93832.88281 | -0.133930424 | 0.548338817 | -0.248512598 | 0.173986552 | -0.413835925 | 0.073664389 |
| 16182.22852 | 16327.0166 | 15381.63184 | -0.134078059 | 0.019930448 | -0.193138364 | 0.023497318 | -0.590800558 | 0.018975627 |
| 29494.4668 | 25805.02344 | 28408.73242 | -0.13422984 | 0.418600089 | 0.937682455 | 0.019141974 | 0.479087279 | 0.051408747 |
| 162515.1875 | 165666.5 | 183774.9531 | -0.135074989 | 0.200137373 | 0.664346209 | 0.01179292 | 0.157550115 | 0.189363824 |
| 120795.1875 | 127734.2422 | 127335.1875 | -0.135308432 | 0.259437282 | 0.020861305 | 0.850477666 | -0.092666023 | 0.445250651 |
| NA | NA | NA | -0.135619354 | 0.576784015 | -0.725029265 | 0.101316342 | NA | NA |
| 37491.83203 | 40637.375 | 41690.09766 | -0.135816532 | 0.820329617 | -0.57340138 | 0.443367839 | 0.023177084 | 0.936804651 |
| 35504.97656 | 32450.55859 | 29976.16211 | -0.135817082 | 0.39188184 | 0.134238909 | 0.426150538 | 0.201024655 | 0.281910706 |
| 9539.220703 | 9934.422852 | 9668.837891 | -0.13601897 | 0.719961518 | 0.527469216 | 0.228822955 | 0.804018431 | 0.122001398 |
| 583538.8125 | 546838.0625 | 414357.7188 | -0.136215748 | 0.229185272 | -0.067371293 | 0.573808561 | -0.28638226 | 0.242076208 |

| | | | | | | | | |
|---|---|---|---|---|---|---|---|---|
| 25831.86719 | 28542.23633 | 27323.61523 | -0.136254386 | 0.597050377 | -0.784616655 | 0.156320824 | -1.338741404 | 0.097319049 |
| 1123556.875 | 1175393.125 | 1222531.625 | -0.136533547 | 0.353923902 | -0.202633839 | 0.15358024 | -0.274946392 | 0.03405215 |
| 11597.9541 | 9607.714844 | 9074.919922 | -0.136682013 | 0.668552376 | -0.492990903 | 0.289832997 | -0.461501772 | 0.291909198 |
| 214018.2031 | 208338.125 | 225987.9844 | -0.137266126 | 0.571651851 | 0.0903295 | 0.732060293 | 0.971863024 | 0.042737273 |
| 1234517.375 | 1520599.625 | 1177702.75 | -0.137319925 | 0.295293871 | 0.224775332 | 0.18586548 | 0.173549827 | 0.421252263 |
| 28298.30273 | 29220.86133 | 25230.07227 | -0.138460997 | 0.125383484 | 0.021729025 | 0.775422699 | -0.315294025 | 0.069955133 |
| 1182034.625 | 1238728.375 | 1261850.25 | -0.138467137 | 0.302262319 | -0.0904866 | 0.417275225 | -0.311267112 | 0.062345205 |
| 190015.5 | 194480.6719 | 166713.1406 | -0.138559141 | 0.217792037 | -0.216182585 | 0.128166006 | -0.413190366 | 0.041088687 |
| 5520.851074 | 7313.13623 | 6387.880371 | -0.138824244 | 0.497232795 | -0.036442027 | 0.711001084 | -0.136187714 | 0.471985509 |
| 130323.2969 | 131409.5625 | 139033.2813 | -0.139132846 | 0.212274306 | 0.040149177 | 0.722366426 | -0.174446853 | 0.181707424 |
| 68739.29688 | 73887.79688 | 51111.625 | -0.13922797 | 0.530888065 | 0.071346055 | 0.760913073 | -0.420897837 | 0.22934165 |
| 26378.58008 | 27178.96875 | 29109.72266 | -0.139284587 | 0.268099056 | -0.170256041 | 0.20225589 | -0.200660242 | 0.137812617 |
| 7630.778809 | 8775.064453 | 11288.11719 | -0.13931188 | 0.81532173 | 1.182054791 | 0.438779269 | 0.336251352 | 0.513062691 |
| 132380.5625 | 126946.7266 | 120017.3125 | -0.139440896 | 0.046319754 | -0.31390471 | 0.047792402 | -0.285991099 | 0.062318926 |
| 14828.20117 | 11550.02539 | 18109.62695 | -0.139483145 | 0.338105128 | -0.059015354 | 0.604317147 | -0.16074429 | 0.563981705 |
| 14486.88379 | 18712.01367 | 16898.60156 | -0.139717469 | 0.164693062 | 0.04176825 | 0.563576269 | -0.104431962 | 0.524048725 |
| 332834.4688 | 332885.8125 | 349167.2188 | -0.140070159 | 0.094486667 | -0.121549251 | 0.15750793 | 0.046852451 | 0.458182767 |
| 5190.867676 | 5270.545898 | 4064.256348 | -0.140146753 | 0.457574834 | -0.490467217 | 0.070723902 | -0.245530552 | 0.245828736 |
| 186465.7813 | 178948.8281 | 184972.1875 | -0.140183758 | 0.023724131 | 0.098953905 | 0.32242293 | 0.078514063 | 0.095057794 |
| 111451.5391 | 113869.7734 | 100772.7344 | -0.140306164 | 0.087075969 | 1.075498038 | 0.014861552 | 0.638012892 | 0.034122066 |
| 275772.625 | 310479.5625 | 328772.5313 | -0.140511978 | 0.527167215 | -0.178083088 | 0.221183294 | -0.160577439 | 0.248833486 |
| 210597.627 | 216815.3555 | 218519.8828 | -0.140571702 | 0.051381763 | -0.378325842 | 0.023005377 | -0.511392309 | 0.022340012 |
| 19927.94727 | 15486.32422 | 18264.41992 | -0.140600399 | 0.07153575 | 0.13454053 | 0.256145059 | -0.583198002 | 0.063886777 |
| 70955.02344 | 74887.96875 | 65769.3125 | -0.141023216 | 0.426576337 | -0.549850289 | 0.105107021 | -0.449013057 | 0.134825131 |
| 10622.46973 | 9950.336914 | 10476.36719 | -0.141064849 | 0.512667848 | 0.467672892 | 0.123184236 | 0.476008872 | 0.120255983 |
| 2279851.5 | 2153711.25 | 2116970 | -0.14128857 | 0.504450935 | -0.024756038 | 0.612877707 | -0.224122815 | 0.049889904 |
| 19356.34961 | 15809.73242 | 12354.77051 | -0.141400023 | 0.415955323 | 0.103046459 | 0.096457635 | -0.031914553 | 0.912221036 |
| 15280.75879 | 14152.46289 | 11380.48047 | -0.141500694 | 0.263777163 | 0.338858519 | 0.096649815 | 0.076362815 | 0.715540356 |
| 147315.4063 | 154776.7813 | 141912.5313 | -0.141596339 | 0.250607381 | 0.326308129 | 0.081503088 | -0.099811768 | 0.432177454 |
| 3251106 | 3442467 | 3284958.5 | -0.141691454 | 0.449917615 | -0.281269393 | 0.088198476 | -0.25800281 | 0.098217186 |
| 48725.98438 | 54348.64844 | 48529.88281 | -0.141971091 | 0.546269698 | 0.060864178 | 0.807058896 | -0.124303715 | 0.639103574 |
| 455631.125 | 779210.625 | 217480.4063 | -0.142055202 | 0.926116021 | -1.158456529 | 0.454547212 | -1.503045819 | 0.382150932 |
| 53092.40625 | 58434.25391 | 57562.53906 | -0.14212577 | 0.619463698 | -0.05263089 | 0.788148361 | -0.074292337 | 0.430149409 |

| | | | | | | | | |
|---|---|---|---|---|---|---|---|---|
| 182395.3906 | 192894.7813 | 168996.4063 | -0.142221945 | 0.218218223 | -1.21526153 | 0.039557441 | -0.909808989 | 0.032218853 |
| 816668.25 | 898775.5625 | 861934.75 | -0.142267134 | 0.425653141 | -0.176522387 | 0.100538317 | -0.241079986 | 0.05183905 |
| 27365.33008 | 24943.16797 | 24364.04102 | -0.14252879 | 0.273969933 | 0.243441724 | 0.052817461 | -0.111475409 | 0.254952757 |
| 416341.2813 | 404139.5625 | 378235.7188 | -0.142646838 | 0.232095578 | 0.344315073 | 0.034038555 | 0.440570485 | 0.026993109 |
| 8362.472656 | 8191.258789 | 6481.230957 | -0.1426935 | 0.398233046 | -0.660189673 | 0.102526216 | -0.58306087 | 0.083206978 |
| 653296.8125 | 711605.125 | 562907.375 | -0.142893108 | 0.307058839 | -0.182889606 | 0.06919378 | -0.309353987 | 0.125291384 |
| 1672126.25 | 1663825.875 | 1608635.75 | -0.142904345 | 0.293152108 | -0.254068405 | 0.144217925 | -0.27712706 | 0.036257728 |
| 6308.609863 | 4313.397461 | 2814.224365 | -0.143024956 | 0.672468558 | 0.317926554 | 0.335331407 | -0.670088991 | 0.222187378 |
| 298283.4375 | 217697.6563 | 289939.2813 | -0.143432516 | 0.20574985 | 0.237562762 | 0.105107021 | 0.097980138 | 0.668657953 |
| 309091.2188 | 281339.5625 | 272416.1563 | -0.143457708 | 0.309938239 | 0.527426151 | 0.063274608 | 0.700608301 | 0.024052583 |
| 209670.1563 | 210957.9688 | 199437.3594 | -0.143478688 | 0.020006619 | 0.14113249 | 0.053543594 | -0.107324252 | 0.083459966 |
| 1147655.75 | 1193791.875 | 1055466.625 | -0.143540646 | 0.370998695 | -0.347339781 | 0.093561793 | -0.093945027 | 0.448958831 |
| 521832.5 | 551497.9375 | 612842.5625 | -0.143583182 | 0.123485909 | -0.284937436 | 0.060352568 | -0.103830293 | 0.385275383 |
| 23038.71875 | 19131.84961 | 10881.9043 | -0.143777919 | 0.472179863 | -0.197088301 | 0.428256136 | -0.658042323 | 0.169861917 |
| 34615.52734 | 35380.35156 | 34762.61328 | -0.144148688 | 0.594194983 | 0.276851424 | 0.35139128 | 0.530905985 | 0.147866982 |
| 23665.97852 | 45414.94141 | 24848.125 | -0.144626057 | 0.812567176 | 0.042697925 | 0.955134985 | 0.146793872 | 0.833116553 |
| 7731.389648 | 8978.994141 | 3383.692871 | -0.144883037 | 0.900501778 | 0.814172757 | 0.54034971 | -0.063236312 | 0.928142182 |
| 6907.018555 | 7662.324707 | 5988.841797 | -0.144911986 | 0.279633164 | -0.175093012 | 0.052437153 | -0.298878589 | 0.148333026 |
| 174114.5938 | 142517.4219 | 157099.1719 | -0.145029218 | 0.223184099 | -0.047279356 | 0.265715011 | -0.137192783 | 0.334732175 |
| 101561.1719 | 94121.42188 | 97871.35938 | -0.145159473 | 0.730519751 | 1.505003155 | 0.200677134 | 2.080571101 | 0.01858328 |
| 27116.08594 | 41354.73438 | 44851.46094 | -0.145269941 | 0.765004512 | -0.258122854 | 0.656902454 | -0.035237001 | 0.954810237 |
| 2710499 | 2775054 | 2540575.25 | -0.145299894 | 0.336682884 | -0.158303416 | 0.228822955 | -0.257703465 | 0.072736493 |
| 48077.55859 | 41725.80078 | 39423.05469 | -0.145548438 | 0.541690293 | -0.214424724 | 0.192313296 | -0.437429619 | 0.066134523 |
| 144988.0313 | 144437.0313 | 138216.9063 | -0.145630965 | 0.207266359 | 0.28135229 | 0.05467585 | 0.127603822 | 0.110032459 |
| 1188653.625 | 1231856.625 | 1271442.25 | -0.14583357 | 0.373059822 | 0.174158495 | 0.31156031 | 0.515741238 | 0.056398361 |
| 47172.74609 | 45635.24609 | 41668.43359 | -0.145867226 | 0.344320467 | -0.500385145 | 0.082096908 | -1.056446292 | 0.00797582 |
| 23641.59375 | 22797.89453 | 21085.71875 | -0.146247508 | 0.226907185 | -0.044701063 | 0.604528039 | -0.298674078 | 0.046487363 |
| 54080.94141 | 52827.90234 | 45691.83203 | -0.146291717 | 0.489383561 | -0.575396797 | 0.142097036 | -0.095361148 | 0.675250595 |
| 151926.6094 | 165032.5156 | 159766.3906 | -0.146351305 | 0.209930575 | -0.054911867 | 0.456419987 | -0.628081848 | 0.016808364 |
| 617144.8125 | 600444.5625 | 569294.125 | -0.146848438 | 0.256554127 | -0.195126668 | 0.087630196 | -0.297786702 | 0.041664392 |
| 55736.58984 | 56349.36328 | 51319.22266 | -0.146958681 | 0.425006009 | -0.413027288 | 0.135015805 | -0.769797405 | 0.072139147 |
| 467744.75 | 477711.0313 | 450052 | -0.147347384 | 0.167150416 | -0.268265137 | 0.118379622 | -0.19441212 | 0.148597902 |
| 26568.28516 | 28332.5 | 27381.17383 | -0.147514224 | 0.058516905 | -0.158882338 | 0.083800868 | 0.289042334 | 0.02527614 |

| | | | | | | | | |
|---|---|---|---|---|---|---|---|---|
| 48507.38281 | 45313.78516 | 22575.18555 | -0.14780534 | 0.485660332 | -0.50692945 | 0.034480785 | -1.488923146 | 0.042737273 |
| 37519.39844 | 37558.86719 | 33144.35938 | -0.147885594 | 0.240263529 | -0.620501692 | 0.074419062 | -0.836234594 | 0.037962113 |
| 25529.30859 | 23005.25977 | 23844.41211 | -0.147943541 | 0.594535741 | 0.096820587 | 0.336933122 | -0.354830529 | 0.045383045 |
| 193215.5781 | 209824.8359 | 215636.0977 | -0.147948355 | 0.598139599 | 0.189203843 | 0.534862174 | 0.126894451 | 0.668752886 |
| 44592.71094 | 53126.82031 | 41730.50781 | -0.148149045 | 0.510558024 | 0.248839866 | 0.240361908 | 0.017616818 | 0.935152011 |
| 12149.20605 | 12605.17188 | 12705.34082 | -0.148345406 | 0.478432564 | -0.135750373 | 0.414751124 | -0.324319919 | 0.16248539 |
| 47279.83203 | 70349.28125 | 49894.66016 | -0.148626924 | 0.737915376 | 0.218579485 | 0.639796921 | 0.080613532 | 0.847639555 |
| 3647.907471 | 4061.415039 | 2899.512939 | -0.148939983 | 0.639033033 | 0.802608942 | 0.048221719 | 0.642876485 | 0.108339667 |
| 35203.39844 | 12452.41504 | 37566.58203 | -0.149055352 | 0.408751663 | 0.662234863 | 0.328806215 | 0.840159302 | 0.366192785 |
| 155941.8281 | 105863.8203 | 55789.77734 | -0.149494298 | 0.846936401 | -0.128463756 | 0.728045745 | -0.972884489 | 0.133339878 |
| 44215.21875 | 28057.35938 | 41023.10938 | -0.149497945 | 0.599725849 | 0.83610017 | 0.065601851 | 0.752777002 | 0.146004119 |
| 64890.42188 | 73335.08594 | 74175.07031 | -0.149574008 | 0.658831436 | 0.524260175 | 0.102454913 | 0.448292588 | 0.117034613 |
| 100115.25 | 103617.6875 | 97930.47656 | -0.149838404 | 0.070027298 | -1.028809041 | 0.031946838 | -0.665394647 | 0.010654184 |
| 31195.61133 | 29450.31641 | 25690.16602 | -0.150119776 | 0.295346854 | -0.404544328 | 0.129412395 | -0.564530773 | 0.063818875 |
| 1090328.375 | 977209.875 | 869115.5625 | -0.150290811 | 0.385226149 | -0.198314404 | 0.14231796 | -0.359170205 | 0.084784384 |
| 15634.2627 | 15817.98242 | 22187.49414 | -0.15116016 | 0.653200607 | -1.469420788 | 0.137745823 | -0.736164653 | 0.215864193 |
| 1270878.625 | 1349732.125 | 1291047.5 | -0.151284474 | 0.011814081 | -0.195048196 | 0.021622074 | -0.265818179 | 0.027629097 |
| 153265.8125 | 214104.8438 | 160734.2188 | -0.151329401 | 0.580758957 | 0.406043146 | 0.192808829 | -0.126134257 | 0.70709851 |
| 47310.44531 | 58460.77344 | 50418.39063 | -0.151834195 | 0.248680662 | 0.515815988 | 0.041910161 | 0.274712313 | 0.181668605 |
| 3846.931885 | 3579.564941 | 3123.932861 | -0.151867462 | 0.482908271 | 1.002770615 | 0.044124904 | 0.232938742 | 0.342106462 |
| 16466.47852 | 22696.42773 | 17334.8418 | -0.151929504 | 0.628378978 | 0.016365038 | 0.953723766 | 0.142456426 | 0.621533646 |
| 189895.7188 | 185687.7188 | 187540.2188 | -0.151937962 | 0.029016212 | -0.001691055 | 0.964457282 | 0.029217901 | 0.225939894 |
| 183169.7656 | 173970.3125 | 193236.2656 | -0.151965511 | 0.16126803 | -0.493706478 | 0.037014778 | 0.119058916 | 0.227026159 |
| 29865.27148 | 31346.39844 | 15205.3457 | -0.151982345 | 0.618124568 | -0.529864087 | 0.287681705 | -0.819470313 | 0.167298956 |
| 90787.45313 | 88691.04688 | 89645.26563 | -0.152075458 | 0.136416657 | 0.773667888 | 0.015625705 | 0.595383232 | 0.021320773 |
| 111649.7188 | 111406.1875 | 103637.4453 | -0.152264812 | 0.173510342 | 0.302067187 | 0.074416318 | 0.159665712 | 0.136713204 |
| 39979.94922 | 42223.87891 | 37730.14844 | -0.15233974 | 0.35633845 | 0.422724997 | 0.090033554 | 0.105497904 | 0.529445312 |
| 26612.33984 | 22769.70898 | 26017.68164 | -0.152340911 | 0.273267781 | -0.027869813 | 0.856414131 | -0.117006313 | 0.453348884 |
| 6552.331055 | 4890.969727 | 6352.532715 | -0.153034129 | 0.749455721 | 0.233446798 | 0.659249191 | 0.420720266 | 0.350816507 |
| 83513.75 | 88140.71875 | 92081.42969 | -0.153197767 | 0.151189017 | 0.110952641 | 0.285390631 | -0.142352271 | 0.218872239 |
| 5715.756348 | 5279.672363 | 3226.291748 | -0.153265092 | 0.701855384 | 0.922462874 | 0.084035804 | 0.485643054 | 0.344155352 |
| 155290.4844 | 162917.0156 | 171954.7188 | -0.15334875 | 0.229185272 | 0.463162257 | 0.113470983 | 0.90547242 | 0.02146481 |
| 25439.32813 | 24335.28516 | 19239.08789 | -0.153416988 | 0.270935076 | -0.241339332 | 0.100046608 | -0.544546709 | 0.060016882 |

| | | | | | | | | |
|---|---|---|---|---|---|---|---|---|
| 623993.375 | 688227 | 633588.75 | -0.154288613 | 0.205630433 | -0.28724427 | 0.053890972 | -0.287069605 | 0.047806561 |
| 11764.71484 | 12816.23242 | 9728.21582 | -0.15432647 | 0.449957266 | -0.769859934 | 0.094689804 | -1.034187163 | 0.057068534 |
| 74743.04688 | 72704.00781 | 68022.16406 | -0.154682467 | 0.35633845 | -0.405918389 | 0.13134246 | -0.339463076 | 0.139075051 |
| 44159.37109 | 39158.42578 | 34201.82031 | -0.154872515 | 0.188470879 | 0.821535349 | 0.063325993 | 0.196705868 | 0.329172546 |
| 44233.72656 | 72417.52344 | 71739.75 | -0.154939218 | 0.896724734 | 0.648081136 | 0.557434542 | 0.233776432 | 0.849688248 |
| 42127.07422 | 45310.16406 | 31037.54688 | -0.155314901 | 0.300368132 | 0.245864307 | 0.165197811 | -0.324585449 | 0.229454106 |
| 3954.463135 | 3429.361328 | 4807.078125 | -0.155374936 | 0.425653141 | 0.107741203 | 0.788148361 | -0.584151532 | 0.068838627 |
| 35828.38672 | 29284.87109 | 23838.52734 | -0.155854452 | 0.348780804 | 0.331851964 | 0.058077004 | 0.36281498 | 0.300973363 |
| 85556.67188 | 84619.03906 | 76922.85156 | -0.156050863 | 0.100751002 | -0.018839648 | 0.809765769 | -0.084517233 | 0.351210463 |
| 96621.10938 | 95149.96094 | 89324.94531 | -0.156224005 | 0.070357633 | -0.122330661 | 0.162505567 | -0.25781835 | 0.039636154 |
| 45881.57813 | 47488.02344 | 42535.21094 | -0.156364113 | 0.101773345 | -0.278318527 | 0.047212306 | -0.441915449 | 0.02183874 |
| 25574.60547 | 23710.57031 | 21618.31055 | -0.156564134 | 0.457079322 | 0.320940532 | 0.125646125 | 0.421002527 | 0.066610782 |
| 209456.3438 | 231132.4688 | 199633.4531 | -0.156863552 | 0.163416464 | -0.243940516 | 0.130198127 | -0.726243947 | 0.026764821 |
| 78354.58594 | 76742.84375 | 65085.39063 | -0.156871437 | 0.267418718 | 0.109671216 | 0.414751124 | 0.159901915 | 0.341019132 |
| 132066.8281 | 125715.125 | 143450.3281 | -0.157654174 | 0.182944903 | 0.622319299 | 0.022035738 | 0.031667038 | 0.79481908 |
| 3350.731934 | 4411.301758 | 4164.992188 | -0.157717751 | 0.161072919 | -0.090372118 | 0.692014727 | -0.424941972 | 0.08767208 |
| 83990.98438 | 89508.63281 | 109163.9609 | -0.157735682 | 0.47099193 | 0.013546318 | 0.961166689 | 1.029447032 | 0.049415261 |
| 76510.66406 | 74550.33594 | 80275.89844 | -0.157784553 | 0.232373511 | -0.016432501 | 0.908389695 | -0.392183746 | 0.041664392 |
| 12965.49707 | 12853.40137 | 13340.47852 | -0.157869273 | 0.278303711 | 0.080045311 | 0.553393539 | 0.216850171 | 0.14336531 |
| 33021.42188 | 30118.26758 | 35373.33984 | -0.157988869 | 0.44400291 | 0.659880212 | 0.024921531 | 0.979182972 | 0.046487363 |
| 4140.566406 | 3948.674072 | 3301.759033 | -0.158300299 | 0.173810033 | 0.266688064 | 0.065871863 | -0.159435274 | 0.329562454 |
| 103901.2734 | 87751.10156 | 89451.44531 | -0.158361977 | 0.171662309 | -0.408477687 | 0.077511889 | -1.136324451 | 0.009824429 |
| 5806.938477 | 7301.213379 | 5901.611328 | -0.158423861 | 0.905320604 | 0.615239544 | 0.616257631 | 0.542078694 | 0.659530356 |
| 11215965.56 | 12875673.56 | 12186454.41 | -0.158693233 | 0.059269869 | -0.364847309 | 0.039320532 | -0.304584179 | 0.052349495 |
| 130387.6875 | 110210.7734 | 136165.5156 | -0.158699009 | 0.343206941 | 0.141603166 | 0.497872234 | 0.066321127 | 0.732453934 |
| 51135.99609 | 55279.97656 | 54103.25391 | -0.158706357 | 0.11973218 | 0.032217493 | 0.691754954 | -0.293633595 | 0.044163517 |
| 90450.9375 | 96054.41797 | 88748.96484 | -0.15902106 | 0.178014687 | -0.149842412 | 0.185362734 | -0.258767139 | 0.070564403 |
| 199888.6563 | 176873.0625 | 171706.3281 | -0.159026794 | 0.277145558 | -0.148138196 | 0.352254195 | -0.29014965 | 0.078304895 |
| 99413.88281 | 104149.2344 | 110357.1328 | -0.159079636 | 0.029906057 | 0.10391267 | 0.094955363 | 0.288868597 | 0.061031191 |
| 38515.625 | 43831.55078 | 57818.20313 | -0.159601321 | 0.825031307 | 0.763027738 | 0.267225595 | 0.795490449 | 0.241451511 |
| 659731.9375 | 661517 | 674396.5625 | -0.15991671 | 0.33278804 | -0.065640339 | 0.730297798 | -0.27751113 | 0.205658715 |
| 1757337.75 | 1674077.375 | 1737129.125 | -0.159919879 | 0.325150883 | -0.106795216 | 0.189272555 | -0.214734622 | 0.041771828 |
| 4969.568848 | 4972.929688 | 4024.855469 | -0.159948065 | 0.420601191 | -0.00189883 | 0.99201113 | -0.403549063 | 0.179416082 |

| | | | | | | | | |
|---|---|---|---|---|---|---|---|---|
| 5459.648438 | 3763.455811 | 7334.086914 | -0.160234534 | 0.716159442 | 0.567198563 | 0.244556831 | 0.13786658 | 0.801376074 |
| 40560.04297 | 40599.92969 | 40011.38281 | -0.160497232 | 0.045302846 | -0.103223384 | 0.100528183 | -0.011789291 | 0.335573935 |
| 52173.11719 | 44314.30859 | 44120.56641 | -0.161619266 | 0.345950084 | 0.035807157 | 0.842416661 | -0.188502514 | 0.370013071 |
| 51589.30078 | 53471.32031 | 36732.70703 | -0.161736133 | 0.696706503 | -0.093067797 | 0.856770955 | 0.431656269 | 0.36278401 |
| 4804.774902 | 4304.119141 | 3754.673584 | -0.161816649 | 0.471667974 | 0.11504774 | 0.631635066 | -0.251448236 | 0.270937884 |
| 315691.7188 | 328055.625 | 311274.5625 | -0.161944716 | 0.191047498 | -0.150773811 | 0.244990627 | -0.150614193 | 0.231806748 |
| 100319.0156 | 110133.4844 | 55792.32422 | -0.162193065 | 0.618124568 | -0.256638003 | 0.463235529 | -0.405025614 | 0.386649359 |
| 26818.91016 | 27025.57617 | 28477.76758 | -0.162206325 | 0.37756775 | -0.700004784 | 0.015625705 | 0.057157684 | 0.276169806 |
| 20563.23633 | 21748.99023 | 19071.32422 | -0.162379016 | 0.365446213 | -0.279845841 | 0.105604495 | -0.569867774 | 0.037582639 |
| 111068.3203 | 95492.32031 | 109700.875 | -0.162394801 | 0.138465855 | 0.185688154 | 0.159217919 | 0.047998364 | 0.703419925 |
| 156844.2344 | 155175.7344 | 152306.125 | -0.16261721 | 0.160731205 | 0.418390243 | 0.064458068 | 0.524893333 | 0.037405698 |
| 16649.40039 | 24710.67578 | 11208.6416 | -0.162757213 | 0.661967827 | 0.309869626 | 0.362329238 | 0.039039085 | 0.948264111 |
| 57873.35547 | 62919.22266 | 66030.33594 | -0.162968869 | 0.425653141 | -0.015571055 | 0.933858968 | 0.15650677 | 0.41499819 |
| 186621.375 | 196192.1563 | 151086.4219 | -0.163013823 | 0.544081454 | 0.245544324 | 0.401539423 | 0.096013916 | 0.752143833 |
| 95063.07031 | 95703.30469 | 91747.61719 | -0.163052977 | 0.334314771 | -0.018862134 | 0.912282749 | -0.133024645 | 0.44852733 |
| 33874.34375 | 30856.45313 | 33981.98828 | -0.163121085 | 0.269574529 | -0.494203689 | 0.114739915 | -1.025099866 | 0.051510832 |
| 436156.875 | 415928.0625 | 429151.25 | -0.163682107 | 0.542409959 | 0.296987838 | 0.333313485 | 0.518101019 | 0.153886288 |
| 18932.4082 | 26610.41406 | 17762.79102 | -0.164242297 | 0.545665828 | 0.097697339 | 0.813471835 | -0.299839566 | 0.32247649 |
| 90005.53125 | 107081.1328 | 109227.8125 | -0.164408377 | 0.615871093 | 0.480405576 | 0.175968875 | 0.258194164 | 0.381068951 |
| 128909.2266 | 146177.9375 | 139811.2031 | -0.164415464 | 0.167779614 | 0.133357284 | 0.264344226 | 0.269880661 | 0.088892469 |
| 702148.125 | 734661.4375 | 748494.875 | -0.164653939 | 0.346314225 | -0.164385119 | 0.205304317 | -0.190553359 | 0.144807901 |
| 61902.00781 | 66331.6875 | 66076.46094 | -0.164847879 | 0.127714789 | -0.010434617 | 0.917056233 | 0.094987576 | 0.313892614 |
| 3644.755127 | 3252.4375 | 3053.726807 | -0.164913497 | 0.464841982 | -0.098479913 | 0.691582507 | -0.21203358 | 0.36727379 |
| 97983.11719 | 84973.07031 | 85512.11719 | -0.164955145 | 0.110302818 | 0.004225286 | 0.966354559 | 0.003163092 | 0.980712992 |
| 207733.3281 | 234878.3281 | 221536.6094 | -0.165043712 | 0.146203541 | -0.079234091 | 0.20864513 | -0.165684819 | 0.123718324 |
| 519945.1563 | 614863.125 | 558679.5625 | -0.165169409 | 0.209563031 | 0.245323585 | 0.132054277 | 0.342387003 | 0.087514859 |
| 6025.224609 | 3748.749023 | 5066.716797 | -0.165188526 | 0.824285485 | 0.838548742 | 0.27329409 | 1.361604127 | 0.095273455 |
| 38922.39063 | 40160.45703 | 44231.97656 | -0.165320805 | 0.06703077 | 0.592130754 | 0.019141974 | 0.255931935 | 0.091619439 |
| 831534.4375 | 797133.25 | 806337.5 | -0.16534986 | 0.040854235 | 0.045526804 | 0.378983428 | -0.170113419 | 0.055019141 |
| 30633.95703 | 27626.46875 | 25744.14453 | -0.165496804 | 0.206331283 | -0.580562193 | 0.05467585 | -0.431527505 | 0.068838627 |
| 194044.7188 | 187629.9688 | 182064.8906 | -0.165631632 | 0.032172412 | 0.032522967 | 0.568203171 | -0.15951359 | 0.047847186 |
| 17625.51172 | 18658.51758 | 17845.11133 | -0.165956994 | 0.079968566 | 0.255804477 | 0.081028372 | 0.796908665 | 0.009014131 |
| 6896.05957 | 5850.599609 | 6063.509277 | -0.166063387 | 0.427440307 | -0.000717315 | 0.996223855 | 0.094745606 | 0.447946456 |

| | | | | | | | | |
|---|---|---|---|---|---|---|---|---|
| 53439.01172 | 49513.35938 | 40728.23438 | -0.166085631 | 0.742611895 | -0.49259309 | 0.052338435 | -0.414444096 | 0.088625686 |
| 50230.92188 | 50179.17969 | 46160.00391 | -0.166204968 | 0.307058839 | 0.497077926 | 0.015639733 | 0.123582483 | 0.167338875 |
| 26001.9707 | 35034.21094 | 42537.78125 | -0.166499837 | 0.811188364 | -1.004150084 | 0.161989613 | -1.20328302 | 0.11327823 |
| 3603.851563 | 6488.061523 | 8081.539551 | -0.166529604 | 0.807202342 | 1.348000853 | 0.167467663 | 1.5213263 | 0.15166044 |
| 281674.0313 | 306839.875 | 291512.5625 | -0.166583796 | 0.098021382 | 0.115261809 | 0.223159833 | 0.054022488 | 0.526508697 |
| 10782124 | 12015159 | 11348021 | -0.166593451 | 0.13878271 | -0.206663613 | 0.167314796 | -0.052045187 | 0.621973931 |
| 33493.14844 | 29635.32617 | 29190.4082 | -0.166612672 | 0.591514625 | 0.375193088 | 0.297241396 | 0.407682189 | 0.246127568 |
| 131060.9219 | 128726.8672 | 129154.1406 | -0.166652584 | 0.250607381 | 0.339694564 | 0.109314009 | 0.290399565 | 0.11743518 |
| 9259.554688 | 7100.754883 | 5064.21875 | -0.167486069 | 0.509688722 | 0.401347243 | 0.170317392 | 0.406264087 | 0.39661161 |
| 14585.63281 | 13490.70801 | 11572.39941 | -0.168108944 | 0.428708346 | 0.01518437 | 0.962732724 | -0.876961425 | 0.049341299 |
| 132461.4219 | 152612.7031 | 148833.0156 | -0.168189134 | 0.146530199 | 0.110960071 | 0.384609578 | 0.307095452 | 0.080462751 |
| 288629.6563 | 295439.1875 | 271459.9688 | -0.168245707 | 0.068444011 | -0.507550405 | 0.020268365 | -0.046751363 | 0.493707745 |
| 67128.63281 | 110461.0547 | 101409.2422 | -0.168354522 | 0.783418048 | 0.543837222 | 0.354607038 | 0.664173433 | 0.196718271 |
| 216141.125 | 217478.375 | 204513.1719 | -0.168464697 | 0.169155397 | -0.20168654 | 0.200654181 | -0.400051134 | 0.081022357 |
| 18048.51563 | 14070.25293 | 13448.75098 | -0.168779942 | 0.450107761 | 0.131214488 | 0.555712947 | -0.403344785 | 0.218790688 |
| 7317.813965 | 6383.903809 | 3572.830811 | -0.168880081 | 0.672802027 | -0.797597585 | 0.276885933 | -0.378159369 | 0.51858543 |
| 6837.563965 | 6321.230469 | 6635.766113 | -0.168894181 | 0.483009333 | 0.020940369 | 0.94481242 | -1.211846241 | 0.083362426 |
| 119265.7813 | 125616.9844 | 134015.8125 | -0.169277087 | 0.555655237 | 0.290548375 | 0.366494549 | 0.0405364 | 0.897135275 |
| 284782.7813 | 290685.4063 | 347853.1875 | -0.169592533 | 0.313971512 | -0.180975429 | 0.336127197 | 0.296074984 | 0.182333546 |
| 90934.63281 | 92906.78125 | 83412.57031 | -0.169671443 | 0.582599493 | 0.522094681 | 0.193402284 | 0.377909186 | 0.276465277 |
| 74480.00781 | 67836.69531 | 67017.44531 | -0.169864797 | 0.192655263 | -0.671264376 | 0.024656861 | -0.920235598 | 0.009026966 |
| 62005.42578 | 75574.57031 | 53372.69531 | -0.169910309 | 0.271759161 | 0.707206849 | 0.203813506 | 0.667434595 | 0.121741414 |
| 91713.4375 | 81445.38281 | 76419.27344 | -0.170291305 | 0.136100201 | -0.301799629 | 0.114739915 | -0.171083874 | 0.249954705 |
| 98603.5 | 95873.72656 | 91114.04688 | -0.170374318 | 0.169155397 | -0.23688127 | 0.019456985 | -0.325042654 | 0.027868741 |
| 62447.06641 | 66094.00781 | 53604.66406 | -0.170472471 | 0.496529012 | 0.499845605 | 0.142579096 | -0.005640028 | 0.986487983 |
| 35864.69141 | 35505.98828 | 35935.90625 | -0.170804341 | 0.575855762 | 0.027214549 | 0.93298262 | -0.378999481 | 0.355778725 |
| 38962.03516 | 39621.14844 | 36416.72266 | -0.170950279 | 0.109633261 | -0.299186155 | 0.105893922 | 0.125575129 | 0.205514355 |
| 149708.7969 | 158720.4375 | 170816.0156 | -0.170985994 | 0.363352652 | -0.039597976 | 0.84398735 | 0.821761712 | 0.027391709 |
| 75387.74219 | 72647.09375 | 63541.71094 | -0.17112749 | 0.173993181 | -0.105869665 | 0.400065651 | -0.274699422 | 0.123837484 |
| 130383.8125 | 148524.3594 | 138472.4688 | -0.17114391 | 0.0885422 | -0.035211711 | 0.702380822 | -0.057440636 | 0.566086724 |
| 32322.97656 | 35715.94531 | 28865.25586 | -0.171308992 | 0.19253889 | 0.268395915 | 0.131263401 | -0.217234804 | 0.202324618 |
| 102110.9375 | 101663.3281 | 95586.46094 | -0.171363987 | 0.204753953 | 0.2927362 | 0.129928099 | 0.319952162 | 0.082507796 |
| 1982454.375 | 2042600 | 1944566.5 | -0.171813684 | 0.326112011 | -0.251572299 | 0.094437122 | -0.28766509 | 0.084613954 |

| | | | | | | | | |
|---|---|---|---|---|---|---|---|---|
| 2785143.25 | 2883606.25 | 2810725.5 | -0.172012048 | 0.337270865 | -0.232741398 | 0.090276742 | -0.215313057 | 0.108589973 |
| 4991.946777 | 9660.172852 | 10101.93262 | -0.172134579 | 0.189747121 | -0.586464891 | 0.073324814 | -0.237952646 | 0.563099327 |
| 38960.05469 | 47363.60156 | 48640.28125 | -0.17226025 | 0.634561612 | 0.5555414 | 0.213309848 | 0.320609757 | 0.413244913 |
| 599785.5625 | 541924.375 | 527382.25 | -0.1725018 | 0.122286474 | 0.283212809 | 0.10267159 | 0.06930029 | 0.523786482 |
| 35037.36328 | 39350.61328 | 38388.24609 | -0.172806892 | 0.201158113 | 0.279909943 | 0.319523239 | 0.015171853 | 0.911447185 |
| 314725.8125 | 328562.625 | 296878.3438 | -0.173574832 | 0.191834923 | 0.091994422 | 0.38622281 | -0.269576884 | 0.079165593 |
| 151188.6563 | 159437.625 | 172038.2969 | -0.173599033 | 0.066425026 | -0.021473946 | 0.674526148 | 0.146812566 | 0.197875324 |
| 506857.5938 | 596584.75 | 539705.6875 | -0.173919649 | 0.515727508 | -0.26484231 | 0.28983684 | -0.217649909 | 0.302853132 |
| 19616.26367 | 10792.93066 | 19500.33789 | -0.174142229 | NA | 0.442662229 | 0.452768719 | 0.786975774 | 0.22434269 |
| 172001.0781 | 163123.9219 | 173761.625 | -0.174235093 | 0.160623983 | 0.227782668 | 0.041058966 | 0.317594271 | 0.02354608 |
| 243641.2813 | 255445 | 259252.25 | -0.174425806 | 0.034151807 | 0.070575555 | 0.182060286 | -0.025099925 | 0.626495487 |
| 146893.875 | 120722.6797 | 123406.4922 | -0.174434666 | 0.49850761 | 0.617679884 | 0.056098132 | 0.308316781 | 0.15573211 |
| 619920.5625 | 679812.875 | 572104.5625 | -0.1746402 | 0.706689159 | -0.355895857 | 0.430750734 | 0.154199449 | 0.660449752 |
| 210687.6094 | 208508.2188 | 216180.5469 | -0.1749984 | 0.121071049 | 0.873075666 | 0.035946996 | 0.629048706 | 0.020337832 |
| 474431.8438 | 658002.3125 | 306370.9375 | -0.175252741 | 0.766532028 | 0.241940033 | 0.775185663 | -0.301501359 | 0.558686385 |
| 135361.9063 | 156937.0469 | 164751.9219 | -0.176081963 | 0.211846788 | 0.033627345 | 0.816812492 | 0.25842241 | 0.168305791 |
| 38785.66797 | 31565.83203 | 35844.51953 | -0.176458967 | 0.55218283 | -0.029022656 | 0.933858968 | -0.016275347 | 0.963827209 |
| 143141.4141 | 146452.3242 | 188048.2617 | -0.176488001 | 0.44400291 | 0.164821446 | 0.395202582 | 0.454938334 | 0.154013101 |
| 16700.00977 | 24064.54102 | 19513.55469 | -0.176679935 | 0.205630433 | -0.319371408 | 0.05610328 | -0.219244944 | 0.370482883 |
| 264728.4375 | 235223.9531 | 237419.25 | -0.176860972 | 0.072606338 | 0.06784637 | 0.428256136 | 0.113770761 | 0.292955254 |
| 35059.61328 | 40363.60156 | 31394.36719 | -0.176951703 | 0.344247041 | -0.446977005 | 0.027730297 | -0.78348898 | 0.027800907 |
| 341466.3125 | 300712.7813 | 334601.8125 | -0.177480195 | 0.26557175 | 0.168917261 | 0.269046934 | 0.058937751 | 0.68269007 |
| 2442.90625 | 2747.923584 | 422.3220825 | -0.177511896 | 0.878992455 | 1.694571097 | 0.143717276 | 0.124633847 | 0.93432622 |
| 21574.4043 | 14046.45215 | 20857.8457 | -0.17757113 | 0.817428054 | 0.83673158 | 0.242349552 | 0.485852465 | 0.486026097 |
| 22899.56641 | 20673.02734 | 22157.75391 | -0.177798001 | 0.17823104 | 0.004202919 | 0.971348973 | 0.053235501 | 0.66247174 |
| 19857.29297 | 16411.62109 | 20223.10742 | -0.177840073 | 0.087656352 | 0.735222387 | 0.021906439 | 0.121252353 | 0.453224015 |
| 44286.84766 | 47188.04297 | 43074.3125 | -0.178440154 | 0.425653141 | -0.042208241 | 0.861195218 | 0.013483363 | 0.957177369 |
| 42463.60547 | 41123.96875 | 37275.09766 | -0.178458401 | 0.040854235 | -0.027252814 | 0.653650751 | -0.181901099 | 0.12131068 |
| 229747.0156 | 274054.5625 | 276157.5 | -0.178615488 | 0.519382561 | 0.799264204 | 0.068499936 | 0.25433196 | 0.370433388 |
| 68787.52344 | 66022.25 | 61284.42578 | -0.178698295 | 0.169155397 | -0.694476714 | 0.054768492 | -0.683597388 | 0.044039968 |
| 27013.17188 | 25212.63281 | 24014.20508 | -0.178710476 | 0.518558018 | 0.128314142 | 0.648824794 | 0.002671697 | 0.988723353 |
| 767474.0625 | 800755.125 | 777206.1875 | -0.17879348 | 0.25396141 | -0.029433329 | 0.788148361 | -0.208054983 | 0.129572086 |
| 20126.56445 | 21944.97461 | 20772.46875 | -0.178808647 | 0.219449972 | 0.117076883 | 0.431158225 | 0.038223325 | 0.733336166 |

| | | | | | | | | |
|---|---|---|---|---|---|---|---|---|
| 2337852.5 | 2505979.75 | 2139968.25 | -0.179028671 | 0.329857534 | 0.312197775 | 0.162317825 | 0.304745823 | 0.148145575 |
| 2566119.75 | 2605139.25 | 2092384.75 | -0.179597982 | 0.392836619 | -0.07156359 | 0.29362323 | -0.384331142 | 0.094349963 |
| 4513.556641 | 3792.537842 | 2866.505371 | -0.179610315 | 0.269696513 | -0.05487998 | 0.687498922 | -0.444512262 | 0.168528497 |
| 15853.72559 | 19797.57227 | 18341.22461 | -0.179833529 | 0.491905235 | -0.092077438 | 0.750780433 | 0.264110627 | 0.368053981 |
| 46990.25781 | 45191.41016 | 46783.70703 | -0.179948916 | 0.097612498 | -0.205267386 | 0.131070383 | -0.140465286 | 0.18892579 |
| 66249.82813 | 75130.125 | 71002.15625 | -0.180595814 | 0.076517108 | 0.183956564 | 0.095325195 | 0.237224768 | 0.089861415 |
| 20473.18359 | 22287.4668 | 21475.8418 | -0.18062928 | 0.458709838 | 0.265380337 | 0.359822419 | -0.311329296 | 0.330416896 |
| 1749442 | 1809211.5 | 1767970.25 | -0.18071529 | 0.379011751 | -0.128943062 | 0.284636711 | -0.249458474 | 0.132019413 |
| 8932.211914 | 12569.62695 | 9989.404297 | -0.180736635 | 0.375208532 | -0.129857682 | 0.532402501 | -0.25434215 | 0.323234781 |
| 87946.59375 | 83976.92969 | 88433.30469 | -0.181103726 | 0.110365189 | 0.037309623 | 0.472254292 | 0.028105336 | 0.5195709 |
| 72054.89063 | 74605.20313 | 65463.05469 | -0.18174339 | 0.086199451 | -0.338935409 | 0.132066965 | -0.158874818 | 0.173821986 |
| 208304.1406 | 190086.4688 | 192761.8438 | -0.181752148 | 0.11900804 | -0.143094338 | 0.256553278 | -0.745239171 | 0.033948107 |
| 92478.90625 | 97607.82031 | 88606.36719 | -0.182525003 | 0.058516905 | -0.184976806 | 0.061296684 | -0.086616754 | 0.258592693 |
| 1353332.586 | 1528067.678 | 1252842.646 | -0.182651109 | 0.433035708 | 0.716882017 | 0.093825034 | 0.68049887 | 0.060352444 |
| 53788.24609 | 53454.53906 | 50079.83594 | -0.183159827 | 0.105014553 | 0.008655646 | 0.935549076 | -0.363682202 | 0.060035964 |
| 33409.97656 | 35208.34375 | 32187.17383 | -0.183216826 | 0.02749963 | 0.077129265 | 0.118990174 | 0.055223323 | 0.395364978 |
| 79804.65625 | 85301.24219 | 82472.3125 | -0.183541809 | 0.17764559 | -1.253067702 | 0.019456985 | -0.719062018 | 0.011546635 |
| 80147.125 | 75808.3125 | 74280.85938 | -0.183936144 | 0.216558281 | -0.331264612 | 0.146003276 | -0.143492667 | 0.317865083 |
| 14510.65234 | 13366.52344 | 11345.61914 | -0.18402442 | 0.213648703 | 0.108288622 | 0.61359516 | -0.579103638 | 0.045353072 |
| 1843371.25 | 1887101.625 | 1951366.125 | -0.184388543 | 0.348637918 | -0.161883552 | 0.137806142 | -0.1546661 | 0.13404473 |
| 89280.03125 | 96791.33594 | 81683.63281 | -0.184846848 | 0.450123638 | 0.520108784 | 0.137284361 | 0.442683301 | 0.060539652 |
| 58834.9375 | 56826.14844 | 54770.35156 | -0.184861476 | 0.151906049 | -0.35102725 | 0.112543356 | -0.46222245 | 0.07008325 |
| 32610.88672 | 33141.18359 | 30000.89648 | -0.185172002 | 0.511583597 | -0.182280782 | 0.139565179 | -0.140455793 | 0.144211186 |
| 13733.47852 | 13517.61035 | 13702.32617 | -0.185355141 | 0.185933663 | 0.789681172 | 0.04632615 | 0.380117435 | 0.018947576 |
| 55374.14844 | 49550.17188 | 49506.14453 | -0.185599383 | 0.155221958 | -0.215576128 | 0.343908073 | 0.54533678 | 0.031006277 |
| 41365.53516 | 39530.55078 | 34713.28125 | -0.185851193 | 0.198998542 | -0.487975011 | 0.031437224 | -0.654531951 | 0.036848746 |
| 41406.56641 | 41369.57031 | 39803.12891 | -0.185872178 | 0.076320591 | 0.134778854 | 0.17063479 | 0.023526555 | 0.752524997 |
| 5359174 | 6159942 | 6164809.5 | -0.186254535 | 0.037600114 | -0.352895665 | 0.052381108 | -0.260377913 | 0.085486138 |
| 39484.85938 | 53756.31641 | 50053.20703 | -0.18628852 | 0.32088092 | 0.546881511 | 0.108508712 | 0.654897771 | 0.10732392 |
| 47188.48828 | 43523.22656 | 44972.35938 | -0.186343083 | 0.09975396 | -0.074484953 | 0.399976168 | -0.353199297 | 0.051942155 |
| 59047.17578 | 52533.66406 | 55096.92578 | -0.186672743 | 0.203813206 | 0.315212704 | 0.105321264 | 0.243585088 | 0.143937923 |
| 8774.707031 | 11255.65527 | 8843.482422 | -0.186783944 | 0.639116447 | 0.090400597 | 0.837478979 | 0.234864645 | 0.573204391 |
| 382829.3438 | 450577.0313 | 419816.0313 | -0.186811209 | 0.139325834 | 0.114339131 | 0.281480596 | 0.295313366 | 0.099088078 |

| | | | | | | | | |
|---|---|---|---|---|---|---|---|---|
| 212057.9844 | 223565.8281 | 195592.9844 | -0.187100489 | 0.088487242 | 0.11541831 | 0.362329238 | -0.145273025 | 0.215760001 |
| 59926.80078 | 64099.99609 | 67892.53125 | -0.187555418 | 0.422079833 | 0.861877902 | 0.0155094 | 1.027159231 | 0.017602156 |
| 40015.78125 | 45556.00781 | 42087.24609 | -0.188082073 | 0.084007167 | 0.123924797 | 0.179553613 | -0.124862995 | 0.24120794 |
| 10515.98828 | 10011.07617 | 12344.87598 | -0.18817657 | 0.593204455 | -0.569907817 | 0.312038217 | -1.00297461 | 0.180002421 |
| 126333.0156 | 134860.5 | 137488.2031 | -0.188684536 | 0.160992897 | 0.188451181 | 0.202293969 | 0.270790109 | 0.094300908 |
| 2401438.25 | 2378056.5 | 2268382.75 | -0.188779107 | 0.222484189 | -0.115613971 | 0.213397043 | -0.243368534 | 0.034933712 |
| 104049.8438 | 110112.0625 | 95792.23438 | -0.188958973 | 0.181247944 | -0.174009261 | 0.119752963 | -0.17399817 | 0.138383382 |
| 73773.60156 | 60418.17969 | 55483.73438 | -0.189549766 | 0.287214289 | 0.308159604 | 0.054768492 | -0.070007426 | 0.720107197 |
| NA | 3081.830078 | 1917.890503 | -0.189550581 | NA | 0.928584811 | NA | -0.293694985 | NA |
| 15227.42383 | 13315.57715 | 8660.035156 | -0.189899933 | 0.139016894 | -0.399630653 | 0.068043648 | -0.484111552 | 0.201758531 |
| 1108168 | 1173414.25 | 1050070.75 | -0.19020225 | 0.282923851 | 0.135294914 | 0.440308034 | -0.122719892 | 0.50761246 |
| 32730.5625 | 25839.29102 | 26969.14258 | -0.190336557 | 0.363436072 | 0.281683086 | 0.38067054 | 0.864841632 | 0.05170067 |
| 13509.65137 | NA | 7969.057617 | -0.190898239 | NA | 0.253706542 | NA | 0.119349996 | NA |
| 68918.57813 | 62721.88672 | 61873.26953 | -0.190918856 | 0.18862527 | 0.284272564 | 0.636663202 | -0.382835621 | 0.047120928 |
| 81781.8125 | 42000.57813 | 55398.02344 | -0.191199047 | 0.49449467 | -0.118118846 | 0.685920313 | -0.378518886 | 0.410415738 |
| 25949.82422 | 28773.40039 | 23525.0625 | -0.191297074 | 0.147715547 | -0.740270099 | 0.03968601 | -0.746255045 | 0.027974321 |
| 2106997 | 2265551.5 | 2144137.5 | -0.191619993 | 0.372499466 | -0.205283103 | 0.114706646 | -0.166633242 | 0.118870803 |
| 98079.25781 | 88415.76563 | 95731.67969 | -0.191709893 | 0.076923702 | -0.141753093 | 0.274788966 | 0.260343844 | 0.064726471 |
| NA | 2185.021729 | NA | -0.192030361 | NA | 0.413331504 | NA | -0.521372612 | NA |
| 1462433.945 | 1642404.484 | 1514621.492 | -0.192210322 | 0.117342707 | 0.047371321 | 0.698341031 | 0.162986452 | 0.19608824 |
| 37413.21094 | 40512.26172 | 33513.66797 | -0.192320407 | 0.355407665 | -0.346477167 | 0.255836673 | -0.630361625 | 0.118033234 |
| 62947.61328 | 61018.72266 | 61532.89844 | -0.192496844 | 0.281702994 | 0.69210349 | 0.079086322 | 0.523769066 | 0.023619478 |
| 50232.08984 | 41300.29297 | 36354.12109 | -0.19255988 | 0.585026867 | -0.414497836 | 0.399249134 | -0.217947107 | 0.609208541 |
| 53782.37109 | 56536.53516 | 42049.39063 | -0.192957733 | 0.233167433 | -0.307754525 | 0.128915752 | -0.256382199 | 0.251283986 |
| 173878.0313 | 171087.5 | 179159.9844 | -0.193077957 | 0.036566441 | 0.190106265 | 0.058077004 | 0.164796221 | 0.060161253 |
| 1110039.375 | 1139500.75 | 1155257.375 | -0.193189706 | 0.282301731 | 0.578399796 | 0.031437224 | 0.878118185 | 0.016795677 |
| 65631.41406 | 58382.73438 | 61653.76953 | -0.193193728 | 0.099307987 | 0.063387047 | 0.336862505 | -0.01884882 | 0.804302828 |
| 102313.4766 | 103465.1797 | 97816.96094 | -0.193529251 | 0.143772566 | 0.060486402 | 0.577045972 | -0.278061574 | 0.128529572 |
| 33580.41406 | 26676.13477 | 31718.80469 | -0.193581396 | 0.56717271 | 0.113582096 | 0.742577273 | -0.205193211 | 0.231478811 |
| 23044.24609 | 24233.26367 | 12303.6875 | -0.193582198 | 0.210059063 | -0.608674458 | 0.08642513 | -0.673780064 | 0.148445737 |
| 196412.3906 | 188481.2656 | 205260.4219 | -0.193749463 | 0.201158113 | 0.068382007 | 0.611432555 | 0.90500211 | 0.015916823 |
| 32737.08984 | 33606.32422 | 39328.74609 | -0.193751179 | 0.058955365 | 0.071563219 | 0.534862174 | 0.395146195 | 0.106029703 |
| 4235.061523 | 5522.055176 | 3428.281494 | -0.19375282 | 0.73774346 | 0.413301819 | 0.466150869 | 0.167729502 | 0.786883312 |

| | | | | | | | | |
|---|---|---|---|---|---|---|---|---|
| 65019.44531 | 60183.82813 | 61490.22656 | -0.193805016 | 0.037492489 | 0.124816809 | 0.217458834 | -0.113386666 | 0.119708622 |
| 522297.875 | 561382.4375 | 556913.5 | -0.194500098 | 0.048963509 | -0.246341592 | 0.015007028 | -0.149715205 | 0.085169708 |
| 12474.20313 | 11739.87402 | 9780.476563 | -0.194612207 | 0.809643539 | 0.134899215 | 0.878943291 | 0.18530929 | 0.829646836 |
| 1635.116699 | 2137.369873 | 988.3890991 | -0.194700387 | 0.661967827 | 0.219877368 | 0.606467062 | -0.51543363 | 0.405062284 |
| 350338.125 | 365711.2188 | 374765.4688 | -0.194722826 | 0.047621263 | -0.104984237 | 0.34315805 | 0.071212766 | 0.298365177 |
| 207562.7031 | 214503.1406 | 224683.75 | -0.194773469 | 0.138598181 | 0.023785624 | 0.821333494 | 0.220668661 | 0.112528358 |
| 111066.7188 | 103110.0625 | 103393.1094 | -0.195041104 | 0.131501491 | -0.622931541 | 0.041554876 | -0.901037759 | 0.025120016 |
| 61521.82813 | 62480.52734 | 50927.28125 | -0.195247702 | 0.085283871 | 0.072894343 | 0.423507188 | -0.18350589 | 0.253685901 |
| 585511.8125 | 632151.0625 | 578681.9375 | -0.195325139 | 0.077183684 | -0.724838008 | 0.032113568 | -0.39669501 | 0.030727383 |
| 17175.87891 | 16669.4707 | 17067.85742 | -0.195832862 | 0.066639151 | -0.56814103 | 0.038609344 | -0.769972419 | 0.002683747 |
| 207059.9219 | 212419.9531 | 186290.5938 | -0.19586998 | 0.023401535 | -0.012328206 | 0.903524394 | -0.054270097 | 0.544786561 |
| 1720037.375 | 1224844.25 | 727275.375 | -0.196281308 | 0.404242885 | 1.572417182 | 0.541442962 | -0.481631855 | 0.335259007 |
| 182750.8594 | 169531.2969 | 173287.1563 | -0.196409445 | 0.025968675 | -0.0280766 | 0.642450324 | -0.230130268 | 0.034567592 |
| 78106.53125 | 78576.2832 | 80755.15234 | -0.196867595 | 0.249779609 | 0.172133338 | 0.334251946 | 0.017282196 | 0.916018251 |
| 1672518.5 | 1755544 | 1676515.375 | -0.197442314 | 0.379283654 | -0.122280661 | 0.582697111 | 0.107828383 | 0.573204391 |
| 2835450 | 2797151.75 | 2801227.5 | -0.197489311 | 0.088240896 | -0.018418681 | 0.62182213 | 0.482869124 | 0.016395134 |
| 7659.774902 | 8439.402344 | 8247.057617 | -0.197578737 | 0.498323734 | 0.013810734 | 0.967452872 | 0.301893356 | 0.311546035 |
| 58066.42578 | 57728.77734 | 58774.91406 | -0.19781733 | 0.080487571 | 0.285299796 | 0.089400459 | 0.006421016 | 0.862115276 |
| 4089.009033 | 4874.198242 | 3336.345947 | -0.197822361 | 0.313971512 | -0.082221042 | 0.715505426 | -0.205517427 | 0.467315264 |
| 238181.4375 | 199786.8906 | 217131.4219 | -0.197835888 | 0.282301731 | -0.116971286 | 0.310169864 | 0.078534915 | 0.528935183 |
| 45849.89844 | 31334.12305 | 37389.03906 | -0.19825183 | 0.536107129 | 0.413098879 | 0.195850577 | 0.175223443 | 0.586713414 |
| 60803.17969 | 59300.58984 | 60593.66016 | -0.198368717 | 0.031790847 | -0.089604583 | 0.280400081 | 0.096203645 | 0.107554917 |
| 49521.32813 | 47369.99219 | 47966.25391 | -0.198376454 | 0.025960199 | -0.070642228 | 0.506362982 | -0.378632888 | 0.01858328 |
| 250255.7344 | 262551.75 | 266118.0625 | -0.19856295 | 0.069242208 | -0.489319847 | 0.012254503 | -0.525855436 | 0.009026966 |
| 80834.32031 | 77277.28906 | 72571.59375 | -0.198882726 | 0.113193912 | -0.314629939 | 0.111083106 | -0.436849324 | 0.055298525 |
| 45444.69531 | 38599.45313 | 41454.10547 | -0.199021956 | 0.20270675 | -0.213349095 | 0.258368613 | -1.041090234 | 0.045495106 |
| 641110.6934 | 792201.166 | 692597.2676 | -0.199232515 | 0.649999812 | -0.282167193 | 0.467378475 | -0.034836665 | 0.916489217 |
| 15504.4043 | 15272.1582 | 10243.71777 | -0.199404669 | 0.208223043 | -0.145984828 | 0.37194326 | -0.342241357 | 0.255236048 |
| 66319.52344 | 59503.79688 | 61876.85938 | -0.199419141 | 0.482181614 | -0.248715452 | 0.422801 | -0.171279006 | 0.52834849 |
| 10756.57813 | 11313.81152 | 10776.22949 | -0.199449101 | 0.556071856 | 0.388771463 | 0.319523239 | 0.005604731 | 0.989076637 |
| 3636.314453 | 4024.518799 | 3520.952148 | -0.199969384 | 0.403414633 | -0.215094217 | 0.457214055 | -0.30016381 | 0.335259007 |
| 40657.28516 | 41011.85938 | 40280.89063 | -0.20002683 | 0.14905608 | -0.374308144 | 0.103986734 | -0.226464982 | 0.185694198 |
| 725458.125 | 709110.875 | 664997.6875 | -0.20010348 | 0.538777426 | -0.011579524 | 0.941386226 | -0.011502305 | 0.93888999 |

| | | | | | | | | |
|---|---|---|---|---|---|---|---|---|
| 166579.3906 | 137318.7031 | 127264.0156 | -0.200242513 | 0.271312801 | 0.673639193 | 0.017037525 | 0.372037259 | 0.179875711 |
| 25058.31445 | 30979.60156 | 26323.02539 | -0.200615437 | 0.308512841 | 0.032114615 | 0.88001899 | -0.07591819 | 0.680795082 |
| 49131.28125 | 53148.42188 | 43555.0625 | -0.200684269 | 0.178859684 | -0.227989349 | 0.238316696 | -0.742014592 | 0.043034683 |
| 20840.85547 | 22796.64258 | 19132.80078 | -0.201139205 | 0.427658072 | -1.15464017 | 0.082380201 | -0.485219446 | 0.136837137 |
| 33662.72266 | 33181.02344 | 23789.25195 | -0.201147537 | 0.132178167 | 0.186428923 | 0.179171247 | -0.167135003 | 0.489401961 |
| 365302.625 | 403858 | 410882.4063 | -0.201204281 | 0.059431981 | -0.027379758 | 0.716062229 | -0.023334924 | 0.795695086 |
| 11340.57129 | 13034.3457 | 11383.2334 | -0.201754113 | 0.332739002 | -0.397178604 | 0.108760394 | -2.158383483 | 0.035414333 |
| 646550.75 | 674037.75 | 593235.0625 | -0.201829983 | 0.207247298 | -0.144864089 | 0.343628392 | -0.2853238 | 0.053012031 |
| 91887.30469 | 85584.41406 | 73902.16406 | -0.201919318 | 0.060903498 | -0.166907446 | 0.159703167 | -0.025899311 | 0.861343486 |
| 235760.7813 | 211973.0469 | 167791.4531 | -0.201934941 | 0.66215868 | 0.272309936 | 0.725384008 | -0.347869896 | 0.481097195 |
| 1308685 | 1263307.375 | 1205095.75 | -0.202032323 | 0.32746691 | -0.072943632 | 0.60094054 | -0.227528285 | 0.178411577 |
| 76945.28906 | 72999.26563 | 64809.77734 | -0.202232748 | 0.142913384 | 0.067681859 | 0.511095278 | -0.262214188 | 0.109624066 |
| 4982.445313 | 6965.514648 | 6924.37207 | -0.202238519 | 0.316823258 | -0.40646342 | 0.09742076 | -0.681085132 | 0.055868946 |
| 72740.21875 | 77683.10938 | 75904.41406 | -0.202308997 | 0.181549839 | 0.398996066 | 0.152134335 | -0.316092022 | 0.027949491 |
| 137938.1563 | 137087.9688 | 119644.2344 | -0.202362671 | 0.103153657 | 0.002190313 | 0.9694773 | -0.503826015 | 0.029219266 |
| 37663.24219 | 45016.77734 | 43394.83203 | -0.202375663 | 0.125639042 | 0.020859483 | 0.881155563 | -0.097754734 | 0.489464179 |
| 23783.98438 | 23290.74023 | 26814.46484 | -0.202407361 | 0.434950944 | 0.074556086 | 0.747804994 | -0.3119356 | 0.290630021 |
| 105259.4844 | 113757.4609 | 112038.9766 | -0.202722538 | 0.133097116 | 0.213745933 | 0.13524054 | -0.037746142 | 0.731910614 |
| 270426.3438 | 260156.375 | 262313.5 | -0.202782659 | 0.57007183 | 0.053242171 | 0.893894927 | -0.261535751 | 0.54164829 |
| 119177.6875 | 132889.3438 | 107252.4297 | -0.202858153 | 0.079342211 | -0.360850208 | 0.050288747 | -0.622072897 | 0.029057465 |
| 10653.76465 | 11055.90918 | 5341.547363 | -0.203116949 | 0.553857303 | -0.089941864 | 0.806649717 | -0.4712941 | 0.371690724 |
| 526370.9375 | 561499.3125 | 540625.8125 | -0.203280036 | 0.062445147 | 0.078115685 | 0.205952446 | 0.10805325 | 0.121525007 |
| 67024.39844 | 68440.17969 | 67191.84375 | -0.203336399 | 0.028903558 | -0.122703712 | 0.066504374 | -0.173140465 | 0.047387434 |
| 860300.625 | 858557.125 | 851193.5 | -0.203803465 | 0.172104341 | -0.307398038 | 0.138103908 | -0.282411641 | 0.169075369 |
| 207931.7031 | 238880.0781 | 212035.125 | -0.204392583 | 0.07362475 | -0.067508668 | 0.428011055 | -0.295842151 | 0.069955133 |
| 21676.9375 | 30614.43359 | 25073.7168 | -0.204445586 | 0.530730634 | 1.317998188 | 0.114455867 | 0.983668375 | 0.069955133 |
| 40062.67578 | 33887.22656 | 38457.33594 | -0.204632955 | 0.157585334 | 0.082571899 | 0.572760293 | -0.279429945 | 0.133712681 |
| 176500.625 | 171991.5625 | 175396.1719 | -0.204639709 | 0.097443336 | 0.058159632 | 0.087781599 | -0.006046151 | 0.796833165 |
| 589748.3242 | 573681.4863 | 623416.4629 | -0.204828999 | 0.583542363 | 0.192917979 | 0.628277537 | 0.087433241 | 0.827301412 |
| 380507.7188 | 375764.8125 | 379218.6875 | -0.204906127 | 0.025818052 | -0.152720345 | 0.08165067 | 0.123301903 | 0.099555681 |
| 13431.77734 | 17193.4375 | 18974.89844 | -0.205353524 | 0.272917388 | 0.560272663 | 0.104928475 | 0.735665978 | 0.100069721 |
| 26982.81445 | 29822.93555 | 28760.63086 | -0.205617788 | 0.561642224 | 0.344456265 | 0.371275269 | 0.251161723 | 0.492365146 |
| 29569.35547 | 29937.4707 | 24270.96484 | -0.205688781 | 0.260535434 | 0.036342794 | 0.834435409 | 0.127641333 | 0.537160122 |

| | | | | | | | | |
|---|---|---|---|---|---|---|---|---|
| 351324.3125 | 332391.4063 | 296885.0313 | -0.205703327 | 0.285678451 | 0.21757366 | 0.254631118 | 0.186959591 | 0.317312079 |
| 106856.6484 | 115681.2031 | 91712.77344 | -0.205835001 | 0.39571103 | 0.614935359 | 0.096721331 | -0.037012865 | 0.872413573 |
| 14576.90332 | 12615.64063 | 12038.62109 | -0.205881906 | 0.102924346 | -0.313712656 | 0.052702293 | -0.312609727 | 0.09843293 |
| 206595.8125 | 197757.2656 | 179989.9844 | -0.205969774 | 0.131546744 | 0.468615963 | 0.066380875 | 0.37698574 | 0.05747179 |
| 21282.72656 | 17285.43164 | 16495.56641 | -0.206188256 | 0.369140023 | 0.187559694 | 0.417577766 | -0.183119216 | 0.500573086 |
| 12861.51074 | 9373.984375 | 11009.89844 | -0.206366771 | 0.506889993 | -0.171730986 | 0.634960279 | 0.312793407 | 0.249456378 |
| 63018.93359 | 58205.62109 | 56813.80469 | -0.206387513 | 0.118081327 | -0.477061348 | 0.041554876 | -0.671382847 | 0.010654184 |
| 1184166.25 | 1165810.875 | 1382116.125 | -0.206512246 | 0.199046058 | -1.352477271 | 0.051593867 | -0.633068952 | 0.056151195 |
| 18691.51758 | 21357.43555 | 12803.83691 | -0.206785729 | 0.436807126 | -0.588687313 | 0.215436149 | -0.848144924 | 0.118298395 |
| 3799.115479 | 3402.1604 | 3662.357666 | -0.206829109 | 0.224955599 | -0.064157389 | 0.703106698 | -0.233345304 | 0.254919095 |
| NA | 5372.925781 | 3194.958984 | -0.206853535 | NA | -0.473653584 | NA | 0.037370767 | NA |
| 27013.59375 | 34170.28516 | 43103.93359 | -0.206996843 | 0.128742961 | 0.279403864 | 0.096609145 | 0.134147272 | 0.671185498 |
| 105635.6563 | 100900.9766 | 70612.64844 | -0.20725761 | 0.096367624 | -0.228399247 | 0.050817911 | -0.473857945 | 0.151776851 |
| 24518.76953 | 26203.36133 | 26749.125 | -0.207514299 | 0.085294501 | -0.012634232 | 0.903478941 | -0.160537799 | 0.149964889 |
| 232824.4844 | 223504.75 | 230984.4063 | -0.207890876 | 0.019286562 | -0.010099454 | 0.904355047 | -0.237922913 | 0.017279709 |
| 175551.2813 | 188258.3906 | 195752.2188 | -0.208242648 | 0.256206147 | 0.172262581 | 0.371192256 | -0.018024895 | 0.923992665 |
| 6122.986328 | 6791.675781 | 14494.43457 | -0.208352785 | 0.595858443 | -0.400480692 | 0.279661196 | 0.555284754 | 0.503055359 |
| 61138.67969 | 62615.10547 | 50216.45313 | -0.20893355 | 0.066179601 | -0.147291277 | 0.143786228 | -0.696119159 | 0.027800907 |
| 20373.45313 | 16386.625 | 17149.5625 | -0.20923411 | 0.517401645 | -0.091986658 | 0.814025375 | -0.242030201 | 0.54168178 |
| 36909.98047 | 39861.07813 | 33575.19141 | -0.209596589 | 0.176121329 | 0.184496145 | 0.189753768 | -0.223884143 | 0.178181856 |
| 8183.918457 | 8053.722168 | 7266.336914 | -0.209672432 | 0.149076522 | -0.227348676 | 0.202852155 | -0.107694369 | 0.443995111 |
| 74529.70313 | 107256.5234 | 86085.92188 | -0.210067985 | 0.675025166 | -0.784629108 | 0.147508999 | -0.692442087 | 0.154835571 |
| 254404.4844 | 236211.6719 | 242534.0156 | -0.210082974 | 0.088903515 | -0.284065594 | 0.091622784 | -0.154802263 | 0.178198454 |
| 7927.30957 | 9686.004883 | 7083.256348 | -0.210122899 | 0.279048922 | 0.299335277 | 0.229462977 | -0.41529084 | 0.167212148 |
| 50035.55078 | 54904.44141 | 46385.51953 | -0.210193843 | 0.197255448 | 0.065118083 | 0.669480124 | -0.029233183 | 0.863987358 |
| 670890.8125 | 674167.5 | 653372.5625 | -0.210210828 | 0.078894657 | 0.30054464 | 0.051630099 | 0.383235033 | 0.041609196 |
| 8588.401367 | 11035.80957 | 7012.837891 | -0.21049845 | 0.374464563 | 0.163776276 | 0.480676341 | -0.034941188 | 0.917547988 |
| 12633.12695 | 8505.088867 | 7515.449707 | -0.210804625 | 0.176820059 | 0.56949588 | 0.08303186 | -0.268468835 | 0.453772169 |
| 22468.07813 | 23671.56641 | 7755.14209 | -0.211189043 | 0.312774598 | -0.387665081 | 0.07692885 | -0.743125527 | 0.233022905 |
| 42336.74609 | 44825.05078 | 44384 | -0.211356407 | 0.191047498 | 0.576246819 | 0.046285147 | 0.328982408 | 0.060939295 |
| 121083.4922 | 130154.8281 | 130315.4766 | -0.211746582 | 0.35077204 | -0.13381485 | 0.591184368 | -0.793233891 | 0.120554018 |
| 162717.7969 | 149609.3594 | 144154.6406 | -0.211946034 | 0.085313321 | 0.065031834 | 0.483916014 | -0.186751299 | 0.153274064 |
| 44145.17969 | 43072.8125 | 47112.08984 | -0.212081051 | 0.111508367 | 0.118102843 | 0.136651367 | 0.143898652 | 0.126741146 |

| | | | | | | | | |
|---|---|---|---|---|---|---|---|---|
| 536149.6875 | 704344.1875 | 595377.5625 | -0.212117073 | 0.566478626 | 0.040243367 | 0.930769393 | -0.141905221 | 0.654410188 |
| 264347.0313 | 262262.625 | 274655.0313 | -0.21213776 | 0.293892903 | 0.07651316 | 0.702808705 | 0.207390203 | 0.328978319 |
| 79496.85938 | 86192.64844 | 83309.01563 | -0.212248441 | 0.063885064 | 0.009107715 | 0.907792135 | -0.15744953 | 0.08376499 |
| 780871.25 | 803542.25 | 639945.25 | -0.21245822 | 0.295023996 | -0.075687949 | 0.612877707 | -0.256970308 | 0.171126239 |
| 735.8850098 | NA | 436.4188538 | -0.21251725 | NA | 0.551785478 | NA | -1.344103821 | NA |
| 374890.1953 | 402001.8672 | 330886.0625 | -0.212564861 | 0.64719338 | 0.117995512 | 0.630524877 | 0.131989586 | 0.546828747 |
| 6526.976563 | 6227.495605 | 9049.803711 | -0.212883143 | 0.366847091 | 0.120873155 | 0.644844761 | -0.789424026 | 0.070883677 |
| 49814.95703 | 44451.53516 | 47645.98828 | -0.213085237 | 0.066932774 | -0.062470082 | 0.380869244 | -0.293011704 | 0.070422955 |
| 459579.0625 | 410403.0313 | 366967.5625 | -0.21339681 | 0.14678023 | -1.046986879 | 0.023005377 | -0.511185134 | 0.046624125 |
| 43794.67969 | 45949.91797 | 38651.40234 | -0.213740307 | 0.036566441 | 0.281717732 | 0.029364116 | -0.021061608 | 0.85680206 |
| 10141.43457 | 10825.73633 | 9516.026367 | -0.214030862 | 0.268547366 | 0.409541821 | 0.109653398 | -0.064559352 | 0.73671998 |
| 67579.10156 | 62064.67969 | 64702.62109 | -0.214221458 | 0.178783758 | -0.260136648 | 0.166004104 | -0.34961947 | 0.108225817 |
| 227391 | 234045.6875 | 212575.2656 | -0.214390246 | 0.231197087 | -0.241364682 | 0.274179898 | -0.653615507 | 0.091523165 |
| 184281.9844 | 178454.1406 | 164164.0469 | -0.214497861 | 0.108697689 | -0.345611913 | 0.01915096 | -0.015536576 | 0.840966234 |
| 44449.19531 | 43494.52734 | 38995.72266 | -0.214644241 | 0.040928921 | -0.064799967 | 0.364459731 | -0.365664398 | 0.038826476 |
| 56864.375 | 56427.39063 | 54375.32031 | -0.214911538 | 0.160229633 | 0.542732261 | 0.046373997 | 0.531614139 | 0.028162772 |
| 2874.533447 | 4728.338867 | 3502.163574 | -0.214957154 | 0.607604225 | -1.110510097 | 0.204401609 | -1.24747826 | 0.167751522 |
| 19826.55859 | 25057.62891 | 25853.65234 | -0.215603168 | 0.137237024 | 0.176882141 | 0.220100738 | 0.078344115 | 0.696855946 |
| 570407.1914 | 639645.9375 | 625381.7656 | -0.215665351 | 0.118973844 | 0.248005713 | 0.080473926 | 0.058485766 | 0.547919535 |
| 463294.75 | 521788.5938 | 495412.1563 | -0.215671267 | 0.038502747 | -0.344669244 | 0.041391338 | -0.232949108 | 0.067864002 |
| 17839.33203 | 23582.90039 | 26735.93359 | -0.216209785 | 0.293345911 | -0.272446777 | 0.191836121 | 0.1240303 | 0.660836387 |
| 170011.375 | 159480.3281 | 162686.125 | -0.216646009 | 0.122673099 | -0.142378757 | 0.318095673 | -0.270929908 | 0.137952392 |
| 101729.8516 | 98005.88281 | 106875.2813 | -0.217136671 | 0.110302818 | -0.81062475 | 0.033121608 | -0.243781355 | 0.062261743 |
| 32105.9043 | 34137.62891 | 32687.84766 | -0.217609229 | 0.26296677 | -0.050190347 | 0.804097821 | -0.305205081 | 0.237901119 |
| 55407.85547 | 56896.94922 | 49372.71875 | -0.217754574 | 0.46081773 | 0.616256182 | 0.039856414 | 0.723973042 | 0.028311992 |
| 399686.1563 | 366158.75 | 399681.0938 | -0.217816607 | 0.232373511 | 0.676435887 | 0.054857325 | 0.293343201 | 0.124444567 |
| 4428.971191 | 2969.024658 | 2511.138916 | -0.217910481 | 0.437231181 | 0.145697012 | 0.588676506 | 0.175600218 | 0.688854424 |
| 294030.875 | 400292.8125 | 338946.4375 | -0.218010527 | 0.32400401 | -0.394612055 | 0.035895043 | -0.10494072 | 0.591863123 |
| 5523.379395 | 6088.356934 | 4924.213379 | -0.218023492 | 0.643599796 | 0.578824885 | 0.279342103 | -0.007162093 | 0.989480447 |
| 95854.83594 | 85334.99219 | 80889.78906 | -0.218430788 | 0.066545395 | -0.211807768 | 0.123694369 | -0.199128251 | 0.167225632 |
| 17775.6582 | 25098.60547 | 12102.36719 | -0.218481007 | 0.644698558 | -0.735642536 | 0.32238411 | -0.958421587 | 0.234646777 |
| 342968.5195 | 445417.1016 | 298095.5469 | -0.218588662 | 0.573636658 | 0.052503743 | 0.885440998 | -0.261972969 | 0.44621387 |
| 17892.20508 | 19072.04492 | 18959.38086 | -0.218756692 | 0.145258682 | -0.305456659 | 0.117428051 | 0.065909353 | 0.553786226 |

| | | | | | | | | |
|---|---|---|---|---|---|---|---|---|
| 62927.05469 | 57602.59375 | 60965.25781 | -0.218851858 | 0.040012377 | 0.366824916 | 0.121785014 | 0.407188512 | 0.025666912 |
| 6032.317383 | 7050.502441 | 1809.254028 | -0.218914997 | NA | -0.23244134 | NA | 1.476155435 | NA |
| 3015588.25 | 2655862.25 | 2274020 | -0.219349731 | 0.168226751 | -0.1184866 | 0.479482194 | -0.433007937 | 0.102230154 |
| 969039.4375 | 1021172.438 | 1011542.375 | -0.219857724 | 0.299113805 | -0.171549084 | 0.078311743 | -0.220992631 | 0.041088687 |
| 197467.5 | 224953.7813 | 234340.3438 | -0.219941381 | 0.173573283 | 0.126556163 | 0.381306421 | 0.245364802 | 0.170776308 |
| 10259.94336 | 11600.28906 | 7456.916504 | -0.220261319 | 0.250607381 | -0.663750321 | 0.098228912 | -0.9332871 | 0.048613954 |
| 52362.77344 | 52858.84375 | 45969.625 | -0.220350684 | 0.15673669 | 0.519599737 | 0.0395519 | 0.091779206 | 0.512645993 |
| 152139.2969 | 158012.9844 | 143026.0625 | -0.220438552 | 0.078758366 | 0.266590041 | 0.229545075 | -0.385699153 | 0.056784127 |
| 4938.495117 | 2018.245117 | 4389.086426 | -0.221454707 | 0.301405522 | 0.410981122 | 0.294325744 | 0.237687135 | 0.680027467 |
| 86886.89844 | 106326.1484 | 82023.24219 | -0.221547258 | 0.53534498 | 0.228434856 | 0.511953424 | -0.393129727 | 0.379731306 |
| 58071.74219 | 61396.9375 | 55356.25 | -0.221587462 | 0.029571758 | -0.650248179 | 0.024404888 | -0.348297479 | 0.028275238 |
| 685316.5 | 807183.625 | 737769.8125 | -0.221642914 | 0.318687987 | -0.246416852 | 0.179460228 | 0.050467978 | 0.731636574 |
| 34941.31641 | 33344.08203 | 31169.43945 | -0.221659992 | 0.110726532 | 0.383610082 | 0.048952672 | -0.145177643 | 0.222819528 |
| 59667.94531 | 55508.99219 | 62576.62109 | -0.221712752 | 0.160583048 | -0.318987977 | 0.147941386 | -0.132129729 | 0.399002054 |
| 12127694 | 14277598 | 13218781 | -0.221778043 | 0.134341857 | -0.117532996 | 0.365797071 | 0.053162734 | 0.700196087 |
| 3990.709473 | 3951.621094 | 3822.170166 | -0.221871423 | 0.456092971 | 0.534026694 | 0.068436129 | 0.222602306 | 0.277551095 |
| 82262.6875 | 75592.15625 | 85229.66406 | -0.222402133 | 0.159989357 | -0.037381066 | 0.61104577 | -0.000869857 | 0.992689539 |
| 49116.23828 | 46191.91016 | 52978.59766 | -0.222486853 | 0.155102978 | 0.672810171 | 0.097043528 | 0.942116847 | 0.020008995 |
| 17676.51953 | 17439.55859 | 17298.93164 | -0.222591182 | 0.206331283 | 0.283192994 | 0.053827713 | 0.380377537 | 0.034567592 |
| 2299991.75 | 2633147 | 2478961.75 | -0.222815647 | 0.273267781 | -0.17961636 | 0.221012263 | -0.188244662 | 0.192622934 |
| 24712.89063 | 21527.54102 | 24937.00195 | -0.222864534 | 0.100743514 | 0.312681605 | 0.091981596 | 0.134448913 | 0.338432733 |
| 376222.875 | 632041.4375 | 486735.9688 | -0.223060332 | 0.052850388 | -0.117070284 | 0.287066192 | -0.055814318 | 0.863987358 |
| 49456.03125 | 45542.0625 | 47032.16797 | -0.223124017 | 0.142526436 | -0.081723801 | 0.52544407 | -0.093240146 | 0.469205269 |
| 67781.34375 | 66315.66406 | 61796.17578 | -0.223203124 | 0.361920409 | 0.56755632 | 0.078617016 | 0.163963519 | 0.384463529 |
| 130352.7656 | 139903.2813 | 197728.6563 | -0.223534982 | 0.512681345 | 0.037951398 | 0.930297693 | -0.083381621 | 0.852856834 |
| 57133.19141 | 52656.32813 | 54204.53906 | -0.224121553 | 0.127230164 | 0.047631618 | 0.660062477 | -0.16557307 | 0.231918617 |
| 767081.5 | 774117.5625 | 900992.375 | -0.224442835 | 0.118081327 | 0.359687995 | 0.05917214 | 0.531371896 | 0.05611292 |
| 12348.88281 | 20105.07227 | 14546.25 | -0.22463396 | 0.767960924 | 0.289812487 | 0.71517609 | 0.320250579 | 0.69482592 |
| 384222.6563 | 282260 | 250505.75 | -0.22498041 | 0.029659089 | 0.386804325 | 0.06919378 | 0.153667602 | 0.621198226 |
| 6703.018066 | 8286.272461 | 4278.177246 | -0.225825528 | 0.061954848 | 0.009429102 | 0.935126799 | -0.310782332 | 0.426923454 |
| 58540.29688 | 59874.73828 | 56419.00391 | -0.225996489 | 0.056529322 | 0.159628803 | 0.124641637 | -0.001427531 | 0.98733347 |
| 11543.77051 | 10204.39453 | 9280.957031 | -0.226273548 | 0.438905184 | 0.231513729 | 0.45827938 | -0.343243141 | 0.370397499 |
| 38665.44922 | 43096.82422 | 40236.19922 | -0.226326844 | 0.231588073 | -0.327323634 | 0.057189248 | 0.250168439 | 0.070587266 |

| | | | | | | | | |
|---|---|---|---|---|---|---|---|---|
| 789100.1875 | 788794.3125 | 779865.3125 | -0.226335225 | 0.065355995 | 0.04866544 | 0.486895725 | -0.284418107 | 0.049415261 |
| 233453.6563 | 233541.8125 | 236205.875 | -0.226425349 | 0.209084852 | -0.107616522 | 0.569398849 | 0.403388618 | 0.110388055 |
| 31216.32617 | 33377.98438 | 31941.24023 | -0.226443544 | 0.010893509 | -0.152478012 | 0.030863246 | -0.10435835 | 0.113248184 |
| 72070.9375 | 65661.42969 | 78214.94531 | -0.226525571 | 0.600673217 | 0.628670682 | 0.178579756 | 0.132435102 | 0.620264646 |
| 572354.8125 | 609773.75 | 556869.375 | -0.22677651 | 0.042260333 | -0.570605268 | 0.018539883 | -0.311833901 | 0.029653052 |
| 10877.12109 | 10357.19434 | 6147.032715 | -0.227102473 | 0.21684615 | 0.315386818 | 0.318820359 | -0.09629487 | 0.791930121 |
| 239024.5625 | 263256.1875 | 265246.3125 | -0.227453324 | 0.085283871 | 0.230176685 | 0.088587863 | -0.233249255 | 0.08720771 |
| 213176.9844 | 215423 | 195936.5469 | -0.227541196 | 0.06561751 | 0.17633747 | 0.212397647 | -0.352854746 | 0.054887084 |
| 82032.03125 | 81228.9375 | 70933.88281 | -0.227572696 | 0.047058169 | -0.015468828 | 0.869754532 | -0.398186327 | 0.040786554 |
| 102954.5 | 105125.1016 | 103174.3594 | -0.227847794 | 0.036071458 | -0.023338287 | 0.250296981 | -0.297485679 | 0.008614663 |
| 87900.67188 | 90804.17969 | 91058.35156 | -0.22797089 | 0.041353498 | 0.045998753 | 0.50345588 | -0.110060143 | 0.135230932 |
| 70222.74219 | 72454.19531 | 69685.41406 | -0.228730501 | 0.098201119 | -0.043917889 | 0.062911841 | -0.123579769 | 0.04728688 |
| 2870.156738 | 3584.892578 | 1726.219116 | -0.228866532 | 0.489291353 | -0.854926536 | 0.183896983 | -1.360219731 | 0.092375143 |
| 36956.91406 | 38929.80469 | 35654.71094 | -0.229300812 | 0.522531517 | 0.020145082 | 0.833630996 | -0.242634609 | 0.08853367 |
| 155372.4219 | 124865.7656 | 133590.5938 | -0.229565373 | 0.171600786 | -1.285530208 | 0.011710842 | -0.632785288 | 0.04014804 |
| 7509.171875 | 6927.737305 | 8575.306641 | -0.230428076 | 0.407278175 | -0.342310621 | 0.093818896 | 0.132433405 | 0.432375389 |
| 79291.92188 | 91673.50781 | 85315.0625 | -0.230616815 | 0.131380074 | 0.045403033 | 0.488412384 | -0.03019534 | 0.765845177 |
| 74526.36719 | 68121.24219 | 70206.94531 | -0.230906481 | 0.145796786 | 0.620051512 | 0.091622784 | 0.316437244 | 0.036834039 |
| 51334.17188 | 38688.71875 | 29345.14258 | -0.230912938 | 0.374464563 | 0.255779652 | 0.314954562 | -0.039939113 | 0.923534132 |
| 143158.5313 | 152748.25 | 150616.7656 | -0.230937221 | 0.182007594 | 0.517303694 | 0.065516721 | 0.778905915 | 0.02527614 |
| 17488.29492 | 16031.28906 | 15265.76758 | -0.231749399 | 0.293152108 | 0.619397276 | 0.040974624 | 0.274395196 | 0.119386672 |
| 6489.453125 | 5165.647461 | 4034.732422 | -0.231831541 | 0.411547567 | 0.33197164 | 0.24184923 | -0.143411496 | 0.672091123 |
| 16625.49023 | 17220.97266 | 17217.16211 | -0.232177539 | 0.200146696 | 0.02975026 | 0.85722804 | -0.070493365 | 0.674614707 |
| 561.8756104 | 619.4343262 | 237.4462585 | -0.23218337 | 0.298673579 | 0.648236969 | 0.111801734 | -0.458703601 | 0.389796538 |
| 1161613 | 1202979.5 | 1068410.875 | -0.232365751 | 0.135340524 | -0.141947766 | 0.098031162 | -0.302913944 | 0.042860672 |
| 129785.3174 | 128424.2488 | 131800.6277 | -0.232611417 | 0.031883977 | -0.571461082 | 0.030767136 | -0.731906506 | 0.011733679 |
| 129466.0391 | 128251.375 | 134810.4219 | -0.232848593 | 0.138906239 | 0.22980892 | 0.092617517 | 0.649808006 | 0.017279709 |
| 12853.87793 | 13364.90137 | 14415.3125 | -0.232880186 | 0.645989146 | 0.83696896 | 0.165197811 | 1.133132291 | 0.095785034 |
| 104800.7656 | 101919.0781 | 95971.58594 | -0.232914893 | 0.120063669 | -0.251712492 | 0.162505567 | -0.545008215 | 0.06710618 |
| 57522.90234 | 63459.45313 | 61137.625 | -0.233040381 | 0.04950513 | -0.101698752 | 0.162317825 | 0.293585635 | 0.047110629 |
| 28570.94531 | 26064.77539 | 24415.95508 | -0.233090535 | 0.105481009 | -0.183324595 | 0.243784441 | -0.269337856 | 0.125742768 |
| 890499.8594 | 869796.6016 | 882195.5391 | -0.233613379 | 0.293152108 | -0.107835898 | 0.639466845 | 0.142846084 | 0.508589617 |
| 317016.6563 | 342729 | 358911.8438 | -0.233891697 | 0.032929334 | 0.577554107 | 0.013427556 | 0.233877933 | 0.090965556 |

| | | | | | | | | |
|---|---|---|---|---|---|---|---|---|
| 1349215.875 | 1382616.5 | 1357544.125 | -0.234173668 | 0.103160411 | 0.309249713 | 0.084714823 | 0.151657635 | 0.246773877 |
| 136731.3438 | 141118.25 | 129838.9453 | -0.234229573 | 0.06853067 | -0.027711802 | 0.6139644 | 0.238544935 | 0.047760658 |
| 658602.75 | 761868.1875 | 656860.4375 | -0.234308746 | 0.15673669 | -0.087708884 | 0.642450324 | 0.083488666 | 0.575112472 |
| 61646.78516 | 53121.67969 | 50996.09766 | -0.234372243 | 0.095292766 | -0.045558938 | 0.680816532 | -0.336663282 | 0.084126995 |
| 132622.5156 | 131821.2969 | 136276.6563 | -0.234419421 | 0.116159237 | -0.310800481 | 0.051593867 | 0.236923825 | 0.032187838 |
| 23627.23438 | 26944.80078 | 25684.43164 | -0.234678371 | 0.366951692 | 0.591368418 | 0.098031162 | 0.312120071 | 0.248833486 |
| 1306937.375 | 1470714.25 | 1299300.625 | -0.234705272 | 0.596932465 | 0.29440773 | 0.512809933 | 0.537392032 | 0.231024496 |
| 32508.01367 | 34116.75 | 30908.86328 | -0.23477825 | 0.109154846 | 0.060046014 | 0.504715483 | -0.160773285 | 0.101487706 |
| 494112.5938 | 461215.9063 | 460266.875 | -0.234794076 | 0.073146457 | -0.331536585 | 0.045329621 | -0.348682598 | 0.030007011 |
| 26264.07422 | 26892.44141 | 33426.91797 | -0.234910098 | 0.096750655 | -0.279799941 | 0.101048965 | -0.012634686 | 0.950160109 |
| 127607.5313 | 113836.3203 | 120118.0156 | -0.23516791 | 0.337114746 | 0.474109246 | 0.234571805 | 0.360857583 | 0.061706361 |
| 880993.0625 | 952102.3125 | 901833.3125 | -0.23644573 | 0.181549839 | -0.116265315 | 0.321230521 | -0.323393274 | 0.082184581 |
| 137582.2813 | 133763.0781 | 148257.9063 | -0.236629469 | 0.028138974 | 0.185069042 | 0.22425544 | 0.206549253 | 0.089383058 |
| 246964.0469 | 238965.4375 | 95030.92969 | -0.236981442 | 0.412298442 | -0.091469661 | 0.61125836 | -0.277225507 | 0.598883912 |
| 310877.1563 | 306255.875 | 324396.125 | -0.237126163 | 0.024409946 | -0.00349035 | 0.949204616 | 0.072800469 | 0.225939894 |
| 17151.67969 | 8197.501953 | 12278.15918 | -0.237160391 | 0.248932045 | 0.067895797 | 0.751895789 | 0.242911106 | 0.631690532 |
| 19339.23438 | 22354.60547 | 23555.53906 | -0.237276322 | 0.140783489 | 0.290384218 | 0.118955051 | 0.360136516 | 0.107554917 |
| 27762.69531 | 28189.58594 | 22432.60352 | -0.237464708 | 0.216922584 | 0.215595625 | 0.280400081 | -0.052878522 | 0.807833796 |
| 108797.1953 | 123899.1875 | 107004.9844 | -0.237677146 | 0.588172074 | 0.176458715 | 0.355229313 | 0.123287116 | 0.377533593 |
| 110599.1953 | 115388.1875 | 58708.96094 | -0.237755531 | 0.743069818 | 0.449834291 | 0.583822478 | -0.376493554 | 0.419154696 |
| 29985.21289 | 27152.38867 | 26367.06641 | -0.237878946 | 0.366967951 | -0.330506567 | 0.351392721 | -0.651855461 | 0.171907805 |
| 185933.6406 | 200413.5 | 210296.7031 | -0.237897933 | 0.060865769 | 0.154362133 | 0.127902424 | 0.178950864 | 0.134102681 |
| 20864.50195 | 18821.84375 | 15617.33691 | -0.238278791 | 0.146753968 | -0.072235876 | 0.340354928 | -0.059166919 | 0.749389169 |
| 24899.67773 | 13483.90234 | 20563.23828 | -0.23834242 | 0.17113456 | -0.210024871 | 0.265989276 | -0.427659234 | 0.279591992 |
| 121248.5234 | 115372.6641 | 109540.8594 | -0.23839823 | 0.067030699 | 0.090146509 | 0.11146465 | -0.139312279 | 0.122439097 |
| 12066.83691 | 12854.36426 | 11733.11328 | -0.238460087 | 0.016471012 | -0.26781173 | 0.052036261 | -0.166043344 | 0.080462751 |
| 20953.30859 | 19469.33594 | 21249.55664 | -0.238559383 | 0.118081327 | 0.005644085 | 0.979550955 | 0.046197499 | 0.713487699 |
| 4430.416992 | 4711.487305 | 2229.029053 | -0.238651576 | 0.69044279 | 1.609965685 | 0.322217412 | 0.233820216 | 0.703811131 |
| 407030.375 | 403974.5625 | 375672.9688 | -0.239456593 | 0.55770286 | 0.337890751 | 0.447621797 | 0.050593977 | 0.898117002 |
| 332669.375 | 348202.625 | 339085.0625 | -0.239755842 | 0.055951111 | -0.276294046 | 0.081994347 | -0.500620804 | 0.048118422 |
| 48576.18359 | 42458.05078 | 42655.88672 | -0.239793366 | 0.052921297 | -0.274017268 | 0.055080662 | -0.234075836 | 0.101012637 |
| 25276.2168 | 33786.03516 | 33502.26953 | -0.239960013 | 0.418600089 | 0.161332659 | 0.649274833 | 0.279284971 | 0.287182177 |
| 1330531.25 | 1442472 | 1349628.5 | -0.240465345 | 0.200171429 | 0.039164383 | 0.797128652 | 0.083997677 | 0.573238987 |

| | | | | | | | | |
|---|---|---|---|---|---|---|---|---|
| 466722.3438 | 462526.0313 | 444587.1563 | -0.24094436 | 0.113205271 | 0.502546892 | 0.069086457 | 0.710379735 | 0.02183874 |
| 7218.041992 | 7774.682617 | 178.9876404 | -0.241271966 | 0.246138931 | -1.457176519 | NA | -1.367447008 | 0.137062098 |
| 121805.3594 | 124178.6953 | 123587.3359 | -0.241411996 | 0.015685639 | -0.067920994 | 0.127892721 | -0.10196934 | 0.063283918 |
| 2896.547119 | NA | 1819.428223 | -0.241426584 | 0.598058346 | -0.339987345 | 0.531589481 | -1.24600385 | NA |
| 12389.83008 | 14345.75 | 11948.57324 | -0.241707524 | 0.229992532 | 0.251353731 | 0.26312545 | -0.198428341 | 0.386628982 |
| 24423.52539 | 25699.20117 | 24496.67773 | -0.241817479 | 0.112787536 | 0.68458903 | 0.027730297 | 0.280826136 | 0.107077213 |
| 31146.40625 | 29844.71875 | 29055.79492 | -0.241952726 | 0.178228644 | 0.29030302 | 0.046568753 | -0.141051039 | 0.137229118 |
| 232730.25 | 245607.1719 | 231226.25 | -0.2420266 | 0.540500239 | 0.534840261 | 0.242042707 | 0.536465905 | 0.239787303 |
| 28278.06055 | 30412.04297 | 35371.65234 | -0.242105798 | 0.250607381 | 0.42959039 | 0.06015933 | -0.31207058 | 0.128425855 |
| 8383.850586 | 8503.612305 | 5718.358398 | -0.242431266 | 0.275379351 | -0.733221438 | 0.045744926 | -0.368205399 | 0.23061805 |
| 1883808.625 | 2021721.125 | 1635632.125 | -0.242925326 | 0.297902313 | -0.207468894 | 0.108875676 | -0.375484083 | 0.070298355 |
| 1336268.75 | 1344659.25 | 1312665.125 | -0.243048352 | 0.036590352 | 0.300213321 | 0.065986072 | 0.485293355 | 0.021134419 |
| 86715.5 | 88400.72656 | 79597.92969 | -0.24316798 | 0.194423598 | -0.081445672 | 0.623797104 | -0.30506573 | 0.185596413 |
| 454634.4688 | 469972.0625 | 395130.6875 | -0.243350927 | 0.538373638 | -0.28925047 | 0.094800581 | -0.138006818 | 0.318972136 |
| 102432.9844 | 99161.97656 | 105004.7734 | -0.243497402 | 0.093909622 | 0.506097779 | 0.048820023 | 0.691551038 | 0.02183874 |
| 36071.72656 | 39181.69141 | 38860.3125 | -0.243943597 | 0.088231376 | -0.035537515 | 0.724710676 | -0.129103954 | 0.287586066 |
| 2537.038574 | 3125.064697 | 4152.509277 | -0.244034974 | NA | -0.460291896 | 0.330821995 | 0.224801396 | 0.596651466 |
| 13872.78711 | 12396.13672 | 12567.40234 | -0.244232434 | 0.066744288 | -0.604863648 | 0.024656861 | -0.636445233 | 0.016395134 |
| 10769.9209 | 11850.0332 | 8814.146484 | -0.244339628 | 0.139542297 | -0.608312721 | 0.047212306 | -0.806524351 | 0.029120377 |
| 102084.1875 | 101248.6641 | 89546.29688 | -0.244408293 | 0.072654291 | 0.366550649 | 0.055605274 | 0.246791705 | 0.112639617 |
| 51926.26953 | 60062.63281 | 49697.60156 | -0.244844719 | 0.302378035 | 0.157945532 | 0.192808829 | -0.364158619 | 0.085047269 |
| 139222.2656 | 133891.0781 | 133142.3438 | -0.244885227 | 0.074349462 | 0.688655639 | 0.047984032 | 0.311061268 | 0.068838627 |
| 24165.25391 | 21450 | 19527.61133 | -0.24490661 | 0.092522848 | 0.007362243 | 0.933440383 | -0.013271003 | 0.931040293 |
| 534591.8125 | 540423.7031 | 553883.9063 | -0.244952942 | 0.036590352 | 0.032465087 | 0.539179679 | 0.081357323 | 0.198621682 |
| 5081.567871 | 8656.047852 | 6479.870605 | -0.245163009 | NA | -0.18658853 | 0.769923035 | -0.175623534 | 0.782285106 |
| 449390.0625 | 448739.25 | 421426.0938 | -0.245356565 | 0.061707718 | 0.418358045 | 0.04183385 | -0.006023588 | 0.950211597 |
| 610208.375 | 604677.875 | 589707.5 | -0.245488156 | 0.288353039 | 0.057622437 | 0.626415094 | -0.032536533 | 0.758543242 |
| 12390.47852 | 12430.53906 | 9895.956055 | -0.245735629 | 0.263608427 | -0.15519049 | 0.531076327 | -0.230606378 | 0.381968648 |
| 148695.9844 | 141616.7344 | 146726.2344 | -0.246066031 | 0.026259973 | -0.166510884 | 0.104557189 | -0.141850097 | 0.098742683 |
| 63126.93359 | 61928 | 58199.42188 | -0.246233616 | 0.359435166 | -0.036792645 | 0.89718243 | 0.120528099 | 0.634317573 |
| 10061.50391 | 8982.975586 | 11447.79492 | -0.246269788 | 0.15260666 | -0.186140705 | 0.355286655 | 0.079768481 | 0.686180113 |
| 184103.0938 | 187748.2656 | 174456.8906 | -0.24652346 | 0.040946717 | 0.011522921 | 0.737531604 | -0.190545202 | 0.05183905 |
| 42363.47266 | 39019.82813 | 37766.32422 | -0.246642231 | 0.060865769 | 0.021004916 | 0.808781142 | -0.342331165 | 0.040202519 |

| | | | | | | | | |
|---|---|---|---|---|---|---|---|---|
| 108874.9688 | 97295.82813 | 97958.85938 | -0.247162463 | 0.200189061 | -0.126995817 | 0.343878142 | 0.162938701 | 0.231126418 |
| 1415987.82 | 1423045.164 | 1414123.219 | -0.247163824 | 0.155524283 | 0.490545617 | 0.052015428 | 0.32779879 | 0.045887602 |
| 23961.375 | 24318.71289 | 22328.42969 | -0.247170502 | 0.109833334 | 0.012061639 | 0.918393626 | -0.254288897 | 0.129496281 |
| 69049.92969 | 63030.39844 | 55828.50781 | -0.247387112 | 0.239569883 | -1.100456886 | 0.071542761 | -0.80303826 | 0.061900682 |
| 66551.95313 | 66140.17188 | 71398.94531 | -0.24751661 | 0.117342707 | 0.19264362 | 0.059562109 | 0.175395361 | 0.080462751 |
| 77093.46875 | 67731.83594 | 72567.17188 | -0.247662262 | 0.086970437 | -0.095218508 | 0.428256136 | -0.128837425 | 0.331779341 |
| 25368.86523 | 28780.48828 | 24358.94922 | -0.247666158 | 0.36684241 | -0.335719012 | 0.350889153 | -0.115087896 | 0.695364867 |
| 230078.8594 | 237063.5938 | 248768.6875 | -0.248154325 | 0.224585525 | 0.26231212 | 0.244072263 | 0.261941499 | 0.22860933 |
| 3657.624268 | 2233.125732 | 3255.689209 | -0.24828208 | 0.340758192 | 0.782880387 | 0.015625705 | -0.480424231 | 0.170158914 |
| 14912.48535 | 16943.89844 | 16590.9043 | -0.248524298 | 0.200881846 | 0.25128423 | 0.228751244 | 0.204238814 | 0.289235979 |
| 60242.73438 | 54676.71094 | 62039.55469 | -0.248651143 | 0.232509252 | -0.045917607 | 0.821991128 | 0.413180175 | 0.105718826 |
| 46231.47656 | 46458.67969 | 41160.30078 | -0.248742331 | 0.038502747 | 0.166195986 | 0.108793898 | -0.318180623 | 0.046745056 |
| 15586.12402 | 13277.93457 | 11694.60742 | -0.249012788 | 0.56477738 | -0.309668882 | 0.31462727 | -0.597316109 | 0.130422289 |
| 56832.39844 | 59940.65625 | 49357.28516 | -0.249323692 | 0.080448585 | 0.291952855 | 0.080227549 | -0.320726996 | 0.095208361 |
| 38198.79199 | 34915.39355 | 43379.9873 | -0.249336482 | 0.13220281 | 0.617314116 | 0.04128058 | 0.528042883 | 0.072903408 |
| 71299.84375 | 78294.72656 | 83188.55469 | -0.249459867 | 0.179094466 | 0.21208602 | 0.214021621 | 0.236340902 | 0.137554403 |
| 2996.825684 | 4735.873535 | 4074.781738 | -0.249599519 | 0.175753206 | 0.611834747 | 0.31156031 | -1.139576372 | 0.026525799 |
| 25984.25781 | 24840.53906 | 22947.45898 | -0.24959965 | 0.070426297 | 0.010270839 | 0.951517411 | -0.257248944 | 0.096439544 |
| 14736.95703 | 18542.07813 | 11945.03125 | -0.249641411 | 0.557161628 | -0.299166189 | 0.559568799 | -1.73531765 | 0.110128352 |
| 4735.230957 | 4467.482422 | 3439.743164 | -0.249752718 | 0.43923187 | 0.182361314 | 0.550128659 | -1.104198346 | 0.032510751 |
| 109741.1406 | 115233.75 | 102226.1875 | -0.250016055 | 0.195683117 | 0.050253534 | 0.767672979 | -0.174167199 | 0.379385206 |
| 311445.0313 | 308761.875 | 307039.3125 | -0.250383556 | 0.062999512 | -0.165677967 | 0.189599836 | 0.104533563 | 0.301801908 |
| 137840.5781 | 133907.4219 | 131788.8438 | -0.250654939 | 0.104597563 | 0.257738589 | 0.132973237 | 0.154040597 | 0.240008368 |
| 26799.79883 | 24099.47461 | 21590.10938 | -0.250706319 | 0.422996137 | -0.005709697 | 0.986379847 | -0.043423181 | 0.896281252 |
| 215427.4063 | 226146.8438 | 173106.8438 | -0.250738963 | 0.356124734 | 0.229094086 | 0.419859836 | 0.009728 | 0.977605725 |
| 1924703.5 | 1910250.875 | 1799138.625 | -0.250739849 | 0.168043685 | -0.16663855 | 0.152314783 | -0.276905805 | 0.0236713 |
| 293254.8623 | 297070.6084 | 294398.4609 | -0.250741826 | 0.095278437 | -0.019939737 | 0.762408602 | -0.362736927 | 0.011394883 |
| 38666.17188 | 38621.73047 | 42392.24219 | -0.251050383 | 0.127811678 | 0.13777277 | 0.407857231 | 0.601806666 | 0.02925669 |
| 1624207.875 | 1917286.375 | 1808934.625 | -0.251220427 | 0.142526436 | -0.130761658 | 0.410302997 | -0.078432261 | 0.617419313 |
| 247284.1563 | 264303 | 223789.1406 | -0.25128856 | 0.443831691 | 0.353099272 | 0.104714354 | -0.050396795 | 0.760384076 |
| 67571.09375 | 71268.34375 | 69945.78125 | -0.251344788 | 0.06941189 | -0.106950613 | 0.296076847 | -0.113053961 | 0.260059332 |
| 19285.55859 | 20695.67578 | 19873.74219 | -0.251664107 | 0.088240896 | -0.393925448 | 0.093751816 | -0.977963824 | 0.032577228 |
| 81191.46094 | 74660.04688 | 74420.23438 | -0.252490297 | 0.178834579 | -0.544020598 | 0.127083816 | -0.39003316 | 0.153803943 |

| | | | | | | | | |
|---|---|---|---|---|---|---|---|---|
| 34508.91016 | 33257.35547 | 30146.17578 | -0.252757301 | 0.195683117 | 0.067620909 | 0.360638437 | -0.177747641 | 0.145702448 |
| 23024.80469 | 18117.55078 | NA | -0.253077454 | 0.428496066 | -0.425390201 | 0.60094054 | -0.127903574 | NA |
| 19291.9707 | 16328.19043 | 13521.80273 | -0.253106737 | 0.02548578 | 0.074431461 | 0.552697805 | 0.120833739 | 0.61132384 |
| 276739.75 | 267380.6875 | 275851.8125 | -0.253182175 | 0.071114381 | -0.194736406 | 0.203650142 | -0.398439198 | 0.050138206 |
| 88334.78906 | 70479.05469 | 83420.32813 | -0.253562712 | 0.151101744 | -0.083971276 | 0.195590467 | -0.713917427 | 0.030633392 |
| 7664.42627 | 9797.950195 | 7827.557617 | -0.253926279 | 0.182007594 | -0.333289887 | 0.135908153 | -0.574868278 | 0.059847517 |
| 5859.319336 | 6033.684082 | 4695.342773 | -0.254142646 | 0.334444005 | 0.35279437 | 0.02887273 | -0.346940891 | 0.137581058 |
| 13460.74219 | 10456.81055 | 9279.859375 | -0.254471601 | 0.06853067 | 1.021801569 | 0.089652719 | 0.915891094 | 0.115401194 |
| 153675.6563 | 129789.7188 | 142425.8438 | -0.254928123 | 0.050437184 | 0.088423072 | 0.168802204 | -0.208417517 | 0.140342819 |
| 177829 | 168713.2813 | 196912.0156 | -0.255028671 | 0.136100201 | -0.083281749 | 0.586104878 | -0.313191479 | 0.145702448 |
| 101400.8359 | 108293.4688 | 119681.9609 | -0.255213994 | 0.358433598 | 0.433342171 | 0.198924075 | 0.115291452 | 0.677060914 |
| 2270.367676 | 2257.739746 | 2000.651978 | -0.255234545 | 0.625931855 | 0.059487193 | 0.922225068 | -0.135386317 | 0.827943664 |
| 1313060.125 | 1211006 | 1244875.625 | -0.255377122 | 0.203513529 | -0.077065321 | 0.415649015 | -0.264642451 | 0.082022058 |
| 411472.0938 | 420455.1875 | 442854.5625 | -0.25539525 | 0.019930448 | -0.190129078 | 0.065601851 | -0.322739147 | 0.023570253 |
| 609163.1875 | 694968.3125 | 728206.4375 | -0.255397214 | 0.120148037 | -0.482163311 | 0.08026683 | 0.170822001 | 0.265568379 |
| 184385.375 | 203481.2656 | 160091.4688 | -0.255627535 | 0.198572024 | 0.445909735 | 0.097708125 | 0.118846017 | 0.578829911 |
| 818281.8125 | 834133.0625 | 865861.3125 | -0.255674809 | 0.103296092 | -0.111485664 | 0.192352326 | 0.005451128 | 0.938779736 |
| 223903.6406 | 223698.2188 | 219413.125 | -0.256054518 | 0.060976526 | 0.457999201 | 0.058877608 | 0.354270829 | 0.051509147 |
| 15665.24316 | 14615.92676 | 13105.46484 | -0.256076098 | 0.086199451 | 0.15779954 | 0.279104324 | -0.268511916 | 0.101487706 |
| 1792228.125 | 1802380.125 | 1346383.375 | -0.256365987 | 0.262015127 | 0.575200556 | 0.046782366 | 0.476988393 | 0.147259899 |
| 31338.68945 | 40479.42578 | 38357.8125 | -0.256872581 | 0.637942139 | 0.573292605 | 0.300186184 | -0.043688387 | 0.942609678 |
| 76607.67188 | 76766.24219 | 74623.89063 | -0.256995664 | 0.046600136 | -0.338582319 | 0.058077004 | 0.265268943 | 0.06969244 |
| 36456.86719 | 39018.44141 | 51515.28516 | -0.257210458 | 0.125874965 | 0.184607854 | 0.246365413 | 0.556347181 | 0.17086882 |
| 71367.65625 | 76347.3125 | 76562.71875 | -0.257253918 | 0.08534895 | -0.01261763 | 0.756936542 | 0.013724916 | 0.785956549 |
| 267822.9688 | 269223.1875 | 270276.0938 | -0.257450776 | 0.051882811 | 0.432492611 | 0.129861867 | 0.182562518 | 0.138522639 |
| 142318.4531 | 148231.5625 | 147452.0781 | -0.25787457 | 0.13478072 | -0.599377656 | 0.033121608 | -0.162687685 | 0.069955133 |
| 41505.51172 | 33321.80859 | 33824.01172 | -0.258063592 | 0.194423598 | 0.08566292 | 0.623543083 | -0.196378273 | 0.386578339 |
| 324312.8125 | 429584.5313 | 331429.125 | -0.258461062 | 0.288733547 | -0.036053512 | 0.496299916 | 0.170561188 | 0.466799273 |
| 16344.67578 | 11077.7959 | 18190.39063 | -0.258678946 | 0.045740682 | 0.019610872 | 0.843432332 | -0.091474617 | 0.761894315 |
| 21285.66211 | 23482.15625 | 18536.56445 | -0.259099318 | 0.255475247 | 0.426064416 | 0.149058018 | 0.273826028 | 0.279150231 |
| 82709.96875 | 77123.19531 | 77129.39063 | -0.259202649 | 0.017824776 | -0.112690972 | 0.154336346 | -0.400736418 | 0.018257145 |
| 30481.70313 | 35898.53516 | 37081.3125 | -0.259335606 | 0.050002651 | -0.02137111 | 0.816502262 | 0.033622074 | 0.811089737 |
| 6038.803223 | 4506.016113 | 5403.275879 | -0.259356733 | 0.096697208 | -0.034887141 | 0.877867274 | 0.097257944 | 0.641750326 |

| | | | | | | | | |
|---|---|---|---|---|---|---|---|---|
| 33070.03516 | 28663.53125 | 29703.76172 | -0.25944002 | 0.072387792 | -0.129118107 | 0.087919389 | -0.006921116 | 0.948264111 |
| 322700.5625 | 248514.0938 | 208429.3438 | -0.259695877 | 0.499844166 | -0.205360387 | 0.708396864 | -0.397139348 | 0.442381195 |
| 8070.390625 | 9907.103516 | 10025.45703 | -0.259740604 | 0.360101307 | -0.086589937 | 0.82763601 | 0.143989089 | 0.519508074 |
| 2600.493408 | 2385.14624 | 2017.566528 | -0.259824093 | 0.538853002 | 0.417756248 | 0.346969068 | -0.284439004 | 0.565837212 |
| 98595.5 | 79258.67188 | 71533.91406 | -0.260023393 | 0.043719896 | 1.222109373 | 0.023005377 | 0.816277076 | 0.106930847 |
| 285293.625 | 311344.4063 | 310806.3438 | -0.260419481 | 0.178228644 | 0.083996704 | 0.269521931 | 0.249737908 | 0.063440273 |
| 10594.41309 | 11757.75684 | 7607.685059 | -0.260588988 | 0.302378035 | 0.185571221 | 0.466161254 | 0.347961987 | 0.340162365 |
| 33335.5 | 25980.86523 | 29223.51563 | -0.260698165 | 0.044763187 | -0.138813017 | 0.52544407 | -0.794236578 | 0.023471308 |
| 132232.4375 | 132297.2969 | 118097.1484 | -0.260742692 | 0.060136197 | -0.245444065 | 0.052036261 | -0.293803481 | 0.080288762 |
| 8858.882813 | 7263.792969 | 5077.98584 | -0.260966257 | 0.139920717 | 0.081906782 | 0.484425023 | -0.500414467 | 0.185585346 |
| 13619.11328 | 12627.53906 | 8255.529297 | -0.261005559 | 0.077857993 | -0.084829007 | 0.364020618 | -0.185365309 | 0.546266077 |
| 1110075.625 | 1126810.625 | 882755.5 | -0.261938825 | 0.148947782 | 0.026401093 | 0.875616587 | 0.043396936 | 0.83795406 |
| 364380.8438 | 392633.125 | 390682.125 | -0.262205581 | 0.16393534 | 0.088283029 | 0.591184368 | 0.171484421 | 0.326296322 |
| 71167.40625 | 50787.74219 | 58608.91016 | -0.262547121 | 0.200542122 | 0.266651972 | 0.198890201 | -0.090321932 | 0.672255917 |
| 1583258.5 | 1726471.875 | 1647519.125 | -0.262895511 | 0.051166099 | 0.112542276 | 0.082161171 | 0.018207987 | 0.771355563 |
| 4992.89209 | 4194.596191 | 4370.663574 | -0.263507169 | 0.546847336 | 0.307281445 | 0.188682494 | -0.423515989 | 0.146451238 |
| 31076.43359 | 38959.07422 | 44830.81641 | -0.263564933 | 0.598139599 | -0.2668753 | 0.674926802 | -0.327322575 | 0.60486123 |
| 723881.625 | 701577.375 | 716905.0625 | -0.263602436 | 0.081686863 | 0.073920279 | 0.500361309 | -0.25923652 | 0.140951005 |
| 2012.125244 | 2446.075684 | 2304.700928 | -0.263678361 | 0.670916189 | 0.316879548 | 0.620856133 | 0.135993752 | 0.835774433 |
| 11416.65723 | 12672.71484 | 13126.22266 | -0.263808963 | 0.407059023 | 0.383271241 | 0.204377767 | 0.307431844 | 0.096285234 |
| 87701.67188 | 87949.55469 | 54207.17969 | -0.263825778 | 0.218876728 | -0.32333361 | 0.107845807 | -0.483769011 | 0.17839917 |
| 115384.7891 | 96511.25781 | 109979.2969 | -0.2640173 | 0.25252148 | 0.030028072 | 0.483003566 | 0.20192839 | 0.214658701 |
| 7953.399902 | 9816.926758 | 6297.750977 | -0.264188225 | 0.644633407 | -0.008258292 | 0.988898404 | -0.688107745 | 0.179047372 |
| 72868.86719 | 73466.10938 | 71296.03125 | -0.264783411 | 0.10227856 | 0.149882536 | 0.52156931 | 1.249848789 | 0.015632484 |
| 96666.54688 | 100498.5547 | 99228.05469 | -0.265090252 | 0.068674829 | 0.062155598 | 0.80599196 | 0.642073205 | 0.026764821 |
| 36145.36719 | 34520.88281 | 32024.26758 | -0.265624915 | 0.26956076 | 0.055408024 | 0.654658287 | -0.146488784 | 0.315114193 |
| 38972.66406 | 33449.5625 | 31788.33203 | -0.265994608 | 0.169680194 | -0.401352553 | 0.116818656 | -0.352162731 | 0.128326365 |
| 35758.94531 | 39951.40625 | 44940.88281 | -0.266095912 | 0.11322605 | 0.368864315 | 0.074416318 | 0.32469645 | 0.149630842 |
| 2833812.75 | 2918286.75 | 2871835.25 | -0.266578798 | 0.058429627 | 0.290656613 | 0.065557261 | 0.28504277 | 0.06688778 |
| 54936.69922 | 59643.48438 | 55739.47656 | -0.266645807 | 0.018999567 | -0.153237256 | 0.094078555 | -0.218684349 | 0.047526425 |
| 1280275.625 | 1405601 | 1266859.875 | -0.266776106 | 0.0992732 | 0.331867921 | 0.110122293 | 0.099060171 | 0.423206002 |
| 8328.824219 | 6898.505371 | 8344.470703 | -0.267478619 | 0.212496914 | 0.531991399 | 0.21691719 | 0.354451452 | 0.136000191 |
| 11470.7793 | 12140.03125 | 9097.307617 | -0.267633256 | 0.038651604 | 0.180189454 | 0.192808829 | -0.254704241 | 0.235404176 |

| | | | | | | | | |
|---|---|---|---|---|---|---|---|---|
| 22680.09961 | 22664.71484 | 16924.83398 | -0.267646982 | 0.420496465 | 0.32159056 | 0.382319312 | -0.160750018 | 0.687054116 |
| 5349149.5 | 6516925 | 6178658 | -0.267863269 | 0.104433134 | 0.491026517 | 0.055241445 | 0.619788896 | 0.051570925 |
| 203598.1094 | 192153.1563 | 201926.8438 | -0.267971134 | 0.037492489 | -0.079204698 | 0.599435333 | -0.445636773 | 0.043404927 |
| 186297.1875 | 189964.1875 | 181308.4531 | -0.26806094 | 0.013585622 | -0.391967148 | 0.01179292 | -0.238625942 | 0.017279709 |
| 68453.61719 | 69302.36719 | 66395.03125 | -0.268171843 | 0.069372981 | 0.214756914 | 0.148589158 | 0.144720522 | 0.229590822 |
| 12108.37109 | 12553.11816 | 10470.87207 | -0.268232064 | 0.119451239 | -0.194434857 | 0.175315476 | -0.103947399 | 0.458205509 |
| 50596.51172 | 51368.30859 | 49535.65625 | -0.268639484 | 0.14249829 | 0.031600899 | 0.834946164 | 0.088205434 | 0.553786226 |
| 244598.7188 | 272198 | 259295.7188 | -0.26927455 | 0.265542969 | 0.299567897 | 0.199074765 | 0.513244457 | 0.060161253 |
| 150453.4375 | 134140.2813 | 142227.9688 | -0.269377298 | 0.191047498 | -0.241210674 | 0.246423036 | -0.211289144 | 0.255838317 |
| 39177.58203 | 27987.01563 | 28850.51953 | -0.269404018 | 0.170436627 | 0.302974323 | 0.195044682 | -0.14571615 | 0.588970577 |
| 3581.692871 | 3854.094971 | 323.5848083 | -0.269417027 | 0.645700069 | 1.48593243 | 0.045329621 | 0.362327369 | 0.75209716 |
| 36291.95703 | 34918.3125 | 29938.81445 | -0.269468651 | 0.067908927 | 0.050618129 | 0.502039149 | -0.496551444 | 0.067125324 |
| 246487.6094 | 271817.5 | 267508.5 | -0.26955998 | 0.186682517 | 0.214986637 | 0.052965881 | 0.24391342 | 0.063440273 |
| 481519.5313 | 470026.125 | 439144.0938 | -0.269561718 | 0.024701492 | -0.149978937 | 0.120506435 | -0.459644245 | 0.017279709 |
| 20007.12305 | 15581.42578 | 14206.26855 | -0.269946914 | 0.207266359 | 0.224635371 | 0.077476727 | -0.462725099 | 0.125291384 |
| 41360.69922 | 41591.18359 | 45123.21875 | -0.270032125 | 0.189698514 | 0.11597017 | 0.532402501 | 0.110841741 | 0.52607612 |
| 22517.92188 | 21716.56055 | 18403.66016 | -0.270319459 | 0.090992566 | -0.181707467 | 0.087342285 | -0.622458615 | 0.029820656 |
| 317078.5 | 276482.5938 | 264263.1875 | -0.270429388 | 0.15559033 | -0.20769727 | 0.15465818 | -0.530559632 | 0.038170182 |
| 11661.89551 | 10655.80859 | 12260.7959 | -0.270734951 | 0.275080134 | 0.77549206 | 0.428474841 | 0.230369851 | 0.138166222 |
| 766830.625 | 731770 | 797555.75 | -0.270956318 | 0.170807941 | -0.173251887 | 0.045588559 | -0.462149569 | 0.014702473 |
| 13135.98242 | 11997.50195 | 10803.91699 | -0.272066716 | 0.256635939 | 0.704004208 | 0.025989516 | 0.164388749 | 0.311264669 |
| 894142.625 | 1065734.875 | 1035477.938 | -0.272237077 | 0.168487308 | -0.115588901 | 0.417759893 | -0.095114102 | 0.538581833 |
| 67870 | 71600.55469 | 74202.125 | -0.272365891 | 0.177998285 | 0.028683708 | 0.750780433 | 0.036881175 | 0.663092542 |
| 475643.25 | 487982.0938 | 446757.1563 | -0.272511635 | 0.02237899 | -0.030352486 | 0.623543083 | 0.210644864 | 0.063283918 |
| 103330.1484 | 108946.7656 | 98694.03906 | -0.272528766 | 0.176857048 | 0.289195485 | 0.197542159 | 0.219579416 | 0.260165601 |
| 39771.07813 | 31896.64453 | 25805.12109 | -0.272589144 | 0.243770538 | 0.017010276 | 0.954173999 | -0.486569603 | 0.183356293 |
| 12397.27148 | 13163.44531 | 11303.84375 | -0.272711124 | 0.144653292 | 0.742383524 | 0.05253541 | -0.043619404 | 0.801376074 |
| 25532.60547 | 20907.82031 | 20687.91602 | -0.272775673 | 0.200146696 | 0.161434986 | 0.433819762 | -0.578635562 | 0.10786231 |
| 179179.1875 | 189680.1719 | 168312.8281 | -0.273013913 | 0.200813951 | 0.366192656 | 0.156712278 | 0.509478946 | 0.074184841 |
| 25883.75586 | 20958.01758 | 24319.30078 | -0.27316775 | 0.096750655 | -0.772948375 | 0.036730784 | -0.256606651 | 0.151930474 |
| 103693.9375 | 91741.10156 | 88692.10156 | -0.273460597 | 0.110708378 | -0.472224264 | 0.085927719 | -0.480321476 | 0.067300418 |
| 12127.62793 | 12702.76172 | 14160.47266 | -0.273581209 | 0.329857534 | 0.401263548 | 0.215436149 | 0.154595147 | 0.578503308 |
| 34247.44531 | 39944.47656 | 37614.56641 | -0.273736646 | 0.135465479 | 0.127270465 | 0.440401575 | 0.035133217 | 0.833250417 |

| | | | | | | | | |
|---|---|---|---|---|---|---|---|---|
| 22318.90625 | 29712.20508 | 33741.08203 | -0.27473197 | 0.746281035 | 0.26595882 | 0.772142973 | 0.192807436 | 0.838930418 |
| 8376.929688 | 9455.087891 | 5836.517578 | -0.274836607 | 0.374517432 | 0.653693486 | 0.053336389 | 0.233224505 | 0.507049834 |
| 27282.23242 | 24656.28711 | 23178.82617 | -0.274839849 | 0.222040127 | 0.461684585 | 0.119643706 | 0.038089316 | 0.863280807 |
| 55048.39063 | 56428.27344 | 56575.71875 | -0.275010512 | 0.057948579 | -0.556042301 | 0.065601851 | -0.348615631 | 0.086670386 |
| 46530.58594 | 46325.35547 | 51483.83594 | -0.275191435 | 0.221052997 | 0.098368786 | 0.663386446 | 1.029695981 | 0.033053706 |
| 216487.7344 | 221523 | 217216.5 | -0.2754901 | 0.116674977 | -0.052525067 | 0.433190294 | -0.088360196 | 0.236290138 |
| 196021.6406 | 189144.5469 | 181521.7188 | -0.275712476 | 0.0424199 | 0.334551075 | 0.039887225 | 0.414960304 | 0.021984636 |
| 345441.7813 | 336373.3438 | 322222.6563 | -0.275721194 | 0.103527693 | 0.294794779 | 0.057189248 | -0.086602158 | 0.225497653 |
| 3331.841553 | 8136.095215 | 4946.60791 | -0.276394038 | 0.513194681 | -0.196602028 | 0.693736734 | -0.517418373 | 0.440512744 |
| 95135.17188 | 92821.69531 | 99173.67969 | -0.2767523 | 0.065343643 | 0.263840723 | 0.094623224 | -0.050150048 | 0.557774325 |
| 1099.365845 | 2093.06665 | 762.4043579 | -0.276789933 | 0.607074449 | -0.938154652 | 0.329328421 | -1.776801763 | 0.189515197 |
| 19140.63867 | 18922.83789 | 17884.09961 | -0.276897047 | 0.251410421 | 0.784304926 | 0.013885891 | 0.037599063 | 0.616686362 |
| 20507.77734 | 21878.0957 | 19974.09375 | -0.277098954 | 0.205586415 | -0.02243009 | 0.778831195 | 0.383210689 | 0.029771119 |
| 242627.7969 | 251500.4531 | 233627.3281 | -0.277185947 | 0.028922381 | 0.325374548 | 0.044099436 | 0.128200044 | 0.140674741 |
| 135199.375 | 146307.5938 | 142861.1719 | -0.277256935 | 0.036547632 | 0.088756844 | 0.355229313 | 0.023158472 | 0.748376124 |
| 81999.67969 | 73966.04688 | 87795.07813 | -0.277377321 | 0.414002282 | -0.650888604 | 0.133753732 | -0.574419916 | 0.118373568 |
| 33700.89063 | 29880.79492 | 25914.48828 | -0.277779882 | 0.146820844 | -0.033482567 | 0.823694131 | 0.005809091 | 0.979696342 |
| 228728.3438 | 256285.9219 | 231903.5469 | -0.27803935 | 0.088240896 | 0.305310444 | 0.079406664 | 0.15209187 | 0.248290955 |
| 109301.4063 | 107323.2031 | 112850.2969 | -0.278534642 | 0.013354673 | -0.191998462 | 0.032675377 | -0.164562563 | 0.032932243 |
| 60578.80859 | 65452.27344 | 62570.27734 | -0.278763391 | 0.18705416 | 0.453915067 | 0.06327555 | 0.34557122 | 0.026764821 |
| 26402.33203 | 27346.33789 | 22508.33594 | -0.278887501 | 0.076923702 | -0.707754352 | 0.062630775 | -0.278870588 | 0.137726349 |
| 4410.59082 | 5840.890137 | 4436.688965 | -0.279706204 | 0.192504708 | -0.245353601 | 0.158923089 | -0.569002846 | 0.085098887 |
| 108159.6953 | 99763.09375 | 104138.9141 | -0.280399731 | 0.072051826 | 0.237088416 | 0.122344477 | 0.714701796 | 0.017279709 |
| 12885.50781 | 11842.7627 | 11366.72461 | -0.280593826 | 0.070275485 | 0.28861757 | 0.117377154 | -0.247633825 | 0.135241092 |
| 53505.27734 | 58989.16016 | 61401.16016 | -0.280680821 | 0.056964374 | 0.326661971 | 0.065601851 | 0.661416553 | 0.029219266 |
| 43576.81641 | 44950.86719 | 44070.94531 | -0.281262329 | 0.12524354 | -0.338191293 | 0.082612243 | -0.65296766 | 0.055516192 |
| 25159.27539 | 21175.00391 | 24223.01953 | -0.281336771 | 0.114778686 | -0.307920258 | 0.360382142 | -0.48700942 | 0.080867847 |
| 696761.625 | 897840.8125 | 704806 | -0.281411214 | 0.244992051 | -0.028568156 | 0.885818377 | 0.048235018 | 0.814604012 |
| 12984.60254 | 19881.66406 | 14509.25098 | -0.281497106 | 0.098618413 | 0.364037334 | 0.199074765 | 0.120084999 | 0.701899856 |
| 820258.5625 | 1367986.25 | 977824.375 | -0.281734972 | 0.401089173 | -0.032762928 | 0.933858968 | -0.208664244 | 0.646958584 |
| 35359.74609 | 42904.24219 | 39866.66406 | -0.28174981 | 0.478874032 | 0.248195473 | 0.553204641 | 0.10725821 | 0.801698552 |
| 102122.0781 | 104865.0469 | 110354.7969 | -0.281903518 | 0.062357731 | -0.000189061 | 0.996796859 | 0.386789011 | 0.04292928 |
| 62619.16406 | 57897.92969 | 60860.72266 | -0.28204488 | 0.039762681 | 0.054641398 | 0.576783001 | 0.18660952 | 0.102620951 |

| | | | | | | | | |
|---|---|---|---|---|---|---|---|---|
| 1826.144043 | 9189.006836 | 4565.219238 | -0.282659287 | NA | 1.147661911 | NA | 0.461291377 | NA |
| 21658.91406 | 22284.77539 | 18731.70703 | -0.282779045 | 0.095446028 | 0.33280521 | 0.021465426 | -0.240286401 | 0.130194761 |
| 14813.87109 | 11506.77734 | 11643.95215 | -0.282799278 | 0.251205996 | 0.144527773 | 0.727709362 | -0.479640051 | 0.180643461 |
| 732168.875 | 714127.75 | 666903.875 | -0.282835061 | 0.63242956 | 0.557934891 | 0.366494549 | 0.280658104 | 0.63905709 |
| 8865.513672 | 10984.95898 | 6588.968262 | -0.28375039 | 0.42956317 | -0.310665771 | 0.280916524 | -0.610913826 | 0.126272884 |
| 103494.7109 | 98112.98438 | 94125.5625 | -0.283826528 | 0.038217062 | -0.478804449 | 0.014951154 | -0.429064534 | 0.01858328 |
| 27021.4668 | 23930.67773 | 22056.08984 | -0.283878898 | 0.458148137 | 0.54901215 | 0.068180046 | 0.498404214 | 0.092384624 |
| 1169267.75 | 1113875.75 | 1132167.75 | -0.284342929 | 0.08771097 | -0.37019815 | 0.051593867 | -0.644408014 | 0.006394096 |
| 1552.768677 | 909.7476807 | 183.6109009 | -0.284516546 | 0.448907681 | 0.469456529 | 0.161959189 | -0.858983838 | 0.312834606 |
| 25688.47266 | 24930.66602 | 17261.04102 | -0.284679729 | 0.050556436 | -0.083359363 | 0.414380687 | -0.671049231 | 0.073130408 |
| 4636.412109 | 6092.828613 | 5637.368164 | -0.285135318 | 0.294814724 | -0.748786074 | 0.05467585 | -0.687072287 | 0.061045405 |
| 389817.375 | 455425.0313 | 544219.75 | -0.285288759 | 0.032963794 | 0.018245859 | 0.81508428 | 0.209000867 | 0.405343283 |
| 1456228.5 | 1560942 | 1513350.375 | -0.28541527 | 0.037600114 | 0.005973796 | 0.947346059 | -0.186453055 | 0.106930847 |
| 18606.03711 | 18709.15039 | 15239.49316 | -0.285679827 | 0.159397666 | 0.035060293 | 0.711115835 | -0.026262488 | 0.872130886 |
| 64799.66797 | 61546.22266 | 66231.46094 | -0.285956926 | 0.258459268 | 0.704851801 | 0.086540831 | 0.541224373 | 0.112266171 |
| 69476.73438 | 76710.82031 | 78566.36719 | -0.285963415 | 0.186686406 | 0.070854172 | 0.577242402 | 0.0120412 | 0.935031407 |
| 554894.625 | 445520.9688 | 459828.2813 | -0.286287957 | 0.067908927 | 0.090326652 | 0.51425117 | -0.095731504 | 0.543664393 |
| 63021.94141 | 65215.91406 | 63164.58203 | -0.286486359 | 0.045092515 | -0.085397145 | 0.28939927 | -0.199666535 | 0.074531134 |
| 2605.365967 | 3930.377441 | 3532.140625 | -0.286663358 | NA | 2.481100535 | 0.198890201 | 2.299655403 | 0.043404927 |
| 19344.56641 | 20115.42578 | 19791.33789 | -0.286767197 | 0.374464563 | -0.045056641 | 0.899806127 | -0.30299326 | 0.450252535 |
| 144206.2031 | 131558.7031 | 143194.7188 | -0.286955715 | 0.087565113 | 0.057721366 | 0.241519943 | 0.258688974 | 0.062063322 |
| 362234.4063 | 376484.2188 | 357398.3438 | -0.287341809 | 0.248195549 | 0.360025062 | 0.200504592 | 0.069886305 | 0.764731388 |
| 8529.789063 | 15835.57422 | 8699.473633 | -0.287645216 | 0.214920041 | 0.012796513 | 0.951848655 | 0.230989364 | 0.669503576 |
| 11610.83008 | 9282.975586 | 9198.5625 | -0.287891108 | 0.371658988 | -0.002421132 | 0.992236761 | -0.124226477 | 0.681870325 |
| 9723.639648 | 8806.578125 | 9220.37793 | -0.287894888 | 0.050002651 | -0.20529443 | 0.102854593 | -0.072944948 | 0.342355842 |
| 202732.3438 | 221802.8281 | 222654.3438 | -0.288352753 | 0.171503557 | 0.064054617 | 0.706776442 | -0.188010596 | 0.361302058 |
| 14291.18066 | 13475.45508 | 8783.364258 | -0.288411193 | 0.278628279 | -0.293655739 | 0.150726837 | -0.538373671 | 0.160431331 |
| 47233.74512 | 42207.81445 | 40557.08398 | -0.288420257 | 0.134212026 | 0.317203411 | 0.110122293 | 0.157918534 | 0.244290663 |
| 24147.62109 | 21953.92578 | 22131.85156 | -0.28854312 | 0.089964482 | -0.227666301 | 0.181752348 | -0.215642239 | 0.178198454 |
| 49154.17188 | 43154.91797 | 34167.35547 | -0.288560039 | 0.106074437 | -0.218688266 | 0.068436129 | -0.349662995 | 0.194351963 |
| 110384.0781 | 115820.2656 | 88475.20313 | -0.288753883 | 0.530526722 | -0.071317118 | 0.734733431 | -0.072320985 | 0.746954593 |
| 39318.07813 | 40834.07813 | 37358.50391 | -0.289028857 | 0.052738763 | -0.000843884 | 0.99247981 | 0.175400153 | 0.152597158 |
| 29561.80273 | 29865.62305 | 31087.20508 | -0.289074046 | 0.052356162 | -0.333158868 | 0.089716327 | 0.383181526 | 0.047134496 |

| | | | | | | | | |
|---|---|---|---|---|---|---|---|---|
| 59339.04297 | 57307.44141 | 59029.08594 | -0.289089073 | 0.345950084 | -0.295508476 | 0.318509566 | 0.290042011 | 0.247217109 |
| 152642.25 | 171890.2344 | 153420.3281 | -0.289327617 | 0.066448651 | 0.148214439 | 0.184317721 | 0.116835559 | 0.29550852 |
| 259763.7656 | 270117.1563 | 257219.625 | -0.289340716 | 0.061354438 | -0.221510616 | 0.055241445 | 0.084253099 | 0.140758419 |
| 392410.9688 | 389418.5313 | 429716.4375 | -0.289736857 | 0.116764456 | 0.476237862 | 0.018195523 | 0.284860233 | 0.05752489 |
| 31071.01758 | 38218.6875 | 44692.61328 | -0.289831726 | 0.795729729 | 0.33311744 | 0.770096621 | -0.093400675 | 0.947307151 |
| 3383.095215 | 3407.776855 | 3022.418457 | -0.289858037 | 0.139278926 | -0.06284345 | 0.632281284 | -0.538681696 | 0.062848011 |
| 3035.154297 | 3332.66626 | 2126.641357 | -0.289930124 | 0.154507223 | 0.098131975 | 0.597251041 | -0.332657547 | 0.292537906 |
| 174023.8125 | 186558.8594 | 169063.6094 | -0.290009234 | 0.238040125 | 0.392989335 | 0.165555532 | 0.158056119 | 0.486378719 |
| 85478.24219 | 98752.72656 | 88897.76563 | -0.290189795 | 0.415273342 | 0.554851608 | 0.208267253 | 0.662957917 | 0.129491158 |
| 159651.8594 | 191479.3438 | 190339.1719 | -0.290208197 | 0.096761263 | 0.08021765 | 0.540771148 | 0.27006525 | 0.164256752 |
| 8476.966797 | 6041.497559 | 10454.71484 | -0.290694185 | 0.639880745 | 0.425315878 | 0.613040078 | 0.015571791 | 0.984470463 |
| 604351.3125 | 547674.1875 | 624337.1875 | -0.29109104 | 0.203029059 | 0.373173032 | 0.168359532 | 0.359665685 | 0.153754411 |
| 207566.2031 | 196826.7344 | 178846.8438 | -0.291474918 | 0.109761076 | -0.306800642 | 0.159703167 | -0.478982496 | 0.071696088 |
| 792610.1875 | 774729.125 | 790885.125 | -0.291515635 | 0.020740891 | 0.359907891 | 0.027373868 | 0.248003449 | 0.049004518 |
| 124877.8828 | 125491.7266 | 145044.0781 | -0.292157904 | 0.045160985 | 0.573608317 | 0.023592908 | 0.564723441 | 0.050810933 |
| 43764.63281 | 45908.17188 | 39243.28906 | -0.292620774 | 0.498334318 | -0.66450707 | 0.191852289 | 0.740304412 | 0.065056294 |
| 160964.6719 | 160061.1094 | 147907.5938 | -0.292630298 | 0.058059531 | 0.346085204 | 0.058877608 | 0.099252971 | 0.329050274 |
| 482540.125 | 452559.875 | 452170.7188 | -0.292724329 | 0.104246701 | 0.238692685 | 0.152415276 | -0.111327499 | 0.428096482 |
| 220153.375 | 225844.25 | 208315.3125 | -0.293095623 | 0.027277127 | -0.195465898 | 0.052702293 | -0.134895853 | 0.09149674 |
| 186995.4688 | 188388.9063 | 181951.625 | -0.293229683 | 0.00799298 | 0.128873015 | 0.174231909 | -0.006014492 | 0.852856834 |
| 222201.6875 | 213117.75 | 193843.8906 | -0.293331917 | 0.346510839 | 0.503017078 | 0.175868431 | 0.172225708 | 0.577906494 |
| 21963.70508 | 22359.53711 | 23849.06836 | -0.293952126 | 0.038502747 | 0.040933405 | 0.614330411 | -0.203839339 | 0.112052905 |
| 41489.52734 | 46096.65625 | 37930.85156 | -0.294066706 | 0.100973803 | -0.149826647 | 0.406071842 | -1.002207317 | 0.040202519 |
| 306596.4355 | 314204.9463 | 295503.8125 | -0.294135196 | 0.012750417 | -0.131159946 | 0.065415915 | -0.39258206 | 0.01123893 |
| 18932.69727 | 13620.71191 | 22387.32031 | -0.29426569 | 0.371838207 | 0.37648065 | 0.316595052 | 0.630857171 | 0.205451497 |
| 50915.42188 | 49013.5625 | 40906.34375 | -0.29512018 | 0.019078369 | 0.148214293 | 0.085196118 | -0.05711637 | 0.695793958 |
| 13782.45215 | 13408.3252 | 13738.52441 | -0.295200272 | 0.332623284 | 0.104717396 | 0.753235332 | 0.282012029 | 0.363709214 |
| 127865.0898 | 136243.0938 | 118741.6055 | -0.295496073 | 0.113472458 | 0.645788668 | 0.04542656 | 0.492163342 | 0.04854063 |
| 33329.98438 | 22026.64063 | 25131.82422 | -0.295768548 | 0.42822149 | -0.232263487 | 0.294543232 | -1.444376244 | 0.02146481 |
| 1411.561768 | 6331.310059 | 5084.921875 | -0.296049636 | 0.741388161 | 0.008828609 | 0.992136662 | -0.423633384 | 0.72699212 |
| 13549.63867 | 14621.28516 | 13875.06348 | -0.296062584 | 0.142526436 | 0.198086043 | 0.297911261 | 0.147274401 | 0.394034633 |
| 194667.9375 | 229518.7031 | 111820.0234 | -0.296195268 | 0.350129741 | -0.91431215 | 0.154025091 | -1.242720102 | 0.089218964 |
| 1284028.625 | 1318800.875 | 1243326.75 | -0.296293836 | 0.107338684 | 0.432783174 | 0.015625705 | 0.419904689 | 0.015233843 |

| | | | | | | | | |
|---|---|---|---|---|---|---|---|---|
| 570859.5625 | 672572.375 | 533848.5625 | -0.296669271 | 0.379283654 | 0.324690791 | 0.130087098 | 0.222765871 | 0.266521029 |
| 66703.28906 | 123997.0703 | 107812.9844 | -0.296860815 | 0.261387089 | -0.197821509 | 0.504460238 | -0.427060392 | 0.31862804 |
| 26410.21289 | 28479.09766 | 26457.96875 | -0.297022151 | 0.049192971 | -0.371406543 | 0.023005377 | 0.108784962 | 0.16200644 |
| 6121.727539 | 4178.128906 | 12267.46387 | -0.297132969 | 0.643797985 | -0.149420462 | 0.492292239 | -0.385416668 | 0.552272008 |
| 42266.22266 | 39240.10938 | 43027.25391 | -0.297140635 | 0.087043567 | 0.029689357 | 0.294272595 | -0.215817226 | 0.063474265 |
| 3809.176025 | 5003.57666 | 4482.897949 | -0.298576473 | 0.192973731 | -0.142268411 | 0.521101932 | -0.612150443 | 0.047120928 |
| 111027.2891 | 103940.0469 | 89055.10156 | -0.298642523 | 0.122520185 | 0.286380092 | 0.127225665 | -0.240426037 | 0.201736266 |
| 919750.1875 | 1155871 | 1237327.625 | -0.298944278 | 0.079968566 | 0.310086875 | 0.05744085 | 0.592332763 | 0.118023574 |
| 236685.7031 | 267606.1563 | 246760.0781 | -0.298989833 | 0.077186454 | -0.207575394 | 0.190057963 | -0.231386175 | 0.134445271 |
| 249607 | 247652.125 | 244702.375 | -0.299069926 | 0.141905535 | -0.311482837 | 0.224248912 | -0.386797927 | 0.176276737 |
| 1146496.75 | 1154736.5 | 1061284 | -0.299435753 | 0.108442252 | 0.106877861 | 0.460748219 | -0.283079147 | 0.16847165 |
| 98913.65625 | 79168.34375 | 76487.77344 | -0.299706943 | 0.110302818 | -0.22922943 | 0.251091228 | -0.216672905 | 0.332898966 |
| 79766.24219 | 80553.74219 | 78325.48438 | -0.299956963 | 0.019286562 | -0.123544034 | 0.172806689 | -0.231574979 | 0.066959828 |
| 42644.26563 | 39005.69531 | 42397.59375 | -0.300116963 | 0.042621515 | -0.116701188 | 0.363939878 | 0.0633528 | 0.528938224 |
| 2866.175049 | 2723.494385 | NA | -0.30031812 | 0.663001353 | -0.036080245 | 0.950464931 | -0.428678615 | NA |
| 102252.75 | 98128.41406 | 88128.84375 | -0.300492429 | 0.165438813 | -0.075848102 | 0.708396864 | -0.131202572 | 0.535435703 |
| 8834.632813 | 8844.644531 | 8873.532227 | -0.301235942 | 0.173221038 | -0.081372357 | 0.790098584 | -0.004407635 | 0.98246789 |
| 64460.50781 | 62965.05859 | 66688.875 | -0.301311624 | 0.108968105 | 0.481599846 | 0.079287615 | 0.038130333 | 0.561579182 |
| 31258.14844 | 33691.47656 | 36564.22656 | -0.301815883 | 0.114205402 | 0.133858761 | 0.392329965 | 0.319665682 | 0.112700927 |
| 5237.205566 | 5096.813477 | 3459.465088 | -0.301828441 | 0.173483544 | -0.353670616 | 0.184506133 | -0.858772404 | 0.049651271 |
| 76369.28906 | 71630.39063 | 84893.65625 | -0.302329283 | 0.036480793 | -0.289171724 | 0.047452686 | 0.352206695 | 0.103879505 |
| 405097 | 352393.5625 | 362641.8438 | -0.302336963 | 0.109594531 | -0.603189616 | 0.100538317 | -0.128406483 | 0.436019867 |
| 26493.77539 | 42908.13281 | 27462.83203 | -0.302911467 | 0.267418718 | -0.277908367 | 0.432802496 | -0.304970973 | 0.402826429 |
| 234856.5 | 237652.875 | 220991.875 | -0.303089547 | 0.146800875 | 0.64767864 | 0.046369992 | 0.585800035 | 0.048839627 |
| 39144.27734 | 32345.7168 | 25223.20117 | -0.303126521 | 0.302635975 | 0.218920204 | 0.205390089 | 0.330979392 | 0.345709627 |
| 157370.9531 | 180080.9219 | 200585.5 | -0.303350335 | 0.100026503 | 0.100586825 | 0.527850691 | 0.218870936 | 0.277551095 |
| 27300.74023 | 29602.43945 | 33078.67188 | -0.303355764 | 0.164604942 | 0.472168286 | 0.029180754 | 0.145752552 | 0.336295567 |
| 24837.48242 | 26779.98438 | 25808.24609 | -0.303492706 | 0.176268956 | 0.393396837 | 0.149709854 | -0.081322567 | 0.192741849 |
| 116877.1133 | 114366.2813 | 105242.2305 | -0.303513672 | 0.034240061 | 0.28102575 | 0.109899281 | -0.228571321 | 0.059847517 |
| 169702.3906 | 161334.7813 | 164388.4688 | -0.303910898 | 0.078894657 | 0.209885422 | 0.058641336 | 0.111204862 | 0.149087782 |
| 49772.01953 | 47534.17969 | 44052.96875 | -0.30409317 | 0.028255845 | -0.714834826 | 0.021408573 | -1.094235569 | 0.013271537 |
| 16077.91113 | 19898.58789 | 18776.91016 | -0.304289263 | 0.53534498 | 0.507272725 | 0.324535676 | 0.083507368 | 0.86993911 |
| 17000.98438 | 18177.60938 | 15322.52637 | -0.304381503 | 0.145925272 | 0.106007464 | 0.534271251 | 0.193326704 | 0.210758021 |

| | | | | | | | | |
|---|---|---|---|---|---|---|---|---|
| 251055.4219 | 277296.625 | 290541.7813 | -0.304436024 | 0.116497357 | 0.100526417 | 0.128476695 | 0.293193088 | 0.097178029 |
| 6824.931641 | 4520.713379 | 3431.701416 | -0.304669796 | 0.237418408 | 0.127333781 | 0.63348405 | -0.542253205 | 0.255592007 |
| 69784.95313 | 70761.33594 | 62672.71484 | -0.304850792 | 0.025587445 | -0.271382837 | 0.078996408 | -0.293157225 | 0.052786471 |
| 52032.68359 | 53435.38281 | 44544.37109 | -0.304865378 | 0.081090664 | -0.148228553 | 0.327201979 | -0.380697491 | 0.089218964 |
| 875.3198853 | 850.6691284 | 401.2958984 | -0.304972306 | 0.212615135 | -0.159470213 | 0.534693749 | -0.463366143 | 0.338685457 |
| 130927.7734 | 160526.4531 | 137006.9688 | -0.305211831 | 0.036275447 | 0.177031555 | 0.147695831 | 0.122419237 | 0.438703042 |
| 16377.86328 | 18694.76953 | 14491.95703 | -0.305325802 | 0.082624617 | -0.245435803 | 0.20617006 | -0.55974751 | 0.060542864 |
| 35857.71875 | 37344.83984 | 34037.32813 | -0.305457663 | 0.02187351 | 0.184540281 | 0.064458068 | 0.010201913 | 0.893534893 |
| 686855.25 | 667771.9375 | 661149.5 | -0.305604139 | 0.029016212 | 0.420424369 | 0.031122153 | 0.416420167 | 0.02909479 |
| 66109.00781 | 68815.89063 | 65395.56641 | -0.305681239 | 0.042134056 | 0.082792073 | 0.349131522 | 0.02498996 | 0.758771158 |
| 61330.68359 | 61940.00781 | 54753.79297 | -0.305873596 | 0.01332549 | 0.049578036 | 0.467062508 | -0.073214867 | 0.423557711 |
| 5936.917969 | 7877.196289 | 4901.613281 | -0.306024863 | 0.774198519 | -0.36368419 | 0.535392728 | -0.12739444 | 0.81895658 |
| 138613.8281 | 129633.1719 | 121297.8516 | -0.306173279 | 0.272383086 | -0.255362774 | 0.192808829 | -0.305128897 | 0.130595595 |
| 49500.32422 | 58702.09375 | 56316.33984 | -0.306517772 | 0.044665667 | 0.099868469 | 0.376711856 | 0.076072595 | 0.551480236 |
| 4840.199219 | 8173.195801 | 7670.996094 | -0.307032916 | 0.369074013 | 0.67914728 | 0.065601851 | 0.664726652 | 0.209524848 |
| 173961.1563 | 158215.2188 | 175670.4844 | -0.307448498 | 0.038502747 | 0.140019255 | 0.224463272 | -0.030088933 | 0.763765593 |
| 3595.804199 | 5241.321289 | NA | -0.307524159 | 0.520142615 | 0.361098621 | 0.423580857 | -0.130153556 | NA |
| 83216.1875 | 80497.39063 | 82143.61719 | -0.30755464 | 0.019930448 | 0.05229171 | 0.539179679 | -0.199408032 | 0.080462751 |
| 155993.0781 | 188115.625 | 141632.8438 | -0.307718265 | 0.337452634 | 0.399870472 | 0.179476554 | 0.383109989 | 0.188422821 |
| 407536.25 | 520628.5 | 241537.0313 | -0.307829575 | 0.234047674 | -0.293772493 | 0.352759971 | -0.491911553 | 0.302401454 |
| 886536.25 | 926711.3125 | 918799.75 | -0.307938262 | 0.180746458 | -0.017204972 | 0.885440998 | -0.14281076 | 0.313944854 |
| 56561.0625 | 58364.99609 | 56124.92188 | -0.308360926 | 0.087565113 | -0.083869542 | 0.319259968 | -0.172547722 | 0.031785503 |
| 268625.3438 | 249857.375 | 200714.4844 | -0.308385198 | 0.187369902 | 0.200234442 | 0.366735281 | -0.349226879 | 0.235870699 |
| 9269.466797 | 5062.750977 | 6077.157227 | -0.308655323 | 0.225237475 | -0.227821075 | 0.336933122 | -0.670876438 | 0.14379126 |
| 760609.5 | 714186.25 | 752884.0625 | -0.309417108 | 0.025086961 | 0.041295988 | 0.582455701 | 0.127580218 | 0.143689091 |
| 14314.03516 | 14284.37988 | 12882.19141 | -0.309685213 | 0.077105145 | -0.076920755 | 0.536218169 | 0.076716137 | 0.536701705 |
| 739058.3047 | 696798.6992 | 653862.5723 | -0.310058038 | 0.040223584 | 0.024886519 | 0.816723703 | 0.152164974 | 0.176254612 |
| 99340.01563 | 95560.09375 | 107797.8047 | -0.310080895 | 0.027930748 | 0.191933904 | 0.230350006 | 0.225606889 | 0.095057794 |
| 190311.5156 | 205469.3125 | 187471.9063 | -0.310188764 | 0.058516905 | -0.248462892 | 0.056194202 | -0.351812133 | 0.029710029 |
| 6875485 | 7978181.5 | 7132952.5 | -0.310342198 | 0.103502264 | 0.437602524 | 0.096710716 | 0.567219745 | 0.042919102 |
| 25739.02734 | 26064.30078 | 24847.20117 | -0.311471727 | 0.122499451 | -0.434179609 | 0.155243406 | -0.417431055 | 0.145662926 |
| 48579.39063 | 46810.97656 | 34211.82031 | -0.311618857 | 0.096461916 | -0.779630155 | 0.029180754 | -0.774895676 | 0.065714742 |
| 39935.19531 | 40703.38672 | 38592.38281 | -0.311839882 | 0.148078807 | -0.175225516 | 0.204833051 | -0.10787515 | 0.344199475 |

| | | | | | | | | |
|---|---|---|---|---|---|---|---|---|
| 7883.802734 | NA | NA | -0.311840413 | NA | -0.173604232 | NA | -0.796698594 | NA |
| 16863.8418 | 16391.38281 | 11845.77539 | -0.311962929 | 0.688585426 | -0.158753019 | 0.775164931 | -0.03391385 | 0.950164303 |
| 54179.57422 | 55098.89844 | 52375.10938 | -0.312066873 | 0.031440016 | -0.227583657 | 0.039808229 | -0.268520603 | 0.026263251 |
| 95576.72656 | 113730.8594 | 85053.1875 | -0.312248399 | 0.163416464 | -0.104839677 | 0.541058779 | -0.128177488 | 0.550609087 |
| 65089.13672 | 79509.07813 | 90606.69531 | -0.312256937 | 0.232183802 | -0.321468354 | 0.331502317 | -0.058335713 | 0.848089906 |
| 36236.71094 | 47074.91016 | 35279.93359 | -0.312337132 | 0.327647019 | -0.065324228 | 0.832929946 | -0.304103966 | 0.252822179 |
| 312225.9375 | 292680.7188 | 280442.875 | -0.312446392 | 0.022440374 | -0.345096546 | 0.045481011 | -0.281118973 | 0.047120928 |
| 21826.68164 | 23823.27344 | 18896.70508 | -0.312794907 | 0.158308661 | -0.452332371 | 0.162564211 | -0.412683094 | 0.148421794 |
| 13308.09668 | 17493.75781 | 16010.25488 | -0.313051675 | 0.140717943 | -0.835685719 | 0.038037652 | -0.815698614 | 0.02909479 |
| 41851.59766 | 47699.28516 | 37993.16797 | -0.313063634 | 0.350711493 | 0.215505627 | 0.34766869 | 0.281255316 | 0.183050335 |
| 20818.22461 | 25311.3457 | 30298.23828 | -0.313566138 | 0.423961947 | 0.691636471 | 0.151849296 | 0.457138666 | 0.282508948 |
| 119.5614243 | 197.4251556 | 163.8505249 | -0.313624026 | 0.618635039 | -1.118311903 | NA | -0.308784452 | 0.701341354 |
| 11423.26074 | 9821.447266 | 9184.669922 | -0.313644917 | 0.077857993 | 0.446306349 | 0.065813387 | 0.243246203 | 0.225165761 |
| 60149.47266 | 61498.69141 | 57544.65234 | -0.31377833 | 0.038008847 | -0.099341497 | 0.300418041 | -0.627005798 | 0.035559086 |
| 374831.8438 | 422239.7188 | 406170.8125 | -0.313878773 | 0.081995281 | 0.14292477 | 0.048593497 | -0.076187566 | 0.365046345 |
| 92997.46094 | 88650.63281 | 78895.59375 | -0.314015148 | 0.509831804 | 0.155519509 | 0.52156931 | -0.072864068 | 0.755944 |
| 61491.91016 | 56846.6875 | 61141.86328 | -0.314111777 | 0.132676546 | -0.232944031 | 0.260555973 | 0.159517044 | 0.329050274 |
| 14395.35352 | 15028.70605 | 8278.219727 | -0.314172349 | 0.148202641 | -0.883005069 | 0.075603505 | -1.17578541 | 0.037571191 |
| 11156.55371 | 11057.21191 | 9763.875 | -0.314195895 | 0.128747675 | -0.187118363 | 0.200210626 | -0.403279481 | 0.060012809 |
| 13162.55469 | 17556.41406 | 17252.37305 | -0.314358572 | 0.195180132 | 0.880183231 | 0.05467585 | 0.938776095 | 0.06688778 |
| 327056.1563 | 370965.5 | 301060.0313 | -0.31443695 | 0.1346049 | 0.297269034 | 0.174475307 | -0.120015247 | 0.549054792 |
| 88238.60156 | 93903.63281 | 86486.39844 | -0.314445237 | 0.095292766 | -0.017326774 | 0.839908525 | -0.321324641 | 0.044994481 |
| 28783.78906 | 40055.93945 | 36469.71777 | -0.31508726 | 0.147754486 | -1.315588732 | 0.021906439 | -0.41174855 | 0.146875127 |
| 154255.2656 | 149404.4531 | 125252.0313 | -0.31529589 | 0.067188303 | -0.181837947 | 0.195013961 | -0.397493224 | 0.071353321 |
| 55016.57617 | 59108.55908 | 61907.61426 | -0.315489897 | 0.060388606 | 0.164630629 | 0.167753597 | 0.215042645 | 0.114246364 |
| 6069.487305 | 5534.225586 | 6120.691406 | -0.315732393 | 0.155747436 | -0.112464818 | 0.673113251 | -0.206065637 | 0.333735773 |
| 285136.1875 | 259961.5156 | 302376.0938 | -0.315788611 | 0.576547365 | 0.588144236 | 0.160918214 | 1.263581394 | 0.031136253 |
| 18252.46484 | 15440.09082 | 16840.93164 | -0.315848984 | 0.171490772 | -0.106777239 | 0.66458436 | -0.197350482 | 0.381068951 |
| 129632.5469 | 116419.7031 | 107537.4688 | -0.3159733 | 0.063350133 | 0.70203842 | 0.015639733 | 0.441429292 | 0.08152982 |
| 38581.98828 | 48036.375 | 63051.02344 | -0.316199299 | 0.218843086 | 0.189046715 | 0.434082643 | 0.06946223 | 0.851823372 |
| 13900.88672 | 14839.63184 | 9711.629883 | -0.31628593 | 0.118300349 | 0.545443884 | 0.143256542 | 0.183472665 | 0.550690788 |
| 204692.4063 | 212101.5625 | 216894.9844 | -0.316395069 | 0.042621515 | 0.068048027 | 0.426846112 | 0.029194193 | 0.722176291 |
| 25062.84766 | 26435.10156 | 30169.24609 | -0.316547865 | 0.077066654 | 0.073587136 | 0.596480117 | 0.156788138 | 0.362075745 |

| | | | | | | | | |
|---|---|---|---|---|---|---|---|---|
| 162014.2188 | 155617.9063 | 170868 | -0.317354467 | 0.029183044 | -0.061282758 | 0.459072637 | -0.24950281 | 0.072903408 |
| 173964.125 | 156167.4844 | 180184.25 | -0.318455581 | 0.119064113 | -0.019836487 | 0.885440998 | 0.01370583 | 0.891581356 |
| 17317.25195 | 19167.1543 | 16826.38281 | -0.318631614 | 0.162830006 | -0.276641063 | 0.081173768 | -0.334532883 | 0.047255794 |
| 366783.8438 | 371274.5313 | 337787.1875 | -0.318981103 | 0.026113102 | 0.047151123 | 0.513083308 | -0.241482351 | 0.068496057 |
| 135474.5 | 145374.3281 | 125307.5 | -0.31904938 | 0.404498232 | 0.69348478 | 0.056103327 | 0.133575601 | 0.371124353 |
| 340710.9375 | 435666.25 | 369541.3125 | -0.319236649 | 0.403630684 | -0.083543486 | 0.660718628 | -0.02885919 | 0.886601496 |
| 13796.07813 | 11390.75684 | 11299.5293 | -0.319267071 | 0.172685916 | 0.227835127 | 0.102454913 | 0.036887192 | 0.807484317 |
| 16649.21289 | 17375.54102 | NA | -0.319896811 | 0.381822856 | 0.184678298 | 0.60094054 | -0.287416612 | NA |
| 83910.125 | 99693.22656 | 87341.82031 | -0.320089352 | 0.282890531 | -0.536916561 | 0.243612448 | -0.913938503 | 0.142772717 |
| 762.5810547 | 956.6744385 | 878.888855 | -0.320122451 | 0.138806746 | 0.405659592 | 0.105893922 | -0.344806849 | 0.18019177 |
| 27829.0332 | 34016.13281 | 29309.02539 | -0.320746594 | 0.028932331 | -0.235098192 | 0.11146465 | -0.835196815 | 0.01875548 |
| 437705.8984 | 375179.6426 | 344090.6094 | -0.320810793 | 0.566541202 | -0.449653574 | 0.562301807 | -0.207199995 | 0.763776161 |
| 19816.88086 | 17440.62109 | 17822.00977 | -0.320880751 | 0.116159237 | 0.338927717 | 0.138141237 | 0.400964047 | 0.084223187 |
| 53095.65625 | 39735.22656 | 35476.80859 | -0.320888368 | 0.064613771 | 0.277267053 | 0.14612569 | -0.025740373 | 0.929478549 |
| 57999.92969 | 58250.58984 | 57349.08984 | -0.32105941 | 0.059440088 | -0.132783859 | 0.303685582 | -0.090235636 | 0.416566132 |
| 290116.0625 | 295122.6875 | 288831.7813 | -0.32132221 | 0.038217062 | 0.452539087 | 0.032159738 | 0.437969861 | 0.006394096 |
| 426179.3438 | 468578.9375 | 460581.625 | -0.321799651 | 0.087237737 | -0.221684308 | 0.184657727 | -0.346746725 | 0.096058082 |
| 2820227.25 | 2682211 | 2565086.75 | -0.322039971 | 0.28847371 | 0.167051467 | 0.130589029 | 0.089380299 | 0.242530269 |
| 186813.9531 | 187313.1406 | 196701.8281 | -0.322076279 | 0.055768726 | 0.279604729 | 0.091478417 | 0.261187695 | 0.093054769 |
| 315779.3438 | 331093.125 | 329031.2188 | -0.322197536 | 0.0424199 | -0.244878846 | 0.112714218 | -0.166656161 | 0.15966324 |
| 65917.85156 | 64714.86719 | 53887.18359 | -0.322214111 | 0.078998007 | 0.072950886 | 0.197056982 | -0.136449885 | 0.358282985 |
| 48585.88281 | 48890.21875 | 40128.13672 | -0.322297892 | 0.028138974 | -0.011636504 | 0.88053234 | -0.264615036 | 0.137062098 |
| 45465.83594 | 49481.01172 | 43986.72266 | -0.322596505 | 0.081004874 | 0.253137186 | 0.172411368 | 0.246740406 | 0.135536925 |
| 61625.82422 | 62314.58203 | 48600.95313 | -0.322818204 | 0.026880281 | -0.350096412 | 0.058011247 | -0.45524653 | 0.07580998 |
| 31303.58984 | 25389.33008 | 33018.69922 | -0.323554076 | 0.309737132 | 0.071827707 | 0.831472283 | 0.237786011 | 0.478935691 |
| 8505.272461 | 8629.671875 | 5832.216797 | -0.323740112 | 0.11060902 | -0.358772662 | 0.163834679 | -0.456532819 | 0.149843463 |
| 33122.10938 | 42295.19922 | 16439.99805 | -0.324078361 | 0.265527675 | -1.091228387 | 0.065260161 | -1.505439407 | 0.036818836 |
| 275677.5 | 259913.5781 | 269133.9688 | -0.324104096 | 0.056093567 | 0.120723151 | 0.275840689 | 0.068062054 | 0.249456378 |
| 305381.3125 | 335231.125 | 277035.75 | -0.324216314 | 0.437509858 | -0.098123936 | 0.644354586 | -0.09370085 | 0.644909155 |
| 17897.53516 | 18239.07031 | 15766.07813 | -0.324245866 | 0.127230164 | -0.301513711 | 0.077476727 | -0.578893739 | 0.02786639 |
| 77457.16406 | 83998.00781 | 89159.26563 | -0.324422709 | 0.027612915 | 0.097814577 | 0.327559963 | 0.184765197 | 0.156978352 |
| 53478.17188 | 55338.98438 | 49113.04297 | -0.324624906 | 0.136100201 | -0.51076508 | 0.101045598 | -0.341673582 | 0.143571995 |
| 99345.63281 | 99439.91406 | 105682.9688 | -0.324634448 | 0.019930448 | -0.103356568 | 0.221717993 | -0.159244481 | 0.100630717 |

| | | | | | | | | |
|---|---|---|---|---|---|---|---|---|
| 468942.25 | 431595.5313 | 473292.75 | -0.32472444 | 0.29342279 | 0.159010964 | 0.593974342 | 0.276276919 | 0.367550122 |
| 145072.0156 | 143982.6563 | 134926 | -0.32476826 | 0.021027835 | -0.135943952 | 0.102497957 | -0.295228183 | 0.029771119 |
| 47831.69141 | 47491.70313 | 49146.69531 | -0.325058935 | 0.286654231 | 0.46945372 | 0.090139838 | 0.435572746 | 0.048582087 |
| 13291.35645 | 14311.78809 | 16148.62402 | -0.325154265 | 0.187635714 | -0.313707324 | 0.275021879 | -0.093207111 | 0.690582254 |
| 279240.5938 | 413554.5 | 340276.875 | -0.325530465 | 0.273682083 | 0.0488555 | 0.858230595 | 0.159332588 | 0.622745767 |
| 71937.14844 | 78853.47656 | 79834.88281 | -0.325703824 | 0.056529322 | 0.349568988 | 0.080440179 | 0.484309252 | 0.033486824 |
| 1638602.375 | 1794581.75 | 1948955.375 | -0.325851374 | 0.182751618 | -0.16124217 | 0.362287701 | 0.47299659 | 0.059258927 |
| 83920.42969 | 84336.23438 | 83728.99219 | -0.326754662 | 0.607487192 | 1.39737504 | 0.06863053 | 1.101479644 | 0.134304215 |
| 80224.79688 | 79941.72656 | 75314.53125 | -0.326771996 | 0.096461916 | -0.384455221 | 0.119711295 | -0.481944731 | 0.097707274 |
| 12121.00684 | 9309.692383 | 9156.418945 | -0.326852312 | 0.18705416 | 0.745754976 | 0.052965881 | -0.168066732 | 0.557456388 |
| 4705.759766 | 5264.306152 | 1795.727905 | -0.326873189 | 0.300849654 | 0.065337138 | 0.675172962 | -0.733144591 | 0.230308748 |
| 26461.32617 | 23268.99414 | 15421.08496 | -0.326912096 | 0.14678023 | -0.485604821 | 0.161939384 | -0.962979731 | 0.060174773 |
| 9773.364258 | 7472.020508 | 6499.574707 | -0.327361666 | 0.40453073 | 0.093390013 | 0.828619605 | 0.1700626 | 0.710545806 |
| 39497.11719 | 43864.67188 | 38215.99609 | -0.328047512 | 0.027405706 | 0.249264038 | 0.058456539 | -0.577650734 | 0.022925497 |
| 36200.22266 | 36474.28516 | 35636.97266 | -0.328296385 | 0.031442732 | -0.046108725 | 0.57675299 | -0.31169269 | 0.08504597 |
| 9983.348633 | 11554.92871 | 19347.125 | -0.328381405 | 0.490885072 | 0.388236855 | 0.198917031 | 1.158145032 | 0.20724059 |
| 133384.5625 | 150263.5313 | 124146.9688 | -0.32870129 | 0.174784378 | 0.648630857 | 0.065266848 | 0.275186155 | 0.244006157 |
| 10244.96973 | 10238.33594 | 9333.876953 | -0.328926816 | 0.079009944 | 0.121275078 | 0.392889701 | -0.312349701 | 0.136887167 |
| 10743.78516 | 5761.703125 | 17769.23047 | -0.329067441 | 0.45968104 | 0.033088499 | NA | 0.201326759 | 0.797672854 |
| 4348.705566 | 4384.776367 | 4010.97583 | -0.329101402 | 0.20354889 | 0.5061621 | 0.105114411 | -0.08590654 | 0.716525566 |
| 31674.43359 | 33347.76563 | 31467.93359 | -0.329290152 | 0.035776373 | -0.319401466 | 0.094128959 | -0.066905916 | 0.481648842 |
| 13544.10449 | 14690.30664 | 11367.50684 | -0.329758295 | 0.08771097 | -0.342270574 | 0.21691719 | -0.670000265 | 0.035377452 |
| 13924.71582 | 11658.9668 | 11597.41992 | -0.330525906 | 0.111266593 | 0.138951545 | 0.430961626 | -0.273594965 | 0.235404176 |
| 297653.375 | 362906.1875 | 334971.375 | -0.330688093 | 0.204963542 | -0.041994114 | 0.831472283 | 0.072530874 | 0.67863809 |
| 12127.88965 | 13752.64844 | 11902.47266 | -0.330820253 | 0.161932491 | -0.281681216 | 0.135981012 | -0.630650019 | 0.044156984 |
| 11287.6543 | 11405.42188 | 12697.42285 | -0.331044393 | 0.182074985 | 0.292391237 | 0.08093378 | -0.127828723 | 0.317865083 |
| 518206.7578 | 513368.9922 | 455295.3203 | -0.331328807 | 0.042146623 | 0.087810872 | 0.400736085 | 0.017465084 | 0.884769453 |
| 244701.8906 | 263893.25 | 261201.1406 | -0.331977723 | 0.036765012 | -0.346852792 | 0.080440179 | -0.253181138 | 0.066667224 |
| 2022.671387 | 2047.660767 | 1350.397339 | -0.331979976 | 0.376428671 | 1.076091858 | 0.090435995 | -0.081713019 | 0.836533248 |
| 1246.920288 | 3409.898438 | NA | -0.332207675 | 0.274068475 | -0.408300839 | 0.224271577 | -1.122070272 | NA |
| 13555.48828 | 14524.35059 | 10732.4707 | -0.332236056 | 0.137725162 | -0.584139236 | 0.109509233 | -1.51608104 | 0.047850077 |
| 2184123.5 | 2649811.75 | 2608945.5 | -0.332590689 | 0.09684552 | 0.245394051 | 0.051861383 | 0.176903907 | 0.2881913 |
| 71848.26563 | 76564.95313 | 80343.58594 | -0.332735827 | 0.04508378 | -0.235617913 | 0.111349456 | -0.407225164 | 0.043897759 |

| | | | | | | | | |
|---|---|---|---|---|---|---|---|---|
| 447861.9063 | 525755.3125 | 487510.8438 | -0.333062023 | 0.097194367 | -0.037319158 | 0.688328439 | 0.178069791 | 0.200720922 |
| 13834.94629 | 12725.27637 | 6580.271484 | -0.33309216 | 0.107464969 | -0.558964438 | 0.076036407 | -0.630952548 | 0.186616329 |
| 59844.26953 | 61983.76172 | 61944.10156 | -0.333229949 | 0.015391678 | 0.023683137 | 0.714831692 | -0.065558291 | 0.270937884 |
| 70713.34375 | 64174.24219 | 65037.53125 | -0.334131836 | 0.09200896 | 0.636804965 | 0.031437224 | 0.30105315 | 0.093361286 |
| 83949.94531 | 87000.02344 | 78692.49219 | -0.335535991 | 0.100934147 | 0.307372683 | 0.09037921 | 0.123423797 | 0.283755733 |
| 84247.57813 | 84171.42188 | 48335.88672 | -0.335600125 | 0.042134056 | -1.326530247 | 0.039887225 | -1.129427864 | 0.041771828 |
| 165294.5781 | 175381.9375 | 169342.0625 | -0.335680863 | 0.021494917 | -0.374655261 | 0.050817911 | -0.277422371 | 0.043976385 |
| 3095820.25 | 3093677.5 | 3294484.75 | -0.335891036 | 0.029183044 | 0.002437048 | 0.974822088 | -0.034741191 | 0.673787999 |
| 10544.95215 | 13063.77832 | 7756.155762 | -0.335946949 | 0.188824978 | 0.122880218 | 0.591860699 | 0.013649537 | 0.973658284 |
| 30525.50781 | 29558.8457 | 25454.67773 | -0.336108479 | 0.047647106 | -0.094601109 | 0.4202179 | -0.126017589 | 0.394984688 |
| 210316.0469 | 233958.25 | 211747.75 | -0.33646192 | 0.009974872 | 0.099390631 | 0.092018139 | -0.037931135 | 0.632470042 |
| 1852.260742 | 1322.855835 | 361.2938538 | -0.336665808 | 0.268867993 | -0.541701442 | 0.229698337 | -1.195269265 | 0.127037403 |
| 534008.9248 | 546920.8994 | 539087.1592 | -0.336746923 | 0.116516598 | 0.314442203 | 0.147323941 | 0.886909268 | 0.037772514 |
| 3916.574219 | 5010.90918 | 3442.345215 | -0.337120377 | 0.056714022 | 0.19990414 | NA | -0.090648371 | 0.72266742 |
| 7965.669434 | 10779.49902 | 7188.761719 | -0.337148341 | 0.119438563 | -0.310327769 | 0.171390921 | -0.48217233 | 0.141846462 |
| 133126.4375 | 136920.5469 | 124892.0078 | -0.337257256 | 0.0640041 | -0.228800674 | 0.14541426 | -0.321895977 | 0.07561358 |
| 6631.529785 | 6940.900391 | 5849.541016 | -0.337272039 | 0.146853115 | -0.420594456 | 0.183896983 | -0.598109368 | 0.119221254 |
| 71443.61719 | 66995.45313 | 58509.49609 | -0.337316959 | 0.067459302 | 0.51702733 | 0.172671211 | 0.434542786 | 0.091109394 |
| 986296.5 | 1018379.375 | 1013355.563 | -0.337683806 | 0.20354889 | 0.098412374 | 0.691779308 | -0.062284274 | 0.806396591 |
| 429217.0313 | 427159.5 | 394150.0313 | -0.337685709 | 0.046600136 | 0.115603933 | 0.265049654 | 0.180511175 | 0.107554917 |
| 175183.7031 | 167307.9844 | 163193.5156 | -0.337708258 | 0.092503073 | 0.723997394 | 0.05087265 | 0.759055418 | 0.029492107 |
| 192342.6094 | 165899.9688 | 153878.5 | -0.338108605 | 0.252969162 | 0.225207607 | 0.178204855 | 0.844548858 | 0.060161253 |
| 63572.54492 | 66993.35352 | 60441.80664 | -0.338247052 | 0.227207392 | -0.34695038 | 0.339683948 | -0.645404762 | 0.177627983 |
| 52501.03906 | 38992.95703 | 60751.14063 | -0.338326685 | 0.165760942 | -0.43329449 | 0.100528183 | -0.347050614 | 0.244446425 |
| 315192.2813 | 379703.6953 | 360517.8164 | -0.338911815 | 0.171503557 | 0.854506303 | 0.273114108 | 1.829295313 | 0.026263251 |
| 1309965 | 1169900.25 | 1072194.5 | -0.338998314 | 0.142870238 | 0.076369019 | 0.568703657 | -0.204971097 | 0.263528267 |
| 42578.27734 | 78801.03125 | 41459.86328 | -0.339075102 | 0.307501176 | 0.564753524 | 0.222489287 | -0.121886798 | 0.815558126 |
| 1093398.281 | 1560669.25 | 1233382.875 | -0.339557483 | 0.109041324 | 1.037841902 | 0.034038555 | 0.343118883 | 0.280119454 |
| 72649.08594 | 71464.10938 | 68043.05469 | -0.339748176 | 0.057253208 | -0.148459277 | 0.105321264 | -0.176588144 | 0.061382227 |
| 42342.71094 | 37191.01953 | 44575.05078 | -0.340484972 | 0.222265151 | 0.620973623 | 0.102454913 | 0.251266605 | 0.365046345 |
| 45349.92188 | 35356.85547 | 39567.47656 | -0.340557876 | 0.103296092 | 0.530576956 | 0.077209499 | 0.250264692 | 0.27322162 |
| 37071.69922 | 36721.73828 | 44997.85938 | -0.340574863 | 0.485679981 | 0.132294448 | 0.805890965 | -0.281636526 | 0.645907062 |
| 399043.3438 | 439146.125 | 437729.375 | -0.340935447 | 0.028196123 | -0.012281805 | 0.852417959 | -0.280174095 | 0.042737273 |

| | | | | | | | | |
|---|---|---|---|---|---|---|---|---|
| 156643.5625 | 154312.1094 | 161182.7969 | -0.341022645 | 0.054152412 | 0.557059818 | 0.046114261 | 0.216944713 | 0.156588849 |
| 10861.58984 | 12682.37305 | 11451.63574 | -0.341314905 | 0.314273917 | 0.063843986 | 0.850403567 | 0.111268449 | 0.737743931 |
| 84244.85156 | 80459.08594 | 74111.97656 | -0.341571434 | 0.147450707 | -0.073302432 | 0.722910121 | -0.091212852 | 0.668912516 |
| 24432.8125 | 29141.84766 | 23479.0918 | -0.342355797 | 0.364472828 | -0.271459912 | 0.553204641 | 0.256998999 | 0.510020897 |
| 35111.01172 | 35471.05859 | 33562.66797 | -0.342449507 | 0.097194367 | 0.445314639 | 0.09273472 | 0.458279159 | 0.07856704 |
| 35952.82031 | 34466.87109 | 32884.05469 | -0.343237289 | 0.036547632 | 0.119534067 | 0.331502317 | 0.305396293 | 0.050976089 |
| 421550.9375 | 524983.3125 | 451515.7188 | -0.343622369 | 0.210383422 | -0.245240722 | 0.097722369 | -0.094142361 | 0.536092868 |
| 224146.0469 | 247033.8281 | 240536.25 | -0.343960735 | 0.053959867 | 0.253481984 | 0.012254503 | 0.221724494 | 0.089805965 |
| 139860.7188 | 208355.5156 | 123568.1406 | -0.344161767 | 0.535155129 | 0.094473545 | 0.808694324 | 0.119080989 | 0.778343026 |
| 86870.20313 | 105968.0391 | 86884.14844 | -0.344231687 | 0.069867455 | 0.650064917 | 0.082833891 | -0.139998053 | 0.455335597 |
| 172158.5781 | 158133.4844 | 151258.7188 | -0.344307782 | 0.09684552 | -0.036859686 | 0.712899971 | 0.062814941 | 0.587981638 |
| 57715.66016 | 58063.0625 | 62182.17578 | -0.344885877 | 0.138650351 | 0.781739289 | 0.044605958 | 0.898237965 | 0.030633392 |
| 8135.07373 | 6651.296875 | 9784.59375 | -0.345461878 | 0.149067264 | 0.619586358 | 0.071811131 | 0.434631343 | 0.223059038 |
| 32857.32813 | 35030.48438 | 31622.31641 | -0.345693213 | 0.062680034 | -0.126001663 | 0.098005404 | -0.220396559 | 0.059417728 |
| 394600.9375 | 312396.7188 | 328099.75 | -0.34579436 | 0.182582097 | -0.209260149 | 0.240174102 | 0.195227537 | 0.327187467 |
| 9686.464844 | 11512.92383 | 10622.95508 | -0.345861233 | 0.086055451 | -0.540392555 | 0.100538317 | -0.801020395 | 0.056482153 |
| 3078.099854 | 4438.512695 | 2620.885254 | -0.345893337 | 0.211452184 | -0.759135872 | 0.136752641 | -1.230890922 | 0.066959828 |
| 80875.39063 | 86938.89063 | 81097.78906 | -0.345925859 | 0.064026439 | 0.17729685 | 0.132067536 | -1.157092212 | 0.027533174 |
| 70913.07813 | 67827.53125 | 64404.29297 | -0.346005648 | 0.050654092 | 0.252201857 | 0.114596423 | 0.17233733 | 0.159743169 |
| 40455.35547 | 36971.04297 | 38233.30078 | -0.346009155 | 0.096694236 | -0.299704239 | 0.079287989 | -0.322474424 | 0.061455564 |
| 128428.0859 | 129230.0703 | 115116.6953 | -0.346170968 | 0.042134056 | 0.34168489 | 0.079580647 | 0.323488746 | 0.0667896 |
| 8467.560547 | 7213.302734 | NA | -0.346402324 | 0.289195889 | -0.392439058 | 0.398087952 | -0.115343546 | NA |
| 9587.703125 | 10617.15137 | 8516.373047 | -0.346493454 | 0.097443336 | 0.344213561 | 0.069777099 | -0.004685202 | 0.978263952 |
| 78683.4375 | 82751.48438 | 76809.11719 | -0.34655838 | 0.088231376 | -0.021331056 | 0.798228787 | -0.237233745 | 0.068265317 |
| 5368.927246 | 6435.611816 | 4943.710449 | -0.346796905 | 0.524060283 | 0.590695491 | 0.191184742 | 0.126810821 | 0.758771158 |
| 7993.394531 | 7102.979004 | 7215.48877 | -0.346892985 | 0.079495636 | -0.172858934 | 0.538096429 | -0.167370428 | 0.149087782 |
| 552785.4375 | 432987.4063 | 575295.625 | -0.347670921 | 0.559171823 | -0.335793941 | 0.516815532 | -0.456574011 | 0.406218733 |
| 54252.47266 | 50839.07031 | 55322.64844 | -0.347955575 | 0.032155886 | -0.160613253 | 0.193488746 | -0.102975039 | 0.317316467 |
| 37048.90234 | 29240.67578 | 24808.01758 | -0.348106986 | 0.053542507 | -0.721651707 | 0.039808229 | -0.884800369 | 0.039446558 |
| 43196.90625 | 46924.53516 | 39519.23828 | -0.348128168 | 0.017355582 | 0.350327641 | 0.024018676 | -0.074832655 | 0.510742601 |
| 25450.01563 | 26542.2168 | 22978.39453 | -0.348249216 | 0.0667907 | 0.011404974 | 0.924647274 | -0.141892324 | 0.339103506 |
| 19289.49023 | 19654.85352 | 29352.66602 | -0.348345261 | 0.13878271 | 0.262160867 | 0.34708379 | 0.603653103 | 0.22095921 |
| 26628.90234 | 33193.64453 | 45534.44922 | -0.34852866 | 0.046568492 | 0.114851047 | 0.550464936 | 0.149981608 | 0.687539514 |

| | | | | | | | | |
|---|---|---|---|---|---|---|---|---|
| 37863.67578 | 46081.08984 | 40116.59766 | -0.348646718 | 0.267129155 | -0.29504657 | 0.023497318 | -0.170853615 | 0.258758163 |
| 48594.35938 | 43895.96484 | 46205.99219 | -0.3490204 | 0.062373059 | 0.368280045 | 0.022410713 | 0.031459623 | 0.687590757 |
| 23783.94922 | 25024.29492 | 17536.75586 | -0.349463426 | 0.275807707 | 0.649632972 | 0.108992465 | -0.486537299 | 0.246773877 |
| 31487.55664 | 29225.90039 | 34003.94141 | -0.349532475 | 0.144732815 | 0.537641254 | 0.031422588 | 0.569548043 | 0.036848746 |
| 74303.05469 | 79839.40625 | 99517.70313 | -0.349652956 | 0.073735071 | 0.047694493 | 0.74357075 | 0.626432474 | 0.107874808 |
| 121592.2969 | 150957.6875 | 132644.0313 | -0.350058102 | 0.691392083 | 0.835719082 | 0.356068496 | 0.515267494 | 0.563084657 |
| 62213.26953 | 61087.3125 | 63047.71484 | -0.350484912 | 0.024554363 | -0.115051985 | 0.300186184 | 0.255063976 | 0.062345205 |
| 128340.4766 | 127840.6563 | 123817.3828 | -0.351099305 | 0.144580907 | 0.022837358 | 0.734697104 | 0.051970986 | 0.460908588 |
| 8374.774414 | 10921.94434 | 4003.5979 | -0.351233627 | 0.104002895 | 0.036036266 | 0.844905766 | -0.595336593 | 0.28013775 |
| 647768.2891 | 636784.3906 | 620322.7969 | -0.351347578 | 0.105495503 | 0.262356454 | 0.216004498 | 0.173148274 | 0.337637057 |
| 15089.25977 | 17888.2793 | 15285.91406 | -0.35135207 | 0.410359728 | -0.132260411 | 0.790517549 | -0.359077391 | 0.513281452 |
| 60815.98828 | 66050.10938 | 60953 | -0.351562025 | 0.039443869 | -0.204198924 | 0.097708125 | -0.244404682 | 0.046581887 |
| 213181.1875 | 230047.2031 | 219146.6563 | -0.351667992 | 0.136865016 | 0.028287997 | 0.602172086 | -0.018672169 | 0.777319405 |
| 363710.0313 | 393918.9688 | 357375.9688 | -0.352196446 | 0.016471012 | -0.505586026 | 0.02118733 | -0.337756975 | 0.029324108 |
| 41615.41016 | 46778.6875 | 37361.89453 | -0.352320998 | 0.078894657 | -0.185740178 | 0.322217412 | -0.360208499 | 0.134536614 |
| 403983.2813 | 426666.3125 | 444214.9375 | -0.352677283 | 0.015732345 | -0.178999433 | 0.079859428 | -0.347058083 | 0.043108871 |
| 147257.3438 | 151348.3438 | 151687.4531 | -0.353727673 | 0.058516905 | 0.325878404 | 0.087018661 | 0.354917111 | 0.076969062 |
| 50516.91406 | 46704.00781 | 51199.04297 | -0.353785669 | 0.061134086 | 0.40170536 | 0.085565133 | 0.084617814 | 0.52834849 |
| 1525.580322 | 1248.029541 | 1061.523315 | -0.35379006 | 0.177971041 | -0.261562571 | 0.358946075 | -1.34569248 | 0.062985995 |
| 163797.375 | 183456.0781 | 196437.7344 | -0.353791572 | 0.184517222 | 0.449817749 | 0.01179292 | 0.552941016 | 0.077094395 |
| 28106.08398 | 27838.79883 | 23050.27539 | -0.354046752 | 0.052520339 | 0.228698553 | 0.149709854 | 0.058298128 | 0.727321619 |
| 22583.91211 | 24823.16406 | 28984.7832 | -0.354314755 | 0.075807406 | -0.06973209 | 0.59763359 | 0.250248034 | 0.251555607 |
| 131838.25 | 128596.7734 | 130374.0313 | -0.354378896 | 0.11255422 | 0.229668254 | 0.363532071 | 1.09608783 | 0.00797582 |
| 72362.875 | 73548.77344 | 77478.85156 | -0.354481224 | 0.110302818 | 0.666871364 | 0.012254503 | 0.919042126 | 0.009587646 |
| 99443.70313 | 86608.96875 | 83548.45313 | -0.354905247 | 0.116546 | 0.487791409 | 0.194188982 | 0.471579387 | 0.082016867 |
| 25309.41797 | 25883.68164 | 26469.70508 | -0.354946268 | 0.111508367 | 0.001955262 | 0.991529214 | 0.126472886 | 0.445250651 |
| 338834.2813 | 323298.2813 | 345344.4063 | -0.355244889 | 0.033091329 | -0.028546136 | 0.717241613 | -0.163119887 | 0.112514654 |
| 6375.262695 | 4943.412109 | 5145.164063 | -0.355300552 | 0.029838966 | -0.157116982 | 0.185649792 | -0.373701327 | 0.114669448 |
| 22430.08789 | 22685.18555 | 22365.88672 | -0.355511496 | 0.308539067 | 0.038396894 | 0.905873111 | 0.275037467 | 0.345554668 |
| 30617.98633 | 51062.25391 | 24609.98438 | -0.355656459 | 0.52413751 | -0.42161973 | 0.511634098 | -0.526651018 | 0.35787308 |
| 105503.9453 | 119983.2969 | 127775.6875 | -0.355997777 | 0.048782796 | 0.271813543 | 0.11151684 | 1.040417796 | 0.035612683 |
| 28690.32422 | 28690.94531 | 24529.93555 | -0.356113889 | 0.094925671 | 0.313272592 | 0.047347558 | 0.282299577 | 0.123718324 |
| 894260.25 | 963347.9375 | 923087.25 | -0.356142881 | 0.082806274 | 0.06381248 | 0.516207413 | -0.107584632 | 0.314191738 |

| | | | | | | | | |
|---|---|---|---|---|---|---|---|---|
| 91804.15625 | 90675.94531 | 84510.625 | -0.356389053 | 0.055041746 | 0.333006652 | 0.026250916 | 0.111480028 | 0.178198454 |
| 687189.125 | 666648.4375 | 675525.6875 | -0.356413442 | 0.131380074 | 0.163900756 | 0.390894216 | -0.095119596 | 0.6063018 |
| 65226.30078 | 67542.58594 | 70073.19531 | -0.356481758 | 0.024810615 | -0.207335577 | 0.11104965 | -0.160499492 | 0.145840147 |
| 10611.89746 | 11335.625 | 8981.454102 | -0.357073224 | 0.051158864 | 0.184113142 | 0.184447357 | -0.113885125 | 0.503444621 |
| 22010.33594 | 21129.60547 | 20675.0918 | -0.357168859 | 0.028238951 | -0.414853638 | 0.045536435 | -0.352107556 | 0.054887084 |
| 386271.9375 | 397952.8438 | 376782.3438 | -0.358075181 | 0.145925272 | 0.204996925 | 0.108795517 | 0.413608879 | 0.028738995 |
| 8648.65625 | 10835.04102 | 9463.707031 | -0.358429305 | 0.121235391 | -0.275761466 | 0.284889693 | -0.527156878 | 0.119188149 |
| 26084.01563 | 25776.16602 | 21283.11523 | -0.358942308 | 0.168558068 | 0.433915384 | 0.129506032 | 0.294475542 | 0.239967727 |
| 454993.6875 | 479482.0313 | 483779.625 | -0.359257233 | 0.046970661 | -0.058425674 | 0.596108635 | -0.743537708 | 0.04923471 |
| 183363.7031 | 157591.625 | 123781.8906 | -0.359457579 | 0.19253889 | -0.002015132 | 0.9901874 | -0.387159753 | 0.193067253 |
| 410968.8125 | 386734.6875 | 385685.5 | -0.359468376 | 0.036160457 | 0.098492165 | 0.368600434 | -0.145217573 | 0.2429355 |
| 157762.5 | 164950.9063 | 167457.6406 | -0.359620163 | 0.265572358 | 0.439103507 | 0.220100738 | 0.699800137 | 0.110895176 |
| 7611.465332 | 6511.165039 | 2186.868652 | -0.359626782 | 0.621182888 | 0.837891652 | 0.280688037 | 0.374807855 | 0.700016179 |
| 113002.0156 | 107204.9766 | 110825.1563 | -0.36076929 | 0.012414001 | 0.140503819 | 0.095219221 | -0.069109496 | 0.256000151 |
| 196843.0625 | 176584.6406 | 171462.5469 | -0.360833704 | 0.026113102 | -0.866426828 | 0.01179292 | -0.415951239 | 0.037907761 |
| 2116956.25 | 2106857.75 | 1772847.375 | -0.361111746 | 0.14275971 | -0.012492501 | 0.912016384 | -0.267286374 | 0.140631421 |
| 4244.241211 | 4834.760742 | 3881.47876 | -0.361128969 | 0.03500296 | -0.168106056 | 0.137058989 | -0.639266259 | 0.030633392 |
| 19774.89063 | 15369.85254 | 17545.89258 | -0.36127508 | 0.111176969 | -0.226519853 | 0.526766343 | 0.049288353 | 0.827903135 |
| 28006.70508 | 27042.3125 | 26660.35547 | -0.36158896 | 0.066276059 | 1.09475623 | 0.030506702 | 1.072308897 | 0.006394096 |
| 21684.93945 | 27450.04102 | 19419.77344 | -0.362020607 | 0.074012043 | 0.533723801 | 0.063325993 | -0.009259034 | 0.973660041 |
| 93032.00781 | 84783.10156 | 83188.35156 | -0.362212961 | 0.093016235 | 0.72304245 | 0.107969678 | 0.40699822 | 0.040202519 |
| 225459.2969 | 228819.5156 | 189579.9844 | -0.362450451 | 0.070570116 | 0.242262845 | 0.137931056 | -0.278244061 | 0.131190542 |
| 83346.25 | 87071.32031 | 93440.8125 | -0.363306618 | 0.270935076 | 0.199060207 | 0.559679082 | 0.187902481 | 0.556912126 |
| 835258.375 | 949838.9375 | 908305.0625 | -0.36334407 | 0.202791888 | -0.097862115 | 0.386997353 | 0.810569736 | 0.029219266 |
| 317067.6563 | 333424.0313 | 302875.2188 | -0.36340614 | 0.070959976 | -0.164748812 | 0.312686937 | -0.331573354 | 0.122834452 |
| 192667.125 | 209245.6094 | 217596.4219 | -0.36346866 | 0.051855812 | 0.05588412 | 0.551083277 | 0.324118338 | 0.058100863 |
| 11530.41406 | 13896.99414 | 3337.544434 | -0.363691159 | 0.16126803 | -0.996380868 | 0.081324743 | -1.387061429 | 0.07175538 |
| 2249.015137 | 1995.90271 | 1182.04187 | -0.363948434 | 0.146466811 | -1.012838212 | 0.10108711 | -1.176239632 | 0.055355517 |
| 1078433.5 | 1060464.5 | 980503.3125 | -0.364644533 | 0.10213643 | -0.389116791 | 0.161933659 | -0.679307499 | 0.082586658 |
| 146642.1563 | 135692.2656 | 131136.4688 | -0.364728992 | 0.097194367 | 0.377742839 | 0.03129572 | 0.019985624 | 0.831540545 |
| 18220.6543 | 21009.60742 | 16646.76172 | -0.364736015 | 0.042340539 | 0.175637792 | 0.204851511 | 0.039957042 | 0.81836013 |
| 590704.25 | 602509 | 620989.4375 | -0.364872128 | 0.026769495 | 0.028841201 | 0.6115573 | -0.174345255 | 0.079169623 |
| 454106.1875 | 410090 | 376744.4375 | -0.364876243 | 0.047406642 | 0.576436412 | 0.058877608 | -0.503470403 | 0.054369794 |

| | | | | | | | | |
|---|---|---|---|---|---|---|---|---|
| 1474897.5 | 1551941.5 | 1893801 | -0.364984196 | 0.0424199 | 0.663174461 | 0.03412759 | 0.35437495 | 0.173509851 |
| 51171.36719 | 61148.99609 | 56791.40625 | -0.365244857 | 0.153779868 | 0.571486994 | 0.098459633 | 1.071200355 | 0.023792795 |
| 68507.25781 | 63169.47656 | 69693.5 | -0.365379336 | 0.037492489 | -0.006021697 | 0.918953337 | 0.010117229 | 0.902744456 |
| 7871.136719 | 7142.280762 | 8485.624023 | -0.365772428 | 0.045230483 | -0.339751926 | 0.407206729 | 0.083920793 | 0.528158253 |
| 6519.757813 | 7763.175293 | 6936.218262 | -0.365875751 | 0.675681446 | 0.832003063 | 0.322202084 | 0.828965029 | 0.311382514 |
| 311408.5938 | 341286.625 | 323353.4688 | -0.365925639 | 0.091607129 | 0.134399678 | 0.134028069 | -0.237791641 | 0.042919102 |
| 573635.1875 | 593842.5 | 620610.375 | -0.366119738 | 0.056659836 | 0.208414011 | 0.147323941 | -0.054069169 | 0.339663786 |
| 10318.35742 | 10261.30762 | 9646.652344 | -0.366527913 | 0.031606514 | -0.063082221 | 0.391326232 | -0.304754153 | 0.04296087 |
| 29542.84375 | 27324.02539 | 27314.28516 | -0.366596546 | 0.546911852 | 0.01972816 | 0.783136193 | -0.113989239 | 0.228116839 |
| 82462.01563 | 77728.71094 | 65271.36328 | -0.366745984 | 0.059269869 | 0.536919552 | 0.055083913 | 0.656447754 | 0.064289264 |
| 71676.33594 | 70859.35938 | 63853.23047 | -0.367161983 | 0.018311389 | 0.056696859 | 0.393711547 | -0.05100304 | 0.576226388 |
| 11926.30957 | 12587.0127 | 12032.43945 | -0.367166358 | 0.032652748 | 0.225843895 | 0.102526216 | 0.114284414 | 0.248754576 |
| 75768.54688 | 86857.48438 | 90633.14844 | -0.367538363 | 0.15346698 | 0.495470515 | 0.05467585 | 0.30262471 | 0.129008887 |
| 343462.125 | 351377.4063 | 337552.9375 | -0.367801659 | 0.004709568 | 0.171705074 | 0.108625401 | -0.151685187 | 0.027087229 |
| 168972.2969 | 162686.25 | 179232.4844 | -0.368114434 | 0.329618532 | 0.670904123 | 0.109579479 | 0.943919777 | 0.062838928 |
| 28477.52344 | 26824.94141 | 23847.92578 | -0.368236422 | 0.196988669 | 0.607054309 | 0.108962882 | 0.592764966 | 0.089381209 |
| 30606.95117 | 28379.03711 | 28869.65625 | -0.369261706 | 0.048963509 | 0.025299045 | 0.61192528 | -0.177576901 | 0.055868946 |
| 48674.76953 | 47486.41406 | 36964.78125 | -0.369315213 | 0.246338956 | -0.15153392 | 0.272826246 | -0.108613534 | 0.576226388 |
| 34025.70703 | 41683.62109 | 29849.85938 | -0.369442121 | 0.066305135 | 0.210547462 | 0.235403731 | -0.038854273 | 0.874328917 |
| 10857.9541 | 15363.8252 | 10887.52734 | -0.369621328 | 0.071306984 | -0.024376593 | 0.808553621 | -0.097293904 | 0.716525566 |
| 4059.89624 | 2974.77002 | 2294.820313 | -0.369637257 | 0.142741867 | 0.845368497 | 0.045481011 | -0.205839059 | 0.557774325 |
| 460569.4063 | 484882.5938 | 456636.3438 | -0.369736043 | 0.030838583 | -0.450558956 | 0.07841071 | -0.31349136 | 0.081870265 |
| 9172.636719 | 8825.631836 | 7431.681152 | -0.369997668 | 0.156220502 | -0.068015224 | 0.636784361 | -0.069243999 | 0.668657953 |
| 4456.714844 | 10128.10156 | 10648.63281 | -0.370351322 | 0.355826028 | 1.17849059 | 0.048024604 | 0.629896056 | 0.37957979 |
| 44488.89063 | 46267.17969 | 38024.69531 | -0.370538669 | 0.06578064 | -0.133440064 | 0.403148694 | -0.166314943 | 0.356805712 |
| 92787.01563 | 98111.42969 | 88807.92188 | -0.370639896 | 0.086929104 | -0.580362939 | 0.052965881 | -0.616353854 | 0.029820656 |
| 34916.17969 | 37629.89063 | 33511.71094 | -0.371643017 | 0.008543222 | 0.025180856 | 0.795244019 | -0.251670325 | 0.063438208 |
| 222908.75 | 238841.0625 | 189514.1406 | -0.371909048 | 0.029258207 | 0.077569968 | 0.420340567 | -0.609943331 | 0.034567592 |
| 2671862.5 | 3033361.75 | 3076248.25 | -0.371942293 | 0.037492489 | 0.142467265 | 0.076036407 | 0.100208328 | 0.375139877 |
| 71383.96875 | 62368.45703 | 53660.17578 | -0.372487987 | 0.056769135 | -0.954581461 | 0.058077004 | -0.481420167 | 0.074131758 |
| 20269.28516 | 20248.48438 | 14592.21973 | -0.372717722 | 0.065637909 | 0.008486737 | 0.876704025 | -0.310991014 | 0.229556812 |
| 10809.88281 | 12444.65039 | 15178.18555 | -0.372751334 | 0.090517853 | 0.168971213 | 0.374414438 | 0.015672039 | 0.955025524 |
| 6421.321777 | 3795.190674 | 3459.981201 | -0.372786316 | 0.040079376 | 0.109812682 | 0.537530415 | -0.18167141 | 0.674280128 |

| | | | | | | | | |
|---|---|---|---|---|---|---|---|---|
| 223585.3125 | 234574.1094 | 222556.0625 | -0.37396821 | 0.042916723 | 0.177456059 | 0.163729974 | -0.154340515 | 0.207881414 |
| 8142.59082 | 17471.69336 | 15014.85254 | -0.374143081 | 0.236422852 | -0.03509176 | 0.919703825 | -0.073316275 | 0.88793846 |
| 28023.50781 | 25030.44531 | 20633.27734 | -0.37434773 | 0.245606762 | 0.842377732 | 0.029364116 | 0.740686421 | 0.081870265 |
| 20047.08398 | 19786.79883 | 13113.60742 | -0.375480896 | 0.063885064 | 0.750144566 | 0.048024604 | 0.494118413 | 0.236563713 |
| 1405762 | 5225760 | 1583413.875 | -0.376234958 | 0.782395042 | -0.600828481 | 0.660062477 | -0.646673864 | 0.593204536 |
| 96729.28125 | 91268.58594 | 84435.53906 | -0.376316209 | 0.139366912 | -0.10215627 | 0.634882033 | -0.037601138 | 0.860757824 |
| 34182.07422 | 34036.64063 | 29477.0957 | -0.376377112 | 0.109774427 | -0.266633223 | 0.251035871 | -0.386639727 | 0.141501171 |
| 4765.504883 | 5140.339844 | 3297.649414 | -0.37639011 | NA | 0.207259336 | NA | -0.080322602 | NA |
| 724729.0625 | 1574304.625 | 844125.4375 | -0.376445087 | 0.585026867 | 0.20221147 | 0.792608337 | 0.40826155 | 0.582450918 |
| 241185.8594 | 219790.0313 | 227922.625 | -0.376464451 | 0.022703538 | 0.219295351 | 0.095410346 | 0.137949207 | 0.143234366 |
| 525048.8125 | 421733.5938 | 380855.5 | -0.376723467 | 0.037913948 | 0.324035616 | 0.279104324 | 0.145785355 | 0.538573538 |
| 39732.5625 | 40765.74609 | 39183.20313 | -0.376861712 | 0.036830197 | -0.058180321 | 0.667331576 | -0.542294796 | 0.06858506 |
| 5909.765137 | 4615.288086 | 3024.755859 | -0.377107396 | 0.108542219 | -0.440651563 | 0.177893197 | -1.538195763 | 0.033994245 |
| 1116852.375 | 1120727.125 | 1064892.75 | -0.37713033 | 0.052052972 | -0.001690465 | 0.991037626 | -0.222771342 | 0.2081433 |
| 3809.93457 | 4959.794922 | 3714.502686 | -0.377495129 | 0.037600114 | -0.252334614 | 0.058077004 | -0.713670309 | 0.052340195 |
| 12010.94922 | 13651.22949 | 12098.87695 | -0.377554403 | 0.20812859 | 0.118585679 | 0.572766243 | -0.726909347 | 0.105718826 |
| 418577.9375 | 392786.5 | 410743.5 | -0.377719872 | 0.080491971 | 0.314463943 | 0.106499477 | 0.278360585 | 0.128960705 |
| 52909.38672 | 54348.81641 | 47726.40625 | -0.378414318 | 0.024701492 | 0.311040298 | 0.05467585 | -0.0376042 | 0.730529882 |
| 163090.0156 | 183325.5313 | 167722.8906 | -0.379232115 | 0.037492489 | -0.348242861 | 0.114884739 | -0.391038611 | 0.068496057 |
| 17108.20508 | 16454.09766 | 11430.05859 | -0.379573579 | 0.013660857 | 0.766078387 | 0.102883408 | 0.811002404 | 0.142342008 |
| 198891.7656 | 219484.9688 | 185979.375 | -0.380282755 | 0.027969454 | 0.038740182 | 0.715505426 | -0.273509761 | 0.098453999 |
| 52961.73438 | 114953.5625 | 93083.08594 | -0.380479823 | 0.11742195 | 0.332825282 | 0.409333864 | 0.463368451 | 0.428577571 |
| 33993.94531 | 32112.47656 | 27457.37109 | -0.380824342 | 0.035474135 | -0.794625915 | 0.013885891 | -0.453294283 | 0.081022357 |
| 54100.42969 | 58842.80078 | 56444.34375 | -0.381208608 | 0.035680768 | -0.008967354 | 0.93847846 | -0.030575427 | 0.745602951 |
| 139262.75 | 140791.9063 | 130902.6953 | -0.38132775 | 0.01802855 | 0.369772166 | 0.032675377 | 0.060618475 | 0.433285959 |
| 71018.70313 | 122033.4688 | 80805.45313 | -0.381730858 | 0.121165219 | 0.105560091 | 0.580508796 | 0.63724096 | 0.262335514 |
| 23710.61523 | 22371.17188 | 20463.62695 | -0.382231752 | 0.059863719 | 0.806419766 | 0.033121608 | 0.409065362 | 0.062822 |
| 8473.460938 | 9100.886719 | 6206.053223 | -0.382430926 | 0.238301994 | -0.267939897 | 0.458532796 | -0.555707017 | 0.21738258 |
| 138624.8594 | 154122.8438 | 151399.8281 | -0.382561213 | 0.047647106 | -0.081836951 | 0.41896461 | -0.172130614 | 0.119608231 |
| 100730.3594 | 102530.6797 | 104582.1328 | -0.382927438 | 0.092143075 | 0.167193135 | 0.281730085 | 0.068438579 | 0.601247781 |
| 38977.22266 | 40546.75 | 40933.69141 | -0.383505248 | 0.337539485 | 1.179877006 | 0.123296741 | 0.051592961 | 0.901897565 |
| 87585.58594 | 80130.17188 | 74498.29688 | -0.3836963 | 0.058429627 | 0.370819693 | 0.119643706 | 0.395532997 | 0.07008325 |
| 970811.3125 | 962609.375 | 713192.5625 | -0.383794237 | 0.218653229 | 0.434551317 | 0.081028372 | 0.235581525 | 0.36267558 |

| | | | | | | | | |
|---|---|---|---|---|---|---|---|---|
| 1524354.25 | 2879204 | 1103586.875 | -0.383984549 | 0.804392171 | -1.195933743 | 0.417490892 | -1.460305306 | 0.359015506 |
| 425.8248596 | 576.3446045 | NA | -0.384433039 | 0.169155397 | 0.121524311 | 0.6777223 | -0.439928508 | NA |
| 68090.64063 | 61114.3125 | 67282.67188 | -0.384447273 | 0.055791428 | 0.110671543 | 0.424281202 | 0.138199834 | 0.249332771 |
| 14684.04004 | 15966.61523 | 13426.57617 | -0.384561365 | 0.047540015 | -0.270774207 | 0.10262657 | -0.250273482 | 0.118033234 |
| 4289.455078 | 3645.4729 | 2754.864014 | -0.385103959 | 0.061674089 | 0.377266085 | 0.150324399 | -0.201291429 | 0.478450914 |
| 84869.03906 | 86490.03125 | 83318.54688 | -0.385184467 | 0.009905914 | 0.102793546 | 0.245062116 | 0.081150278 | 0.077005785 |
| 45921.80469 | 41566.73047 | 39958.38281 | -0.38526103 | 0.019694688 | -0.435690378 | 0.023005377 | -0.602783617 | 0.019880539 |
| 14766.97559 | 10074.44238 | 11659.63672 | -0.385697727 | 0.129051952 | 0.665647841 | 0.052892568 | 0.039651057 | 0.878629235 |
| 508134.9375 | 531287.8125 | 484050.5313 | -0.385875825 | 0.04599437 | 0.092034965 | 0.311954306 | 0.382336964 | 0.029215308 |
| 3994.074707 | 5457.799805 | 2639.391113 | -0.385916915 | 0.22827254 | 0.175566611 | 0.313982423 | -0.914087757 | 0.086851601 |
| 37306.25 | 37584.68359 | 40537.63672 | -0.385941272 | 0.194809208 | -0.150749674 | 0.242349552 | -0.058329357 | 0.537160122 |
| 88607.71094 | 94372.34375 | 80507.67188 | -0.38632617 | 0.263045302 | 0.348903532 | 0.066380875 | 0.192960356 | 0.208408498 |
| 127800.7109 | 120782.1563 | 114895.4141 | -0.386395906 | 0.019930448 | 0.163788183 | 0.159908538 | -0.315715367 | 0.047120928 |
| 56100.73438 | 57974.86719 | 36383.78516 | -0.386641558 | 0.148656306 | -0.611504141 | 0.122344477 | -0.509164297 | 0.16522197 |
| 9145.477539 | 8211.414063 | 8033.363281 | -0.386833283 | 0.076604576 | 0.171562172 | 0.339659199 | -0.119054254 | 0.515909789 |
| 1735240.75 | 1790519.75 | 1667834.375 | -0.387625551 | 0.187040833 | 0.128897803 | 0.52544407 | -0.141238979 | 0.517915901 |
| 88188.35156 | 97515.33594 | 83224.39844 | -0.388009349 | 0.023527743 | -0.258291379 | 0.035493707 | -0.37142879 | 0.053053903 |
| 15998.81543 | 16725.89063 | 11125.42676 | -0.388141196 | 0.24464973 | -0.73808109 | 0.020431201 | -0.679416989 | 0.094664375 |
| 1387649 | 1398759.125 | 1269590.875 | -0.388517812 | 0.068786749 | 0.147119088 | 0.432898177 | -0.229475726 | 0.240274412 |
| 11726.61523 | 12994.09863 | 9338.455078 | -0.388599813 | 0.088231376 | -0.282146771 | 0.259477021 | -0.424973522 | 0.151995 |
| 81293.89063 | 96751.05469 | 88731.8125 | -0.38870196 | 0.115288524 | 0.432314578 | 0.11332333 | 0.185356367 | 0.387879439 |
| 50331.84375 | 51182.75195 | 46766.57617 | -0.389075147 | 0.045235089 | -0.222462656 | 0.076036407 | -0.349395273 | 0.027629097 |
| 37560.41406 | 37240.01563 | 36164.67188 | -0.389231433 | 0.063885064 | 0.991466767 | 0.021204995 | 0.774184947 | 0.008614663 |
| 115805.8438 | 129950.3438 | 122893.2734 | -0.389781751 | 0.427440307 | -0.056398033 | 0.783217426 | 0.222650167 | 0.213371407 |
| 1619922.875 | 1643970.375 | 1512517.25 | -0.390723988 | 0.216818598 | 0.049431353 | 0.61730772 | -0.164326152 | 0.206215456 |
| 62192.86328 | 66743.82813 | 56552.16016 | -0.391003477 | 0.095292766 | -0.136647626 | 0.429618137 | -0.447615023 | 0.095209744 |
| 61025.12109 | 56443.36328 | 58914.37891 | -0.391052948 | 0.070477311 | 0.255431988 | 0.189616058 | -0.02384794 | 0.882286856 |
| 10208.19434 | 10301.74316 | 11238.875 | -0.391095264 | 0.041542612 | 0.724093871 | 0.027730297 | 0.327191774 | 0.067123192 |
| 32508.8457 | 23204.93164 | 24979.01953 | -0.391234226 | 0.209930575 | 0.339140639 | 0.09227179 | 0.320874849 | 0.296894328 |
| 8317.918945 | 10482.65527 | 10516.41406 | -0.39137001 | 0.167028974 | 0.051539868 | 0.842124822 | -0.508263246 | 0.197748841 |
| 140456.5981 | 142947.2061 | 148356.3184 | -0.391419594 | 0.222905479 | 0.153751478 | 0.615862301 | -0.000869887 | 0.997993879 |
| 20604.50586 | 17351.43359 | 17845.72461 | -0.391424051 | 0.211214576 | 0.195022526 | 0.06863053 | 0.149623176 | 0.311287625 |
| 418046.75 | 405949.9063 | 384953.2188 | -0.391533473 | 0.018805749 | 0.093162891 | 0.121033071 | -0.197631632 | 0.050501801 |

| | | | | | | | | |
|---|---|---|---|---|---|---|---|---|
| 259566.3438 | 286383.6875 | 292896.9375 | -0.391652555 | 0.097443336 | 0.136247932 | 0.253840892 | 0.2679014 | 0.083375759 |
| 2277802.75 | 2234053.5 | 2246145.5 | -0.39177817 | 0.010484689 | -0.184274125 | 0.054857325 | -0.474426873 | 0.030633392 |
| 8566.324219 | 8341.105469 | 7328.618164 | -0.39240043 | 0.051502878 | 0.113424199 | 0.427225198 | 0.564606609 | 0.063440273 |
| 39271.32031 | 45253.4375 | 50659.69141 | -0.392559763 | 0.395173171 | 0.659109749 | 0.222516879 | 0.631492714 | 0.213371407 |
| 6133.580078 | 5895.251953 | 4666.485352 | -0.392607316 | 0.291066025 | 0.100239069 | 0.589689805 | -1.147697189 | 0.03161115 |
| 18460.89453 | 11259.31934 | 13051.63184 | -0.392819844 | 0.035879745 | 0.121913908 | 0.419136477 | -0.461470346 | 0.215760001 |
| 11682.02637 | 2872.061035 | NA | -0.392873687 | 0.338882582 | 0.306778307 | 0.399962117 | -0.299732392 | NA |
| 40053.09766 | 38060.4375 | 26962.0957 | -0.393251943 | 0.182007594 | 0.58212631 | 0.03968601 | 0.301676938 | 0.351174019 |
| 1257181.125 | 1231361 | 1282426.375 | -0.393365595 | 0.048657867 | 0.048550177 | 0.749640075 | 0.256741872 | 0.090232821 |
| 207787.75 | 216000.9688 | 193671.7969 | -0.393476532 | 0.113223175 | 0.516883254 | 0.063663792 | 0.504576443 | 0.060858857 |
| 69188.49219 | 78517.83594 | 75115.82031 | -0.393548164 | 0.439514374 | 0.492678805 | 0.37394121 | 1.724295276 | 0.035750975 |
| 323388.7813 | 341616.4375 | 369451.2813 | -0.393632895 | 0.016885305 | 0.097841957 | 0.213397043 | 0.179150242 | 0.151995 |
| 18815.34375 | 22086.65039 | 19717.42578 | -0.394079806 | 0.005752145 | -0.146760135 | 0.087974015 | -0.545356742 | 0.039602697 |
| 71585.375 | 73301.89844 | 76189.45313 | -0.394156285 | 0.019286562 | 0.097662763 | 0.211440521 | -0.067660831 | 0.33287633 |
| 30079.01953 | 28831.45117 | 27593.72266 | -0.394371625 | 0.017507272 | 0.230747613 | 0.10725287 | 0.255101162 | 0.045861991 |
| 93438.64063 | 87548.46875 | 96407.57813 | -0.394493673 | 0.138906239 | 0.25594863 | 0.047180214 | 0.050027914 | 0.505671265 |
| 144749.0313 | 144556.2031 | 148234.9688 | -0.394671644 | 0.122673099 | -0.033930325 | 0.860538167 | -0.096319274 | 0.588958618 |
| 92103.42969 | 86380.92188 | 82397.04688 | -0.39485596 | 0.02749963 | 0.251098756 | 0.065678888 | 0.120583563 | 0.224071828 |
| 4896.975098 | 5239.997559 | 4334.074219 | -0.395197194 | 0.218843086 | 0.228197375 | 0.496299916 | 0.420633481 | 0.205451497 |
| 151831.2031 | 158510.5 | 137466.9844 | -0.395308399 | 0.14905608 | -0.198835096 | 0.292501538 | -0.286008443 | 0.093070634 |
| 265183 | 281772.9688 | 275842.25 | -0.395356941 | 0.064600103 | 0.571307017 | 0.038121589 | 0.077082812 | 0.159215918 |
| 4437.478027 | 5254.3125 | 3931.147949 | -0.395426952 | 0.118526303 | -0.318365387 | 0.266880013 | -0.394195264 | 0.200720922 |
| 74824.52344 | 84288.625 | 64974.30078 | -0.396292429 | 0.270847227 | -0.251405302 | 0.330992339 | -0.301177903 | 0.268750571 |
| 28467.35547 | 30304.67969 | 27578.53906 | -0.396566058 | 0.019316731 | 0.003692874 | 0.955379211 | -0.173976317 | 0.095599918 |
| 244944.6094 | 243025.3281 | 239746.8125 | -0.396581704 | 0.099983289 | -0.320805462 | 0.102526216 | -0.228887673 | 0.062835547 |
| 221528.9844 | 210729.125 | 215588.4375 | -0.396666125 | 0.01150099 | 0.166129959 | 0.052817461 | 0.06278398 | 0.228883111 |
| 4877.663574 | 3777.792236 | 5378.788086 | -0.39678619 | 0.173500617 | 0.763176168 | 0.059340088 | 0.181938053 | 0.558333765 |
| 29220.71094 | 29408.31445 | 28467.46484 | -0.397130007 | 0.019286562 | 0.41403246 | 0.026826422 | 0.691267629 | 0.015632484 |
| 68483.82813 | 72225.60156 | 76459.25781 | -0.397231843 | 0.051425002 | -0.019793062 | 0.747450252 | 0.201410008 | 0.092375143 |
| 873339.8125 | 827507.125 | 508185.1875 | -0.397263595 | 0.152253903 | 0.381048995 | 0.192295741 | 0.070347494 | 0.859576357 |
| 171941.6719 | 178528.9219 | 180015.8125 | -0.397381796 | 0.013964852 | 0.35768882 | 0.025196036 | 0.419862768 | 0.017279709 |
| 316735.4688 | 339366.9688 | 314663.5938 | -0.397596507 | 0.09684552 | 0.026244531 | 0.776241637 | -0.037863759 | 0.665829523 |
| 12983.81543 | 14725.68555 | 13177.12695 | -0.398396402 | 0.363215631 | 0.34832739 | 0.450480395 | 0.069575251 | 0.882846687 |

| | | | | | | | | |
|---|---|---|---|---|---|---|---|---|
| 54616.85156 | 57537.71094 | 48294.88281 | -0.398560123 | 0.053893741 | -0.458403593 | 0.079312553 | -0.73628061 | 0.041176746 |
| 271472.0938 | 288881.625 | 285361.25 | -0.398805588 | 0.157678205 | 0.223922977 | 0.046782366 | 0.073585423 | 0.260904046 |
| 4046.753174 | 4834.485352 | 2652.720703 | -0.399212116 | 0.453634157 | -0.524154324 | 0.506362982 | -0.69112035 | 0.399002054 |
| 101379.7734 | 100868.9922 | 113456.9844 | -0.399387542 | 0.029514505 | -0.055952759 | 0.521536521 | 0.389339896 | 0.078538417 |
| 1287.085815 | 925.8080444 | 5118.768066 | -0.399888976 | 0.35552667 | -0.204561254 | 0.686661605 | -0.076497394 | 0.953592692 |
| 69891.57813 | 89291.21094 | 75327.83594 | -0.399954199 | 0.118293642 | -0.029696321 | 0.895586983 | 0.07653261 | 0.746954593 |
| 182082.7813 | 187055.9531 | 152746.6094 | -0.400071365 | 0.09809023 | -0.273719249 | 0.28126844 | -0.831178759 | 0.069232216 |
| 27162.22852 | 40354.40625 | 30532.87891 | -0.400542591 | 0.469719585 | 0.516263344 | 0.102916055 | 0.05018964 | 0.857787595 |
| 511251.5625 | 507598.875 | 486198.5938 | -0.400756167 | 0.039781174 | 0.075578266 | 0.079641904 | -0.145646269 | 0.04174272 |
| 197905.4063 | 207943.9063 | 197287.7656 | -0.400878639 | 0.01317261 | 0.034450397 | 0.626170555 | -0.083124569 | 0.24326618 |
| 94377.10156 | 83205.08594 | 91519.69531 | -0.401125543 | 0.108721666 | 0.291961727 | 0.230974993 | 0.826316303 | 0.030633392 |
| 19269.43359 | 18729.39063 | 18191.70508 | -0.401341049 | 0.022724126 | -0.044168237 | 0.608397014 | -0.095054968 | 0.312053461 |
| 505484.25 | 531856.375 | 553104.625 | -0.401376329 | 0.25462554 | 0.10041599 | 0.596340933 | 0.185904273 | 0.344337372 |
| 28578.80469 | 27676.27734 | 24413.48047 | -0.401575361 | 0.029390153 | -0.118591078 | 0.287681705 | -0.139451995 | 0.291532902 |
| 24418.32617 | 26844.37109 | 28332.17188 | -0.401940258 | 0.32827513 | 0.450983775 | 0.308741463 | 0.707715022 | 0.145046754 |
| 6071.51416 | 8229.641602 | 7331.532715 | -0.402223107 | 0.489346028 | 0.503914393 | 0.41861873 | 0.449903181 | 0.458870452 |
| 237888.4844 | 261182.167 | 250684.3906 | -0.402966114 | 0.173573283 | 0.285954337 | 0.331489808 | 0.266264649 | 0.343177751 |
| 4848.411621 | NA | 5854.69043 | -0.403100084 | NA | -0.400435783 | NA | -0.394543522 | NA |
| 23494.23242 | 32703.77148 | 28475.82031 | -0.403518814 | 0.117038402 | 0.240237698 | 0.321892407 | 0.166978425 | 0.545099828 |
| 3136.193848 | 2865.912842 | 1735.279419 | -0.40354977 | 0.081995281 | 1.011627678 | 0.026024574 | -0.307540175 | 0.409889954 |
| 195198.9844 | 154686.3594 | 156594.9375 | -0.40396917 | 0.09943294 | -0.106159404 | 0.463028085 | -0.432443547 | 0.085236358 |
| 15256.25781 | 24188.4082 | 20655.40625 | -0.404263018 | 0.030049926 | -0.090022086 | 0.630482257 | -0.045657543 | 0.872743168 |
| 27565.125 | 24053.82422 | 28092.34766 | -0.405204029 | 0.042878535 | 0.740963826 | 0.020431201 | 0.150803889 | 0.307121273 |
| 88760.48438 | 78216.10156 | 67981.41406 | -0.405542044 | 0.03228159 | -0.668835918 | 0.051861383 | -0.296996311 | 0.157967349 |
| 92041.60938 | 97171.64063 | 83745.14844 | -0.405880933 | 0.036566441 | -0.457048571 | 0.015497619 | -0.399780363 | 0.056784127 |
| 98461.625 | 70948.95313 | 131560.4219 | -0.405911914 | 0.034569907 | 0.077875042 | 0.497840929 | -0.196220672 | 0.595317341 |
| 49607.04297 | 47817.38672 | 46757.91406 | -0.405987018 | 0.065566406 | 0.175896971 | 0.161989613 | 0.0773676 | 0.252058261 |
| 133754.4531 | 138946.0313 | 144187.1563 | -0.406284262 | 0.034320767 | -0.017253585 | 0.847145011 | -0.082743552 | 0.402147464 |
| 267331.1875 | 271568.2813 | 258291.8438 | -0.406311418 | 0.068441045 | 0.134933659 | 0.112545088 | -0.145749205 | 0.107077213 |
| 4231.579102 | 4021.973389 | 3179.616211 | -0.406395953 | 0.058310315 | -0.159960505 | 0.325495749 | -0.265168853 | 0.219471057 |
| 810041.25 | 629805.875 | 643004.75 | -0.406490429 | 0.140338324 | 0.401787125 | 0.162035692 | 0.069344152 | 0.801698552 |
| 18338.77344 | 23445.96484 | 17846.12305 | -0.406751229 | 0.081936227 | 0.165204291 | 0.340489335 | -0.145367915 | 0.531405981 |
| 49214.52246 | 43894.88574 | 35778.27344 | -0.406824354 | 0.026586316 | 0.272494841 | 0.046114261 | 0.094908159 | 0.654410188 |

| | | | | | | | | |
|---|---|---|---|---|---|---|---|---|
| 189981.2656 | 198457.8438 | 188537.6875 | -0.40709278 | 0.018999567 | -0.471252117 | 0.058077004 | -0.616675684 | 0.034457303 |
| 5789.583984 | 6939.546387 | 6825.862305 | -0.407172328 | 0.165522295 | 0.520485419 | 0.168700158 | 0.07333554 | 0.786883312 |
| 161214.8281 | 176396.625 | 163424.4219 | -0.407250679 | 0.040287826 | -0.127549055 | 0.130087098 | -0.668316988 | 0.014388274 |
| 92308.32031 | 132297.2344 | 83137.97656 | -0.407939351 | 0.156741193 | -0.185996709 | 0.491954038 | -0.370084488 | 0.283693593 |
| 9241.504883 | 9669.826172 | 9897.052734 | -0.408130171 | 0.20862342 | 0.108521575 | 0.723850432 | -0.032892955 | 0.920227312 |
| 137644.9219 | 144171.8906 | 142512.125 | -0.408223462 | 0.054479995 | 0.225787236 | 0.01179292 | 0.261139066 | 0.030813706 |
| 183759.4824 | 179565.6914 | 191516.0566 | -0.408389526 | 0.026631215 | 0.253683572 | 0.04128058 | 0.458889932 | 0.021984636 |
| 201096.4531 | 143171.4219 | 127646.7344 | -0.408731649 | 0.491789048 | 0.54227236 | 0.645244823 | -0.758854791 | 0.07008325 |
| 40268.80078 | 44419.57227 | 42092.19336 | -0.408907812 | 0.275794055 | 0.712475869 | 0.150649728 | 0.021454795 | 0.958657593 |
| 32348.2168 | 34351.53516 | 31641.16992 | -0.409045908 | 0.029183044 | 0.028945274 | 0.773489083 | 0.036235972 | 0.731684806 |
| 1261083.289 | 1184198.401 | 1022978.193 | -0.409310518 | 0.095703193 | 0.001626895 | 0.992236761 | -0.042000331 | 0.843349698 |
| 619872.125 | 592185.8125 | 592130.1875 | -0.409884023 | 0.073778017 | 0.707508312 | 0.021906439 | -0.117757508 | 0.316209448 |
| 260547.1563 | 294989.125 | 199167.0469 | -0.410320285 | 0.359445722 | 0.781849652 | 0.119439839 | 0.422811951 | 0.347215504 |
| 36605.34766 | 38046.09375 | 36789.61328 | -0.411074391 | 0.035399497 | -0.274201543 | 0.094078555 | -0.256996423 | 0.105824879 |
| 30594.91602 | 25964.81641 | 28244.24609 | -0.411080663 | 0.042404592 | 0.041981651 | 0.750780433 | -0.546359345 | 0.043034683 |
| 13497.29785 | 11593.88281 | 7987.559082 | -0.411111591 | 0.018220349 | -0.271767399 | 0.07245119 | -0.810615962 | 0.076836055 |
| 67477.19531 | 72278.97656 | 45683.17578 | -0.411250234 | 0.058516905 | -0.062673155 | 0.79527904 | -0.245783495 | 0.408044366 |
| 260532.3203 | 292482.4238 | 294196.7148 | -0.411292362 | 0.046747318 | 1.028725952 | 0.015395012 | 0.658005805 | 0.026263251 |
| 7814.244629 | 7280.728516 | 5605.215332 | -0.411467324 | 0.309938239 | -0.729620549 | 0.244556831 | -0.827938977 | 0.194263423 |
| 520188.4375 | 547358.1875 | 555883.3125 | -0.412069086 | 0.02548578 | 0.005947556 | 0.93298262 | -0.140036137 | 0.067893499 |
| 13519.08594 | 8378.480469 | 12962.2998 | -0.412226722 | 0.120468071 | 0.783767718 | 0.102883408 | 0.618407583 | 0.21819369 |
| 28471.64844 | 22991.09766 | 20673.90625 | -0.412508645 | 0.065089491 | 0.072426687 | 0.635139401 | -0.173417285 | 0.458870452 |
| 12653.12207 | 17785.03125 | 5406.172363 | -0.413464817 | 0.099542225 | -0.701055932 | 0.102963557 | -1.080348969 | 0.110105852 |
| 84248.58594 | 81808.46875 | 77945.50781 | -0.413647427 | 0.04424731 | 0.073204548 | 0.552558639 | -0.065009584 | 0.609208541 |
| 84992.92188 | 80968.25 | 83564.11719 | -0.413858871 | 0.042146623 | -0.03765011 | 0.732942123 | -0.113814362 | 0.354443352 |
| 1528762 | 1790028.875 | 1445208.75 | -0.413976063 | 0.224955599 | -0.121872025 | 0.578166827 | -0.14187775 | 0.515665974 |
| 52419.77344 | 48467.41406 | 47553.25781 | -0.4141472 | 0.074012043 | -0.466202928 | 0.153662285 | -0.913675825 | 0.074385174 |
| 55457.05469 | 53354.21094 | 49656.00781 | -0.414337278 | 0.038217062 | 0.219537339 | 0.11417961 | 0.091266579 | 0.346395659 |
| 42938.05469 | 44343.36719 | 36819.63672 | -0.414376946 | 0.093428026 | 0.019911908 | 0.925262844 | -0.129806245 | 0.548141599 |
| 112902.0078 | 114256.6953 | 93166.96875 | -0.414736604 | 0.020823572 | -0.006245957 | 0.885440998 | -0.176818154 | 0.268490418 |
| 345536.75 | 343110.1875 | 347807.7813 | -0.414737781 | 0.019286562 | -0.352615811 | 0.047069121 | -0.346925047 | 0.066274975 |
| 69925.35938 | 75691.78125 | 70089.46094 | -0.414850909 | 0.016764271 | 0.072984103 | 0.147710173 | -0.321384229 | 0.027331143 |
| 97726.76563 | 84551.15625 | 67477.26563 | -0.414970164 | 0.102272218 | 0.152345988 | 0.467062508 | -0.184418762 | 0.51695586 |

| | | | | | | | | |
|---|---|---|---|---|---|---|---|---|
| 35358.65625 | 32390.81641 | 37286.82813 | -0.415534218 | 0.047406642 | -0.023107596 | 0.816076757 | -0.30011392 | 0.057068534 |
| 164249.8281 | 148777.2656 | 145813.3281 | -0.41604329 | 0.592240482 | 0.639803034 | 0.431424722 | 0.525609044 | 0.512102022 |
| 39335.30469 | 40689.84375 | 32269.10547 | -0.416568952 | 0.027690255 | 0.485556781 | 0.06015933 | -0.128806615 | 0.448929881 |
| 35258.33594 | 39965.37891 | 24248.92383 | -0.416588638 | 0.158308661 | -0.064033641 | 0.850198859 | -0.450650775 | 0.25973778 |
| 40682.19922 | 42298.70703 | 57255.11719 | -0.416968163 | 0.073661708 | 0.077093378 | 0.617383969 | 0.20759543 | 0.478832582 |
| 169677.0469 | 185841.2656 | 168060.2344 | -0.417963605 | 0.053861475 | 0.619862422 | 0.029180754 | 0.606075009 | 0.025517732 |
| 28040.46289 | 22313.2793 | 25577.68359 | -0.418148635 | 0.045092515 | -0.014660597 | 0.910648359 | -0.30496361 | 0.14131016 |
| 30160.53711 | 31603.25781 | 25524.42969 | -0.418187201 | 0.170113636 | 0.799506791 | 0.048271656 | 0.381338145 | 0.147031451 |
| 5208.520996 | 5749.366699 | 4206.245605 | -0.418236627 | 0.100529206 | 0.535011876 | 0.03809973 | 0.071860169 | 0.738904113 |
| 91128.25781 | 92666.5 | 83409.89844 | -0.418663997 | 0.02194812 | 0.242458929 | 0.142713982 | 0.119739212 | 0.254146446 |
| 10441.86816 | 7016.42041 | 10228.00586 | -0.418708156 | 0.106137102 | -0.180026737 | 0.47714217 | -0.352216812 | 0.282822905 |
| 13897.47852 | 12500.40137 | 11674.4082 | -0.418806013 | 0.342592569 | 0.63517329 | 0.181757098 | 0.16819309 | 0.663092542 |
| 58.53773117 | NA | NA | -0.418988348 | 0.117038402 | NA | NA | -2.633452267 | NA |
| 2458.041016 | 2382.325684 | 1267.893188 | -0.419053145 | 0.077013222 | -0.672786005 | 0.097722369 | -1.204971973 | 0.044885273 |
| 99521.83594 | 103129.4063 | 108018.6016 | -0.419079642 | 0.018311389 | 0.170542413 | 0.05818214 | 0.378810143 | 0.028275238 |
| 29556.83984 | 28247.66211 | 30748.04297 | -0.420033379 | 0.06487731 | 0.379299028 | 0.168464234 | 0.222846435 | 0.138173555 |
| 59722.22266 | 45929.44141 | 48691.44531 | -0.420335861 | 0.077066654 | -0.03003678 | 0.852417959 | 0.029037566 | 0.886748012 |
| 77508.05469 | 77367.60938 | 79418.47656 | -0.420574358 | 0.016105265 | 0.261035812 | 0.112996996 | -0.343470556 | 0.062345205 |
| 11331.67773 | 9944.291016 | 9177.000977 | -0.420599359 | 0.037492489 | -0.385960031 | 0.11146465 | -0.550016762 | 0.052091631 |
| 367931.4063 | 406931.375 | 362933.125 | -0.420659699 | 0.355016641 | 0.021702415 | 0.90147703 | 0.005510424 | 0.960764822 |
| 42368.72656 | 45009.90234 | 43855.99219 | -0.421191302 | 0.135239277 | 0.294790996 | 0.07036809 | 0.175696171 | 0.145071579 |
| 175378.2188 | 174670.8438 | 163886.3906 | -0.42142676 | 0.069981647 | 0.204032906 | 0.046114261 | 0.083446689 | 0.194692917 |
| 245692.6875 | 282403.7188 | 252770.2969 | -0.421697688 | 0.069692211 | 0.027795673 | 0.8496215 | -0.293216106 | 0.149748939 |
| 223797.0313 | 241943.0781 | 260936.5781 | -0.42174339 | 0.077013222 | -0.047385517 | 0.501016819 | 0.625255793 | 0.045311345 |
| 162012.6719 | 23037.85938 | 133027.9063 | -0.421830489 | 0.740331859 | 1.705939746 | 0.291036386 | 2.384801724 | 0.27757357 |
| 125907.5938 | 133622.0547 | 126890.5195 | -0.421844051 | 0.032963794 | -0.109970496 | 0.312686937 | -0.138208315 | 0.226153996 |
| 61511.11719 | 61734.39844 | 58064.91406 | -0.421889402 | 0.027158619 | 0.048159277 | 0.645770679 | -0.240386269 | 0.120554018 |
| 56352.19922 | 106131.5 | 60007.64844 | -0.421927116 | 0.308539067 | -0.071491097 | 0.877927692 | -0.20738759 | 0.699603473 |
| 65169.02734 | 49629.22266 | 64250.90625 | -0.422110717 | 0.021498678 | 0.380349208 | 0.05744085 | 0.208727806 | 0.356805712 |
| 79051 | 76955.83594 | 78464.94531 | -0.422159721 | 0.033123824 | -0.379113259 | 0.045064018 | -0.625529359 | 0.034567592 |
| 30975228 | 32382768 | 32143256 | -0.42307069 | 0.043802925 | -0.085282708 | 0.249697654 | -0.146680468 | 0.106029703 |
| 5325.121094 | 4449.537598 | 3627.895996 | -0.423306375 | 0.142526436 | -0.239750098 | 0.430961626 | -1.171659799 | 0.077081711 |
| 28778.43555 | 26582.2168 | 25521.07227 | -0.423350267 | 0.036566441 | -0.036215313 | 0.767078045 | -0.496385813 | 0.064571615 |

| | | | | | | | | |
|---|---|---|---|---|---|---|---|---|
| 120794.3359 | 113099.8438 | 112589.8125 | -0.423731937 | 0.019286562 | -0.324498156 | 0.025196036 | -0.334748424 | 0.021564465 |
| 102191.7656 | 114749.0781 | 104379.875 | -0.424075415 | 0.06257795 | 0.089775657 | 0.583013317 | -0.043846797 | 0.801045589 |
| 177107.3906 | 171996.2969 | 164677.5313 | -0.424186353 | 0.010893509 | -0.147907383 | 0.022410713 | -0.36387226 | 0.021984636 |
| 11708.15234 | 12440.95801 | 10809.72266 | -0.424478142 | 0.046970661 | 0.008172267 | 0.945953011 | -0.475182427 | 0.051065331 |
| 60584.33984 | 87243.08594 | 86181.17969 | -0.424848566 | 0.449984317 | 0.77630084 | 0.088308366 | 1.361033924 | 0.06414243 |
| 159340.4063 | 294578.1875 | 139701.2656 | -0.424981372 | 0.616535572 | 0.318681661 | 0.663006884 | 0.30343184 | 0.629623612 |
| 100265.2969 | 103858.4297 | 92164.89063 | -0.425373192 | 0.444866261 | 0.075900699 | 0.752789866 | 0.047750238 | 0.763765593 |
| 85835.125 | 102372.8984 | 81393.07813 | -0.425389994 | 0.170677244 | -0.316669752 | 0.243813899 | -0.192387099 | 0.374127623 |
| 6834.298828 | 6951.616211 | 7636.471191 | -0.425824423 | 0.590378147 | 1.038094168 | 0.286605361 | 0.347794481 | 0.508977597 |
| 118622.6484 | 122807.0078 | 131760.2188 | -0.426063554 | 0.050055523 | 0.497862359 | 0.052756858 | 0.260327905 | 0.142358981 |
| 19065.92969 | 18035.99609 | 15857.99121 | -0.426263131 | 0.027231414 | 0.119682266 | 0.147323941 | -0.315468035 | 0.079785507 |
| 125713.6094 | 146536.6719 | 127264.9219 | -0.426339214 | 0.049203263 | -0.012891803 | 0.909428871 | -0.053891676 | 0.681798536 |
| 3533.168945 | 5785.519043 | 7053.462402 | -0.426427928 | 0.619239039 | 0.698658775 | 0.418601481 | 0.538846523 | 0.549312085 |
| 25757.00195 | 24806.81055 | 22877.57227 | -0.426792578 | 0.066713713 | 0.090103185 | 0.562301807 | 0.029813712 | 0.851983677 |
| 10638.63184 | 10296.17969 | 11525.41406 | -0.427332121 | 0.553857303 | 0.949085896 | 0.228114774 | 0.619995509 | 0.408792062 |
| 34602.84766 | 36349.36719 | 37435.41406 | -0.427512821 | 0.038648015 | 0.103141434 | 0.228997603 | -0.088434325 | 0.192235011 |
| 7783.581543 | 7783.423828 | 5712.806641 | -0.42873911 | 0.107191715 | 0.527585758 | 0.027730297 | -0.10985748 | 0.604419494 |
| 42114.60938 | 38065.23828 | 34412.19922 | -0.429390537 | 0.079968566 | 0.506426418 | 0.111778257 | -0.183594131 | 0.223412133 |
| 21553.66797 | 25545.03516 | 6441.646973 | -0.430188504 | 0.036480793 | 0.586818849 | 0.395857551 | -0.485117527 | 0.454451558 |
| 168723.875 | 165359.4063 | 175660.8438 | -0.430295068 | 0.160518598 | 0.242931953 | 0.099165657 | -0.067871332 | 0.377810872 |
| 22100.54102 | 24788.22266 | 25392.05078 | -0.430509189 | 0.078894657 | 0.210504258 | 0.433819762 | 0.292712564 | 0.167225632 |
| 11054.58984 | 11303.03906 | 11891.29883 | -0.430912939 | 0.221775028 | 0.060477998 | 0.875290527 | -0.066625834 | 0.851983677 |
| 134803.2656 | 140494.625 | 129932.2656 | -0.430943596 | 0.036218901 | 0.30688406 | 0.073326865 | 0.391304398 | 0.022968762 |
| 165472.5 | 165851.6719 | 195469.3438 | -0.430955314 | 0.096166178 | 0.313888522 | 0.042887894 | 0.044361282 | 0.746954593 |
| 12587.14551 | 11286.74805 | 8096.604004 | -0.431074083 | 0.077955129 | -0.164859677 | 0.412941247 | -0.635384343 | 0.082189314 |
| 154069.9531 | 158218.625 | 156819.6406 | -0.431167157 | 0.023591212 | -0.194553071 | 0.106684108 | -0.11863234 | 0.206265186 |
| 8119.598145 | 8081.509277 | 3526.121582 | -0.431422262 | 0.030838583 | -0.425681746 | 0.092617517 | -0.751427705 | 0.166657728 |
| 41626.58984 | 37216.63281 | 37744.47266 | -0.431506513 | 0.026698267 | 0.585643269 | 0.055241445 | 0.185406893 | 0.166657728 |
| 67522.61719 | 62605.41016 | 59633.40234 | -0.431576557 | 0.066276059 | 0.233830932 | 0.149671955 | 0.274918426 | 0.106413249 |
| 575524.6875 | 622687.375 | 809638.875 | -0.431639629 | 0.086055451 | -0.979923651 | 0.082612243 | -0.939340845 | 0.059641593 |
| 45079.88281 | 45512.16016 | 41703.44922 | -0.432253461 | 0.010297098 | 0.522515192 | 0.131196907 | 0.116711206 | 0.166745689 |
| 30092.24219 | 29026.49023 | 26008.74219 | -0.432357478 | 0.010297098 | 0.085942651 | 0.101031629 | 0.341947922 | 0.09056925 |
| 17573.93945 | 14367.29297 | 15832.20508 | -0.43280749 | 0.073031078 | -0.056541358 | 0.758682909 | 0.104322893 | 0.592878834 |

| | | | | | | | | |
|---|---|---|---|---|---|---|---|---|
| 117737.0703 | 117374.0781 | 107169.5078 | -0.432873811 | 0.012041023 | -0.072245996 | 0.258199098 | -0.304636884 | 0.034677634 |
| 4856.885254 | 4833.130371 | 4302.896484 | -0.433980975 | 0.150766851 | 0.252159954 | 0.18325637 | 0.022652152 | 0.833250417 |
| 226878.7695 | 247587.0313 | 259479.0469 | -0.434278466 | 0.196263279 | 0.340120701 | 0.305853052 | 0.270899959 | 0.385444173 |
| 10131.29883 | 7939.280273 | 7933.375488 | -0.434900578 | 0.142694155 | 2.759417045 | 0.023005377 | 2.284502031 | 0.052503815 |
| 23090.83789 | 22767.4375 | 17123.37891 | -0.435791526 | 0.403414633 | 0.483344323 | 0.155305603 | 0.631735221 | 0.112423117 |
| 122775.5625 | 126598.8438 | 141717.0938 | -0.435875243 | 0.580848626 | 1.086923083 | 0.097735536 | 1.319453853 | 0.039508617 |
| 131491.1563 | 137109.5313 | 137099.6094 | -0.436154043 | 0.005752145 | 0.149452204 | 0.154908183 | 0.038184844 | 0.344839158 |
| 112023.6797 | 115695.1172 | 106856.75 | -0.436779446 | 0.189533944 | -0.204597819 | 0.077996184 | -0.265376517 | 0.040202519 |
| 52147.05469 | 53126.27734 | 62913.78906 | -0.437268324 | 0.04179011 | 0.053226424 | 0.723122465 | -0.053288363 | 0.762071822 |
| 107577.5234 | 104381.1172 | 95121.32813 | -0.437882569 | 0.062325508 | -0.160659341 | 0.364459731 | -0.152577877 | 0.376493076 |
| 35314.42578 | 38693.21094 | 44855.09766 | -0.438425246 | 0.1051608 | 0.239754902 | 0.297997596 | 0.185736612 | 0.408044366 |
| 113072.9141 | 108523.1875 | 116688.5234 | -0.438659794 | 0.019463657 | 0.10330106 | 0.275753538 | 0.253466519 | 0.061202655 |
| 97043.85156 | 106438.7188 | 120843.9922 | -0.438787093 | 0.041709412 | -0.244394731 | 0.122107272 | -0.268901226 | 0.143173978 |
| 28002.41797 | 29010.94922 | 25861.36914 | -0.43886228 | 0.03740048 | 0.658779248 | 0.021408573 | 0.237478793 | 0.082507796 |
| 59572.10156 | 57138.93359 | 53696.25781 | -0.439077679 | 0.013660857 | 0.419488633 | 0.024814918 | 0.180564985 | 0.101483387 |
| 184424.2344 | 189469.3594 | 193320.9375 | -0.439168858 | 0.055757407 | 0.177327769 | 0.087781599 | 0.230246946 | 0.046487363 |
| 38250.32813 | 40540.92578 | 36994.99219 | -0.439609587 | 0.032802153 | -0.145178702 | 0.313230636 | -0.355096639 | 0.085307756 |
| 1192.03064 | 1802.055176 | 443.7984924 | -0.439881864 | NA | -0.177767282 | NA | -0.509113847 | NA |
| 7588.614258 | 9359.109375 | 7056.28125 | -0.440312563 | 0.486426842 | 0.716708493 | 0.256103156 | -0.032346028 | 0.964243952 |
| 79016.91406 | 72033.95313 | 74832.90625 | -0.440592193 | 0.037492489 | -0.111740414 | 0.275278246 | -0.121729975 | 0.21738258 |
| 129186.7578 | 123369.2813 | 130864.3125 | -0.440616818 | 0.006395507 | 0.086482022 | 0.306017296 | -0.065227469 | 0.20646932 |
| 23463.26563 | 22893.98438 | 11760.4043 | -0.440925502 | 0.094057963 | -0.791310704 | 0.101856807 | -1.212412385 | 0.045311345 |
| 376850.25 | 392968.4375 | 373651.5938 | -0.441419017 | 0.060948791 | 0.079616903 | 0.332892691 | 0.011071876 | 0.829744218 |
| 37088.81641 | 38629.91016 | 37728.89844 | -0.441878132 | 0.053424232 | 0.414216664 | 0.024939682 | 0.419326695 | 0.015712916 |
| 16400.20117 | 25112.15625 | 22172.8457 | -0.442594272 | 0.141668804 | 0.508159319 | 0.111349456 | 0.354849642 | 0.328604461 |
| 11610.58984 | 5407.4375 | 7657.184082 | -0.443024323 | 0.250607381 | -0.122428801 | 0.562301807 | -0.191431019 | 0.689648336 |
| 17211.14844 | 16811.16016 | 17184.97852 | -0.443142098 | 0.028143177 | -0.0361722 | 0.773489083 | 0.302905097 | 0.030727383 |
| 5637.899902 | 5229.682129 | 5861.214355 | -0.443301095 | 0.106525325 | 0.035435583 | 0.825034469 | 0.443183025 | 0.036016754 |
| 16602.93359 | 13091.62891 | 11875.56152 | -0.444306751 | 0.035146788 | 0.032899141 | 0.812563709 | -0.632972054 | 0.060937162 |
| 78851.03125 | 76440.69531 | 61975.44141 | -0.444559307 | 0.016105265 | 0.035897012 | 0.706396106 | -0.087937056 | 0.588834899 |
| 36380.75098 | 37160.48755 | 36678.99487 | -0.444574704 | 0.028368096 | 0.173291076 | 0.179978582 | 0.186018355 | 0.169021874 |
| 15058988 | 18450338 | 19393520 | -0.445000504 | 0.010893509 | 0.058132485 | 0.436404714 | 0.151289367 | 0.423263948 |
| 7212.092285 | 7517.988281 | 5633.63623 | -0.445012301 | 0.06543655 | 0.064269202 | 0.713346788 | -0.203378102 | 0.320774355 |

| | | | | | | | | |
|---|---|---|---|---|---|---|---|---|
| 50592.58984 | 52418.38672 | 52113.23828 | -0.445716301 | 0.10042667 | 0.050688646 | 0.798421669 | -0.080086816 | 0.692301013 |
| 34441.9375 | 32344.0332 | 36030.92969 | -0.445722162 | 0.037562551 | 0.276341473 | 0.119711295 | 0.862981695 | 0.016683666 |
| 1558384.875 | 5015086.5 | 1448372.5 | -0.446020918 | 0.736518539 | -0.522119268 | 0.697931686 | -0.750652607 | 0.531726747 |
| 377216.7188 | 385485.375 | 373661.1875 | -0.44626438 | 0.008009891 | 0.071617809 | 0.234875537 | 0.074707445 | 0.12440873 |
| 9390.208008 | 6095.332031 | 23621.21094 | -0.446273319 | 0.344215704 | -0.365280275 | 0.596219861 | -0.251406373 | 0.763776161 |
| 257498 | 273587.75 | 253384.875 | -0.446322678 | 0.007485535 | -0.058404356 | 0.102350969 | -0.091711844 | 0.17538628 |
| 85630.42969 | 87028.32031 | 78362.77344 | -0.446696404 | 0.069867455 | 0.080546394 | 0.279028781 | -0.06454624 | 0.399380775 |
| 265818.125 | 266408.5 | 260617.8594 | -0.446972872 | 0.136265249 | 0.421389487 | 0.191226807 | 0.451631025 | 0.169799715 |
| 56609.30469 | 57711.52734 | 54763.48047 | -0.447118491 | 0.18607177 | 0.056167035 | 0.69690851 | 0.178106275 | 0.065367829 |
| 8709.370117 | 8767.333984 | 8908.364258 | -0.447195524 | 0.073155658 | 0.218036961 | 0.094078555 | -0.035601626 | 0.349400005 |
| 38432.46094 | 39684.64063 | 38235.39453 | -0.447341156 | 0.048502917 | 0.787183304 | 0.046487658 | -0.105790151 | 0.512774056 |
| 8278.860352 | 9512.962891 | 10523.91211 | -0.447523042 | 0.111176969 | 0.228158086 | 0.137745823 | 0.276725211 | 0.201634825 |
| 17430.10352 | 17555.63086 | 13986.48145 | -0.448268417 | 0.059460749 | 0.281430489 | 0.17813923 | 0.146978368 | 0.464975591 |
| 17033.375 | 16834.05859 | 13942.25781 | -0.448359847 | 0.038942433 | 0.075223048 | 0.535392728 | -0.031563442 | 0.836718763 |
| 108272.9766 | 123758.2266 | 110410.3359 | -0.449319718 | 0.131122522 | -0.388451663 | 0.110122293 | -0.267704019 | 0.152372879 |
| 38953.79688 | 38161.46094 | 36489.83594 | -0.449368791 | 0.064133462 | -0.25717637 | 0.205468655 | -1.029681797 | 0.057713404 |
| 80641.35938 | 75195.88281 | 77410.32031 | -0.449832522 | 0.121782631 | 0.163340159 | 0.313413007 | 0.266617242 | 0.041258108 |
| 6069.958496 | 4936.692383 | 3856.119141 | -0.450410464 | 0.134042679 | 0.321140343 | 0.532402501 | -0.406517247 | 0.277551095 |
| 42857.61719 | 40833.30078 | 45871.28516 | -0.450552509 | 0.252906284 | 0.964063338 | 0.059093142 | 0.967812008 | 0.034677634 |
| 14536.08984 | 15390.24219 | 12620.18359 | -0.450947073 | 0.039409697 | 0.22883101 | 0.13125649 | -0.049170839 | 0.74659203 |
| 12013.84961 | 10186.92969 | 6213.495605 | -0.450958754 | 0.227207392 | -0.072301227 | 0.805534524 | -0.238106116 | 0.559687295 |
| 2509.320801 | 3031.64502 | 1749.606812 | -0.450999663 | 0.221774211 | -0.05521486 | 0.890279825 | -0.521148866 | 0.30758607 |
| 524216.4375 | 769954.75 | 468305.7813 | -0.451026656 | 0.440692775 | 0.381705504 | 0.432898177 | 0.430156869 | 0.34800693 |
| 7127.080078 | 1721.64563 | 6275.126465 | -0.451087322 | 0.707492306 | -0.093491151 | 0.926456674 | -0.756970101 | 0.382150932 |
| 98286.57031 | 100071.1094 | 106247.8438 | -0.451565516 | 0.037389698 | 0.073226344 | 0.232060257 | 0.303754808 | 0.032840382 |
| 4026.639648 | 4098.334961 | 4255.946777 | -0.451590664 | NA | 0.147508953 | 0.613687235 | -0.536676001 | 0.159389389 |
| 105756.6875 | 115920.1016 | 56032.50781 | -0.451874868 | 0.186886504 | -0.911171919 | 0.030053634 | -0.976510597 | 0.088892469 |
| 402589.4063 | 355277.5938 | 301207.1563 | -0.451876311 | 0.155839329 | 0.879710025 | 0.052946166 | 0.52598114 | 0.13403167 |
| 22865.73438 | 19754.08398 | 23784.80273 | -0.452179409 | 0.240333703 | 0.632629757 | 0.049160495 | 0.474770345 | 0.08420993 |
| 7122.253906 | 6296.541992 | 5899.672852 | -0.453330835 | 0.084766561 | -0.214093014 | 0.343878142 | -0.075602806 | 0.713487699 |
| 2057.855957 | 2042.437134 | 1251.171509 | -0.453705312 | 0.08116125 | 0.661933573 | 0.239014656 | -0.573333288 | 0.136049841 |
| 310140.6875 | 272645.2813 | 290608.4063 | -0.453926237 | 0.012774217 | 0.176550379 | 0.079406664 | 0.559470541 | 0.03884284 |
| 376909.0625 | 553343.0625 | 579014 | -0.454977838 | 0.473151187 | 0.834245865 | 0.039887225 | 1.71028055 | 0.084223187 |

| | | | | | | | | |
|---|---|---|---|---|---|---|---|---|
| 40664.08594 | 34811.35547 | 38494.375 | -0.455478811 | 0.20799233 | 1.383508111 | 0.024018676 | 1.822573984 | 0.029710029 |
| 9970.519531 | 9157.547852 | 6505.951172 | -0.4556396 | 0.02753514 | 0.552305255 | 0.056328031 | -0.113732537 | 0.67224785 |
| 13894.55273 | 19571.83008 | 14588.42188 | -0.45585025 | 0.181071128 | -0.052364645 | 0.540113573 | -0.275322797 | 0.296309595 |
| 7575.978027 | 7882.902832 | 5415.492676 | -0.455886033 | 0.028446198 | 0.394924894 | 0.056328031 | -0.07920708 | 0.747716162 |
| 16642.0918 | 16715.07422 | 13741.4502 | -0.456183021 | 0.121422892 | 0.276871222 | 0.156712278 | -0.012633418 | 0.950211597 |
| 12956.84766 | 14282.43359 | 12108.18164 | -0.456379029 | 0.019694688 | 0.457906522 | 0.046899287 | 0.10684692 | 0.414665837 |
| 1739750.375 | 1808086 | 1689762.125 | -0.456650059 | 0.029183044 | 0.241028216 | 0.102526216 | 0.077712674 | 0.428343498 |
| 34412.87109 | 64638.75 | 30640.77734 | -0.45718493 | 0.261850791 | -0.016583911 | 0.976816962 | -0.197919097 | 0.737230437 |
| 23055.23828 | 24413.92969 | 22643.70508 | -0.457225128 | 0.044205353 | -0.025187801 | 0.728877771 | -0.284957845 | 0.026764821 |
| 14434.84961 | 16985.72852 | 13887.35742 | -0.457813773 | 0.038651604 | 0.09642602 | 0.471790095 | -0.515772968 | 0.053375696 |
| 5905.50293 | 3059.478027 | 4323.471191 | -0.458047746 | 0.216922584 | 1.324291629 | 0.116818656 | 0.285642113 | 0.543407482 |
| 8340.18457 | 7496.597168 | 5938.542969 | -0.458107774 | 0.037647614 | -0.332012475 | 0.114167274 | -0.579549287 | 0.066292531 |
| 122423.4766 | 111826.6797 | 116100.1094 | -0.45811275 | 0.137237024 | 0.862934015 | 0.043784011 | 1.578512008 | 0.019022762 |
| 81325.74219 | 93129.83594 | 89405.85156 | -0.458163831 | 0.136416657 | 0.38568007 | 0.186393782 | 0.142948133 | 0.549614221 |
| 134711.7656 | 131306.2031 | 80796.90625 | -0.458216255 | 0.05112493 | 0.17470128 | 0.399976168 | -0.630899574 | 0.142328609 |
| 143915.7344 | 129068.8125 | 132202.8438 | -0.458561252 | 0.02059262 | 0.537075352 | 0.023068087 | 0.416635352 | 0.036013501 |
| 27130.76367 | 41782.74219 | 13916.17969 | -0.458942598 | 0.799151672 | -1.216414773 | 0.512809933 | -1.604543571 | 0.436830073 |
| 15088.59473 | 14247.93359 | 12077.6416 | -0.459125588 | 0.040854235 | -0.059193339 | 0.663386446 | -0.363765371 | 0.107874808 |
| 162072.9043 | 170961.6355 | 167164.5237 | -0.459746967 | 0.039978912 | -0.008196153 | 0.938617908 | -0.109117144 | 0.345320824 |
| 5593945 | 5688600 | 5310614.5 | -0.459824886 | 0.013660857 | 0.29632715 | 0.028019426 | 0.033645741 | 0.574749876 |
| 195308.7813 | 195670.3125 | 201573.1719 | -0.460403403 | 0.268867993 | 0.678640742 | 0.081994347 | 0.714752947 | 0.0890834 |
| 5088.937988 | 9291.280273 | 8316.860352 | -0.461108861 | 0.055791428 | -0.534759659 | 0.241155984 | -0.284122491 | 0.433777033 |
| 124351.0547 | 121044.4766 | 119052.3906 | -0.46112785 | 0.036448621 | -0.299134789 | 0.050410119 | -0.338551217 | 0.029219266 |
| 65248.66016 | 75323.46094 | 65043.40234 | -0.46124896 | 0.068715601 | 0.135753643 | 0.270113498 | 0.187162904 | 0.22024877 |
| 233990.7344 | 220058.1563 | 194456.6719 | -0.462008406 | 0.05579064 | 0.047195233 | 0.776741539 | -0.053908459 | 0.780623364 |
| 717202.625 | 637989.3125 | 719748.8125 | -0.462388089 | 0.067008234 | 0.085127823 | 0.305630144 | -0.277129871 | 0.065074878 |
| 114898.8125 | 106832.9297 | 95068.4375 | -0.46246974 | 0.015991503 | 0.034882578 | 0.475179451 | -0.142453491 | 0.294777228 |
| 101742.0234 | 79567.57813 | 90333.61719 | -0.462692836 | 0.009974872 | 0.255061877 | 0.081451772 | -0.161072039 | 0.340144283 |
| 26970.46289 | 29224.35938 | 27768.26953 | -0.462720704 | 0.047058169 | 0.248165904 | 0.10262657 | 0.285741127 | 0.07008325 |
| 98267.58594 | 103678.7109 | 104557.7422 | -0.462750952 | 0.035776373 | 0.100533289 | 0.374394756 | 0.14564207 | 0.178198454 |
| 54347.47656 | 60797.64453 | 60146.02344 | -0.463020583 | 0.024200411 | 0.375429515 | 0.05283098 | 0.872473761 | 0.020045376 |
| 11456.8457 | 14567.22363 | 12703.62109 | -0.463709482 | 0.172685916 | 0.122418098 | NA | 0.388950285 | 0.231509452 |
| 9225.520508 | 9168.767578 | 8299.72168 | -0.463797793 | 0.188632173 | 0.921467608 | 0.107845807 | 0.592992469 | 0.118373568 |

| | | | | | | | | |
|---|---|---|---|---|---|---|---|---|
| 9742.618164 | 10006.0791 | 10528.50586 | -0.463958472 | 0.203641334 | 0.230238768 | 0.179324339 | -0.206663513 | 0.095599918 |
| 14551.33398 | 14704.45996 | 15111.98926 | -0.464456062 | 0.097993976 | -0.64733279 | 0.021969813 | -0.042645592 | 0.512102022 |
| 121305.5234 | 113266.8906 | 126994.2578 | -0.464510709 | 0.105790758 | 0.294411492 | 0.142833812 | 0.364076362 | 0.080072763 |
| 5220.517578 | 6487.488281 | 6062.186035 | -0.464725138 | 0.086199451 | 0.557432451 | 0.268940886 | 0.114512875 | 0.479397694 |
| 44213.80664 | 41242.40527 | 41986.52637 | -0.464839345 | 0.022557251 | -0.024828241 | 0.844314089 | -0.26175554 | 0.09056925 |
| 27294.07227 | 33012.87109 | 38948.77734 | -0.464889999 | 0.275423604 | 0.270985253 | 0.319259968 | 0.158172408 | 0.558743536 |
| 8345.375977 | 15382.37207 | 11038.74023 | -0.465601479 | 0.416948441 | -0.028937881 | 0.955923696 | 0.337102848 | 0.506039655 |
| 177728.8438 | 167102.9688 | 173809.0469 | -0.465730309 | 0.022628223 | 0.275645563 | 0.08026683 | 0.438982131 | 0.024293977 |
| 76458.32813 | 79438.98438 | 82614.36719 | -0.465819562 | 0.025740944 | 0.082760215 | 0.316240692 | 0.361139938 | 0.027982615 |
| 400949.75 | 388238.4375 | 380005.7188 | -0.465873888 | 0.053542507 | 0.466116654 | 0.03412547 | 0.364212131 | 0.049004012 |
| 43488.92969 | 43940.64844 | 44585.04688 | -0.466338863 | 0.086699923 | 0.291131503 | 0.184506133 | 0.202074012 | 0.296064354 |
| 5414.044922 | 6538.772949 | 6115.44043 | -0.466375537 | 0.16947575 | -0.385352506 | 0.285225778 | -0.182455499 | 0.309046955 |
| 951711.5 | 1080592.75 | 1059275.25 | -0.466668247 | 0.033164349 | 0.015468131 | 0.84622109 | -0.145986332 | 0.183050335 |
| 113444.6094 | 117113.1875 | 119142.6953 | -0.466699181 | 0.01111345 | 0.48763091 | 0.032675377 | 0.931646784 | 0.006394096 |
| 1511088.375 | 1792644.125 | 1660508.75 | -0.466700135 | 0.021681865 | 0.387943744 | 0.019141974 | 0.200520501 | 0.190976702 |
| 318137.9063 | 333662.9688 | 353507 | -0.466880804 | 0.066333303 | -0.160974406 | 0.065601851 | 0.612174926 | 0.041161261 |
| 65201.24609 | 74170.73438 | 75880.19531 | -0.467448472 | 0.029603305 | 0.257991703 | 0.122334368 | 0.374619006 | 0.072139147 |
| 10425.78516 | 10150.20508 | 9657.145508 | -0.467487193 | 0.118081327 | 0.230975569 | 0.338407158 | 0.280447876 | 0.255006067 |
| 48285.29297 | 51306.26953 | 58454.74219 | -0.467670562 | 0.029297283 | -0.515882513 | 0.083732049 | -0.377616585 | 0.076235519 |
| 182223.6094 | 184511.7656 | 198842.9688 | -0.467828519 | 0.08302054 | 0.031790957 | 0.528875081 | 0.405373775 | 0.054109248 |
| 20554.99023 | 18672.02148 | 17301.17578 | -0.468047097 | 0.008854154 | 0.014557564 | 0.825529139 | 0.10600556 | 0.401325961 |
| 76509.25 | 76203.52344 | 71268.17969 | -0.468304153 | 0.10610949 | 0.035844939 | 0.458093846 | 0.353917877 | 0.026764821 |
| 104812.5234 | 105204.3203 | 106646.1719 | -0.468619967 | 0.012414001 | 0.022294104 | 0.821547091 | 0.225414464 | 0.077005785 |
| 52091.77734 | 32125.12695 | 46710.11328 | -0.46884227 | 0.171609668 | -0.197377834 | 0.163136465 | -0.694704444 | 0.084223187 |
| 47669.57422 | 48041.22656 | 44063.21875 | -0.469474577 | 0.061766802 | -0.138141068 | 0.462804615 | -0.244352997 | 0.253716363 |
| 26880.75 | 25241.03906 | 19715.99609 | -0.469858198 | 0.026565565 | 0.995516496 | 0.044605958 | 0.318348732 | 0.247325674 |
| 32622.97656 | 37728.65234 | 41924.74219 | -0.470028092 | 0.466667935 | 1.383720453 | 0.114968148 | 2.727370101 | 0.020157267 |
| 22940.04688 | 23105.38086 | 12906.90625 | -0.47002853 | 0.214706043 | -0.859249304 | 0.196359715 | -1.013908061 | 0.133042137 |
| 5853.819824 | 6316.179688 | 4524.041992 | -0.470069642 | 0.037314581 | 0.004620574 | 0.952846884 | -0.322317159 | 0.208408498 |
| 4652886.5 | 4828030 | 4395528.5 | -0.470174172 | 0.019930448 | 0.092333799 | 0.363955689 | -0.201140407 | 0.126675389 |
| 21938.78516 | 19625.58008 | 18719.625 | -0.47019106 | 0.252787526 | 0.627964668 | 0.175175027 | 0.583409918 | 0.191339598 |
| 156674.4219 | 153413.1719 | 138533.6406 | -0.470322436 | 0.05267493 | 0.145331285 | 0.309474627 | 0.453415065 | 0.04617854 |
| 3895.404541 | 3241.527588 | 1521.219849 | -0.470482682 | 0.16063032 | -1.30340243 | 0.099830362 | -1.032290179 | 0.120695883 |

| | | | | | | | | |
|---|---|---|---|---|---|---|---|---|
| 19143.16602 | 19082.7793 | 19437.13086 | -0.470638984 | 0.050618264 | 0.049053165 | 0.654658287 | -0.06232073 | 0.405672325 |
| 10032.87207 | 10538.74707 | 7303.842773 | -0.470645126 | 0.408467937 | -0.34953737 | 0.321888877 | -0.584446257 | 0.155706725 |
| 49353.10938 | 13177.88867 | 27381.71875 | -0.471014737 | 0.158538614 | 0.753743314 | 0.428256136 | -0.265642518 | 0.721615814 |
| 50760.21484 | 48828.82813 | 40875.05078 | -0.471203571 | 0.047406642 | 0.076061695 | 0.521659639 | -0.434275517 | 0.071457666 |
| 1404.480347 | 1378.043091 | 1611.976074 | -0.471255591 | 0.200294988 | -0.000321623 | 0.999222442 | -0.431198797 | 0.295151813 |
| 479672.1563 | 490647.25 | 499230.625 | -0.473125816 | 0.081289628 | 0.841374878 | 0.065122057 | 1.115912937 | 0.01054584 |
| 82175.36719 | 71693.92969 | 95779.3125 | -0.473470375 | 0.051365679 | 0.579913355 | 0.051888756 | 1.098825604 | 0.076193308 |
| 21456.25391 | 30108.54688 | 33303.19141 | -0.473789406 | 0.421606192 | 1.246883369 | 0.116818656 | 1.070923144 | 0.122834452 |
| 310725.125 | 381204.9688 | 295090.4063 | -0.473815687 | 0.125585919 | -0.056794469 | 0.721103647 | -0.208138748 | 0.302062013 |
| 381332.9688 | 343020.8438 | 444558.0938 | -0.474537828 | 0.081861007 | 0.867134536 | 0.14150374 | 1.356028522 | 0.04296087 |
| 101825.3047 | 120457.8516 | 123975.2969 | -0.47508796 | 0.013585622 | 0.179416664 | 0.023005377 | 0.246112615 | 0.207777277 |
| 49886.09375 | 44656.69141 | 48789.94922 | -0.475868782 | 0.036590352 | 0.013797152 | 0.922008276 | 0.003416355 | 0.983784117 |
| 1171867.75 | 1134756 | 921257.5625 | -0.4761568 | 0.216822285 | -0.408788659 | 0.260639434 | -0.687415729 | 0.138383382 |
| 518383.875 | 516812.3438 | 479556.5625 | -0.476895677 | 0.032101298 | -0.200150211 | 0.102916055 | -0.27928732 | 0.034655694 |
| 35363.73047 | 39041.83203 | 33219.33984 | -0.477615717 | 0.040928921 | 0.129478377 | 0.397903448 | -0.159115965 | 0.34800693 |
| 79032.10156 | 84435.17969 | 90531.52344 | -0.477833294 | 0.036480793 | 0.607466399 | 0.01915096 | 0.068508741 | 0.532718231 |
| 15183.97656 | 19056.82227 | 12761.0791 | -0.477930001 | 0.146203541 | -0.345761121 | 0.418214561 | 0.154393739 | 0.669450319 |
| 15040.91699 | 16238.99121 | 13026.90332 | -0.478060121 | 0.020852448 | 0.582516176 | 0.139513199 | 0.266984128 | 0.193898229 |
| 4931.623047 | 5700.875977 | 4014.34082 | -0.478081108 | 0.299350044 | -0.543307074 | 0.416045961 | -0.729415317 | 0.319455937 |
| 5257.272461 | 6107.199219 | 4743.230469 | -0.47813039 | 0.109220627 | -0.510537325 | 0.205564527 | -0.1570805 | 0.584717012 |
| 12011.78809 | 12434.50879 | 14476.7002 | -0.478212451 | 0.196367569 | 0.340581816 | 0.348195132 | 0.46677248 | 0.211941572 |
| 17381.27539 | 18087.9668 | 17395.38477 | -0.478353107 | 0.024701492 | -0.165024628 | 0.244921988 | -0.259414303 | 0.140469801 |
| 27469.88867 | 27741.37695 | 28686.10742 | -0.479091843 | 0.152253903 | 0.35128234 | 0.337374554 | 0.923100478 | 0.069782688 |
| 90589.82031 | 87927.16406 | 74830.58594 | -0.479870612 | 0.058566116 | -0.069403504 | 0.653854214 | 0.23427557 | 0.19608824 |
| 161897.6563 | 157922.3438 | 173335.2813 | -0.47989967 | 0.015044187 | 0.38225296 | 0.060234163 | 0.429831426 | 0.027834206 |
| 19520.09961 | 13501.39746 | 11323.66406 | -0.480030981 | 0.194423598 | 0.049957055 | 0.84058234 | -0.028735787 | 0.942571287 |
| 34652.94922 | 38677.57813 | 33353.84375 | -0.480799702 | 0.008070298 | -0.050763394 | 0.414751124 | -0.154325371 | 0.201736266 |
| 47155.4375 | 48148.23047 | 43458.14063 | -0.480851521 | 0.044857631 | -0.153289312 | 0.388564687 | -0.317301244 | 0.161733344 |
| 232732.1563 | 353039.7813 | 213137.7344 | -0.480979012 | 0.288497712 | 0.046005297 | 0.90544874 | -0.314745575 | 0.404560212 |
| 271877.2314 | 309422.1104 | 323769.3408 | -0.481487693 | 0.307746917 | 2.122634481 | 0.241244667 | 1.815461746 | 0.017015337 |
| 766136.3125 | 801423.3125 | 818127.625 | -0.48191346 | 0.0424199 | 0.528553667 | 0.06345934 | 0.342097862 | 0.095017861 |
| 126611.3828 | 168814.9219 | 156310.6719 | -0.482009873 | 0.135465479 | 0.077627538 | 0.683204566 | 0.142551131 | 0.520080533 |
| 430219 | 502584.7188 | 493039.7813 | -0.482094291 | 0.155874305 | 0.176374633 | 0.540113573 | 0.121950547 | 0.669450319 |

| | | | | | | | | |
|---|---|---|---|---|---|---|---|---|
| 34418.80078 | 36631.52734 | 26777.00195 | -0.48279718 | 0.01983805 | -0.185809735 | 0.185362734 | -0.479908613 | 0.08581567 |
| 16326.7207 | 14681.17383 | 12792.23633 | -0.484009829 | 0.111266593 | -0.129341561 | 0.640731417 | -0.403889114 | 0.256745248 |
| 25678.9707 | 25659.08789 | 24107.6582 | -0.484188092 | 0.026060442 | 0.044004491 | 0.503628106 | -0.241185366 | 0.047847186 |
| 176057.3594 | 176160.9063 | 196793.9688 | -0.48431058 | 0.047323083 | 0.155541697 | 0.283989936 | 0.308797769 | 0.06509693 |
| 278358.625 | 284964.4375 | 272005.0625 | -0.48442668 | 0.020651493 | -0.057327999 | 0.421213795 | -0.144319354 | 0.134102681 |
| 227085.2344 | 228567.8281 | 213237.0781 | -0.484606882 | 0.031284382 | -0.271553935 | 0.080680546 | -0.248355991 | 0.082507796 |
| 42235.96094 | 44862.55859 | 45912.40234 | -0.484691353 | 0.054968615 | -0.207796738 | 0.076036407 | -0.387699083 | 0.020008995 |
| 112707.7422 | 117274.8594 | 88951.34375 | -0.484863732 | 0.116475008 | 0.314206272 | 0.142833812 | 0.252311476 | 0.305073527 |
| 7406.606934 | 9159.852539 | 4358.51709 | -0.484984012 | 0.127797647 | -0.528796203 | 0.20617006 | -1.260474392 | 0.063474265 |
| 8649.669922 | 9894.376953 | 7295.834961 | -0.485367437 | 0.137182236 | 0.031019914 | 0.922759718 | -0.004705035 | 0.988723353 |
| 5815.675293 | 3575.660156 | 7435.961914 | -0.485370895 | 0.211214576 | -0.182523738 | 0.674604697 | 0.019073619 | 0.975148062 |
| 139757.625 | 134488.6406 | 138491.7031 | -0.485561697 | 0.026565565 | 0.451964674 | 0.045064797 | 0.6058332 | 0.029820656 |
| 114636.6563 | 111228.3594 | 121082.4609 | -0.48563812 | 0.054968615 | 0.245042174 | 0.243784441 | 0.005739393 | 0.977247866 |
| 524334.5625 | 514444.25 | 454137.6875 | -0.486065312 | 0.038671807 | -0.19733339 | 0.24779163 | -0.486270049 | 0.069955133 |
| 58188.32031 | 66334.90625 | 68344.32031 | -0.486309048 | 0.062989607 | 0.071563348 | 0.205232896 | 0.242285942 | 0.15573211 |
| 114378.4297 | 113638.7578 | 119988.2188 | -0.486947665 | 0.016938156 | 0.105695001 | 0.223722442 | 0.188129347 | 0.072336923 |
| 10260.06152 | 11283.22852 | 10423.81445 | -0.487028642 | 0.209865964 | 0.971760678 | 0.102026311 | 0.502089554 | 0.034722024 |
| 45264.69922 | 44014.72266 | 30351.51172 | -0.488055537 | 0.112434322 | -0.679782092 | 0.021906439 | -0.776845133 | 0.06280986 |
| 15751.43457 | 12839.40918 | 12328.11328 | -0.488210103 | 0.010484689 | 0.224348849 | 0.288975313 | -0.010494295 | 0.954795145 |
| 121820.4688 | 121910.875 | 110375.2578 | -0.488441419 | 0.020082924 | -0.034984808 | 0.563797862 | -0.394667014 | 0.027868741 |
| 42106.32031 | 36225.47266 | 36731.87109 | -0.488478618 | 0.032290095 | 0.932735364 | 0.020651809 | 0.23701158 | 0.145687527 |
| NA | NA | NA | -0.488712968 | NA | 0.096053942 | NA | NA | NA |
| 28153.37891 | 27485.11523 | 20714.33008 | -0.488843009 | 0.123070256 | 0.726294203 | 0.060921412 | 0.940531468 | 0.083375759 |
| 10297.84082 | 8731.084961 | 10260.8584 | -0.48894732 | 0.20270675 | 0.322804582 | 0.384749949 | -0.236796858 | 0.558864682 |
| 34168.75 | 25659.43164 | 29455.29102 | -0.489009324 | 0.112988822 | 0.735735328 | 0.058077004 | -0.032208263 | 0.914576246 |
| 36770 | 35514.55469 | 32917.10156 | -0.489258316 | 0.0424199 | 0.261101139 | 0.083757952 | 0.30998224 | 0.053337315 |
| 48138.00781 | 55409.61328 | 49602.36328 | -0.489628483 | 0.06500624 | -0.301668148 | 0.05818214 | -0.541706629 | 0.045311345 |
| 121558.1484 | 116051.2266 | 125857.1328 | -0.489718124 | 0.024262842 | -0.025304352 | 0.542644692 | 0.291286145 | 0.036848746 |
| 292.7205811 | 423.1038818 | 291.7931519 | -0.489952121 | NA | 0.752440001 | NA | 0.231222353 | NA |
| 92960.11719 | 90884.3125 | 97560.82813 | -0.490215708 | 0.175573104 | -0.415314666 | 0.14231796 | -0.288307927 | 0.200261065 |
| 23799.20703 | 22261.74805 | 25023.6543 | -0.490320042 | 0.037380825 | -0.061374168 | 0.593433268 | -0.46791031 | 0.054887084 |
| 32796.96094 | 37732.69141 | 33886.19141 | -0.490673149 | 0.028903558 | -0.099508756 | 0.294259446 | -0.236649906 | 0.101462623 |
| 17217.55859 | 24545.71289 | 17908.66602 | -0.491233036 | 0.274371306 | 0.473740844 | 0.210152001 | 0.28802568 | 0.458699163 |

| | | | | | | | | |
|---|---|---|---|---|---|---|---|---|
| 126794.3125 | 133522.9375 | 143660.4375 | -0.491270524 | 0.01964718 | 0.13990212 | 0.234049008 | 0.293870588 | 0.072416132 |
| 110288.7891 | 116498.8281 | 103816.1953 | -0.491334793 | 0.033353006 | -0.222575726 | 0.05467585 | -0.378350426 | 0.027868741 |
| 271747.625 | 237590.125 | 241958.4844 | -0.492028832 | 0.016189975 | -0.019214906 | 0.796507507 | -0.139406371 | 0.218219327 |
| 816308.5 | 813482.3125 | 733170 | -0.492398123 | 0.042404592 | 0.641283031 | 0.045064018 | -0.091320428 | 0.582972866 |
| 148831.25 | 147457.8438 | 141883.0469 | -0.492940967 | 0.018311389 | 0.321325821 | 0.096169757 | 0.138462983 | 0.099555681 |
| 22565.33594 | 20424.24805 | 22750.09375 | -0.492959202 | 0.134341857 | 0.797401517 | 0.05467585 | 0.924371943 | 0.019816721 |
| 17831.2793 | 43337.42578 | 20632.33789 | -0.493343654 | 0.318687987 | -0.015196175 | 0.981156167 | -0.034873238 | 0.968145079 |
| 11389.73535 | 9382.883789 | 12968.55664 | -0.493556625 | 0.069002061 | 0.007927707 | 0.977833898 | -0.323278639 | 0.25191185 |
| 67554 | 61801.57422 | 59656.71484 | -0.494160731 | 0.030344162 | 0.349983475 | 0.026826422 | 0.265063691 | 0.084288402 |
| 8765.924805 | 7649.099609 | 7557.483887 | -0.494855436 | 0.024508165 | -0.073121408 | 0.639049031 | -0.473578703 | 0.044743801 |
| 141409.6875 | 125558.0156 | 142316.7188 | -0.495258857 | 0.041945948 | 0.548779879 | 0.045588559 | 0.908054506 | 0.020008995 |
| 53399.77734 | 43700.26563 | 47231.76563 | -0.495332695 | 0.212467151 | 0.404526133 | 0.234571805 | 0.279795092 | 0.165312588 |
| 2039458.5 | 2434837 | 2340579.25 | -0.495691971 | 0.071472027 | 0.055148466 | 0.629908746 | -0.120738018 | 0.343177751 |
| 108535.7969 | 85306.61719 | 99948.70313 | -0.495728721 | 0.138039095 | -0.115026697 | 0.618374257 | -0.585100541 | 0.124211802 |
| 3745.119141 | 8286.912109 | 12125.68945 | -0.496185551 | 0.48077548 | 1.91665379 | 0.173072655 | 1.739832042 | 0.228883111 |
| 34882.86719 | 33381.69531 | 29605.0918 | -0.496294969 | 0.110302818 | -0.027990263 | 0.644374085 | -0.096207606 | 0.408380417 |
| 47651.16406 | 49510.27344 | 45068.49609 | -0.496480155 | 0.237633931 | 1.246225436 | 0.048593497 | 1.503898687 | 0.034722024 |
| 345383.7188 | 294380.875 | 320270.5313 | -0.497046438 | 0.044763187 | 0.127780529 | 0.406076773 | -0.102219315 | 0.535817134 |
| 1552.431641 | 1264.263184 | 1094.456909 | -0.497634637 | 0.050873498 | -0.183067231 | 0.393523298 | -0.608264219 | 0.086930144 |
| 115812.8203 | 114422.4531 | 112272.3281 | -0.497661679 | 0.008503176 | 0.184597933 | 0.051888756 | 0.056855829 | 0.274556211 |
| 134182 | 154127.0156 | 126319.5859 | -0.498416875 | 0.076958005 | 0.32461994 | 0.048024604 | 0.218190116 | 0.221863877 |
| 2502112.5 | 2258281.5 | 2136635.5 | -0.498823015 | 0.300286839 | 0.09298096 | 0.238020687 | -0.07805333 | 0.491638437 |
| 251342.5 | 255786.4063 | 245528.1406 | -0.498918772 | 0.022709737 | 0.41135208 | 0.037725739 | 0.255057365 | 0.047850077 |
| 30909.75586 | 26835.08594 | 25816.81445 | -0.499198189 | 0.074071792 | -0.224148403 | 0.364883875 | -0.623462368 | 0.107557275 |
| 113140.25 | 112243.4063 | 125051.3672 | -0.499651157 | 0.010297098 | 0.136020725 | 0.11629032 | 0.399921316 | 0.046487363 |
| 1377736.25 | 1597470.125 | 1395103.875 | -0.499865993 | 0.064978842 | -0.332543129 | 0.212262475 | -0.029537253 | 0.882940754 |
| 39710.83594 | 41682.92578 | 42340.81641 | -0.500047387 | 0.037142448 | -0.406701812 | 0.083732049 | -1.105693403 | 0.042737273 |
| 16611.36719 | 17334.5293 | 12473.64941 | -0.500147308 | 0.178228644 | -0.014227302 | 0.922008276 | -0.155309972 | 0.50437262 |
| 23700.65234 | 24974.80859 | 25917.9375 | -0.500294597 | 0.038058333 | 0.238698723 | 0.232975249 | 0.517664668 | 0.029820656 |
| 52291.50391 | 53830.77734 | 55547.05469 | -0.500791727 | 0.182007594 | 0.611973378 | 0.137136478 | 0.452907839 | 0.225167244 |
| 51855.22266 | 46137.89844 | 46456.35938 | -0.500889104 | 0.030588379 | -0.145104549 | 0.244990627 | -0.162633131 | 0.186378157 |
| 60097.64063 | 61650.60156 | 46110.19531 | -0.50103383 | 0.028910784 | -0.031329322 | 0.828575501 | -0.214000186 | 0.331779341 |
| 25498.10547 | 25189.95703 | 21829.44336 | -0.501127449 | 0.038502747 | 0.413611719 | 0.054829172 | 0.34577129 | 0.086177515 |

| | | | | | | | | |
|---|---|---|---|---|---|---|---|---|
| 153634.2969 | 150131.7188 | 158714.0156 | -0.501190994 | 0.034518918 | 0.121433362 | 0.082329519 | 0.268548836 | 0.022925497 |
| 4321.080078 | 5634.607422 | 4567.079102 | -0.501665603 | 0.115723457 | 0.914275767 | 0.021217492 | 0.470077334 | 0.130712196 |
| 118545.0547 | 115969.3594 | 136277.6094 | -0.501787387 | 0.016471012 | -0.045665062 | 0.655353432 | 0.236329743 | 0.159338304 |
| 164194.1094 | 165672.6563 | 157773.2656 | -0.502269491 | 0.031140524 | -0.162024452 | 0.033804616 | -0.21395898 | 0.021863066 |
| 14517.14258 | 16532.37305 | 15574.09961 | -0.502349814 | 0.12643787 | 0.256720295 | 0.688650291 | -0.255029005 | 0.395955794 |
| 290513.25 | 305967.4063 | 303029.6563 | -0.502563579 | 0.048312249 | 0.416362946 | 0.072725977 | 0.243884099 | 0.133959444 |
| 115994.6328 | 115102.3516 | 121295.4375 | -0.503038454 | 0.049249627 | 0.032929427 | 0.739638889 | 0.23099065 | 0.103328406 |
| 78098.75781 | 90640.625 | 98185.04688 | -0.503143327 | 0.170436627 | -0.347857015 | 0.427472799 | -0.007210327 | 0.986487983 |
| 10709.60547 | 12396.37207 | 12215.0332 | -0.50347365 | 0.03699257 | 0.406019471 | 0.109438848 | -0.002907809 | 0.984804767 |
| 15120.10449 | 14750.02734 | 20416.99023 | -0.503645273 | 0.404501862 | 0.210384131 | 0.746355251 | 0.814093401 | 0.212263796 |
| 77260.54688 | 77840 | 76531.46875 | -0.503783332 | 0.063399586 | 0.118832341 | 0.163729974 | 0.219207987 | 0.076783998 |
| 110378.7891 | 117429.1641 | 117621.7422 | -0.503874662 | 0.052848772 | 0.103531251 | 0.218322637 | 0.330587105 | 0.027331143 |
| 21699.91992 | 42542.99609 | 29460.94531 | -0.504294471 | 0.046697667 | -0.397480245 | 0.150761343 | -0.333017742 | 0.429961702 |
| 32460.53125 | 31108.61523 | 31408.83203 | -0.504467413 | 0.008009891 | 0.189417798 | 0.03968601 | -0.072939633 | 0.134102681 |
| 5558.235352 | 4768.458984 | 2688.534668 | -0.504549657 | 0.024409946 | -0.078967444 | 0.57025817 | -0.690809882 | 0.149748939 |
| 9501.825195 | 4033.976563 | 3579.522705 | -0.504928019 | 0.597550855 | 2.002936188 | 0.516940522 | 0.690779103 | 0.534352678 |
| 14679777 | 15232577 | 14948994 | -0.50506324 | 0.019286562 | 0.183674406 | 0.049927795 | 0.122998826 | 0.063440273 |
| 15888.67969 | 16845.47266 | 14077.36523 | -0.505187782 | 0.029811229 | -0.077199292 | 0.533779394 | -0.18217066 | 0.257876793 |
| 3376.052979 | 5144.351074 | 4131.01416 | -0.505752816 | 0.142870238 | 0.099485117 | 0.599664693 | -0.143161279 | 0.596749137 |
| 6527.960938 | 9772.154297 | 10310.63086 | -0.505796249 | 0.2756587 | 0.180369034 | 0.640908913 | -0.052887313 | 0.868574715 |
| 10415.84961 | 8845.257813 | 8387.469727 | -0.506047924 | 0.159512994 | 0.105879071 | 0.735373025 | -0.138629979 | 0.686180113 |
| 88556.78125 | 106827.0859 | 105479.6641 | -0.506095633 | 0.035078404 | 0.459394417 | 0.053890972 | 0.4002396 | 0.092063331 |
| 358451.5444 | 486870.6094 | 412076.8564 | -0.506816403 | 0.218376737 | -0.151833995 | 0.685354976 | -0.450386368 | 0.215760001 |
| 56668.31641 | 54451.24219 | 56140.67578 | -0.506988063 | 0.08028391 | 0.271154023 | 0.227543266 | 0.268107168 | 0.225533768 |
| 230794.6875 | 251989.125 | 248130.8594 | -0.507179997 | 0.032345916 | 0.080581583 | 0.215727673 | 0.345650996 | 0.033948107 |
| 446598.3125 | 505977.0938 | 479377.1875 | -0.507902733 | 0.058992808 | -0.541244384 | 0.013938321 | -0.883807064 | 0.010236624 |
| 3816.105713 | 5358.811035 | 1737.430176 | -0.507959869 | 0.122673099 | -0.470591279 | 0.147741919 | -1.546842628 | 0.054887084 |
| 13246.8457 | 12710.94922 | 12161.17383 | -0.508051776 | 0.006735623 | -0.053819032 | 0.321962613 | -0.212550757 | 0.055835745 |
| 70437.67188 | 51519.37109 | 62866.12109 | -0.50833879 | 0.074085004 | 0.484354728 | 0.124127483 | 0.098957399 | 0.713705355 |
| 608126 | 545512.625 | 612945.375 | -0.508510361 | 0.011814081 | 0.132763868 | 0.191377912 | -0.175905724 | 0.129065826 |
| 15231.53125 | 18878.62109 | 6869.430664 | -0.508567863 | 0.066139345 | -0.963542591 | 0.069318558 | -1.232226529 | 0.063937169 |
| 10701.08691 | 11027.63867 | 10018.90918 | -0.508881015 | 0.015175173 | 0.049699907 | 0.537359718 | -0.344470879 | 0.045861991 |
| 42426.85547 | 46833.27734 | 42838.26953 | -0.508883716 | 0.054037379 | -0.634530845 | 0.086540831 | -0.746755466 | 0.061395833 |

| | | | | | | | | |
|---|---|---|---|---|---|---|---|---|
| 316876.1875 | 330797.7188 | 332596.625 | -0.509101845 | 0.008685066 | 0.190877525 | 0.097735536 | -0.104575314 | 0.105453219 |
| 177790.0469 | 124687.75 | 131110.3906 | -0.509165783 | 0.102861044 | 1.075542064 | 0.257322951 | 1.349741422 | 0.074674553 |
| 370196.7813 | 362932.375 | 375376.7188 | -0.509333817 | 0.045302846 | -0.242553165 | 0.077648523 | -0.350443921 | 0.060937162 |
| 21972.58789 | 22448.56641 | 16554.24219 | -0.509446506 | 0.105883354 | 0.36221421 | 0.197991611 | -0.022036933 | 0.942386927 |
| 516.6091919 | 2536.723877 | NA | -0.509665801 | 0.546731421 | -0.02553257 | 0.987735084 | 0.415245492 | NA |
| 111933.25 | 101547.0938 | 100236.9063 | -0.509863218 | 0.031396254 | 0.229025107 | 0.037760167 | 0.302959946 | 0.07008325 |
| 164957.6563 | 155557.5313 | 169437.1875 | -0.510016385 | 0.029239337 | -0.088240419 | 0.379832076 | -0.154786422 | 0.18596657 |
| 1173595.375 | 1167634.625 | 1057783.125 | -0.510628869 | 0.036480793 | 0.104336947 | 0.259097064 | -0.247875834 | 0.077710677 |
| 7875.881348 | 7641.44043 | 6025.206543 | -0.511107056 | 0.10030267 | -0.645827478 | 0.023796962 | -1.052190205 | 0.017279709 |
| 11641.75879 | 18165.17578 | 34632.86719 | -0.512642865 | 0.470665764 | 0.732142919 | 0.335837303 | 0.2366329 | 0.767865119 |
| 74210.99219 | 66194.50781 | 62114.78906 | -0.512669654 | 0.017112862 | 0.328342691 | 0.050817911 | 0.427327428 | 0.074385174 |
| 33251.92969 | 33893.51172 | 33359.95703 | -0.512884501 | 0.050093742 | 0.030287299 | 0.866635988 | -0.443562143 | 0.160431331 |
| 21038.01953 | 17371.33984 | 13440.65723 | -0.51315986 | 0.023724131 | -0.03470348 | 0.720955012 | 0.057770478 | 0.844045457 |
| 1907049.016 | 2550120.969 | 2521739.125 | -0.513396535 | 0.046600136 | 0.061277884 | 0.704789036 | 0.479560106 | 0.142342008 |
| 101056.5547 | 43464.875 | 22406.92383 | -0.513448577 | 0.154341803 | 1.377008192 | 0.114739915 | 0.953604652 | 0.481006414 |
| 96532.29688 | 101478.2891 | 108893.1719 | -0.513486886 | 0.03766688 | 0.075707785 | 0.189627399 | 0.307576404 | 0.068414793 |
| 45980.28906 | 41801.04688 | 38088.66406 | -0.513982259 | 0.068152067 | 0.099669983 | 0.782462825 | 0.419381243 | 0.116918544 |
| 36343.10938 | 36818.625 | 38547.07422 | -0.514060715 | 0.03500296 | 0.013954383 | 0.909428871 | -0.169989938 | 0.230231868 |
| 1165854.25 | 1081846.625 | 1021029.375 | -0.514105156 | 0.062785336 | -0.627128328 | 0.09742076 | -0.348237886 | 0.147031451 |
| 9107.75293 | 9907.087891 | 2908.015625 | -0.514133347 | 0.077426021 | -0.329363393 | 0.241244667 | -0.5869897 | 0.343143637 |
| 54148.14844 | 51491.90234 | 48050.33594 | -0.514140646 | 0.033529204 | 0.271572927 | 0.150798451 | 0.07449951 | 0.611365676 |
| 22123612 | 12734712 | 20882660 | -0.514466213 | 0.53413865 | 0.798202255 | 0.176310792 | 0.861664074 | 0.168496789 |
| 98086.30469 | 100897.4609 | 97871.375 | -0.514653041 | 0.024681796 | -0.130487431 | 0.323345452 | -0.287655063 | 0.132636805 |
| 72706.53906 | 67622.39844 | 66321.64844 | -0.514746097 | 0.034713383 | 0.271311101 | 0.134444814 | 0.372717937 | 0.045009854 |
| 39542.84375 | 44551.07422 | 41419.80859 | -0.514801186 | 0.159539615 | 0.110834042 | 0.714090192 | 0.379203745 | 0.25220534 |
| 21197.55078 | 30998.64453 | 28022.98047 | -0.515181547 | 0.275788376 | 0.664349925 | 0.217532051 | 0.62954286 | 0.225935566 |
| 386313.4375 | 472210.5625 | 416627.5 | -0.515402584 | 0.221587537 | 0.501705279 | 0.06919378 | 0.49977285 | 0.071122316 |
| 46261.17188 | 46367.63672 | 48487.08203 | -0.515650736 | 0.164371576 | -0.007134245 | 0.953027781 | 0.156156461 | 0.064464306 |
| 6167.014648 | 4914.895996 | 5427.887695 | -0.515696477 | 0.134574193 | 0.171110326 | 0.574712761 | 0.674459367 | 0.086914819 |
| 53010.57813 | 51123.10156 | 52551.74219 | -0.515887314 | 0.017507272 | 0.16927228 | 0.05468055 | 0.230890777 | 0.030321148 |
| 20452.32227 | 20335.59766 | 19266.26953 | -0.516371302 | 0.110178191 | 0.555860093 | 0.092974661 | 0.333402287 | 0.215760001 |
| 49251.77734 | 52311.73438 | 30656.62109 | -0.516602735 | 0.044053484 | -0.04434459 | 0.509478737 | -0.027494123 | 0.938451831 |
| 204376.0625 | 201104.7969 | 235588.3125 | -0.517343125 | 0.010893509 | 0.246853736 | 0.0280168 | 0.23964246 | 0.170423153 |

| | | | | | | | | |
|---|---|---|---|---|---|---|---|---|
| 2861.3396 | 3213.009521 | 2268.647461 | -0.517667468 | 0.047406642 | 0.744070319 | 0.034038555 | 0.158434072 | 0.544636732 |
| 34291.55859 | 36254.05469 | 39690.96094 | -0.518642022 | 0.054926056 | 0.247832248 | 0.25116103 | 0.355887836 | 0.128960705 |
| 8829.831055 | 8666.943359 | 10202.15723 | -0.518798366 | 0.038306202 | -0.210853929 | 0.238316696 | -0.935973916 | 0.040591316 |
| 37977.85156 | 41188.98828 | 32143.32031 | -0.518895356 | 0.117342707 | 0.292088184 | 0.160878456 | 0.337591905 | 0.159582522 |
| 195388.2656 | 195770.3906 | 207646.4375 | -0.519016646 | 0.005752145 | 0.15648417 | 0.076036407 | 0.365033895 | 0.029710029 |
| 8191.445313 | 6388.555176 | 6795.833984 | -0.5190356 | 0.181440382 | 0.827977377 | 0.083773082 | 0.967592634 | 0.055868946 |
| 1579.392944 | 3190.409424 | 1697.13855 | -0.519262231 | 0.250895179 | -0.187573804 | 0.714090192 | -1.261800592 | 0.174355634 |
| 302182.375 | 310202.0938 | 301495 | -0.519421625 | 0.029801769 | 0.282941715 | 0.01179292 | -0.184801055 | 0.01858328 |
| 625070.375 | 716093.0625 | 623310.0625 | -0.519722507 | 0.08344219 | -0.011265381 | 0.934248513 | -0.020525894 | 0.889982534 |
| 28067.48242 | 23190.26758 | 27860 | -0.519848178 | 0.156854993 | 0.034906752 | 0.799141879 | 0.119402396 | 0.452083839 |
| 3910.084473 | 3710.022705 | 3072.867432 | -0.51993893 | 0.039520533 | -0.46633899 | 0.041260113 | -0.513228031 | 0.053785506 |
| 80084.05469 | 89874.46875 | 77211.21875 | -0.520555694 | 0.01031238 | 0.083610024 | 0.188018725 | -0.139444892 | 0.244006157 |
| 601.9941406 | 1147.304077 | 779.6897583 | -0.520795823 | 0.206326402 | 0.264413241 | 0.506573912 | 0.127954388 | 0.81578493 |
| 1661687.125 | 1773967.375 | 1542079.875 | -0.521157489 | 0.010228174 | -0.17363297 | 0.105321264 | -0.326271581 | 0.047387434 |
| 19491.79102 | 18815.64648 | 7878.589844 | -0.521377929 | 0.10498796 | 0.29898977 | 0.269570571 | -0.201773862 | 0.706396046 |
| 6418.480957 | 2288.73291 | 2752.171631 | -0.521818642 | 0.188824978 | 0.257348641 | 0.598335471 | -0.947555542 | 0.213371407 |
| 257008.75 | 237833.5625 | 252676.2969 | -0.522460365 | 0.048069896 | 0.31789649 | 0.033121608 | 0.000310881 | 0.997178143 |
| 107317.0156 | 107300.375 | 107587.5391 | -0.522517751 | 0.040854235 | 0.339023919 | 0.096721331 | 0.232790691 | 0.189363824 |
| 158154.2969 | 157094.8438 | 165566.9688 | -0.522605381 | 0.088487242 | 0.547073599 | 0.021906439 | 0.533600641 | 0.020256835 |
| 21432.76563 | 16265.39551 | 25466.69141 | -0.523006653 | 0.028901981 | -0.658303478 | 0.111882773 | -0.059081707 | 0.834063546 |
| 6525.907715 | 6766.520996 | 5514.492676 | -0.523081972 | 0.029906057 | 0.201480287 | 0.154164226 | -0.241467917 | 0.17565098 |
| 7643.730957 | 7391.763184 | 5973.807617 | -0.523226226 | 0.130427426 | -0.427118383 | 0.197991611 | -0.163845157 | 0.352334437 |
| 799079.6875 | 781323.125 | 840803.1875 | -0.523449531 | 0.02930047 | -0.043851534 | 0.587707401 | 0.141041102 | 0.147937915 |
| 26859.29297 | 31055.98242 | 25337.125 | -0.523971044 | 0.17764559 | 0.270000487 | 0.073326865 | 0.133892214 | 0.397364957 |
| 860305.875 | 898988.9375 | 891135.625 | -0.523980941 | 0.019930448 | -0.00109056 | 0.99201113 | -0.195174111 | 0.188422821 |
| 94234.84375 | 96328.01563 | 97153.10156 | -0.524361263 | 0.011063961 | 0.128759135 | 0.173986552 | 0.139699571 | 0.136000191 |
| 35030.47656 | 27509.19922 | 26832.78906 | -0.524479715 | 0.047035938 | -0.553253414 | 0.048024604 | -0.599944529 | 0.076783998 |
| 165626.9688 | 163790.7031 | 176725.8281 | -0.524725518 | 0.054728089 | 0.124145463 | 0.088308366 | 0.300732753 | 0.034722024 |
| 141650.4219 | 139843.8906 | 140219.9375 | -0.525115382 | 0.058238988 | 0.360725244 | 0.152460574 | 0.456094379 | 0.110105852 |
| 1745.512939 | 1672.440552 | 1889.921753 | -0.525368754 | 0.367630259 | -0.465717078 | 0.142682072 | -1.06991148 | 0.068366118 |
| 106994.1484 | 106974.9063 | 108083.3516 | -0.525386465 | 0.022233058 | 0.178058546 | 0.110122293 | 0.054552822 | 0.293986304 |
| 585990.875 | 550311.375 | 526102.5625 | -0.525568475 | 0.096471873 | 0.7815132 | 0.042608596 | 0.414969647 | 0.103046053 |
| 10398.125 | 11675.57324 | 8511.775391 | -0.525774862 | 0.045302846 | 0.226061409 | 0.353224762 | -0.416469015 | 0.118373568 |

| | | | | | | | | |
|---|---|---|---|---|---|---|---|---|
| 81045.1875 | 101450.7578 | 89424.44531 | -0.525823268 | 0.081289628 | -0.058893363 | 0.647976234 | 0.046781507 | 0.771355563 |
| 28934.50781 | 28466.55664 | 28615.32617 | -0.526128716 | 0.133330992 | 0.428982367 | 0.043644056 | 0.196544931 | 0.16055099 |
| 10719.01465 | 11567.00195 | 10957.98438 | -0.52614885 | 0.064124584 | -0.530505515 | 0.126731731 | -0.799010602 | 0.079165593 |
| 13535.67188 | 15710.26563 | 13529.02148 | -0.527198389 | 0.070959976 | 0.908504775 | 0.028434447 | 0.160889202 | 0.267519066 |
| 161255.0313 | 145371.3281 | 133180.1875 | -0.527208141 | 0.050541732 | 0.653667783 | 0.088308366 | 0.340231842 | 0.144501069 |
| 52807.39063 | 53532.32422 | 53547.50781 | -0.527233991 | 0.025437157 | 0.566611466 | 0.035318687 | 0.388078747 | 0.063474265 |
| 134344.2656 | 133884.7813 | 140969.0938 | -0.527376195 | 0.042392829 | 0.170171235 | 0.152134335 | 0.221980308 | 0.088642968 |
| 21297.16016 | 21076.80078 | 15600.10547 | -0.527446057 | 0.012364839 | -0.21819198 | 0.06106734 | -0.413920053 | 0.139434127 |
| 74412.10156 | 70126.47656 | 75820.4375 | -0.527476802 | 0.011979502 | 0.450244999 | 0.092761595 | 0.30539053 | 0.035887862 |
| 12668.66992 | 33090.16406 | 9098.720703 | -0.528066351 | 0.542216889 | 1.298218481 | 0.178660809 | 0.443166632 | 0.709579616 |
| 38366.69141 | 42038.74609 | 42791.5 | -0.528185014 | 0.227260721 | 0.349151958 | 0.425452152 | 0.53610525 | 0.24207605 |
| 46582.55469 | 50593.35938 | 56194.47656 | -0.528305941 | 0.024701492 | 0.063745399 | 0.308832804 | 0.308093393 | 0.138724083 |
| 4173.040039 | 5763.631348 | 2154.281494 | -0.528364428 | 0.388565287 | 0.214589021 | 0.742577273 | -0.475411033 | 0.536592057 |
| 180028.9688 | 183098.8438 | 183950.2813 | -0.529128687 | 0.016105265 | -0.085701628 | 0.21914142 | -0.063222097 | 0.088388282 |
| 34448.5 | 34341.26172 | 33202.09766 | -0.530809217 | 0.053424232 | -0.029340217 | 0.802718747 | -0.128979286 | 0.244315159 |
| 5355.108887 | 3566.966309 | 3386.57373 | -0.530911388 | 0.091315146 | 0.673311904 | 0.089335127 | 0.377326954 | 0.389456703 |
| 2131.416504 | 3790.045898 | 1872.947998 | -0.531128703 | 0.340812741 | -0.355365345 | 0.61104577 | -0.384272617 | 0.61110795 |
| 945840.375 | 726897.6875 | 628708 | -0.531398784 | 0.157842529 | 1.064289142 | 0.068255066 | 0.598257665 | 0.188843753 |
| 12084.64453 | 12987.73926 | 11312.65137 | -0.532253223 | 0.026744661 | 0.190257659 | 0.292396668 | 0.057915375 | 0.646302665 |
| 47718.84766 | 46872.92969 | 48400.14844 | -0.532414408 | 0.117038402 | 0.685337914 | 0.152415276 | 0.636252943 | 0.046092295 |
| 9394.349609 | 11061.62793 | 11299.24023 | -0.532880333 | 0.158538614 | 0.648032846 | 0.079859428 | 0.470655021 | 0.075839948 |
| 11207.13184 | 14672.61914 | 10328.36621 | -0.533129342 | 0.171512612 | 0.368812413 | 0.048024604 | 0.124811893 | 0.632802777 |
| 17304.44727 | 19733.00391 | 13459.8291 | -0.533167693 | 0.01111345 | -0.030152334 | 0.687639724 | -0.350300267 | 0.210234556 |
| 3970.147949 | 4498.591797 | 3029.220459 | -0.533935821 | 0.255838638 | 0.600856306 | 0.22425544 | 0.345816967 | 0.295259111 |
| NA | NA | NA | -0.534008629 | 0.030661161 | -0.168425403 | 0.516940522 | NA | NA |
| 501456.1875 | 458494.125 | 538861.625 | -0.53423408 | 0.014306629 | 0.144657526 | 0.165855735 | -0.082094529 | 0.485667129 |
| 98452.61719 | 85326.32813 | 76296.97656 | -0.535364757 | 0.056932452 | 0.387040801 | 0.115989509 | 0.012638036 | 0.957199444 |
| 954659.0625 | 987617.25 | 913803.875 | -0.535451368 | 0.02930047 | -0.194156922 | 0.096073212 | -0.147093344 | 0.124069253 |
| 35923.90234 | 32647.83594 | 38711.26172 | -0.535673649 | 0.094031642 | 0.932886671 | 0.308118796 | 2.51433344 | 0.021984636 |
| 24340.98047 | 13871.19824 | 17430.10156 | -0.535968948 | 0.055042495 | 0.674394626 | 0.050016319 | 0.325061058 | 0.461591343 |
| 20361.70898 | 20798.4668 | 22226.15039 | -0.53632869 | 0.210218452 | 0.272341945 | 0.514648743 | 0.036351218 | 0.934597273 |
| 69685.39844 | 69881.30469 | 64220.04297 | -0.536572571 | 0.071158565 | -0.37989942 | 0.199435309 | -0.682469979 | 0.101895754 |
| 32714.03516 | 27883.80664 | 28503.16602 | -0.536856105 | 0.054037379 | -0.509148703 | 0.14010832 | -1.268633816 | 0.065710106 |

| | | | | | | | | |
|---|---|---|---|---|---|---|---|---|
| 453947.3125 | 532266.3125 | 430737.875 | -0.537346379 | 0.061073891 | 0.113083064 | 0.297951899 | 0.02341461 | 0.885245898 |
| 55142.36328 | 60139.26563 | 45858.15234 | -0.537592922 | 0.140276781 | 0.410084911 | 0.259408195 | 0.582950426 | 0.135715333 |
| 21483.26563 | 14772.21484 | 12148.81836 | -0.537601155 | 0.039126605 | 0.235811502 | 0.255118722 | -0.271623731 | 0.474198763 |
| 66625.36719 | 65750.04688 | 63104.25781 | -0.537768113 | 0.019286562 | 0.092674314 | 0.160099354 | 0.079210746 | 0.14559756 |
| 21065.25195 | 27469.71484 | 29027.89063 | -0.5378067 | 0.22713782 | 0.297705274 | 0.505056955 | 0.493946901 | 0.288104776 |
| 128725.9531 | 118882.7813 | 101808.4297 | -0.537949167 | 0.023770785 | 0.45545512 | 0.174475307 | 0.538083748 | 0.08023139 |
| 6404.599609 | 6688.959961 | 7801.726074 | -0.537983434 | 0.134212026 | -0.274650312 | 0.238302636 | -0.730816929 | 0.062345205 |
| 36414.45313 | 32962.54688 | 19599.625 | -0.538194233 | 0.034518918 | 0.50889839 | 0.05467585 | -0.274262677 | 0.461172821 |
| 44112.01563 | 42082.58984 | 39744.64063 | -0.538354694 | 0.047142848 | -0.031413552 | 0.85987839 | -0.182029901 | 0.370576128 |
| 289272.6875 | 247921.2969 | 231861.6406 | -0.538819055 | 0.079009944 | 0.507197475 | 0.106558364 | 0.815926817 | 0.046581887 |
| 128517.4844 | 147438.5313 | 148832.7188 | -0.540047772 | 0.0424199 | 0.019620292 | 0.54508114 | 0.249036059 | 0.142328609 |
| 146134.4531 | 126483.2344 | 117173.9219 | -0.540183136 | 0.023724131 | -0.162678269 | 0.018539883 | -0.44599333 | 0.087190025 |
| 28631.58398 | 28616.77148 | 20828.72852 | -0.540392313 | 0.056869835 | 0.312789151 | 0.19256535 | 0.014974474 | 0.959722655 |
| 9966.287109 | 10079.08691 | 7616.79541 | -0.540450798 | 0.143738987 | 0.445160138 | 0.243646891 | 0.390852503 | 0.289842489 |
| 10761.12891 | 14313.03809 | 12749.98047 | -0.540468755 | 0.158200263 | 0.946596659 | 0.289867338 | 0.783255151 | 0.068838627 |
| 68759.94531 | 72811.39063 | 79630.90625 | -0.541087213 | 0.063046822 | -0.128913677 | 0.557806724 | 0.399621409 | 0.118373568 |
| 108051.5313 | 106827.1016 | 73248.92969 | -0.541481493 | 0.034953337 | 0.039261753 | 0.643665701 | -0.326848816 | 0.263528267 |
| 96313.375 | 105296.4766 | 101458.2109 | -0.541767311 | 0.023527743 | 0.112357101 | 0.190466684 | 0.222465185 | 0.063283918 |
| 112328.9063 | 116932.7422 | 119868.75 | -0.541884378 | 0.032646789 | 0.007931129 | 0.961166689 | -0.133019699 | 0.397676581 |
| 46319.4375 | 49556.85156 | 43594.07031 | -0.542219128 | 0.029599705 | -0.140386296 | 0.069257825 | -0.061574527 | 0.479299319 |
| 8508.660156 | 7982.518555 | 6380.147949 | -0.542512226 | 0.064591518 | -0.078033667 | 0.70052948 | -0.461552564 | 0.137062098 |
| 7305.883789 | 7974.529785 | 6231.503418 | -0.5428887 | 0.03211333 | 0.169347352 | 0.306401385 | 0.052565563 | 0.790483404 |
| 330766.0938 | 333150.9688 | 322671.1875 | -0.542943609 | 0.035907705 | 0.477300935 | 0.063274608 | 0.295508496 | 0.022340012 |
| 17062.02539 | 15587.49121 | 19549.99805 | -0.543031442 | 0.537689662 | 0.866808955 | 0.341435242 | 0.193996696 | 0.845545984 |
| 31191.72656 | 34822.96094 | 35790.96484 | -0.544370825 | 0.092516251 | 0.001104927 | 0.996723914 | -0.527869184 | 0.145702448 |
| 45523.13281 | 45419.33594 | 42298.40234 | -0.544775229 | 0.066158993 | 0.234989546 | 0.050817911 | 0.0885637 | 0.210903657 |
| 97656.73438 | 103032.5703 | 113060 | -0.546036212 | 0.032587088 | 0.217280269 | 0.162505567 | 0.386066147 | 0.063076534 |
| 165489.5156 | 169837.5469 | 172084.8906 | -0.546191757 | 0.027671686 | 0.093850352 | 0.176864019 | 0.178612555 | 0.060539652 |
| 140060.3594 | 156506.6563 | 142987.7656 | -0.546272808 | 0.013354673 | -0.045378438 | 0.486829517 | -0.375391971 | 0.031136253 |
| 5447.336426 | 5465.65625 | 6106.193359 | -0.546348919 | 0.034144692 | 0.241661052 | 0.14612569 | 0.039669183 | 0.689076806 |
| 9361.902344 | 9265.47168 | 5368.078125 | -0.54659116 | 0.238472464 | 0.409660228 | 0.407540737 | 0.149809341 | 0.789032408 |
| 137421.5625 | 140185.4375 | 118270.5391 | -0.547062784 | 0.39188184 | 0.150589479 | 0.487853215 | 0.034461606 | 0.844045457 |
| 202157.9219 | 208997.9531 | 220127.4844 | -0.547206614 | 0.086909654 | -0.225681125 | 0.425499456 | -0.159521648 | 0.534750592 |

| | | | | | | | | |
|---|---|---|---|---|---|---|---|---|
| 3072409.5 | 3254954 | 2856886.75 | -0.547438142 | 0.005790576 | 0.067600966 | 0.191852289 | -0.319736615 | 0.062835547 |
| 181343.4844 | 186101 | 193910.9063 | -0.547462983 | 0.009194956 | 0.086773139 | 0.397807267 | 0.19183244 | 0.06065526 |
| 146893.5938 | 144527.7031 | 136405.5156 | -0.547766264 | 0.191685231 | 0.045784163 | 0.604317147 | -0.183359254 | 0.051065331 |
| 696019.5625 | 1840874.75 | 756712.6875 | -0.547786681 | 0.450433079 | -0.43560247 | 0.622913596 | -0.074186454 | 0.935152011 |
| 74463.65625 | 71640.39063 | 76975.25781 | -0.547909099 | 0.025025466 | 0.273412905 | 0.090012942 | -0.237827631 | 0.131961597 |
| 11944.45996 | 12106.08691 | 13083.81445 | -0.547948219 | 0.027671686 | 0.445901679 | 0.068436129 | 0.222278334 | 0.075384367 |
| 45990.59375 | 46443.02734 | 48578.90625 | -0.549235339 | 0.198743433 | -0.066717032 | 0.875616587 | 0.364901652 | 0.366285302 |
| 54249.60547 | 46970.83203 | 55571.92578 | -0.549277768 | 0.043773448 | -0.108834962 | 0.555712947 | -0.127880591 | 0.516522435 |
| 10362.10352 | 12951.98145 | 8144.967285 | -0.54929828 | 0.113673327 | 0.671701187 | 0.100046608 | 0.485892008 | 0.262538234 |
| 55876.48828 | 52410.71484 | 51019.63672 | -0.549417311 | 0.040854235 | 0.723201567 | 0.011710842 | 0.381206872 | 0.030894309 |
| 9959.977539 | 9944.147461 | 9377.717773 | -0.549571406 | 0.019930448 | 0.059560514 | 0.470177395 | -0.100657297 | 0.269529321 |
| NA | 227505.375 | 97493.60156 | -0.549820373 | 0.202243414 | -0.874451434 | 0.226930121 | 1.435915383 | NA |
| 279184.4688 | 264907.9375 | 282528.2813 | -0.551525647 | 0.009008041 | 0.273579797 | 0.065601851 | 0.284443319 | 0.032583622 |
| 119614.3984 | 125853.1016 | 133123.3594 | -0.551747876 | 0.025437157 | 0.137321727 | 0.236607128 | 0.395844019 | 0.050570269 |
| 94834.57813 | 108671.0938 | 100628.2344 | -0.552306094 | 0.202476876 | 0.253323027 | 0.066380875 | 0.491297241 | 0.042737273 |
| 52983.66406 | 50365.12891 | 49570.8125 | -0.552650321 | 0.006735623 | 0.082207114 | 0.371290006 | -0.327354744 | 0.023619478 |
| 234408.6563 | 246547.5781 | 265065.7188 | -0.552717342 | 0.034240061 | 0.16561107 | 0.099450521 | 0.356359437 | 0.054870367 |
| 1169.84021 | 838.6275024 | NA | -0.553321032 | 0.189500928 | -0.370728791 | 0.539433554 | -0.597420273 | NA |
| 147442.8594 | 147355.0313 | 161051.2188 | -0.553580923 | 0.007109595 | 0.076929006 | 0.348633551 | 0.332642134 | 0.063818875 |
| 164145.375 | 161566.3438 | 150514.375 | -0.553668246 | 0.021326467 | 0.416735635 | 0.01179292 | 0.083082804 | 0.252413552 |
| 20916.82813 | 19491.98047 | 19453.81445 | -0.554081359 | 0.100858145 | 0.008116686 | 0.962732724 | 0.22740909 | 0.050570269 |
| 121594.8672 | 127949.8516 | 143028.1406 | -0.554442953 | 0.016471012 | 0.140761006 | 0.133201112 | 0.326956122 | 0.115929208 |
| 12777.2627 | 16508.38477 | 9090.350586 | -0.555328425 | 0.252787526 | 0.306136037 | 0.471943698 | -0.963288875 | 0.207383673 |
| 7779.598145 | 7089.418457 | 7672.633301 | -0.555791963 | 0.01332549 | -0.009379854 | 0.909219658 | -0.154113733 | 0.167225632 |
| 15843.29785 | 15207.64063 | 15889.52637 | -0.555968752 | 0.021027835 | -0.581264814 | 0.059340088 | -0.358043677 | 0.108600169 |
| 145499.9531 | 137306.1719 | 144526.4844 | -0.557153804 | 0.024701492 | 0.135690084 | 0.228751244 | 0.264778037 | 0.050665122 |
| 23706.96875 | 28653.83203 | 18684.31445 | -0.557675423 | 0.08662098 | 0.394468891 | 0.075404888 | 0.181995906 | 0.544452231 |
| 124061.1797 | 135918.5156 | 139355.625 | -0.558151019 | 0.03777619 | 0.251311268 | 0.09163831 | -0.066024297 | 0.515420217 |
| 20653.39844 | 23441.18555 | 21269.07227 | -0.558295559 | 0.034061991 | -0.217528435 | 0.244612494 | -0.751788629 | 0.015233843 |
| 65174.46094 | 70745.9375 | 56686.86328 | -0.558735849 | 0.056813934 | 0.007289347 | 0.953027781 | -0.121493943 | 0.477536926 |
| 27019.5332 | 28967.62305 | 30946.72461 | -0.559376036 | 0.249522174 | 0.754500985 | 0.132066965 | 0.244552689 | 0.114499805 |
| 60439.73047 | 54098.08594 | 53390.50781 | -0.559642882 | 0.022489083 | 0.644347786 | 0.050958871 | 0.547068434 | 0.033709967 |
| 32895.85156 | 38919.42969 | 34607.71094 | -0.559900266 | 0.167506197 | 0.288978662 | 0.08165067 | 0.143339951 | 0.321796756 |

| | | | | | | | | |
|---|---|---|---|---|---|---|---|---|
| 3613.797363 | 3964.116943 | 2425.02832 | -0.560056498 | 0.042404592 | 0.862117031 | 0.038037652 | 1.02029339 | 0.131502216 |
| 272933.875 | 286578.1875 | 287962.0938 | -0.560493193 | 0.004709568 | 0.148509747 | 0.041058966 | 0.247994814 | 0.027868741 |
| 4355.555176 | 4923.299316 | 4851.733887 | -0.560829098 | 0.045092515 | 0.375598926 | 0.148283241 | 0.128478339 | 0.315565256 |
| 114630.8984 | 115416.7734 | 119305.8438 | -0.561345398 | 0.027375615 | -0.056451941 | 0.591668834 | -0.416027734 | 0.062822 |
| 2195.98291 | 4408.109375 | 3860.302002 | -0.561821451 | 0.16947575 | -0.215278015 | 0.508503755 | -0.727878004 | 0.136049841 |
| 16757.49023 | 19047.39063 | 18395.89453 | -0.562916983 | 0.037314581 | -0.222471129 | 0.265104001 | -0.185497554 | 0.117034613 |
| 13672.20801 | 14671.6709 | 12083.4082 | -0.56297589 | 0.019286562 | -0.132355094 | 0.3150978 | -0.713255857 | 0.031136253 |
| 40338.43359 | 45451.42578 | 39073.85547 | -0.563282209 | 0.017507272 | 0.012176223 | 0.909761575 | -0.153601948 | 0.277166323 |
| 108173.5391 | 106172.4844 | 118344.4453 | -0.563534447 | 0.036899714 | 1.109464286 | 0.016310878 | 0.885304973 | 0.020159082 |
| 6544.773438 | 6651.510254 | 7176.734375 | -0.565081929 | 0.105631439 | -0.224719069 | 0.479088235 | -0.090791315 | 0.747860314 |
| 39728.62109 | 44292.35938 | 44091.125 | -0.56512649 | 0.22690371 | 0.585755513 | 0.142285137 | 0.767325179 | 0.08420993 |
| 96064.78125 | 81005.34375 | 83694.88281 | -0.565472494 | 0.068432057 | 0.252407786 | 0.193121972 | 0.157718732 | 0.39271806 |
| 6208.92627 | 8056.713867 | 4812.675781 | -0.565533151 | 0.104350517 | -0.008232343 | 0.967452872 | -0.641638739 | 0.118472908 |
| 8409.921875 | 9974.010742 | 5658.375977 | -0.565568122 | 0.114478366 | -0.485427552 | 0.258746601 | -0.77777225 | 0.134304215 |
| 109519.2656 | 107128.9375 | 109957.875 | -0.565660841 | 0.03510946 | 0.047131058 | 0.562301807 | 0.224307654 | 0.020008995 |
| 153254.8438 | 166285.9531 | 152054.4688 | -0.566068827 | 0.014306629 | 0.769182323 | 0.042000044 | 0.28407486 | 0.060012809 |
| 9155.796875 | 7498.907227 | 6284.317383 | -0.566402504 | 0.054037379 | 0.268749465 | 0.149078465 | -0.478588306 | 0.118023574 |
| 592352.25 | 624508.3125 | 661900.875 | -0.56648661 | 0.021027835 | 0.452371451 | 0.106670824 | 0.740890867 | 0.030690989 |
| 17611.35156 | 17826.56055 | 14495.62207 | -0.566750703 | 0.034514668 | 0.042277767 | 0.781053515 | 0.037045317 | 0.847737136 |
| 47601.97656 | 58531.44922 | 38943.41406 | -0.567506992 | 0.391855884 | -0.011713469 | 0.979876653 | -0.097168259 | 0.778018685 |
| 34423.69922 | 33079.31641 | 27212.99414 | -0.567536466 | 0.247825184 | 0.11890984 | 0.811209306 | -0.179398638 | 0.742102915 |
| 3673.946533 | 3233.801514 | 2614.512695 | -0.567538376 | 0.213145643 | 0.447739451 | 0.306000099 | -0.272673291 | 0.569184065 |
| 542726.5625 | 557802.9375 | 553906.5625 | -0.567841275 | 0.016859654 | -0.161067099 | 0.131030789 | -0.406606609 | 0.051133451 |
| 73229.58594 | 73818 | 62888.32031 | -0.568155116 | 0.012925173 | 0.537555954 | 0.099100403 | 1.121973127 | 0.037571191 |
| 986516.25 | 1039614.375 | 943815 | -0.568806723 | 0.056696683 | -0.168185082 | 0.169675649 | 0.44142834 | 0.049415261 |
| 1101.647339 | NA | NA | -0.5693139 | NA | -0.928002055 | NA | -0.697875992 | NA |
| 13918.25684 | 12078.2041 | 12182.02246 | -0.569490106 | 0.421615231 | 0.234613318 | 0.765565426 | -0.19833148 | 0.82622823 |
| 41175.25 | 40150.53516 | 43632.19531 | -0.570082976 | 0.045554362 | 0.022535908 | 0.753193033 | -0.055907715 | 0.466490285 |
| 93979.14844 | 92530.8125 | 95095.11719 | -0.570361743 | 0.027375615 | -0.146500922 | 0.162505567 | -0.152798634 | 0.155548876 |
| 439703.9688 | 430516.3125 | 396843.0625 | -0.570604038 | 0.029933445 | 0.296084135 | 0.105216387 | 0.297067164 | 0.079082931 |
| 105806.5156 | 113207.1172 | 102360.2656 | -0.570843563 | 0.078335617 | 0.451931927 | 0.125646125 | 0.356643128 | 0.180594841 |
| 1682543.375 | 2716986.578 | 1895112.406 | -0.571086972 | 0.379357439 | 0.031414554 | 0.947443559 | 0.070500714 | 0.861875296 |
| 81106.57813 | 78631.46875 | 88712.07031 | -0.571122348 | 0.009008041 | 0.290163293 | 0.059562109 | 0.388029851 | 0.070873774 |

| | | | | | | | | |
|---|---|---|---|---|---|---|---|---|
| 16321.70801 | 20592.86133 | 11912.84082 | -0.571776358 | 0.46508728 | -0.077316082 | 0.873585467 | 0.086961294 | 0.823907108 |
| 693953.625 | 766659.1875 | 743470.9375 | -0.572011895 | 0.10995741 | -0.642338079 | 0.020431201 | 0.136639409 | 0.152372879 |
| 360003.5938 | 370631.125 | 316331.8438 | -0.572051497 | 0.223354704 | -0.273952152 | 0.343660317 | -0.224542577 | 0.397275308 |
| 51845.74609 | 52973.04297 | 44499.31641 | -0.57210718 | 0.06853067 | -0.322536581 | 0.312233367 | -0.276951703 | 0.341447574 |
| 82704.14063 | 90445.71094 | 88250.89063 | -0.572198536 | 0.100529206 | 0.589183049 | 0.117302538 | 0.357995645 | 0.230606211 |
| 41841.97656 | 41123.35156 | 28710.55664 | -0.572292812 | 0.05114005 | 0.181036603 | 0.406503625 | -0.120054385 | 0.687757656 |
| 131320.6406 | 152408.5938 | 135182.1563 | -0.572361773 | 0.028238951 | -0.205653512 | 0.315239445 | -1.649543814 | 0.036848746 |
| 54659.85547 | 52161.04297 | 38887.12109 | -0.572657262 | 0.027671686 | -0.198403291 | 0.280688037 | -0.5917273 | 0.071644236 |
| 50333.29297 | 44690.53125 | 39455.88672 | -0.572701738 | 0.029906057 | -0.127677755 | 0.094690058 | -0.486261135 | 0.067682423 |
| 338295.1875 | 328290.4063 | 313643.2188 | -0.572918541 | 0.039647218 | 0.144260953 | 0.28533223 | -0.027142311 | 0.824693806 |
| 685982.9375 | 626820.5 | 642749.875 | -0.572927712 | 0.090273155 | -0.153719816 | 0.384749949 | -0.00780753 | 0.935152011 |
| 23983.65625 | 23536.04102 | 22096.49414 | -0.572937619 | 0.047710339 | -0.102359831 | 0.15090616 | -0.119348612 | 0.119231025 |
| 46781.1875 | 50432.73438 | 48126.96484 | -0.57327777 | 0.031356602 | 0.159292595 | 0.282993231 | 0.201224586 | 0.192079081 |
| 19883.05859 | 23891.2168 | 24398.21289 | -0.573336597 | 0.050873498 | -0.065724068 | 0.736018403 | -0.020053397 | 0.931604241 |
| 200686.9688 | 199980.1563 | 216112.8125 | -0.573429054 | 0.042292137 | 0.128905722 | 0.124478178 | 0.303351672 | 0.060774742 |
| 11370.55273 | 11008.84961 | 11808.6875 | -0.574334508 | NA | 0.682645783 | 0.212815319 | 0.231430655 | 0.557774325 |
| 140917.9063 | 142258.4688 | 133167.3438 | -0.574936757 | 0.046681455 | 0.199762102 | 0.116818656 | 0.322095535 | 0.024862205 |
| 38026.26172 | 42995.28906 | 42108.92969 | -0.575118514 | 0.02187351 | -0.054518762 | 0.649577647 | -0.246190491 | 0.125718789 |
| 8961.134766 | 9952.542969 | 7950.125488 | -0.575255784 | 0.079503363 | 0.054896187 | 0.635624309 | -0.266139132 | 0.167717489 |
| 18875.37891 | 36620.54297 | 35904.26172 | -0.575279493 | 0.327070857 | -1.353838623 | 0.281024002 | 0.033504135 | 0.966538084 |
| 9653.291992 | 9648.728516 | 7149.204102 | -0.575848829 | 0.086055451 | 0.607333453 | 0.032229098 | 0.587328002 | 0.1278105 |
| 61453.30078 | 92902.14063 | 51857.87891 | -0.575937304 | 0.340055181 | -0.000411067 | 0.999222442 | 0.02237791 | 0.963218346 |
| 5145.274414 | 3933.662598 | 2548.053711 | -0.576243954 | 0.104002895 | -0.898476255 | 0.024018676 | -1.128993101 | 0.059641593 |
| 73511.26563 | 78260.29688 | 67836.0625 | -0.576314853 | 0.082384942 | 0.154725609 | 0.187574879 | -0.594003439 | 0.032502 |
| 157426.4688 | 145608.0313 | 141602.75 | -0.576401309 | 0.117230721 | -0.23224378 | 0.098688136 | -0.311962878 | 0.056784127 |
| 889714.8369 | 963561.8877 | 914295.7339 | -0.576581236 | 0.037496289 | 0.391978196 | 0.102026311 | -0.150512429 | 0.080867847 |
| 2850.056152 | 2031.249268 | 1266.374023 | -0.576906619 | 0.013964852 | 0.331112739 | 0.398018503 | 0.02920996 | 0.956668754 |
| 14706.49023 | 12812.52051 | 11256.19043 | -0.577055151 | 0.030417399 | 0.154269906 | 0.52210667 | 0.075659835 | 0.672052206 |
| 293281.3438 | 298349.0625 | 292065.25 | -0.577588334 | 0.148814893 | -0.206230516 | 0.22111982 | -0.498591079 | 0.084288402 |
| 200724.7813 | 229891.4688 | 192285.7969 | -0.577713313 | 0.138429888 | -0.283154226 | 0.220633017 | -0.126841409 | 0.341019132 |
| 78715.9375 | 65258.66797 | 74134.45313 | -0.578638608 | 0.012414001 | 0.286614496 | 0.113912361 | 0.27615235 | 0.139758304 |
| 206192.625 | 209164.5625 | 209754.2813 | -0.578645763 | 0.029603305 | 0.072309708 | 0.72552298 | 0.008702349 | 0.954795145 |
| 999.9839478 | 1570.226318 | 1333.339966 | -0.579038547 | 0.087897715 | 0.18012786 | 0.25339535 | -0.468765551 | 0.174887902 |

| | | | | | | | | |
|---|---|---|---|---|---|---|---|---|
| 110716.7031 | 107986.6953 | 90281.75781 | -0.579186092 | 0.048836369 | 0.180508204 | 0.090033554 | 0.045839024 | 0.749389169 |
| 55065.02344 | 43592.83594 | 29361.56836 | -0.579881765 | 0.211452184 | 0.434821741 | 0.157734764 | 0.046886684 | 0.917097845 |
| 128770.25 | 119238.7656 | 107947.2891 | -0.580124757 | 0.062609 | 0.650011454 | 0.140225049 | 0.411695159 | 0.128960705 |
| 9172.670898 | 8147.681152 | 6780.30127 | -0.580256524 | 0.06906613 | 0.282213878 | 0.355229313 | 0.243460814 | 0.3964865 |
| 1636.692749 | 854.7792969 | NA | -0.581124669 | 0.148078396 | -1.025253 | 0.178486735 | -2.319031407 | NA |
| 129910.9688 | 201426.4531 | 147725.2188 | -0.581300969 | 0.174564561 | -0.102939526 | 0.809940067 | -0.018497272 | 0.962025175 |
| 43908.21094 | 35551.91016 | 38548.79688 | -0.581723636 | 0.084757955 | 0.399746198 | 0.054829172 | 0.348557882 | 0.124736435 |
| 58967.07031 | 53680.92969 | 48884.67188 | -0.582558692 | 0.062357731 | 0.243168796 | 0.418488198 | 0.032295658 | 0.878850339 |
| 72541.39063 | 66905.4375 | 78075.74219 | -0.583474888 | 0.116159237 | 0.03454149 | 0.32371757 | 0.05146734 | 0.61132384 |
| 38371.01172 | 44164.28516 | 44595.21094 | -0.583485586 | 0.071158565 | 0.274073464 | 0.083732049 | 0.380322098 | 0.089805965 |
| 2692110.75 | 2835073 | 3140378.25 | -0.583555631 | 0.038502747 | 0.187924082 | 0.217926074 | 0.086480843 | 0.555018201 |
| 14150.73438 | 9033.530273 | 11182.89844 | -0.584728695 | 0.076438569 | -0.197679453 | 0.254161368 | -0.703899244 | 0.071151453 |
| 155913.1719 | 159935.125 | 161867.375 | -0.58477495 | 0.006395507 | 0.188013281 | 0.030241535 | 0.271970779 | 0.015632484 |
| 34923.14063 | 30648.30273 | 26460.79883 | -0.585486343 | 0.053893741 | 1.201644276 | 0.05467585 | 0.313631461 | 0.223194941 |
| 15862.9082 | 20099.25586 | 36585.63672 | -0.585934516 | 0.017143851 | 0.250488387 | 0.137136478 | 0.262026919 | 0.680808788 |
| 46143.92969 | 52113.52344 | 54471.79297 | -0.586111451 | 0.097443336 | 0.313366552 | 0.388254654 | 0.760832622 | 0.066335881 |
| 59687.6875 | 66410.85156 | 71788.73438 | -0.586206399 | 0.187781848 | 0.474937424 | 0.280546656 | 1.019055215 | 0.056809612 |
| 2128.642578 | 8913.972656 | 6206.311523 | -0.586240558 | 0.295381969 | 0.416286652 | 0.47004298 | 0.218289166 | 0.815961723 |
| 157282.5938 | 155158.7969 | 155080.4688 | -0.586259428 | 0.038942433 | 0.139859504 | 0.013885891 | 0.2384276 | 0.006394096 |
| 34864.11328 | 42391.04688 | 36828.71484 | -0.587120928 | 0.097194367 | 0.211736614 | 0.467054186 | 0.918423971 | 0.042175609 |
| 146398.3906 | 144808.5781 | 160351.5781 | -0.588284686 | 0.012637622 | 0.173302199 | 0.099808559 | 0.288725637 | 0.063474265 |
| 23753.11719 | 15667.12305 | 14511.68457 | -0.588312551 | 0.138598181 | 0.668705761 | 0.138224464 | 0.166776391 | 0.713487699 |
| 35833.84766 | 37282.97266 | 34886.33594 | -0.588721172 | 0.013618049 | 0.416945285 | 0.045481011 | -0.359528993 | 0.046178241 |
| 149904.2813 | 167678.5469 | 132942.4063 | -0.589308098 | 0.146753968 | 0.062048608 | 0.41941562 | 0.195555549 | 0.293140864 |
| 10910.17285 | 10922.50293 | 10323.56836 | -0.589321743 | 0.013723547 | 0.403427908 | 0.022904665 | -0.041792025 | 0.477695497 |
| 11512.99219 | 8400.431641 | 9342.951172 | -0.589415171 | 0.021745834 | 0.013612658 | 0.909761575 | -0.480748657 | 0.089541557 |
| 39861.42578 | 74966.47656 | 47483.73438 | -0.589514447 | 0.15142156 | -0.028121694 | 0.947346059 | 0.158785135 | 0.764791569 |
| 10453.52246 | 10007.58398 | 3560.770264 | -0.589539955 | NA | 0.916047726 | NA | 0.677684032 | NA |
| 21427.0918 | 20316.96484 | 16429.04297 | -0.589618608 | 0.073792129 | -0.081739401 | 0.723713085 | -0.431449107 | 0.194140122 |
| 16790.98828 | 16080.85059 | 14333.49219 | -0.590517875 | 0.070422084 | -0.188328844 | 0.353137578 | -0.373562451 | 0.062942605 |
| 5505.73291 | 5282.89209 | 4326.34082 | -0.590572701 | 0.160003781 | 1.016143036 | 0.081080964 | 0.830326369 | 0.077902102 |
| 6568.735352 | 7882.39209 | 4061.407227 | -0.590967854 | 0.100135417 | -0.485865866 | 0.147741919 | -0.760367772 | 0.106375556 |
| 36493.28125 | 29314.81641 | 34016.62891 | -0.591039958 | 0.046112206 | -0.070646549 | 0.462179438 | -0.318251518 | 0.101462623 |

| | | | | | | | | |
|---|---|---|---|---|---|---|---|---|
| 60315.41016 | 71668.97656 | 82211.03906 | -0.591219269 | 0.191383717 | 0.355410488 | 0.416953273 | 0.316084614 | 0.457083172 |
| 9622.418945 | 10500.19434 | 12858.96777 | -0.592014543 | 0.214914807 | 0.783934225 | 0.167607585 | 0.356333164 | 0.455428652 |
| 187728.2188 | 177878.5313 | 195482.25 | -0.59208277 | 0.036534375 | 0.144275249 | 0.041850551 | 0.274246302 | 0.059641593 |
| 31469.13281 | 40864.53906 | 24349.84375 | -0.592192941 | 0.288353039 | -0.154803413 | 0.678734633 | -0.405054271 | 0.262168702 |
| 48579.92188 | 52479.56641 | 50779.08203 | -0.592329673 | 0.076331689 | 0.326291083 | 0.27203717 | -0.28849706 | 0.374308247 |
| 180048.9844 | 198049.9375 | 181456.9063 | -0.592349708 | 0.045233346 | -0.412225894 | 0.020268365 | -0.745907338 | 0.022638191 |
| 734781.25 | 702694.1875 | 615260.125 | -0.592525621 | 0.041945948 | -0.045432763 | 0.735029422 | -0.352534543 | 0.111102977 |
| 6282.468262 | 6394.444824 | 7158.141113 | -0.592687737 | 0.042292137 | 0.077485766 | 0.65108636 | 0.049475206 | 0.780636174 |
| 43748.13672 | 50857.72266 | 48007.10156 | -0.592849826 | 0.032310891 | 0.002070352 | 0.99120847 | 0.233997192 | 0.164457546 |
| 5930.611328 | 6384.887207 | 4447.519043 | -0.592957792 | 0.025086961 | -0.31028608 | 0.097033041 | -0.57200441 | 0.07832051 |
| 67061.17188 | 64273.19531 | 55611.76953 | -0.59301599 | 0.050437184 | 0.224820894 | 0.092617517 | 0.105068803 | 0.445699998 |
| 20339.48438 | 17870.55664 | 15615.70605 | -0.594010572 | 0.091842354 | 0.173594846 | 0.266303698 | 0.020982143 | 0.92031567 |
| 446998.8359 | 471270.6953 | 451267.7813 | -0.594573581 | 0.035025385 | 0.609413709 | 0.07036809 | 0.551917188 | 0.010236624 |
| 73775.96875 | 75117.85156 | 69412.125 | -0.595321755 | 0.068352646 | 0.053883796 | 0.806856566 | -0.490530142 | 0.169075369 |
| 9352.124023 | 11037.1416 | 9001.290039 | -0.595512639 | NA | 3.171719572 | NA | 2.649335416 | NA |
| 1451314.25 | 1583299.875 | 1547480.75 | -0.595815046 | 0.005648534 | 0.006369409 | 0.933858968 | -0.097097296 | 0.179985011 |
| 30200.41602 | 30892.83203 | 31268.25977 | -0.596377848 | 0.110302818 | 0.216921358 | 0.456419987 | 0.104305472 | 0.706933965 |
| 16533.75391 | 17396.01367 | 16966.02344 | -0.596523965 | 0.031790847 | -0.24791557 | 0.198890201 | -0.854063006 | 0.061670859 |
| 69781.41406 | 59182.94141 | 57791.53906 | -0.596530319 | 0.077955129 | -0.123956206 | 0.25405646 | -0.735716042 | 0.022280208 |
| 339188.8438 | 307554.9375 | 307596.625 | -0.597057045 | 0.009093304 | 0.336582995 | 0.277641591 | 0.328296632 | 0.049082028 |
| 21240064 | 20329132 | 20831560 | -0.597077531 | 0.037600114 | 1.668513983 | 0.046285147 | 1.146674791 | 0.023619478 |
| 134845.5313 | 1094302.625 | 186763.4219 | -0.597282855 | 0.612958191 | -0.18662542 | 0.911760787 | -0.081844351 | 0.961294722 |
| 54345.72656 | 52861.69922 | 52488.97656 | -0.597598123 | 0.030959545 | -0.236476874 | 0.183896983 | -0.1587051 | 0.036230872 |
| 20574.58398 | 17512.76758 | 18045.6582 | -0.59768274 | 0.179079166 | 0.204782849 | 0.647292503 | -0.02786643 | 0.953276013 |
| 24881.79883 | 26634.64063 | 29868.69531 | -0.599013834 | 0.106835217 | 0.431483891 | 0.32242293 | -0.024666843 | 0.946096095 |
| 20548.66016 | 19549.54688 | 16045.6709 | -0.599191453 | 0.08771097 | 0.461050299 | 0.075810296 | -0.249485877 | 0.193326667 |
| 319105.1563 | 333285.0625 | 321391.8438 | -0.599510275 | 0.018112028 | 0.151912408 | 0.066508521 | 0.260855894 | 0.020470112 |
| 19860.69727 | 18212.64258 | 21286.81641 | -0.600032568 | 0.029906057 | 0.104857089 | 0.41941562 | 0.047727862 | 0.742102915 |
| 8679.217773 | 10488.34473 | 5961.916992 | -0.600046619 | 0.073155658 | -0.244062581 | 0.437151431 | -0.211722653 | 0.5849965 |
| 11197.10645 | 14542.96387 | 10935.1084 | -0.60019482 | 0.18862527 | 0.179992567 | 0.66660509 | 0.123442147 | 0.783657196 |
| 39927.84375 | 46380.19922 | 33618.90625 | -0.600738776 | 0.036971527 | -0.474667943 | 0.052036261 | -0.742484991 | 0.040516462 |
| 1233230.375 | 1160451.625 | 1284164.875 | -0.600876671 | 0.04977426 | 0.495924358 | 0.141416656 | 0.786062294 | 0.030417592 |
| 167094.5313 | 163794.8438 | 191605.625 | -0.601233882 | 0.01802855 | 0.163626672 | 0.18915961 | 0.314461537 | 0.101567963 |

| | | | | | | | | |
|---|---|---|---|---|---|---|---|---|
| 1557.958862 | 788.8487549 | NA | -0.601775732 | 0.044763187 | 0.33793831 | 0.152452638 | -1.059941046 | NA |
| 200418.2617 | 197269.9766 | 234041.4102 | -0.602454054 | 0.007142399 | 0.018761547 | 0.841230106 | -0.125277804 | 0.352925267 |
| 138783.1406 | 166119.1563 | 123473.3359 | -0.602456618 | 0.061766802 | 0.428497407 | 0.119480602 | 0.210475743 | 0.426701953 |
| 213044.2969 | 315820.875 | 173979.6563 | -0.602805695 | 0.384607321 | 0.316182014 | 0.613423653 | 0.297088919 | 0.529658074 |
| 36753.53906 | 40011.10156 | 38397.68359 | -0.604023072 | 0.029016212 | -0.254919617 | 0.163729974 | -0.67663729 | 0.056570421 |
| 33471.76367 | 28627.01514 | 35058.25684 | -0.604341558 | 0.080149199 | 0.933273196 | 0.054857325 | -0.264018773 | 0.378492763 |
| 323067.875 | 364920.9688 | 307163.1563 | -0.604754404 | 0.042916723 | 0.11799077 | 0.458532796 | 0.03048448 | 0.816697307 |
| 54873.19141 | 57727.03125 | 52155.31641 | -0.604887758 | 0.04765314 | 0.186658003 | 0.268940886 | 0.080801463 | 0.586390561 |
| 187889.5938 | 187416.9688 | 195475.9219 | -0.605038177 | 0.004709568 | 0.174561905 | 0.023005377 | 0.319463309 | 0.017279709 |
| 39556.55078 | 35865.40234 | 35694.55469 | -0.605255179 | 0.01391113 | 0.279822725 | 0.052305796 | -0.036050978 | 0.636715098 |
| 125612.7275 | 121495.9019 | 111147.1836 | -0.605681203 | 0.025086961 | -0.110206566 | 0.312686937 | 0.070570725 | 0.519989522 |
| 375422.625 | 428307.8125 | 337912.5625 | -0.607155834 | 0.170413823 | -0.552782853 | 0.059615832 | -0.890451448 | 0.029219266 |
| 25436.07813 | 23848.75195 | 25367.33008 | -0.608506546 | 0.134486207 | 0.883729701 | 0.097877924 | 1.226212464 | 0.046624125 |
| 89079.875 | 94156.16406 | 91801.55469 | -0.608516745 | 0.010893509 | 0.326014454 | 0.039184987 | 0.336742868 | 0.029219266 |
| 46833.41797 | 43059.52539 | 42573.72266 | -0.609001843 | 0.046410702 | -0.086393781 | 0.430916401 | -0.173366617 | 0.190517393 |
| 61885.26953 | 43159.33984 | 54639.55469 | -0.609154324 | 0.071921738 | -0.141628115 | 0.6777223 | -0.193764068 | 0.4289352 |
| 120753.4922 | 118232.7969 | 125904.5469 | -0.609184648 | 0.026670912 | 0.135306724 | 0.184945314 | 0.293243645 | 0.045887602 |
| 10821.5918 | 8633.988281 | 4466.303711 | -0.609380651 | 0.019536332 | 0.236256051 | 0.147248245 | -0.298882384 | 0.536804361 |
| 109235.875 | 111518.6641 | 115074.8516 | -0.609693668 | 0.162395091 | 0.591981713 | 0.189895512 | 0.467671085 | 0.10119601 |
| 61646.17578 | 58555.46094 | 50347.15625 | -0.609791572 | 0.12468265 | 0.820446758 | 0.081324743 | -0.021223451 | 0.872236202 |
| 3482313.5 | 3697498.25 | 3533046.5 | -0.60984704 | 0.005752145 | 0.949666544 | 0.024656861 | 0.41929528 | 0.017015337 |
| 127212.2188 | 137772.4688 | 136914.4219 | -0.610103342 | 0.021857251 | 0.42696692 | 0.034488442 | 0.494408943 | 0.02146481 |
| 41980.00391 | 40620.65625 | 37984.625 | -0.610388235 | 0.014943375 | -0.404220386 | 0.011710842 | -0.703440603 | 0.015111332 |
| 8847.431641 | 8505.78418 | 7297.74707 | -0.611002651 | 0.023510048 | -0.110486975 | 0.392264454 | -0.178545928 | 0.235870699 |
| 43590.09375 | 45603.375 | 38847.31641 | -0.611288478 | 0.011814081 | -0.187623464 | 0.058077004 | -0.103536453 | 0.360575146 |
| 4072.723877 | 4376.73584 | 3313.323486 | -0.611432266 | 0.010297098 | 0.268642492 | 0.22112427 | -0.2282585 | 0.261164148 |
| 73516.10938 | 70656.36719 | 70704.14063 | -0.611641624 | 0.017254559 | -0.271761208 | 0.021969813 | -0.445805654 | 0.010308904 |
| 1977817 | 2039334 | 2123731 | -0.612390224 | 0.065091163 | -0.315491878 | 0.02118733 | -0.314939563 | 0.020008995 |
| 39184.70703 | 40220.875 | 42369.60938 | -0.613174111 | 0.018311389 | 0.2457987 | 0.144097346 | -0.019321657 | 0.82622823 |
| 171909.6719 | 211541.9688 | 168731.5 | -0.613383188 | 0.055345971 | -0.532637373 | 0.110237532 | -0.54326461 | 0.073672213 |
| 137182.1719 | 136375.6719 | 135460.4219 | -0.614687069 | 0.061786866 | 0.646029145 | 0.038037652 | 0.543221054 | 0.027868741 |
| 326739.3125 | 217281 | 186812.7031 | -0.615221607 | 0.041635583 | 1.653054462 | 0.180997148 | 1.758406922 | 0.126766337 |
| 11332.87793 | 10806.99512 | 11880.4541 | -0.615337367 | 0.029811229 | -0.179830851 | 0.332429777 | -0.202196778 | 0.276951269 |

| | | | | | | | | |
|---|---|---|---|---|---|---|---|---|
| 156576.1719 | 159693.125 | 162486.6719 | -0.615491614 | 0.006351124 | 0.109052029 | 0.109509233 | 0.188612367 | 0.040133731 |
| 41545.71094 | 42508.09375 | 34145.86719 | -0.61549643 | 0.02194812 | -0.169760929 | 0.193121972 | -0.556369584 | 0.041386295 |
| 19648.25391 | 20048.10742 | 10697.79102 | -0.616201042 | 0.077108326 | -0.512313309 | 0.111349456 | -0.934335319 | 0.067414905 |
| 7299.432617 | 7251.586426 | 7112.358398 | -0.616718797 | 0.080487571 | -0.42957622 | 0.288306358 | -0.171871055 | 0.573635205 |
| 12402.30273 | 13859.53809 | 5916.827637 | -0.61696711 | 0.042134056 | -0.516956053 | 0.089948012 | -0.862005146 | 0.118023574 |
| 1930.766602 | 2295.583496 | 1385.60376 | -0.617334245 | 0.028255845 | 0.421351469 | 0.252424496 | -0.594595553 | 0.135721265 |
| 54687.13672 | 55399.50781 | 56648.70313 | -0.61733532 | 0.061344134 | -0.203279578 | 0.065130124 | -0.215349891 | 0.042414121 |
| 6991.580078 | 8430.953125 | 8162.581055 | -0.617444803 | 0.048201324 | -0.191448771 | 0.481415807 | -0.064556367 | 0.786883312 |
| 42152.64453 | 43706.71484 | 51247.41016 | -0.61766123 | 0.020626238 | 0.01619021 | 0.904939768 | 0.370551198 | 0.119166108 |
| 11276.9668 | 8217.138672 | 8956.258789 | -0.618394131 | 0.07465848 | 0.243873316 | 0.402164209 | -0.069669771 | 0.82870971 |
| 70892.1875 | 100142.875 | 71714.96094 | -0.618425777 | 0.046945973 | -0.21640235 | 0.48473721 | -0.085796527 | 0.785580722 |
| 21758.52734 | 19939.16406 | 19427.14258 | -0.619644359 | 0.010228174 | 0.324264371 | 0.043784011 | -0.063492391 | 0.513062691 |
| 17192.46484 | 17909.60938 | 16169.5293 | -0.619852261 | 0.051555299 | -0.3906347 | 0.055241445 | -0.215692483 | 0.111095796 |
| 50293.63672 | 48331.72266 | 34527.67969 | -0.620215051 | 0.146530199 | 1.156781077 | 0.111308998 | -0.024101801 | 0.926180903 |
| 52985.40625 | 59246.33984 | 54067.54688 | -0.620788507 | 0.019930448 | 0.068294229 | 0.382105006 | -0.089415256 | 0.366898574 |
| 65096.57813 | 69847.98438 | 78048.02344 | -0.6214362 | 0.050928677 | 0.172765863 | 0.071544796 | 0.491532773 | 0.07832051 |
| 43084.625 | 48761.77734 | 50146.80469 | -0.621846299 | 0.037718146 | 0.206249355 | 0.043360794 | 0.434557519 | 0.074385174 |
| 7598.695313 | 6470.829102 | 7091.835449 | -0.622779034 | 0.063843436 | -0.290014563 | 0.109314009 | -0.305954543 | 0.070298132 |
| NA | 1921.274658 | 419.0759888 | -0.623075909 | 0.388688484 | 0.439263162 | 0.637841781 | -0.396839032 | NA |
| 6083.186523 | 5457.347656 | 3427.121094 | -0.623532025 | 0.115723457 | 0.636533988 | 0.138525684 | -0.307843561 | 0.504929127 |
| 282472.3438 | 535862.25 | 256914.1094 | -0.623651046 | 0.346157494 | 0.21063973 | 0.755290951 | 0.203765023 | 0.742793738 |
| 318176.375 | 338136.9688 | 350548.9375 | -0.624041303 | 0.037883744 | 0.094464832 | 0.374500771 | 0.532705016 | 0.02146481 |
| 50288.29688 | 54566.4375 | 52248.74609 | -0.624721145 | 0.034514668 | -0.103678308 | 0.276405455 | -0.223718271 | 0.068221145 |
| 180204.5625 | 180273.2344 | 201526.2656 | -0.625213533 | 0.015157115 | 0.158989862 | 0.233271513 | 0.307877884 | 0.076508873 |
| 30350.46094 | 32457.79883 | 25813.03516 | -0.625730617 | 0.291292002 | -0.131212737 | 0.639049031 | -0.149144873 | 0.549537719 |
| 10837.04199 | 10575.24121 | 11830.28516 | -0.626064106 | 0.034151807 | 0.698401897 | 0.029468331 | 0.376496468 | 0.069955133 |
| 5241.645508 | 6219.59082 | 4305.088867 | -0.62758996 | 0.115723457 | 0.516551308 | 0.188920591 | 0.15433485 | 0.685677252 |
| 199985.8281 | 208199.1094 | 184498.9375 | -0.627714824 | 0.019286562 | 0.426025413 | 0.042753324 | -0.047586626 | 0.68342299 |
| 31067.04297 | 33084.99219 | 31198.69336 | -0.627845413 | 0.186381308 | 0.574563835 | 0.255136723 | 0.338836404 | 0.459479226 |
| 133615.9531 | 138561.3594 | 132918.7188 | -0.628719706 | 0.011206547 | 0.097145404 | 0.052036261 | 0.214647131 | 0.03161115 |
| 129198.2788 | 127479.6021 | 129360.3789 | -0.629279091 | 0.017507272 | -0.082716508 | 0.362329238 | -0.30326712 | 0.049410629 |
| 11692.4668 | 10439.86426 | 11228.12598 | -0.629501641 | 0.082806274 | 1.378136761 | 0.093818896 | 0.064992633 | 0.528606397 |
| 145104.75 | 164903.7656 | 169382.7813 | -0.629639508 | 0.156615262 | 0.021616119 | 0.956152889 | 0.105096593 | 0.787435595 |

| | | | | | | | | |
|---|---|---|---|---|---|---|---|---|
| 17318.73242 | 18948.81055 | 17758.66797 | -0.629946518 | 0.020662324 | 0.099022704 | 0.173549208 | -0.076414003 | 0.325682638 |
| 96691.36719 | 98373.09375 | 93898.07813 | -0.630187613 | 0.013618049 | 0.328290933 | 0.050410119 | 0.257525764 | 0.069955133 |
| 7900.219727 | 28757.38867 | 7498.860352 | -0.630198815 | 0.279738117 | 1.339715826 | 0.33801894 | 1.200545462 | 0.467886278 |
| 39634.00781 | 36145.36328 | 36610.51563 | -0.631044695 | 0.025300184 | -0.369225666 | 0.162505567 | -0.427088593 | 0.117397927 |
| 21532.52734 | 22471.94531 | 15423.49023 | -0.631376416 | 0.01964718 | 0.361492473 | 0.198890201 | 0.24042884 | 0.413538473 |
| 108185.4844 | 179666.4375 | 100371.7891 | -0.632242744 | 0.128106288 | -0.32065874 | 0.425452152 | -0.941238838 | 0.118785258 |
| 57491.96875 | 77102.77344 | 60989.57031 | -0.632475167 | 0.303535621 | -0.604984535 | 0.28983684 | -0.532562274 | 0.309468368 |
| 1011044.188 | 1049421 | 1144829.625 | -0.632562208 | 0.028767912 | -0.141020996 | 0.229545075 | 0.043247039 | 0.643137528 |
| 77664.96875 | 62935.27734 | 69505.35938 | -0.632790416 | 0.037729073 | 0.969019379 | 0.05975961 | 0.417504591 | 0.109643103 |
| 134120.875 | 137432.8594 | 132505.4063 | -0.63452465 | 0.009974872 | 0.230101879 | 0.048024604 | -0.259091612 | 0.046092295 |
| 541.8858643 | 480.9403992 | NA | -0.635236144 | 0.072163652 | 0.451692489 | 0.165757248 | -0.605336521 | NA |
| 1002.24646 | 1725.84082 | 1234.658081 | -0.635486122 | 0.137764527 | 1.475321045 | 0.124035194 | 0.687874523 | 0.221497805 |
| 7476.598145 | 6418.138184 | 6886.911133 | -0.636013521 | 0.22827254 | -0.620711308 | 0.265347062 | -0.388491937 | 0.395364978 |
| 50930.50781 | 45466.13281 | 44768.25781 | -0.637383981 | 0.055510745 | -0.355465095 | 0.148613888 | -0.504729957 | 0.071102463 |
| 23578.26172 | 23992.30859 | 22489.26172 | -0.637440211 | 0.07153575 | -0.262573567 | 0.406536014 | -0.185655261 | 0.515022525 |
| 81172.72656 | 85218 | 83048.5625 | -0.637845763 | 0.074117769 | 0.222014944 | 0.387717813 | 0.039112217 | 0.870796801 |
| 1160523.375 | 1065919.75 | 990319.875 | -0.638491566 | 0.005752145 | -0.036646924 | 0.555994171 | 0.367530599 | 0.088296374 |
| 8463.951172 | 5997.472168 | 9774.295898 | -0.638655736 | 0.197302391 | 0.592849208 | 0.25313315 | 0.189341653 | 0.721666648 |
| 45081.44531 | 59519.41797 | 66312.65625 | -0.639101502 | 0.517914463 | 0.850411135 | 0.407350368 | 0.681119431 | 0.505116773 |
| 7452.693848 | 3476.896484 | 2393.539795 | -0.639274738 | 0.343477561 | 0.710841337 | 0.323323292 | -0.700537569 | 0.522523924 |
| 12332.22559 | 13535.14453 | 10636.7959 | -0.639303479 | 0.102272218 | -0.528878467 | 0.28225379 | -0.458122532 | 0.302053963 |
| 16074.45313 | 14688.42773 | 12236.18457 | -0.639966112 | 0.057465973 | 0.485399735 | 0.086634371 | -0.108057749 | 0.618961985 |
| 6574.317383 | 7620.511719 | 5252.334473 | -0.640231859 | 0.256635939 | 0.029082824 | 0.964257738 | 0.071372247 | 0.91311394 |
| 861267.4375 | 1622515.375 | 1095272.375 | -0.640299117 | 0.217707799 | -0.123097259 | 0.849641769 | -0.584659703 | 0.404479667 |
| 22457.71289 | 20441.49219 | 20280.69141 | -0.640509202 | 0.10498796 | 0.460540387 | 0.054829172 | 0.169295049 | 0.266989013 |
| 32329.32422 | 34013.46484 | 46709.53516 | -0.640555028 | 0.161959121 | 0.42546681 | 0.336127197 | 0.04125041 | 0.934434551 |
| 11502.3584 | 13419.82422 | 7708.163574 | -0.640726138 | 0.042427338 | 0.098187223 | 0.741595046 | -0.432674328 | 0.248318189 |
| 11388.24609 | 16062.20801 | 14269.05176 | -0.640797626 | 0.02237899 | 0.592031675 | 0.039887225 | -0.663147798 | 0.059258927 |
| 32145.32617 | 30605.38086 | 38444.91016 | -0.64191428 | 0.082811166 | 0.212903795 | 0.432686549 | 0.001048846 | 0.997782424 |
| 30298.33984 | 41960.84375 | 29518.72656 | -0.64219757 | 0.051033437 | -0.26702407 | 0.231386286 | -0.508936505 | 0.118023574 |
| 11889.12012 | 10514.32422 | 10779.30762 | -0.642686641 | 0.019610279 | 0.570516047 | 0.029435257 | 0.303929805 | 0.077440496 |
| 9683.740234 | 8398.910156 | 9110.888672 | -0.643097625 | 0.089177752 | 1.107630743 | 0.039567204 | 0.196810584 | 0.189363824 |
| 281157.8438 | 354313.3125 | 297084.0625 | -0.646334642 | 0.10036677 | 0.111048313 | 0.45111432 | 0.102585908 | 0.584717012 |

| | | | | | | | | |
|---|---|---|---|---|---|---|---|---|
| 2177.993408 | 2825.185303 | 2348.591553 | -0.646471263 | 0.087043567 | 0.123083569 | 0.68693 | 0.431919182 | 0.146392846 |
| 100653.3359 | 100348.7813 | 102756.4688 | -0.646637486 | 0.025025466 | 0.197528769 | 0.166408179 | 0.304590058 | 0.091810798 |
| 3751.518311 | 5619.544922 | 4356.574707 | -0.646701281 | 0.113705211 | -0.684487437 | 0.233498164 | -0.735324478 | 0.202642834 |
| 40668.92578 | 46095.95313 | 36835.33594 | -0.647114115 | 0.166231401 | -0.700581643 | 0.287681705 | -1.149835316 | 0.186529757 |
| 34799.72656 | 48420.18359 | 38242.30469 | -0.647154525 | 0.219397202 | 0.327855309 | 0.150271527 | 0.096309002 | 0.713487699 |
| 356852.5313 | 258719.9688 | 198311.4375 | -0.647600955 | 0.448907681 | 0.551038672 | 0.328095647 | -0.399433678 | 0.298633387 |
| 107008.1719 | 119258.1484 | 116918.8203 | -0.647952875 | 0.022960243 | -0.407518919 | 0.142439766 | 0.030674533 | 0.840720144 |
| 136781.4844 | 117938.3984 | 89492.02344 | -0.648495387 | 0.063544728 | -0.609265325 | 0.027859864 | -0.908911002 | 0.044156984 |
| 87449.82813 | 85196.28906 | 85505.91406 | -0.649299068 | 0.004709568 | -0.272454973 | 0.068436129 | 0.061408115 | 0.142091675 |
| 16896.30273 | 16603.82617 | 13284.15625 | -0.649502662 | 0.042916723 | -0.160161513 | 0.517413777 | 0.269213077 | 0.302401454 |
| 117087.2188 | 122381.2656 | 103005.4375 | -0.649626335 | 0.173343608 | 0.721595439 | 0.193209925 | 0.338873639 | 0.317506407 |
| 25374.64844 | 30899.43359 | 22871.58008 | -0.649667034 | 0.358576039 | -0.027962277 | 0.935209325 | -0.159838426 | 0.594883754 |
| 33509.56641 | 33701.76563 | 41272.03516 | -0.649815686 | 0.032652748 | 0.735134718 | 0.05818214 | 1.190792619 | 0.057068534 |
| 92009.70313 | 95871.36719 | 86362.8125 | -0.65002534 | 0.009008041 | 1.014997774 | 0.033121608 | 0.843063886 | 0.029820656 |
| 23855.7793 | 23055.43555 | 13113.9834 | -0.651284947 | 0.010893509 | -1.197582056 | 0.018793019 | -0.90807752 | 0.082507796 |
| 1251.686646 | 1346.841187 | 1253.712891 | -0.651619371 | 0.319383004 | 0.251496204 | 0.732942123 | -0.911930861 | 0.414468675 |
| 7917.709473 | 9457.008789 | 5562.432617 | -0.651670501 | 0.058516905 | -1.026511165 | 0.03268871 | -0.824370326 | 0.06578999 |
| 2046023.25 | 1481712.25 | 1716287.625 | -0.651871335 | 0.022292412 | 0.338524667 | 0.090012942 | -0.026651813 | 0.910443707 |
| 380648.3438 | 394386.4375 | 398729.5625 | -0.652064752 | 0.178550789 | -0.21150612 | 0.106499477 | 0.028878944 | 0.701495778 |
| 348605.6563 | 381903.9375 | 367596.0313 | -0.652226217 | 0.065343643 | 0.285022024 | 0.149058018 | 0.127600008 | 0.354260455 |
| 47094.78516 | 52740 | 58003.28516 | -0.653390309 | 0.031057643 | 0.159446566 | 0.26604978 | 0.409257279 | 0.091523165 |
| 18323.24609 | 19437.79102 | 17987.59375 | -0.653535953 | 0.023401535 | -0.796642391 | 0.04939273 | -0.431025469 | 0.057068534 |
| 147911.3281 | 170054.5625 | 146137.3594 | -0.654009102 | 0.024962646 | -0.303866903 | 0.06919378 | 0.177442231 | 0.224000425 |
| 179830.3906 | 160970.3438 | 172874.0938 | -0.654357513 | 0.018311389 | 0.156040643 | 0.055241445 | 0.179992435 | 0.116287044 |
| 10419.91504 | 5317.069824 | 6095.419922 | -0.655296478 | 0.404223207 | -0.805484813 | 0.398870378 | 1.041953022 | 0.22355782 |
| 24079.29297 | 40351.86328 | 25036.67383 | -0.656138748 | 0.161916175 | -0.025372621 | 0.94724497 | -0.334274826 | 0.414213561 |
| 7725.036133 | 8661.709961 | 7044.170898 | -0.657085703 | 0.167094844 | 0.427251582 | 0.241244667 | 0.11882334 | 0.71939873 |
| 90453.74219 | 111107.3359 | 32197.98828 | -0.657986019 | 0.110708378 | -1.225276418 | 0.117377154 | -0.941065774 | 0.167338875 |
| 10947.07227 | 8895.282227 | 10418.66602 | -0.658793488 | 0.058516905 | 0.342780428 | 0.228952165 | 0.428874981 | 0.148459899 |
| 5274.057617 | 9160.304688 | 3039.562012 | -0.65896629 | 0.155863438 | 0.095436598 | 0.893275827 | -0.863873758 | 0.210234556 |
| 16051.50488 | 17132.8457 | 15627.35645 | -0.659712733 | 0.015746372 | 0.449255104 | 0.12145287 | 0.566369783 | 0.020008995 |
| 33603.86328 | 35411.09766 | 35264.85156 | -0.6600055 | 0.031883977 | 0.128578774 | 0.307431149 | -0.168471419 | 0.235399699 |
| 85895.875 | 85302.29688 | 67347.03906 | -0.660342773 | 0.104538446 | 0.551620021 | 0.341247073 | 0.609702165 | 0.081899796 |

| | | | | | | | | |
|---|---|---|---|---|---|---|---|---|
| 875.6879883 | 1334.178345 | 1845.611694 | -0.660960863 | 0.118613952 | 0.28435001 | 0.301633849 | 0.465195988 | 0.424862908 |
| 9695.025391 | 9777.628906 | 8134.85498 | -0.661110849 | 0.01434847 | 0.052197591 | 0.675235839 | -0.315152687 | 0.091797709 |
| 153486.1563 | 156737.4531 | 73487.14844 | -0.661729806 | 0.081995281 | -0.745231667 | 0.124287645 | -0.918138563 | 0.096744551 |
| 45970.04688 | 46817.58203 | 36685.75 | -0.662049155 | 0.058516905 | 0.227809125 | 0.326116631 | -0.182529266 | 0.481225203 |
| 10697.32813 | 10354.60059 | 8154.255859 | -0.66266651 | 0.066425026 | 0.184402189 | 0.353953112 | -0.079541817 | 0.707564204 |
| 33464.69141 | 27655.86133 | 30310.25 | -0.662738283 | 0.018050472 | 0.250173405 | 0.134252261 | -1.018145545 | 0.014388274 |
| 210277.9688 | 210922.875 | 225415.9375 | -0.663064053 | 0.013619747 | 0.219315532 | 0.130039916 | 0.277097238 | 0.064887728 |
| 36377.875 | 36906.77344 | 43397.73438 | -0.663362575 | 0.030009295 | 0.792741442 | 0.429618137 | -0.217936707 | 0.256562304 |
| 10740.74902 | 10673.16016 | 9588.448242 | -0.663728957 | 0.061148248 | -0.054420021 | 0.691978994 | -0.413458785 | 0.036229875 |
| 58113.29688 | 61853.88281 | 62567.44922 | -0.664728714 | 0.049051285 | 0.679721514 | 0.043418125 | 0.678097574 | 0.020849231 |
| 317802.6875 | 307035.7188 | 327031.7188 | -0.665199936 | 0.018479329 | 0.162491337 | 0.119711295 | 0.352031743 | 0.028449017 |
| 5192.760254 | 6200.029297 | 5403.876465 | -0.665403203 | 0.169155397 | 0.36466967 | 0.409475152 | 0.215328483 | 0.610411174 |
| 30736.31641 | 33888.85938 | 26459.86523 | -0.666364637 | 0.30022468 | -0.104639304 | 0.695245169 | -0.091762324 | 0.71939873 |
| 348525.875 | 338555.0938 | 307084.5 | -0.666657968 | 0.139278926 | 0.077821233 | 0.563676672 | 0.142426568 | 0.343566025 |
| 14346.08984 | 13738.19141 | 10782.69141 | -0.667495481 | 0.037600114 | 0.183113559 | 0.321892407 | -0.344623617 | 0.140761434 |
| 64331.21875 | 70653.98438 | 71017.96094 | -0.667795995 | 0.037238828 | 0.15629318 | 0.241244667 | -0.242997973 | 0.06479708 |
| 12764.57227 | 13617.97461 | 10839.30859 | -0.66821548 | 0.031638928 | -0.44162597 | 0.091622784 | -0.480135498 | 0.059641593 |
| 41302.98828 | 43230.99609 | 42179.00391 | -0.669206472 | 0.019930448 | 0.157026493 | 0.256229464 | 0.324858758 | 0.086952277 |
| 1870.367798 | 1623.903076 | NA | -0.669246077 | 0.12780052 | -0.227177556 | 0.682050046 | -0.083589715 | NA |
| 4248.392578 | 5311.96875 | 2579.008545 | -0.669411486 | 0.116475008 | -0.905867648 | 0.213397043 | -0.733279972 | 0.237683126 |
| 319787.875 | 321299.1875 | 334318.4063 | -0.669490839 | 0.025818052 | 0.51600279 | 0.081503088 | 0.222795244 | 0.213227295 |
| 2980.143799 | NA | 2948.507568 | -0.669807656 | 0.28128155 | 1.107705702 | 0.094211938 | 0.812519547 | NA |
| 25880.00977 | 14863.57031 | 34644.32031 | -0.670228924 | 0.178859946 | -0.211215514 | 0.639324884 | -0.309707041 | 0.557769502 |
| 9687.630859 | 7372.770508 | 5248.900879 | -0.670604025 | 0.036566441 | 0.068945978 | 0.80843958 | 0.052233173 | 0.901062666 |
| 598168.1055 | 635206.9063 | 707026.625 | -0.671201476 | 0.038502747 | 0.51682726 | 0.03968601 | 0.816561136 | 0.034567592 |
| 7286.155762 | 7409.417969 | 4979.061523 | -0.672075088 | 0.066713713 | 0.253247255 | 0.380869244 | 0.205160258 | 0.558655773 |
| 1906.693848 | 2294.998047 | 1969.115479 | -0.672222284 | 0.01332549 | 0.155696075 | 0.577348282 | -0.008475047 | 0.95773598 |
| 24686.49609 | 39354.35547 | 37995.84766 | -0.672276725 | 0.19187233 | 0.599325414 | 0.234571805 | 0.438721754 | 0.397327146 |
| 68701.32813 | 89617.28125 | 57543.60156 | -0.672387904 | 0.07827194 | -1.053546888 | 0.11104965 | -0.744760451 | 0.130708956 |
| 39813.03516 | 31402.82031 | 527.9295654 | -0.672468634 | 0.036566441 | 0.195406647 | 0.247821361 | -0.093090682 | 0.934434551 |
| 580887 | 615922.9375 | 711785.1875 | -0.672640282 | 0.013964852 | 0.546918029 | 0.050817911 | 1.277507299 | 0.040349459 |
| 5241.35498 | 4429.445313 | 2889.945068 | -0.673935324 | 0.091607129 | -0.070301837 | 0.873764511 | -1.006446308 | 0.054887084 |
| 29502.71289 | 27635.9082 | 25151.53125 | -0.674191071 | 0.011914125 | 0.597247088 | 0.057105325 | 1.096064131 | 0.028161099 |

| | | | | | | | | |
|---|---|---|---|---|---|---|---|---|
| 244047.2031 | 234156.9375 | 251985.5625 | -0.674664257 | 0.017533506 | 0.183737895 | 0.121388364 | 0.133375521 | 0.187797081 |
| 257812.3438 | 243952.7656 | 227003.7813 | -0.675218836 | 0.178701914 | 0.330977075 | 0.478999939 | 0.299428578 | 0.511855731 |
| 29735.78125 | 30189.70313 | 28424.6875 | -0.675262954 | 0.017729013 | -0.311313746 | 0.131196907 | -0.184524613 | 0.249396712 |
| 55679.51172 | 63808.33203 | 48943.36328 | -0.676179306 | 0.058153443 | 0.048580989 | 0.60188291 | -0.241741346 | 0.234416337 |
| 22952.71484 | 21969.78516 | 19338.125 | -0.676376344 | 0.0140944 | -0.109358226 | 0.508533269 | -0.304446123 | 0.101462623 |
| 55577.60156 | 53143.98438 | 55826.48047 | -0.676635209 | 0.05946073 | 1.189149082 | 0.05467585 | 1.550187657 | 0.005647029 |
| 125777.9297 | 145507.6875 | 128235.5234 | -0.676666399 | 0.023840475 | 0.148539624 | 0.203650142 | 0.100350204 | 0.442716166 |
| 9937.625 | 13868.19141 | 11098.40918 | -0.676867583 | 0.337591889 | 0.045286206 | 0.925262844 | -0.214280571 | 0.479079389 |
| 123105.7188 | 284885.2813 | 80125.74219 | -0.677287271 | 0.639880745 | -0.365511401 | 0.797975895 | -1.155556782 | 0.368053981 |
| 41146.94141 | 42713.81641 | 38758.77344 | -0.677372461 | 0.009974872 | 0.513263663 | 0.045483405 | 0.215049766 | 0.082896489 |
| 4393.746094 | 5498.887695 | 4617.69043 | -0.67855493 | 0.022677015 | -0.476428056 | 0.083115167 | -0.618374012 | 0.051065331 |
| 192607.4375 | 183000.3594 | 180005.2656 | -0.678951016 | 0.029390153 | 0.581863911 | 0.052457323 | 0.132564585 | 0.360870004 |
| 19550.98047 | 31764.87305 | 20133.1582 | -0.678962477 | 0.055366286 | 0.515059878 | 0.525110952 | 0.059680978 | 0.886748012 |
| 39905.79688 | 38826.65234 | 43153.11328 | -0.679917519 | 0.011471576 | -0.17378775 | 0.181752348 | -0.089092916 | 0.413448538 |
| 87878.29688 | 91561.35938 | 69300.25 | -0.680187276 | 0.153722588 | 1.940568956 | 0.038159713 | 2.058055894 | 0.029370864 |
| 31906.74023 | 38301.94141 | 37093.25781 | -0.680631315 | 0.154530346 | 0.409919998 | 0.31156031 | 0.289841957 | 0.443505091 |
| 34250.61719 | 31828.26953 | 31018.3418 | -0.681063544 | 0.042260333 | 0.054956093 | 0.603759375 | -0.142586909 | 0.177868363 |
| 26040.67969 | 59464.78906 | 23464.79102 | -0.681272504 | 0.3918194 | -0.19049147 | 0.798645885 | -0.30987315 | 0.678055454 |
| 9233.191406 | 7933.44873 | 7288.999023 | -0.682144412 | 0.194014788 | 0.307502962 | 0.087919389 | 0.291643201 | 0.190240855 |
| 33744.45703 | 34660.67188 | 27678.33008 | -0.682694711 | 0.046747318 | -0.007960831 | 0.952244753 | -0.243322537 | 0.181045061 |
| 1025105.438 | 1065975.375 | 979857.3125 | -0.683356792 | 0.04646045 | 0.232017372 | 0.094230752 | -0.305746199 | 0.069955133 |
| 174674.375 | 184700.8906 | 201170.5156 | -0.683544549 | 0.078030759 | -0.148489456 | 0.160878456 | -0.543747262 | 0.024764165 |
| 159516.2969 | 153694.7813 | 162223.875 | -0.684366439 | 0.026206894 | 0.148964425 | 0.150271527 | 0.284648886 | 0.02183874 |
| 9963.772461 | 11512.56348 | 10086.43066 | -0.685174027 | 0.081090664 | -0.068739209 | 0.844905766 | 0.151671436 | 0.639103574 |
| 13466.58496 | 13171.44922 | 14445.20605 | -0.685533429 | 0.089820226 | 0.824350188 | 0.098616588 | 0.272807471 | 0.084288402 |
| 21330.59961 | 21548.58008 | 18071.64844 | -0.685779721 | 0.158308661 | 0.492073399 | 0.116818656 | 0.383537971 | 0.121306313 |
| 2635.211914 | 1145.076416 | 983.5717773 | -0.685941698 | 0.068463493 | 0.662429246 | 0.099835584 | 0.049672806 | 0.952338298 |
| 5078.515625 | 5246.847656 | 4177.945313 | -0.686084158 | 0.202939587 | -0.510764357 | 0.087781599 | -0.358087836 | 0.109020087 |
| 46543.41406 | 53711.77344 | 37118.84375 | -0.686097881 | 0.10217612 | 0.054248129 | 0.885440998 | -0.422134827 | 0.35911176 |
| 12314.93359 | 14190.86816 | 13002.09961 | -0.686262496 | 0.048292506 | -0.795351914 | 0.137806142 | -1.004320174 | 0.107874808 |
| 13635.27344 | 12251.93652 | 5637.013184 | -0.686820311 | 0.01964718 | -0.588869978 | 0.097735536 | -1.060488377 | 0.080327609 |
| 59664.39063 | 55529.73828 | 63985.36328 | -0.68688596 | 0.055228595 | 0.235016239 | 0.198421115 | 0.413450997 | 0.069537512 |
| 28194.41211 | 20281.07031 | 19828.03516 | -0.686889338 | 0.2756587 | 2.360089205 | 0.228114774 | 2.596260421 | 0.069207996 |

| | | | | | | | | |
|---|---|---|---|---|---|---|---|---|
| 68509.96094 | 65165.17578 | 65653.15625 | -0.687432847 | 0.012182163 | 0.244568826 | 0.077996184 | 0.233974078 | 0.071696088 |
| 42724.92969 | 37244.97656 | 38051.46484 | -0.687507725 | 0.040802701 | 0.34713147 | 0.089652719 | 0.559968903 | 0.063474265 |
| 950.8185425 | 1590.205444 | NA | -0.687866848 | 0.425399143 | -0.475623933 | 0.334528184 | -0.970278402 | NA |
| 467637.9375 | 525438.375 | 511477.5313 | -0.688635941 | 0.011814081 | 0.024402632 | 0.777841261 | 0.126532843 | 0.266563506 |
| 74774.86719 | 81635.42188 | 67027.55469 | -0.689005608 | 0.27752677 | -0.056388632 | 0.842386539 | -0.129826434 | 0.572297095 |
| 204387.1719 | 195568.5469 | 205241.6719 | -0.690104408 | 0.060650522 | 0.224432115 | 0.061296684 | 0.421596617 | 0.017015337 |
| 196169.2969 | 184250.4531 | 69673.03125 | -0.690382161 | 0.246087007 | 1.075620361 | 0.509030551 | 0.116079601 | 0.861196469 |
| 9256.007813 | 11232.60352 | 11583.28809 | -0.690507419 | 0.013585622 | 0.246828283 | 0.167314796 | 0.193142532 | 0.302724499 |
| 37101.00781 | 40510.86719 | 41439.71484 | -0.690552676 | 0.035879745 | 0.216731733 | 0.30772129 | 0.015119525 | 0.923522245 |
| 3445.927246 | 6377.184082 | 4833.363281 | -0.690624858 | 0.045212188 | 0.019938073 | 0.955457512 | -0.985988124 | 0.079129451 |
| 191298.5156 | 181423.5781 | 217986.875 | -0.691547815 | 0.100274678 | 0.114672157 | 0.367508475 | 0.34896861 | 0.110787119 |
| 20011.30273 | 17370.90039 | 11105.11914 | -0.69214031 | 0.011914125 | 0.416669629 | 0.04939273 | -0.610483301 | 0.146265302 |
| 109666.9297 | 102255.2891 | 103996.9141 | -0.692389405 | 0.045092515 | 0.222070893 | 0.356068496 | -0.246470068 | 0.369290322 |
| 7070.985352 | 5569.722168 | 8185.020508 | -0.693862911 | 0.10217612 | 0.528512748 | 0.185493647 | 0.231956801 | 0.547207255 |
| 41259.12891 | 40662.79688 | 32608.18555 | -0.694328763 | 0.060886955 | -0.425604233 | 0.054285958 | -0.744494576 | 0.027087229 |
| 533038.625 | 577898.5 | 529585.9375 | -0.694459934 | 0.095292766 | 0.470367446 | 0.234860891 | 0.18307306 | 0.587664227 |
| 113447.3125 | 113118.5234 | 108405.8438 | -0.695206569 | 0.013618049 | 0.053606997 | 0.649577647 | -0.179157573 | 0.215760001 |
| 9320.302734 | 9617.194336 | 7269.413574 | -0.695820842 | 0.030009295 | -0.01532152 | 0.834180529 | -0.562442517 | 0.062345205 |
| 12737.01172 | 10896.94141 | 10753.14941 | -0.696176635 | 0.02552716 | 0.861728183 | 0.086089321 | 0.410726276 | 0.113248184 |
| 3392.881592 | 3347.73584 | 2422.905518 | -0.697177444 | 0.064886852 | 0.355236912 | 0.34166974 | -0.529041946 | 0.226542002 |
| 5195.020996 | 5947.560547 | 2117.331543 | -0.697221784 | 0.264058004 | 0.595716418 | 0.075608836 | 0.500106524 | 0.495174091 |
| 34536.72656 | 34708.21484 | 45477.83203 | -0.697367961 | 0.039588532 | 1.208777976 | 0.048655973 | 2.033137562 | 0.060580787 |
| 14030.33789 | 13327.01758 | 11365.96289 | -0.698837118 | 0.027969454 | 0.555672189 | 0.059937144 | 0.282673886 | 0.174083052 |
| 5351.843262 | 7891.651367 | 4017.909912 | -0.700135378 | 0.141259086 | 1.163877137 | 0.115267536 | -2.498586207 | 0.026263251 |
| 203078.0938 | 218915.3438 | 208067.5469 | -0.700418333 | 0.029508458 | 0.136368547 | 0.156320824 | 0.220494324 | 0.06365738 |
| 17694.20508 | 13624.07813 | 12086.82031 | -0.700498483 | 0.071859228 | -0.479702055 | 0.059340088 | -0.850343996 | 0.041009821 |
| 114633.2969 | 134202.1875 | 136864.3594 | -0.700995474 | 0.02318259 | 0.961914747 | 0.034480785 | 1.1166725 | 0.035626762 |
| 3205.567383 | 4444.185059 | 3734.600098 | -0.701525552 | 0.147715547 | -0.116736276 | 0.433819762 | 0.08524435 | 0.697701928 |
| 138187.293 | 151738.8984 | 139053.6875 | -0.701784922 | 0.076320591 | 0.722594196 | 0.118137091 | 1.059082752 | 0.017279709 |
| 131758.5938 | 119216.7813 | 130913.0078 | -0.702137219 | 0.03807887 | 0.052697451 | 0.388058343 | -0.622324228 | 0.013258914 |
| 15391.19629 | 16041.19531 | 15396.02734 | -0.702616641 | 0.061674089 | 0.015523812 | 0.930307469 | -0.105611393 | 0.433601327 |
| 587716.8125 | 606109.4375 | 601280.5625 | -0.70369297 | 0.038541031 | 0.086260034 | 0.635177865 | 0.467166389 | 0.09039268 |
| 18808.77539 | 17333.66602 | 15265.37012 | -0.703708107 | 0.056093567 | -0.099537024 | 0.60218417 | 0.088457108 | 0.533751353 |

| | | | | | | | | |
|---|---|---|---|---|---|---|---|---|
| 31142.26367 | 30381.09961 | 37488.44141 | -0.704409461 | 0.061707718 | 0.275315288 | 0.162317825 | -0.095325441 | 0.538909094 |
| 1269.681274 | 1260.150024 | 29331.24805 | -0.704670965 | 0.252282983 | 0.256246501 | 0.669562874 | 2.657013794 | 0.544636732 |
| 7521.853516 | 6304.023926 | 7297.99707 | -0.704858569 | 0.048312249 | 0.369913664 | 0.115283074 | -0.518427814 | 0.071043799 |
| 11920.84277 | 17653.19141 | 9056.007813 | -0.705434034 | 0.333748899 | 0.724704446 | 0.313230636 | 0.408929701 | 0.446063092 |
| 10061.13477 | 10294.5459 | 9836.038086 | -0.706018012 | 0.271312801 | 1.162785601 | 0.125467937 | 0.654219839 | 0.009587646 |
| 8082.424805 | 5319.376465 | 7215.533691 | -0.706854233 | 0.089968483 | -0.566692873 | 0.313673617 | -0.555638336 | 0.284828458 |
| 43491.73828 | 40357.53516 | 37667.63281 | -0.707668484 | 0.027325542 | -0.032144555 | 0.736903884 | -0.379826853 | 0.063236537 |
| 27934.34375 | 25862.01758 | 12915.13184 | -0.708884037 | 0.353604572 | 0.120944564 | 0.847462119 | -0.088210101 | 0.88423328 |
| 134293.3438 | 135344.4063 | 143929.2188 | -0.709347055 | 0.015248463 | 0.165560787 | 0.045481011 | 0.322687523 | 0.031906483 |
| 8576.32135 | 15095.37598 | 6983.750977 | -0.709517352 | 0.245606762 | 0.90887663 | 0.05975961 | 0.071481703 | 0.905430917 |
| 69291.26563 | 72323.52344 | 66641.36719 | -0.709685361 | 0.056932452 | 0.053088288 | 0.501164698 | -0.251924166 | 0.04753509 |
| 28376.29492 | 24908.62109 | 23974.42969 | -0.709848765 | 0.019536332 | 0.263393417 | 0.112848816 | 0.212952945 | 0.201736266 |
| 51721.91406 | 50458.49219 | 51693.75781 | -0.711085371 | 0.045302846 | -0.292158027 | 0.232151348 | -0.417432211 | 0.151865906 |
| 6243.194824 | 4424.964355 | 4300.515137 | -0.711123135 | 0.086168506 | -0.16334677 | 0.513385141 | -0.47385737 | 0.160431331 |
| 69208.24219 | 69673.20313 | 74720.90625 | -0.711364724 | 0.030063148 | 0.87250416 | 0.029690454 | 0.407383142 | 0.086670386 |
| 20909.88477 | 18577.95117 | 18987.97266 | -0.711427165 | 0.018093601 | 0.090285535 | 0.515273002 | 0.042438199 | 0.771442892 |
| 5145.176758 | 5659.363281 | 2136.760986 | -0.71201941 | 0.182953348 | -0.365195447 | 0.123008764 | -0.783801981 | 0.173821986 |
| 6391.117188 | 8495.320313 | 8351.889648 | -0.712580178 | 0.038185326 | -0.891859555 | 0.073324814 | -0.345099162 | 0.215864193 |
| 13994.30469 | 9138.674805 | 11593.69824 | -0.712653309 | 0.010297098 | 0.204229468 | 0.102454913 | -0.419622129 | 0.176448901 |
| 73850.9375 | 60463.76172 | 58248.05469 | -0.71311019 | 0.021708974 | 0.698363985 | 0.023578081 | 0.203515825 | 0.336101329 |
| 522863.0938 | 505886.7188 | 492502.5313 | -0.714671725 | 0.049206079 | 0.614974867 | 0.05913715 | 0.761548735 | 0.044743801 |
| 723043.5 | 755742.5625 | 781595.75 | -0.714773659 | 0.016048565 | 0.256867108 | 0.048520879 | 0.091939927 | 0.176248484 |
| 43323.28125 | 47388.75391 | 58547.28125 | -0.714932055 | 0.011979502 | -0.211659953 | 0.170700977 | 0.134385107 | 0.548024371 |
| 235538.0938 | 255876.7813 | 239594.7656 | -0.715470533 | 0.006936499 | 0.028596961 | 0.646592818 | 0.072554344 | 0.362406407 |
| 10816.14551 | 13174.61719 | 13670.29199 | -0.716553463 | 0.030396703 | 0.248111226 | 0.265715011 | 0.042710302 | 0.854395797 |
| 50880.37891 | 53587.14453 | 55483.27734 | -0.716588763 | 0.079009944 | 0.60633736 | 0.137745823 | 0.433887811 | 0.23557113 |
| 14272.73535 | 13532.36719 | 10285.61816 | -0.716844698 | 0.042260333 | 0.313987997 | 0.193741352 | 0.518883374 | 0.150131299 |
| 11612.43555 | 10971.94629 | 10630.08789 | -0.717239141 | 0.023124008 | -0.089522669 | 0.669338991 | -0.238622749 | 0.269669899 |
| 236567.7734 | 245567.4258 | 252242 | -0.71750471 | 0.116159237 | 1.614155043 | 0.137951093 | 2.165182086 | 0.019231091 |
| 8115.883301 | 8387.376953 | 6255.875977 | -0.717709586 | 0.327856987 | 1.897818132 | 0.063274608 | 1.409519727 | 0.09056925 |
| 74718.38281 | 78902.89063 | 68384.32031 | -0.717795123 | 0.025939018 | 0.06412574 | 0.549134414 | 0.164778221 | 0.234647912 |
| 102404.8984 | 92714.16406 | 92731.125 | -0.718102328 | 0.019286562 | 0.472706601 | 0.149229274 | 0.835159956 | 0.020008995 |
| 128002.6406 | 126301.5703 | 130733.1953 | -0.718376864 | 0.043222991 | 0.360225567 | 0.046755278 | 0.405669943 | 0.009026966 |

| | | | | | | | | |
|---|---|---|---|---|---|---|---|---|
| 26873.31445 | 36696.05078 | 31333.4043 | -0.718884077 | 0.138101644 | 0.43663098 | 0.230859884 | 0.247818229 | 0.339148153 |
| 7722.415527 | 6946.134277 | 6996.483887 | -0.720596131 | 0.034151807 | -0.008124652 | 0.925623784 | -0.428102077 | 0.039408515 |
| 2526.18042 | 3158.706299 | 2407.736328 | -0.720834659 | 0.132174911 | 0.223879663 | 0.584503222 | 0.227969932 | 0.578503308 |
| 13114.58984 | 12836.86523 | 11819.10449 | -0.721014313 | 0.121041004 | 0.656839541 | 0.031861061 | 0.026974507 | 0.805082256 |
| 11990.33789 | 14685.08887 | 12371.44629 | -0.721090689 | 0.042979653 | 0.574266512 | 0.159258737 | 0.879781234 | 0.03977281 |
| 19001.47852 | 25288.6875 | 11553.99219 | -0.721163319 | 0.018311389 | -0.967055498 | 0.043983378 | -1.381144284 | 0.039446558 |
| 233457.1875 | 249869.6719 | 312176.9688 | -0.72246426 | 0.056620854 | 0.314936522 | 0.082612243 | 0.45535576 | 0.16248539 |
| 14005.78027 | 10658.29004 | 9610.621094 | -0.723386135 | 0.064328842 | 0.576097355 | 0.340114217 | -0.196246501 | 0.514699959 |
| 39344.67969 | 40577.88672 | 37445.19141 | -0.724381468 | 0.015175173 | -0.254320576 | 0.150727713 | -0.748021669 | 0.052124898 |
| 21621.91602 | 13374.93262 | 16111.07617 | -0.725036648 | 0.049895713 | -0.005232078 | 0.930333414 | 0.313832258 | 0.412469761 |
| 124805.7969 | 131626.0781 | 135043.5781 | -0.725273936 | 0.0360176 | 0.527248435 | 0.032675377 | 0.376935408 | 0.041477764 |
| 41713.10938 | 25049.76953 | 33143.29688 | -0.725880373 | 0.115609849 | 0.926110345 | 0.162681498 | 0.573507215 | 0.244446425 |
| 147031.2969 | 155465.8281 | 169855.9375 | -0.726628589 | 0.045092515 | -0.448326019 | 0.038022436 | 0.106355566 | 0.338848169 |
| 155269.0625 | 144530.1719 | 157538.8594 | -0.727863712 | 0.157842529 | 0.779706482 | 0.179476554 | 0.863742066 | 0.135824856 |
| 17376.88281 | 13730.67871 | 13090.60059 | -0.728034762 | NA | 1.914634155 | NA | 3.661735268 | NA |
| 3483.908203 | 2832.413574 | 2355.529785 | -0.728050342 | 0.155196387 | 0.58319801 | 0.27504497 | -0.332368994 | 0.578085212 |
| 915105.4375 | 878812.5625 | 1015312.313 | -0.72822671 | 0.018310596 | 0.628046903 | 0.05382061 | 0.035360605 | 0.720582737 |
| 118994.4844 | 107751.2344 | 96507.59375 | -0.729173645 | 0.01434847 | 1.342307933 | 0.072641707 | 0.396499848 | 0.114860889 |
| 558356.125 | 581548.5625 | 599217.1875 | -0.729209178 | 0.027405706 | 0.108831617 | 0.204964225 | 0.039890513 | 0.572303677 |
| 136171.1875 | 140254.8594 | 176459.125 | -0.72921568 | 0.059908598 | 1.094255325 | 0.031422588 | 3.301070475 | 0.047120928 |
| 118578.3594 | 105494.9766 | 143476.2188 | -0.730849982 | 0.392211726 | -0.047228866 | 0.957138366 | 1.027192598 | 0.075782294 |
| 260330.8281 | 277352.6797 | 222131.7109 | -0.731753874 | 0.059547685 | -0.163217173 | 0.134872788 | -0.289703212 | 0.150000617 |
| 124397.9375 | 124256.3203 | 128218.4844 | -0.732138681 | 0.012952445 | 0.644010954 | 0.028096606 | 0.616866866 | 0.015632484 |
| 136566.2344 | 231274.1719 | 147516.6875 | -0.732278123 | 0.161609672 | 0.018743334 | 0.961183343 | 0.006212384 | 0.989213317 |
| 1187968.25 | 1164454.25 | 1406155.375 | -0.732959531 | 0.038299729 | 0.958289809 | 0.142579096 | 1.243210395 | 0.031136253 |
| 11846.36035 | 10469.63379 | 9276.728516 | -0.733571183 | 0.104222252 | -0.056830802 | 0.71242314 | 0.112246946 | 0.529273607 |
| 24661.18555 | 29332.97852 | 32082.91406 | -0.733970852 | 0.071922239 | 0.419746886 | 0.159258737 | 0.268863816 | 0.332165755 |
| 22414.49805 | 20612.7793 | 22618.47852 | -0.733991256 | 0.173500617 | 0.969302425 | 0.130198127 | 0.672826749 | 0.222120646 |
| 40120.05859 | 41589.86719 | 40837.93359 | -0.735019112 | 0.062398014 | 1.575725674 | 0.018539883 | 2.244989662 | 0.011038949 |
| 193444.6094 | 121061.6094 | 211022.8281 | -0.735302952 | 0.276307466 | 0.818030734 | 0.345433593 | 1.045723567 | 0.1666149 |
| 4410.13916 | 5055.757324 | 5679.321289 | -0.735402374 | 0.064437538 | -0.87884529 | 0.164449495 | -0.654599713 | 0.202826206 |
| 50774.57031 | 53765.31641 | 50216.38281 | -0.735508292 | 0.035591101 | -0.106286626 | 0.295943747 | -0.425104013 | 0.019577244 |
| 12341.59277 | 14741.03711 | 13308.43848 | -0.735879157 | 0.036547632 | 0.20920208 | 0.107963035 | -0.190190048 | 0.195513278 |

| | | | | | | | | |
|---|---|---|---|---|---|---|---|---|
| 108869.1094 | 108657.3672 | 76241.02344 | -0.736115298 | 0.034131624 | 0.004471246 | 0.954690016 | -0.132306187 | 0.583408989 |
| 5754.103516 | 5054.661133 | 5095.442871 | -0.736165941 | 0.102673034 | -0.249656567 | 0.233817631 | -0.497688153 | 0.085245136 |
| 84391.53125 | 86855.92969 | 80172.63281 | -0.736294947 | 0.031790847 | -0.922426741 | 0.038454242 | -0.321390327 | 0.085333617 |
| 492527.9375 | 504695.0625 | 511755.1875 | -0.736388872 | 0.020318767 | 0.022745934 | 0.794724285 | 0.042509731 | 0.620264646 |
| 10448.64355 | 7810.86377 | 8190.555664 | -0.736988925 | 0.041987159 | -0.365632961 | 0.265924368 | -0.801108022 | 0.094277833 |
| 305470.8125 | 290100.0625 | 267466.875 | -0.737338347 | 0.026011511 | 0.053153312 | 0.768741797 | 0.378241171 | 0.095385921 |
| 23310.97852 | 26540.82031 | 27785.41992 | -0.73823587 | 0.053015916 | 0.317079301 | 0.098459633 | 0.580246302 | 0.045887602 |
| 29785.64453 | 28727.2168 | 24988.7832 | -0.738960322 | 0.127443855 | 0.066603334 | 0.553733446 | 0.841737077 | 0.041664392 |
| 12755.20801 | 10008.02148 | 8675.085938 | -0.741253934 | 0.019694688 | -0.211006031 | 0.131070383 | -0.534250748 | 0.105824879 |
| 4668.042969 | 6399.79541 | 2099.999756 | -0.742876018 | 0.160731205 | -1.390422403 | 0.205952446 | -1.001881121 | 0.251406907 |
| 32752.13477 | 29162.89844 | 30137.60352 | -0.743872024 | 0.029838966 | 0.394561736 | 0.091622784 | 0.62958761 | 0.027800907 |
| 96445.67188 | 106028.25 | 95724.00781 | -0.743897977 | 0.054479995 | -0.042138549 | 0.832010656 | -0.114337166 | 0.346395659 |
| 2443.052734 | 1411.488525 | 802.9898682 | -0.744247874 | 0.043150956 | -0.866800186 | 0.172405242 | -1.71376328 | 0.046581887 |
| 7296.576172 | 7378.877441 | 7791.053711 | -0.74481056 | 0.079112163 | 0.606451661 | 0.287272611 | -0.903295609 | 0.16733016 |
| 403627.1875 | 428129.5 | 378538.9688 | -0.745355545 | 0.018002294 | -0.060054646 | 0.708875993 | 0.56540846 | 0.034457303 |
| 60611.69531 | 53617.71484 | 60722.07422 | -0.745463222 | 0.022677015 | 0.245181929 | 0.123694369 | -0.187179763 | 0.207832079 |
| 22562.24609 | 24203.75195 | 19334.45508 | -0.746148035 | 0.131541836 | 0.835430788 | 0.127993682 | 0.243159616 | 0.577277473 |
| 172218.7656 | 182804.3906 | 186038.2813 | -0.746978088 | 0.047279091 | -0.054084708 | 0.585220616 | 0.128652191 | 0.230308748 |
| 292976.7813 | 332583.875 | 290939.0938 | -0.747021805 | 0.022729015 | 0.354592275 | 0.0480335 | -0.205961848 | 0.128326365 |
| 35308.82813 | 39121.46094 | 37445.01172 | -0.747249761 | 0.090517853 | -0.10635494 | 0.701829892 | -0.315694462 | 0.317922761 |
| 70217.26563 | 108390.0859 | 110732.8906 | -0.748108899 | 0.249779609 | 0.187398095 | 0.638775352 | 0.268591326 | 0.448315524 |
| 7914.955078 | 6254.367188 | 2701.8479 | -0.748136417 | 0.136409711 | -0.262318032 | 0.64741062 | -0.017495047 | 0.983784117 |
| 65582.29688 | 52502.52734 | 48444.34766 | -0.749592335 | 0.072220768 | 0.166940455 | 0.687866158 | 0.255603529 | 0.459294132 |
| 5030.999512 | 4785.810059 | 3050.680664 | -0.749592383 | 0.053170792 | -0.013836554 | 0.941574529 | -0.333043539 | 0.316440408 |
| 62345.43945 | 61547.42188 | 59261.82227 | -0.75066265 | 0.087656352 | 0.477686663 | 0.208565384 | 0.634388488 | 0.136000191 |
| 62133.64844 | 62360.95313 | 54549.83984 | -0.751058263 | 0.030839901 | -0.078034934 | 0.208500335 | -0.202756376 | 0.132698576 |
| 9867.601563 | 11720.9043 | NA | -0.751753241 | 0.023724131 | -0.195886818 | 0.344272662 | -0.627882019 | NA |
| 11478.96289 | 11453.03223 | 13455.23828 | -0.752396041 | 0.08292593 | 0.486312747 | 0.316476374 | -0.182860634 | 0.62573426 |
| 5652.125488 | 7158.974121 | 5802.412109 | -0.753022653 | 0.047406642 | 0.098770826 | 0.528259024 | 0.068475365 | 0.717621825 |
| 21308.93555 | 19046.13086 | 14933.41309 | -0.753345091 | 0.01255155 | 0.630430481 | 0.050288747 | -0.035515162 | 0.879611016 |
| 76974.59375 | 99964.95313 | 61370.75391 | -0.753639638 | 0.142751092 | 0.058873778 | 0.873764511 | -0.395717244 | 0.260266111 |
| 47841.44531 | 14472.22363 | 29412.66016 | -0.75379469 | 0.177638439 | 0.990072038 | 0.336933122 | 0.723482976 | 0.448159735 |
| 22394.12891 | 24105.39063 | 12129.72852 | -0.75453839 | 0.1152822 | -0.582011776 | 0.054219446 | -0.901077622 | 0.089275366 |

| | | | | | | | | |
|---|---|---|---|---|---|---|---|---|
| 505622.3438 | 542168 | 592057.5625 | -0.755144861 | 0.009194956 | -0.096613166 | 0.366953709 | 0.079826573 | 0.52055086 |
| 8388.603516 | 10696.64844 | 7690.380859 | -0.755912651 | 0.084948587 | 0.115341237 | 0.292501538 | 0.069081385 | 0.772377085 |
| 4409.617676 | 2883.630859 | 1794.426147 | -0.756381653 | 0.082411613 | 0.798977431 | 0.105893922 | -0.146857441 | 0.785068679 |
| 32166.42383 | 28406.45898 | NA | -0.756690688 | 0.039600055 | 0.17549766 | 0.43398523 | -0.242908731 | NA |
| 31543.14258 | 31160.23633 | 31098.24414 | -0.757430226 | 0.008009891 | 0.06576 | 0.355035916 | 0.048750409 | 0.443629763 |
| 30767.07031 | 55637.75 | 30197.08594 | -0.757983718 | 0.271405489 | 0.640685486 | 0.209926022 | 0.548362838 | 0.391808699 |
| 3994.606445 | 6060.422363 | 6664.025391 | -0.759703027 | 0.415536302 | 0.994687724 | 0.140225049 | 1.19127384 | 0.111095796 |
| 1732314.75 | 1636260.75 | 1450397.25 | -0.759929405 | 0.045484912 | 0.598256734 | 0.17813923 | 0.378336202 | 0.122550756 |
| 3816.406494 | 3970.430664 | 3496.008545 | -0.762068207 | 0.340890742 | 1.30942604 | 0.147811629 | 0.9151559 | 0.270797304 |
| 15677.39453 | 14780.3916 | 12667.82715 | -0.762351037 | 0.248375454 | -0.199802595 | 0.558732642 | -0.06757038 | 0.669988616 |
| 28617.39844 | 110117.2656 | 18432.37305 | -0.76254091 | 0.634429464 | -0.223859802 | 0.886810149 | -0.693501084 | 0.591060803 |
| 259975.5781 | 279099.4063 | 298184.2188 | -0.762584131 | 0.086055451 | 0.380239048 | 0.283496174 | 0.284973127 | 0.38215706 |
| 20511.65625 | 20649.86719 | 20268.02539 | -0.763157809 | 0.019930448 | 0.627938854 | 0.023005377 | -0.01550452 | 0.881655423 |
| 19280.79297 | 15956.01172 | 16393.2168 | -0.763253325 | 0.12468265 | 0.696326585 | 0.171025806 | -0.222501094 | 0.633703609 |
| 2623.54126 | 3440.519287 | 2069.797852 | -0.763458633 | 0.058310315 | 0.25101739 | 0.504348764 | -0.77216784 | 0.142837957 |
| 48626.81641 | 42213.08203 | 45963.56641 | -0.763836995 | 0.008262405 | 0.065879734 | 0.414751124 | -0.189092067 | 0.139285214 |
| 129430.2578 | 134124.75 | 117395.8281 | -0.76490882 | 0.029390153 | 0.362982865 | 0.098459633 | 0.115133962 | 0.393173467 |
| 95437.34375 | 94119.02344 | 90416.35938 | -0.765172202 | 0.028368096 | 1.007748086 | 0.015993065 | 0.409555691 | 0.02354608 |
| 9727.563477 | 11309.51367 | 8872.604492 | -0.765185645 | 0.184095742 | -0.772459755 | 0.112848816 | -0.573705017 | 0.138668568 |
| 98005.35938 | 93161.69531 | 76572.09375 | -0.765538765 | 0.049995393 | 0.28066662 | 0.336933122 | -0.086575728 | 0.77583338 |
| 6013.02002 | 5918.200684 | 3974.51001 | -0.765562369 | 0.022703538 | 0.038596624 | 0.795956929 | -0.173278698 | 0.544636732 |
| 46585.44141 | 46193.8125 | 43809.74609 | -0.765589292 | 0.006395507 | 0.037529761 | 0.82536887 | -0.461435684 | 0.036848746 |
| 1244.513306 | 962.4783325 | NA | -0.765693755 | NA | -0.381295175 | NA | -0.201422432 | NA |
| 8071.089355 | 4512.924805 | 6046.82666 | -0.765859547 | 0.081936227 | 0.199826954 | 0.509387885 | -0.676919615 | 0.12523068 |
| 52900.90234 | 53377.84766 | 52037.26172 | -0.766955911 | 0.037600114 | 0.17120518 | 0.107063979 | 0.052072531 | 0.303258583 |
| 68912.07813 | 69082.03906 | 78236.83594 | -0.767231551 | 0.013723547 | 0.18611526 | 0.219332589 | 0.572409829 | 0.035349276 |
| 40674.55078 | 51830.53906 | 37908.10938 | -0.76747263 | 0.102334557 | 0.062423495 | 0.834128891 | -0.090050071 | 0.710507923 |
| 111962.625 | 121531.2031 | 111650.3125 | -0.767803509 | 0.037314581 | 0.632114798 | 0.068834111 | 0.197137097 | 0.382150932 |
| 2800828.5 | 2588433.75 | 2621984.5 | -0.767970584 | 0.146800875 | 0.276680489 | 0.271287773 | 0.760625152 | 0.042660494 |
| 895880.5625 | 980509.9375 | 927876.375 | -0.768123871 | 0.013857751 | 0.179509046 | 0.135823019 | 0.036613939 | 0.687682325 |
| 1138.373047 | 2318.769287 | 1256.163818 | -0.768410576 | 0.06611715 | -0.798982762 | 0.129861867 | -1.058301696 | 0.079165593 |
| 32326.52539 | 31791.74414 | 31132.85547 | -0.768519937 | 0.005783136 | 0.042690923 | 0.484425023 | -0.508569657 | 0.030633392 |
| 3189888 | 3166488.25 | 3202615.25 | -0.769208661 | 0.013953967 | 0.162547764 | 0.246470557 | -0.160617766 | 0.274971315 |

| | | | | | | | | |
|---|---|---|---|---|---|---|---|---|
| 26703.39648 | 53532.40234 | 29826.76563 | -0.769734426 | 0.377135523 | 1.105716557 | 0.162505567 | 1.346539937 | 0.187964955 |
| 27973.78516 | 22987.05469 | 24800.72266 | -0.770099785 | 0.064338292 | 0.056153876 | 0.415767255 | -0.253396353 | 0.144611007 |
| 481289.9063 | 506813.875 | 472607.375 | -0.770787525 | 0.00744439 | -0.16624552 | 0.152444893 | 0.033233839 | 0.680261203 |
| 25658.20508 | 27785.23438 | 27132.41016 | -0.771426611 | 0.025587445 | -0.736785221 | 0.093662394 | 0.139492982 | 0.478866737 |
| 1025.463135 | 1592.612183 | NA | -0.771641324 | 0.11900804 | -0.166778184 | 0.519411653 | -0.424815593 | NA |
| 2222.833008 | 2365.159912 | 1528.261475 | -0.772978122 | 0.075761347 | 0.604047542 | 0.129928099 | -0.172197346 | 0.670107333 |
| 132317.3438 | 143592.3125 | 151674.2656 | -0.774993004 | 0.151178293 | 0.395832464 | 0.130757652 | 0.237569277 | 0.170833198 |
| 19765.29297 | 21391.58008 | 19445.4375 | -0.77562139 | 0.097957545 | 0.595570903 | 0.164062441 | 0.420644746 | 0.279338111 |
| 92187.9375 | 96718.45313 | 86365.67188 | -0.775740862 | 0.010893509 | -0.121499156 | 0.399976168 | -0.227939707 | 0.0741034 |
| 32089.08984 | 30630.14648 | 28039.09766 | -0.776374356 | 0.020732204 | 0.729279141 | 0.018539883 | -0.037746147 | 0.686261238 |
| 95893.85938 | 84539.79688 | 81955.08594 | -0.778267429 | 0.044225724 | 0.437310293 | 0.122107272 | 0.237528513 | 0.300806788 |
| 51851.01172 | 45345.08203 | 34913.37109 | -0.778320765 | 0.11816999 | 0.292552459 | 0.699189397 | 0.604315314 | 0.159261086 |
| 31115.54688 | 34454.99219 | 31742.55273 | -0.778348177 | 0.0436254 | -0.298572367 | 0.123694369 | -0.433817761 | 0.073108053 |
| 39523.87891 | 27970.76172 | 33594.35547 | -0.779119233 | 0.032155886 | 0.174312945 | 0.440017251 | 0.162477562 | 0.567552768 |
| 6279.461426 | 8650.649414 | 7862.070313 | -0.779791726 | 0.206031511 | 0.481199616 | 0.320417864 | 0.2936691 | 0.27322162 |
| 18785.01563 | 18566.68359 | 17059.28906 | -0.780074352 | 0.055571373 | 0.198918057 | 0.455479964 | -0.016639672 | 0.953117728 |
| 69790.92969 | 72878.76563 | 57733.97656 | -0.781182573 | 0.015732345 | -0.165799249 | 0.312015025 | -0.13000792 | 0.461591343 |
| 71065.69531 | 79247.32813 | 72768.39844 | -0.782982582 | 0.024701492 | 0.417100681 | 0.045230118 | 0.390102278 | 0.039602697 |
| 11078.53613 | 11672.28613 | 10733.42773 | -0.783022238 | 0.005752145 | 0.24768637 | 0.056841901 | 0.140296092 | 0.118373568 |
| 66657.38281 | 59847.11328 | 57196.54297 | -0.783692797 | 0.060816981 | -0.170834926 | 0.161933659 | -0.363297905 | 0.076362612 |
| 27359.93945 | 30005.50781 | 10280.50098 | -0.784066539 | 0.143726661 | 0.545422128 | 0.236617457 | 0.499902054 | 0.511062465 |
| 15872.78809 | 13935.27051 | 14886.77441 | -0.785146428 | 0.197563952 | 0.92922936 | 0.190217492 | 1.109399081 | 0.121741414 |
| 373140.375 | 324210.1563 | 328275.7188 | -0.785174569 | 0.014665071 | -0.282045115 | 0.165197811 | 0.09081256 | 0.536804361 |
| 1203149.25 | 1287751.75 | 1217166.125 | -0.785196022 | 0.023227019 | 0.090346917 | 0.470469741 | 0.065939113 | 0.584137475 |
| 3264.218262 | 4100.565918 | 2705.687256 | -0.785262694 | 0.027158619 | 0.572880088 | 0.057533032 | -0.0949846 | 0.730529882 |
| 25599.26367 | 24460.80615 | 20519.54443 | -0.785740148 | 0.021667504 | -0.570824486 | 0.126359156 | -0.206490004 | 0.359569317 |
| 11612.32422 | 11602.67578 | 8354.762695 | -0.786544576 | 0.022885276 | 0.291246981 | 0.122435857 | -0.478532609 | 0.106930847 |
| 7161.769043 | 7034.950195 | 8579.667969 | -0.786676028 | 0.052356162 | -0.370025063 | 0.280688037 | -0.217293353 | 0.454451558 |
| 49160.91406 | 47703.81641 | 43675.18359 | -0.788608836 | 0.0140944 | -0.126090121 | 0.093561793 | -0.206028417 | 0.094664375 |
| 9989.597656 | 8031.762695 | 6860.650879 | -0.78989572 | NA | 2.236166038 | NA | 1.458537632 | NA |
| 292091.75 | 327647.0313 | 328150.6875 | -0.790323642 | 0.034061991 | 0.088559894 | 0.334528184 | 0.386104379 | 0.04816468 |
| 150049.0938 | 157954.0313 | 154526.9531 | -0.790821089 | 0.019930448 | 0.422543708 | 0.006408045 | 0.228863059 | 0.037176621 |
| 582938.5452 | 810373.0044 | 650872.3354 | -0.792799337 | 0.155203679 | 0.124358319 | 0.539004608 | 0.318853024 | 0.283248296 |

| | | | | | | | | |
|---|---|---|---|---|---|---|---|---|
| 38719.49219 | 35518.68359 | 22773.04492 | -0.794428826 | 0.058578037 | -0.526161002 | 0.142833812 | -1.339081234 | 0.034677634 |
| 6239.128418 | 6882.645508 | 7100.081055 | -0.794658028 | 0.04219674 | 0.14959839 | 0.513385141 | -0.241652677 | 0.206265186 |
| 7886.879395 | 7968.679199 | 6771.085938 | -0.795187595 | 0.211214576 | 1.826217387 | 0.048024604 | -1.633273622 | 0.268750571 |
| 8615.669922 | 8329.02832 | 5149.263672 | -0.795212325 | 0.009072032 | -0.002280244 | 0.990876631 | -0.618946985 | 0.128242084 |
| 125722.3594 | 115524.3594 | 128863.9063 | -0.796840915 | 0.024701492 | 0.076497063 | 0.265813401 | 0.163020696 | 0.134445271 |
| 9255.584961 | 9443.146484 | 8962.129883 | -0.796860084 | 0.04428557 | 0.152288835 | 0.440030647 | 0.111522897 | 0.548308998 |
| 18975.19531 | 24223.39063 | 19939.42383 | -0.797913702 | 0.033164349 | 0.550213408 | 0.125004947 | -0.206106196 | 0.473264096 |
| 16857.88867 | 18251.16406 | 16692.54883 | -0.798623047 | 0.025307586 | -0.29887939 | 0.089248231 | -0.363245898 | 0.062838928 |
| 230470.4844 | 229637.0469 | 222161.1719 | -0.799605167 | 0.025955593 | 1.210365905 | 0.048024604 | 0.410610944 | 0.053570169 |
| 13326.25488 | 12978.30957 | 10180.75 | -0.799618914 | 0.048416918 | 0.259438409 | 0.406744405 | 0.420668352 | 0.22024877 |
| 61631.64063 | 61890.87109 | 60304.45703 | -0.800360357 | 0.01837679 | 0.193856495 | 0.246470557 | 0.198140839 | 0.231505615 |
| 16318.20117 | 13691.03418 | 13666.99316 | -0.801284917 | 0.077638666 | 2.07800109 | 0.220633017 | 1.747197271 | 0.020008995 |
| 81703.70313 | 78966.29688 | 80067.71875 | -0.801613153 | 0.036359027 | 0.290658721 | 0.233439789 | 0.240154853 | 0.293140864 |
| 4358.158691 | 3849.461914 | 3136.892822 | -0.802484492 | 0.029514505 | 0.031998227 | 0.903519658 | -0.452993695 | 0.185258988 |
| 110141.6563 | 103880.8359 | 131494.1563 | -0.803447357 | 0.008009891 | 0.158846216 | 0.175466445 | -0.212248505 | 0.249109031 |
| 202196.7969 | 191850.9063 | 182853.8594 | -0.803629013 | 0.009974872 | -0.087331632 | 0.321888877 | -0.279164515 | 0.048032607 |
| 75840.22656 | 70870.57031 | 75883.28125 | -0.805211886 | 0.00953954 | -0.043075752 | 0.660718628 | 0.143824057 | 0.192001691 |
| 40454.14063 | 43883.12891 | 38496.05469 | -0.805374804 | 0.01391113 | -0.231419615 | 0.157837558 | -0.059856303 | 0.625379855 |
| 16810.22852 | 18843.15234 | 16633.83594 | -0.805412285 | 0.039878354 | 0.641456531 | 0.113692461 | 0.619374015 | 0.089724983 |
| 1681764.875 | 1743807.75 | 1541876.375 | -0.805481608 | 0.004709568 | 0.191197314 | 0.110498088 | -0.05087085 | 0.544452231 |
| 4719.01123 | 4480.092285 | 5137.589844 | -0.805824549 | 0.156018312 | -0.610121013 | 0.437151431 | 0.95489152 | 0.132347246 |
| 9556.40625 | 18617.2793 | 8253.675781 | -0.808466034 | 0.302378035 | 0.096610681 | 0.923246587 | -0.11828945 | 0.885220307 |
| 44472.53906 | 15116.47852 | 28962.16992 | -0.808888592 | 0.404501862 | 0.821527017 | 0.431639915 | -0.058421143 | 0.948114044 |
| 1449.487305 | 3498.335938 | 2421.762451 | -0.808939475 | 0.090710537 | -0.14506037 | 0.764863365 | -0.236802716 | 0.69422618 |
| 85331.36719 | 98823.98438 | 115080.4922 | -0.809493169 | 0.086929104 | 0.554121088 | 0.236313717 | 1.148405412 | 0.048032607 |
| 10301.90137 | 13598.22168 | 3211.530273 | -0.809975976 | 0.065816378 | -0.683168342 | 0.039582042 | -1.079833927 | 0.148405728 |
| 985211.3125 | 874885.1875 | 766282.6875 | -0.810065509 | 0.062989607 | 1.272842275 | 0.08807311 | 0.2596038 | 0.380780911 |
| 72216.32813 | 74541.63281 | 71268.10938 | -0.811133798 | 0.054037379 | 0.306393571 | 0.033121608 | -0.047094652 | 0.263528267 |
| 27685.94531 | 15730.35547 | 16388.58789 | -0.811416361 | 0.032802153 | 0.33503088 | 0.111682839 | -0.364215126 | 0.385259401 |
| 34265.11328 | 38987.9375 | 26260.46875 | -0.812427788 | 0.139521033 | 0.065048206 | 0.675235839 | -0.320427358 | 0.244446425 |
| 31773.41992 | 45974.75781 | 31855.58398 | -0.812510934 | 0.300318309 | -0.059456919 | 0.913882409 | 0.635633925 | 0.194351963 |
| 34591.77734 | 43907.34375 | 42683.56641 | -0.812581445 | 0.013529896 | 0.337025474 | 0.042884533 | 0.161069715 | 0.389456703 |
| 82809.32813 | 79143.9375 | 70377.1875 | -0.812939669 | 0.035058209 | 0.29136308 | 0.234161898 | 1.081059956 | 0.02146481 |

| | | | | | | | | |
|---|---|---|---|---|---|---|---|---|
| 242626.4844 | 233491.2344 | 275424.5938 | -0.81312718 | 0.009011817 | -0.076518134 | 0.481934316 | 0.187921116 | 0.223168984 |
| 4960.828613 | 5778.520508 | 8421.988281 | -0.813791192 | 0.076923702 | 0.392770257 | 0.098459633 | 0.337286196 | 0.441150671 |
| 97237.80469 | 180613.6719 | 144964.8281 | -0.813822911 | 0.230359299 | 0.601010461 | 0.28299945 | 0.728076137 | 0.225750747 |
| 479536.0313 | 486564.7188 | 491730.0938 | -0.815032103 | 0.006735623 | 0.237468983 | 0.07158592 | 0.187067032 | 0.097707274 |
| 19303.86523 | 18555.33594 | 21524.75781 | -0.815054528 | 0.019853827 | -0.226109953 | 0.38067054 | 0.425348668 | 0.063440273 |
| 165423.5469 | 157831.9531 | 147641.9375 | -0.81747791 | 0.038299729 | 0.366734044 | 0.142630817 | 0.312958756 | 0.173821986 |
| 14869.29102 | 14080.33887 | 17408.45508 | -0.818669066 | 0.104002895 | 0.782664623 | 0.057189248 | 0.658739357 | 0.060937162 |
| 30382.12695 | 36136.66406 | 35433.60156 | -0.818818804 | 0.142871999 | 0.393654141 | 0.226925343 | 0.50151107 | 0.128326365 |
| 491950.9375 | 498086.1875 | 483190.2188 | -0.81905924 | 0.013723547 | 0.356943249 | 0.024676726 | 0.09863418 | 0.14379126 |
| 19086.76758 | 20758.99023 | 20786.33594 | -0.820419609 | 0.020740891 | 0.730175328 | 0.025444899 | 0.438335611 | 0.036624925 |
| NA | NA | NA | -0.820580143 | NA | -0.21807827 | NA | NA | NA |
| 20497.41992 | NA | 30456.50586 | -0.821146014 | NA | 1.51048576 | NA | 1.054188271 | NA |
| 3668.798584 | 2754.725342 | 2677.734619 | -0.823463988 | 0.046088702 | -0.546212079 | 0.110712084 | -0.365873102 | 0.211105363 |
| 5249.196289 | 5679.711914 | 5056.20166 | -0.824271422 | 0.023105653 | 0.091614534 | 0.734889757 | -0.508561801 | 0.14559756 |
| 5500.846191 | 4530.129883 | NA | -0.82444539 | 0.238040125 | 1.421131941 | 0.256838624 | 0.290012511 | NA |
| 21565.15234 | 19727.94531 | 19287.91406 | -0.826556008 | 0.012364839 | 0.289687687 | 0.054382553 | -0.095476674 | 0.320074674 |
| 88002.32031 | 88033.42188 | 78667.94531 | -0.826895104 | 0.012750417 | 0.322277996 | 0.123184236 | -0.317303333 | 0.04292928 |
| 21306.34375 | 26405.02539 | 19225.48438 | -0.827791526 | 0.036480793 | -0.094180105 | 0.370196377 | 0.117568152 | 0.60604939 |
| 25289.44922 | 23130.48242 | 24731.37695 | -0.827996059 | 0.018161103 | 0.416680672 | 0.05253541 | 0.145084629 | 0.223102126 |
| 13663.47461 | 24089.66406 | 11009.72168 | -0.828559189 | 0.389396389 | 0.885866208 | 0.215799184 | 0.893893024 | 0.299598031 |
| 65665.53125 | 60182.77344 | 63313.39063 | -0.82960428 | 0.027930748 | -0.050745129 | 0.803978674 | -0.399949278 | 0.192489677 |
| 755875.9375 | 781889.5625 | 818150.8125 | -0.831121323 | 0.034127829 | 0.301288009 | 0.066380875 | 0.232293595 | 0.057202212 |
| 24625.30664 | 19924.92773 | 16939.41797 | -0.832129065 | 0.024259746 | 0.147444978 | 0.094508793 | -0.510120444 | 0.121383661 |
| 10226.04297 | 21540.94727 | 13165.04199 | -0.83319025 | 0.255300969 | 0.492741973 | 0.430750734 | 0.791422834 | 0.299674772 |
| 309943.6563 | 314565.5 | 364157.3125 | -0.83331401 | 0.333000288 | -0.185205281 | 0.64877337 | 0.103775728 | 0.778661035 |
| 30263.53125 | 27980.98242 | 25606.79102 | -0.833835476 | 0.05114005 | 0.257798231 | 0.029364116 | -0.214530002 | 0.150748843 |
| 106588.3203 | 95059.35938 | 105001.0156 | -0.834829566 | 0.049775794 | 0.094803346 | 0.108875676 | -0.148346497 | 0.152319293 |
| 17452.04883 | 16654.73633 | 15498.09277 | -0.835318148 | 0.011954463 | 0.277953664 | 0.054811357 | -0.243626149 | 0.060937162 |
| 28644.18555 | 24480.86133 | 26890.35352 | -0.835697981 | 0.009316552 | 0.938370473 | 0.011710842 | 0.046870506 | 0.682933573 |
| 22111.08398 | 22269.79688 | 21148.5 | -0.835840411 | 0.027790926 | 0.282001122 | 0.167314796 | 0.18628385 | 0.035986915 |
| 28555.55273 | 37070.63281 | 26276.09766 | -0.835883867 | 0.236422852 | -0.172642589 | 0.687706188 | -0.238752784 | 0.428546505 |
| 88516.8125 | 85669.33594 | 83809.04688 | -0.836569044 | 0.005752145 | -0.027181036 | 0.563589094 | -0.150266615 | 0.049829884 |
| 898018.9375 | 1090156.875 | 1096521.5 | -0.83746971 | 0.071655577 | 0.87384043 | 0.094689804 | 0.709484178 | 0.103046053 |

| | | | | | | | | |
|---|---|---|---|---|---|---|---|---|
| 13473.15527 | 13041.5918 | 11966.67773 | -0.837904303 | 0.024111712 | 0.109411234 | 0.32306131 | -0.10013326 | 0.400632465 |
| 645480.3125 | 698166.75 | 700687.8125 | -0.838281328 | 0.042146623 | -0.417627709 | 0.11151684 | 0.50076483 | 0.034150933 |
| 138639.3438 | 125038.2344 | 143341.3281 | -0.838369926 | 0.014563672 | 1.064119738 | 0.047601616 | 0.791213165 | 0.026263251 |
| 40367.38281 | 44481.07422 | 39612.83594 | -0.839890198 | 0.004709568 | -0.441091958 | 0.020431201 | -0.453683465 | 0.023920487 |
| 24681.36133 | 22456.93359 | 21713.03711 | -0.840299854 | 0.016471012 | 0.257302136 | 0.121847149 | -0.246080819 | 0.134304215 |
| 94837.00781 | 98824.04688 | 102186.7656 | -0.840899062 | 0.017507272 | -0.289422602 | 0.168359532 | -0.415792822 | 0.103046053 |
| 6850.71875 | 5353.206055 | 1353.725952 | -0.841004311 | 0.021595434 | 0.073592245 | 0.693736734 | -0.560347658 | 0.431199487 |
| 18049.36328 | 15857.0498 | 15532.20117 | -0.842721304 | 0.022480145 | -0.002302444 | 0.987735084 | -0.501019857 | 0.047847186 |
| 2527.700928 | 1966.53894 | 1139.117798 | -0.842761222 | 0.038176133 | 0.104965469 | 0.714090192 | -0.250057199 | 0.607360809 |
| 35618.36719 | 28929.17188 | 29173.76172 | -0.843268135 | 0.018311389 | -0.192655878 | 0.374780416 | 0.297448304 | 0.214396489 |
| 11625.0498 | 14034.19141 | 9652.438477 | -0.843525237 | 0.229185272 | -0.158061593 | 0.454547212 | -0.045996142 | 0.844714145 |
| 63712.71875 | 55043.32422 | 49510.79688 | -0.844631545 | 0.010893509 | -0.099475228 | 0.466656023 | 0.36238805 | 0.151192408 |
| 51514.10156 | 62814.22266 | 60340.36328 | -0.845411959 | 0.005752145 | 0.029490407 | 0.777841261 | 0.049647342 | 0.721357803 |
| 3693.73877 | 5707.017578 | 4053.087646 | -0.846018107 | 0.249155595 | 1.343501772 | 0.07468596 | 0.434341773 | 0.341867323 |
| 48395.65234 | 46591.26563 | 46536.95313 | -0.846187947 | 0.111432182 | -0.338666102 | 0.254014221 | -0.164461305 | 0.464552866 |
| NA | NA | NA | -0.846265192 | 0.04251435 | -0.014726885 | NA | NA | NA |
| 38607.75781 | 122039.4453 | 36273.02344 | -0.846268719 | 0.240263529 | -0.036809362 | 0.966354559 | -0.064347141 | 0.954810237 |
| 45341.91406 | 45347.69336 | 44409.74023 | -0.846817717 | 0.015716954 | -0.098040881 | 0.317178531 | -0.349792778 | 0.038799047 |
| 26828.45703 | 23306.38867 | 16391.61328 | -0.847256391 | 0.014813633 | -0.057545324 | 0.711282743 | -0.743023819 | 0.070968049 |
| 2954.633057 | 2774.03125 | 2730.782227 | -0.847448542 | 0.174262283 | 0.80167276 | 0.121179966 | -0.307146528 | 0.032116314 |
| 20520.43359 | 23057.92188 | 18935.00977 | -0.847526113 | 0.029390153 | 0.472547536 | 0.072424745 | -0.058439506 | 0.762717395 |
| 158605.2969 | 151969.8906 | 161288.0938 | -0.848523538 | 0.024408916 | 0.201791705 | 0.093367083 | 0.759112314 | 0.01054584 |
| 7297.416992 | 7171.180176 | 7921.92334 | -0.848533949 | 0.015638289 | -0.409794259 | 0.082612243 | -0.813237418 | 0.043437905 |
| 13190.64844 | 10925.33398 | 5106.415527 | -0.849796972 | 0.150232054 | 1.432883386 | 0.052702293 | 0.654310223 | 0.40585939 |
| 52487.64063 | 56270.36719 | 51439.35938 | -0.850173038 | 0.053863679 | 0.518003998 | 0.024921531 | -0.416576044 | 0.026764821 |
| 12005.91211 | 9667.168945 | 7445.460938 | -0.850513678 | 0.12468265 | 0.323219201 | 0.082612243 | -0.060093936 | 0.838339899 |
| NA | NA | NA | -0.851240588 | NA | 1.852049686 | NA | NA | NA |
| 526356.875 | 460344.9688 | 436756.1563 | -0.851595855 | 0.029088124 | 0.371709631 | 0.284636711 | -1.134911997 | 0.081747287 |
| 62139.94141 | 69316.17969 | 76321.71094 | -0.852712164 | 0.08662098 | 0.520949396 | 0.187490341 | 1.329321167 | 0.023619478 |
| 7942.661621 | 7626.742188 | 2430.426758 | -0.852947198 | 0.048340163 | 0.688365075 | 0.127902424 | 0.302931963 | 0.689054401 |
| 18521.70313 | 15774.18164 | 15111.39355 | -0.854200255 | 0.010603888 | 0.353509372 | 0.080440542 | -0.104004514 | 0.489401961 |
| 50270.10156 | 53598.9375 | 52371.01563 | -0.854250783 | 0.044614292 | 0.099516929 | 0.30354249 | 0.018827839 | 0.654381087 |
| 9013.109375 | 11255.87891 | 7788.410156 | -0.854520901 | 0.044080868 | 1.668114257 | 0.05467585 | 0.610035 | 0.145687527 |

| | | | | | | | | |
|---|---|---|---|---|---|---|---|---|
| 90783.89063 | 75687.78125 | 82233.74219 | -0.85465425 | 0.019286562 | 0.330655631 | 0.160918214 | 0.253721612 | 0.216424718 |
| 13334.85449 | 8396.508789 | 8636.876953 | -0.855097922 | 0.034352005 | -0.345176908 | 0.15465818 | -0.440721618 | 0.234646777 |
| 10546.90137 | 12401.28613 | 10353.04102 | -0.855250804 | 0.04646045 | -0.98016 | 0.178109487 | 0.894434568 | 0.039195867 |
| 15993.58105 | 19969.22461 | 14575.29883 | -0.856862513 | 0.068077455 | 0.343321531 | 0.137540222 | -0.214564579 | 0.34800693 |
| 105122.25 | 111840.125 | 96619.8125 | -0.857001338 | 0.027230193 | 0.088898909 | 0.623543083 | 0.325286147 | 0.125718789 |
| 38066.19141 | 31141.12109 | 30608.38672 | -0.857020552 | 0.026154212 | 0.648014134 | 0.087593657 | 0.677892455 | 0.064631313 |
| 2500.479248 | 3207.917236 | 2057.10498 | -0.857585664 | 0.0424199 | -1.11441178 | 0.131843649 | -0.917786108 | 0.130708956 |
| 27447.66211 | 22559.82227 | 24645.40234 | -0.857828629 | 0.035760906 | -0.137520427 | 0.686521476 | -0.216414801 | 0.496031681 |
| 13549.14648 | 9261.703125 | 8018.90332 | -0.858214136 | 0.131546744 | 0.463676871 | 0.400065651 | -0.571468185 | 0.447116057 |
| 9936.416016 | 10562.09473 | 7974.09082 | -0.858231235 | 0.02883182 | 0.5492963 | 0.042884533 | -0.030645375 | 0.869150523 |
| 43401.44141 | 48981.16016 | 54211.17578 | -0.858879501 | 0.131546744 | 0.214678252 | 0.215837791 | 0.349817519 | 0.132183117 |
| 68654.38281 | 70357.17188 | 56812.47266 | -0.859686888 | 0.025587445 | 0.962181726 | 0.034480785 | -0.073038533 | 0.772996243 |
| 43346.53906 | 46586.59375 | 46697.04688 | -0.861792223 | 0.027671686 | 0.793409009 | 0.103892924 | 1.212915202 | 0.009461368 |
| 529148.5625 | 501657.6875 | 491571.6563 | -0.862741553 | 0.011814081 | 0.232366682 | 0.157684191 | 0.414420616 | 0.057477654 |
| 19272.1582 | 22075.60352 | 21458.35352 | -0.863156979 | 0.015598284 | 0.141406273 | 0.372701329 | 0.183609608 | 0.274556211 |
| 12880.43359 | 8878.805664 | 11705.73242 | -0.863924234 | 0.039781174 | -0.645252384 | 0.199074765 | -0.237985892 | 0.5189513 |
| 6969.879883 | 7192.118652 | 7385.790039 | -0.86410762 | 0.193297417 | 0.676942092 | 0.190057963 | 0.130307803 | 0.682933573 |
| 18081.34375 | 18561.86133 | 13760.88672 | -0.86417915 | 0.066355015 | 0.73819995 | 0.092429302 | 0.452814624 | 0.225833549 |
| 14063.12207 | 17291.41602 | 16693.24023 | -0.864996947 | 0.038620049 | 0.683226982 | 0.089652719 | -0.16216339 | 0.549861553 |
| 120361.3438 | 125988.4844 | 119529.6328 | -0.865250087 | 0.010881569 | 0.317337439 | 0.058070756 | -0.175071416 | 0.15696864 |
| 106722.9766 | 107241.2344 | 106189.8672 | -0.865465317 | 0.033688384 | 0.987940019 | 0.052447104 | 0.988935645 | 0.010236624 |
| 652488.0625 | 659035 | 686642.5 | -0.866581667 | 0.009974872 | 0.916277032 | 0.015625705 | 1.052223867 | 0.00995521 |
| 8405.360352 | 11089.6543 | 6057.887695 | -0.867056204 | 0.026675593 | -0.90441014 | 0.081663959 | -1.119378823 | 0.055259337 |
| 15794.61133 | 14136.94531 | 11129.83594 | -0.867216422 | 0.029258207 | 0.303299667 | 0.276647436 | 0.169866137 | 0.573238987 |
| 6330.378906 | 5868.837402 | 5653.3125 | -0.868418638 | 0.029390153 | 0.517682674 | 0.282731898 | -0.275304211 | 0.361411188 |
| 46472.91406 | 48194.93359 | 46761.8125 | -0.868710178 | 0.011866927 | 0.14662731 | 0.30106838 | 0.215760799 | 0.173821986 |
| 16938.51563 | 19076.32227 | 13126.71191 | -0.869442636 | 0.027671686 | -0.142938481 | 0.562301807 | -0.577183797 | 0.118373568 |
| 38399.51953 | 45383.07813 | 39889.27344 | -0.870096548 | 0.087043567 | 0.327930155 | 0.467718612 | 0.236594205 | 0.58875957 |
| 41720.58203 | 48791.875 | 19423.9043 | -0.870180288 | 0.035262392 | -0.863030508 | 0.05468055 | -1.145078523 | 0.069955133 |
| 44655.28516 | 41799.3125 | 49734.46484 | -0.870565852 | 0.029051003 | 0.923711079 | 0.048024604 | 0.079818021 | 0.732715412 |
| 70896.46094 | 67606.25781 | 56993.21875 | -0.870826016 | 0.010297098 | 0.086042451 | 0.269570571 | -0.676024469 | 0.027331143 |
| 307375.5938 | 332718.9375 | 324288.7188 | -0.871282386 | 0.025855185 | 0.82539009 | 0.045588559 | 0.966279727 | 0.034567592 |
| 11304.82227 | 10428.12402 | 7314.616699 | -0.872479172 | 0.120016833 | 0.017453621 | 0.645544015 | -0.156021857 | 0.558686385 |

| | | | | | | | | |
|---|---|---|---|---|---|---|---|---|
| 5907.358398 | 6245.439453 | 4970.974609 | -0.872698798 | 0.147715547 | -0.336068974 | 0.10262657 | -1.211304412 | 0.02183874 |
| 20114.91797 | 21277.75977 | 16611.08594 | -0.873796394 | 0.06270748 | -0.850678768 | 0.23288998 | -1.069116282 | 0.174083052 |
| 44051.89063 | 40609.10156 | 26737.99023 | -0.874825365 | 0.037752026 | 0.334004218 | 0.262642953 | 0.165529162 | 0.624656745 |
| 53836.92188 | 52266.10938 | 61244.63281 | -0.875003847 | 0.016861009 | 0.024093153 | 0.853299816 | -0.473667808 | 0.056784127 |
| 26511.14453 | 21125.30273 | 21473.13086 | -0.875087143 | 0.019930448 | 0.451579782 | 0.093708586 | 0.566672764 | 0.086201739 |
| 35861.05078 | 41779.5 | 34926.04688 | -0.876390419 | 0.031890241 | 0.045711048 | 0.863198658 | -0.433504921 | 0.215760001 |
| 322756.6936 | 316274.9375 | 318915.4688 | -0.876794357 | 0.009008041 | 0.095874961 | 0.348524737 | 0.195090957 | 0.142342008 |
| 253048.5625 | 267573 | 274536.8125 | -0.876912768 | 0.037238828 | -0.08846392 | 0.228822955 | -0.029340775 | 0.676744371 |
| 19932.58789 | 17658.76172 | 11905.00098 | -0.877116819 | 0.030009295 | 0.453697384 | 0.172669944 | 0.123546545 | 0.750765983 |
| 49487.24609 | 51704.90234 | 59586.70703 | -0.877160941 | 0.047458 | 0.203599349 | 0.053446178 | 0.13899827 | 0.354370577 |
| 29345.89453 | 31739.73047 | 26268.79492 | -0.880576574 | 0.005752145 | -0.168796928 | 0.11288505 | -0.404249289 | 0.051977257 |
| 7873.791992 | 12628.92188 | 6227.336426 | -0.881999069 | 0.40645545 | 0.024354224 | 0.975992768 | -0.235473864 | 0.6857936 |
| 2886.820313 | 3872.07373 | NA | -0.883395409 | 0.074826564 | 1.283067365 | 0.161747209 | 0.186398666 | NA |
| 3793.964355 | 4540.731445 | 3402.880615 | -0.88353226 | 0.137237024 | 0.580325107 | 0.344413701 | 0.199168952 | 0.714462192 |
| 142733.6875 | 140643.625 | 149108.5938 | -0.884576388 | 0.022489083 | -0.929905667 | 0.098036283 | -0.693936714 | 0.124182636 |
| 42855.46875 | 45472.78516 | 35858.0625 | -0.884726921 | 0.031140524 | -0.055202182 | 0.643779366 | 0.003195448 | 0.985804247 |
| 4492.898926 | 6627.229004 | 4385.466797 | -0.884805933 | 0.020807424 | 0.136122096 | 0.41861873 | -0.419608996 | 0.217152139 |
| 24905.56836 | 24710.46484 | 19785.07813 | -0.886005726 | 0.14548836 | -0.200204988 | 0.213120095 | -0.472612145 | 0.066089234 |
| 9637.135742 | 8778.123047 | 8394.797852 | -0.886174117 | 0.018311389 | -0.079411409 | 0.445857703 | -0.020094294 | 0.854933382 |
| 28729.68164 | 31227.93164 | 31185.87695 | -0.886947069 | 0.015169362 | 0.206586314 | 0.09803344 | 0.377388742 | 0.031906483 |
| 16869.99609 | 15427.91602 | 17919.36523 | -0.88697003 | 0.056373954 | -0.758600173 | 0.068436129 | 0.07487998 | 0.644909155 |
| 21232.56445 | 23025.18555 | 16140.38281 | -0.887529294 | 0.154959228 | 0.107580939 | 0.777099212 | 0.07094954 | 0.794565284 |
| 213171.5625 | 200341.4531 | 211581.625 | -0.887683848 | 0.035199008 | -0.443575403 | 0.27866616 | -0.745980838 | 0.166745689 |
| 35090.47266 | 36239.9375 | 36360.89063 | -0.890401157 | 0.016471012 | -0.790412711 | 0.052457323 | 0.542013343 | 0.048428024 |
| 48057.50781 | 45097.78516 | 54659.75781 | -0.890995664 | 0.078489996 | 1.190747954 | 0.054285958 | 1.996243848 | 0.030482431 |
| 23977.08594 | 26379.02148 | 27053.83203 | -0.891581967 | 0.045092515 | 0.487995112 | 0.031234888 | 0.326523613 | 0.083570684 |
| 30880.77539 | 35621.60547 | 32954.33203 | -0.891806957 | 0.224955599 | 0.724660011 | 0.34872471 | 0.47609337 | 0.524925524 |
| 7876.835938 | 7525.881836 | 7595.854004 | -0.893578779 | 0.153142215 | 0.818826733 | 0.073324814 | 1.430071681 | 0.03434959 |
| 6653.218262 | 6154.845703 | 6134.614746 | -0.893834106 | 0.166292161 | 0.924968947 | 0.185173357 | -0.508889108 | 0.544452231 |
| 127144.3516 | 162109.7344 | 136235.8594 | -0.898310344 | 0.139135078 | 0.481900634 | 0.256838624 | 0.582119659 | 0.176540122 |
| 2698.96582 | 3794.247803 | 2576.969727 | -0.899107994 | 0.256719915 | 1.660823018 | 0.055241445 | 0.735437143 | 0.263528267 |
| 15720.4248 | 17017.02148 | 25605.18945 | -0.899788996 | 0.072654291 | 0.656399823 | 0.175682075 | 0.077545269 | 0.884436094 |
| 193503.9219 | 194000.7031 | 166650.5781 | -0.900986518 | 0.034144692 | 0.138413438 | 0.60094054 | -0.56996894 | 0.168323482 |

| | | | | | | | | |
|---|---|---|---|---|---|---|---|---|
| 16047.44043 | 23055.60352 | 19439.87305 | -0.901737073 | 0.092593056 | -0.571982756 | 0.352759971 | 0.384568478 | 0.386649359 |
| 5283.184082 | 3630.931641 | NA | -0.904227672 | 0.252787526 | 0.291405767 | 0.532317074 | -0.404973174 | NA |
| 6258.363281 | 6206.258789 | 6711.812988 | -0.904637734 | 0.027846024 | -0.496338332 | 0.218601949 | -0.223860617 | 0.430145932 |
| 40447.29688 | 48203.42188 | 32853.82031 | -0.905085769 | 0.137336288 | -0.555875263 | 0.248010278 | -0.220534904 | 0.385275383 |
| 29151.67773 | 20788.71875 | 23971.82422 | -0.906498922 | 0.113223175 | 0.458734983 | 0.393273993 | 2.416627715 | 0.055868946 |
| 32749.77539 | 45168.89453 | 28358.29688 | -0.907362773 | 0.078924537 | 0.013934134 | 0.967378164 | -0.401601744 | 0.30332534 |
| 6454.047852 | 6313.968262 | 12052.09961 | -0.907849771 | 0.206331283 | 1.814610972 | 0.165197811 | 2.775953974 | 0.136474059 |
| 12060.40625 | 11544.54297 | 10094.02441 | -0.908315907 | 0.016132048 | -0.625034018 | 0.100528183 | -0.322526216 | 0.162869873 |
| 246849.2656 | 275746.9063 | 310364.5313 | -0.9084764 | 0.09943294 | 0.937048444 | 0.012254503 | 0.896456112 | 0.055516192 |
| 17475.11133 | 17167.84766 | 17839.82617 | -0.909242652 | 0.007536593 | 0.000877662 | 0.991322905 | -0.41187715 | 0.00797582 |
| 408.5338745 | 612.1486816 | NA | -0.909838401 | NA | NA | NA | 0.371167427 | NA |
| 40454.125 | 33024.08203 | 34906.36719 | -0.910495999 | 0.023177803 | -0.031312432 | 0.863910543 | -0.390101561 | 0.152269127 |
| 5874.055664 | 9681.852539 | 3972.02832 | -0.911495737 | 0.137345712 | 0.686070011 | 0.250511865 | 0.169522966 | 0.827659891 |
| 3896.142334 | 3491.790771 | 2397.534668 | -0.911543012 | 0.024408916 | -0.250891404 | 0.331607261 | -0.506511167 | 0.161137259 |
| 1255.928223 | 1430.873779 | 884.9938965 | -0.912659184 | 0.422468945 | 0.823183658 | 0.485037816 | -0.482533665 | 0.778013904 |
| 76725.33594 | 80489.03125 | 70823.44531 | -0.912713289 | 0.124546636 | 0.83027748 | 0.162724701 | 0.378616608 | 0.448315524 |
| 2116.006836 | 2389.510742 | 1263.911865 | -0.915089174 | 0.059872767 | 0.063373936 | 0.805890965 | -0.762054069 | 0.12996555 |
| 118961.4375 | 123752.7578 | 119991.9688 | -0.915604812 | 0.02237899 | 0.599883256 | 0.051139483 | 0.954046054 | 0.027331143 |
| 3597.467529 | 2605.639893 | 4180.150391 | -0.915804678 | 0.032032289 | 0.102708308 | 0.576955884 | -0.178223488 | 0.56267449 |
| 7645.685547 | 6520.339844 | 4034.751709 | -0.91669501 | 0.014038682 | 0.835266898 | 0.021906439 | 0.396243156 | 0.413244913 |
| 2689899 | 2693851.5 | 2769065.5 | -0.917288145 | 0.004709568 | -0.251344206 | 0.051097951 | -0.214473976 | 0.06365738 |
| 11166.78027 | 9630.644531 | 13344.5625 | -0.918087973 | 0.021416605 | -0.144917682 | 0.485773334 | -0.452124898 | 0.106930847 |
| 31342.75977 | 30289.70117 | 28383.66211 | -0.918989791 | 0.056155267 | 0.786075091 | 0.086909686 | 1.14194972 | 0.045096398 |
| 36967.21875 | 33361.91797 | 34070.23047 | -0.919336304 | 0.048752402 | -0.507363622 | 0.077576966 | 0.20405816 | 0.093609212 |
| 19298.61914 | 16756.05273 | 18946.70313 | -0.920347404 | 0.018311389 | 0.65934089 | 0.046782366 | 0.472082116 | 0.054109248 |
| 121677.1328 | 112805.3125 | 121780.125 | -0.920748411 | 0.033865703 | 0.551421846 | 0.049480102 | 0.564127692 | 0.021984636 |
| 49947.10938 | 52939.52344 | 40598.84766 | -0.920780945 | 0.100751002 | -0.218654643 | 0.227092221 | -0.316038216 | 0.143427363 |
| 6783.708984 | 6984.986328 | 5572.921875 | -0.921907135 | 0.016603853 | -0.569462993 | 0.100046608 | -1.302426624 | 0.044802319 |
| 114525.2813 | 112308.0234 | 106575.7734 | -0.922667787 | 0.021020174 | 0.022737766 | 0.75018573 | -0.272744749 | 0.05752489 |
| 220466.5625 | 245872.9531 | 207804.4531 | -0.923933686 | 0.008009891 | -0.161702147 | 0.221894535 | -0.252611247 | 0.122834452 |
| 35524.36719 | 37649.13672 | 34493.75 | -0.924022241 | 0.011206547 | -0.317917537 | 0.051112404 | -0.069451509 | 0.397327146 |
| 25214.05664 | 22869.31836 | 18851.82617 | -0.924805766 | 0.076613377 | 0.520161188 | 0.336705725 | 0.474148036 | 0.148421794 |
| 152972.9531 | 165462.0781 | 158089.8594 | -0.924977113 | 0.029906057 | 0.262536964 | 0.187490341 | 0.223616434 | 0.049415261 |

| | | | | | | | | |
|---|---|---|---|---|---|---|---|---|
| 290086.9531 | 289383.3438 | 318601.4375 | -0.927254559 | 0.008593584 | 0.55332552 | 0.056841901 | 0.625737673 | 0.021984636 |
| 3552.381104 | 3616.599365 | 3115.039307 | -0.930241696 | NA | -0.106879288 | 0.604373663 | -0.200482216 | 0.135721265 |
| 210026.0625 | 243074.0313 | 234785.1719 | -0.930513906 | 0.113922974 | 0.653044493 | 0.27343485 | 0.257004202 | 0.641018025 |
| 31265.60352 | 40613.14453 | 34295.90625 | -0.930965634 | 0.155490733 | -0.288487295 | 0.498770255 | -0.199754431 | 0.597655284 |
| 13190.99512 | 11023.5293 | 11252.78906 | -0.937396217 | 0.017952706 | -0.425677161 | 0.132364679 | -0.313834332 | 0.185378125 |
| 7136.581543 | 6587.61377 | 6445.392578 | -0.93906973 | 0.020626238 | -0.327439791 | 0.272117866 | -0.433828448 | 0.194351963 |
| 10643.41309 | 10191.5957 | 7276.604004 | -0.942641088 | 0.017713193 | -0.424631374 | 0.119032068 | -1.006938909 | 0.036066713 |
| 3415.187744 | 2231.497559 | 3292.28833 | -0.944255396 | 0.125515217 | 0.026445868 | NA | 0.187836324 | 0.764204423 |
| 20897.8125 | 5772.32959 | 8413.625977 | -0.944845216 | 0.043150549 | 0.567230287 | 0.11146465 | -0.299632363 | 0.713487699 |
| 43996.16016 | 47094.02344 | 51597.53125 | -0.945365768 | 0.018002294 | 0.628346713 | 0.024676726 | 0.216630787 | 0.169381511 |
| 2096.565918 | 797.3242188 | 4574.415039 | -0.945474877 | NA | 1.741064453 | NA | 0.011034002 | NA |
| 146638.375 | 133974.2656 | 143792.8438 | -0.945583258 | 0.018999567 | 0.442293523 | 0.116818656 | 0.131783242 | 0.528680834 |
| 169718.0313 | 167731.3438 | 176410.4844 | -0.946063327 | 0.005790576 | 0.11244381 | 0.280427668 | -0.008502481 | 0.916719934 |
| 413274.7188 | 401276.2188 | 390558.9375 | -0.946998523 | 0.027556237 | 0.1827913 | 0.21418297 | -0.369872014 | 0.102372782 |
| 1256.195923 | 1335.303833 | 562.1189575 | -0.948529937 | 0.067161873 | 0.791333826 | 0.052437153 | -0.761653386 | 0.164342717 |
| 92009.86719 | 104771.9141 | 107569.4141 | -0.948999816 | 0.080663793 | -0.261958573 | 0.088308366 | 0.14193894 | 0.28320943 |
| 14525.91406 | 16627.49219 | 13490.1709 | -0.950175641 | 0.045092515 | 0.111481387 | 0.458532796 | -0.415856587 | 0.091109394 |
| 98359.04688 | 91512.69531 | 89708.23438 | -0.950682697 | 0.021462764 | 0.410304485 | 0.163676753 | 0.287591907 | 0.247523967 |
| 2455.308594 | 2379.480469 | 2155.533203 | -0.95170426 | 0.066355015 | -0.196614985 | 0.625452135 | -0.531505606 | 0.297583227 |
| 29806.56055 | 25066.99805 | 23996.07617 | -0.952763869 | 0.03211333 | 0.270333534 | 0.228114774 | -0.039714239 | 0.860403762 |
| 6082.944824 | NA | NA | -0.953663633 | 0.203513529 | 0.059035502 | 0.885440998 | -0.299275909 | NA |
| 1679.933594 | 1927.930298 | 1470.929565 | -0.954720896 | 0.021947631 | 0.200143378 | 0.275888795 | -0.457670701 | 0.084223187 |
| 5891.236816 | 6939.020508 | 5548.752441 | -0.954818146 | 0.052321316 | -0.525539422 | 0.30642692 | -1.301392825 | 0.131502216 |
| 20434.42773 | 21450.4668 | NA | -0.955466732 | NA | 0.575803692 | NA | -0.024657721 | NA |
| 68714.09375 | 83453.60156 | 74426.17969 | -0.957619195 | 0.135044793 | 0.196590159 | 0.546823528 | 0.250580083 | 0.228109691 |
| 25889.22852 | 26859.10547 | 22488.99023 | -0.958690466 | 0.034381342 | 0.334107558 | 0.110237532 | -0.063351051 | 0.717271407 |
| 221784.7656 | 281113.9063 | 226582.7969 | -0.959303189 | 0.018432262 | -0.206364174 | 0.160970091 | 0.225795329 | 0.302010752 |
| 11511.71777 | 10468.50293 | 11590.71387 | -0.959847456 | 0.024701492 | 0.492850744 | 0.131789366 | 0.209053433 | 0.414598632 |
| 4365.773926 | 8594.102539 | 8790.56543 | -0.961909525 | 0.058516905 | -0.269183809 | 0.330637946 | -0.840377706 | 0.103046053 |
| 39552.41797 | 42150.93359 | 27698.34375 | -0.961985088 | 0.17437849 | -0.34982304 | 0.274420464 | -0.59819393 | 0.091920126 |
| 61664.47266 | 55801.67578 | 61352.3125 | -0.965102134 | 0.015638289 | 1.627627394 | 0.042753324 | 2.504350964 | 0.017983315 |
| 1844.580933 | 2227.171387 | 1199.289795 | -0.968133804 | 0.025050105 | -0.321674647 | 0.444175174 | -0.699564794 | 0.112700927 |
| 18985.69336 | 23860.12305 | 18235.48047 | -0.969812856 | 0.008543222 | -0.729747531 | 0.04667516 | -0.891131614 | 0.029324108 |

| | | | | | | | | |
|---|---|---|---|---|---|---|---|---|
| 1358.14978 | 2571.928467 | 1513.837158 | -0.970546343 | 0.156563923 | 0.909844005 | 0.074416318 | -1.628139969 | 0.024052583 |
| 115630.2891 | 118591.5391 | 133242.8438 | -0.970767062 | 0.09222768 | 0.42481947 | 0.433283386 | 0.532750102 | 0.321284513 |
| 25155.34766 | 23972.15625 | 23711.01953 | -0.972527466 | 0.024259746 | -0.168666059 | 0.094128959 | 0.296919011 | 0.023786578 |
| 41362.46875 | 39397.58203 | 35241.93359 | -0.975729448 | 0.073567321 | -0.108574017 | 0.517215148 | -0.053893574 | 0.753922782 |
| 599.2550049 | 857.4698486 | 459.6673584 | -0.976400195 | 0.264210628 | 1.049450765 | 0.285423852 | 0.020811269 | 0.963218346 |
| 49707.41406 | 55583.89453 | 55346.3125 | -0.977861225 | 0.042392829 | 0.441381003 | 0.074724231 | 0.188091785 | 0.215503868 |
| 133906.5156 | 125479.4453 | 111955.3828 | -0.978009196 | 0.016471012 | 0.280351051 | 0.116818656 | 0.091456185 | 0.548845982 |
| 16200.8457 | 30737.16211 | 13969.96973 | -0.979293661 | 0.333309126 | 0.443127467 | 0.626134586 | 0.027903587 | 0.971950381 |
| 42927.52344 | 42246.89844 | 43474.61328 | -0.980411056 | 0.047406642 | 2.245522698 | 0.023068087 | 1.694827983 | 0.007057904 |
| 3953.8396 | NA | 1952.413818 | -0.981121192 | NA | 0.384616434 | NA | -0.910367499 | NA |
| 36441.24219 | 31247.18945 | 28282.31445 | -0.981896583 | 0.005752145 | 0.79388459 | 0.047137518 | -0.446170193 | 0.088642968 |
| 3649.632813 | 3215.454346 | NA | -0.982033377 | 0.048340163 | 0.070350808 | 0.907258259 | -0.463687473 | NA |
| 27310.09766 | 19463.33984 | 22647.31836 | -0.983226323 | 0.123415711 | 0.207901958 | 0.711282743 | 1.059238697 | 0.081394533 |
| 39167.94531 | 42565.64844 | 45921.89453 | -0.983558264 | 0.007142399 | 0.063109368 | 0.706396106 | 0.246312023 | 0.133712681 |
| 16829.17969 | 12346.6543 | 10983.39063 | -0.983683795 | 0.106137102 | 0.830005132 | 0.051924975 | 0.256022225 | 0.473709978 |
| 32435.26563 | 34933.86328 | 32830.66797 | -0.985501742 | 0.027375615 | 0.209029947 | 0.220100738 | -0.192873469 | 0.238635537 |
| 3560.508789 | 2968.771729 | 3907.265137 | -0.98576323 | 0.0830364 | -0.103399681 | 0.781102846 | -0.368106852 | 0.391347168 |
| 3079.811279 | 3469.236084 | 1989.635864 | -0.986700197 | 0.006564049 | -0.234757306 | 0.1787498 | -0.921216263 | 0.063419062 |
| 30138.65039 | 31062.45508 | 22791.18945 | -0.986702881 | 0.138213581 | -0.266731856 | 0.164062441 | -0.43611194 | 0.135536925 |
| 15196.30859 | 27481.02539 | 18038.31055 | -0.988461718 | 0.077426021 | 0.198306453 | 0.610889349 | -0.262014977 | 0.535773657 |
| 73166.66406 | 87864.67969 | 97318.34375 | -0.988627564 | 0.012620338 | 0.216517388 | 0.232151348 | 1.219750594 | 0.055096798 |
| 891179.3125 | 959734.9375 | 901699.9375 | -0.988671298 | 0.012414001 | 0.195759529 | 0.300186184 | -0.178748301 | 0.366486609 |
| 12629.63184 | 13188.74805 | 9834.336914 | -0.989047562 | 0.04113039 | -0.0333234 | 0.936042943 | 0.916483405 | 0.067123192 |
| 7267.695313 | 9998.505859 | 8785.802734 | -0.991813973 | 0.137661186 | 0.082388052 | 0.885440998 | 0.181060122 | 0.744940584 |
| 25185.58203 | 23254.91016 | 26420.7793 | -0.995647754 | 0.040104768 | -0.147693565 | 0.585714677 | -0.432008679 | 0.233347919 |
| 251131.4063 | 299494.0625 | 307574.6563 | -0.996008595 | 0.0885422 | 1.219373682 | 0.090139838 | 0.997337226 | 0.099088078 |
| NA | NA | NA | -0.996137634 | 0.016105265 | 1.001747808 | 0.312153356 | NA | NA |
| 7386.444336 | 7632.999023 | 6721.162598 | -0.996680764 | 0.04950513 | 0.52700135 | 0.13681424 | 0.322368377 | 0.146875127 |
| 4966.190918 | 6039.904297 | 6505.486816 | -0.997274691 | 0.019930448 | 0.345789711 | 0.223404971 | 0.253916224 | 0.350904531 |
| 24840.12305 | 21289.54688 | 21726.93555 | -0.9982773 | 0.011946285 | 0.810075968 | 0.02118733 | 0.453002874 | 0.059847517 |
| 118412.8594 | 144862.2813 | 115585.8906 | -0.998300474 | 0.046600136 | 0.86856563 | 0.082612243 | 0.461052113 | 0.185585346 |
| 87685.11719 | 81130.72656 | 75567.04688 | -0.998321684 | 0.105079901 | 1.534782553 | 0.123404644 | 1.905116107 | 0.020337832 |
| 13944.31836 | 14241.12207 | 10079.46777 | -0.999025758 | 0.018311389 | 0.136002381 | 0.180354112 | -0.507012811 | 0.107790858 |

| | | | | | | | | |
|---|---|---|---|---|---|---|---|---|
| 6460.241699 | 19318.00781 | 14648.22949 | -1.000605383 | 0.154891947 | 0.319162565 | 0.560671011 | 0.055268695 | 0.948264111 |
| 72196.10938 | 130469.2422 | 66852.89844 | -1.002805586 | 0.156631718 | 0.043295542 | 0.936763383 | -0.141190097 | 0.797329975 |
| 13521.46777 | 10662.60938 | 6308.67334 | -1.005433519 | 0.013618049 | 0.324609448 | 0.16494964 | -0.687751194 | 0.156112176 |
| 12547.79785 | 10665.24121 | 10644.02734 | -1.006122558 | 0.061707718 | 1.353001615 | 0.074618176 | 0.886220783 | 0.032304888 |
| 3853.968018 | 5209.258301 | 5725.317871 | -1.006273395 | 0.005752145 | 0.115725057 | 0.352029077 | -0.894902602 | 0.038799047 |
| 16337.78711 | 14566.12402 | 15814.9873 | -1.006983104 | 0.029750807 | 0.3824451 | 0.036339782 | 0.031218172 | 0.755295455 |
| 206260.0625 | 202869.1094 | 202928.6719 | -1.007057788 | 0.03474627 | 0.994376036 | 0.045536435 | 1.194906119 | 0.050623317 |
| 3345.862061 | 3601.493164 | 2603.304688 | -1.008968068 | 0.089267599 | 0.953116671 | 0.130087098 | 0.249309389 | 0.626462088 |
| 17184.51367 | 18792.72461 | 21949.76563 | -1.009885192 | 0.013596234 | -0.200575467 | 0.364883875 | 0.101660951 | 0.64498426 |
| 12348.50391 | 11891.49609 | 11031.47949 | -1.010595273 | 0.052969391 | 0.186667771 | 0.224845397 | 0.054591914 | 0.49125304 |
| 54563.76563 | 57939.37891 | 53680.70313 | -1.010729594 | 0.01391113 | -0.025787764 | 0.892168845 | 0.648978402 | 0.01858328 |
| NA | NA | NA | -1.010842737 | 0.053991433 | -0.672697397 | NA | NA | NA |
| 2105.49585 | 3696.646973 | 1464.53833 | -1.011624451 | 0.256095131 | 2.002451792 | 0.039887225 | 0.316811234 | 0.717270871 |
| 10865.55566 | 11436.54102 | 11364.96191 | -1.012361459 | 0.041859693 | 0.110072347 | 0.474591418 | -0.128418811 | 0.159646627 |
| 7704.472168 | 9752.599609 | 4016.126465 | -1.012834682 | 0.063167428 | -0.740546458 | 0.033121608 | -1.265427665 | 0.06065526 |
| NA | 3795.361084 | 2528.046875 | -1.012858566 | NA | 0.572239049 | NA | 0.000166611 | NA |
| 265866.4688 | 239211.4219 | 254702.6719 | -1.012913687 | 0.030417399 | 0.257673792 | 0.138540382 | 0.703739355 | 0.021723356 |
| 73216.64844 | 82308.96094 | 83004.21875 | -1.014468165 | 0.028452905 | 0.015110509 | 0.952506289 | -0.343154544 | 0.233299785 |
| 42336.86328 | 51962.05859 | 35599.62891 | -1.014748766 | 0.104607763 | -0.668096282 | 0.163722766 | -0.423183095 | 0.150131299 |
| 54253.9375 | 55085.57422 | 51939.10938 | -1.0154695 | 0.014033928 | -0.012443265 | 0.926005043 | -0.491038203 | 0.08674947 |
| 90831.875 | 94285.70313 | 46456.01172 | -1.015511685 | 0.033164349 | -0.591549359 | 0.043830603 | -1.36471802 | 0.060171057 |
| 2825.350098 | 3140.346436 | 2260.865967 | -1.01587655 | 0.031442732 | 0.267709043 | 0.258384367 | -0.570630318 | 0.071891702 |
| 6044.152344 | 5986.061523 | 3313.651367 | -1.018034505 | 0.017824776 | 0.260605082 | 0.278495147 | -0.753222175 | 0.102620951 |
| 32727.5293 | 46613.28516 | 37094.74219 | -1.019707656 | 0.045212188 | 0.71985201 | 0.125736204 | 0.206823637 | 0.543506636 |
| 38189.81641 | 35899.17188 | 33556.74219 | -1.019766975 | 0.16932956 | 0.148742076 | 0.761868395 | -0.595588687 | 0.070298132 |
| 4417.194824 | 3478.77002 | 5429.845215 | -1.01977348 | 0.248212377 | -1.001914068 | NA | 0.144567714 | 0.643933046 |
| 64045.40625 | 69817.76563 | 72031.58203 | -1.01998794 | 0.035947727 | -0.168502852 | 0.323565507 | -0.074877105 | 0.510430611 |
| 16918.16602 | 14151.49902 | 15915.27246 | -1.023364076 | 0.009583258 | 0.710964855 | 0.058077004 | 0.202256827 | 0.215349457 |
| 106469.1328 | 93660.07031 | 86055.77344 | -1.02342436 | 0.029801769 | 0.418949762 | 0.203813506 | 0.203429181 | 0.487511859 |
| 31083.49414 | 32190.60938 | 31608.20898 | -1.02489633 | 0.004709568 | 0.253272819 | 0.333634157 | 0.29891337 | 0.023319813 |
| 10904.44141 | 9104.357422 | 9450.44043 | -1.025011204 | 0.0140944 | 0.071826446 | 0.667649144 | -0.269629761 | 0.175730736 |
| 15545.07422 | 15494.87109 | 13777.45508 | -1.02761628 | 0.010228174 | 0.452399846 | 0.065369282 | -0.638980339 | 0.062318926 |
| 3515.847412 | 3547.980469 | 4434.100098 | -1.029571584 | 0.041945948 | -0.329187653 | 0.428011055 | -0.563637104 | 0.235404176 |

| | | | | | | | | |
|---|---|---|---|---|---|---|---|---|
| 56180.96875 | 58773.64844 | 64504.67188 | -1.030881767 | 0.011946285 | 0.010264402 | 0.959223724 | 0.337539554 | 0.077903915 |
| 54206.22266 | 53602.94141 | 55007.73047 | -1.031291418 | 0.024262842 | 0.255086064 | 0.055241445 | 0.198579251 | 0.092731087 |
| 989381.3125 | 1062502.125 | 898924.8125 | -1.03395972 | 0.058893147 | 0.833377169 | 0.091622784 | 0.222025751 | 0.218250475 |
| 14910.00879 | 16286.0332 | 15730.02246 | -1.035855336 | 0.220923499 | 1.246880991 | 0.193772919 | 0.817736074 | 0.354260455 |
| 23529.75391 | 14738.89258 | 15594.87305 | -1.036134148 | 0.113223175 | 1.034913469 | 0.227071428 | 1.028316652 | 0.157128166 |
| 46844.57031 | 44674.22656 | 32842.75391 | -1.039245726 | 0.018497551 | 0.483313586 | 0.088308366 | -0.290259798 | 0.261699898 |
| 3008.039795 | 3449.060303 | 6324.227539 | -1.04381714 | 0.161897705 | 1.06050348 | 0.094344084 | 0.394867514 | 0.535783088 |
| 11666.77441 | 8720.476563 | 7696.42334 | -1.044713434 | 0.06440189 | 1.033877373 | 0.102454913 | 0.911597491 | 0.124509143 |
| 10262.87109 | 10664.79102 | 8975.725586 | -1.045909765 | 0.044665667 | 0.667243906 | 0.157633835 | 0.844687757 | 0.073672213 |
| 4442.351074 | 5461.31543 | 4849.530273 | -1.046226133 | 0.051358293 | -0.337736192 | 0.579358181 | 0.165110882 | 0.318716777 |
| 29283.92578 | 30185.20117 | 27237.15039 | -1.047977098 | 0.005752145 | 0.73617784 | 0.073853963 | 0.762932101 | 0.029120377 |
| 126772.7656 | 142830.2969 | 111564.2891 | -1.050455378 | 0.099983289 | 0.188320499 | 0.311954306 | -0.152937866 | 0.364586325 |
| 1225.868042 | 2163.029053 | 2837.18042 | -1.051642505 | 0.037492489 | -1.052748472 | 0.139103828 | 0.018979971 | 0.976070752 |
| 15392.21387 | 13884.24902 | 13039.89648 | -1.054147279 | 0.036547632 | 0.157053572 | 0.393273993 | -0.066768594 | 0.592495191 |
| 160100.2813 | 162735.3125 | 172674.0469 | -1.054795361 | 0.017117516 | -0.095854657 | 0.537692101 | -0.081501905 | 0.395555054 |
| 12871.41211 | 9237.799805 | 9769.599609 | -1.056979783 | 0.019930448 | 0.4529519 | 0.038647166 | -0.154066784 | 0.519426302 |
| 31070.81445 | 25765.36328 | 26049.79102 | -1.059816156 | 0.016830771 | -0.186789284 | 0.101612528 | -0.759092901 | 0.021195162 |
| 25912.04688 | 23906.31836 | 21464.86133 | -1.063952169 | 0.060864381 | 0.293111017 | 0.474591418 | 0.505940502 | 0.220869174 |
| 53113.03125 | 77140 | 60919.58594 | -1.064939047 | 0.086307687 | 0.301161685 | 0.402480724 | 0.124107756 | 0.695406237 |
| 138531.3906 | 137836.7813 | 133513.0781 | -1.068337708 | 0.009011817 | 0.070796013 | 0.27866616 | -0.159283911 | 0.096058082 |
| 133616.7031 | 134400.626 | 141999.3115 | -1.070101224 | 0.03211333 | 0.384015164 | 0.233503043 | 0.104517668 | 0.695643018 |
| 74877.875 | 62147.84766 | 64555.54297 | -1.071391338 | 0.01508889 | -0.379270735 | 0.03968601 | -0.126028526 | 0.362743641 |
| 16703.42969 | 18220.40039 | 19881.91602 | -1.071495504 | 0.015175173 | 0.603616286 | 0.043921768 | 0.640030979 | 0.039195867 |
| 17923.99414 | 14323.19043 | 19026.21094 | -1.072799042 | 0.019610279 | 0.410174606 | 0.104959595 | 0.10333952 | 0.63605603 |
| 39037 | 52839.29297 | 47643.92578 | -1.0728392 | 0.029432233 | 0.340769512 | 0.310739135 | -0.081973309 | 0.817331351 |
| 125731.2969 | 146442.1563 | 120334.2422 | -1.07360433 | 0.051607729 | 0.209411852 | 0.297020932 | 0.227074175 | 0.235870699 |
| 490896.0625 | 523565 | 306310.4063 | -1.076935509 | 0.009008041 | -0.450121216 | 0.107207509 | -0.795858876 | 0.071891702 |
| 25920.10352 | 25800.91797 | 24410.38672 | -1.077524537 | 0.054920992 | 1.018645498 | 0.104315579 | 1.409110315 | 0.009587646 |
| 119364.375 | 158848.6563 | 101479.0859 | -1.083755941 | 0.16335134 | -0.122355584 | 0.816076757 | -0.05718179 | 0.879190302 |
| 86778.55469 | 90790.26563 | 88970.92969 | -1.084102392 | 0.034212471 | 0.58886281 | 0.083437934 | 0.322626833 | 0.067112928 |
| 44394.17188 | 66124.875 | 87939.80469 | -1.084840223 | 0.175388722 | 0.396331783 | 0.236672563 | -0.004049977 | 0.993663084 |
| 41847.26953 | 43393.5 | 39953.91797 | -1.085788893 | 0.010603888 | 0.539104963 | 0.165757248 | 0.872616893 | 0.017279709 |
| 203000.2813 | 194226.1875 | 177261.2656 | -1.086552869 | 0.013723547 | -0.049627907 | 0.77405498 | -0.288035675 | 0.057068534 |

| | | | | | | | | |
|---|---|---|---|---|---|---|---|---|
| 8365.441406 | 8037.71875 | 7280.830566 | -1.087077861 | 0.024059634 | 0.29141522 | 0.319523239 | -0.060732885 | 0.831156656 |
| 121395.2578 | 163558.125 | 135918.0156 | -1.08743951 | 0.072756623 | -0.217771337 | 0.59453601 | -0.025835085 | 0.933264795 |
| 5101.82373 | 7673.655273 | 5102.585449 | -1.088004476 | 0.125515217 | 0.664885471 | 0.131196907 | 0.757319275 | 0.191055359 |
| 4341.796875 | 6512.051758 | 4736.003906 | -1.088644175 | 0.167266141 | 0.667276017 | 0.18136293 | 1.269738869 | 0.079217245 |
| 9505.511719 | 9699.144531 | 10267.40039 | -1.090018023 | 0.039647218 | 0.542345248 | 0.198202277 | 0.831941081 | 0.010654184 |
| 195850.7402 | 198140.3066 | 189974.7388 | -1.092018558 | 0.00953954 | -0.001992514 | 0.979876653 | 0.001198689 | 0.989056083 |
| 4019.782959 | 3807.527344 | 3884.723877 | -1.092974739 | 0.032101455 | -0.214394436 | 0.537753876 | 0.200363529 | 0.477266228 |
| 25459.44336 | 28513.08594 | 18772.51367 | -1.093154035 | 0.020740891 | -0.130745964 | 0.205568001 | -0.273371224 | 0.321023436 |
| 10333.49609 | 18756.63086 | 17620.70313 | -1.098381753 | NA | 0.096457697 | NA | 0.351534705 | NA |
| 37198.60547 | 33936.02344 | 28124.05078 | -1.099380098 | 0.01656229 | 1.224181577 | 0.021485221 | 0.283549609 | 0.276498861 |
| 1189627.25 | 1206649 | 1143359.5 | -1.100140651 | 0.032538953 | 0.850599058 | 0.013427556 | 0.147879267 | 0.085245136 |
| 40578.53906 | 47607.20703 | 42026.80469 | -1.10143606 | 0.025818052 | 0.579458993 | 0.138536351 | 0.462885587 | 0.187623713 |
| 32322.75195 | 31825.30664 | 28243.58789 | -1.102694841 | 0.026784983 | -0.11791269 | 0.729173554 | -0.309667789 | 0.428160396 |
| 201528.0894 | 276810.1987 | 280571.9233 | -1.10278198 | 0.207266359 | 1.347090833 | 0.16693375 | 0.962845543 | 0.290161902 |
| 122662.4141 | 130213.0313 | 123121.5625 | -1.106306332 | 0.008009891 | 0.010380678 | 0.921325037 | -0.131226864 | 0.196983665 |
| 45033.89453 | 56161.78906 | 56375.91406 | -1.10749105 | 0.025773059 | 1.114542302 | 0.021622074 | 0.552352729 | 0.081899796 |
| NA | NA | NA | -1.110247269 | NA | NA | NA | NA | NA |
| 1444.723267 | 1739.522461 | 1975.664307 | -1.11323868 | 0.089388811 | 0.305167311 | 0.616392146 | -0.025643358 | 0.972311744 |
| 9145.710938 | 9592.105469 | 8086.074707 | -1.117429235 | 0.016471012 | -0.279927442 | 0.167899496 | -0.744569904 | 0.048431709 |
| 37562.68018 | 42034.93506 | 31075.78589 | -1.118966885 | 0.031567192 | 0.066847517 | 0.82536887 | -0.427347434 | 0.299167804 |
| 35541.80469 | 55231.67578 | 35500.91406 | -1.11930623 | 0.200968261 | -0.095630237 | 0.87503879 | 0.181156224 | 0.682718378 |
| 38185.11719 | 37739.33594 | 41023.34766 | -1.119352581 | 0.068152067 | 0.360150594 | 0.195109477 | 1.390971585 | 0.016402675 |
| 15724.58301 | 35070.03125 | 9777.696289 | -1.119360625 | 0.308721026 | 0.299932842 | 0.740759582 | -0.149321802 | 0.863987358 |
| 134693.8691 | 139289.8789 | 130100.7188 | -1.119753614 | 0.039443869 | 0.869432034 | 0.098459633 | -0.246432754 | 0.597655284 |
| 272038.2813 | 215076.375 | 193977.375 | -1.120533169 | 0.026060442 | 0.155603852 | 0.524069349 | 0.189476118 | 0.533485192 |
| 14537.10742 | 12495.00488 | 14811.23828 | -1.1228057 | 0.226456157 | -0.062673699 | 0.92625747 | 1.651373623 | 0.034567592 |
| 56417.74609 | 65798.85156 | 58615.50781 | -1.124642884 | 0.032595469 | 0.178054005 | 0.494663499 | -0.085339116 | 0.670209128 |
| 5331.812012 | 7891.006348 | 4392.90332 | -1.128466685 | 0.039911484 | -0.424247971 | 0.353953112 | -0.759840083 | 0.188422821 |
| 12216.08984 | 11812.69727 | 12778.7666 | -1.129816205 | 0.219303383 | 0.376690657 | 0.667526405 | 0.667897187 | 0.074131758 |
| 14896.25684 | 12492.42383 | 11167.27051 | -1.130909994 | 0.175388722 | -0.112847215 | 0.577348282 | -0.319212354 | 0.22024877 |
| 12260.25 | 12323.41797 | 11315.54785 | -1.132529429 | 0.013676424 | -0.290848367 | 0.189805528 | -0.407820868 | 0.118461316 |
| 21335.98047 | 22835.46875 | 20173.18359 | -1.133852091 | 0.023105653 | 0.600189514 | 0.133753732 | 0.37920571 | 0.246406034 |
| 6391.165527 | 4735.919434 | 3557.00415 | -1.134818321 | 0.368127801 | 1.274226766 | 0.514818315 | 0.787493558 | 0.395364978 |

| | | | | | | | | |
|---|---|---|---|---|---|---|---|---|
| 11310.7168 | 11090.2041 | 11626.02441 | -1.137923252 | 0.00953954 | 0.267965219 | 0.198421115 | 0.310909358 | 0.063440273 |
| 18059.0918 | 22172.22461 | 21221.86133 | -1.139459003 | 0.023619216 | 0.200196446 | 0.255621964 | -0.94893061 | 0.015905912 |
| 32681.5 | 32391.97461 | 30036.9082 | -1.139604736 | 0.200756526 | 1.95553387 | 0.174871312 | 3.229137034 | 0.006394096 |
| 35158.49609 | 41015.33984 | 26862.27344 | -1.139635074 | 0.115877831 | 0.074943377 | 0.876454953 | -0.205936116 | 0.495216499 |
| 10199.1084 | 8200.705078 | 7590.500488 | -1.13979438 | 0.025181767 | 0.722574592 | 0.123269689 | 0.47143806 | 0.160292276 |
| 10451.92676 | 10901.48828 | 9067.232422 | -1.140194524 | 0.052584221 | 0.688978731 | 0.034488442 | -0.068332735 | 0.574930993 |
| 8055.807617 | 6465.324707 | 5126.244629 | -1.141298697 | 0.016189975 | 0.593943482 | 0.117092557 | -0.266842958 | 0.363680339 |
| 32386.86719 | 31146.27539 | 24911.91602 | -1.141611958 | 0.009974872 | -0.170862972 | 0.291253414 | -0.429052218 | 0.090965556 |
| 42088.94141 | 79521.35156 | 39743.21875 | -1.141622334 | 0.218843086 | 0.091473867 | 0.903989395 | 0.137250045 | 0.827301412 |
| 17271.46094 | 15406.3252 | 11272.50488 | -1.141705354 | 0.181464022 | -0.173097066 | 0.345466606 | -0.186054729 | 0.505012783 |
| NA | 4546.726074 | 2027.293213 | -1.147471521 | 0.045092515 | 0.377486449 | 0.530439879 | 0.516852745 | NA |
| 51100.98828 | 53056.0625 | 41556.65625 | -1.151601067 | 0.025307586 | 0.157442088 | 0.205952446 | -0.319547994 | 0.129872829 |
| 15571.05566 | 15955.21094 | 12546.80957 | -1.152434579 | 0.012333246 | 0.571342999 | 0.115391022 | -0.336553306 | 0.215760001 |
| 16728.52148 | 16640.6875 | 14288.42188 | -1.154575822 | 0.016105265 | -0.104410347 | 0.749163925 | -0.128131528 | 0.445473288 |
| 227092.7344 | 282452.1563 | 258508.5 | -1.161345659 | 0.00744439 | 0.306023342 | 0.121247389 | 0.379705257 | 0.113248184 |
| 10867.4707 | 8679.387695 | 8613.680664 | -1.164679847 | 0.03900016 | 0.55727742 | 0.16693375 | -0.203593163 | 0.570887829 |
| 5753.972656 | 7139.963379 | 6975.486328 | -1.166423565 | 0.029390153 | -0.669914026 | 0.111682839 | -0.409744305 | 0.16889005 |
| 112250.5625 | 113477.5391 | 119709.2188 | -1.16688604 | 0.041436302 | 0.385108007 | 0.521465225 | 2.142872965 | 0.01858328 |
| 29197.29102 | 27680.66602 | 20538.49219 | -1.166964473 | 0.013723547 | 1.066862421 | 0.058877608 | 0.897766671 | 0.084288402 |
| 231651.4375 | 310537.375 | 251620.5469 | -1.167127308 | 0.005904605 | -0.070756067 | 0.610552707 | -0.00526477 | 0.982643219 |
| 24450.26953 | 24168.0293 | 21377.31055 | -1.16793507 | 0.025944205 | -0.573804843 | 0.13347815 | -0.648732092 | 0.101555637 |
| 158335.9375 | 173889.6875 | 153188.8438 | -1.170597173 | 0.038526889 | 1.843973819 | 2.91E-05 | 2.567236616 | 0.025930246 |
| 29769.78125 | 27758.85547 | 20472.31641 | -1.170677117 | 0.02624826 | 1.036331652 | 0.052579611 | 0.204400725 | 0.593379512 |
| 16040.23926 | 12461.96484 | 13194.71094 | -1.171573461 | NA | 1.773728821 | NA | 2.042102285 | NA |
| 228242.1406 | 213513.0625 | 214972.8125 | -1.171774609 | 0.087167557 | -0.189883999 | 0.095410346 | 0.282280824 | 0.033948107 |
| 1784.627441 | 1076.133301 | 2319.165527 | -1.177163596 | 0.089040397 | -0.643100181 | NA | -0.907688472 | 0.111750711 |
| 448711.3125 | 560592.8125 | 619424.3125 | -1.178867769 | 0.046600136 | -1.072778322 | 0.031122153 | 1.570331869 | 0.063437047 |
| 8327.358398 | 11240.26465 | 13562.97852 | -1.181175052 | 0.134574193 | 0.281847192 | 0.565383328 | 0.837665627 | 0.149640099 |
| 71951.23438 | 60416.04688 | 63071.73438 | -1.185011558 | 0.071550701 | 0.300114449 | 0.418003637 | -0.929225336 | 0.137062098 |
| 4860.121582 | 5111.057617 | 3346.409912 | -1.186091039 | 0.08771097 | 0.493478945 | 0.166408179 | 0.512453827 | 0.2258352 |
| 459893.9688 | 872022.9375 | 351972.9688 | -1.187123482 | 0.185922642 | 0.003429665 | 0.995740584 | -0.556080705 | 0.383863356 |
| 54027.11328 | 58601.43359 | 65415.81641 | -1.187198306 | 0.030855213 | -0.030746387 | 0.732004961 | 0.416738393 | 0.08901833 |
| 27614.0625 | 25839.60938 | 20575.04492 | -1.188231766 | 0.006564049 | 0.346311083 | 0.052756858 | -0.690865576 | 0.038799047 |

| | | | | | | | | |
|---|---|---|---|---|---|---|---|---|
| 14476.74805 | 3399.046631 | 20084.68359 | -1.188247625 | 0.106456043 | 2.112795779 | 0.417803644 | 0.956312013 | 0.454451558 |
| 186021.6094 | 183065.8281 | 179888.6875 | -1.191902853 | 0.075250823 | 0.762738412 | 0.352309135 | 0.354699371 | 0.074184841 |
| NA | 650.1316528 | NA | -1.193488136 | 0.006395507 | 0.02754671 | 0.939293277 | -2.595139434 | NA |
| 59832.17188 | 93554.82813 | 53397.33203 | -1.194596746 | 0.101686005 | 0.074511707 | 0.88886483 | -0.188808288 | 0.668752886 |
| 1828.054565 | 1887.17749 | 474.056488 | -1.195446491 | 0.224229053 | 0.296346398 | 0.431724911 | -0.487390403 | 0.487511859 |
| 49970.88281 | 48628.87109 | 52167.83594 | -1.196951022 | 0.027696987 | 1.20396715 | 0.067394457 | 1.38893036 | 0.009703417 |
| 24920.02344 | 20025.78516 | 18260.35352 | -1.200523521 | 0.071056032 | -0.019974772 | 0.94549331 | 0.458561285 | 0.168872074 |
| 51595.76953 | 46896.80078 | 44075.69141 | -1.202432616 | 0.030269876 | 0.649785662 | 0.051097951 | 0.180637672 | 0.354977058 |
| 4216.710938 | 3453.528809 | 1509.3396 | -1.202459553 | NA | 5.549964735 | NA | 1.747467396 | NA |
| 11501.42285 | 8402.077148 | 4568.081543 | -1.202494662 | 0.032490498 | -0.167388873 | 0.737531604 | -1.465391354 | 0.12298361 |
| 19348.03906 | 24795.42969 | 18384.88281 | -1.203090856 | 0.015175173 | 0.222192964 | 0.462700416 | -0.248571516 | 0.389796538 |
| 8724.411133 | 8922.075195 | 7048.321289 | -1.209084143 | 0.056696683 | 1.300857524 | 0.107969678 | -0.118816131 | 0.682933573 |
| 5619.316895 | 5165.060059 | 3596.549805 | -1.20931024 | 0.189289028 | 1.228382389 | 0.221634958 | 0.644969738 | 0.168323482 |
| 31578.84766 | 59843.48438 | 25501.51953 | -1.217077788 | 0.101183195 | 0.132484053 | 0.84622109 | -0.073368928 | 0.914576246 |
| 106609.1563 | 127542.5234 | 76111.73438 | -1.218971063 | 0.080233873 | -1.002439299 | 0.14612569 | -1.188039884 | 0.118163173 |
| 5306.825195 | 3894.110352 | 4695.041504 | -1.222129073 | 0.052584604 | 0.204930015 | 0.681063929 | 0.206457977 | 0.481431043 |
| 3262.246338 | 3233.19165 | 3876.260254 | -1.223515392 | 0.053424232 | 0.280984049 | 0.533779394 | 0.26287295 | 0.552026106 |
| NA | 2133.432617 | 10314.91992 | -1.227573799 | NA | 1.689705078 | NA | 0.885620619 | NA |
| 16963.57422 | 22127.05859 | 15664.36719 | -1.230838789 | 0.139278926 | 0.467035752 | 0.194396063 | 0.10871367 | 0.720635723 |
| 145814.25 | 131160.2031 | 146974.5 | -1.232473358 | 0.032484233 | 0.556657983 | 0.140576585 | 1.15463976 | 0.027868741 |
| 12943.83887 | 11937.67578 | 9935.319336 | -1.232699475 | 0.030588379 | 1.208271004 | 0.059029787 | -0.390855919 | 0.448447963 |
| NA | 3167.955566 | 2535.991699 | -1.23809532 | 0.052321316 | -0.045878966 | 0.888375183 | 0.130305809 | NA |
| NA | NA | NA | -1.238565738 | NA | -1.974724735 | NA | NA | NA |
| 130485.3672 | 125292.2578 | 173159.2344 | -1.238608295 | 0.009337528 | 0.012193695 | 0.955923696 | 0.6391582 | 0.134445271 |
| 1534.673096 | 1225.118408 | 591.4165649 | -1.240568703 | 0.047142848 | 0.516374632 | 0.327611438 | 0.075239397 | 0.900311098 |
| 33648.97656 | 29033.67578 | 46514.75781 | -1.244982344 | 0.263383916 | 1.613450679 | 0.256103156 | 1.21648351 | 0.118023574 |
| 104527.1172 | 126762.6172 | 55190.23438 | -1.246298458 | 0.039721652 | -0.136242359 | 0.374951761 | -0.736213297 | 0.156978352 |
| 20294.53711 | 27153.76953 | 18626.31836 | -1.249812923 | 0.147088229 | 0.153142791 | 0.666618357 | 0.023089228 | 0.948081141 |
| 13089.87012 | 13257.97168 | 10414.40332 | -1.252861985 | 0.037600114 | -0.503401594 | 0.091622784 | -0.766332112 | 0.040786554 |
| 53926.83984 | 63304.13281 | 54495.08984 | -1.25436938 | 0.008009891 | 0.051909074 | 0.765250128 | 0.031890428 | 0.861343486 |
| 17729.83789 | 18086.10156 | 17397.12695 | -1.256208335 | 0.027930748 | 0.320322687 | 0.06494295 | 0.193957035 | 0.022924177 |
| 83878.90625 | 63955.82813 | 84347.32031 | -1.26044441 | 0.012367336 | -0.051191789 | 0.553204641 | -0.075374032 | 0.695643018 |
| 18951.24414 | 24512.84375 | 14534.20898 | -1.260821602 | 0.194842086 | 0.122385087 | 0.816632277 | 0.196929452 | 0.647709869 |

| | | | | | | | | |
|---|---|---|---|---|---|---|---|---|
| 24428.14453 | 28751.64063 | 22780.94727 | -1.26762477 | 0.015044187 | -0.077908221 | 0.747060046 | -0.41651811 | 0.215213867 |
| 15418.56836 | 12995.8252 | 15033.73926 | -1.267649437 | 0.036590352 | 0.035582417 | 0.87026131 | -0.334102326 | 0.173821986 |
| 7196.814941 | 7344.299805 | 3718.067871 | -1.271910603 | 0.016136356 | 0.346530269 | 0.240374068 | -0.364130058 | 0.385275383 |
| 21672.4668 | 16623.51563 | 16631.83594 | -1.273617686 | 0.027035662 | 1.21368588 | 0.023005377 | 0.520425323 | 0.136000191 |
| 324931.4063 | 344033.5 | 330933.8125 | -1.274370741 | 0.004709568 | -0.08746352 | 0.366273258 | 0.594879587 | 0.017279709 |
| 2923.078857 | 2941.515625 | 2174.071777 | -1.280436249 | 0.006735623 | 0.380382847 | 0.098408786 | -0.068254427 | 0.766301418 |
| 94176.85938 | 74659.35156 | 440252.5625 | -1.280775478 | 0.167266141 | 1.603377616 | 0.518528002 | 0.918265181 | 0.604954789 |
| 6362.000488 | 6415.1875 | 5533.209473 | -1.28248194 | 0.087897715 | 1.398879453 | 0.088308366 | 2.433860705 | 0.014448708 |
| 34886.52734 | 39819.91016 | 29475.75195 | -1.289113989 | 0.003898264 | 0.685457593 | 0.054209378 | -0.401679149 | 0.123837484 |
| 65815.52344 | 83757.13281 | 120110.0391 | -1.292539646 | 0.017824776 | 1.387782115 | 0.018539883 | 1.980903941 | 0.1187593 |
| NA | 1126.642212 | NA | -1.296652875 | 0.06866705 | 0.789886809 | 0.219649135 | 0.48083186 | NA |
| 20912.82227 | 16675.70313 | 12396.68457 | -1.29974895 | 0.034240061 | -0.302318046 | 0.297911261 | -1.039195162 | 0.061670859 |
| 20918.9043 | 20627.77148 | 21872.70117 | -1.303451779 | 0.027470745 | 0.579882364 | 0.046373997 | 0.516966427 | 0.015233843 |
| 11239.14063 | 11721.59961 | 10566.59668 | -1.306551725 | 0.152253903 | 0.893591564 | 0.455732934 | 1.747485359 | 0.017279709 |
| 3132.484619 | NA | 8591.589844 | -1.31002211 | 0.033464629 | -0.625333681 | NA | 0.289522786 | NA |
| 47049.03516 | 45194.10938 | 46019.57813 | -1.311758842 | 0.004709568 | 0.075514547 | 0.327735 | 0.187589832 | 0.069553713 |
| 20680.03516 | 19427.44531 | 21654.10742 | -1.314458238 | 0.028452905 | -0.566093525 | 0.160878456 | -0.802860067 | 0.12131068 |
| 479325.8438 | 496513.25 | 558233.125 | -1.315398474 | 0.015030322 | 1.638038476 | 0.089248231 | 1.603863586 | 0.030832901 |
| 27648.08398 | 25253.63086 | 22901.99414 | -1.316125758 | 0.049572442 | -0.177153647 | 0.24146716 | -0.427047875 | 0.066335881 |
| 2150508.5 | 2510698 | 3235874 | -1.319614769 | 0.039911484 | 0.271652497 | 0.45527367 | -0.1592839 | 0.713313063 |
| 14887.646 | 11331.17902 | 12350.34766 | -1.31971813 | 0.011817481 | 0.442668251 | 0.131269584 | 0.149531515 | 0.557774325 |
| 77334.20313 | 67299.17188 | 60106.72266 | -1.320709102 | 0.02930047 | -0.129182235 | 0.393390579 | -1.034348574 | 0.016834605 |
| NA | NA | 8112.352051 | -1.325436207 | NA | 0.037656788 | NA | -0.391847255 | NA |
| 268843.625 | 271271.7813 | 262081.6406 | -1.326633191 | 0.014955435 | 0.618724238 | 0.035356834 | 0.222170459 | 0.118785258 |
| 1108.124878 | NA | NA | -1.33244064 | 0.135239277 | NA | NA | -1.338609087 | NA |
| 175399.75 | 174508.0469 | 162526.3281 | -1.333662609 | 0.009974872 | 0.682908675 | 0.082833891 | 0.210959581 | 0.056962705 |
| 25815.6543 | 39027.59375 | 24364.83398 | -1.33445828 | 0.076775419 | 0.209792777 | 0.693736734 | 0.0474019 | 0.901062666 |
| 12900.47559 | 10640.90723 | 9253.089844 | -1.338086334 | 0.104512489 | 1.653567624 | 0.094777667 | 2.42220639 | 0.031136253 |
| 19293.97852 | 20775.07031 | 19997.09961 | -1.33974506 | 0.066639151 | 1.012399008 | 0.415338319 | 2.012755786 | 0.00797582 |
| 10550.72363 | 9151.892578 | 7605.170898 | -1.341691916 | 0.005752145 | -0.222725388 | 0.157377951 | -0.772906538 | 0.035612683 |
| 243792.3906 | 221242.2188 | 254907.4688 | -1.343149875 | 0.020651493 | -0.614747622 | 0.012254503 | -0.102252668 | 0.316709896 |
| 3112.800293 | 3627.876953 | 1735.556152 | -1.345451259 | 0.300318309 | 0.480450555 | 0.541557838 | 1.073513582 | 0.221229381 |
| 12444.96582 | 9842.620117 | 10377.67383 | -1.348892472 | 0.145015199 | -0.06294833 | 0.731605116 | -0.279526194 | 0.253731741 |

| | | | | | | | | |
|---|---|---|---|---|---|---|---|---|
| 3777.072998 | 4038.446533 | 3999.604248 | -1.352791293 | 0.005752145 | 0.899614834 | 0.018539883 | 0.476001466 | 0.051693365 |
| 3360.470703 | 1985.362305 | 1437.919312 | -1.363338132 | 0.031097865 | 0.232286758 | 0.572580393 | -0.23548744 | 0.713313063 |
| 16496.17383 | 21826.93945 | 15599.23438 | -1.366569034 | 0.012364839 | -0.25275707 | 0.265670827 | 0.187649096 | 0.487850018 |
| 17847.92773 | 17693.125 | 12232.56934 | -1.366639176 | 0.004709568 | -0.076352955 | 0.546574023 | -0.427333424 | 0.15573211 |
| 61057.91797 | 64905.0625 | 68787.61719 | -1.367163239 | 0.00744439 | 1.060385744 | 0.038407677 | 0.682556284 | 0.023619478 |
| 100365.1328 | 124464.6484 | 130733.8594 | -1.368235367 | 0.023401535 | 1.295052978 | 0.029180754 | 1.025776175 | 0.047120928 |
| 24893.43359 | 25975.42969 | 25759.53516 | -1.379591709 | 0.025187448 | 0.318314411 | 0.183085834 | 0.229061497 | 0.225533768 |
| 156391.625 | 149874.875 | 143011 | -1.379995481 | 0.057336343 | 0.410261447 | 0.134845721 | 0.114975296 | 0.20861125 |
| 148108.5625 | 131749.5625 | 113249.3281 | -1.384742811 | 0.012750417 | 0.268204324 | 0.067039668 | 0.16313315 | 0.408925574 |
| 3760.731445 | 3647.783691 | 1756.52002 | -1.389376479 | 0.042292137 | 0.061134299 | 0.63755951 | -0.459682753 | 0.31960533 |
| 18855.81116 | 28179.04688 | 28096.64941 | -1.390343849 | 0.009072032 | -1.636672193 | 0.046739326 | -1.735947157 | 0.021984636 |
| 21338.45703 | 33022.52344 | 15093.13672 | -1.393569497 | 0.164028708 | 0.40232327 | 0.573219709 | 0.065253159 | 0.920227312 |
| 18985.99414 | 18646.26758 | 17153.50977 | -1.395207018 | 0.055037162 | 0.459670188 | 0.240555748 | 0.132797995 | 0.607583742 |
| 5177.566406 | 3904.053223 | 4095.79834 | -1.39761018 | 0.038106838 | 0.442863814 | 0.080723218 | -0.129427426 | 0.528354562 |
| 200050.8594 | 211446.9844 | 194874.375 | -1.40310683 | 0.039126605 | 0.908527843 | 0.012254503 | 0.799976288 | 0.011723127 |
| 13925.46582 | 9162.130859 | 12982.87695 | -1.404103115 | 0.033270591 | 0.180937447 | 0.671961254 | 0.043227617 | 0.904535319 |
| 4231.537109 | 3455.120605 | 3184.57959 | -1.40510181 | 0.061453753 | 0.343546358 | 0.124972519 | -0.157463631 | 0.500325294 |
| 30051.19141 | 30036.58203 | 26323.24219 | -1.408541507 | 0.028047191 | 1.188380027 | 0.028434447 | 0.736469699 | 0.054887084 |
| 10220.43164 | 15517.4375 | 27953.73242 | -1.409874909 | 0.171174352 | -0.475370916 | 0.604209303 | -0.422397032 | 0.663201766 |
| 4536.182129 | 4574.772949 | 4900.469238 | -1.410382763 | 0.012414001 | -0.089104936 | 0.675235839 | -0.670814439 | 0.117593322 |
| 966.0788574 | 1128.103638 | NA | -1.411296381 | 0.017077173 | 0.472762764 | 0.176015705 | -0.782706682 | NA |
| 63742.61719 | 15033.41113 | 3715.450684 | -1.414123796 | 0.316898283 | 1.173939751 | 0.105107021 | 1.77332272 | 0.511169671 |
| 44612.65625 | 54894.26123 | 51212.89844 | -1.41648661 | 0.006710768 | 0.929916537 | 0.016400229 | 0.766597538 | 0.045009854 |
| 49227.25 | 47767.55469 | 45598.33203 | -1.417596931 | 0.021667504 | 0.405360865 | 0.056128038 | 0.038288402 | 0.577343498 |
| 2160.388916 | 1591.930542 | 1241.486084 | -1.42162001 | 0.070727326 | 0.529150181 | 0.390625946 | 0.007879779 | 0.991353382 |
| 28449.83789 | 32838.37109 | 24278.22461 | -1.422814521 | 0.042840863 | 1.116464851 | 0.125350518 | 0.347582532 | 0.545099828 |
| 152925.5938 | 146638.9844 | 158493.1406 | -1.425664906 | 0.032646789 | 0.527272277 | 0.027730297 | 0.809773017 | 0.010623938 |
| 43686.13672 | 19543.63086 | 125555.9922 | -1.426655272 | 0.119247856 | 1.499075639 | 0.534693749 | 0.957924782 | 0.546266077 |
| NA | 341.3919373 | NA | -1.427132368 | 0.081995281 | -0.088350474 | 0.903989395 | -1.916998666 | NA |
| 126285.4609 | 135717.8906 | 144884.5156 | -1.427302229 | 0.038502747 | 0.710992078 | 0.087342285 | 1.323121745 | 0.017972665 |
| 8326.439453 | 8495.25 | 8832.43457 | -1.427829952 | 0.024037284 | 0.55461664 | 0.078072776 | 0.623061373 | 0.017511333 |
| 4630.996094 | NA | NA | -1.431039611 | NA | 0.131653622 | NA | -0.331238841 | NA |
| 15149.21973 | 13820.33594 | 13419.86328 | -1.434327104 | 0.004709568 | 0.892333821 | 0.036723028 | 0.197903127 | 0.129496281 |

| | | | | | | | | |
|---|---|---|---|---|---|---|---|---|
| 2425.568848 | 2874.67749 | 1391.834595 | -1.43449246 | 0.052356162 | -0.133877172 | 0.610552707 | -1.006457395 | 0.071851903 |
| NA | NA | NA | -1.437312366 | NA | -0.155653461 | NA | NA | NA |
| 4138.882324 | 4878.102539 | 3734.633545 | -1.438575278 | 0.025086961 | 0.358397402 | 0.130039916 | -1.287457795 | 0.047847186 |
| 34143.11719 | 20814.17773 | 26616.97266 | -1.439162423 | 0.201598743 | 0.745698659 | 0.066365764 | 0.208962894 | 0.548141599 |
| 61516.40234 | 54458.29297 | 58351.01563 | -1.440076661 | 0.009072032 | 0.885107119 | 0.065871863 | 0.861399966 | 0.019799501 |
| 102531.0469 | 85082.35156 | 72362.41406 | -1.442798928 | 0.156655374 | 1.000717467 | 0.414380687 | 2.163745491 | 0.060309077 |
| 7661.525391 | NA | 29478.93945 | -1.457588444 | NA | 1.423229297 | NA | 1.466859545 | NA |
| 23039.85156 | 24804.98242 | 23477.15625 | -1.459585095 | 0.038502747 | -0.498611842 | 0.034840225 | -0.295162875 | 0.052505925 |
| 3275.835693 | 3929.820801 | 3654.559814 | -1.459857534 | 0.07287987 | -0.181446459 | 0.527065457 | -0.392770367 | 0.18657303 |
| NA | 144.3400726 | NA | -1.461121954 | NA | NA | NA | -0.149240959 | NA |
| 4333.73877 | 5030.205566 | 5545.667969 | -1.4675697 | 0.029906057 | -0.123827649 | 0.834946164 | -0.67796537 | 0.355531806 |
| 4113.684082 | NA | 2177.077393 | -1.467797008 | 0.029391485 | -0.42200196 | NA | -0.519653722 | NA |
| 2531.373291 | 2177.190186 | 2020.131104 | -1.469648329 | 0.030049926 | -0.240903657 | 0.479790187 | -0.594202059 | 0.198576209 |
| 19926.94727 | 19690.42969 | 17118.63672 | -1.471785222 | 0.015991503 | 1.269968691 | 0.015625705 | 0.789116238 | 0.03161115 |
| 8335.378906 | 8902.412109 | 7199.774414 | -1.476641028 | 0.015685639 | -0.074623917 | 0.705913085 | 0.639274384 | 0.053570169 |
| 42515.57031 | 55484.32422 | 56973.98828 | -1.477333388 | 0.040256153 | -1.712203271 | 0.046369992 | -1.106140606 | 0.04016603 |
| 8502.963867 | 11717.61133 | 12913.15723 | -1.482983537 | 0.025818052 | -0.398951829 | 0.28570212 | 1.721736663 | 0.079165593 |
| 13353.97559 | 4670.680176 | NA | -1.48598219 | 0.025453132 | 0.873697598 | 0.122107272 | 0.612977947 | NA |
| 213386.0996 | 211295.5225 | 178292.9146 | -1.488907418 | 0.005790576 | -0.576886119 | 0.099830362 | -1.379844309 | 0.02163333 |
| 1212.284668 | 331.1016846 | 2447.869385 | -1.496674335 | 0.295346854 | 3.156185288 | 0.387750232 | -0.051570825 | 0.980712992 |
| 4512.910645 | 6090.053711 | 3248.437744 | -1.497172057 | 0.135044793 | 0.032569127 | 0.943661372 | -0.783707702 | 0.26766605 |
| 73958.58594 | 67926.10938 | 61192.21094 | -1.505156197 | 0.084819075 | 1.099979171 | 0.504715483 | 2.223746736 | 0.025666912 |
| 1024.52063 | 627.2304077 | 562.6016235 | -1.506153631 | 0.034918419 | -0.397670901 | NA | -0.647510907 | 0.197103593 |
| 10914.13281 | 9923.828125 | 9147.330078 | -1.508265795 | 0.005944135 | 0.317380736 | 0.02383791 | -0.352868725 | 0.065303359 |
| 18093.90039 | 5141.344238 | 52570.52734 | -1.508862817 | 0.120050919 | 1.780861456 | 0.41703591 | 1.00436813 | 0.566209266 |
| 7186.300293 | 13888.08594 | 8332.515625 | -1.510354983 | 0.074826564 | 0.95185813 | 0.291036386 | 0.76675307 | 0.376019906 |
| 30550.93359 | 26397.875 | 16759.54883 | -1.515218804 | 0.013317304 | 0.297475251 | 0.395202582 | -0.176116687 | 0.695549453 |
| 41336.89063 | 41546.00781 | 36799.72656 | -1.529392818 | 0.005790576 | -0.07023261 | 0.62182213 | 0.04182779 | 0.765741675 |
| 1590433 | 1713574.125 | 1657203 | -1.53726169 | 0.063328311 | 1.233740037 | 0.052817461 | 0.38957726 | 0.073420995 |
| 51665.57813 | 80876.17969 | 55561.41016 | -1.537483106 | 0.101951046 | 0.155390663 | 0.745775286 | 0.076073345 | 0.84431148 |
| 17178.4375 | 7903.651855 | 11007.73047 | -1.548283965 | 0.019126217 | 0.160555544 | 0.861576827 | -0.160349391 | 0.79356962 |
| 20956.38867 | 17959.55078 | 16185.56836 | -1.549747689 | 0.107619023 | -0.156126714 | 0.612877707 | -0.398924989 | 0.290771554 |
| 167759.5938 | 166752.2969 | 170845.9531 | -1.552703209 | 0.10870824 | 1.939701301 | 0.012690505 | 3.303144091 | 0.000739989 |

| | | | | | | | | |
|---|---|---|---|---|---|---|---|---|
| 84727.90625 | 124034.1641 | 64978.6875 | -1.557307111 | 0.057253208 | 0.304373002 | 0.557007626 | -0.147784507 | 0.765741675 |
| 62890.98828 | 105651.5547 | 79500.57031 | -1.563067543 | 0.081796003 | 0.307306214 | 0.60094054 | 0.967969514 | 0.135684801 |
| 3641.683105 | 2304.955811 | 2662.13916 | -1.566507483 | 0.02194812 | -0.065120746 | 0.914856491 | -0.20593583 | 0.716051255 |
| NA | NA | NA | -1.575762985 | NA | 0.527188049 | NA | NA | NA |
| 18422.12012 | 19919.91309 | 16546.42725 | -1.576302647 | 0.138728728 | 0.471016932 | 0.634593889 | 0.227219125 | 0.690244068 |
| 5499.871094 | 8040.120605 | 8328.064453 | -1.577221471 | 0.014328894 | 0.105267924 | 0.785487872 | 0.760794053 | 0.136837137 |
| NA | 17738.64063 | NA | -1.57737566 | 0.029750807 | 0.40196289 | NA | 0.91776056 | NA |
| 19543.65625 | 20274.58789 | 24118.70313 | -1.578260094 | 0.066744288 | -0.069978688 | 0.946377163 | 0.662125828 | 0.426273569 |
| 7737.474121 | 4167.876465 | NA | -1.579001152 | NA | 0.668305757 | NA | 0.875194578 | NA |
| 12483.20313 | 11987.32324 | 11026.37891 | -1.591575395 | 0.032969545 | 1.925491605 | 0.114167274 | 1.384789684 | 0.012916212 |
| 35431.25 | 25783.33398 | 183021.7344 | -1.593182191 | 0.081289628 | 1.825869826 | 0.419290117 | 1.18055617 | 0.566251903 |
| 283546.5938 | 288102.0625 | 229706.1094 | -1.593754602 | 0.064602236 | 0.912081144 | 0.048024604 | -0.346102349 | 0.260534667 |
| 174779.3281 | 49802.95703 | 541392.5625 | -1.595497903 | 0.077105145 | 1.962507706 | 0.340513249 | 1.12014692 | 0.557774325 |
| 40075.08203 | 40191.61328 | 39907.03125 | -1.598005683 | 0.02237899 | 0.469950753 | 0.197534048 | 1.269319407 | 0.049004518 |
| 60165.46094 | 74510.17969 | 80869.01563 | -1.607371976 | 0.017524813 | -0.246283437 | 0.098031162 | -0.261609765 | 0.223102126 |
| 44086.85938 | 43832.60156 | 39543.98828 | -1.617579439 | 0.009657125 | 0.060331567 | 0.525306265 | -0.538689805 | 0.027979863 |
| 87953.82813 | 77939.01563 | 64460.40625 | -1.617863969 | 0.04443276 | 1.259002128 | 0.048593497 | 0.534372152 | 0.24207605 |
| 229175.9219 | 25255.83203 | 387412.5313 | -1.632677113 | 0.11060902 | 2.682836324 | 0.242699931 | 1.150177062 | 0.497782932 |
| 2869.325928 | 914.6201172 | 2830.593994 | -1.632692416 | 0.354269327 | 0.880380235 | 0.174475307 | -0.519043346 | 0.54796267 |
| 7225.155273 | 6427.133789 | 4027.644775 | -1.644077578 | NA | 1.088027696 | NA | 0.948029517 | NA |
| 7750.061035 | 6133.901367 | 5453.961426 | -1.64443593 | 0.056893333 | 1.537634823 | 0.100538317 | 0.228399049 | 0.573238987 |
| 76251.57031 | 67727.36719 | 68940.51563 | -1.648099442 | 0.036506605 | 0.355589729 | 0.075842188 | -0.103285601 | 0.436906116 |
| 31833.27734 | 35851.28516 | 44915.94922 | -1.649489262 | 0.010463664 | -0.560662968 | 0.149299933 | 1.614221953 | 0.062345205 |
| 35942.16016 | 43034.64844 | 34460.00781 | -1.65415231 | 0.04816919 | -0.017295818 | 0.955166858 | -0.313755972 | 0.193350927 |
| 121732.7813 | 121420.6953 | 88163.07813 | -1.65719712 | 0.138429888 | 0.196635013 | 0.537721043 | 0.332993691 | 0.289284863 |
| 83382.77344 | 83148.34375 | 76597.88281 | -1.667140138 | 0.019202395 | -0.03051758 | 0.777841261 | 0.083562221 | 0.436264113 |
| 14791.09766 | 6724.997559 | 12558.08887 | -1.670161444 | 0.100789035 | 0.433292102 | 0.359867822 | 1.392282957 | 0.175598599 |
| 31577.99414 | 36592.57813 | 29681.20703 | -1.675167241 | 0.005752145 | -0.645412809 | 0.06824263 | -0.452301153 | 0.085236358 |
| 4135.234375 | 3939.18042 | 3870.51709 | -1.675183097 | 0.011814081 | 0.425046799 | 0.062726463 | 0.28159796 | 0.035953811 |
| 50441.16016 | 12094.14453 | 119898.7188 | -1.675187968 | 0.171832714 | 2.02511341 | 0.286186534 | 0.971266052 | 0.563099327 |
| 4248.090332 | NA | 3884.90625 | -1.688691633 | 0.072654291 | 0.990462106 | 0.112543356 | -0.345932285 | NA |
| 22998.69727 | 26527.68945 | 30491.64258 | -1.691502722 | 0.025960199 | 1.371565614 | 0.159908538 | 3.098356581 | 0.042737273 |
| 26763.49805 | 21852.52734 | 19883.95508 | -1.692035696 | 0.008012423 | 0.928860036 | 0.184506133 | 1.109333894 | 0.057068534 |

| | | | | | | | | |
|---|---|---|---|---|---|---|---|---|
| 6967.993652 | 8723.498047 | 7205.352539 | -1.692340497 | 0.071940398 | 1.302415288 | 0.065601851 | -0.628648903 | 0.413241482 |
| 437267.4688 | 497524.75 | 541126.4375 | -1.7016167 | 0.029750807 | 1.055915776 | 0.147741919 | 1.69823763 | 0.022638191 |
| 16313.54199 | 17777.81836 | 12197.47363 | -1.707596677 | 0.010893509 | -0.242672026 | 0.312416108 | -0.578936788 | 0.083213718 |
| 49526.09766 | 60538.10547 | 55512.14453 | -1.711318146 | 0.077066654 | 1.125304879 | 0.026826422 | 1.003112903 | 0.030044909 |
| 30619.05078 | 31601.96875 | 26378.04492 | -1.714409507 | 0.081861007 | 0.384283289 | 0.524701907 | 1.825622374 | 0.019816721 |
| 182139.7031 | 144397.0469 | 138398.1094 | -1.72059658 | 0.095307674 | -1.525125041 | 0.157797838 | -0.122659715 | 0.790450343 |
| 62027.48828 | 47995.90625 | 43305.34766 | -1.727274066 | 0.009974872 | 1.475031177 | 0.042884533 | 0.719178004 | 0.12131068 |
| 10240.70703 | 11881.05859 | 12264.65723 | -1.728528098 | 0.026113102 | 0.787525776 | 0.078283911 | 0.650818883 | 0.087278746 |
| 51221.92969 | 47475.08984 | 49033.72266 | -1.730554017 | 0.006351124 | 0.054569825 | 0.817133443 | 0.556219991 | 0.089464864 |
| 10668.36914 | 3907.606201 | 5924.37793 | -1.731416189 | 0.047406642 | -0.068524491 | 0.812069222 | -0.099682479 | 0.881517111 |
| 15512.42773 | 16677.07813 | 4960.175781 | -1.731493045 | NA | 1.14328636 | NA | 0.27914423 | NA |
| 30713.49609 | 30351.79102 | 28831.80469 | -1.737081817 | 0.013585622 | 0.784888683 | 0.077511889 | 0.485114496 | 0.178911811 |
| 73228.65625 | 77474.48438 | 90110.78906 | -1.738948796 | 0.020740891 | -0.039566045 | 0.836170745 | 0.515630267 | 0.074889461 |
| 27904.02734 | 28618.5625 | 17605.35938 | -1.739380191 | 0.034890445 | 1.238001495 | 0.03588199 | 0.401922002 | 0.339006931 |
| NA | NA | NA | -1.755481439 | NA | NA | NA | NA | NA |
| 709677.125 | 503860.375 | 496345.0625 | -1.762257604 | 0.03777619 | 0.270298911 | 0.817213245 | 3.736011504 | 0.062835547 |
| 11076.70996 | 13763.11328 | 13280.60645 | -1.7752988 | 0.018310596 | 0.321115425 | 0.338904732 | 0.099669299 | 0.745242088 |
| 20298.41992 | 17933.67383 | 21546.88867 | -1.776775121 | 0.009194956 | -0.113260195 | 0.324535676 | 0.756501455 | 0.043970318 |
| 41594.10938 | 44232.04688 | 45348.125 | -1.801864666 | 0.004709568 | -0.388605228 | 0.24186282 | 0.007933423 | 0.963218346 |
| 2909.436768 | 3052.467285 | 2420.814697 | -1.804234557 | NA | -0.434333824 | NA | -0.258754848 | NA |
| 2280.050781 | 3988.936035 | 2323.512207 | -1.809680794 | 0.069311915 | -2.481552202 | 0.04654357 | -2.934909823 | 0.031136253 |
| 5730.135742 | 5494.9375 | 6423.566406 | -1.813265451 | 0.022288357 | 0.525994101 | 0.278495147 | 0.805255921 | 0.124562645 |
| 9916.585938 | 10749.29297 | 4154.89502 | -1.821062481 | 0.063885064 | 0.184475078 | 0.688199662 | 0.008974031 | 0.989480447 |
| 408828.918 | 120877.1934 | 279568.4258 | -1.830515826 | 0.146746483 | -2.730758899 | 0.237779438 | -1.177324254 | 0.355778725 |
| NA | 861.2233276 | NA | -1.863673673 | 0.020651493 | -0.512849772 | NA | 0.471756956 | NA |
| 24244.43555 | 22269.66797 | 23252.51953 | -1.902544615 | 0.029061881 | 0.282697854 | 0.230614464 | 0.462994887 | 0.097278088 |
| 33000.57813 | 34029.97656 | 62676.375 | -1.919610125 | 0.089412683 | 0.582063746 | 0.365007609 | 1.926743367 | 0.154835571 |
| 56835.01563 | 50352.86328 | 40704.41797 | -1.923031199 | 0.079968566 | 0.711402381 | 0.406071842 | 0.004124421 | 0.988723353 |
| 23652.08984 | 23631.28711 | 24460.70703 | -1.933541615 | 0.004709568 | -0.375033857 | 0.159573401 | -0.442872811 | 0.143807259 |
| 31425.10938 | 41983.95313 | 41276.62891 | -1.937500651 | 0.062989607 | 0.126540897 | 0.563537425 | -0.325762462 | 0.249456378 |
| 29744.44141 | 39211.36328 | 42813 | -1.94000375 | 0.033675729 | -0.704224985 | 0.20617006 | -0.125020648 | 0.743582483 |
| 15210.46387 | 16066.97852 | 15994.44141 | -1.954013188 | 0.01904327 | 0.863707305 | 0.197242348 | 1.894121289 | 0.029710029 |
| 15940.96094 | 12043.31543 | 13287.88867 | -1.966075342 | 0.010484689 | 1.127273398 | 0.020431201 | 0.562970884 | 0.106413249 |

| | | | | | | | | |
|---|---|---|---|---|---|---|---|---|
| 10613.10547 | 7714.940918 | 12066.02734 | -1.967978149 | 0.023298213 | 0.738555457 | 0.197998947 | 0.46884672 | 0.374268561 |
| 1231356 | 655131.1875 | 1912629.625 | -1.985956223 | 0.169115491 | 1.238812547 | 0.414751124 | 0.57281937 | 0.481758088 |
| 70429.125 | 69615.34375 | 55000.57031 | -1.990027217 | 0.022444016 | 0.251447209 | 0.11146465 | 0.501384213 | 0.113248184 |
| 2958.260742 | 1856.585693 | 1989.661499 | -1.991446459 | 0.025680323 | 0.280497453 | 0.399209403 | -0.922980273 | 0.07498931 |
| 2394.861816 | 3767.108398 | 2885.956299 | -1.993692048 | 0.018310596 | 0.150986287 | 0.582455701 | -0.317945695 | 0.327784837 |
| 2004.829834 | 3446.220947 | 1880.651245 | -2.00443374 | 0.06572043 | 0.144073329 | 0.697931686 | -0.541412706 | 0.249456378 |
| 93042.61914 | 101703.7617 | 90387.73901 | -2.025353257 | 0.024701492 | -0.094900183 | 0.909761575 | -0.098350683 | 0.9064439 |
| 11522.98242 | 6248.359863 | 8730.307617 | -2.043159026 | 0.02753514 | 1.848955837 | 0.052691197 | 1.326961216 | 0.144898741 |
| 3464.21167 | 4530.785156 | 2894.397705 | -2.073501487 | 0.027405706 | 0.614304312 | 0.068812228 | -0.183237915 | 0.57264617 |
| 5524.154785 | 9630.93457 | 8136.009277 | -2.083620651 | 0.029514505 | 0.308946134 | 0.660549779 | 0.47056533 | 0.370433388 |
| 970012.9785 | 965099.6221 | 840439.6997 | -2.091929524 | 0.012414001 | 0.607318823 | 0.167467663 | -1.144147669 | 0.009587646 |
| 17652.22266 | 23505.29688 | 19958.22461 | -2.094589469 | 0.012750417 | 0.407157576 | 0.19074133 | 0.119051959 | 0.67072268 |
| NA | 5788.032227 | 3153.595459 | -2.105457678 | NA | 1.497808448 | NA | 0.814850038 | NA |
| 17486.43555 | 18125.81055 | 18959.45117 | -2.119955632 | 0.047364099 | -0.13026137 | 0.710906719 | -0.643534888 | 0.208350613 |
| 8329.625977 | 5738.331543 | 10825.97754 | -2.132800674 | 0.019286562 | 0.331758395 | 0.171581754 | 0.085522015 | 0.841259847 |
| NA | 1721.691528 | NA | -2.144581887 | NA | -0.537149525 | NA | -0.093205476 | NA |
| 31452.90234 | 41663.41406 | 49298.81641 | -2.15585032 | 0.01446452 | -0.095555816 | 0.85722804 | 0.737458968 | 0.173157377 |
| 6201.301758 | 4988.960938 | 9659.449219 | -2.167111874 | 0.207480826 | 0.630370986 | 0.320787116 | 0.223395714 | 0.713487699 |
| 199264.9063 | 183447.4063 | 194044.875 | -2.169658723 | 0.039443869 | 0.639177812 | 0.237987125 | -1.290304699 | 0.050810933 |
| 11908.38965 | 11064.28711 | 9466.121094 | -2.186225475 | 0.015732345 | 2.311259261 | 0.104692976 | 0.884851356 | 0.057202212 |
| 1195.625977 | 2620.227539 | 587.2550659 | -2.186239534 | 0.01031238 | 1.46624367 | 0.353137578 | 0.753206695 | 0.536319749 |
| 48794.67407 | 42440.61914 | 41758.60645 | -2.190234407 | 0.066744288 | 0.373511077 | 0.037014778 | -0.186948728 | 0.178181856 |
| 170360.5625 | 150490.1719 | 149523.6719 | -2.20673646 | 0.025960199 | 1.638987672 | 0.085927719 | 2.92870689 | 0.020008995 |
| 3593.927734 | 2867.455078 | 1686.931763 | -2.221399491 | NA | 1.044268384 | NA | 1.248428143 | NA |
| 21680.83008 | 25032.51367 | 24938.67383 | -2.230180275 | 0.010893509 | 1.400861246 | 0.071050221 | 0.953511454 | 0.02527614 |
| 208464.3594 | 166313.6719 | 142030.8281 | -2.238700437 | 0.006395507 | 1.736775217 | 0.086089321 | 1.824829304 | 0.057544683 |
| 3357.619141 | 3870.780029 | 2219.219482 | -2.252075941 | 0.075463513 | 0.275138653 | 0.300186184 | -0.596919538 | 0.158083869 |
| 83522.51563 | 89384.36719 | 73079.67969 | -2.252337854 | 0.015288185 | -0.405750215 | 0.344912699 | -0.192916197 | 0.582706431 |
| 423326.9063 | 449428.9688 | 412413.0625 | -2.257079005 | 0.013660857 | 0.861887751 | 0.105107021 | -0.148250278 | 0.512544319 |
| NA | NA | 245065.6563 | -2.27116921 | NA | -0.403838482 | NA | 5.292428563 | NA |
| 21373.76367 | 15937.87988 | 13330.36035 | -2.27973581 | 0.235000038 | 2.849709711 | 0.353604166 | 2.740084233 | 0.076783998 |
| 18703.94336 | 16489.92969 | 14057.77246 | -2.288876255 | 0.175952077 | 2.176432319 | 0.363287481 | 2.620305386 | 0.041088687 |
| 138697.8906 | 113775.5156 | 89571.52344 | -2.313444034 | 0.173323809 | 2.405954174 | 0.328806215 | 2.508456653 | 0.070968049 |

| | | | | | | | | |
|---|---|---|---|---|---|---|---|---|
| NA | NA | NA | -2.317491582 | NA | -0.257072948 | NA | NA | NA |
| 41090.37109 | 43339.39063 | 53347.58984 | -2.341421036 | 0.020082924 | -0.119711905 | 0.689502716 | 0.120505471 | 0.620956029 |
| 32566.4043 | 28281.16406 | 22615.98047 | -2.344847703 | 0.14905608 | 1.542928158 | 0.422801 | 2.213016714 | 0.041258108 |
| 5883.041504 | 3866.151855 | 2789.9021 | -2.348921972 | 0.049763403 | -0.578852678 | 0.27504497 | -3.153858606 | 0.082507796 |
| NA | 417.4758911 | NA | -2.353554042 | NA | -0.403755662 | NA | 0.866197158 | NA |
| NA | 2436.73999 | 3865.247559 | -2.36194754 | 0.008731287 | -0.303431564 | 0.655094166 | 0.292642399 | NA |
| 516.2731323 | 306.6769409 | NA | -2.367057161 | 0.203407262 | -0.155510405 | NA | -1.402825201 | NA |
| 5347.838379 | NA | NA | -2.371327077 | 0.056034046 | 0.26702255 | 0.540404601 | 0.772688709 | NA |
| 71932.85156 | 71098.82031 | 45913.71875 | -2.37158523 | 0.032427351 | 0.322691664 | 0.736903884 | 1.183828868 | 0.111080976 |
| 2749.380859 | 3362.34668 | 3987.463623 | -2.379403925 | 0.025818052 | -0.059899131 | 0.760205385 | -0.849099992 | 0.045254155 |
| 40373.375 | 42254.60938 | 40138.69922 | -2.382591472 | 0.031288094 | 0.105779205 | 0.793826848 | 0.697576298 | 0.149239217 |
| 3535.796143 | 2961.994385 | 2488.808105 | -2.39097376 | NA | 2.918576734 | NA | 0.038359401 | NA |
| NA | NA | NA | -2.410835482 | NA | 4.832615436 | NA | NA | NA |
| NA | NA | 4091.661377 | -2.436591267 | NA | NA | NA | -0.431366273 | NA |
| 486.9956055 | 1122.130615 | NA | -2.443167516 | 0.066814763 | -1.914762002 | NA | -1.015707479 | NA |
| 9616.939453 | 12035.81152 | 10587.22852 | -2.449061521 | 0.036971527 | 2.081361432 | 0.053960002 | -0.419469692 | 0.070298132 |
| 13080.75391 | 11396.90332 | 12184.29102 | -2.471387241 | 0.046628852 | 0.102920981 | 0.54034971 | -0.592065814 | 0.035626762 |
| 19652.75391 | 17635.21094 | 13335.64355 | -2.48357377 | 0.005752145 | 1.152108448 | 0.050472986 | 0.82728134 | 0.100662842 |
| 110535.9688 | 88122.6875 | 88645.5625 | -2.484496657 | 0.086055451 | 1.205475776 | 0.253646775 | 3.500590122 | 0.037571191 |
| 4376.556641 | 4186.726074 | 3839.357666 | -2.537941642 | NA | 1.330545085 | NA | 1.906843809 | NA |
| NA | NA | 45.94494629 | -2.551275961 | NA | 2.242221973 | NA | -4.287519763 | NA |
| NA | NA | NA | -2.579960724 | 0.004709568 | NA | NA | NA | NA |
| 62344.84375 | 70140.60938 | 88092.32813 | -2.582186099 | 0.018310596 | -1.369401677 | 0.444176279 | -1.569746345 | 0.405884448 |
| 2480.580566 | 4526.077637 | 5987.490234 | -2.592934714 | 0.036817204 | -0.353999939 | NA | 1.798001356 | 0.152597158 |
| 573.0206909 | 776.427063 | NA | -2.596713582 | 0.053789965 | -0.528353604 | NA | -0.902019634 | NA |
| 111227.1875 | 97408.92188 | 102285.8125 | -2.617376917 | 0.006395507 | 0.895804963 | 0.065827569 | 2.15224625 | 0.010178497 |
| 8538.089844 | 7149.818848 | 14911.25586 | -2.675516894 | 0.168887111 | 1.700834413 | 0.146984394 | 1.500814152 | 0.194092071 |
| 13530140 | 7999361 | 14205338 | -2.677627872 | 0.083280787 | 1.692645755 | 0.103044963 | 0.189603718 | 0.674026884 |
| 36112.80078 | 43925.42969 | 34741.70313 | -2.730346613 | 0.05239238 | 0.595421216 | 0.187643348 | 0.895618934 | 0.069207996 |
| NA | 4606.909668 | 1560.352173 | -2.751152128 | 0.017324375 | NA | NA | -0.529850471 | NA |
| 18545.17969 | 5979.238281 | 10707.4707 | -2.79375314 | 0.01964718 | -0.470095634 | 0.222412514 | 1.400614341 | 0.286537586 |
| 2011.270264 | 2199.127686 | NA | -2.796129342 | 0.098361037 | 1.565141118 | 0.051888756 | 1.118838567 | NA |
| 21486.20508 | 21164.21875 | 20753.20117 | -2.8294635 | 0.01964718 | 0.930377079 | 0.07468596 | 0.807506676 | 0.071891702 |

| | | | | | | | | |
|---|---|---|---|---|---|---|---|---|
| 36735.00781 | 35347.61328 | 19495.59766 | -2.832935129 | 0.039413354 | 0.345655899 | 0.482872626 | 0.985550011 | 0.177001447 |
| 46218.82813 | 25712.92773 | 35658.12109 | -2.835754997 | 0.42956317 | 0.450891917 | 0.628879959 | 0.231913583 | 0.613727616 |
| 15698.48633 | 18160.52148 | 18798.65625 | -2.878063477 | 0.003149457 | -0.245575557 | 0.208500335 | 0.224418088 | 0.210335561 |
| 488896.75 | 448574.8125 | 456870.4063 | -2.885057019 | 0.004709568 | 0.248999967 | 0.509884399 | 0.216953774 | 0.266829778 |
| 14615.52148 | 15980 | 15730.85449 | -2.951116723 | 0.07799635 | 0.216726149 | 0.431639915 | 0.766256192 | 0.052091631 |
| NA | NA | NA | -2.957823772 | 0.025783969 | -0.486393816 | NA | NA | NA |
| 2476.481201 | 1430.147705 | 8701.625977 | -2.960917068 | 0.184955964 | 1.961817715 | 0.398870378 | 1.119723336 | 0.529505889 |
| 1867.593384 | 2036.228394 | 2393.80249 | -2.974257495 | 0.075287012 | 0.413100824 | 0.500849741 | -0.323032484 | 0.66403022 |
| NA | 8100.45752 | 12261.08203 | -2.97920525 | 0.032850107 | 0.011353323 | 0.97978487 | 1.444742218 | NA |
| 57704.59375 | 70693.88281 | 67603.5918 | -3.019762301 | 0.067495684 | 0.218216626 | 0.218573457 | 0.322395625 | 0.139434127 |
| 819445.375 | 382392.625 | 1147011.75 | -3.04313242 | 0.041200447 | 1.400309192 | 0.24706799 | 1.139816355 | 0.294384476 |
| 39385.46484 | 31104.21289 | 32594.39063 | -3.076762398 | 0.048937447 | 0.420223554 | 0.228822955 | 0.54264568 | 0.0890834 |
| 65735.21094 | 30374.51758 | 48637.83203 | -3.113704763 | 0.089446936 | 1.709531908 | 0.179324339 | 1.840245181 | 0.135524989 |
| 39397.64063 | 40300.01172 | 39770.9375 | -3.127566246 | 0.011954463 | 1.5232595 | 0.024656861 | 1.183665839 | 0.008614663 |
| 6140.084961 | 16332.01855 | 9405.226563 | -3.169107812 | 0.099406137 | 1.144219331 | 0.114022318 | -0.2499136 | 0.680988887 |
| 108282.9004 | 104409.749 | 107412.0488 | -3.213368532 | 0.031288094 | 0.34190807 | 0.090666606 | 0.288161783 | 0.063830205 |
| 8512.178711 | 6271.660156 | 5820.926758 | -3.286360134 | 0.081374079 | 2.153019518 | 0.354648933 | 0.996543472 | 0.117547897 |
| 30468.33398 | 43454.06641 | 38037.95313 | -3.319874325 | 0.028027914 | 3.053106212 | 0.084356576 | 2.281889684 | 0.059847517 |
| 3586.816406 | 4940.02002 | 5039.874512 | -3.367264232 | 0.092623368 | 2.473656586 | 0.024921531 | 1.341024946 | 0.109006643 |
| 2925.263916 | 3385.269531 | 4475.580566 | -3.381166454 | NA | 2.373800619 | NA | 2.963864949 | NA |
| 3160.128906 | 4214.197266 | 3214.4021 | -3.405010463 | 0.028138974 | 1.774212697 | 0.055241445 | 0.209368823 | 0.406371799 |
| 245165.8906 | 198815.1094 | 142894.3594 | -3.410447482 | 0.10217612 | 1.465014207 | 0.408992835 | 2.42199067 | 0.089949026 |
| 490.350708 | NA | NA | -3.421878498 | 0.035068902 | -0.01540915 | 0.973748201 | -0.901058388 | NA |
| 5487.073242 | 4586.611328 | NA | -3.445846396 | 0.092740746 | -0.860893588 | 0.127611794 | -0.597975986 | NA |
| NA | NA | NA | -3.483678695 | NA | 2.29489493 | NA | NA | NA |
| 11127.68848 | 13384.43066 | 11731.44824 | -3.49170058 | 0.025097519 | 1.811757833 | 0.013427556 | 1.047459493 | 0.028107172 |
| 5034.223633 | 4298.788574 | 3284.63623 | -3.616653243 | NA | 3.647442849 | NA | 2.362469948 | NA |
| 6062.11377 | 2026.395996 | 4695.383789 | -3.618318735 | NA | 1.635117911 | NA | 1.316640991 | NA |
| 3888.407715 | 4289.97998 | 3920.38208 | -3.635993184 | NA | 3.225269679 | NA | 2.84211637 | NA |
| 10429.86816 | 13295.56738 | 16150.10059 | -3.651910187 | 0.013277203 | -0.409836713 | 0.374048576 | -0.592930491 | 0.247217109 |
| 11270.04492 | 13719.08887 | 13225.64844 | -3.664656367 | 0.026060442 | 1.517463421 | 0.073036563 | -0.643428026 | 0.301769552 |
| 10549.94434 | 6902.84375 | 7508.144531 | -3.75639706 | NA | 0.165416008 | NA | 0.347193315 | NA |
| 32017.01758 | 37030.48828 | 34817.22656 | -3.7621218 | 0.064602538 | 2.66119858 | 0.048593497 | 2.922915538 | 0.022312939 |

| | | | | | | | | |
|---|---|---|---|---|---|---|---|---|
| 6081.275391 | NA | 6169.848145 | -3.824485184 | NA | -0.233764882 | NA | 0.083337269 | NA |
| 96203.03125 | 77471.65625 | 70723.75781 | -4.001443719 | NA | 3.487957797 | NA | 4.964888515 | NA |
| 23096.81445 | 22332.89063 | 19958.5957 | -4.019607733 | 0.049982407 | 0.3283738 | 0.263772792 | 0.183932359 | 0.173821986 |
| NA | NA | 2781.094238 | -4.052934017 | 0.084307838 | -0.170766397 | 0.786124415 | -2.183665092 | NA |
| 162921.1406 | 195000.6406 | 250131.8281 | -4.20285868 | 0.055768726 | 2.261346532 | 0.095325195 | 3.730355158 | 0.064122807 |
| NA | NA | NA | -4.222540728 | 0.061350067 | 2.126540996 | 0.017037525 | NA | NA |
| 2069.142822 | 1391.570679 | 1434.557861 | -4.31611367 | NA | 4.433937129 | NA | 3.681296984 | NA |
| 2543.948975 | 5253.241211 | 4326.470703 | -4.383994692 | 0.065774506 | -0.045149959 | 0.833023999 | -0.230026705 | 0.583048837 |
| NA | NA | 154.7383575 | -4.577949319 | NA | -0.016520487 | NA | 0.523117755 | NA |
| 7561.730957 | 7934.213379 | 9194.517578 | -4.788037455 | 0.021027835 | 0.280698302 | 0.630482257 | 0.253356257 | 0.504418582 |
| NA | 1144.937988 | NA | -4.880990996 | NA | NA | NA | -0.08979184 | NA |
| NA | 403.256073 | NA | -5.265537945 | NA | 0.609390734 | NA | -0.879898314 | NA |
| 12342.58203 | 9664.641602 | 16531.91992 | -6.134363072 | 0.04977426 | 0.925108577 | 0.168942668 | -0.139314482 | 0.749715423 |

LTED
