## Supplementary Table 8 for "GLYATL1 is associated with metabolic and epigenetic changes and with endocrine resistance in luminal breast cancer"

| HMDB_ID | Metabolite | C_parental_1 | C_parental_2 | C_parental_3 | C_parental_4 | C_parental_5 | Mean_parent | C_LTED_1 | C_LTED_2 | C_LTED_3 | C_LTED_4 |
|---|---|---|---|---|---|---|---|---|---|---|---|
| #NV | 2-Hydroxybut | 33498.7254 | 28451.5128 | 57509.3567 | 15364.4981 | 32691.2794 | 33503.0745 | 32126.0384 | 35803.5878 | 33034.5474 | 21670.2233 |
| #NV | 2-Hydroxyglu | 2.81 | 1.96 | 2.45 | 2.19 | 1.6 | 2.202 | 1.73 | 1.68 | 1.49 | 1.71 |
| HMDB005965 | 3-Hydroxybut | 244482.906 | 233344.555 | 291402.227 | 148630.705 | 153166.368 | 214205.352 | 151244.383 | 274762.535 | 179968.196 | 98984.1039 |
| HMDB005965 | 3-Phosphogly | 743406.901 | 627003.299 | 764969.287 | 576512.467 | 595218.332 | 661422.057 | 255328.805 | 34015.7691 | 230444.754 | 222623.029 |
| HMDB000080 | 5-Oxoproline | 83551382.8 | 87026522.3 | 99229098 | 100716167 | 85501413.9 | 91204916.8 | 63177797.3 | 60332672.3 | 55248307.5 | 67055407.6 |
| HMDB000026 | 6-Phosphoglu | 225445.657 | 219872.848 | 262758.545 | 244079.021 | 243379.335 | 239107.081 | 43989.2542 | 4471.27486 | 37299.2292 | 44365.8889 |
| HMDB000131 | Acetylcarnitin | 1604766.96 | 2020380.78 | 1861329.9 | 2446548.65 | 2359736.07 | 2058552.47 | 712520.38 | 552641.754 | 557538.828 | 595040.441 |
| HMDB000020 | Adenosine2,2 | 1681700.48 | 1629454.52 | 2376104.47 | 2191046.08 | 1078530.17 | 1791367.14 | 169837.351 | 187067.169 | 178698.132 | 302268.387 |
| HMDB000005 | ADP6,968neg | 2113360.15 | 1880471.48 | 2370568.79 | 2809730.84 | 2645476.42 | 2363921.54 | 1197201.34 | 128235.652 | 1103964.42 | 1293562.57 |
| HMDB000134 | AMP6,601neg | 339949.194 | 387497.933 | 426303.831 | 546267.734 | 614772.014 | 462958.141 | 173239.511 | 82797.4646 | 149528.961 | 140622.93 |
| HMDB000004 | ATP6,964neg | 5016397.24 | 4205677.42 | 5130318.31 | 7322000.02 | 5037131.85 | 5342304.97 | 2533965.18 | 129987.523 | 2261622.69 | 3970552.51 |
| HMDB000053 | Betaine6,134 | 274330537 | 268296298 | 294051722 | 283029779 | 241316189 | 272204905 | 228320656 | 231308289 | 189772499 | 231392722 |
| HMDB000004 | Butanoyl-Car | 151249674 | 173307314 | 186535709 | 199715065 | 154811427 | 173123838 | 102397191 | 98093215 | 89740502.7 | 96582496.2 |
| HMDB000201 | CDP7,01neg | 204594.693 | 195584.786 | 273215.191 | 266415.364 | 222566.27 | 232475.261 | 131851.95 | 8588.10484 | 110167.116 | 156698.979 |
| HMDB000154 | CDP-choline6 | 3.25E+05 | 337786.081 | 358760.566 | 465846.281 | 459589.494 | 389373.509 | 397616.058 | 358285.655 | 395297.698 | 421010.159 |
| HMDB000141 | Choline10,23 | 74263434.4 | 71851277.8 | 73610364.7 | 72513154.3 | 55659723.8 | 69579591 | 49997977.9 | 50475841.2 | 38726312.2 | 55827058.7 |
| HMDB000009 | cis-Aconitic a | 5943219.42 | 4975918.61 | 5817105.92 | 6908525.37 | 5944734.32 | 5917900.73 | 3843561.19 | 2202965.67 | 2945745.02 | 3743719.76 |
| HMDB000080 | Citric acid7,58 | 70.7088808 | 68.2255852 | 75.630885 | 83.9213481 | 68.1242349 | 73.3221868 | 54.9 | 68 | 44.4 | 54.8 |
| HMDB000009 | Citrulline6,85 | 473205.366 | 400946.333 | 482490.312 | 383735.567 | 282812.586 | 404638.033 | 224874.768 | 276964.302 | 191410.528 | 216501.119 |
| HMDB000090 | CMP6,758pos | 2.75E+05 | 295576.476 | 369961.672 | 415513.337 | 330126.379 | 337229.588 | 166458.511 | 103437.199 | 148587.353 | 179157.284 |
| HMDB000009 | Creatine6,508 | 110.412733 | 109.458486 | 129.172247 | 127.568902 | 104.431095 | 116.208693 | 25.9 | 23.8 | 23.7 | 27.5 |
| HMDB000006 | Creatinine3,1 | 37274635.9 | 38533664.1 | 44794950.1 | 36534872.7 | 42876376.1 | 40002899.8 | 3226788.28 | 2414501.81 | 2742286.62 | 2025259.62 |
| HMDB000056 | CTP7,094pos | 430439.465 | 402904.944 | 450323.411 | 406477.324 | 338196.651 | 405668.359 | 282358.563 | 13223.9151 | 233937.632 | 365274.96 |
| HMDB000008 | Cytosine3,831 | 716510.037 | 1386709.77 | 1162735.23 | 1735967.98 | 2498904.3 | 1500165.46 | 818437.377 | 499553.723 | 654981.855 | 647792.749 |
| HMDB000063 | Dihydroxyace | 183280.512 | 168675.579 | 185166.324 | 160676.025 | 102281.608 | 160016.01 | 129045.969 | 52203.9897 | 114085.534 | 223384.591 |
| HMDB000147 | Dimethylglyci | 2983589.03 | 2325627.98 | 2441233.43 | 2879338.48 | 2451777.86 | 2616313.36 | 1092242.78 | 749726.152 | 763087.044 | 826357.106 |
| HMDB000009 | Fructose5,579 | 4894298.95 | 4642239.68 | 5364976.59 | 5088187.94 | 4597905.87 | 4917521.8 | 3307888.73 | 4577292.8 | 3230679.99 | 4710716.19 |
| HMDB000066 | Fumaric acid6 | 73.5231014 | 86.0918204 | 107.194733 | 153.291926 | 135.426773 | 111.105671 | 25.6 | 16.8 | 19.5 | 26.3 |
| HMDB000013 | gamma-Glu-a | 53364.8797 | 61052.812 | 78819.5684 | 66970.1241 | 87107.1841 | 69462.9137 | 90991.196 | 68954.3779 | 93467.3163 | 102270.845 |
| HMDB000576 | GDP7,117neg | 268627.927 | 251312.34 | 289401.911 | 343799.157 | 291593.908 | 288947.049 | 152406.238 | 20749.2108 | 128816.304 | 208923.944 |
| HMDB000120 | Glucose5,964 | 44285656.6 | 31836311.5 | 34629234.8 | 29489053.6 | 20098952.9 | 32067841.9 | 34975952.3 | 36522300.2 | 25797261 | 31982299.3 |
| HMDB000122 | Glucose-6-Ph | 1442076.52 | 1524781.47 | 1496309.89 | 2181533.04 | 1758455.35 | 1680631.25 | 502582.863 | 127284.529 | 445876.884 | 556463.153 |

| | | | | | | | | | | | |
|---|---|---|---|---|---|---|---|---|---|---|---|
| HMDB00014( | Glutathione6 | 14881643.1 | 16706168.2 | 16350379.6 | 21406748.4 | 19241856.2 | 17717359.1 | 7858141.68 | 3535456.51 | 6071139.47 | 6526946.08 |
| HMDB00012 | Glycine6,748 | 240447.816 | 202285.305 | 183109.965 | 107029.7 | 88275.1948 | 164229.596 | 288468.861 | 180266.851 | 179705.029 | 144882.092 |
| HMDB00012 | GMP6,78neg | 634402.391 | 687223.985 | 707345.898 | 958464.702 | 829774.388 | 763442.273 | 269674.979 | 77329.9962 | 278004.866 | 282971.763 |
| HMDB02405 | GTP7,044neg | 330152.585 | 295489.292 | 322760.277 | 427623.031 | 341301.805 | 343465.398 | 181724.723 | 20532.5044 | 164798.351 | 294234.422 |
| HMDB00000 | Guanine4,53 | 2575414.45 | 2320863.63 | 2692089.15 | 1392354.25 | 1267999.34 | 2049744.16 | 979057.335 | 1171838.93 | 788452.655 | 868964.502 |
| HMDB00013 | Hypoxanthine | 5561893.71 | 3715234.71 | 3878280.13 | 2664042.35 | 2612429.76 | 3686376.13 | 349634.84 | 299688.9 | 261844.188 | 382816.651 |
| HMDB00015 | IMP6,656neg | 233491.652 | 198497.619 | 222441.328 | 256116.979 | 199515.407 | 222012.597 | 13951.9558 | 0 | 23472.0537 | 23792.2114 |
| HMDB00017 | Inosine4,105 | 247861.22 | 218828.689 | 309707.812 | 242124.03 | 249329.412 | 253570.233 | 32456.4592 | 30843.6532 | 31188.2718 | 32206.2199 |
| HMDB00019 | L-Alanine6,46 | 236.58778 | 293.070013 | 252.360338 | 359.375252 | 288.466688 | 285.972014 | 128.366526 | 120.712098 | 89.1826767 | 166.638064 |
| HMDB00596 | L-Arginine11, | 258.541462 | 221.331986 | 236.140694 | 223.672098 | 167.504661 | 221.43818 | 201.033428 | 227.296624 | 160.077426 | 220.235428 |
| HMDB00051 | L-Asparagine | 25.8 | 19.9 | 23.7 | 27.9 | 17.8 | 23.02 | 21.3 | 20.2 | 20.3 | 22.5 |
| HMDB00051 | L-Aspartic aci | 279.415365 | 307.590617 | 355.106308 | 359.2194 | 290.104856 | 318.287309 | 136.709493 | 109.484529 | 120.929934 | 128.659159 |
| HMDB00019 | L-Carnitine7, | 314799112 | 370728662 | 396167291 | 355360083 | 360142049 | 359439439 | 147129426 | 123246263 | 127286787 | 128849162 |
| HMDB00006 | L-Cystathioni | 2230785.59 | 2309347.33 | 2502797.35 | 2856756.31 | 2260411.43 | 2432019.6 | 1181242.72 | 690616.524 | 877469.782 | 891258.999 |
| HMDB00009 | L-Glutamic ac | 926.255907 | 908.973817 | 977.113834 | 1139.6197 | 907.463818 | 971.885416 | 452.91238 | 385.159438 | 356.34992 | 433.46207 |
| HMDB00014 | L-Glutamine6 | 2954.80693 | 2862.76528 | 2720.90187 | 2990.31279 | 2015.34595 | 2708.82656 | 1419.57768 | 1344.85888 | 1173.82164 | 1509.5936 |
| HMDB00064 | L-Histidine10 | 165.357003 | 138.13364 | 166.069444 | 155.133758 | 123.888536 | 149.716476 | 133.391764 | 142.656332 | 105.621499 | 141.258618 |
| HMDB00017 | L-Isoleucine5 | 594.486732 | 474.147568 | 579.802895 | 521.49372 | 457.372664 | 525.460716 | 467.358129 | 480.208071 | 384.145534 | 427.01468 |
| HMDB00017 | L-Lactic acid5 | 2420.95511 | 2210.28703 | 2518.50643 | 2239.35567 | 1600.01104 | 2197.82306 | 1607.90231 | 1544.55628 | 1156.80667 | 1508.54384 |
| HMDB00019 | L-Lysine12,34 | 541.219646 | 476.168768 | 528.800052 | 467.763818 | 338.636768 | 470.51781 | 428.723292 | 472.853354 | 344.878422 | 454.193462 |
| HMDB00018 | L-Methionine | 167.938916 | 170.044938 | 154.242894 | 156.758418 | 111.721631 | 152.141359 | 123.104369 | 124.72209 | 104.573885 | 126.035053 |
| HMDB00069 | L-Phenylalani | 340.654081 | 283.010237 | 343.74818 | 329.597014 | 240.777441 | 307.557391 | 283.905967 | 273.628982 | 202.7655 | 285.229927 |
| HMDB00015 | L-Proline6,36 | 1.78E+02 | 179.276242 | 199.057216 | 220.321277 | 184.645704 | 192.265345 | 81.2557547 | 68.3 | 63.0232695 | 74.1382042 |
| HMDB00016 | L-Serine6,85 | 386.271854 | 368.088156 | 288.029104 | 412.096594 | 292.780649 | 349.453271 | 301.273974 | 257.752681 | 246.694076 | 359.059422 |
| HMDB00016 | L-Threonine6 | 967.588796 | 775.336976 | 837.061643 | 891.566251 | 608.176778 | 815.946089 | 709.124065 | 752.689204 | 575.288815 | 795.48716 |
| HMDB00016 | L-Tryptophan | 70.3206786 | 57.0600858 | 63.3462893 | 61.5724464 | 49.4 | 60.3399 | 46.3 | 49.7 | 41.6 | 54.3 |
| HMDB00092 | L-Tyrosine5,9 | 227.22026 | 190.036527 | 254.008125 | 248.06912 | 179.851642 | 219.837135 | 187.983276 | 187.629991 | 161.964487 | 195.656549 |
| HMDB00015 | L-Valine6,179 | 706.763388 | 470.979767 | 550.594254 | 737.486064 | 450.705609 | 583.305817 | 459.347317 | 437.865222 | 339.607049 | 493.8183 |
| HMDB00008 | Malic acid6,5 | 64.9157966 | 62.4863643 | 66.1791161 | 73.8203312 | 63.6762808 | 66.2155778 | 32.2 | 36.8 | 32.3 | 37.3 |
| HMDB00015 | myo-Inositol6 | 6377885.13 | 5278585.04 | 6388022.75 | 6553734.55 | 5307037.37 | 5981052.97 | 3558055.74 | 3874692.14 | 2979379.58 | 3763663.3 |
| HMDB00002 | N-Acetyl-L-as | 117677997 | 122228394 | 137456528 | 136935896 | 138405650 | 130540893 | 82004657.8 | 82706101.1 | 60210686.5 | 79265532.3 |
| HMDB00008 | NAD6,613pos | 2744439.12 | 3257483.93 | 3377060.94 | 4083306.75 | 3532961.41 | 3399050.43 | 2219383.23 | 1779476.24 | 2084642.37 | 2378820.22 |
| HMDB00009 | NADH6,626p | 105460.017 | 98889.399 | 104050.273 | 116490.552 | 96856.0573 | 104349.26 | 78178.2701 | 69808.7896 | 69782.2346 | 87301.3194 |

| | | | | | | | | | | | |
|---|---|---|---|---|---|---|---|---|---|---|---|
| HMDB00014{ | NADP6,954pc | 124890.861 | 143339.613 | 106459.603 | 152047.864 | 98933.6 | 125134.308 | 151741.842 | 32747.554 | 98517.2511 | 97138.3801 |
| HMDB00002: | Ornithine12,4 | 217.819085 | 173.714035 | 247.224949 | 213.726561 | 167.174188 | 203.931764 | 172.944173 | 189.641977 | 144.380057 | 181.470457 |
| HMDB00002: | Oxidized glut | 7239536.55 | 6999236.88 | 8107112.2 | 10960297.2 | 10293297.4 | 8719896.04 | 5359976.51 | 4089325.61 | 4518801.61 | 5485221.79 |
| HMDB00033: | Oxoglutaric a | 53.9120489 | 62.9757331 | 61.9205124 | 61.8684253 | 42.6 | 56.6553439 | 40.2 | 32.6 | 28.1 | 39.7 |
| HMDB0002( | pa_Hypotaur | 2175698.06 | 2269959.42 | 2428906.71 | 1973101.28 | 1567881.81 | 2083109.46 | 1028156.72 | 640048.885 | 746786.968 | 697490.454 |
| HMDB0009( | pa_Itaconic a | 3576635.27 | 3177260.55 | 4264042.47 | 5546666.42 | 5128806.39 | 4338682.22 | 2451755.16 | 1463422.19 | 2027780.67 | 3165531.4 |
| HMDB00209 | pa_Sedohept | 256242.103 | 251910.126 | 268743.274 | 358330.708 | 276751.353 | 282395.513 | 81475.2156 | 11630.4844 | 70752.633 | 90980.4917 |
| HMDB0000( | Pantothenic a | 131469571 | 97660131.7 | 119471856 | 127798456 | 93000327.3 | 113880068 | 47557316 | 68939256.7 | 35891981.8 | 51898149.3 |
| HMDB00002: | Phosphocrea | 8489903.61 | 8146633.32 | 10309722.5 | 10649551.3 | 9343579.72 | 9387878.1 | 2909439.11 | 965844.36 | 2253537.35 | 2457685.98 |
| HMDB00015: | Phosphoenol | 997394.516 | 1018769.38 | 902584.52 | 736800.853 | 1031840.76 | 937478.005 | 417675.273 | 93585.8303 | 259184.671 | 312073.589 |
| HMDB0002( | Propionyl-Ca | 1.22E+08 | 134500640 | 151327404 | 141001489 | 138247541 | 137347239 | 119469453 | 96819841.3 | 112086913 | 106569732 |
| HMDB0008: | Pyridoxine1,6 | 38888802.2 | 38723321.9 | 37429328.4 | 26536004.2 | 25619655.5 | 33439422.4 | 30521577.4 | 35332549.5 | 23827229 | 27587439.5 |
| HMDB0002: | Pyruvic acid1 | 7320.7608 | 4089.36879 | 5350.95201 | 5496.13793 | 6366.84249 | 5724.8124 | 4014.82074 | 4314.84359 | 4220.69929 | 8669.16693 |
| HMDB0002: | Succinic acid6 | 63.6180801 | 53.1946636 | 59.7765662 | 55.5 | 40.7 | 54.557862 | 164.061055 | 128.442792 | 116.588196 | 137.276518 |
| HMDB0002! | Taurine5,991 | 21194298.7 | 24112522.9 | 24192171.4 | 29535930 | 27144679.3 | 25235920.4 | 18314070.3 | 13393381.6 | 15868000.7 | 16077805.8 |
| HMDB0002! | Thiamine10,7 | 867206.96 | 889645.517 | 933485.185 | 972281.779 | 818676.576 | 896259.204 | 663993.582 | 675538.818 | 560578.489 | 796419.754 |
| HMDB0002: | UDP6,939neg | 1127552.86 | 1131722.85 | 1342400.83 | 1869853.05 | 1430525.74 | 1380411.07 | 833209.918 | 413850.892 | 767331.313 | 1088728.43 |
| HMDB00029 | UDP-glucoror | 3623856.44 | 3984200.33 | 4800046.47 | 5696451.99 | 4831236.5 | 4587158.35 | 4191416.13 | 3321104.31 | 3755708.19 | 4627596.52 |
| HMDB00093 | UDP-glucose6 | 1924663.2 | 1794196.57 | 2444293.64 | 3034670.2 | 2725120.02 | 2384588.73 | 1519588.35 | 1214627.46 | 1409289.54 | 1578652.53 |
| HMDB0002{ | UDP-N-acetyl | 37437000.1 | 38727230.7 | 44455226.1 | 55538646 | 53617262.9 | 45955073.1 | 39683087.9 | 38297753.4 | 36356473.1 | 42222850.1 |
| HMDB00029 | UMP6,625ne | 2097265.91 | 2083246.49 | 2020327.41 | 2828354.24 | 2431062.7 | 2292051.35 | 929966.257 | 627024.373 | 863597.846 | 931212.534 |
| HMDB0002{ | Uracil2,29pos | 158653.777 | 169633.166 | 157678.859 | 163806.668 | 211327.747 | 172220.043 | 135121.165 | 186828.346 | 126482.958 | 215955.591 |
| HMDB0003( | Uridine2,274 | 644299.667 | 593264.417 | 720771.508 | 595356.978 | 612013.897 | 633141.293 | 516938.4 | 706673.863 | 550521.3 | 585242.295 |
| HMDB00029 | UTP6,992neg | 1748021.93 | 1445721.14 | 1836591.02 | 2963641.34 | 1862455.79 | 1971286.24 | 923292.633 | 56258.2868 | 904312.437 | 2127282.14 |
| HMDB0002{ | Xanthine3,19 | 4783431.24 | 4125161.07 | 5062962.02 | 3130117.12 | 2845474.02 | 3989429.09 | 31959.3301 | 41541.0531 | 38967.3274 | 46649.2952 |
| HMDB00029 | Xanthosine4, | 88879.9884 | 69853.7359 | 83030.7531 | 99390.2704 | 224991.749 | 113229.299 | 21186.1675 | 10147.2139 | 9778.70479 | 5842.17773 |

| C_LTED_5 | Mean_LTED | C_KO1_1 | C_KO1_2 | C_KO1_3 | C_KO1_4 | C_KO1_5 | Mean_KO1 | C_KO2_1 | C_KO2_2 | C_KO2_3 | C_KO2_4 |
|---|---|---|---|---|---|---|---|---|---|---|---|
| 21609.2716 | 27110.0201 | 23141.0829 | 29690.7582 | 34625.3 | 20121.4228 | 49045.2605 | 31324.7649 | 21754.7623 | 27368.4671 | 19361.9512 | 18779.0789 |
| 1.78 | 1.6775 | 1.75 | 1.85 | 2.16 | 3.12 | 2.31 | 2.238 | 1.69 | 1.78 | 1.7 | 1.73 |
| 162048.608 | 148061.323 | 160766.443 | 239651.116 | 242731.742 | 161190.069 | 224982.522 | 205864.378 | 150024.005 | 209879.984 | 160316.281 | 100552.053 |
| 286919.816 | 248829.101 | 442835.852 | 390832.625 | 315189.145 | 373372.448 | 313288.561 | 367103.726 | 300802.356 | 512778.868 | 536459.673 | 432495.422 |
| 75498907.7 | 65245105 | 53233360.6 | 47493803.9 | 54145331.7 | 61823955.4 | 76031705.3 | 58545631.4 | 70518501.4 | 78124046.4 | 72551638.7 | 69941014.6 |
| 37311.0581 | 40741.3576 | 53392.6503 | 63117.1922 | 55967.9007 | 72306.0393 | 39693.4959 | 56895.4557 | 28393.2457 | 62369.6228 | 50446.7693 | 47347.8339 |
| 741991.571 | 651772.805 | 714922.765 | 479106.605 | 630939.895 | 545645.409 | 1088434.41 | 691809.817 | 438114.509 | 461454.634 | 418176.552 | 334174.828 |
| 316669.007 | 241868.219 | 200786.377 | 279386 | 254147.054 | 666546.249 | 552751.44 | 390723.424 | 302732.422 | 406784.485 | 273473.016 | 344923.814 |
| 1201777.49 | 1199126.46 | 1535993.86 | 1560278.07 | 1392763 | 1870898.27 | 1346658.13 | 1541318.26 | 2342461.54 | 4152719.69 | 3512838.76 | 3729859.21 |
| 171490.18 | 158720.395 | 308942.251 | 207403.291 | 240893.62 | 258380.897 | 228538.424 | 248831.696 | 413667.167 | 526611.912 | 507507.385 | 608675.122 |
| 3062288.72 | 2957107.27 | 3021453.97 | 3658311.34 | 2392512.83 | 4749040.51 | 3505216.13 | 3465306.96 | 5205995.63 | 8707875.76 | 8052969.81 | 9220041.14 |
| 223163594 | 218162368 | 205598382 | 218160739 | 217817297 | 247728918 | 276578168 | 233176701 | 231566808 | 215907460 | 207859947 | 183903832 |
| 101081092 | 97450320.3 | 110826900 | 108313013 | 96481798.1 | 81762763.4 | 121201795 | 103717254 | 69806019.9 | 79899722.1 | 69530294.1 | 72190556.7 |
| 122956.884 | 130418.732 | 168155.316 | 166768.048 | 121909.037 | 198415.037 | 161047.689 | 163259.025 | 259520.957 | 407071.289 | 339281.603 | 354300.454 |
| 482680.433 | 424151.087 | 1088713.45 | 983352.149 | 985797.88 | 1214956.15 | 1222560.76 | 1099076.08 | 1051139.86 | 1062390.37 | 778045.448 | 1048364.16 |
| 58247209.6 | 50699639.6 | 48190039.6 | 47362616.6 | 54058128.4 | 68006347.8 | 59038725 | 55331171.5 | 57866321.1 | 62326080.1 | 50587833.9 | 58030636 |
| 3525376.96 | 3514600.73 | 3274658.81 | 2955495.13 | 2788339.01 | 2865427.59 | 4375190.9 | 3251822.29 | 3323825.66 | 3277880.18 | 2652520.01 | 2750291.05 |
| 51 | 51.275 | 45.019714 | 44.4938982 | 42.832129 | 51.9681621 | 56.8 | 48.2227807 | 49.0124364 | 50.4995889 | 43.6610017 | 46.3505822 |
| 207344.643 | 210032.765 | 140576.364 | 178673.18 | 194001.208 | 209318.586 | 215612.575 | 187636.382 | 145573.086 | 149177.116 | 139462.887 | 125600.004 |
| 194738.828 | 172235.494 | 761780.692 | 744748.951 | 706422.734 | 942575.849 | 945273.346 | 820160.315 | 1084073.7 | 1156170.62 | 1004069.74 | 1095139.96 |
| 26.4 | 25.875 | 32.1 | 32.1 | 31.1 | 34.3 | 34.6 | 32.84 | 27.3 | 31.6 | 25.7 | 27.1 |
| 2260289.29 | 2563655.95 | 4288756.03 | 2131956.65 | 4186511.45 | 3337967.06 | 2335087.01 | 3256055.64 | 3251246.22 | 4546963.3 | 2940516.08 | 3651071.99 |
| 271950.412 | 288380.392 | 323619.569 | 366082.579 | 276168.996 | 438241.685 | 310878.532 | 342998.272 | 516879.675 | 736294.665 | 595642.926 | 598778.106 |
| 547808.258 | 667255.06 | 517558.698 | 396891.86 | 331614.839 | 234838.788 | 726993.43 | 441579.523 | 438823.233 | 526009.988 | 282945.506 | 240890.209 |
| 174227.654 | 160185.937 | 188406.673 | 167605.091 | 159234.629 | 132249.382 | 203886.977 | 170276.55 | 102100.432 | 82729.1521 | 83664.8381 | 79431.3632 |
| 1022541.43 | 926057.091 | 1104054.76 | 922897.904 | 797214.342 | 1192746.67 | 1100894.13 | 1023561.56 | 1897362.17 | 2131000.39 | 1735709.52 | 1789907.43 |
| 5786894.81 | 4259044.93 | 3731881.99 | 4582901.94 | 6329481.26 | 8662329.18 | 7444098.57 | 6150138.59 | 3272557.06 | 4309723.01 | 3208563.86 | 3574535.55 |
| 34.9 | 26.575 | 50.2653119 | 46.4529699 | 46.2293412 | 71.0304307 | 78.8662956 | 58.5688699 | 57.0766874 | 77.2485595 | 68.275506 | 80.4471167 |
| 67928.2114 | 88664.3922 | 59475.6632 | 60456.8261 | 65302.6893 | 57597.0884 | 116082.492 | 71782.9519 | 307361.808 | 246394.889 | 227158.661 | 265709.114 |
| 142542.265 | 158172.188 | 178355.411 | 192768.068 | 170605.115 | 254683.208 | 170172.707 | 193316.902 | 242531.177 | 424759.229 | 363759.6 | 359313.791 |
| 28526649.1 | 30320540.4 | 31221429.2 | 31894939.1 | 29428682.1 | 28797437.4 | 34805170.5 | 31229531.7 | 27788434.9 | 25579930.5 | 18815553.6 | 18522183.7 |
| 532358.956 | 509320.464 | 682490.379 | 668991.852 | 563944.52 | 819738.543 | 863401.854 | 719713.43 | 926926.336 | 975201.7 | 904549.212 | 829587.36 |

| | | | | | | | | | | | |
|---|---|---|---|---|---|---|---|---|---|---|---|
| 8223535.5 | 7169940.68 | 9352574.7 | 9207941.04 | 7611844.8 | 9376810.76 | 8669810.97 | 8843796.45 | 16802855.3 | 17691979.1 | 19523997.9 | 17771370.1 |
| 109047.075 | 180525.764 | 339745.906 | 323301.709 | 223379.624 | 186839.188 | 124521.691 | 239557.623 | 418947.119 | 348322.905 | 241340.604 | 187186.265 |
| 343898.832 | 293637.61 | 552711.571 | 498674.326 | 411290.069 | 590260.665 | 463834.819 | 503354.29 | 622328.799 | 914097.986 | 714022.081 | 851777.072 |
| 195209.2 | 208991.674 | 179974.373 | 229704.229 | 140524.566 | 330406.841 | 270717.181 | 230265.438 | 291266.092 | 496621.932 | 427983.23 | 495886.361 |
| 1136023.29 | 943124.445 | 1336147.21 | 1464644.27 | 1613545.3 | 1104623.63 | 1279616.01 | 1359715.28 | 1542826.69 | 2309396.79 | 1510330.56 | 1806794.22 |
| 402142.002 | 349109.42 | 174191.296 | 225166.955 | 199285.12 | 261919.502 | 372421.765 | 246596.927 | 158609.992 | 266601.838 | 143605.96 | 253905.354 |
| 22658.2122 | 20968.6083 | 6447.87757 | 15085.2957 | 7534.53406 | 9044.27681 | 12267.5546 | 10075.9077 | 0 | 19785.7165 | 0 | 12979.9985 |
| 39434.5943 | 33821.3863 | 49481.0818 | 51431.6486 | 53698.3231 | 51520.2625 | 100740.969 | 61374.4569 | 36536.2115 | 55243.2583 | 43164.5025 | 43846.9639 |
| 139.802853 | 130.99753 | 186.407431 | 152.198502 | 116.447112 | 166.881 | 365.215074 | 197.429824 | 196.396279 | 199.4558 | 145.171043 | 171.617761 |
| 221.328733 | 200.668754 | 166.369793 | 163.369676 | 177.633541 | 213.317117 | 230.736498 | 190.285325 | 154.304759 | 163.354394 | 130.272134 | 151.741323 |
| 26.9 | 22.75 | 26.7 | 18.5 | 19.6 | 24.2 | 27.3 | 23.26 | 21.2 | 26.2 | 21.4 | 21.4 |
| 131.46404 | 129.440657 | 176.867072 | 199.286601 | 156.113571 | 186.030353 | 178.525616 | 179.364643 | 358.931929 | 373.318831 | 312.789977 | 306.083795 |
| 138650501 | 135478969 | 170444078 | 161105787 | 163509195 | 176842684 | 184166578 | 171213664 | 189637374 | 206577775 | 160616776 | 140843834 |
| 1062565.5 | 1003134.25 | 974677.939 | 944030.07 | 824175.307 | 966103.713 | 806298.094 | 903057.025 | 2994079.03 | 3343219.92 | 2596865.13 | 2731060.52 |
| 393.284865 | 409.002309 | 543.340153 | 536.339875 | 450.760902 | 531.462699 | 492.61741 | 510.904208 | 726.927038 | 816.077722 | 722.975187 | 733.392798 |
| 1473.49007 | 1394.12075 | 1126.9671 | 928.374664 | 1144.71586 | 1175.86047 | 1231.37742 | 1121.4591 | 1522.48346 | 1791.28594 | 1531.97838 | 1369.88928 |
| 133.633289 | 128.476293 | 101.686483 | 108.317004 | 110.492846 | 130.591637 | 133.861741 | 116.989942 | 104.234957 | 115.421723 | 99.7070365 | 103.531862 |
| 489.070472 | 441.897204 | 375.527504 | 388.002276 | 410.837319 | 496.941117 | 504.797378 | 435.221119 | 372.409686 | 392.565206 | 336.069813 | 373.076165 |
| 1739.43184 | 1503.17116 | 1360.81631 | 1728.93832 | 1511.83112 | 1748.35234 | 1847.76273 | 1639.54016 | 977.024542 | 988.909037 | 1138.21654 | 1038.82453 |
| 447.953578 | 418.937189 | 354.623806 | 365.10592 | 378.872514 | 511.251206 | 471.594588 | 416.289607 | 311.354836 | 317.224358 | 255.400176 | 295.404983 |
| 137.15415 | 122.716864 | 98.491446 | 109.477258 | 108.584226 | 127.744087 | 126.511213 | 114.161646 | 112.148178 | 131.505631 | 99.7092709 | 113.533114 |
| 293.729105 | 266.407625 | 218.986477 | 211.852165 | 219.097701 | 265.37366 | 285.925048 | 240.24701 | 206.287369 | 298.307817 | 186.448021 | 208.298086 |
| 77.7272503 | 74.0361197 | 120.723176 | 114.175238 | 113.889799 | 122.680764 | 128.604904 | 120.014776 | 147.655786 | 185.418021 | 172.086867 | 162.861206 |
| 318.422589 | 306.362515 | 325.700735 | 227.539398 | 259.551576 | 226.723794 | 317.183808 | 271.339862 | 311.543325 | 362.502644 | 340.729845 | 291.925455 |
| 663.655218 | 685.888814 | 603.72294 | 576.465507 | 552.514453 | 733.648752 | 657.713264 | 624.812983 | 895.578482 | 943.863509 | 756.454483 | 782.431209 |
| 51.1 | 48.325 | 36.8 | 40.4456384 | 40.9310506 | 51.9223695 | 50.7 | 44.1598117 | 40.6 | 43.5935466 | 35.951967 | 41.1542438 |
| 194.310359 | 184.978668 | 152.363154 | 162.7489 | 160.917637 | 126.549869 | 135.802132 | 147.676339 | 150.964465 | 171.168567 | 148.053179 | 151.859043 |
| 539.366078 | 458.034686 | 440.35625 | 425.364498 | 426.538433 | 492.842151 | 616.567219 | 480.33371 | 297.4089 | 366.160322 | 287.867016 | 437.393216 |
| 35.8 | 34.4 | 38.2 | 38.8 | 36.4 | 35.6 | 40.7 | 37.94 | 46.1807156 | 52.8236296 | 58.7702146 | 49.0511786 |
| 3792334.69 | 3523358.33 | 4230427.35 | 3801051.07 | 4013753.45 | 4075887.48 | 4999955.81 | 4224215.03 | 3351552.74 | 3522821.99 | 3009981.15 | 2812891.23 |
| 75019144.6 | 74125005.3 | 115178806 | 102896523 | 104906780 | 89828583.2 | 165174042 | 115596947 | 99339169.4 | 103550931 | 93099740 | 82537601.8 |
| 2293501.32 | 2244086.79 | 3059500.07 | 2829034.62 | 2655991.58 | 3036456.5 | 2878598.21 | 2891916.19 | 3936883.15 | 3633190.66 | 3260484.17 | 3731518.16 |
| 101733.788 | 84248.903 | 108202.874 | 90292.3311 | 74606.5659 | 120066.453 | 111261.947 | 100886.034 | 127383.093 | 132892.675 | 103985.847 | 154458.292 |

| | | | | | | | | | | | |
|---|---|---|---|---|---|---|---|---|---|---|---|
| 150114.635 | 124378.027 | 124619.461 | 142239.013 | 110593.197 | 146950.445 | 127669.101 | 130414.243 | 206024.564 | 242113.966 | 210690.109 | 209023.619 |
| 207.091327 | 176.471504 | 131.818352 | 154.202005 | 154.547977 | 163.281274 | 218.22032 | 164.413985 | 120.847121 | 133.580824 | 110.514249 | 126.91514 |
| 6420708.76 | 5446177.16 | 5709537.87 | 5391415.83 | 4921345.17 | 7430321.98 | 8208949.02 | 6332313.97 | 6296895.98 | 8046171.3 | 5927957.97 | 7534820.85 |
| 40.6 | 37.15 | 46.9789908 | 55.0984971 | 31.1 | 48.3474581 | 81.7543358 | 52.6558564 | 80.1993464 | 74.2597324 | 60.9426481 | 56.0713595 |
| 809902.34 | 820584.121 | 998712.319 | 927226.204 | 779886.649 | 973435.224 | 548164.293 | 845484.938 | 1038915.51 | 1173249.64 | 860905.44 | 874141.985 |
| 2975614.99 | 2655170.56 | 2135833.45 | 2047336.8 | 2052778.08 | 2391541.97 | 3915612.43 | 2508620.55 | 2145971.74 | 2142303.8 | 1935470.11 | 2386176.98 |
| 59646.4161 | 75713.6891 | 121916.891 | 129655.395 | 101808.747 | 125393.413 | 150791.419 | 125913.173 | 98523.2261 | 117017.704 | 117150.099 | 103779.107 |
| 52974933 | 47080595 | 31595103.5 | 29018187.6 | 15173893.6 | 15996236.8 | 47337899.2 | 27824264.1 | 27344536.1 | 50673977.1 | 44306528.3 | 54823606 |
| 2736707.2 | 2589342.41 | 3388350.93 | 3045257.67 | 2991967.92 | 4055735.91 | 3484140.07 | 3393090.5 | 2916329.9 | 3131891.02 | 3067374.89 | 2619784.19 |
| 436763.691 | 356424.306 | 522619.77 | 646815.626 | 509590.062 | 611452.624 | 723203.182 | 602736.253 | 450565.195 | 437205.238 | 671387.656 | 299082.918 |
| 109265336 | 111847858 | 180216370 | 183413583 | 190279742 | 189344287 | 195559134 | 187762623 | 203843113 | 237085757 | 187306416 | 156091921 |
| 31545043.2 | 28370322.3 | 25009451.2 | 28345014.3 | 24538437.8 | 26261590.8 | 31058833.9 | 27042665.6 | 24970624.8 | 26870799 | 16319973.2 | 22083400.3 |
| 4891.69187 | 5449.0947 | 5923.44162 | 2873.99703 | 5642.2498 | 6524.96523 | 6278.49608 | 5448.62995 | 3272.74653 | 3443.56399 | 3103.45506 | 2749.72382 |
| 145.456926 | 140.845674 | 53.5677622 | 50.8228342 | 49.2710242 | 61.5155664 | 72.2540238 | 57.4862422 | 44.8581624 | 45.7955788 | 39.3108706 | 43.4986253 |
| 17937604.5 | 17049370.3 | 23564885.8 | 21614827.2 | 19739422.5 | 23037935 | 22403742.2 | 22072162.5 | 30139289.5 | 35369918.7 | 27946767.1 | 28930299.5 |
| 960746.75 | 745434.644 | 587822.112 | 606642.628 | 576325.634 | 813374.145 | 885579.084 | 693948.721 | 533630.201 | 517858.992 | 468653.318 | 552797.67 |
| 899894.161 | 897290.955 | 833009.336 | 935816.009 | 750851.149 | 1048191.99 | 1009007 | 915375.097 | 1331035.72 | 2385750.05 | 1999343.33 | 1922386.71 |
| 4401968.11 | 4244172.24 | 3144019.09 | 2949246.49 | 2306157.02 | 2855883.07 | 3769170.14 | 3004895.16 | 1390627.1 | 1643971.29 | 1580932.1 | 1420039.36 |
| 1759389.29 | 1566729.93 | 1667902.45 | 1730051.47 | 1450965.34 | 1773296.94 | 2093572.74 | 1743157.79 | 3183580 | 3000641.36 | 2877995.95 | 3072656.71 |
| 42823617.5 | 40271507.1 | 30539663.1 | 28368085.1 | 30602561.7 | 25794536.2 | 36655099.8 | 30391989.2 | 33995215.3 | 38439301.8 | 36639422.7 | 34554157.4 |
| 1002083.92 | 931715.139 | 1433821.46 | 1004796.99 | 985216.001 | 1526608.49 | 1109707.61 | 1212030.11 | 2233080.49 | 2534304.98 | 2534481.33 | 2726778.42 |
| 311152.178 | 197177.973 | 87813.2638 | 79950.6362 | 87815.0926 | 137144.919 | 140331.762 | 106611.135 | 123037.728 | 179739.316 | 111381.329 | 227072.064 |
| 779526.022 | 608057.004 | 297617.142 | 329750.679 | 376956.251 | 388640.424 | 474376.462 | 373468.192 | 496058.528 | 658916.901 | 554982.468 | 611052.583 |
| 1315819.86 | 1317676.77 | 1059610.72 | 1411072.57 | 828662.168 | 1579932.75 | 1505522.44 | 1276960.13 | 2110197.79 | 3388790.54 | 3189376.28 | 3369101.2 |
| 43685.8747 | 40315.4568 | 47963.5104 | 51761.4355 | 70257.4831 | 66532.2281 | 50467.0391 | 57396.3392 | 38258.584 | 53754.475 | 53065.0531 | 46602.5927 |
| 10802.3697 | 11902.3549 | 18307.4698 | 12983.2806 | 16158.3422 | 6438.001 | 15775.3194 | 13932.4826 | 2535.52102 | 7393.97827 | 8812.0542 | 3560.37245 |

| C_KO2_5 | Mean_KO2 | FC_LTED_vs_ | log2FC_LTED_ | pcal_LTED_vs | padj_LTED_vs | FC_KO1_vs_L | log2FC_KO1_ | pcal_KO1_vs_LTE | padj_KO1_vs_ | FC_KO2_vs_L |
|---|---|---|---|---|---|---|---|---|---|---|
| 15188.2151 | 20490.4949 | 0.80918007 | -0.3054673 | 0.4623008 | 0.5079017 | 1.15546815 | 0.2084775 | 0.26609034868 | 0.3341661 | 0.75582736 |
| 1.44 | 1.668 | 0.76180745 | -0.3925017 | 0.06579286 | 0.09354922 | 1.33412817 | 0.41589727 | 0.04310009260 | 0.09338353 | 0.99433681 |
| 101560.463 | 144466.557 | 0.69121206 | -0.5327997 | 0.09956244 | 0.1352266 | 1.39039943 | 0.47549939 | 0.03085418067 | 0.07588461 | 0.9757211 |
| 323386.301 | 421184.524 | 0.37620321 | -1.4104159 | 4.2854E-05 | 0.00025998 | 1.47532473 | 0.56103254 | 0.00302241954 | 0.0144758 | 1.69266586 |
| 67102042.4 | 71647448.7 | 0.71536829 | -0.4832419 | 0.00222752 | 0.00471405 | 0.89731837 | -0.1563081 | 0.17541884711 | 0.259654 | 1.09812757 |
| 47416.5483 | 47194.804 | 0.17038959 | -2.5530909 | 8.5987E-08 | 7.82E-06 | 1.39650368 | 0.48181937 | 0.01952268120 | 0.05551762 | 1.15840038 |
| 305306.773 | 391445.459 | 0.31661705 | -1.6591892 | 0.00011082 | 0.00048021 | 1.06142787 | 0.08600633 | 0.38115133220 | 0.4129139 | 0.60058575 |
| 382268.713 | 342036.49 | 0.13501879 | -2.8887679 | 0.00058881 | 0.00167443 | 1.61543929 | 0.69192653 | 0.10883933170 | 0.2021302 | 1.414144 |
| 3362632.84 | 3420102.41 | 0.50726153 | -0.9791983 | 0.00055988 | 0.00164351 | 1.28536757 | 0.36218098 | 0.00840601416 | 0.02942105 | 2.85216158 |
| 478913.185 | 507074.954 | 0.34283962 | -1.5443942 | 0.00123257 | 0.00295169 | 1.56773612 | 0.64868275 | 0.00168160677 | 0.01177125 | 3.19476872 |
| 8809157.19 | 7999207.91 | 0.55352648 | -0.8532758 | 0.00977218 | 0.01778536 | 1.17185703 | 0.22879657 | 0.19382905733 | 0.2686205 | 2.7050787 |
| 166604605 | 201168530 | 0.80146376 | -0.3192908 | 0.00446443 | 0.0088318 | 1.06882183 | 0.09602138 | 0.20149652387 | 0.2736744 | 0.92210463 |
| 63803746.8 | 71046067.9 | 0.56289372 | -0.8290656 | 0.00020466 | 0.00074082 | 1.06430901 | 0.08991708 | 0.23105062708 | 0.3003658 | 0.7290491 |
| 323903.157 | 336815.492 | 0.56100048 | -0.8339261 | 0.0013852 | 0.00323213 | 1.25180656 | 0.32401165 | 0.04207711763 | 0.09338353 | 2.5825699 |
| 1042845.4 | 996557.048 | 1.08931676 | 0.12342353 | 0.40070521 | 0.450175 | 2.59123721 | 1.37364109 | 0.00000615974 | 0.00018685 | 2.34953317 |
| 50977985.3 | 55957771.3 | 0.72865676 | -0.4566887 | 0.01106792 | 0.01974865 | 1.09135236 | 0.12611698 | 0.22421726557 | 0.2957068 | 1.10371142 |
| 2812982.01 | 2963499.78 | 0.59389315 | -0.7517247 | 0.00046112 | 0.0014329 | 0.92523235 | -0.1121124 | 0.25357172874 | 0.3250004 | 0.84319671 |
| 41.3884059 | 46.182403 | 0.69931084 | -0.5159942 | 0.00091125 | 0.00227307 | 0.94047354 | -0.0885407 | 0.21870610788 | 0.2926802 | 0.9006807 |
| 111601.71 | 134282.961 | 0.51906333 | -0.9460175 | 0.00220899 | 0.00471405 | 0.8933672 | -0.1626748 | 0.10765863562 | 0.2021302 | 0.63934292 |
| 1121803.3 | 1092251.46 | 0.5107366 | -0.9693486 | 0.0009083 | 0.00227307 | 4.76185422 | 2.25152345 | 0.00000563470 | 0.00018685 | 6.34161656 |
| 26.3 | 27.6 | 0.22265976 | -2.1670872 | 1.0827E-06 | 2.46E-05 | 1.26917874 | 0.34389526 | 0.00014357674 | 0.00217758 | 1.06666667 |
| 3477900.36 | 3573539.59 | 0.06408675 | -3.96383 | 1.8569E-07 | 8.45E-06 | 1.27008292 | 0.34492269 | 0.12866178909 | 0.2209099 | 1.39392323 |
| 615526.787 | 612624.432 | 0.71087721 | -0.4923277 | 0.00853757 | 0.01585549 | 1.18939527 | 0.25022824 | 0.10624366927 | 0.2021302 | 2.1243623 |
| 663645.014 | 430462.79 | 0.44478764 | -1.1688114 | 0.04559925 | 0.06802512 | 0.6617852 | -0.5955651 | 0.03759643957 | 0.0855319 | 0.6451248 |
| 87574.9235 | 87100.1418 | 1.00106194 | 0.00153124 | 0.99526321 | 0.9952632 | 1.06299313 | 0.08813227 | 0.35344196268 | 0.3922344 | 0.543744 |
| 1666656.12 | 1844127.13 | 0.35395496 | -1.4983623 | 1.7714E-05 | 0.00014655 | 1.10528991 | 0.14442483 | 0.19482367006 | 0.2686205 | 1.9913752 |
| 3721947.02 | 3617465.3 | 0.86609579 | -0.2074015 | 0.2794489 | 0.3260237 | 1.44401825 | 0.53008897 | 0.07308266514 | 0.1546633 | 0.84936068 |
| 84.2654206 | 73.462658 | 0.23918671 | -2.0637909 | 0.00166518 | 0.00378827 | 2.20390856 | 1.14006437 | 0.00301209429 | 0.0144758 | 2.76435214 |
| 226136.41 | 254552.176 | 1.27642777 | 0.35211191 | 0.08074998 | 0.11305 | 0.80960293 | -0.3047136 | 0.13636765174 | 0.2298047 | 2.87096285 |
| 338034.5 | 345679.659 | 0.54740891 | -0.8693092 | 0.00083785 | 0.00227307 | 1.22219275 | 0.28947183 | 0.09104160323 | 0.180104 | 2.18546424 |
| 14730066.1 | 21087233.7 | 0.94551235 | -0.0808318 | 0.72494402 | 0.7670919 | 1.02997939 | 0.04261547 | 0.34110812755 | 0.3922344 | 0.69547684 |
| 899597.093 | 907172.34 | 0.30305307 | -1.7223576 | 0.00013679 | 0.0005412 | 1.41308563 | 0.49884889 | 0.00715192245 | 0.02711771 | 1.78114253 |

| | | | | | | | | | | |
|---|---|---|---|---|---|---|---|---|---|---|
| 13595474.3 | 17077135.4 | 0.4046845 | -1.3051305 | 0.00012778 | 0.00052854 | 1.23345462 | 0.30270464 | 0.01263754108 | 0.03965573 | 2.38176801 |
| 148038.956 | 268767.17 | 1.09922796 | 0.13649061 | 0.74015611 | 0.7741863 | 1.32699964 | 0.40816798 | 0.16921473689 | 0.2566424 | 1.48880228 |
| 760579.433 | 772561.074 | 0.3846232 | -1.3784823 | 0.00022527 | 0.00075923 | 1.71420238 | 0.77753744 | 0.00050704447 | 0.00434538 | 2.63100178 |
| 508140.25 | 443979.573 | 0.60847956 | -0.7167193 | 0.00729595 | 0.0138319 | 1.1017924 | 0.13985242 | 0.32763804865 | 0.3822444 | 2.12438881 |
| 1455645.68 | 1724998.79 | 0.46011813 | -1.1199238 | 0.01530696 | 0.02579506 | 1.44171354 | 0.52778454 | 0.00473393442 | 0.01936563 | 1.82902563 |
| 286440.118 | 221832.652 | 0.0947026 | -3.4004521 | 0.00092422 | 0.00227307 | 0.70635999 | -0.5015245 | 0.03462559316 | 0.08079305 | 0.63542442 |
| 16545.9368 | 9862.33035 | 0.09444783 | -3.4043385 | 8.6069E-07 | 2.46E-05 | 0.48052344 | -1.0573213 | 0.00268482688 | 0.0144758 | 0.47033786 |
| 45903.238 | 44938.8348 | 0.13338074 | -2.9063777 | 4.113E-06 | 6.02E-05 | 1.81466414 | 0.85970256 | 0.02248883054 | 0.06019069 | 1.32871061 |
| 226.925489 | 187.913274 | 0.45807815 | -1.1263344 | 0.00085733 | 0.00227307 | 1.50712631 | 0.59180033 | 0.11813994662 | 0.2067449 | 1.43447952 |
| 115.485439 | 143.03161 | 0.90620666 | -0.142088 | 0.3589985 | 0.4083608 | 0.94825588 | -0.0766517 | 0.30783889407 | 0.3685966 | 0.71277469 |
| 21 | 22.24 | 0.98827107 | -0.0170213 | 0.91579077 | 0.9363703 | 1.02241758 | 0.03198455 | 0.41927585149 | 0.4401693 | 0.97758242 |
| 321.647596 | 334.554426 | 0.40667866 | -1.2980388 | 2.1819E-05 | 0.00016546 | 1.38569015 | 0.47060469 | 0.00030381239 | 0.00345587 | 2.58461626 |
| 157433104 | 171021773 | 0.37691737 | -1.4076798 | 1.8225E-06 | 3.32E-05 | 1.26376563 | 0.33772893 | 0.00038030554 | 0.00384531 | 1.26234923 |
| 2828211.05 | 2898687.13 | 0.41246964 | -1.2776402 | 2.5242E-05 | 0.00017669 | 0.90023546 | -0.1516257 | 0.11478172333 | 0.2048066 | 2.8896303 |
| 658.003657 | 731.47528 | 0.42083388 | -1.2486772 | 1.4589E-05 | 0.00013276 | 1.24914749 | 0.32094383 | 0.00368637120 | 0.01597428 | 1.78843802 |
| 1161.81582 | 1475.49057 | 0.51465855 | -0.9583125 | 0.00047239 | 0.0014329 | 0.80442035 | -0.3139785 | 0.00887603317 | 0.02991552 | 1.05836641 |
| 79.6812692 | 100.515369 | 0.85813062 | -0.2207308 | 0.10878776 | 0.1434737 | 0.91059557 | -0.1351177 | 0.14437728675 | 0.2388788 | 0.78236511 |
| 273.980526 | 349.620279 | 0.84097096 | -0.2498721 | 0.05909049 | 0.0867296 | 0.98489222 | -0.0219622 | 0.43123648253 | 0.4459377 | 0.79118011 |
| 455.92458 | 919.779846 | 0.68393639 | -0.5480659 | 0.01352923 | 0.02322943 | 1.09072087 | 0.12528195 | 0.19480641966 | 0.2686205 | 0.61189296 |
| 238.99037 | 283.674944 | 0.89037477 | -0.1675154 | 0.3030223 | 0.349051 | 0.99368024 | -0.0091464 | 0.47580402640 | 0.4810907 | 0.67713001 |
| 97.1294185 | 110.805123 | 0.80659766 | -0.3100789 | 0.06357131 | 0.09182522 | 0.9302849 | -0.1042555 | 0.17976047933 | 0.259654 | 0.90293313 |
| 178.791874 | 215.626633 | 0.86620459 | -0.2072203 | 0.20381305 | 0.2540683 | 0.90180231 | -0.1491169 | 0.16689483985 | 0.2566424 | 0.80938612 |
| 171.800188 | 167.964414 | 0.38507262 | -1.3767976 | 5.5469E-06 | 6.31E-05 | 1.62103007 | 0.69691085 | 0.00001194501 | 0.00027175 | 2.26868202 |
| 265.50659 | 314.441572 | 0.87669093 | -0.1898598 | 0.25875289 | 0.3057989 | 0.88568232 | -0.1751388 | 0.15275074133 | 0.2396606 | 1.0263709 |
| 616.562261 | 798.977989 | 0.84060555 | -0.2504991 | 0.14751267 | 0.1867274 | 0.91095374 | -0.1345503 | 0.14972932464 | 0.2396606 | 1.16487975 |
| 33.1 | 38.8799515 | 0.80087968 | -0.3203426 | 0.0356585 | 0.05408206 | 0.91380883 | -0.1300357 | 0.17708197046 | 0.259654 | 0.8045515 |
| 125.884285 | 149.585908 | 0.84143504 | -0.2490762 | 0.0994697 | 0.1352266 | 0.79834254 | -0.3249202 | 0.00489461005 | 0.01936563 | 0.80866572 |
| 308.234426 | 339.412776 | 0.78523936 | -0.3487956 | 0.14774039 | 0.1867274 | 1.04868414 | 0.06858021 | 0.35026134510 | 0.3922344 | 0.74101981 |
| 43.5303316 | 50.071214 | 0.51951521 | -0.9447621 | 4.6296E-06 | 6.02E-05 | 1.10290698 | 0.14131111 | 0.02628798680 | 0.06834877 | 1.45555855 |
| 2764971.45 | 3092443.71 | 0.58908663 | -0.7634483 | 0.00025092 | 0.00081548 | 1.19891724 | 0.26173207 | 0.02209831223 | 0.06019069 | 0.87769776 |
| 86757039.6 | 93056896.3 | 0.56782977 | -0.8164696 | 5.6903E-05 | 0.0003046 | 1.55948652 | 0.64107108 | 0.01540575662 | 0.0467308 | 1.25540492 |
| 3318595.58 | 3576134.34 | 0.66020991 | -0.5990033 | 0.00249228 | 0.0051545 | 1.28868287 | 0.36589727 | 0.00016998223 | 0.00220977 | 1.59358112 |
| 119266.476 | 127597.277 | 0.80737423 | -0.3086906 | 0.02614113 | 0.04031938 | 1.19747594 | 0.25999667 | 0.08762984933 | 0.177207 | 1.51452745 |

| | | | | | | | | | | |
|---|---|---|---|---|---|---|---|---|---|---|
| 213052.646 | 216180.981 | 0.99395624 | -0.0087458 | 0.96724446 | 0.9779916 | 1.04853121 | 0.06836981 | 0.35299711959 | 0.3922344 | 1.73809624 |
| 106.515356 | 119.674538 | 0.86534584 | -0.2086513 | 0.21891035 | 0.2656112 | 0.93167442 | -0.1021022 | 0.28193044158 | 0.3466982 | 0.6781522 |
| 8570664.47 | 7275302.11 | 0.62456905 | -0.679067 | 0.01222452 | 0.02139291 | 1.16270804 | 0.21748888 | 0.15065954052 | 0.2396606 | 1.33585484 |
| 65.8795264 | 67.4705226 | 0.65571926 | -0.6088498 | 0.00670194 | 0.01297611 | 1.4173851 | 0.50323179 | 0.07783057196 | 0.1609678 | 1.81616481 |
| 823199.526 | 954082.422 | 0.39392271 | -1.3440155 | 0.00020948 | 0.00074082 | 1.03034523 | 0.04312782 | 0.41683959048 | 0.4401693 | 1.16268692 |
| 2567956.69 | 2235575.86 | 0.61197627 | -0.7084524 | 0.01924018 | 0.0312653 | 0.9448058 | -0.0819103 | 0.38070006249 | 0.4129139 | 0.84197072 |
| 124751.491 | 112244.326 | 0.26811222 | -1.8990911 | 4.1703E-05 | 0.00025998 | 1.66301728 | 0.73380316 | 0.00111228450 | 0.00843482 | 1.48248391 |
| 41832435.8 | 43796216.6 | 0.41342261 | -1.2743108 | 0.00021166 | 0.00074082 | 0.59099219 | -0.758789 | 0.01875920437 | 0.05506734 | 0.93023923 |
| 2419422.3 | 2830960.46 | 0.27581764 | -1.8582134 | 6.5832E-06 | 6.66E-05 | 1.31040626 | 0.39001415 | 0.00757084562 | 0.02755788 | 1.09331252 |
| 418678.28 | 455383.857 | 0.38019485 | -1.3951891 | 9.1714E-05 | 0.0004582 | 1.69106383 | 0.75793112 | 0.00199366423 | 0.01295882 | 1.27764535 |
| 174712121 | 191807866 | 0.8143437 | -0.2962903 | 0.0037317 | 0.00754633 | 1.67873239 | 0.74737227 | 0.00000012050 | 1.10E-05 | 1.71489976 |
| 17968799.6 | 21642719.4 | 0.84840946 | -0.2371674 | 0.2180774 | 0.2656112 | 0.95320262 | -0.0691452 | 0.26806727248 | 0.3341661 | 0.76286477 |
| 5496.23107 | 3613.14409 | 0.95183813 | -0.0712119 | 0.81517651 | 0.8429666 | 0.99991471 | -0.0001231 | 0.49985280352 | 0.4998528 | 0.66307236 |
| 42.8347052 | 43.2595884 | 2.58158345 | 1.36825623 | 4.6207E-05 | 0.0002628 | 0.40815057 | -1.2928266 | 0.00003270790 | 0.00059528 | 0.30714176 |
| 31776474.5 | 30832549.9 | 0.6755993 | -0.5657603 | 0.00200248 | 0.00444453 | 1.2946028 | 0.37250953 | 0.00052526556 | 0.00434538 | 1.80842748 |
| 557858.044 | 526159.645 | 0.8317177 | -0.2658342 | 0.1073884 | 0.1434737 | 0.93093167 | -0.1032528 | 0.32038243933 | 0.3786338 | 0.70584276 |
| 1876307.46 | 1902964.66 | 0.65001721 | -0.6214502 | 0.02246228 | 0.03524254 | 1.02015416 | 0.02878717 | 0.42082117355 | 0.4401693 | 2.12078885 |
| 1572850.58 | 1521684.08 | 0.92522907 | -0.1121175 | 0.46325104 | 0.5079017 | 0.708005 | -0.4981685 | 0.00276196570 | 0.0144758 | 0.35853495 |
| 3136418.28 | 3054258.46 | 0.65702312 | -0.605984 | 0.02032349 | 0.03244627 | 1.11260898 | 0.15394666 | 0.11466497855 | 0.2048066 | 1.94944796 |
| 35765764.7 | 35878772.4 | 0.87632343 | -0.1904647 | 0.23917519 | 0.2863808 | 0.75467722 | -0.4060684 | 0.00229953461 | 0.01395051 | 0.89092202 |
| 2614022.46 | 2528533.54 | 0.40649837 | -1.2986785 | 0.00010623 | 0.00048021 | 1.30085909 | 0.3794647 | 0.03393250470 | 0.08079305 | 2.71384829 |
| 184646.158 | 165175.319 | 1.14491884 | 0.19524534 | 0.54694308 | 0.5925217 | 0.54068481 | -0.8871403 | 0.03052626132 | 0.07588461 | 0.83769661 |
| 495988.81 | 563399.858 | 0.96038121 | -0.0583209 | 0.67974671 | 0.7277288 | 0.61419931 | -0.7032212 | 0.00340538882 | 0.01549452 | 0.92655763 |
| 3842936.99 | 3180080.56 | 0.66843502 | -0.5811408 | 0.13458631 | 0.1749622 | 0.96909968 | -0.045283 | 0.44778433636 | 0.4578469 | 2.41339958 |
| 49066.2099 | 48149.3829 | 0.01010557 | -6.6287054 | 9.5669E-05 | 0.0004582 | 1.42368074 | 0.50962565 | 0.01154105778 | 0.03750844 | 1.1943157 |
| 7181.78534 | 5896.74226 | 0.10511727 | -3.2499284 | 0.01637637 | 0.02709545 | 1.17056521 | 0.22720531 | 0.30023859663 | 0.3642895 | 0.49542652 |

| log2FC_KO2_ | pcal_KO2_vs_ | padj_KO2_vs_LTED |
|---|---|---|
| -0.4038714 | 0.10834785 | 0.1493887 |
| -0.0081935 | 0.9166788 | 0.9166788 |
| -0.0354593 | 0.90050753 | 0.9105132 |
| 0.75929721 | 0.01755367 | 0.03549742 |
| 0.13504566 | 0.17518759 | 0.2214177 |
| 0.21213398 | 0.34847503 | 0.382063 |
| -0.7355578 | 0.00159193 | 0.00467308 |
| 0.49992903 | 0.05810553 | 0.08528392 |
| 1.51205571 | 0.00033851 | 0.00128351 |
| 1.67571149 | 3.0406E-05 | 0.00026059 |
| 1.43567057 | 0.00073225 | 0.00256287 |
| -0.1169976 | 0.31221915 | 0.3464871 |
| -0.4559121 | 0.00025085 | 0.00099251 |
| 1.3688074 | 0.00016742 | 0.00072548 |
| 1.23237414 | 4.7071E-05 | 0.0003295 |
| 0.14236301 | 0.29064748 | 0.3306115 |
| -0.2460589 | 0.05341141 | 0.07967931 |
| -0.1509123 | 0.1202195 | 0.160882 |
| -0.6453382 | 0.00013602 | 0.0006189 |
| 2.66485065 | 1.0009E-08 | 9.11E-07 |
| 0.0931094 | 0.24901548 | 0.2905181 |
| 0.47915111 | 0.0346852 | 0.05636345 |
| 1.08702983 | 0.00022682 | 0.0009382 |
| -0.6323498 | 0.0516405 | 0.07832143 |
| -0.8790005 | 0.01300832 | 0.02752924 |
| 0.99376507 | 9.0988E-05 | 0.00046498 |
| -0.2355508 | 0.307163 | 0.3450844 |
| 1.46694141 | 0.00012868 | 0.00061632 |
| 1.52153466 | 3.9999E-05 | 0.00030332 |
| 1.12793977 | 0.00142842 | 0.0045629 |
| -0.5239256 | 0.02533433 | 0.04520439 |
| 0.83280297 | 7.4489E-06 | 8.47E-05 |

| | | |
|---|---|---|
| 1.2520329 | 7.29E-05 | 0.00041462 |
| 0.57415217 | 0.22650014 | 0.2748202 |
| 1.39561222 | 9.1973E-05 | 0.00046498 |
| 1.08704783 | 0.00295981 | 0.00727953 |
| 0.87107529 | 0.00462004 | 0.01106377 |
| -0.6542075 | 0.02130106 | 0.04128889 |
| -1.0882306 | 0.06825423 | 0.09858944 |
| 0.41002692 | 0.02223248 | 0.04128889 |
| 0.52052737 | 0.0307297 | 0.05084369 |
| -0.488482 | 0.0087802 | 0.01997496 |
| -0.0327098 | 0.7732467 | 0.7996074 |
| 1.3699501 | 3.1161E-06 | 5.67E-05 |
| 0.33611109 | 0.03926093 | 0.06267974 |
| 1.53088493 | 6.7105E-06 | 8.47E-05 |
| 0.83870012 | 3.1499E-05 | 0.00026059 |
| 0.08183918 | 0.56567749 | 0.5985657 |
| -0.3540861 | 0.02201499 | 0.04128889 |
| -0.3379219 | 0.02149136 | 0.04128889 |
| -0.7086488 | 0.01225876 | 0.02720848 |
| -0.5624952 | 0.00202787 | 0.005592 |
| -0.1473089 | 0.23325444 | 0.2792915 |
| -0.3051 | 0.14186875 | 0.1844294 |
| 1.18185442 | 6.5577E-06 | 8.47E-05 |
| 0.03755218 | 0.78321285 | 0.8008131 |
| 0.22018103 | 0.1826991 | 0.2277482 |
| -0.3137433 | 0.02300093 | 0.0418617 |
| -0.3063846 | 0.01291268 | 0.02752924 |
| -0.432416 | 0.04665313 | 0.07195652 |
| 0.54157287 | 0.00183793 | 0.0052266 |
| -0.1882039 | 0.111525 | 0.1514743 |
| 0.32815277 | 0.01749755 | 0.03549742 |
| 0.67227246 | 5.5774E-05 | 0.00033836 |
| 0.59886773 | 0.00602213 | 0.01405164 |

| | | |
|---|---|---|
| 0.79750797 | 0.00056351 | 0.00205116 |
| -0.560319 | 0.0028938 | 0.00727953 |
| 0.41776324 | 0.02890712 | 0.04871385 |
| 0.86089513 | 0.00102276 | 0.00344708 |
| 0.21746267 | 0.21784113 | 0.2678857 |
| -0.248158 | 0.14835035 | 0.1901392 |
| 0.56801644 | 0.00270449 | 0.00713524 |
| -0.1043263 | 0.62088356 | 0.6494299 |
| 0.12870584 | 0.26615115 | 0.3065792 |
| 0.35348743 | 0.2445994 | 0.289072 |
| 0.77812425 | 0.00145411 | 0.0045629 |
| -0.3905008 | 0.04349818 | 0.06824714 |
| -0.5927618 | 0.1401048 | 0.1844294 |
| -1.7030234 | 1.024E-05 | 0.00010353 |
| 0.85473575 | 5.1368E-05 | 0.00033389 |
| -0.5025813 | 0.02625027 | 0.04592229 |
| 1.08460099 | 0.00155765 | 0.00467308 |
| -1.4798143 | 9.8606E-07 | 2.24E-05 |
| 0.96306564 | 6.4919E-07 | 2.11E-05 |
| -0.1666289 | 0.02674595 | 0.04592229 |
| 1.44034007 | 6.9702E-07 | 2.11E-05 |
| -0.2555003 | 0.49855289 | 0.5379414 |
| -0.1100474 | 0.50247274 | 0.5379414 |
| 1.2710668 | 0.00274432 | 0.00713524 |
| 0.25618424 | 0.10719702 | 0.1493887 |
| -1.013257 | 0.10131565 | 0.1440582 |
